# Exhaustive identification of conserved upstream open reading frames with potential translational regulatory functions from animal genomes

**DOI:** 10.1101/672840

**Authors:** Hiro Takahashi, Shido Miyaki, Hitoshi Onouchi, Taichiro Motomura, Nobuo Idesako, Anna Takahashi, Masataka Murase, Shuichi Fukuyoshi, Toshinori Endo, Kenji Satou, Satoshi Naito, Motoyuki Itoh

**Author notes:** Correspondence., Correspondence may also be addressed to Motoyuki Itoh. Joint first authors (Hiro Takahashi, Shido Miyaki, Hitoshi Onouchi).

## Abstract

Upstream open reading frames (uORFs) are present in the 5’-untranslated regions of many eukaryotic mRNAs, and some peptides encoded by these regions play important regulatory roles in controlling main ORF (mORF) translation. We previously developed a novel pipeline, ESUCA, to comprehensively identify plant uORFs encoding functional peptides, based on genome-wide identification of uORFs with conserved peptide sequences (CPuORFs). Here, we applied ESUCA to diverse animal genomes, because animal CPuORFs have been identified only by comparing uORF sequences between a limited number of species, and how many previously identified CPuORFs encode regulatory peptides is unclear. By using ESUCA, 1,517 (1,373 novel and 144 known) CPuORFs were extracted from four evolutionarily divergent animal genomes. We examined the effects of 17 human CPuORFs on mORF translation using transient expression assays. Through these analyses, we identified seven novel regulatory CPuORFs that repressed mORF translation in a sequence-dependent manner, including one conserved only among Eutheria. We discovered a much higher number of animal CPuORFs than previously identified. Since most human CPuORFs identified in this study are conserved across a wide range of Eutheria or a wider taxonomic range, many CPuORFs encoding regulatory peptides are expected to be found in the identified CPuORFs.

## Introduction

The human genome contains many regions encoding potential functional small peptides outside the canonical protein-coding regions ^1^. Some upstream open reading frames (uORFs), which are located in the 5’-untranslated regions (5’-UTRs) of mRNAs, have been shown to encode such functional small peptides ^2–5^. uORFs are cis-acting regulatory elements that control the translation of protein-coding main ORFs (mORFs) in various ways ^6,7^. In eukaryotes, 43S pre-initiation complexes (PICs) scan for a start codon along an mRNA from the 5’ end. Therefore, PICs can recognize the start codon of a uORF and translate the uORF before reaching the downstream mORF. In many cases, after translating a uORF, ribosomes dissociate from the mRNA or small ribosomal subunits remain bound to the mRNA and resume scanning. When ribosomes dissociate from the mRNA after uORF translation, ribosomes that have translated the uORF do not translate the downstream mORF. Therefore, if the translation initiation efficiency of the uORF is high, the uORF exerts a substantial repressive effect on mORF translation ^6,7^. When a small ribosomal subunit resumes scanning after uORF translation, the ribosomes can reinitiate translation at a downstream AUG codon. However, the reinitiation efficiency depends on the time needed for the uORF translation and the distance between the uORF stop codon and the downstream start codon ^8–11^. The intercistronic distance required for efficient reinitiation depends on cellular availability of the ternary complex that comprises eukaryotic initiation factor 2 (eIF2), GTP, and Met-tRNA ^Met^, and the level of the available ternary complex is reduced under starvation or stress conditions ^12^. These properties are utilized for the translational regulation of yeast *GCN4* and mammalian *ATF4* and *ATF5* mRNAs ^13–18^. In these mRNAs, there is an inhibitory uORF downstream of the uORF that allows reinitiation. Under normal conditions, reinitiation preferentially occurs at the start codon of the inhibitory uORF, and therefore, mORF translation is repressed. In contrast, under starvation or stress conditions, reinitiation is delayed due to the reduced availability of the ternary complex, and therefore, ribosomes more frequently bypass the inhibitory uORF and reinitiate translation at the start codon of the mORF, resulting in enhanced mORF translation. Apart from mORF translation control, uORFs can affect mRNA stability through the nonsense-mediated RNA decay pathway ^19^. While the effects of most uORFs on the expression of the mORF-encoded proteins are independent of the uORF-encoded sequences, certain uORFs repress mORF translation in a peptide sequence-dependent manner. Most of these uORFs encode regulatory peptides that cause ribosome stalling by interacting with components of the ribosomal exit tunnel during uORF translation ^4^. Ribosome stalling on a uORF results in translational repression of the downstream mORF because the stalled ribosomes block the scanning of subsequently loaded PICs and prevent them from reaching the start codon of the mORF ^20^. In some genes, uORF-encoded peptides are involved in translational regulation in response to metabolites or environmental stresses, whereby the uORF translation initiation efficiency or the efficiency of ribosome stalling is regulated in a condition-dependent manner ^4,21^. In the sequence-dependent regulatory uORF of the mouse antizyme inhibitor (*AZIN1*) gene, which begins with a non-canonical start codon ^22^, polyamine induces ribosome stalling, and the stalled ribosome causes ribosome queuing by blocking the scanning of PICs ^23^. This ribosome queuing promotes translation initiation at the non-canonical start codon of the uORF by positioning PICs near the start codon, thereby enhancing the repressive effect of the uORF on mORF translation. Apart from uORFs encoding regulatory peptides, some uORFs have been reported to code for proteins with functions independent of the control of the downstream mORF ^24–26^.

To comprehensively identify uORFs encoding functional peptides or proteins, genome-wide searches for uORFs with conserved peptide sequences (CPuORFs) have been conducted using comparative genomic approaches in plants ^27–32^. To date, 157 CPuORF families have been identified by comparing 5’-UTR sequences among plant species. Of these, 101 families were identified in our previous studies by applying our original methods, BAIUCAS ^29^ and ESUCA (an advanced version of BAIUCAS) ^32^ to the genomes of *Arabidopsis*, rice, tomato, poplar, and grape.

ESUCA has many unique functions ^32^, such as efficient comparison of uORF sequences among an unlimited number of species using BLAST, automatic determination of taxonomic ranges of CPuORF sequence conservation, systematic calculation of *K*_a_/*K*_s_ ratios of CPuORF sequences, and wide compatibility with any eukaryotic genome whose sequence database is registered in ENSEMBL ^33^. By comparing uORF sequences from certain species and those from many other species whose transcript sequence databases are available, ESUCA enables more comprehensive identification of CPuORFs conserved in various taxonomic ranges than conventional comparative genomic approaches, in which uORF sequences are compared among limited numbers of selected species. In addition, to distinguish between “spurious” CPuORFs conserved because they encode parts of mORF-encoded proteins and “true” CPuORFs conserved because of the functional constraints of their encoded small peptides, ESUCA assesses whether a transcript containing a fusion of a uORF and an mORF is a major or minor form among homologous transcripts ^32^. By using these functions, ESUCA is able to efficiently identify CPuORFs likely to encode functional small peptides. In fact, our recent study demonstrated that poplar CPuORFs encoding regulatory peptides were efficiently identified using ESUCA by selecting ones conserved across diverse eudicots ^32^.

Several studies on genome-wide identification of animal CPuORFs have been reported. By comparing uORF sequences between human and mouse, 204 and 198 CPuORFs have been identified in human and mouse, respectively ^34^. In addition, by comparing uORF sequences among several species in dipteran, 44 CPuORFs have been identified in fruit fly ^35^. More recently, among translatable uORFs identified by ribosome profiling studies, 118, 80, 13, 50, and 37 CPuORFs in human, mouse, zebrafish, fruit fly, and nematode, respectively, have been identified by Mackowiak *et al*. ^36^, and 97 CPuORFs in human have been identified by Samandi *et al*. ^26^. In these previous studies, uORF sequences were compared between a limited number of species. Therefore, further comprehensive identification of animal CPuORFs was expected by applying the approach using ESUCA to animal genomes. In addition, the relationships between the taxonomic ranges of CPuORF conservation and the likelihood of having a regulatory function have not been studied in animals.

Accordingly, in this study, we applied ESUCA to the genomes of fruit fly, zebrafish, chicken, and human to exhaustively identify animal CPuORFs and to determine the taxonomic range of their sequence conservation. Using ESUCA, we identified 1,517 animal (1,373 novel and 144 known) CPuORFs belonging to 1,430 CPuORF families. Using transient expression assays, we examined the effects of 17 CPuORFs conserved in various taxonomic ranges on mORF translation. Through this analysis, we identified seven novel regulatory CPuORFs that repress mORF translation in a sequence-dependent manner.

## Results

### Genome-wide search for animal CPuORFs using ESUCA

Prior to ESUCA application (Supplementary Fig. S1a and S1b), we counted the number of protein-coding genes in four species, i.e., fruit fly, zebrafish, chicken, and human. The genes whose mORF-encoded amino acid sequences were available in ENSEMBL (http://www.ensembl.org) were defined as protein-encoding genes in the present study. As shown in Table 1, 13,938, 25,206, 14,697, and 19,956 genes were extracted for fruit fly, zebrafish, chicken, and human, respectively. In step 1 of ESUCA, we extracted uORF sequences from the 5’-UTR sequence of these genes, using the transcript sequence datasets described in the Methods section. In these datasets, different transcript IDs are assigned to each splice variant from the same gene. To extract sequences of uORFs and their downstream mORFs from all splice variants, we extracted uORF and mORF sequences from each of the transcripts with different transcript IDs. The uORFs were extracted by searching the 5’-UTR sequence of each transcript for an ATG codon and its nearest downstream in-frame stop codon (Supplementary Fig. S2a). As shown in Table 1, 17,035, 39,616, 8,929, and 44,085 uORFs were extracted from 7,066, 14,453, 3,535, and 12,321 genes of fruit fly, zebrafish, chicken, and human genomes, respectively. In this analysis, when multiple uORFs from splice variants of a gene shared the same stop or start codon, they were counted as one, but all uORFs in splice variants were retained for further analyses. In step 2, we calculated the uORF–mORF fusion ratio for each of the extracted uORFs. To assess whether transcripts bearing a uORF–mORF fusion are minor or major forms among homologous transcripts, the ratio of the NCBI reference sequence (RefSeq) RNAs with a uORF–mORF fusion to all RefSeq RNAs with both sequences similar to the uORF and its downstream mORF was calculated as the uORF–mORF fusion ratio (Supplementary Fig. S2b). We discarded uORFs with uORF–mORF fusion ratios equal to or greater than 0.3 (Supplementary Table S1). As shown in Table 1, the numbers of uORFs were dramatically reduced after this step, suggesting that this step effectively excluded “spurious” CPuORFs that were conserved because they encode parts of mORF-encoded proteins. In step 3.1, we performed homology searches of the uORF amino acid sequences, using tBLASTn with an *E*-value cutoff of 2,000 (uORF-tBLASTn analysis). In this search, the amino acid sequence of each uORF was queried against an animal transcript sequence database that contained contigs of assembled expressed sequence tags (ESTs) and transcriptome shotgun assemblies (TSAs), singleton EST/TSA sequences, and RefSeq RNAs (See the Methods for details). The uORFs with tBLASTn hits from other species were selected in this step. In step 3.2, an ORF containing the amino acid sequence similar to the original uORF sequence was extracted from each of the uORF-tBLASTn hits as putative uORFs (Supplementary Fig. S3a). In step 4.1, to confirm whether the suORF-tBLASTn hits were derived from homologs of the original uORF-containing gene, the downstream sequences of putative uORFs in the uORF-tBLASTn hits were subjected to another tBLASTn analysis (Supplementary Fig. S3b and S3c). In this analysis, the amino acid sequence of the mORF in each original uORF-containing transcript was used as a query, and the uORF-tBLASTn hits matching the mORF with an *E*-value of less than 10^−1^ were extracted (mORF-tBLASTn analysis) (Supplementary Fig. S3c). If a uORF-tBLASTn hit contained a partial or intact ORF sequence similar to the original mORF amino acid sequence downstream of the putative uORF, it was considered to be derived from a homolog of the original uORF-containing gene. In step 4.2, potential contaminant sequences derived from contaminating organisms, such as parasites and infectious microorganisms, were excluded from the mORF-tBLASTn hits. In step 4.3, we selected uORFs conserved in mORF homologs from at least two orders other than the order of the original uORF; for example, in the search for human CPuORFs, we selected uORFs conserved in mORF homologs from at least two orders other than Primates. When uORFs with identical sequences were extracted from different splice variants of the same gene, we selected the one with the lowest median *E*-value in uORF-tBLASTn analysis. In step 5, we calculated *K*_a_/*K*_s_ ratios of the uORFs to assess whether these uORF sequences were conserved at the nucleotide or amino acid level. A *K*_a_/*K*_s_ ratio close to 1 indicates neutral evolution, whereas a *K*_a_/*K*_s_ ratio close to 0 suggests that purifying selection acted on the amino acid sequences ^32^. For each of the uORFs extracted in step 4.3, one representative mORF-tBLASTn hit was selected from each order in which mORF-tBLASTn hits were identified, and the putative uORFs in the selected mORF-tBLASTn hits and the original uORF sequence were used for pairwise calculations of *K*_a_/*K*_s_ ratios. The uORFs with *K*_a_/*K*_s_ ratios less than 0.5 showing significant differences from those of negative controls (*q* < 0.05) were selected as candidate CPuORFs (Supplementary Table S1). In step 6, we determined the taxonomic range of sequence conservation of the candidate CPuORFs. In this step, the representative mORF-tBLASTn hits selected in step 5 were classified into the 19 taxonomic categories shown in Fig. 1a. On the basis of the presence of the mORF-tBLASTn hits in each taxonomic category, the taxonomic range of sequence conservation was determined for each candidate CPuORF (Supplementary Table S2). After the final step of ESUCA, 49, 192, 261, and 1,495 candidate CPuORFs were extracted from fruit fly, zebrafish, chicken, and human, respectively (Table 1). To validate the sequence conservation of the candidate CPuORFs, we generated multiple amino acid sequence alignment for each candidate CPuORF, using the original uORF sequence and the representative putative uORF sequences used for calculating the *K*_a_/*K*_s_ ratio (Supplementary Figs. S4 and S5). If the amino acid sequence of a uORF is evolutionarily conserved because of functional constraints of the uORF-encoded peptide, the amino acid sequence in the functionally important region of the peptide is expected to be conserved among the uORF and its orthologous uORFs. Therefore, we manually checked whether the amino acid sequences in the same region are conserved among the uORF and putative uORF sequences in the alignment of each candidate CPuORF. Then, we removed sequences that do not share the consensus amino acid sequence in the conserved region. When this change resulted in the number of orders with the mORF-tBLASTn hits becoming less than two, the candidate CPuORFs were discarded. In analyses of the genomes of the four species, 66 candidate CPuORFs were discarded for this reason. In addition, when multiple original uORFs in splice variants from the same gene partially shared amino acid sequences, the one with the longest conserved region was manually selected on the basis of the uORF amino acid sequence alignments, and the others were discarded. In the case where the length of the conserved regions is the same among splice variants, the one with the lowest *K*_a_/*K*_s_ ratio was selected. When multiple original uORFs in splice variants from the same gene overlapped each other in different reading frames, the one with the lowest *K*_a_/*K*_s_ ratio was selected, and the others were discarded. After the manual validation, 1,517 animal CPuORFs (37 for fruit fly, 156 for zebrafish, 230 for chicken, and 1,094 for human) were identified as CPuORFs (Fig. 1). Of these, 1,373 CPuORFs have not been reported and are therefore novel CPuORFs. The amino acid sequence alignments and detailed information of the identified CPuORFs are shown in Supplementary Figs. S4, S5 and Table S1. It should be noted that the sequence alignments shown in Supplementary Figs. S4 and S5 do not contain the putative uORF sequences removed by manual validation. In the cases where one or more putative uORF sequences homologous to a CPuORF were removed by the manual validation, we determined the taxonomic range of sequence conservation of the CPuORF again after excluding the mORF-tBLASTn hits corresponding to the removed putative uORF sequences. Figure 1b shows the numbers of the CPuORFs conserved in each of the 19 taxonomic categories. The results shown in this figure indicate that CPuORFs conserved in various taxonomic ranges were identified by ESUCA analyses of the fruit fly, zebrafish, chicken, and human genomes. The identified CPuORF-containing genes were classified into 1,363 ortholog groups on the basis of similarities of mORF-encoded amino acid sequences using OrthoFinder ^37^. CPuORFs with similar amino acid sequences from the same ortholog groups were categorized as the same CPuORF families (homology groups [HGs]; see the Methods for details). The identified 1,517 CPuORFs were classified into 1,430 HGs. We assigned HG numbers to 1,430 HGs so that CPuORF families conserved across wider taxonomic ranges and in higher numbers of orders could have smaller HG numbers. When multiple CPuORF families were identified in the same ortholog groups, the same HG number with a different subnumber was assigned to each of the families (e.g., HG0004.1 and HG0004.2; Supplementary Table S1). It should be noted that the amino acid sequence alignments for each of the identified CPuORFs were separately generated and are individually shown in Supplementary Figs. S4 and S5, even when multiple homologous CPuORFs belonging to the same HG were identified.

**Table 1.**
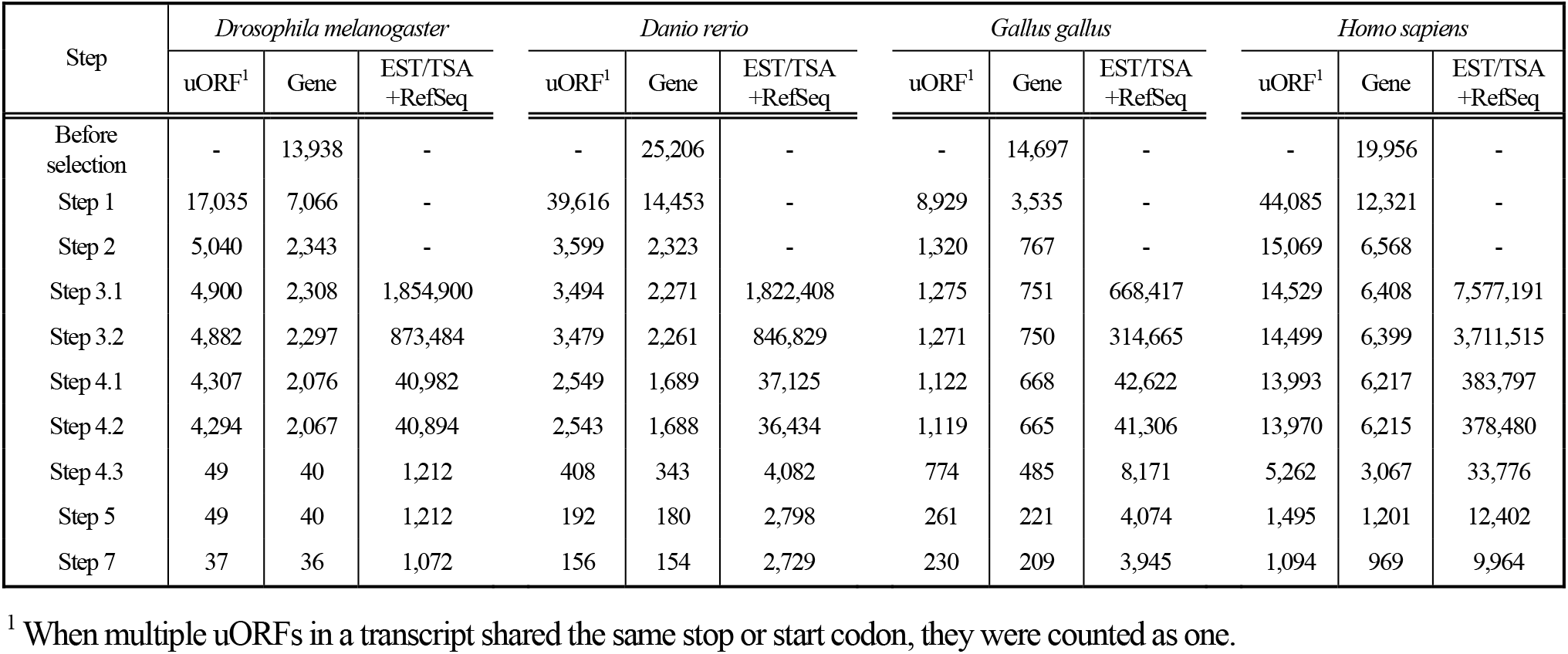
Numbers of uORFs, protein-coding genes, and assembled EST/TSA and RefSeq sequences extracted at each step of ESUCA.

**Figure 1.**
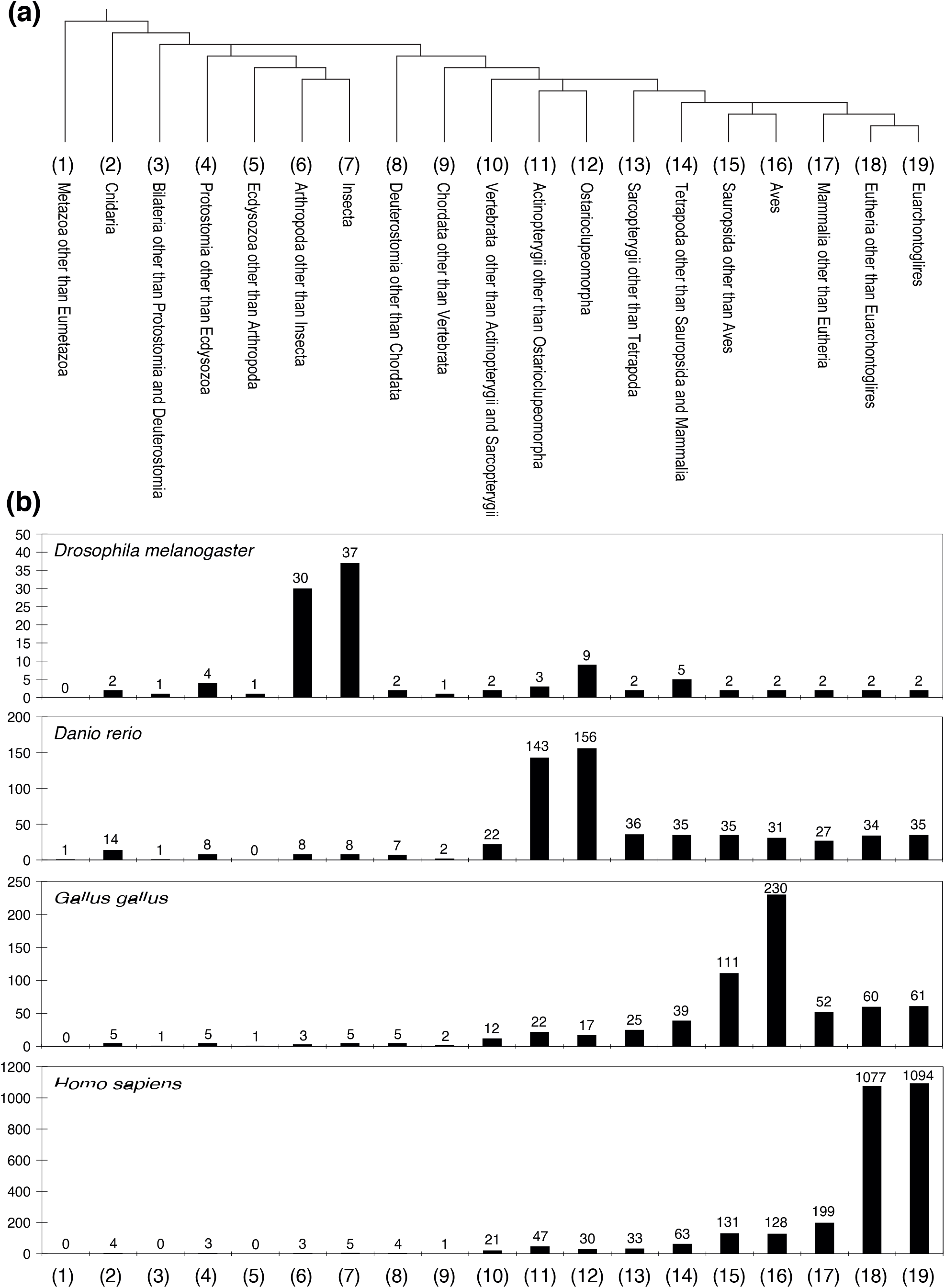
Numbers of CPuORFs extracted by ESUCA in each taxonomic category. (**a**) Cladogram showing the relationship among the 19 taxonomic categories defined in this study. Fruit fly, zebrafish, chicken, and human belong to Diptera, Cypriniformes, Galliformes, and Primates, respectively. Diptera, Cypriniformes, Galliformes, and Primates belong to Insecta, Ostarioclupeomorpha, Aves, and Euarchontoglires, respectively. (**b**) Graphs showing the numbers of CPuORFs extracted by ESUCA analyses of the genomes of the indicated species.

### Sequence-dependent effects of CPuORFs on mORF translation

To address the relationship between taxonomic ranges of CPuORF conservation and the likelihood of having a regulatory function using transient expression assays, we selected 17 human CPuORFs conserved in various taxonomic ranges, including a previously identified sequence-dependent regulatory CPuORF, the *PTP4A1* CPuORF ^38^, as a positive control. Besides the *PTP4A1* CPuORF, we selected CPuORFs with well-conserved C-terminal regions. This is because in many known sequence-dependent regulatory CPuORFs, the C-terminal region of the CPuORF-encoded nascent peptide has been shown to be important to cause ribosome stalling ^4^. Of these 16 selected CPuORFs, those in the genes *eIF5*, *MKKS*, *MIEF1*, and *SLC35A4* have been reported to play roles in regulating mORF translation ^39–41^. However, the sequence dependence of their effects on mORF translation have not been reported; therefore, we included these four CPuORFs. We examined the sequence-dependent effects of these 17 CPuORFs on the expression of the downstream reporter gene using transient expression assays (Fig. 2). Other uORFs overlapping any of the selected CPuORFs were eliminated by introducing mutations that changed the ATG codons of the overlapping uORFs to other codons but did not alter the amino acid sequences of the CPuORFs (Supplementary Fig. S6). The resulting modified CPuORFs were used as CPuORFs bearing the wild-type amino acid sequences (WT-aa CPuORFs) (Fig. 2b). To assess the importance of amino acid sequences for the effects of these CPuORFs on mORF expression, frameshift mutations were introduced into the WT-aa CPuORFs such that the amino acid sequences of their conserved regions could be altered (see the Methods and Supplementary Figure S6 for details). In eight of the 17 CPuORFs including the *PTP4A1* CPuORF, the introduced frameshift mutations significantly upregulated the expression of the reporter gene, indicating that these CPuORFs repressed mORF expression in a sequence-dependent manner (Fig. 2c). One of the eight sequence-dependent regulatory CPuORFs, the *TMEM184C* CPuORF, is conserved only among Eutheria (Fig. 2a). This result suggests that CPuORFs conserved only among Eutheria can have sequence-dependent regulatory effects. This study identified five novel regulatory CPuORFs (in the genes *SLC6A8*, *FAM13B*, *KAT6A*, *LRRC8B*, and *TMEM184C*). In addition, our results suggest that the repressive effects of the *MKKS* and *MIEF1* CPuORFs on mORF expression at least partly depend on their encoding sequences.

**Figure 2.**
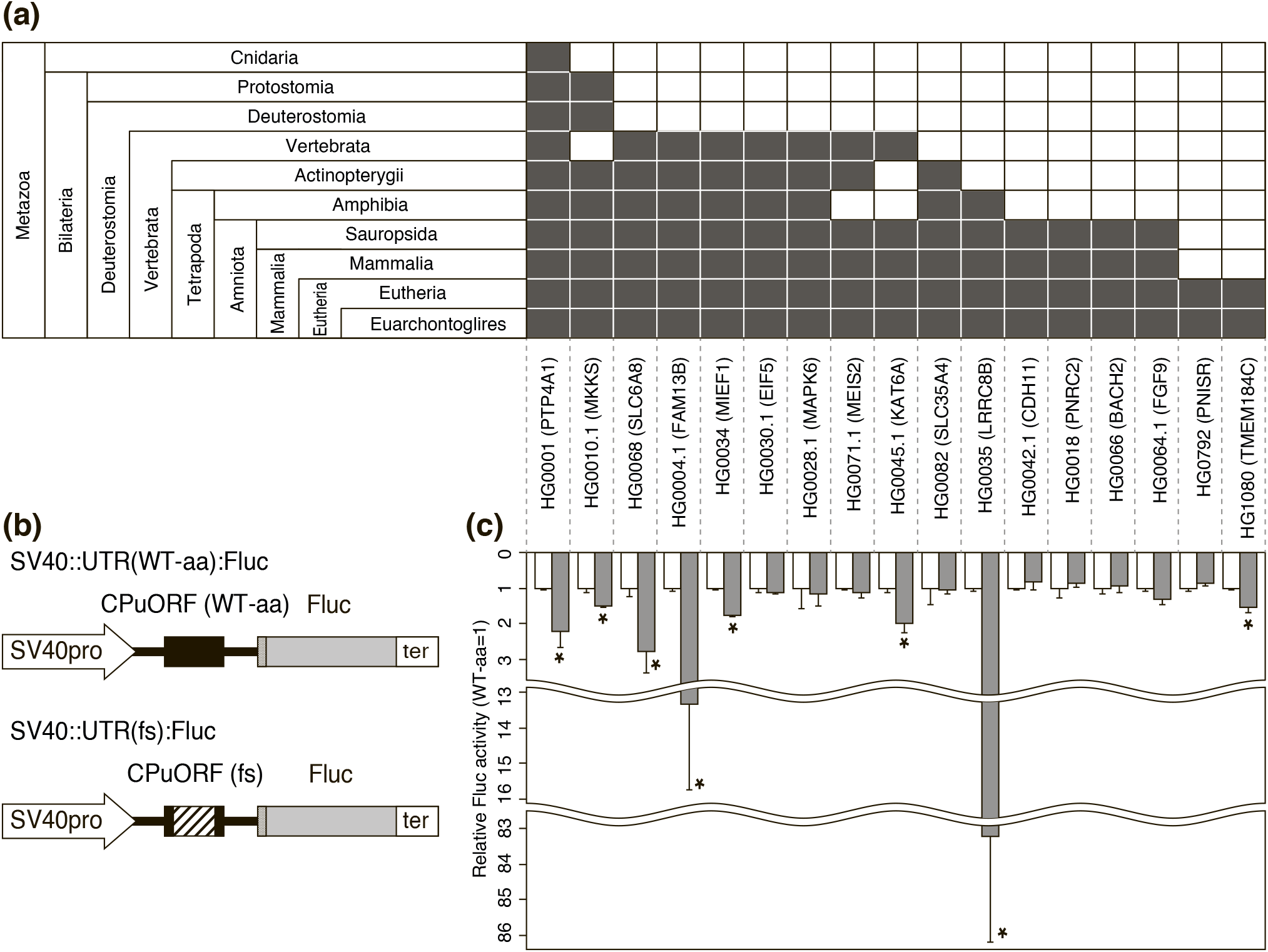
Taxonomic conservation and experimental validation of 17 selected human CPuORFs. (**a**) Taxonomic ranges of conservation of CPuORFs examined in transient assays. Filled cells in each taxonomic category indicate the presence of mORF-tBLASTn hits for CPuORFs of the indicated genes. (**b**) Reporter constructs used for transient assays. The hatched box in the frameshift (fs) mutant CPuORF indicates the frame-shifted region. Dotted boxes represent the first five nucleotides of the mORFs associated with the 17 human CPuORFs. See Supplementary Fig. S6 for the exact position and the length of each CPuORF and the exact frame-shifted region. (**c**) Relative luciferase (Fluc) activities of WT-aa (white) or fs (gray) CPuORF reporter plasmids. Means ± SDs of at least three biological replicates are shown. \**p* < 0.05.

## Discussion

In the current study, we identified 1,517 (1,373 novel and 144 known) CPuORFs belonging to 1,430 HGs by applying ESUCA to four animal genomes and selecting uORFs conserved across more than two orders. This number of identified CPuORFs is much higher than those identified by previous studies on genome-wide identification of CPuORFs ^26,34–36^. Thus, our results demonstrate that the approach using ESUCA, in which uORF sequences from certain species are compared with those from many other species, is highly effective in comprehensively identifying CPuORFs conserved in various taxonomic ranges. Since 1,082 of the 1,094 CPuORFs identified from the human genome are conserved beyond Euarchontoglires (Supplementary Table S2), and our transient expression assays suggested that CPuORFs conserved only among Eutheria can have sequence-dependent regulatory functions, it is likely that many of the human CPuORFs identified in this study are conserved because of functional constraints of their encoded peptides. It remains unknown why a much higher number of CPuORFs was identified in the human genome by our search compared to the other three species. One possible explanation is that in humans compared with the other three species, there are a higher number of transcript isoforms with different 5’-UTR sequences, produced from the same genes via alternative splicing or alternative transcription start site selection. The total numbers of such 5’-UTR variants of protein-coding genes registered in the ENSEMBL transcript databases are 22,352, 32,974, 12,413, and 64,619 for fruit fly, zebrafish, chicken, and human, respectively. The upregulation of mORF translation induced by the removal of an inhibitory uORF via alternative splicing or alternative transcription start site selection has been reported ^42–44^. We speculate that the human genome might need many CPuORFs to differentially control the translation efficiencies of the mORF among the 5’-UTR variants produced from the same gene.

The present study identified seven regulatory CPuORFs; however, the physiological roles of translational regulation mediated by these CPuORFs remain to be elucidated. One of the known physiological roles of CPuORFs is translational repression of the mORF in response to metabolites ^45^. Alternatively, CPuORFs are known to be involved in the promotion of mORF translation under specific conditions. In the latter case, a CPuORF is efficiently translated by ribosomes under normal conditions, and therefore, mORF translation is repressed, whereas ribosomes frequently bypass the CPuORF via leaky scanning under specific conditions such as stress conditions, thereby promoting mORF translation ^21^. The mORF of *SLC6A8*, one of the genes controlled by the CPuORFs identified in this study, codes for a creatine transporter that transports creatine into cells ^46^. Creatine is used as a readily available phosphate pool to regenerate ATP from ADP in cells. Therefore, the CPuORF in *SLC6A8* could mediate feedback regulation of SLC6A8 expression in response to the cellular creatinelevel to maintain creatine homeostasis. Alternatively, the CPuORF in *SLC6A8* could be involved in the upregulation of SLC6A8 expression under ATP-deficient conditions, via the leaky scanning mechanism in which the *SLC6A8* CPuORF is bypassed by ribosomes under these conditions. The *LRRC8B* mORF codes for a Ca^2+^ leak channel localized in the endoplasmic reticulum and participates in intracellular Ca^2+^ homeostasis ^47^. Therefore, the CPuORF in *LRRC8B* could play a role in feedback regulation of LRRC8B expression in response to the cytoplasmic Ca^2+^ level to maintain Ca^2+^ homeostasis. Interestingly, the *MKKS* and *MIEF1* CPuORFs have been reported to code for mitochondrial proteins ^24,26,39^. Therefore, these CPuORFs have been suggested to play two roles, i.e., translational regulation and coding for mitochondrial proteins ^26,39^. Our results additionally suggest that the effects of the *MKKS* and *MIEF1* CPuORFs on mORF translation depend on their amino acid sequences, and therefore, polypeptides encoded by these CPuORFs may have dual functions.

Nine of the 17 CPuORFs tested by transient expression assays exhibited no sequence-dependent effect on mORF expression. One possible reason for this is that these CPuORFs might not have been translated in our experimental conditions. The start codons of two of the nine CPuORFs are in a poor initiation context, with no purine at −3 and no guanine at +4, where the A in ATG is +1. In contrast, the start codons of the remaining seven CPuORFs are in an optimal or sub-optimal context, containing a purine at −3 and/or a guanine at +4 (Supplementary Fig. S6). If these CPuORFs were translated in our experimental conditions, other possible explanations for the lack of sequence-dependent effects of these CPuORFs would be that these CPuORFs might encode peptides with functions other than the control of mORF translation, or they might exert sequence-dependent regulatory effects only under certain conditions.

Chemical screening recently identified a compound that causes nascent peptide-mediated ribosome stalling in the mORF of the human *PCSK9* gene, resulting in specific translational inhibition of *PCSK9* and a reduction in total plasma cholesterol levels ^48^. Nascent peptide-mediated ribosome stalling in some of the previously identified regulatory CPuORFs is promoted by metabolites, such as polyamine, arginine, and sucrose 4,49. Therefore, compounds that promote nascent peptide-mediated ribosome stalling in CPuORFs could be identified by chemical screening through a method similar to that used for screening the stall-inducing compound for *PCSK9*. The data from the current study may be useful for selecting CPuORFs as potential targets for pharmaceuticals and for identifying regulatory CPuORFs.

## Methods

All procedures and protocols were approved by the Institutional Safety Committee for Recombinant DNA Experiments at Chiba University. All methods were carried out in accordance with approved guidelines. All human 5’-UTR sequences containing the CPuORFs used in the present study were artificially synthesized (GenScript, Piscataway, NJ, USA) according to the RefSeq sequences (see “Plasmid construction and transient reporter assays” for details).

### Extraction of CPuORFs using ESUCA

ESUCA was developed as an advanced version of BAIUCAS ^29^ in our previous study ^32^. ESUCA consists of six steps, and some of these steps are divided into substeps, as shown in Supplementary Fig. S1a and S1b. To identify animal CPuORFs using ESUCA, the following eight-step procedures were conducted, including the six ESUCA steps: (0) data preparation for ESUCA, (1) uORF extraction from the 5*’*-UTR (Supplementary Fig. S2a), (2) calculation of uORF–mORF fusion ratios (Supplementary Fig. S2b), (3) uORF-tBLASTn against transcript sequence databases (Supplementary Fig. S3a), (4) mORF-tBLASTn against downstream sequence datasets for each uORF (Supplementary Fig. S3b and S3c), (5) calculation of *K*_a_/*K*_s_ ratios, (6) determination of the taxonomic range of uORF sequence conservation, and (7) manual validation after ESUCA. See the Materials and Methods in our previous study ^32^ for details.

### Transcript dataset construction based on genome information (step 0.1)

To identify animal CPuORFs, data preparation for ESUCA (step 0.1) was conducted as described in our previous study ^32^. We conducted data preparation for ESUCA to identify animal CPuORFs as follows. We used a genome sequence file in FASTA format and a genomic coordinate file in GFF3 format obtained from Ensemble Metazoa Release 33 (https://metazoa.ensembl.org/index.html) ^50^ to extract fruit fly (*Drosophila melanogaster*) uORF sequences. We used genome sequence files in FASTA format and genomic coordinate files in GFF3 format obtained from Ensemble Release 86 (https://metazoa.ensembl.org/index.html) ^50^ for zebrafish (*Danio rerio*), chicken (*Gallus gallus*), and human (*Homo sapiens*). We extracted exon sequences from genome sequences on the basis of genomic coordinate information and constructed transcript sequence datasets by combining exon sequences. On the basis of the transcription start site and the translation initiation codon of each transcript in the genomic coordinate files, we extracted 5’-UTR and mORF RNA sequences from the transcript sequence datasets, as shown in Supplementary Fig. S1a (step 0.1). The 5’-UTR sequences were used at step 1 of ESUCA. The mORF RNA sequences were translated into amino acid sequences (mORF proteins) and were used at step 4.1 of ESUCA.

### Transcript base sequence dataset construction from EST/TSA/RefSeq RNA (step 0.2)

To identify animal CPuORFs, data preparation for ESUCA (step 0.2) was conducted as described in our previous study ^32^. We conducted data preparation for ESUCA to identify animal CPuORFs. As shown in Supplementary Fig. S1b, Metazoa RefSeq RNA sequences were used at steps 2 and 3.1 of ESUCA. Assembled EST/TSA sequences generated by using Velvet ^51^ and Bowtie2 ^52^ were used at step 3.1 of ESUCA. Intact and merged EST/TSA/RefSeq sequences were used at step 4.2 of ESUCA. Taxonomy datasets derived from EST/TSA/RefSeq databases were used at steps 4.3 and 6 of ESUCA. See the Materials and Methods in our previous study ^32^ for details.

### Determination of the taxonomic range of uORF sequence conservation for animal CPuORFs (step 6)

To automatically determine the taxonomic range of the sequence conservation of each CPuORF, we first defined 20 animal taxonomic categories. The 20 defined taxonomic categories were Euarchontoglires, Eutheria other than Euarchontoglires, Mammalia other than Eutheria, Aves, Sauropsida other than Aves, Amphibia (Tetrapoda other than Sauropsida and Mammalia), Sarcopterygii other than Tetrapoda, Ostarioclupeomorpha, Actinopterygii other than Ostarioclupeomorpha, Vertebrata other than Euteleostomi (Actinopterygii and Sarcopterygii), Chordata other than Vertebrata, Deuterostomia other than Chordata, Insecta, Arthropoda other than Insecta, Ecdysozoa other than Arthropoda, Lophotrochozoa (Protostomia other than Ecdysozoa), Bilateria other than Protostomia and Deuterostomia, Cnidaria, Ctenophora (Eumetazoa other than Cnidaria and Bilateria), and Metazoa other than Eumetazoa. Based on taxonomic lineage information of EST, TSA, and RefSeq RNA sequences, which were provided by NCBI Taxonomy, the mORF-tBLASTn hit sequences selected for *K*_a_/*K*_s_ analysis were classified into the 19 taxonomic categories (Fig. 1a and Supplementary Table S2). The category “Ctenophora” was omitted from animal taxonomic categories because no sequences were classified to this category. For each CPuORF, the numbers of transcript sequences classified into each category were counted and are shown in Supplementary Table S2. These numbers represent the number of orders in which the amino acid sequence of each CPuORF is conserved.

### Classification of animal CPuORFs into HGs

Systematic numbering of animal CPuORF families (HGs) has not been reported to date. Here, we defined systematic HG numbers for the identified 1,517 animal CPuORFs. Among these identified CPuORFs, those with both similar uORF and mORF amino acid sequences were classified into the same HGs. We first determined ortholog groups of CPuORF-containing genes, referred to as mORF clusters, based on similarities of mORF-encoded amino acid sequences, using OrthoFinder ^37^. The identified CPuORF-containing genes were classified into 1,194 mORF clusters. CPuORFs contained in each ortholog group (mORF-cluster) were further classified into uORF clusters as follows. We conducted a pairwise comparison of uORF peptide similarity using BLASTp with *E*-values less than 2,000 in each mORF cluster. Binarized distance matrixes consisting of 0 (hit) or 1 (no-hit) were generated by this comparison. Hierarchical clustering with single linkage with the cutoff parameter (*h* = 0.5) was applied to these matrixes for the construction of uORF clusters. In total, 1,336 uORF–mORF clusters were generated automatically. We determined 1,430 clusters by manually checking the alignments of uORFs and mORFs. We assigned HG numbers to the 1,430 clusters so that CPuORF families conserved across wider taxonomic categories could have smaller HG numbers. When there were multiple CPuORF families conserved in the same taxonomic categories, smaller HG numbers were assigned to CPuORF families conserved in higher numbers of orders. The same HG number with a different sub-number was assigned to CPuORFs in genes of the same ortholog group with dissimilar uORF sequences (e.g., HG0004.1 and HG0004.2; Supplementary Table S1).

### Plasmid construction and transient reporter assays

pSV40:Fluc was generated by inserting the SV40 promoter (*Bgl*II/*Hind*III fragment) from pRL-SV40 (Promega, Madison, WI, USA) into the *Kpn*I site of pGL4.10[luc2] (Promega) by blunt-end cloning. The 5’-UTR sequences containing the selected CPuORFs (*Sac*I/*Xho*I fragment) were fused to the Fluc-coding sequence by subcloning the CPuORFs into the *Sac*I/*Xho*I site of pSV40:luc2 to generate the WT-aa reporter construct (pSV40:UTR(WT-aa):Fluc; Fig. 2b, Supplementary Fig. S6). To assess the importance of the amino acid sequences with regard to the effects of these CPuORFs on mORF translation, frameshift mutations were introduced into the CPuORFs so that the amino acid sequences of their conserved regions could be altered. A +1 or −1 frameshift was introduced upstream or within the conserved region of each CPuORF, and another frameshift was introduced before the stop codon to shift the reading frame back to the original frame (pSV40:UTR(fs):Fluc; Fig. 2b, Supplementary Fig. S6). DNA fragments containing the CPuORFs of either WT-aa or fs mutants from the *PTP4A1*, *MKKS*, *SLC6A8*, *FAM13B*, *MIEF1*, *EIF5*, *MAPK6*, *MEIS2*, *KAT6A*, *SLC35A4*, *LRRC8B*, *CDH11*, *PNRC2*, *BACH2*, *FGF9*, *PNISR*, and *TMEM184C* genes were synthesized (GenScript) and subcloned into pSV40:Fluc, as shown in Fig. 2b and Supplementary Table S5. These reporter constructs were each transfected into human HEK293T cells. HEK293T cells (16,000/well) were co-transfected with 80 ng/well of a pSV40:UTR:Fluc reporter plasmid and 1.6 ng/well pGL4.74[hRluc/TK] plasmid (Promega). After 24 h, firefly luciferase (Fluc) and *Renilla* luciferase (Rluc) activities were measured according to the Dual-Luciferase Reporter Assay protocol (Promega) using the GloMaxR-Multi Detection System (Promega). Fluc activity was normalized by Rluc activity to correct for differences in cell viability and transfection efficiency.

### Statistical and informatics analyses

All programs, except for existing stand-alone programs, such as NCBI-BLAST+ ver. 2.6.0 ^53^, Clustal Ω ver. 1.2.2 54, OrthoFinder ver. 1.1.4 ^37^, Velvet ver. 1.2.10 ^51^, Bowtie2 ver. 2.2.9 ^52^, and Jalview ver. 2.10.2 ^55^, were written in R (www.r-project.org). We also used R libraries, GenomicRanges ver. 1.32.7 ^56^, exactRankTests ver. 0.8.30, Biostrings ver. 2.48.0, and seqinr ver. 3.4.5 ^57^. Statistical differences between the control (WT-aa) and fs constructs were determined by Student’s *t*-tests in transient assays.

## Data availability

Individual FASTA-formatted files used in Supplementary Fig. S4 are available from http://www.p.kanazawa-u.ac.jp/~bukka/data/FASTA-Formatted.1517CPuORFs.zip or https://www.takahashi-lab.com/db/FASTA-Formatted.1517CPuORFs.zip. The datasets generated and analyzed during the current study are available from the corresponding author on reasonable request.

## Acknowledgement

This work was supported by the Japan Society for the Promotion of Science (JSPS) KAKENHI (grant nos. JP19H02917 and JP16K07387 to H.O.; JP19K22892 to H.T.; JP18H03330 to H.T., M.I, and H.O.; and JP18H02568 to M.I.); the Ministry of Education, Culture, Sports, Science and Technology (MEXT) KAKENHI (grant nos. JP17H05658 to S.N., JP26114703 to H.T, and JP17H05659 to H.T); the Naito Foundation (to H.O.); and the Research Foundation for the Electrotechnology of Chubu (to H.T.). We would like to thank Editage (www.editage.com) for English language editing. This is a preprint of an article published in *Scientific Reports*. The final authenticated version is available online at: https://doi.org/10.1038/s41598-020-73307-6.

## Author contributions

H.T., H.O., and M.I. designed the study. H.T. and S.M. performed experiments and analyzed the data under the supervision of S.F., T.E., K.S., S.N., and M.I. H.T., M.M., N.I., T.M., and A.T. contributed reagents/materials/analysis tools. H.T., H.O., M.I., and S.M. wrote the article with the contribution of all coauthors.

## Additional information

Supplementary information accompanies this paper.

### Competing financial interests

The authors declare no competing financial interests.

## Notes

### Competing Interest Statement

The authors have declared no competing interest.

