## Supplementary Figure S1 for "Exhaustive identification of conserved upstream open reading frames with potential translational regulatory functions from animal genomes"

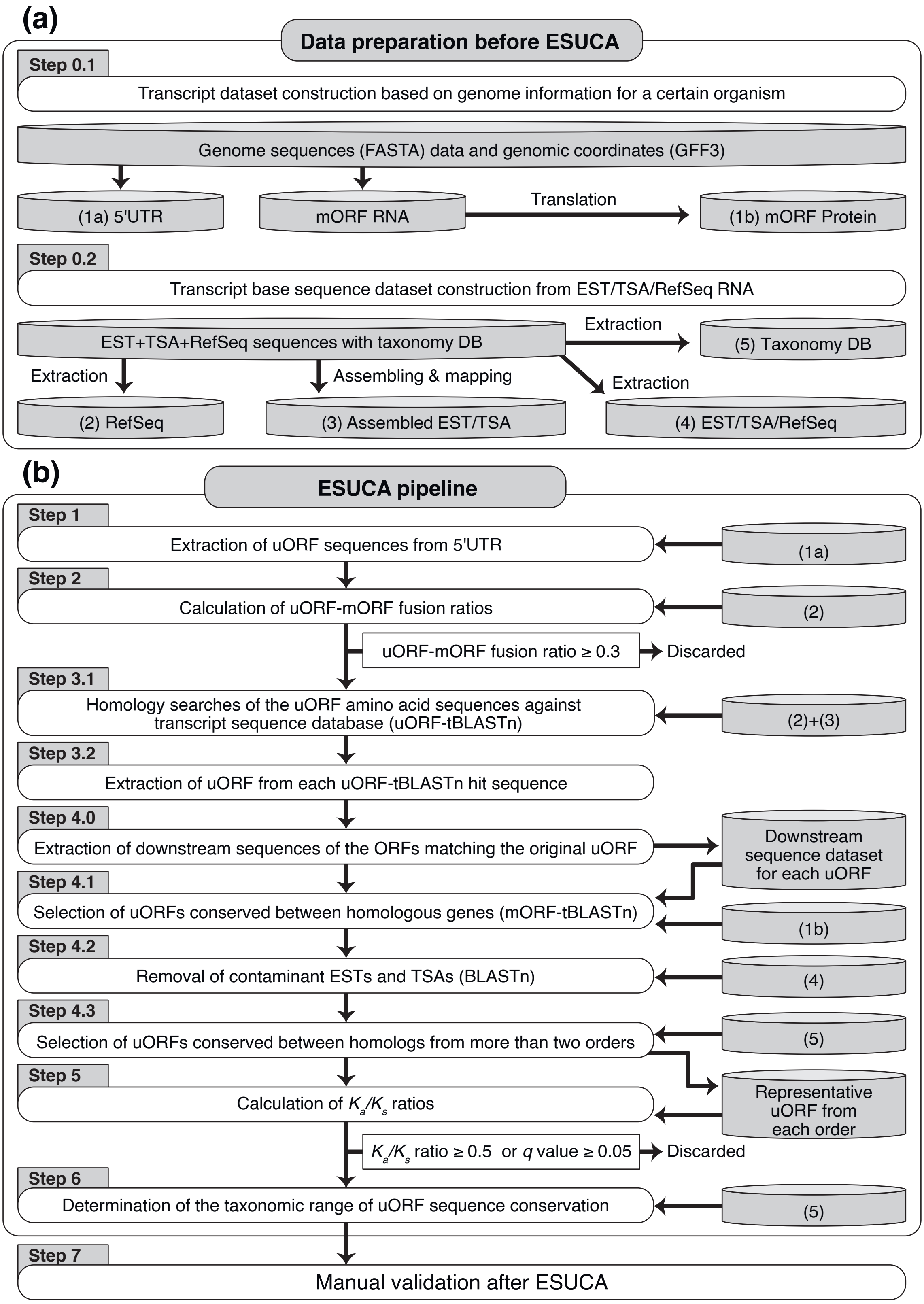


**Supplementary Figure S1.** Identification of animal CPuORFs using ESUCA. (**a**) Data preparation. (**b**) Outline of the ESUCA pipeline. Numbers with parenthesis indicate datasets labeled with the same numbers in (**a**).
