## Supplementary Figure S2 for "Exhaustive identification of conserved upstream open reading frames with potential translational regulatory functions from animal genomes"

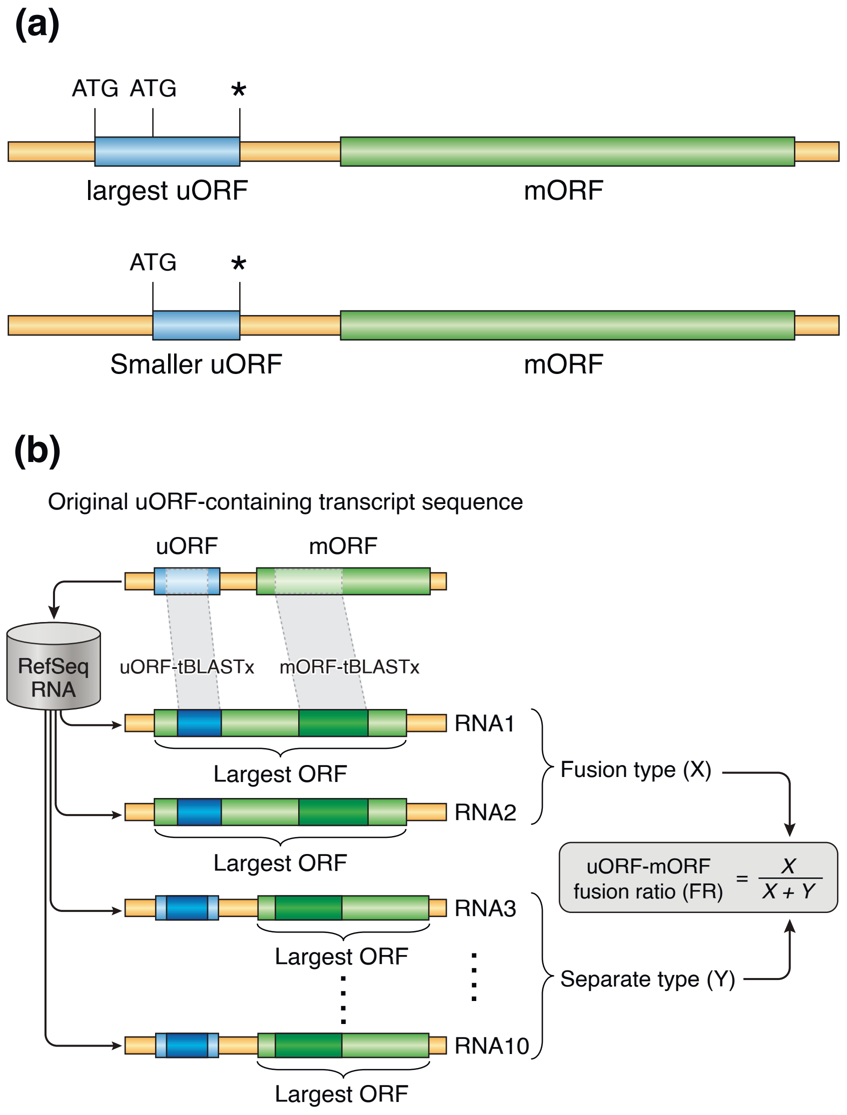


**Supplementary Figure S2.** Extraction of the largest uORF sequences and calculations of uORF-mORF fusion ratio. **(a)** Extraction of the largest uORF sequences from the 5′-UTR. Extraction of the largest uORF sequences from the 5′-UTR. After data preparation for ESUCA (Supplementary Fig. S1b), we conducted the extraction of uORF sequences by searching the 5′-UTR sequences for an ATG codon and its nearest downstream in-frame stop codon at step 1 of ESUCA (Supplementary Fig. S1b). Sequences starting with an ATG codon and ending with the nearest in-frame stop codon were extracted as uORF sequences. When multiple uORFs shared the same stop codon in a transcript, only the longest uORF sequence was used for further analyses. **(b)** Outline for uORF-mORF fusion ratio calculations. For each original uORF-containing transcript sequence, RefSeq RNAs containing both sequences similar to the uORF and the mORF of each uORF-containing transcript were selected using uORF-tBLASTx and mORF-tBLASTx from the RefSeq RNA database (database (2) in Supplementary Fig. S1a). For example, the selected RNA sequences are RNA1, 2, 3...10, as illustrated. Based on whether the uORF-tBLASTx-hit region was included in the largest RefSeq RNA ORF, the selected RefSeq RNAs were classified into two types, namely fusion ($X$) (RNA1 and 2) and separate types ($Y$) (RNA3-10). For each original uORF-containing transcript, the uORF-mORF fusion ratio was calculated as $X/(X+Y)$.
