## Supplementary Figure S3 for "Exhaustive identification of conserved upstream open reading frames with potential translational regulatory functions from animal genomes"

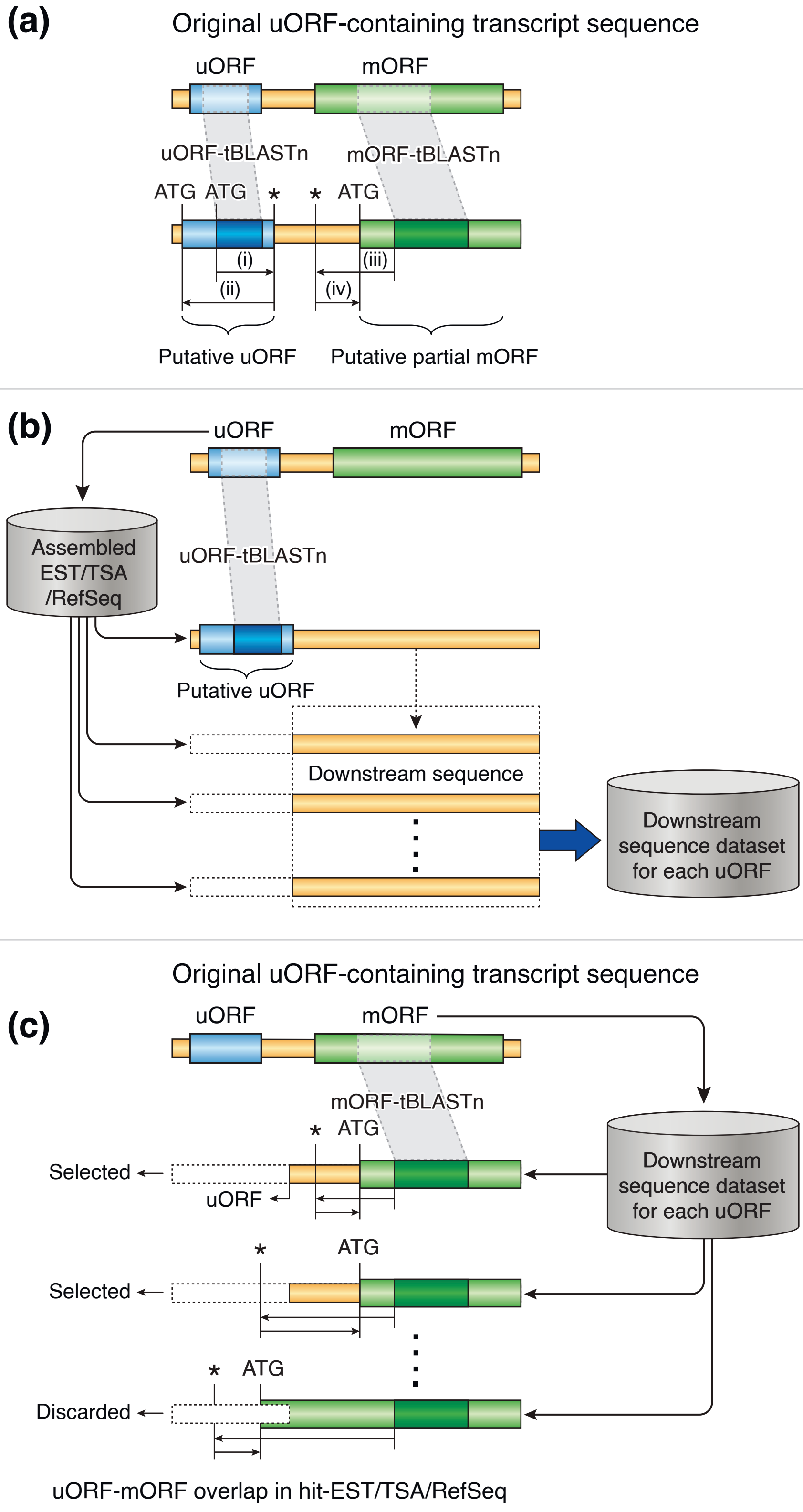


**Supplementary Figure S3.** Outline of homology searches for uORFs with amino acid sequences conserved between homologous genes. **(a)** For each original uORF-containing transcript, sequences containing both similar regions to the uORF and the mORF of uORF-containing transcripts were selected using uORF-tBLASTn (step 3.1 of ESUCA) and mORF-tBLASTn (step 4.1 of ESUCA). A transcript sequence database consisting of RefSeq RNAs (database (2) in Supplementary Fig. S1a) served as data source, while an assembled EST/TSA (database (3) in Supplementary Fig. S1a) was generated at step 0.2 of data preparation for ESUCA. Asterisks represent stop codons. At step 3.2 of ESUCA, the largest tBLASTn-hit region-overlapping uORF was extracted. **(b)** Detailed illustration of step 4.0 of ESUCA. Putative uORF extraction and downstream sequence dataset construction were conducted systematically for each uORF-tBLASTn hit sequence. **(c)** Detailed illustration of step 4.1 of ESUCA. After mORF-tBLASTn, the 5'-most in-frame ATG codon located downstream of the selected stop codon was identified as the initiation codon of the putative partial or intact mORF. uORF-mORF overlaps were discarded as fusion types, according to the positional relationship between them, when found in the hit-assembled EST/TSA+RefSeq sequences.
