## Supplementary Figure S4 for "Exhaustive identification of conserved upstream open reading frames with potential translational regulatory functions from animal genomes"

## HG0001

ENSDARG00000006242 (ptp4a1)

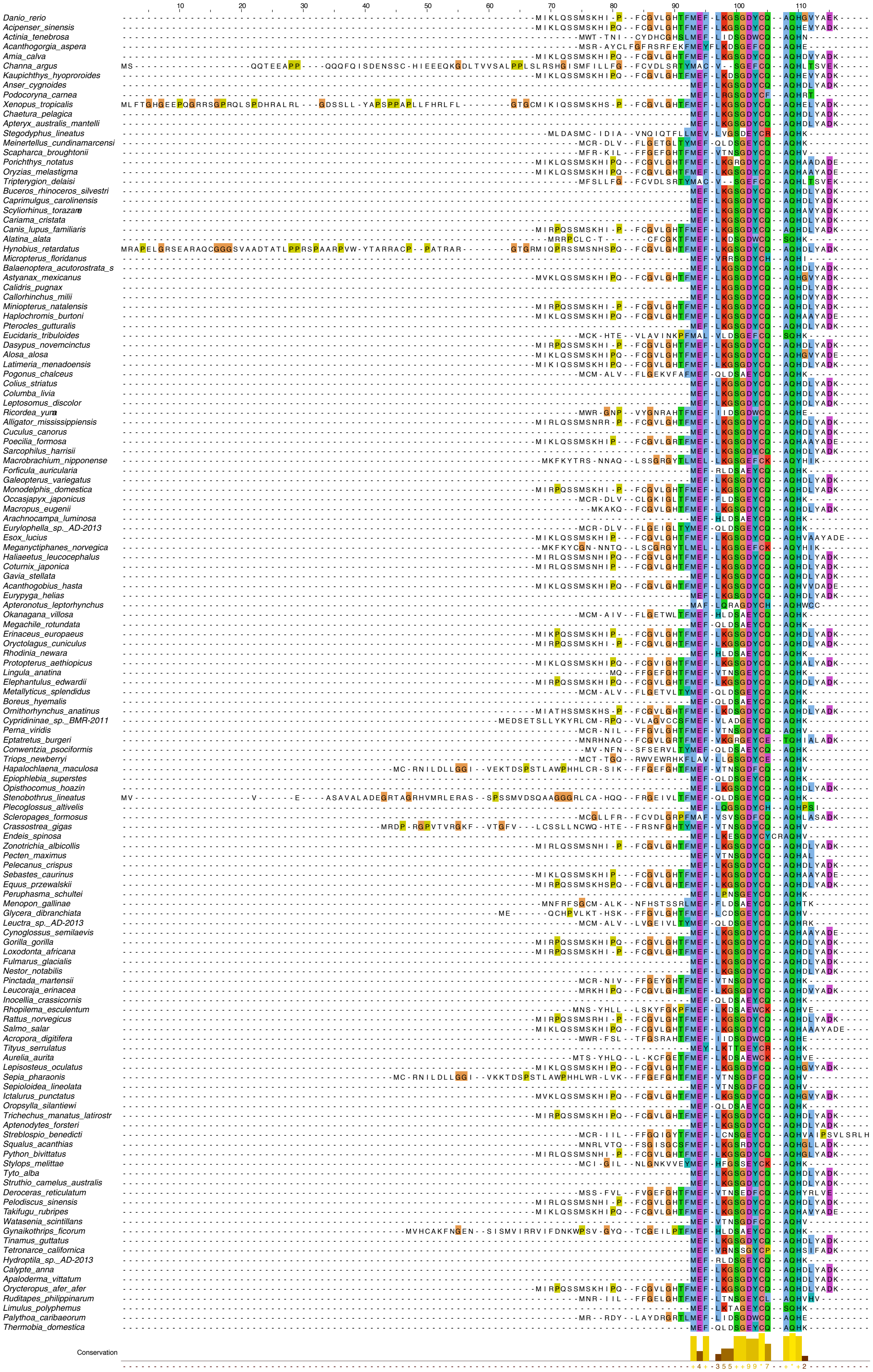

# HG0001

### ENSDARG00000035676 (ptp4a2b)

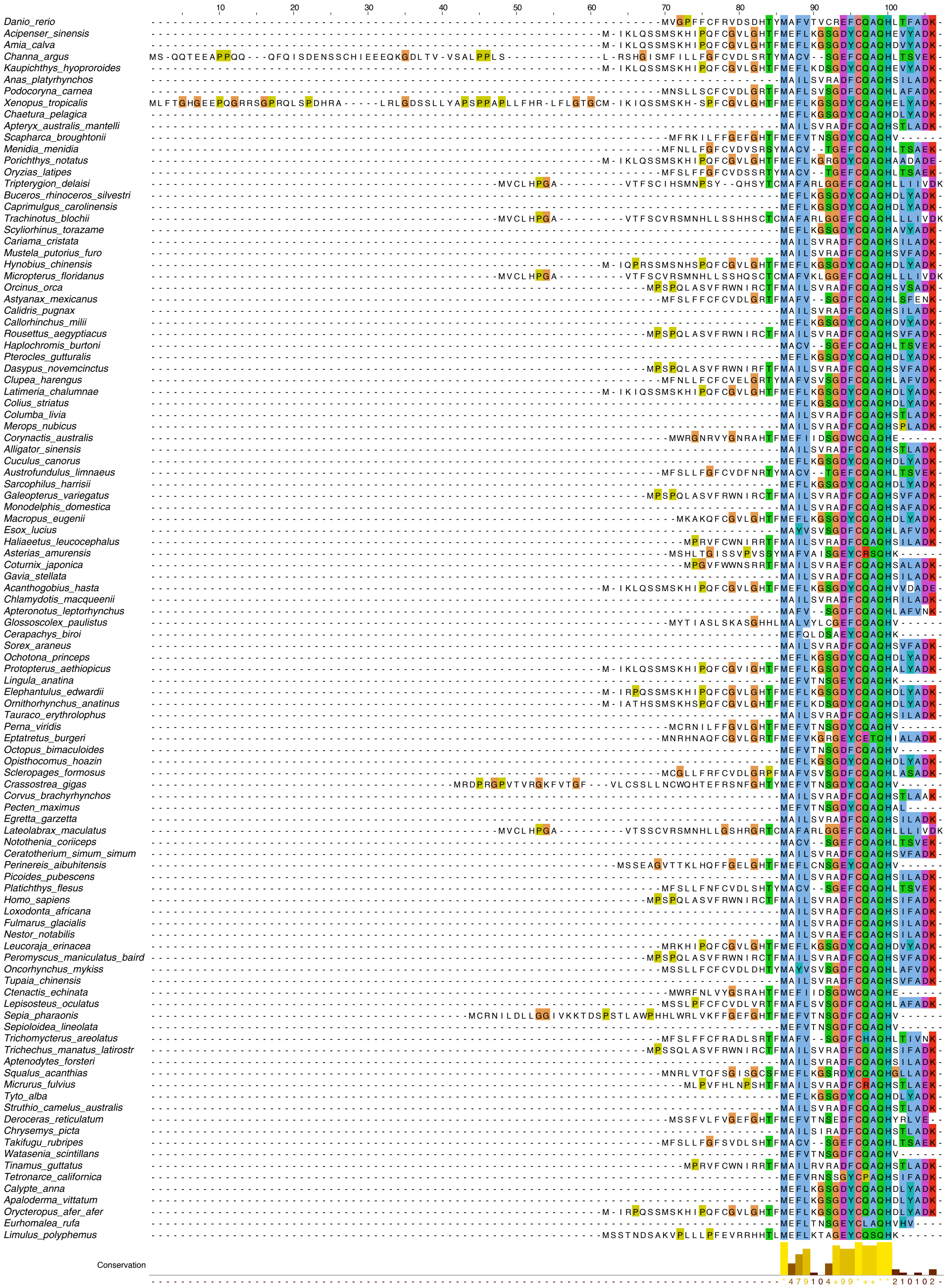

# HG0001

### ENSDARG00000039997 (ptp4a3)

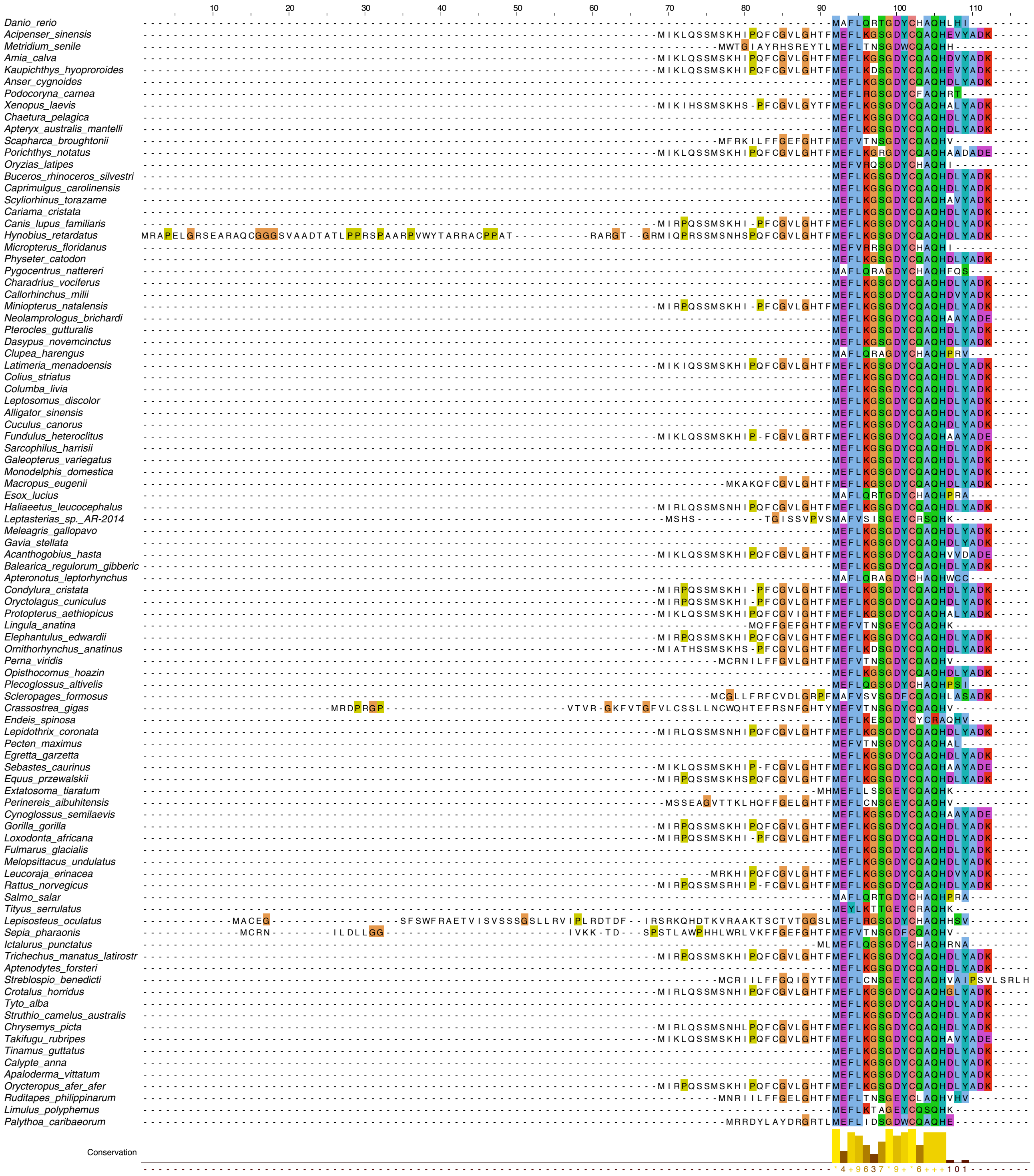

# HG0001

### ENSDARG00000054814 (PTP4A3)

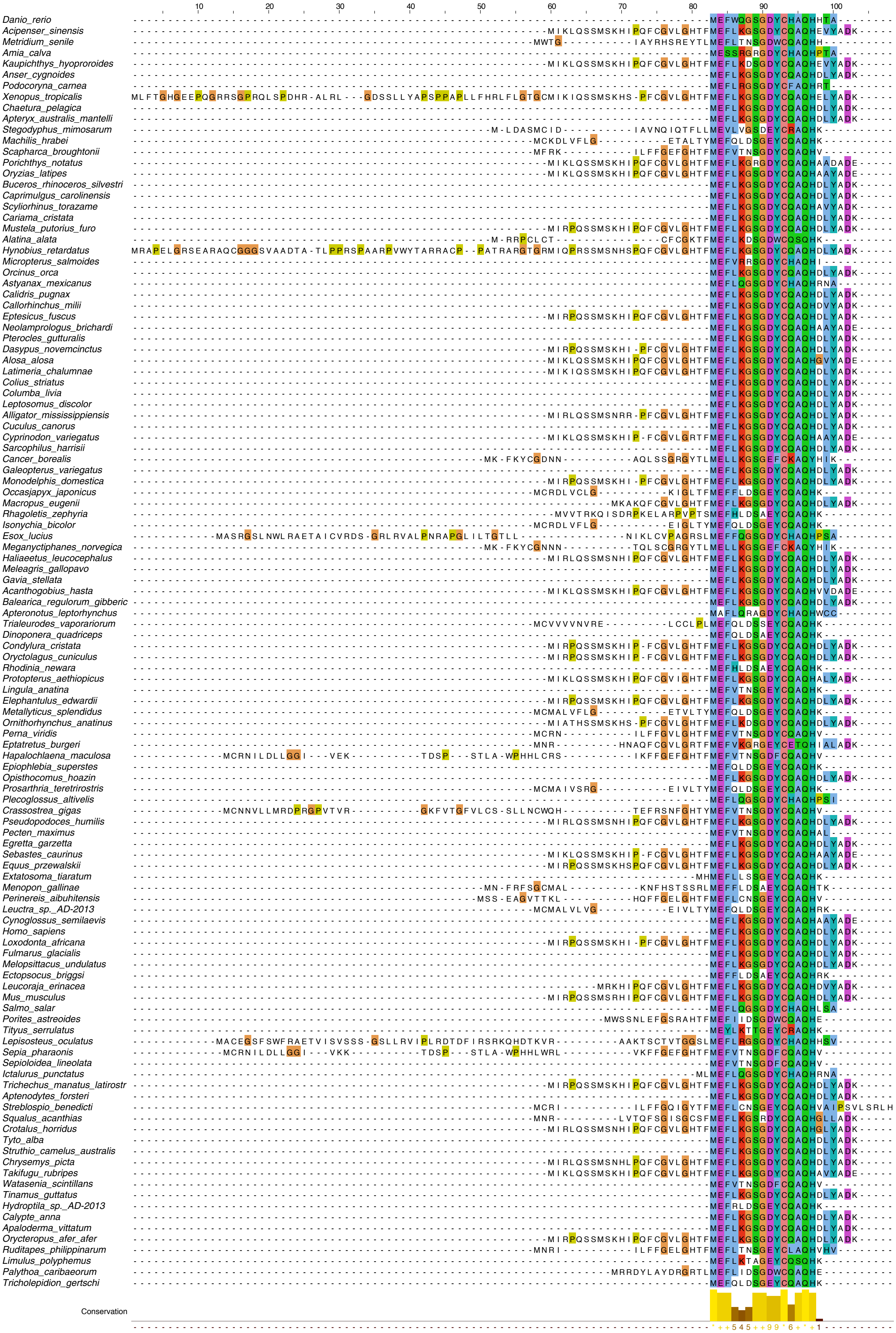

# HG0001

### ENSDARG00000087443 (ptp4a2a)

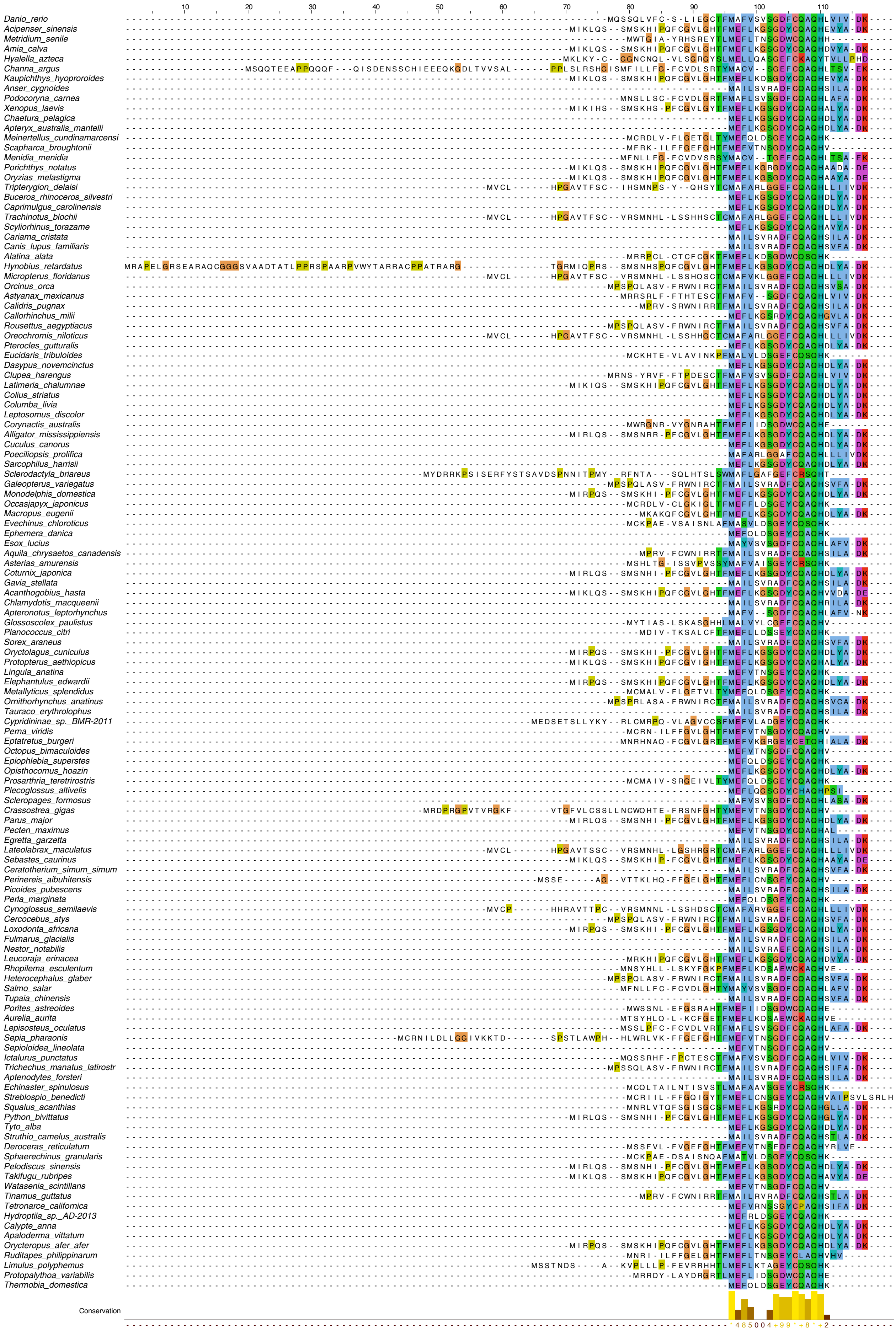

# HG0001

### ENSG00000112245 (PTP4A1)

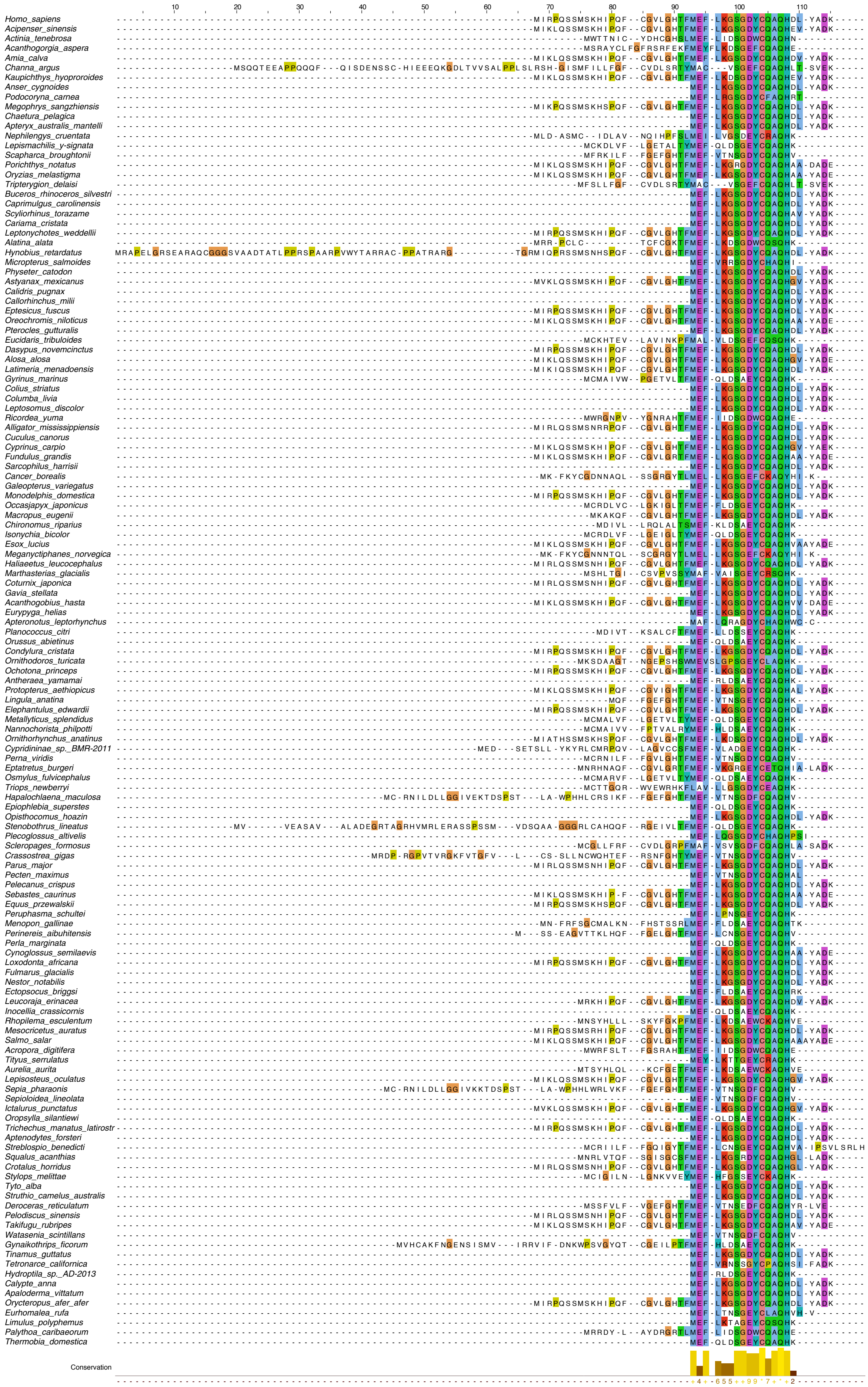

# HG0001

### ENSG00000184007 (PTP4A2)

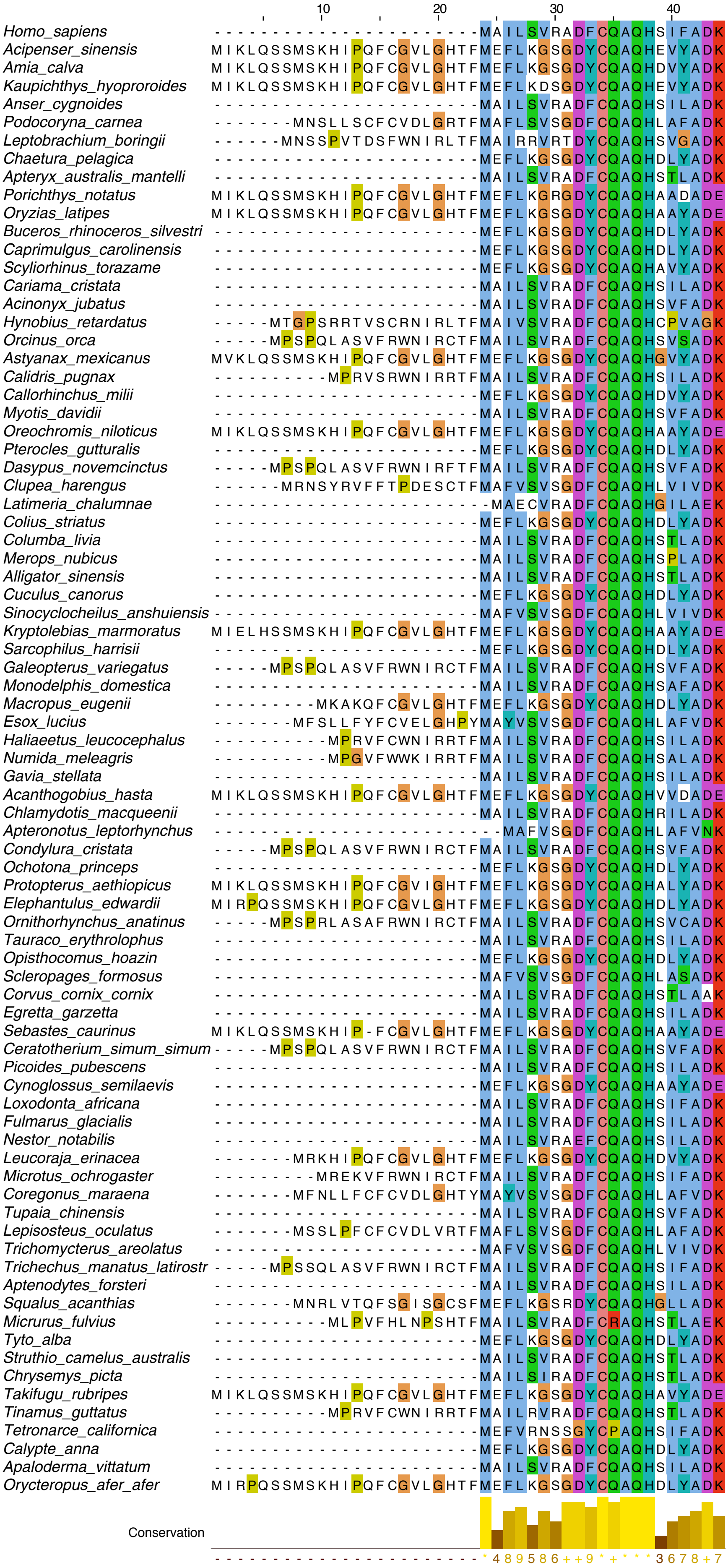

# HG0001

### ENSGALG00000003265 (PTP4A2)

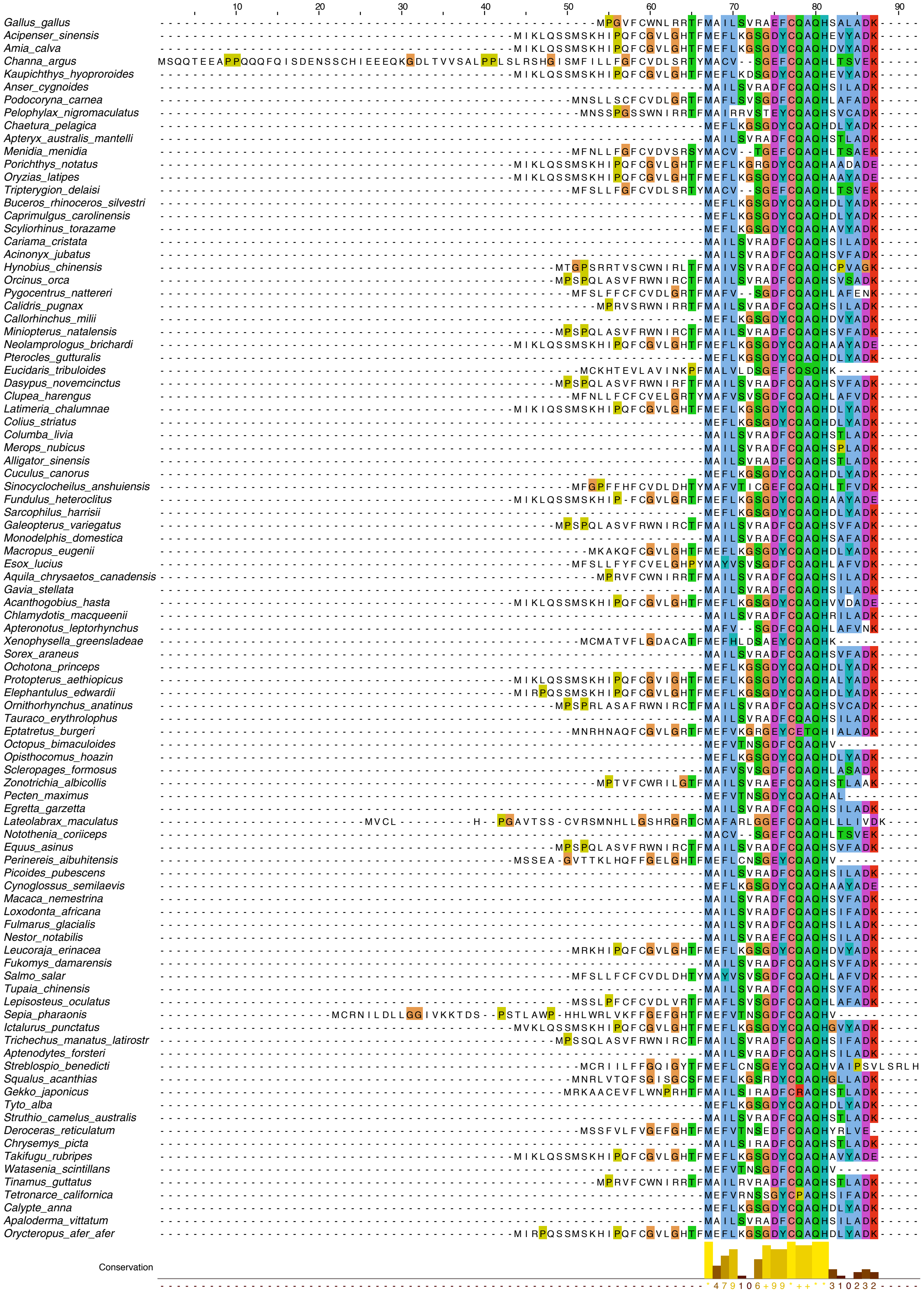

# HG0001

### ENSGALG00000016271 (PTP4A1)

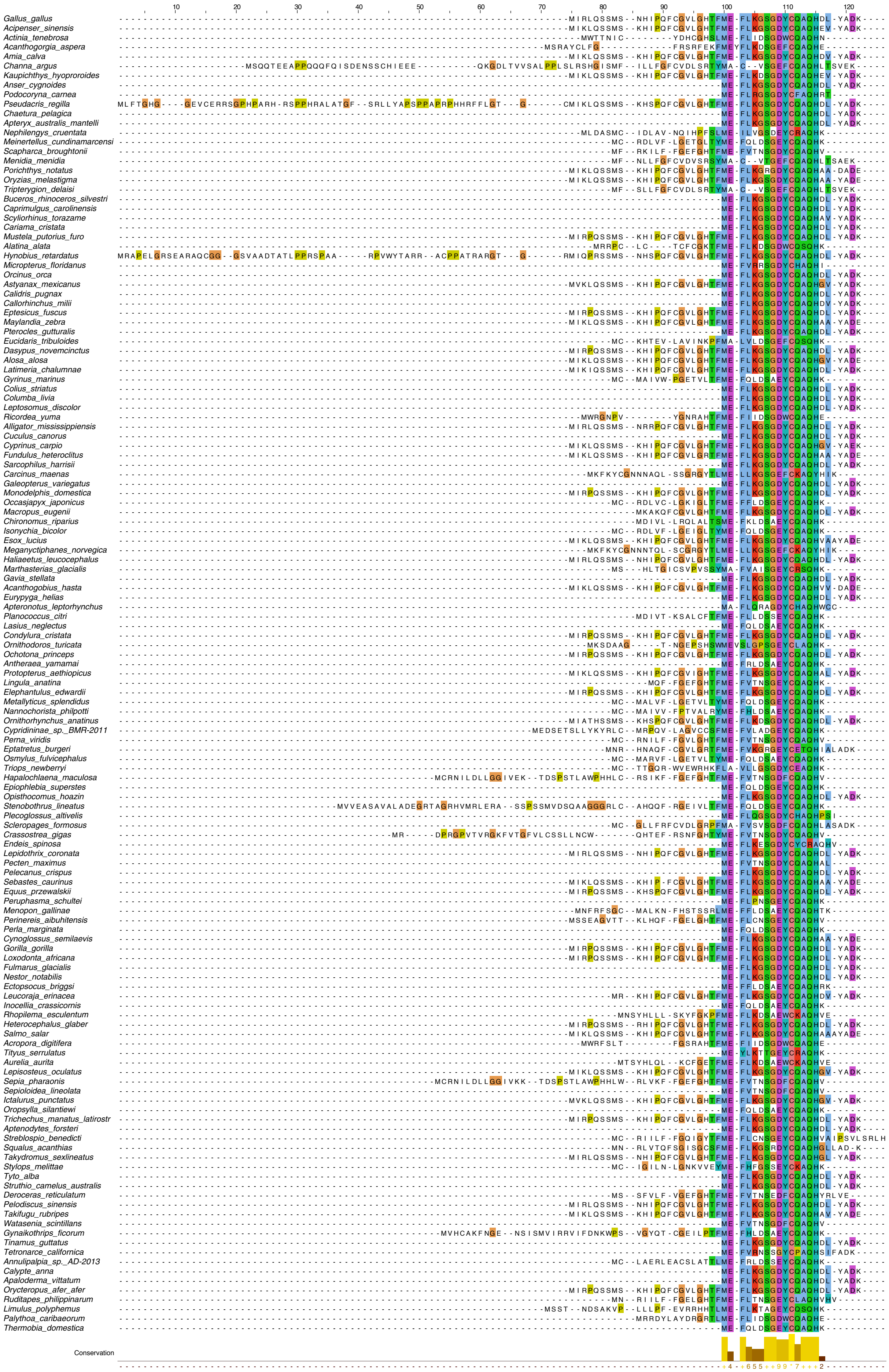

# HG0001

### FBgn0024734 (PRL-1)

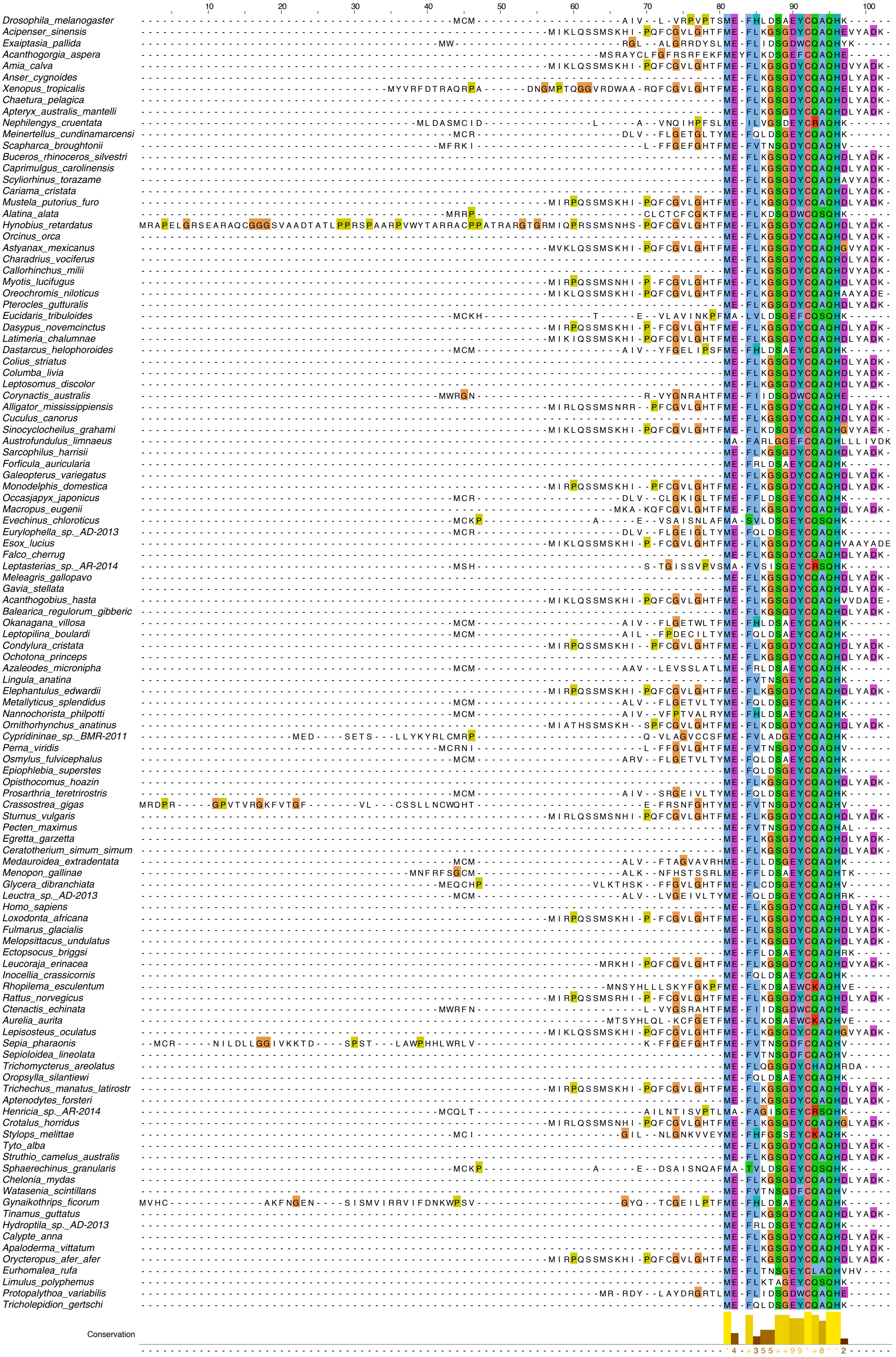

# HG0002

### ENSDARG00000003077

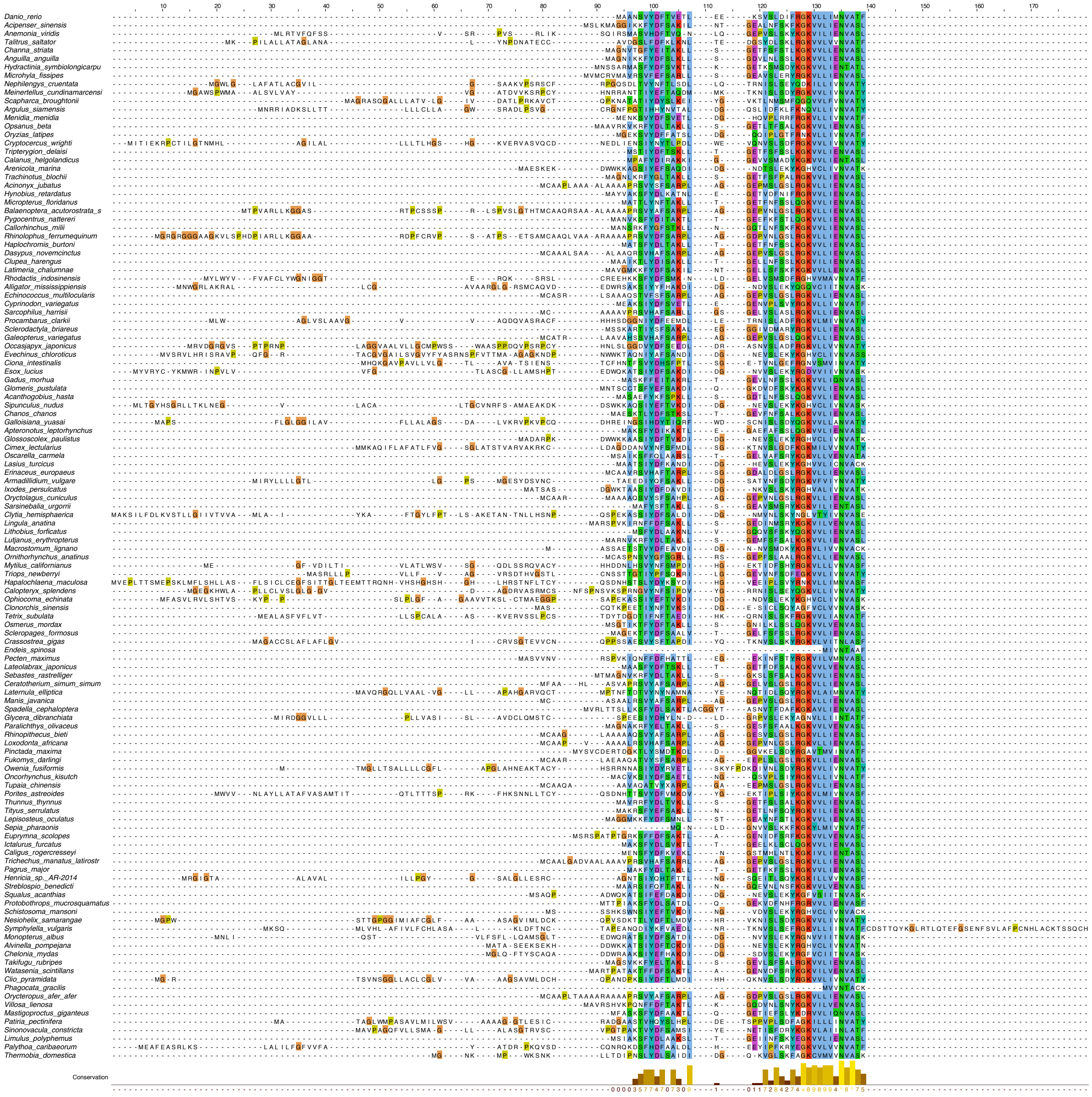

# HG0003

### ENSGALG00000014906 (MOCS2)

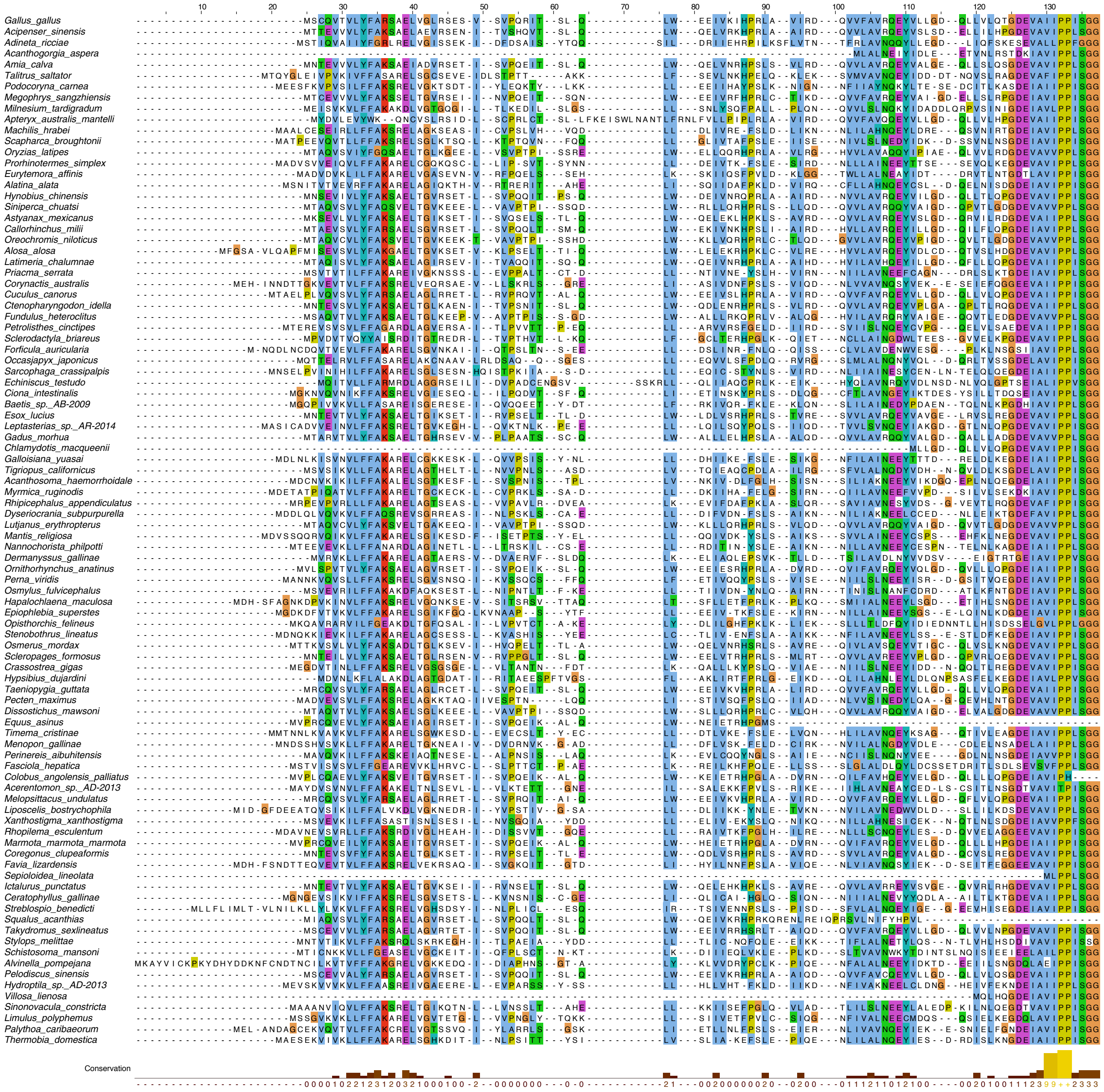

# HG0003

### FBgn0039280 (Mocs2)

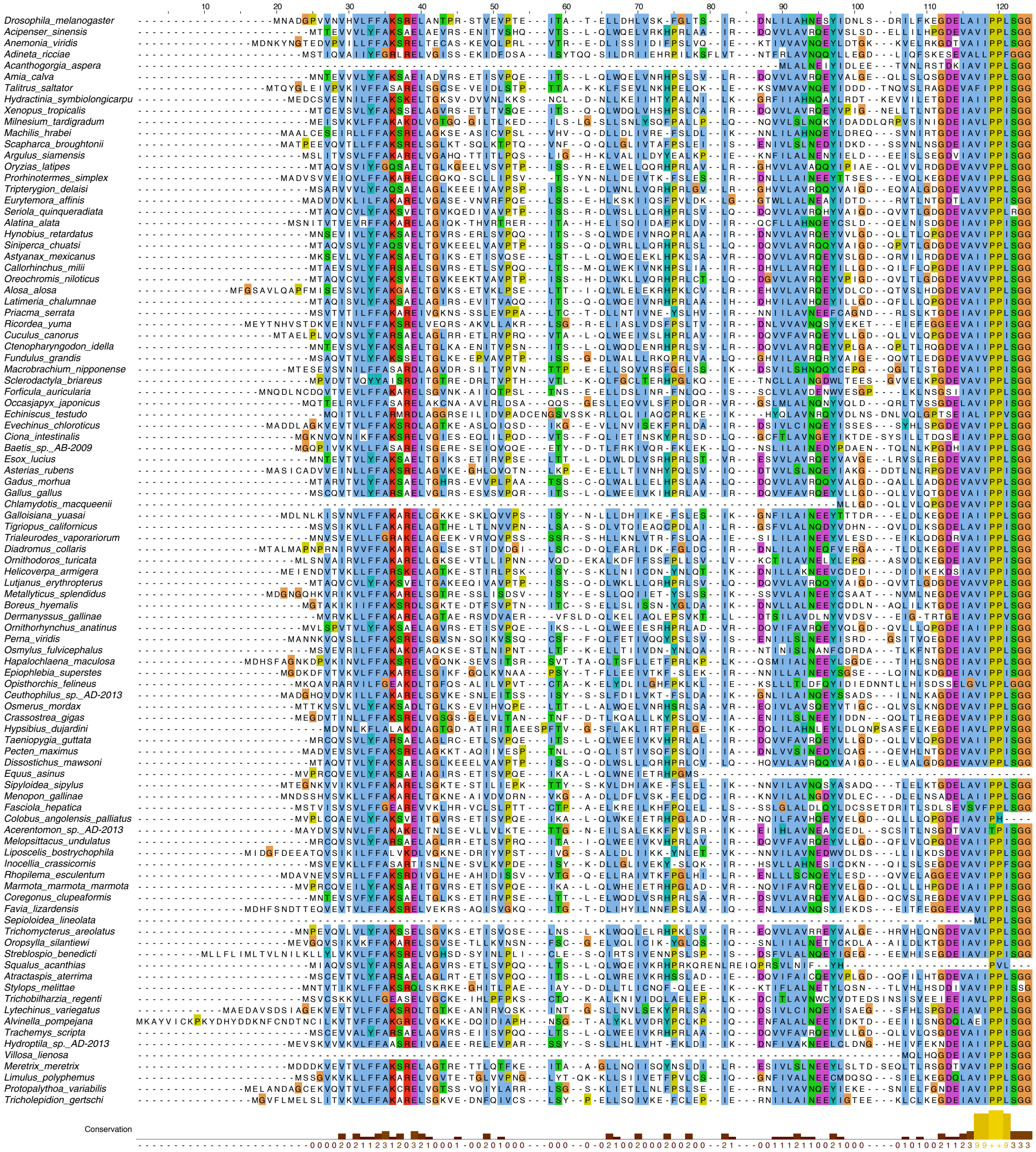

# HG0004.1

### ENSDARG00000076779 (fam13b)

```
                                       10        20        30        40        50        60        70
Danio_rerio                     ---------------------------------------------------------------MRNYRLIFCHNLKQP
Acipenser_sinensis              ---------------------------------------------------------------MRNYRLIFCHNLKQP
Amia_calva                      ---------------------------------------------------------------MRNYRLIFCHNLKQP
Channa_punctata                 ---------------------------------------------------------------MRNYRLIFCHNLKQP
Anas_platyrhynchos              ---------------------------------------------------------------MRNYRLIFCHNLKQP
Xenopus_laevis                  MVTSCSDCGPLAVNAFKCIHKILTERDCWGQRELKGLSKRKPQQNQS-YRHTAEELIELWISSMRNYRLIFCHNLKQP
Chaetura_pelagica               ---------------------------------------------------------------MRNYRLIFCHNLKQP
Menidia_menidia                 ---------------------------------------------------------------MRNYRLIFCHNLKQP
Porichthys_notatus              ---------------------------------------------------------------MRNYRLIFCHNLKQP
Oryzias_latipes                 ---------------------------------------------------------------MRNYRLIFCHNLKQP
Tripterygion_delaisi            ---------------------------------------------------------------MRNYRLIFCHNLKQP
Buceros_rhinoceros_silvestri    ---------------------------------------------------------------MRNYRLIFCHNLKQP
Caprimulgus_carolinensis        ---------------------------------------------------------------MRNYRLIFCHNLKQP
Cariama_cristata                ---------------------------------------------------------------MRNYRLIFCHNLKQP
Canis_lupus_familiaris          ---------------------------------------------------------------MRNYRLIFCHNLKQP
Hynobius_retardatus             ---------------------------------------------------------------MRNYRLIFCHNLKQP
Micropterus_floridanus          ---------------------------------------------------------------MRNYRLIFCHNLKQP
Balaenoptera_acutorostrata_s    ---------------------------------------------------------------MRNYRLIFCHNLKQP
Pygocentrus_nattereri           ---------------------------------------------------------------MRNYRLIFCHNLKQP
Calidris_pugnax                 ---------------------------------------------------------------MRNYRLIFCHNLKQP
Callorhinchus_milii             ---------------------------------------------------------------MRNYRLIFCHNLKQP
Pteropus_vampyrus               ---------------------------------------------------------------MRNYRLIFCHNLKQP
Maylandia_zebra                 ---------------------------------------------------------------MRNYRLIFCHNLKQP
Pterocles_gutturalis            ---------------------------------------------------------------MRNYRLIFCHNLKQP
Clupea_harengus                 --------------------------------------MRKPFQ-------LSEVLHGICGLSMRNYRLIFCHNLKQP
Latimeria_chalumnae             ---------------------------------------------------------------MRNYRLIFCHNLKQP
Colius_striatus                 ---------------------------------------------------------------MRNYRLIFCHNLKQP
Columba_livia                   ---------------------------------------------------------------MRNYRLIFCHNLKQP
Merops_nubicus                  ---------------------------------------------------------------MRNYRLIFCHNLKQP
Alligator_mississippiensis      ---------------------------------------------------------------MRNYRLIFCHNLKQP
Cuculus_canorus                 ---------------------------------------------------------------MRNYRLIFCHNLKQP
Kryptolebias_marmoratus         ---------------------------------------------------------------MRNYRLIFCHNLKQP
Sarcophilus_harrisii            ---------------------------------------------------------------MRNYRLIFCHNLKQP
Galeopterus_variegatus          ---------------------------------------------------------------MRNYRLIFCHNLKQP
Monodelphis_domestica           ---------------------------------------------------------------MRNYRLIFCHNLKQP
Anopheles_funestus              ---------------------------------------------------------------MLNYRLVSTHFLKQP
Esox_lucius                     ---------------------------------------------------------------MRNYRLIFCHNLKQP
Falco_cherrug                   ---------------------------------------------------------------MRNYRLIFCHNLKQP
Gallus_gallus                   ---------------------------------------------------------------MRNYRLIFCHNLKQP
Gavia_stellata                  ---------------------------------------------------------------MRNYRLIFCHNLKQP
Balearica_regulorum_gibberic    ---------------------------------------------------------------MRNYRLIFCHNLKQP
Erinaceus_europaeus             ---------------------------------------------------------------MRNYRLIFCHNLKQP
Oryctolagus_cuniculus           ---------------------------------------------------------------MRNYRLIFCHNLKQP
Eudarcia_simulatricella         ---------------------------------------------------------------MLNYRLVSTHFLKQP
Tauraco_erythrolophus           ---------------------------------------------------------------MRNYRLIFCHNLKQP
Opisthocomus_hoazin             ---------------------------------------------------------------MRNYRLIFCHNLKQP
Plecoglossus_altivelis          ----------------------------------------MLFAPVVFDLLTSEILNRICGLSMRNYRLIFCHNLKQP
Scleropages_formosus            ---------------------------------------------------------------MRNYRLIFCHNLKQP
Lepidothrix_coronata            ---------------------------------------------------------------MRNYRLIFCHNLKQP
Nipponia_nippon                 ---------------------------------------------------------------MRNYRLIFCHNLKQP
Trematomus_bernacchii           ---------------------------------------------------------------MRNYRLIFCHNLKQP
Ceratotherium_simum_simum       ---------------------------------------------------------------MRNYRLIFCHNLKQP
Manis_javanica                  ---------------------------------------------------------------MRNYRLIFCHNLKQP
Glycera_dibranchiata            ---------------------------------------------------------------MHGYRLILCRTLKQP
Picoides_pubescens              ---------------------------------------------------------------MRNYRLIFCHNLKQP
Platichthys_stellatus           ---------------------------------------------------------------MRNYRLIFCHNLKQP
Homo_sapiens                    ---------------------------------------------------------------MRNYRLIFCHNLKQP
Loxodonta_africana              ---------------------------------------------------------------MRNYRLIFCHNLKQP
Fulmarus_glacialis              ---------------------------------------------------------------MRNYRLIFCHNLKQP
Nestor_notabilis                ---------------------------------------------------------------MRNYRLIFCHNLKQP
Mus_musculus                    ---------------------------------------------------------------MRNYRLIFCHNLKQP
Salmo_salar                     ---------------------------------------------------------------MRNYRLIFCHNLKQP
Tupaia_chinensis                ---------------------------------------------------------------MRNYRLIFCHNLKQP
Lepisosteus_oculatus            ---------------------------------------------------------------MRNYRLIFCHNLKQP
Ameiurus_nebulosus              ---------------------------------------------MTVFCLTTDVLYRICGPSMRNYRLIFCHNLKQP
Trichechus_manatus_latirostr    ---------------------------------------------------------------MRNYRLIFCHNLKQP
Aptenodytes_forsteri            ---------------------------------------------------------------MRNYRLIFCHNLKQP
Crotalus_horridus               ---------------------------------------------------------------MRNYRLIFCHNLKQP
Tyto_alba                       ---------------------------------------------------------------MRNYRLIFCHNLKQP
Struthio_camelus_australis      ---------------------------------------------------------------MRNYRLIFCHNLKQP
Chelonia_mydas                  ---------------------------------------------------------------MRNYRLIFCHNLKQP
Takifugu_rubripes               ---------------------------------------------------------------MRNYRLIFCHNLKQP
Tinamus_guttatus                ---------------------------------------------------------------MRNYRLIFCHNLKQP
Calypte_anna                    ---------------------------------------------------------------MRNYRLIFCHNLKQP
Apaloderma_vittatum             ---------------------------------------------------------------MRNYRLIFCHNLKQP
Orycteropus_afer_afer           ---------------------------------------------------------------MRNYRLIFCHNLKQP

Conservation
                                ---------------------------------------------------------------*++***+++++****
```

**HG0004.1**

ENSG00000031003 (FAM13B)

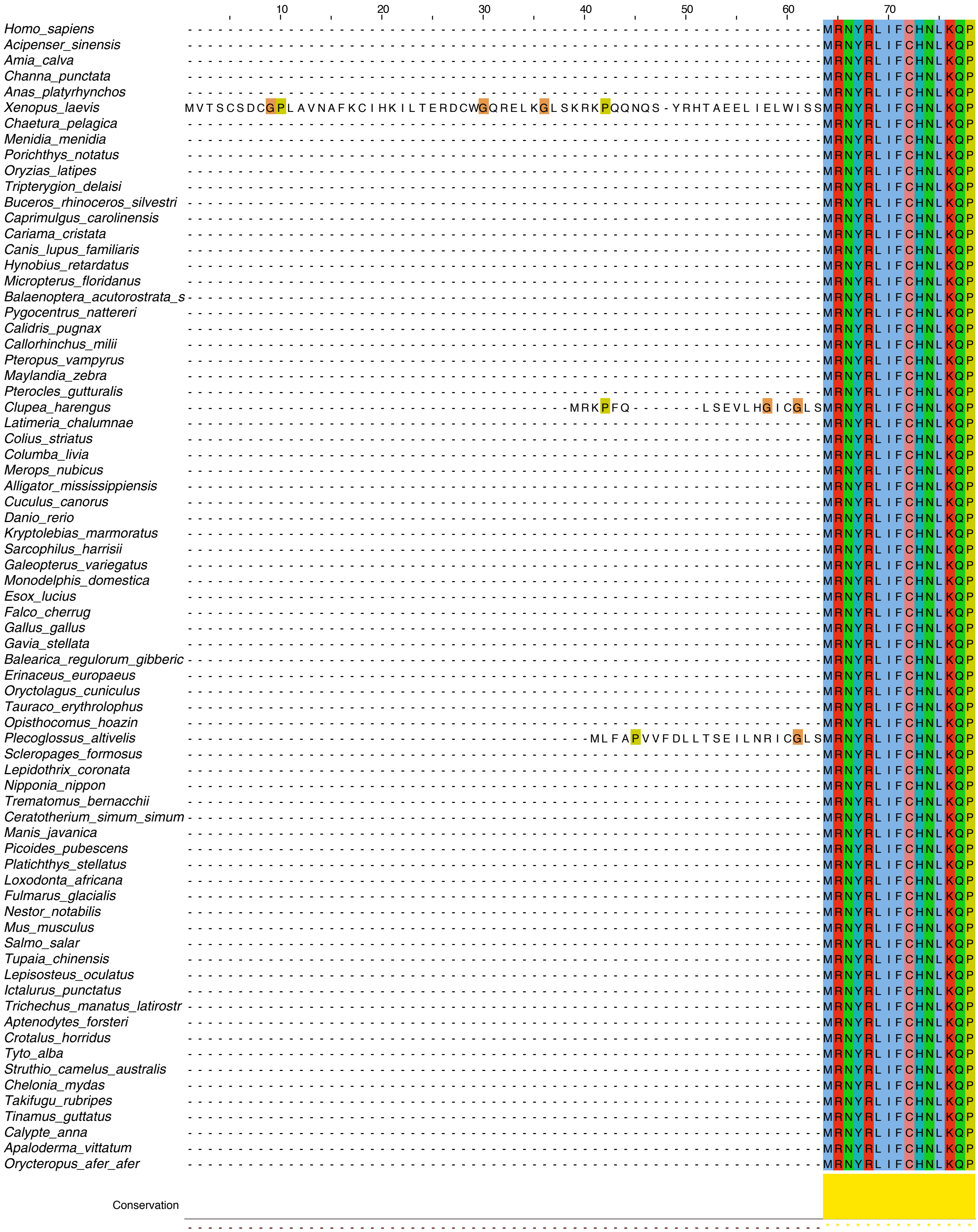

## HG0004.1

ENSGALG00000041706

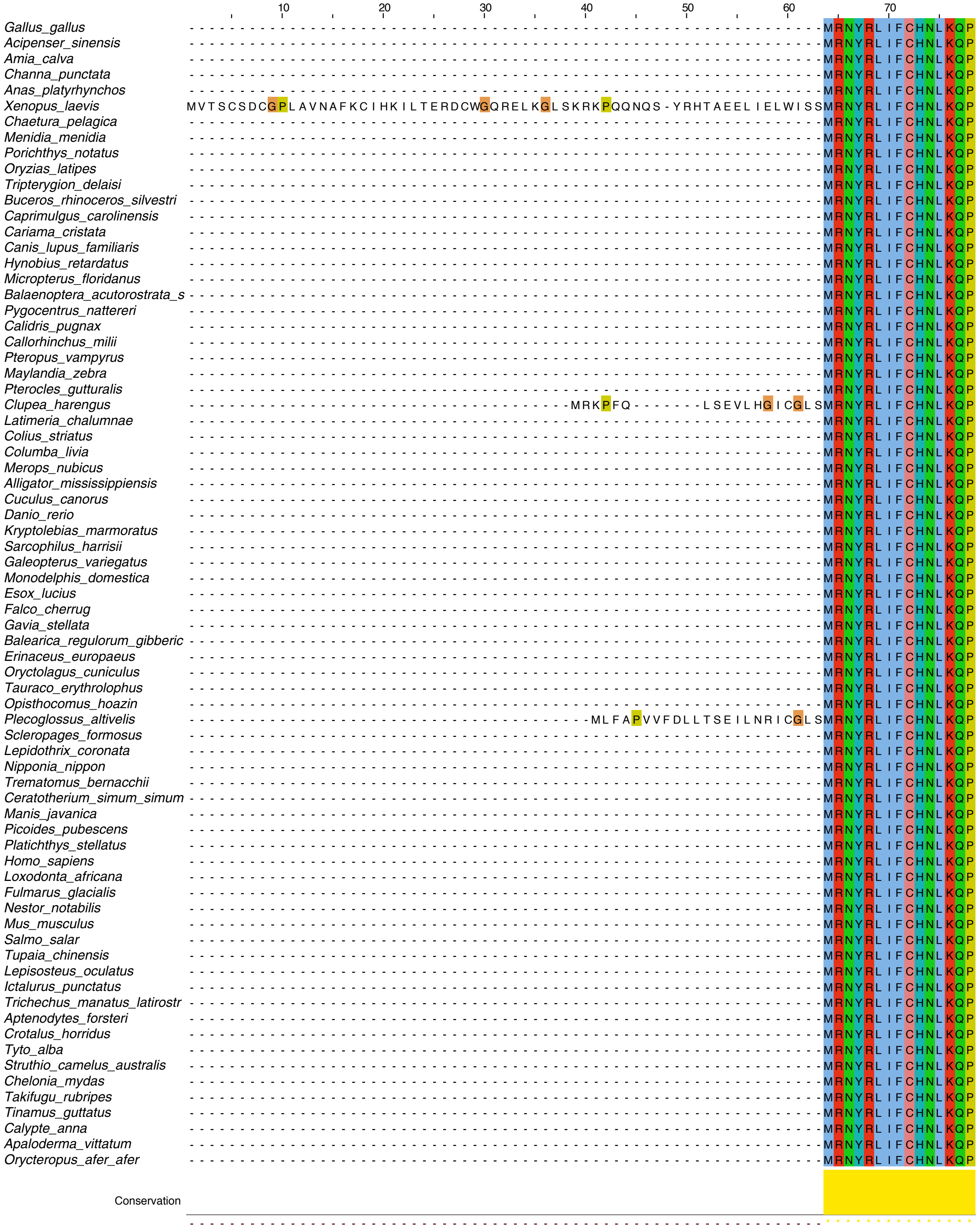

**HG0004.1**

FBgn0028494

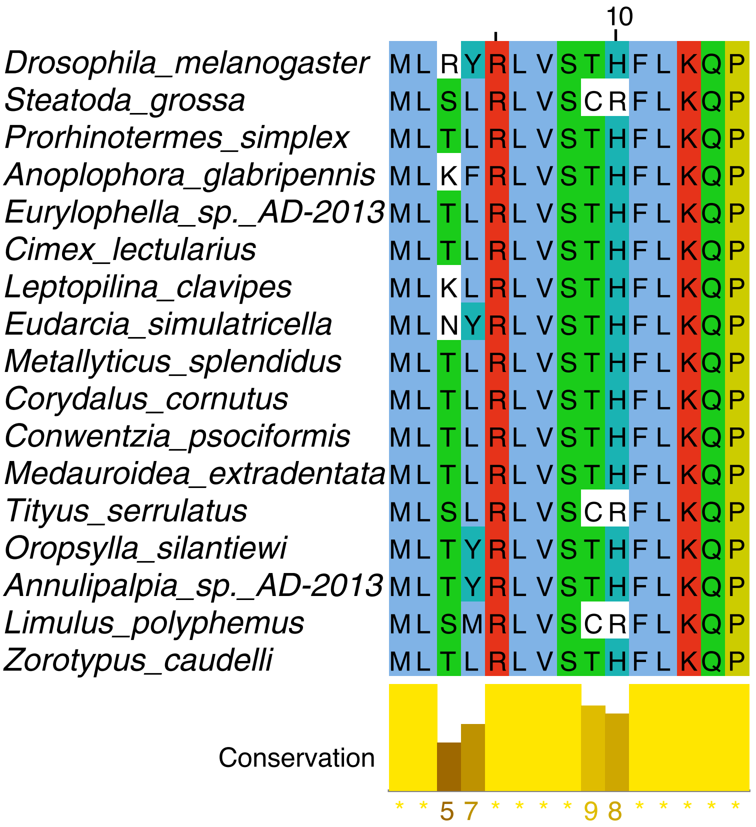

**HG0004.2**

ENSG00000138640 (FAM13A)

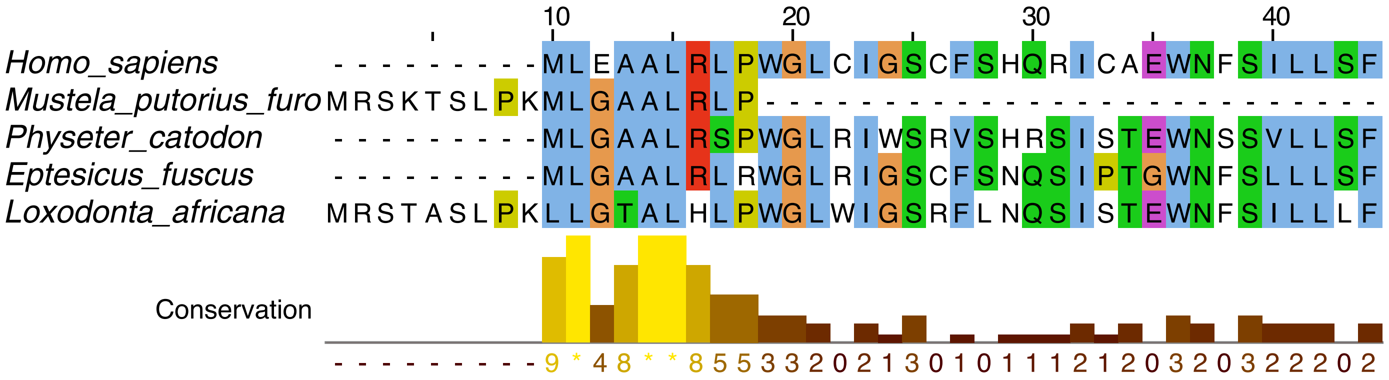

# HG0005

### ENSDARG00000011055 (fbxo9)

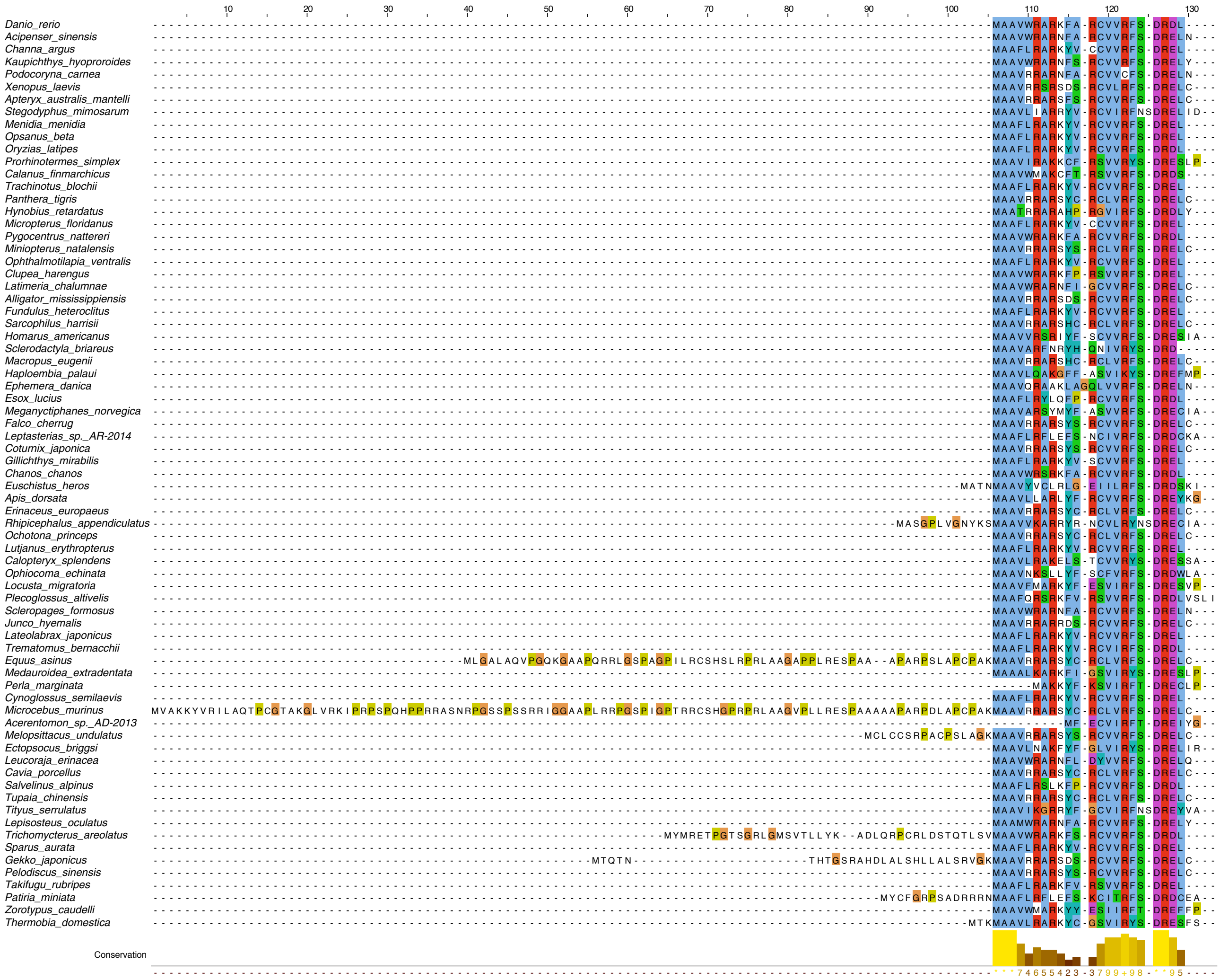

# HG0005

### ENSGALG00000016320

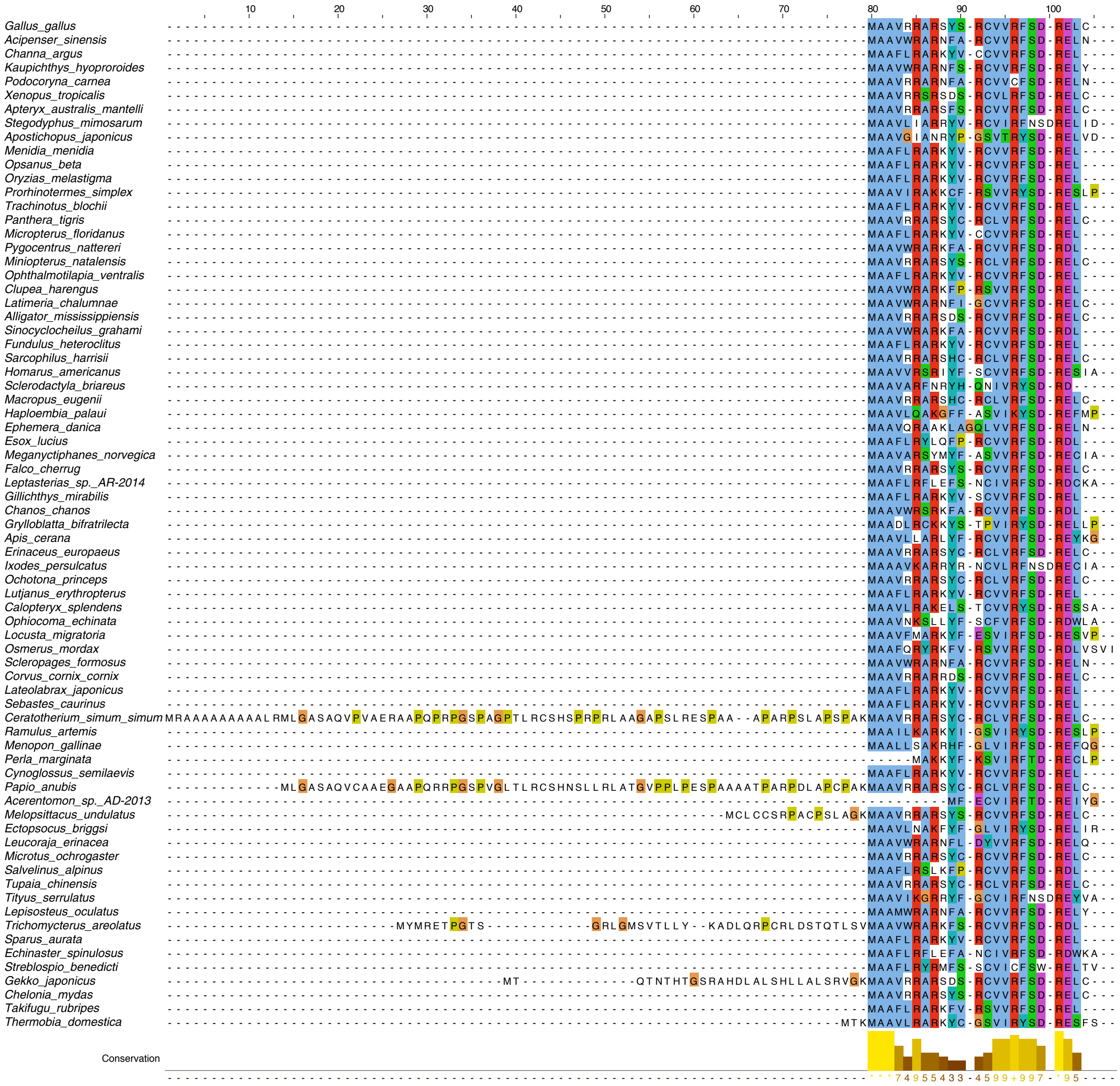

**HG0006**

FBgn0029971 (CG18624)

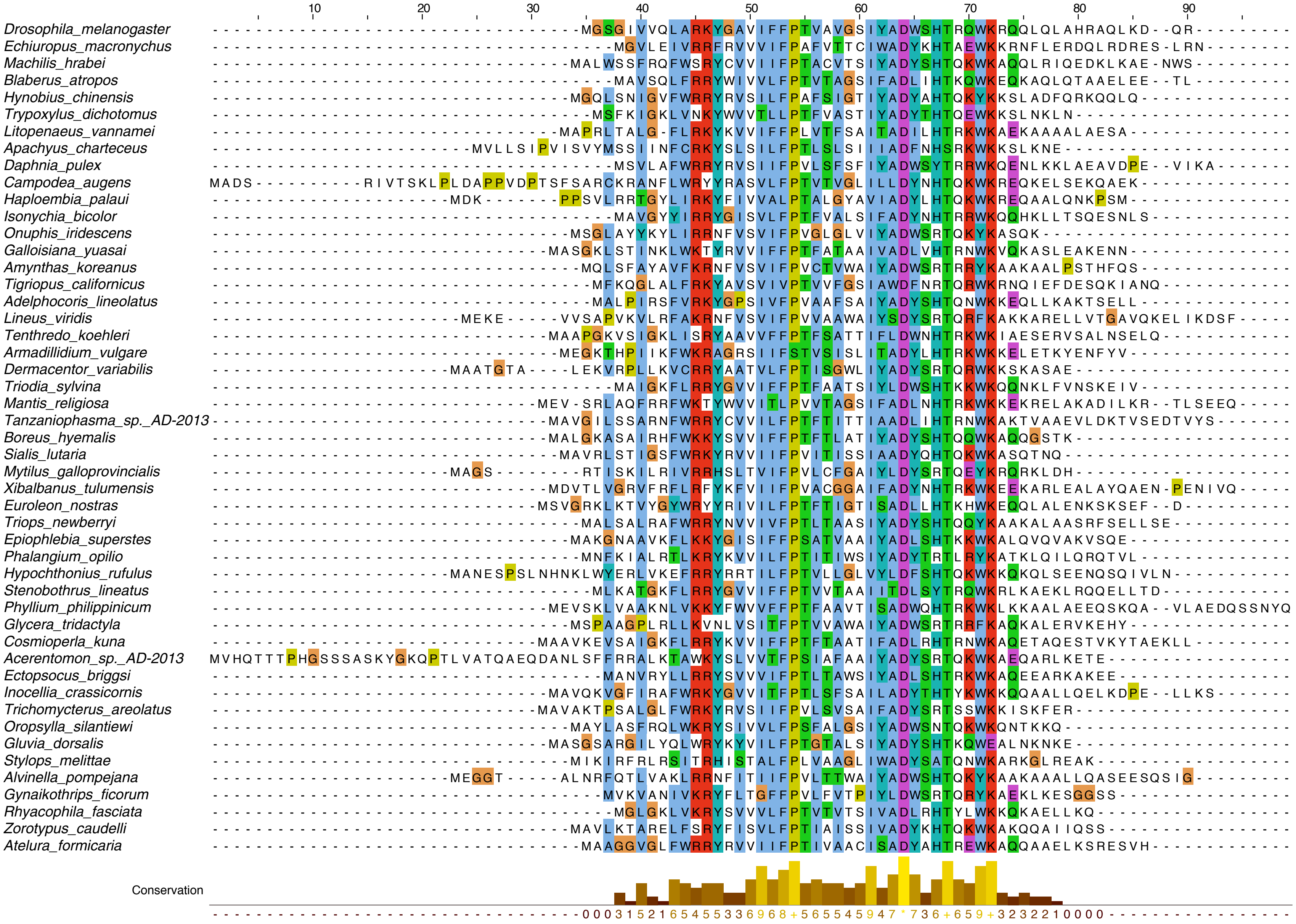

# HG0007

### FBgn0034372 (Gint3)

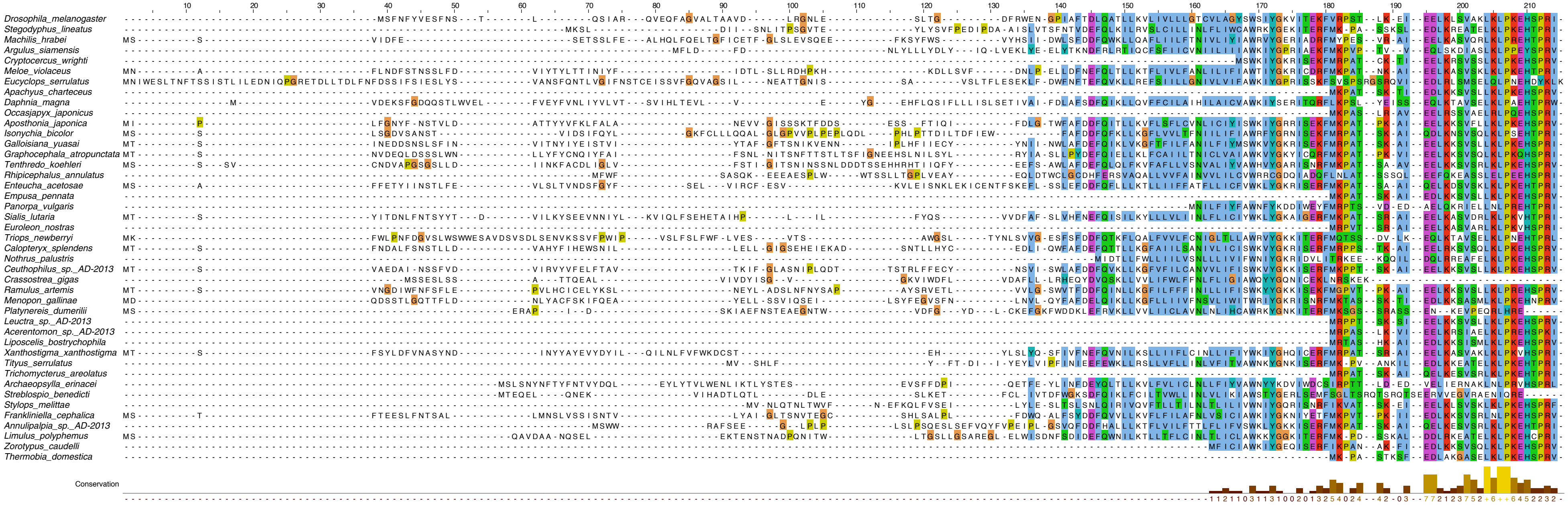

**HG0008**

ENSG00000175567 (UCP2)

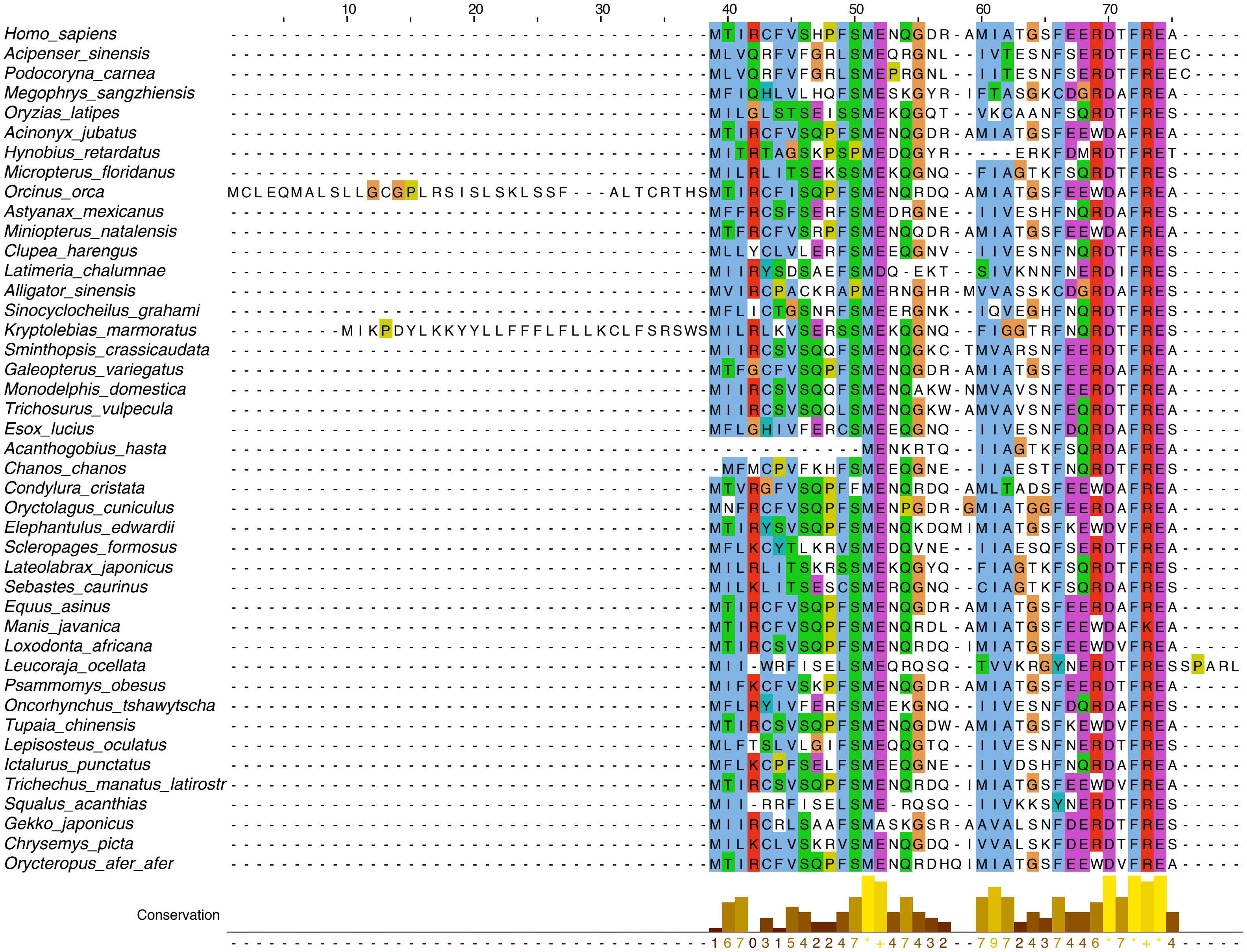

**HG0009**

FBgn0050100 (CG30100)

```
                                       10        20        30        40        50        60        70
Drosophila_melanogaster           -----MKMVQPSRQQILRLYKHLIRYGNHLKLTDKNYFLGRVRHEFRDNRSLTNPVEVEFSFKRGETLLKKGRIL
Talitrus_saltator                 -----------------------------MELSDRRFFEKRIRREFTKQRNS--PEELDTAVQRAEAFLEKGRLL
Latrodectus_geometricus           --------MTVSRQKVLSLYKELLKYSKNLKYTDKSYYLNRLKNEFLENRNLQNASEINFQIQRGKNFLEKKRLL
Meinertellus_cundinamarcensi      -------MQIPNKQQVLCLYKSLLRYGETLRLTDKQYFRTRVRSEFQKNKSSDDSKEIVFQYEKGLKVLNSKRIL
Blattella_germanica               -------MSNPTAREVLKLYRDILKYGRQLQFTDKNYFGQRIRNEFKAGKDLKNQEDINFHFQRGEALLKNKRVV
Calanus_finmarchicus              ------------MSQATLLYKSLLRYGRTLQMTDKDYFYRRVRFEFEKNSVLLEKQEVQFQLDKARQFLKNKRLV
Photinus_pyralis                  --------MHPNAKDILKLYKNLLRYGQELQYTDKNYFARRIKNEFRRNRDITDEADIQFNYKRGISLLEKRAVV
Apachyus_charteceus               -----MASTLPNTRQILSLYKNLMRYGQKLKFTNQSYFYERVRREFENNKTIPED-KREFYYKKGLSLLEKKRVV
Daphnia_pulex                     MNTIFIIMNVPNKTQIFELFRSCIRYSGQLKLSDKRLVSKRIKSEFHKNKTLSNPEEICKAYEKGQSFLSKRRLL
Occasjapyx_japonicus              ---MAAYMGVPSPVQVLRLYRSLVRYGKELKYTDRDFYQRRIRNEFERHRLLTKDEDIRFQYEKGLAFISRKRII
Haploembia_palaui                 -------MSSPSASEVLRLYRELLRFGRQLQFTDQKYFLRRIKTQFQENKHLEDKEQIEFQFNKGQSLLKRKRVV
Baetis_sp._AD-2013                --------MSVARSEVLNLYRSILKYGKDIKLTDKDYFRKRIRQEFRSNQKVTNAEEIQFHVERAKAFLANKRVV
Glomeris_pustulata                --------MTIQRQSVLKLYRELLRYGQTLQYTDKKFYFRRIRKEFEQNRSLTAPADIQYYYEKGLTFLTNKRLI
Galloisiana_yuasai                -------MTNIGGRQIIKLYRDLLRYGKRLEFTDINYFNRRIKTEFLKNKNLASLDEIQFQFERGISLLKNKRVV
Tigriopus_californicus            ---------MSQRSEALRLYKSLLRYGQSLRLTDSAYLHRRCTAEFRTPRST--PAQIERALAQARALLSQRRLL
Cercopis_vulnerata                -------MNKPTVKQIVQLYKDLLRYGNNLKLTDKEYYIDRIHLEFRKNQSLDNPKDIQFCFDRGLALLKNKRVV
Argochrysis_armilla               -----------MRSAILRLYKEILRYGETLKYTDKTYFRERIRTTFRNNKDLTNEKDIDFQFKKGQVFLEKRRVV
Rhipicephalus_appendiculatus      -----MSKPPSKQAVLSLYRSLVRYGHRLELSDKNYYFKRLREEFEARKTLTEPKELQFYFNKGKAFLERKRLI
Papilio_machaon                   --------MSITRSEILNLYKKLMQYSQSLNLTDVQYFRTRVSNEFRRNKTLTKPEDISFAYKKGEALLRKGSVL
Mantis_religiosa                  -------MSIPSPRQVLQLYKSILRYGQELKFTDKEYFYYRINQEFKNNKDIKRTEDIIFNFQRGQALLKNKRVV
Panorpa_vulgaris                  -------MSVPNSSHILRLYKDLLRYGQSLQLTDKLYFRSKIVNEFKKNKTIENHKDISFNYEKGKSLLKKGRVV
Dermanyssus_gallinae              ---MGPVSGAPSVQAMRSLYKQLARYGQQLVLTDRRWFERQLRNEFAKPRVT--AEEQNFAYRKGQRFLELKRLV
Osmylus_fulvicephalus             -------MLIPNNGQVRNLYRHILRYGQTLRYTDKIYFYNRVRNEFKQNKSLTDPADIVFNFKKGEELLRKKTVC
Triops_newberryi                  -------MSGPSPYQVLKLYRSLLRYSSTLQYTDKSFYIKRIRSEFKKKKDLTNEEEIRKAFEKGLSFLSKKRIV
Calopteryx_splendens              -------MCQPSPSQVLSLYRSLLRYGNQLKYTDKKYFIKRIRKEFESNRNLIP-KERGFQYERGLVLLHNKRVI
Hypochthonius_rufulus             --------MSVSREHILRLYKNILRYSNSLVYTDKSYYLQRVKREFRENKQLTDDSDIRFYYEKGITFLNKKRLI
Prosarthria_teretrirostris        -------MNAPTAHEVLRLYRKLLQYGKQLKLTDKKYFFDRIRKEFKTNKTLTNQEDVTFQFKRGLALLENKRVV
Ramulus_artemis                   -------MNTPARVQVLRLYRDLLRYGQNLQLTDKDYFCKRVREEFRNNRVLENKNDLLFHFNKGESLLKTKRVV
Menopon_gallinae                  -------MCVPTTREVLTLYKRLLRYGRQLKYTDQDYYSKRIKQEFQSNKDVKGN-EAIFHYEKGLQLLKLKRVV
Perla_marginata                   -------MSGPTAKQVIQLYRDLLRYSKHLKFTDKNFFTKRVRKEFKTNKRLSNEEDIQFHYQKGLILLKNKSVI
Polydesmus_angustus               -----MSASVVGKSQILHLYRQLLRYGRTLTYTDKNYYYGRIRAEFEKHREAEL-KTQLFQYKKGKALLENGRLI
Acerentomon_sp._AD-2013           --MCWQQMGIPTNGQILKLYRSLLRYGRTLQYTDKDFFKRKVQQEFHRNGAITAPNEIKFQYEKGISFLSRKRLV
Liposcelis_bostrychophila         -------MSAPSRTQVLRLYKDILRYGEELTYTNKTYFKNRVKNEFLSAKSAKDNKEIDNLYQKGVALLMRRRVV
Xanthostigma_xanthostigma         -------MEVPSSTQVLRLYRQLIRYGQKLQYTDKNYYYKRVIAEFKRNRNLEDPKEIKFQFEKGQSLLKRKSVI
Trichomycterus_areolatus          ---------MPSRSEILKLYKQLLRYGQQLRLTDKSYYQERINQEFKQHKNLASNKDITFNFEKGKKLLEKGLVM
Ctenocephalides_felis             -------MNTPSAAQVLRLYKNLLRYGQNLQYTDKDYFCNRIRKEFRNNKTCSRTEDLIFNFEVCI---------
Stylops_melittae                  -----------MRLKVLNLYKSLMRYGKSLRLTDNEYFLERIRSEFRRNKDLYSNQEIDYHFKKGETLLKKKLLV
Symphylella_vulgaris              --------MNPTRVQILSLYKTLLRYADNLKYSDTNYLKQRVRVEFSQHRSLTAKKDIEFYFKRGKSFLERKRLV
Frankliniella_cephalica           ------MSTLINSNQVLQLYRSLIRYGQSLKLTDQKYYLKRVRKEFRANKDLQDPEKIEFMYKKGVSLLDRKRLI
Philopotamus_ludificatus          ---------MPSKREILKLYKQLLRYGQQLQLTDKTYFQQRIVNEFKKNKDLSKSDDITFNFKKGQQLLNKGLVL
Zorotypus_caudelli                --------MNPGKPGILILYKSLIRYGKNLKYTDKKYFIERIRQEFKNRKELENEEDIKFAFQRGVSVLNNKRII
Atelura_formicaria                -------MVAPSSKQVLQLYRNLMRYGRQLKYTGKDYYSKRVREEFKRNKSLENDEKKIFHFEKGQALLSQKRIV

Conservation
```

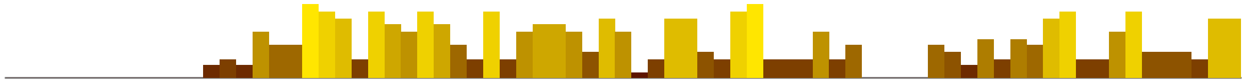

```
--------0000232755*+93+87+853+3788749713994 3+*333735000054263659+337+444399
```

**HG0010.1**

ENSG00000125863 (MKKS)

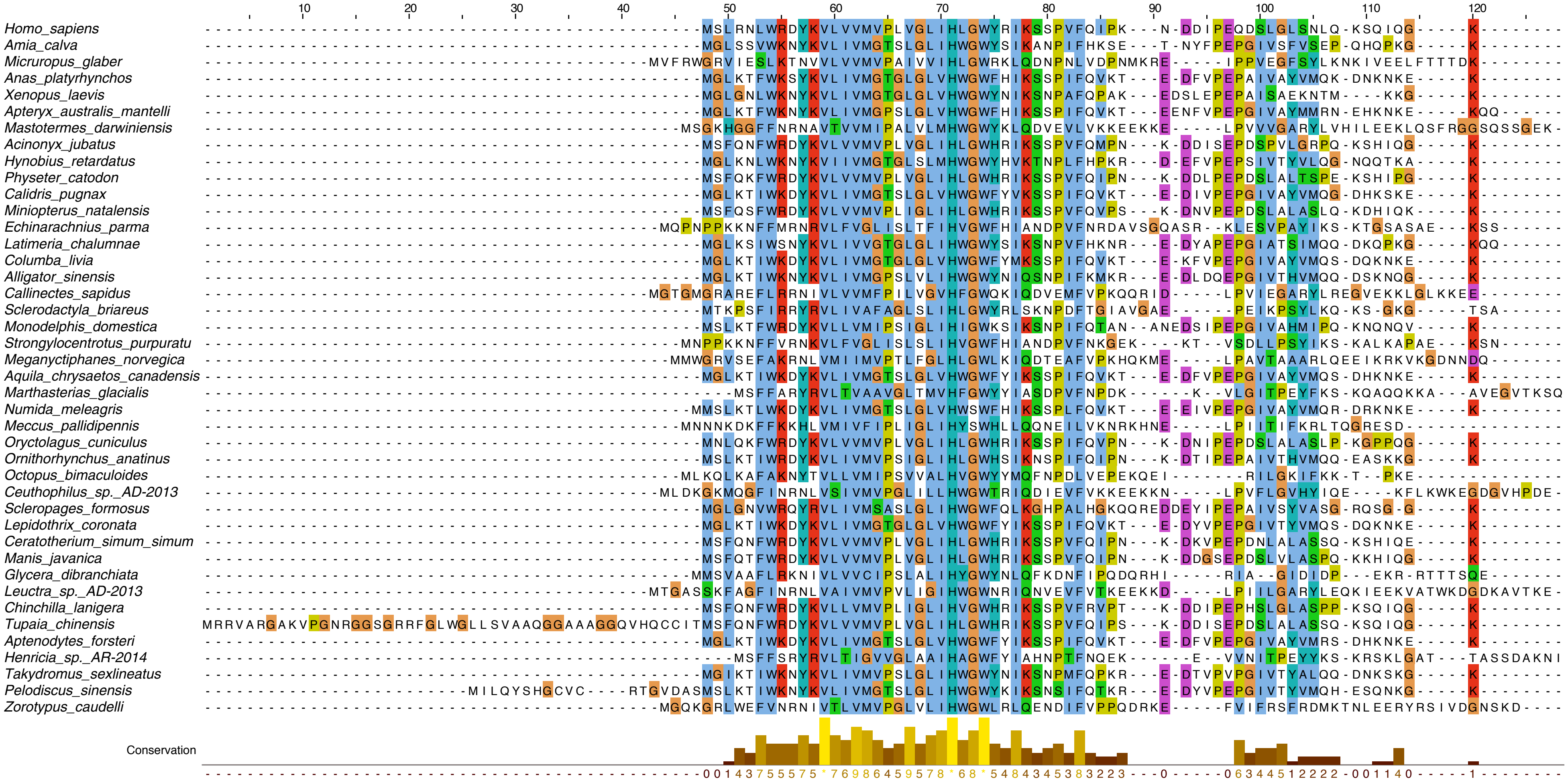

**HG0010.1**

ENSGALG00000009013 (MKKS)

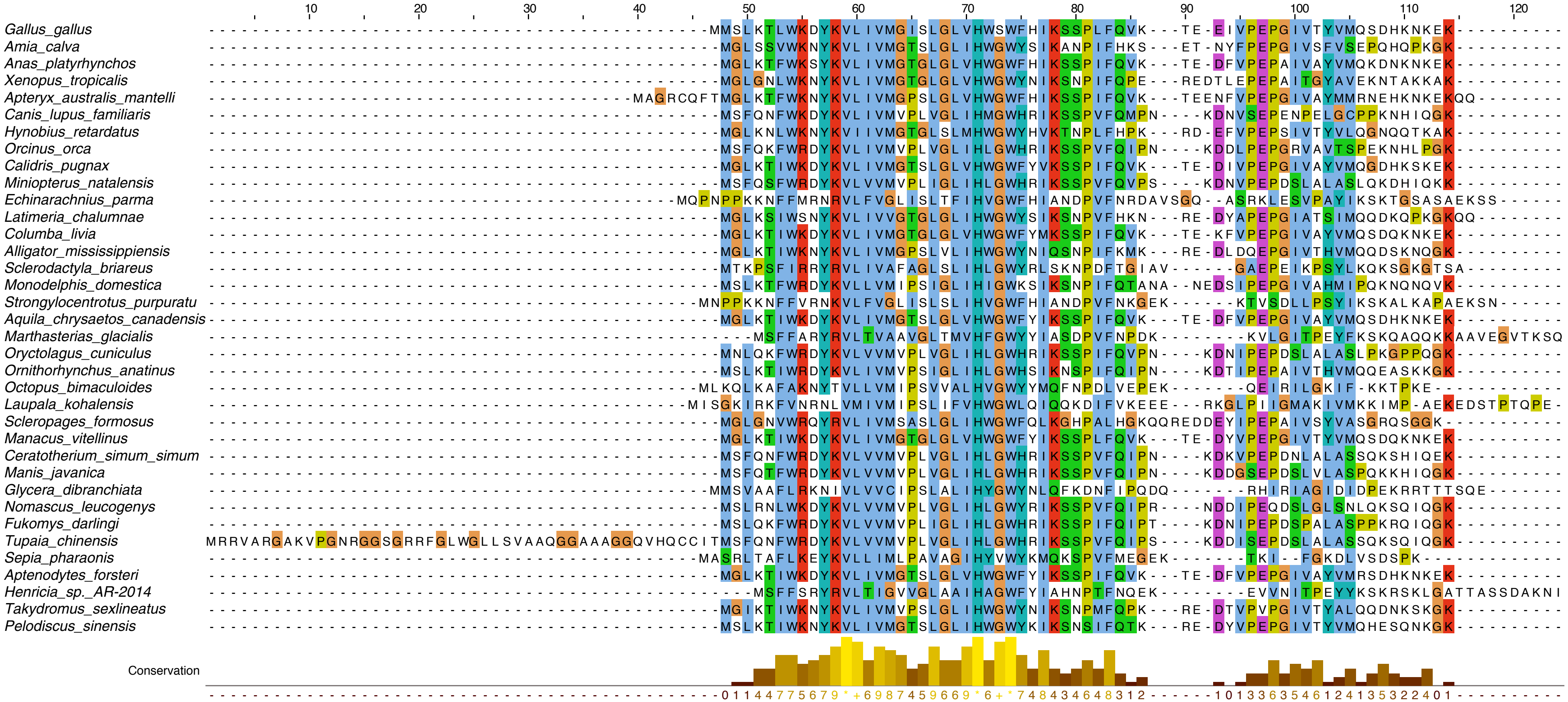

## HG0010.2

ENSG00000125863 (MKKS)

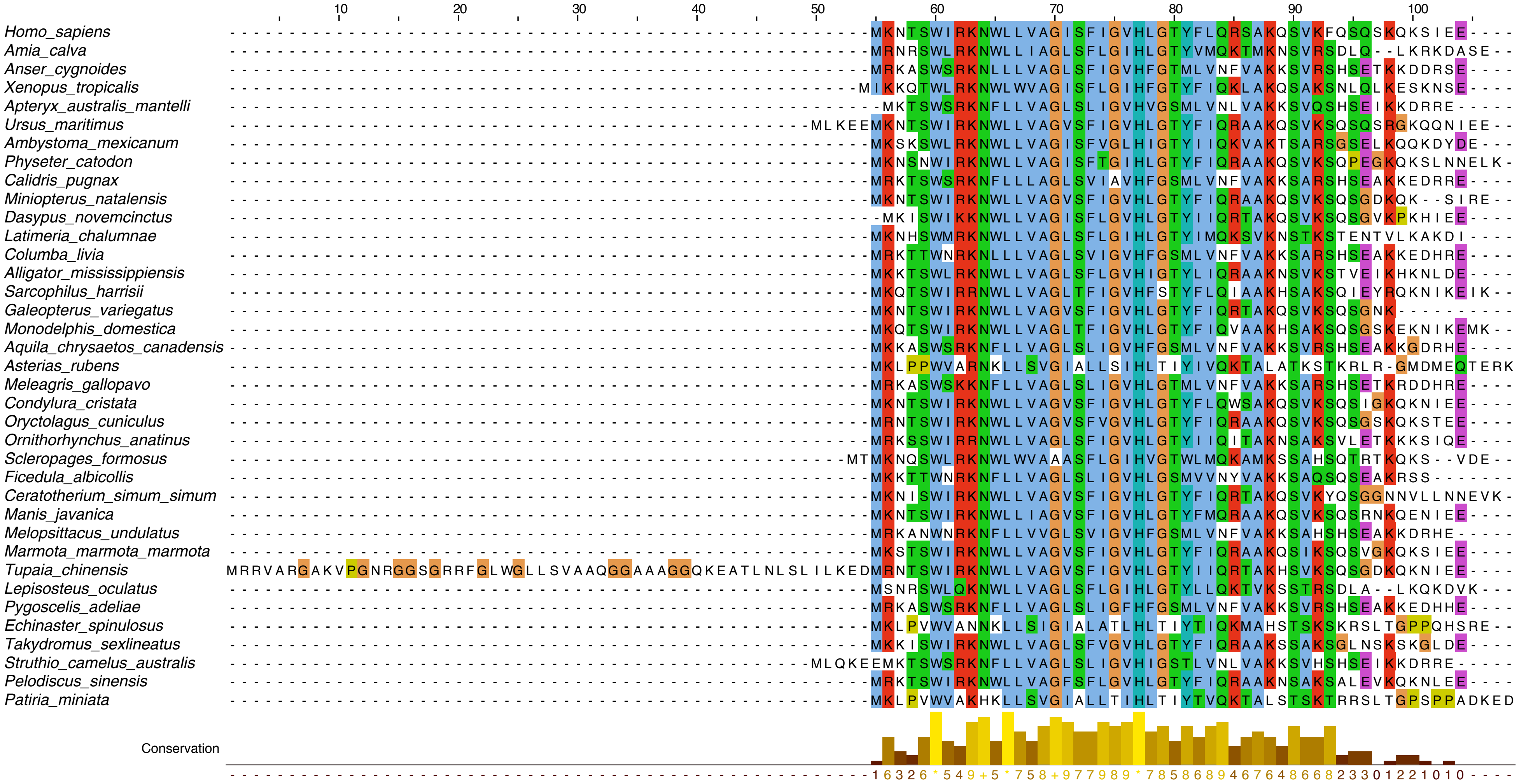

**HG0010.2**

ENSGALG00000009013 (MKKS)

```
                                     10        20        30        40        50
Gallus_gallus                   -----MRKASWSKKNFLLVAGLSLIGVHFGTMLVNFVAKKSARSHSETK--RDNHRE---
Amia_calva                      -----MRNRSWLRKNWLLIAGLSFLGIHLGTYVMQKTMKNSVRSDLQ--LKRKDASE---
Anser_cygnoides                 -----MRKASWSRKNLLLVAGLSFIGVHFGTMLVNFVAKKSVRSHSETK--KDDRSE---
Xenopus_tropicalis              ----MIKKQTWLRKNWLWVAGISFLGIHFGTYFIQKLAKQSAKSNLQLK--ESKNSE---
Apteryx_australis_mantelli      ------MKTSWSRKNFLLVAGLSLIGVHVGSMLVNLVAKKSVQSHSEIK--KDRRE----
Mustela_putorius_furo           -----MKNISWIRKNWLLVAGVSFIGVHLGTYFIQRAAKQSVKSQSTGK--QESIKE---
Ambystoma_mexicanum             -----MKSKSWLRKNWLLVAGISFVGLHIGTYIIQKVAKTSARSGSELK--QQKDYDE--
Balaenoptera_acutorostrata_s----MKNSNWIRKNWLLVAGISFTGIHLGTYLIQRAAKQSVKSQSEGK--QKSLNNELK
Calidris_pugnax                 -----MRKTSWSRKNFLLLAGLSVIAVHFGSMLVNFVAKKSARSHSEAK--KEDRRE---
Eptesicus_fuscus                -----MKTTSWIRKNWLLVAGVSFIGVHLGTYVIQWAAKQSVKSQSGNK--QKSIKE---
Dasypus_novemcinctus            ------MKISWIKKNWLLVAGISFLGVHLGTYIIQRTAKQSVKSQSGVK--PKHIEE---
Latimeria_chalumnae             -----MKNHSWMRKNWLLVAGLSFLGIHLGTYIMQKSVKNSTKSTENTVLKAKDI-----
Columba_livia                   -----MRKTTWNRKNLLLVAGLSVIGVHFGSMLVNFVAKKSARSHSEAK--KEDHRE---
Alligator_mississippiensis      -----MKKTSWLRKNWLLVAGLSFLGVHIGTYLIQRAAKNSVKSTVEIK--HKNLDE---
Sarcophilus_harrisii            -----MKQTSWIRRNWLLVAGLTFIGVHFSTYFLQIAAKHSAKSQIEYR--QKNIKEIK-
Galeopterus_variegatus          -----MKNTSWIRKNWLLVAGVSFIGVHLGTYFIQRTAKQSVKSQSGNK-----------
Monodelphis_domestica           -----MKQTSWIRKNWLLVAGLTFIGVHLGTYFIQVAAKHSAKSQSGSK--EKNIKEMK-
Aquila_chrysaetos_canadensis ---MKKASWSRKNFLLVAGLSLIGVHFGSMLVNFVAKKSVRSHSEAK--KGDRHE---
Condylura_cristata              -----MKNTSWIRKNWLLVAGVSFIGVHLGTYFLQWSAKQSVKSQSIGK--QKNIEE---
Oryctolagus_cuniculus           -----MRNTSWIRKNWLLVAGVSFVGVHLGTYFIQRAAKQSVKSQSGSK--QKSTEE---
Ornithorhynchus_anatinus        -----MRKSSWIRRNWLLVAGLSFIGVHLGTYIIQITAKNSAKSVLETK--KKSIQE---
Scleropages_formosus            ---MTMKNQSWLRKNWLWVAAASFLGIHVGTWLMQKAMKSSAHSQTRTKQKS--VDE---
Corvus_brachyrhynchos           -----MKKTTWSRKNFLLVAGLSLIGVHLGSMLVNFVAKKSAQSHSEAK--RDRHE----
Equus_asinus                    -----MKNISWIRKNWLLVAGVSFLGVHLGTYFIQRAAKQSVKSQSGGN--KVLLNDEVK
Manis_javanica                  -----MKNTSWIRKNWLLIAGVSFIGVHLGTYFMQRAAKQSVKSQSRNK--QENIEE---
Microcebus_murinus              -----MKNTSWVRKNWLLVAGLSLIGVHLGTYFIQRAAKQSVKSQSGAK--QKSIEE---
Melopsittacus_undulatus         -----MRKANWNRKNFLLVVGLSVIGVHFGSMLVNFVAKKSAHSHSEAK--KDRHE----
Dipodomys_ordii                 -----MKNPSWIRKNWLLVAGVSFIGVHLGTYFMQRIAKQSVRSQSADK--QRNVEE---
Tupaia_chinensis                -----MRNTSWIRKNWLLVAGVSFIGVHLGTYIIQRTAKHSVKSQSGDK--QKNIEE---
Lepisosteus_oculatus            -----MSNRSWLQKNWLLVAGLSFLGIHLGTYLLQKTVKSSTRSDLA--LKQKDVK----
Pygoscelis_adeliae              -----MRKASWSRKNFLLVAGLSLIGFHFGSMLVNFVAKKSVRSHSEAK--KEDHHE---
Anolis_carolinensis             -----MKKISWIRKNWLLVAGLTFVGVHLGTYLVQRAAKSSAKSGLDIK--SKGLDE---
Struthio_camelus_australis MLQKEEMKTSWSRKNFLLVAGLSLIGVHIGSTLVNLVAKKSVHSHSEIK--KDRRE----
Trachemys_scripta               -----MRKTSWIKKNWLLVAGLSFLGVHLGTYFIQRAAKNSAKSALEIK--QKDLGE---
```

Conservation

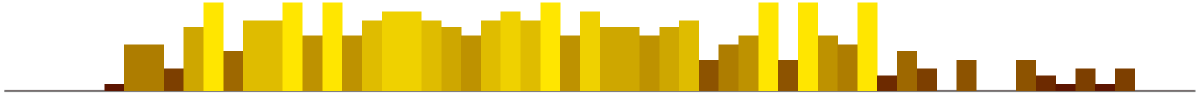

```
-----16638*599*7*79++9879+9*7+88789467*4*76*25304--421313---
```

**HG0010.3**

ENSG00000125863 (MKKS)

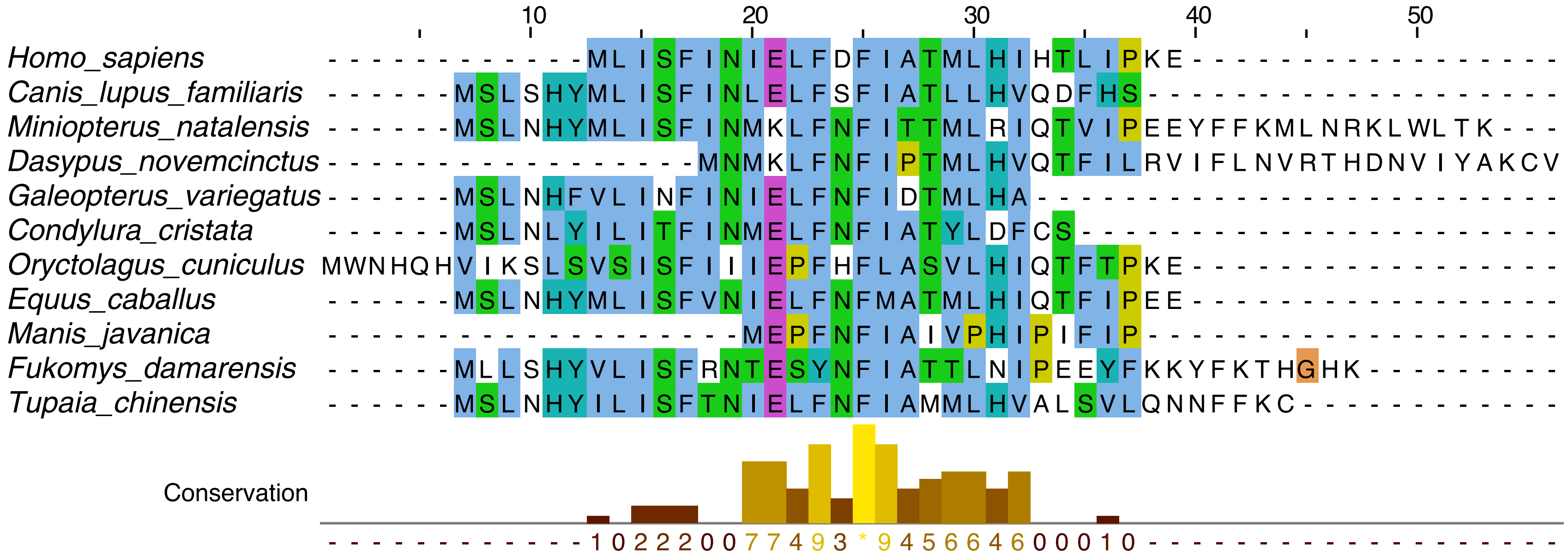

**HG0011**

FBgn0027360 (Tim10)

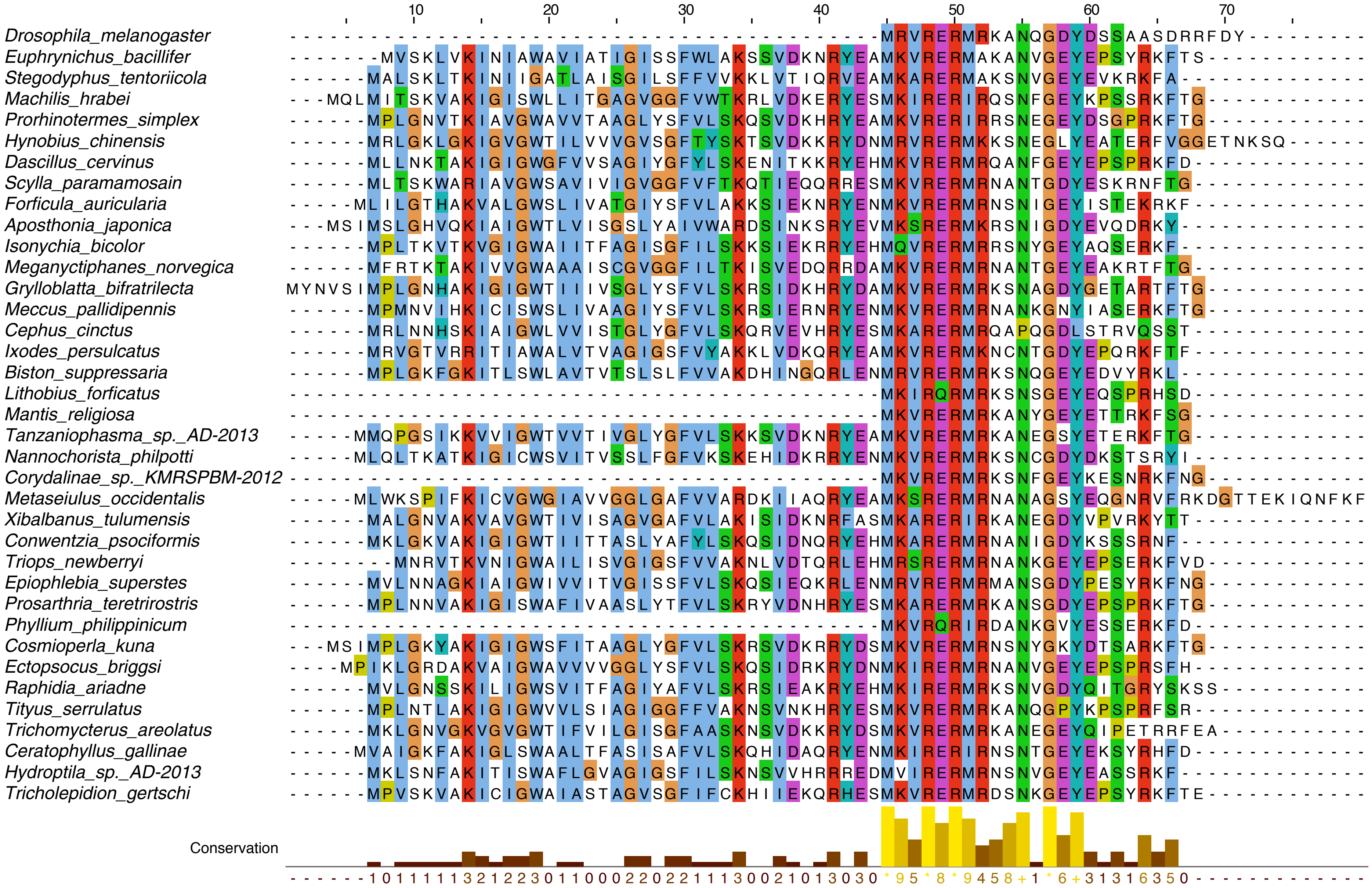

**HG0012**

FBgn0064116 (CG33713)

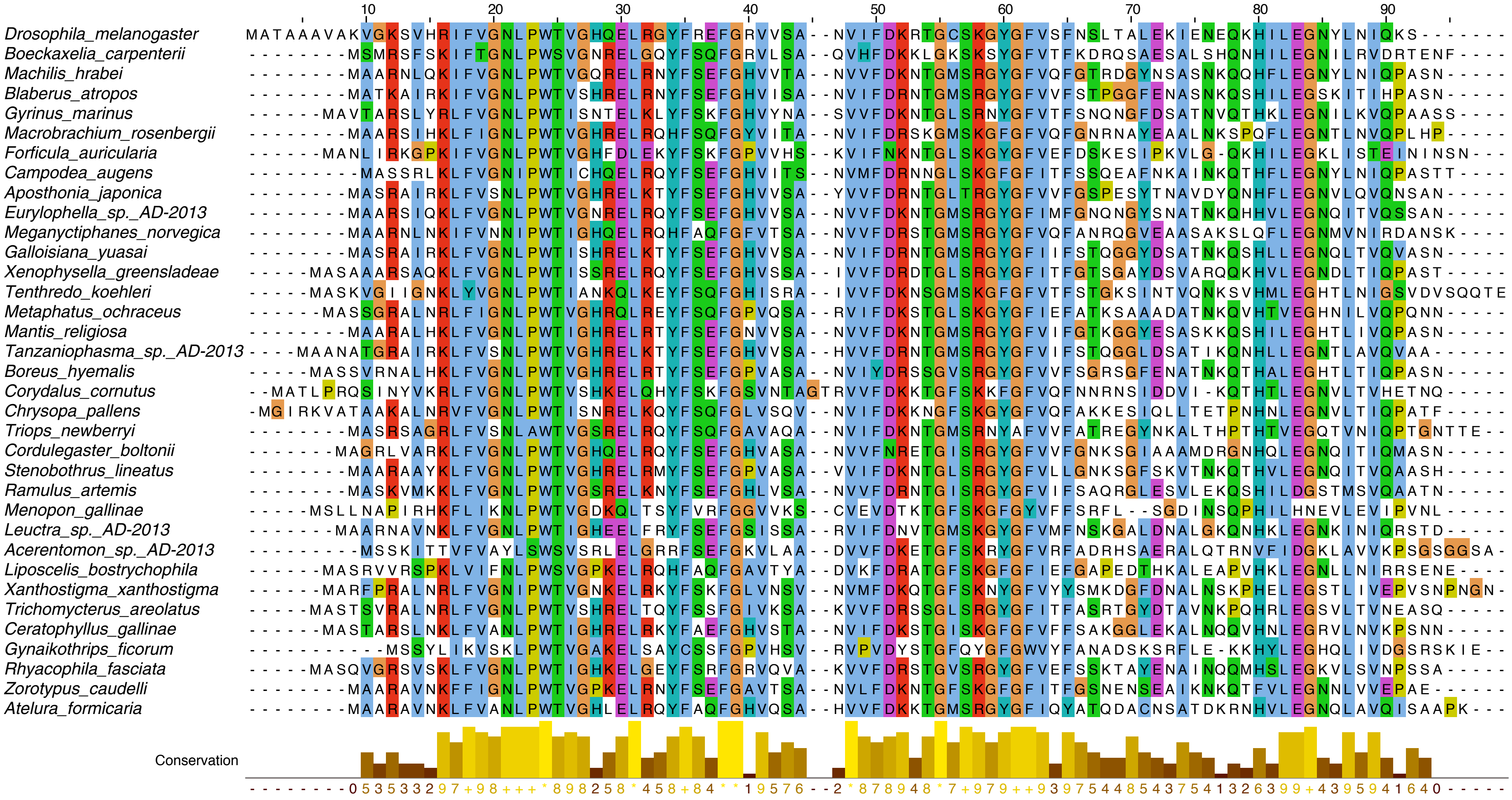

**HG0013**

FBgn0261381 (mtTFB1)

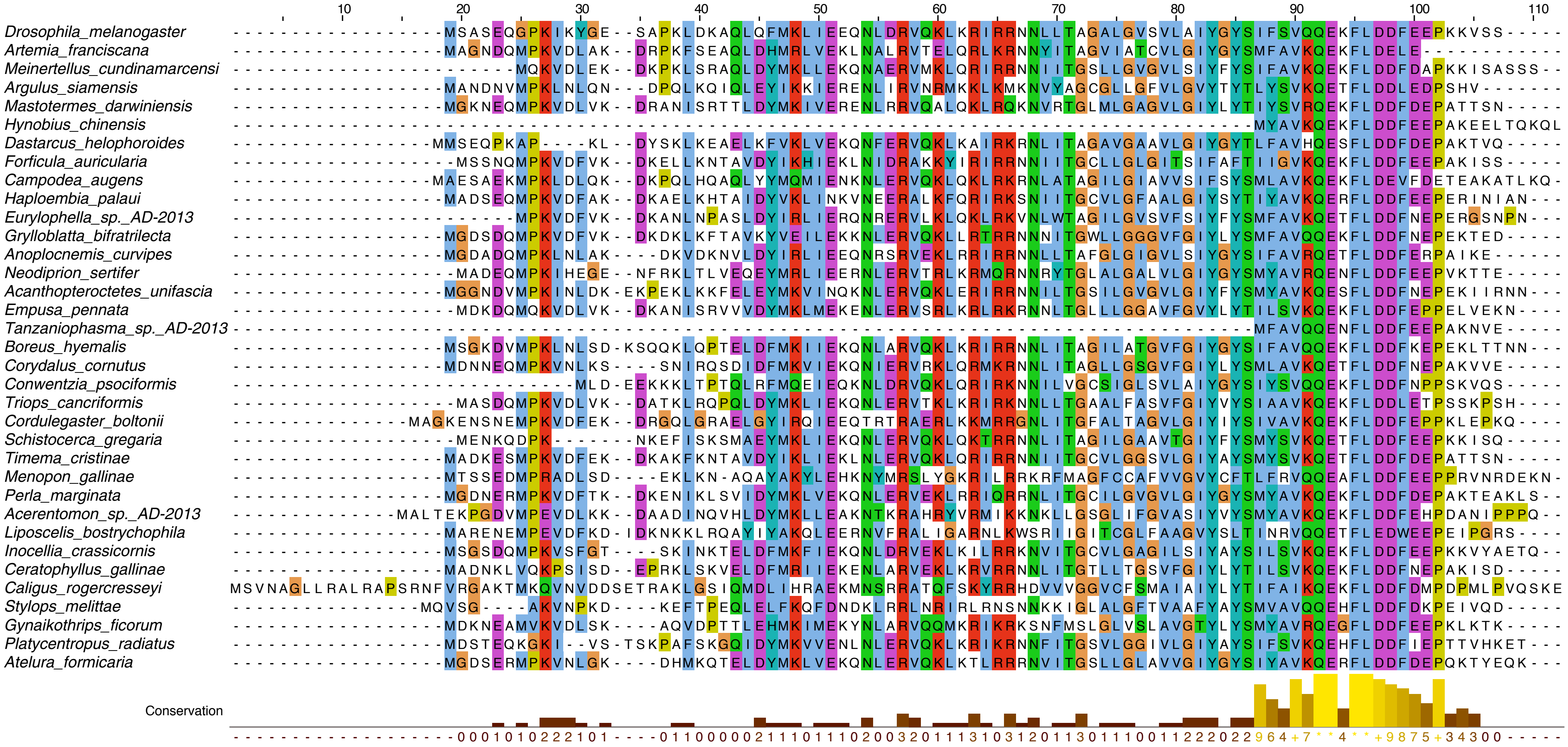

# HG0014

FBgn0260392 (CG42518)

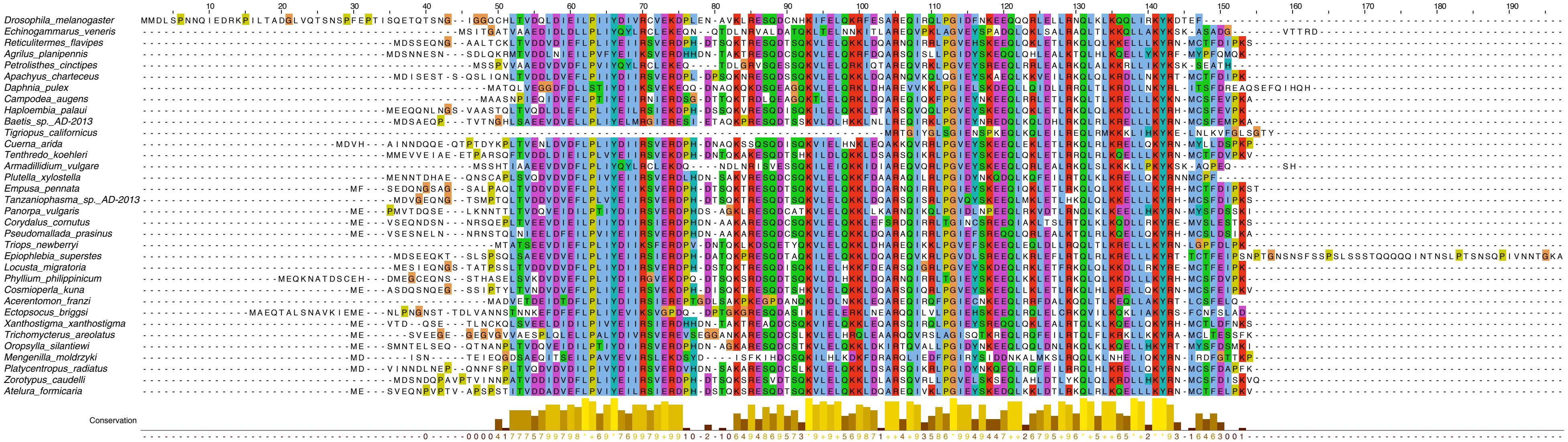

**HG0015**

ENSDARG00000007377 (odc1)

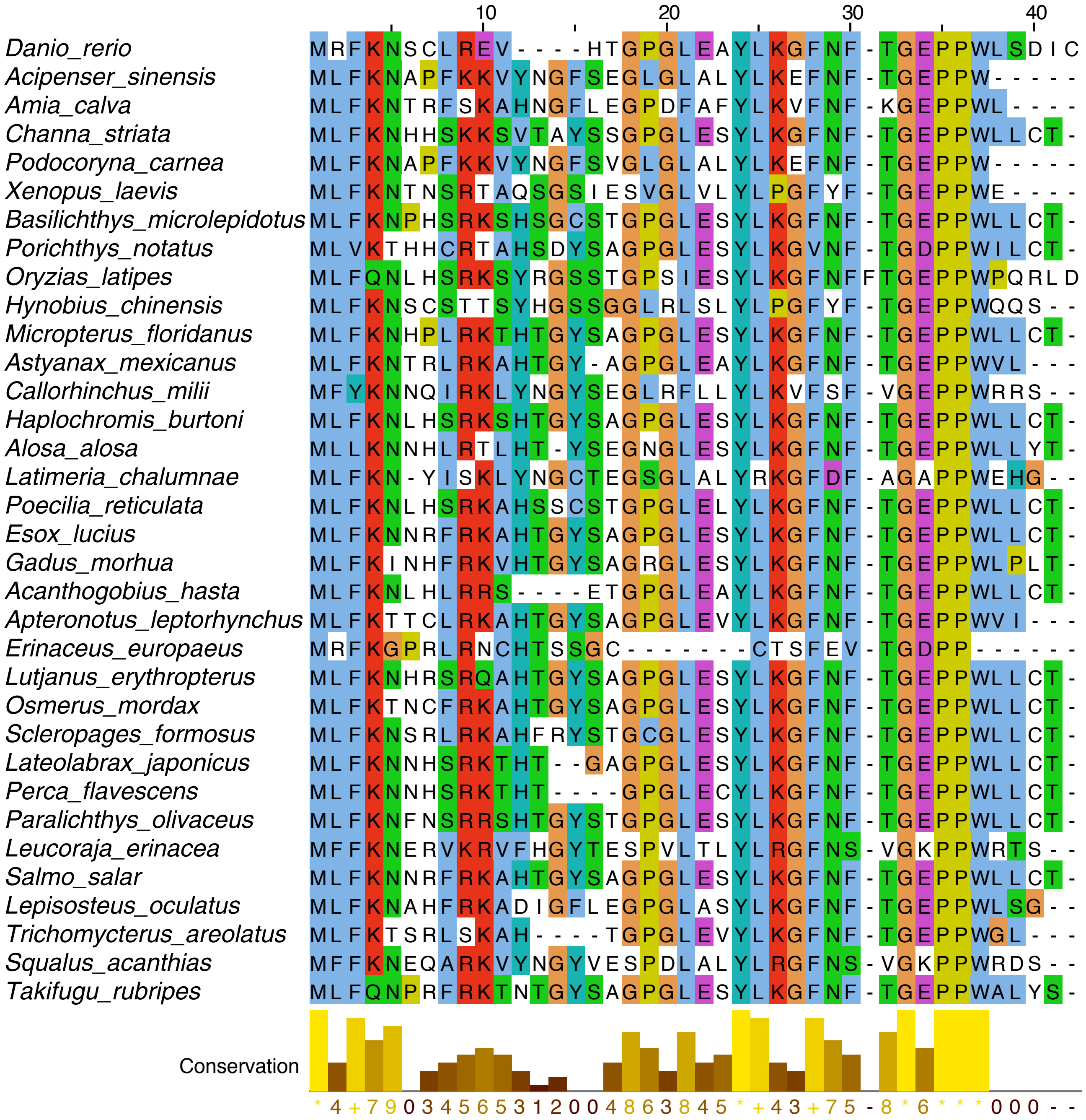

**HG0016**

ENSGALG00000010399 (SELENOT)

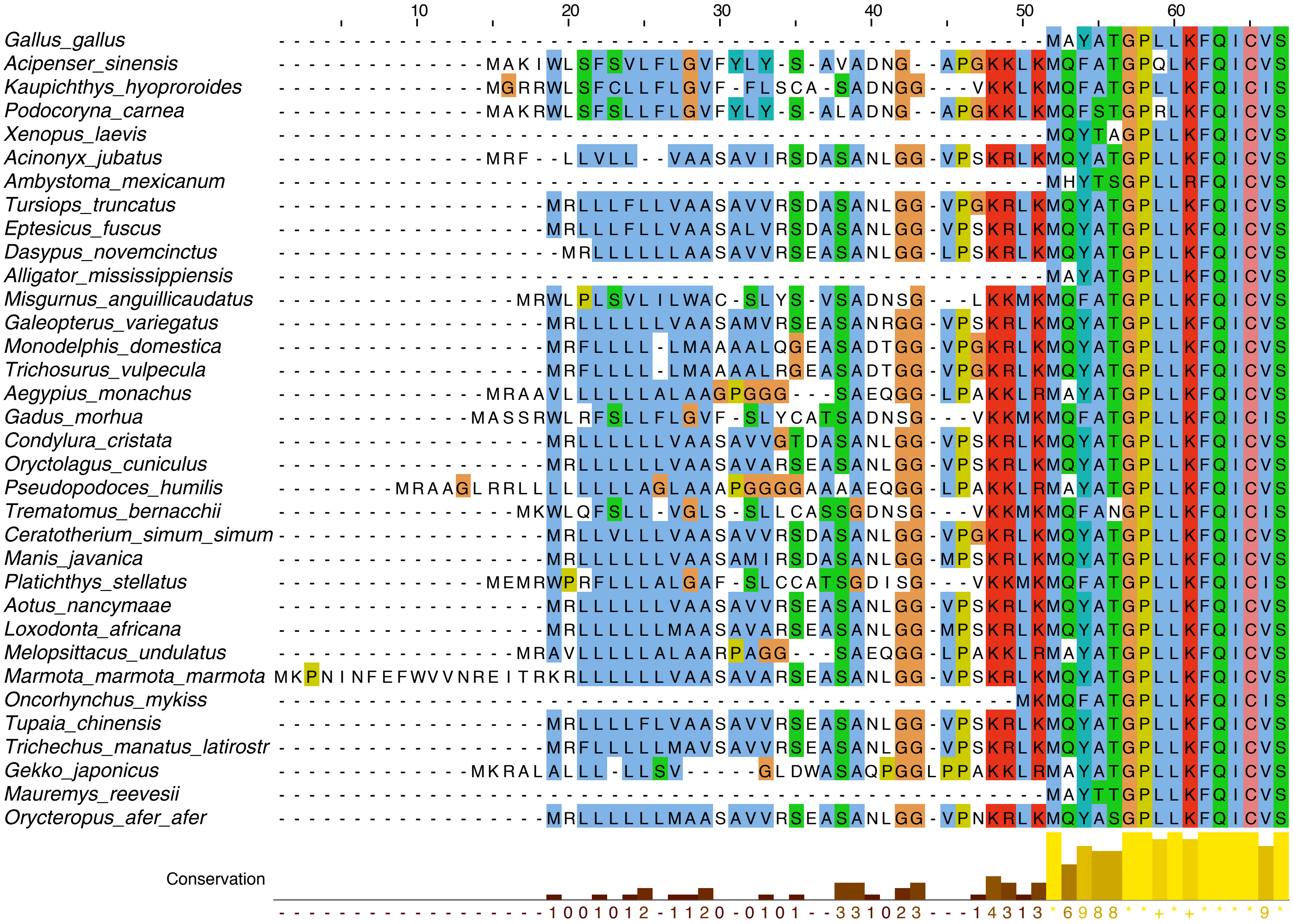

**HG0017**

ENSDARG00000043154 (ucp2)

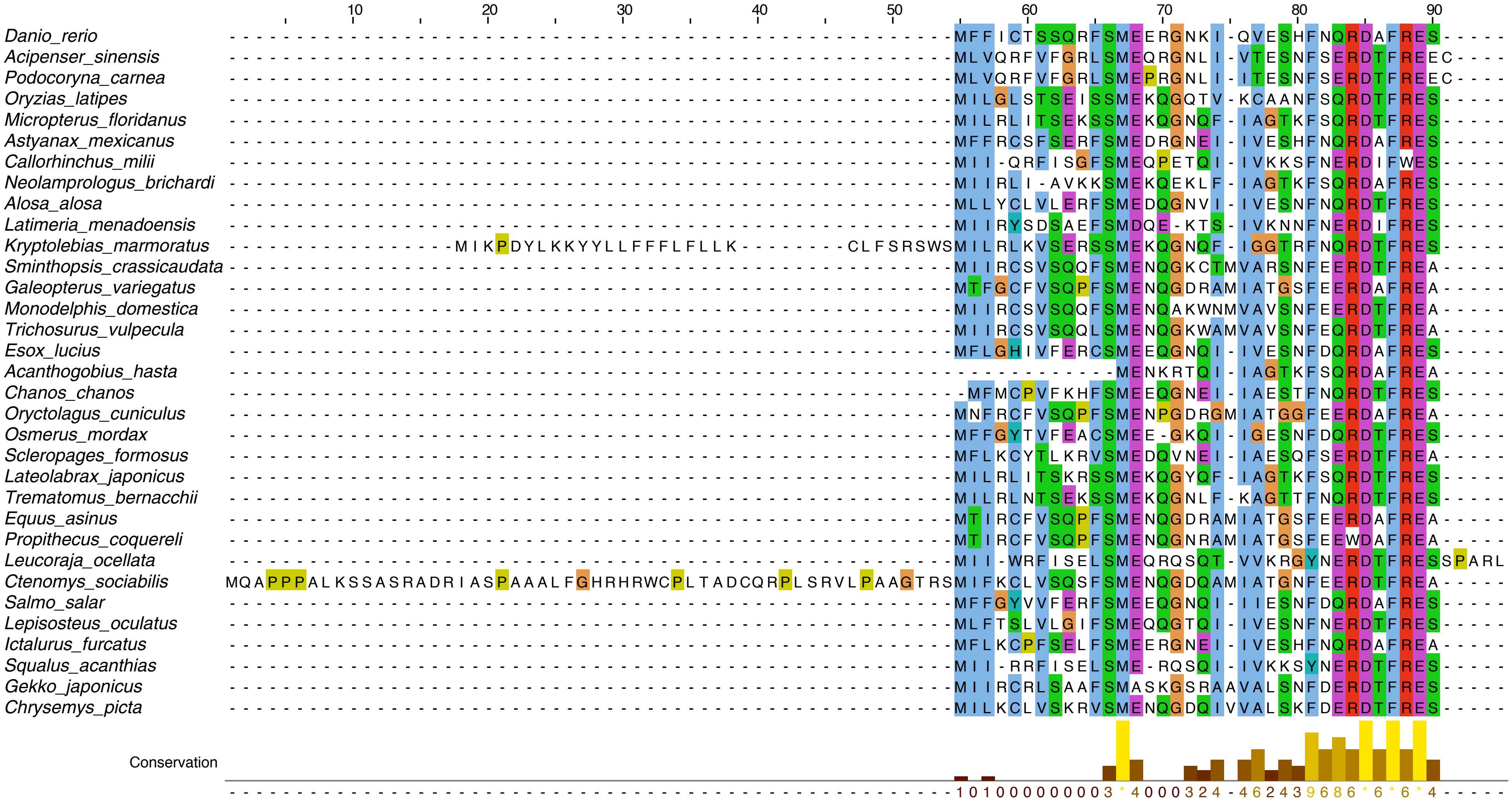

## HG0018

ENSDARG00000053291 (pnrc2)

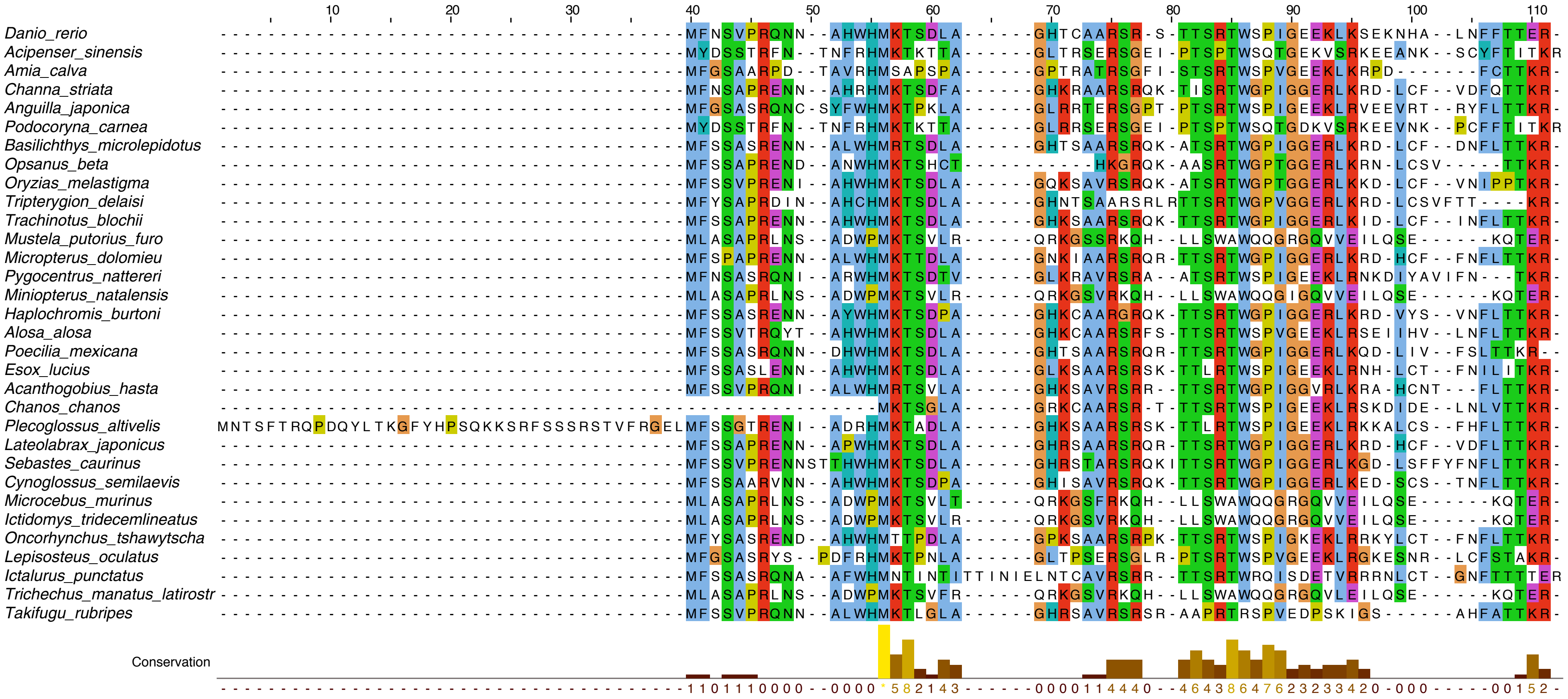

## HG0018

ENSG00000189266 (PNRC2)

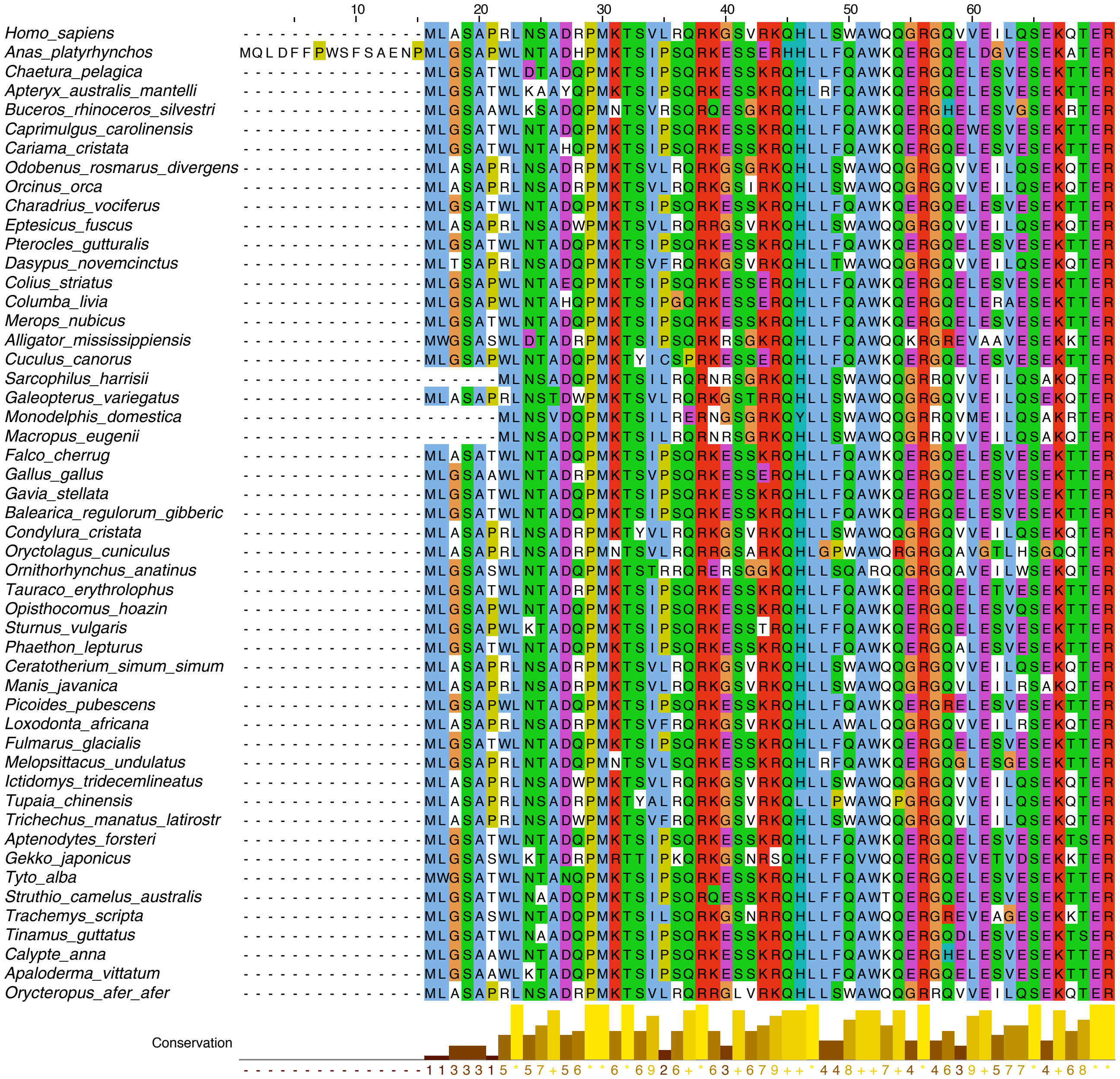

**HG0018**

ENSGALG00000004122 (PNRC2)

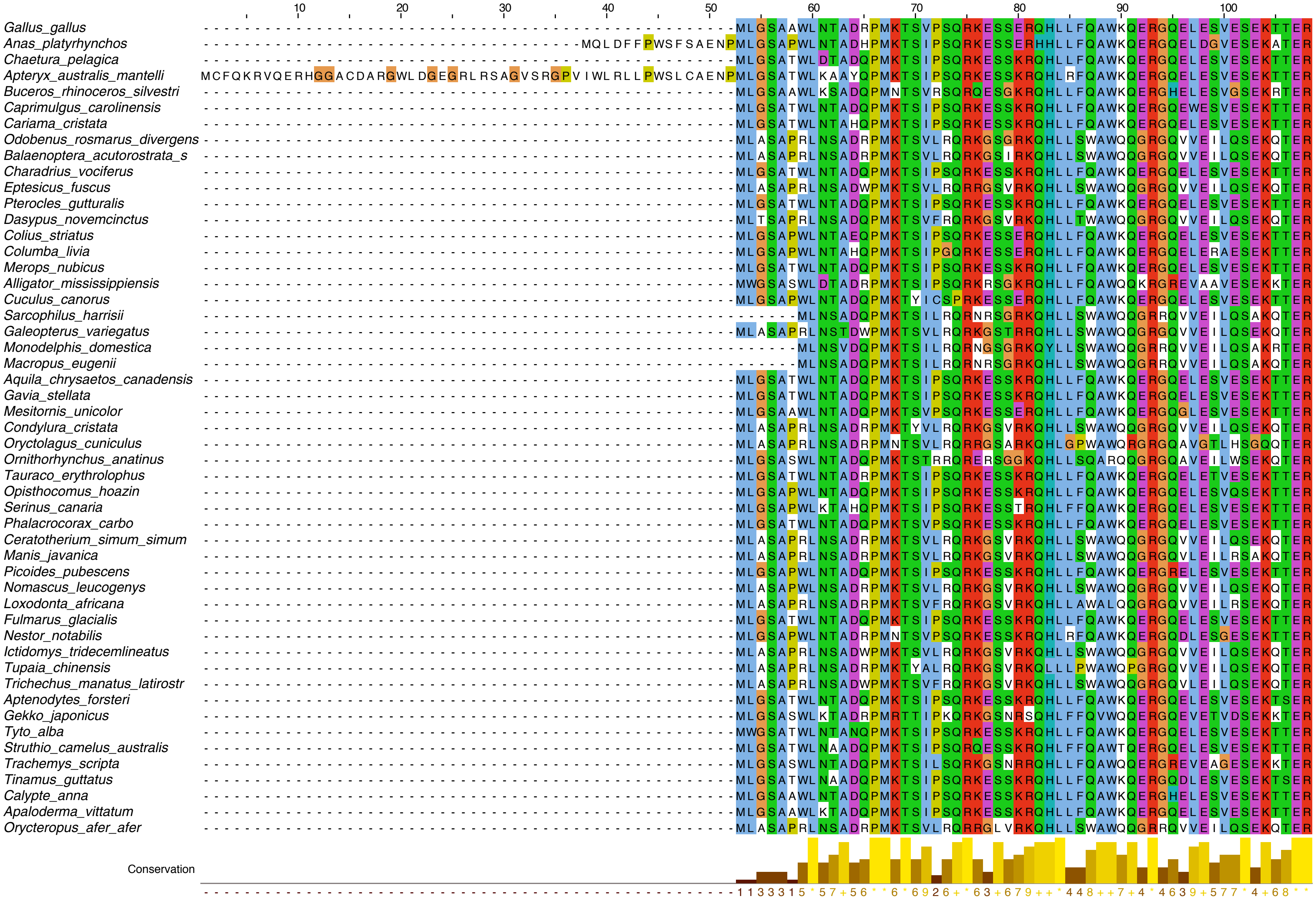

**HG0019**

ENSG00000178397 (FAM220A)

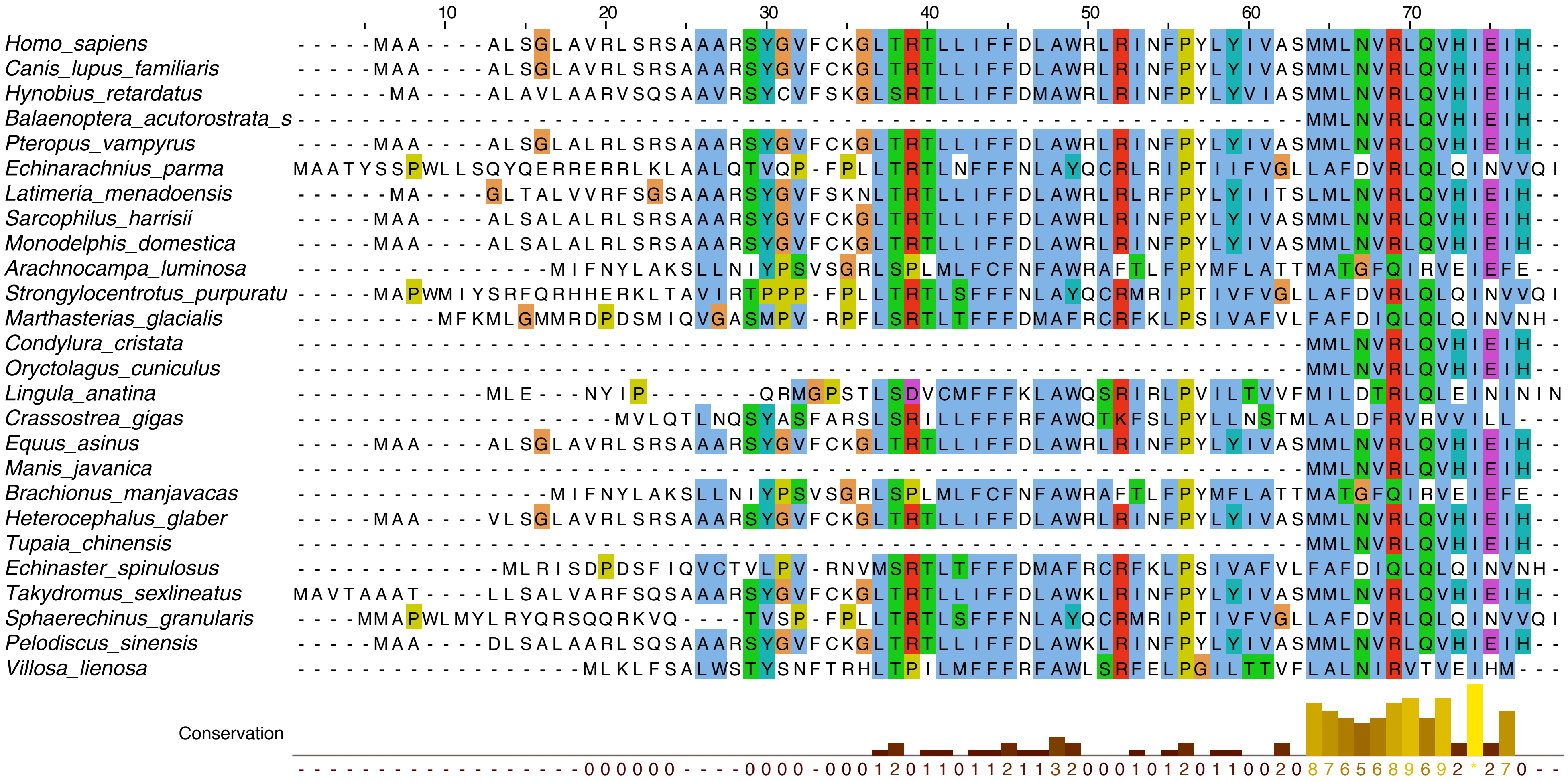

# HG0020

### FBgn0035436 (CG12016)

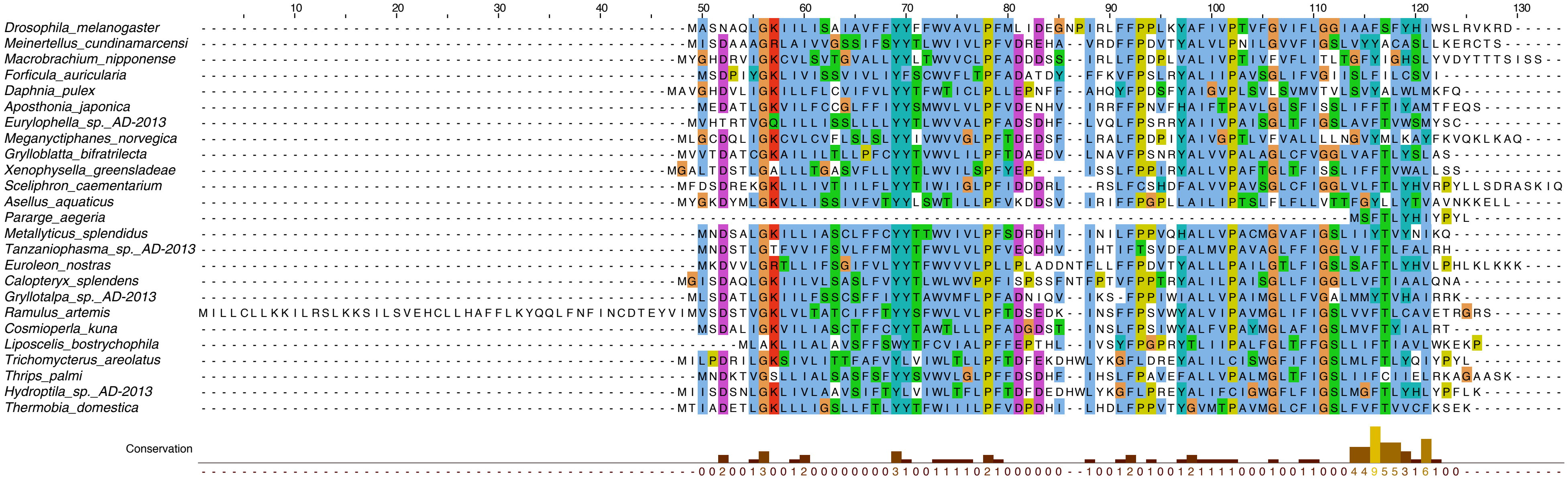

**HG0021.1**

ENSDARG00000068708 (ifrd1)

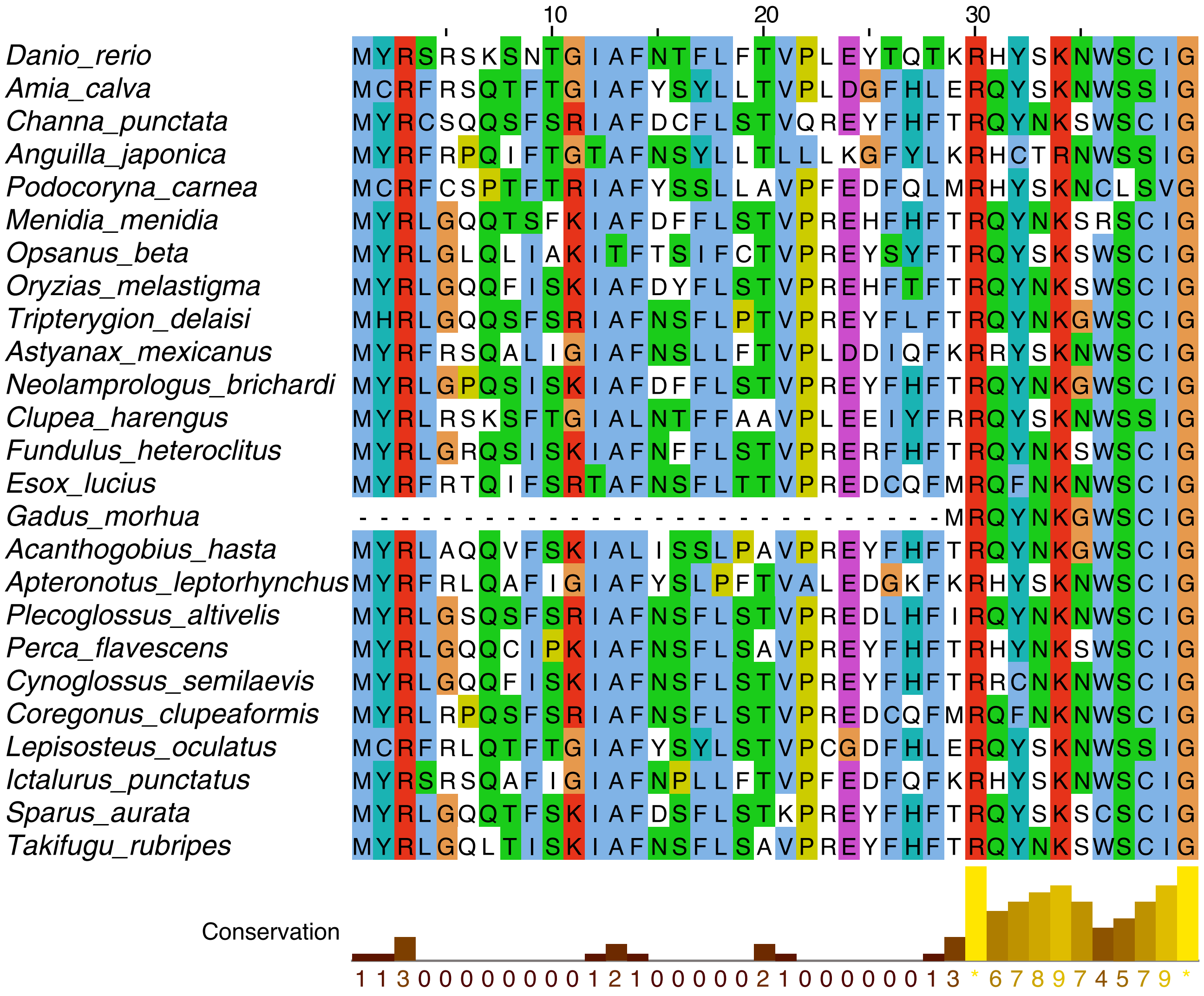

**HG0021.1**

ENSG00000006652 (IFRD1)

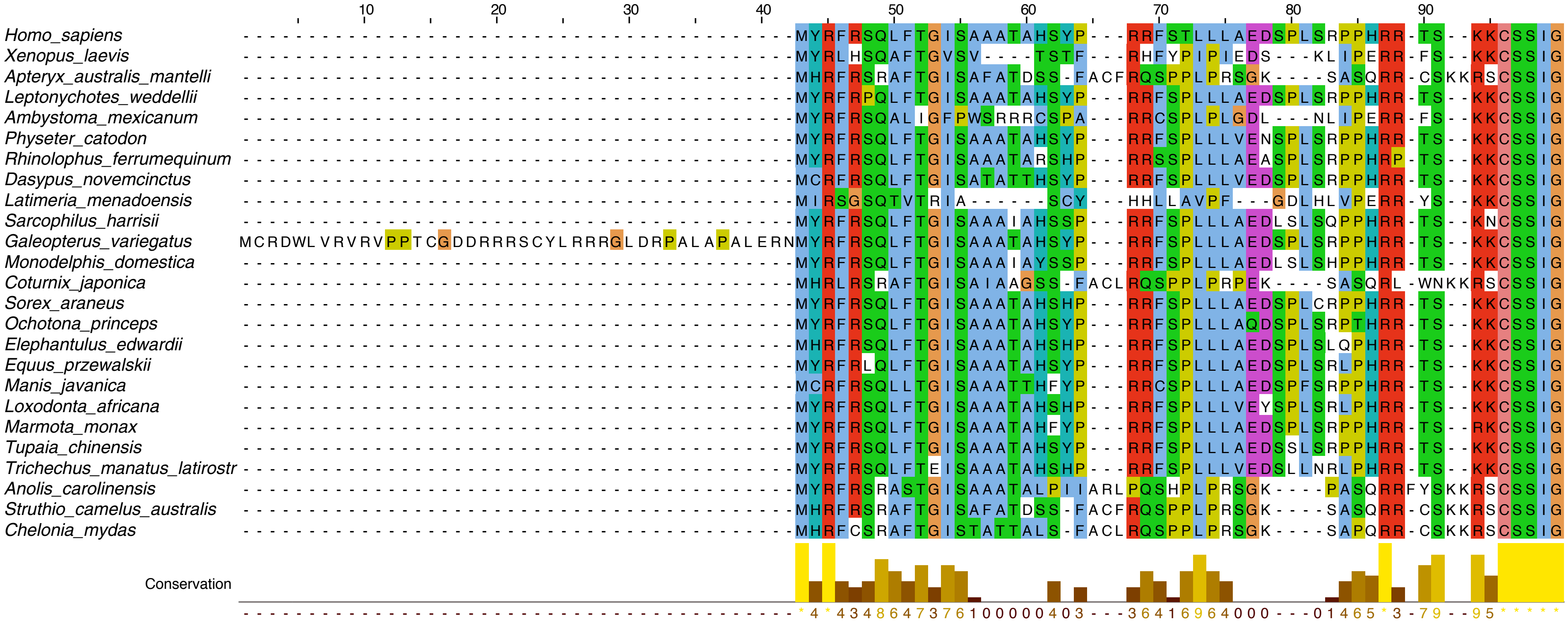

**HG0021.1**

ENSGALG00000009448 (IFRD1)

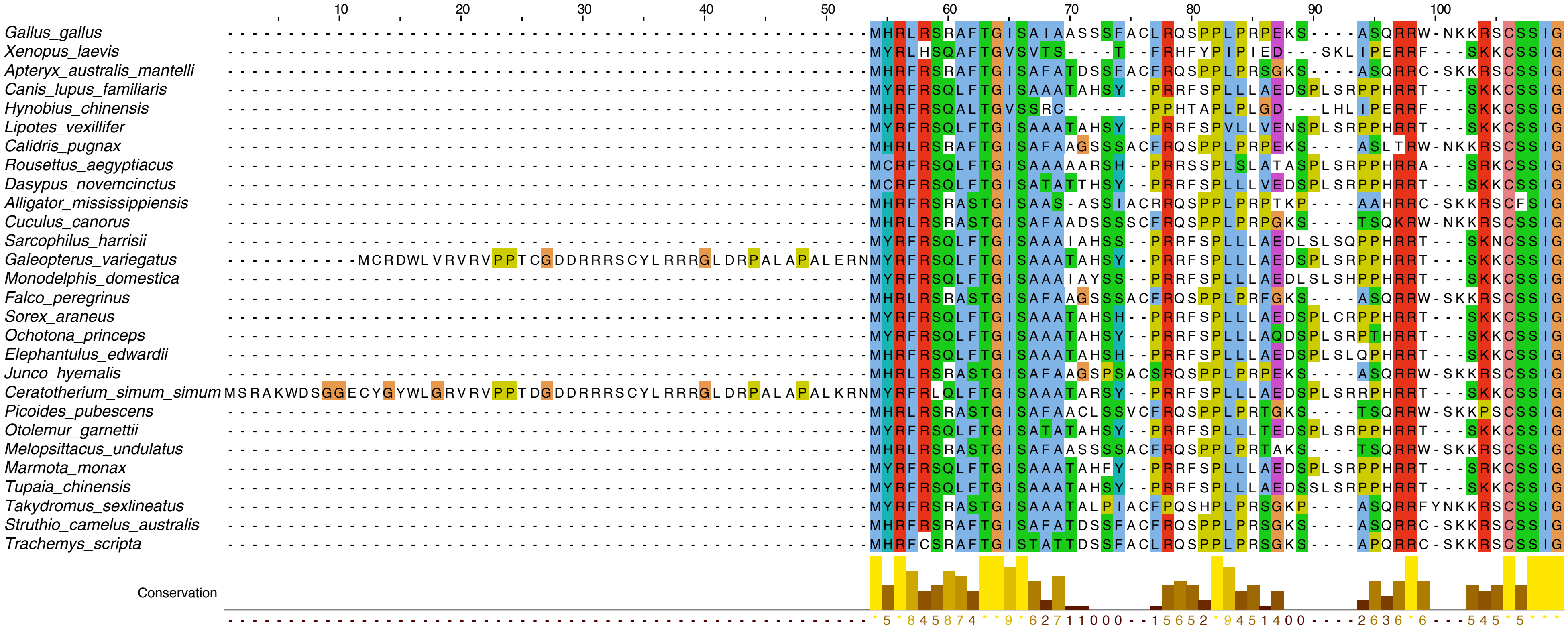

**HG0021.2**

ENSDARG00000036811 (ifrd2)

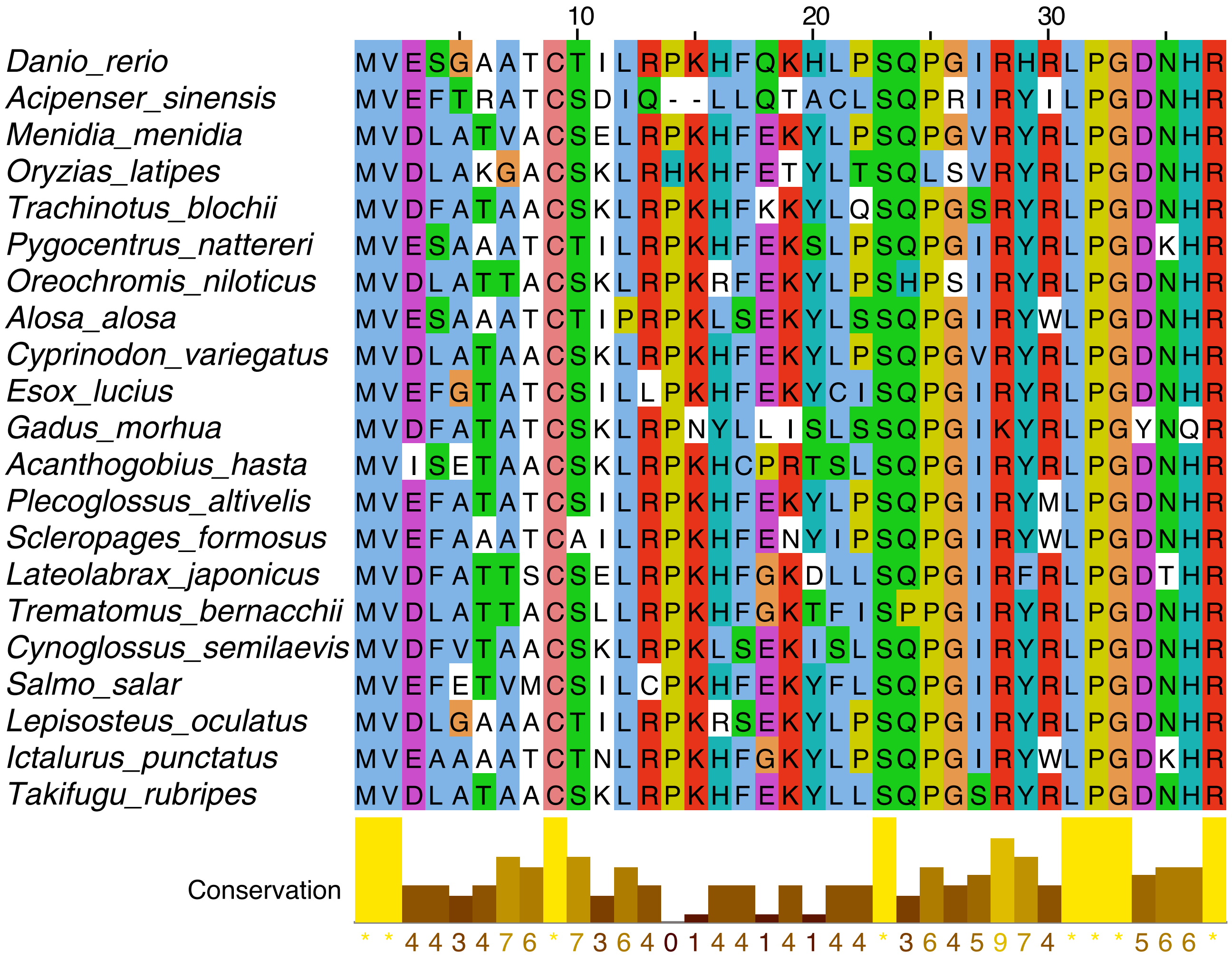

**HG0022**

FBgn0061359 (CG33671)

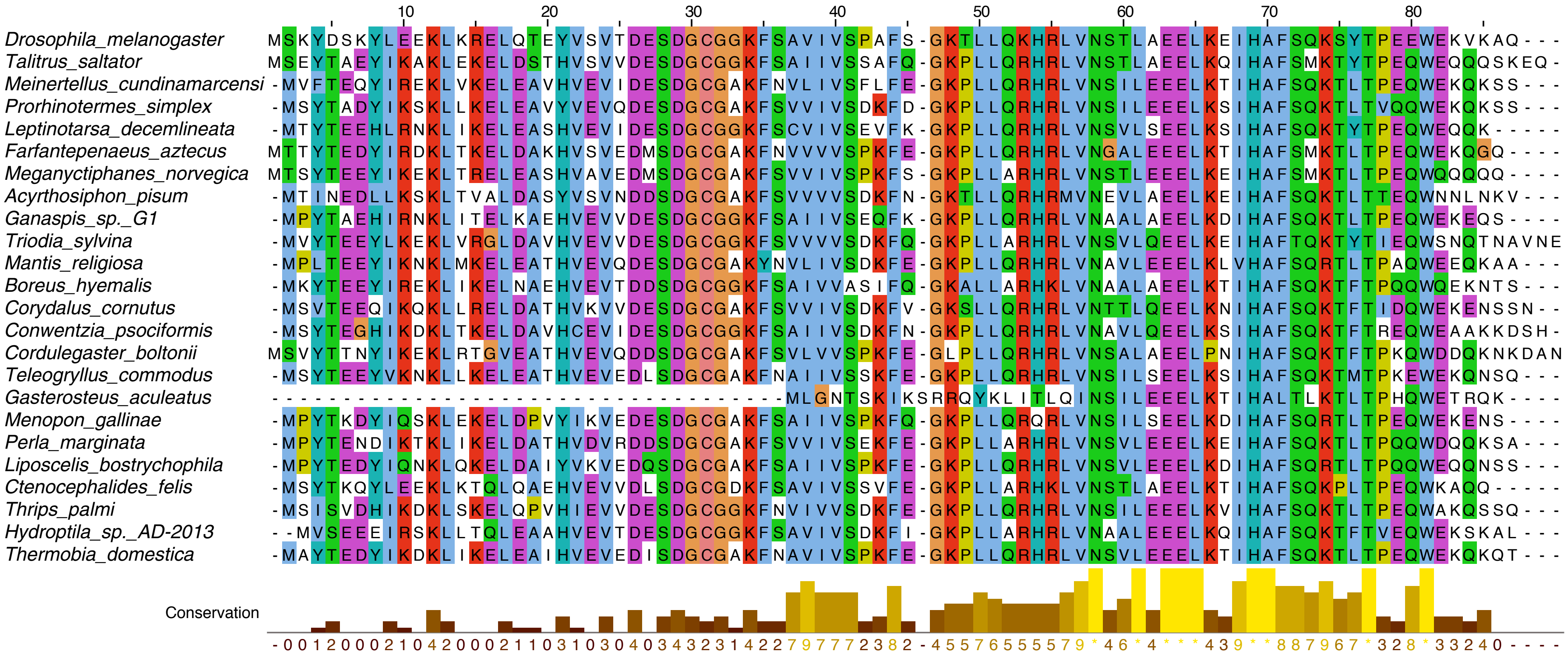

**HG0023.1**

ENSG00000175197 (DDIT3)

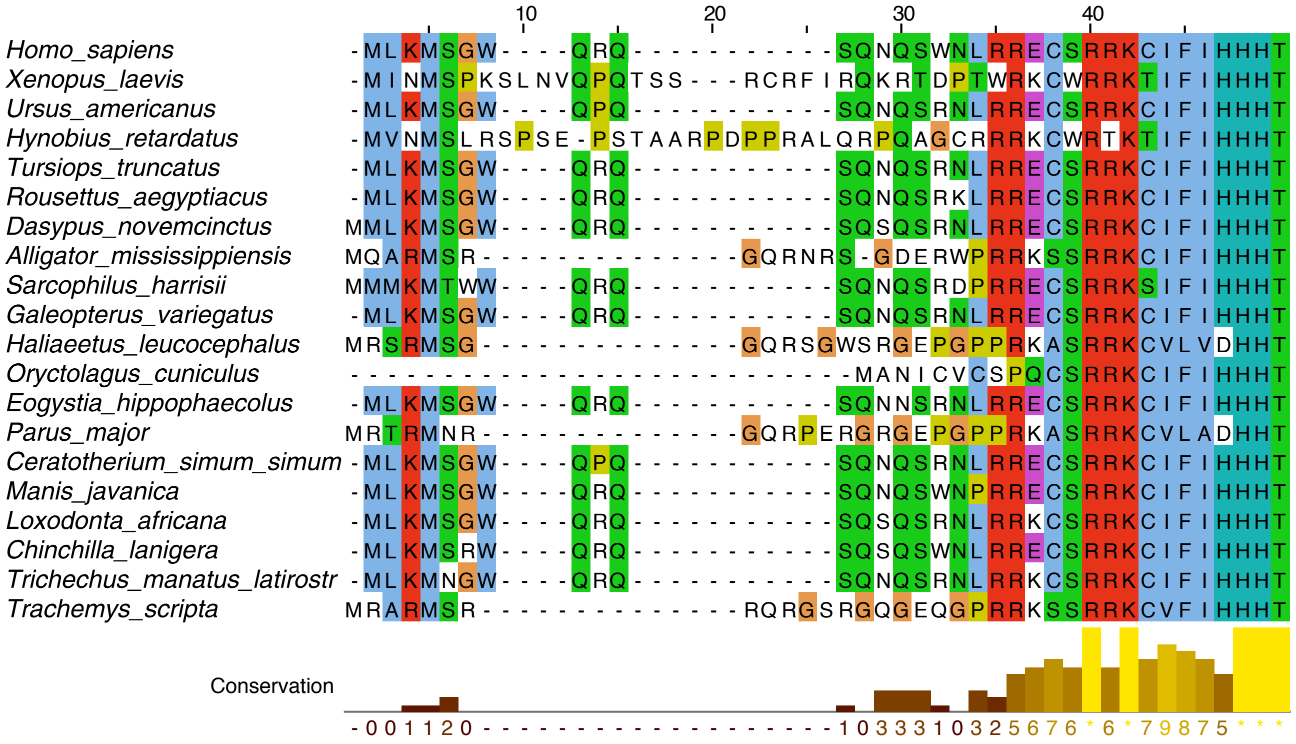

**HG0023.2**

ENSDARG00000059836 (ddit3)

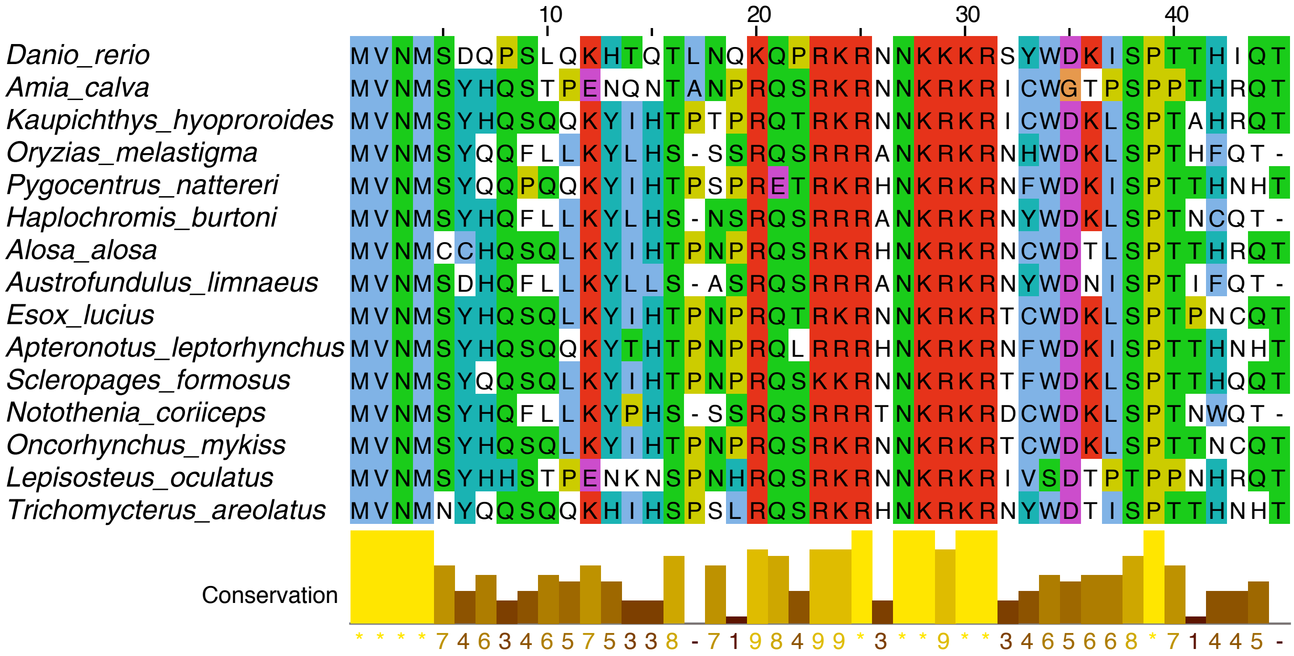

**HG0024.1**

ENSDARG00000093406

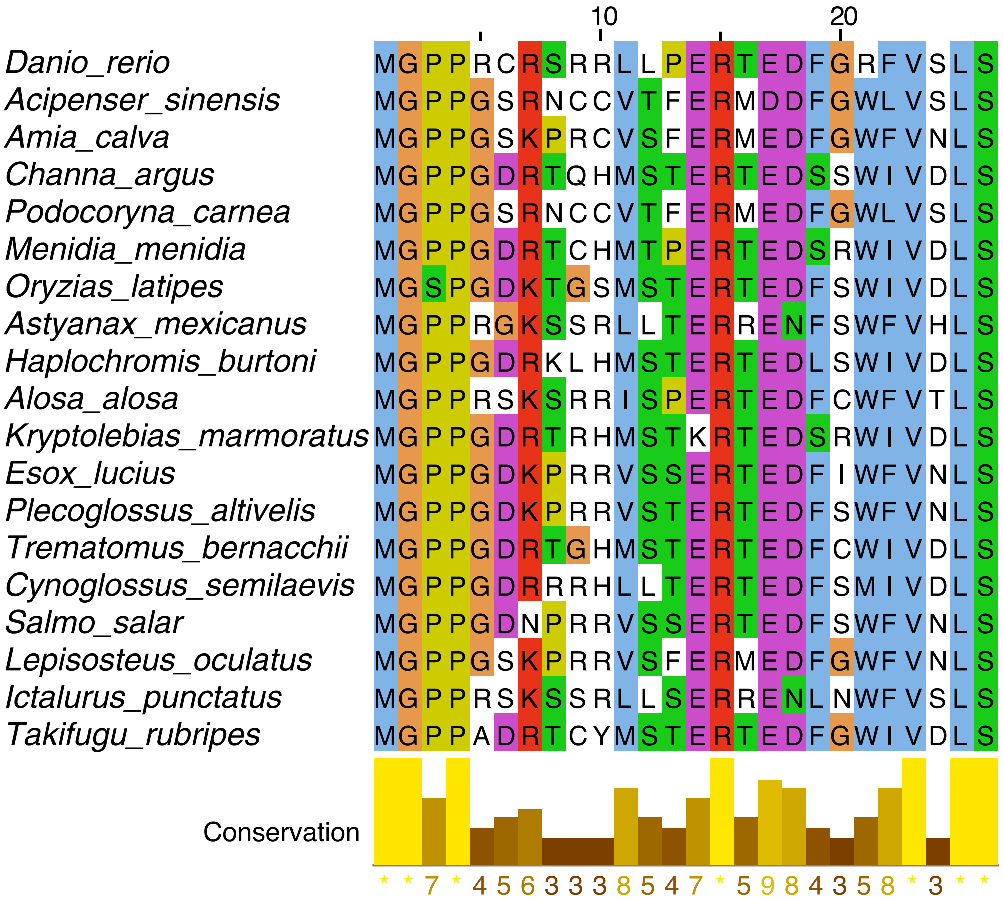

**HG0024.2**

ENSG00000112308 (C6orf62)

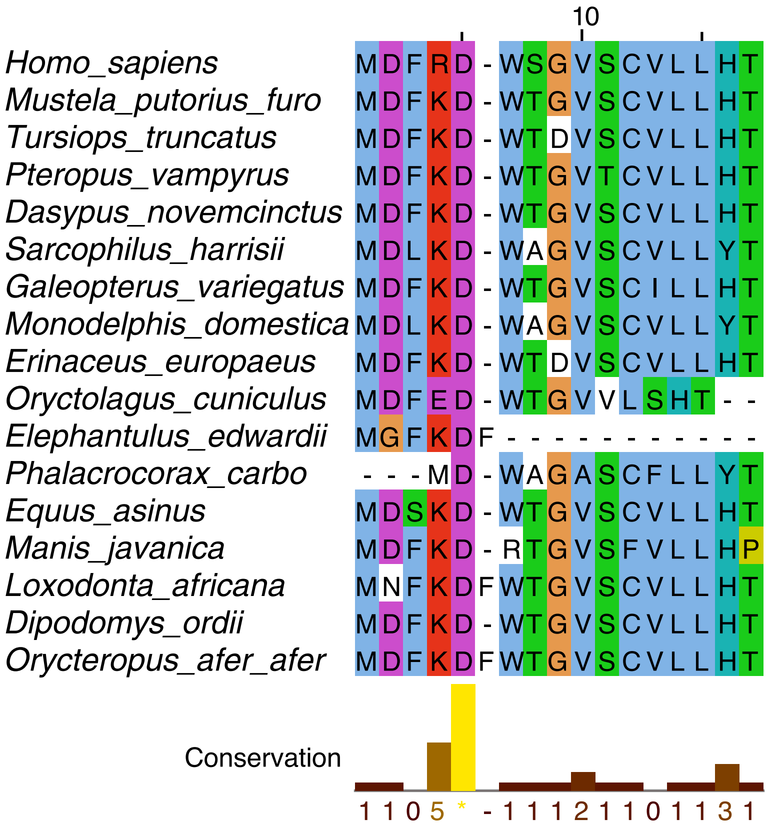

**HG0024.2**

ENSGALG00000013628 (C6orf62)

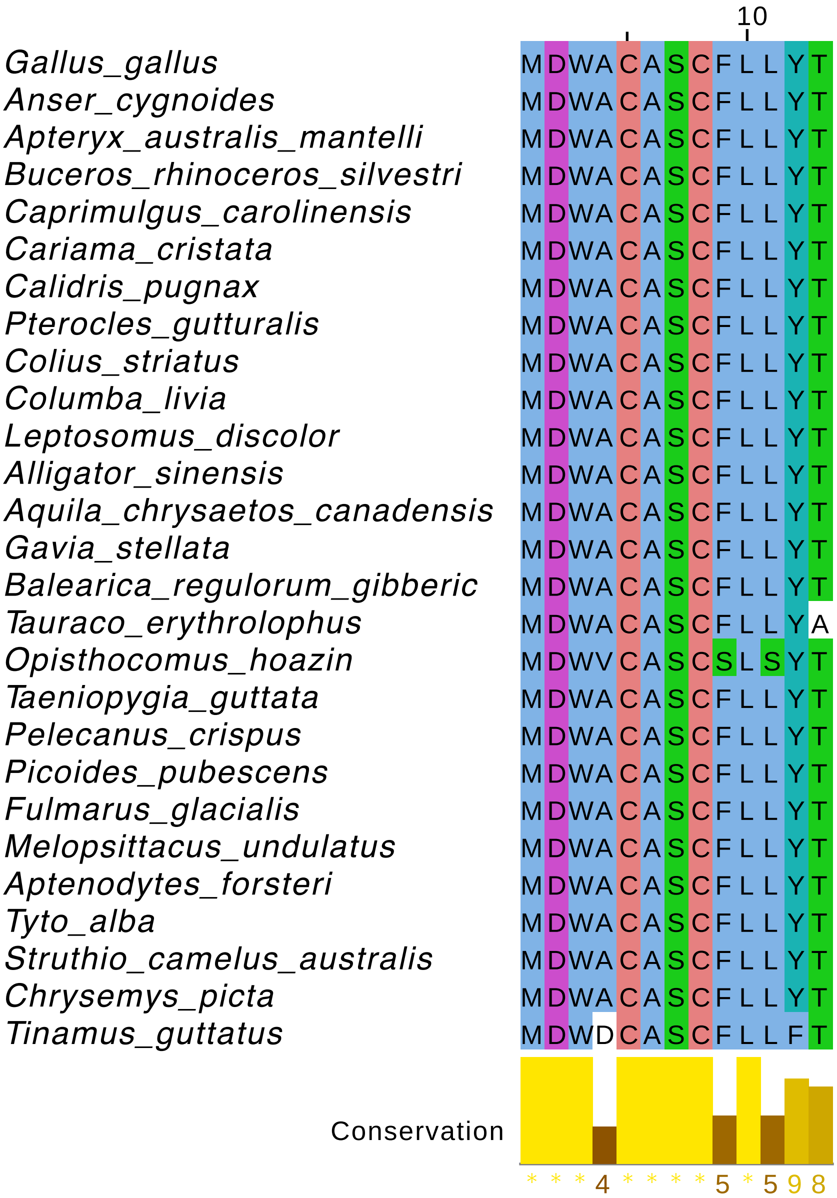

**HG0024.3**

ENSGALG00000013628 (C6orf62)

```
                                       10        20
Gallus_gallus                 ---------MAGQLLILFLDPSNTTFP-
Anser_cygnoides               MVKPVCCPAMAGQLL-LFLDPSDTTFS-
Apteryx_australis_mantelli    ---------MVGQLLILFLDPSNKAFS-
Buceros_rhinoceros_silvestri  ---------MAGQLFSLFLDPSNTTFS-
Caprimulgus_carolinensis      ---------MAGQLLTLVFDPSDTTSS-
Cariama_cristata              ---------MAGQLLRLFLDPSDTTTFS
Calidris_pugnax               ---------MAGQLLTLFLDPSNTTFS-
Pterocles_gutturalis          ---------MAGQLLTLFRDPSDTTFS-
Colius_striatus               ---------MAGQLLILFRGPSDTTVS-
Columba_livia                 ---------MAGQLLTLFLGPSDTTFS-
Leptosomus_discolor           ---------MTGQLLTLFLDPSDTTFS-
Alligator_mississippiensis    ---------MVGQRLILFLDPFNMTVS-
Cuculus_canorus               ---------MAGQLLTLFLDPSNTMFS-
Falco_cherrug                 ---------MAGQLLTLFLDPSNTTFS-
Gavia_stellata                ---------MAGQLLSLFLDPSDTTFS-
Chlamydotis_macqueenii        ---------MAGQLLTLFLDPSDTTFS-
Tauraco_erythrolophus         ---------MAGQLLTLFLDPSDTTFS-
Opisthocomus_hoazin           ---------MAGRLLTLFLDPSNATFS-
Corvus_cornix_cornix          ---------MAGQLLSLFLDPSNTTFS-
Pelecanus_crispus             ---------MAGQLLTLFLDPSNTTFS-
Picoides_pubescens            ---------MAEQLLSWFLDPSDTTFS-
Fulmarus_glacialis            ---------MAGQLLTLFLDPSDTTFS-
Nestor_notabilis              ---------MAGQLLTLFLDPSERPFS-
Aptenodytes_forsteri          ---------MAGQLLTLFLDPSDTTLS-
Anolis_carolinensis           ---------MVGRLLILFYDPSNKTVS-
Tyto_alba                     ---------MAGQLLTLFLDPSDTTFS-
Struthio_camelus_australis    ---------MVGQLLILFLDPSNTTFS-
Chelonia_mydas                ---------MVGQLHSLFLDPSNMIAS-
Tinamus_guttatus              ---------MVGQLLILFLDPSNTTFS-

Conservation
                              ---------*748550774 5*574444-
```

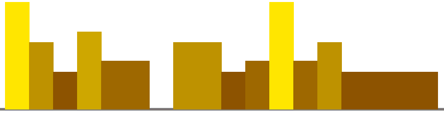

**HG0025**

ENSDARG00000060054 (epc1b)

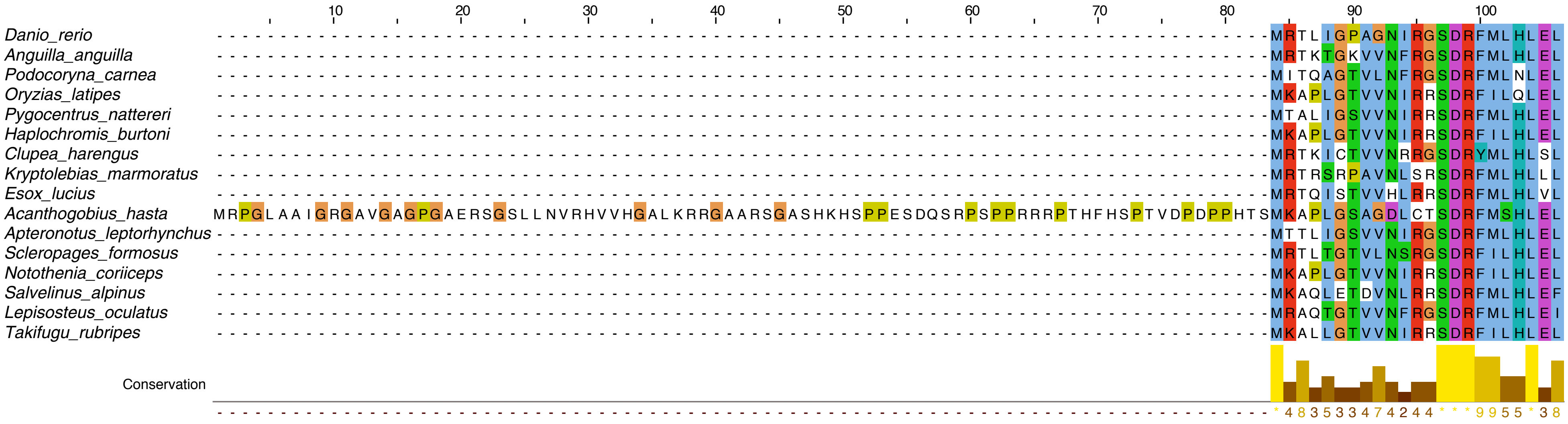

**HG0026**

ENSDARG00000087059 (fam213ab)

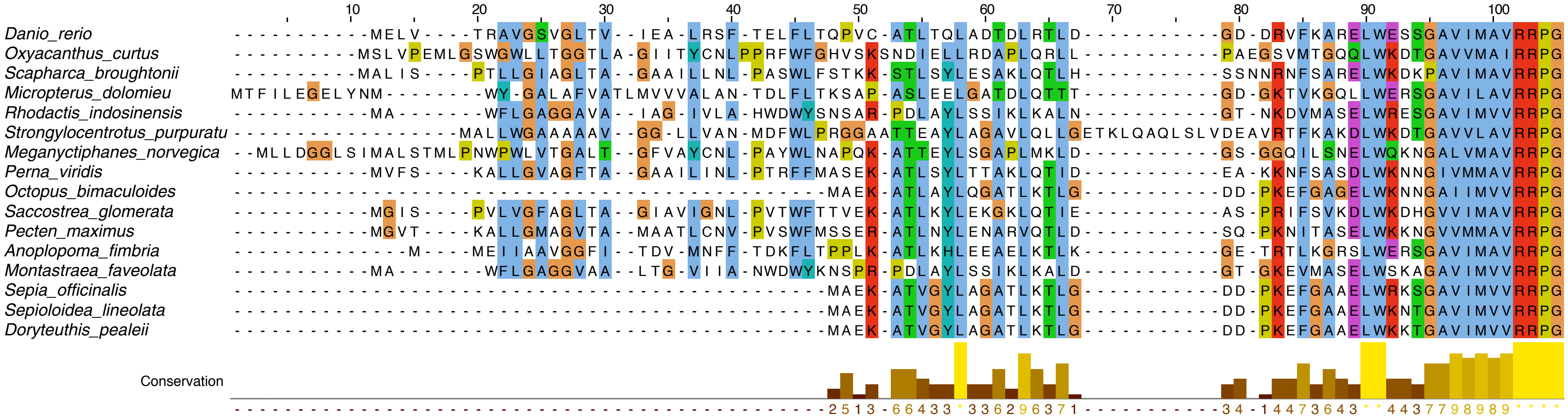

**HG0027**

ENSG00000077254 (USP33)

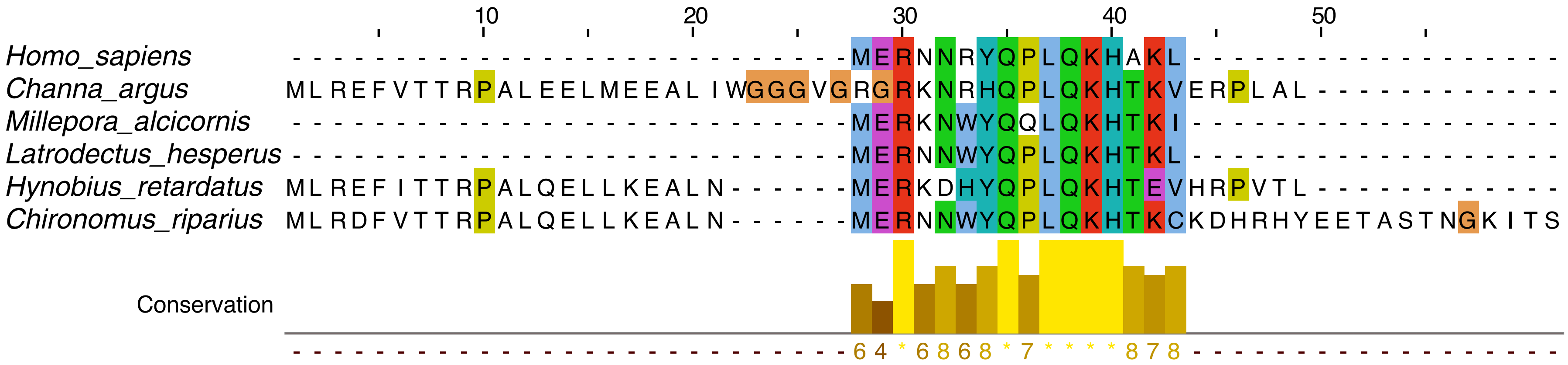

## HG0028.1

ENSDARG00000032103 (mapk6)

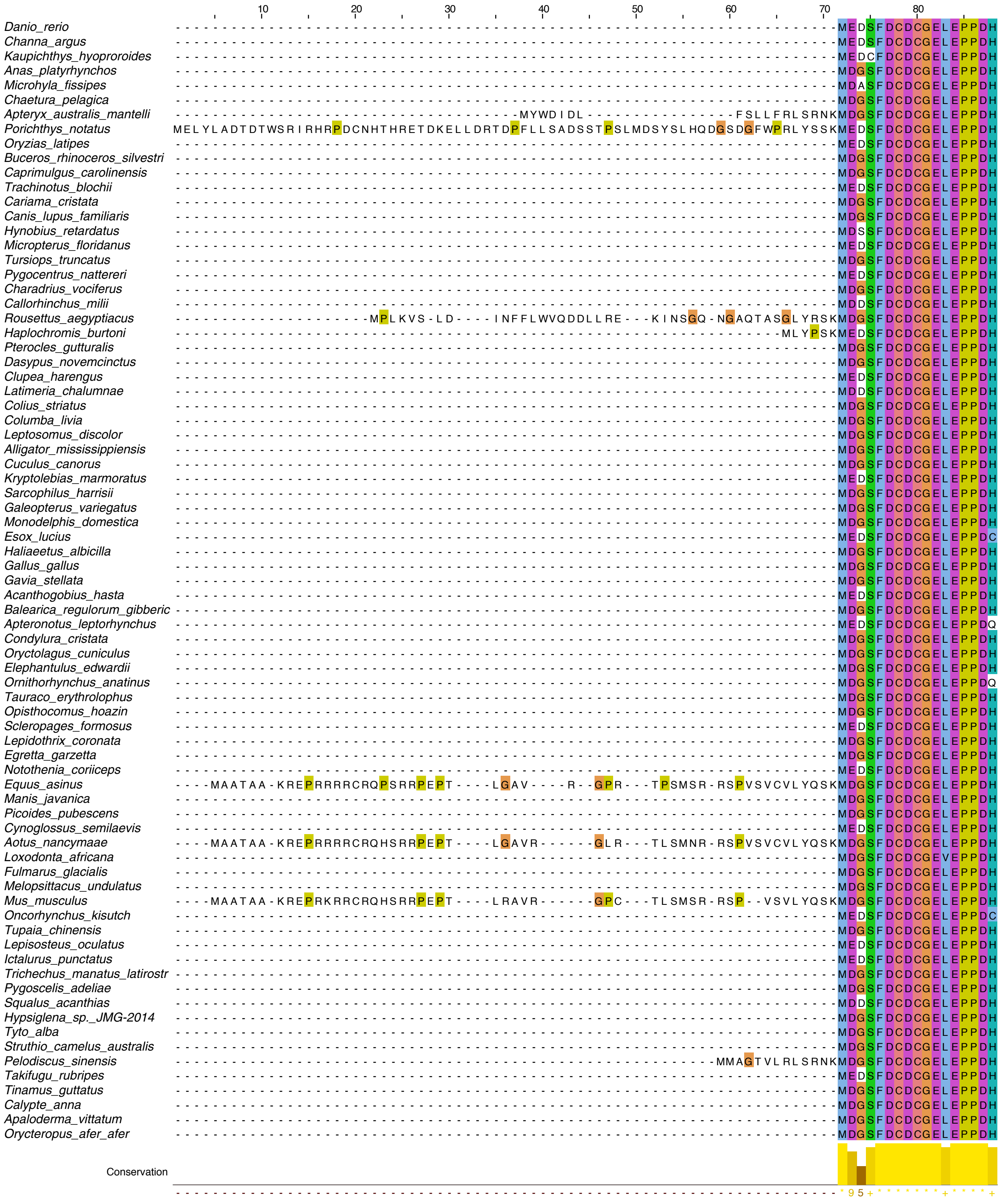

# HG0028.1

### ENSG00000069956 (MAPK6)

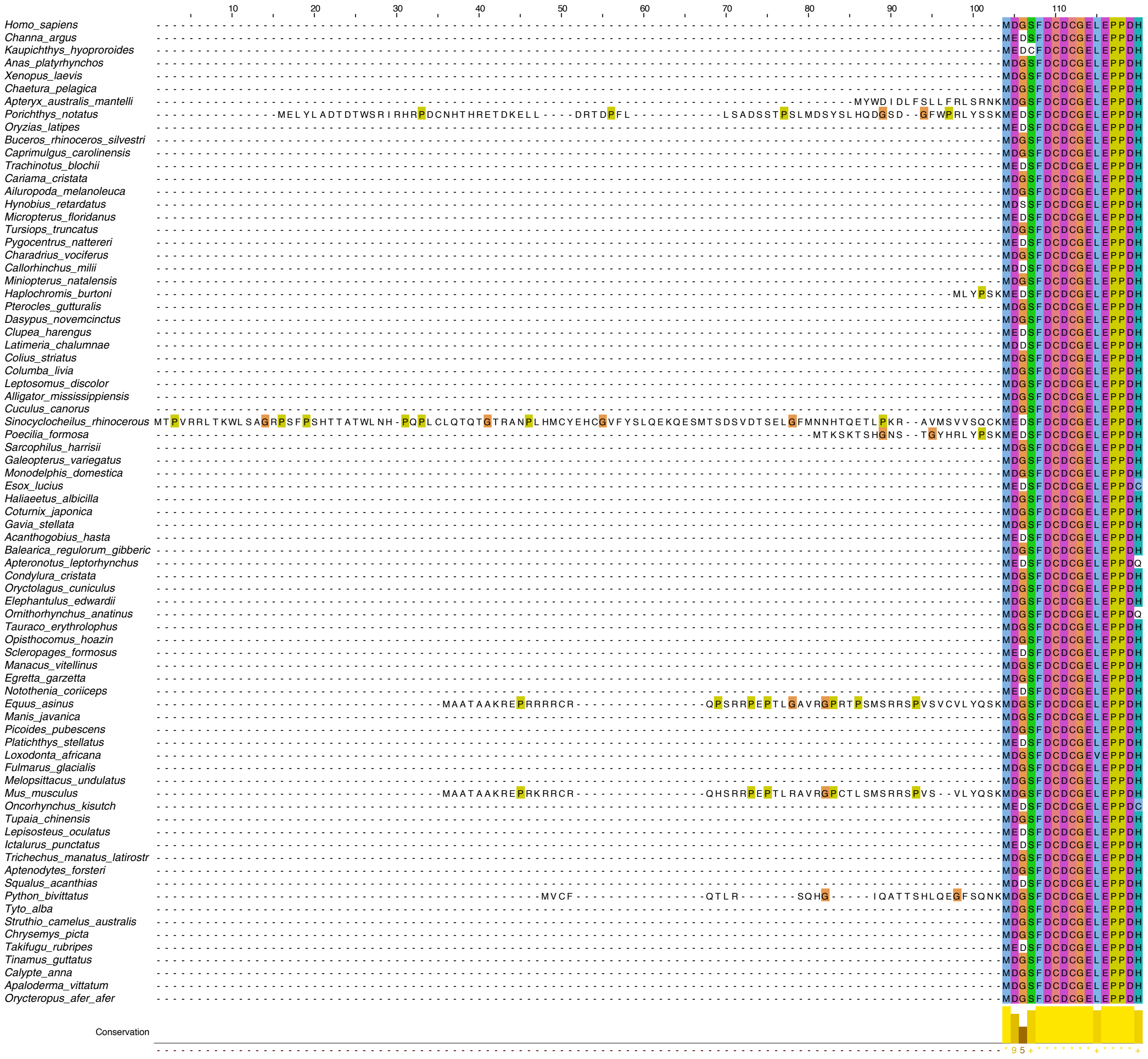

# HG0028.1

### ENSGALG00000031448 (MAPK6)

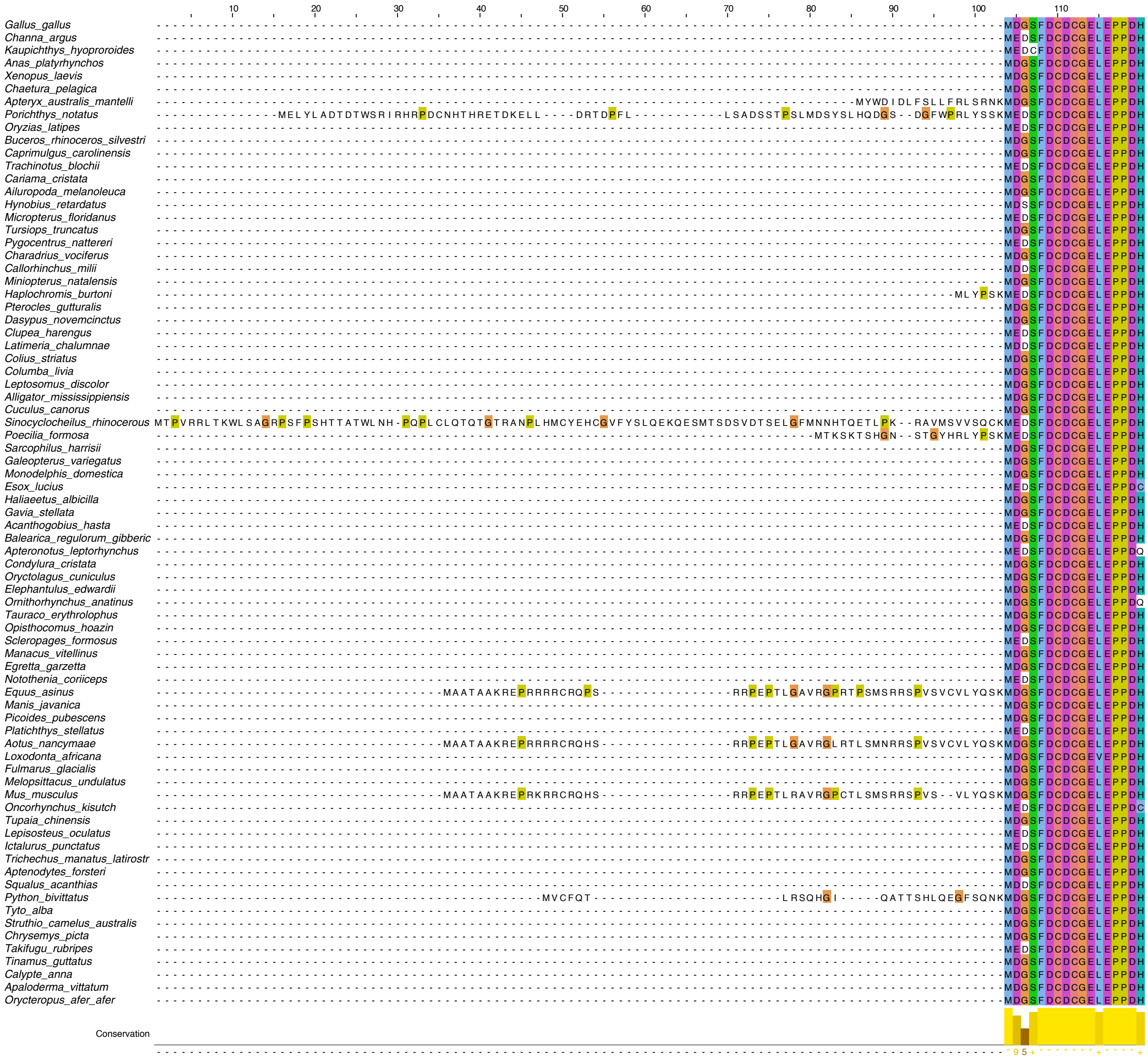

**HG0028.2**

ENSG00000069956 (MAPK6)

| Species | Sequence (positions 1–23) |
|---|---|
| *Homo_sapiens* | MFIEACCVHIYSHFC-------- |
| *Anser_cygnoides* | MLIEAYYVHIYSHFC-------- |
| *Rhacophorus_omeimontis* | MMIEAYILHIYSHFCSMSTCFAH |
| *Chaetura_pelagica* | MLIEAYYVHIYSHFC-------- |
| *Apteryx_australis_mantelli* | MLIEAYYVHIYSHFC-------- |
| *Buceros_rhinoceros_silvestri* | MLIEAYYVHIYSHFC-------- |
| *Caprimulgus_carolinensis* | MLIEAYYVHIYSHFC-------- |
| *Cariama_cristata* | MLIEAYYVHIYSHFC-------- |
| *Acinonyx_jubatus* | MFIEACCVHIYSHFC-------- |
| *Balaenoptera_acutorostrata_s* | MFIEACCVHIYSHFC-------- |
| *Calidris_pugnax* | MLIEAYYVHIYSHFC-------- |
| *Eptesicus_fuscus* | MFIEACCVHIYSHFC-------- |
| *Pterocles_gutturalis* | MLIEAYYVHIYSHFC-------- |
| *Dasypus_novemcinctus* | MFIEACCVHIYSHFC-------- |
| *Colius_striatus* | MLIEAYYVHIYSHFC-------- |
| *Columba_livia* | MLIEAYYVHIYSHFC-------- |
| *Leptosomus_discolor* | MLIEAYYVHIYSHFC-------- |
| *Alligator_sinensis* | MLIEAYYVHIYSYFC-------- |
| *Cuculus_canorus* | MLIEAYYVHIYSHFC-------- |
| *Sarcophilus_harrisii* | MFREAYCVHIYSLFC-------- |
| *Galeopterus_variegatus* | MFIEACCVHIYSHFC-------- |
| *Aquila_chrysaetos_canadensis* | MLIEAYYVHIYSHFC-------- |
| *Coturnix_japonica* | MLIEAYYVHIYSHFC-------- |
| *Gavia_stellata* | MLIEAYYVHIYSHFC-------- |
| *Chlamydotis_macqueenii* | MFIEAYYVHIYSHFC-------- |
| *Condylura_cristata* | MFIEACCVHIYSHFC-------- |
| *Oryctolagus_cuniculus* | MFIEACCVHIYSHFC-------- |
| *Elephantulus_edwardii* | MSIEACCVHIYSHFC-------- |
| *Ornithorhynchus_anatinus* | MLIEACFVHIYSHFC-------- |
| *Tauraco_erythrolophus* | MLKEAYYVHIYSHFC-------- |
| *Opisthocomus_hoazin* | MLIETYYVHIYSHFC-------- |
| *Acanthisitta_chloris* | MLIEAYYVHIYSHFC-------- |
| *Egretta_garzetta* | MLIEAYYVHIYSHFC-------- |
| *Ceratotherium_simum_simum* | MFIEACCVHIYSHFC-------- |
| *Picoides_pubescens* | MLIEAYYVHIYSHFC-------- |
| *Loxodonta_africana* | MFIEACCVHIYSHFC-------- |
| *Fulmarus_glacialis* | MLIEAYYVHIYSHFC-------- |
| *Melopsittacus_undulatus* | MLIEAYYVHIYSHFC-------- |
| *Marmota_marmota_marmota* | MFIEACCVHIYSHFC-------- |
| *Tupaia_chinensis* | MFLEACCVHIYSHFC-------- |
| *Trichechus_manatus_latirostr* | MFIEARCVHIYSRFC-------- |
| *Pygoscelis_adeliae* | MLIEAYYVHIYSHFC-------- |
| *Takydromus_sexlineatus* | MLIEAYYVHIYSHFC-------- |
| *Tyto_alba* | MLIEAYYVHIYSHFC-------- |
| *Struthio_camelus_australis* | MLIEAYYVHIYSHFC-------- |
| *Pelodiscus_sinensis* | MLLEAHYVHIYSHFC-------- |
| *Tinamus_guttatus* | MLIEAYYVHIYSHFC-------- |
| *Calypte_anna* | MLIEAYYVHIYSHFC-------- |
| *Apaloderma_vittatum* | MLIEAYYVHIYSHFC-------- |
| *Orycteropus_afer_afer* | MFIEACCVHIYSHFC-------- |
| Conservation | * 8 + * + 7 7 + * * * * + * * - - - - - - - - |

**HG0029**

ENSDARG00000007523 (kmt2e)

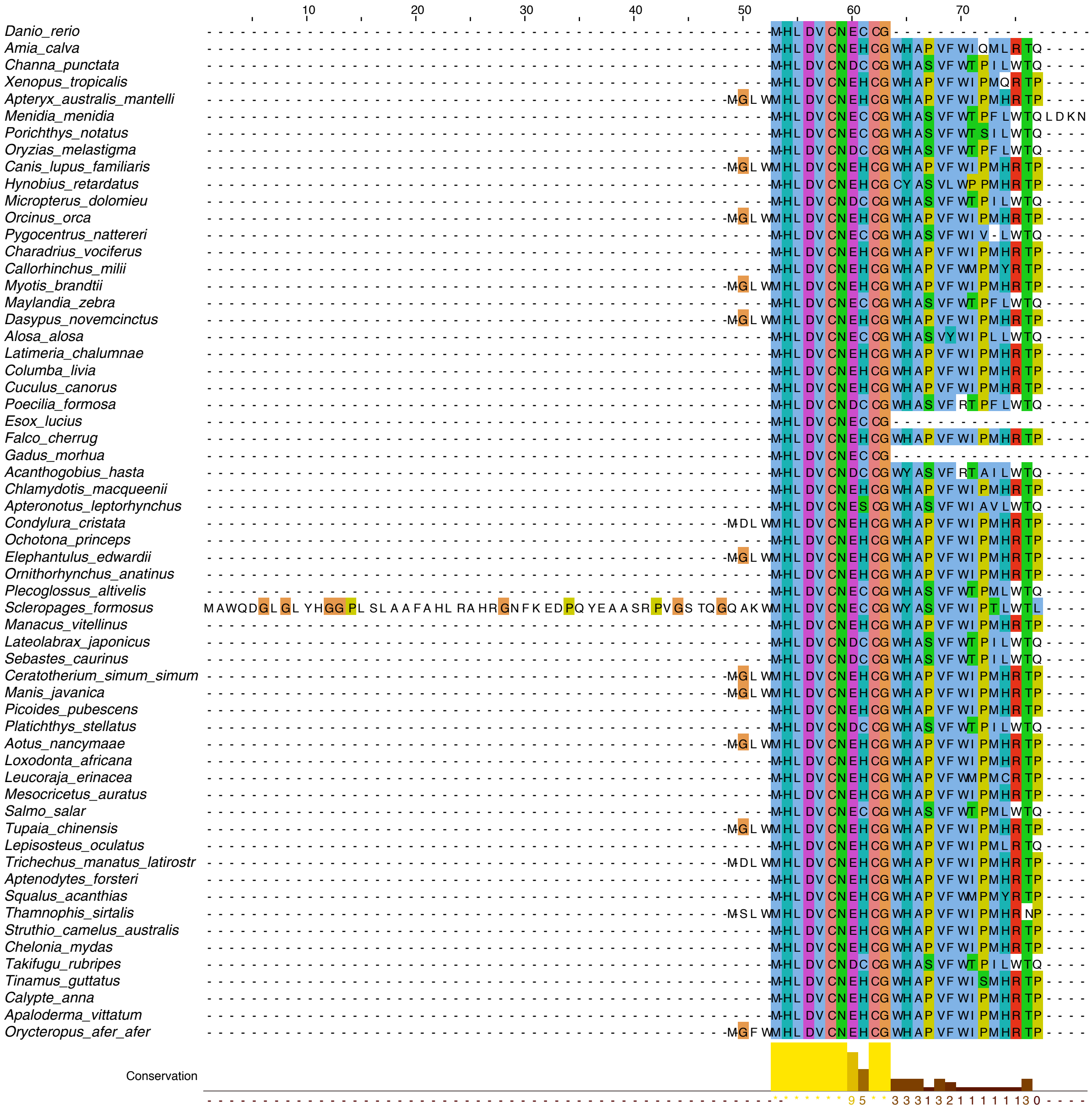

## HG0029

ENSG00000005483 (KMT2E)

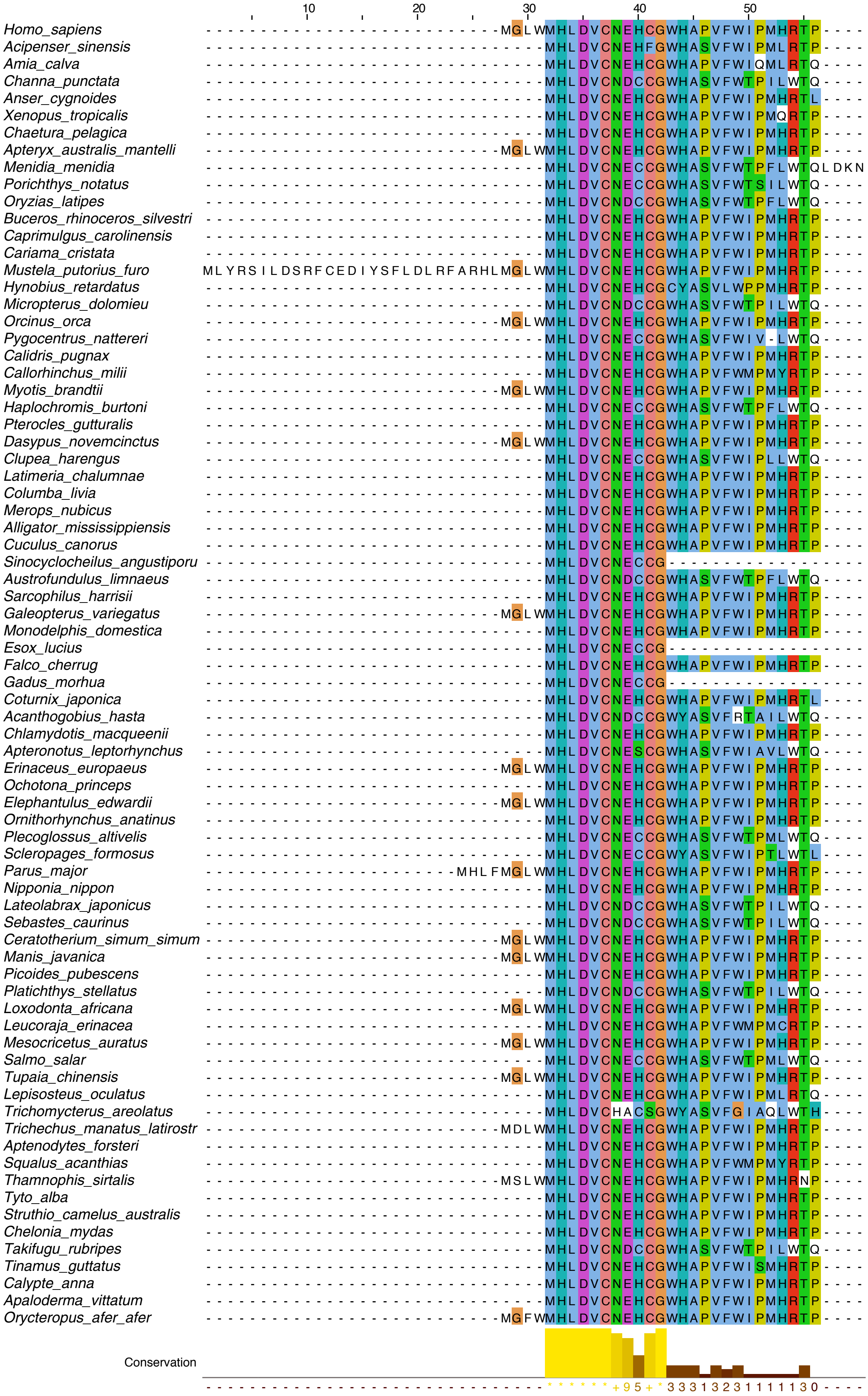

**HG0029**

ENSG00000168137

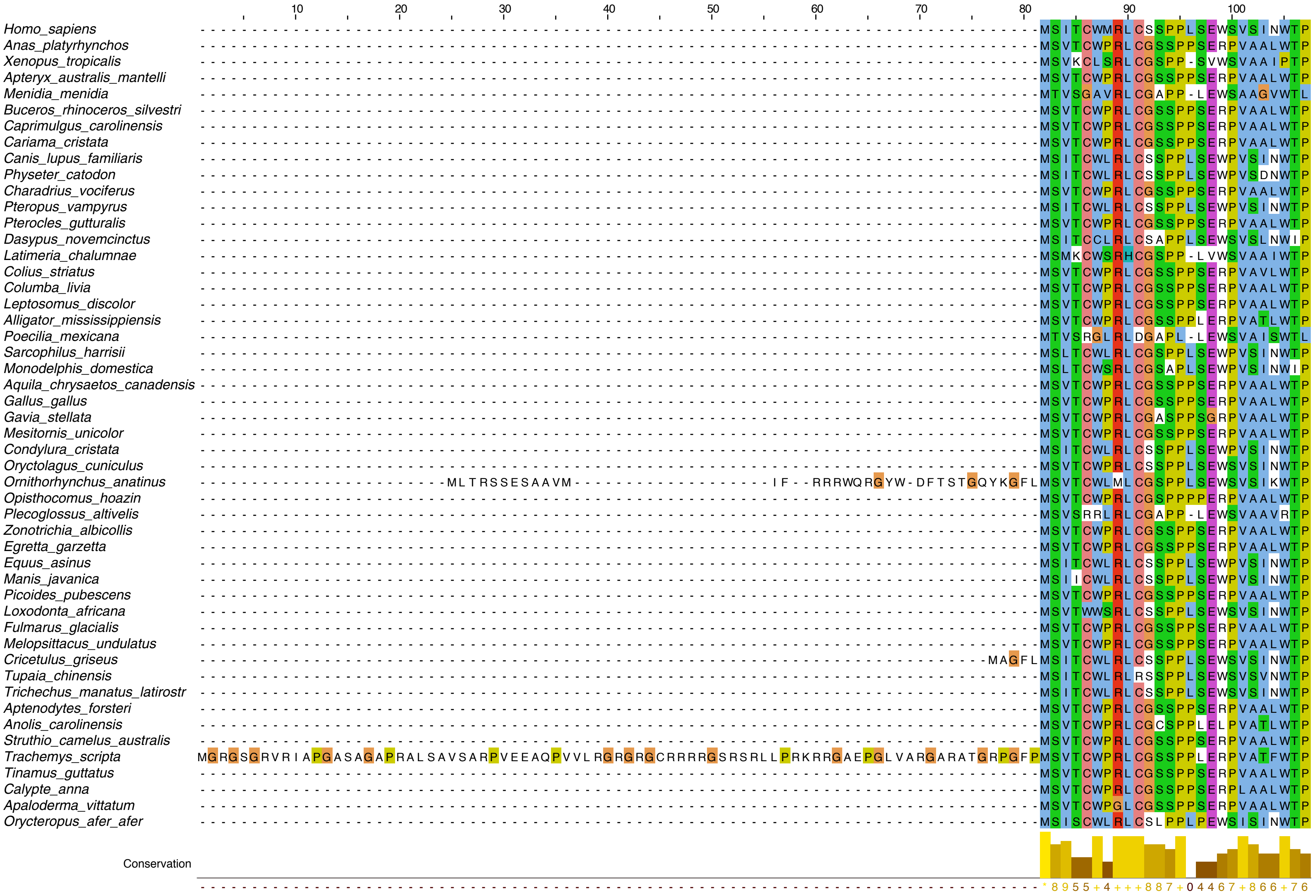

**HG0029**

ENSGALG00000008167

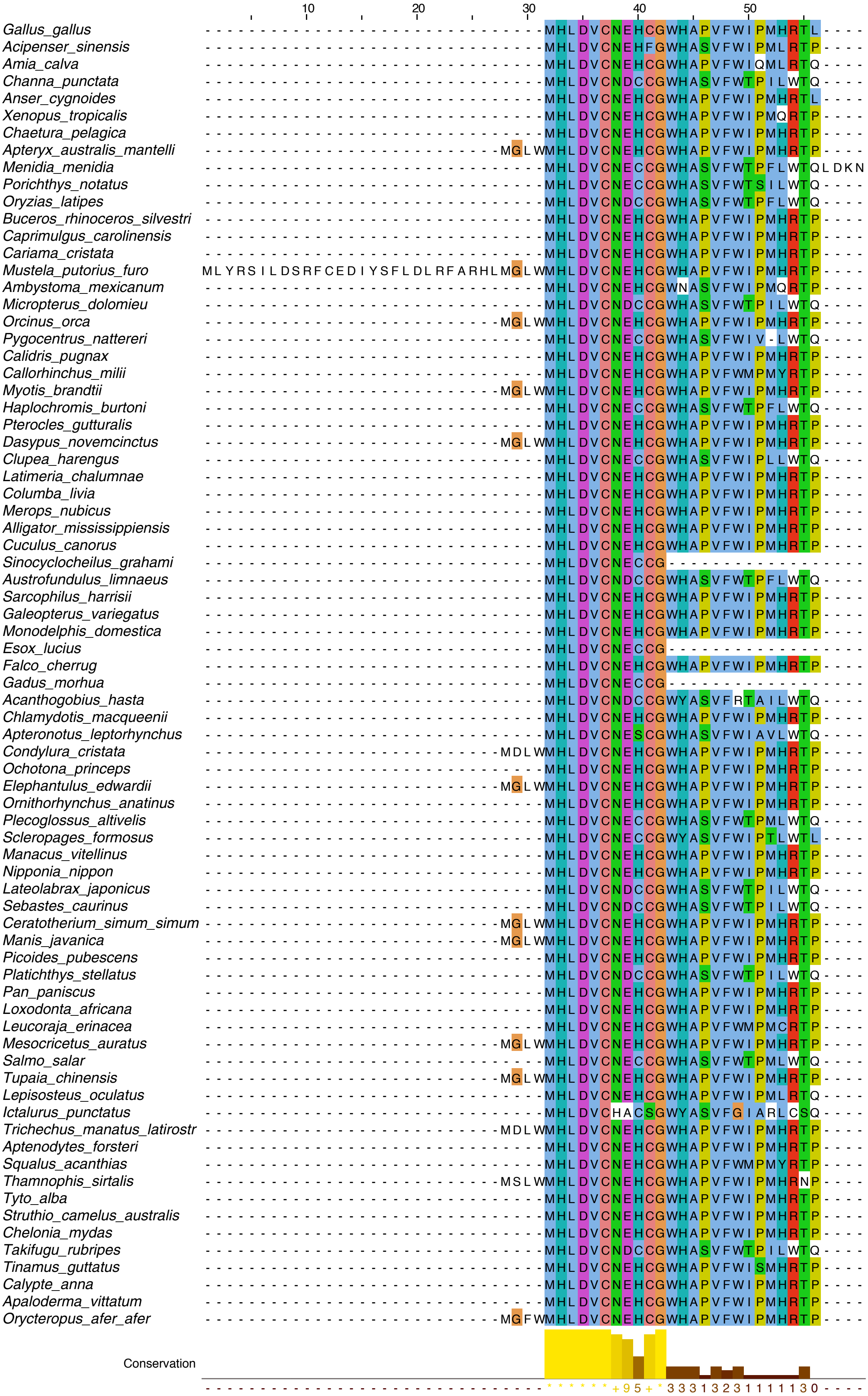

# HG0030.1

### ENSG00000100664 (EIF5)

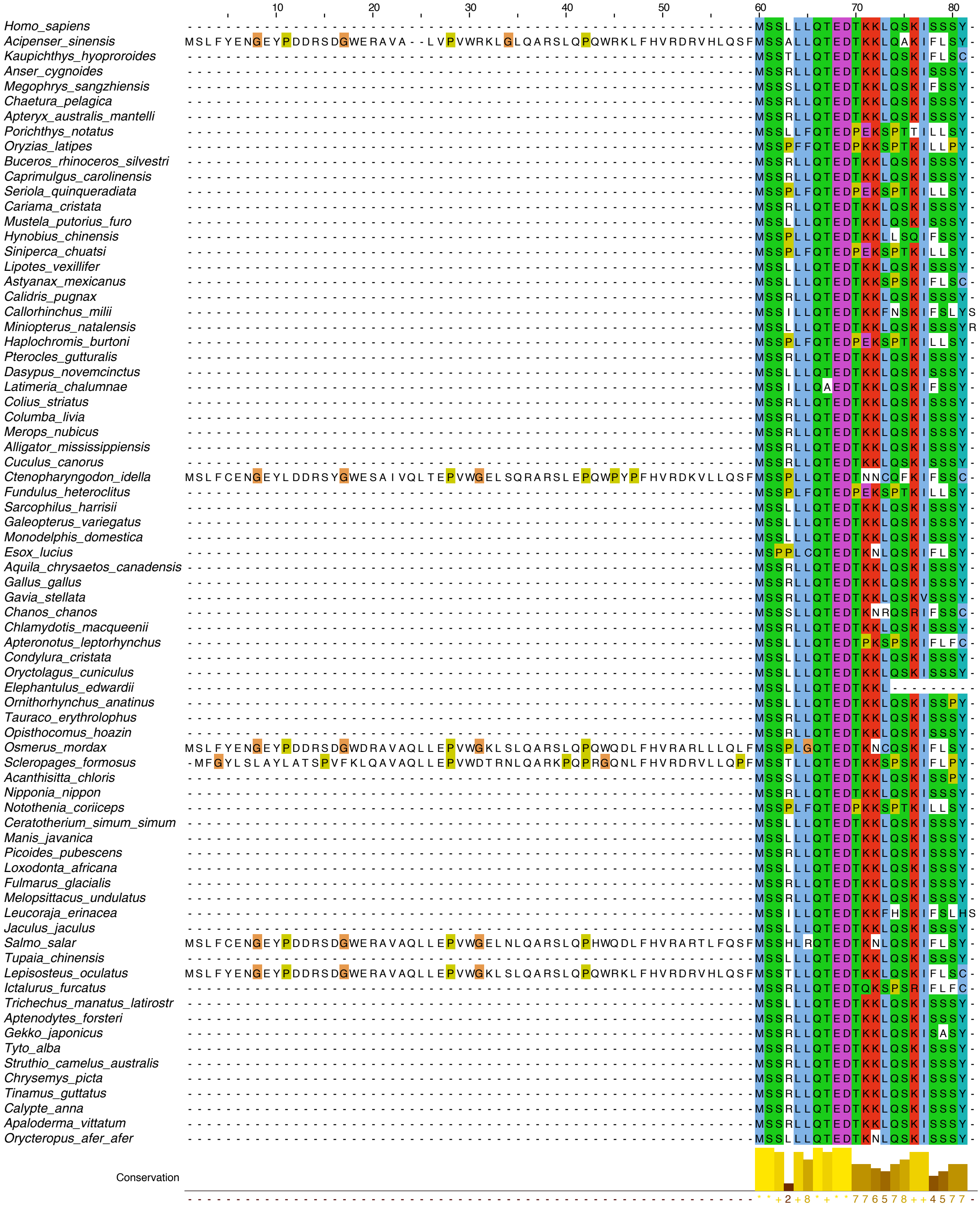

**HG0030.1**

ENSGALG00000011464 (EIF5)

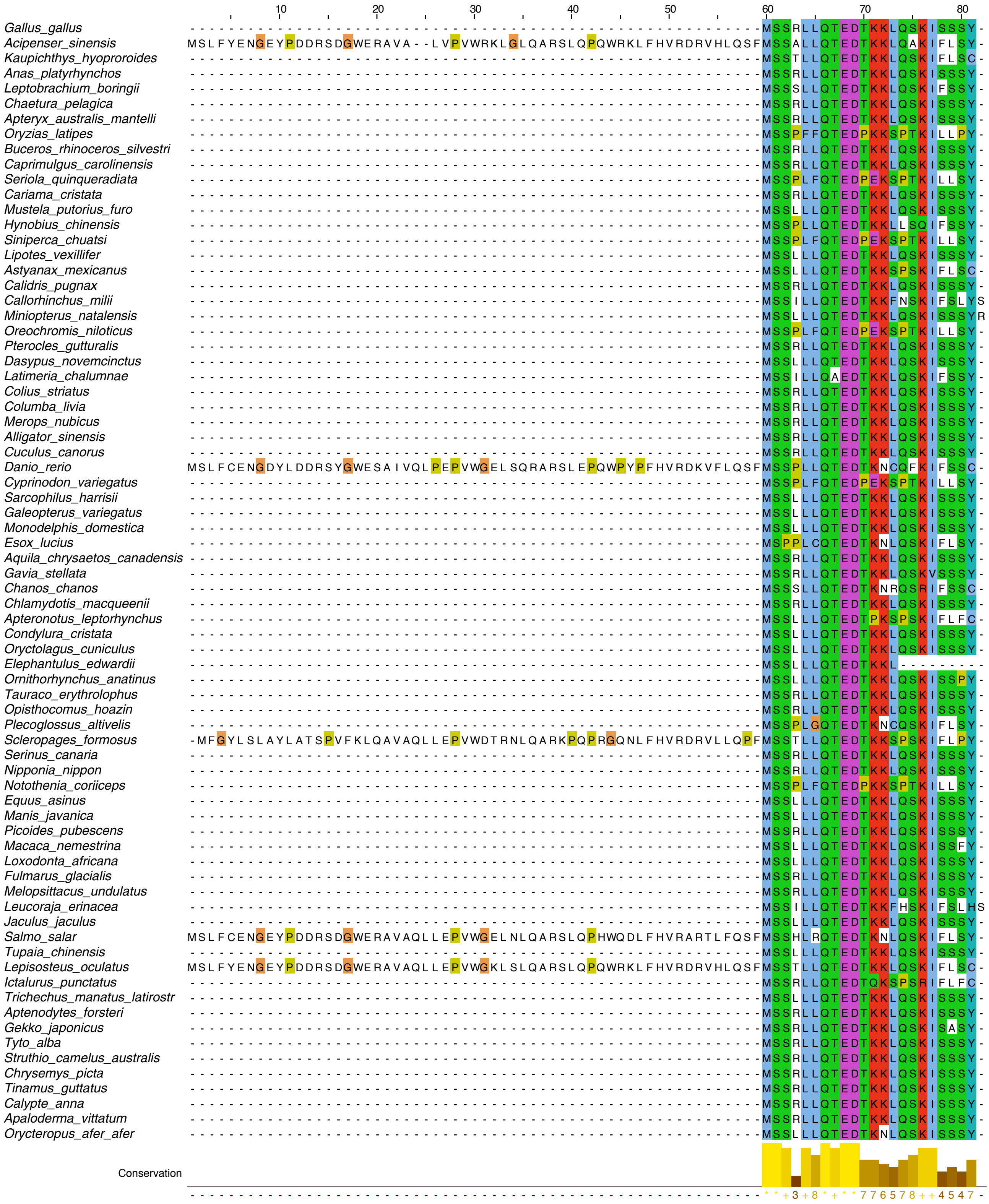

# HG0030.2

FBgn0030719 (eIF5)

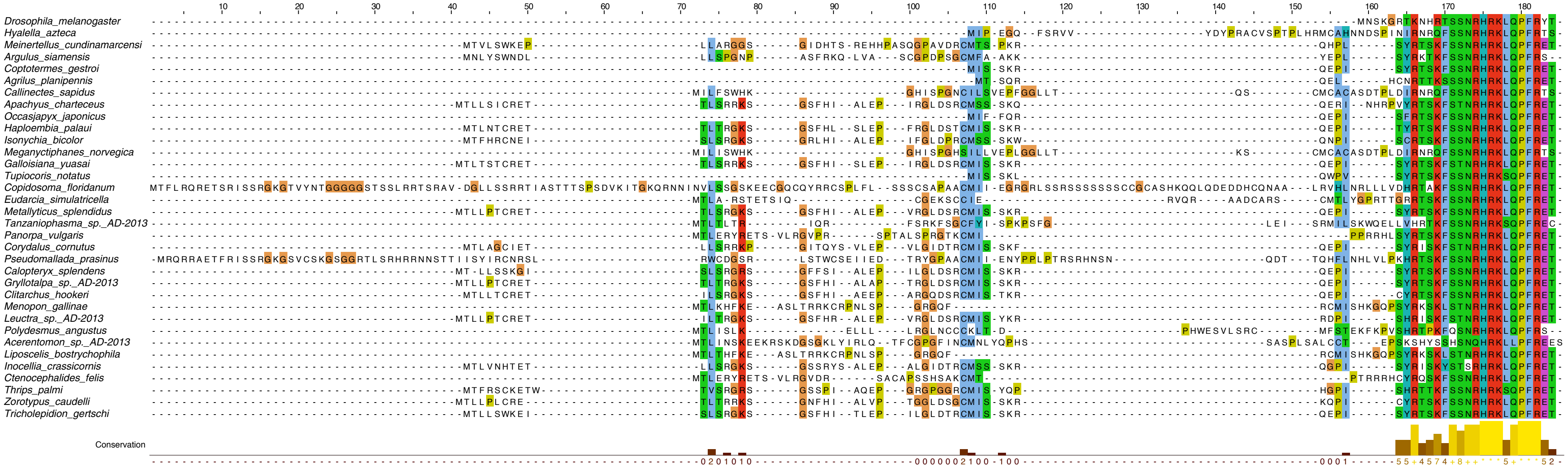

## HG0031

ENSDARG00000000540 (asnsd1)

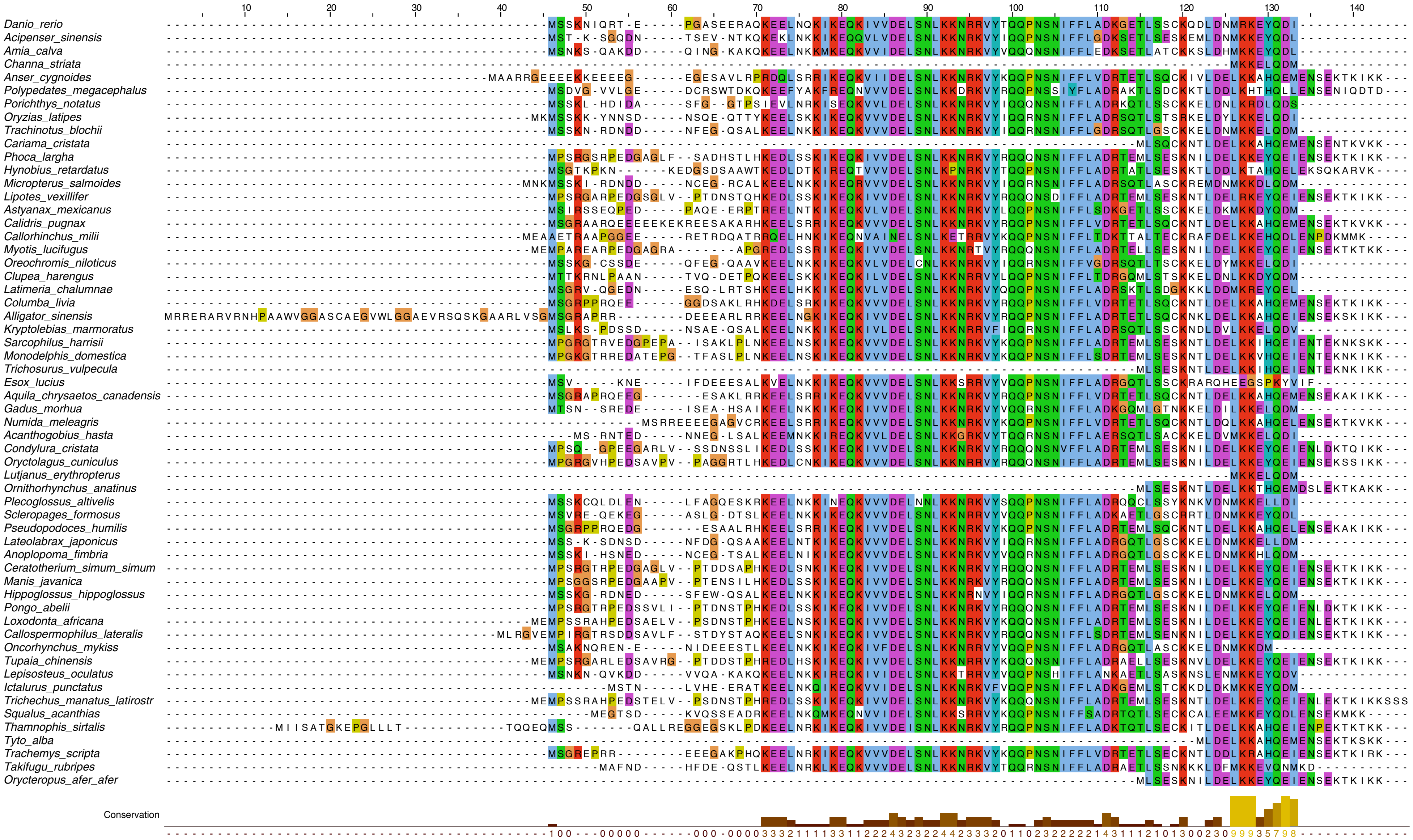

# HG0031

### ENSG00000138381 (ASNSD1)

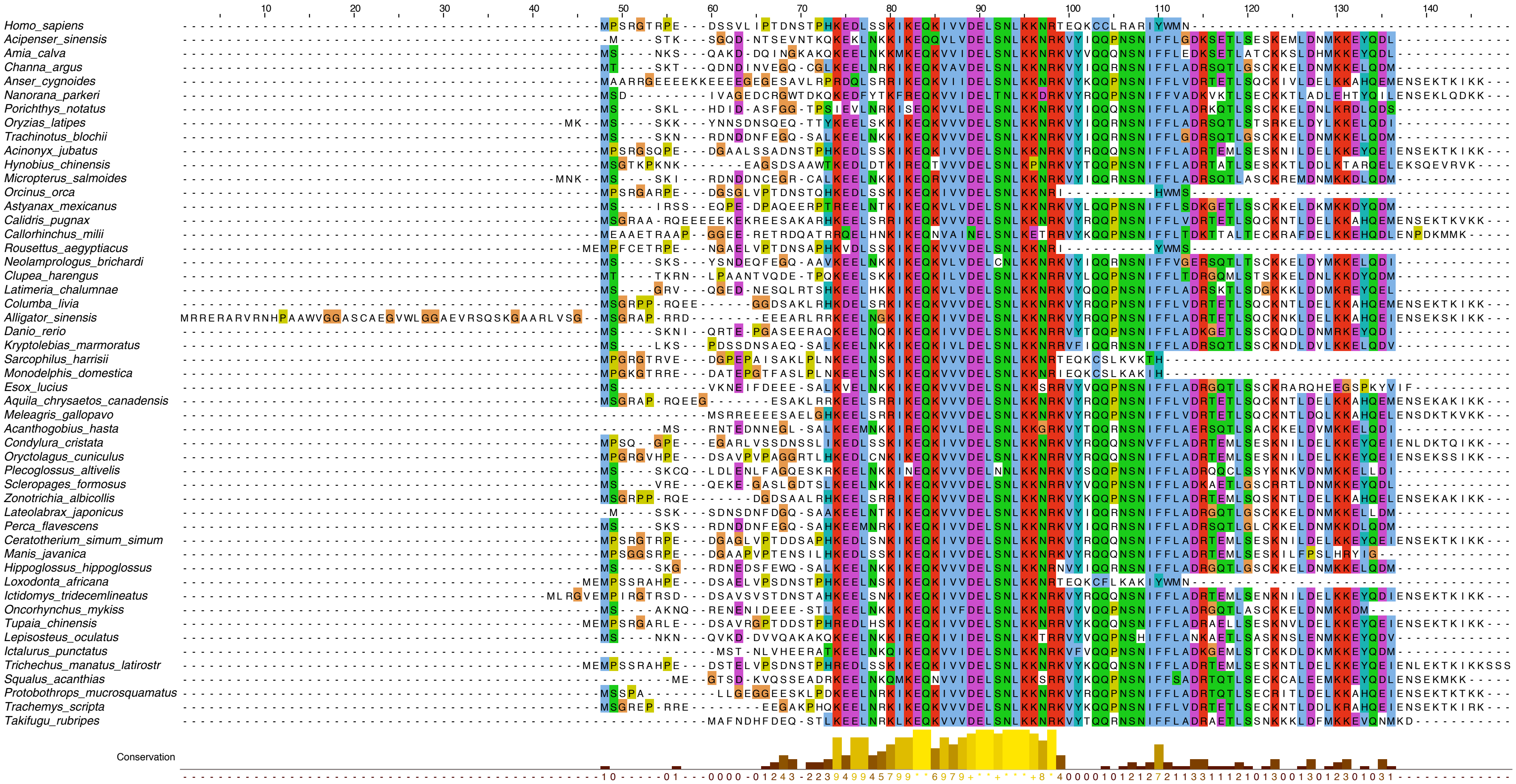

# HG0031

### ENSGALG00000041666

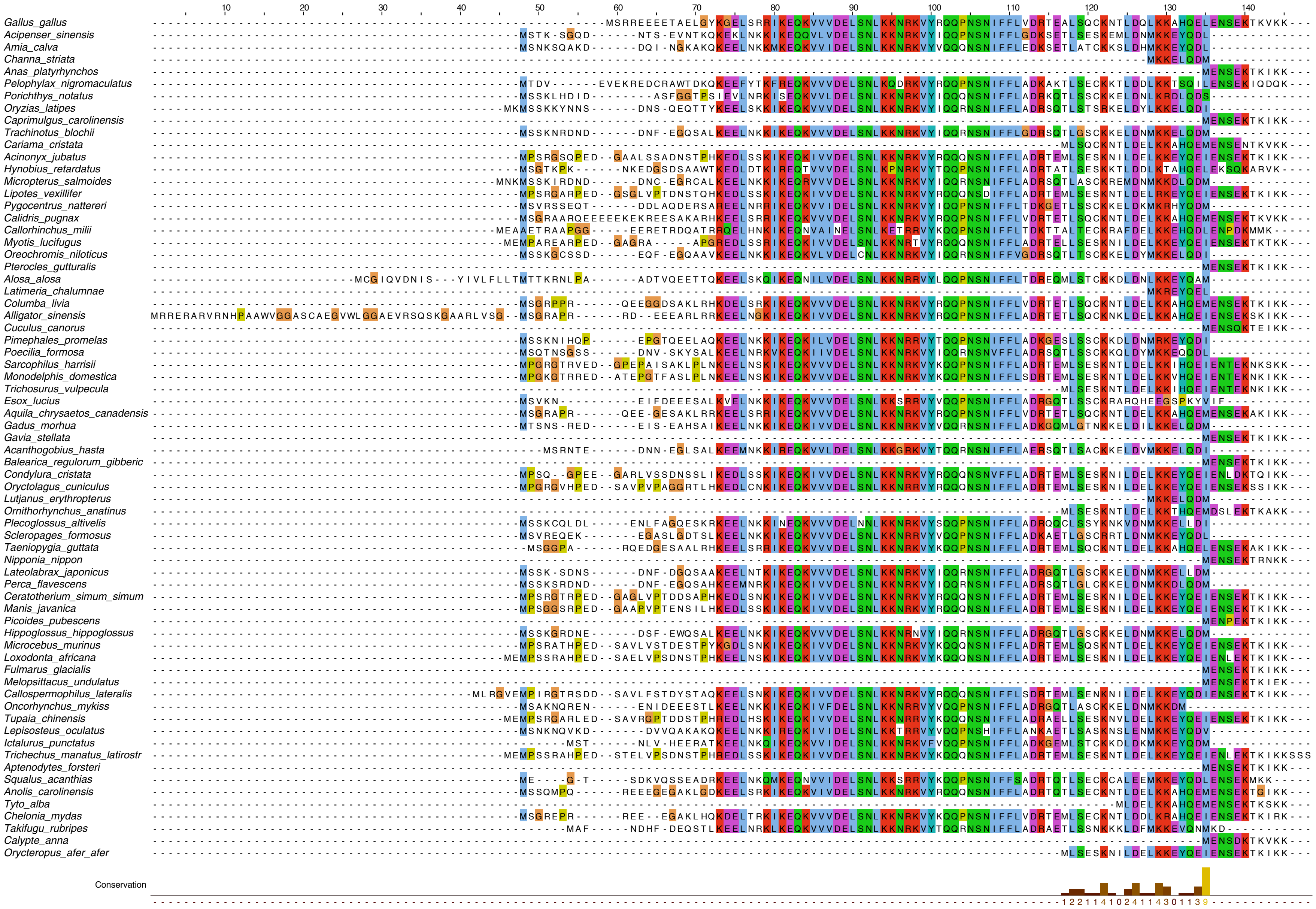

**HG0032**

ENSDARG00000104981 (SLC45A4)

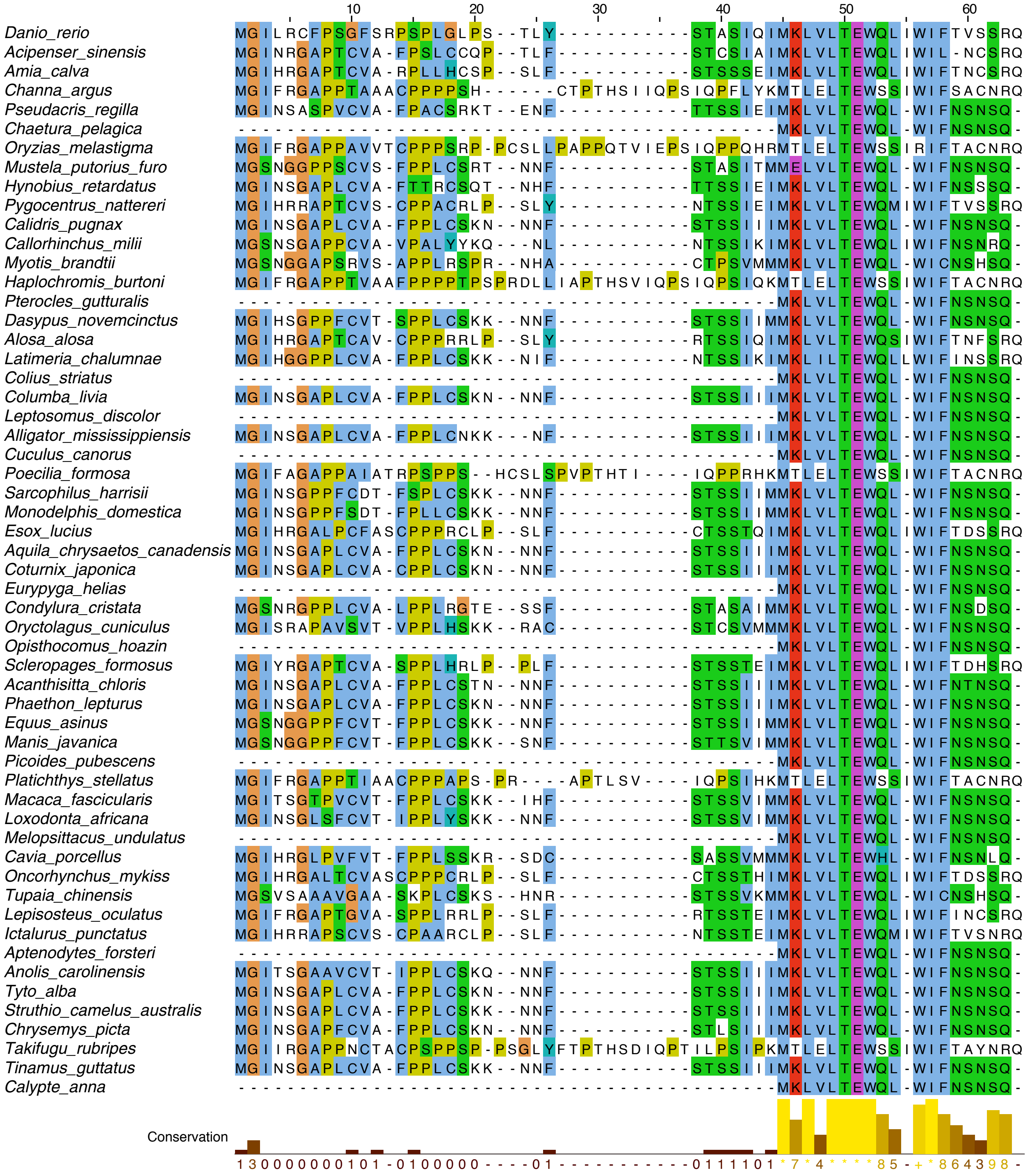

# HG0033

### ENSG00000112144 (ICK)

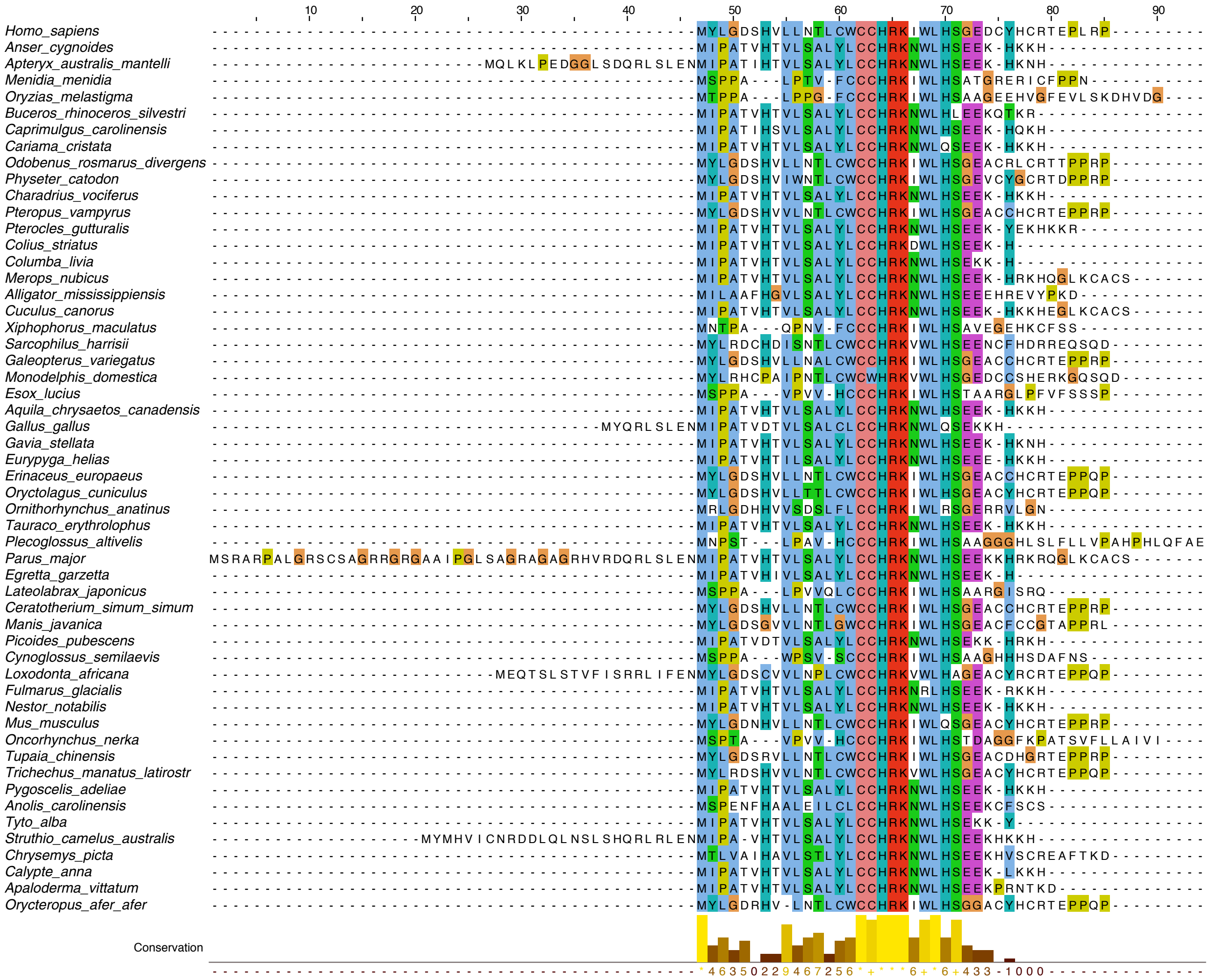

**HG0033**

ENSGALG00000016321 (ICK)

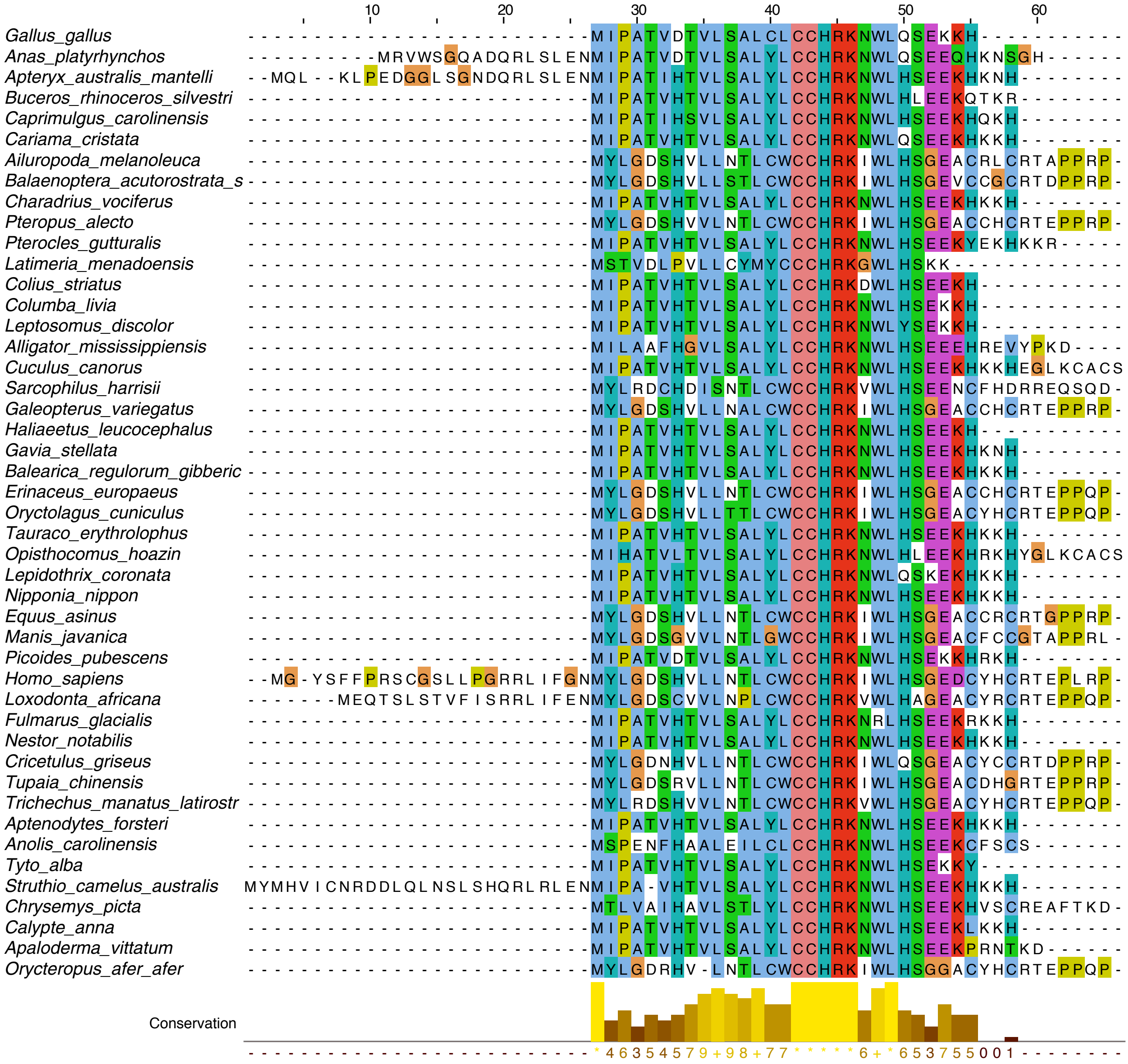

# HG0034

### ENSDARG00000060991 (mief1)

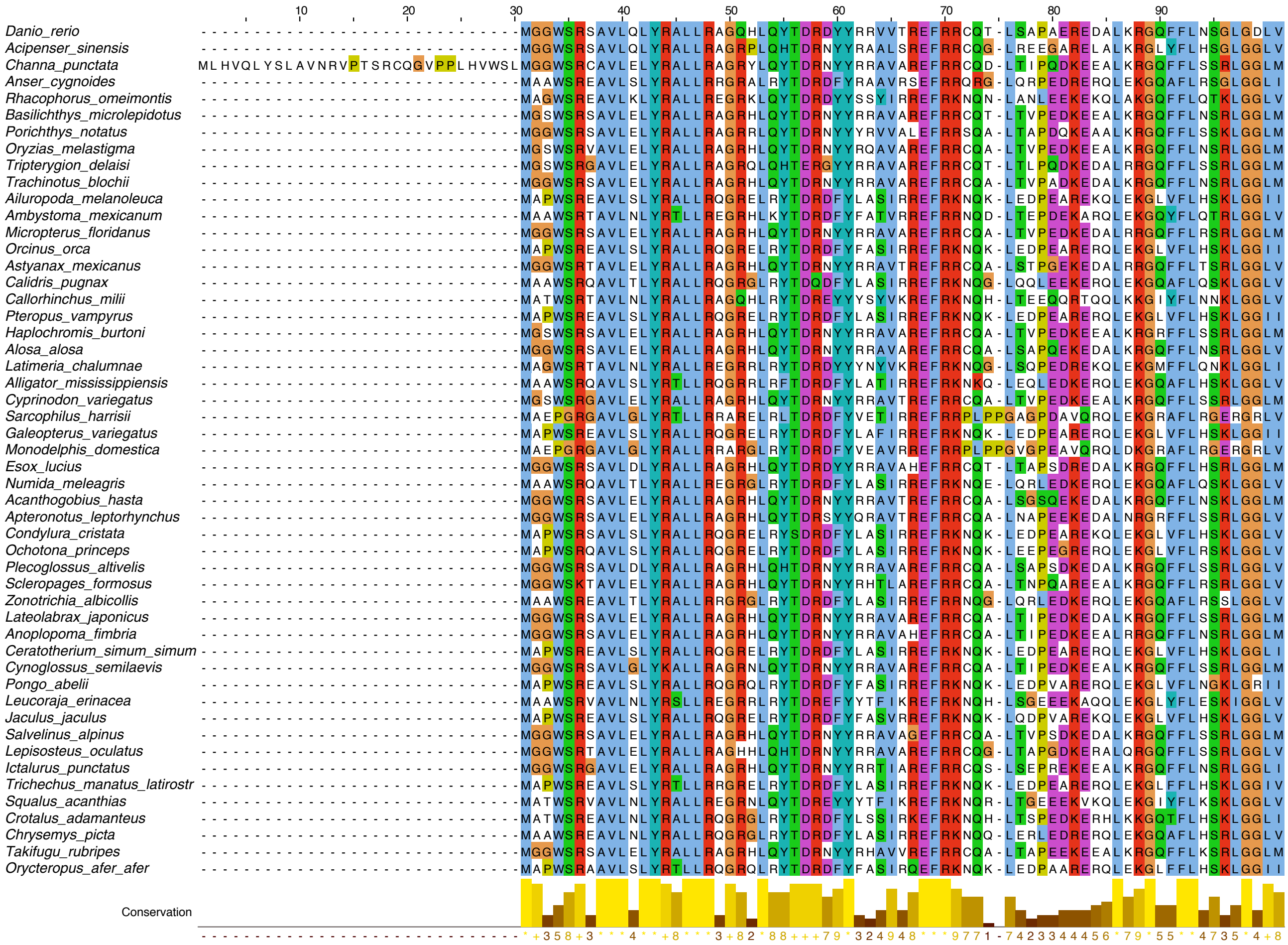

# HG0034

### ENSG00000100335 (MIEF1)

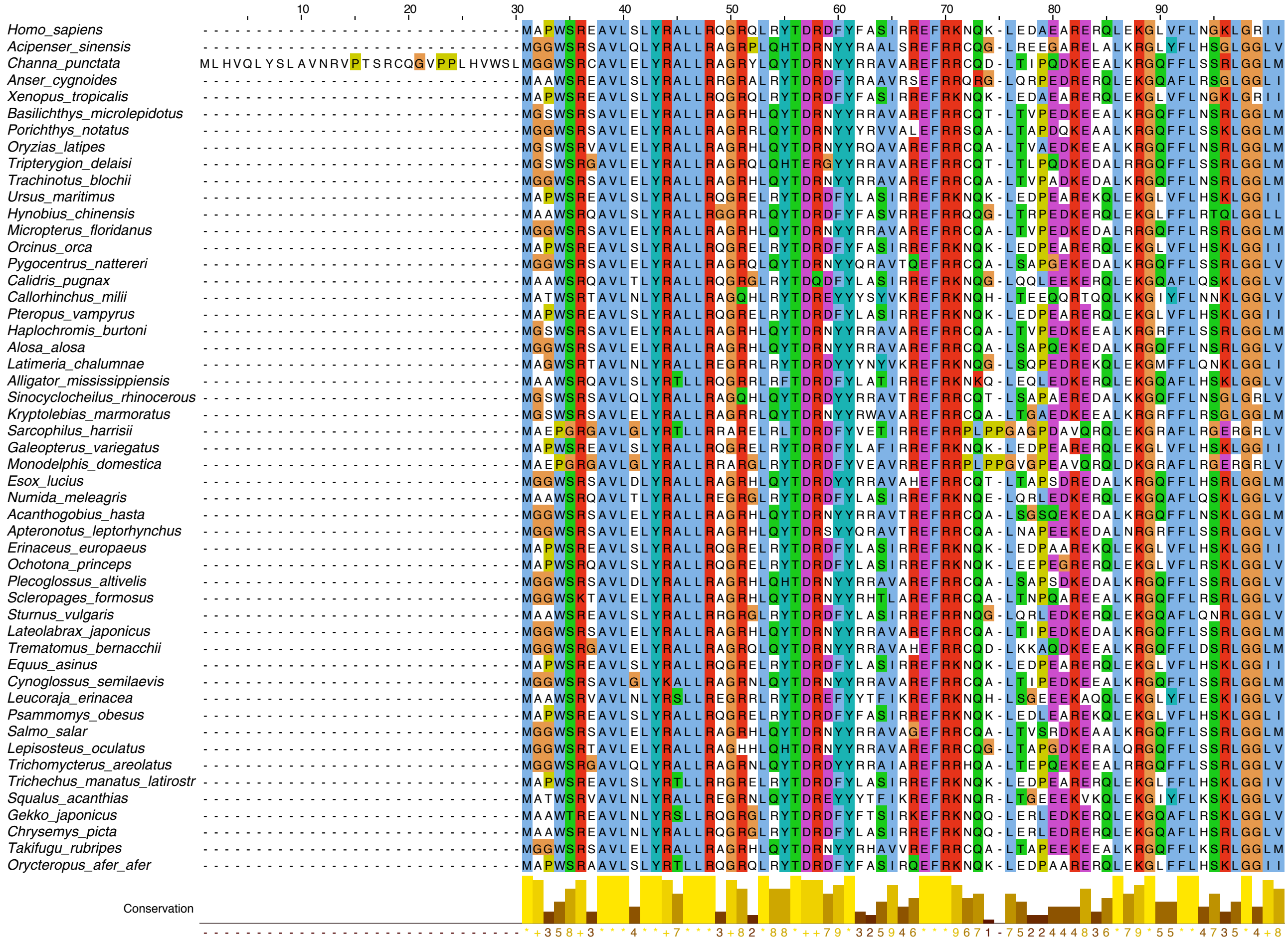

**HG0035**

ENSG00000197147 (LRRC8B)

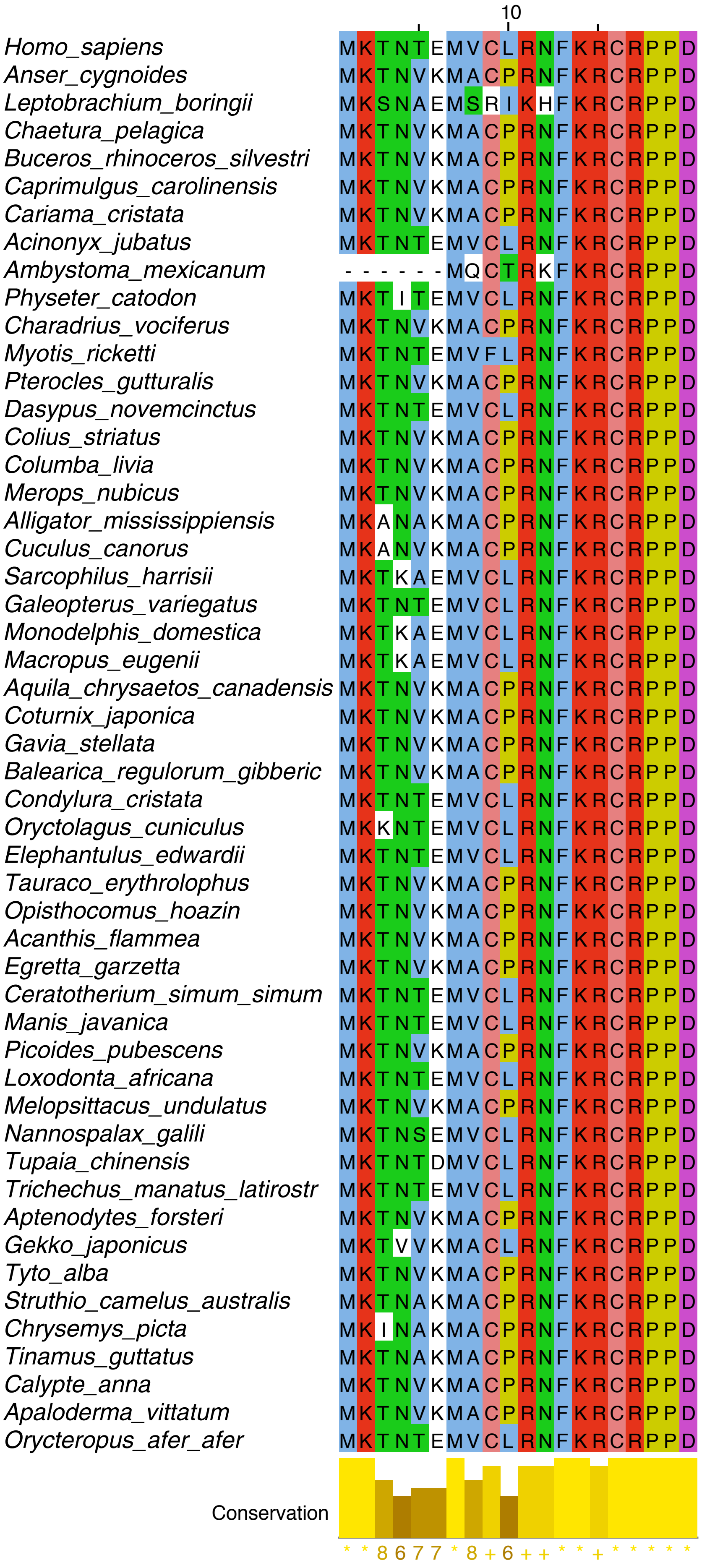

**HG0035**

ENSGALG00000006167 (LRRC8B)

```
                                          10        20        30        40
Gallus_gallus                   ----------------------MKTNVKMACPRNFKRCRPPD
Anser_cygnoides                 ----------------------MKTNVKMACPRNFKRCRPPD
Leptobrachium_boringii          ----------------------MKSNAEMSRIKHFKRCRPPD
Chaetura_pelagica               ----------------------MKTNVKMACPRNFKRCRPPD
Buceros_rhinoceros_silvestri    ----------------------MKTNVKMACPRNFKRCRPPD
Caprimulgus_carolinensis        ----------------------MKTNVKMACPRNFKRCRPPD
Cariama_cristata                ----------------------MKTNVKMACPRNFKRCRPPD
Felis_catus                     --------------MLAAHKCRMKTNTEMVCLRNFKRCRPPD
Ambystoma_mexicanum             ----------------------------MQCTRKFKRCRPPD
Physeter_catodon                ----------------------MKTITEMVCLRNFKRCRPPD
Charadrius_vociferus            ----------------------MKTNVKMACPRNFKRCRPPD
Myotis_ricketti                 ----------------------MKTNTEMVFLRNFKRCRPPD
Pterocles_gutturalis            ----------------------MKTNVKMACPRNFKRCRPPD
Dasypus_novemcinctus            ----------------------MKTNTEMVCLRNFKRCRPPD
Colius_striatus                 ----------------------MKTNVKMACPRNFKRCRPPD
Columba_livia                   ----------------------MKTNVKMACPRNFKRCRPPD
Merops_nubicus                  ----------------------MKTNVKMACPRNFKRCRPPD
Alligator_mississippiensis      ----------------------MKANAKMACPRNFKRCRPPD
Cuculus_canorus                 ----------------------MKANVKMACPRNFKRCRPPD
Sarcophilus_harrisii            ----------------------MKTKAEMVCLRNFKRCRPPD
Galeopterus_variegatus          ----------------------MKTNTEMVCLRNFKRCRPPD
Monodelphis_domestica           ----------------------MKTKAEMVCLRNFKRCRPPD
Macropus_eugenii                ----------------------MKTKAEMVCLRNFKRCRPPD
Aquila_chrysaetos_canadensis    ----------------------MKTNVKMACPRNFKRCRPPD
Gavia_stellata                  ----------------------MKTNVKMACPRNFKRCRPPD
Balearica_regulorum_gibberic    ----------------------MKTNVKMACPRNFKRCRPPD
Condylura_cristata              ----------------------MKTNTEMVCLRNFKRCRPPD
Oryctolagus_cuniculus           ----------------------MKKNTEMVCLRNFKRCRPPD
Elephantulus_edwardii           ----------------------MKTNTEMVCLRNFKRCRPPD
Tauraco_erythrolophus           ----------------------MKTNVKMACPRNFKRCRPPD
Opisthocomus_hoazin             ----------------------MKTNVKMACPRNFKKCRPPD
Acanthis_flammea                ----------------------MKTNVKMACPRNFKRCRPPD
Egretta_garzetta                ----------------------MKTNVKMACPRNFKRCRPPD
Ceratotherium_simum_simum       ----------------------MKTNTEMVCLRNFKRCRPPD
Manis_javanica                  ----------------------MKTNTEMVCLRNFKRCRPPD
Picoides_pubescens              ----------------------MKTNVKMACPRNFKRCRPPD
Homo_sapiens                    ----------------------MKTNTEMVCLRNFKRCRPPD
Loxodonta_africana              ----------------------MKTNTEMVCLRNFKRCRPPD
Melopsittacus_undulatus         ----------------------MKTNVKMACPRNFKRCRPPD
Nannospalax_galili              ----------------------MKTNSEMVCLRNFKRCRPPD
Tupaia_chinensis                ----------------------MKTNTDMVCLRNFKRCRPPD
Trichechus_manatus_latirostr    ----------------------MKTNTEMVCLRNFKRCRPPD
Aptenodytes_forsteri            ----------------------MKTNVKMACPRNFKRCRPPD
Anolis_carolinensis             ----------------------MKTLVRMACPRNFKRCRPPD
Tyto_alba                       ----------------------MKTNVKMACPRNFKRCRPPD
Struthio_camelus_australis      ----------------------MKTNAKMACPRNFKRCRPPD
Chelonia_mydas                  MRVLFWKREPTELESHLKCLSRMKINAKMACPRNFKRCRPPD
Tinamus_guttatus                ----------------------MKTNAKMACPRNFKRCRPPD
Calypte_anna                    ----------------------MKTNVKMACPRNFKRCRPPD
Apaloderma_vittatum             ----------------------MKTNVKMACPRNFKRCRPPD
Orycteropus_afer_afer           ----------------------MKTNTEMVCLRNFKRCRPPD

Conservation                    ----------------------**8677*8+6++**+******
```

**HG0036**

ENSG00000155096 (AZIN1)

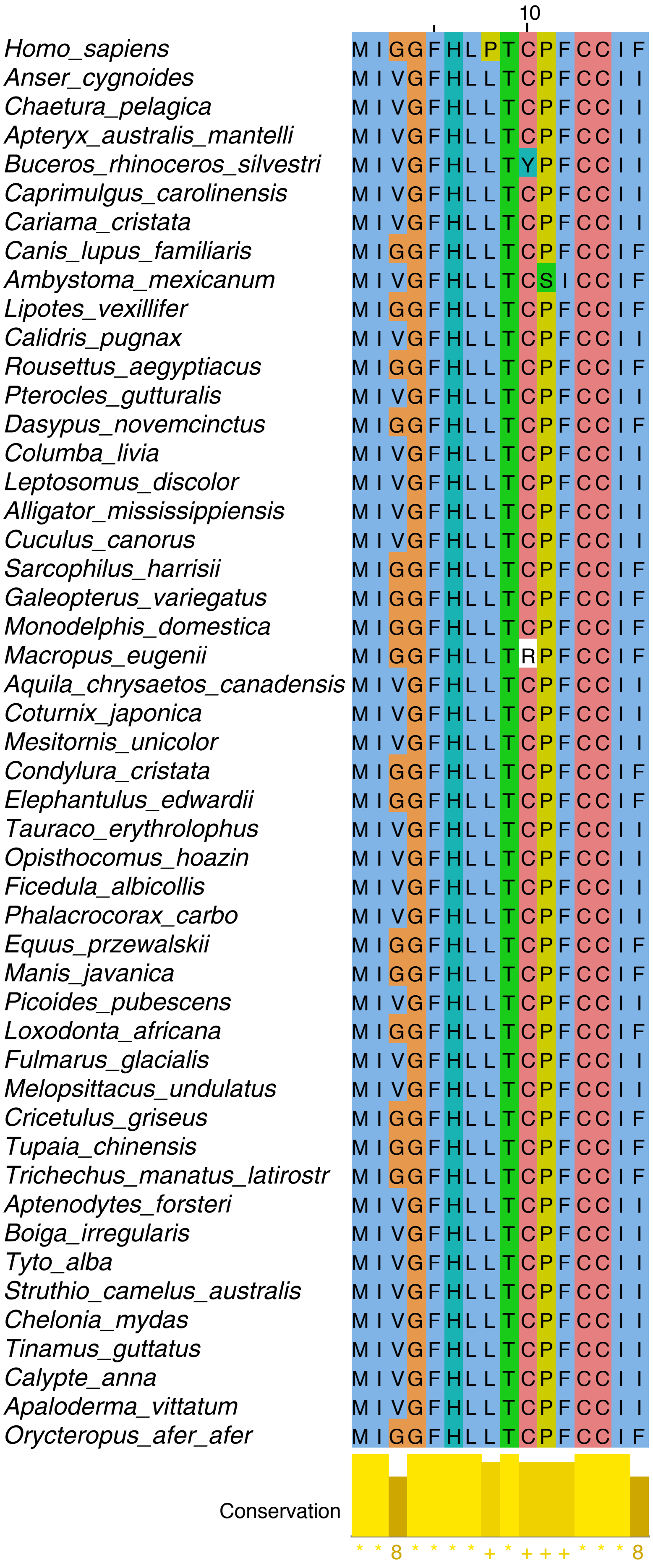

**HG0036**

ENSGALG00000029295

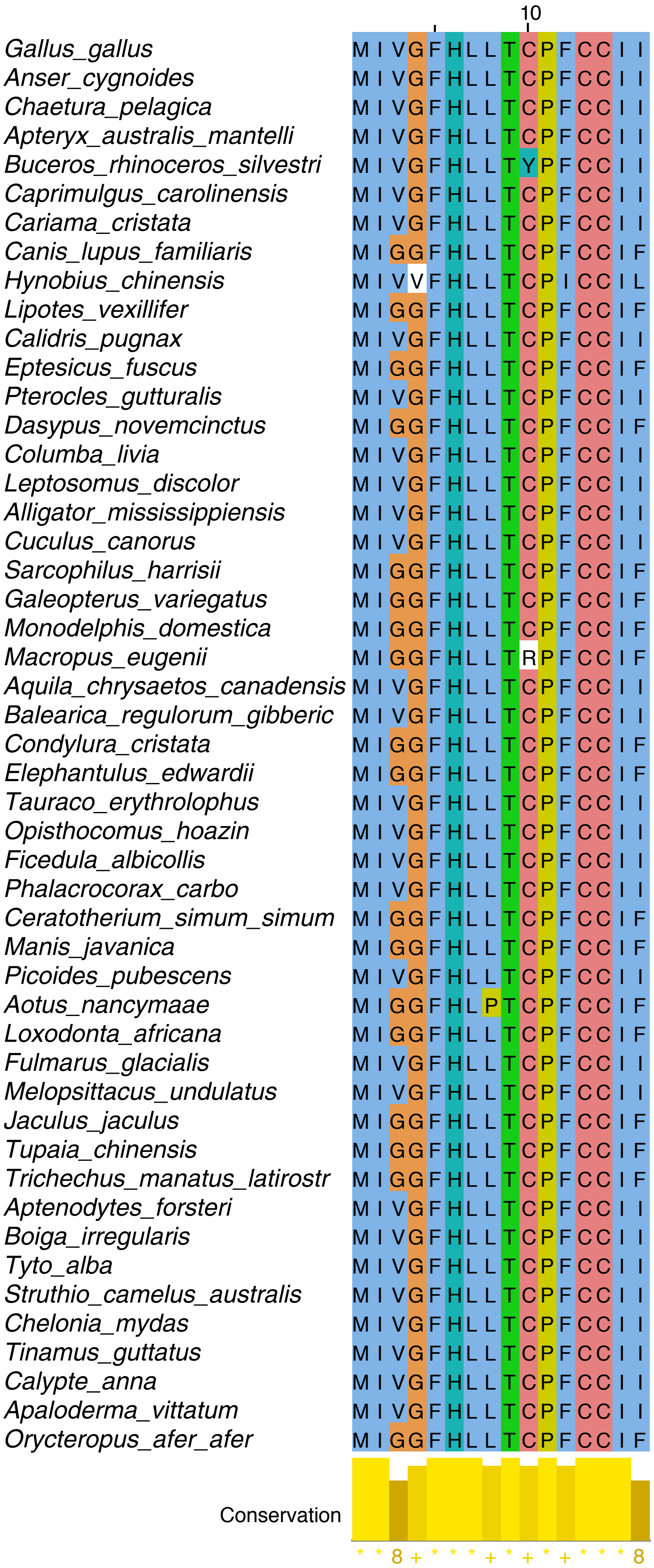

**HG0037**

ENSG00000140092 (FBLN5)

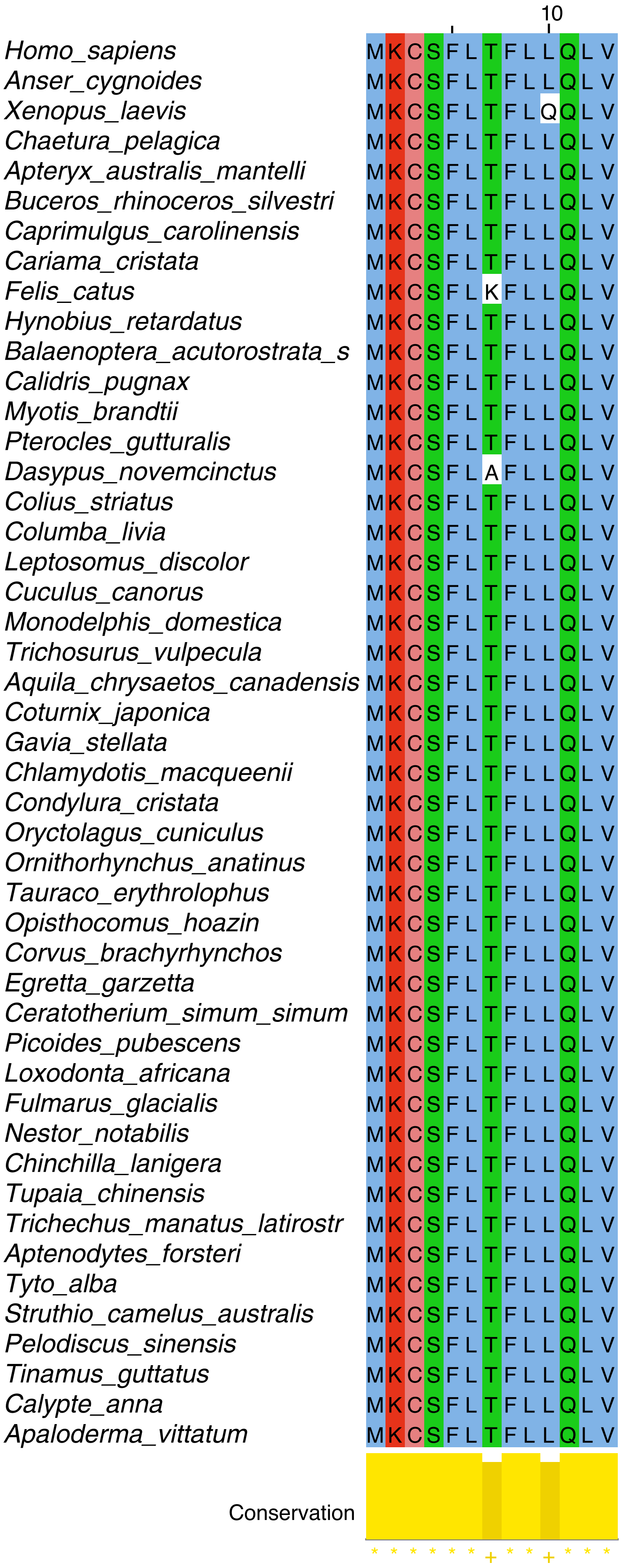

**HG0037**

ENSGALG00000010747 (FBLN5)

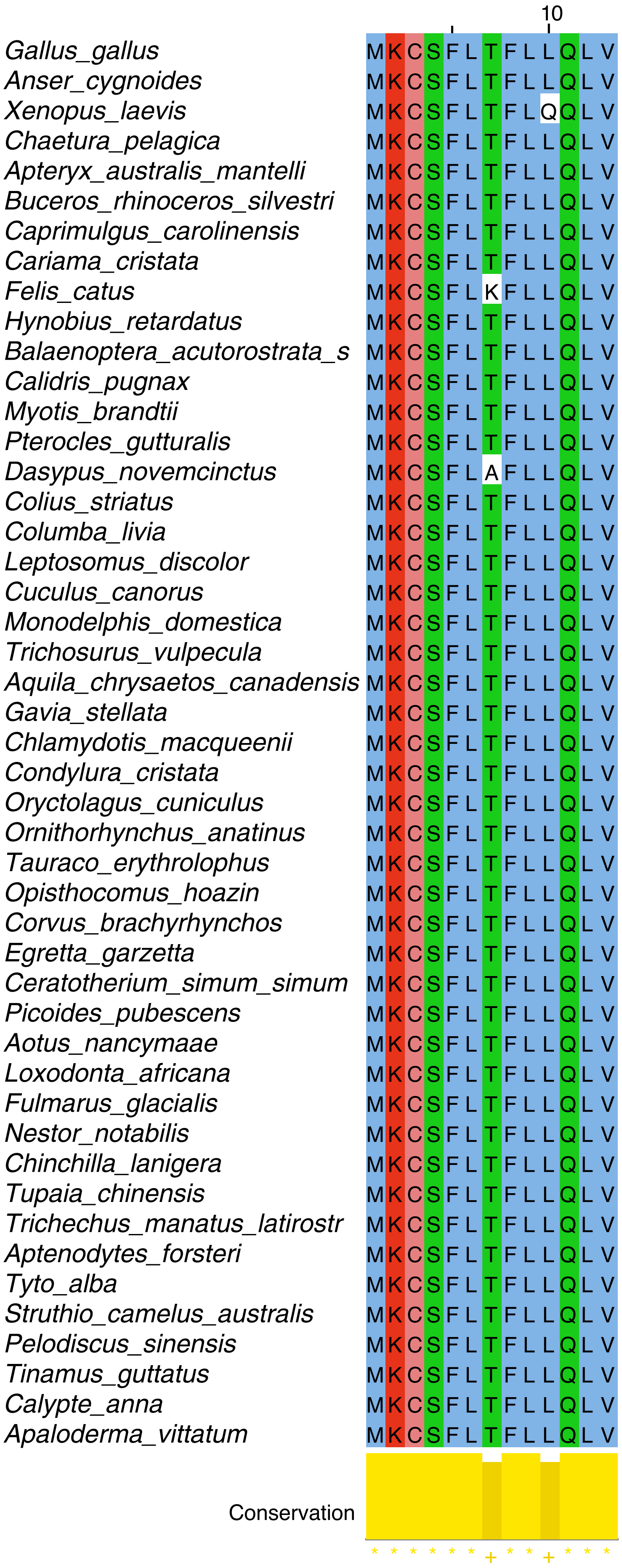

**HG0038**

ENSGALG00000006571

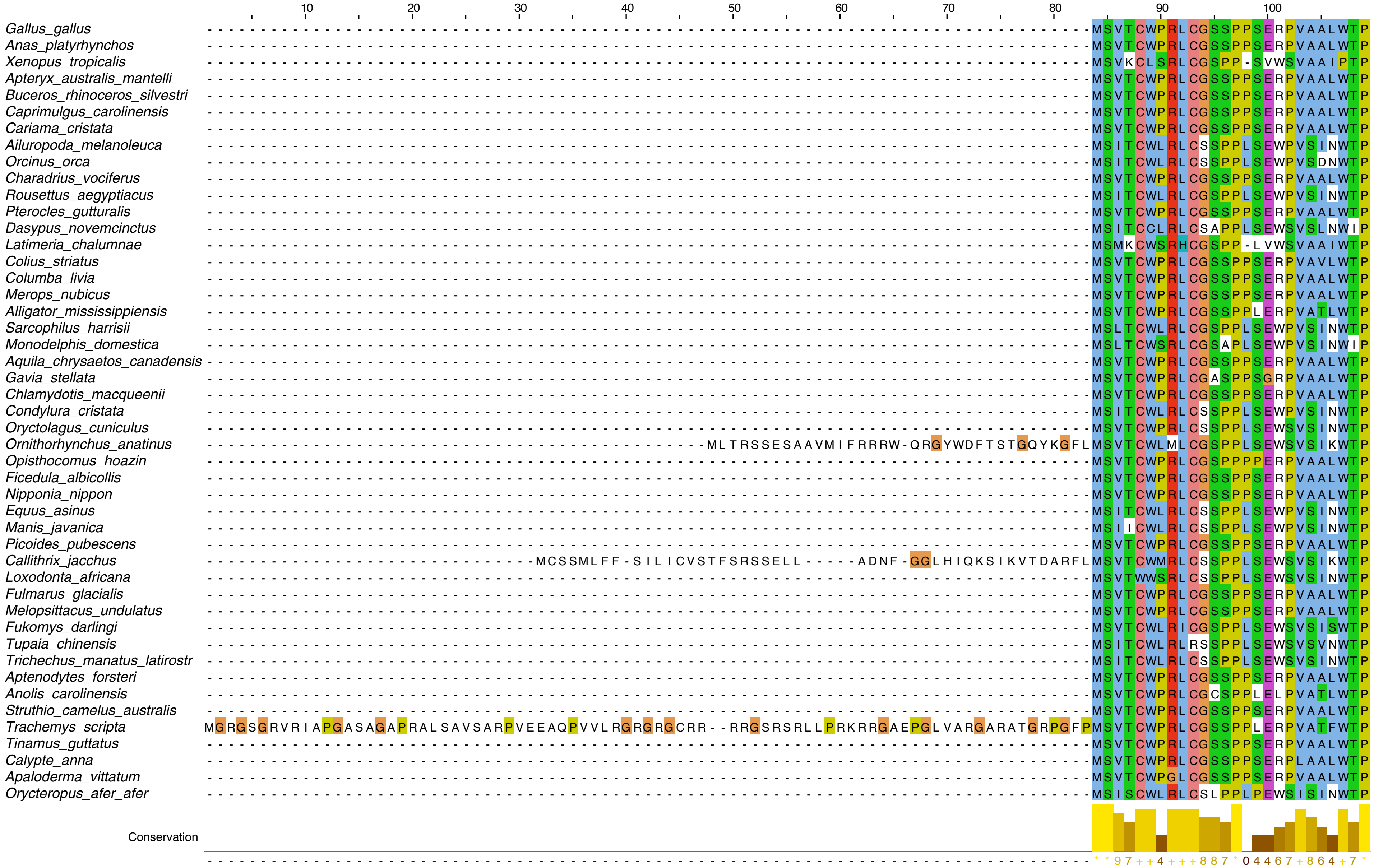

**HG0039.1**

ENSG00000077092 (RARB)

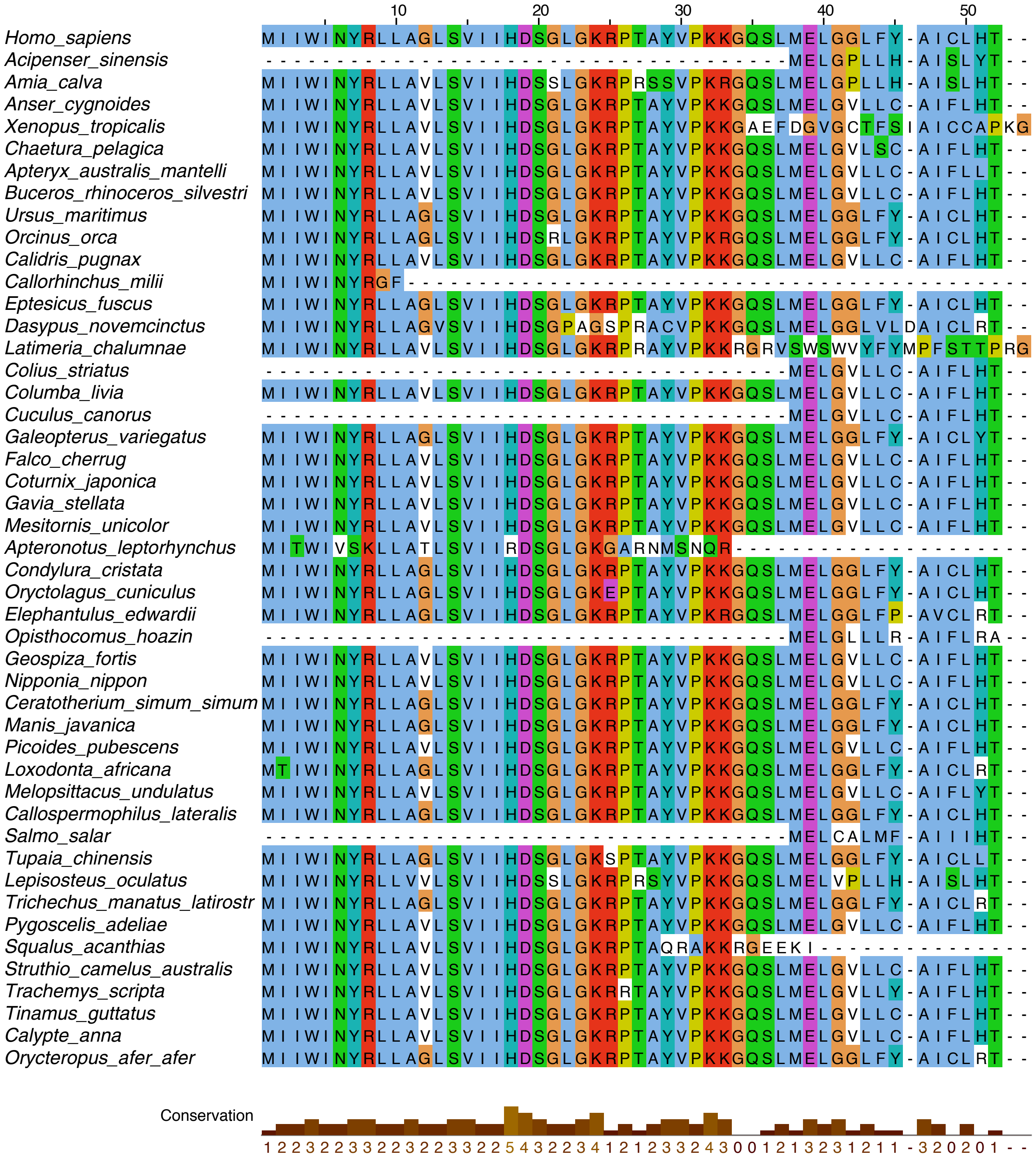

**HG0039.2**

ENSG00000077092 (RARB)

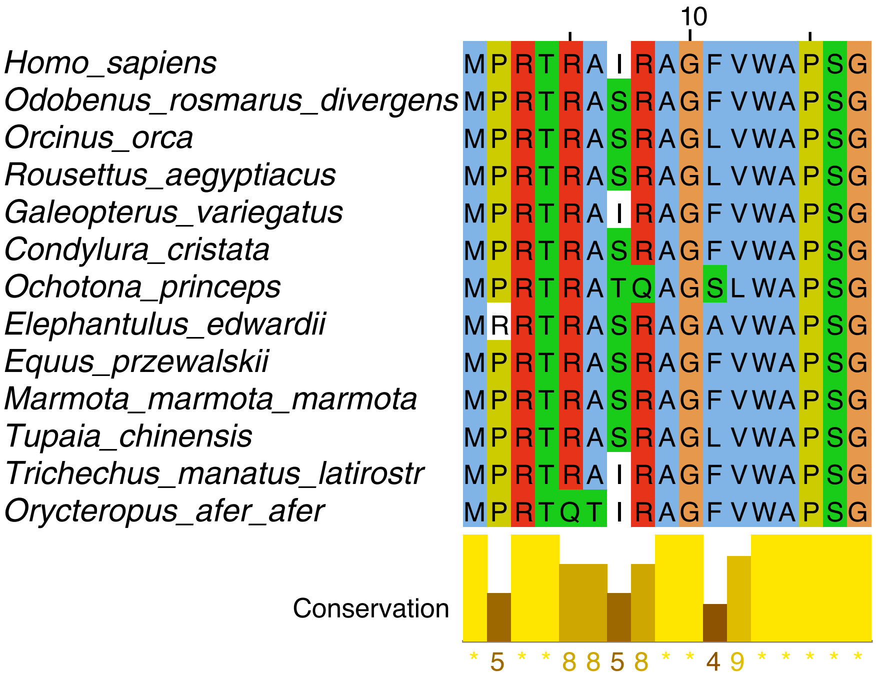

# HG0040.1

### ENSG00000164190 (NIPBL)

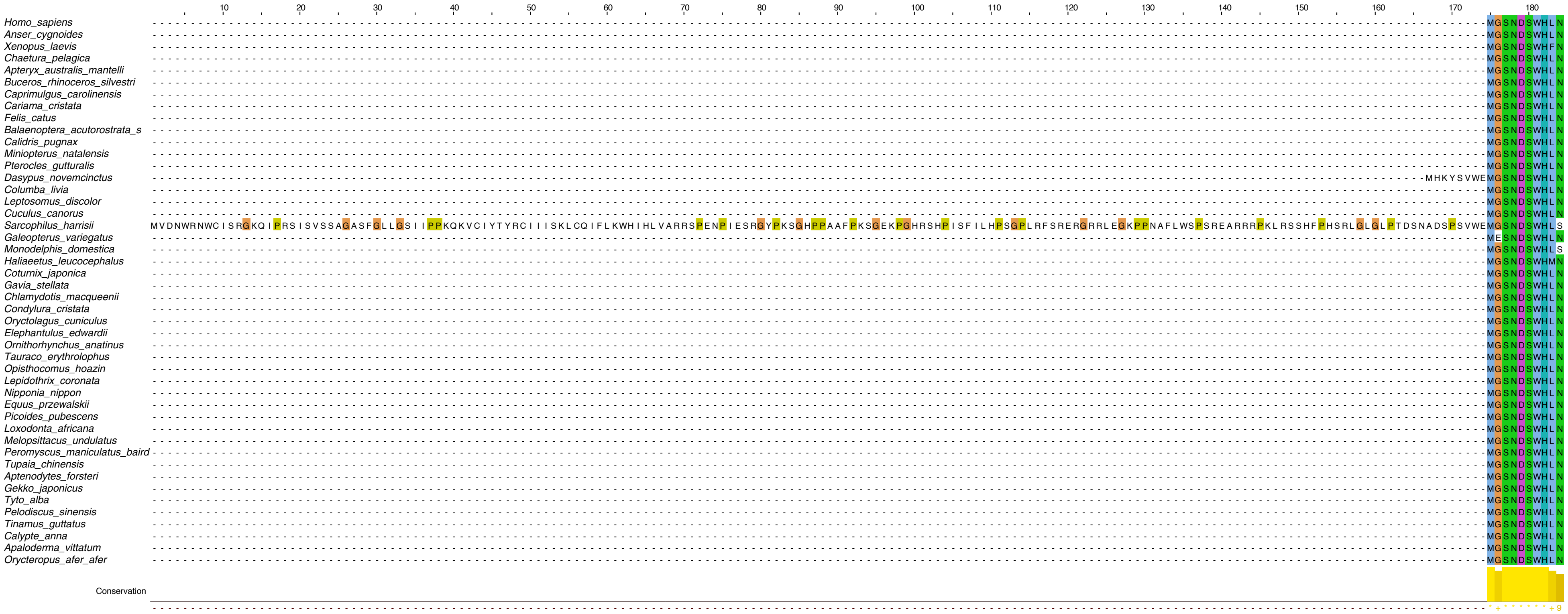

**HG0040.2**

ENSG00000164190 (NIPBL)

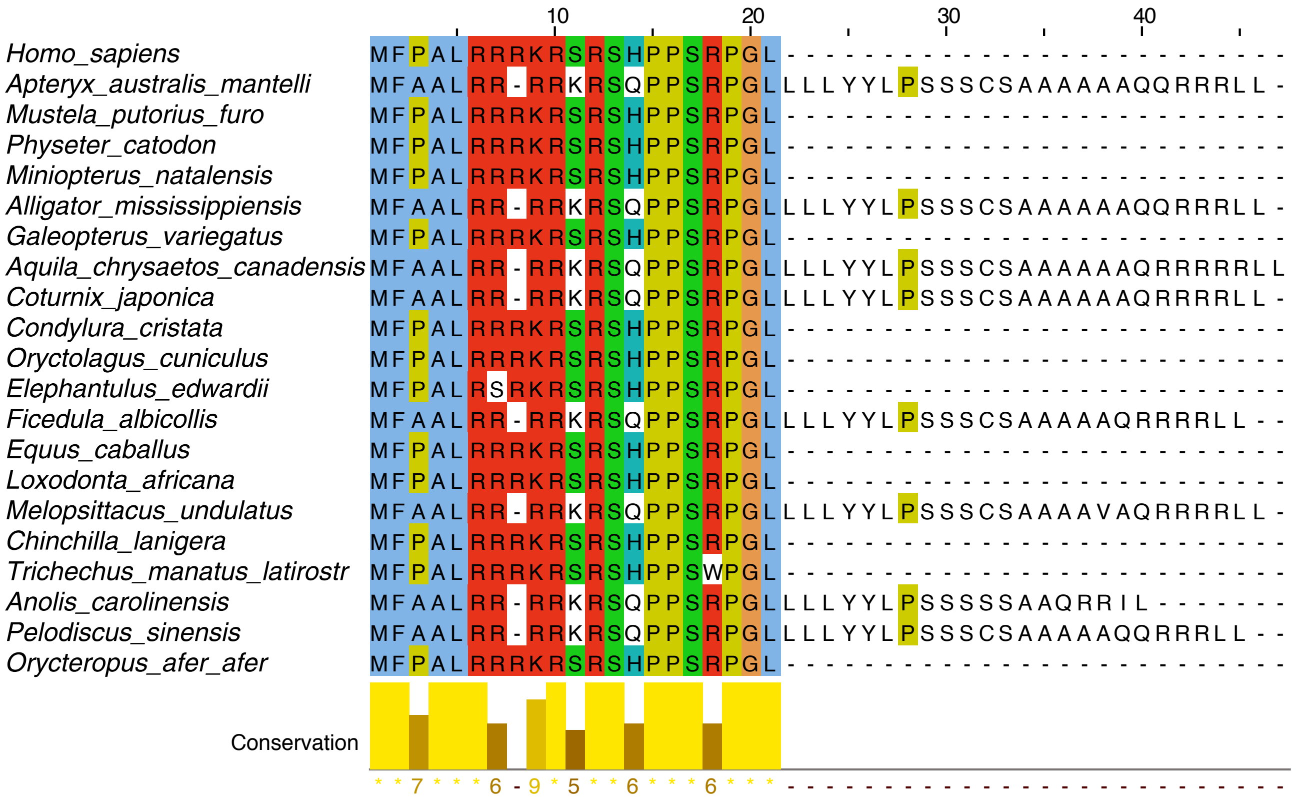

**HG0041**

ENSDARG00000061352 (bcl11aa)

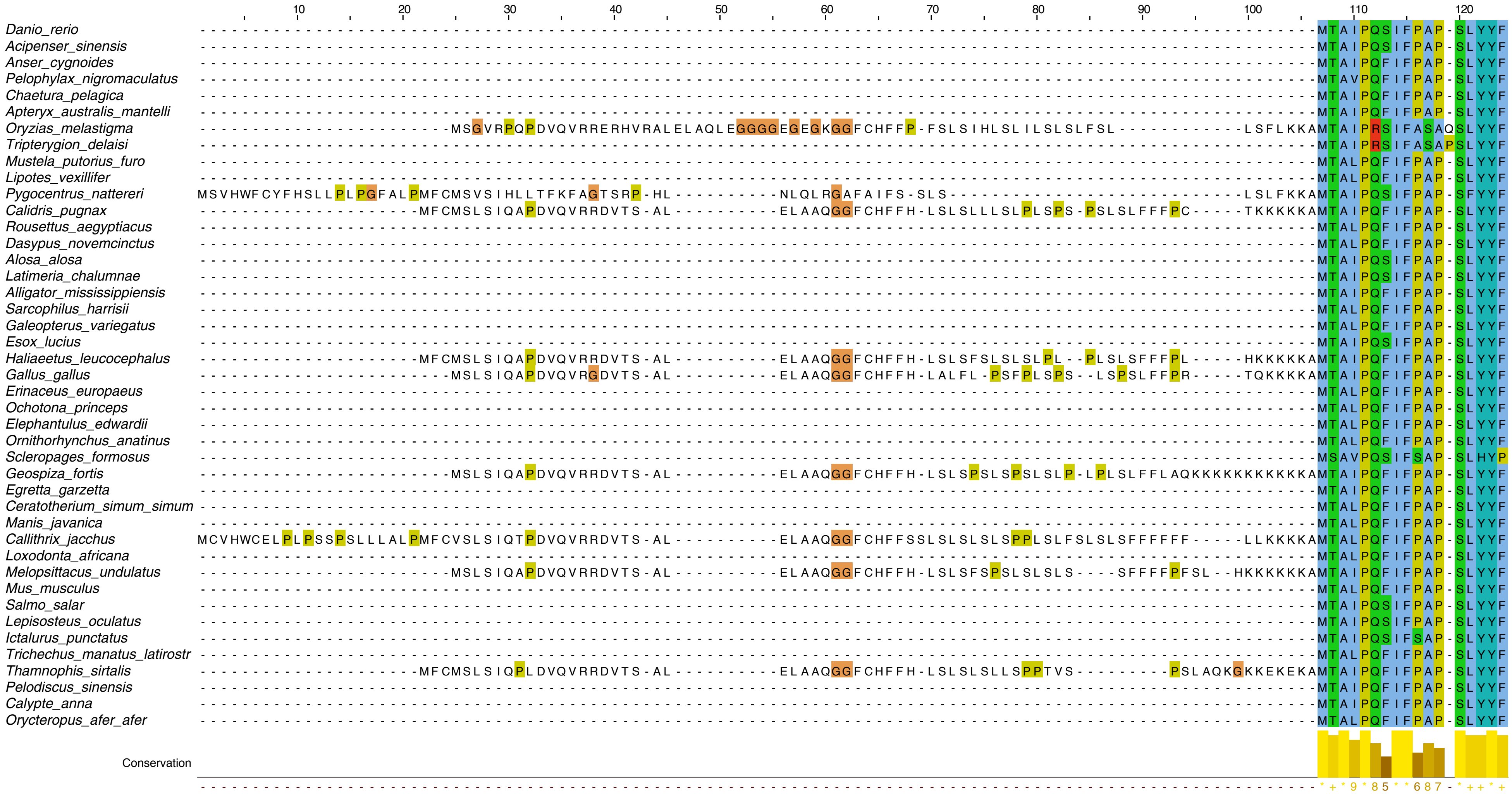

**HG0041**

ENSG00000119866 (BCL11A)

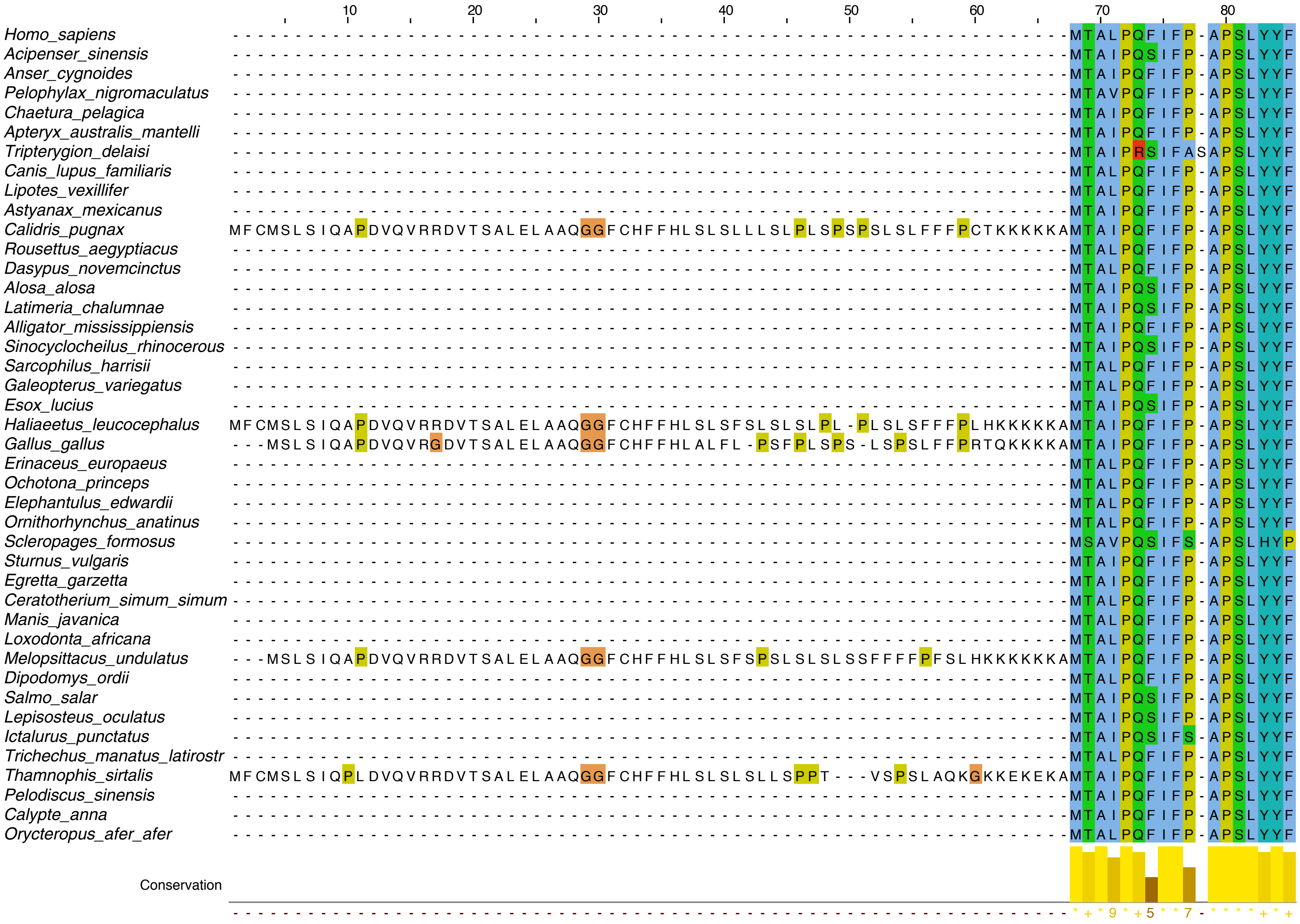

**HG0041**

ENSGALG00000034048

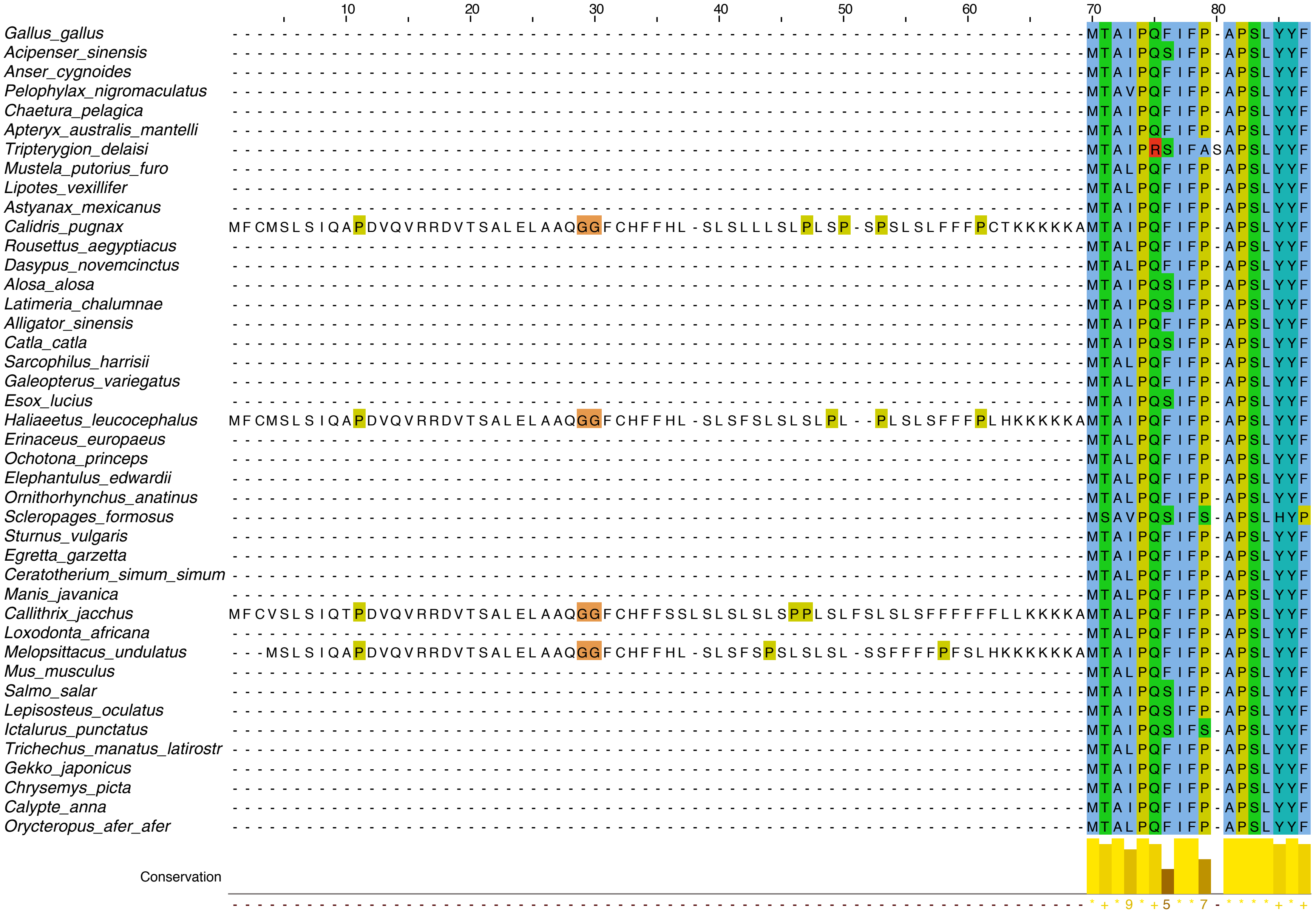

**HG0042.1**

ENSG00000140937 (CDH11)

```
                                            10
Homo_sapiens                       ----MGG-VCYRNWQQVSNG
Anas_platyrhynchos                 ----MCG-ICYVNWQQVSSG
Chaetura_pelagica                  ----MCG-ICYENWQRVSSG
Apteryx_australis_mantelli         ----MCG-ICYVSWQQVSSG
Buceros_rhinoceros_silvestri       ----MCG-ICYINWQQVSSG
Caprimulgus_carolinensis           ----MCG-ICYINWQQVSSG
Acinonyx_jubatus                   ----MGG-VCYRNWQQVSNG
Balaenoptera_acutorostrata_s       MTNQMGGVLRYRNWQQVSNG
Charadrius_vociferus               ----MCG-ICYINWQQVSSG
Eptesicus_fuscus                   ----MGG-VCYRNWQQVSNG
Pterocles_gutturalis               ----MCG-ICYINWQQVSSG
Dasypus_novemcinctus               ----MGE-LSYRNWQQVSNG
Columba_livia                      ----MCG-ICYINWQQVSSG
Leptosomus_discolor                ----MCG-ICYINWQQVSSG
Alligator_mississippiensis         ----MGG-ICYINWQQVTNG
Galeopterus_variegatus             ----MGR-VCYRNWQQVSNG
Monodelphis_domestica              ----MGG-VCYRNWQQVSNG
Falco_cherrug                      ----MCG-ICYINWQQVSSG
Coturnix_japonica                  ----MCG-ICYINWQRVSSG
Gavia_stellata                     ----MCG-ICYINWQQVSSG
Chlamydotis_macqueenii             ----MCG-ICYINWQQVSSG
Sorex_araneus                      ----MGG-VCYRNWQQVSNG
Oryctolagus_cuniculus              ----MGG-VCYRNWQQVSNG
Tauraco_erythrolophus              ----MCG-ICYINWQQVSSG
Opisthocomus_hoazin                ----MCG-ICYINWQQVSSG
Pseudopodoces_humilis              ----MCG-ICYINWQQVSSG
Phaethon_lepturus                  ----MCG-ICYINWQQVSNG
Ceratotherium_simum_simum          ----MGG-VCYRNWQQVSNG
Manis_javanica                     ----MGG-VCYRNWQQVSNG
Picoides_pubescens                 ----MCG-ICYINWQQVPSG
Loxodonta_africana                 ----MGG-VYYRNWQQVSNG
Fulmarus_glacialis                 ----MCG-ICYINWQQVSSG
Melopsittacus_undulatus            ----MCG-ICYINWQQVSSG
Heterocephalus_glaber              ----MGG-VCYRNWQQVSNG
Tupaia_chinensis                   ----MGG-LCYRNWQQVSNG
Aptenodytes_forsteri               ----MSG-ICYITWQQVSSG
Takydromus_sexlineatus             ----MGG-ICYFNWQWVSSG
Struthio_camelus_australis         ----MCG-ICYRSWQQVSSG
Chelonia_mydas                     ----MGG-ICYINWQQVSSG
Tinamus_guttatus                   ----MCG-ICYISWQQVSSG
Calypte_anna                       ----MCG-ICYINWQQVSSG
Apaloderma_vittatum                ----MCG-TCYINWQQVSSG
Orycteropus_afer_afer              MTNQMGR-VCYRNWQQVSNG
```

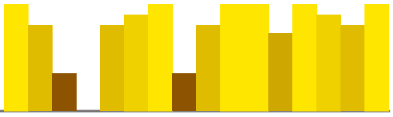

Conservation

```
----*94-9+*49**8*+9*
```

**HG0042.2**

ENSGALG00000039985

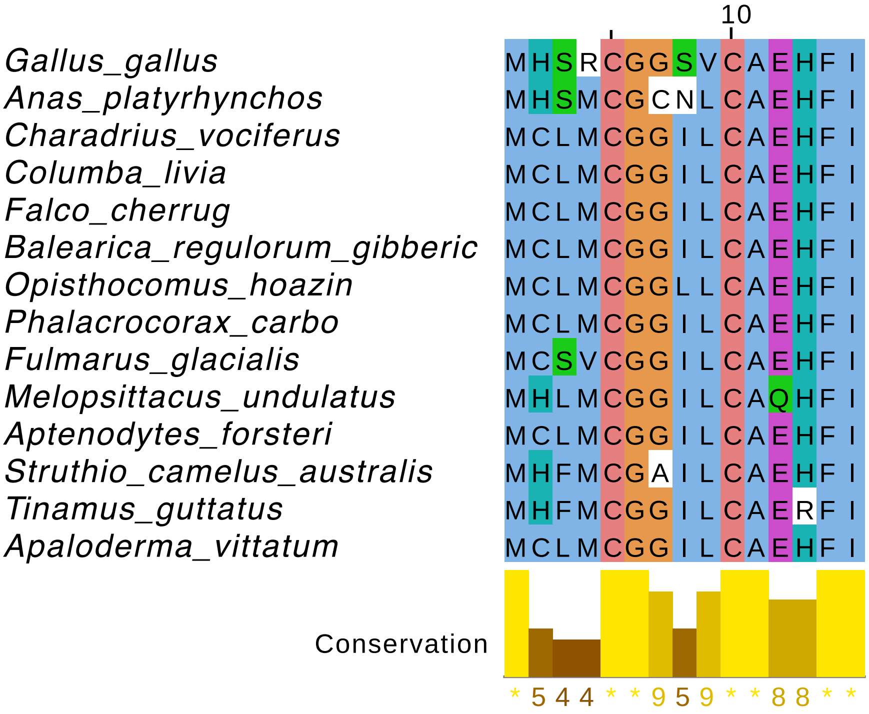

**HG0042.3**

ENSG00000140937 (CDH11)

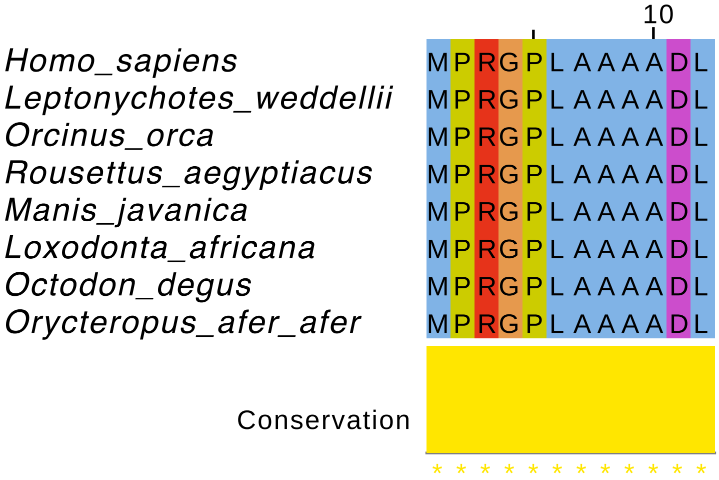

**HG0042.4**

ENSG00000140937 (CDH11)

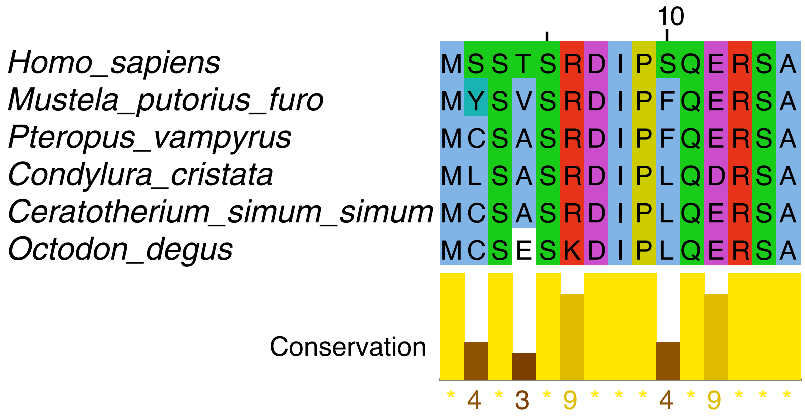

**HG0042.5**

ENSG00000140937 (CDH11)

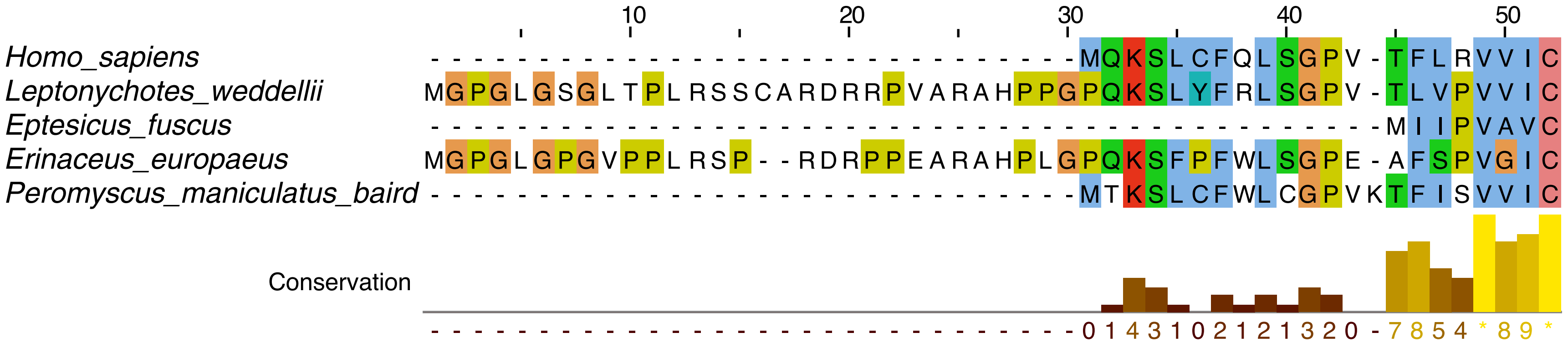

**HG0043**

ENSG00000172995 (ARPP21)

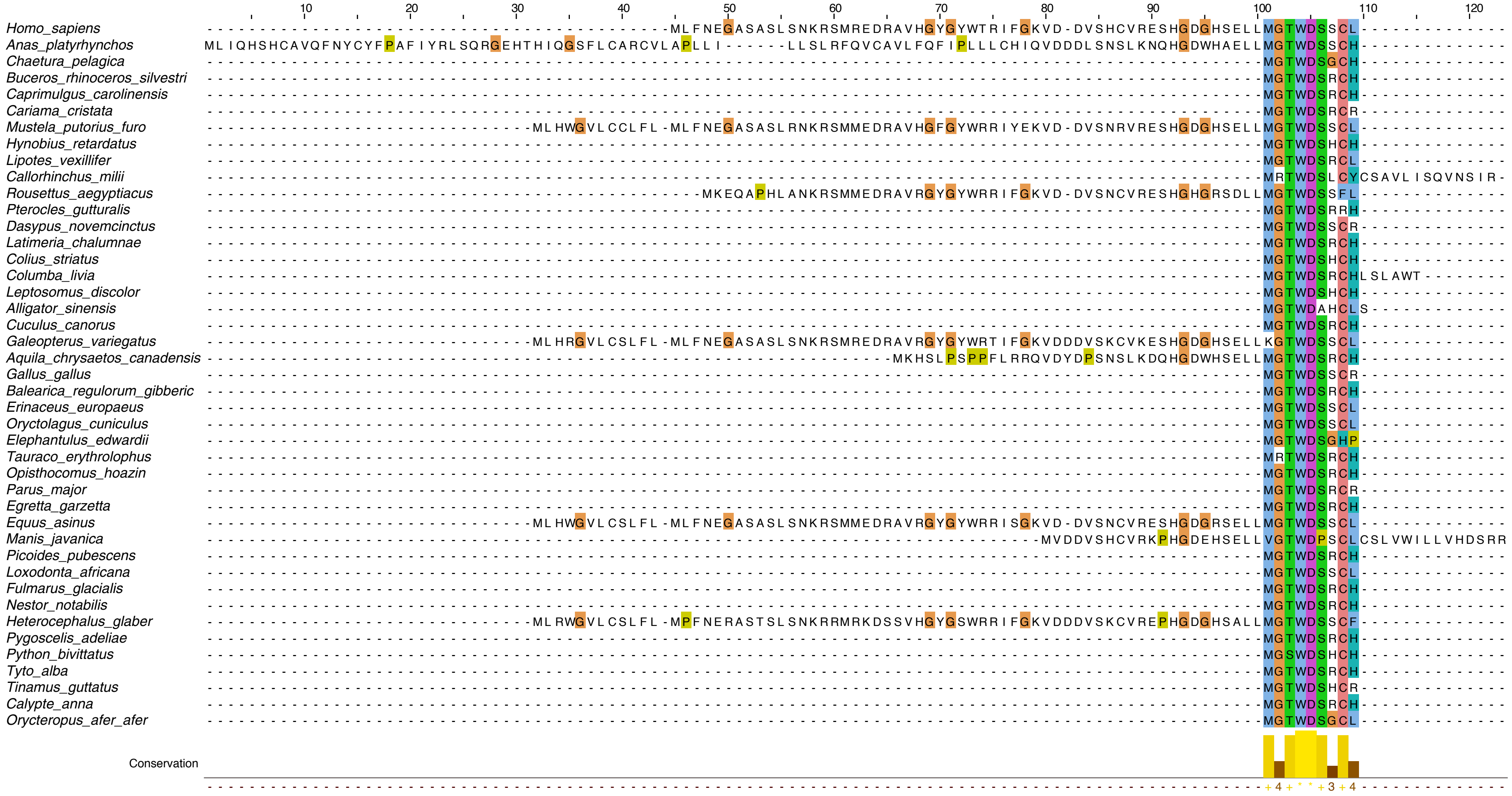

**HG0044**

ENSDARG00000061687 (atxn1a)

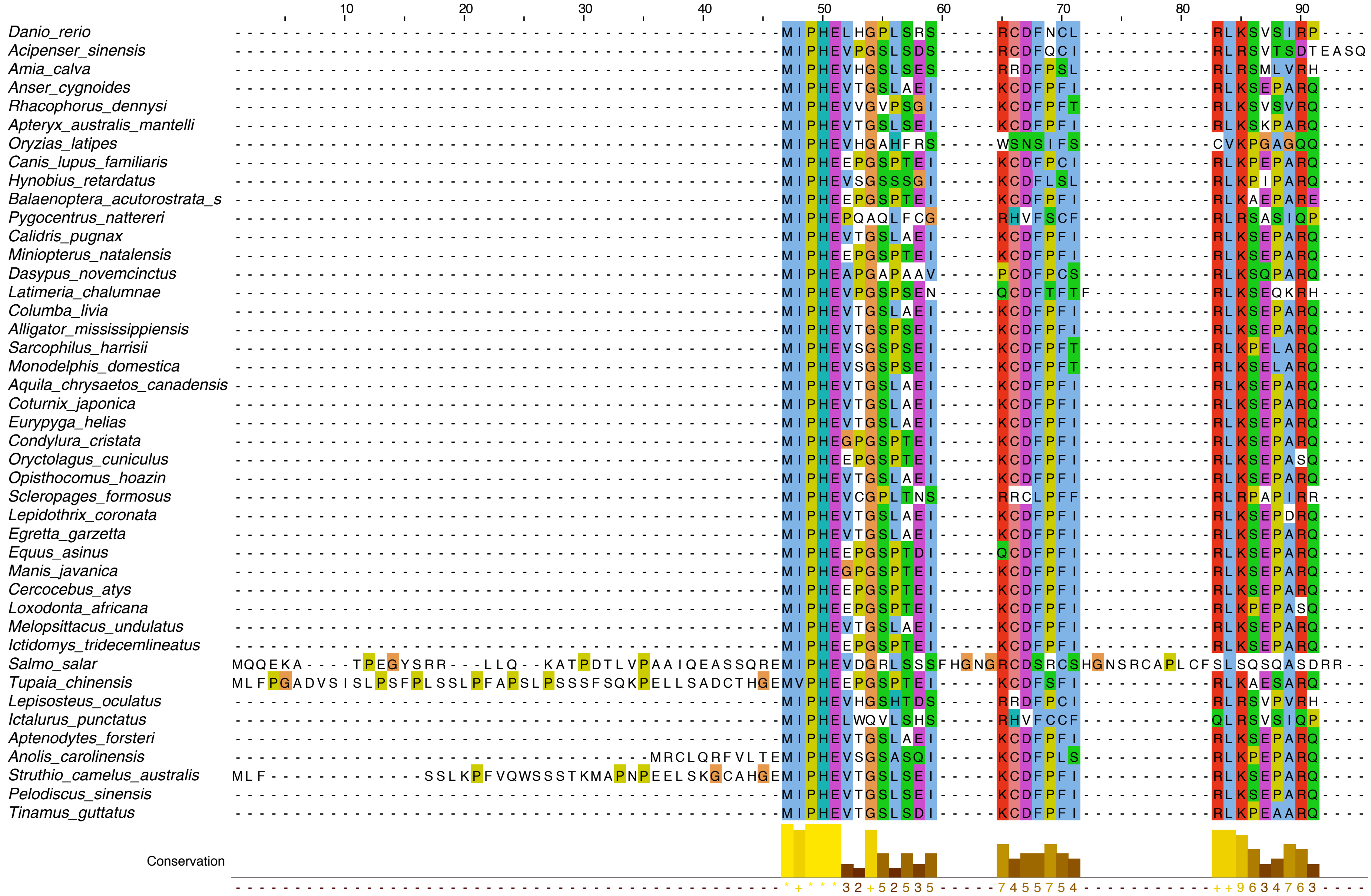

**HG0044**

ENSG00000124788 (ATXN1)

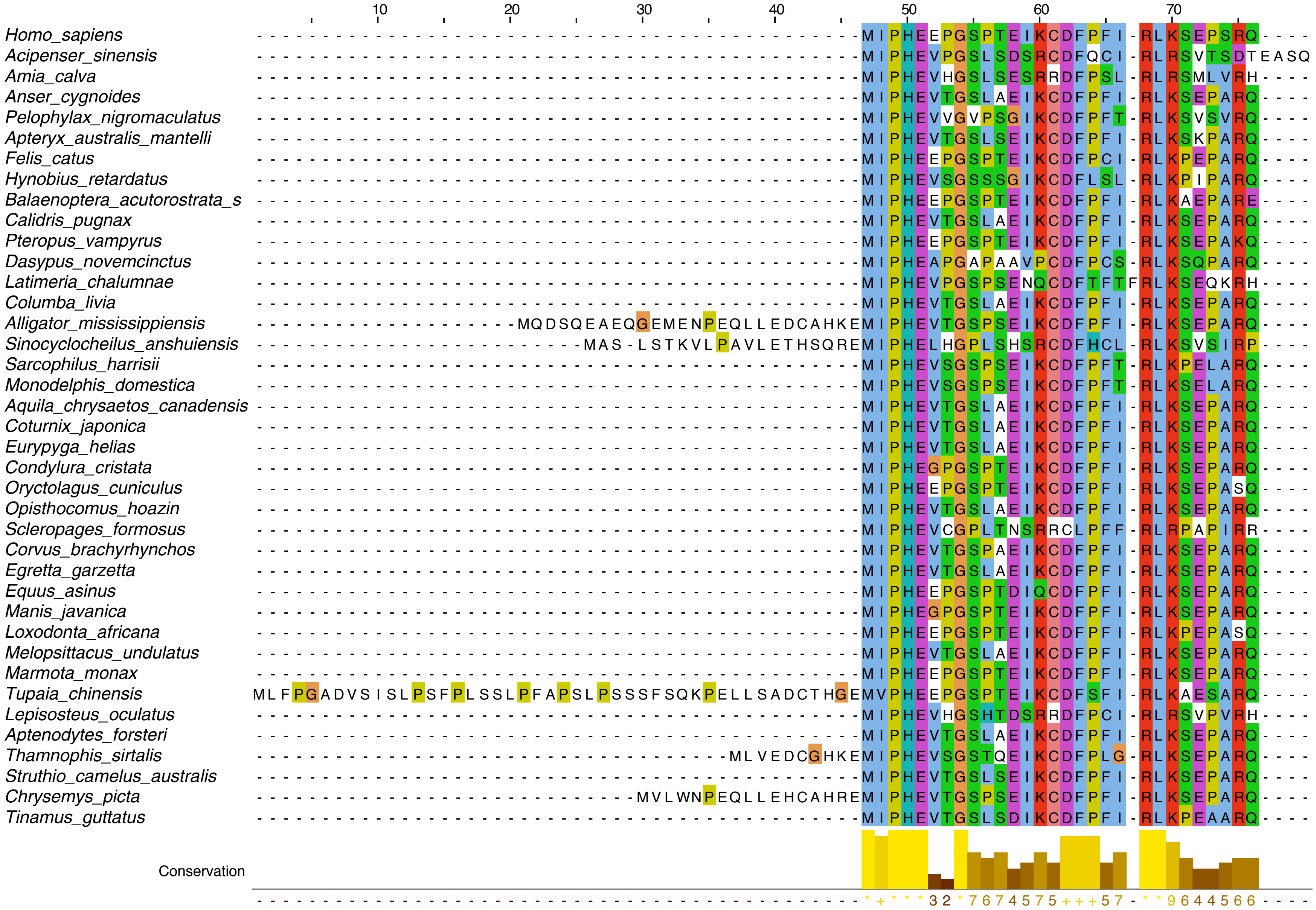

**HG0045.1**

ENSG00000083168 (KAT6A)

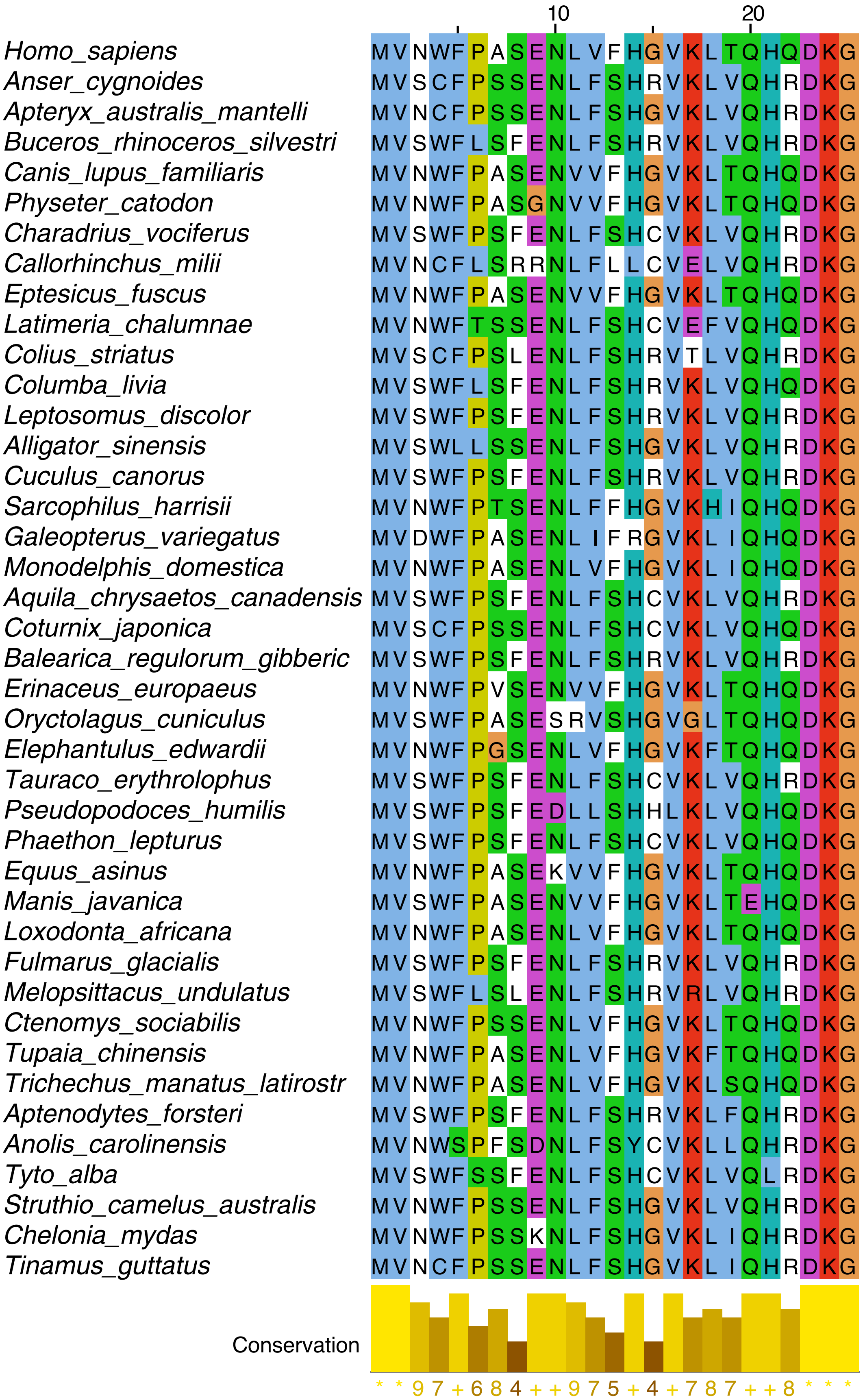

**HG0045.2**

ENSG00000083168 (KAT6A)

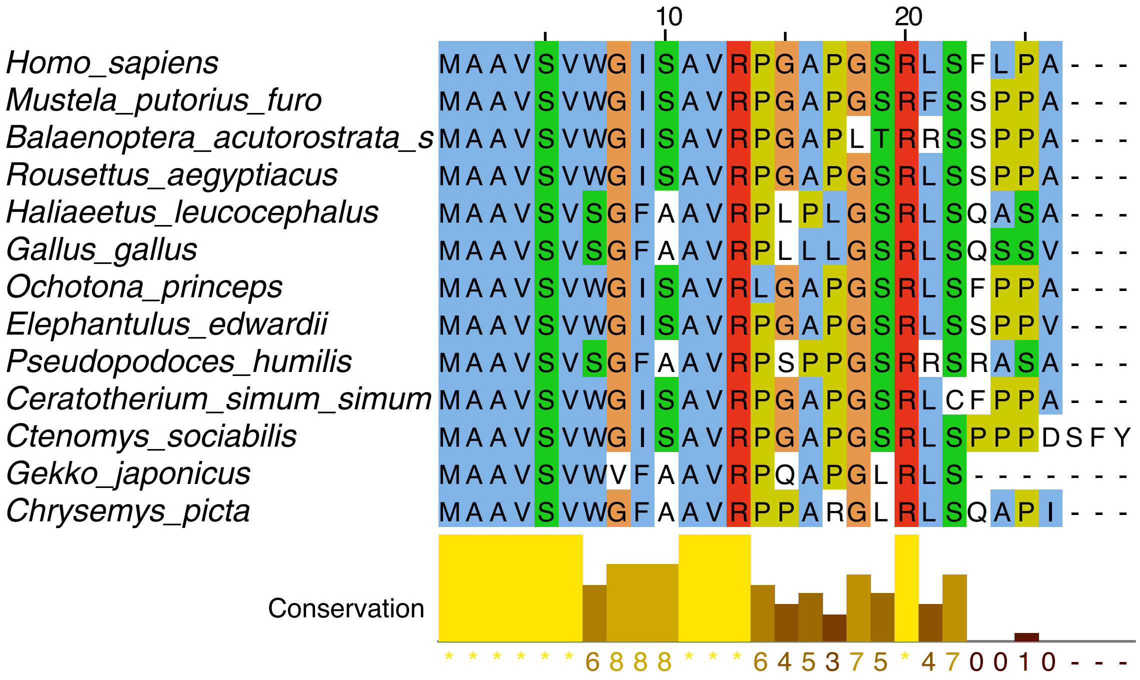

**HG0046**

ENSDARG00000018817 (bdnf)

```
                              10
Danio_rerio                   MMTSNKTHYLPASVGVCGY
Anser_cygnoides               MMTSNKTHYLPASVGETGF
Nanorana_parkeri              MMTSNKRHYLPASVGETGF
Chaetura_pelagica             MMTSNKTHYLPASVGETGF
Canis_lupus_familiaris        MMTSNKTHYLPASVGETGF
Lipotes_vexillifer            MMTSNKTHYLPASVGETGF
Calidris_pugnax               MMTSNKTHYLPASVGETGF
Miniopterus_natalensis        MMTSNKTHYLPASVGETGF
Haplochromis_burtoni          MMTSNKTHYLPASVGVSGY
Columba_livia                 MMTSNKTHYLPASVGETGF
Cuculus_canorus               MMTSNKTHYLPASVGETGF
Austrofundulus_limnaeus       MMTSNKTHYLPASVGVSGY
Galeopterus_variegatus        MMTSNKTHYLPASVGETRF
Monodelphis_domestica         MMTSNKTHYLPASVGETGF
Aquila_chrysaetos_canadensis  MMTSNKTHYLPASVGETGF
Coturnix_japonica             MMTSNKTHYLPASVGETRF
Condylura_cristata            MMTSNKTHYLPASVGETGF
Ochotona_princeps             MMTSNKTHYLPASVGETRF
Elephantulus_edwardii         MMTSNKTHYLPASVGETGF
Opisthocomus_hoazin           MMTSNKTHYLPASVRETGF
Corvus_brachyrhynchos         MMTSNKTHYLPASVGETGF
Egretta_garzetta              MMTSNKTHYLPASVGETGF
Notothenia_coriiceps          MMTSNKTHYLPASVGVSGY
Ceratotherium_simum_simum     MMTSNKTHYLPASVGETGF
Manis_javanica                MMTSNKTHYLPASVGETGF
Cynoglossus_semilaevis        MMTSNKTHYLPASVGVSGY
Aotus_nancymaae               MMTSNKTHYLPASVGETRF
Loxodonta_africana            MMTSNKTHYLPASVGETGF
Melopsittacus_undulatus       MMTSNKTHYLPASVGETGF
Dipodomys_ordii               MMTSNKTHYLPASVGETGF
Salmo_salar                   MMTSNKTHYLPASVGVSGY
Lepisosteus_oculatus          MMTSNKTHYLPASVGESGF
Trichechus_manatus_latirostr  MMTSNKTHYLPASVGETGF
Aptenodytes_forsteri          MMTSNKTHYLPASVGETGF
Anolis_carolinensis           MMTSNETHSLPASGGETGL
Struthio_camelus_australis    MMTSNKTHYLPASVGETGF
Chelonia_mydas                MMTSNKTHYLPASVGETGF
Tinamus_guttatus              MMTSNKTHYLPASVGETGF
Calypte_anna                  MMTSNKTHYLPASVGETGF
Orycteropus_afer_afer         MMTSNKTHYLPASVGETRF
```

Conservation

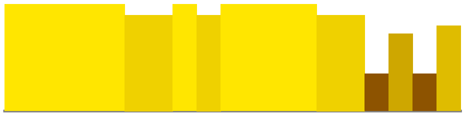

```
* * * * * + + * + * * * * + + 4 8 4 9
```

**HG0046**

ENSG00000176697 (BDNF)

| | |
|---|---|
| *Homo_sapiens* | MMTSNKTHYLPASVGETRF |
| *Anser_cygnoides* | MMTSNKTHYLPASVGETGF |
| *Nanorana_parkeri* | MMTSNKRHYLPASVGETGF |
| *Chaetura_pelagica* | MMTSNKTHYLPASVGETGF |
| *Canis_lupus_familiaris* | MMTSNKTHYLPASVGETGF |
| *Lipotes_vexillifer* | MMTSNKTHYLPASVGETGF |
| *Calidris_pugnax* | MMTSNKTHYLPASVGETGF |
| *Eptesicus_fuscus* | MMTSNKTHYLPASVGETGF |
| *Haplochromis_burtoni* | MMTSNKTHYLPASVGVSGY |
| *Columba_livia* | MMTSNKTHYLPASVGETGF |
| *Cuculus_canorus* | MMTSNKTHYLPASVGETGF |
| *Danio_rerio* | MMTSNKTHYLPASVGVCGY |
| *Austrofundulus_limnaeus* | MMTSNKTHYLPASVGVSGY |
| *Galeopterus_variegatus* | MMTSNKTHYLPASVGETRF |
| *Monodelphis_domestica* | MMTSNKTHYLPASVGETGF |
| *Aquila_chrysaetos_canadensis* | MMTSNKTHYLPASVGETGF |
| *Gallus_gallus* | MMTSNKTHYLPASVGETRF |
| *Condylura_cristata* | MMTSNKTHYLPASVGETGF |
| *Ochotona_princeps* | MMTSNKTHYLPASVGETRF |
| *Elephantulus_edwardii* | MMTSNKTHYLPASVGETGF |
| *Opisthocomus_hoazin* | MMTSNKTHYLPASVRETGF |
| *Acanthis_flammea* | MMTSNKTHYLPASVGETGF |
| *Egretta_garzetta* | MMTSNKTHYLPASVGETGF |
| *Notothenia_coriiceps* | MMTSNKTHYLPASVGVSGY |
| *Ceratotherium_simum_simum* | MMTSNKTHYLPASVGETGF |
| *Manis_javanica* | MMTSNKTHYLPASVGETGF |
| *Cynoglossus_semilaevis* | MMTSNKTHYLPASVGVSGY |
| *Loxodonta_africana* | MMTSNKTHYLPASVGETGF |
| *Melopsittacus_undulatus* | MMTSNKTHYLPASVGETGF |
| *Rattus_norvegicus* | MMTSNKTHYLPASVGETRF |
| *Salmo_salar* | MMTSNKTHYLPASVGVSGY |
| *Lepisosteus_oculatus* | MMTSNKTHYLPASVGESGF |
| *Trichechus_manatus_latirostr* | MMTSNKTHYLPASVGETGF |
| *Aptenodytes_forsteri* | MMTSNKTHYLPASVGETGF |
| *Anolis_carolinensis* | MMTSNETHSLPASGGETGL |
| *Struthio_camelus_australis* | MMTSNKTHYLPASVGETGF |
| *Chelonia_mydas* | MMTSNKTHYLPASVGETGF |
| *Tinamus_guttatus* | MMTSNKTHYLPASVGETGF |
| *Calypte_anna* | MMTSNKTHYLPASVGETGF |
| *Orycteropus_afer_afer* | MMTSNKTHYLPASVGETRF |
| Conservation | * * * * * + + * + * * * * + + 4 8 4 9 |

**HG0047**

ENSGALG00000015422 (NRG1)

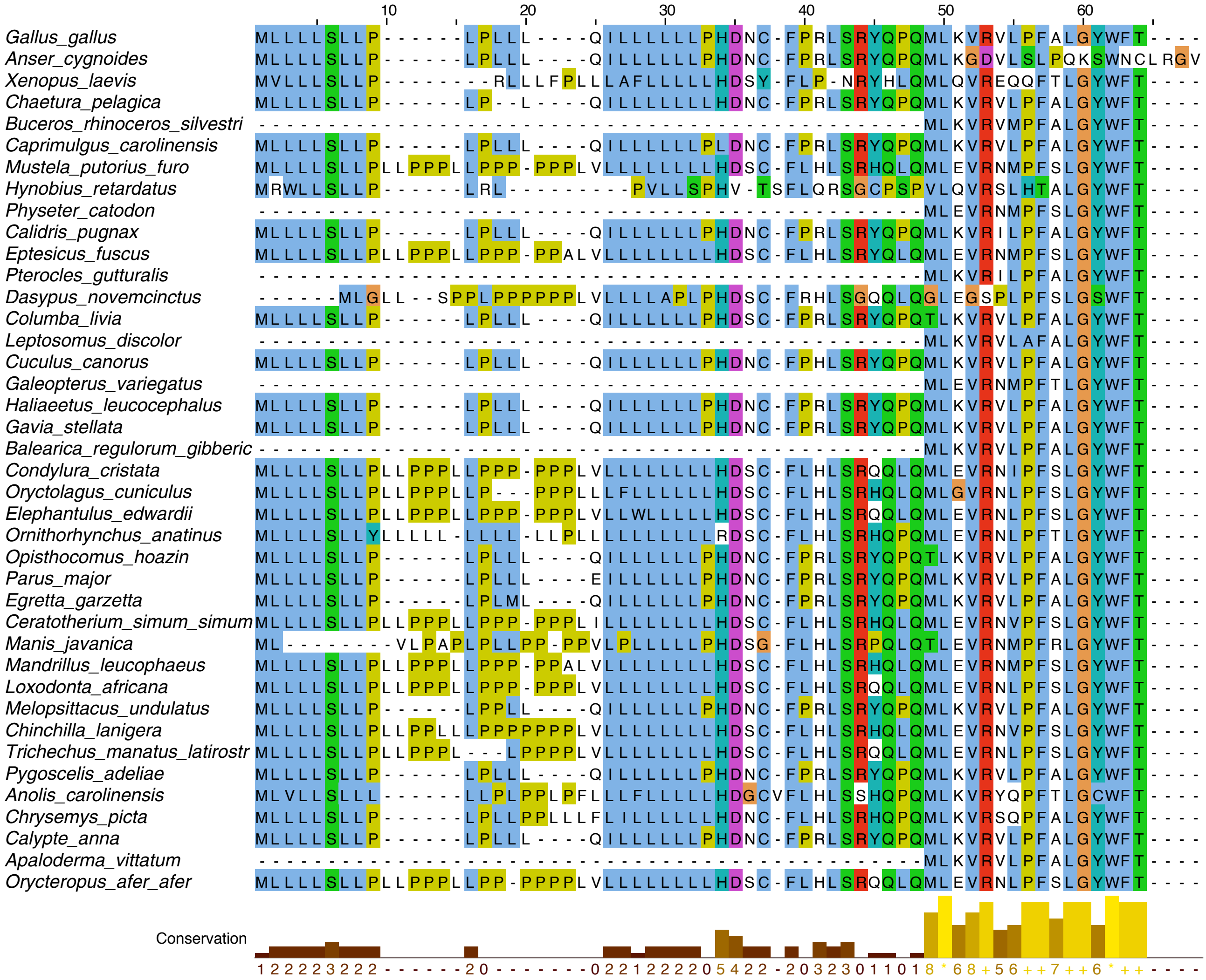

**HG0048**

ENSG00000177565 (TBL1XR1)

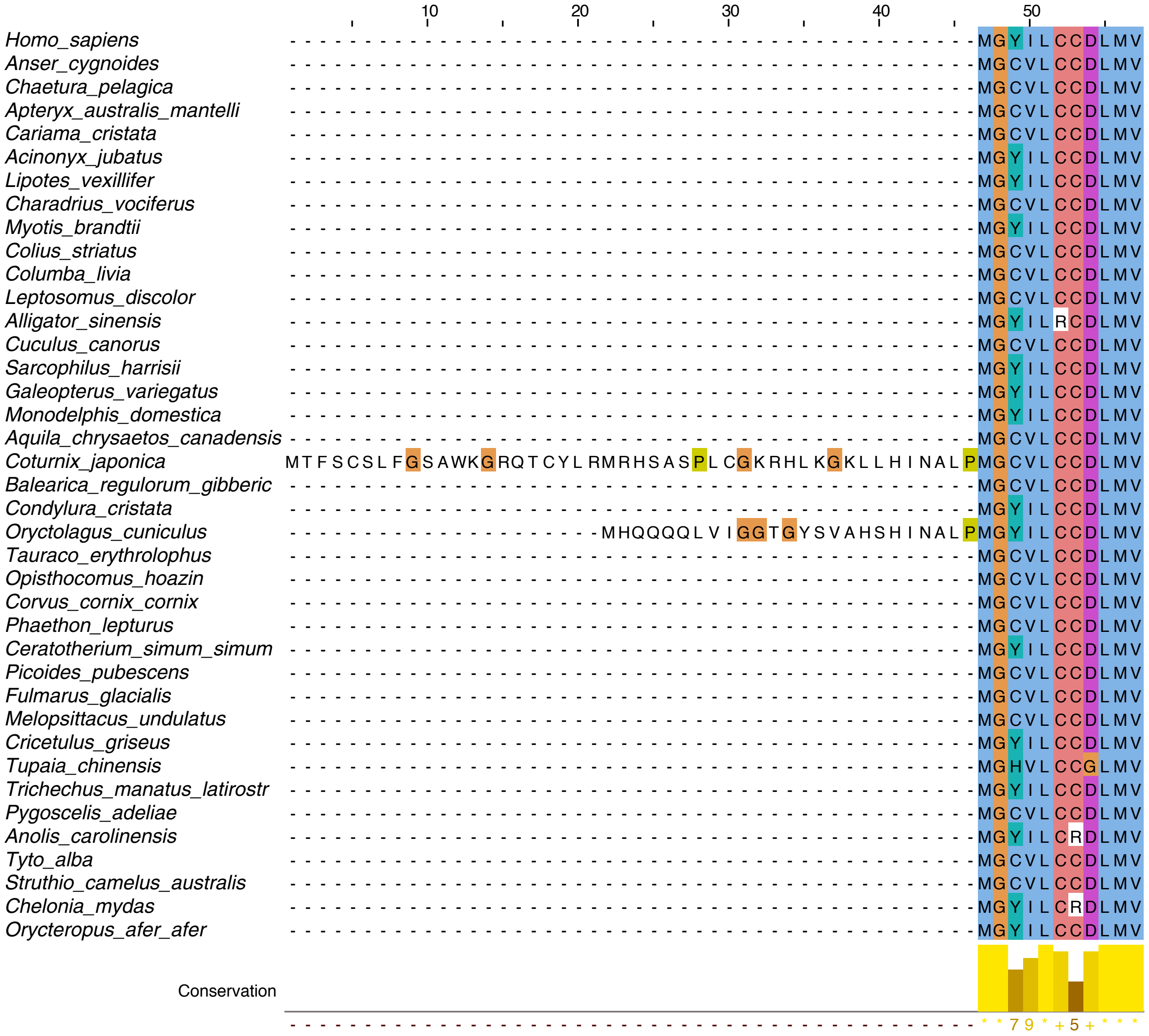

**HG0048**

ENSGALG00000039182

| | |
|---|---|
| *Gallus_gallus* | -------------------------MGCVLCCDLMV |
| *Anser_cygnoides* | -------------------------MGCVLCCDLMV |
| *Chaetura_pelagica* | -------------------------MGCVLCCDLMV |
| *Apteryx_australis_mantelli* | -------------------------MGCVLCCDLMV |
| *Cariama_cristata* | -------------------------MGCVLCCDLMV |
| *Ailuropoda_melanoleuca* | -------------------------MGYILCCDLMV |
| *Lipotes_vexillifer* | -------------------------MGYILCCDLMV |
| *Calidris_pugnax* | -------------------------MGCVLCCDLMV |
| *Rousettus_aegyptiacus* | -------------------------MGYILCCDLMV |
| *Colius_striatus* | -------------------------MGCVLCCDLMV |
| *Columba_livia* | -------------------------MGCVLCCDLMV |
| *Leptosomus_discolor* | -------------------------MGCVLCCDLMV |
| *Cuculus_canorus* | -------------------------MGCVLCCDLMV |
| *Sarcophilus_harrisii* | -------------------------MGYILCCDLMV |
| *Galeopterus_variegatus* | -------------------------MGYILCCDLMV |
| *Monodelphis_domestica* | -------------------------MGYILCCDLMV |
| *Aquila_chrysaetos_canadensis* | -------------------------MGCVLCCDLMV |
| *Balearica_regulorum_gibberic* | -------------------------MGCVLCCDLMV |
| *Condylura_cristata* | -------------------------MGYILCCDLMV |
| *Oryctolagus_cuniculus* | MHQQQQLVIGGTGYSVAHSHINALPMGYILCCDLMV |
| *Tauraco_erythrolophus* | -------------------------MGCVLCCDLMV |
| *Opisthocomus_hoazin* | -------------------------MGCVLCCDLMV |
| *Acanthisitta_chloris* | -------------------------MGCVLCCDLMV |
| *Egretta_garzetta* | -------------------------MGCVLCCDLMV |
| *Equus_asinus* | -------------------------MGFILCCDLMV |
| *Picoides_pubescens* | -------------------------MGCVLCCDLMV |
| *Cercocebus_atys* | -------------------------MGYILCCDLMV |
| *Fulmarus_glacialis* | -------------------------MGCVLCCDLMV |
| *Melopsittacus_undulatus* | -------------------------MGCVLCCDLMV |
| *Chinchilla_lanigera* | -------------------------MGYVLCCDLMV |
| *Tupaia_chinensis* | -------------------------MGHVLCCGLMV |
| *Trichechus_manatus_latirostr* | -------------------------MGYILCCDLMV |
| *Aptenodytes_forsteri* | -------------------------MGCVLCCDLMV |
| *Gekko_japonicus* | -------------------------MGCILRCDLMV |
| *Tyto_alba* | -------------------------MGCVLCCDLMV |
| *Struthio_camelus_australis* | -------------------------MGCVLCCDLMV |
| *Apaloderma_vittatum* | -------------------------MDCVLCCDLMV |
| *Orycteropus_afer_afer* | -------------------------MGYILCCDLMV |
| Conservation | -------------------------*+79*+*+*** |

**HG0049**

ENSDARG00000044485 (sall4)

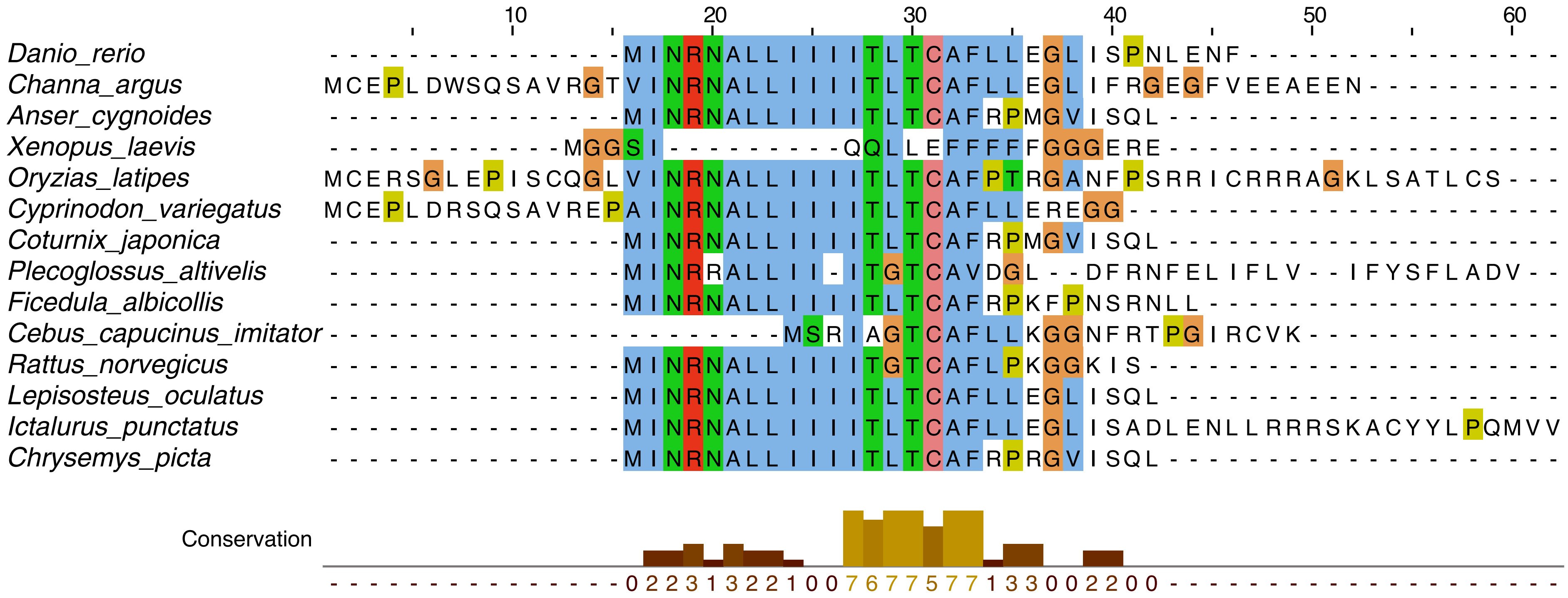

**HG0049**

ENSGALG00000039238

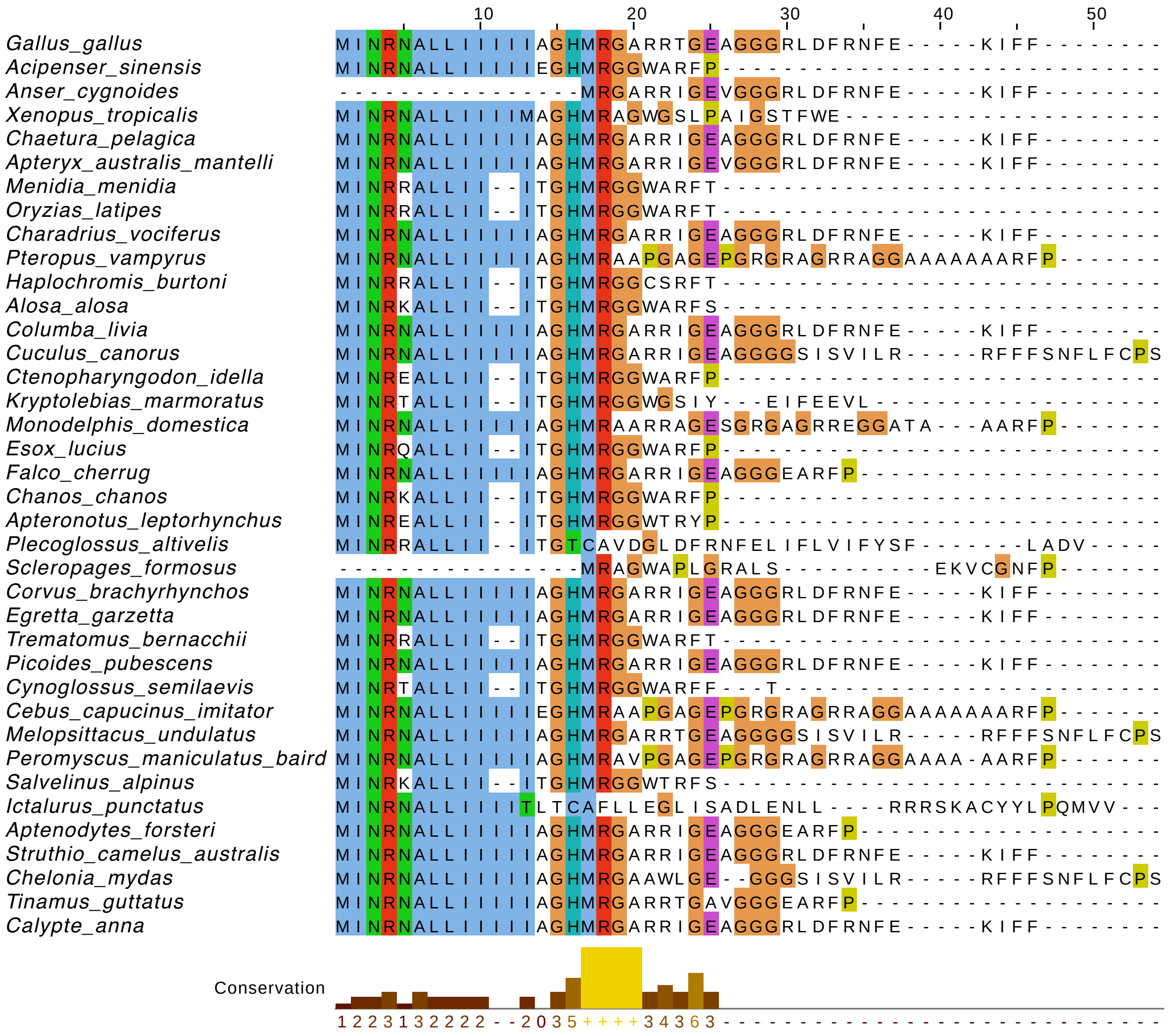

## HG0050.1

ENSG00000074054 (CLASP1)

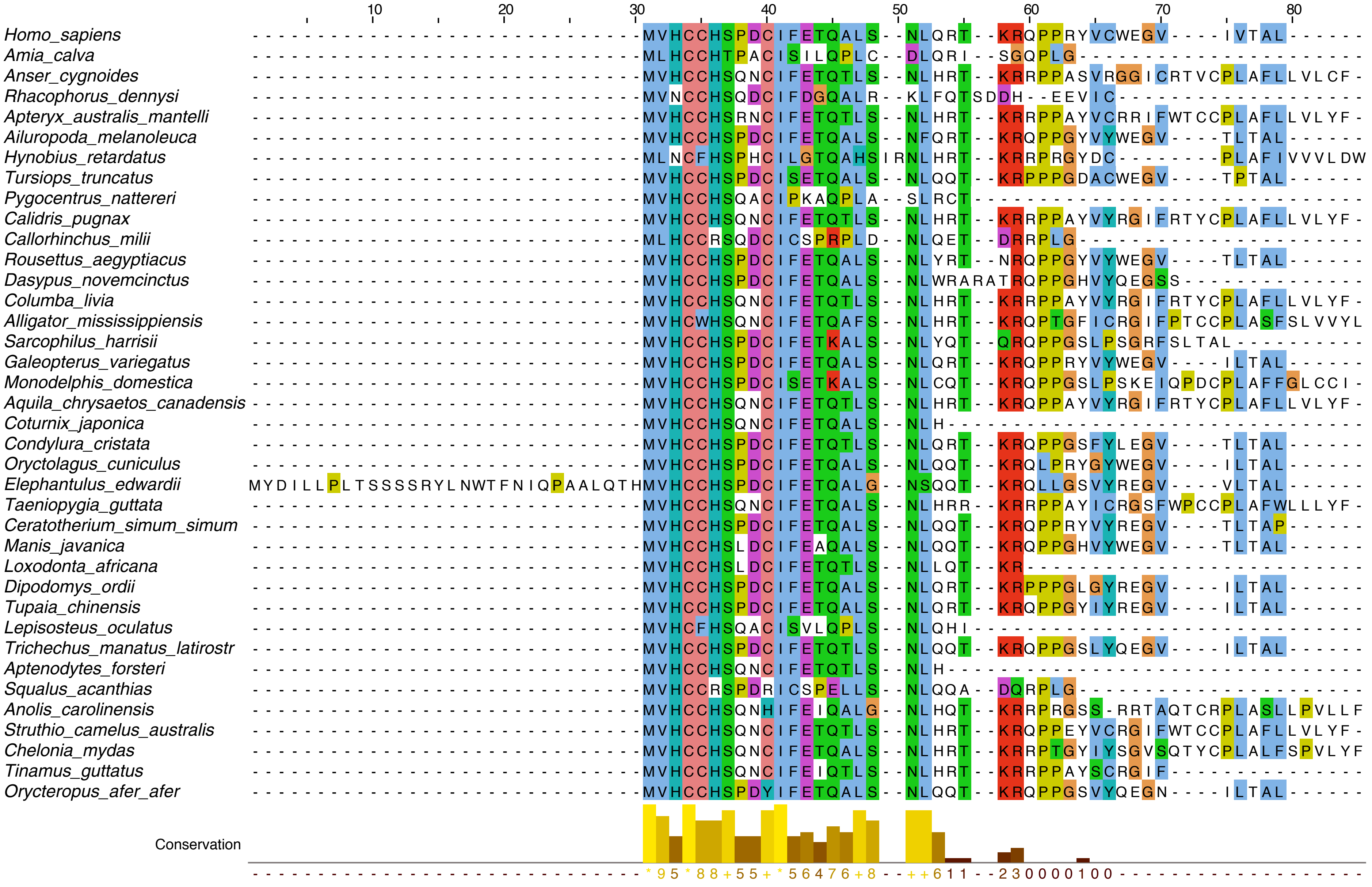

**HG0050.2**

ENSG00000074054 (CLASP1)

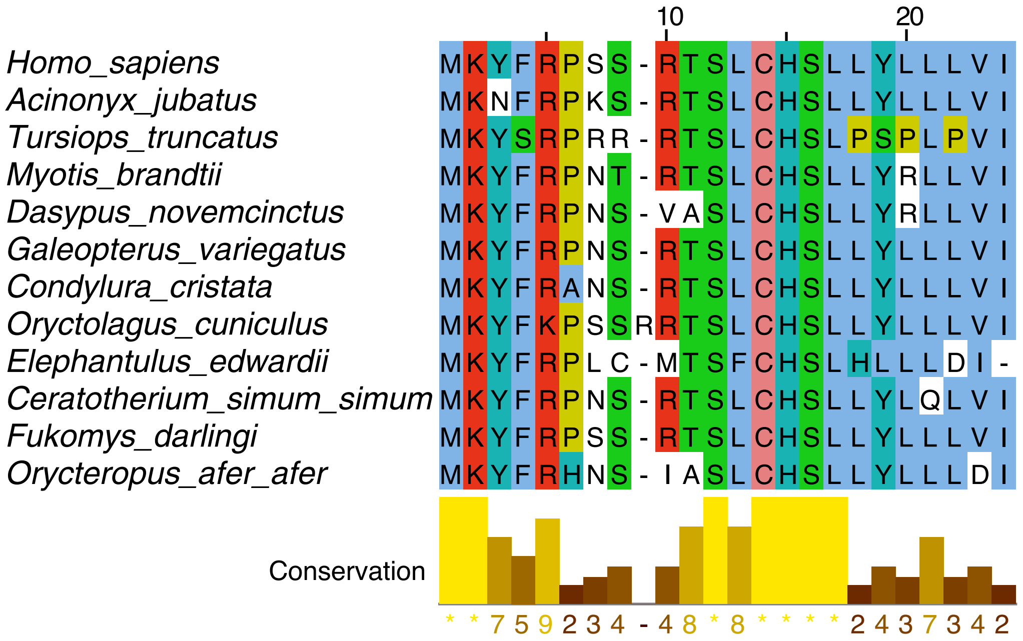

**HG0051**

ENSG00000116679 (IVNS1ABP)

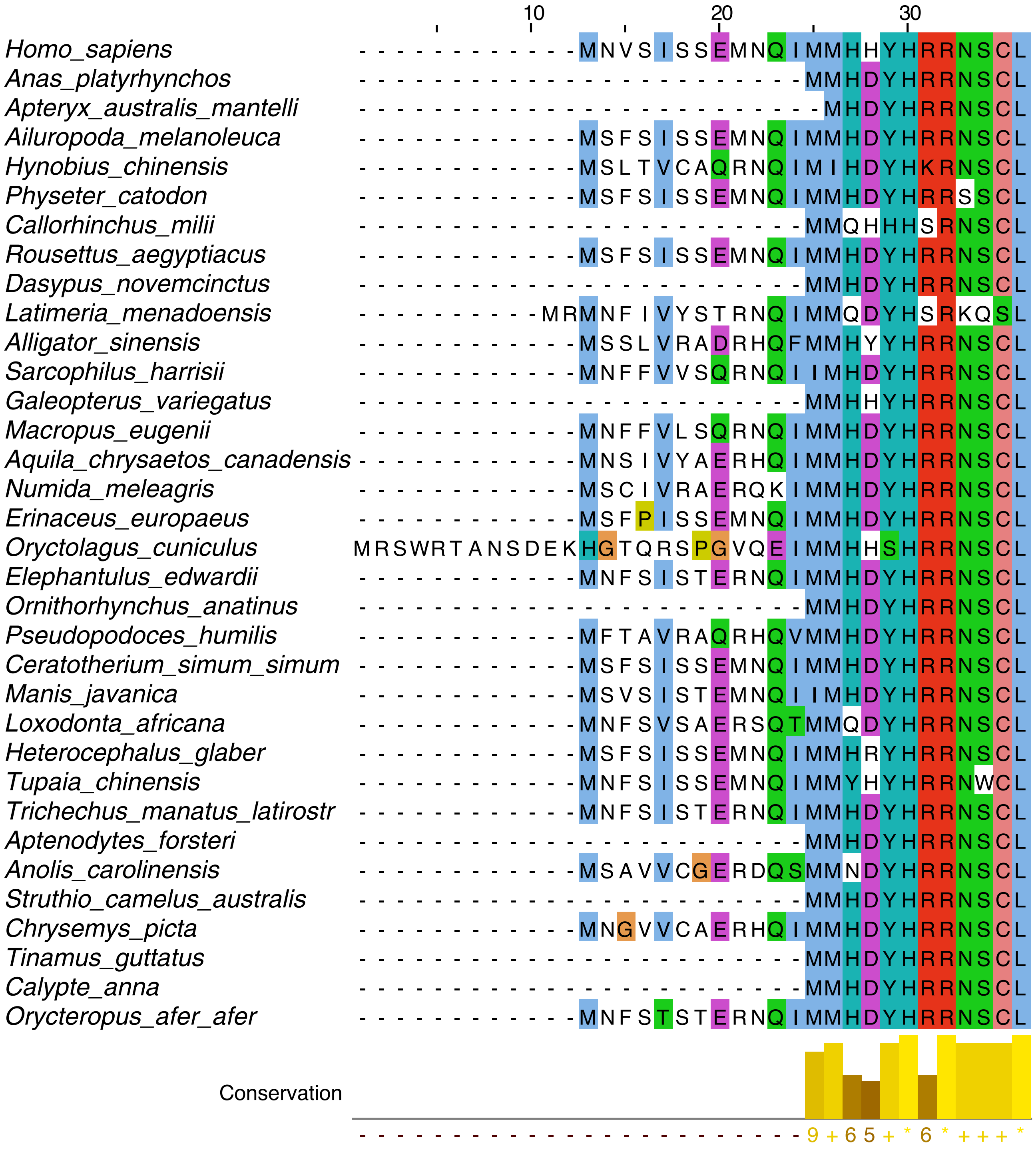

**HG0051**

ENSGALG00000037162

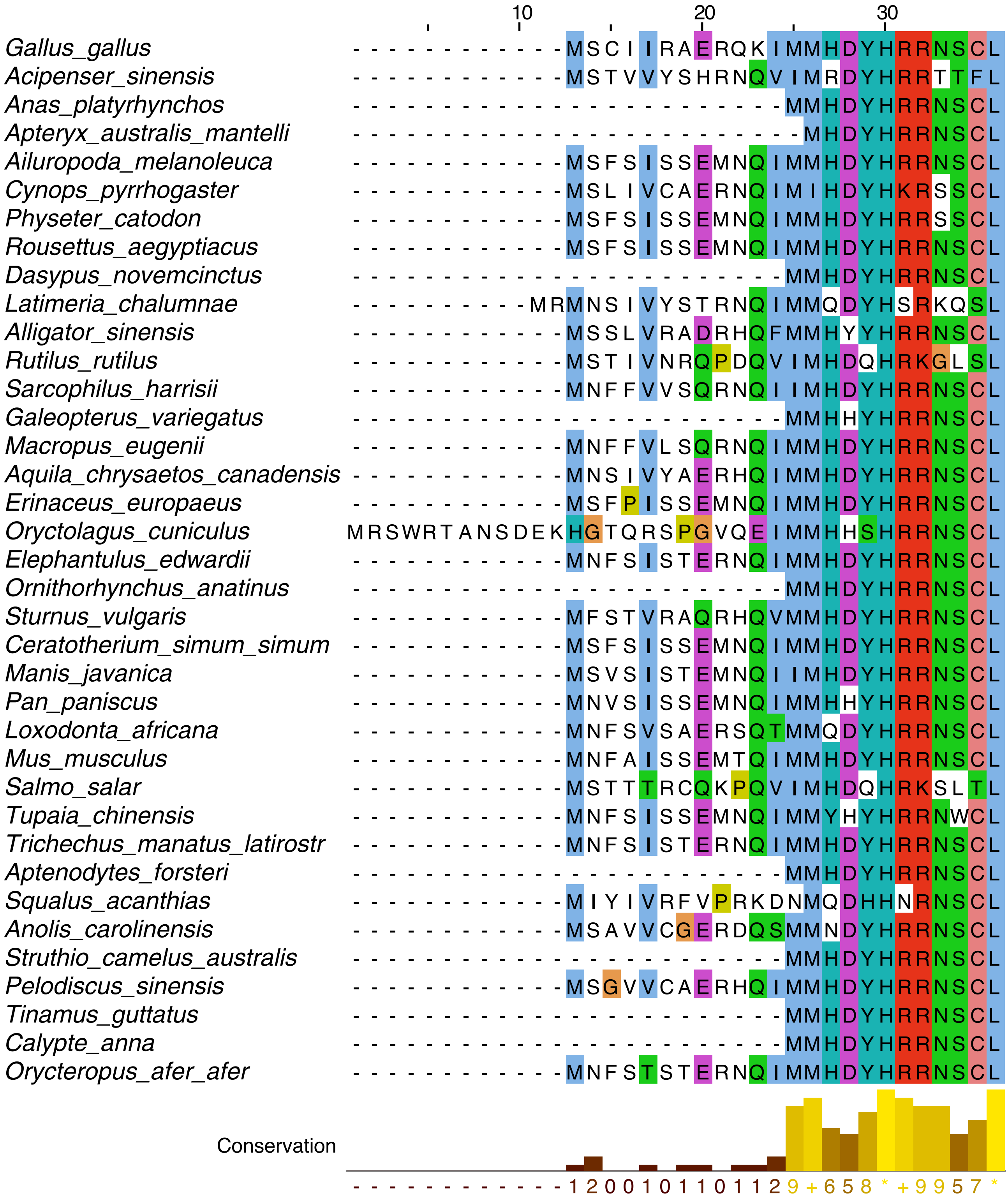

**HG0052**

ENSG00000178235 (SLITRK1)

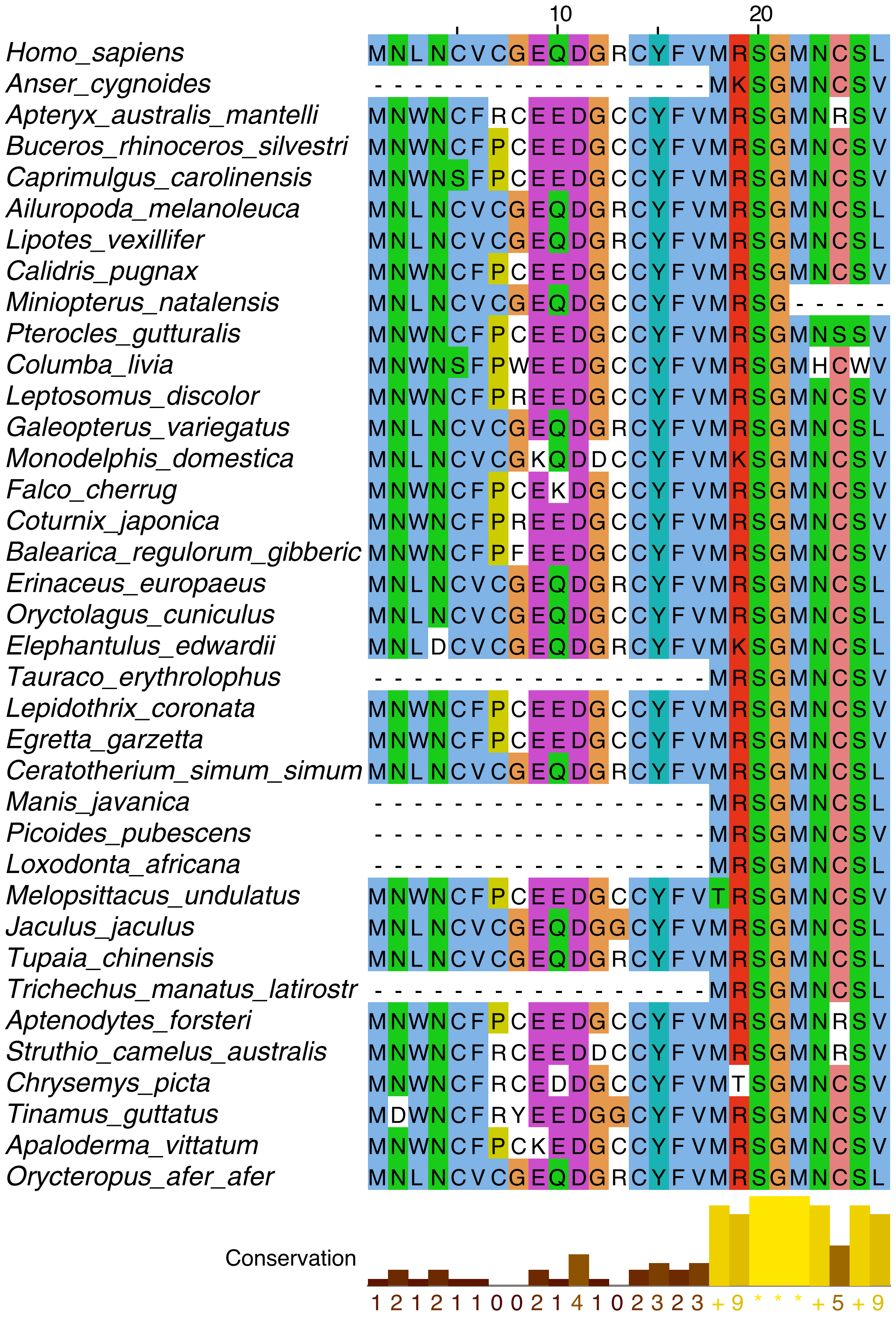

**HG0053.1**

ENSG00000184564 (SLITRK6)

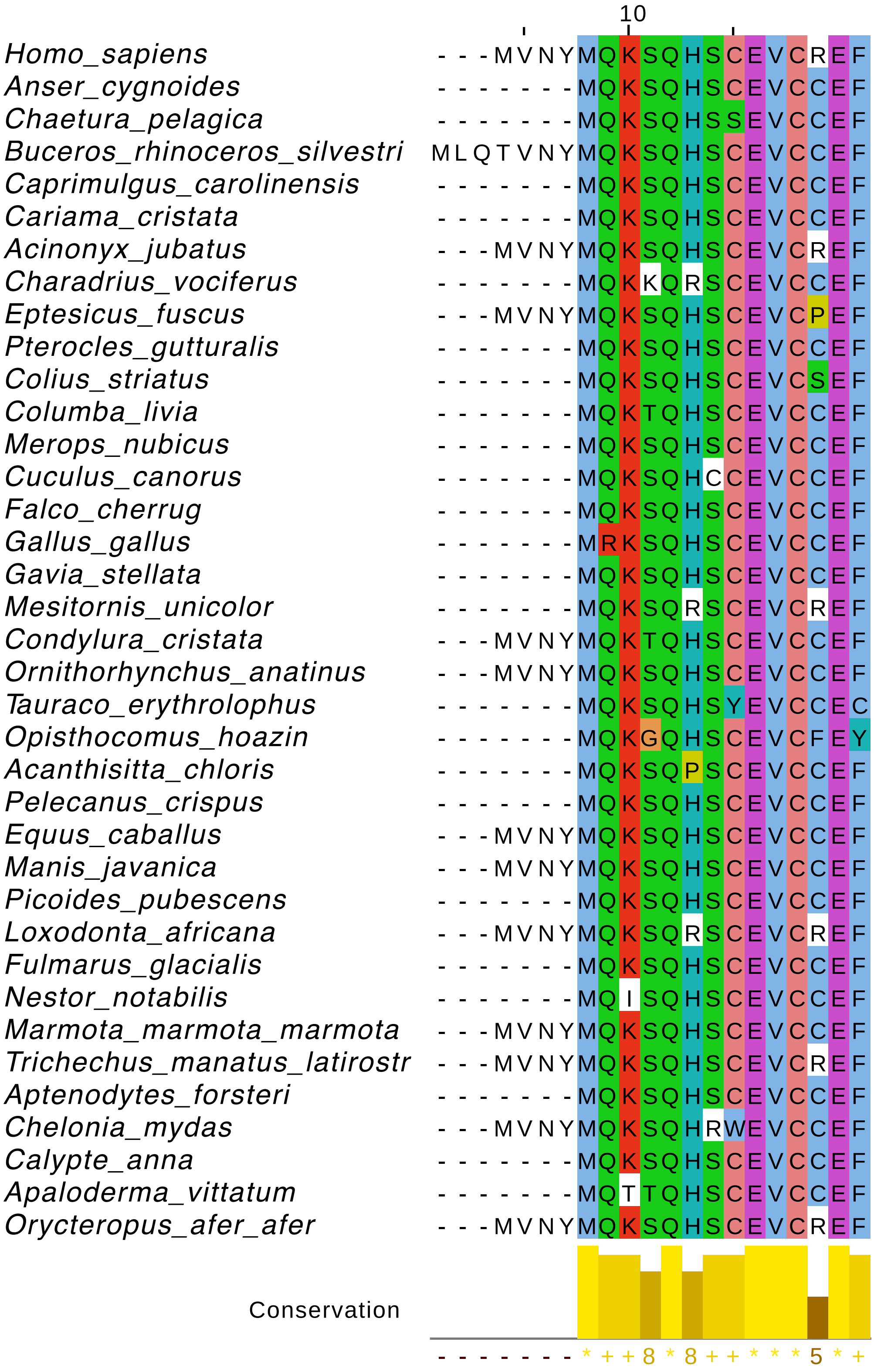

**HG0053.2**

ENSG00000184564 (SLITRK6)

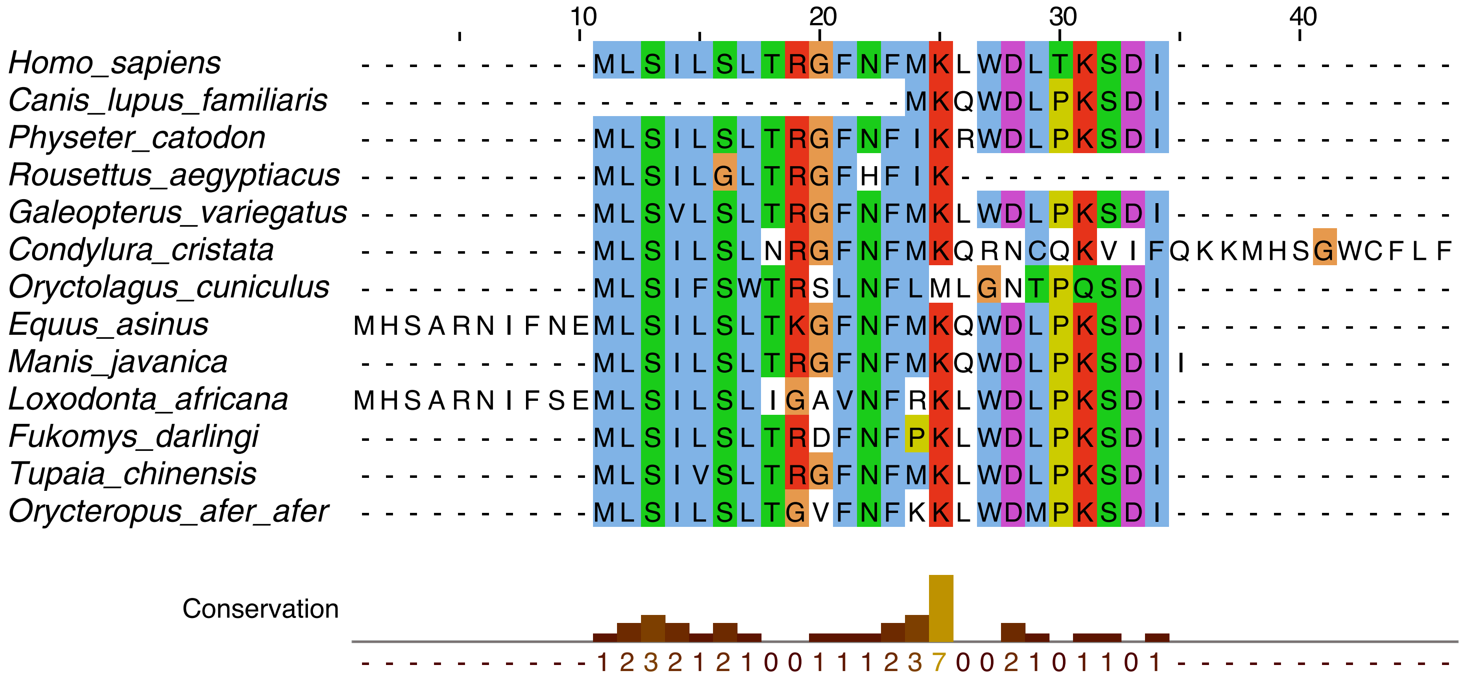

**HG0054**

ENSG00000124479 (NDP)

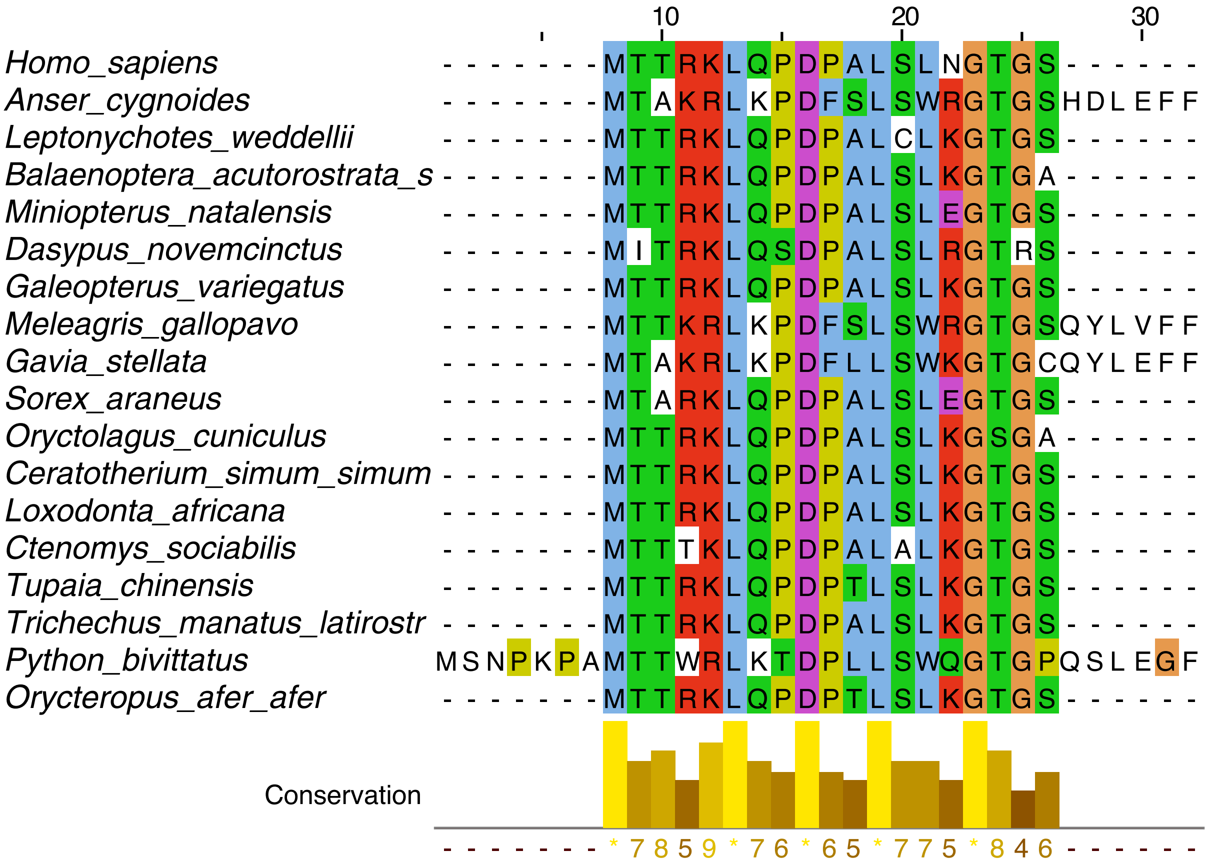

**HG0054**

ENSGALG00000016222 (NDP)

```
                                           10        20        30
                                            |    |    |    |    |
Gallus_gallus                      -----------MTAKRLKPDFSLSWRGTGSQYLEFF
Anser_cygnoides                    -----------MTAKRLKPDFSLSWRGTGSHDLEFF
Apteryx_australis_mantelli         -----------MTAKRPKPDFSPSWKETGSQYLEFF
Cariama_cristata                   -----------MTAKRLKPDFSPSWKGTGCQYLEFF
Felis_catus                        -----------MTTRKLRPDRALCWEGTGS------
Balaenoptera_acutorostrata_s       -----------MTTRKLQPDPALSLKGTGA------
Calidris_pugnax                    -----------MTAKRLKPDFSPSWKGTGCQYLEFF
Pteropus_vampyrus                  -----------MTTRKLQPDPALSLKGTGS------
Pterocles_gutturalis               -----------MTAKRLKPDFSPSWKGTGCQYLEFF
Colius_striatus                    -----------MTSKRLKPVFSPSWEGTGGQYLEFC
Leptosomus_discolor                -----------MTAKRLKPDFSPSWKGTGCQCLEFF
Cuculus_canorus                    -----------MPAKRLKPGFSPSWKGTDCQYLEFF
Galeopterus_variegatus             -----------MTTRKLQPDPALSLKGTGS------
Haliaeetus_albicilla               -----------MTAKRLKPDFSPSWRGTGCQYLEFF
Gavia_stellata                     -----------MTAKRLKPDFLLSWKGTGCQYLEFF
Balearica_regulorum_gibberic       -----------MTAKRLKPDFSPSWKGTGCQYLEFF
Sorex_araneus                      -----------MTARKLQPDPALSLEGTGS------
Oryctolagus_cuniculus              -----------MTTRKLQPDPALSLKGSGA------
Tauraco_erythrolophus              -----------MTAKRLKPDFSPSWKGTGCQDLEFF
Opisthocomus_hoazin                -----------MTAKRLKPDFSPSWKGTGCPYLEFF
Nipponia_nippon                    MISVNSNSRRRMTAKRLKPDFSPSWKGTGCQYLEFF
Ceratotherium_simum_simum          -----------MTTRKLQPDPALSLKGTGS------
Aotus_nancymaae                    -----------MTTRKLQPDPALSLKGTGS------
Loxodonta_africana                 -----------MTTRKLQPDPALSLKGTGS------
Fulmarus_glacialis                 -----------MTAKRLKPDFSPSWKGTGCQYLEFF
Nestor_notabilis                   -----------MTAKALKPDFSPSWKGTGCHYLEFF
Heterocephalus_glaber              -----------MTTKKLRPDPALTLKGTGS------
Tupaia_chinensis                   -----------MTTRKLQPDPTLSLKGTGS------
Trichechus_manatus_latirostr       -----------MTTRKLQPDPALSLKGTGS------
Pygoscelis_adeliae                 -----------MTAKRLKPDFSSSWKGTGCQYLEFF
Gekko_japonicus                    ----MSTPKLAMTTQRLKTDLSLSSKGTGPQSLECF
Struthio_camelus_australis         -----------MTAKRLKPDFSPSWKGTGSRYLEFF
Chrysemys_picta                    -----------MTTKRLKPGFSSSLKGTESQYLEFF
Tinamus_guttatus                   -----------MTVKRTNPDFSLSWKGMGSQYLEFF
Calypte_anna                       -----------MTAKRLKPDSSPSWKGTDCQVLEFF
Orycteropus_afer_afer              -----------MTTRKLQPDPTLSLKGTGS------

Conservation                       -----------*+899+7+5674+77++57------
```

**HG0055.1**

ENSGALG00000010533 (ELAVL4)

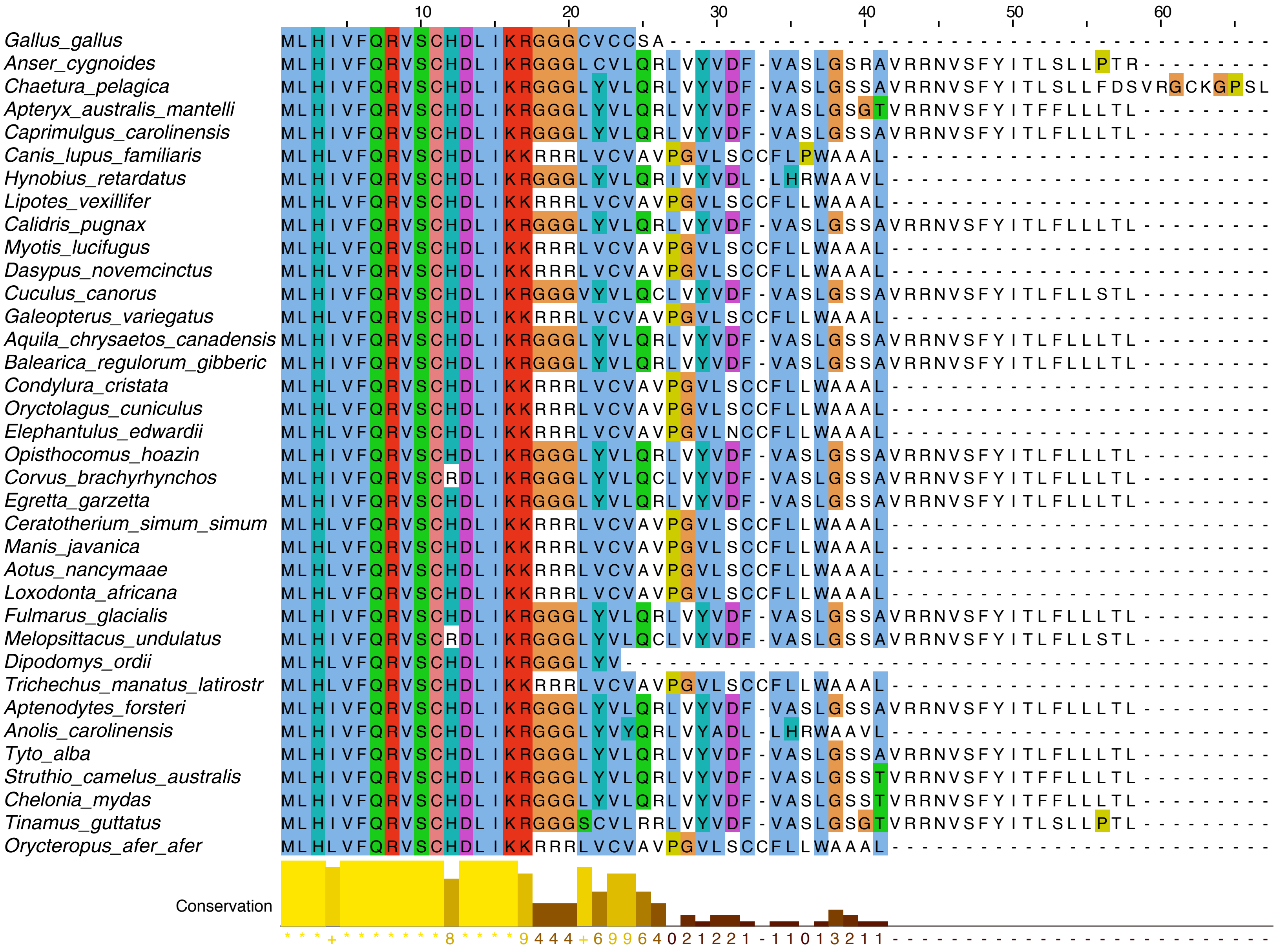

**HG0055.2**

ENSGALG00000010533 (ELAVL4)

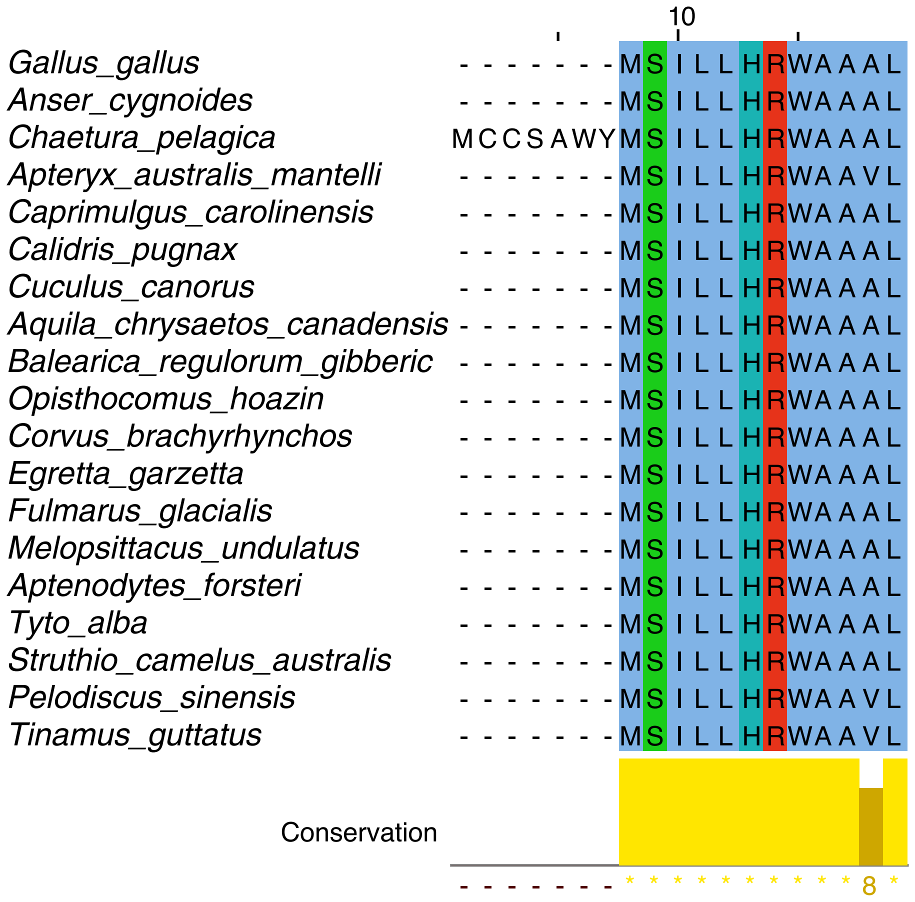

**HG0056**

ENSG00000139329 (LUM)

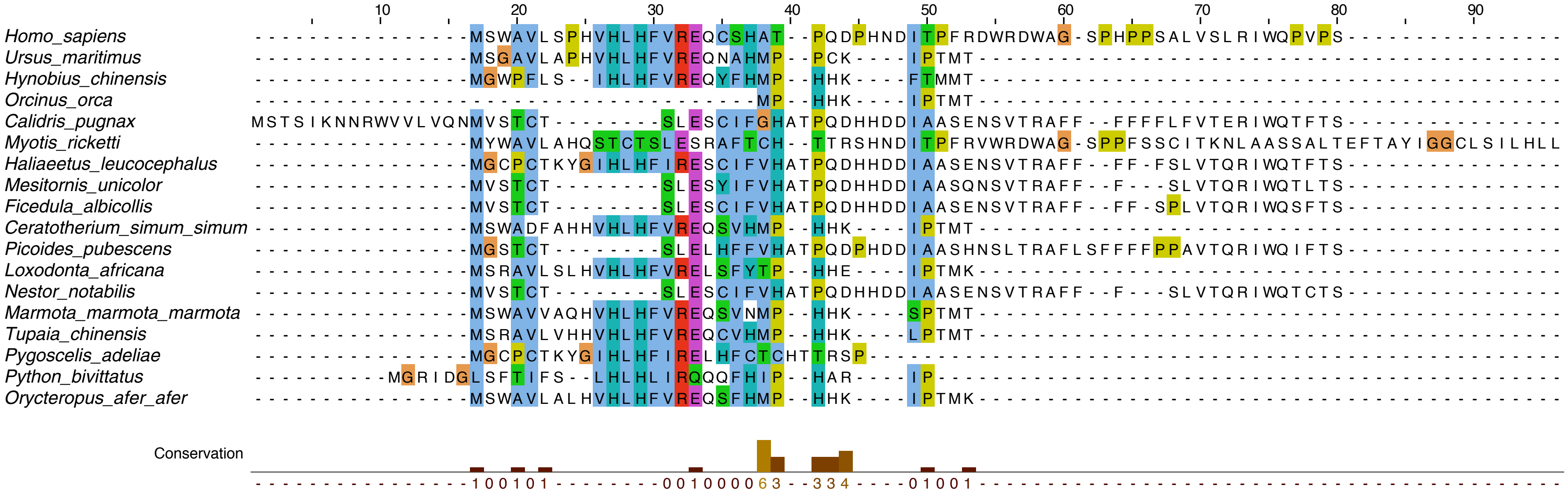

**HG0056**

ENSGALG00000011271 (LUM)

```
                                                  10        20        30
                                         .    |    .    |    .    |
Gallus_gallus                  MGCPCTECGIHLHFIKRAAFSYMPHHKITMMT
Anser_cygnoides                ----------------------MPHHKITMMT
Apteryx_australis_mantelli     ----------------------MPHHKITMMT
Caprimulgus_carolinensis       ----------------------MPHHKITMMT
Cariama_cristata               ----------------------MPHHKITMMT
Hynobius_chinensis             MGWPF--LSIHLHFVRE-QYFHMPHHKFTMMT
Charadrius_vociferus           ----------------------MPHHKITMMT
Myotis_ricketti                ----------------------MPHHKIPQ--
Pterocles_gutturalis           ----------------------MPHHKITMMT
Latimeria_chalumnae            ---------MHLHLVKE-LCFFIPHHKLIMMT
Colius_striatus                MDCPYTKYGIHLHFIRE-LHFCSCHTTISMMT
Columba_livia                  ----------------------MPHHKITMMT
Merops_nubicus                 ----------------------MPHHKITMMT
Alligator_sinensis             ----------------------MPHHKITMMT
Haliaeetus_leucocephalus       MVS-------TCTSLERAAFLYMPHHRITMMT
Gavia_stellata                 ----------------------MPHHKITMMT
Chlamydotis_macqueenii         MGCPCTKCGIHLHFIRE-LHFRTCYTTNSP--
Condylura_cristata             ----------------------MPHHKIPTMT
Ficedula_albicollis            MGCPCTKYGIHLHFIRE-LHFCTCHTTRSP--
Nipponia_nippon                MGCPCTKYGIHLHFIRD-LHFSTCHTTRSP--
Equus_przewalskii              ----------------------MPHHKIPTMT
Picoides_pubescens             MGCPYTKYGIHLHFIRA-AFFCPCHTTRSP--
Propithecus_coquereli          MNWAVLAPHVHLHFVRE-QSVHMPHHKIPTMT
Loxodonta_africana             MSRAVLSLHVHLHFVRE-LSFYTPHHEIPTMK
Fulmarus_glacialis             ----------------------MPHHKITMMT
Nestor_notabilis               ----------------------MPHHKITMMT
Tupaia_chinensis               MSRAVLVHHVHLHFVRE-QCVHMPHHKLPTMT
Pygoscelis_adeliae             ----------------------MPHHKITMMT
Gekko_japonicus                ----------------------MPHHQNPMMT
Tyto_alba                      ----------------------MPHHKITMMT
Struthio_camelus_australis     ----------------------MPHHKITMMT
Pelodiscus_sinensis            MGCPCTVYTVHLHFIRE-QHLYIPHHKIIMMT
Tinamus_guttatus               MGCPCTKYGIHLHFITE-LHFCTYHTTRSL--
Apaloderma_vittatum            ----------------------MPHHKITMMT
Orycteropus_afer_afer          MSWAVLALHVHLHFVRE-QSFHMPHHKIPTMK

Conservation                   ----------------------78+6744612
```

**HG0057**

ENSG00000280987 (MATR3)

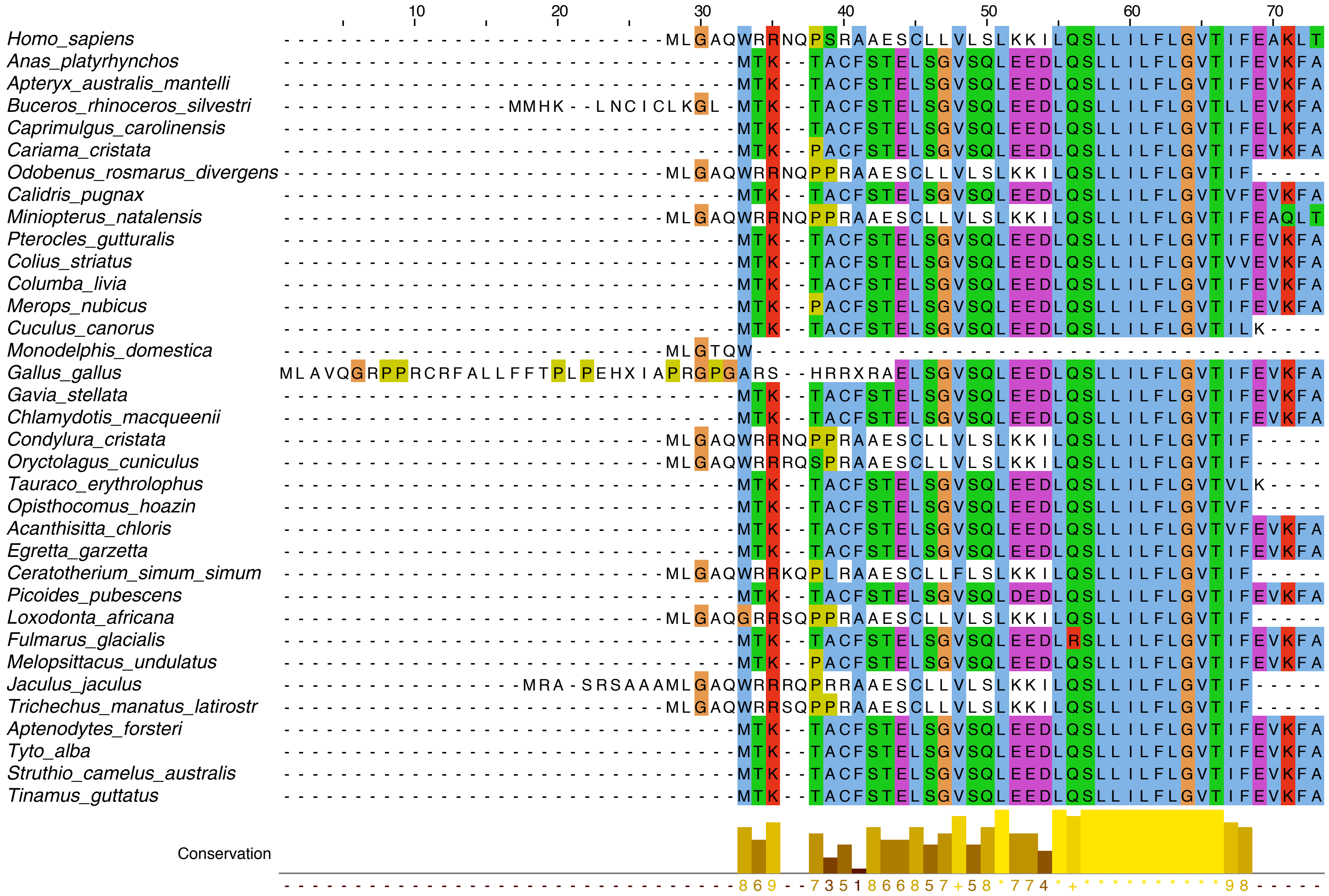

**HG0058**

ENSG00000180332 (KCTD4)

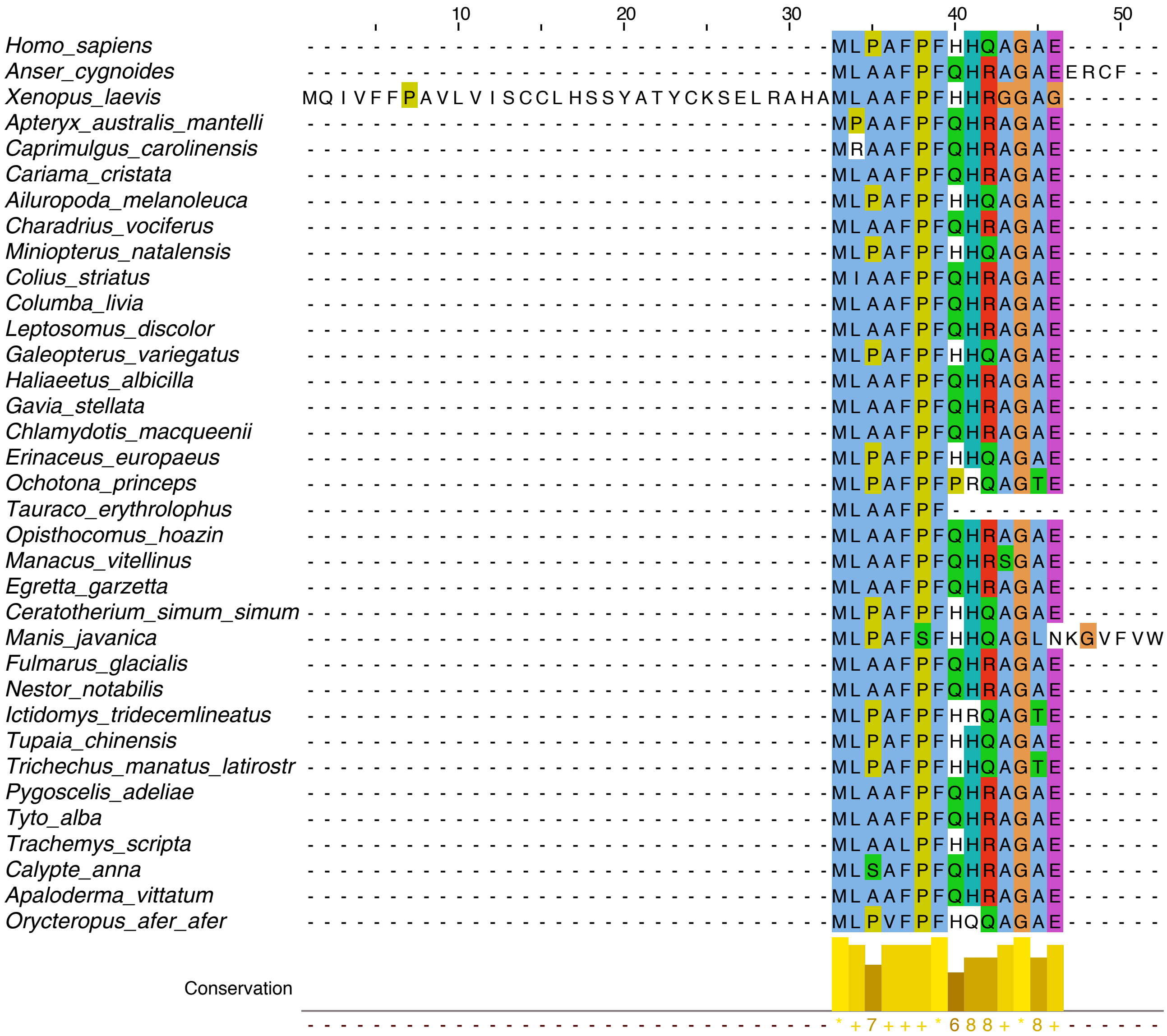

**HG0059.1**

ENSDARG00000018060 (pik3r2)

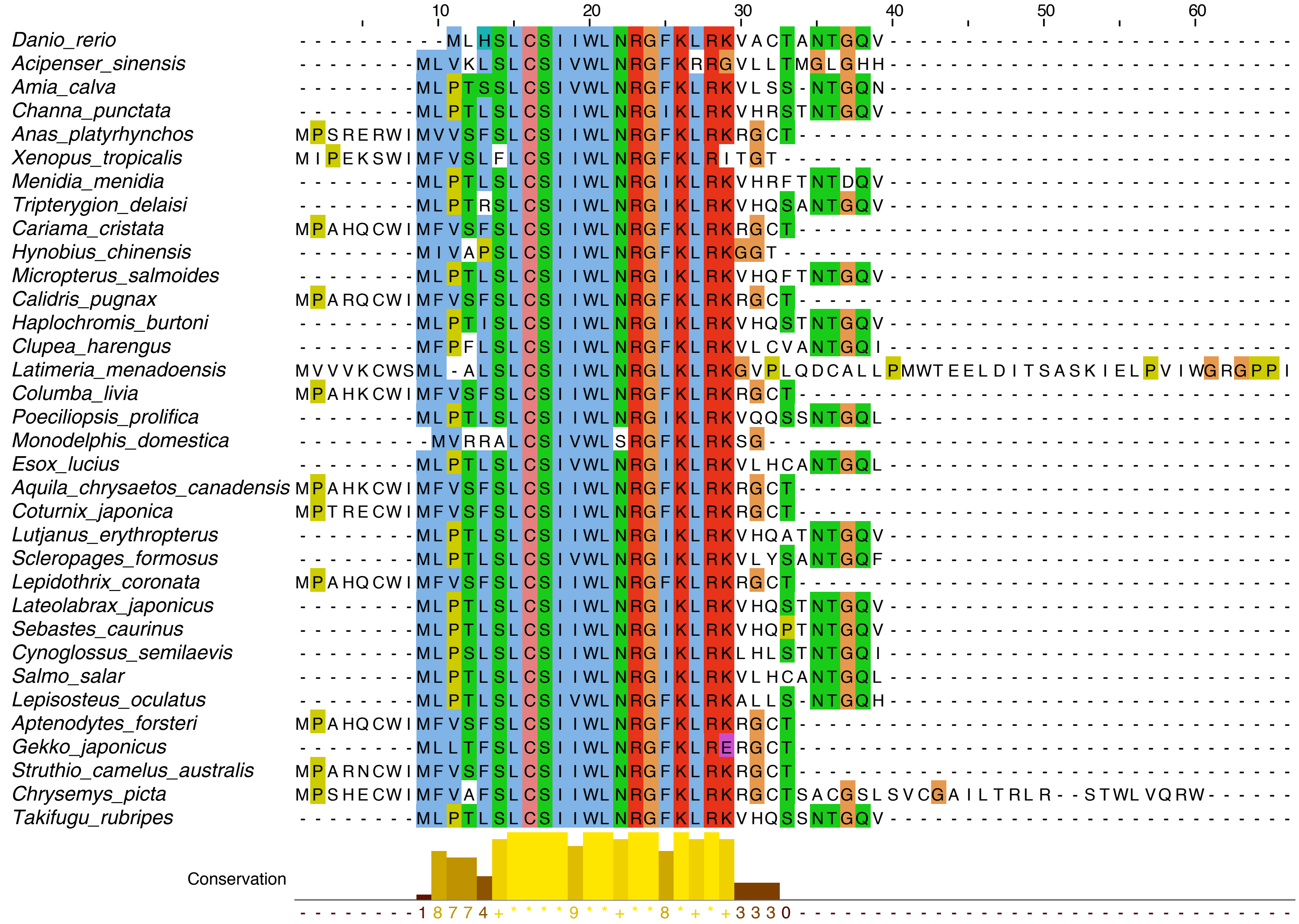

**HG0059.1**

ENSGALG00000003428

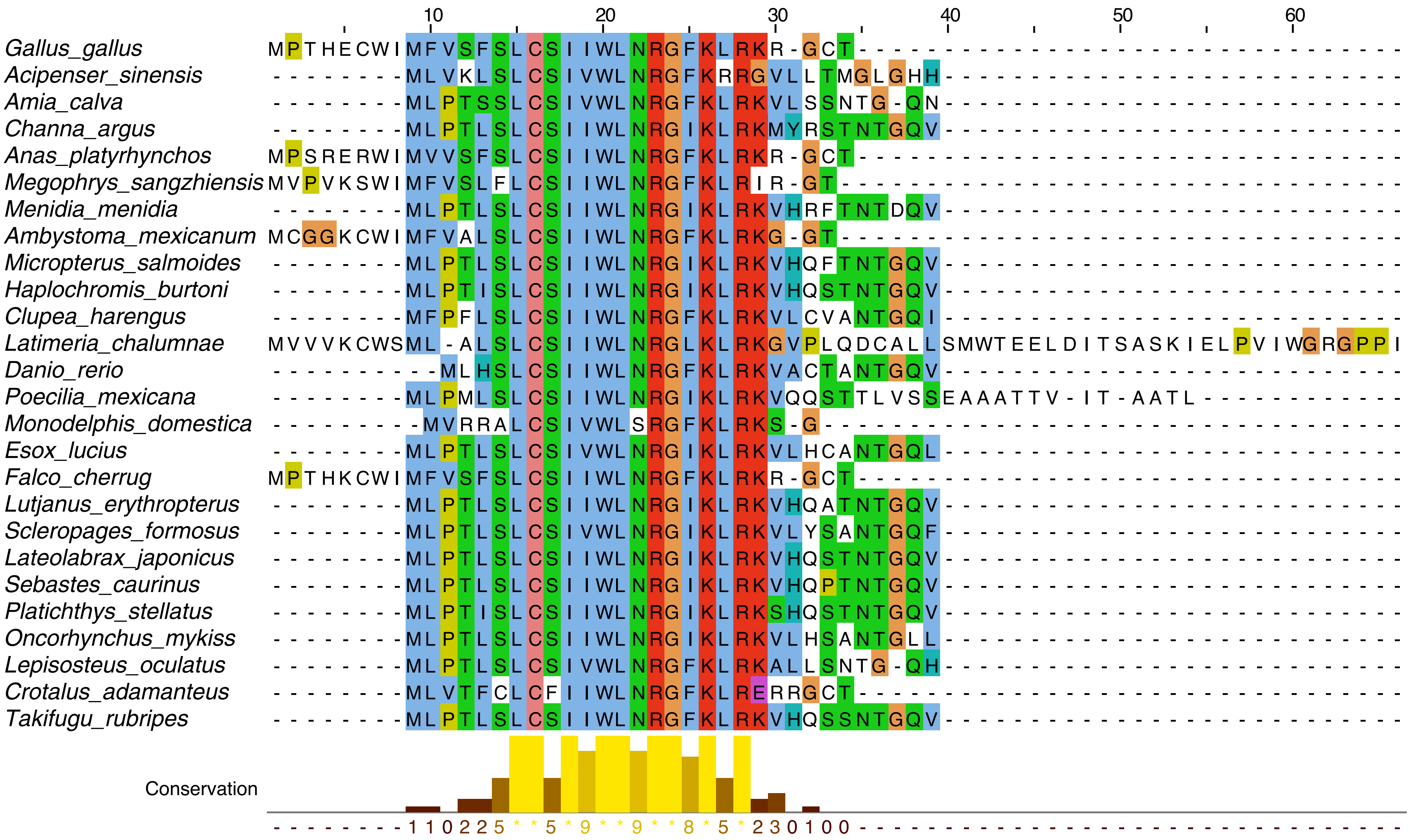

**HG0059.2**

ENSG00000145675 (PIK3R1)

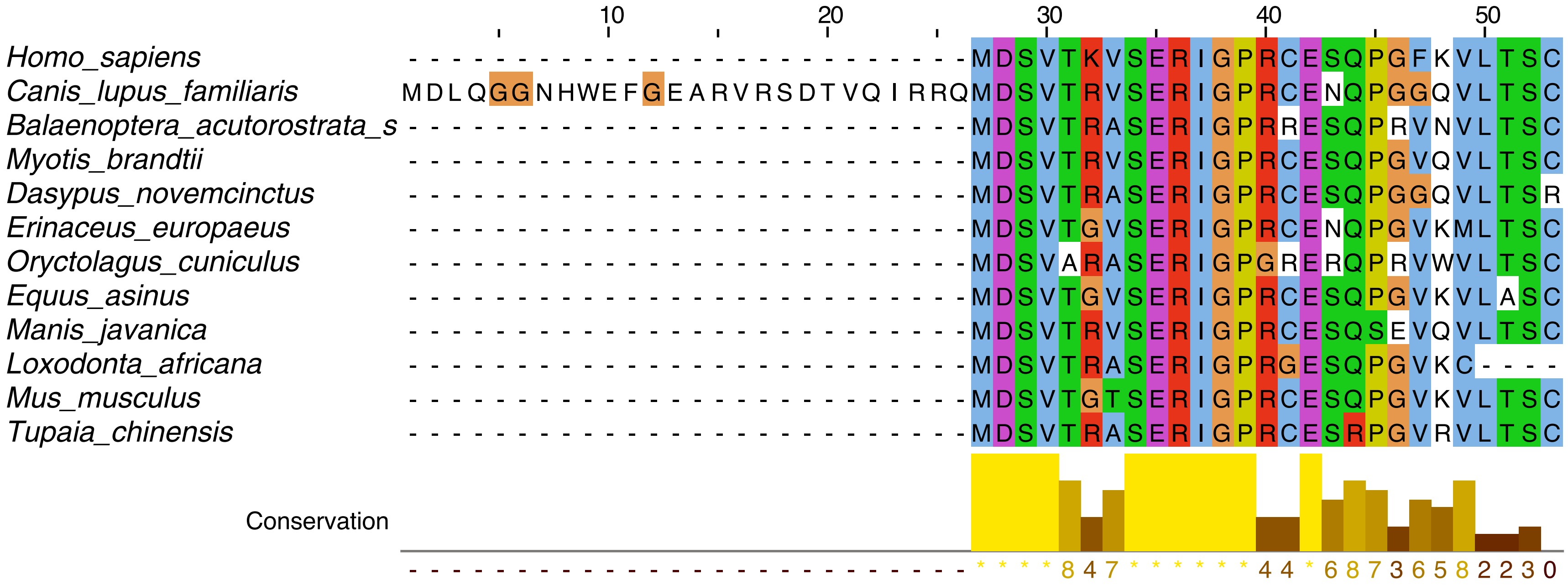

**HG0059.3**

ENSG00000105647 (PIK3R2)

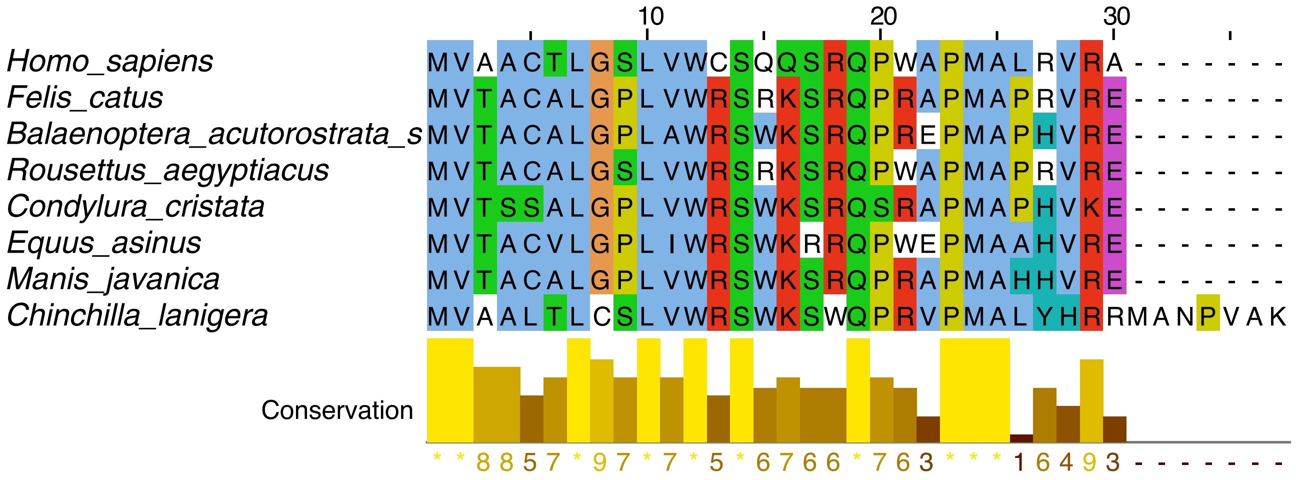

**HG0059.4**

ENSG00000145675 (PIK3R1)

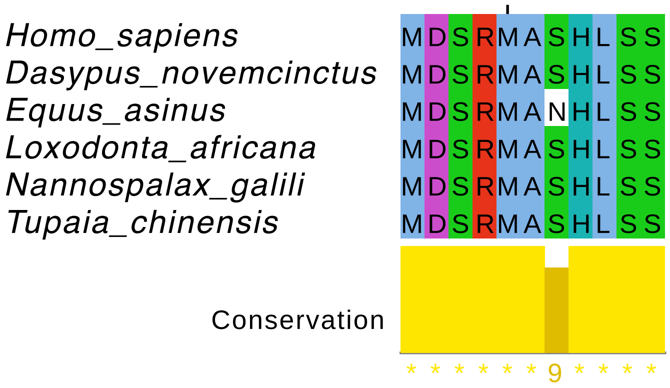

**HG0059.5**

ENSDARG00000038524 (pik3r1)

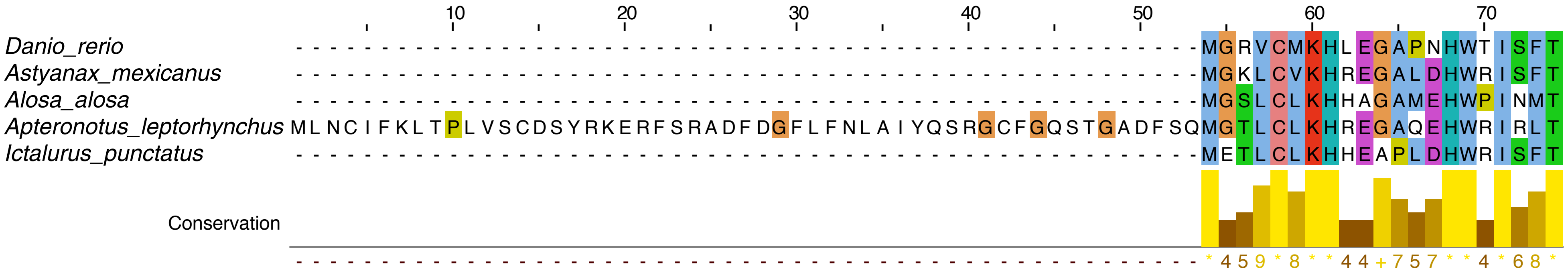

## HG0060.1

ENSG00000049192 (ADAMTS6)

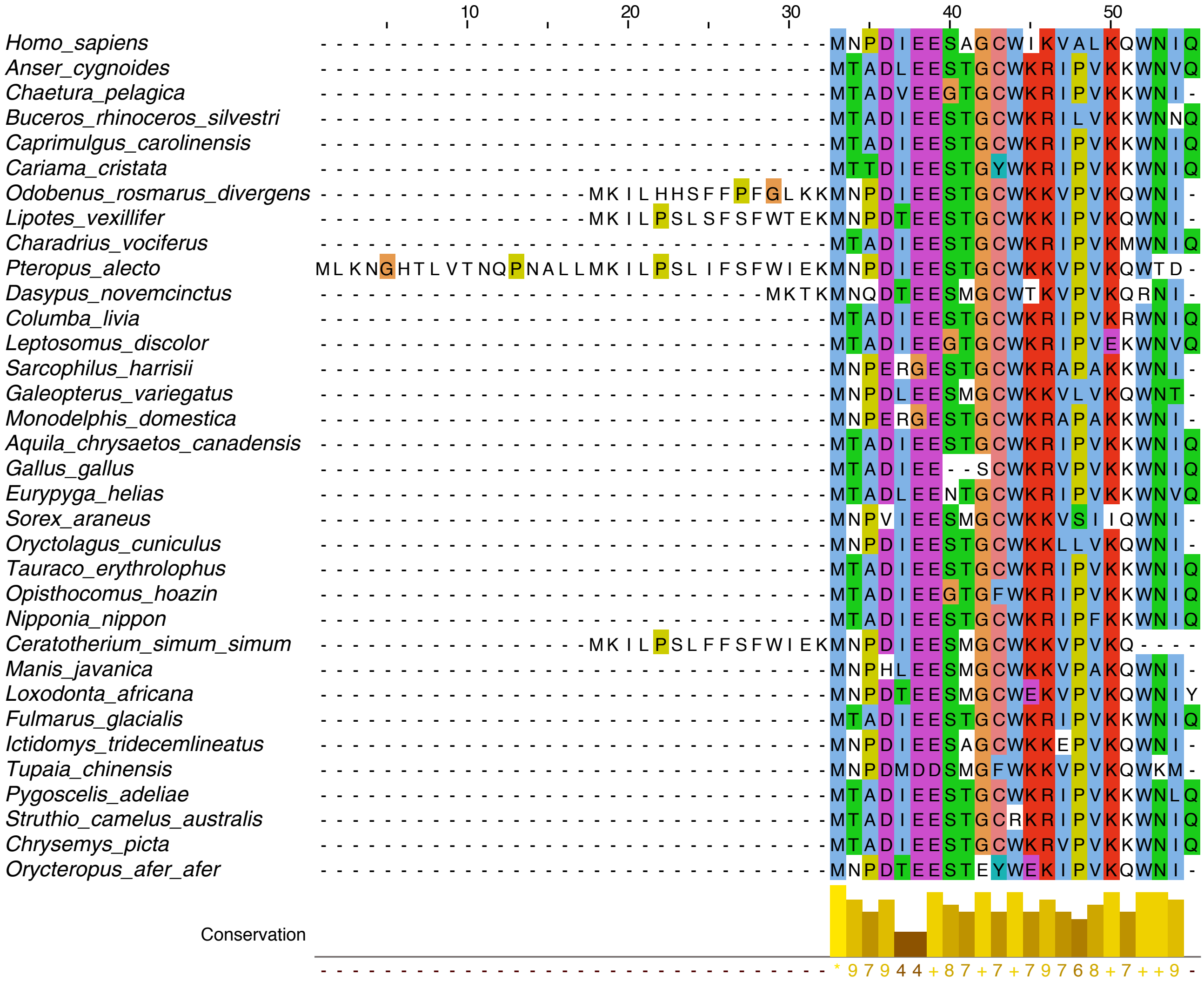

**HG0060.1**

ENSGALG00000014751

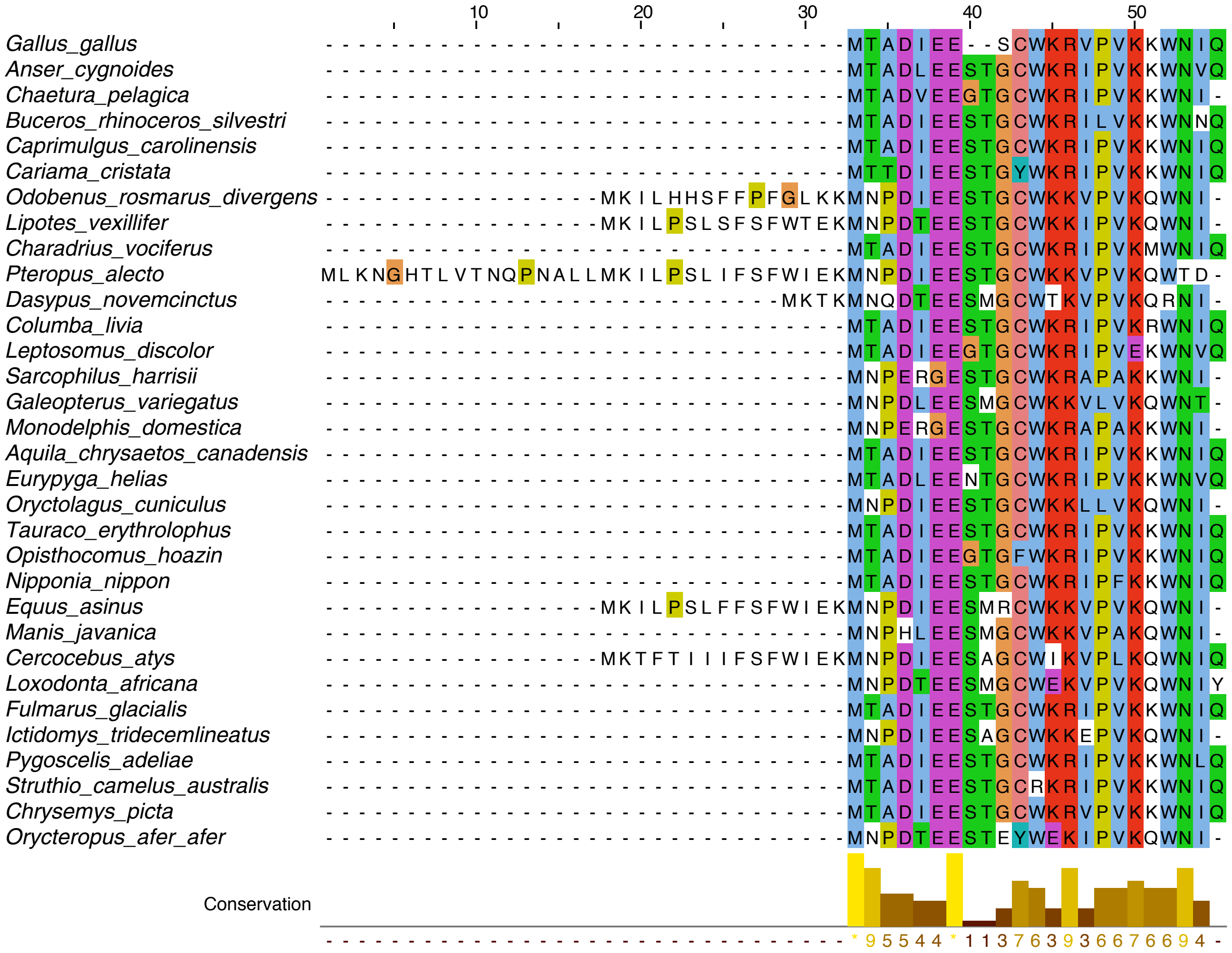

**HG0060.2**

ENSG00000049192 (ADAMTS6)

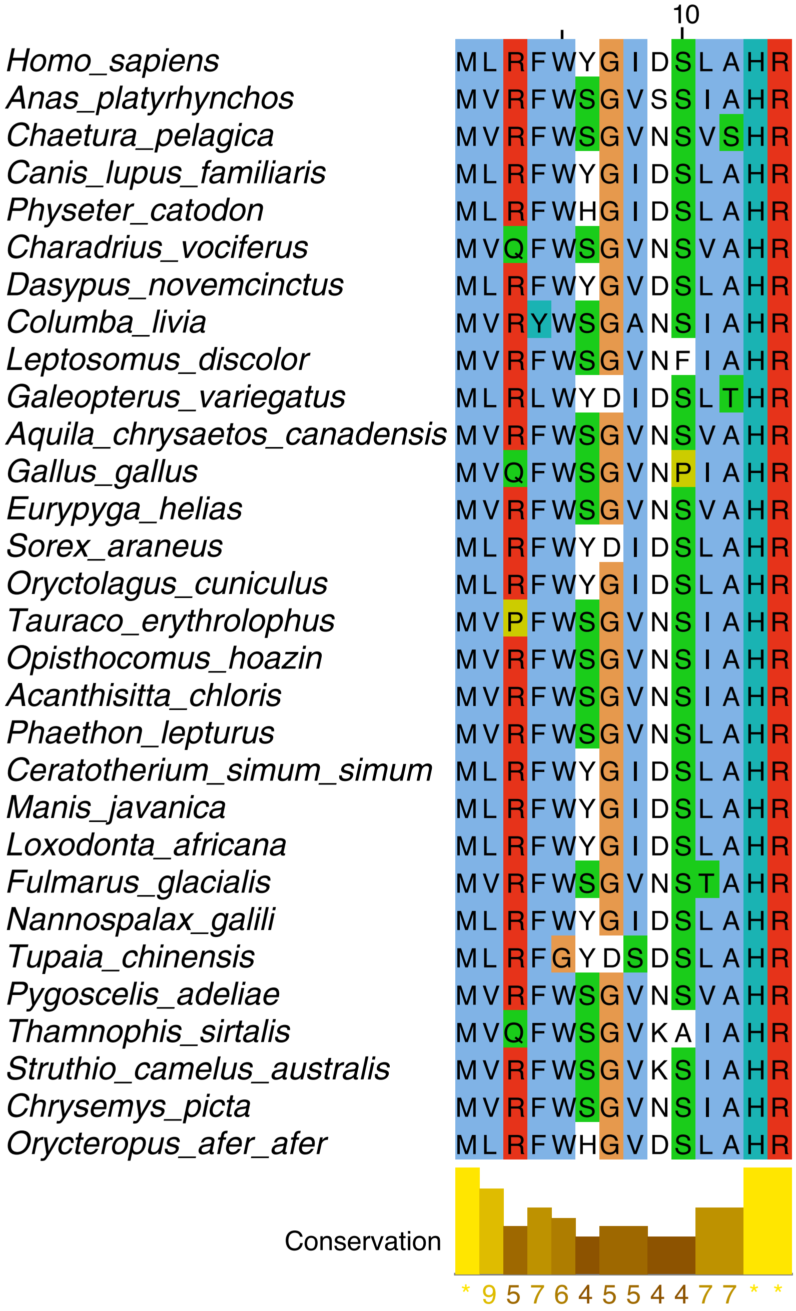

**HG0060.2**

ENSGALG00000014751

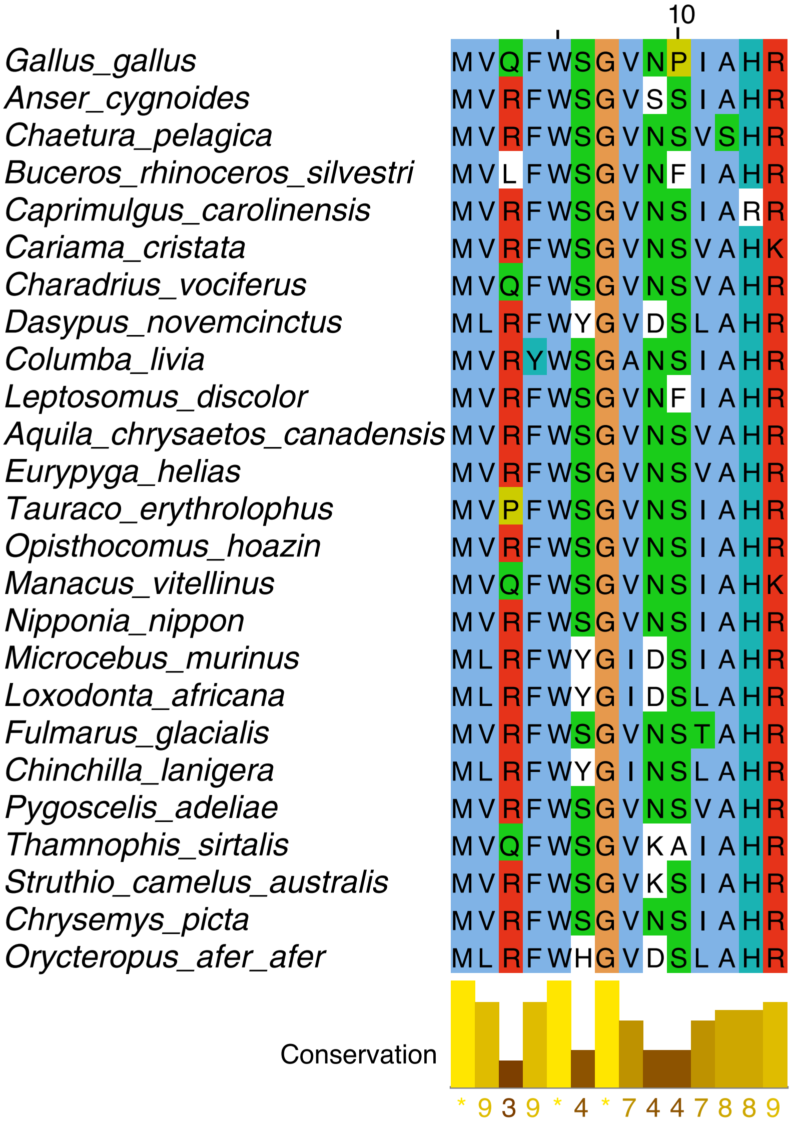

**HG0061.1**

ENSG00000166398

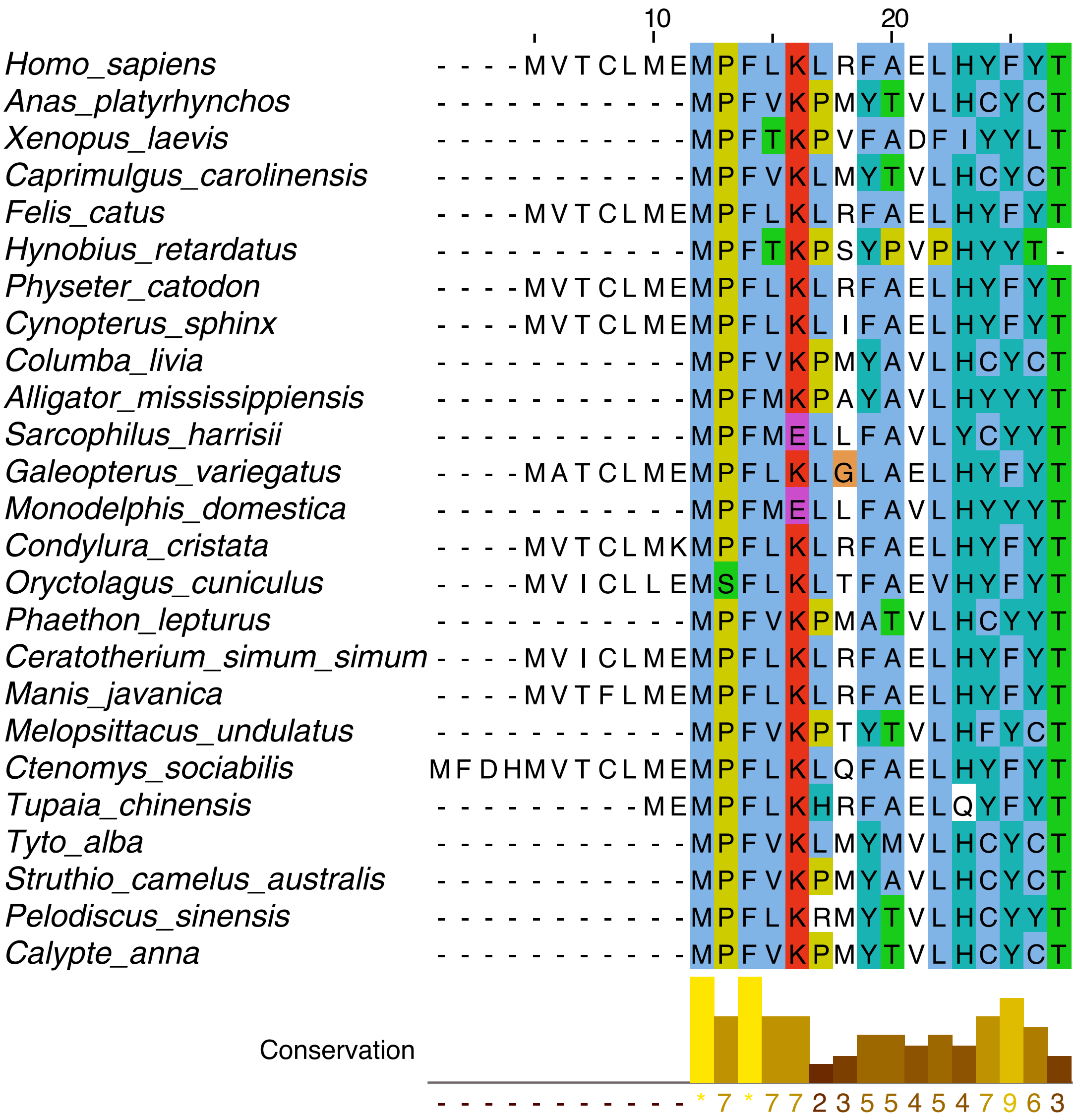

**HG0061.1**

ENSGALG00000004949

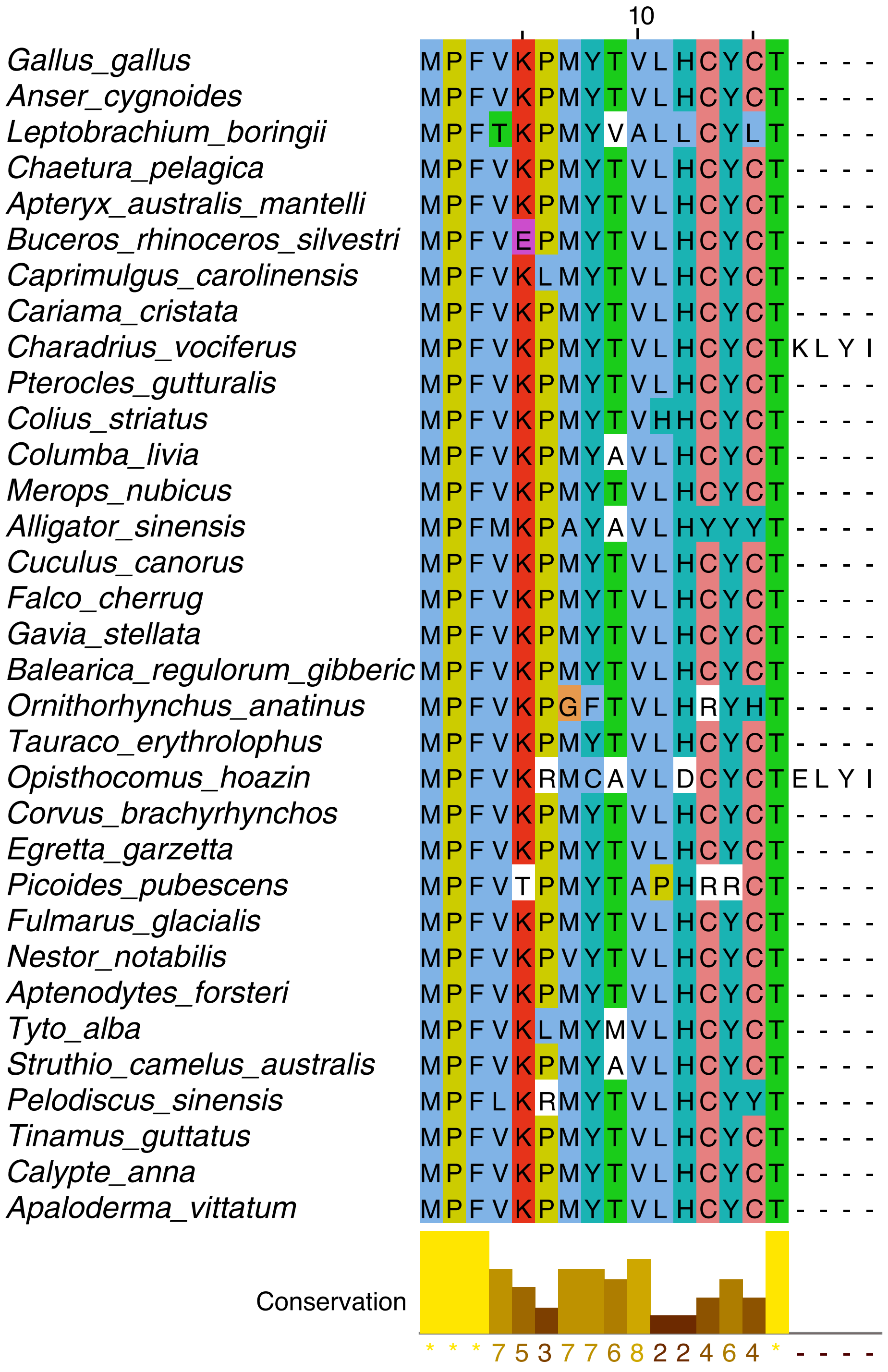

**HG0061.2**

ENSGALG00000004949

```
                                    10        20        30        40        50        60        70        80        90        100
Gallus_gallus                  --------------------------------------------------------MLQRKQLPQQTRKGEEGLQ----------------------------------
Anas_platyrhynchos             --------------------------------------------------------MLQRKQLPQQTRKGEEGLQ----------------------------------
Calidris_pugnax                --------------------------------------------------------MLQRKQLPQQRRKGGKGKKACNNK-HSNF------------------------
Columba_livia                  --------------------------------------------------------MLQRKQLPQQTRKGEEGLQ----------------------------------
Sarcophilus_harrisii           --------------------------------------------------------MLQRKQLPQKTKGKEEAAEQLEITNTPNF------------------------
Monodelphis_domestica          --------------------------------------------------------MLQRKQLPQKTEGKEEAPWQLEITNTPVFNNLTCEKRRHFIFFSCSIKKKHYR
Aquila_chrysaetos_canadensis   --------------------------------------------------------MLQRKQLPQQTRKGEEGLQ----------------------------------
Corvus_brachyrhynchos          --------------------------------------------------------MLQRKQLPQQTGKGEEGLQ----------------------------------
Aptenodytes_forsteri           --------------------------------------------------------MLQRKQLPQQTRKGEEGLQ----------------------------------
Struthio_camelus_australis     MNLLLLKYSKPVNSLESLLTEHRSCGNDLEALQIHLCIFVPSLRLRANAIWKHHFLTLQRKQLPQQTRKGEEGLQ----------------------------------

Conservation                   --------------------------------------------------------8********763554 7+25----------------------------------
```

**HG0061.3**

ENSG00000166398

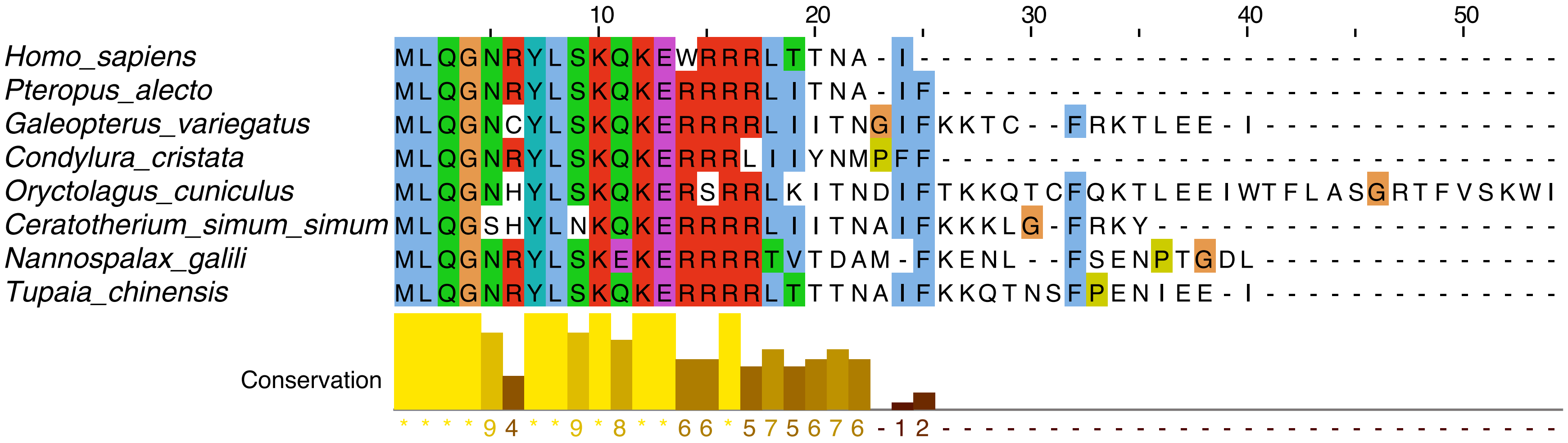

**HG0062.1**

ENSDARG00000075569 (gpbp1l1)

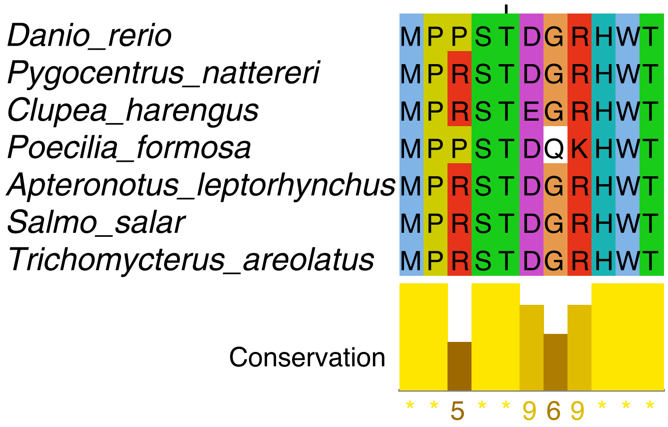

**HG0062.1**

ENSGALG00000010276 (GPBP1L1)

| | |
|---|---|
| *Gallus_gallus* | MPHFIEGRHWT |
| *Anser_cygnoides* | MPHFIEGRHWT |
| *Chaetura_pelagica* | MPHFIEGRHWT |
| *Apteryx_australis_mantelli* | MPHFIEGRHWT |
| *Buceros_rhinoceros_silvestri* | MPHFIEGRHWT |
| *Caprimulgus_carolinensis* | MPHFVEGRHWT |
| *Cariama_cristata* | MPHFIEGRHWT |
| *Physeter_catodon* | MPHLVEKRHWT |
| *Charadrius_vociferus* | MPHFIEGRHWT |
| *Pterocles_gutturalis* | MPHFIEGRHWT |
| *Colius_striatus* | MPHFIEGRHWT |
| *Columba_livia* | MPHFIEGRHWT |
| *Leptosomus_discolor* | MPHFIEGRHWT |
| *Alligator_mississippiensis* | MPHFIEGRHWT |
| *Cuculus_canorus* | MPHFIEGRHWT |
| *Falco_cherrug* | MPHFIEGRHWT |
| *Eurypyga_helias* | MPHFIEGRHWT |
| *Tauraco_erythrolophus* | MPHFIEGRHWT |
| *Opisthocomus_hoazin* | MPHFVEGRHWT |
| *Geospiza_fortis* | MPHFIEGRHWT |
| *Egretta_garzetta* | MPHFIEGRHWT |
| *Picoides_pubescens* | MPHFIEGRHWT |
| *Fulmarus_glacialis* | MPHFIEGRHWT |
| *Melopsittacus_undulatus* | MPHFIEGRHWT |
| *Oncorhynchus_mykiss* | MPHSTDGRHWT |
| *Aptenodytes_forsteri* | MPHFIEGRHWT |
| *Gekko_japonicus* | MPRFIEGRHWT |
| *Tyto_alba* | MPHFIEGRHWT |
| *Struthio_camelus_australis* | MPHFIEGRHWT |
| *Pelodiscus_sinensis* | MPHFIEGRHWT |
| *Tinamus_guttatus* | MPHFIERRHWT |
| *Calypte_anna* | MPHFVEGRHWT |
| *Apaloderma_vittatum* | MPHFIEGRHWT |
| Conservation | * * 8 4 7 9 4 * * * * |

**HG0062.2**

ENSGALG00000010276 (GPBP1L1)

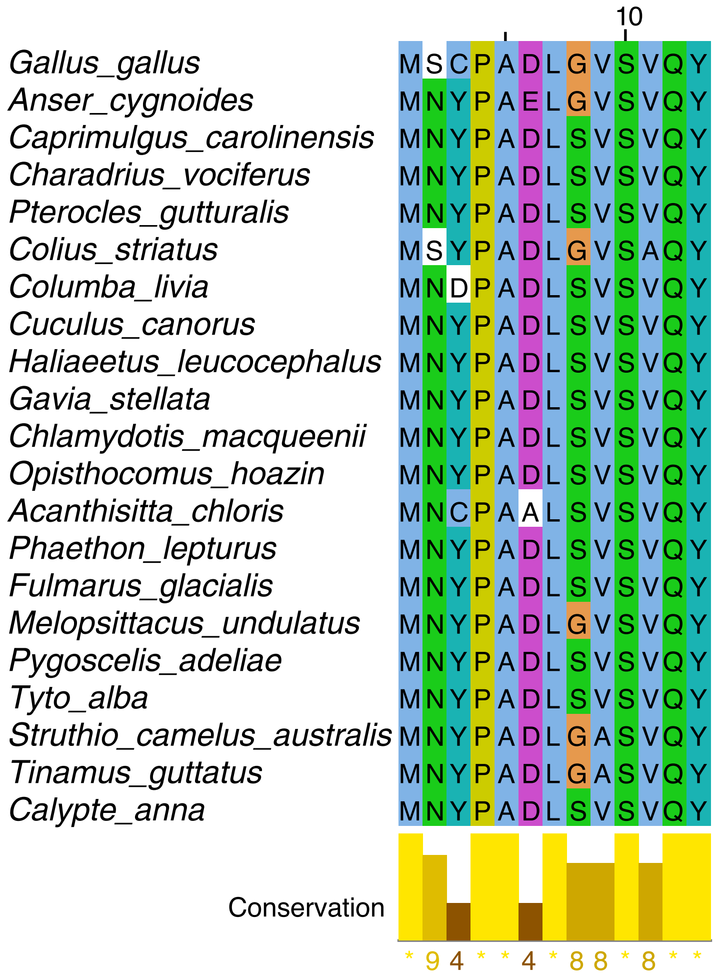

**HG0062.3**

ENSG00000159592 (GPBP1L1)

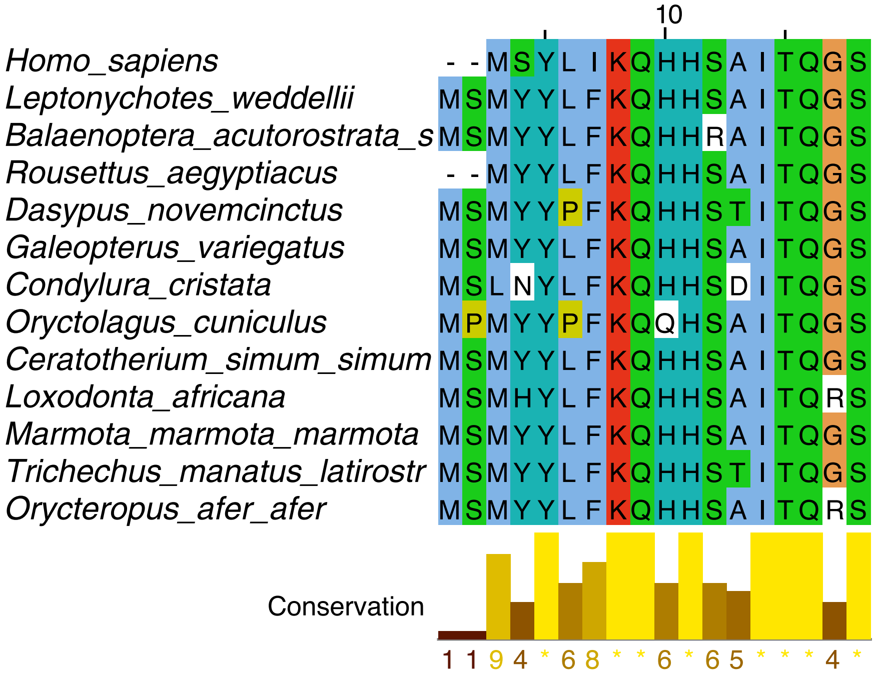

**HG0063.1**

ENSG00000162630 (B3GALT2)

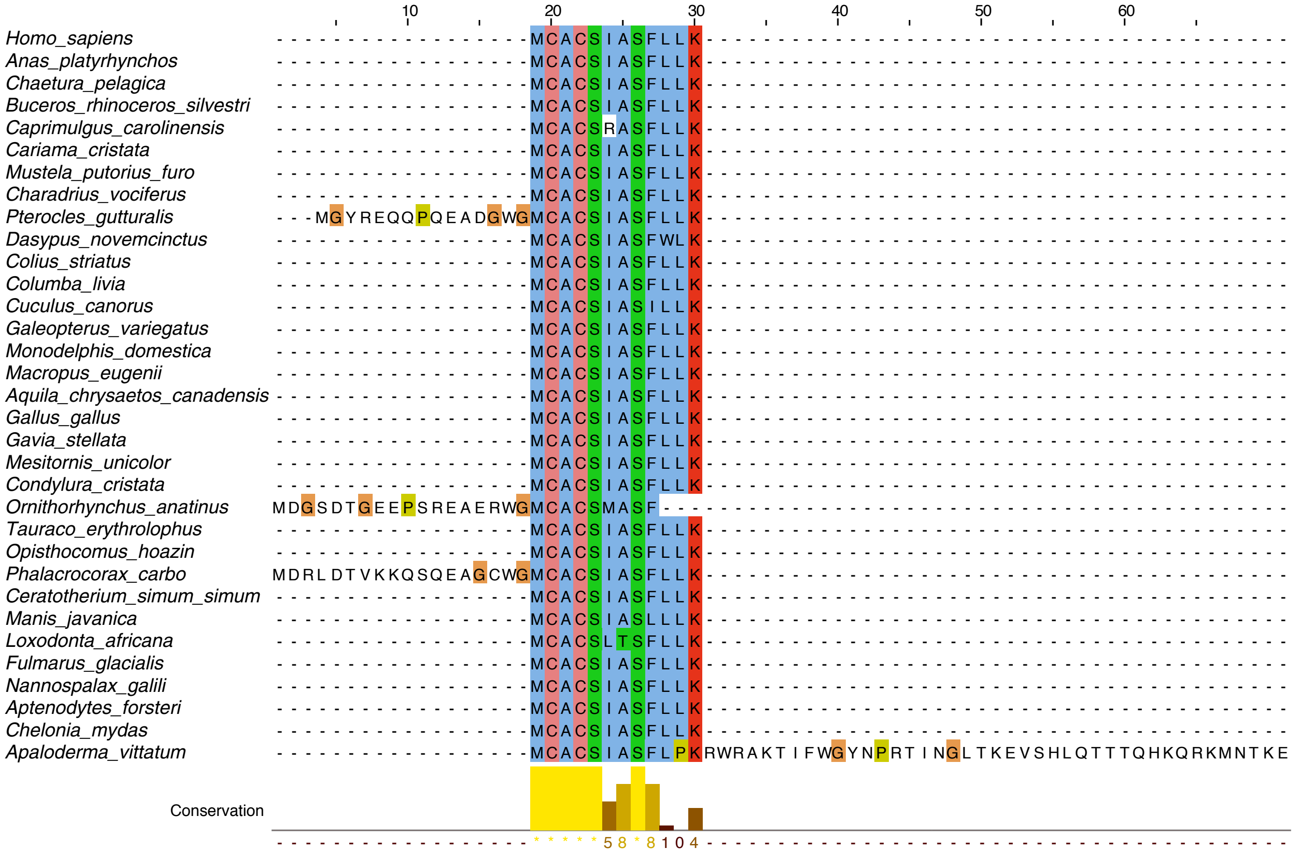

**HG0063.1**

ENSGALG00000002477

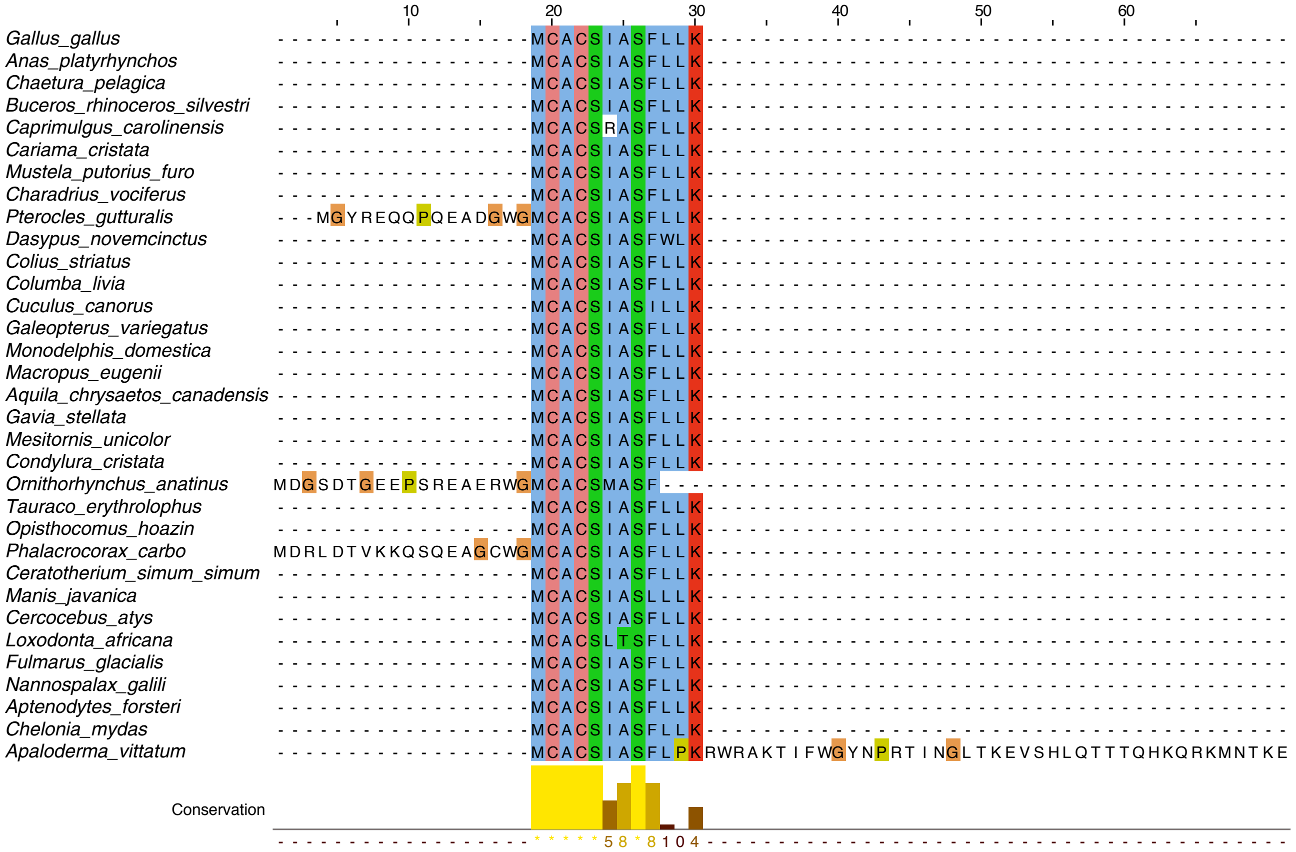

**HG0063.2**

ENSG00000162630 (B3GALT2)

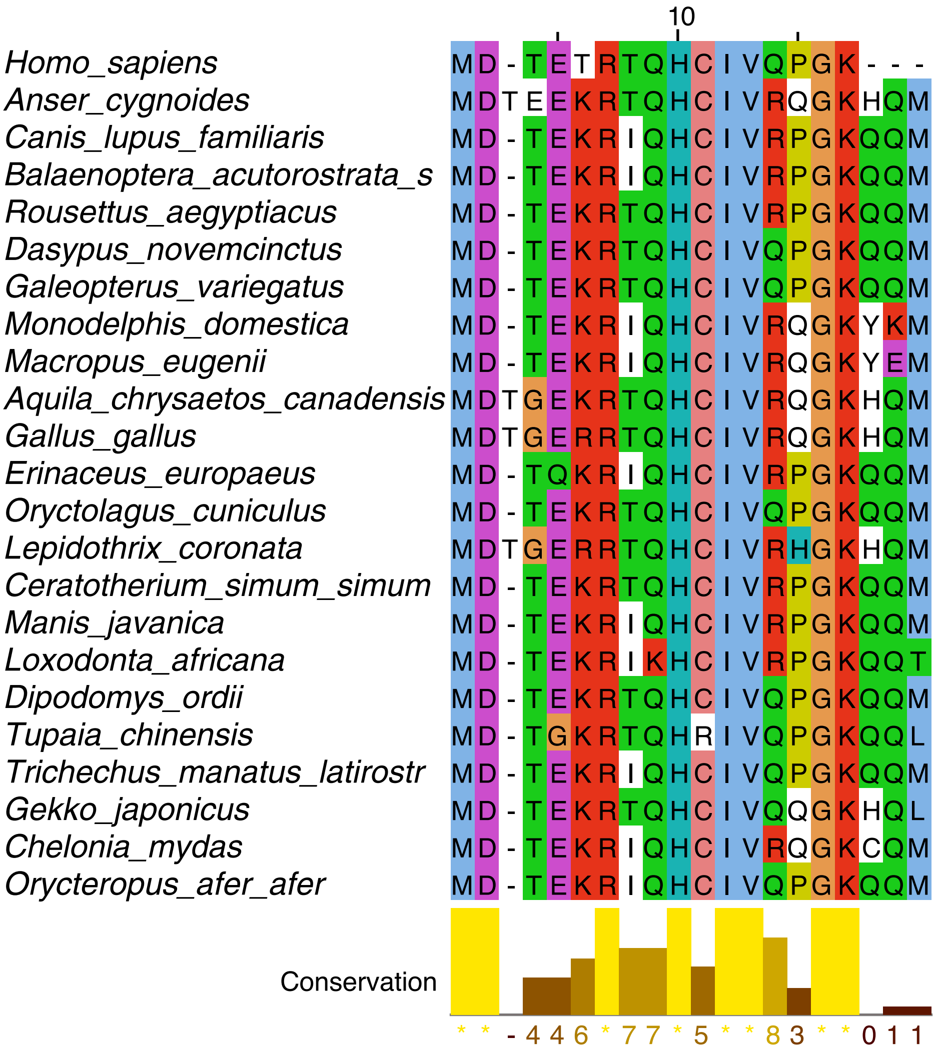

**HG0064.1**

ENSG00000102678 (FGF9)

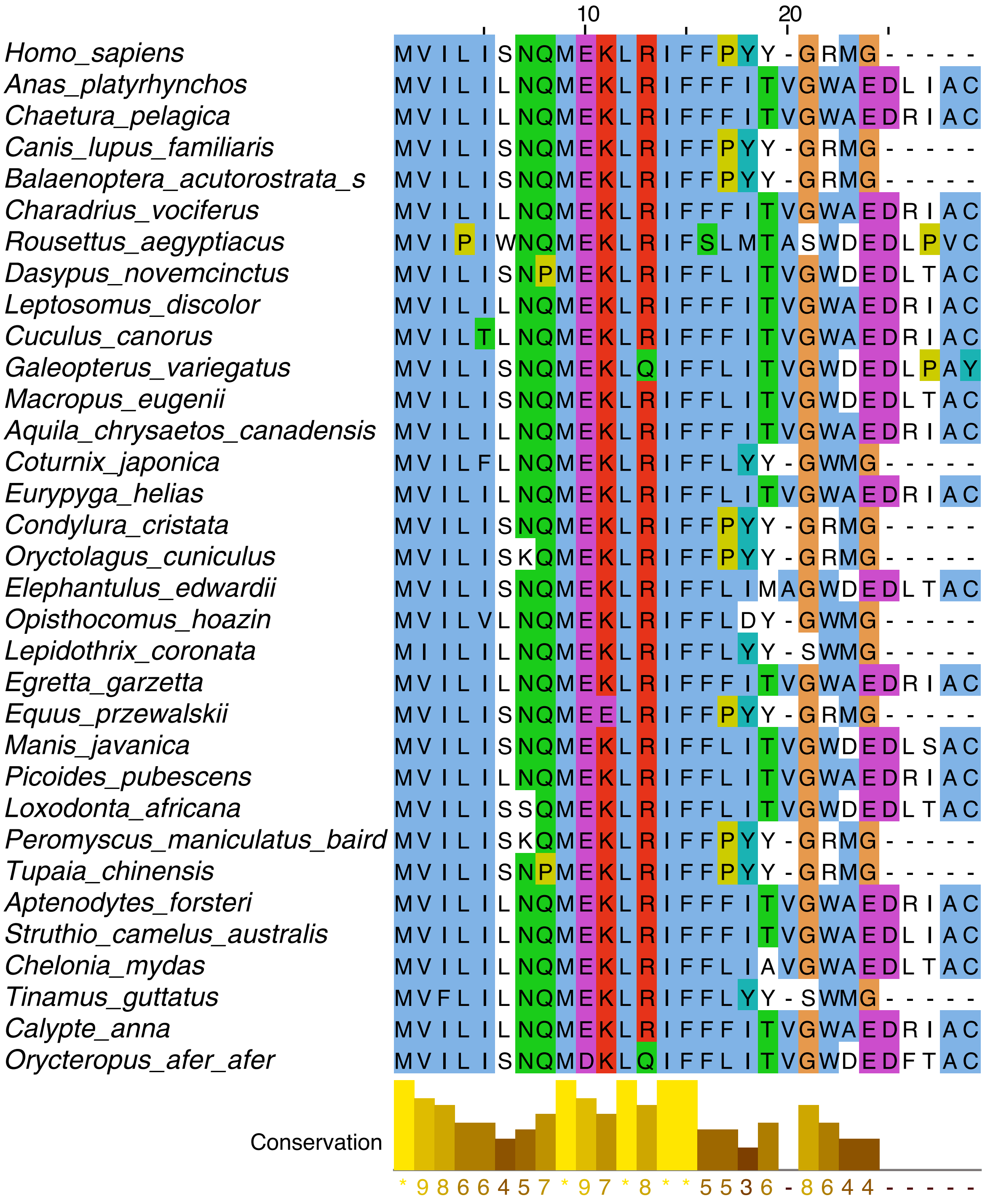

## HG0064.2

ENSG00000113578 (FGF1)

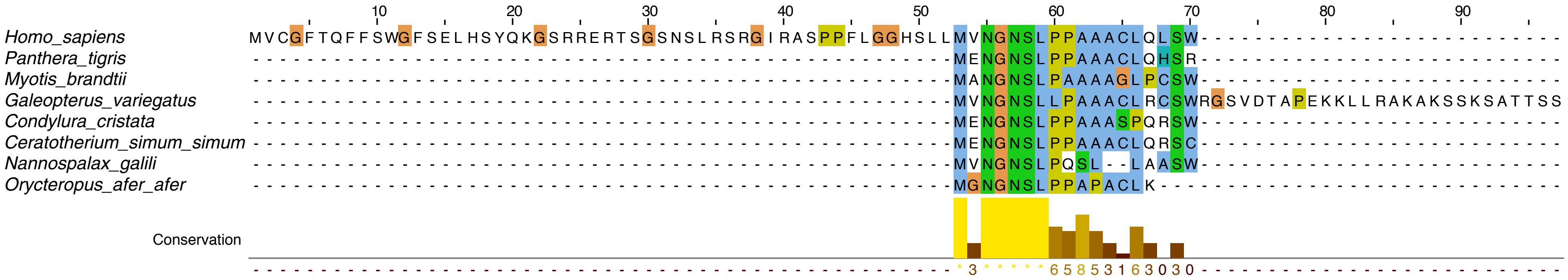

# HG0065

### ENSGALG00000029927 (STRBP)

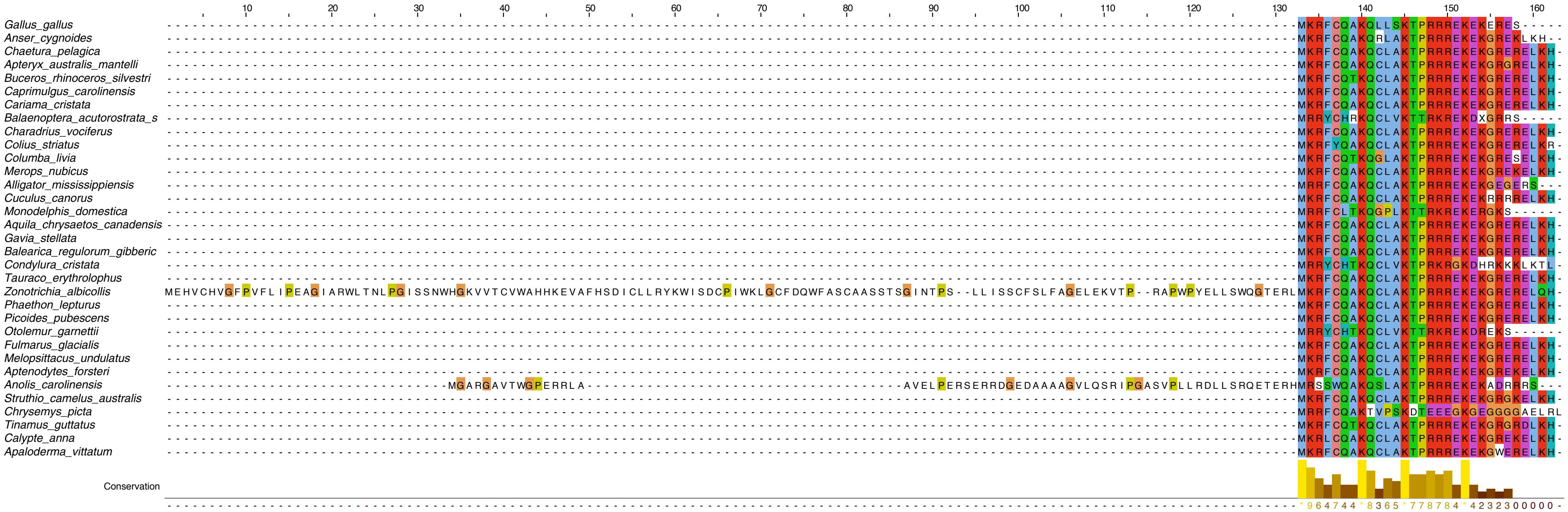

**HG0066**

ENSG00000112182 (BACH2)

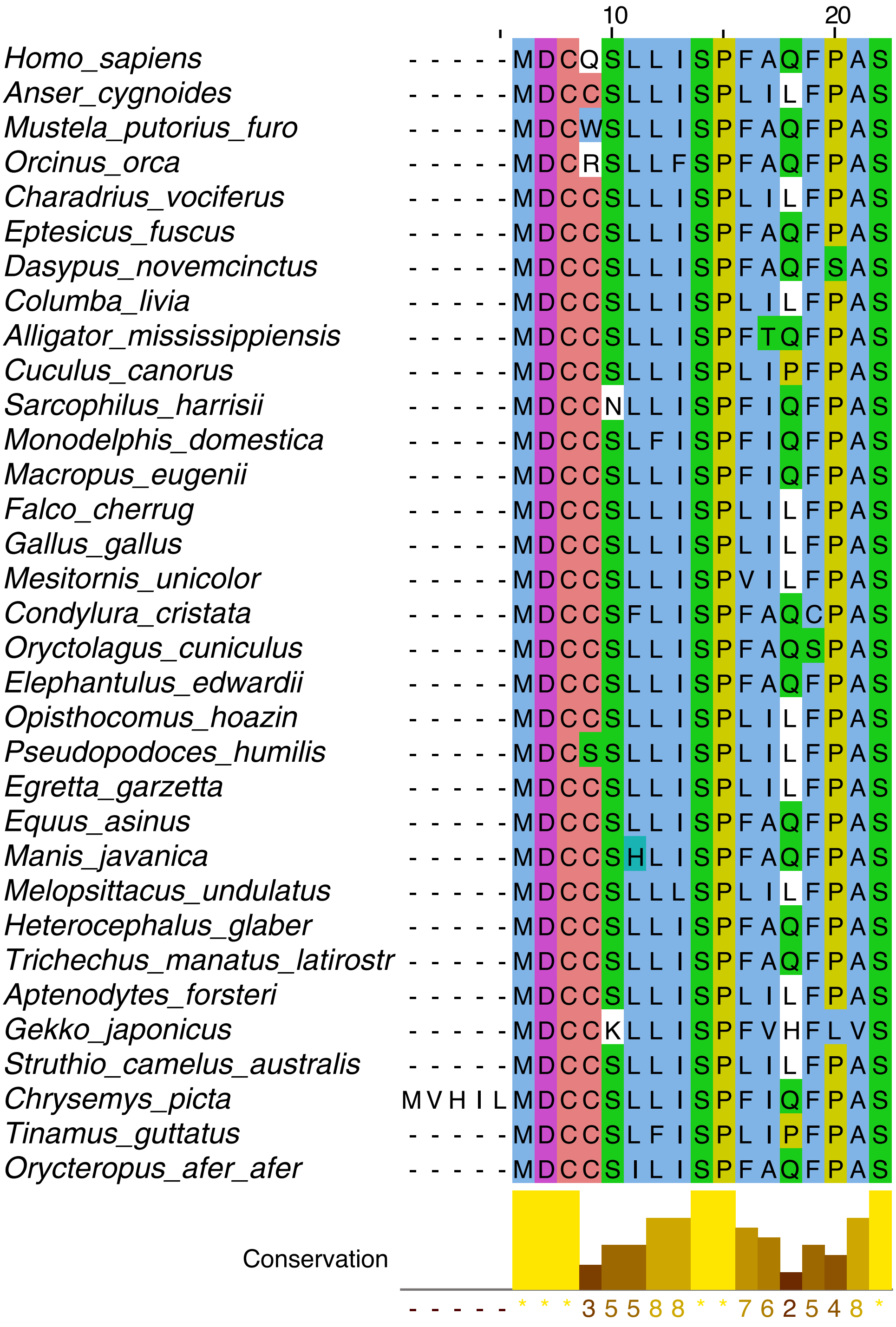

**HG0067**

ENSG00000166862 (CACNG2)

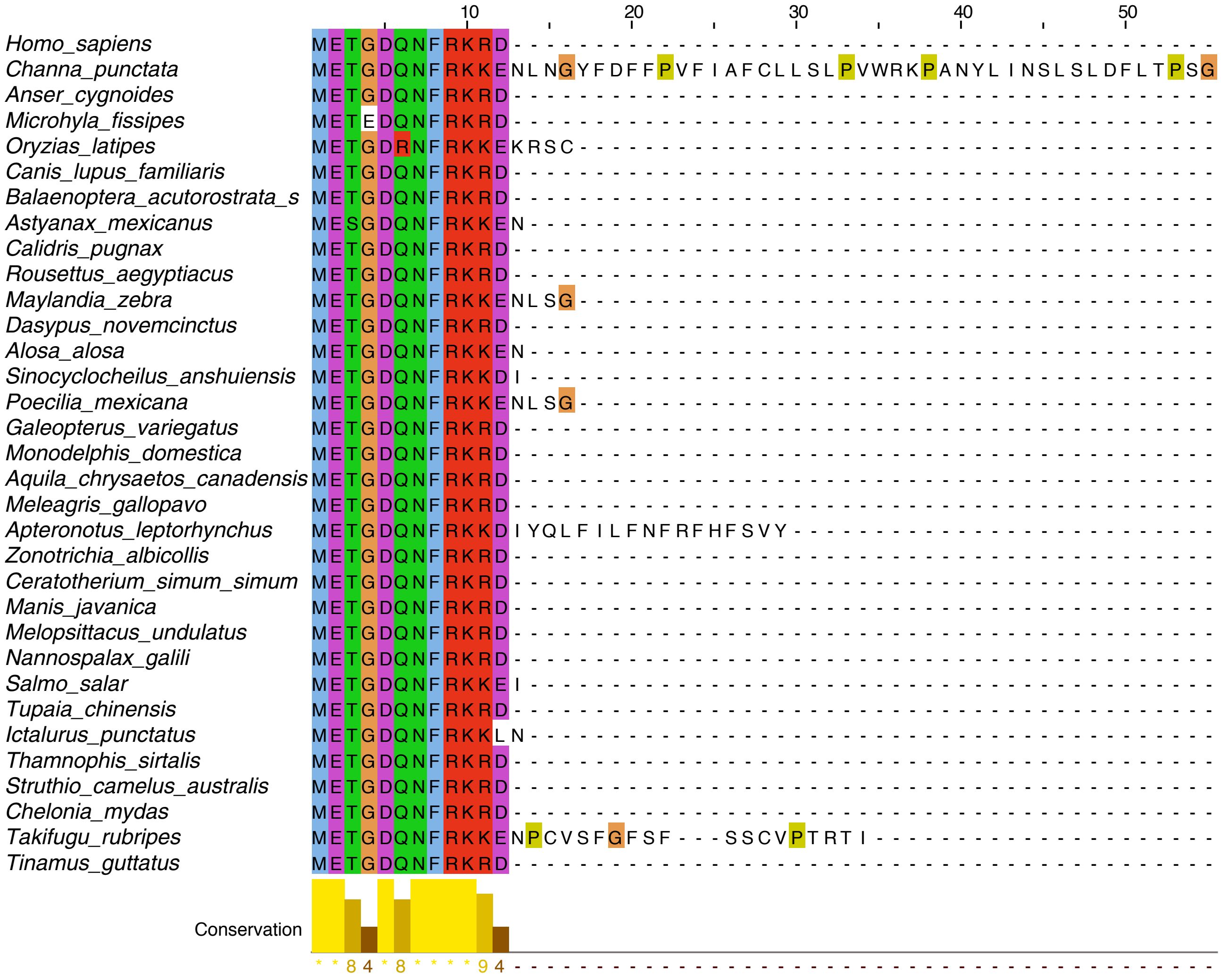

**HG0068**

ENSDARG00000071235

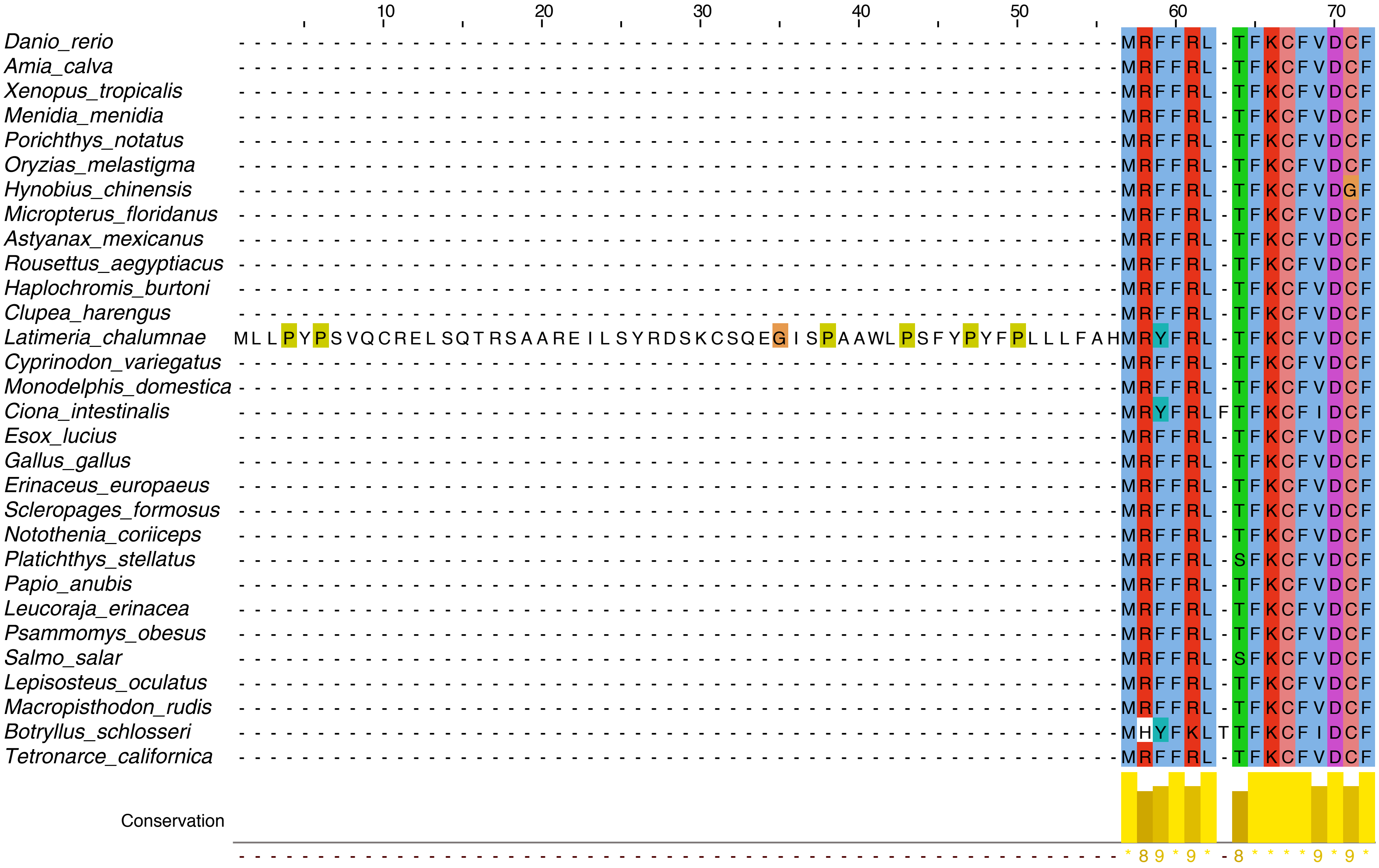

**HG0068**

ENSG00000130821 (SLC6A8)

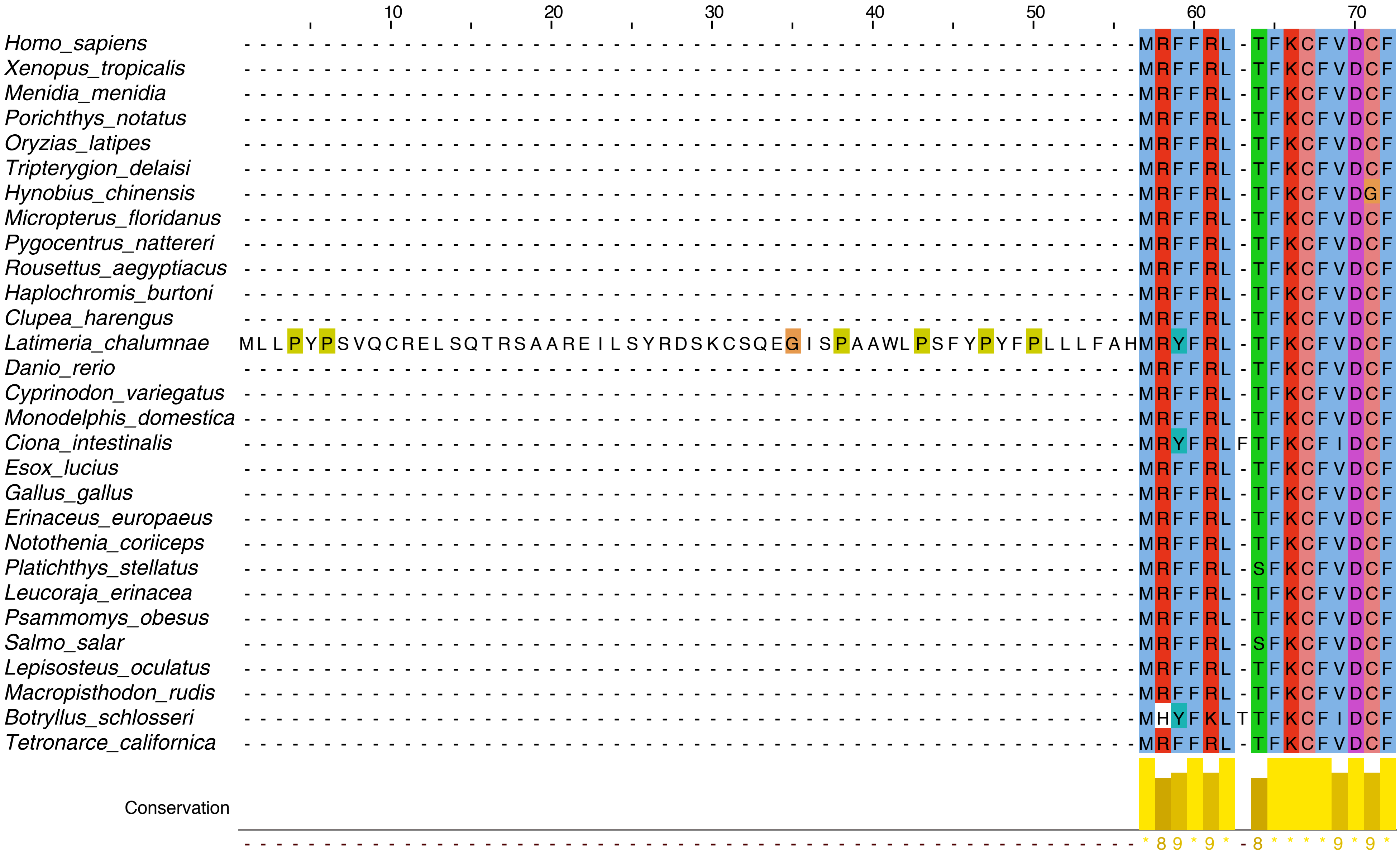

**HG0068**

ENSGALG00000041419

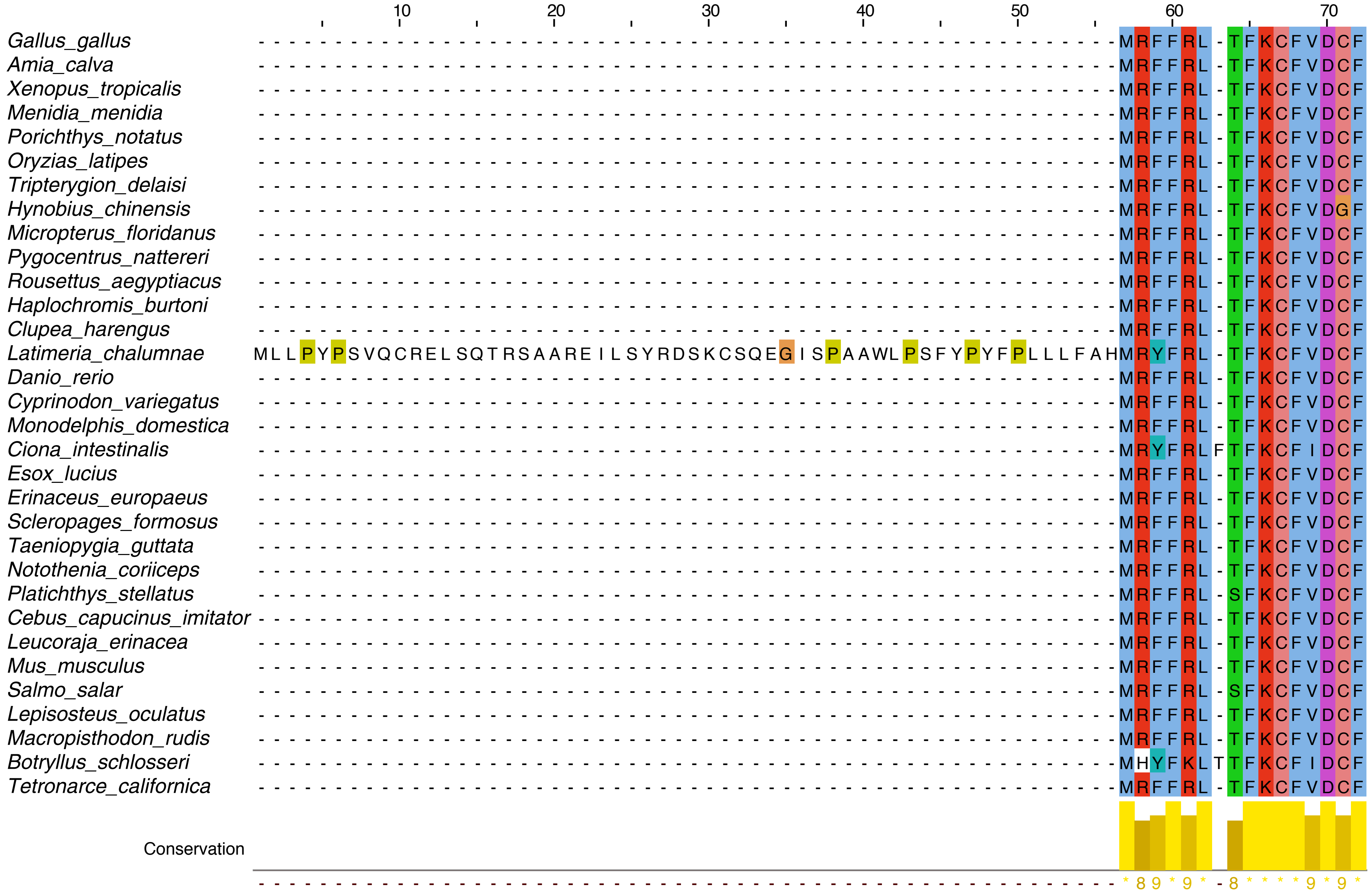

**HG0069**

ENSG00000100697 (DICER1)

| Species | Alignment (position 10 marked) |
|---|---|
| *Homo_sapiens* | -MSKCSSDKSNTEISNIKT |
| *Apteryx_australis_mantelli* | MMSKCSSGSINTEVAEIKT |
| *Cariama_cristata* | MMSKCSSGSINTETAEIKT |
| *Ailuropoda_melanoleuca* | -MSKCSSDNSNTEISNIKT |
| *Balaenoptera_acutorostrata_s* | -MSKCSSDNSNTEISNIKT |
| *Charadrius_vociferus* | MMSKCSSGSINTETAEIKT |
| *Pteropus_alecto* | -MSKCSSDNSNTEISNIKT |
| *Pterocles_gutturalis* | MMSKCSSGSINTETAEIKT |
| *Dasypus_novemcinctus* | -MSKCSSDNSNTEISNIKT |
| *Columba_livia* | MMSKCSSGSINTETAEIKT |
| *Alligator_sinensis* | MMSKCSSENTNTKVSEIKT |
| *Galeopterus_variegatus* | -MSKCSSDNSNTEISNIKT |
| *Aquila_chrysaetos_canadensis* | MMSKCSSGSINTETAEIKT |
| *Gavia_stellata* | MMSKCSSGSINTETAEIKT |
| *Mesitornis_unicolor* | MMSKCSSGSVNTETAEIKT |
| *Erinaceus_europaeus* | MMSKCSSDNSNTEIANIKT |
| *Elephantulus_edwardii* | -MSKCGSDSSNTDIADIKT |
| *Ornithorhynchus_anatinus* | MMSKCSSGNINTEVSEIKT |
| *Ficedula_albicollis* | MMSKCSSGGINTETAEIKT |
| *Phaethon_lepturus* | MMSKCSSGSINTEAAEIKT |
| *Ceratotherium_simum_simum* | -MSKCSSDNSNTEISNIKT |
| *Loxodonta_africana* | -MSKCSSDNSNAEISNIKT |
| *Melopsittacus_undulatus* | MMSKCSSGSINTETAEIKT |
| *Ictidomys_tridecemlineatus* | -MSKCSSDSSNTEISNIKT |
| *Trichechus_manatus_latirostr* | -MSKCSSDNSNTEISNIKT |
| *Aptenodytes_forsteri* | MMSKCSSGSINTETAEIKT |
| *Tyto_alba* | MMSKCSSGSINTETAEIKT |
| *Struthio_camelus_australis* | MMSKCNSGSINTEVAEIKT |
| *Chrysemys_picta* | MMSKCSSKNTNTEISEIKT |
| *Tinamus_guttatus* | MMSKCSSGNINTEVAEIKT |
| *Apaloderma_vittatum* | MMSKCSSGSINTETAEIKT |
| *Orycteropus_afer_afer* | -MSGCSSDGSNTEASHIKT |

Conservation

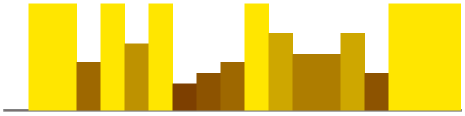

- * * 5 * 7 * 3 4 5 * 8 6 6 8 4 * * *

**HG0069**

ENSGALG00000010999 (DICER1)

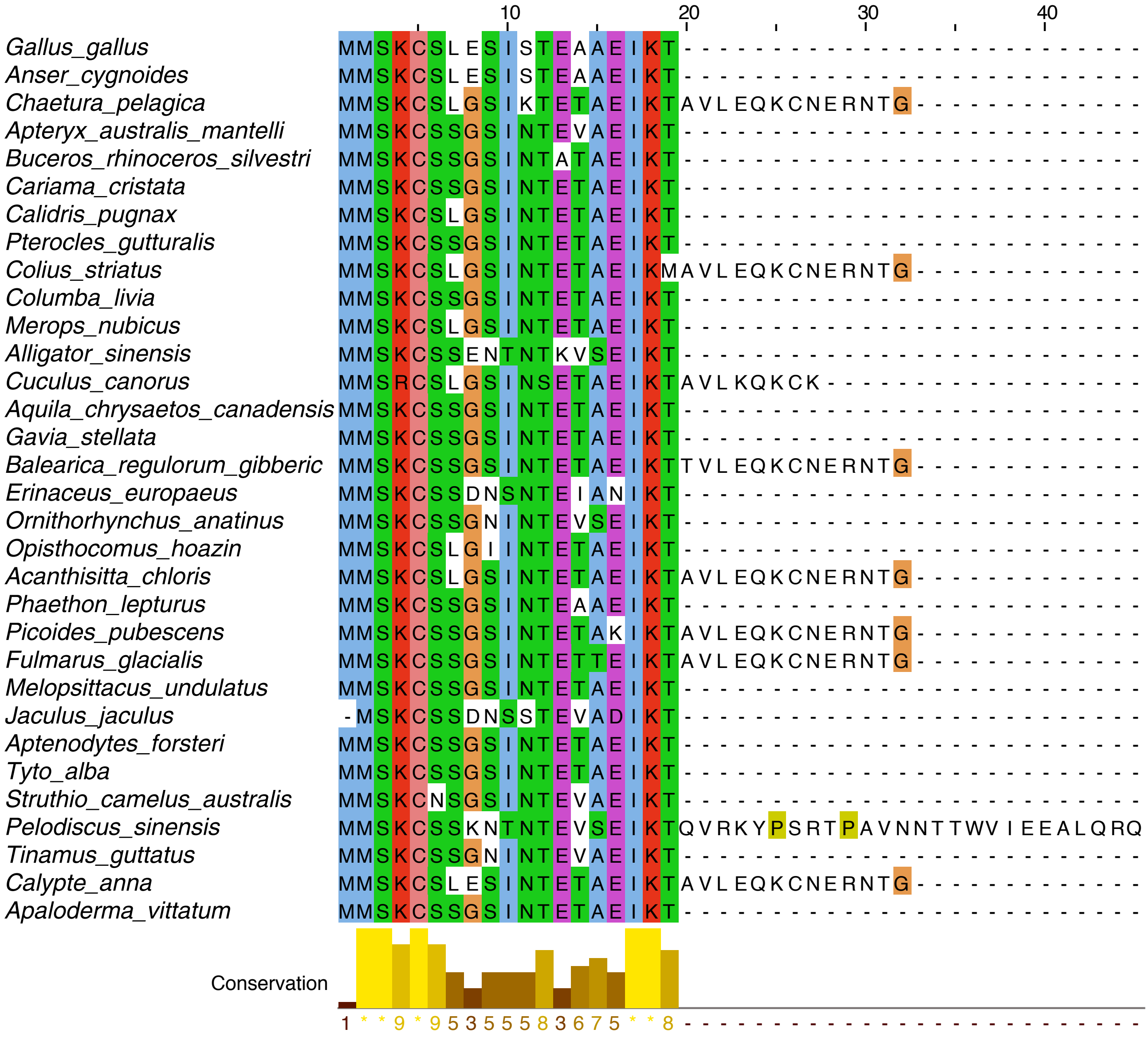

**HG0070**

ENSDARG00000040926 (nr2f2)

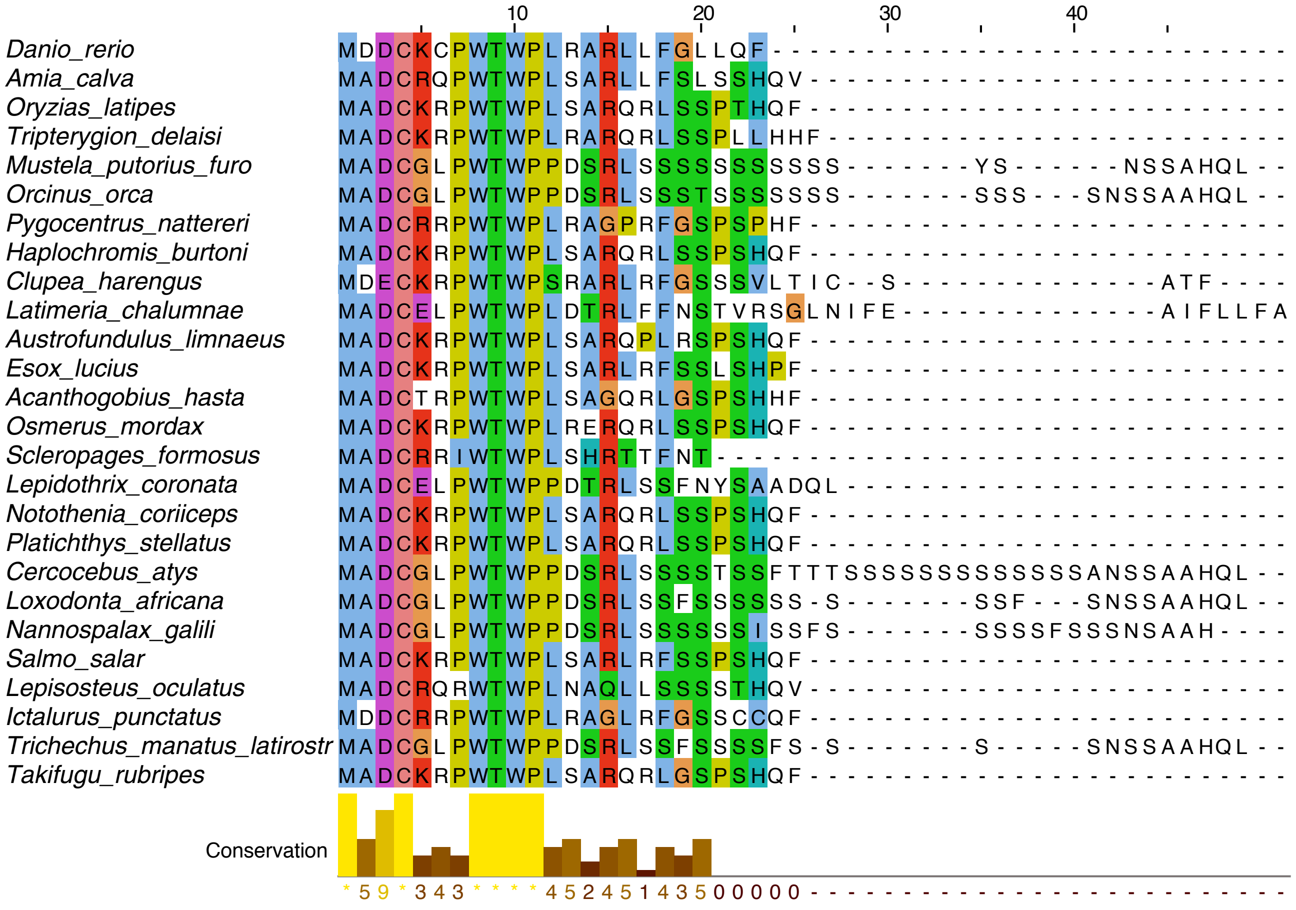

**HG0070**

ENSDARG00000052695 (nr2f1a)

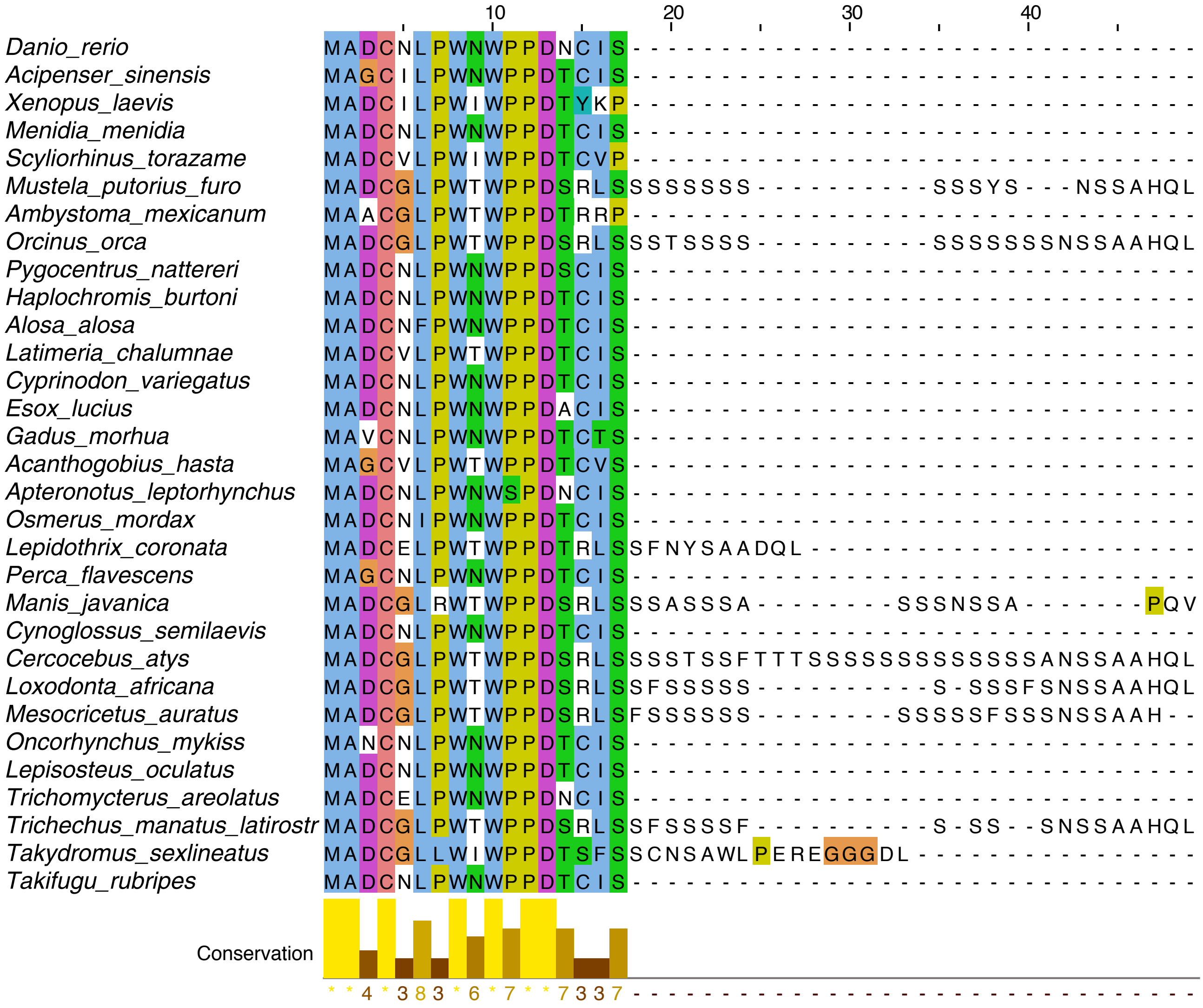

**HG0070**

ENSG00000175745 (NR2F1)

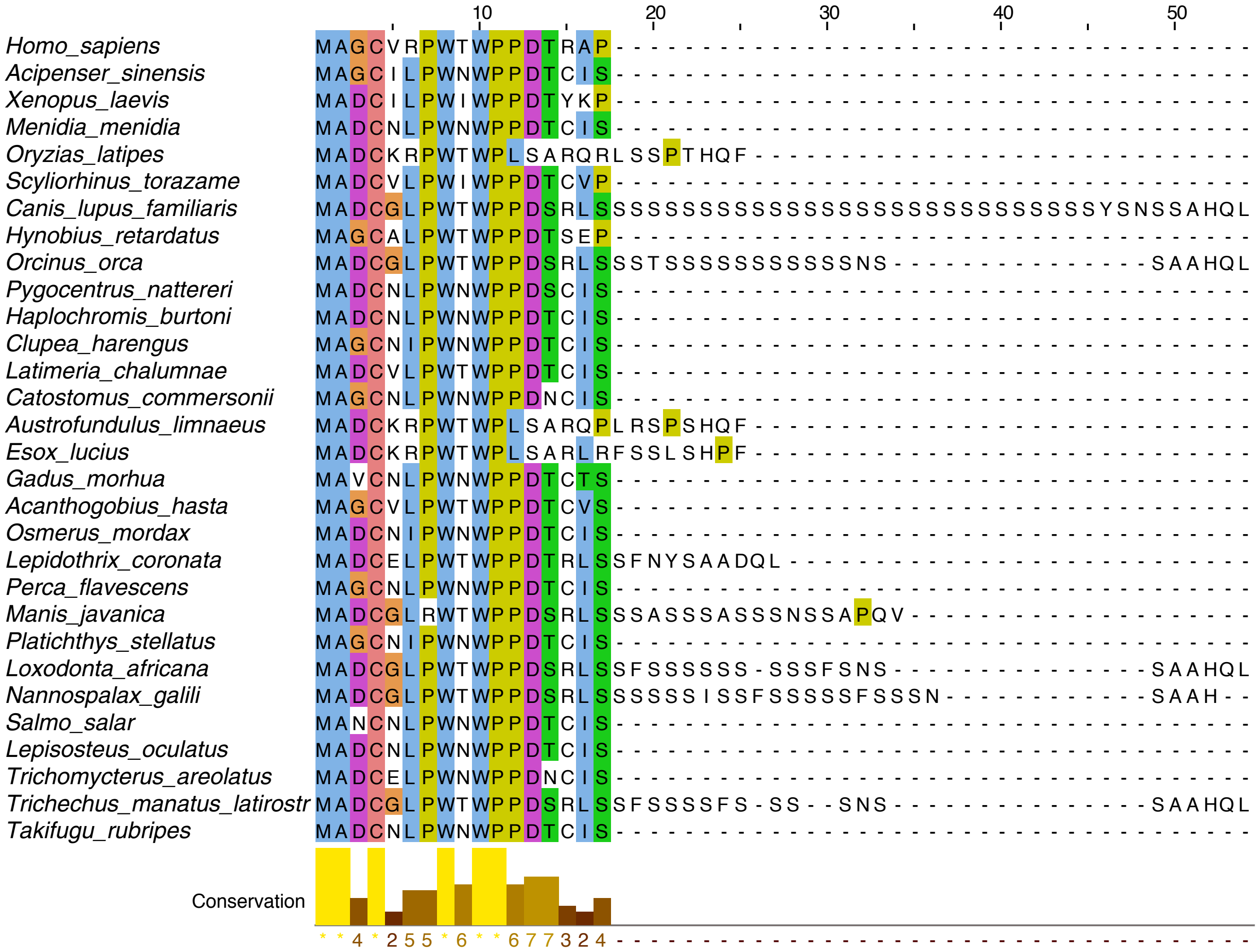

**HG0070**

ENSG00000185551 (NR2F2)

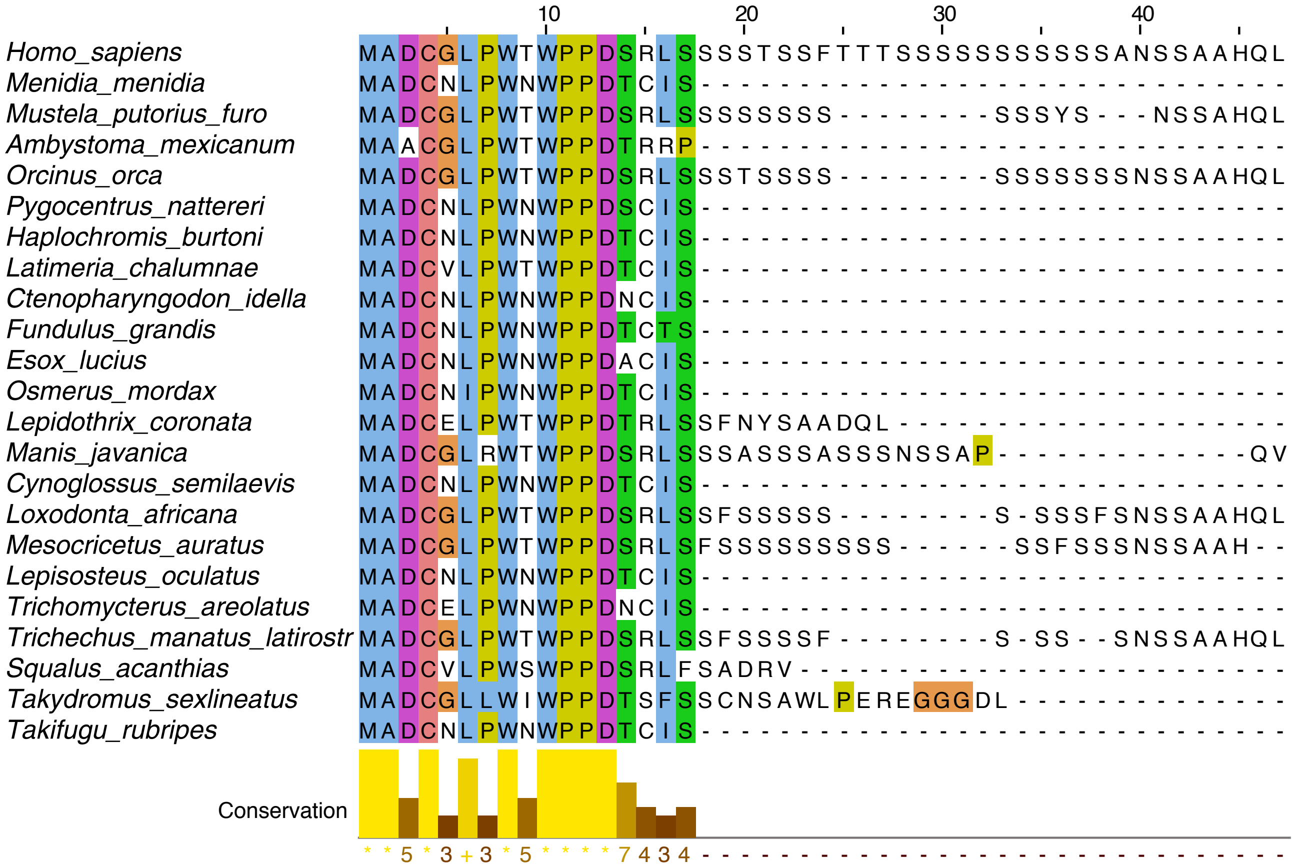

**HG0071.1**

ENSDARG00000098240 (meis2a)

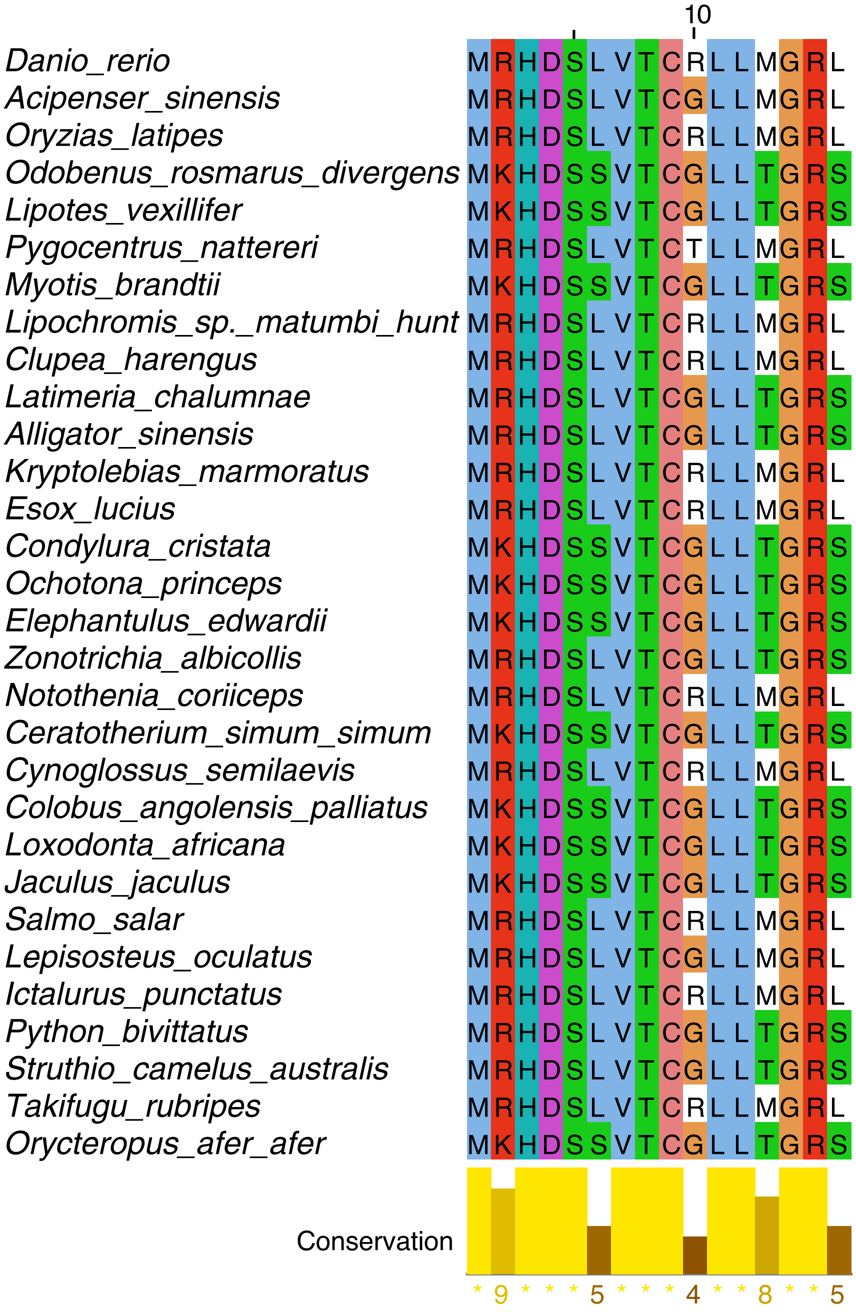

**HG0071.1**

ENSG00000134138 (MEIS2)

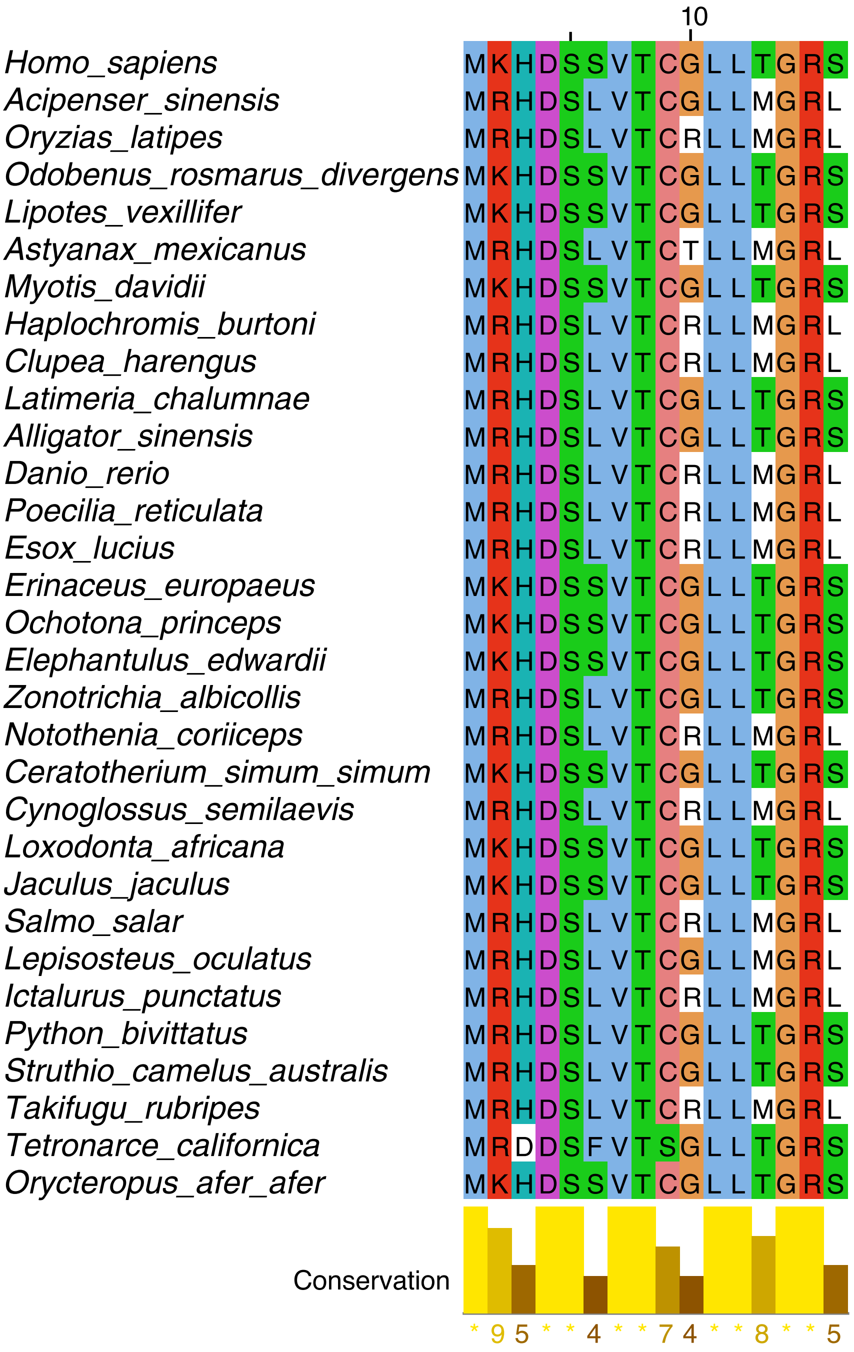

**HG0071.2**

ENSG00000143995 (MEIS1)

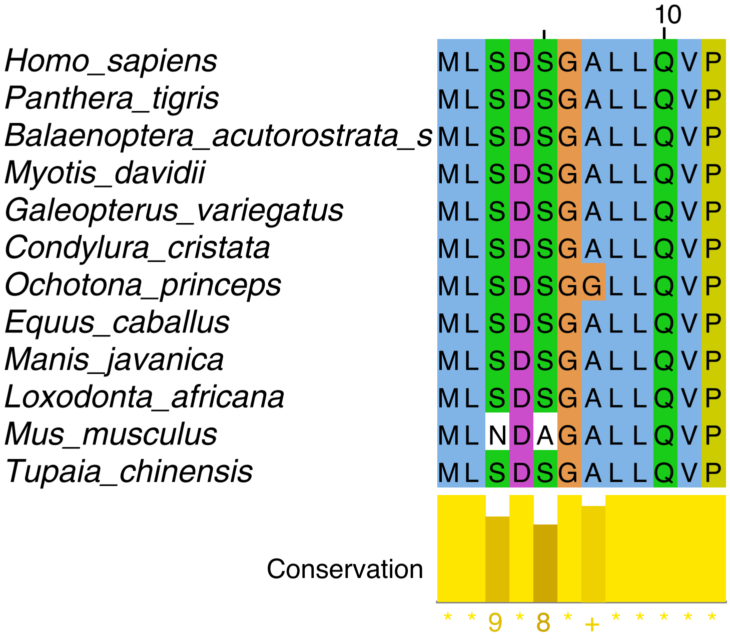

**HG0072**

ENSG00000104435 (STMN2)

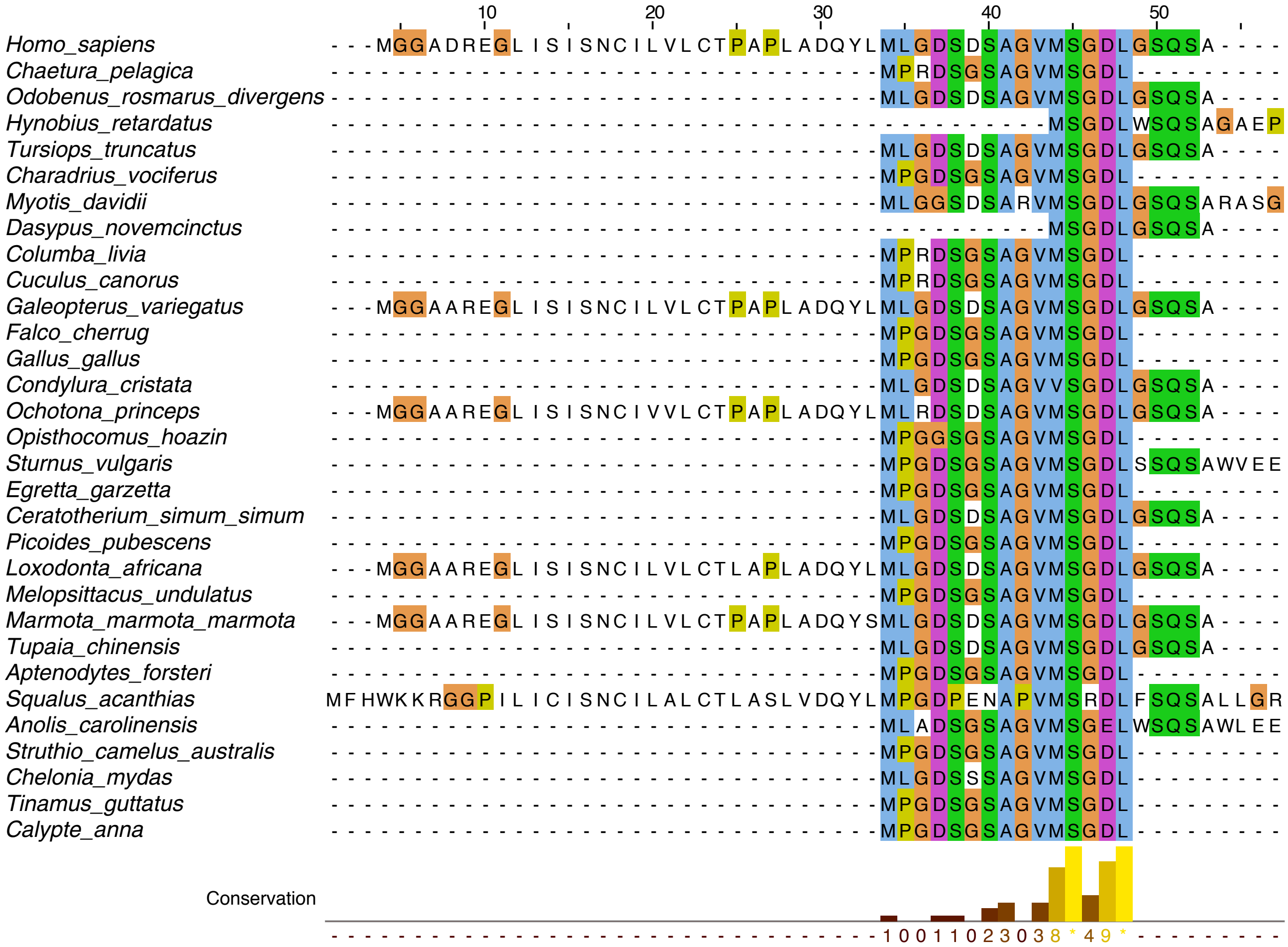

**HG0072**

ENSGALG00000039690 (STMN2)

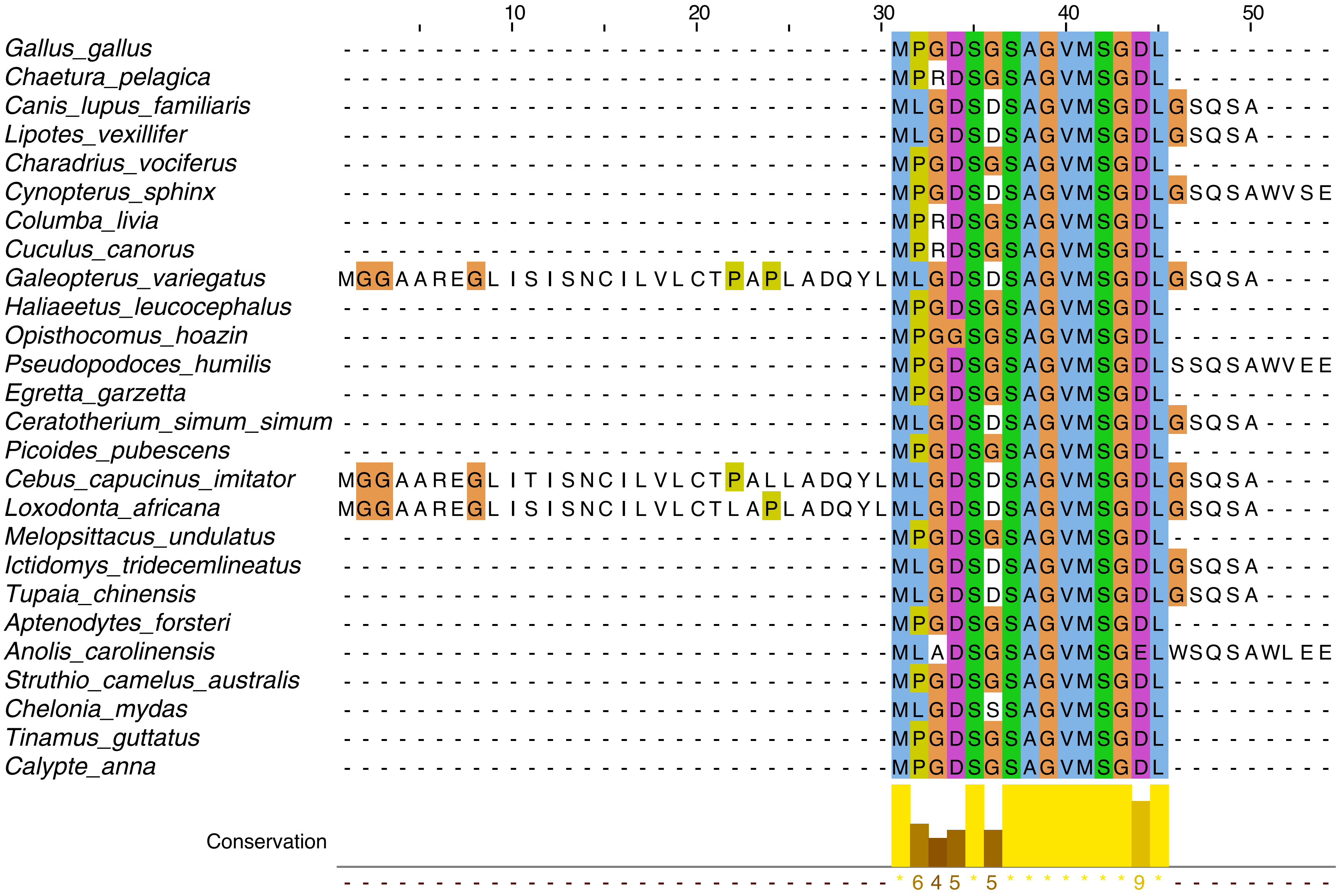

**HG0073**

ENSDARG00000041708 (csrp2bp)

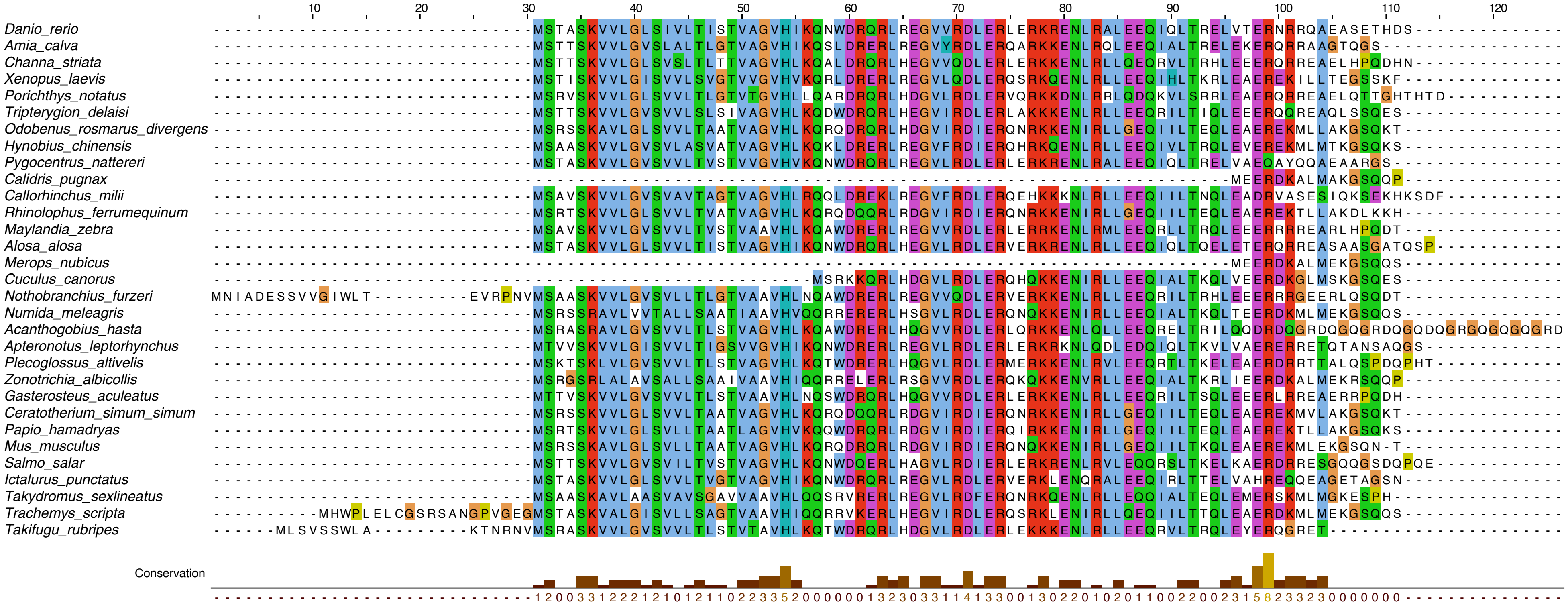

**HG0074**

ENSDARG00000078624 (arhgef9b)

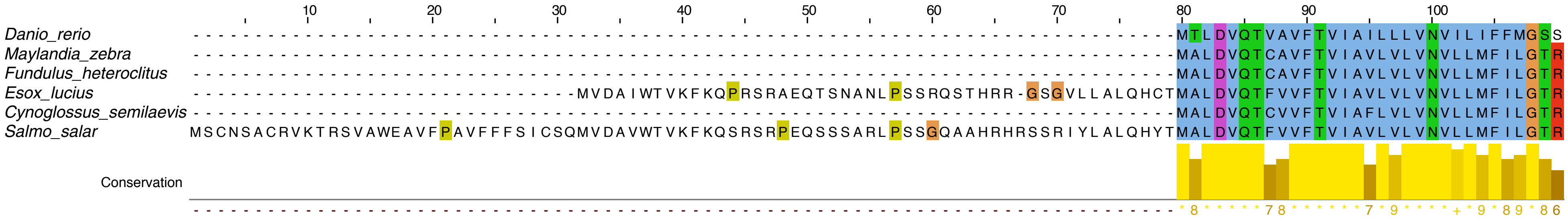

**HG0074**

ENSG00000131089 (ARHGEF9)

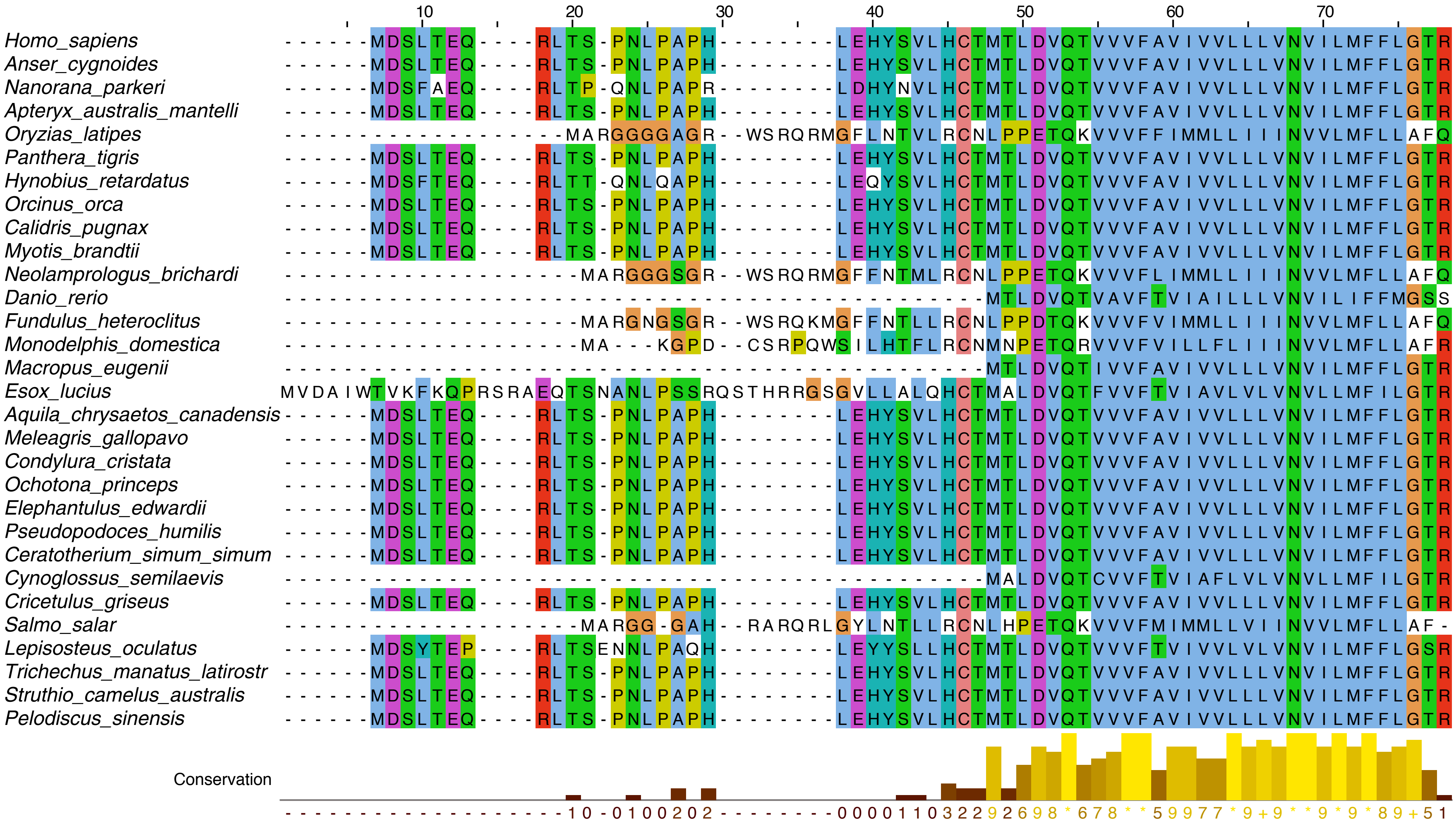

**HG0075**

ENSG00000165699 (TSC1)

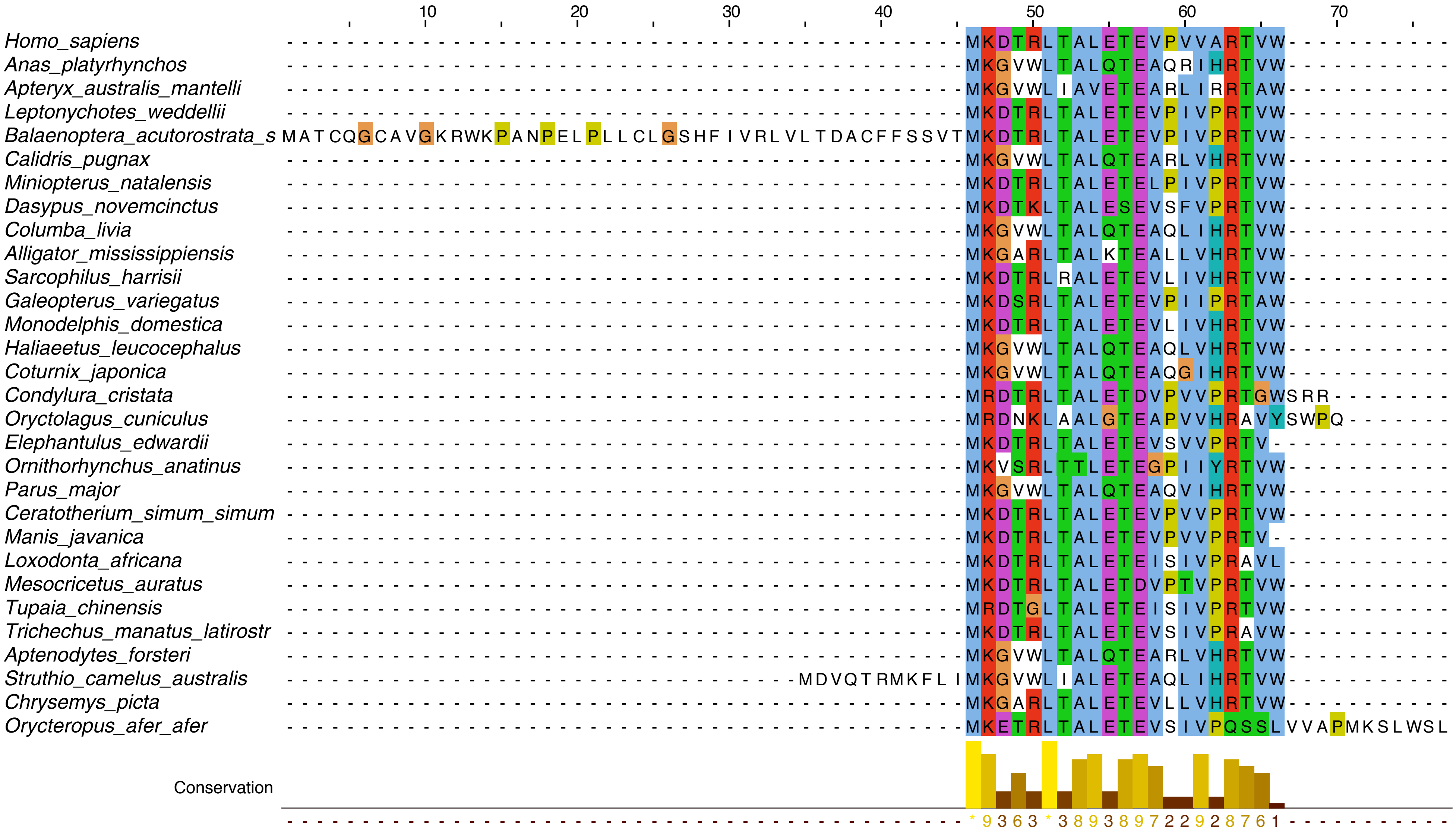

# HG0076

### ENSDARG00000040008 (neurod6a)

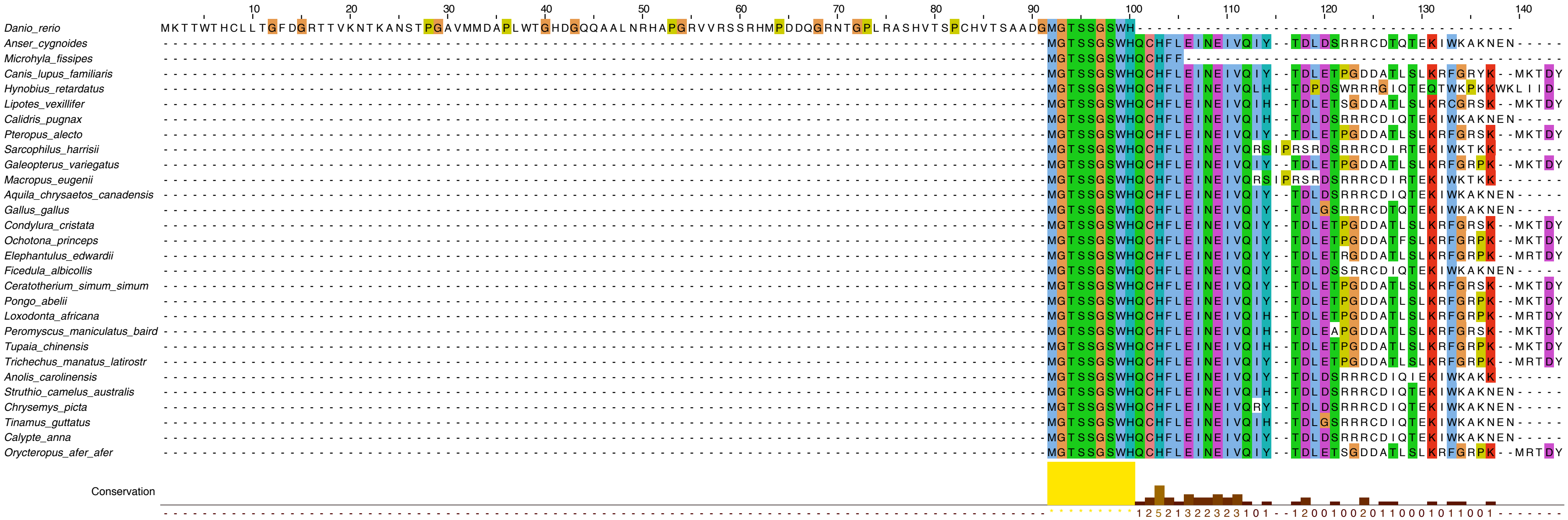

**HG0076**

ENSG00000164600 (NEUROD6)

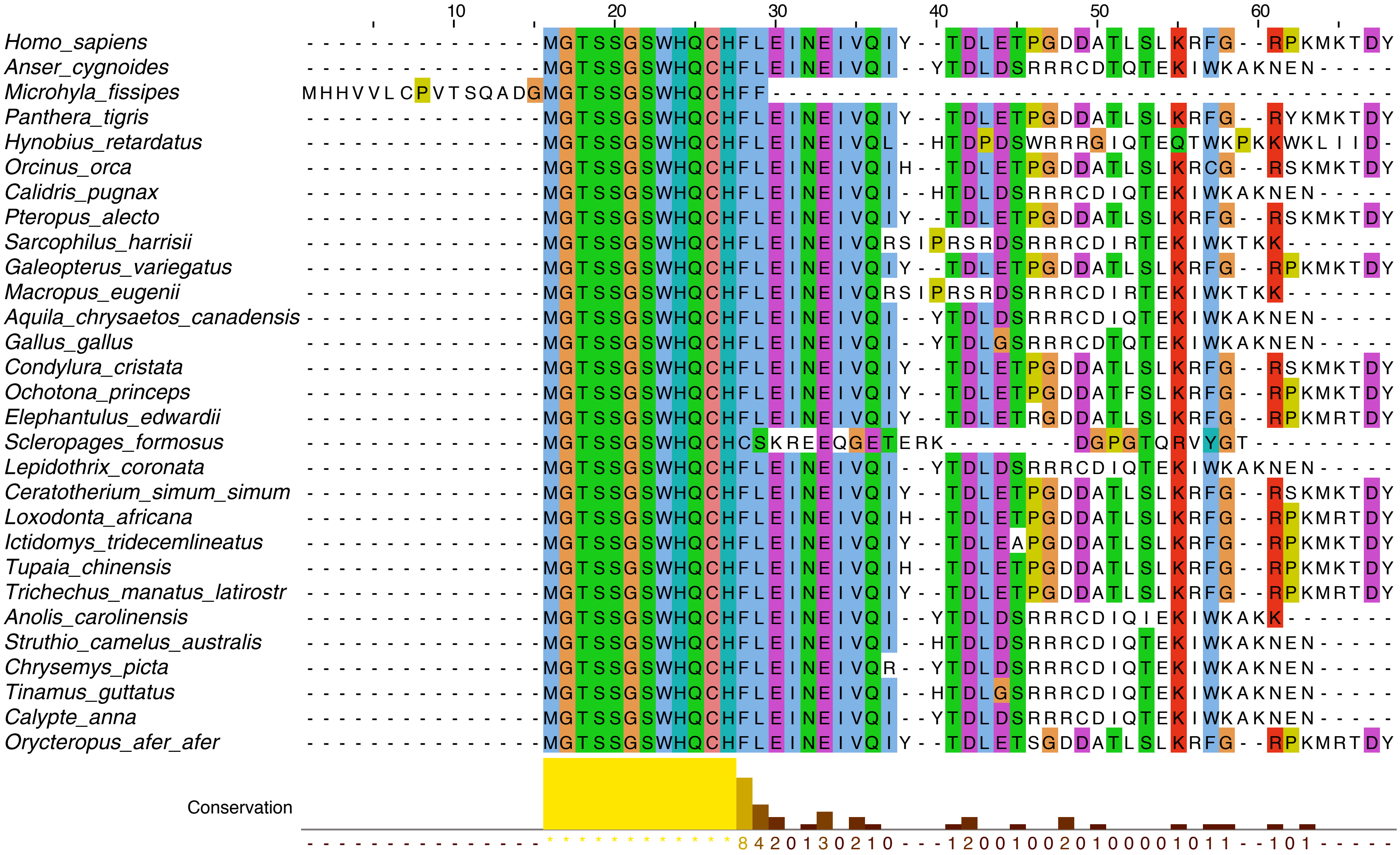

**HG0077.1**

ENSGALG00000005074

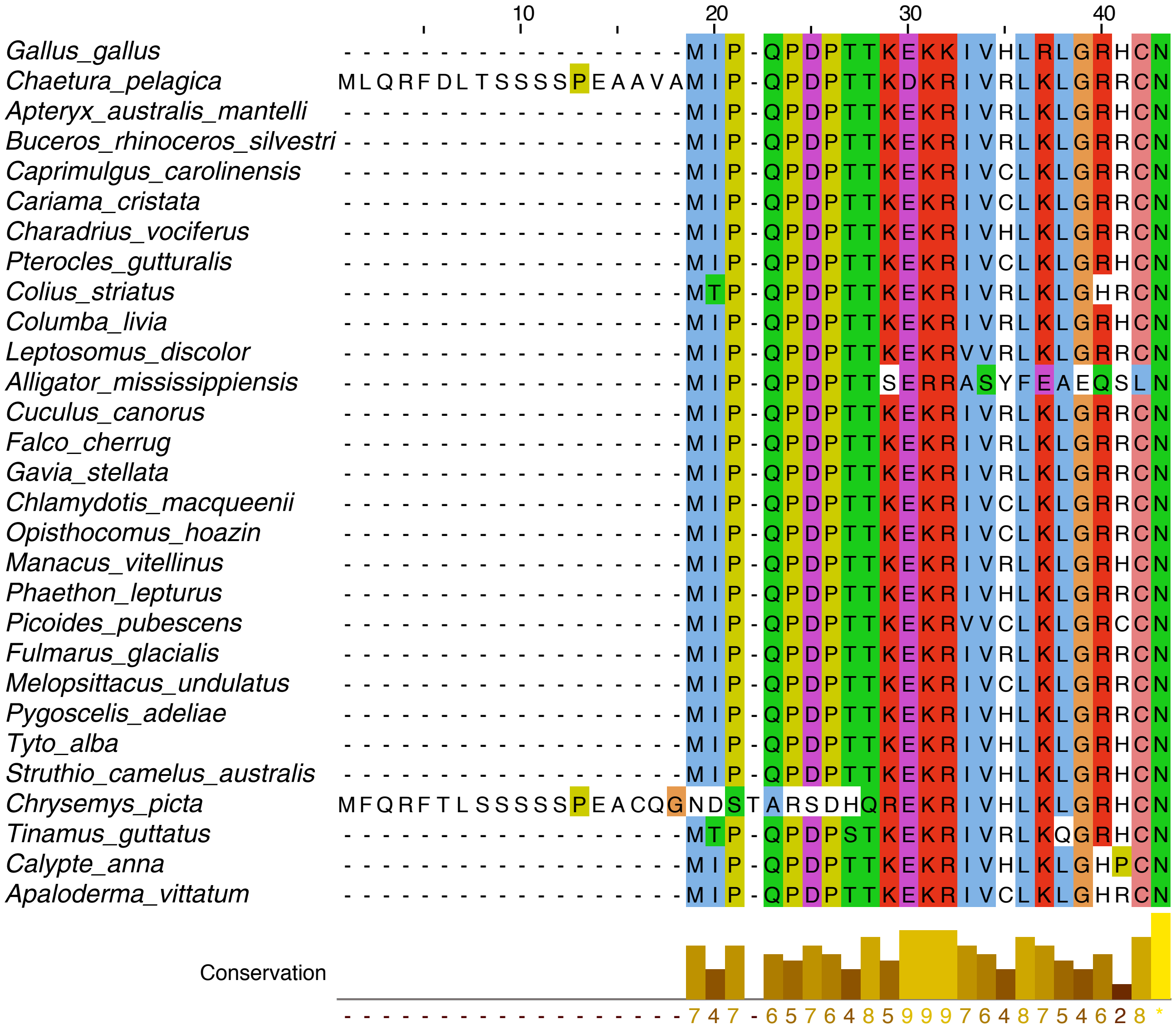

**HG0077.2**

ENSGALG00000005074

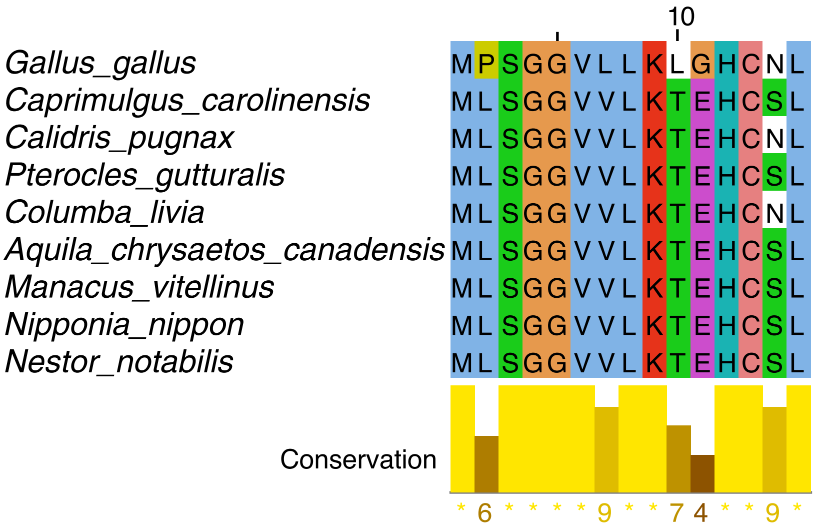

**HG0078.1**

ENSGALG00000016633 (HMBOX1)

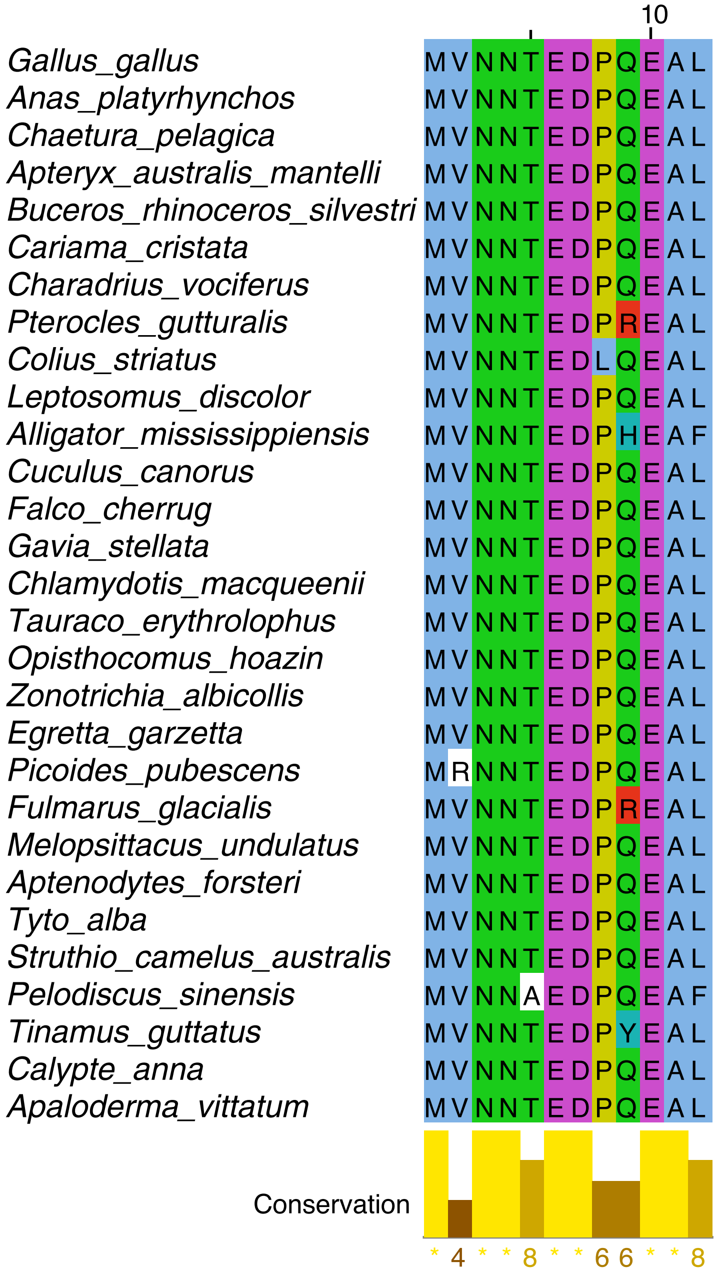

**HG0078.2**

ENSG00000147421 (HMBOX1)

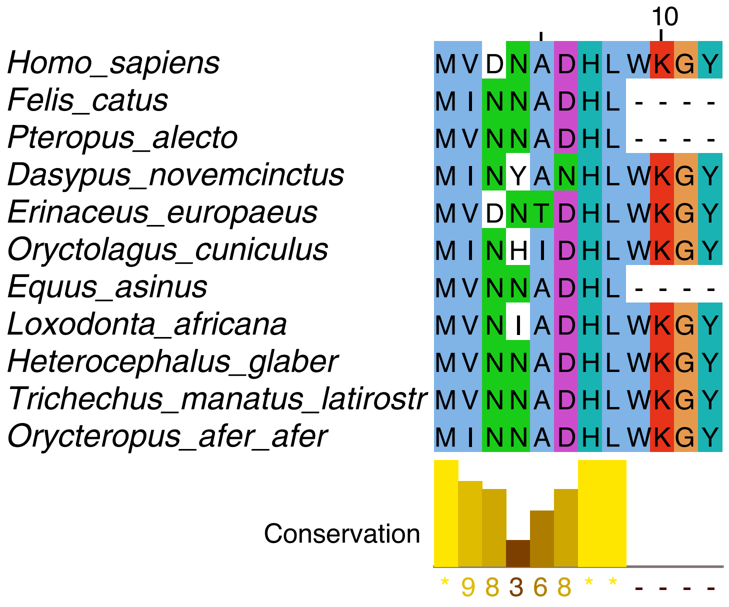

**HG0079**

ENSG00000204406 (MBD5)

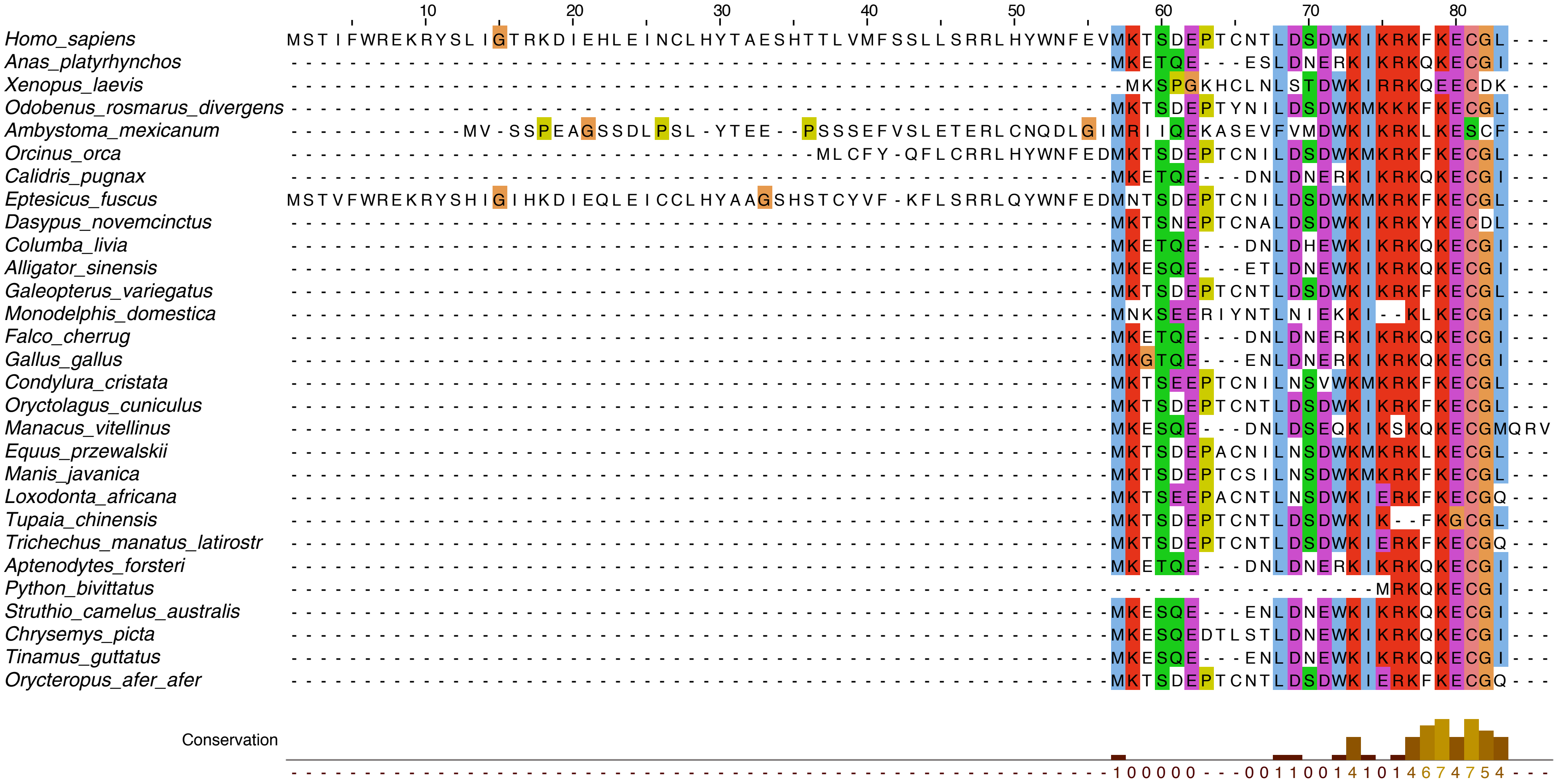

**HG0080.1**

ENSG00000179603 (GRM8)

```
                                     10        20        30        40
Homo_sapiens                  ---------------------------MGPDGSLQKVQPQVVPFLLWQE
Anser_cygnoides               ---------------------MVLYYLLGTDGSLQKLQFQMVNLIL---
Canis_lupus_familiaris        ---------------------------MGPDGSLQKVQPLVVPLLLWQE
Orcinus_orca                  ---------------------------MGPDGSLQKVQPQVVPLLLWQE
Calidris_pugnax               ---------------------MILYYLLGTDGSLQKVQLQMVNLIL---
Myotis_lucifugus              MEIKLNLRGLTVTFQSRKLAEAHRAFLRGPDGSLQKVRPQVVPLLLWQE
Dasypus_novemcinctus          ---------------------------MGPDGSLQKVQPQVVPLLLWRE
Galeopterus_variegatus        ---------------------------MGPDGSLQKEQPQVVPLLLWRE
Monodelphis_domestica         ---------------------------MGPDGSLQKV------------
Aquila_chrysaetos_canadensis  ---------------------MVLYYLLGTDGSLQKVQRQMVNLIL---
Coturnix_japonica             ---------------------MVLYHLLGTDGSLQKVQFQMVNLSL---
Sorex_araneus                 ---------------------------MGPDGRLQTVEPQVVLFLLWQE
Oryctolagus_cuniculus         ---------------------------MGPDGSLQKVQPQAVPLLLWQE
Elephantulus_edwardii         ---------------------------MGPNGGLQQTQPQVVLLLLWQE
Ornithorhynchus_anatinus      ---------------------------MGPDDTRWKVRLQTVPLLLQQE
Lepidothrix_coronata          ---------------------MVLYHLLGTDGSLQKVQLQMVSLIL---
Equus_asinus                  ---------------------------MGPDGSLQKVQPQVVPLLLWQE
Manis_javanica                ---------------------------MGPDGSLQKVQALVVPLLLRQK
Loxodonta_africana            ---------------------------MGPDGSLQKVQPQVVPLFLWQE
Mus_musculus                  ---------------------------MGPDGSLQKVQPQVVPLLLWQE
Tupaia_chinensis              ---------------------------MGPDGSLQKVQAQVVPFLLWQE
Trichechus_manatus_latirostr  ---------------------------MGPDGSLRKVQPQVVPLFLWQE
Anolis_carolinensis           --------------------MVLYSASLGTDGSLQKGKLQIVHFLL---
Struthio_camelus_australis    ---------------------MVLYYLLGTDGSLQKVQLQMVNLIL---
Trachemys_scripta             ----------------------------------------MVHLLF---
Tinamus_guttatus              ---------------------MVLYYLLGTDGSLQKVQLQMVNFTL---
Calypte_anna                  ----------------------------------------MVNLIL---
Orycteropus_afer_afer         ---------------------------MGPDGSLQKVQPQVVPILLWQE

Conservation                  ---------------------------0312100110100130101---
```

**HG0080.1**

ENSGALG00000039403

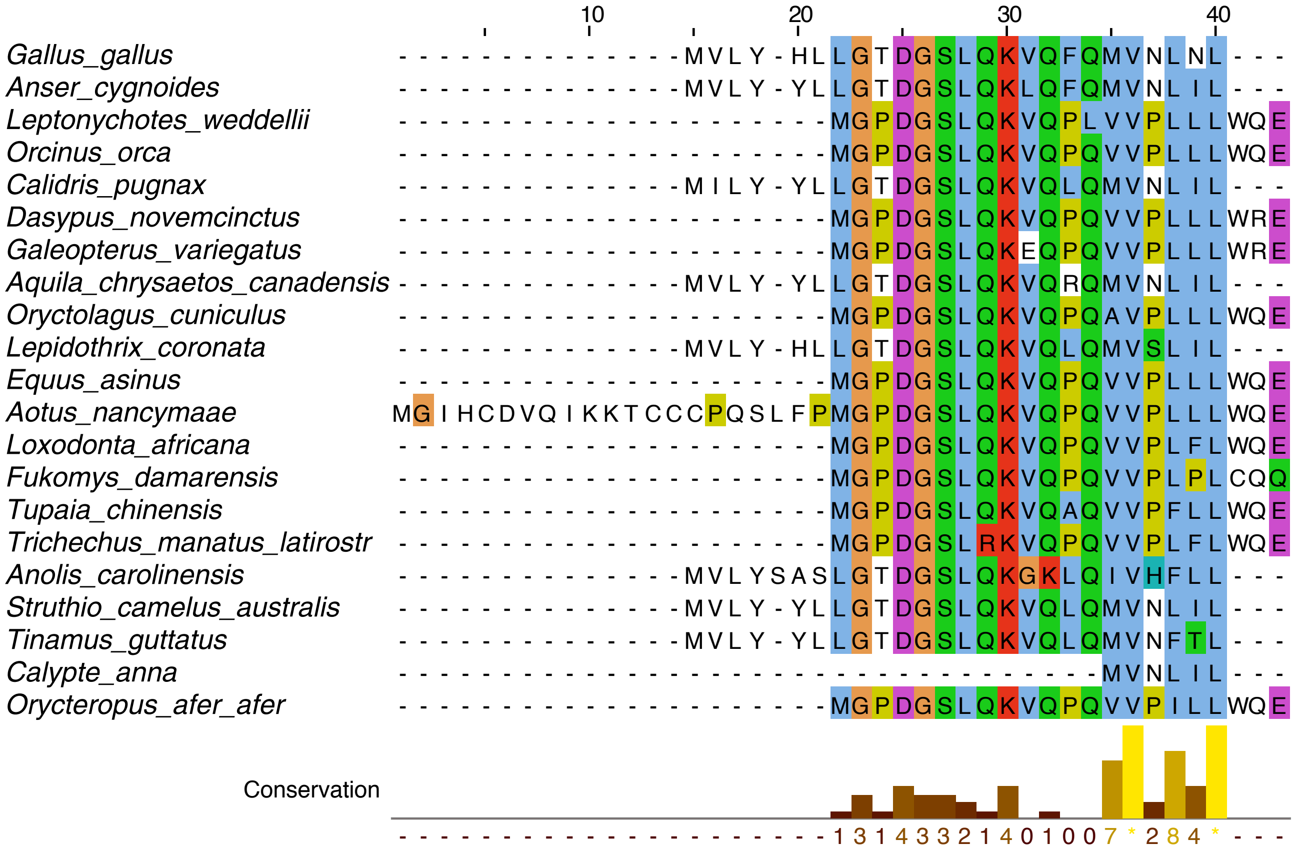

**HG0080.2**

ENSG00000179603 (GRM8)

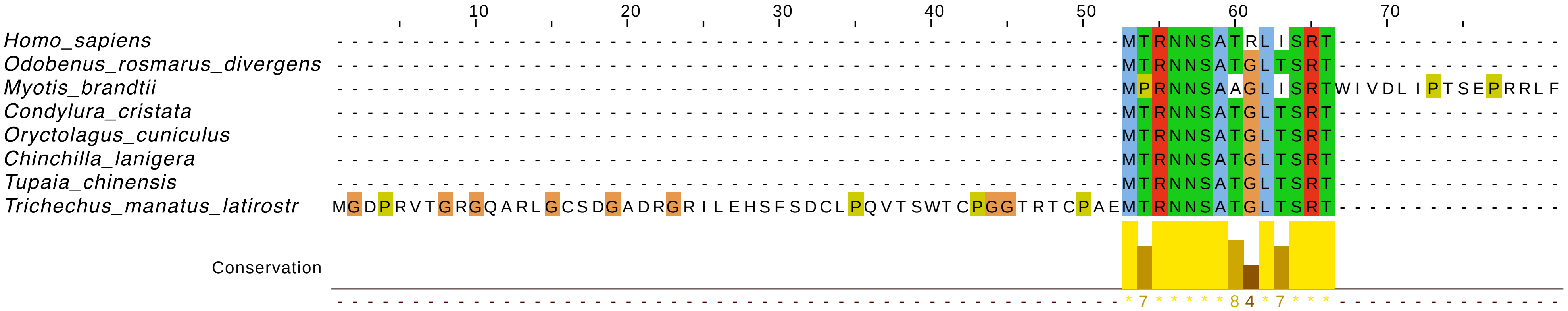

**HG0080.3**

ENSG00000179603 (GRM8)

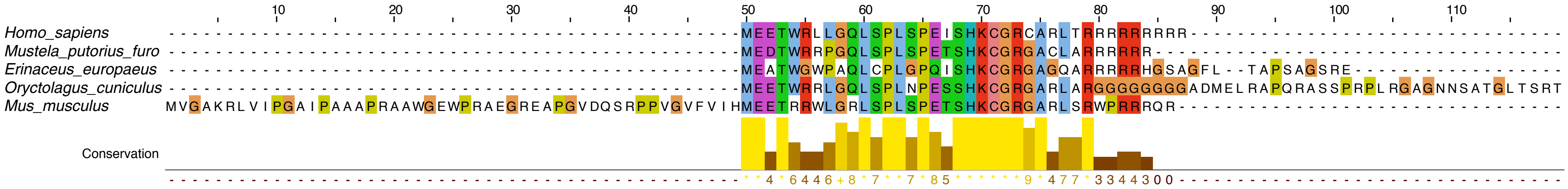

**HG0081.1**

ENSDARG00000028228 (zbtb18)

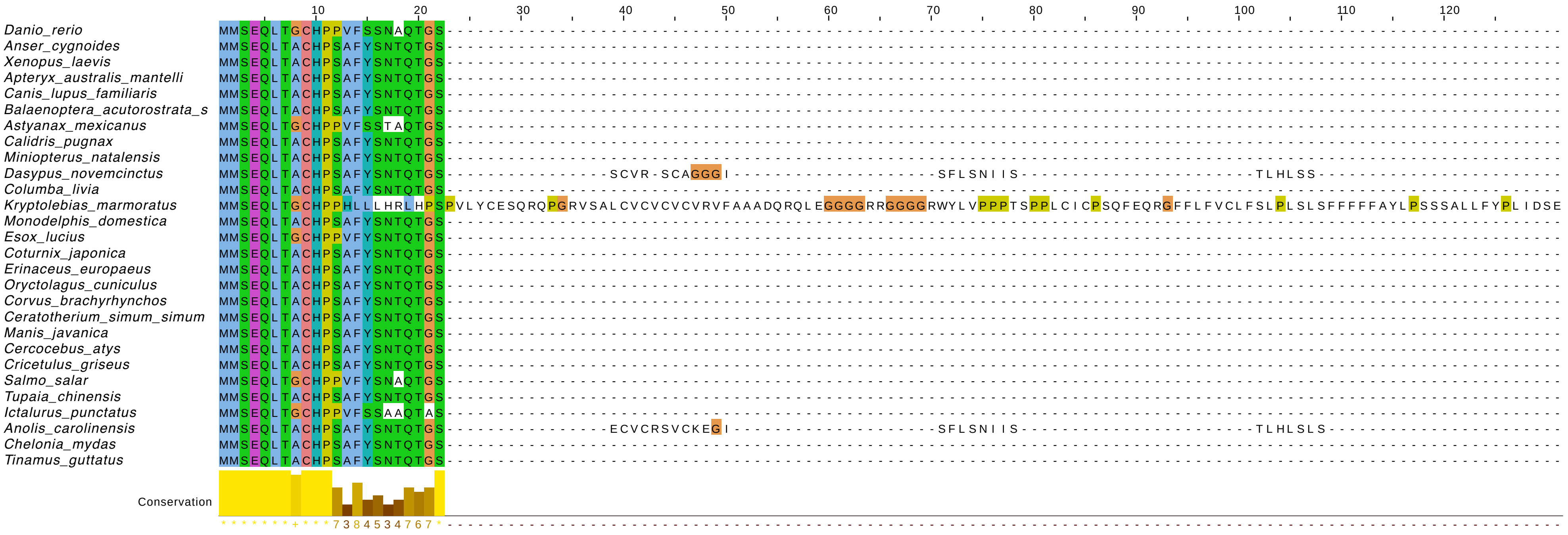

**HG0081.2**

ENSDARG00000028228 (zbtb18)

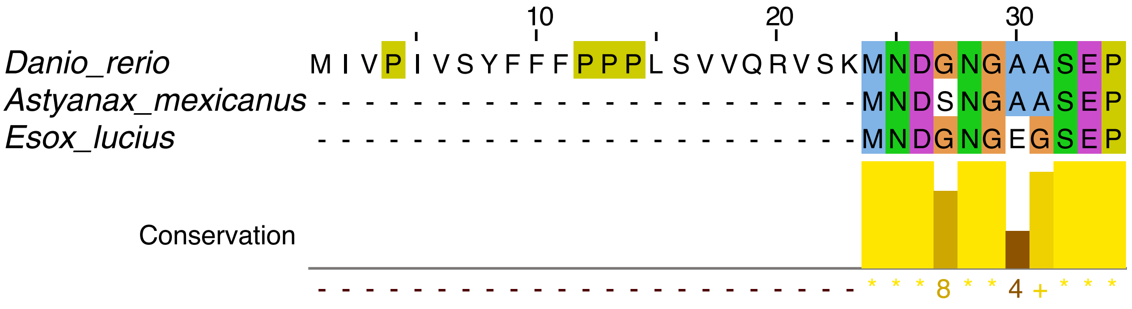

# HG0082

### ENSDARG00000062379 (slc35a4)

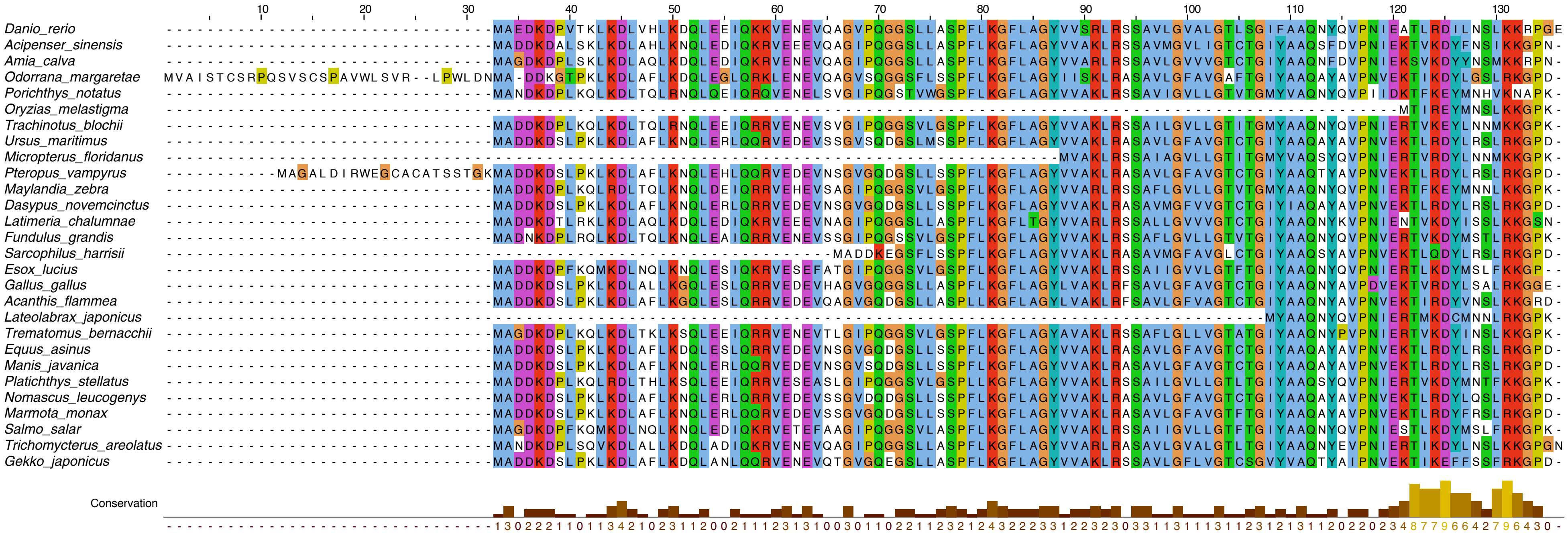

**HG0082**

ENSG00000176087 (SLC35A4)

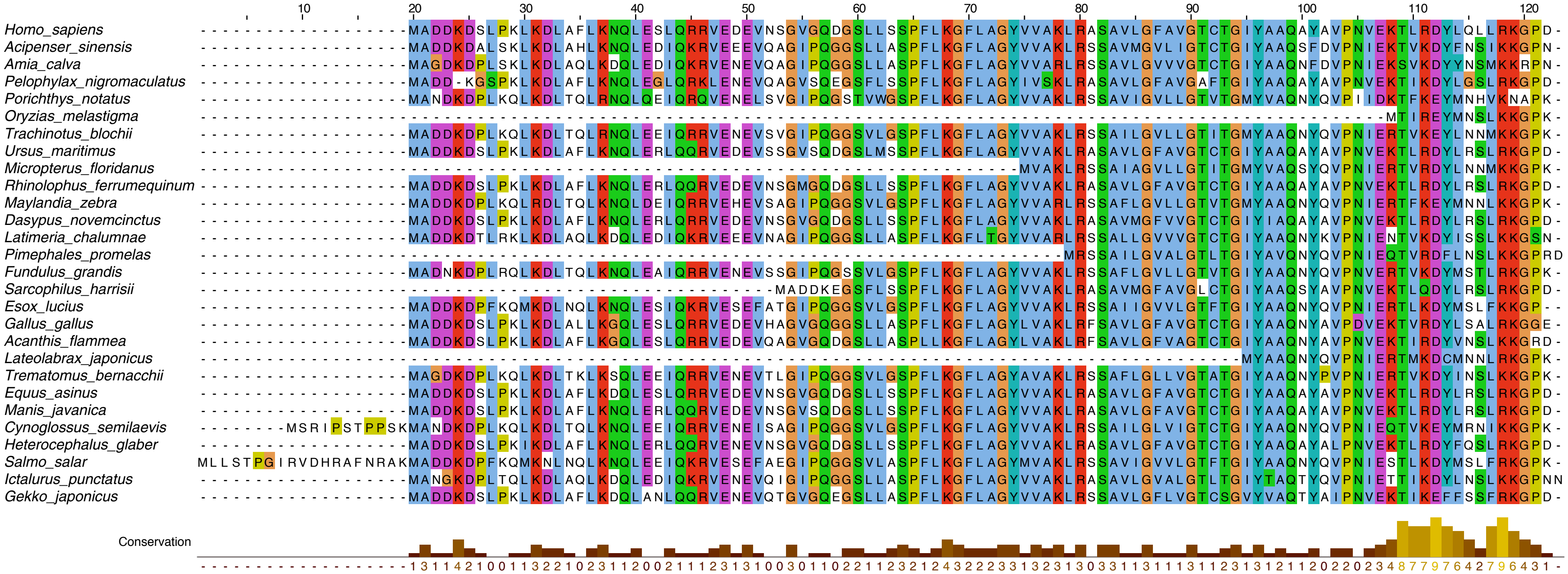

**HG0083.1**

ENSGALG00000008039 (MFSD13A)

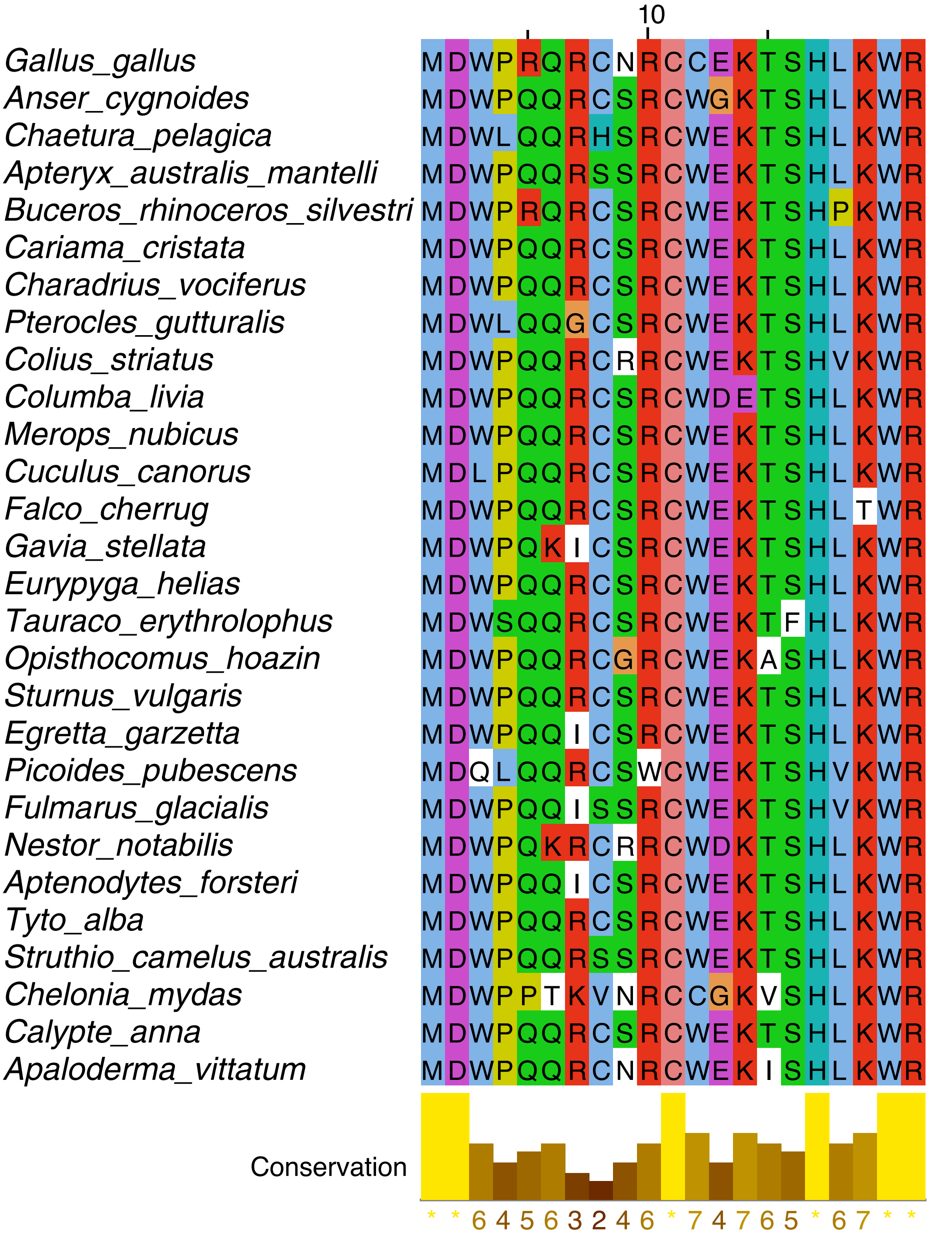

**HG0083.2**

ENSG00000138111 (MFSD13A)

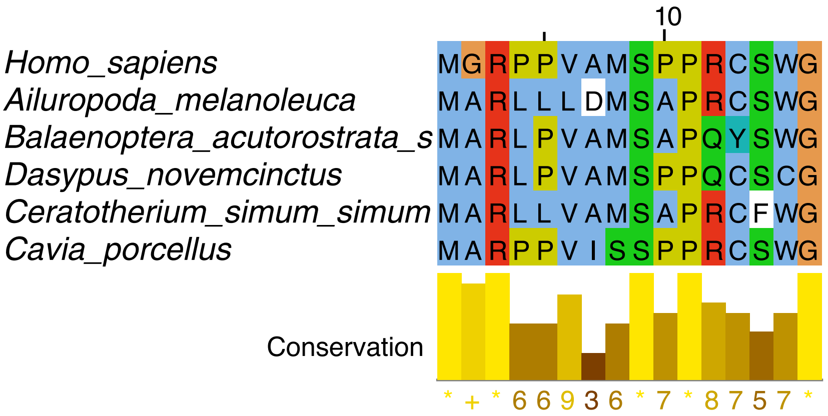

**HG0084.1**

ENSGALG00000009207 (TRIM2)

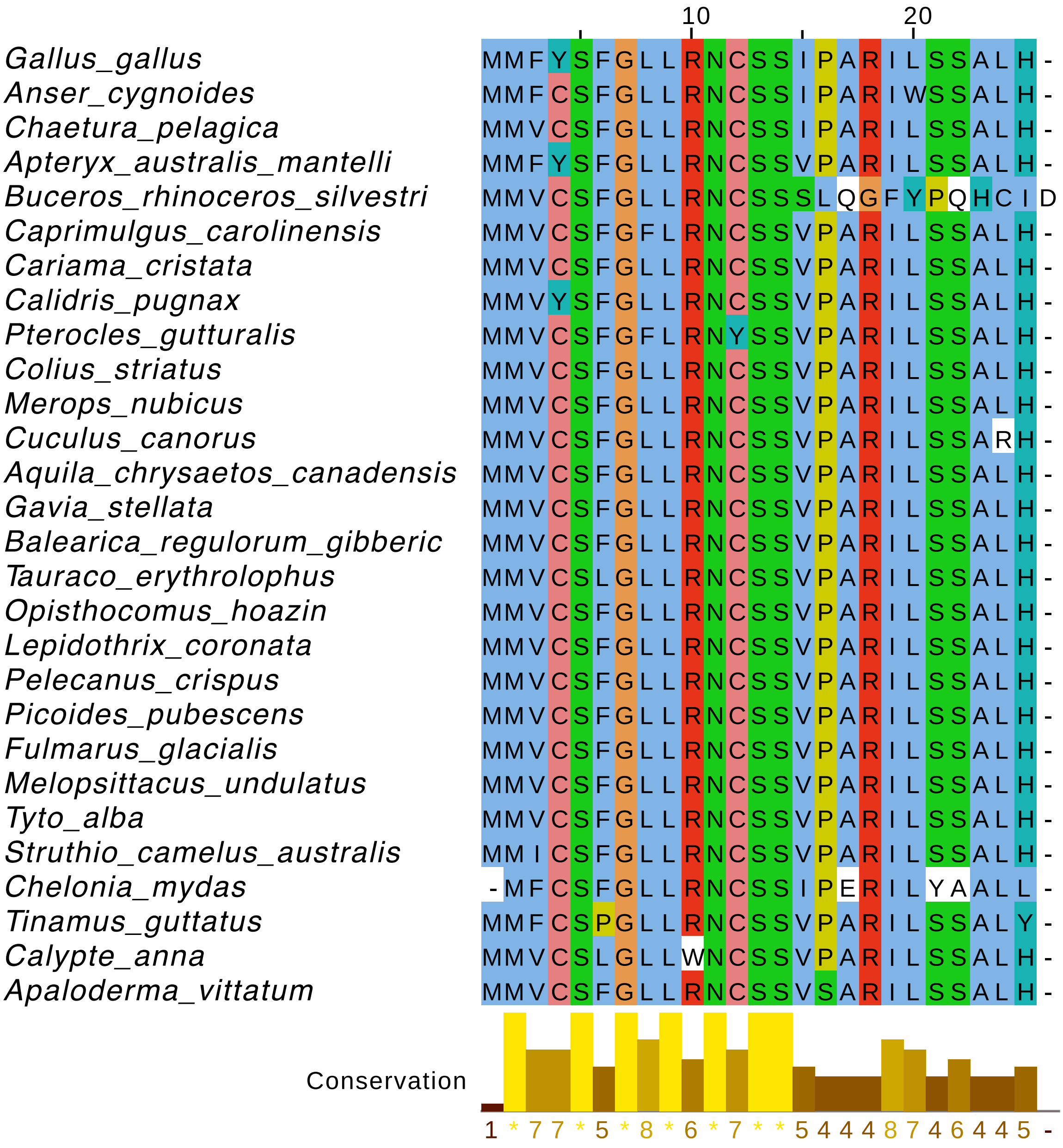

**HG0084.2**

ENSG00000154447 (SH3RF1)

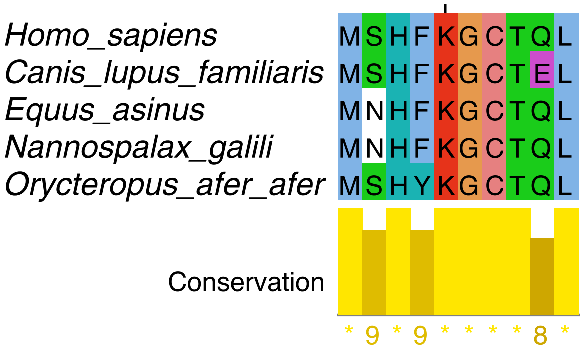

**HG0085**

ENSG00000107779 (BMPR1A)

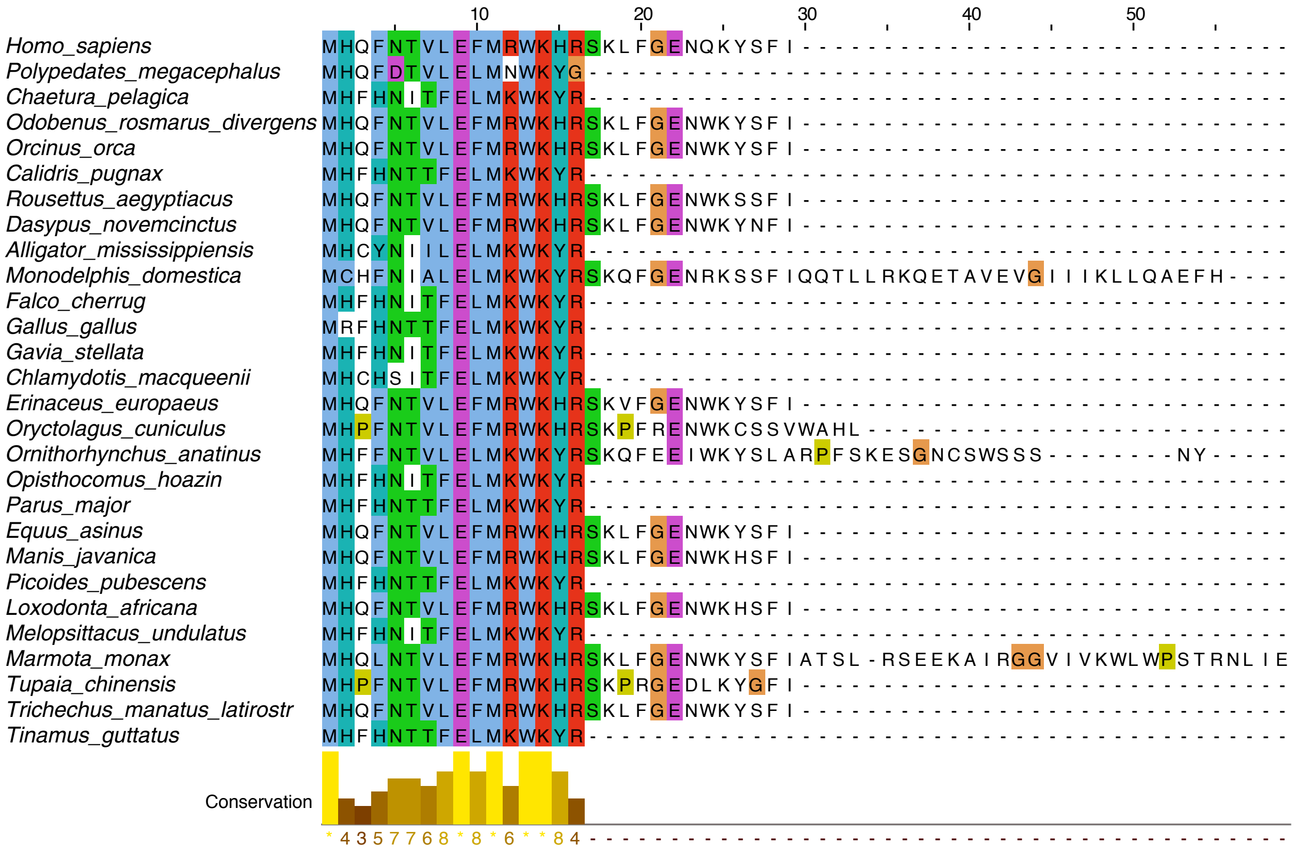

**HG0085**

ENSGALG00000002003 (BMPR1A)

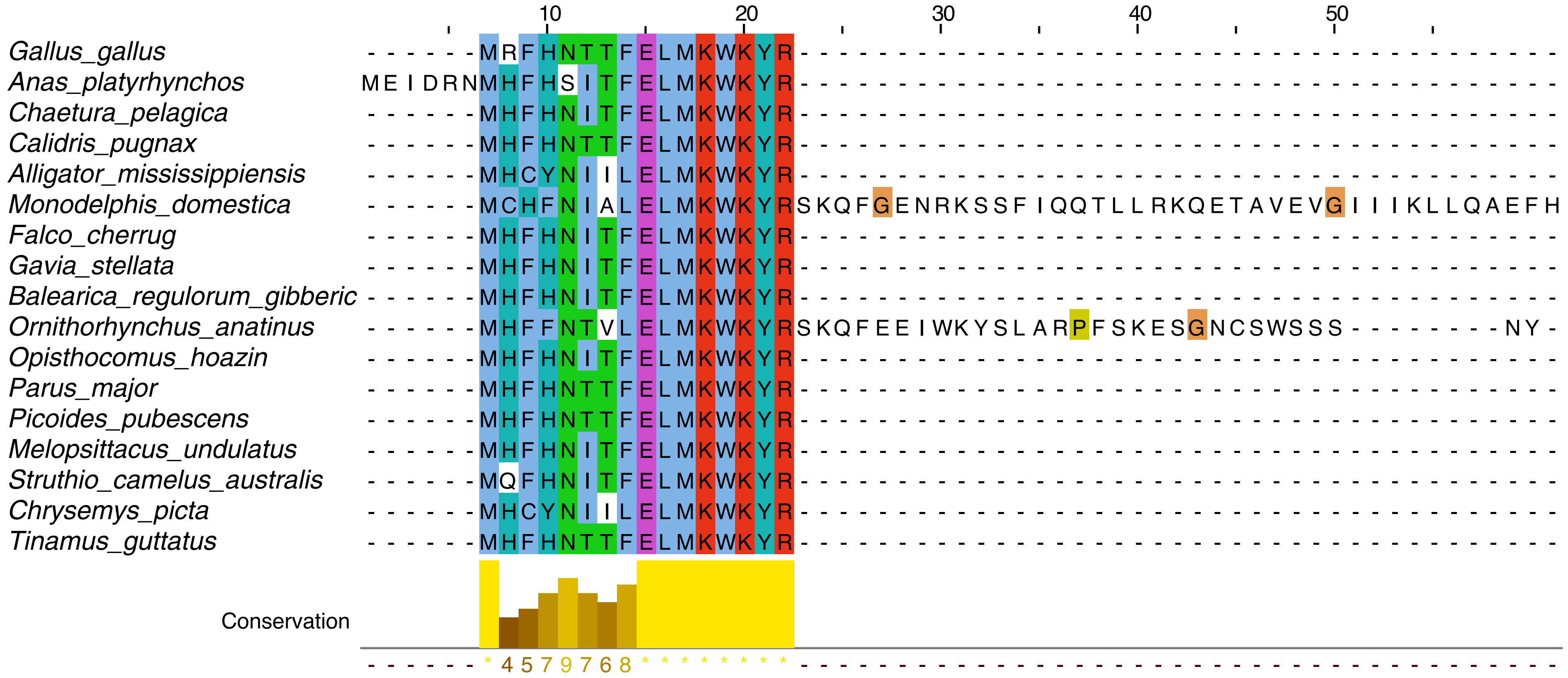

**HG0086**

ENSGALG00000009835 (XPOT)

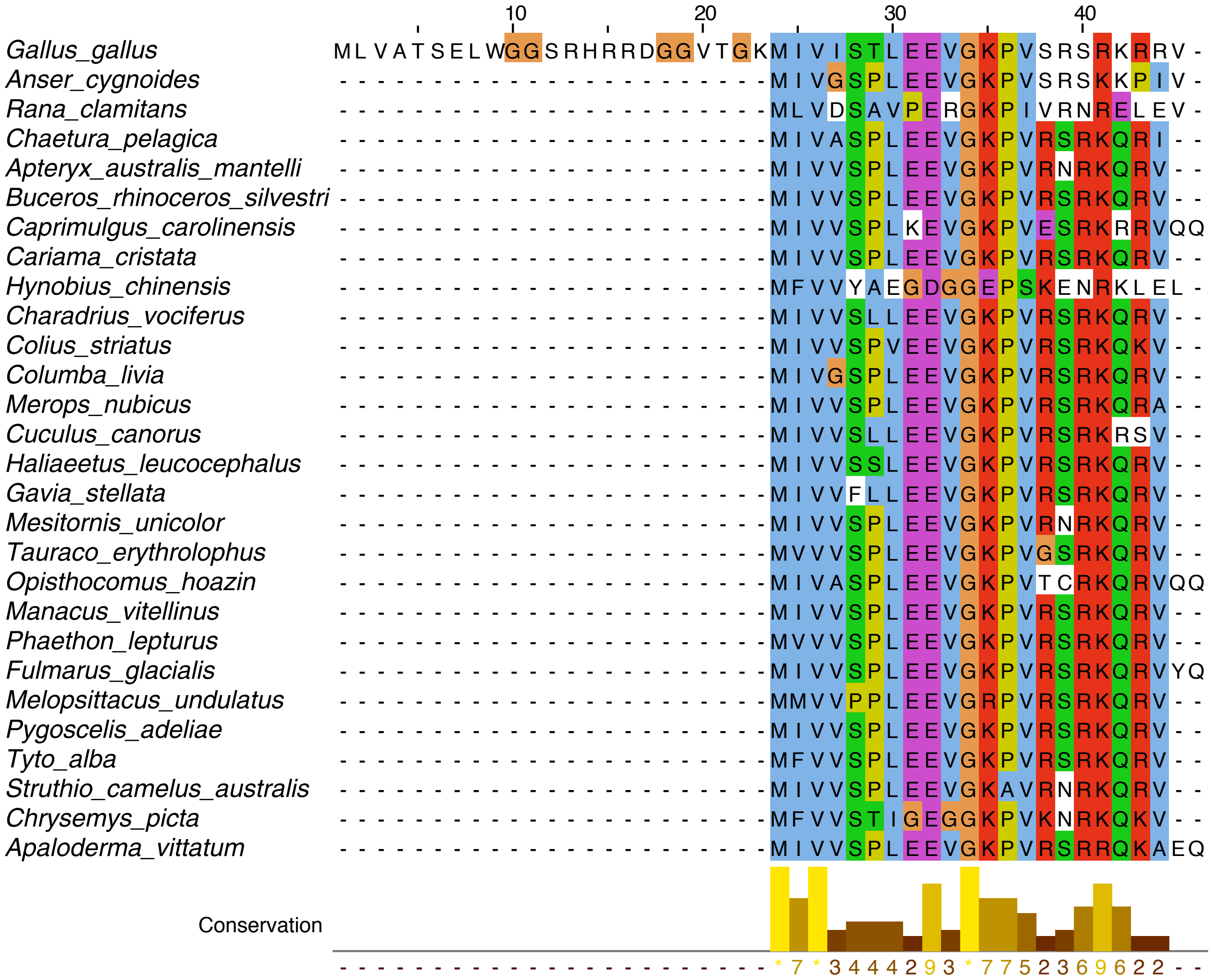

**HG0087**

ENSG00000164548 (TRA2A)

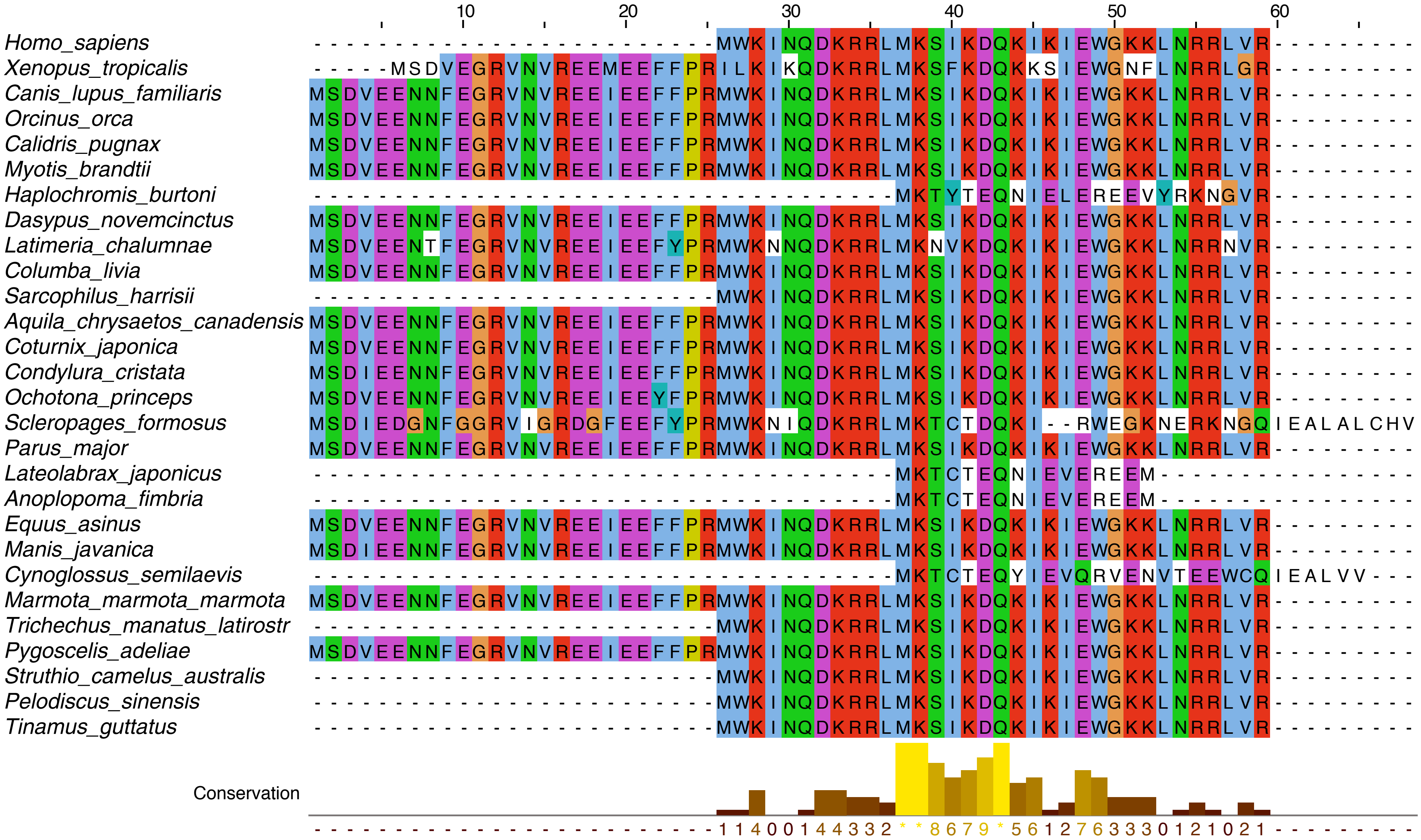

## HG0088

ENSG00000273079 (GRIN2B)

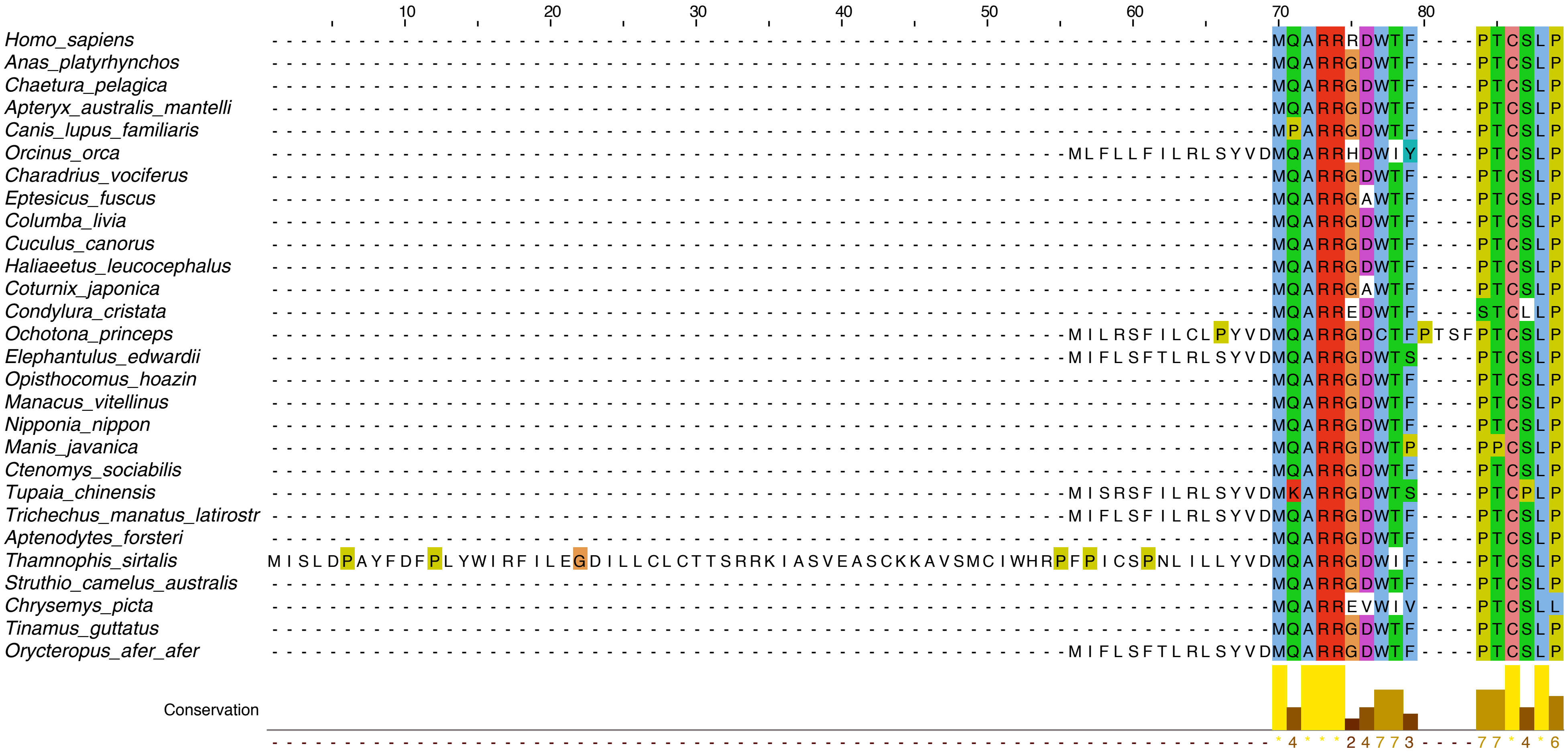

**HG0089.1**

ENSG00000092421 (SEMA6A)

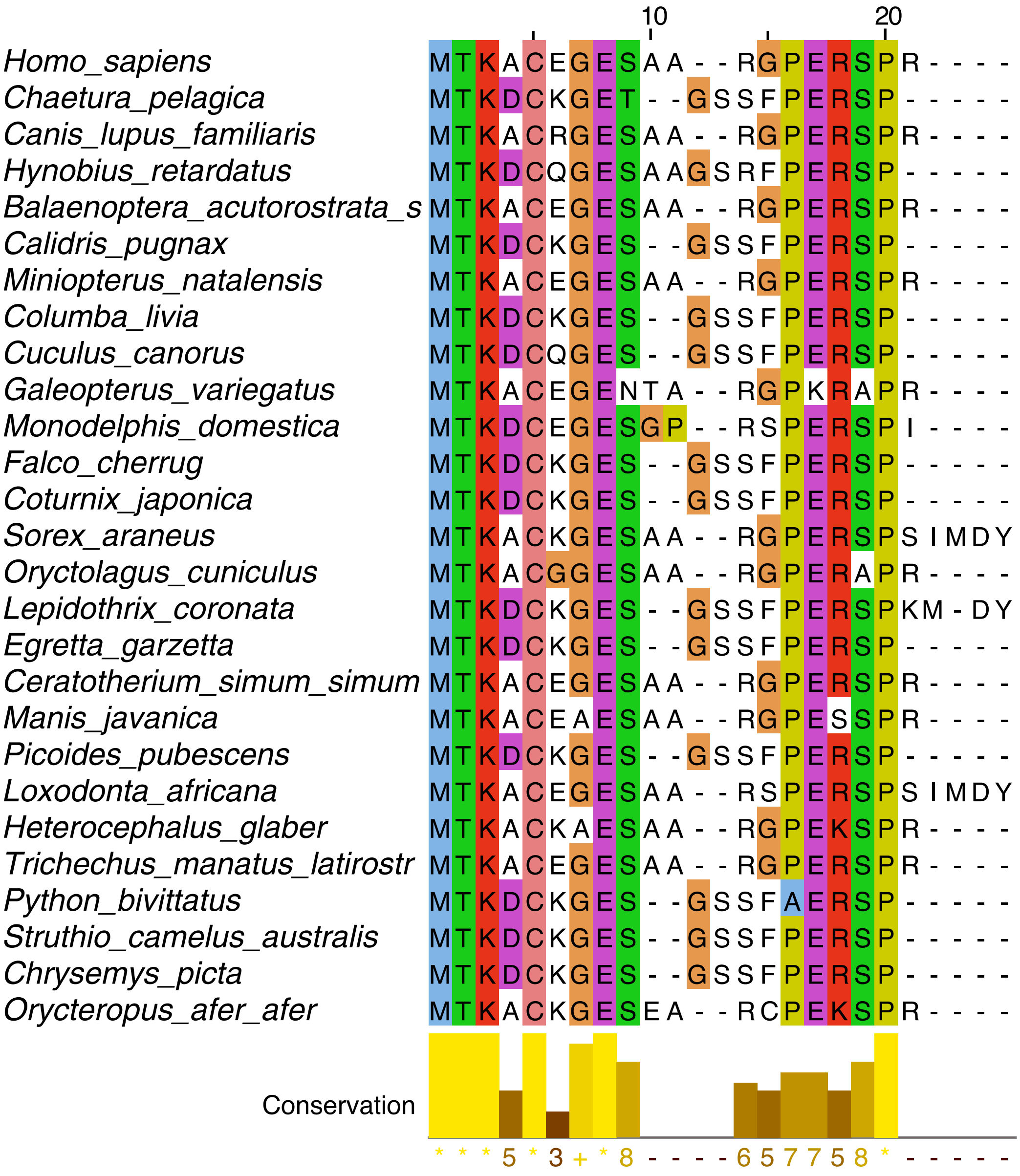

**HG0089.1**

ENSGALG00000002294 (SEMA6A)

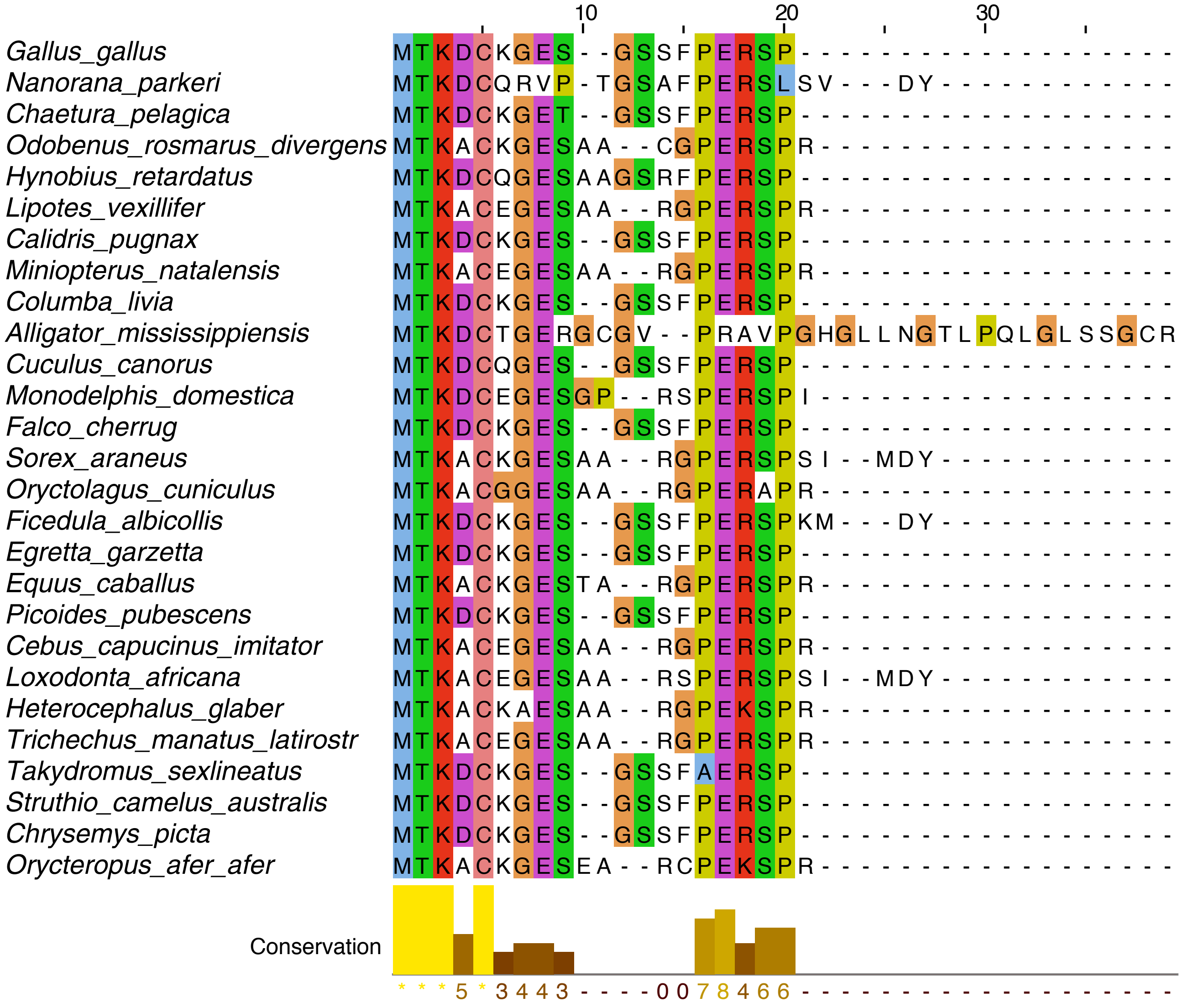

**HG0089.2**

ENSG00000092421 (SEMA6A)

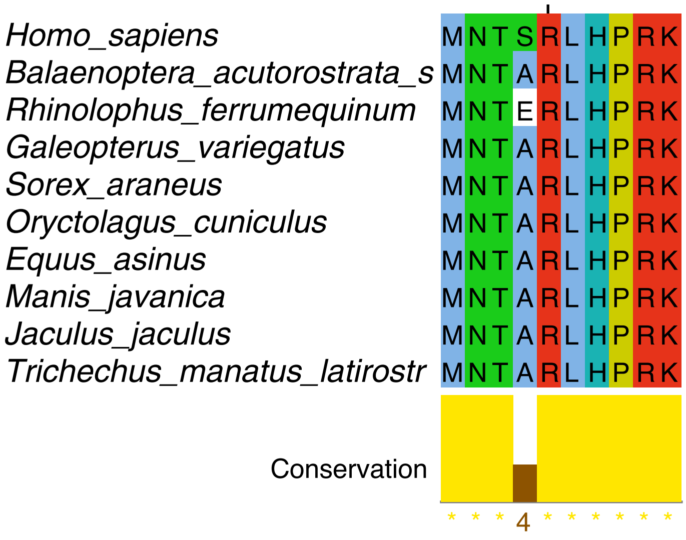

**HG0089.3**

ENSGALG00000002294 (SEMA6A)

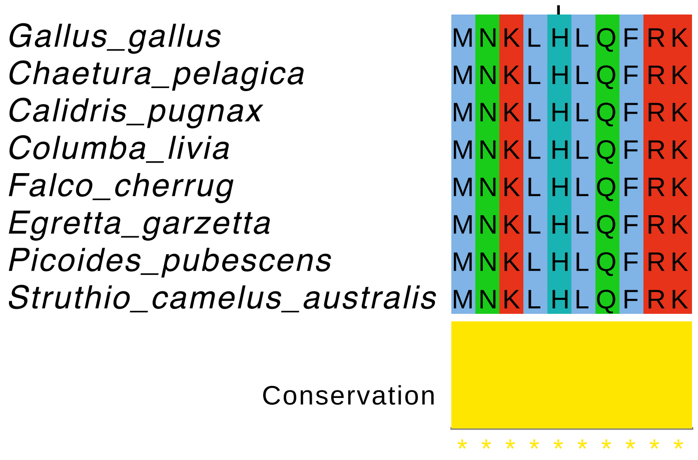

**HG0090.1**

ENSG00000172765 (TMCC1)

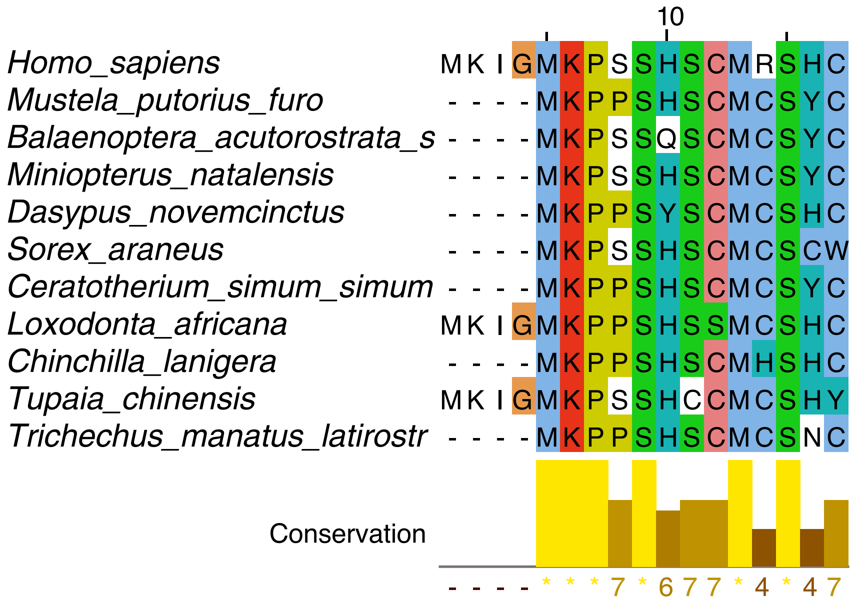

**HG0090.1**

ENSGALG00000008487 (TMCC1)

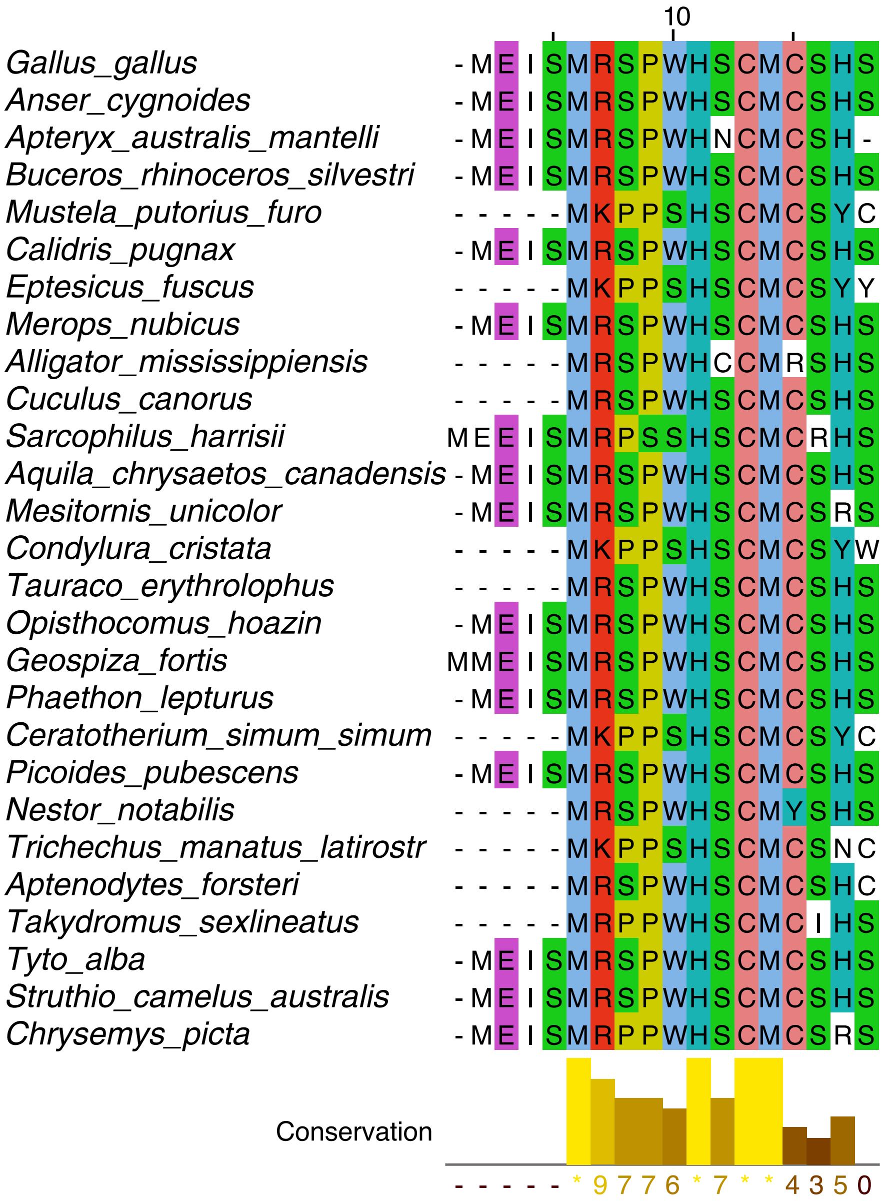

**HG0090.2**

ENSDARG00000060954 (tmcc1b)

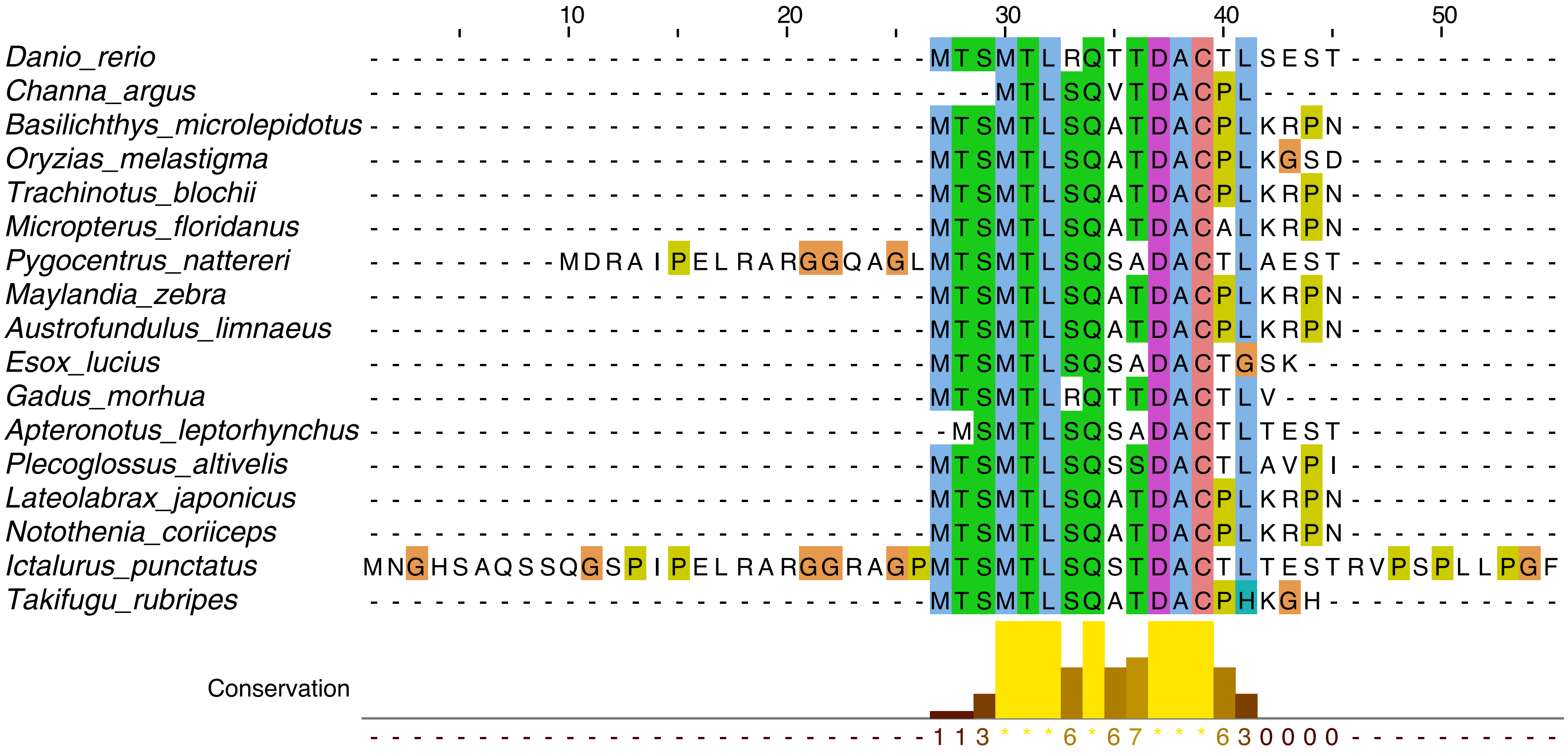

**HG0090.3**

ENSGALG00000008487 (TMCC1)

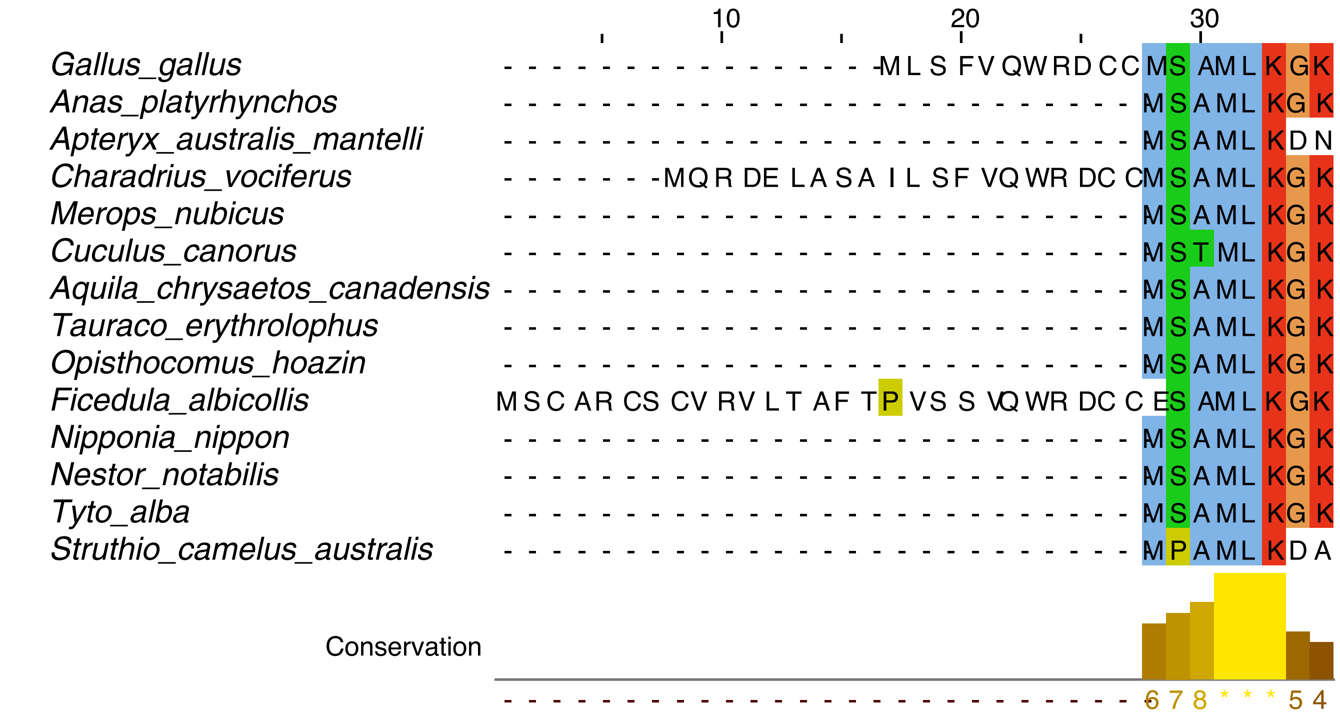

**HG0091.1**

ENSG00000172209 (GPR22)

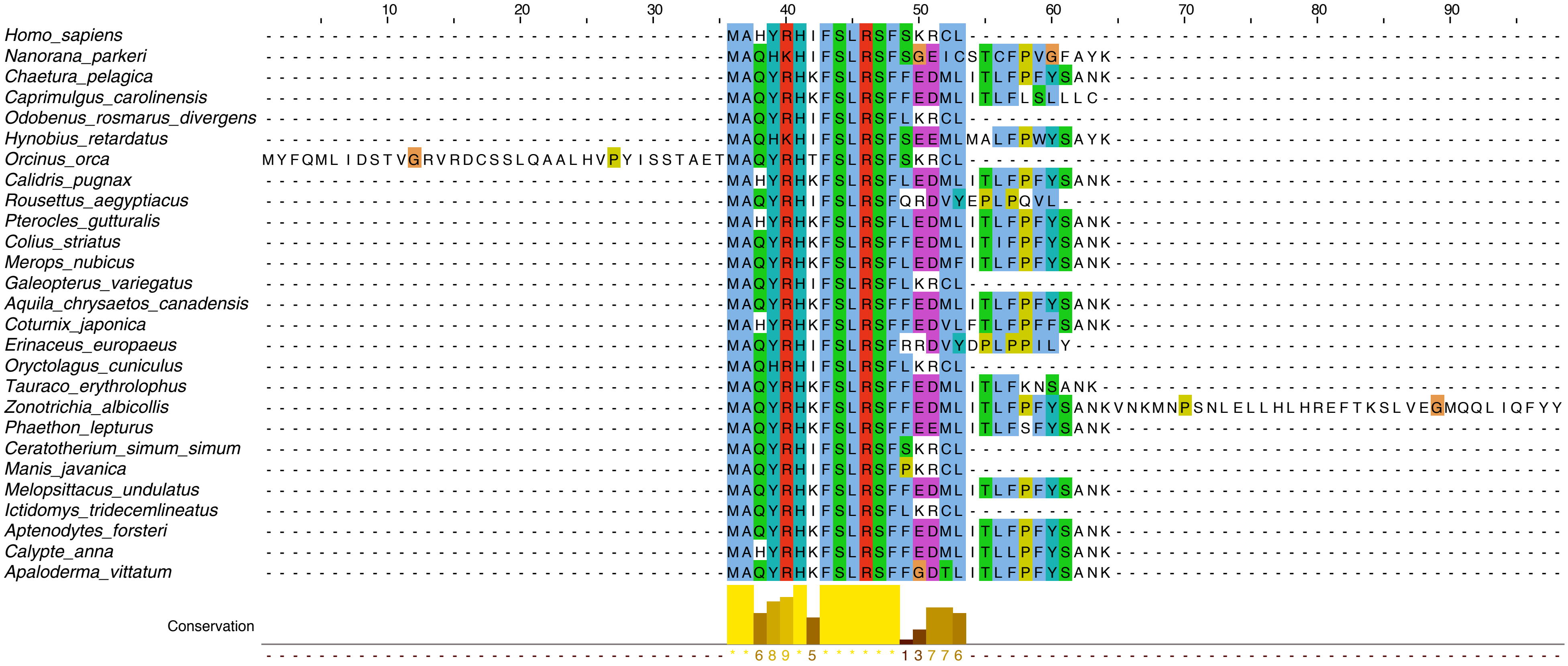

**HG0091.2**

ENSG00000172209 (GPR22)

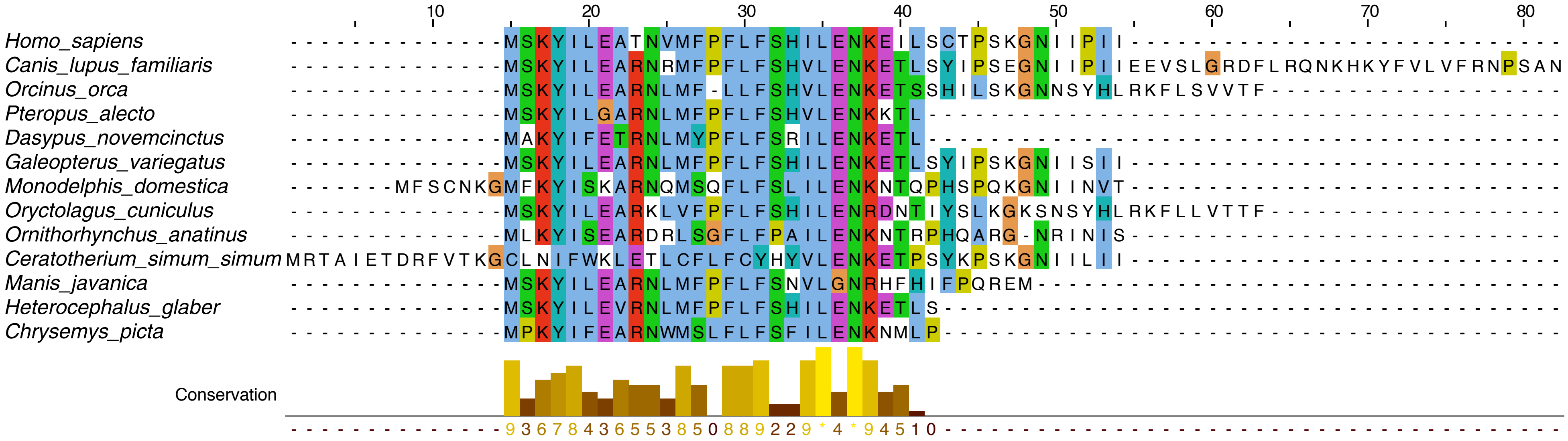

**HG0091.3**

ENSG00000172209 (GPR22)

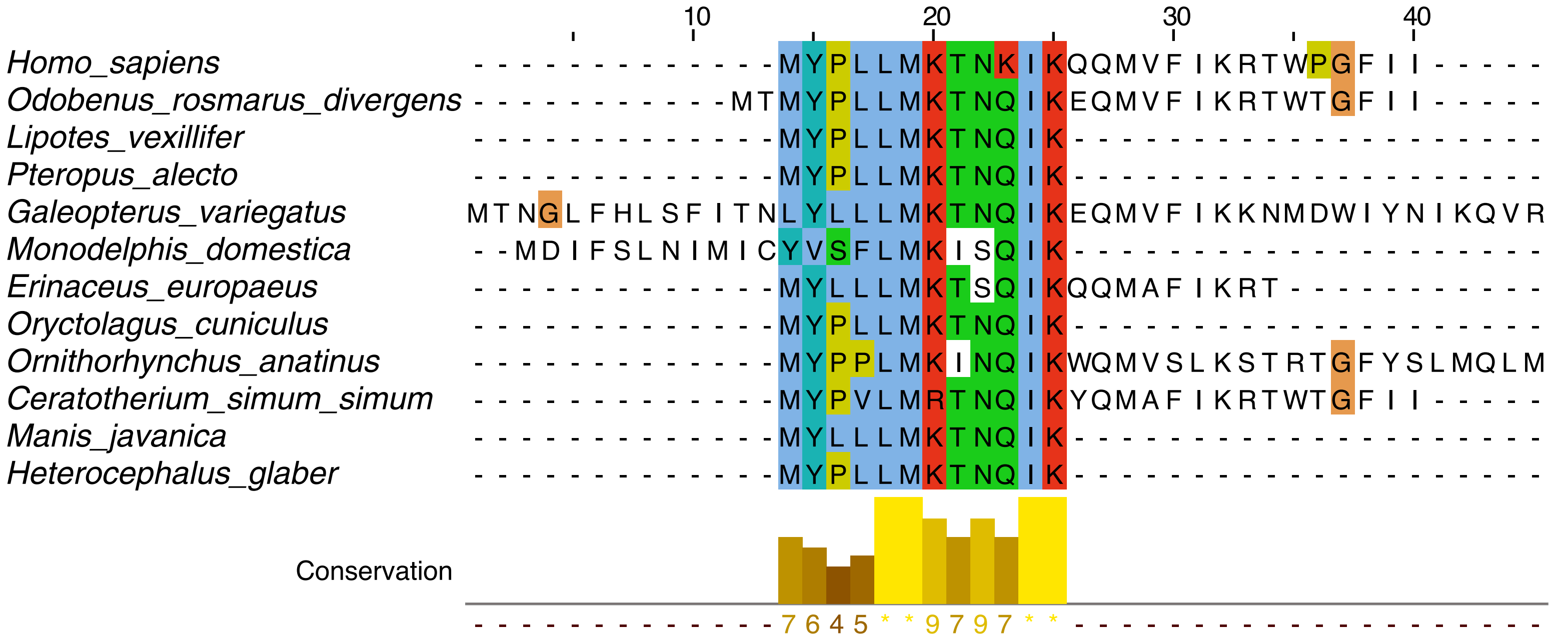

**HG0092.1**

ENSGALG00000009966 (FRS2)

```
                                    10        20        30        40        50        60
Gallus_gallus                   --------------------------------------MISSVTRQHDTLKYDRSPDAATEAGNINGQI
Anser_cygnoides                 --------------MVIKQLLSCSAAVC----LHVLGQMISSVTRQHDTLKYDRTPDAATEAGNINGQI
Xenopus_laevis                  --------------------------------------------MRHVILV------------------
Apteryx_australis_mantelli      ------------MLHGAKCLMLMTLGWK----VTFLDQMITSVTRQHDTLKYDRSPDAAAEAGNINGQI
Buceros_rhinoceros_silvestri    MI-SSVTRQHDTLSMTEALMLQQRLGILMAESRDHMVF-------------------------------
Charadrius_vociferus            MI-SSVTRQHDALSMTEALMLQQRLGILMAKSRDHMVF-------------------------------
Pteropus_alecto                 MRGIWFKLQYREFRLDIKNFQKIKYQILKR-------FVMSSNKKKHGILKYD----------------
Dasypus_novemcinctus            --------------MQLKLFKQIGTKLHLFFSEESSTRLSQQT--KHVILKYD----------------
Latimeria_menadoensis           -------------------------------------------------MKYERGPDVARGIRNLNGQI
Colius_striatus                 MI-SSMTRQHDTLSMTEALMLQQRLGILMAKSRDLMVF-------------------------------
Columba_livia                   --------------------------------------MISSVTRHHDTLKYDRSPDAAAEPGNINGQI
Leptosomus_discolor             MI-SSVTRQHDTLSMTEALMLQQRLGILMAKSRDHMVF-------------------------------
Alligator_mississippiensis      ------------MSPFDDC---SDMGACI---ECILDPVVAHVKRQHDILKYDRSPDAATEARNINGQI
Cuculus_canorus                 MI-SSVTRQHDTSSMTEALMLQQRLGILMAKSRDHMVF-------------------------------
Sarcophilus_harrisii            --------------MQLLLFSHIRRKLHLIFSKEKFARLCKQA--KYGILKYD----------------
Aquila_chrysaetos_canadensis    MI-SSVTRQHDTLSMTEALMLQQRLGILMAKSRDHMVF-------------------------------
Gavia_stellata                  MI-SSVTRQHDPLSMTEALMLQQRLGILMAKSRDHMVF-------------------------------
Balearica_regulorum_gibberic    --------------------------------------MISSVTRQHDTLKYDRSPDAAAEAGNINGQI
Condylura_cristata              --------------MQLWLFNQIRAKLYLFFSEDNSTRISHHTEKSHGILKYD----------------
Tauraco_erythrolophus           MI-SSVTRQHDTLSMTEALMLQQRLGILMAKCRDHMVF-------------------------------
Opisthocomus_hoazin             --------------------------------------MISSVTRQHDTLKYDRSPDAAAEAGNINGQI
Zonotrichia_albicollis          MI-SSVTRQHDTLSMTEALMLQQRLGILMAKSRDQMVF-------------------------------
Phaethon_lepturus               --------------------------------------MISSVTRQHDTLKYDRSPDAAAEAGNINGQI
Fulmarus_glacialis              MI-SSVTRQYDTLSMTEALMLQQRLGILMARSRDHMVF-------------------------------
Aptenodytes_forsteri            MI-SSVTRQHDTLSMTEALMLQQRLGILMAKSRDHMVF-------------------------------
Pelodiscus_sinensis             --------------------------------------MFVYVRRQHAILKYDRSPDAATEARNINGQI
Calypte_anna                    MI-SSVARQHDTLSMTEALMLQQRLGILMAKSRDPMVF-------------------------------

Conservation
                                ---------------------------------------------------------------------
```

**HG0092.2**

ENSG00000166225 (FRS2)

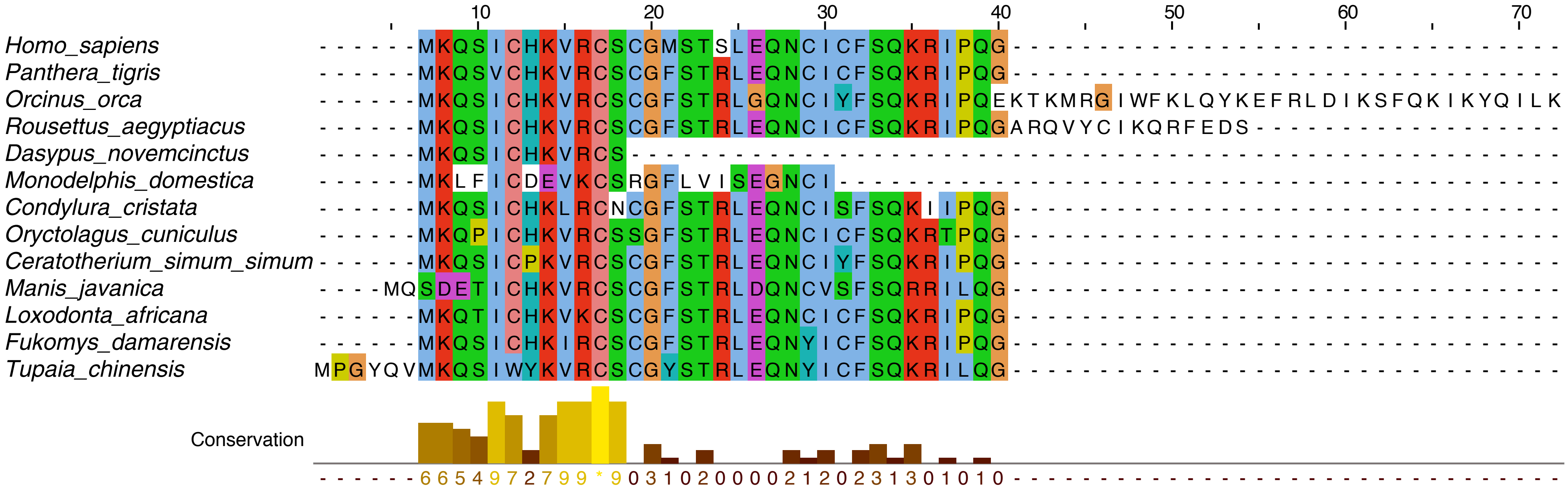

**HG0093**

ENSG00000169554 (ZEB2)

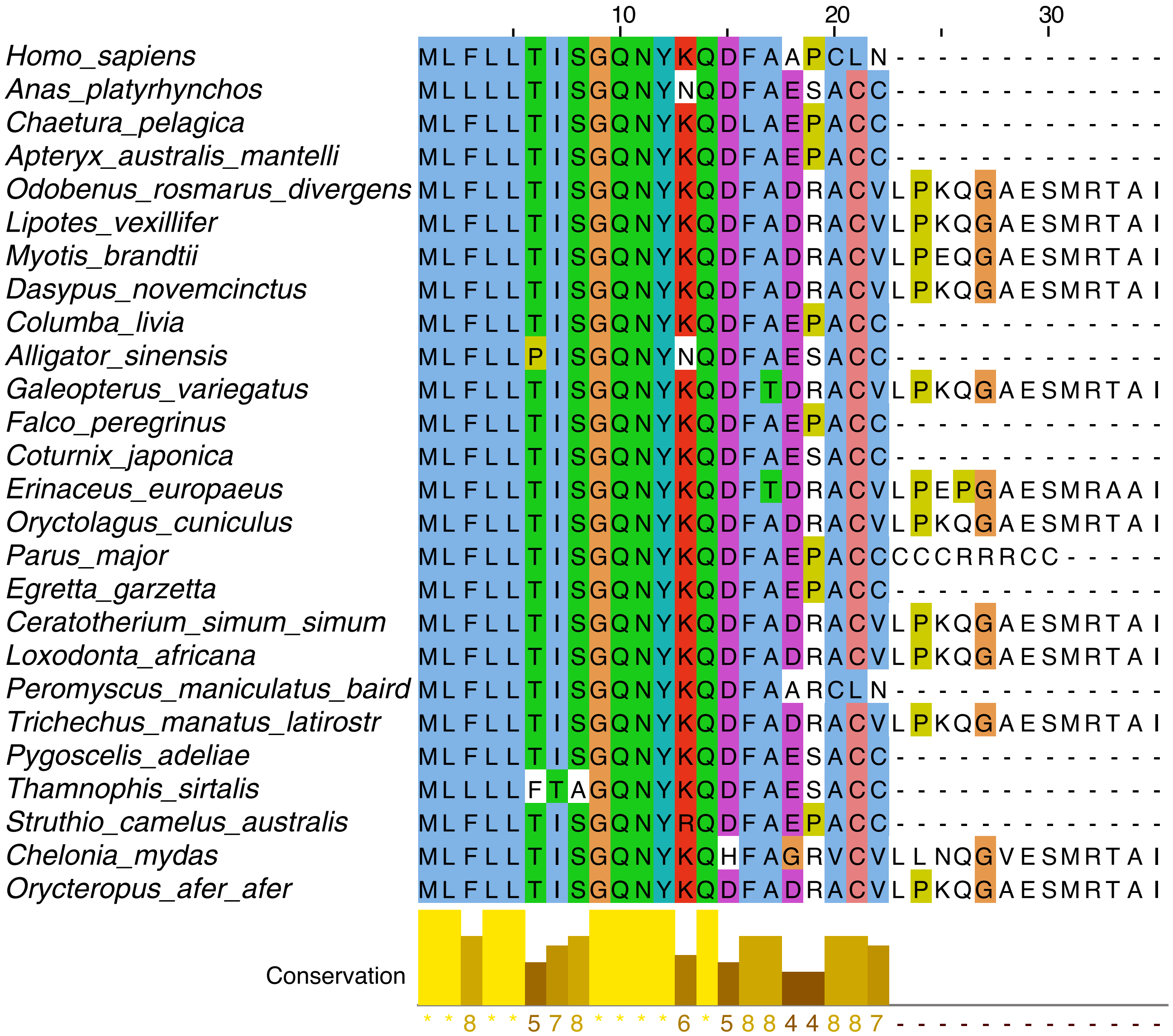

**HG0093**

ENSGALG00000040465 (Sip1)

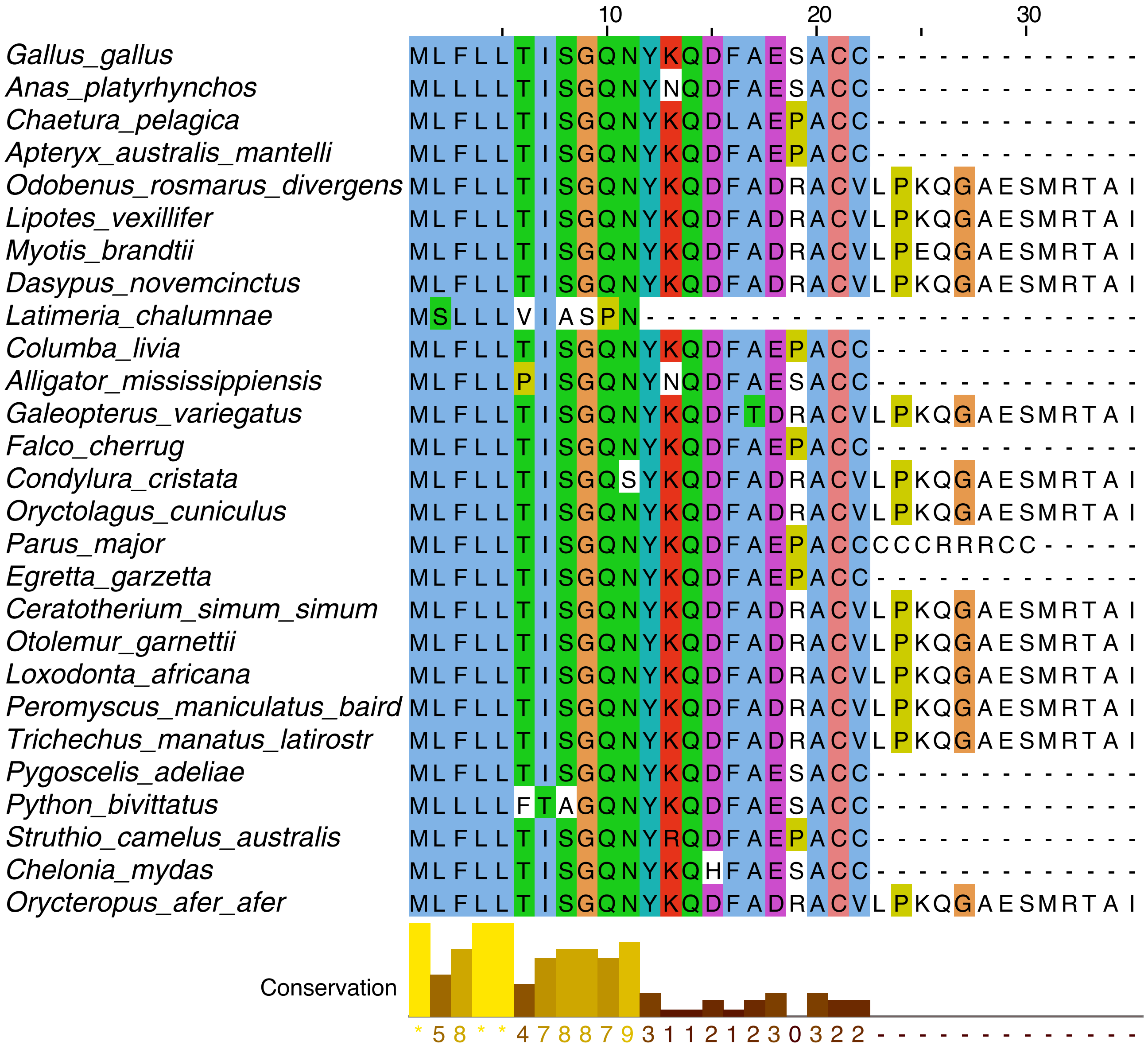

**HG0094.1**

ENSG00000112175 (BMP5)

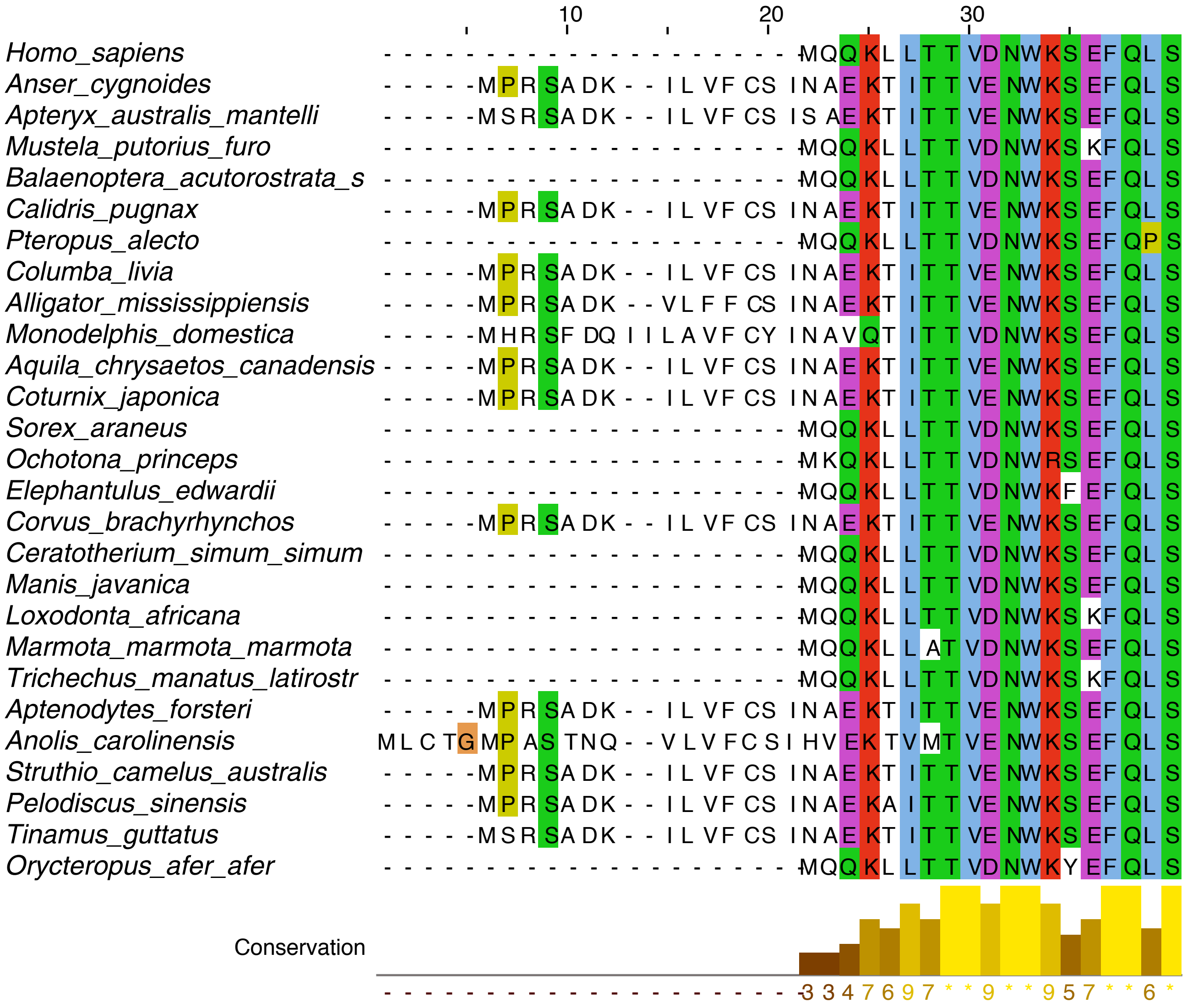

**HG0094.2**

ENSG00000112175 (BMP5)

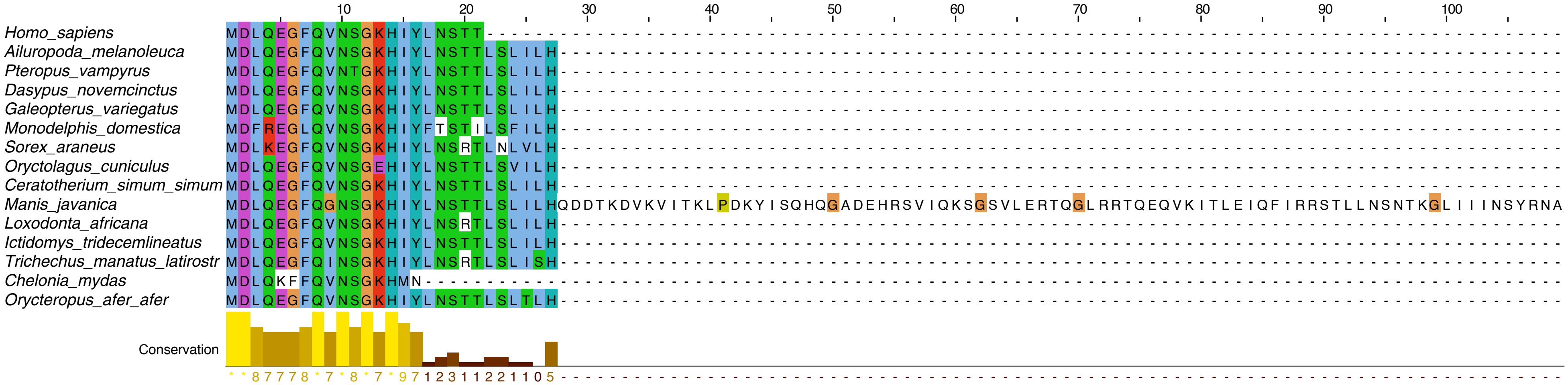

**HG0095**

ENSDARG00000013708 (usp9)

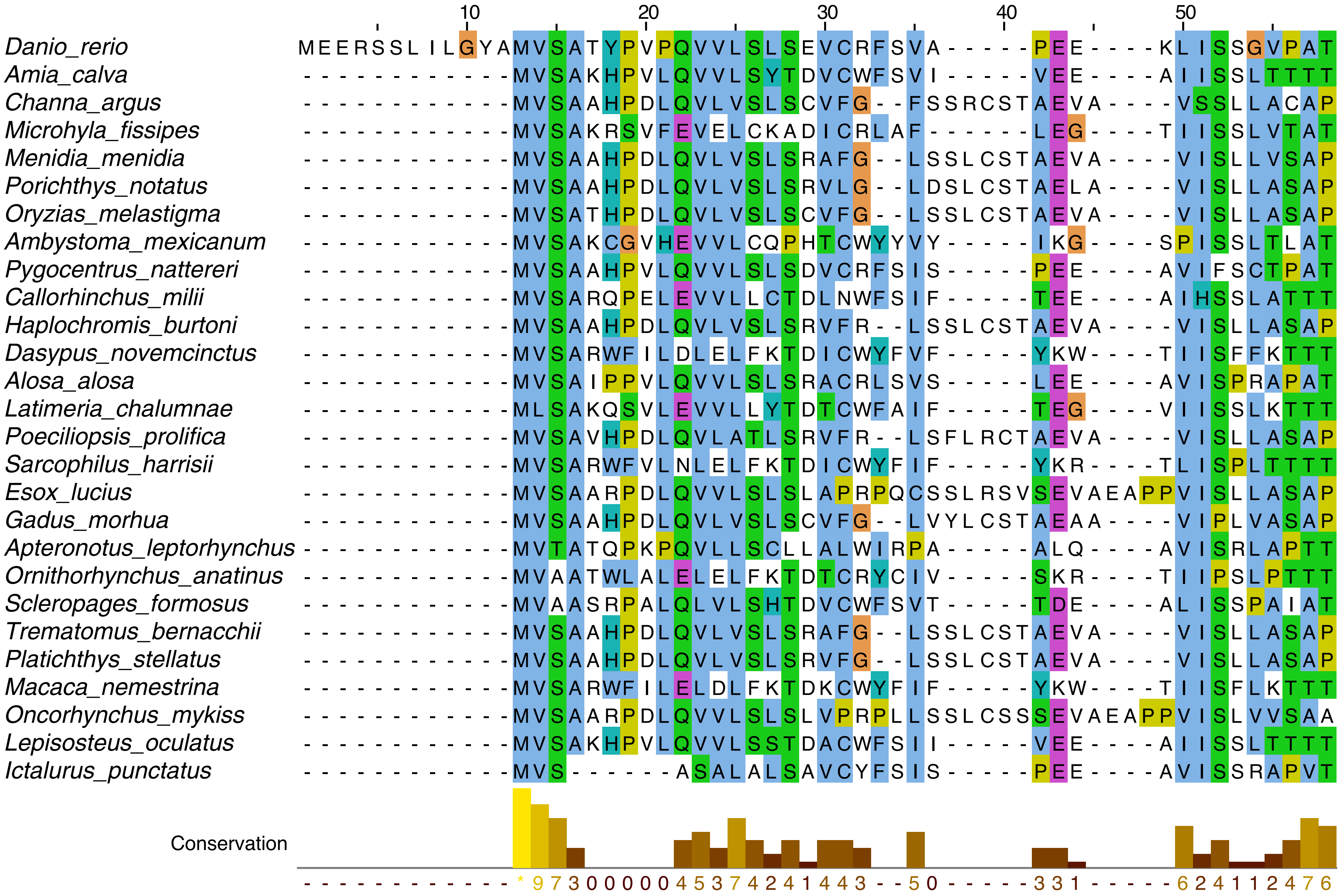

**HG0096**

ENSGALG00000014186 (MPPED1)

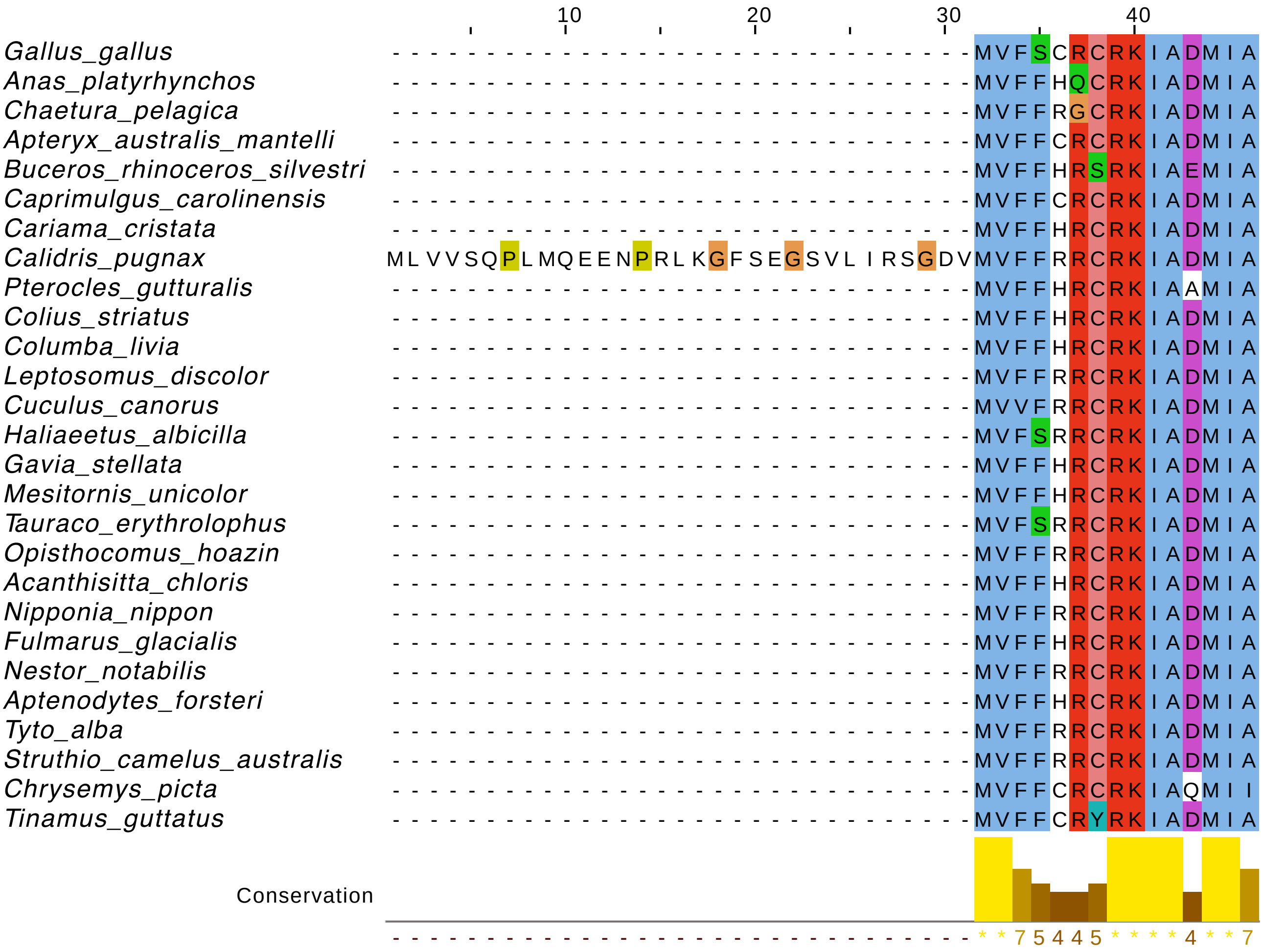

**HG0097**

ENSGALG00000037496

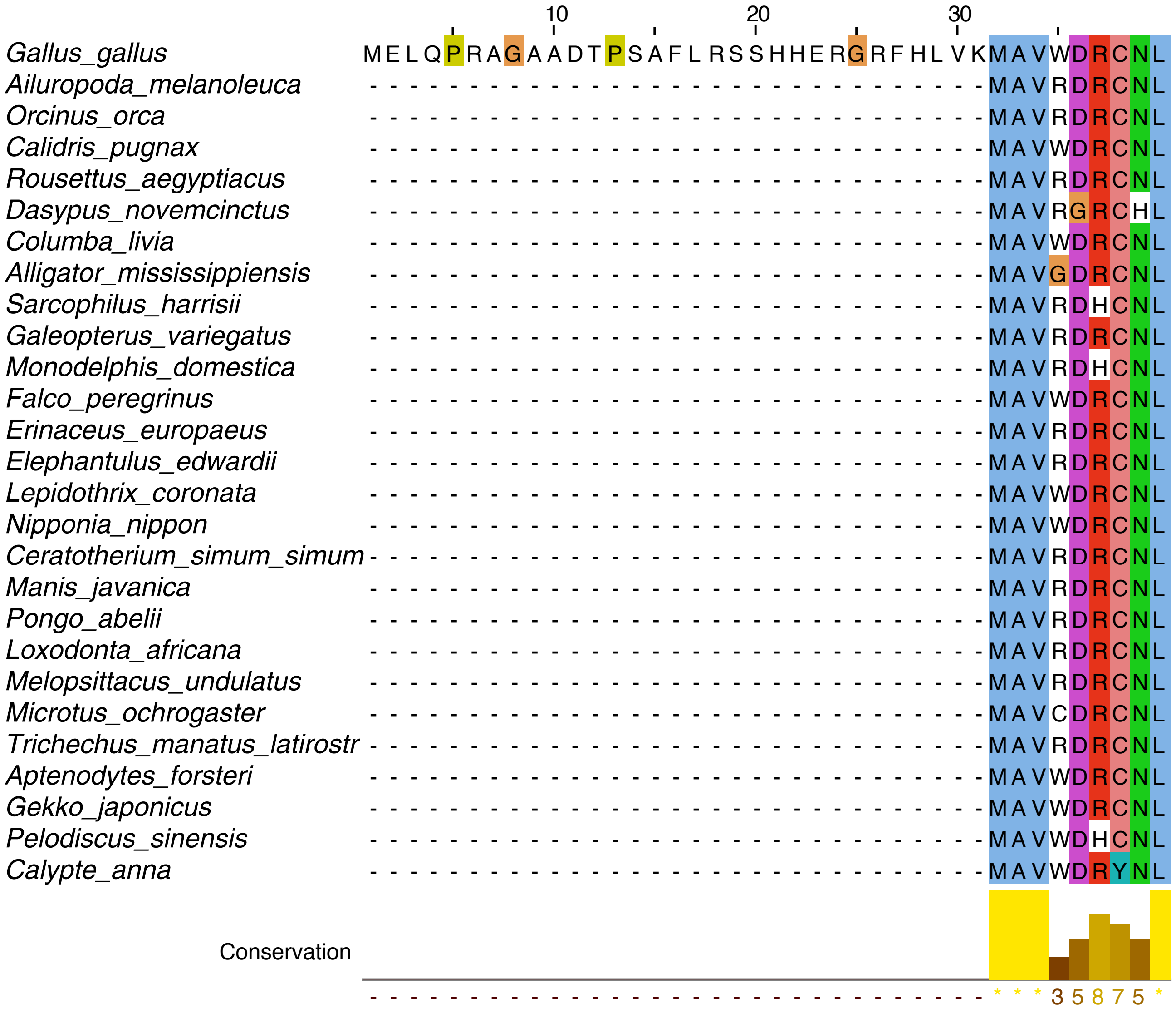

**HG0098**

ENSG00000198739 (LRRTM3)

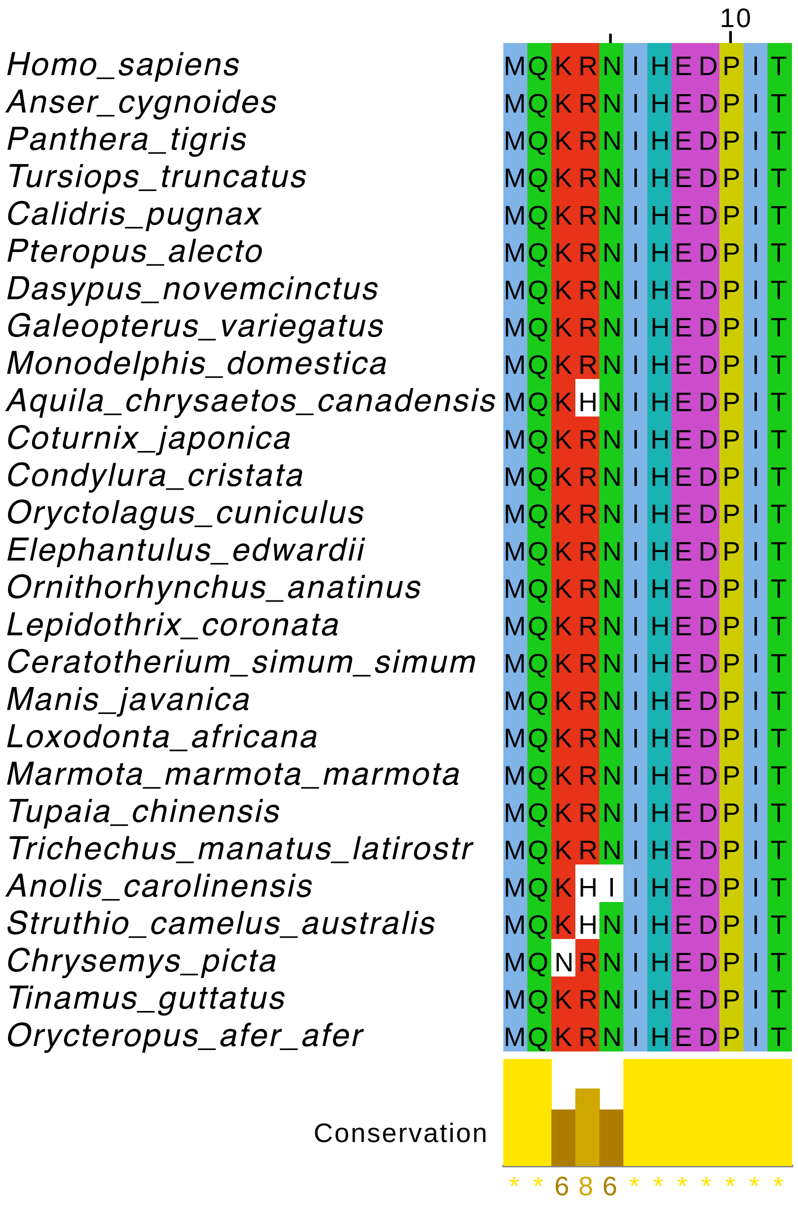

**HG0099.1**

ENSG00000168575 (SLC20A2)

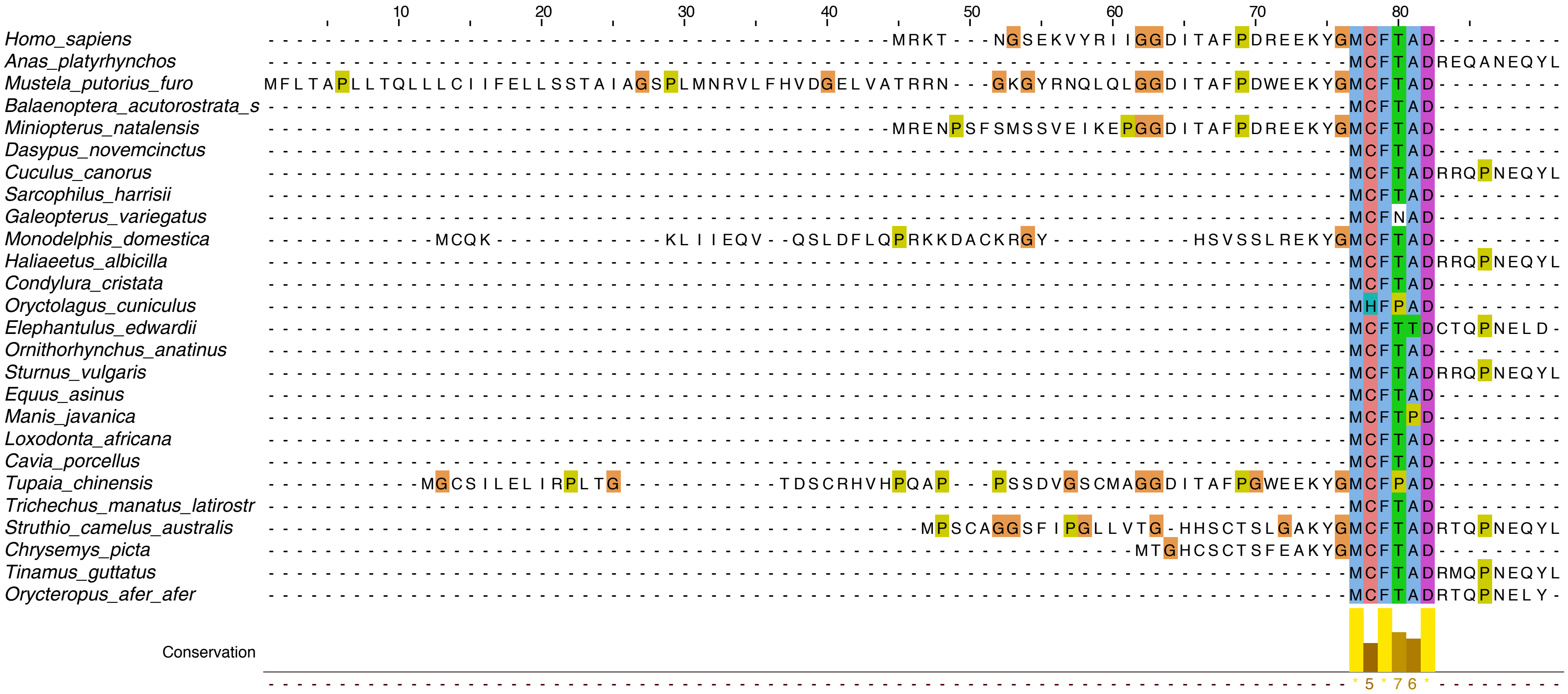

**HG0099.2**

ENSG00000144136 (SLC20A1)

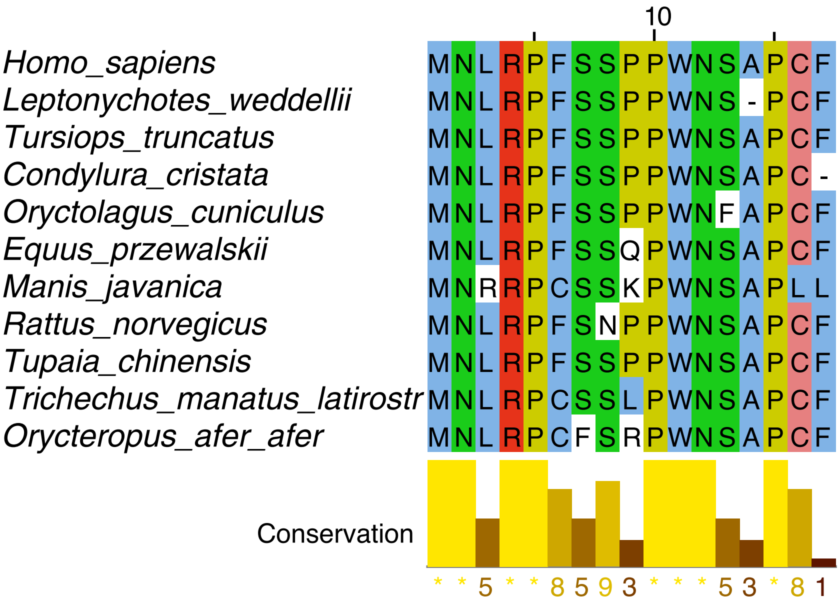

**HG0099.3**

ENSG00000168575 (SLC20A2)

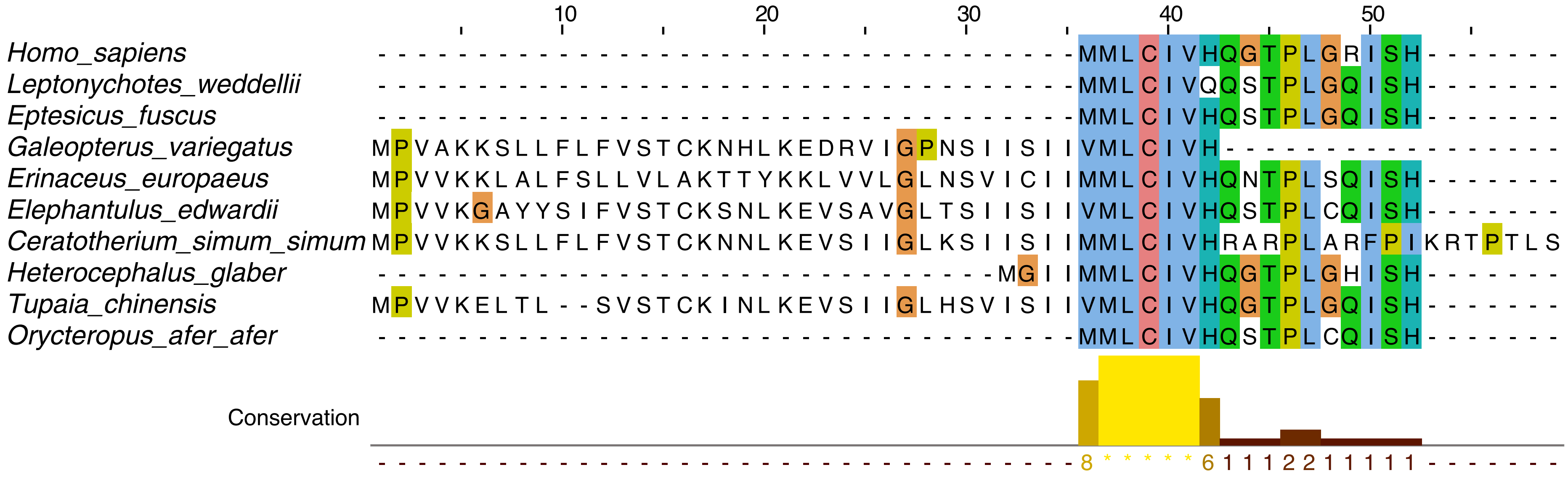

**HG0099.4**

ENSDARG00000010641 (slc20a1b)

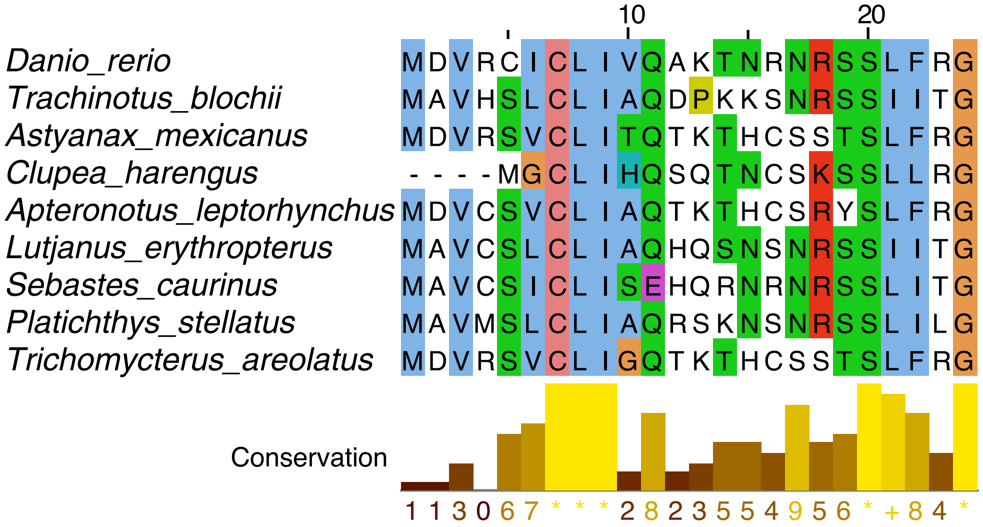

**HG0100.1**

ENSDARG00000104148 (crebbpb)

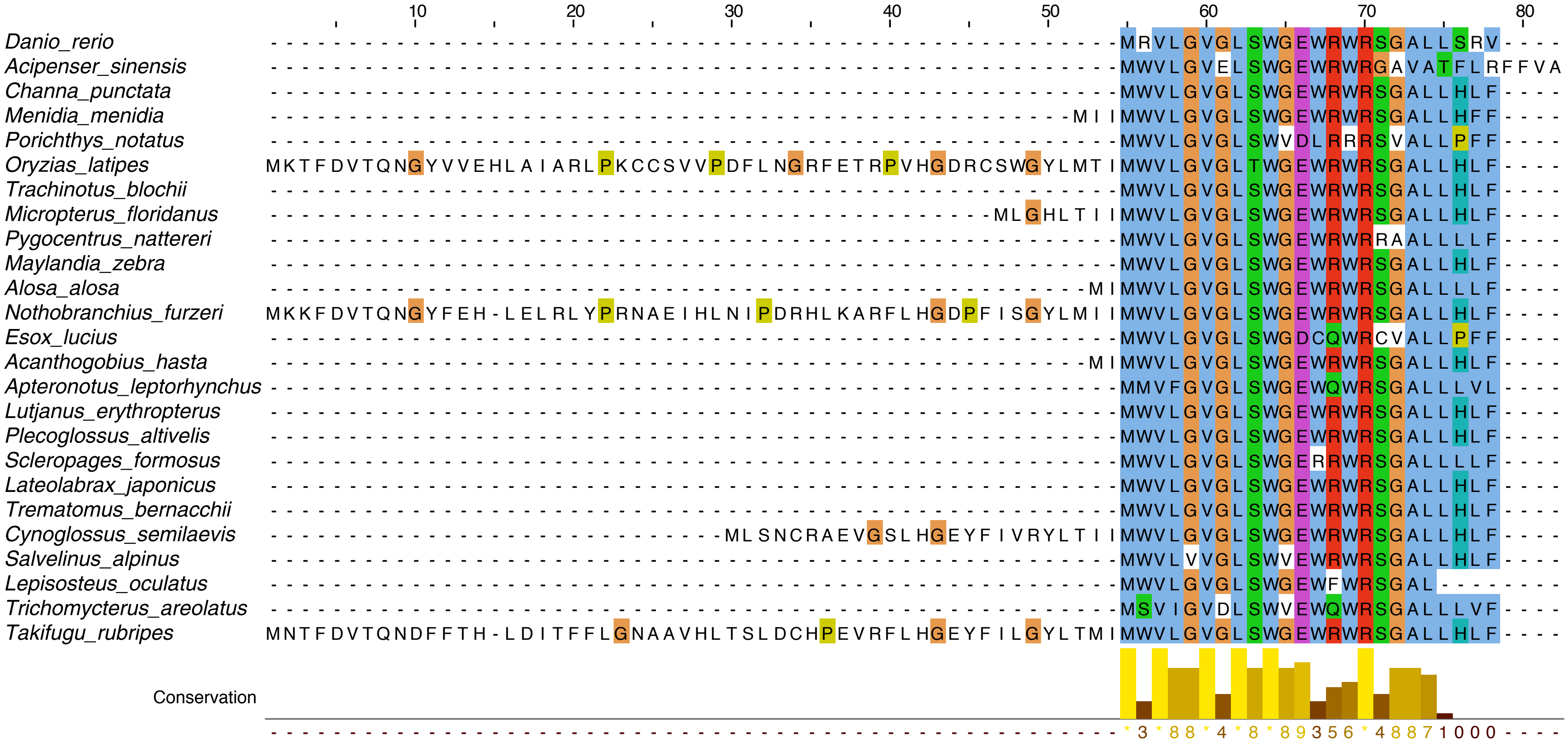

**HG0100.1**

ENSDARG00000104609 (crebbpa)

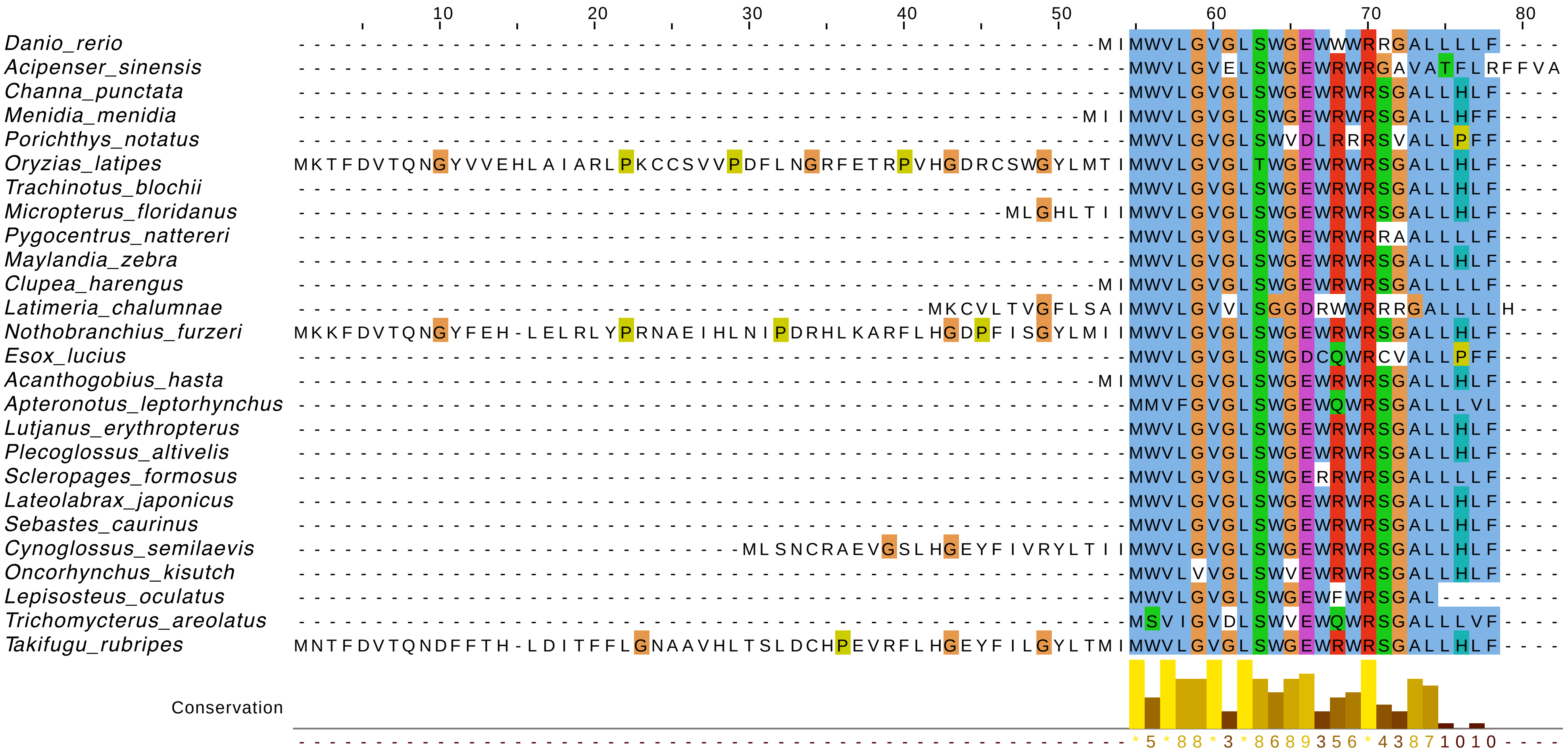

**HG0100.2**

ENSDARG00000061108 (ep300b)

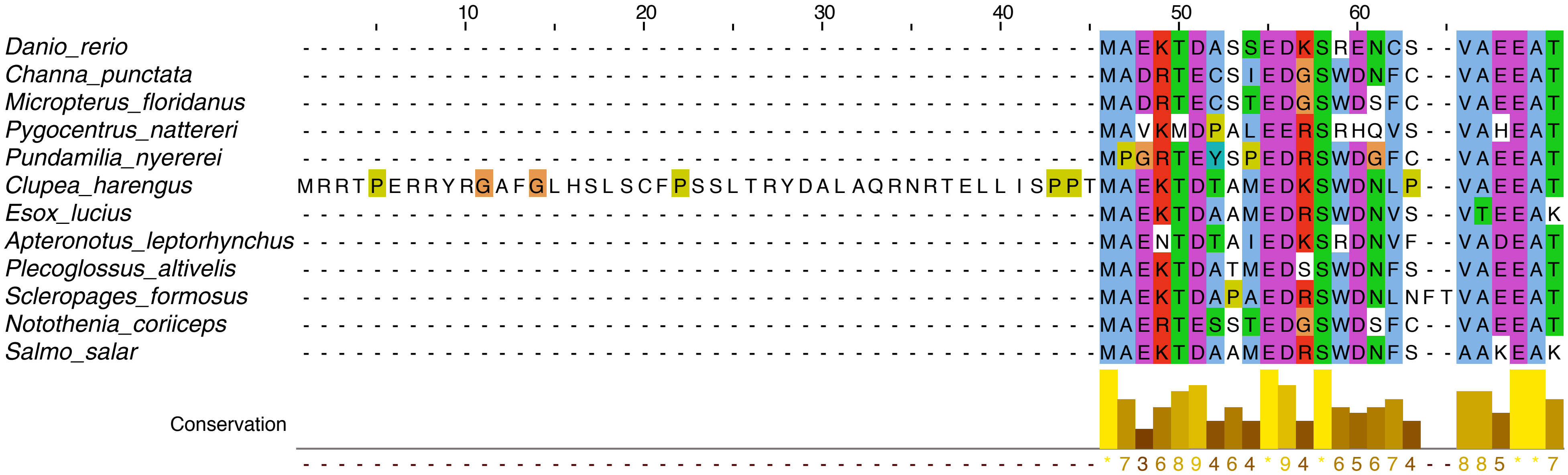

**HG0101**

ENSG00000099250 (NRP1)

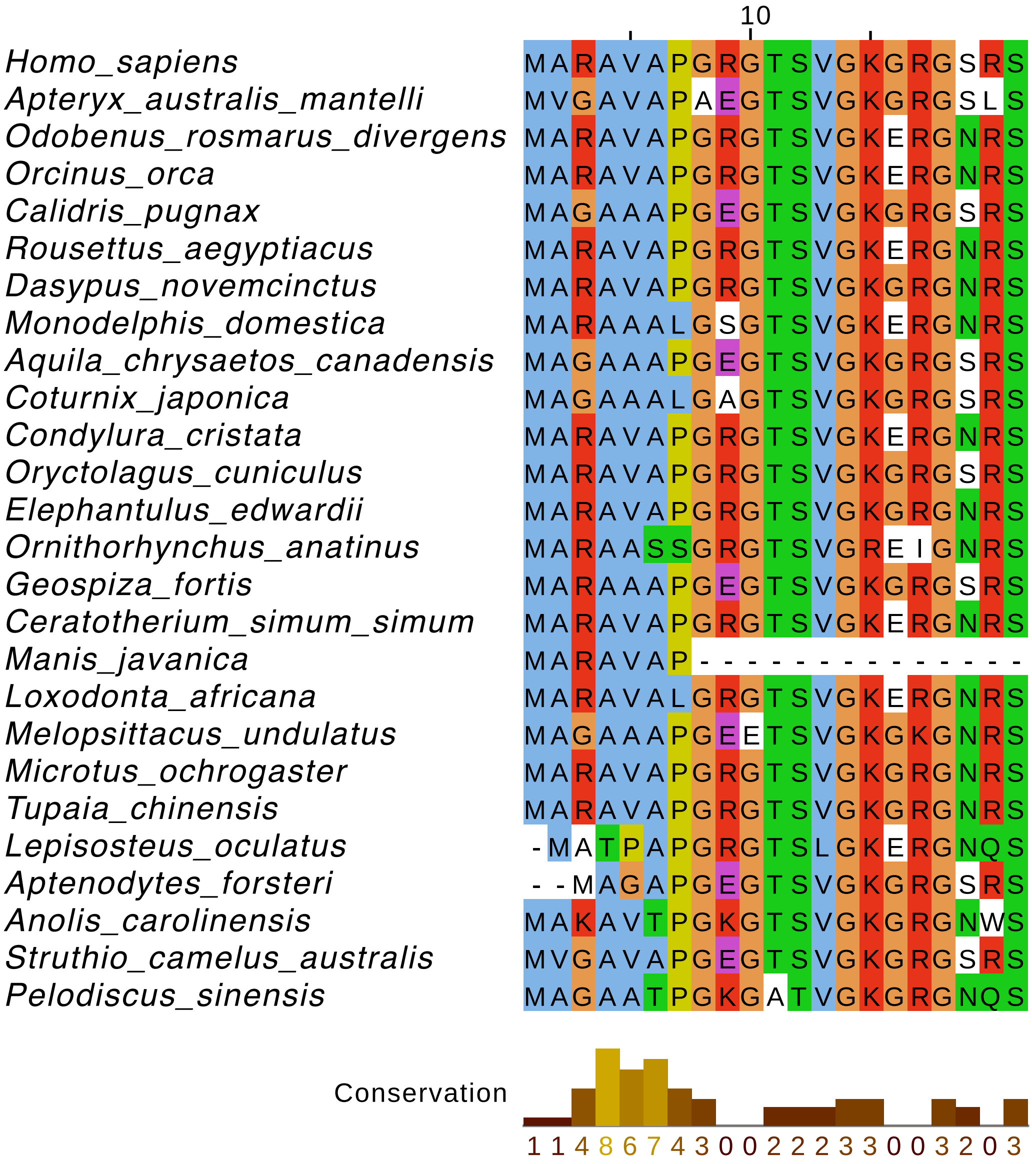

**HG0102**

ENSG00000182263 (FIGN)

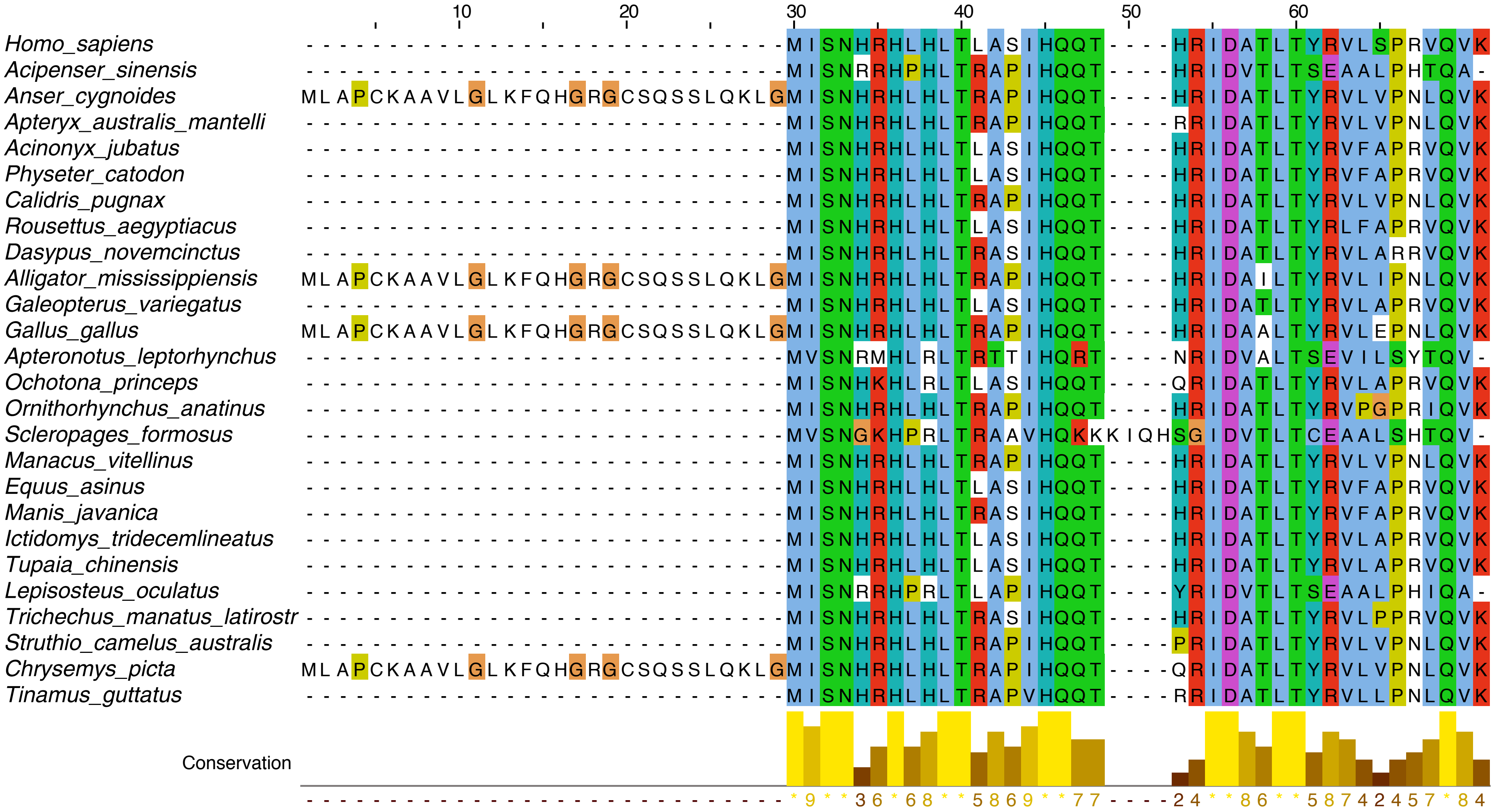

**HG0103**

ENSG00000165197 (VEGFD)

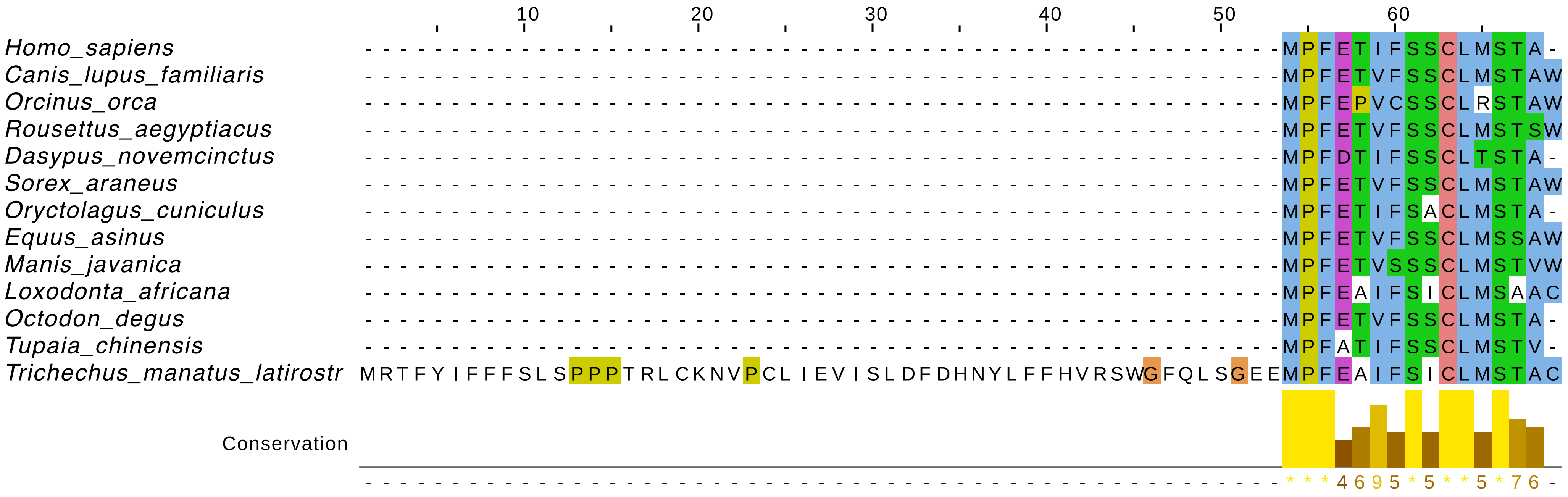

**HG0103**

ENSGALG00000016558 (VEGFD)

```
                                        10        20
Gallus_gallus                   ----------------MHFKRDASSCMTTTA
Anser_cygnoides                 MTEKLESFLVCIGDREMHFKRDASSCVTTTA
Chaetura_pelagica               ----------------MHFETDTSSCMTTTA
Apteryx_australis_mantelli      ----------------MHFKTATSSCMTTTV
Buceros_rhinoceros_silvestri    ----------------MHFETDASICMTTTA
Cariama_cristata                ----------------MHFETDASSCMTTTA
Charadrius_vociferus            ----------------MHFETDASSCMTTTA
Pterocles_gutturalis            ----------------MHFETDASSCMTTTA
Colius_striatus                 ----------------MHFKTDVSSCMTTTA
Columba_livia                   ----------------MHFETDTSSYMTTTA
Merops_nubicus                  ----------------MHFETDASSCMTTTE
Alligator_mississippiensis      ----------------MHFKTDSSSCTISTA
Cuculus_canorus                 ----------------MHFETDASSCVMTTA
Aquila_chrysaetos_canadensis    ----------------MHFETDASSCMTTTA
Mesitornis_unicolor             ----------------MHFKTDASSCMTTTA
Ornithorhynchus_anatinus        ----------------MHFERDSSSCTISTA
Tauraco_erythrolophus           ----------------MHFKTDACSCMTTTA
Opisthocomus_hoazin             ----------------MHFKTDASSCMMTTA
Lepidothrix_coronata            ----------------MHFETDASSCMTTTA
Egretta_garzetta                ----------------MHFETDASSCMTTTA
Melopsittacus_undulatus         ----------------MHFETDASSCMTTSA
Aptenodytes_forsteri            ----------------MHFETDASSCMTTTA
Struthio_camelus_australis      ----------------MHFETDASSCMTITA
Chelonia_mydas                  ----------------MHFETEPSSCMISTA
Apaloderma_vittatum             ----------------MHFKTDASSCMTLTA

Conservation                    ----------------***764575777583
```

**HG0104.1**

ENSGALG00000038097 (EI24)

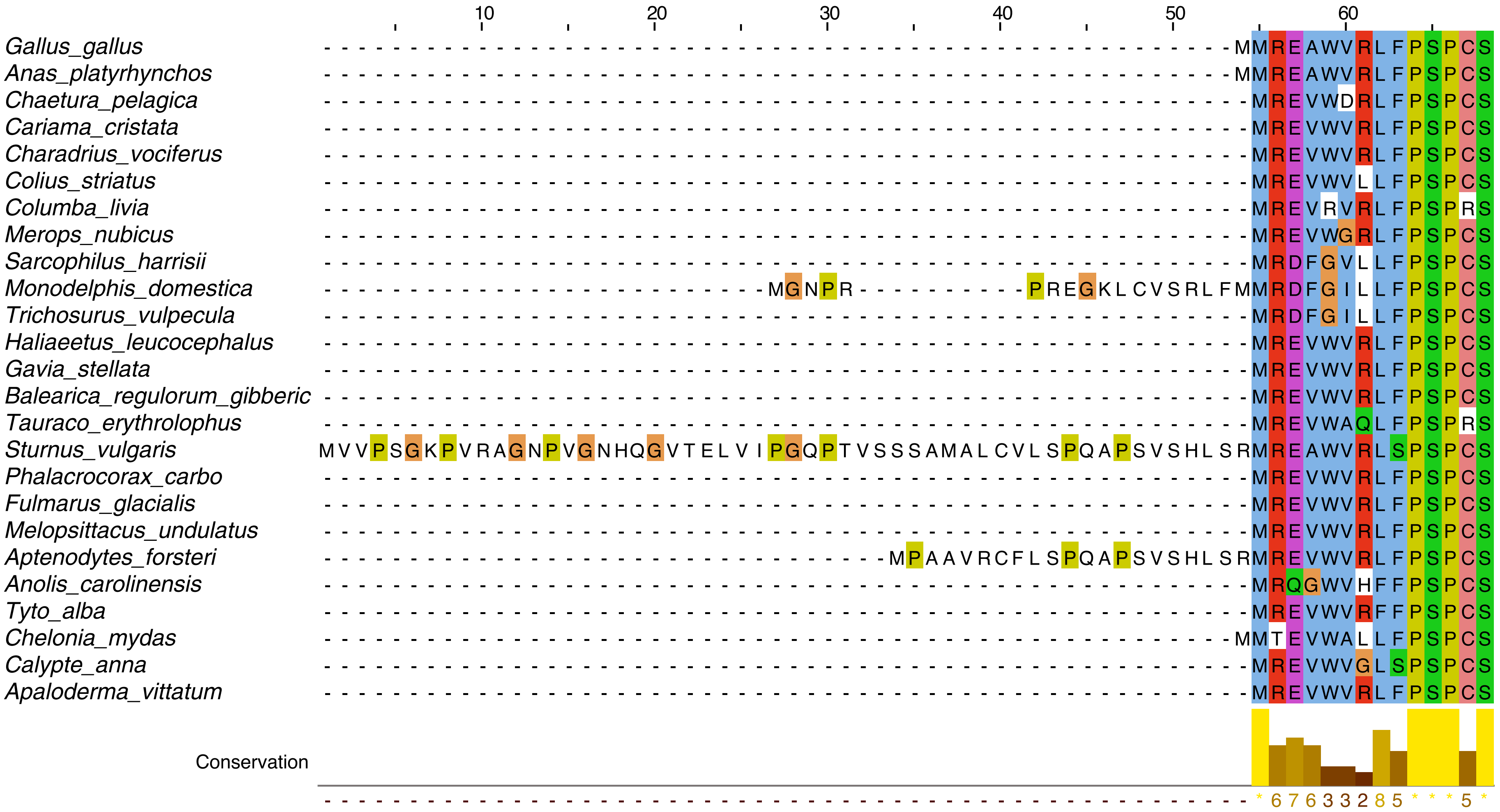

**HG0104.2**

ENSG00000149547 (EI24)

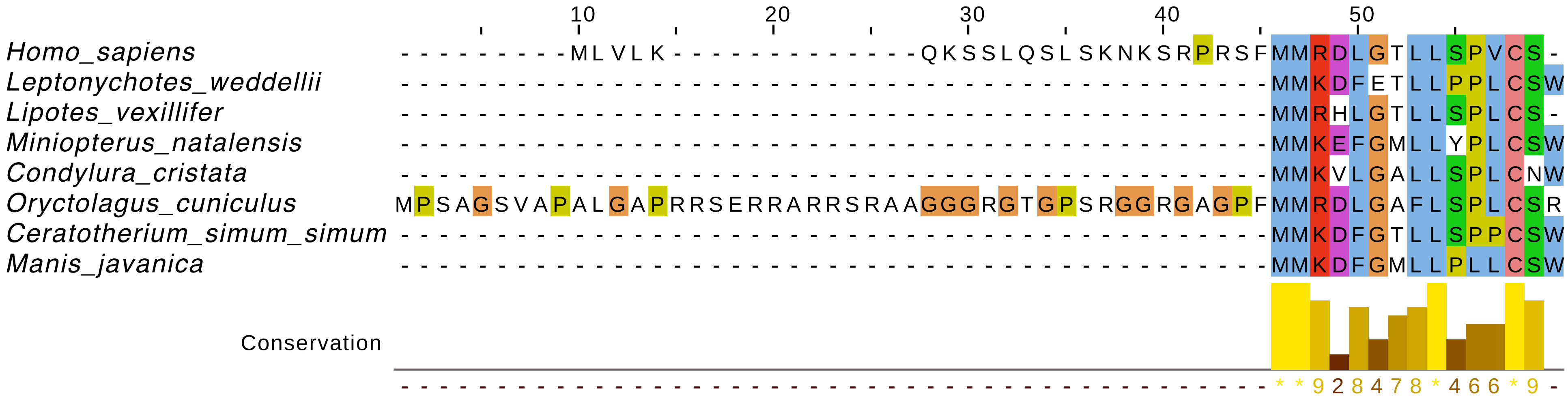

**HG0105**

ENSG00000135298 (ADGRB3)

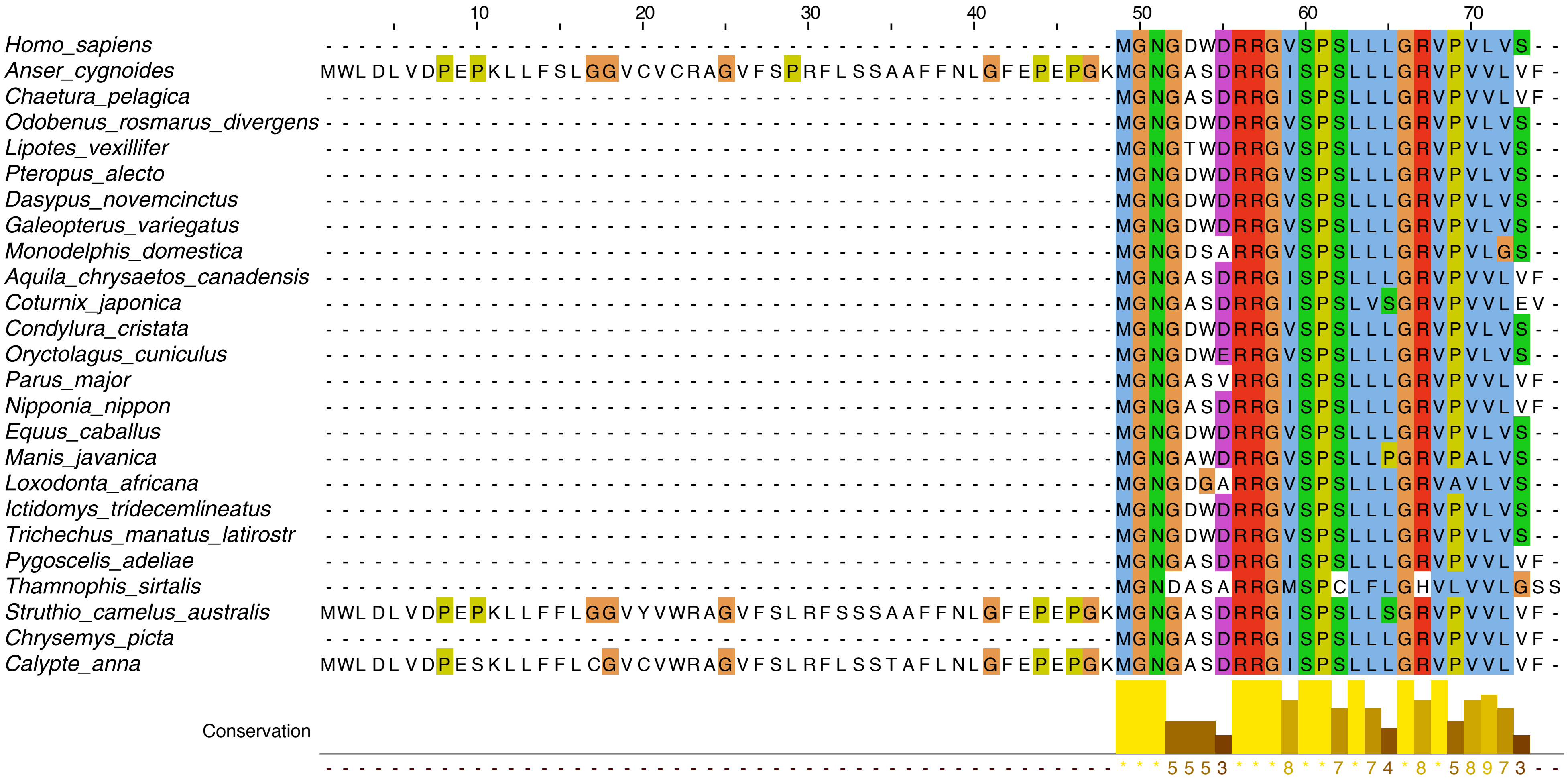

**HG0105**

ENSGALG00000016176

```
                                    10        20        30        40        50        60        70        80
Gallus_gallus                     MWLDLADPEPELLFFRGAVGVREACGGYLAAFFSLLLPFFFFFFFFLVGFEPEPGKMGNGASDRRGISPSLVSGRVPVVLEV-
Anser_cygnoides                   MWLDLVDPEPKLLFSLGGVCVCRAG------VFS--PRFLSSAAFFNLGFEPEPGKMGNGASDRRGISPSLLLGRVPVVLVF-
Chaetura_pelagica                 --------------------------------------------------------MGNGASDRRGISPSLLLGRVPVVLVF-
Leptonychotes_weddellii           --------------------------------------------------------MGNGDWDRRGVSPSLLLGRVPVLVS--
Orcinus_orca                      --------------------------------------------------------MGNGAWDRRGVSPSLLLGRVPVLVS--
Dasypus_novemcinctus              --------------------------------------------------------MGNGDWDRRGVSPSLLLGRVPVLVS--
Galeopterus_variegatus            --------------------------------------------------------MGNGDWDRRGVSPSLLLGRVPVLVS--
Monodelphis_domestica             --------------------------------------------------------MGNGDSARRGVSPSLLLGRVPVLGS--
Aquila_chrysaetos_canadensis      --------------------------------------------------------MGNGASDRRGISPSLLLGRVPVVLVF-
Condylura_cristata                --------------------------------------------------------MGNGDWDRRGVSPSLLLGRVPVLVS--
Ochotona_princeps                 --------------------------------------------------------MGNGGWERRGISRSLLLGRVPVLVS--
Lepidothrix_coronata              MWLDLVDPEPKLLFFLGGVCVWRAG------IFS--LRFLSSAAFFNLGFEPEPGKMGNGASVRRGISPSLLLGRVPVVLVF-
Nipponia_nippon                   --------------------------------------------------------MGNGASDRRGISPSLLLGRVPVVLVF-
Manis_javanica                    --------------------------------------------------------MGNGAWDRRGVSPSLLPGRVPALVS--
Microcebus_murinus                --------------------------------------------------------MGNGDWDRRGVSPSLLLGRVPVLVS--
Loxodonta_africana                --------------------------------------------------------MGNGDGARRGVSPSLLLGRVAVLVS--
Ictidomys_tridecemlineatus        --------------------------------------------------------MGNGDWDRRGVSPSLLLGRVPVLVS--
Trichechus_manatus_latirostr      --------------------------------------------------------MGNGDWDRRGVSPSLLLGRVPVLVS--
Pygoscelis_adeliae                --------------------------------------------------------MGNGASDRRGISPSLLLGRVPVVLVF-
Thamnophis_sirtalis               --------------------------------------------------------MGNDASARRGMSPCLFLGHVLVVLGSS
Struthio_camelus_australis        MWLDLVDPEPKLLFFLGGVYVWRAG------VFS--LRFSSSAAFFNLGFEPEPGKMGNGASDRRGISPSLLSGRVPVVLVF-
Pelodiscus_sinensis               --------------------------------------------------------MGNGASDRRGISPSLVLGRVPVVLVF-
Calypte_anna                      MWLDLVDPESKLLFFLCGVCVWRAG------VFS--LRFLSSTAFLNLGFEPEPGKMGNGASDRRGISPSLLLGRVPVVLVF-
```

Conservation

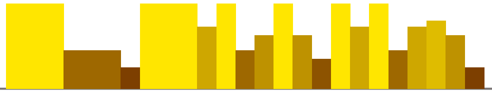

```
--------------------------------------------------------***5553***8*57*74*8*58973--
```

**HG0106**

ENSG00000147548 (NSD3)

```
                                        10        20        30        40        50
Homo_sapiens                   ----------------------------------------------MEQTEGPMRERES
Pseudacris_regilla             ----------------------------------------------MEQTEGPMRERES
Canis_lupus_familiaris         ----------------------------------------------MEQTEGPMRERES
Hynobius_retardatus            ----------------------------------------------MDQTEGPMRERES
Balaenoptera_acutorostrata_s   ----------------------------------------------MEQSEGPMRERES
Calidris_pugnax                ----------------------------------------------MEQTEGPMRRRES
Miniopterus_natalensis         ----------------------------------------------MEQTEGPMRERES
Latimeria_chalumnae            ----------------------------------------------MEQTEGPMRKRES
Sarcophilus_harrisii           ----------------------------------------------MEQTEGPMRERES
Galeopterus_variegatus         ----------------------------------------------MEQTEGPMRERES
Aegypius_monachus              MRRGALRAPLGARRKGSHLPARRGRRRPRRRPLSPLGWVDAAGAVRMEQTEGPMRKRES
Gallus_gallus                  ----------------------------------------------MEQTEGPMRKRES
Condylura_cristata             ----------------------------------------------MEQTEGPMRERES
Oryctolagus_cuniculus          ----------------------------------------------MEQTEGPMRERES
Elephantulus_edwardii          ----------------------------------------------MEQTEVPMREREN
Ornithorhynchus_anatinus       MRLRPAPAPS------PSAPGSSCGPTLLRAPVSLLGSHRHGRGVRMEQTEGPMRERES
Lepidothrix_coronata           ----------------------------------------------MEVTEGPMRKRES
Ceratotherium_simum_simum      ----------------------------------------------MEQTEGPMRERES
Manis_javanica                 ----------------------------------------------MEQTEGPMRERES
Ictidomys_tridecemlineatus     ----------------------------------------------MEQTEGPMRERES
Lepisosteus_oculatus           ----------------------------------------------MEHAEGPVRERES
Trichechus_manatus_latirostr   ----------------------------------------------MEQTEGPMREREN
Takydromus_sexlineatus         ----------------------------------------------MEQTDGPMRKRES
Chrysemys_picta                ----------------------------------------------MEQTDGPMRERES
Orycteropus_afer_afer          ----------------------------------------------MEQTEGPMREREN

Conservation                   ----------------------------------------------*93798*8*7**9
```

**HG0107.1**

ENSG00000058668 (ATP2B4)

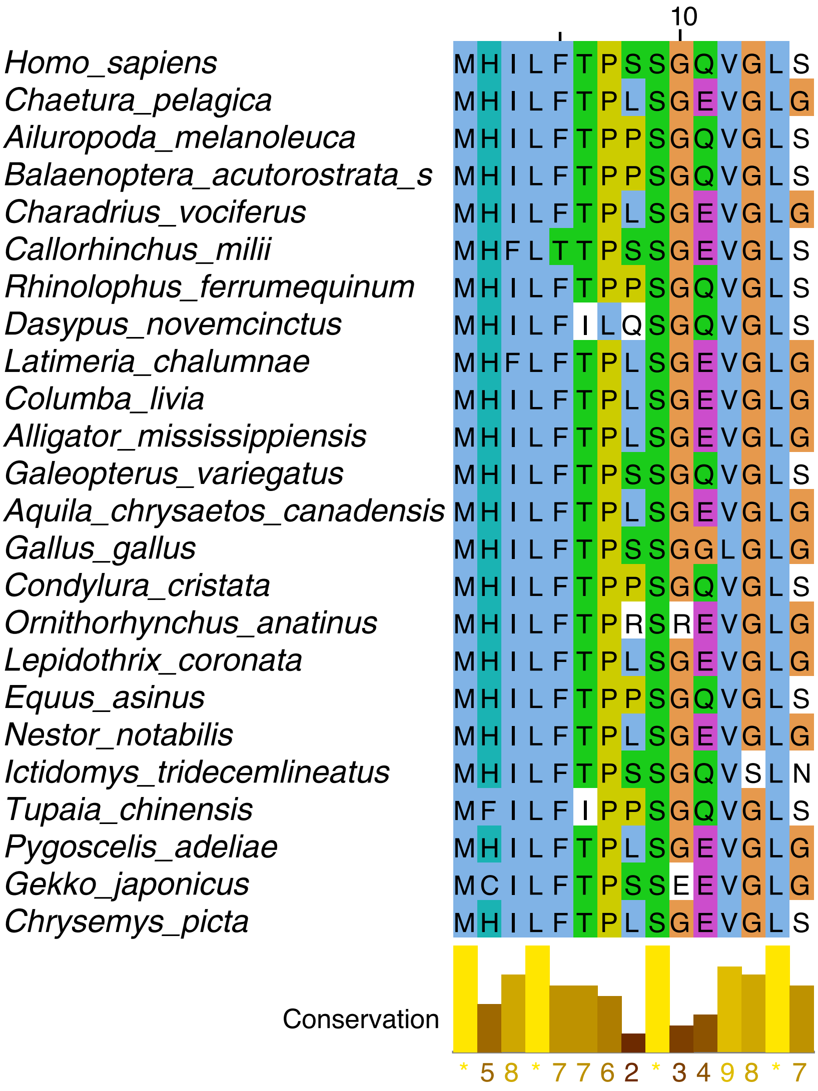

**HG0107.1**

ENSGALG00000003601 (ATP2B4)

| Species | Sequence (ruler mark at 10) |
|---|---|
| *Gallus_gallus* | M H I L F T P S S G G L G L G |
| *Anser_cygnoides* | M R I L F T P A S G G E G L G |
| *Chaetura_pelagica* | M H I L F T P L S G E V G L G |
| *Ailuropoda_melanoleuca* | M H I L F T P P S G Q V G L S |
| *Balaenoptera_acutorostrata_s* | M H I L F T P P S G Q V G L S |
| *Calidris_pugnax* | M H I L F T P L S G E V G L G |
| *Rhinolophus_ferrumequinum* | M H I L F T P P S G Q V G L S |
| *Latimeria_chalumnae* | M H F L F T P L S G E V G L G |
| *Columba_livia* | M H I L F T P L S G E V G L G |
| *Alligator_sinensis* | M H I L F T P L S G E V G L G |
| *Galeopterus_variegatus* | M H I L F T P S S G Q V G L S |
| *Aquila_chrysaetos_canadensis* | M H I L F T P L S G E V G L G |
| *Condylura_cristata* | M H I L F T P P S G Q V G L S |
| *Ornithorhynchus_anatinus* | M H I L F T P R S R E V G L G |
| *Ficedula_albicollis* | M H I L F T P S S P Q L G L G |
| *Equus_asinus* | M H I L F T P P S G Q V G L S |
| *Cebus_capucinus_imitator* | M H I L F T P S S G Q V G L S |
| *Nestor_notabilis* | M H I L F T P L S G E V G L G |
| *Ictidomys_tridecemlineatus* | M H I L F T P S S G Q V S L N |
| *Pygoscelis_adeliae* | M H I L F T P L S G E V G L G |
| *Gekko_japonicus* | M C I L F T P S S E E V G L G |
| *Chrysemys_picta* | M H I L F T P L S G E V G L S |

Conservation

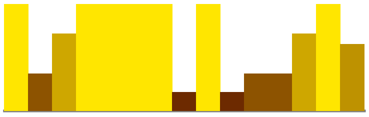

| | |
|---|---|
| Conservation | * 4 8 * * * * 2 * 2 4 4 8 * 7 |

**HG0107.2**

ENSG00000070961 (ATP2B1)

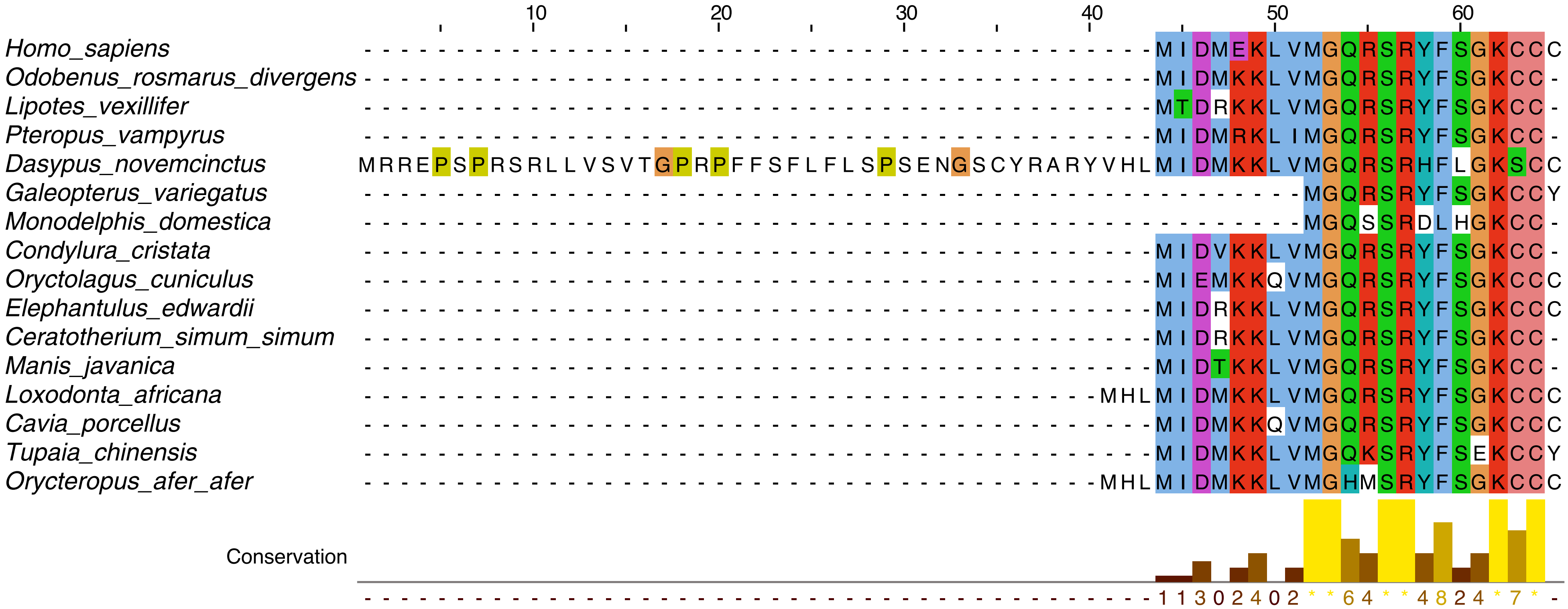

**HG0107.2**

ENSGALG00000030550

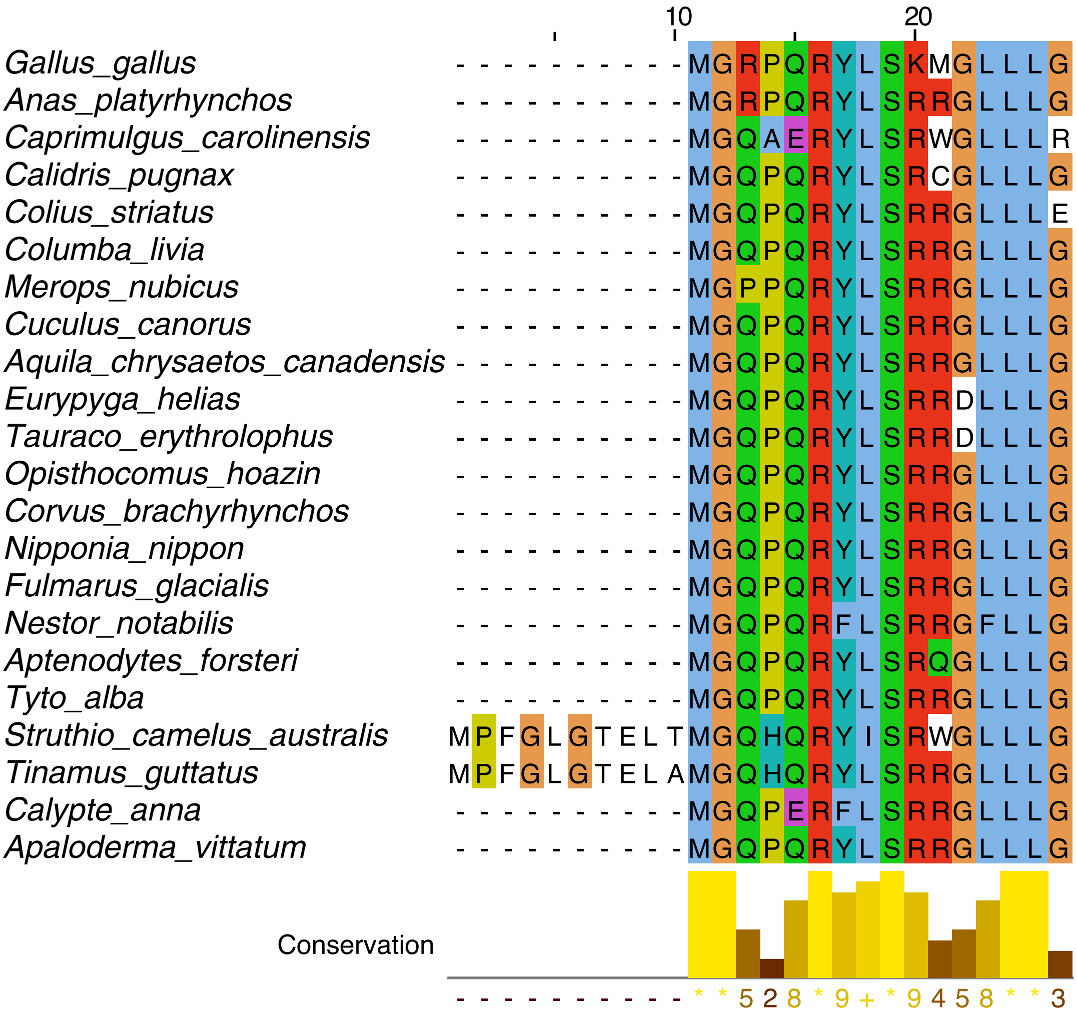

**HG0107.3**

ENSDARG00000063433 (atp2b2)

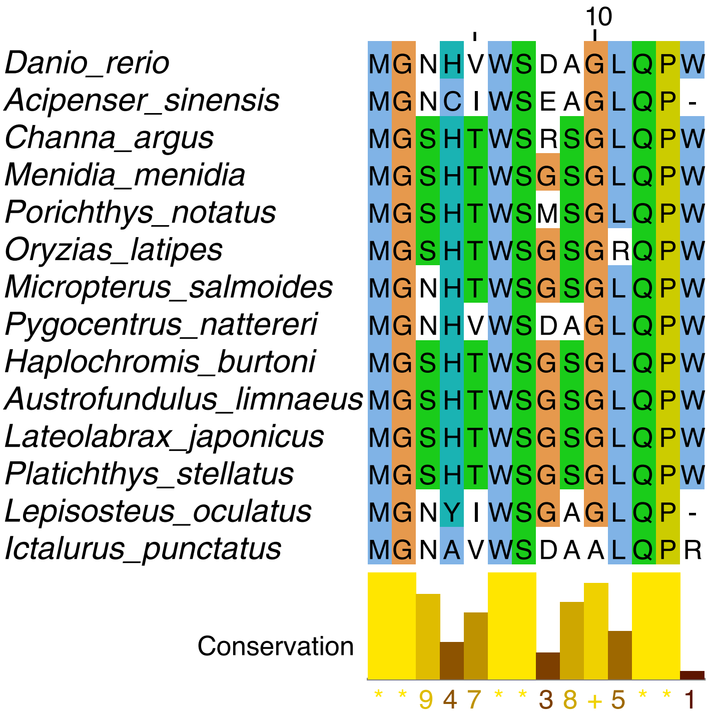

**HG0107.3**

ENSG00000157087 (ATP2B2)

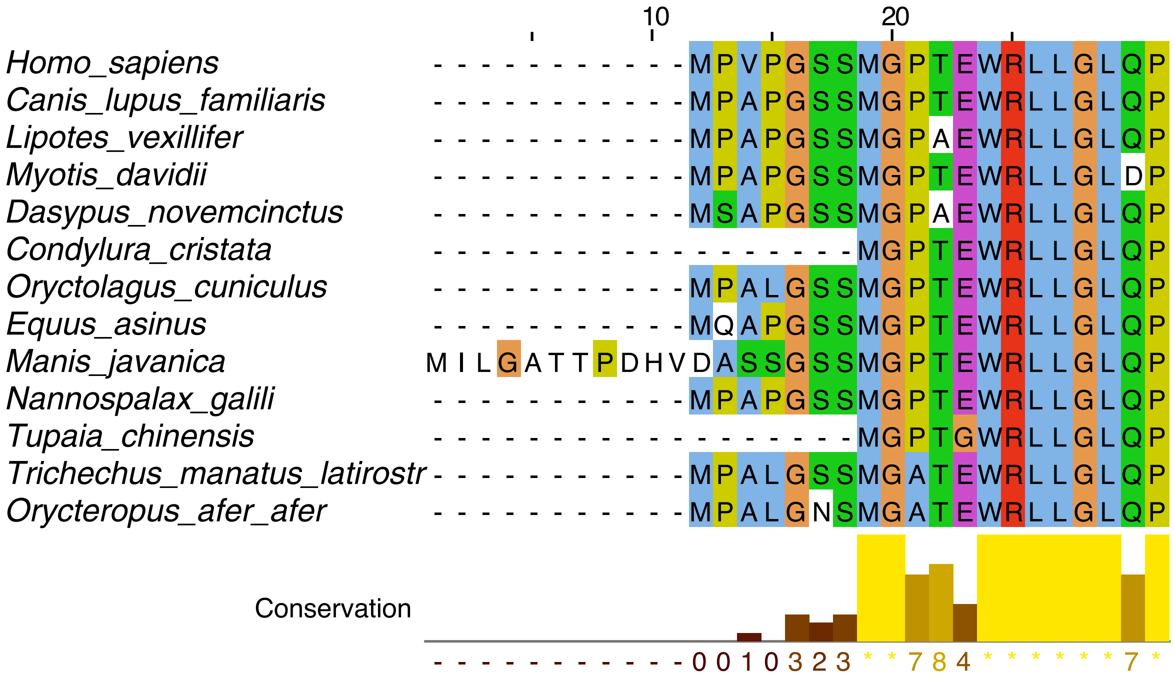

**HG0107.4**

ENSG00000070961 (ATP2B1)

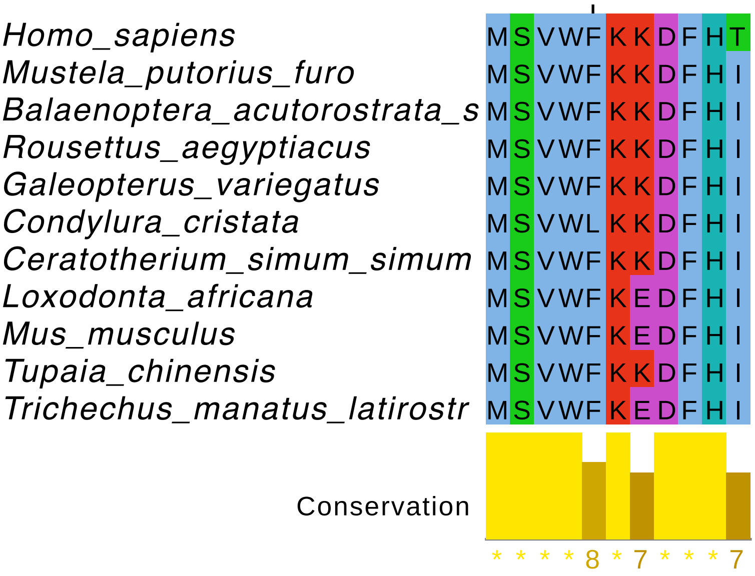

**HG0108.1**

ENSGALG00000007396 (TAOK3)

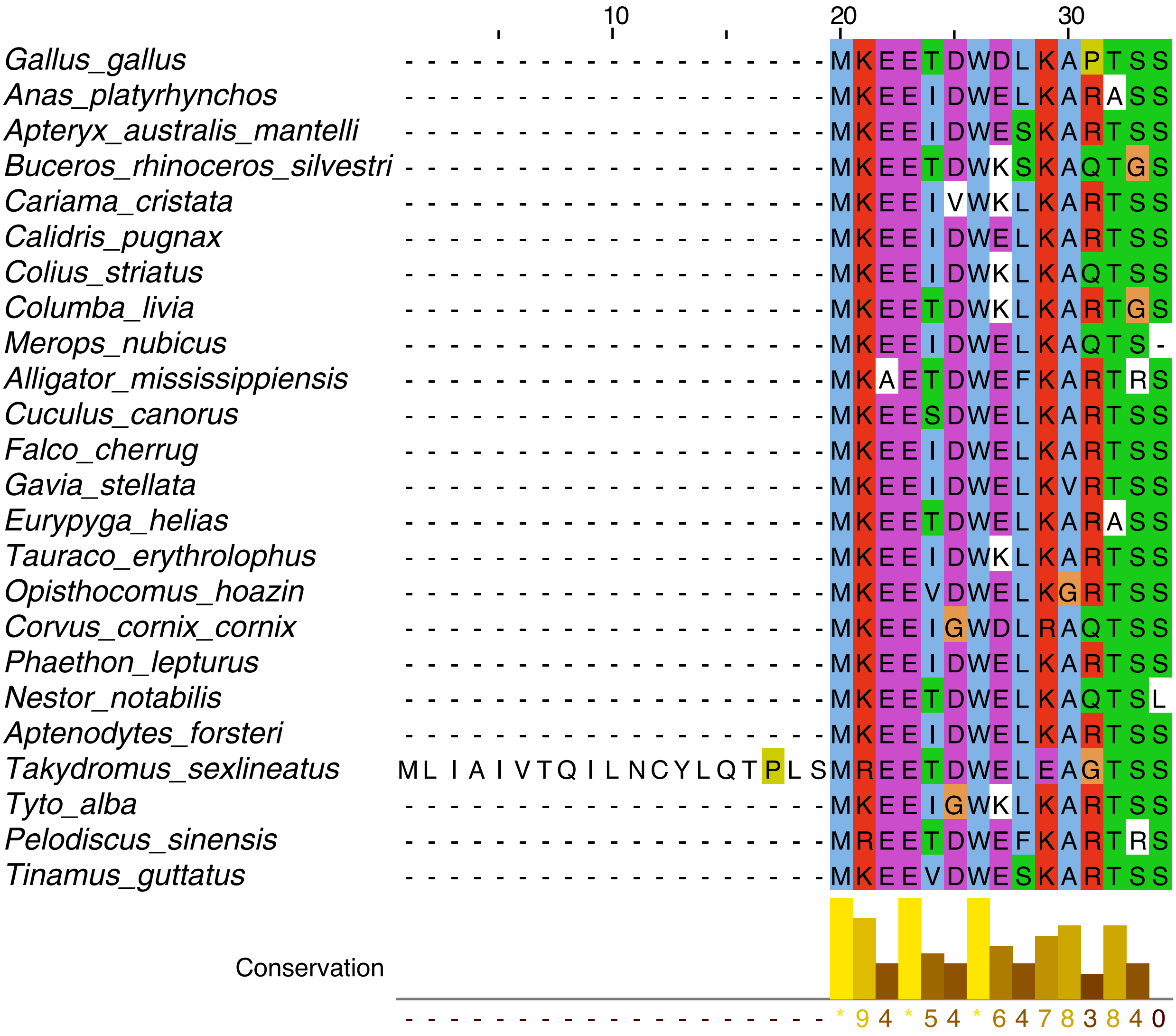

**HG0108.2**

ENSG00000135090 (TAOK3)

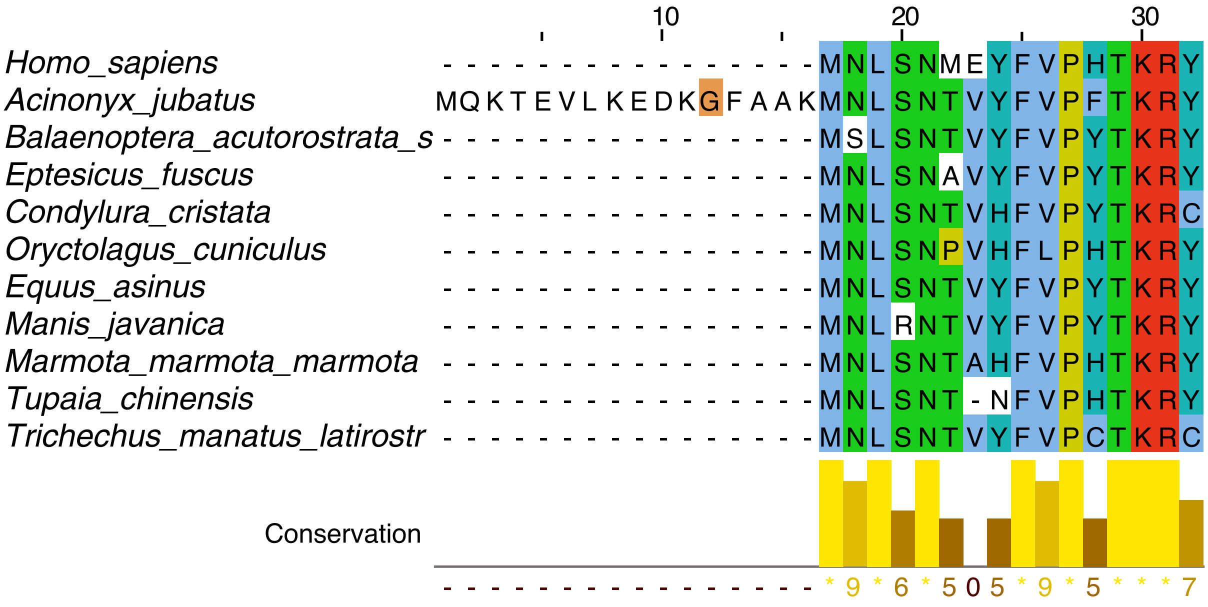

**HG0108.3**

ENSDARG00000098304 (taok1b)

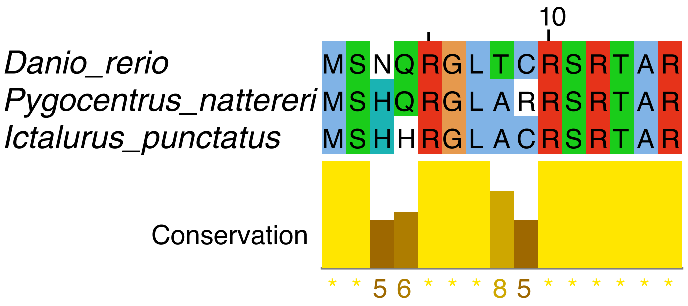

**HG0109.1**

ENSG00000048540 (LMO3)

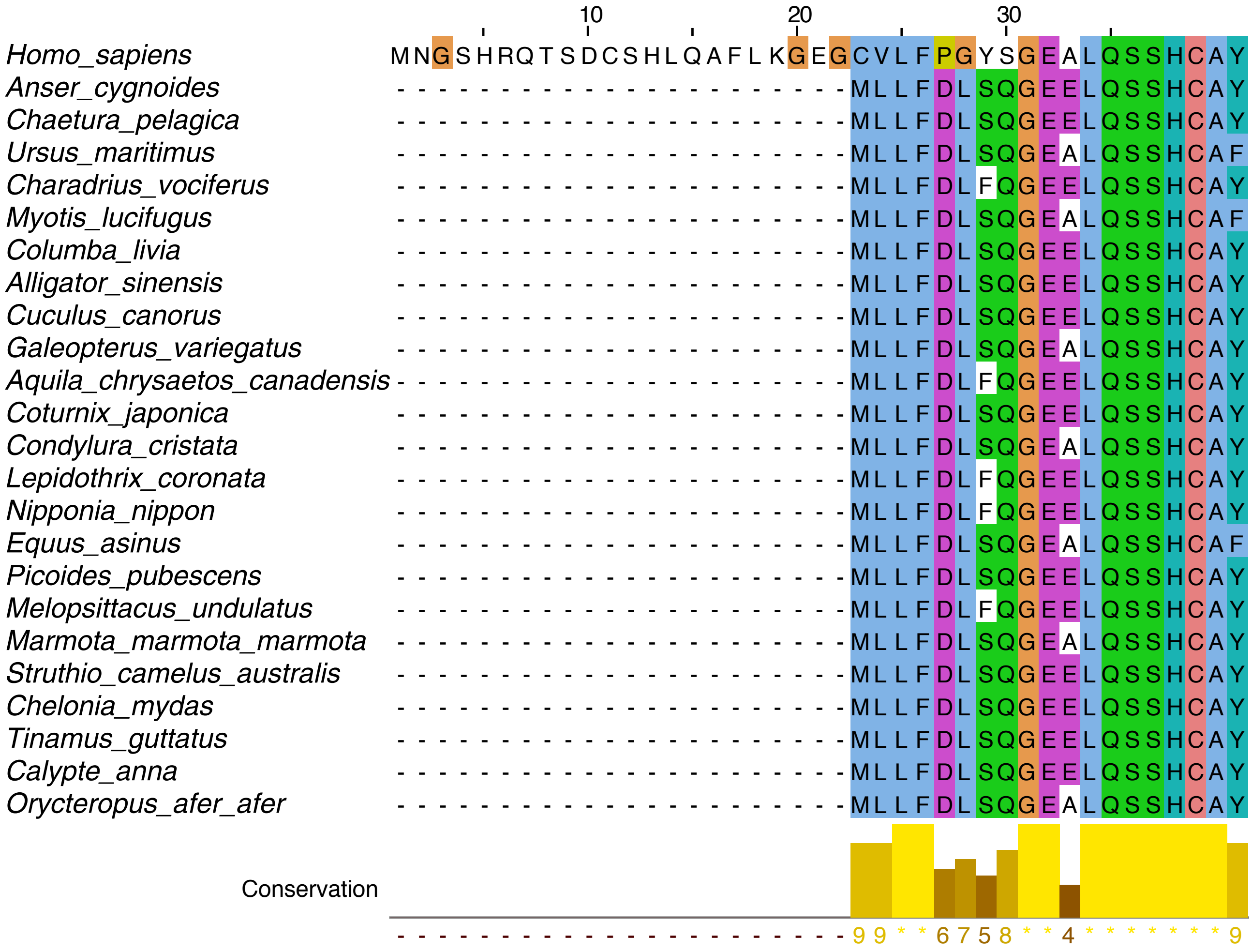

## HG0109.2

ENSG00000048540 (LMO3)

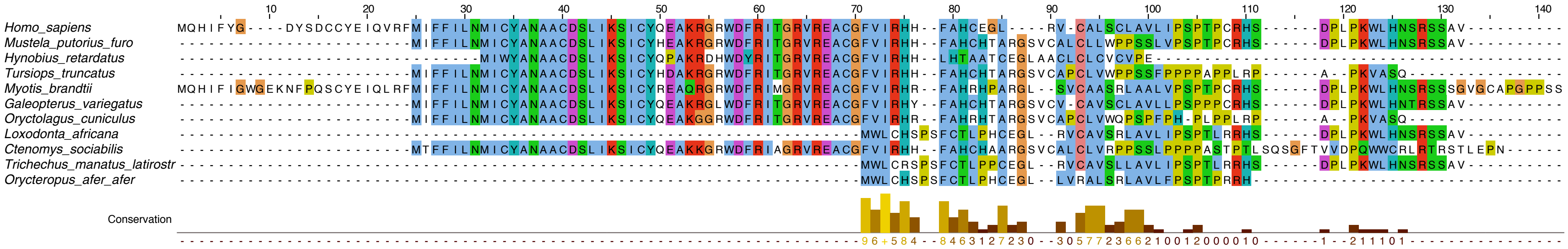

**HG0109.3**

ENSG00000048540 (LMO3)

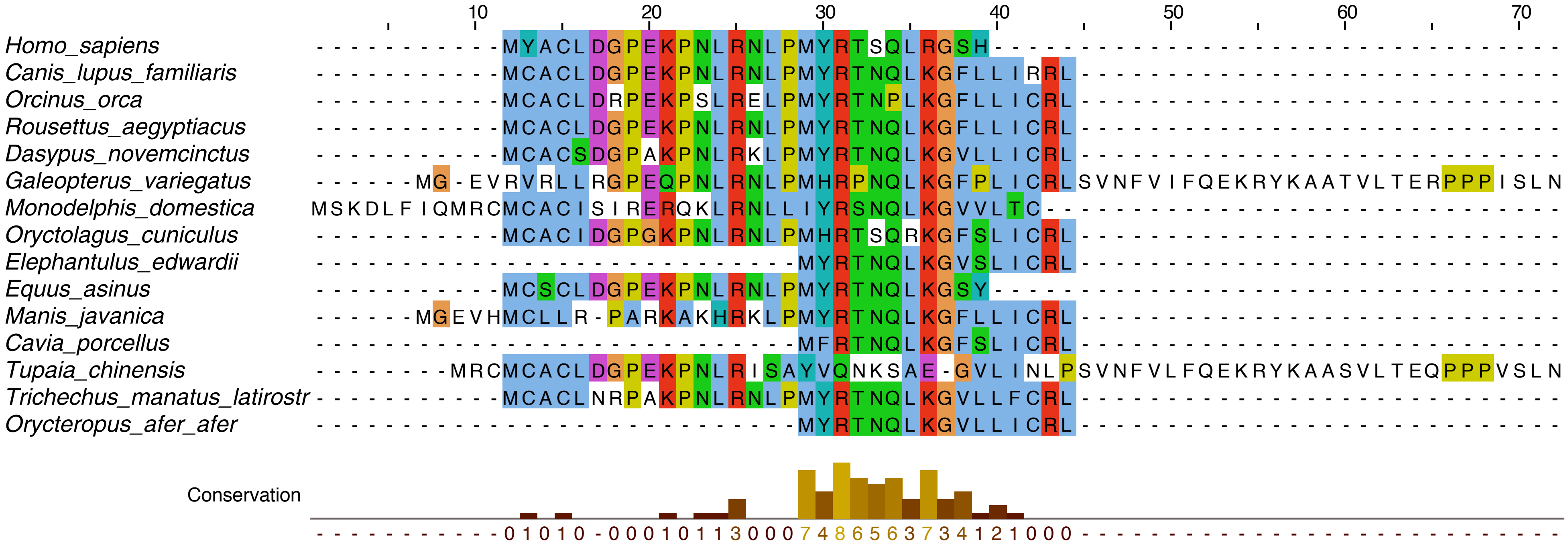

**HG0110.1**

ENSG00000091656 (ZFHX4)

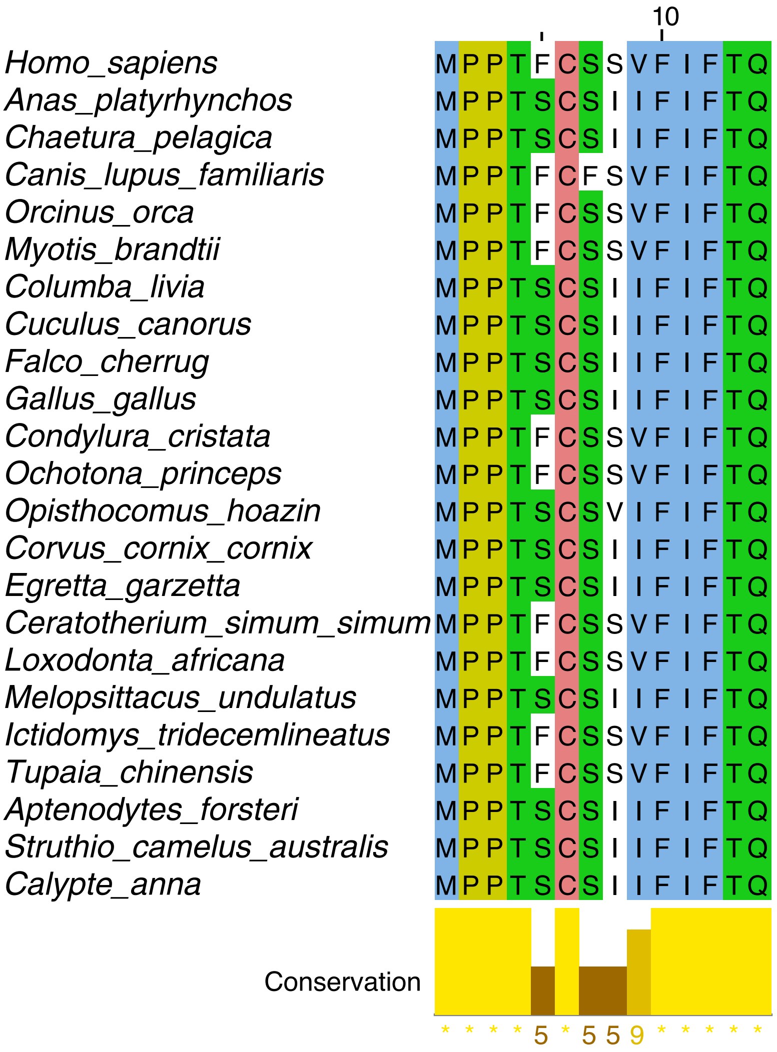

# HG0110.2

### ENSDARG00000073944

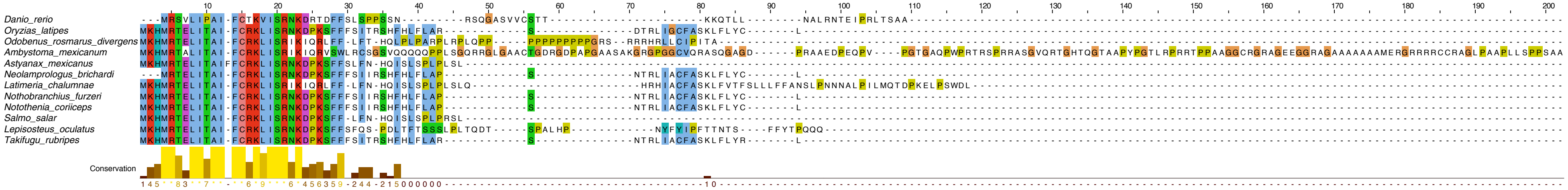

**HG0111**

ENSGALG00000034944

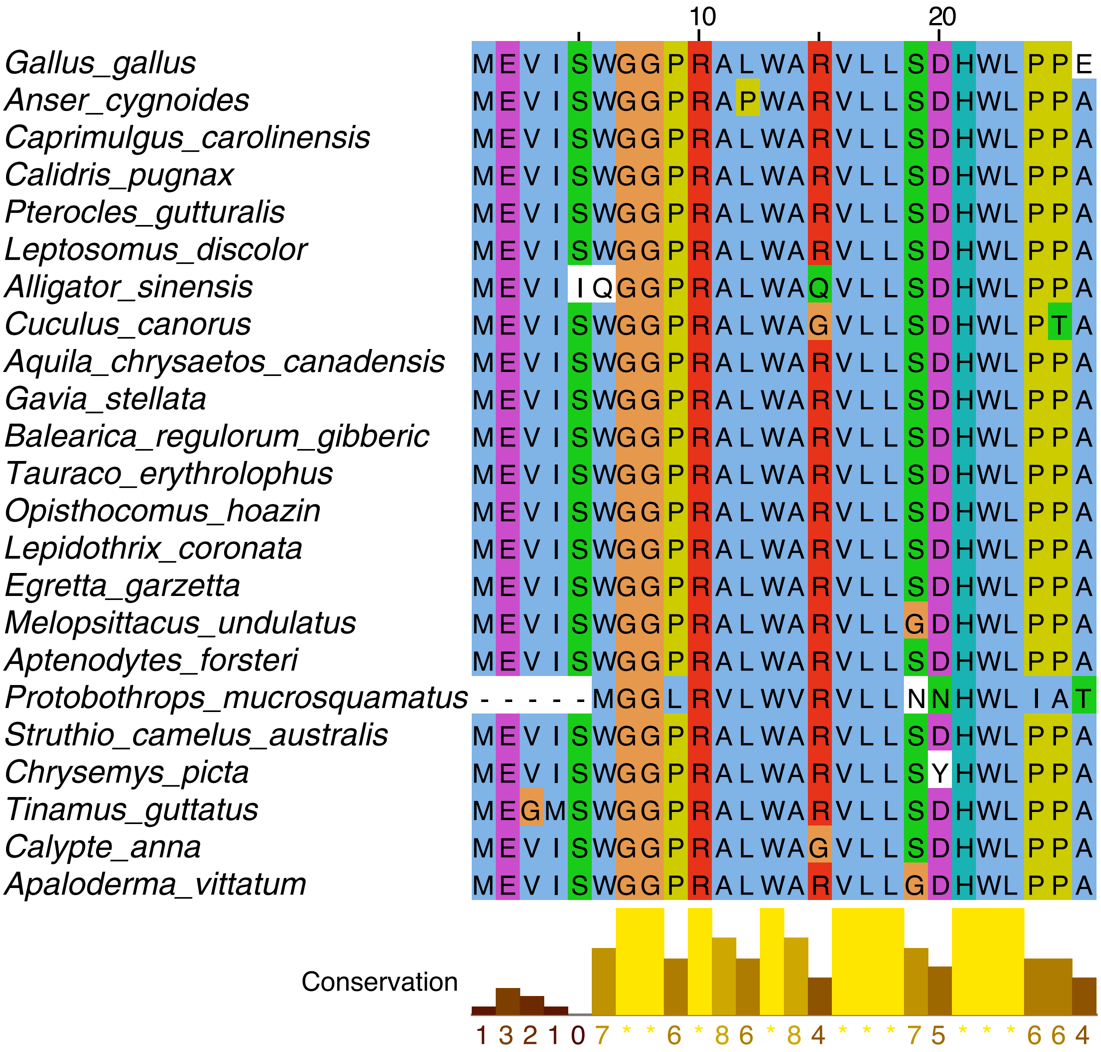

**HG0112**

ENSG00000105997 (HOXA3)

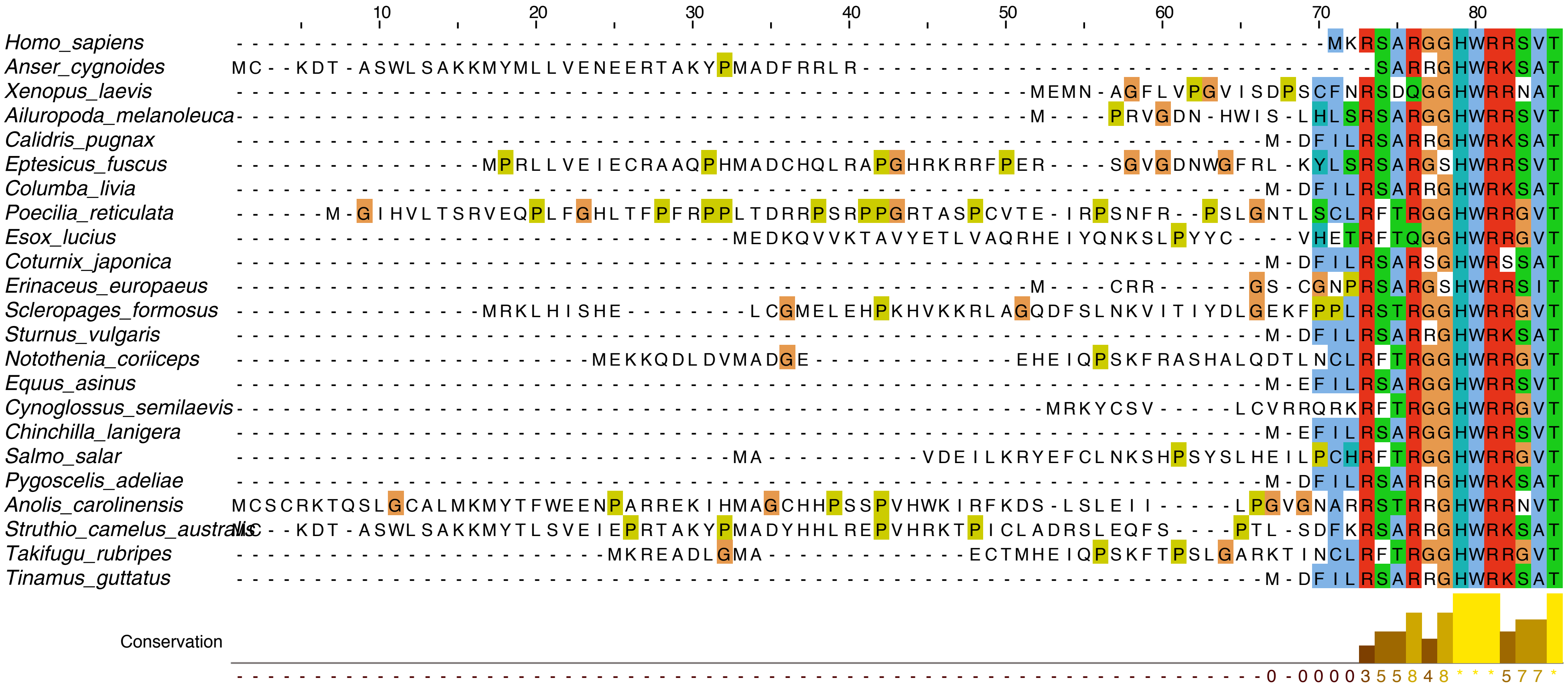

**HG0113**

ENSDARG00000099727 (ssbp2)

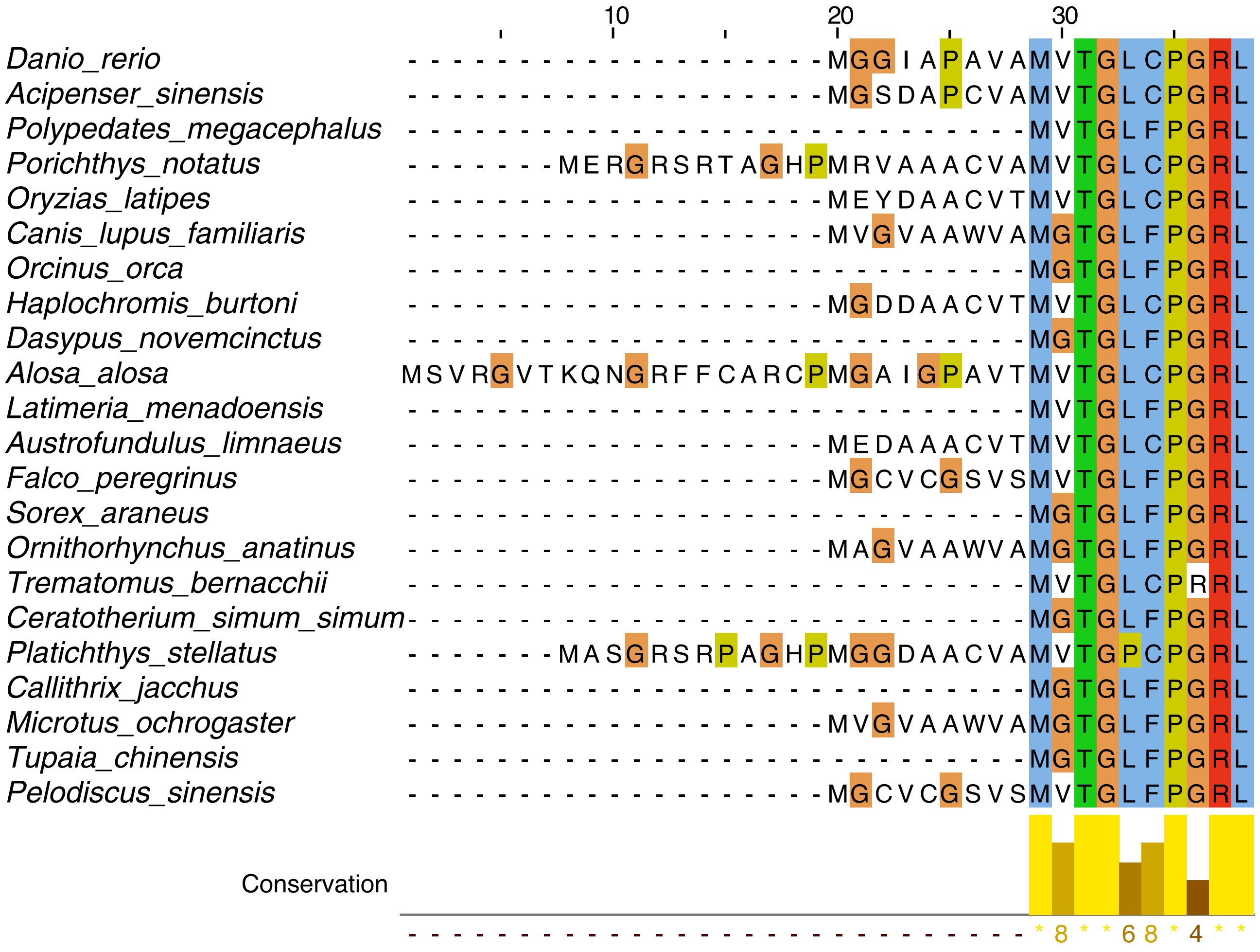

**HG0113**

ENSGALG00000023089 (SSBP2)

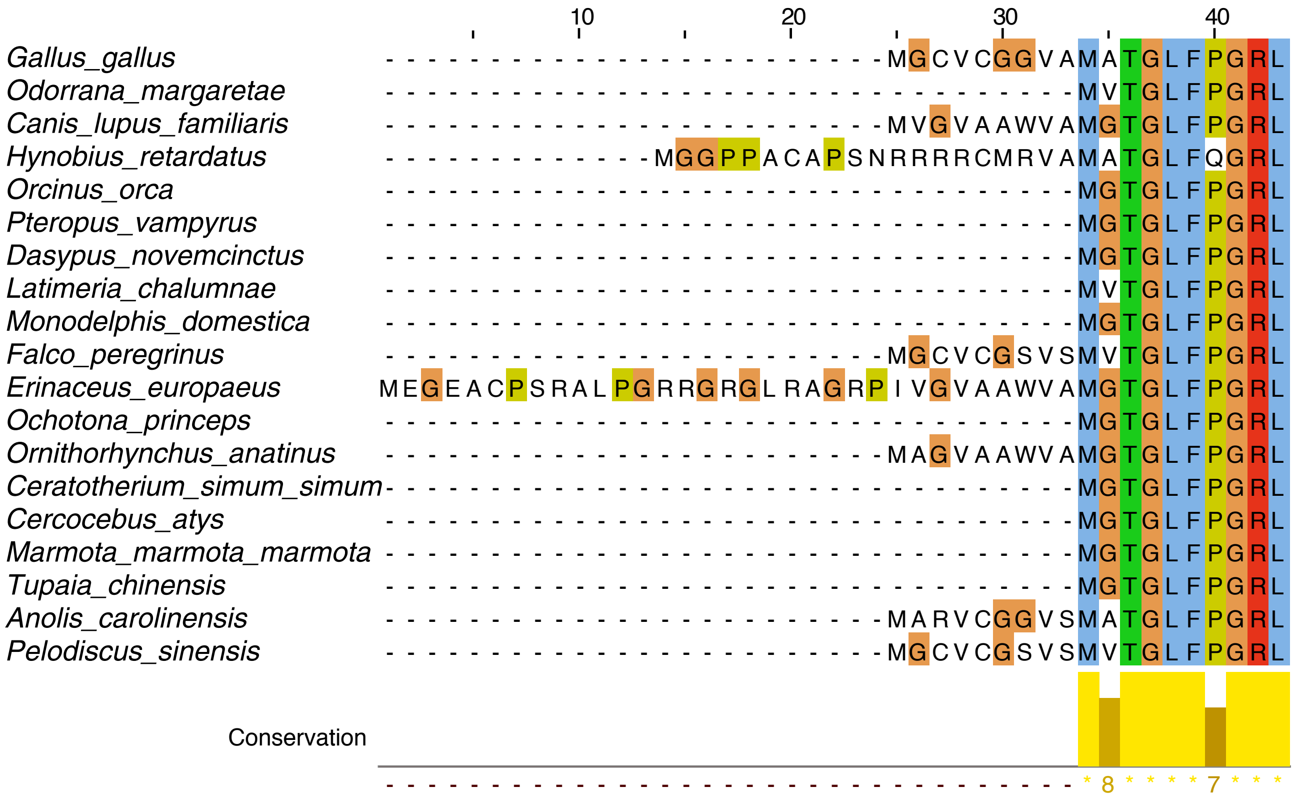

**HG0114.1**

ENSGALG00000003701 (SGMS1)

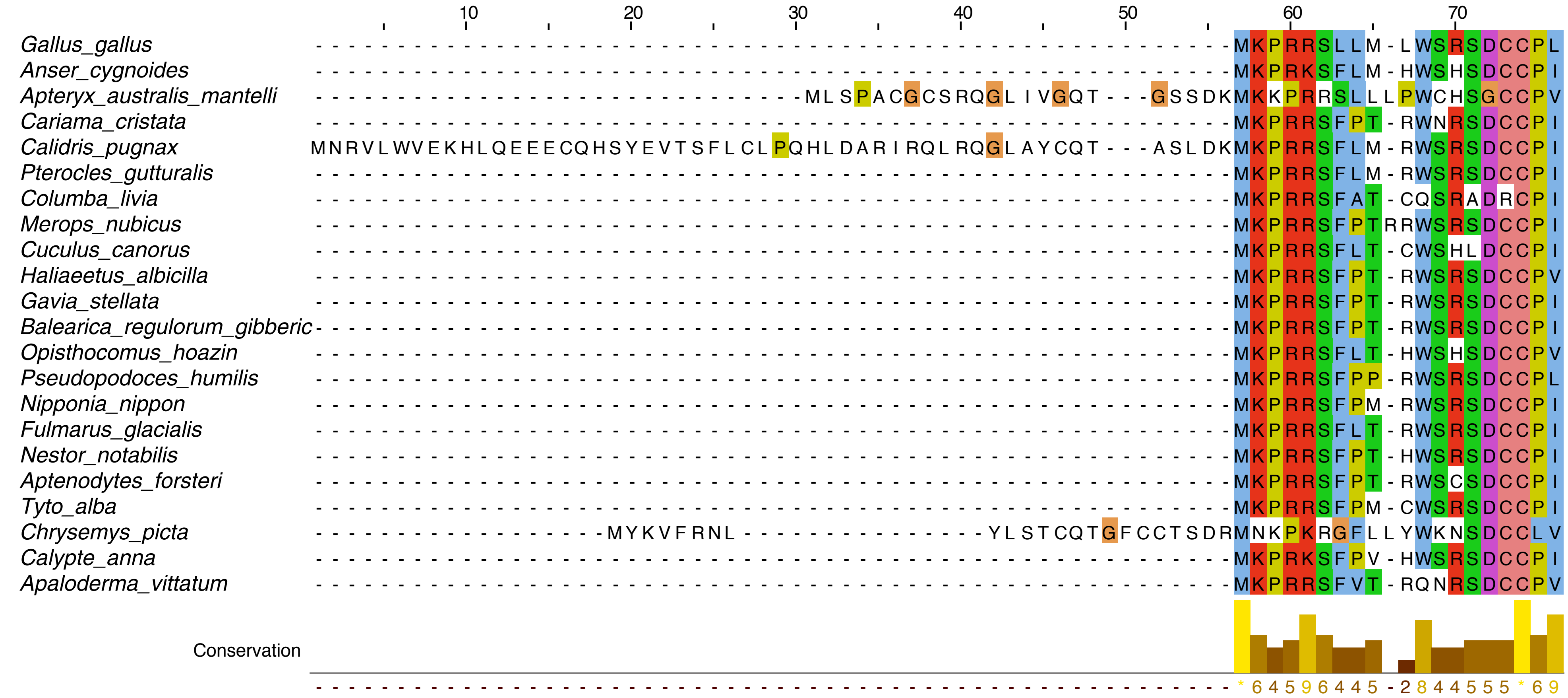

**HG0114.2**

ENSG00000198964 (SGMS1)

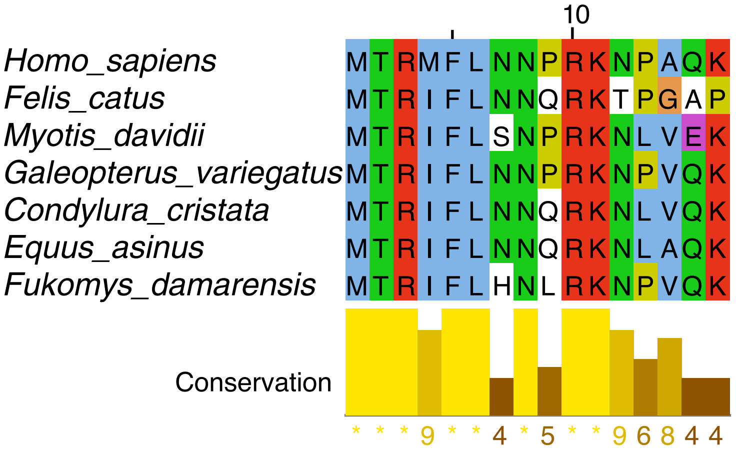

**HG0115.1**

ENSGALG00000010976 (CCDC126)

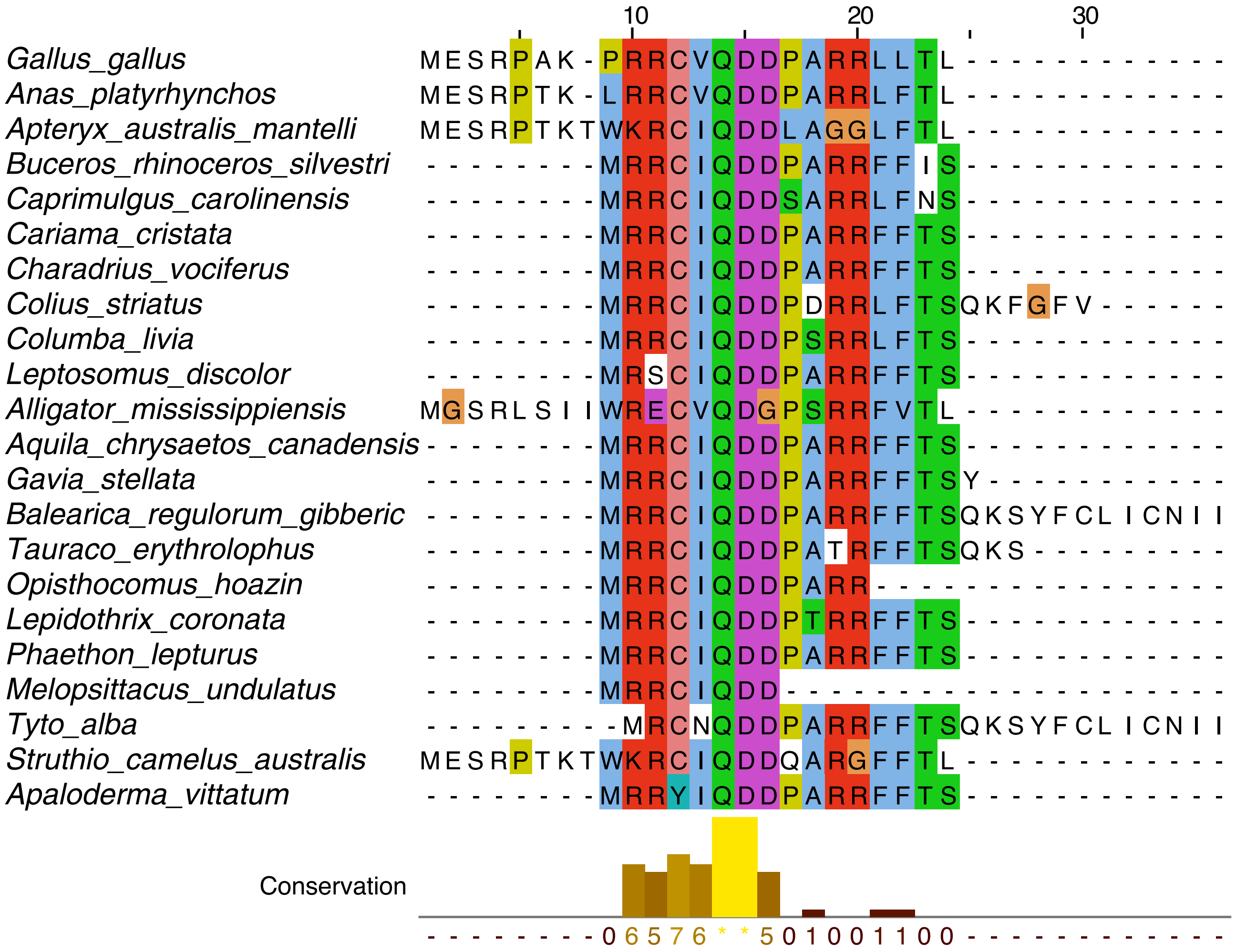

**HG0115.2**

ENSG00000169193 (CCDC126)

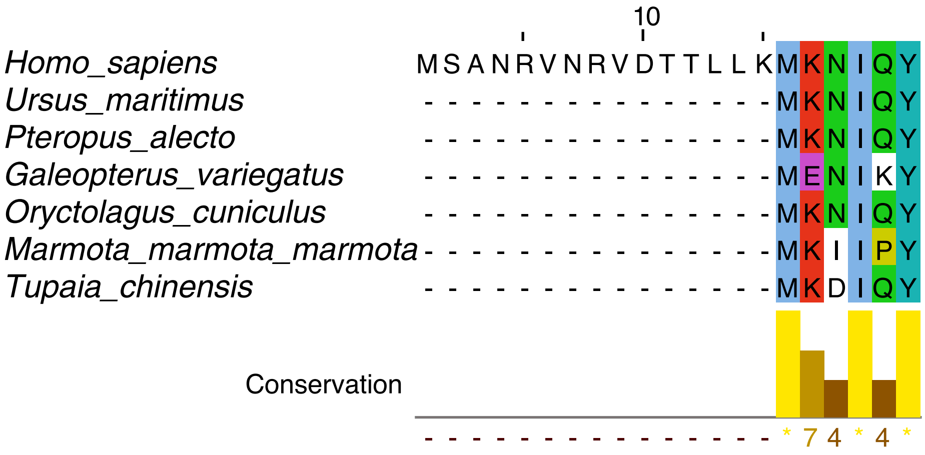

**HG0116**

ENSG00000010270 (STARD3NL)

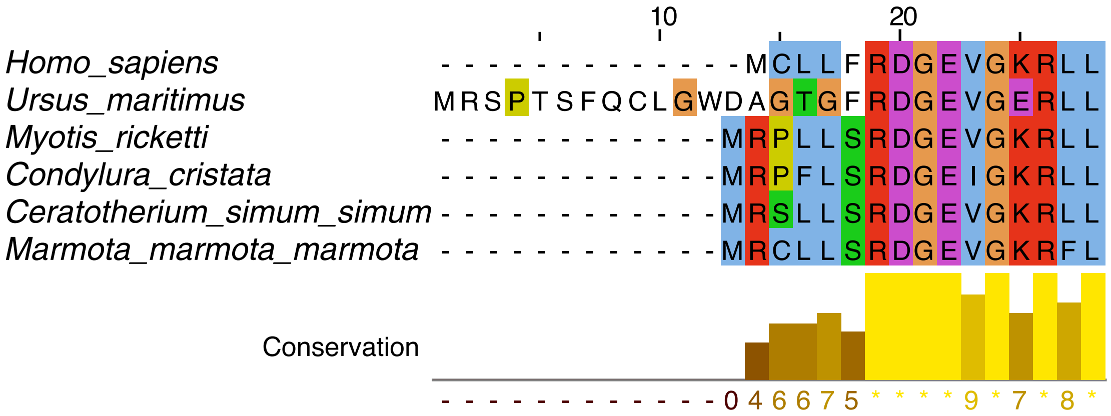

## HG0116

ENSGALG00000033038

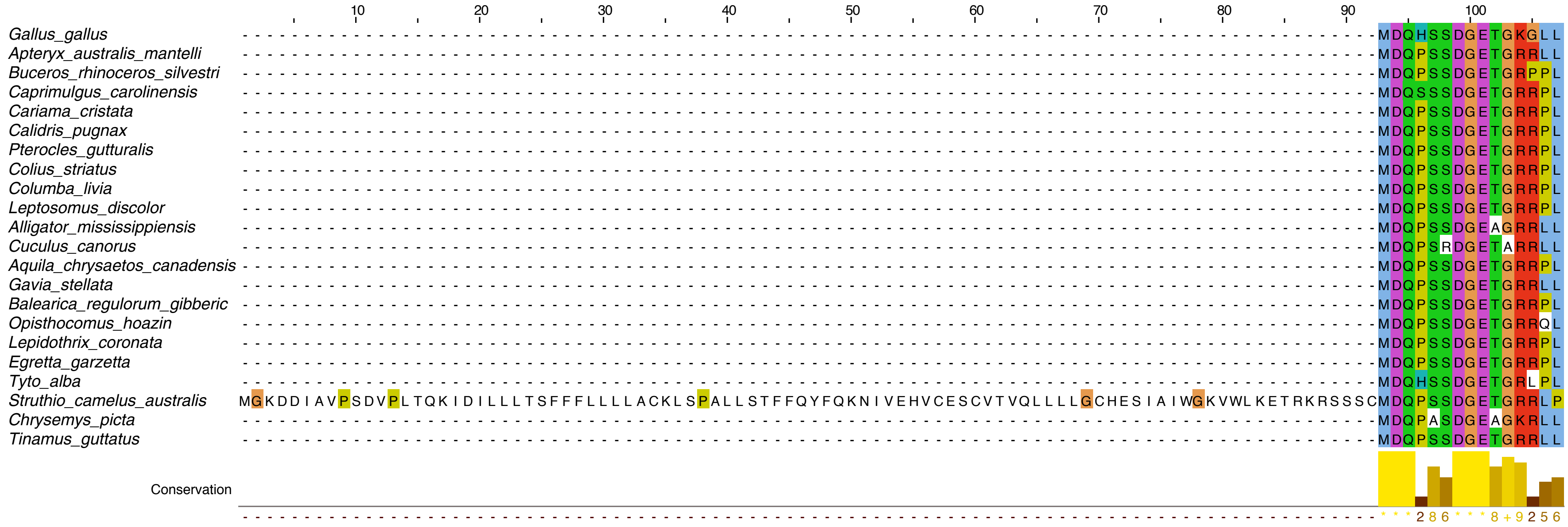

**HG0117.1**

ENSG00000140945 (CDH13)

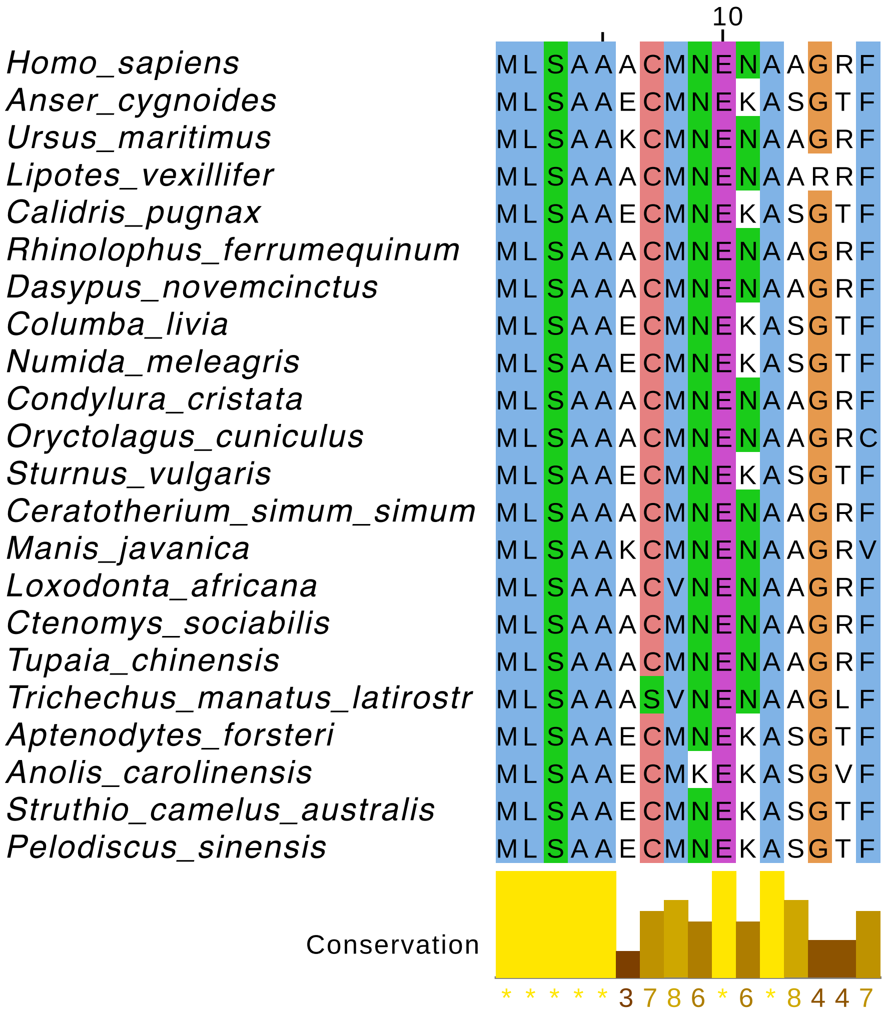

**HG0117.2**

ENSG00000140945 (CDH13)

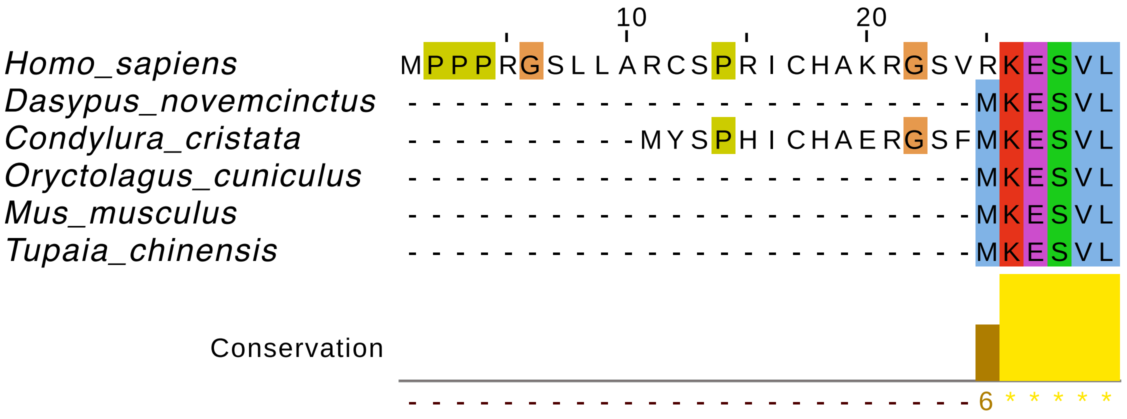

**HG0118.1**

ENSG00000183662 (FAM19A1)

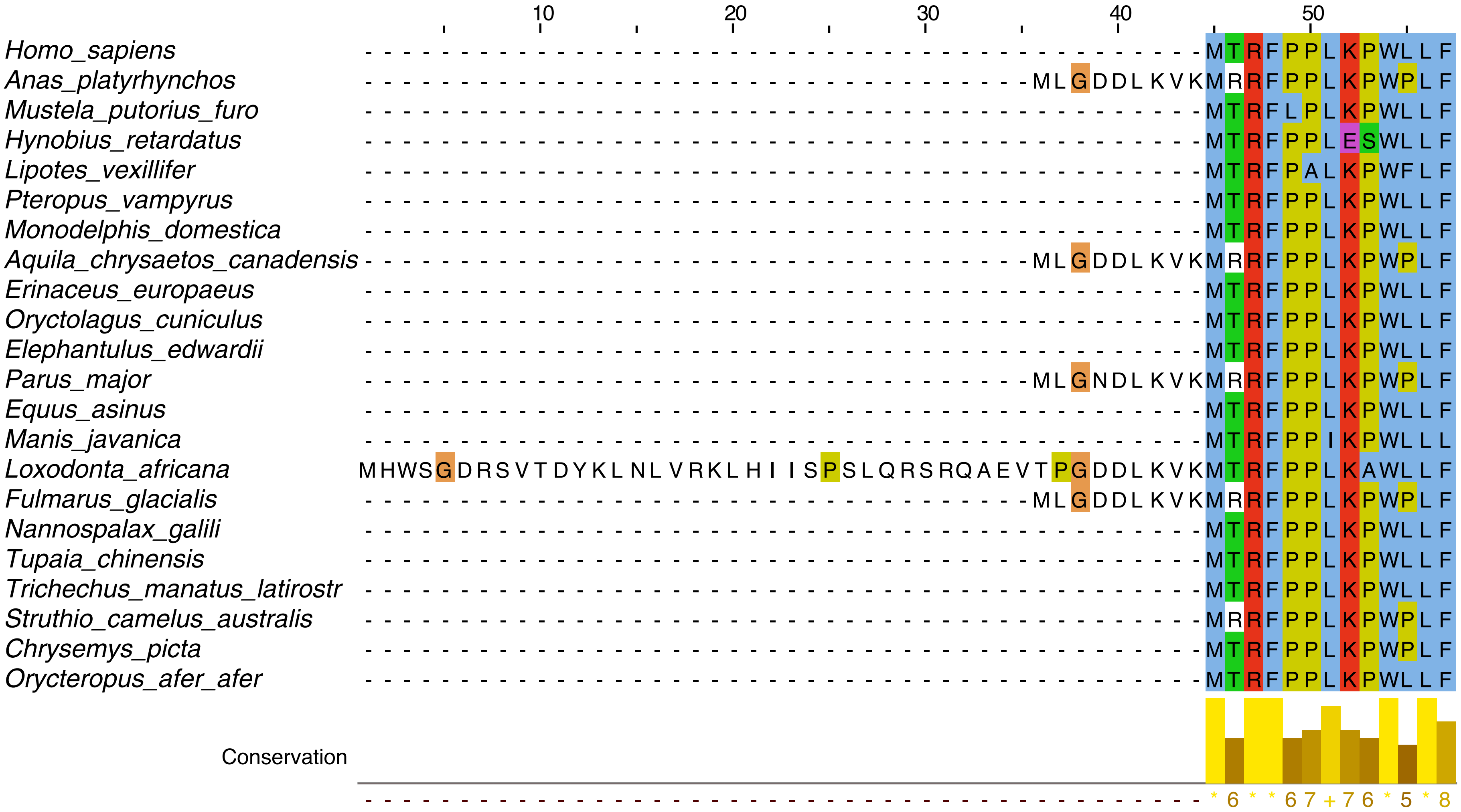

**HG0118.2**

ENSG00000183662 (FAM19A1)

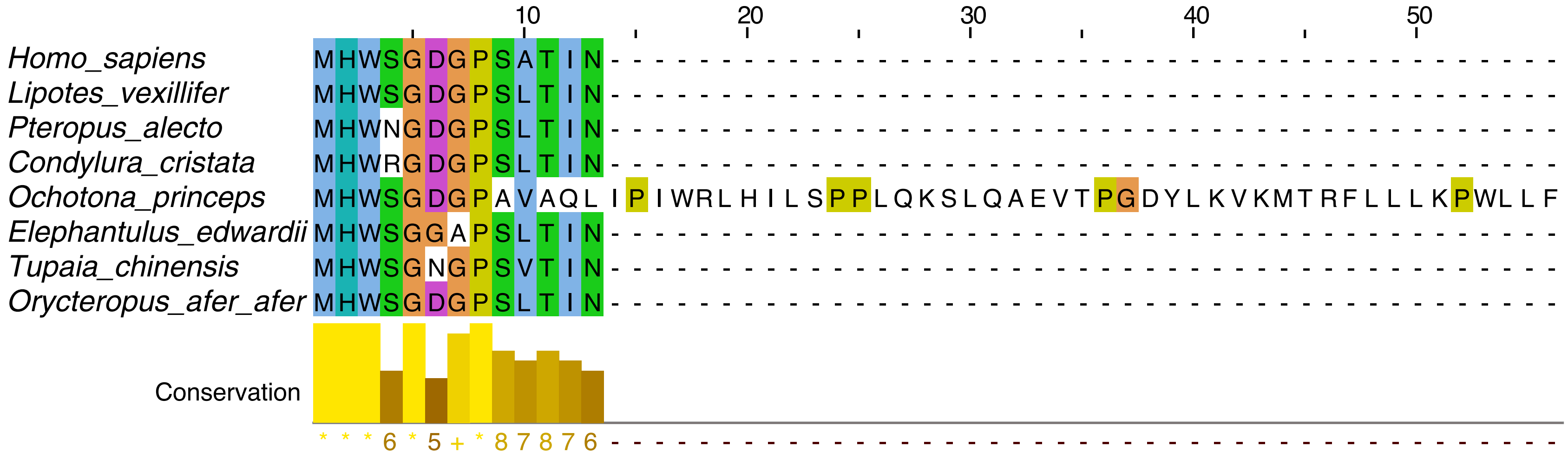

**HG0119**

ENSGALG00000043570 (SGK3)

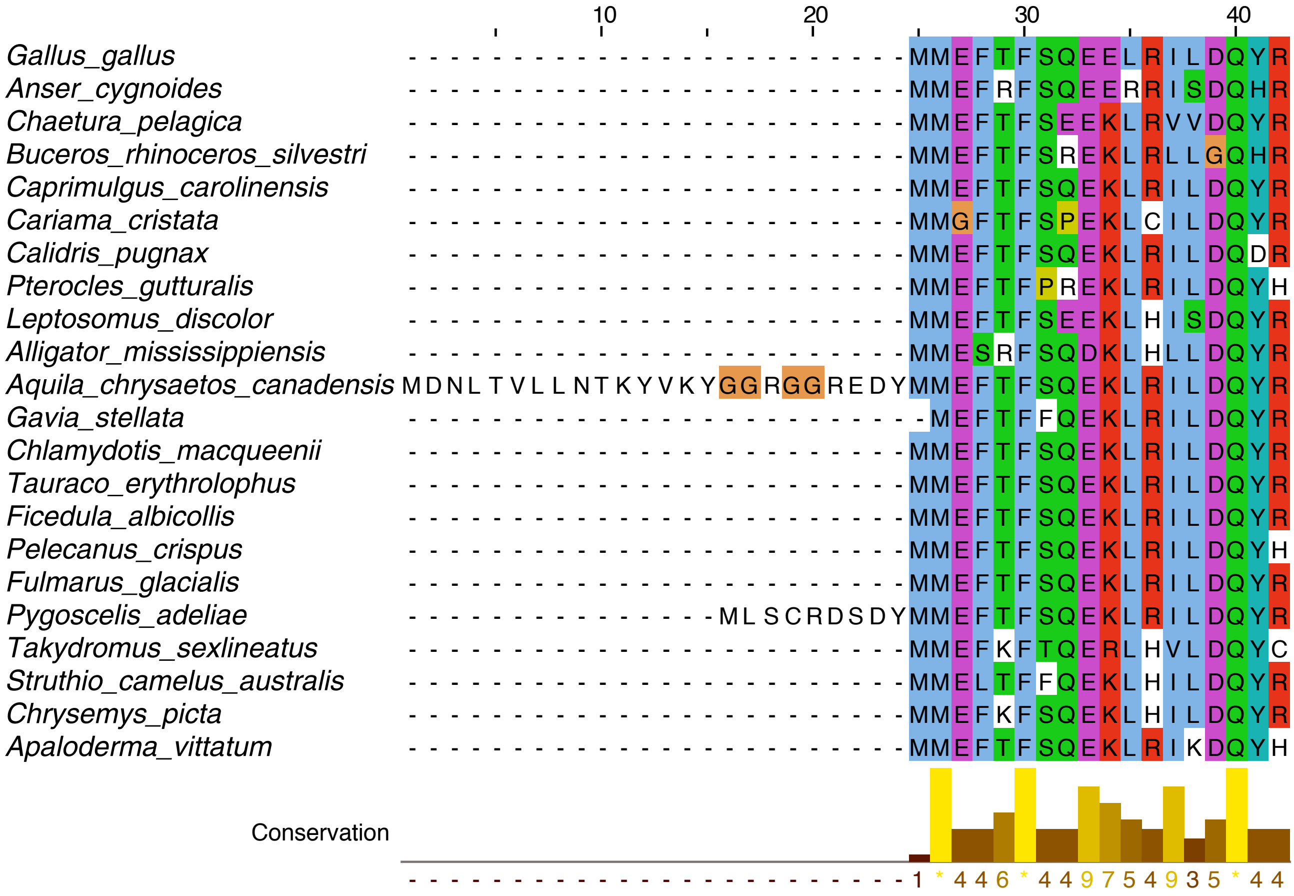

**HG0120**

ENSG00000022355 (GABRA1)

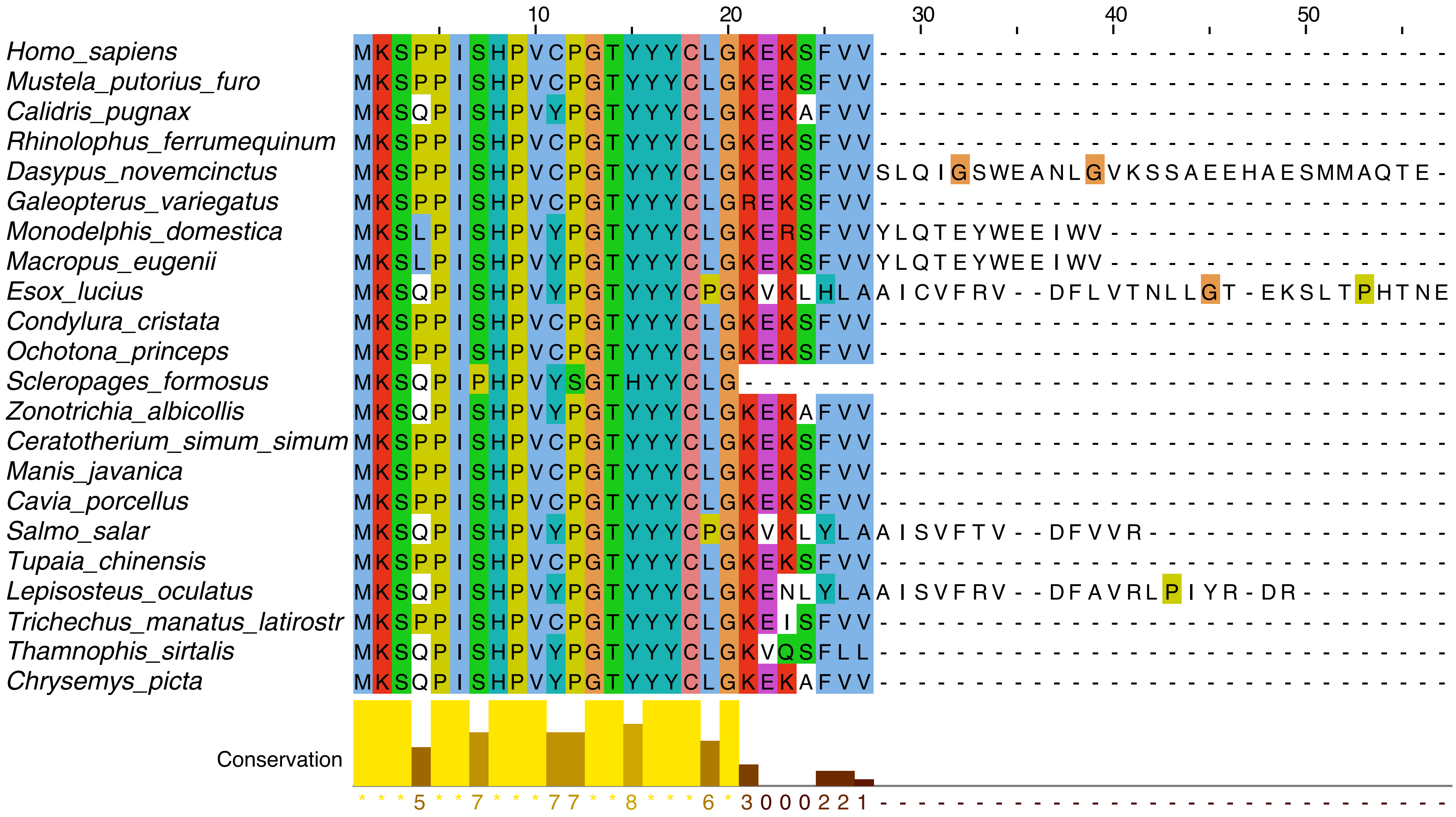

**HG0121**

ENSG00000122786 (CALD1)

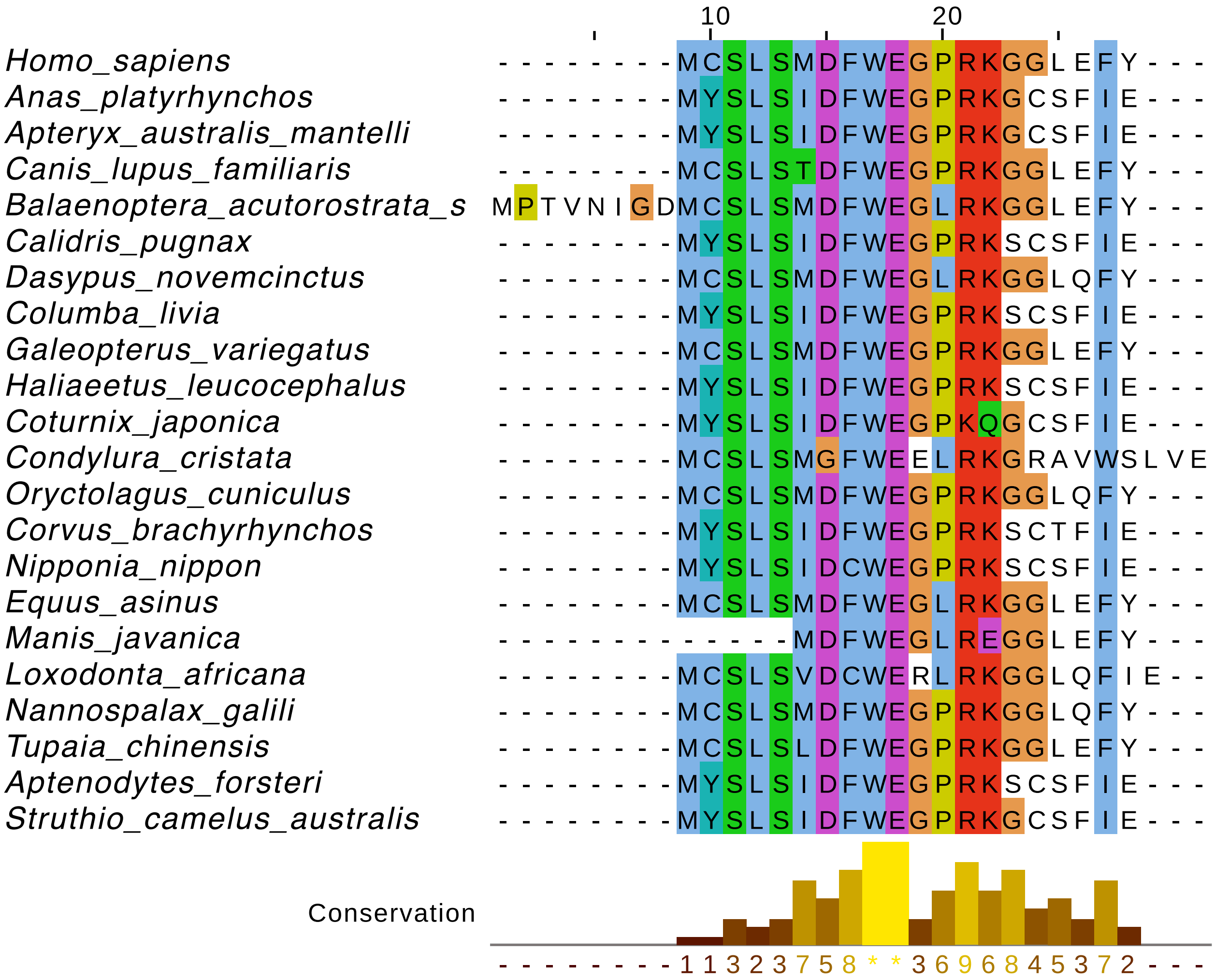

**HG0122**

ENSG00000169925 (BRD3)

```
                                       10        20        30        40        50        60        70        80        90        100
Homo_sapiens                  -----------------------------MLSCDRPSSCLEEAGLQPGSL---------PLCPD-HPHIPKAVAWLGPEMGRRAFSGSVEAARASPSRCCDRK-
Felis_catus                   -------------------------------------------------------------------------------MGRRASERSTVEARASPSRCCDGQ-
Balaenoptera_acutorostrata_s  -------------------------------------------------------------------------------MGCRTSAGSTAEARASPSCCCRDGQ
Calidris_pugnax               -------------------------------------------------------------------------------MGCQSRSGSIKEERKSPSRCCRDSE
Pteropus_alecto               -------------------------------------------------------------------------------MGCHASAGSTEEARAPPSRCCDGQ-
Cuculus_canorus               -------------------------------------------------------------------------------MGCQSRSGNIKEERKSPSRCCCDSE
Sarcophilus_harrisii          -------------------------------------------------------------------------------MGCQSYSGSIKKERESPSCCCDSK-
Galeopterus_variegatus        MQMPRRQTCRAGTTASCLPDWLRASPCHSLLPSQTPEL-LLLVMIEPRSVPCSRAPAPHAVCGSAEVWGAAGVAQLGPEMGCHTSPGSIGAARASRSRCCDRK-
Monodelphis_domestica         -------------------------------------------------------------------------------MGCQSYSGSIKKERESPSCCCDSK-
Macropus_eugenii              -------------------------------------------------------------------------------MGCQSYSGSIKKERESPSCCCDSK-
Balearica_regulorum_gibberic  -------------------------------------------------------------------------------MGCQSHSGSIKEERKSPSCCCCDRE
Erinaceus_europaeus           -------------------------------------------------------------------------------MGCHAPKGSTEHISTSLSRCCDN--
Oryctolagus_cuniculus         -------------------------------------------------------------------------------MGRRASLGSVDAARAPARRRCCGRK
Ornithorhynchus_anatinus      -------------------------------------------------------------------------------MGCQSYSGSIKQVKESPSCCCDDSE
Equus_caballus                -------------------------------------------------------------------------------MGRRASRGSTEELRASPSRCCDGQ-
Manis_javanica                -------------------------------------------------------------------------------MGCHASAGSTEETRASLSRCCDGQ-
Loxodonta_africana            -------------------------------------------------------------------------------MGCRAASGGVEGERAPAGCGCDRK-
Ictidomys_tridecemlineatus    -------------------------------------------------------------------------------MGCQASSGSTEGAGAFWSCSCDGK-
Tupaia_chinensis              -------------------------------------------------------------------------------MGCRGPSGSIAAARAPASPCRDGK-
Trichechus_manatus_latirostr  -------------------------------------------------------------------------------MGCRAASGSLEGARASAGRGCDGK-
Chelonia_mydas                -------------------------------------------------------------------------------MGCQSHSGSIKEDRESPSCCCDSK-
Orycteropus_afer_afer         -------------------------------------------------------------------------------MGCRASSGIVEGTWASLGRGCDE--

Conservation                  -------------------------------------------------------------------------------**5672145722333414445430-
```

**HG0123**

ENSG00000173276 (ZBTB21)

```
                                   10        20
Homo_sapiens               MTSSLVWDDVKFKMDGITAAAAAAARASRV-
Amia_calva                 -------------MDGITTATRSTEF-----
Channa_argus               MTSSLVWDDVKFKMDGITTSAQKSTQH----
Xenopus_laevis             MTSSVVWDDVKFKMDGITAAAARQRLAE---
Menidia_menidia            -------------MDGITISSQKFTQH----
Oryzias_latipes            -------------MDGITTKTQKSCTTVNKV
Hynobius_retardatus        -------------MDGITAAARAGQF-----
Myotis_lucifugus           MTSSLVWDDVKFKMDGITAAAAAAARASRV-
Oreochromis_niloticus      MTSSLVWDDVKFKMDGITTSAEKSTQH----
Pimephales_promelas        MTSSLEWDDVKYKMDGITTASCSTQL-----
Fundulus_heteroclitus      MTSSLVWDDVKFKMDGITTSRLKATQH----
Galeopterus_variegatus     MTSSLVWDDVKFKMDGITAAAAAVARASRV-
Esox_lucius                -------------MDGITTTTHSTEF-----
Coturnix_japonica          MTSSLVWDDVKFKMDGITAAAAARAAEP---
Lateolabrax_japonicus      MTSSLVWDDVKFKMDGITTSAQKSTQH----
Trematomus_bernacchii      ----MVWDDVKFKMDGITTSAQKSTQQH---
Platichthys_stellatus      MTSSLVWDDVKFKMDGITTSAQKAPQQ----
Heterocephalus_glaber      MTSSLVWDDVKFKMDGITAAAAAAARASRV-
Salmo_salar                MTSSLE-------------------------
Thamnophis_sirtalis        MTSSLVWDDVKFKMDGITAVAAARAGES---
Chrysemys_picta            MTSSLVWDDVKFKMDGITAVAAARAGES---
Takifugu_rubripes          MTSSLVWDDVKFKMDGITTAA----------
```

Conservation

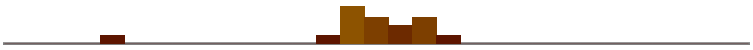

```
----10-------1432310000000-----
```

**HG0124.1**

ENSDARG00000031763 (smad6b)

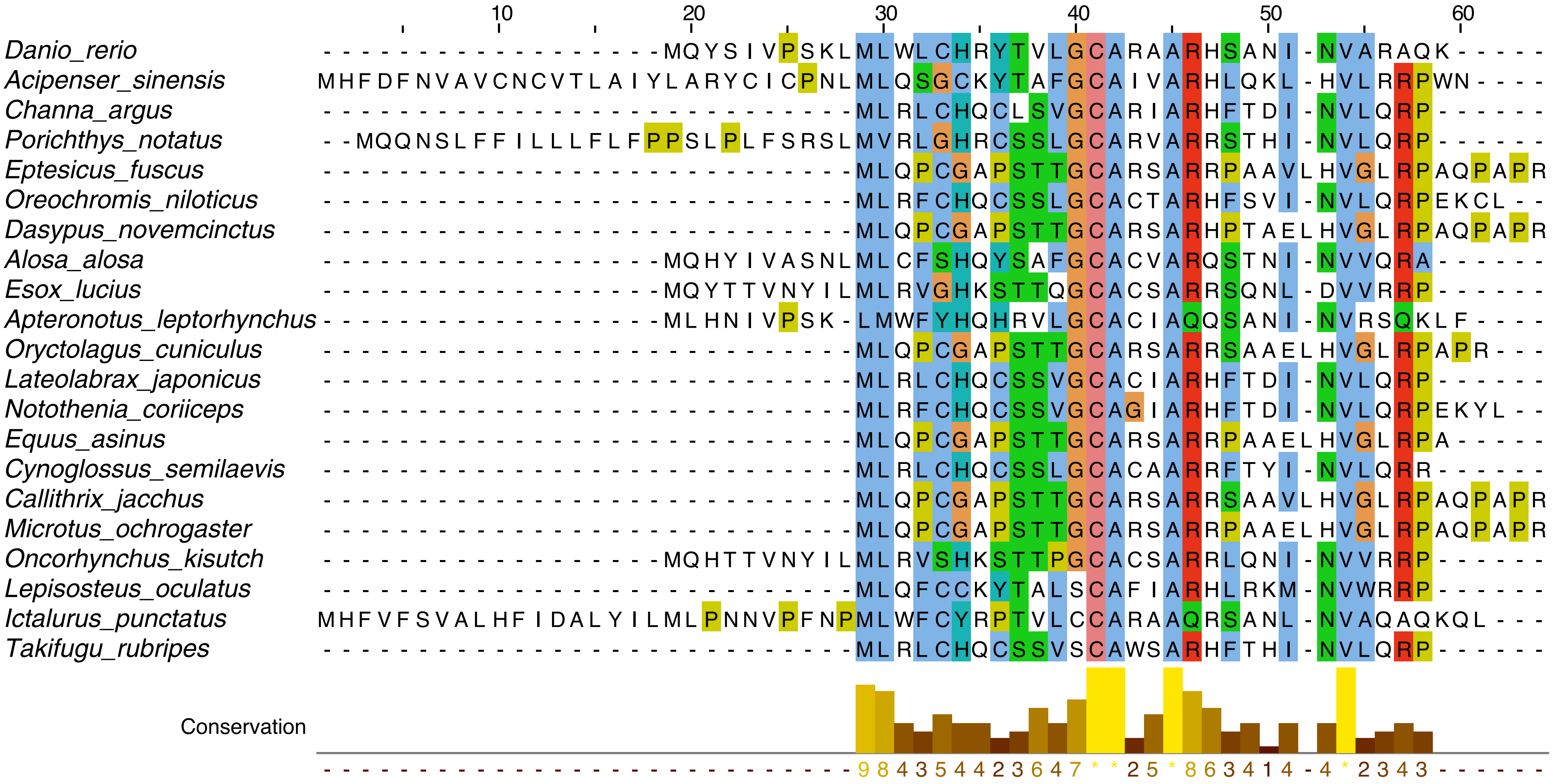

**HG0124.2**

ENSDARG00000053209 (smad6a)

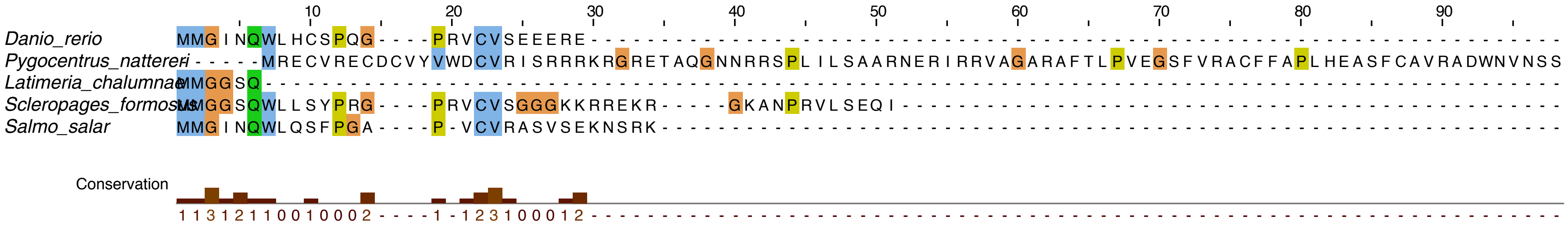

**HG0125.1**

ENSDARG00000051926 (plag1)

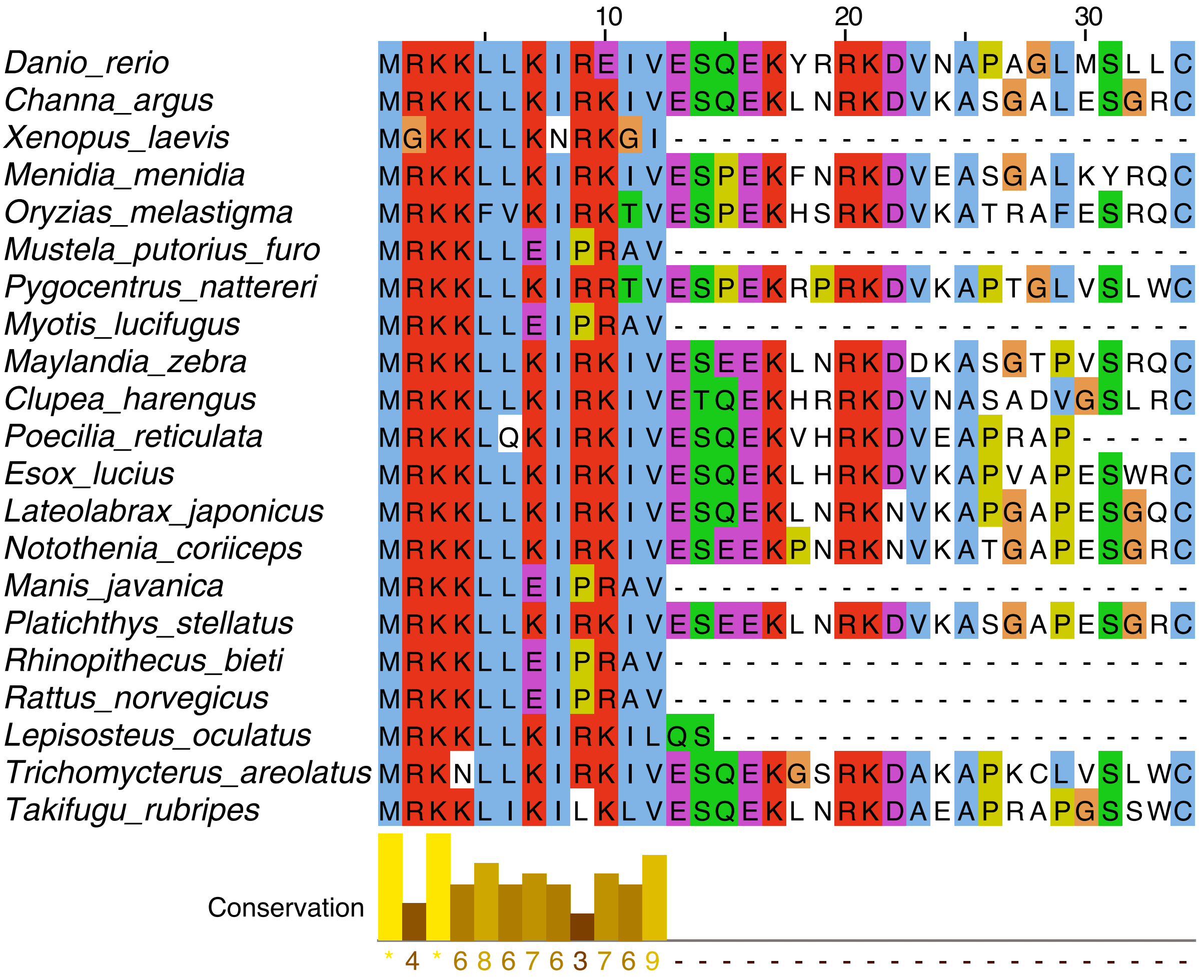

**HG0125.2**

ENSG00000181690 (PLAG1)

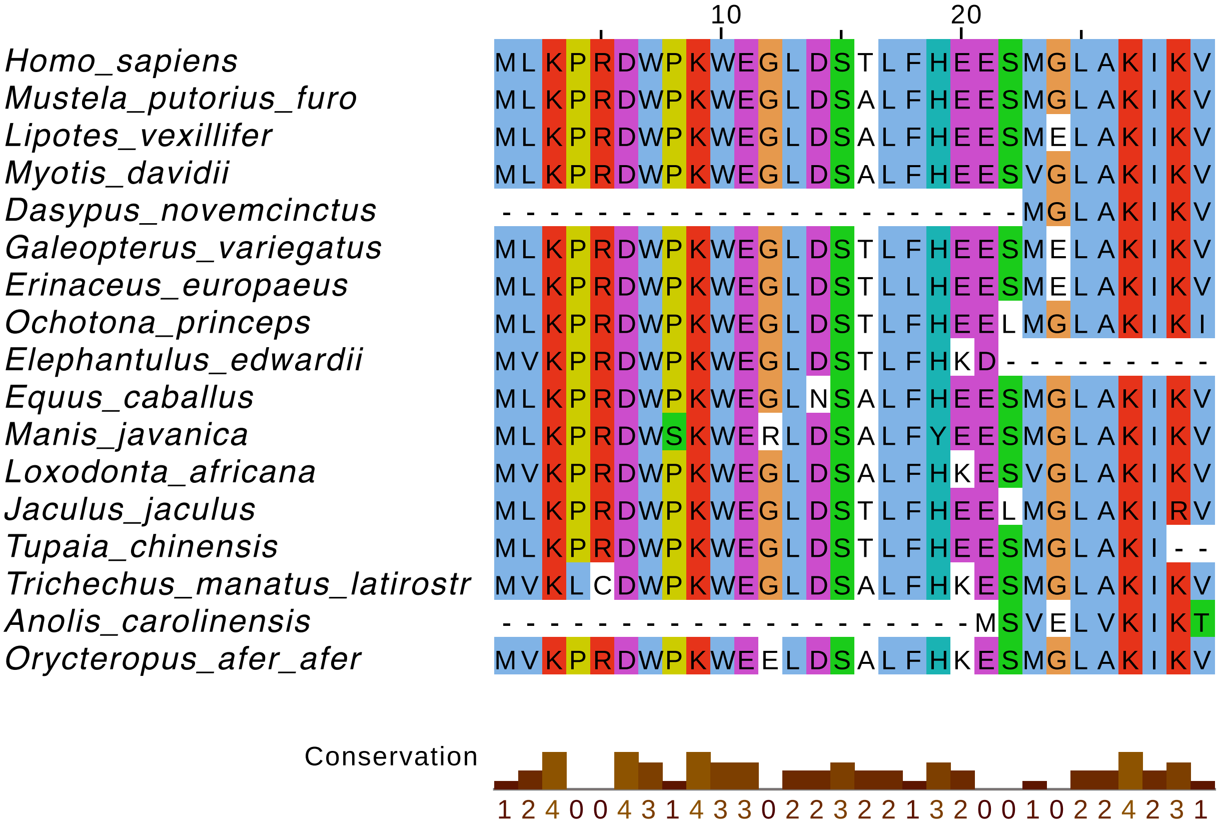

**HG0126.1**

ENSGALG00000010899 (G6PC2)

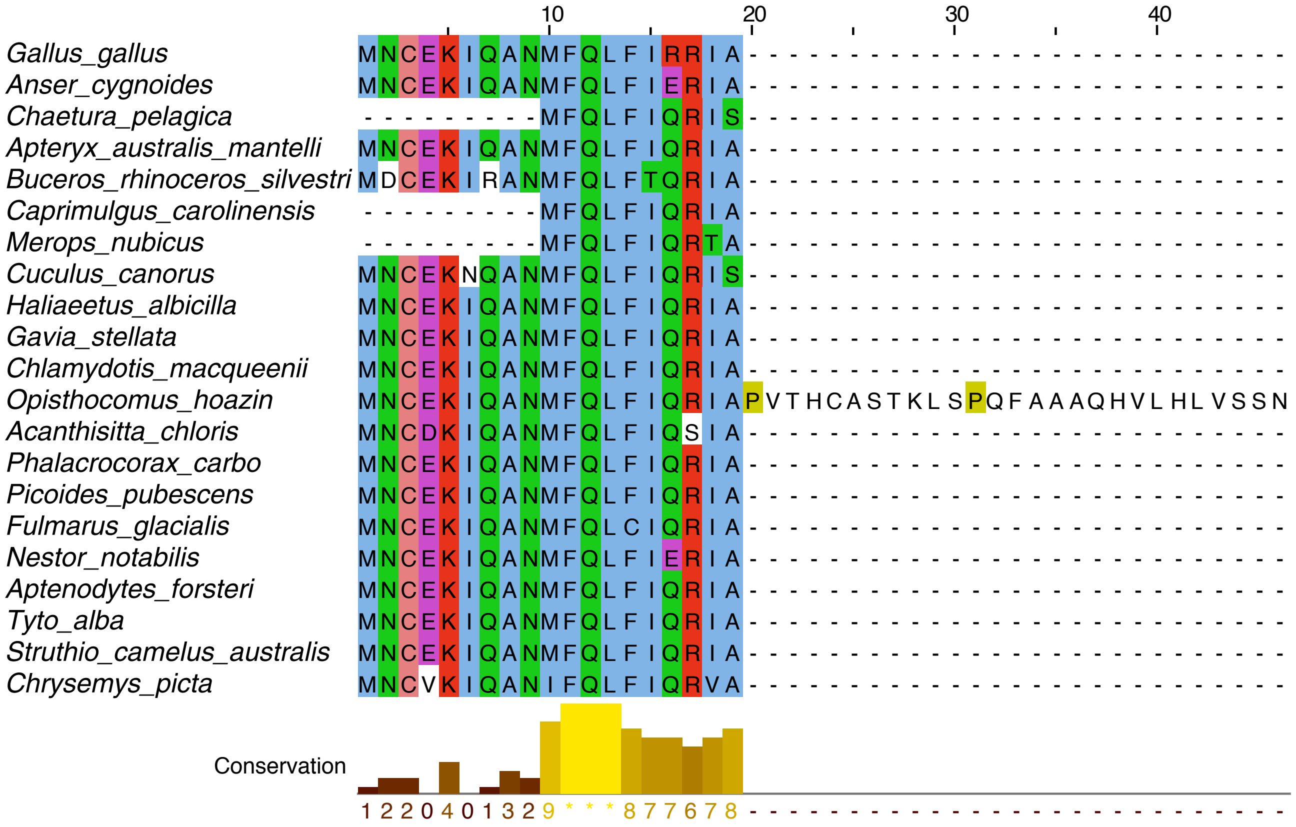

**HG0126.2**

ENSGALG00000010899 (G6PC2)

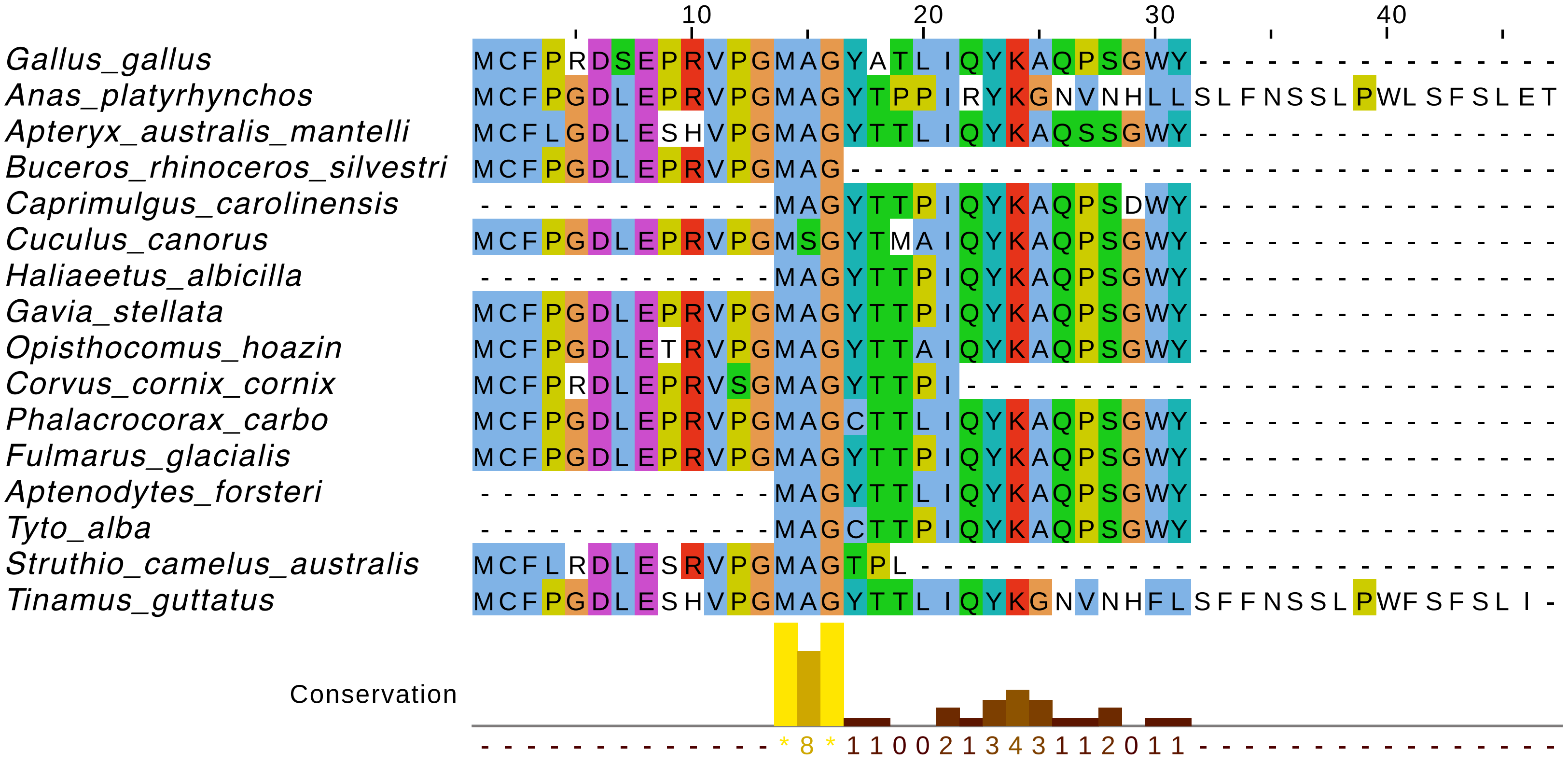

**HG0127.1**

ENSGALG00000029709

```
                                      10        20        30
Gallus_gallus                  MQNQSTH-EEKSPLRGNVRGIAVGDNLLHSPTP-----
Anas_platyrhynchos             MQNPSAH-EEKSPLHGNVRGIALGDNLLHSAAP-----
Buceros_rhinoceros_silvestri   MQNSSTH-EEKSPL--VVMSEE----------------
Caprimulgus_carolinensis       MQNPST--------------------------------
Cariama_cristata               MQNPSTH-EEPSHG--NVRGIAIGDNLLHSATL-----
Calidris_pugnax                MQNPSTH-EEKSPL--IVMLEE----------------
Colius_striatus                MQNPSTH-EEKSSL--VVMLEE----------------
Columba_livia                  -----------MISRGNVRGIAAGDNPLHSAAL-----
Falco_cherrug                  -----------MISRGNVRGIAVGDNPLHSATVQDKHR
Gavia_stellata                 MQNPSTH-EENSPL--VVMLEE----------------
Eurypyga_helias                MQNPSTH-EEKSPL--VVMLEE----------------
Corvus_brachyrhynchos          -----ME-DFEGERHGNVRGIAVGDNLLHSATL-----
Pelecanus_crispus              MQNPSTH-EEKSPL--VVMLEE----------------
Picoides_pubescens             MQNPSSH-EEKSPLL--VTLEE----------------
Fulmarus_glacialis             MQNPSTH-EEESPL--VVMLEE----------------
Pygoscelis_adeliae             MQNPSTH-EEKSPL--VVMLEE----------------
Tyto_alba                      MQNPSTH-EEKSPL--VVMLEE----------------
Struthio_camelus_australis     MQNLFTQ-EERS-----CLGIATGDNLLHSDPLQEKHH
Trachemys_scripta              MQNHTVSQEENSPHYA----------------------
Tinamus_guttatus               MQNPSTQ-EEKSPL------------------------
Apaloderma_vittatum            MQNPSTH-EEKSPL--VIMLKE----------------

Conservation                   1120001-300000--010000----------------
```

**HG0127.2**

ENSG00000109452 (INPP4B)

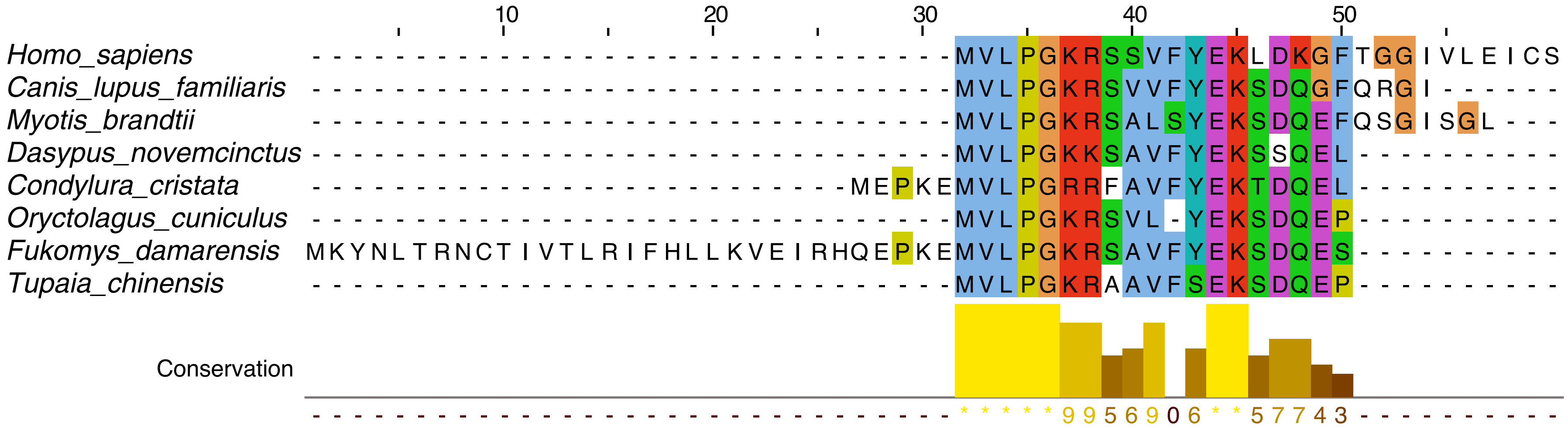

**HG0128**

ENSDARG00000091029 (phox2bb)

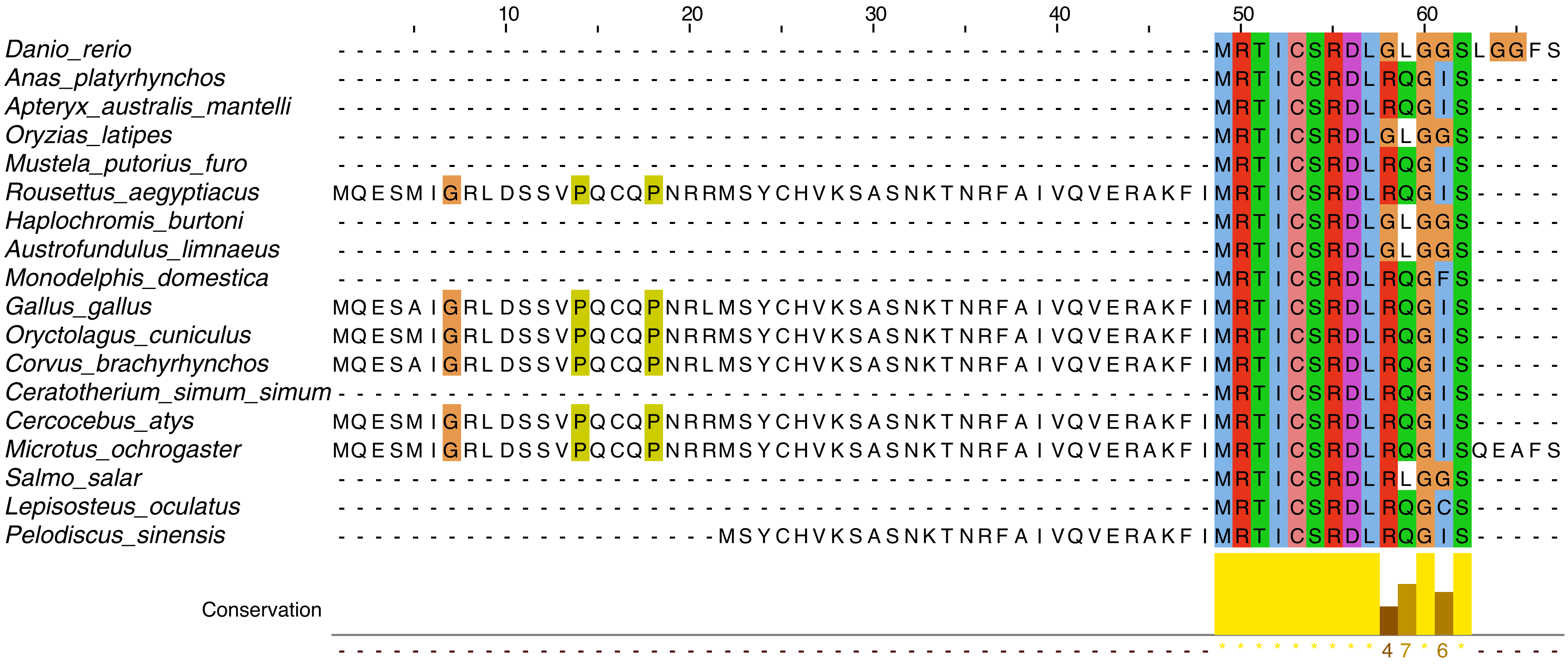

**HG0128**

ENSG00000109132 (PHOX2B)

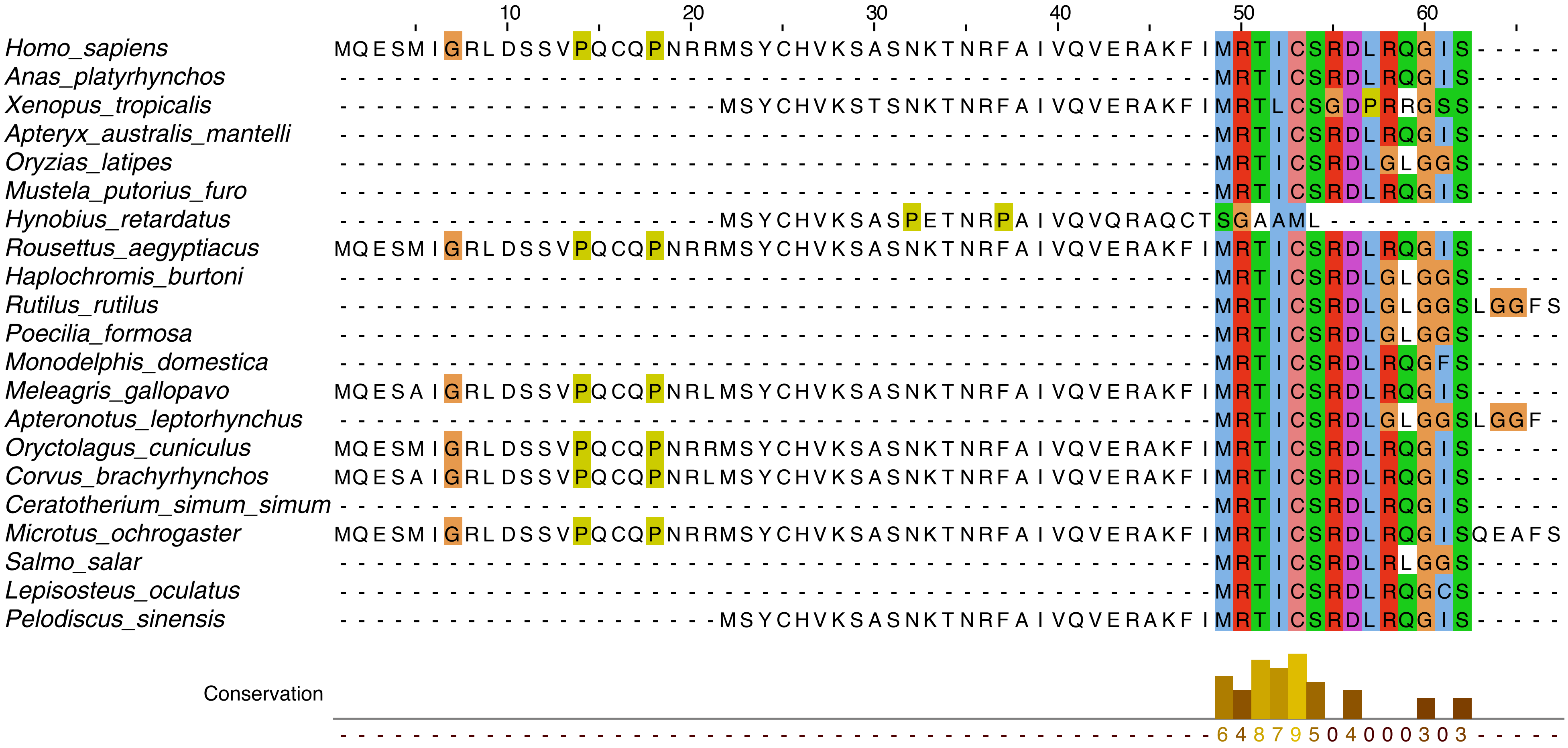

**HG0129**

ENSDARG00000099437

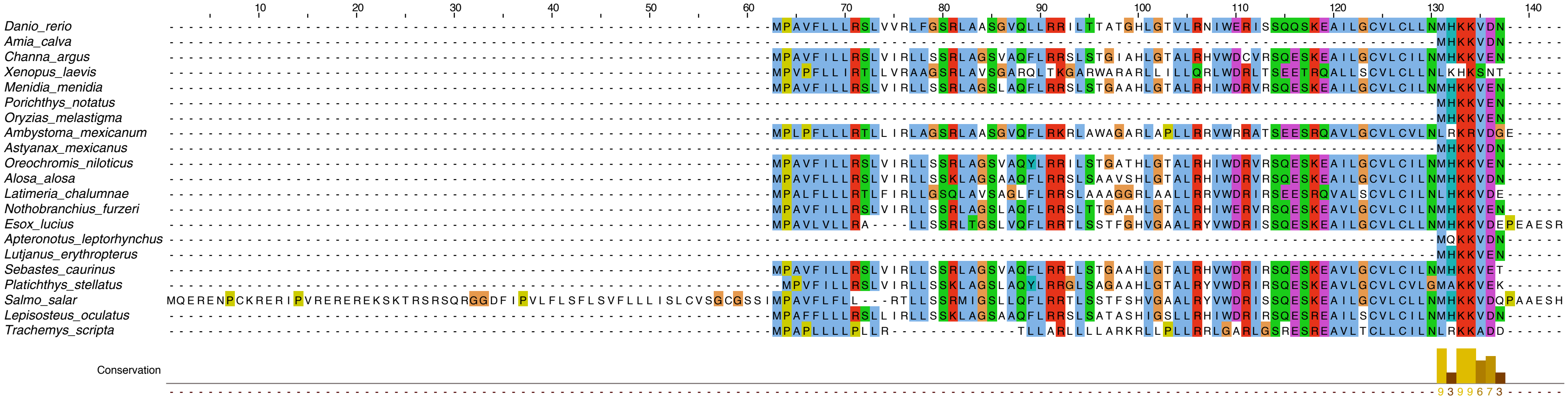

# HG0130

### ENSGALG00000002993

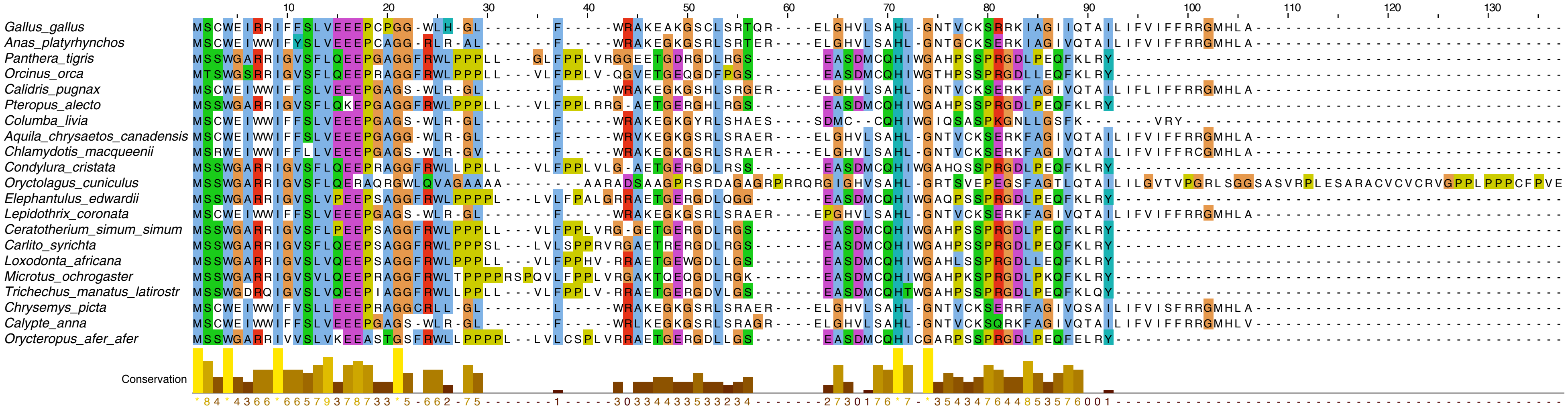

**HG0131**

ENSGALG00000013135 (GALNT1)

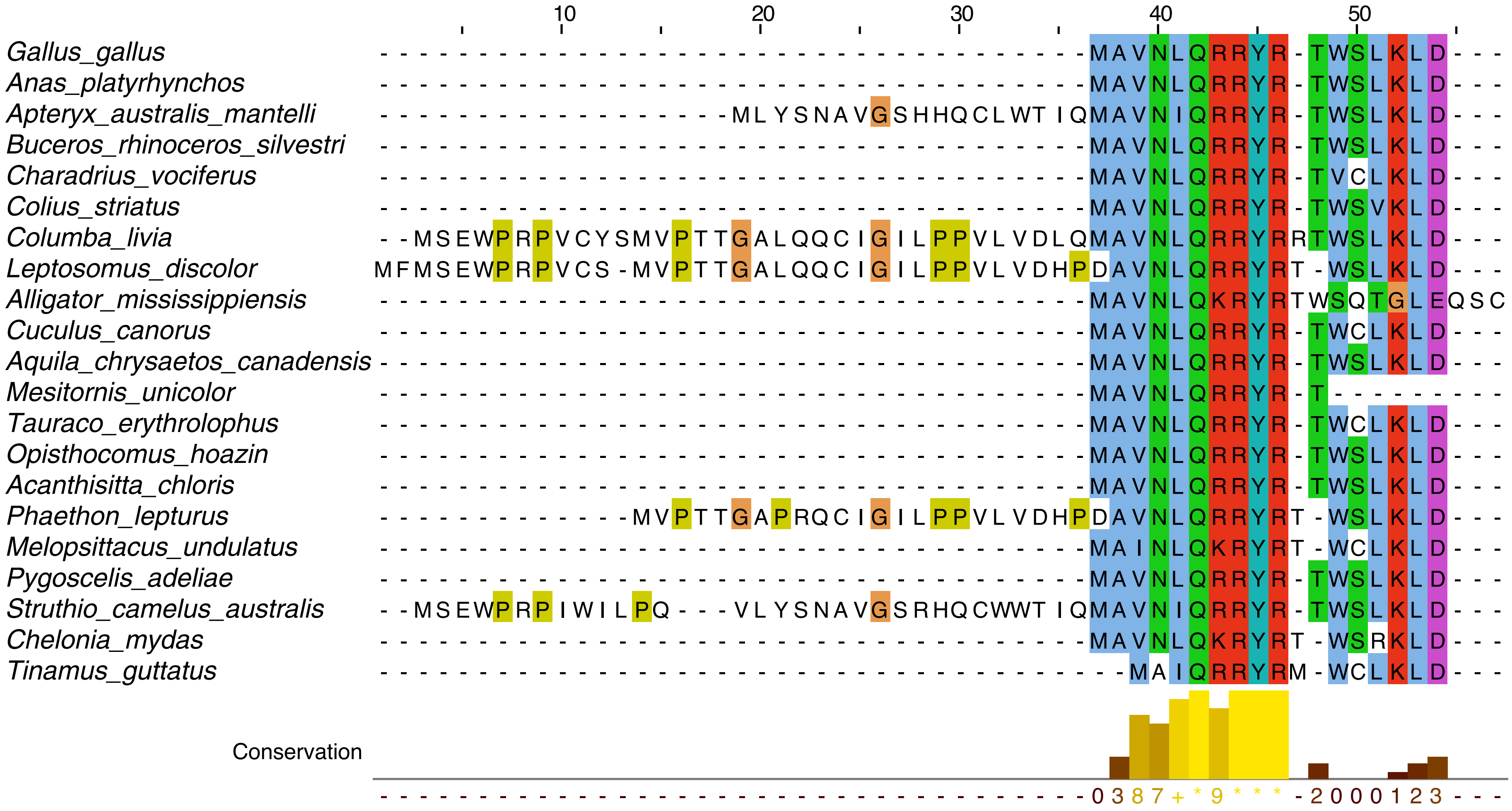

**HG0132**

ENSG00000151067 (CACNA1C)

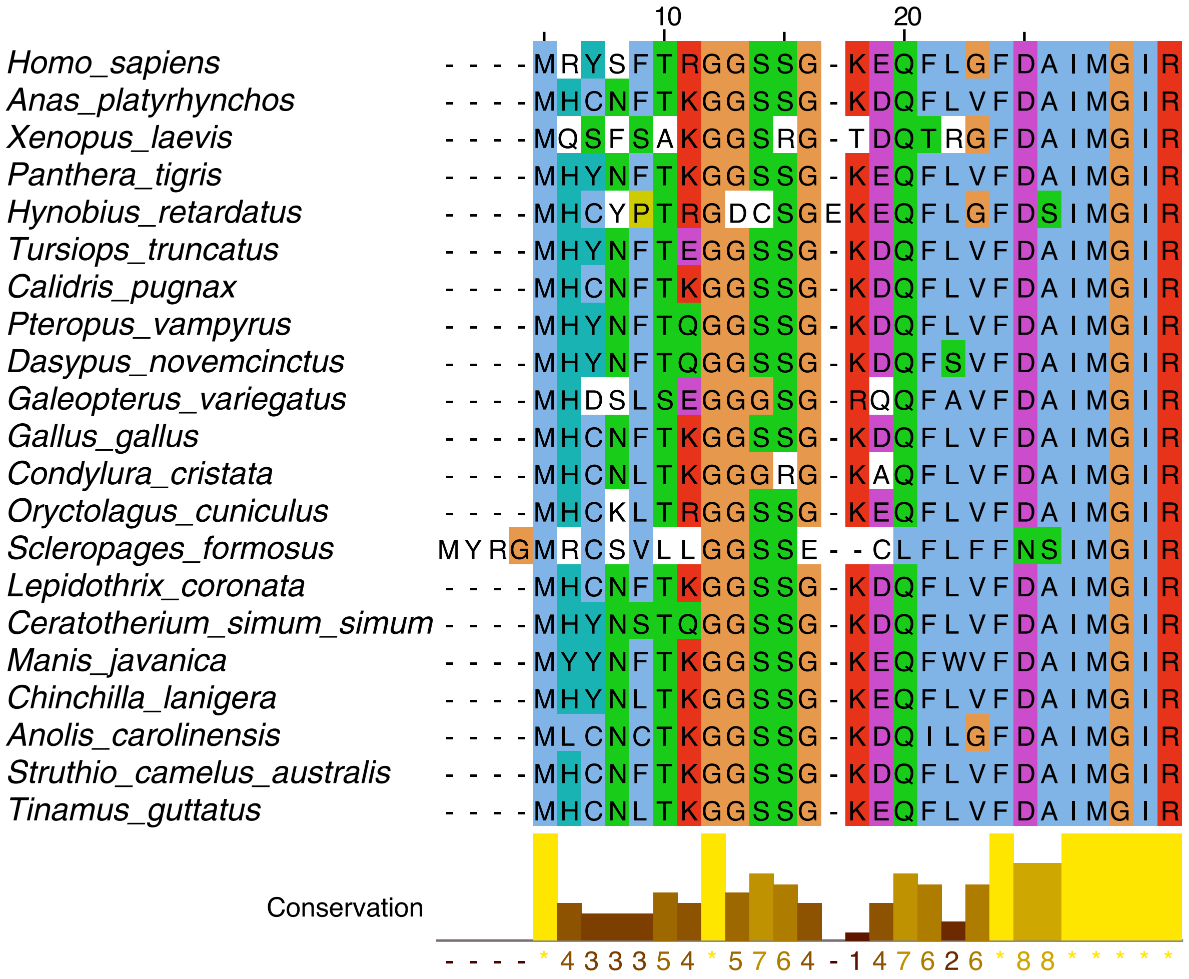

**HG0133.1**

ENSDARG00000051886 (ankrd11)

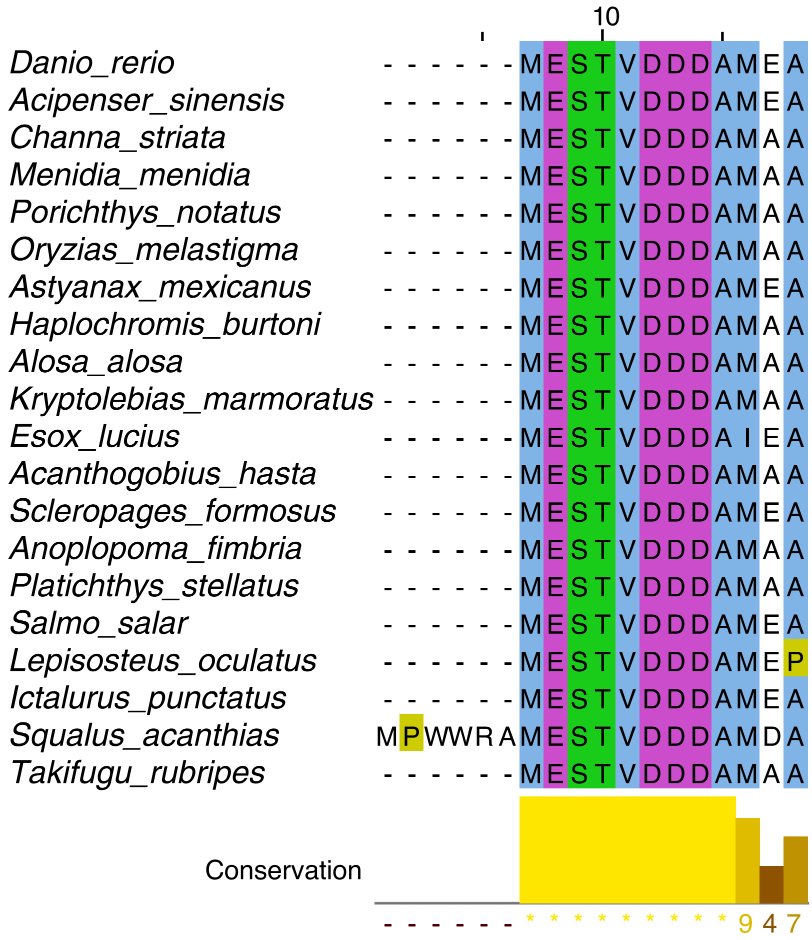

**HG0133.1**

ENSG00000167522 (ANKRD11)

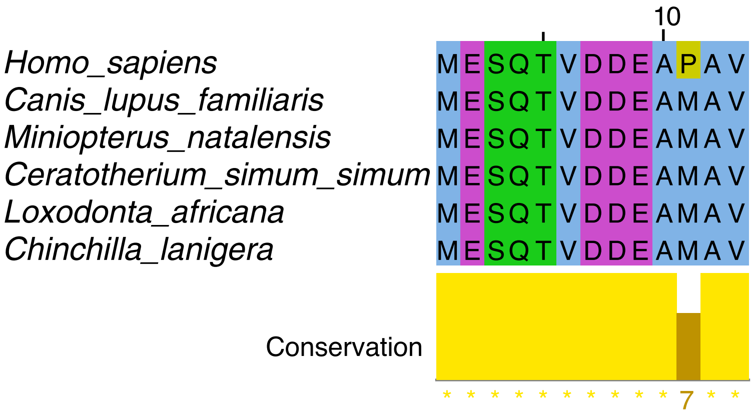

**HG0133.2**

ENSDARG00000051886 (ankrd11)

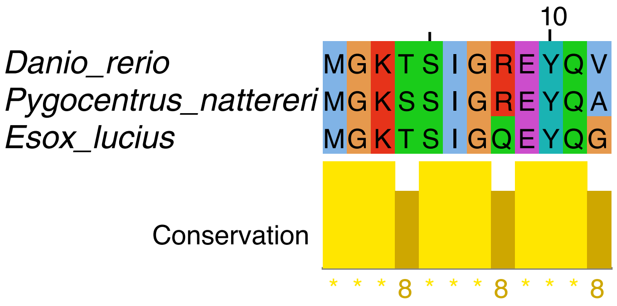

**HG0133.3**

ENSG00000167522 (ANKRD11)

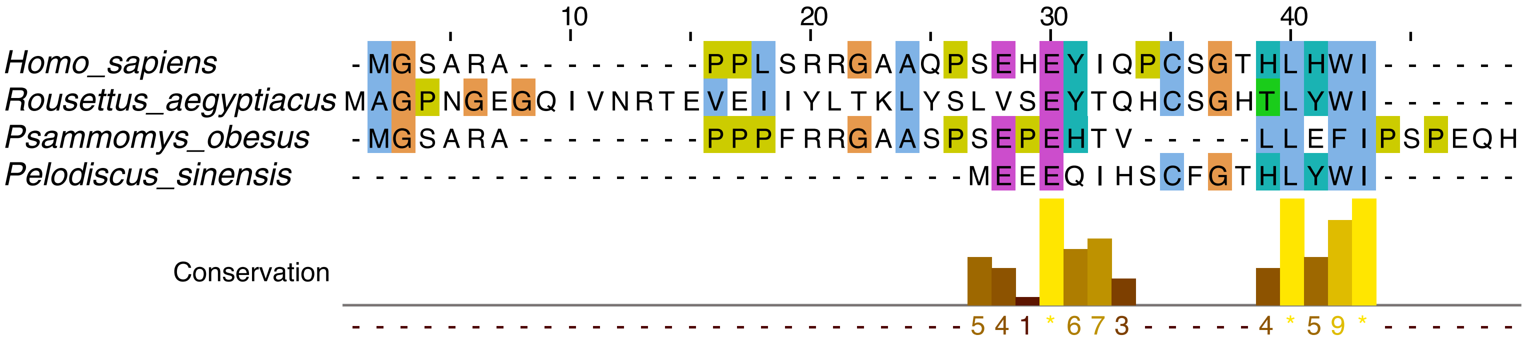

**HG0134.1**

ENSGALG00000041267 (BMI1)

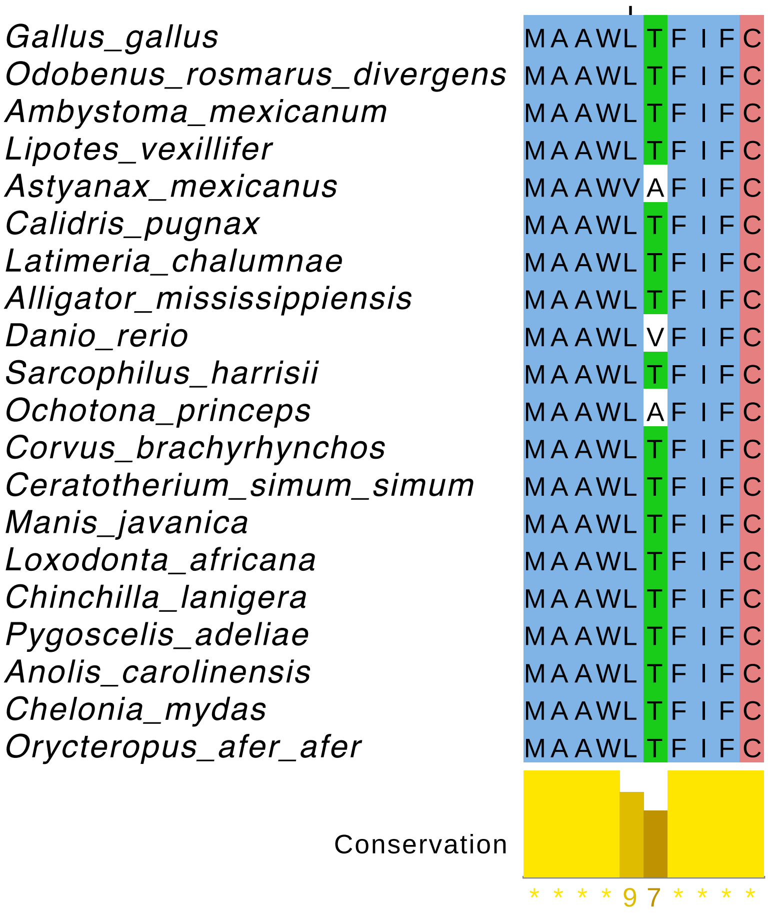

**HG0134.2**

ENSG00000168283 (BMI1)

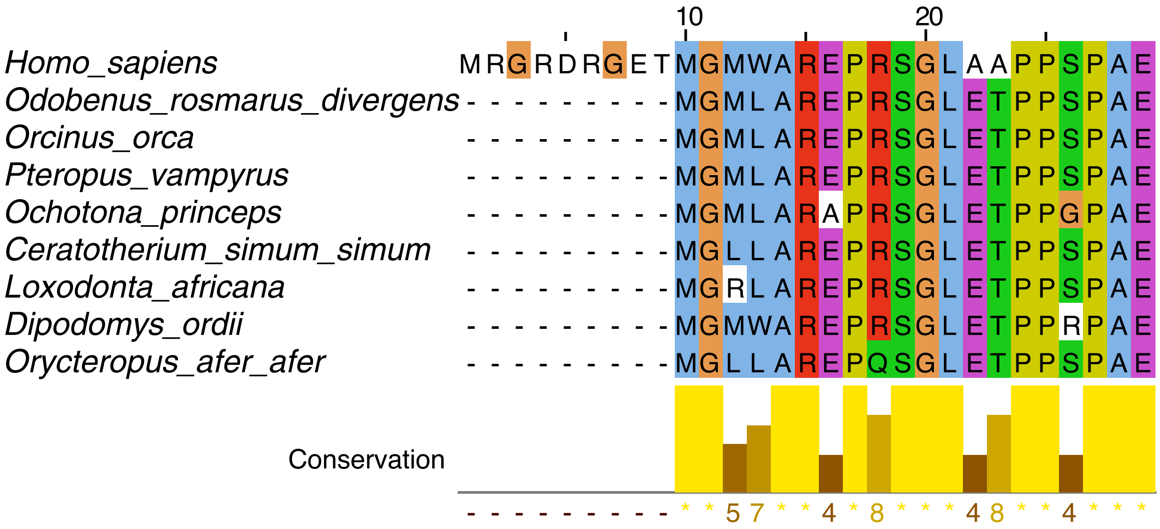

**HG0135**

ENSDARG00000094132 (igf1)

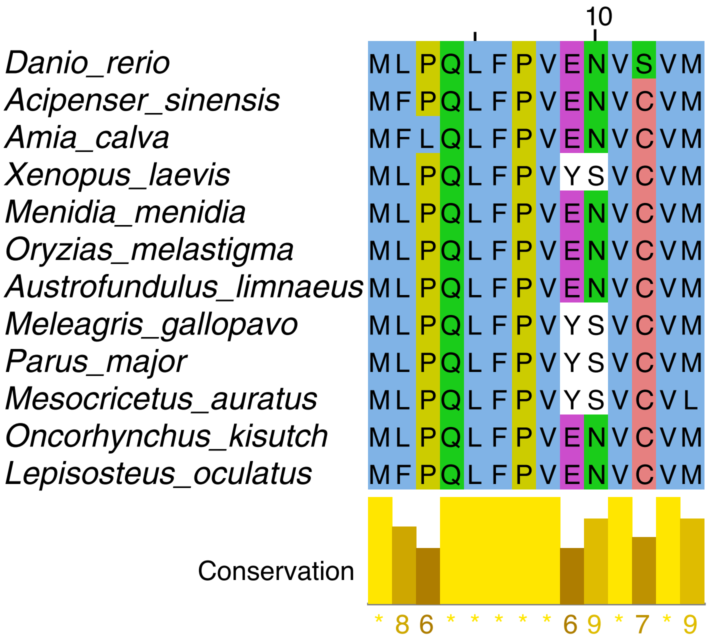

**HG0135**

ENSG00000017427 (IGF1)

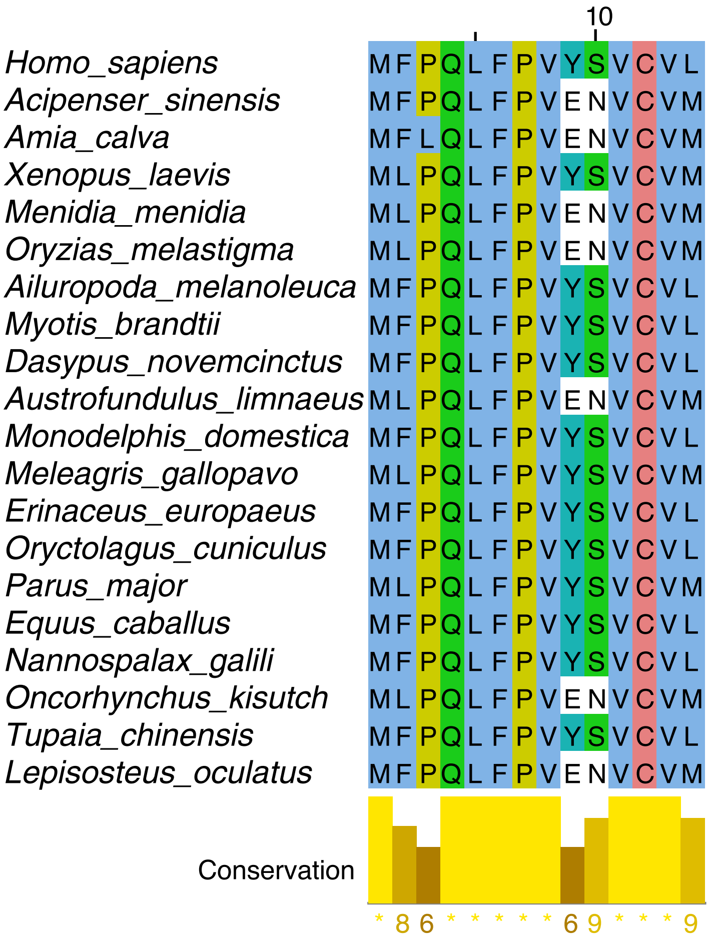

**HG0136.1**

ENSG00000125845 (BMP2)

```
                                    10        20        30        40        50        60        70
Homo_sapiens                 ------------------------------------------------------MWTLFQWTCPRV---LLRRTAVS
Felis_catus                  ------------------------------------------------------MWRLFQWTCPRV---LLRRSAVS
Physeter_catodon             ------------------------------------------------------MWRLFQWTCPRV---LLRRSAVS
Eptesicus_fuscus             ------------------------------------------------------MWRLFQWTCPRV---LLRRSAVS
Dasypus_novemcinctus         ------------------------------------------------------MRRRSRWTCPRV---LPRRSAAS
Latimeria_chalumnae          ------------------------------------------------------MWILFQWTCSSV---LLSQSAAS
Galeopterus_variegatus       ------------------------------------------------------MWRLFQWTCPRV---LLRRTAVS
Monodelphis_domestica        ------------------------------------------------------MWRLFQWTCPRV---LLSQSAVS
Condylura_cristata           ------------------------------------------------------MWRFFQWTCPRV---LLRRSAVS
Oryctolagus_cuniculus        ------------------------------------------------------MWRLFQWTCPRG---LLRRTAVS
Ornithorhynchus_anatinus     MRKKTHREQRRLPQTPAPPPSPSSSRSPSRSPWSPPRPAPDPPAAPVSSARDFSMWRLFRWTCPRV---LLRQSAVS
Ceratotherium_simum_simum    ------------------------------------------------------MWRLFQWTCPRV---LLRRSAVS
Manis_javanica               ------------------------------------------------------MWRLFQWTCPRV---LLRRSAVS
Loxodonta_africana           ------------------------------------------------------MWRLFQWTCPRV---LLRRSAVS
Dipodomys_ordii              ------------------------------------------------------MWRLFQWTCPRV---LLRRTAVS
Tupaia_chinensis             ------------------------------------------------------MWRLFQWTCHRV---LLRRTAVS
Lepisosteus_oculatus         ------------------------------------------------------MWKLFQWTCQRV---LLSRSPLS
Trichechus_manatus_latirostr ------------------------------------------------------MWRLFQWTCLRV---LLRRSAVS
Trachemys_scripta            ------------------------------------------------------MWSPFQWTCPSAPPLLLRRSAGS
Orycteropus_afer_afer        ------------------------------------------------------MWRLFQWTCPRV---LLRRSAVS

Conservation                 ------------------------------------------------------*63258***168---*668877*
```

**HG0136.2**

ENSG00000125845 (BMP2)

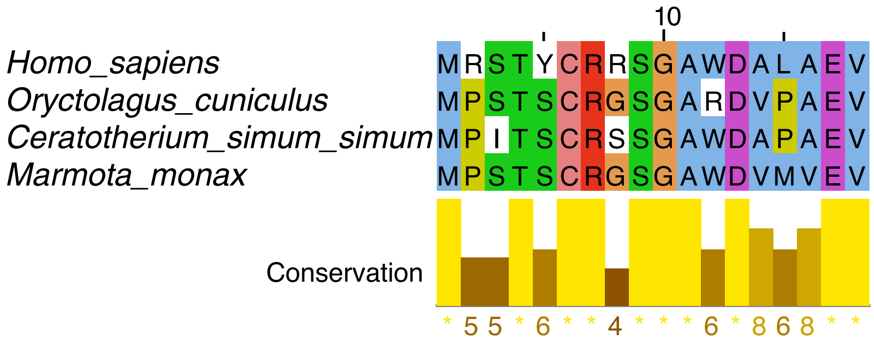

**HG0137.1**

ENSG00000181722 (ZBTB20)

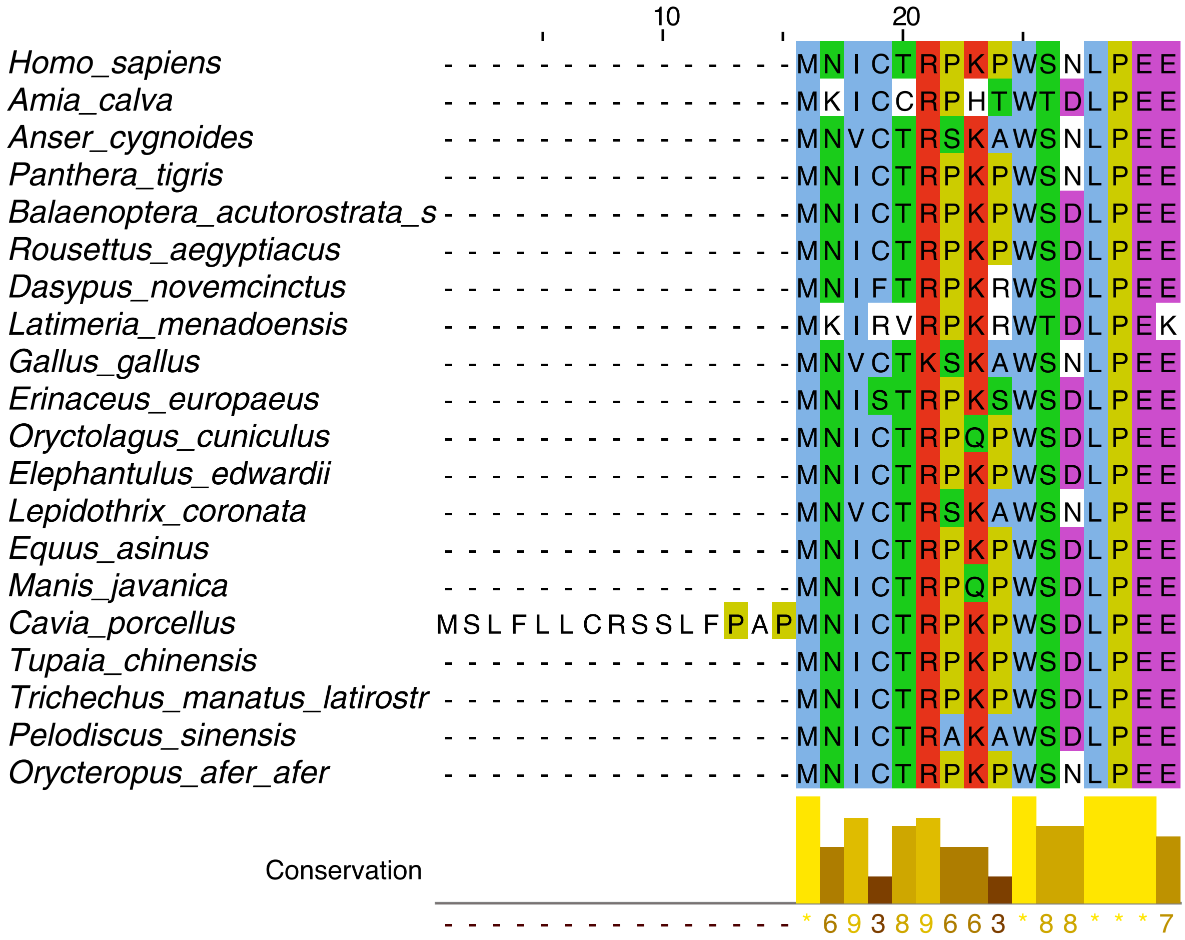

**HG0137.2**

ENSG00000181722 (ZBTB20)

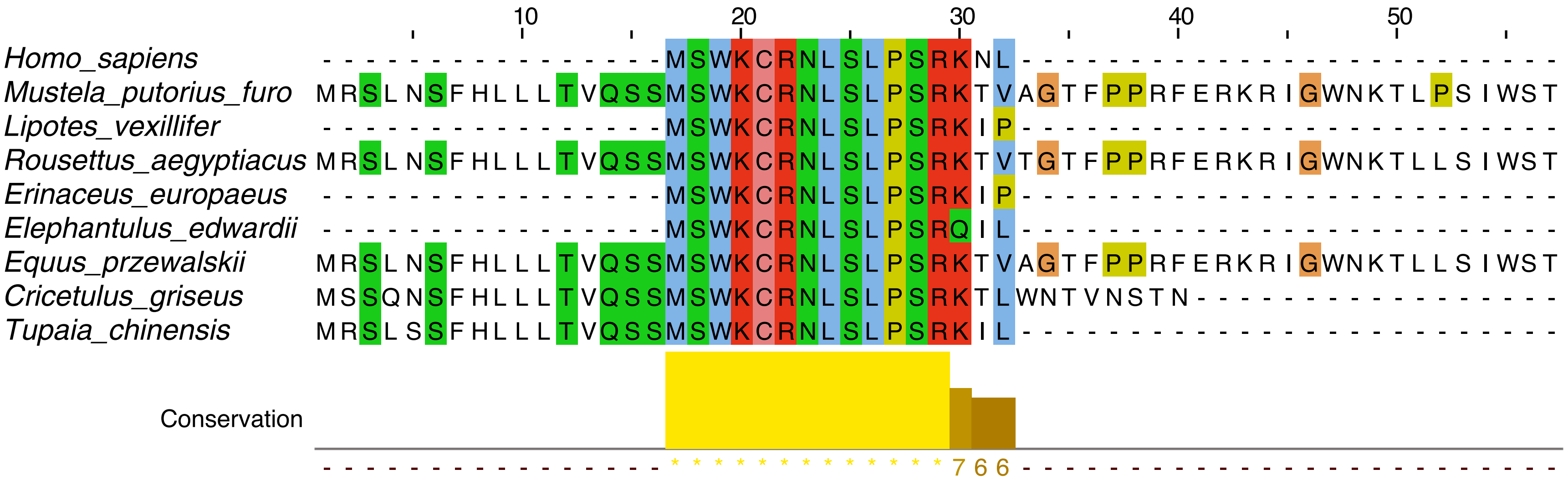

**HG0138.1**

ENSG00000185070 (FLRT2)

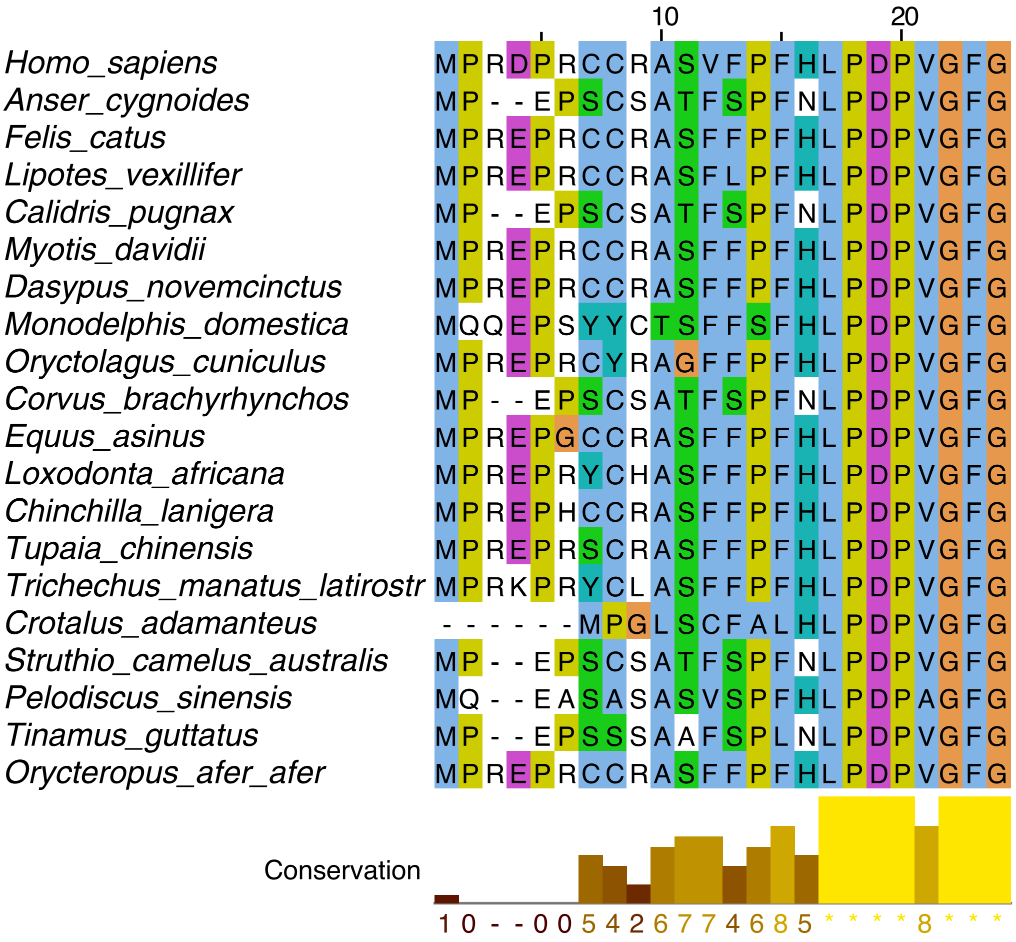

**HG0138.2**

ENSG00000185070 (FLRT2)

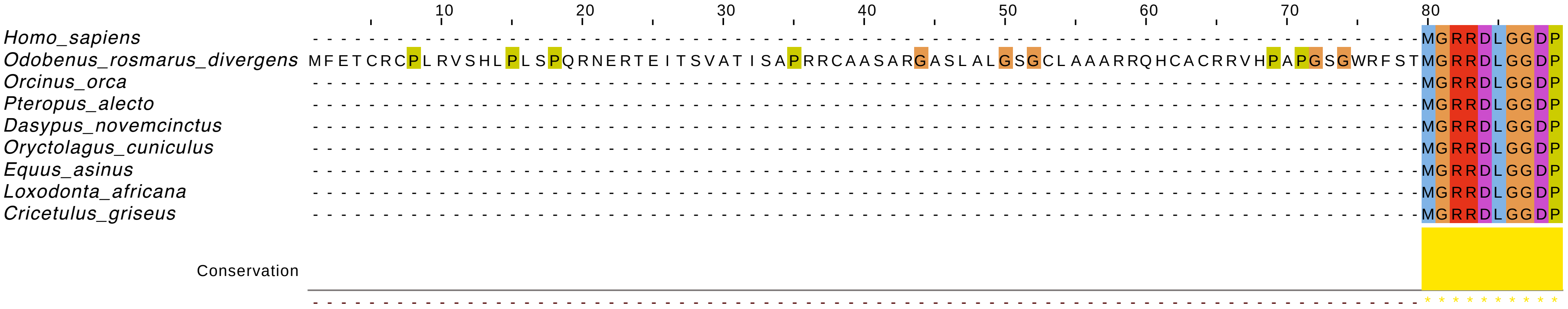

**HG0139**

ENSGALG00000006379 (SHH)

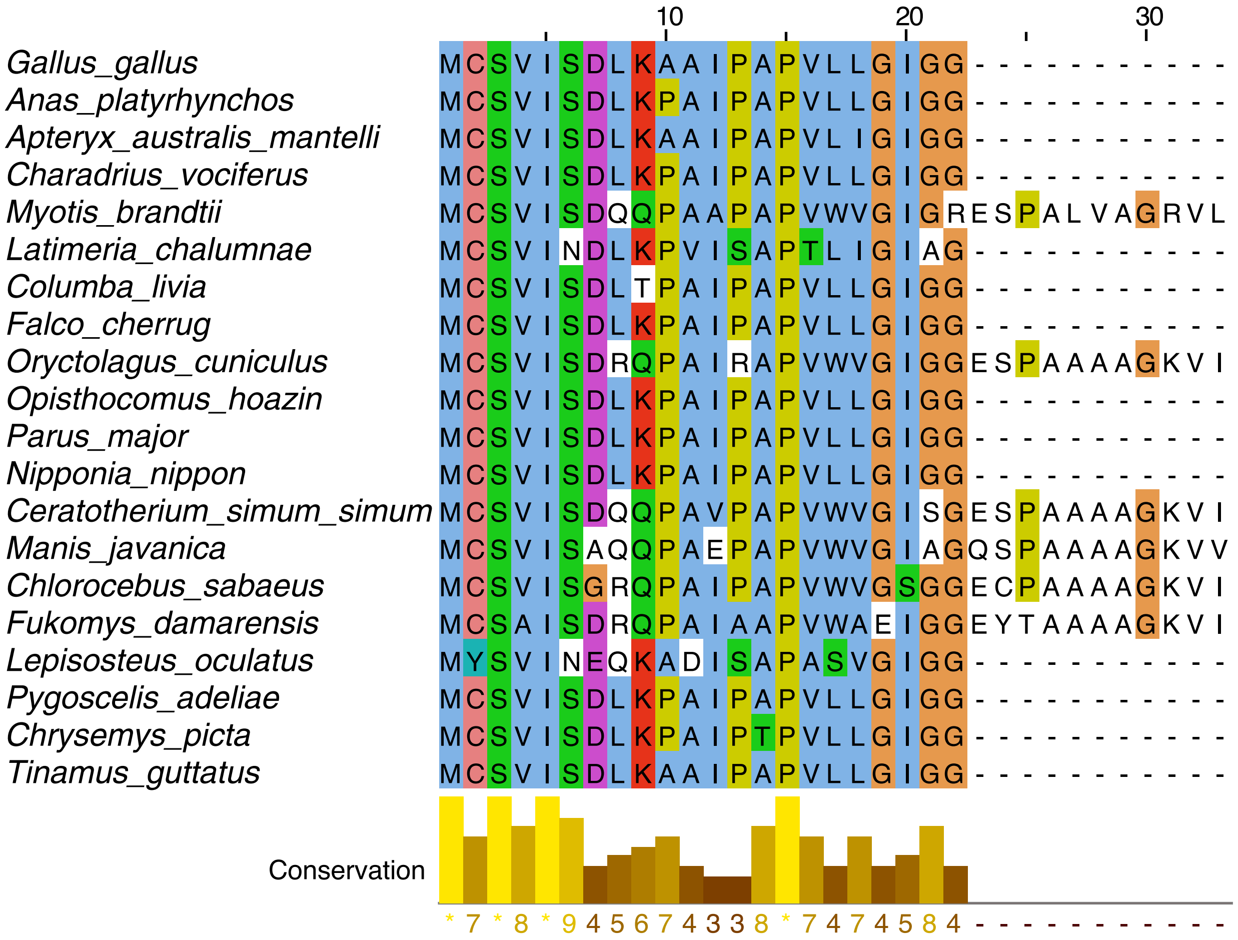

**HG0140**

ENSG00000009307 (CSDE1)

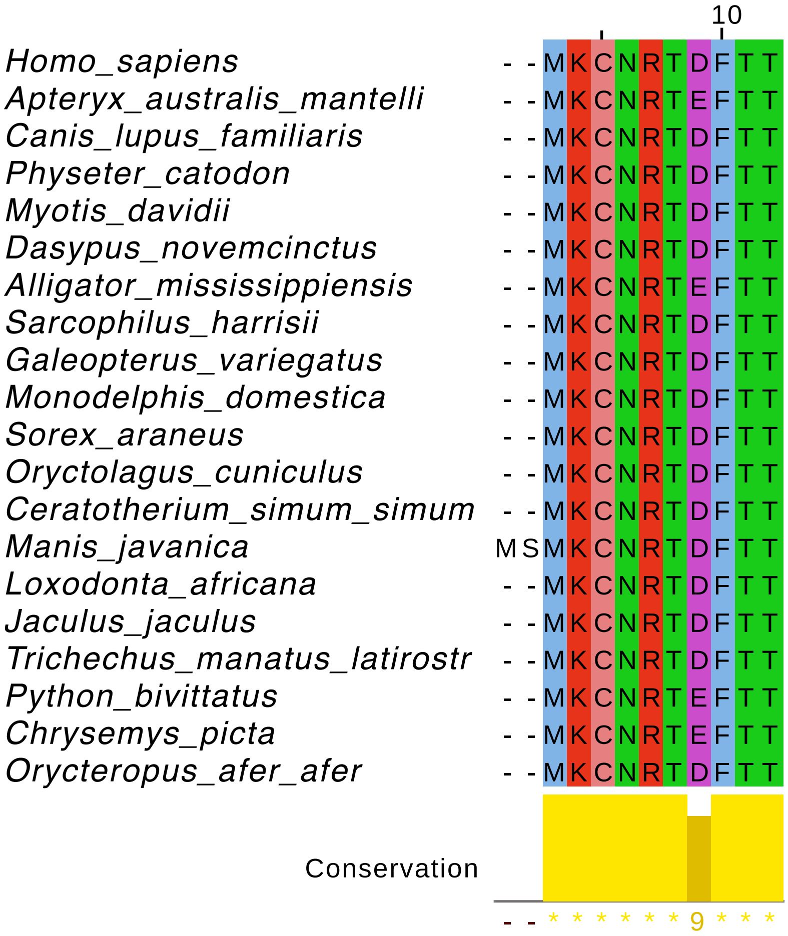

**HG0141.1**

ENSGALG00000010435 (RASGRP3)

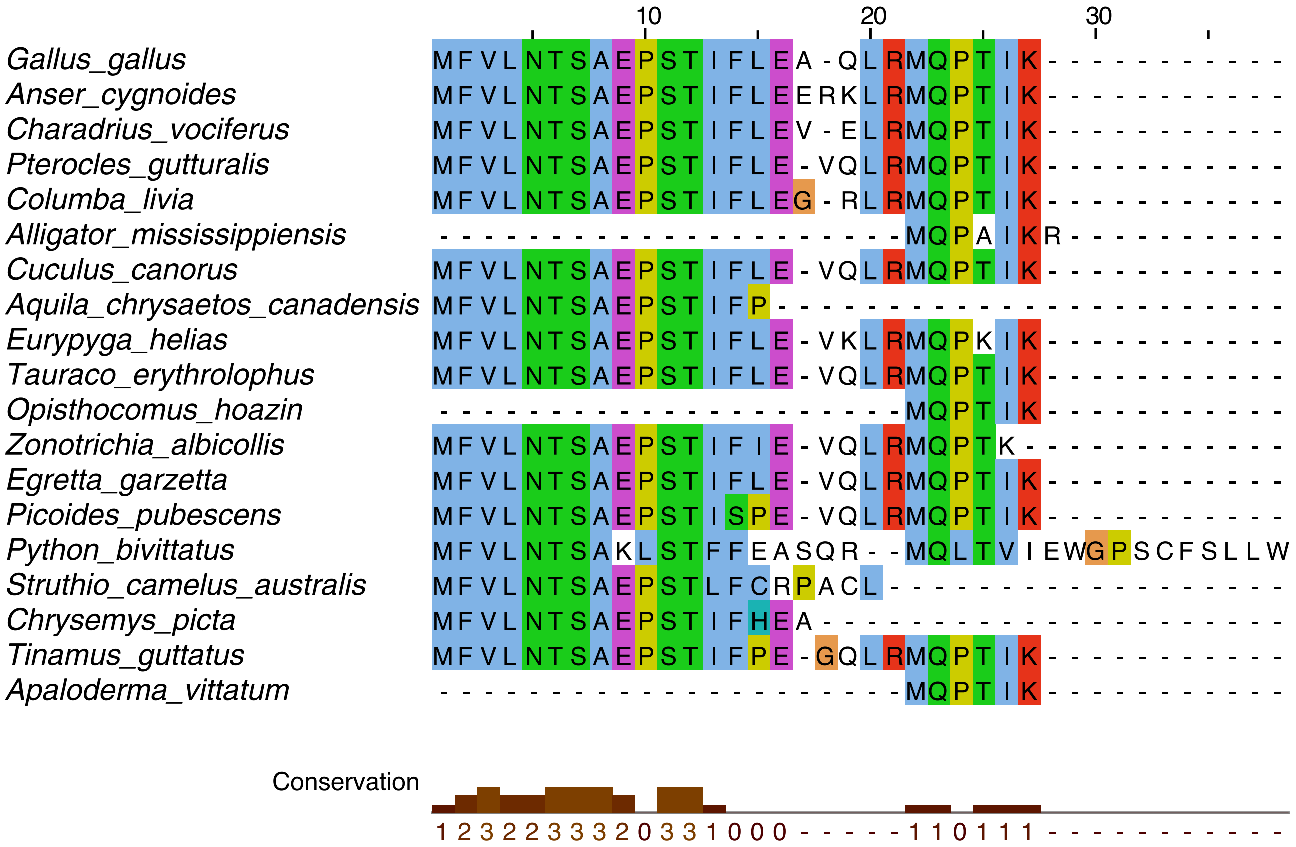

**HG0141.2**

ENSG00000152689 (RASGRP3)

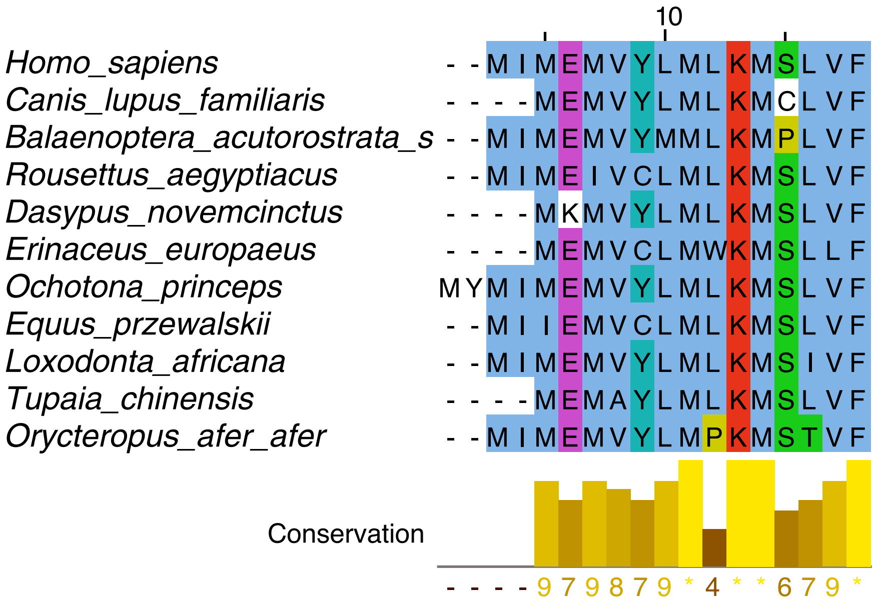

**HG0141.3**

ENSG00000152689 (RASGRP3)

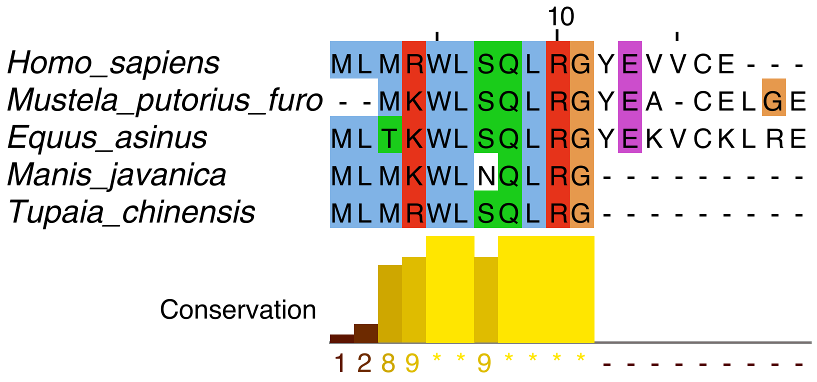

**HG0142.1**

ENSG00000204217 (BMPR2)

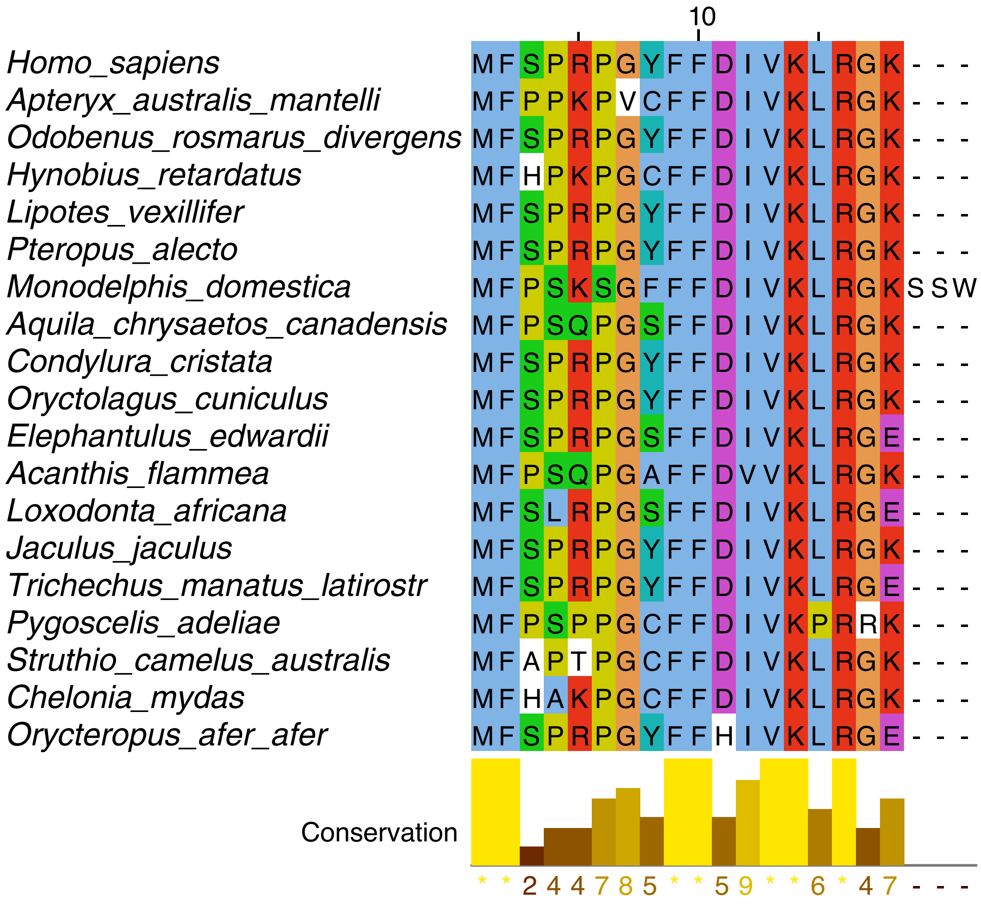

**HG0142.2**

ENSG00000204217 (BMPR2)

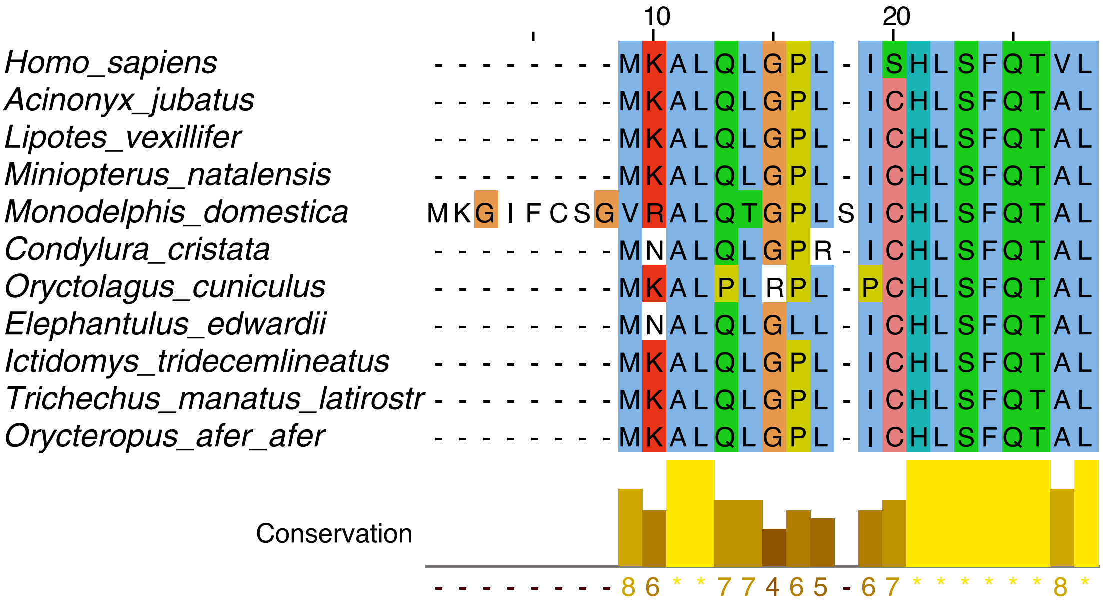

**HG0143**

ENSG00000101958 (GLRA2)

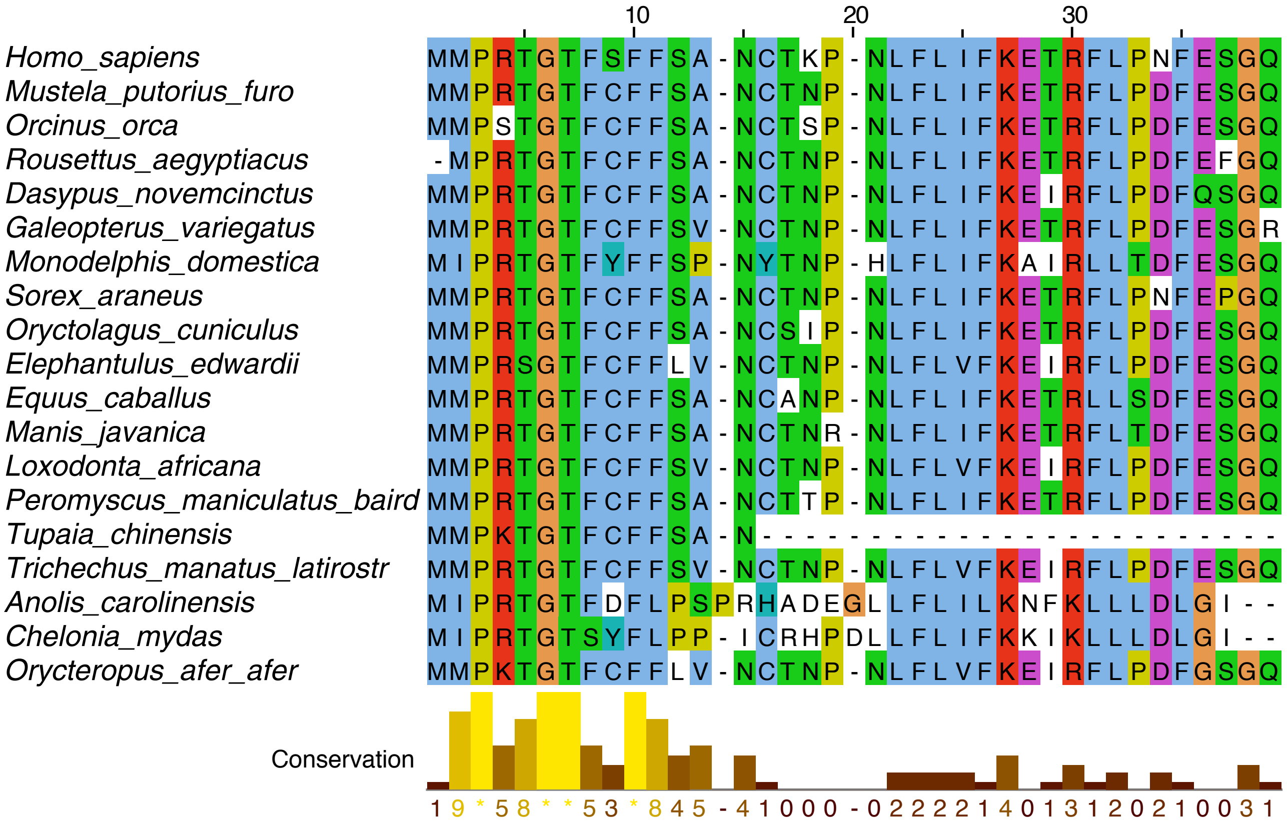

**HG0144**

ENSG00000117114 (ADGRL2)

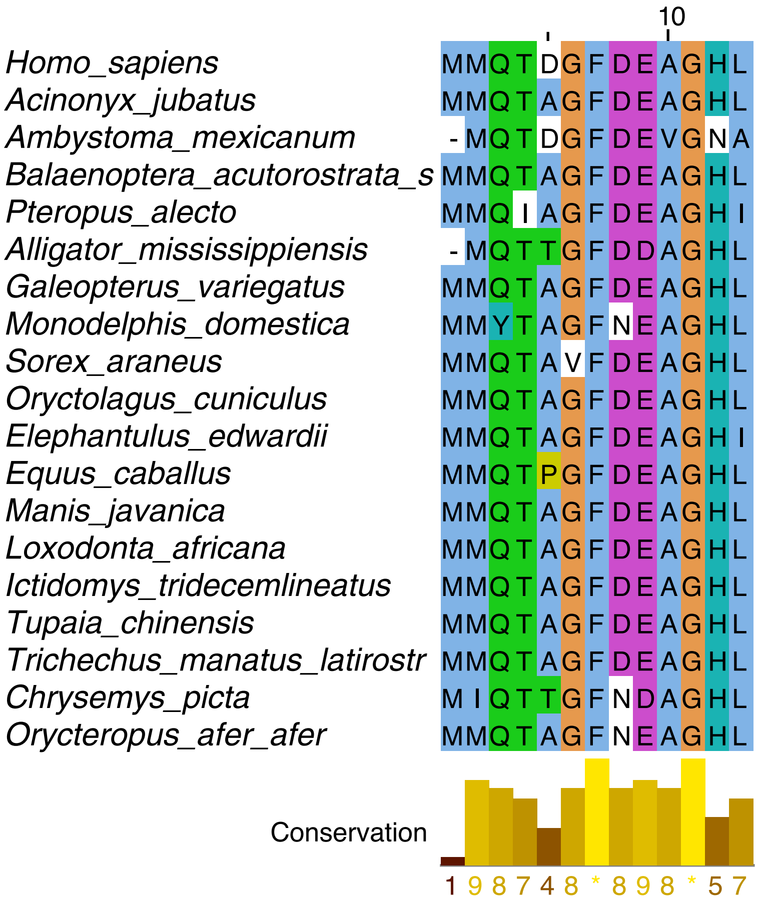

**HG0145**

ENSG00000162599 (NFIA)

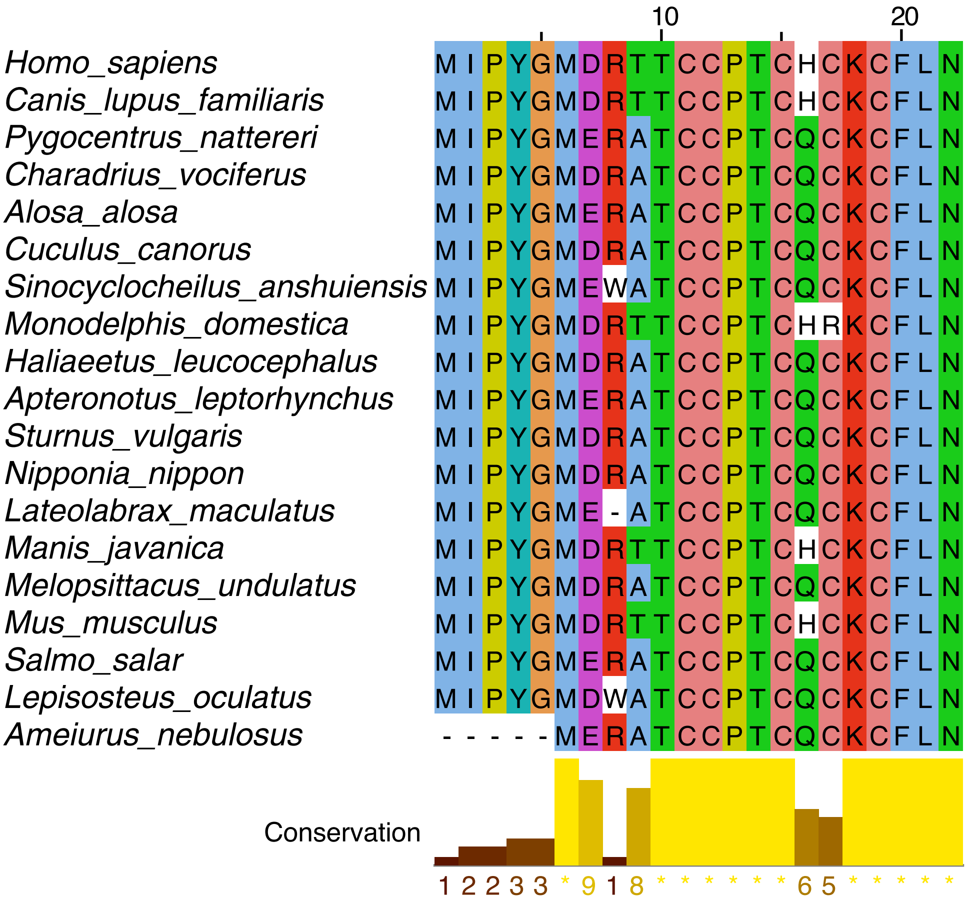

**HG0146.1**

ENSG00000163681 (SLMAP)

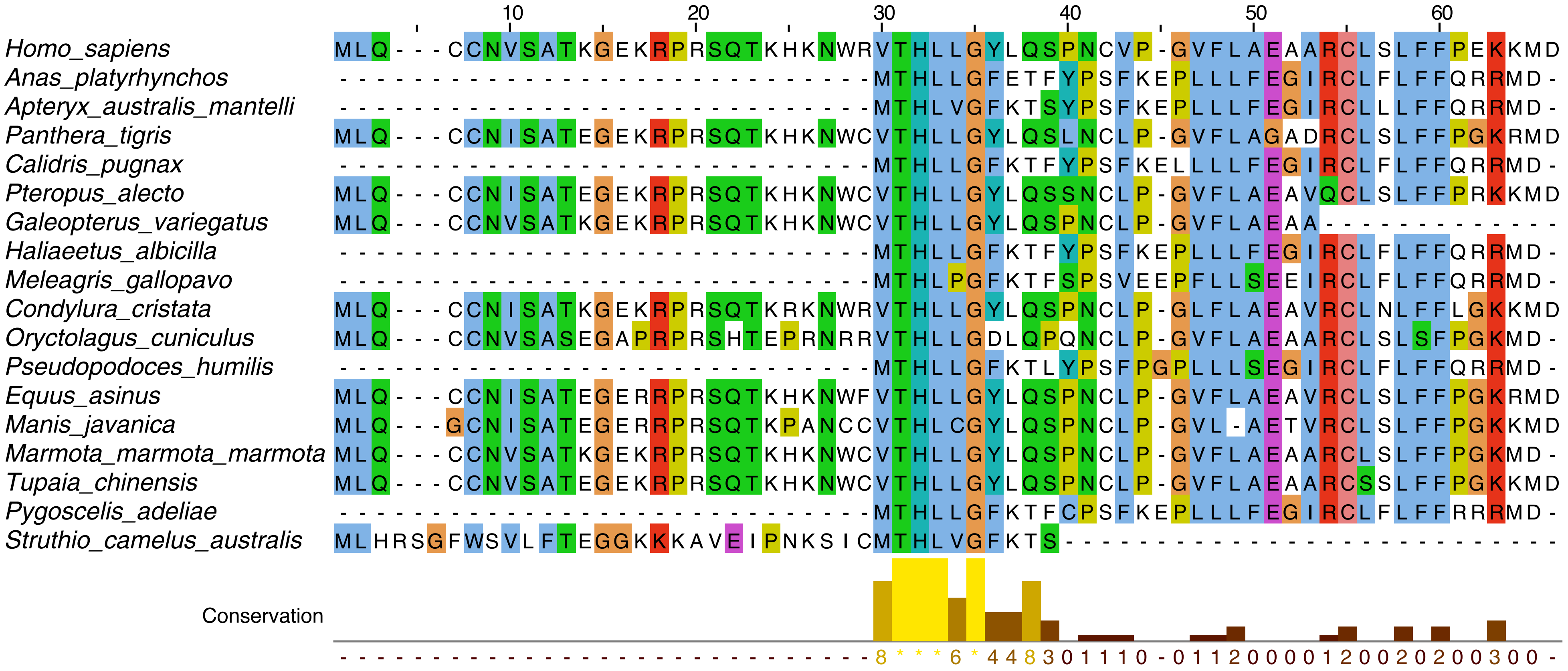

**HG0146.2**

ENSG00000163681 (SLMAP)

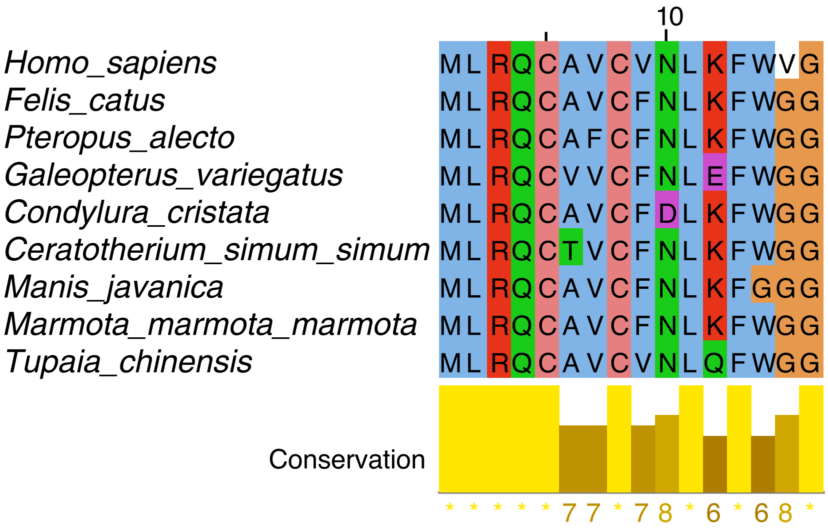

**HG0146.3**

ENSG00000163681 (SLMAP)

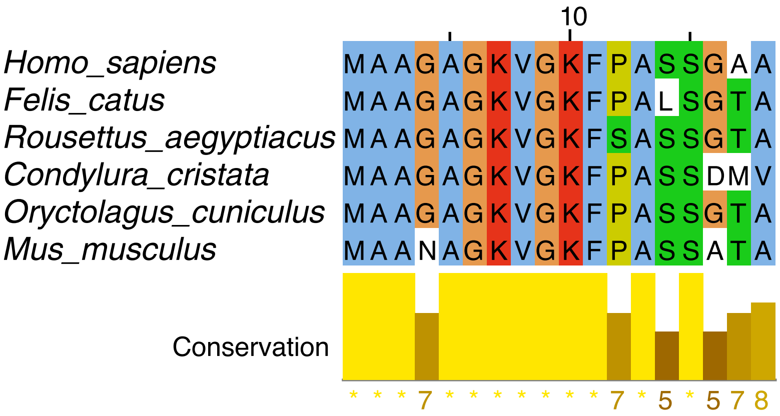

**HG0147**

ENSG00000108984 (MAP2K6)

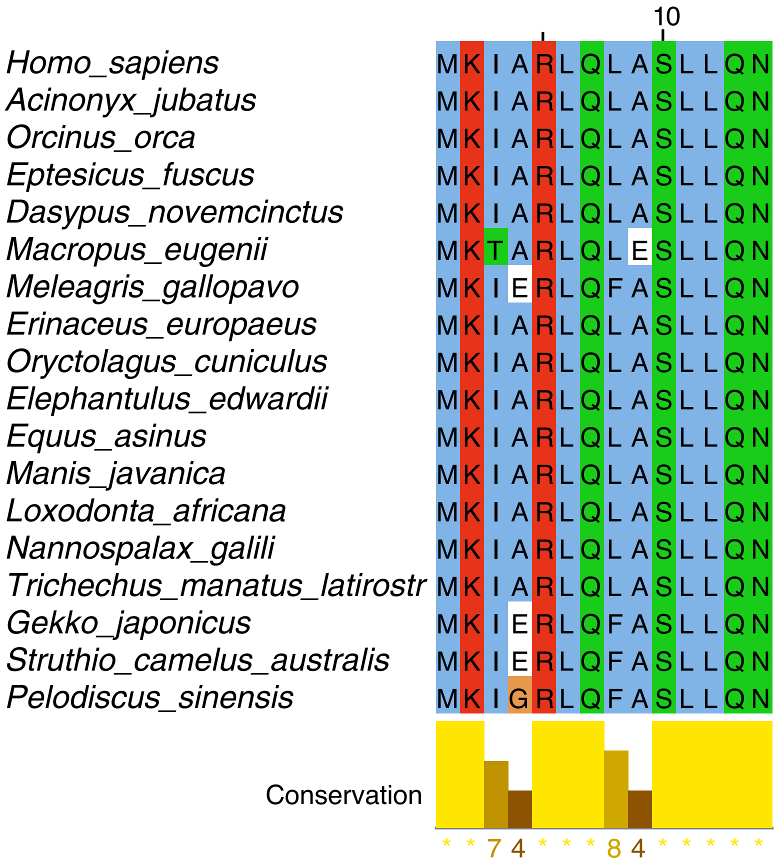

**HG0147**

ENSGALG00000004370 (MAP2K6)

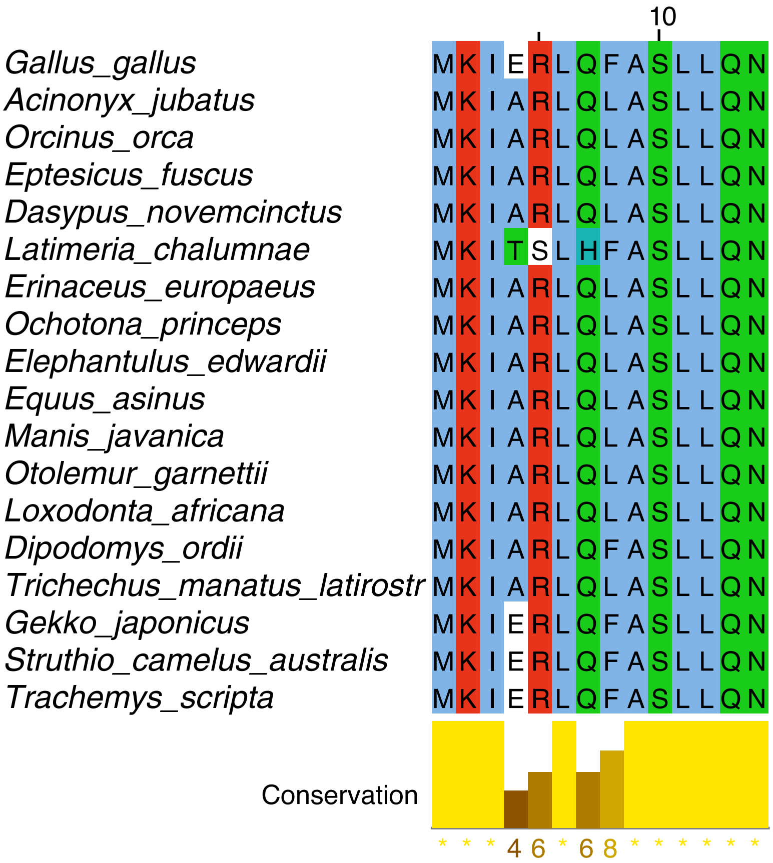

**HG0148.1**

ENSGALG00000011855 (RHOJ)

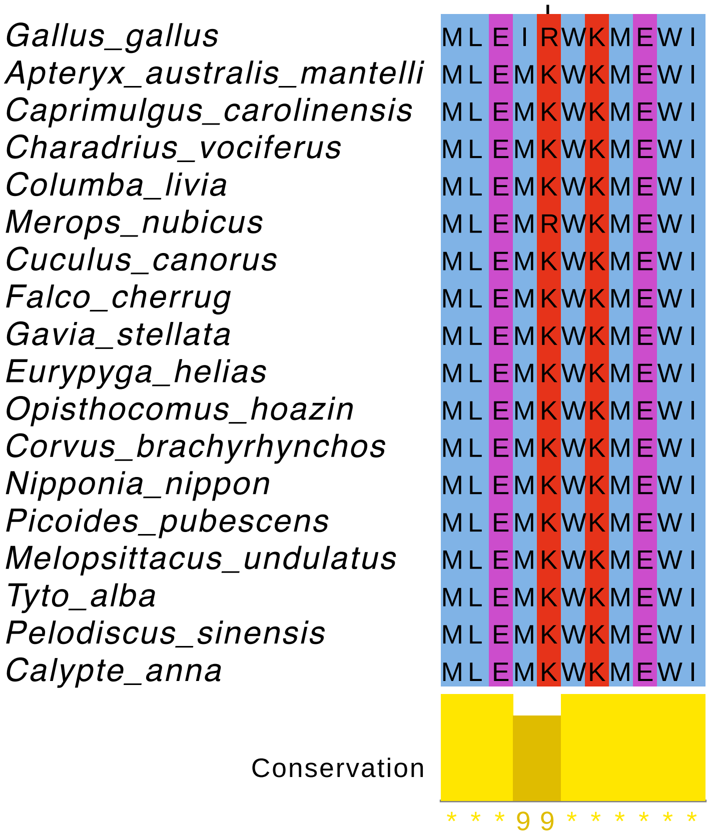

**HG0148.2**

ENSGALG00000011855 (RHOJ)

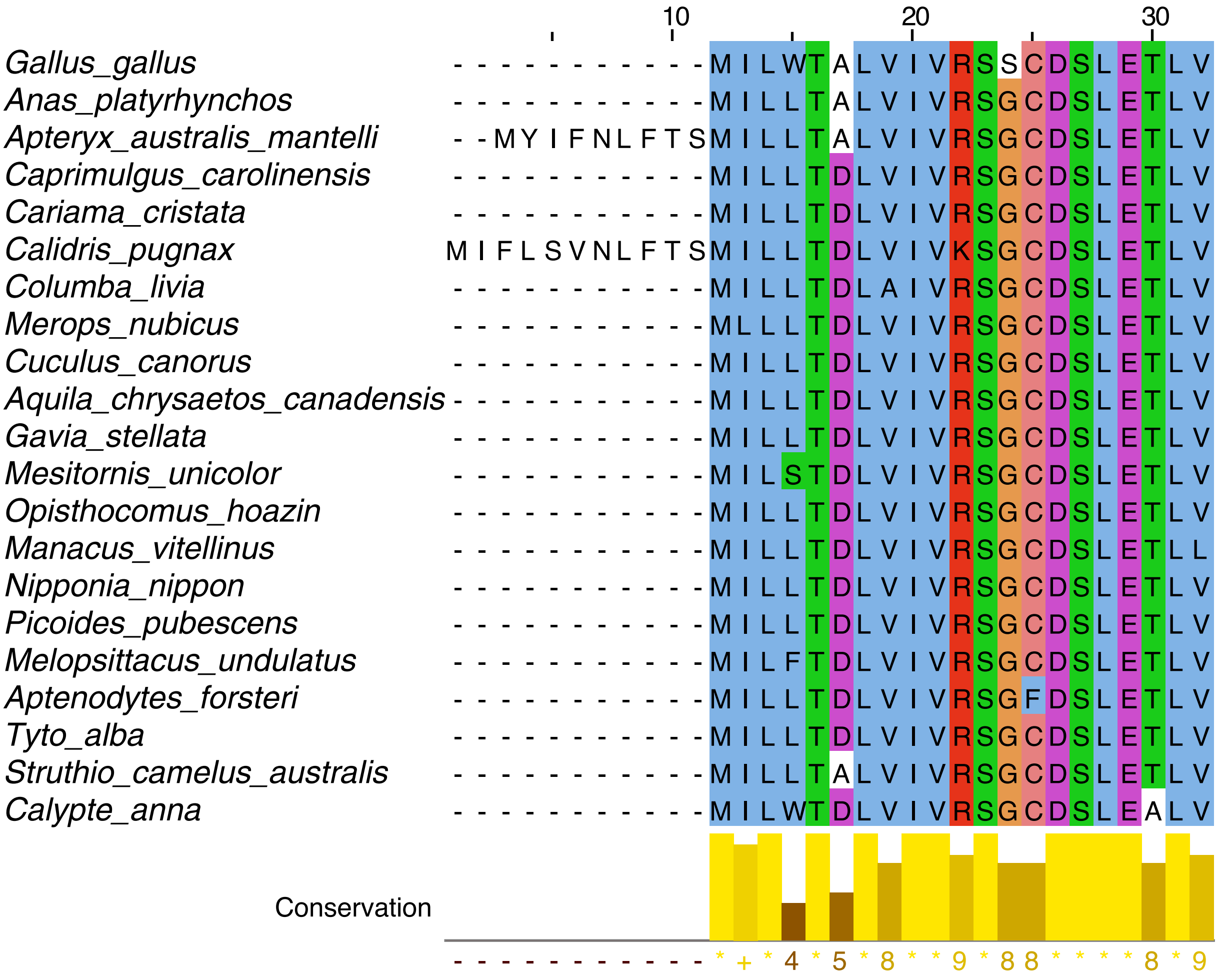

**HG0149.1**

ENSGALG00000014645 (MEF2C)

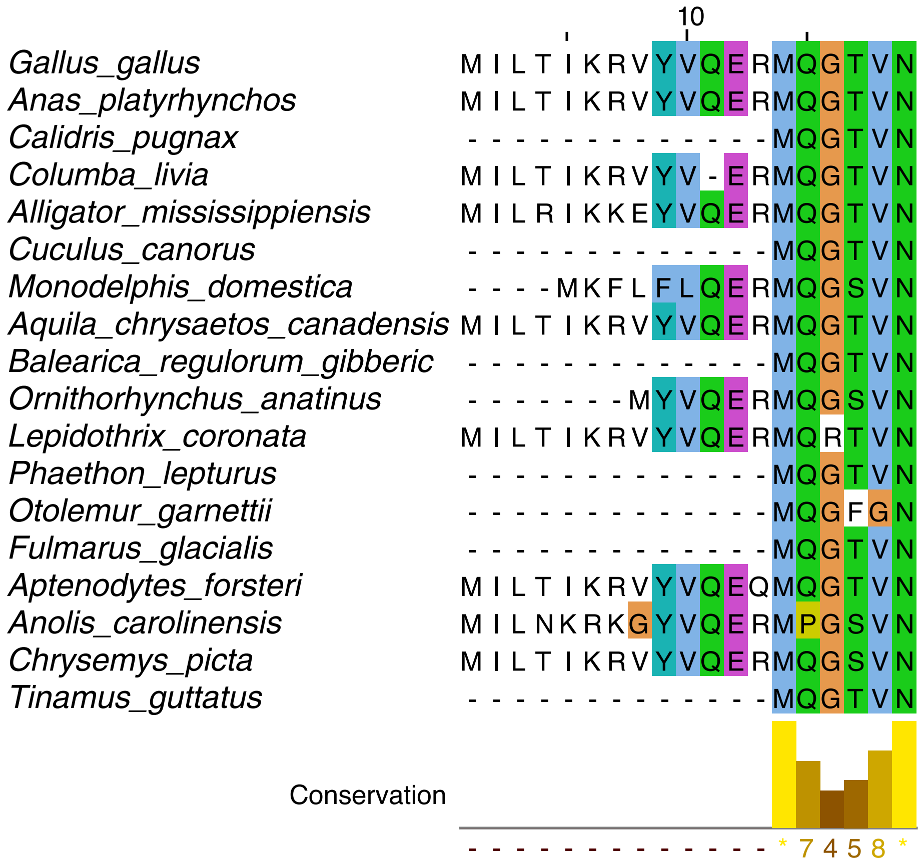

**HG0149.2**

ENSG00000081189 (MEF2C)

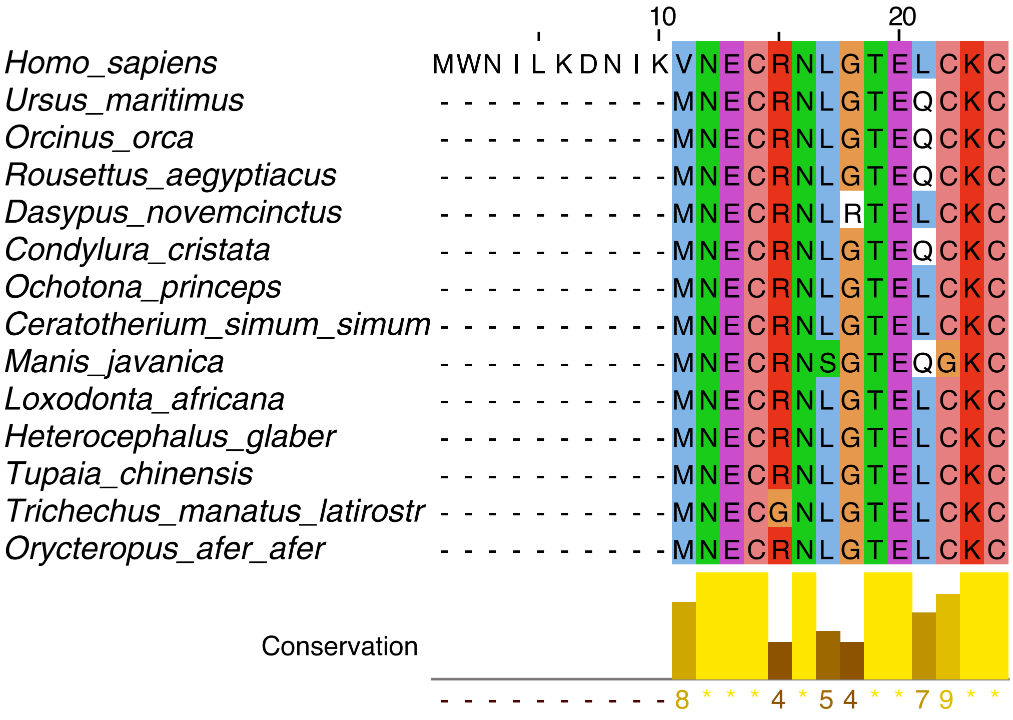

**HG0150.1**

ENSGALG00000032289 (IKZF5)

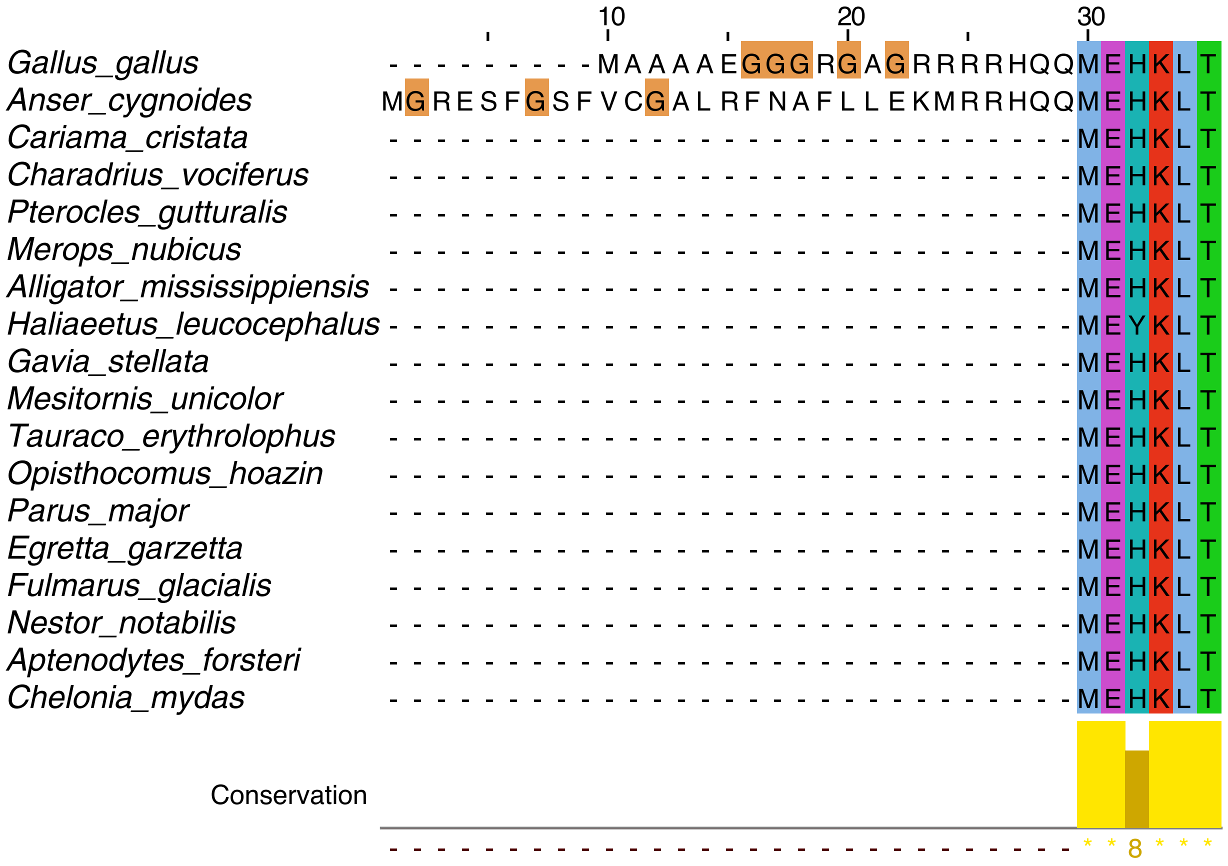

**HG0150.2**

ENSG00000095574 (IKZF5)

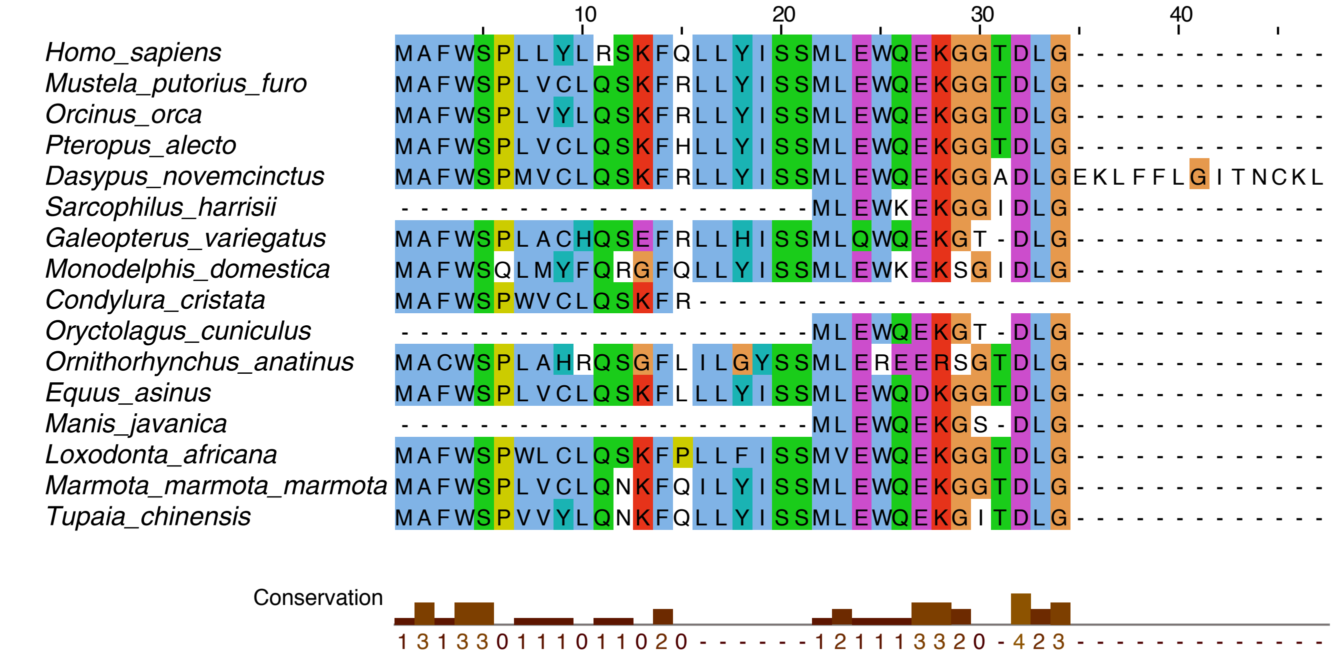

**HG0151**

ENSG00000006468 (ETV1)

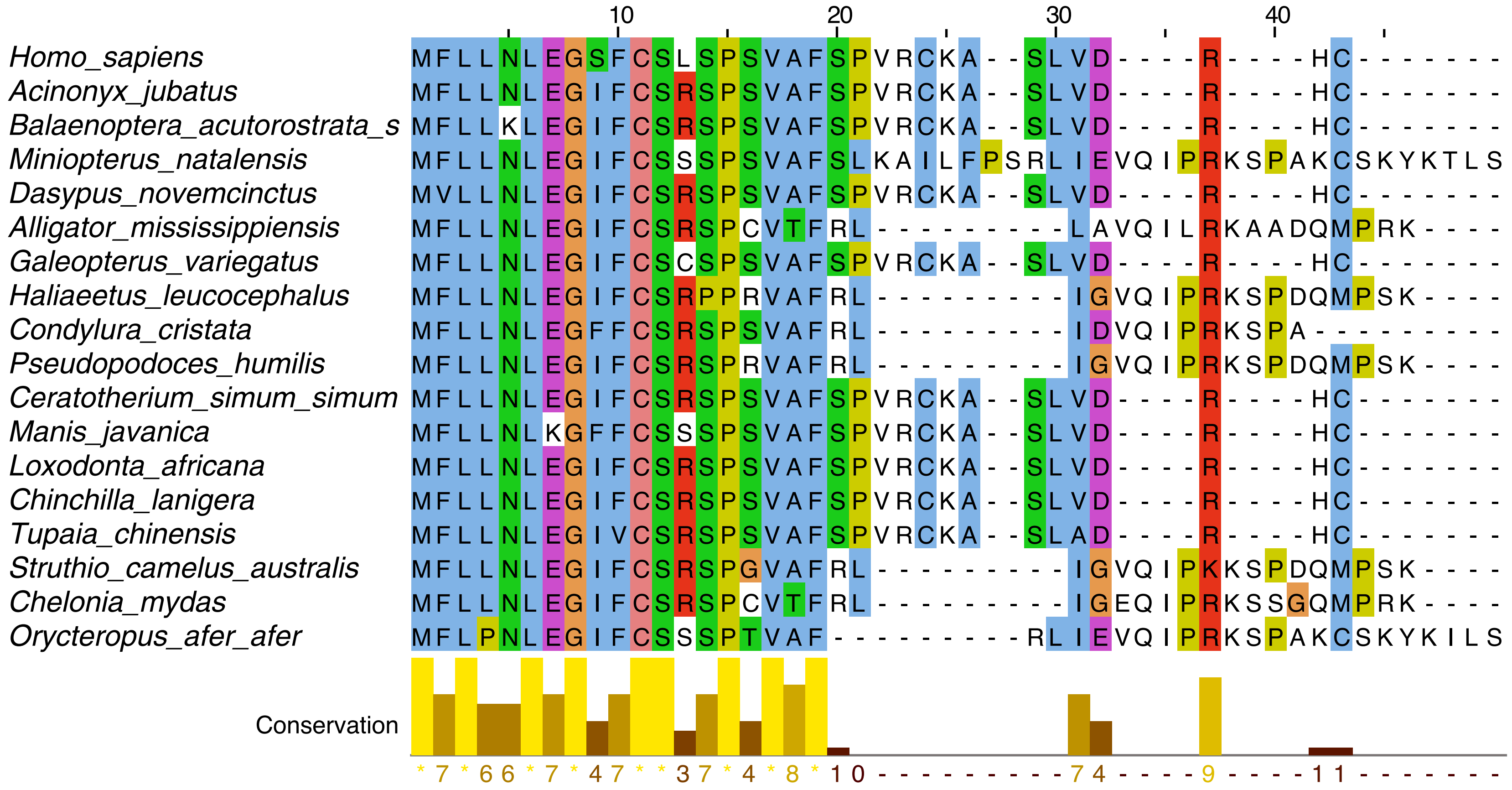

**HG0151**

ENSGALG00000035504 (ER81)

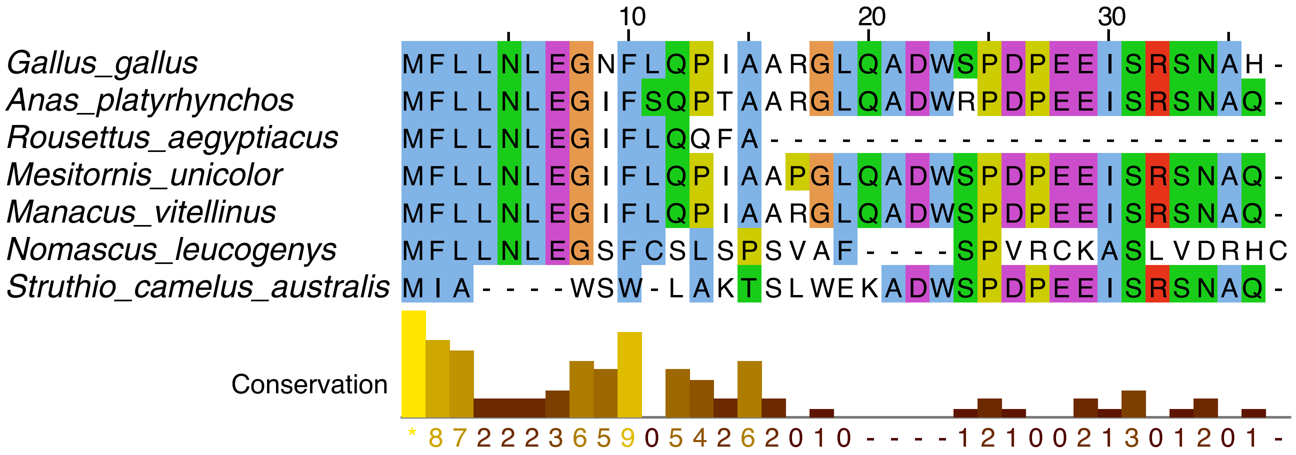

**HG0152**

ENSDARG00000015536 (sox6)

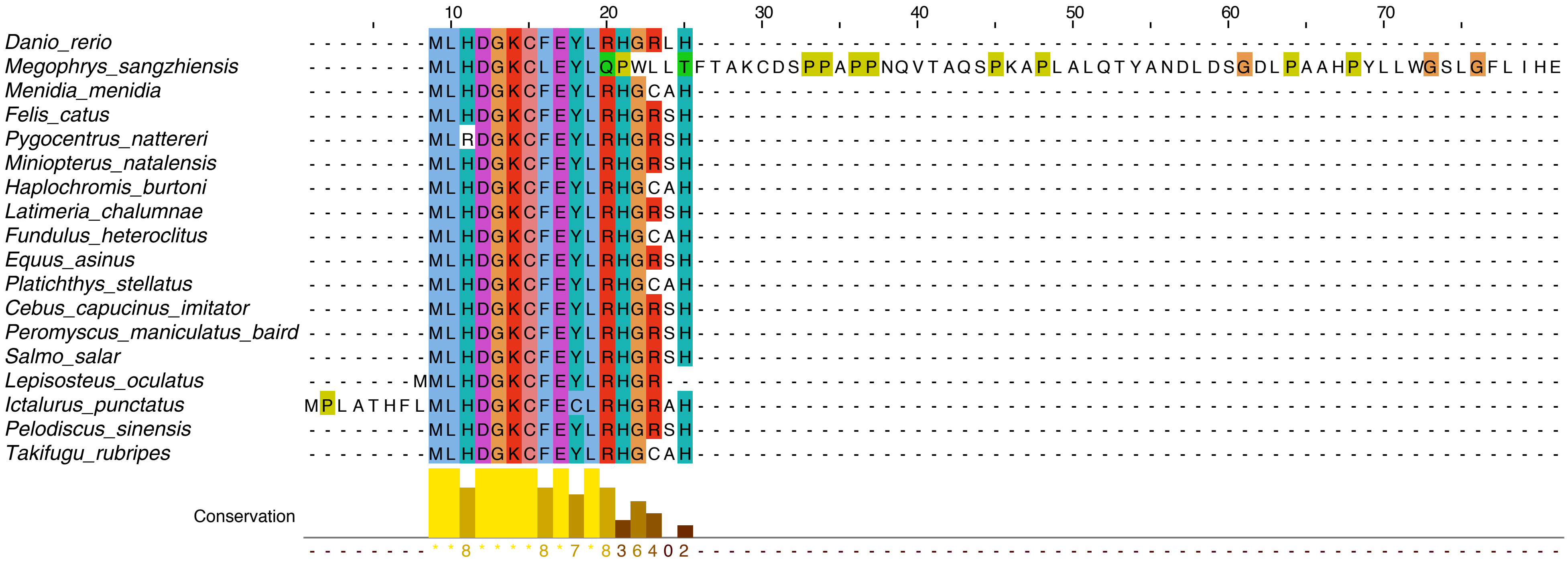

**HG0153**

ENSGALG00000003136 (IKZF2)

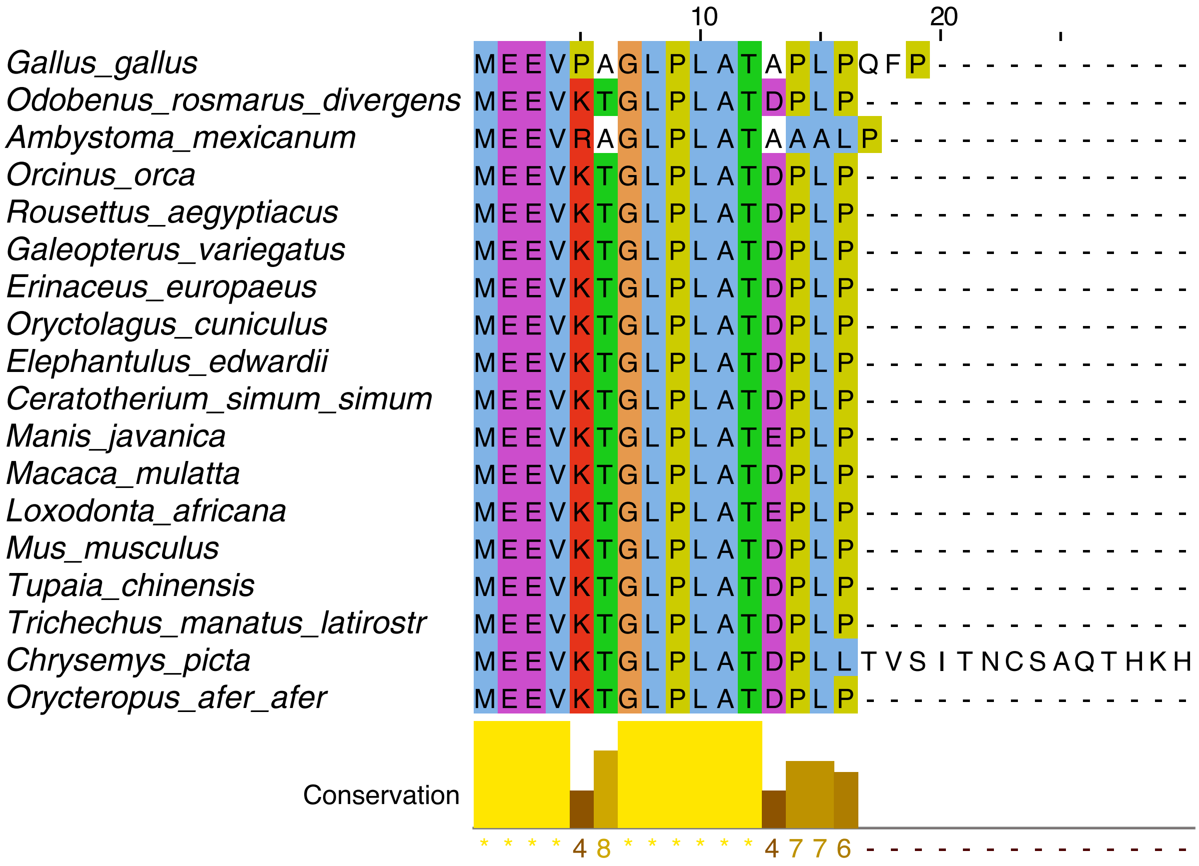

**HG0154**

ENSG00000128573 (FOXP2)

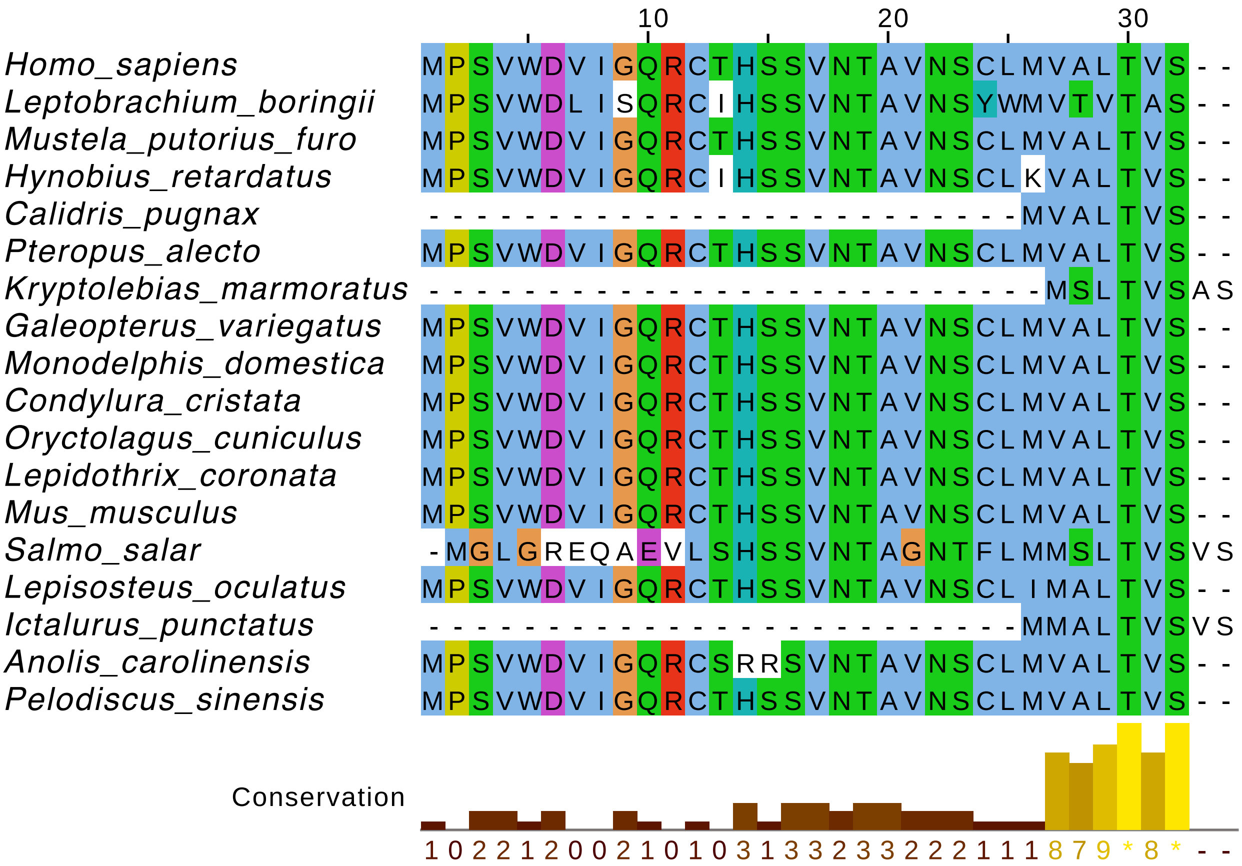

**HG0155**

ENSG00000136535 (TBR1)

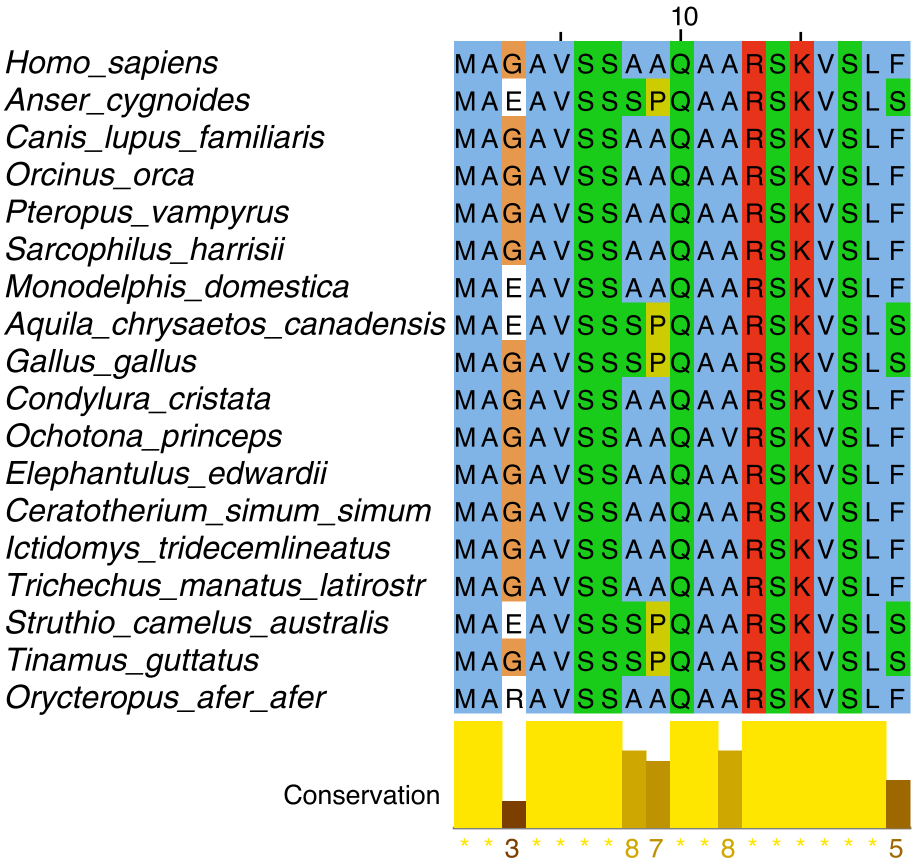

**HG0156**

ENSG00000156113 (KCNMA1)

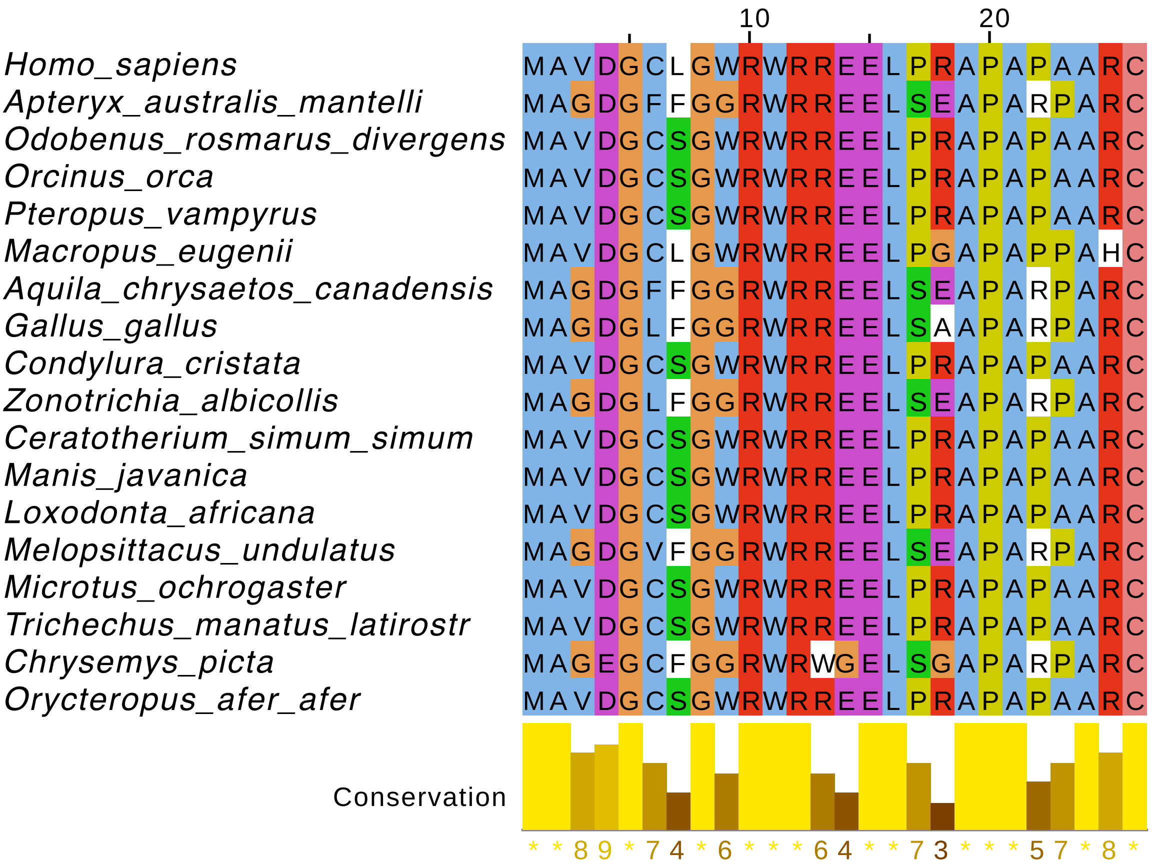

**HG0157**

ENSG00000173926 (3-Mar)

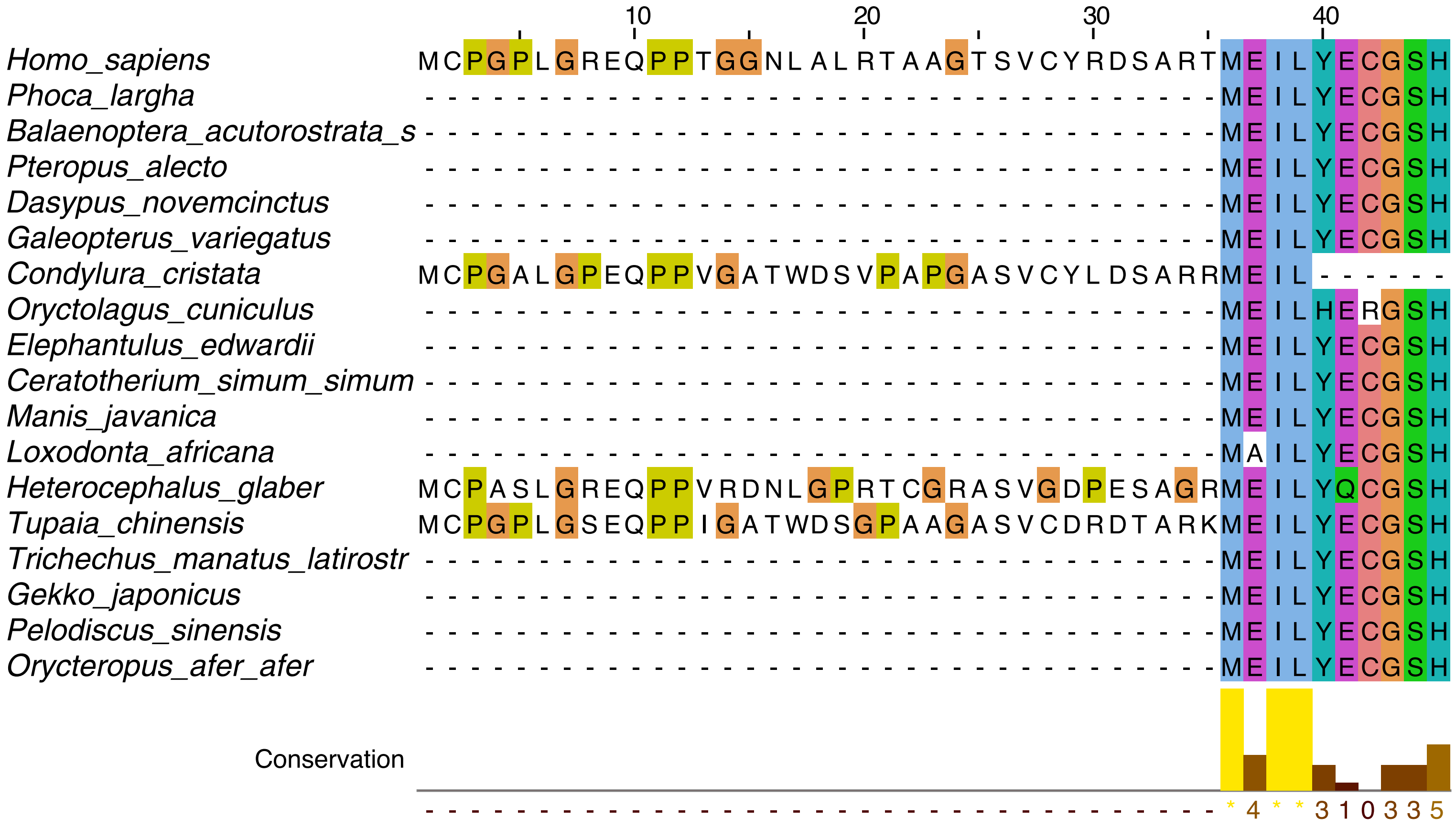

# HG0158

### ENSG00000198939 (ZFP2)

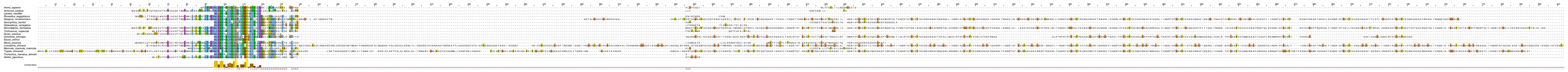

**HG0159.1**

ENSG00000182197 (EXT1)

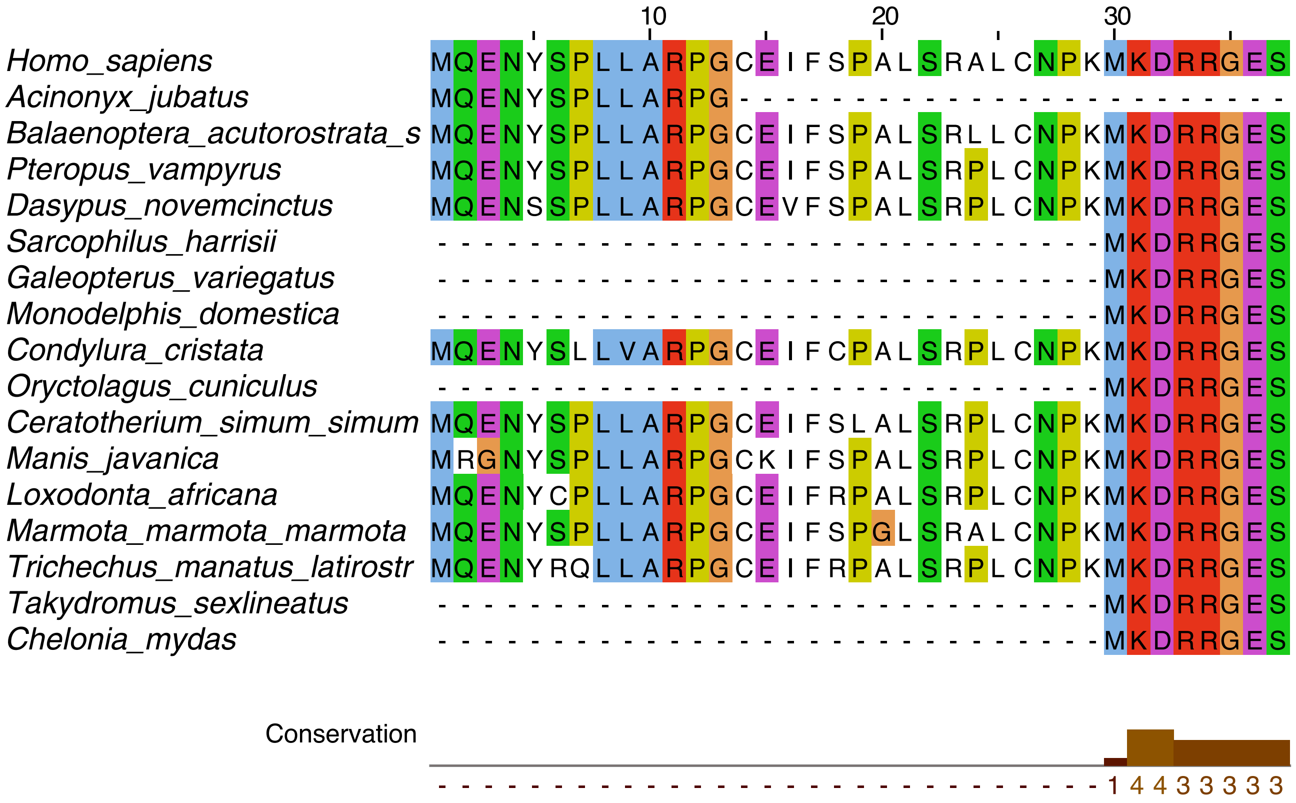

**HG0159.2**

ENSG00000182197 (EXT1)

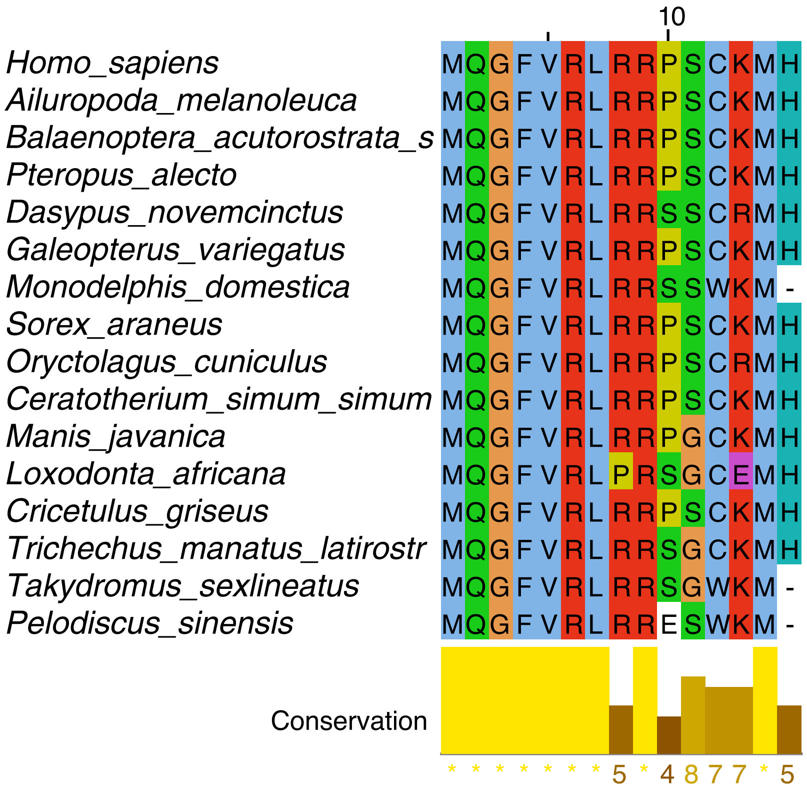

**HG0160**

ENSDARG00000060089 (btaf1)

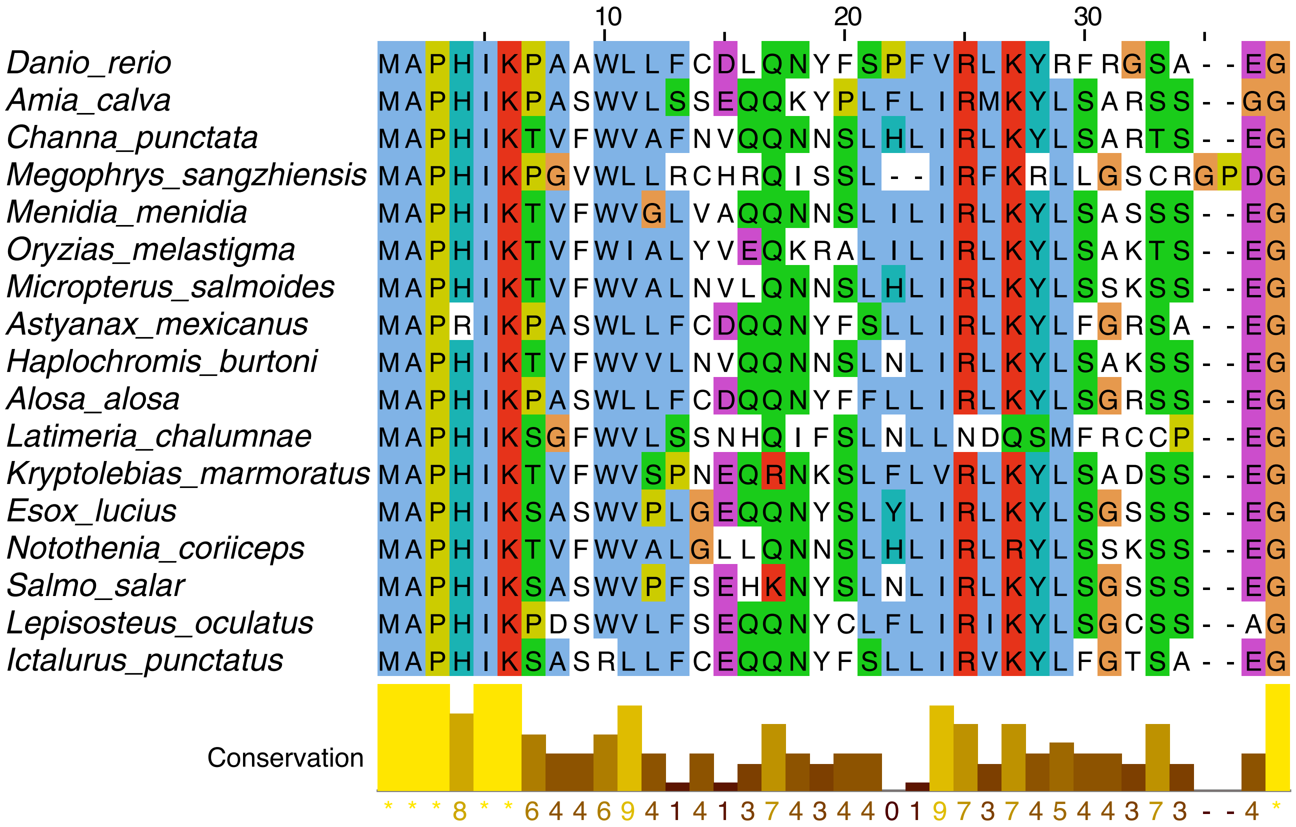

**HG0161**

ENSGALG00000017191

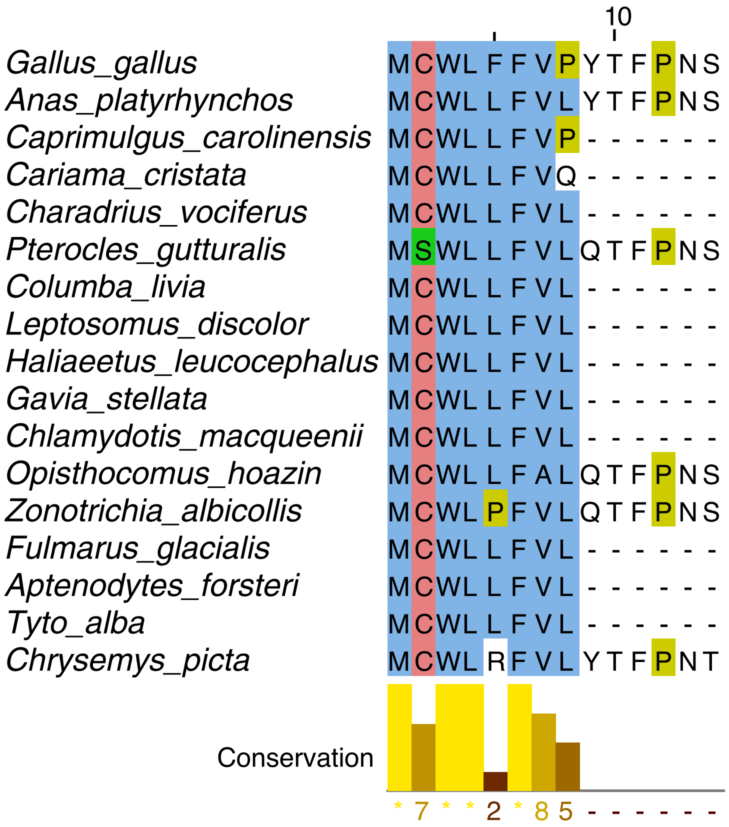

**HG0162**

ENSG00000138347 (MYPN)

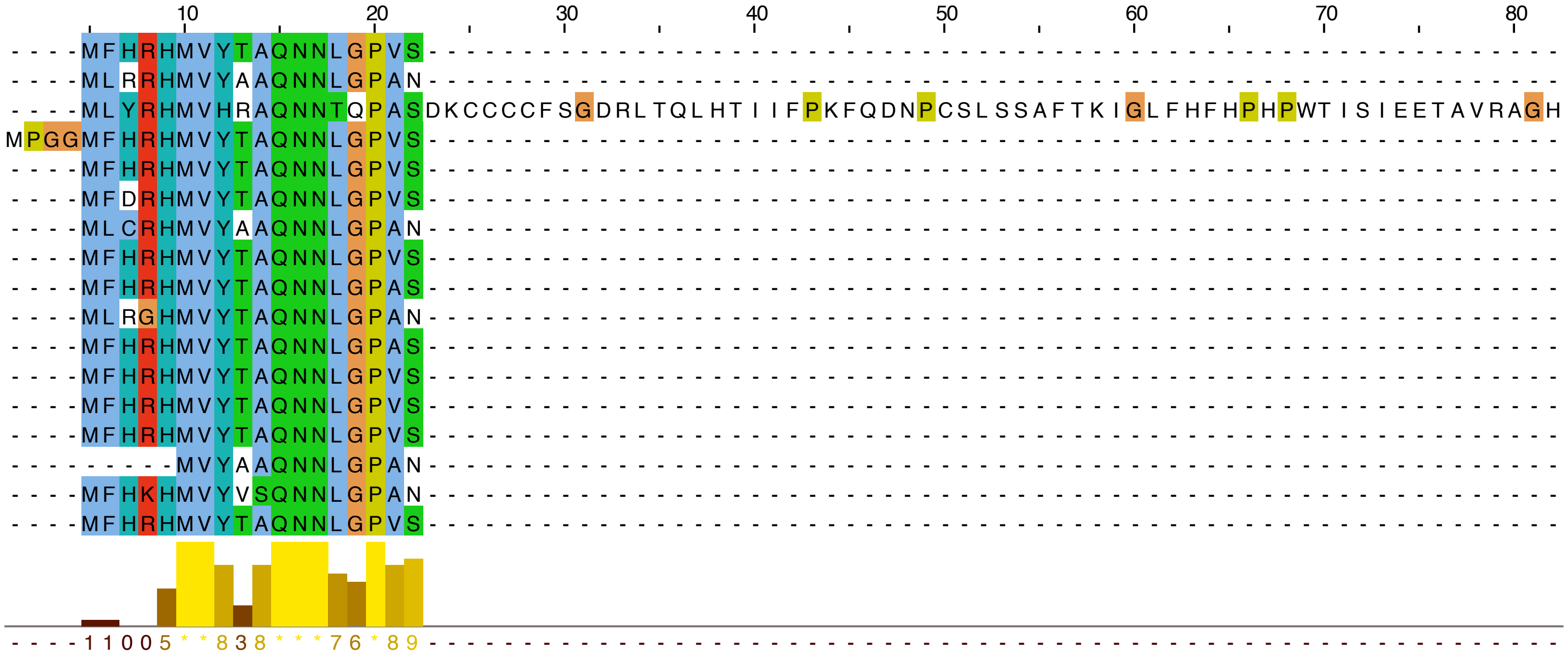

**HG0163**

ENSG00000146285 (SCML4)

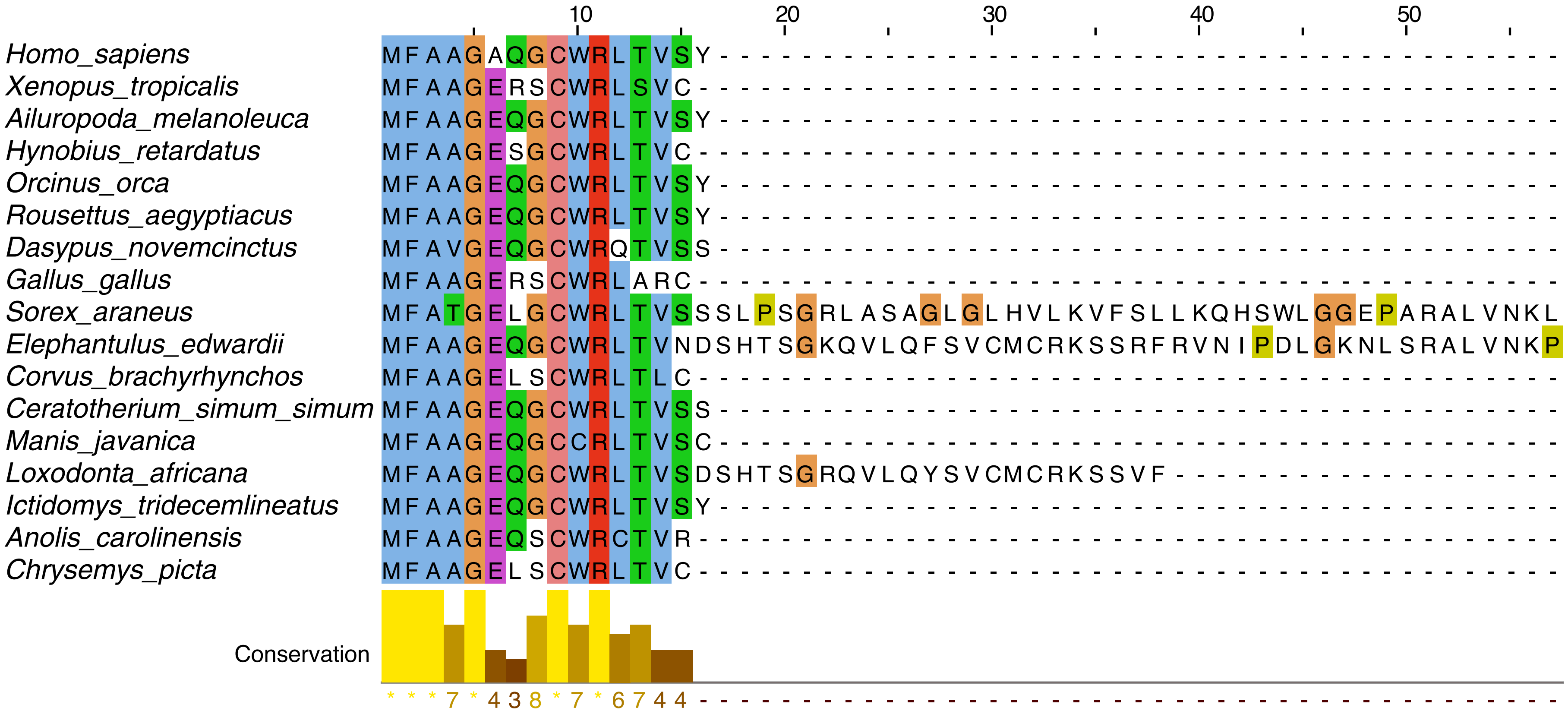

**HG0164**

ENSG00000151292 (CSNK1G3)

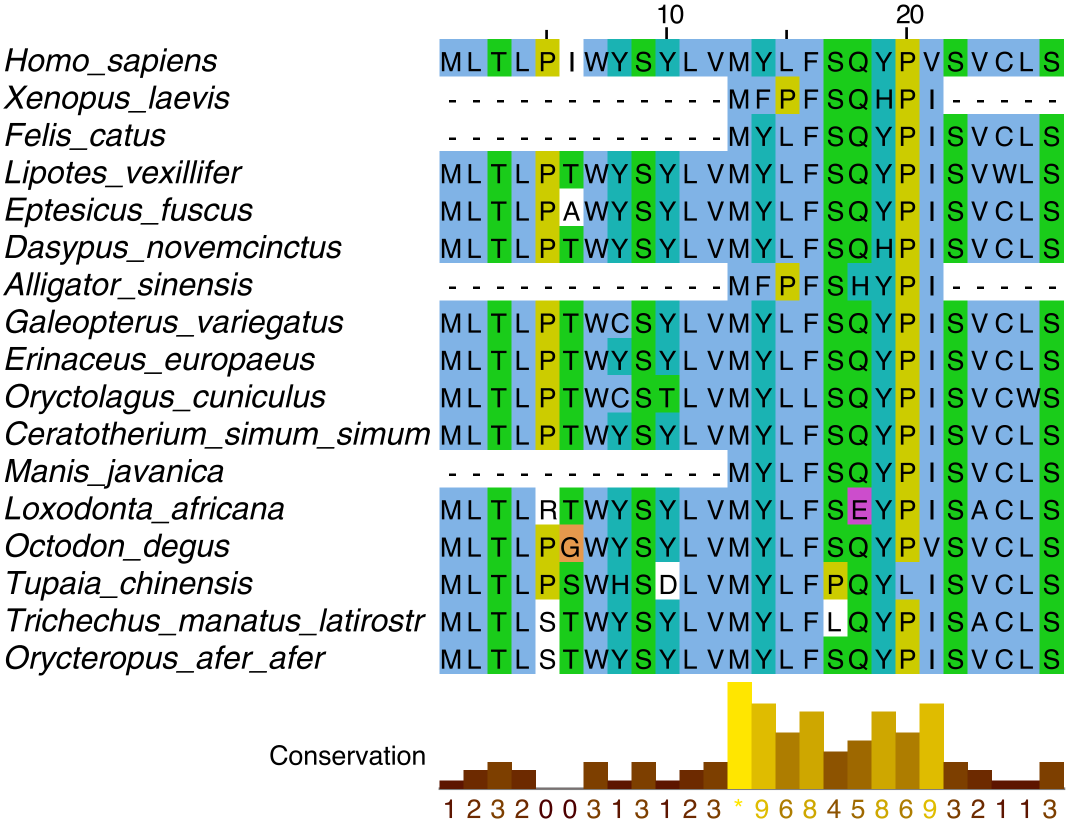

**HG0165**

ENSG00000177508 (IRX3)

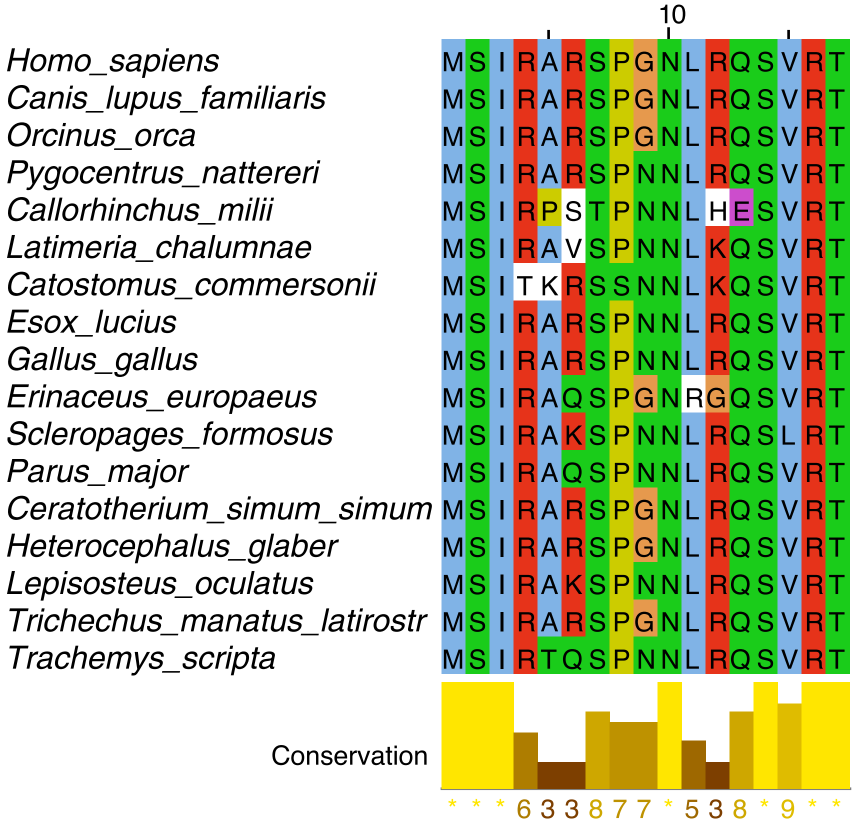

**HG0166.1**

ENSG00000148948 (LRRC4C)

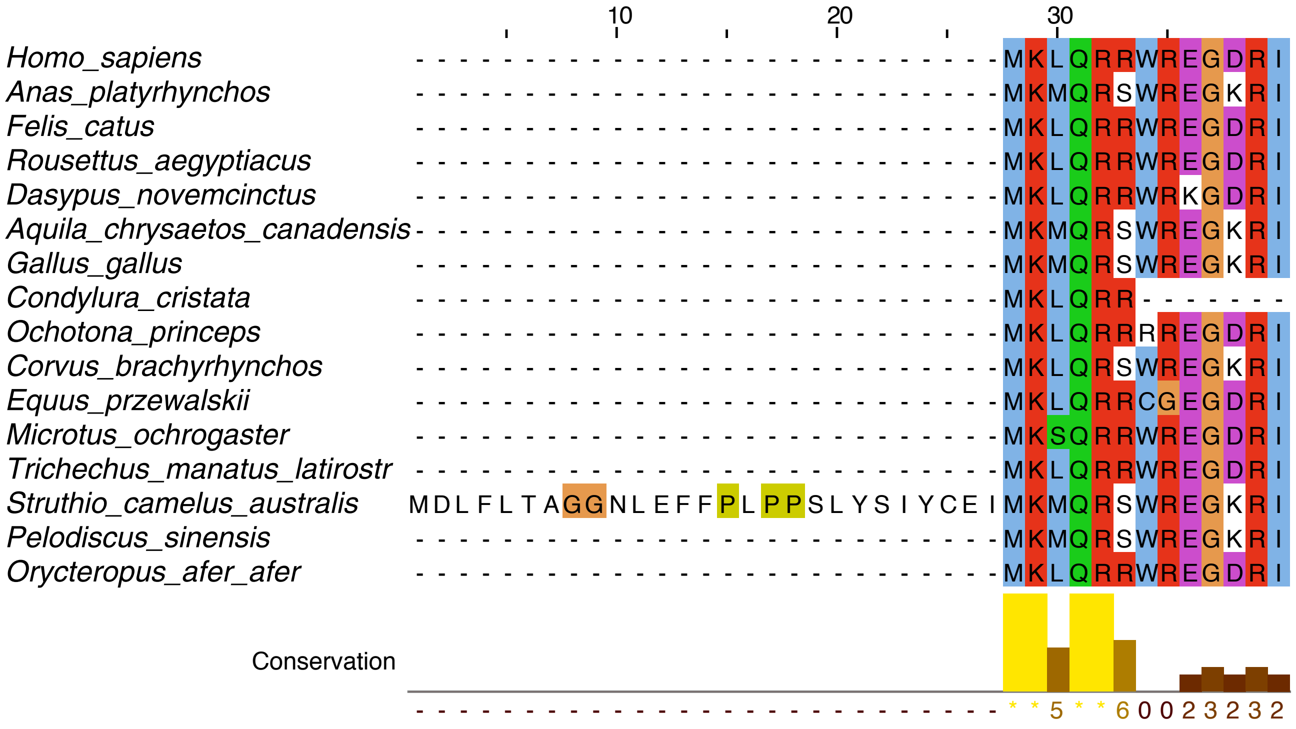

**HG0166.2**

ENSG00000148948 (LRRC4C)

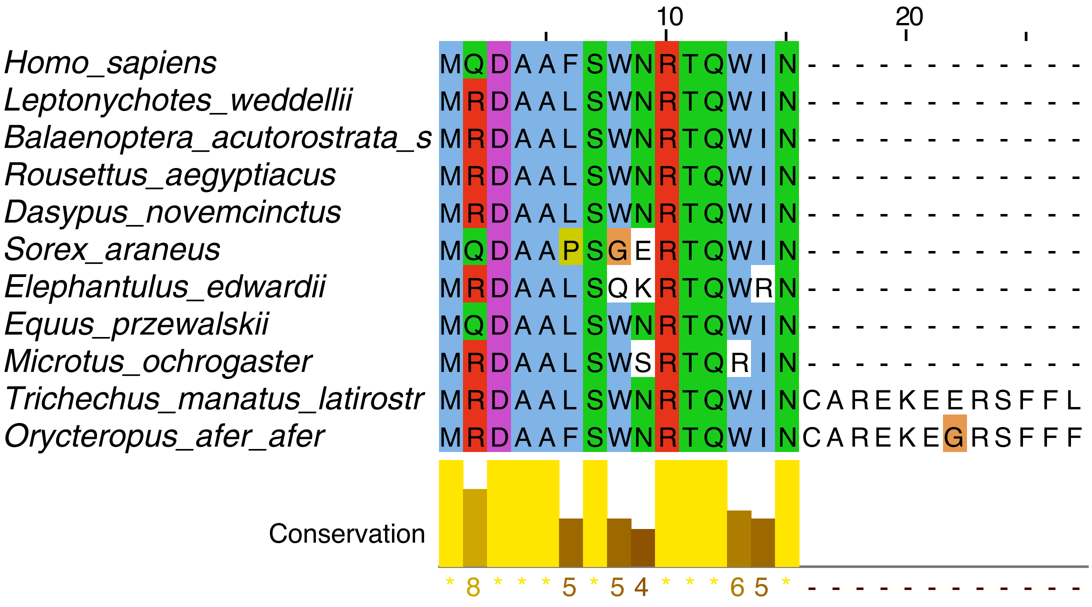

**HG0166.3**

ENSG00000148948 (LRRC4C)

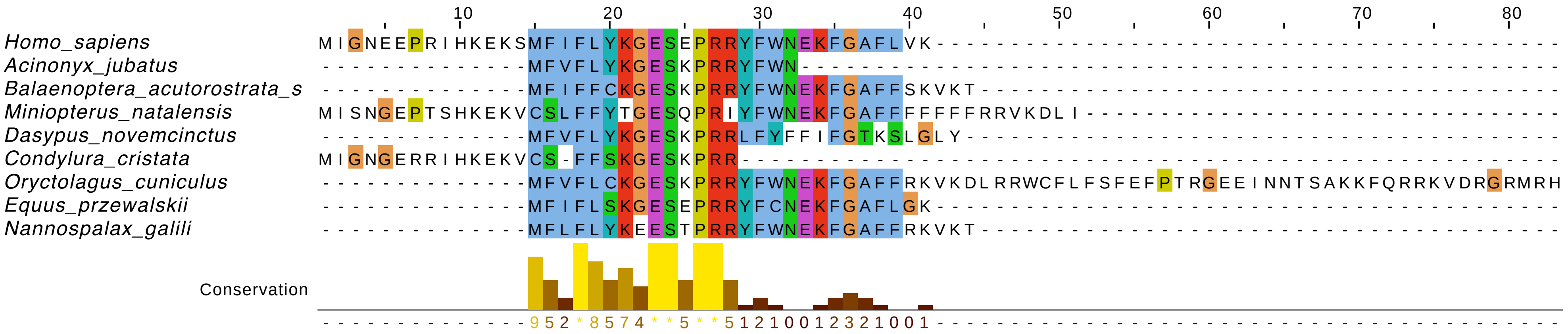

**HG0166.4**

ENSG00000148948 (LRRC4C)

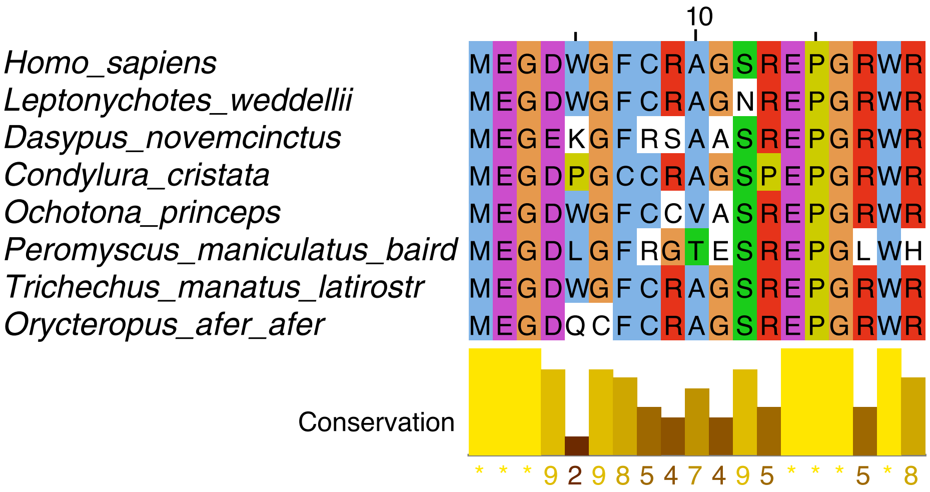

**HG0167**

ENSDARG00000104005 (slc4a3)

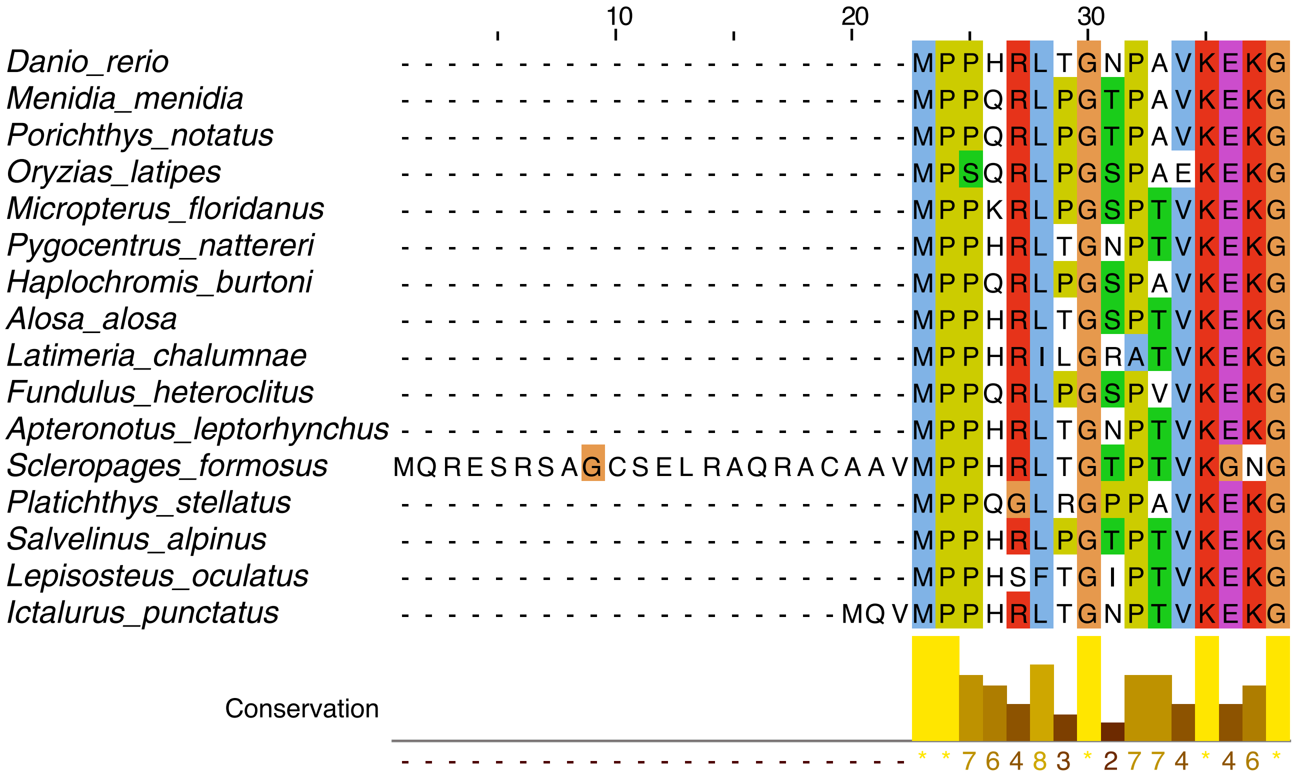

**HG0168**

ENSGALG00000006997 (BRINP1)

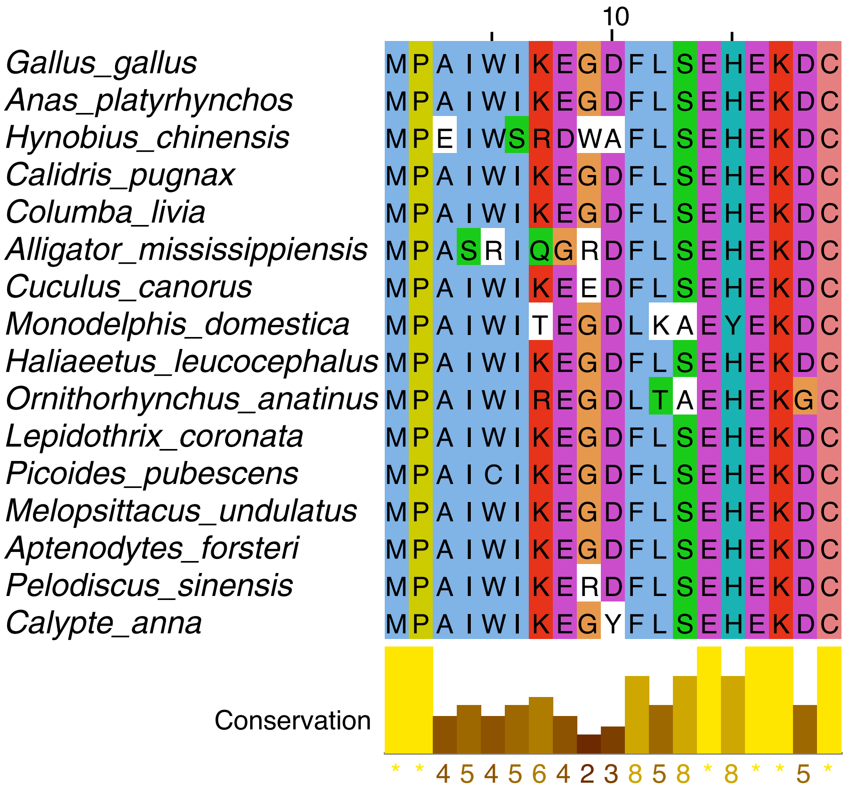

**HG0169**

ENSG00000114933 (INO80D)

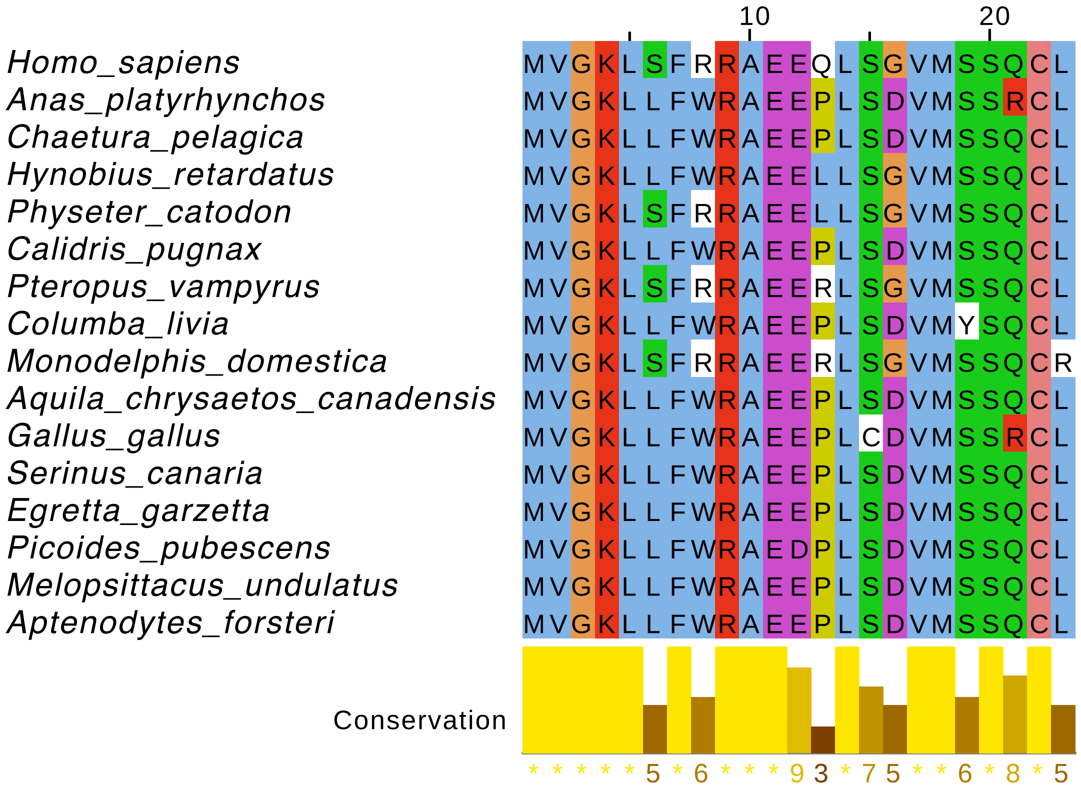

**HG0170**

ENSG00000180530 (NRIP1)

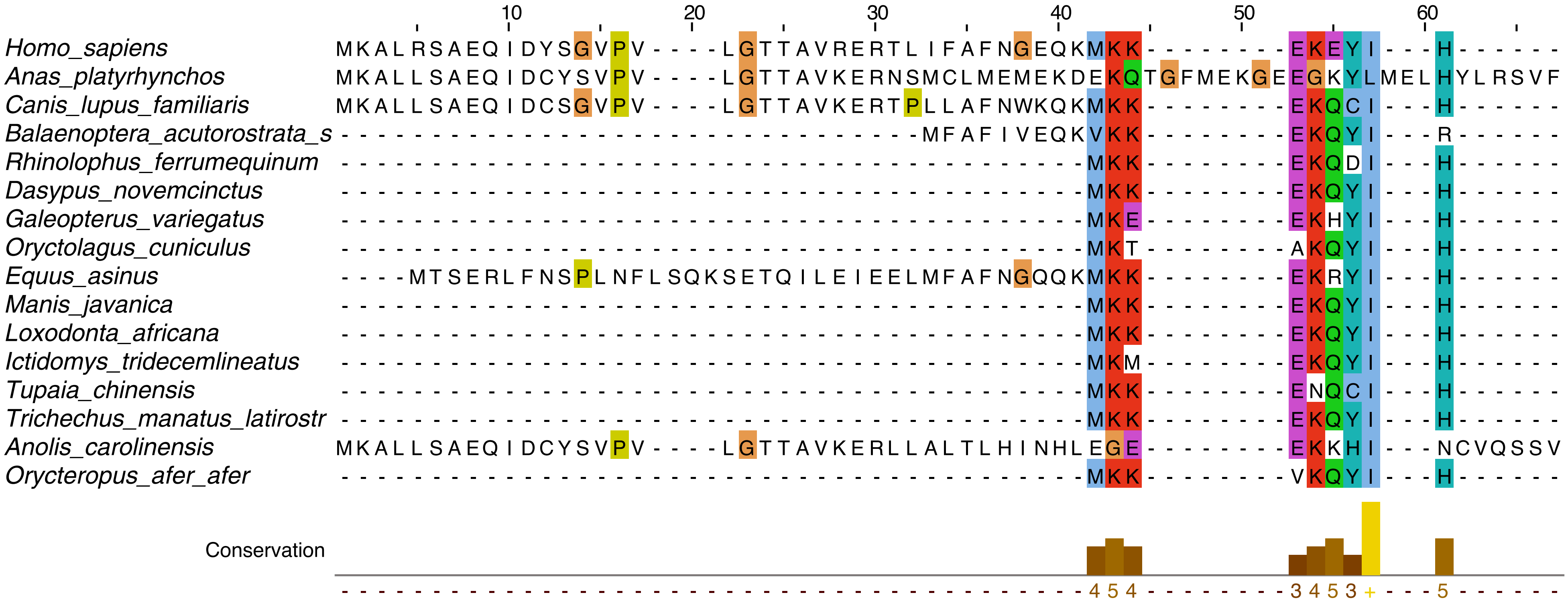

**HG0171.1**

ENSG00000177311 (ZBTB38)

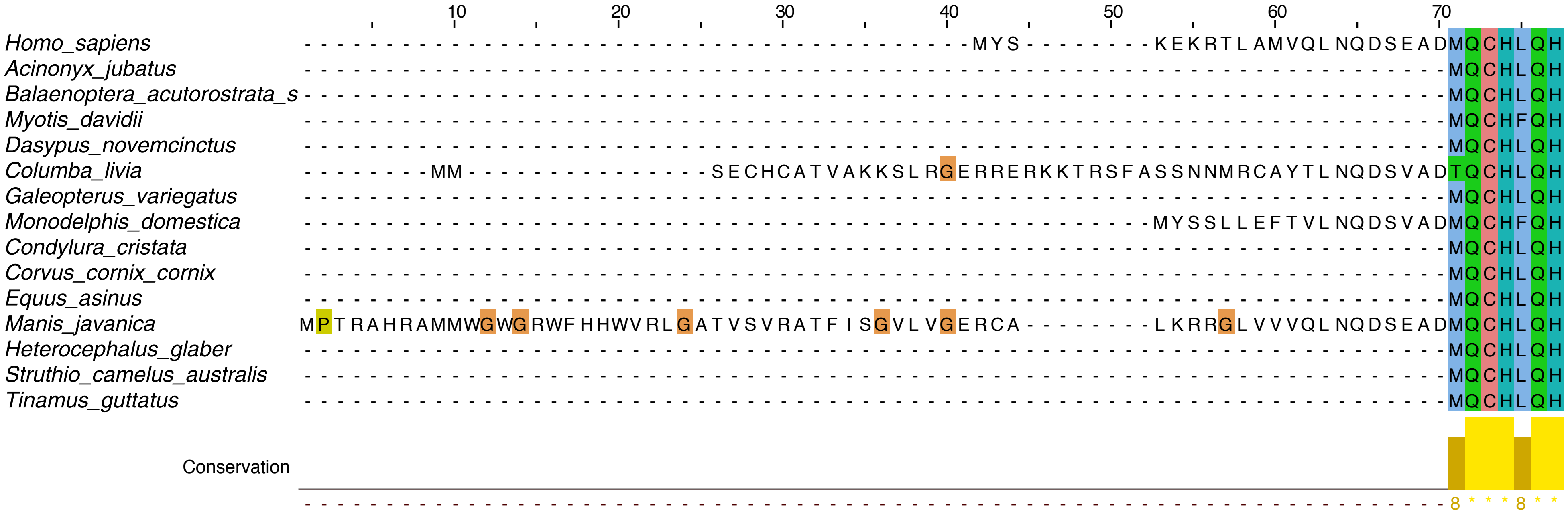

**HG0171.2**

ENSG00000177311 (ZBTB38)

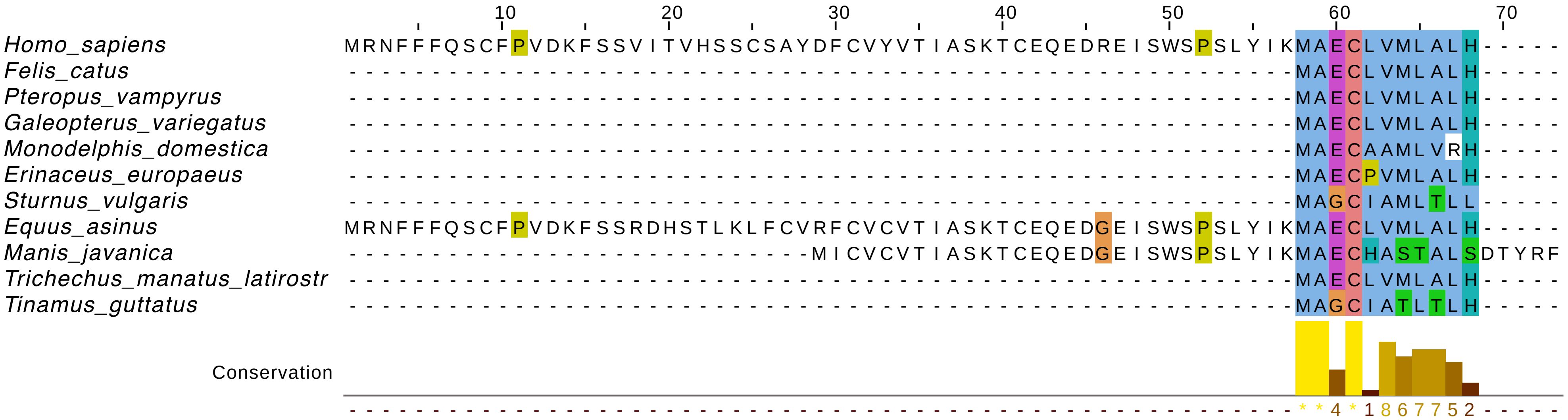

**HG0171.3**

ENSG00000177311 (ZBTB38)

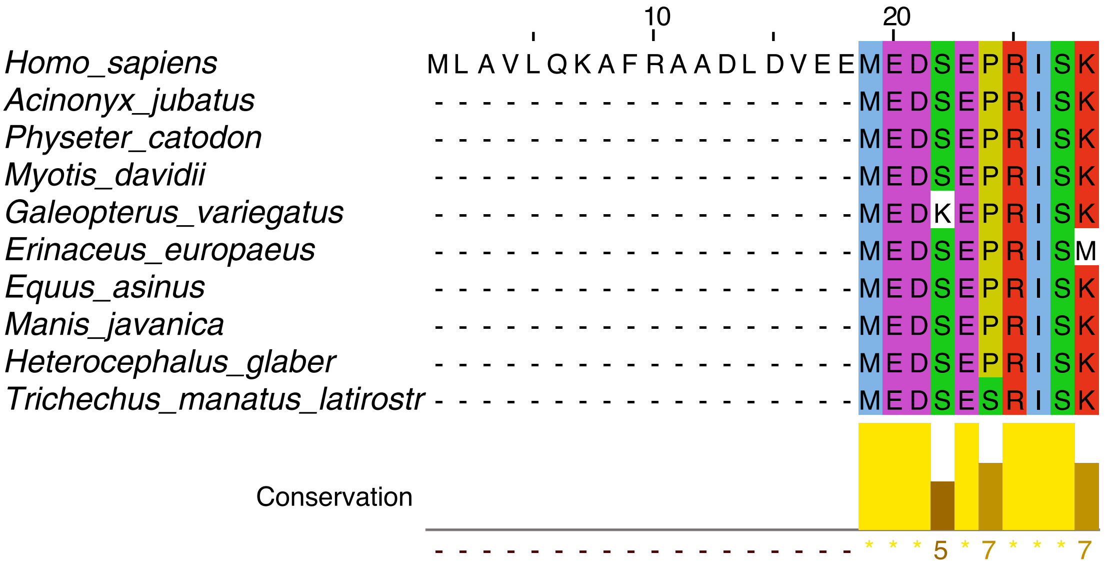

**HG0171.4**

ENSG00000177311 (ZBTB38)

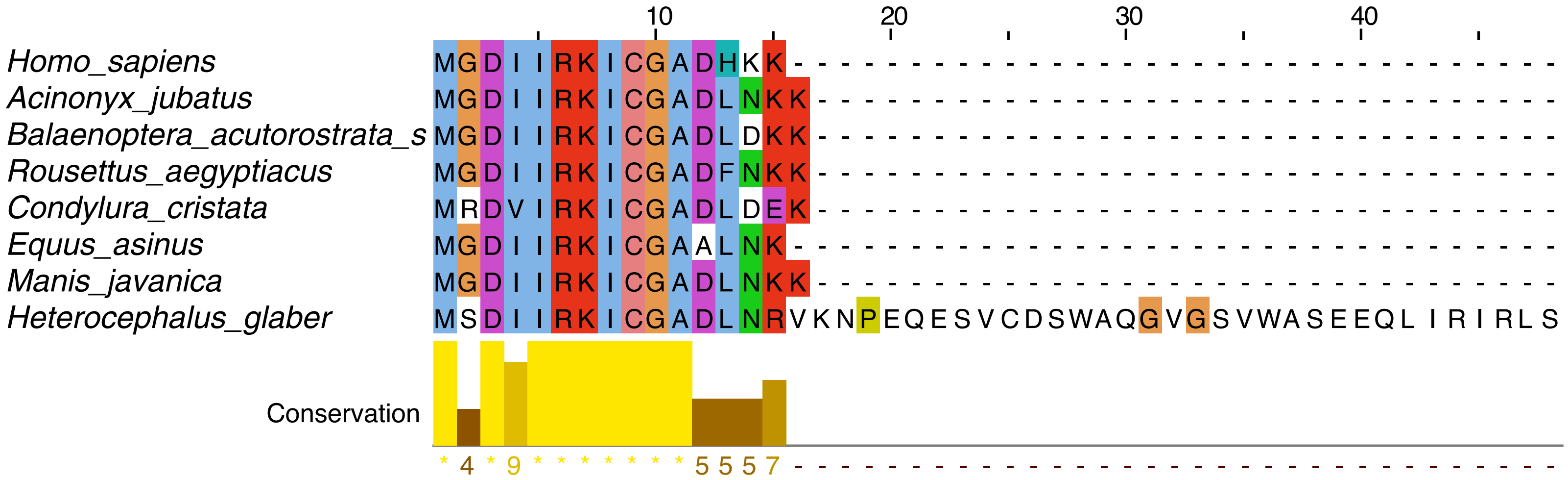

**HG0172.1**

ENSG00000196782 (MAML3)

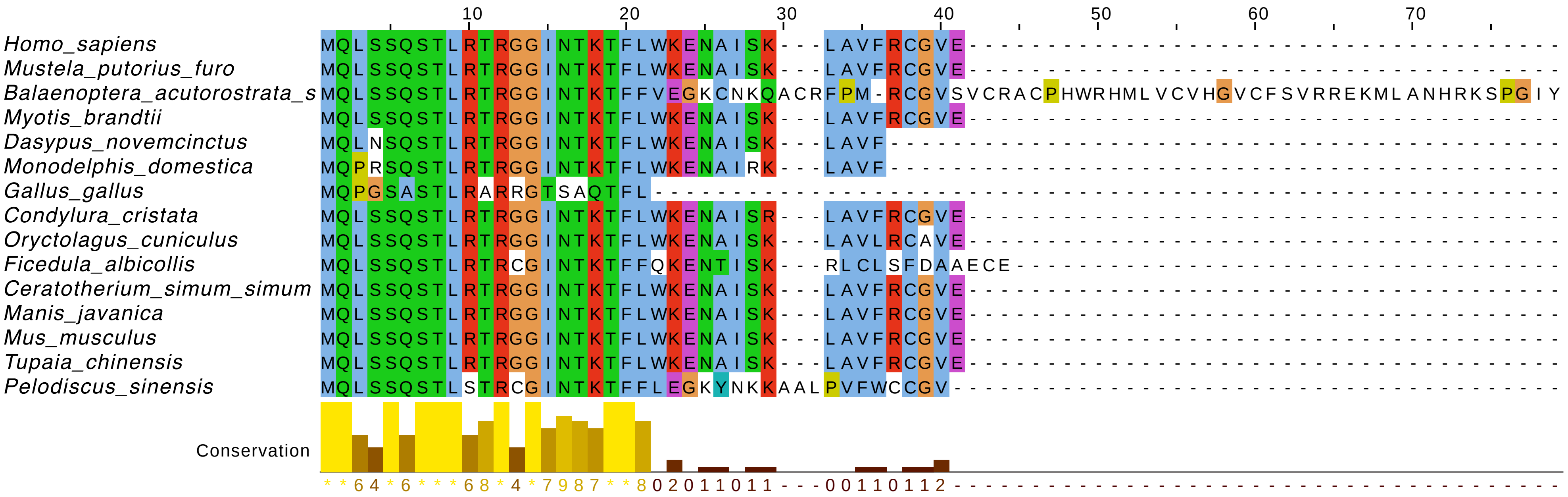

**HG0172.2**

ENSG00000196782 (MAML3)

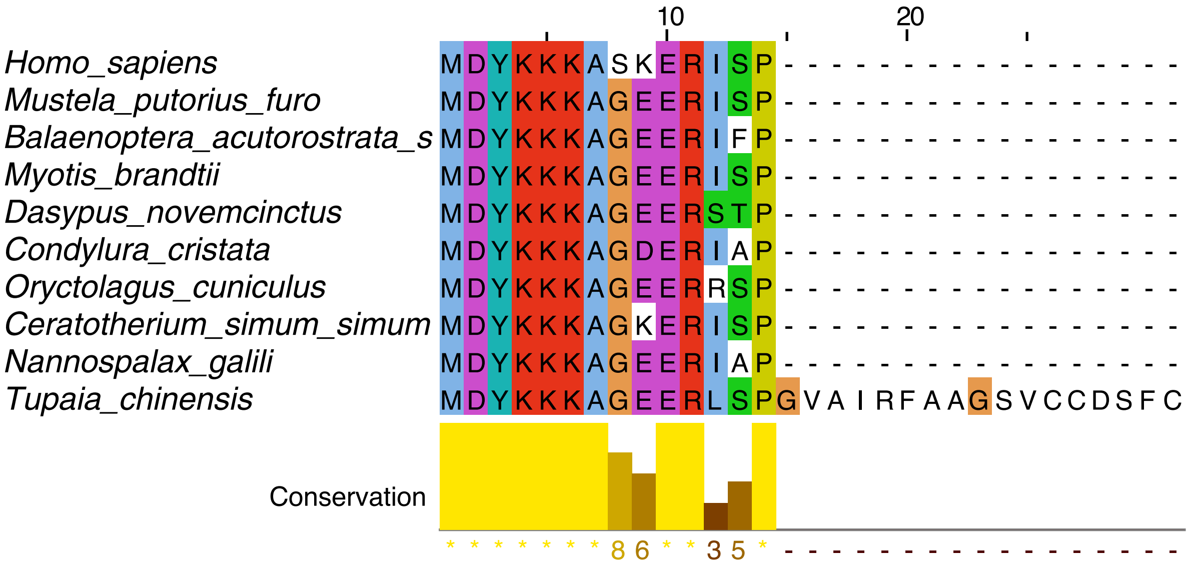

**HG0173**

ENSG00000101746 (NOL4)

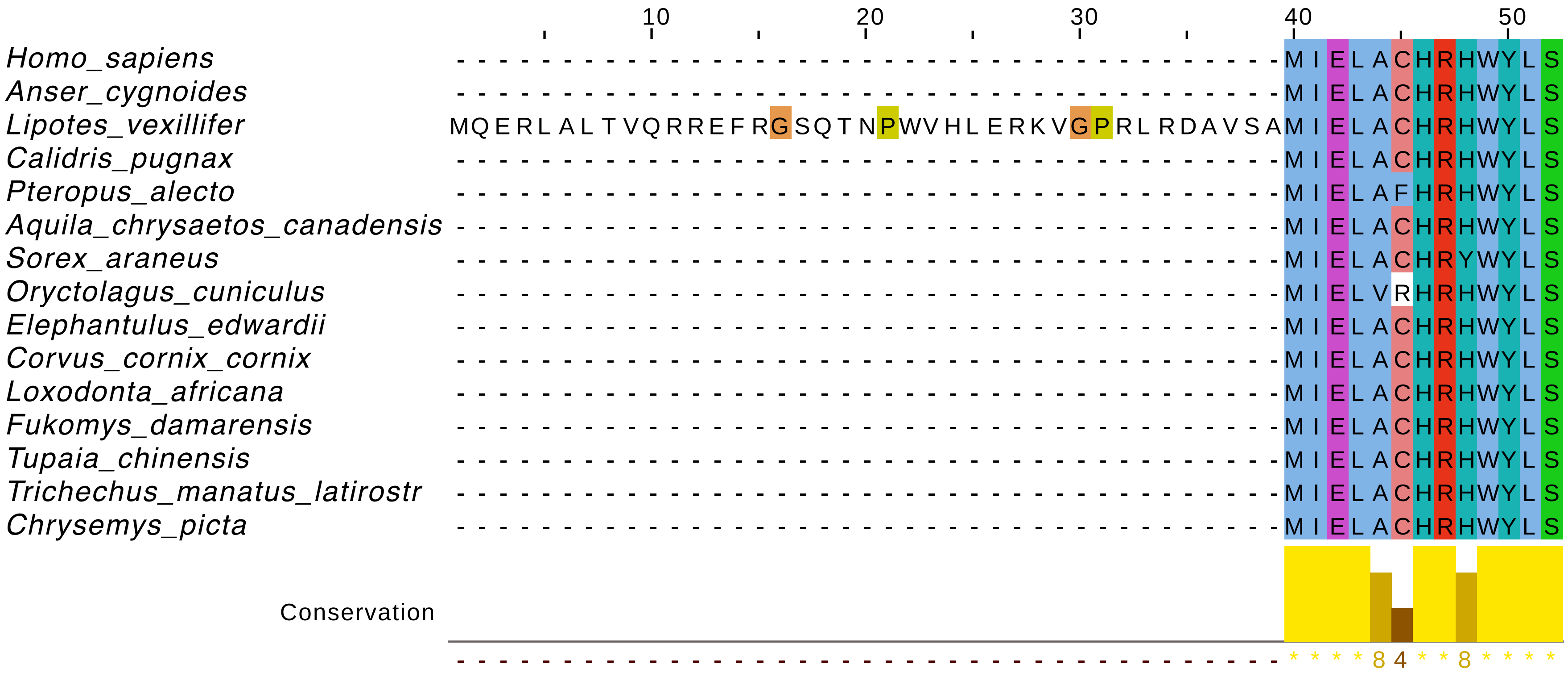

**HG0174**

ENSG00000144355 (DLX1)

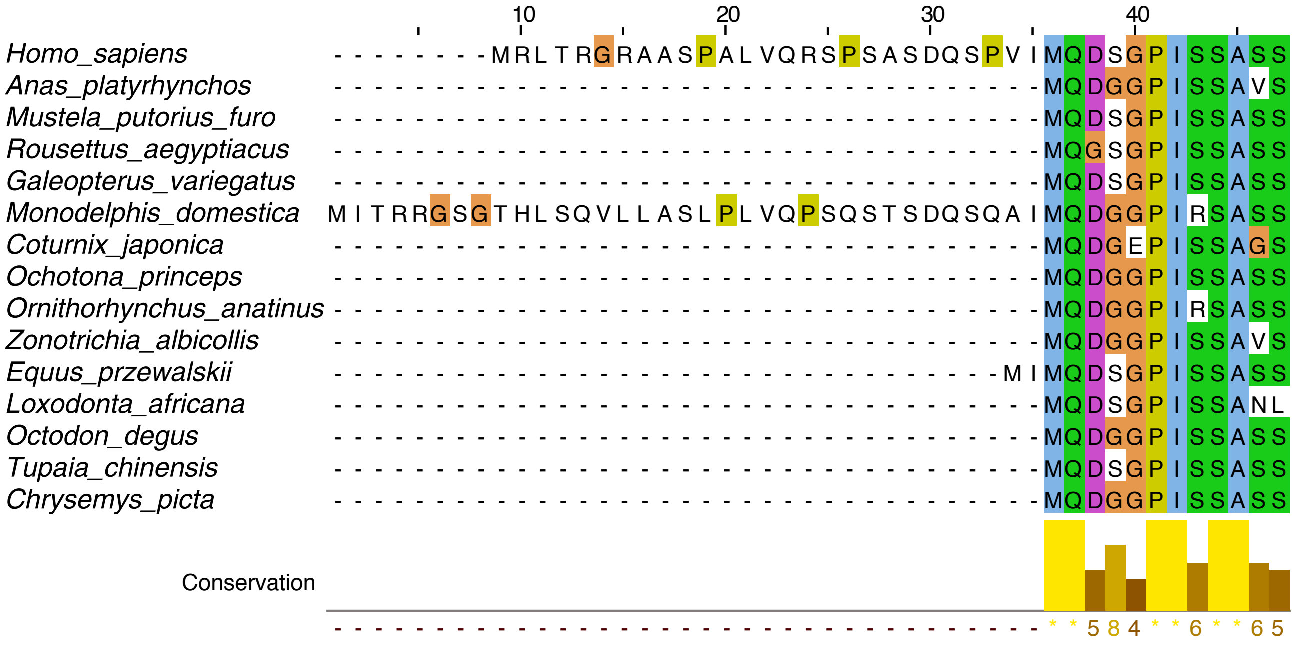

**HG0175**

ENSG00000151320 (AKAP6)

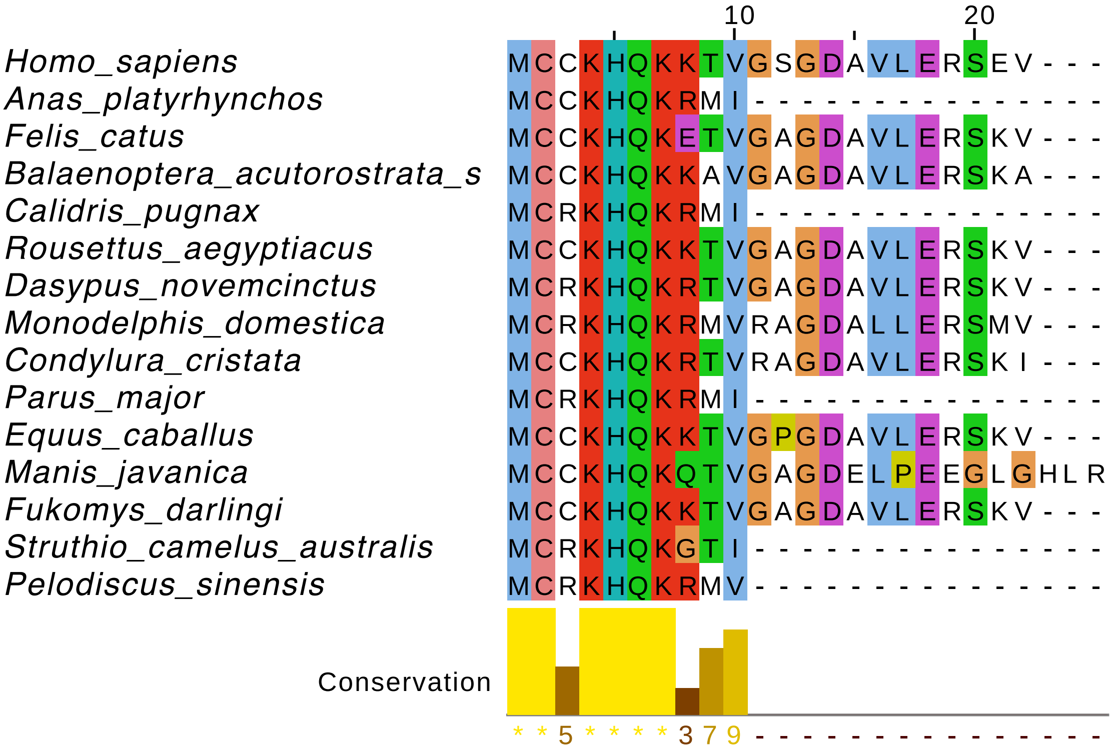

**HG0176**

ENSG00000170962 (PDGFD)

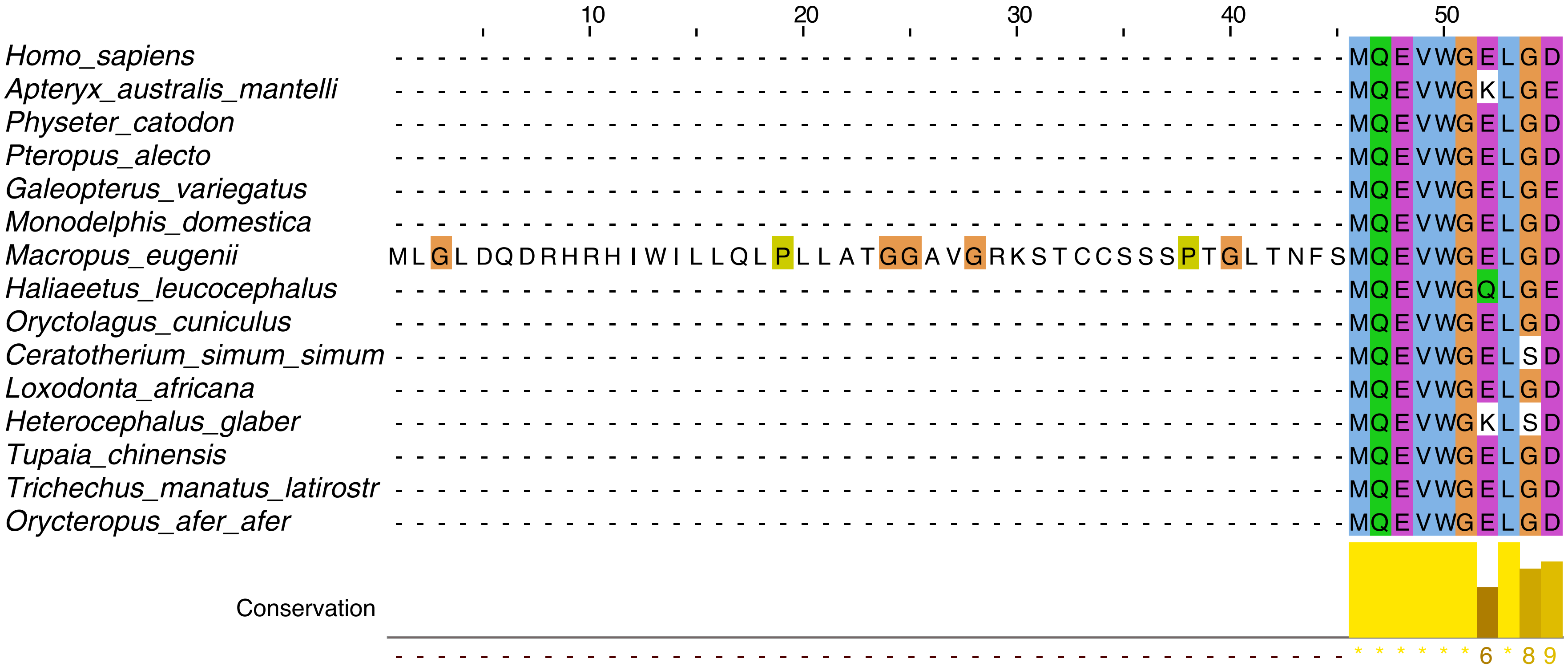

**HG0177**

ENSG00000171843 (MLLT3)

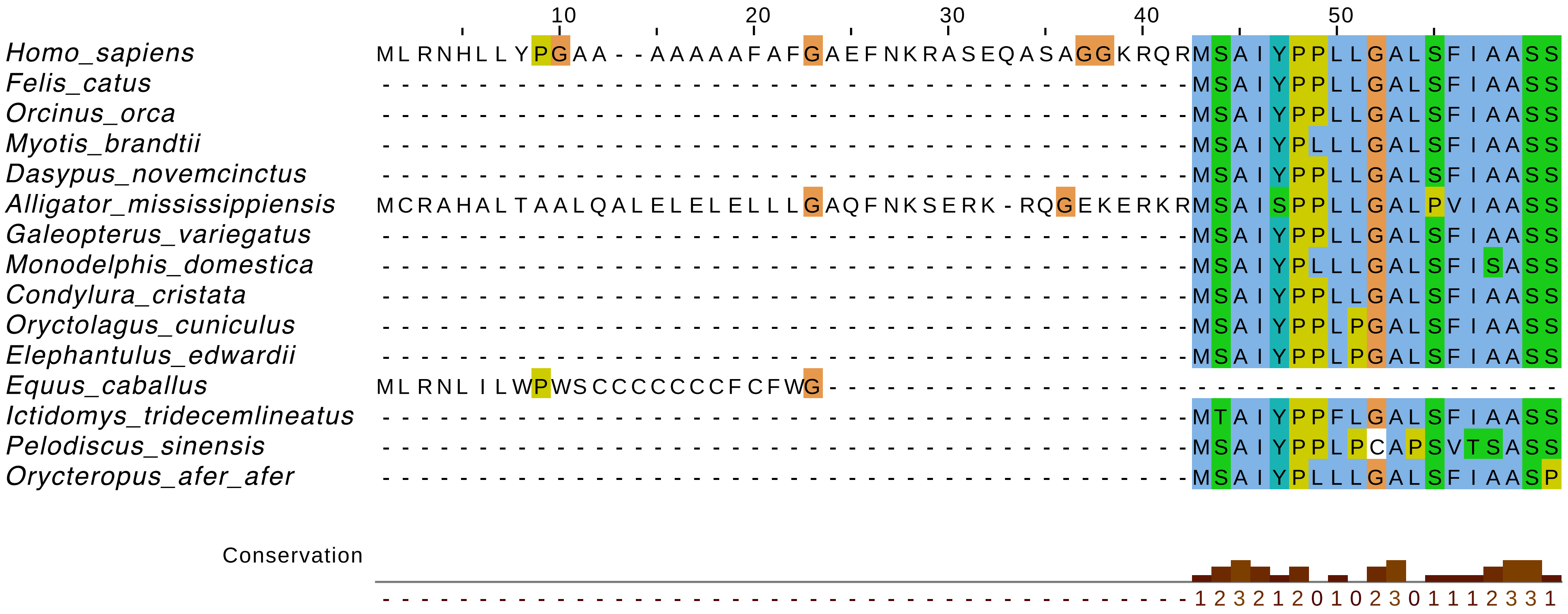

**HG0178.1**

ENSG00000131634 (TMEM204)

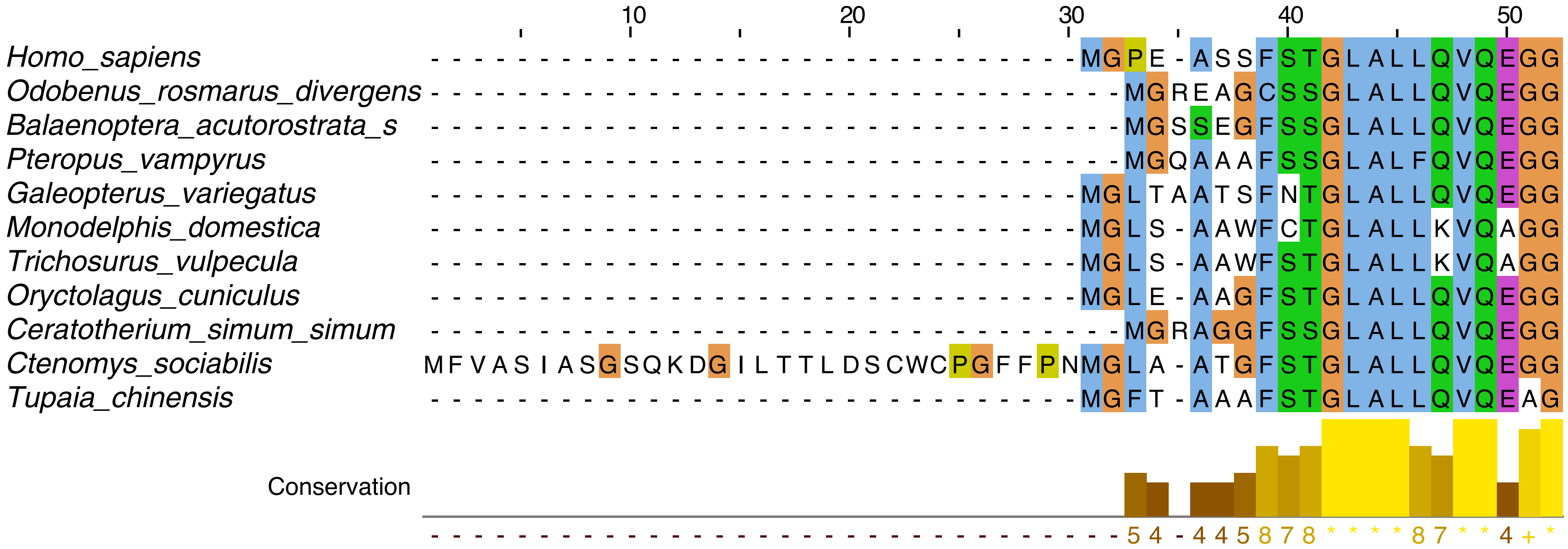

**HG0178.1**

ENSGALG00000029047 (TMEM204)

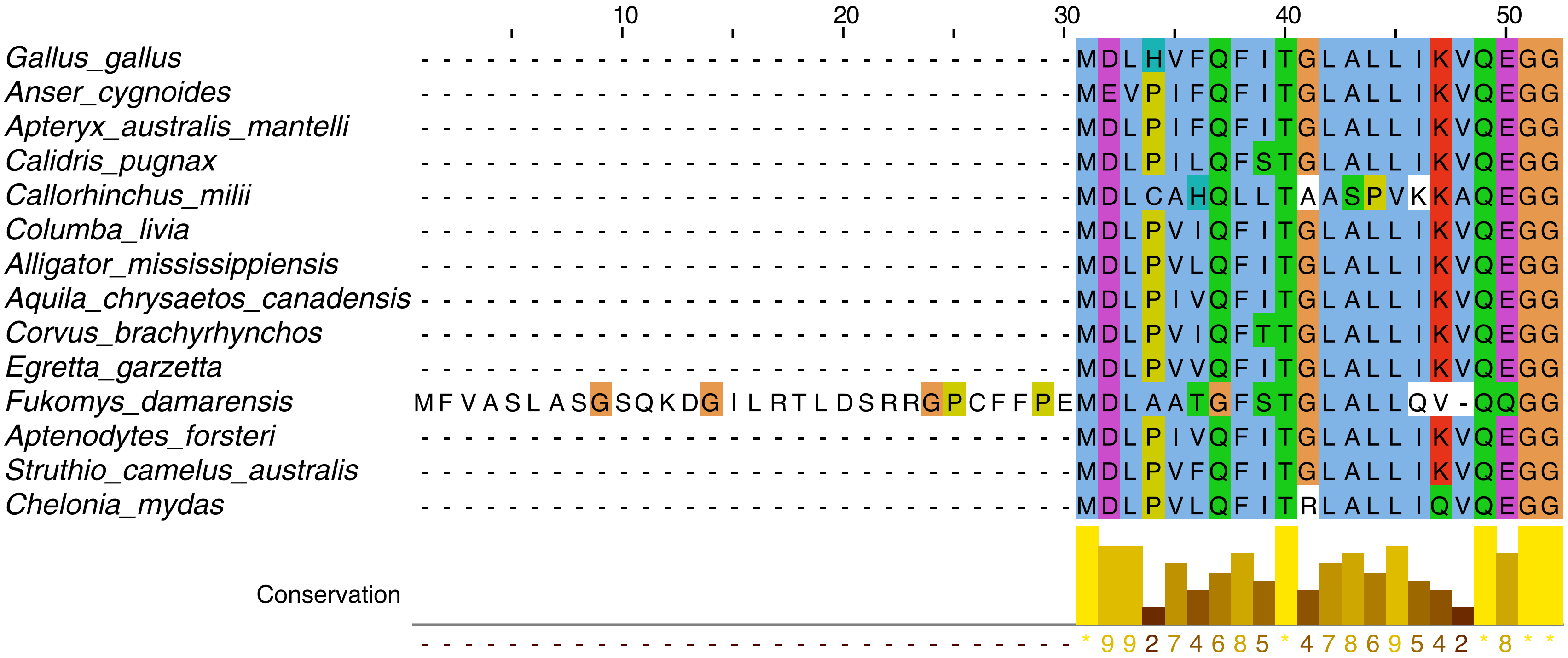

# HG0178.2

### ENSGALG00000029047 (TMEM204)

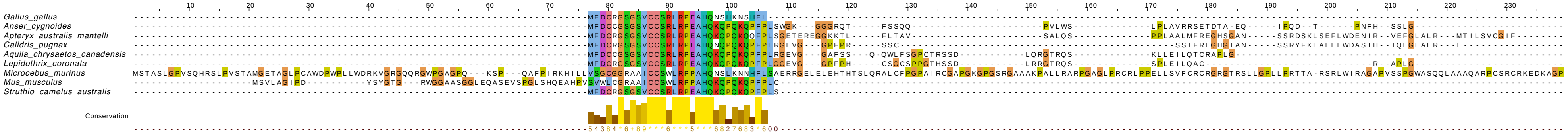

**HG0179**

ENSDARG00000036549 (agpat3)

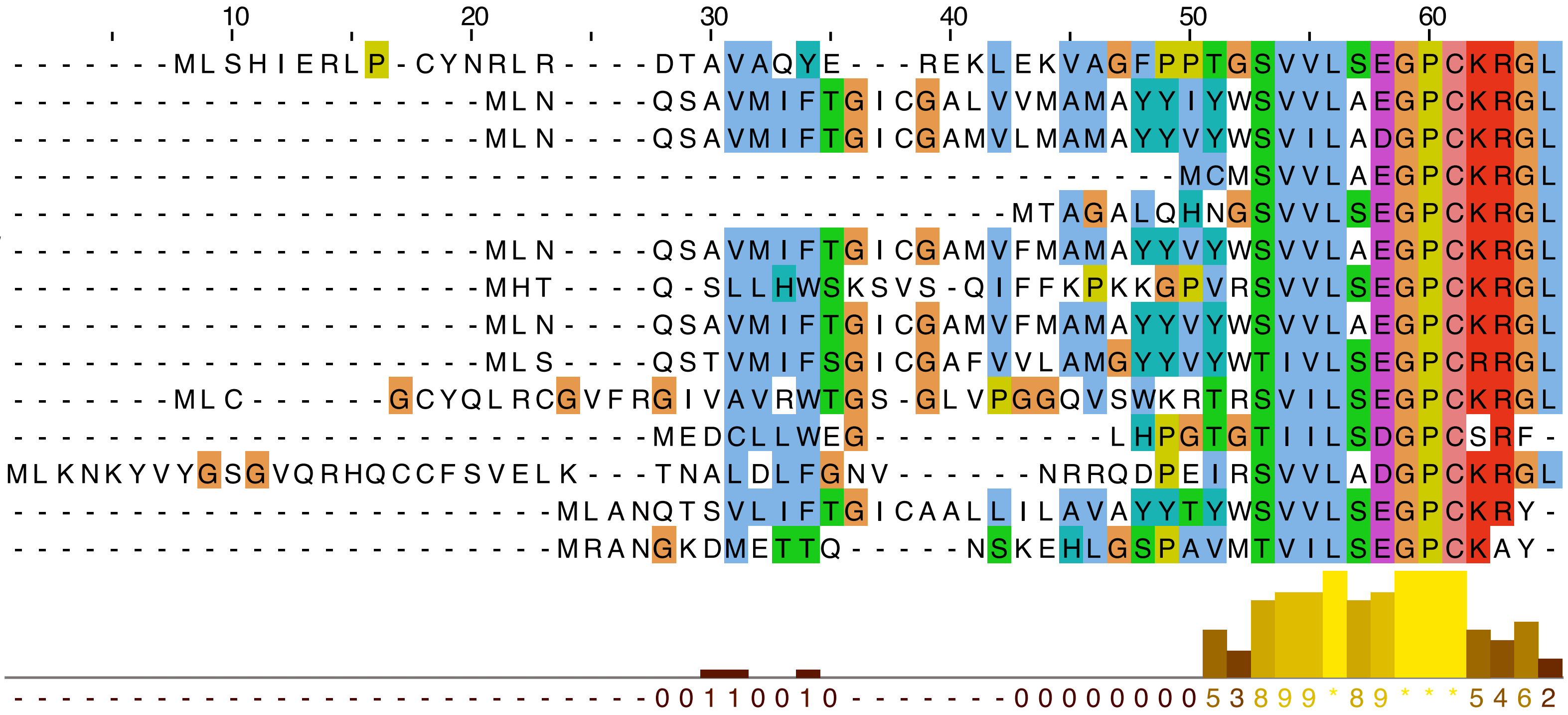

**HG0179**

ENSG00000160216 (AGPAT3)

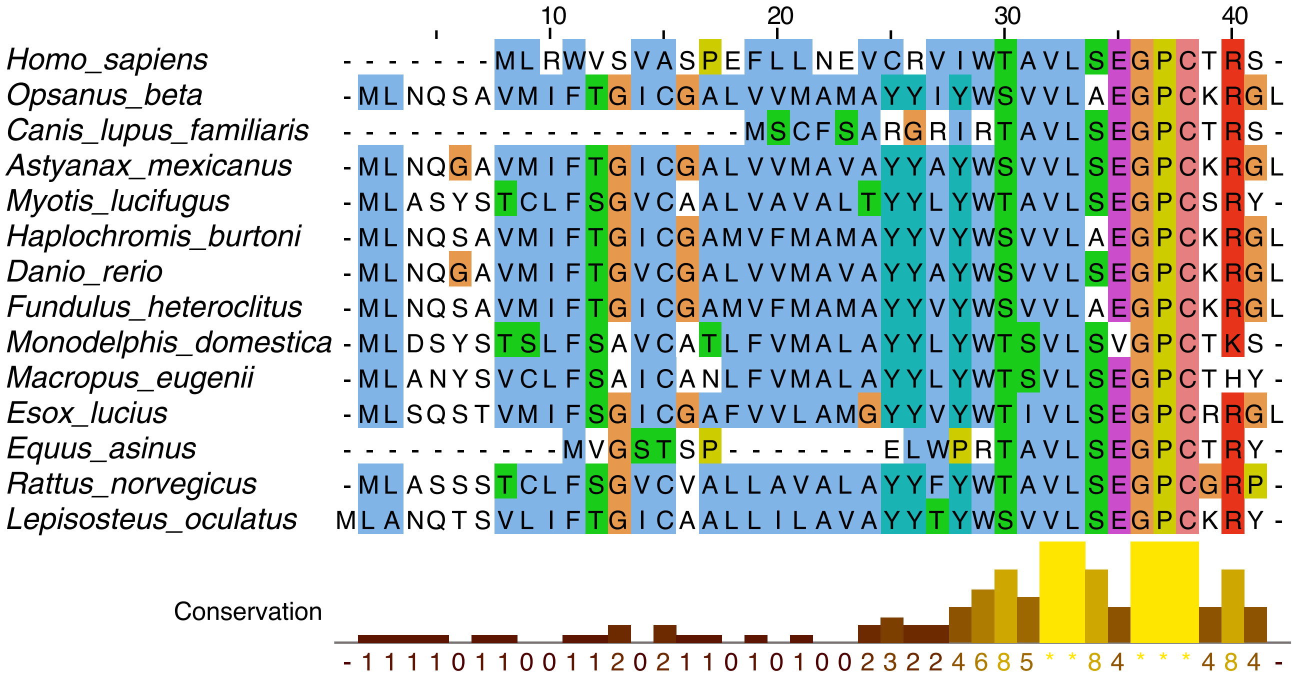

**HG0180.1**

ENSG00000112246 (SIM1)

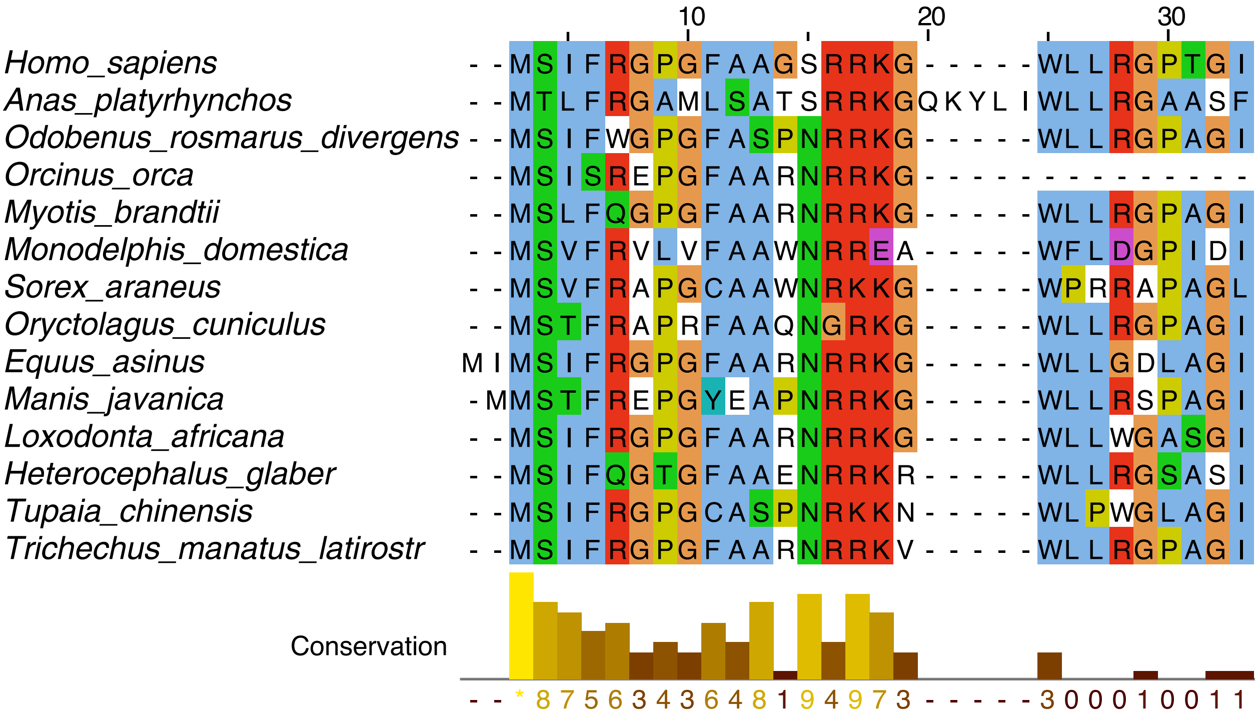

**HG0180.2**

ENSG00000112246 (SIM1)

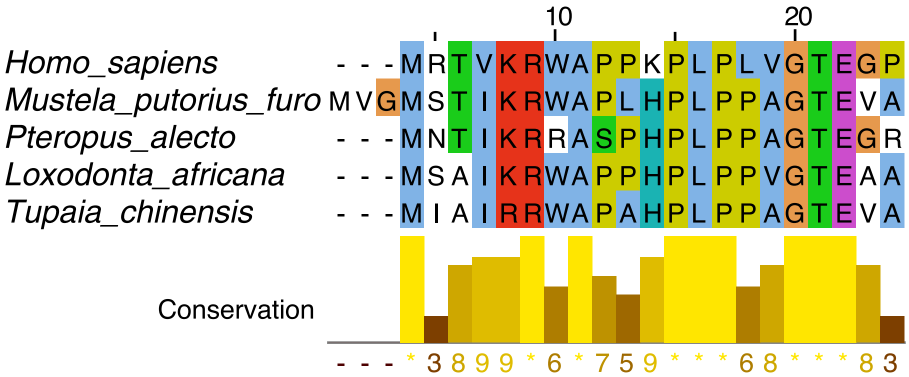

**HG0181.1**

ENSG00000143776 (CDC42BPA)

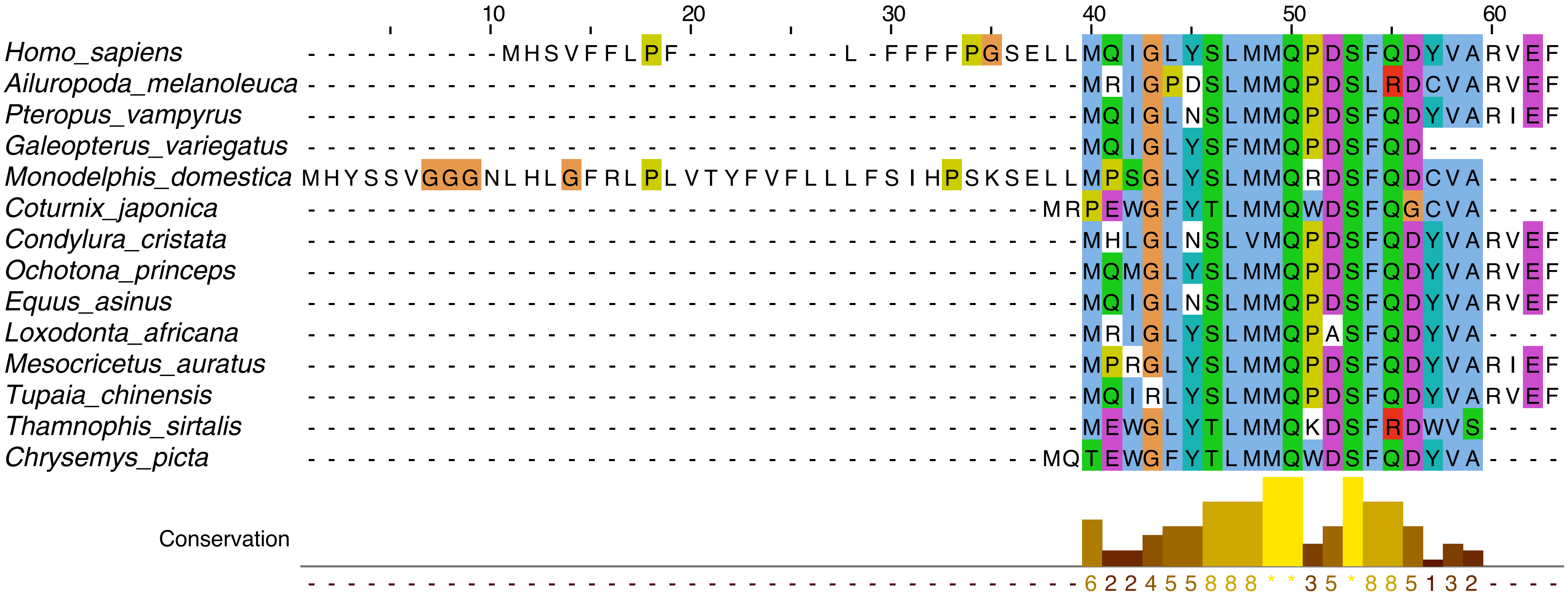

**HG0181.2**

ENSG00000143776 (CDC42BPA)

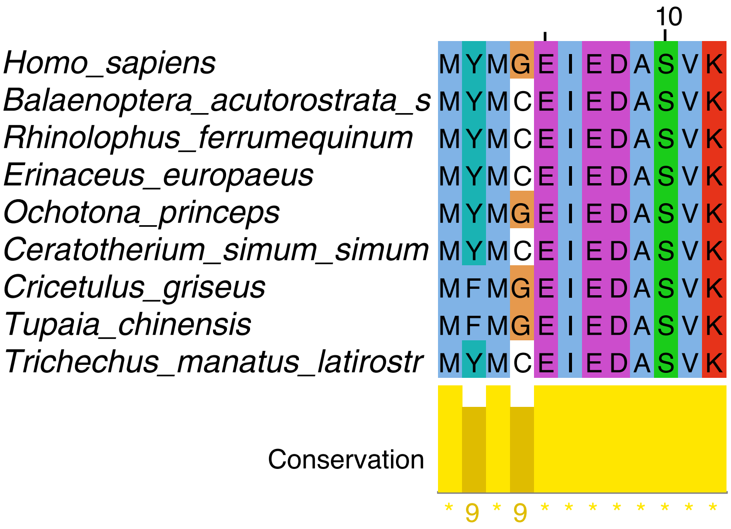

**HG0182**

ENSGALG00000031991

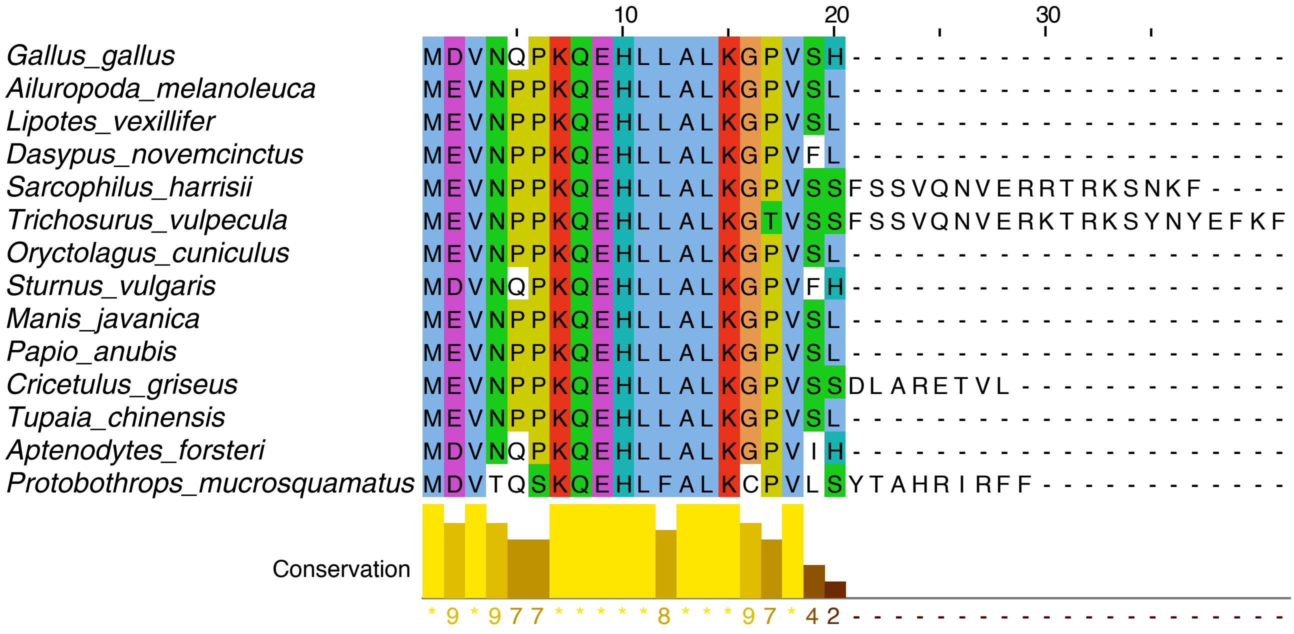

**HG0183**

ENSG00000011465 (DCN)

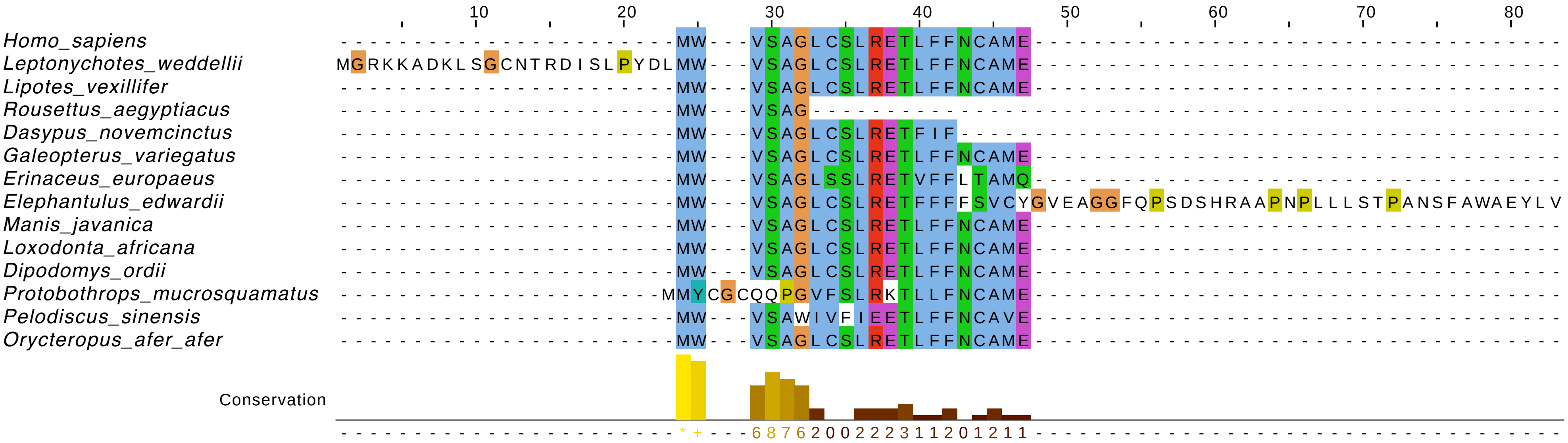

**HG0184**

ENSG00000136542 (GALNT5)

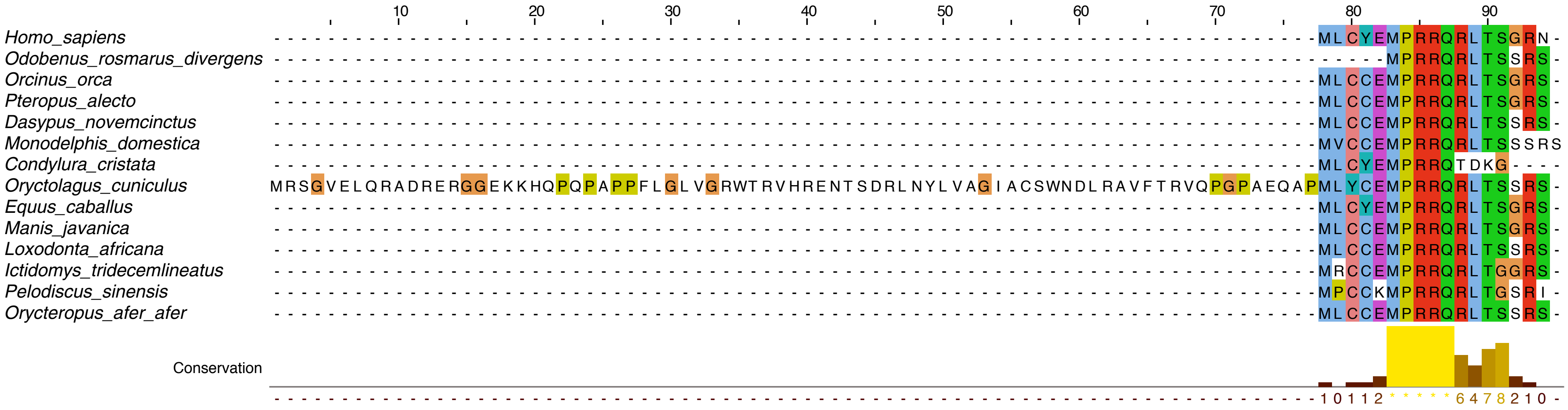

**HG0185**

ENSG00000143033 (MTF2)

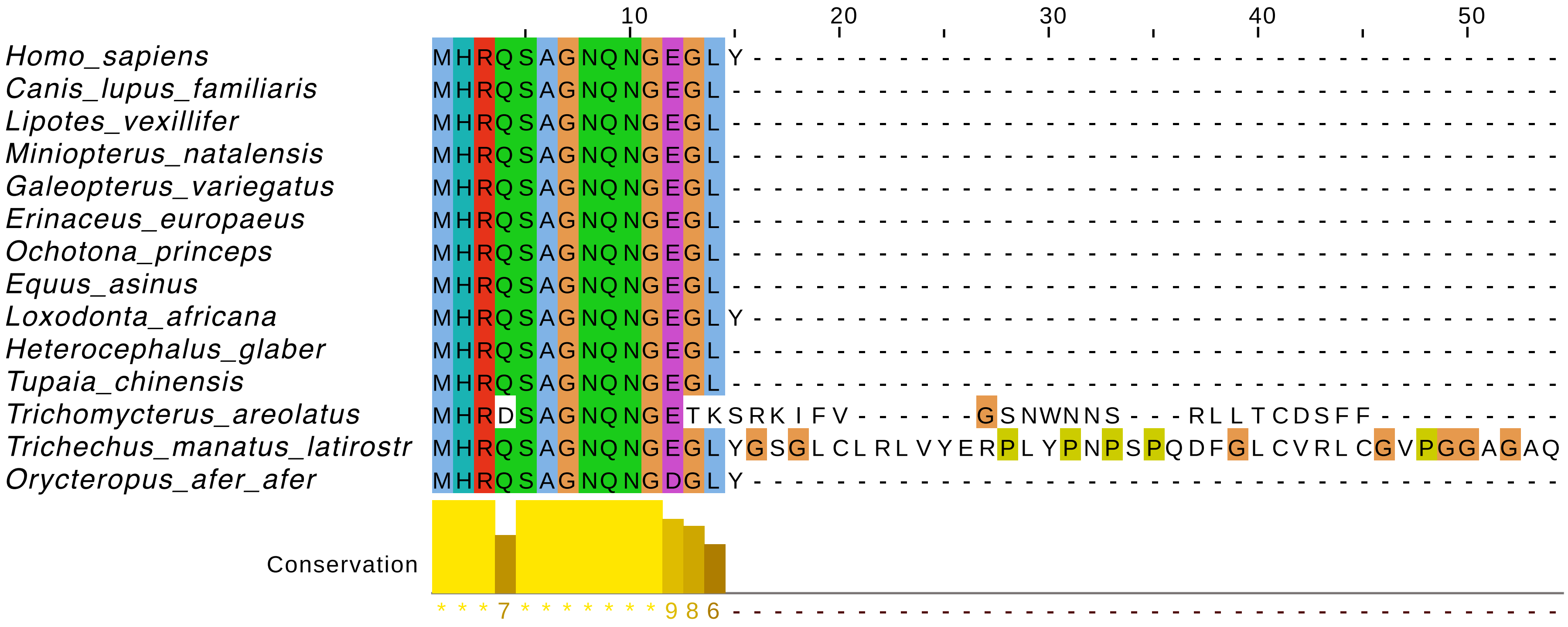

**HG0186**

ENSG00000167552 (TUBA1A)

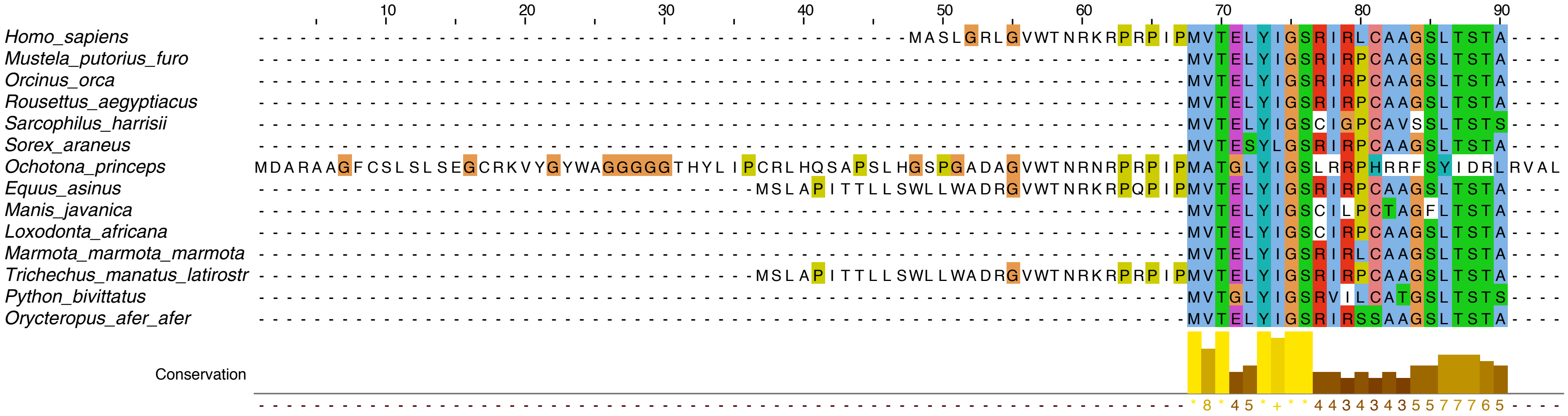

**HG0187.1**

ENSGALG00000004058 (GPR146)

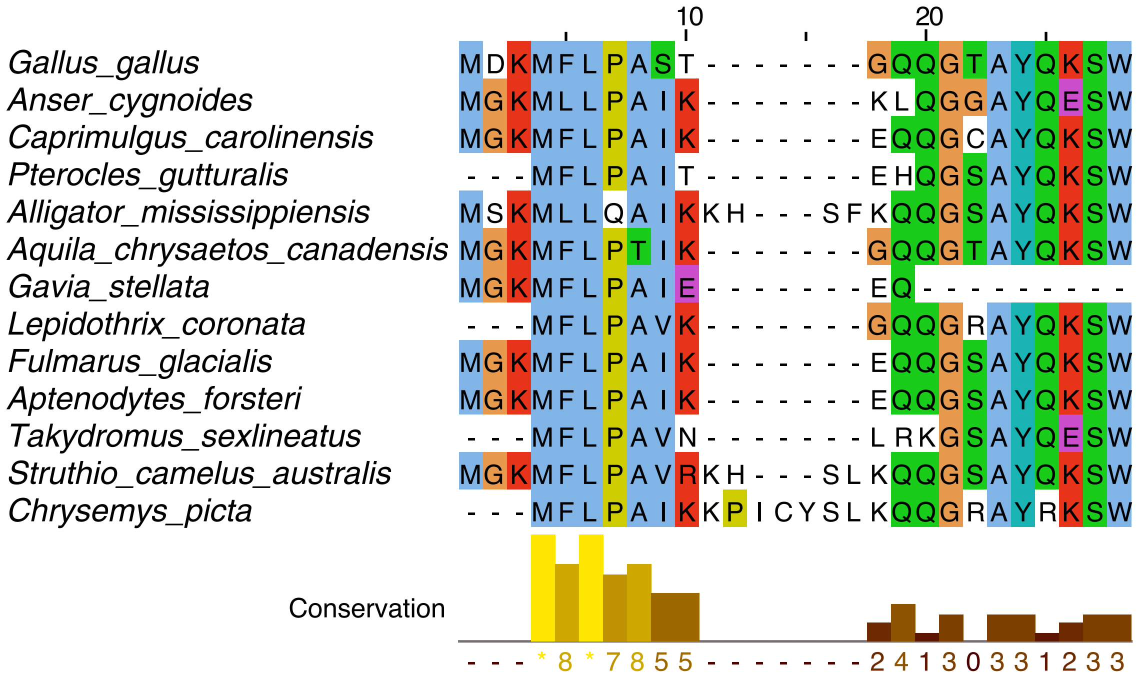

**HG0187.2**

ENSDARG00000059610 (gpr146)

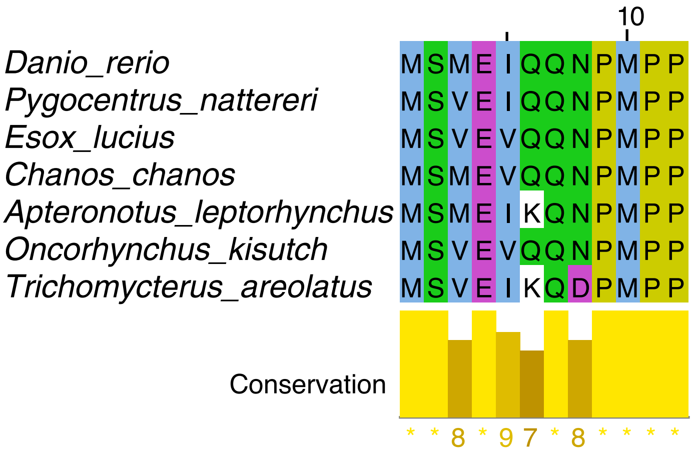

**HG0187.3**

ENSGALG00000004058 (GPR146)

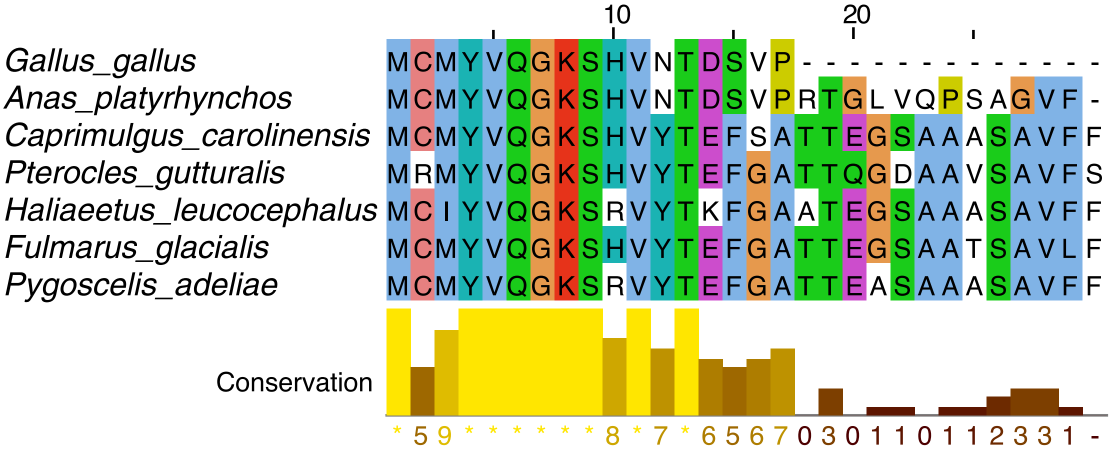

**HG0188**

ENSG00000157625 (TAB3)

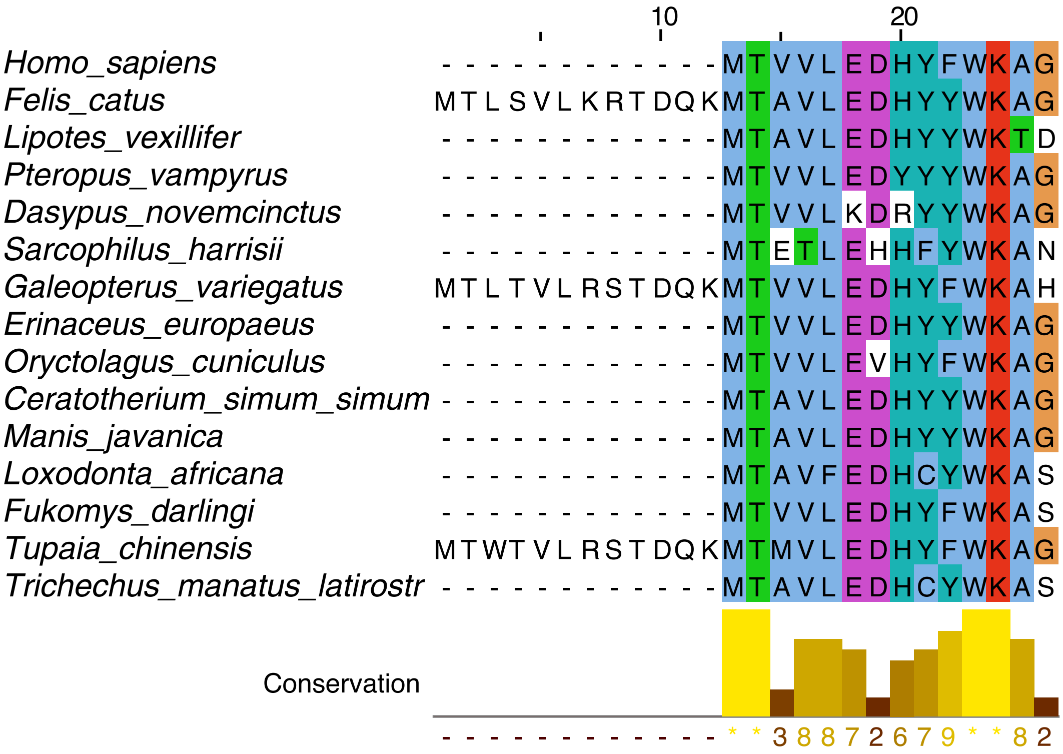

**HG0188**

ENSGALG00000016284 (TAB3)

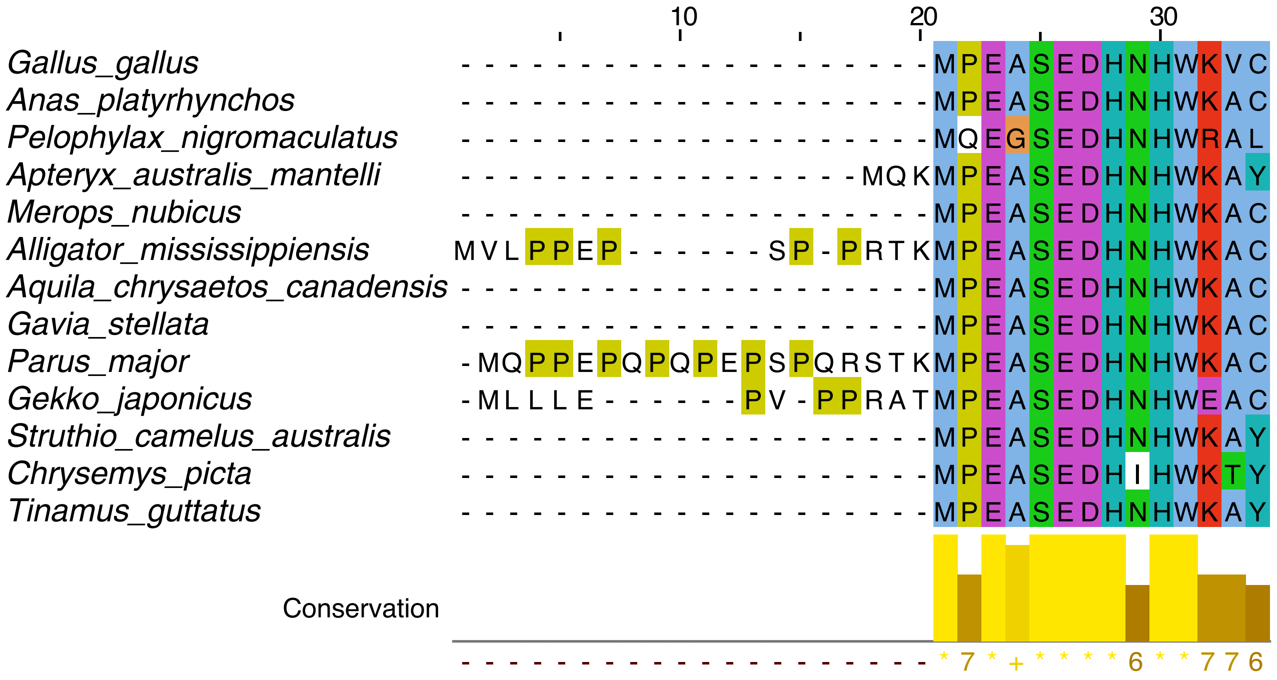

**HG0189**

ENSDARG00000001807 (tnfrsf21)

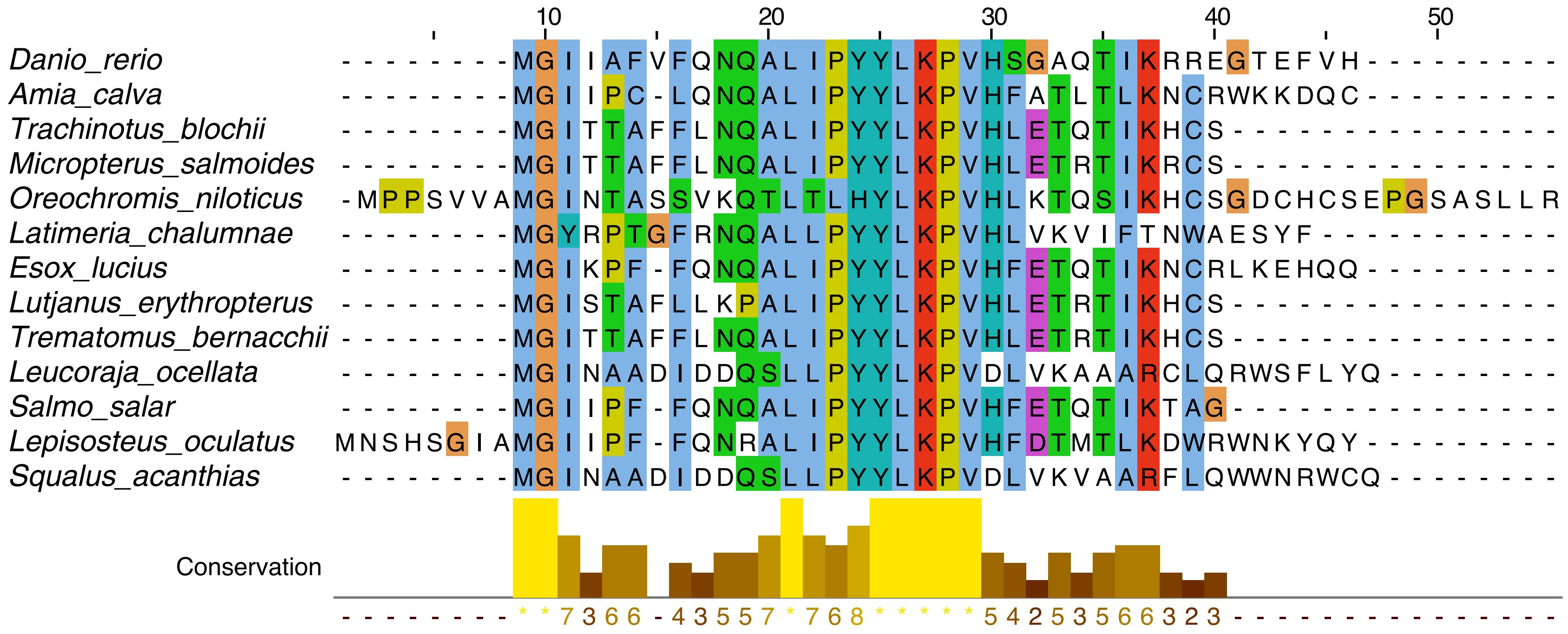

**HG0190**

ENSGALG00000006949 (CHD2)

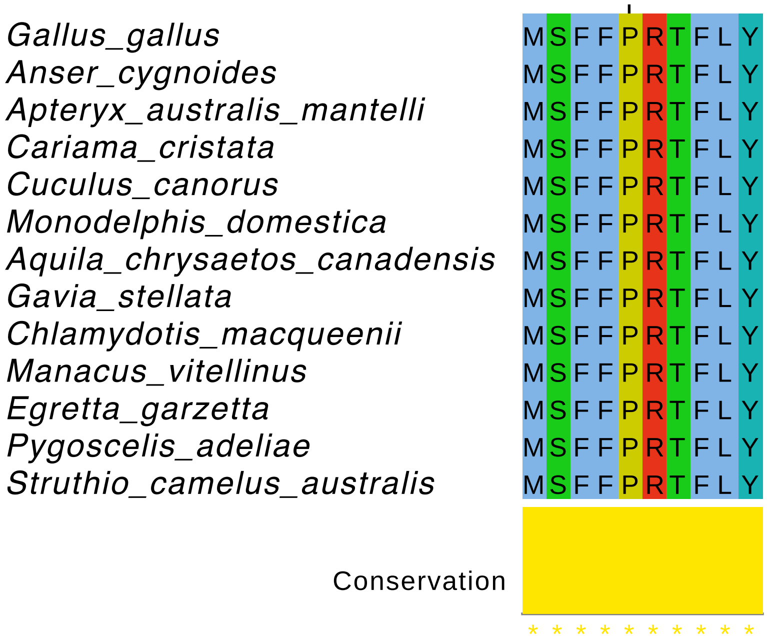

**HG0191**

ENSGALG00000036097

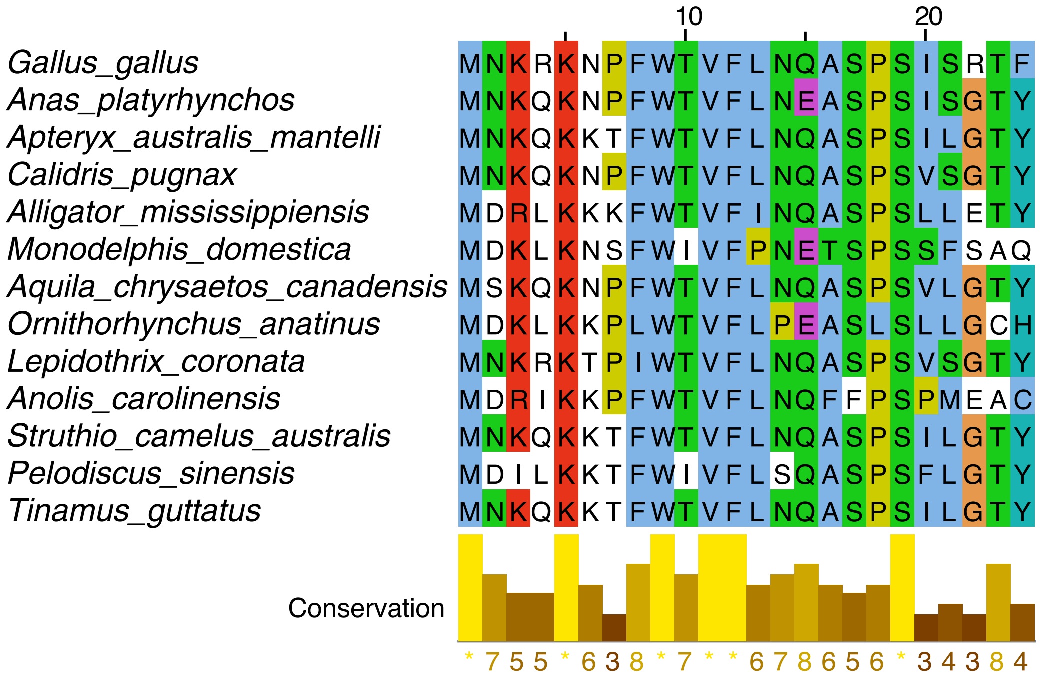

**HG0192**

ENSG00000125107 (CNOT1)

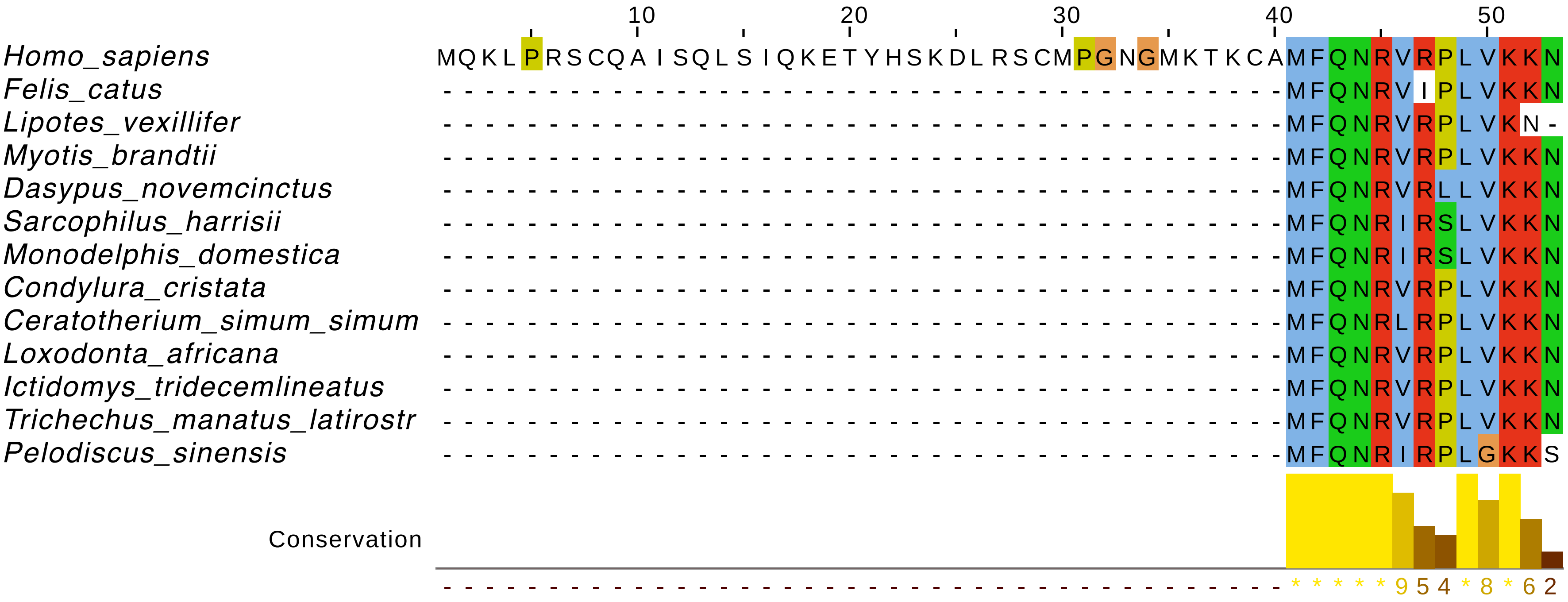

**HG0193**

ENSG00000134853 (PDGFRA)

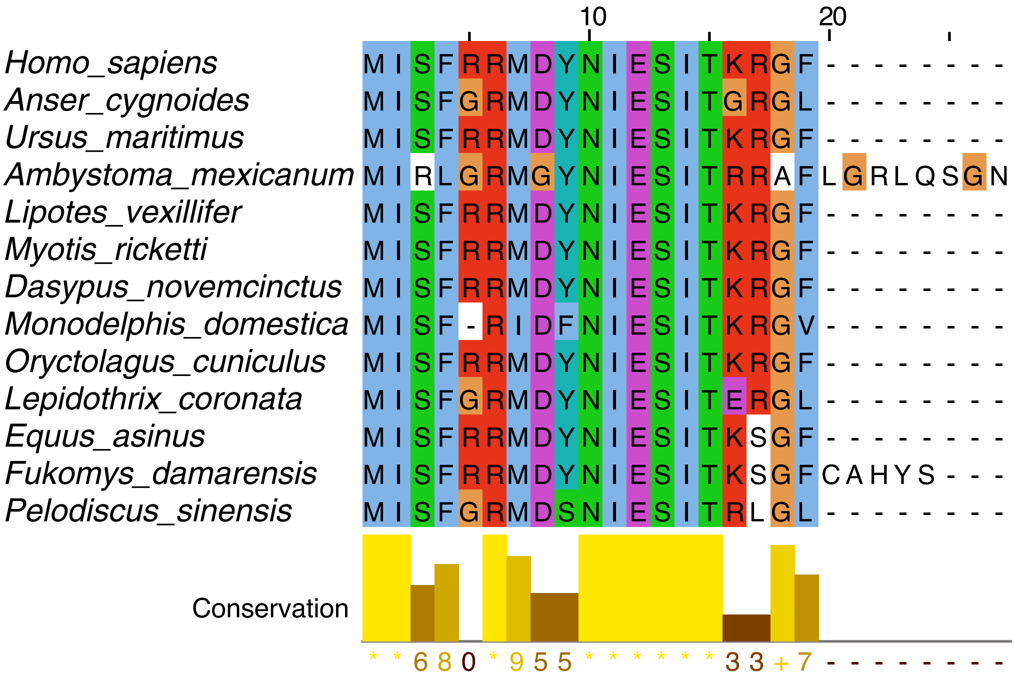

**HG0194**

ENSG00000143507 (DUSP10)

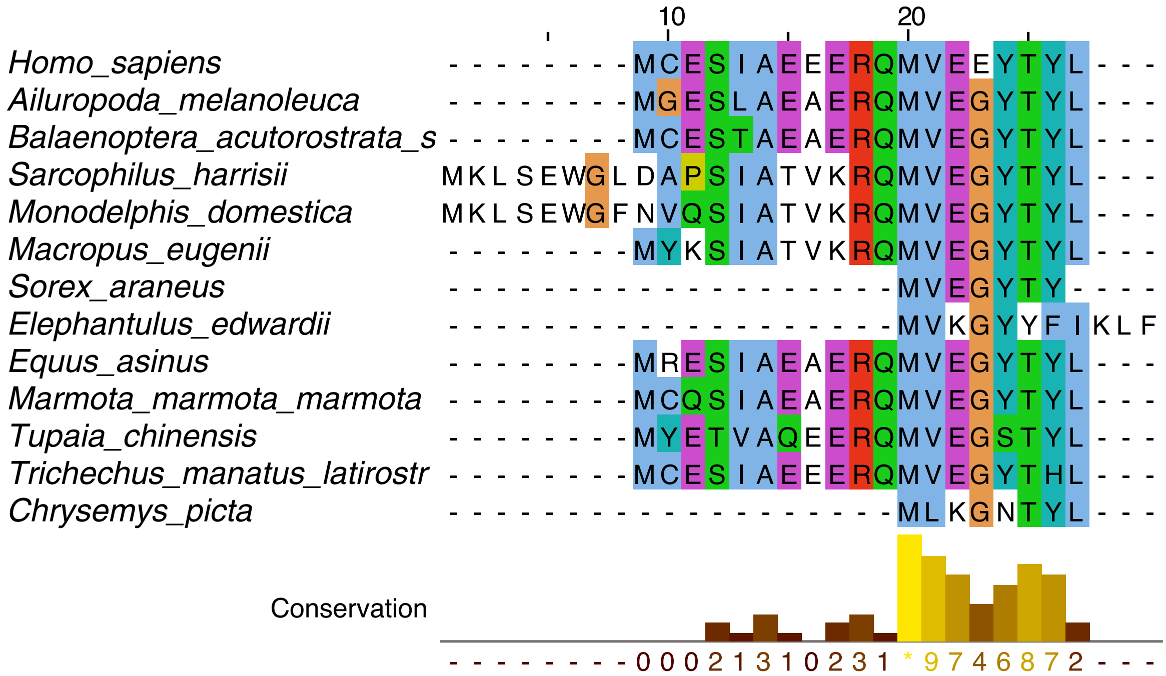

**HG0195**

ENSG00000165966 (PDZRN4)

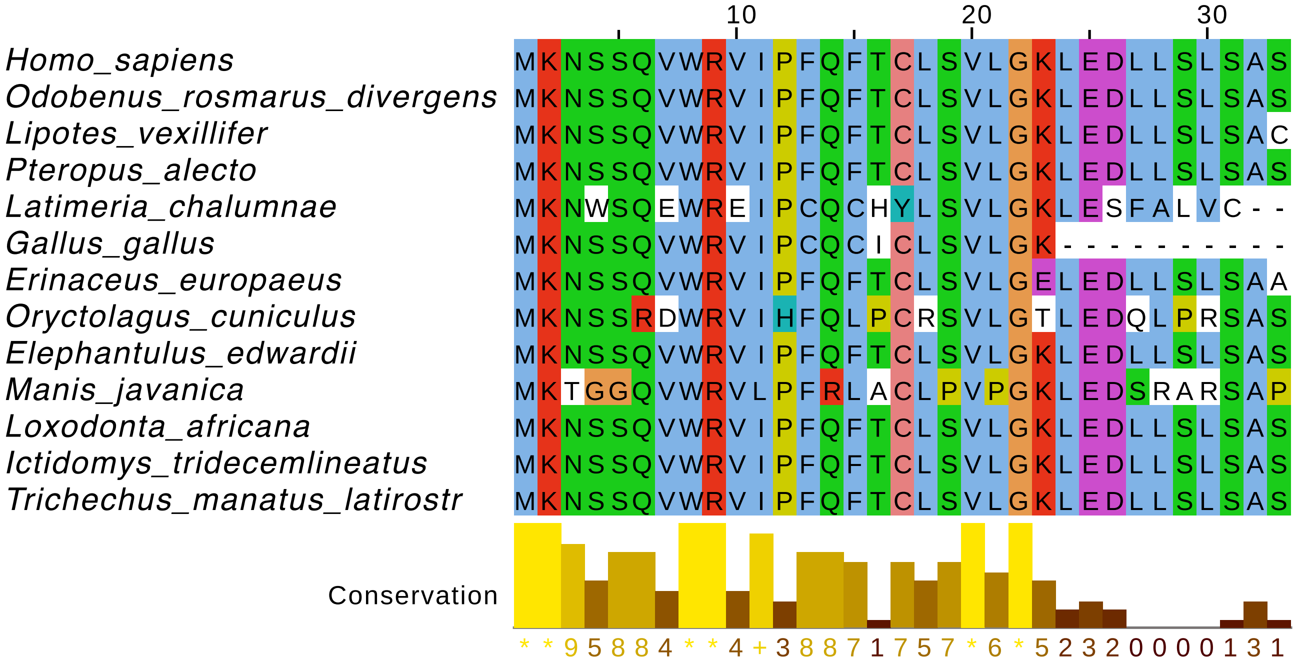

**HG0196**

ENSG00000189403 (HMGB1)

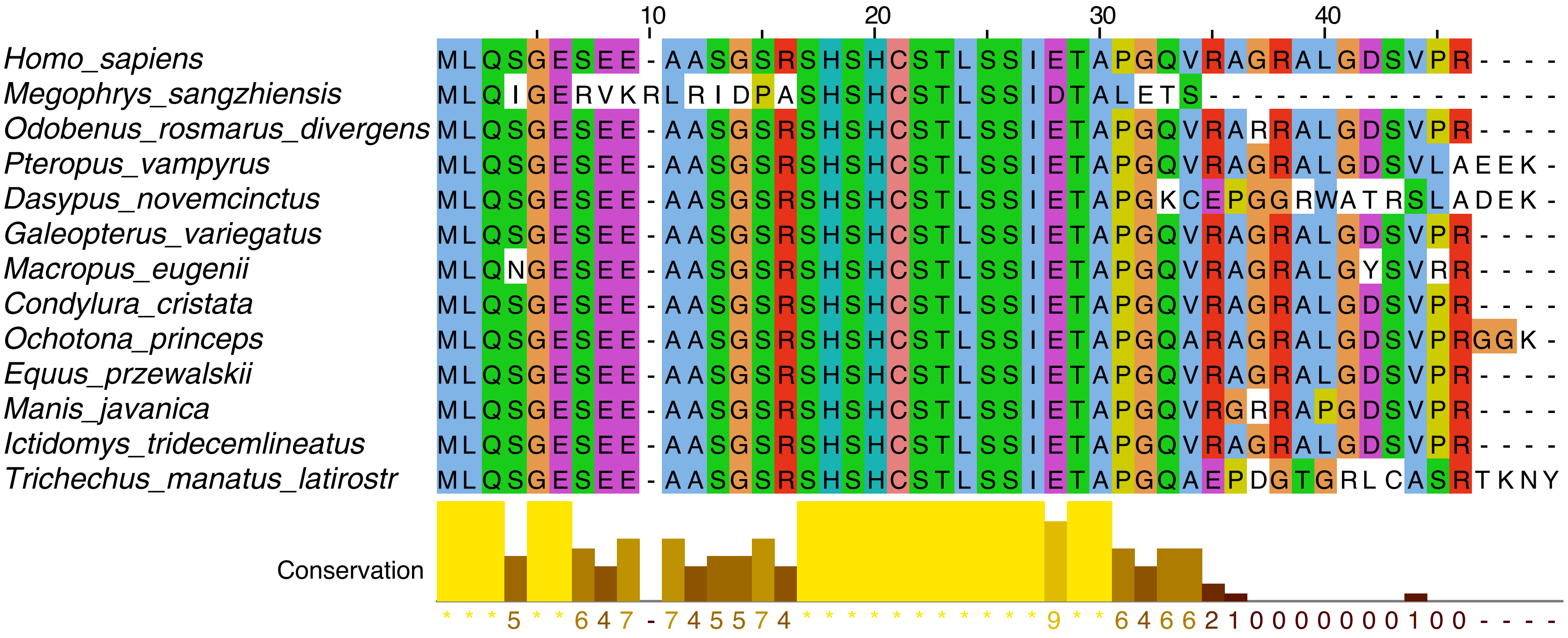

**HG0197**

ENSG00000196132 (MYT1)

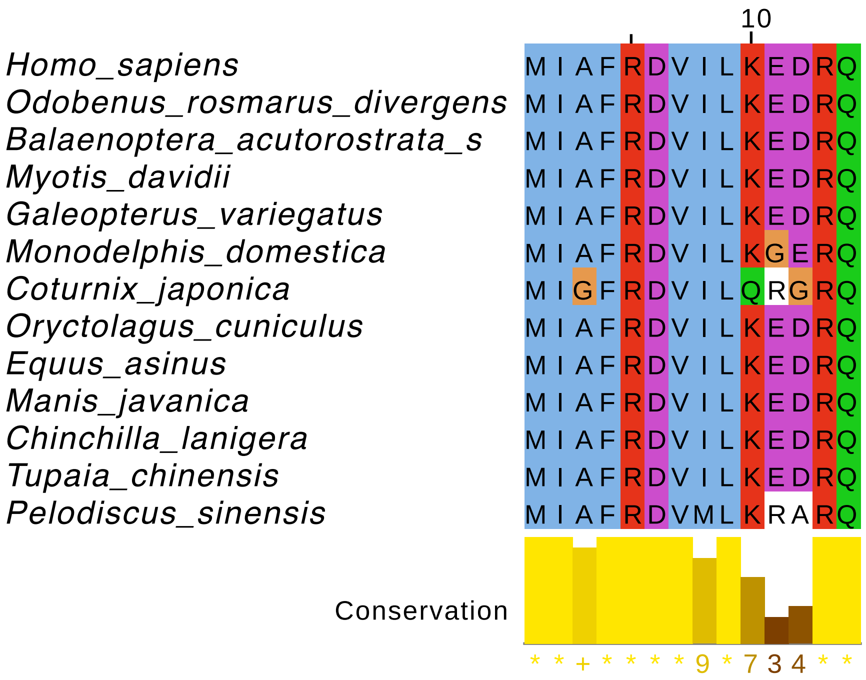

**HG0198.1**

ENSDARG00000018688 (elk3)

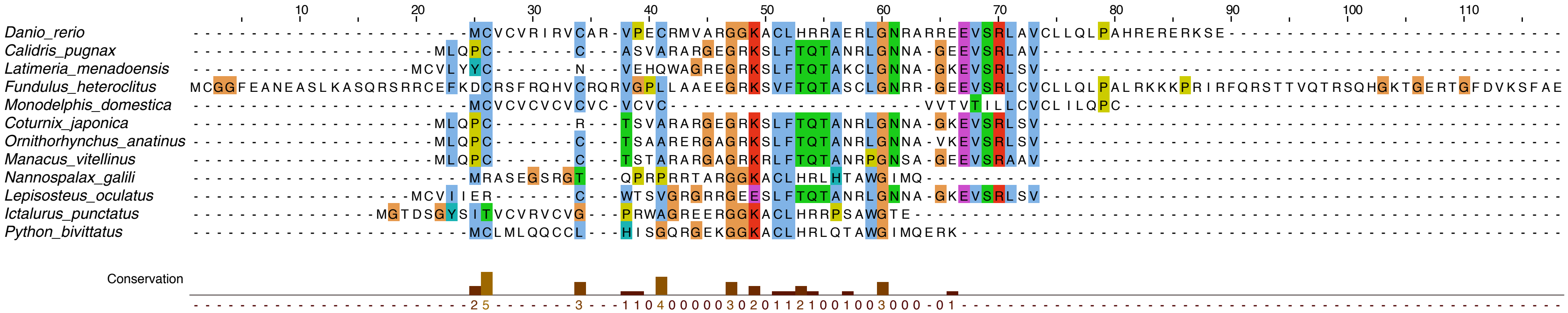

**HG0198.2**

ENSG00000111145 (ELK3)

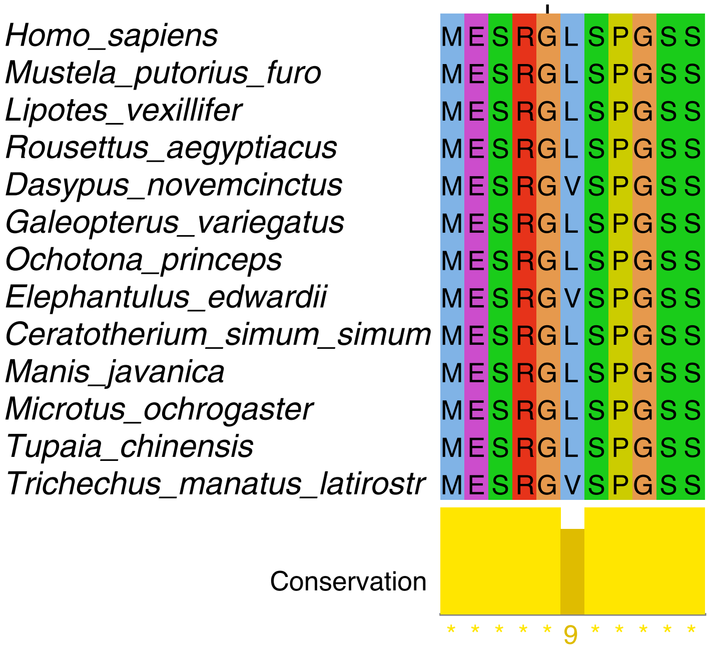

**HG0199**

ENSGALG00000007989 (ADNP)

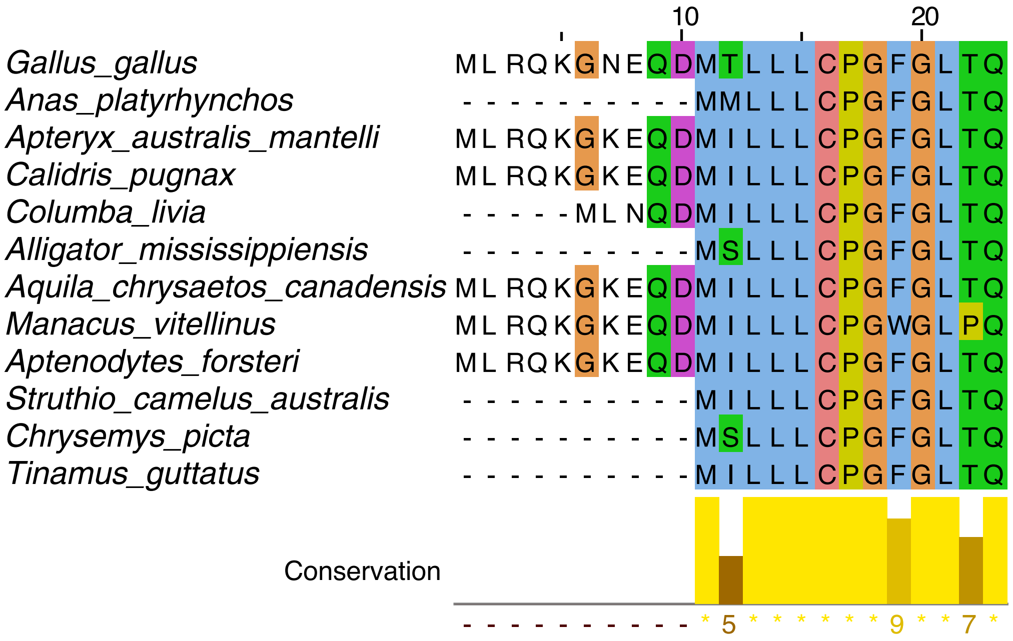

**HG0200**

ENSGALG00000012015 (NDST4)

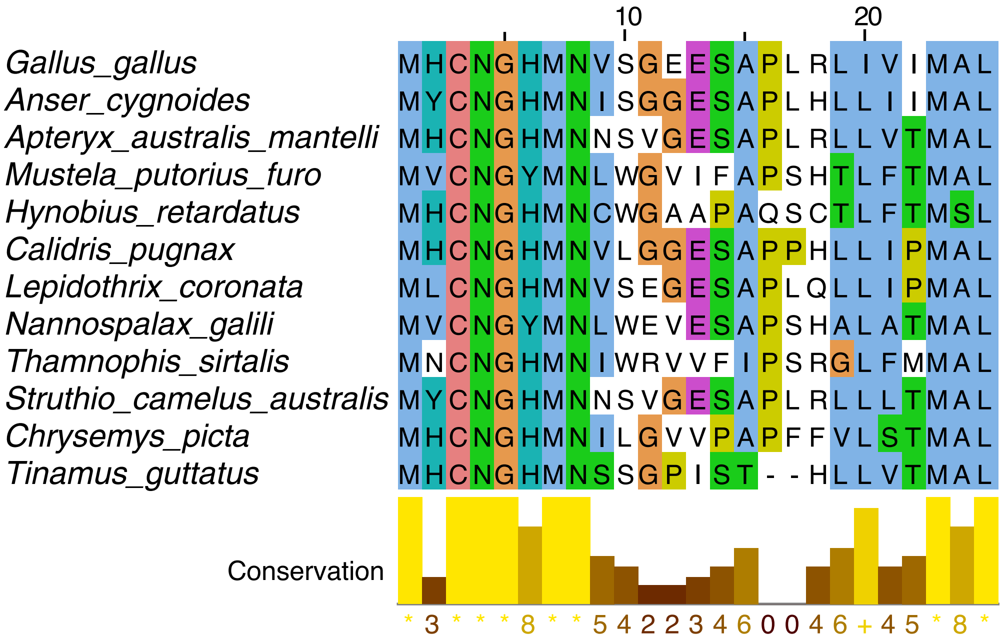

**HG0201**

ENSGALG00000015256 (PAM)

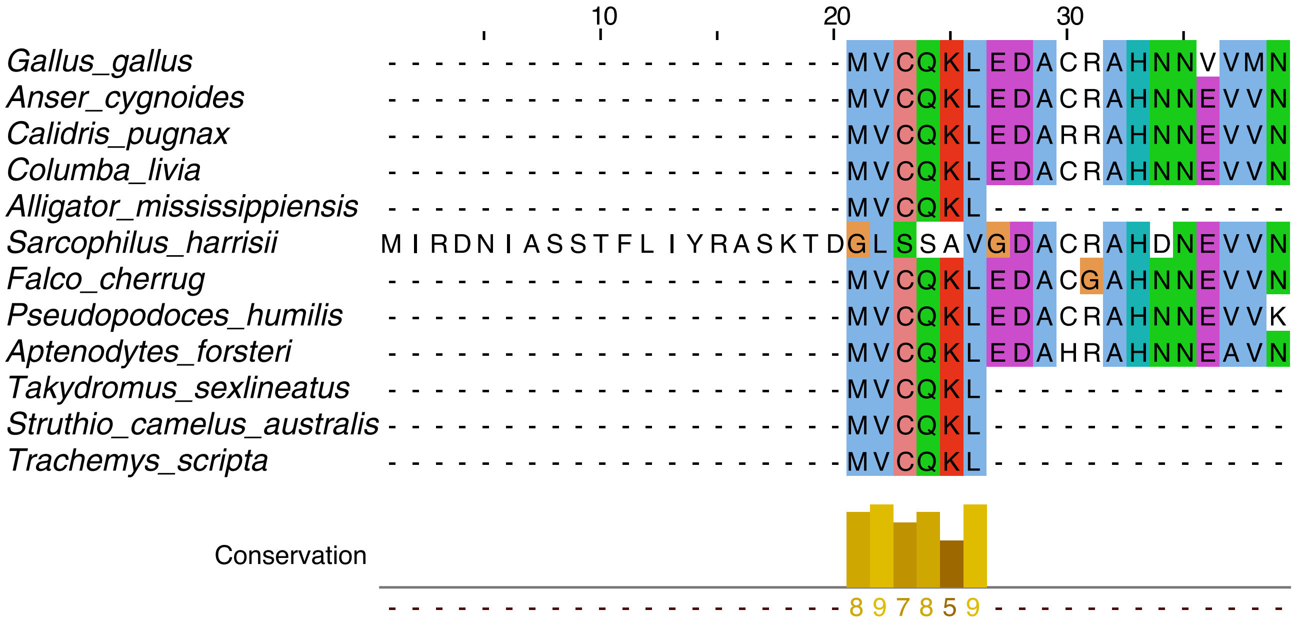

# HG0202

### ENSG00000124766 (SOX4)

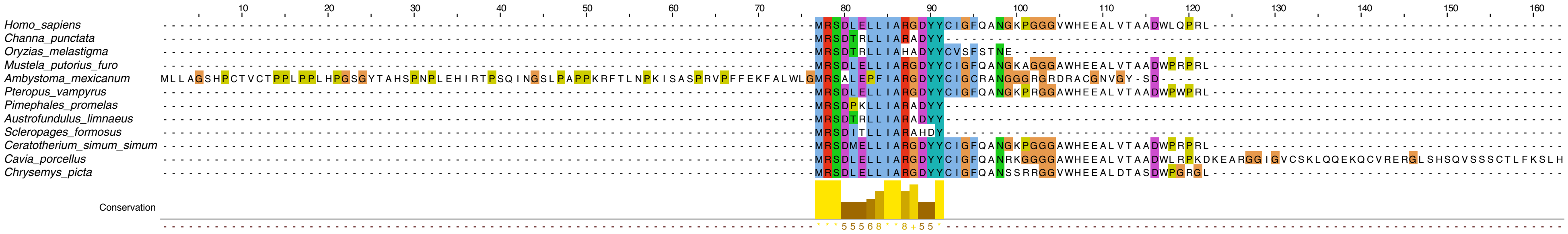

**HG0203**

ENSG00000128606 (LRRC17)

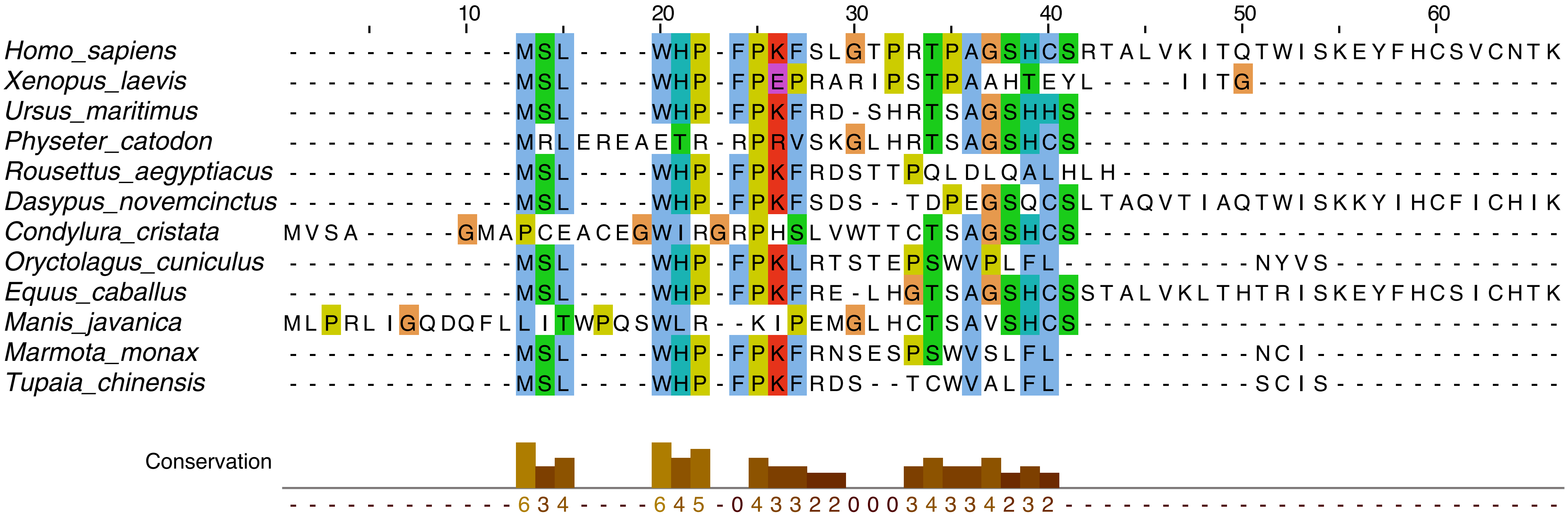

**HG0204**

ENSG00000162599 (NFIA)

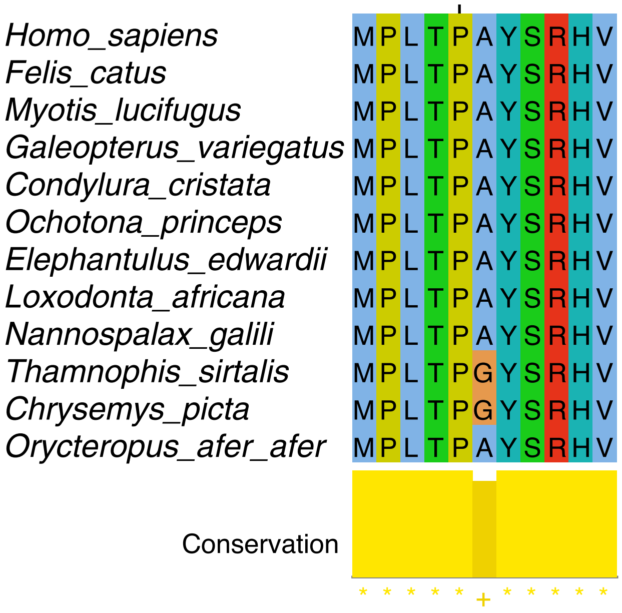

**HG0205**

ENSG00000184611 (KCNH7)

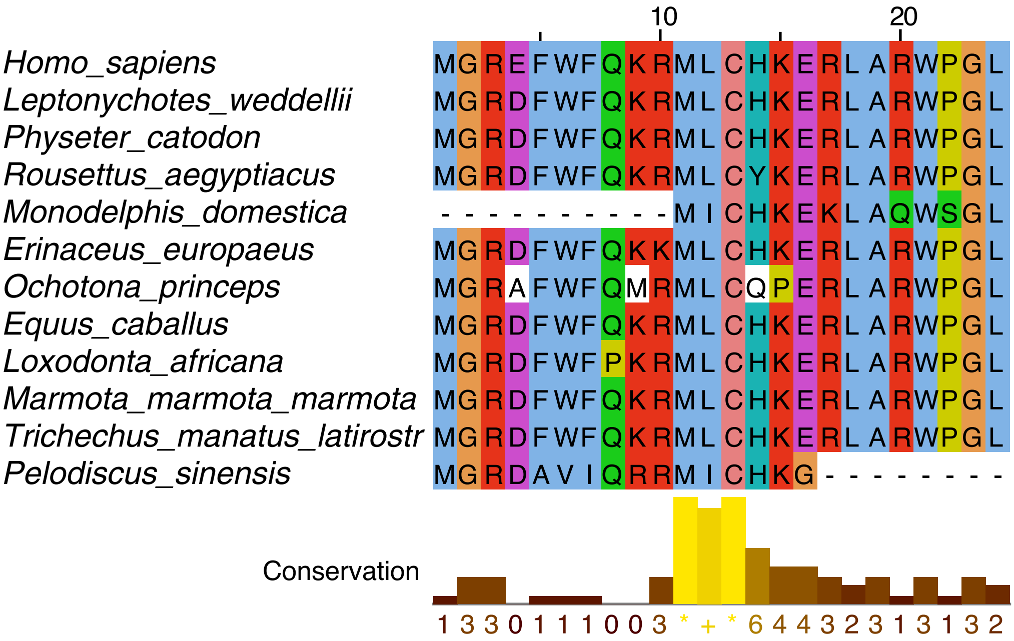

**HG0206.1**

ENSGALG00000002069

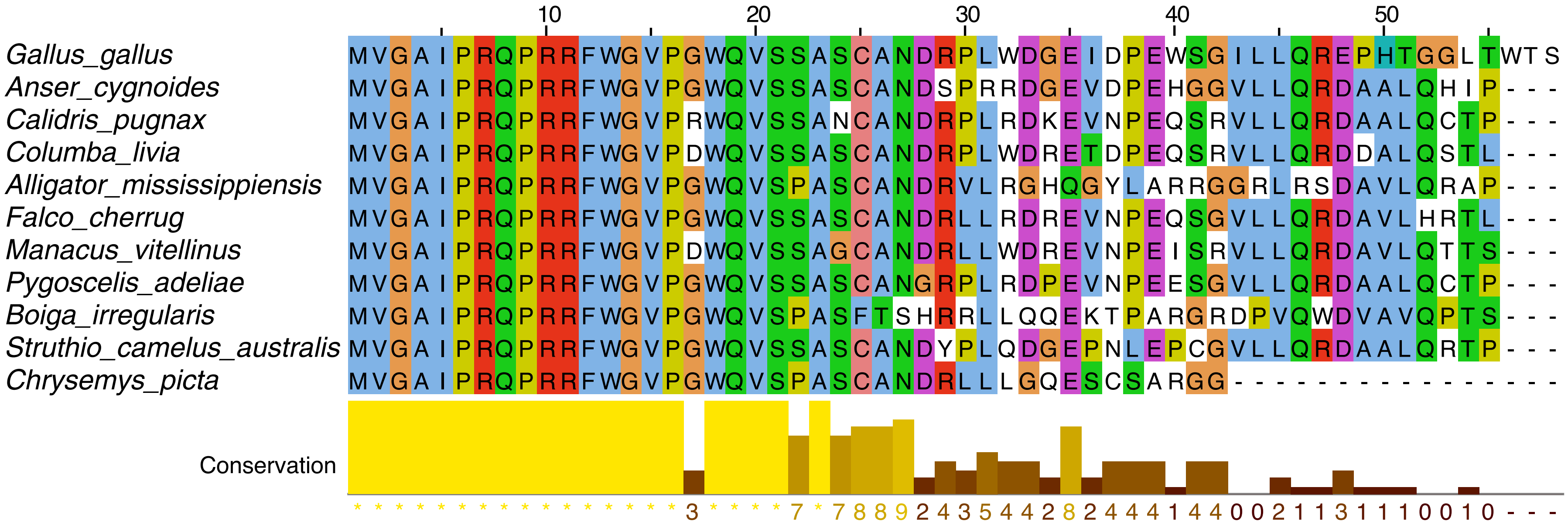

**HG0206.2**

ENSG00000005238 (FAM214B)

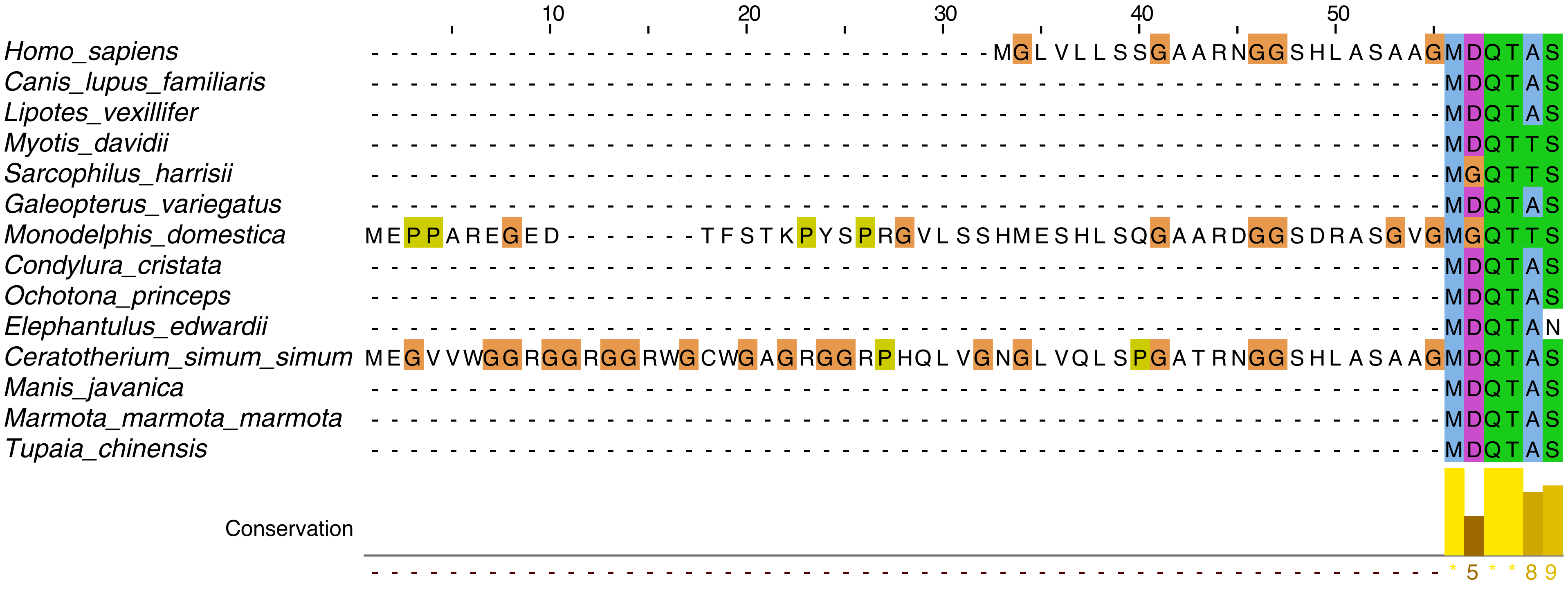

**HG0207.1**

ENSGALG00000007121 (TNFSF8)

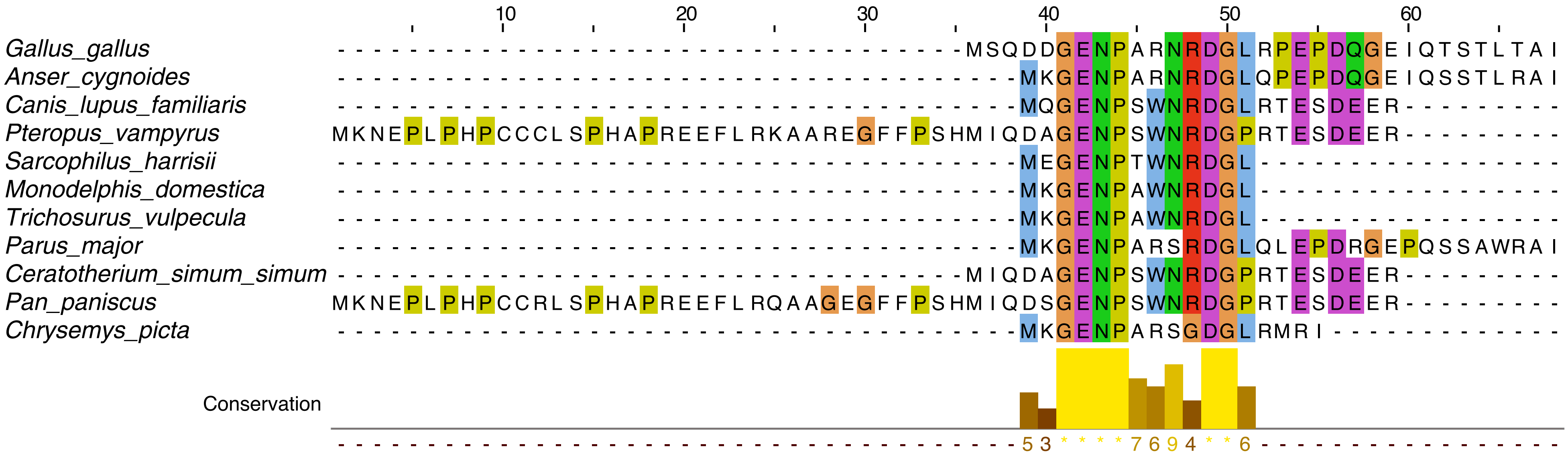

**HG0207.2**

ENSG00000106952 (TNFSF8)

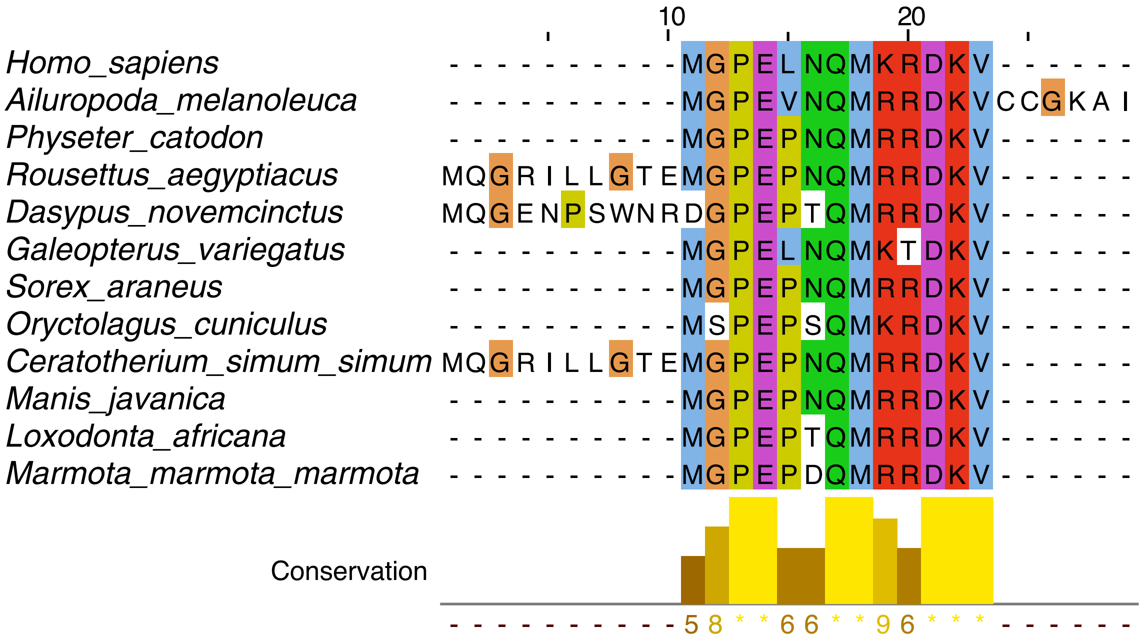

**HG0208**

ENSGALG00000013272 (PRMT8)

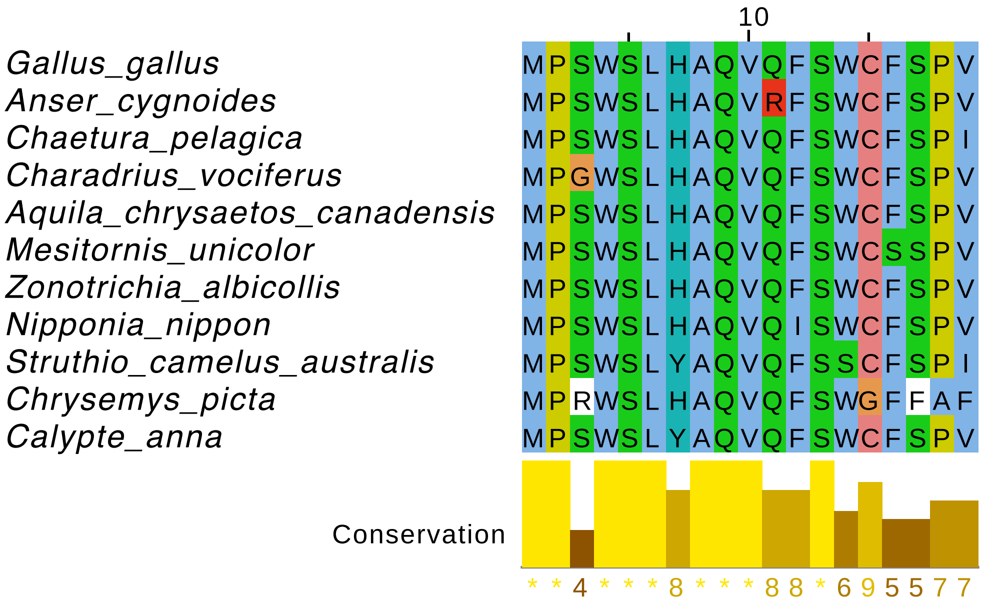

**HG0209**

ENSG00000101040 (ZMYND8)

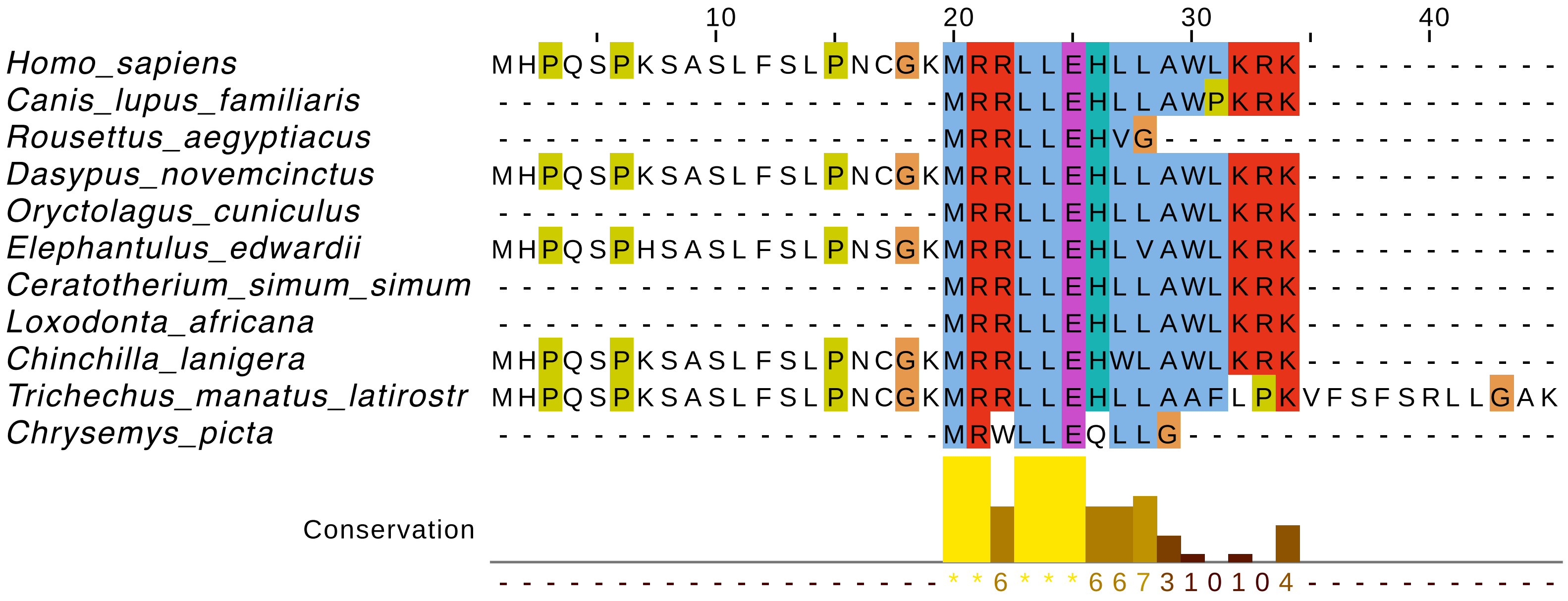

**HG0210**

ENSG00000155849 (ELMO1)

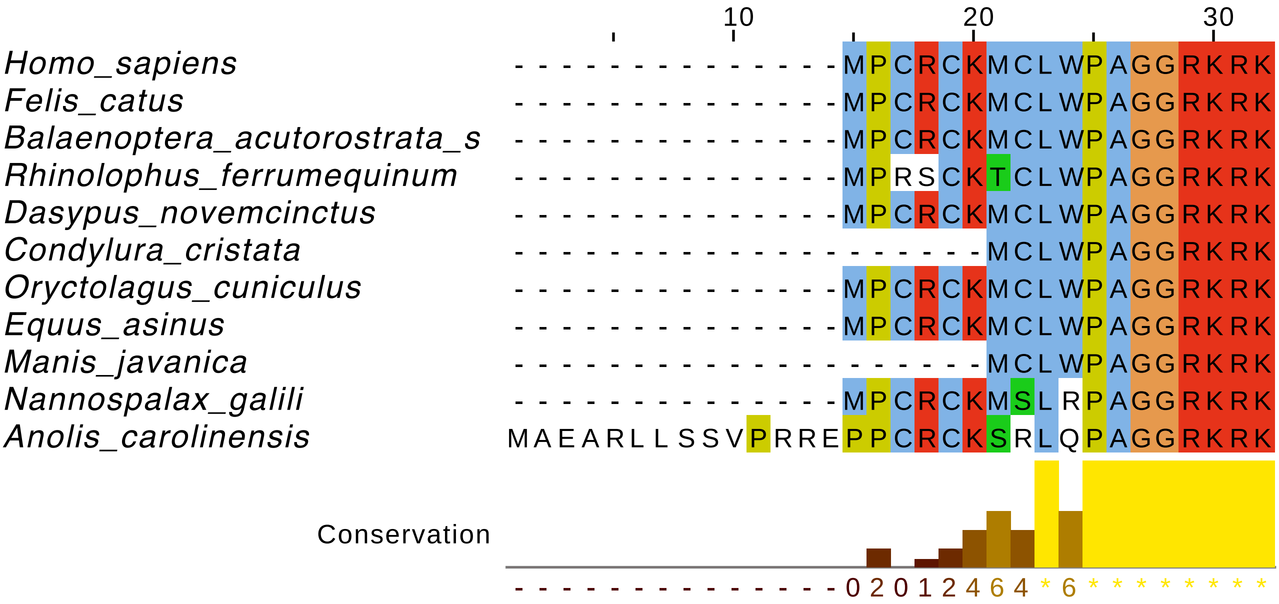

**HG0211**

ENSG00000176204 (LRRTM4)

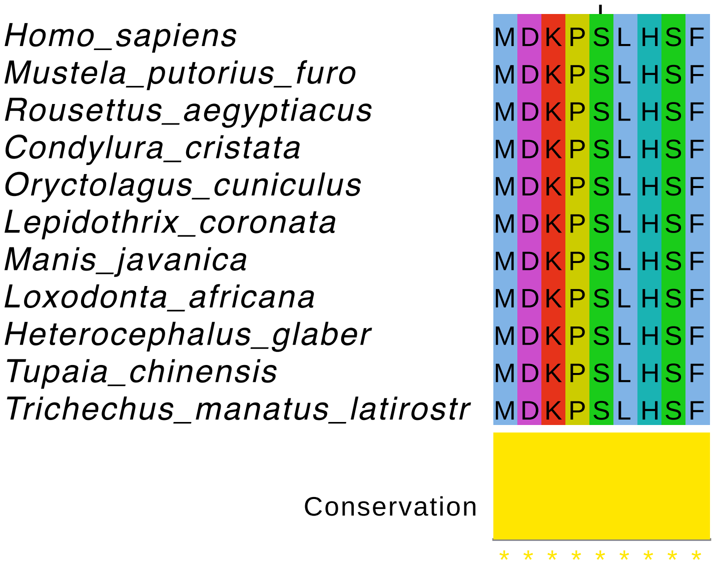

**HG0212**

ENSDARG00000100244 (ebf3a)

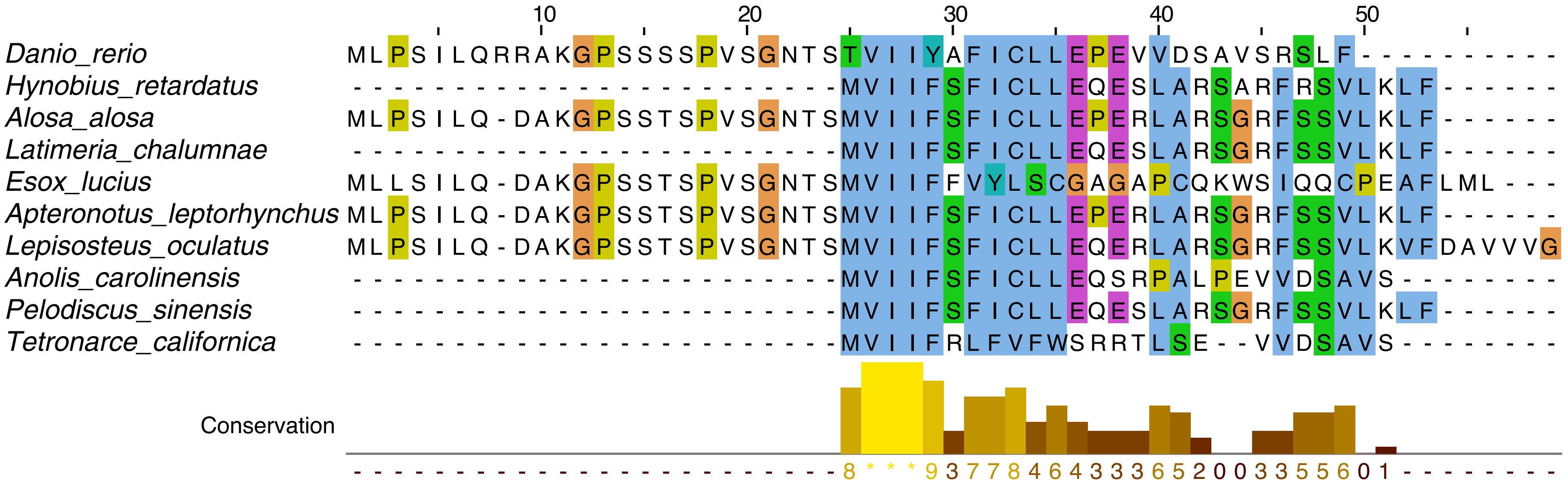

**HG0213.1**

ENSGALG00000001946 (RSPO1)

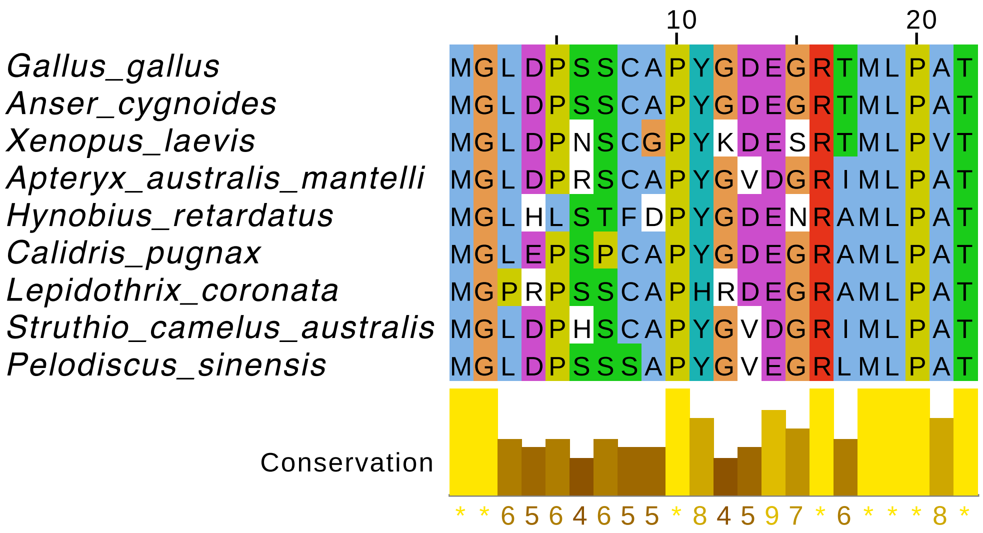

**HG0213.2**

ENSGALG00000001946 (RSPO1)

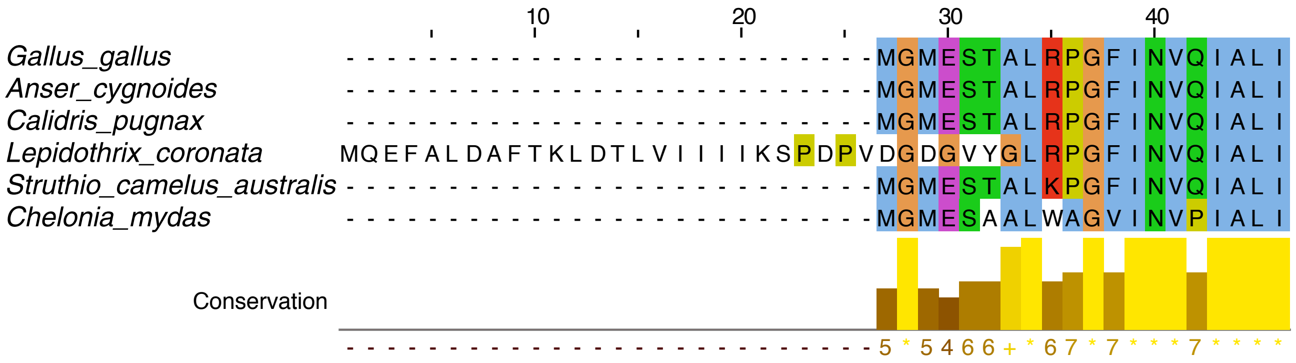

**HG0214.1**

ENSGALG00000012917 (CDH6)

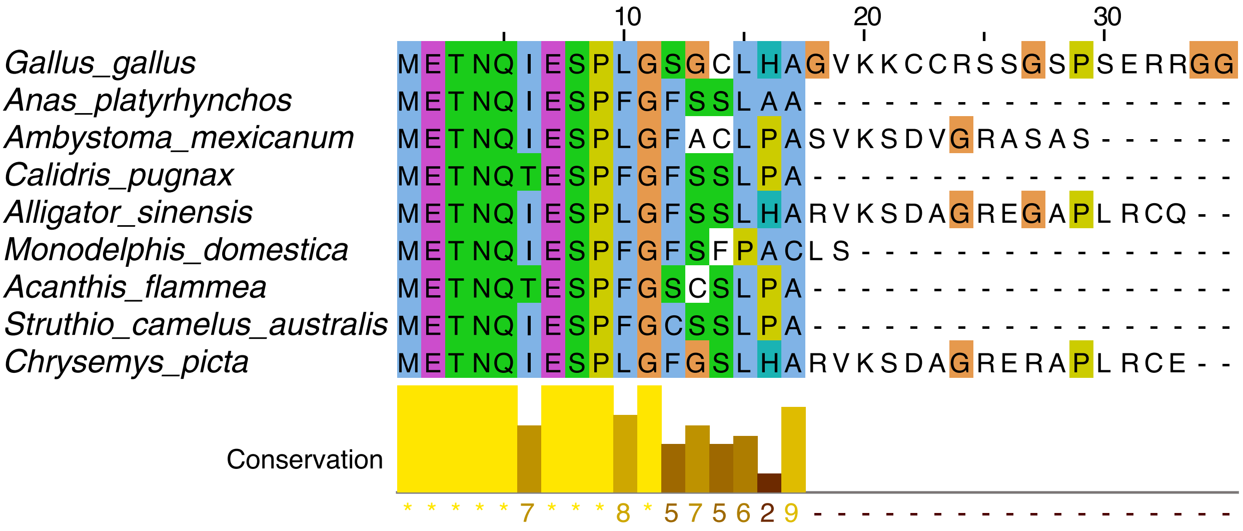

**HG0214.2**

ENSG00000040731 (CDH10)

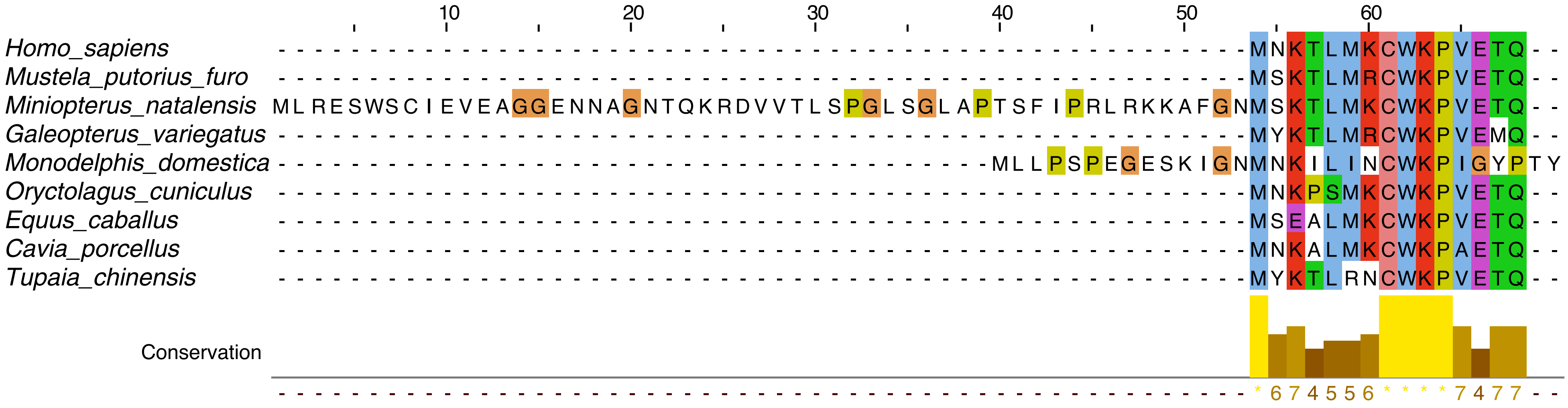

**HG0215**

ENSDARG00000079280 (dhrsx)

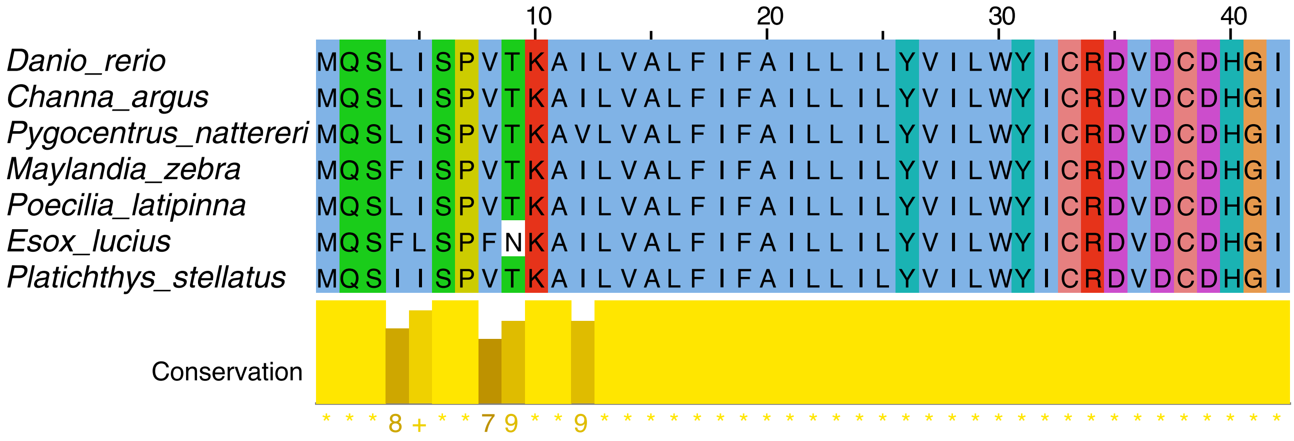

**HG0215**

ENSGALG00000016681 (DHRSX)

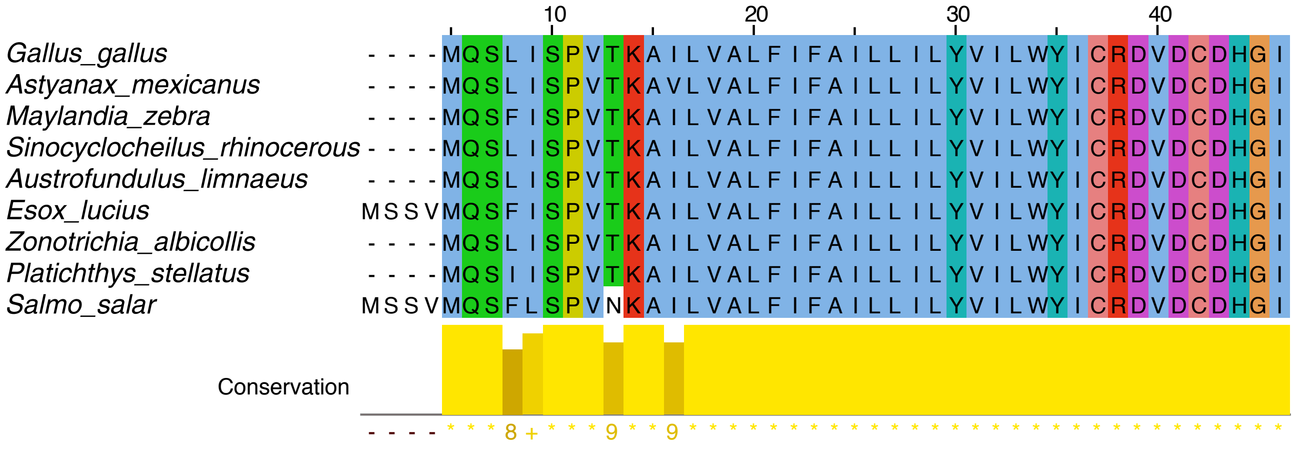

**HG0216.1**

ENSGALG00000020922

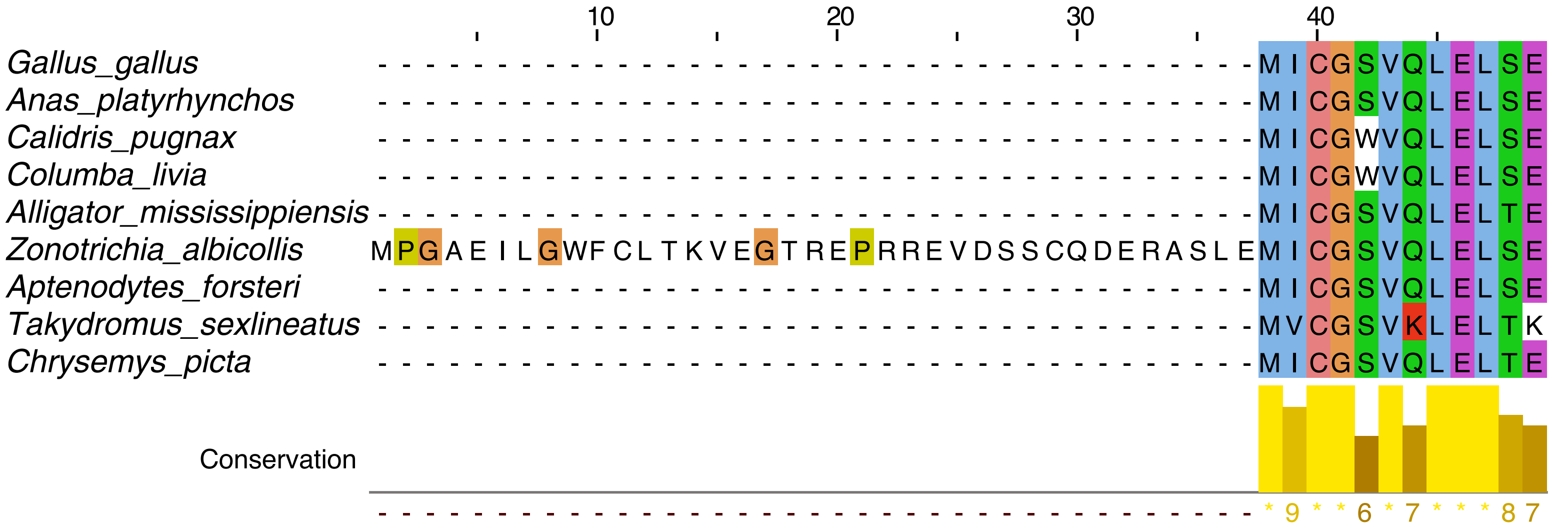

**HG0216.2**

ENSG00000116667 (C1orf21)

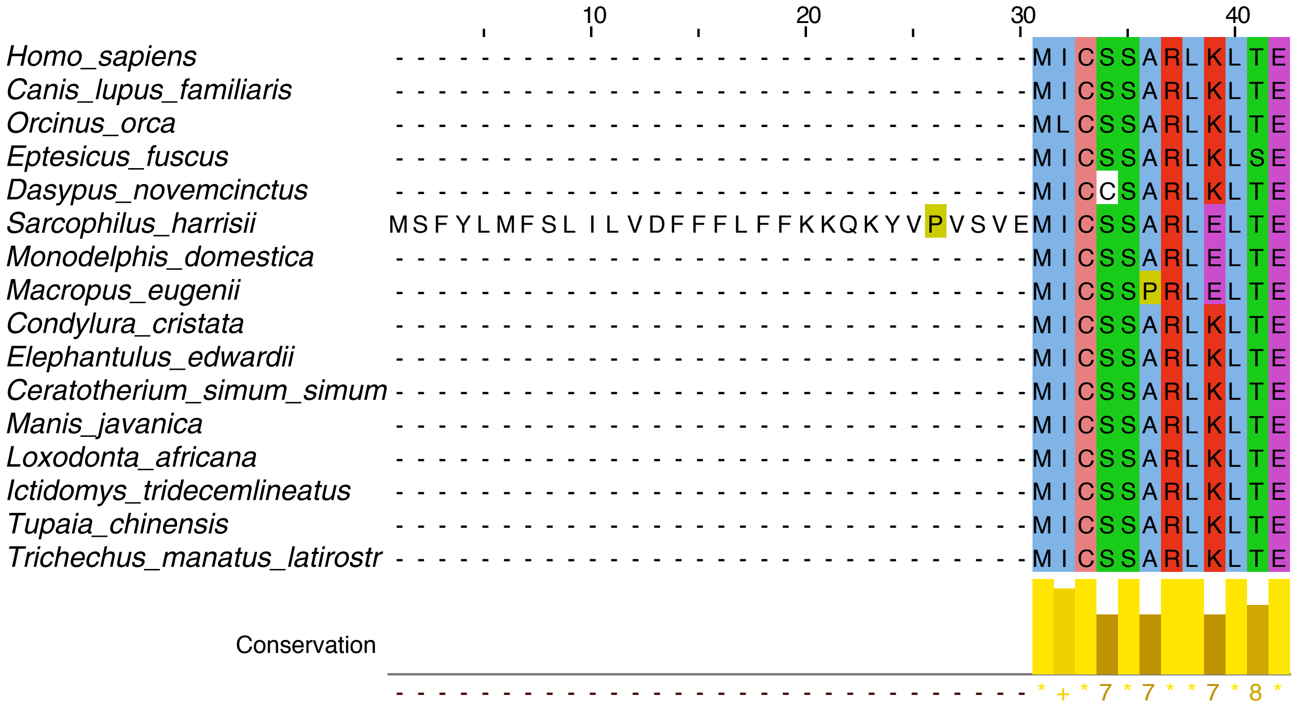

**HG0217.1**

ENSG00000150394 (CDH8)

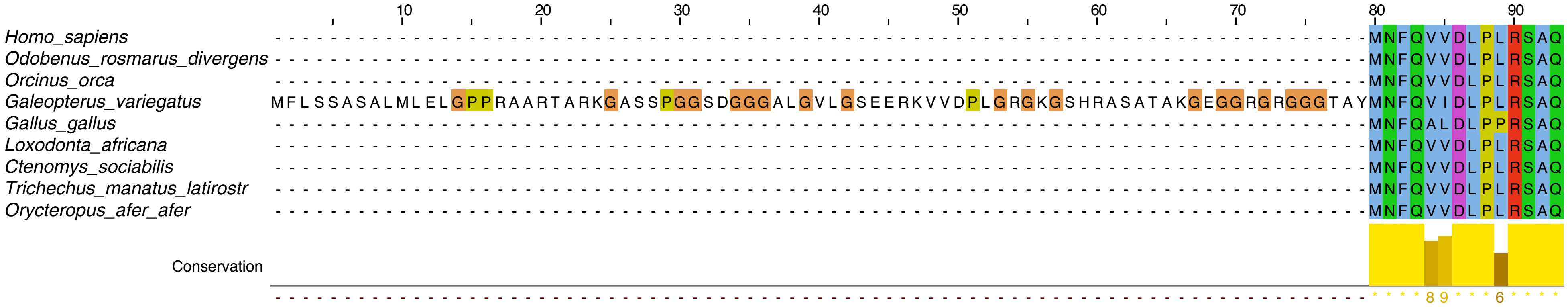

**HG0217.2**

ENSGALG00000005319 (CDH8)

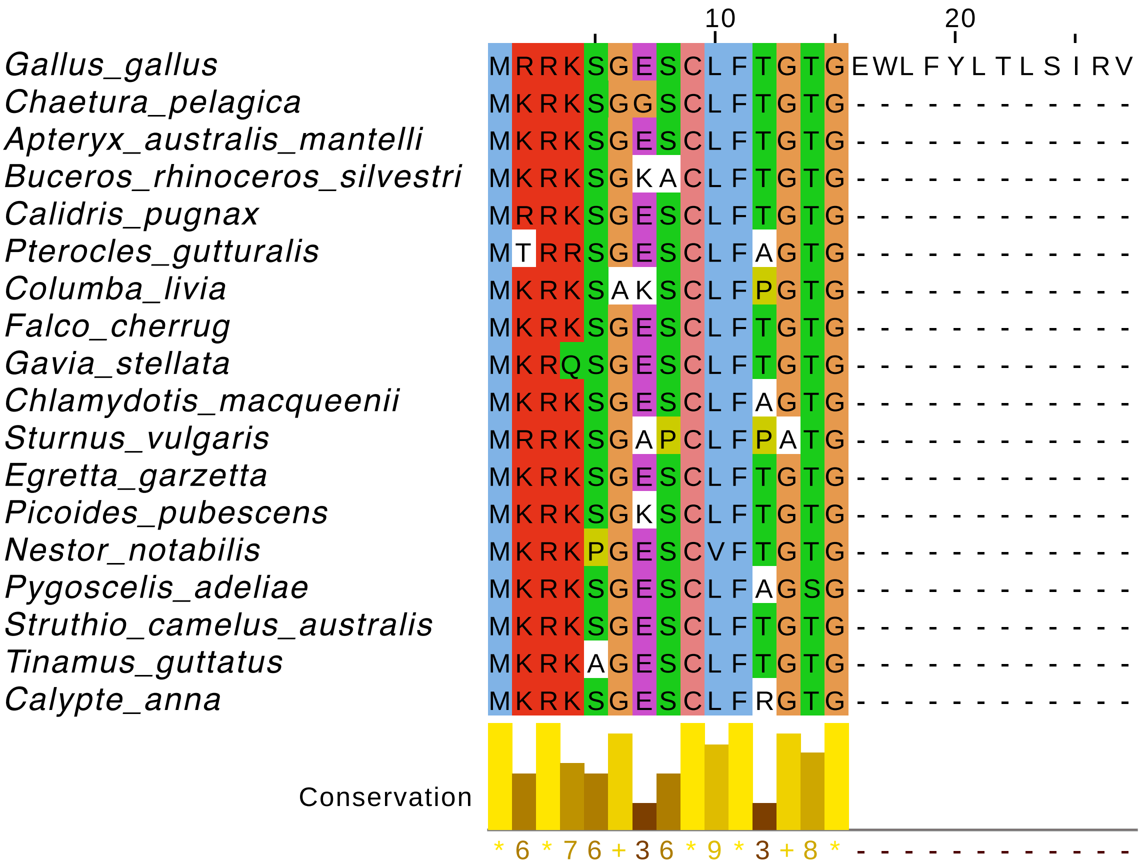

**HG0218**

ENSDARG00000100149 (axin2)

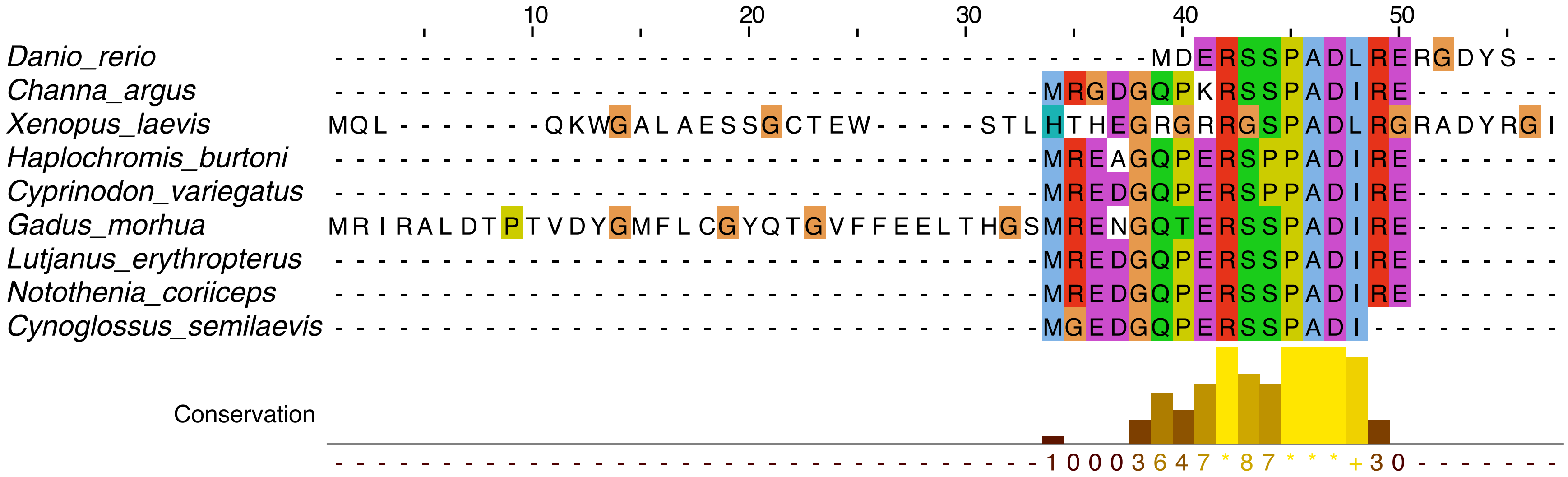

**HG0219**

ENSDARG00000104082 (trps1)

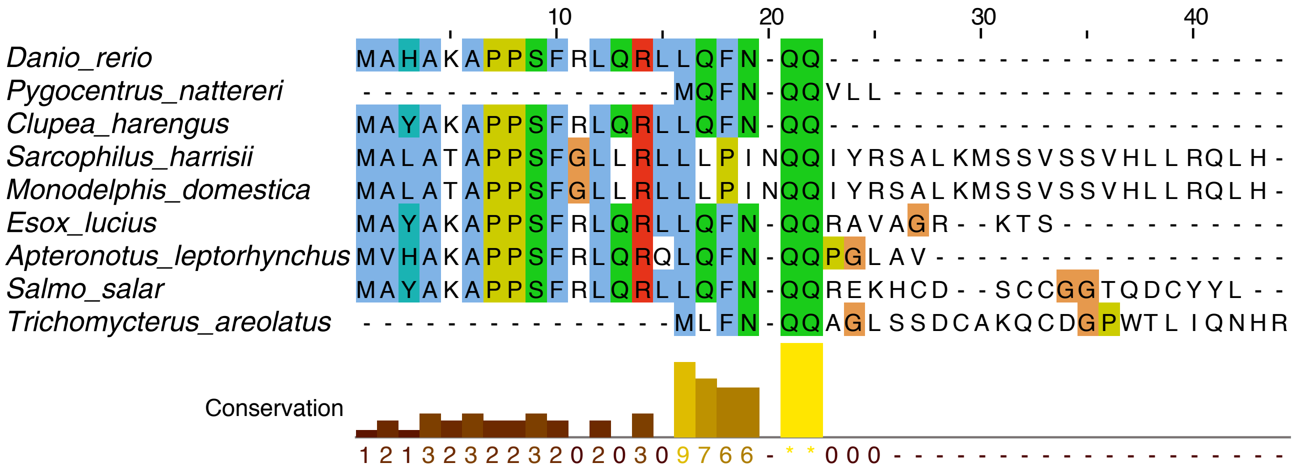

**HG0220**

ENSGALG00000008418 (LARGE2)

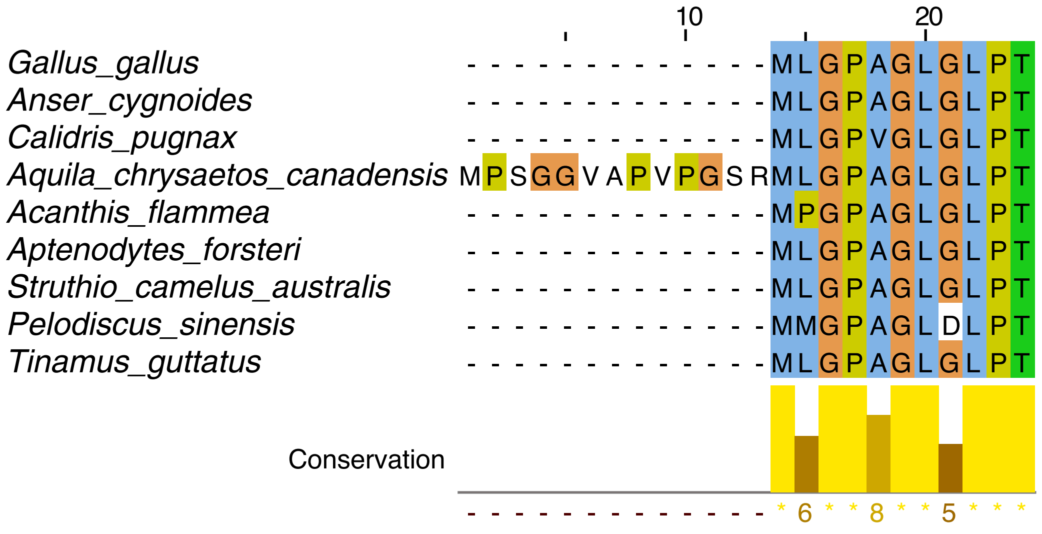

**HG0221**

ENSGALG00000010859 (EPGN)

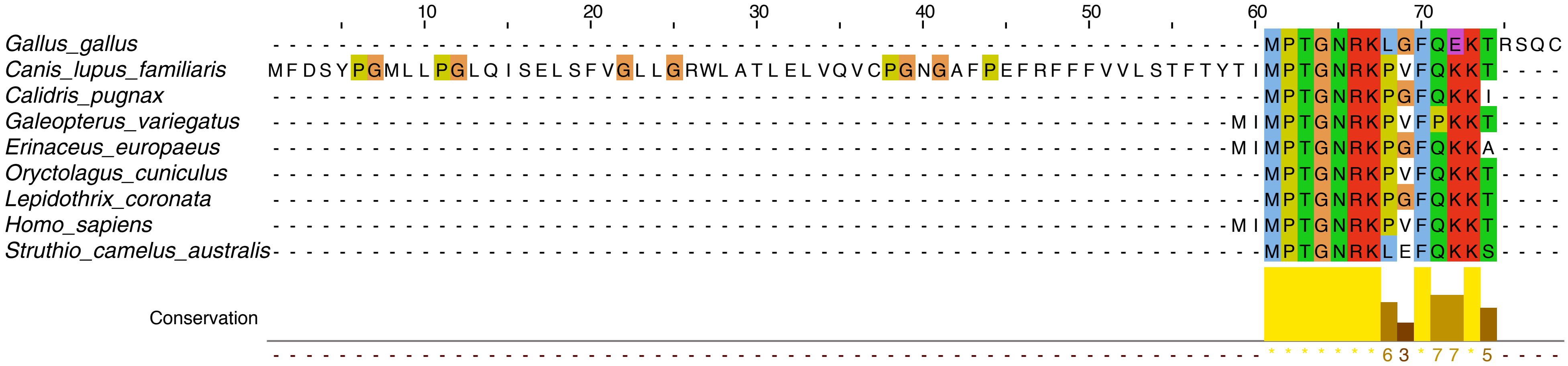

**HG0222**

ENSGALG00000011564 (RUFY3)

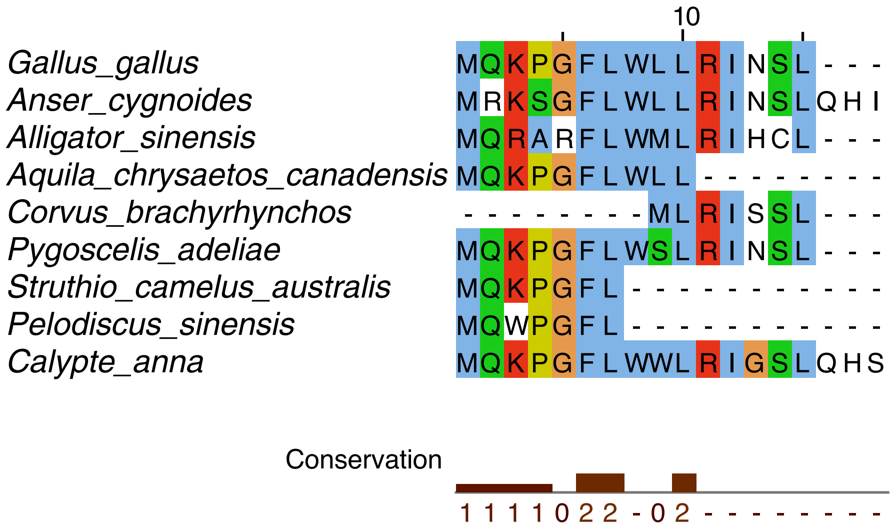

**HG0223**

ENSG00000091656 (ZFHX4)

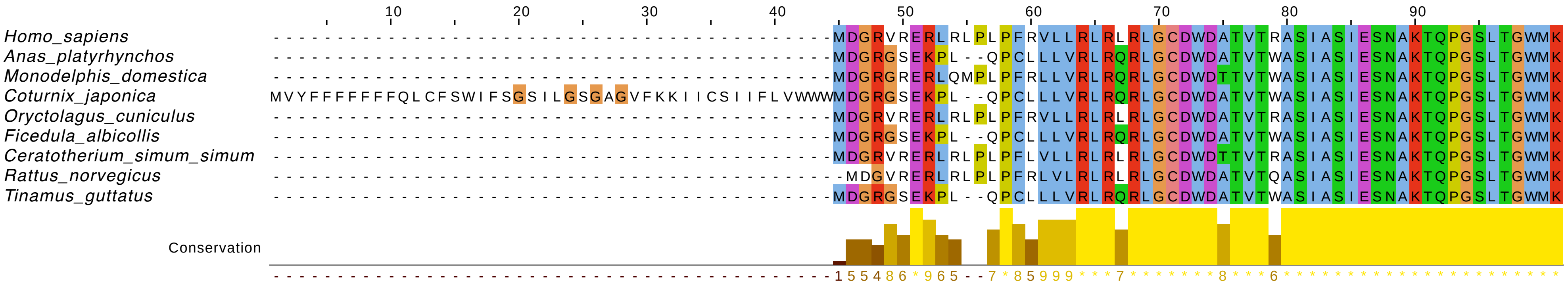

**HG0224**

ENSG00000118007 (STAG1)

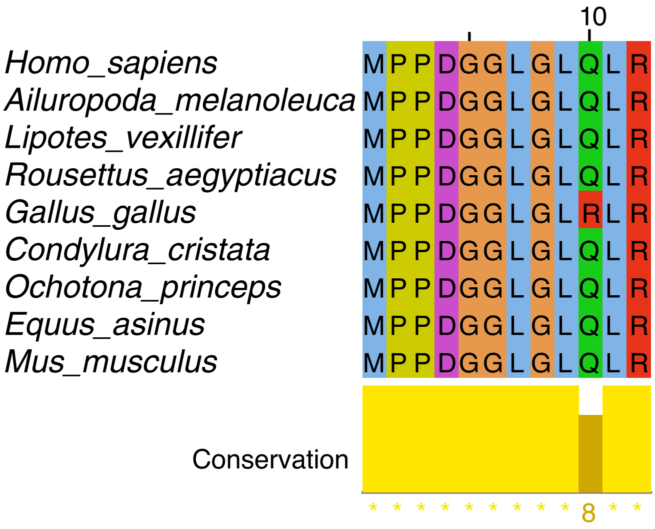

**HG0225**

ENSG00000150051 (MKX)

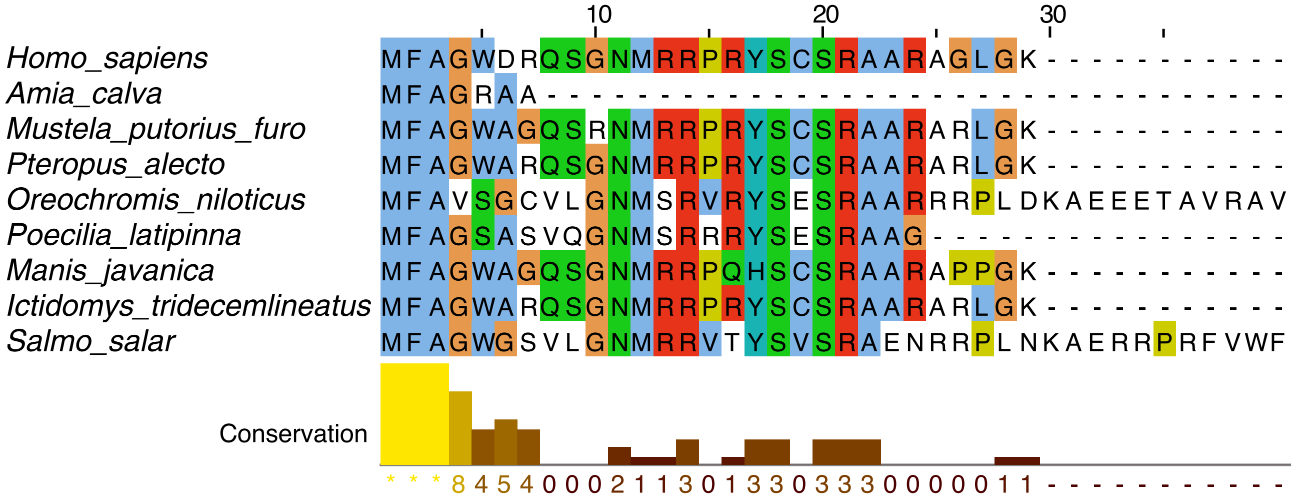

**HG0226.1**

ENSG00000180592

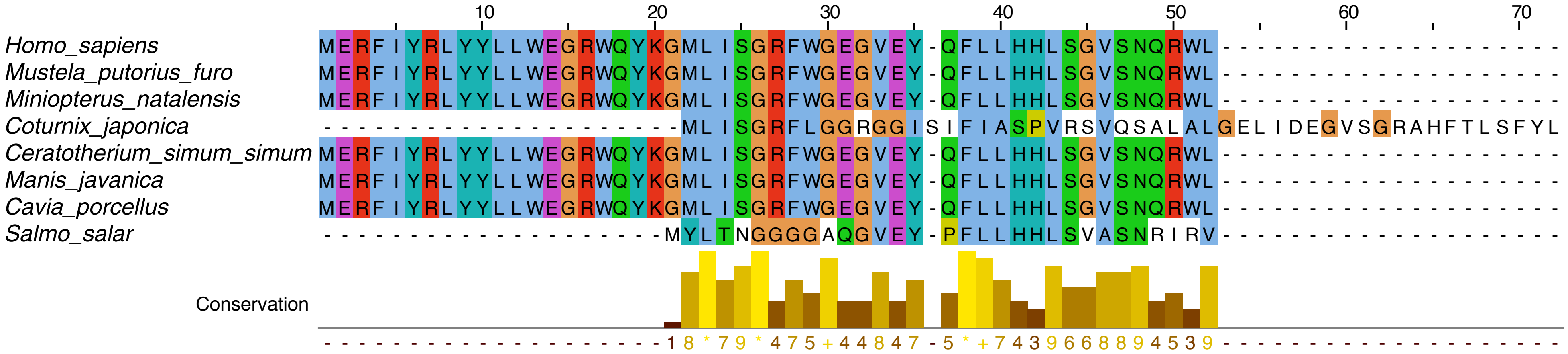

**HG0226.2**

ENSG00000180592

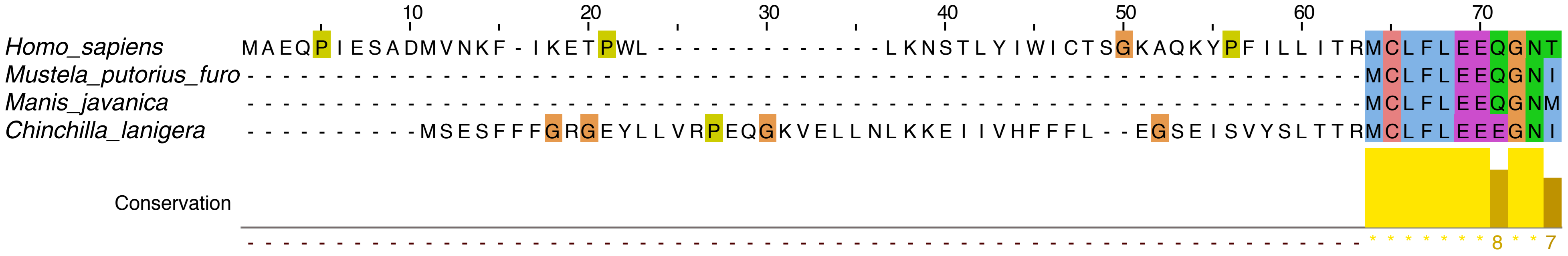

**HG0226.3**

ENSG00000180592

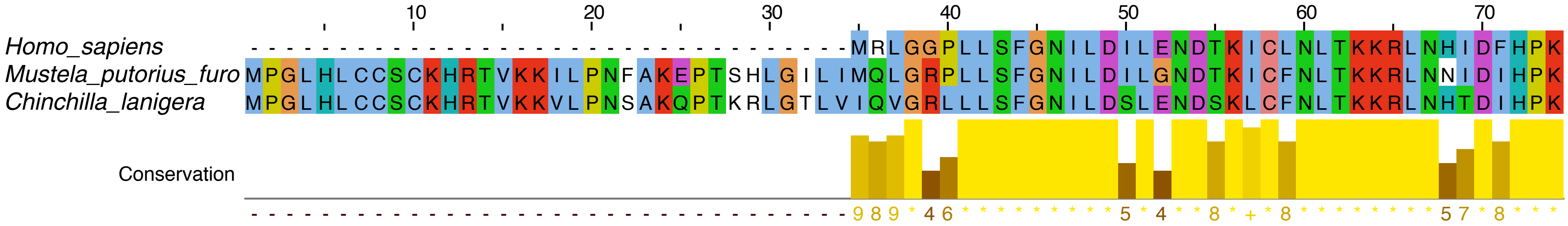

**HG0227.1**

ENSGALG00000031929

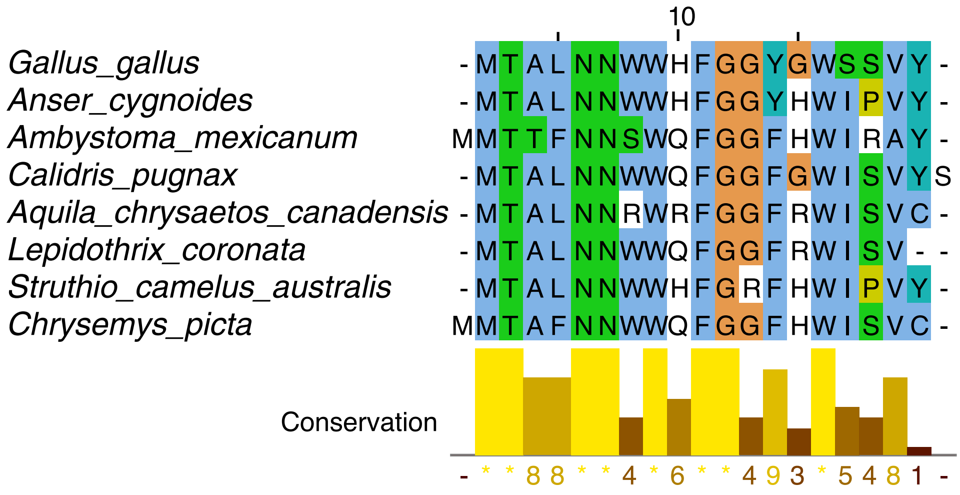

**HG0227.2**

ENSG00000275163

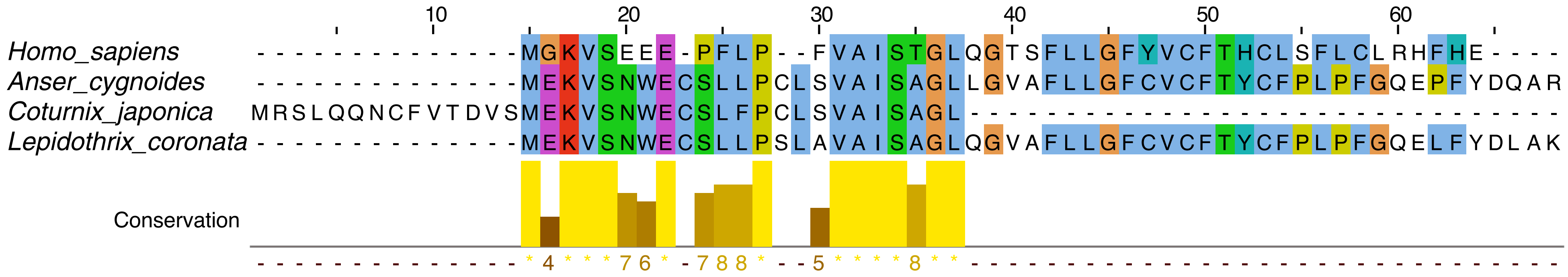

**HG0228**

ENSDARG00000087887 (phf21b)

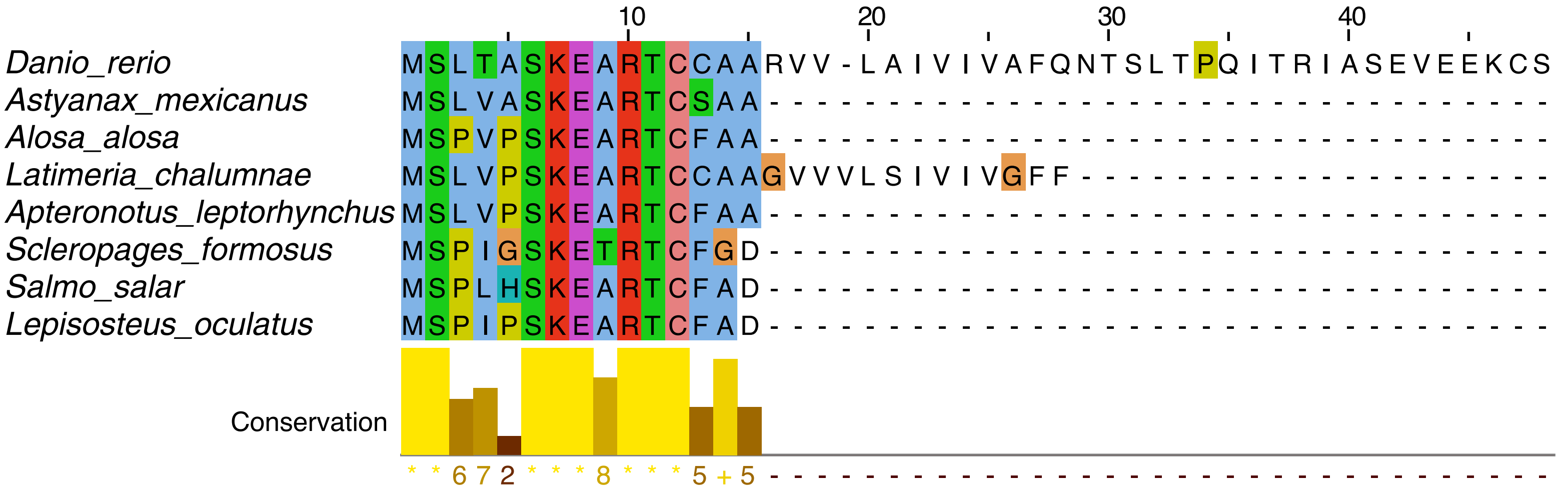

**HG0229**

ENSGALG00000005474

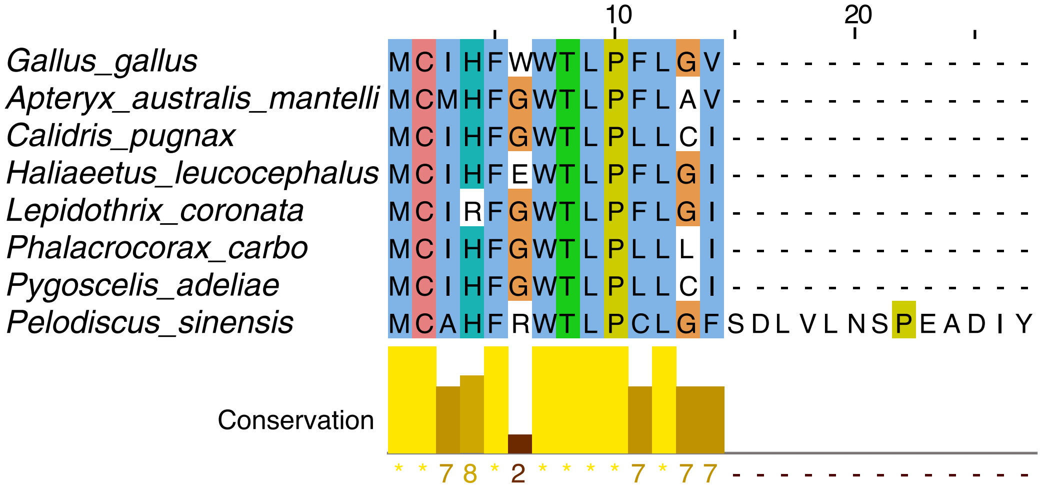

**HG0230**

ENSGALG00000011007 (OSBPL3)

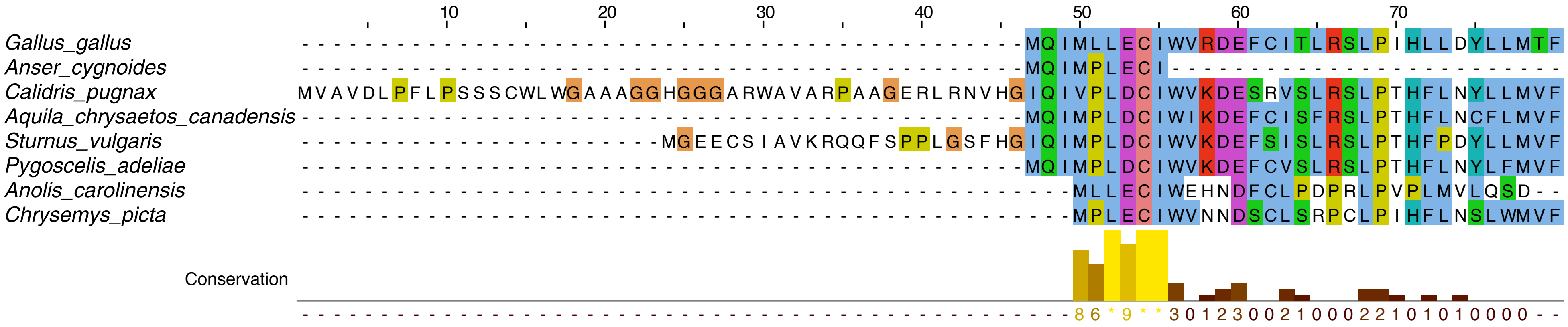

**HG0231**

ENSG00000150636

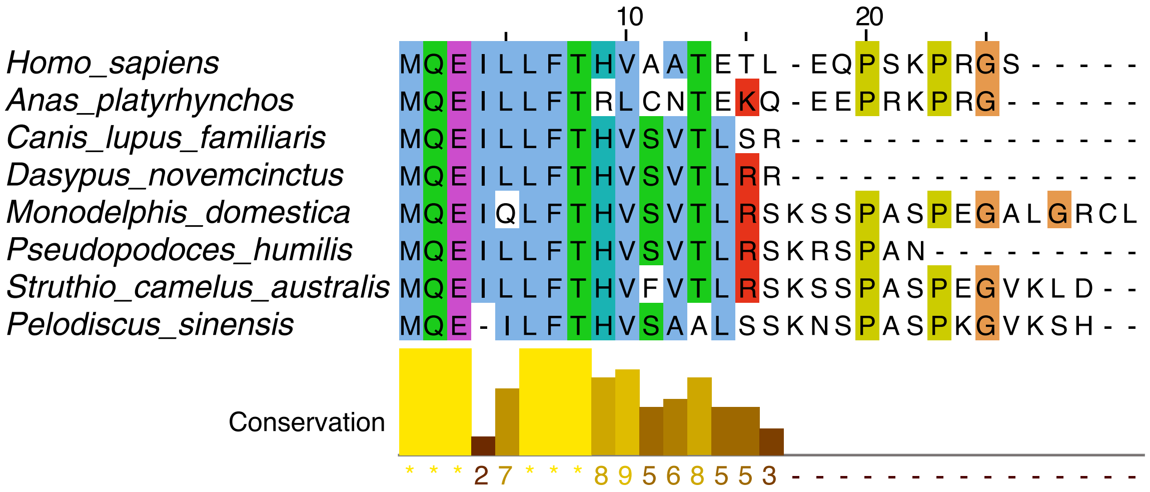

**HG0232**

ENSG00000180875 (GREM2)

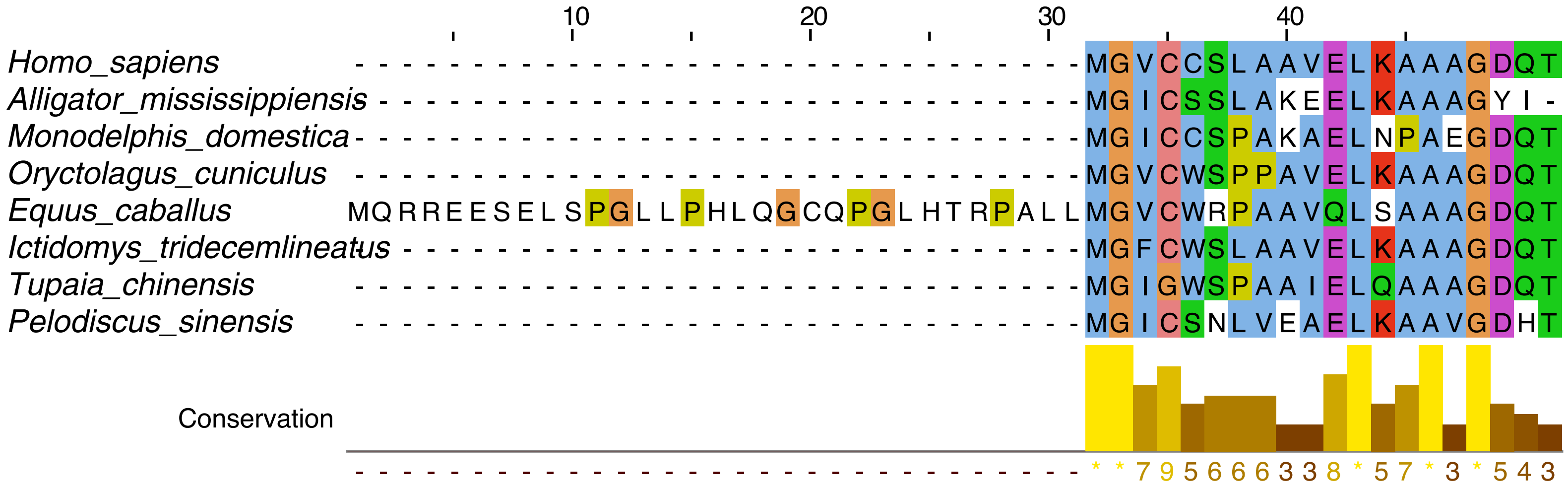

**HG0233**

ENSG00000206432 (TMEM200C)

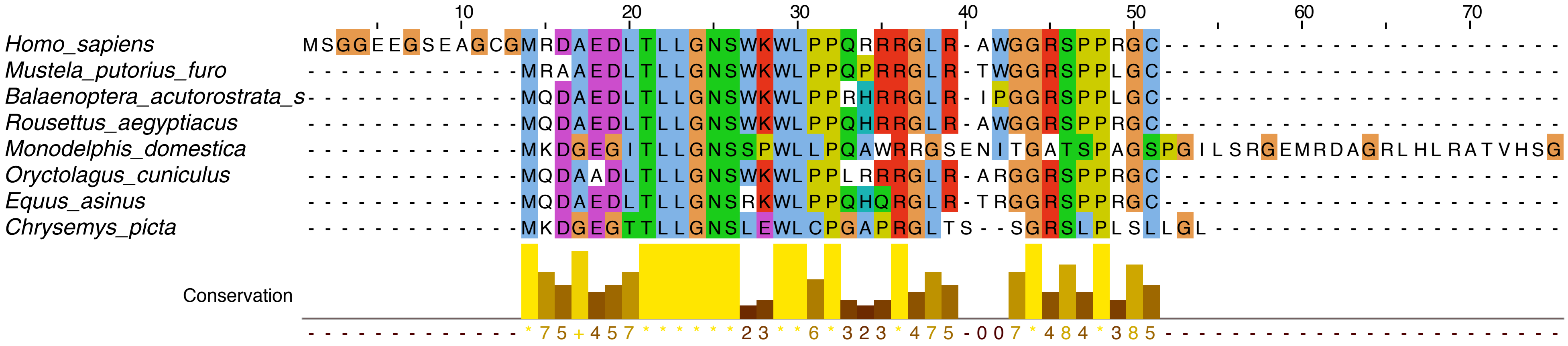

**HG0234.1**

ENSGALG00000011250 (ASIC4)

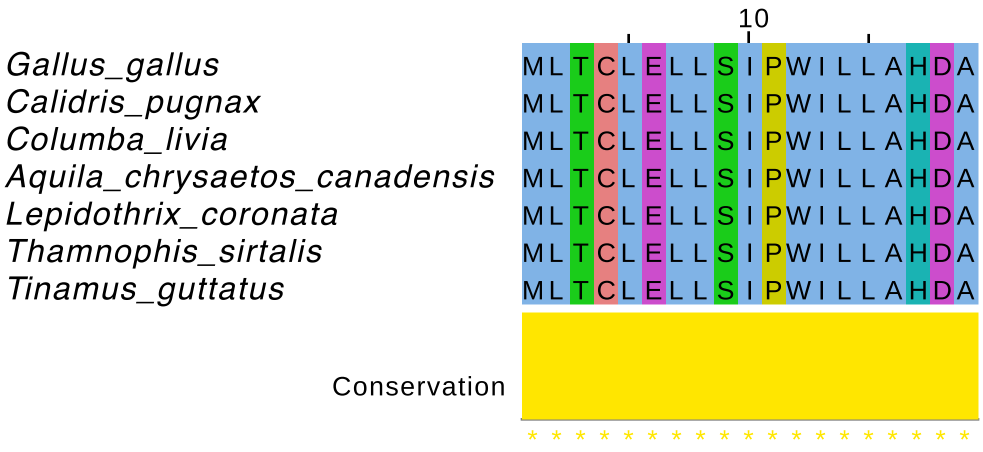

**HG0234.2**

ENSG00000108684 (ASIC2)

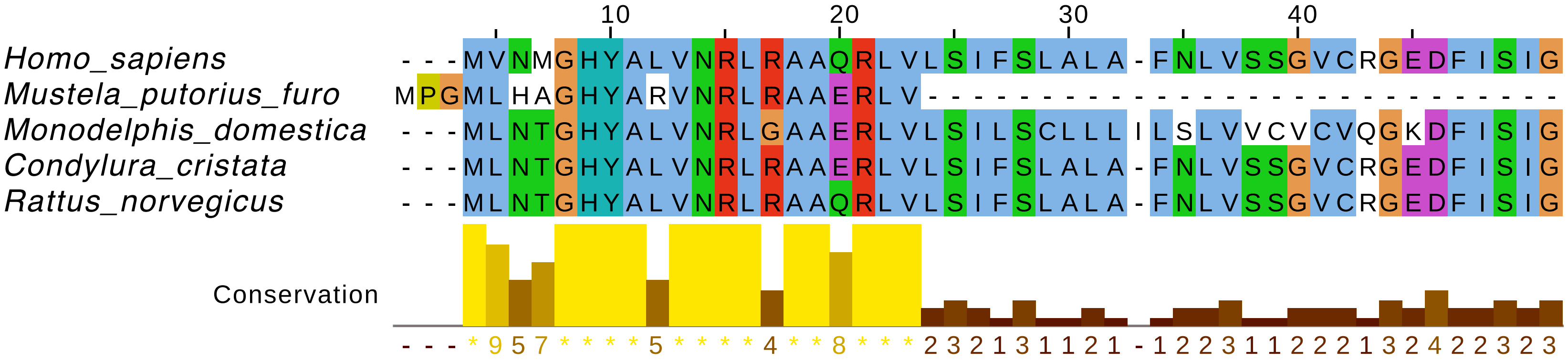

**HG0235.1**

ENSGALG00000035927

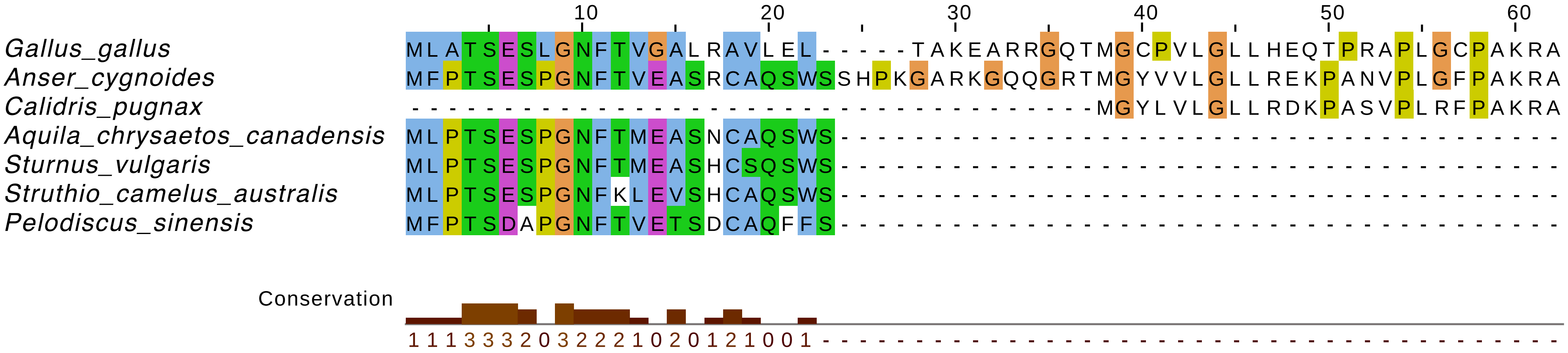

**HG0235.2**

ENSG00000101638 (ST8SIA5)

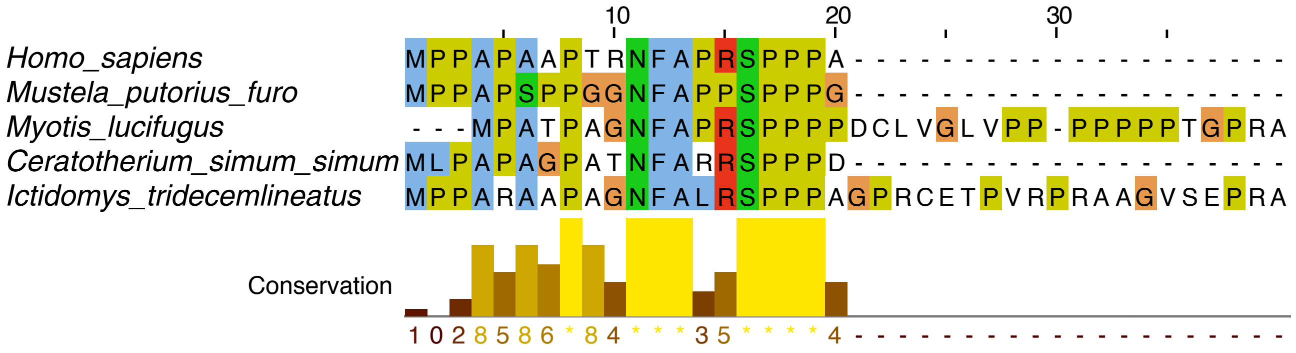

**HG0236**

ENSGALG00000004437 (LHFPL2)

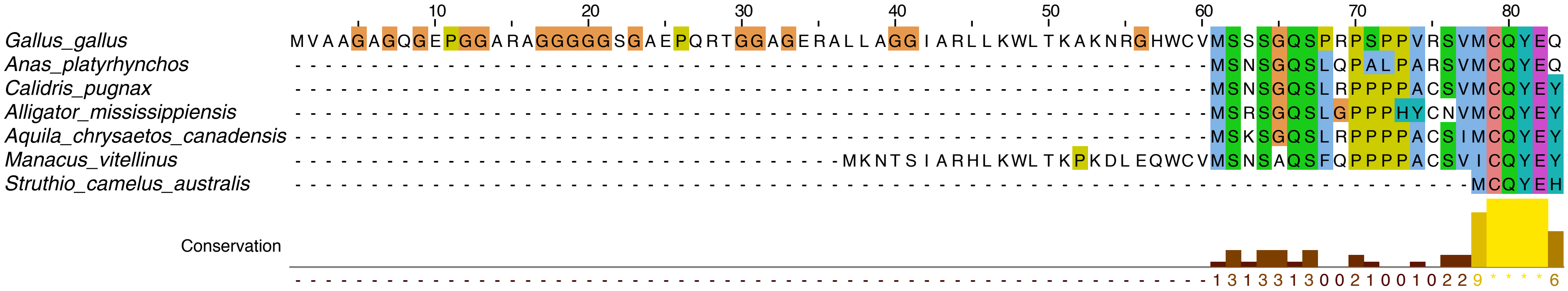

**HG0237**

ENSGALG00000012123 (PAX6)

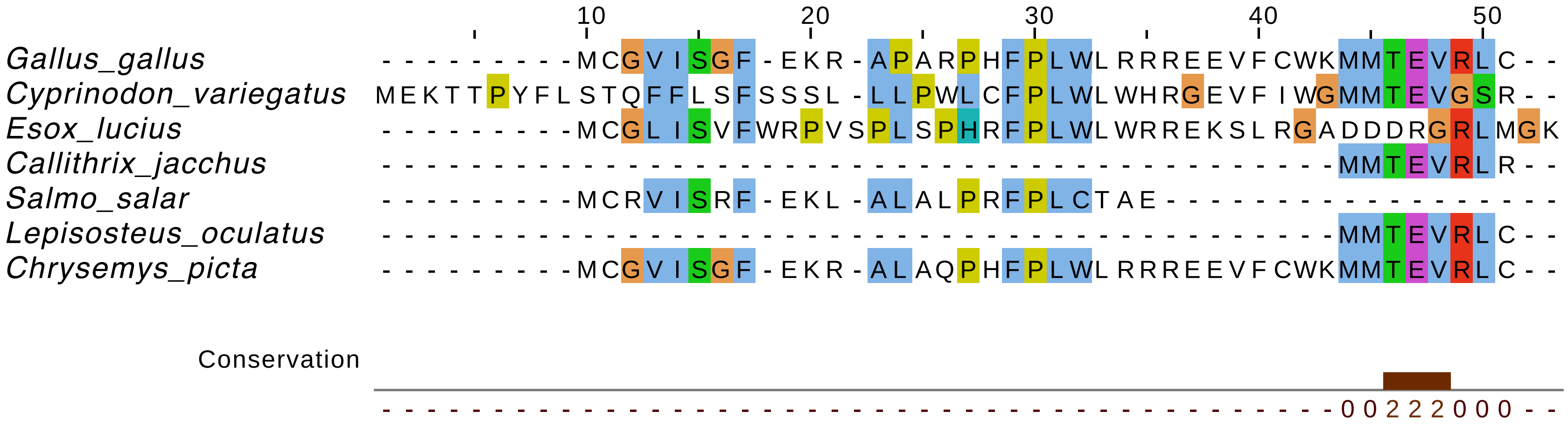

**HG0238**

ENSG00000049618 (ARID1B)

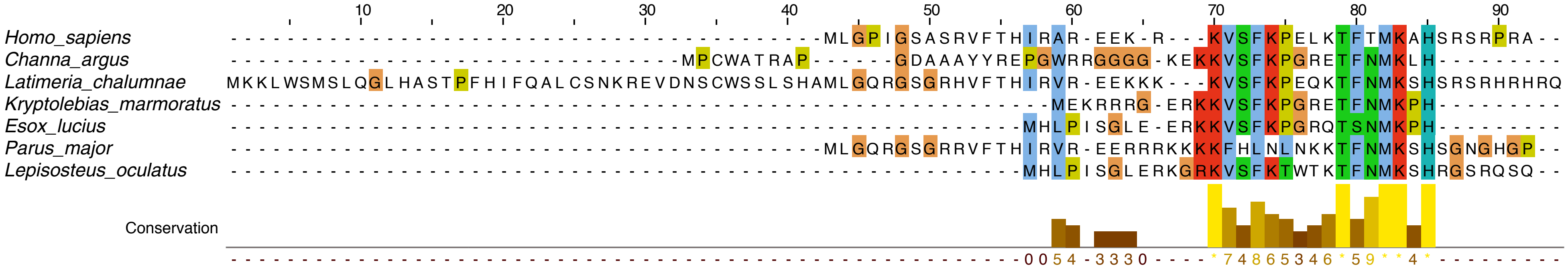

**HG0239**

ENSG00000149256 (TENM4)

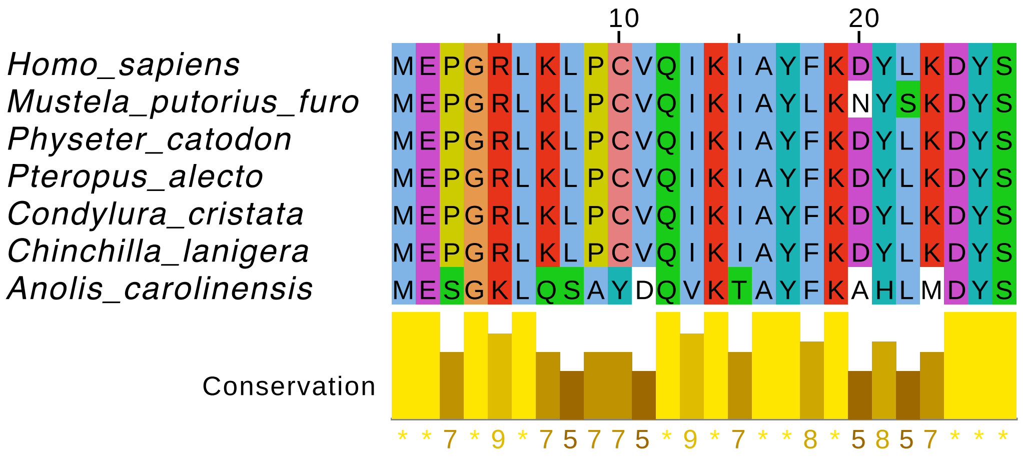

**HG0240**

ENSG00000166342 (NETO1)

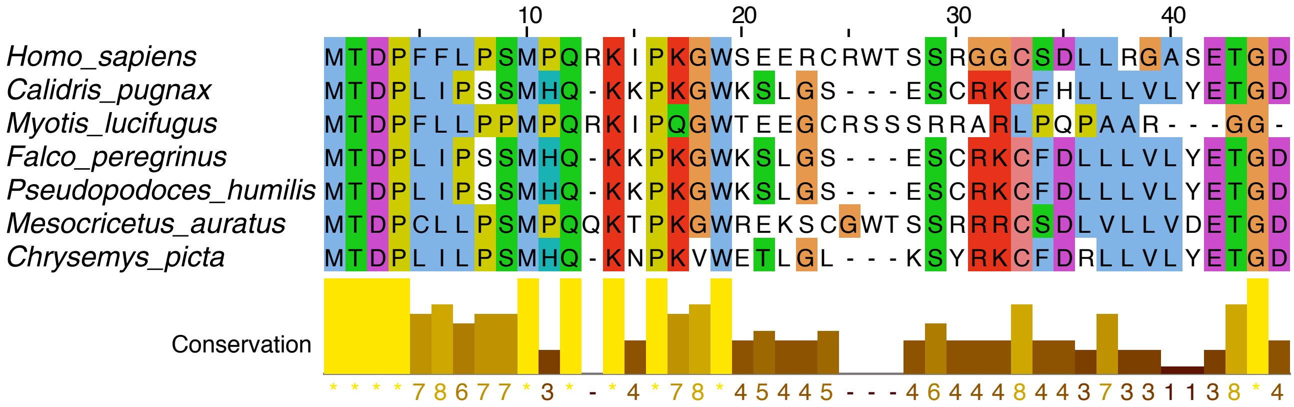

# HG0241

### ENSG00000198561 (CTNND1)

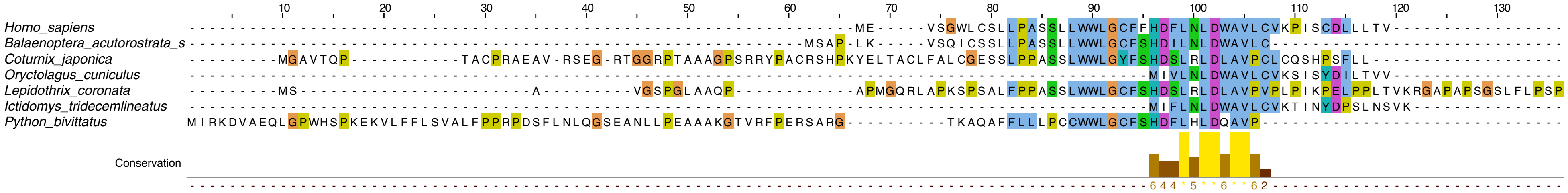

**HG0242.1**

ENSGALG00000007112 (LINS1)

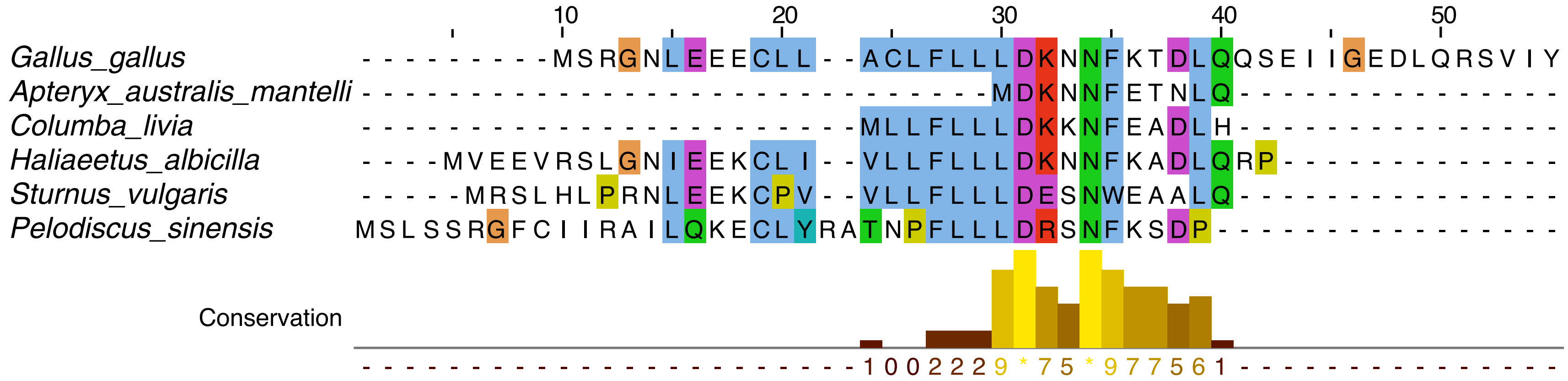

**HG0242.2**

ENSG00000140471 (LINS1)

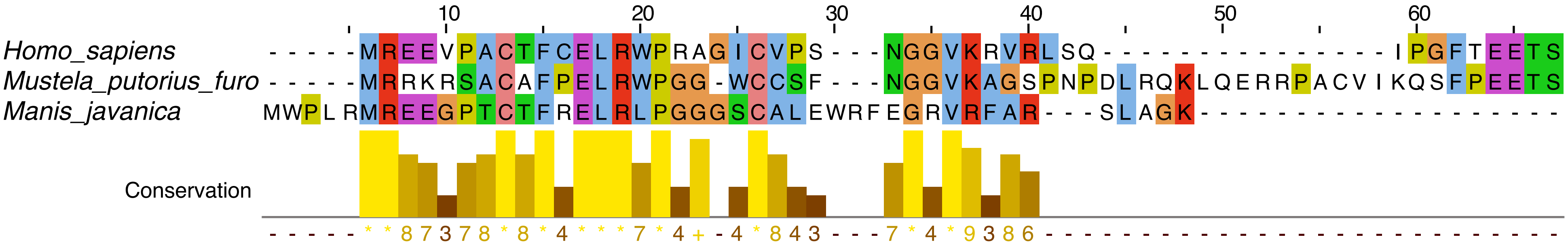

**HG0243.1**

ENSG00000163697 (APBB2)

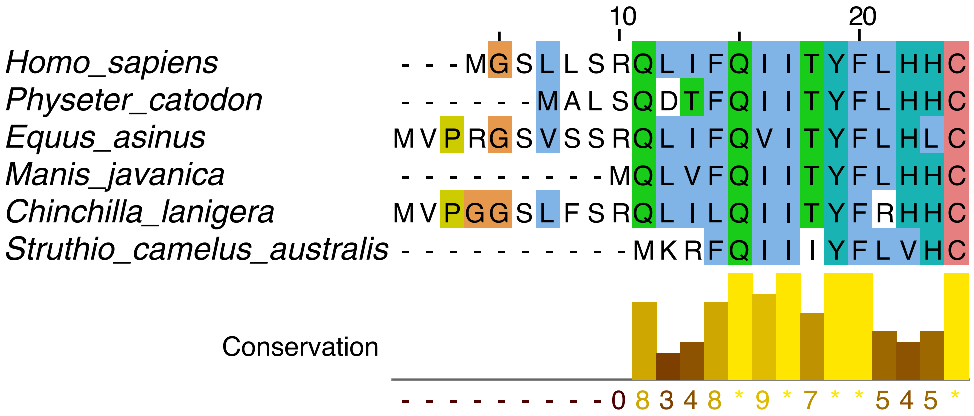

**HG0243.2**

ENSG00000163697 (APBB2)

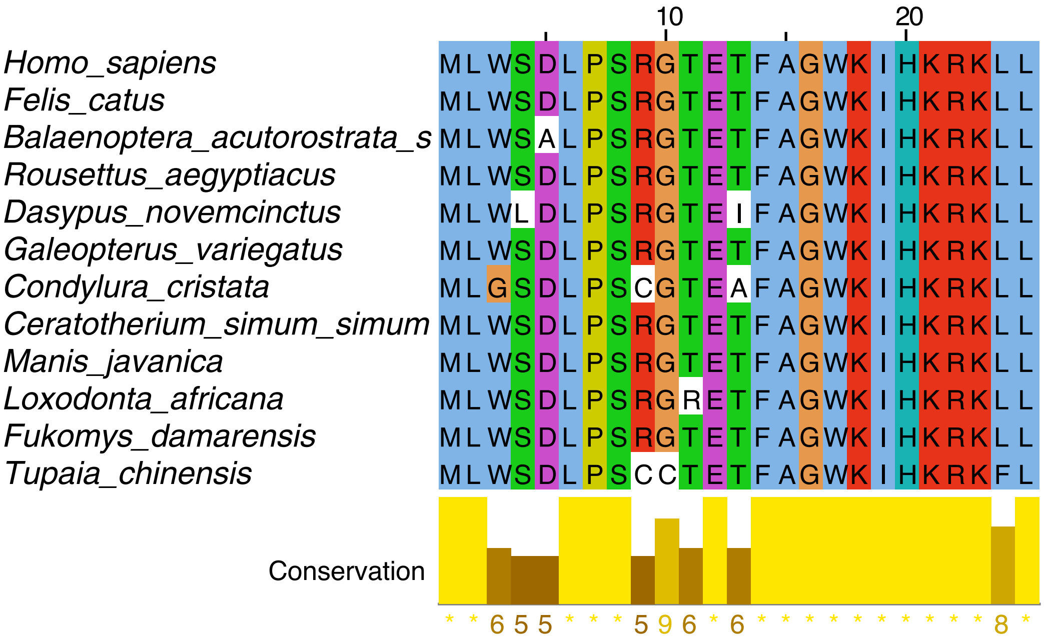

**HG0244**

ENSGALG00000010837 (ASB5)

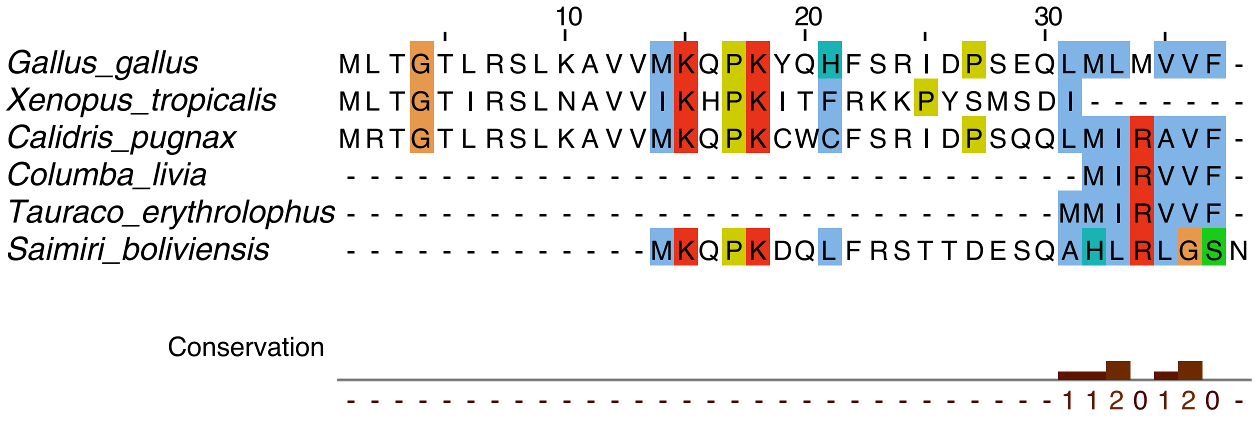

# HG0245

### ENSG00000091831 (ESR1)

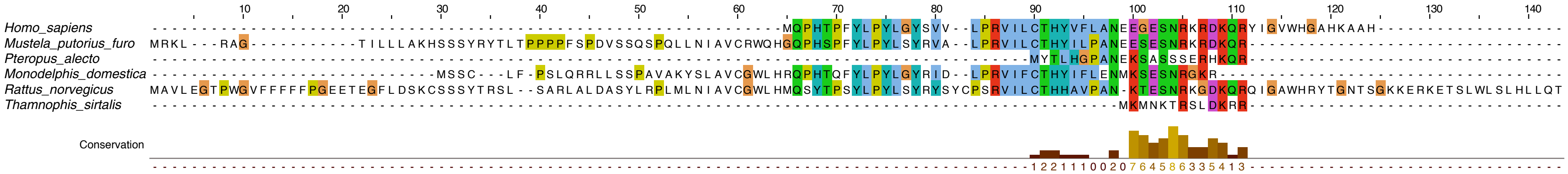

# HG0246

ENSG00000162676 (GFI1)

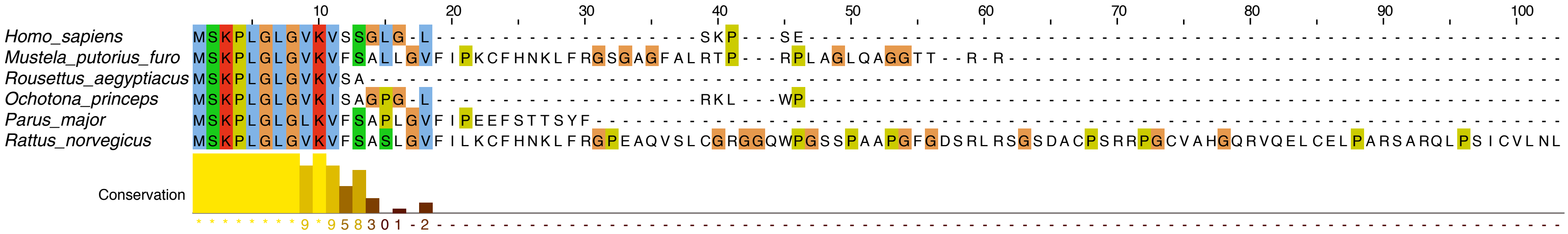

**HG0247.1**

ENSDARG00000014746 (rbfox1)

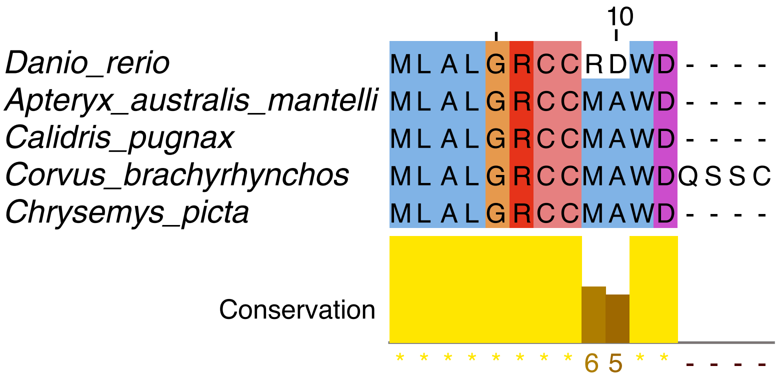

**HG0247.2**

ENSG00000100320 (RBFOX2)

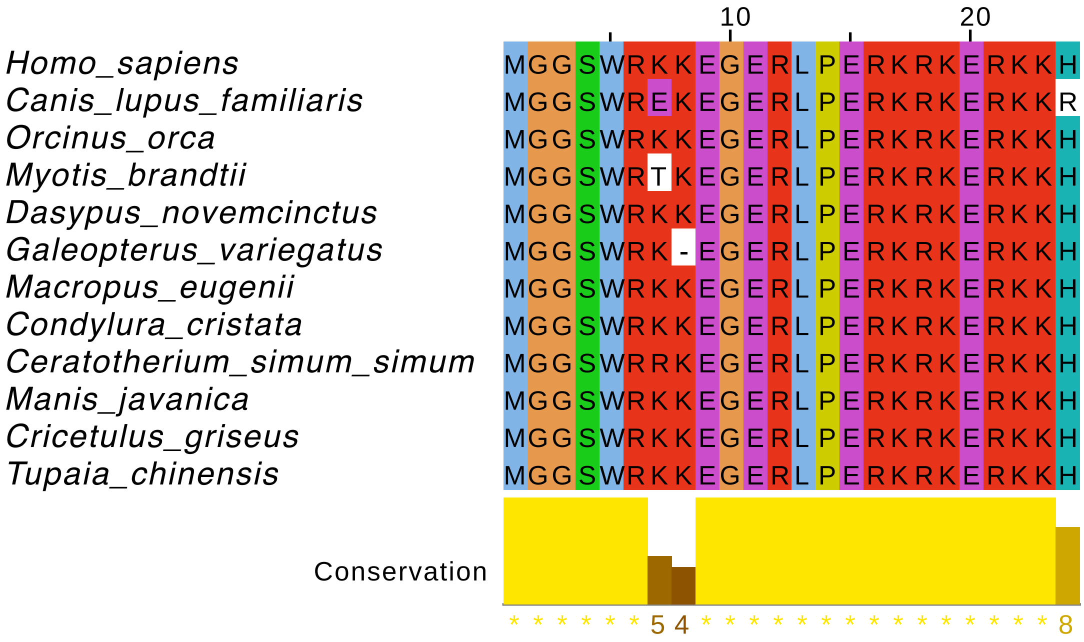

**HG0248.1**

ENSGALG00000004570 (NCOA3)

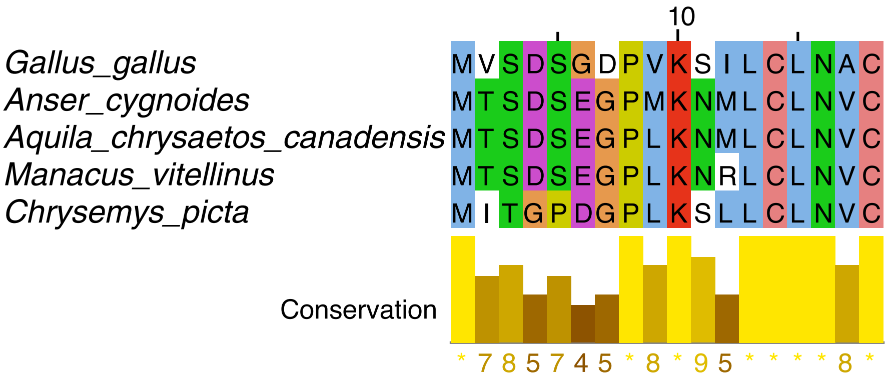

**HG0248.2**

ENSG00000124151 (NCOA3)

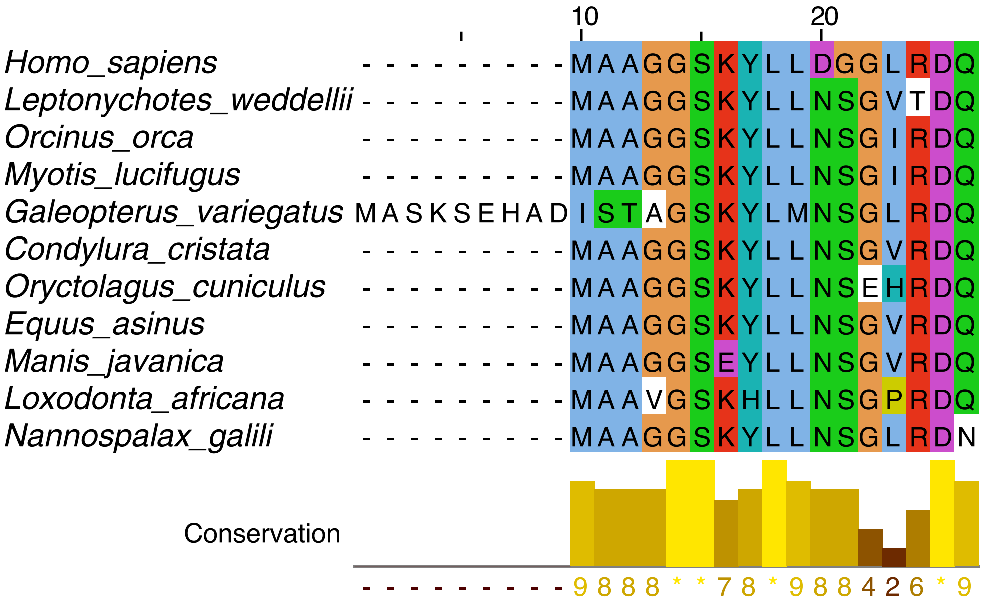

**HG0249**

ENSDARG00000062550 (rc3h1a)

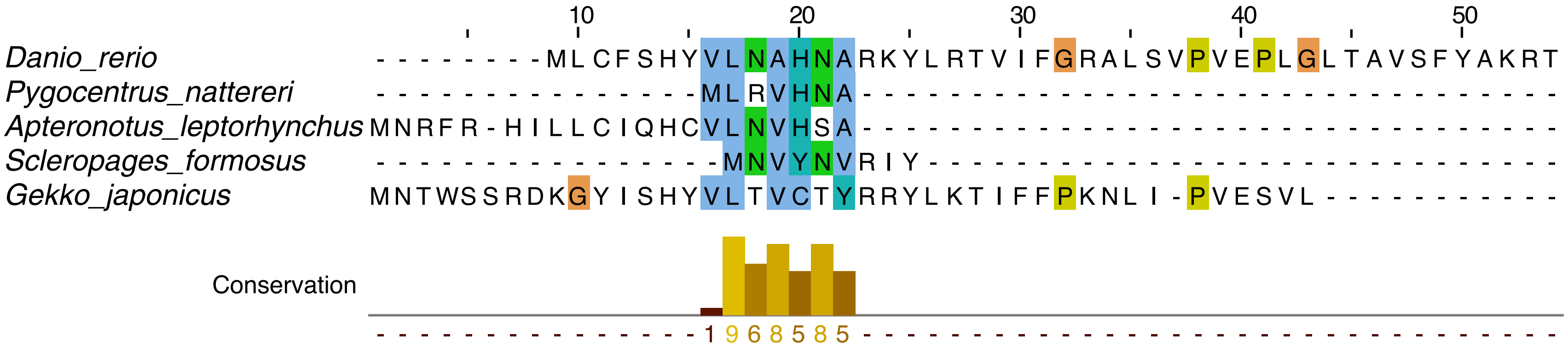

**HG0250**

ENSGALG00000014861

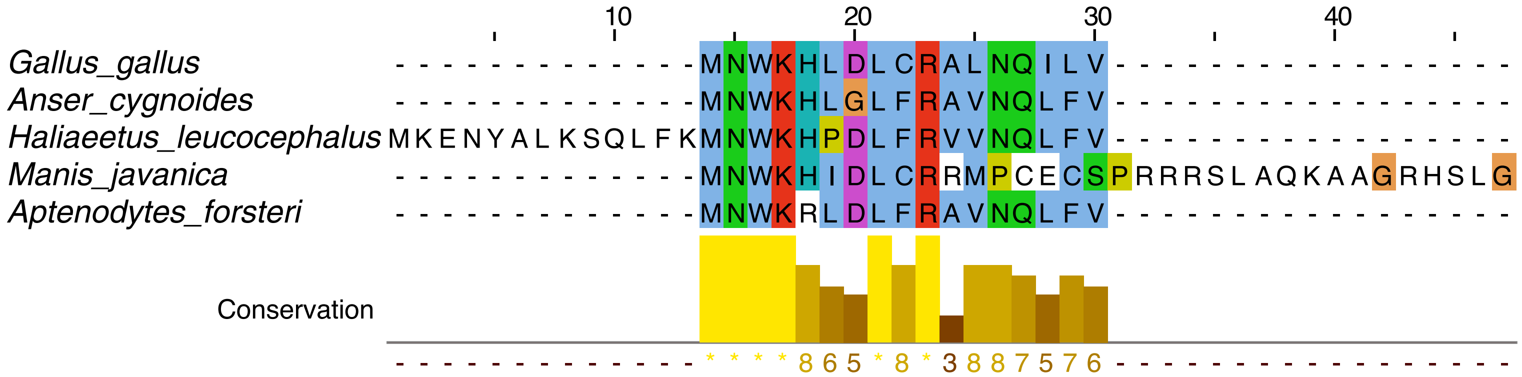

**HG0251**

ENSG00000185053 (SGCZ)

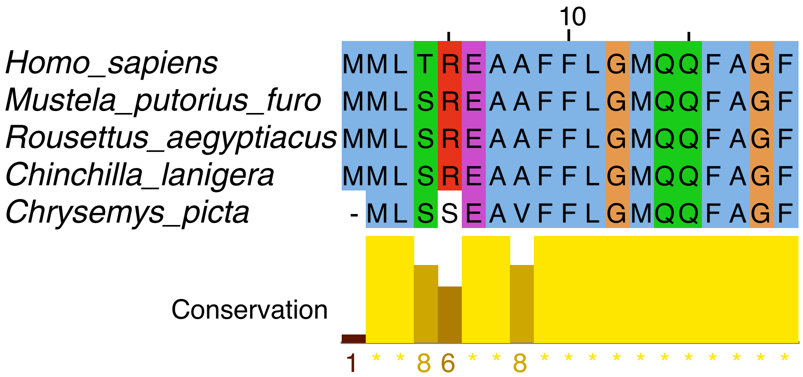

**HG0252.1**

ENSGALG00000009645 (ESRRG)

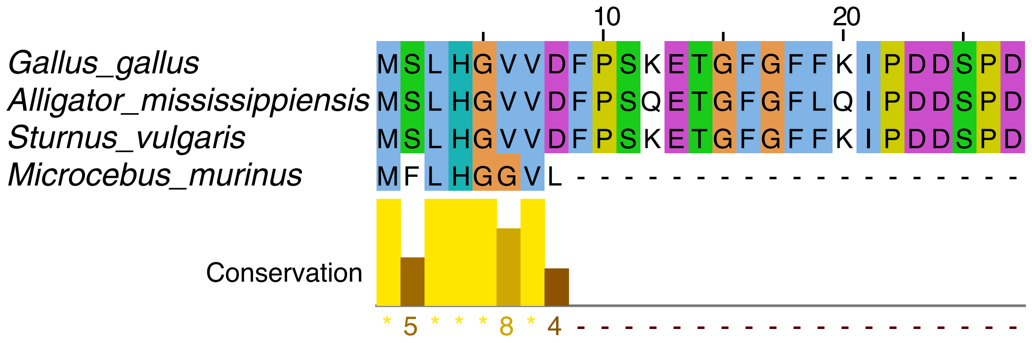

**HG0252.2**

ENSG00000119715 (ESRRB)

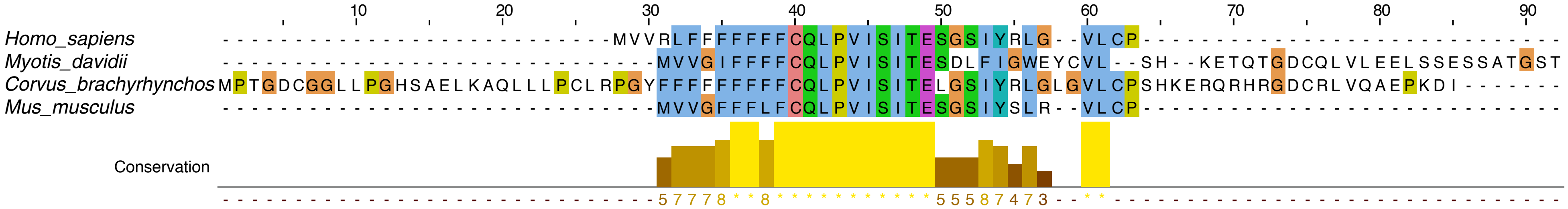

**HG0252.3**

ENSG00000196482 (ESRRG)

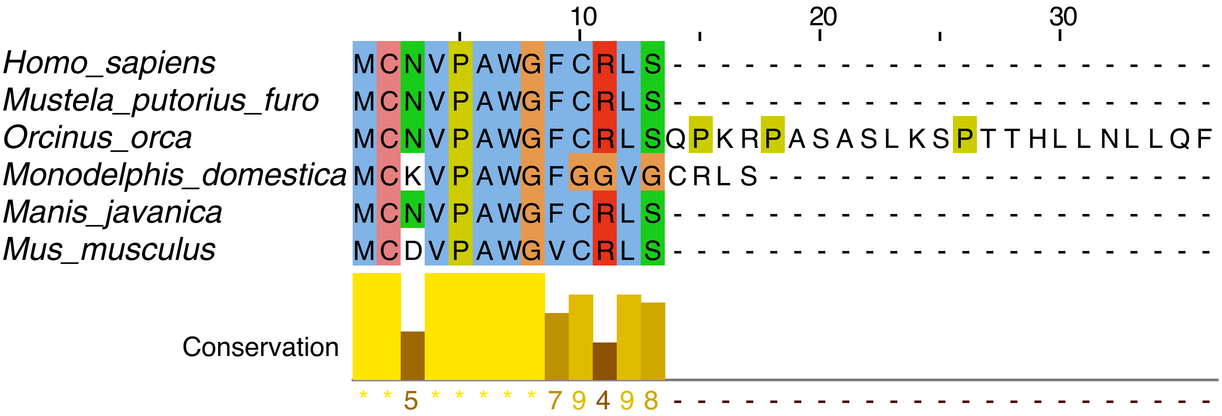

**HG0253.1**

ENSGALG00000015184

```
                            10        20        30        40        50        60        70        80        90
Gallus_gallus          MYQNVNDEALLNLVKGVTSAERAVGAARSGGAGRRAGQGRVGQGGPGRAGHGAPSSAAAASRSLDRQRRARKPNTNKAARTVPSSSSLSLFLIF
Lipotes_vexillifer     MYQNVND---------------------------------------------------------------------------------------
Pteropus_vampyrus      MYQNVND---------------------------------------------------------------------------------------
Anolis_carolinensis    MYQNVNDPPLLNLVKGVTSAESVVRAVPSGGAVPGLPLLLLLPRG--RD---------------------------------------------
Conservation
                       *******---------------------------------------------------------------------------------------
```

**HG0253.2**

ENSG00000140332 (TLE3)

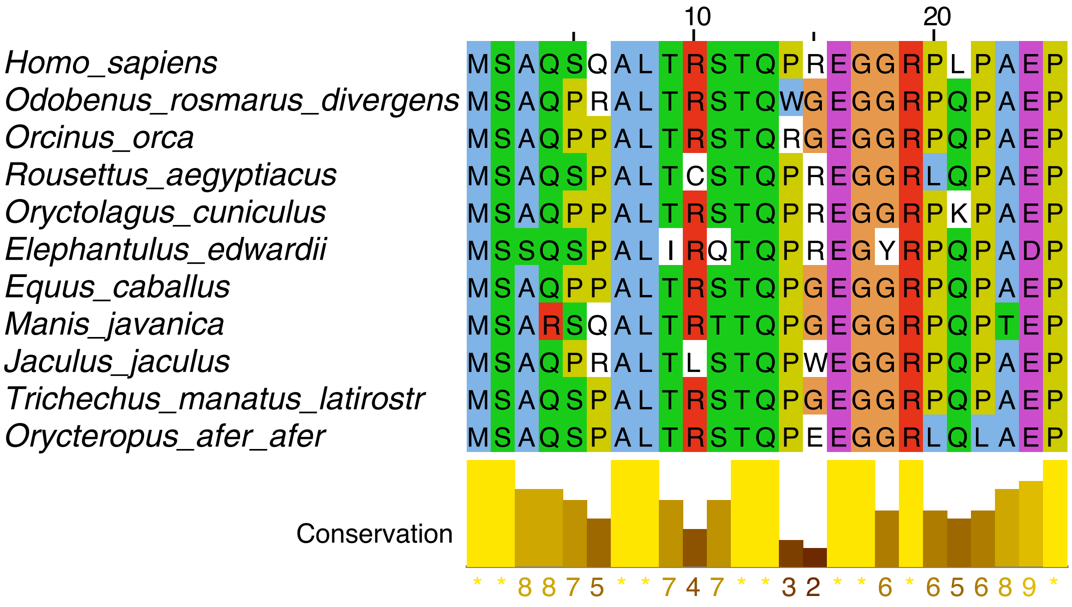

**HG0253.3**

ENSG00000140332 (TLE3)

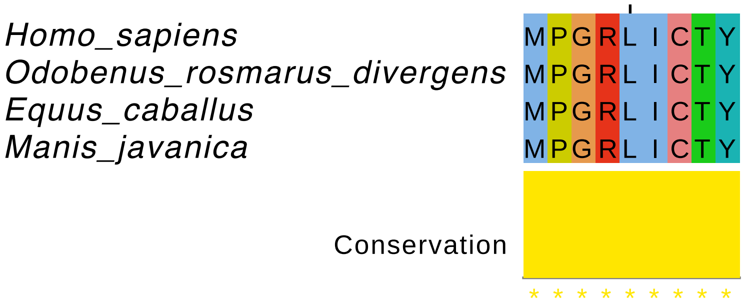

**HG0254.1**

ENSGALG00000035919

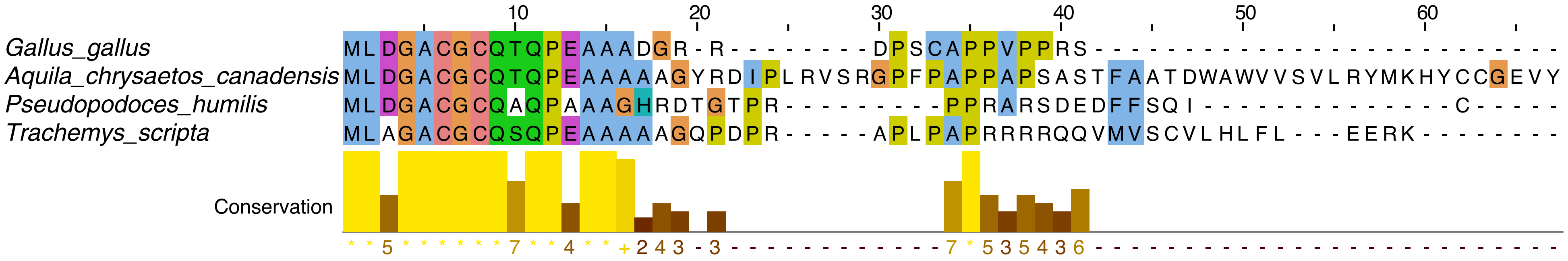

**HG0254.2**

ENSG00000072134 (EPN2)

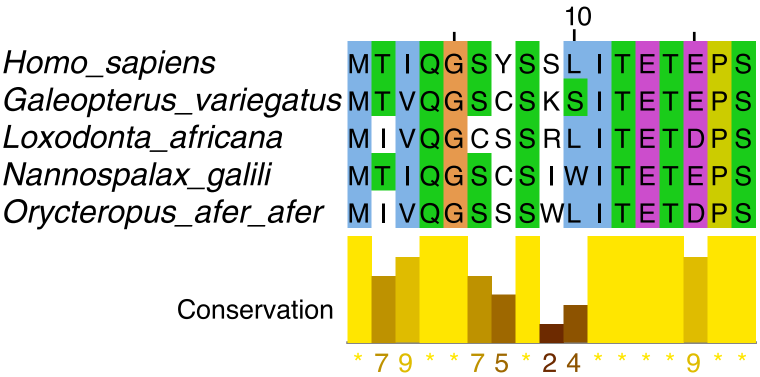

**HG0255**

ENSGALG00000012139 (ARL14EP)

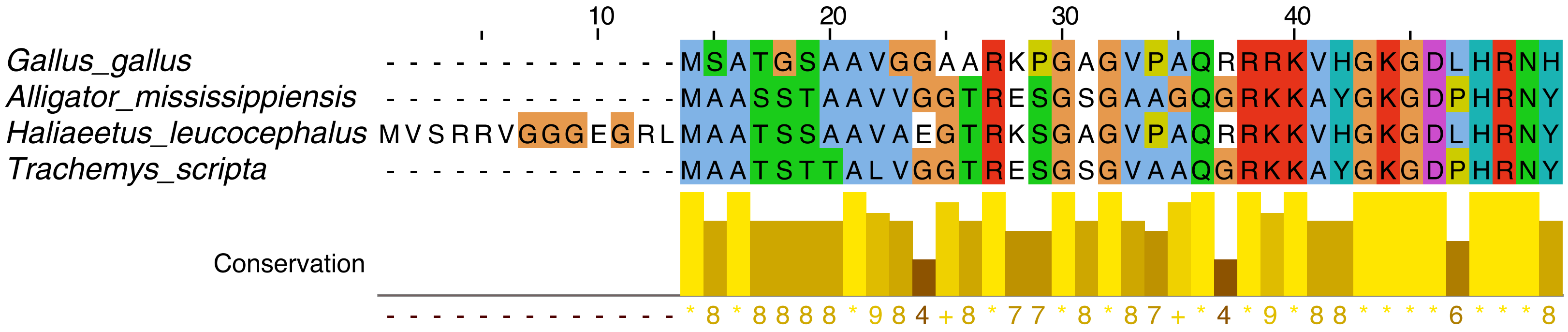

**HG0256**

ENSGALG00000039901

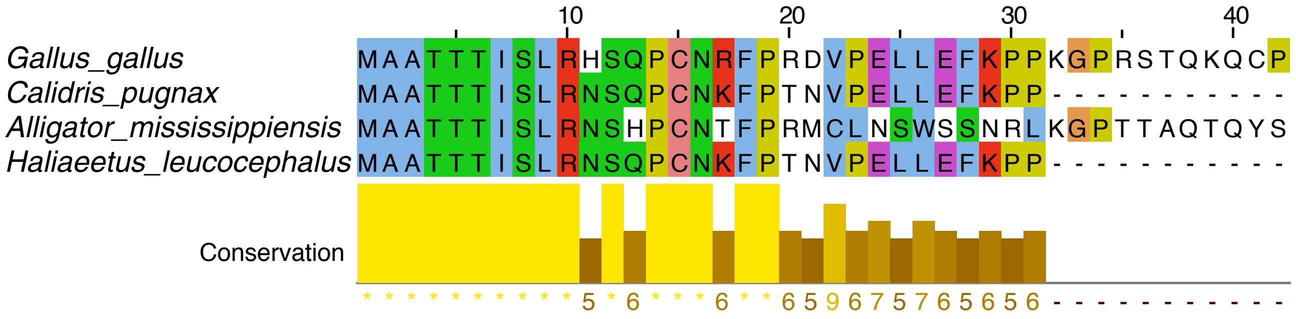

**HG0257**

ENSG00000198791 (CNOT7)

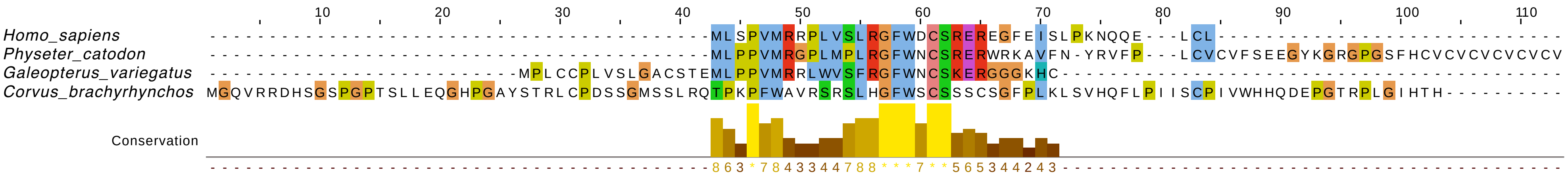

**HG0258.1**

ENSGALG00000016797 (FHL2)

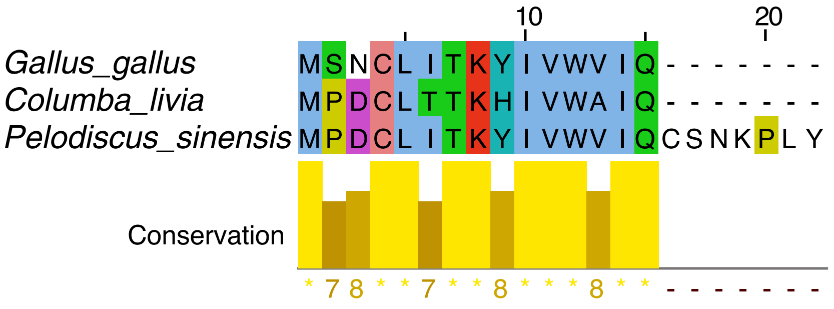

**HG0258.2**

ENSGALG00000016797 (FHL2)

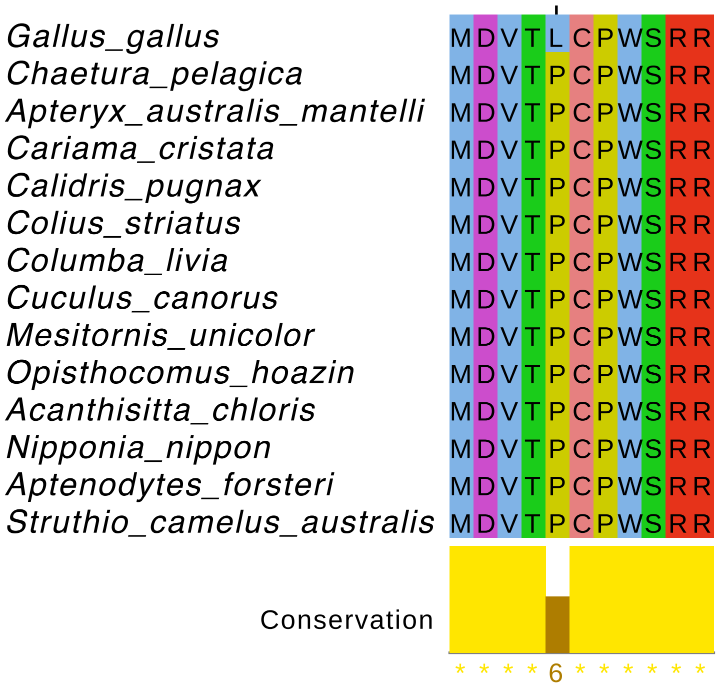

**HG0259**

ENSGALG00000009936 (AP4S1)

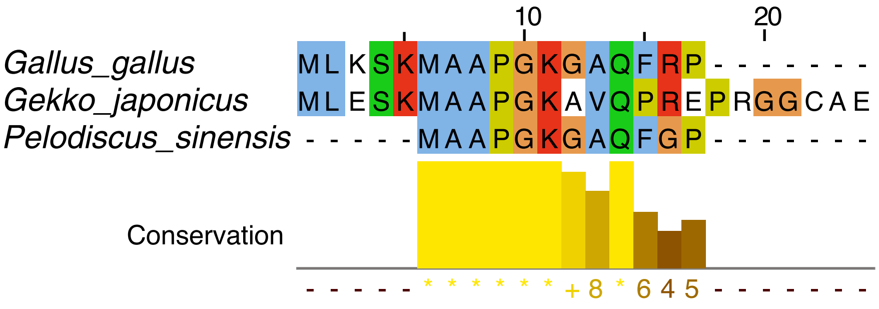

**HG0260.1**

ENSGALG00000016890 (MBNL2)

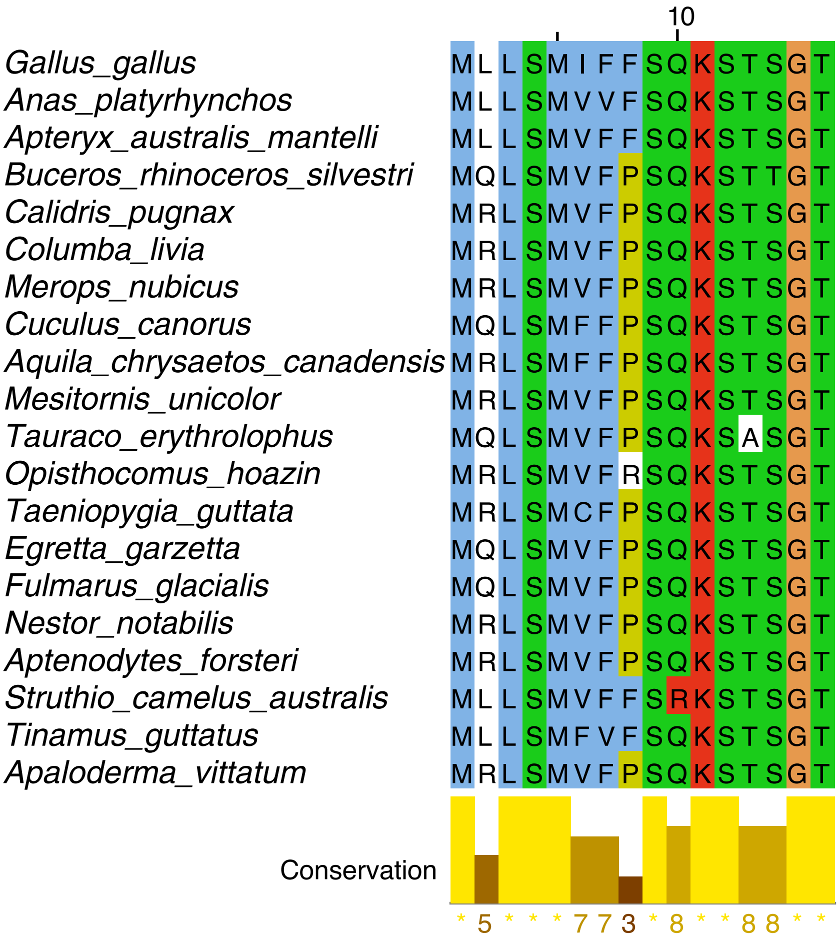

**HG0260.2**

ENSG00000139793 (MBNL2)

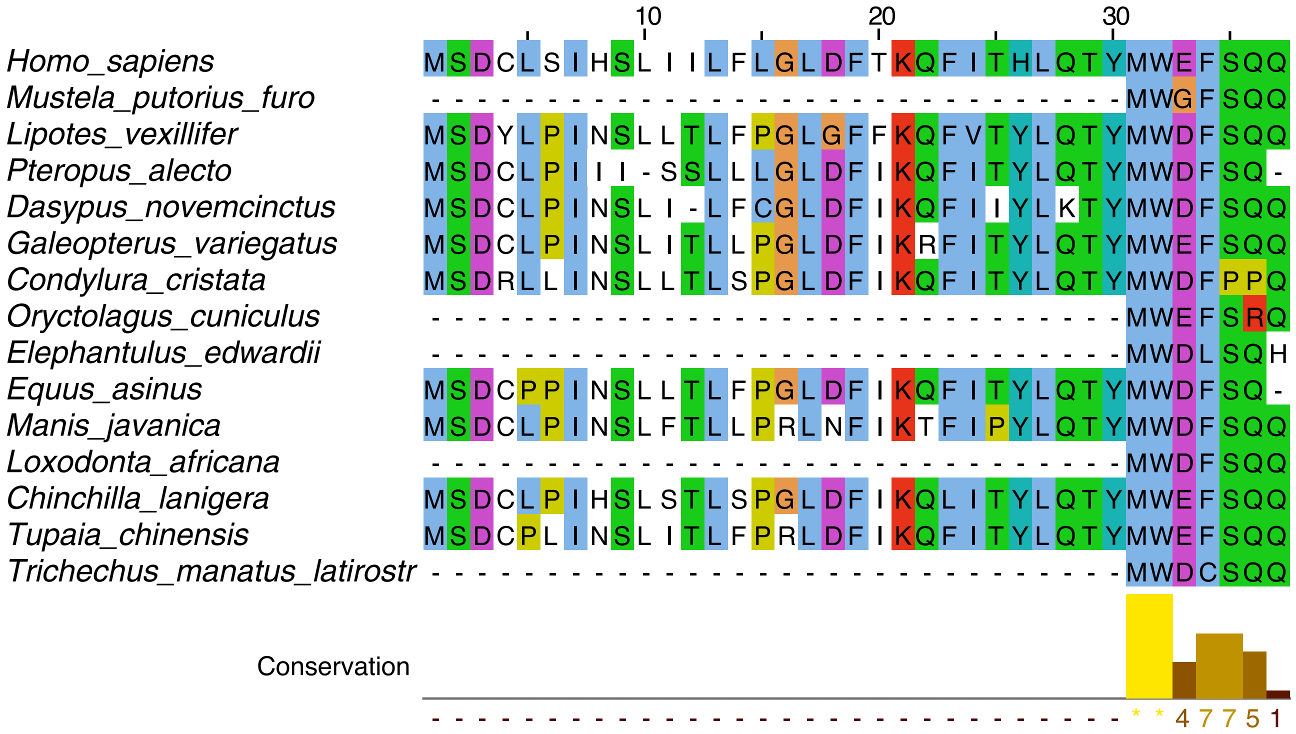

**HG0260.3**

ENSG00000152601 (MBNL1)

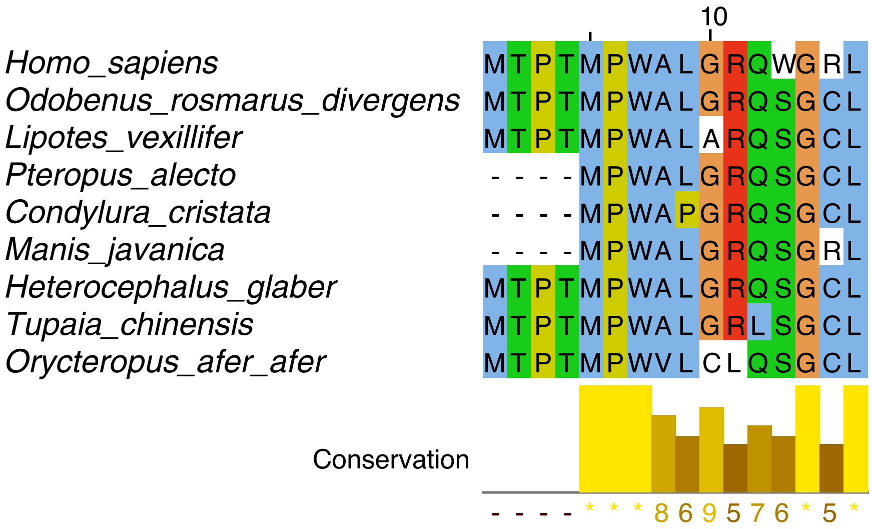

**HG0260.4**

ENSG00000152601 (MBNL1)

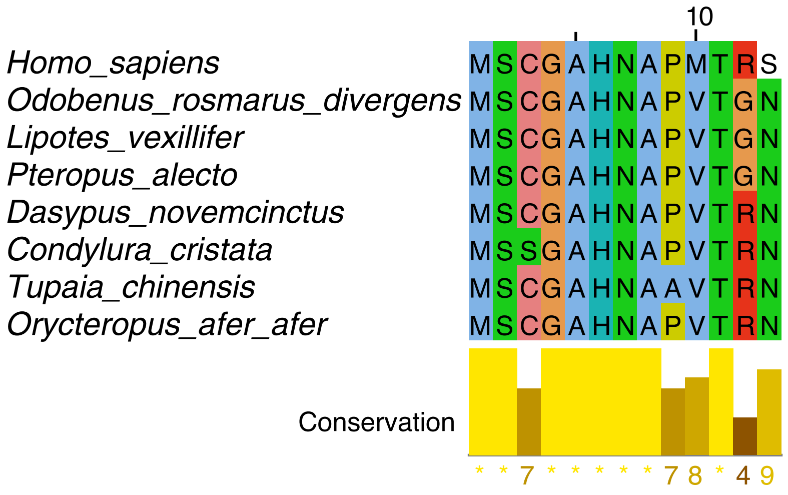

**HG0260.5**

ENSG00000152601 (MBNL1)

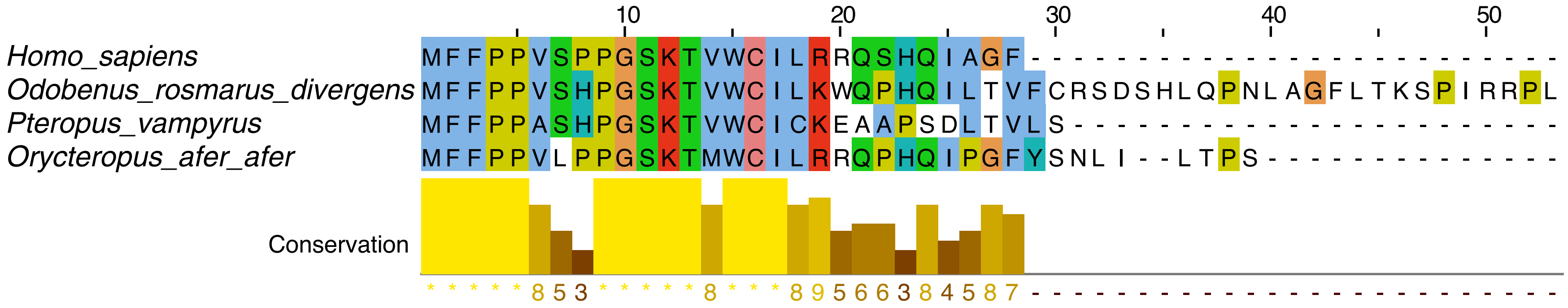

**HG0260.6**

ENSG00000152601 (MBNL1)

```
                                    10        20        30        40        50        60        70
Homo_sapiens                 --MGLRFICSGDNMAYLPWNIPVFQK----------------------------------------------------
Odobenus_rosmarus_divergens  MIWGCGVFCSNNNMT---------------------------------------------------------------
Orcinus_orca                 -------------MTYLPWTITVFKKMNILC-----------------------------------------------
Pteropus_vampyrus            -------------MTYLPWNSPVFQKMVFYFLKRGPLLSINHIMQSWGERGFYISNIIYKLDGKLVCECLCLCDNQFY

Conservation
                             -------------*8---------------------------------------------------------------
```

**HG0261.1**

ENSGALG00000005867 (8-Mar)

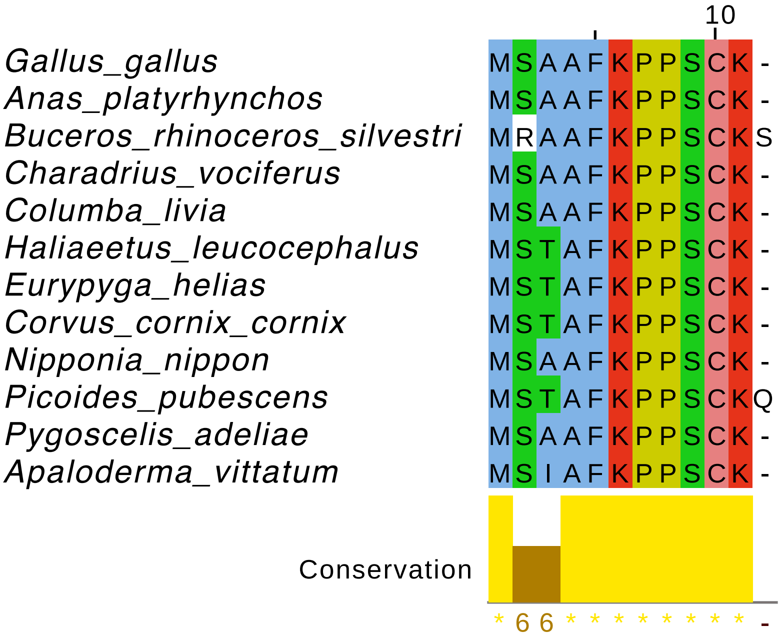

**HG0261.2**

ENSGALG00000009520 (1-Mar)

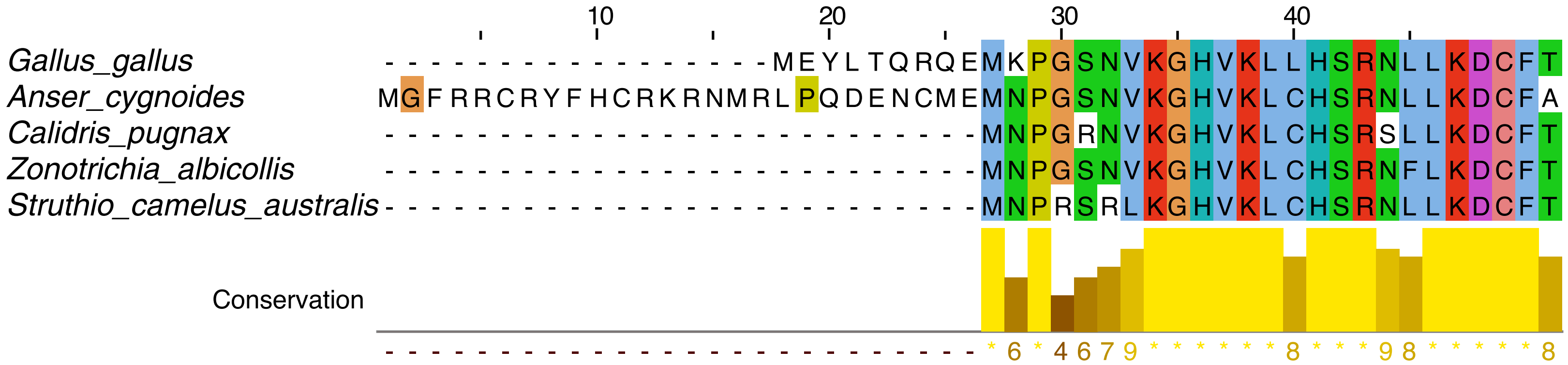

**HG0261.3**

ENSG00000145416 (1-Mar)

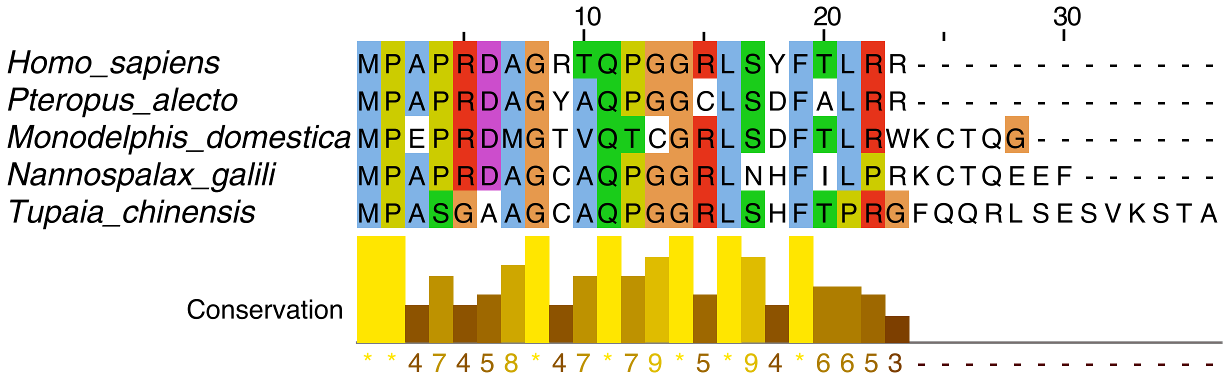

**HG0261.4**

ENSG00000165406 (8-Mar)

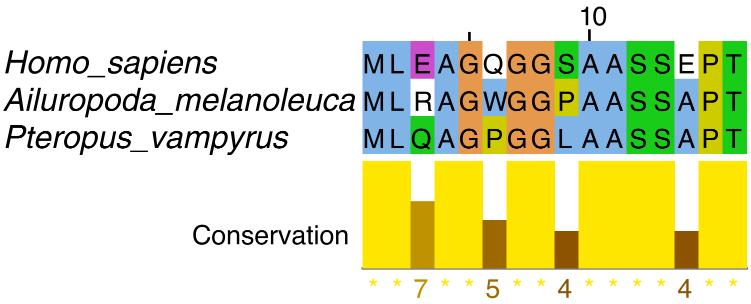

**HG0262.1**

ENSG00000023445 (BIRC3)

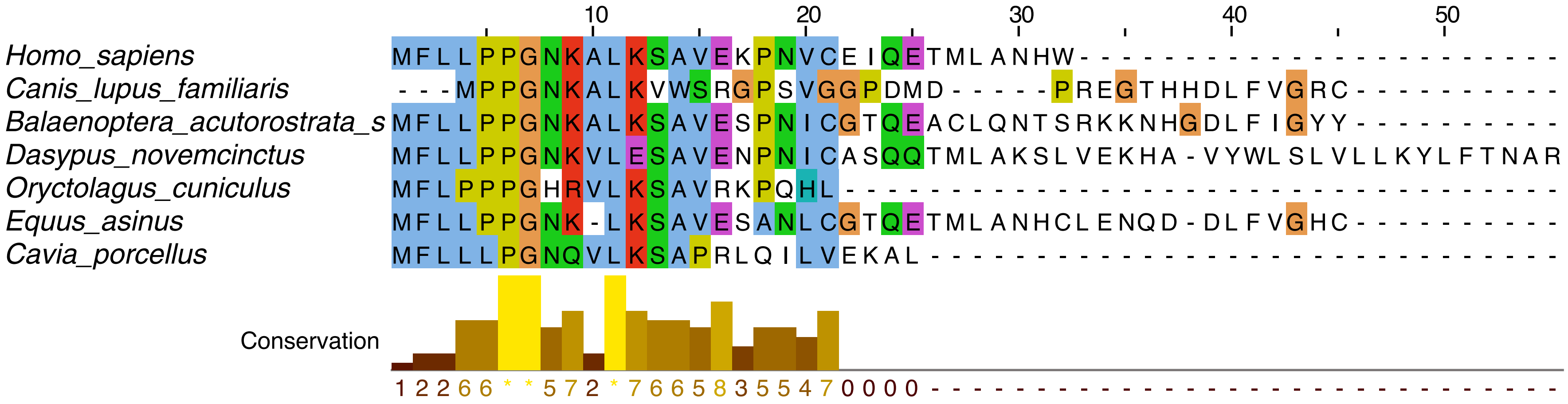

**HG0262.2**

ENSG00000023445 (BIRC3)

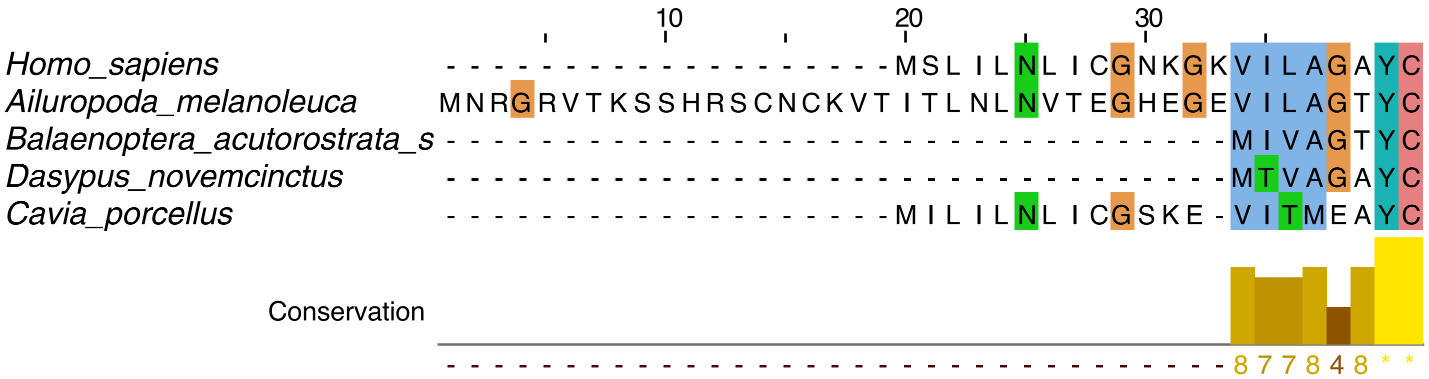

**HG0262.3**

ENSGALG00000017186

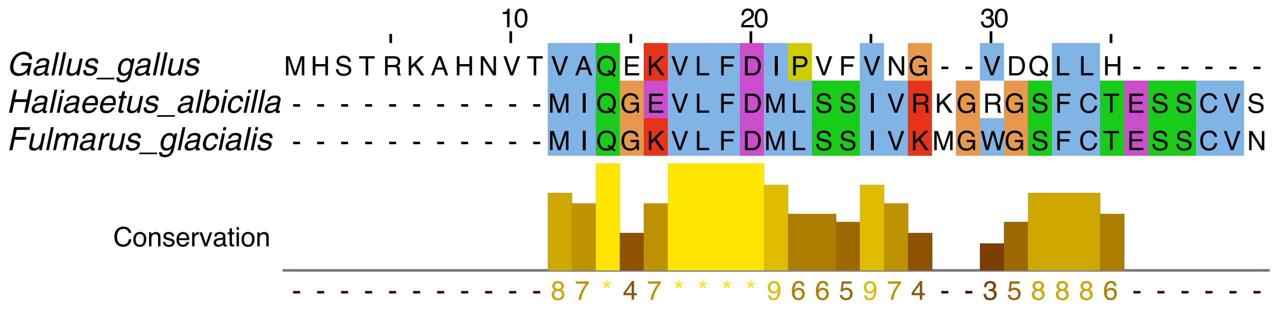

**HG0262.4**

ENSG00000023445 (BIRC3)

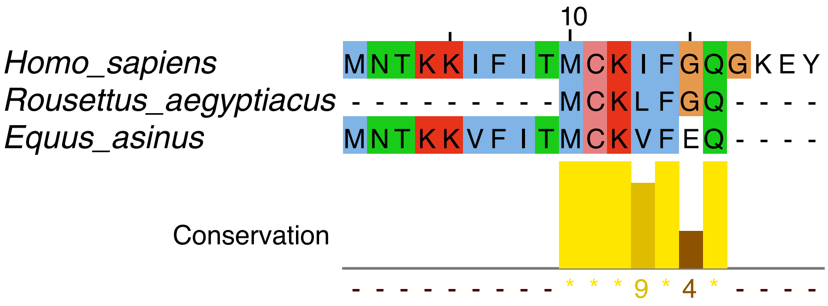

**HG0263.1**

ENSG00000169826 (CSGALNACT2)

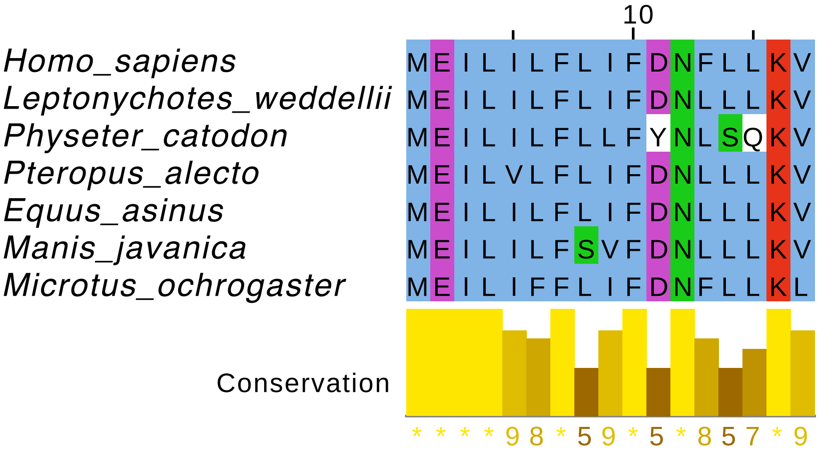

**HG0263.2**

ENSGALG00000002543 (CSGALNACT2)

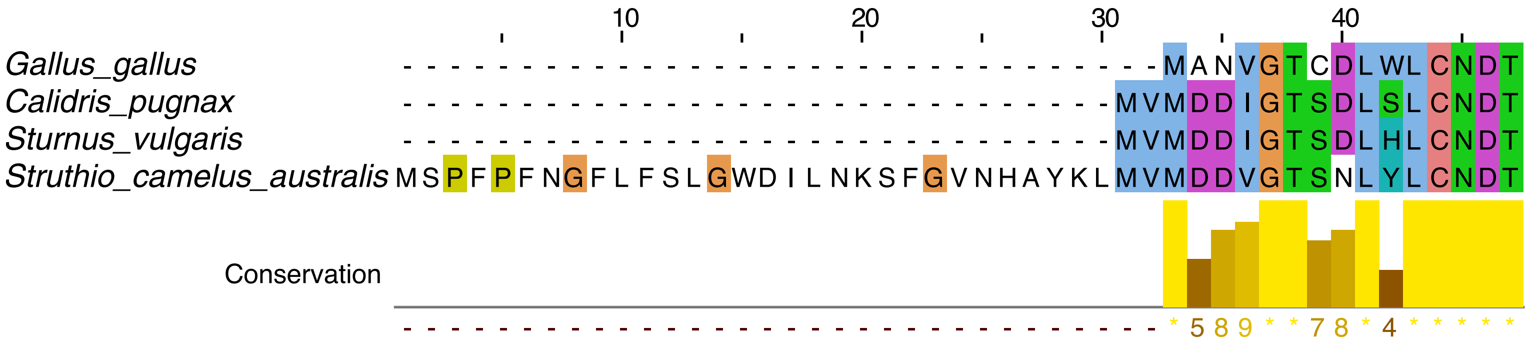

**HG0263.3**

ENSGALG00000002543 (CSGALNACT2)

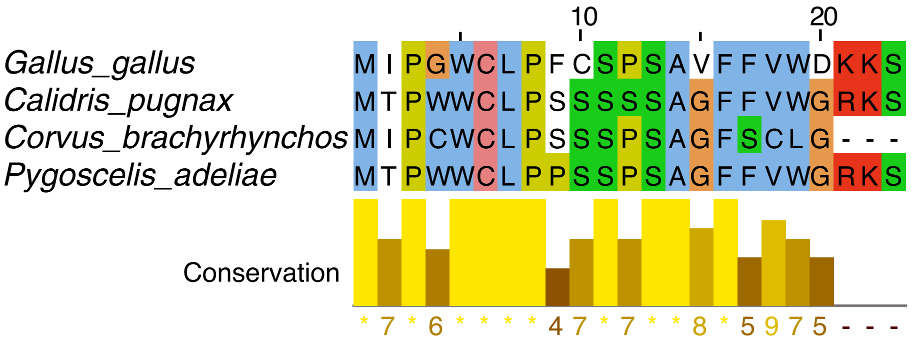

**HG0263.4**

ENSG00000147408 (CSGALNACT1)

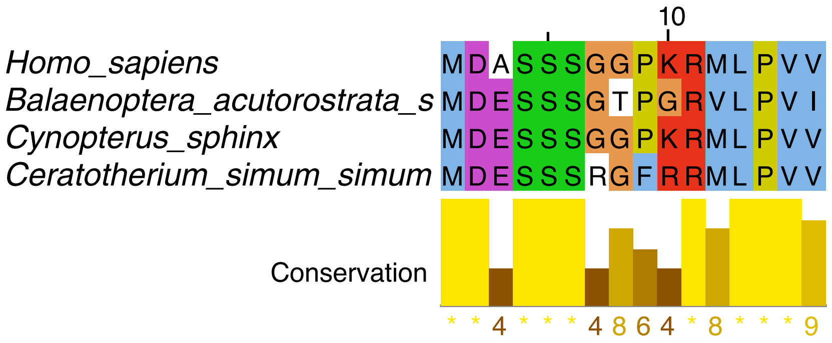

**HG0264.1**

ENSG00000151458 (ANKRD50)

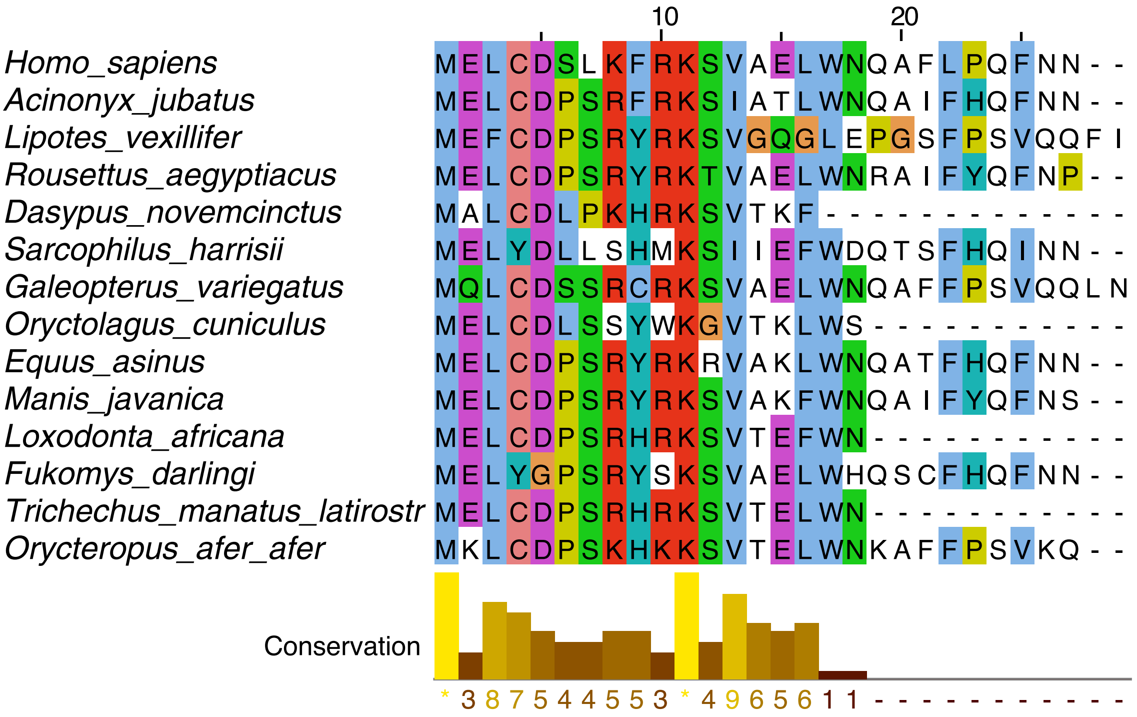

**HG0264.2**

ENSG00000151458 (ANKRD50)

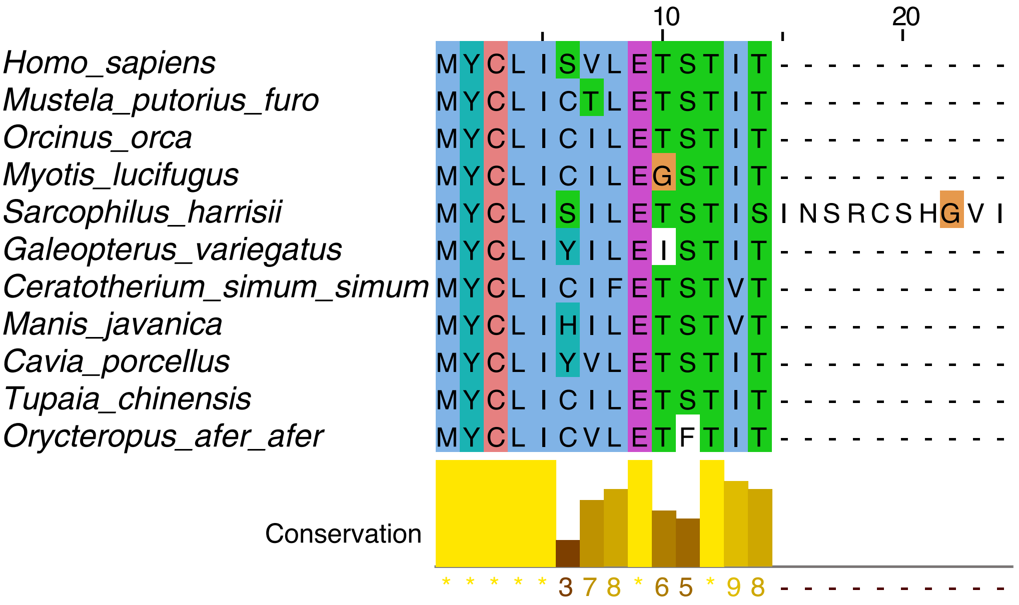

**HG0264.3**

ENSGALG00000011831 (ANKRD50)

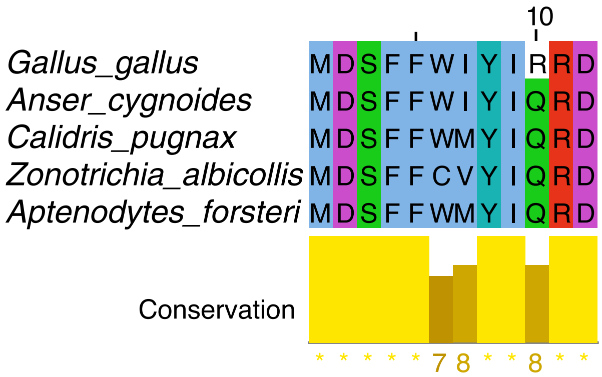

**HG0264.4**

ENSG00000151458 (ANKRD50)

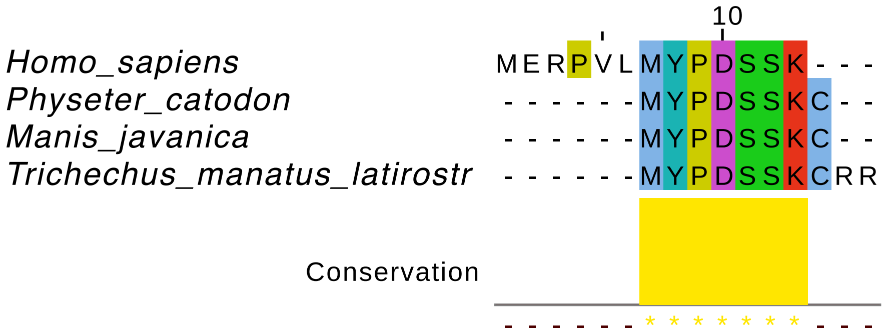

**HG0265.1**

ENSG00000144285 (SCN1A)

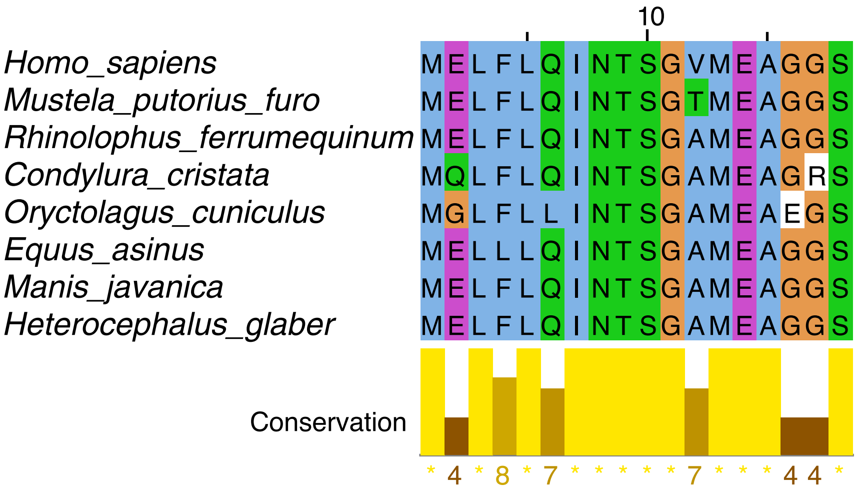

**HG0265.2**

ENSG00000196876 (SCN8A)

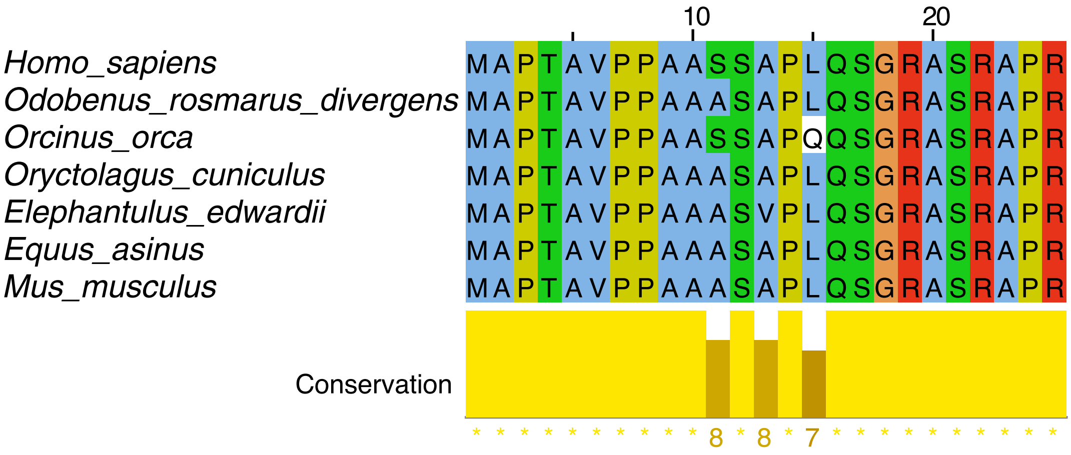

**HG0265.3**

ENSDARG00000018032 (scn8ab)

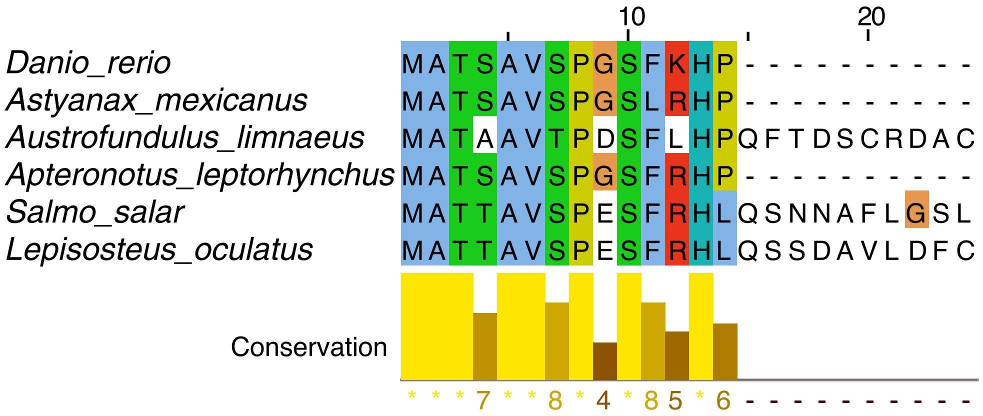

**HG0266.1**

ENSG00000184185 (KCNJ12)

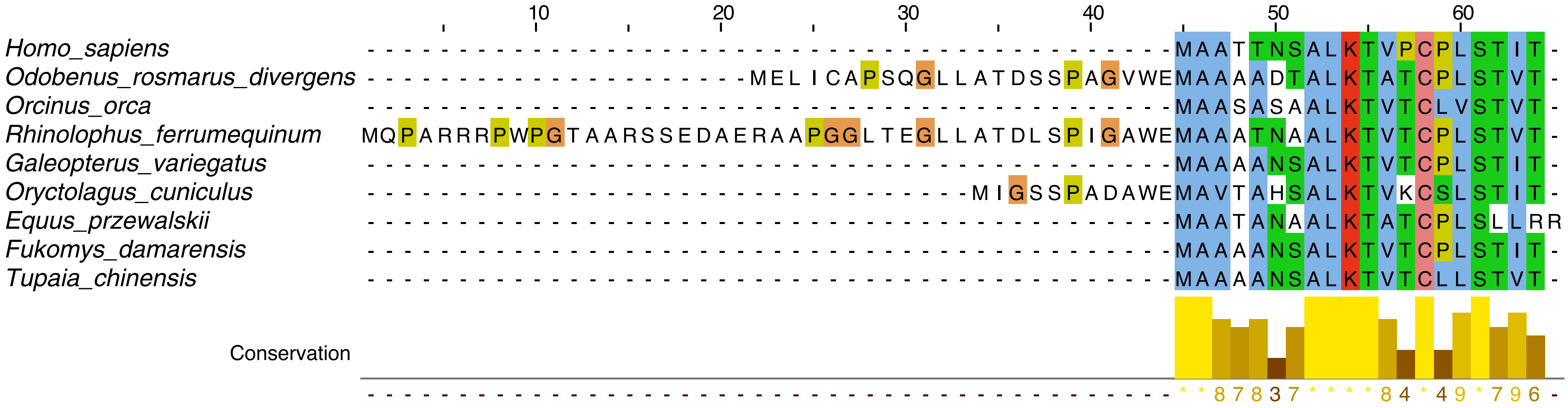

**HG0266.1**

ENSG00000260458 (KCNJ18)

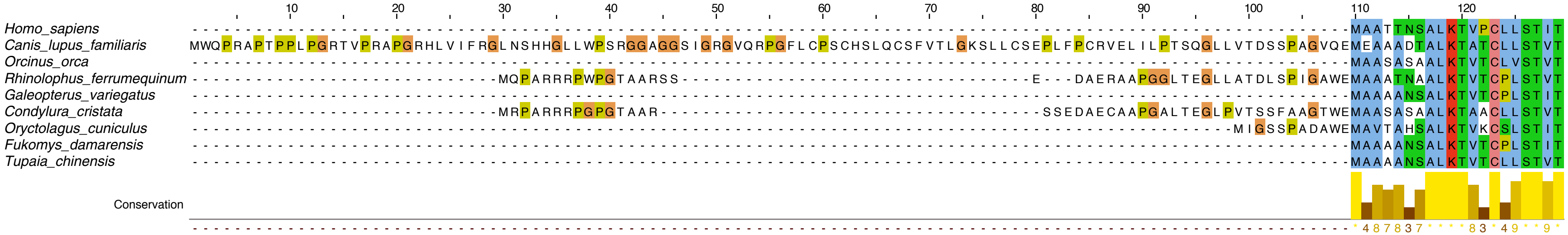

**HG0266.2**

ENSGALG00000001181 (KCNJ5)

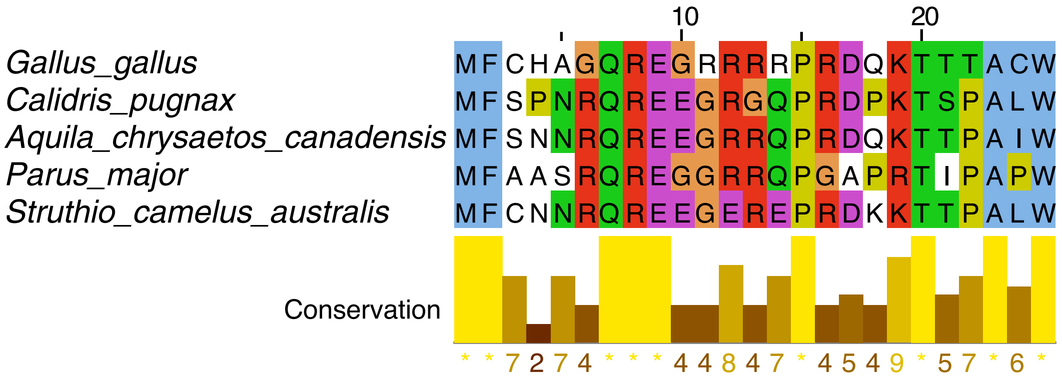

**HG0267**

ENSDARG00000028173 (slc4a2a)

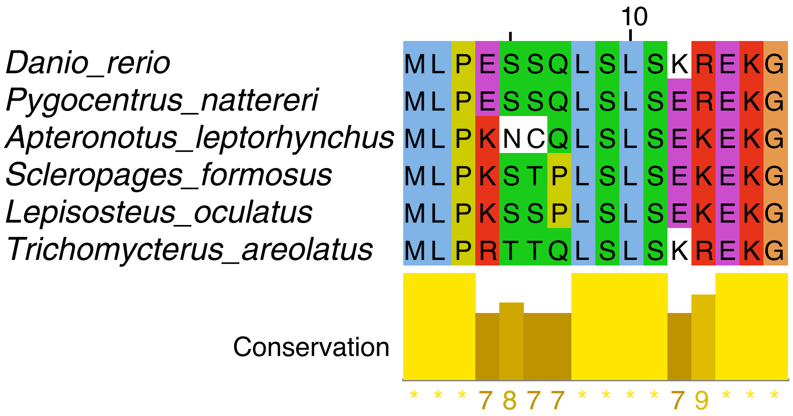

**HG0267**

ENSDARG00000052330 (slc4a2b)

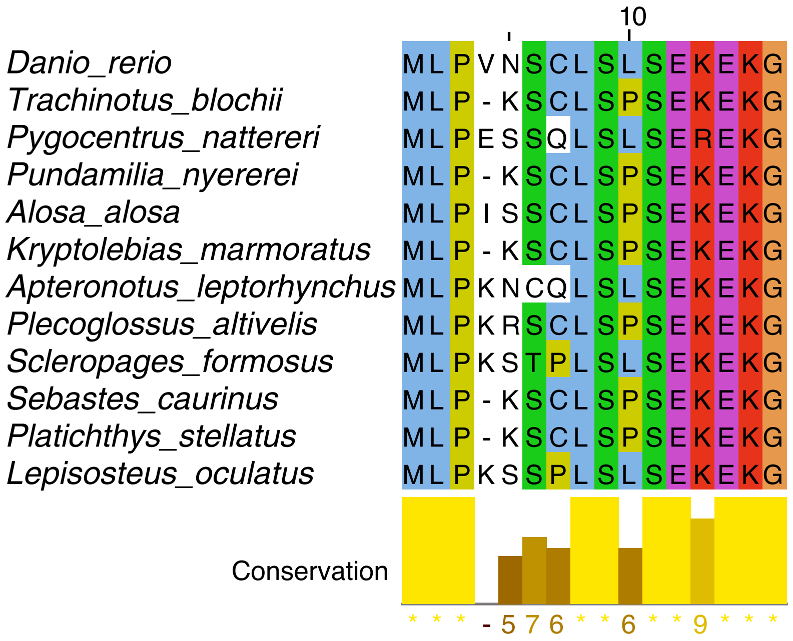

**HG0267**

ENSG00000164889 (SLC4A2)

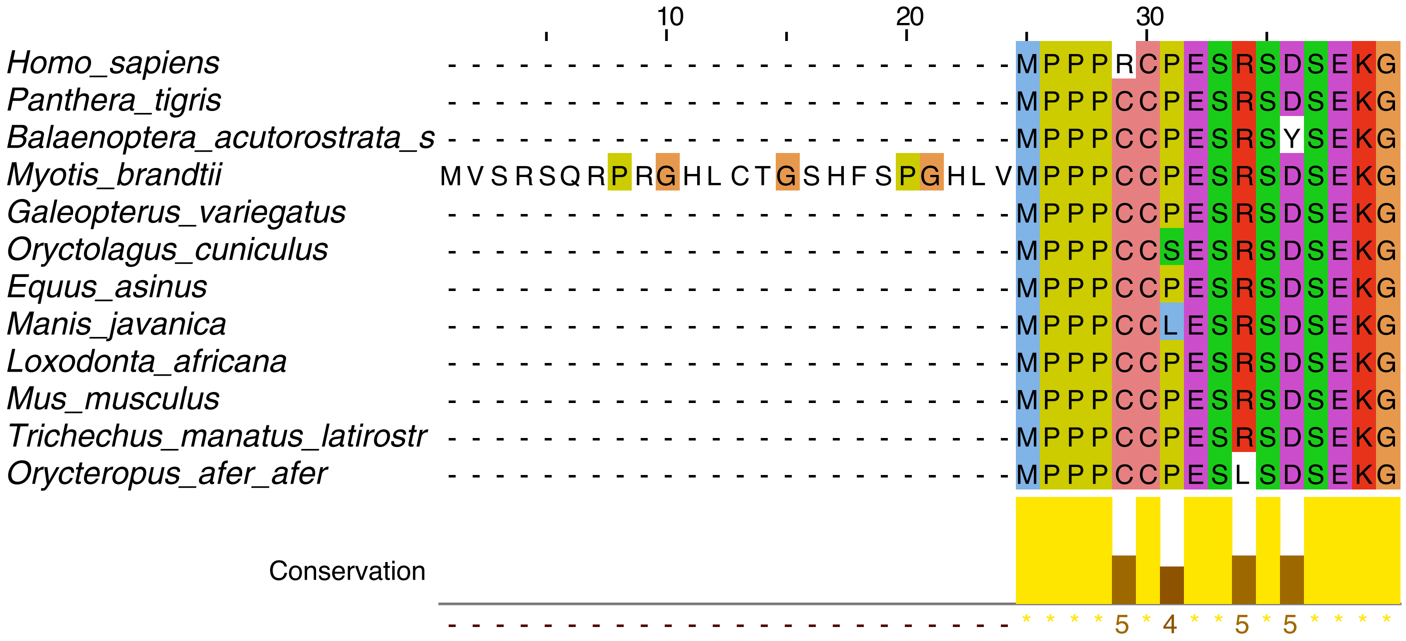

**HG0268.1**

ENSDARG00000011703 (clocka)

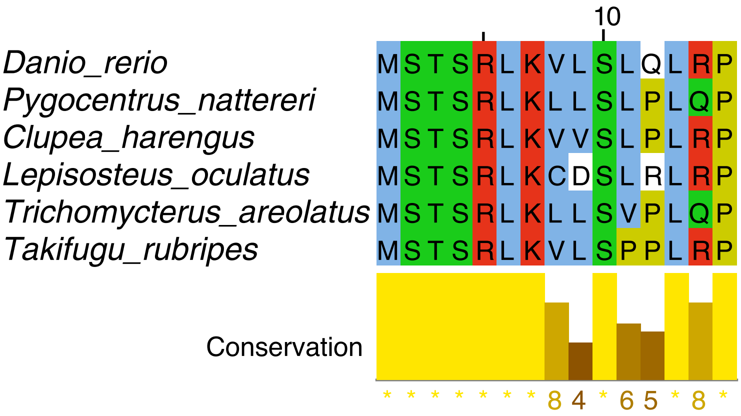

**HG0268.2**

ENSG00000134852 (CLOCK)

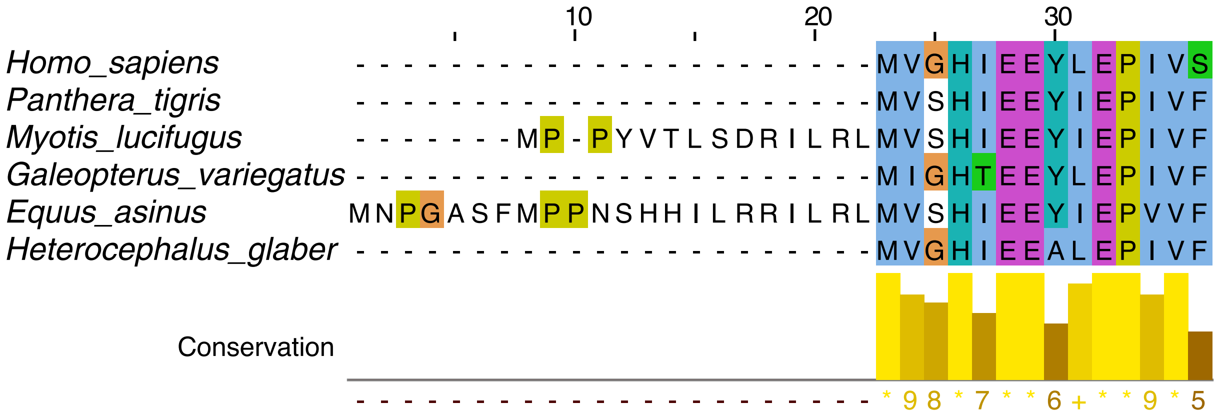

**HG0268.3**

ENSG00000134852 (CLOCK)

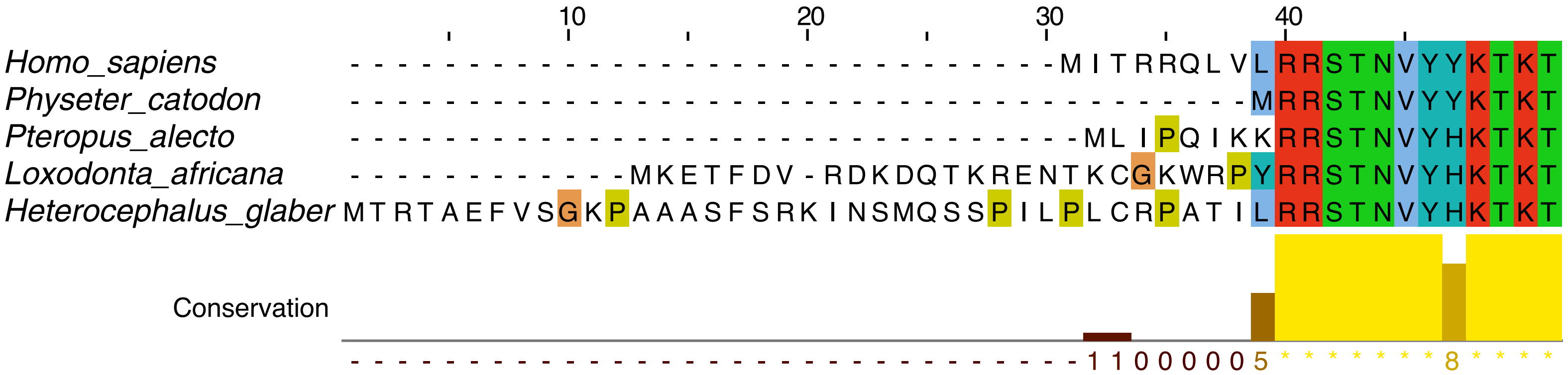

**HG0269.1**

ENSGALG00000028685 (SEMA5A)

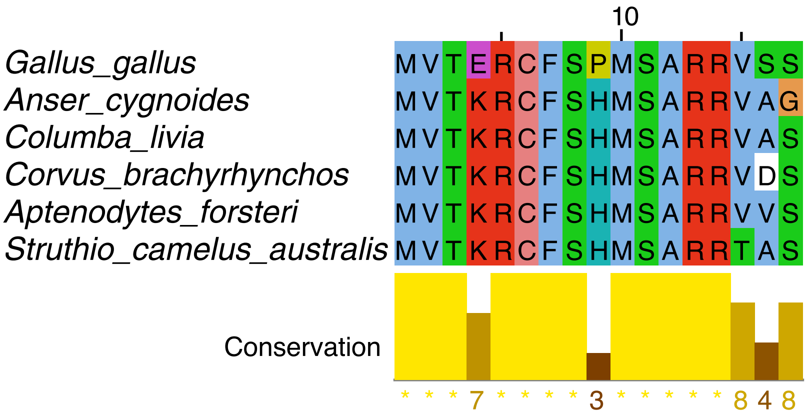

**HG0269.2**

ENSDARG00000061471 (sema5ba)

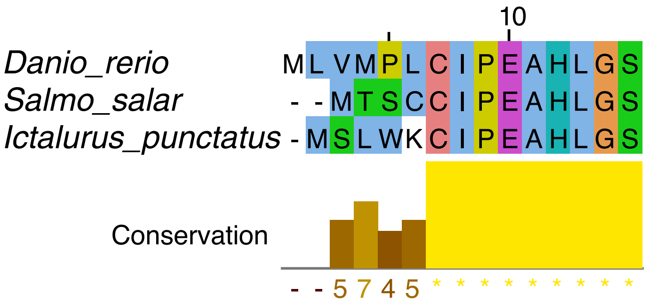

**HG0269.3**

ENSGALG00000011686 (SEMA5B)

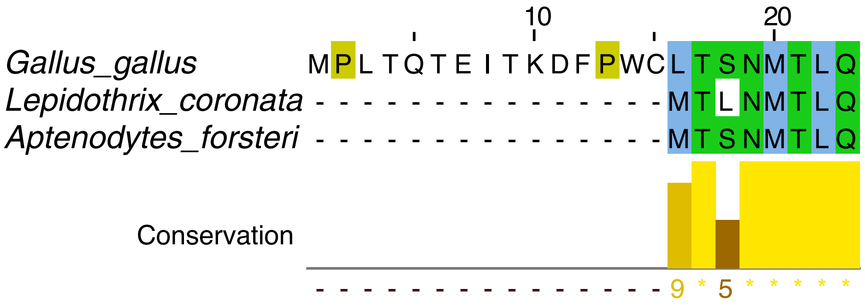

**HG0270.1**

ENSGALG00000015147 (ALDH1A1)

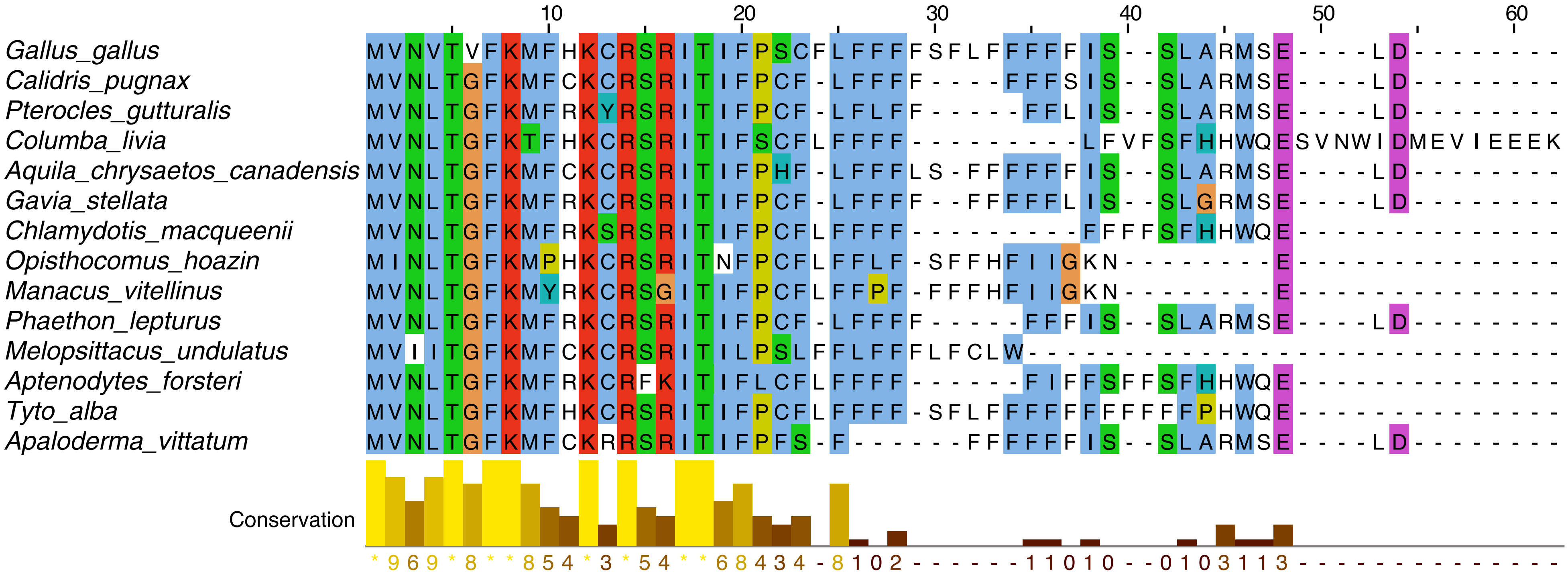

**HG0270.2**

ENSGALG00000015147 (ALDH1A1)

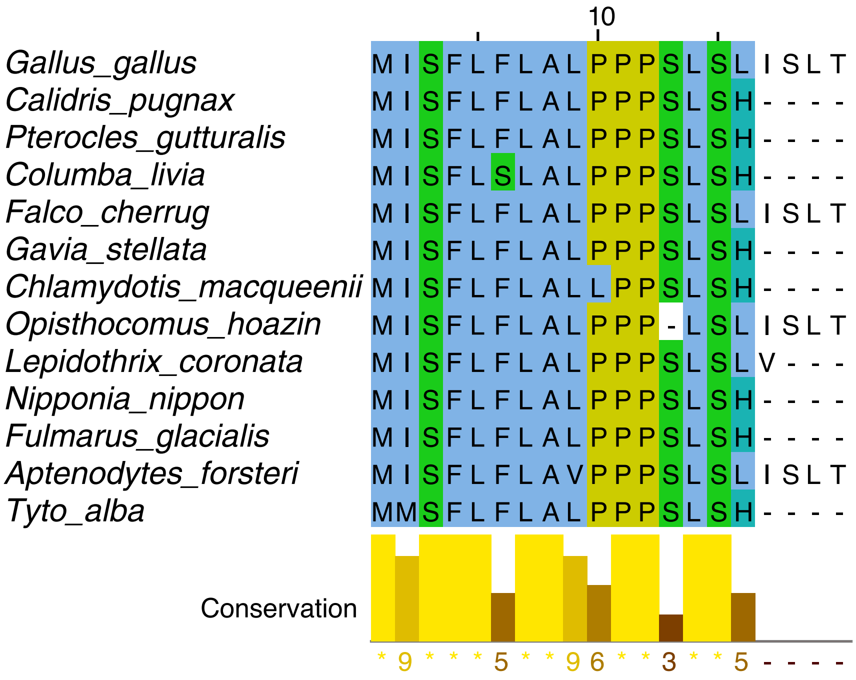

**HG0270.3**

ENSG00000128918 (ALDH1A2)

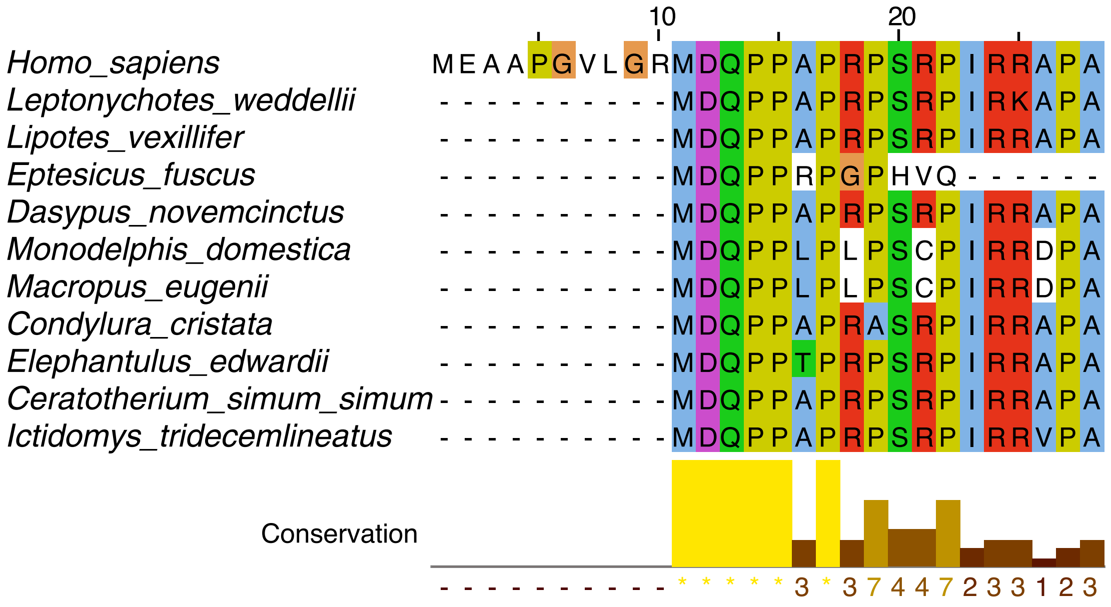

**HG0271.1**

ENSGALG00000004064 (GPER1)

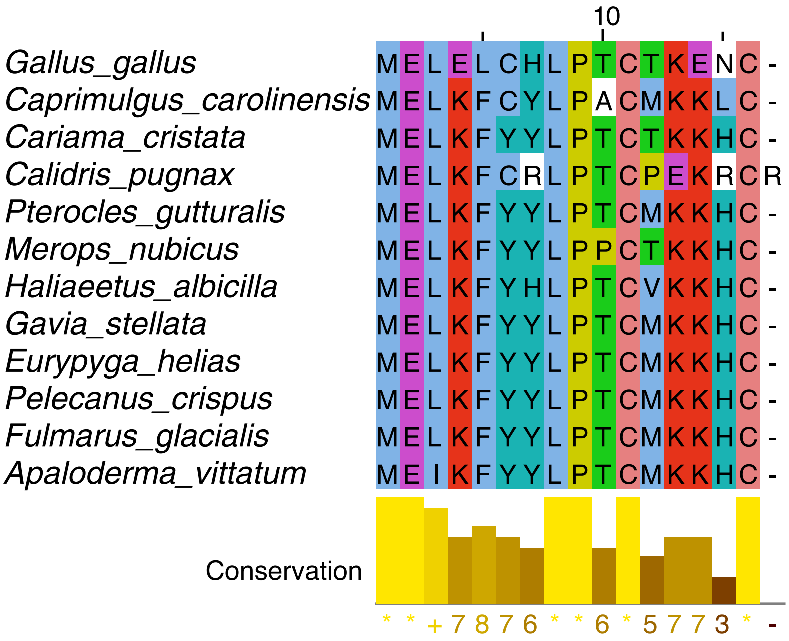

**HG0271.2**

ENSG00000164850 (GPER1)

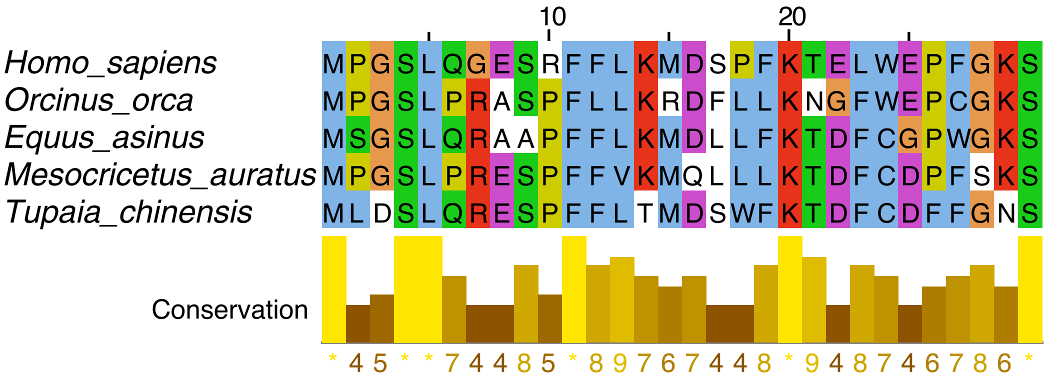

**HG0271.3**

ENSDARG00000074661 (gper1)

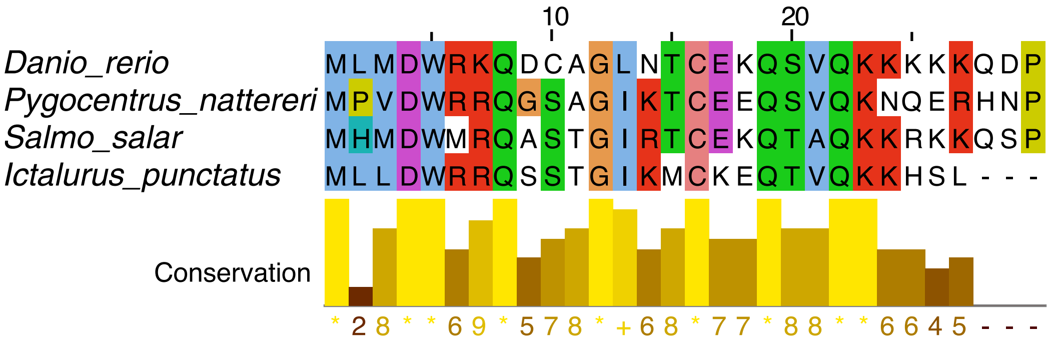

**HG0272.1**

FBgn0034649 (CG9865)

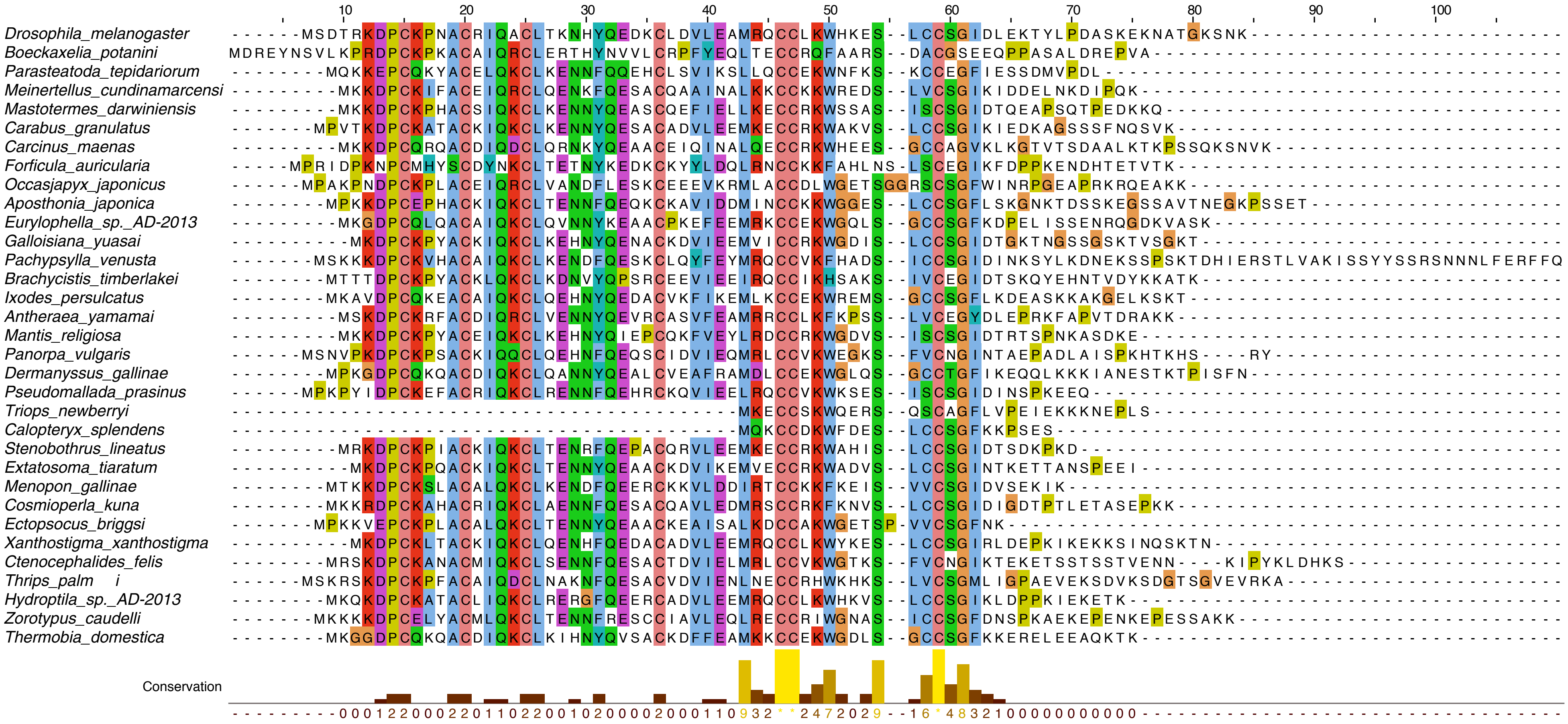

**HG0272.2**

FBgn0034649 (CG9865)

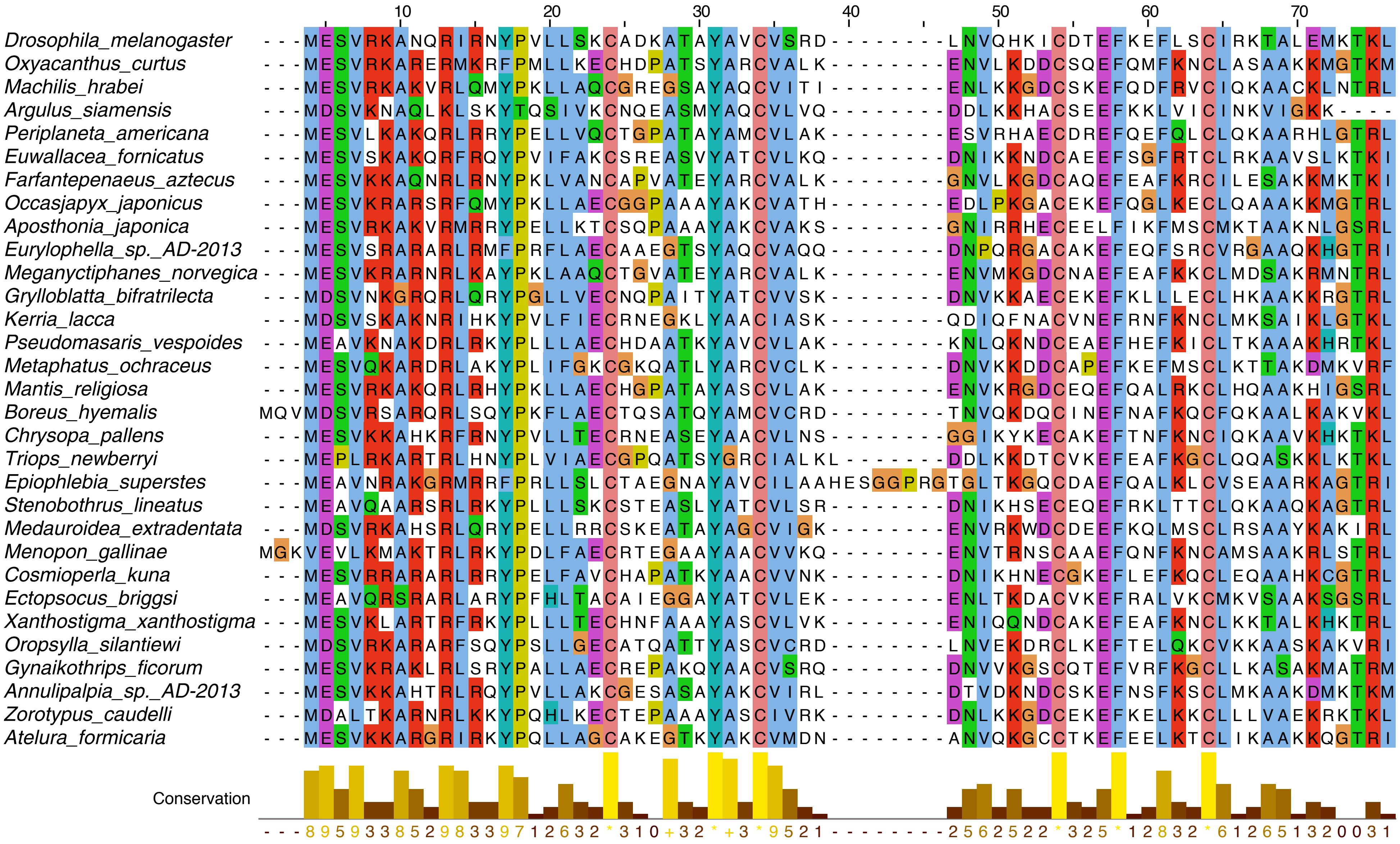

**HG0272.3**

ENSG00000143315 (PIGM)

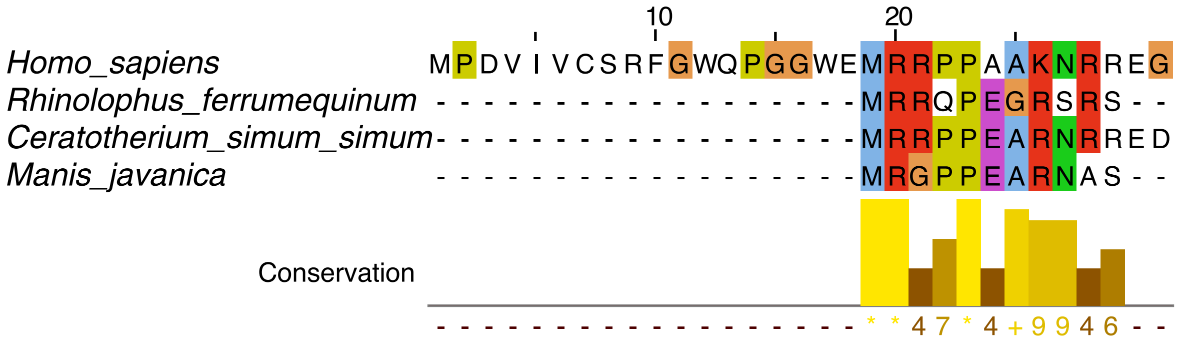

**HG0273.1**

ENSGALG00000003293 (MBTPS1)

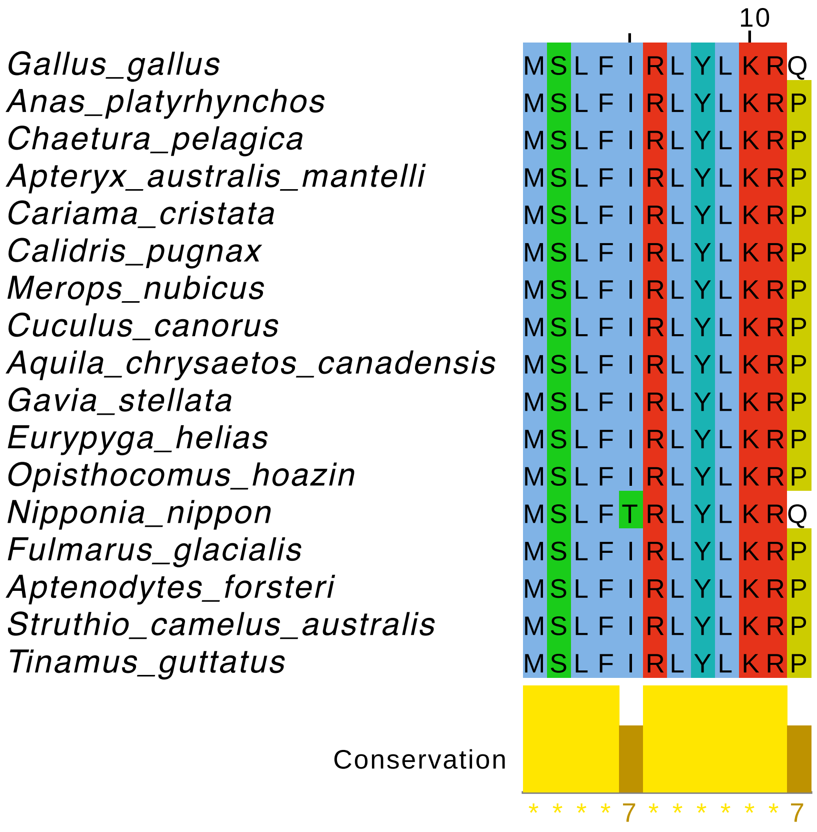

**HG0273.2**

ENSG00000140943 (MBTPS1)

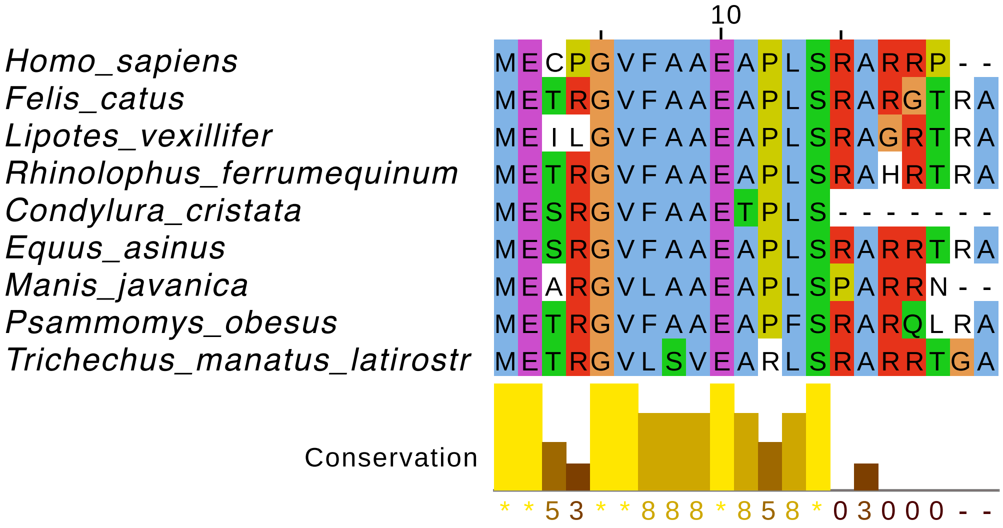

**HG0273.3**

ENSG00000140943 (MBTPS1)

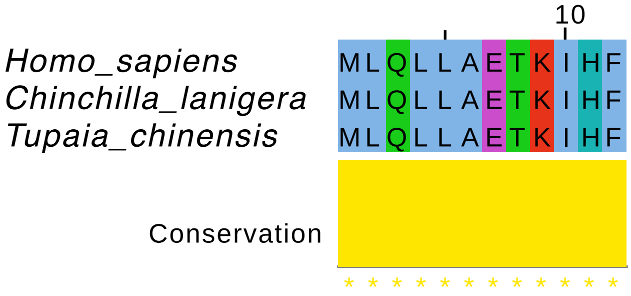

**HG0274.1**

ENSG00000204256 (BRD2)

| Species | Alignment (positions 1–47; scale marks 10, 20, 30, 40) |
|---|---|
| *Homo_sapiens* | MTGPSGQYRPPRSPWSSPFSGPAQSSESVHPLYSALKRISKMEAAAP |
| *Ailuropoda_melanoleuca* | MTGPGGQYRPHRSPRRSPFSGPAQSSESVHPLYSALKRISKMEAAAP |
| *Orcinus_orca* | MTGPGGQYRPHRSPRSSPFSSPAQSSESVHPLYSALKRISKMEAAAP |
| *Myotis_davidii* | MTVPGGQYRPHRSPRSSPFSGPAQSSESVHPLYSALKRISKMEAAAP |
| *Dasypus_novemcinctus* | MTGPGGQY--------------------------------------- |
| *Galeopterus_variegatus* | MTGPCGQCRPHRSPRSSPFSGPAQSSESVHPLYSALKRISKMEAAAP |
| *Condylura_cristata* | -----------------------------------------MEAAAP |
| *Oryctolagus_cuniculus* | MTGPGGQYRPYRSPRSSPFSGPAQSSESVHPLYSALKRISKMEAASP |
| *Elephantulus_edwardii* | MTRPGGQYRPHRSPRSSPFPGPAQSSESIHPLYSALKRISKMEAAAP |
| *Equus_caballus* | MTGPGGQYRPHRSPRSSPFSGPAQSSESIHPLYSALKRISKMEAAAP |
| *Manis_javanica* | MIGPGGQYRPHRSPRSFPFSGPAQSSESVHPLYSALERISKMVAAAP |
| *Loxodonta_africana* | MTDTGGQYRPHRSPRSSPFPGPAQSSESIHPLYSALKRISKMEAAAP |
| *Trichechus_manatus_latirostr* | MTGPGGQYRPHRSPRSSPFPGPAQSSESVRPLYSALKRISKMEAAAP |
| *Orycteropus_afer_afer* | MTGPGGQYRPHRSPRSSPFLGPAQSSESIHPLYSALKRISKMEAAAP |
| Conservation | 1 1 0 1 1 3 1 1 3 2 0 3 3 2 1 1 0 2 2 0 2 2 3 1 3 3 3 3 2 3 2 2 3 3 3 2 2 3 2 3 4 1 0 3 3 2 2 |

**HG0274.2**

ENSDARG00000100129

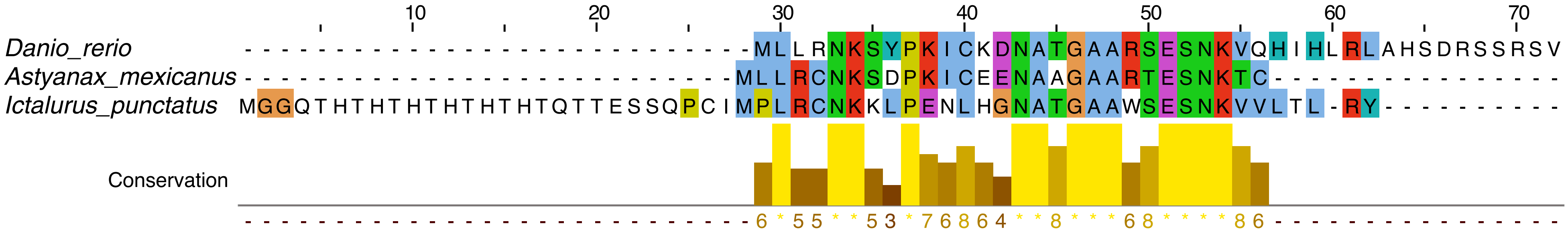

**HG0275.1**

ENSG00000100239 (PPP6R2)

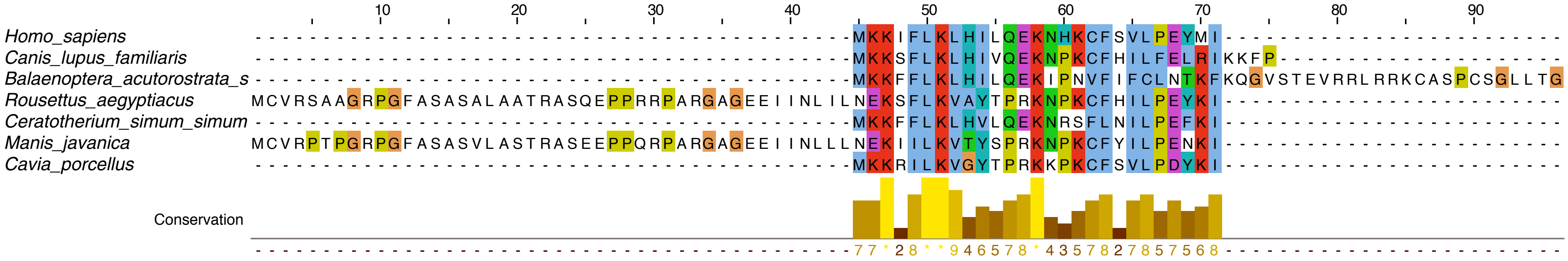

**HG0275.2**

ENSDARG00000045540 (ppp6r2a)

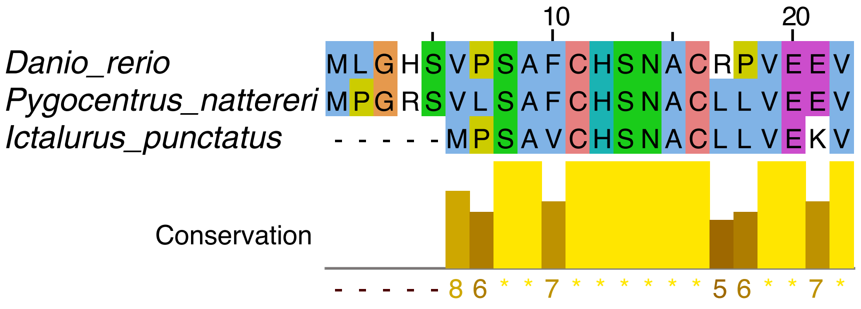

**HG0276.1**

ENSGALG00000008912

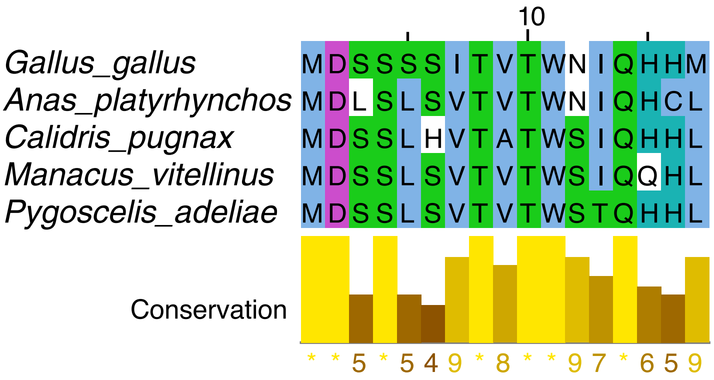

**HG0276.2**

ENSG00000073734 (ABCB11)

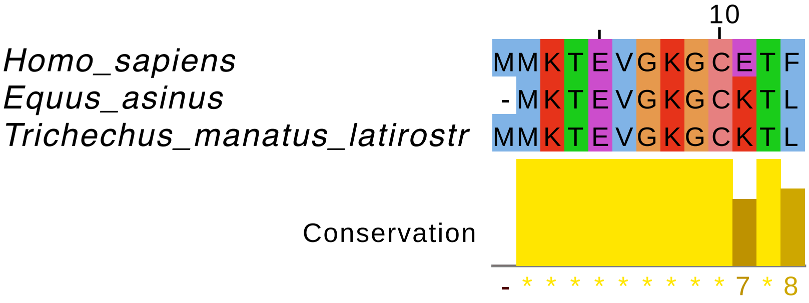

**HG0277.1**

ENSG00000102531 (FNDC3A)

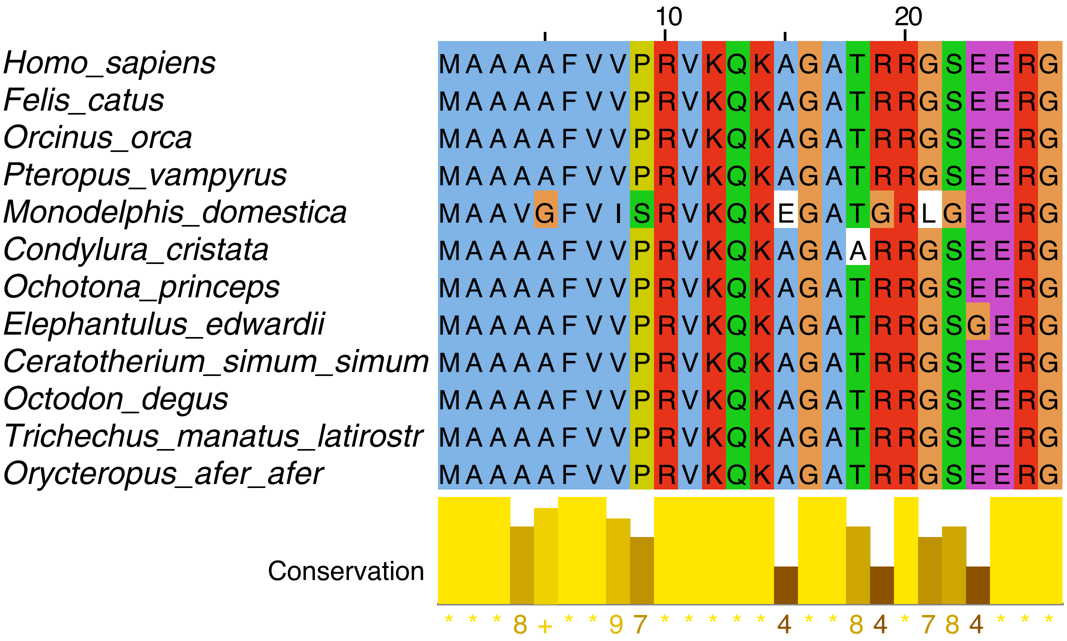

**HG0277.2**

ENSGALG00000004169

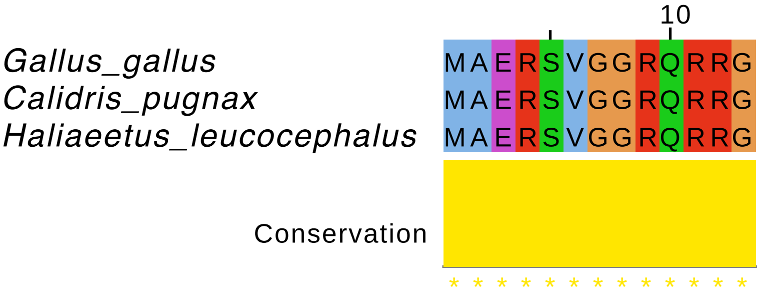

**HG0278.1**

ENSG00000119946 (CNNM1)

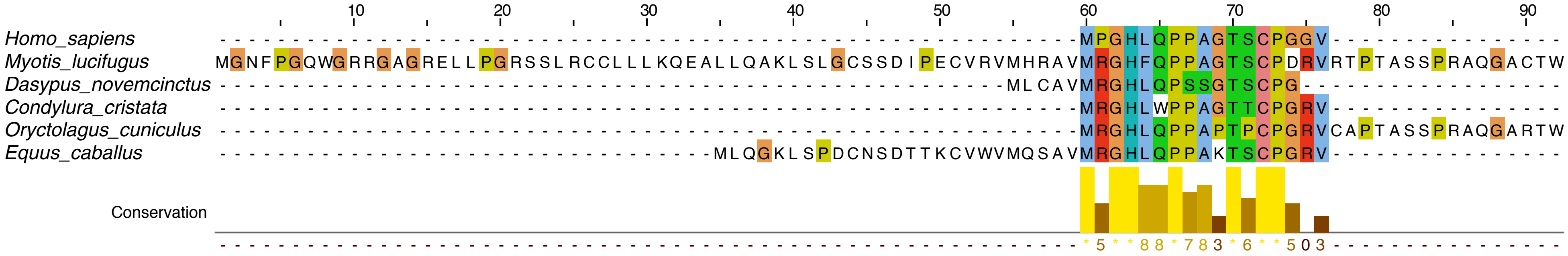

**HG0278.2**

ENSDARG00000078733 (cnnm2b)

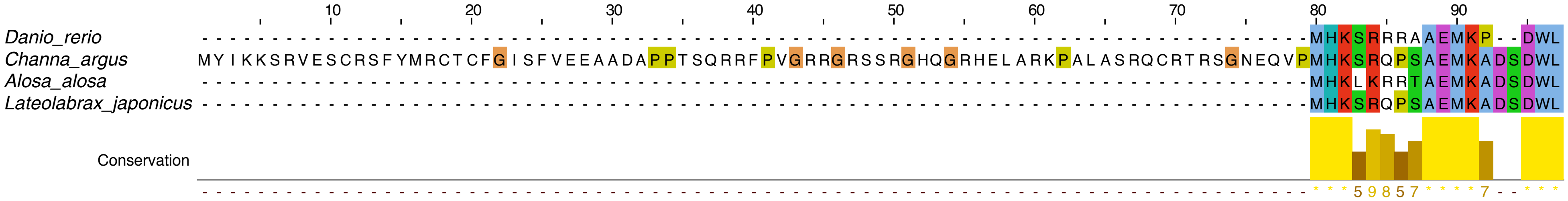

**HG0279.1**

ENSG00000175928 (LRRN1)

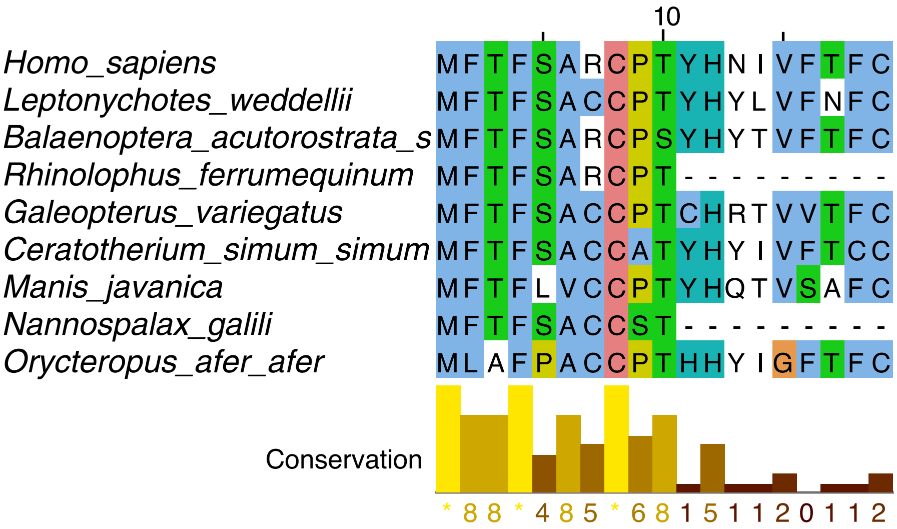

**HG0279.2**

ENSGALG00000035099 (Lrrn1)

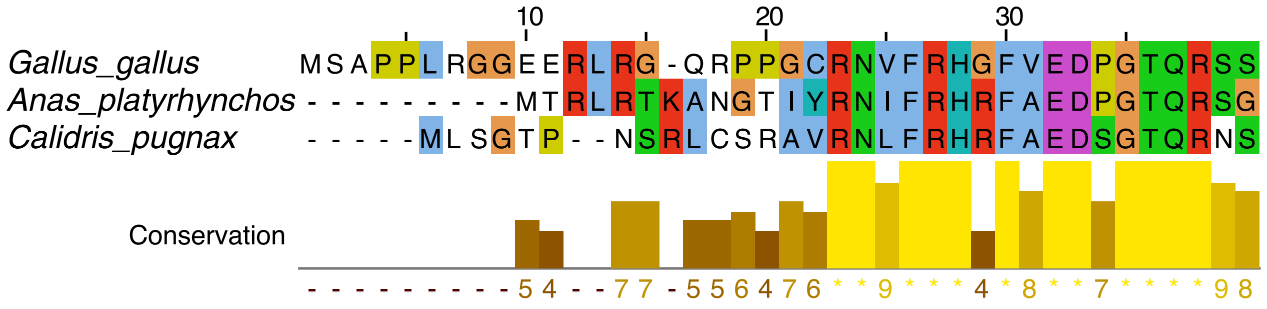

**HG0280.1**

ENSG00000156299 (TIAM1)

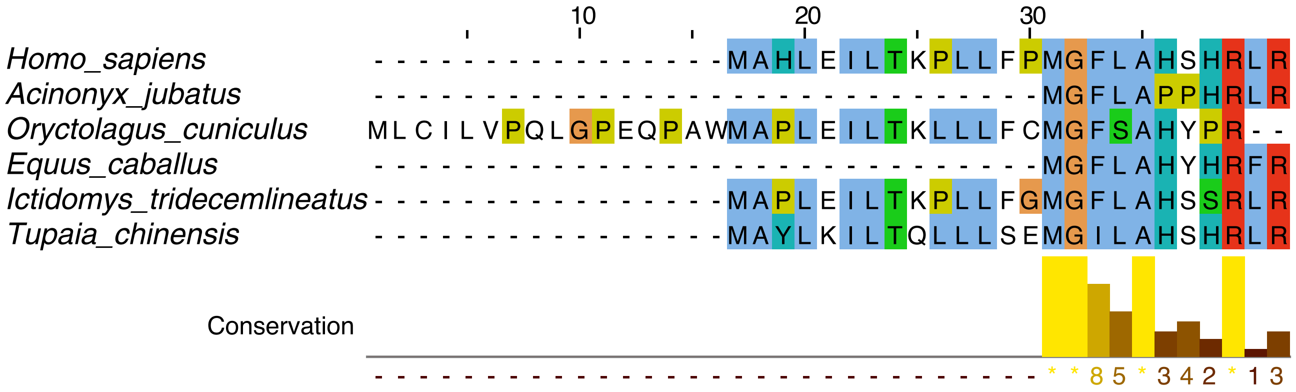

**HG0280.2**

FBgn0085447 (sif)

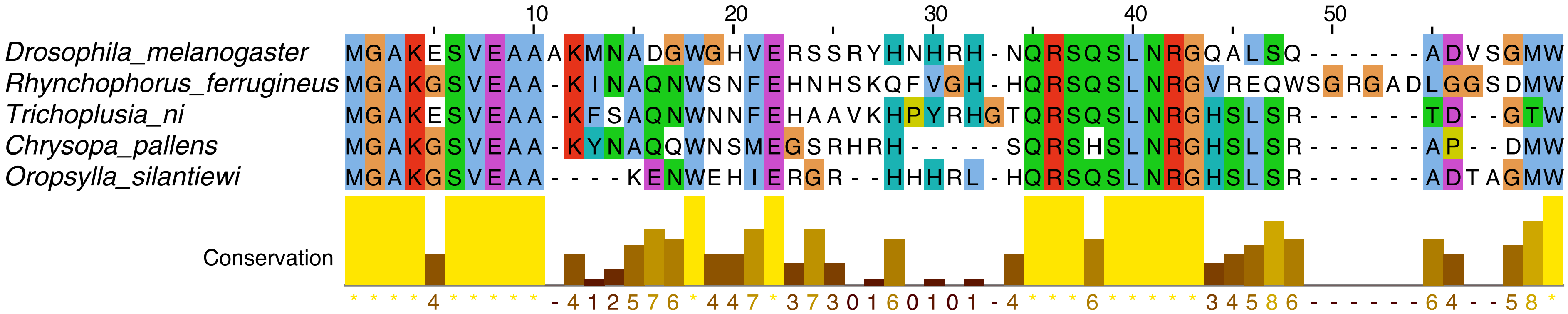

**HG0281.1**

ENSGALG00000006766 (DEPDC5)

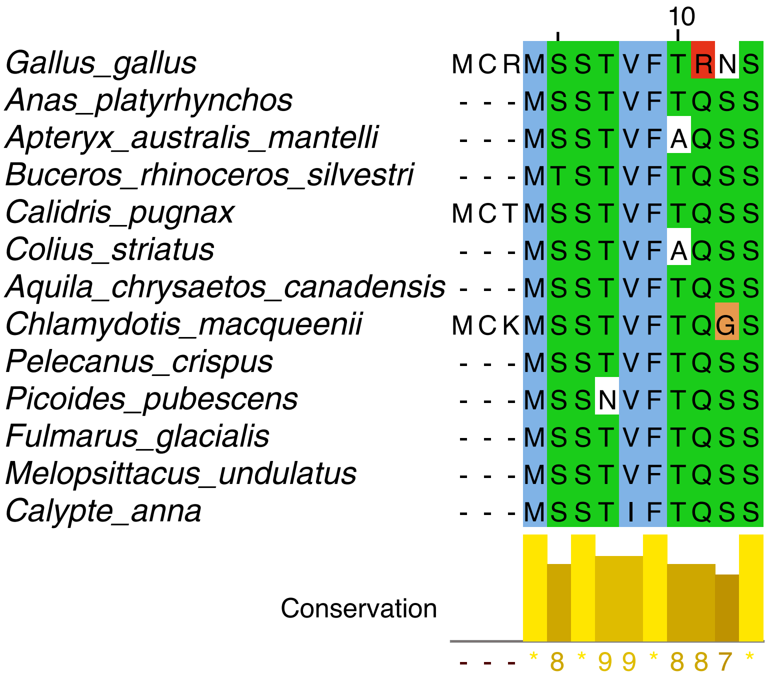

**HG0281.2**

ENSG00000100150 (DEPDC5)

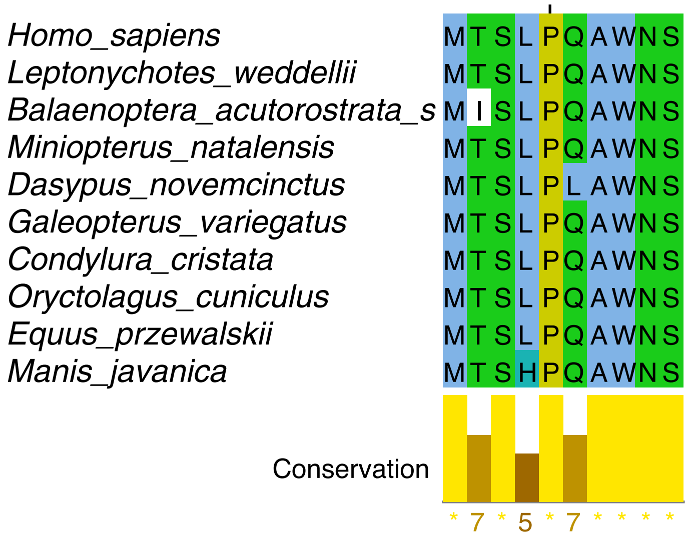

**HG0282.1**

ENSDARG00000061328 (cdon)

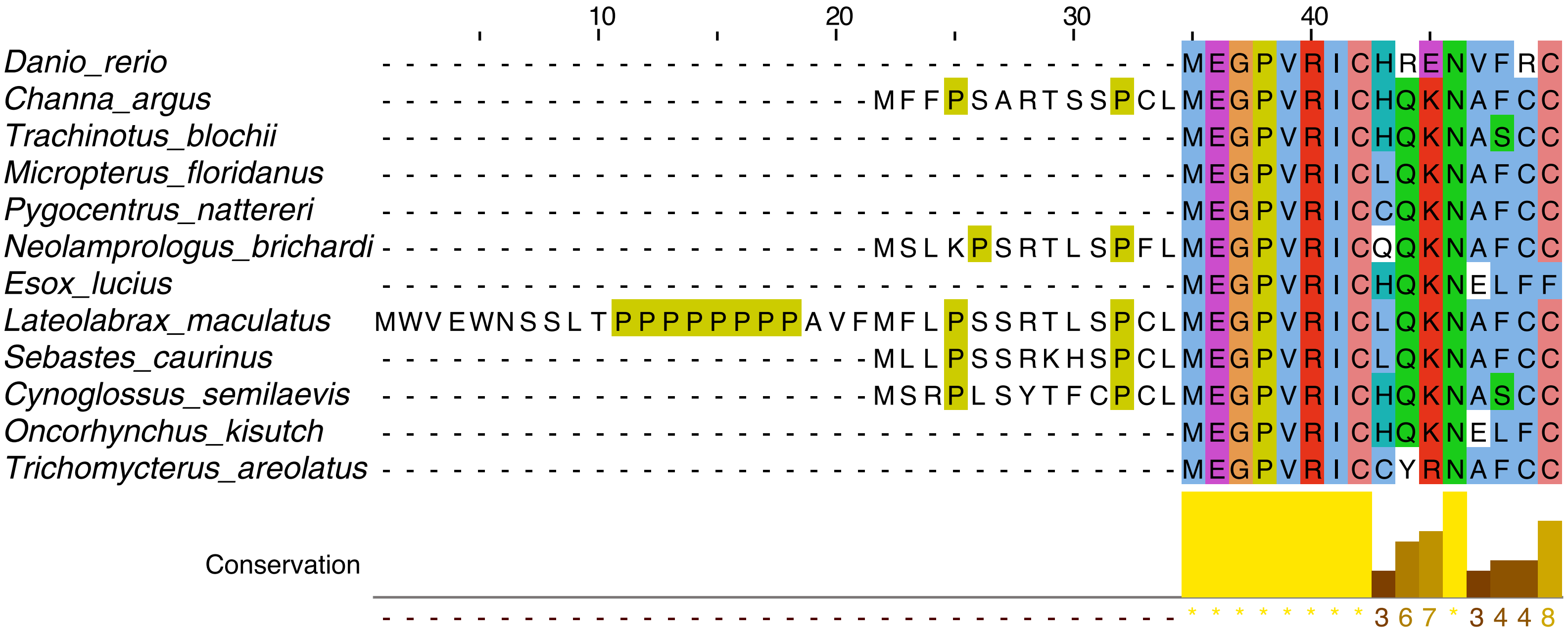

**HG0282.2**

ENSG00000144857 (BOC)

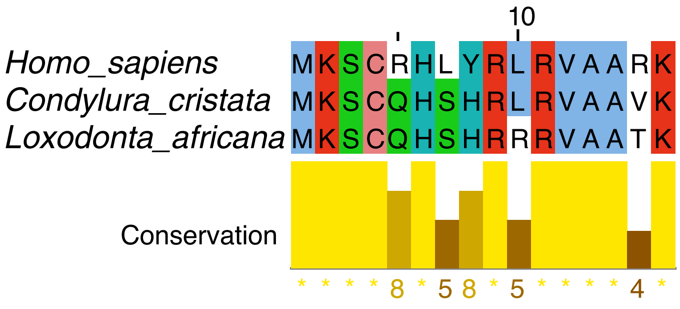

**HG0283.1**

ENSGALG00000010270 (AREL1)

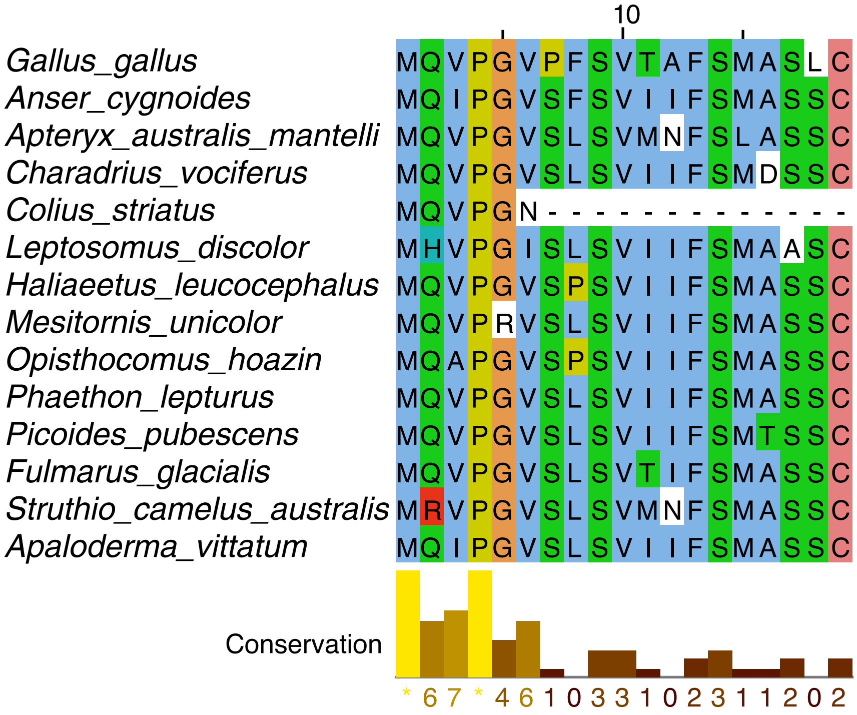

**HG0283.2**

ENSG00000119682 (AREL1)

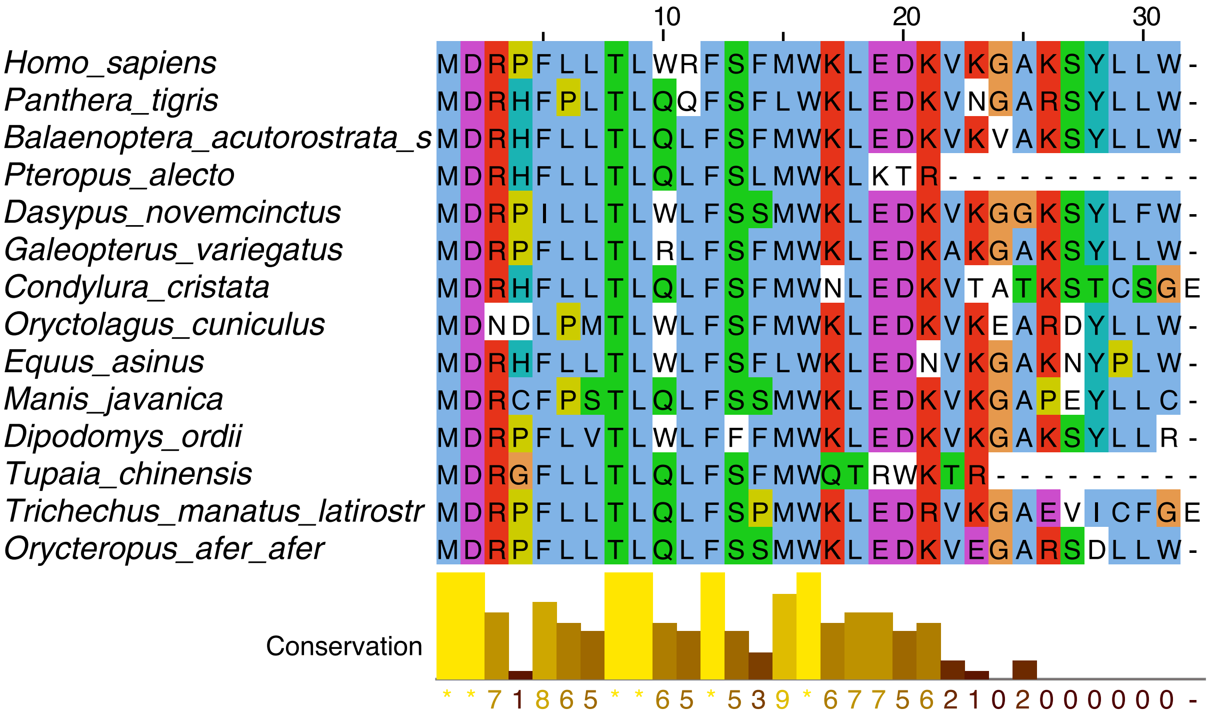

**HG0284.1**

ENSG00000007168 (PAFAH1B1)

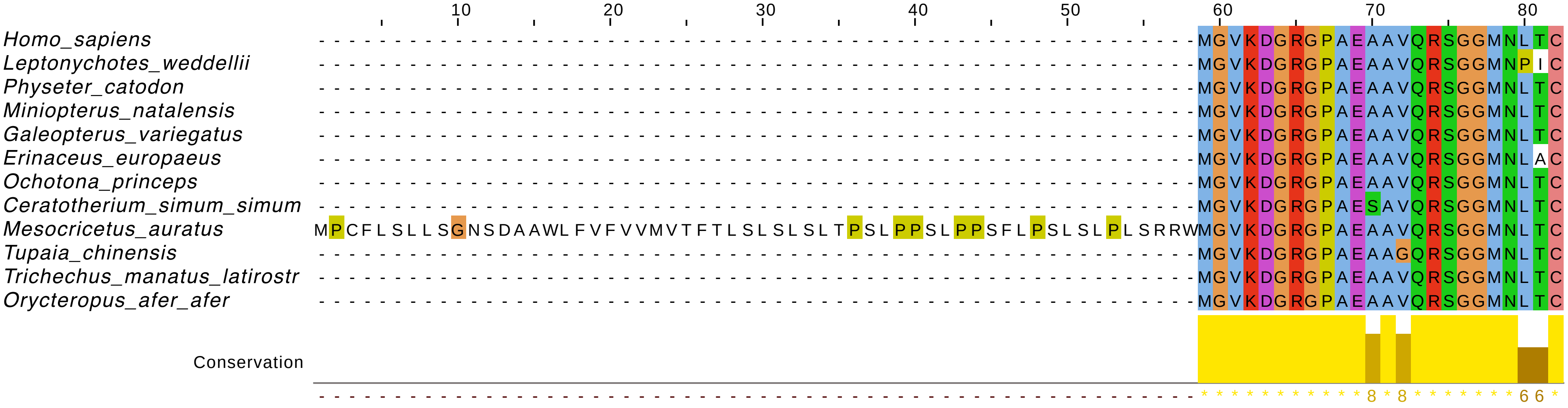

**HG0284.2**

ENSGALG00000005834 (PAFAH1B1)

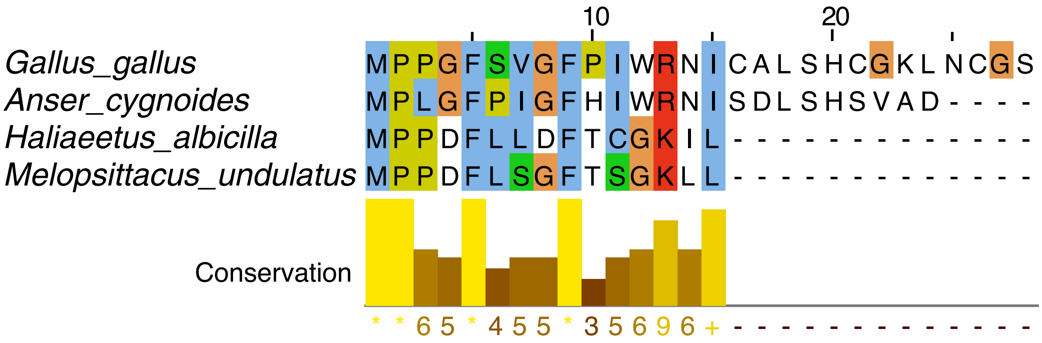

**HG0285.1**

ENSGALG00000012156 (DPP10)

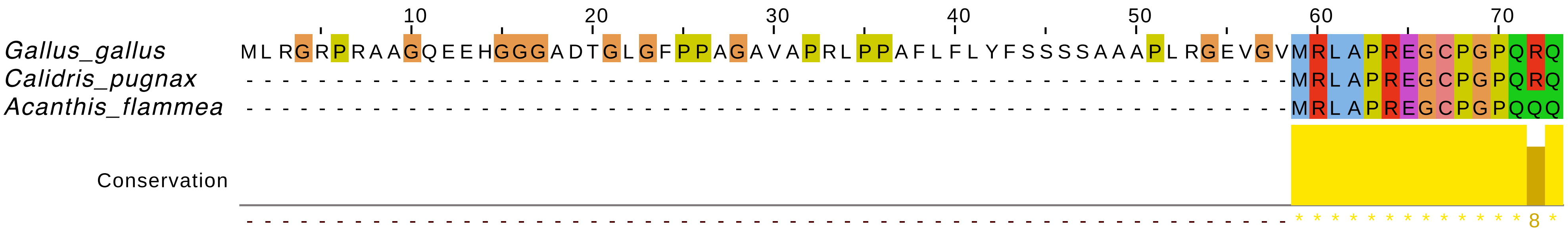

**HG0285.2**

ENSG00000175497 (DPP10)

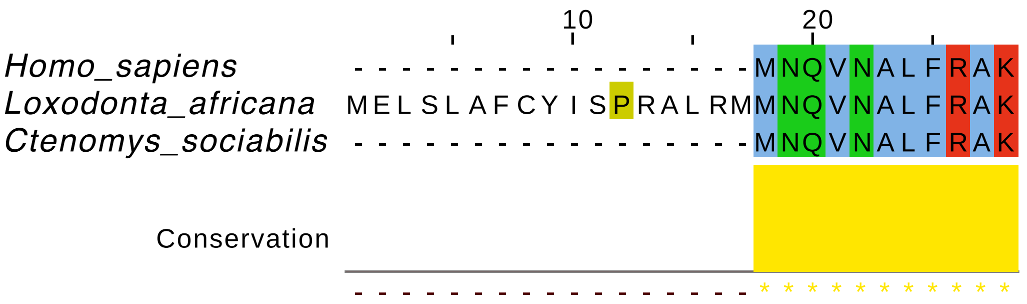

**HG0286.1**

ENSG00000116698 (SMG7)

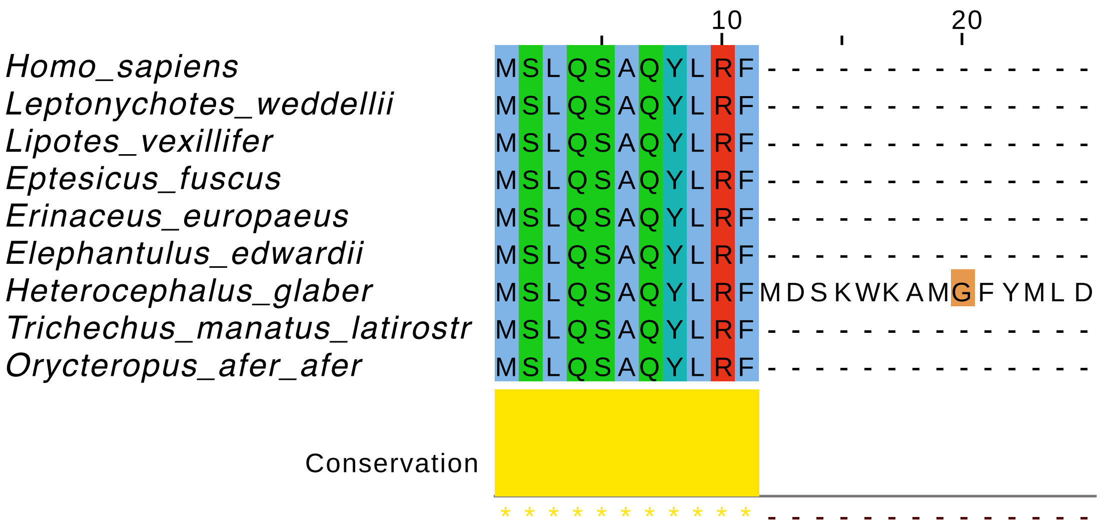

**HG0286.2**

ENSDARG00000060767 (smg7)

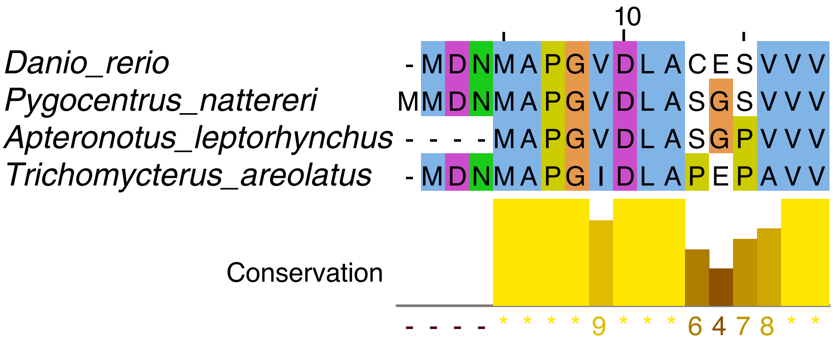

**HG0287.1**

ENSG00000170325 (PRDM10)

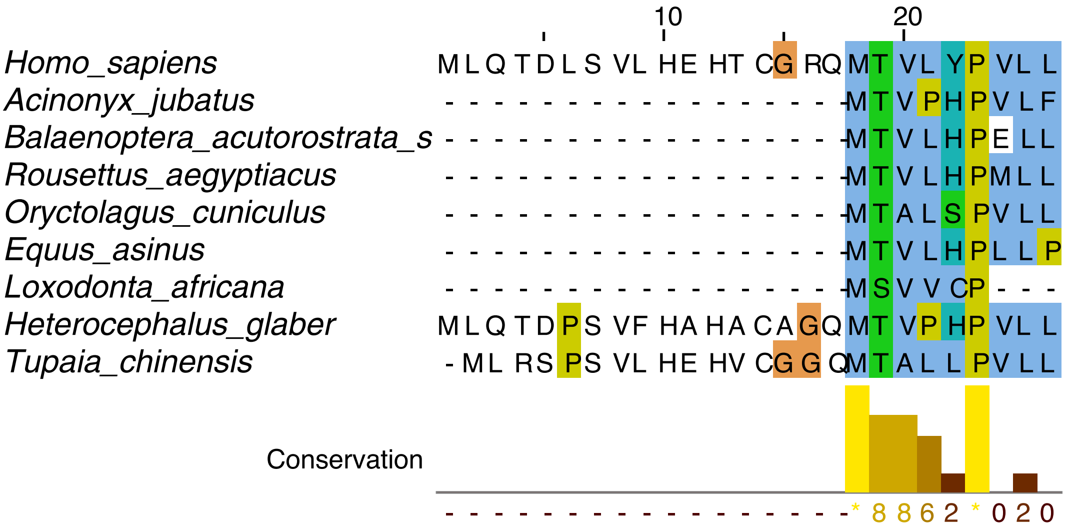

**HG0287.2**

ENSDARG00000104251 (prdm10)

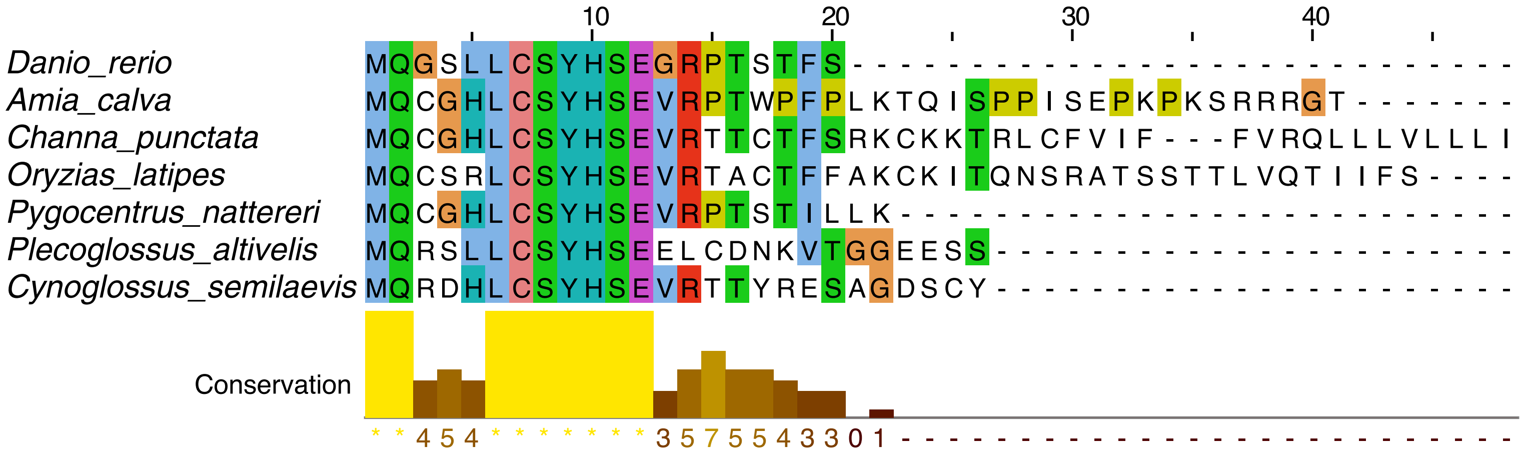

**HG0288.1**

ENSGALG00000006655 (MAP3K13)

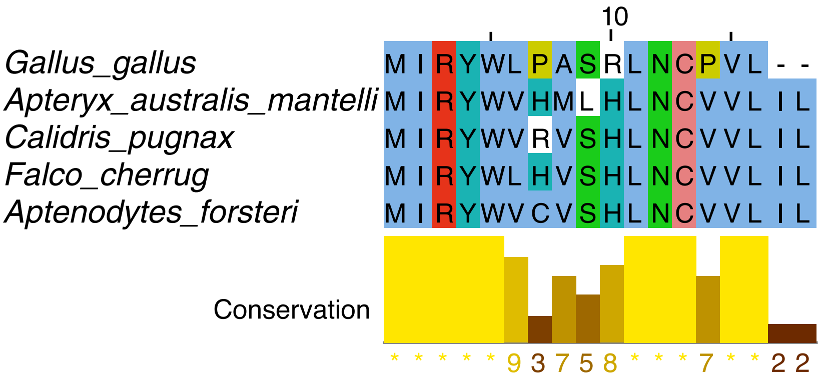

**HG0288.2**

ENSG00000073803 (MAP3K13)

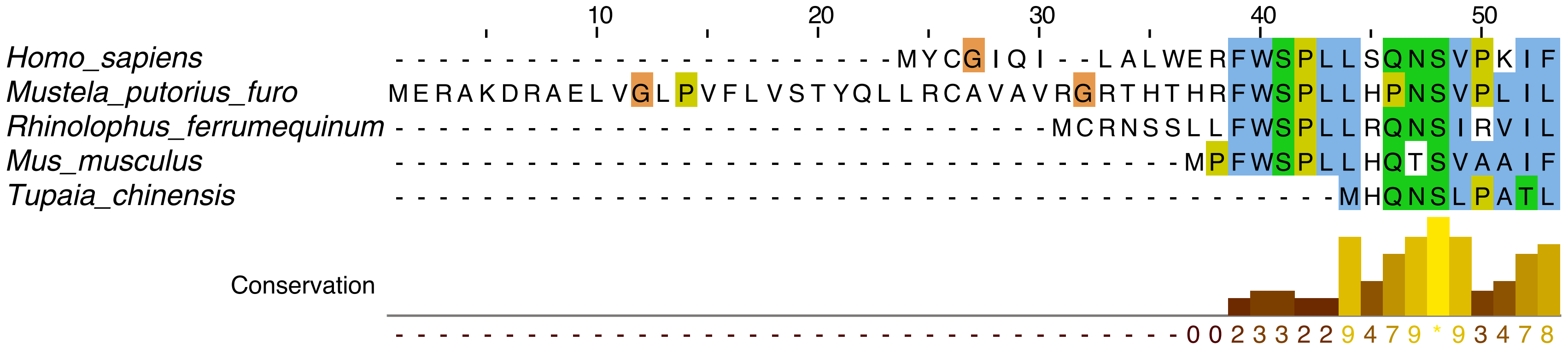

**HG0289**

ENSG00000072415 (MPP5)

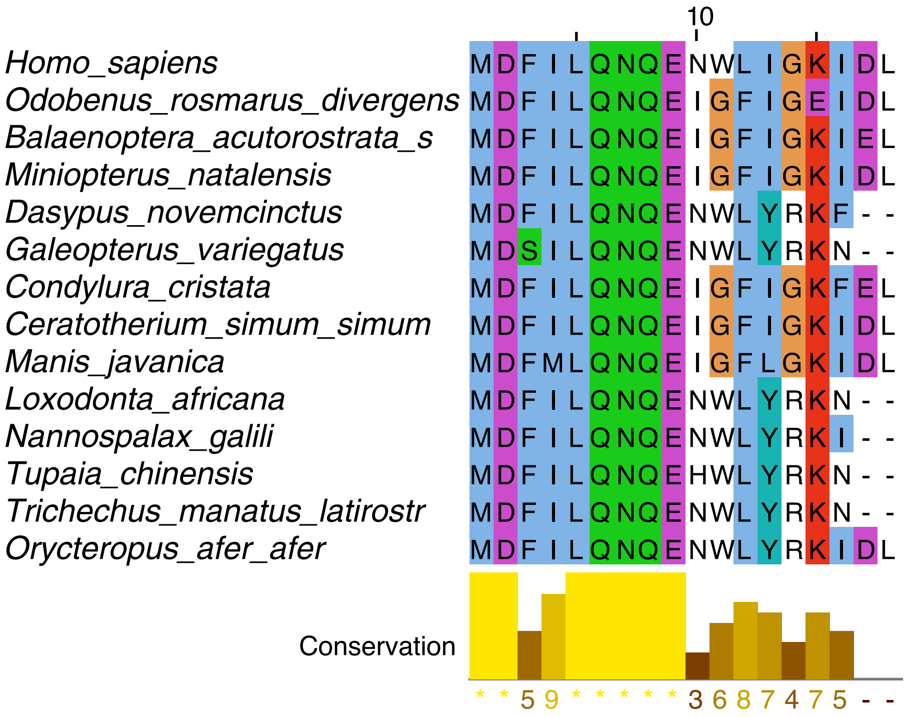

**HG0289**

ENSGALG00000009581 (MPP5)

```
                                      10        20
Gallus_gallus                    --------MDFTQQRNENWPNISISI---
Anser_cygnoides                  --------MDFTQHHNKNWPNSISV----
Chaetura_pelagica                --------MDFTQHHNENWPHSSISV---
Apteryx_australis_mantelli       --------MDFIPHHNKNWPYSSICV---
Buceros_rhinoceros_silvestri     --------MDFTQHHNENWPNSSISVCGE
Caprimulgus_carolinensis         --------MDFTQHHNENWPNRSISV---
Cariama_cristata                 --------MDFTQHRNKNWPNSSISV---
Calidris_pugnax                  --------MDFTQHHNENWPNISISV---
Pterocles_gutturalis             --------MDFTQHHNENWPSSSISV---
Colius_striatus                  --------MDSTQHHNKNCPNSSISV---
Columba_livia                    --------MDFTQHRNKNWPSSSISV---
Leptosomus_discolor              --------MDFMQRHNENWPGSSISV---
Cuculus_canorus                  --------MDFTQHHNENWPNSSISV---
Aquila_chrysaetos_canadensis     --------MDFTQHHNENWPNSSISV---
Gavia_stellata                   --------MDFTQHHNKNWPSSSISV---
Mesitornis_unicolor              --------MDFMQHHNENWPNSSISV---
Tauraco_erythrolophus            --------MDFTQHHNENCPNSSISV---
Opisthocomus_hoazin              --------MDSTQHHNKNWPNSTISV---
Lepidothrix_coronata             -------MMDFTQHHNKNWPNSSICV---
Phalacrocorax_carbo              --------MDFTQHRNENWPNSSISV---
Picoides_pubescens               --------MDFTHHHNENWLNSSIFA---
Fulmarus_glacialis               --------MDSTQQHNKNWPNSSISV---
Melopsittacus_undulatus          --------MDFTQCHSENWPNNSISI---
Aptenodytes_forsteri             --------MDFTQHHNKNWPNSSISV---
Tyto_alba                        --------MDFTQHHNENWPNSSISD---
Struthio_camelus_australis       --------MDFMPHHNENWPYSSICV---
Tinamus_guttatus                 --------MDFIPHHNENWPFSTICV---
Calypte_anna                     MGNIFQVVMDFRQHQNESWPSSSISA---
Apaloderma_vittatum              --------MDFTQHPNENWPNSSFSV---
```

Conservation

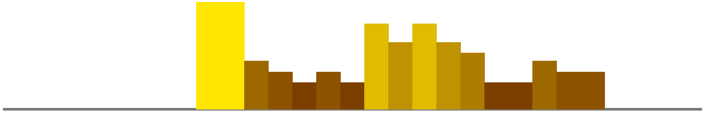

```
--------**5434397976335440---
```

**HG0290.1**

ENSG00000116539 (ASH1L)

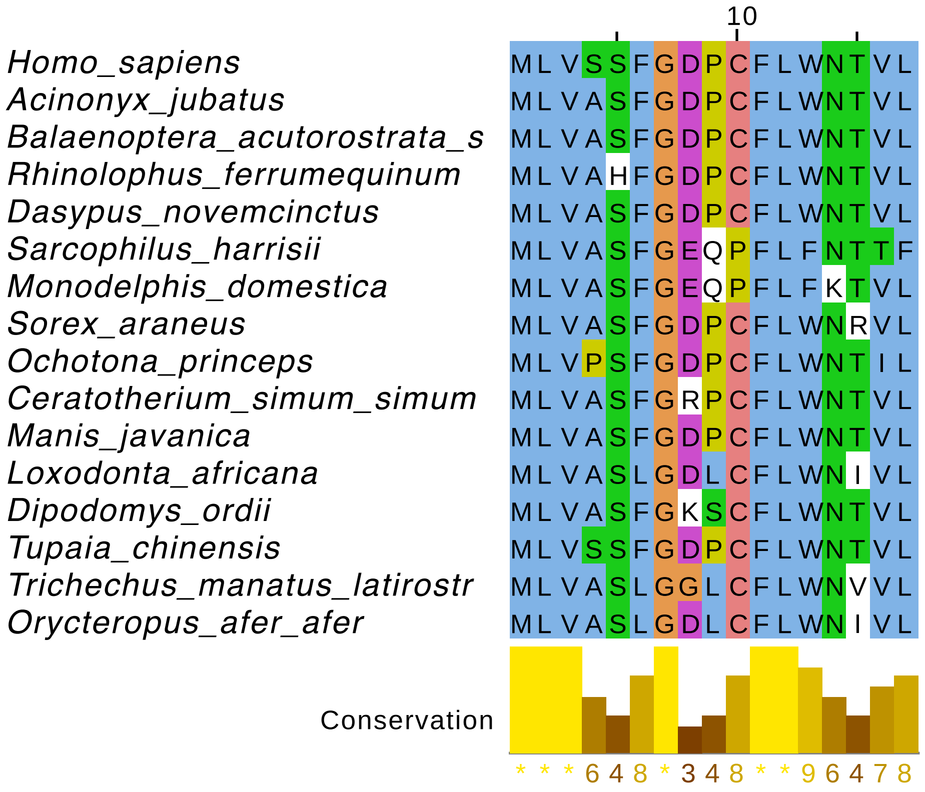

**HG0290.2**

ENSDARG00000070981 (ash1l)

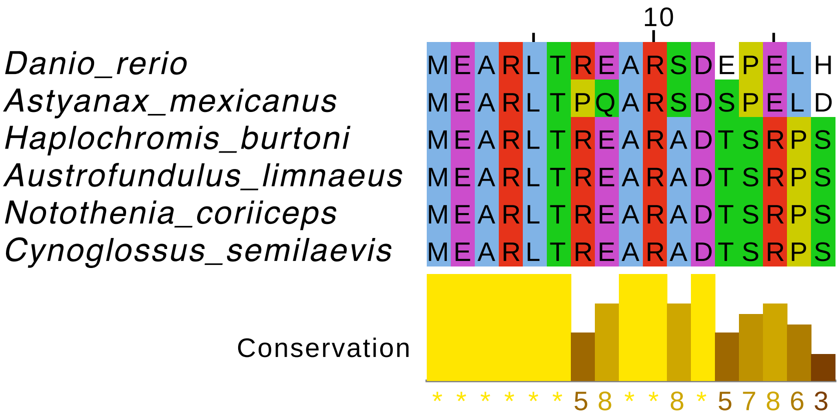

**HG0291.1**

ENSGALG00000009173 (GFRA1)

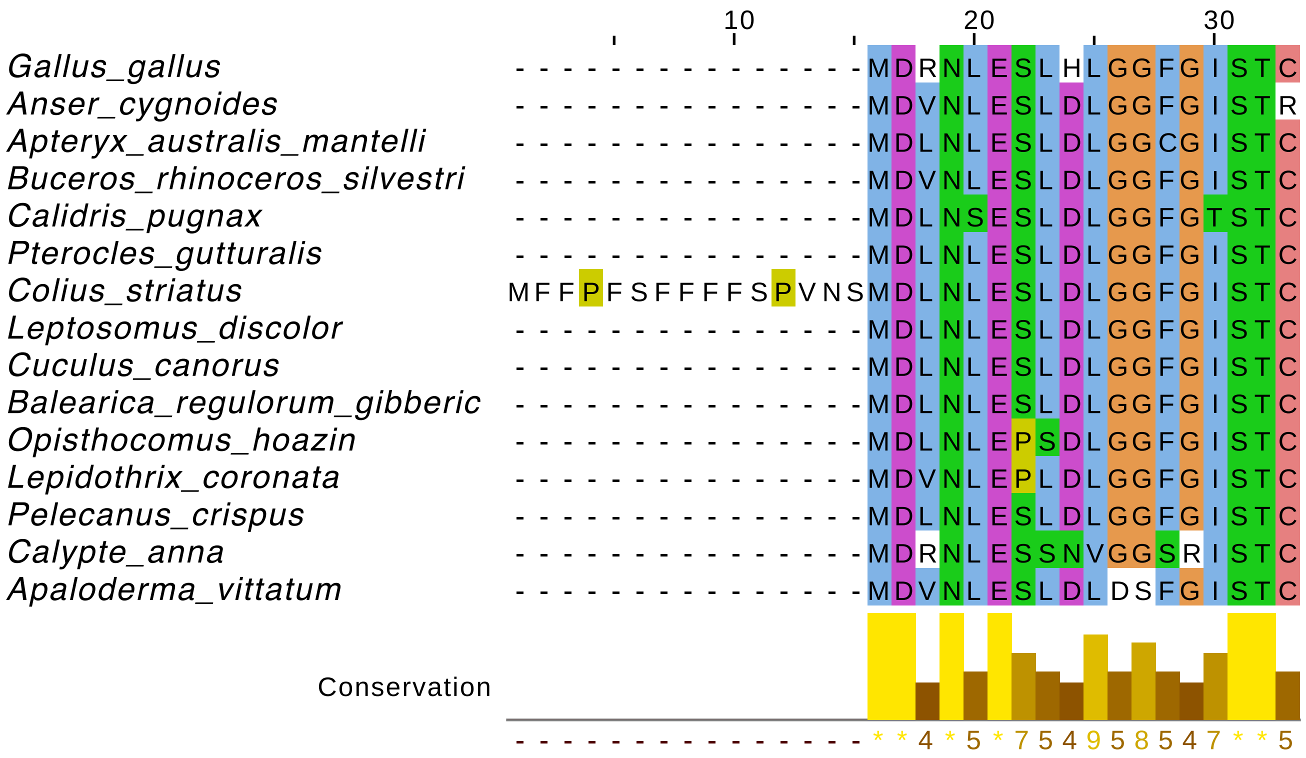

**HG0291.2**

ENSG00000151892 (GFRA1)

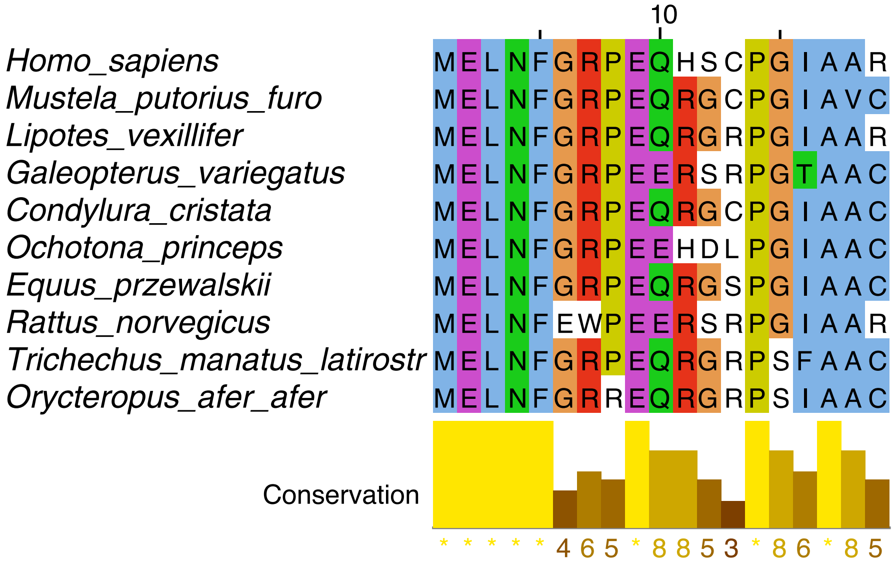

**HG0292.1**

ENSGALG00000006502 (PCGF5)

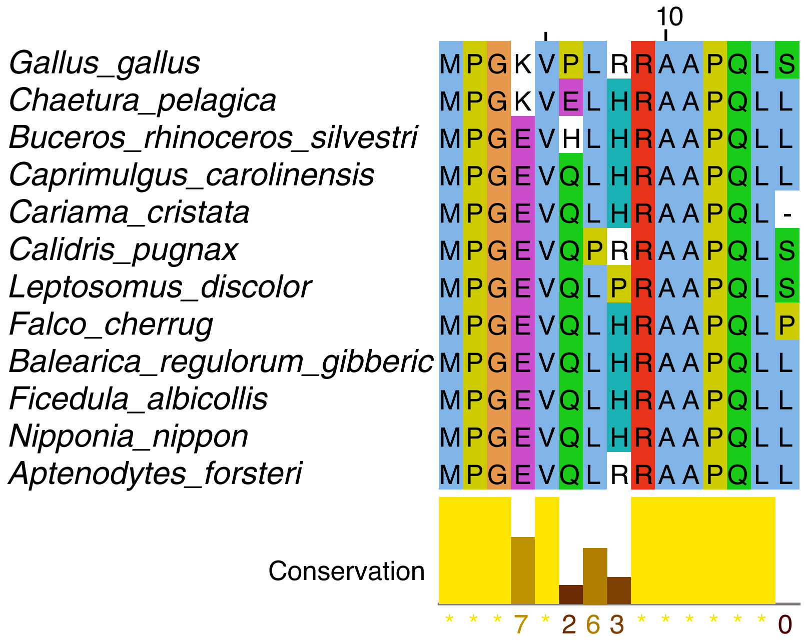

**HG0292.2**

ENSG00000180628 (PCGF5)

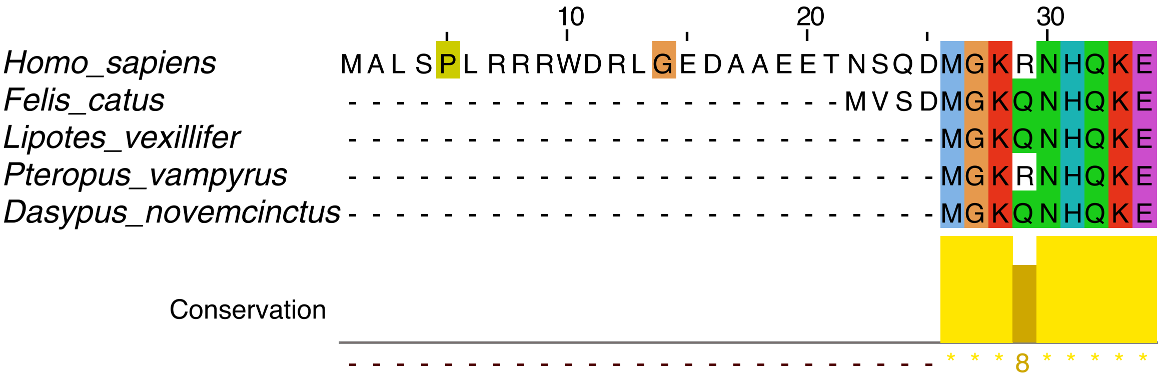

**HG0293**

ENSG00000152127 (MGAT5)

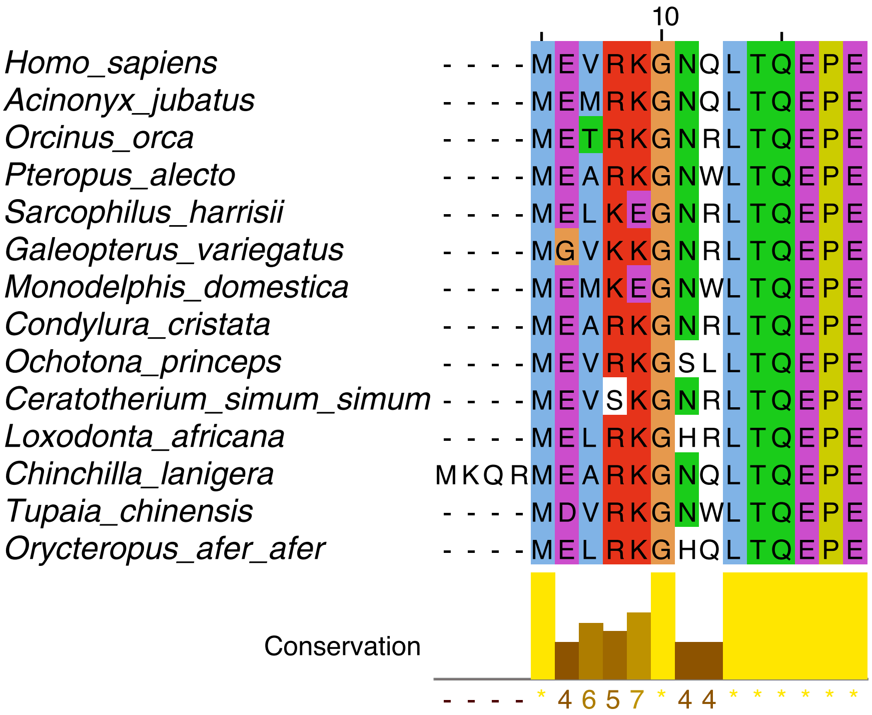

**HG0293**

ENSGALG00000012187 (MGAT5)

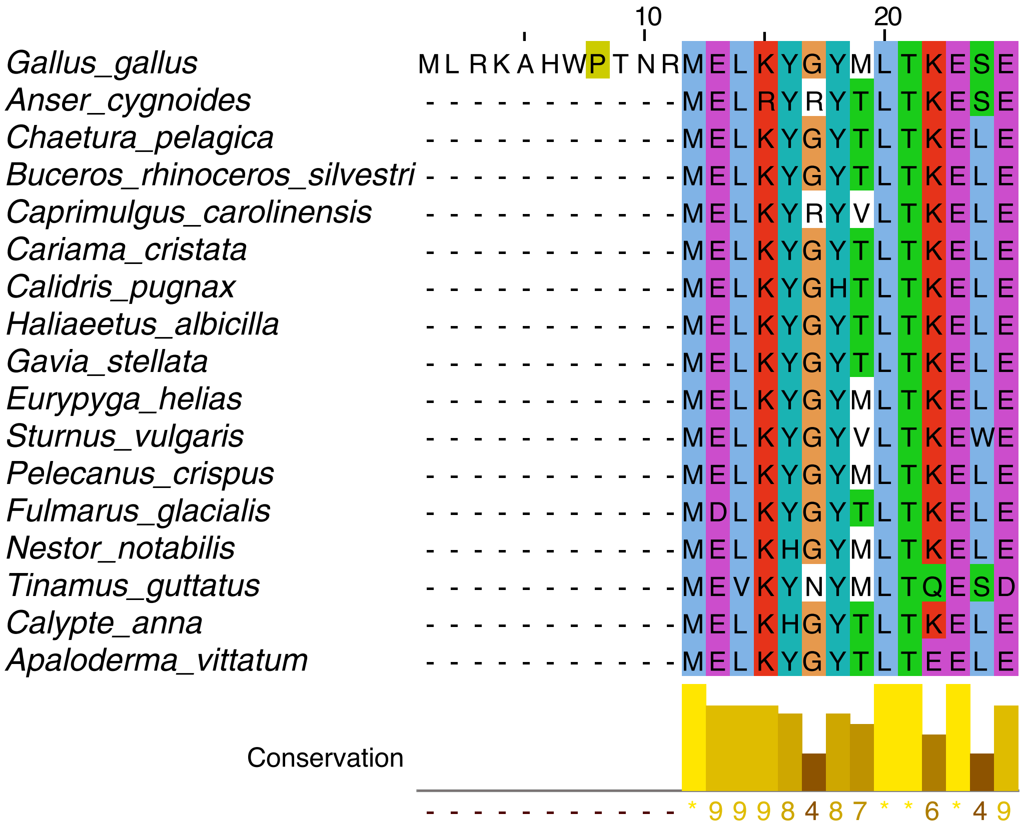

**HG0294.1**

ENSGALG00000017036 (LHFP)

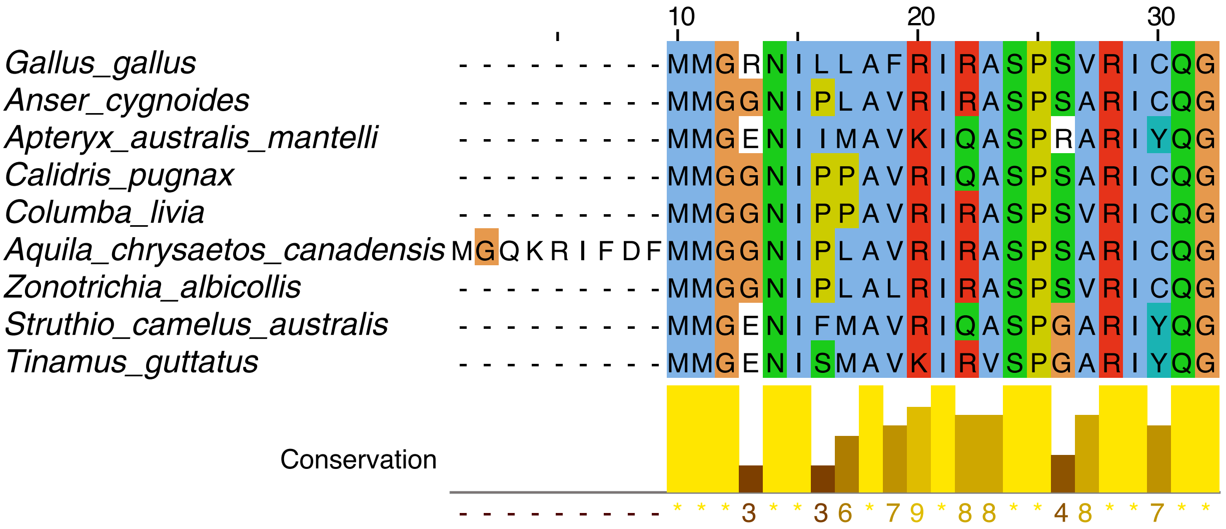

**HG0294.2**

ENSG00000183722 (LHFP)

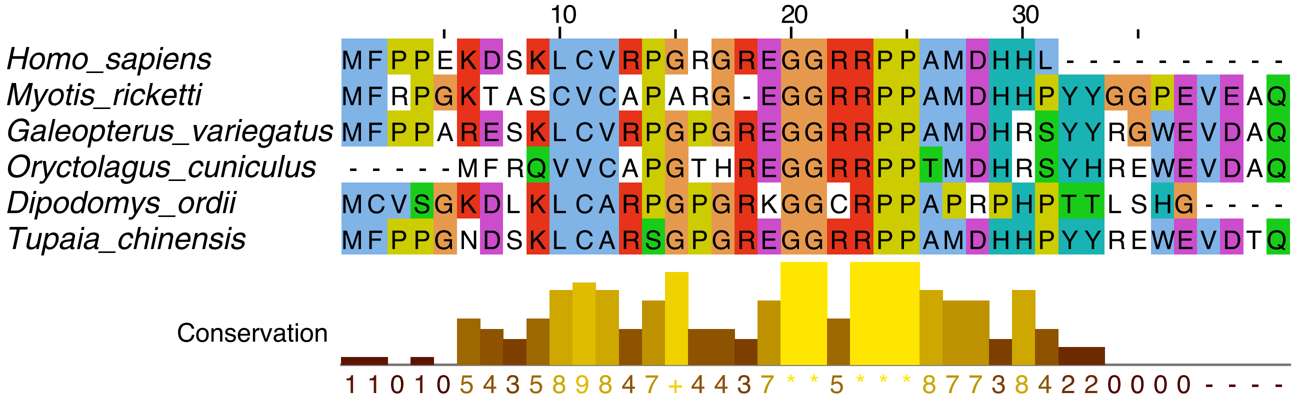

**HG0295.1**

ENSG00000166888 (STAT6)

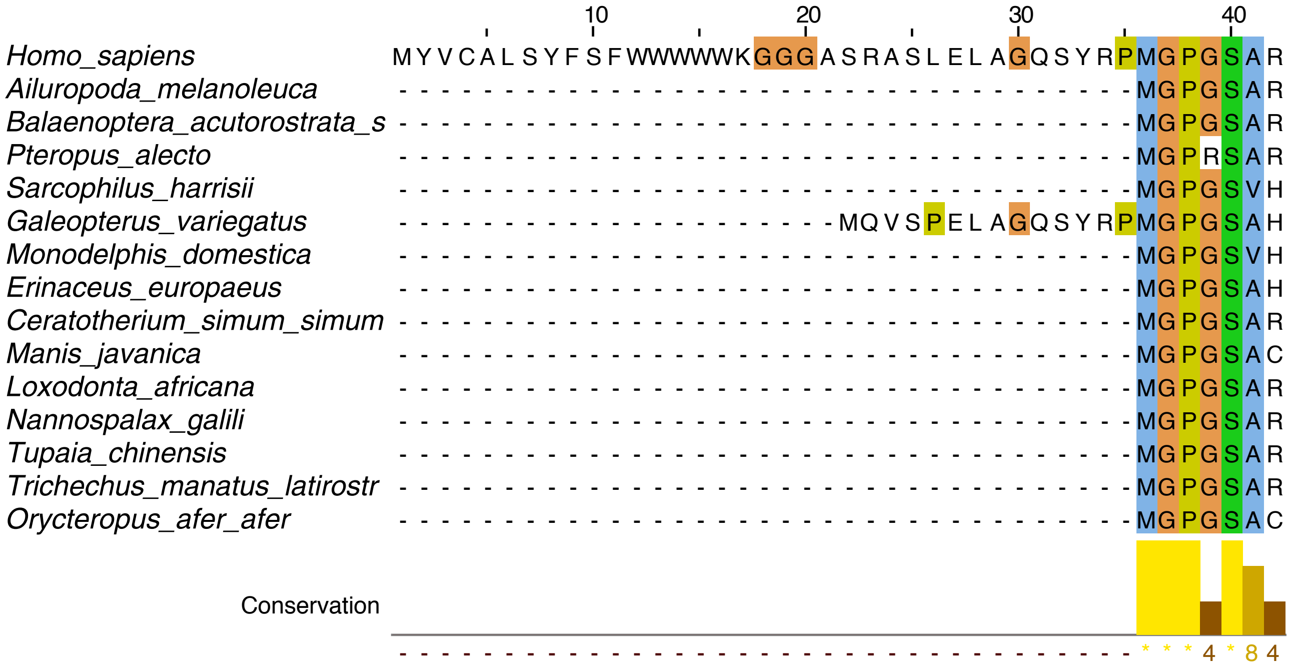

**HG0295.2**

ENSGALG00000003282 (STAT5B)

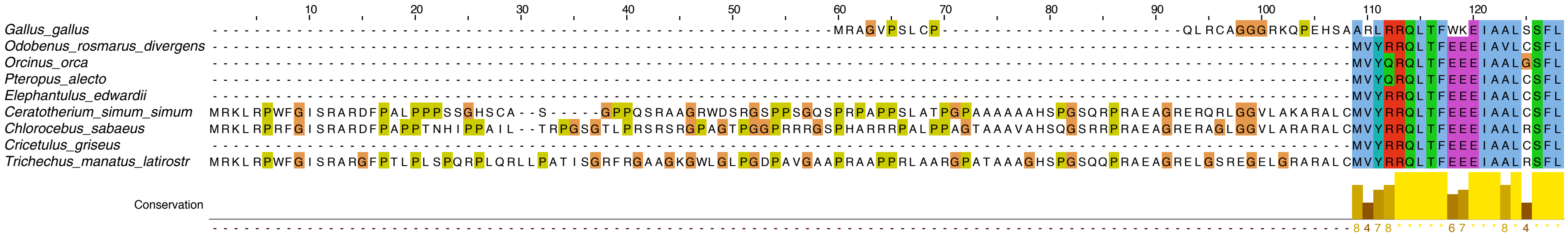

**HG0296**

ENSGALG00000006912 (CPEB3)

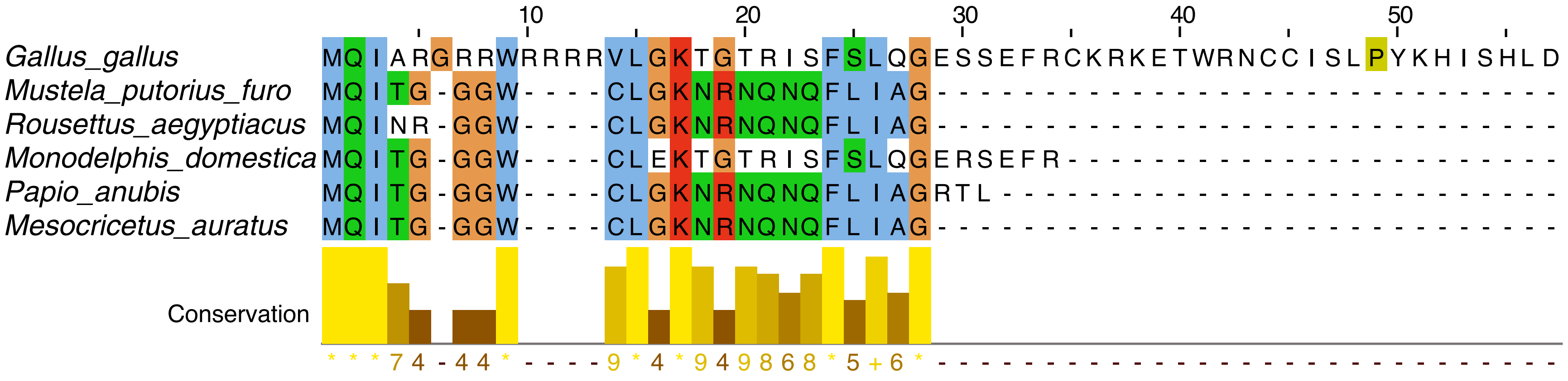

**HG0297**

ENSGALG00000031684

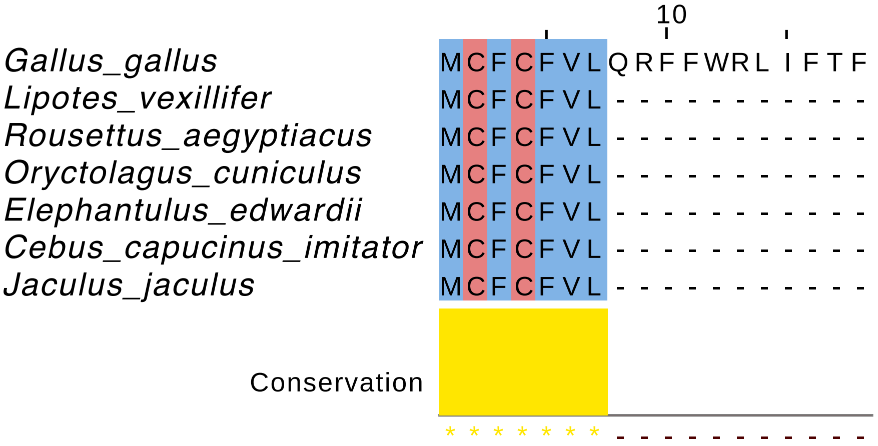

**HG0298**

ENSGALG00000042308

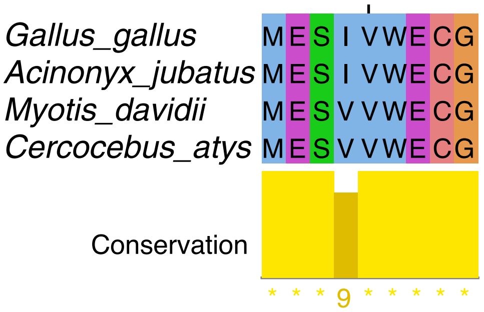

**HG0299**

ENSGALG00000040465 (Sip1)

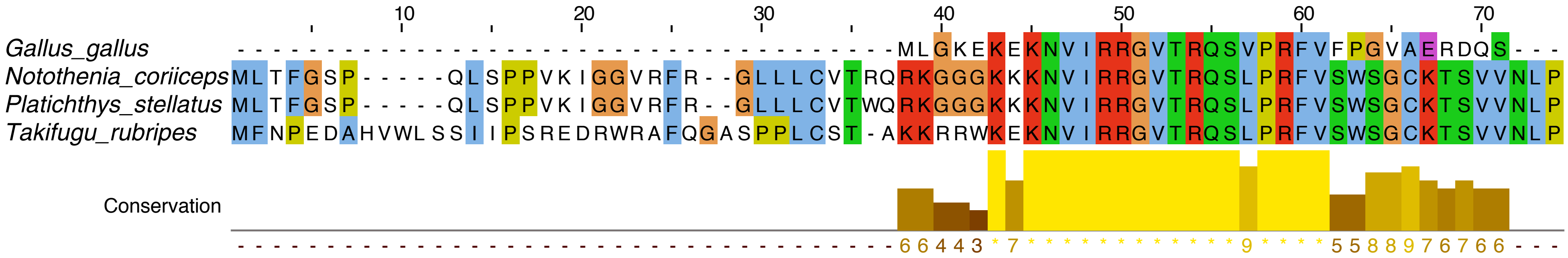

**HG0300**

FBgn0031668 (CG31917)

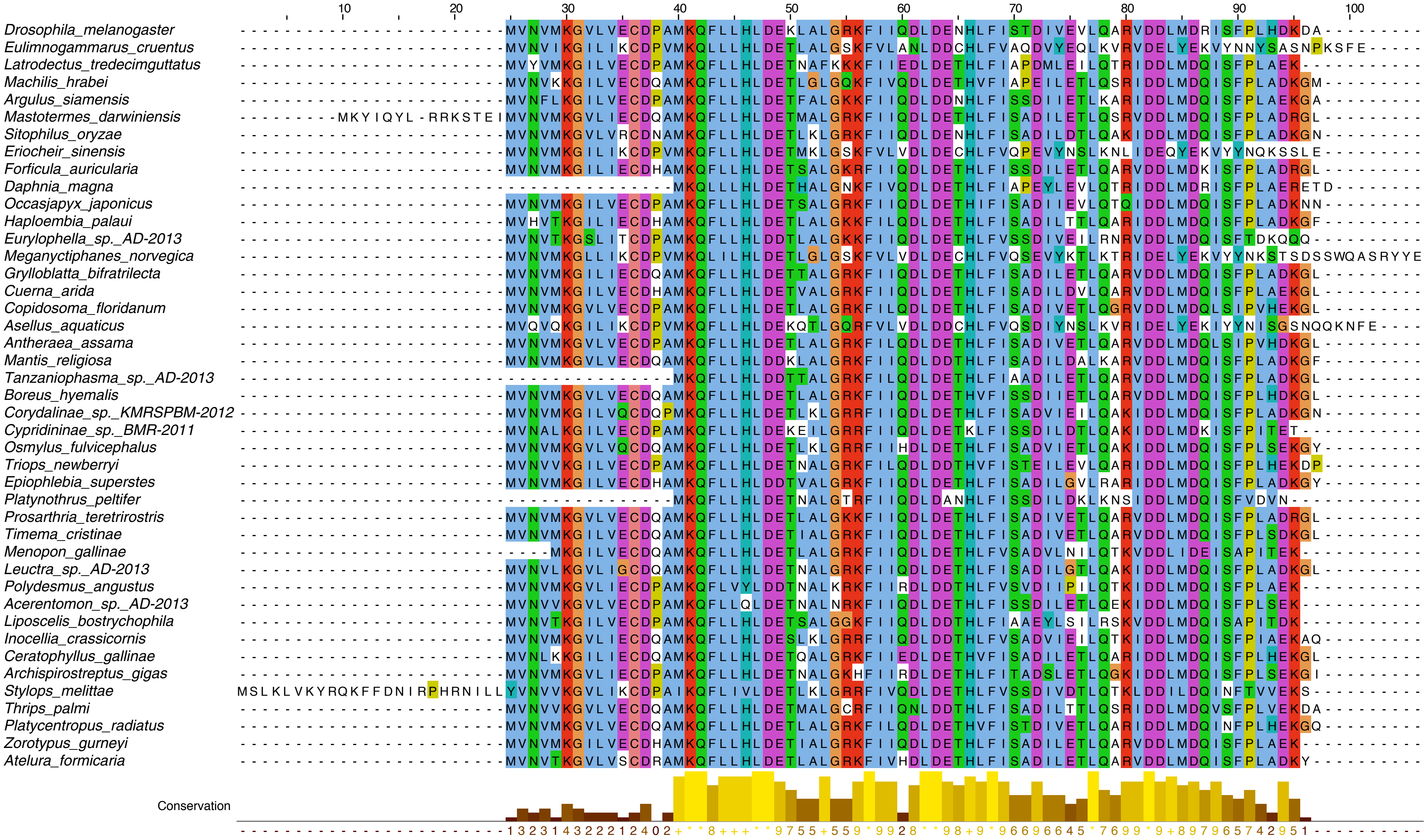

# HG0301

### FBgn0036856 (CG9666)

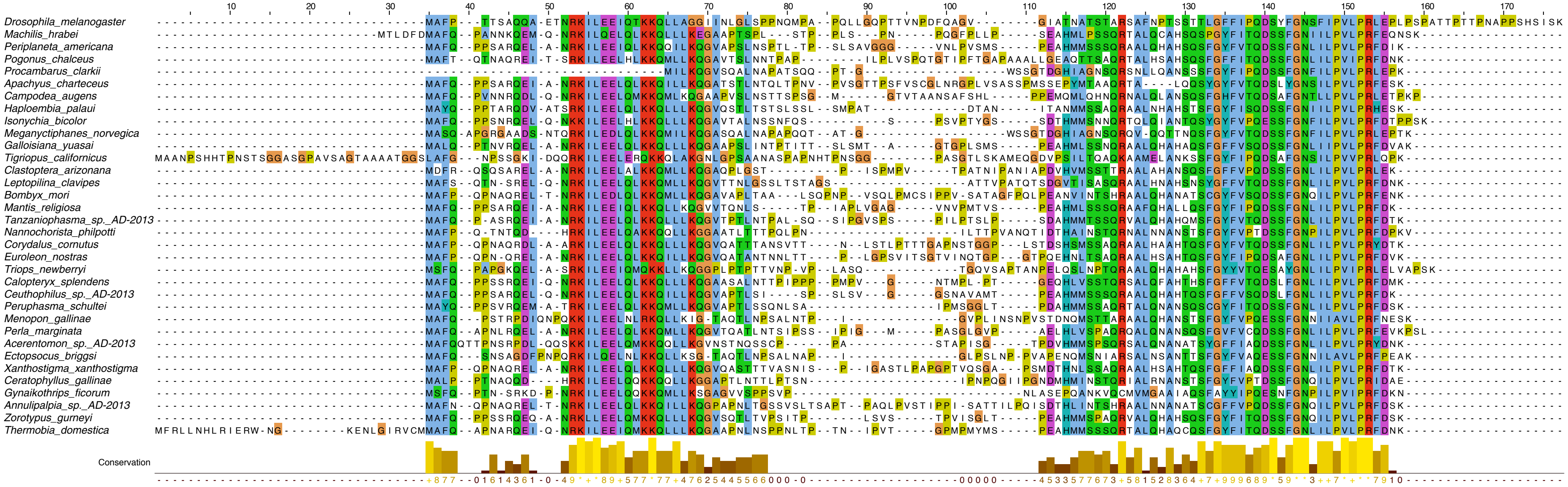

# HG0302

### FBgn0031454 (CG9960)

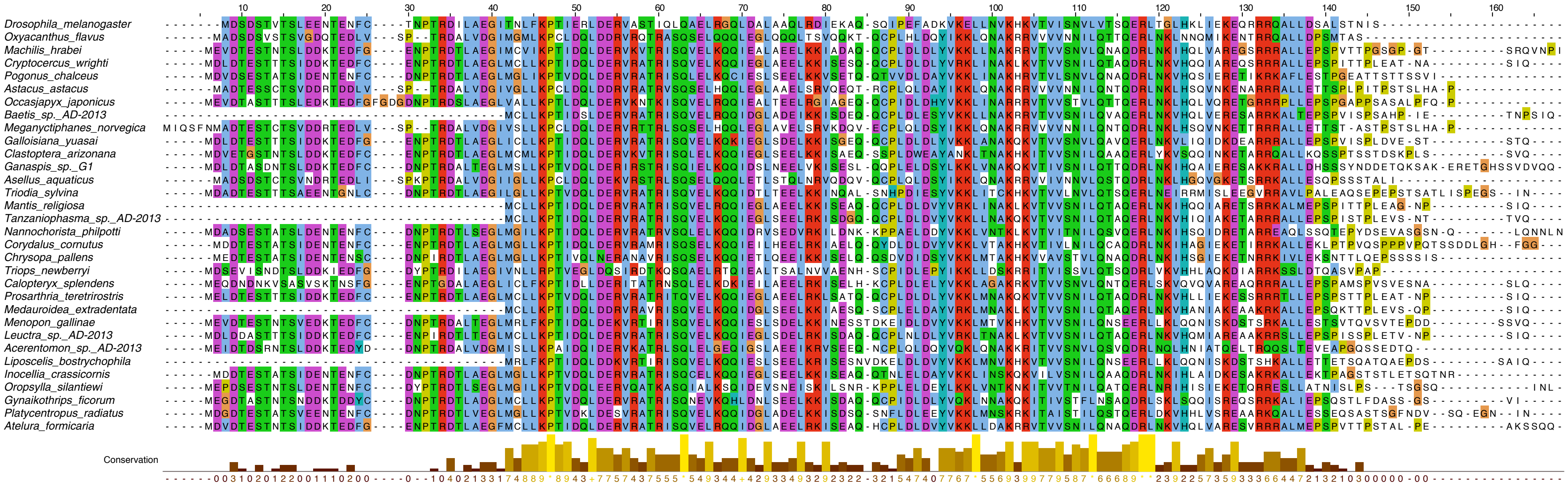

**HG0303**

FBgn0260464

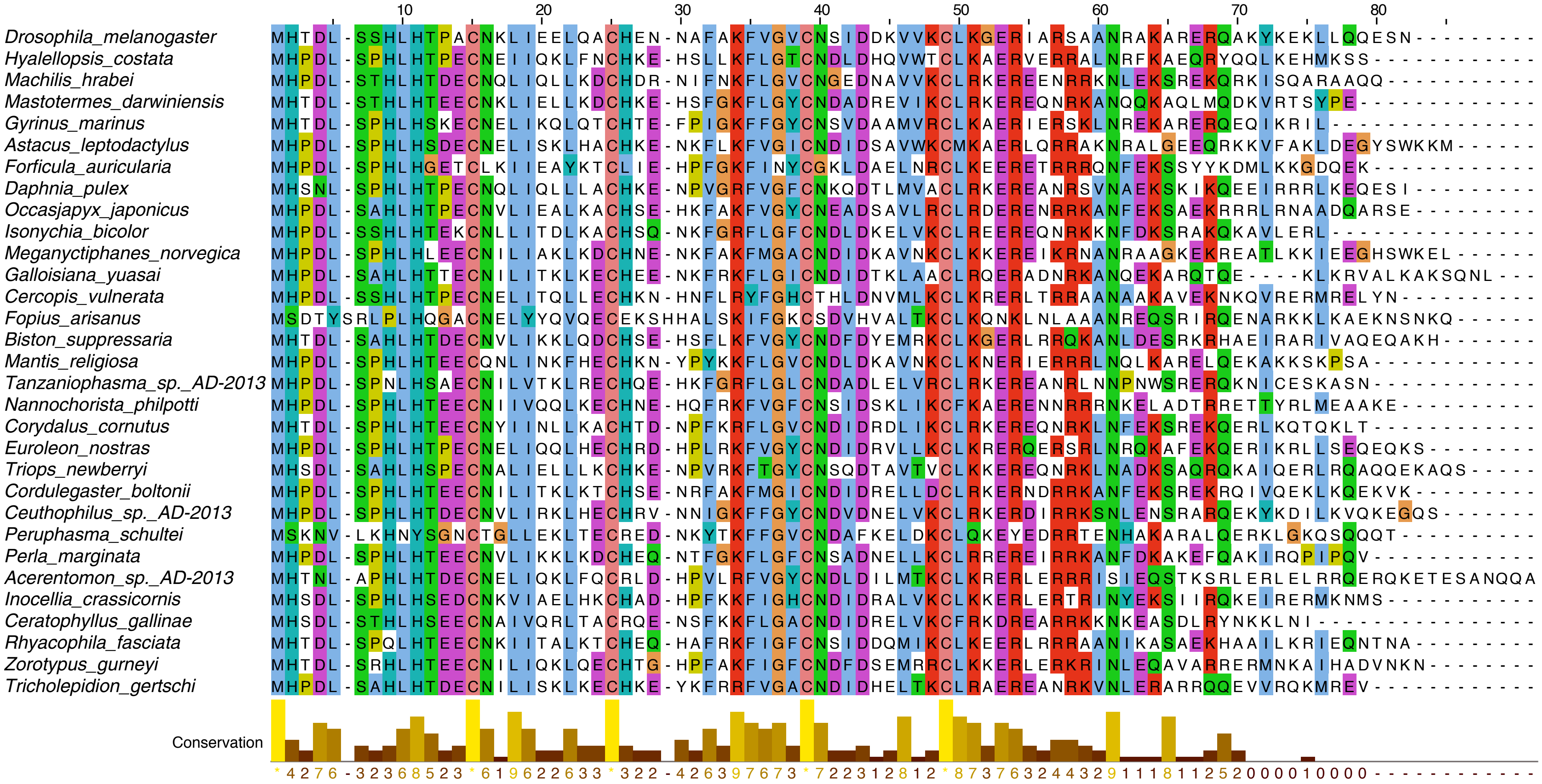

**HG0304**

FBgn0260467 (CG7071)

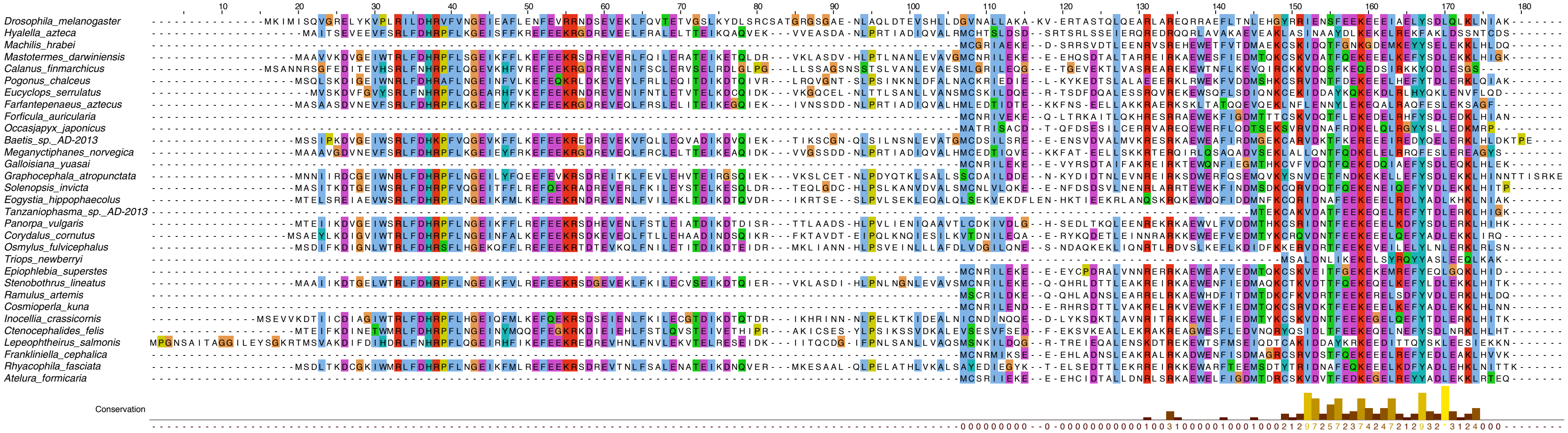

**HG0305**

FBgn0050290 (Ppcdc)

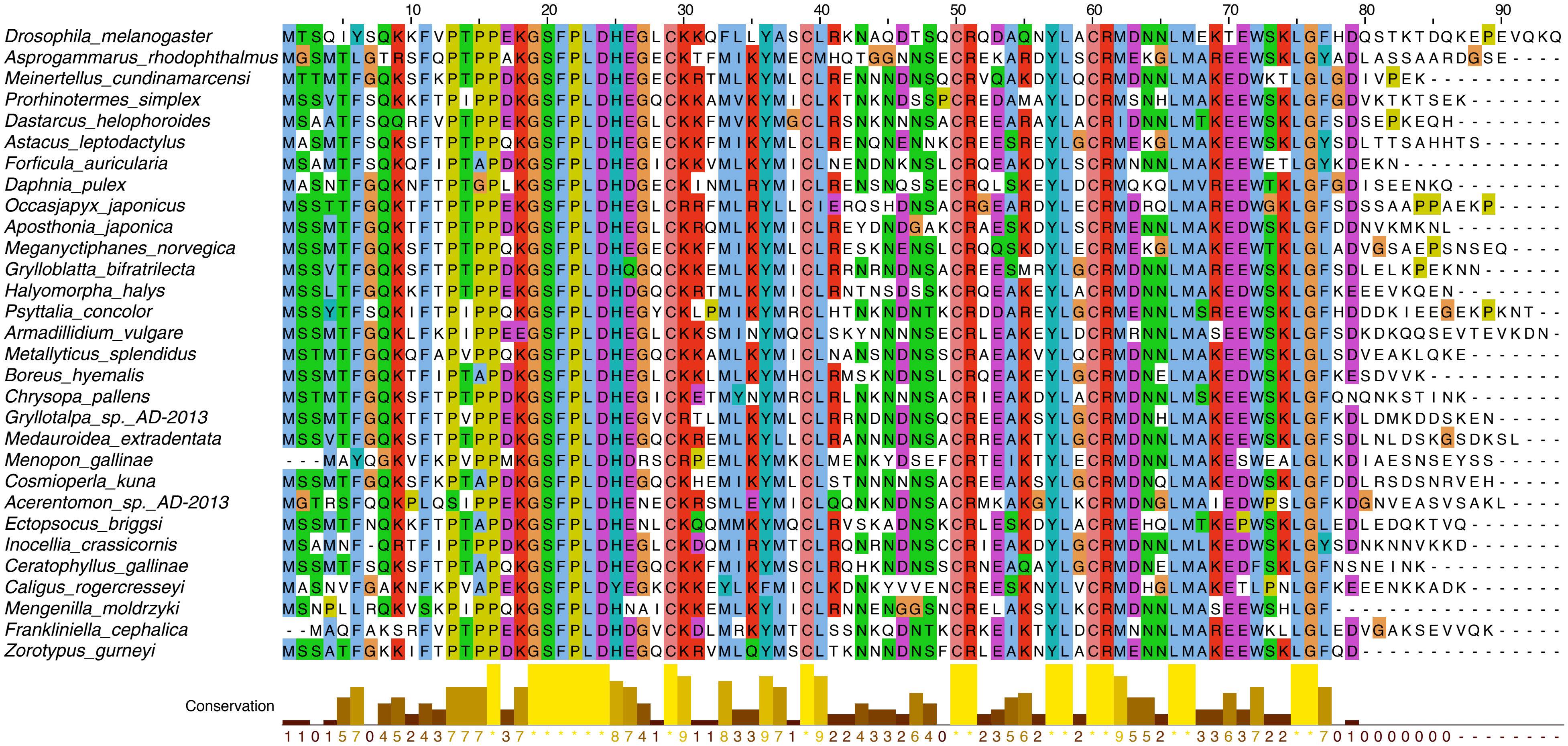

# HG0306

FBgn0259726

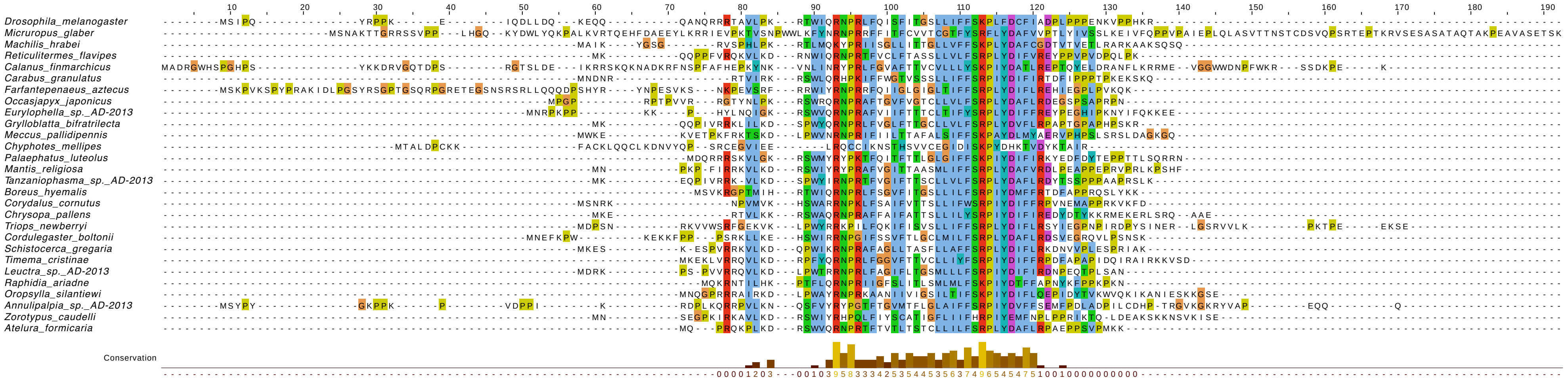

**HG0307**

FBgn0037822 (CG14683)

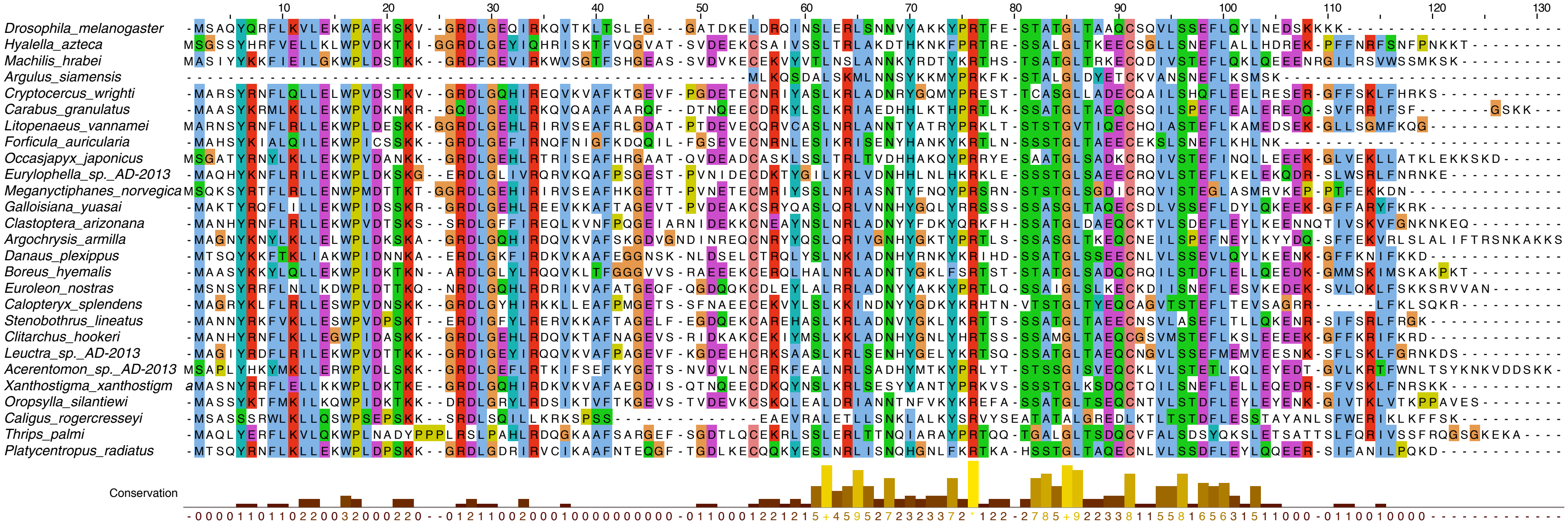

**HG0308**

FBgn0264743 (CG44001)

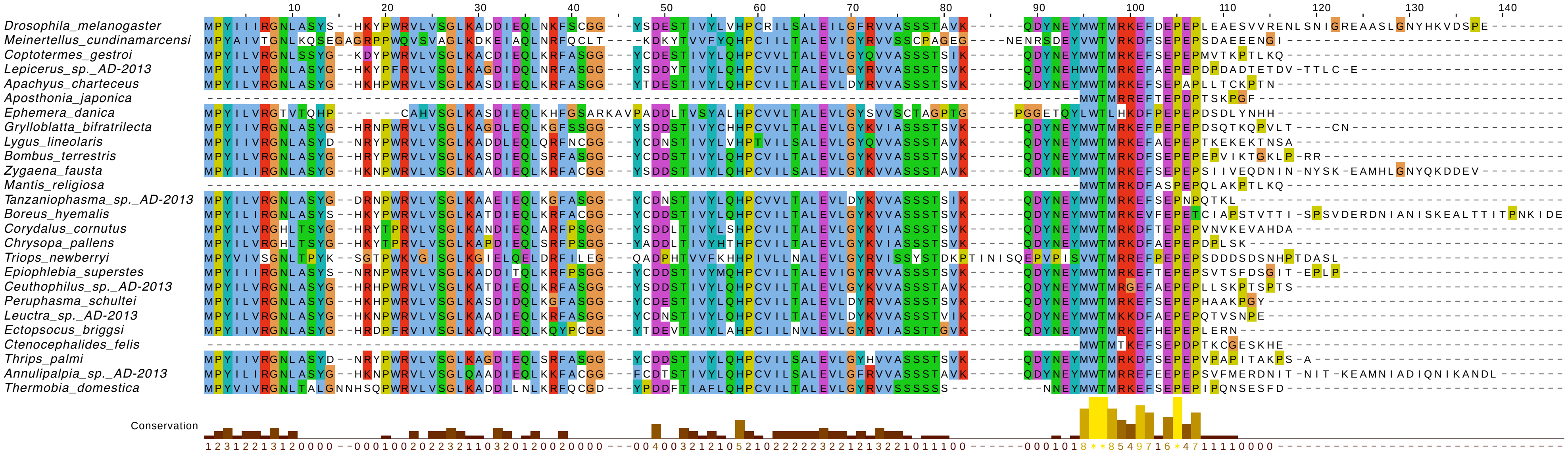

**HG0309**

FBgn0037689 (CG8135)

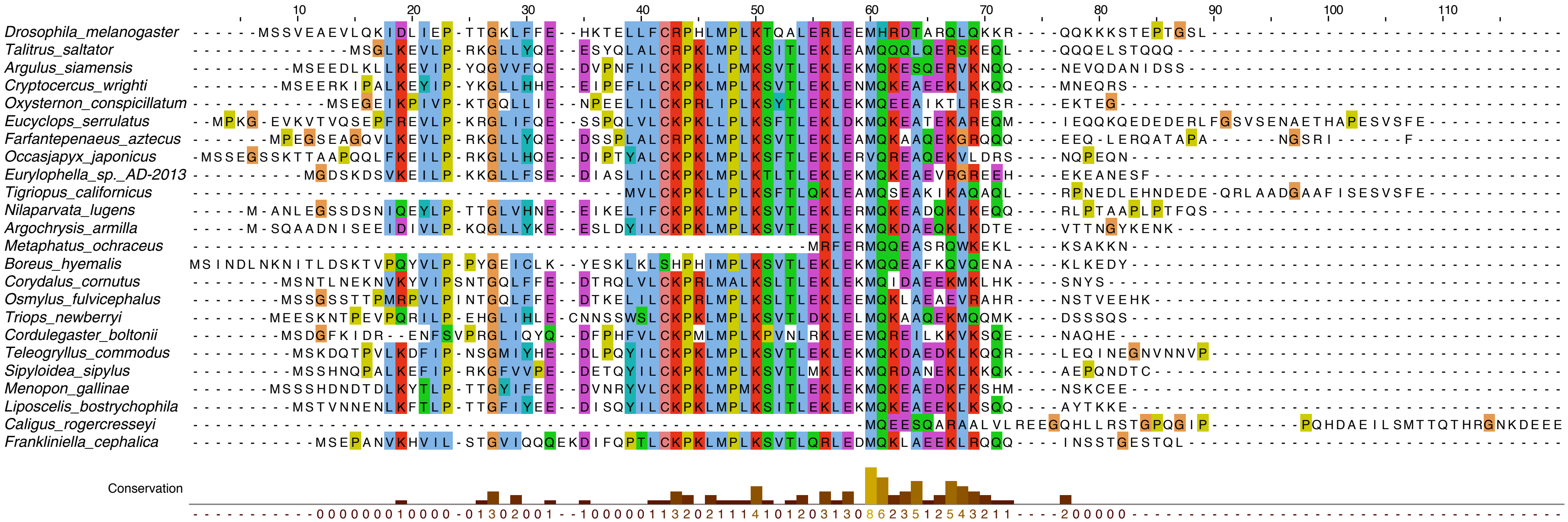

**HG0310**

FBgn0039339 (CG5116)

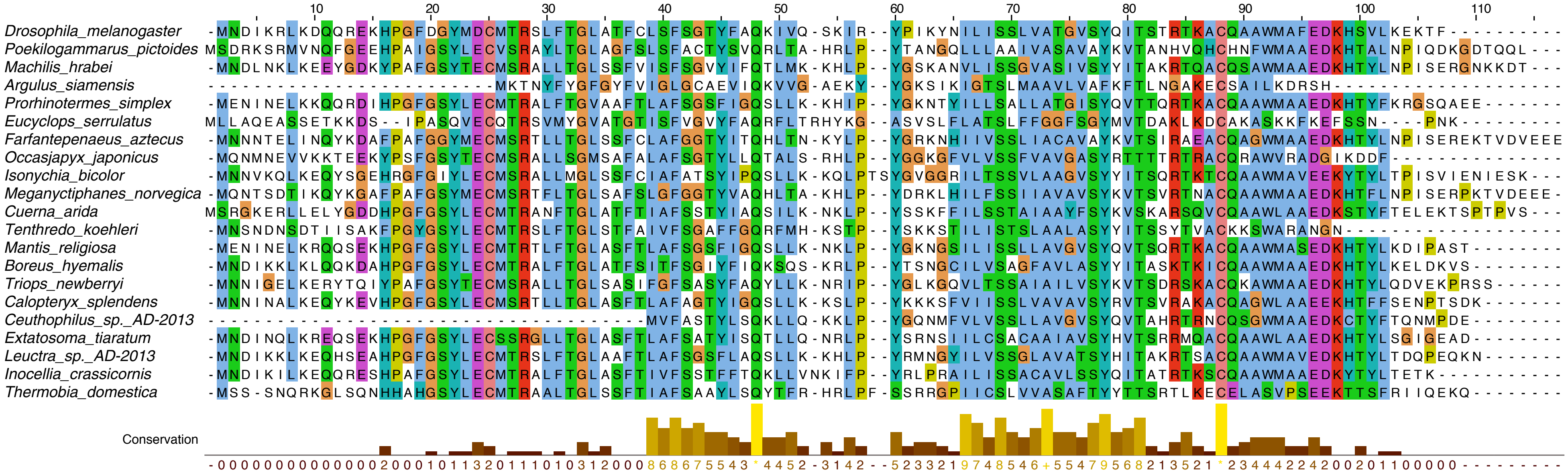

**HG0311**

FBgn0263251 (vnc)

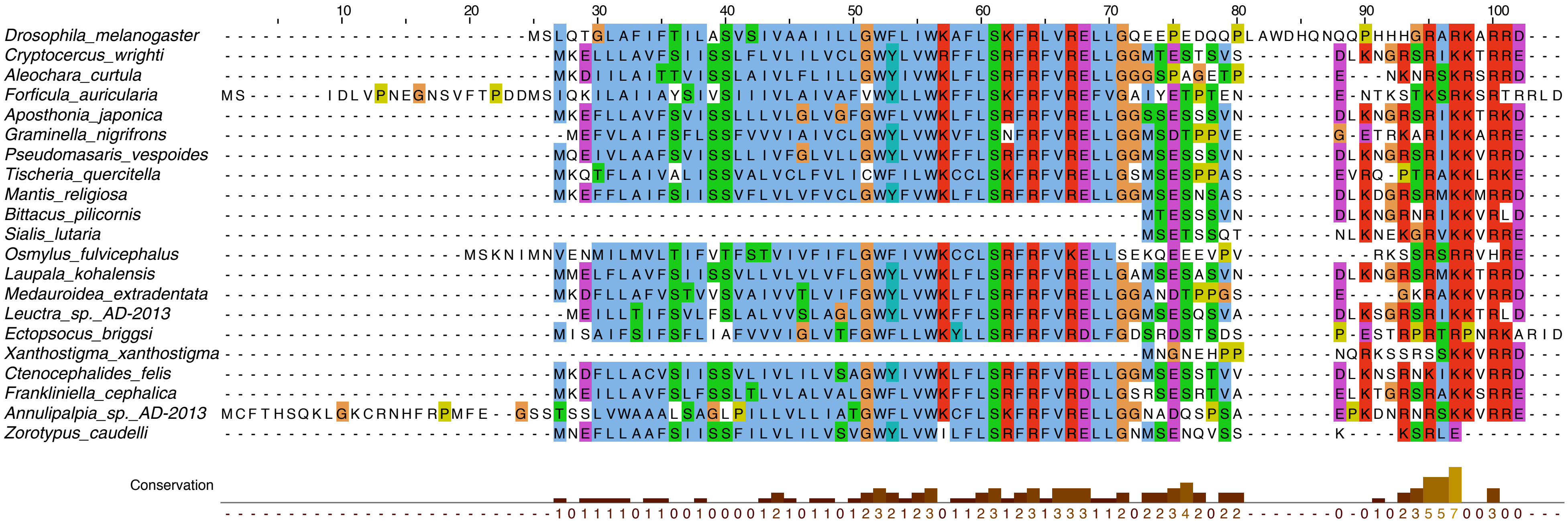

**HG0312**

FBgn0038641 (CG7708)

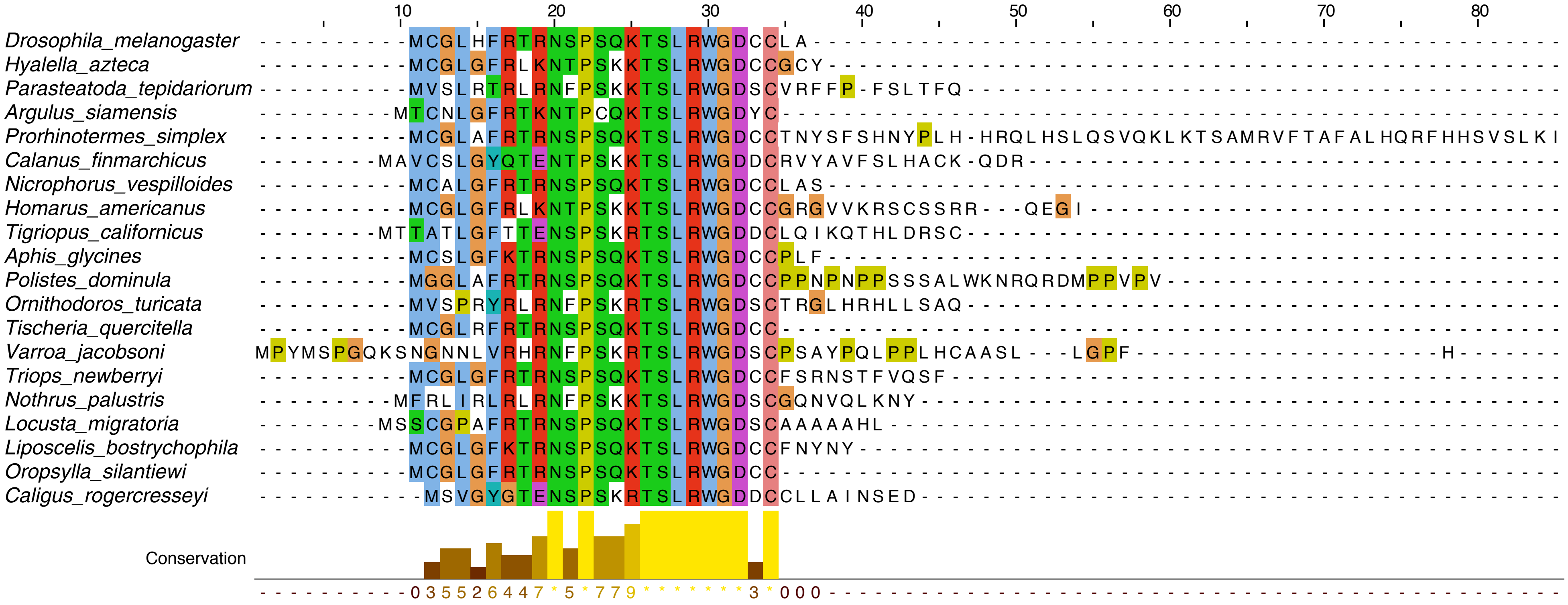

**HG0313**

FBgn0259725

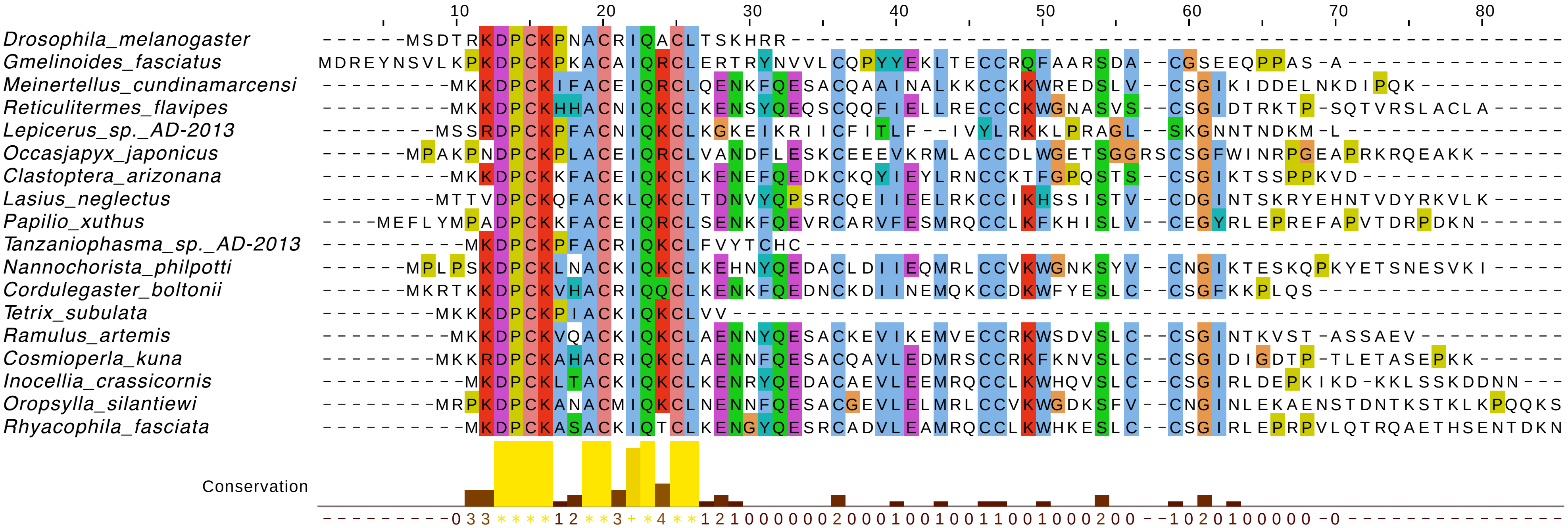

# HG0314

### FBgn0260468 (CG7950)

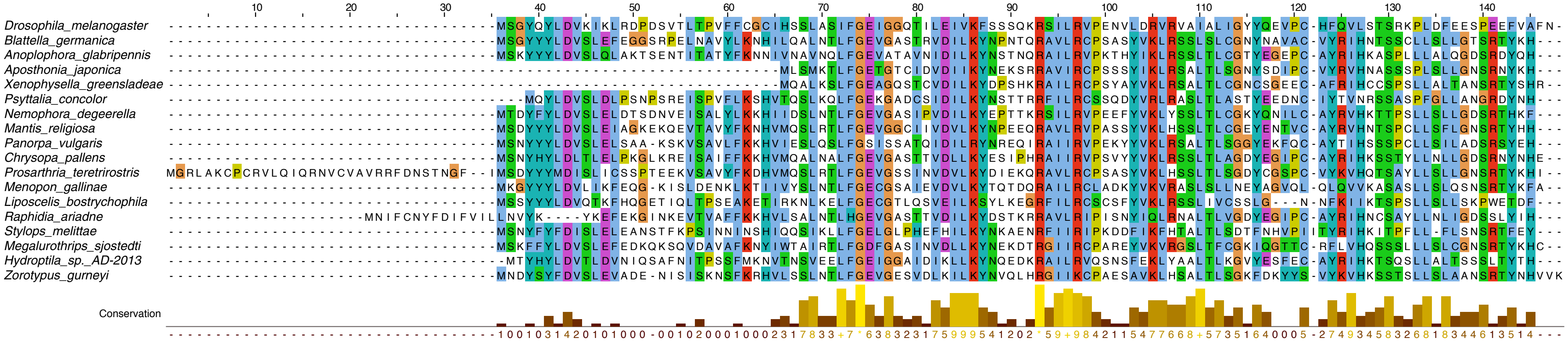

**HG0315**

FBgn0260234 (CG42508)

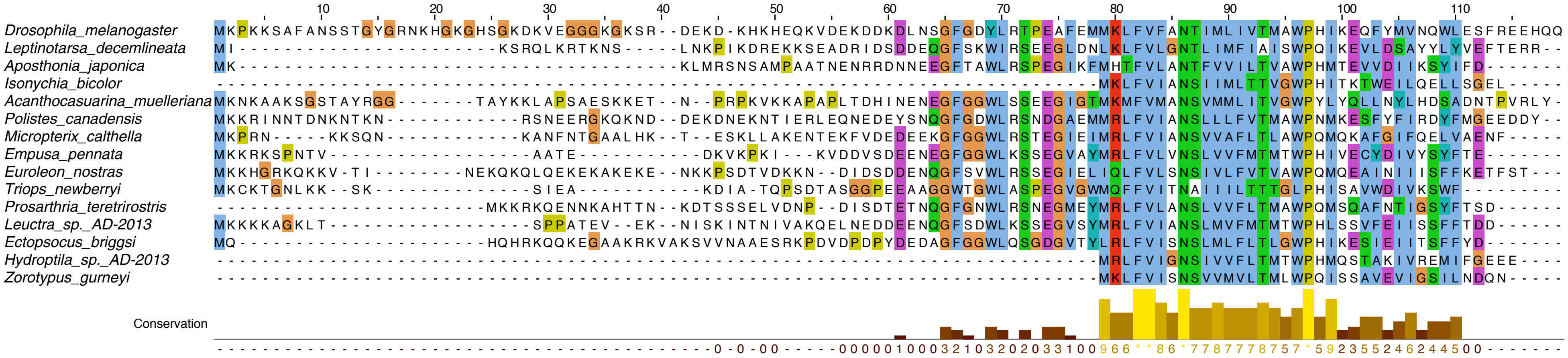

# HG0316

### FBgn0041164 (armi)

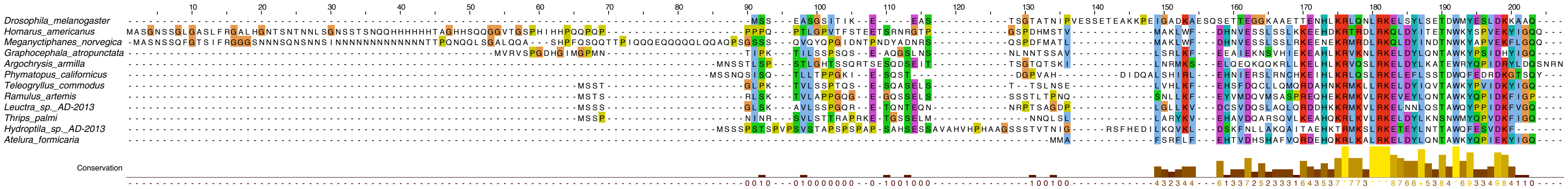

**HG0317**

FBgn0050011 (gem)

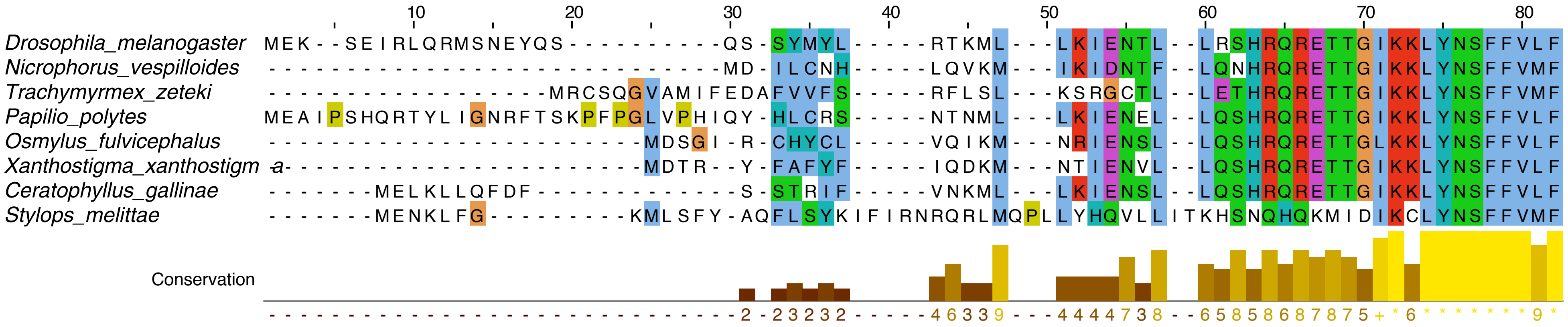

**HG0318**

FBgn0035293 (CG5687)

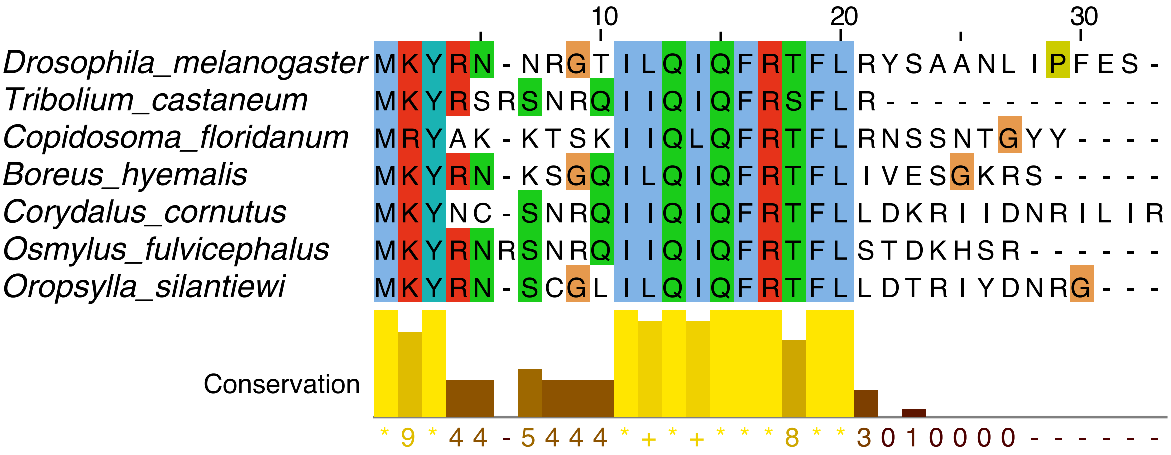

# HG0319

### FBgn0026778 (Rad1)

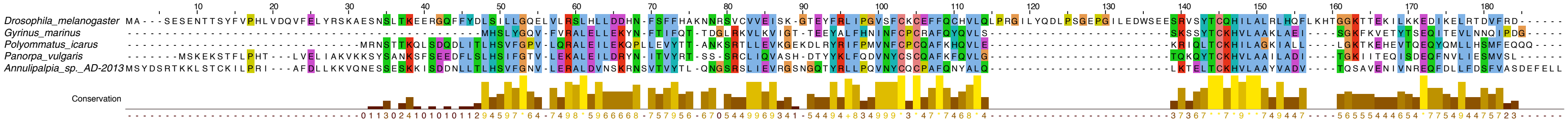

**HG0320**

FBgn0040993

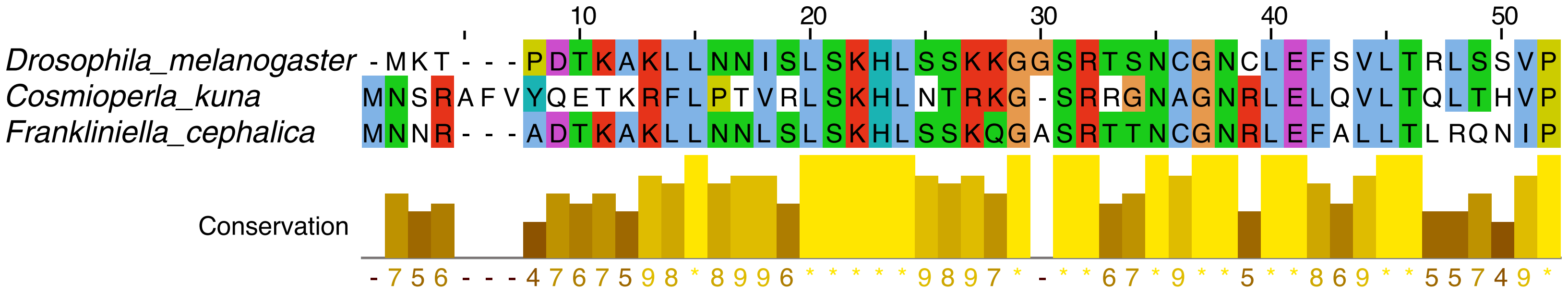

**HG0321**

ENSDARG00000070426 (chac1)

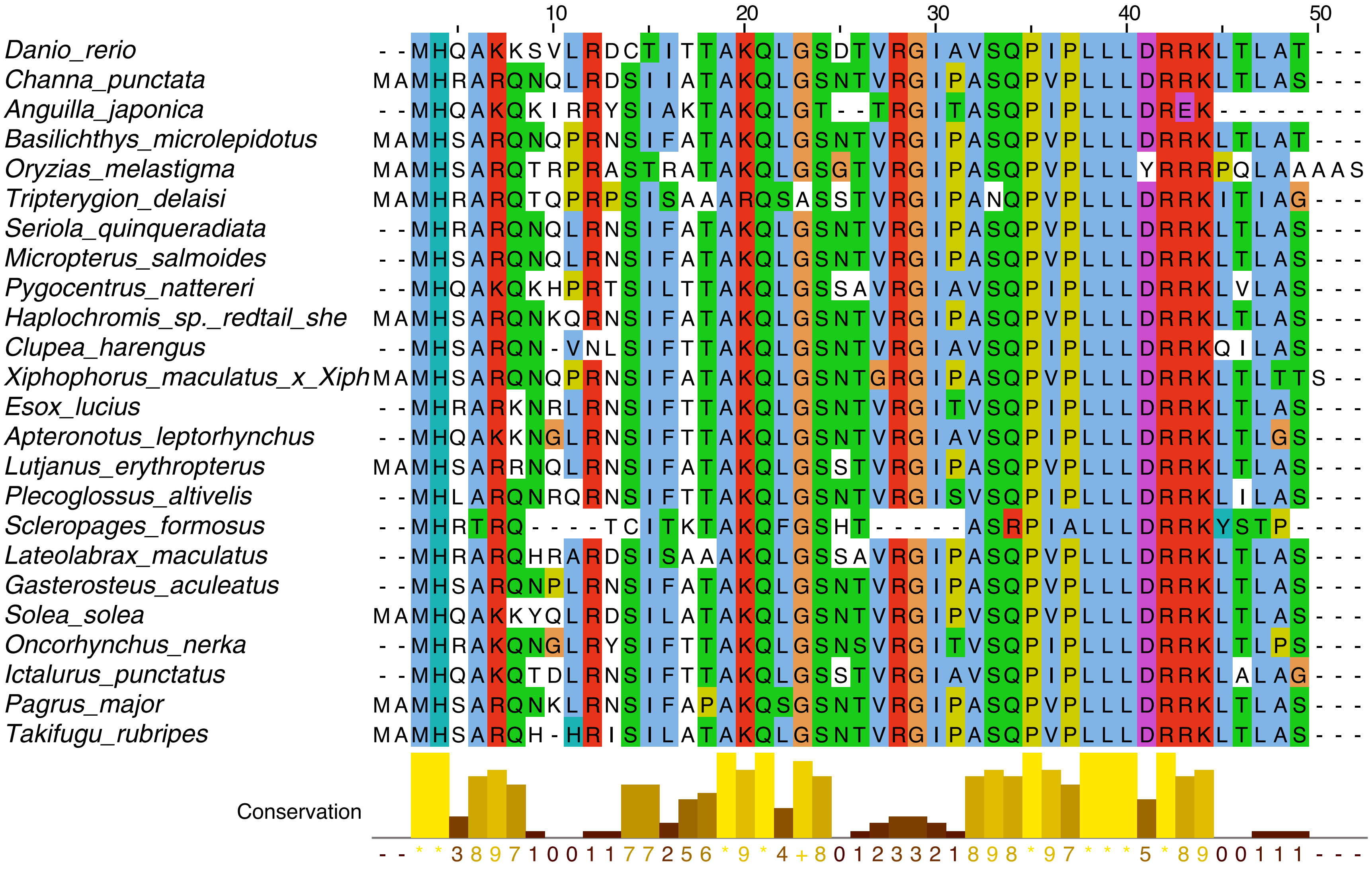

**HG0322**

ENSDARG00000070917 (kitlga)

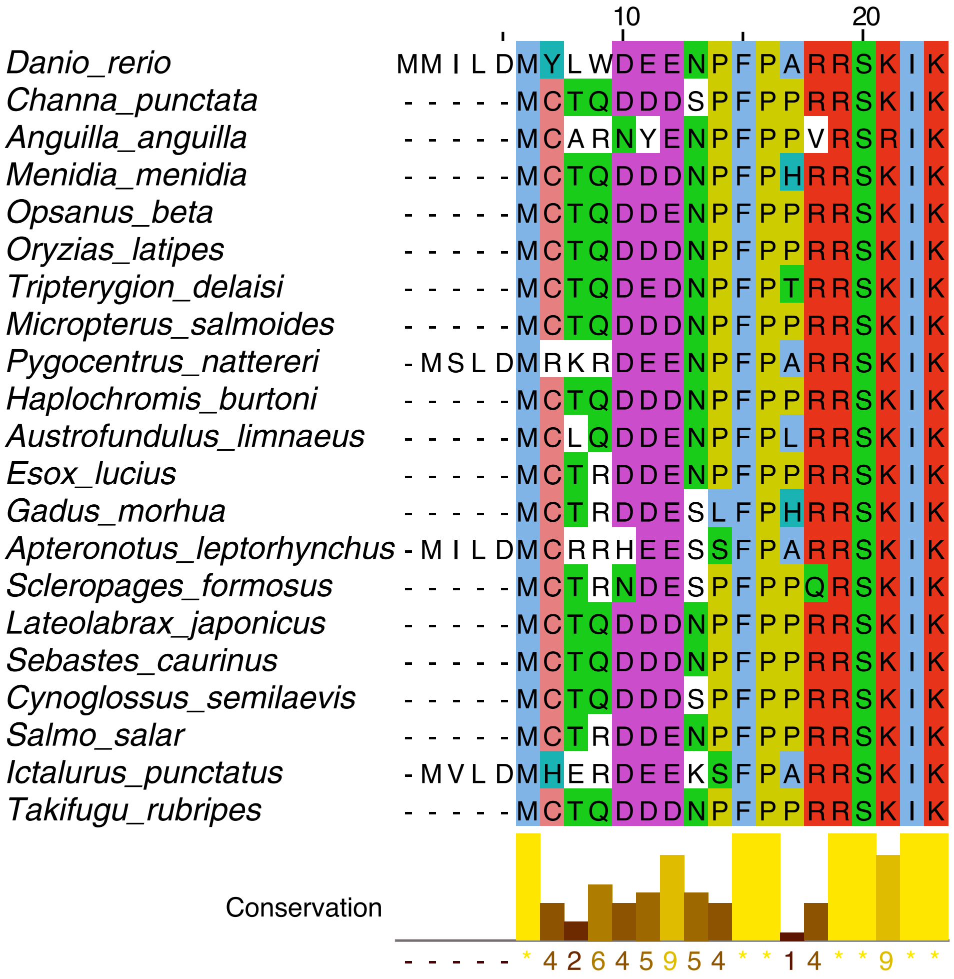

**HG0323**

ENSDARG00000071197 (usp40)

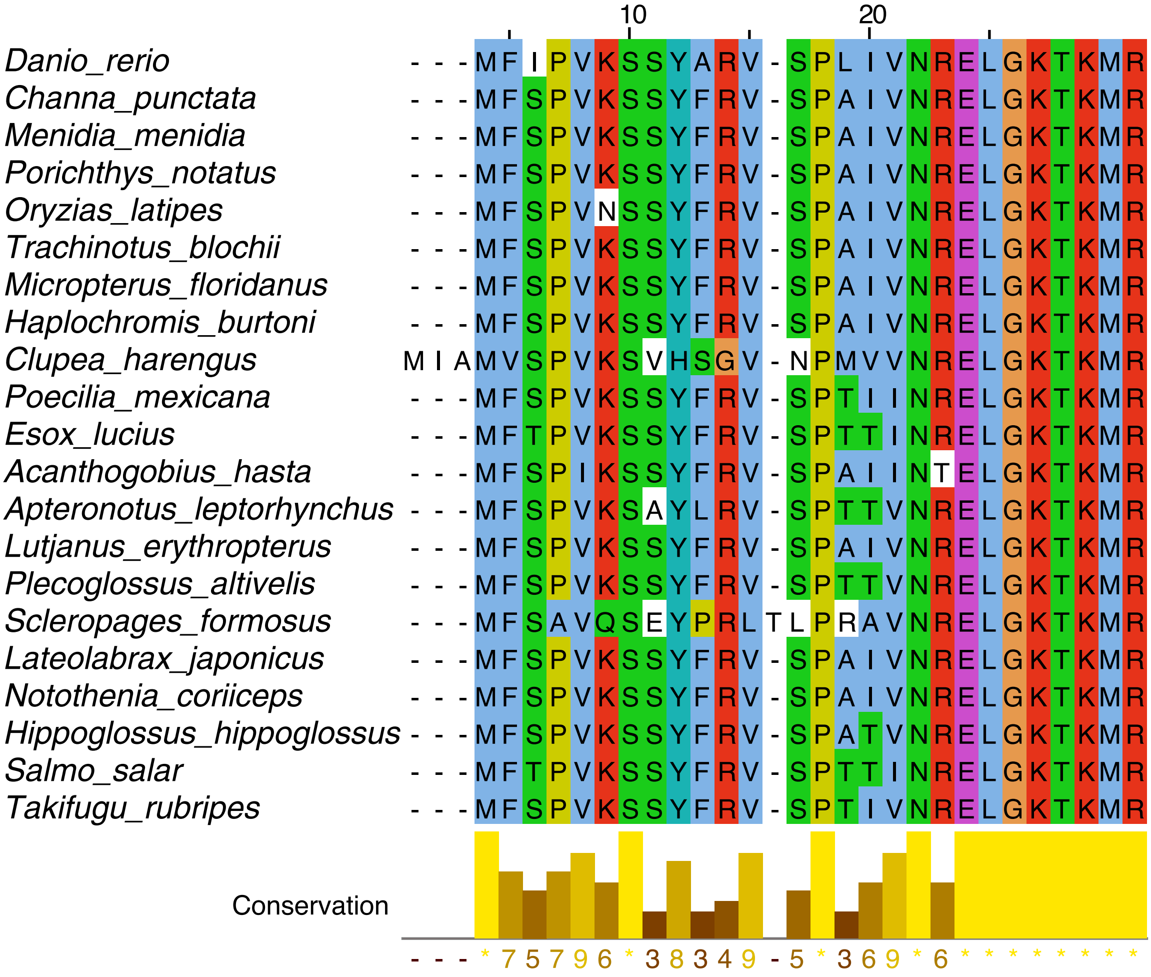

**HG0324**

ENSDARG00000094965 (nfil3-5)

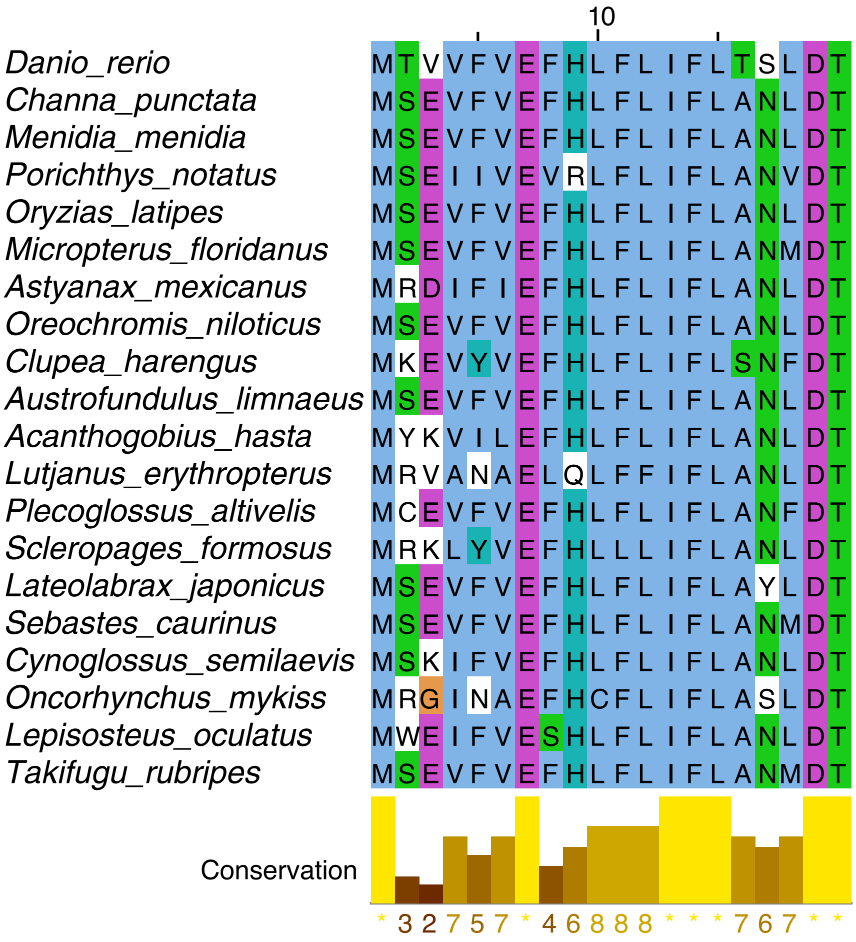

**HG0325**

ENSDARG00000079738 (znf219)

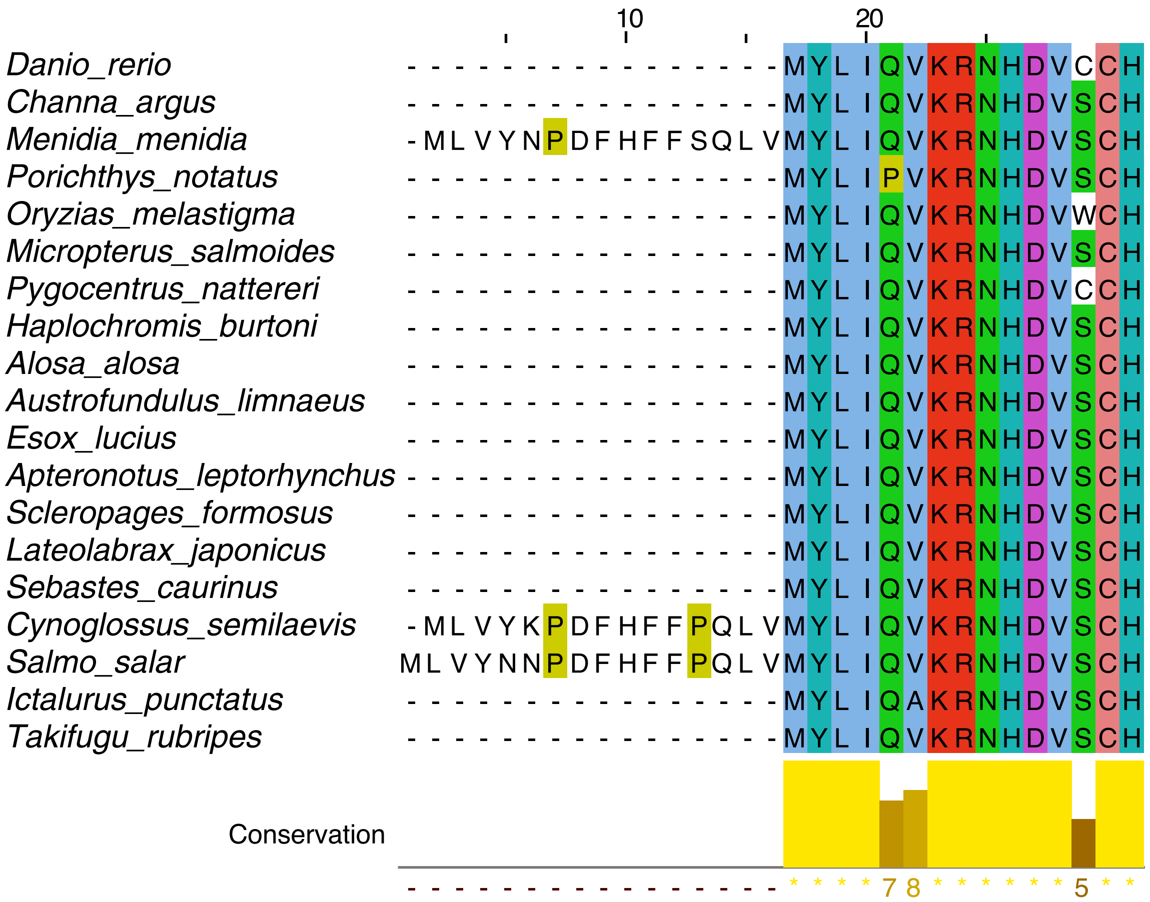

**HG0326**

ENSDARG00000012848 (arih2)

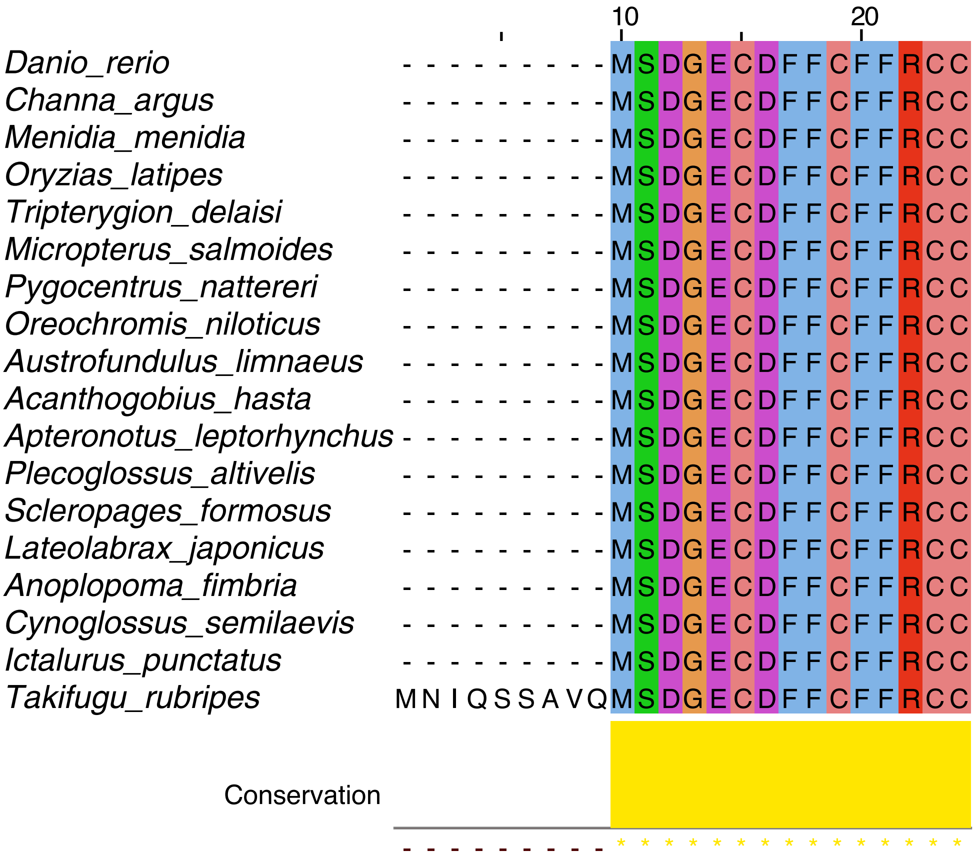

**HG0327**

ENSDARG00000022203 (gtf2ird1)

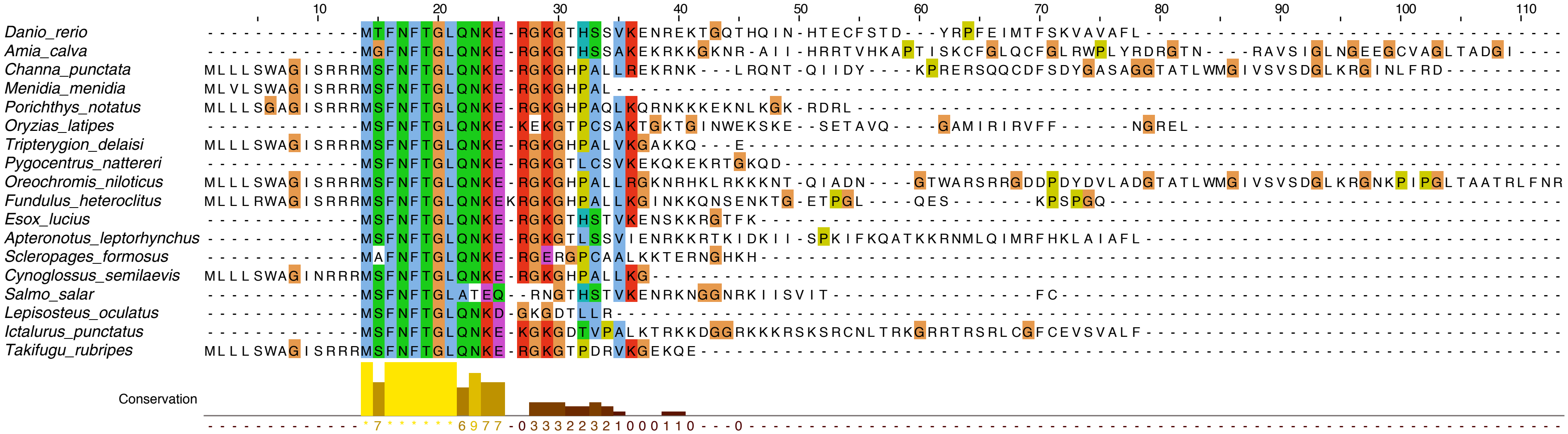

**HG0328**

ENSDARG00000099999 (tcf3b)

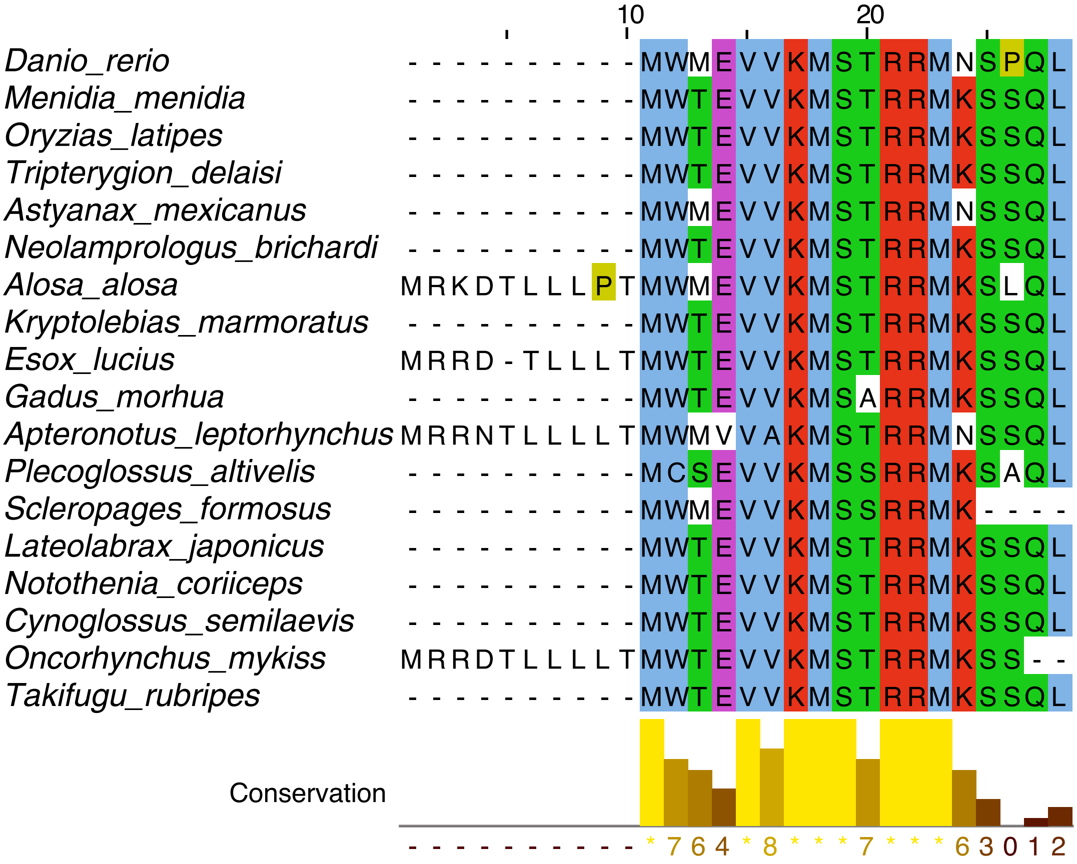

**HG0329**

ENSDARG00000036442 (atp11c)

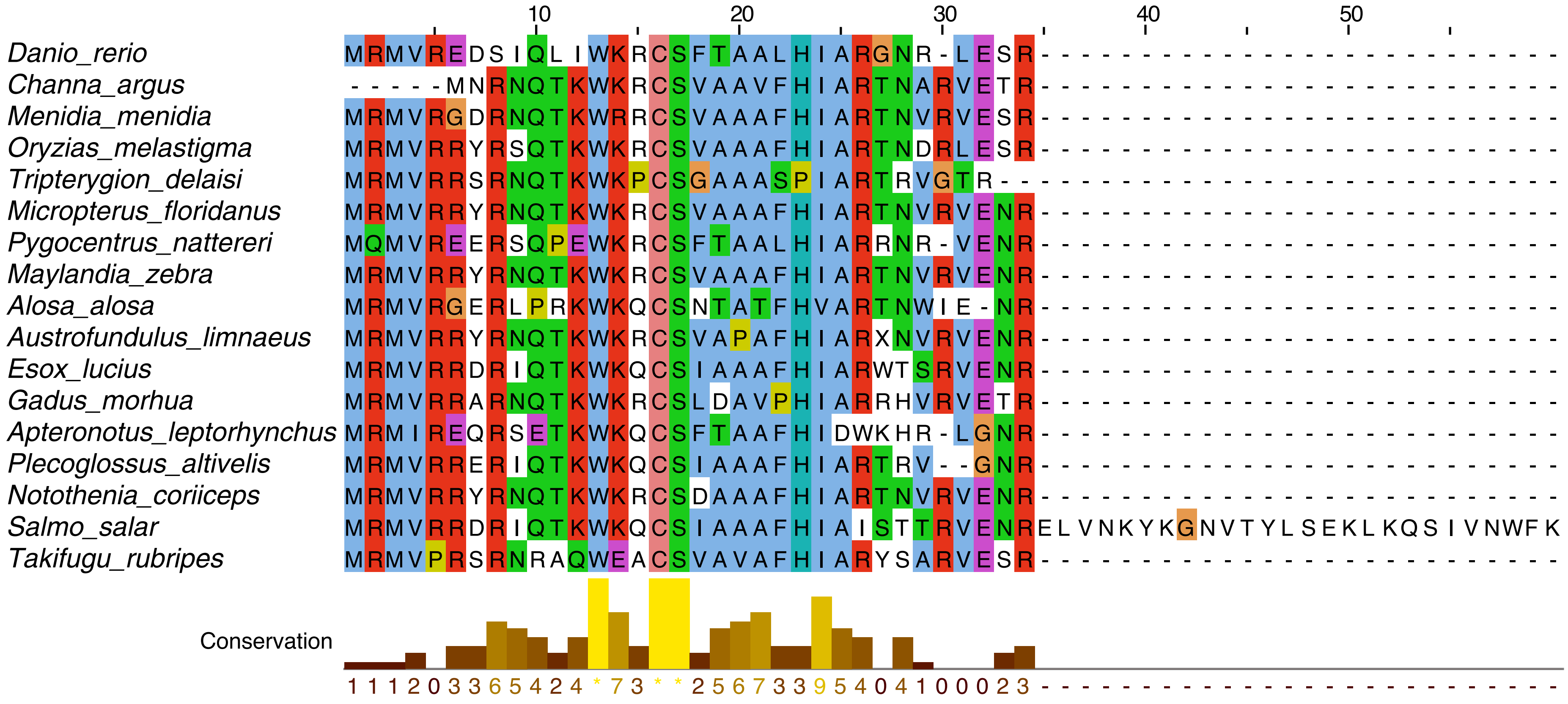

**HG0330**

ENSDARG00000039392 (wnk1b)

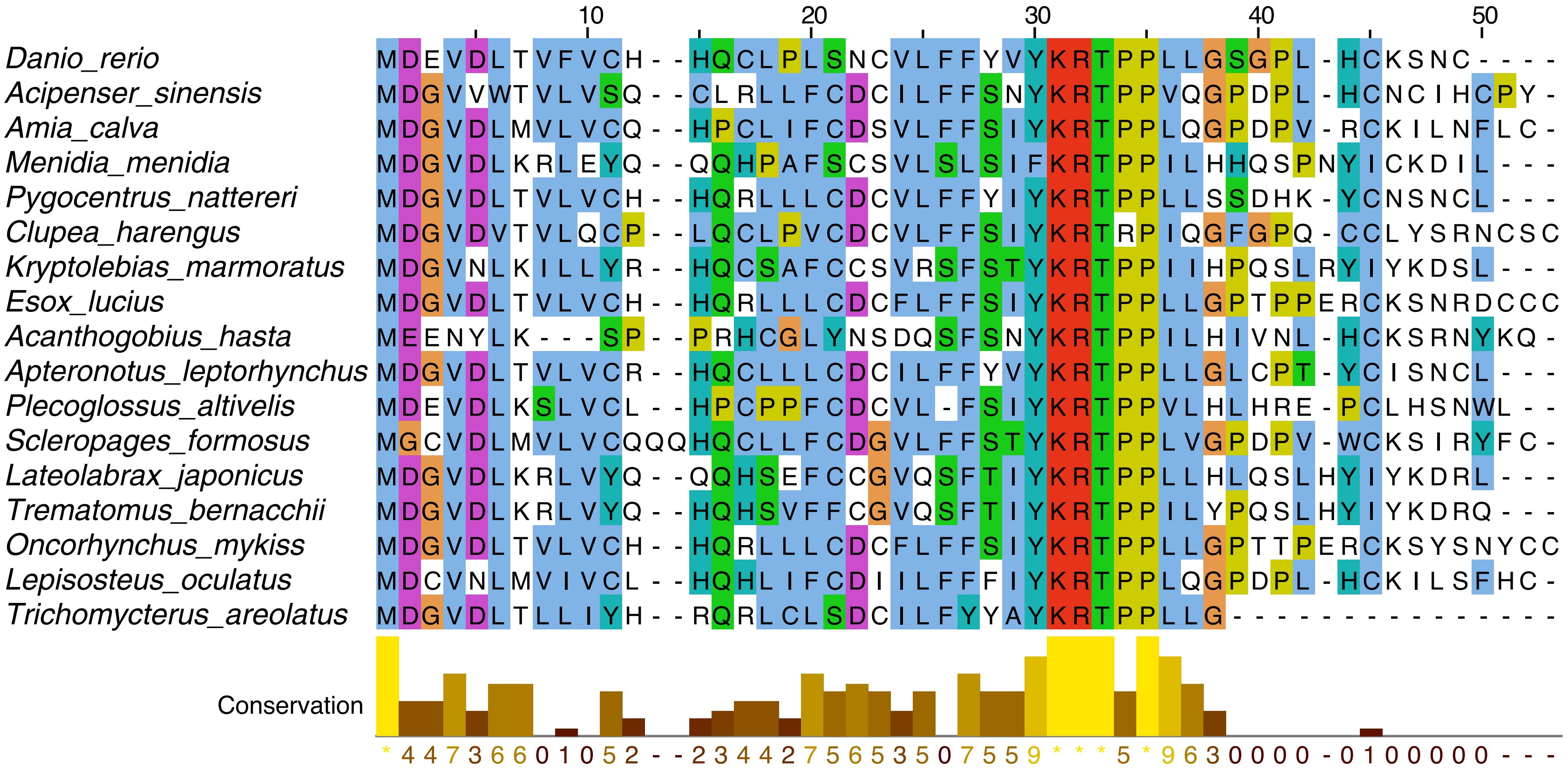

**HG0331**

ENSDARG00000044899 (tmem183a)

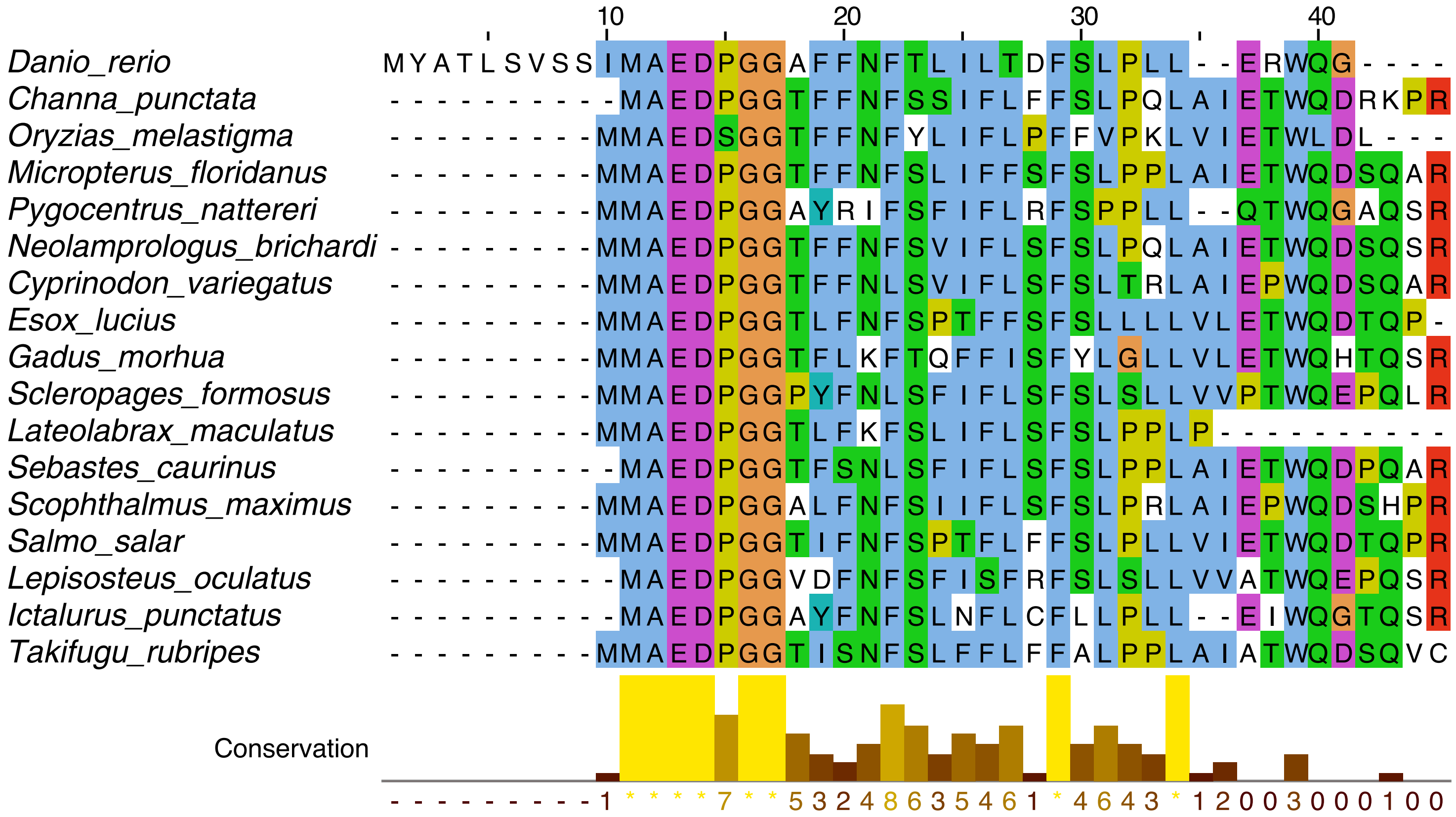

**HG0332**

ENSDARG00000102893 (atg7)

```
                                    10                  20                  30
                             .   |   .    |    .   |    .    |    .   |    .
Danio_rerio                  M K R N S V Y R F L S V F S C L L V C V F L F - F W G F A F Y S M V L A P - -
Amia_calva                   M K R N G V Y Q F F S V F S C L V V F L F L F L F W F F V F Y T M V M T P - -
Channa_argus                 M K R N G A Y R F L S V F S C F V V C A F L F L F W F F I F Y S L V L A P - -
Menidia_menidia              M K R N G A Y R F L S V F S C F I V C V F L F L F W F F V F Y S M M H A P - -
Porichthys_notatus           M K R N G A Y R F L S V F S C F V V C V F L F L F W F F V F Y S L V L A P - -
Oryzias_melastigma           M K R N G A Y R F L S V F S C F I V C A F L F L F W S F V F Y S M L R A P - -
Pygocentrus_nattereri        M K R T G A Y R F L S V F S C L L V C V F L F L F W F F V F Y S M V L A P - -
Haplochromis_burtoni         M K R N G A Y R F L S V F S C F I V C A F L F M F W F F V F Y S M V L A P - -
Poecilia_formosa             M K R N G A Y R F L S V F S C F V V C V F L F L F W F F V F Y S M V L A P - -
Esox_lucius                  - M R S G A Y K F L T V F S C L I V C G F M F M F W F F V F Y S F A P - - - -
Apteronotus_leptorhynchus    M K R D G A Y R L L S V F S C L V I F V F L L F F W V F V F Y S M V L A P - -
Plecoglossus_altivelis       - M K R G A Y K F L S V F S C F V V C A F L F L F W L F I F Y S M M L A P - -
Scleropages_formosus         M K K I C A Y R L L S V L S C V G I C L F L F L F W F F V F Y T L V M A P - -
Lateolabrax_japonicus        M K R N G A Y K F L S V F S C F I V C A F L L L F W F F I F Y S M V L A P - -
Platichthys_stellatus        M K R S G A Y R F L S V L S F F A V C A S L L L F W F I I F Y T M M H V T - -
Trichomycterus_areolatus     M K R N G A Y R F L S V F S C F A V C A F I F I F W F F V F Y S M F L A P E N
Takifugu_rubripes            M K R N G A Y K F L S V F S C F I V C A F L F I F W F F I F Y S M V H A P - -
```

Conservation

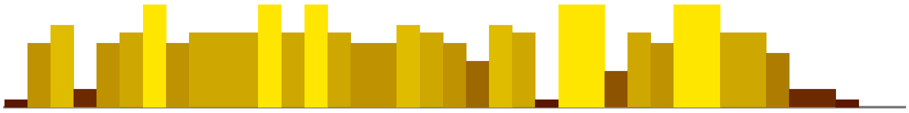

```
1 7 9 2 7 8 * 7 8 8 8 * 8 * 8 7 7 9 8 7 5 9 8 1 * * 4 8 7 * * 8 8 6 2 2 1 - -
```

**HG0333**

ENSDARG00000042518 (hipk2)

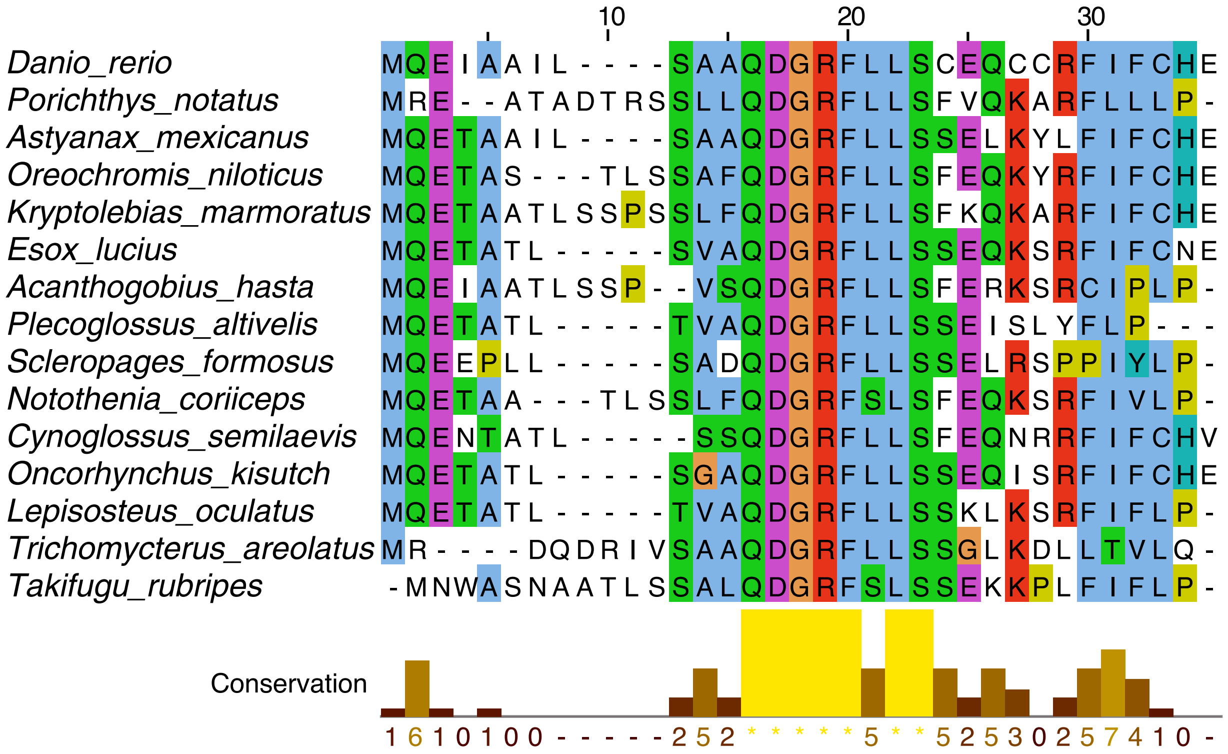

**HG0334**

ENSDARG00000075349

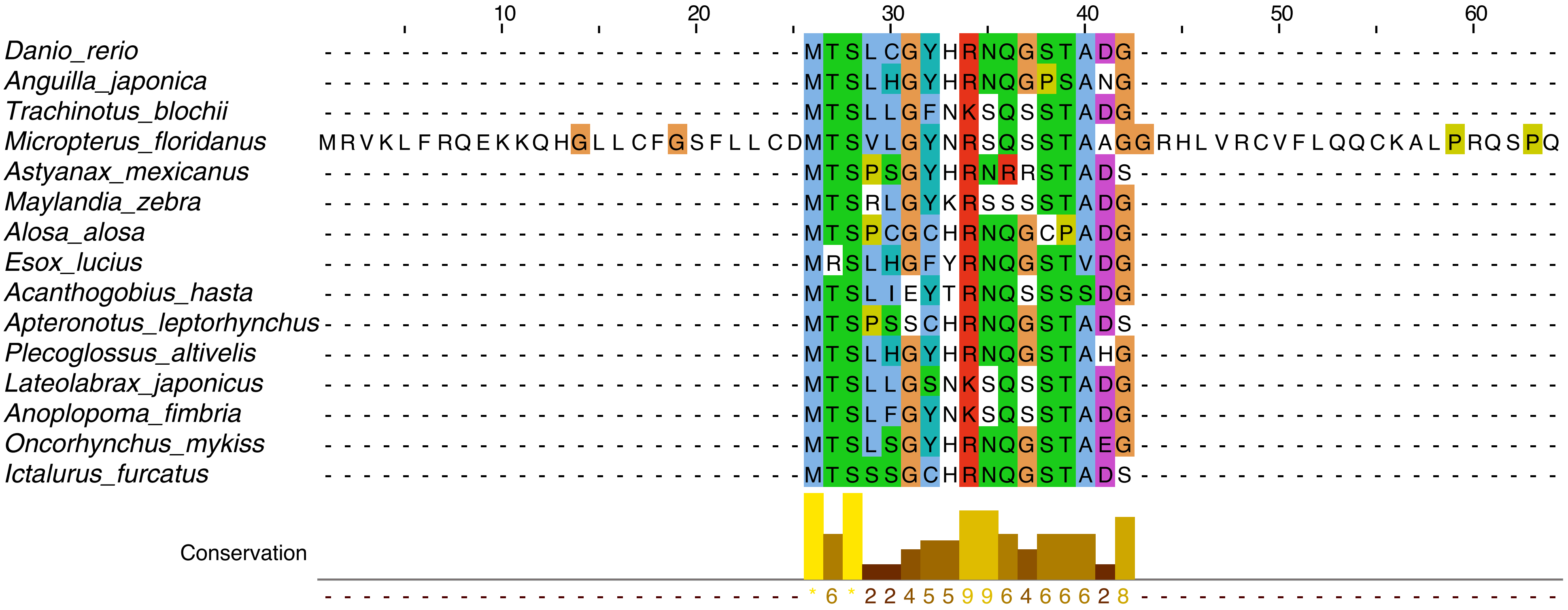

**HG0335**

ENSDARG00000034700 (vegfab)

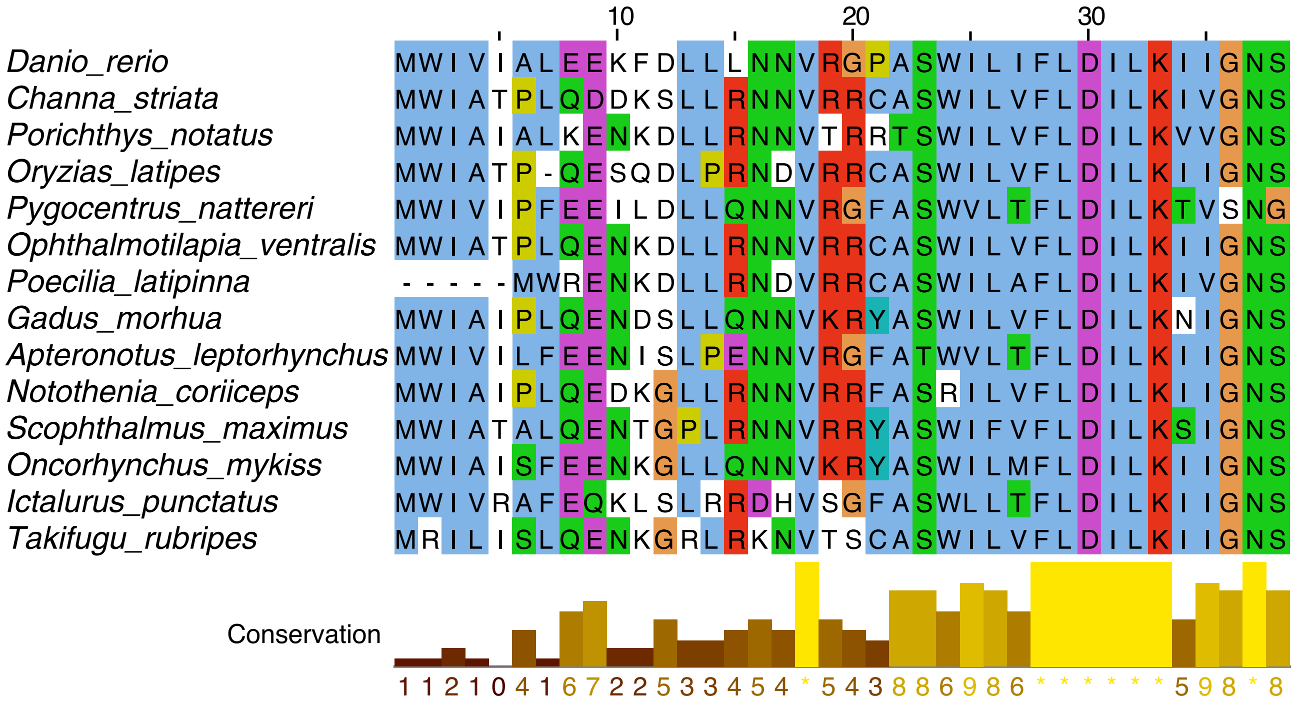

**HG0336**

ENSDARG00000028676 (nuak1b)

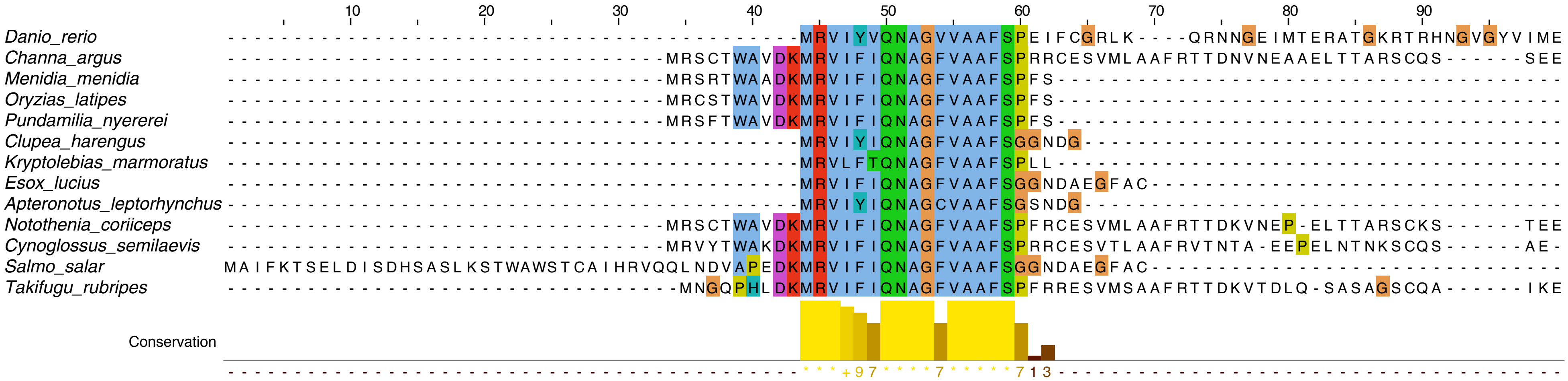

**HG0337**

ENSDARG00000054906

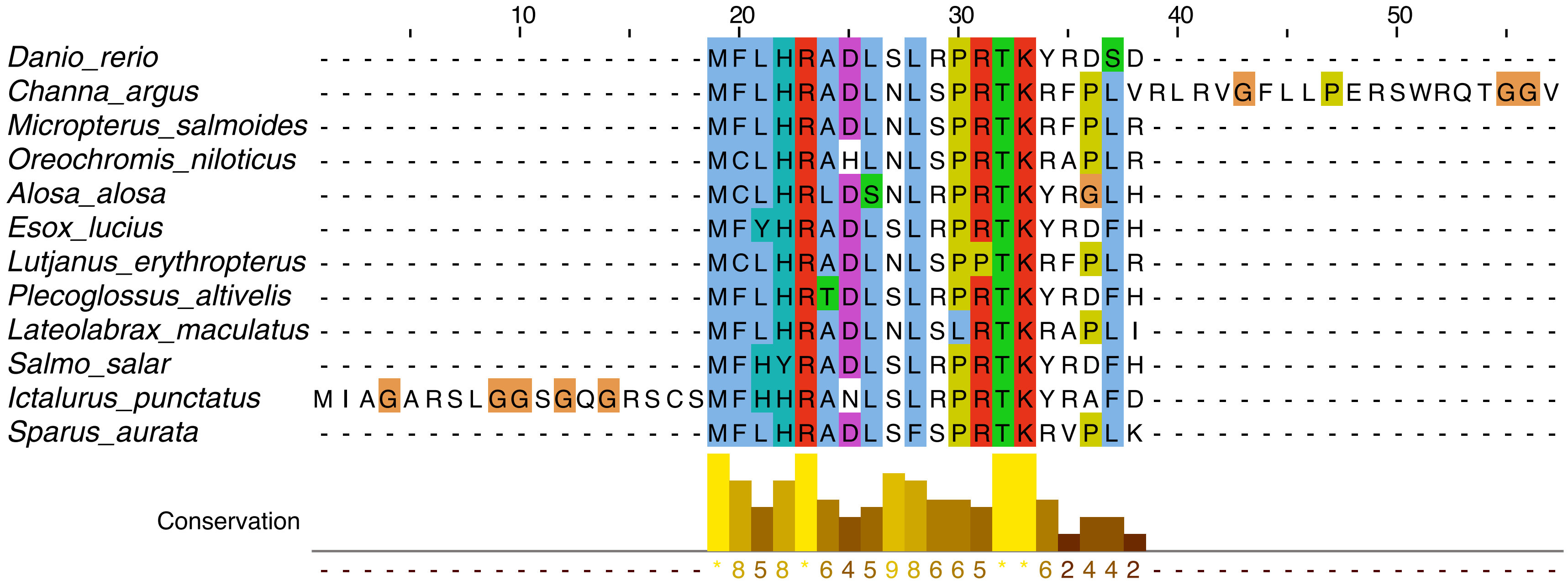

**HG0338**

ENSDARG00000000151 (thraa)

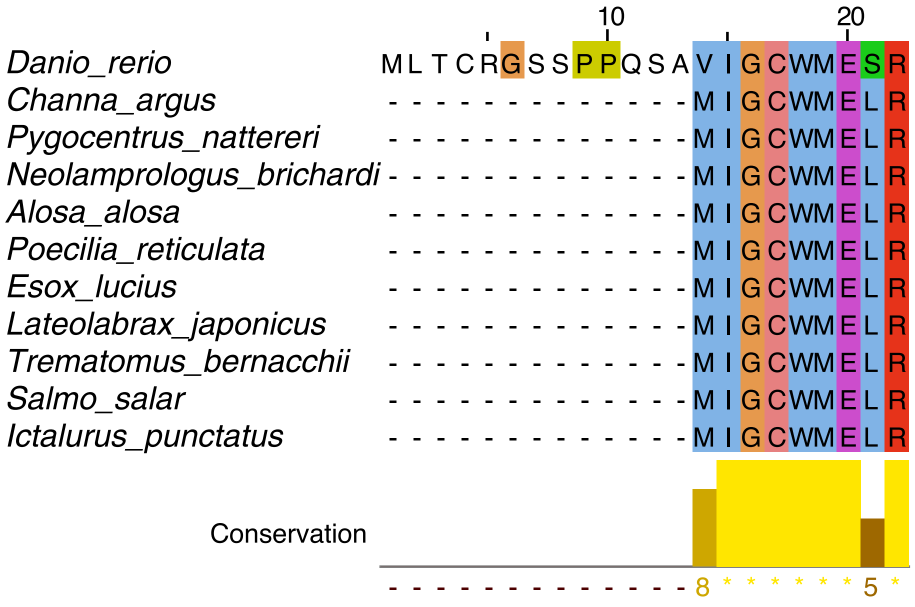

**HG0339**

ENSDARG00000063144 (shisa7b)

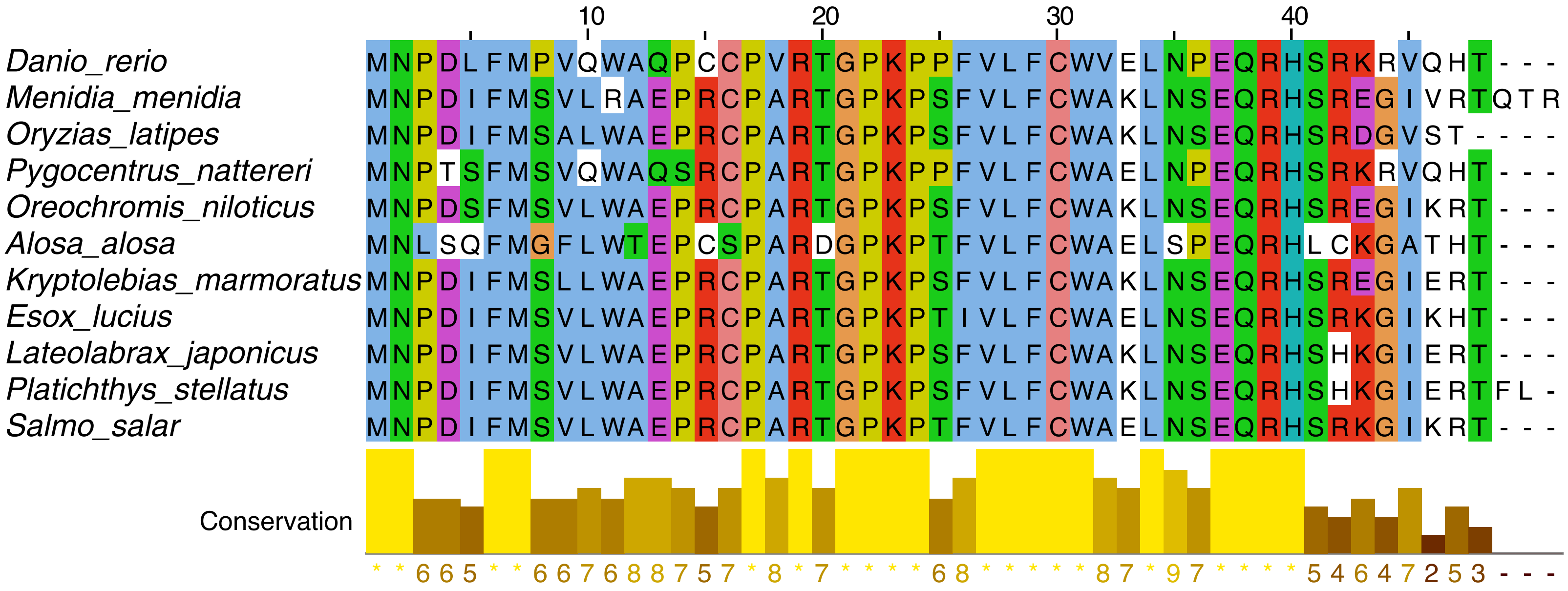

**HG0340**

ENSDARG00000017591 (fat1a)

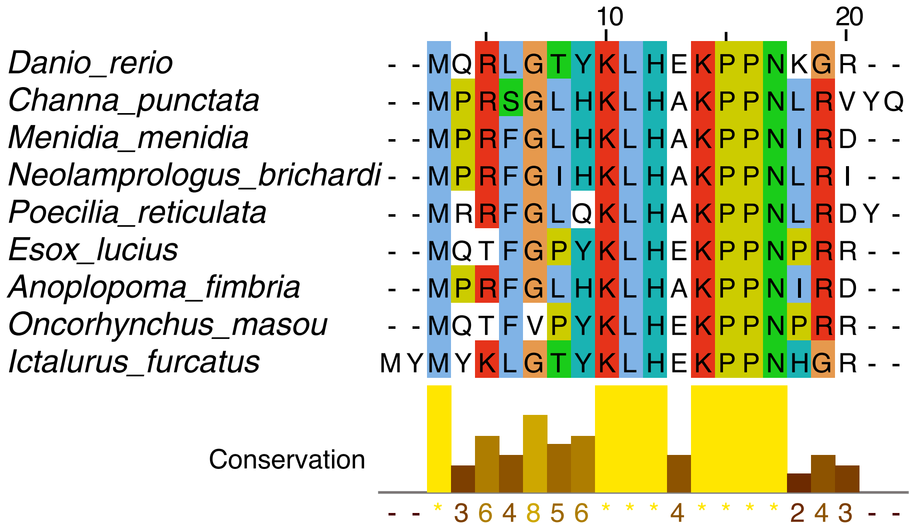

**HG0341**

ENSDARG00000039430 (msl2b)

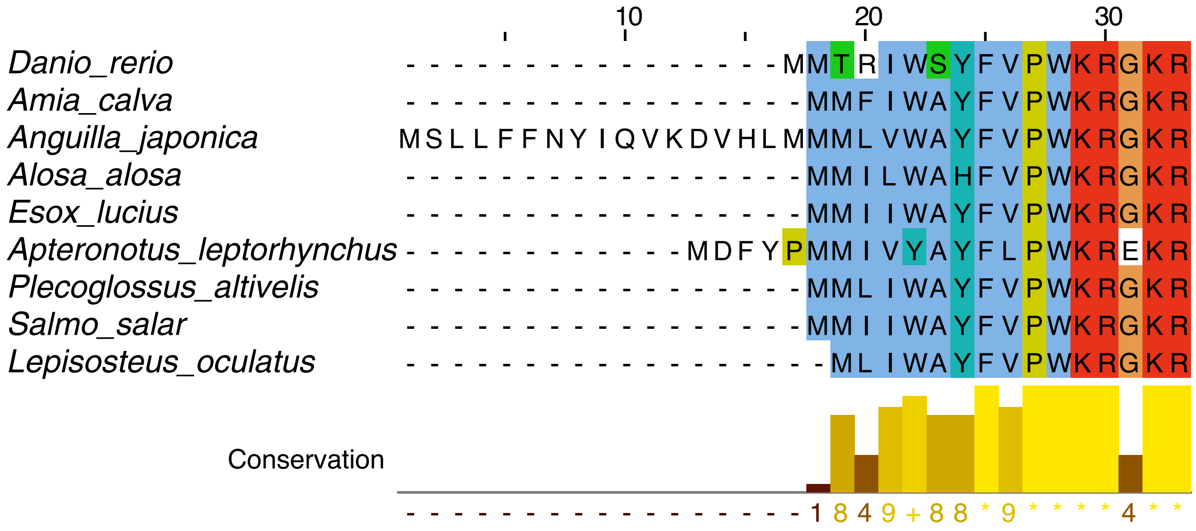

**HG0342**

ENSDARG00000045801 (fbxl14b)

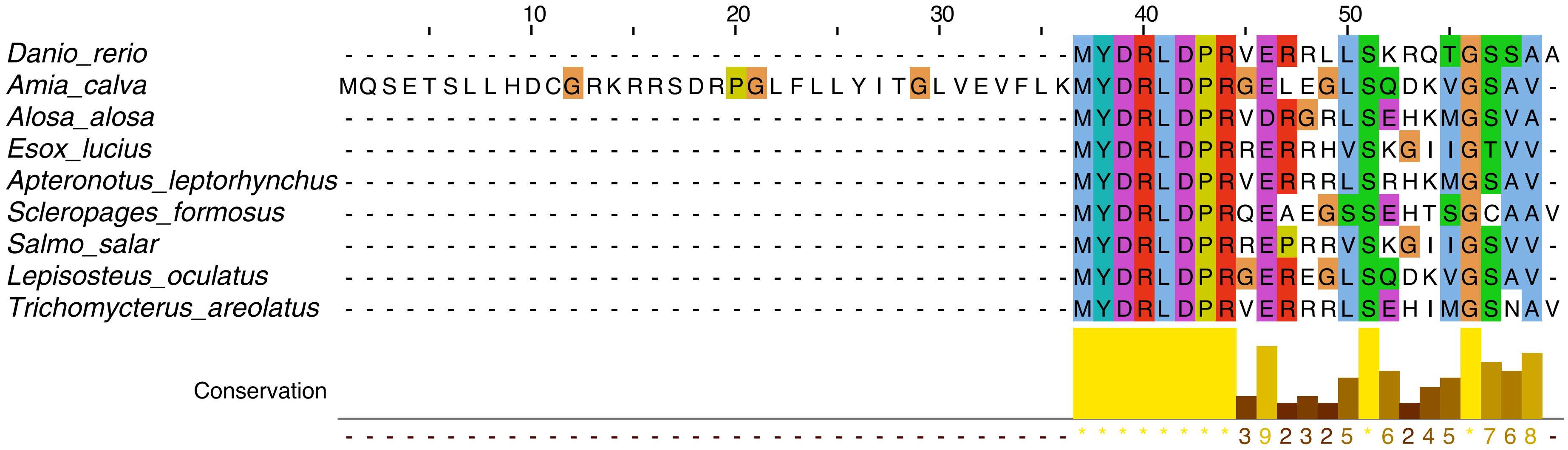

**HG0343**

ENSDARG00000057062

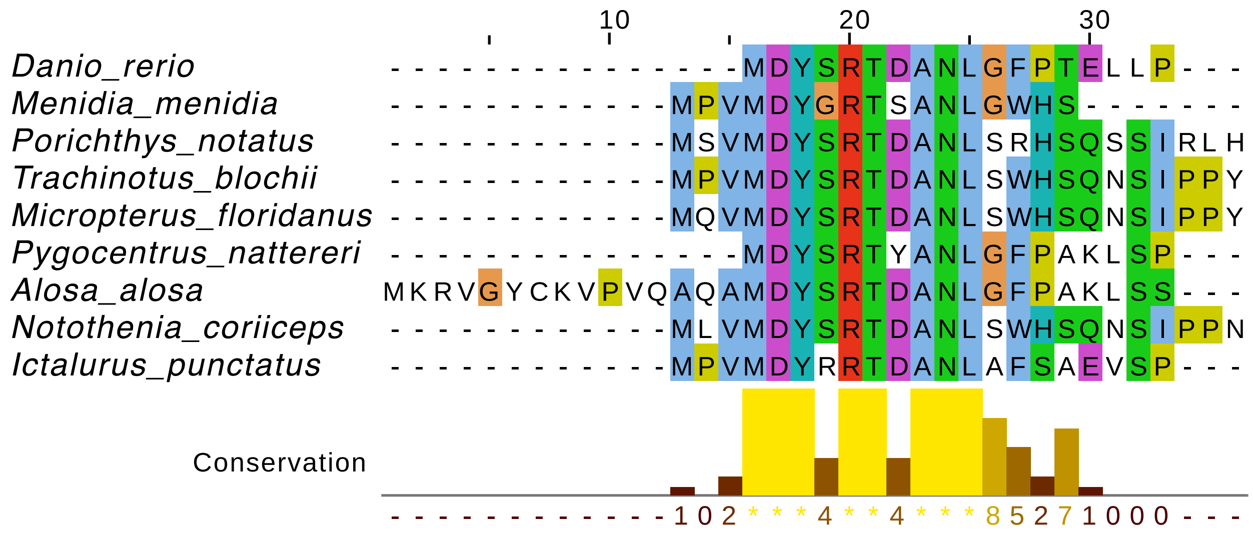

**HG0344**

ENSDARG00000032072 (cdk16)

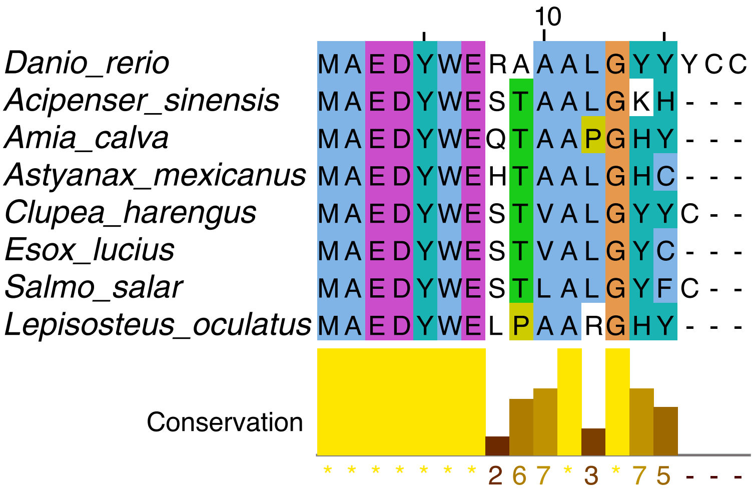

**HG0345**

ENSDARG00000035910 (znf281b)

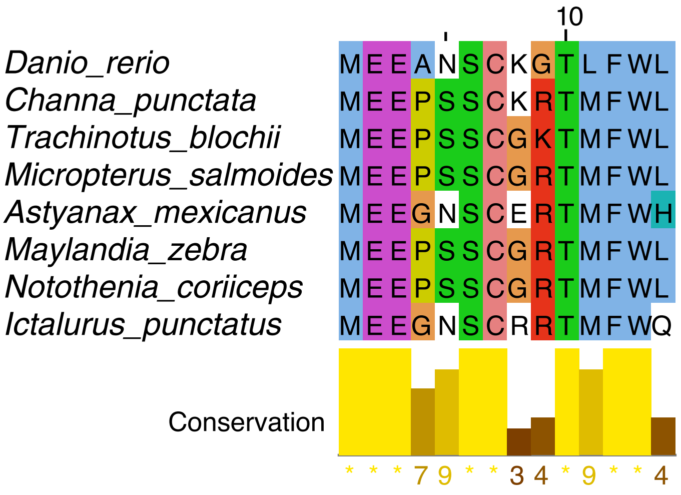

**HG0346**

ENSDARG00000041982 (adamts6)

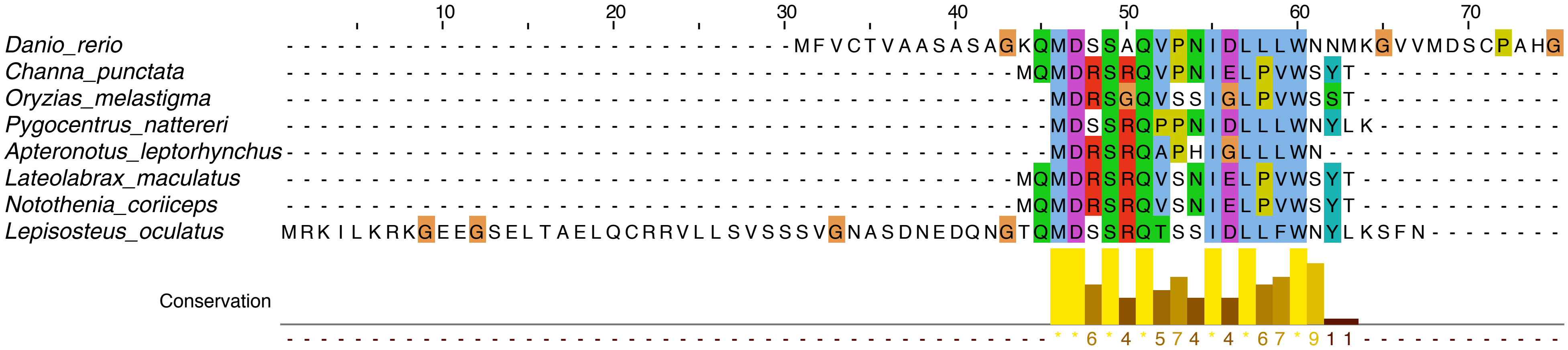

**HG0347**

ENSDARG00000063194

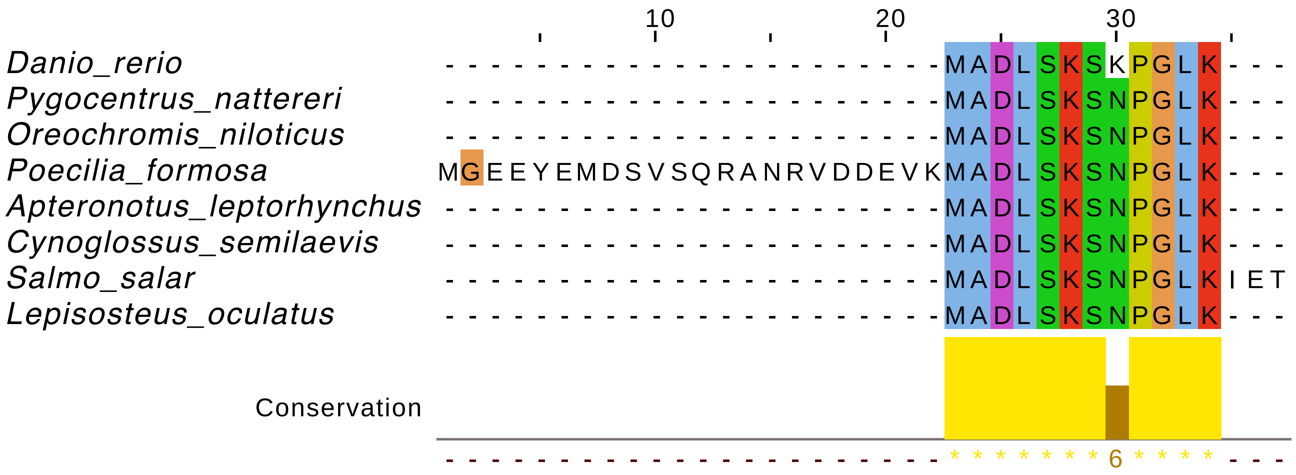

**HG0348**

ENSDARG00000036139 (lctla)

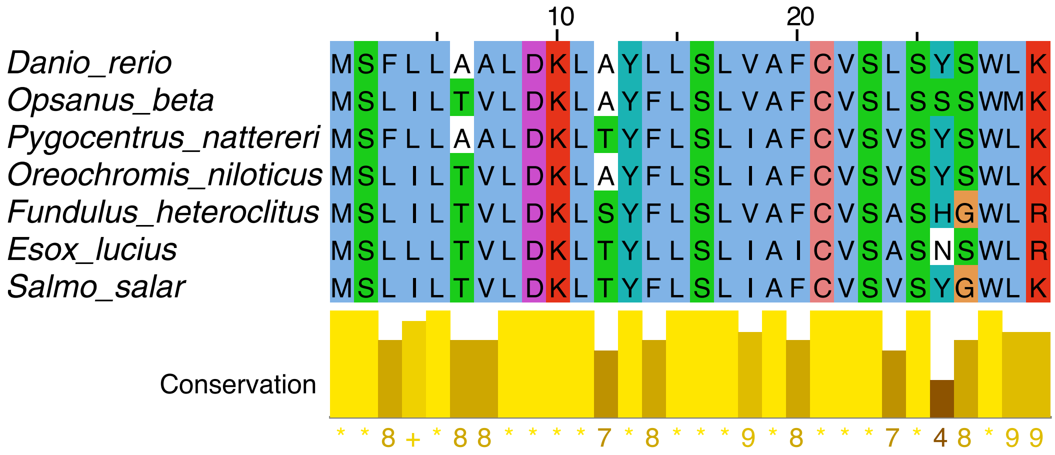

**HG0349**

ENSDARG00000028699 (crkl)

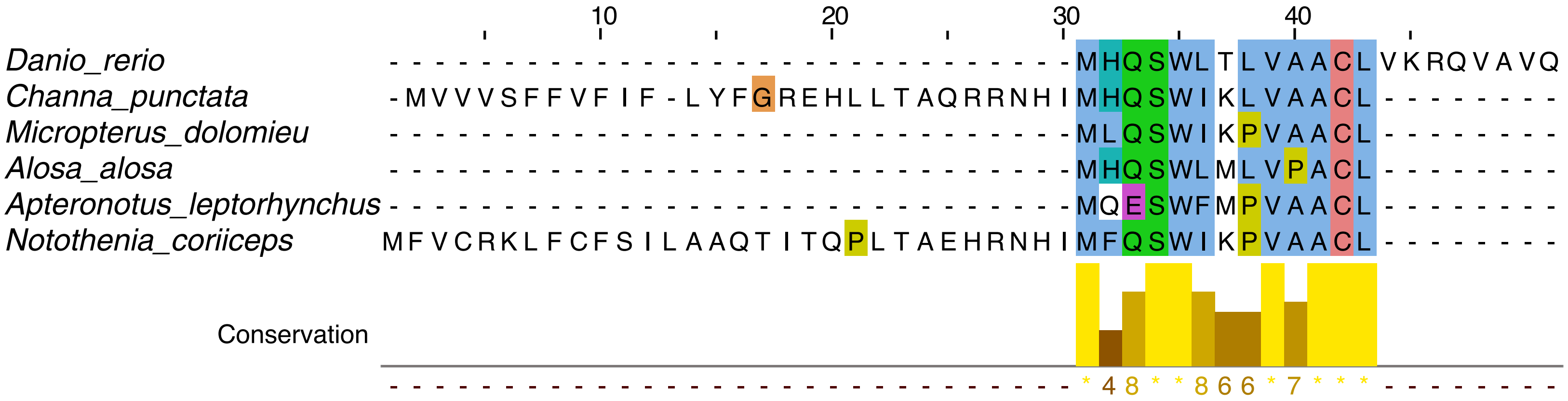

## HG0350

ENSDARG00000036194 (pitx2)

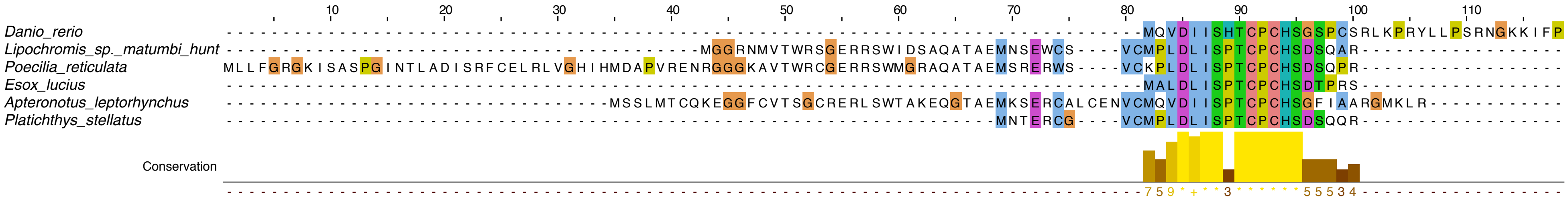

**HG0351**

ENSDARG00000068096 (atf5a)

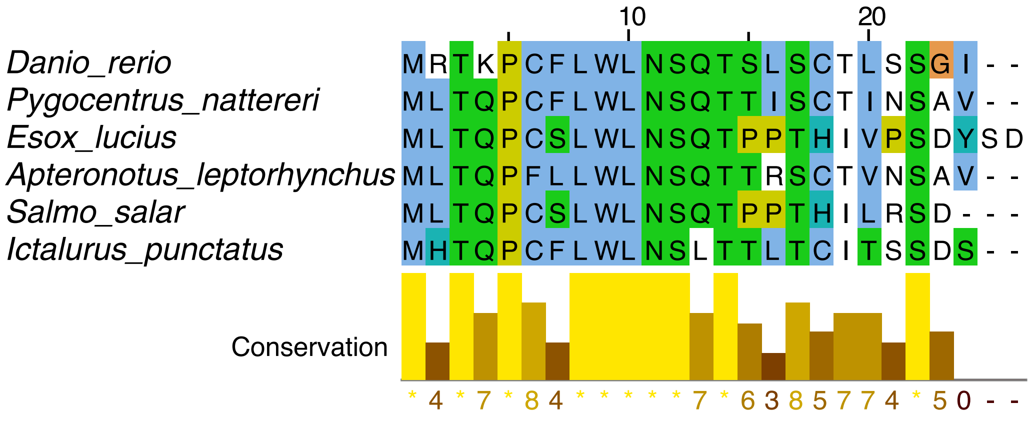

**HG0352**

ENSDARG00000070214 (slc41a1)

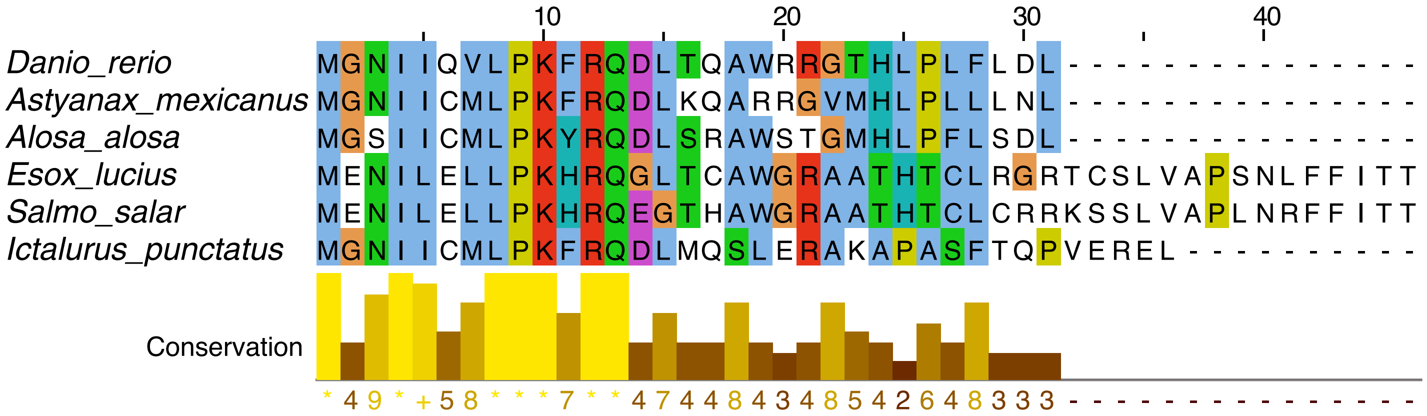

**HG0353**

ENSDARG00000078864 (lats2)

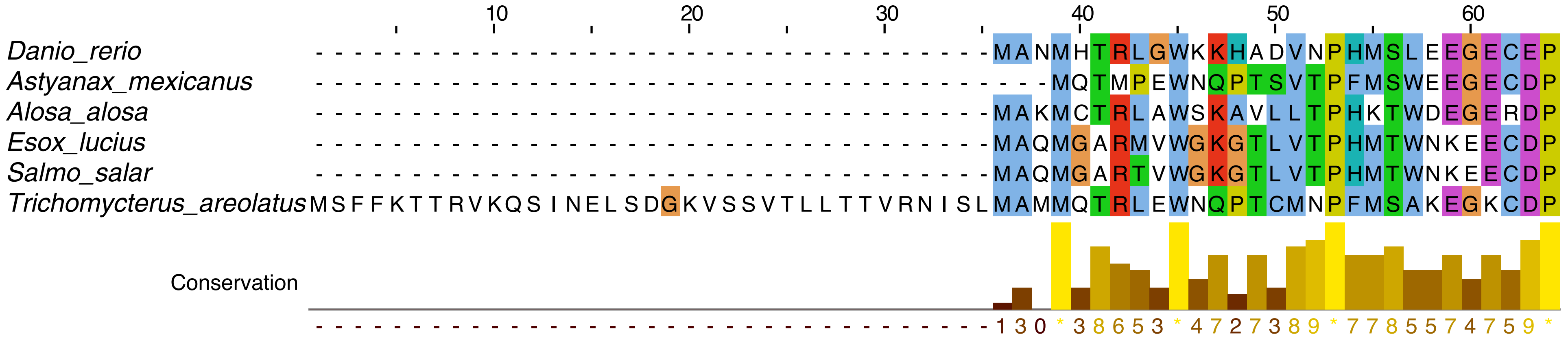

**HG0354**

ENSDARG00000019897 (gpat4)

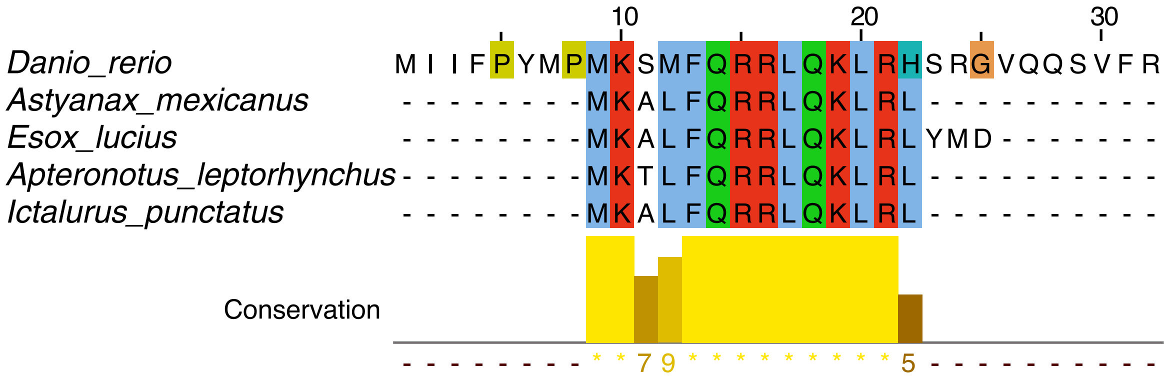

**HG0355**

ENSDARG00000019930 (tal1)

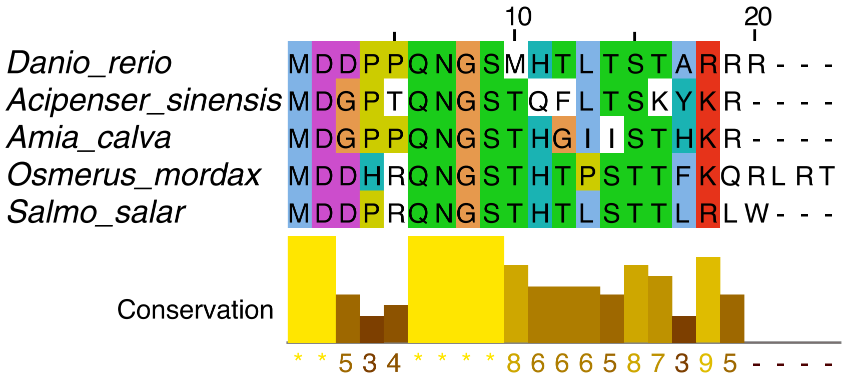

**HG0356**

ENSDARG00000020475 (znrf1)

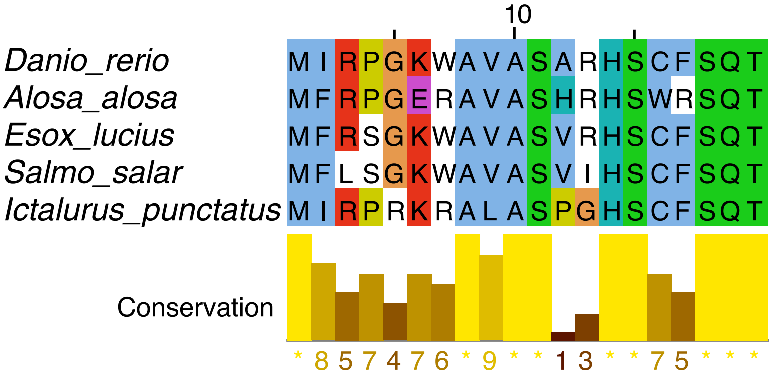

**HG0357**

ENSDARG00000037285 (mipa)

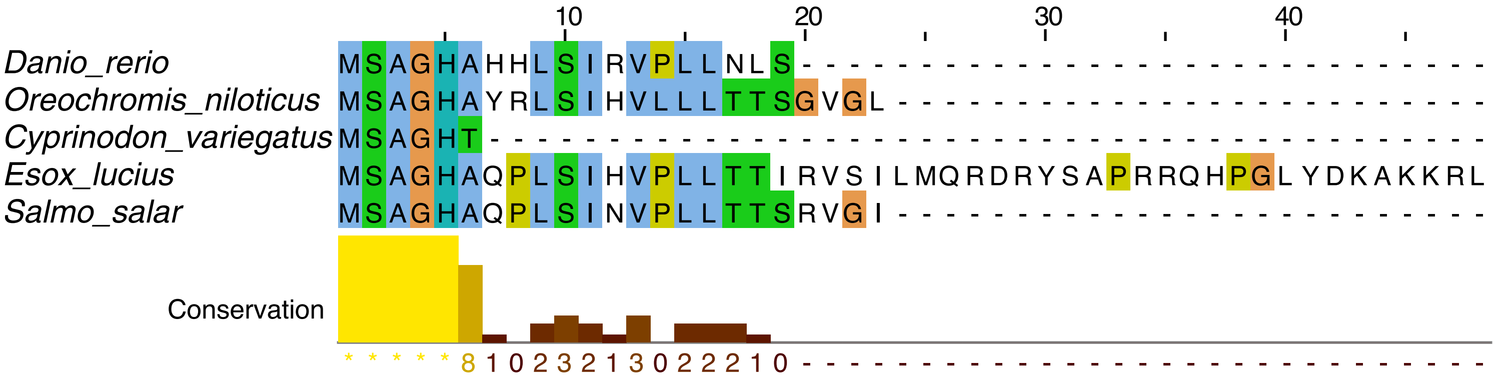

**HG0358**

ENSDARG00000062506 (zc3h18)

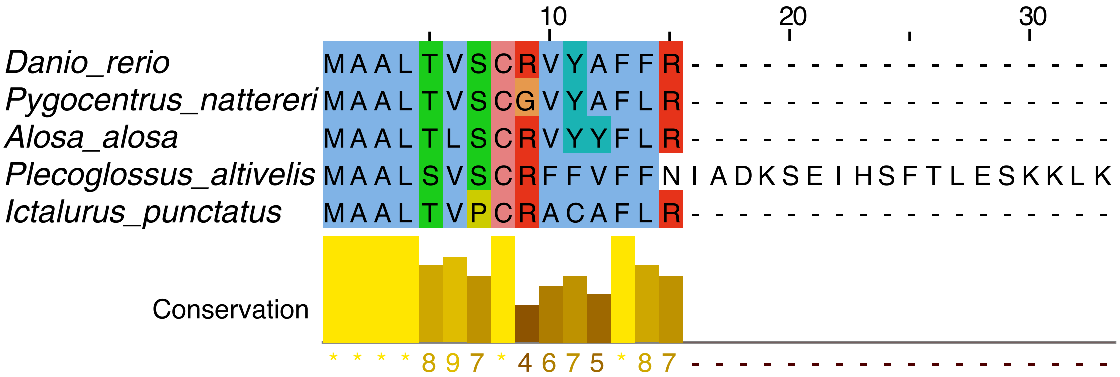

**HG0359**

ENSDARG00000070348 (hoxc10a)

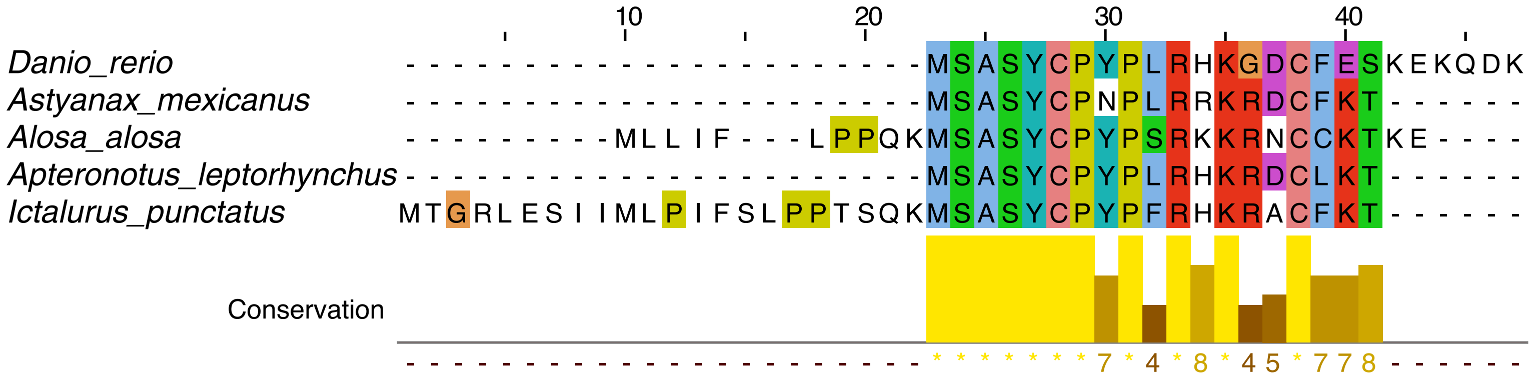

**HG0360**

ENSDARG00000070801 (opa1)

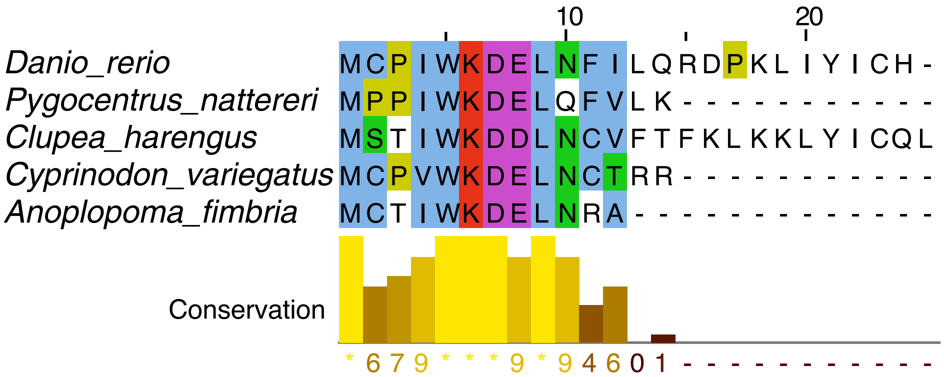

**HG0361**

ENSDARG00000077581 (zzz3)

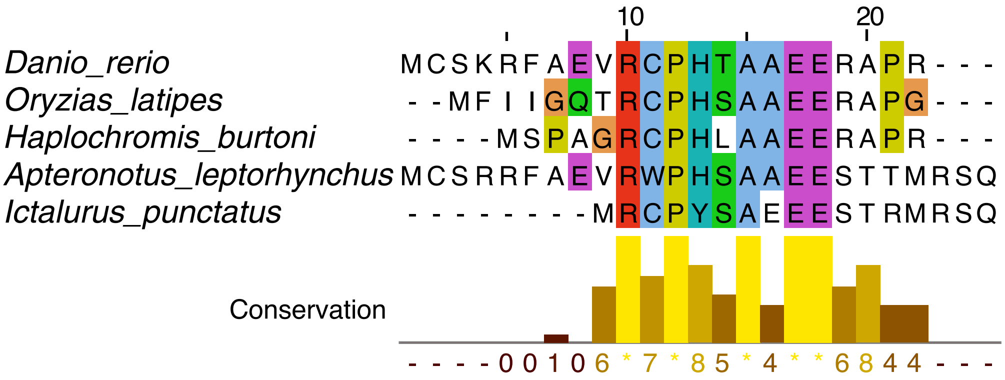

**HG0362**

ENSDARG00000078355 (zc3h4)

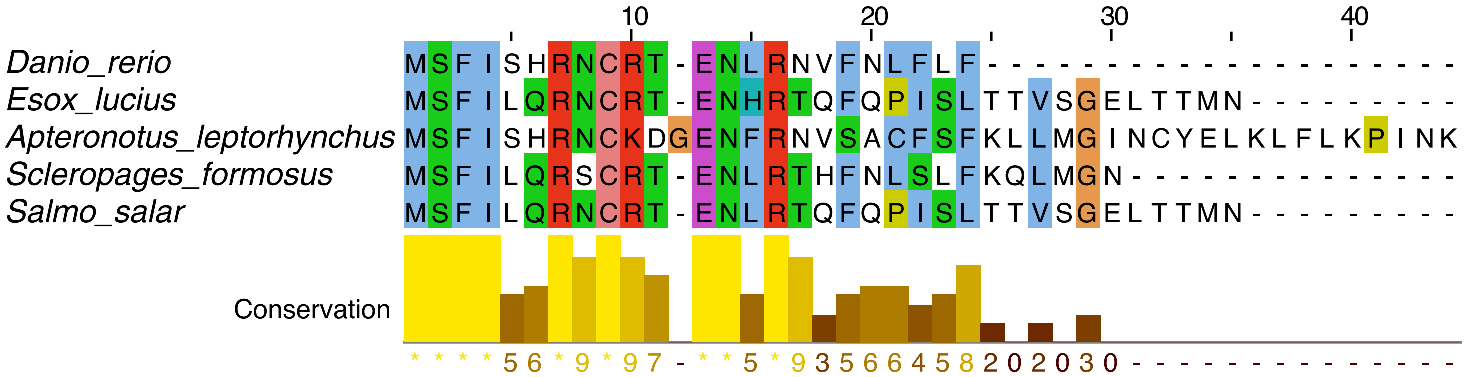

**HG0363**

ENSDARG00000095896 (pou3f3b)

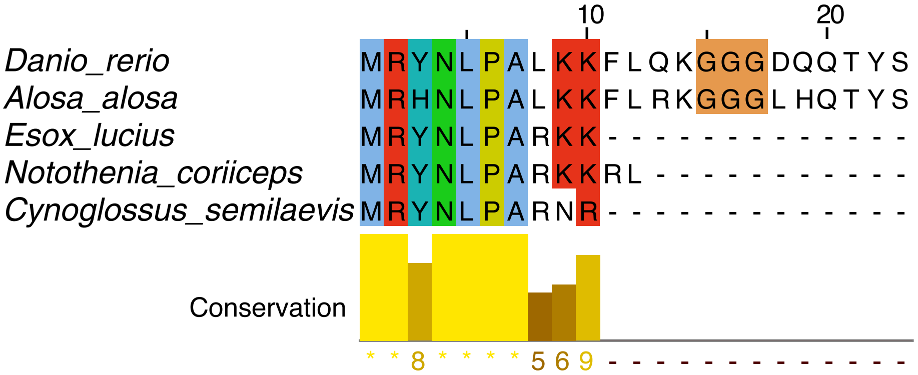

**HG0364**

ENSDARG00000004988 (ppp3ca)

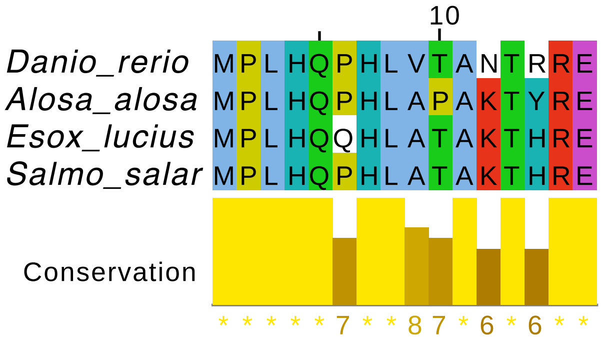

**HG0365**

ENSDARG00000016132 (keap1a)

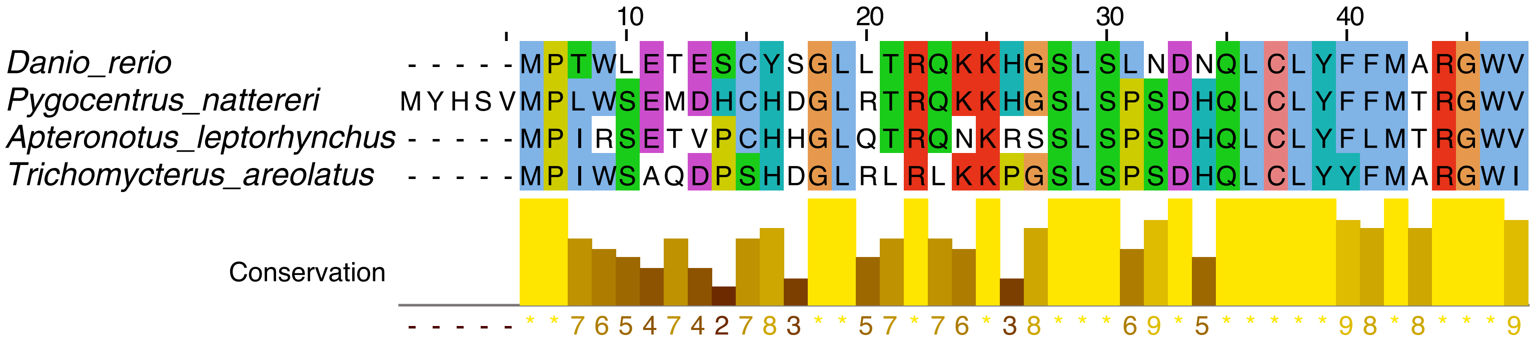

**HG0366**

ENSDARG00000017803 (gsk3b)

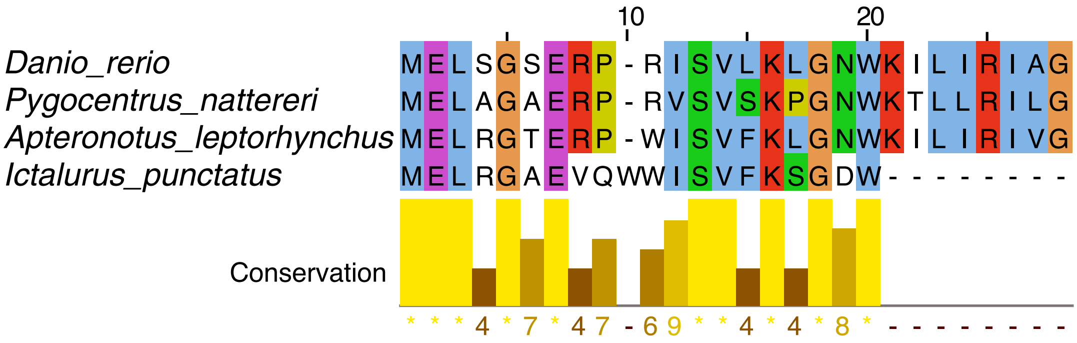

**HG0367**

ENSDARG00000034056 (csnk1g2b)

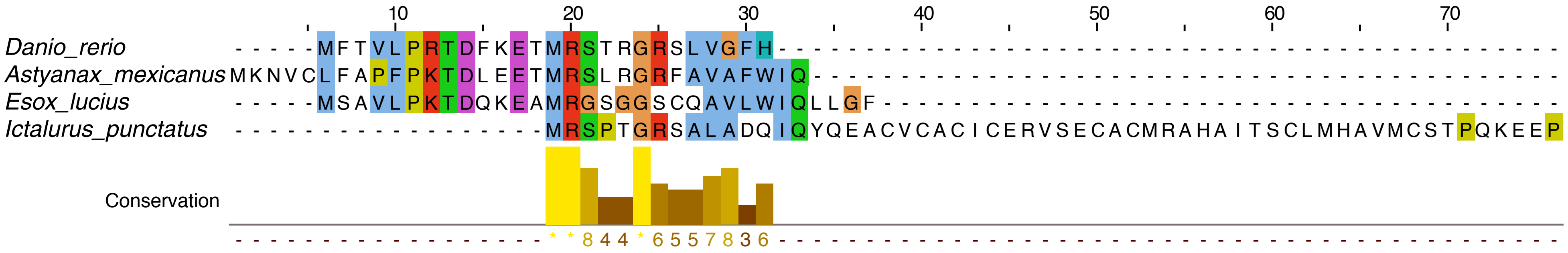

**HG0368**

ENSDARG00000058606 (sik1)

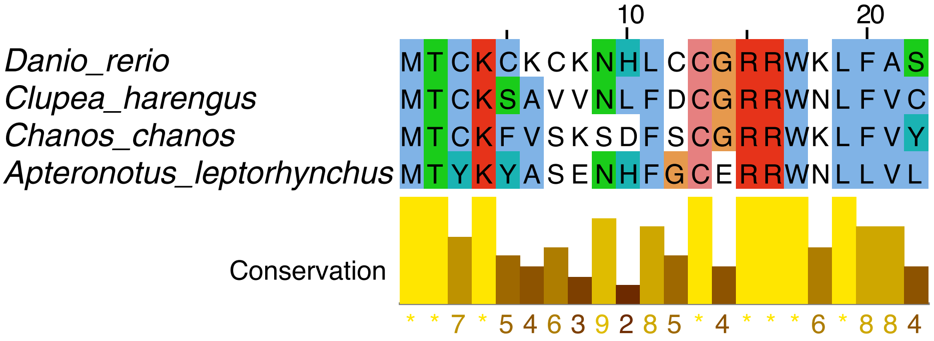

**HG0369**

ENSDARG00000059276 (hoxd4a)

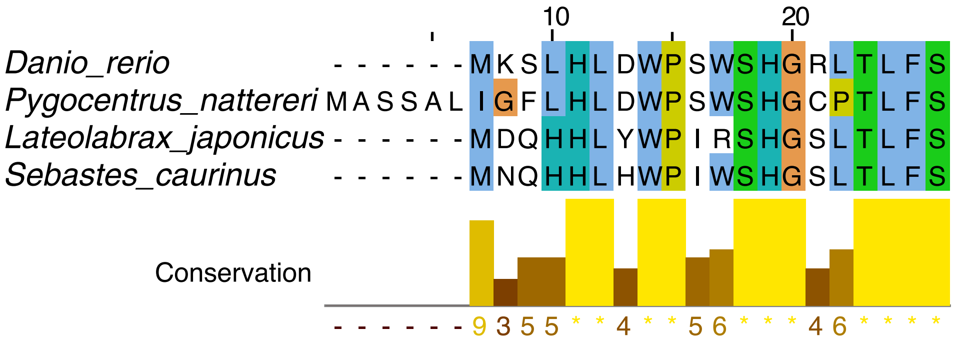

**HG0370**

ENSDARG00000062693 (nrxn3b)

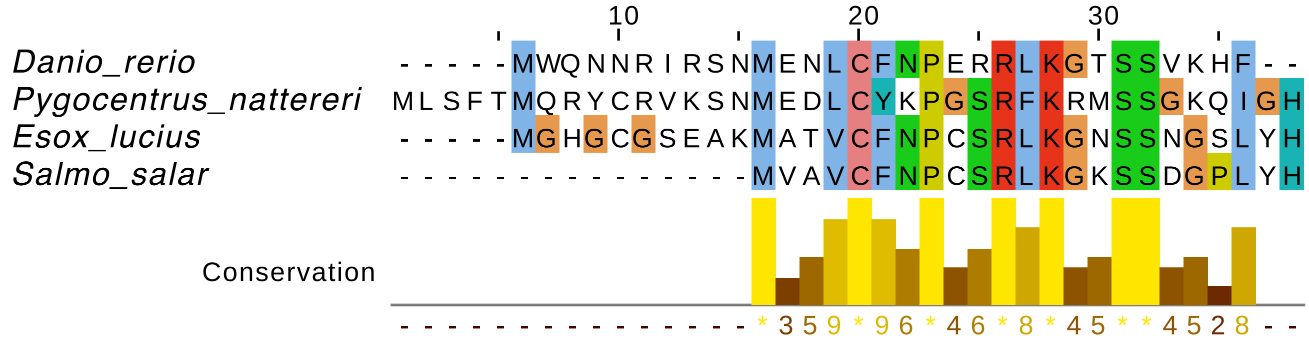

**HG0371**

ENSDARG00000062765 (whsc1l1)

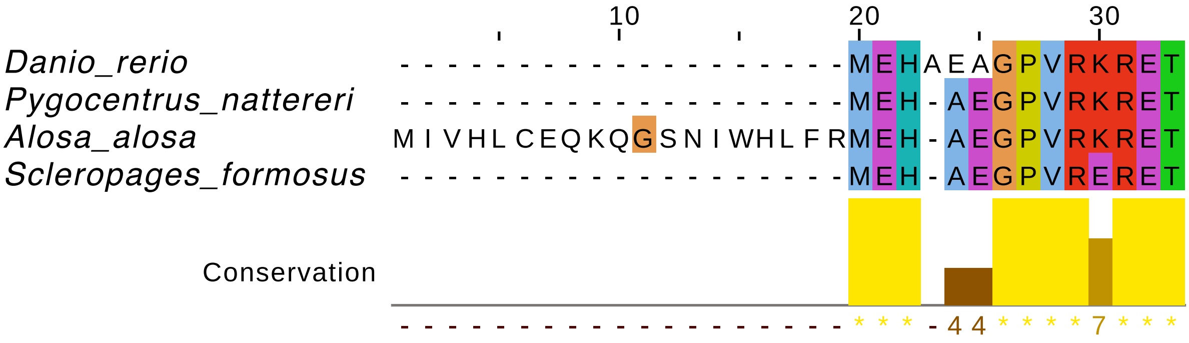

**HG0372**

ENSDARG00000069467 (igsf9bb)

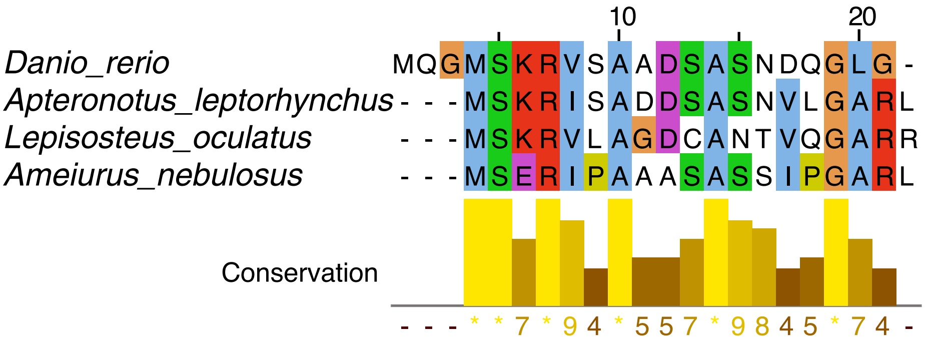

**HG0373**

ENSDARG00000074611 (wdr37)

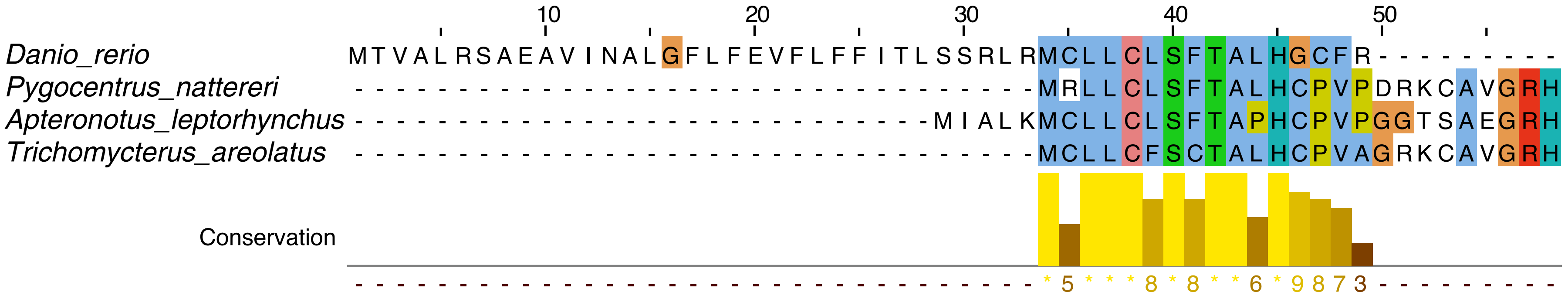

**HG0374**

ENSDARG00000075147 (lrrc38a)

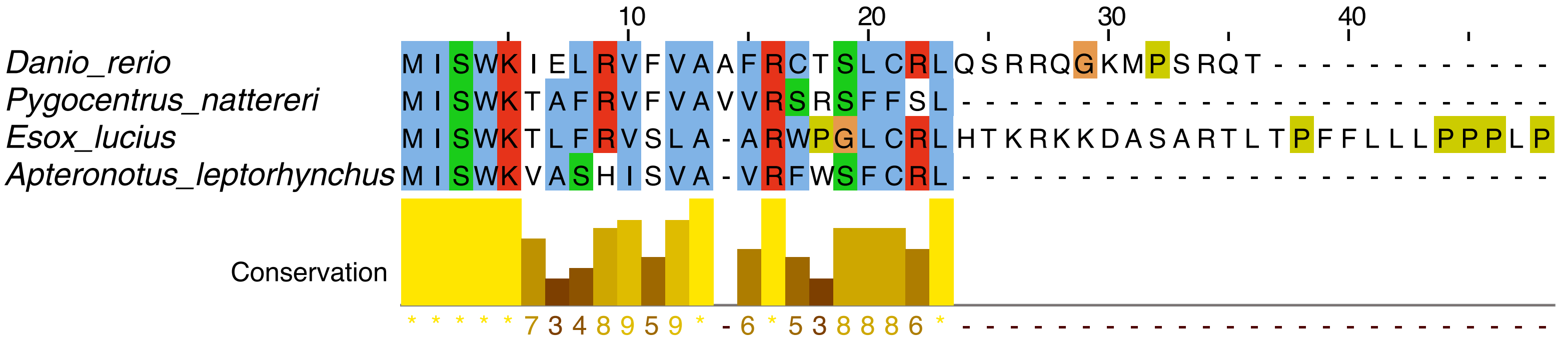

**HG0375**

ENSDARG00000076171 (znf827)

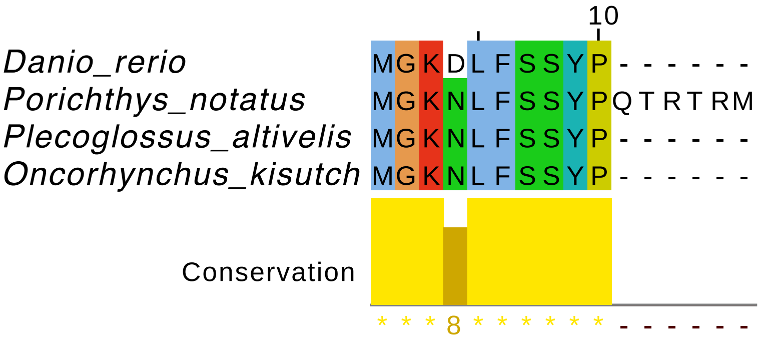

**HG0376**

ENSDARG00000077229 (ano8b)

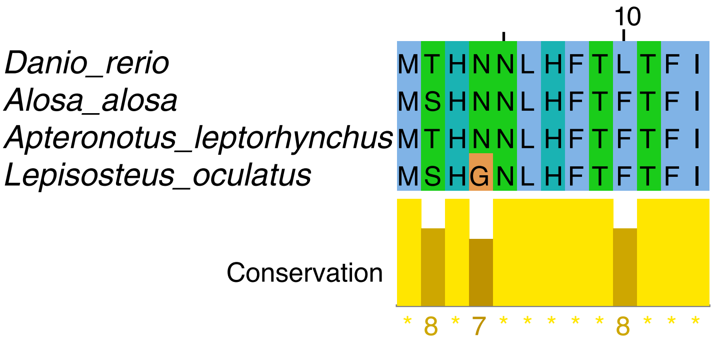

**HG0377**

ENSDARG00000079549 (cdc42ep1a)

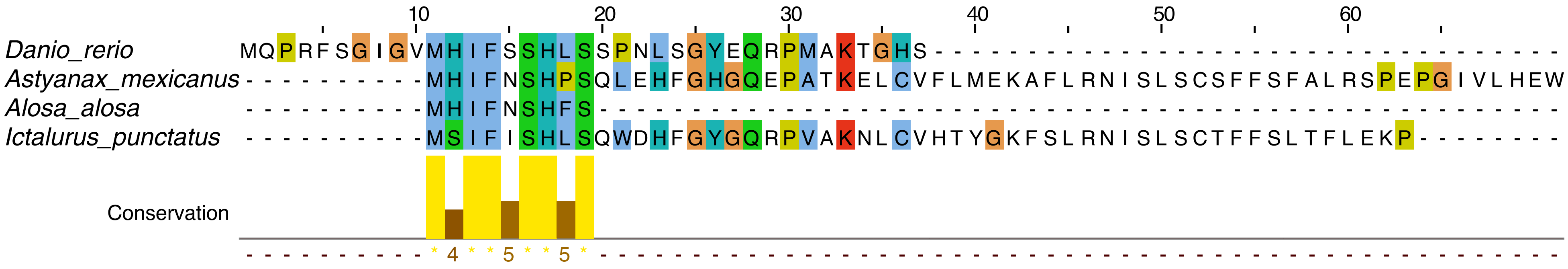

**HG0378**

ENSDARG00000080009 (bahcc1b)

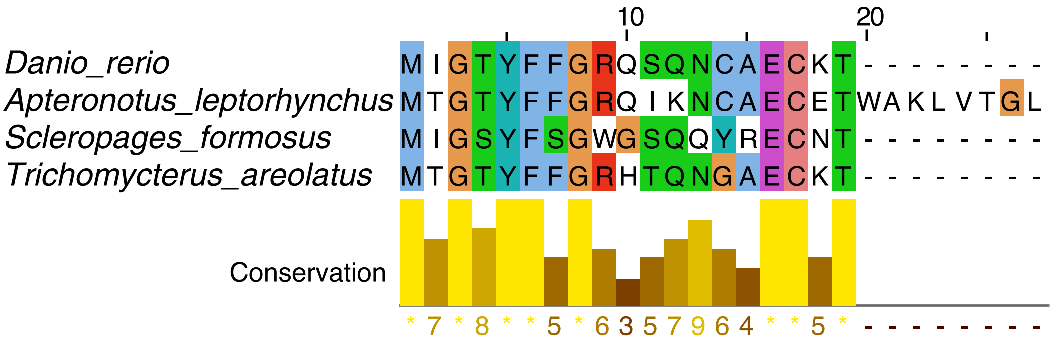

**HG0379**

ENSDARG00000104372 (gnb1b)

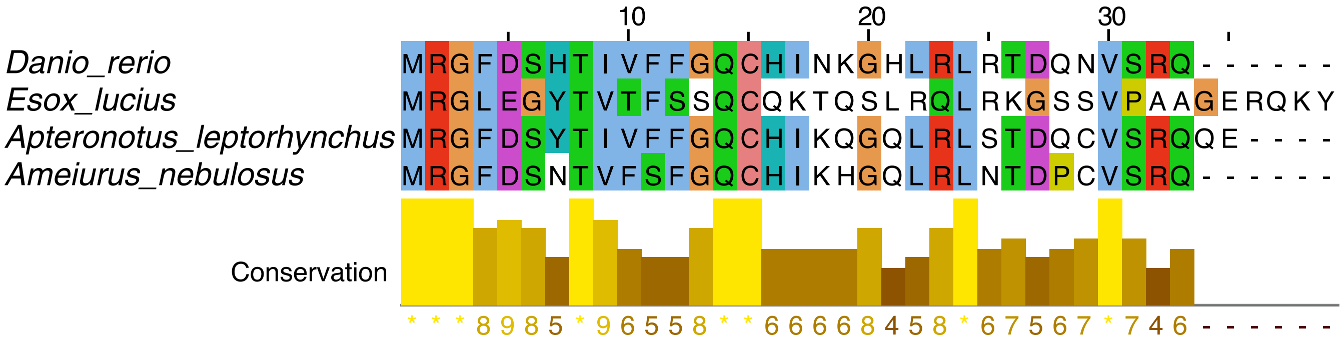

**HG0380**

ENSDARG00000005350 (dip2ba)

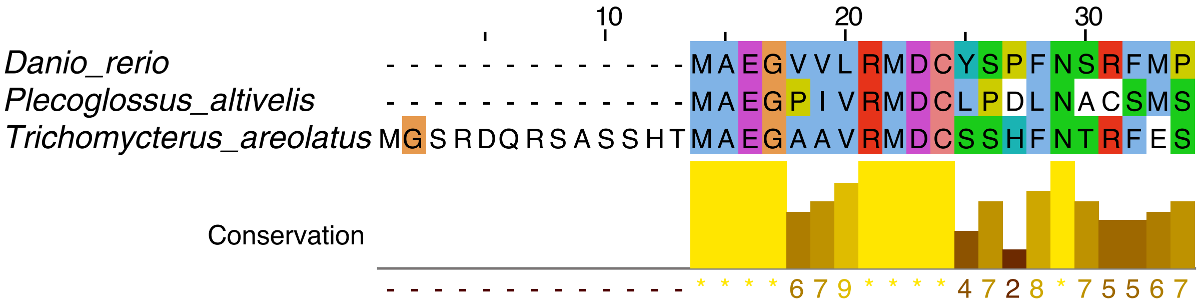

**HG0381**

ENSDARG00000006124 (kdm5c)

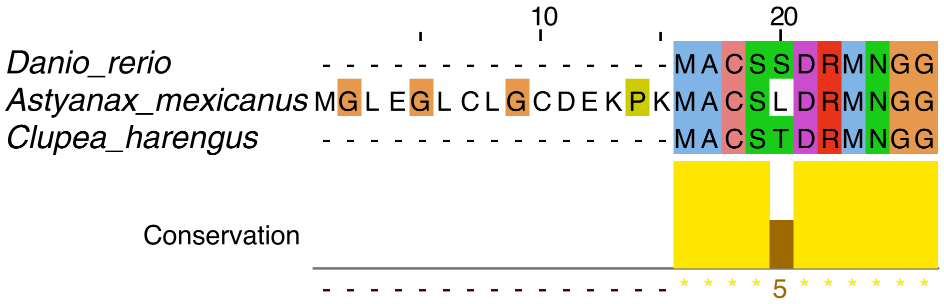

**HG0382**

ENSDARG00000017242

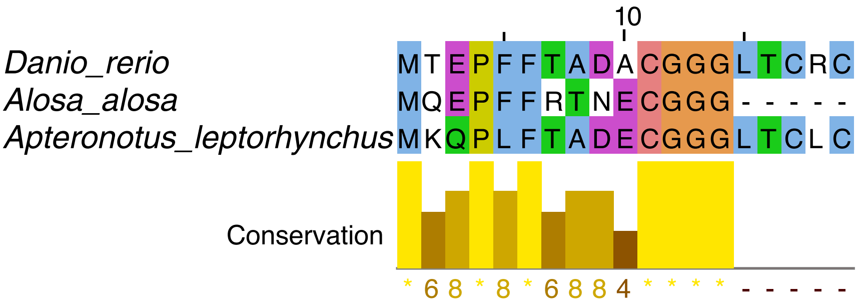

**HG0383**

ENSDARG00000036501 (rab39bb)

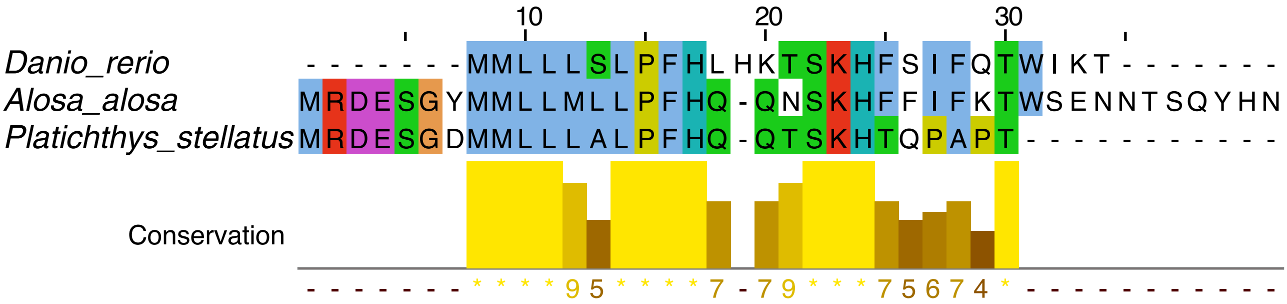

**HG0384**

ENSDARG00000054748 (cuedc1b)

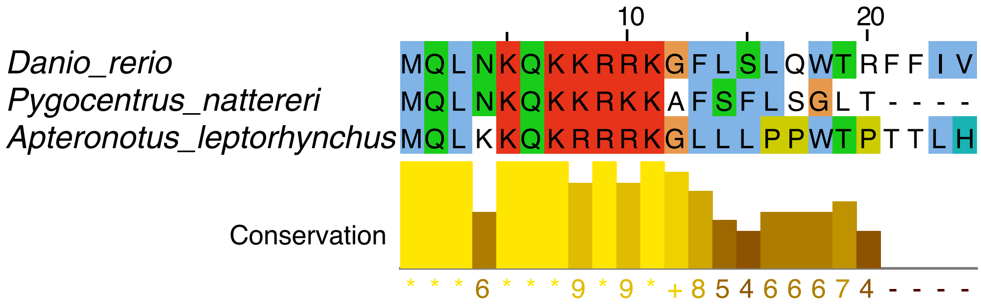

**HG0385**

ENSDARG00000059549 (akap10)

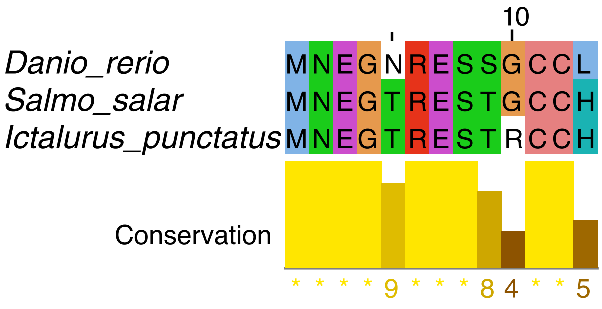

**HG0386**

ENSDARG00000062084 (clcn1a)

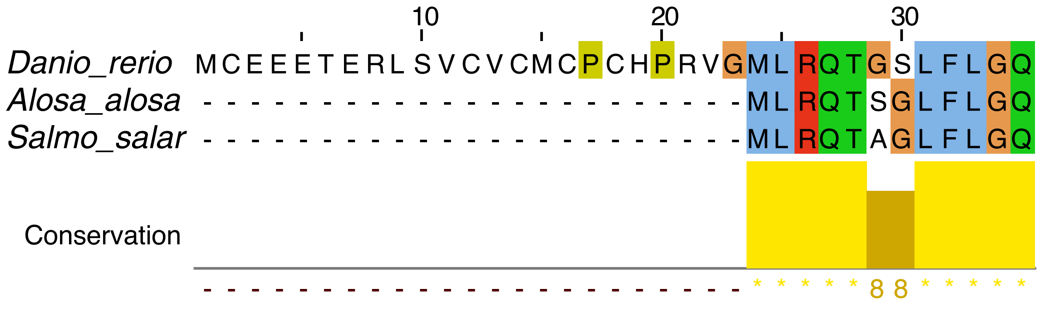

**HG0387**

ENSDARG00000071150 (cica)

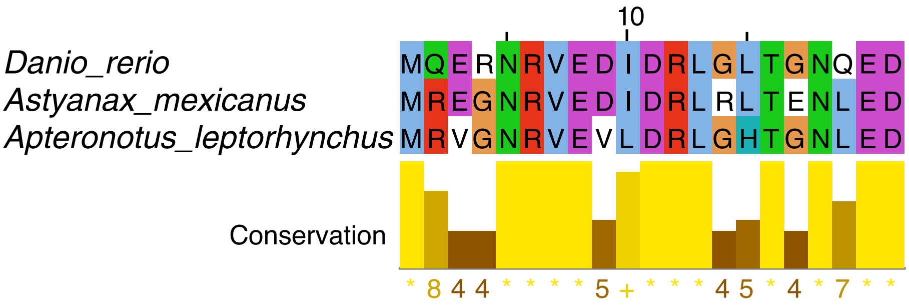

**HG0388**

ENSDARG00000077228 (ntrk3a)

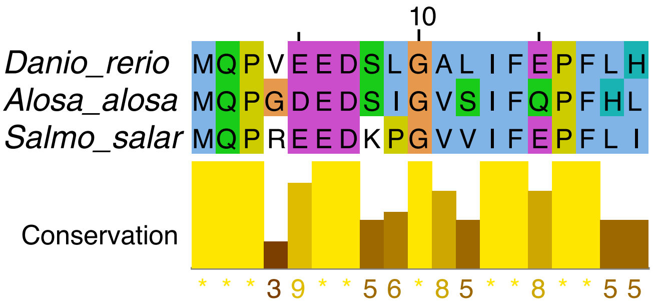

**HG0389**

ENSDARG00000077361 (bptf)

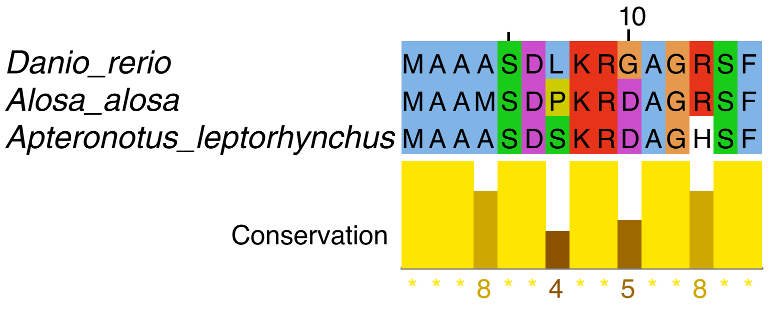

**HG0390**

ENSDARG00000090035 (rtn4r)

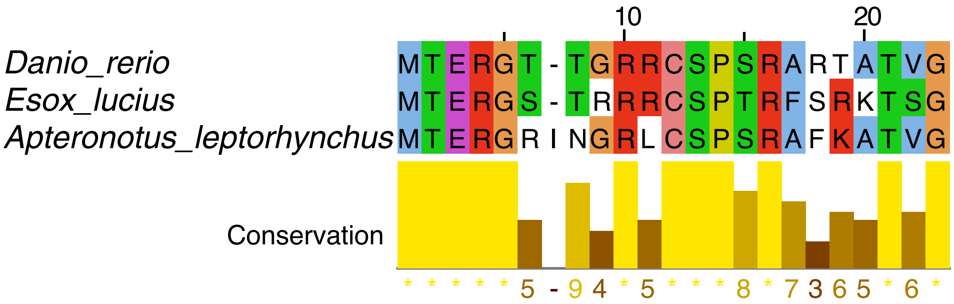

**HG0391**

ENSDARG00000099634 (elna)

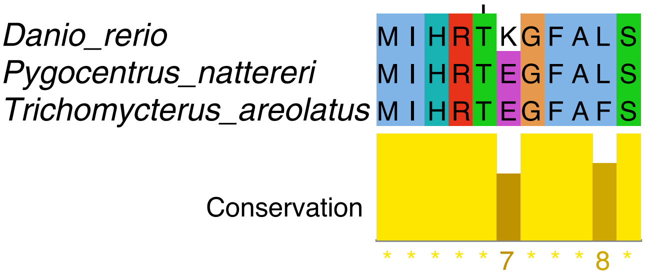

**HG0392**

ENSDARG00000100003 (glulb)

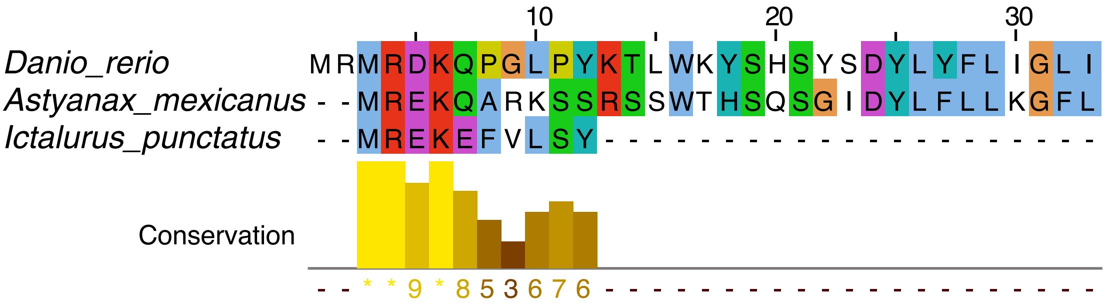

**HG0393**

ENSGALG00000002090 (GPSM2)

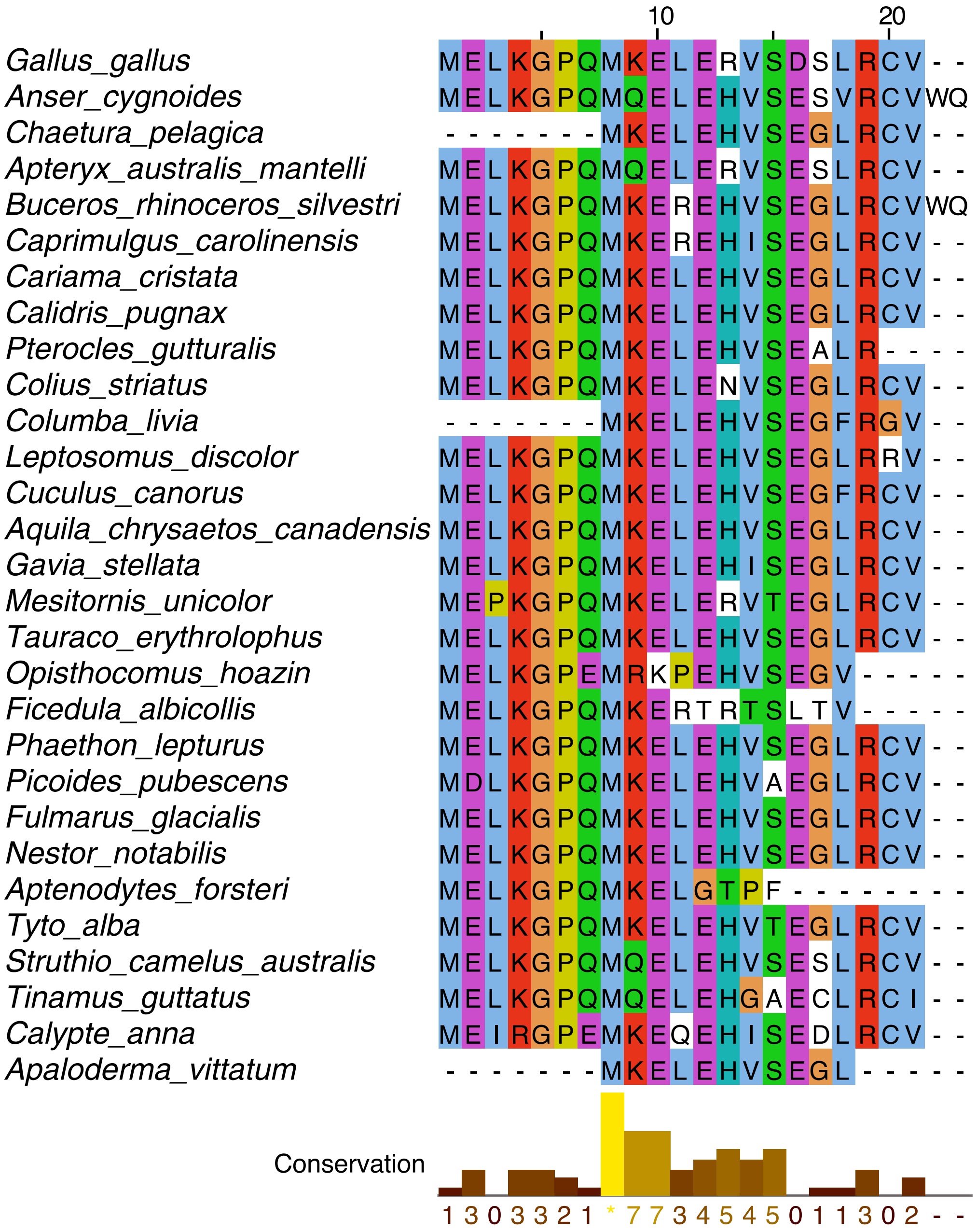

**HG0394**

ENSGALG00000002479 (MAT1A)

```
                                     10        20
Gallus_gallus                MRSPCMHTQIQAHADFTD----
Anser_cygnoides              MHTPCVHAQIQAHAEFTR----
Chaetura_pelagica            MHSPCMCTQMQAHADFTH----
Apteryx_australis_mantelli   MHTPCTHTQIQAHADFTH----
Caprimulgus_carolinensis     MHTPCMRTQMQAHADFTH----
Cariama_cristata             MHAPCMRTQMQAHAEFTH----
Charadrius_vociferus         MHTPCMRTQMQARADFTY----
Pterocles_gutturalis         MHTPCMRTQMQAHADFTH----
Colius_striatus              MHTPCMHAQMQAHAGFTH----
Columba_livia                MHTPCMCTQMQAHADFTH----
Merops_nubicus               ---------MQAHAEFTHSNLS
Aquila_chrysaetos_canadensis MHAPCMHTQMQAHADFTH----
Gavia_stellata               MHTACMRTQMQAHADFTH----
Chlamydotis_macqueenii       MHAPCMRTQMQAHADFTR----
Tauraco_erythrolophus        MHTPCMCTRMQAHADFTH----
Manacus_vitellinus           MRSPCVHAQMQAYAAFTH----
Egretta_garzetta             MHTPCMCTQMQAHADFTH----
Fulmarus_glacialis           MHTPCMRTQMQAHADFTH----
Nestor_notabilis             MHTPCMCTQMQAHAVFTH----
Pygoscelis_adeliae           MHTPCMRTQMQAHADFTH----
Tyto_alba                    MHTPCMRTQMQAHADFTH----
Struthio_camelus_australis   MHAPCTHT--------------
Tinamus_guttatus             MHTACTHSQLQARADFTH----
Calypte_anna                 MHAPCMCTQMKAHADFIH----
Apaloderma_vittatum          MHTPCVCTQMQTHADFTH----

Conservation                 131121011112130211----
```

**HG0395**

ENSGALG00000014936 (ESCO1)

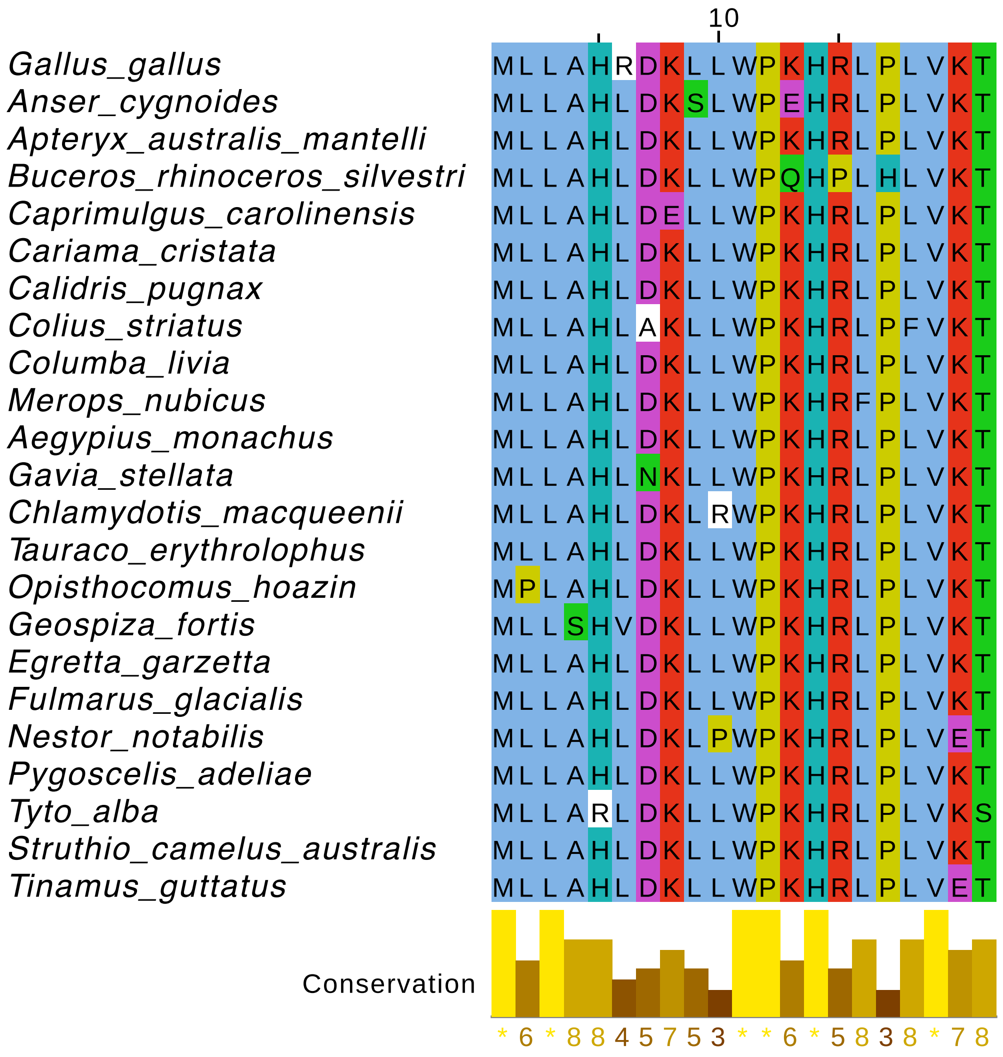

**HG0396.1**

ENSGALG00000034253

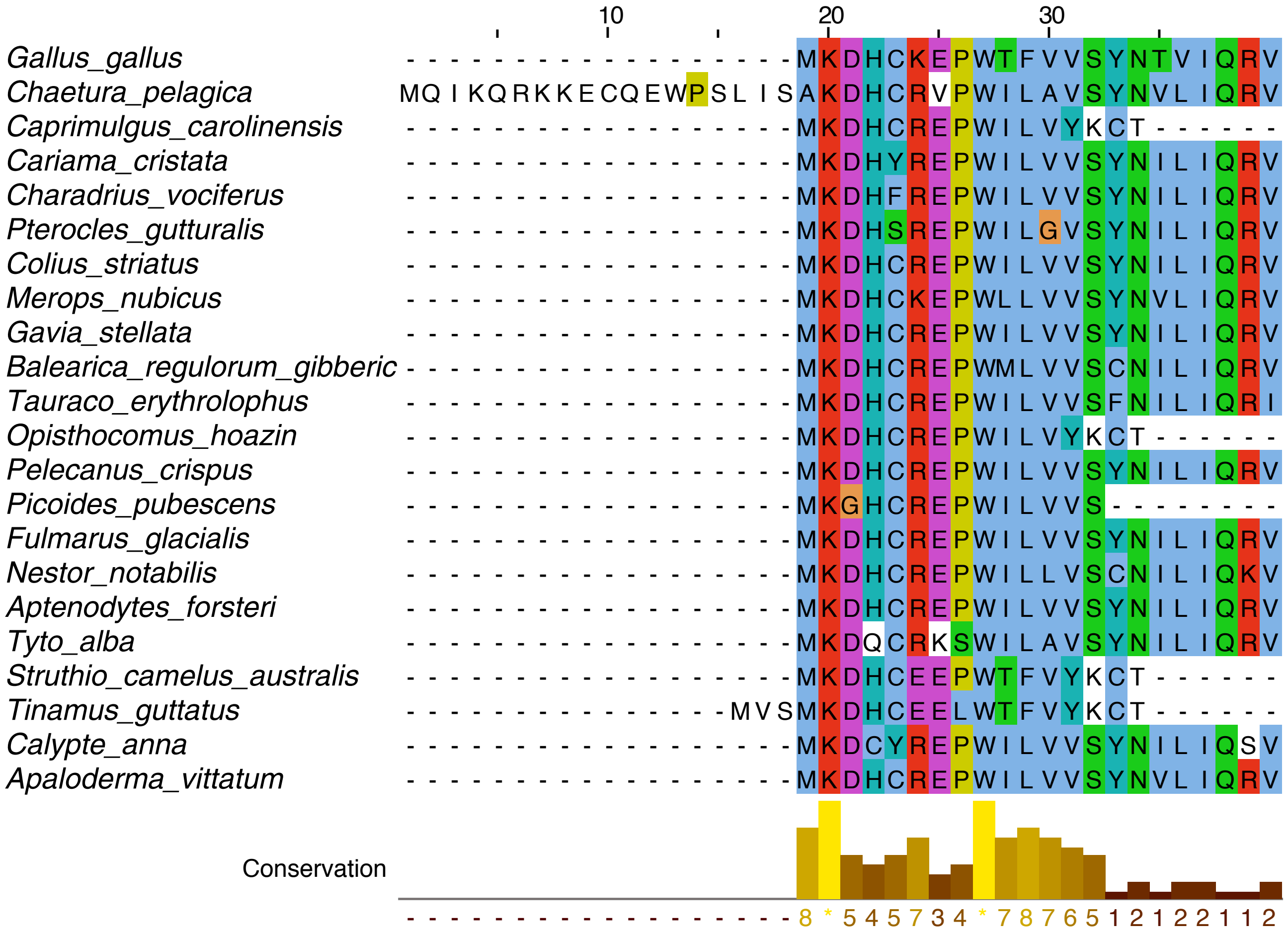

**HG0396.2**

ENSGALG00000034253

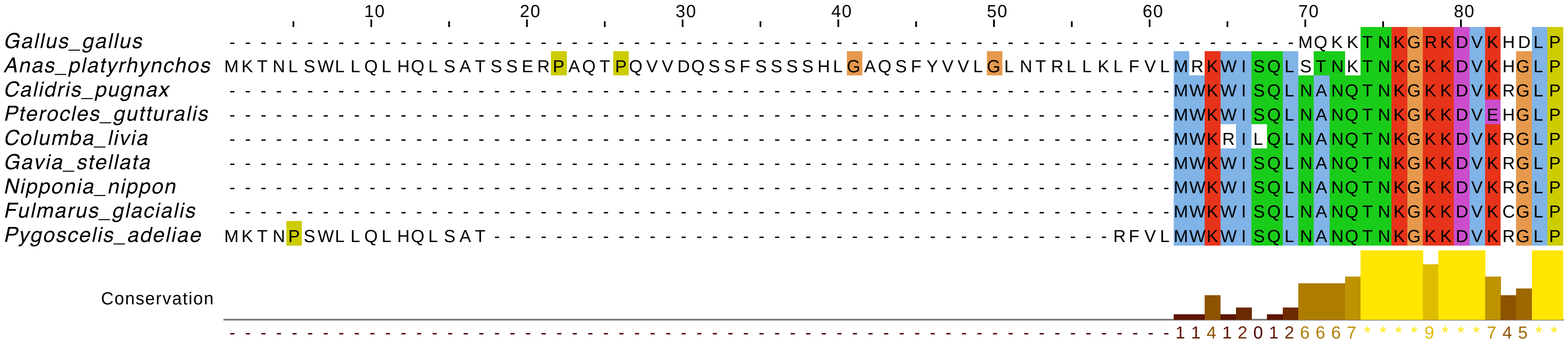

**HG0397**

ENSGALG00000009438 (SLC39A9)

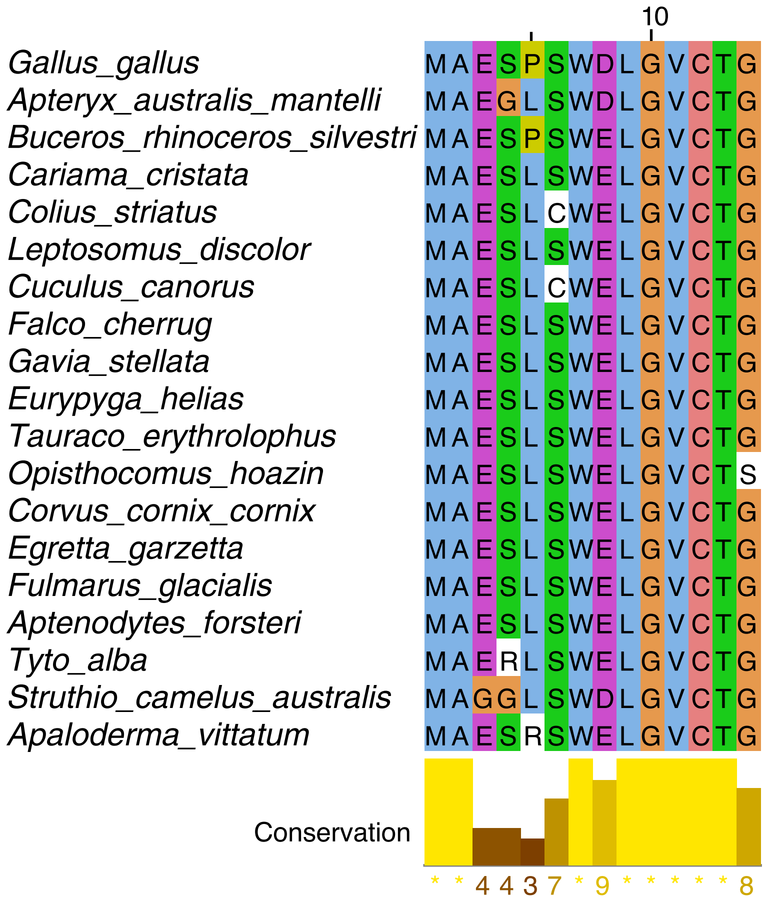

**HG0398**

ENSGALG00000005369 (BTBD10)

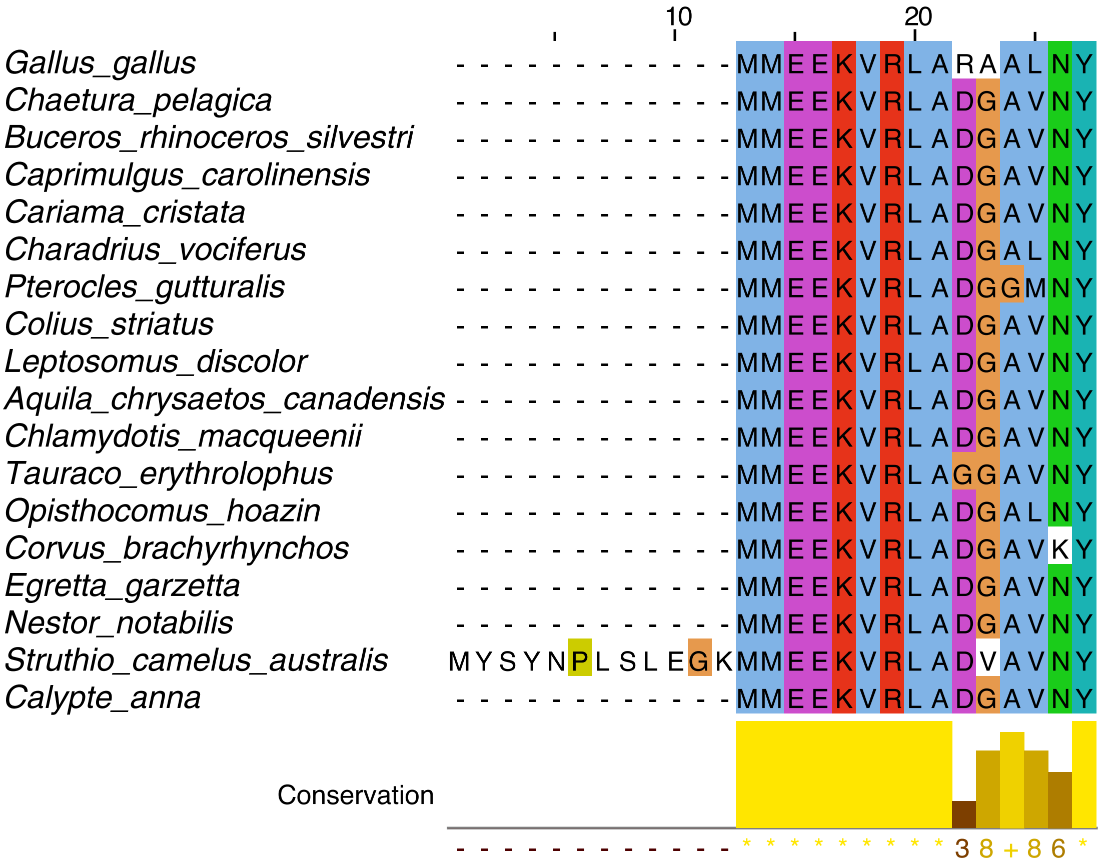

**HG0399**

ENSGALG00000032659

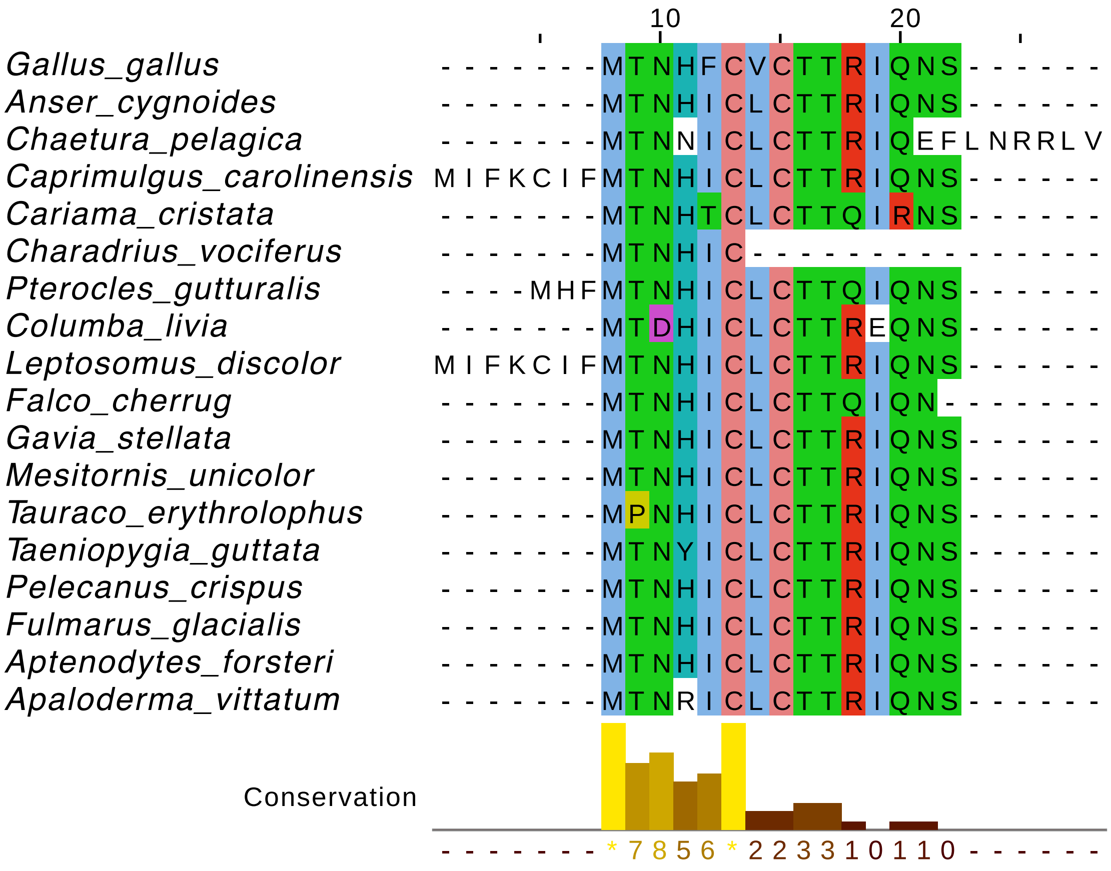

**HG0400**

ENSGALG00000004261 (AQP9)

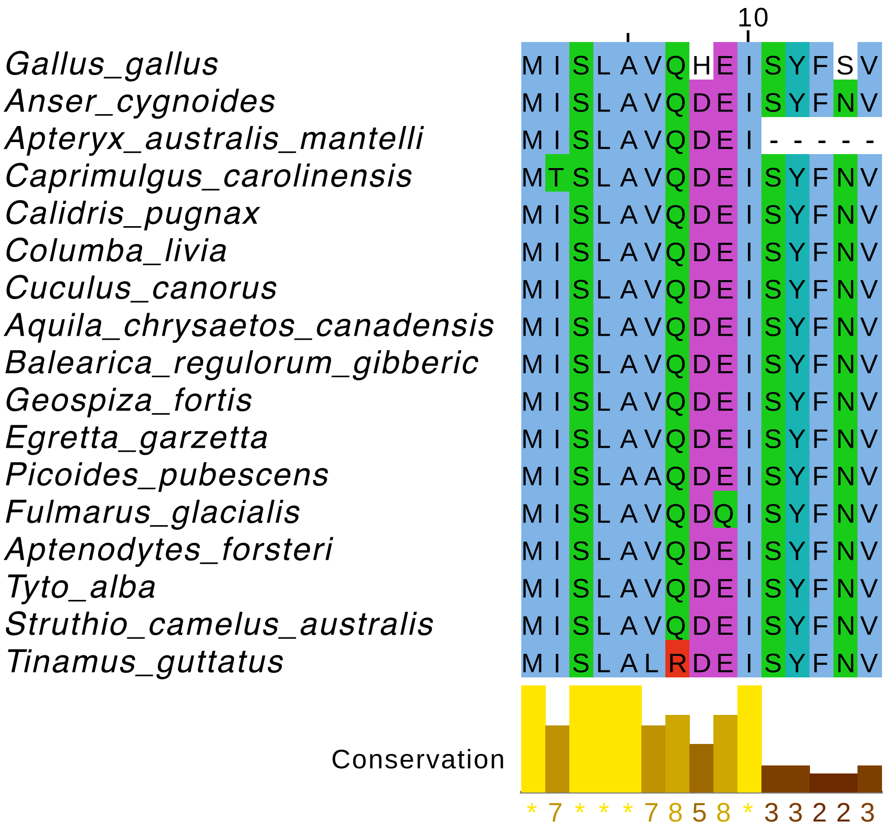

**HG0401**

ENSGALG00000009057 (TASP1)

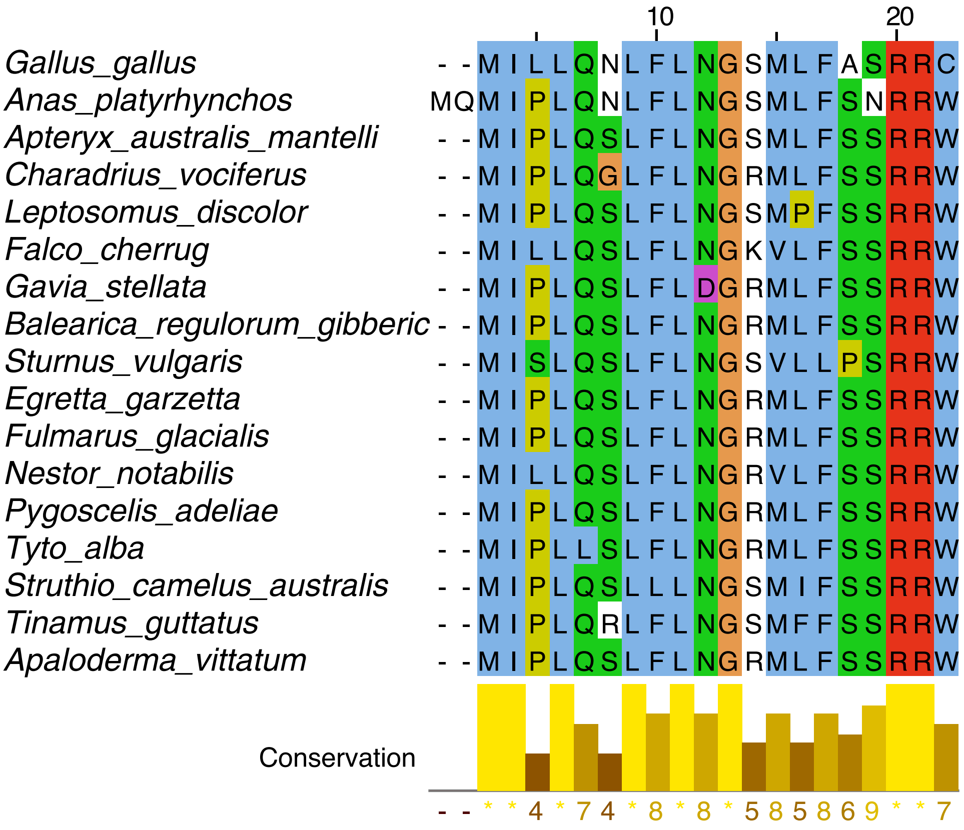

**HG0402**

ENSGALG00000035584

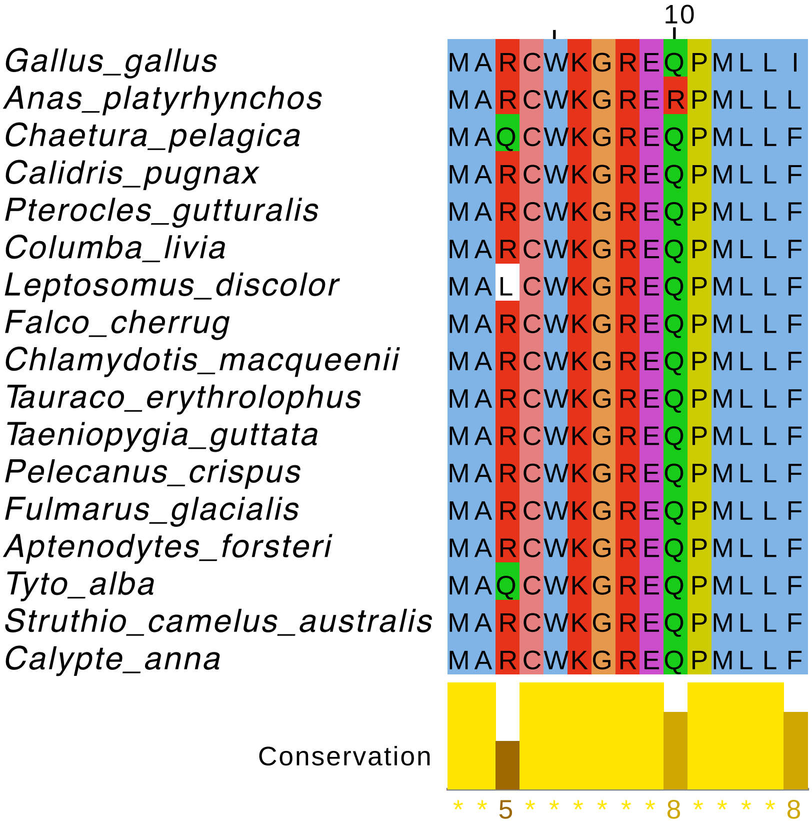

**HG0403.1**

ENSGALG00000007717 (GCNT7)

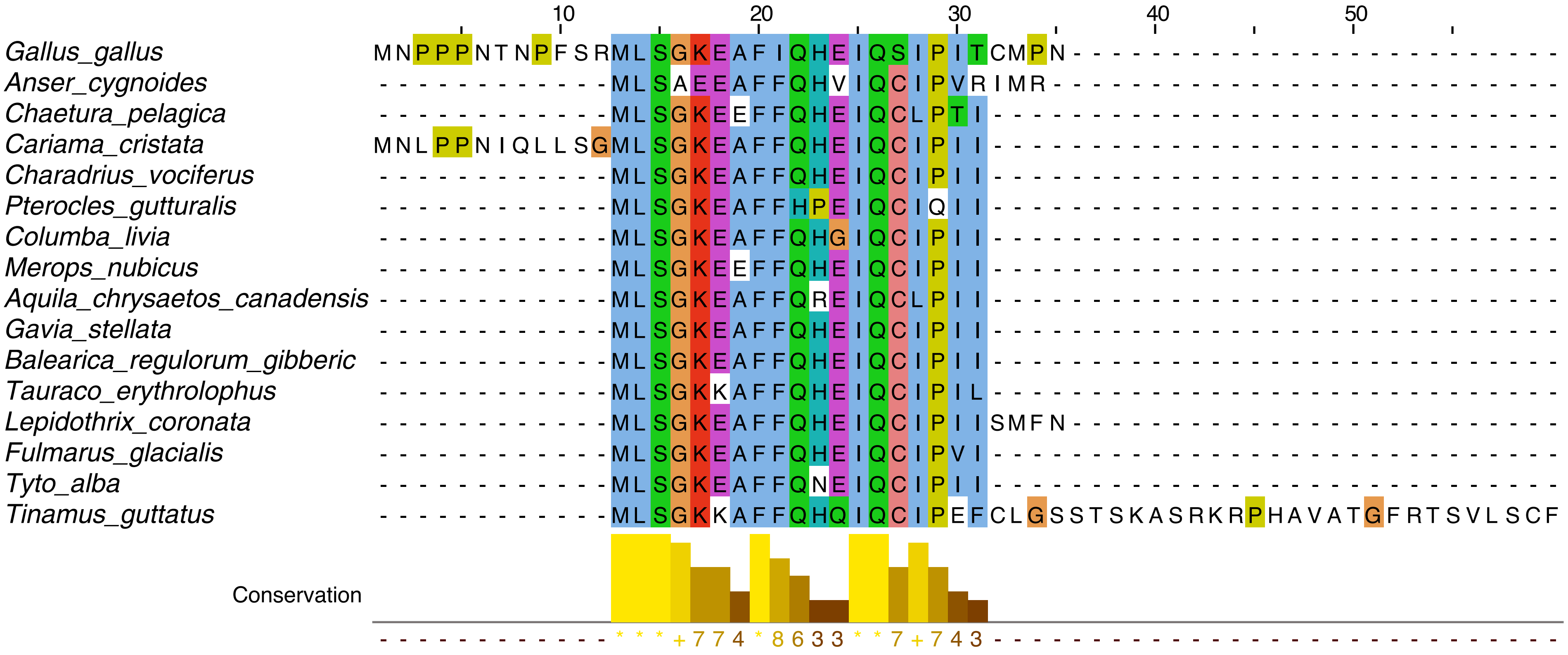

**HG0403.2**

ENSGALG00000007717 (GCNT7)

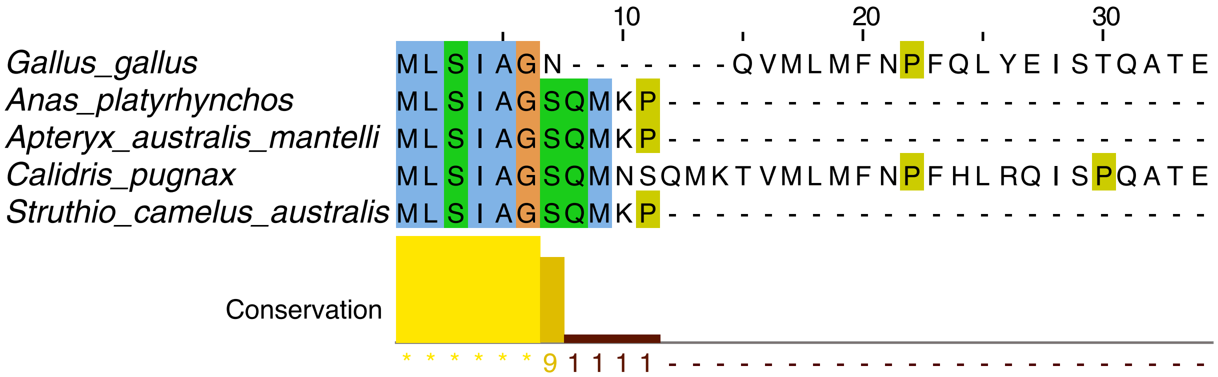

**HG0404**

ENSGALG00000008700 (PDCD4)

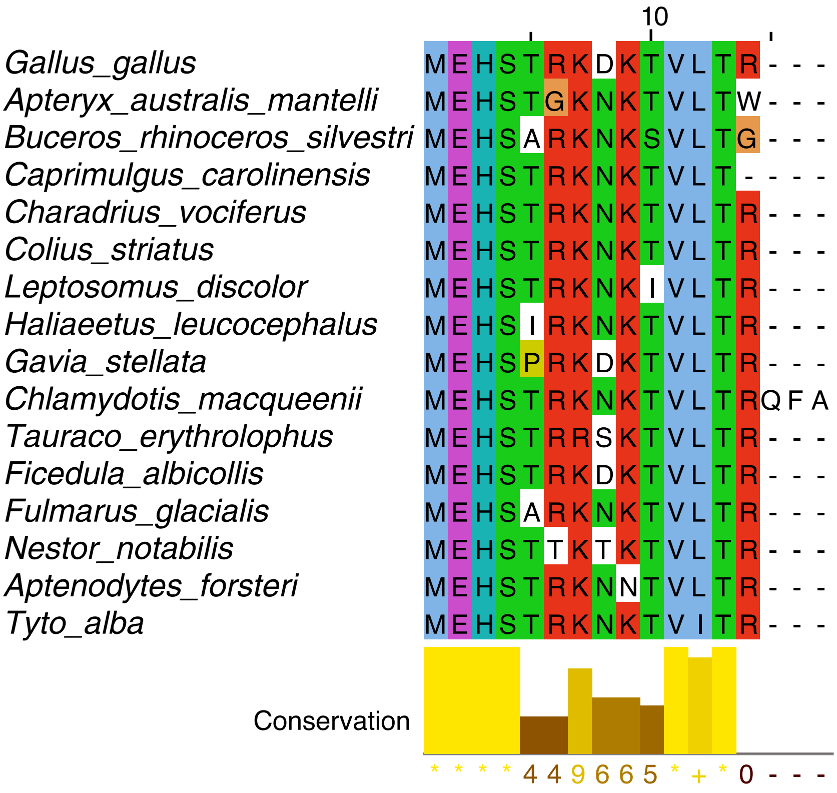

**HG0405.1**

ENSGALG00000009185

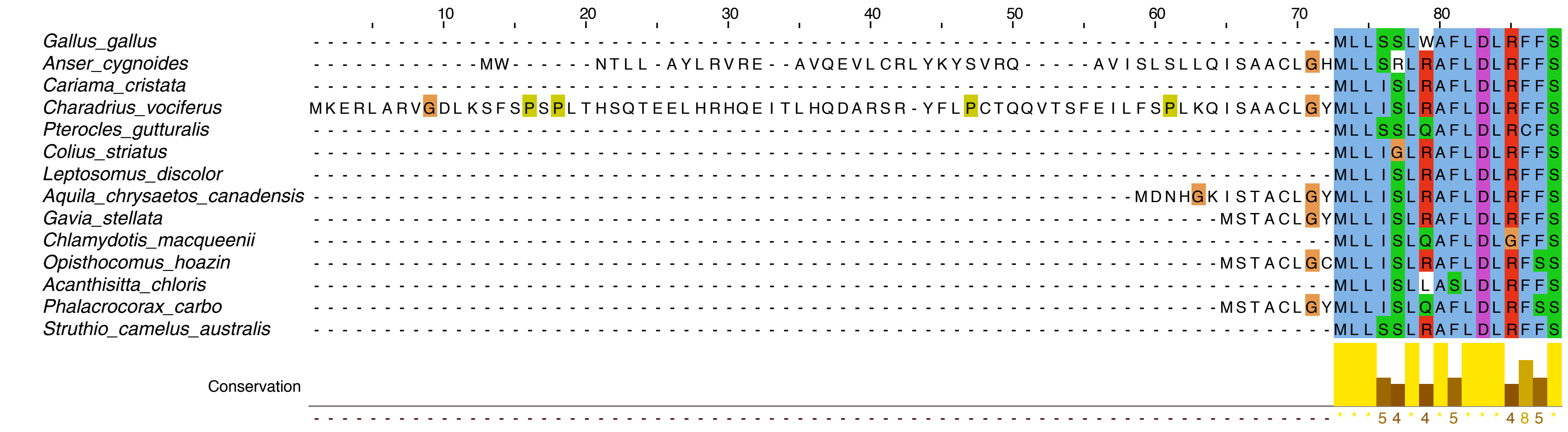

**HG0405.2**

ENSGALG00000009185

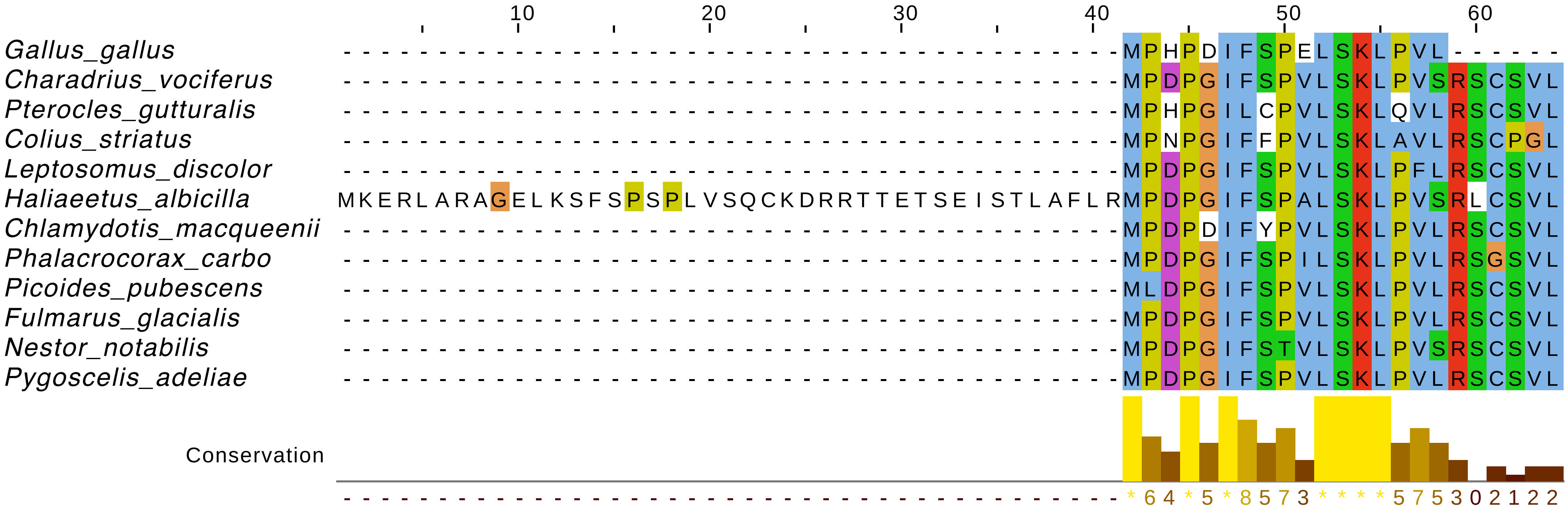

**HG0406**

ENSGALG00000004935 (LSM14A)

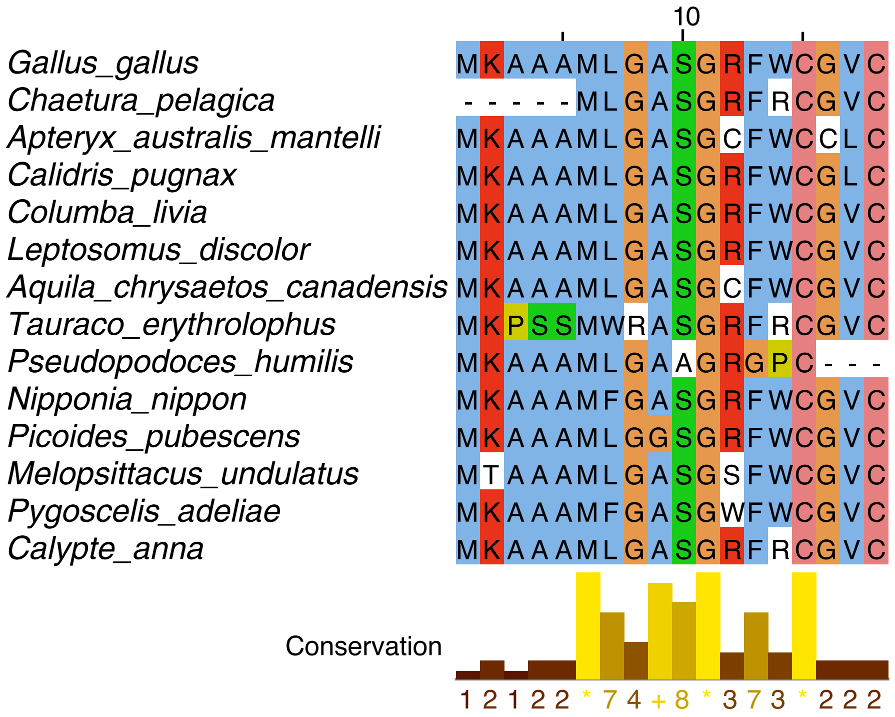

**HG0407**

ENSGALG00000015027 (JAK2)

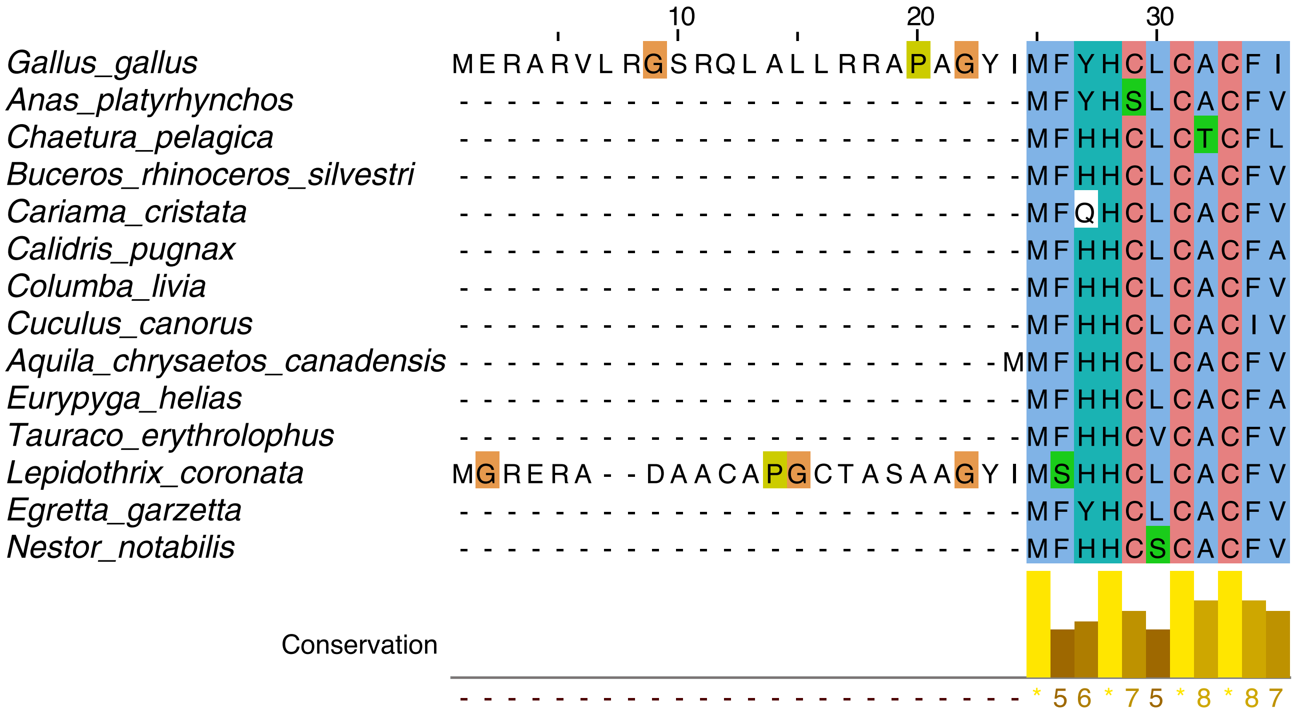

**HG0408**

ENSGALG00000036255 (WNT7B)

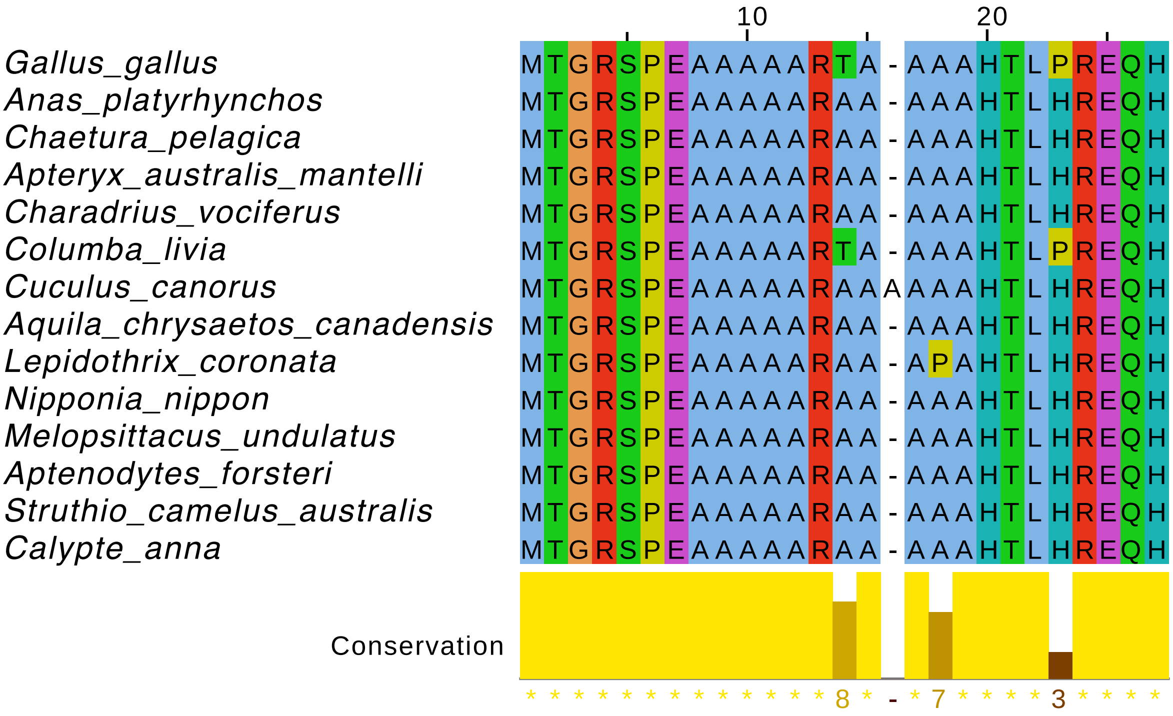

**HG0409**

ENSGALG00000002315

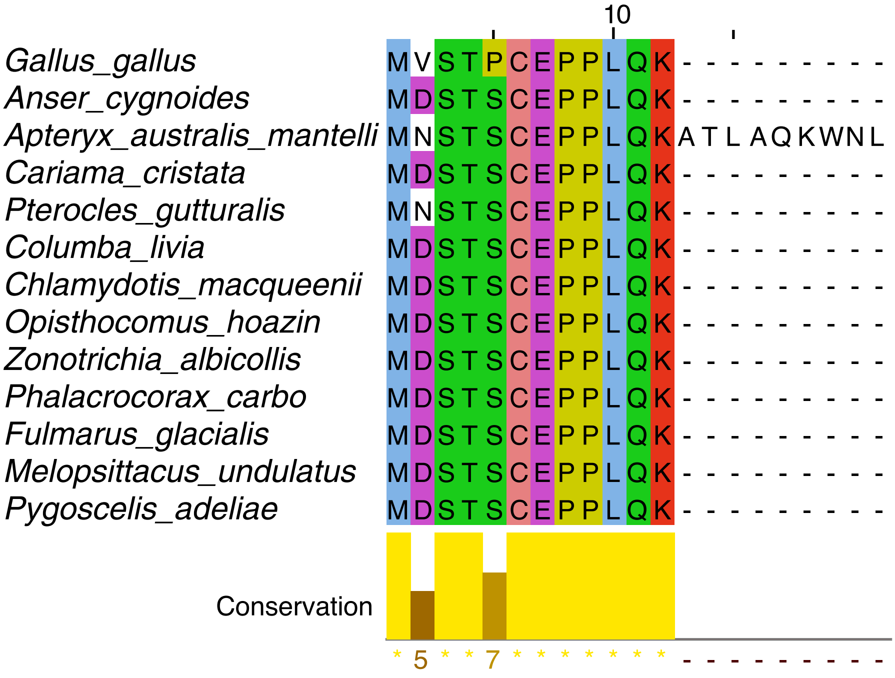

**HG0410**

ENSGALG00000008148 (MYBPC3)

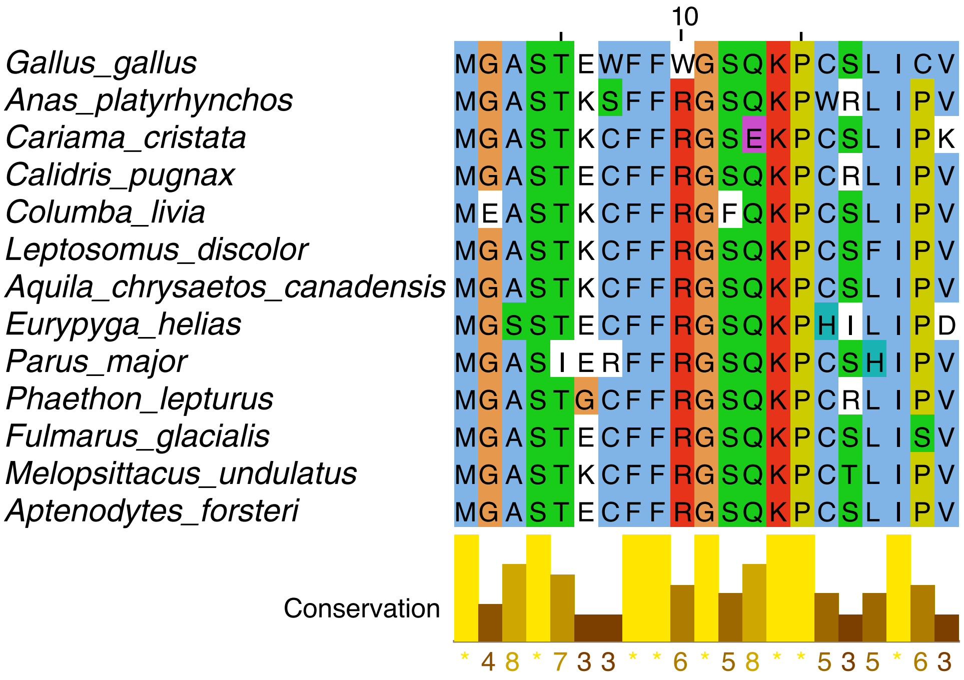

**HG0411**

ENSGALG00000012166 (SLC35F5)

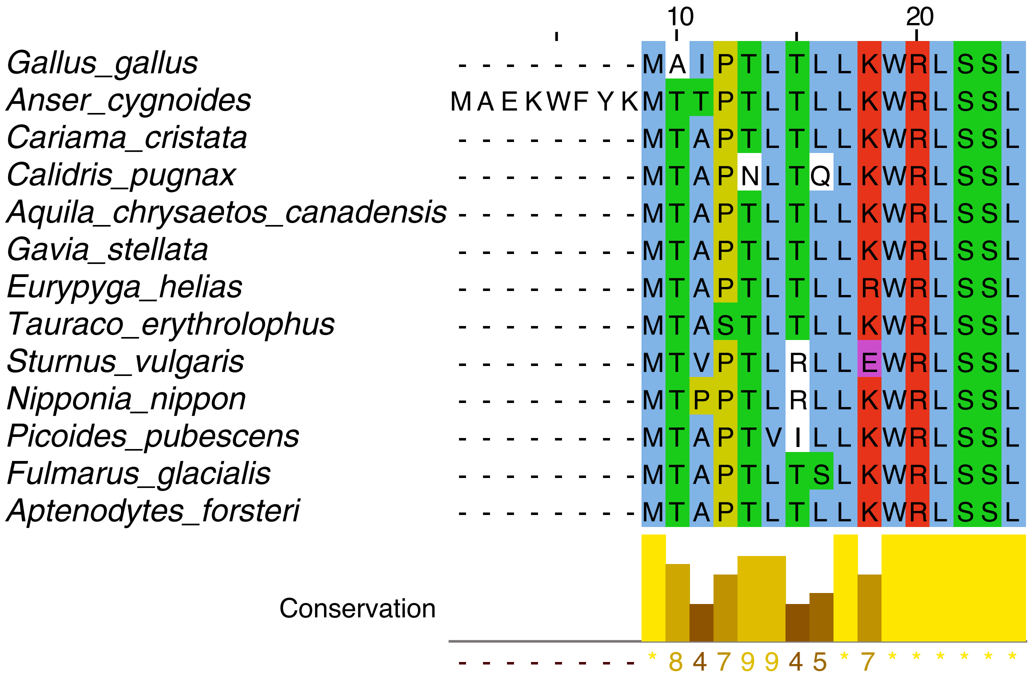

**HG0412**

ENSGALG00000003446 (PRLR)

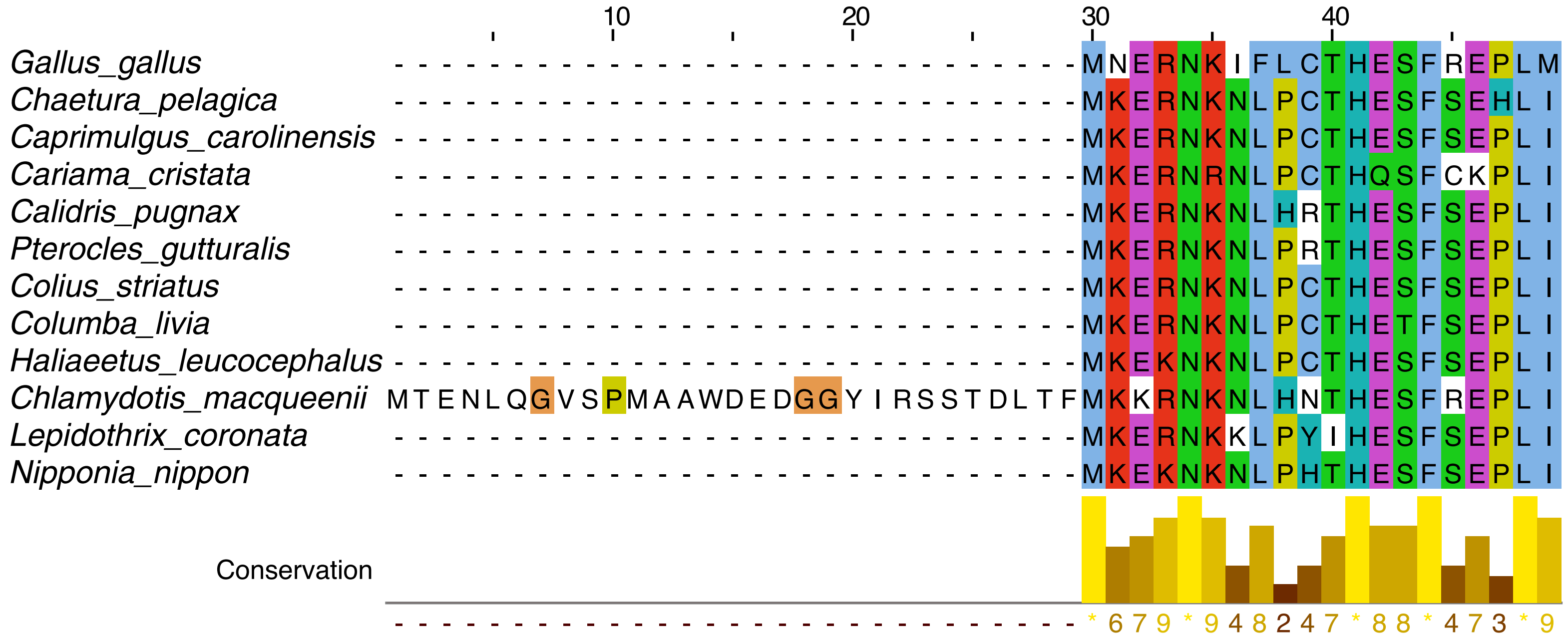

**HG0413**

ENSGALG00000005475

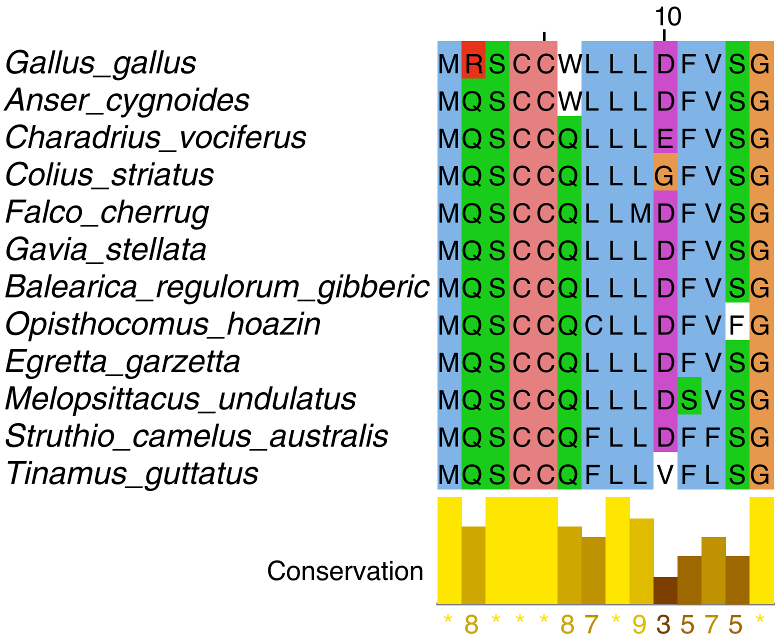

**HG0414**

ENSGALG00000032599

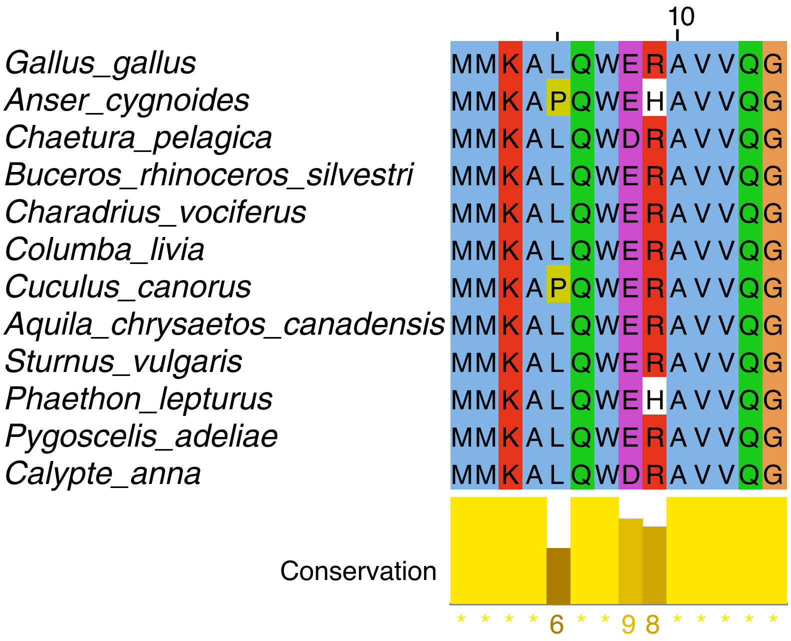

**HG0415**

ENSGALG00000034970

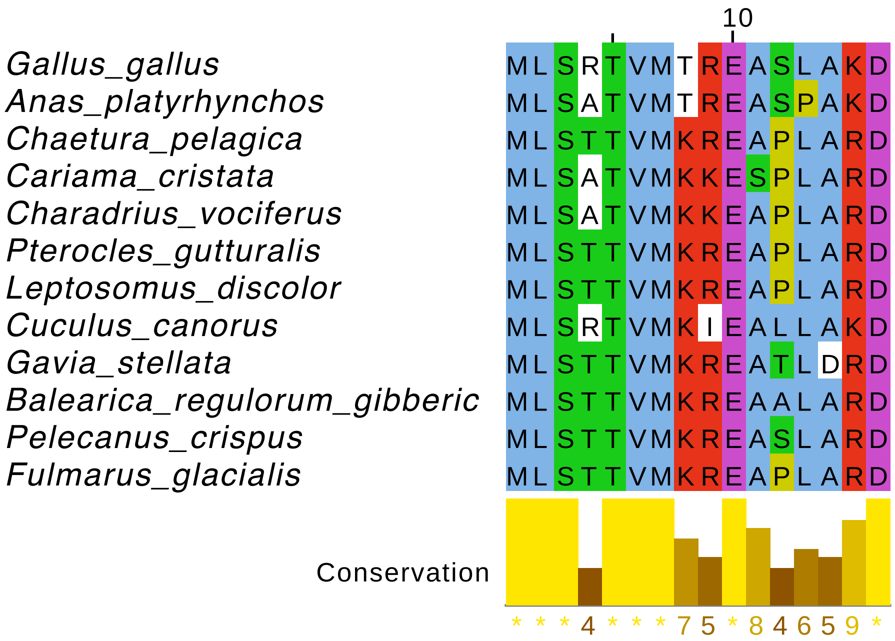

**HG0416**

ENSGALG00000015874 (TTK)

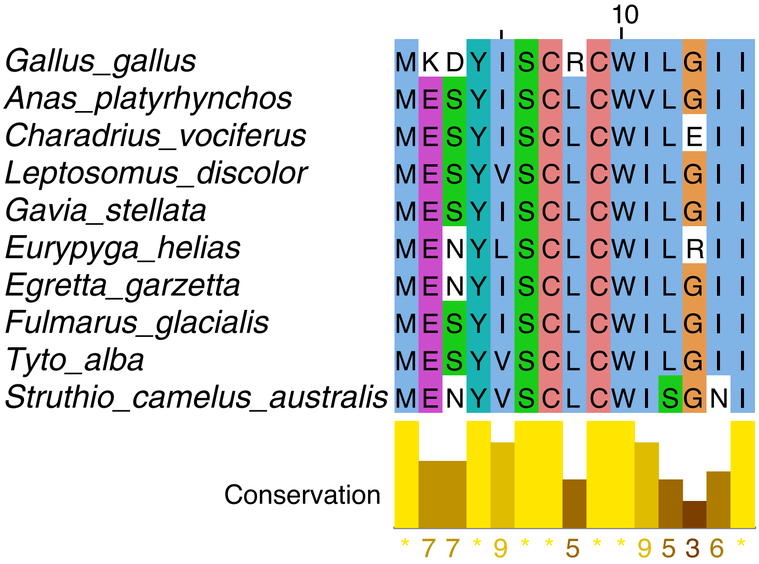

**HG0417**

ENSGALG00000001522 (PHF20)

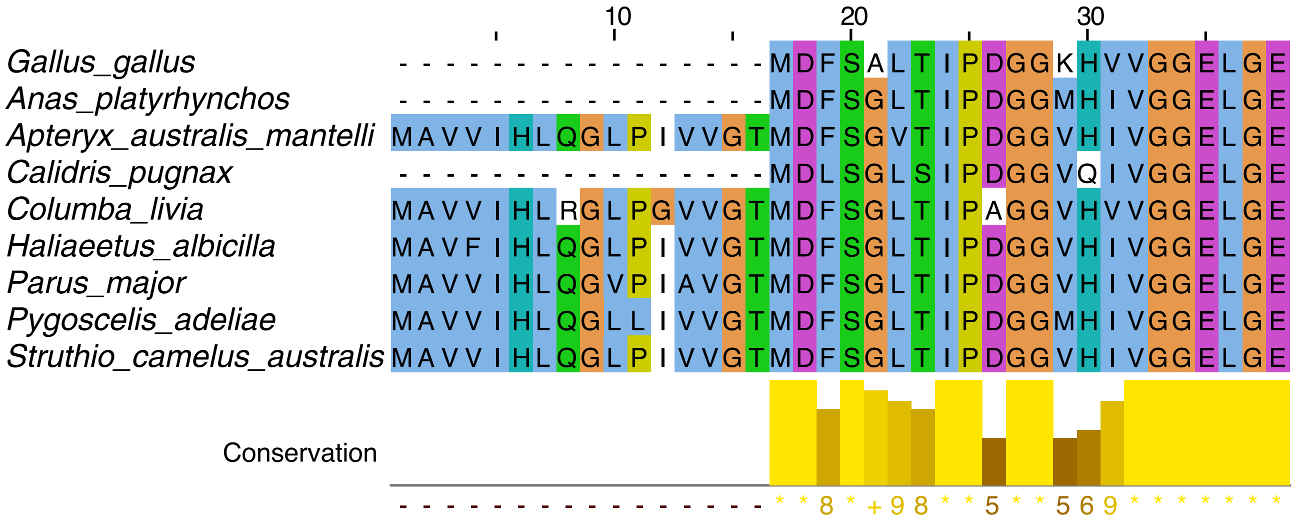

**HG0418**

ENSGALG00000010337 (GPR149)

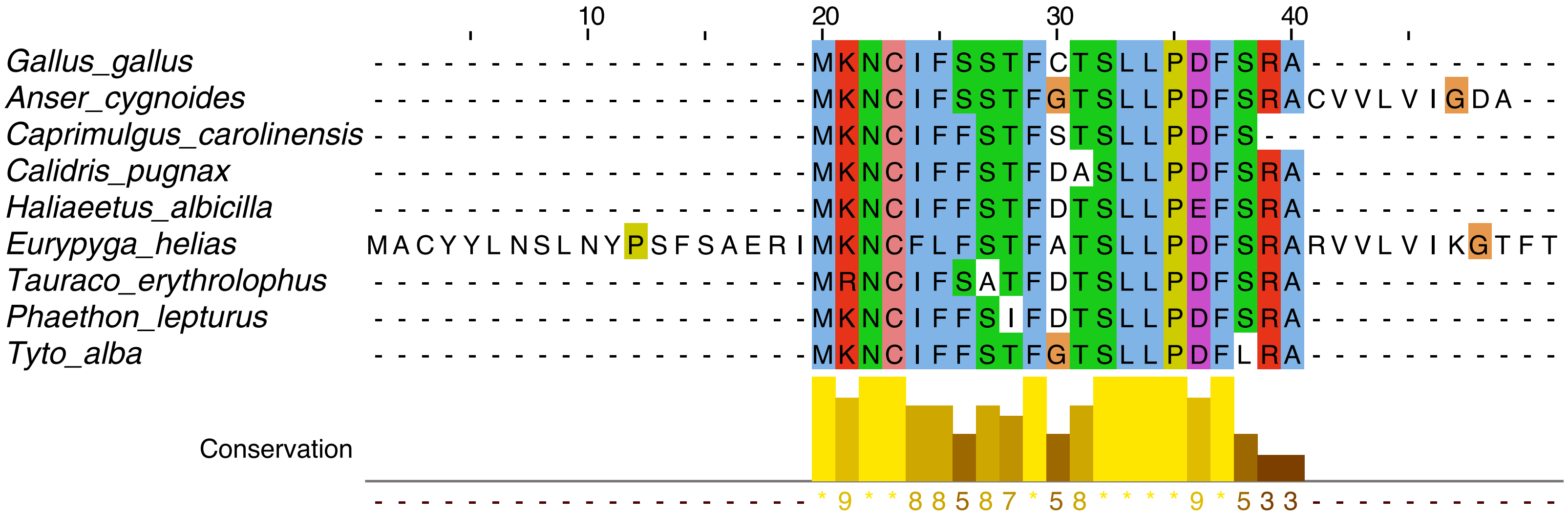

**HG0419**

ENSGALG00000012542 (RASD2)

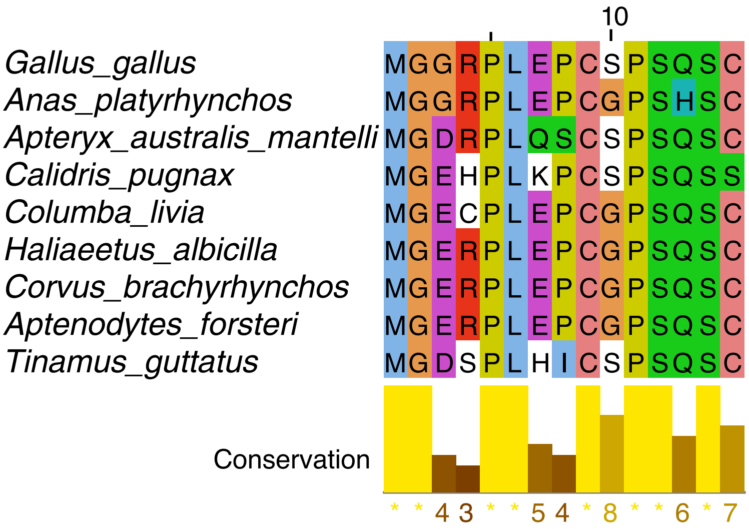

**HG0420**

ENSGALG00000000336 (PPP1R12B)

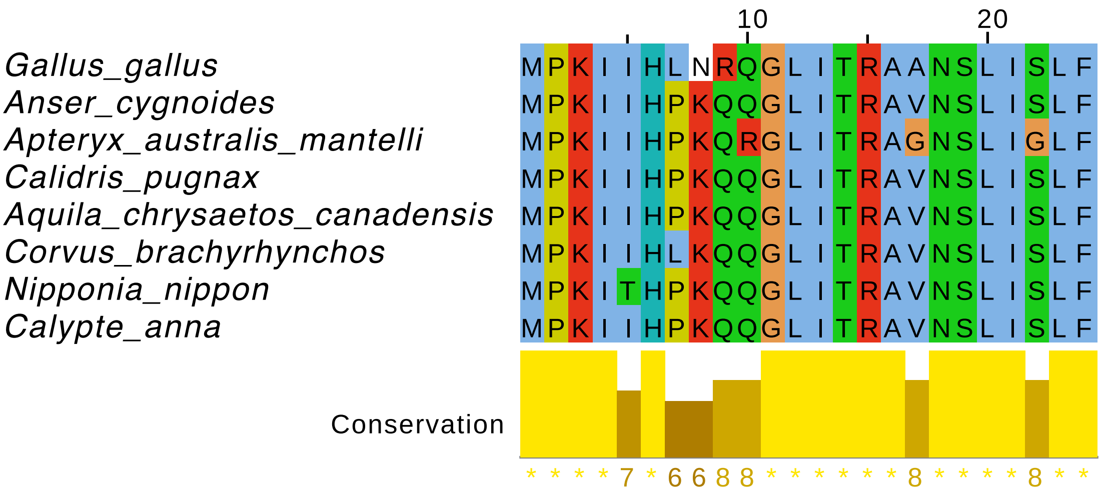

# HG0421

### ENSGALG00000003492

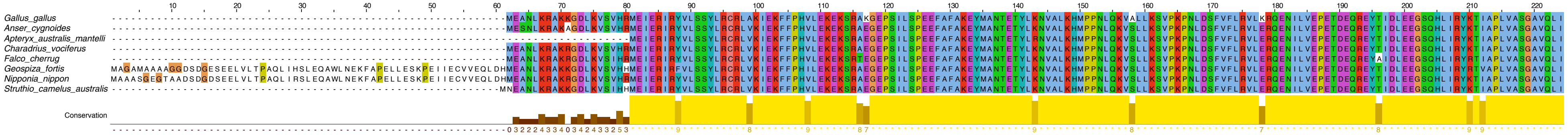

**HG0422**

ENSGALG00000005772 (BOK)

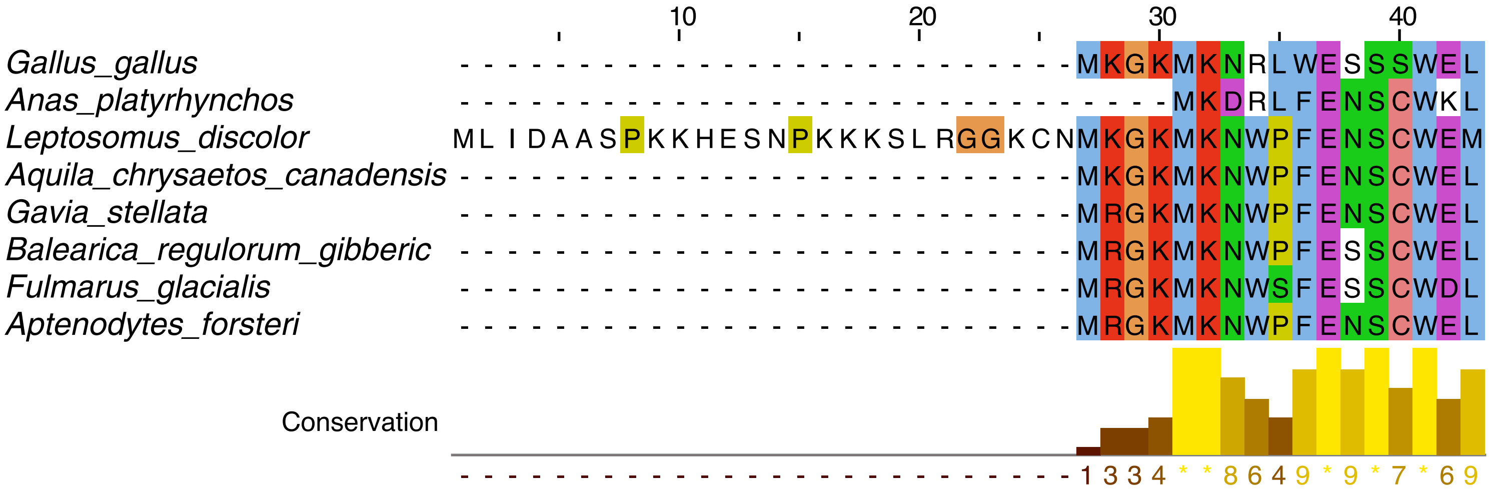

**HG0423**

ENSGALG00000006329

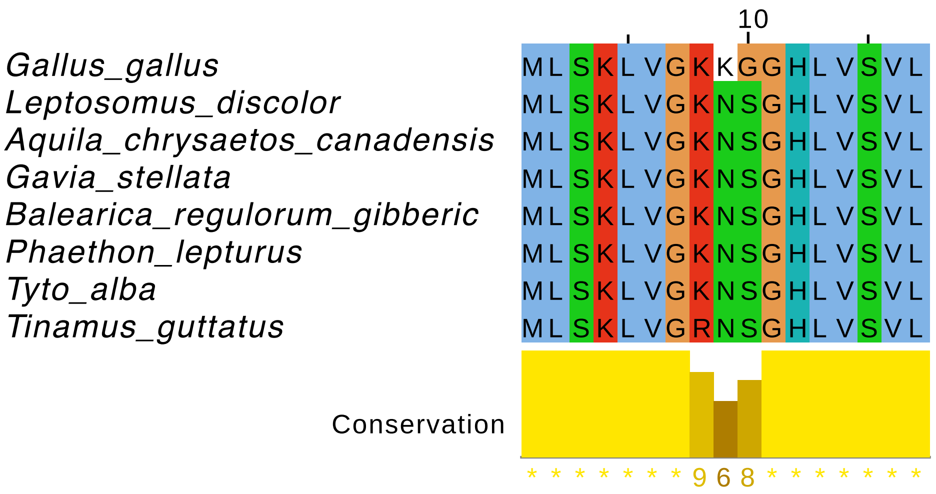

**HG0424**

ENSGALG00000039553

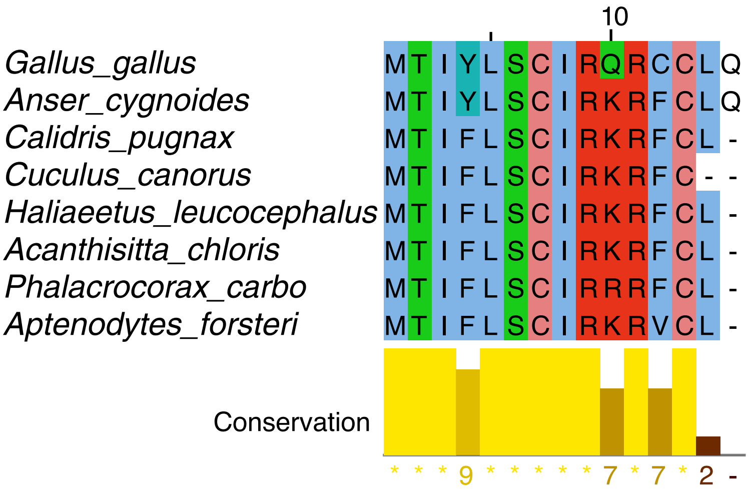

**HG0425**

ENSGALG00000043044

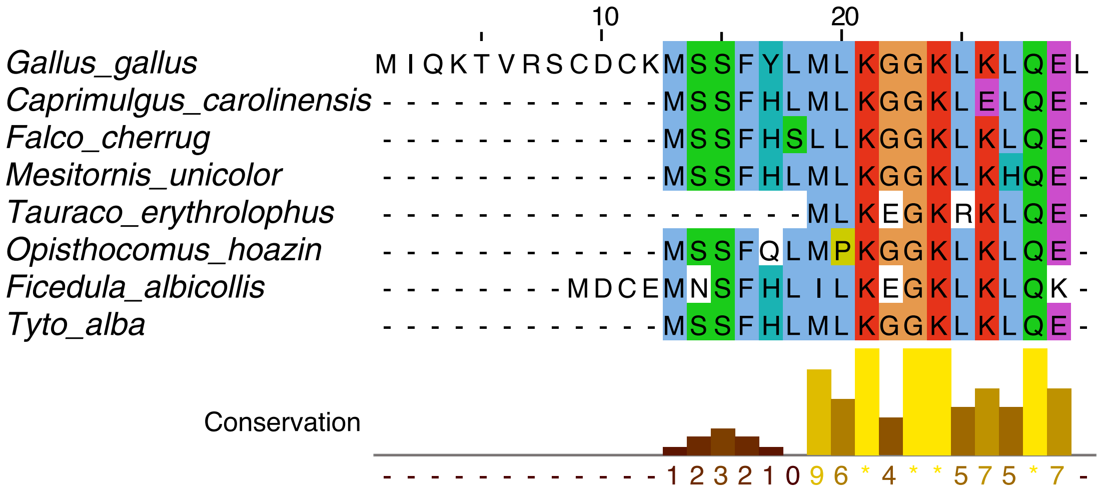

**HG0426**

ENSGALG00000002192 (PTPRC)

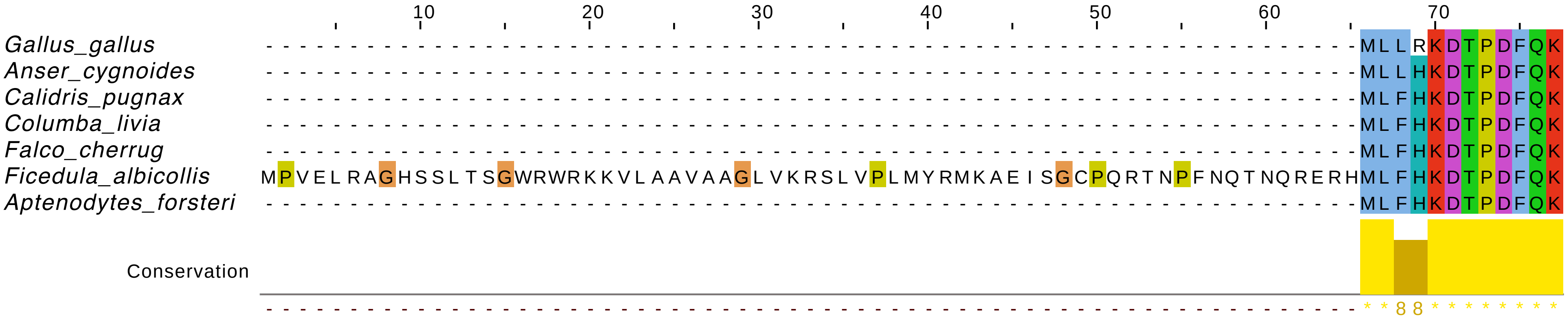

**HG0427**

ENSGALG00000005550 (ST6GAL1)

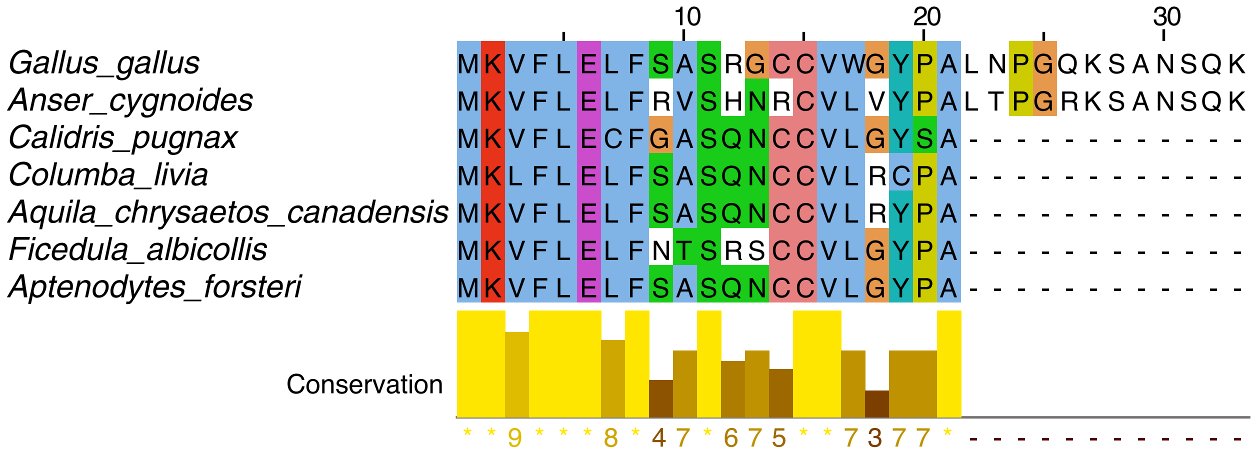

**HG0428**

ENSGALG00000005976 (TCEA2)

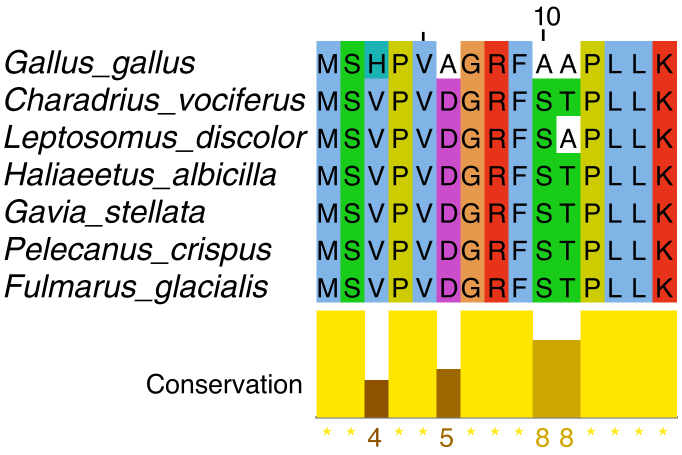

**HG0429**

ENSGALG00000006626 (PDE6C)

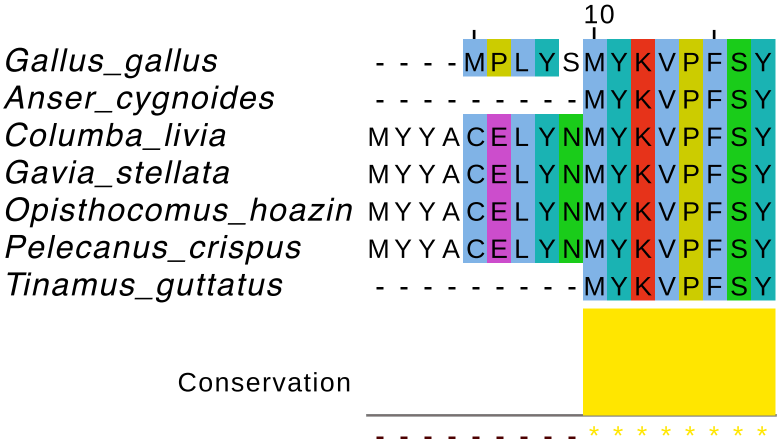

**HG0430**

ENSGALG00000015490

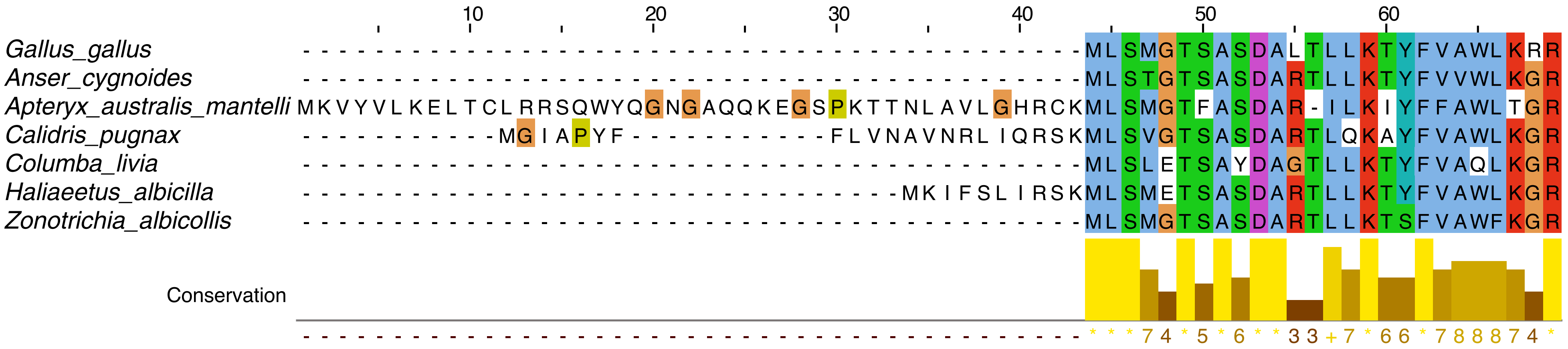

**HG0431**

ENSGALG00000026542 (FAM217B)

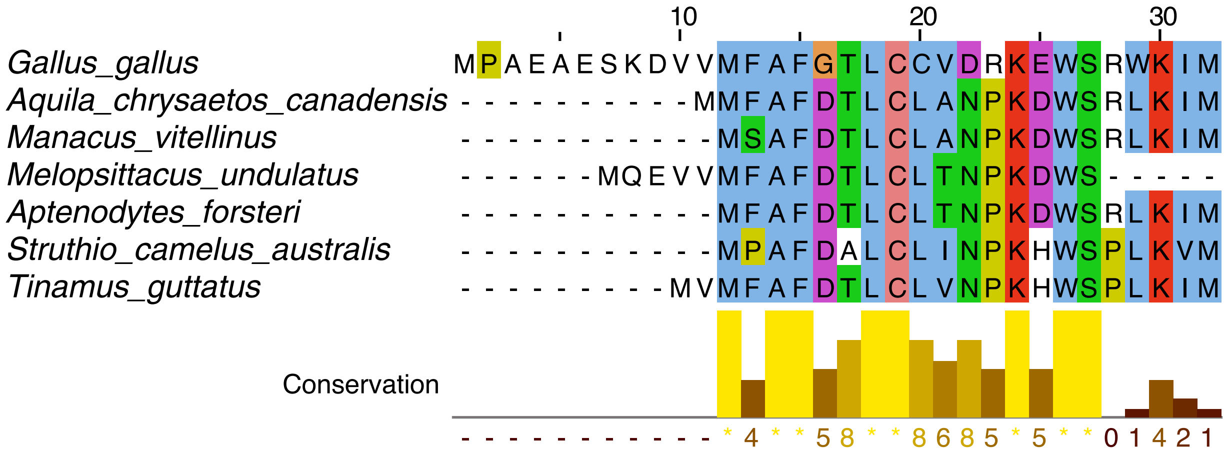

**HG0432**

ENSGALG00000040546

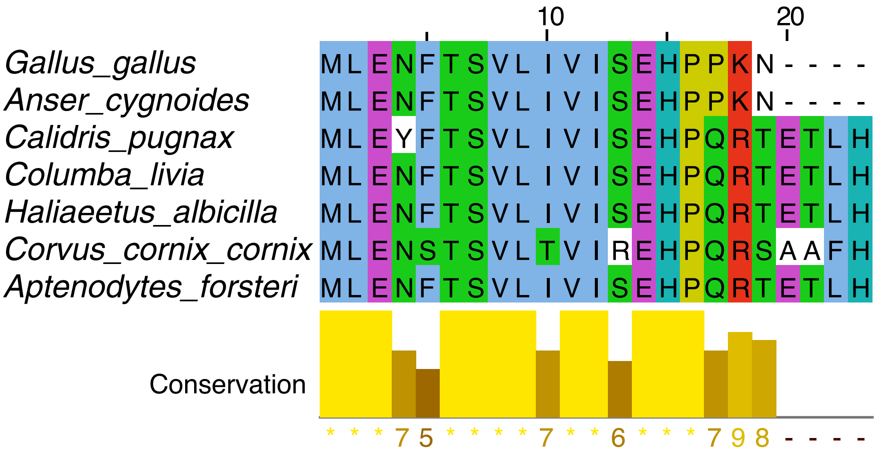

**HG0433**

ENSGALG00000002952 (NRG4)

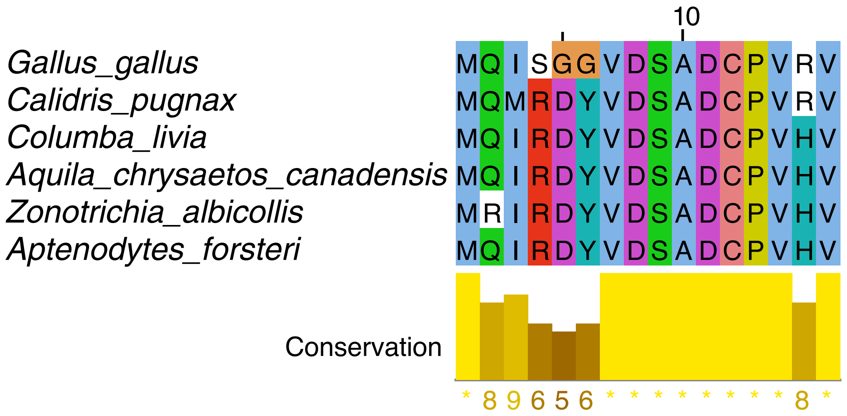

**HG0434**

ENSGALG00000008357 (TPCN1)

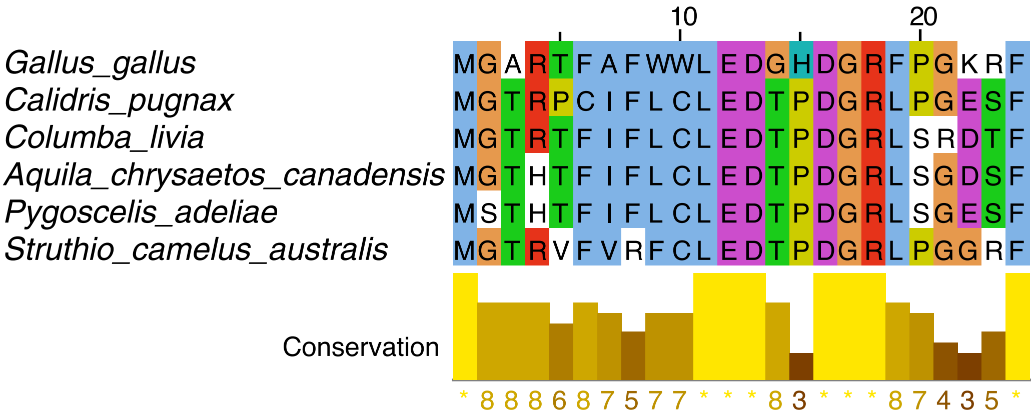

**HG0435**

ENSGALG00000010778 (CHRM3)

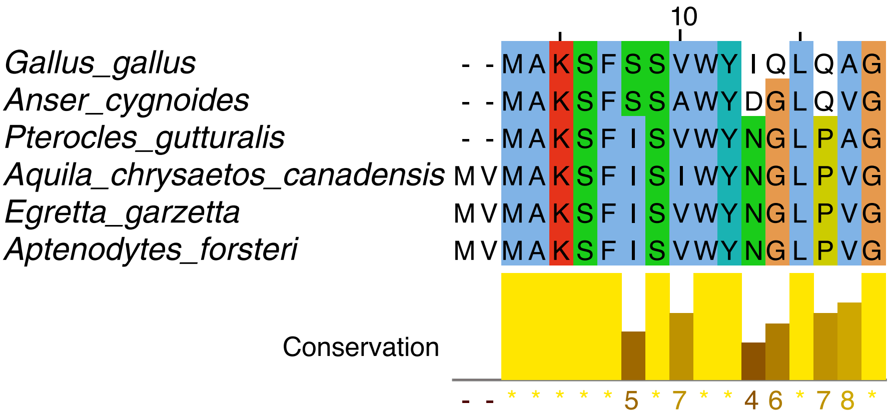

**HG0436**

ENSGALG00000026736 (OGN)

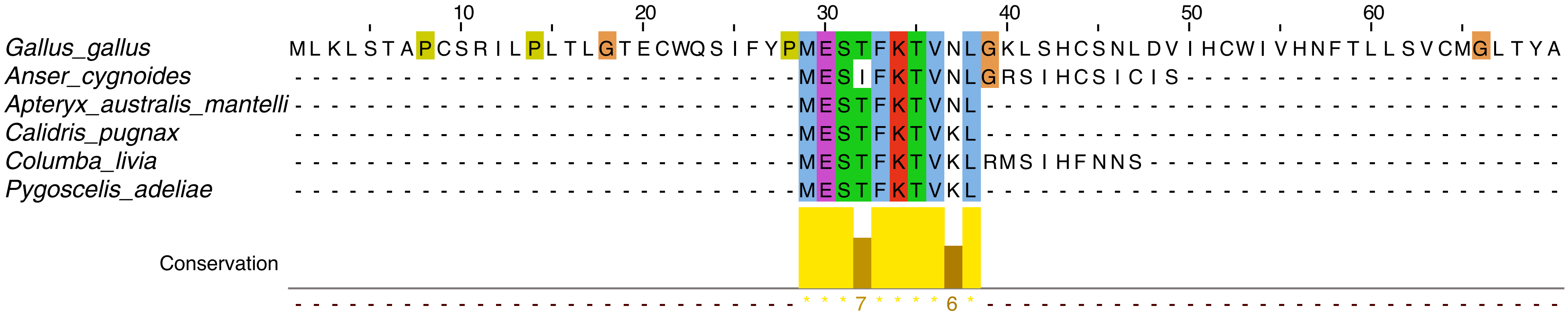

**HG0437**

ENSGALG00000001728 (NACC2)

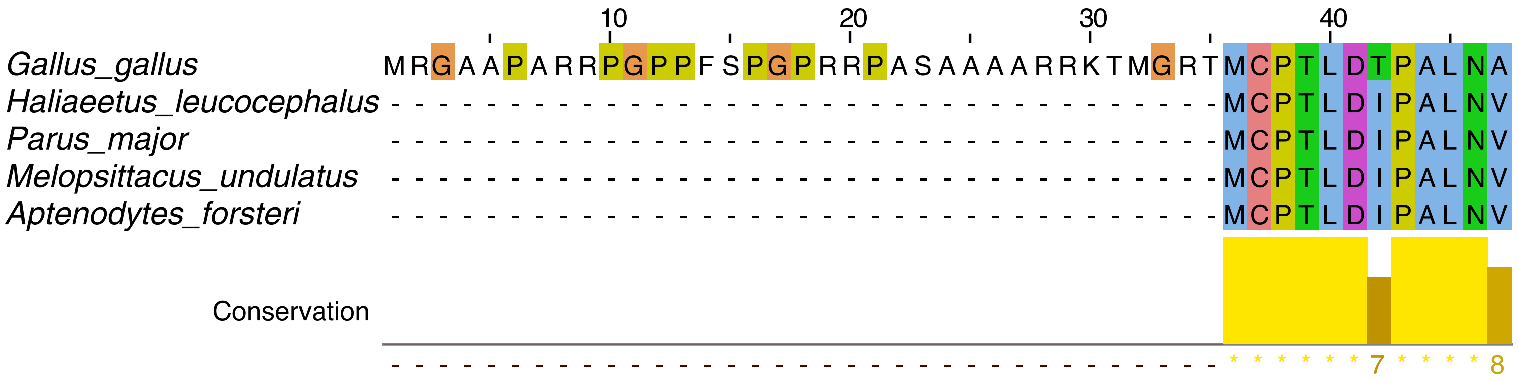

**HG0438**

ENSGALG00000008545 (SENP5)

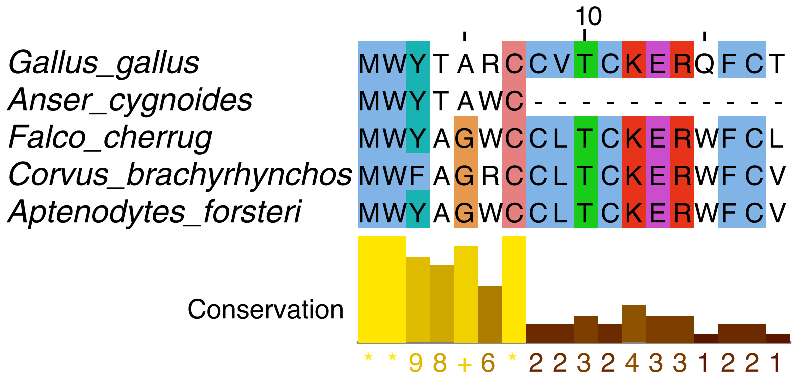

**HG0439**

ENSGALG00000009172 (OSBPL6)

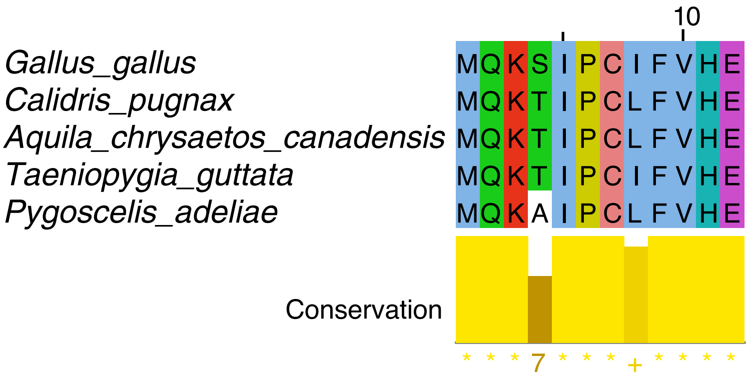

**HG0440**

ENSGALG00000010090

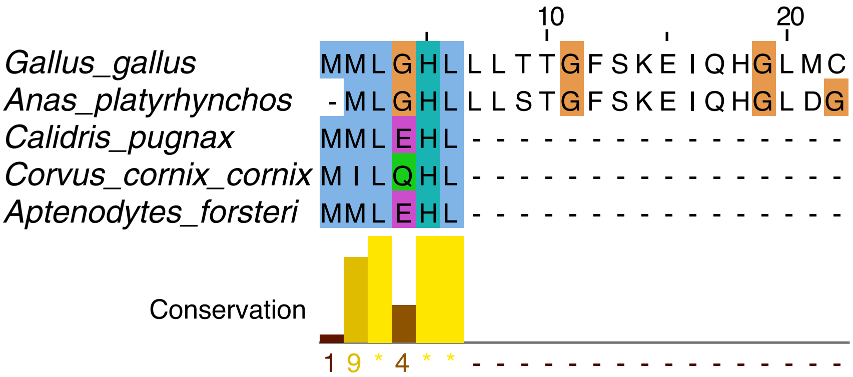

**HG0441**

ENSGALG00000015069 (B4GALT4)

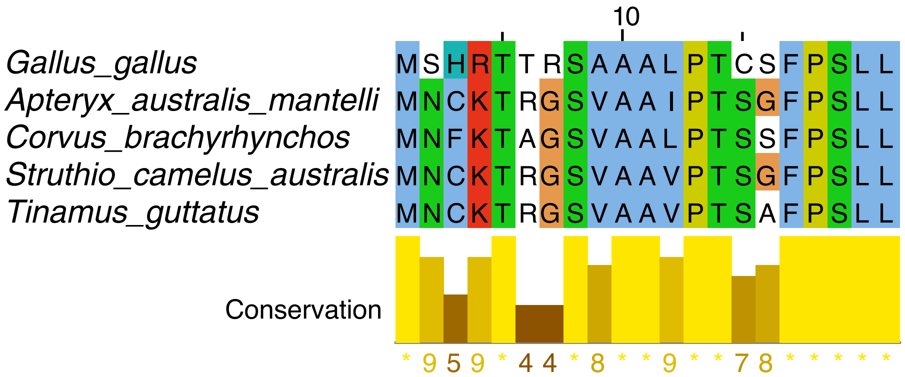

**HG0442**

ENSGALG00000030250

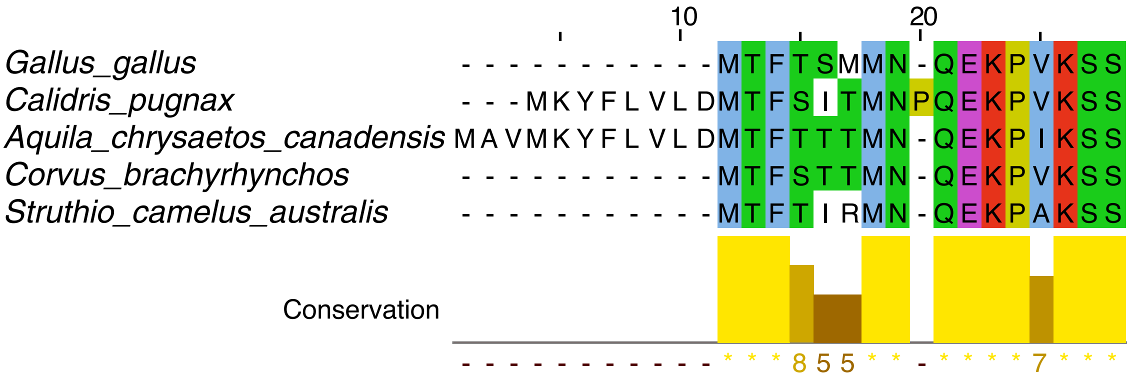

**HG0443**

ENSGALG00000039454

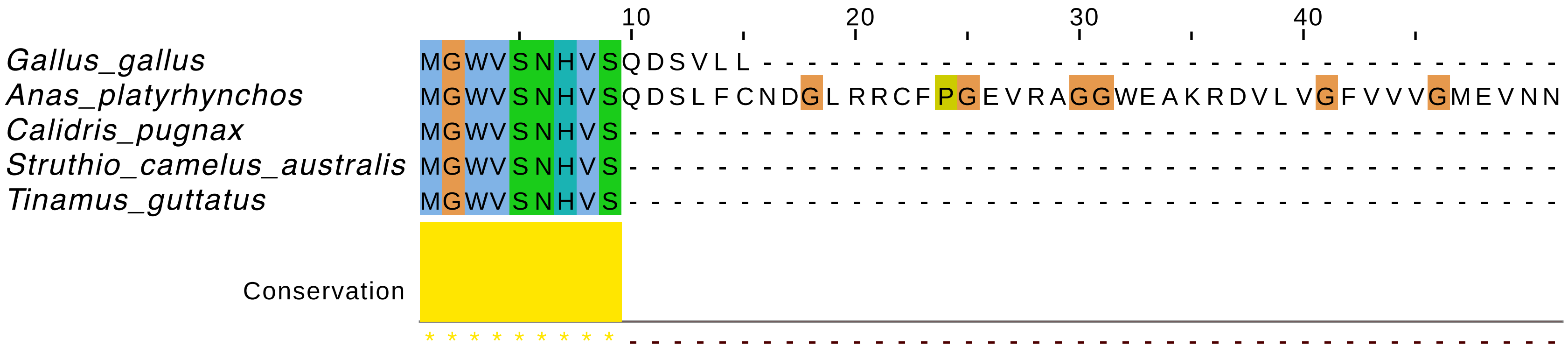

**HG0444**

ENSGALG00000000558 (SLC1A6)

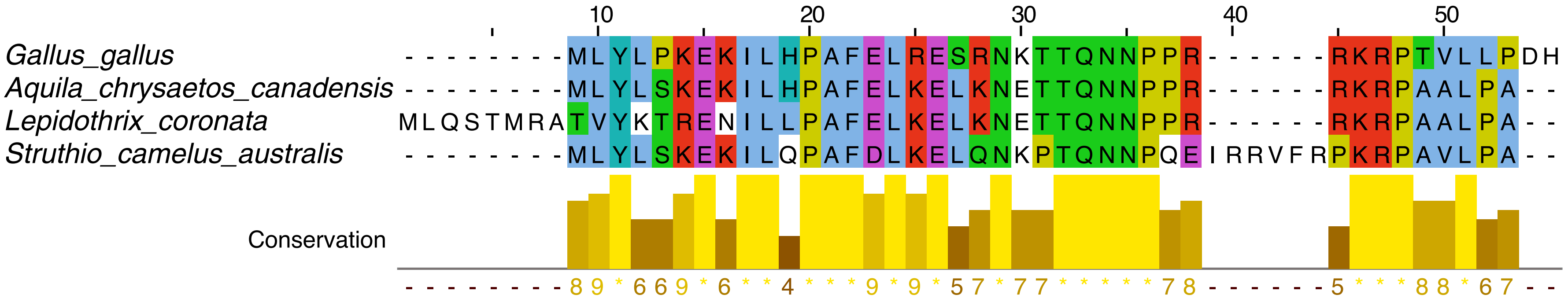

**HG0445**

ENSGALG00000004017

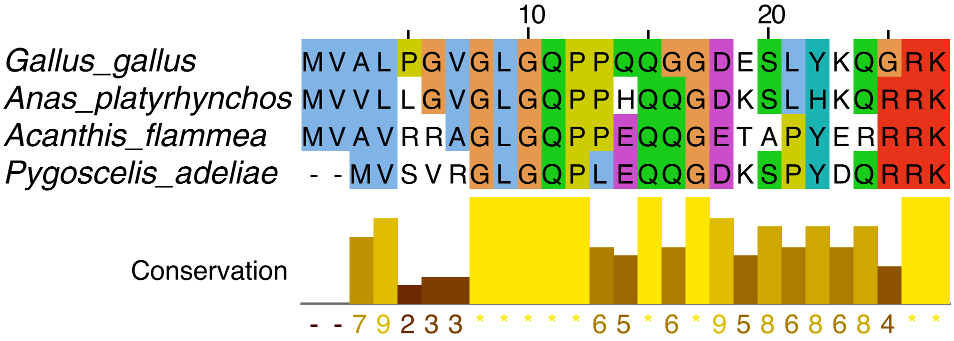

**HG0446**

ENSGALG00000006807 (PLSCR5)

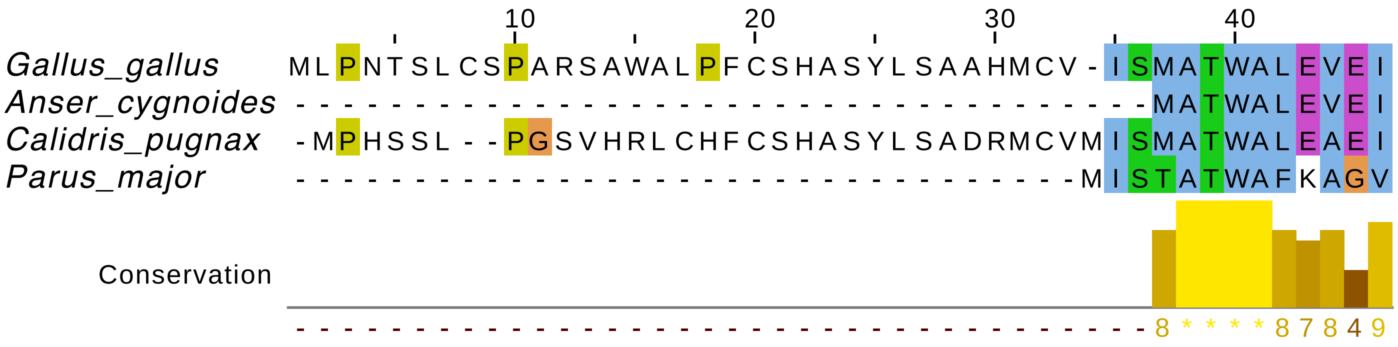

**HG0447**

ENSGALG00000008784

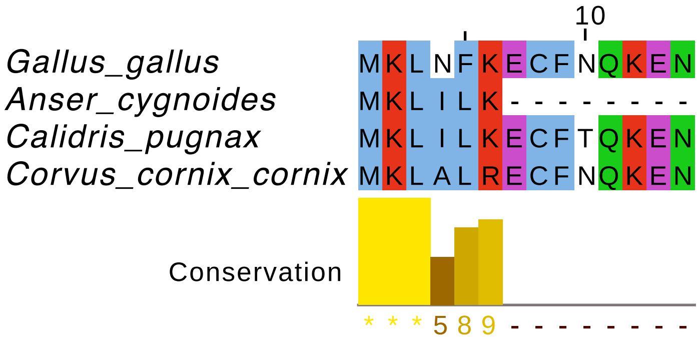

**HG0448**

ENSGALG00000011762 (UBA6)

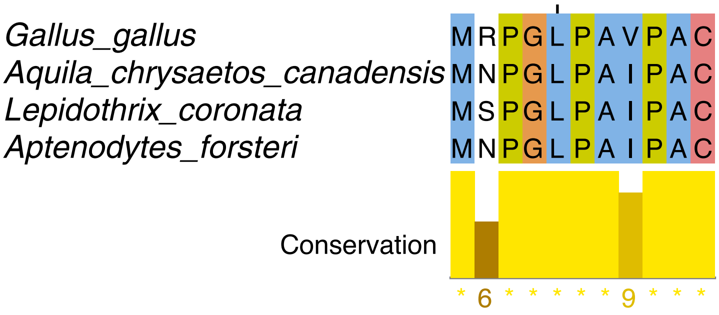

**HG0449**

ENSGALG00000014106 (PPFIBP1)

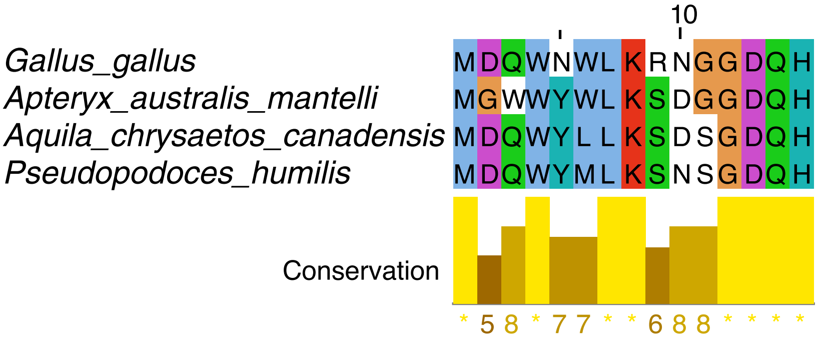

**HG0450**

ENSGALG00000015528 (JAKMIP1)

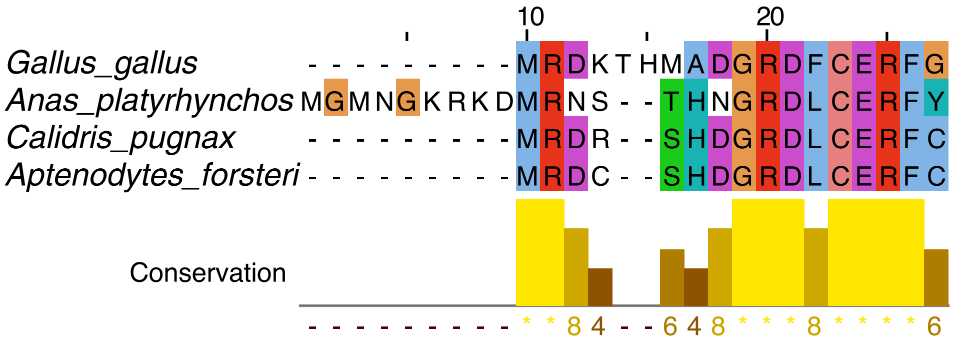

**HG0451**

ENSGALG00000019489 (CHST9)

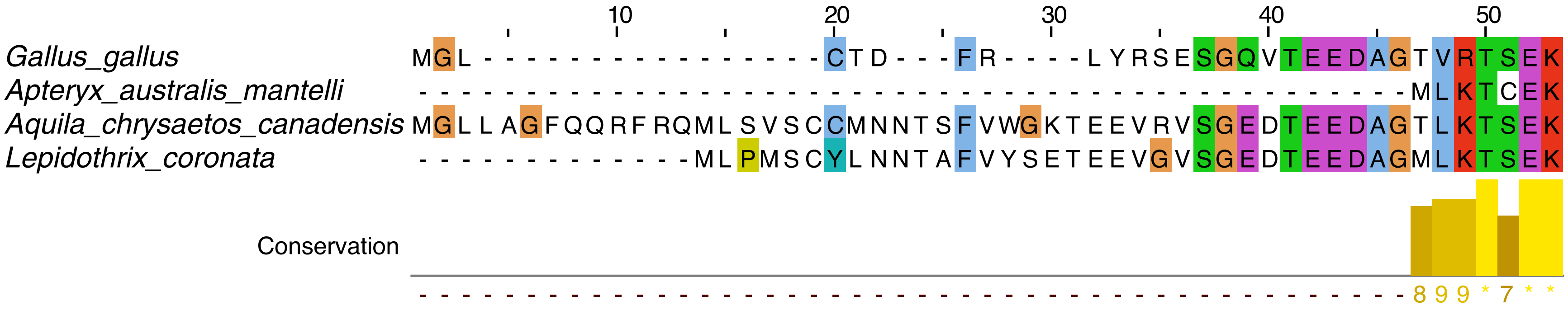

**HG0452**

ENSGALG00000004227 (FAM20B)

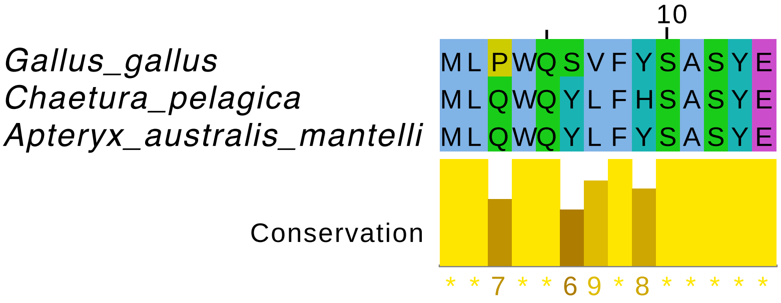

**HG0453**

ENSGALG00000005995 (SLC13A5)

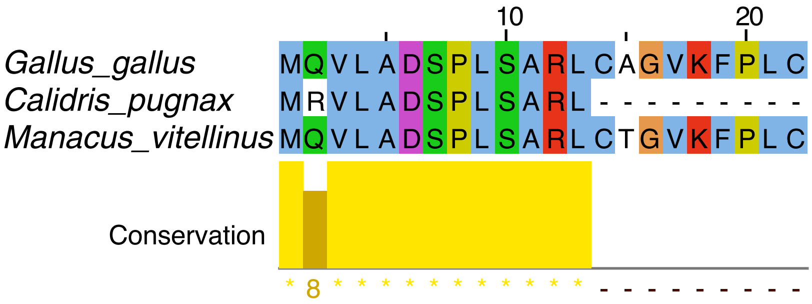

**HG0454**

ENSGALG00000007186 (TMEM268)

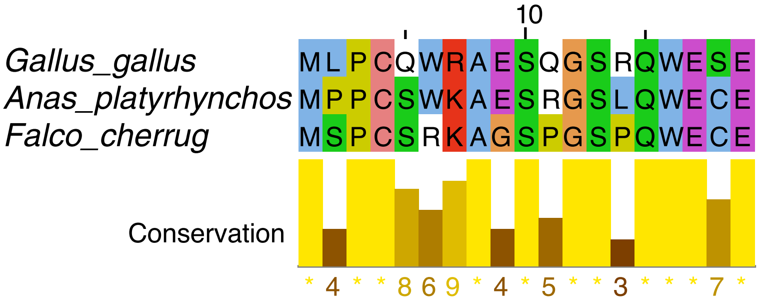

**HG0455**

ENSGALG00000010718 (THSD7A)

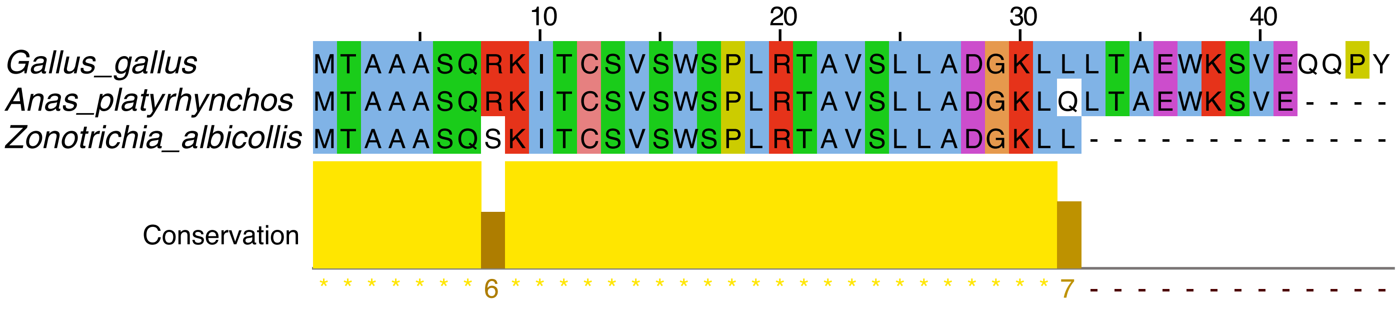

**HG0456**

ENSGALG00000033256

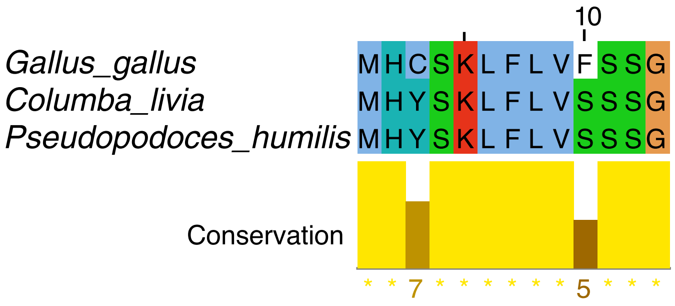

**HG0457**

ENSGALG00000039538 (CLDND1)

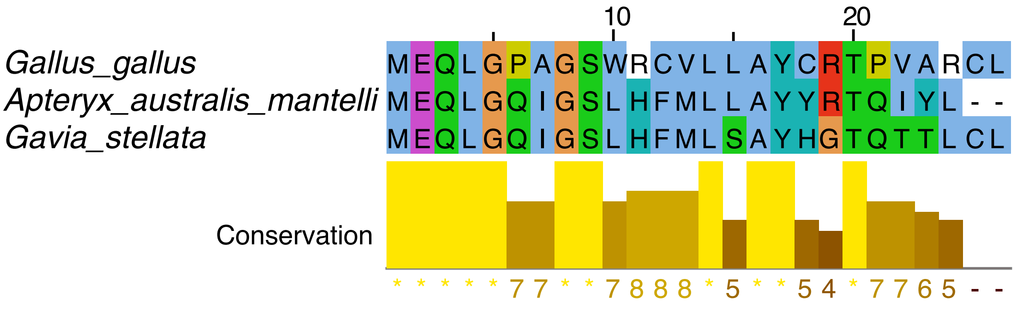

**HG0458**

ENSG00000060339 (CCAR1)

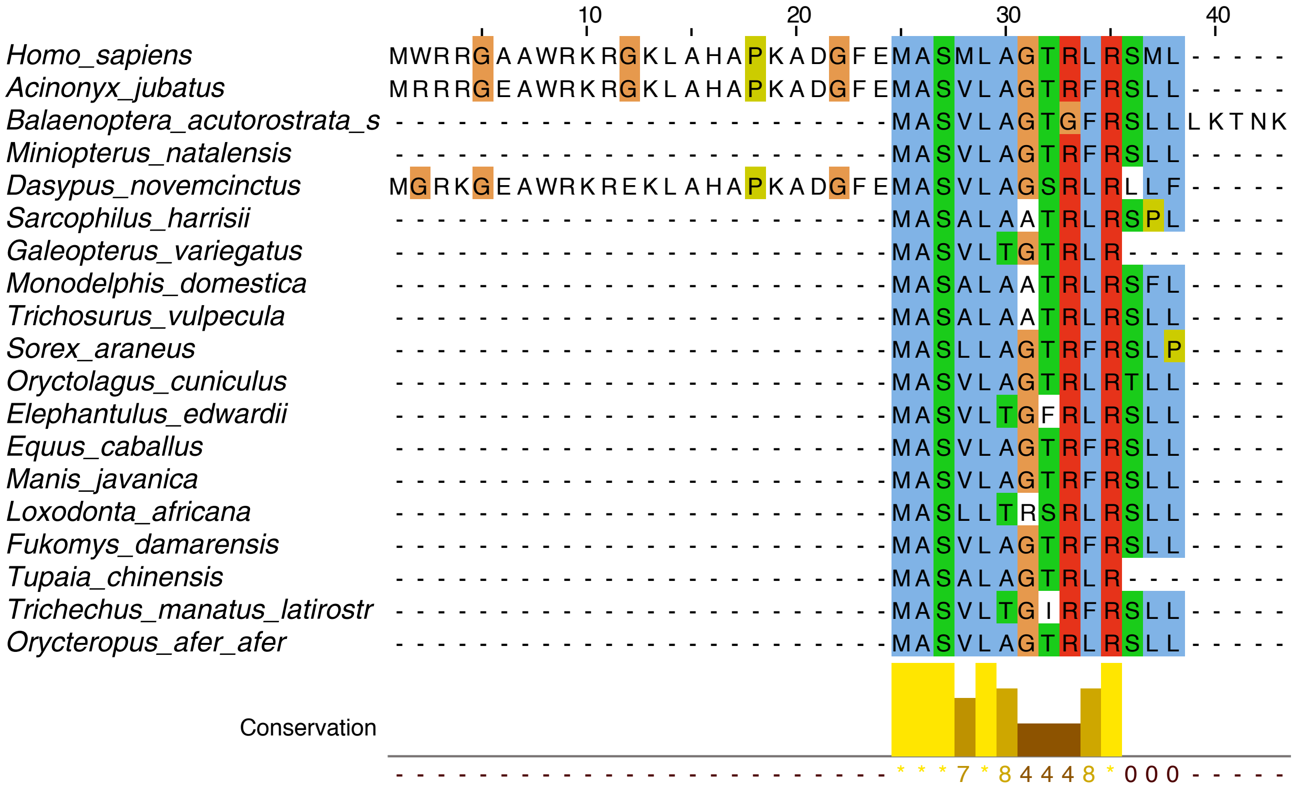

**HG0459.1**

ENSG00000163848 (ZNF148)

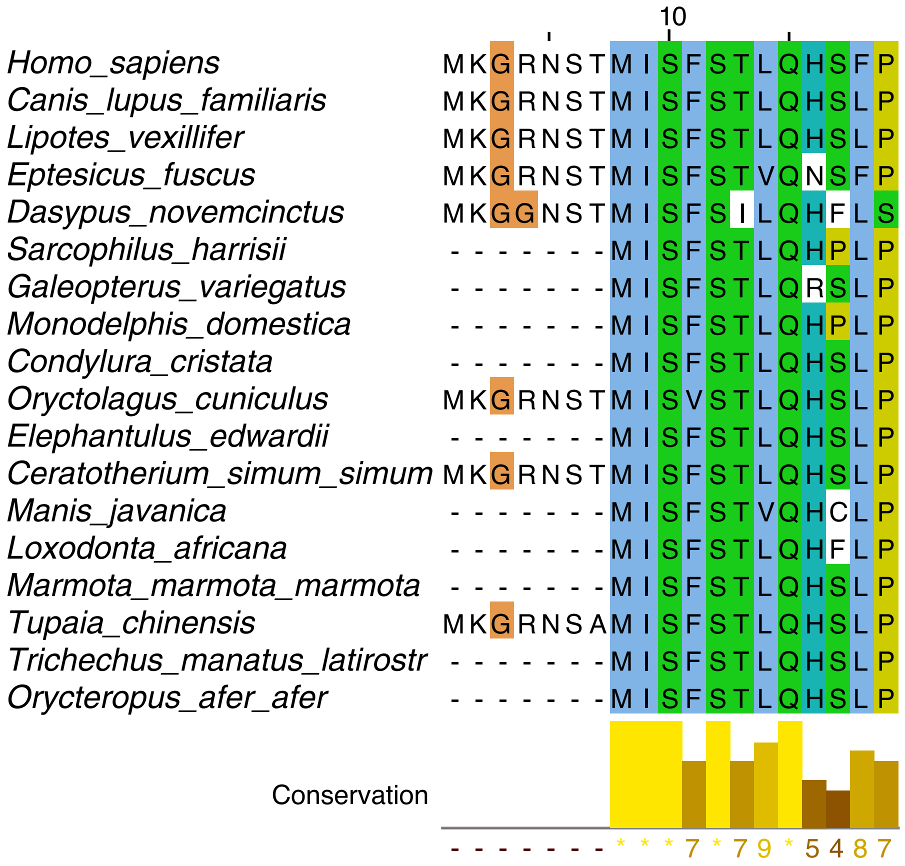

**HG0459.2**

ENSG00000163848 (ZNF148)

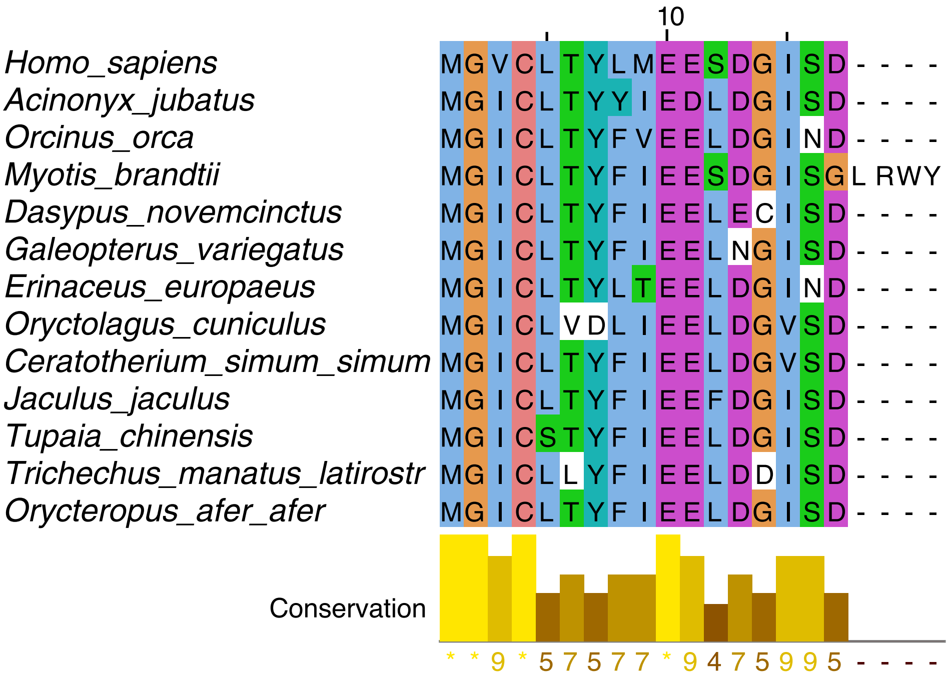

**HG0459.3**

ENSG00000163848 (ZNF148)

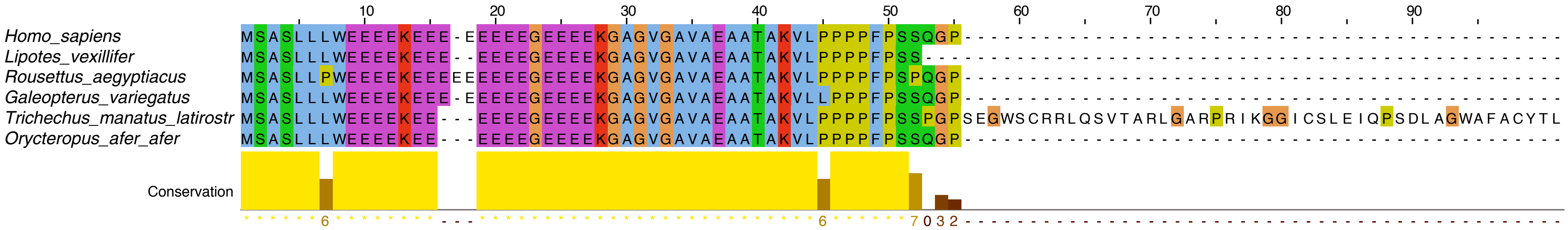

**HG0460**

ENSG00000180008 (SOCS4)

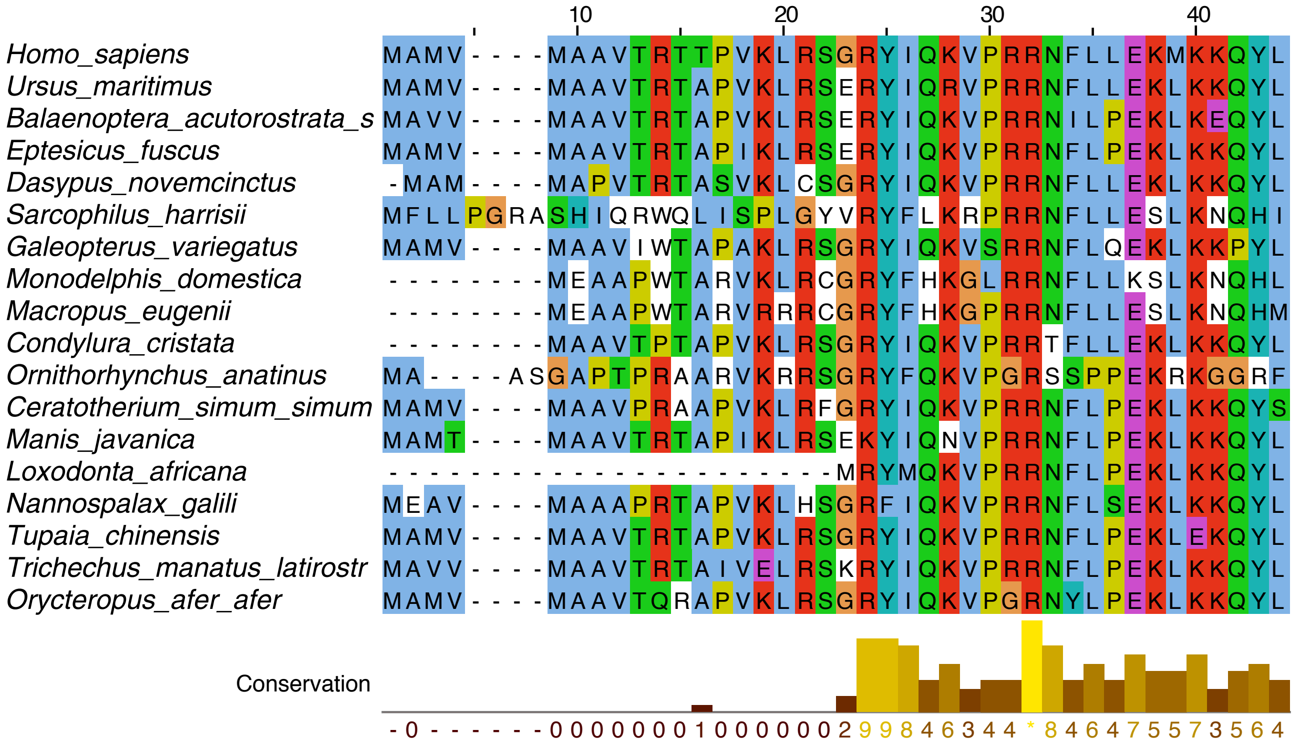

**HG0461**

ENSG00000064419 (TNPO3)

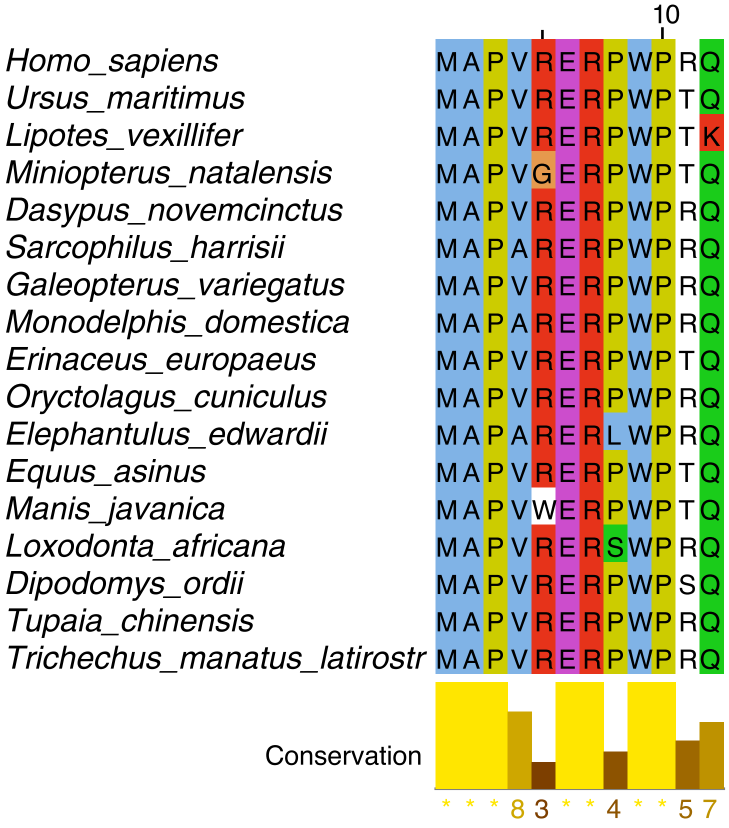

**HG0462**

ENSG00000125686 (MED1)

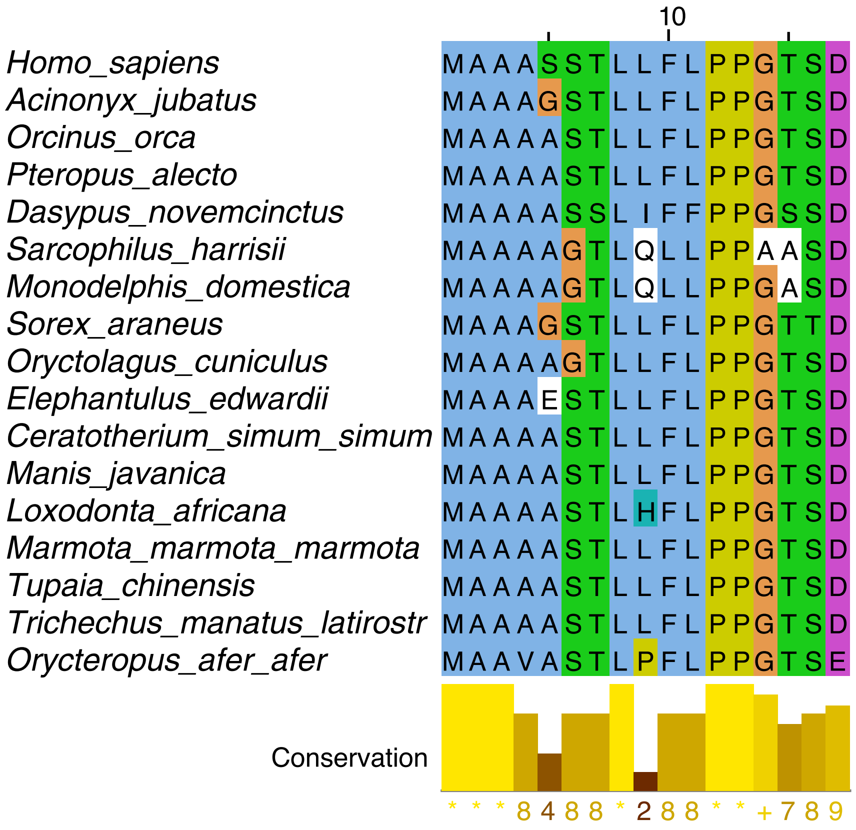

**HG0463**

ENSG00000129473 (BCL2L2)

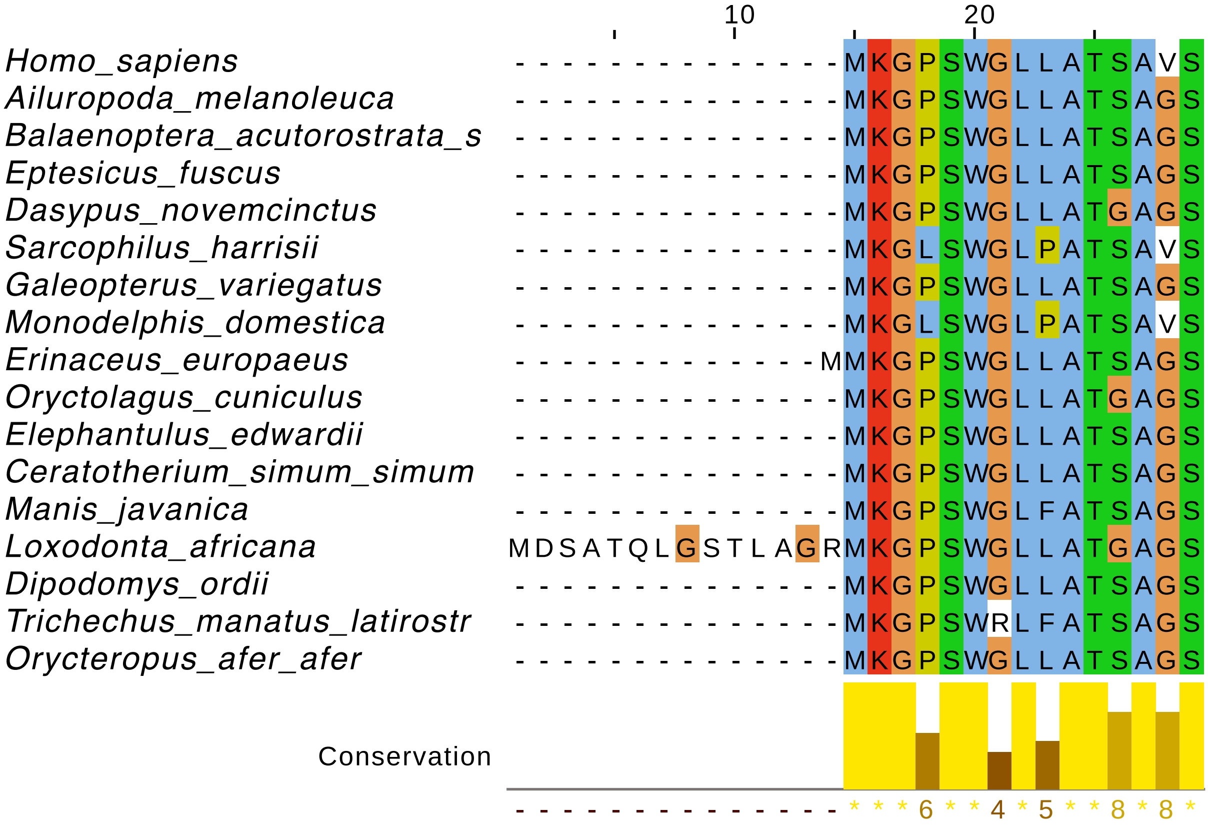

**HG0464**

ENSG00000188215 (DCUN1D3)

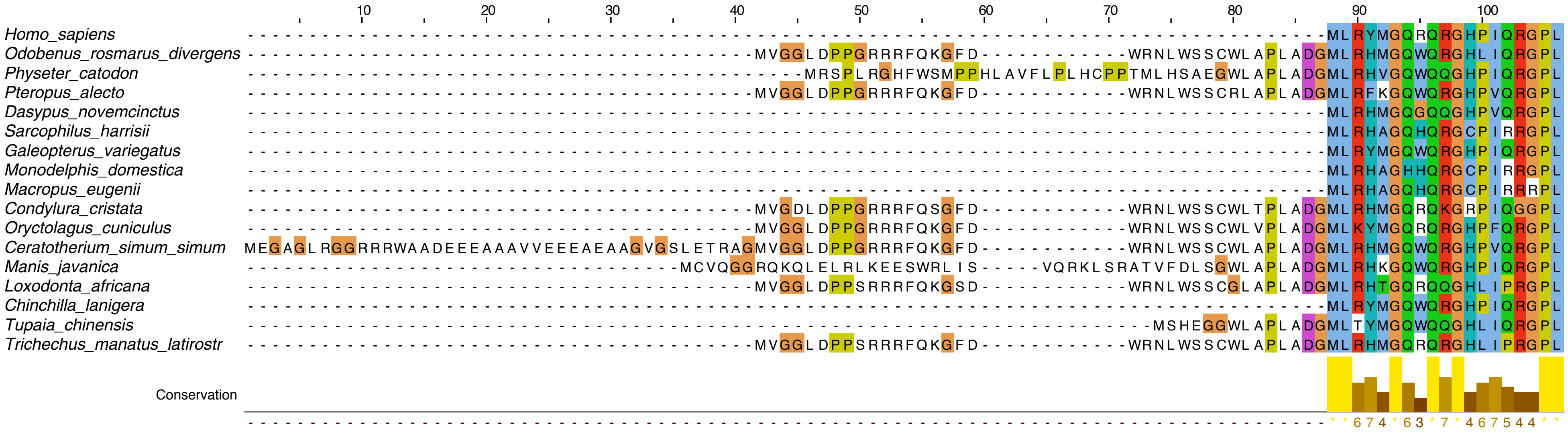

# HG0465

### ENSG00000132153 (DHX30)

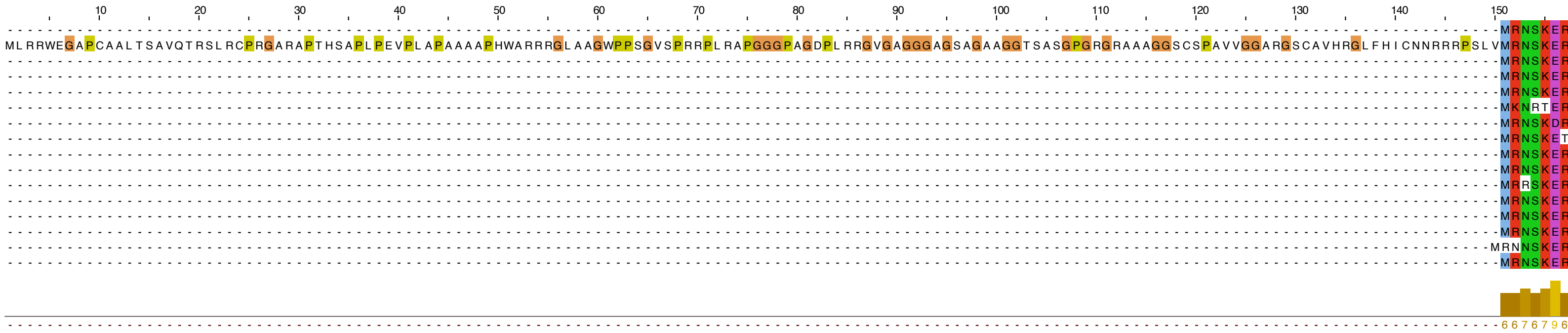

**HG0466**

ENSG00000139679 (LPAR6)

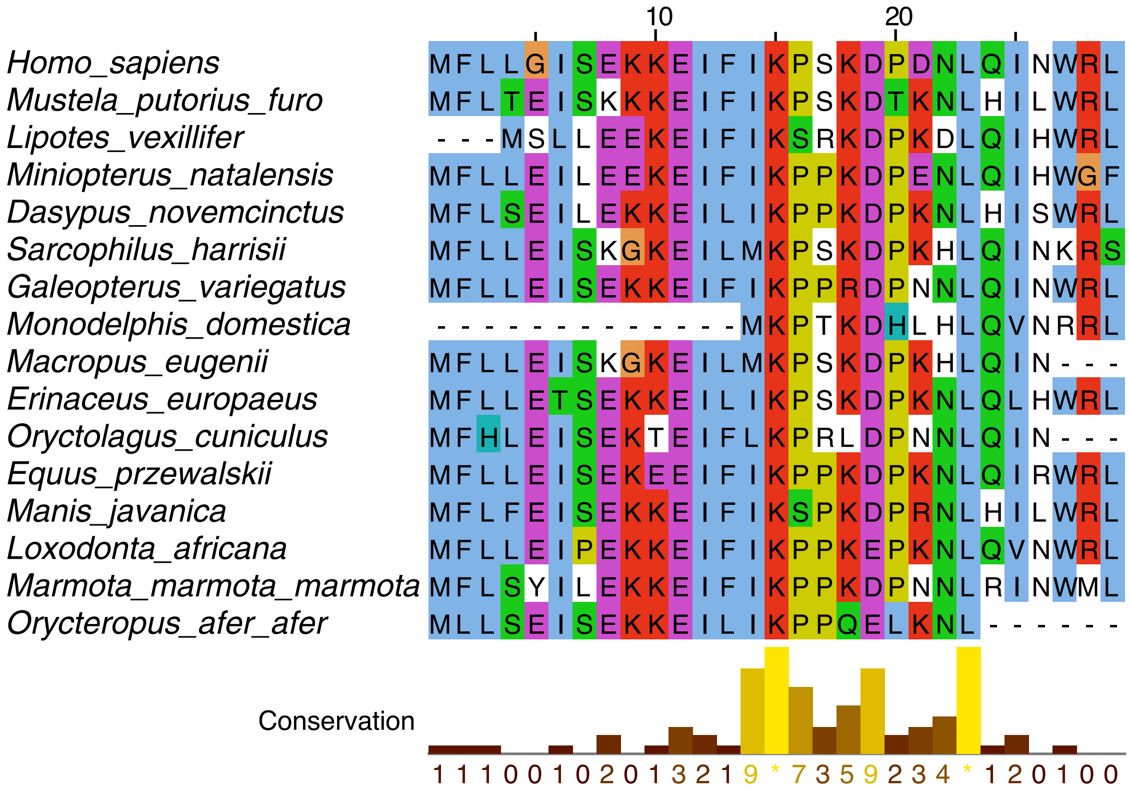

**HG0467.1**

ENSG00000163320 (CGGBP1)

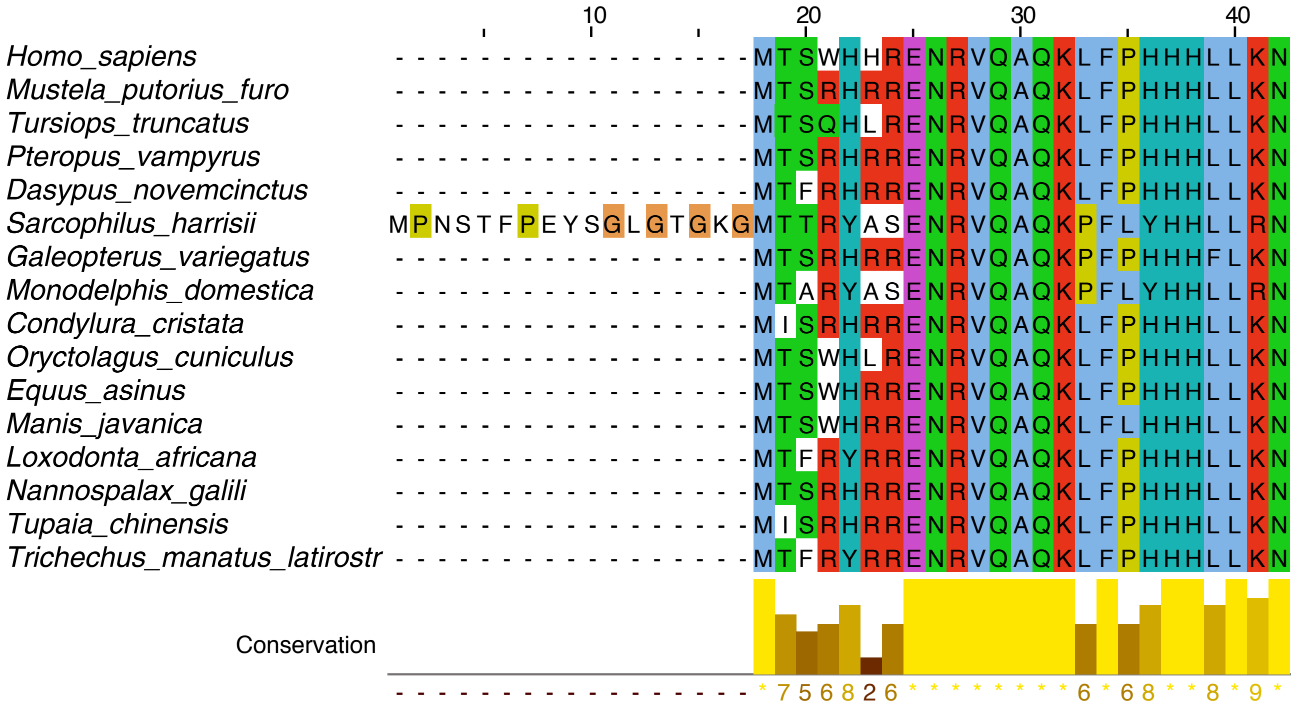

**HG0467.2**

ENSG00000163320 (CGGBP1)

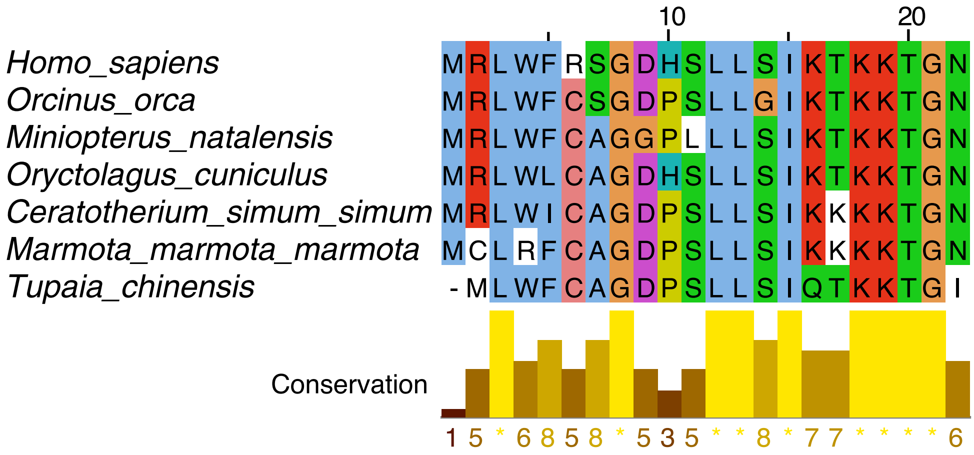

**HG0468**

ENSG00000187778 (MCRS1)

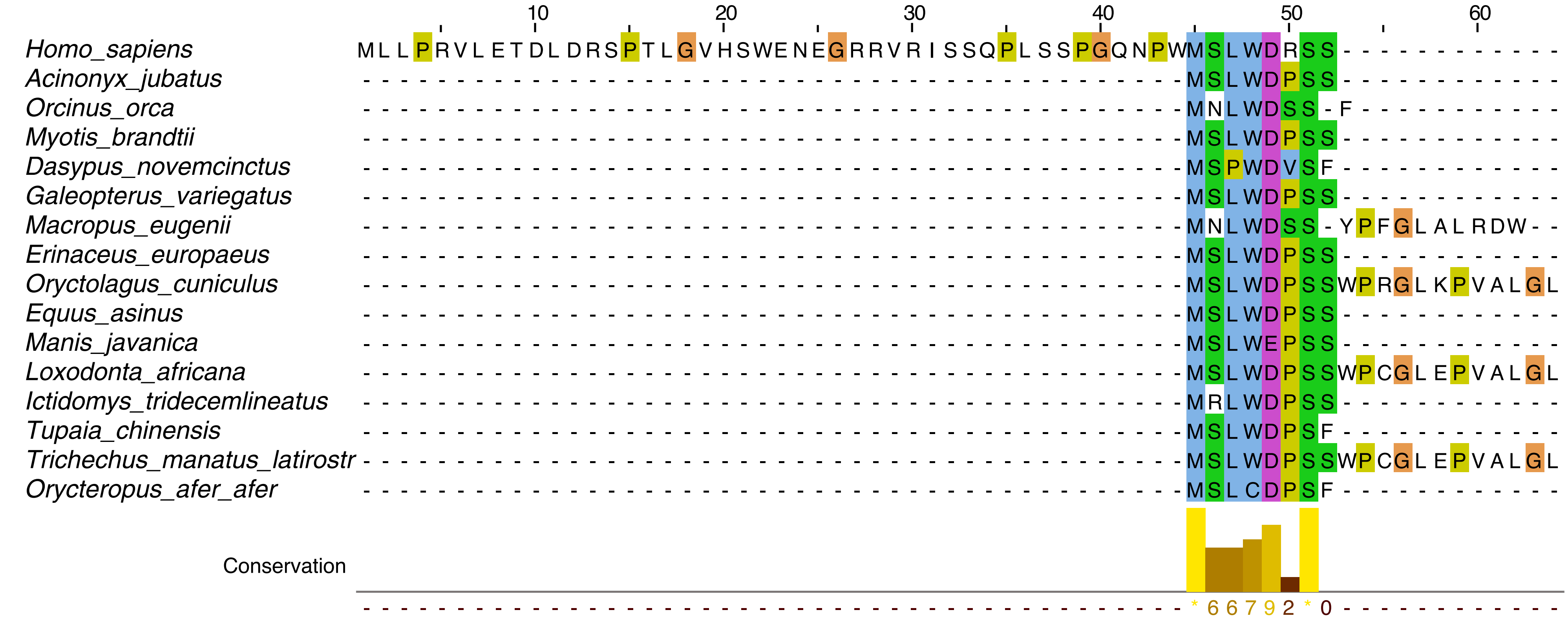

**HG0469**

ENSG00000198369 (SPRED2)

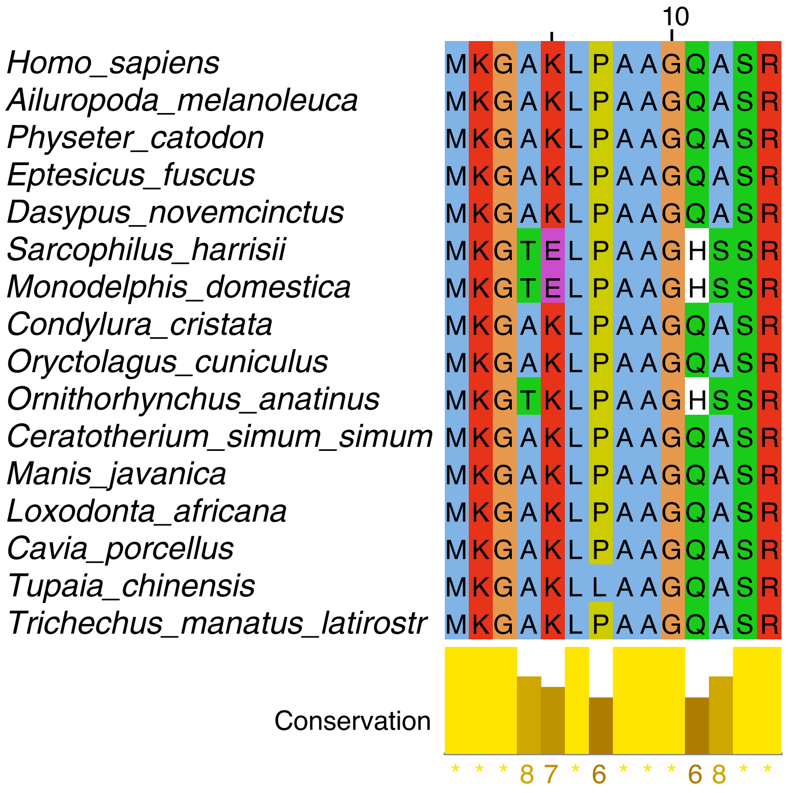

**HG0470**

ENSG00000224470 (ATXN1L)

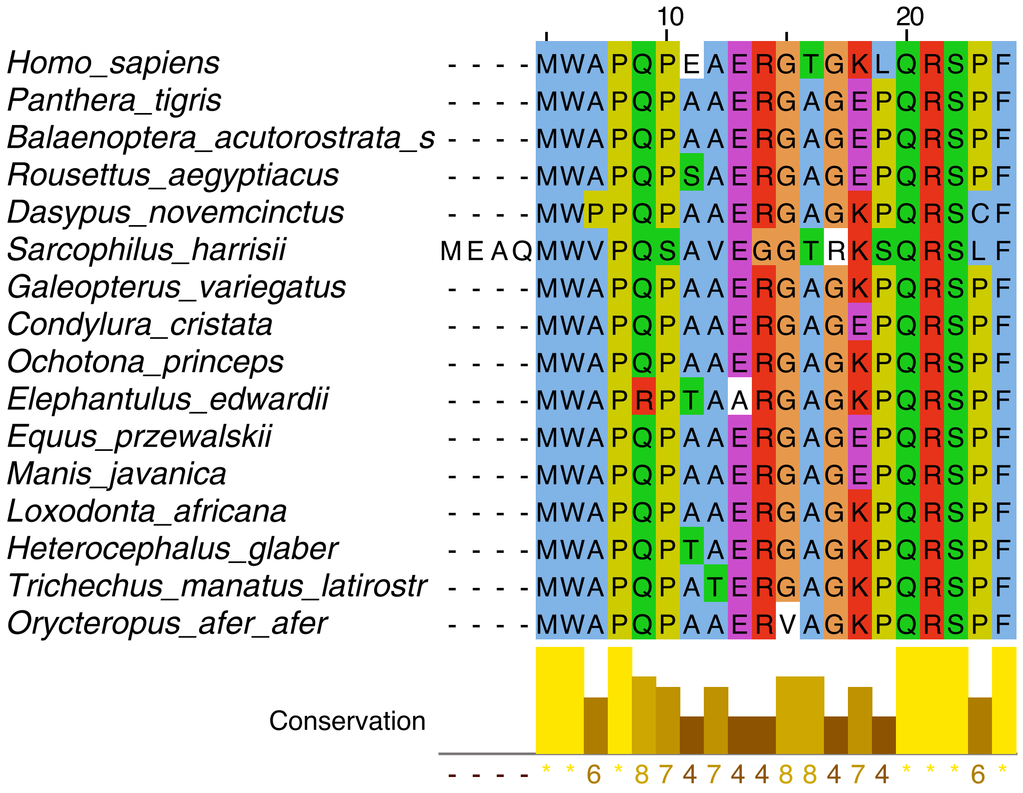

**HG0471**

ENSG00000023041 (ZDHHC6)

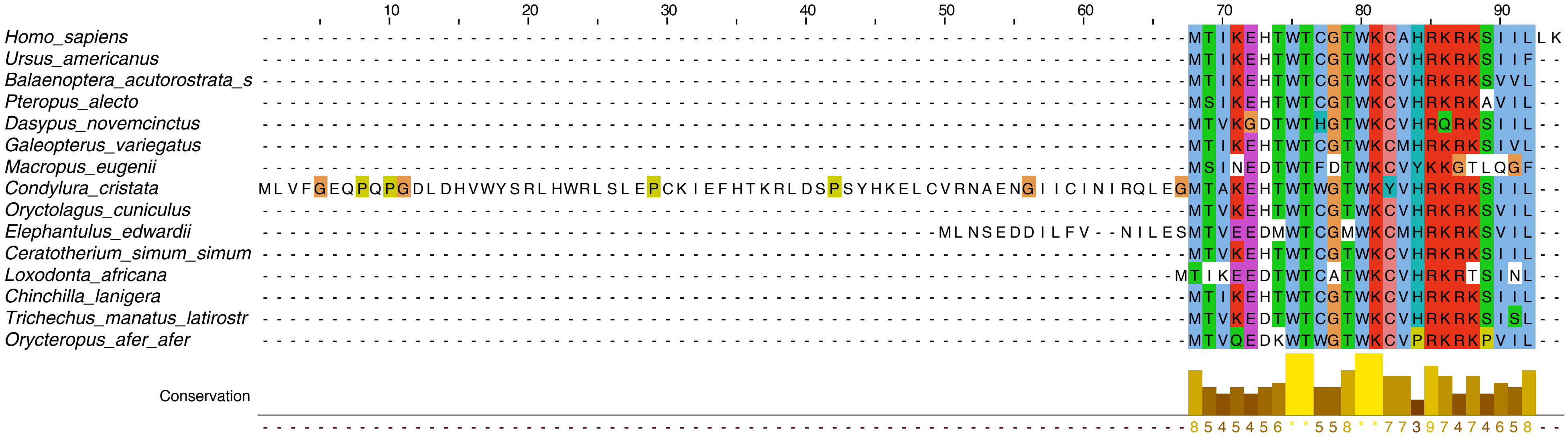

**HG0472**

ENSG00000105991 (HOXA1)

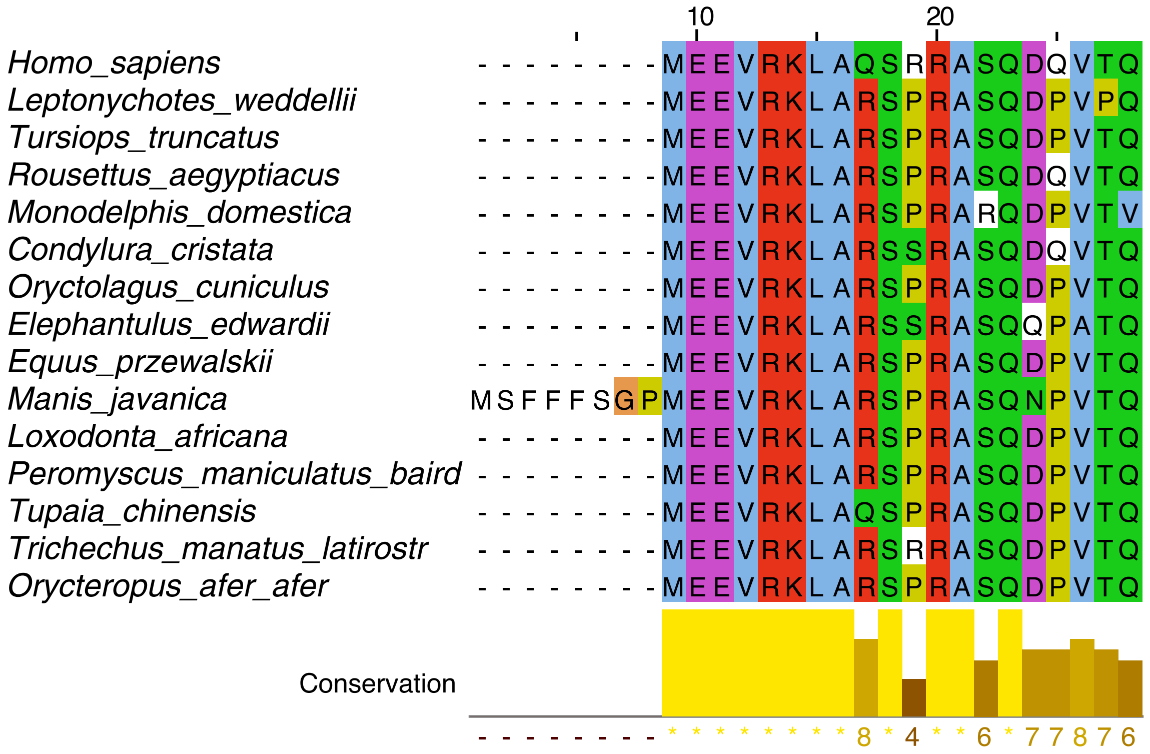

**HG0473.1**

ENSG00000138650 (PCDH10)

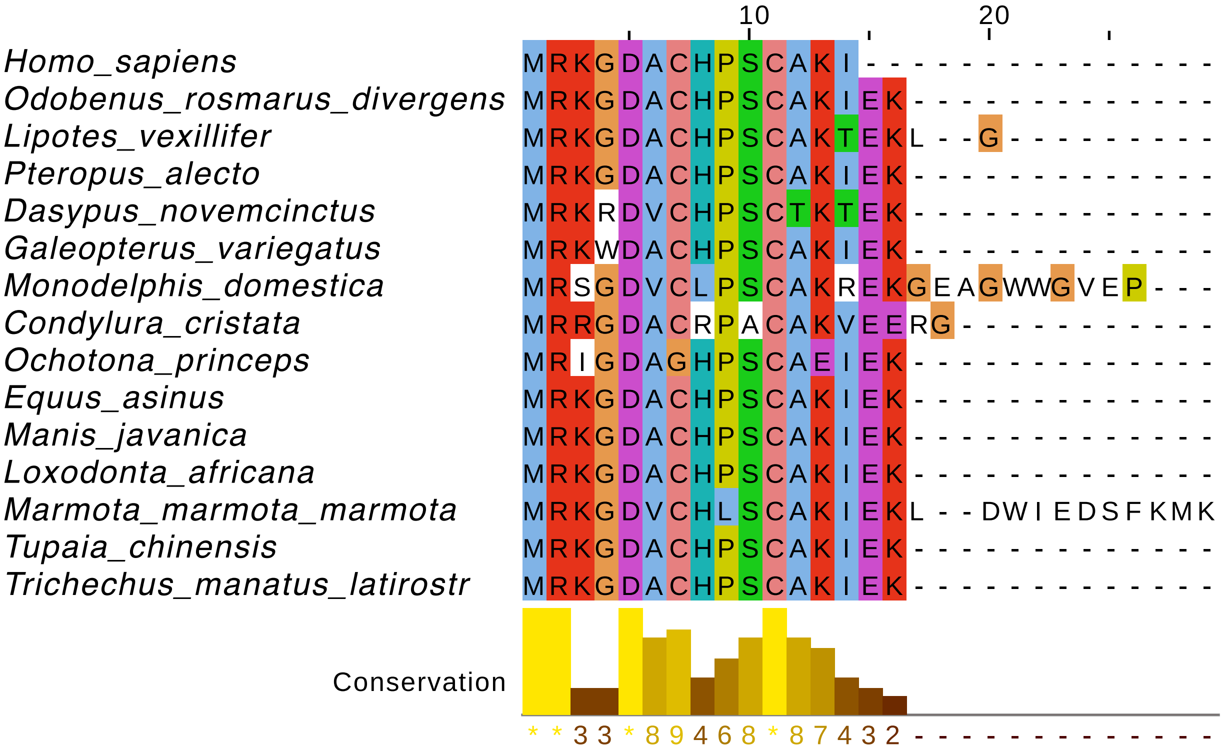

**HG0473.2**

ENSG00000240184 (PCDHGC3)

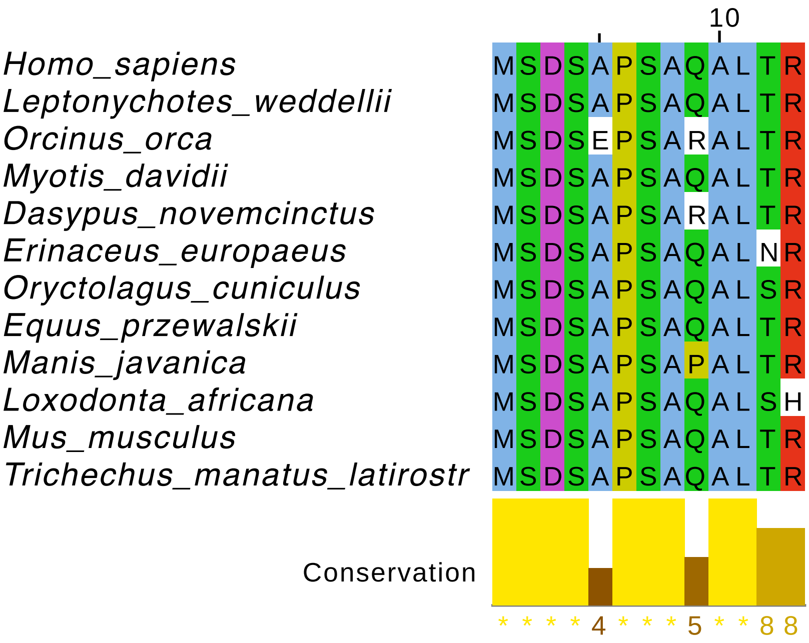

**HG0474**

ENSG00000139083 (ETV6)

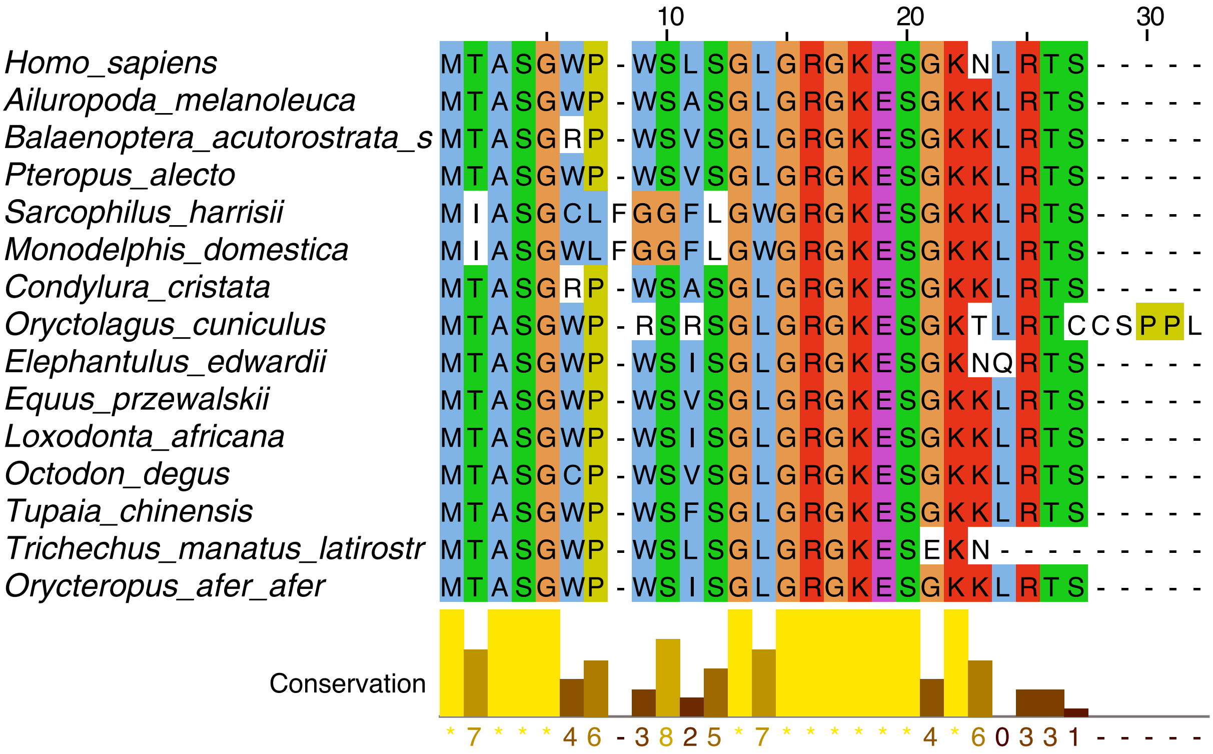

**HG0475**

ENSG00000109118 (PHF12)

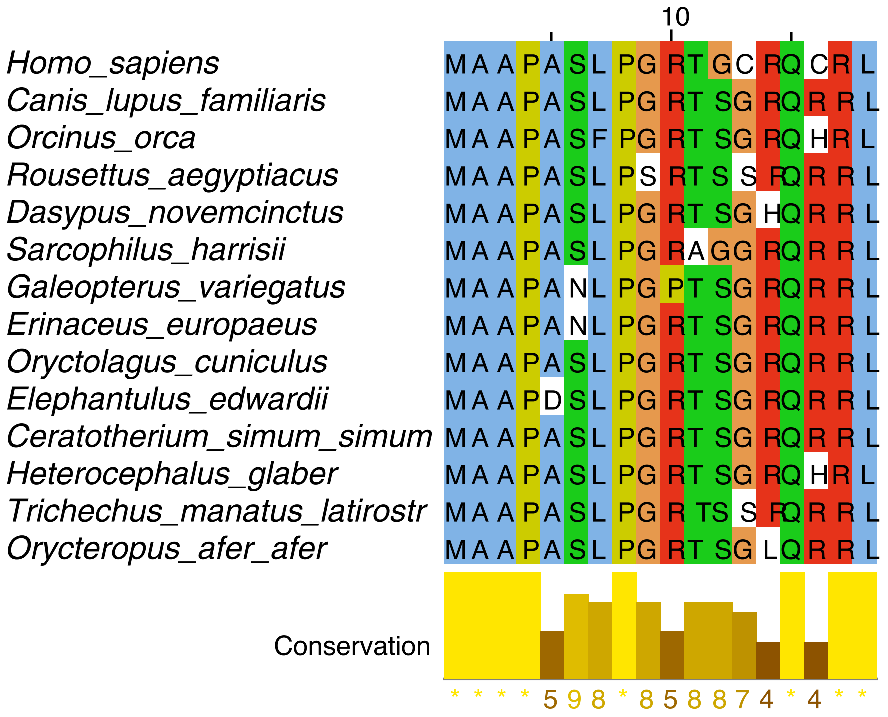

**HG0476**

ENSG00000130939 (UBE4B)

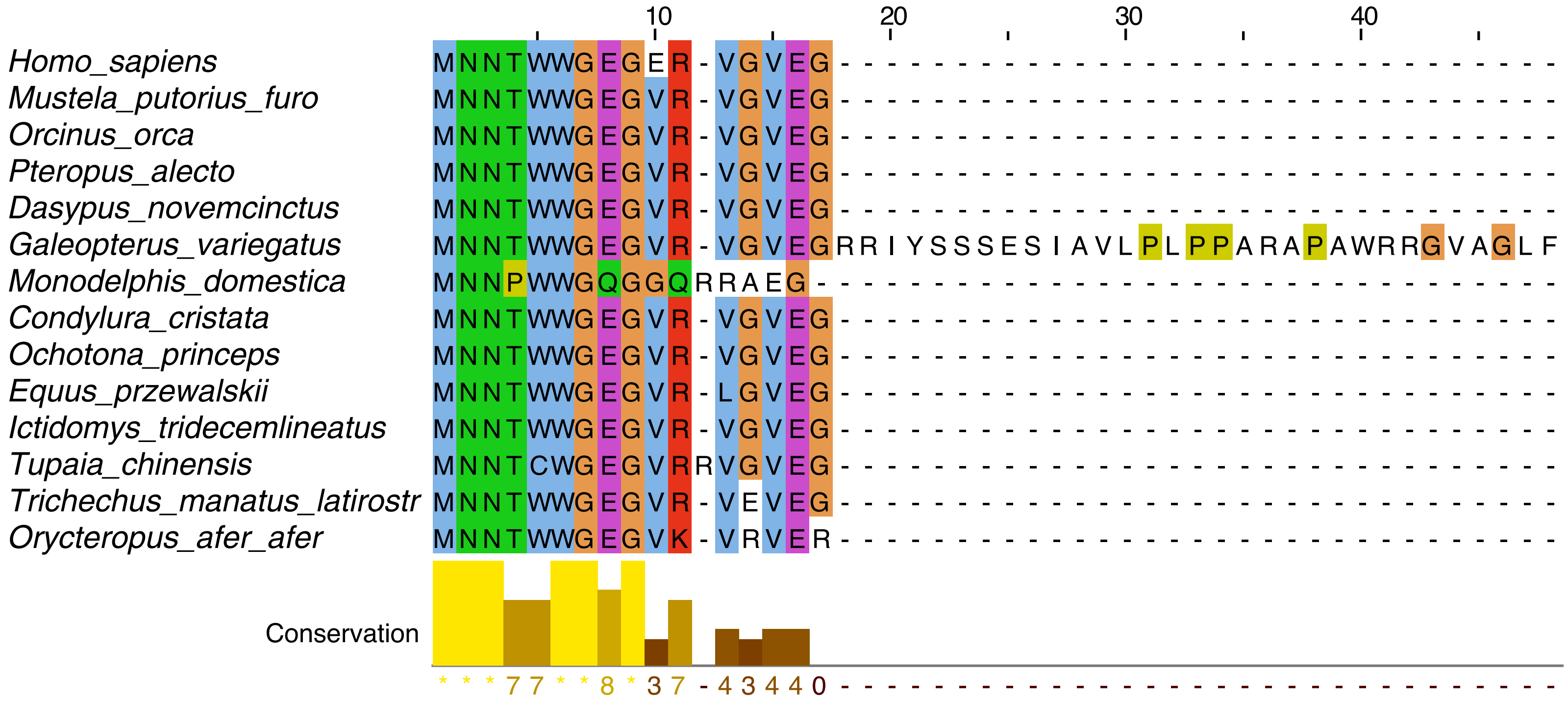

**HG0477**

ENSG00000166925 (TSC22D4)

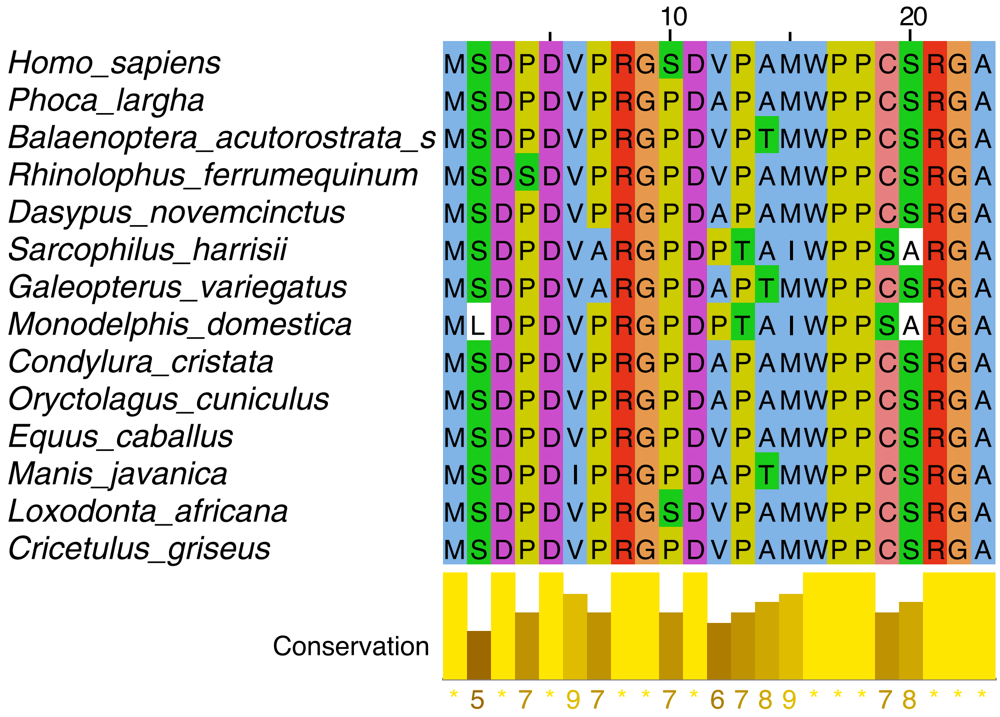

**HG0478**

ENSG00000168214 (RBPJ)

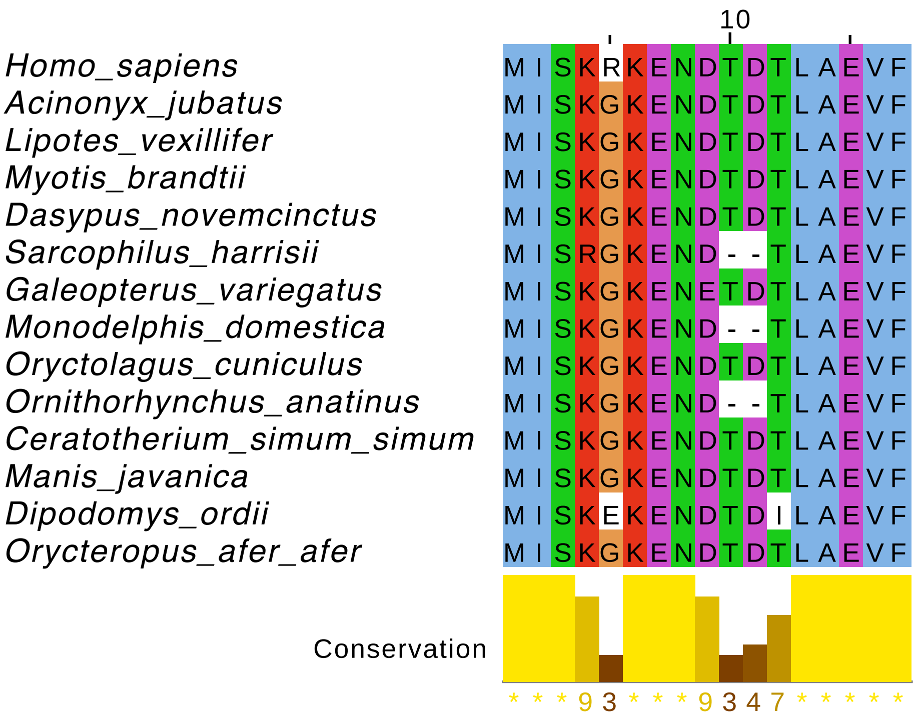

# HG0479.1

### ENSG00000168453 (HR)

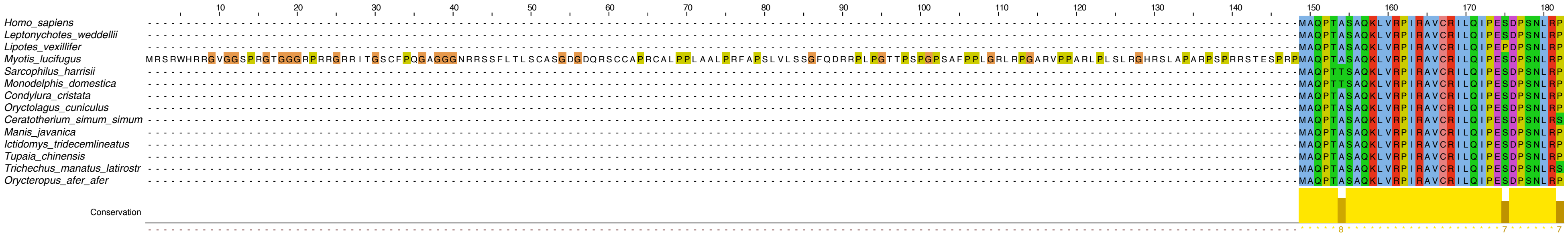

**HG0479.2**

ENSG00000168453 (HR)

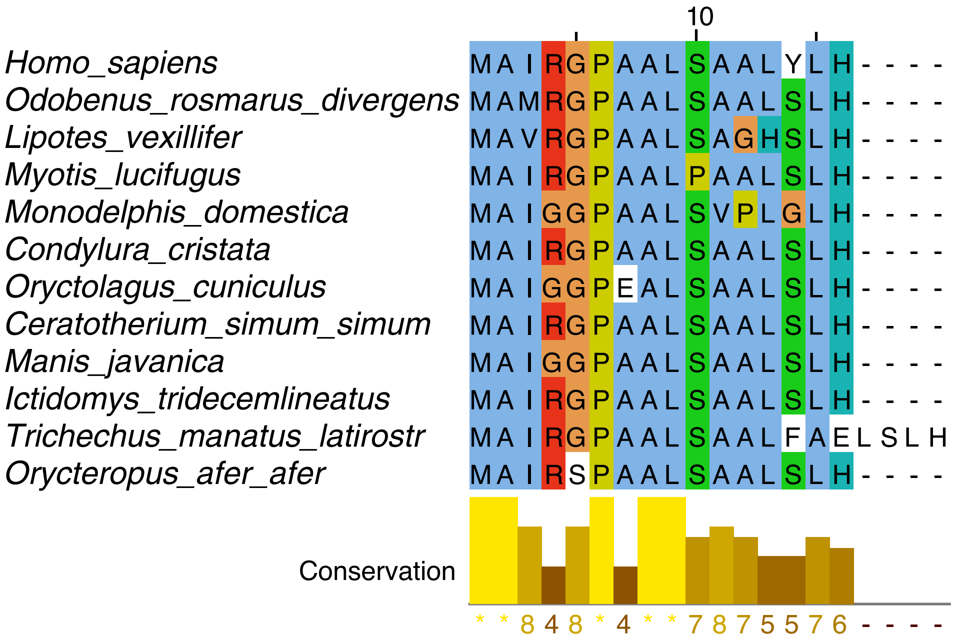

**HG0480**

ENSG00000198018 (ENTPD7)

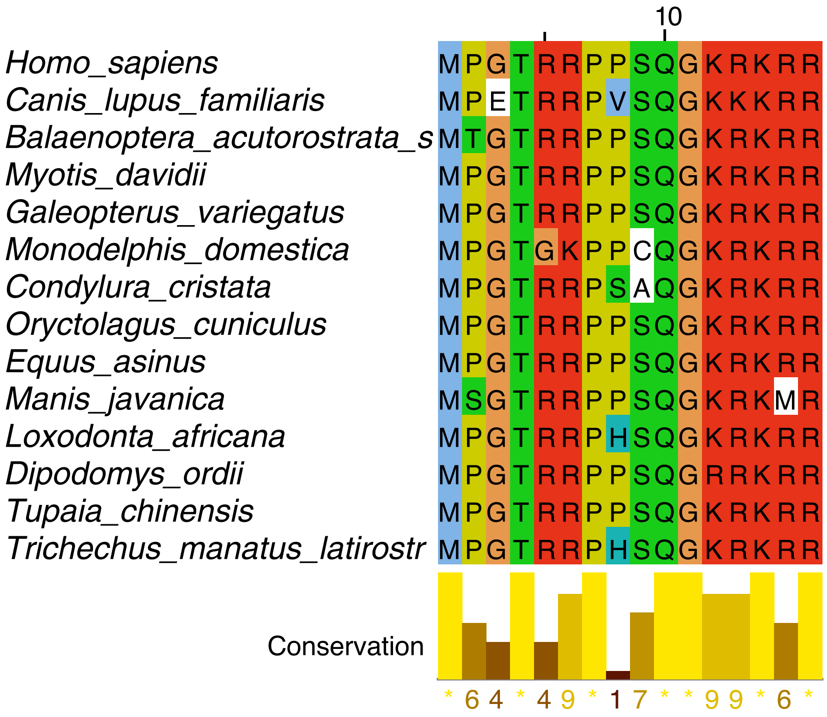

**HG0481**

ENSG00000198963 (RORB)

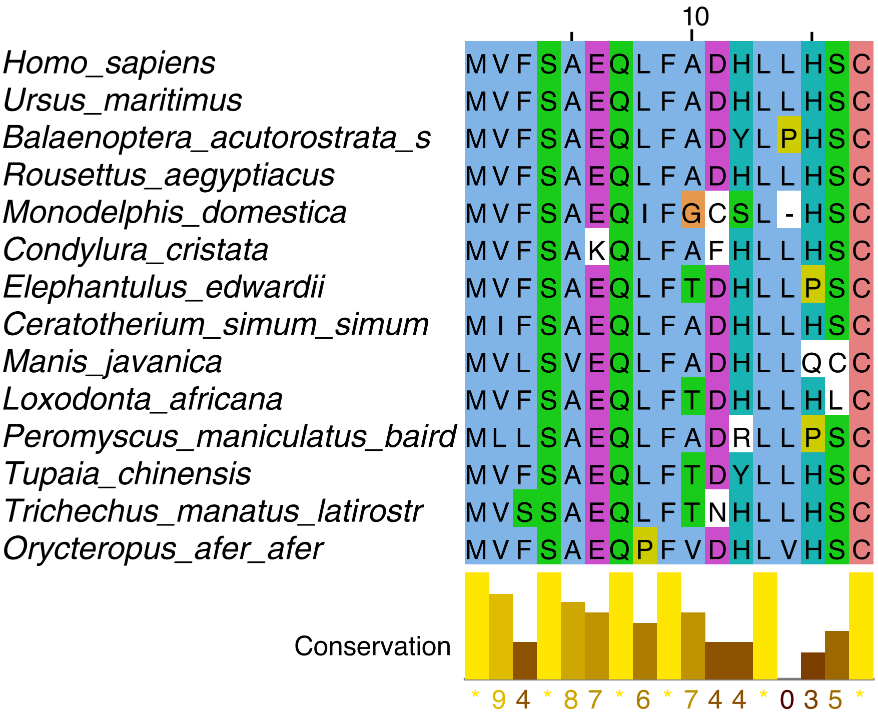

**HG0482**

ENSG00000067167 (TRAM1)

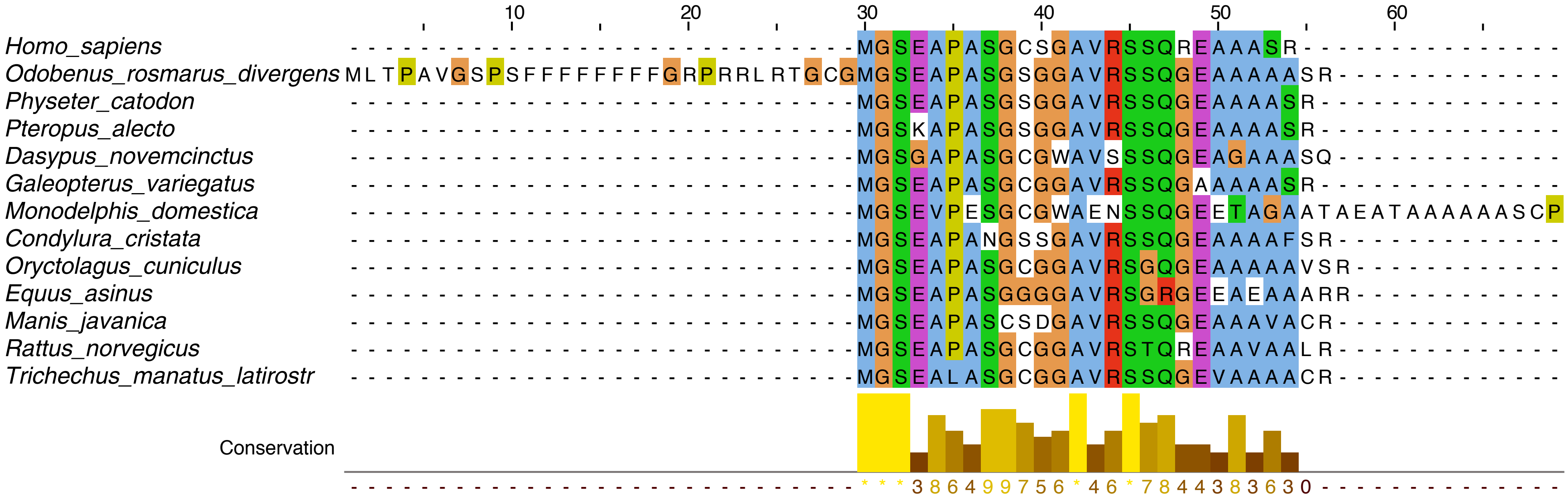

**HG0483**

ENSG00000164463 (CREBRF)

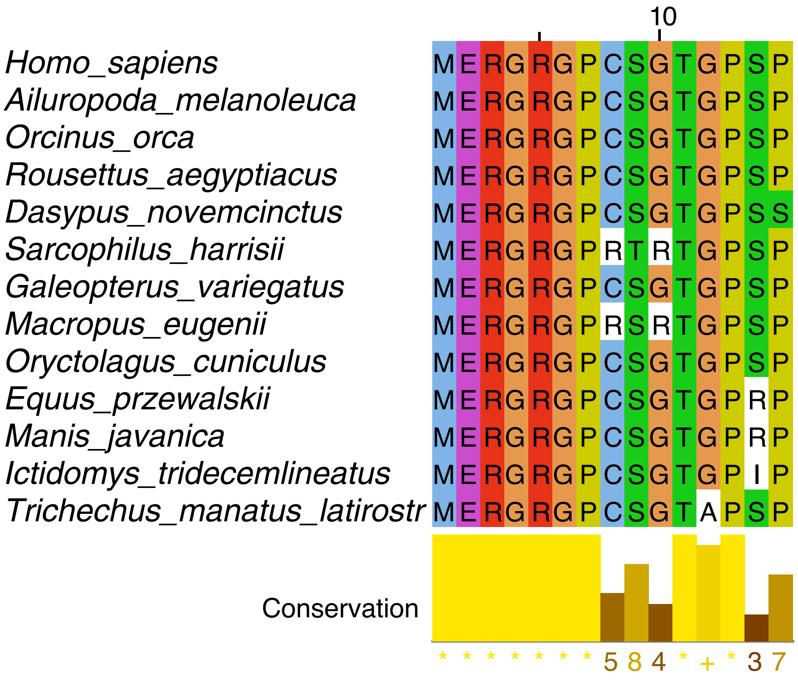

**HG0484**

ENSG00000166444 (ST5)

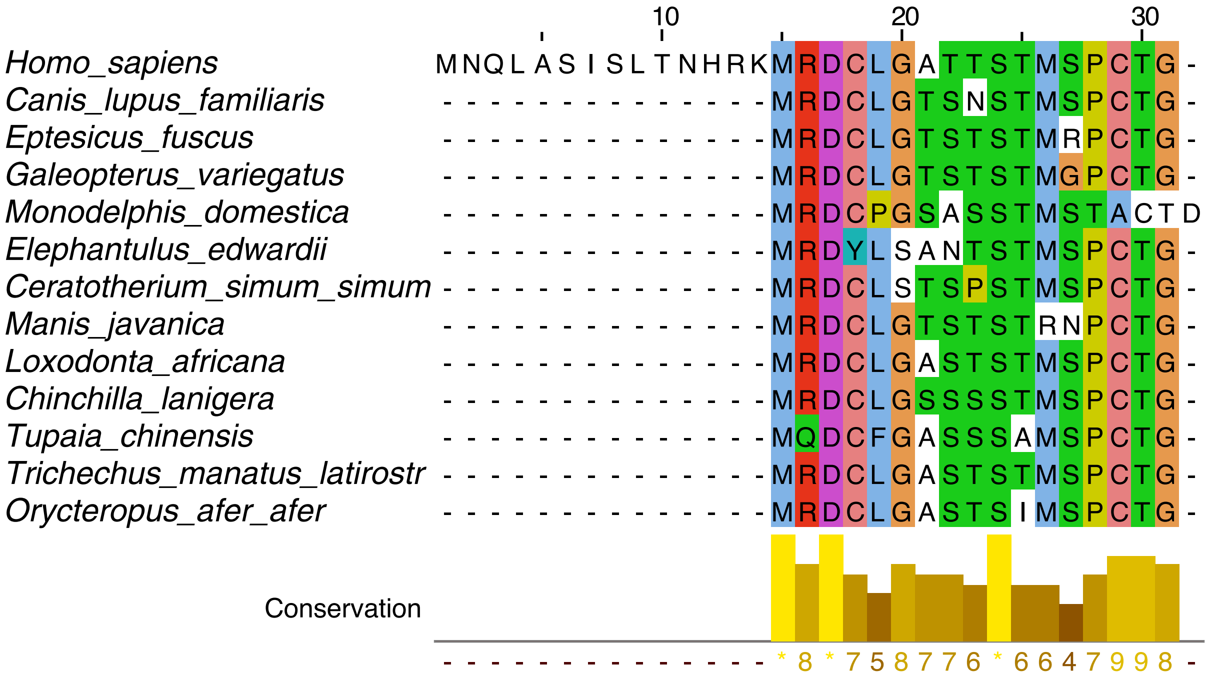

**HG0485.1**

ENSG00000182667 (NTM)

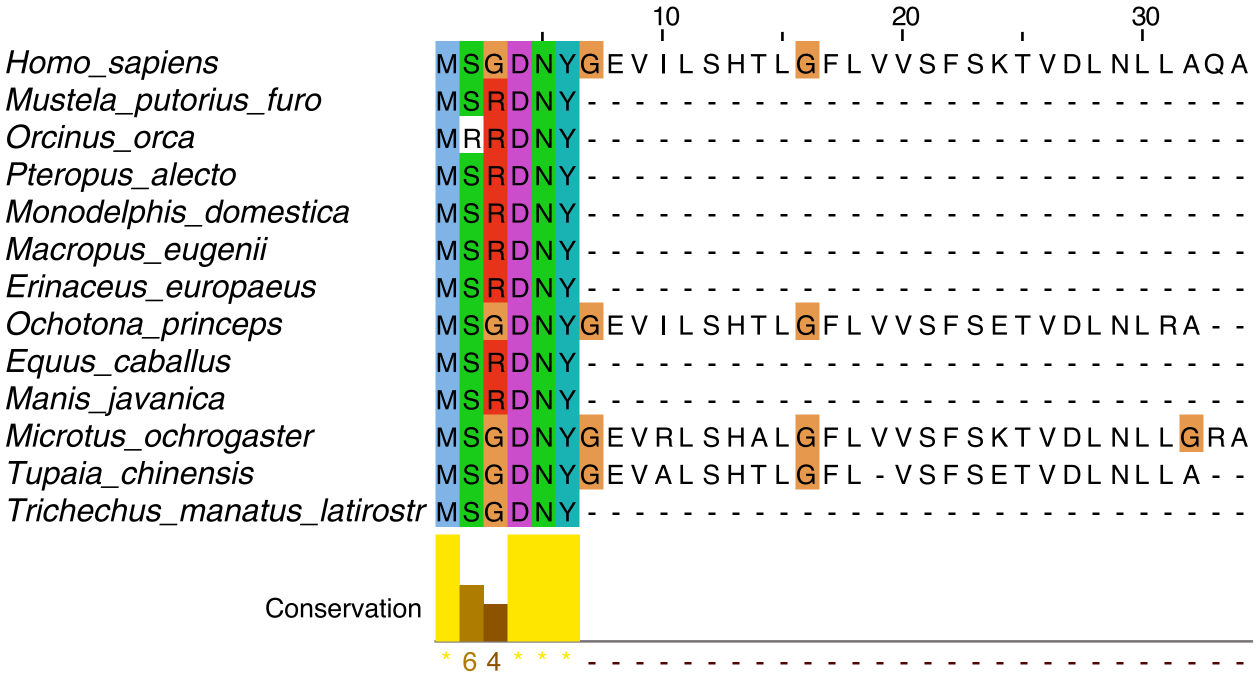

**HG0485.2**

ENSG00000183715 (OPCML)

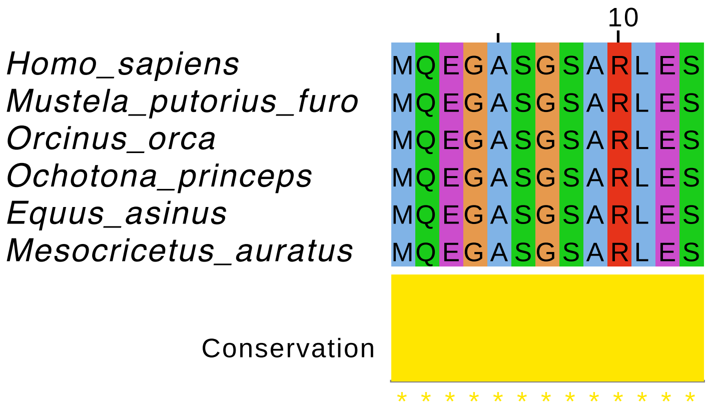

**HG0486**

ENSG00000067900 (ROCK1)

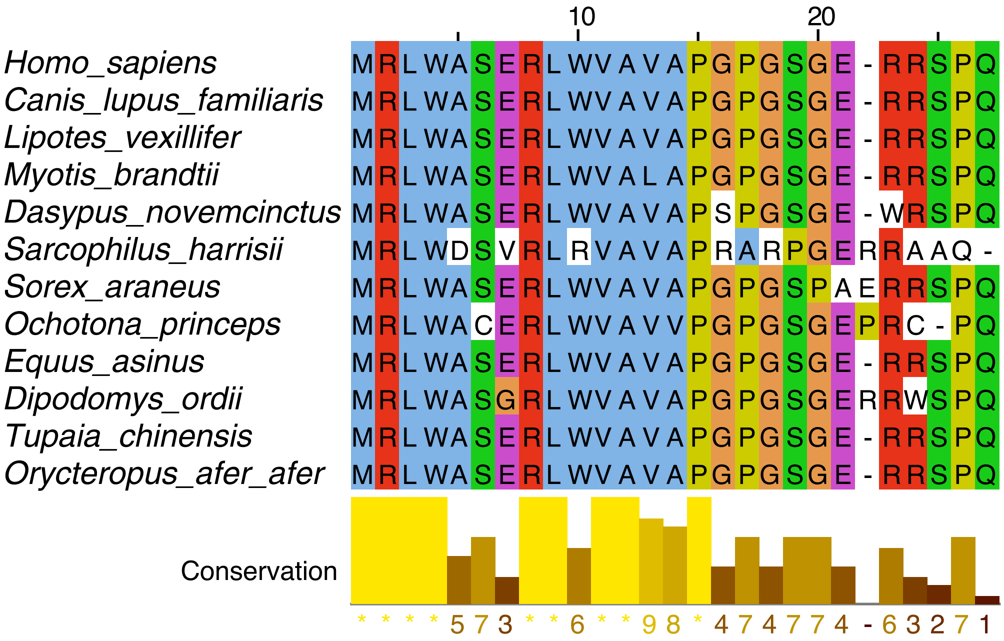

**HG0487**

ENSG00000091831 (ESR1)

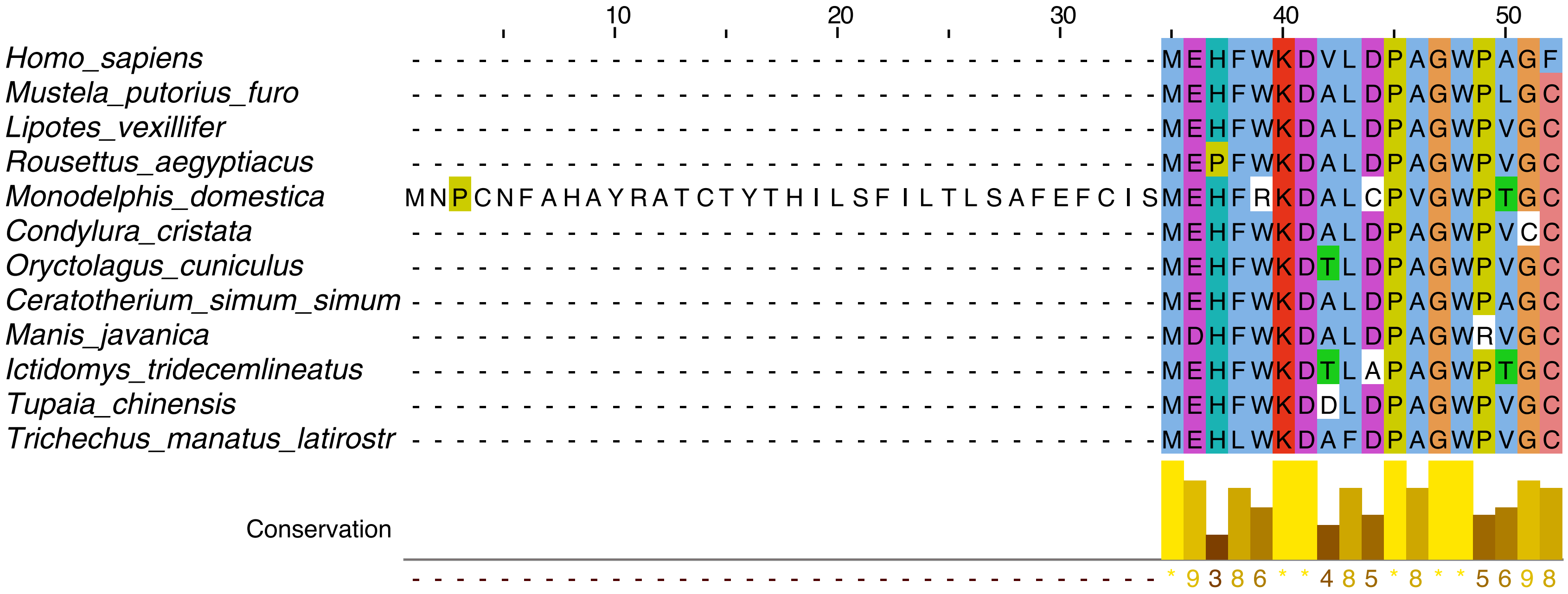

**HG0488.1**

ENSG00000116194 (ANGPTL1)

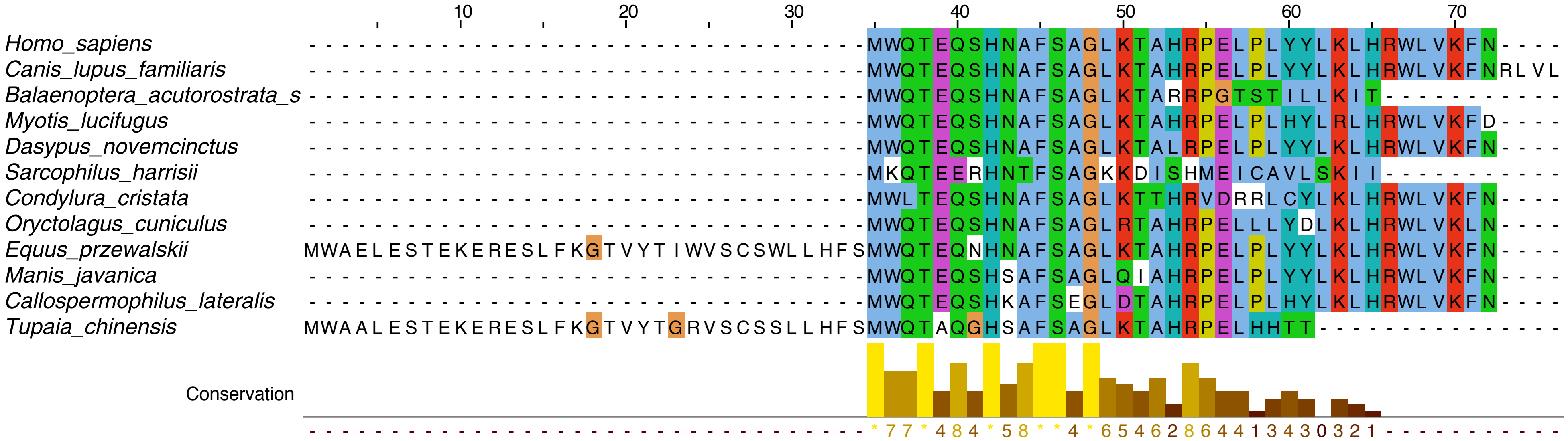

**HG0488.2**

ENSG00000116194 (ANGPTL1)

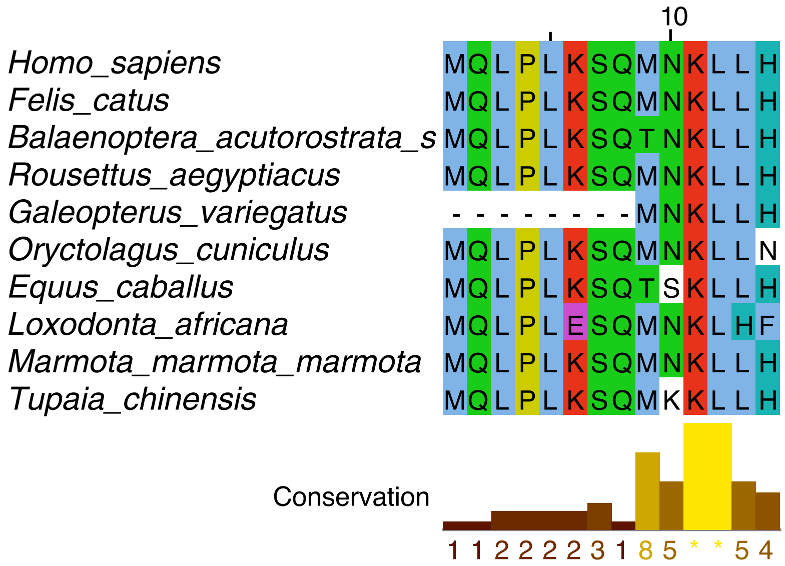

**HG0489**

ENSG00000143847 (PPFIA4)

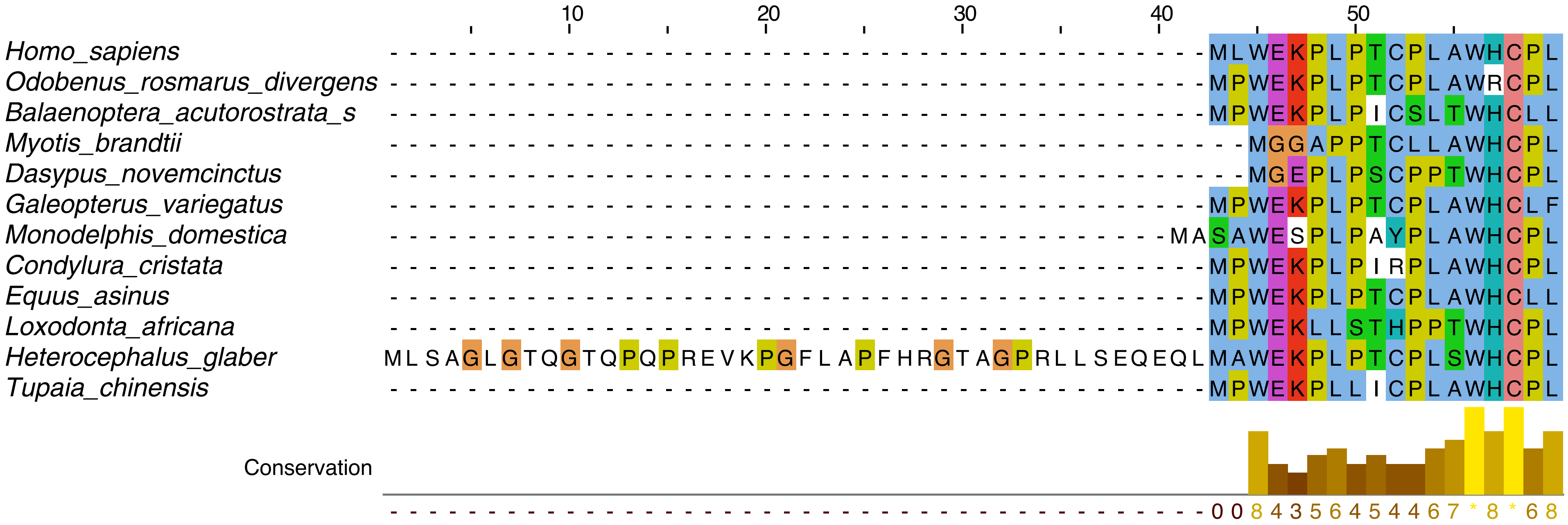

**HG0490.1**

ENSG00000152413 (HOMER1)

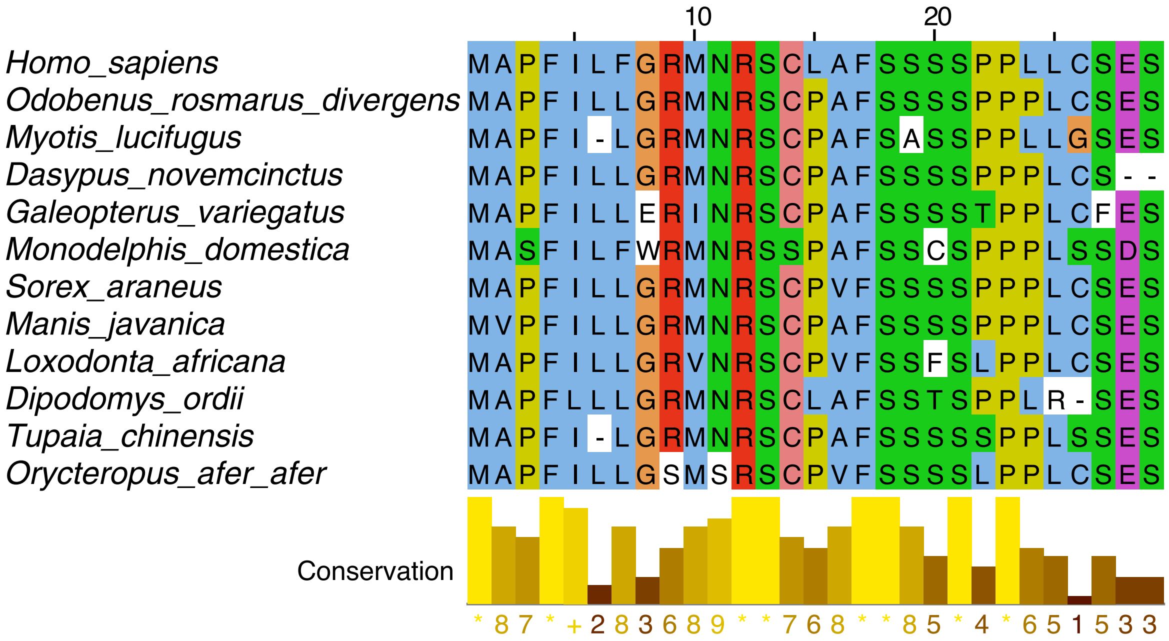

**HG0490.2**

ENSG00000152413 (HOMER1)

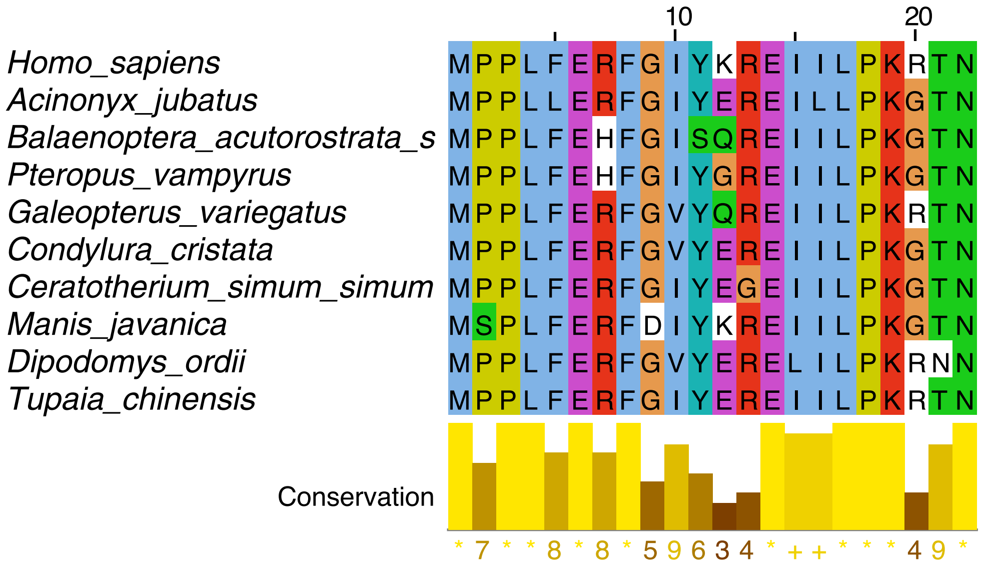

**HG0491**

ENSG00000154114 (TBCEL)

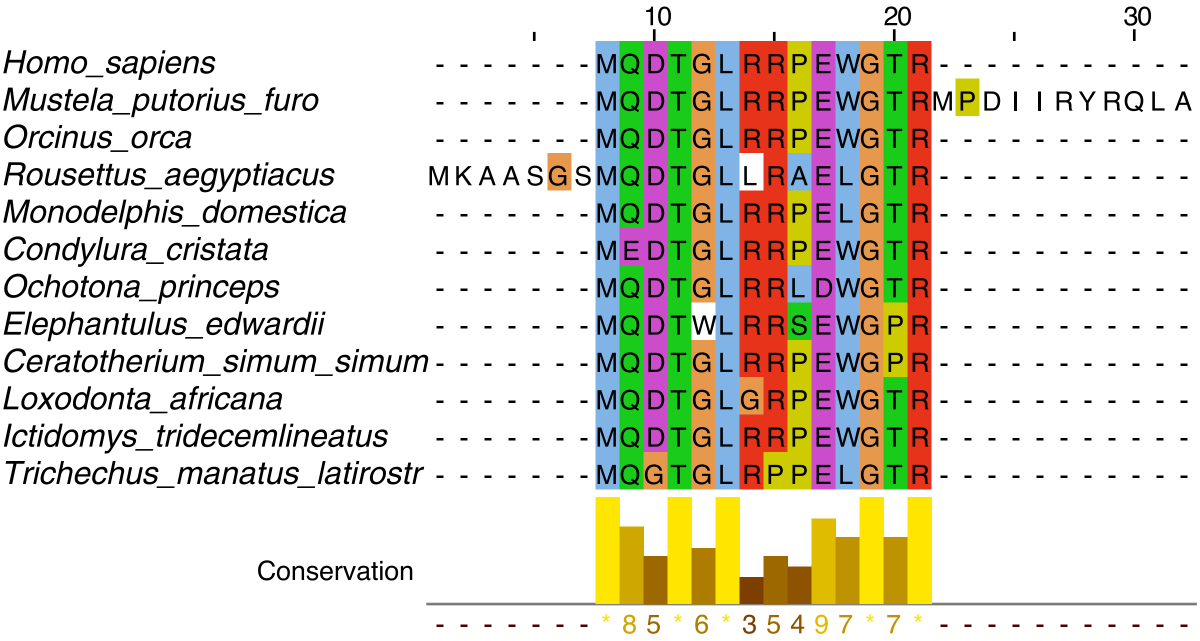

**HG0492**

ENSG00000162670 (BRINP3)

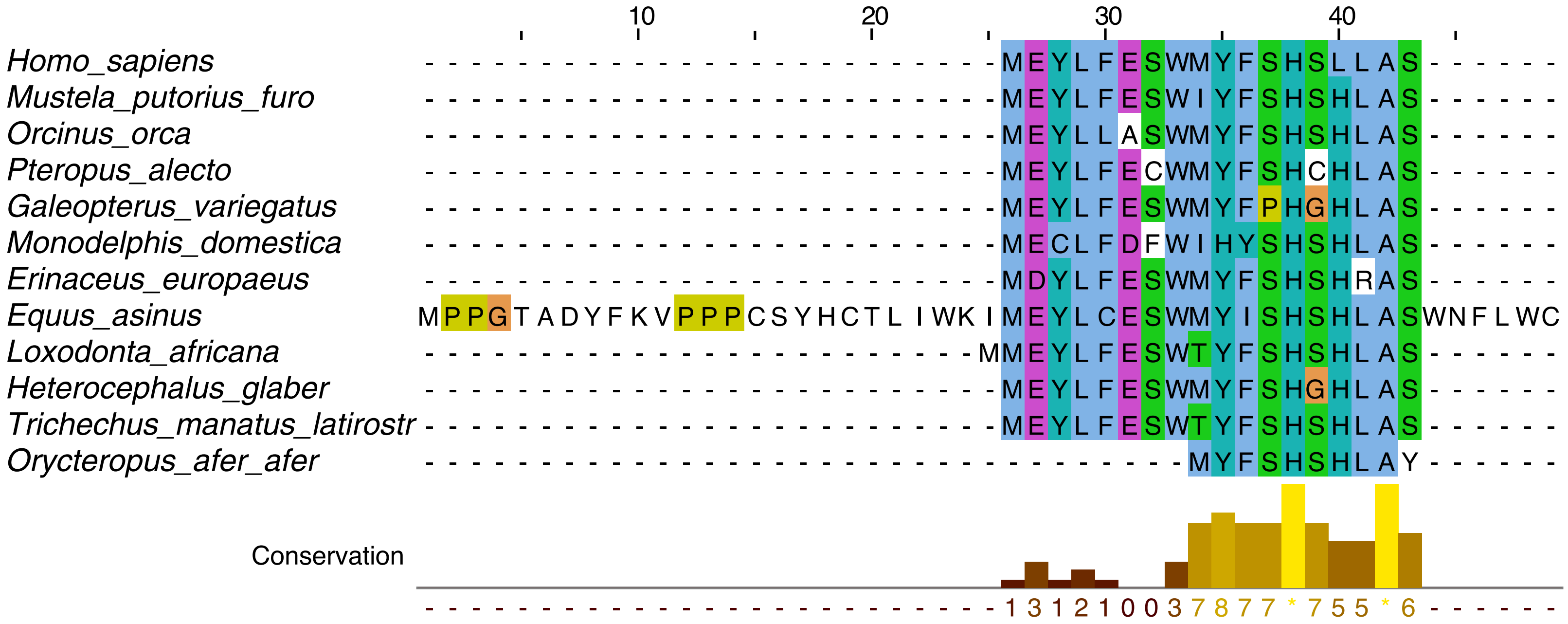

**HG0493**

ENSG00000164707 (SLC13A4)

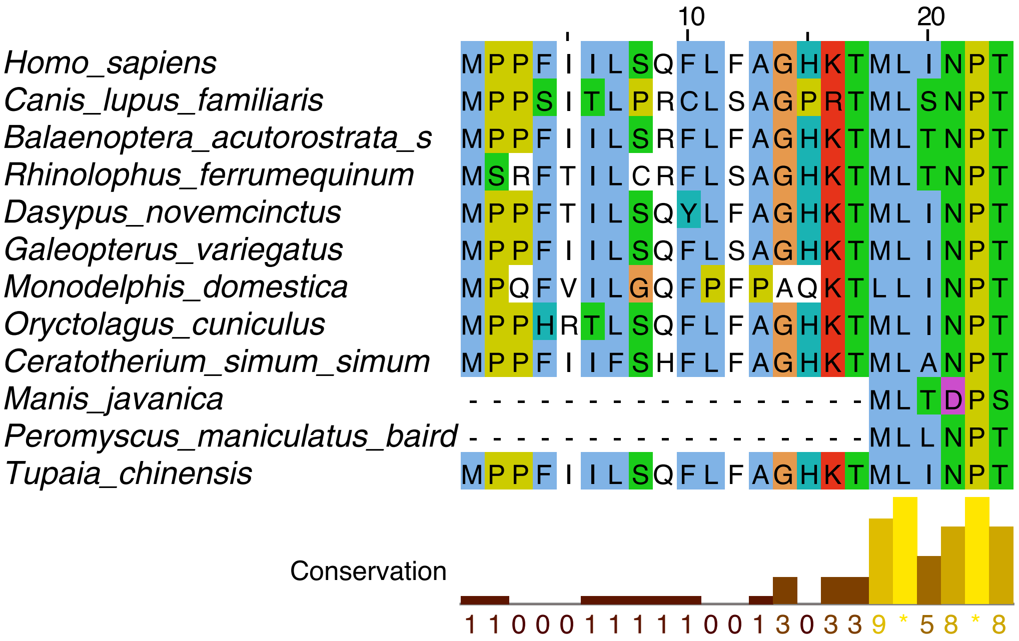

**HG0494**

ENSG00000184226 (PCDH9)

```
                                       10        20        30        40        50        60        70        80
Homo_sapiens                  -----------------------------------------MHAAVENIPELKKCNYVLCVLNAQKN---P---DPVGKRDEMWRE
Mustela_putorius_furo         -----------------------------------------MLDAVENTPELKKCNGMLC--VCVKYQKSP---DPVGNRDEMWRE
Orcinus_orca                  -----------------------------------------MFDAVENTPELKKCNWMLCVR------------------------
Rousettus_aegyptiacus         -----------------------------------------MLDPVENIPELKKCNWMLCVLNTRRAQTSWVIETKCGEND-----
Dasypus_novemcinctus          MRNFAGNCHLGEQPESPHHPNSDVSYQKLETSAVSISDGNRLLDAVGNTPALKKCSWMLCVC------------------------
Monodelphis_domestica         -----------------------------------------MQSTVENIPELKKCNWMLCVCSIQGAPIS------CG--------
Condylura_cristata            -----------------------------------------MLDAVENTPELKKCNWMVCVCVCVKYQKSP---DPMGNKDEMWRE
Ceratotherium_simum_simum     -----------------------------------------MLDAVENTPELKKCNWMLC--VC----------------------
Loxodonta_africana            -----------------------------------------MLDAVENTPELKKCNWMVCVCEIPE---EP---RPSG--------
Nannospalax_galili            -----------------------------------------MLDAVENTPELKKCNWMWCLLNTRS----P---DPVGNRDEMWRE
Trichechus_manatus_latirostr  -----------------------------------------MLDAVENSLELKKCNWMLCVC------------------------
Orycteropus_afer_afer         -----------------------------------------MLGAVENTPEVKKCNWVLCVC------------------------
```

Conservation

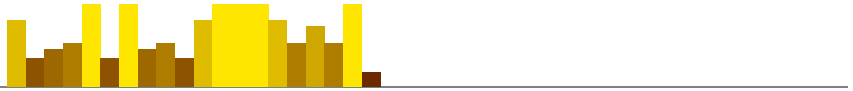

```
-----------------------------------------9456*4*5649***9686*20------------------------
```

**HG0495.1**

ENSG00000196730 (DAPK1)

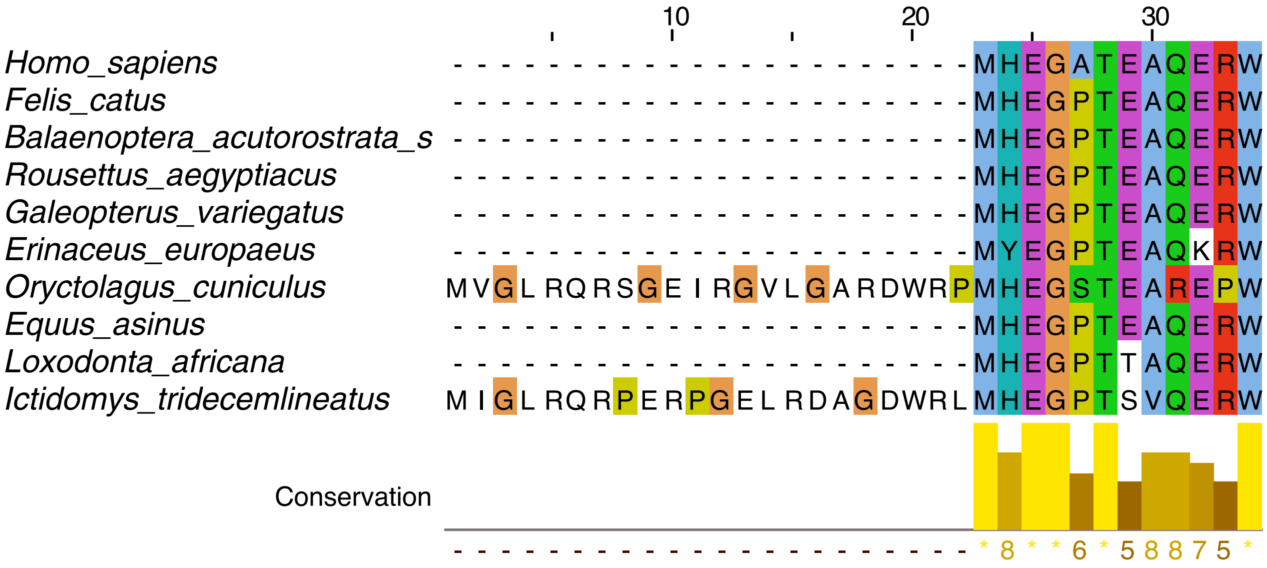

**HG0495.2**

ENSG00000196730 (DAPK1)

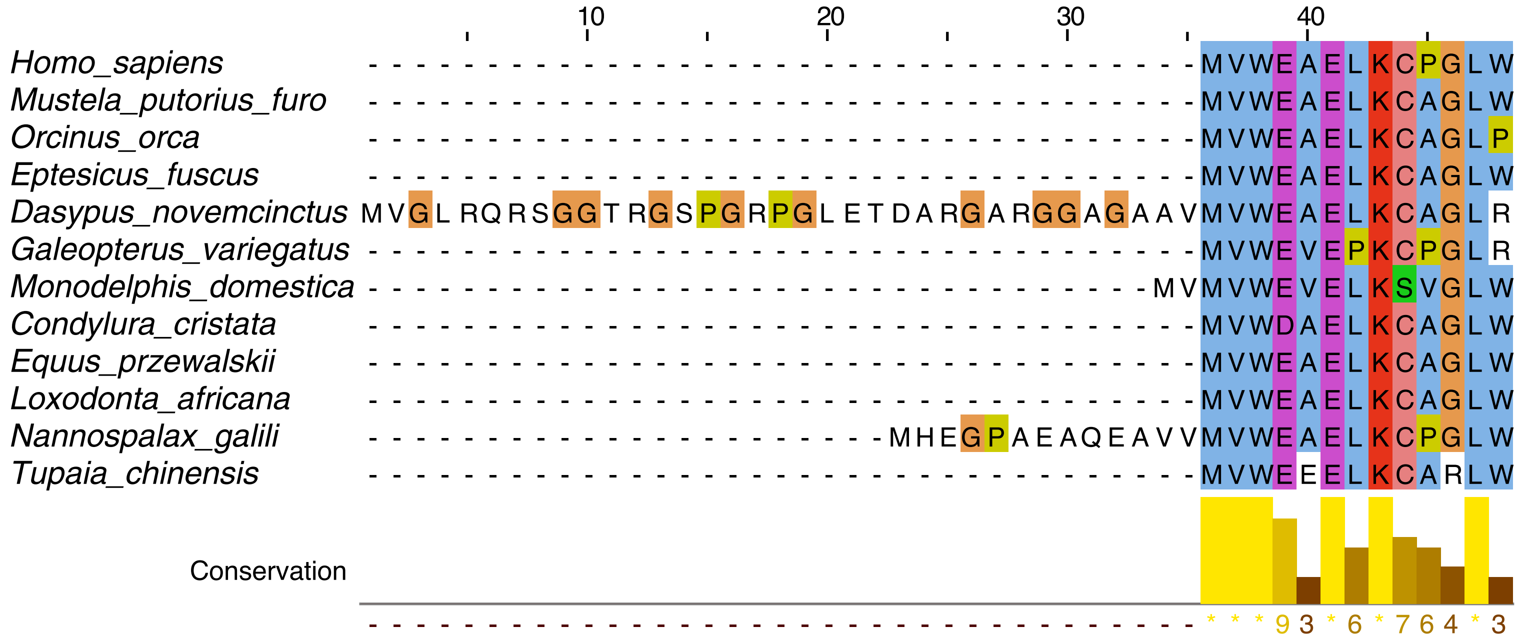

**HG0496**

ENSG00000135913 (USP37)

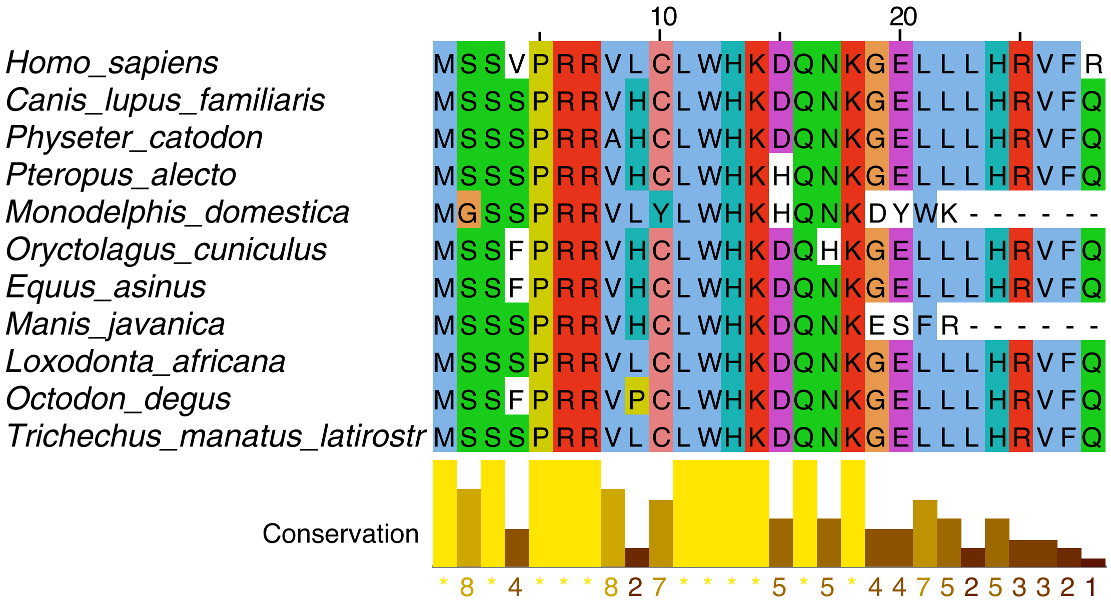

**HG0497**

ENSG00000147571 (CRH)

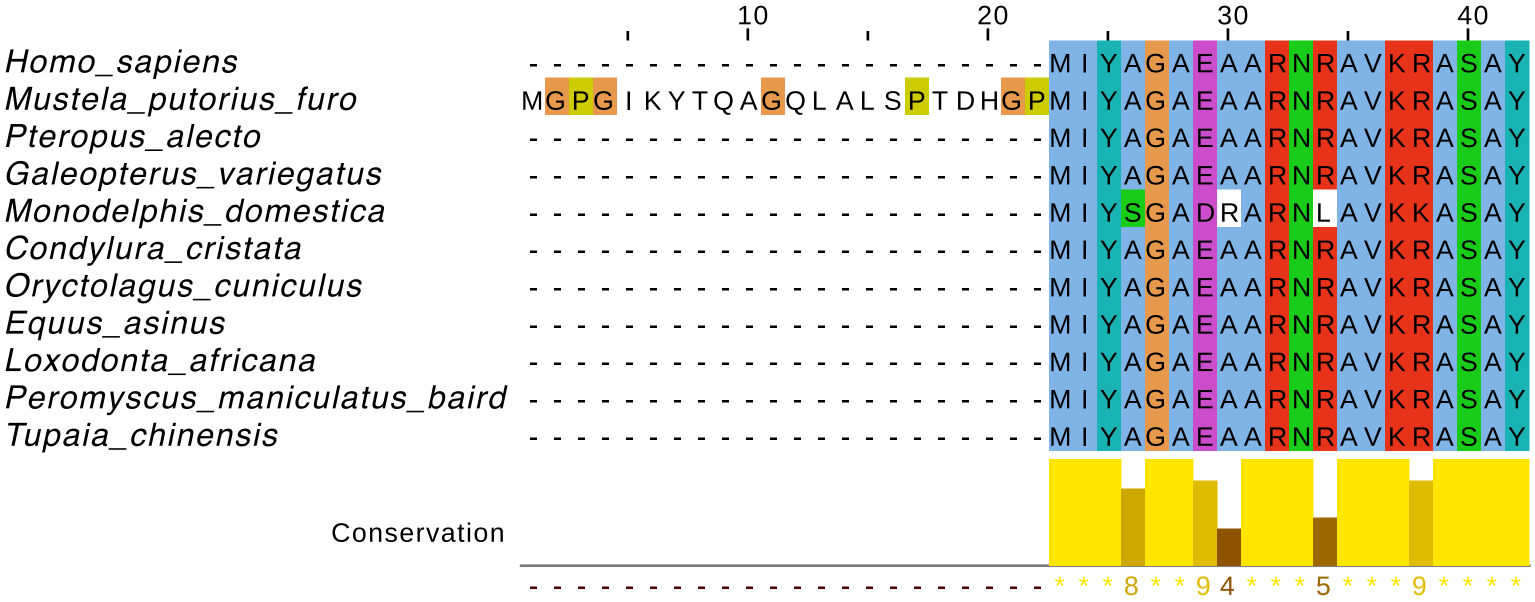

**HG0498**

ENSG00000005379 (TSPOAP1)

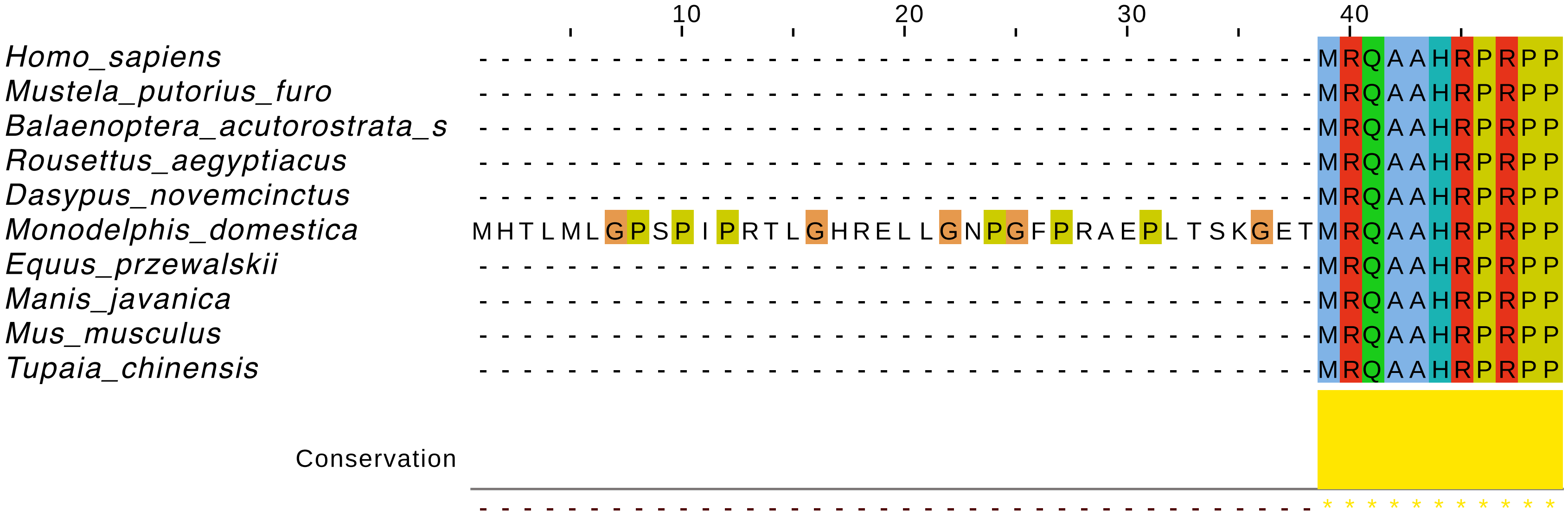

**HG0499**

ENSG00000011258 (MBTD1)

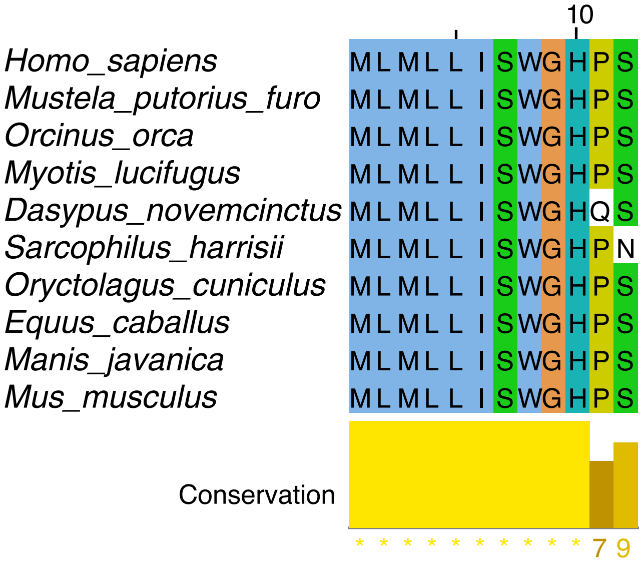

**HG0500**

ENSG00000070778 (PTPN21)

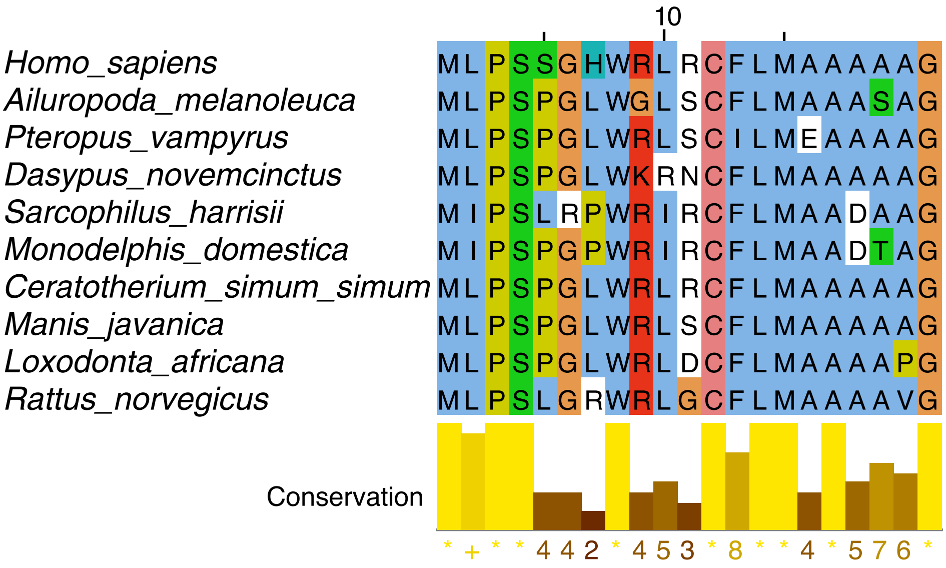

**HG0501**

ENSG00000110400 (NECTIN1)

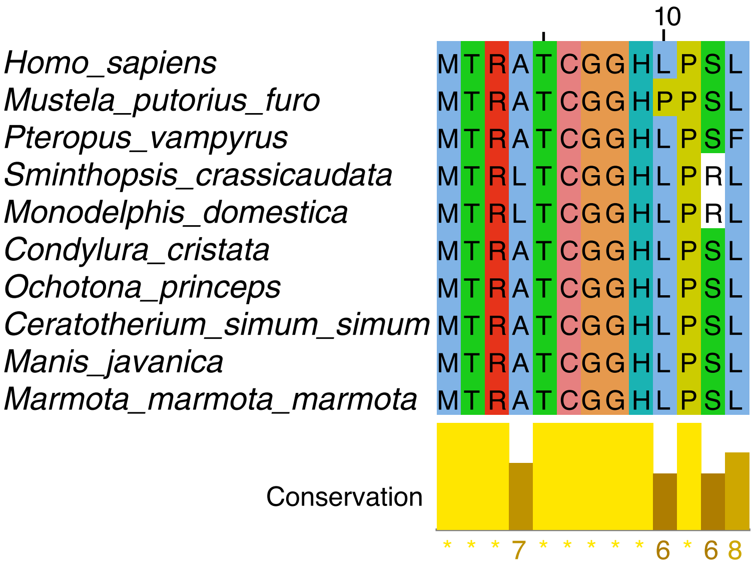

**HG0502**

ENSG00000126882

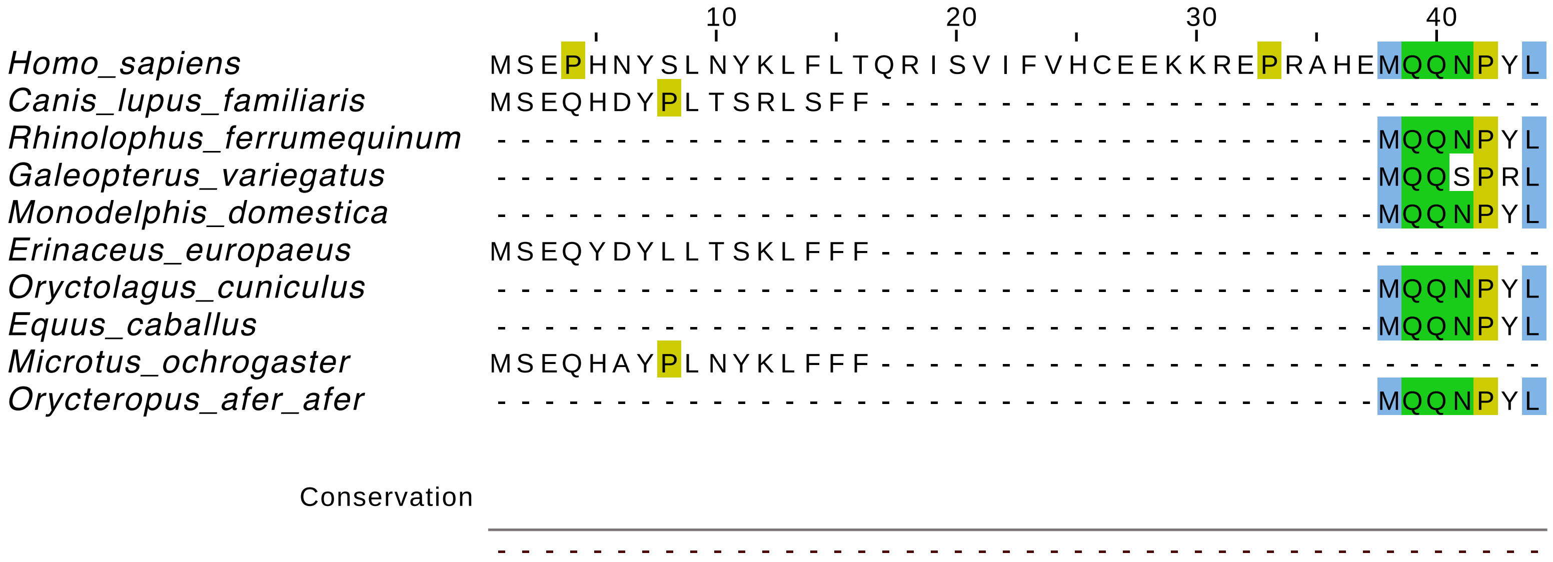

**HG0503**

ENSG00000141564 (RPTOR)

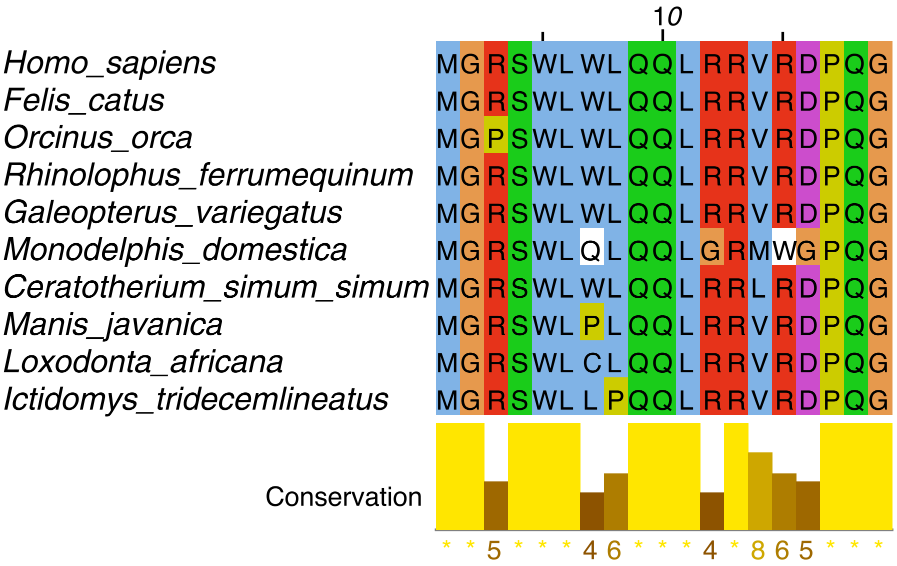

**HG0504**

ENSG00000144460 (NYAP2)

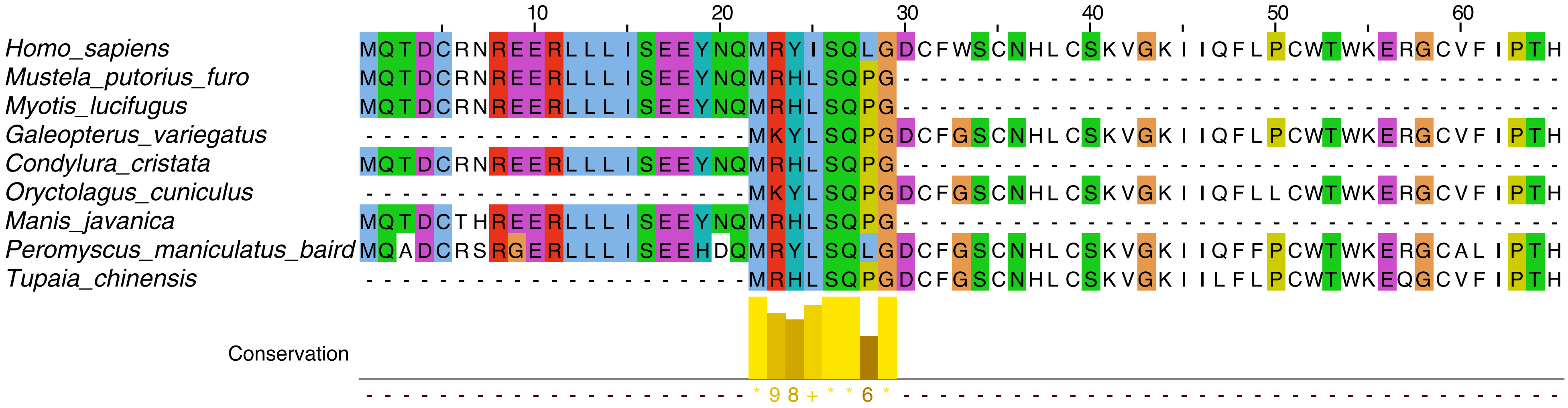

**HG0505**

ENSG00000157470

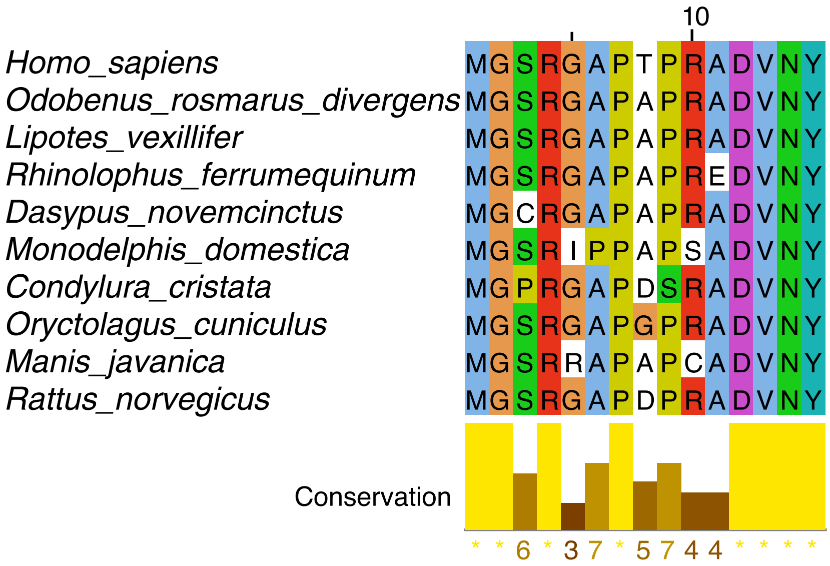

**HG0506.1**

ENSG00000177853 (ZNF518A)

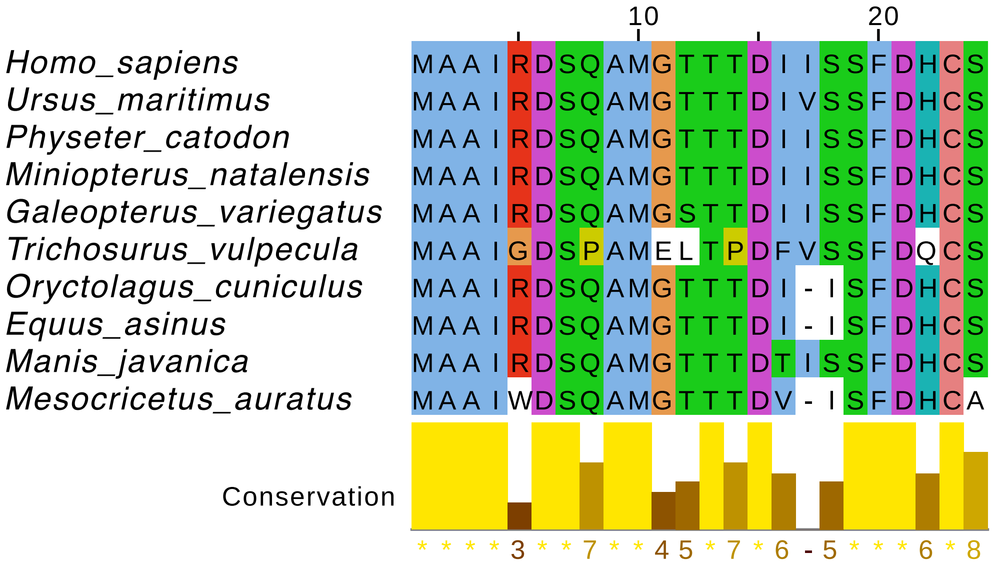

**HG0506.2**

ENSG00000177853 (ZNF518A)

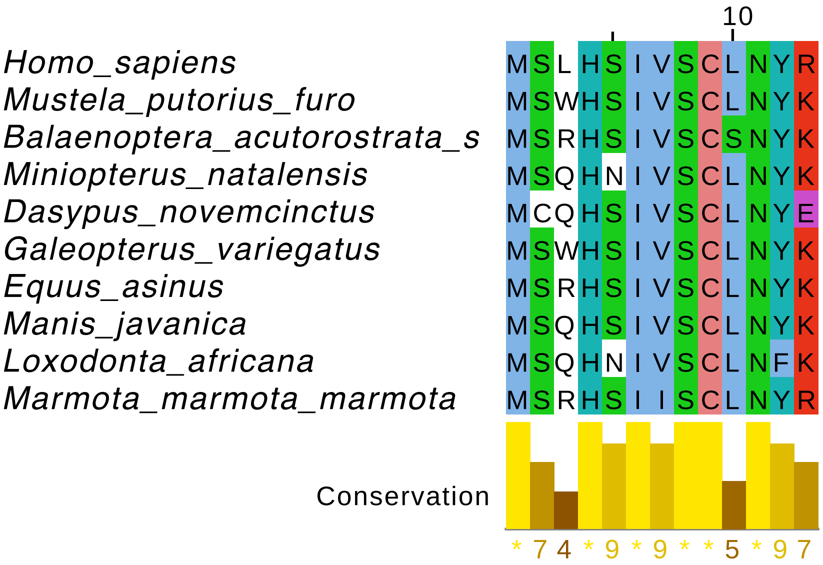

**HG0507**

ENSG00000060237 (WNK1)

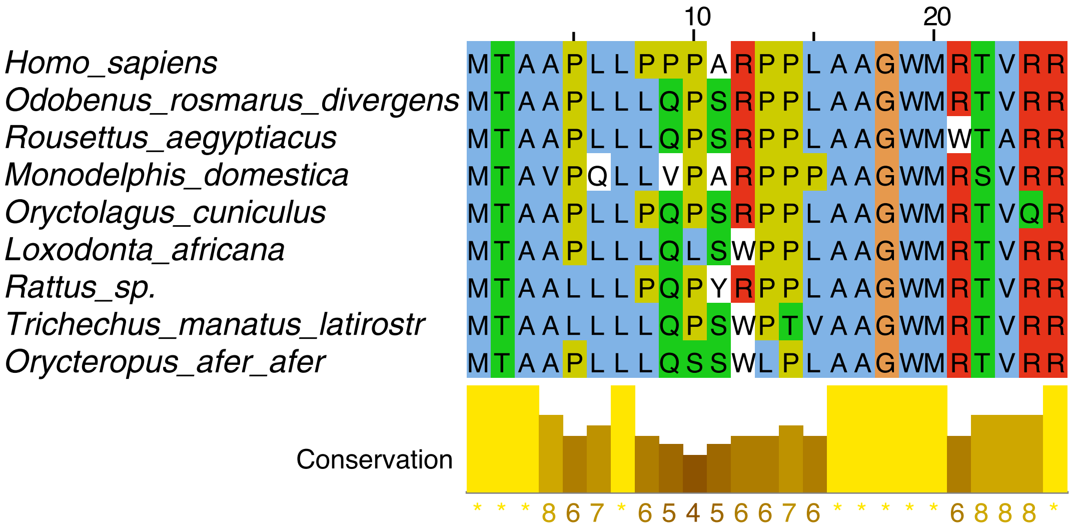

**HG0508**

ENSG00000061936 (SFSWAP)

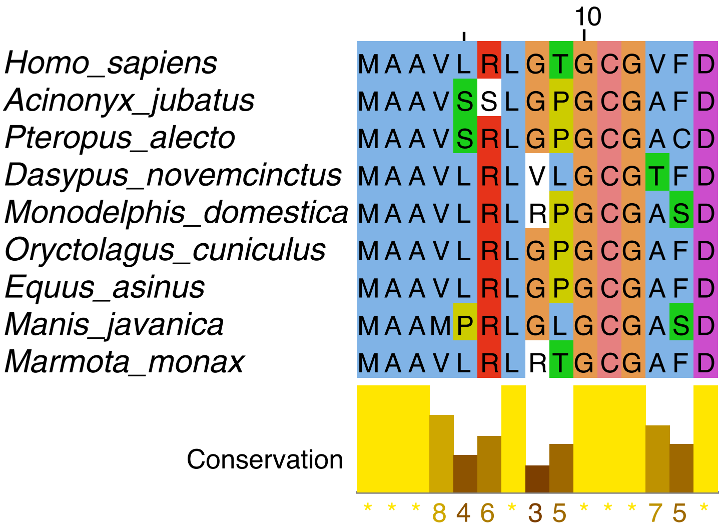

**HG0509**

ENSG00000068024 (HDAC4)

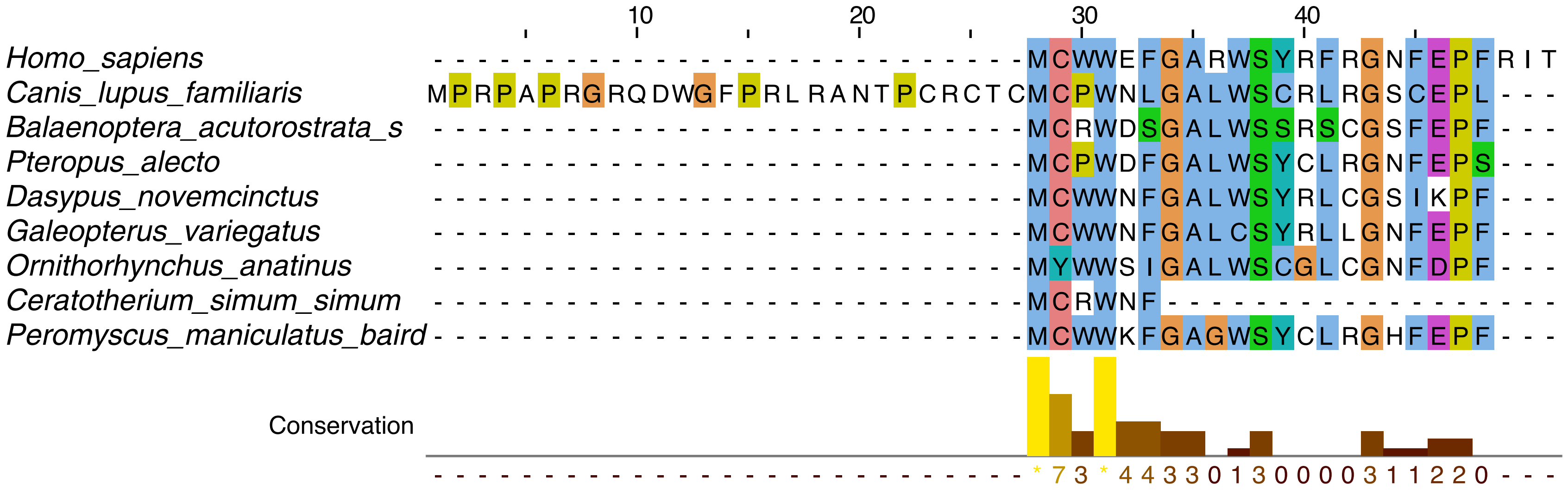

**HG0510**

ENSG00000134769 (DTNA)

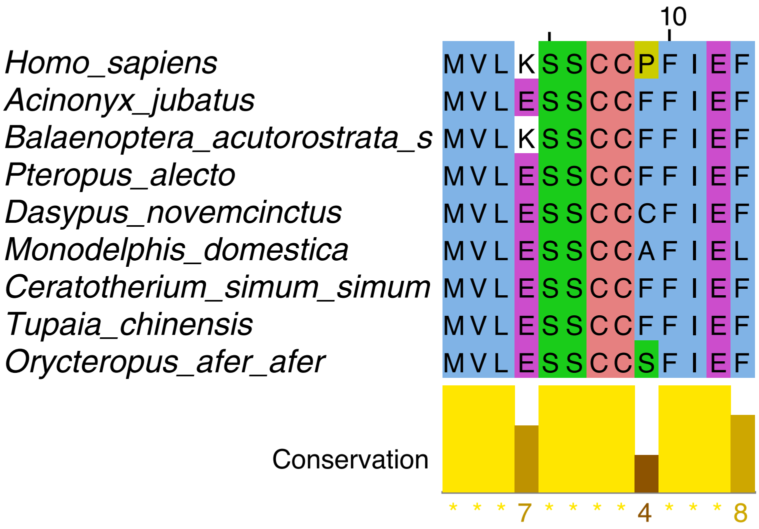

**HG0511**

ENSG00000143337 (TOR1AIP1)

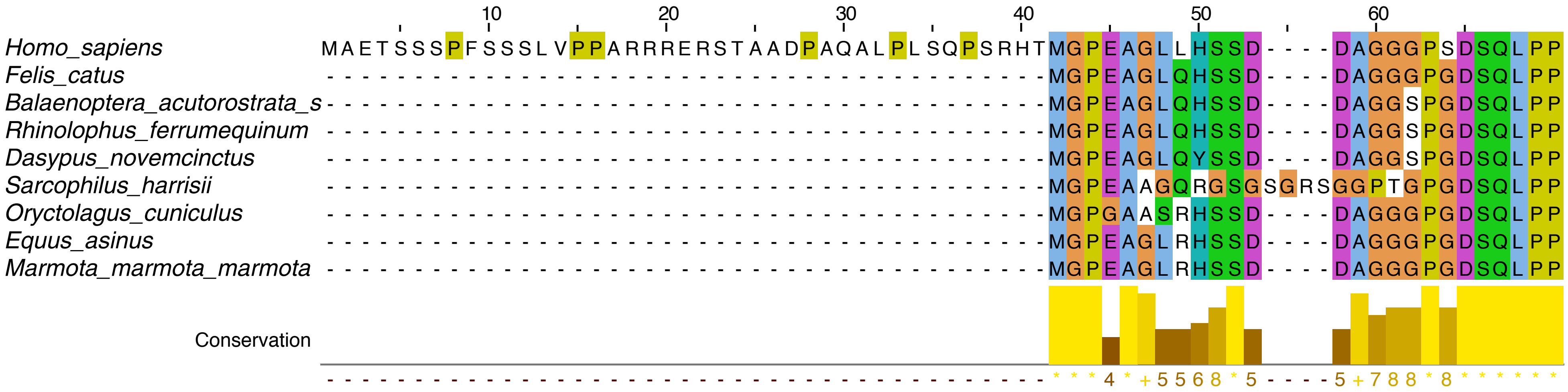

**HG0512.1**

ENSG00000112319 (EYA4)

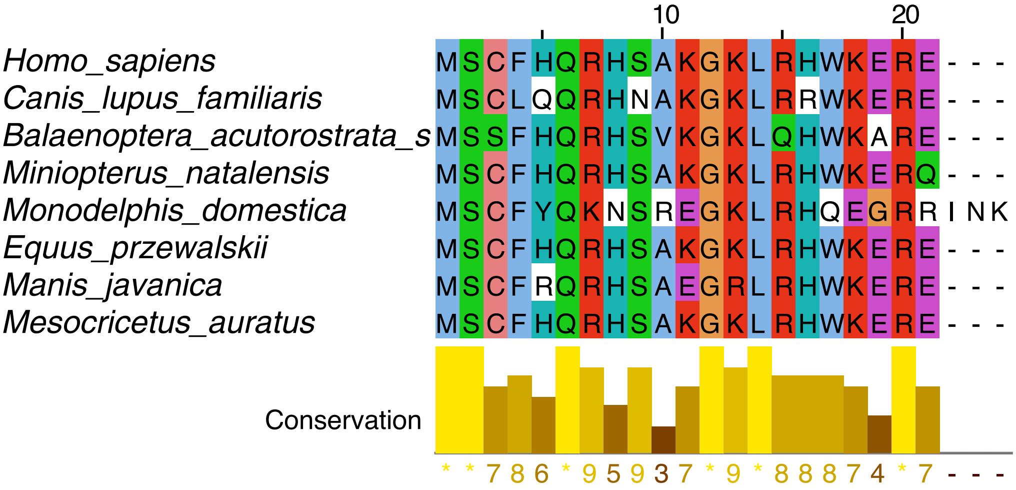

**HG0512.2**

ENSG00000104313 (EYA1)

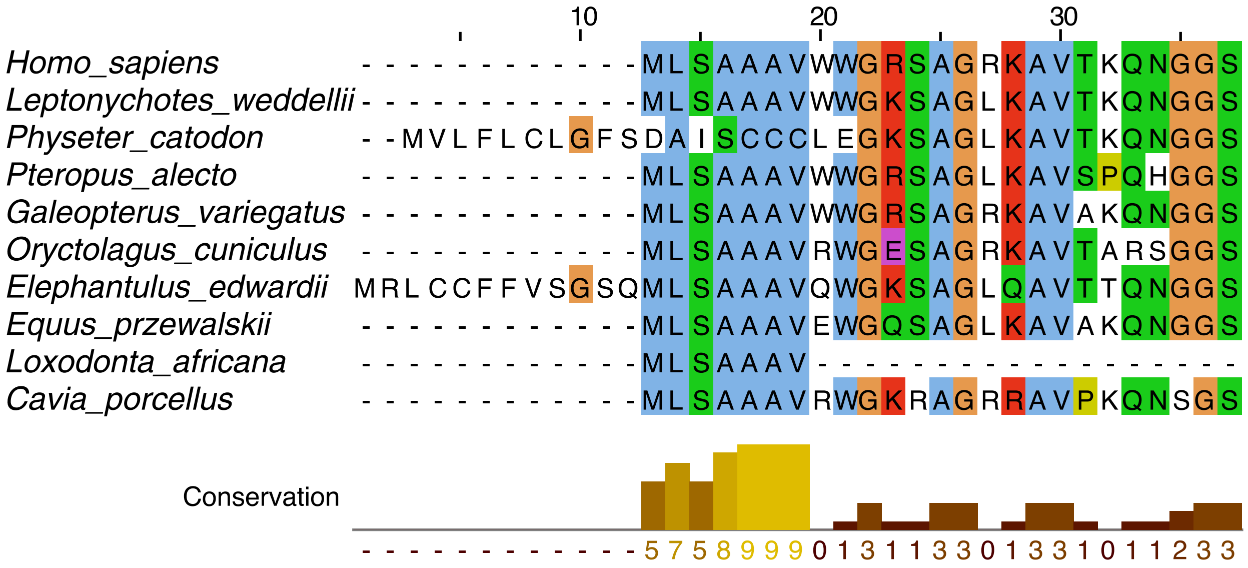

**HG0512.3**

ENSG00000104313 (EYA1)

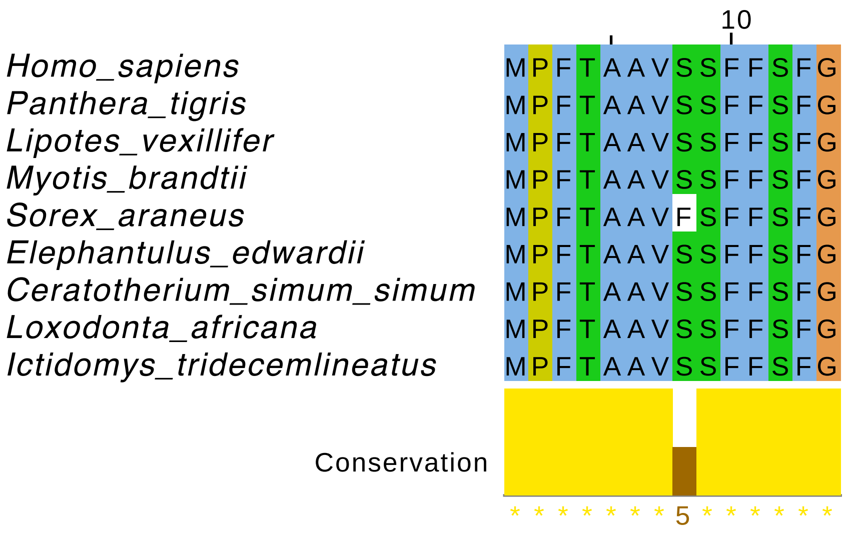

**HG0513**

ENSG00000126860 (EVI2A)

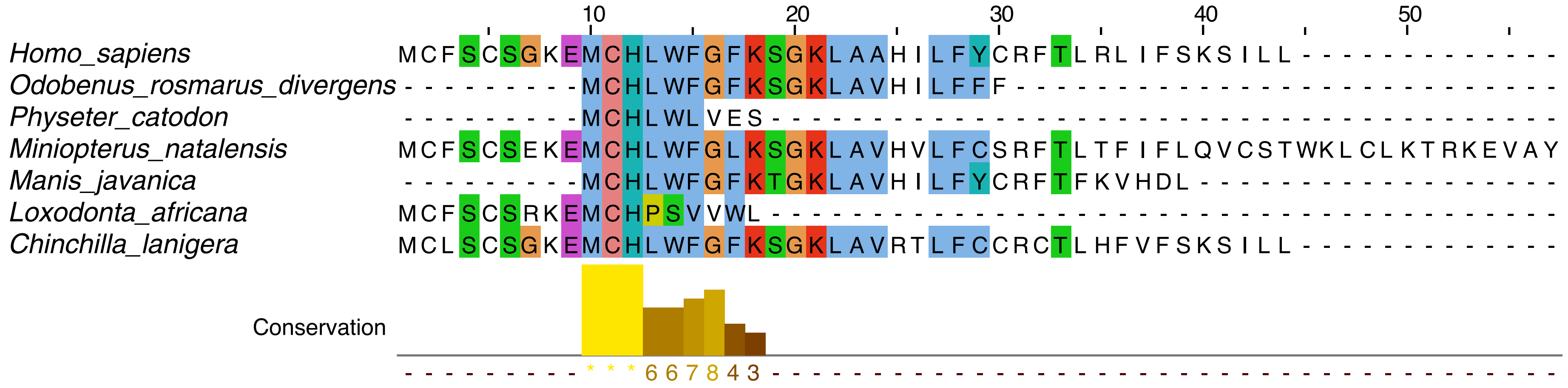

**HG0514**

ENSG00000147570 (DNAJC5B)

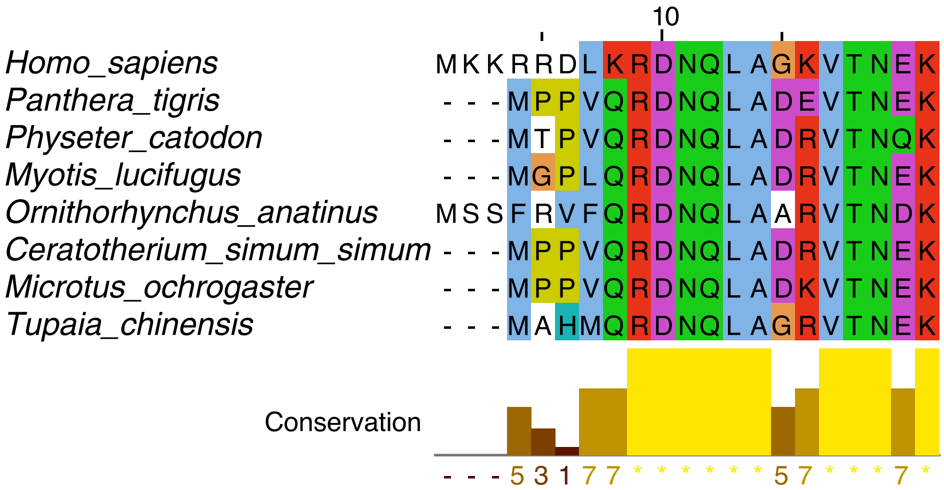

**HG0515**

ENSG00000170471 (RALGAPB)

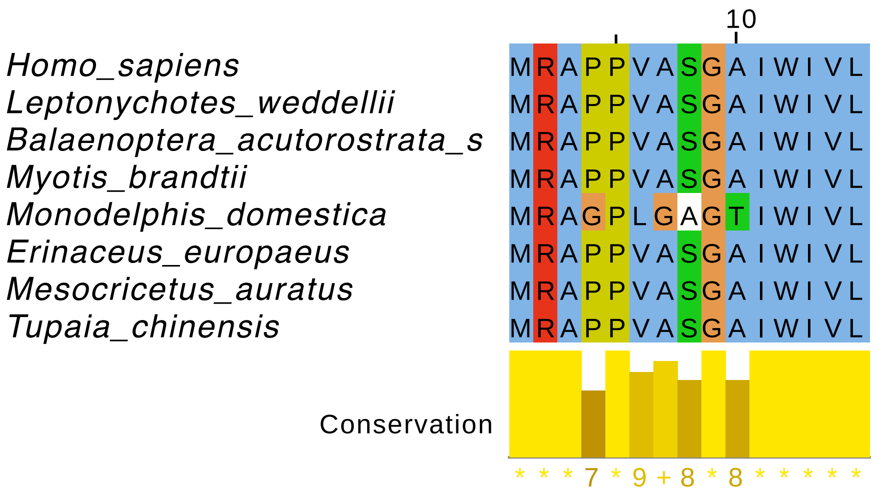

**HG0516**

ENSG00000041515 (MYO16)

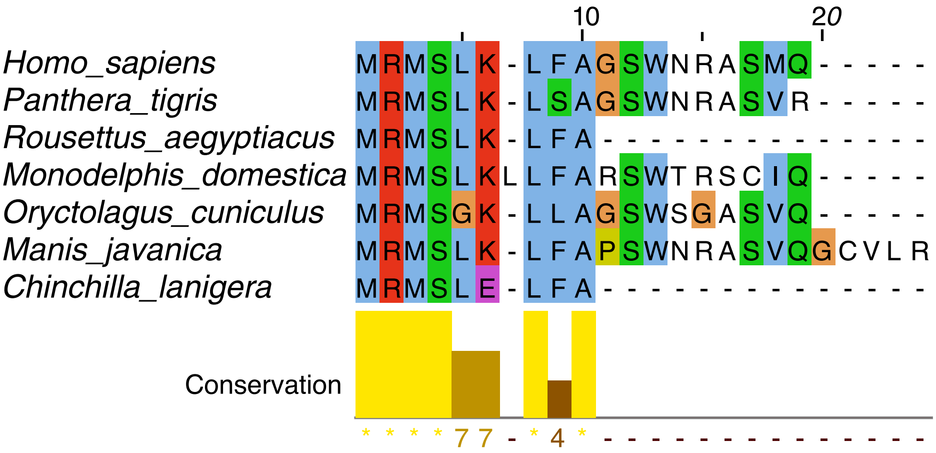

**HG0517**

ENSG00000086717 (PPEF1)

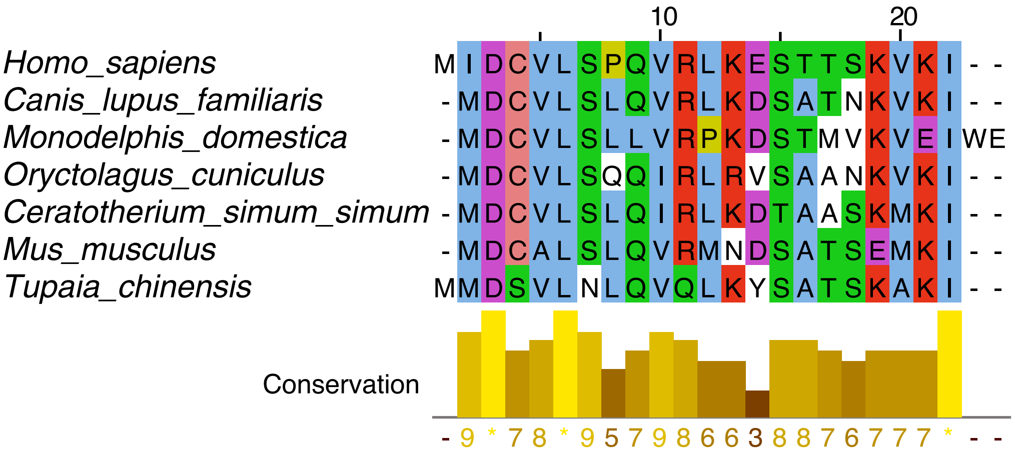

**HG0518**

ENSG00000111254 (AKAP3)

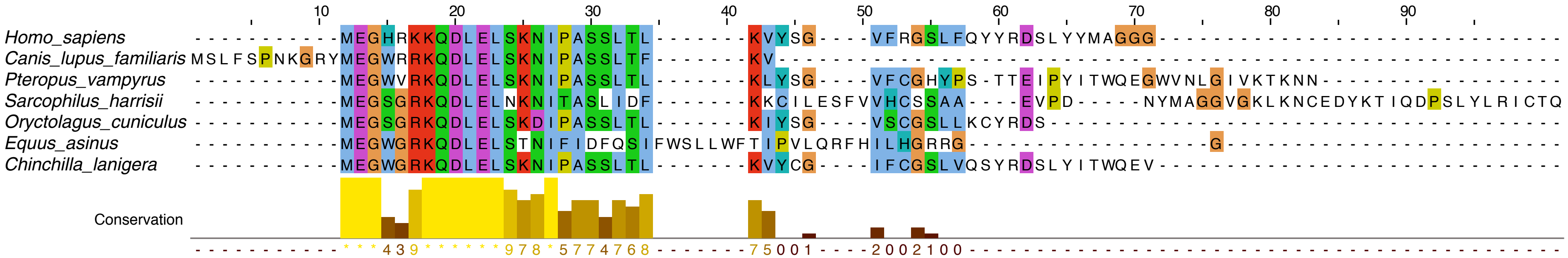

**HG0519**

ENSG00000127074 (RGS13)

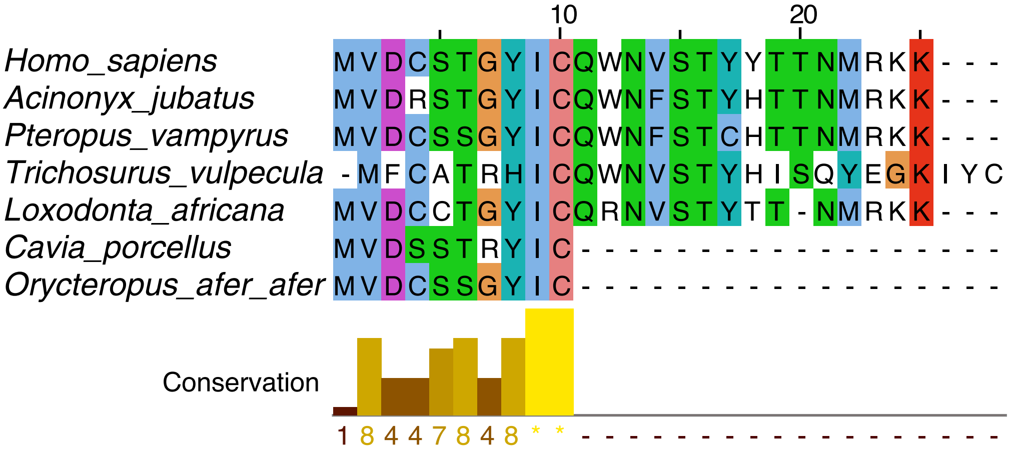

**HG0520**

ENSG00000197415 (VEPH1)

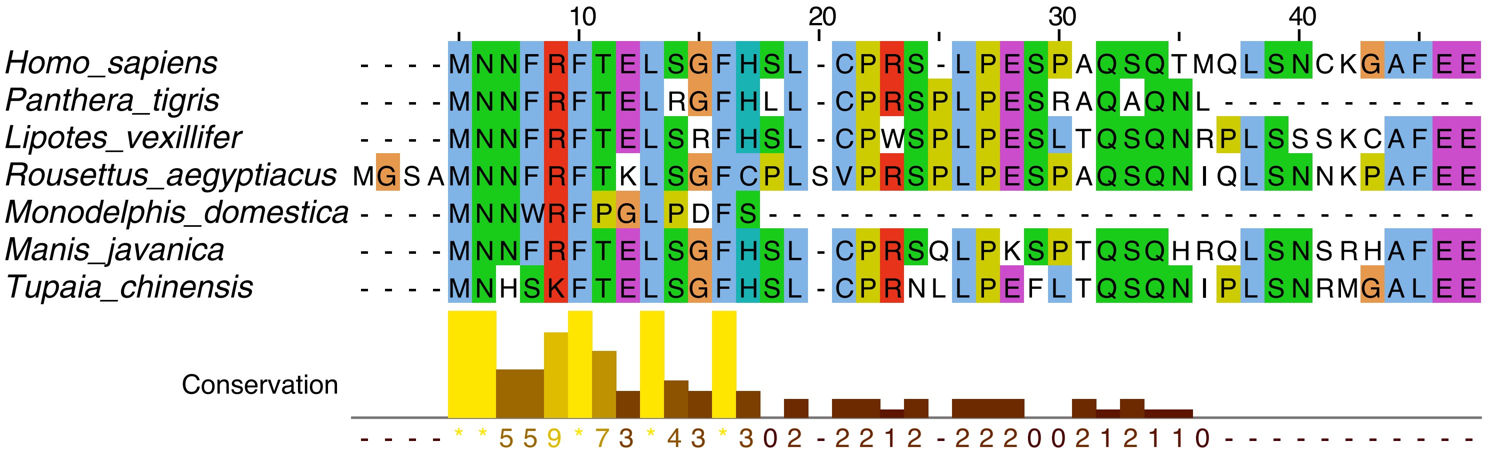

**HG0521**

ENSG00000038274 (MAT2B)

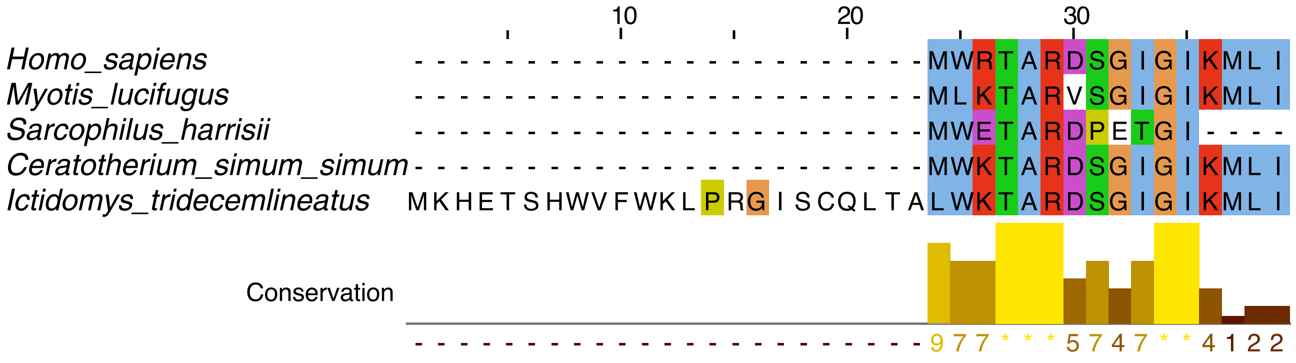

**HG0522**

ENSG00000140396 (NCOA2)

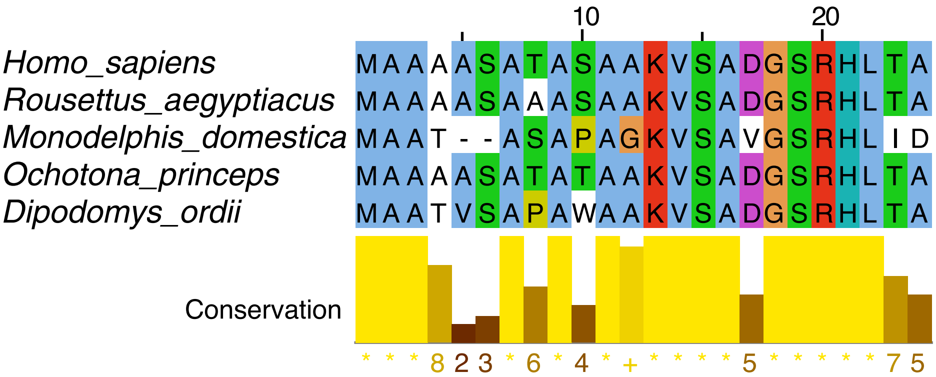

**HG0523**

ENSG00000075213 (SEMA3A)

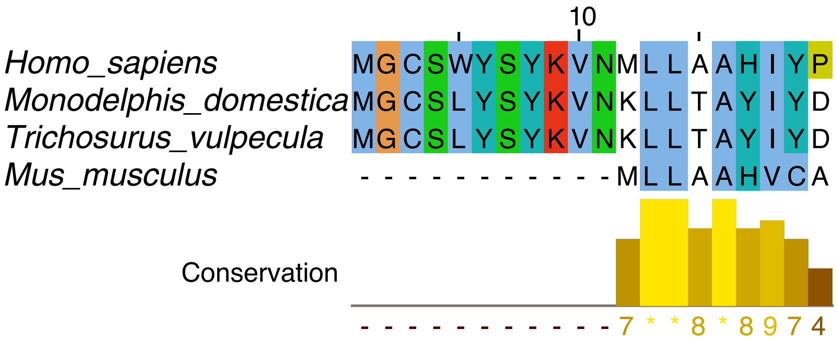

**HG0524**

ENSG00000107929 (LARP4B)

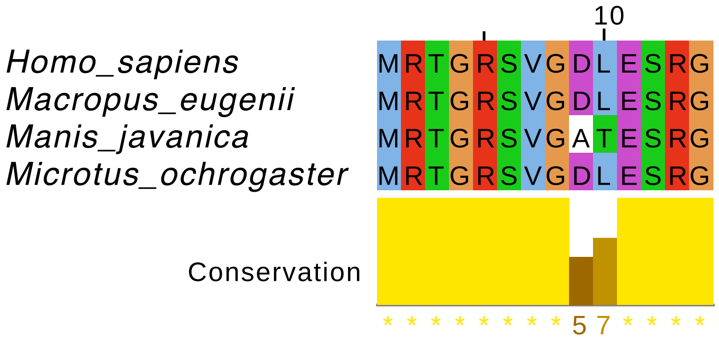

**HG0525**

ENSG00000136167 (LCP1)

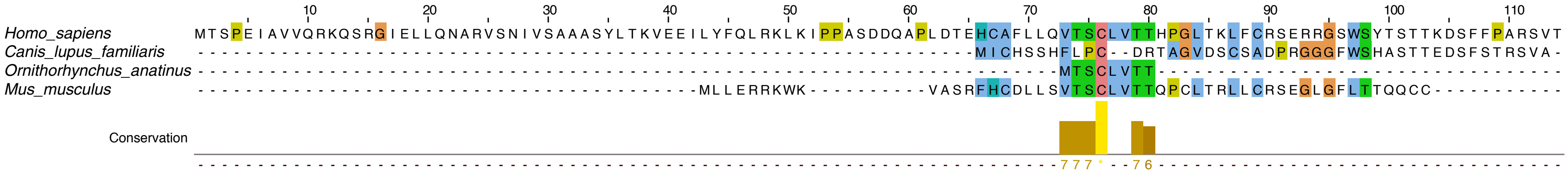

**HG0526**

ENSG00000143569 (UBAP2L)

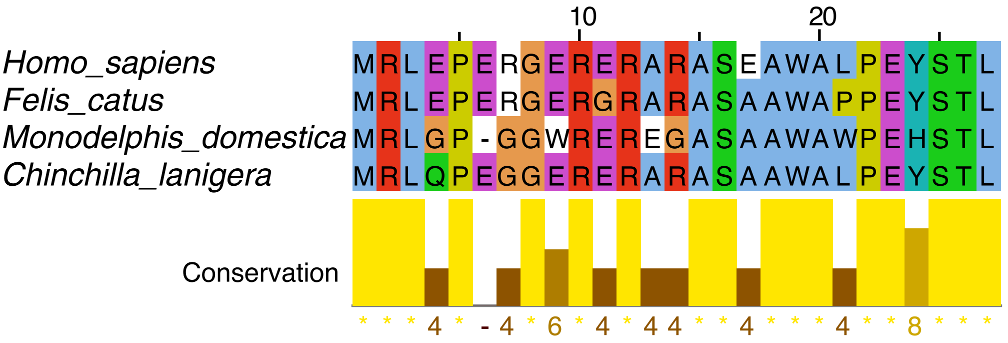

**HG0527**

ENSG00000198382 (UVRAG)

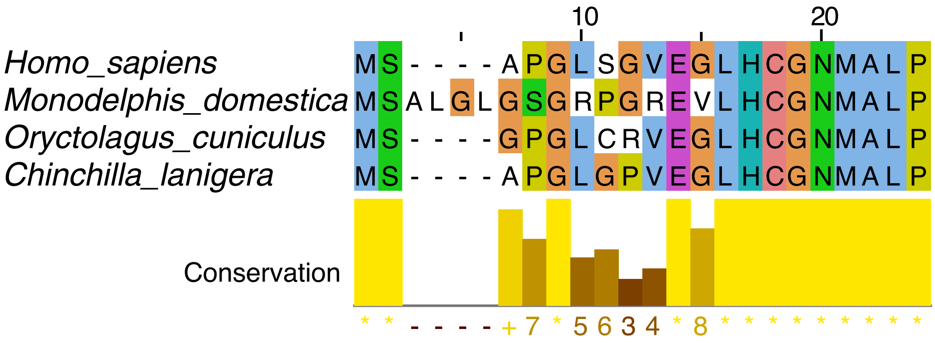

**HG0528**

ENSG00000204767

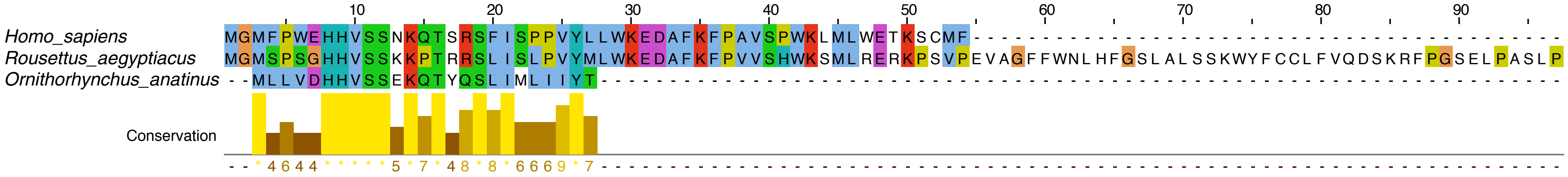

**HG0529**

ENSG00000112851 (ERBIN)

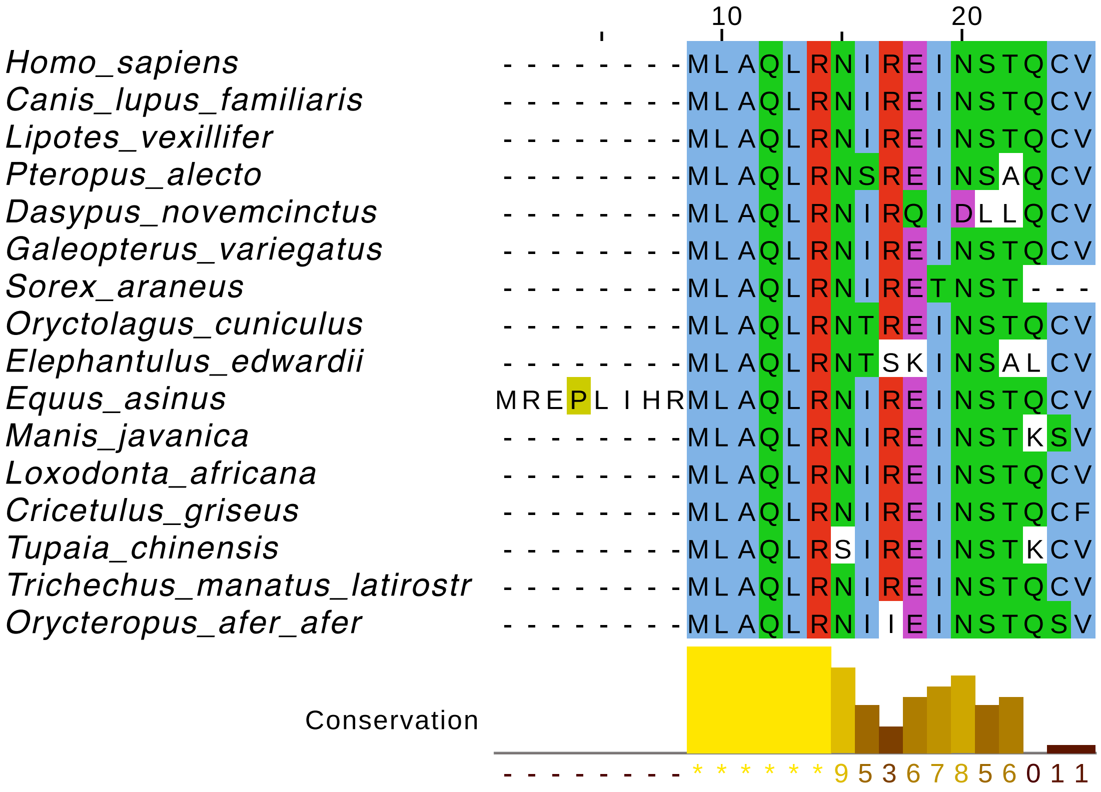

**HG0530**

ENSG00000128594 (LRRC4)

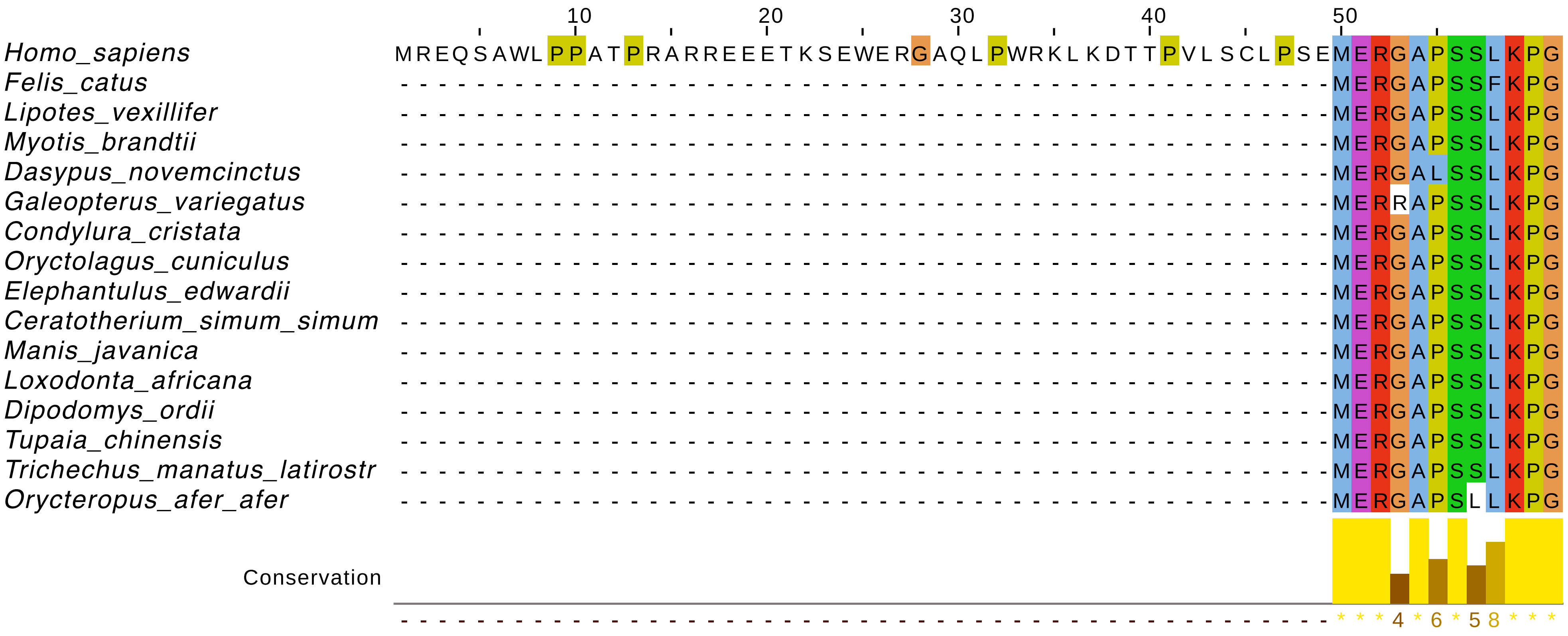

**HG0531**

ENSG00000146834 (MEPCE)

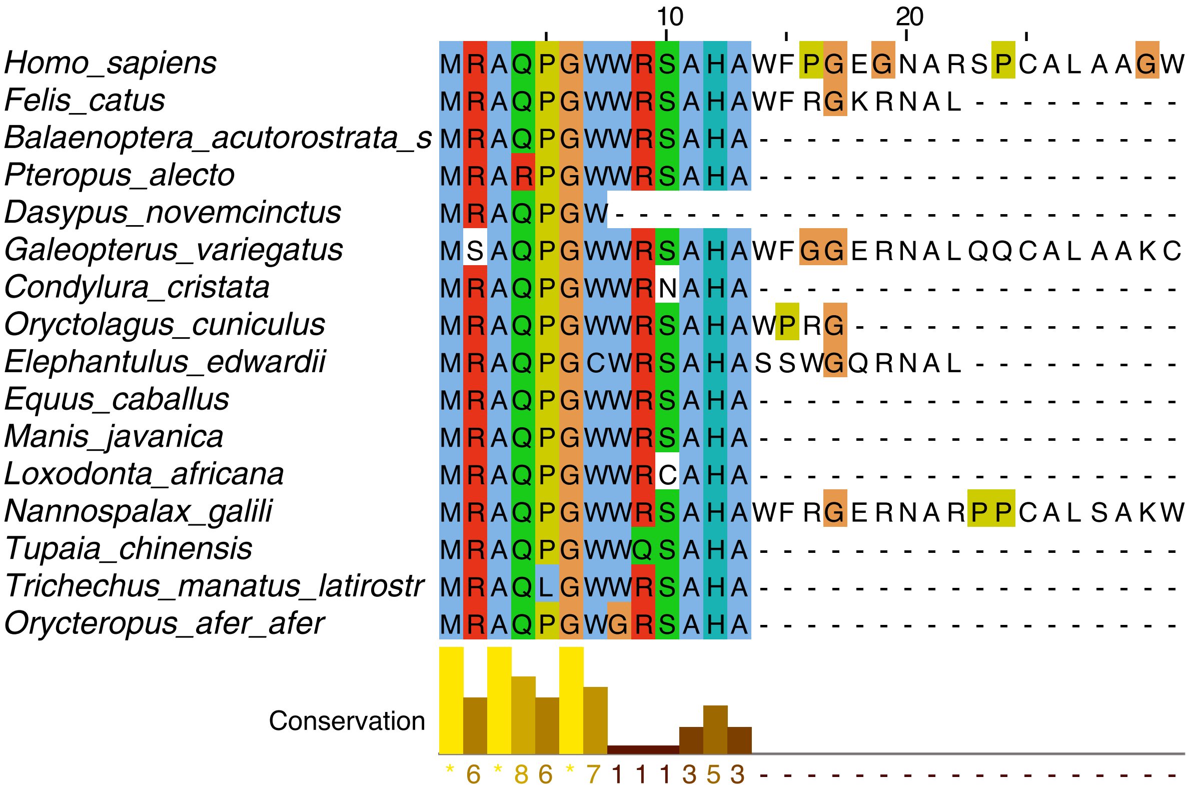

**HG0532**

ENSG00000105221 (AKT2)

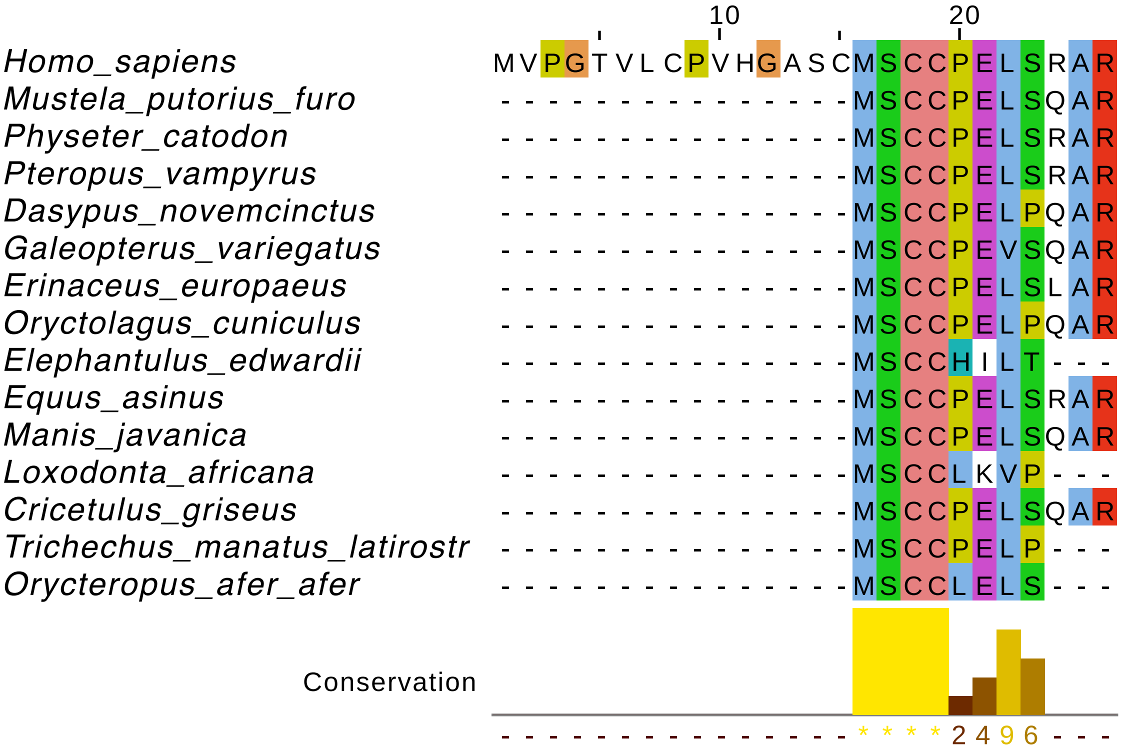

**HG0533**

ENSG00000114209 (PDCD10)

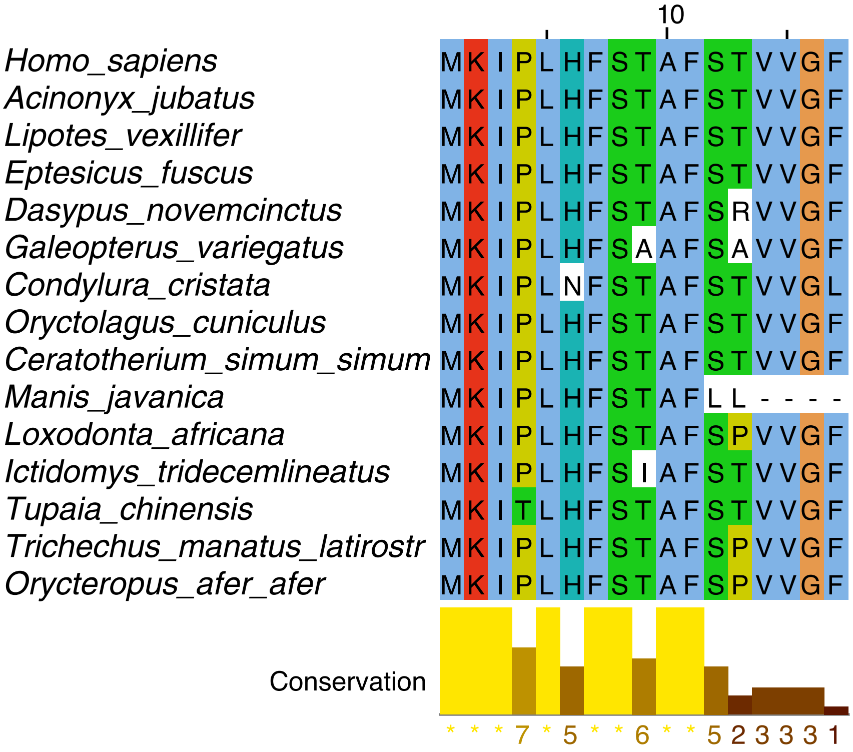

**HG0534**

ENSG00000117408 (IPO13)

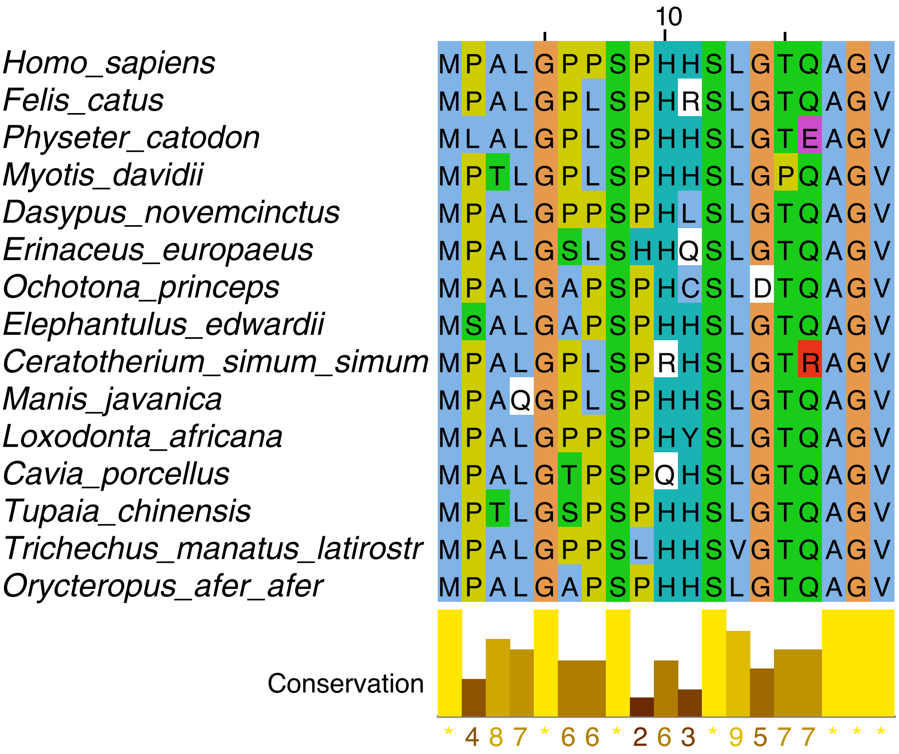

**HG0535**

ENSG00000118407 (FILIP1)

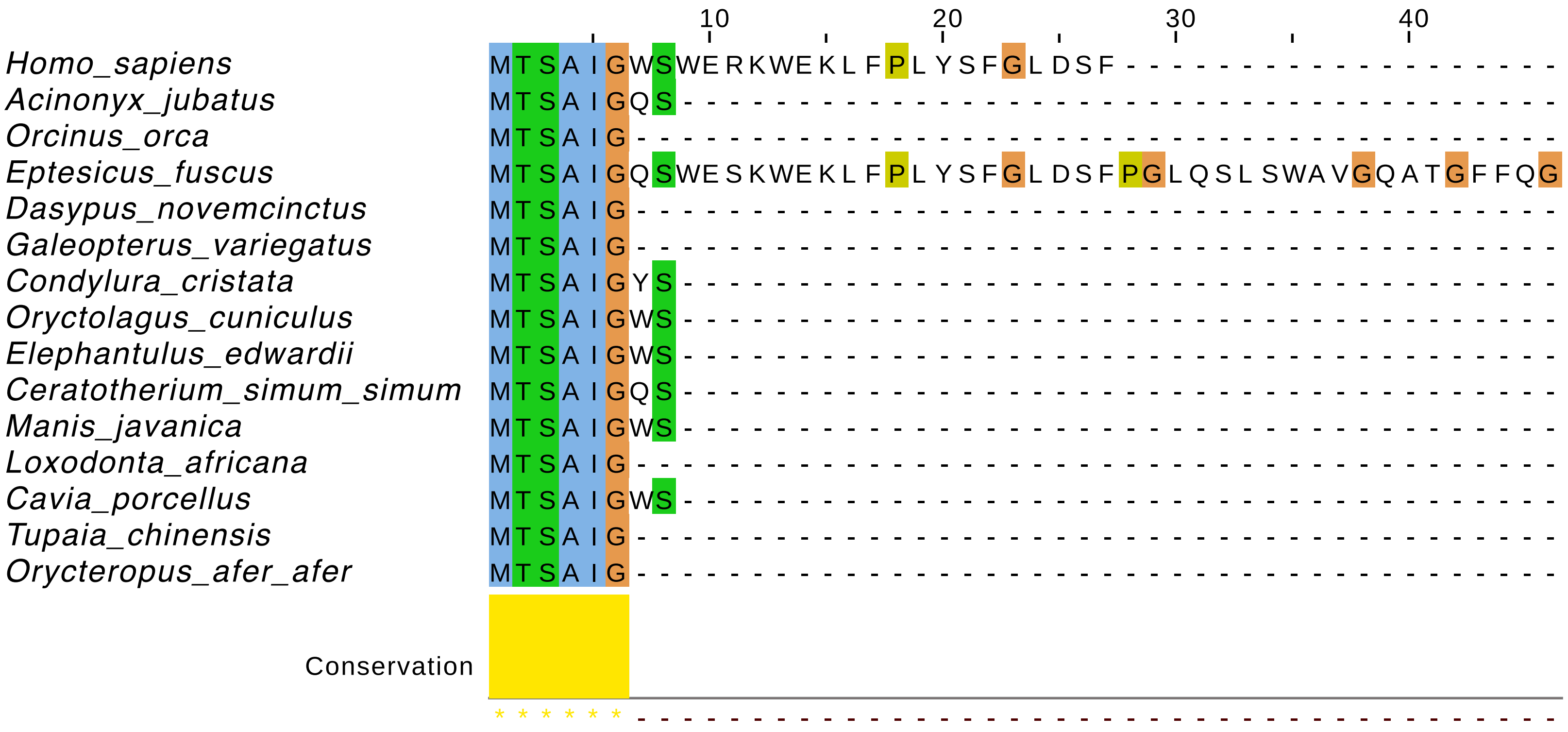

**HG0536**

ENSG00000181751 (C5orf30)

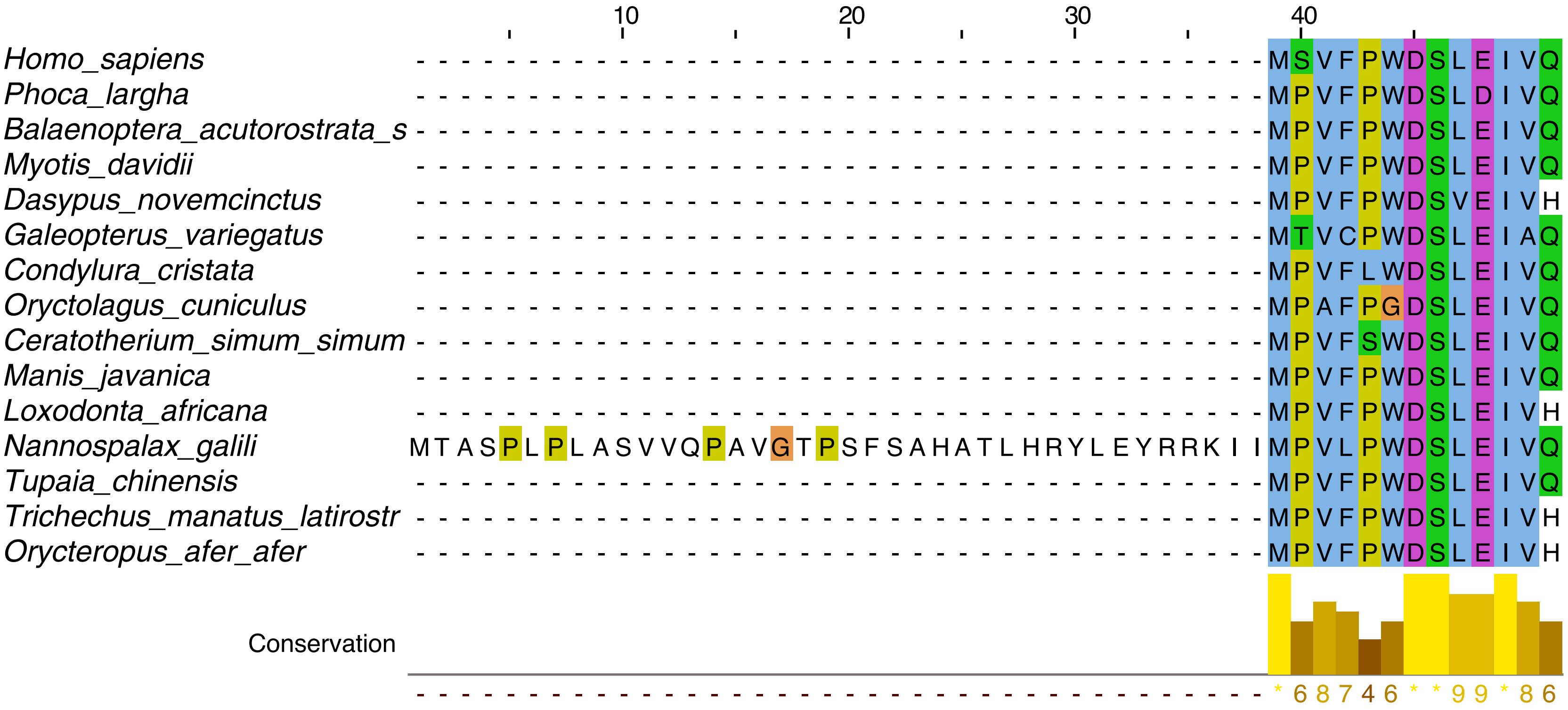

**HG0537**

ENSG00000184481 (FOXO4)

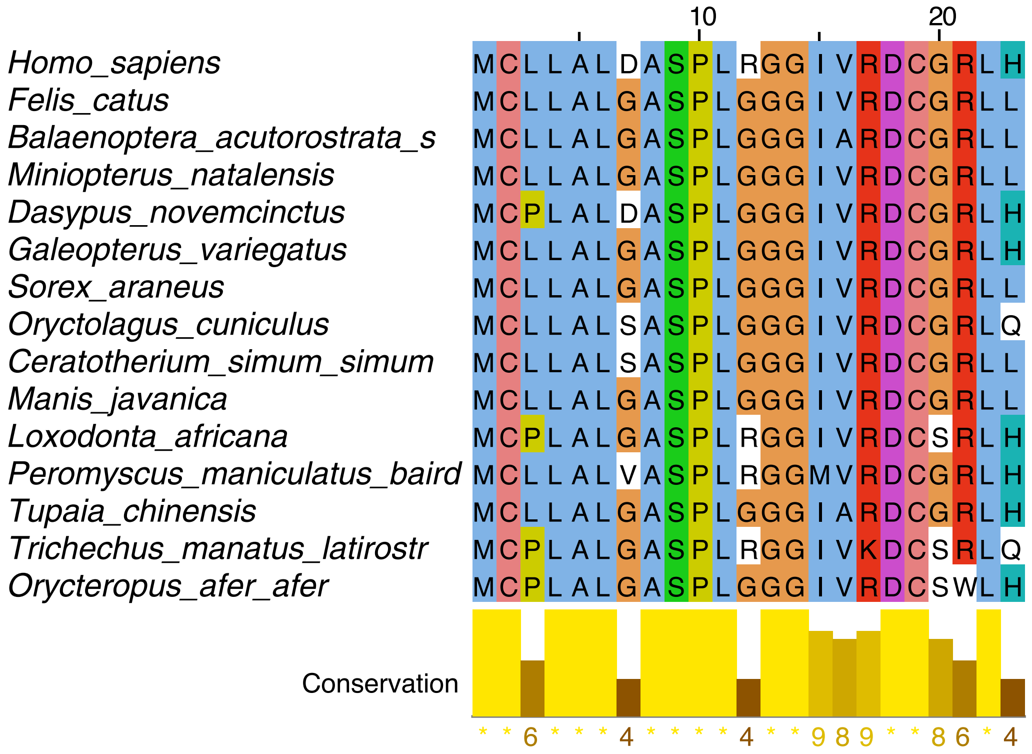

**HG0538**

ENSG00000006116 (CACNG3)

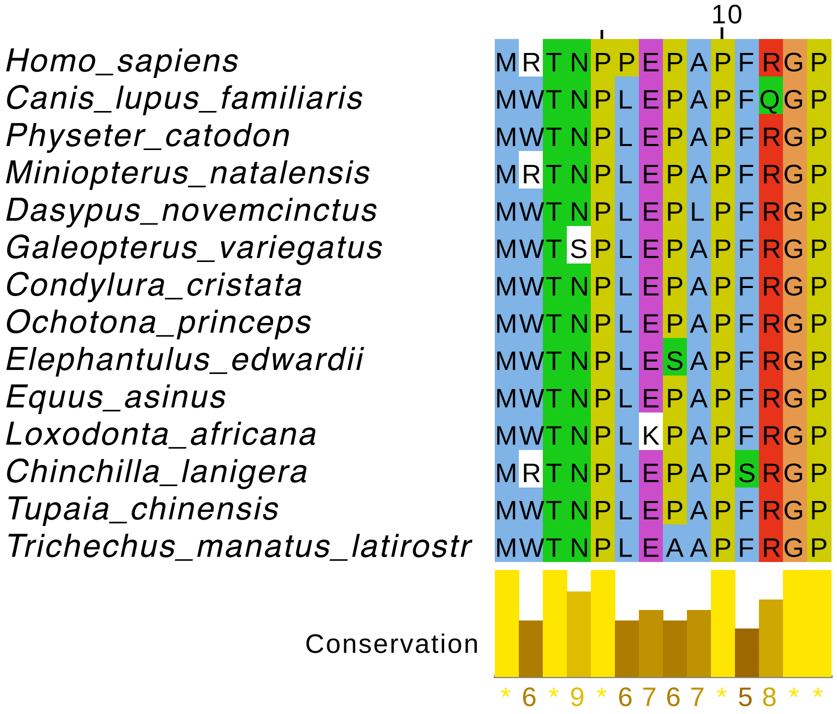

**HG0539**

ENSG00000115966 (ATF2)

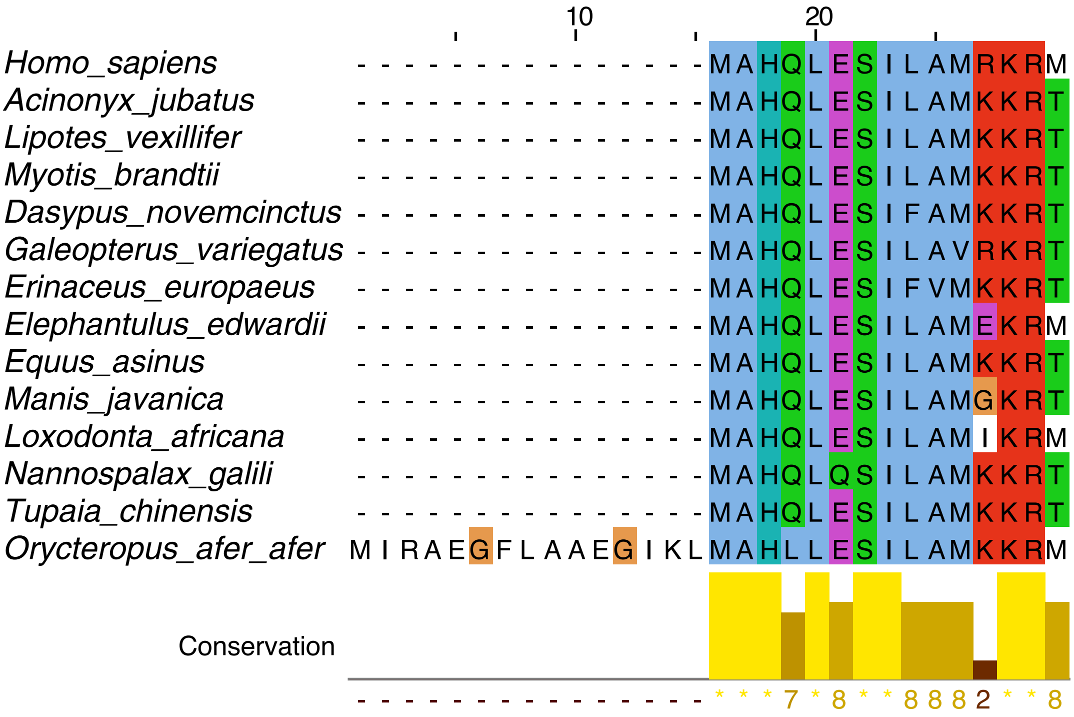

**HG0540**

ENSG00000120519 (SLC10A7)

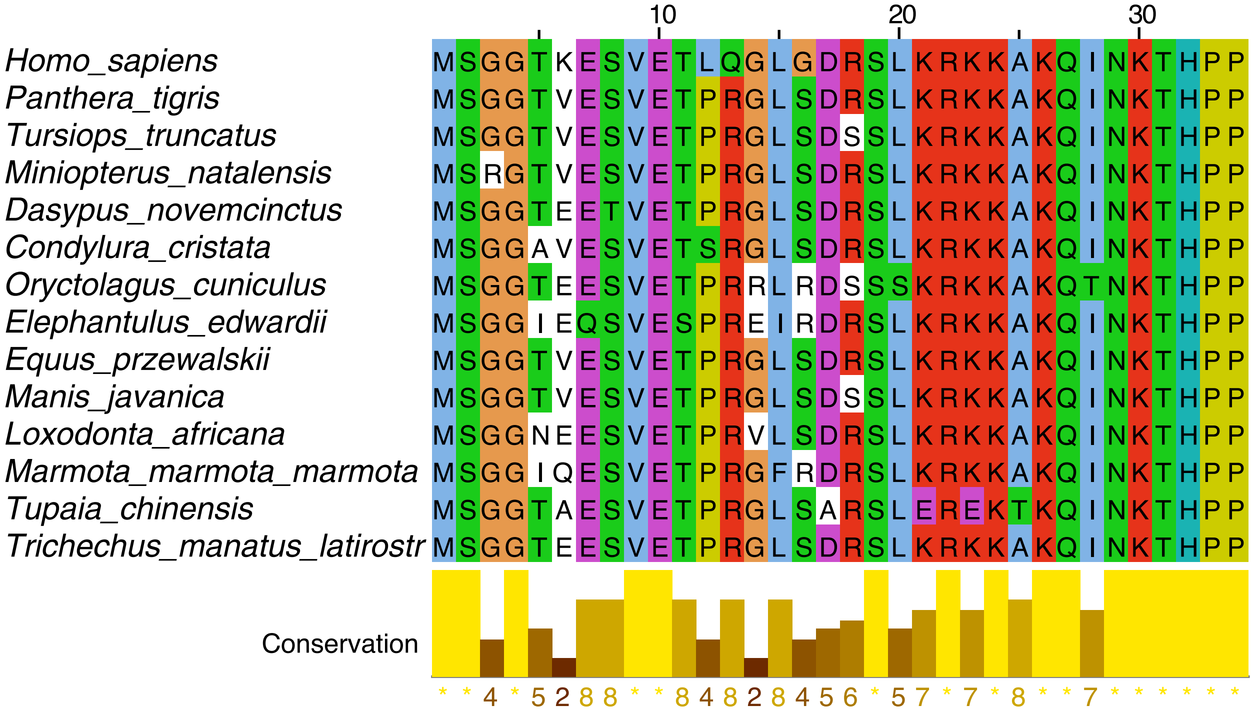

**HG0541.1**

ENSG00000122584 (NXPH1)

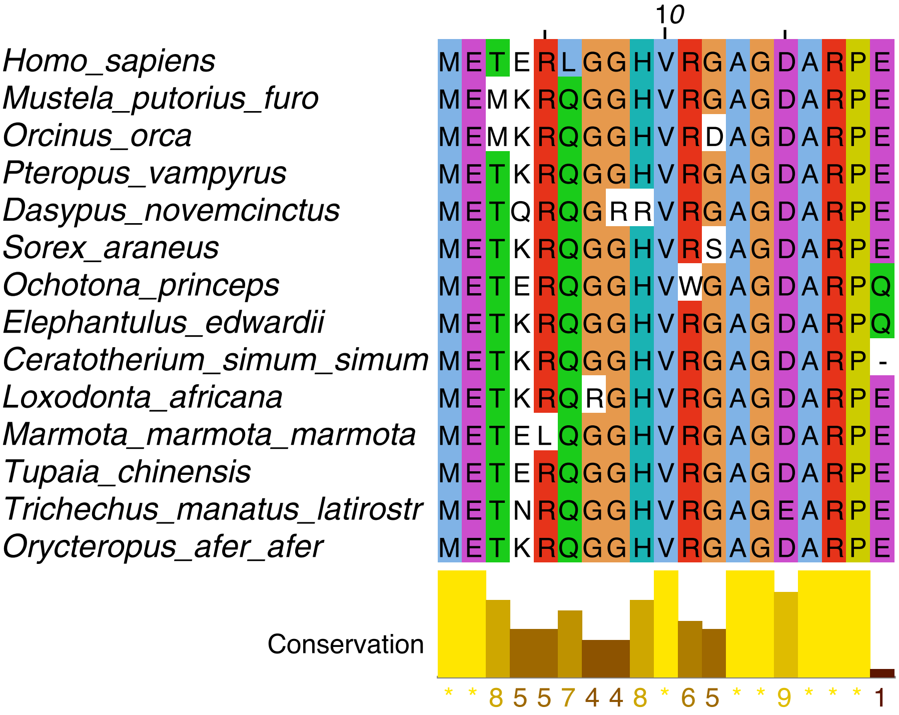

**HG0541.2**

ENSG00000122584 (NXPH1)

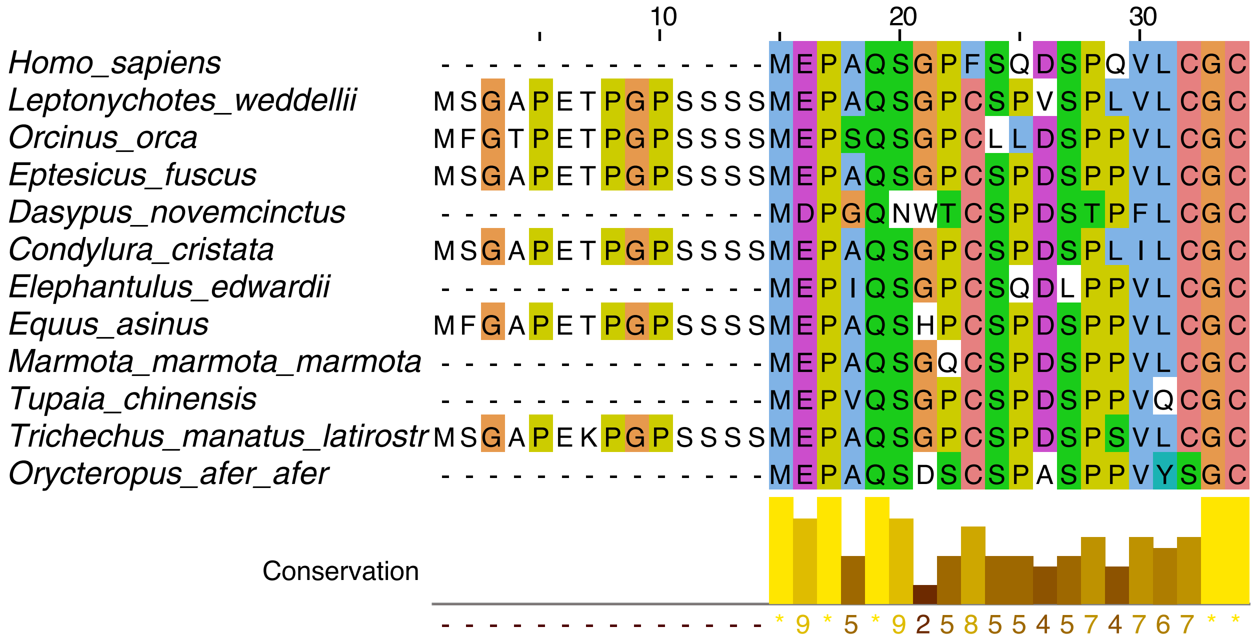

**HG0542**

ENSG00000126581 (BECN1)

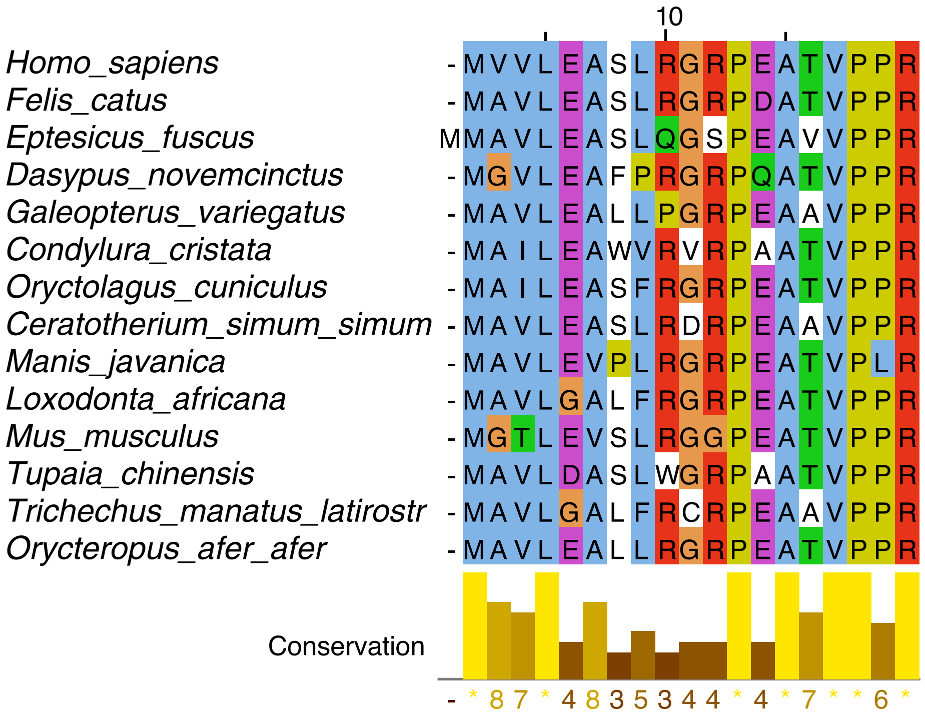

**HG0543**

ENSG00000131931 (THAP1)

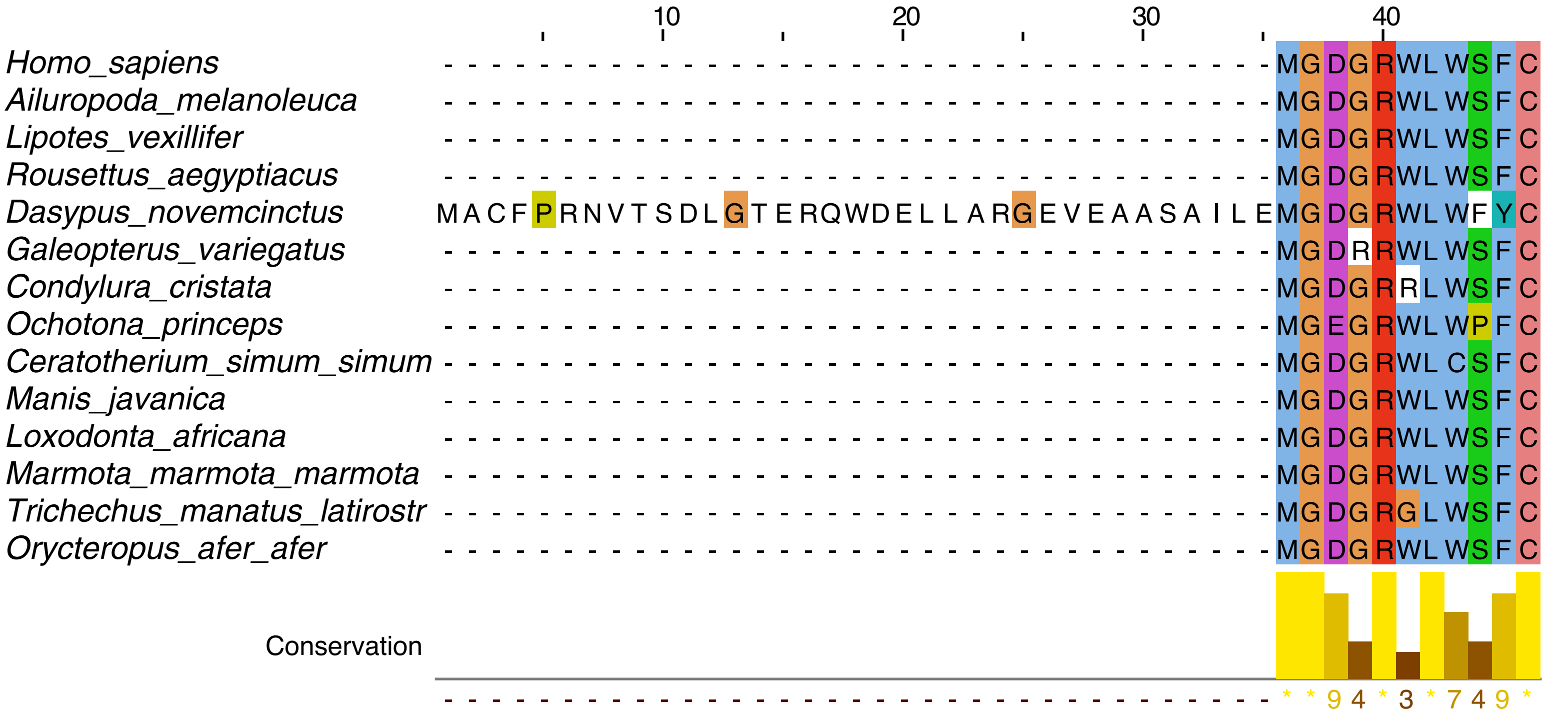

**HG0544**

ENSG00000136802 (LRRC8A)

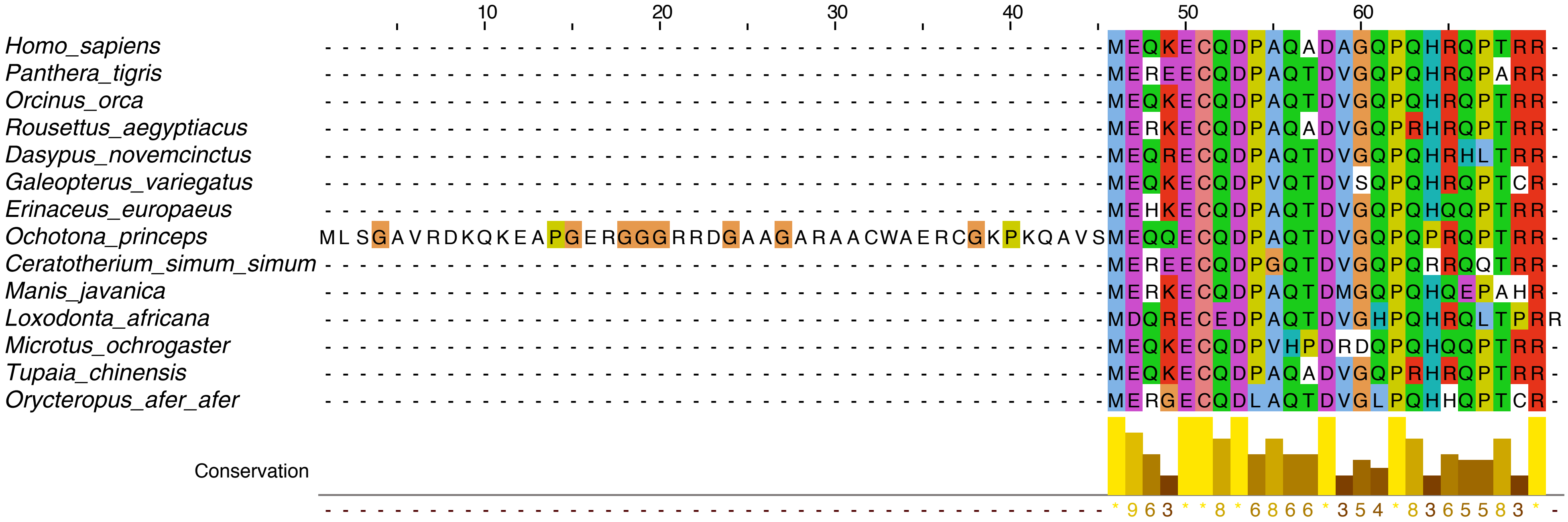

**HG0545**

ENSG00000142784 (WDTC1)

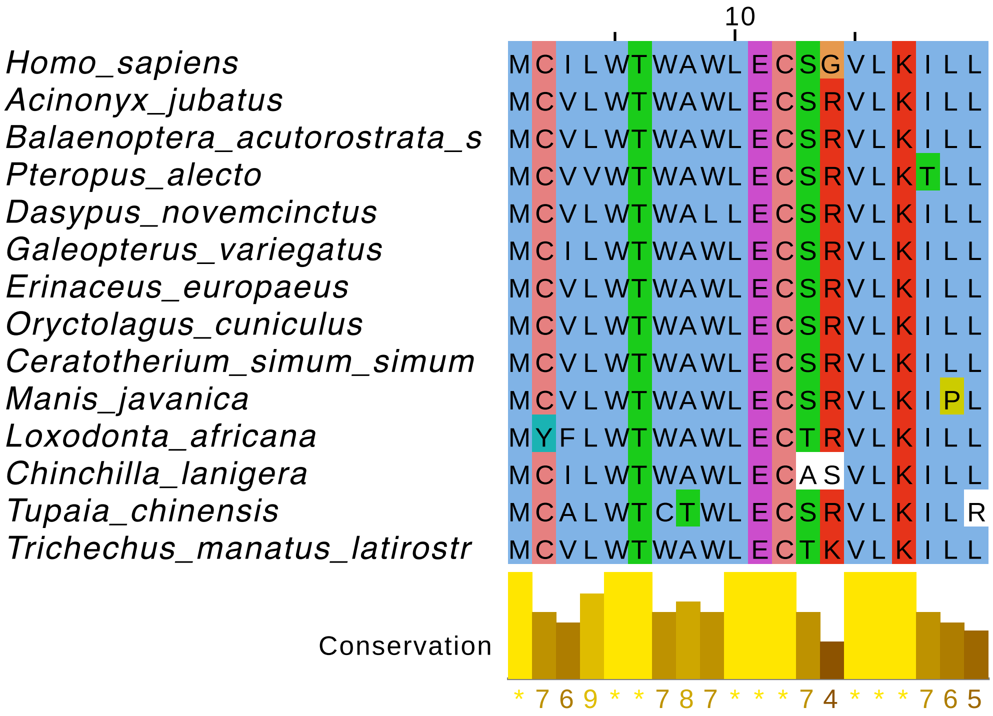

**HG0546**

ENSG00000148143 (ZNF462)

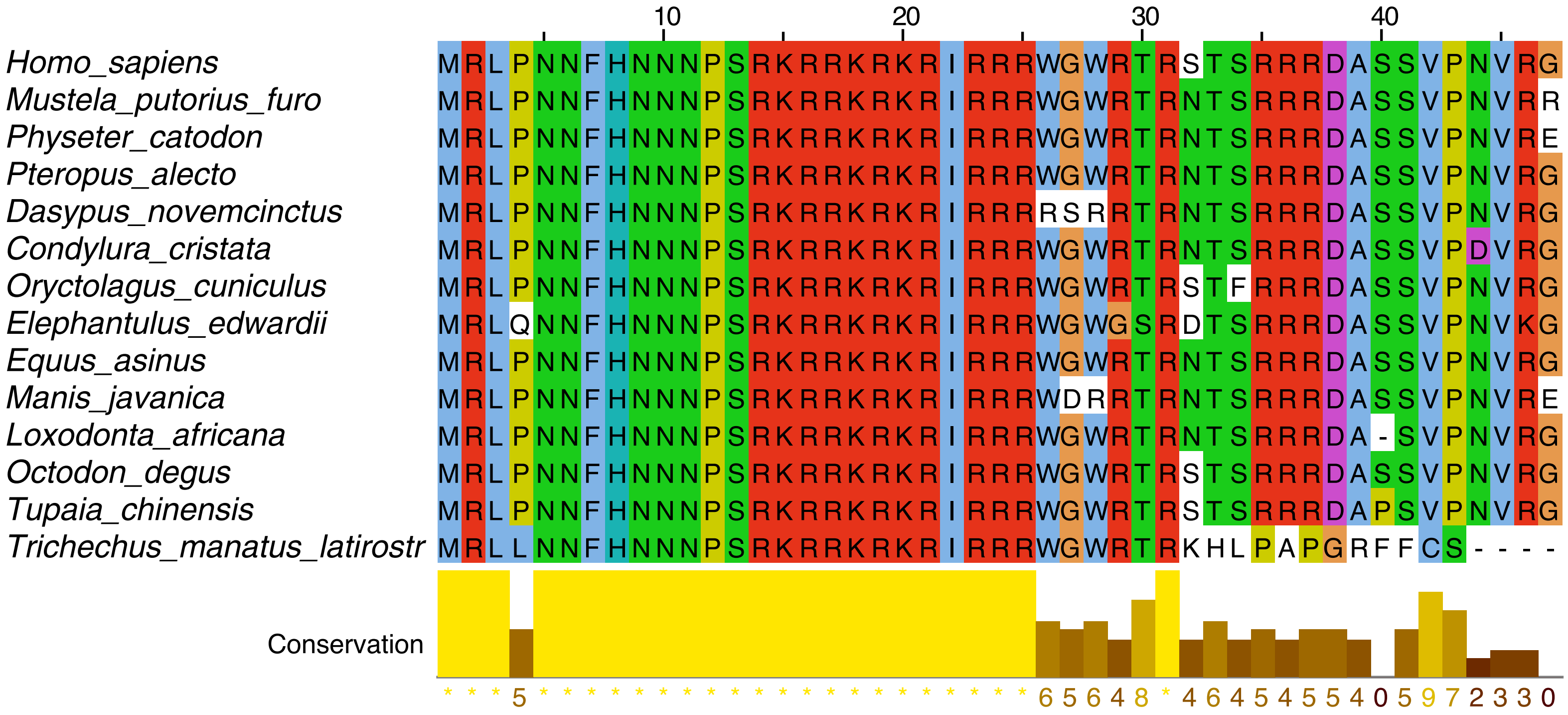

**HG0547**

ENSG00000149596 (JPH2)

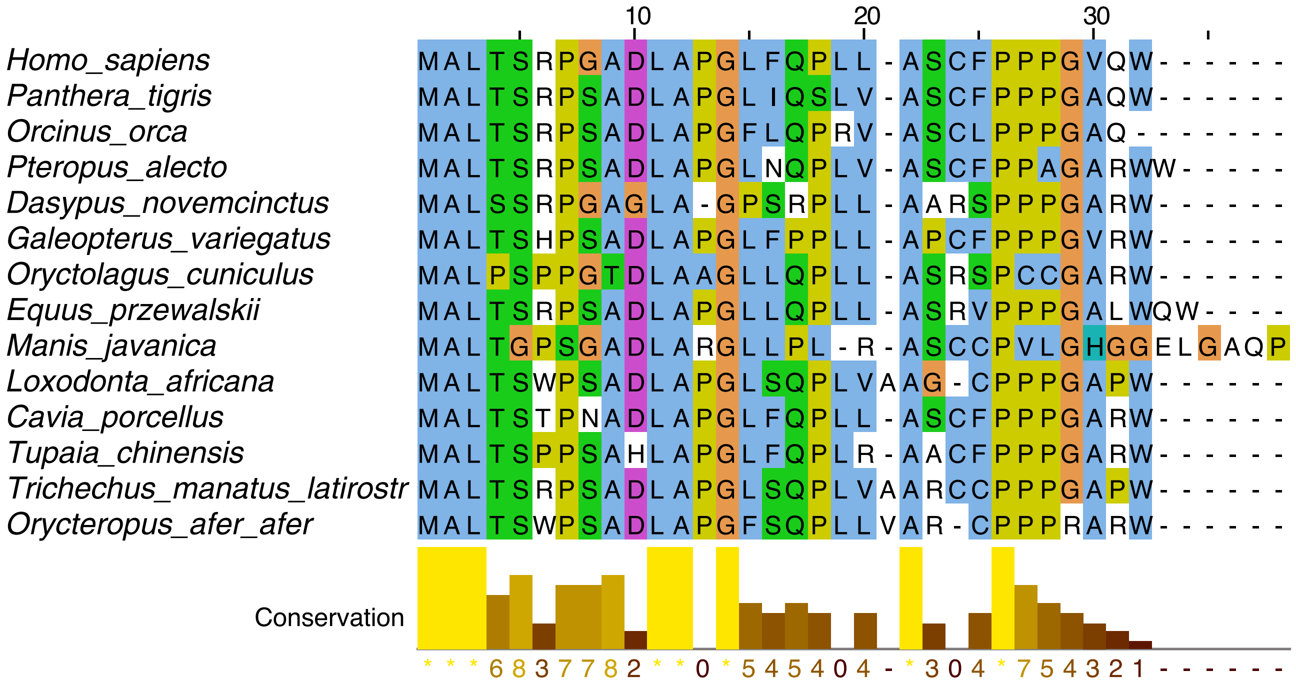

**HG0548**

ENSG00000155744 (FAM126B)

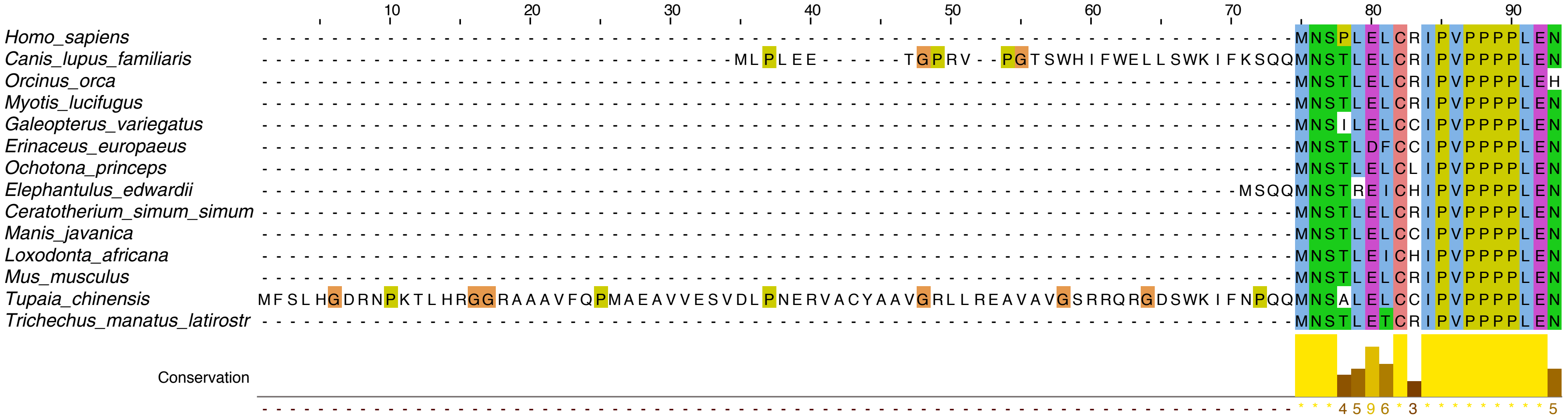

**HG0549**

ENSG00000156256 (USP16)

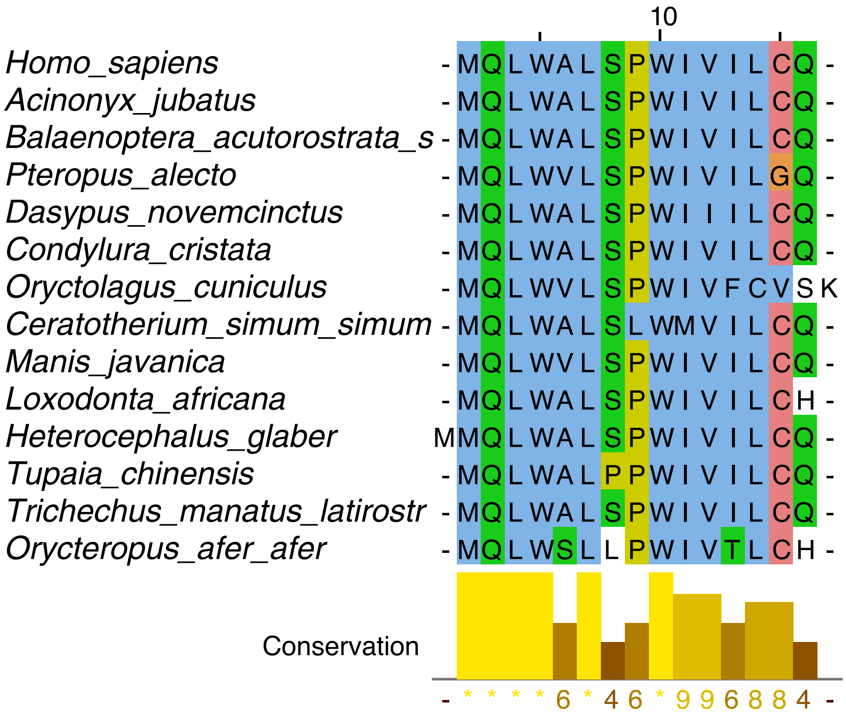

**HG0550.1**

ENSG00000156650 (KAT6B)

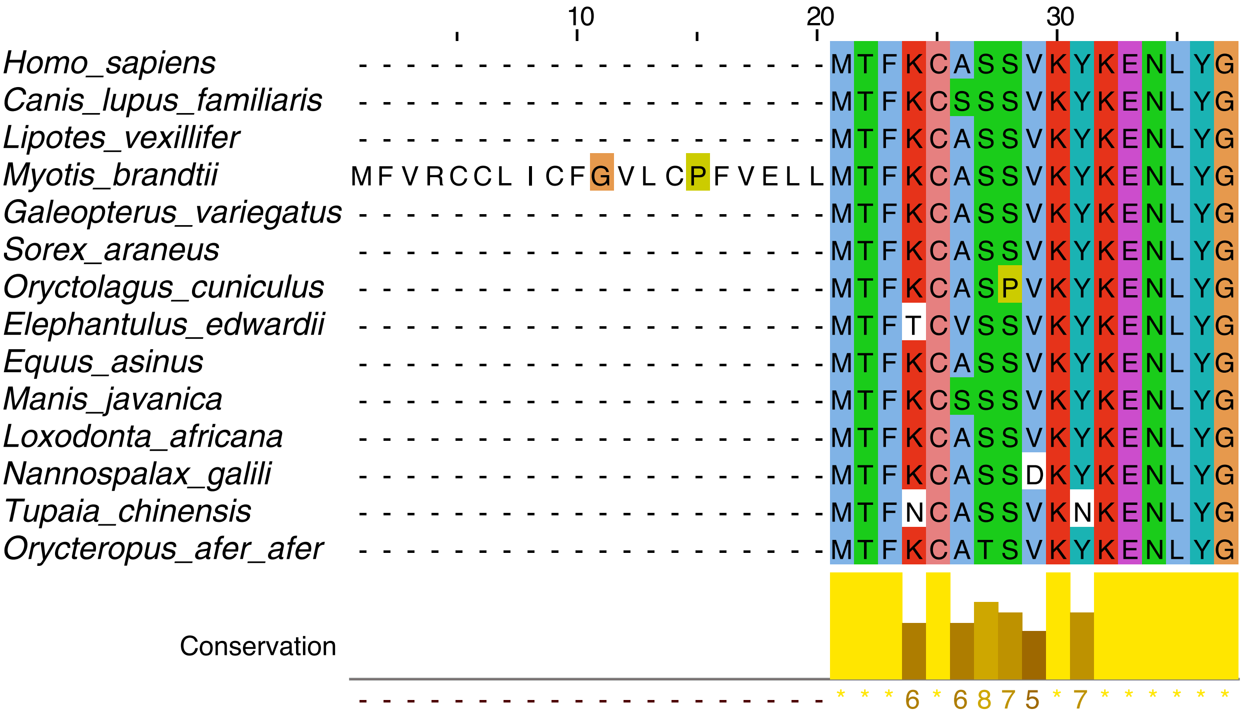

**HG0550.2**

ENSG00000156650 (KAT6B)

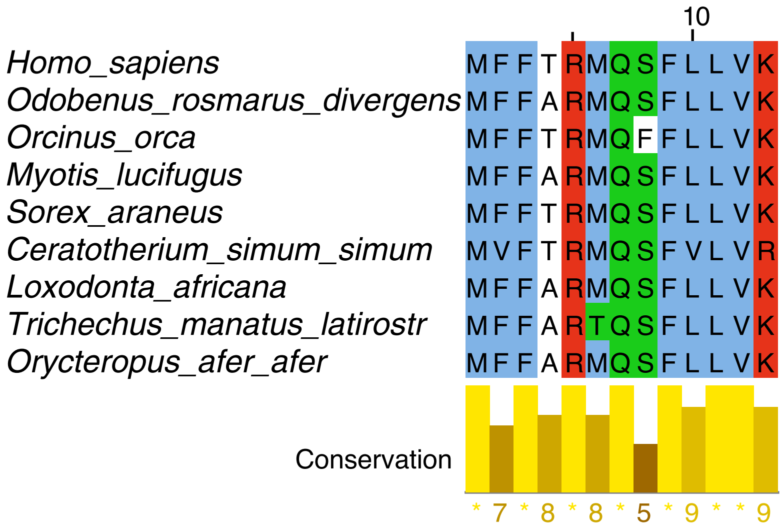

**HG0551**

ENSG00000162951 (LRRTM1)

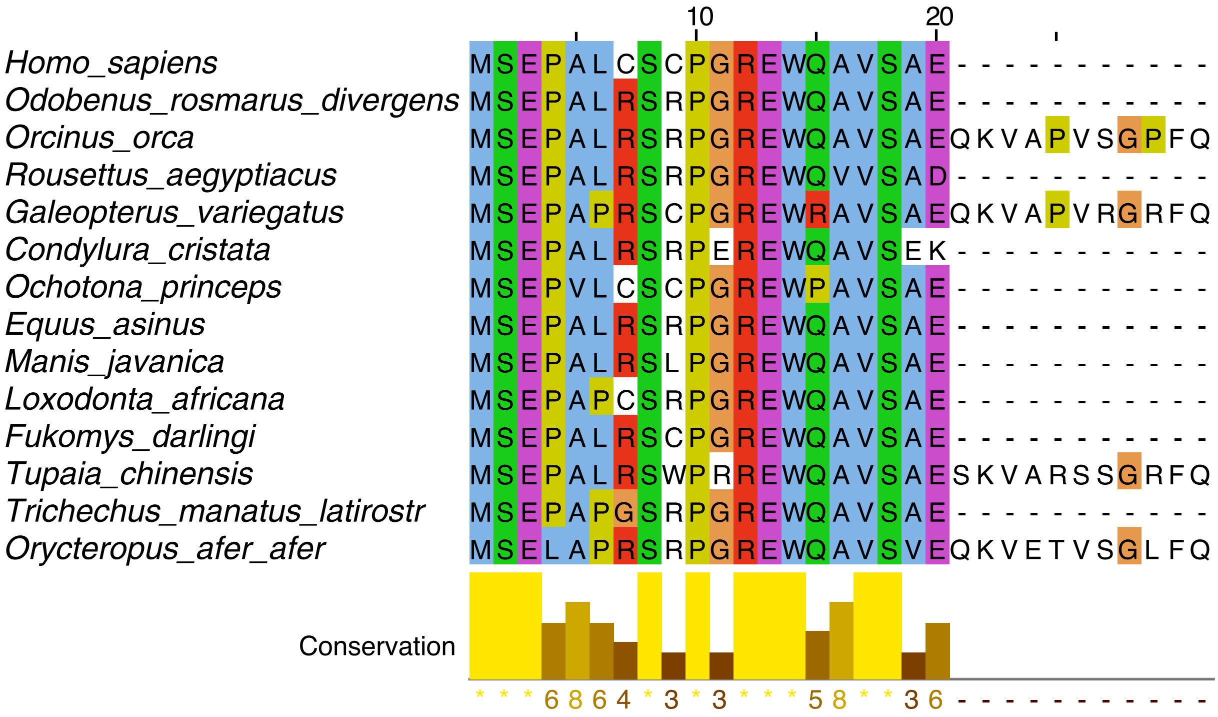

**HG0552**

ENSG00000169641 (LUZP1)

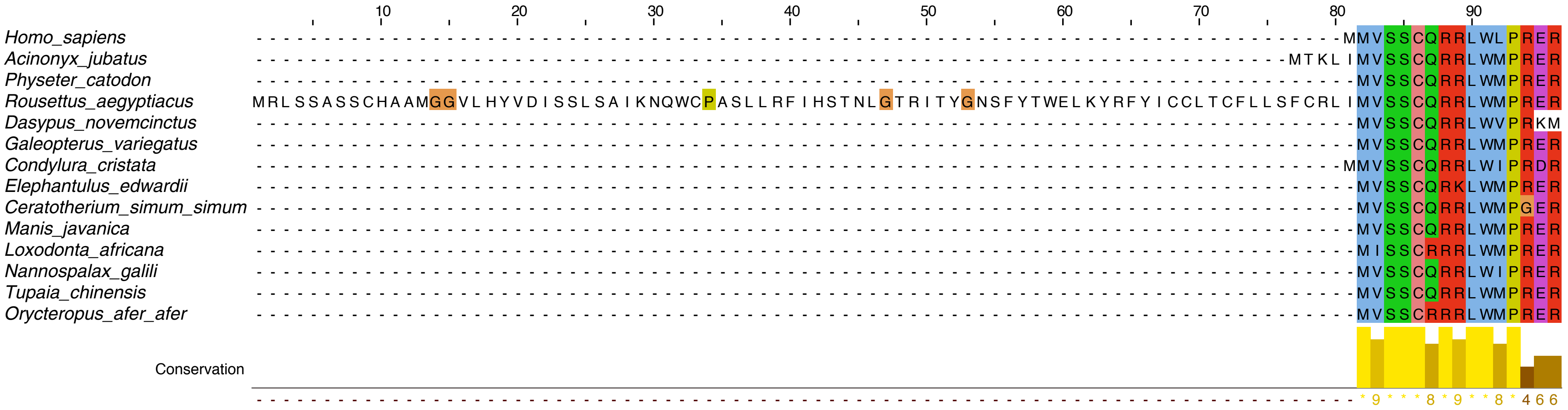

**HG0553**

ENSG00000183475 (ASB7)

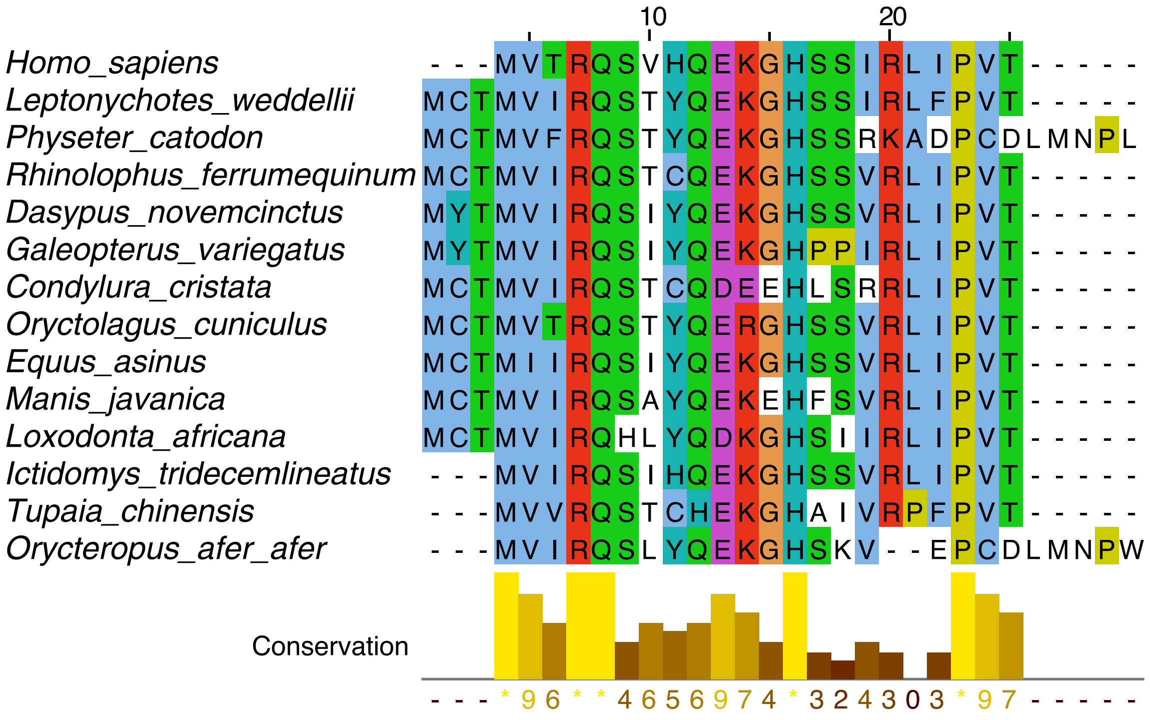

**HG0554**

ENSG00000196914 (ARHGEF12)

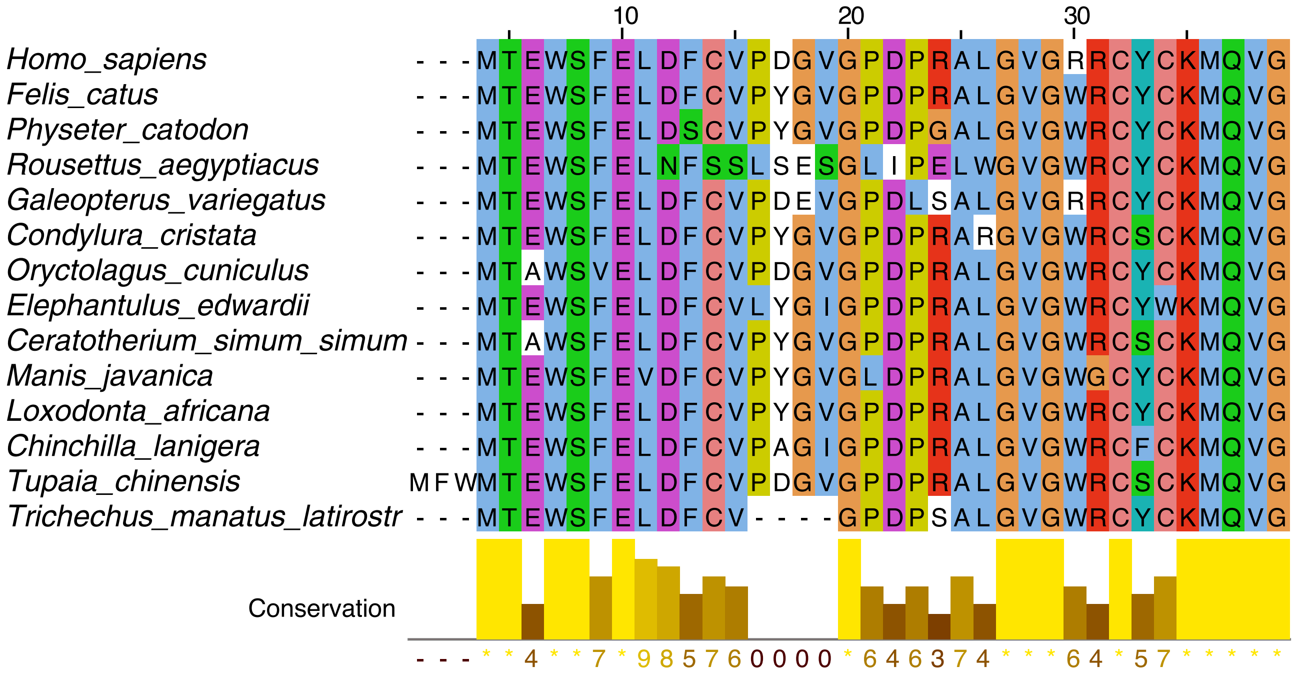

**HG0555**

ENSG00000204310 (AGPAT1)

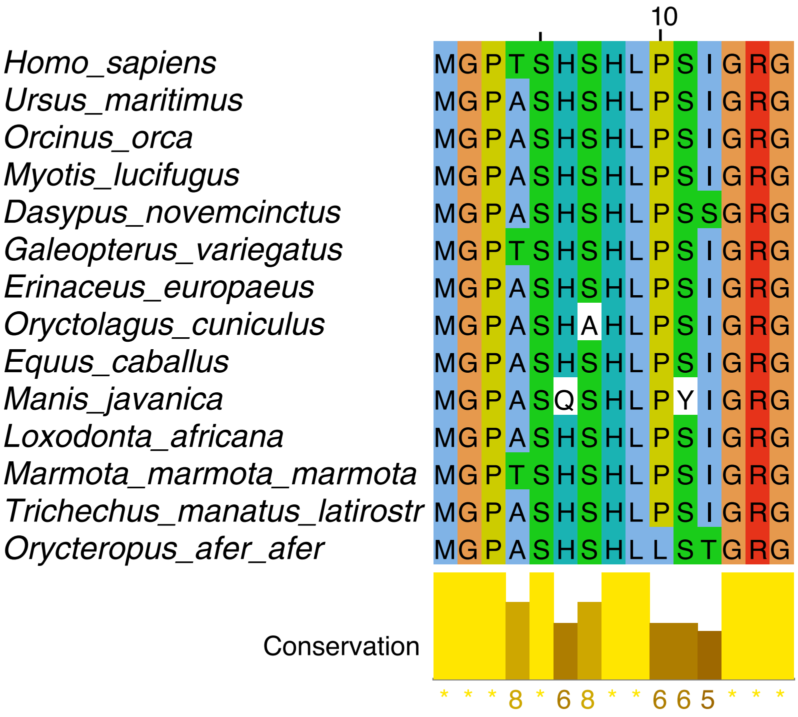

**HG0556**

ENSG00000273841 (TAF9)

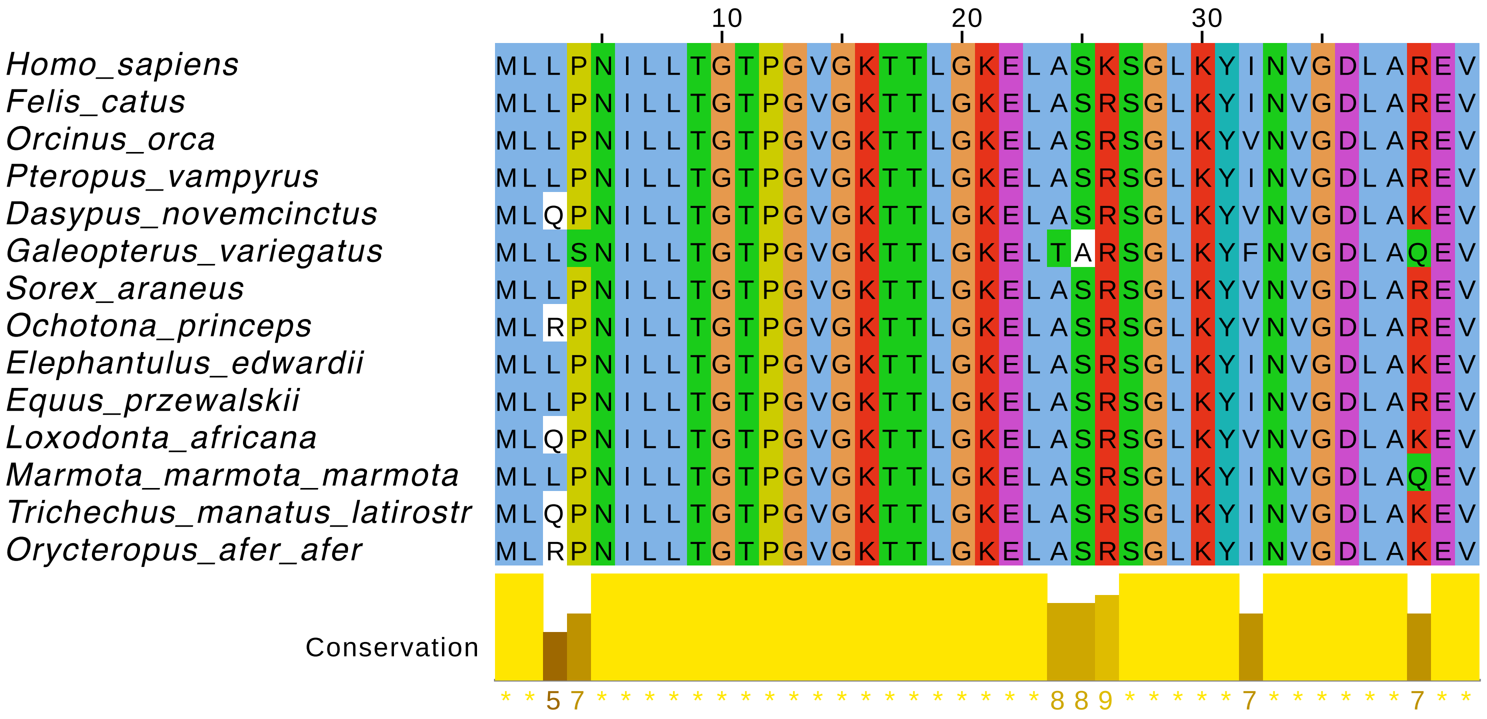

**HG0557**

ENSG00000012232 (EXTL3)

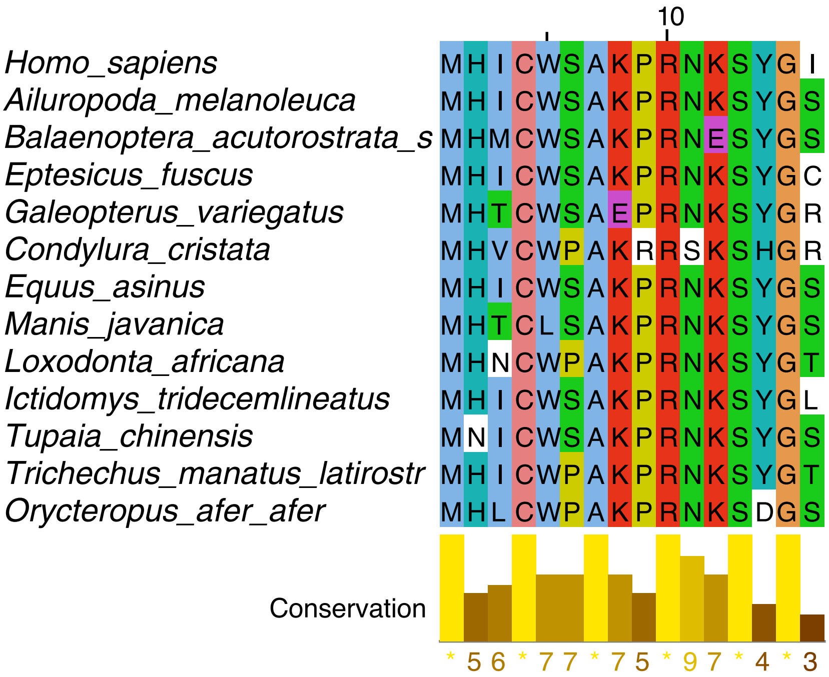

**HG0558**

ENSG00000029363 (BCLAF1)

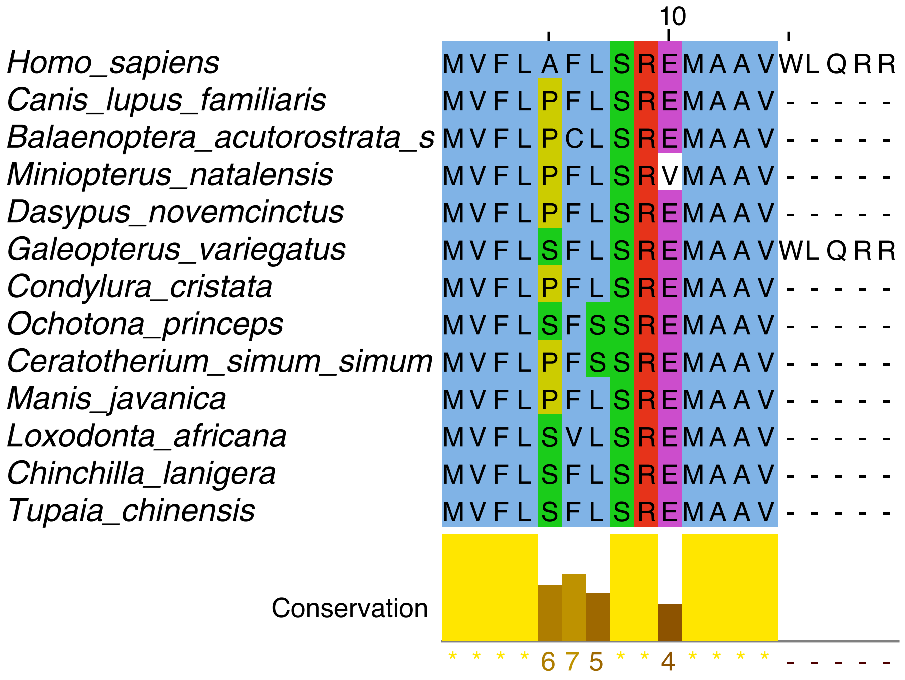

**HG0559**

ENSG00000078747 (ITCH)

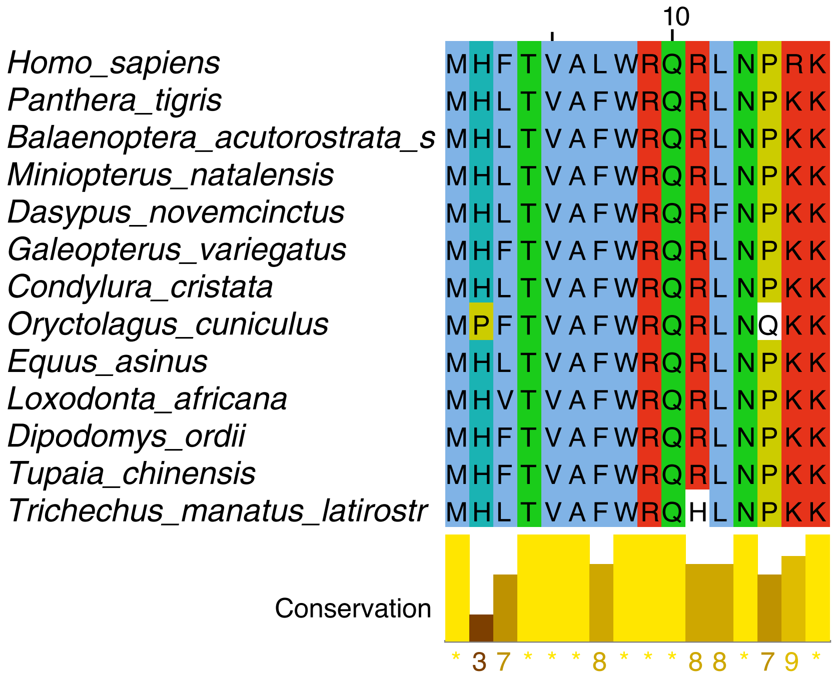

**HG0560.1**

ENSG00000092051 (JPH4)

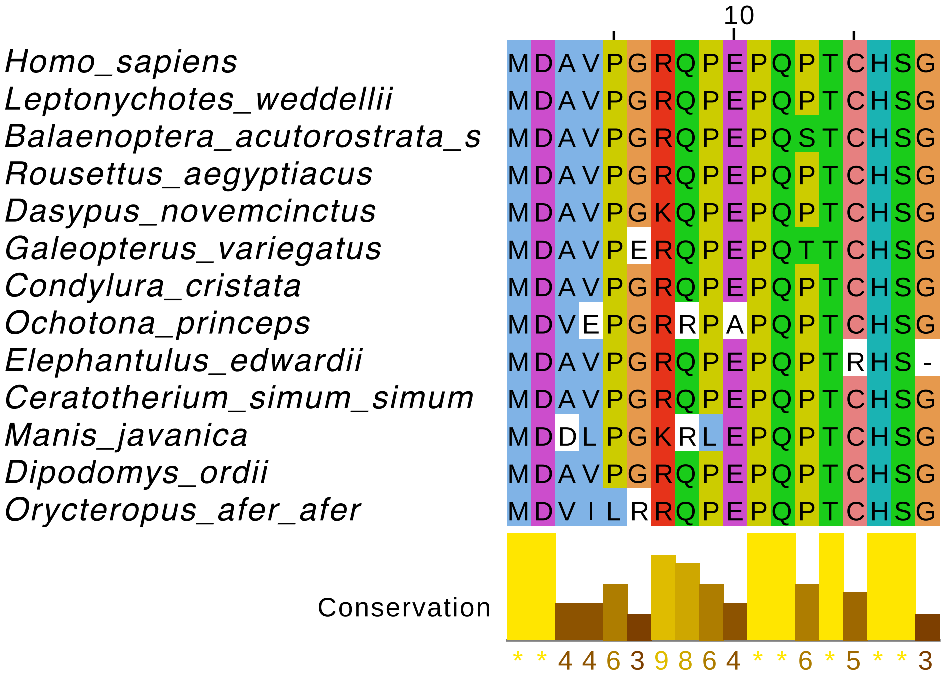

**HG0560.2**

ENSG00000092051 (JPH4)

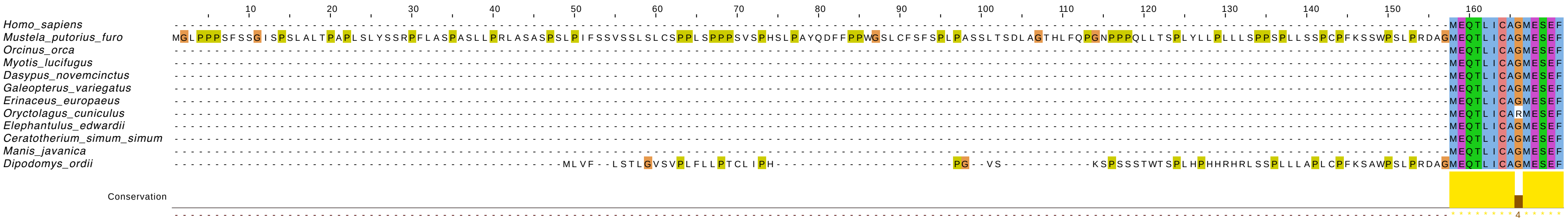

**HG0561**

ENSG00000100393 (EP300)

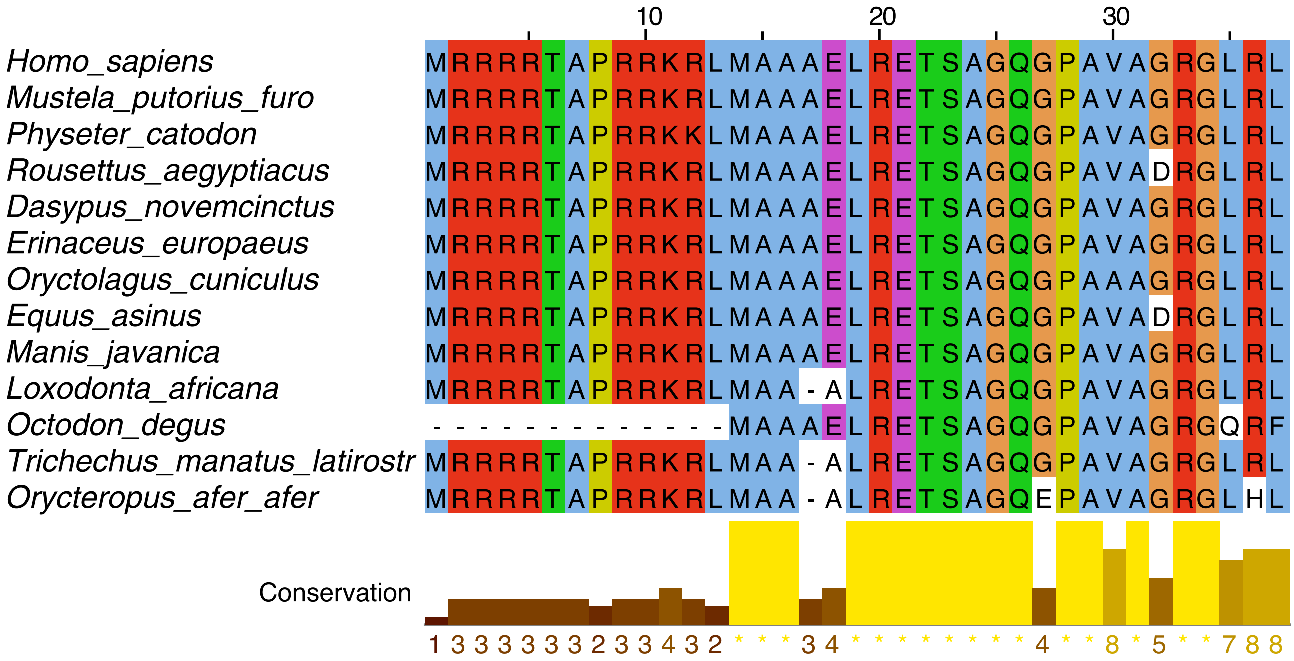

**HG0562**

ENSG00000105821 (DNAJC2)

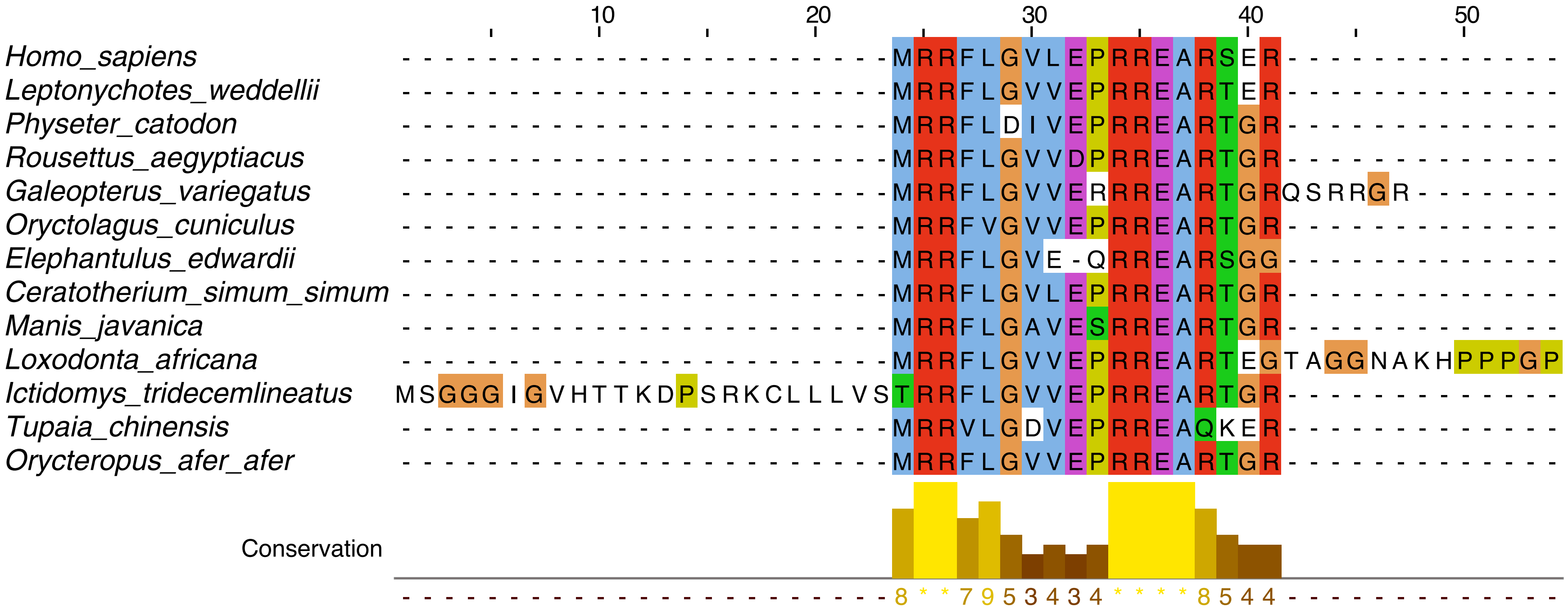

**HG0563**

ENSG00000107249 (GLIS3)

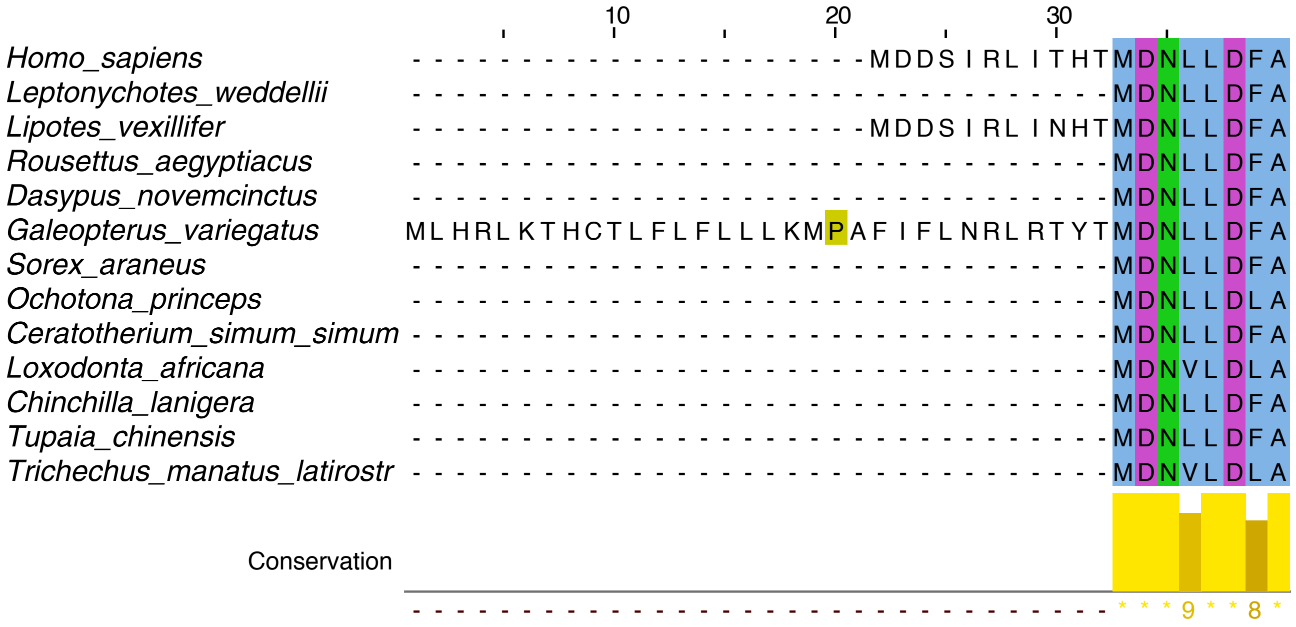

**HG0564**

ENSG00000109171 (SLAIN2)

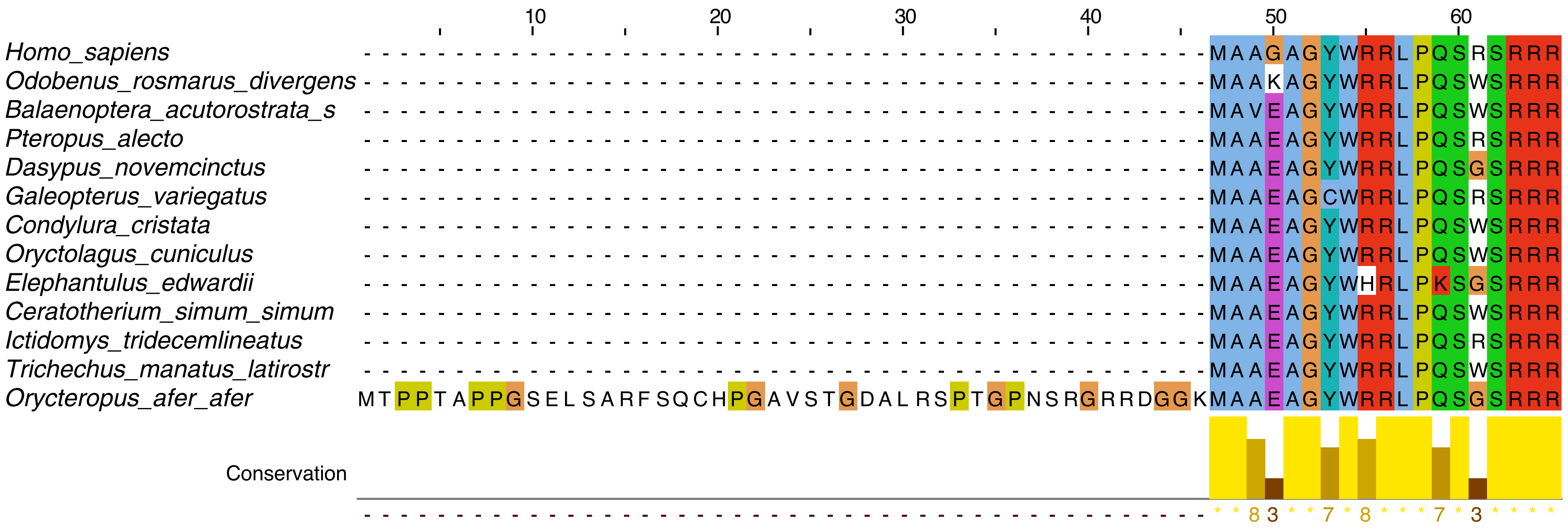

**HG0565**

ENSG00000120533 (ENY2)

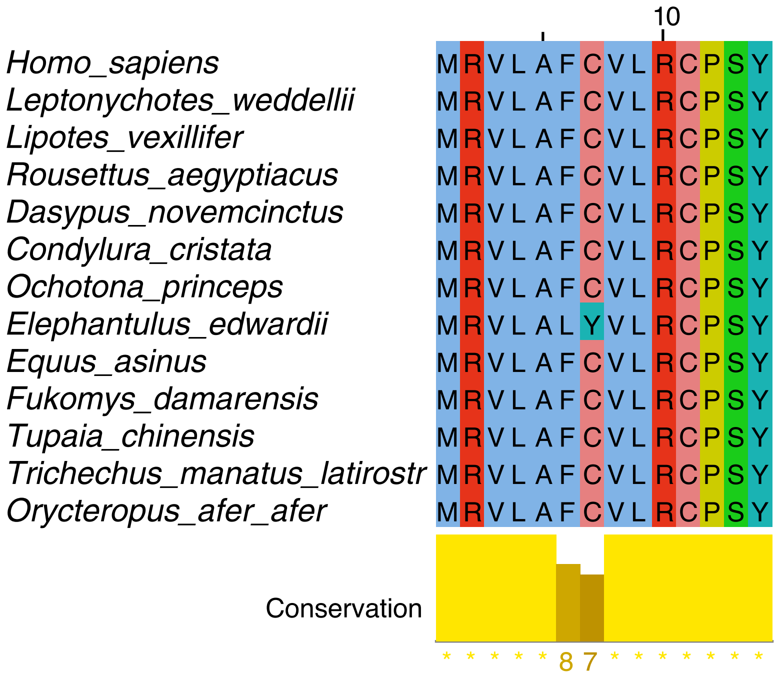

**HG0566**

ENSG00000124496 (TRERF1)

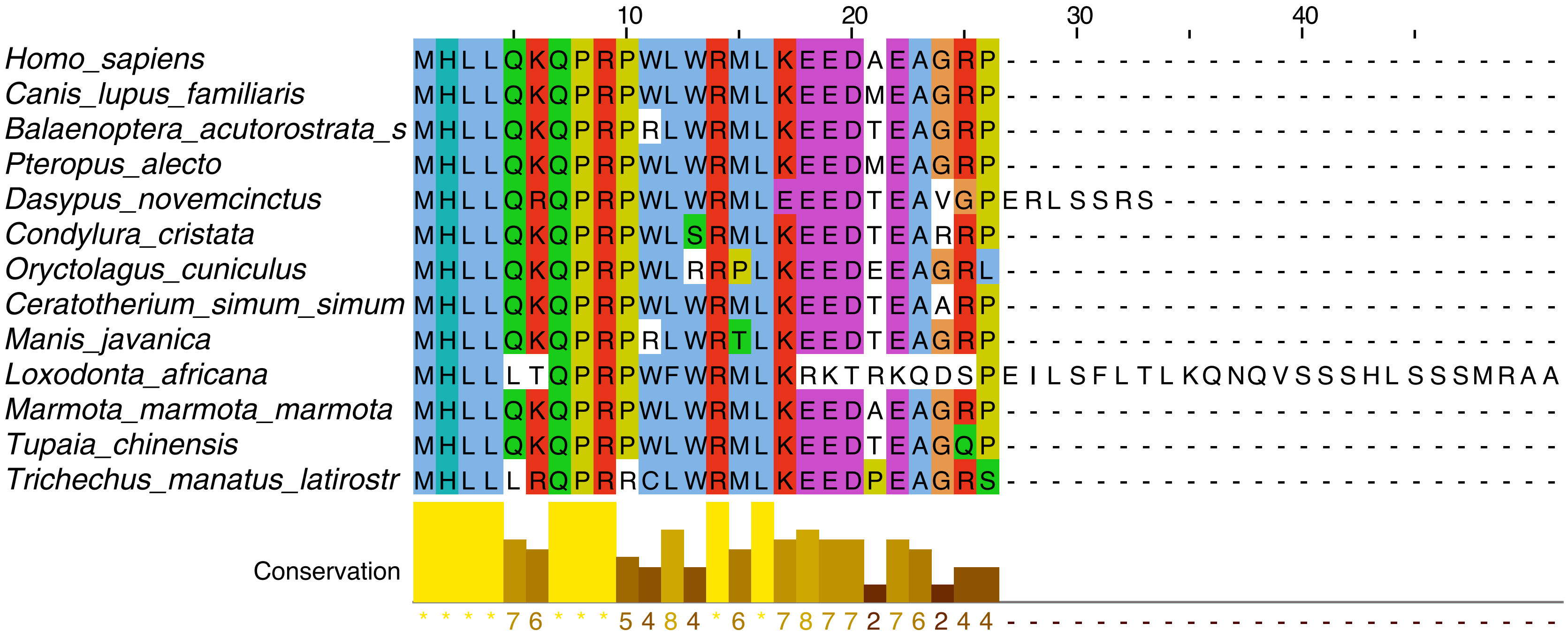

**HG0567**

ENSG00000132155 (RAF1)

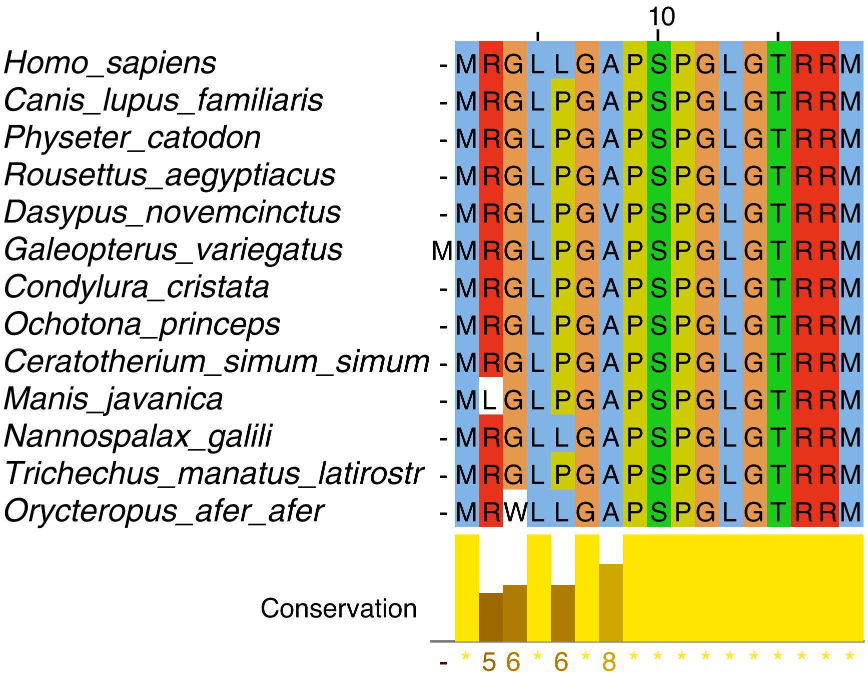

**HG0568.1**

ENSG00000138271 (GPR87)

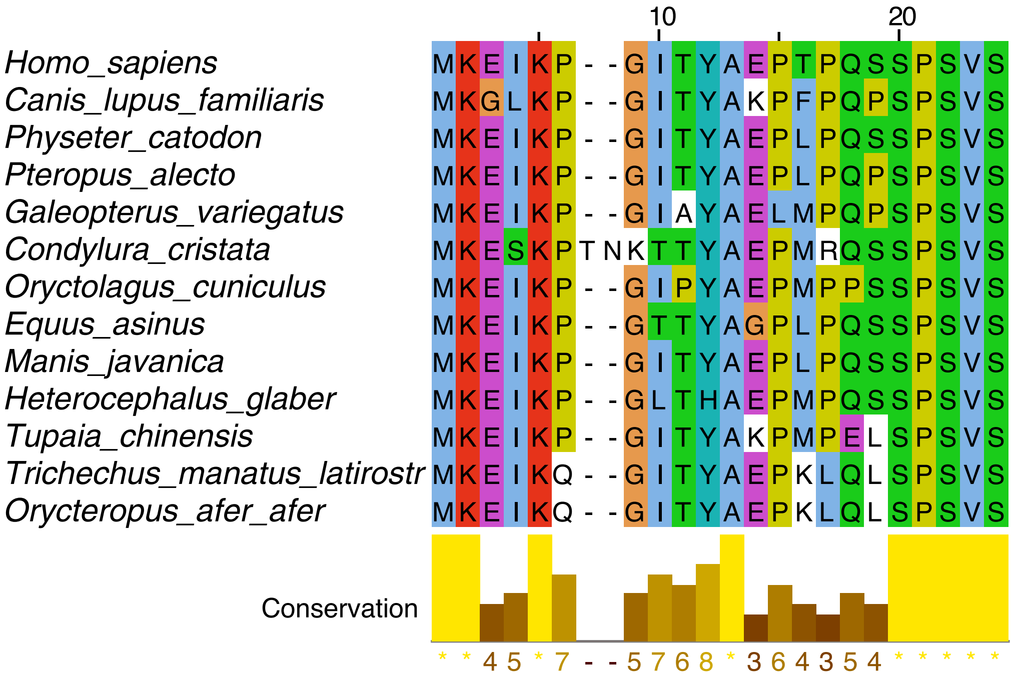

**HG0568.2**

ENSG00000138271 (GPR87)

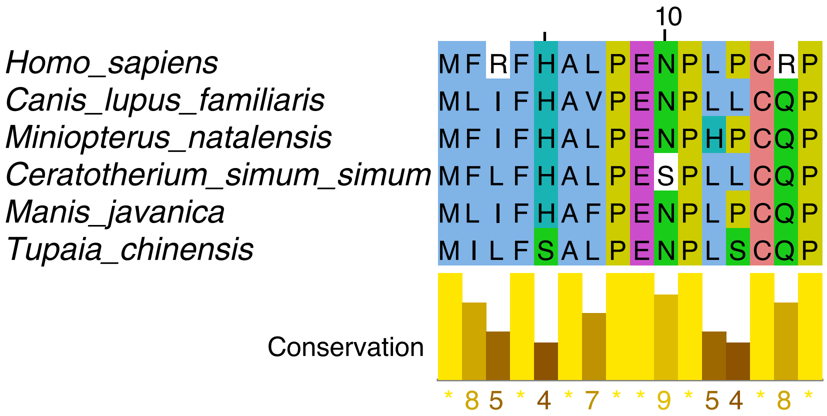

**HG0569**

ENSG00000139651 (ZNF740)

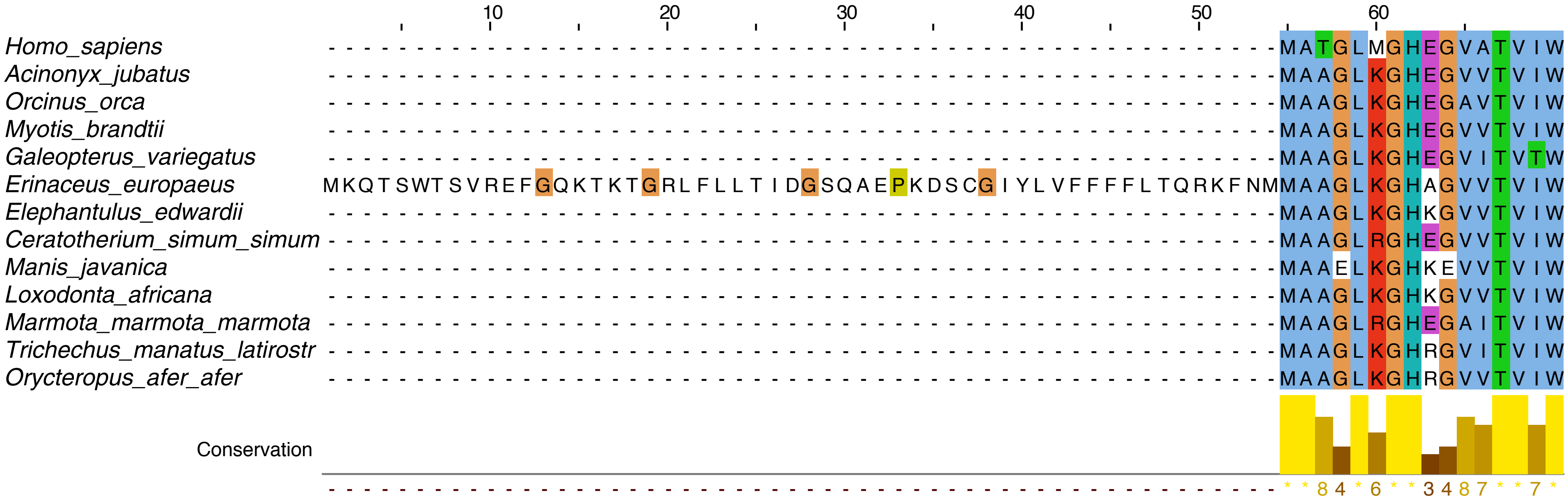

**HG0570**

ENSG00000152214 (RIT2)

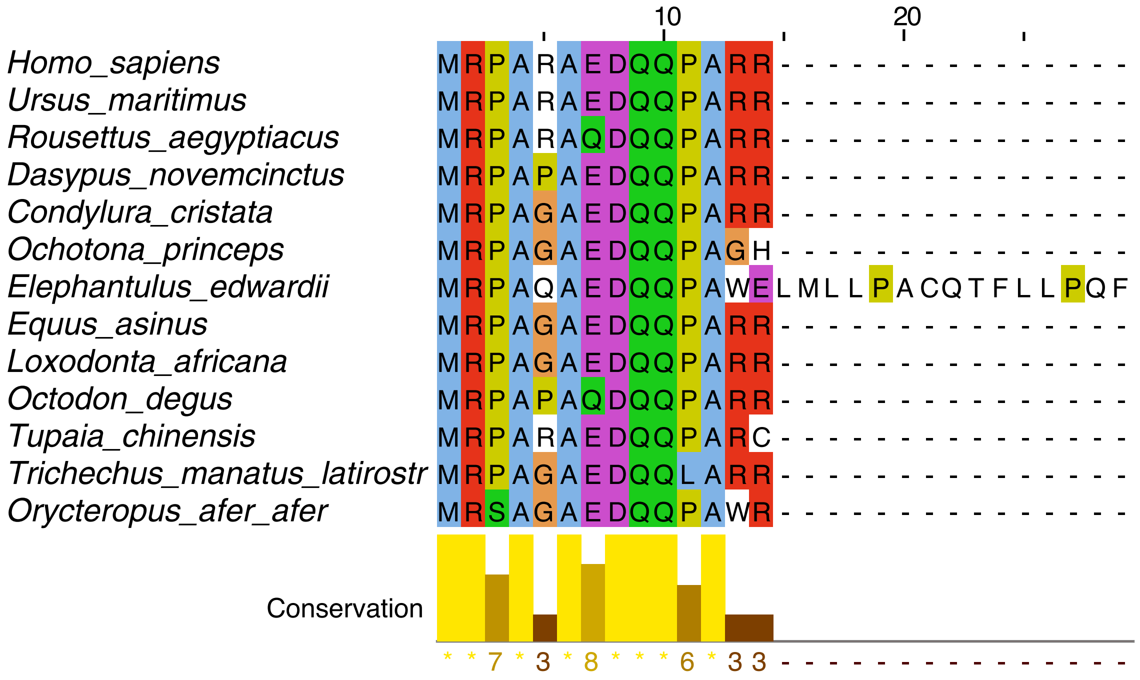

**HG0571**

ENSG00000153266 (FEZF2)

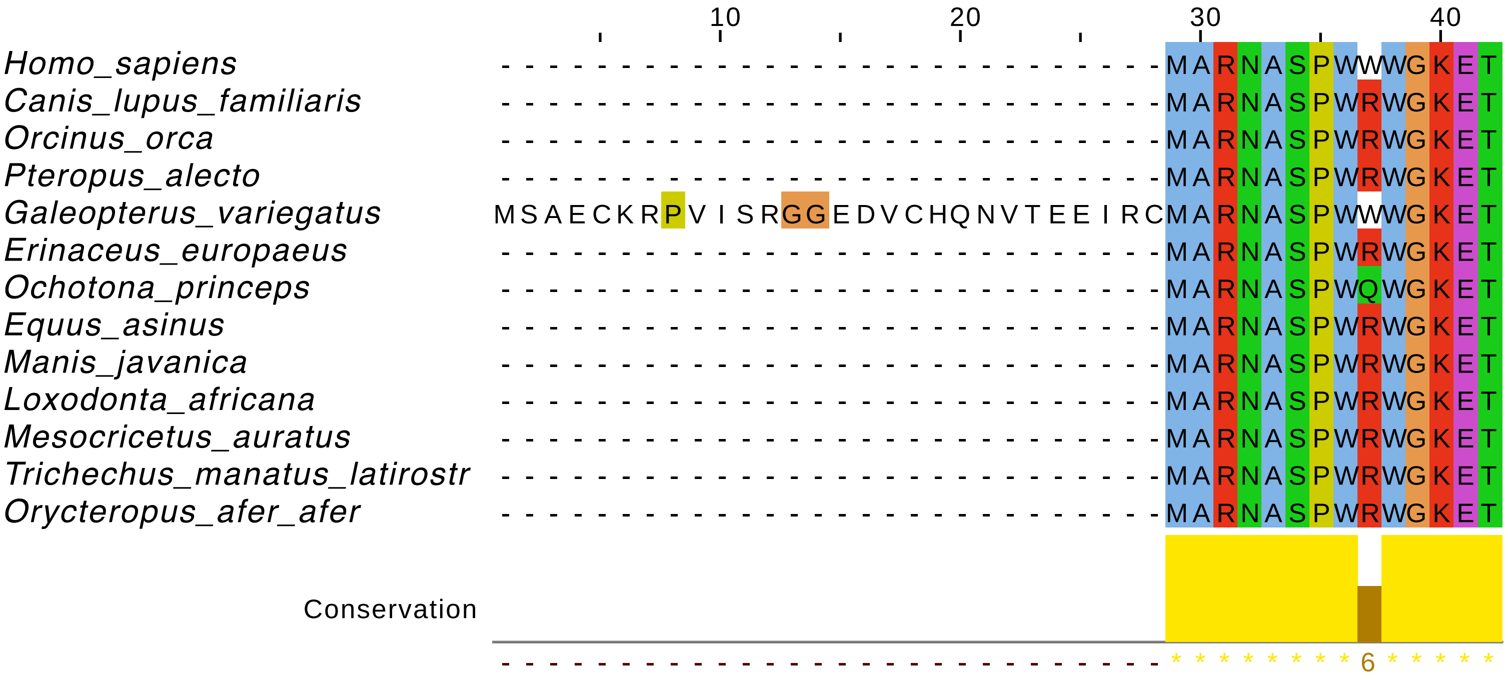

**HG0572**

ENSG00000158161 (EYA3)

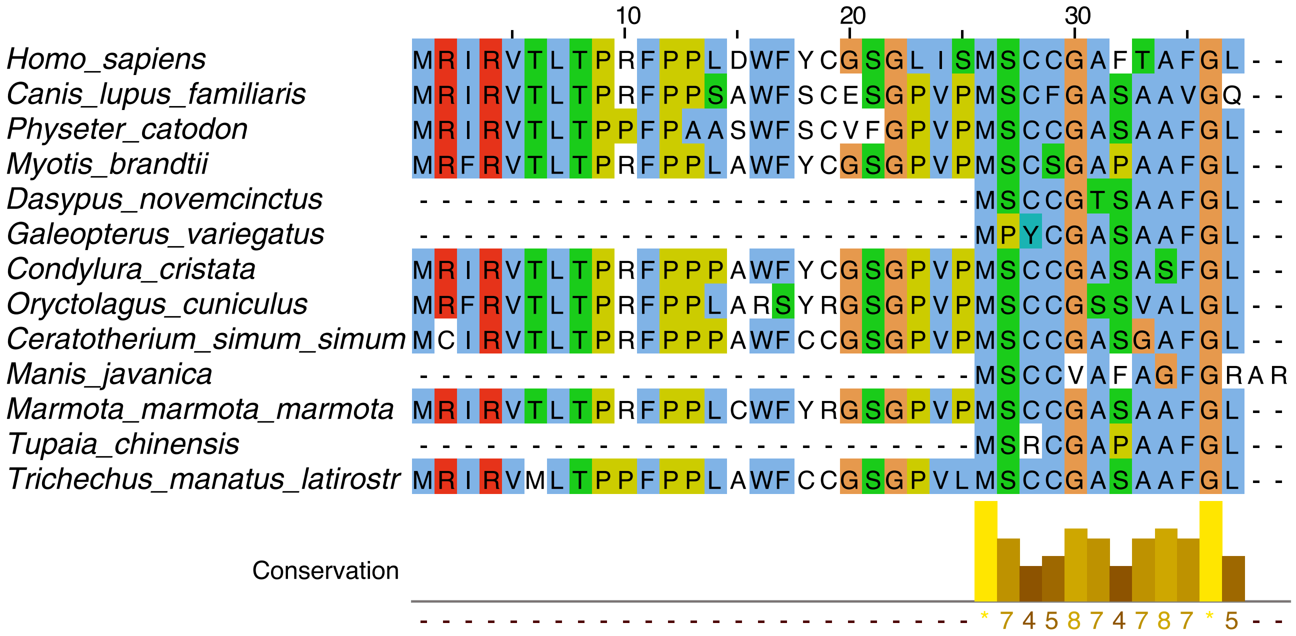

**HG0573.1**

ENSG00000162526 (TSSK3)

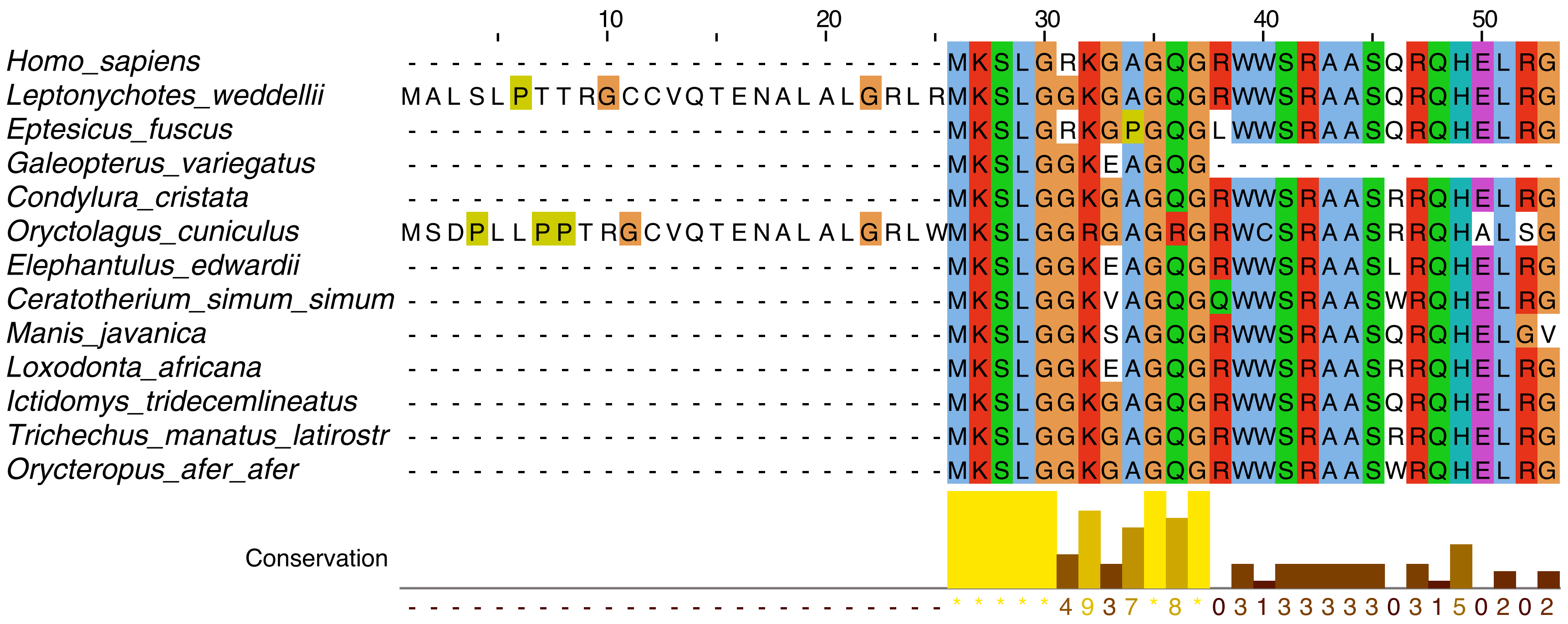

**HG0573.2**

ENSG00000206203 (TSSK2)

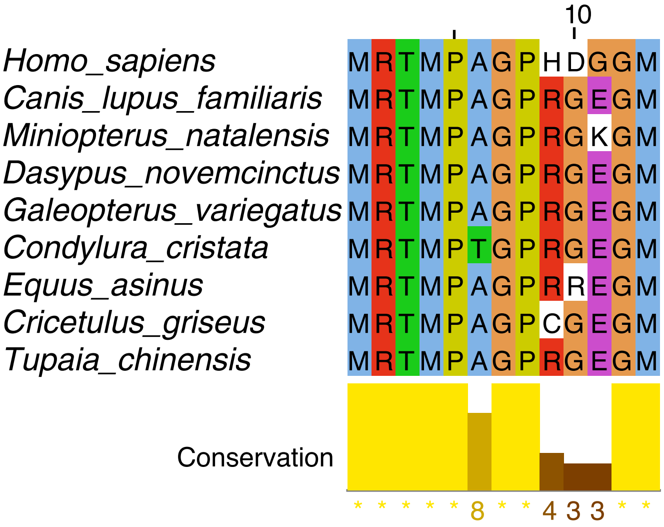

**HG0574**

ENSG00000164663 (USP49)

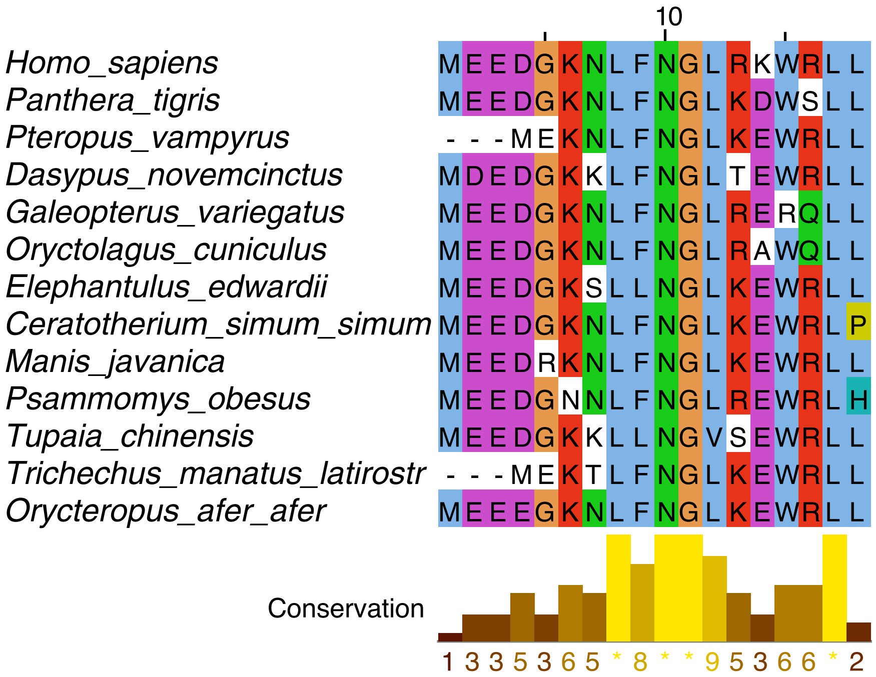

**HG0575**

ENSG00000164663 (USP49)

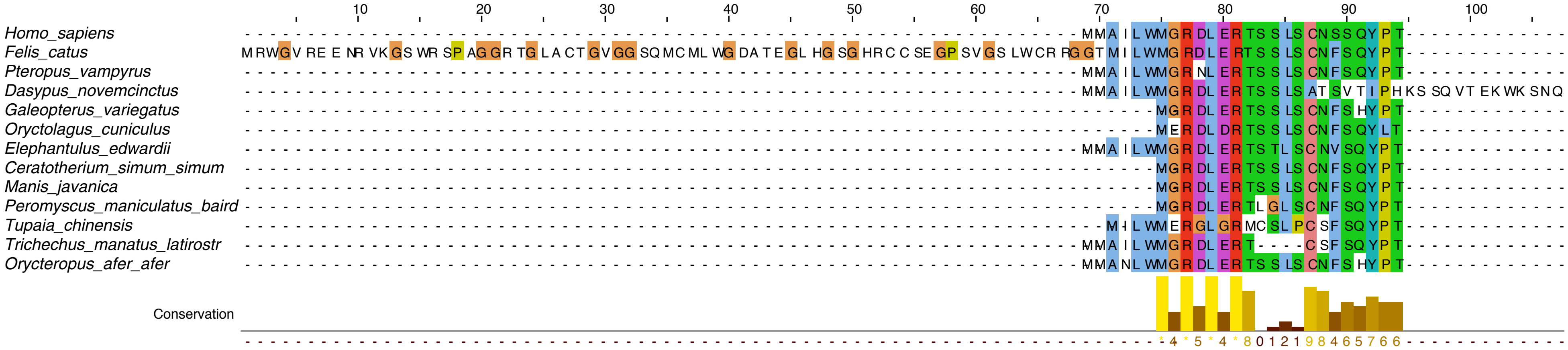

**HG0576**

ENSG00000175029 (CTBP2)

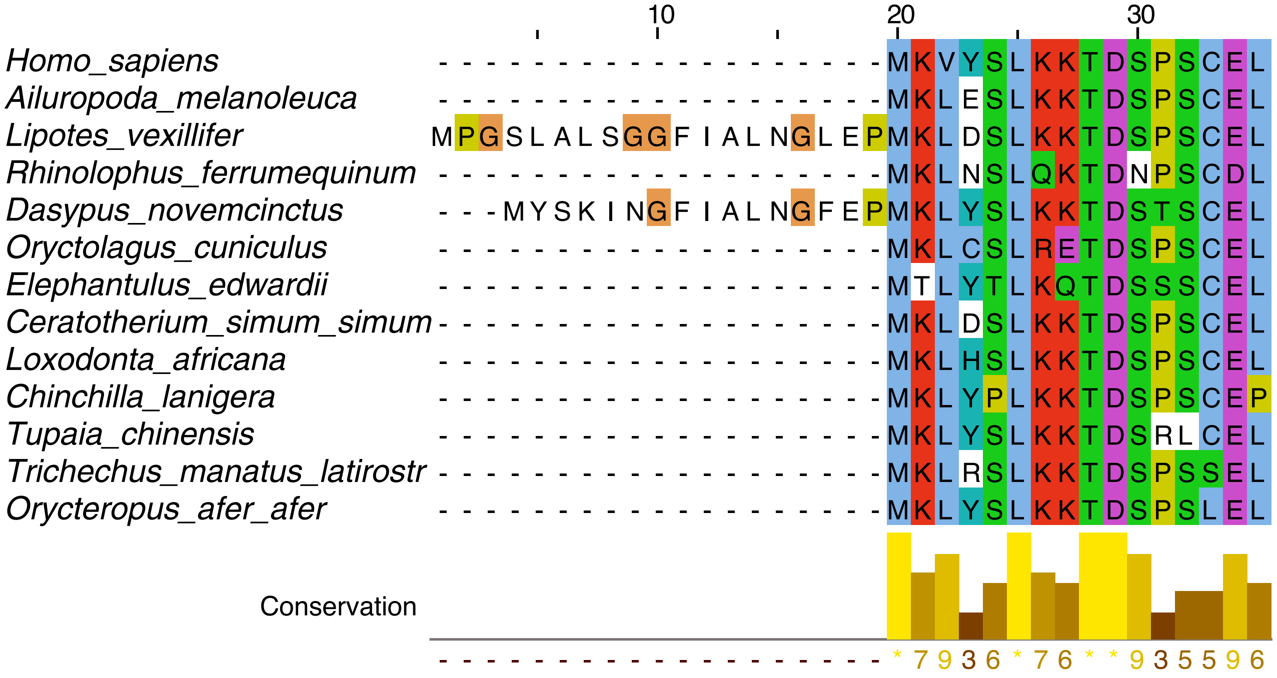

**HG0577**

ENSG00000250510 (GPR162)

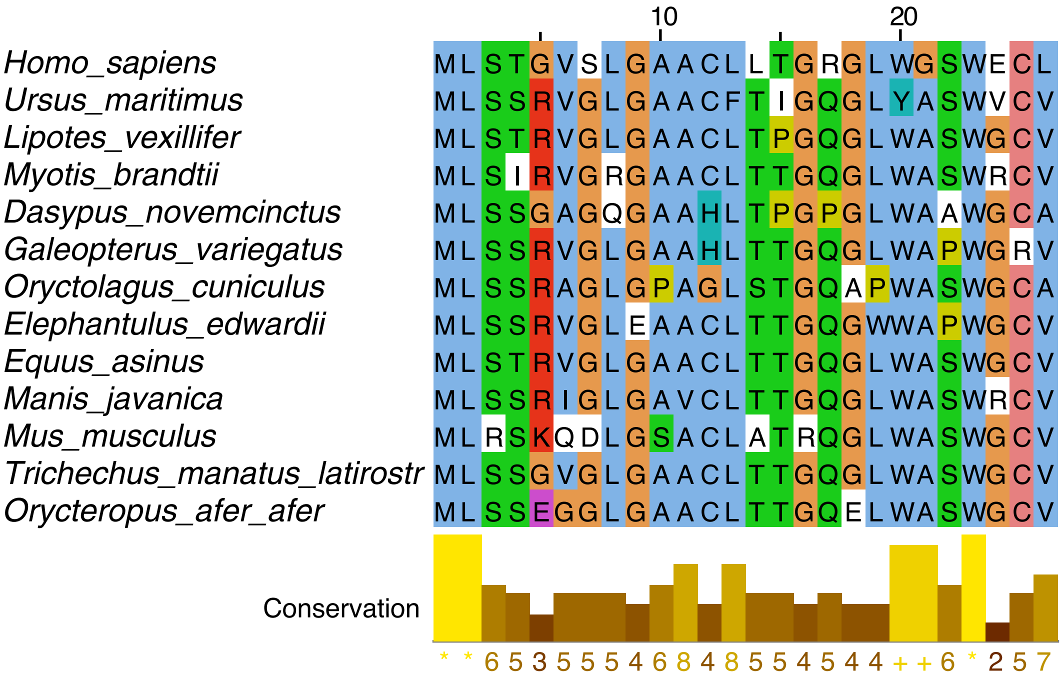

**HG0578**

ENSG00000004897 (CDC27)

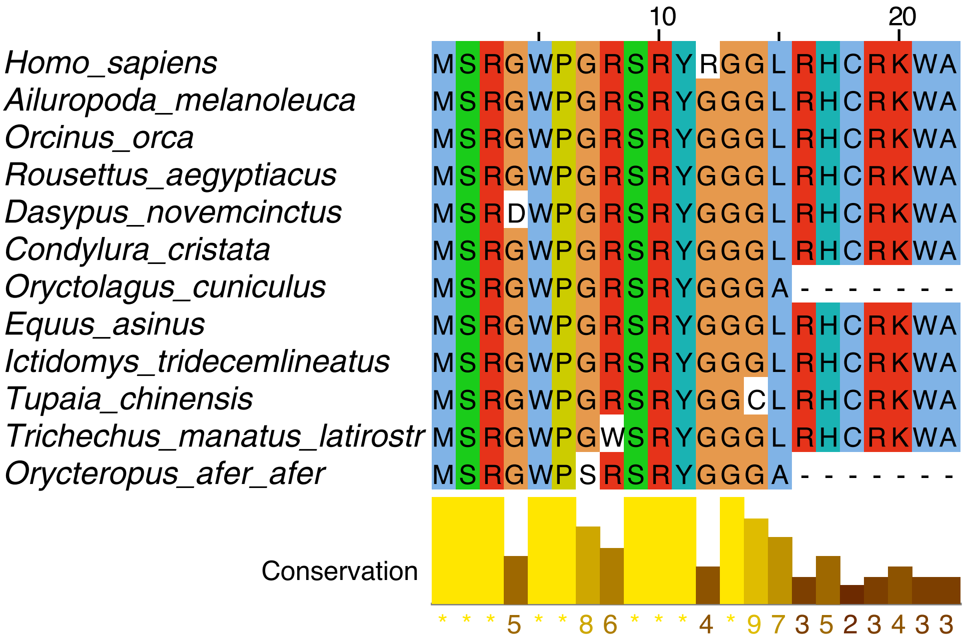

**HG0579**

ENSG00000010818 (HIVEP2)

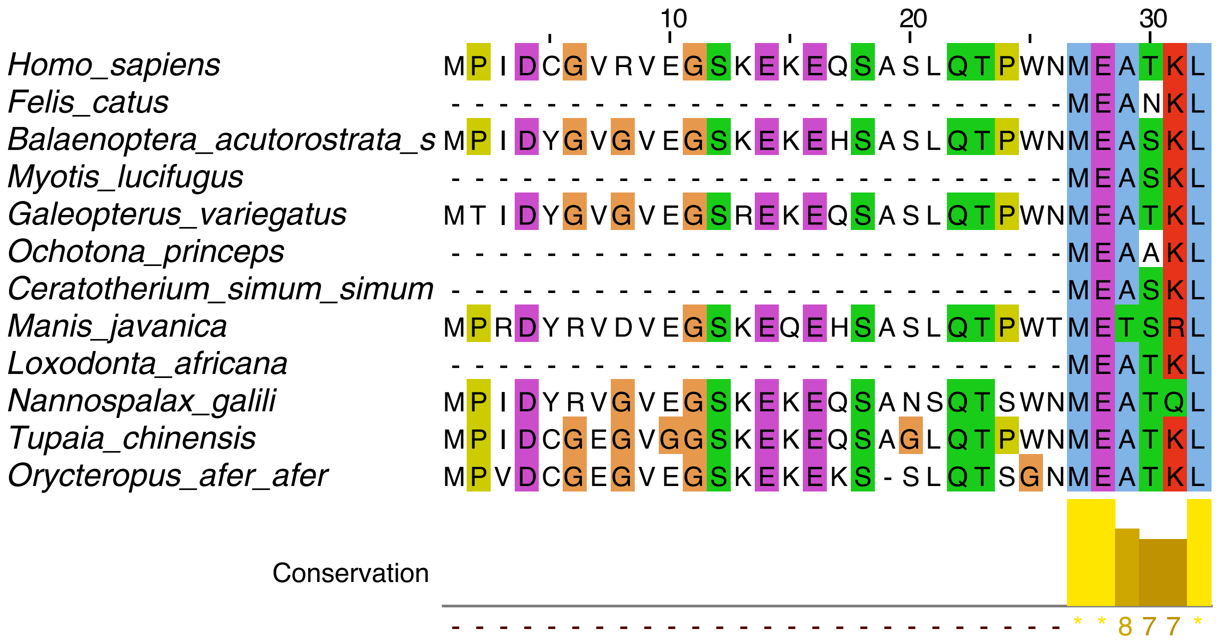

**HG0580**

ENSG00000073711 (PPP2R3A)

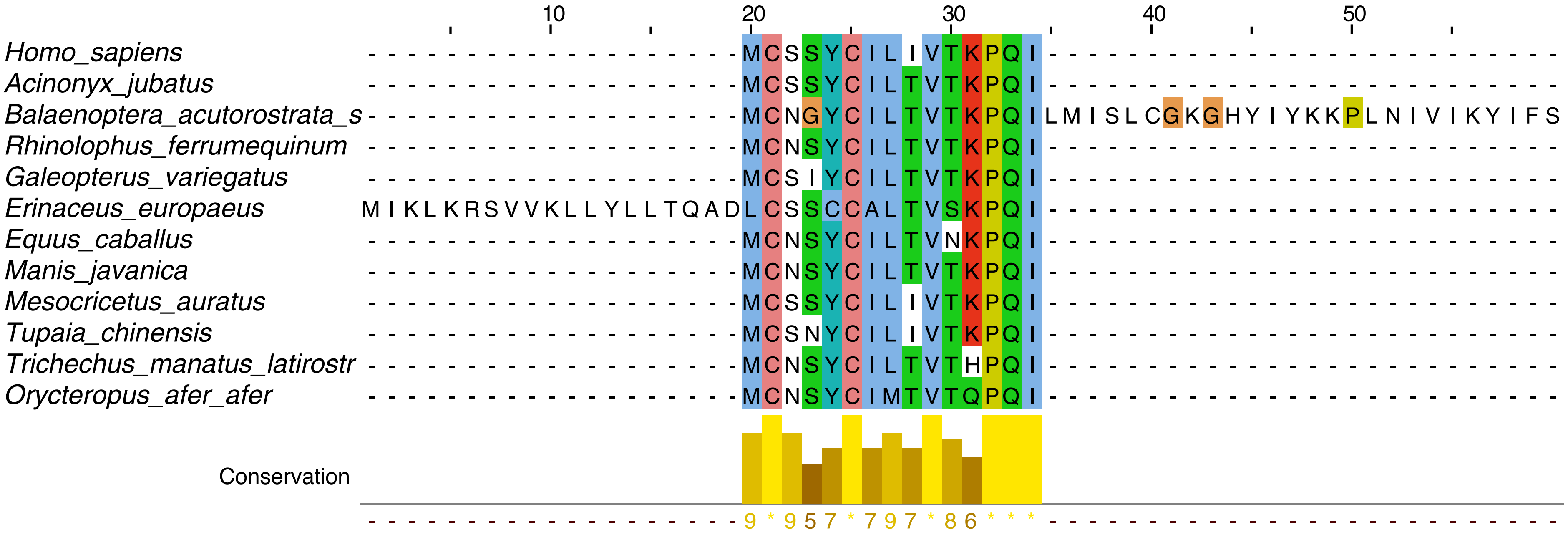

**HG0581**

ENSG00000087074 (PPP1R15A)

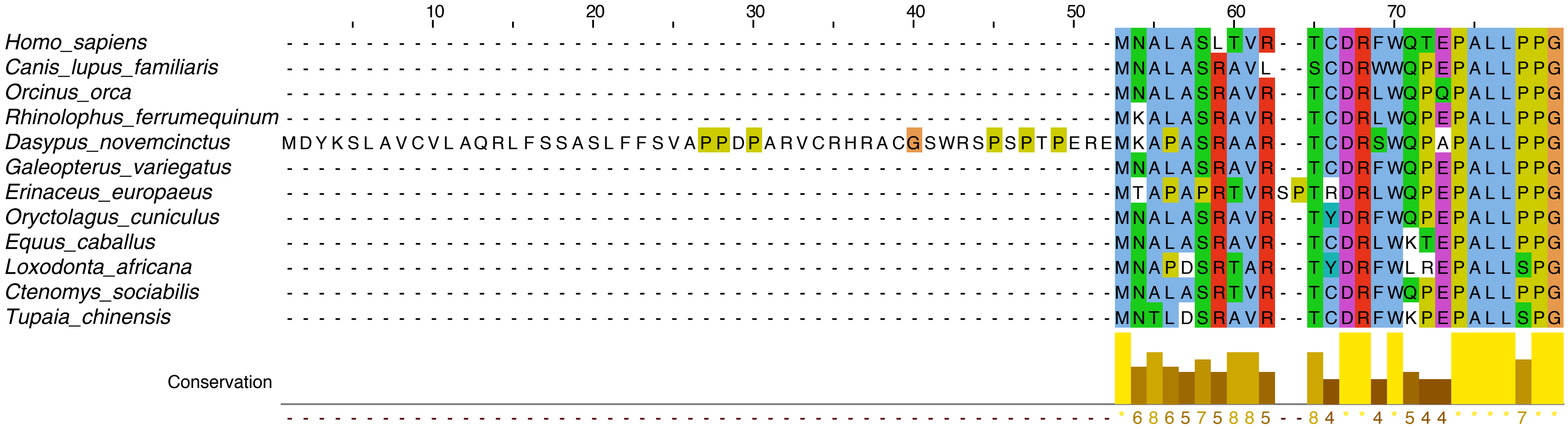

**HG0582**

ENSG00000087338 (GMCL1)

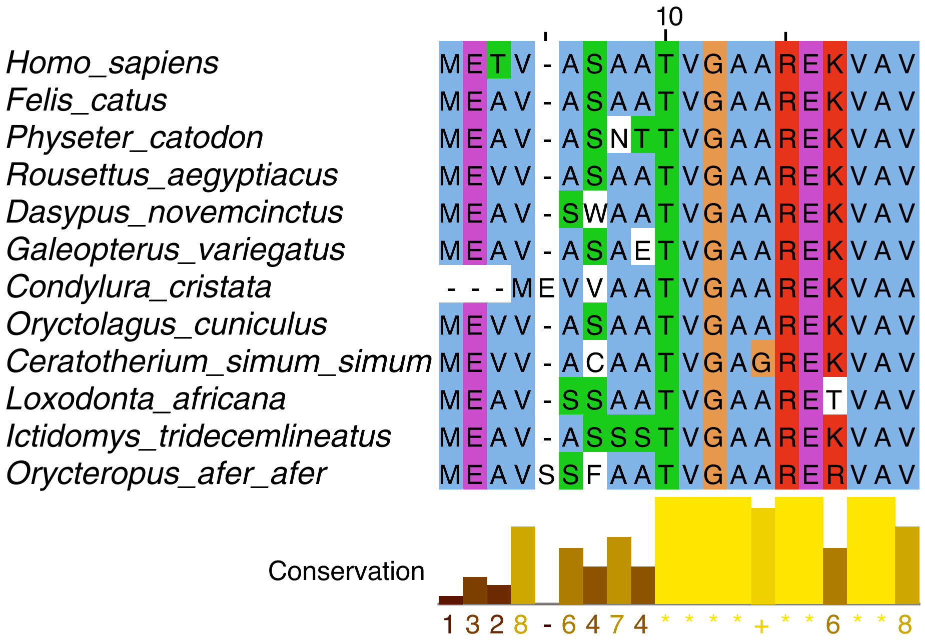

**HG0583**

ENSG00000087502 (ERGIC2)

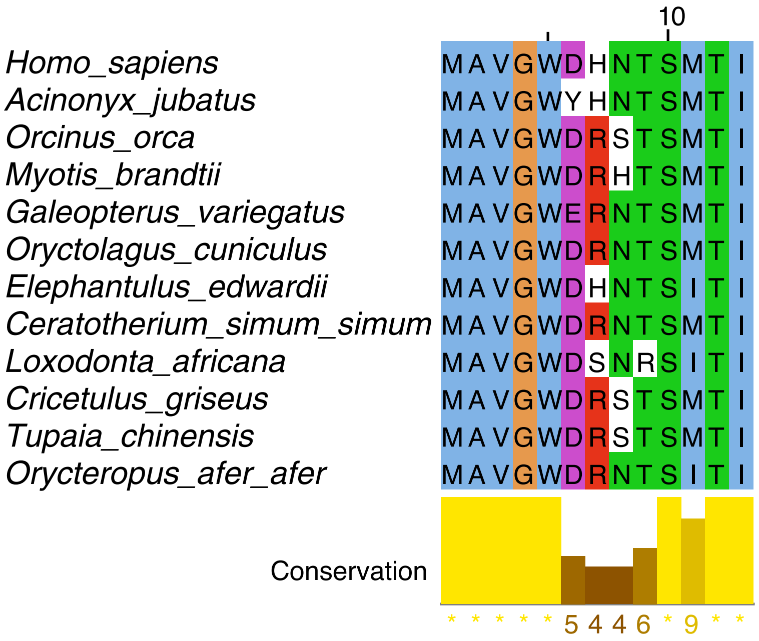

**HG0584**

ENSG00000092051 (JPH4)

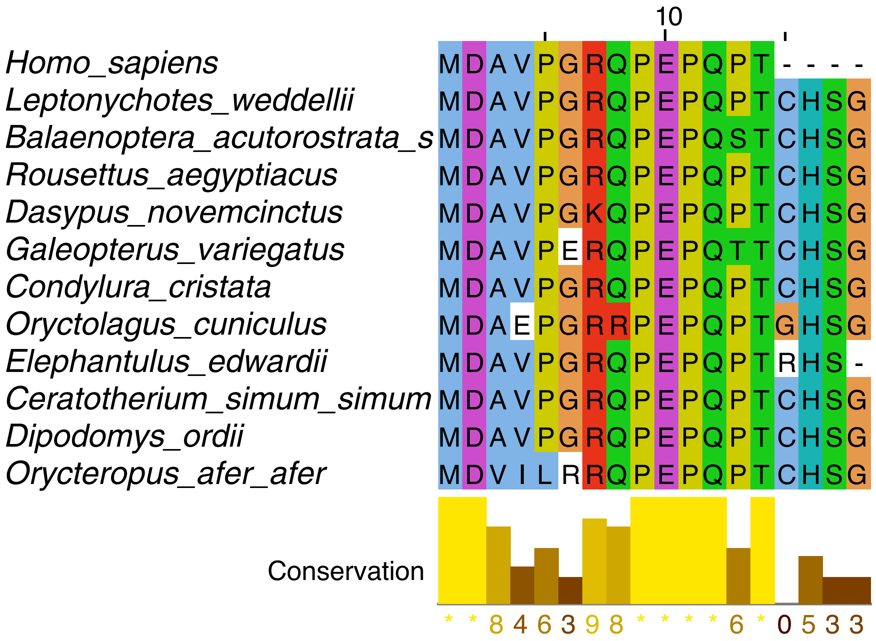

**HG0585**

ENSG00000106038 (EVX1)

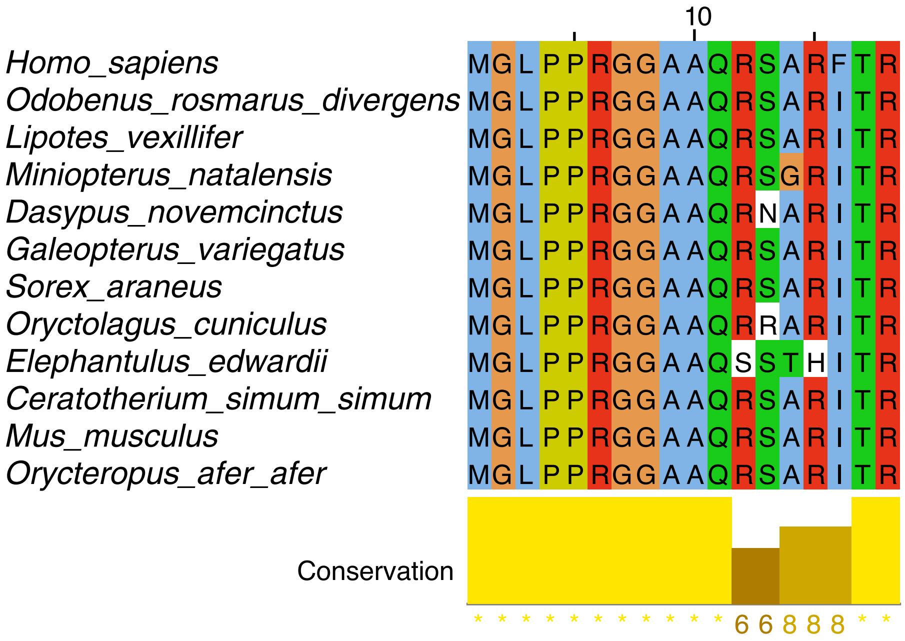

**HG0586.1**

ENSG00000108375 (RNF43)

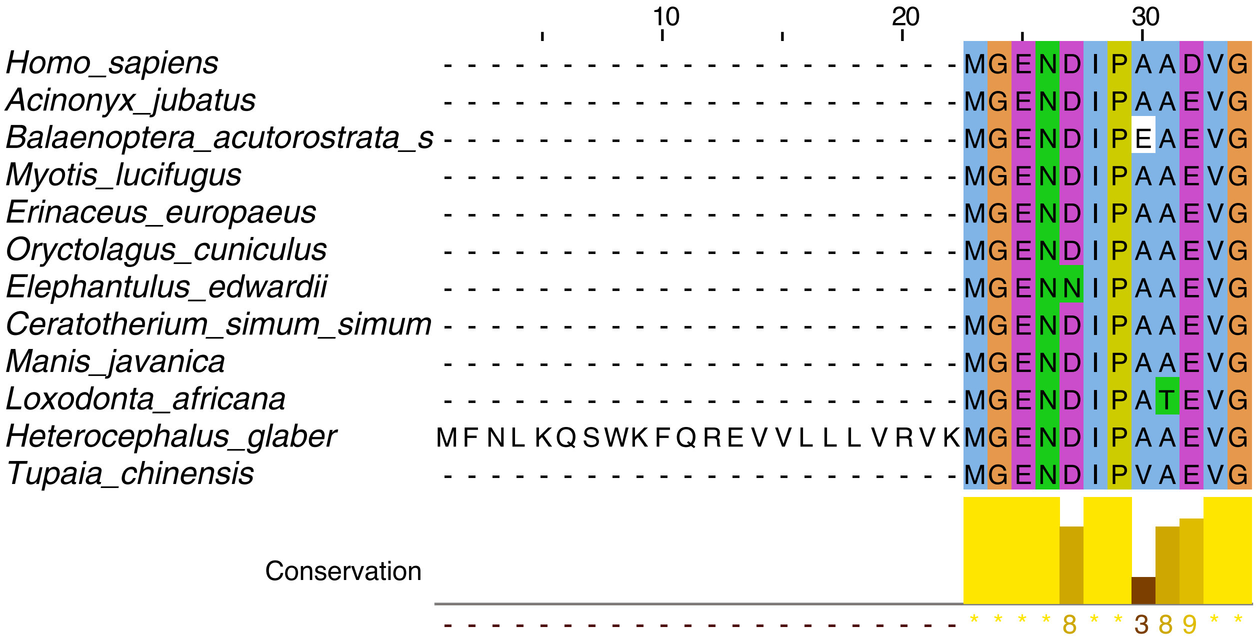

**HG0586.2**

ENSG00000108375 (RNF43)

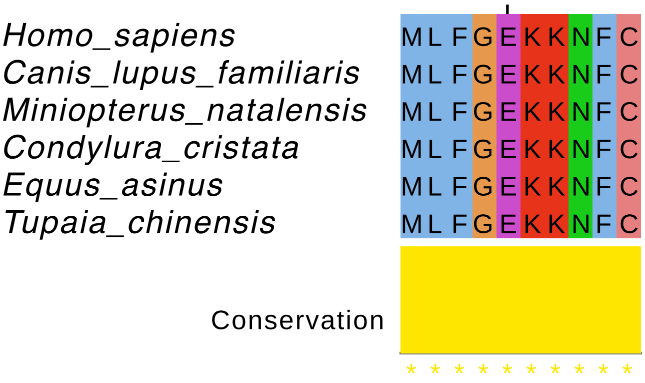

**HG0587**

ENSG00000119138 (KLF9)

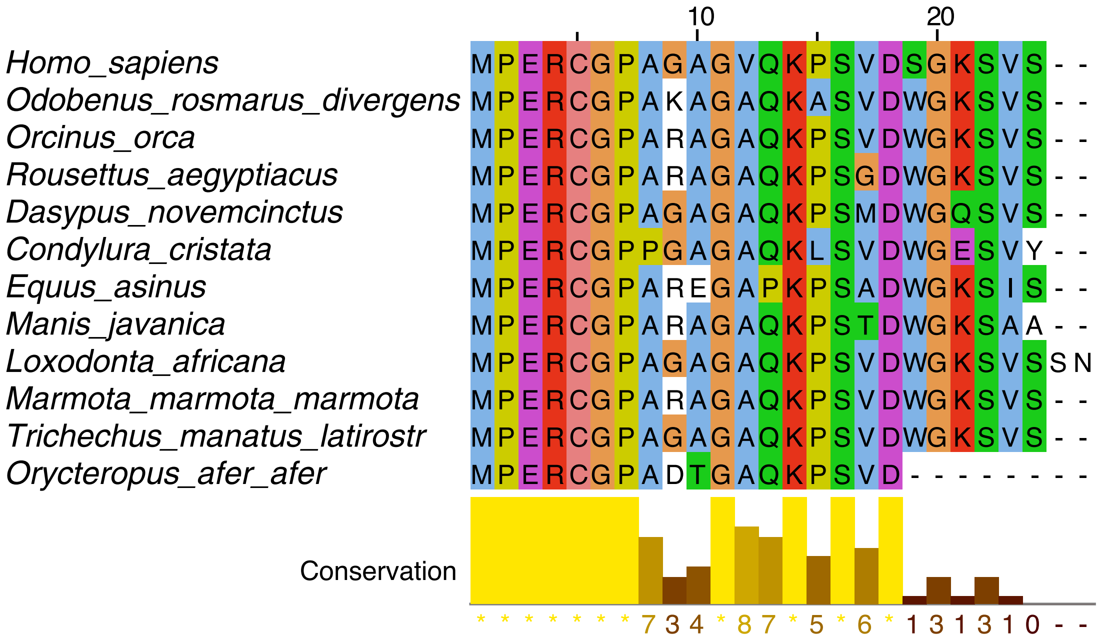

**HG0588**

ENSG00000120162 (MOB3B)

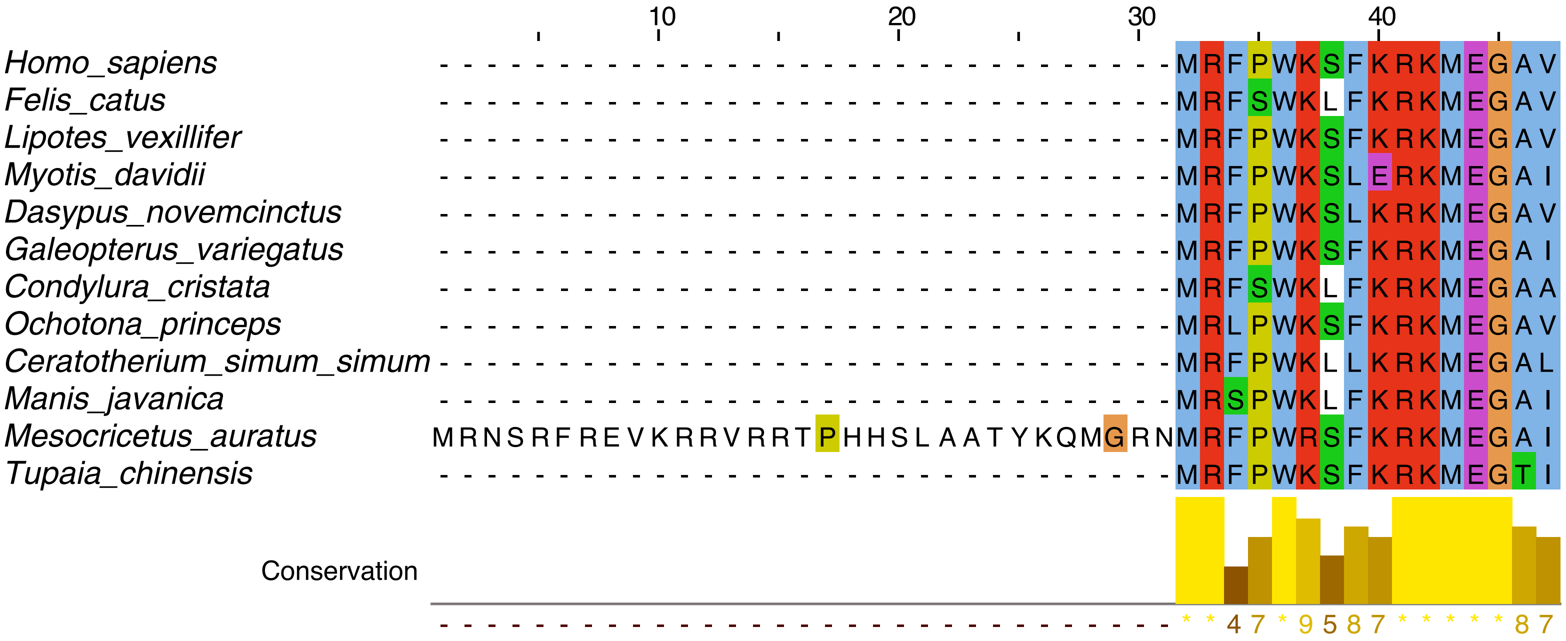

**HG0589.1**

ENSG00000121871 (SLITRK3)

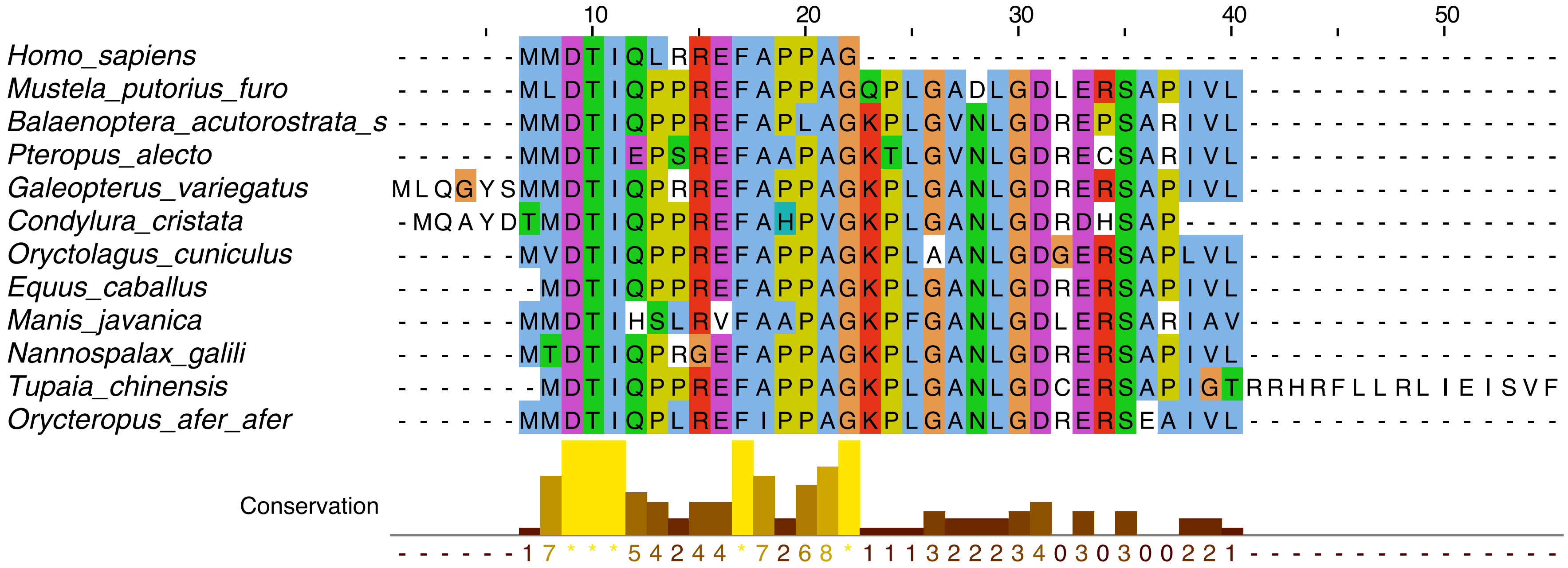

**HG0589.2**

ENSG00000121871 (SLITRK3)

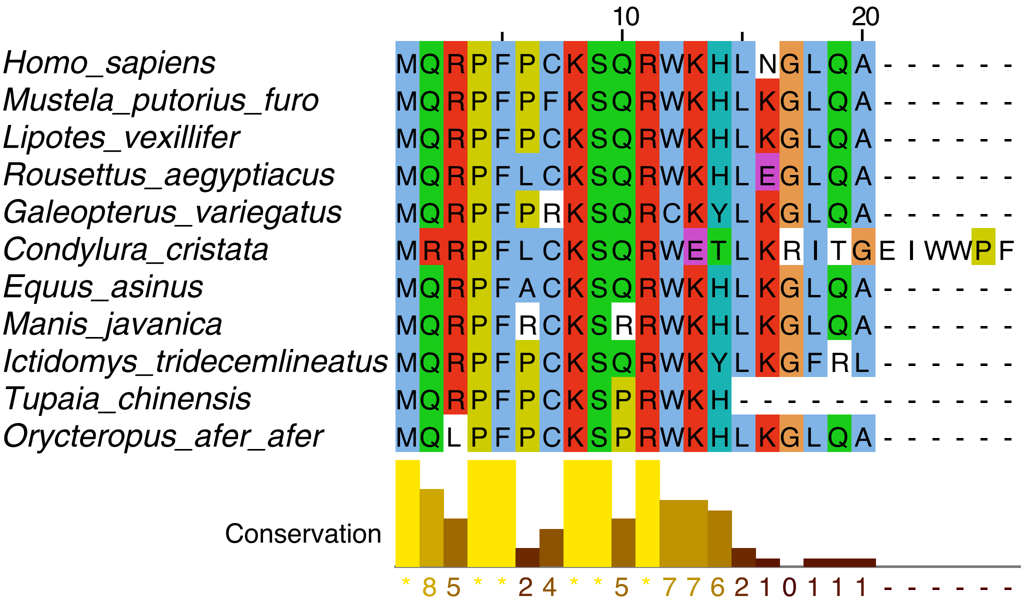

**HG0589.3**

ENSG00000121871 (SLITRK3)

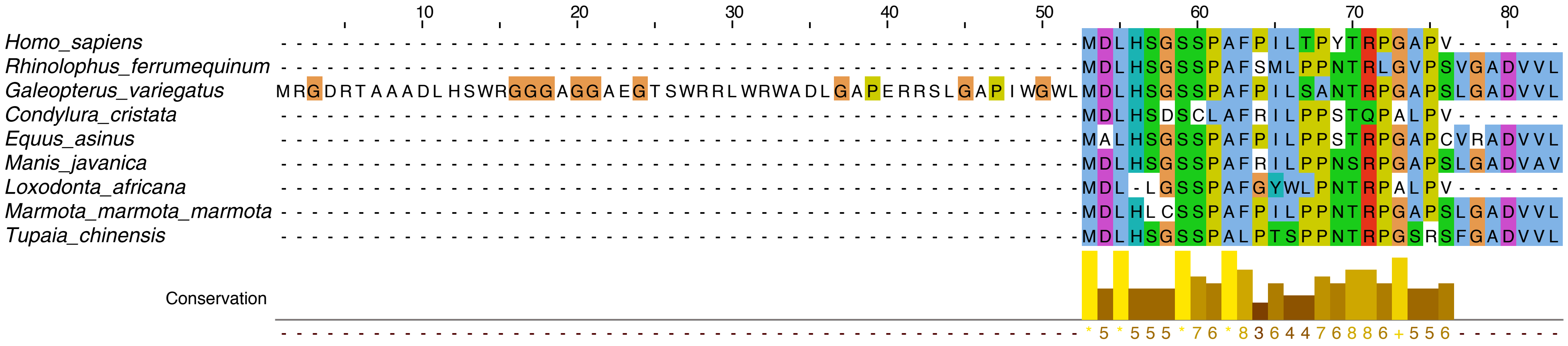

# HG0590

### ENSG00000122557 (HERPUD2)

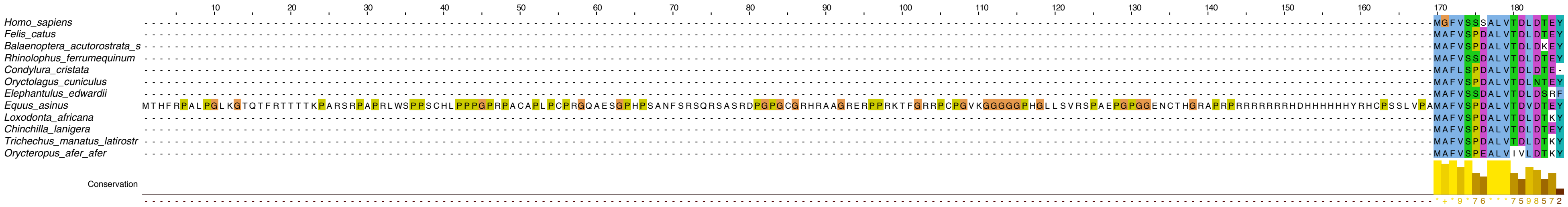

**HG0591.1**

ENSG00000127124 (HIVEP3)

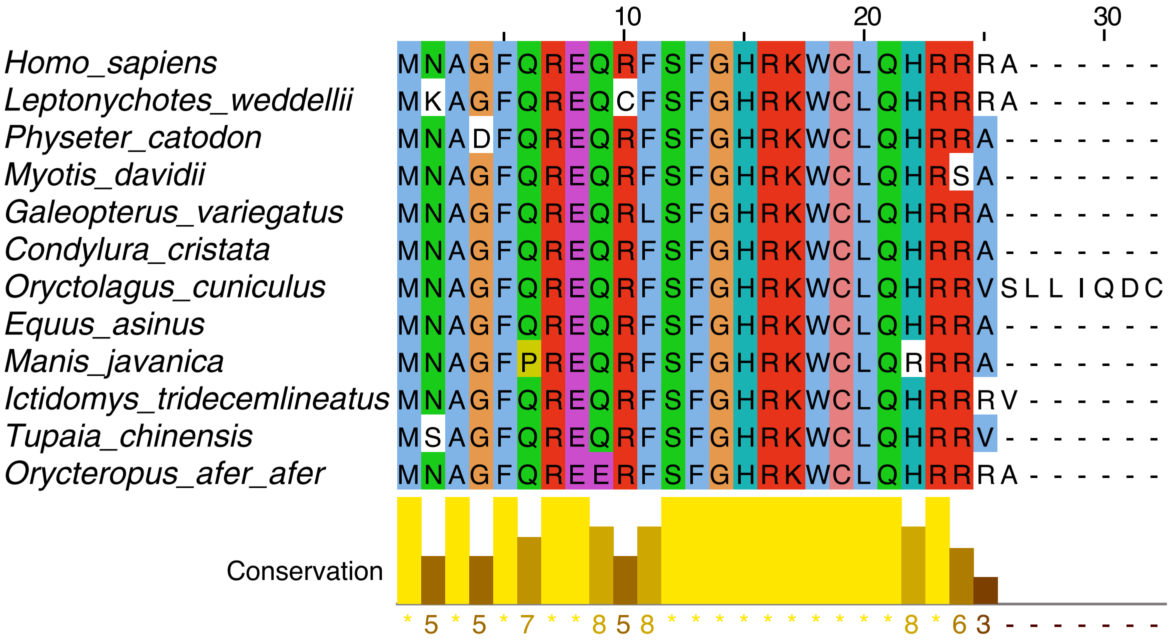

**HG0591.2**

ENSG00000127124 (HIVEP3)

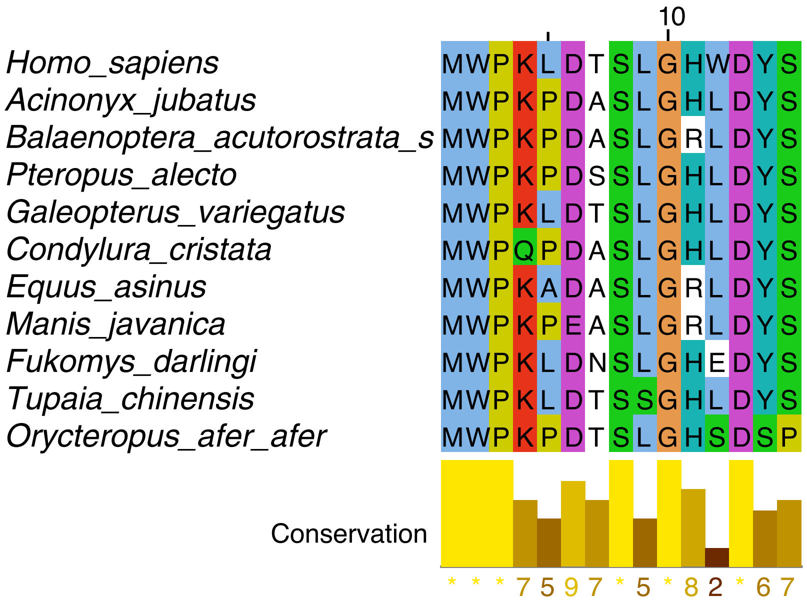

**HG0591.3**

ENSG00000127124 (HIVEP3)

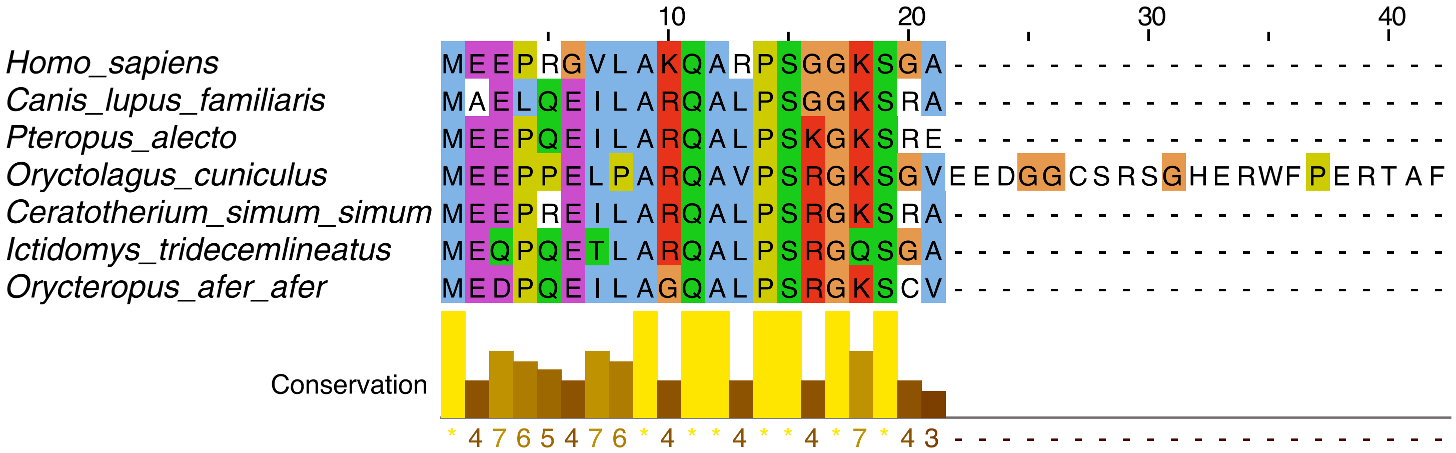

**HG0591.4**

ENSG00000127124 (HIVEP3)

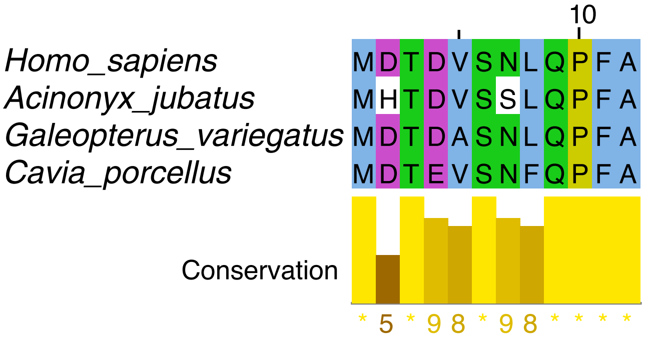

**HG0592**

ENSG00000133401 (PDZD2)

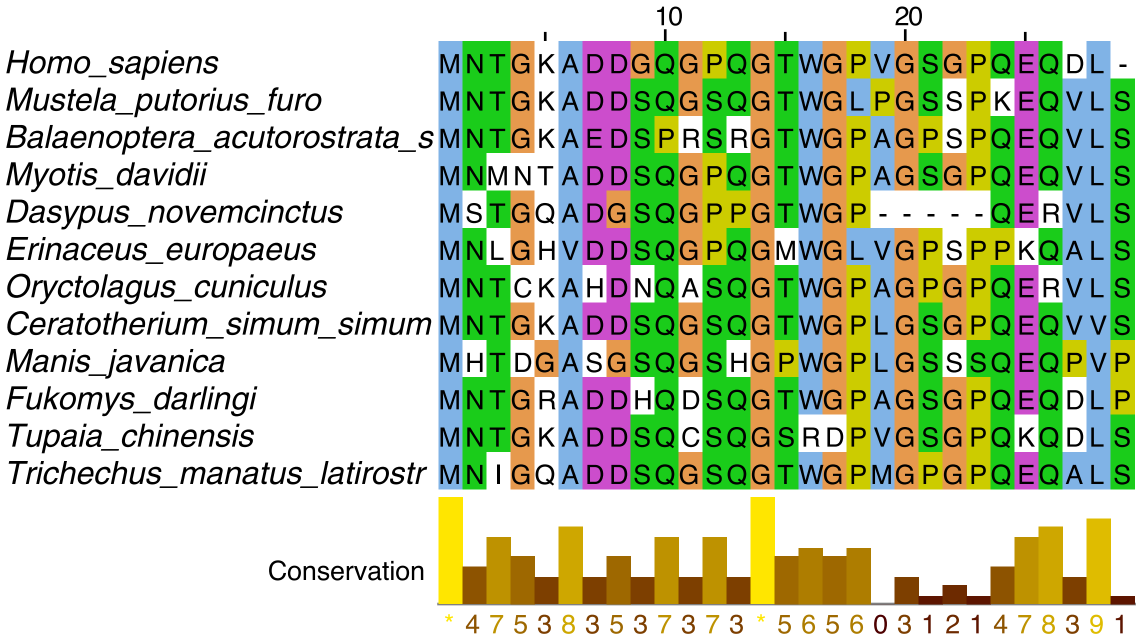

**HG0593**

ENSG00000135048 (TMEM2)

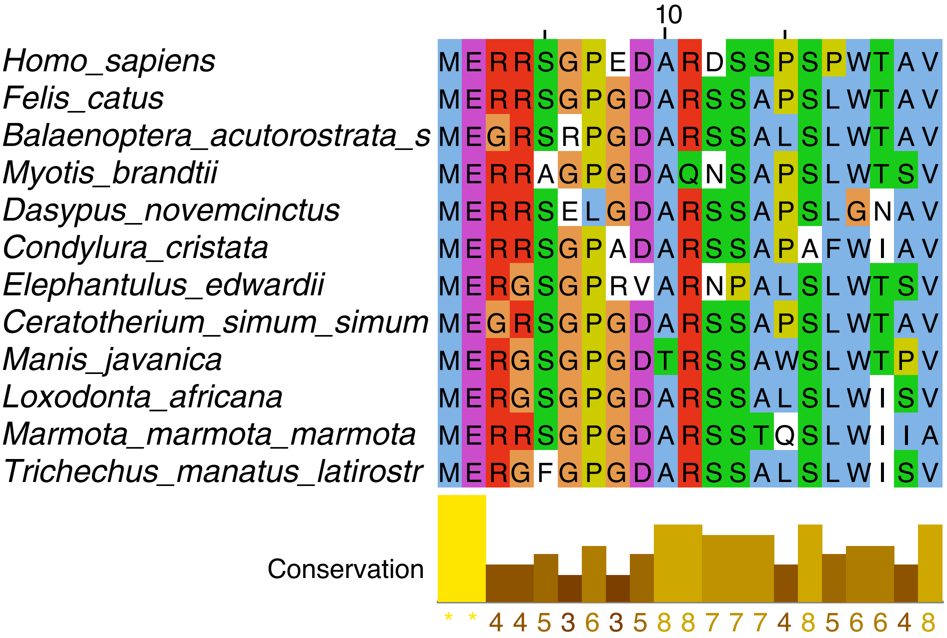

**HG0594**

ENSG00000136169 (SETDB2)

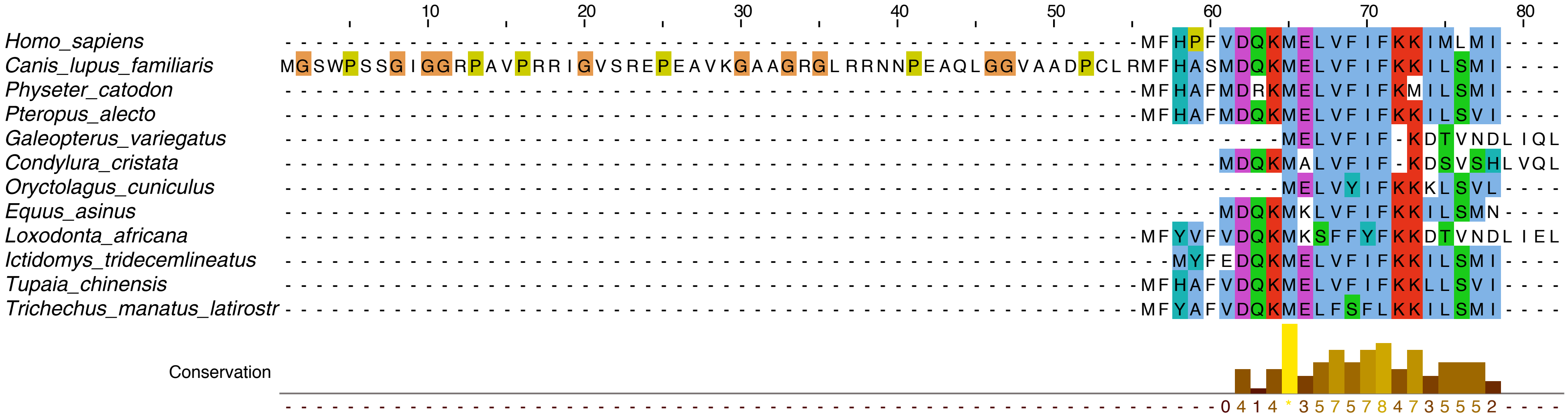

**HG0595**

ENSG00000141098 (GFOD2)

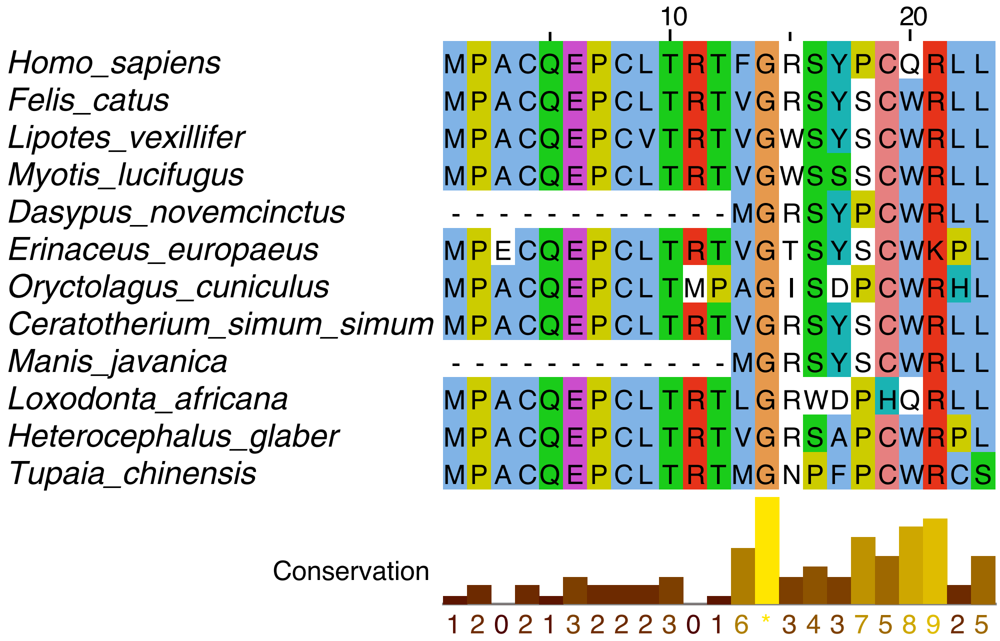

**HG0596**

ENSG00000157483 (MYO1E)

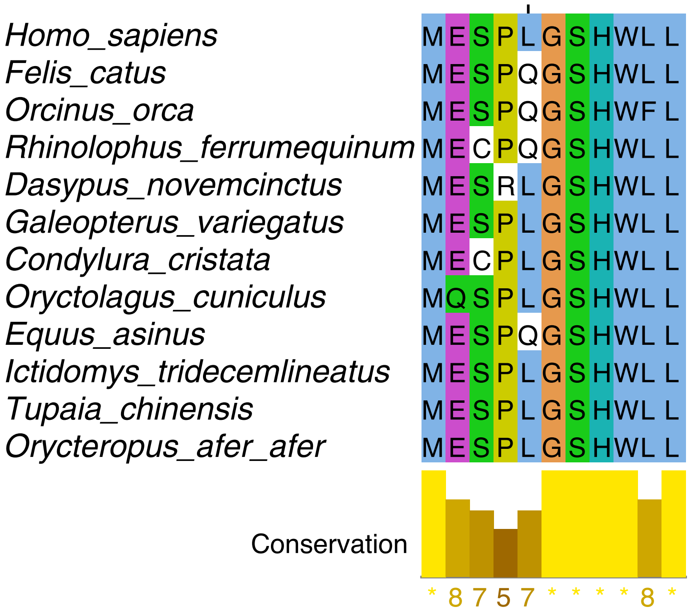

**HG0597**

ENSG00000163032 (VSNL1)

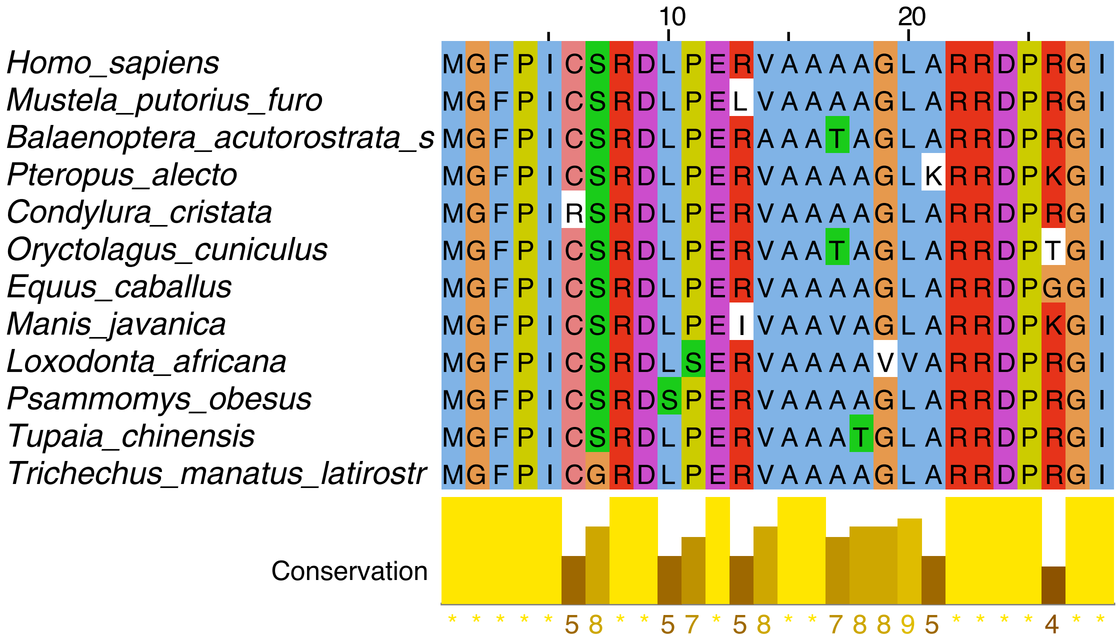

**HG0598**

ENSG00000163788 (SNRK)

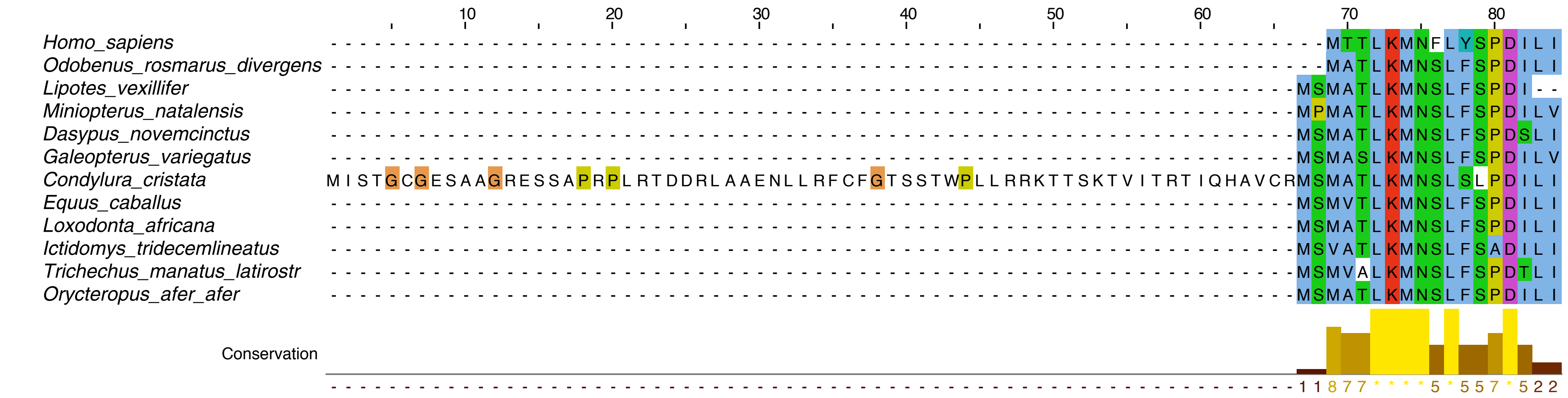

**HG0599**

ENSG00000164930 (FZD6)

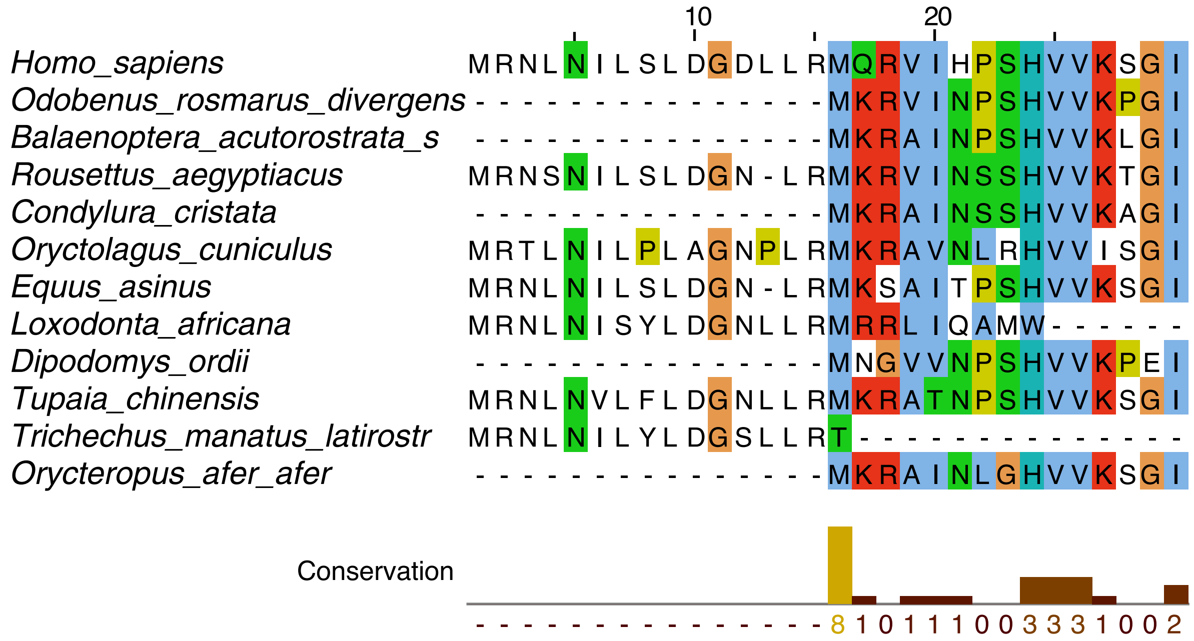

**HG0600**

ENSG00000173065 (FAM222B)

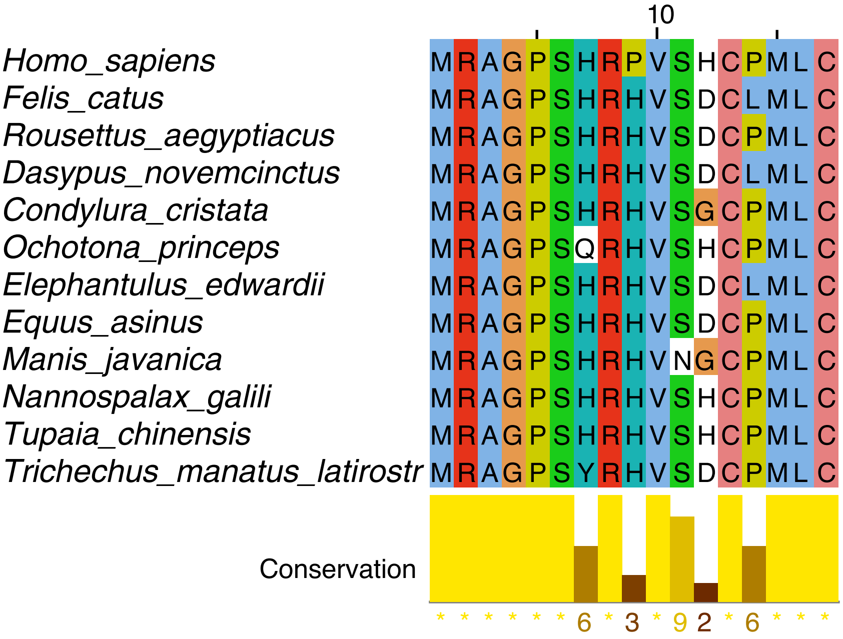

**HG0601**

ENSG00000175893 (ZDHHC21)

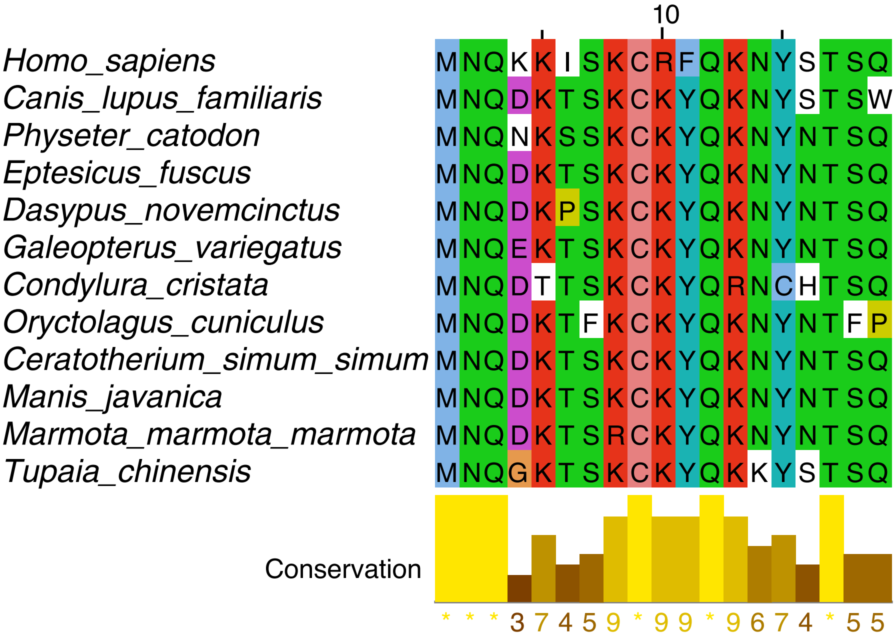

**HG0602**

ENSG00000179195 (ZNF664)

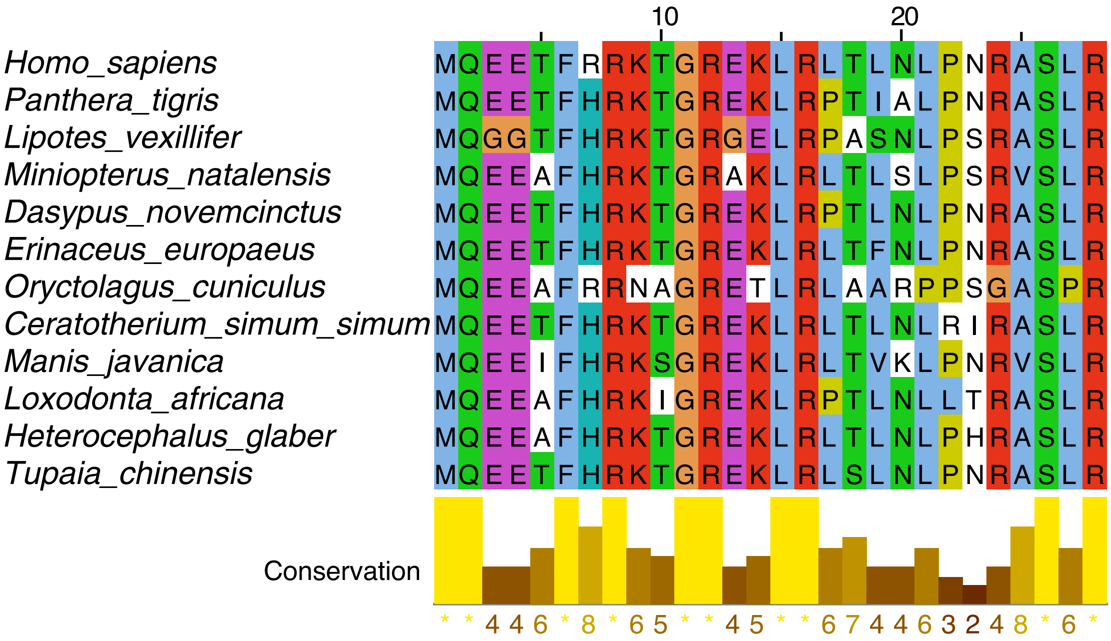

**HG0603**

ENSG00000197329 (PELI1)

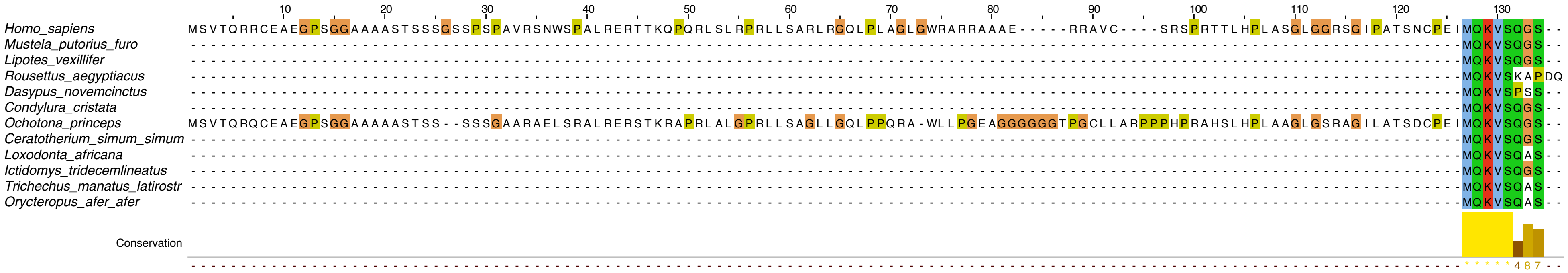

**HG0604.1**

ENSG00000003147 (ICA1)

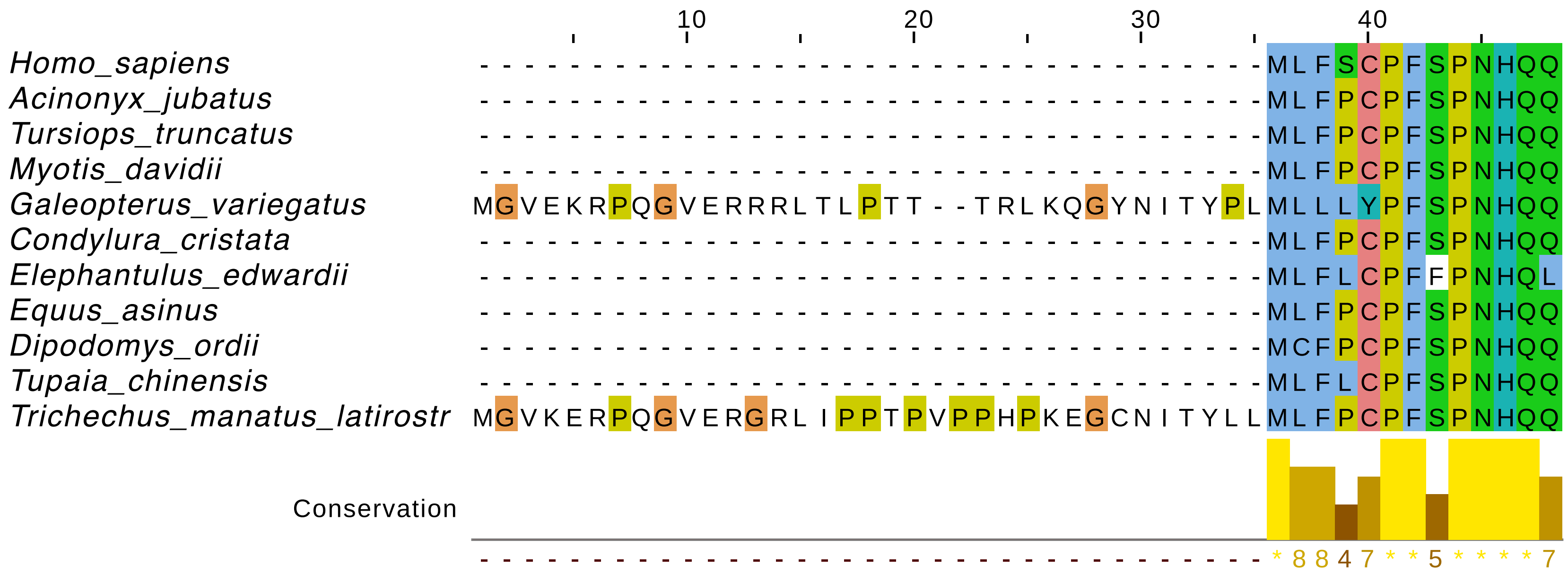

**HG0604.2**

ENSG00000003147 (ICA1)

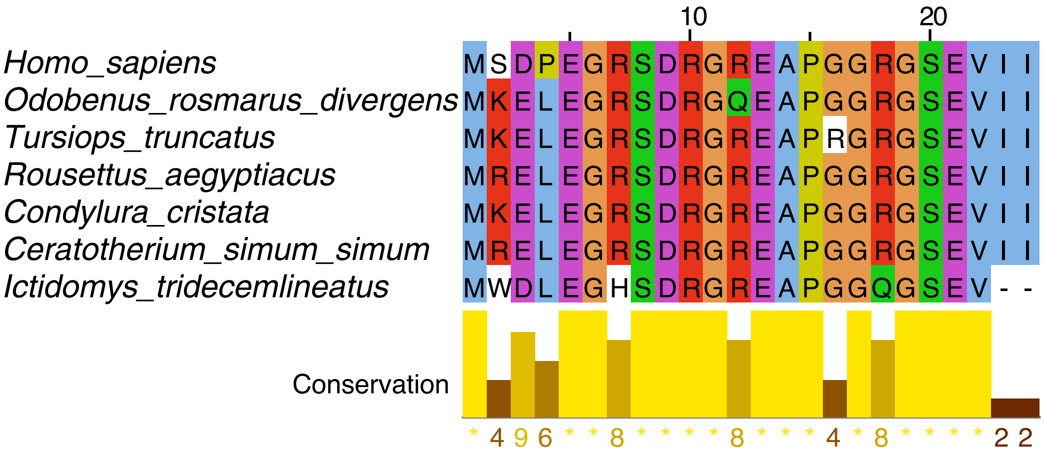

**HG0604.3**

ENSG00000003147 (ICA1)

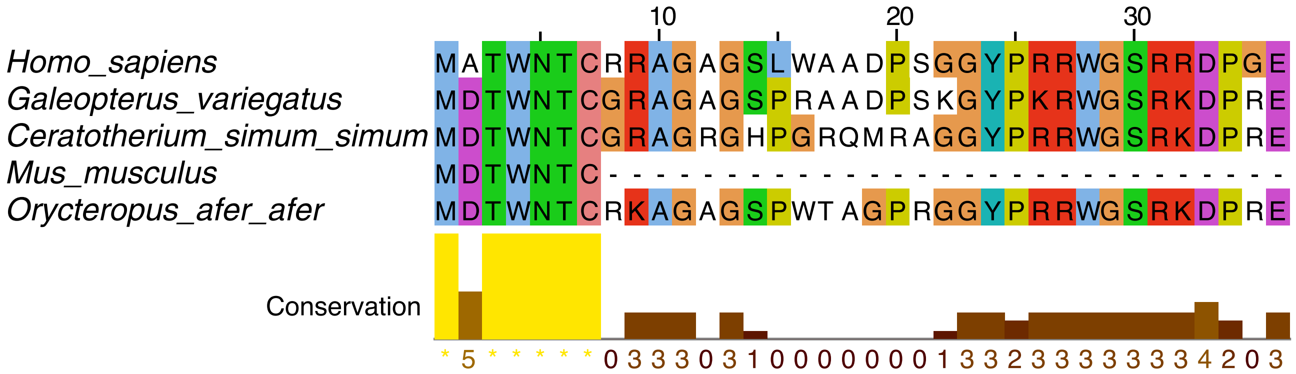

**HG0605**

ENSG00000053254 (FOXN3)

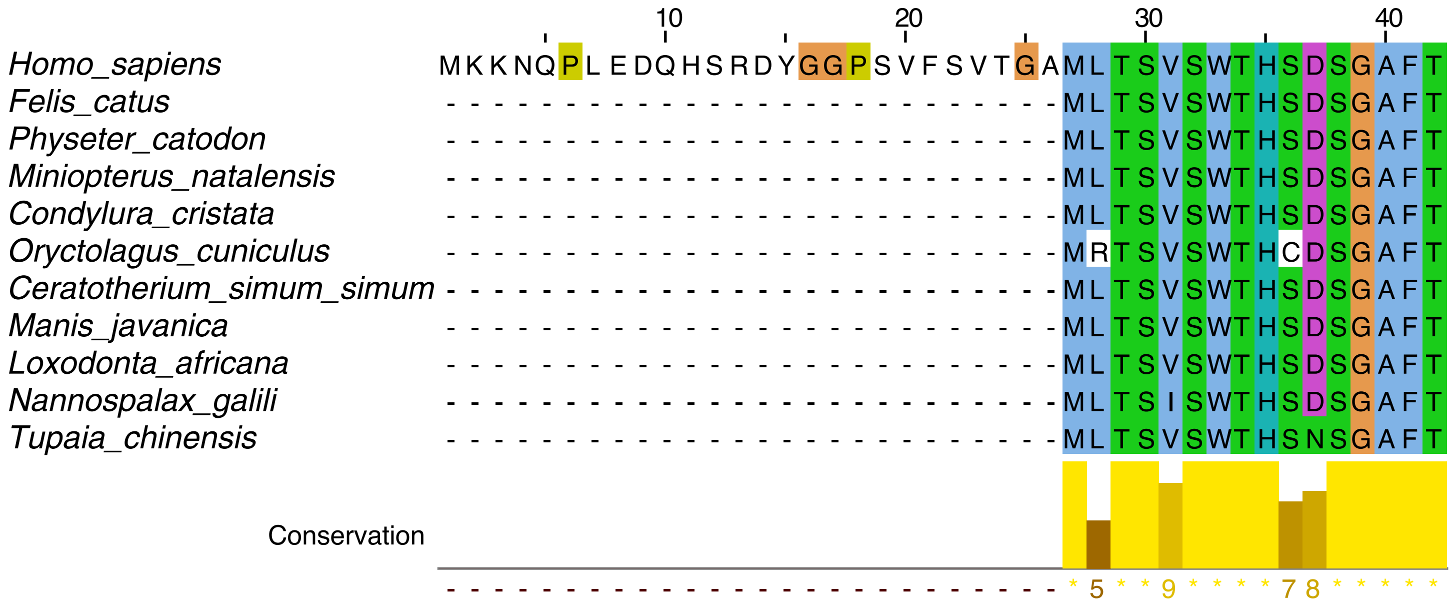

**HG0606**

ENSG00000055130 (CUL1)

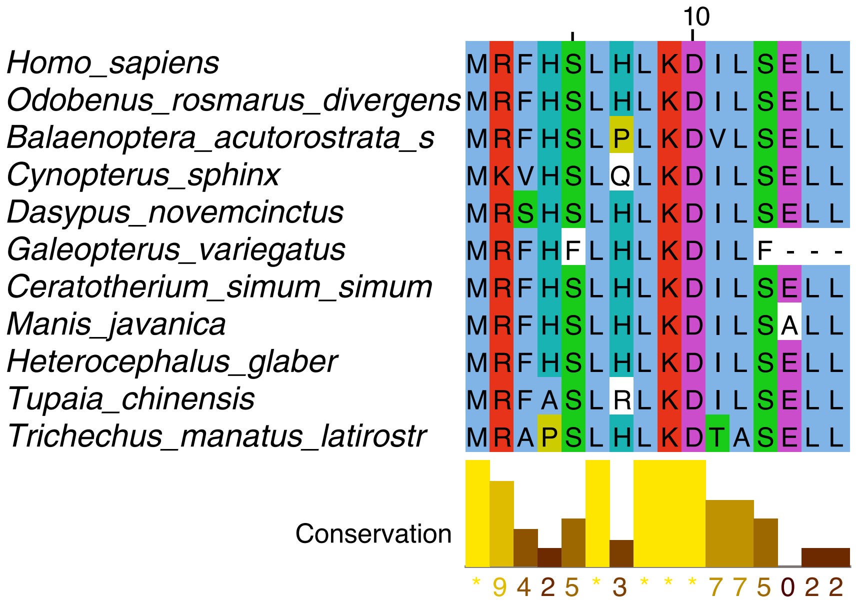

**HG0607**

ENSG00000074657 (ZNF532)

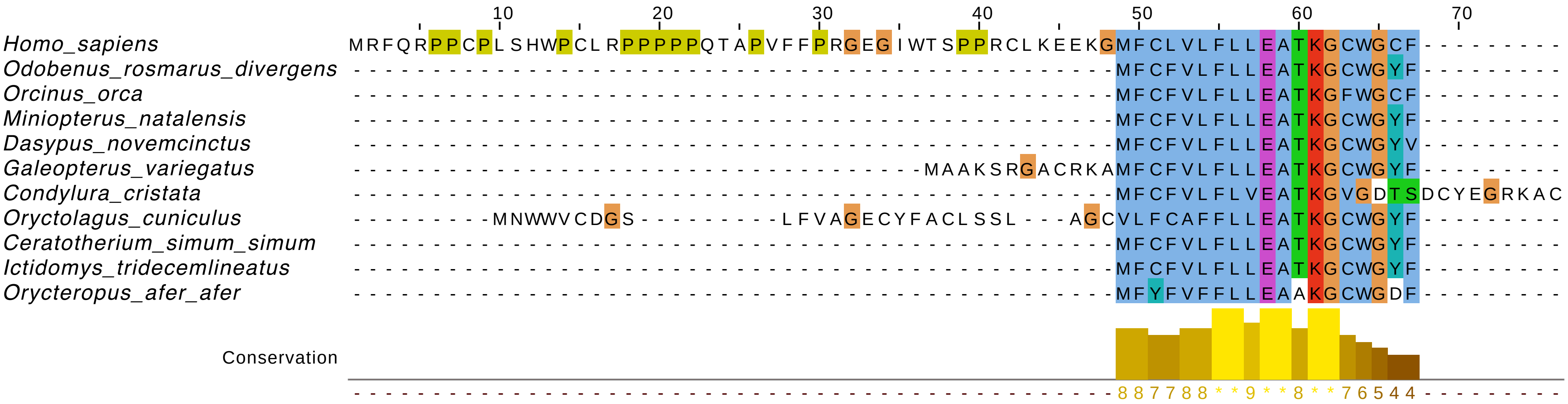

**HG0608**

ENSG00000084731 (KIF3C)

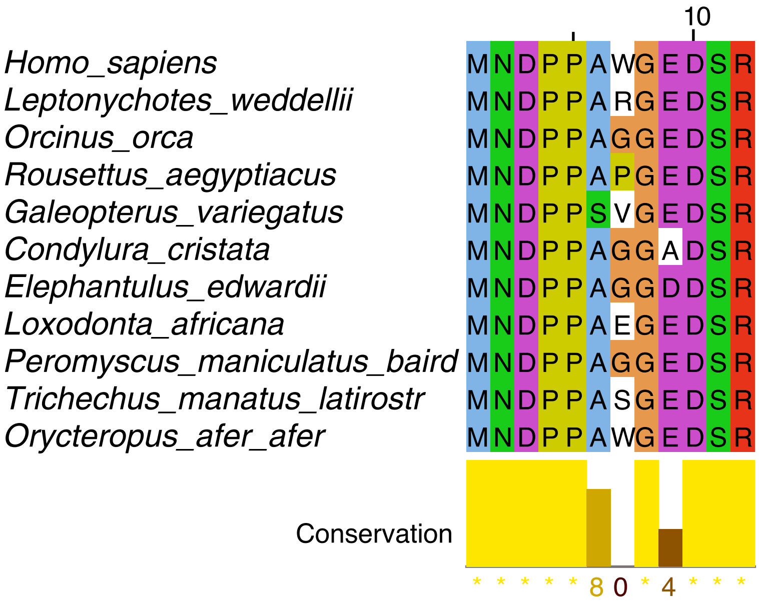

**HG0609**

ENSG00000095539 (SEMA4G)

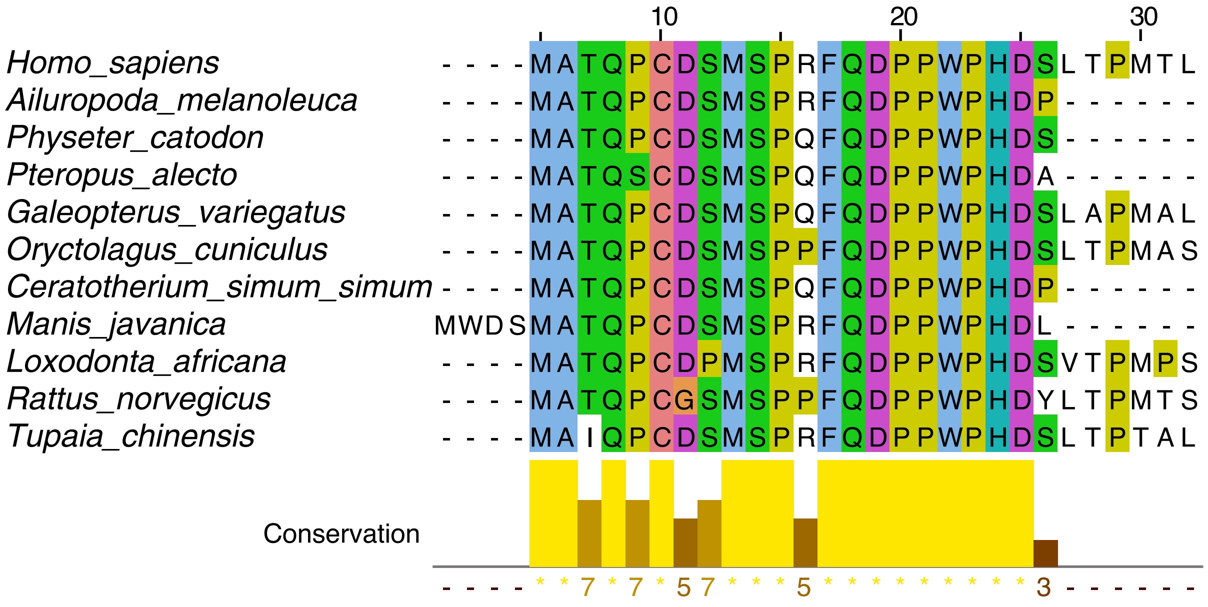

**HG0610**

ENSG00000106069 (CHN2)

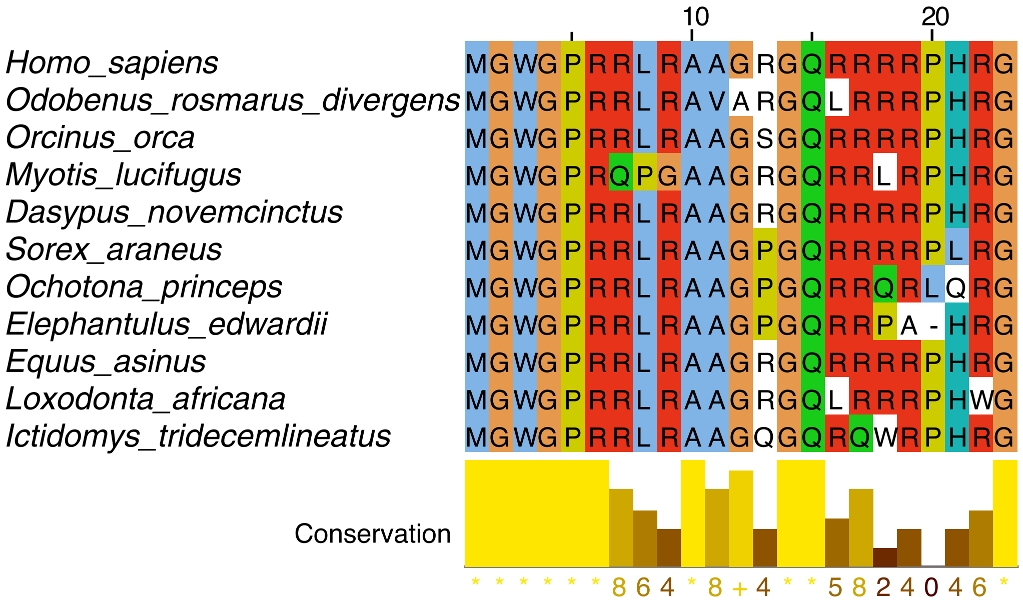

**HG0611**

ENSG00000108309 (RUNDC3A)

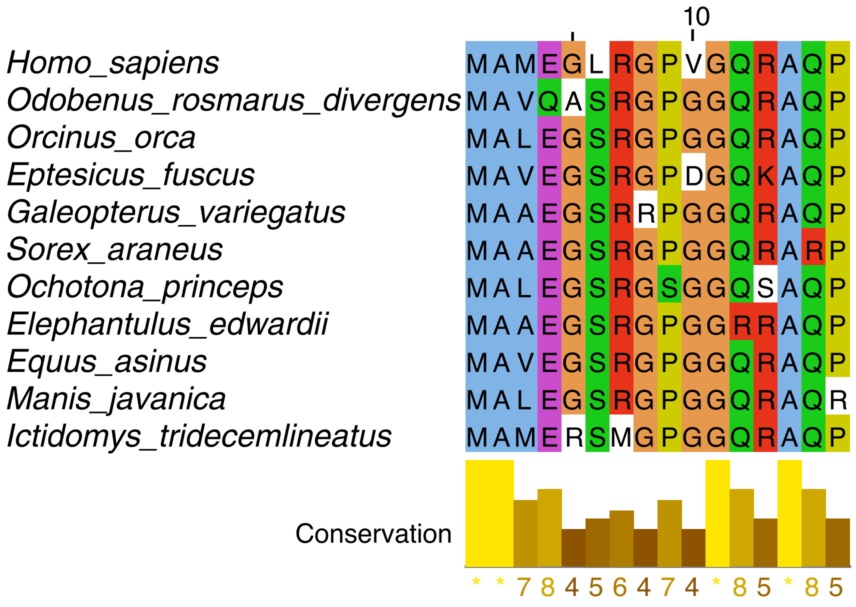

**HG0612.1**

ENSG00000110422 (HIPK3)

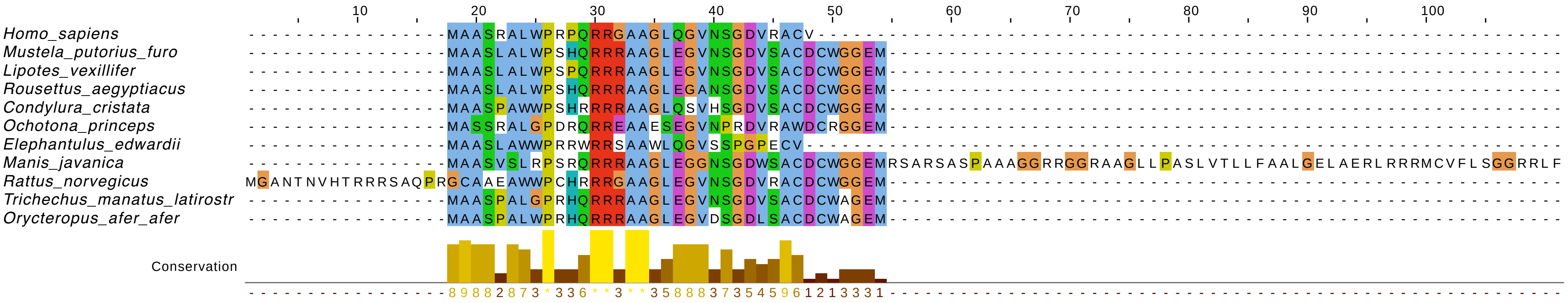

**HG0612.2**

ENSG00000110422 (HIPK3)

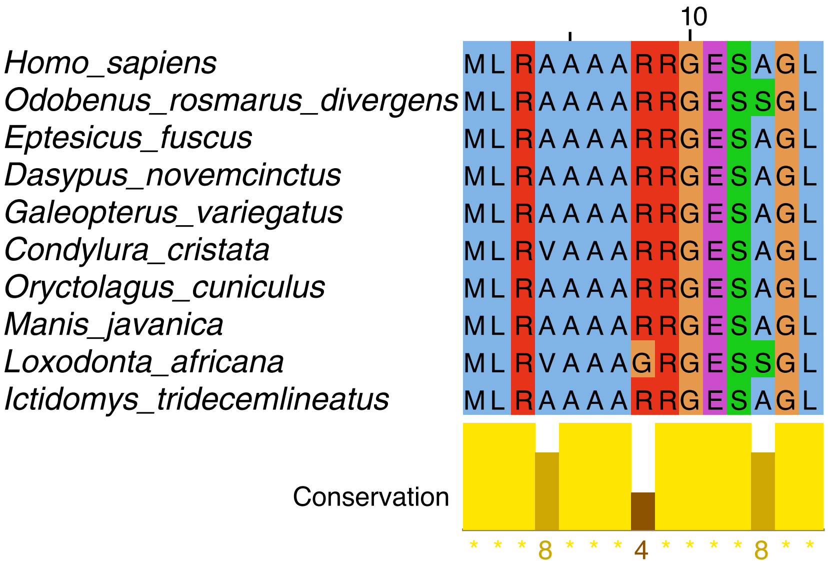

**HG0612.3**

ENSG00000110422 (HIPK3)

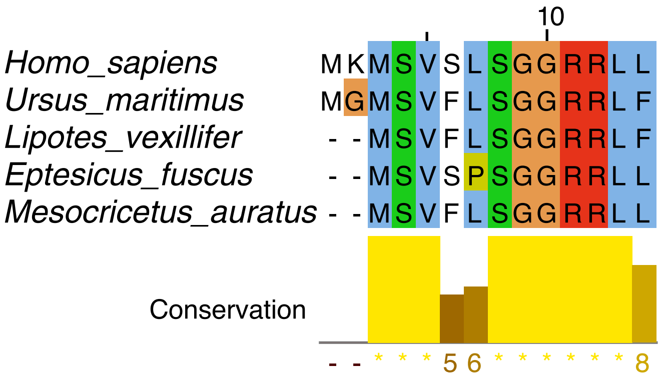

**HG0613**

ENSG00000114861 (FOXP1)

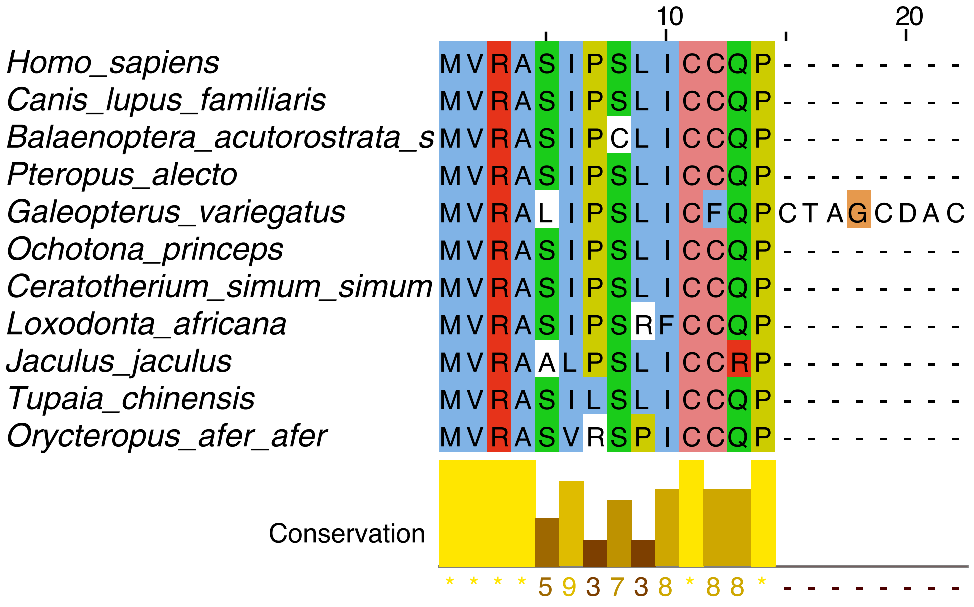

# HG0614

### ENSG00000122203 (KIAA1191)

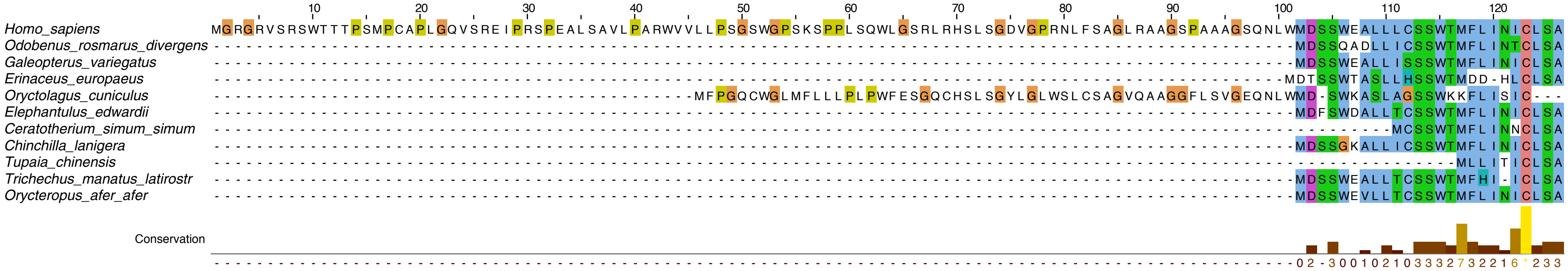

**HG0615**

ENSG00000128191 (DGCR8)

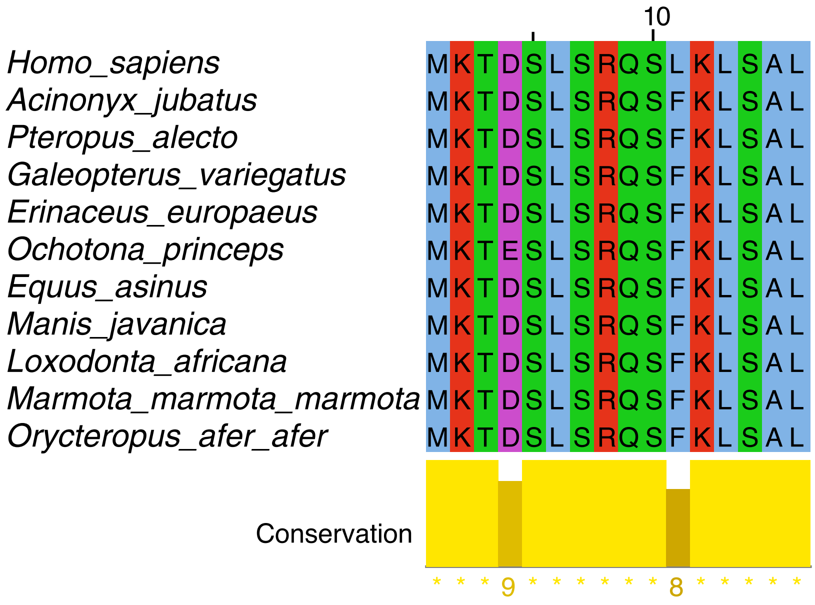

**HG0616**

ENSG00000128739 (SNRPN)

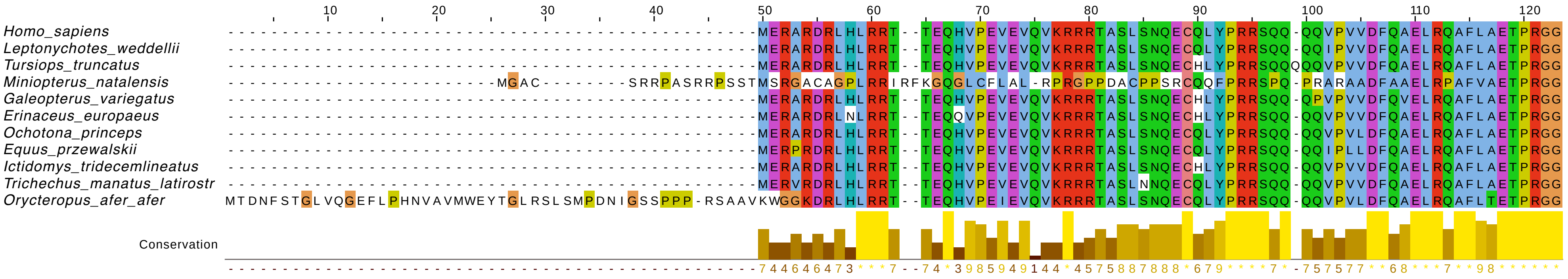

**HG0617**

ENSG00000135164 (DMTF1)

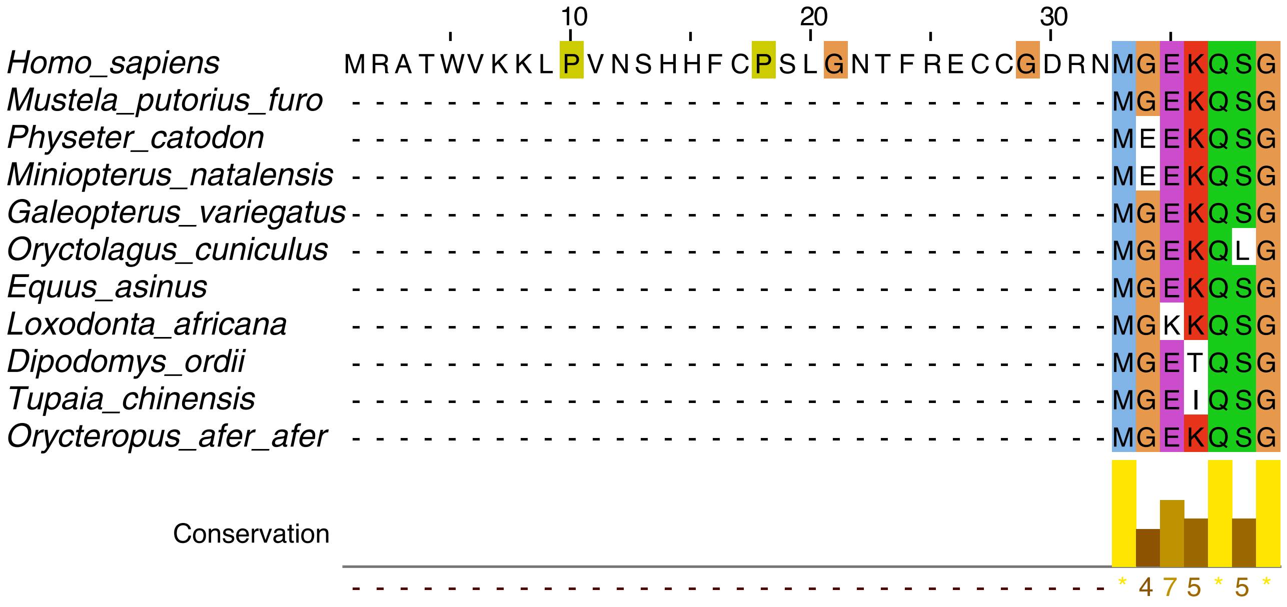

**HG0618**

ENSG00000137817 (PARP6)

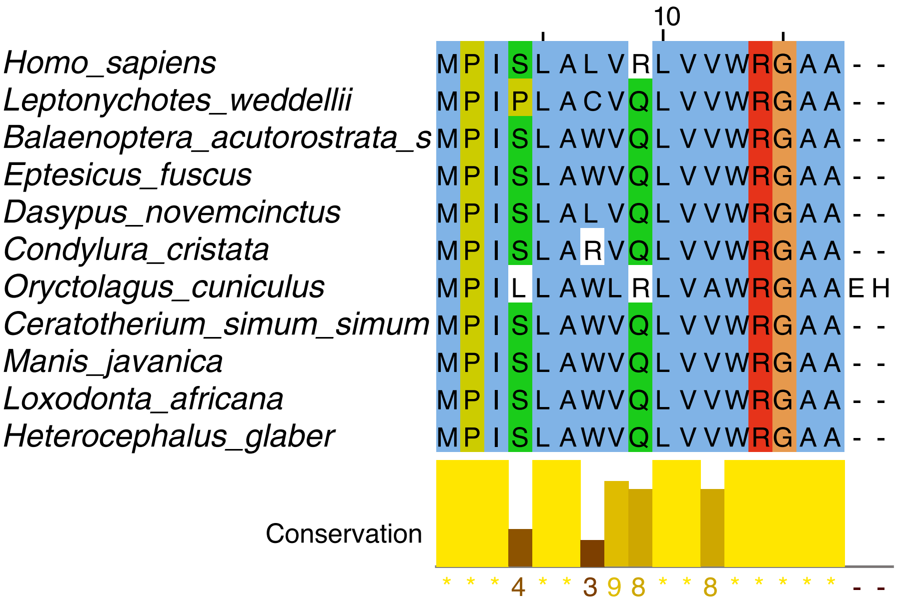

**HG0619**

ENSG00000138593 (SECISBP2L)

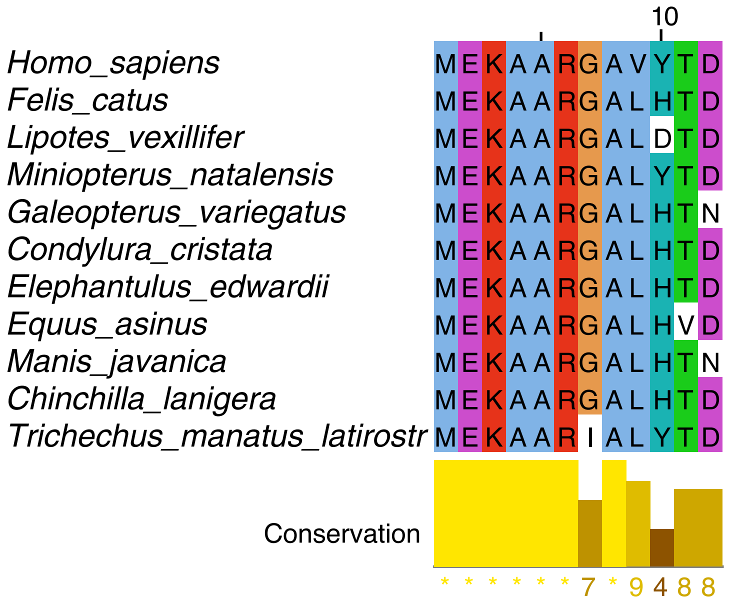

**HG0620**

ENSG00000145555 (MYO10)

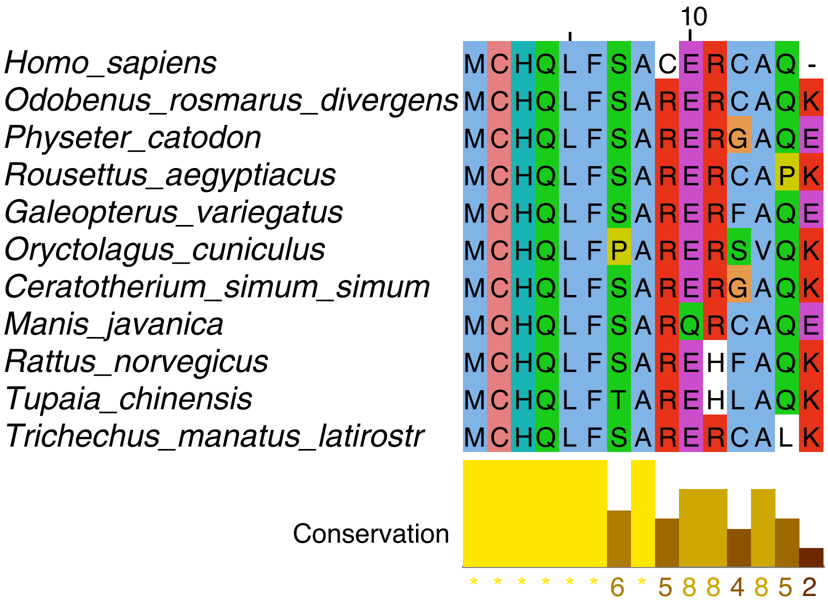

**HG0621.1**

ENSG00000146352 (CLVS2)

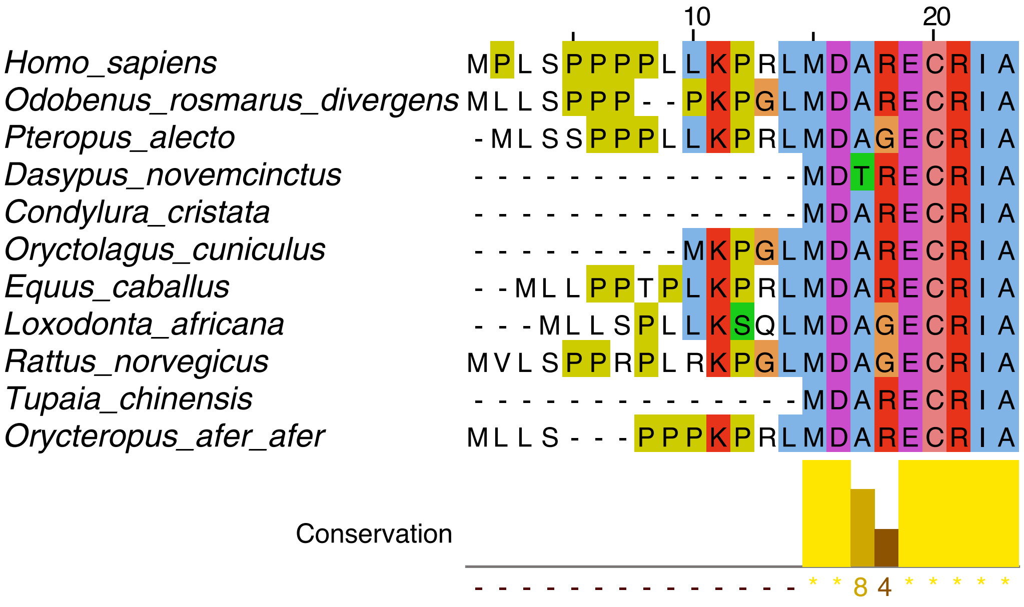

**HG0621.2**

ENSG00000146352 (CLVS2)

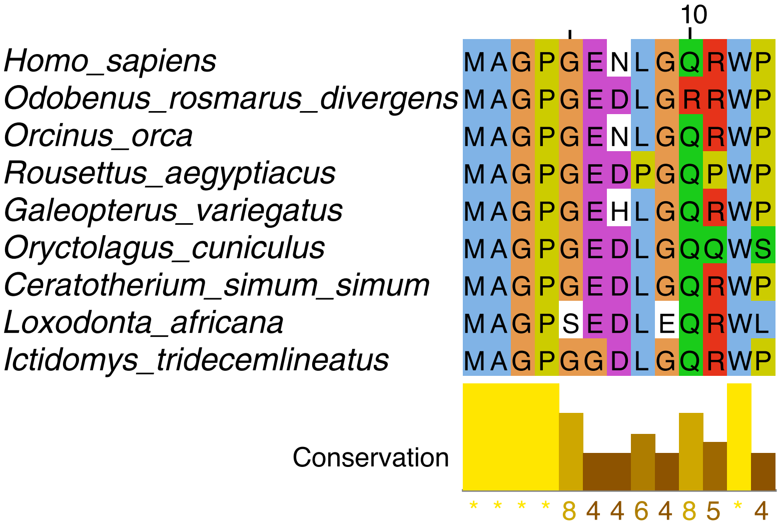

**HG0621.3**

ENSG00000146352 (CLVS2)

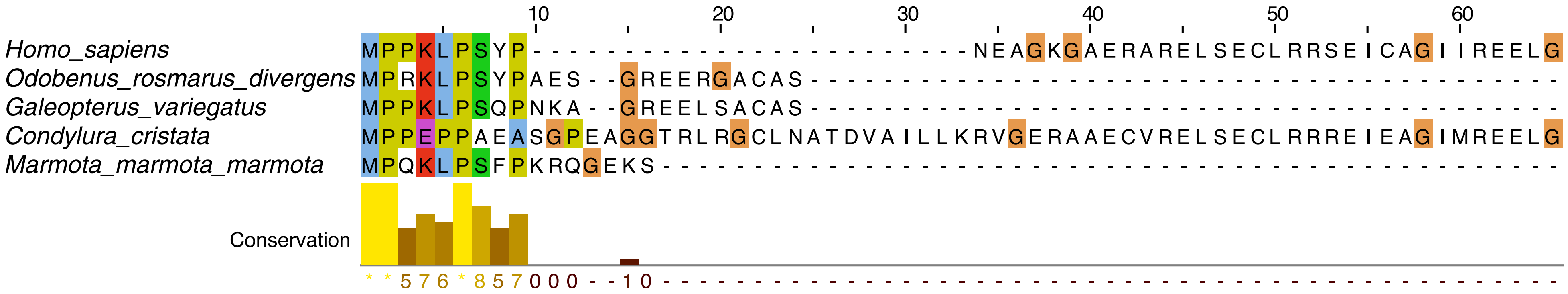

**HG0622**

ENSG00000148677 (ANKRD1)

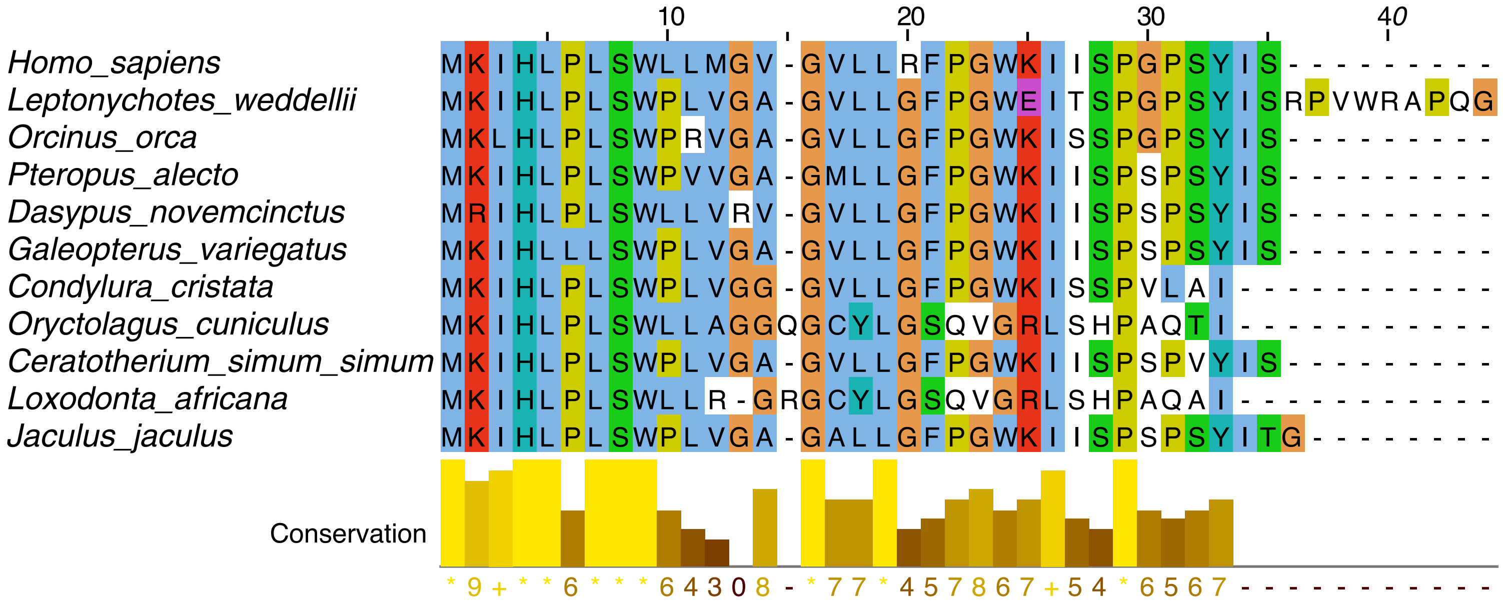

**HG0623**

ENSG00000151422 (FER)

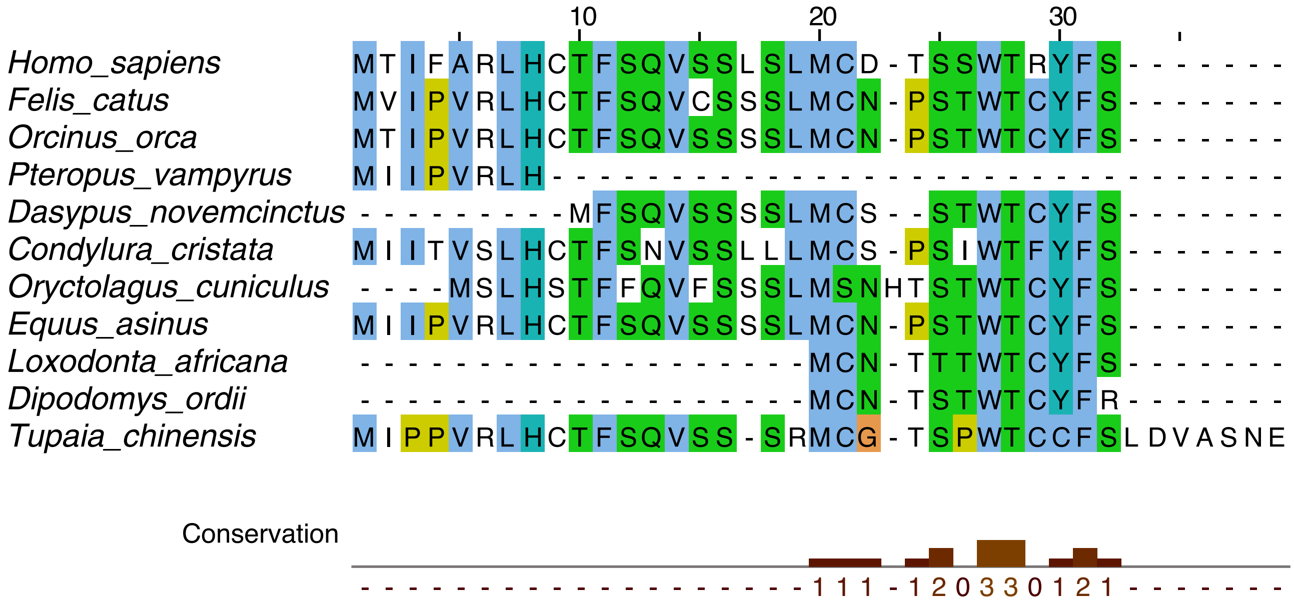

**HG0624**

ENSG00000156853 (ZNF689)

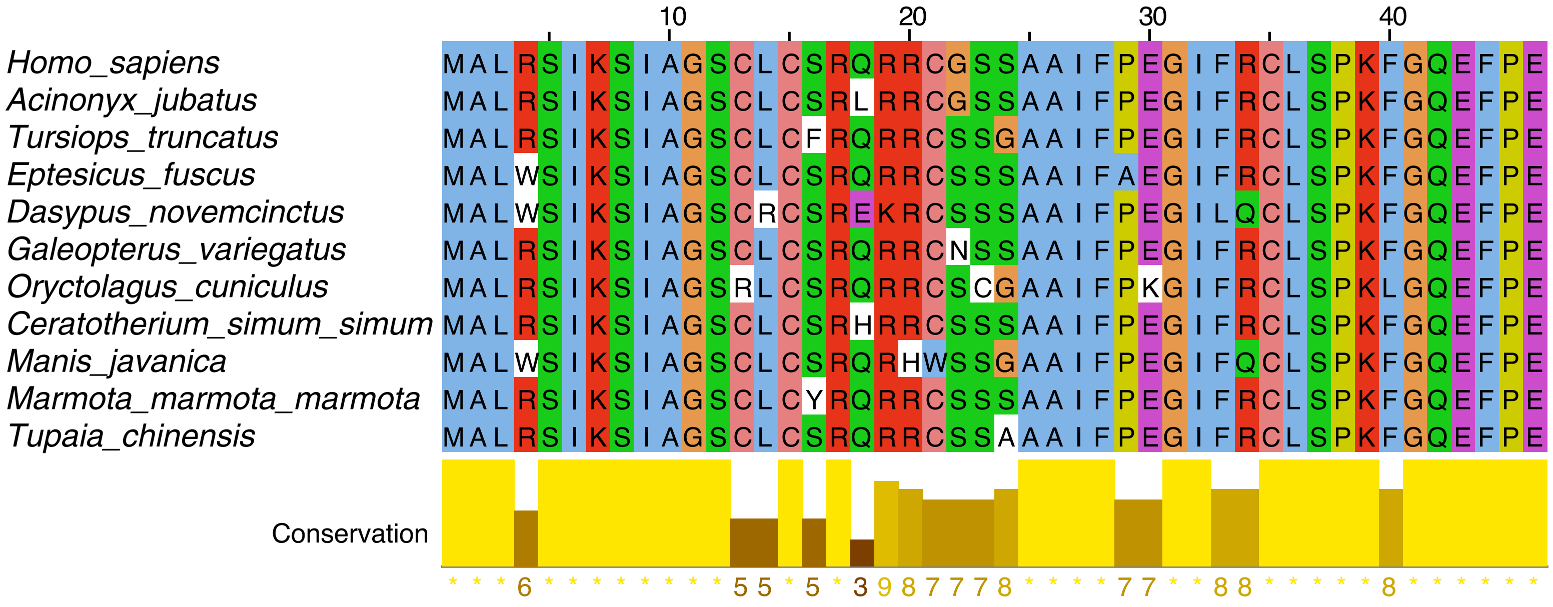

**HG0625**

ENSG00000164050 (PLXNB1)

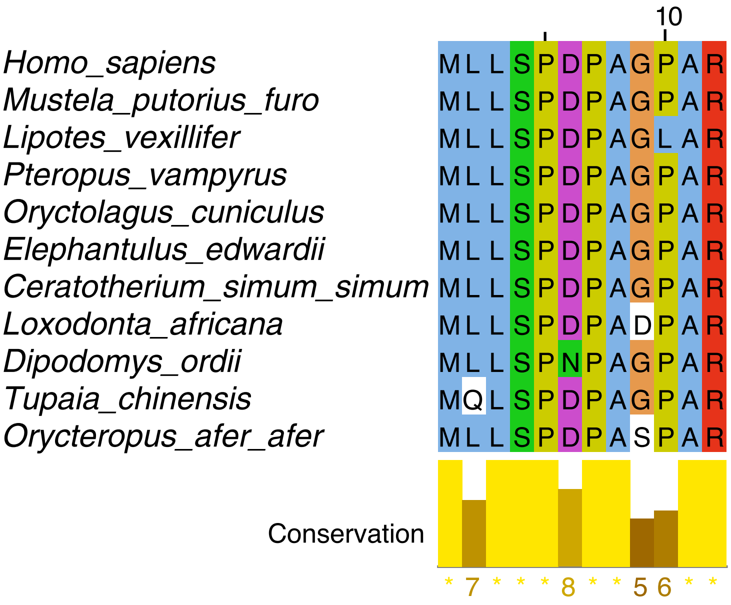

**HG0626**

ENSG00000168939 (SPRY3)

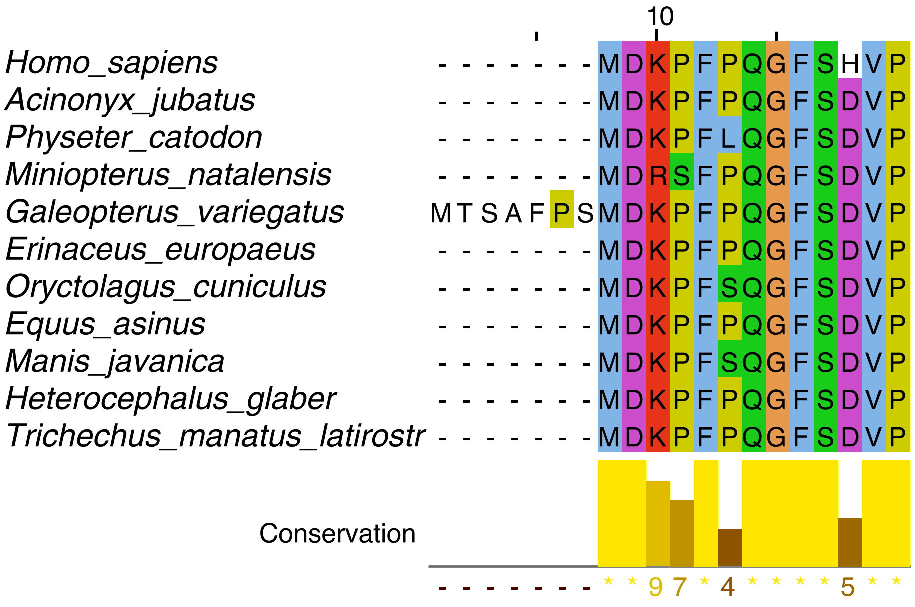

**HG0627**

ENSG00000169760 (NLGN1)

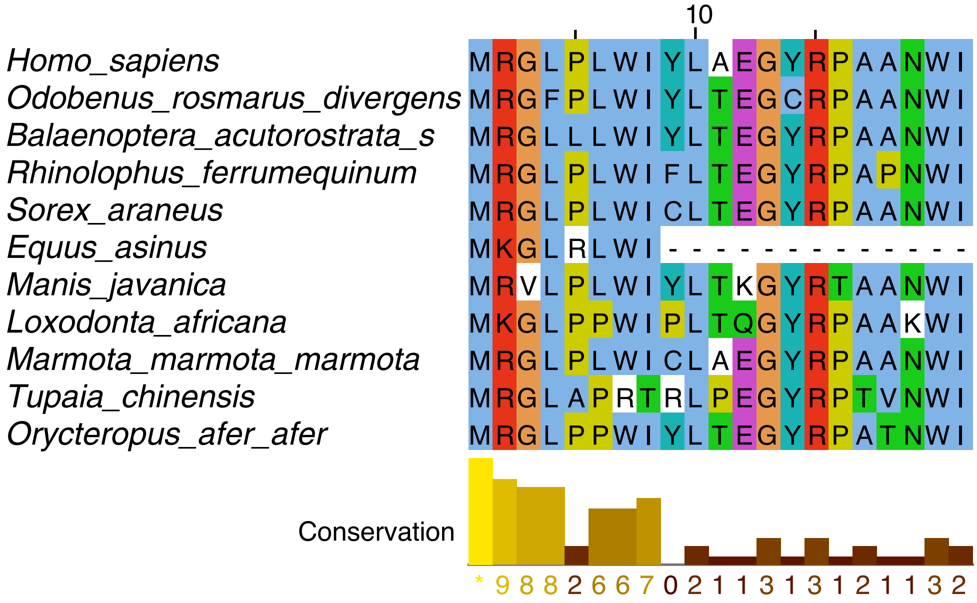

**HG0628**

ENSG00000170677 (SOCS6)

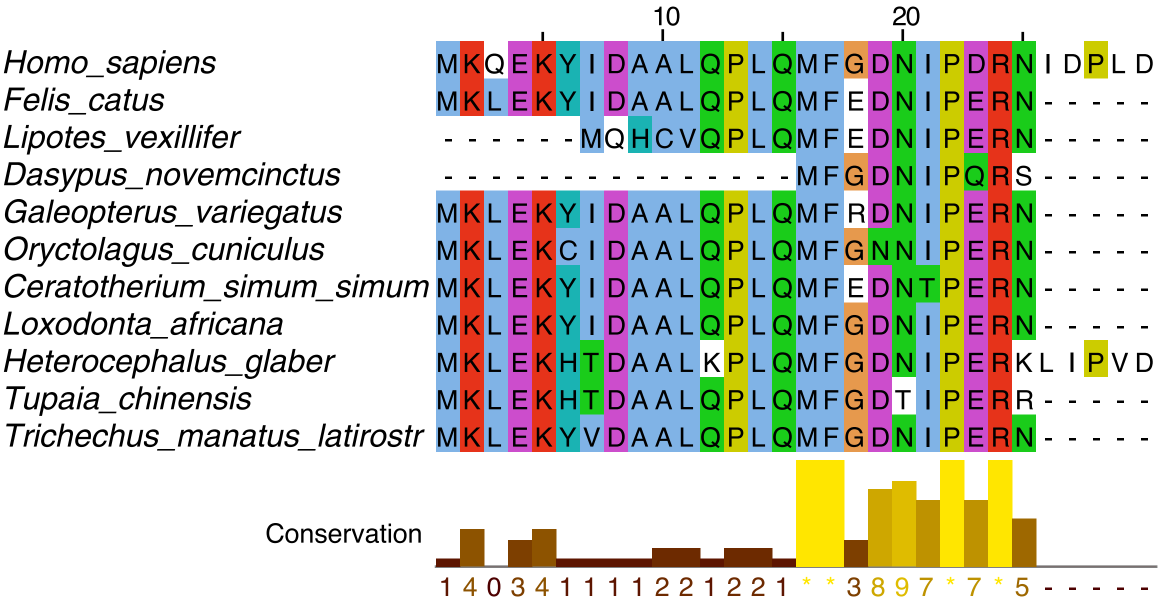

**HG0629**

ENSG00000170961 (HAS2)

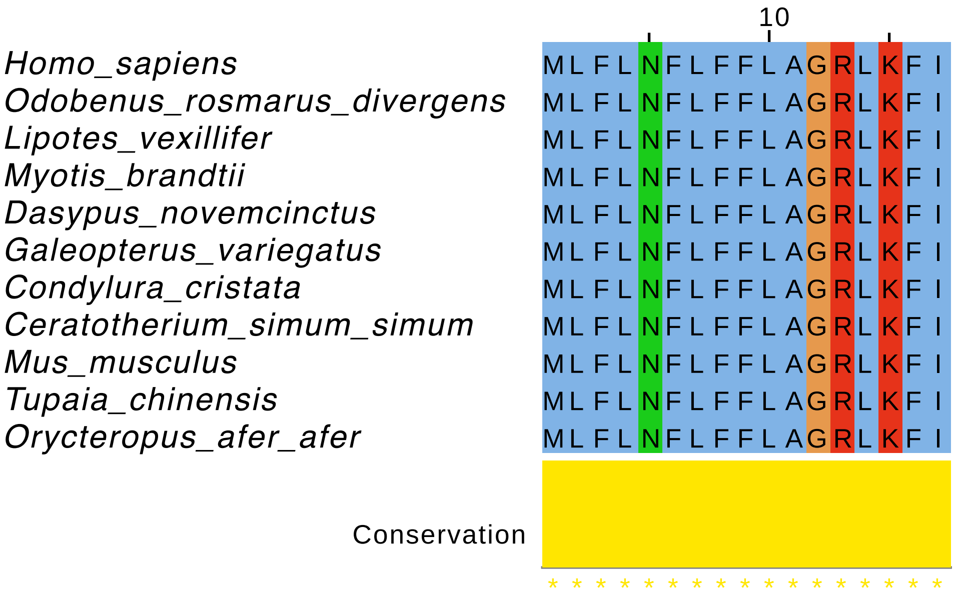

**HG0630**

ENSG00000171552 (BCL2L1)

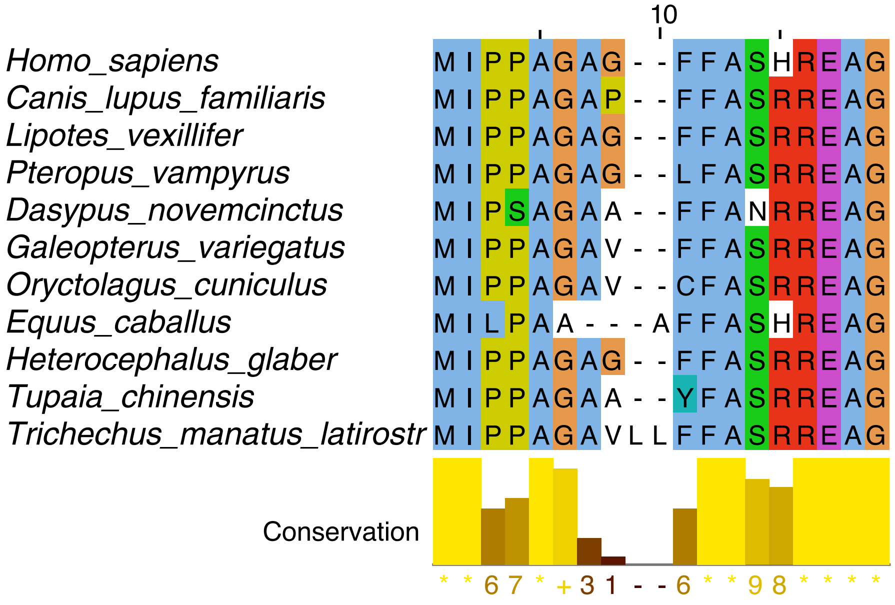

**HG0631**

ENSG00000173548 (SNX33)

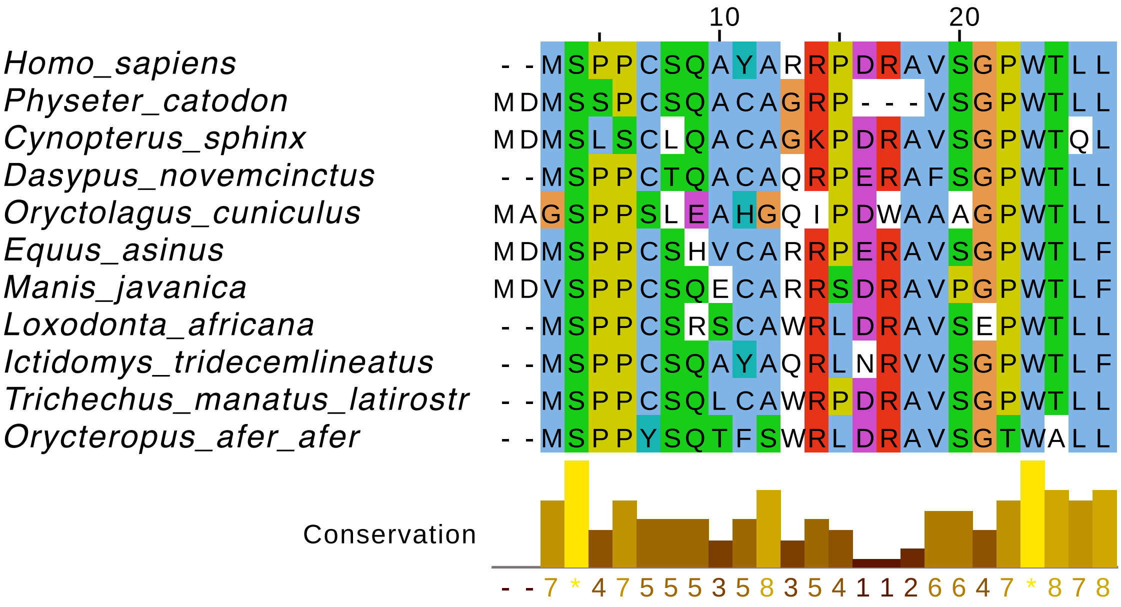

**HG0632.1**

ENSG00000181315 (ZNF322)

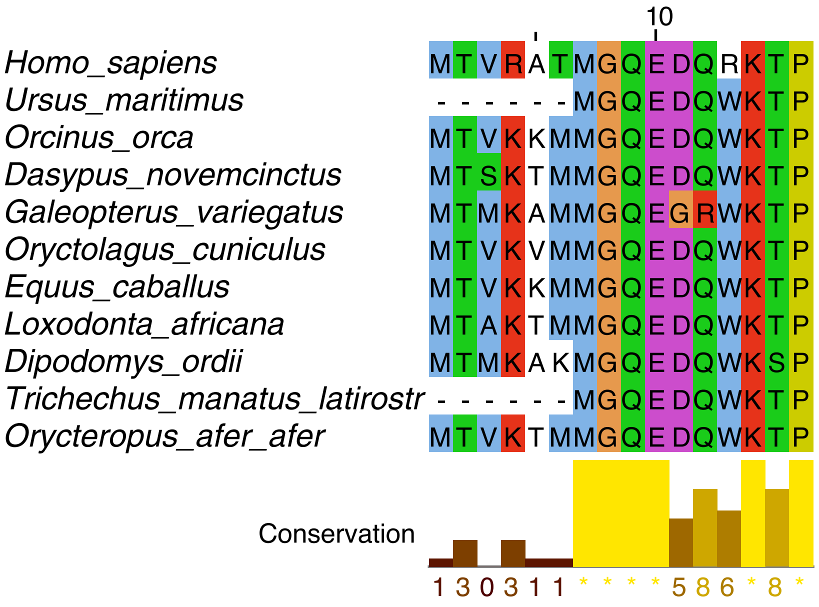

**HG0632.2**

ENSG00000181315 (ZNF322)

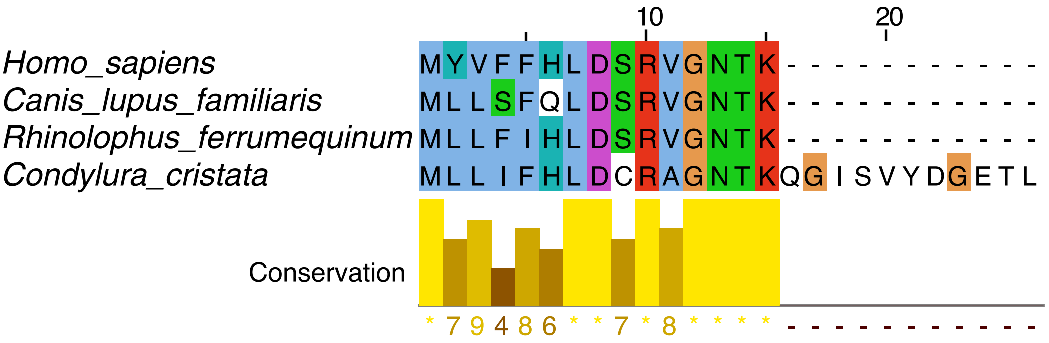

**HG0633**

ENSG00000186174 (BCL9L)

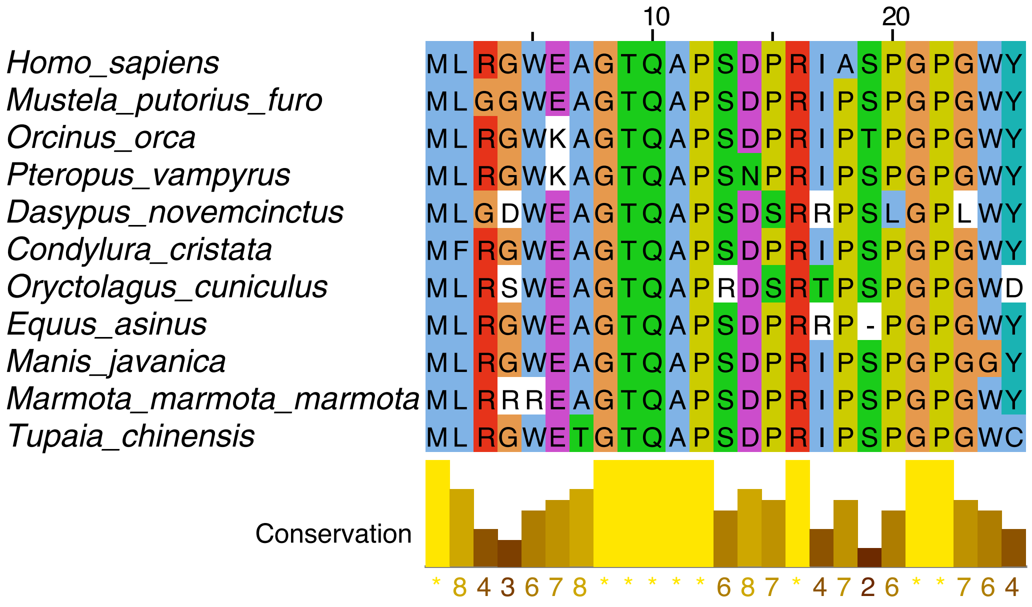

**HG0634**

ENSG00000198026 (ZNF335)

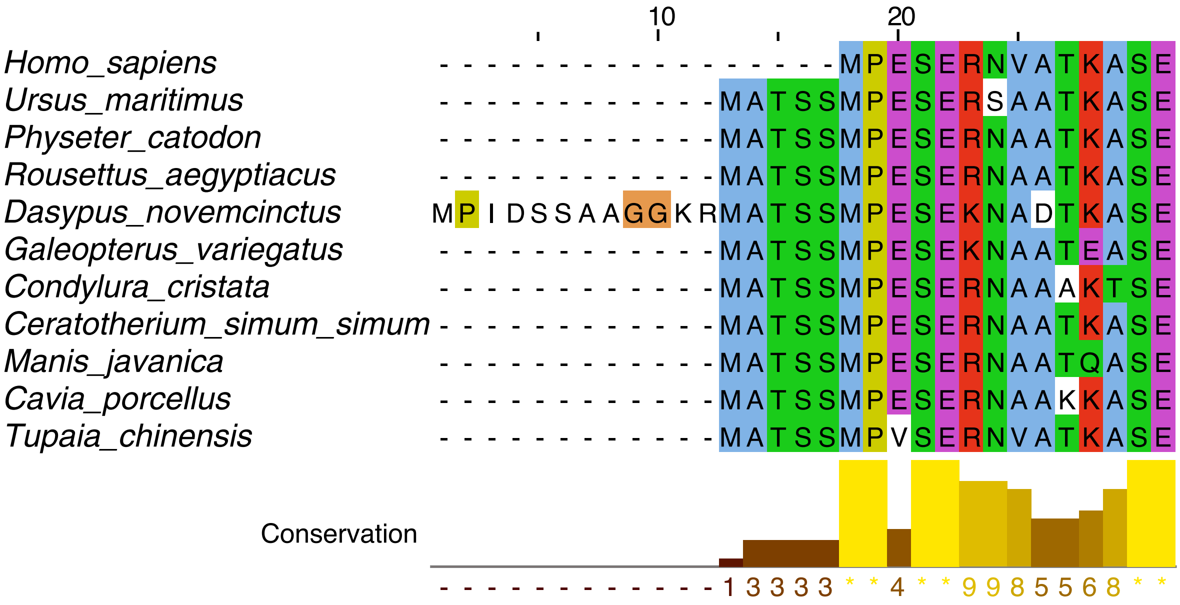

**HG0635**

ENSG00000259332 (ST20-MTHFS)

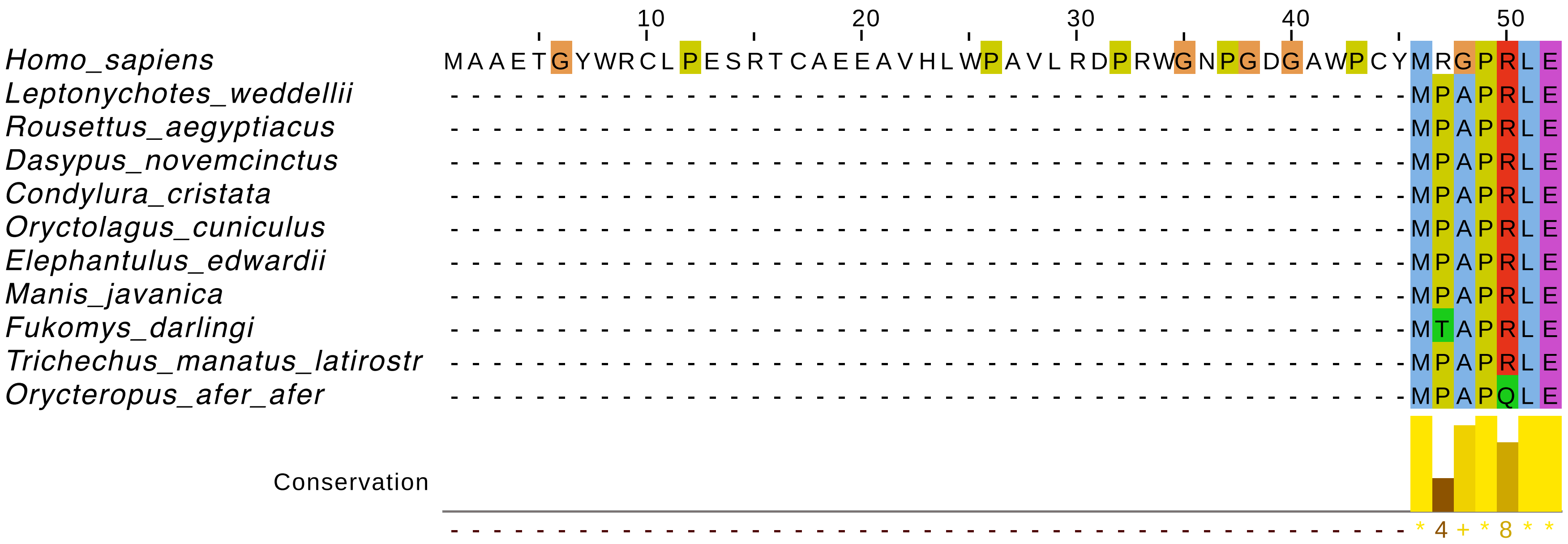

# HG0636

### ENSG00000002587 (HS3ST1)

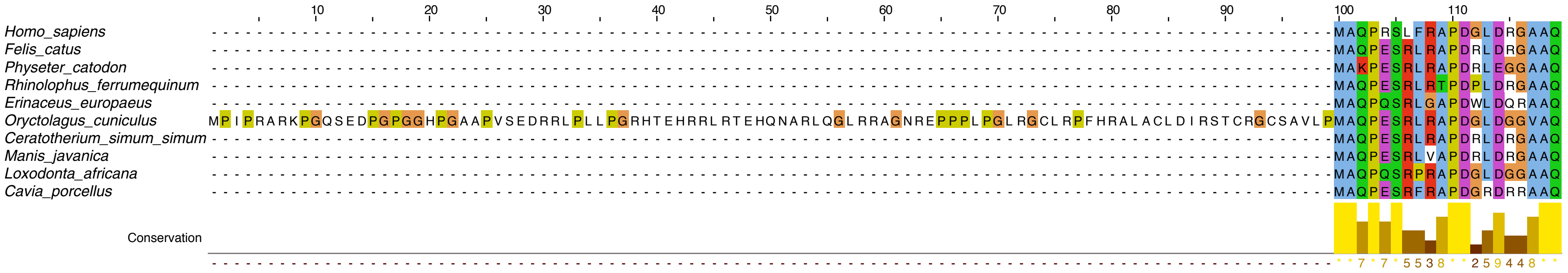

**HG0637**

ENSG00000005102 (MEOX1)

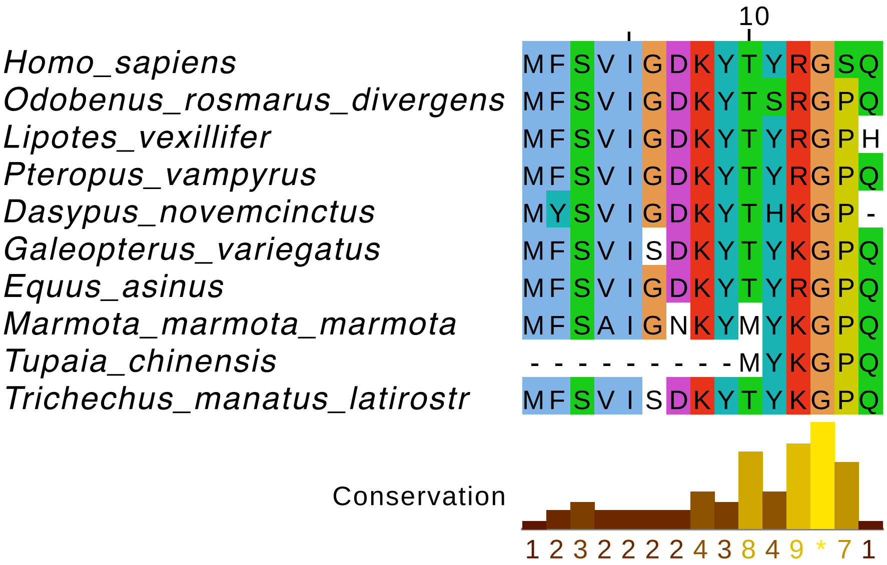

**HG0638**

ENSG00000013561 (RNF14)

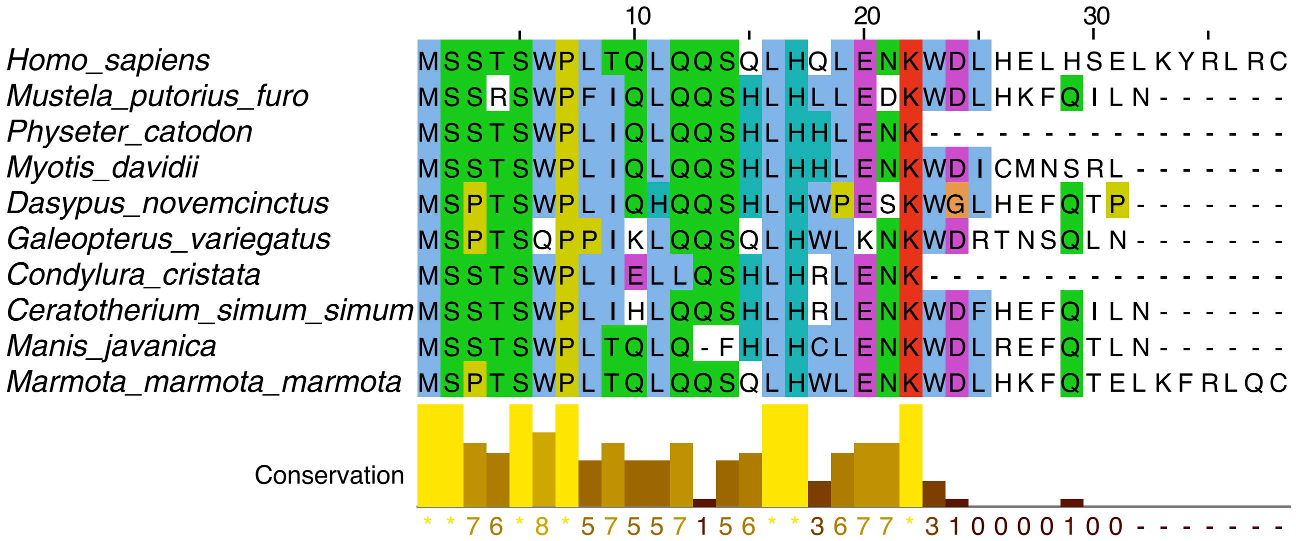

**HG0639**

ENSG00000018236 (CNTN1)

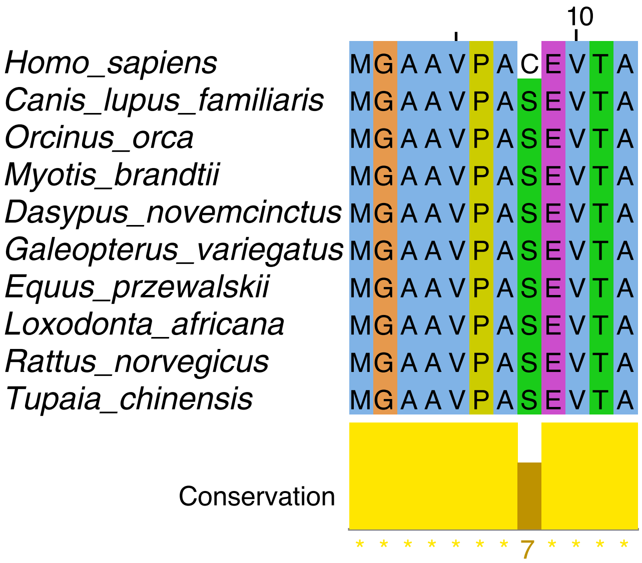

**HG0640**

ENSG00000062194 (GPBP1)

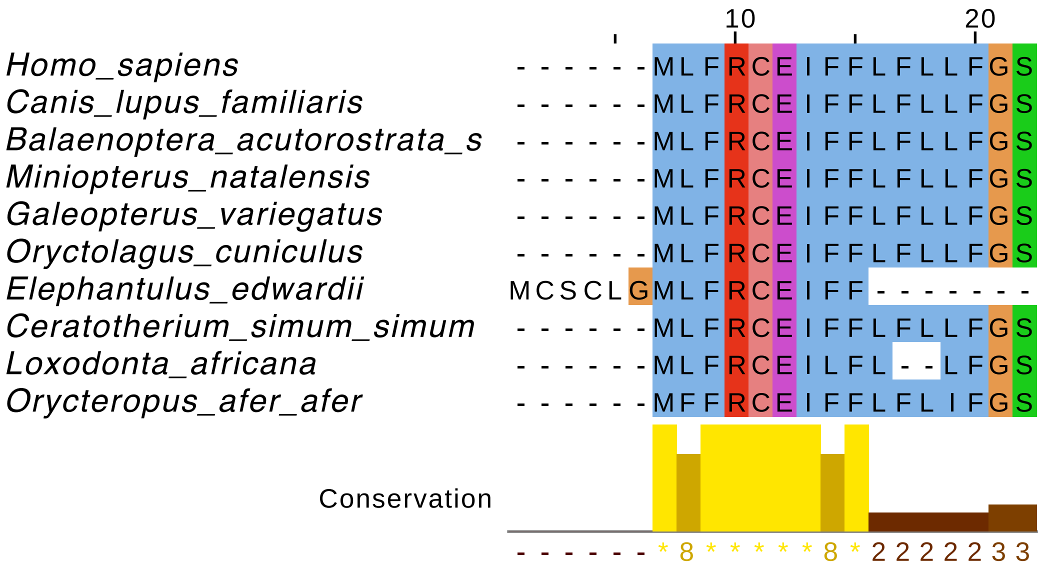

**HG0641**

ENSG00000100359 (SGSM3)

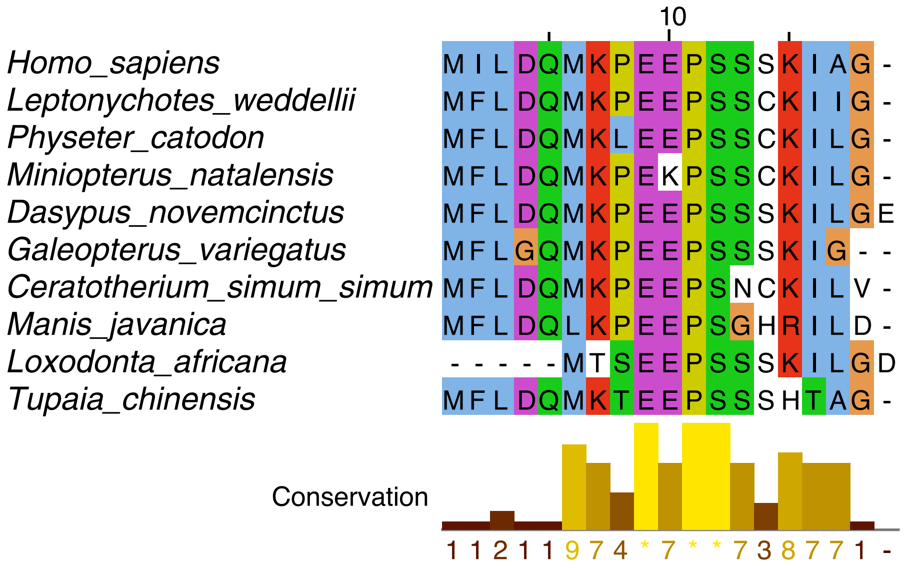

**HG0642**

ENSG00000100815 (TRIP11)

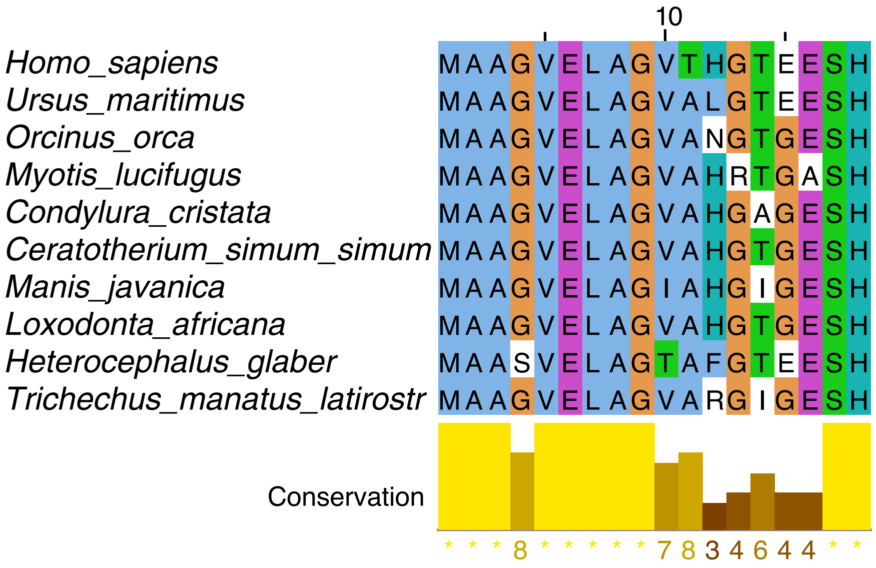

**HG0643**

ENSG00000103653 (CSK)

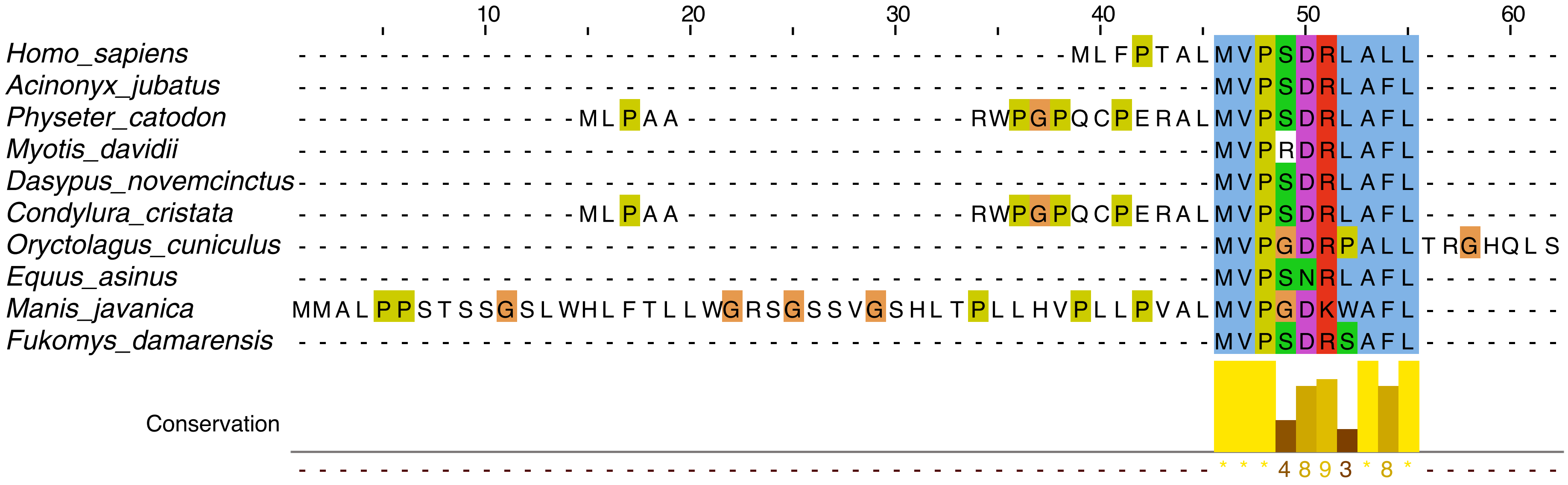

**HG0644**

ENSG00000104299 (INTS9)

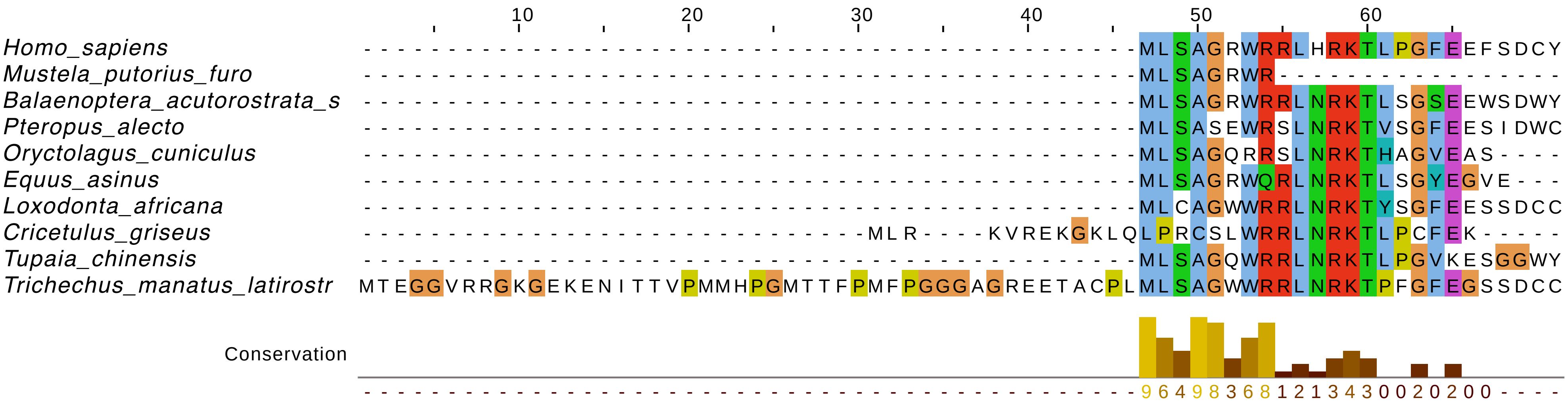

**HG0645**

ENSG00000107282 (APBA1)

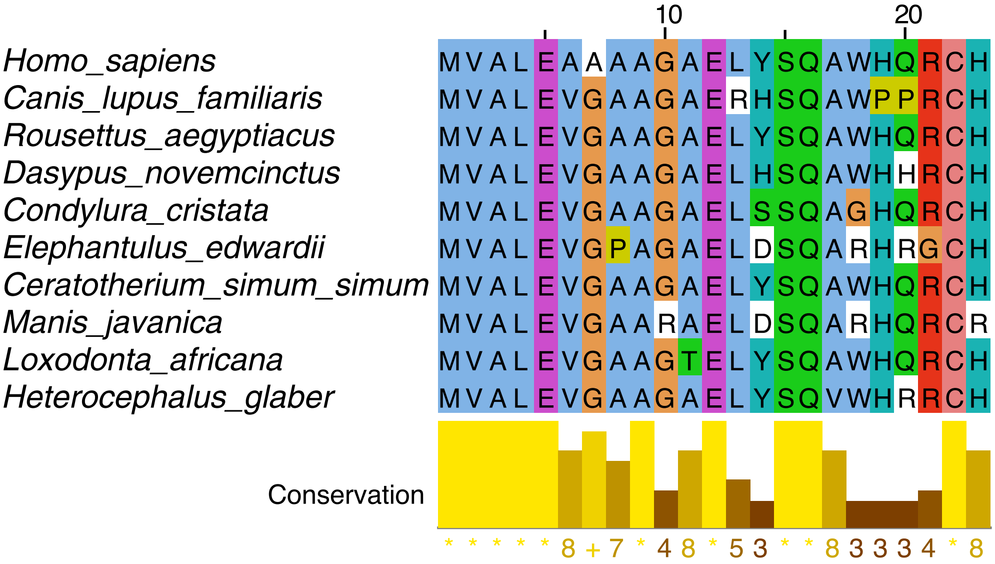

**HG0646**

ENSG00000108312 (UBTF)

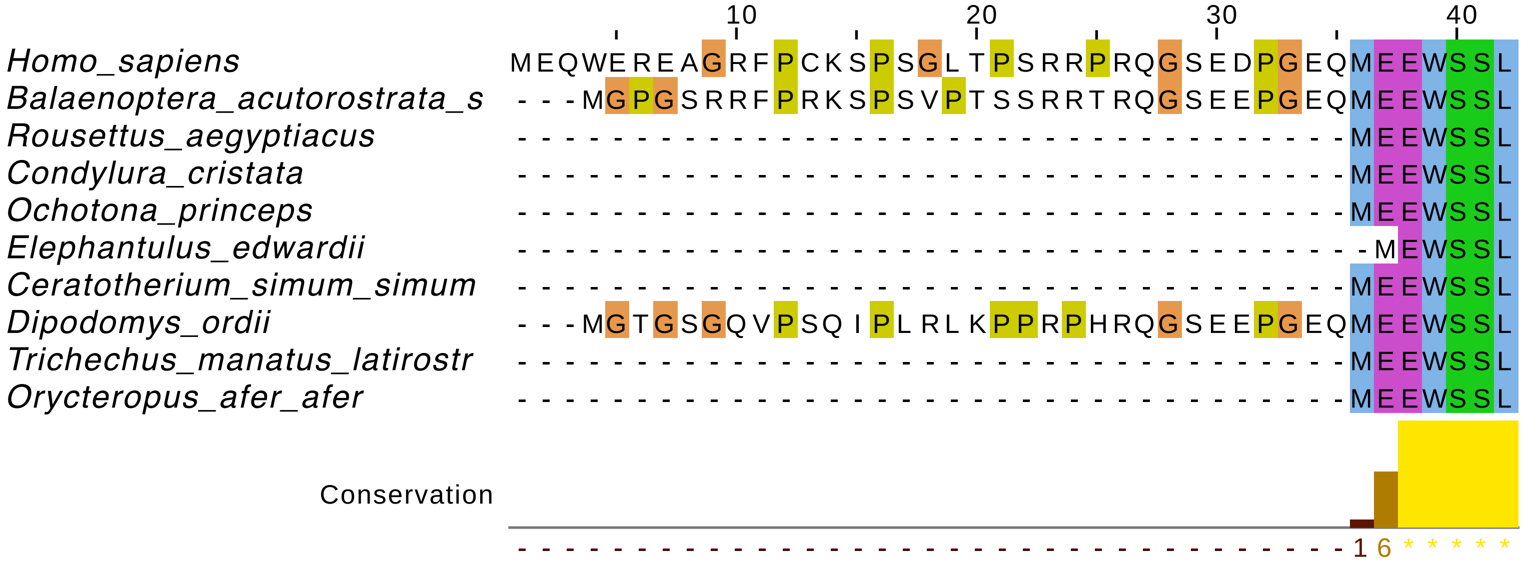

**HG0647**

ENSG00000108395 (TRIM37)

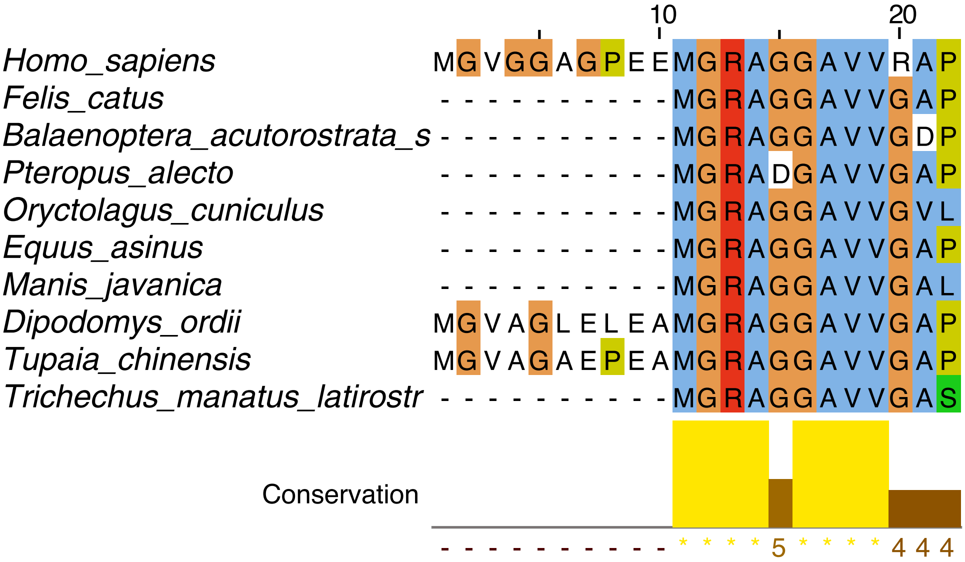

**HG0648**

ENSG00000114331 (ACAP2)

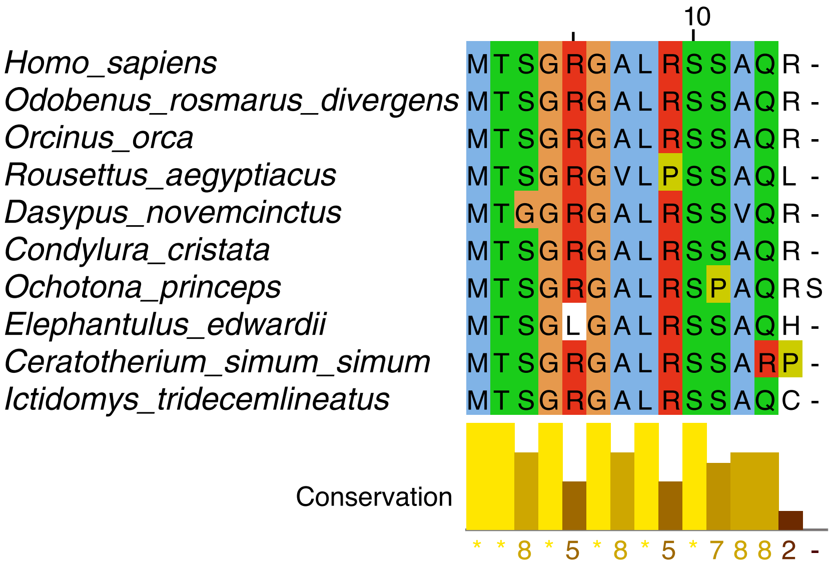

**HG0649.1**

ENSG00000114439 (BBX)

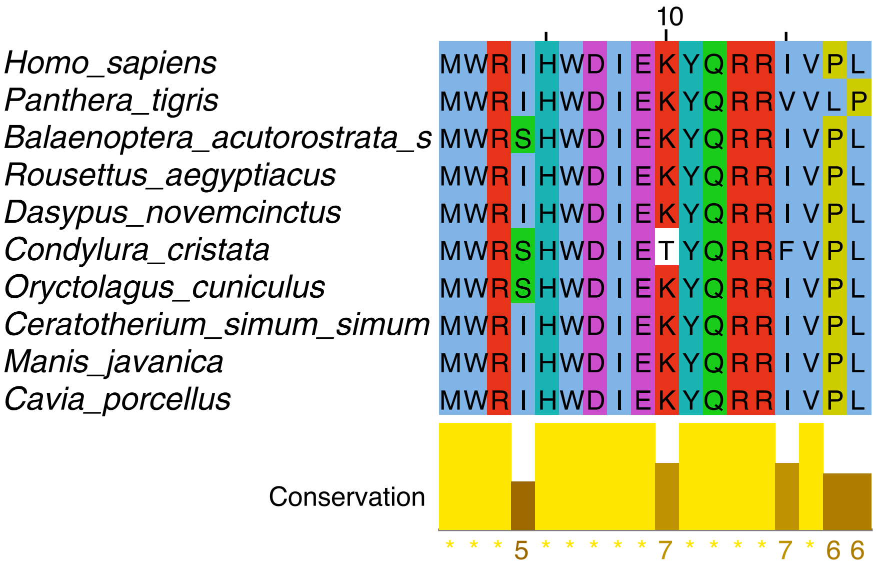

**HG0649.2**

ENSG00000114439 (BBX)

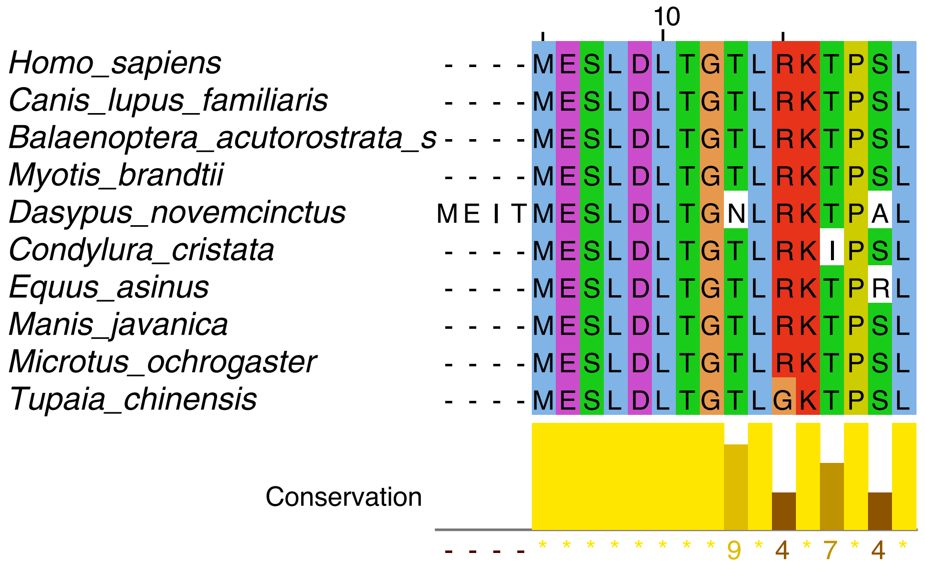

**HG0650**

ENSG00000115226 (FNDC4)

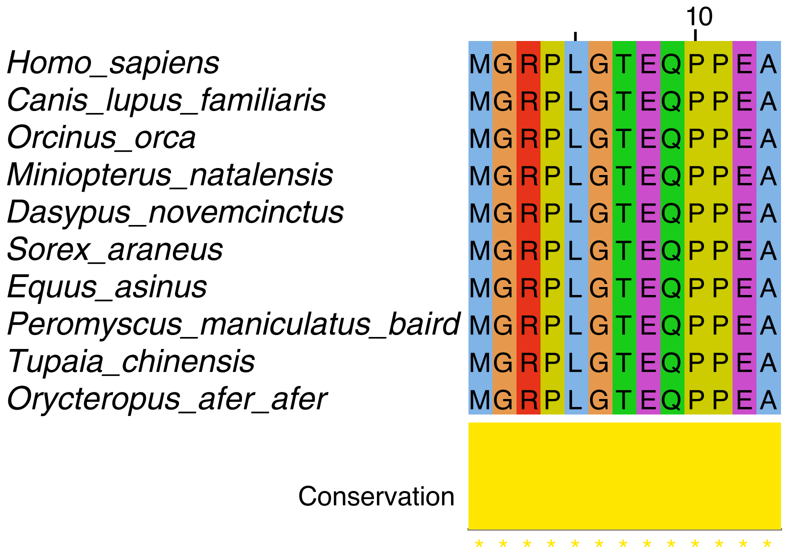

**HG0651**

ENSG00000115935 (WIPF1)

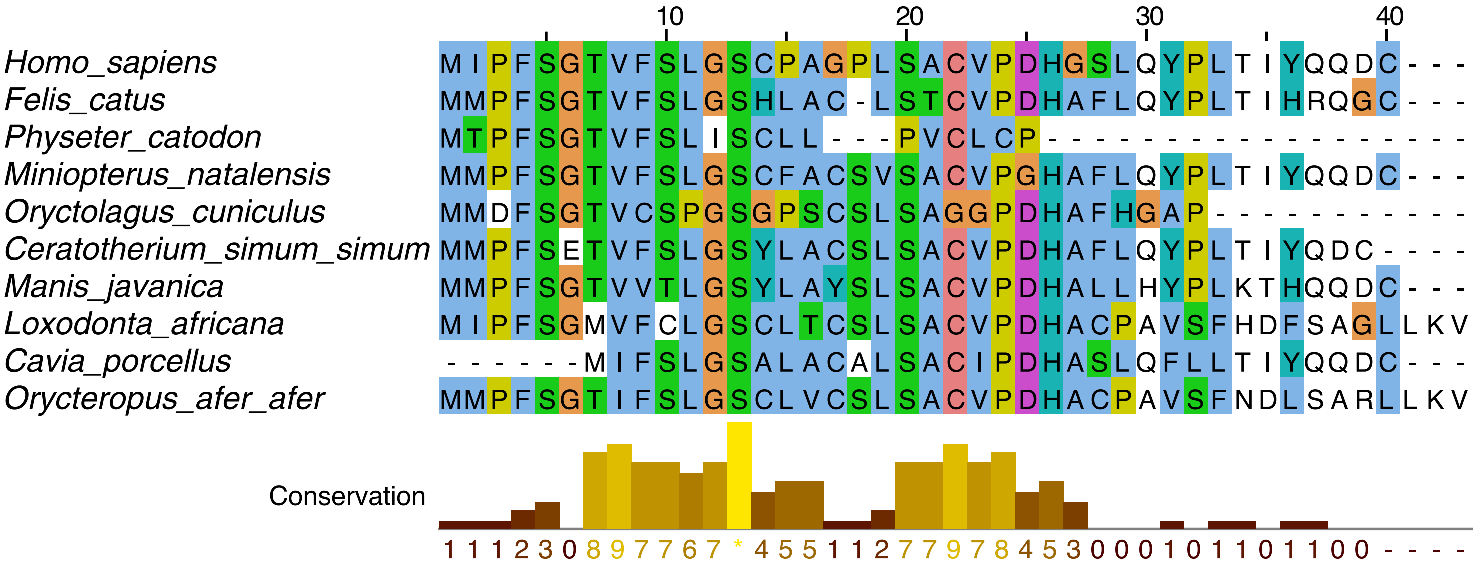

**HG0652**

ENSG00000118515 (SGK1)

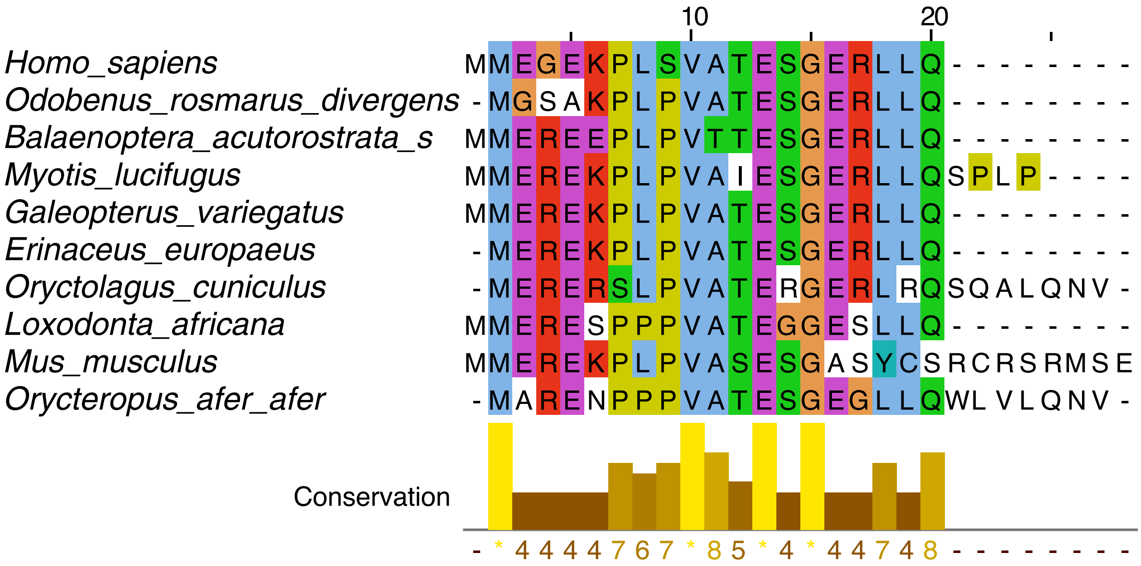

**HG0653**

ENSG00000120334 (CENPL)

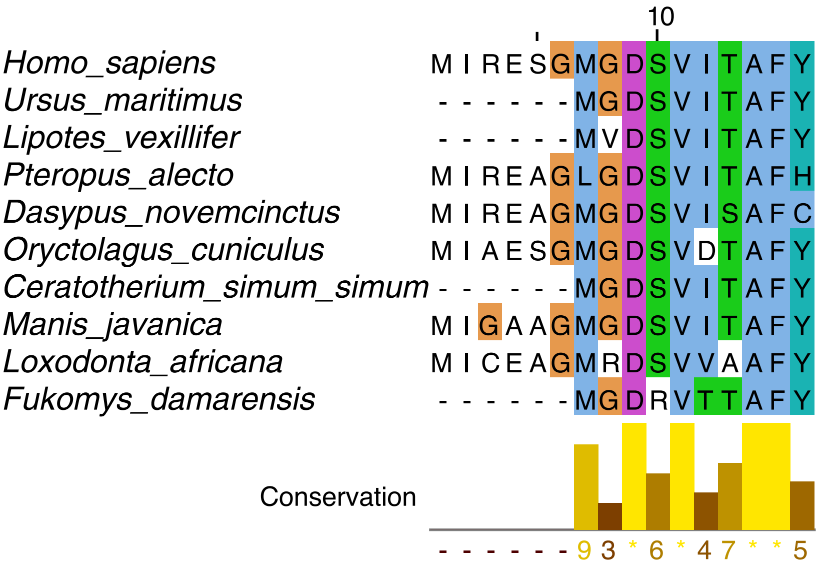

**HG0654**

ENSG00000133794 (ARNTL)

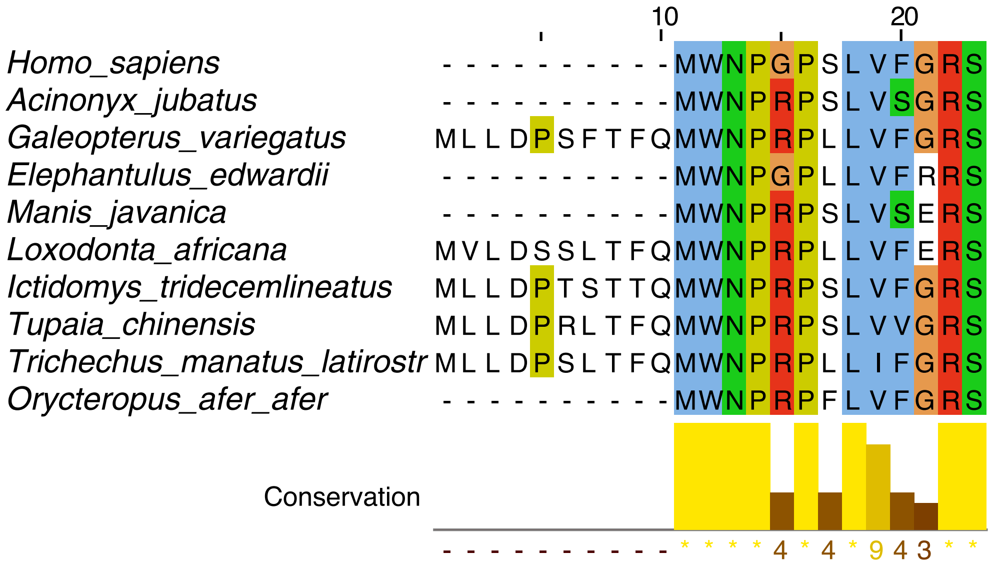

**HG0655**

ENSG00000143061 (IGSF3)

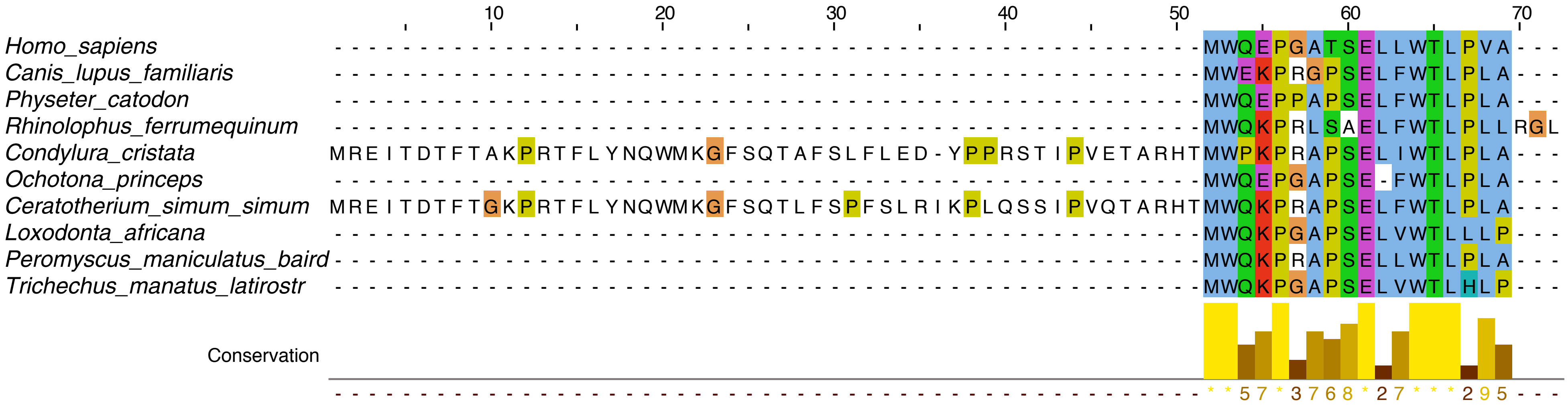

**HG0656**

ENSG00000150776 (C11orf57)

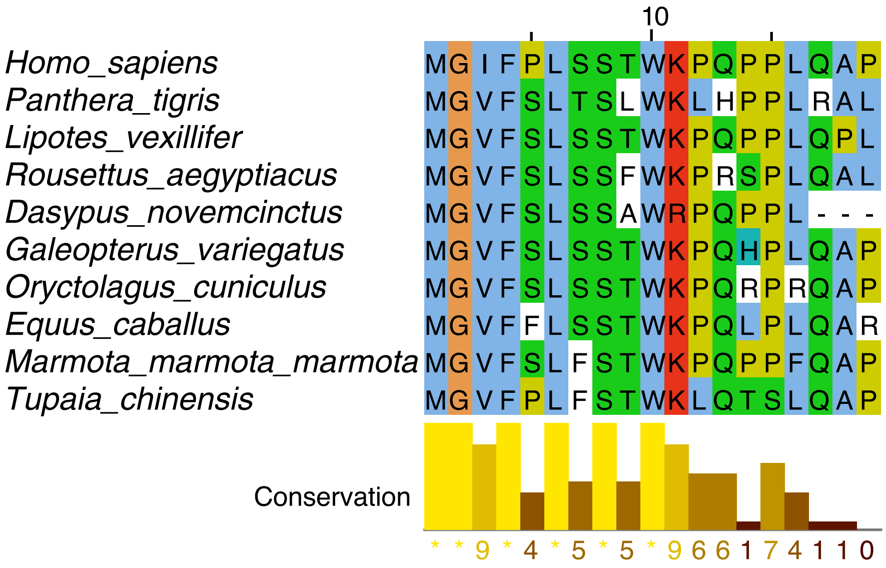

**HG0657**

ENSG00000151276 (MAGI1)

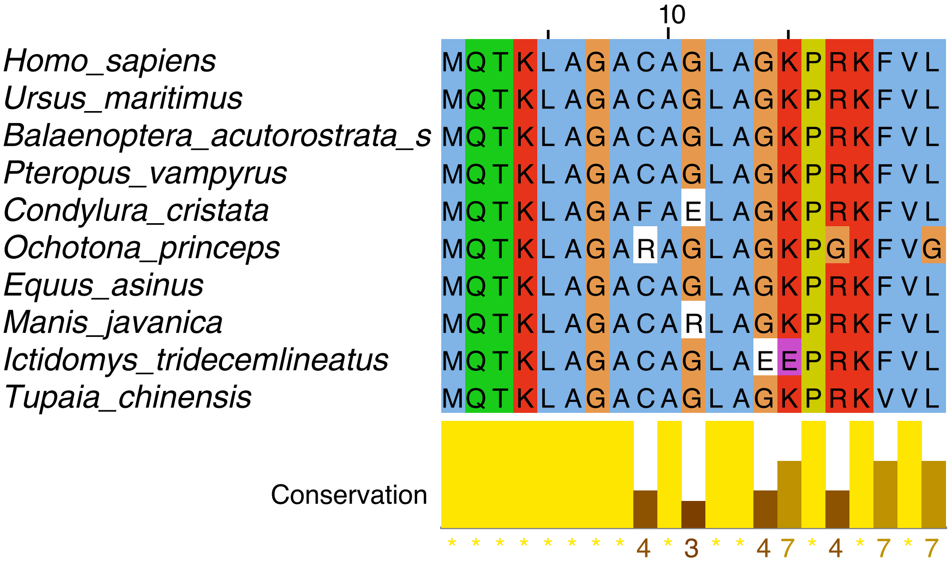

**HG0658**

ENSG00000153786 (ZDHHC7)

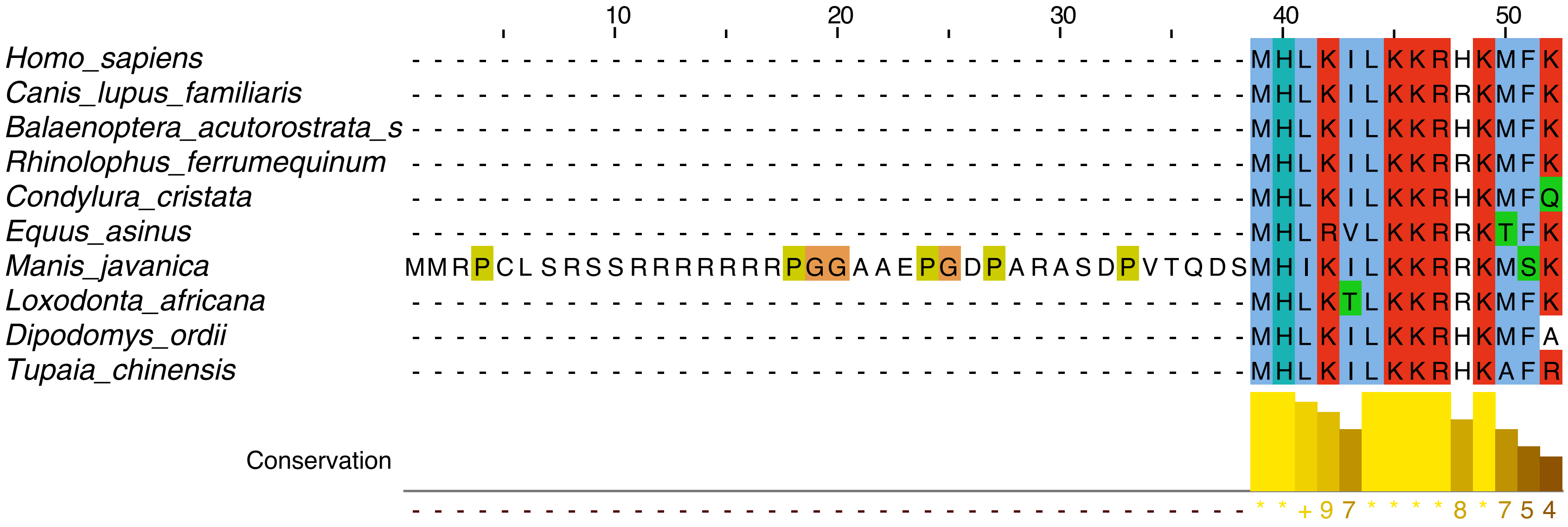

**HG0659.1**

ENSG00000158856 (DMTN)

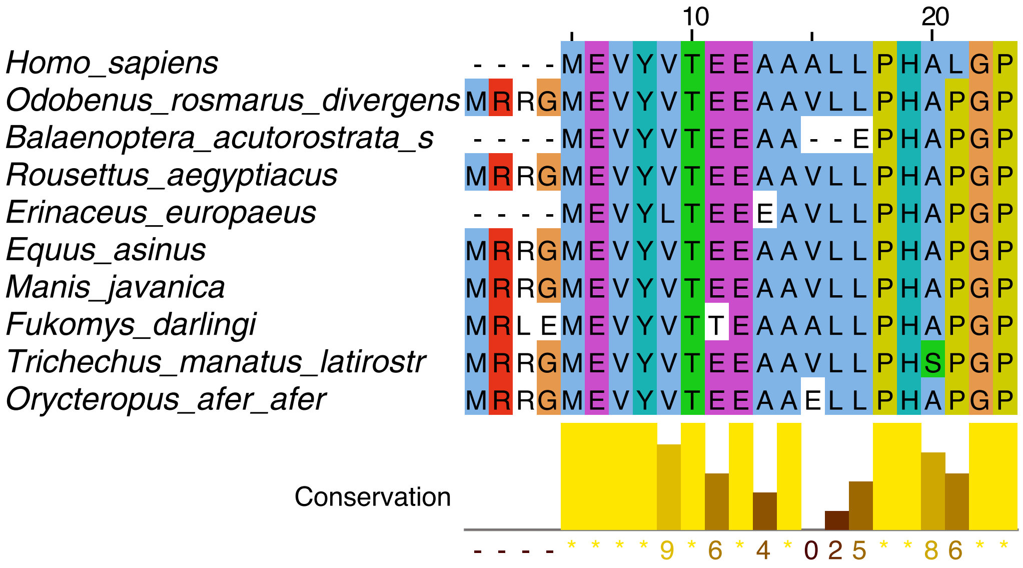

**HG0659.2**

ENSG00000158856 (DMTN)

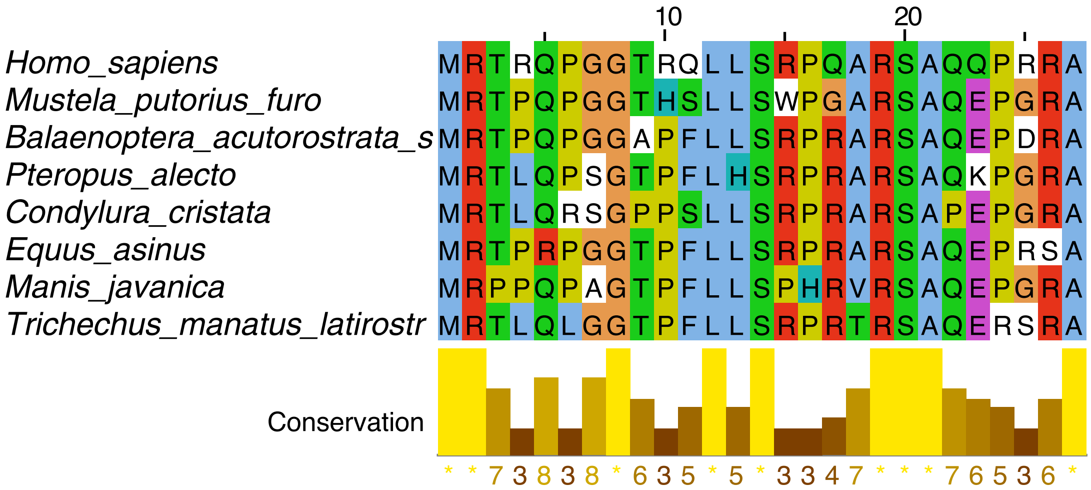

**HG0660**

ENSG00000165861 (ZFYVE1)

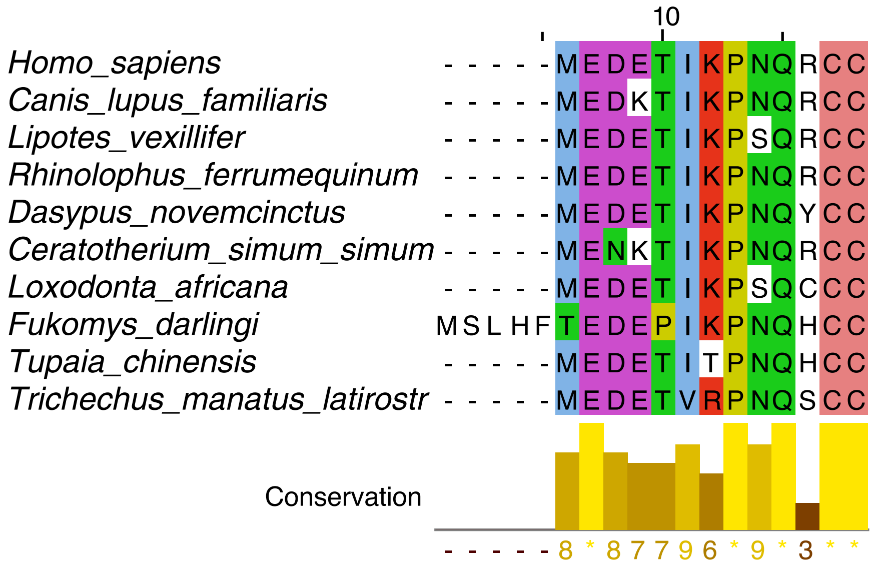

**HG0661**

ENSG00000166845 (C18orf54)

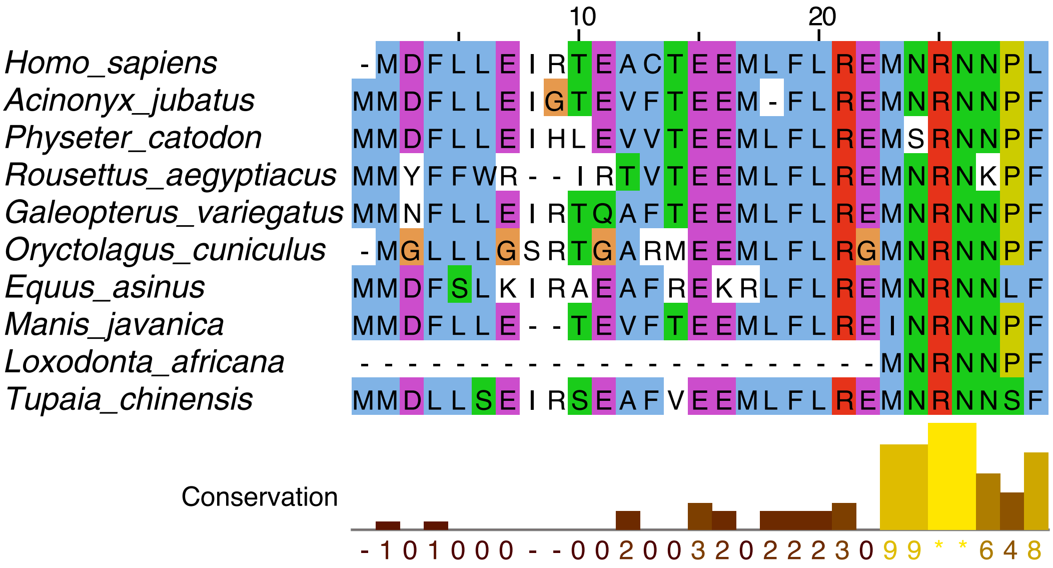

**HG0662**

ENSG00000166897 (ELFN2)

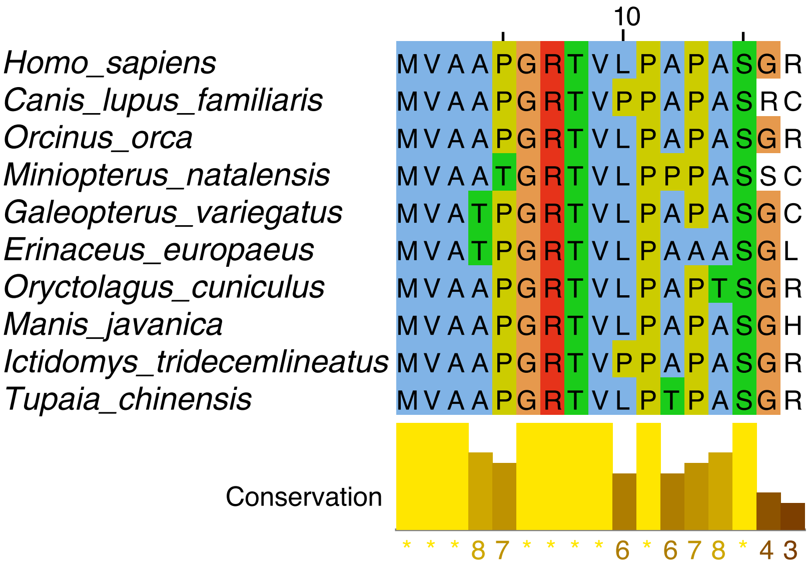

**HG0663**

ENSG00000168615 (ADAM9)

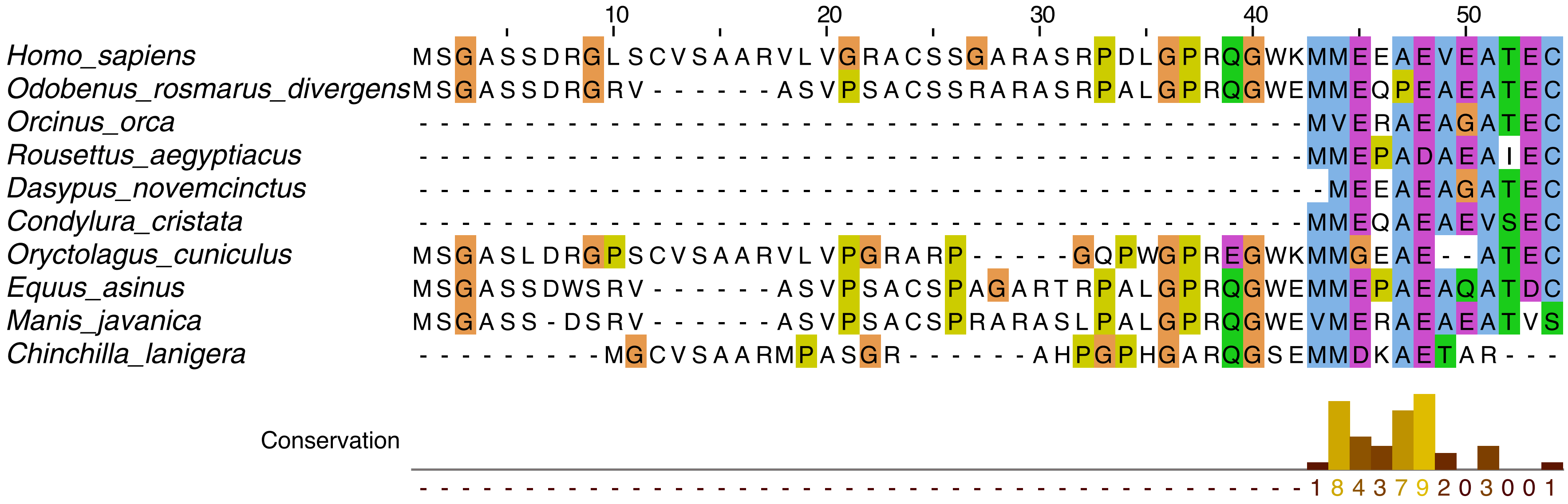

**HG0664**

ENSG00000169085

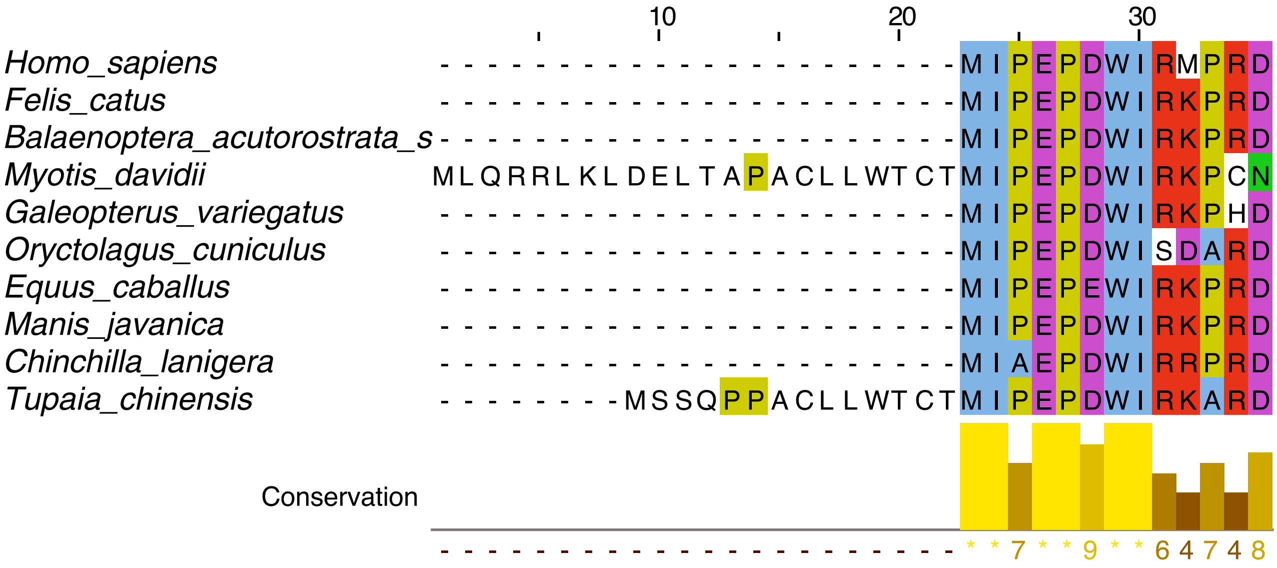

**HG0665**

ENSG00000169515 (CCDC8)

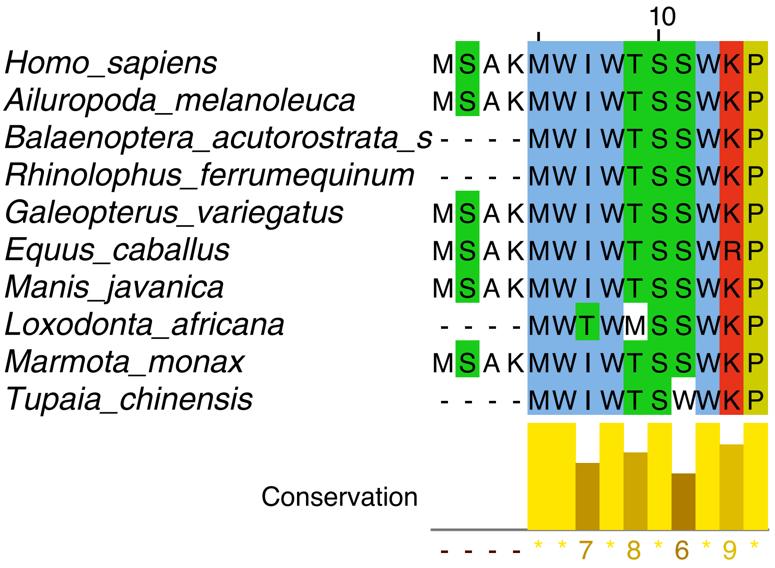

**HG0666**

ENSG00000171943 (SRGAP2C)

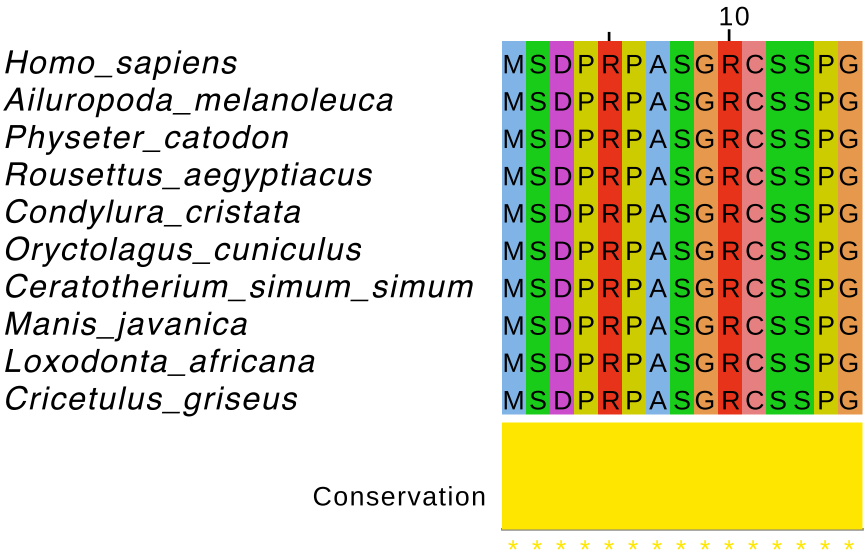

**HG0667.1**

ENSG00000182700 (IGIP)

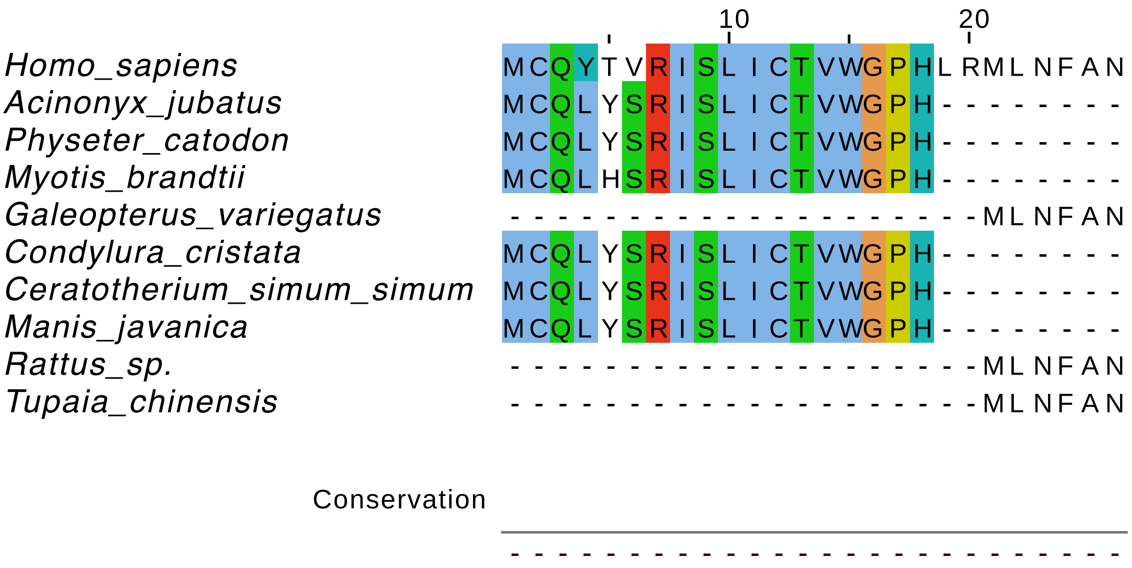

**HG0667.2**

ENSG00000182700 (IGIP)

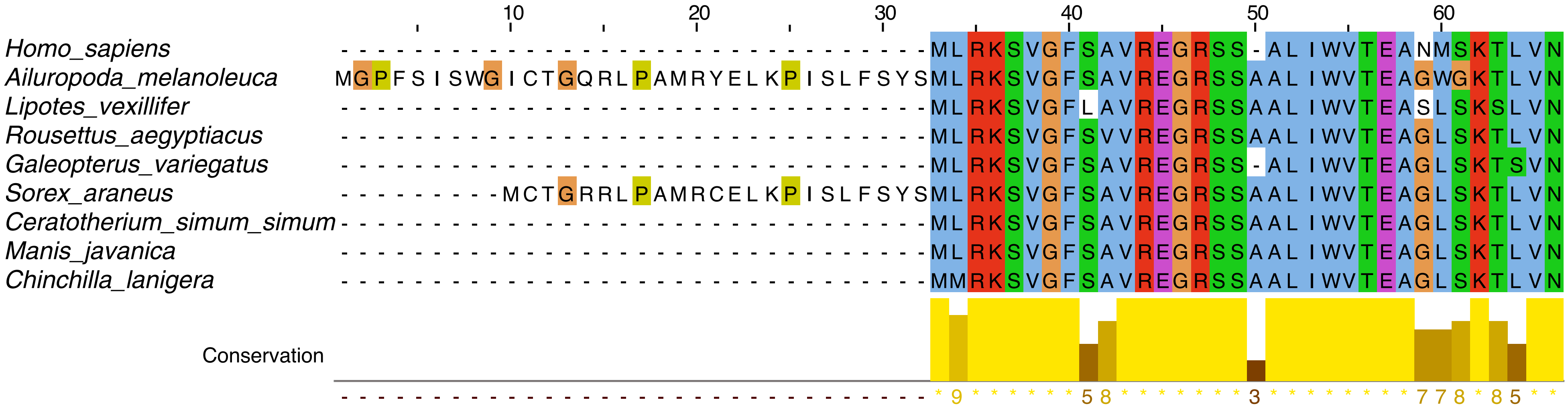

**HG0668**

ENSG00000182944 (EWSR1)

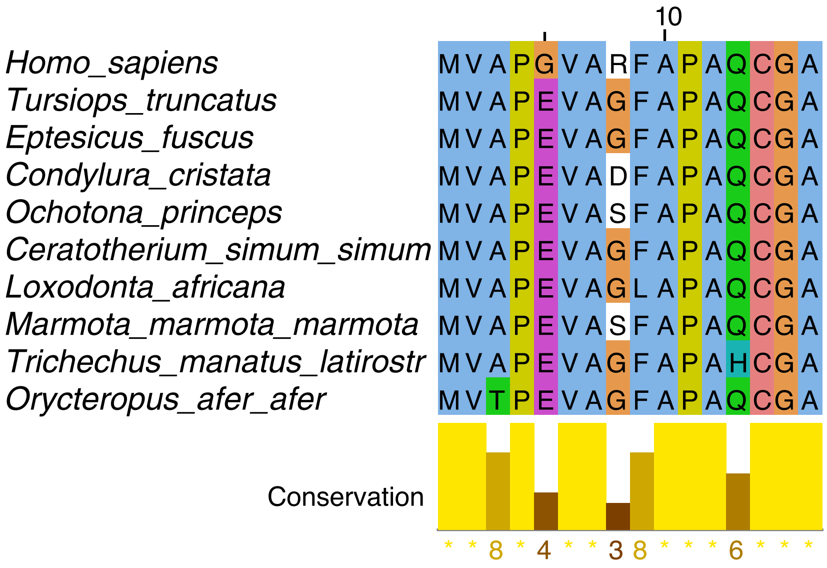

**HG0669**

ENSG00000183876 (ARSI)

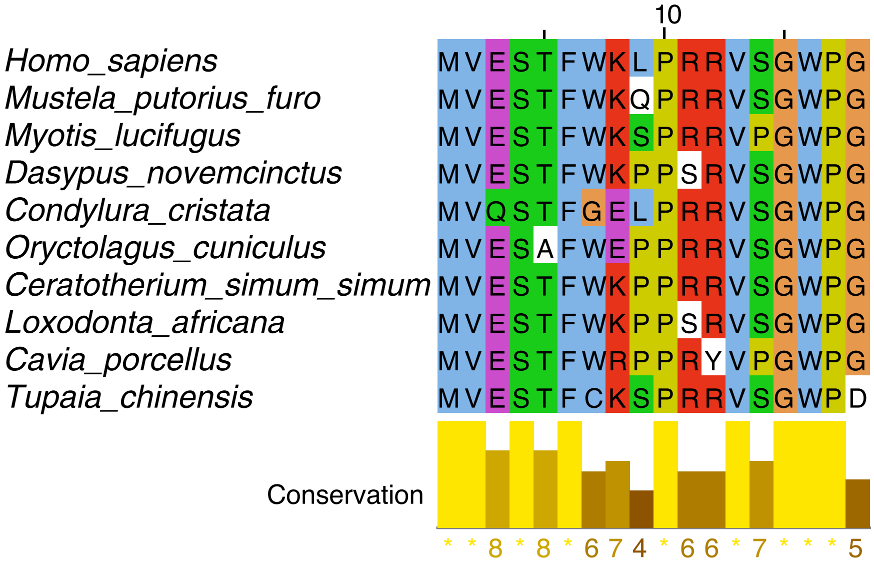

**HG0670**

ENSG00000187164 (SHTN1)

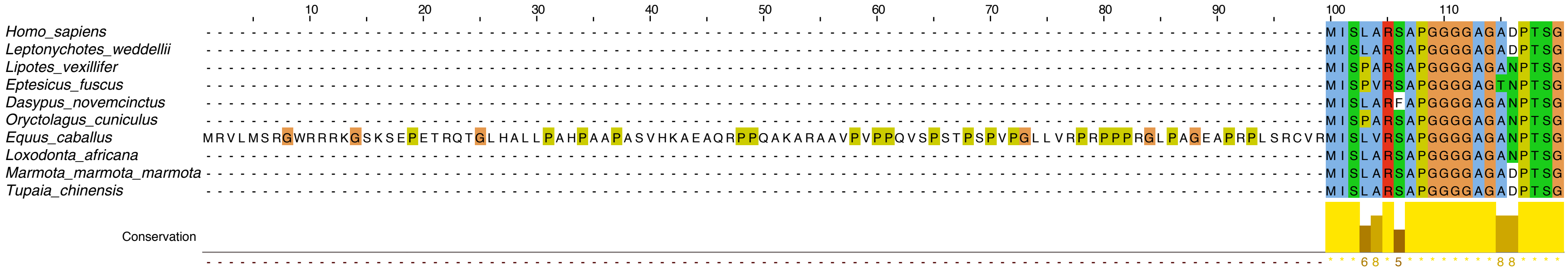

**HG0671**

ENSG00000189184 (PCDH18)

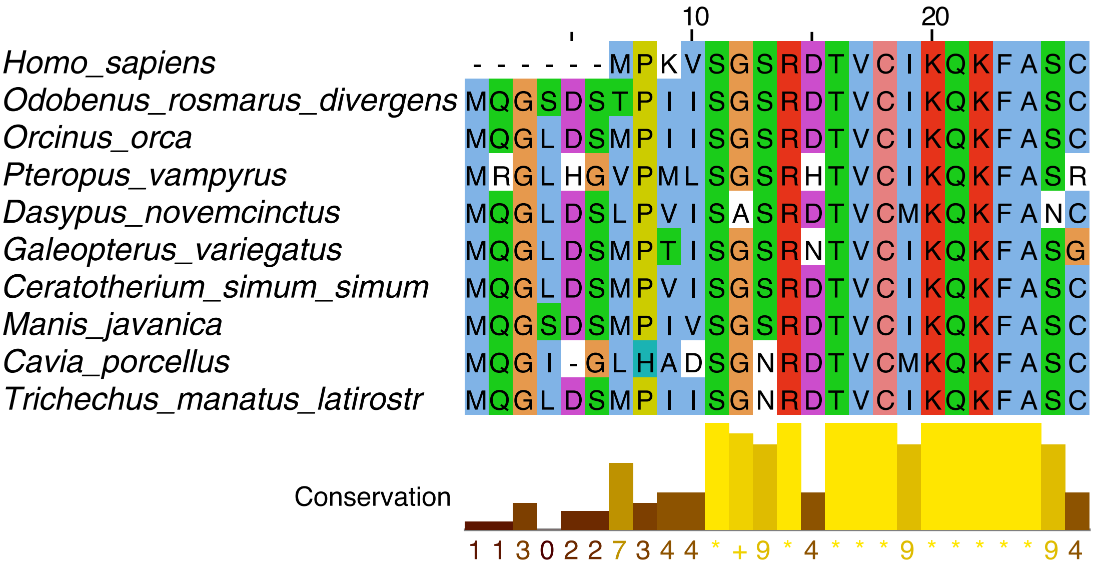

**HG0672**

ENSG00000196369 (SRGAP2B)

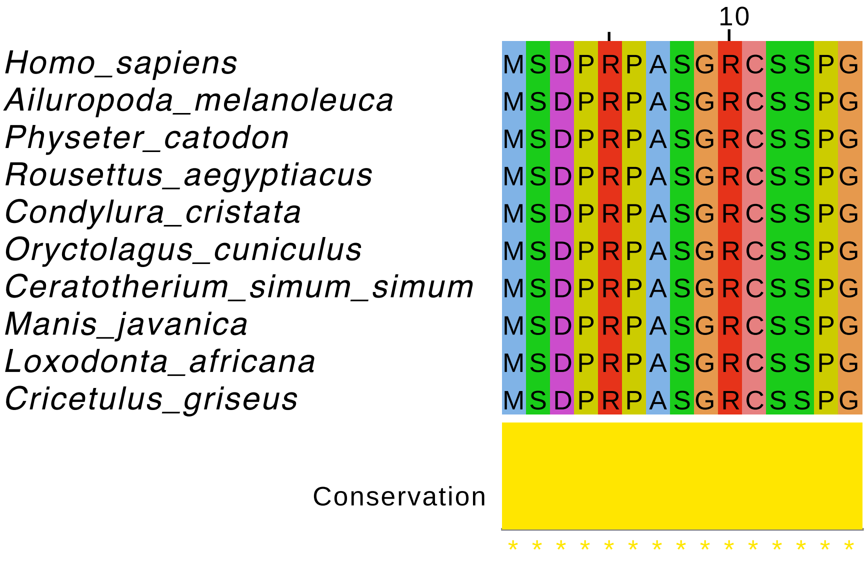

**HG0673**

ENSG00000266028 (SRGAP2)

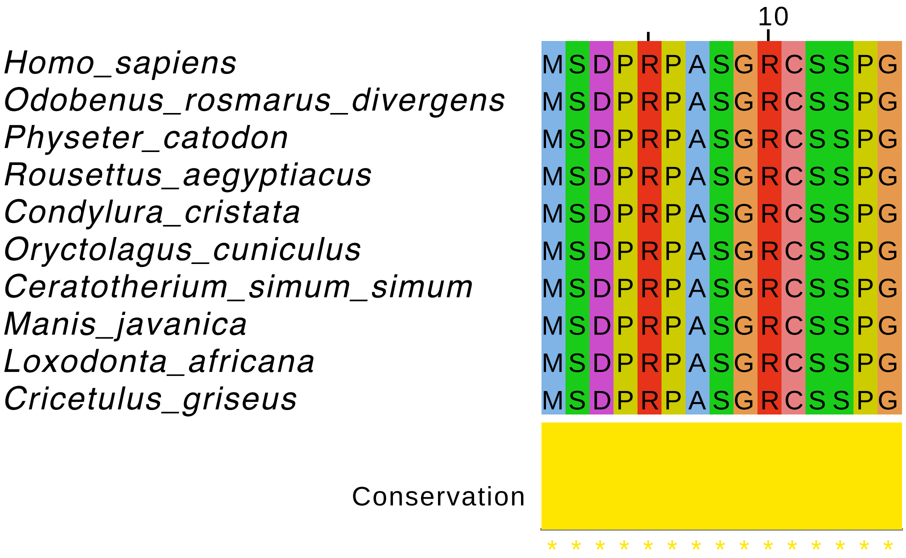

**HG0674.1**

ENSG00000006576 (PHTF2)

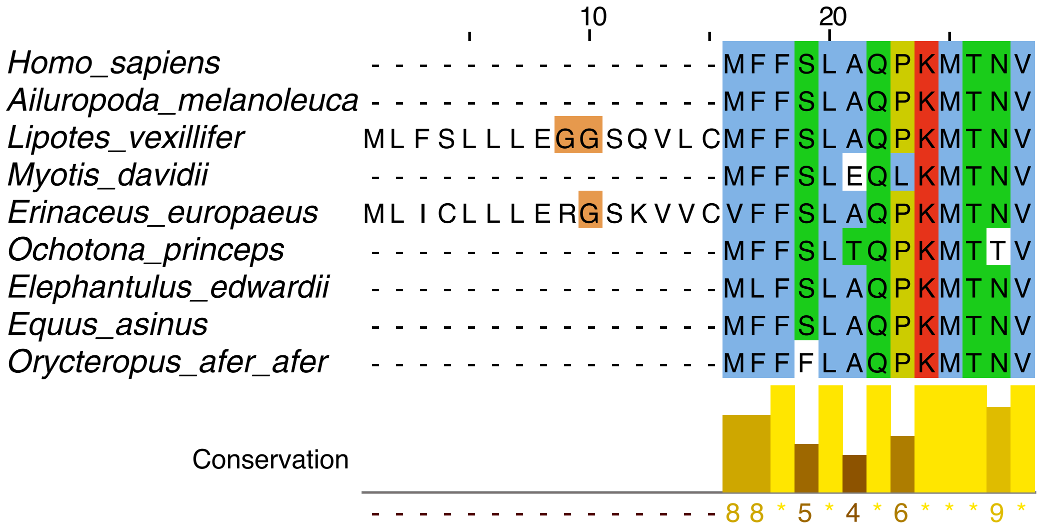

**HG0674.2**

ENSG00000116793 (PHTF1)

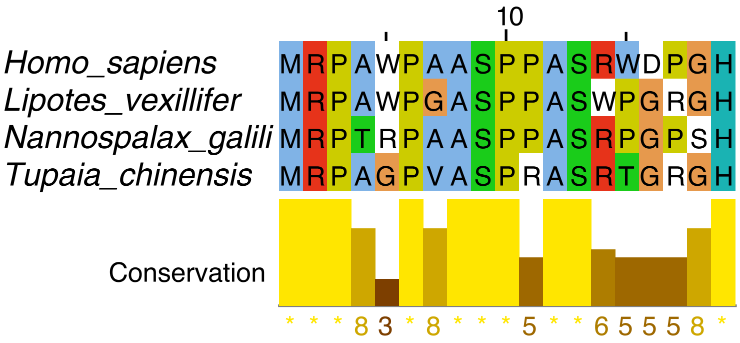

**HG0675**

ENSG00000048991 (R3HDM1)

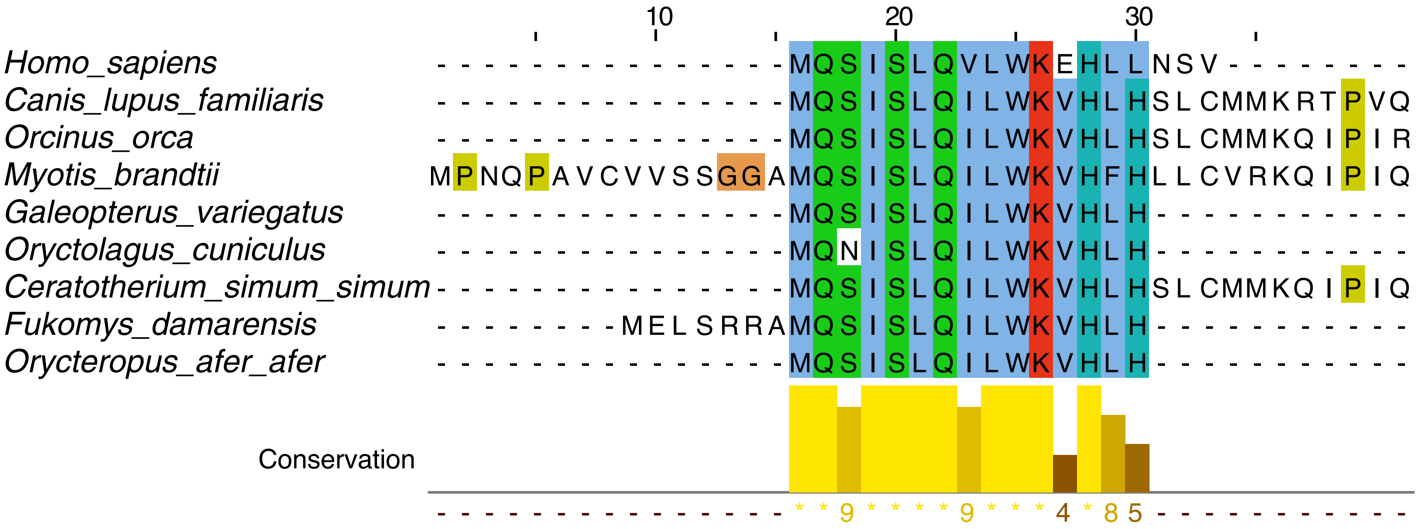

**HG0676**

ENSG00000058673 (ZC3H11A)

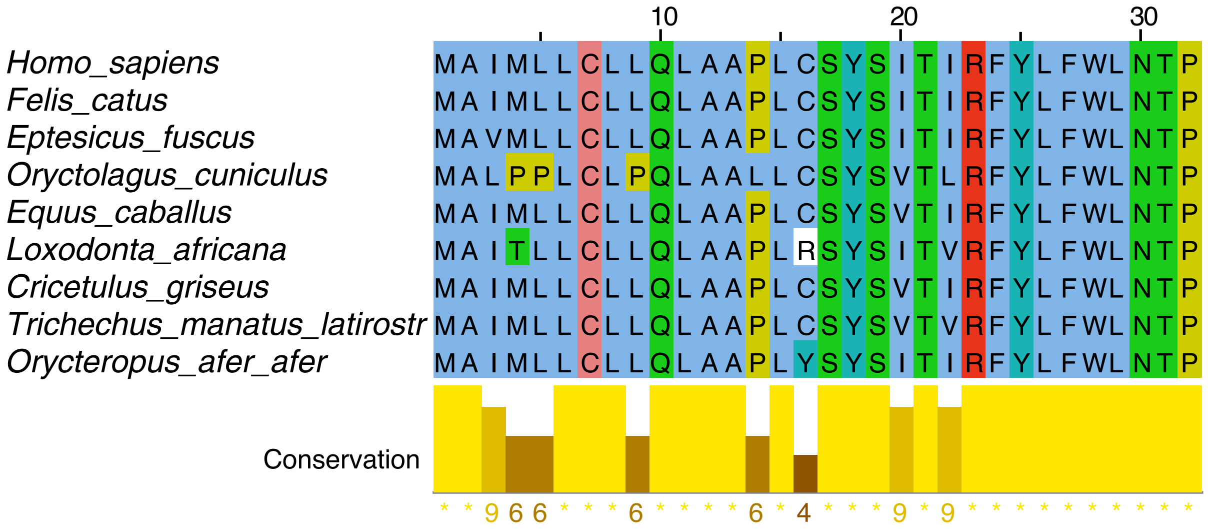

**HG0677**

ENSG00000064989 (CALCRL)

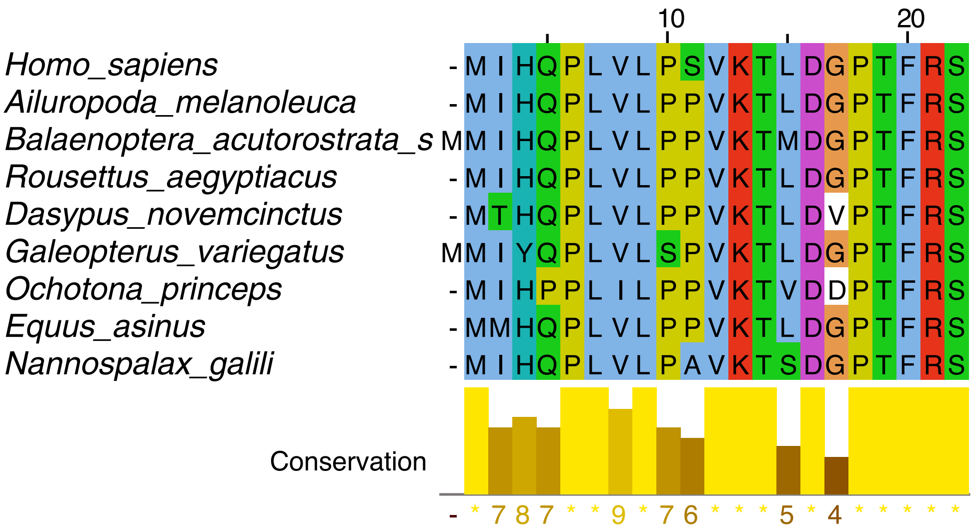

**HG0678.1**

ENSG00000090776 (EFNB1)

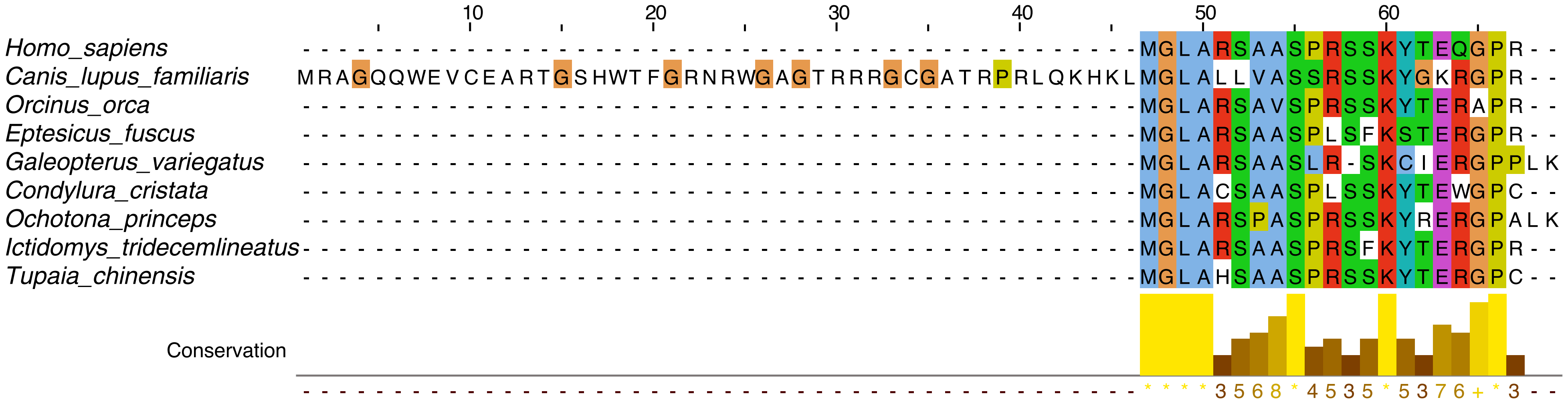

**HG0678.2**

ENSG00000108947 (EFNB3)

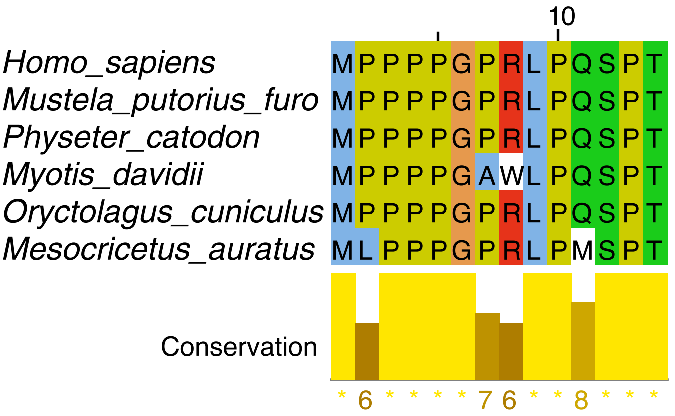

**HG0679**

ENSG00000100281 (HMGXB4)

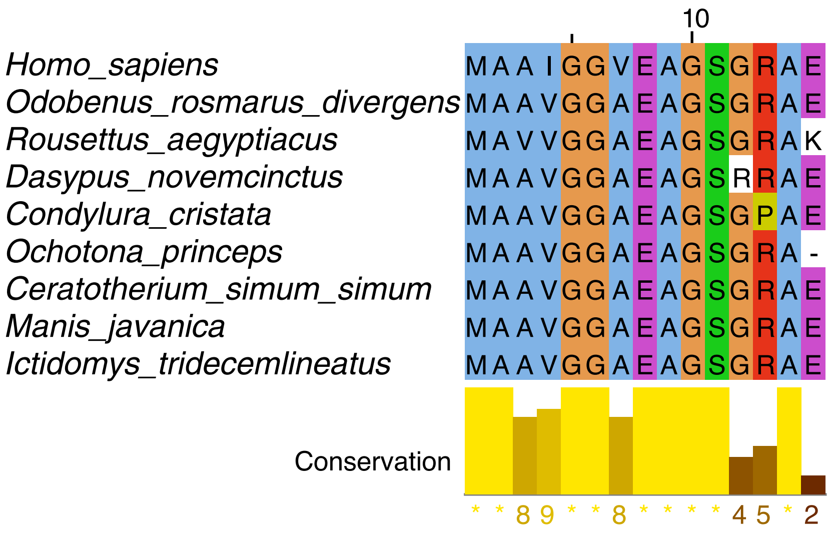

**HG0680**

ENSG00000100678 (SLC8A3)

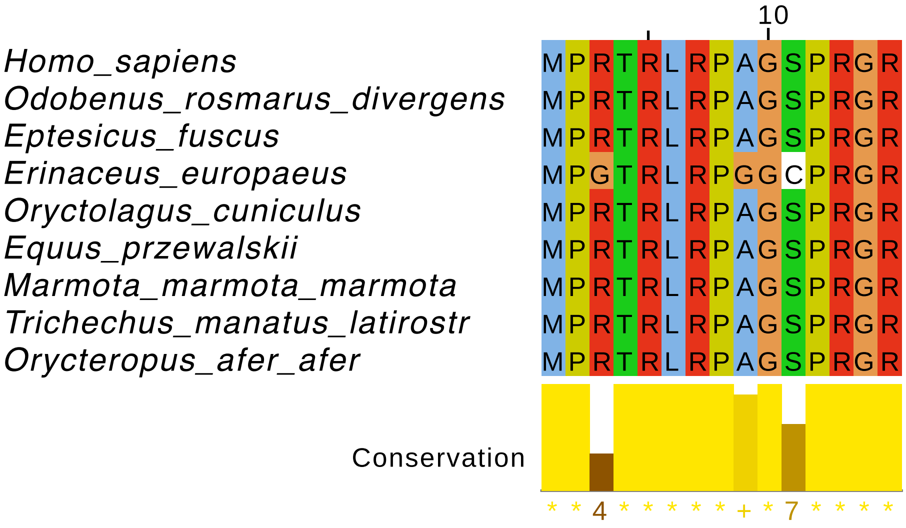

**HG0681**

ENSG00000101349 (PAK5)

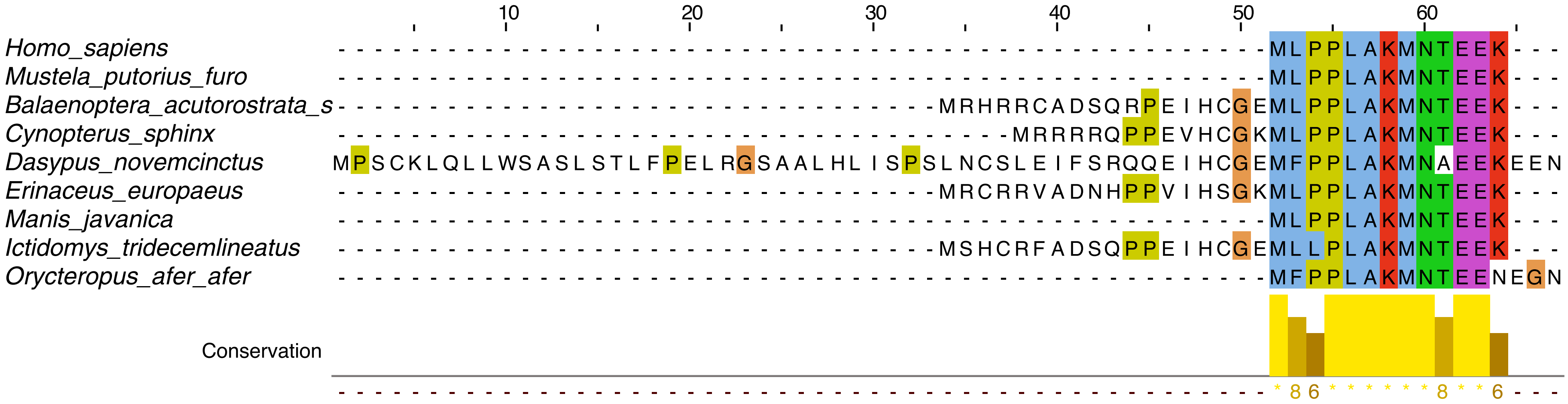

**HG0682**

ENSG00000105829 (BET1)

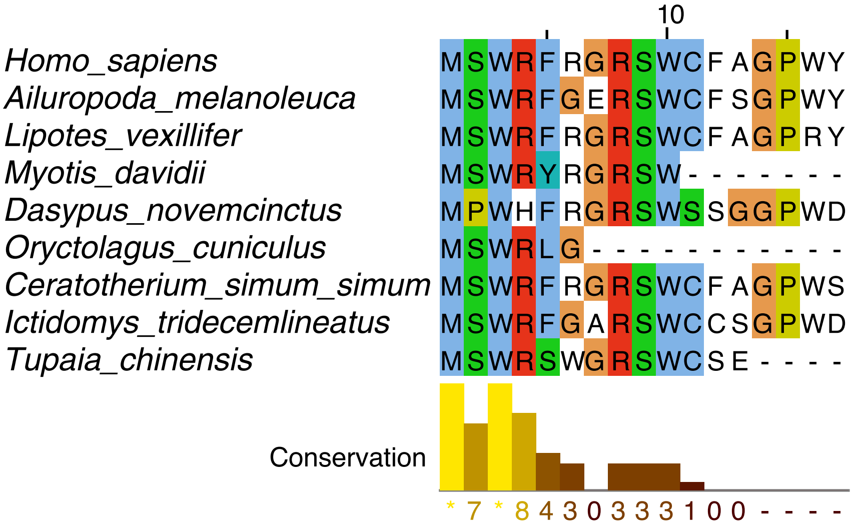

**HG0683**

ENSG00000111837 (MAK)

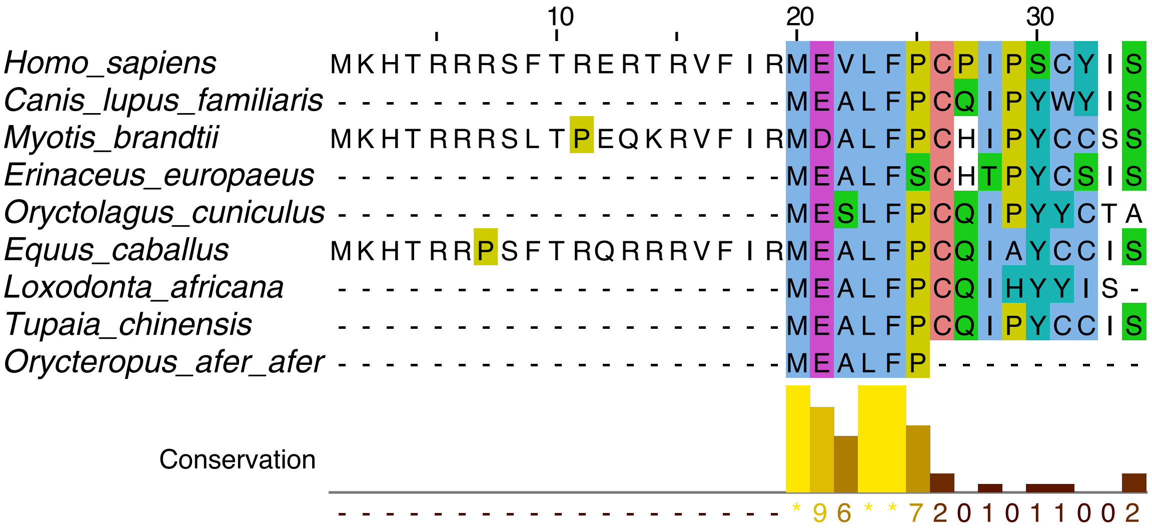

**HG0684**

ENSG00000113916 (BCL6)

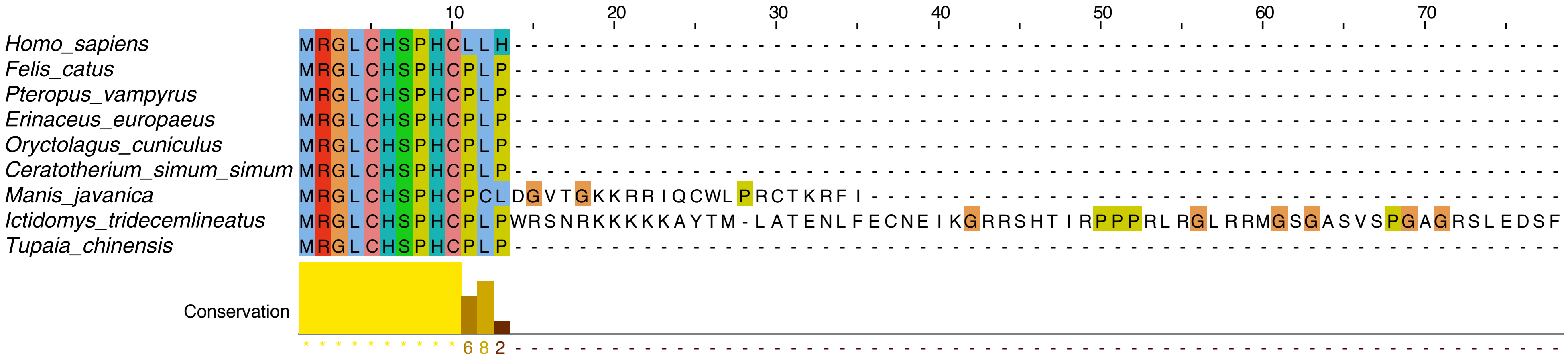

**HG0685**

ENSG00000117262 (GPR89A)

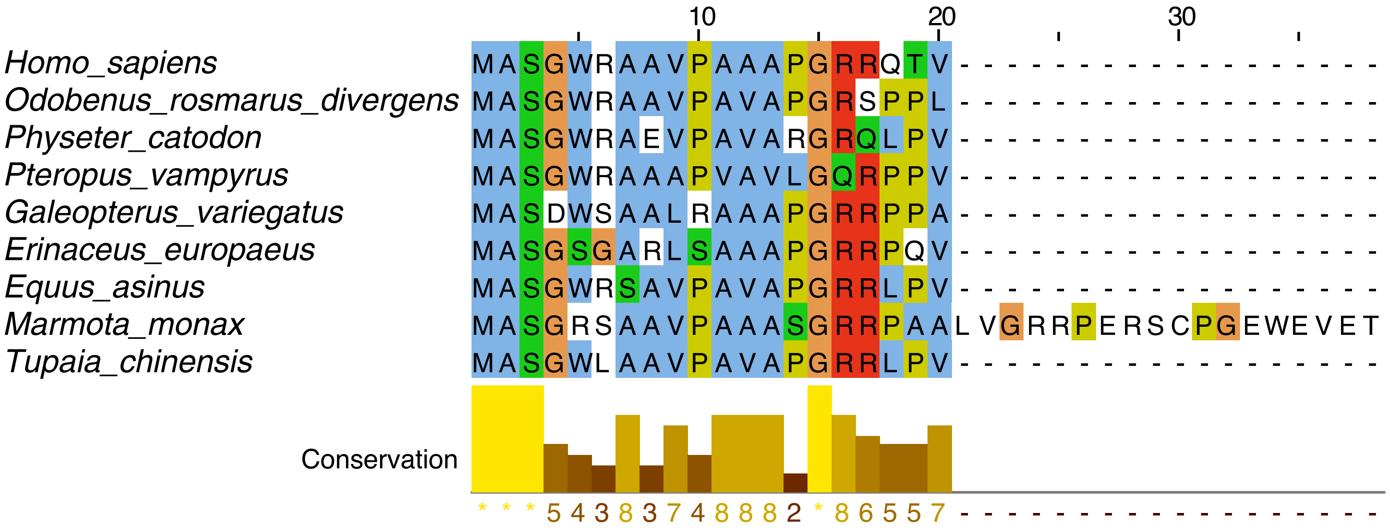

**HG0686**

ENSG00000118985 (ELL2)

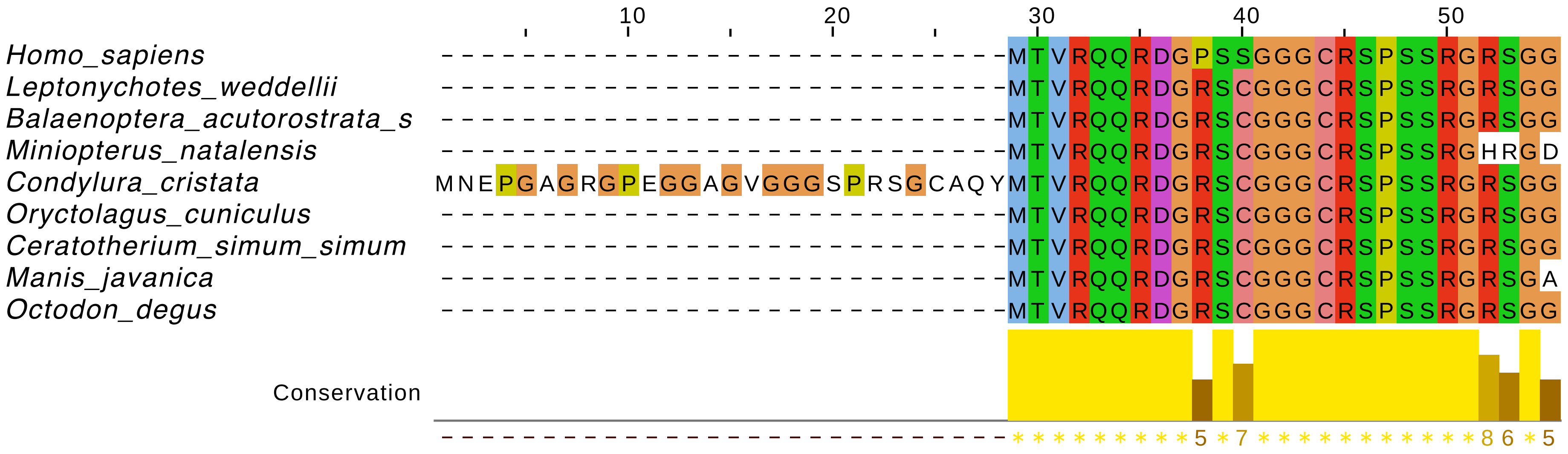

**HG0687.1**

ENSG00000120549 (KIAA1217)

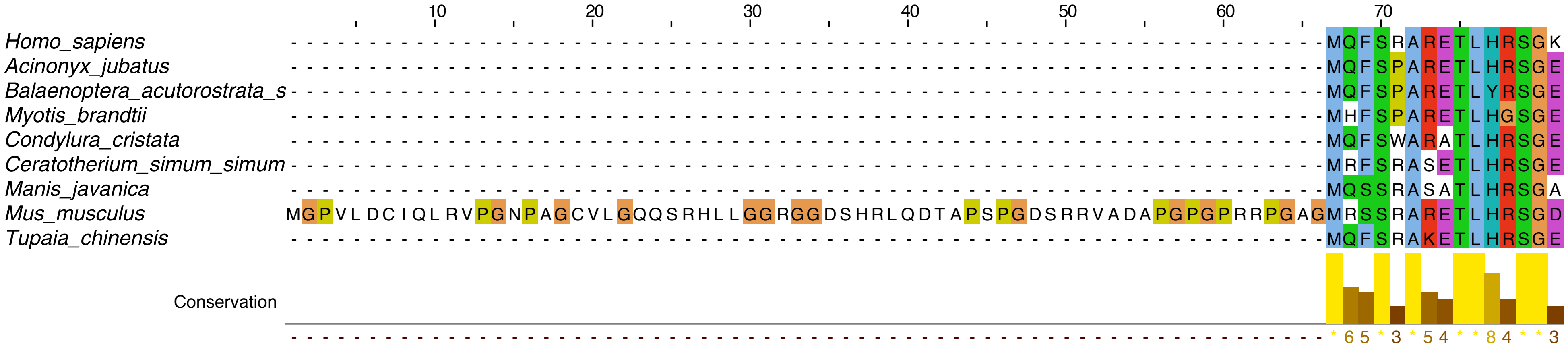

**HG0687.2**

ENSG00000120549 (KIAA1217)

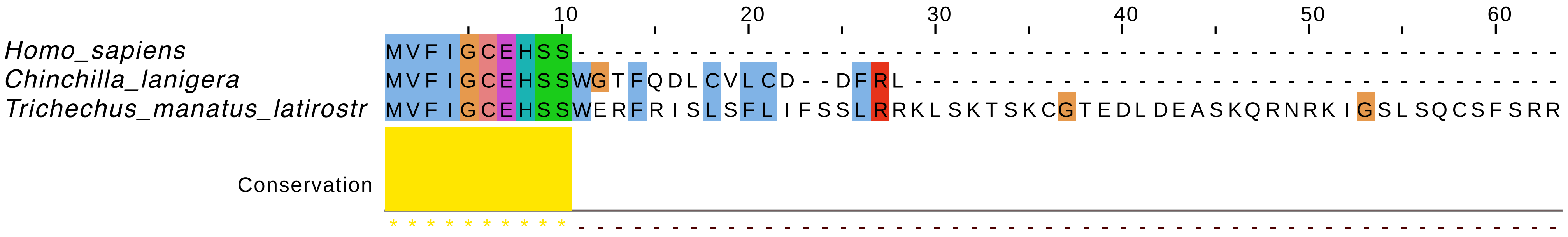

**HG0688**

ENSG00000122420 (PTGFR)

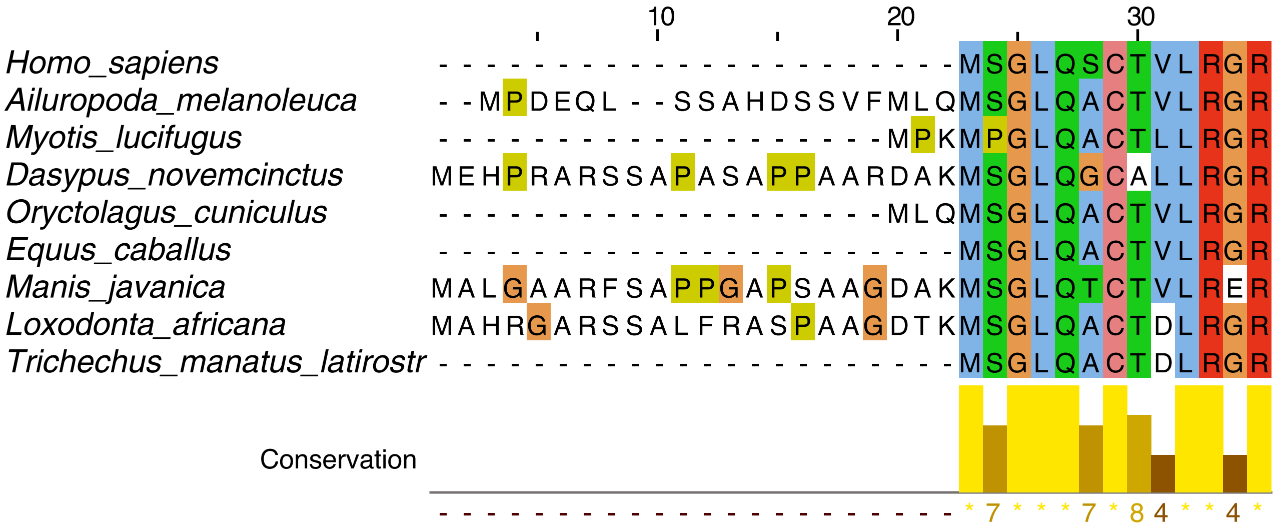

**HG0689**

ENSG00000123352 (SPATS2)

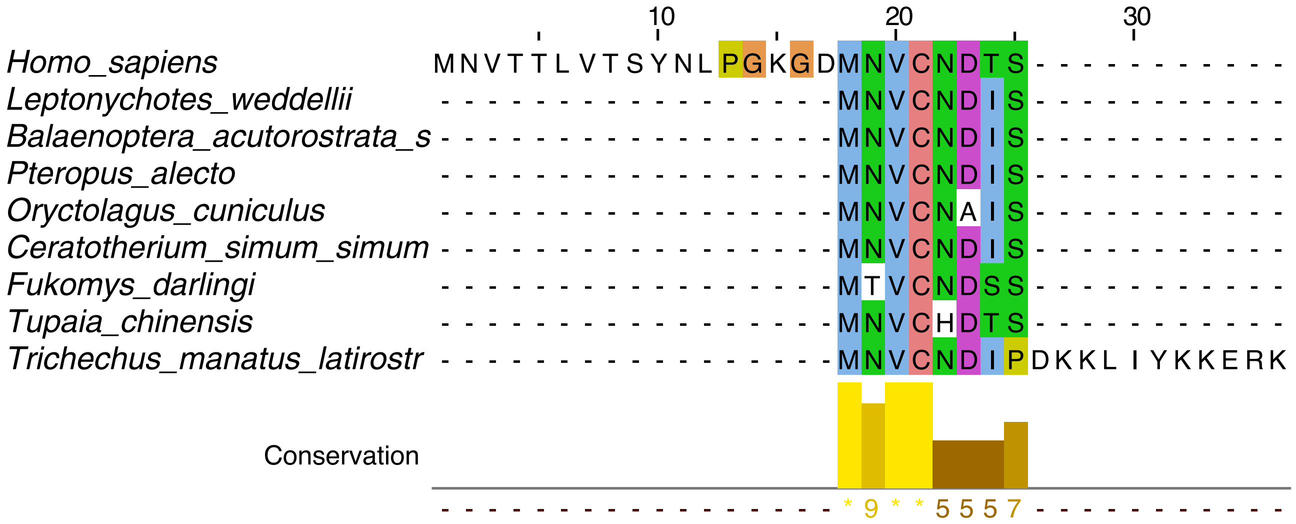

**HG0690**

ENSG00000125931 (CITED1)

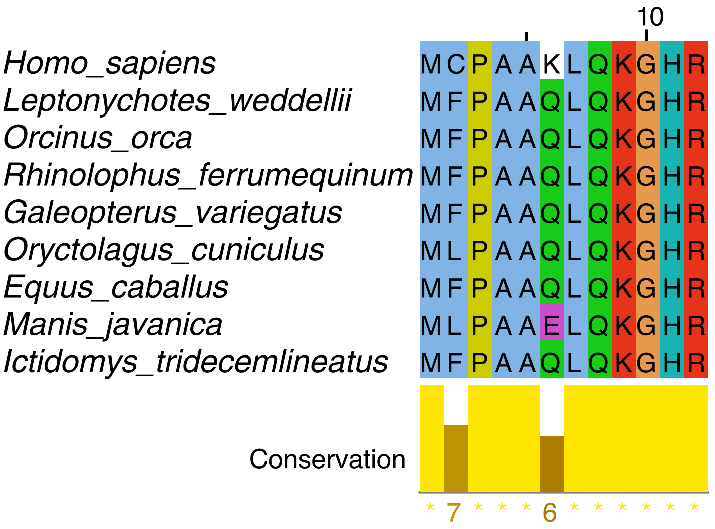

**HG0691**

ENSG00000129535 (NRL)

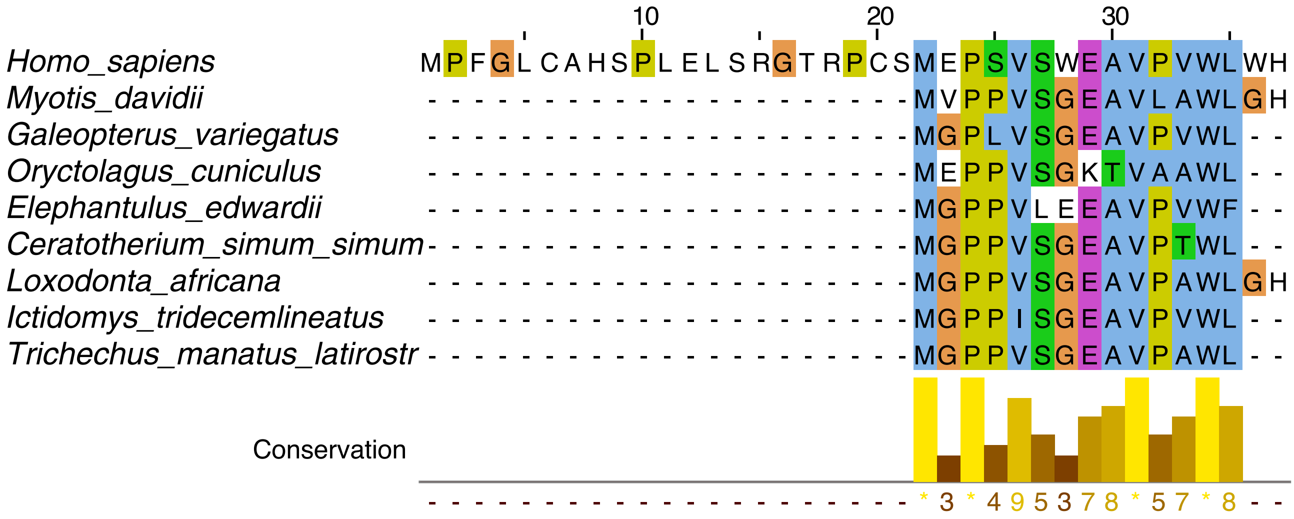

**HG0692.1**

ENSG00000130338 (TULP4)

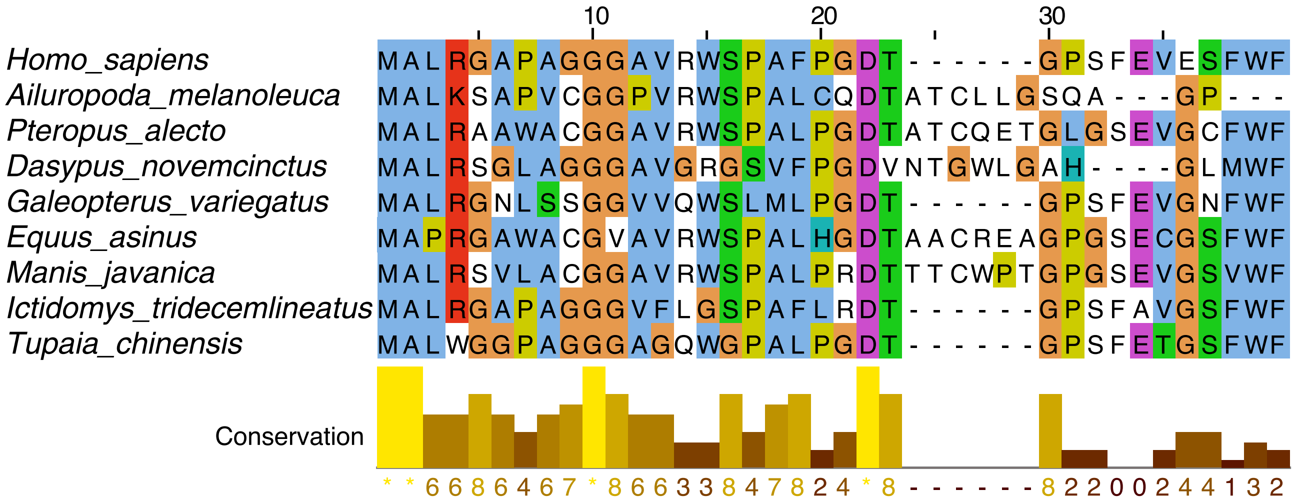

**HG0692.2**

ENSG00000130338 (TULP4)

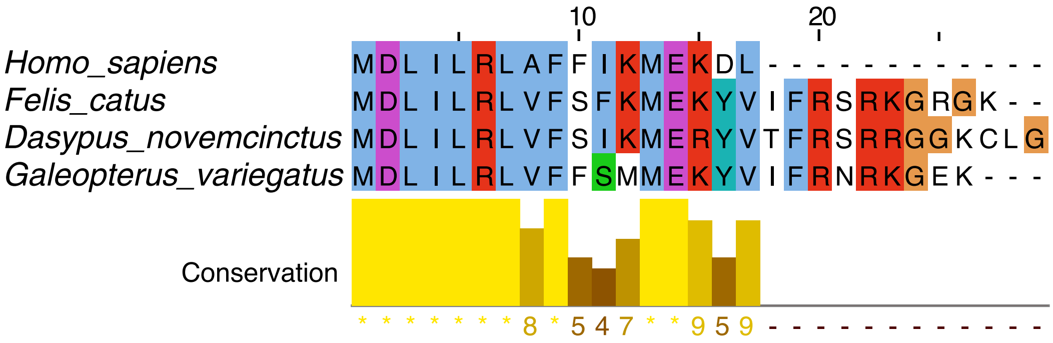

**HG0693**

ENSG00000131653 (TRAF7)

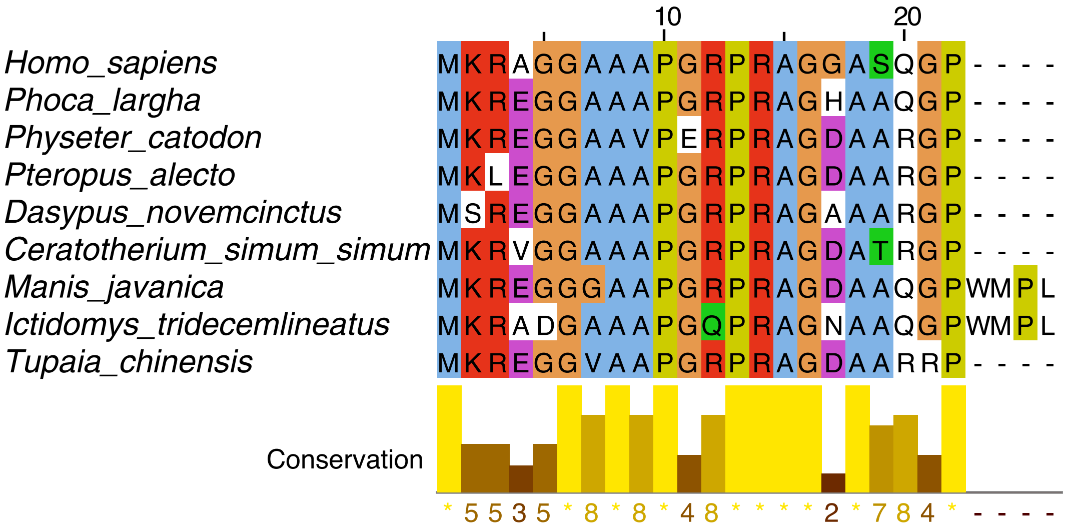

**HG0694**

ENSG00000133265 (HSPBP1)

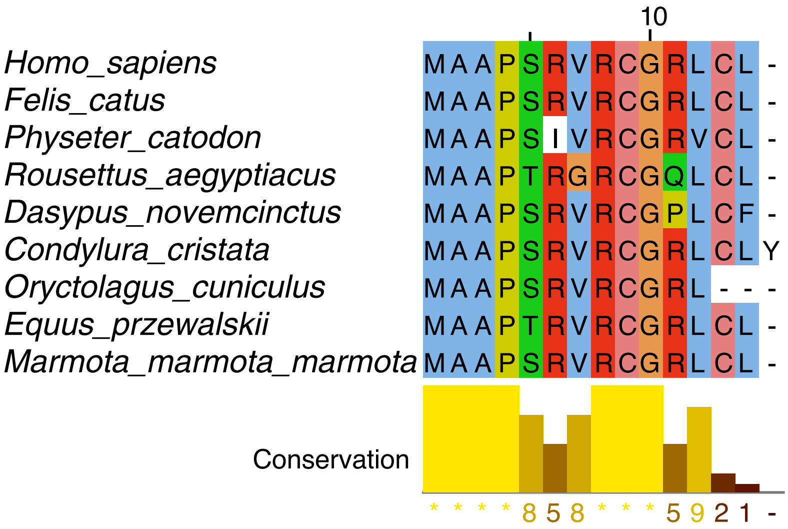

**HG0695**

ENSG00000133704 (IPO8)

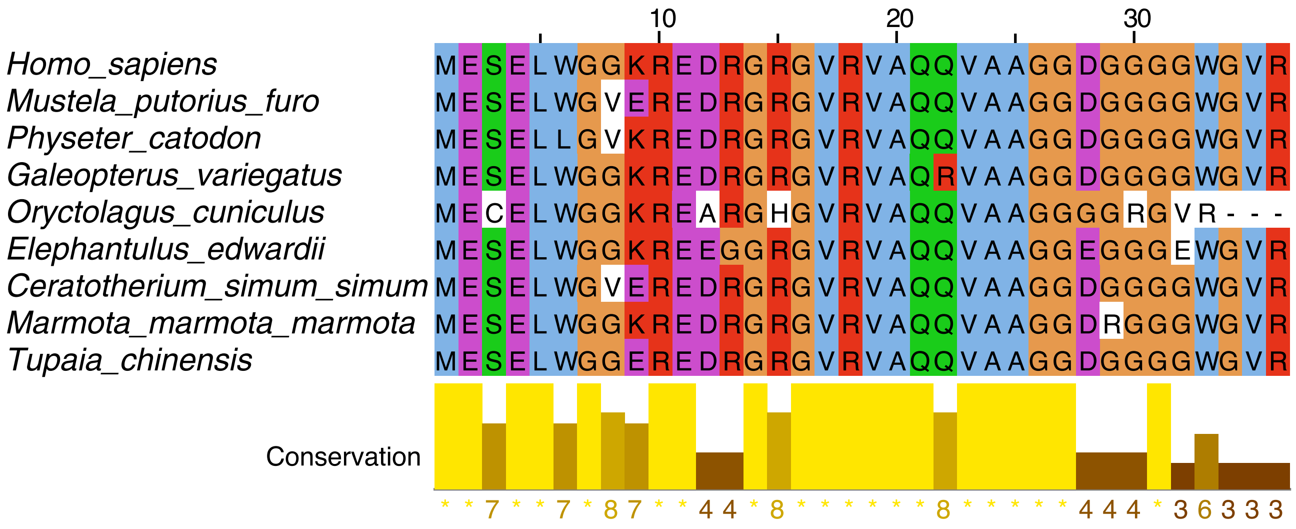

**HG0696**

ENSG00000138380 (CARF)

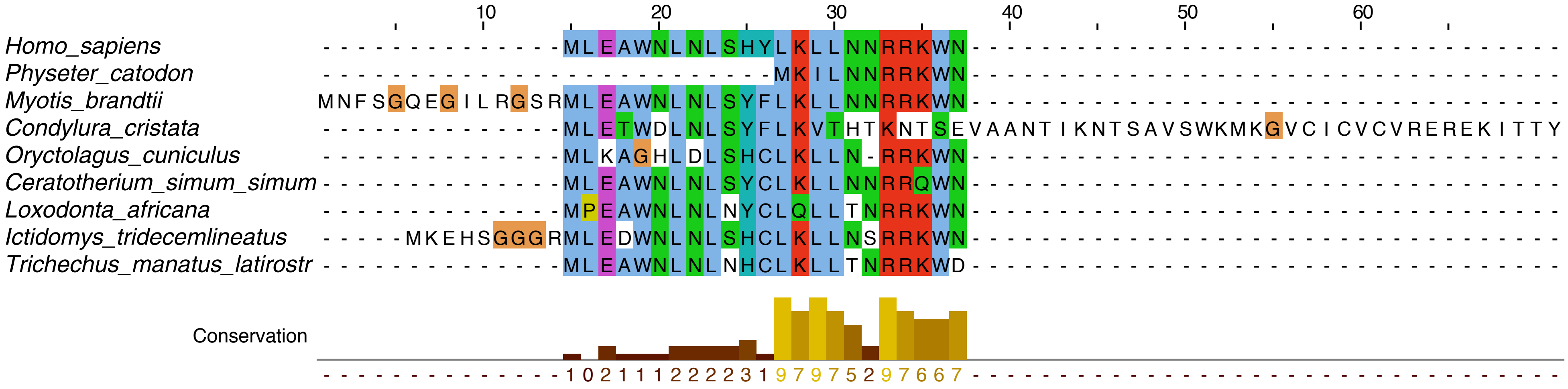

**HG0697**

ENSG00000143344 (RGL1)

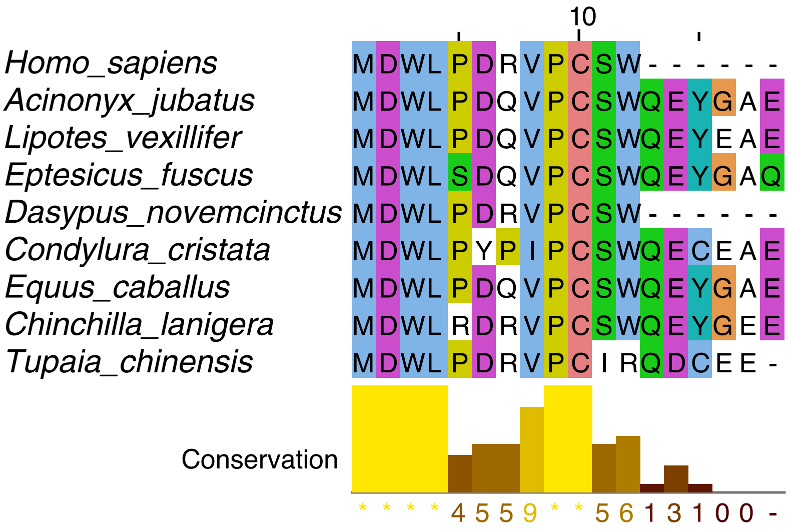

**HG0698**

ENSG00000157554 (ERG)

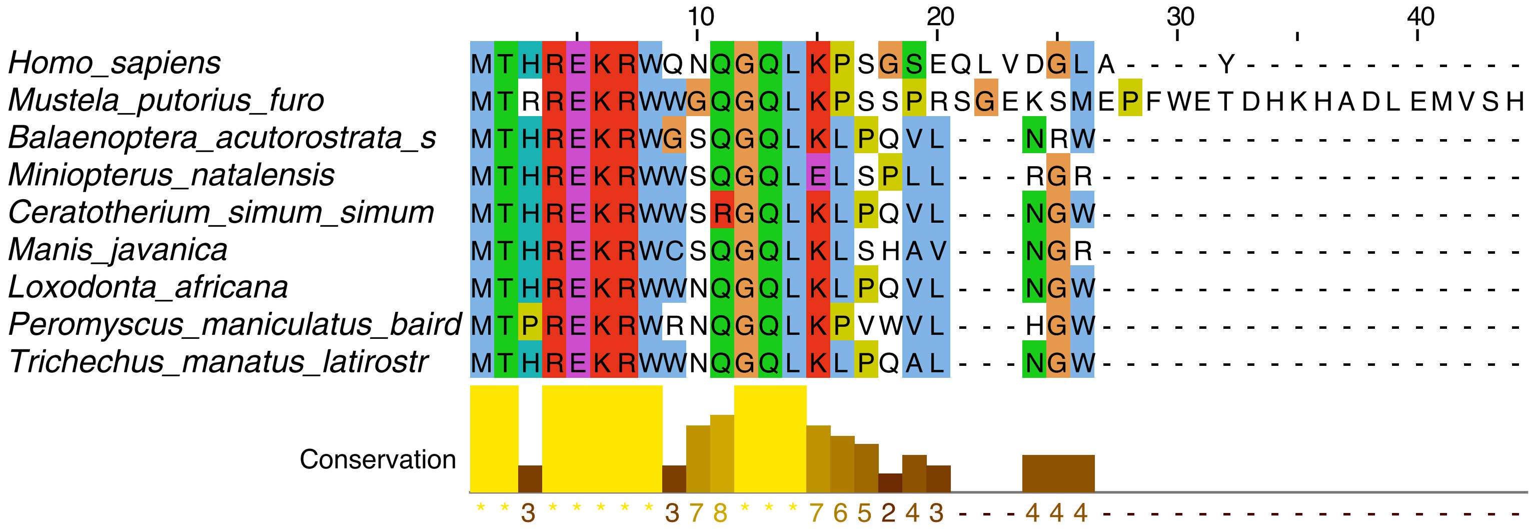

**HG0699**

ENSG00000160785 (SLC25A44)

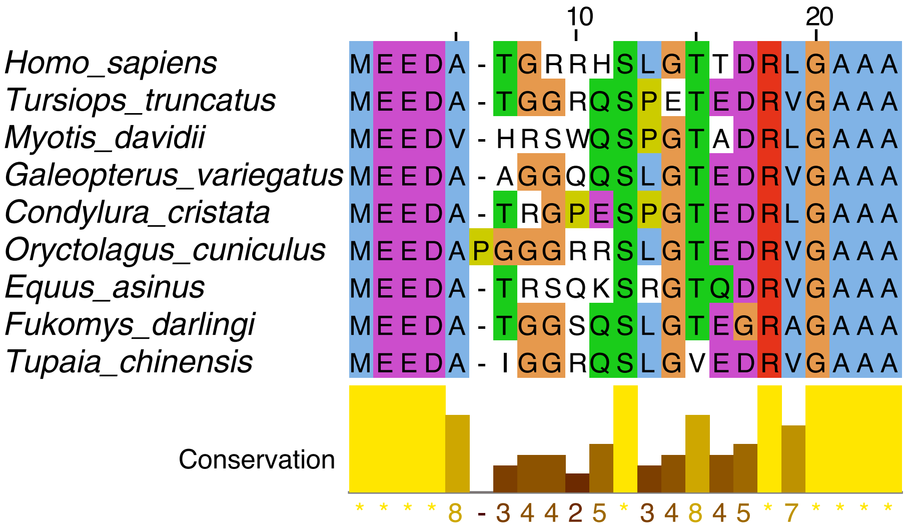

**HG0700**

ENSG00000165156 (ZHX1)

```
                                     10        20        30        40        50        60
Homo_sapiens                         MQDG-GGVALRGAAAPAGSGASRLRDQRARRLWAAGGNWSAESAAGTGKEKTDCCFGPVGNDGLS
Odobenus_rosmarus_divergens          --------------------------------------------------MADRCFRPVGNDGLS
Lipotes_vexillifer                   MQDG-GGGALRGTAAPAGDGR--------------------------------------------
Myotis_ricketti                      --------------------------------------------------MADQCFGPAGNDGLS
Erinaceus_europaeus                  MQDG-GGRALRGAAAPAGHGR--------------------------------------------
Ceratotherium_simum_simum            --------------------------------------------------MADRYFGPVGNDGLS
Manis_javanica                       --------------------------------------------------MADRYFGPVGNDGLS
Ictidomys_tridecemlineatus           MQDG-GGGVLRGAAAPAGED---------------------------------------------
Trichechus_manatus_latirostr         MQDGDGGGALRGAADPAGESR--------------------------------------------

Conservation
                                     -----------------------------------------------------------------
```

**HG0701**

ENSG00000166292 (TMEM100)

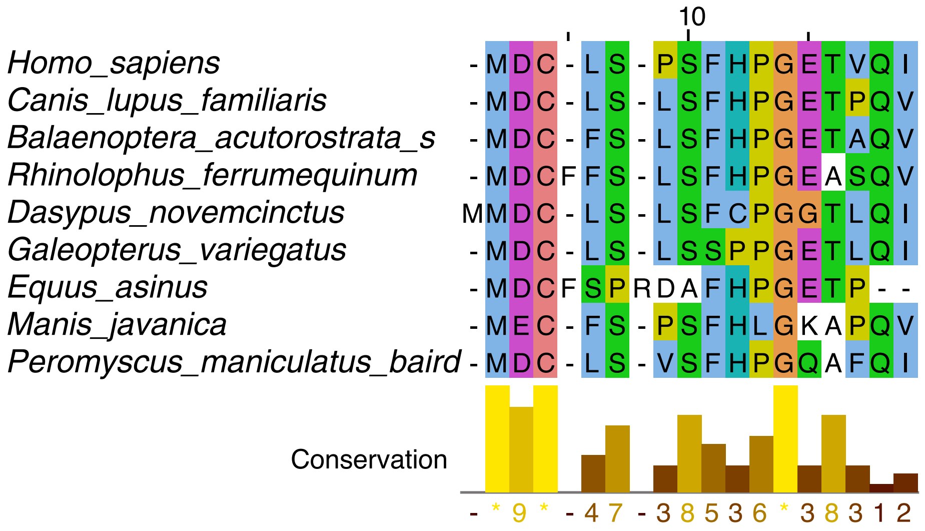

**HG0702**

ENSG00000167377 (ZNF23)

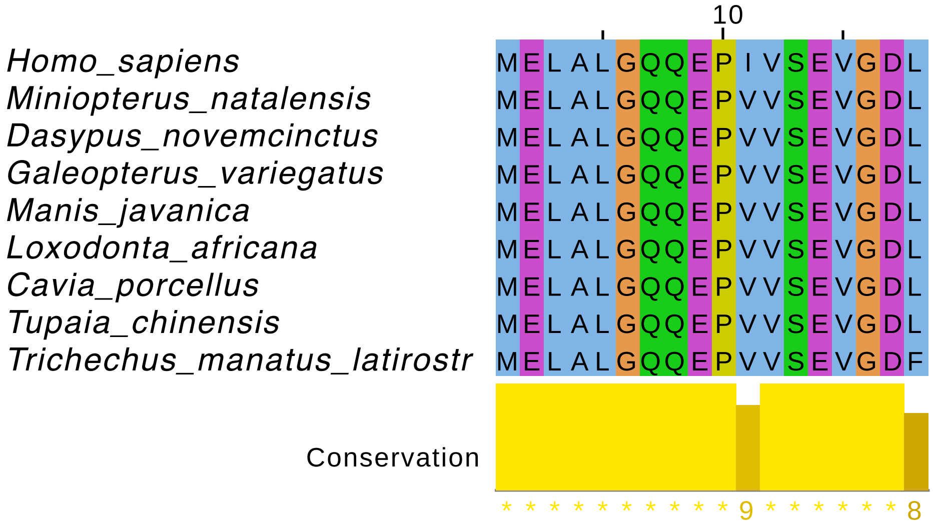

**HG0703**

ENSG00000171862 (PTEN)

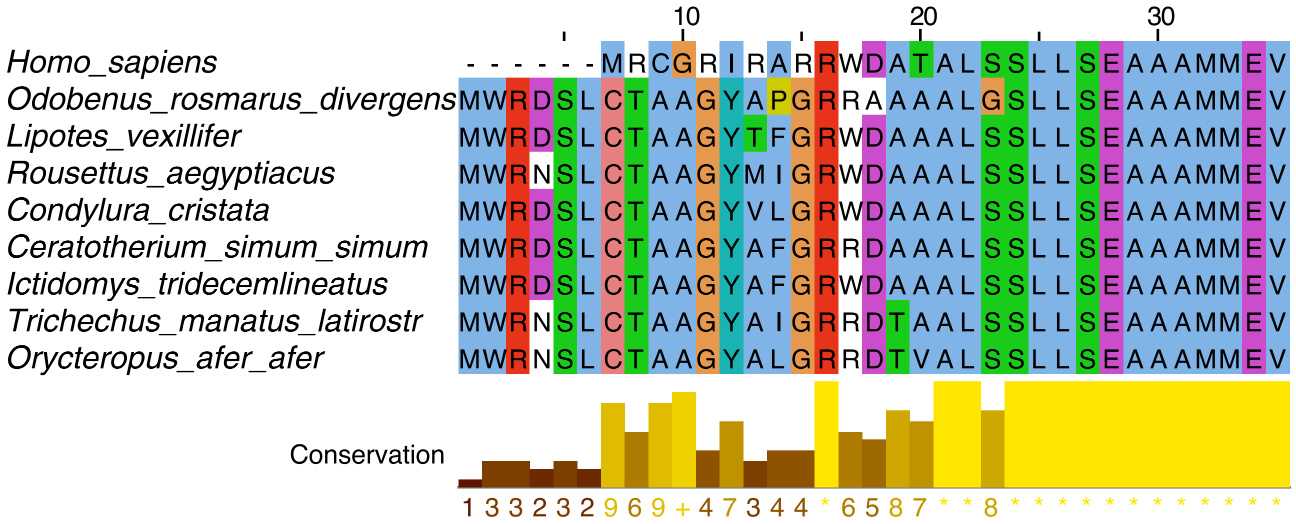

**HG0704.1**

ENSG00000173598 (NUDT4)

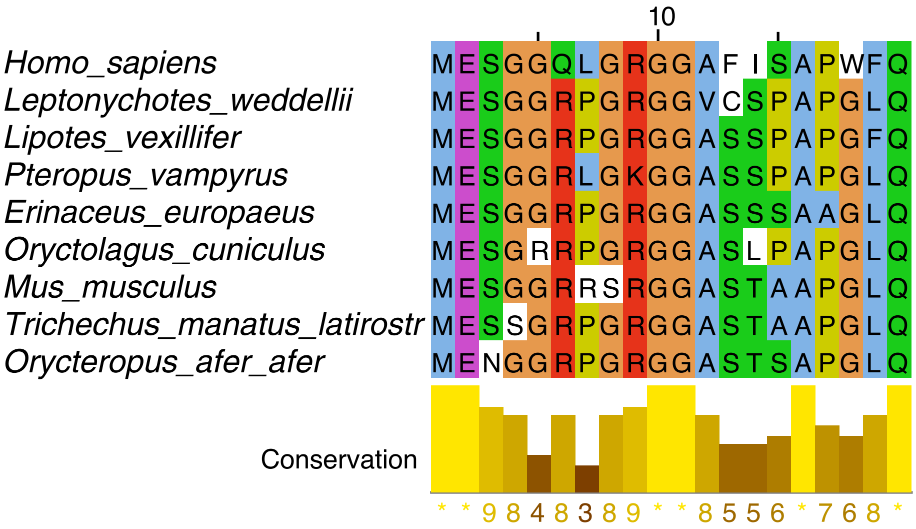

**HG0704.2**

ENSG00000272325 (NUDT3)

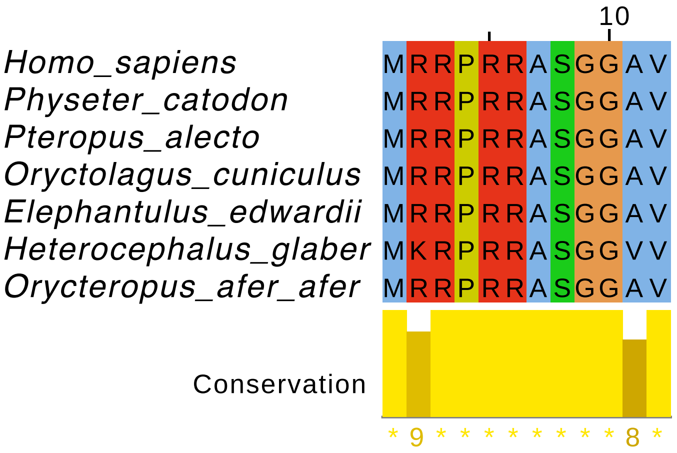

**HG0705.1**

ENSG00000175224 (ATG13)

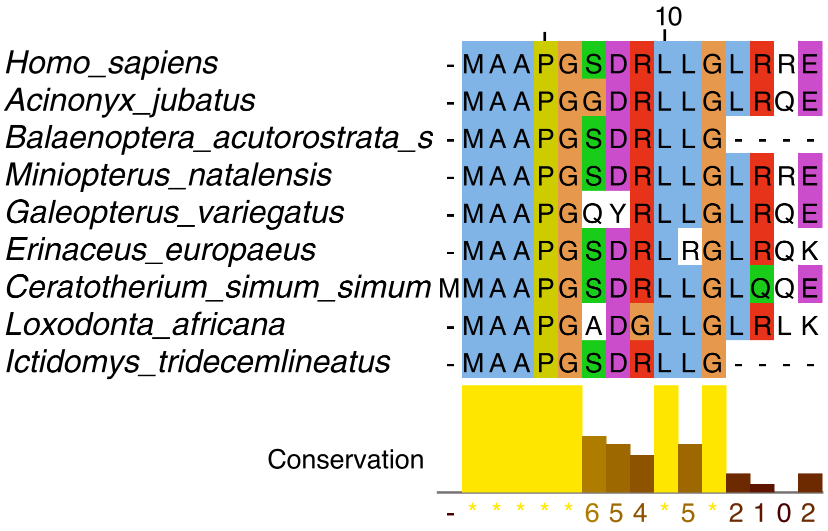

**HG0705.2**

ENSG00000175224 (ATG13)

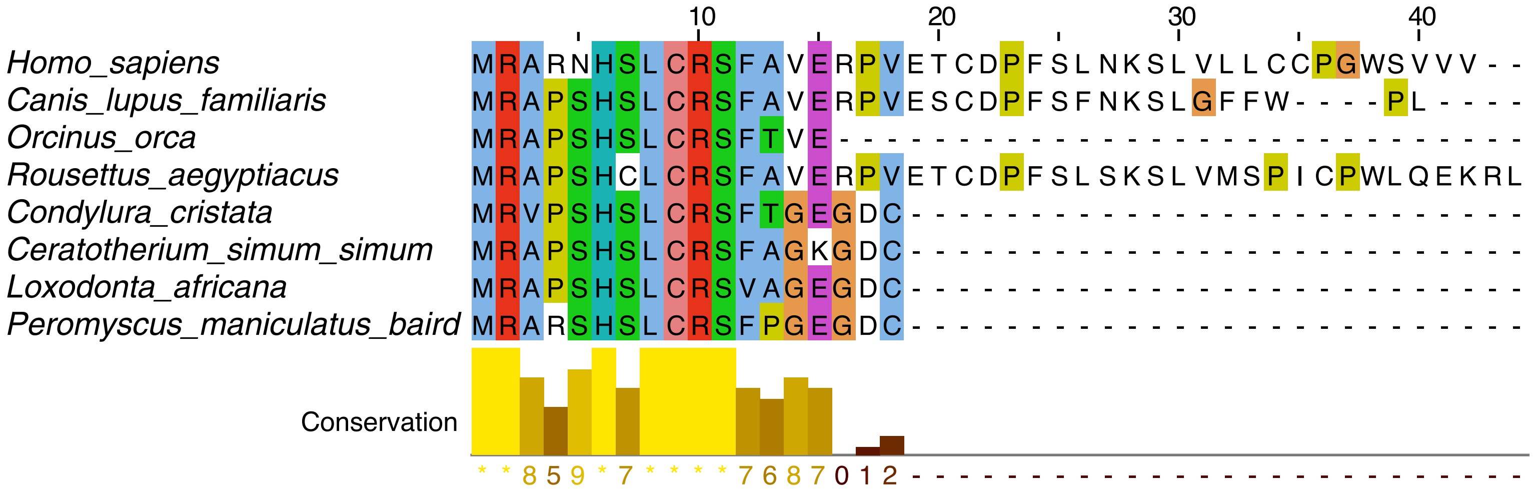

**HG0706**

ENSG00000178498 (DTX3)

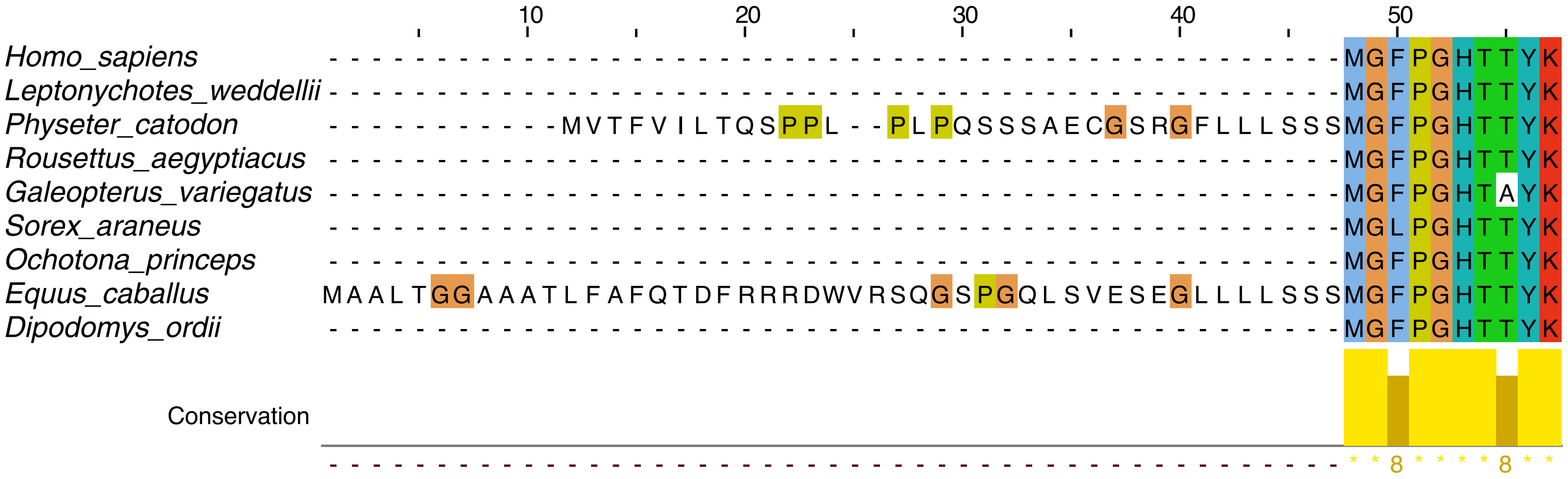

**HG0707**

ENSG00000179796 (LRRC3B)

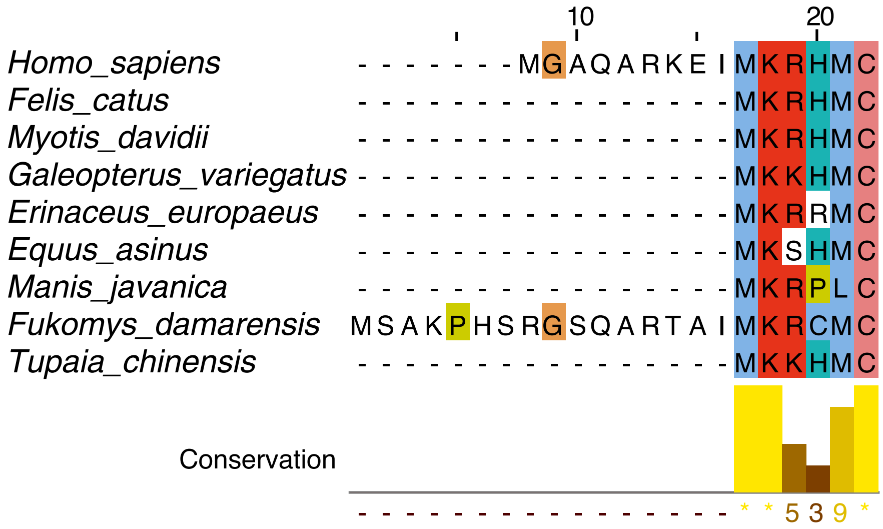

**HG0708**

ENSG00000184613 (NELL2)

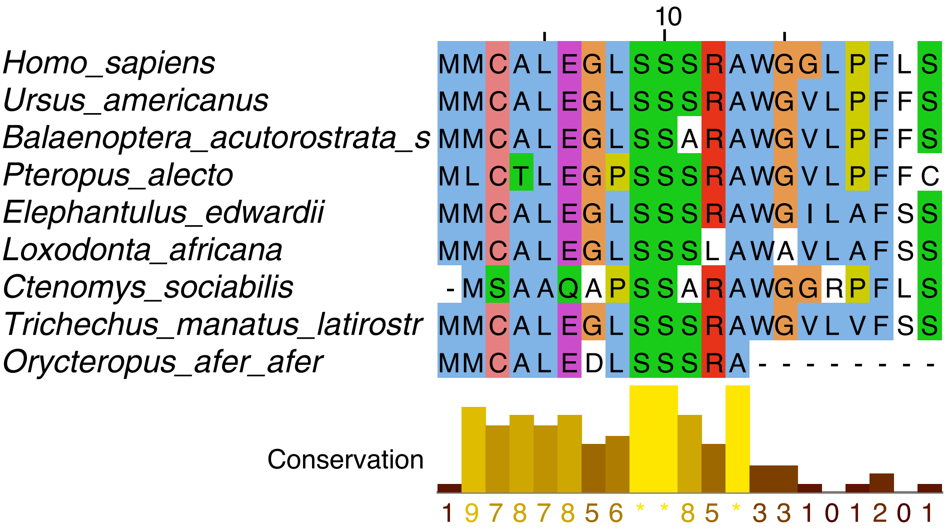

**HG0709**

ENSG00000186439 (TRDN)

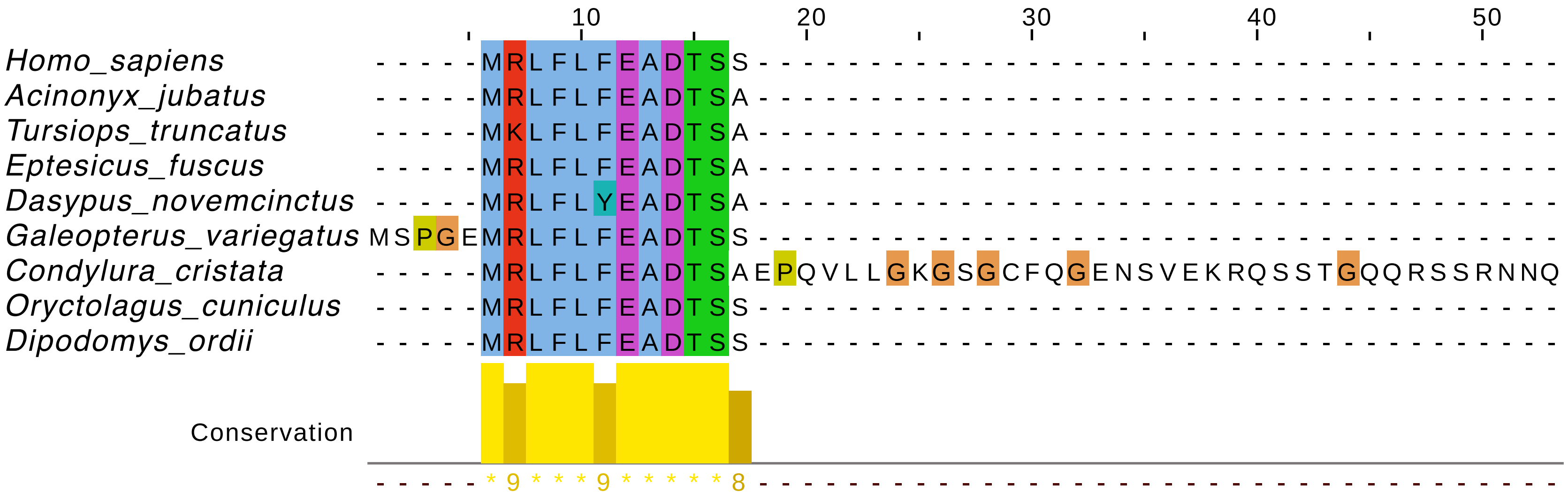

**HG0710**

ENSG00000196505 (GDAP2)

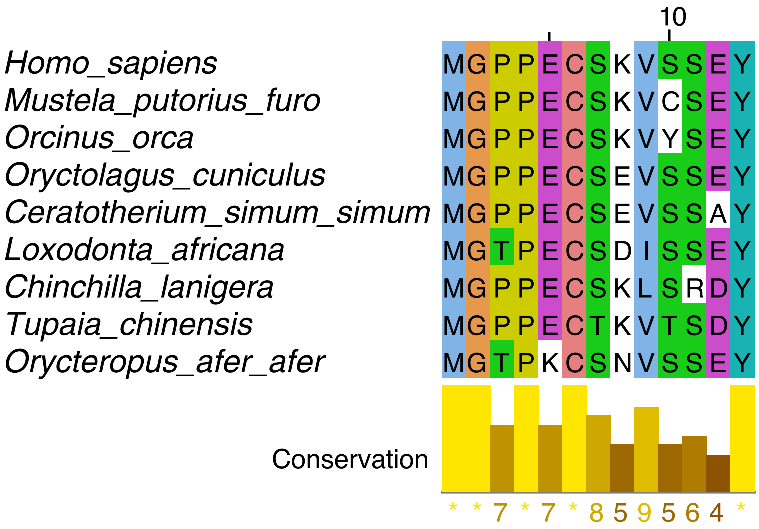

**HG0711**

ENSG00000198300 (PEG3)

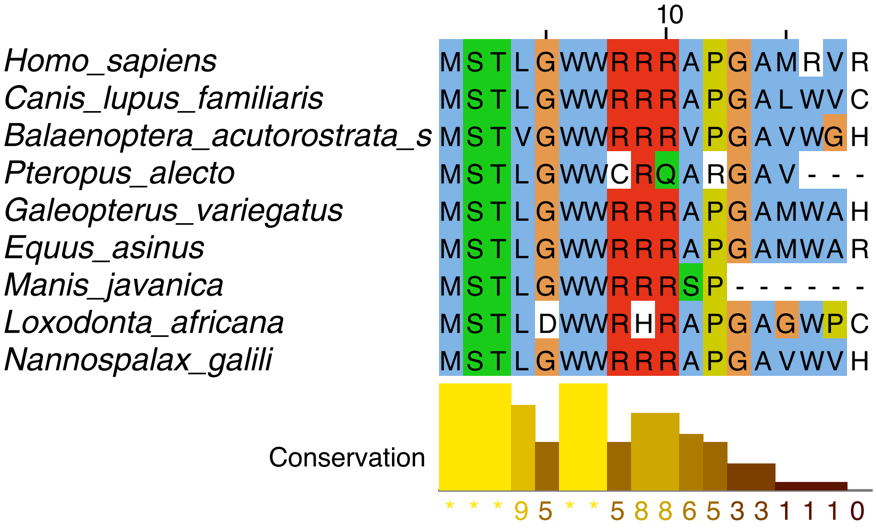

**HG0712**

ENSG00000213949 (ITGA1)

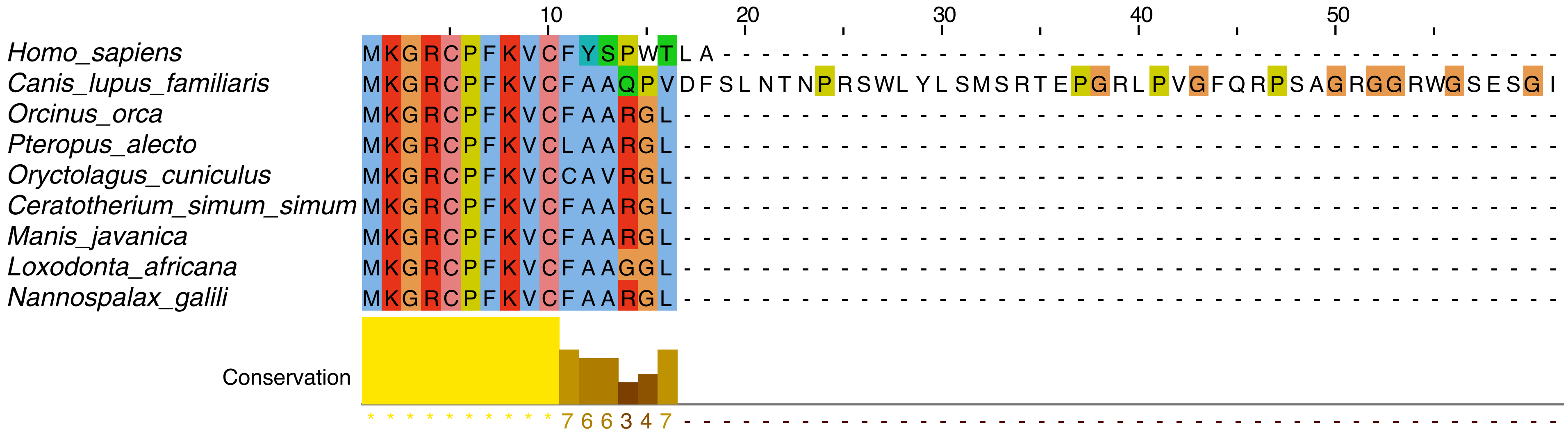

**HG0713**

ENSG00000008513 (ST3GAL1)

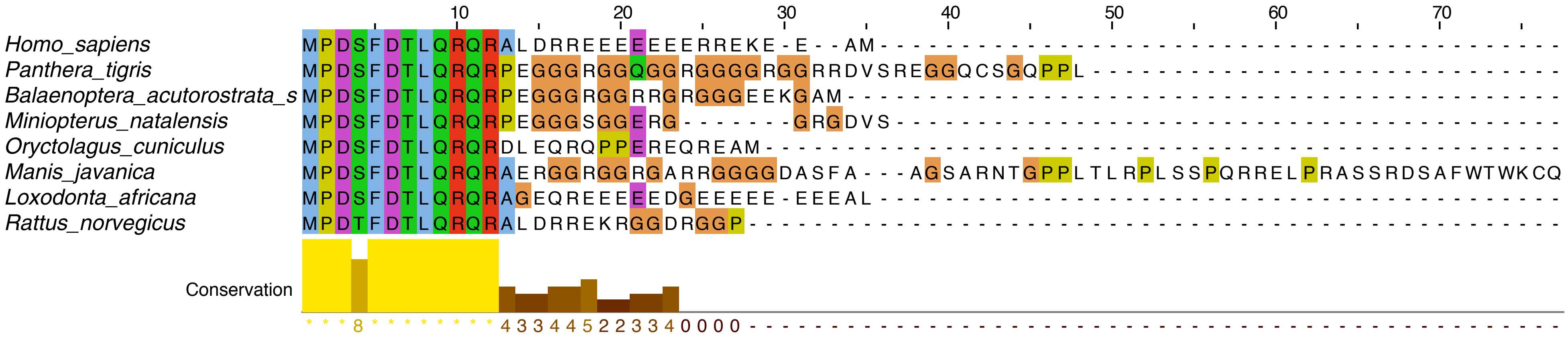

**HG0714**

ENSG00000051825 (MPHOSPH9)

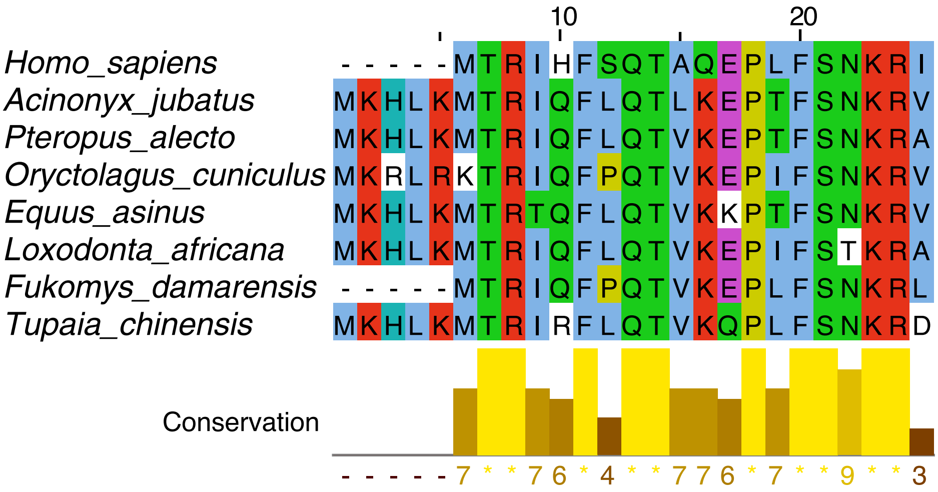

**HG0715**

ENSG00000071246 (VASH1)

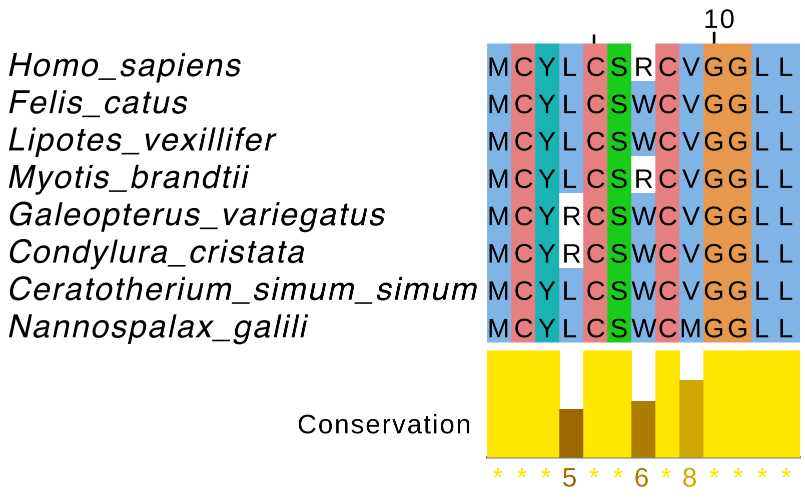

**HG0716**

ENSG00000075413 (MARK3)

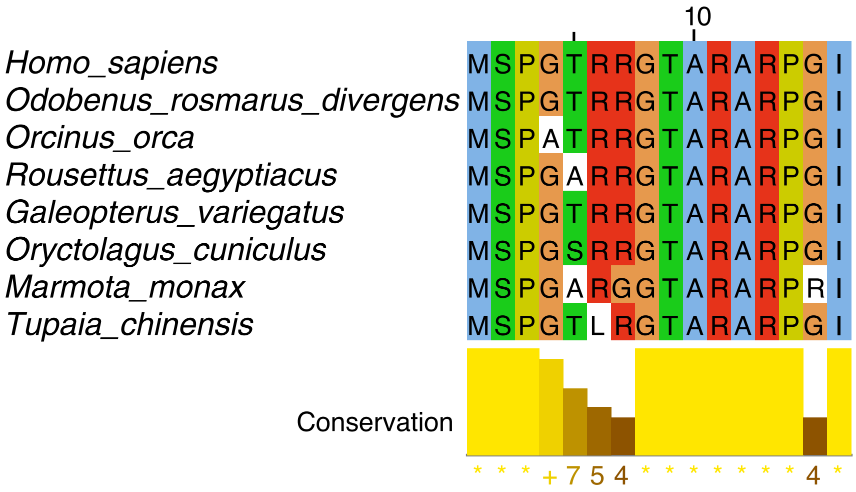

**HG0717**

ENSG00000083857 (FAT1)

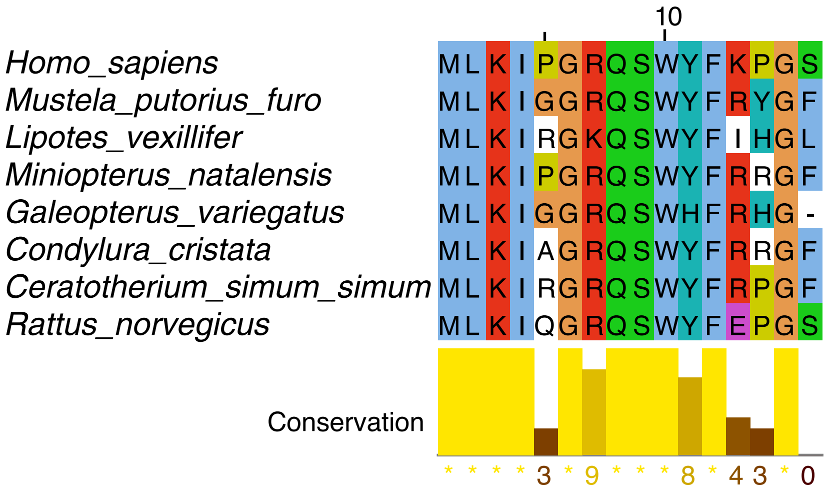

## HG0718

ENSG00000085274 (MYNN)

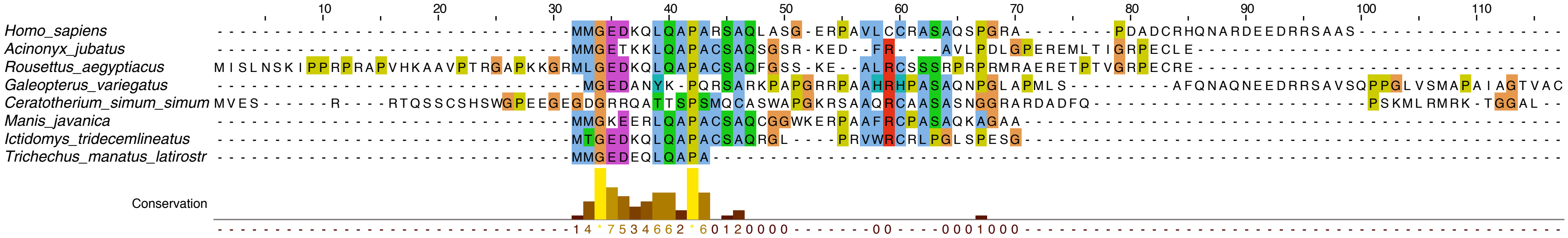

**HG0719**

ENSG00000085998 (POMGNT1)

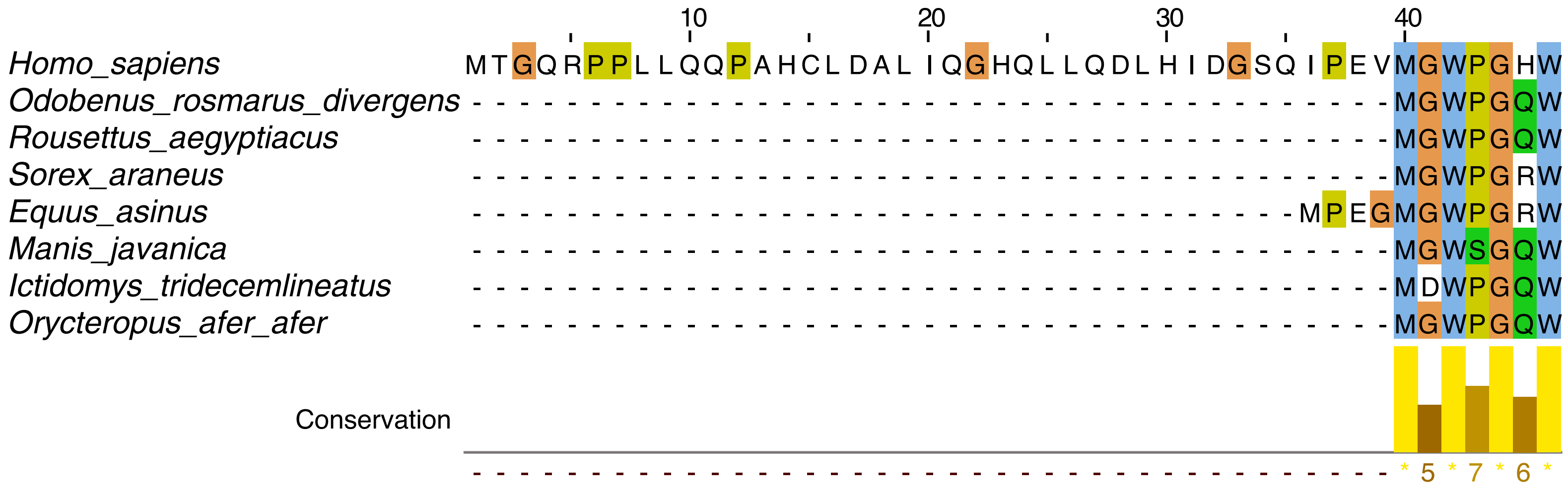

**HG0720**

ENSG00000087053 (MTMR2)

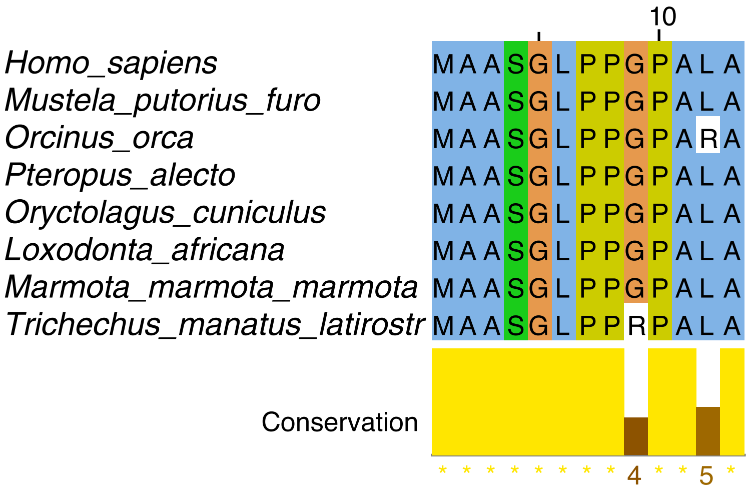

**HG0721**

ENSG00000095951 (HIVEP1)

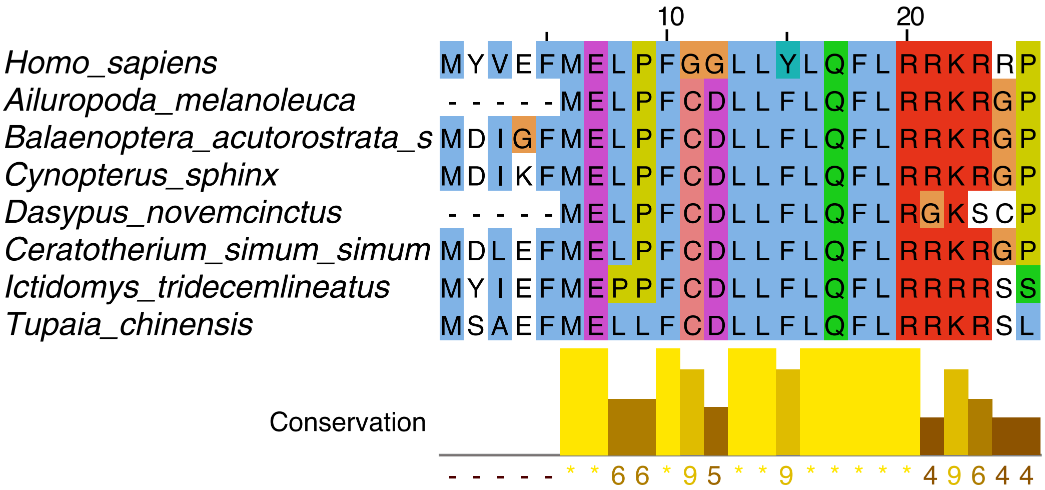

**HG0722**

ENSG00000101445 (PPP1R16B)

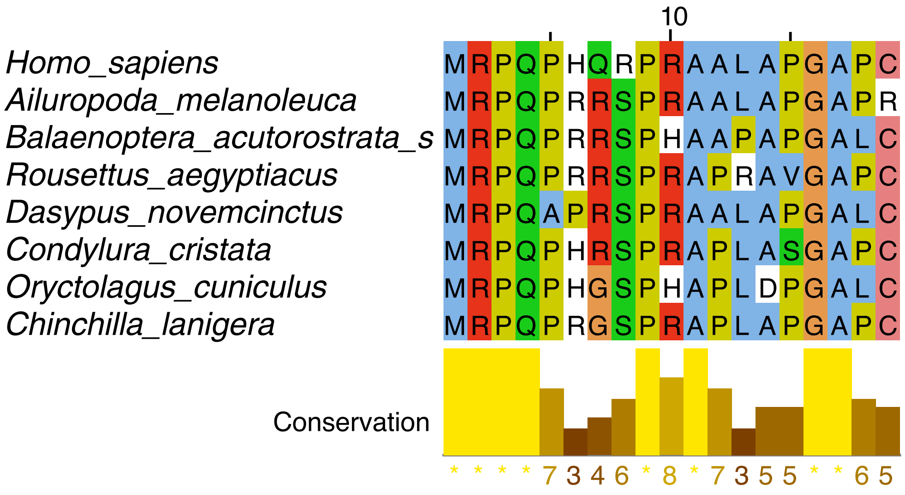

# HG0723

### ENSG00000102974 (CTCF)

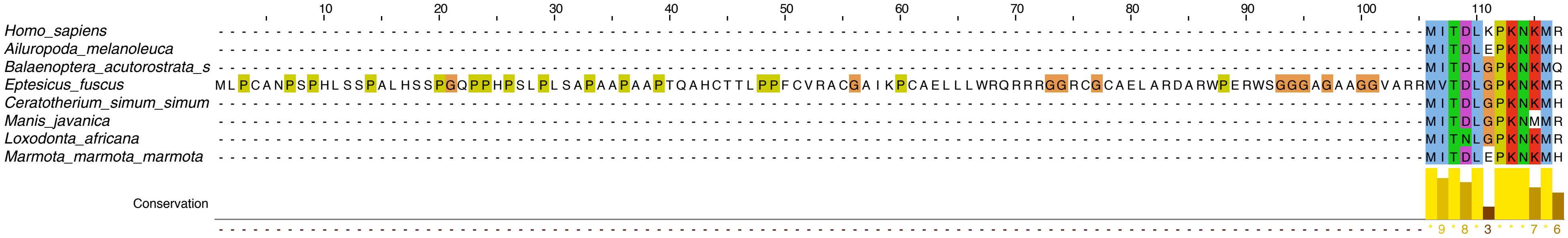

## HG0724

ENSG00000104783 (KCNN4)

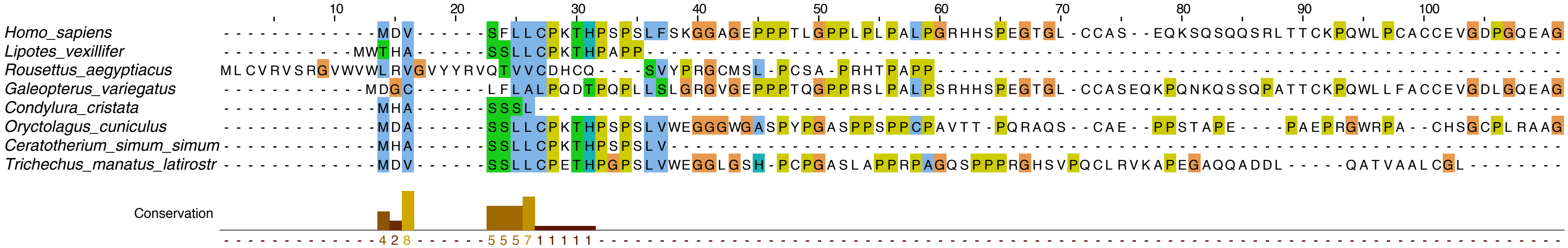

**HG0725**

ENSG00000112218 (GPR63)

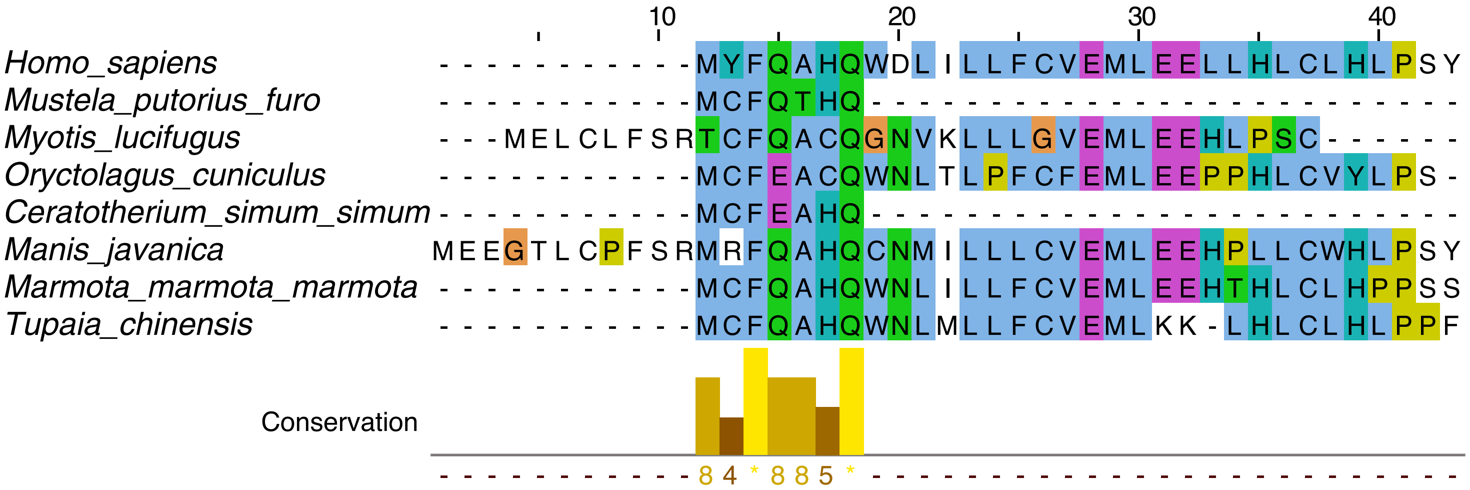

**HG0726.1**

ENSG00000115504 (EHBP1)

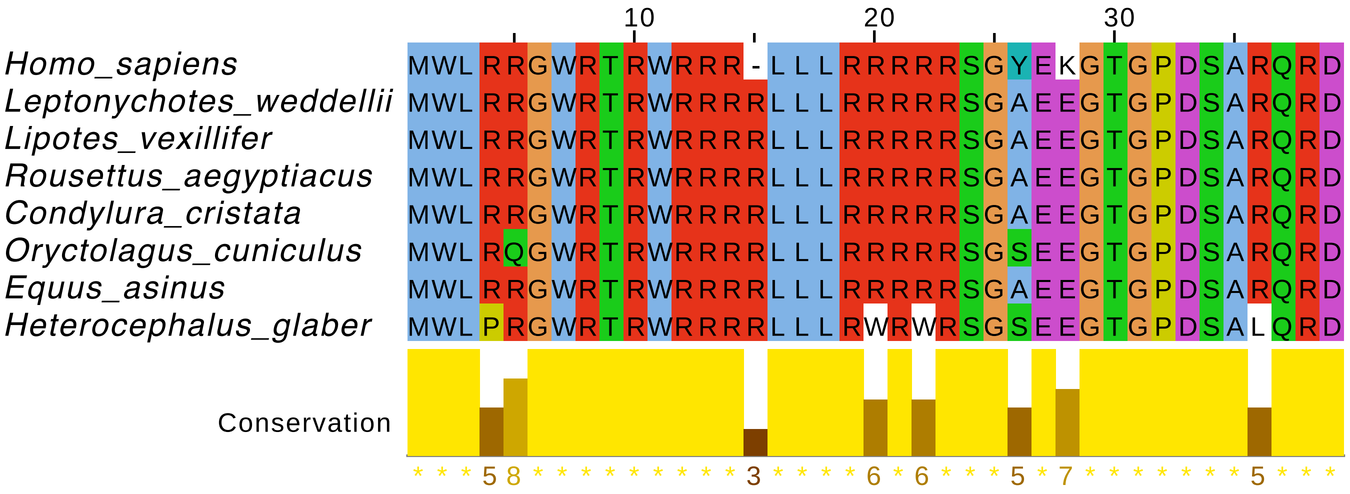

**HG0726.2**

ENSG00000115504 (EHBP1)

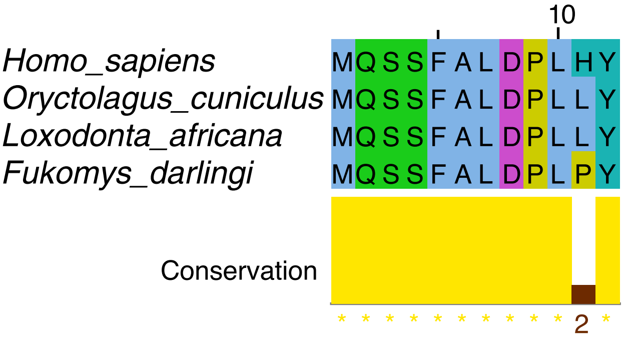

**HG0727**

ENSG00000115825 (PRKD3)

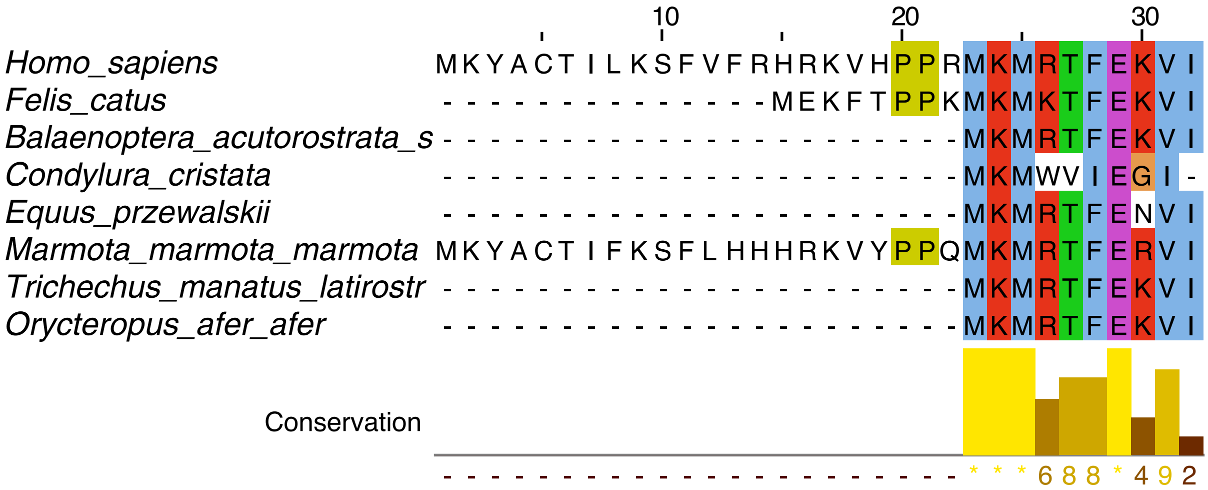

**HG0728**

ENSG00000116991 (SIPA1L2)

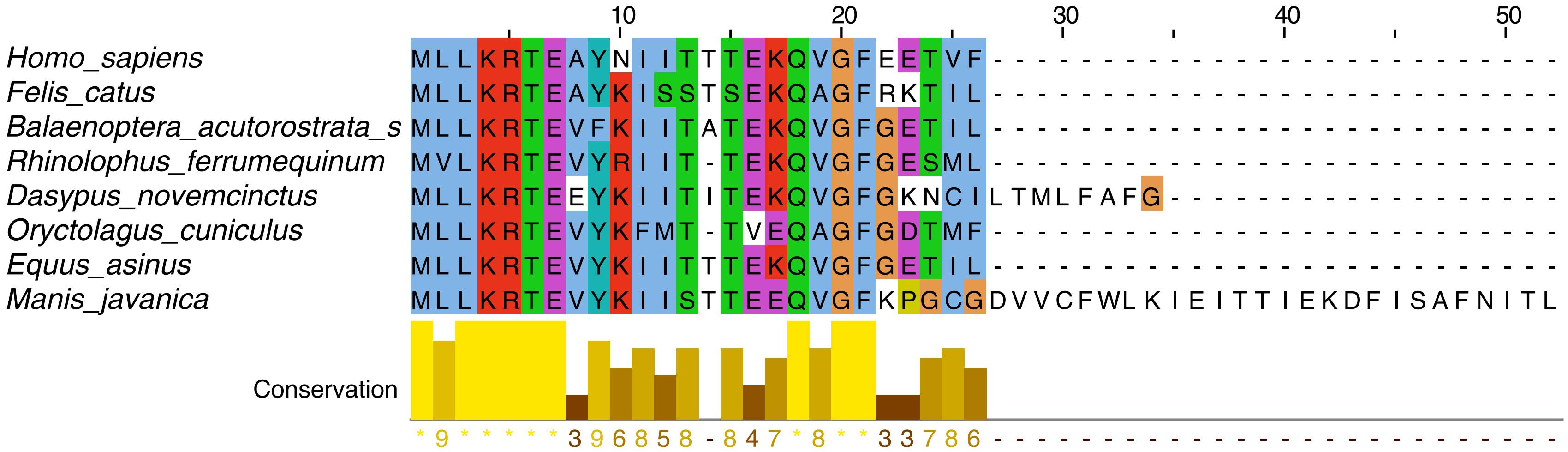

**HG0729**

ENSG00000119383 (PTPA)

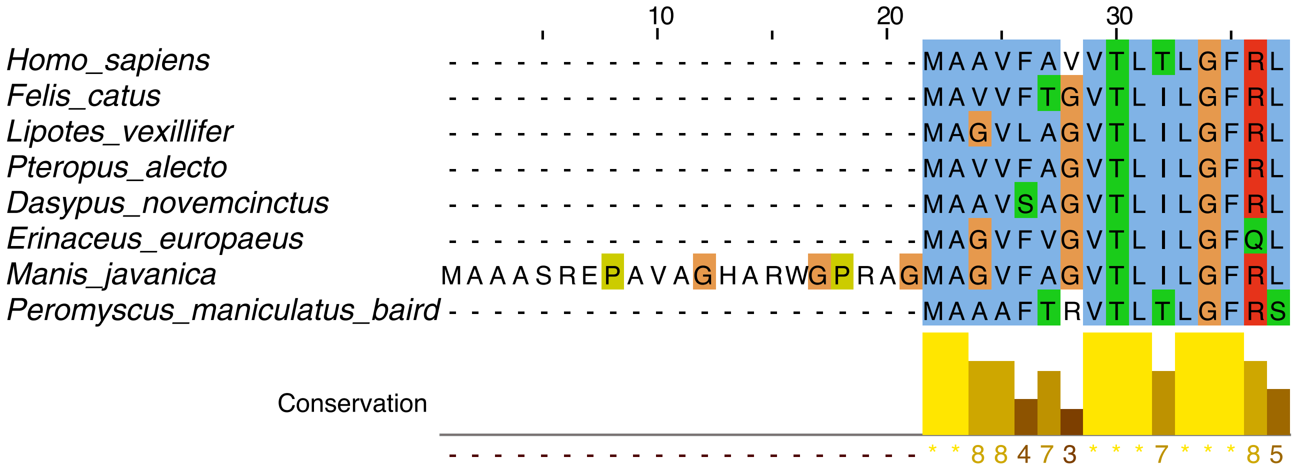

**HG0730**

ENSG00000125170 (DOK4)

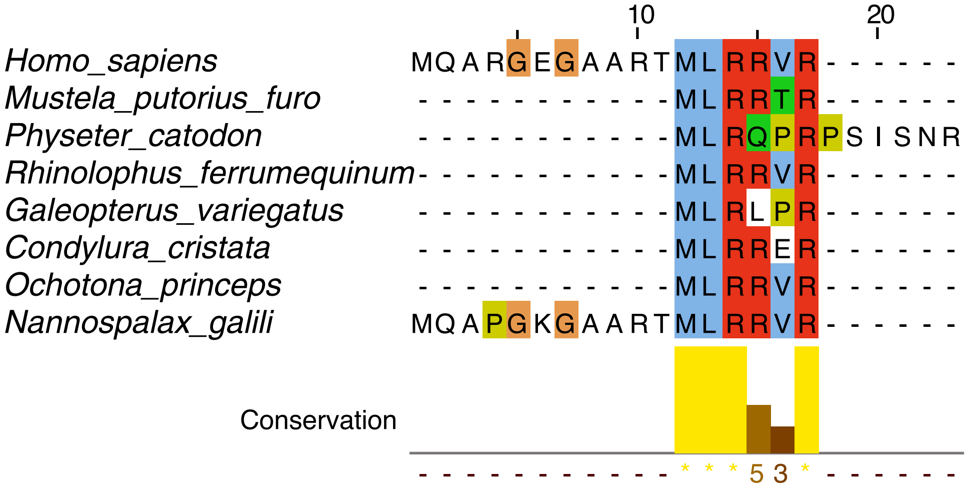

**HG0731**

ENSG00000129255 (MPDU1)

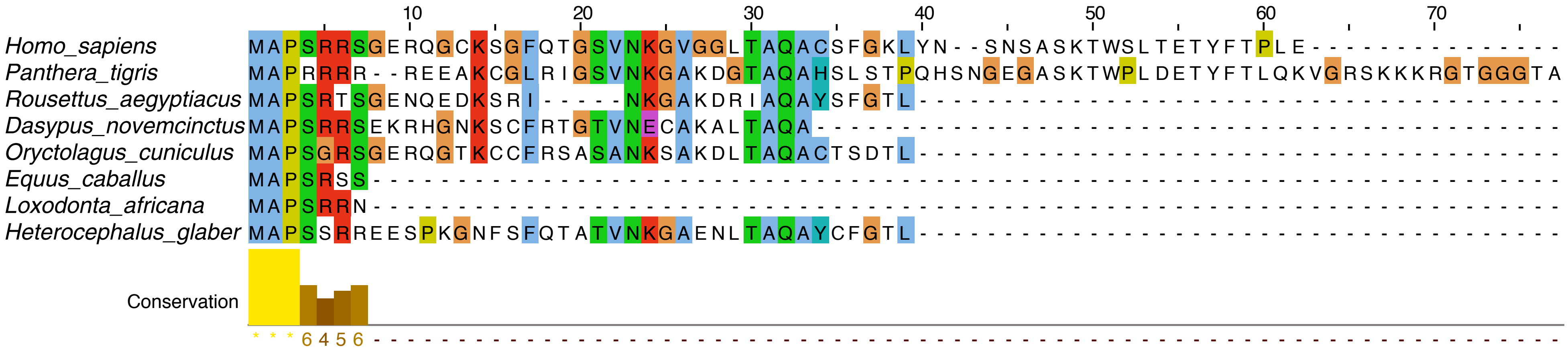

**HG0732**

ENSG00000135914 (HTR2B)

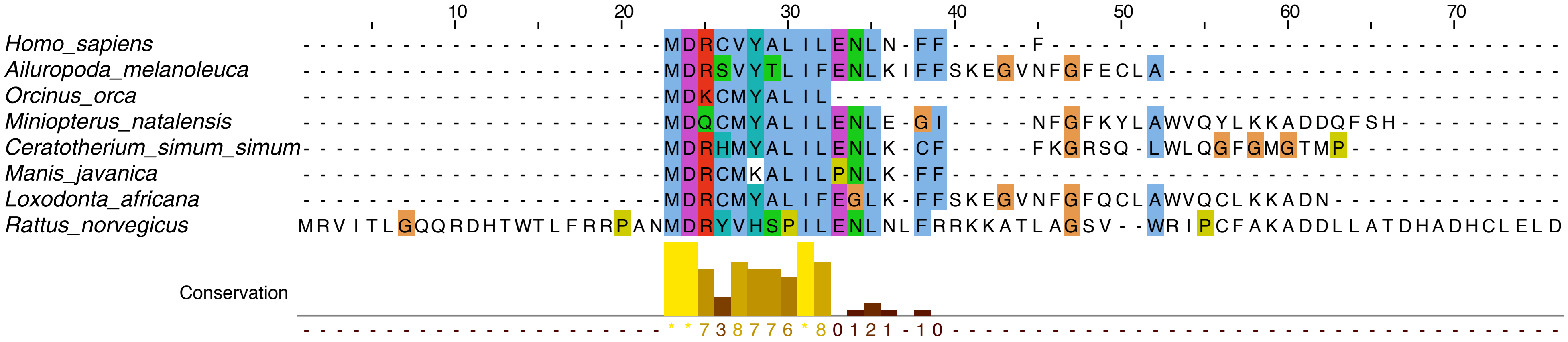

**HG0733**

ENSG00000136247 (ZDHHC4)

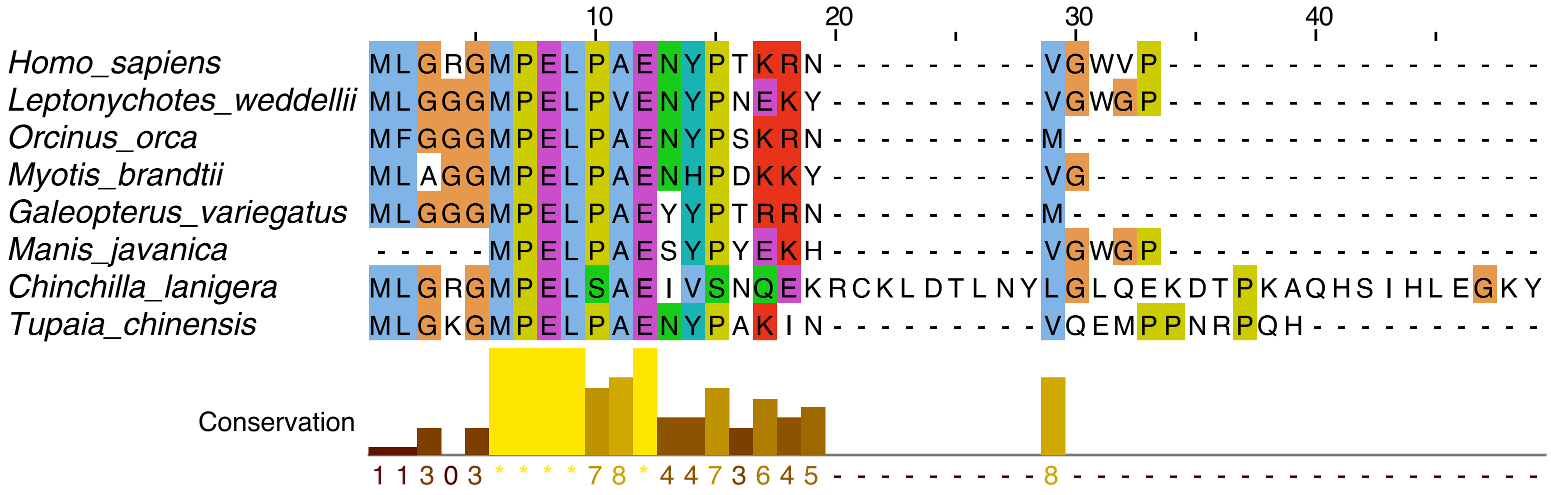

**HG0734**

ENSG00000139318 (DUSP6)

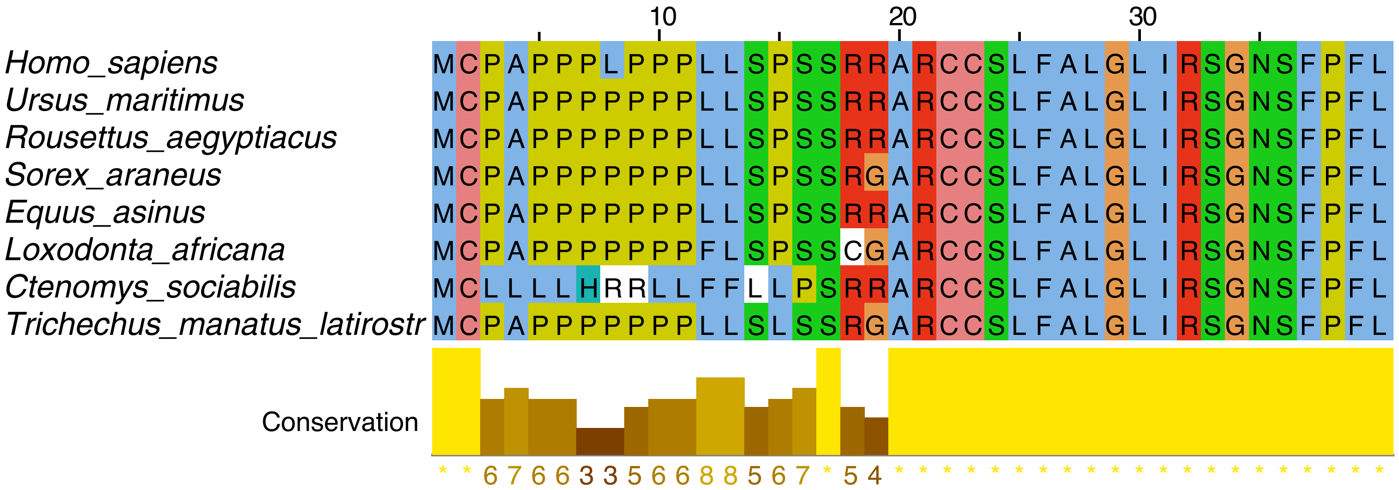

**HG0735**

ENSG00000139579 (NABP2)

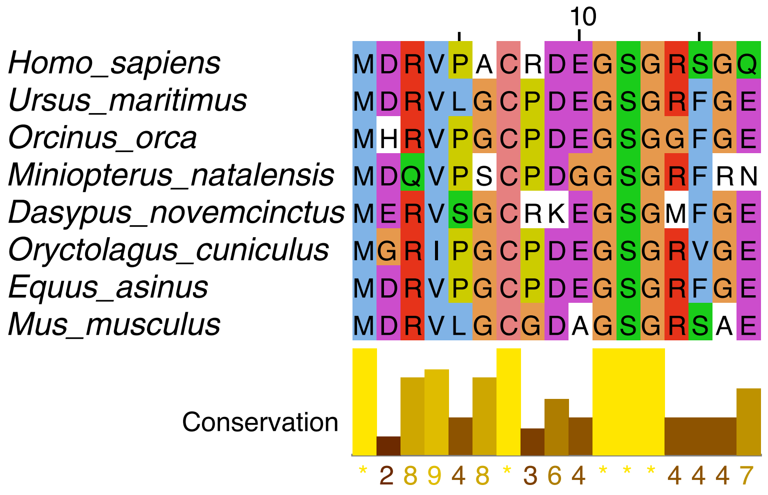

**HG0736**

ENSG00000139826 (ABHD13)

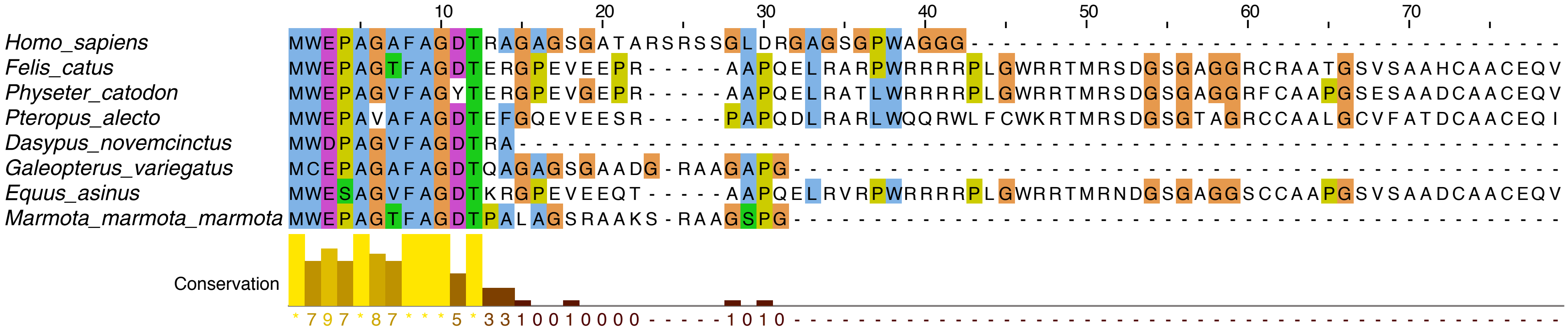

**HG0737**

ENSG00000141639 (MAPK4)

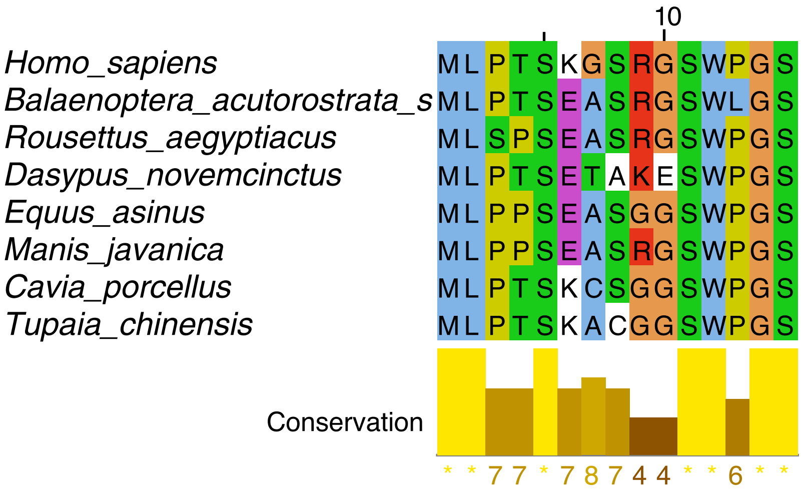

**HG0738**

ENSG00000143393 (PI4KB)

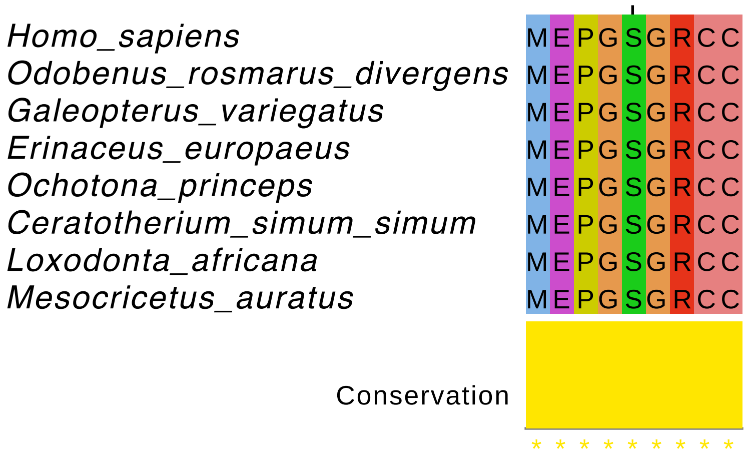

**HG0739**

ENSG00000144331 (ZNF385B)

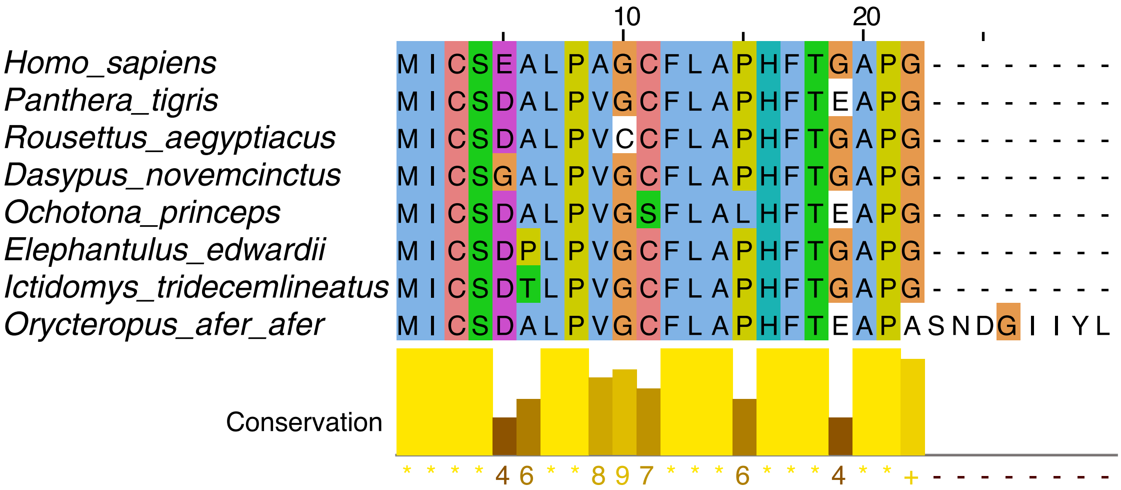

**HG0740**

ENSG00000144331 (ZNF385B)

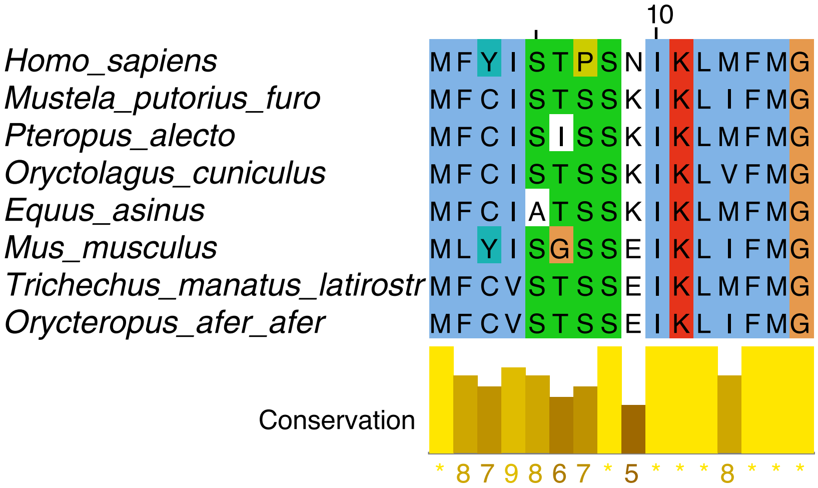

**HG0741**

ENSG00000144834 (TAGLN3)

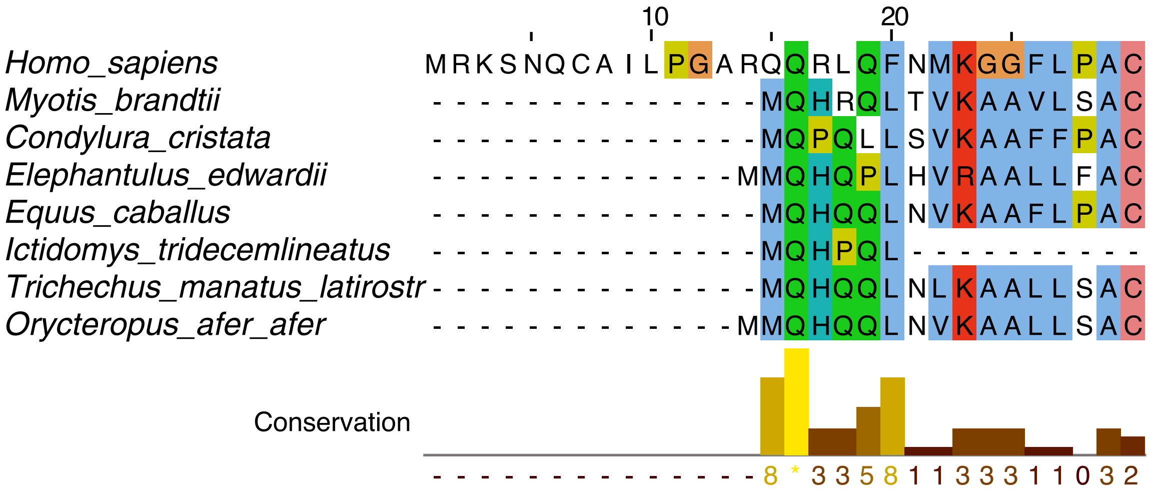

**HG0742**

ENSG00000154556 (SORBS2)

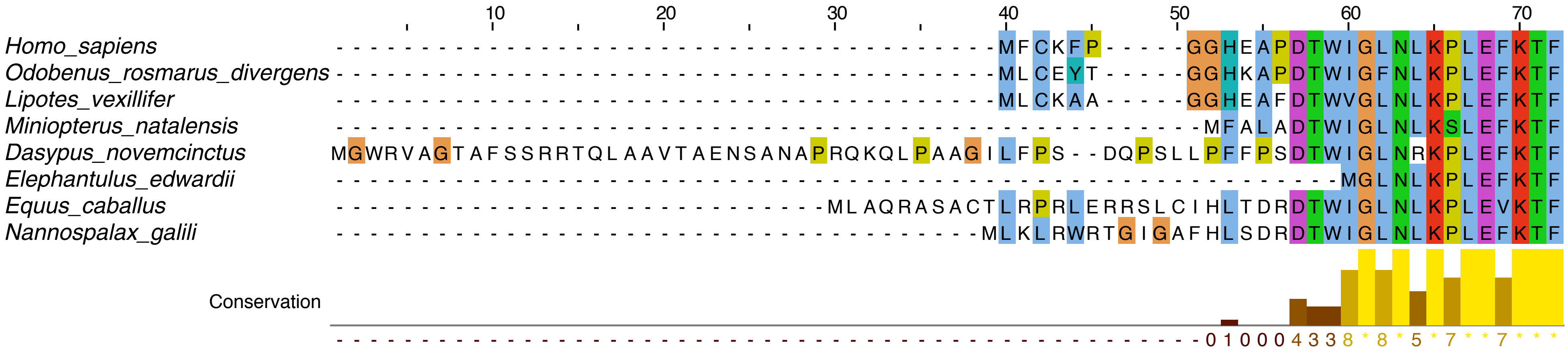

**HG0743**

ENSG00000154814 (OXNAD1)

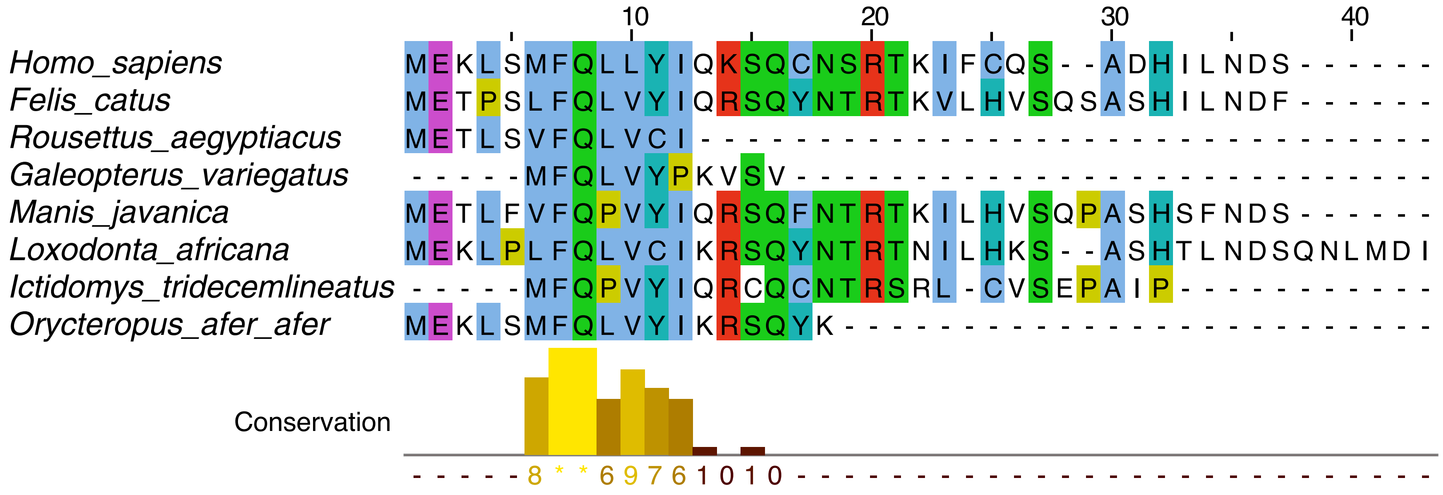

**HG0744**

ENSG00000163053 (SLC16A14)

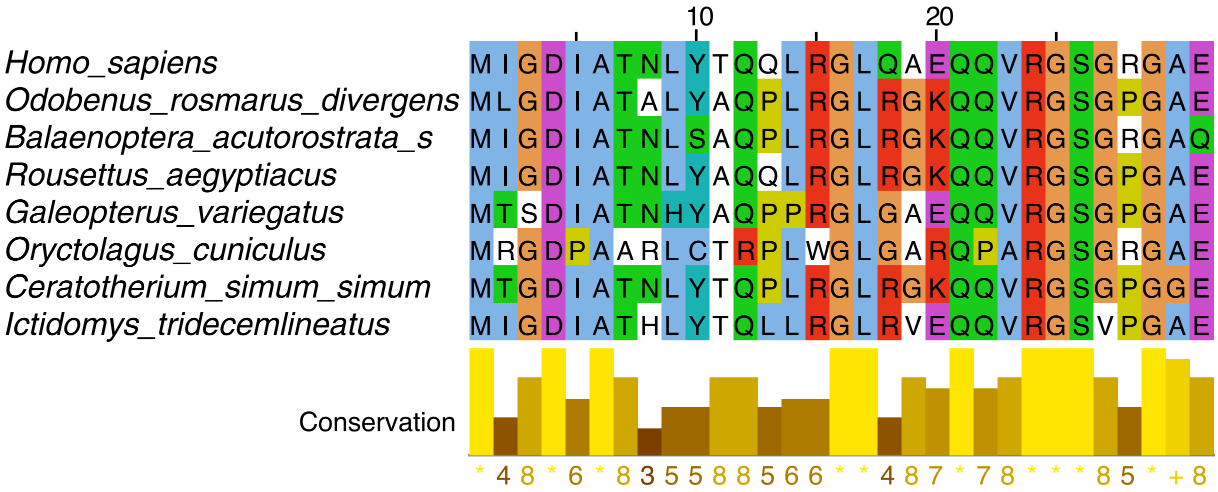

**HG0745**

ENSG00000163145 (C1QTNF7)

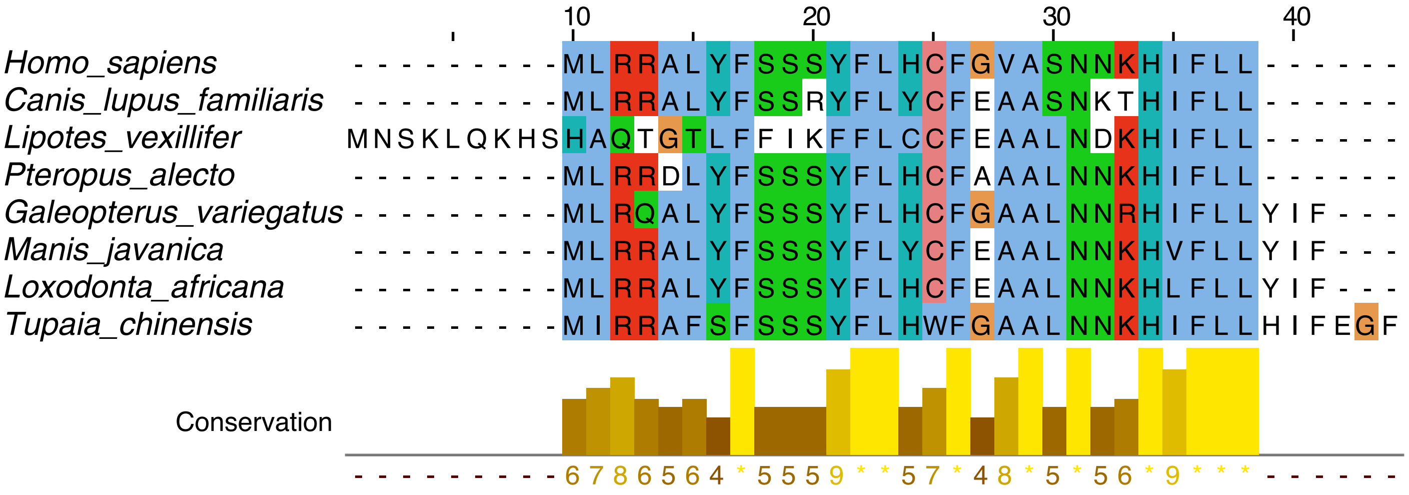

**HG0746**

ENSG00000164741 (DLC1)

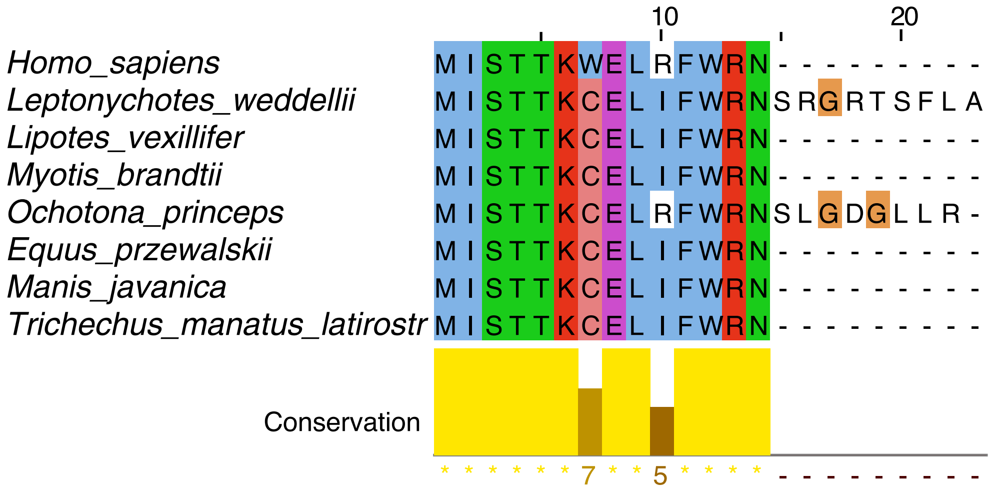

**HG0747**

ENSG00000168036 (CTNNB1)

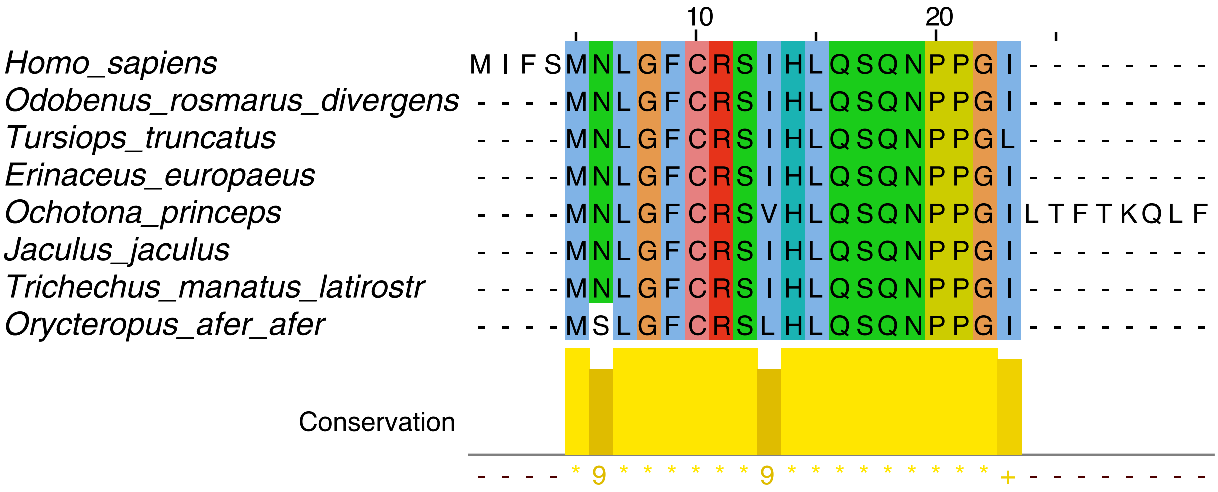

**HG0748**

ENSG00000168830 (HTR1E)

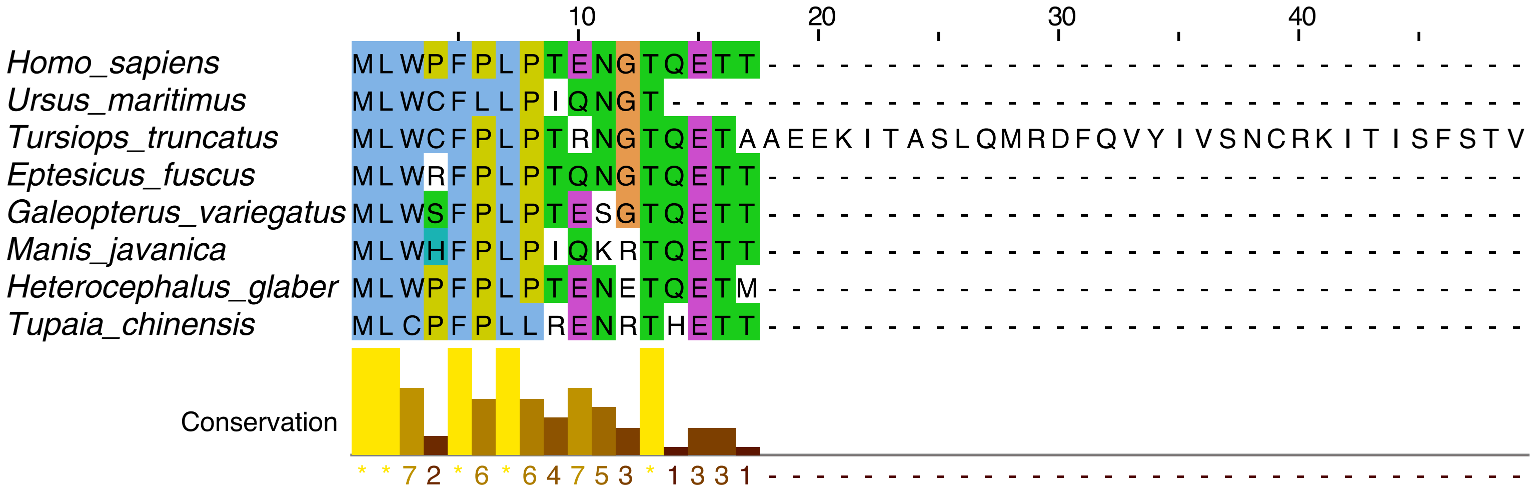

**HG0749**

ENSG00000169760 (NLGN1)

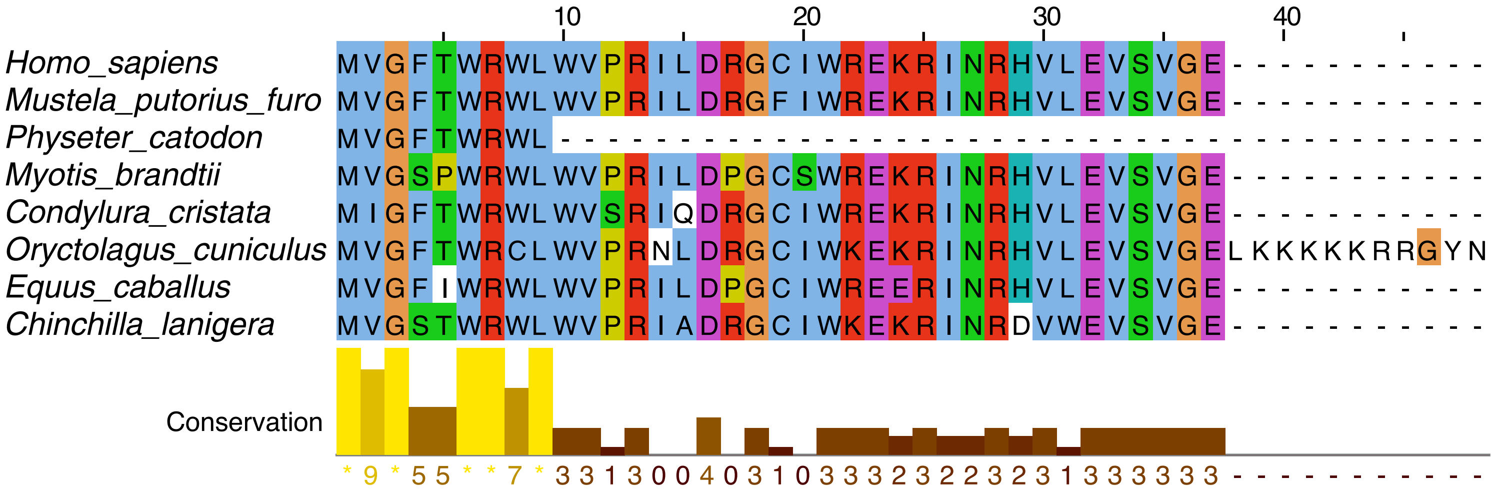

**HG0750**

ENSG00000170881 (RNF139)

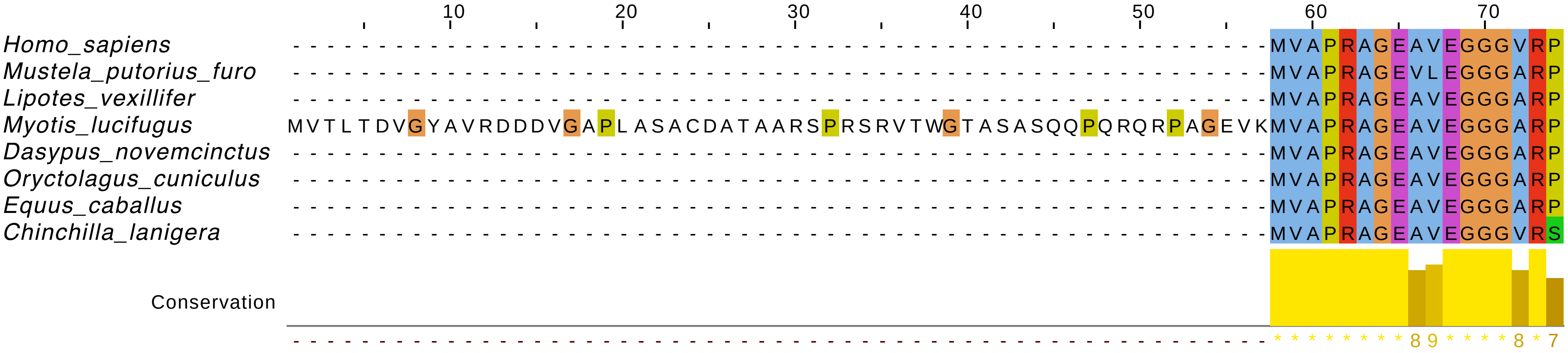

**HG0751**

ENSG00000171791 (BCL2)

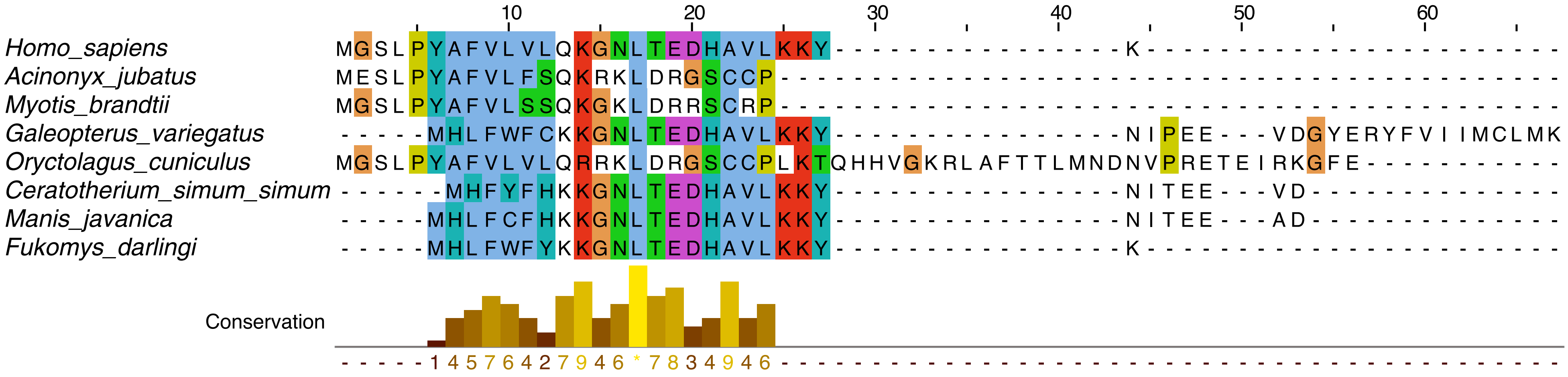

**HG0752**

ENSG00000172269 (DPAGT1)

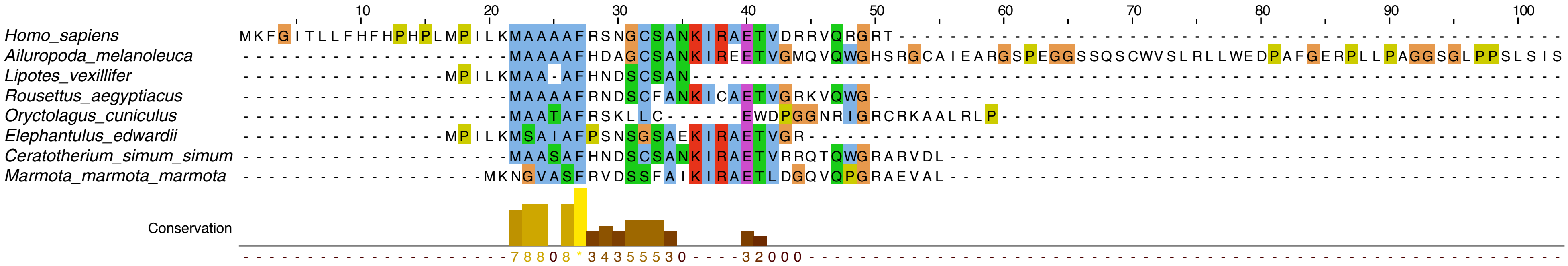

**HG0753**

ENSG00000173406 (DAB1)

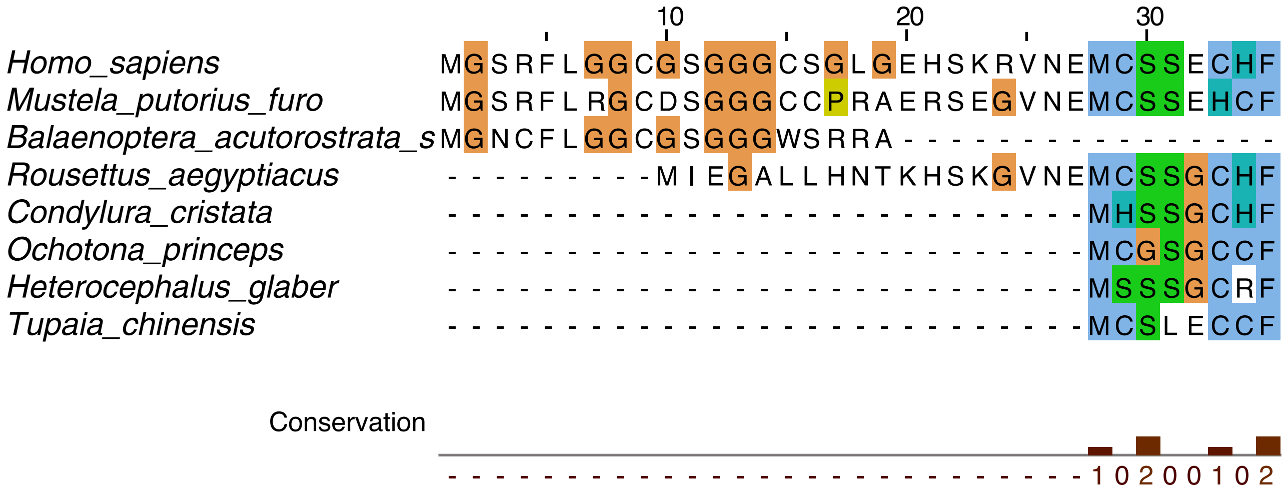

**HG0754**

ENSG00000175182 (FAM131A)

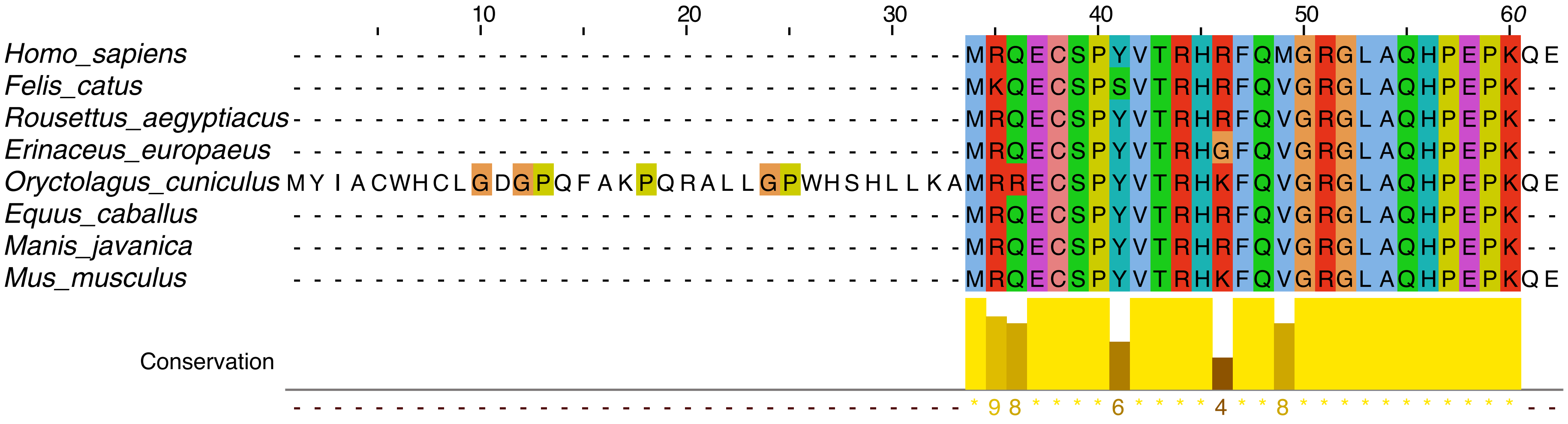

**HG0755**

ENSG00000176597 (B3GNT5)

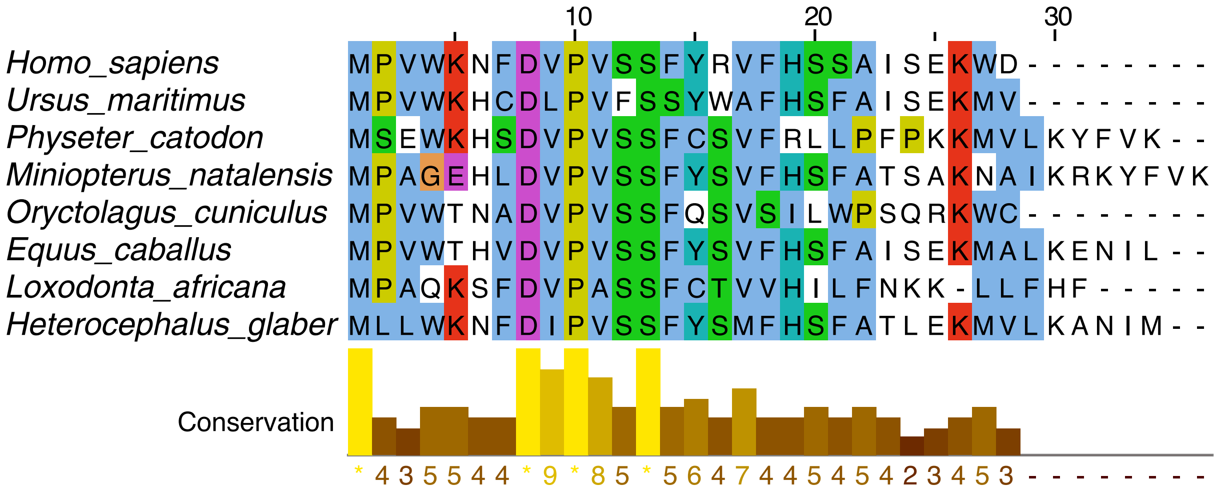

**HG0756**

ENSG00000177051 (FBXO46)

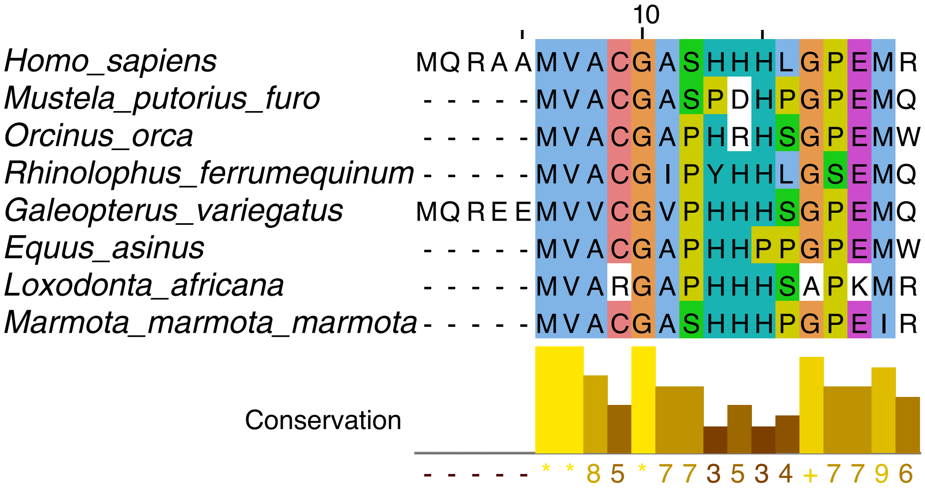

**HG0757.1**

ENSG00000177301 (KCNA2)

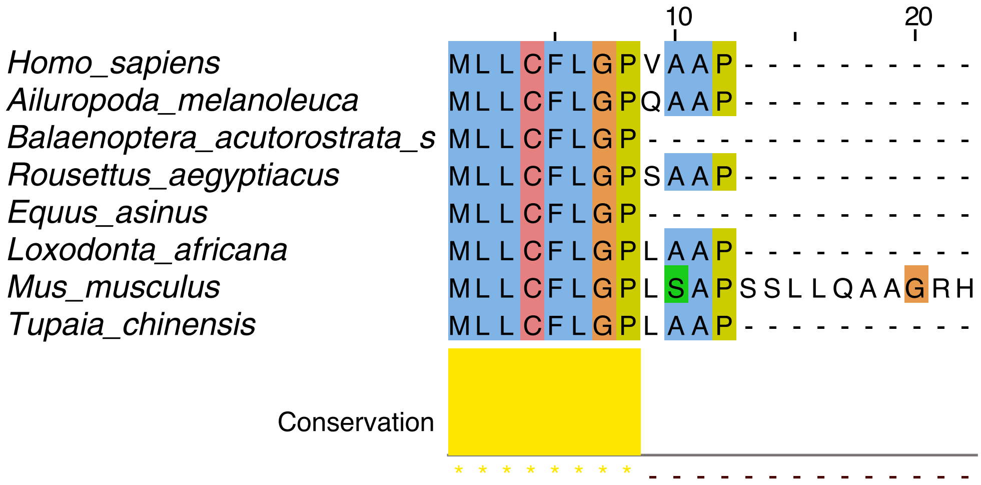

**HG0757.2**

ENSG00000182255 (KCNA4)

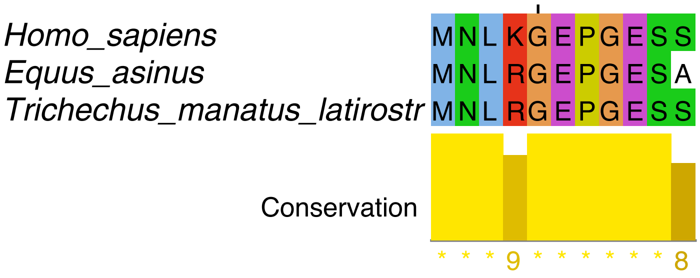

**HG0758**

ENSG00000179299 (NSUN7)

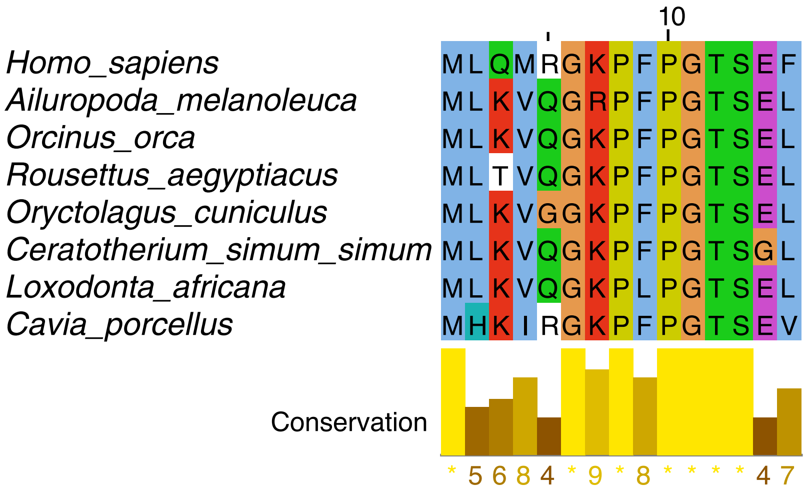

**HG0759**

ENSG00000180929 (GPR62)

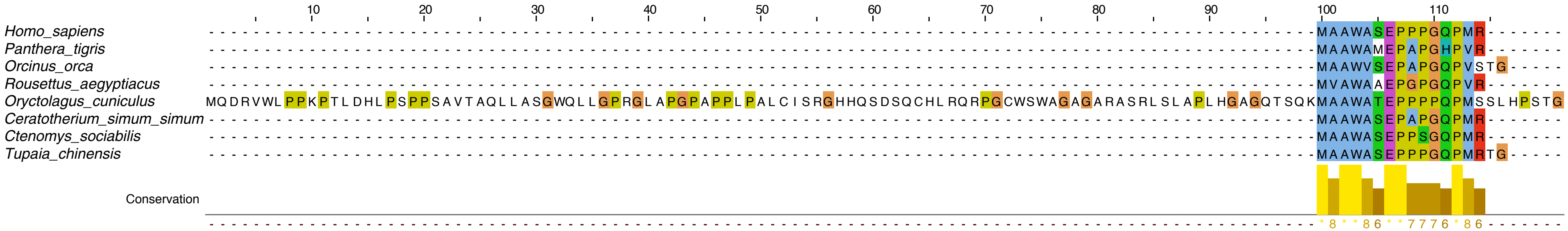

# HG0760

### ENSG00000182463 (TSHZ2)

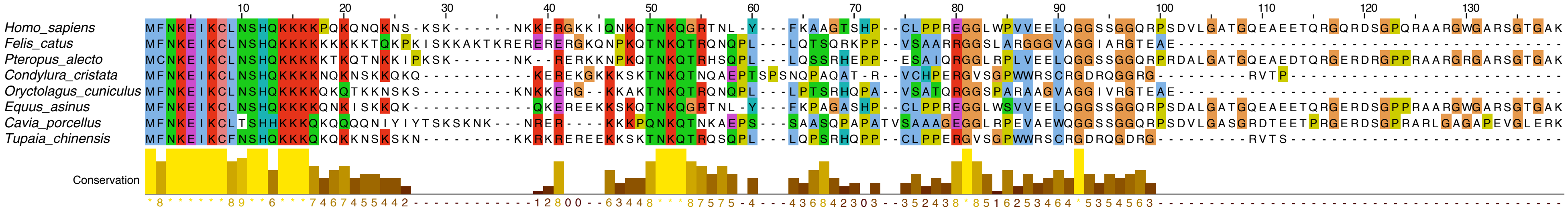

**HG0761**

ENSG00000185985 (SLITRK2)

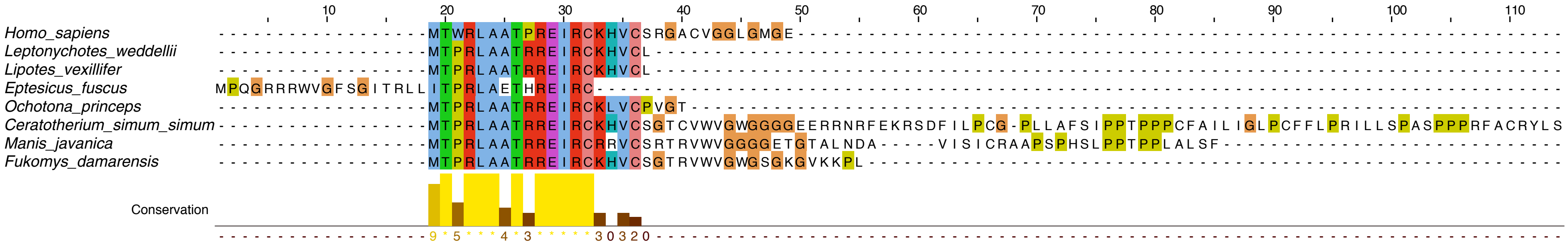

**HG0762**

ENSG00000187147 (RNF220)

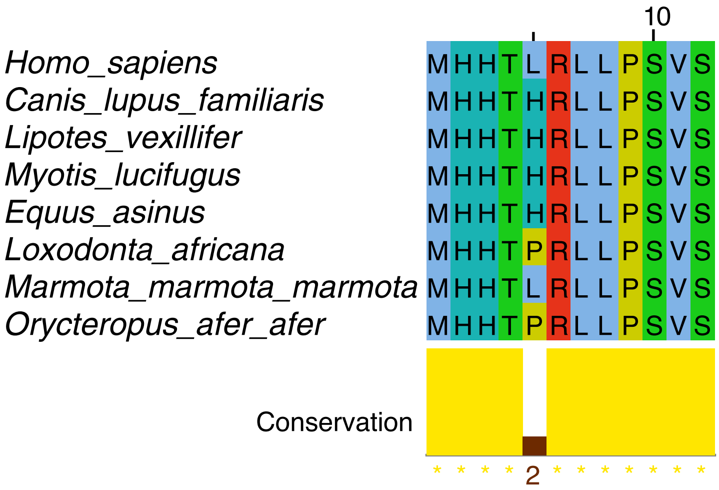

**HG0763**

ENSG00000188050 (RNF133)

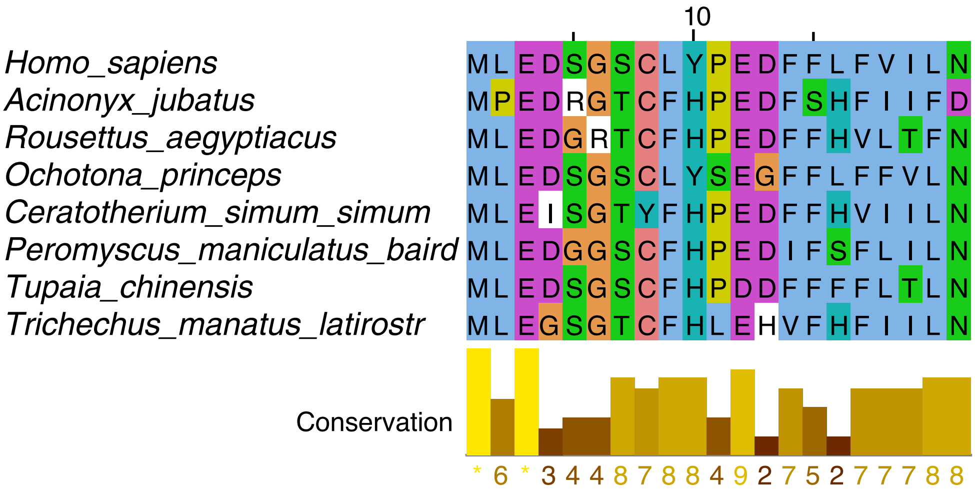

**HG0764**

ENSG00000188725 (SMIM15)

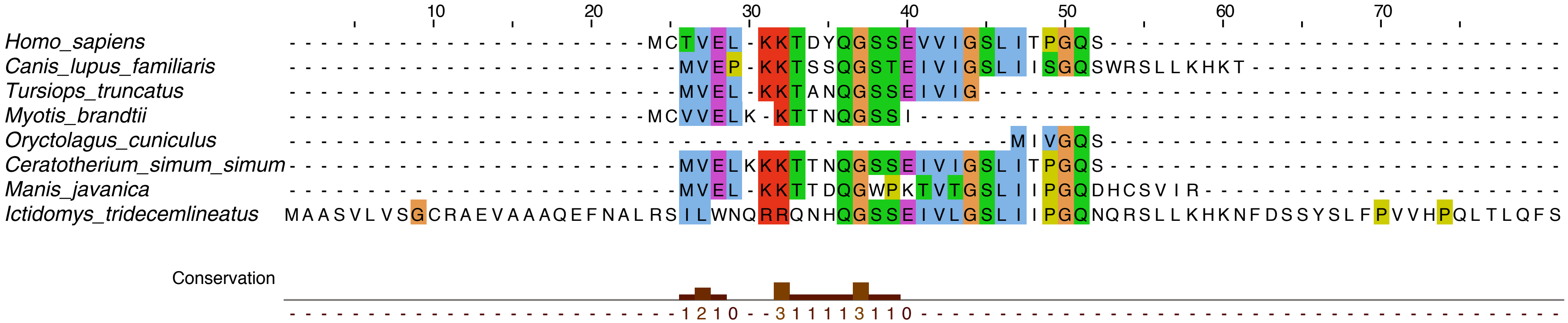

**HG0765**

ENSG00000196123

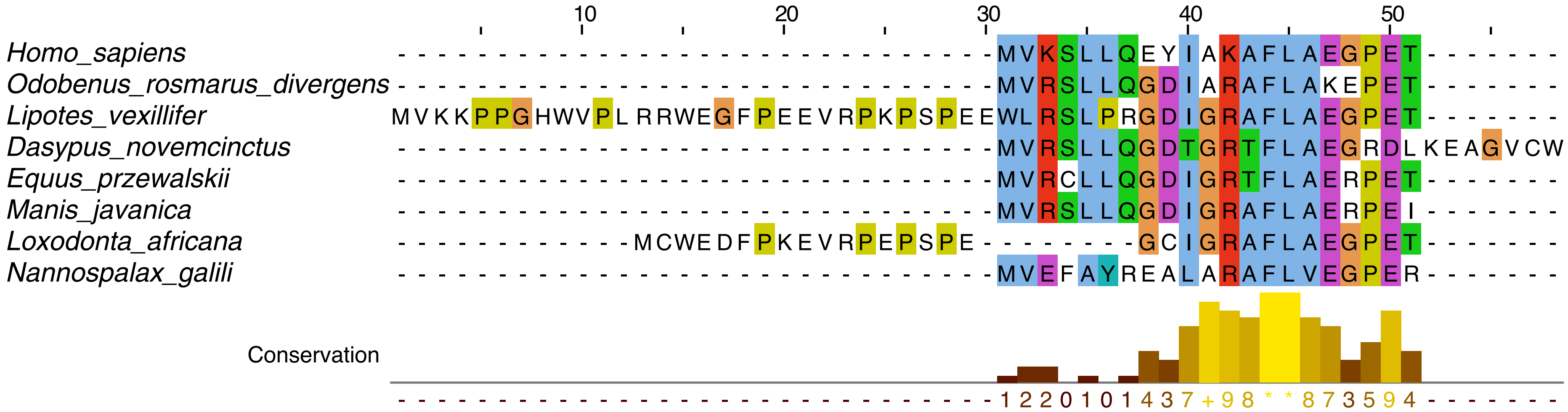

**HG0766**

ENSG00000196189 (SEMA4A)

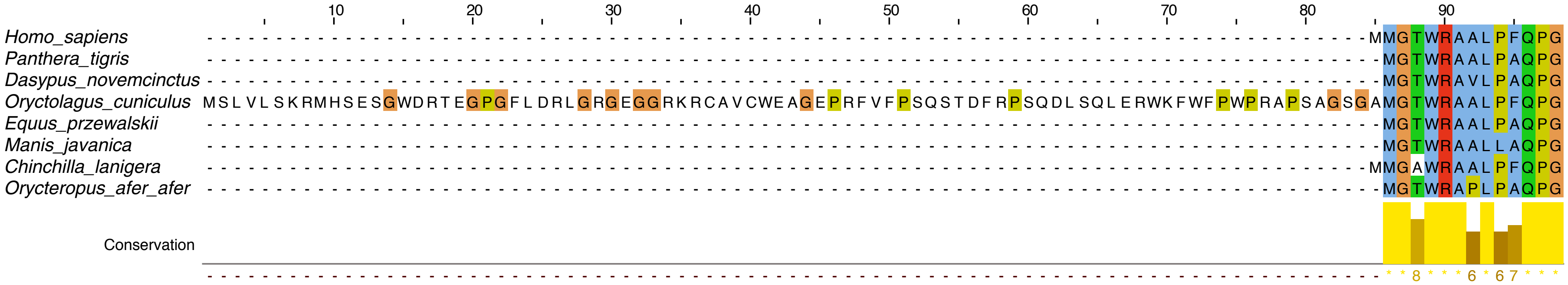

**HG0767.1**

ENSG00000197555 (SIPA1L1)

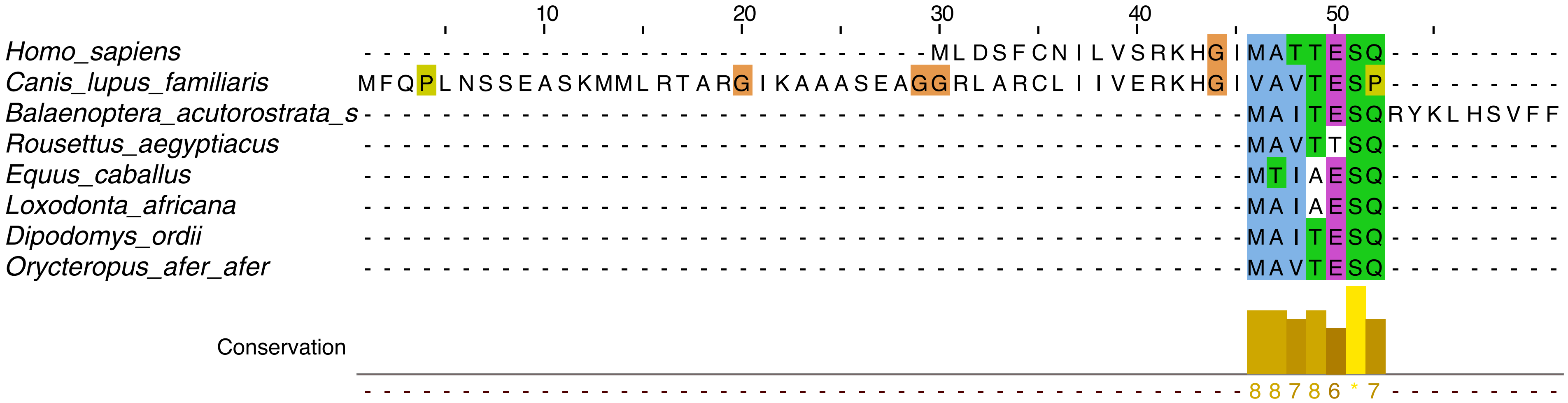

**HG0767.2**

ENSG00000197555 (SIPA1L1)

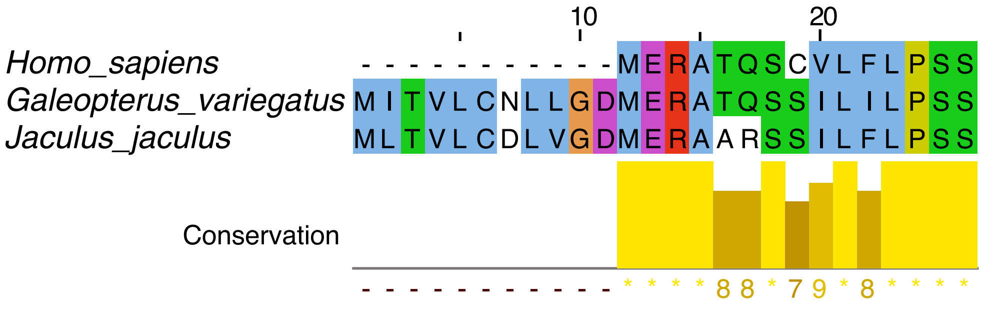

**HG0768**

ENSG00000213023 (SYT3)

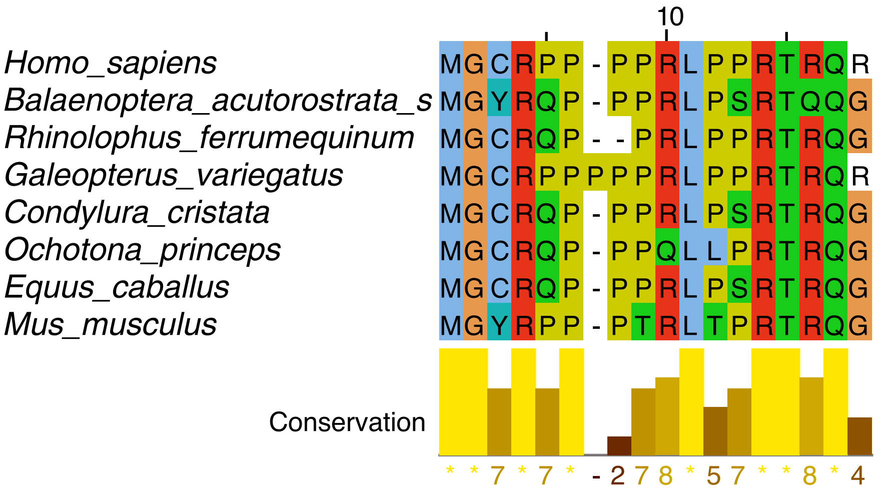

**HG0769.1**

ENSG00000280165 (PCDH20)

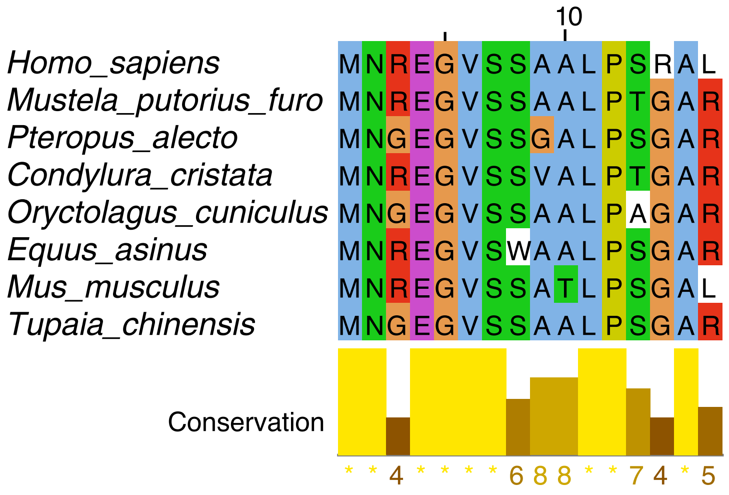

**HG0769.2**

ENSG00000197991

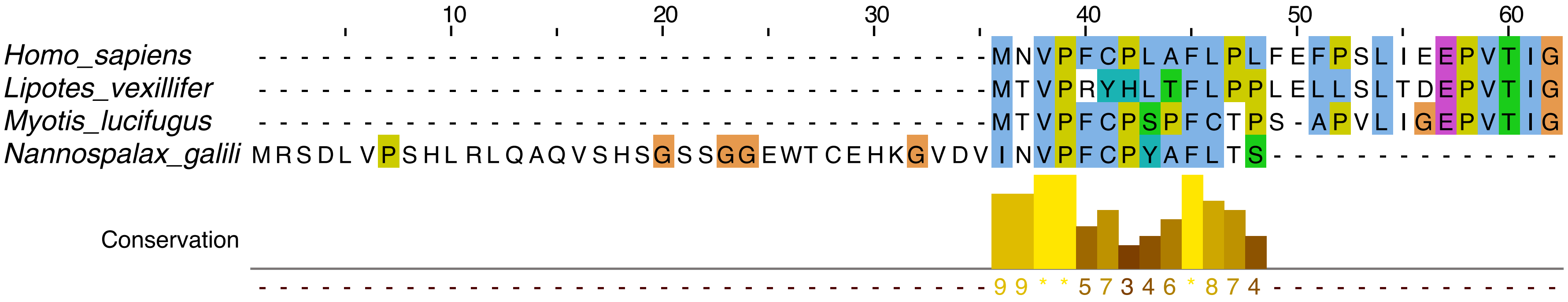

**HG0770**

ENSG00000010165 (METTL13)

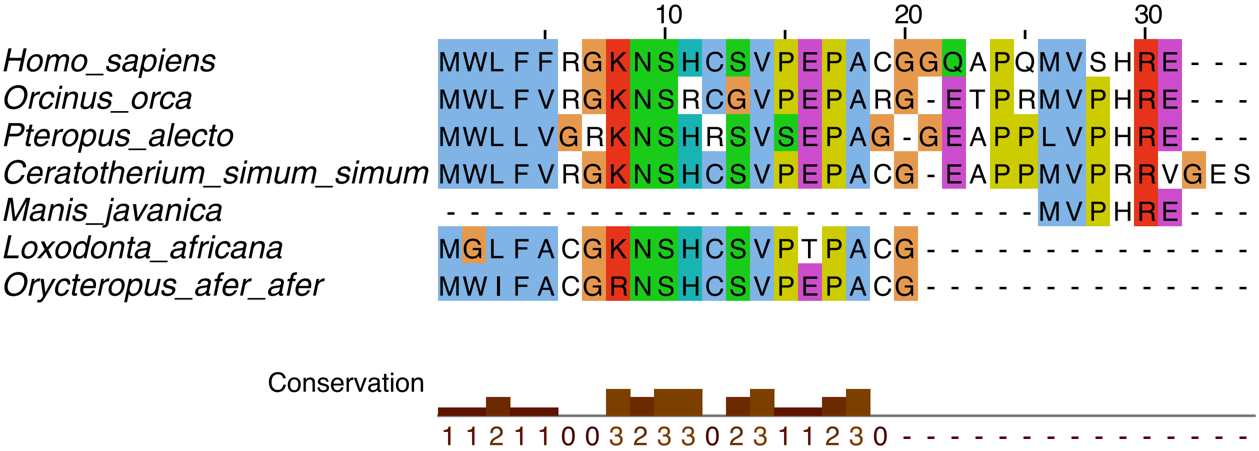

**HG0771**

ENSG00000013306 (SLC25A39)

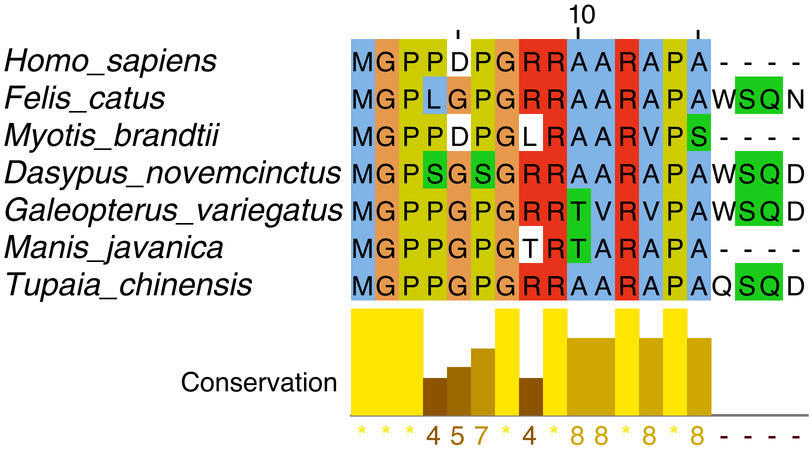

**HG0772**

ENSG00000064309 (CDON)

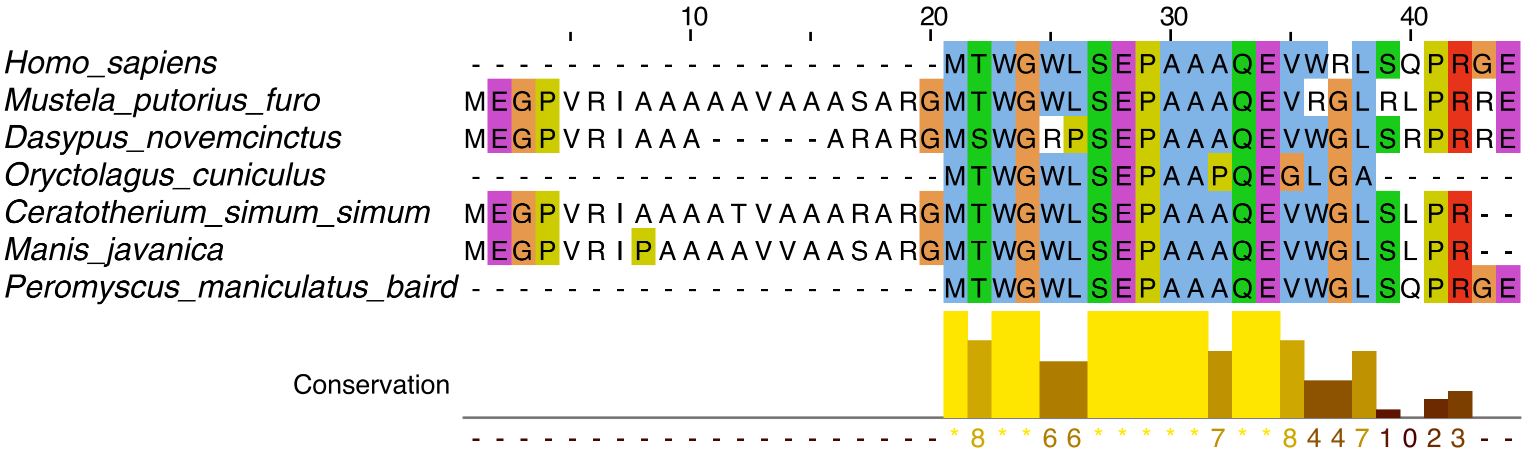

**HG0773**

ENSG00000065548 (ZC3H15)

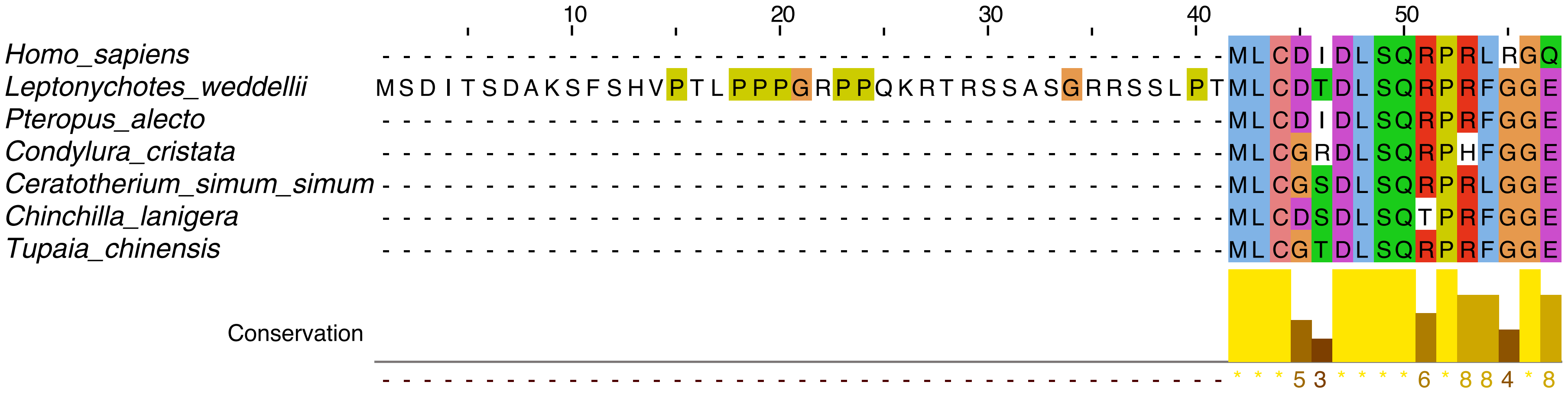

**HG0774**

ENSG00000066933 (MYO9A)

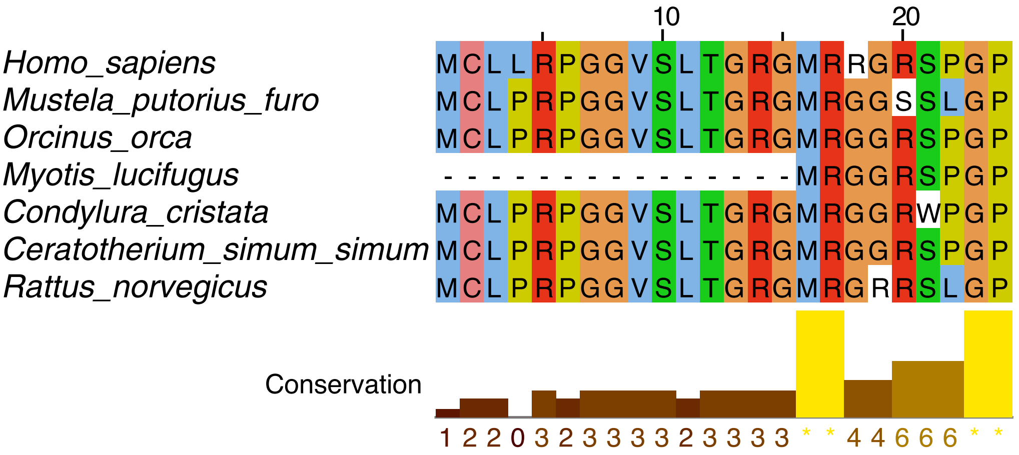

**HG0775**

ENSG00000068001 (HYAL2)

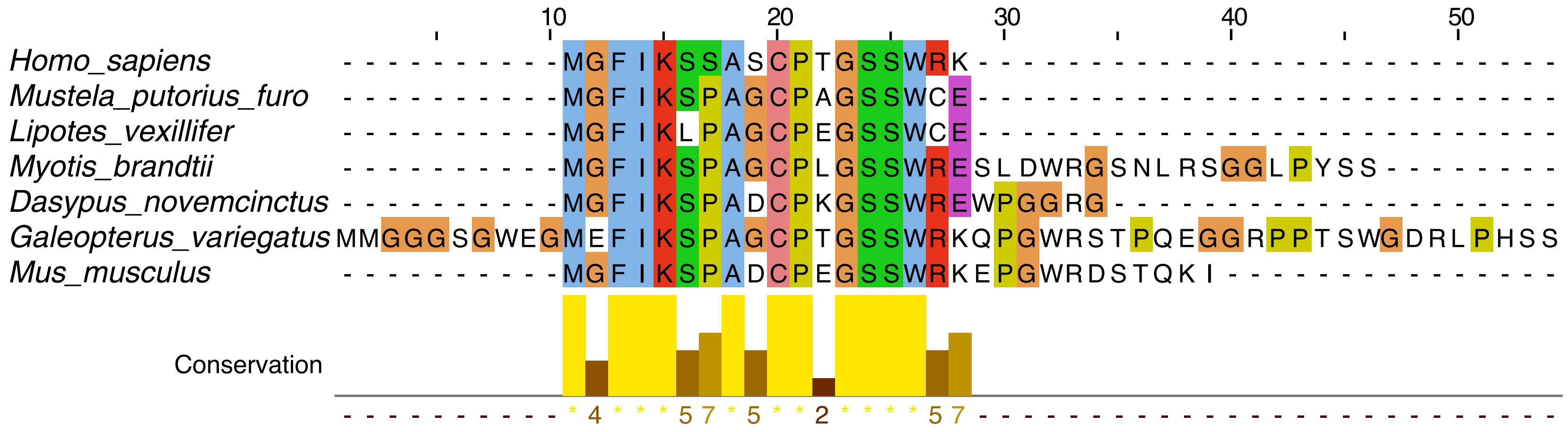

**HG0776**

ENSG00000082014 (SMARCD3)

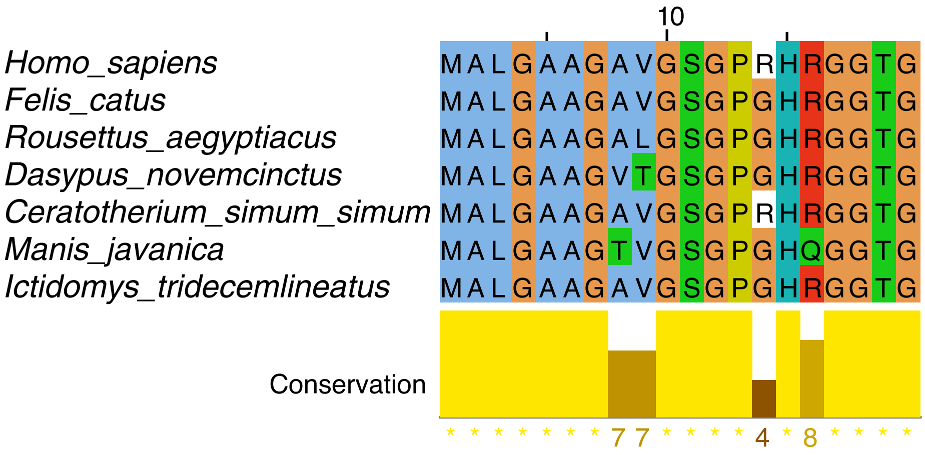

**HG0777**

ENSG00000085644 (ZNF213)

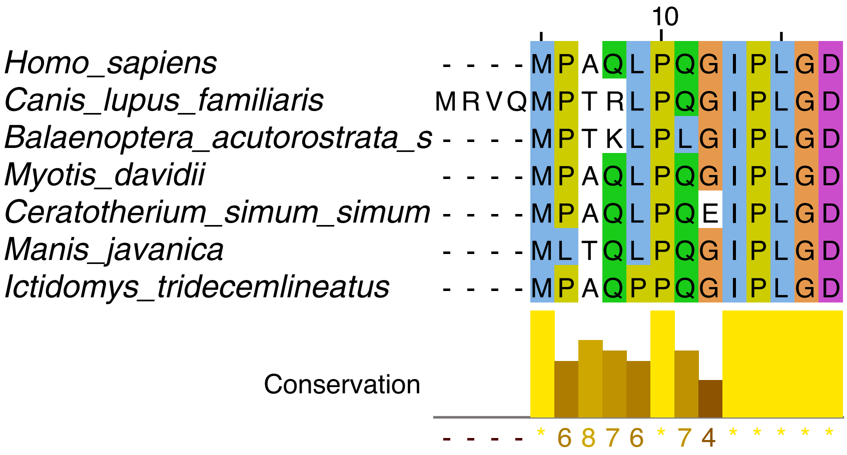

**HG0778**

ENSG00000089116 (LHX5)

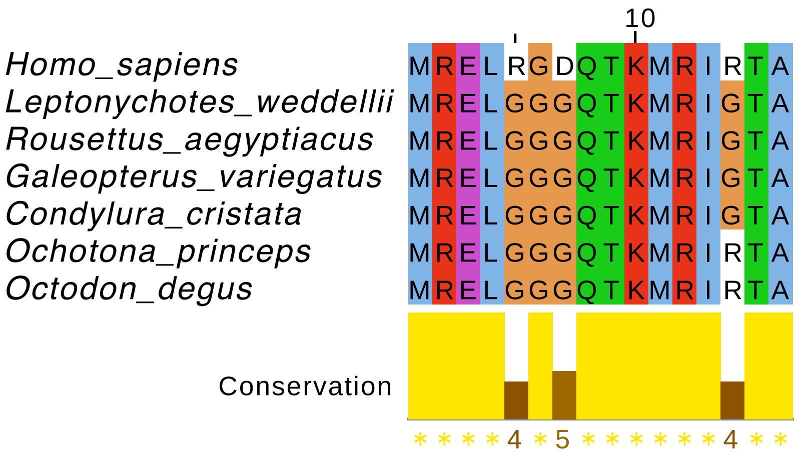

**HG0779**

ENSG00000091140 (DLD)

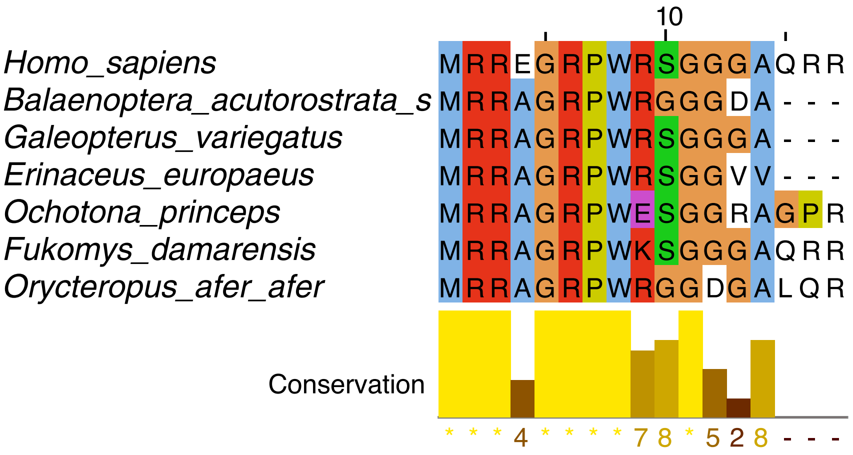

**HG0780**

ENSG00000092199 (HNRNPC)

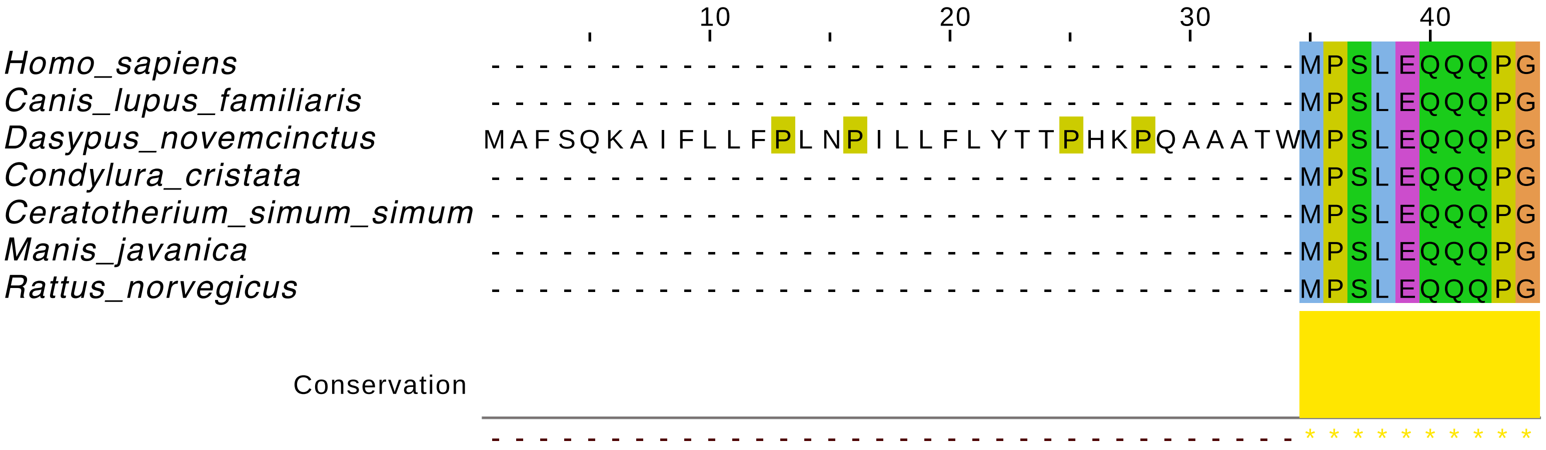

**HG0781**

ENSG00000100099 (HPS4)

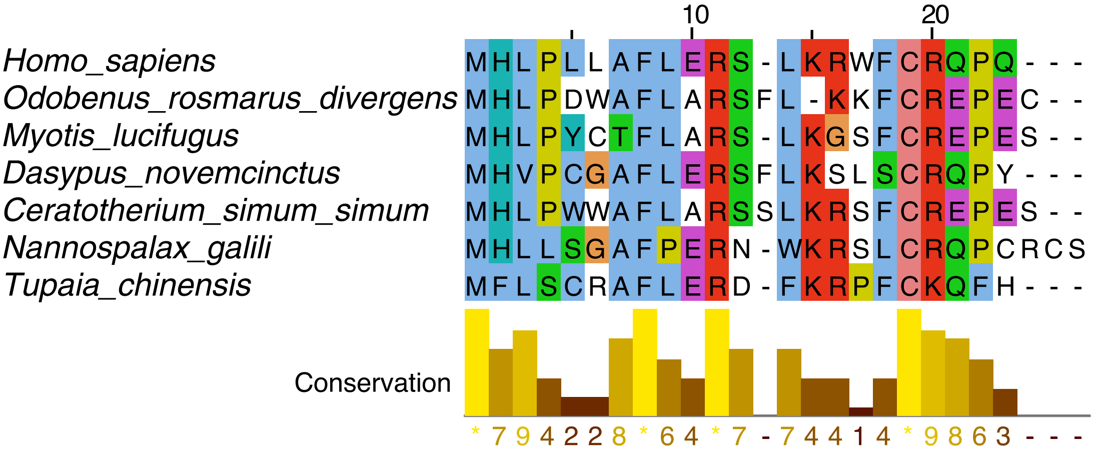

**HG0782**

ENSG00000100368 (CSF2RB)

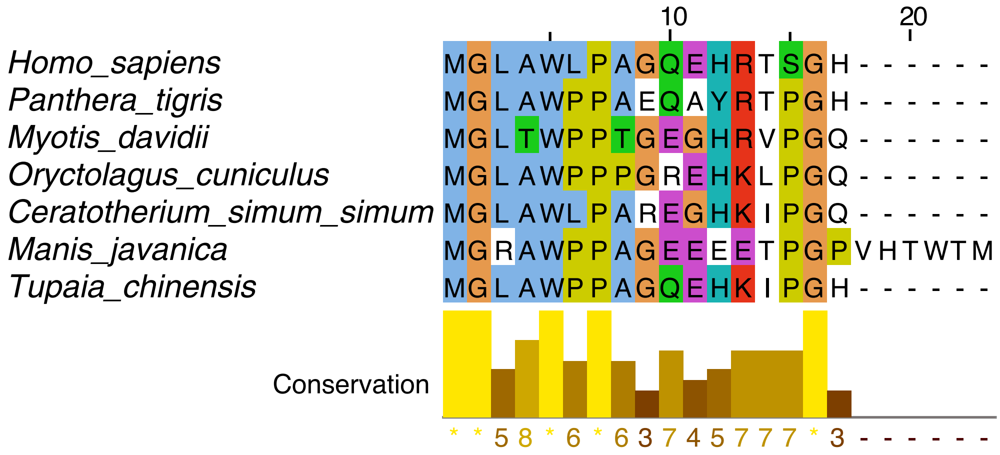

**HG0783**

ENSG00000105053 (VRK3)

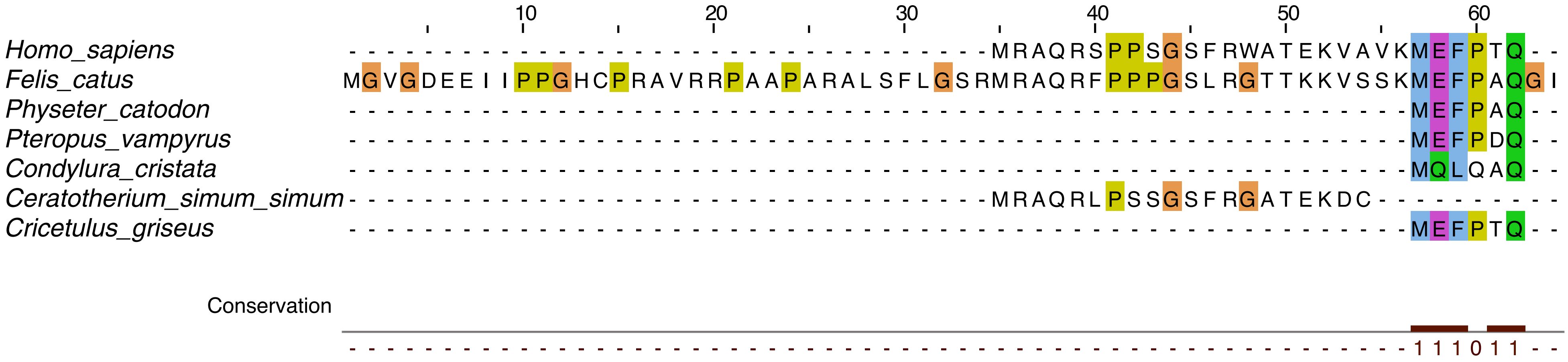

**HG0784**

ENSG00000105851 (PIK3CG)

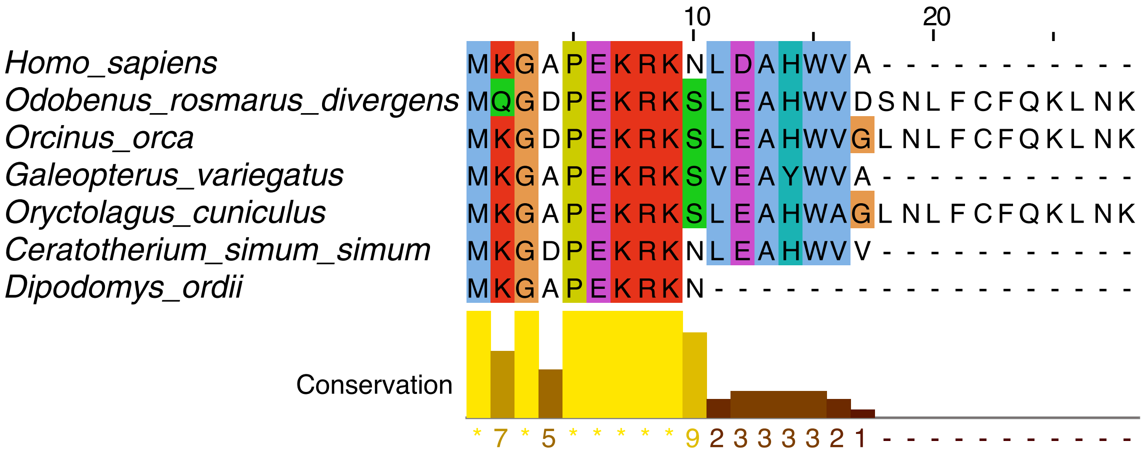

**HG0785**

ENSG00000107242 (PIP5K1B)

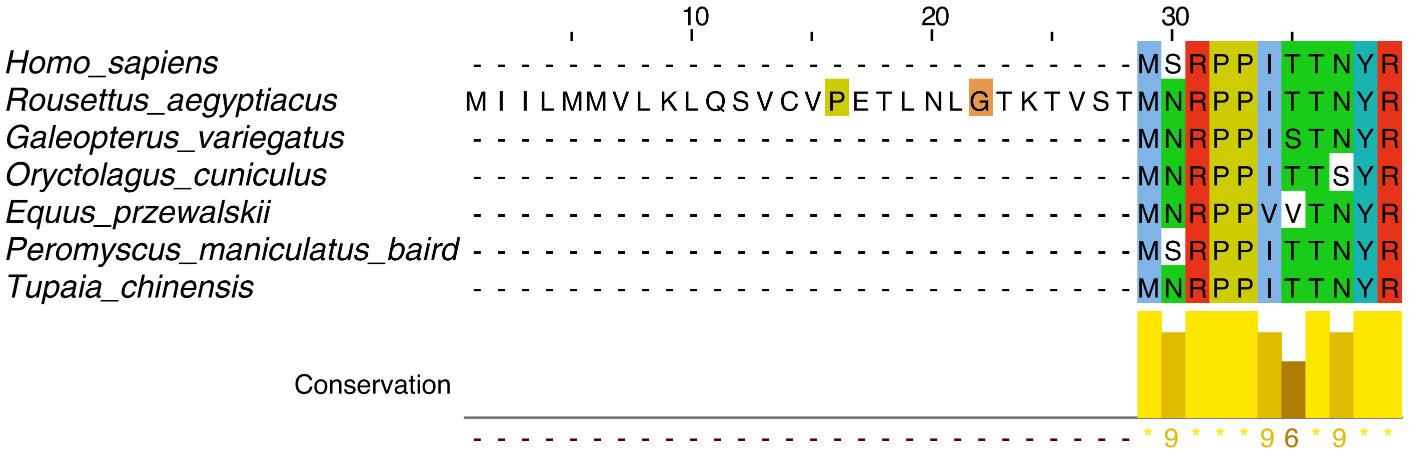

**HG0786**

ENSG00000107862 (GBF1)

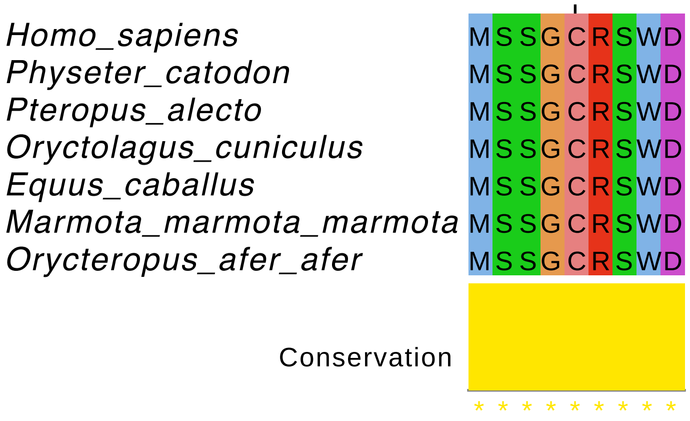

**HG0787**

ENSG00000108592 (FTSJ3)

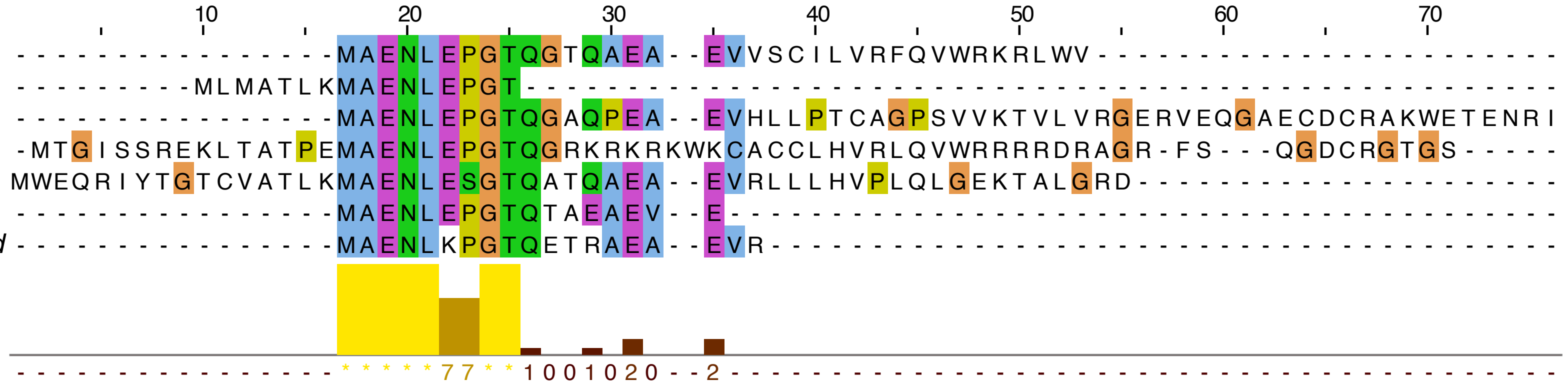

**HG0788**

ENSG00000113739 (STC2)

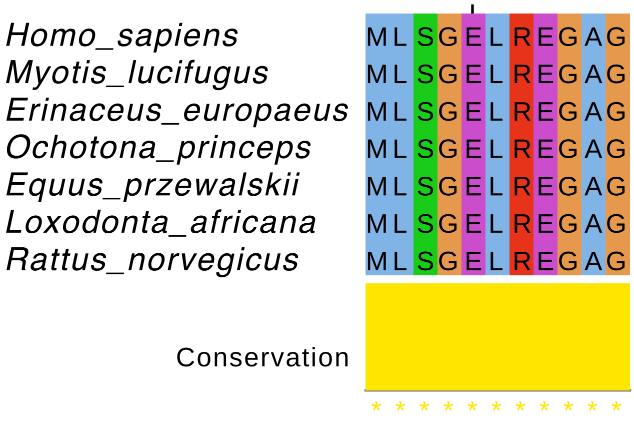

**HG0789**

ENSG00000115355 (CCDC88A)

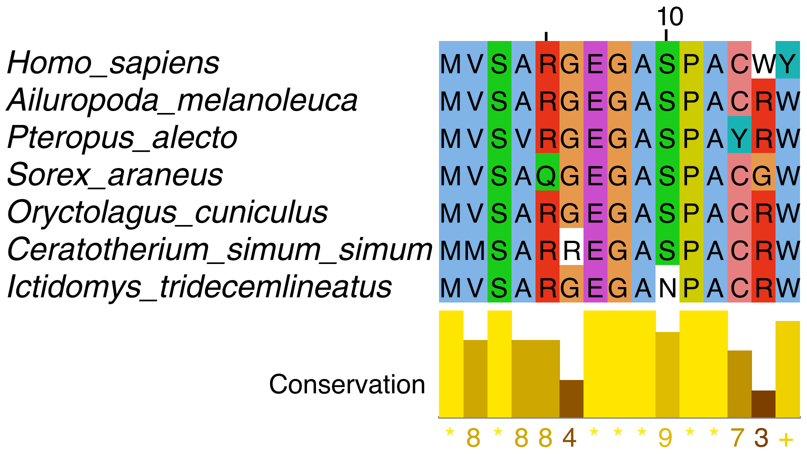

**HG0790.1**

ENSG00000115568 (ZNF142)

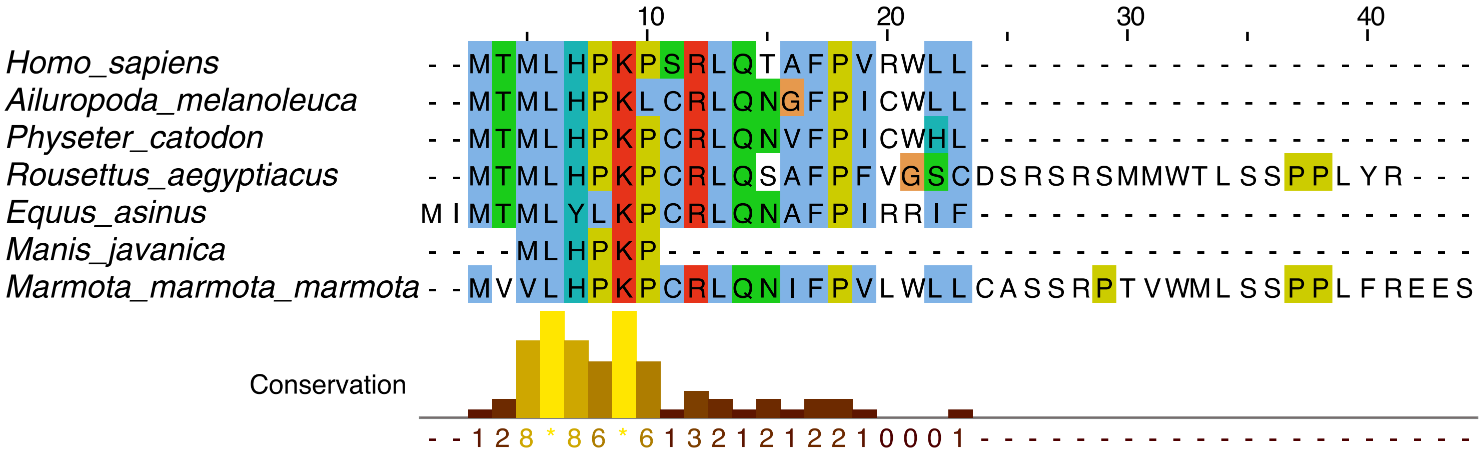

**HG0790.2**

ENSG00000115568 (ZNF142)

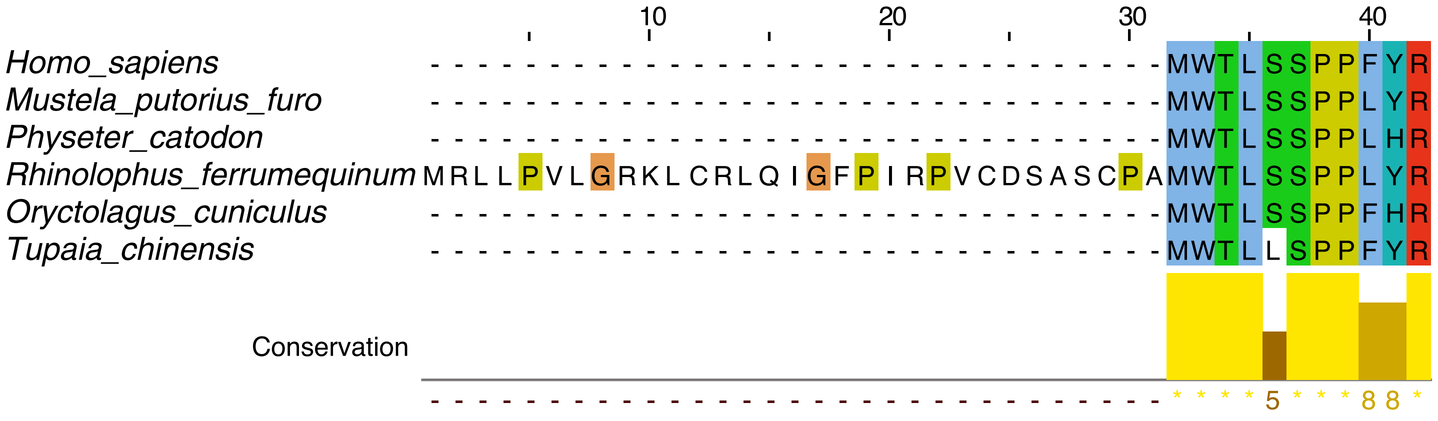

**HG0791**

ENSG00000131323 (TRAF3)

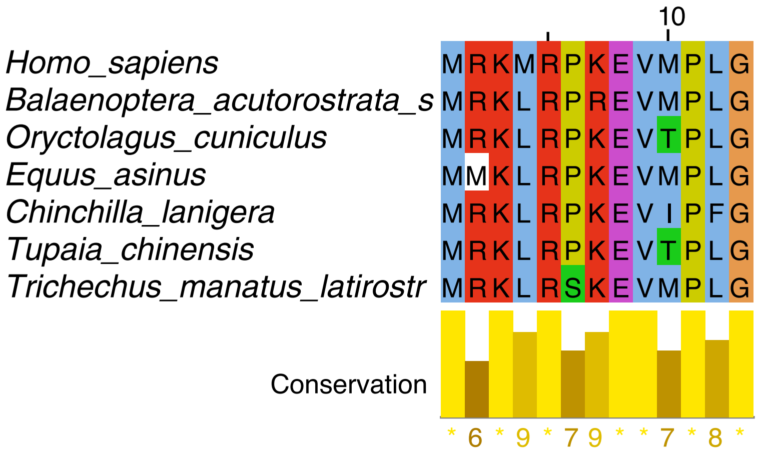

**HG0792**

ENSG00000132424 (PNISR)

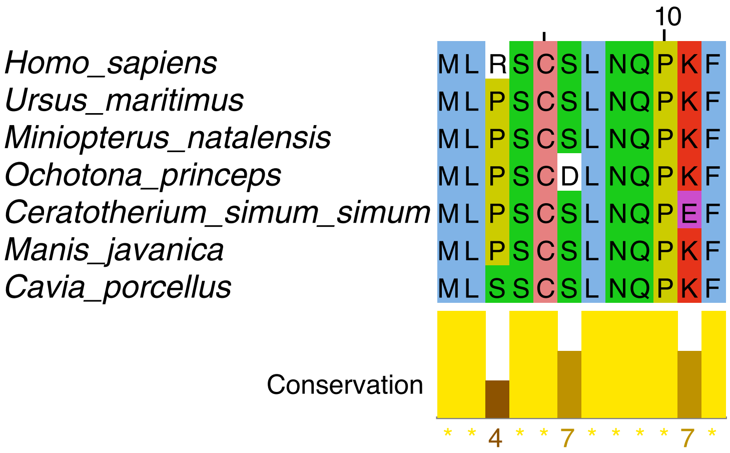

**HG0793**

ENSG00000132698 (RAB25)

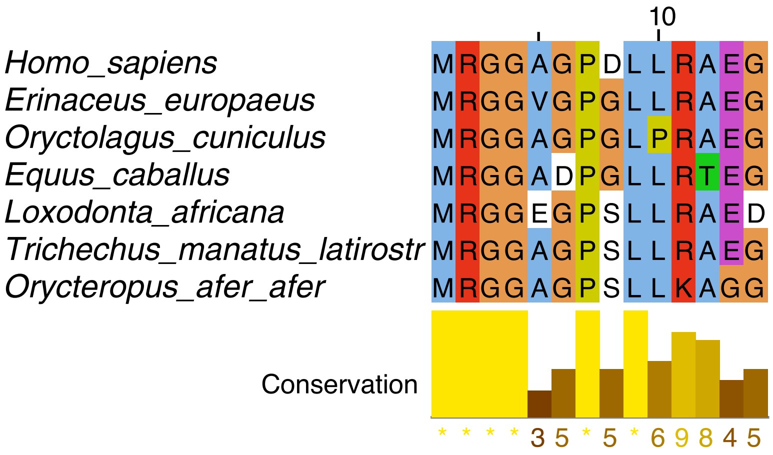

**HG0794.1**

ENSG00000135218 (CD36)

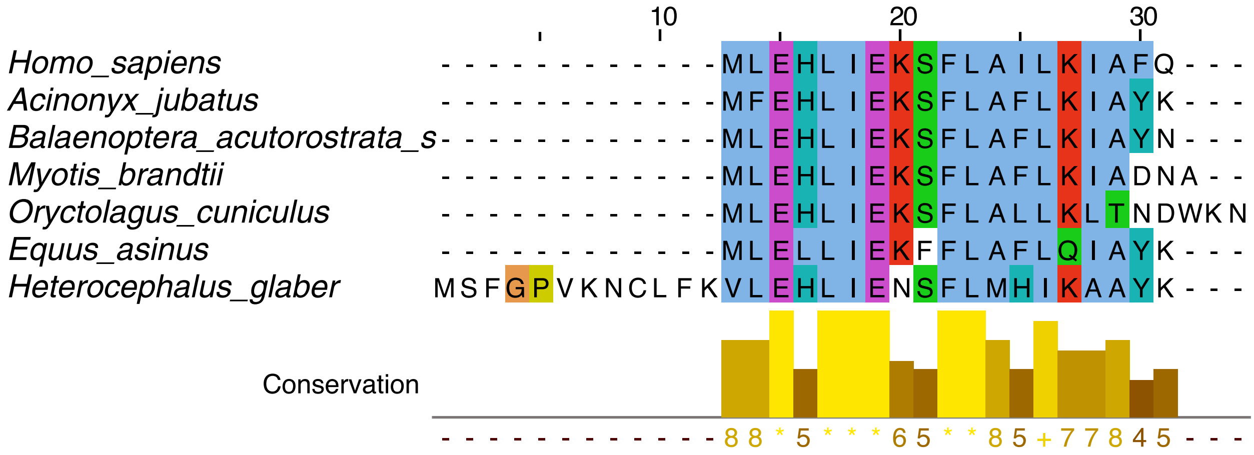

**HG0794.2**

ENSG00000135218 (CD36)

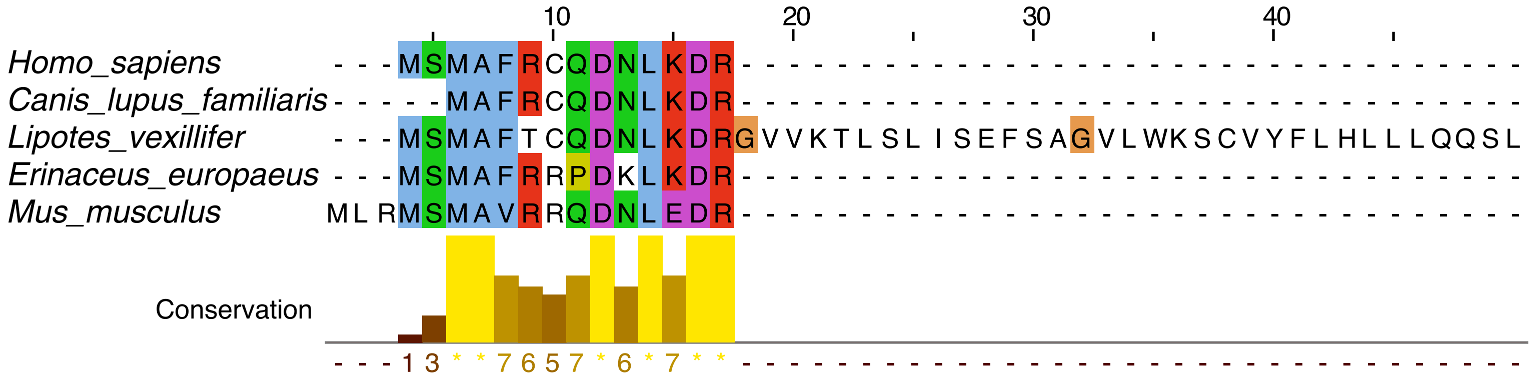

**HG0795**

ENSG00000137601 (NEK1)

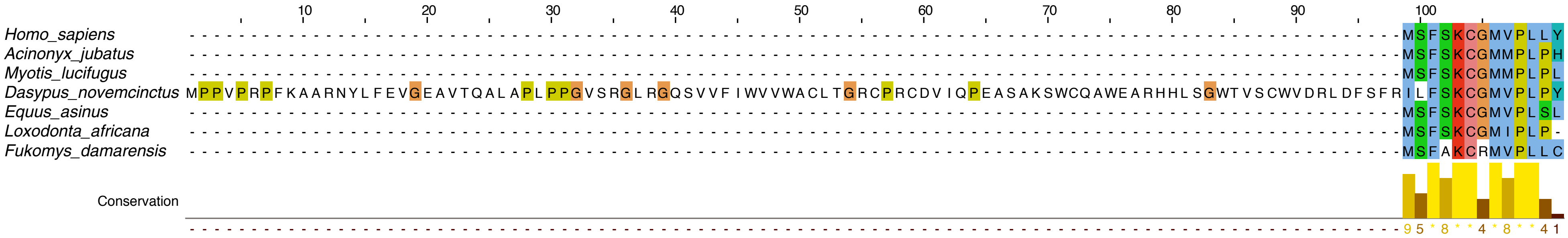

**HG0796**

ENSG00000138074 (SLC5A6)

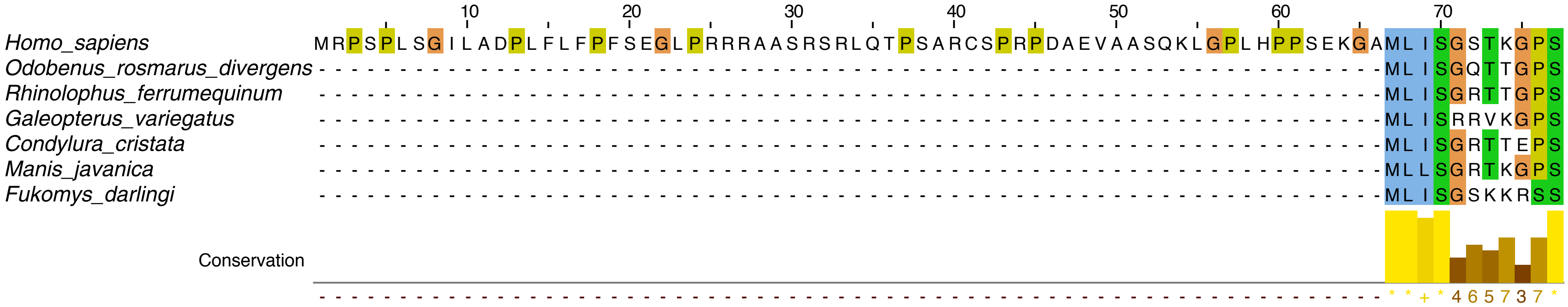

**HG0797**

ENSG00000140526 (ABHD2)

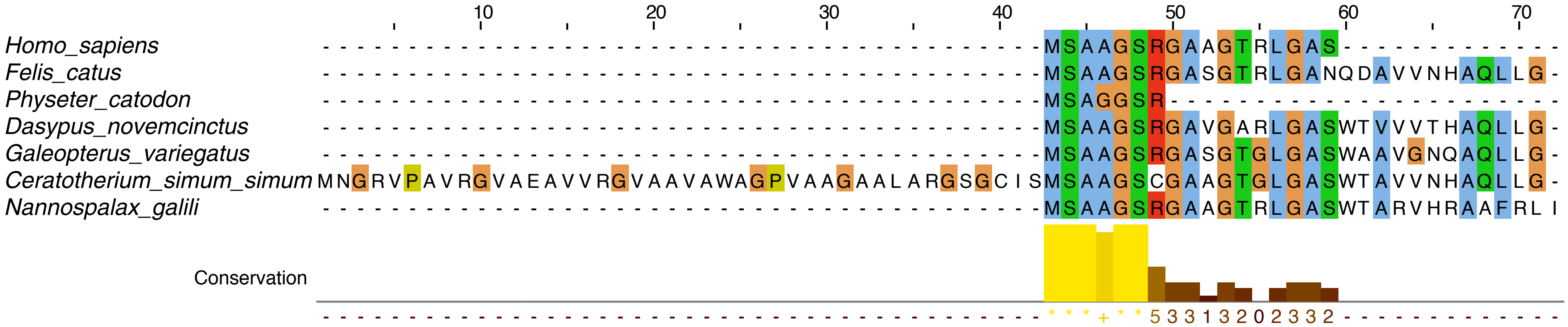

**HG0798.1**

ENSG00000142949 (PTPRF)

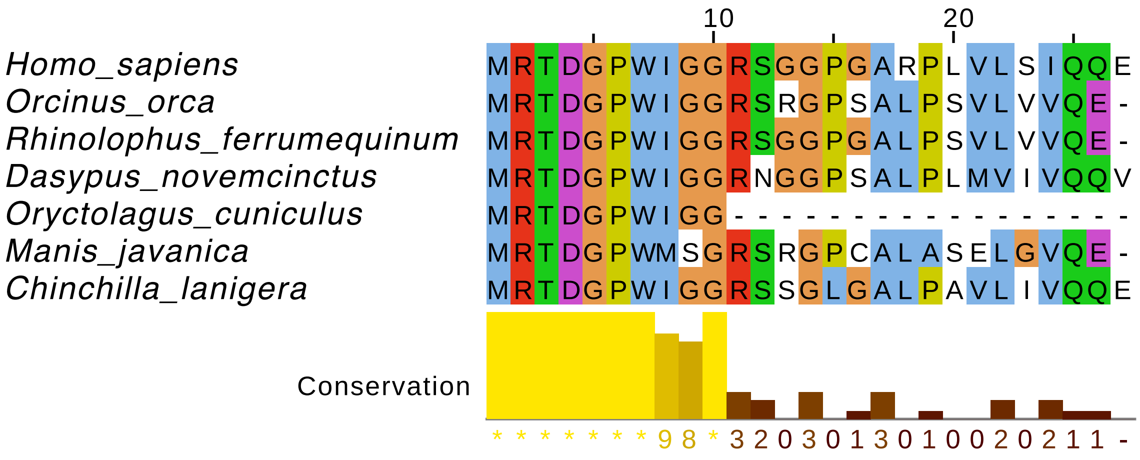

**HG0798.2**

ENSG00000153707 (PTPRD)

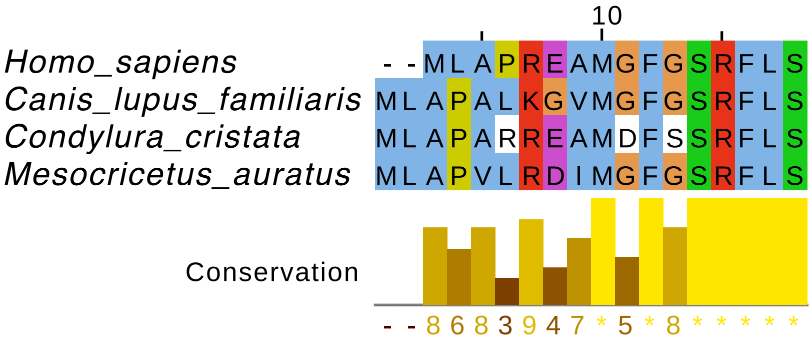

**HG0799**

ENSG00000143398 (PIP5K1A)

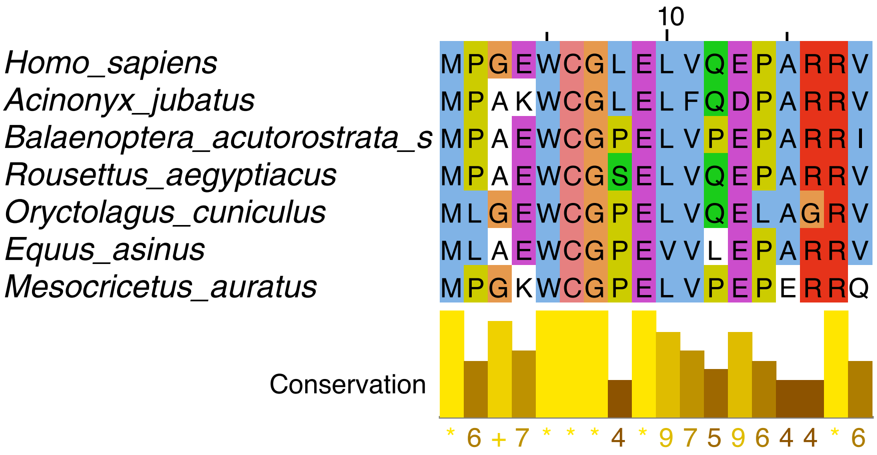

**HG0800**

ENSG00000143612 (C1orf43)

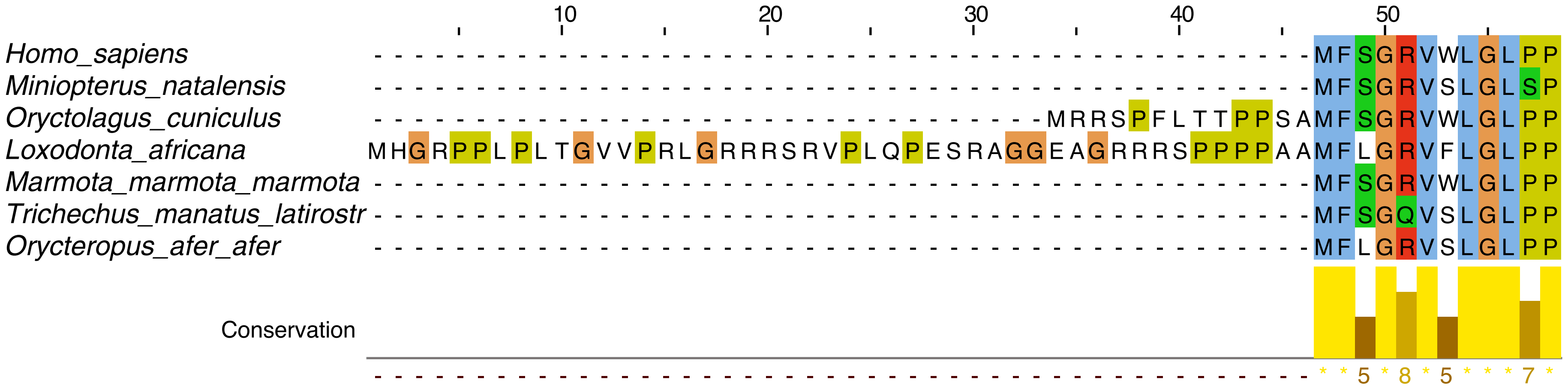

**HG0801**

ENSG00000144278 (GALNT13)

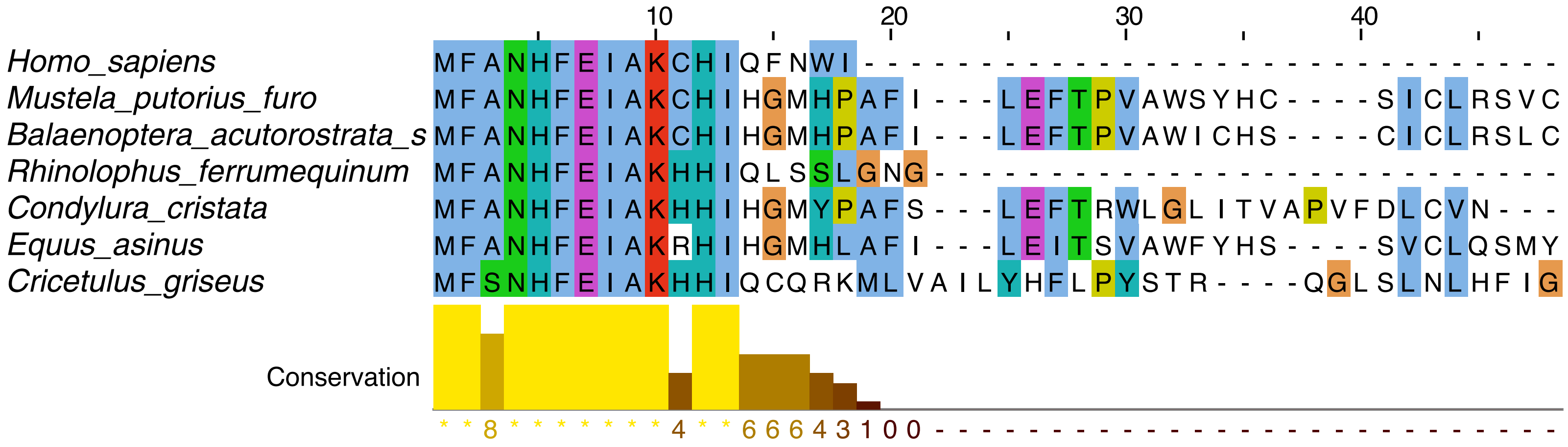

**HG0802**

ENSG00000144283 (PKP4)

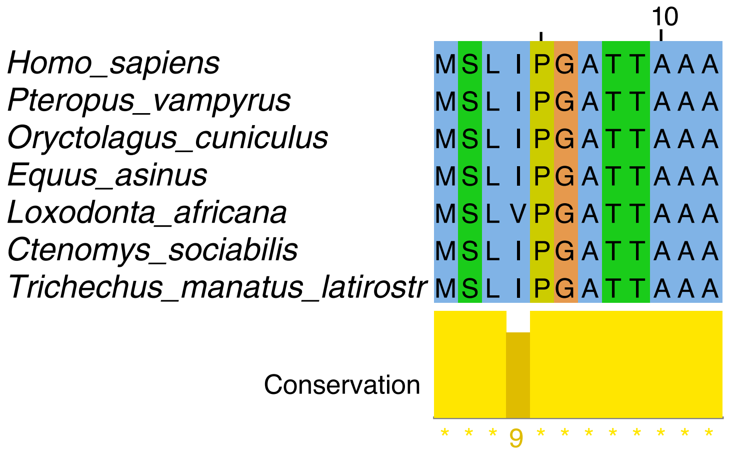

**HG0803**

ENSG00000144802 (NFKBIZ)

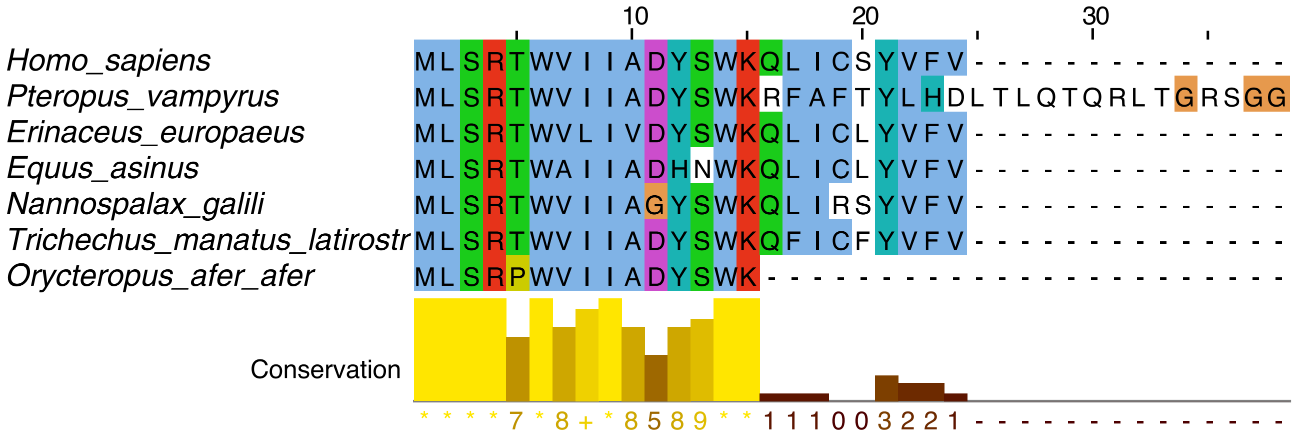

**HG0804**

ENSG00000144815 (NXPE3)

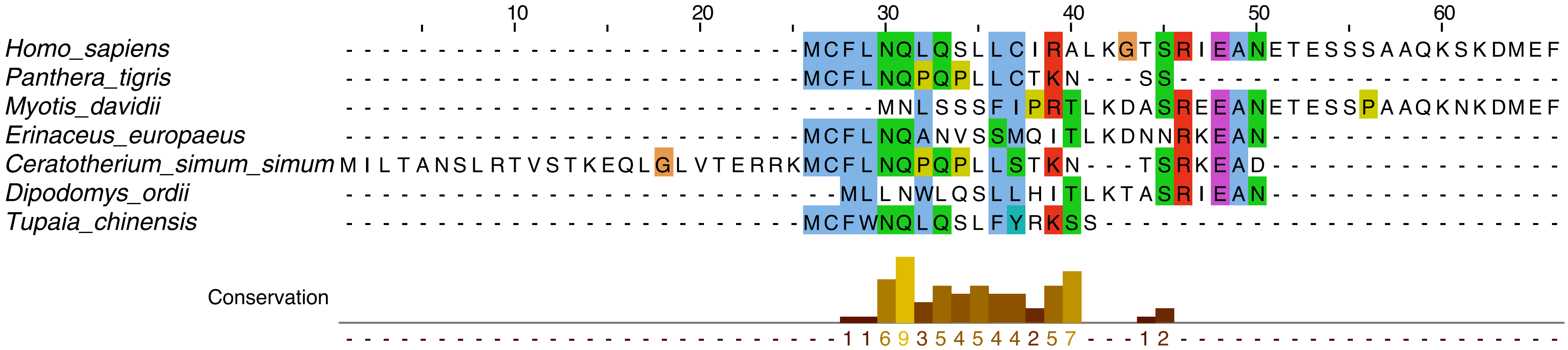

**HG0805.1**

ENSG00000145936 (KCNMB1)

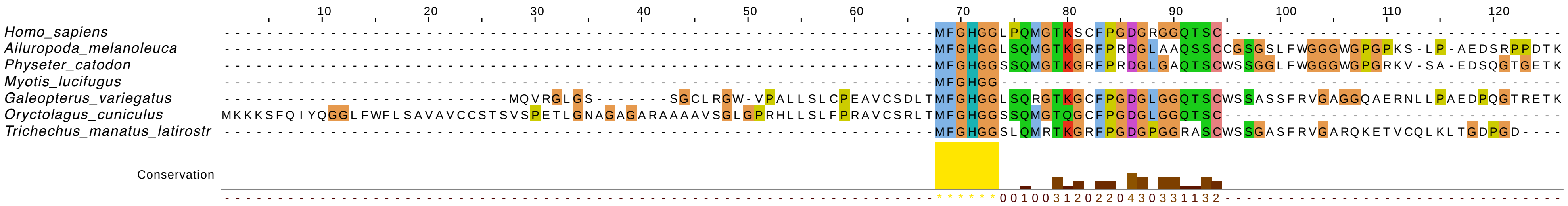

**HG0805.2**

ENSG00000145936 (KCNMB1)

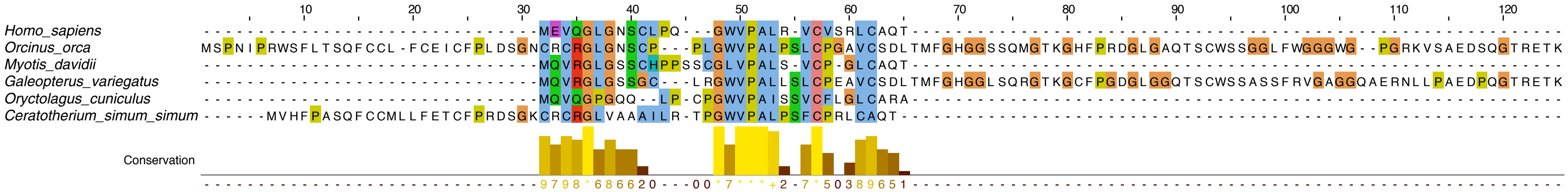

**HG0806**

ENSG00000147481 (SNTG1)

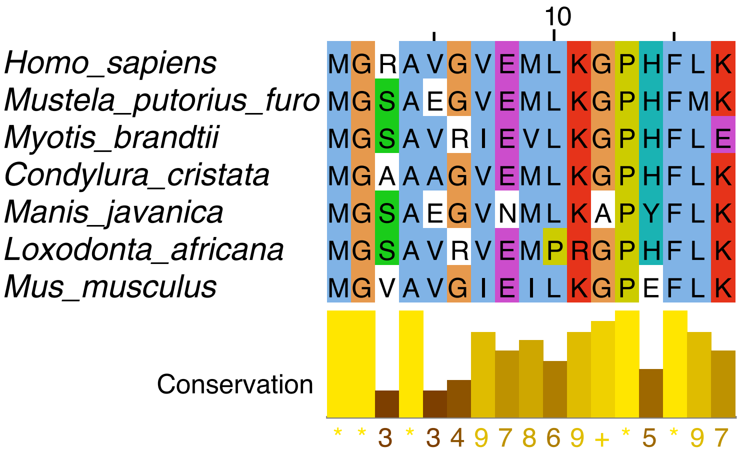

**HG0807.1**

ENSG00000147488 (ST18)

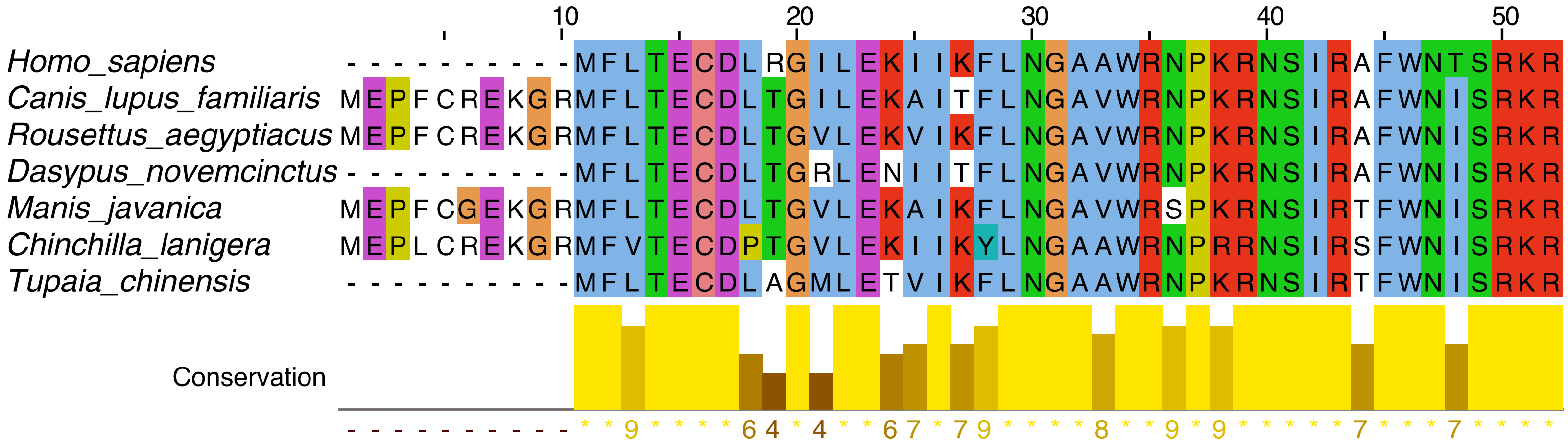

**HG0807.2**

ENSG00000147488 (ST18)

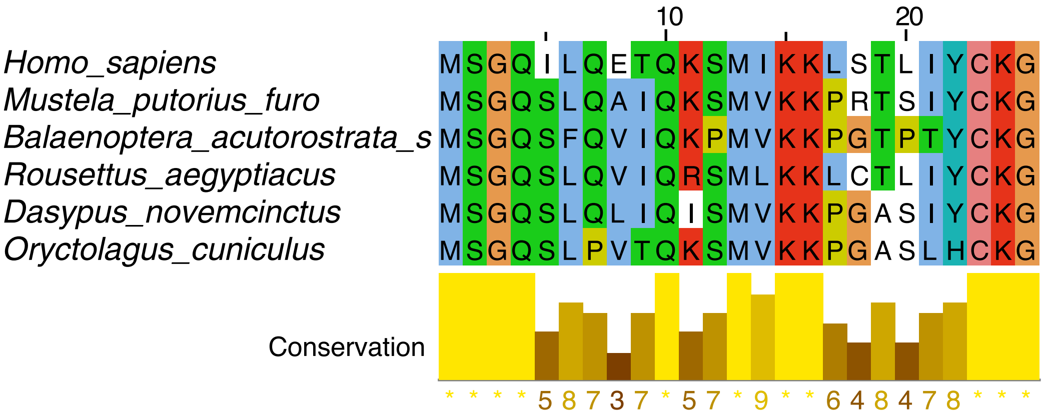

**HG0807.3**

ENSG00000186487 (MYT1L)

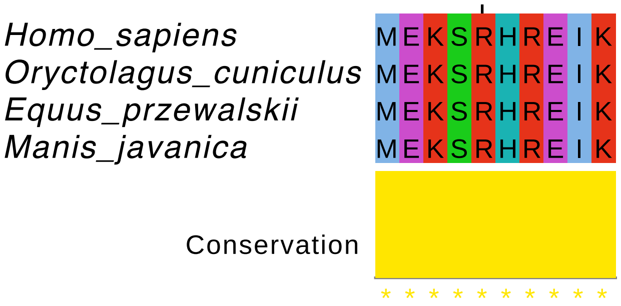

**HG0808**

ENSG00000149269 (PAK1)

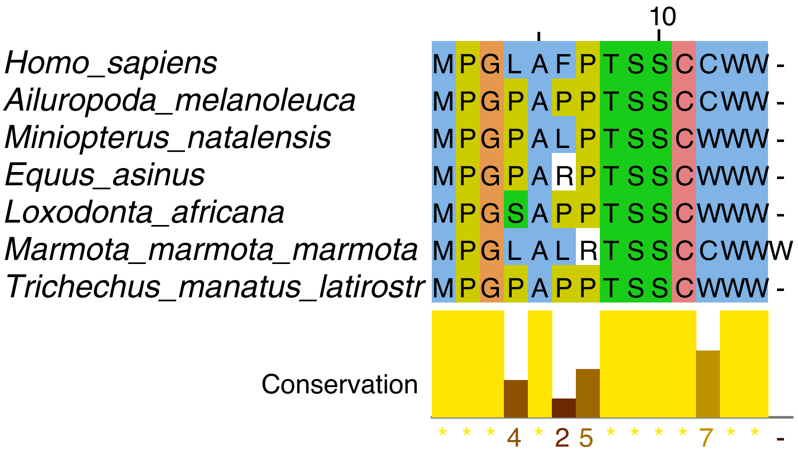

**HG0809**

ENSG00000151287 (TEX30)

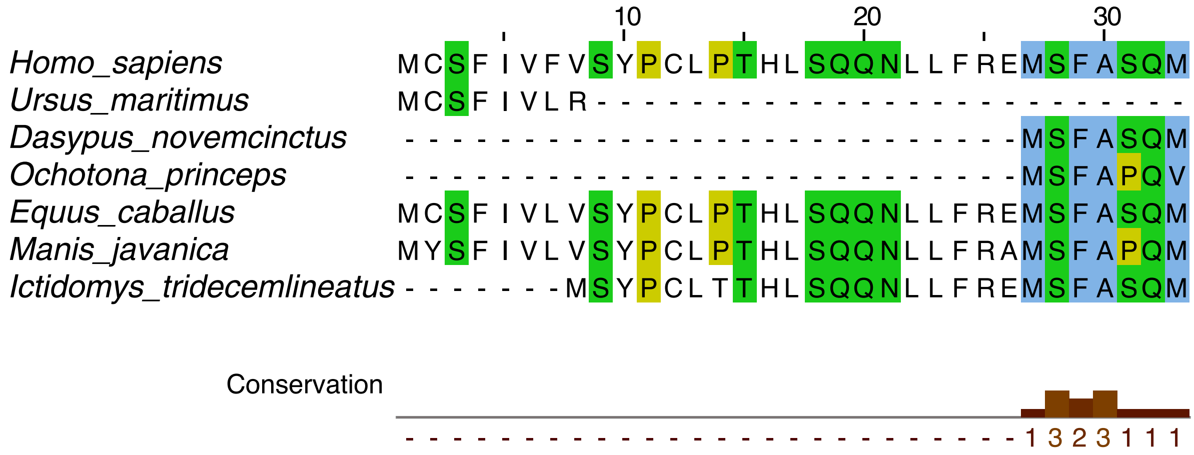

**HG0810**

ENSG00000156504 (FAM122B)

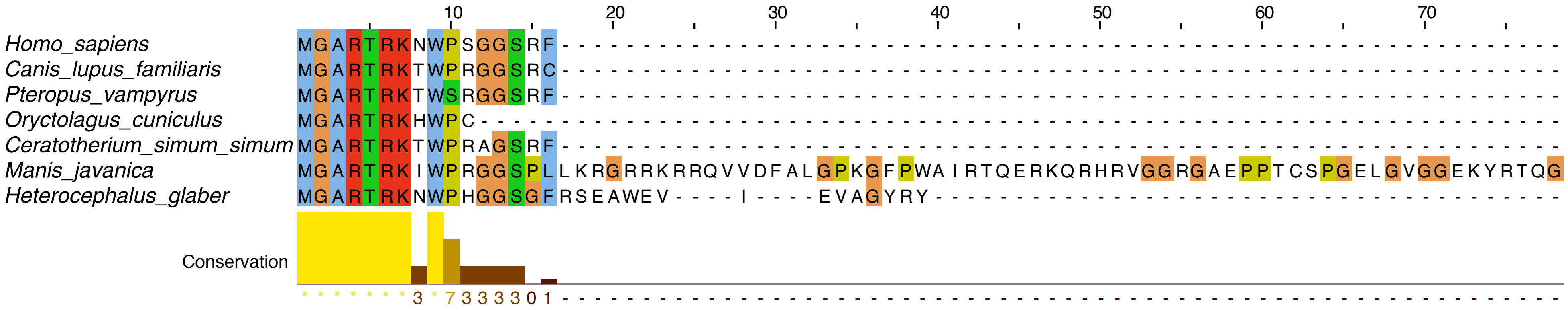

**HG0811**

ENSG00000157350 (ST3GAL2)

```
                                      10        20        30        40        50        60
Homo_sapiens                    MGSRRARPTAERISISAAEFSHTHRASCFCPLRVGVWLPSSGRGFGPGGGTTLAWNPSPLKPLCSLL
Ailuropoda_melanoleuca          MGTRRAEPSARRILVSAAEFSDTHRASWVSSLRTGALFPPLAEGLGQEGQFQWGRLSDGIQAT----
Balaenoptera_acutorostrata_s    MGSRRAEPNARRILVS-EEFSGTHRASWVSSLWIGAWFPFLPWQRGWALRAN---------------
Miniopterus_natalensis          --------------------------------------------------MPLAWNQGCSKPTCSLL
Galeopterus_variegatus          --------------------------------------------------MTLAWNLGPLKPTCSPL
Equus_asinus                    MGRGTAEPNTRRILVSAAEFSDTHRASCVSSLRIGAWFPPLAEGLGQEGQQRWGRLLHGIQAA----
Rattus_norvegicus               -------------------------------------------------MRTRAWDAGLLKPVCSLL

Conservation                    --------------------------------------------------3403100000111----
```

**HG0812**

ENSG00000157368 (IL34)

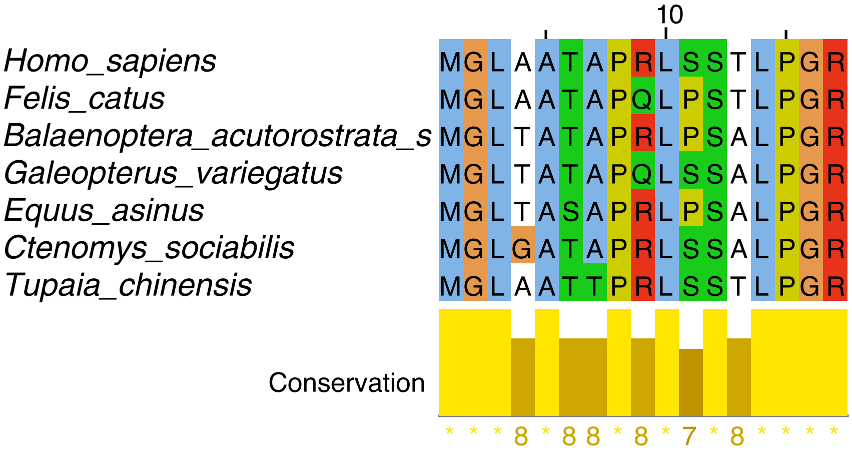

**HG0813**

ENSG00000160199 (PKNOX1)

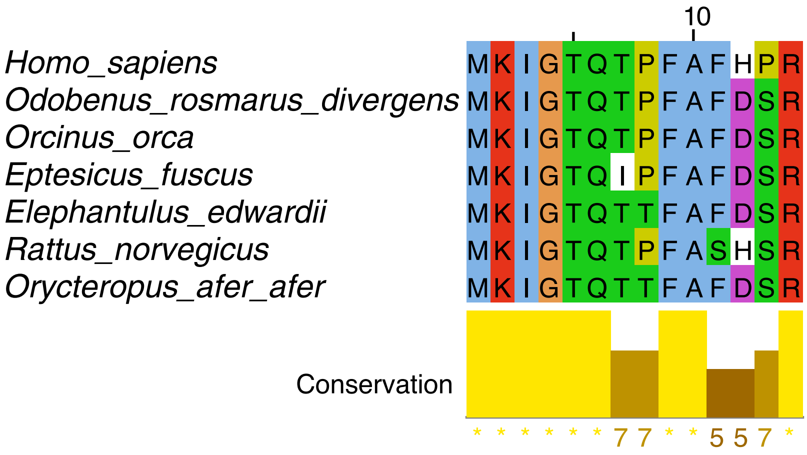

**HG0814**

ENSG00000163071 (SPATA18)

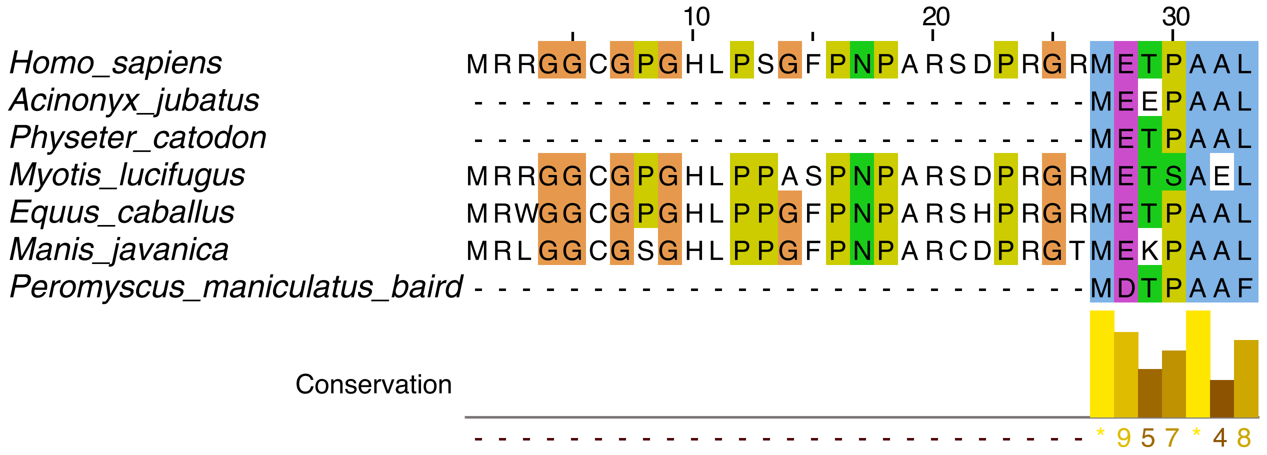

**HG0815**

ENSG00000163072 (NOSTRIN)

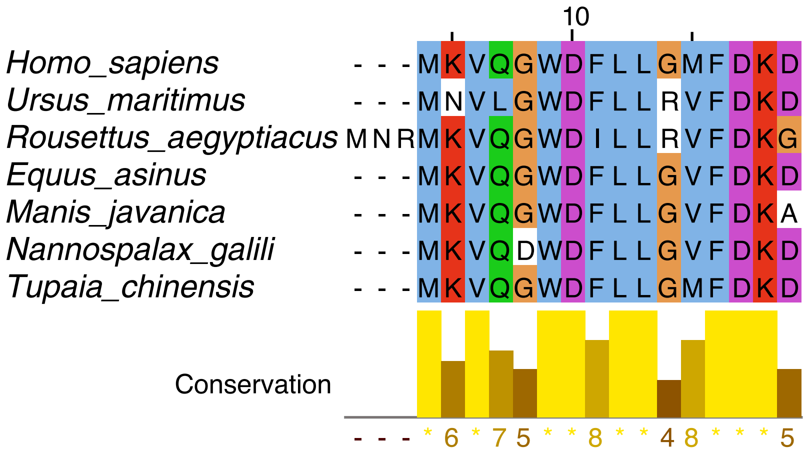

**HG0816**

ENSG00000163794 (UCN)

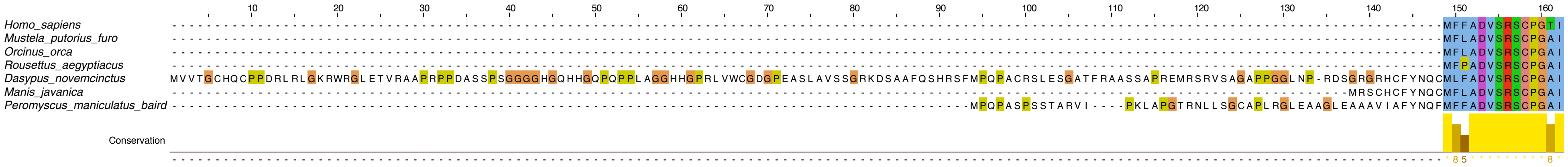

**HG0817**

ENSG00000163888 (CAMK2N2)

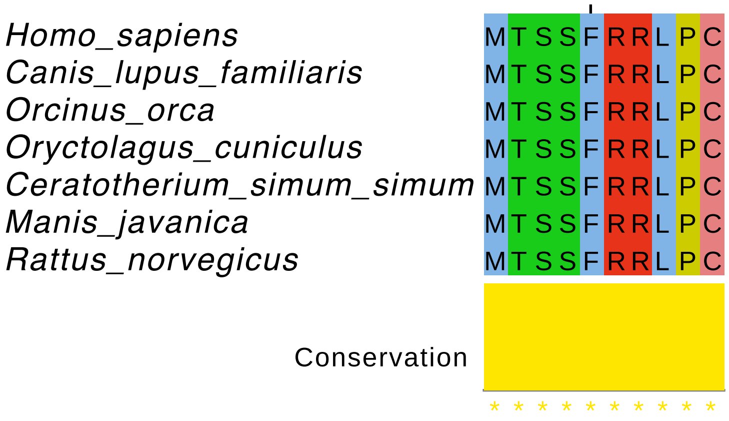

**HG0818**

ENSG00000164125 (FAM198B)

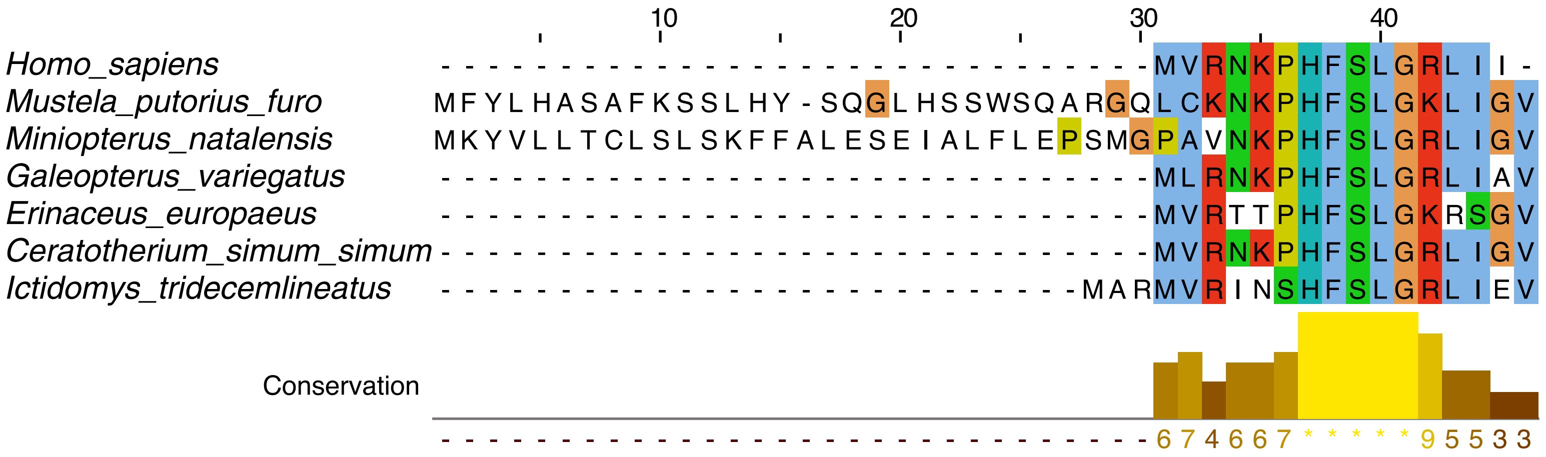

**HG0819**

ENSG00000164188 (RANBP3L)

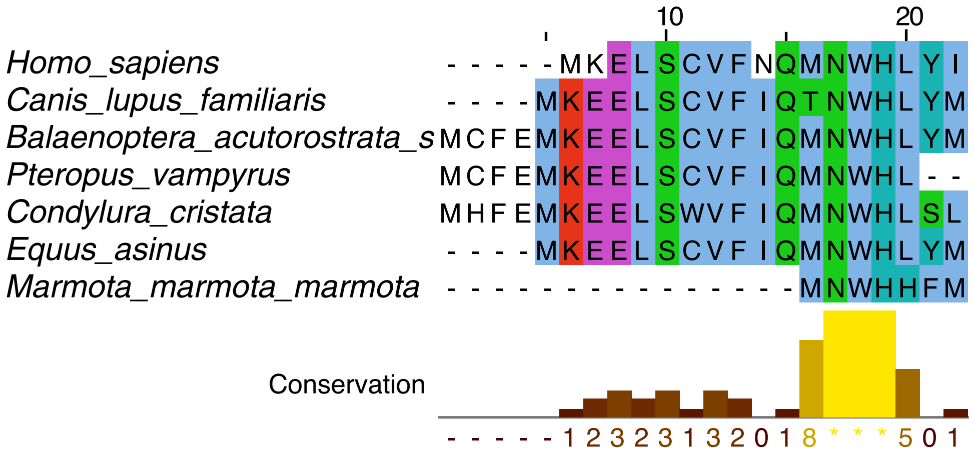

**HG0820**

ENSG00000164946 (FREM1)

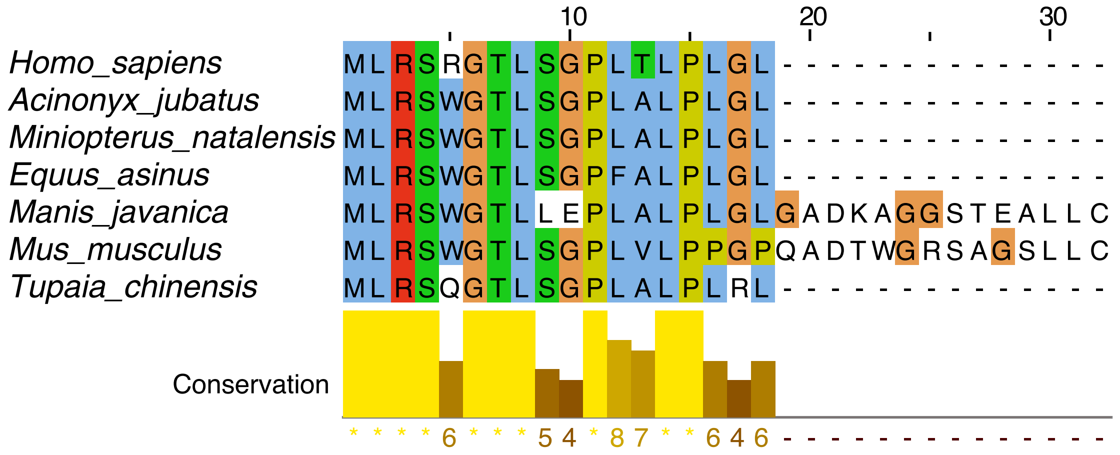

**HG0821**

ENSG00000165300 (SLITRK5)

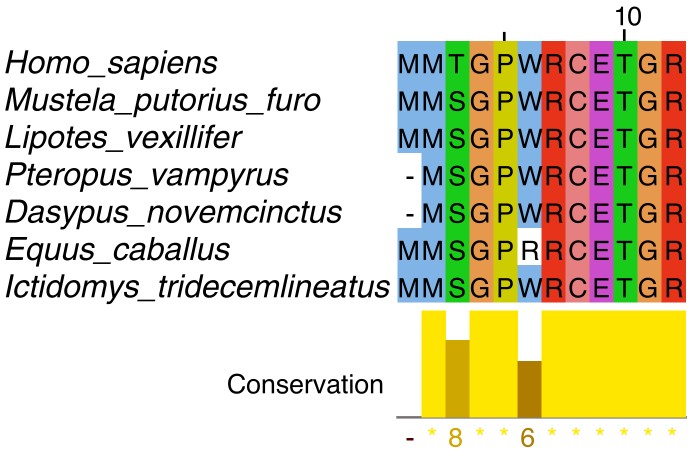

**HG0822**

ENSG00000166170 (BAG5)

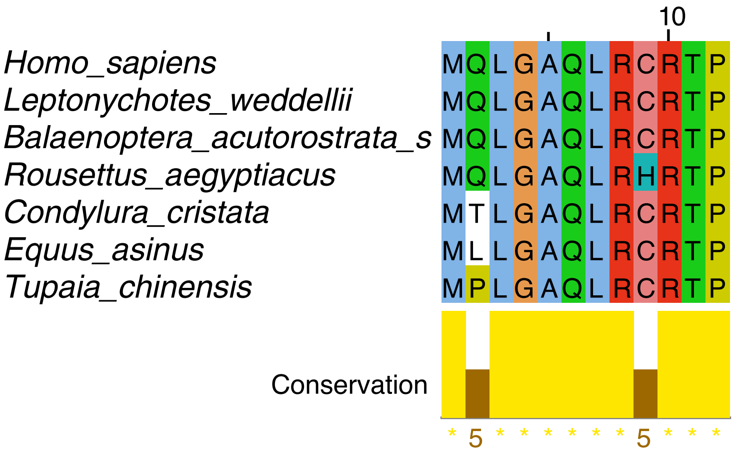

**HG0823**

ENSG00000166268 (MYRFL)

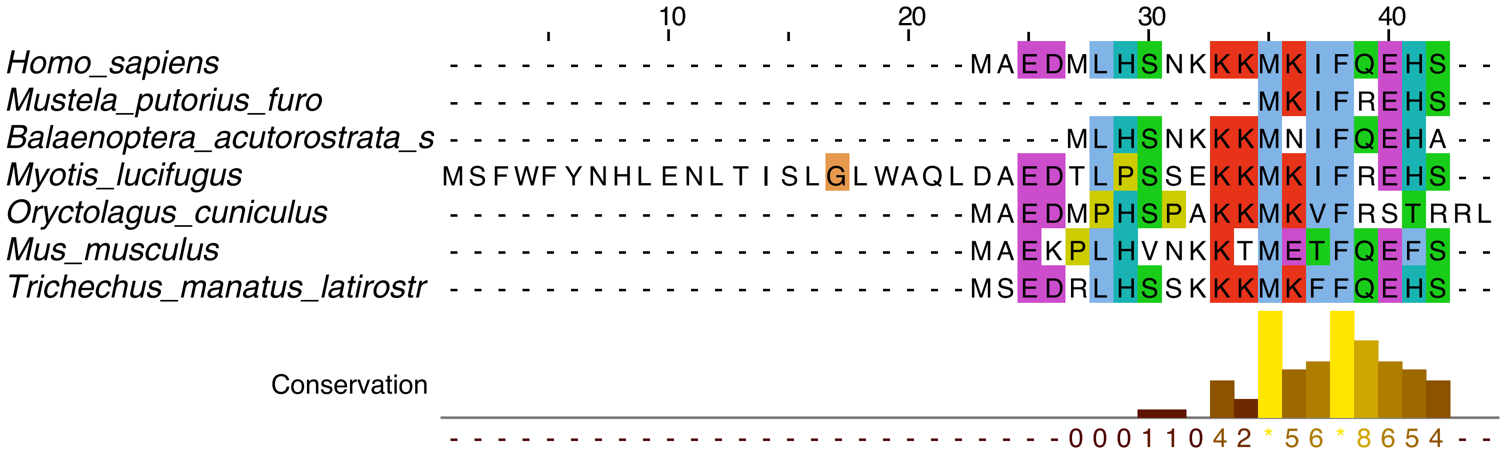

**HG0824**

ENSG00000167100

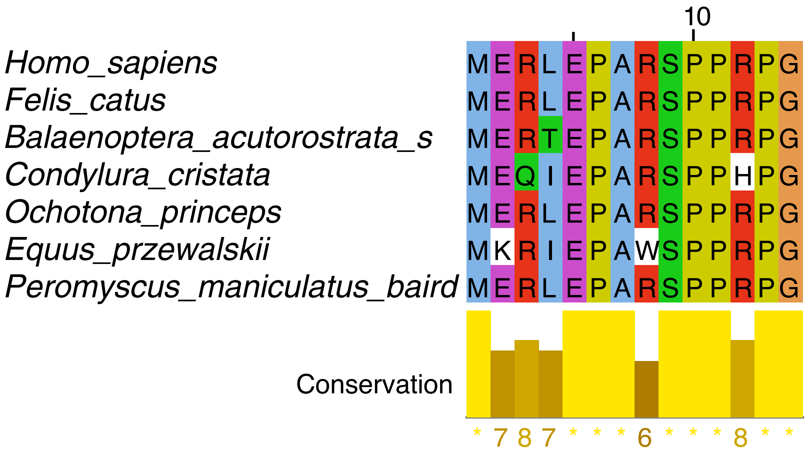

**HG0825**

ENSG00000169180 (XPO6)

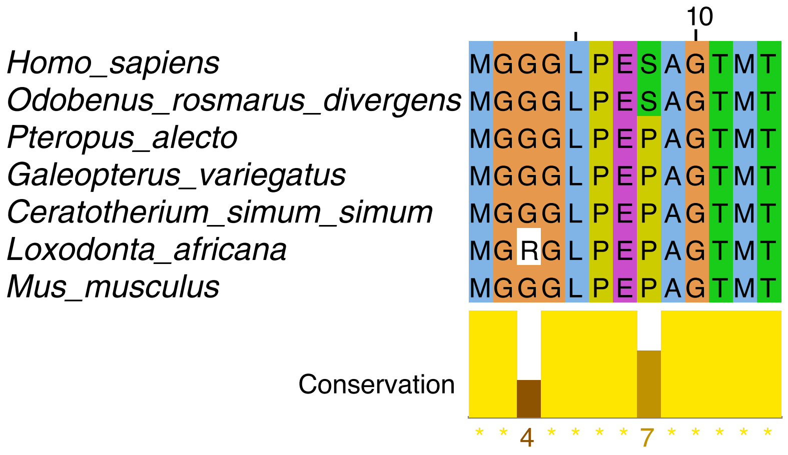

**HG0826**

ENSG00000169851 (PCDH7)

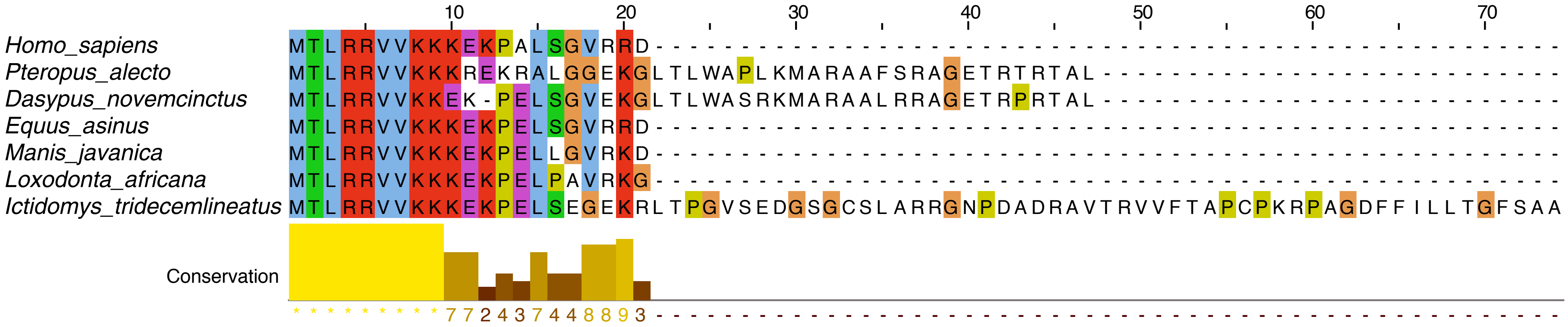

## HG0827

ENSG00000170365 (SMAD1)

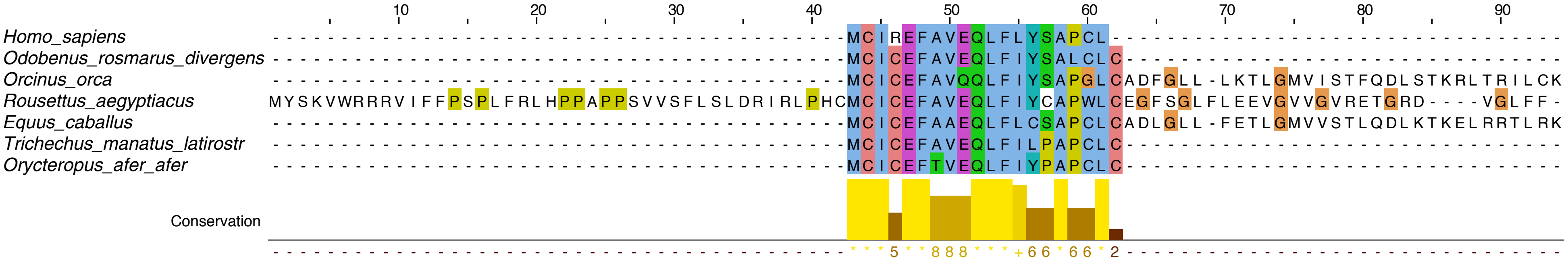

**HG0828**

ENSG00000170390 (DCLK2)

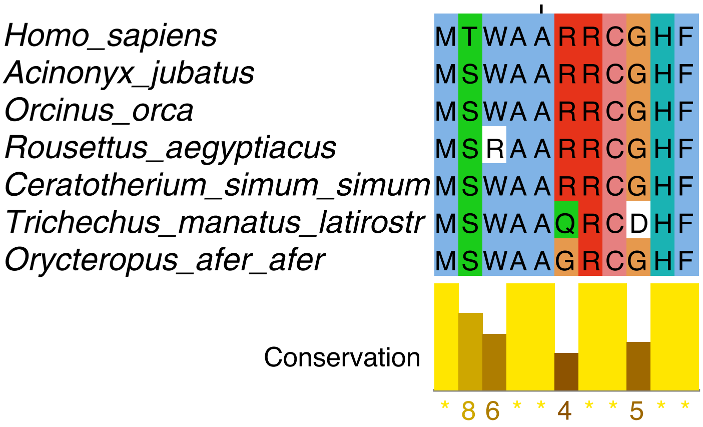

**HG0829**

ENSG00000171502 (COL24A1)

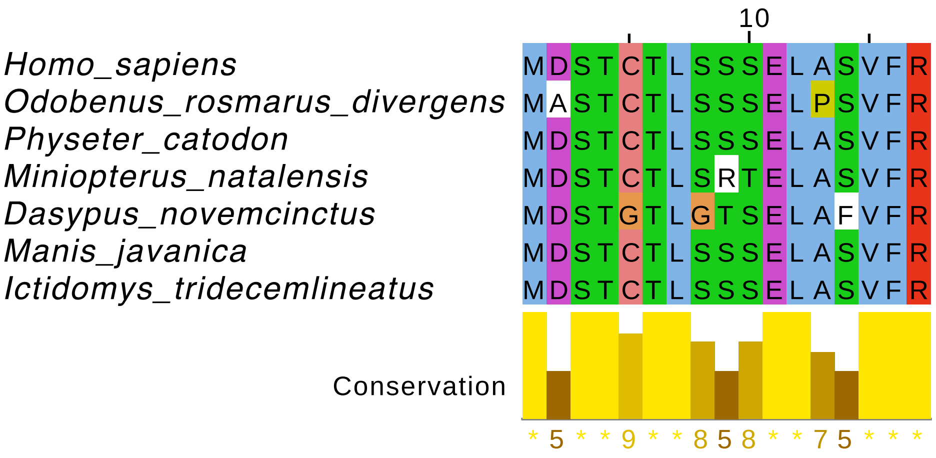

**HG0830**

ENSG00000171631 (P2RY6)

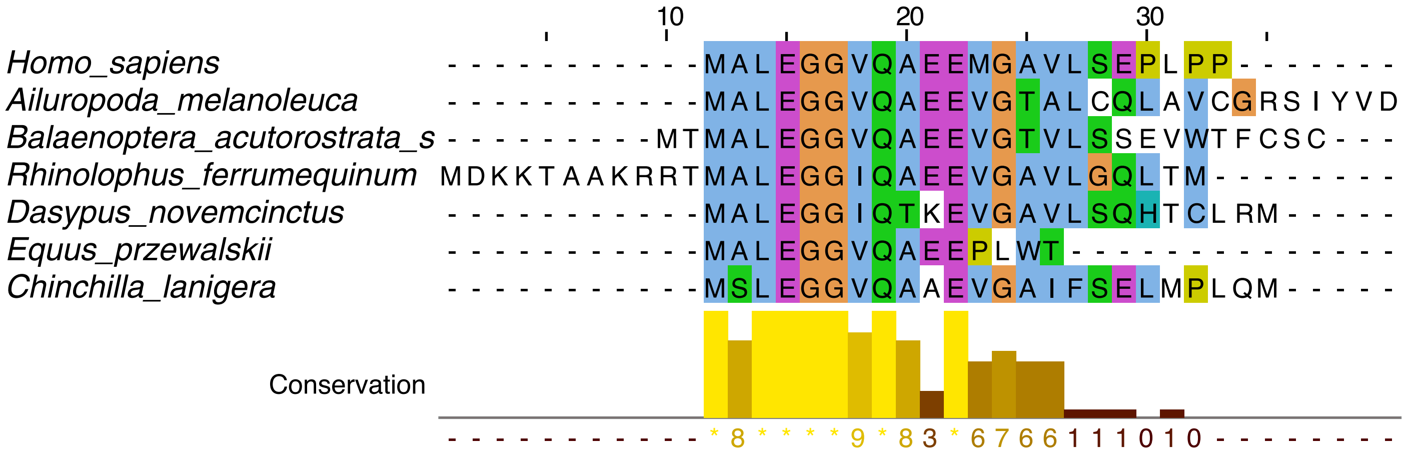

**HG0831.1**

ENSG00000174482 (LINGO2)

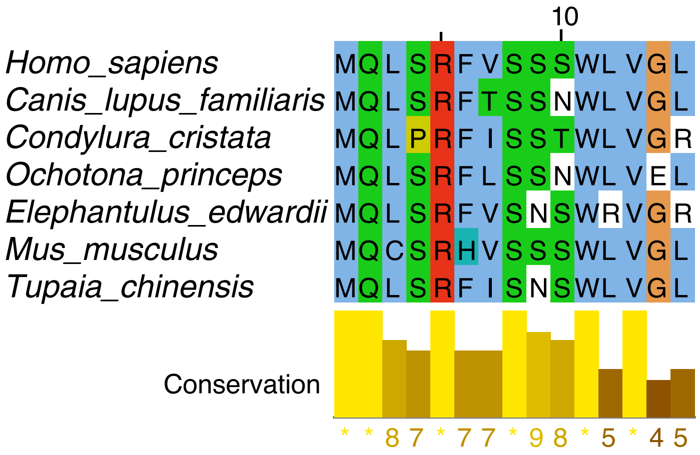

**HG0831.2**

ENSG00000174482 (LINGO2)

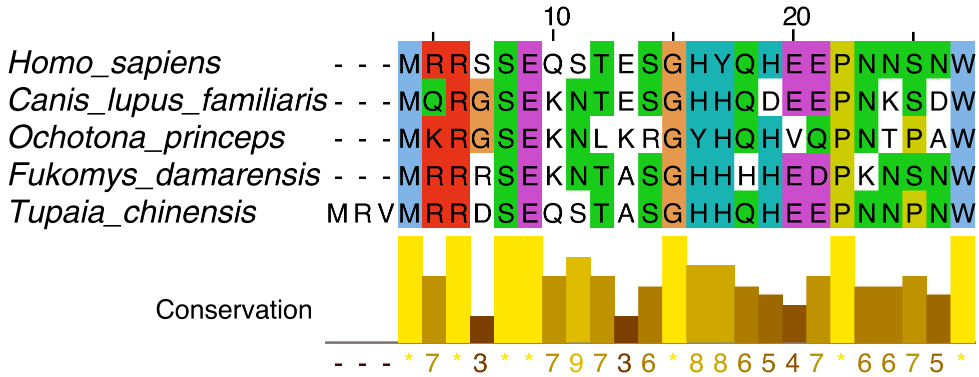

**HG0832**

ENSG00000175048 (ZDHHC14)

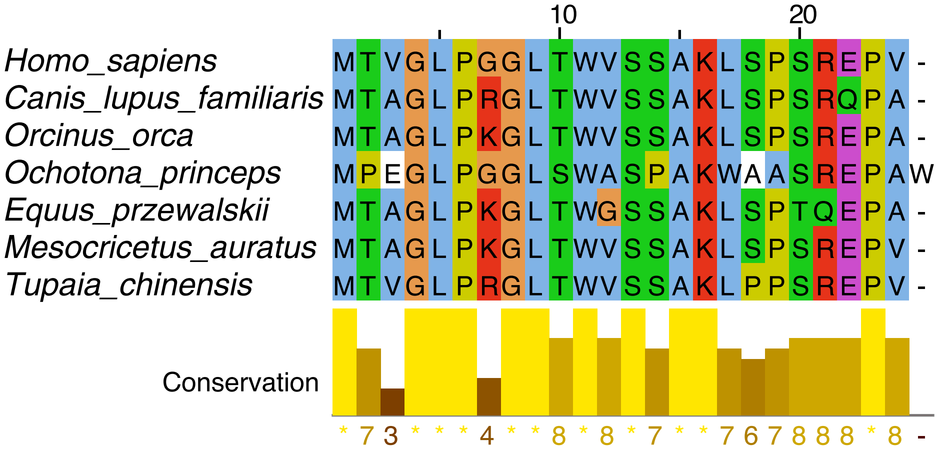

**HG0833**

ENSG00000176641 (RNF152)

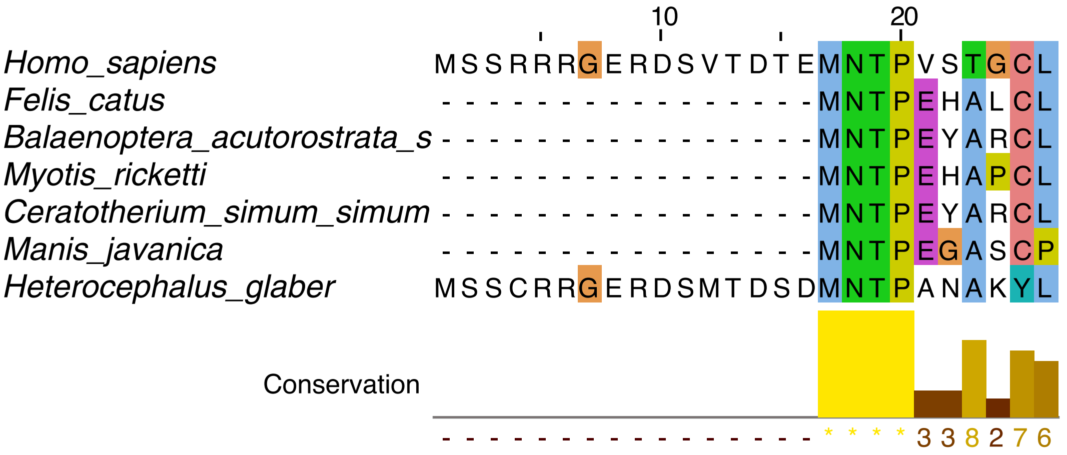

**HG0834**

ENSG00000182489 (XKRX)

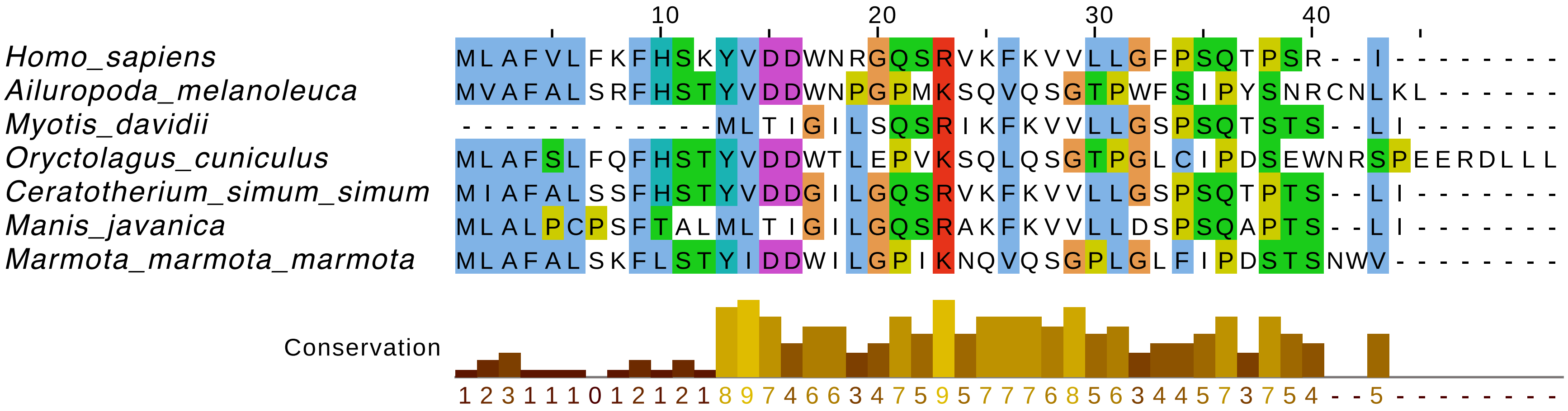

**HG0835**

ENSG00000182901 (RGS7)

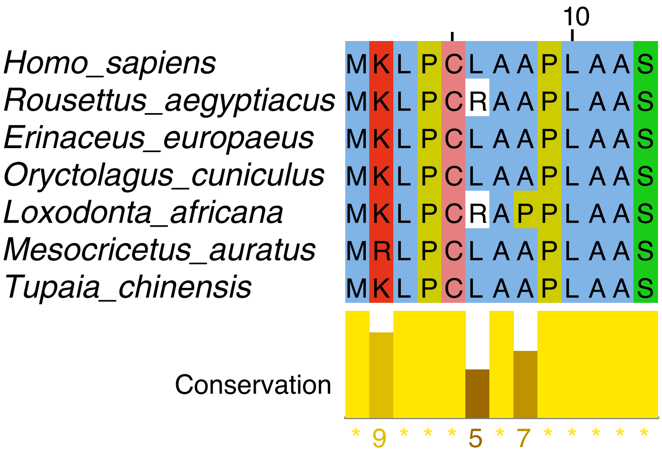

**HG0836**

ENSG00000197461 (PDGFA)

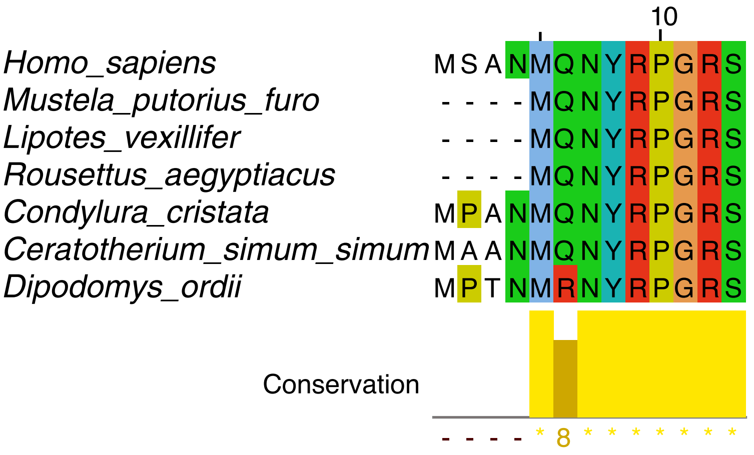

**HG0837**

ENSG00000198690 (FAN1)

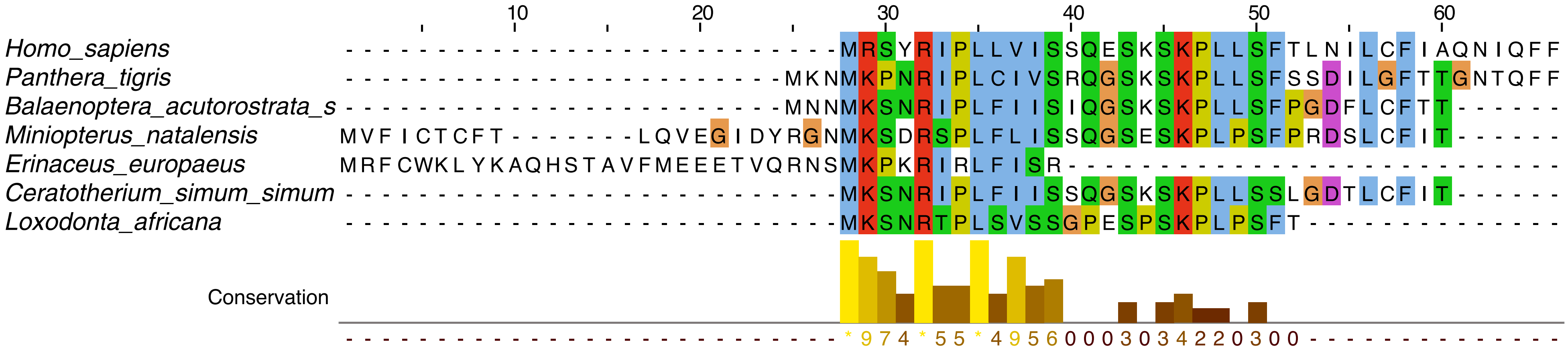

**HG0838**

ENSG00000198707 (CEP290)

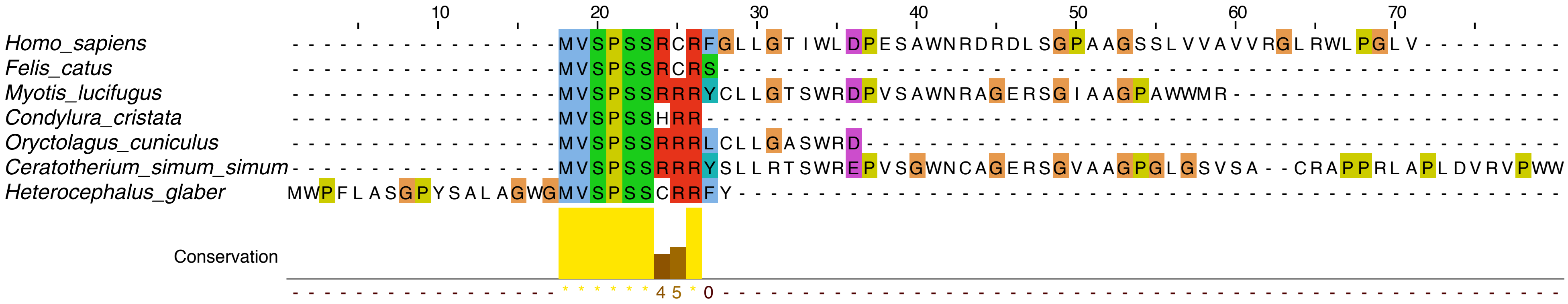

**HG0839**

ENSG00000198791 (CNOT7)

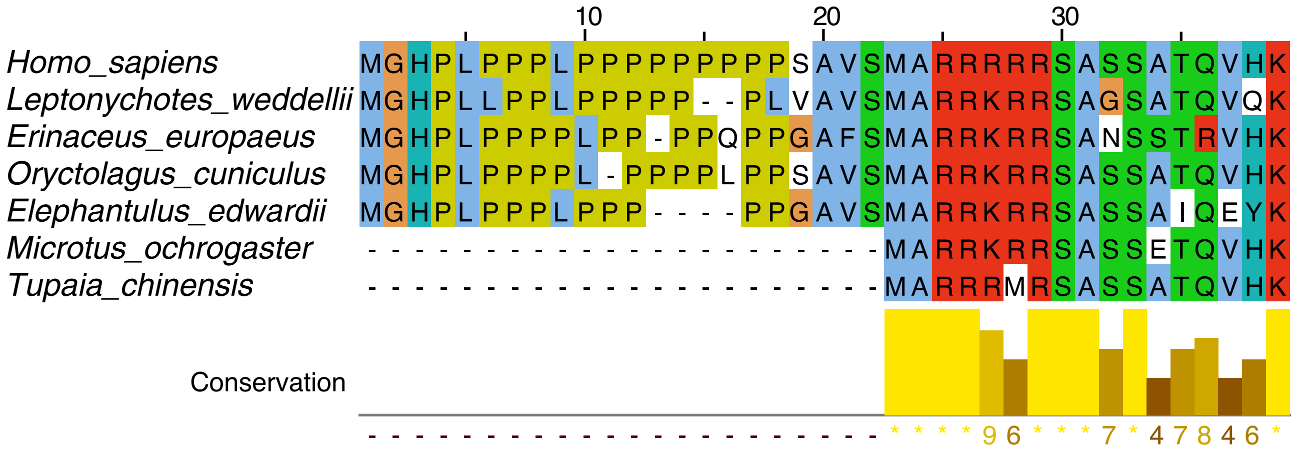

**HG0840**

ENSG00000278195 (SSTR3)

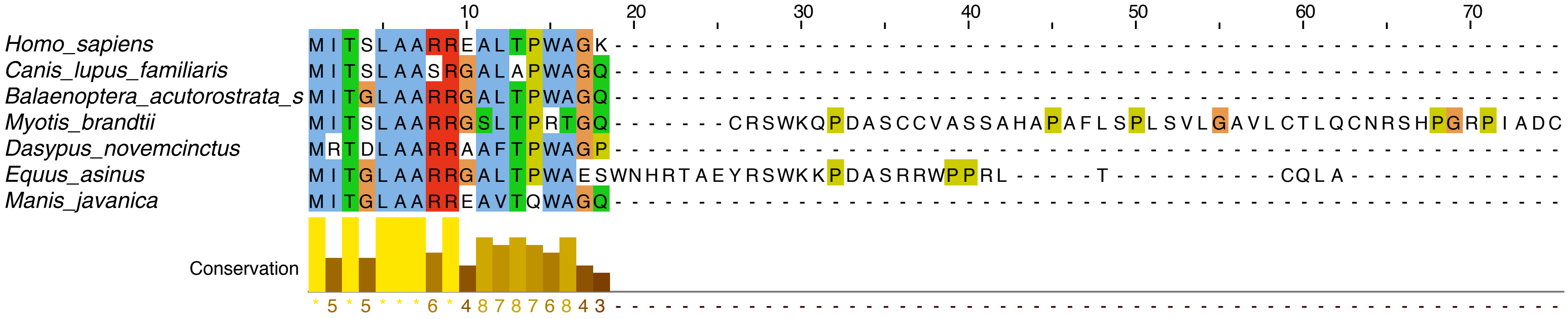

**HG0841**

ENSG00000278259 (MYO19)

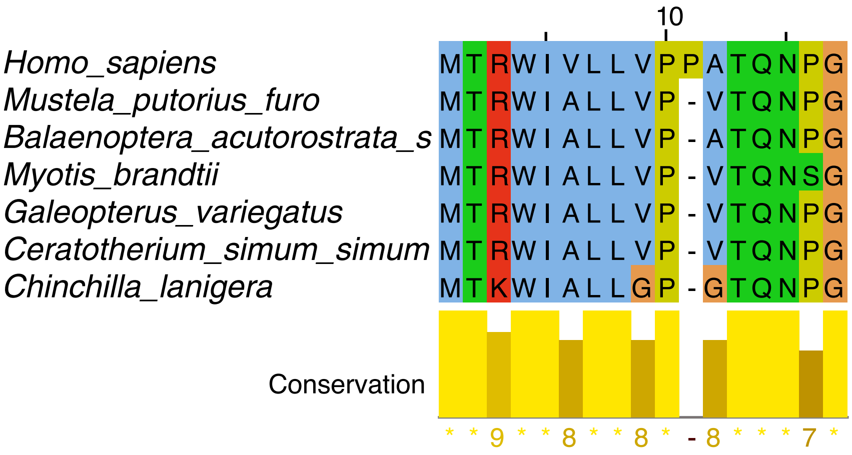

**HG0842**

ENSG00000002746 (HECW1)

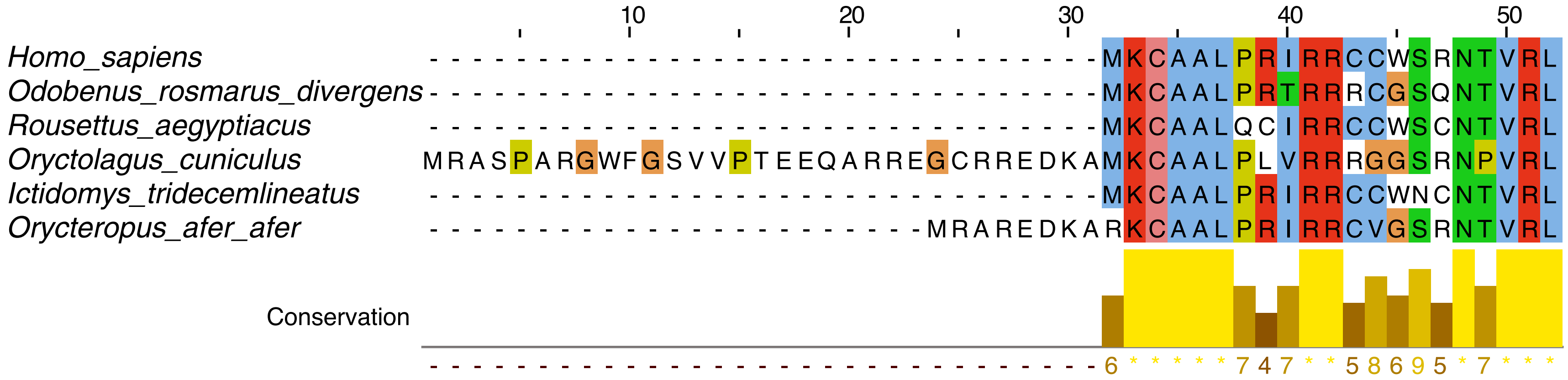

**HG0843**

ENSG00000043093 (DCUN1D1)

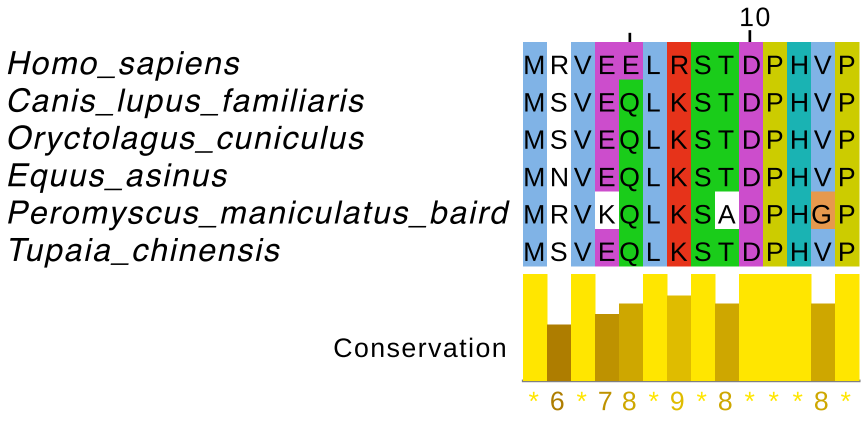

**HG0844**

ENSG00000091009 (RBM27)

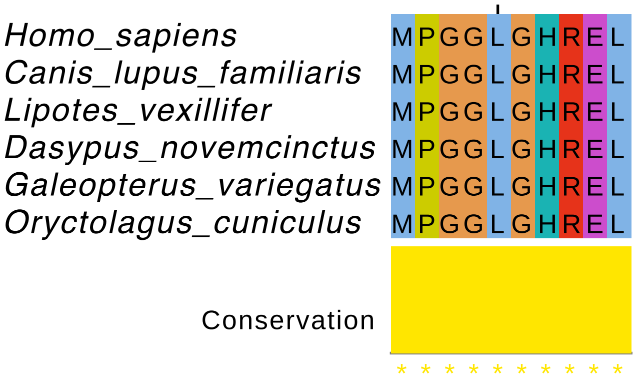

**HG0845**

ENSG00000099260 (PALMD)

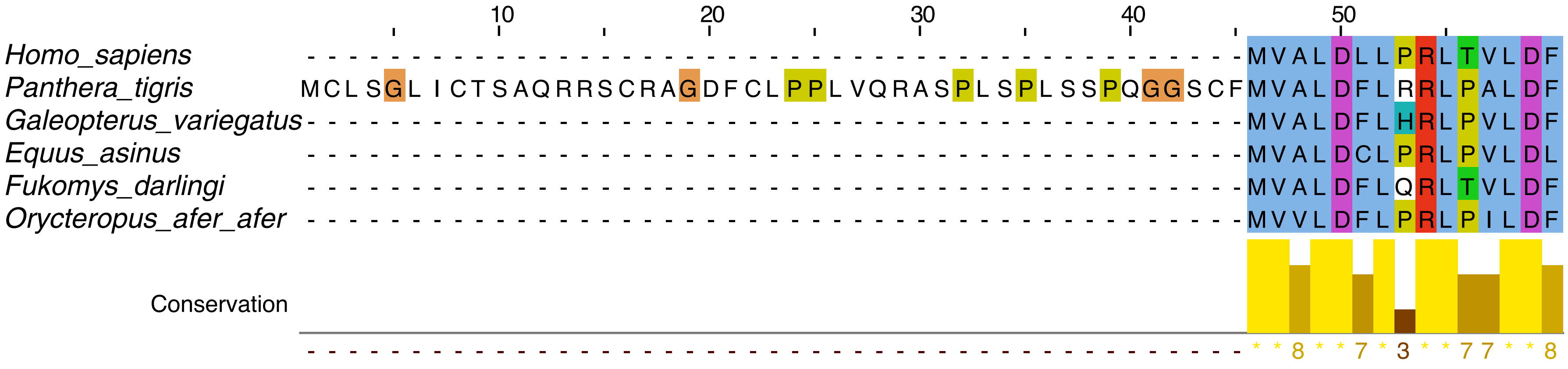

**HG0846**

ENSG00000100852 (ARHGAP5)

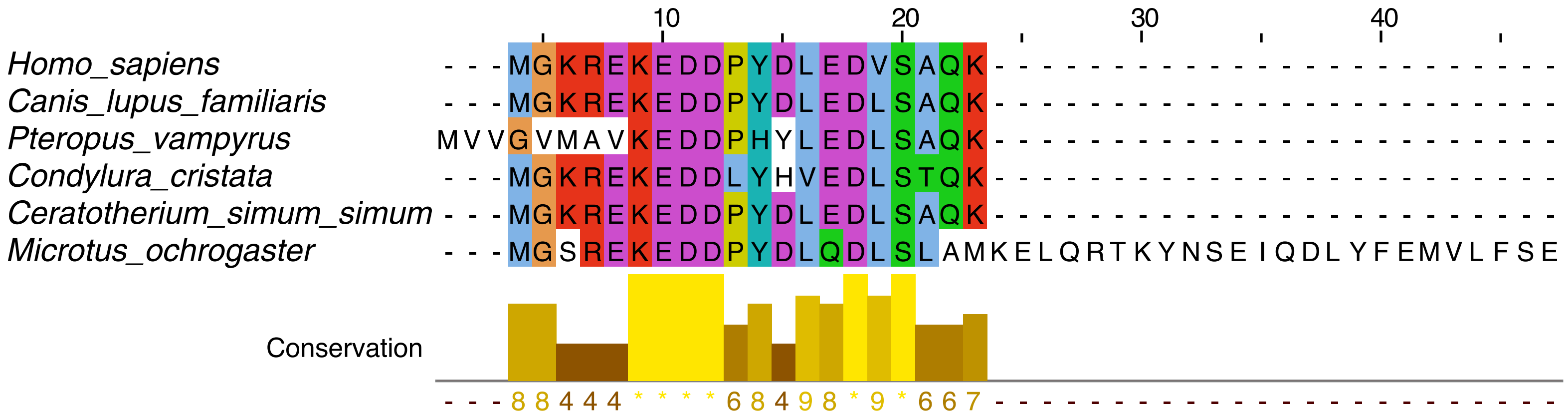

**HG0847**

ENSG00000106086 (PLEKHA8)

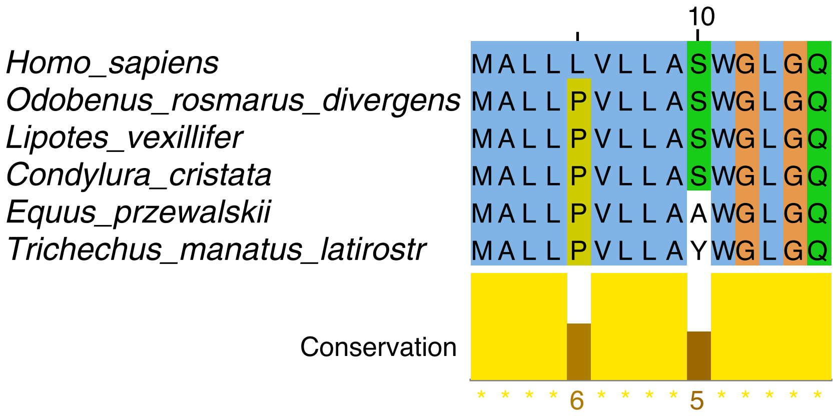

**HG0848**

ENSG00000111817 (DSE)

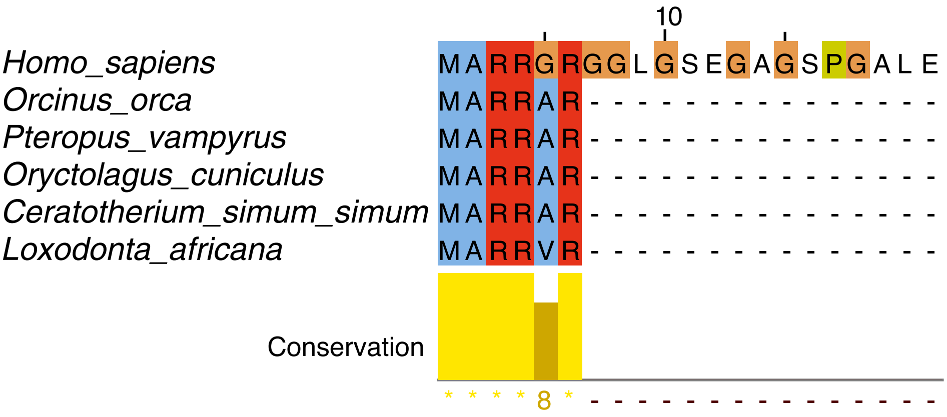

**HG0849**

ENSG00000111846 (GCNT2)

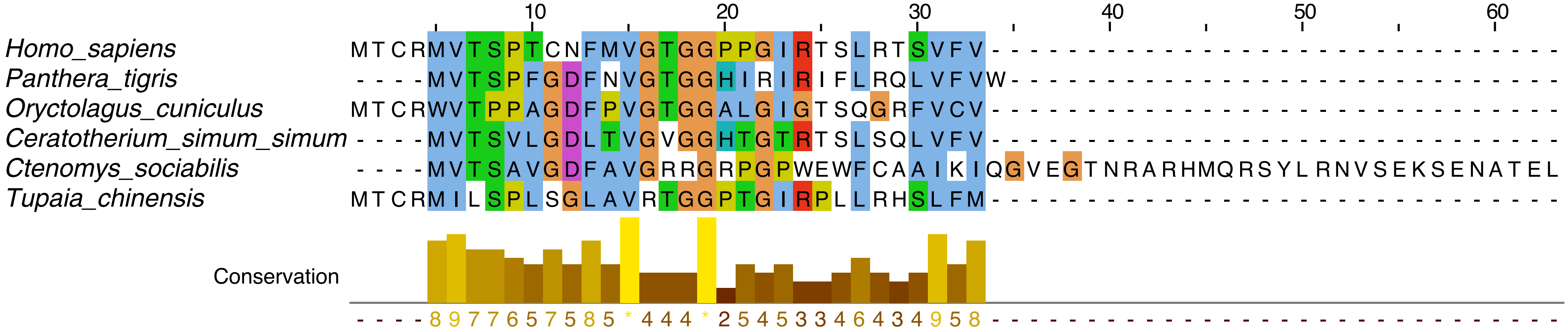

**HG0850**

ENSG00000113916 (BCL6)

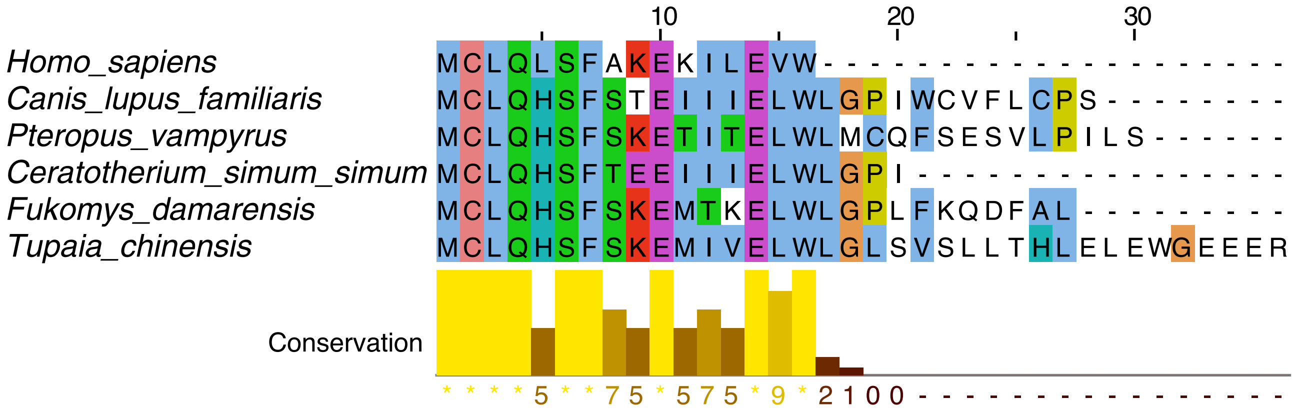

**HG0851**

ENSG00000114279 (FGF12)

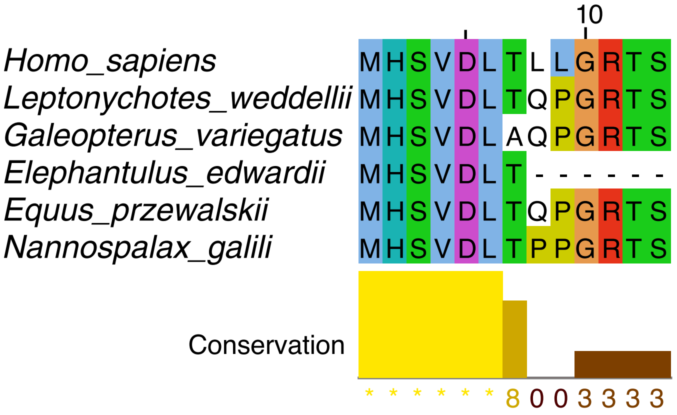

**HG0852**

ENSG00000114942 (EEF1B2)

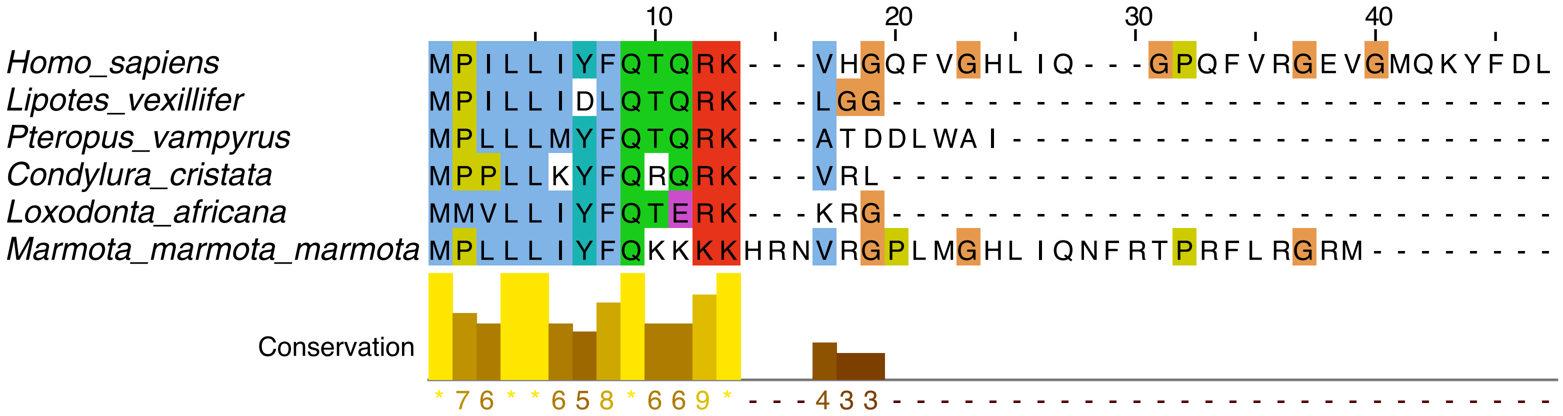

**HG0853**

ENSG00000121807 (CCR2)

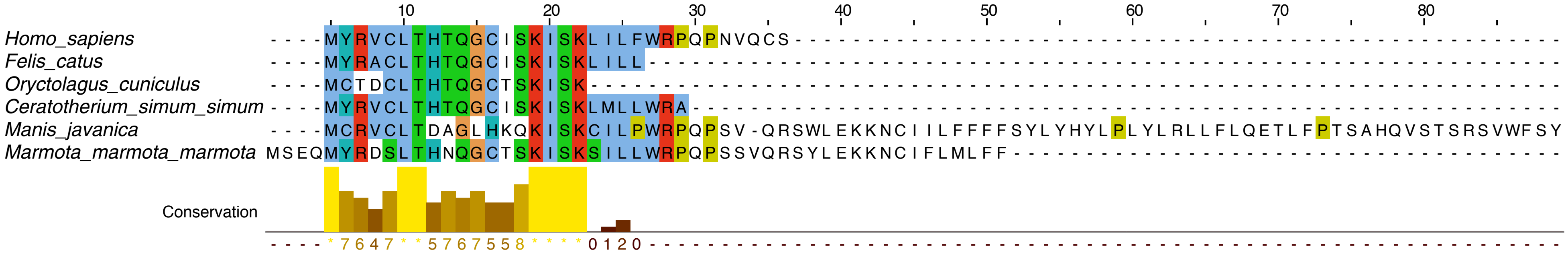

**HG0854**

ENSG00000127249 (ATP13A4)

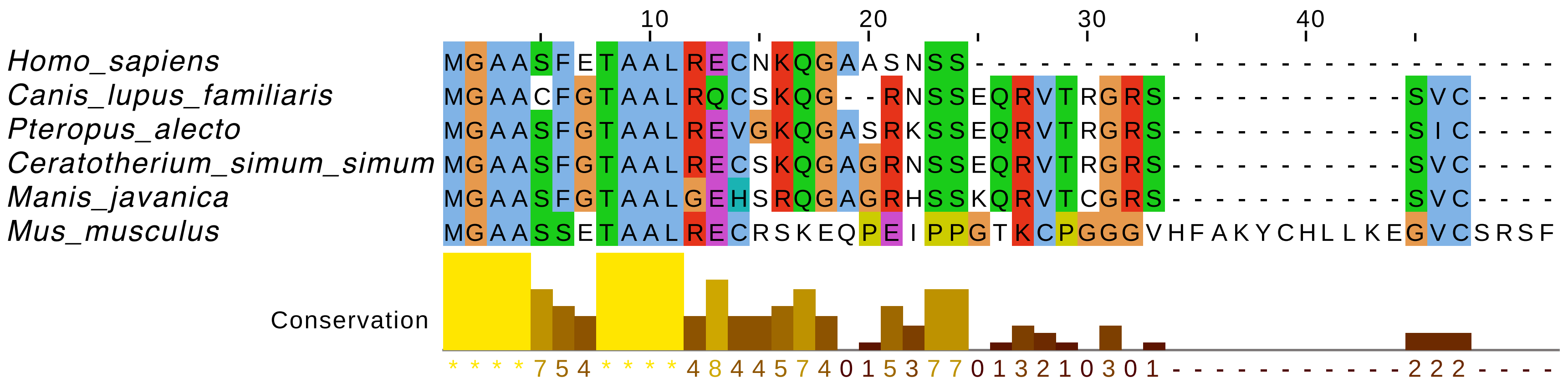

**HG0855**

ENSG00000127804 (METTL16)

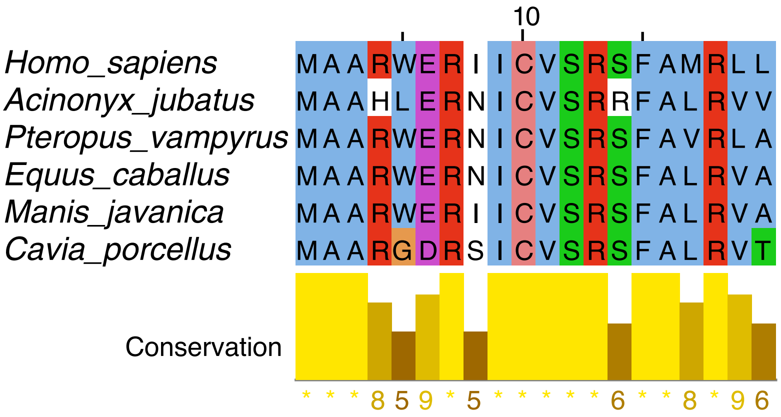

**HG0856**

ENSG00000130055 (GDPD2)

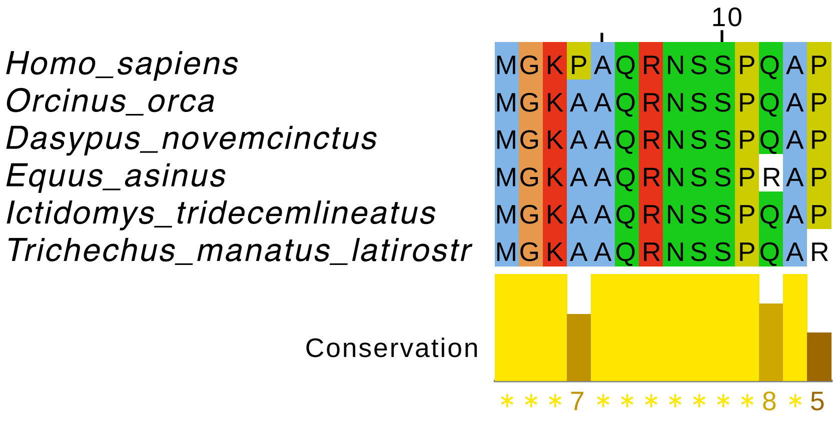

**HG0857**

ENSG00000130347 (RTN4IP1)

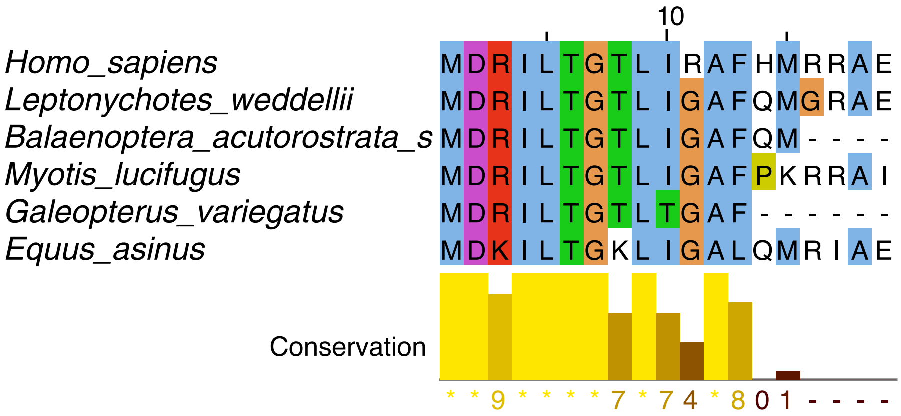

**HG0858**

ENSG00000132702 (HAPLN2)

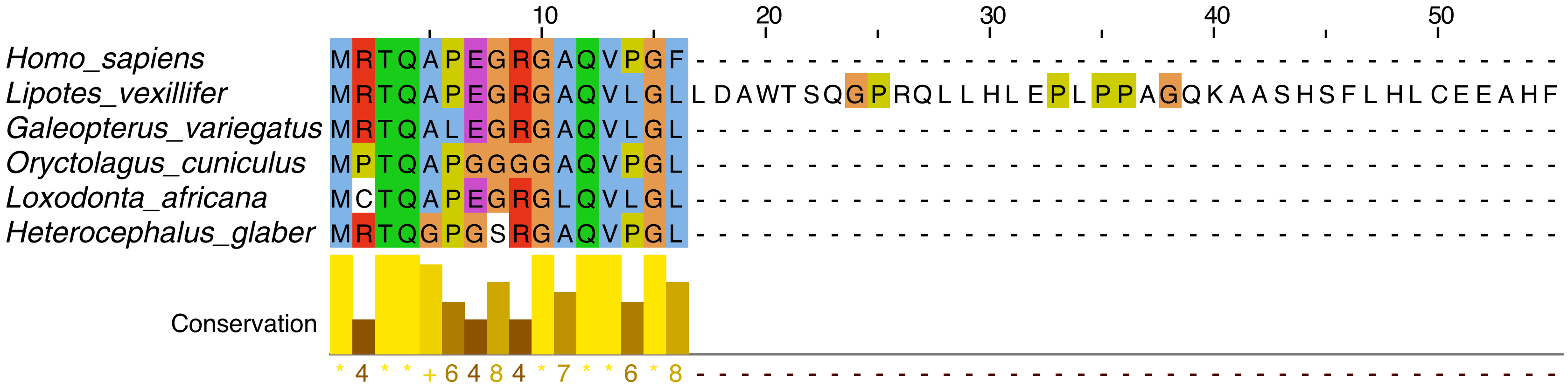

**HG0859**

ENSG00000132842 (AP3B1)

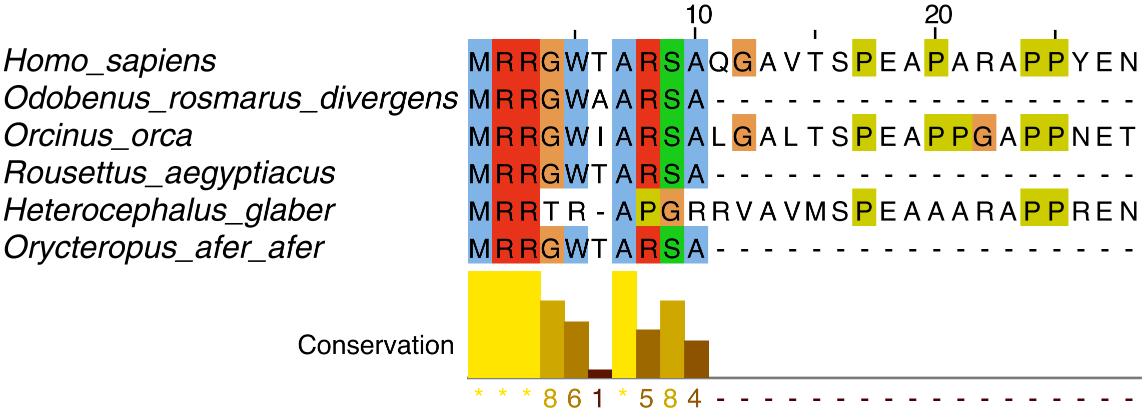

**HG0860**

ENSG00000133056 (PIK3C2B)

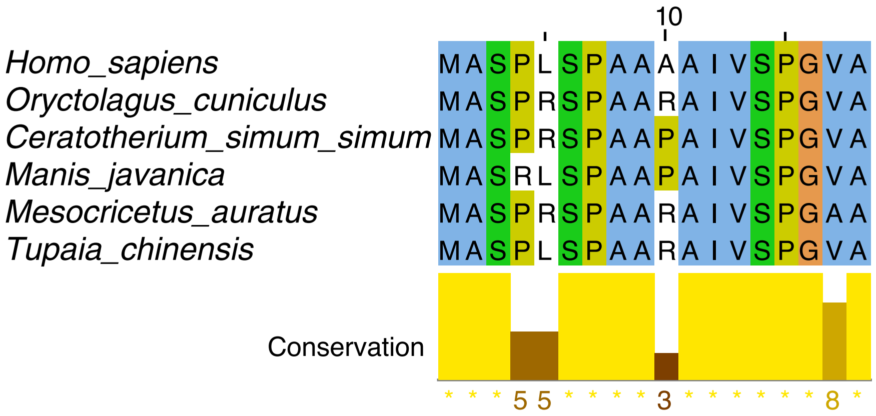

**HG0861**

ENSG00000134001 (EIF2S1)

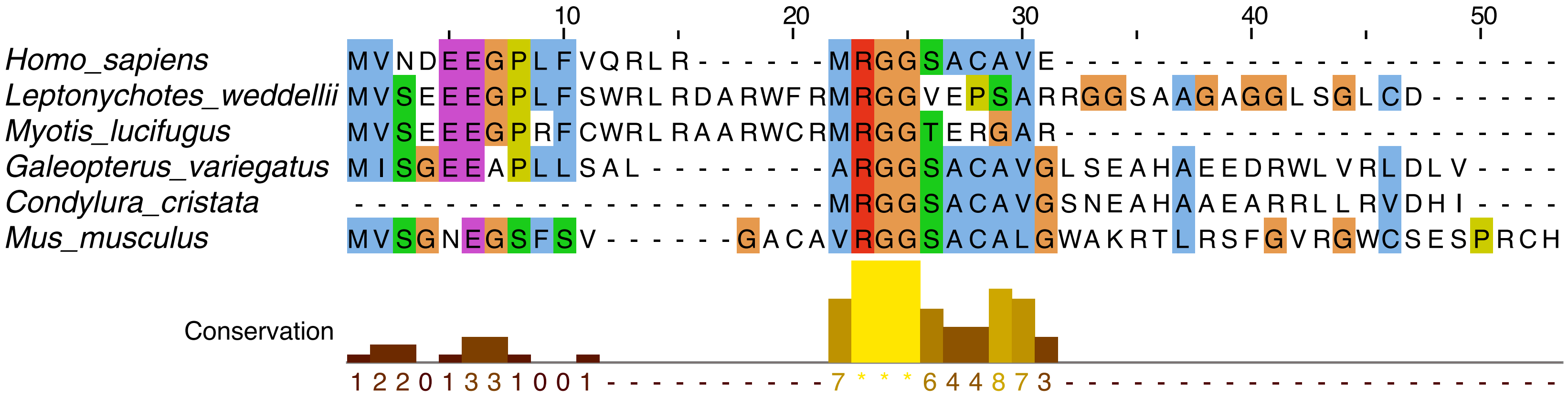

**HG0862**

ENSG00000134245 (WNT2B)

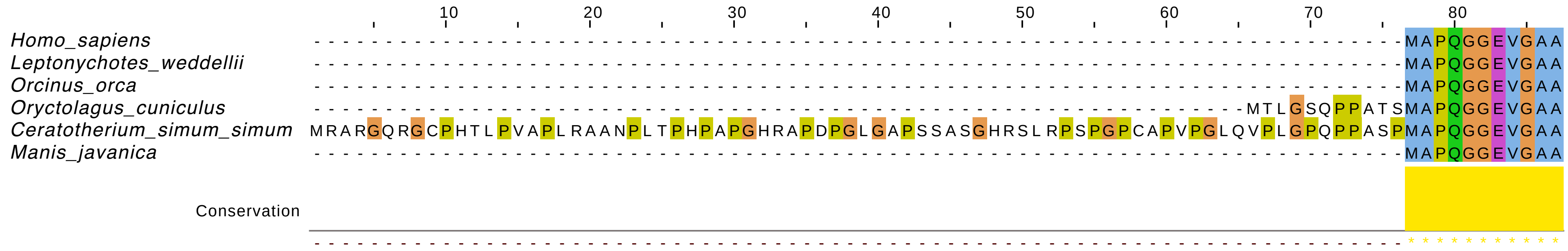

**HG0863**

ENSG00000134313 (KIDINS220)

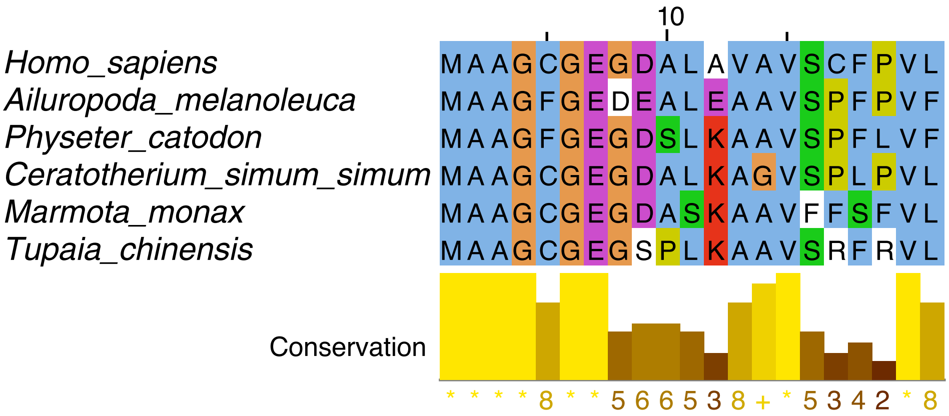

**HG0864**

ENSG00000134533 (RERG)

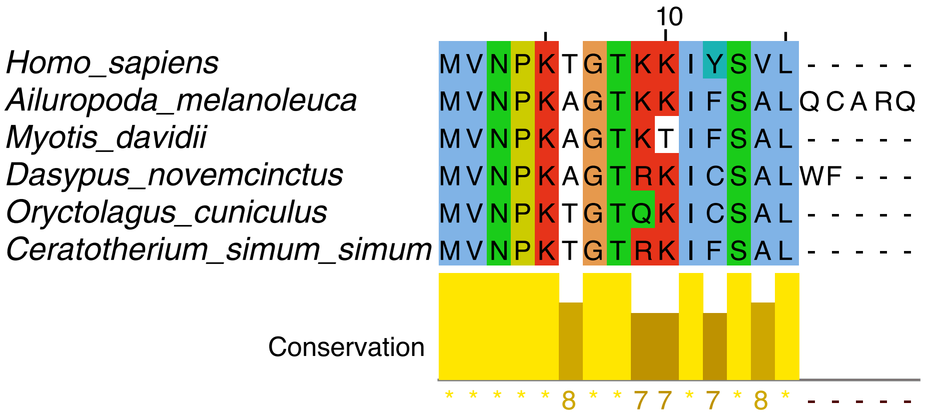

**HG0865**

ENSG00000135540 (NHSL1)

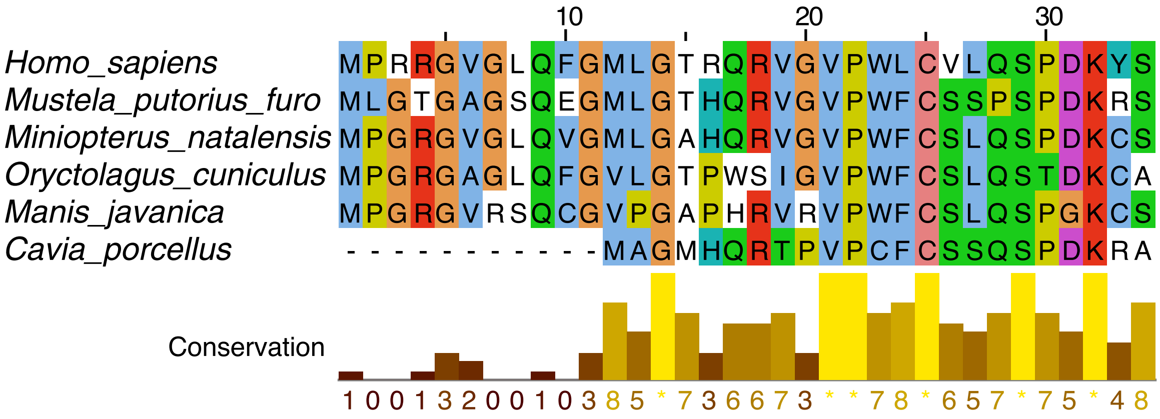

**HG0866**

ENSG00000136011 (STAB2)

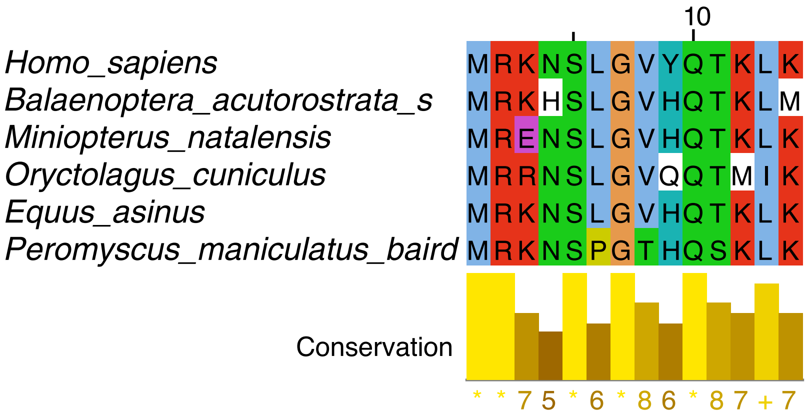

**HG0867**

ENSG00000137203 (TFAP2A)

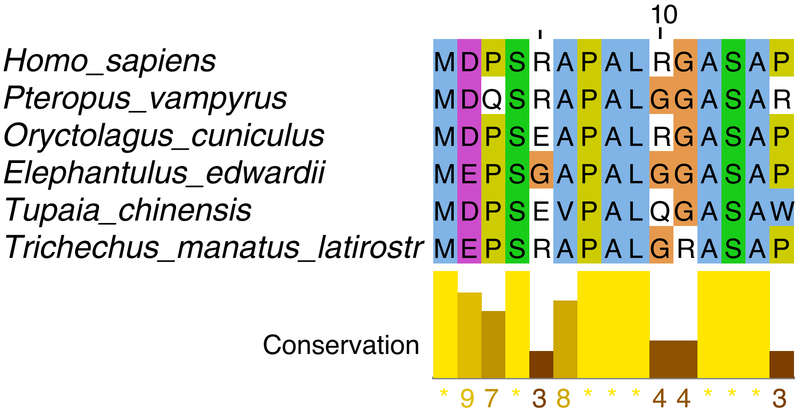

**HG0868**

ENSG00000137843 (PAK6)

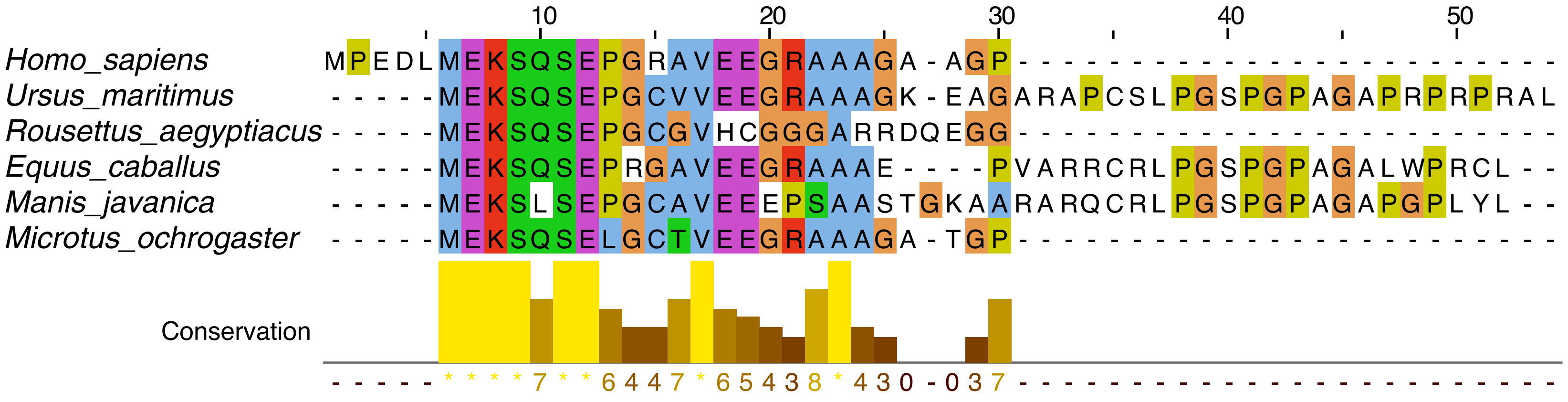

**HG0869**

ENSG00000139910 (NOVA1)

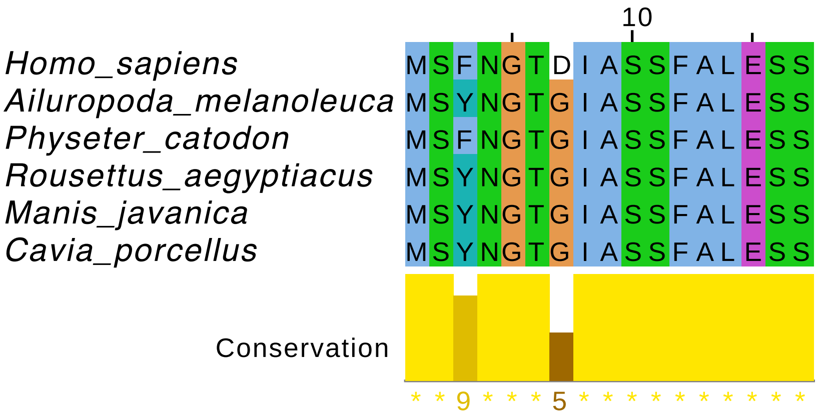

**HG0870**

ENSG00000140993 (TIGD7)

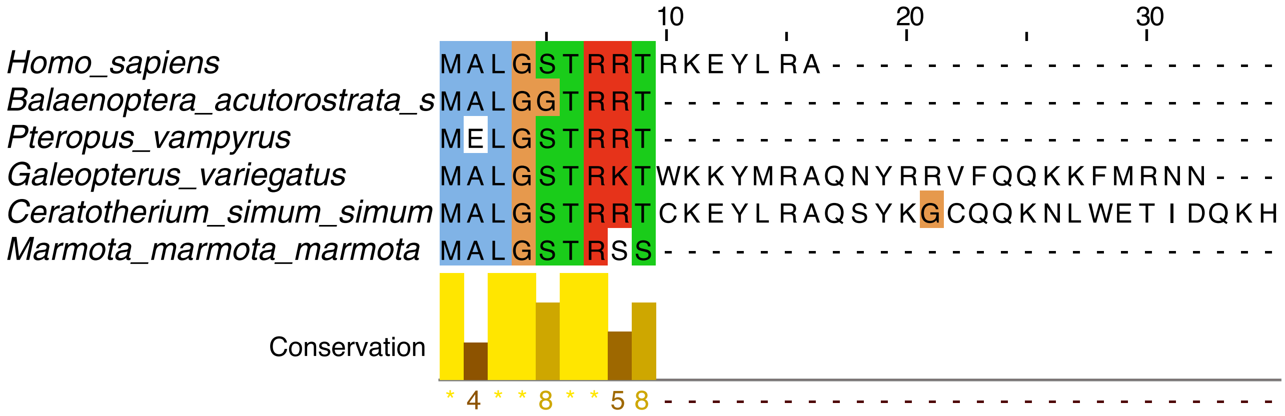

**HG0871**

ENSG00000141127 (PRPSAP2)

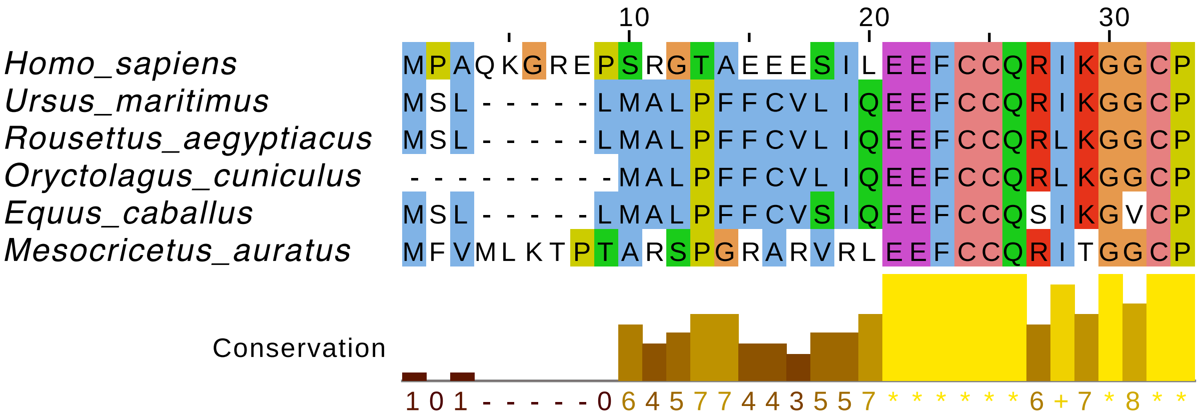

**HG0872**

ENSG00000142583 (SLC2A5)

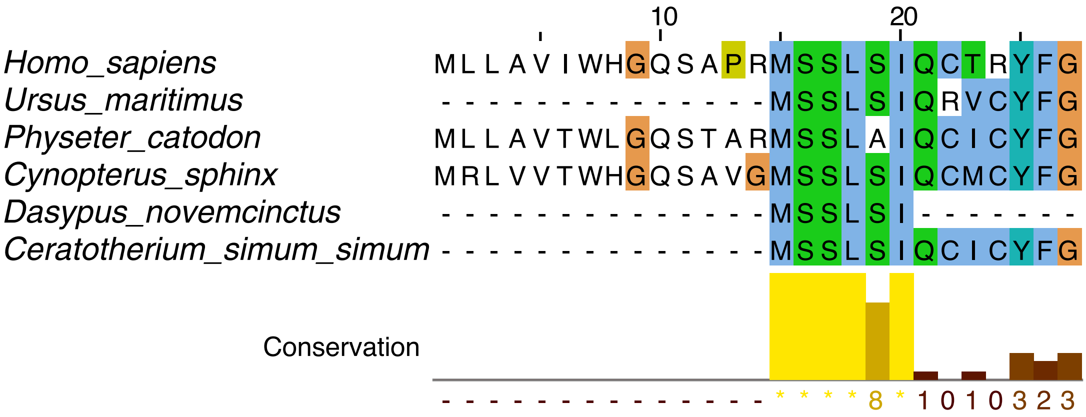

**HG0873**

ENSG00000142875 (PRKACB)

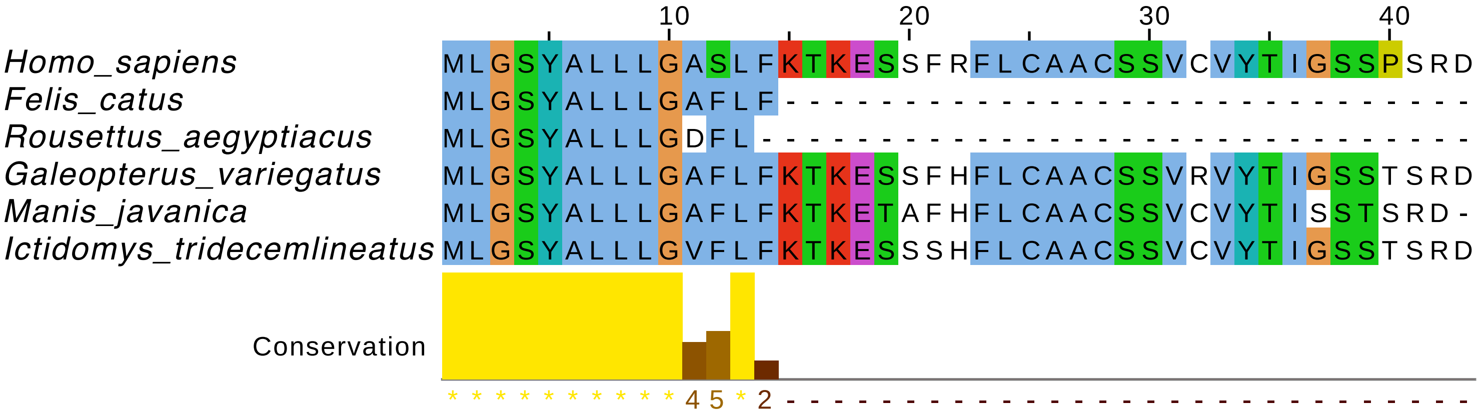

**HG0874**

ENSG00000143515 (ATP8B2)

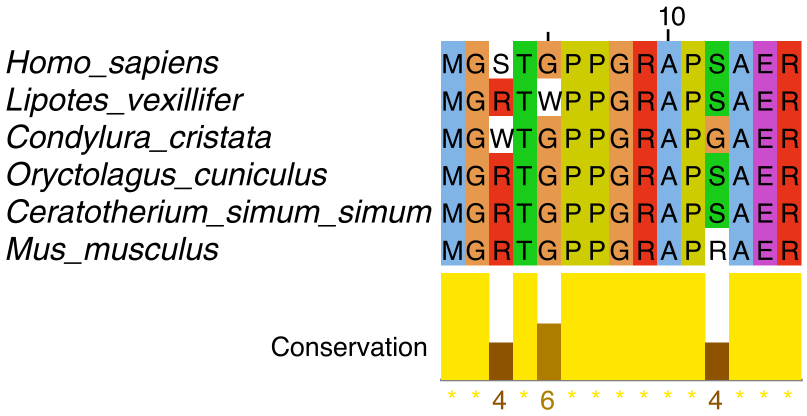

**HG0875**

ENSG00000145284 (SCD5)

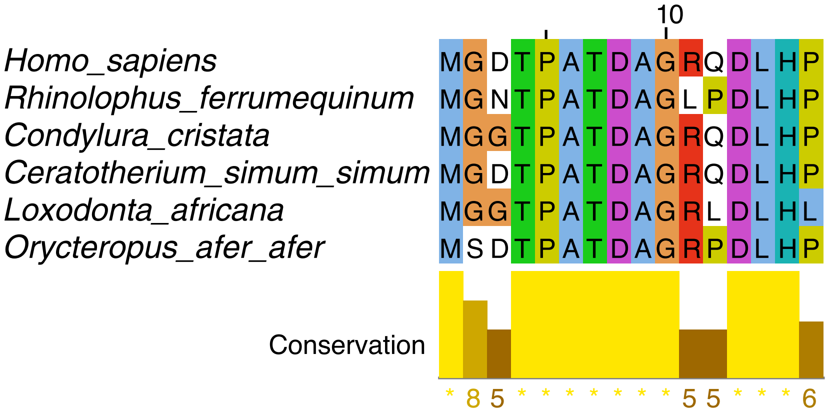

**HG0876**

ENSG00000145908 (ZNF300)

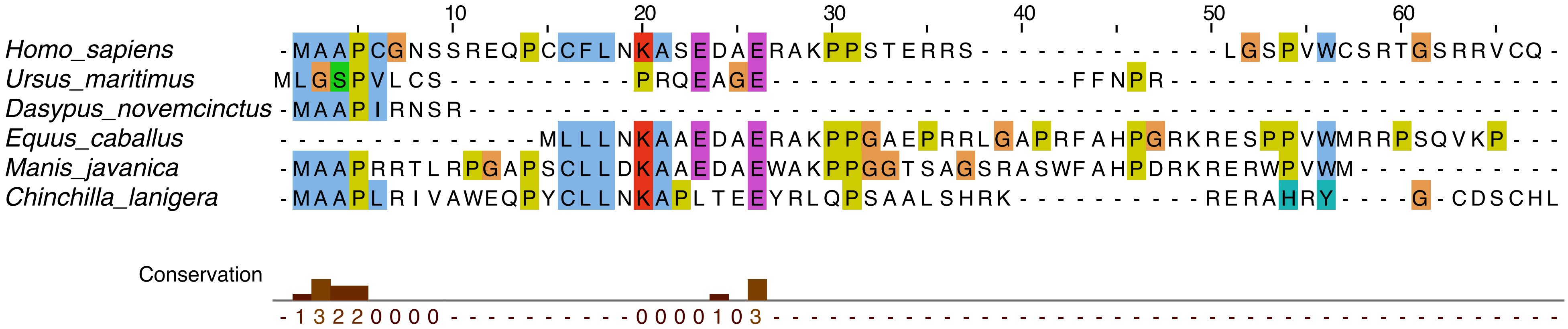

**HG0877**

ENSG00000146426 (TIAM2)

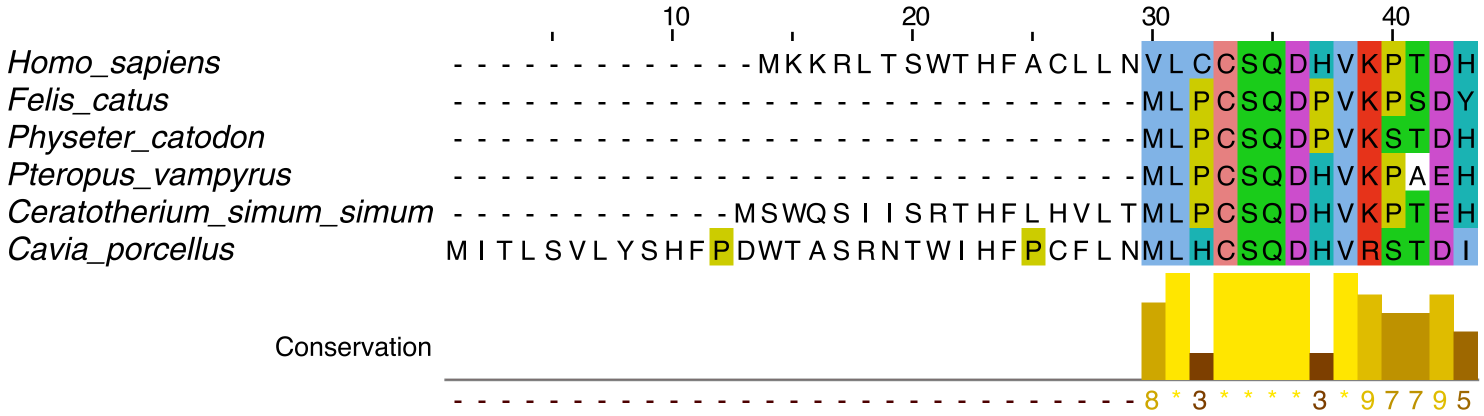

**HG0878**

ENSG00000146802 (TMEM168)

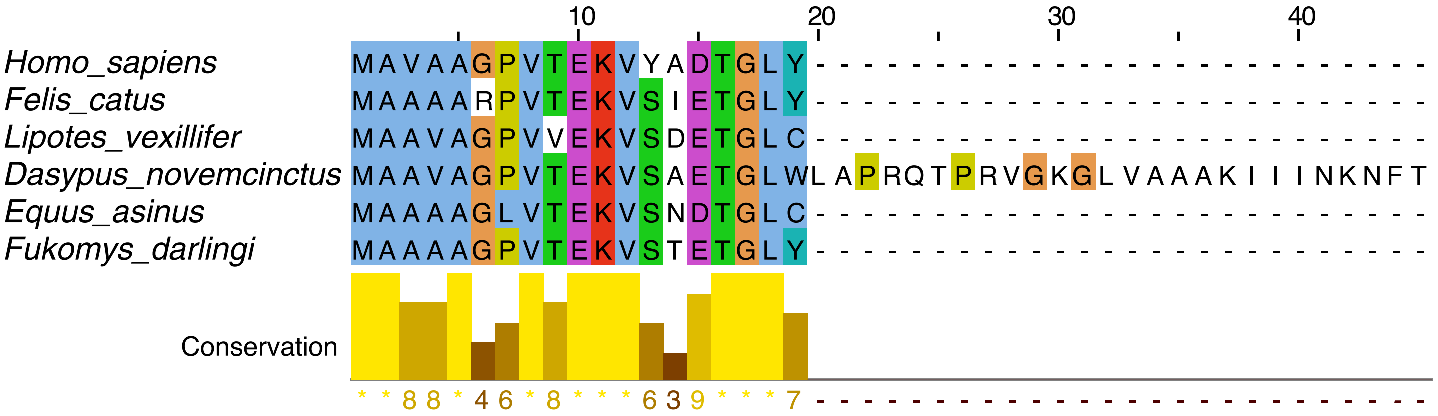

**HG0879**

ENSG00000147145 (LPAR4)

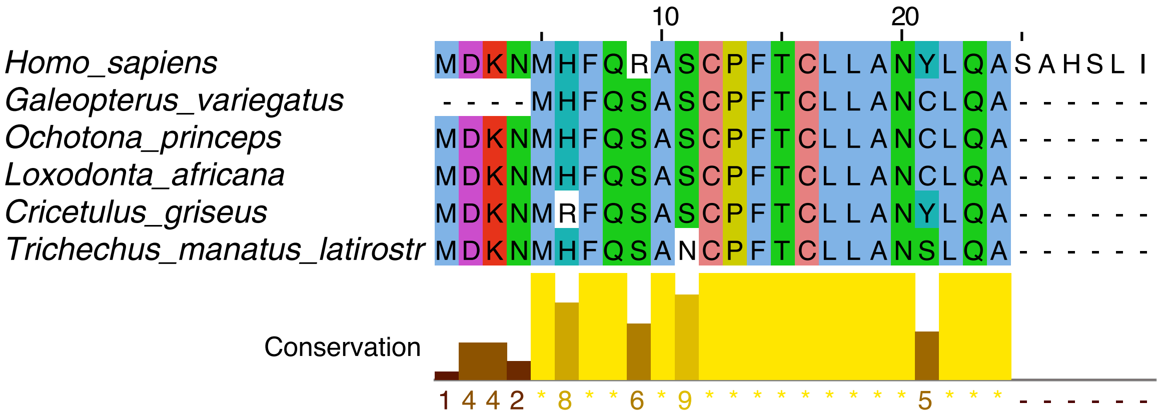

**HG0880**

ENSG00000151789 (ZNF385D)

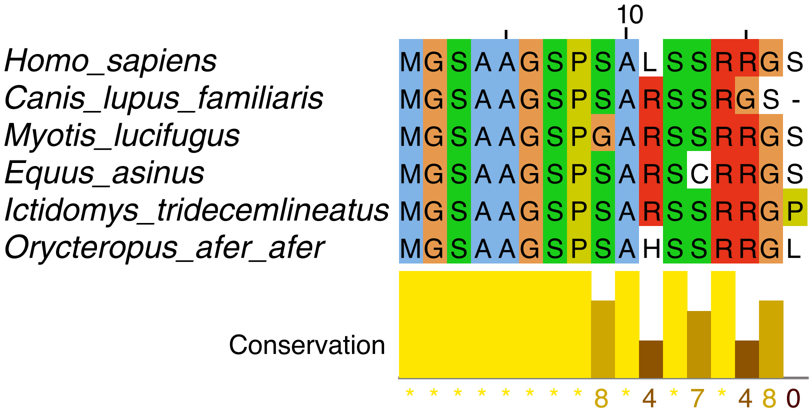

**HG0881**

ENSG00000153822 (KCNJ16)

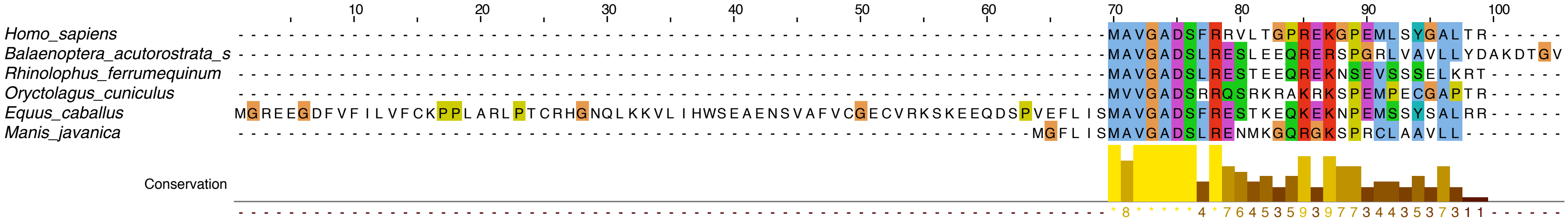

# HG0882.1

### ENSG00000154832 (CXXC1)

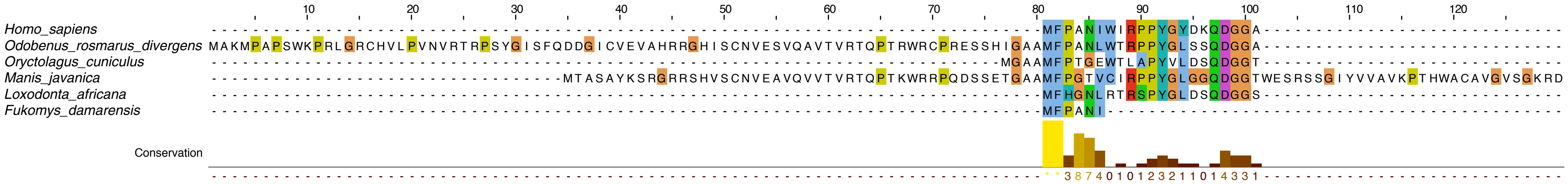

**HG0882.2**

ENSG00000154832 (CXXC1)

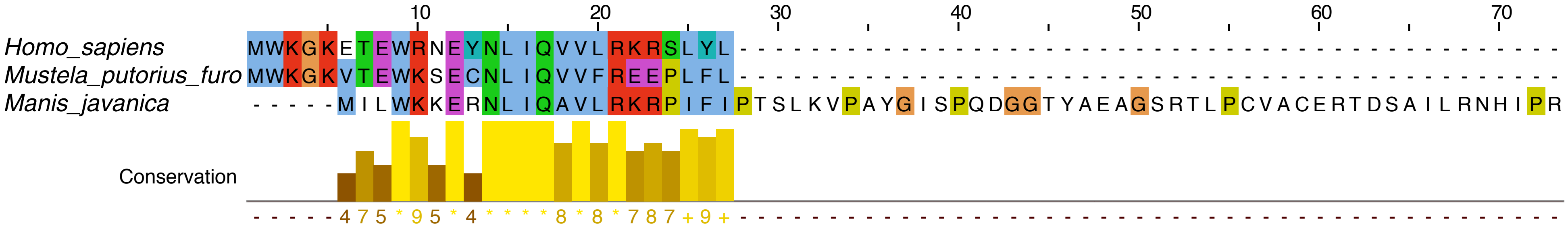

**HG0883**

ENSG00000156011 (PSD3)

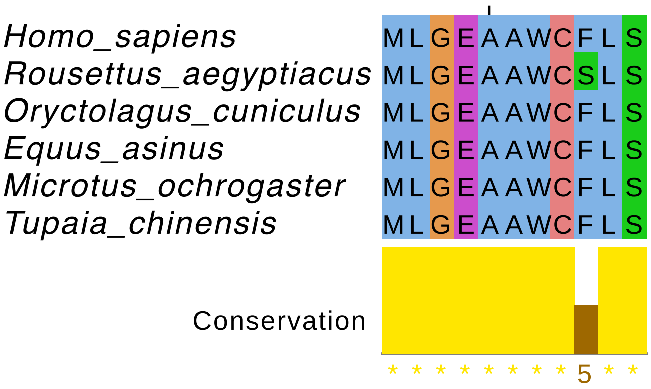

**HG0884**

ENSG00000156463 (SH3RF2)

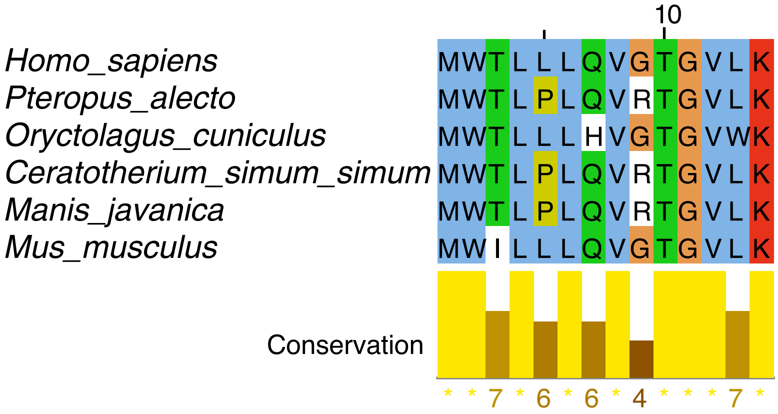

**HG0885.1**

ENSG00000157502 (MUM1L1)

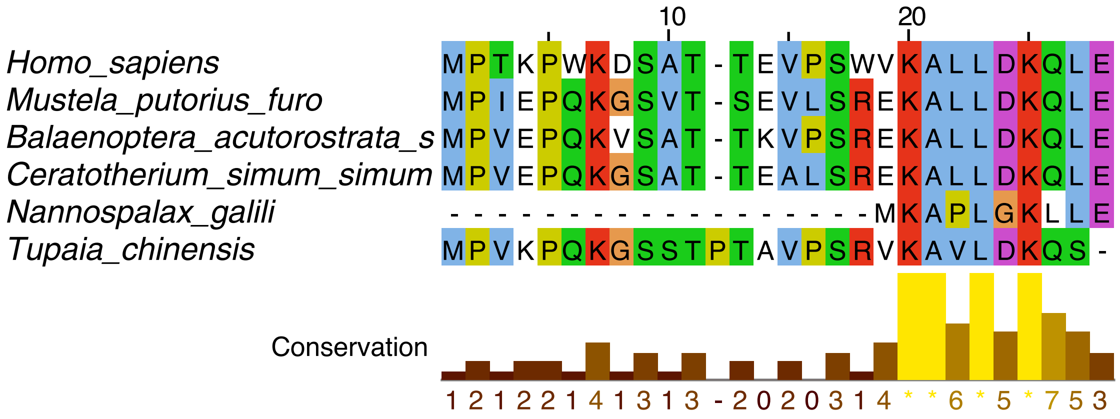

**HG0885.2**

ENSG00000157502 (MUM1L1)

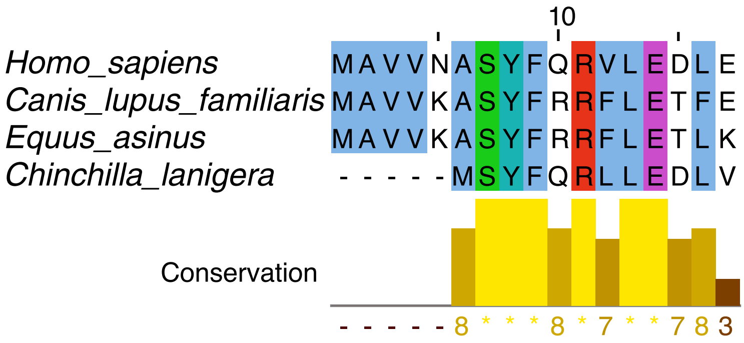

**HG0886**

ENSG00000158850 (B4GALT3)

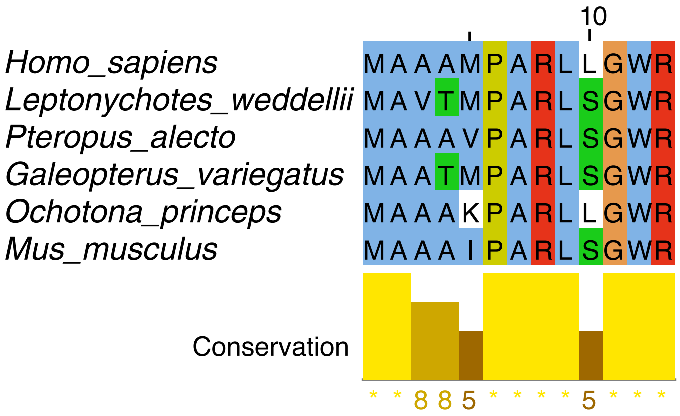

**HG0887**

ENSG00000160959

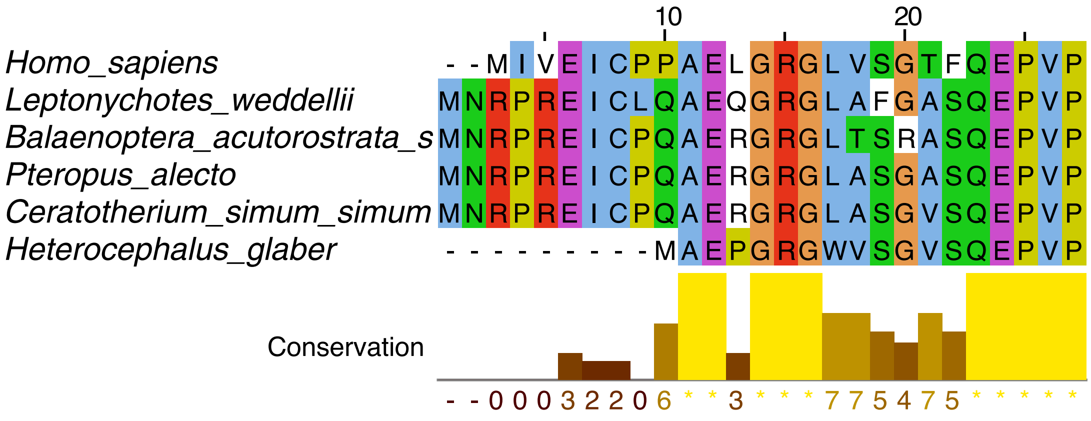

**HG0888**

ENSG00000164023 (SGMS2)

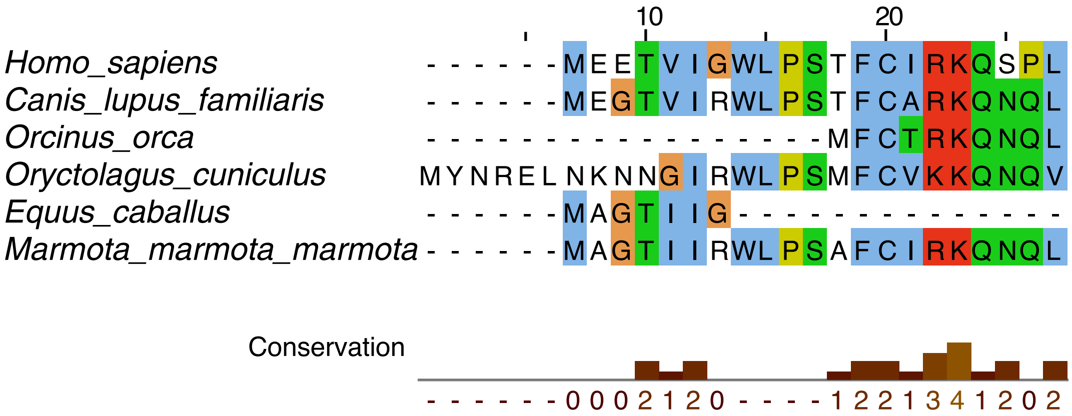

**HG0889**

ENSG00000164091 (WDR82)

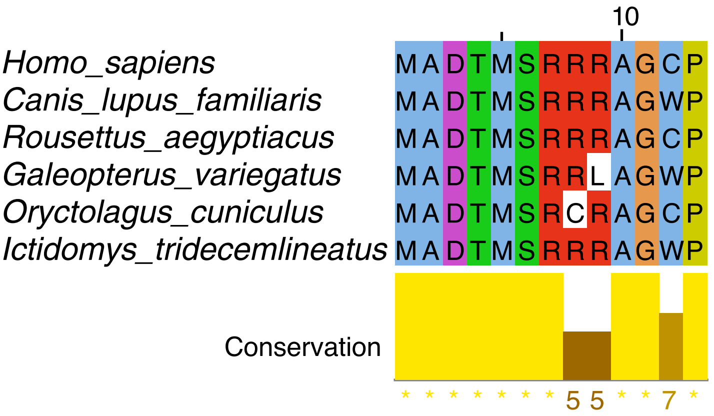

**HG0890**

ENSG00000165813

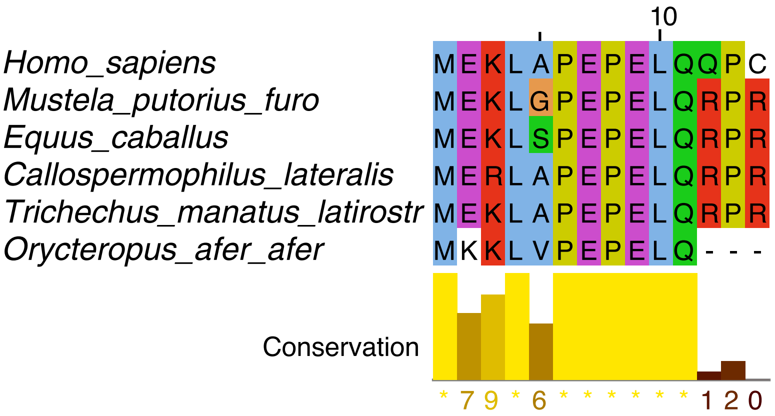

**HG0891**

ENSG00000166211 (SPIC)

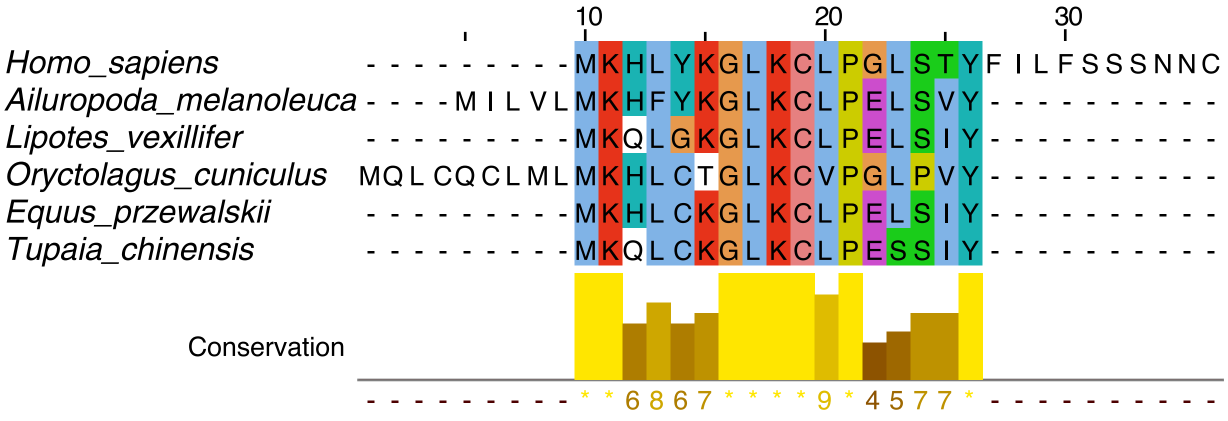

**HG0892**

ENSG00000168418 (KCNG4)

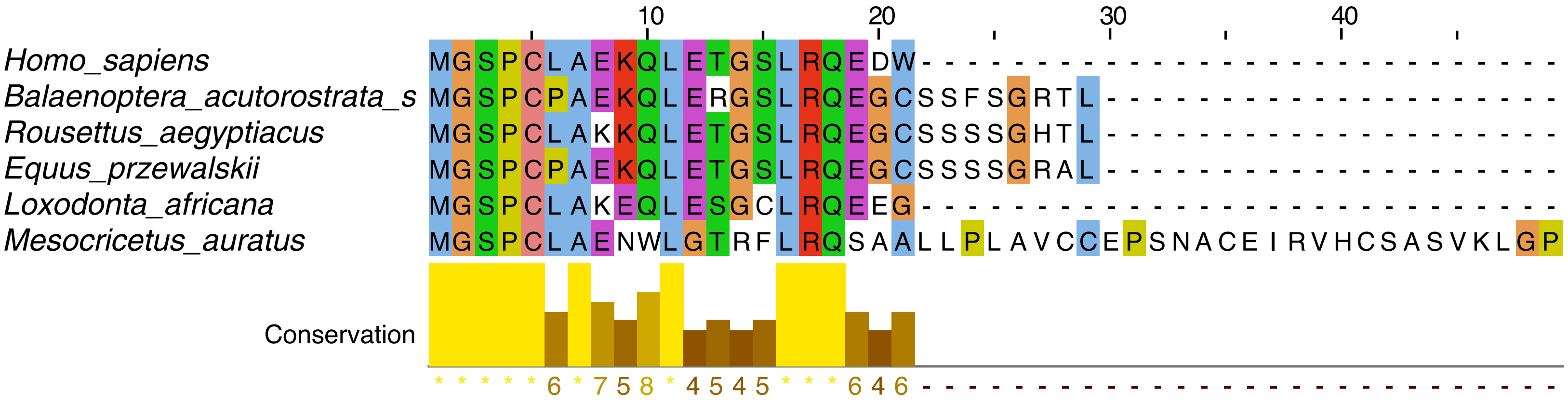

**HG0893**

ENSG00000168769 (TET2)

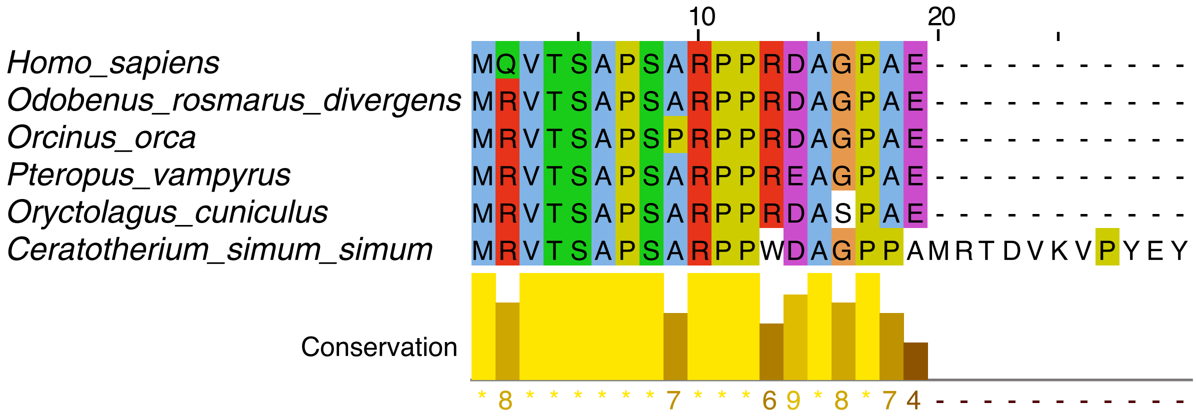

**HG0894.1**

ENSG00000169282 (KCNAB1)

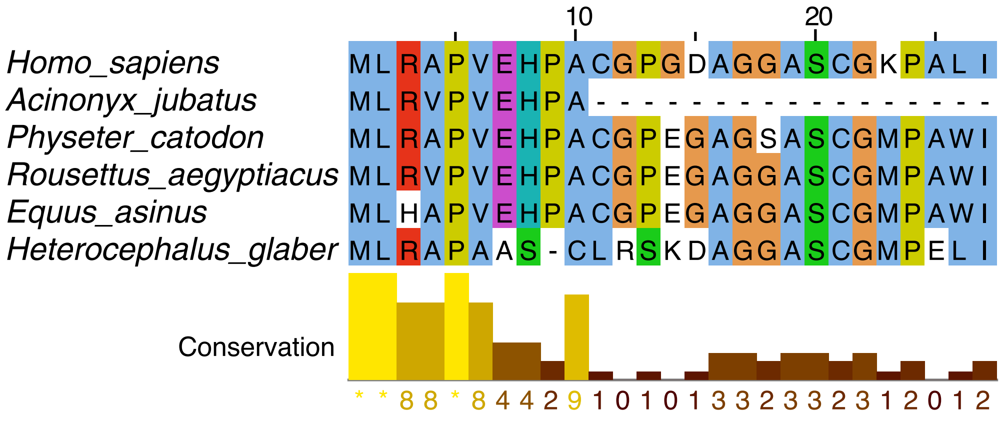

**HG0894.2**

ENSG00000069424 (KCNAB2)

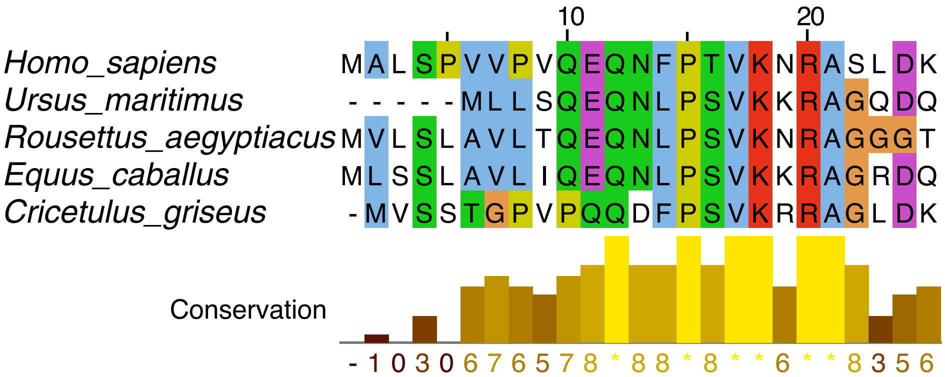

**HG0895**

ENSG00000170035 (UBE2E3)

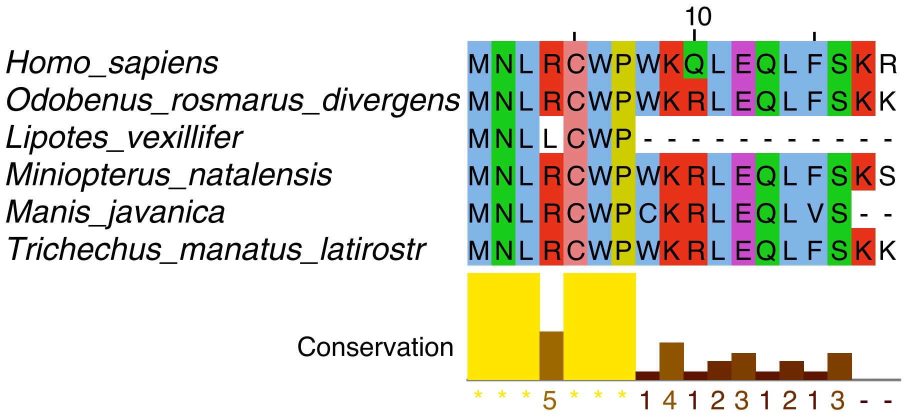

**HG0896**

ENSG00000170365 (SMAD1)

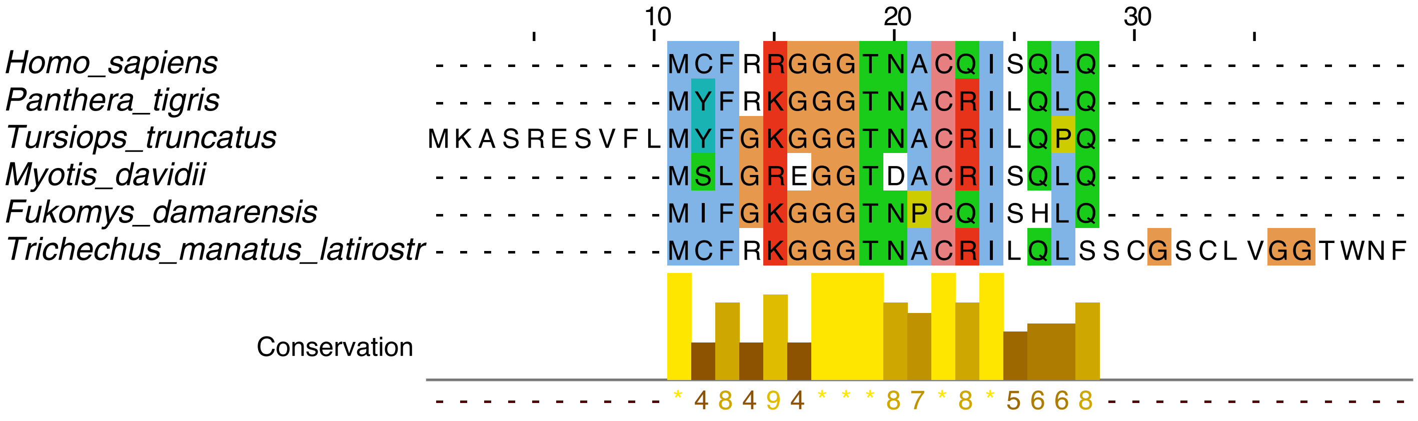

**HG0897**

ENSG00000171435 (KSR2)

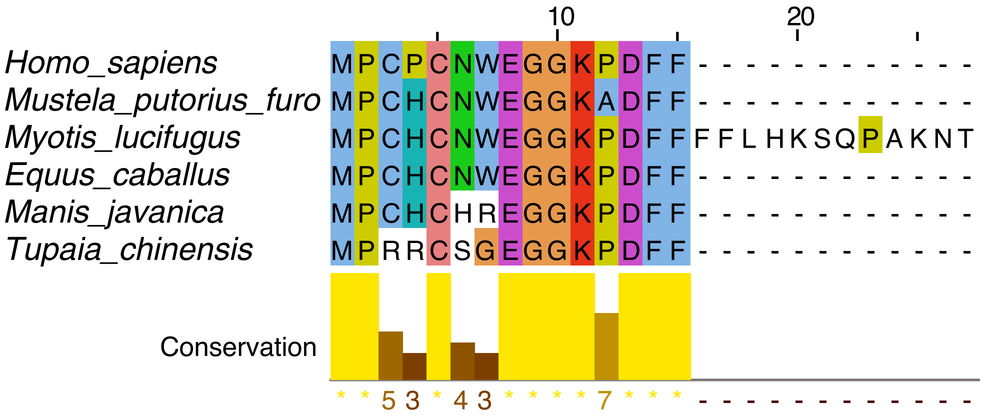

**HG0898**

ENSG00000171729 (TMEM51)

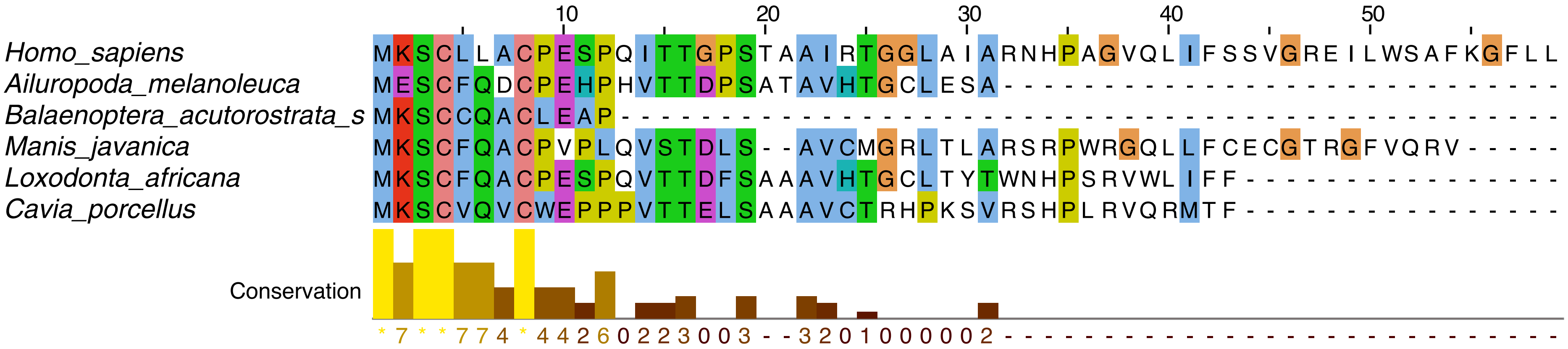

**HG0899**

ENSG00000173320 (STOX2)

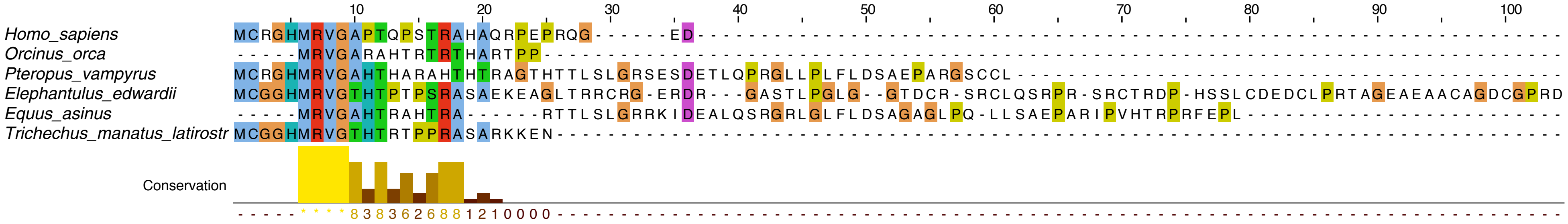

**HG0900**

ENSG00000173517 (PEAK1)

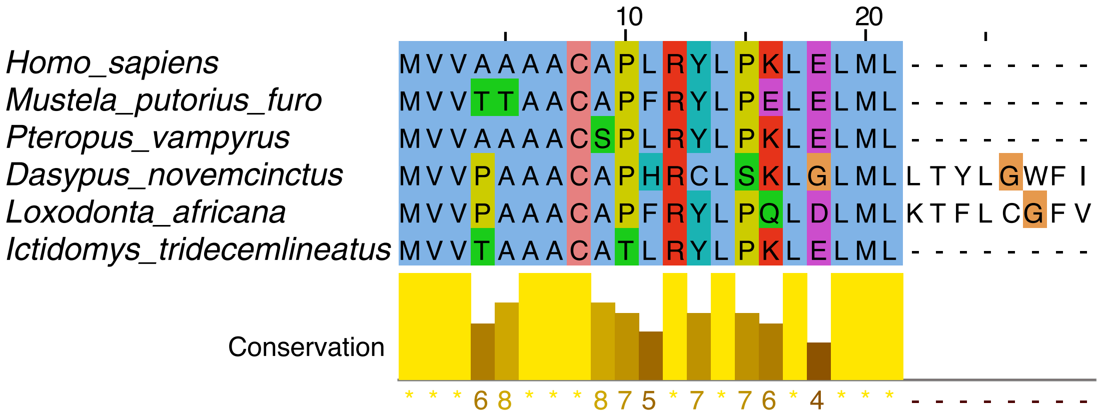

**HG0901**

ENSG00000175697 (GPR156)

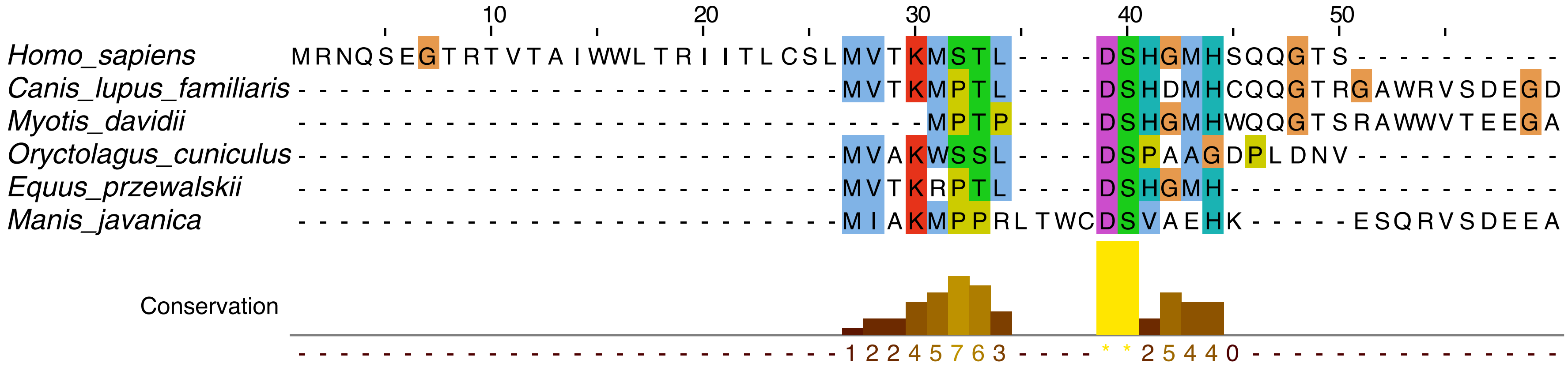

**HG0902**

ENSG00000177182 (CLVS1)

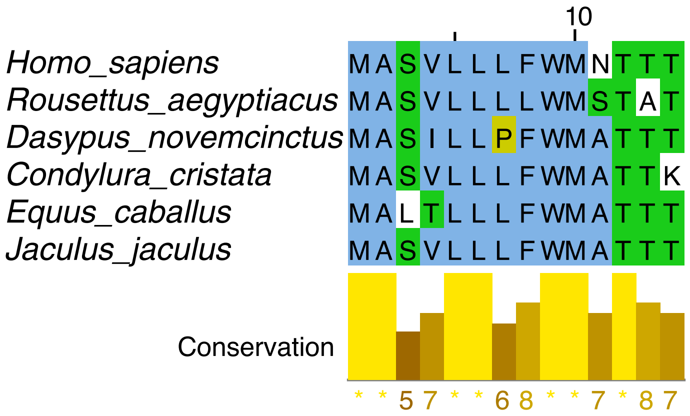

**HG0903**

ENSG00000180616 (SSTR2)

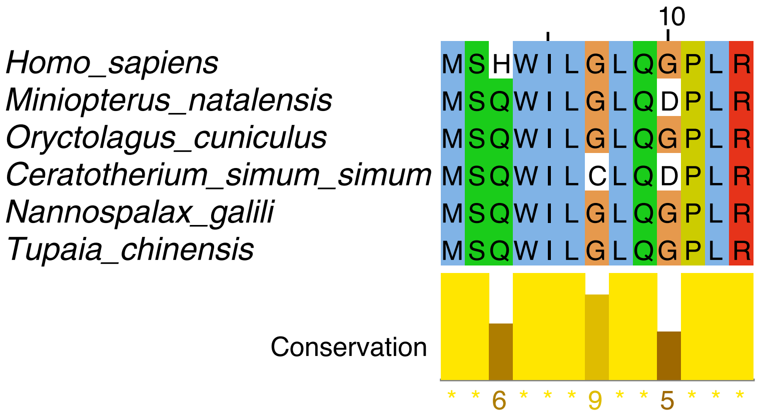

**HG0904**

ENSG00000182329

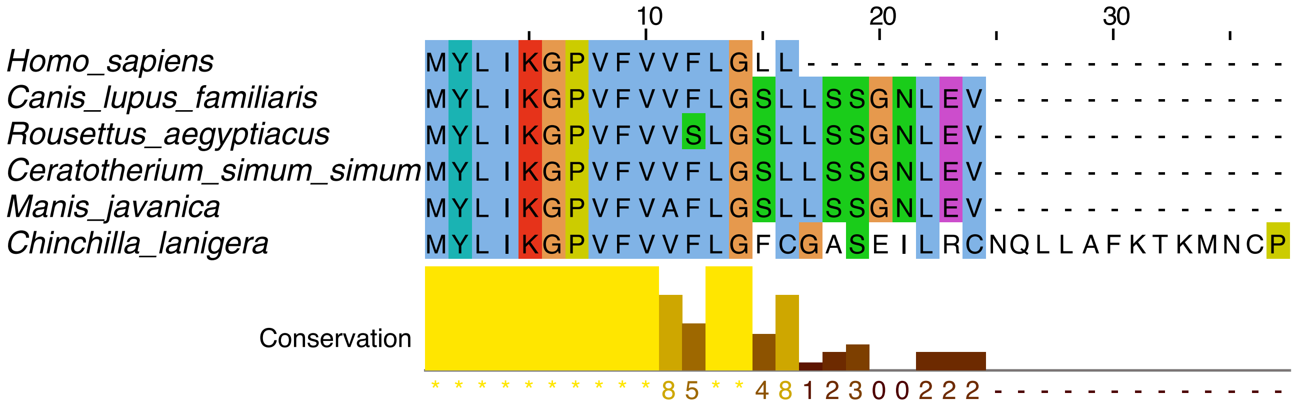

**HG0905**

ENSG00000183337 (BCOR)

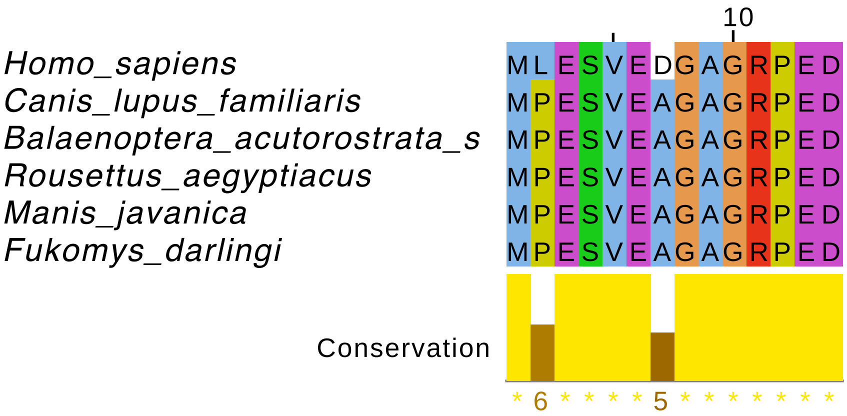

**HG0906**

ENSG00000183773 (AIFM3)

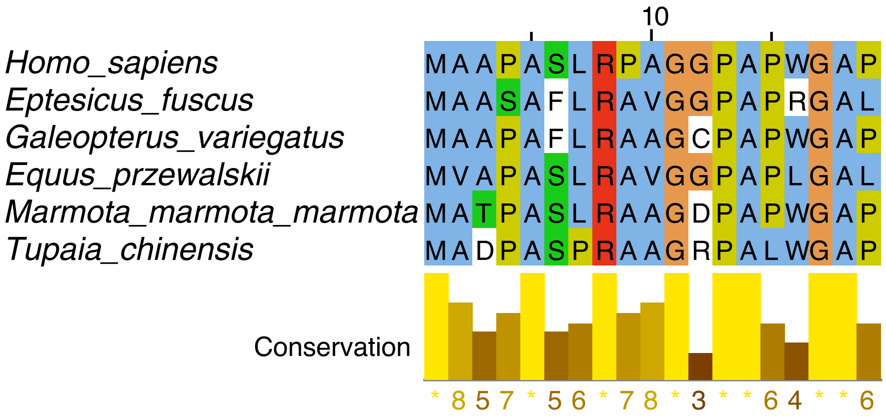

**HG0907**

ENSG00000186862 (PDZD7)

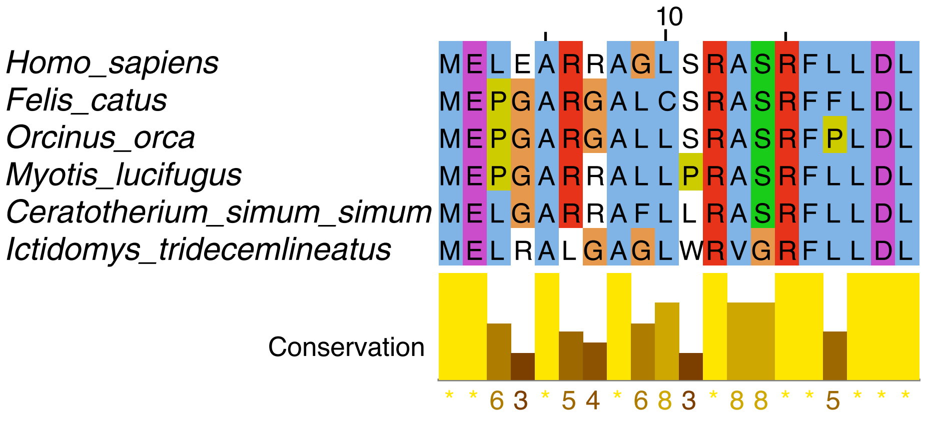

**HG0908.1**

ENSG00000188554 (NBR1)

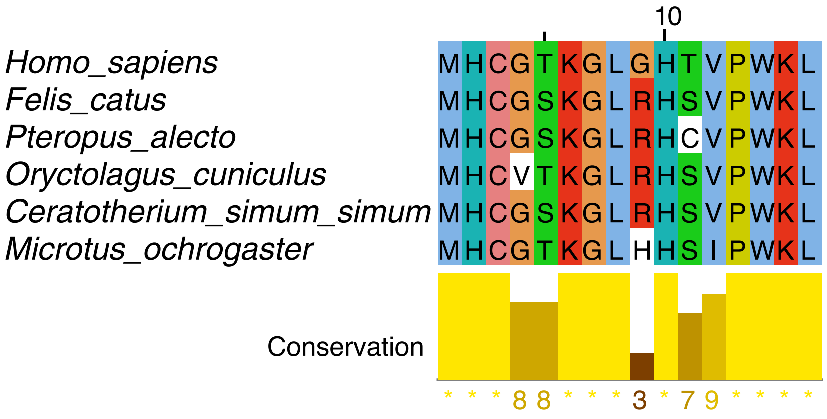

**HG0908.2**

ENSG00000188554 (NBR1)

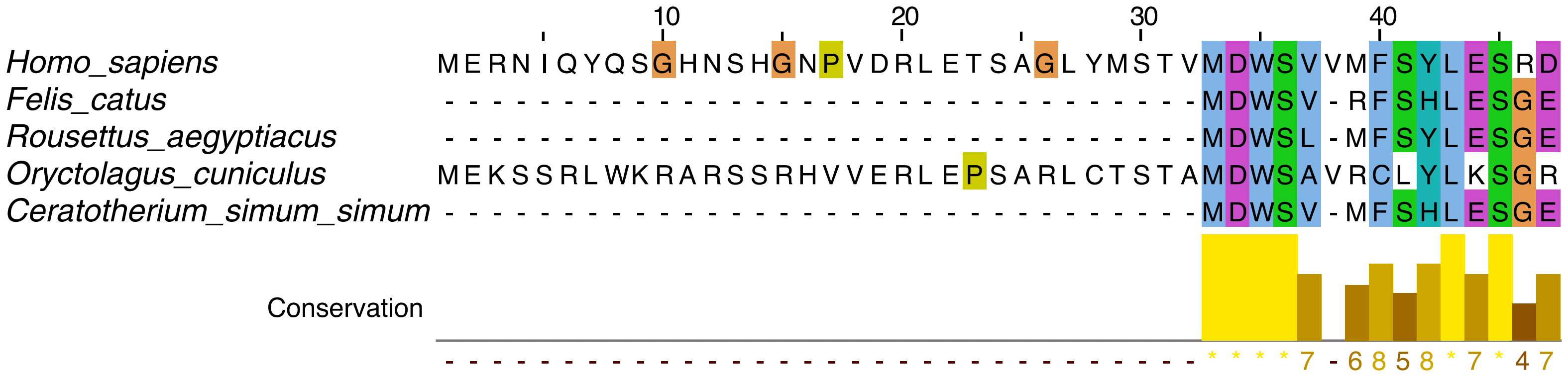

**HG0909**

ENSG00000189079 (ARID2)

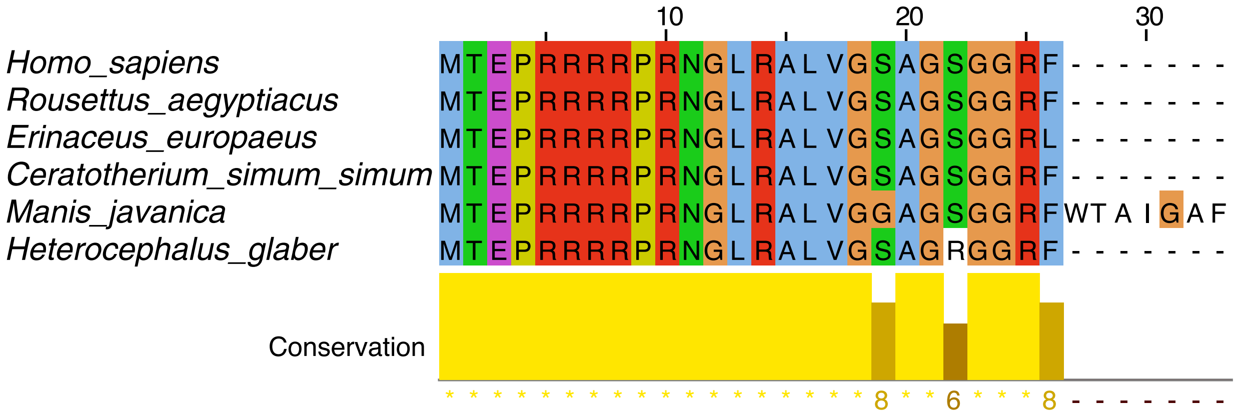

**HG0910**

ENSG00000197935 (ZNF311)

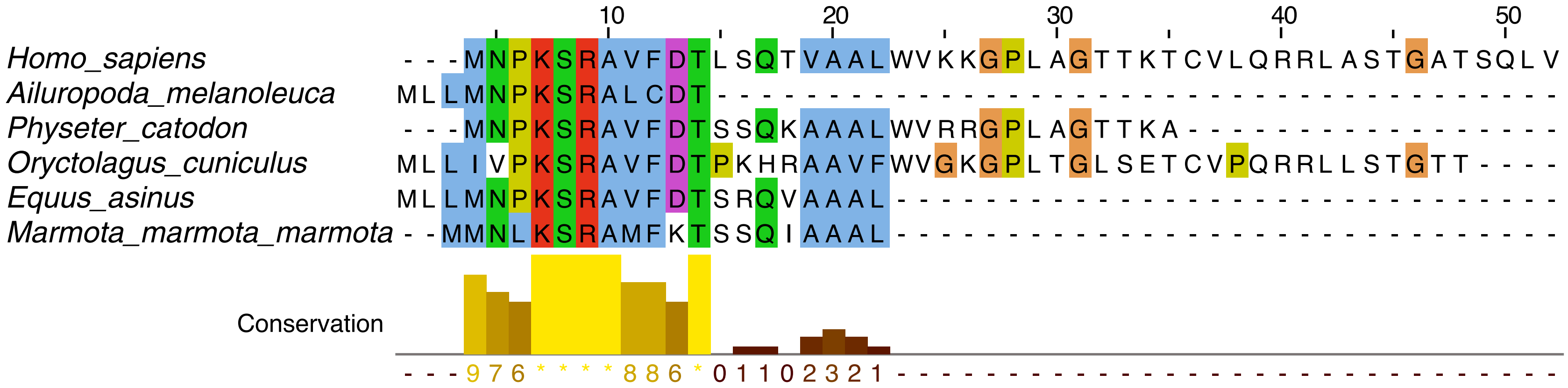

**HG0911**

ENSG00000198673 (FAM19A2)

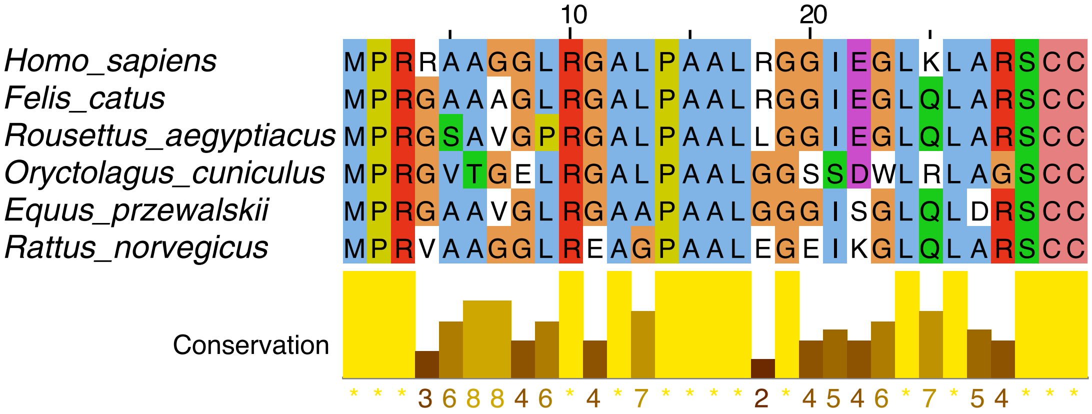

**HG0912**

ENSG00000198822 (GRM3)

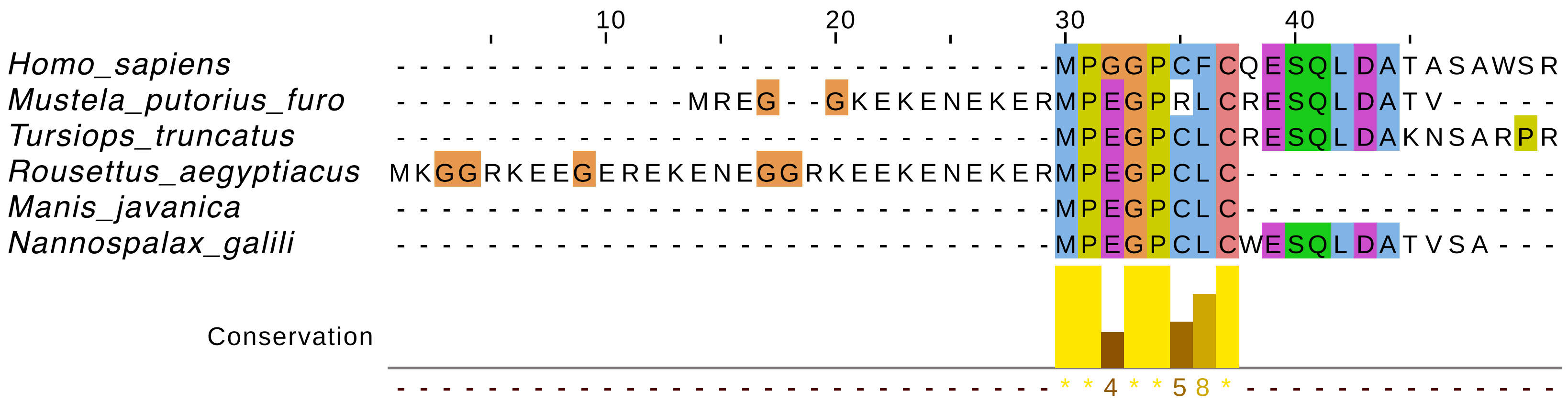

**HG0913.1**

ENSG00000204599 (TRIM39)

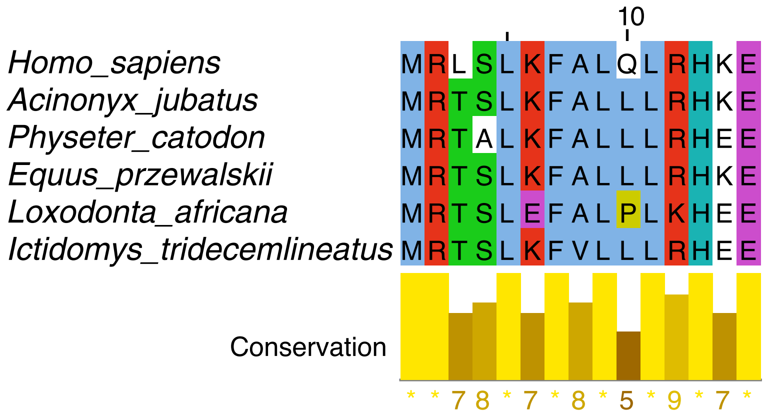

**HG0913.2**

ENSG00000204599 (TRIM39)

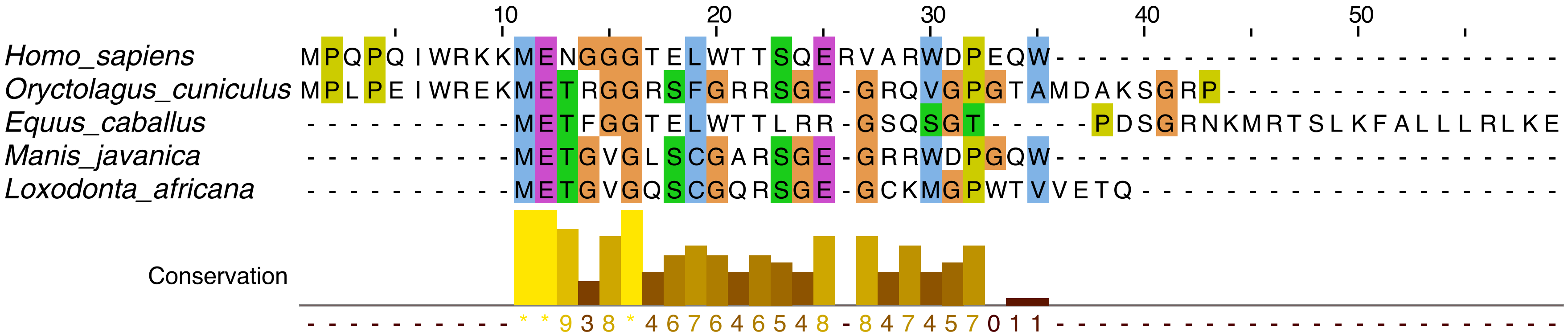

**HG0914**

ENSG00000227877 (MRLN)

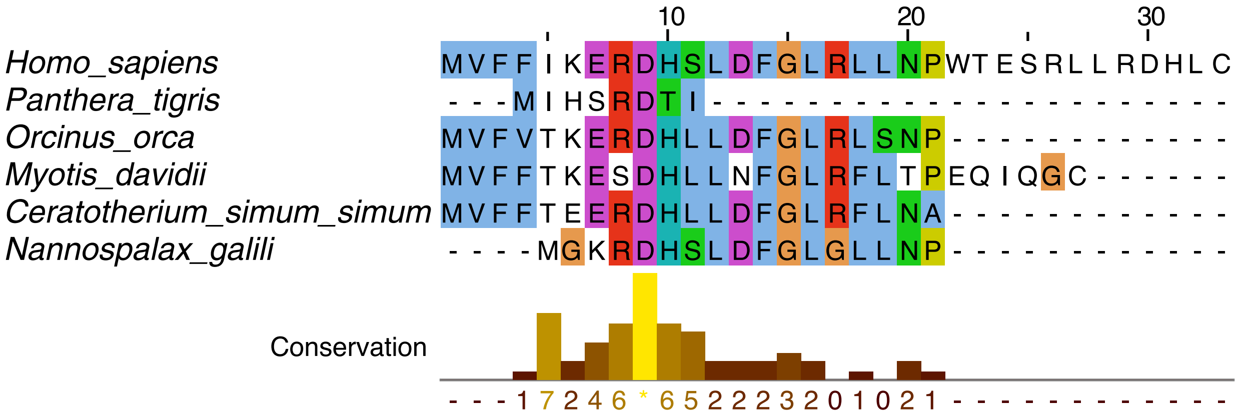

**HG0915**

ENSG00000254469 (AP002495.2)

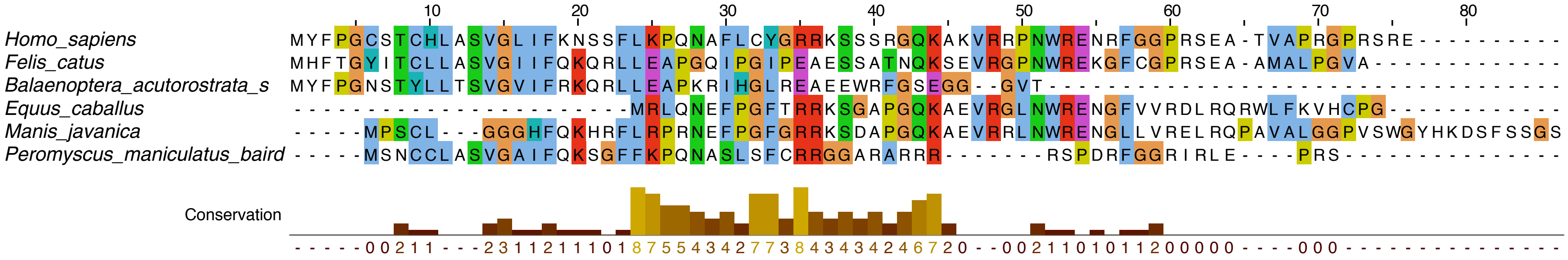

**HG0916**

ENSG00000020633 (RUNX3)

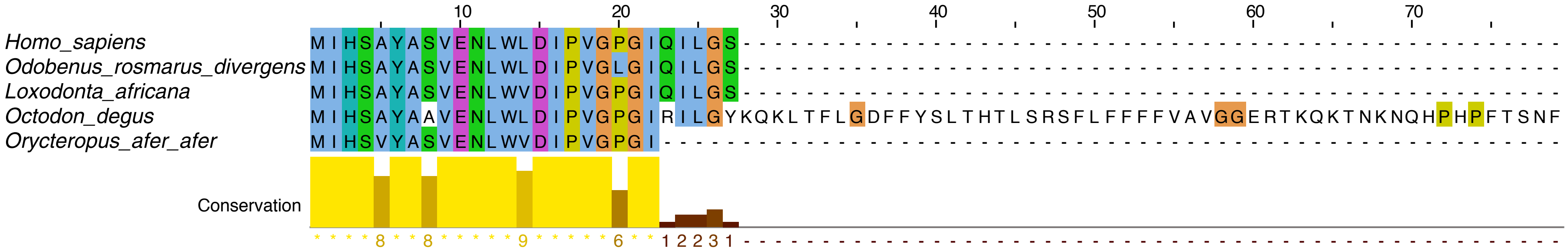

**HG0917**

ENSG00000042980 (ADAM28)

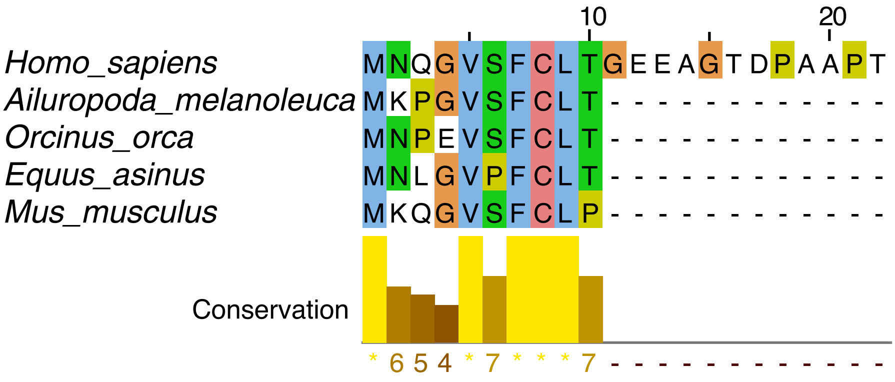

**HG0918**

ENSG00000047648 (ARHGAP6)

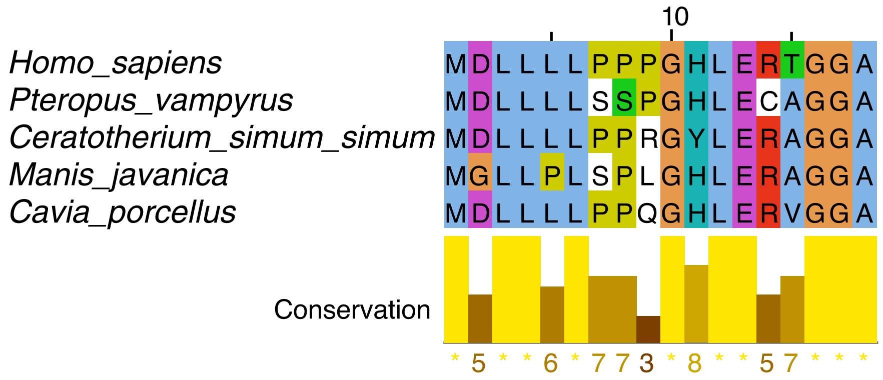

**HG0919**

ENSG00000051009 (FAM160A2)

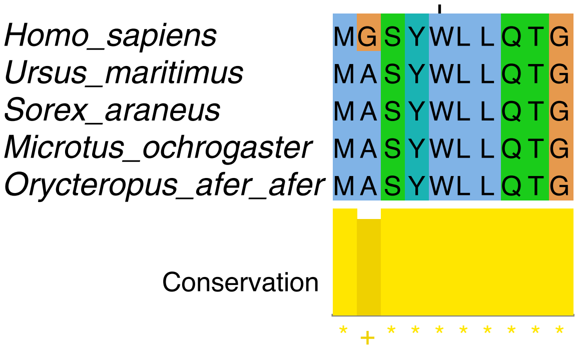

**HG0920**

ENSG00000051382 (PIK3CB)

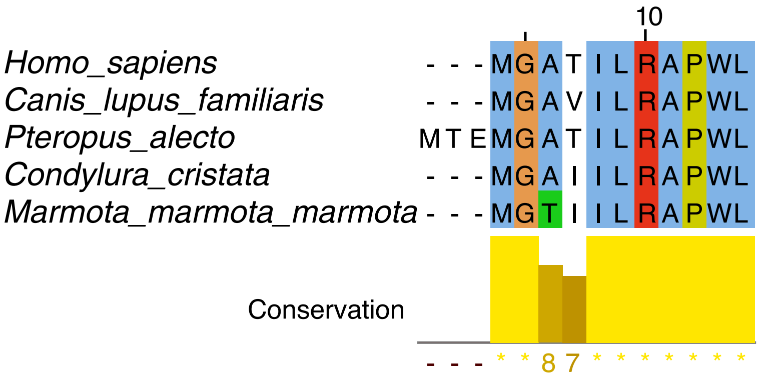

**HG0921**

ENSG00000058799 (YIPF1)

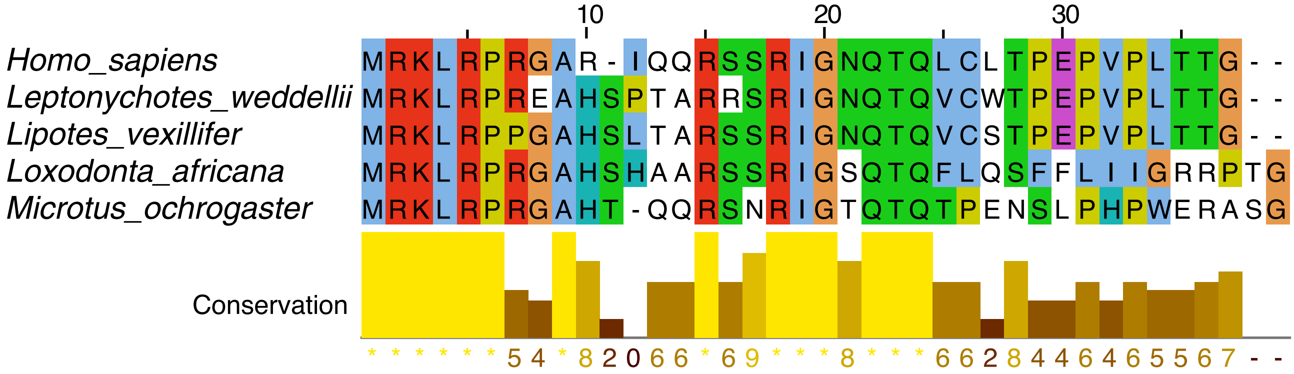

**HG0922**

ENSG00000079841 (RIMS1)

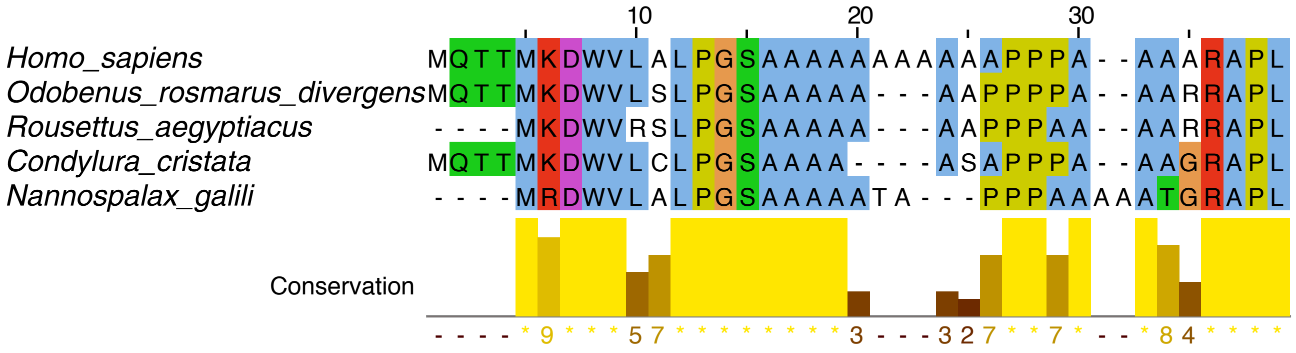

**HG0923**

ENSG00000081138 (CDH7)

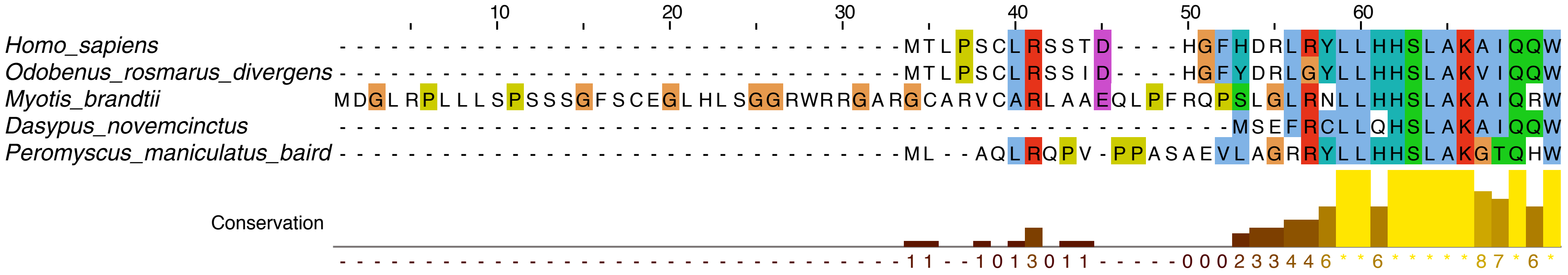

**HG0924**

ENSG00000081307 (UBA5)

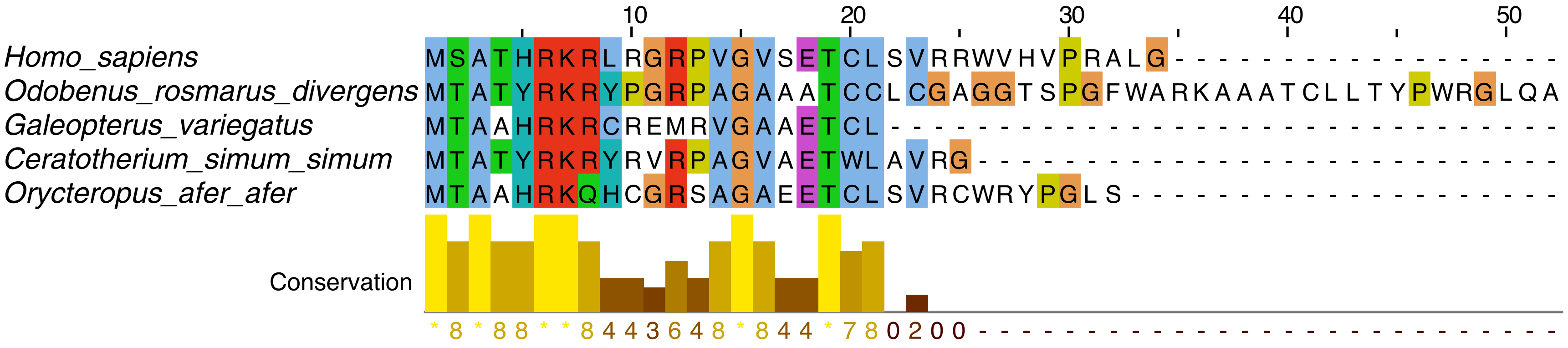

**HG0925**

ENSG00000088766 (CRLS1)

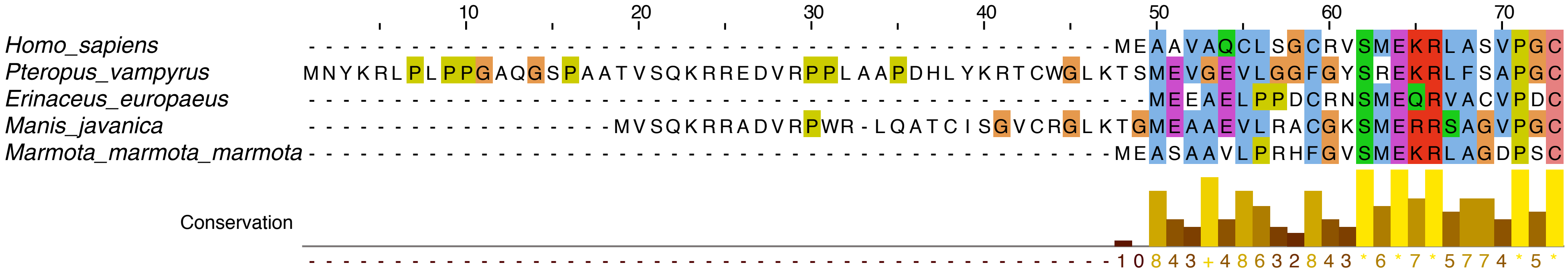

**HG0926**

ENSG00000095397 (WHRN)

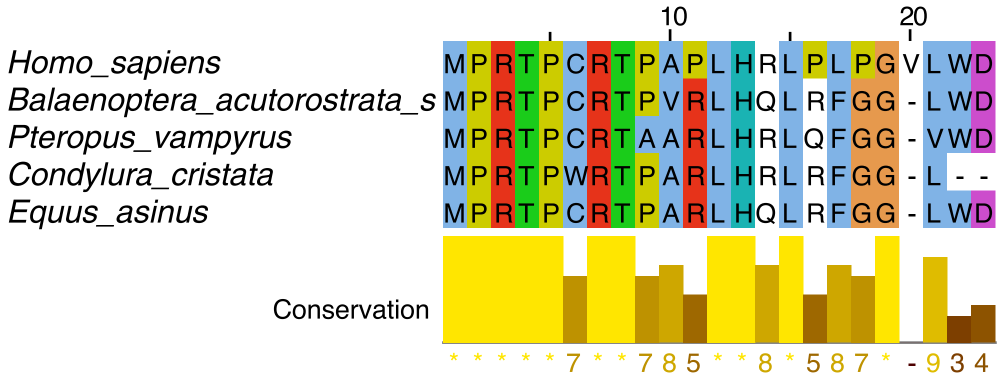

**HG0927**

ENSG00000095951 (HIVEP1)

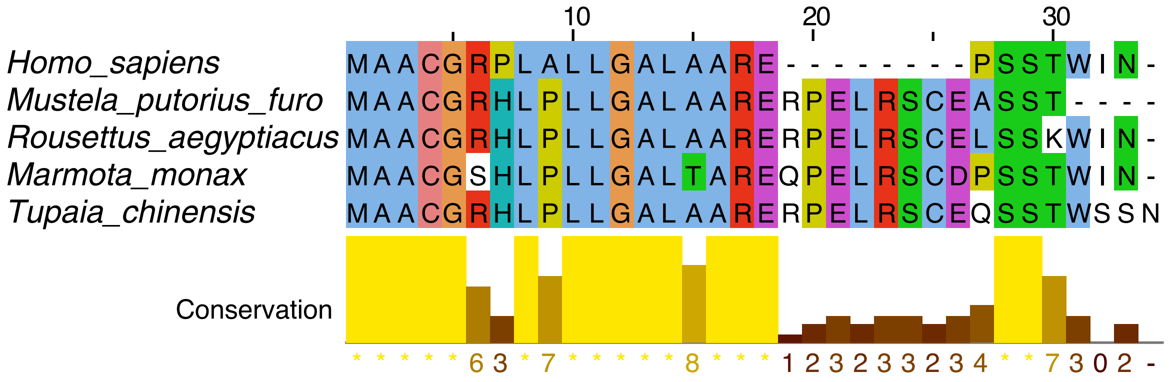

**HG0928**

ENSG00000102057 (KCND1)

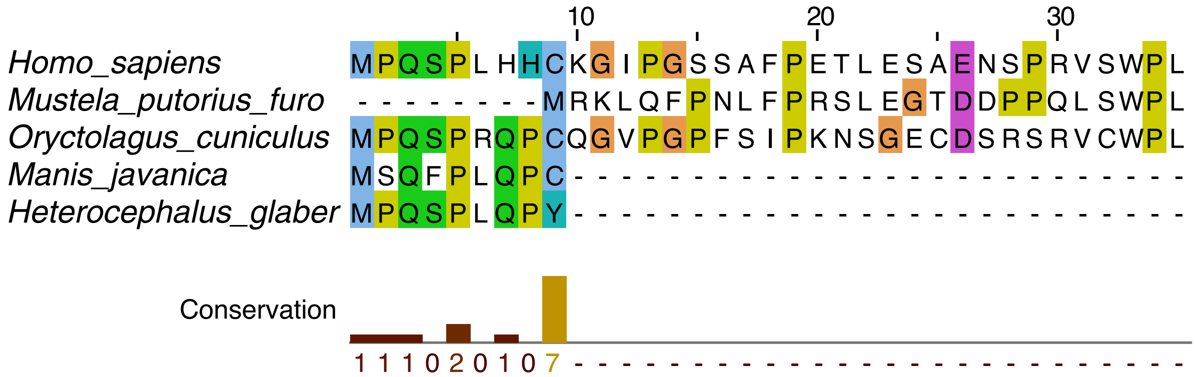

**HG0929**

ENSG00000102239 (BRS3)

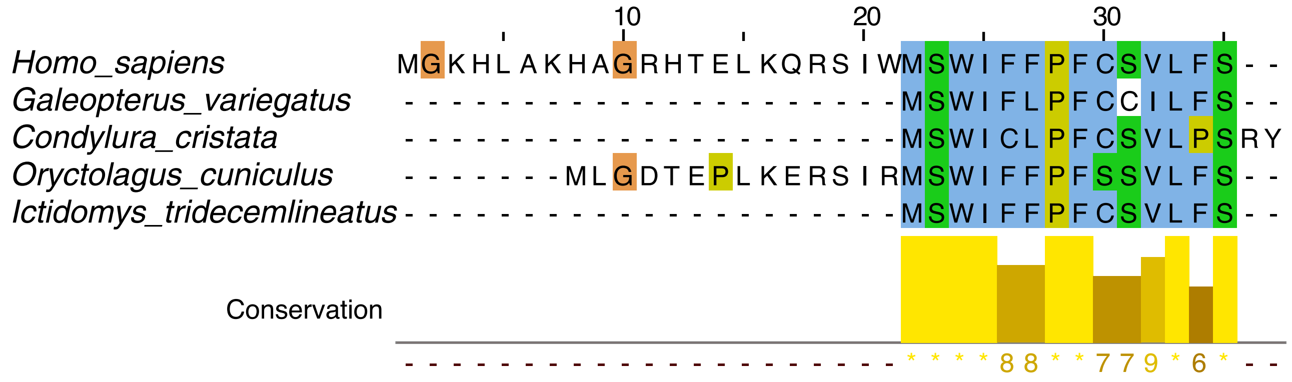

**HG0930**

ENSG00000102524 (TNFSF13B)

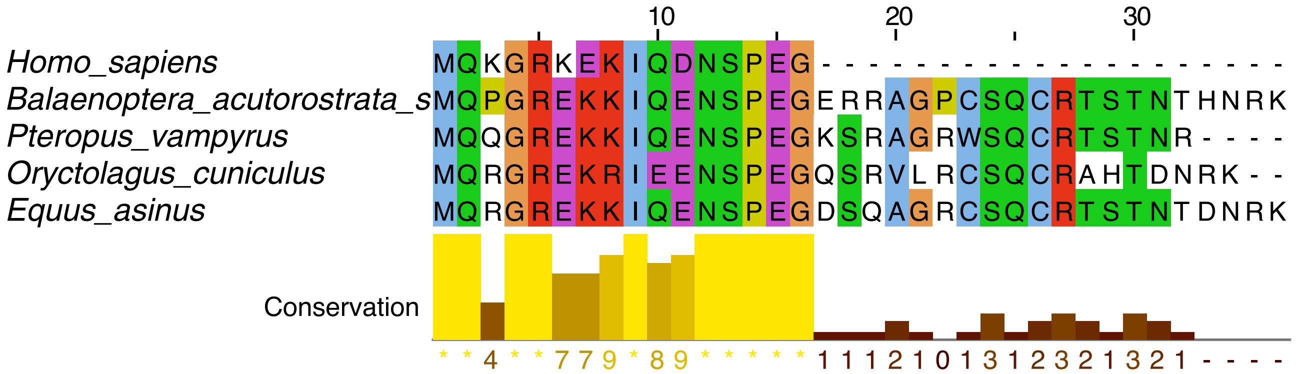

**HG0931**

ENSG00000104951 (IL4I1)

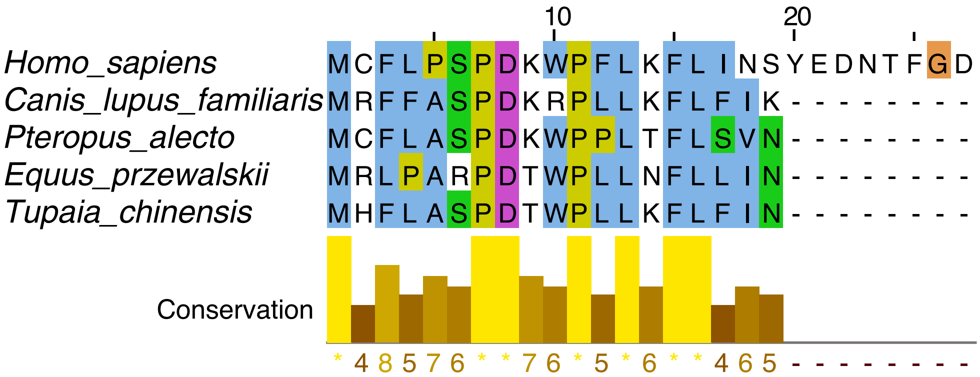

**HG0932**

ENSG00000105997 (HOXA3)

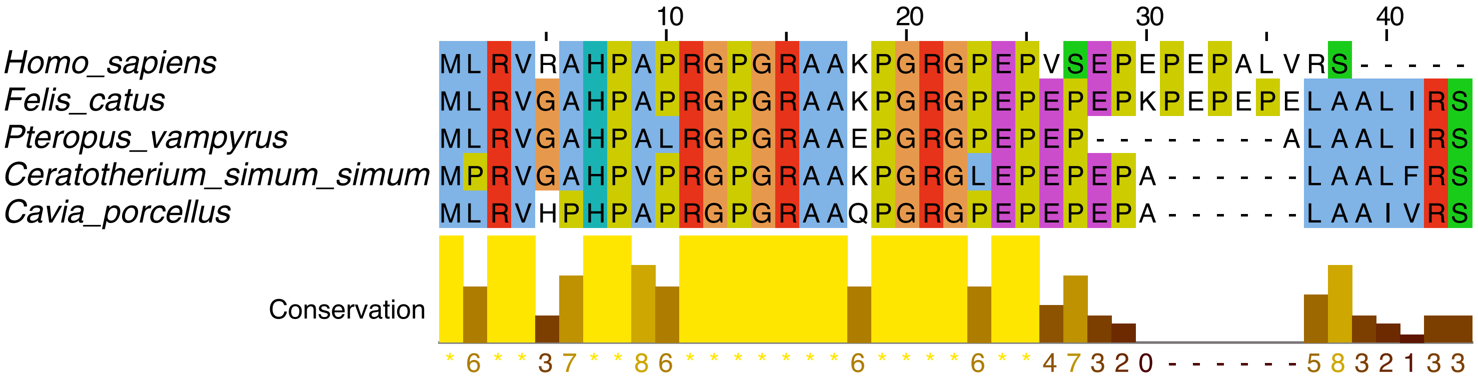

**HG0933**

ENSG00000106330 (MOSPD3)

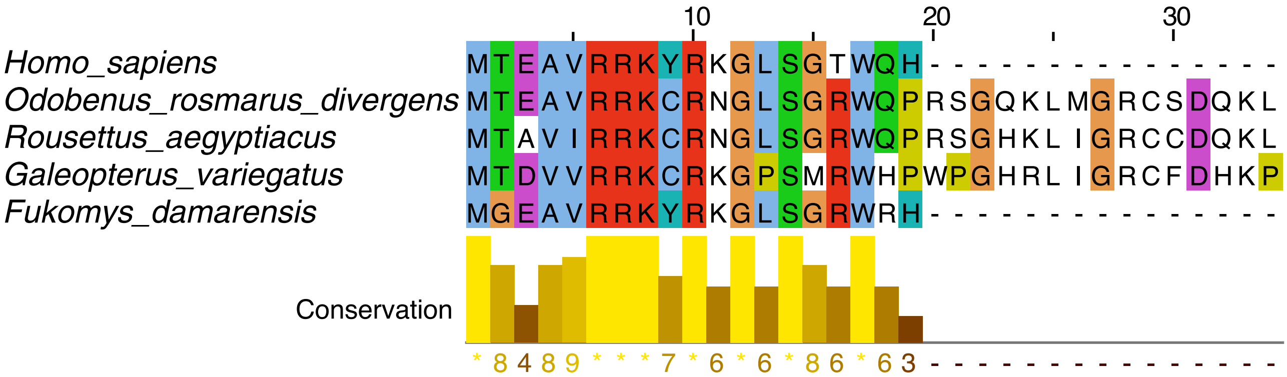

**HG0934**

ENSG00000107443 (CCNJ)

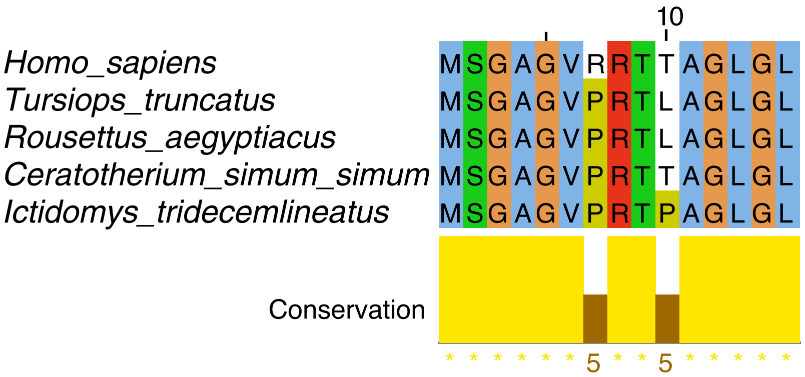

**HG0935**

ENSG00000108518 (PFN1)

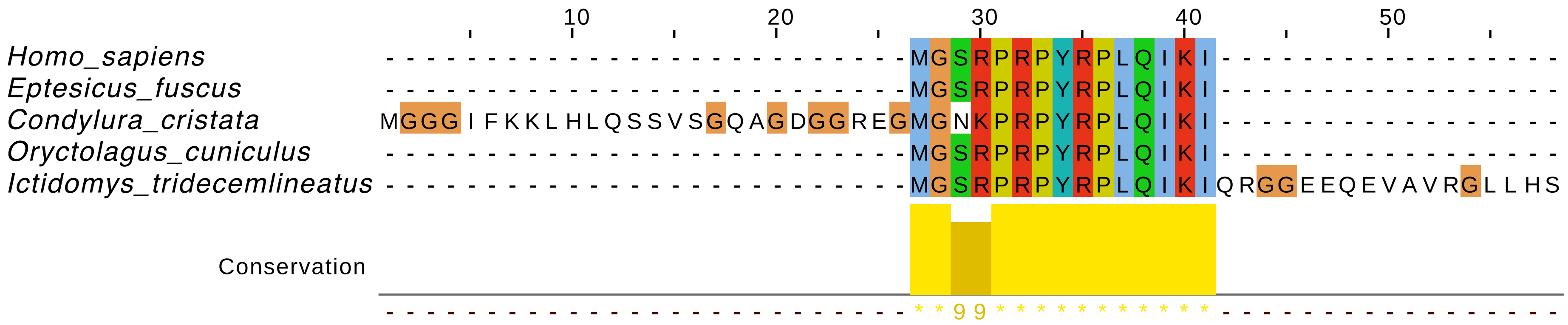

**HG0936**

ENSG00000108799 (EZH1)

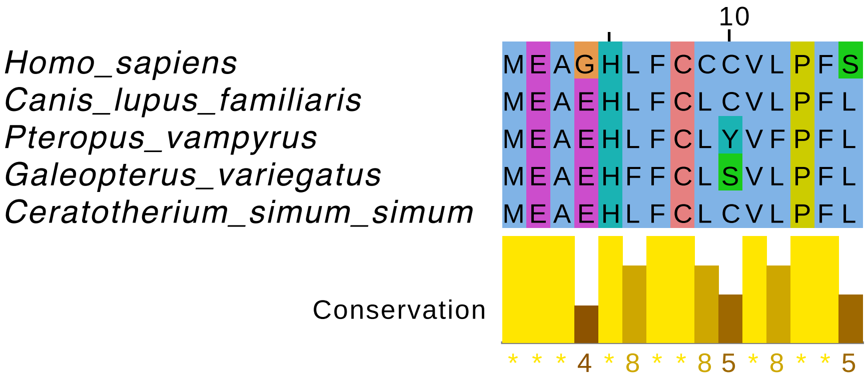

**HG0937**

ENSG00000109846 (CRYAB)

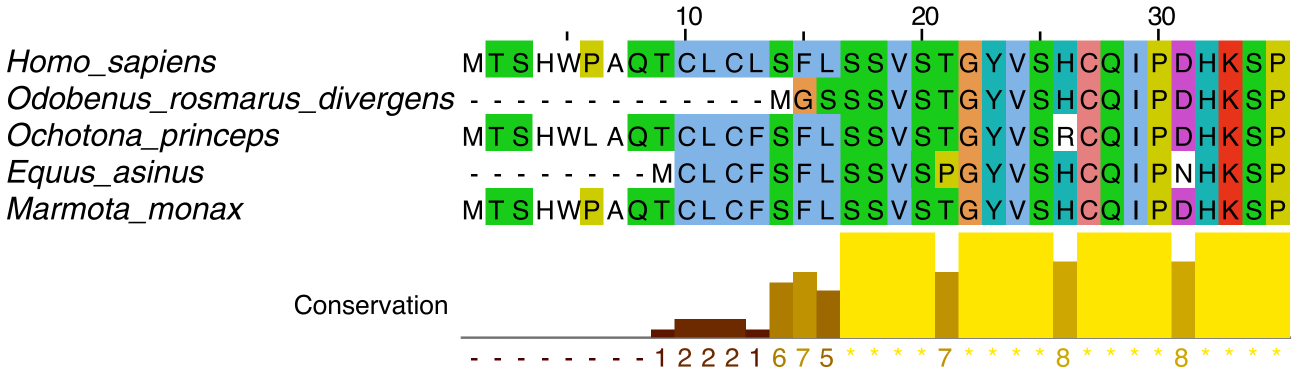

**HG0938**

ENSG00000111665 (CDCA3)

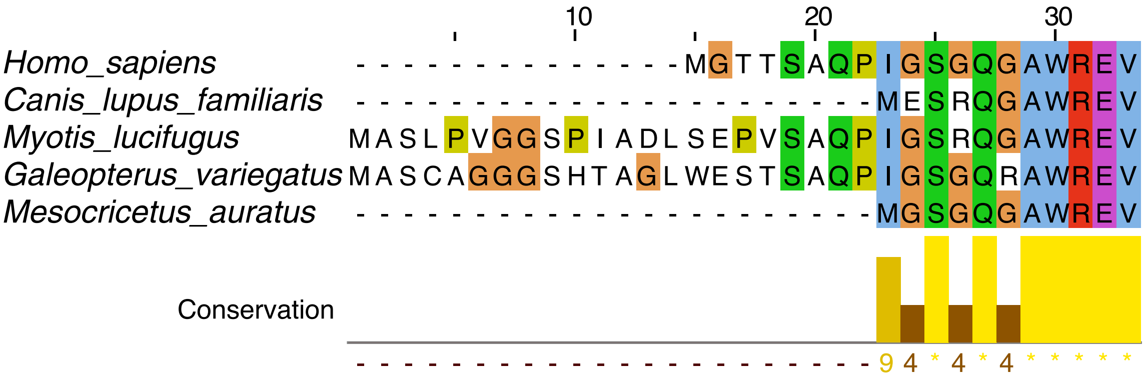

**HG0939**

ENSG00000111674 (ENO2)

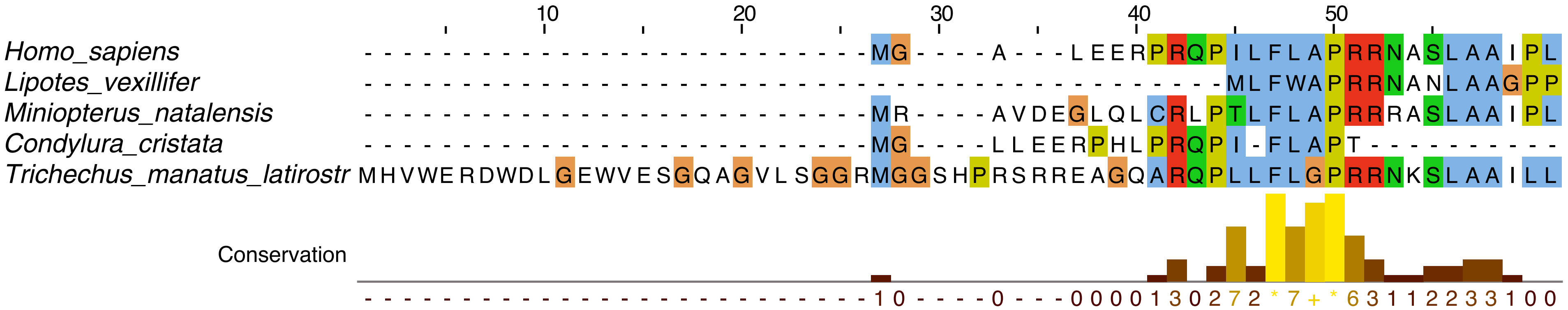

**HG0940**

ENSG00000115556 (PLCD4)

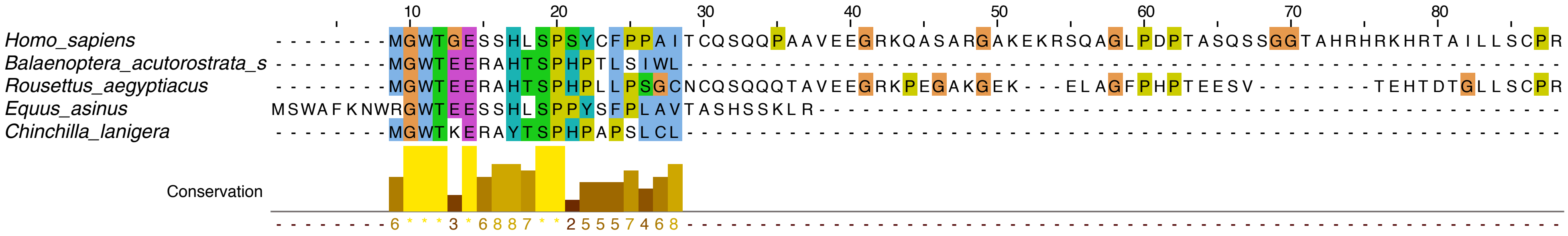

**HG0941**

ENSG00000116285 (ERRFI1)

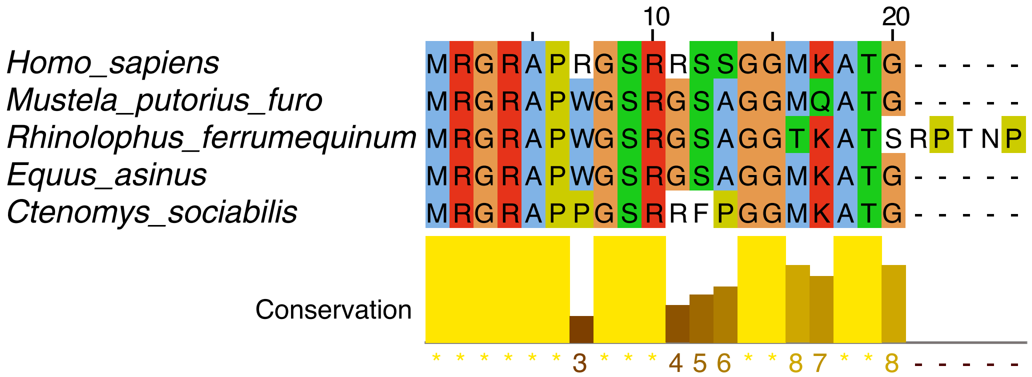

**HG0942**

ENSG00000117477

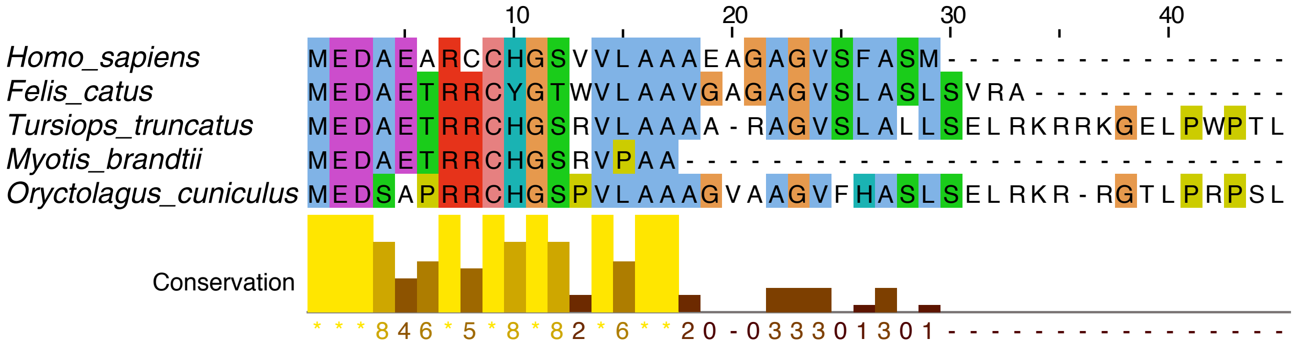

**HG0943**

ENSG00000120690 (ELF1)

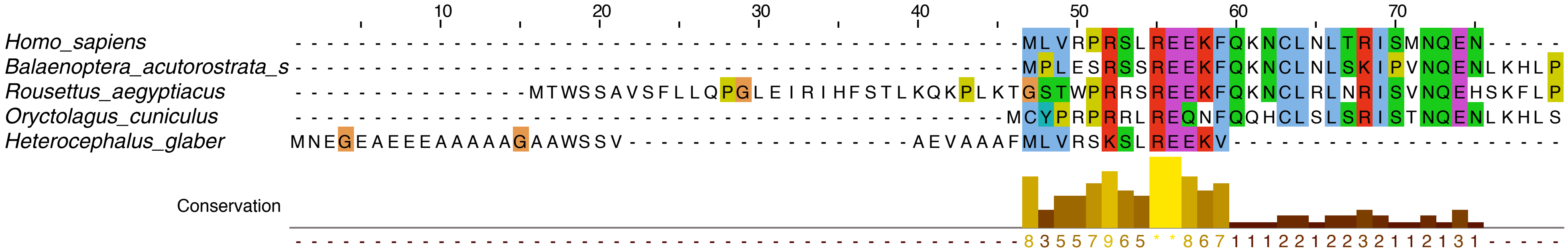

**HG0944**

ENSG00000120833 (SOCS2)

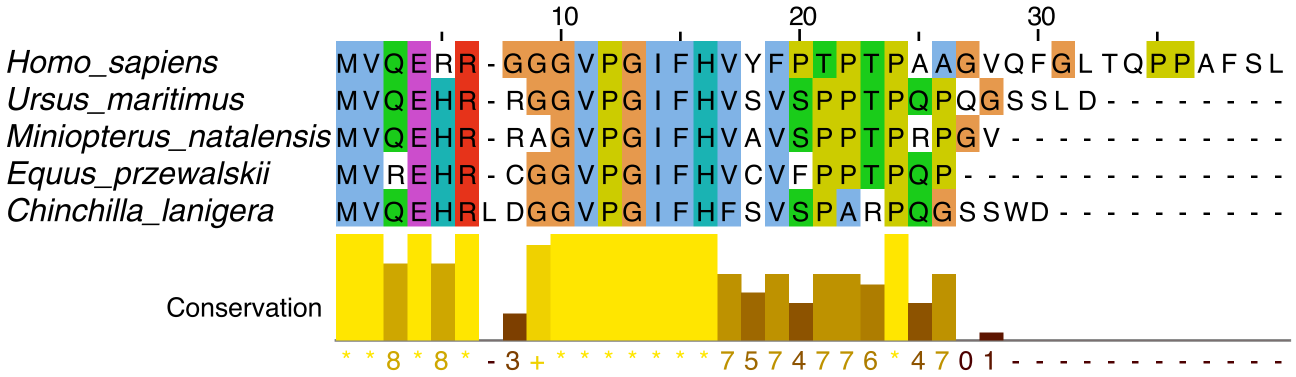

**HG0945**

ENSG00000124782 (RREB1)

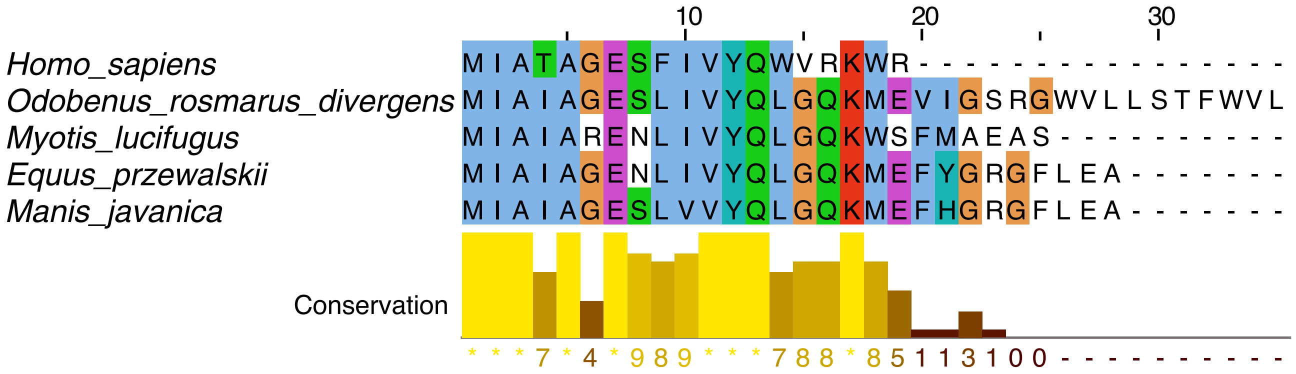

**HG0946.1**

ENSG00000127324 (TSPAN8)

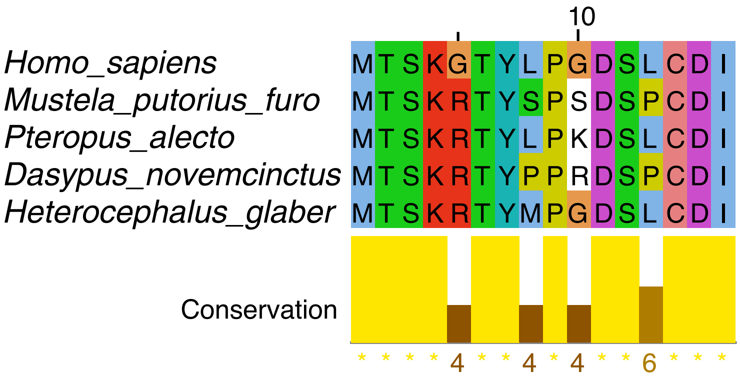

**HG0946.2**

ENSG00000127324 (TSPAN8)

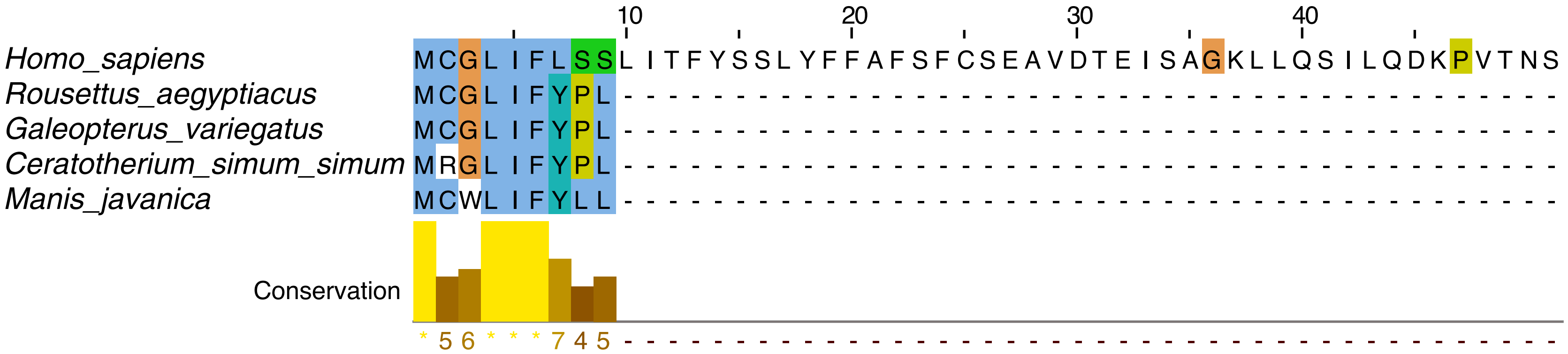

**HG0947**

ENSG00000129226 (CD68)

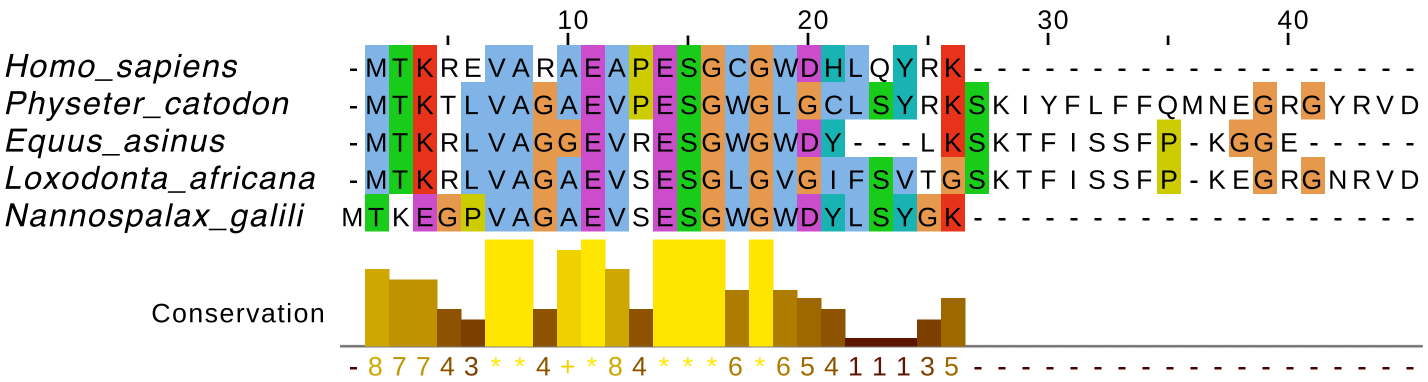

**HG0948**

ENSG00000129595 (EPB41L4A)

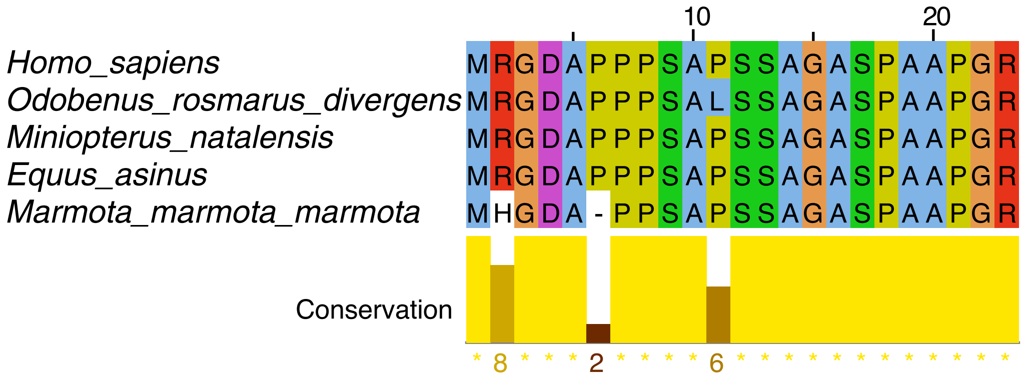

**HG0949**

ENSG00000131381 (RBSN)

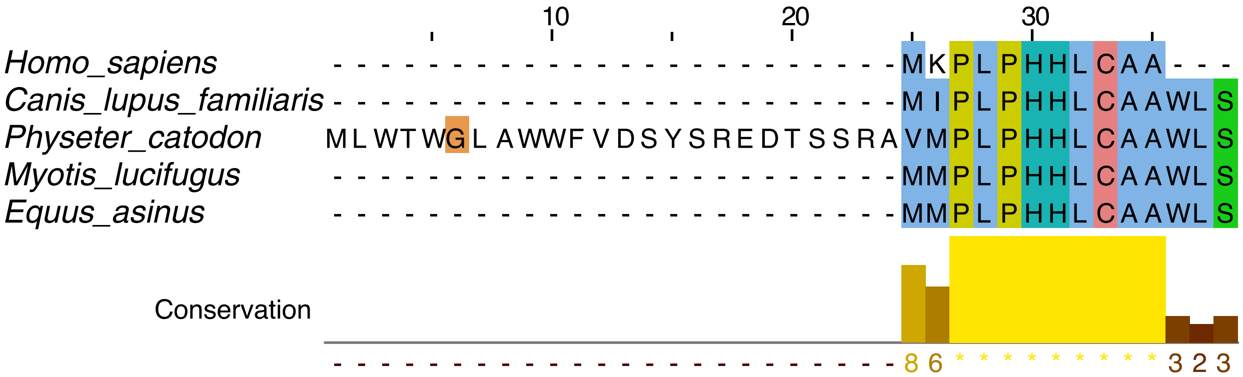

**HG0950**

ENSG00000132964 (CDK8)

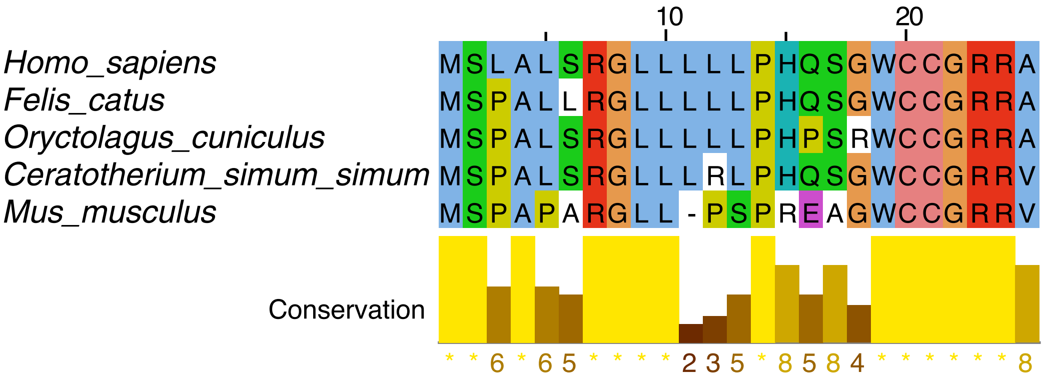

**HG0951**

ENSG00000134061 (CD180)

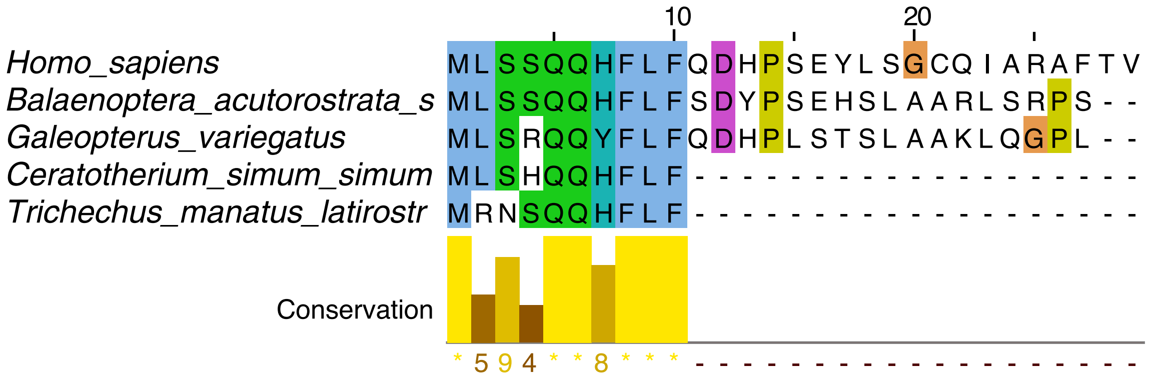

**HG0952**

ENSG00000134453 (RBM17)

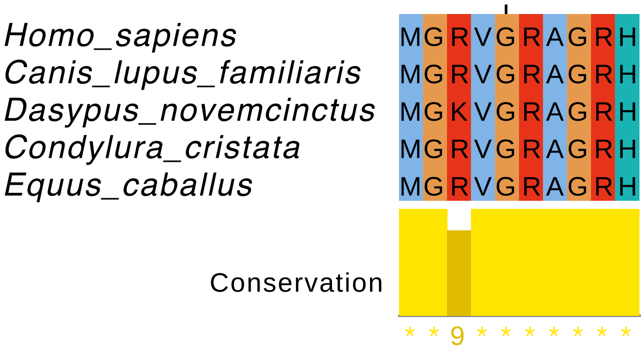

**HG0953**

ENSG00000134815 (DHX34)

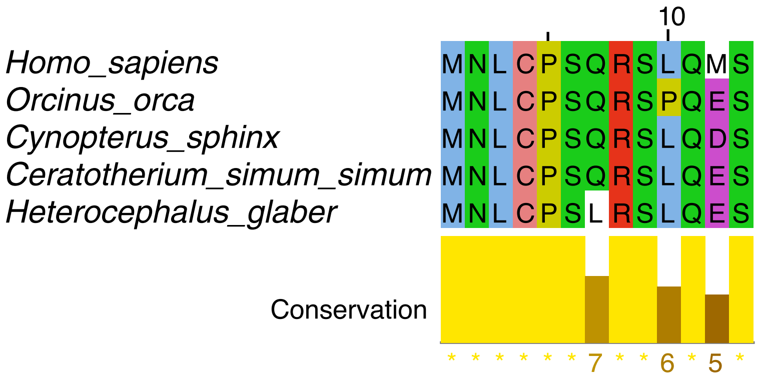

**HG0954**

ENSG00000134874 (DZIP1)

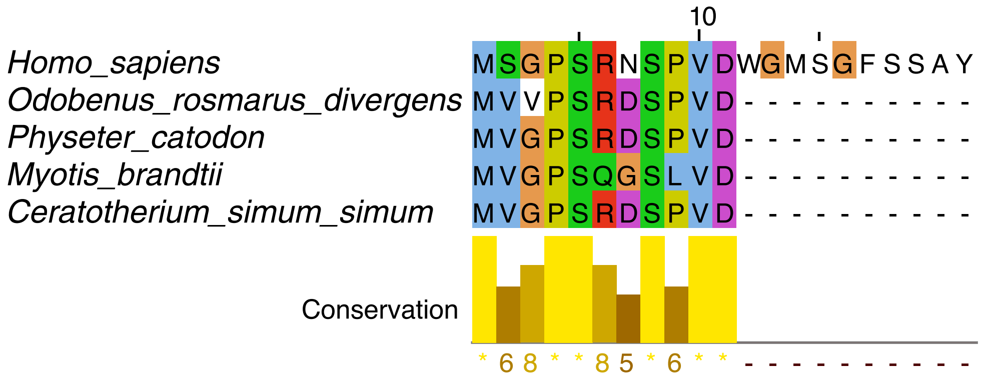

**HG0955**

ENSG00000134982 (APC)

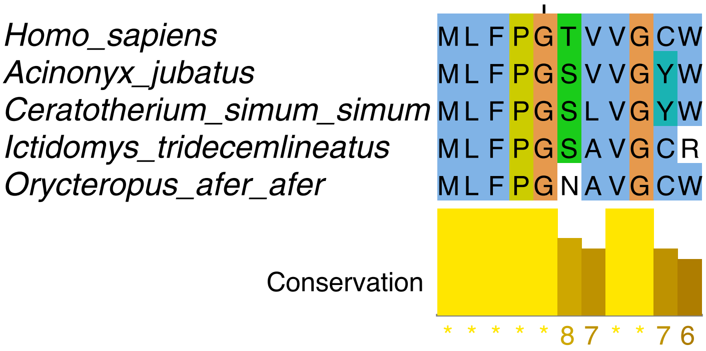

**HG0956**

ENSG00000136235 (GPNMB)

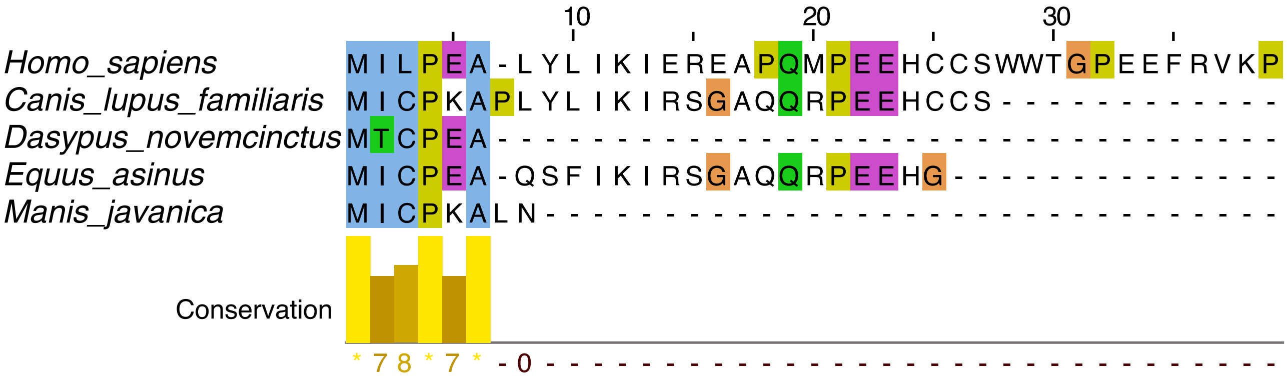

**HG0957**

ENSG00000136874 (STX17)

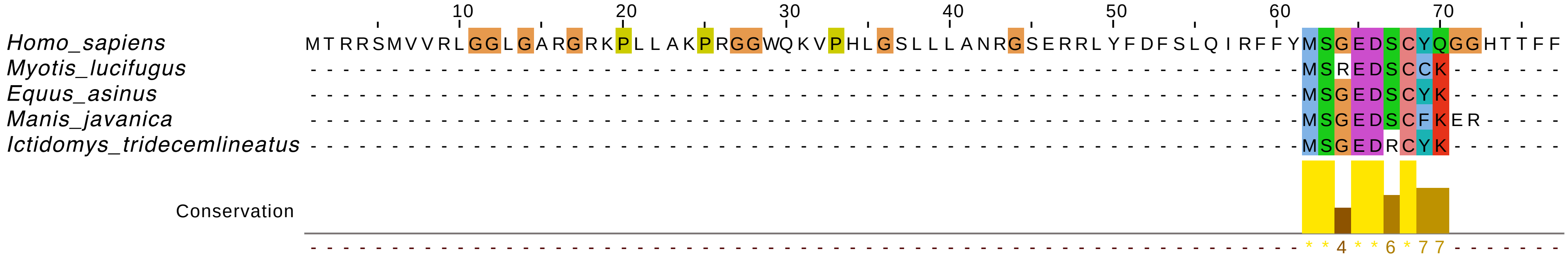

**HG0958**

ENSG00000137497 (NUMA1)

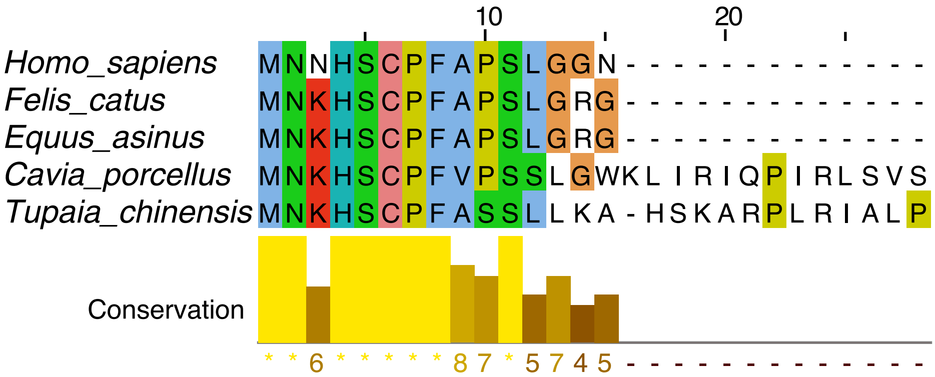

**HG0959**

ENSG00000137558 (PI15)

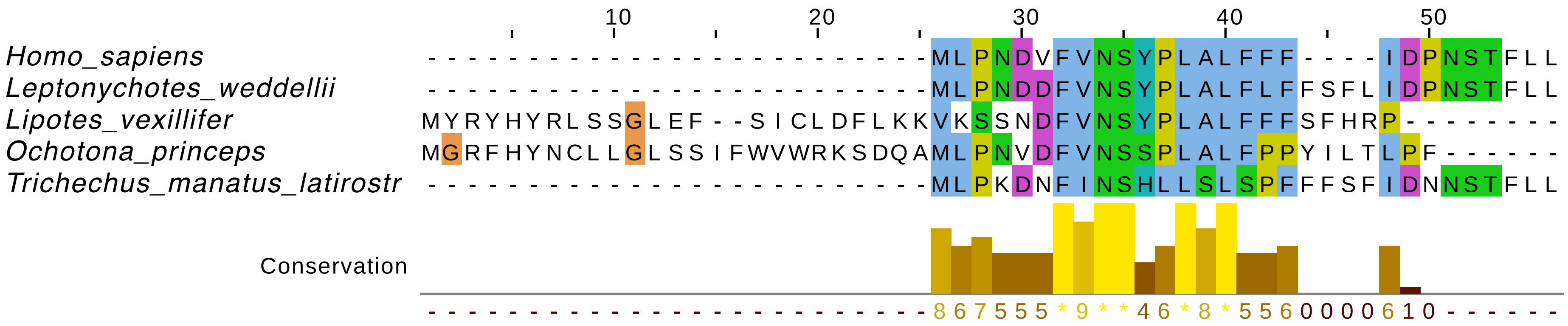

**HG0960**

ENSG00000140297 (GCNT3)

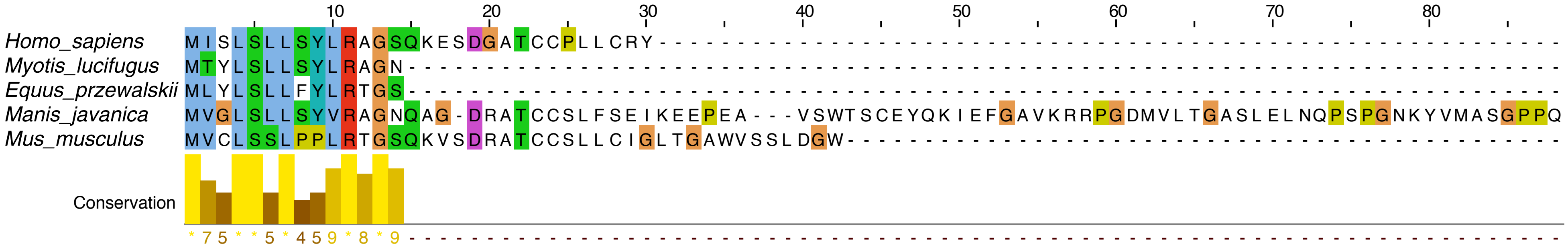

**HG0961**

ENSG00000140853 (NLRC5)

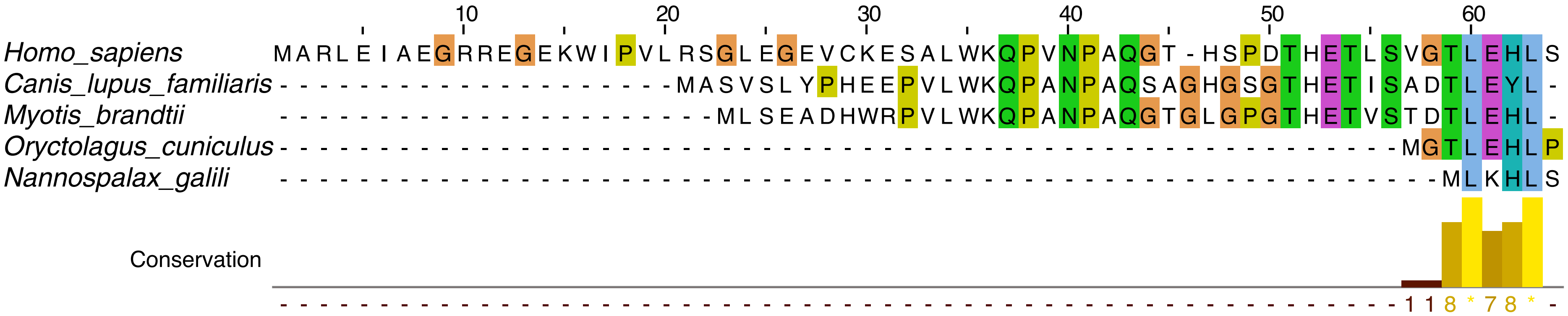

# **HG0962.1**

### ENSG00000143319 (ISG20L2)

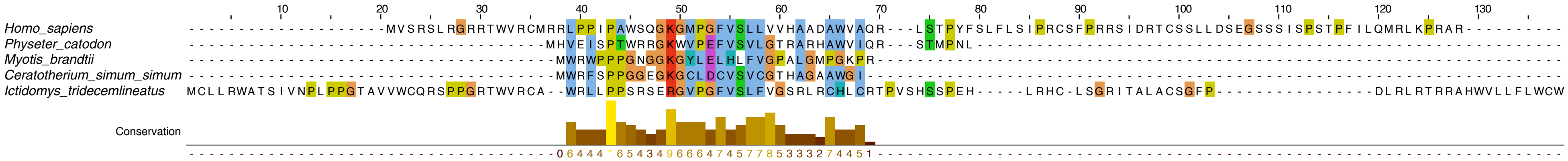

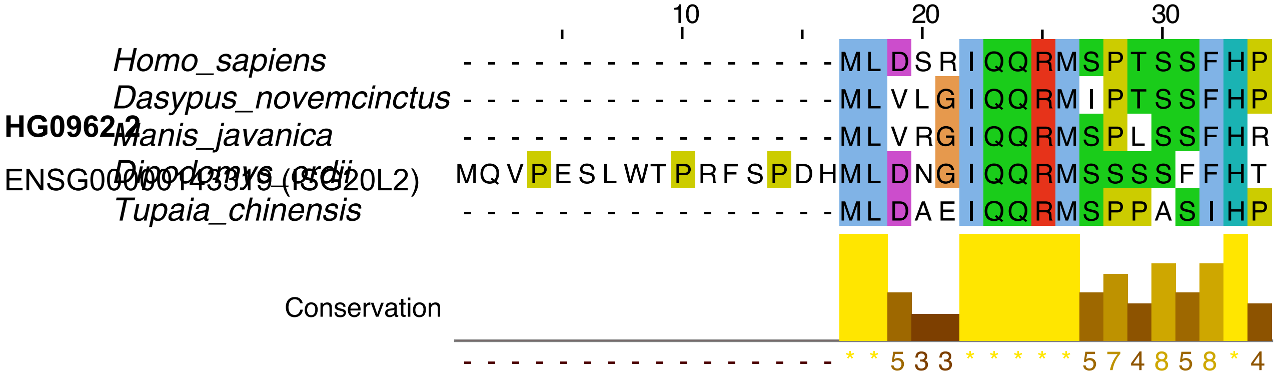
HG0962.2
ENSG00000143319 (ISG20L2)
10
20
30
Homo_sapiens
Dasypus_novemcinctus
Manis_javanica
Dipodomys_ordii
Tupaia_chinensis
Conservation

**HG0963**

ENSG00000143867 (OSR1)

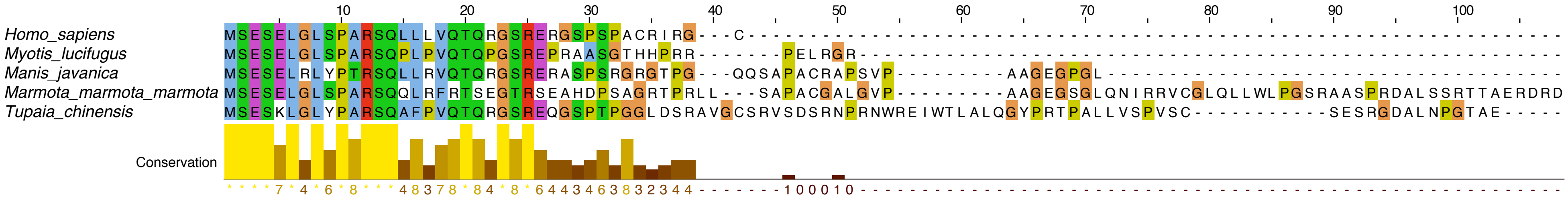

**HG0964**

ENSG00000145246 (ATP10D)

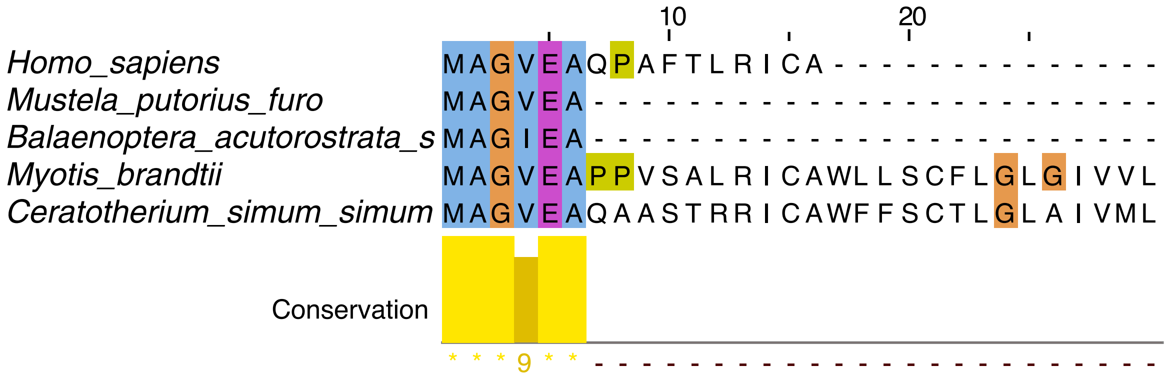

**HG0965**

ENSG00000146006 (LRRTM2)

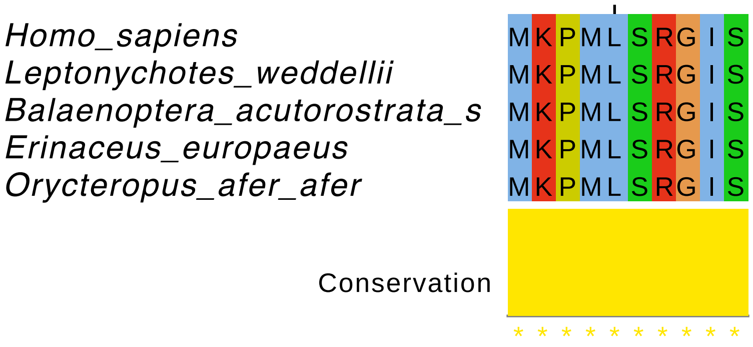

**HG0966**

ENSG00000148356 (LRSAM1)

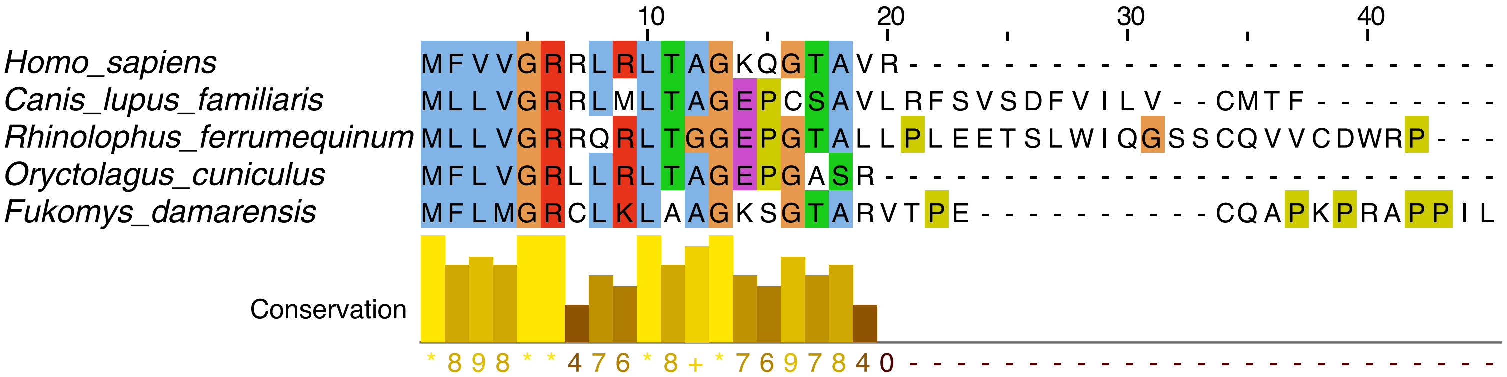

**HG0967**

ENSG00000149925 (ALDOA)

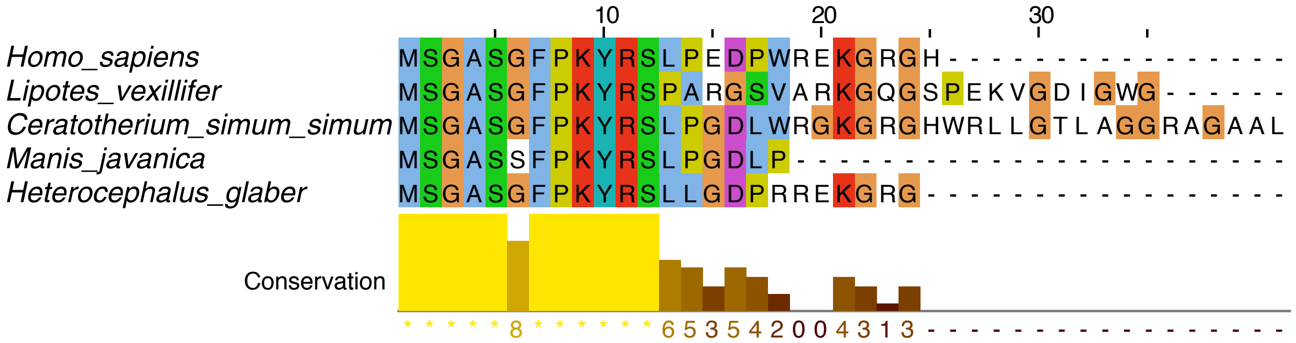

**HG0968**

ENSG00000154319 (FAM167A)

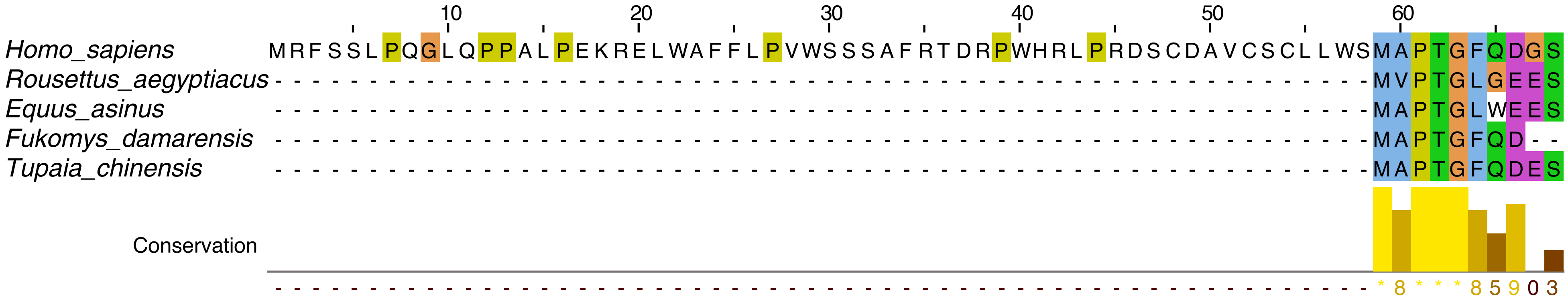

**HG0969**

ENSG00000154645 (CHODL)

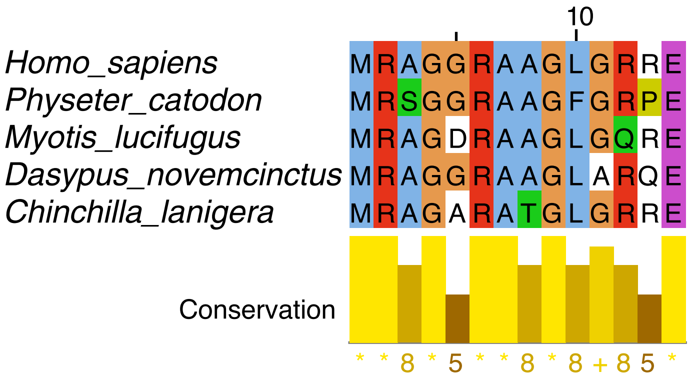

**HG0970**

ENSG00000154727 (GABPA)

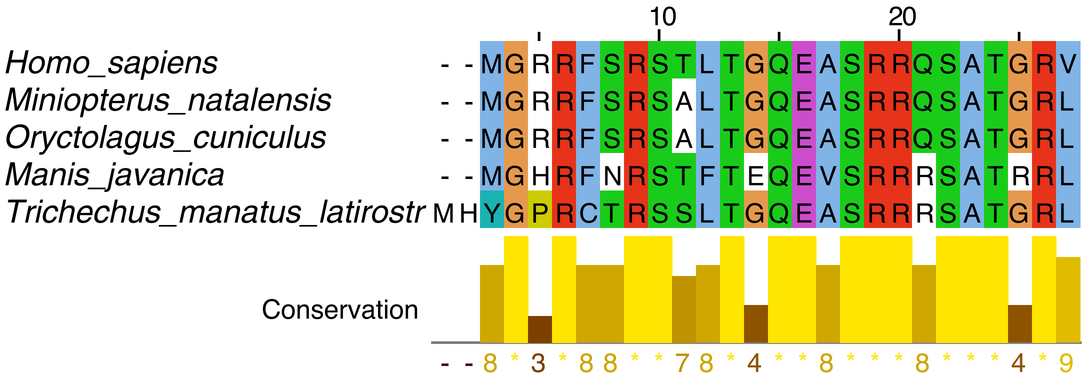

**HG0971**

ENSG00000155592 (ZKSCAN2)

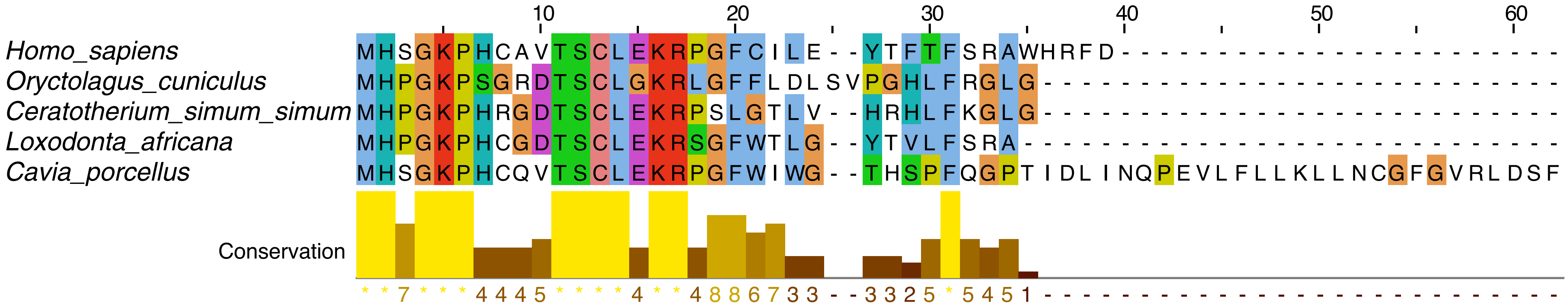

**HG0972**

ENSG00000155926 (SLA)

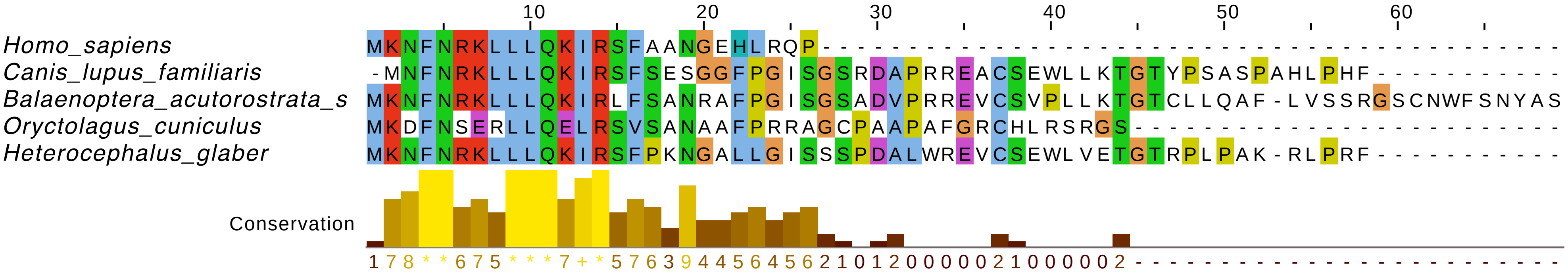

**HG0973**

ENSG00000156535 (CD109)

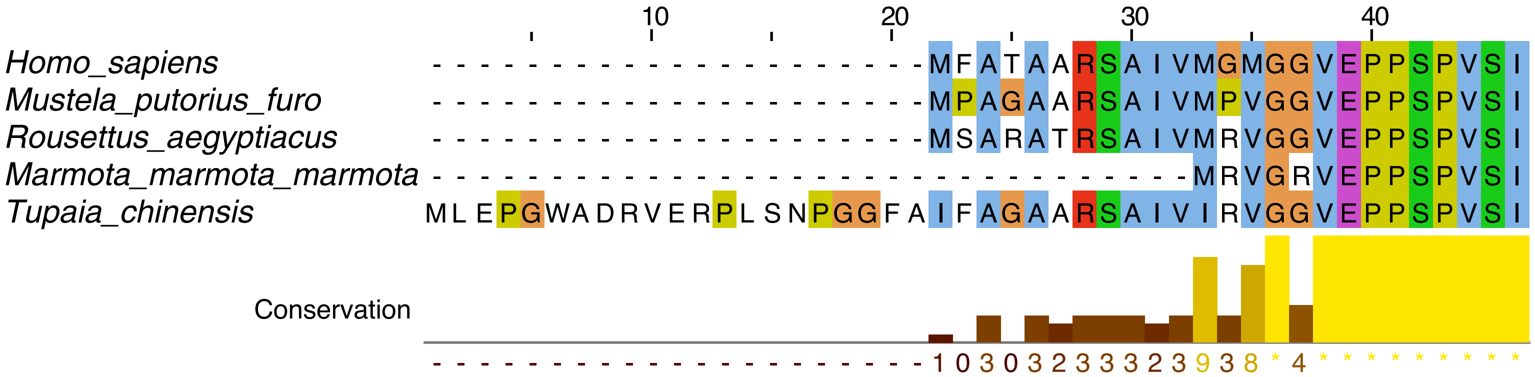

**HG0974**

ENSG00000157578 (LCA5L)

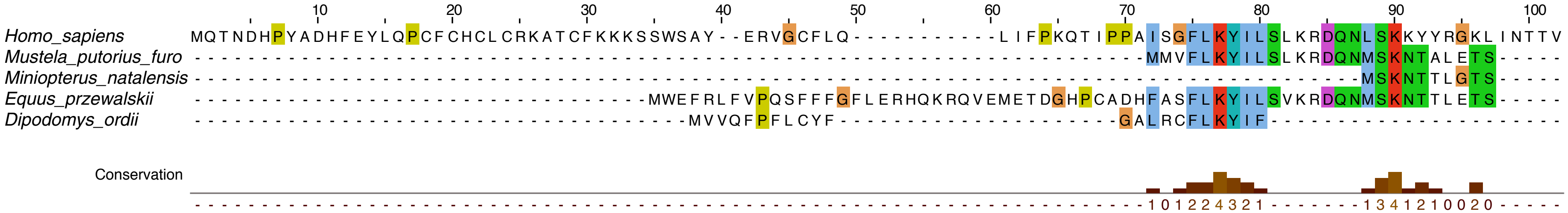

**HG0975.1**

ENSG00000182158 (CREB3L2)

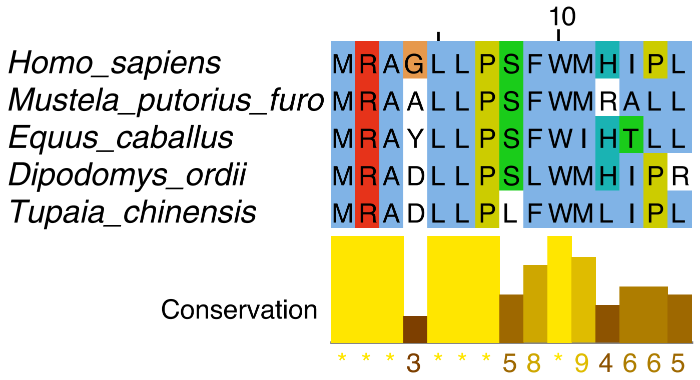

**HG0975.2**

ENSG00000157613 (CREB3L1)

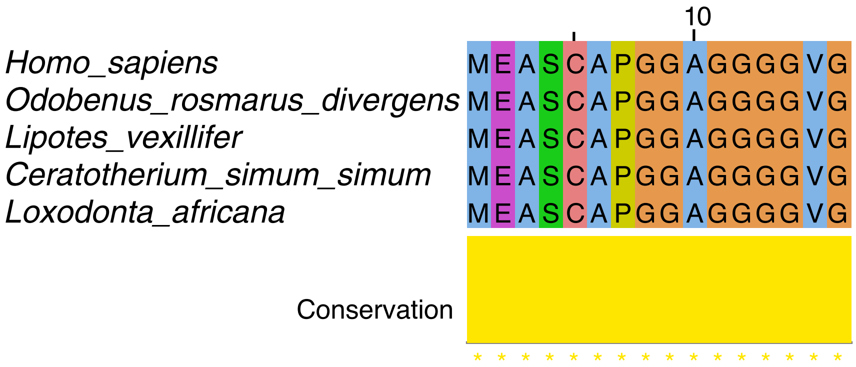

**HG0976**

ENSG00000158008 (EXTL1)

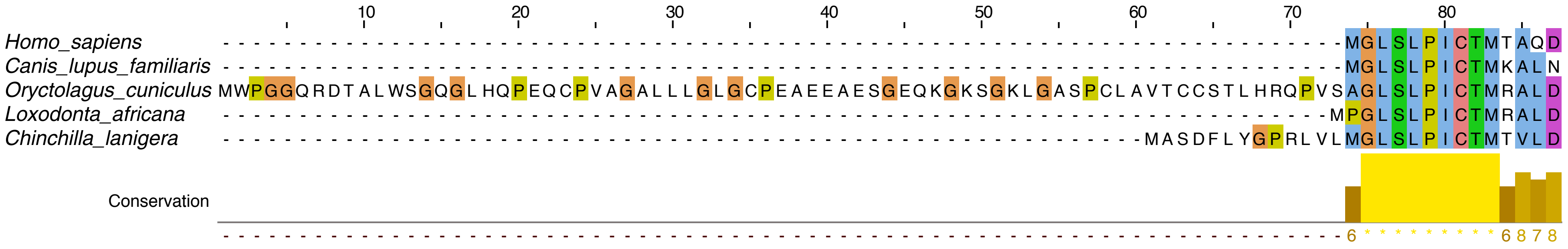

**HG0977**

ENSG00000158321 (AUTS2)

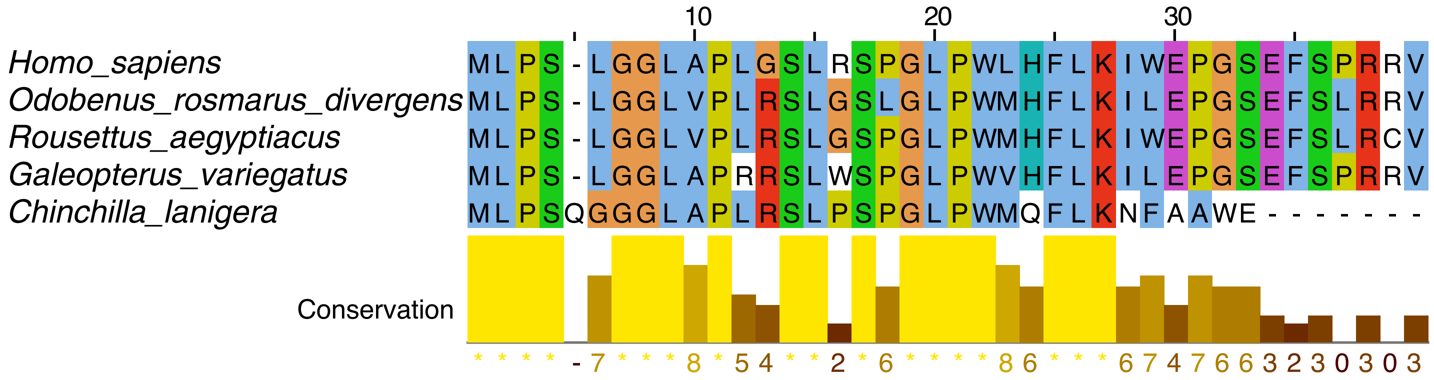

**HG0978**

ENSG00000159023 (EPB41)

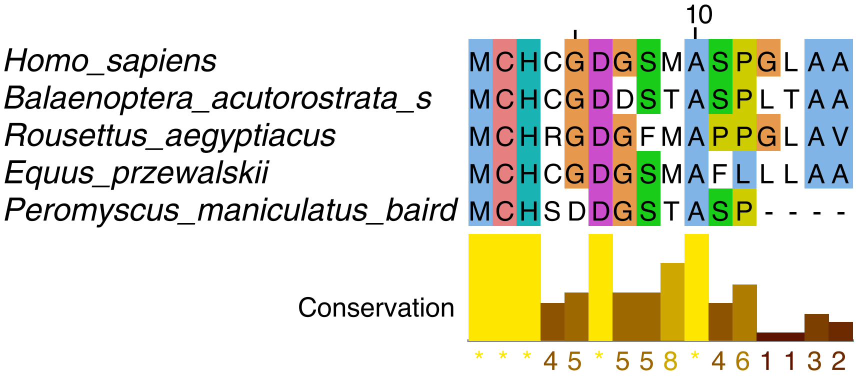

**HG0979**

ENSG00000159202 (UBE2Z)

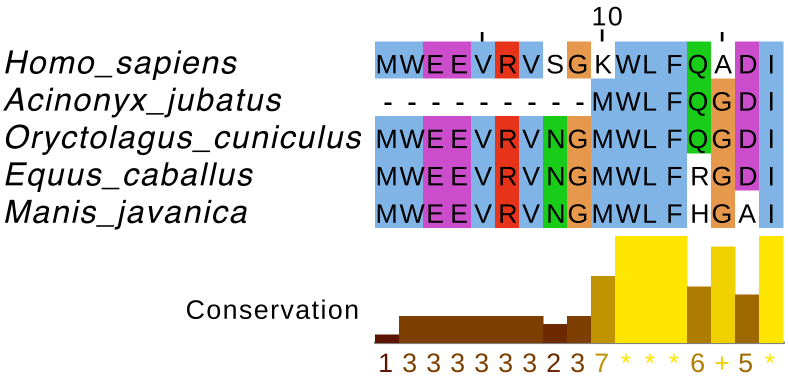

**HG0980**

ENSG00000162670 (BRINP3)

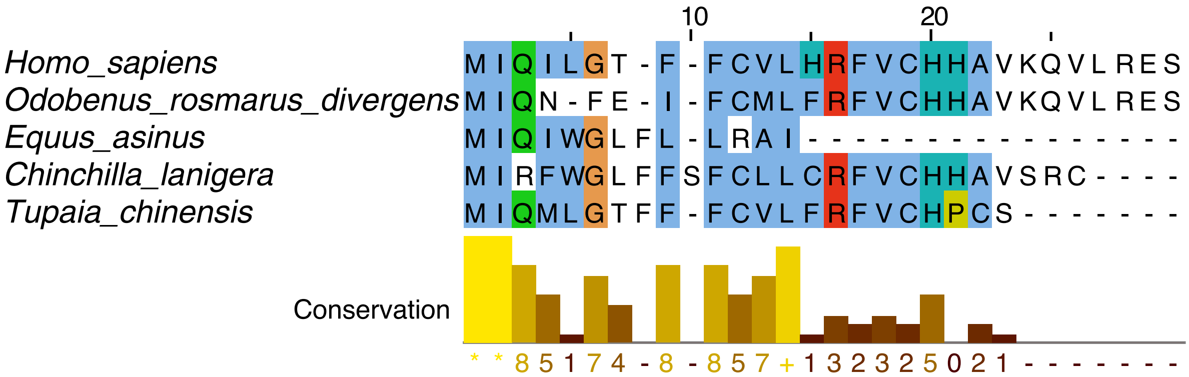

**HG0981**

ENSG00000163629 (PTPN13)

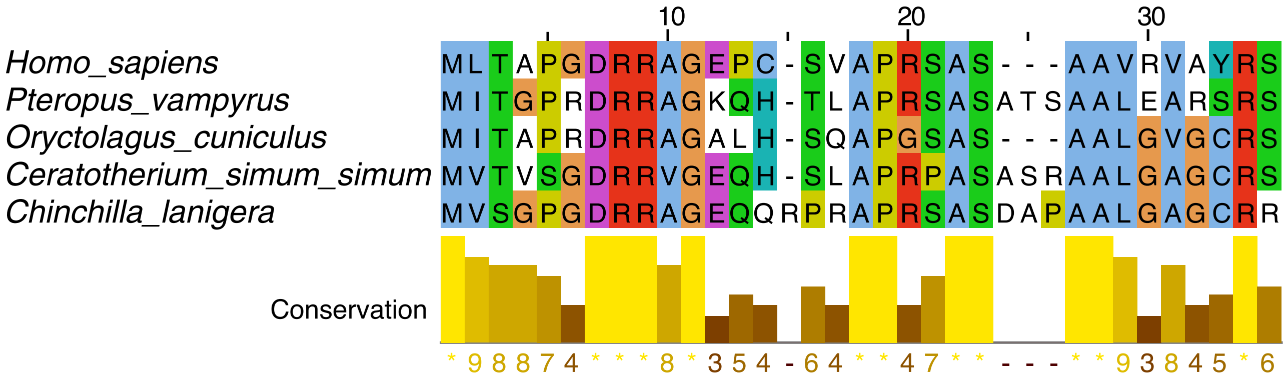

**HG0982**

ENSG00000164116 (GUCY1A3)

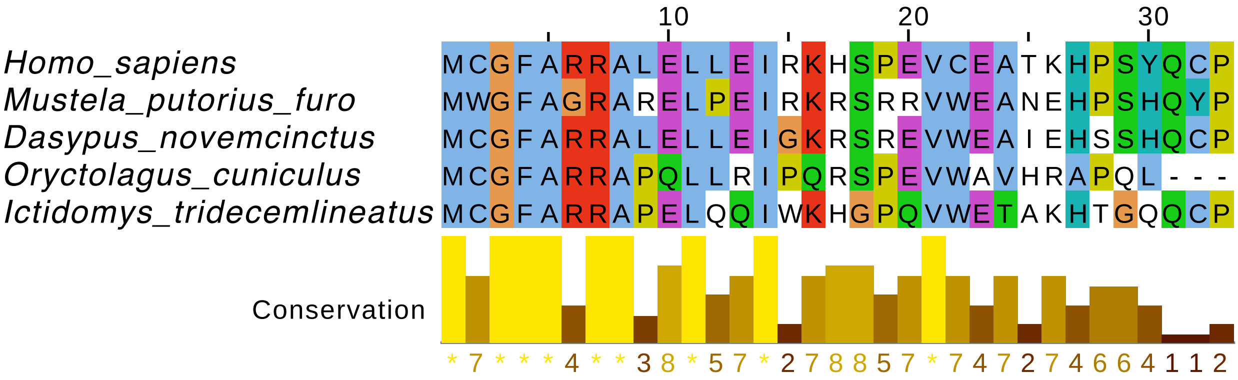

**HG0983**

ENSG00000164128 (NPY1R)

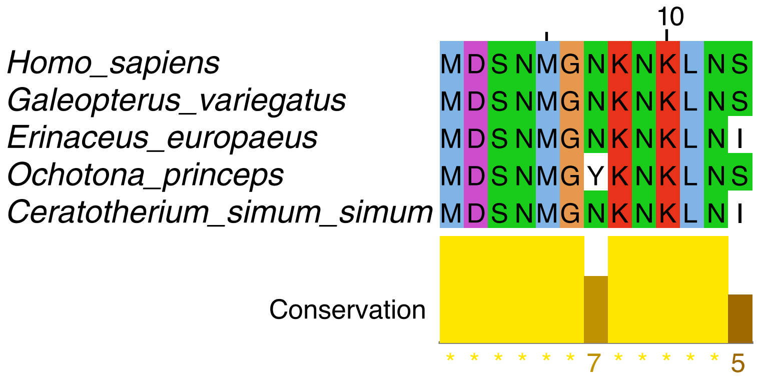

**HG0984**

ENSG00000164185 (ZNF474)

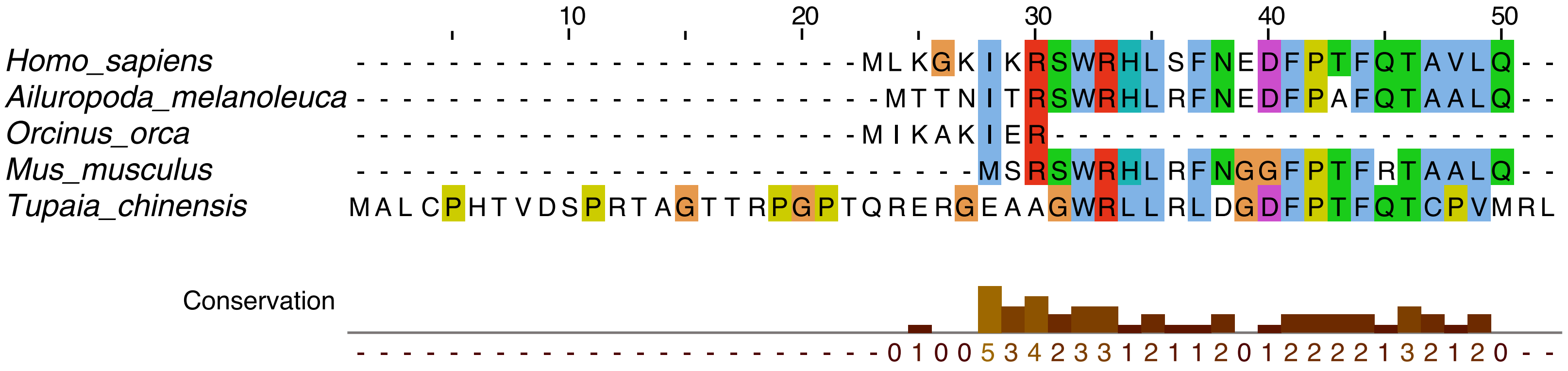

**HG0985**

ENSG00000164604 (GPR85)

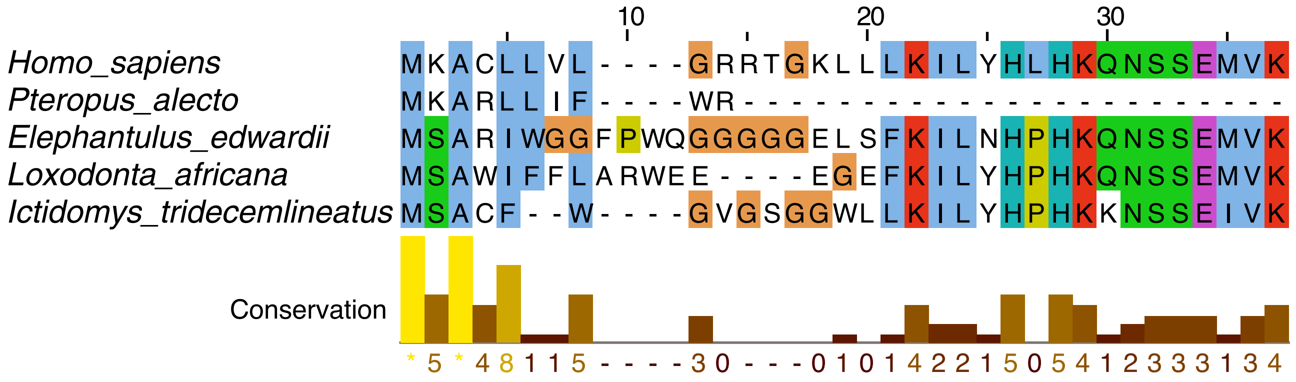

**HG0986**

ENSG00000164663 (USP49)

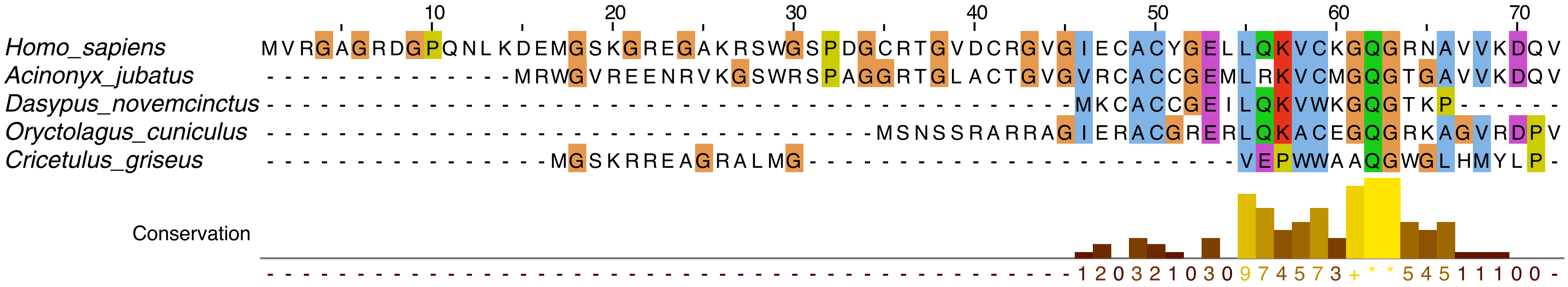

**HG0987**

ENSG00000164932 (CTHRC1)

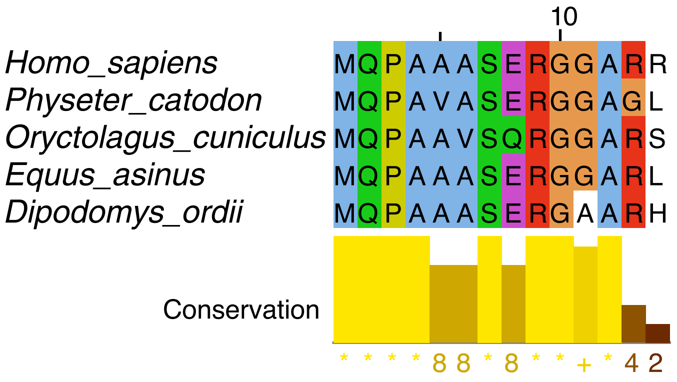

**HG0988**

ENSG00000166130 (IKBIP)

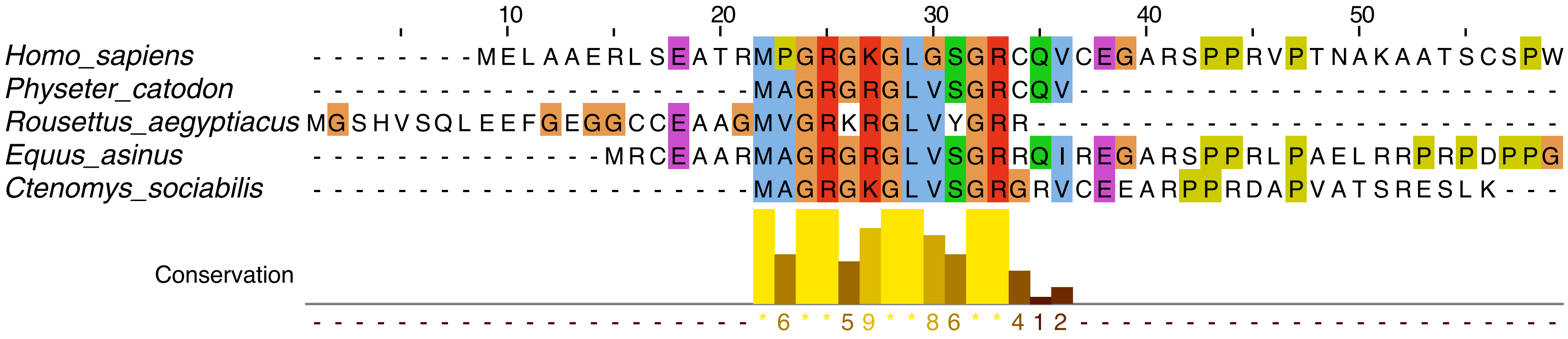

**HG0989**

ENSG00000166603 (MC4R)

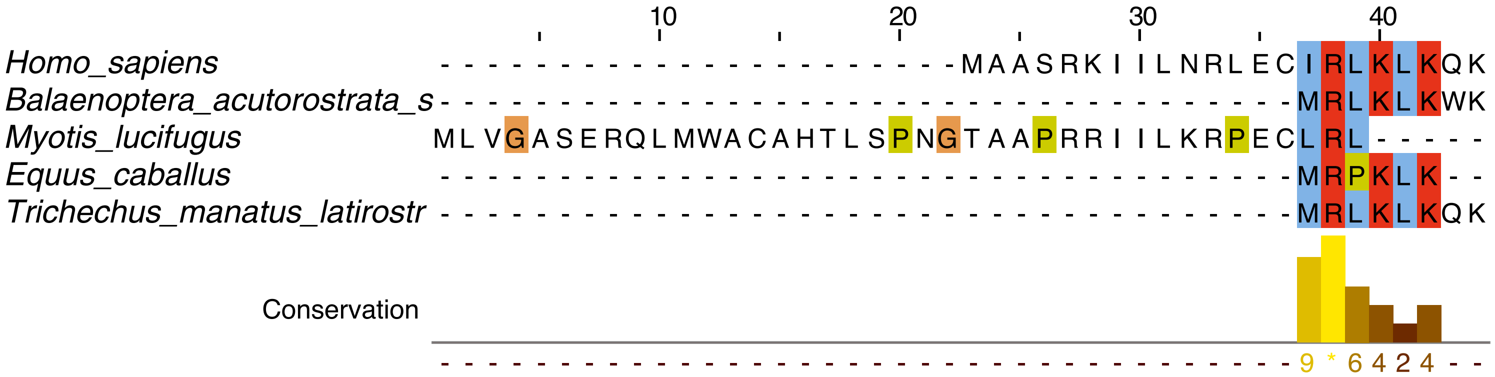

**HG0990**

ENSG00000166866 (MYO1A)

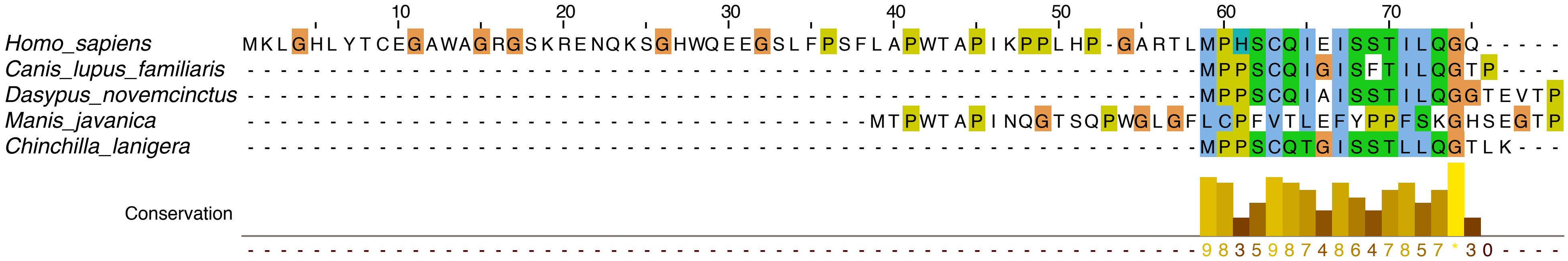

**HG0991**

ENSG00000167654 (ATCAY)

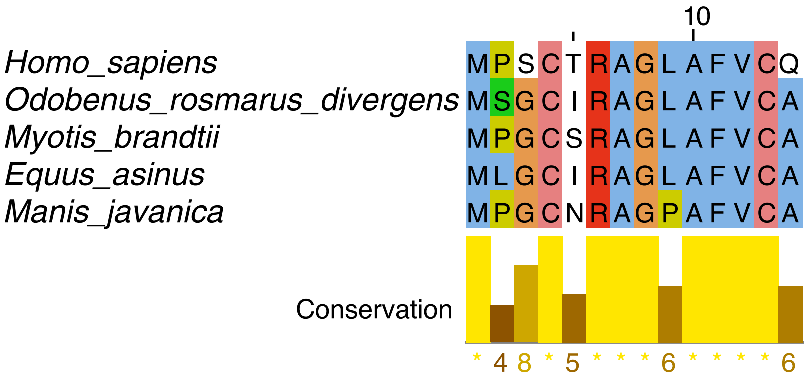

**HG0992**

ENSG00000168000 (BSCL2)

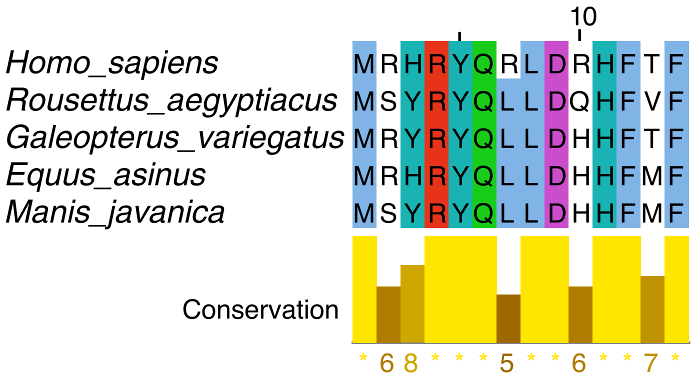

**HG0993**

ENSG00000169306 (IL1RAPL1)

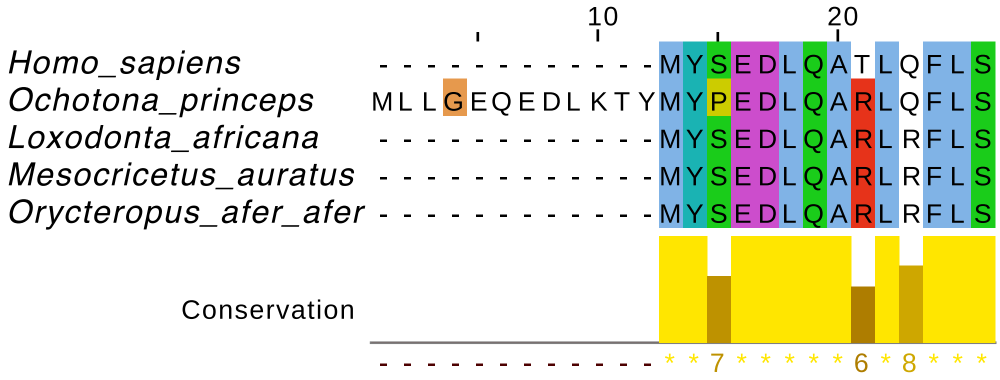

**HG0994**

ENSG00000170852 (KBTBD2)

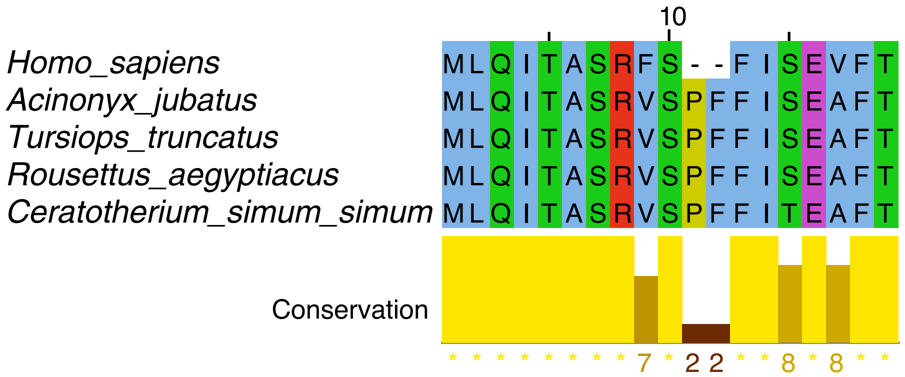

**HG0995**

ENSG00000171307 (ZDHHC16)

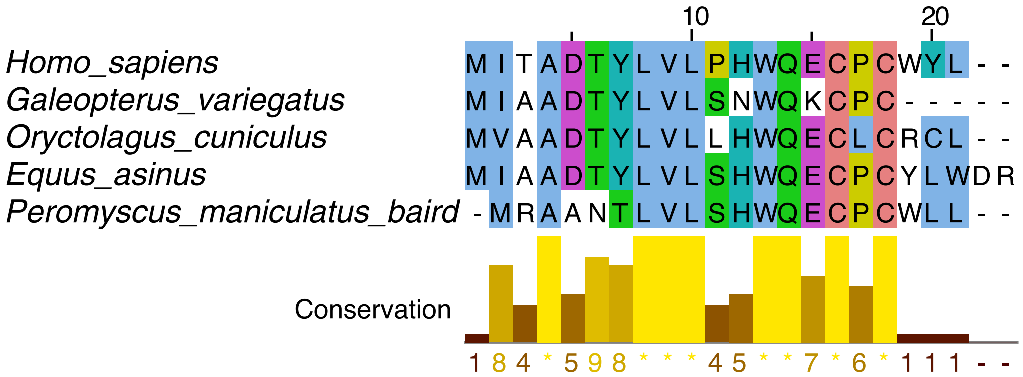

**HG0996**

ENSG00000173890 (GPR160)

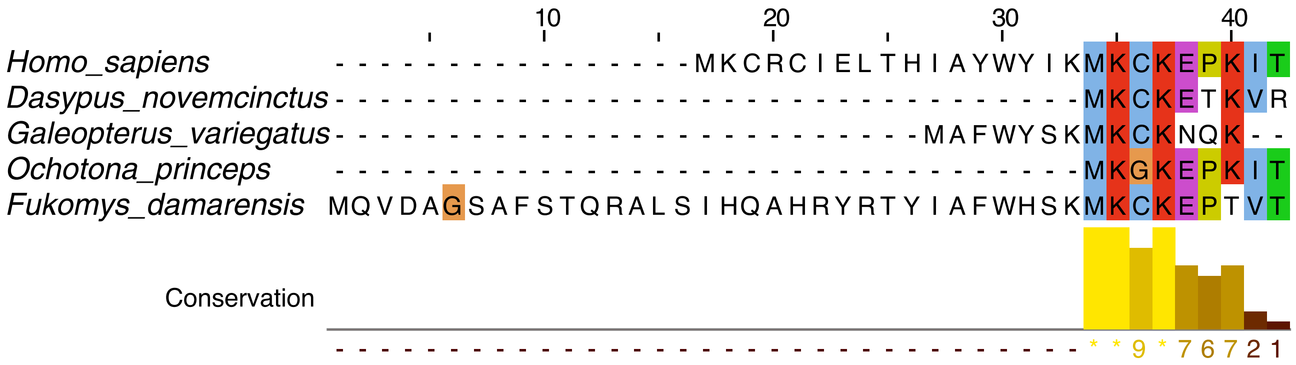

**HG0997**

ENSG00000177728 (TMEM94)

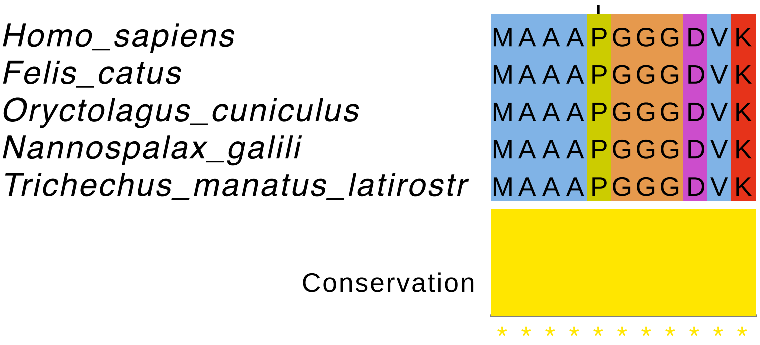

**HG0998**

ENSG00000178172 (SPINK6)

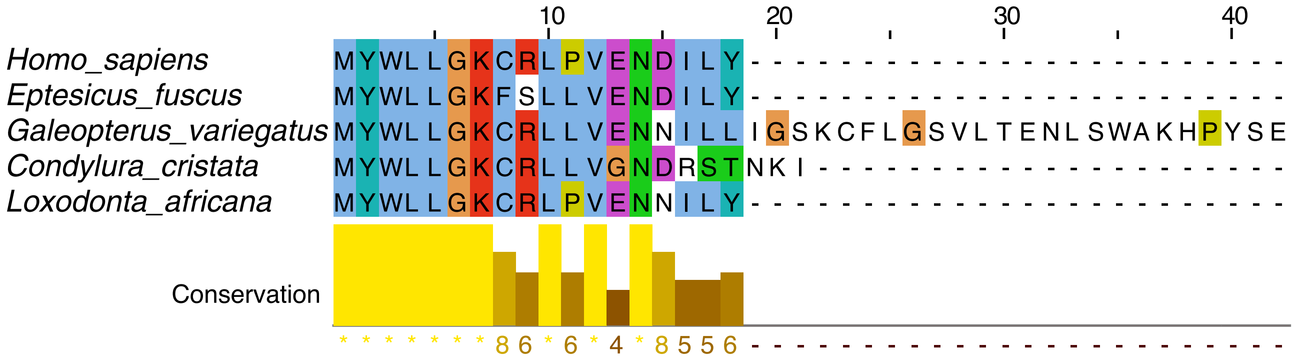

**HG0999**

ENSG00000178401 (DNAJC22)

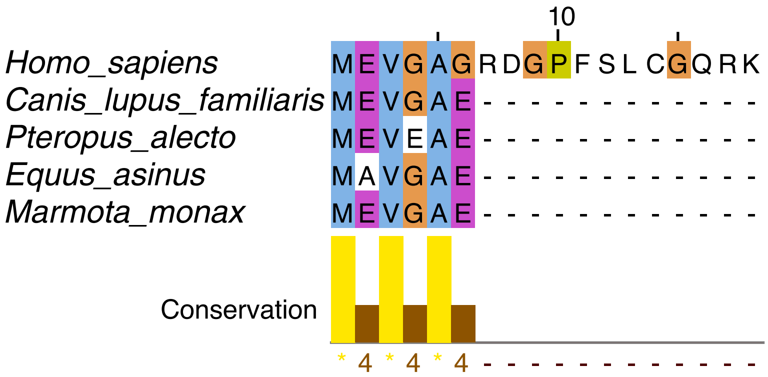

**HG1000**

ENSG00000179083 (FAM133A)

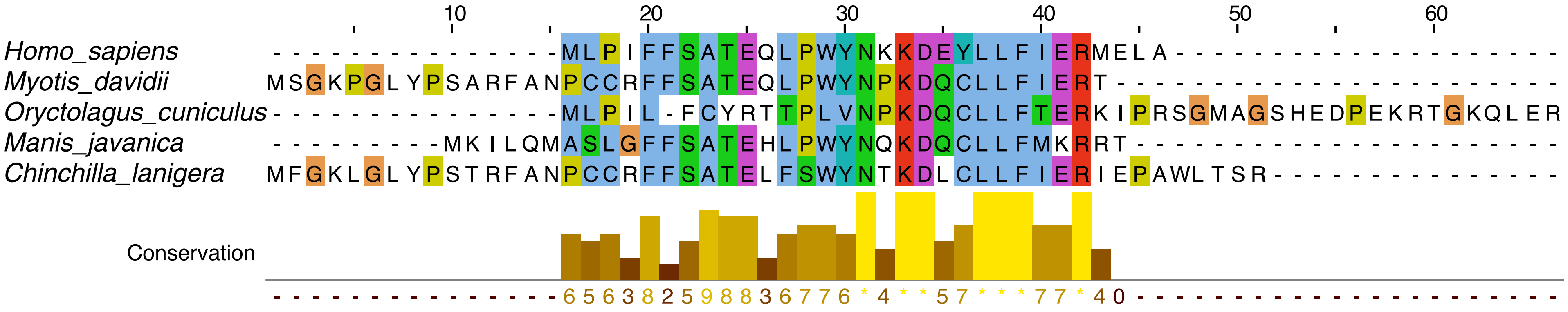

**HG1001**

ENSG00000179761 (PIPOX)

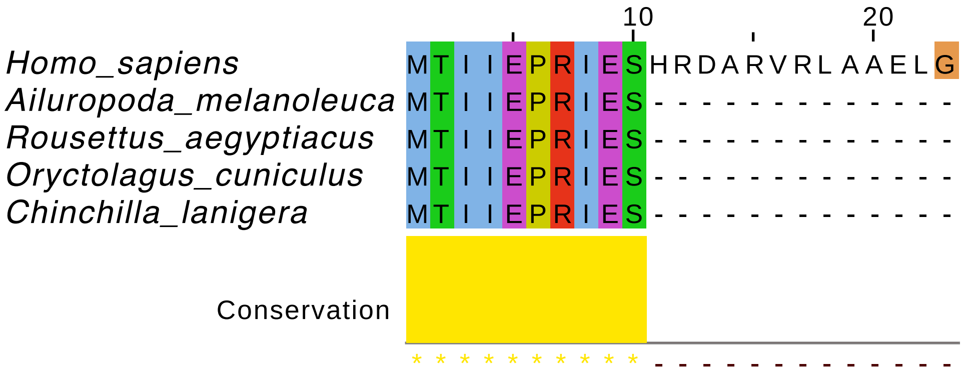

**HG1002**

ENSG00000181027 (FKRP)

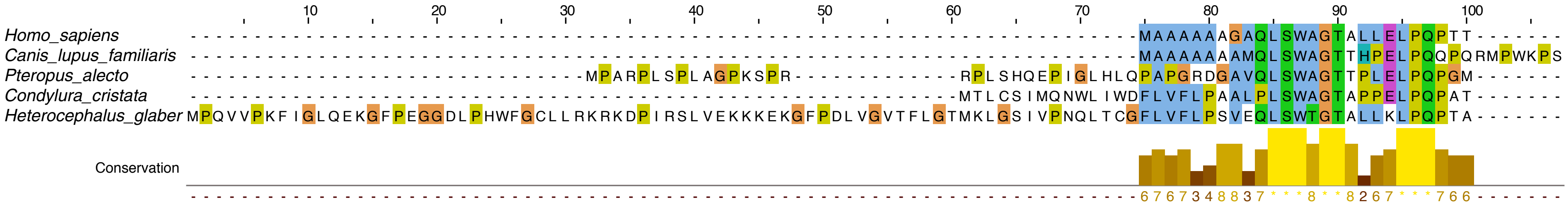

**HG1003**

ENSG00000184867 (ARMCX2)

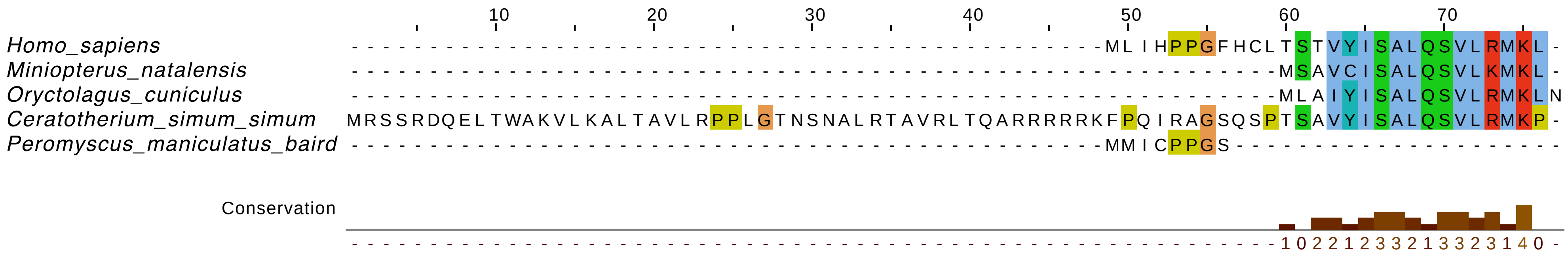

**HG1004**

ENSG00000185875 (THNSL1)

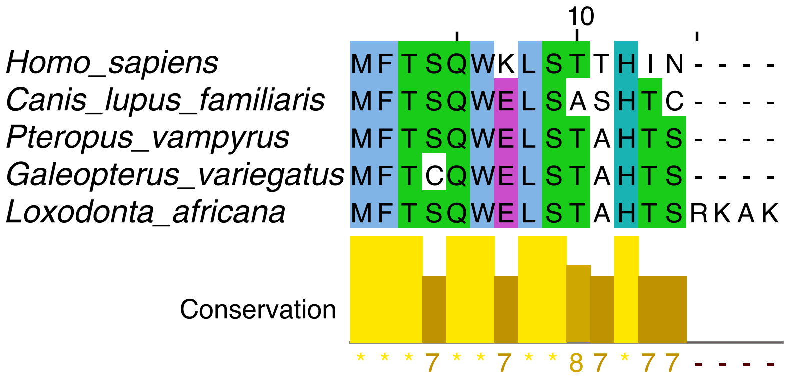

**HG1005**

ENSG00000186472 (PCLO)

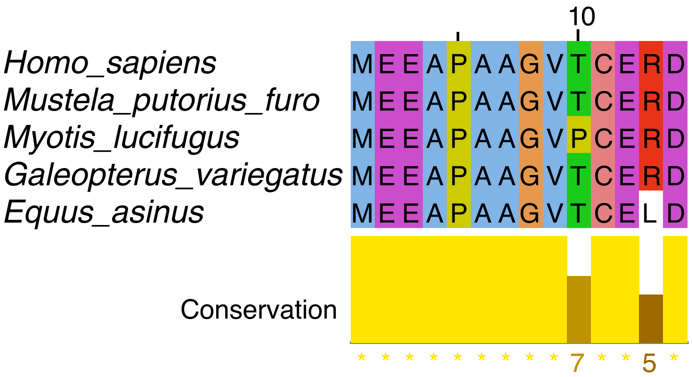

**HG1006.1**

ENSG00000188227 (ZNF793)

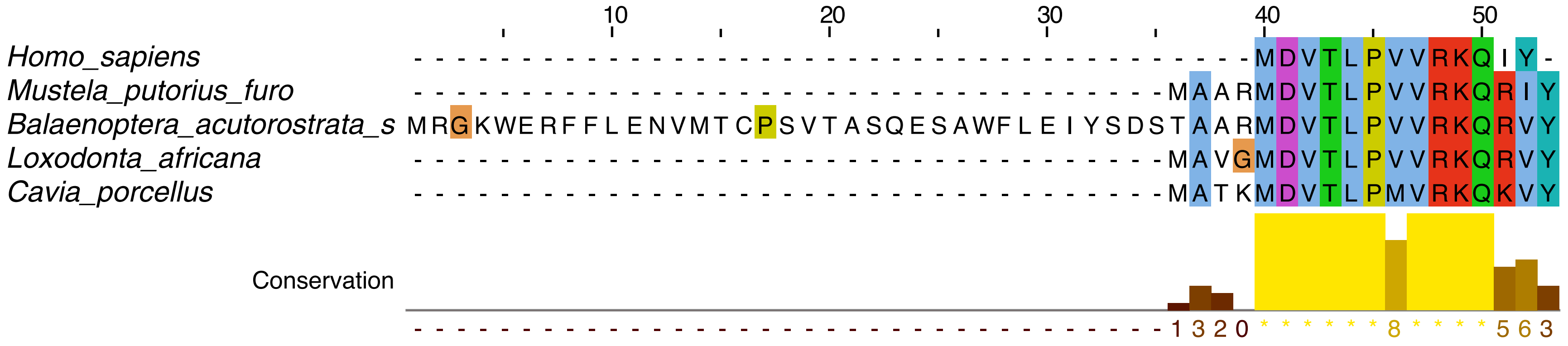

**HG1006.2**

ENSG00000188227 (ZNF793)

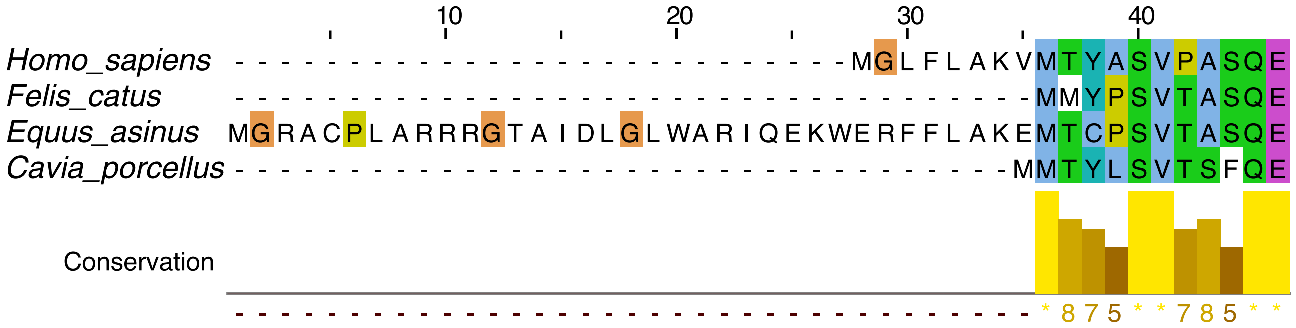

**HG1007**

ENSG00000188910 (GJB3)

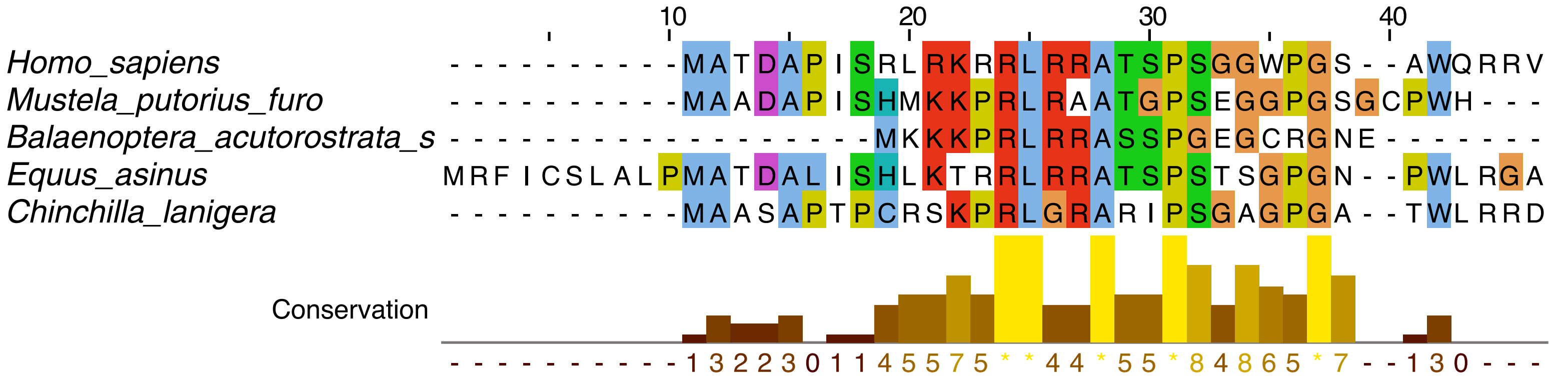

**HG1008**

ENSG00000198252 (STYX)

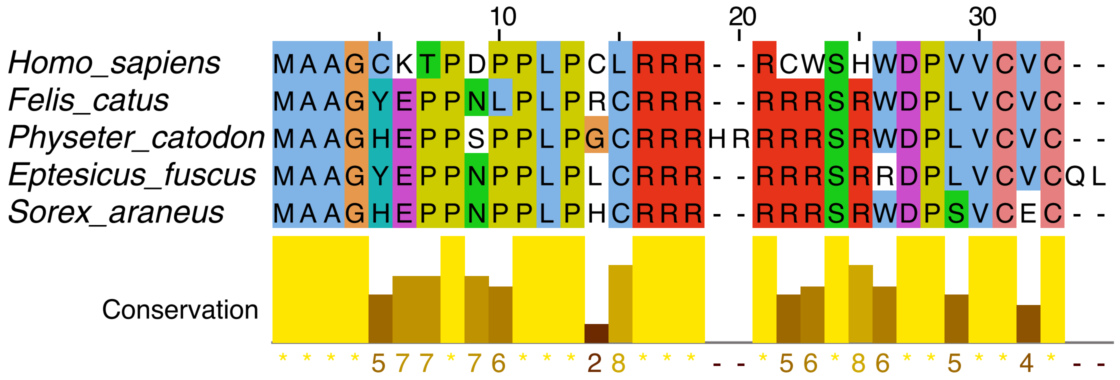

**HG1009**

ENSG00000198909 (MAP3K3)

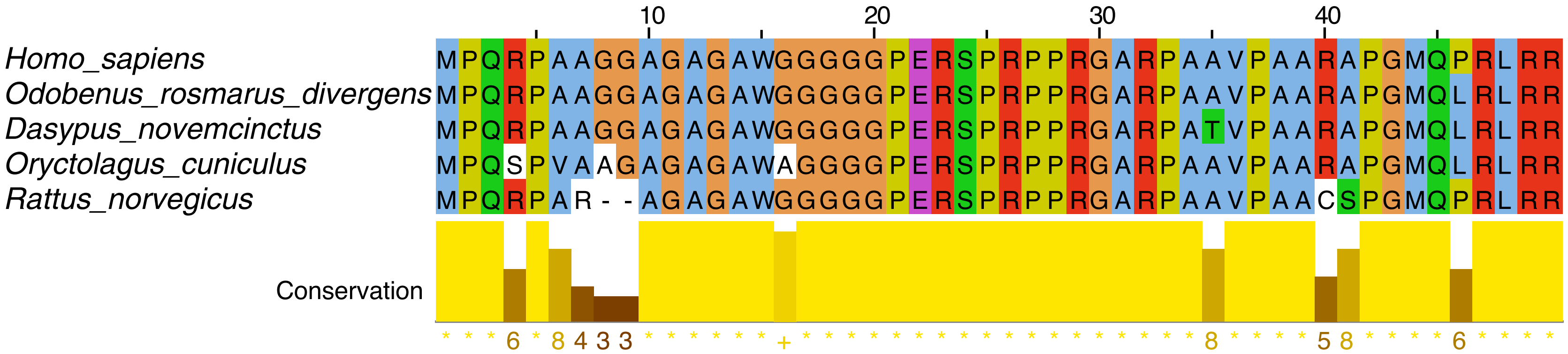

**HG1010**

ENSG00000203740 (METTL11B)

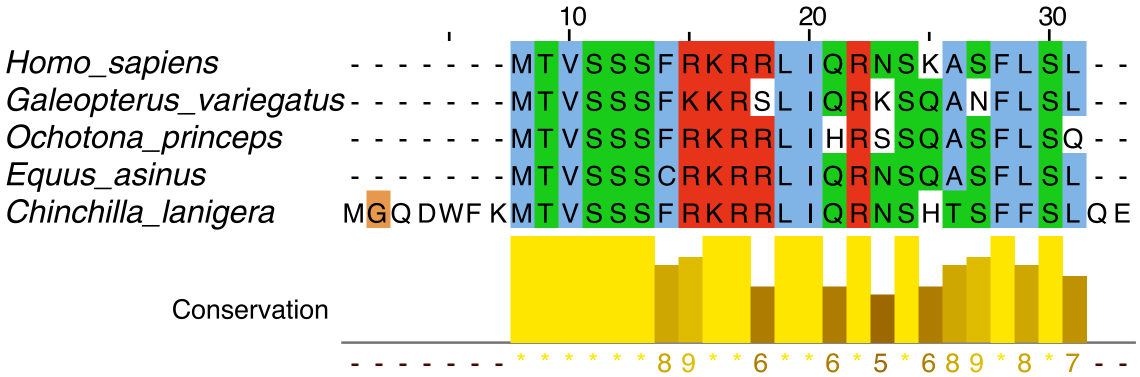

**HG1011**

ENSG00000204439 (C6orf47)

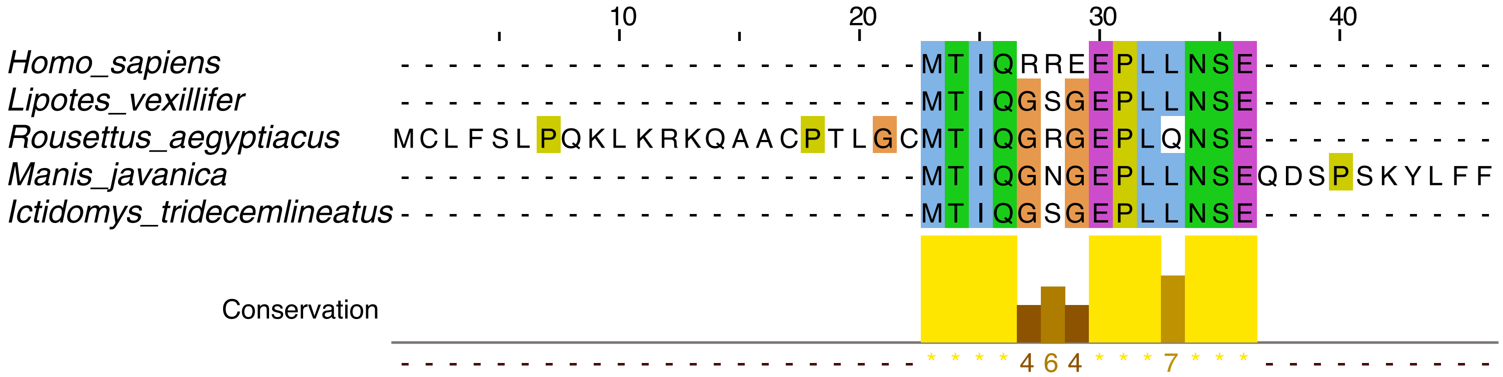

**HG1012**

ENSG00000213694 (S1PR3)

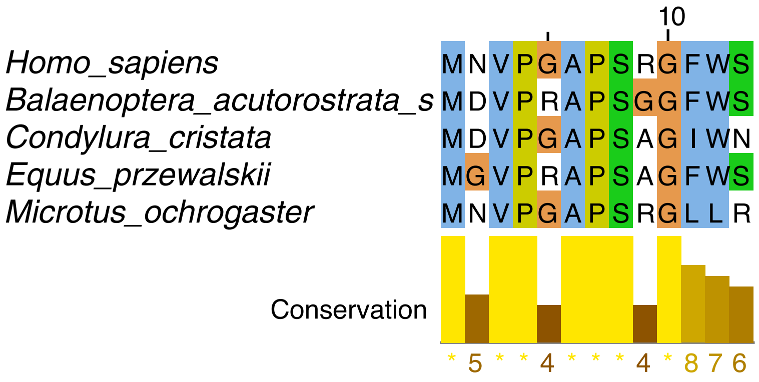

**HG1013**

ENSG00000214022 (REPIN1)

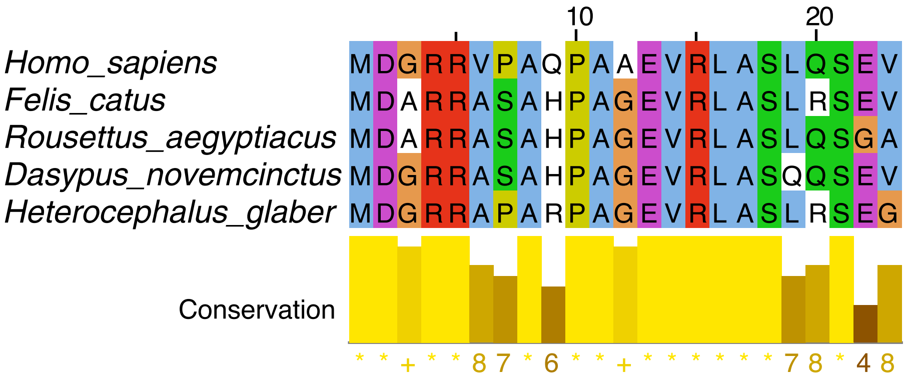

**HG1014.1**

ENSG00000214595 (EML6)

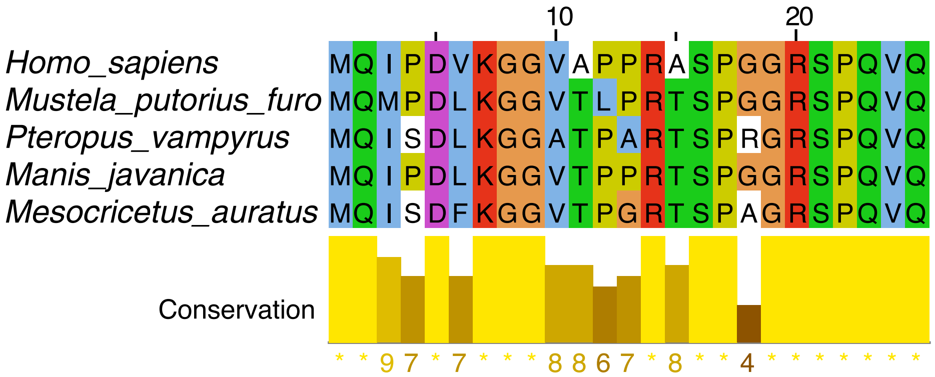

**HG1014.2**

ENSG00000214595 (EML6)

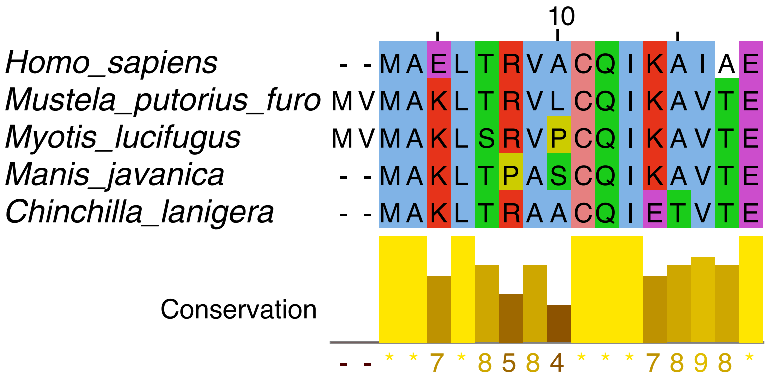

**HG1015**

ENSG00000240720 (LRRD1)

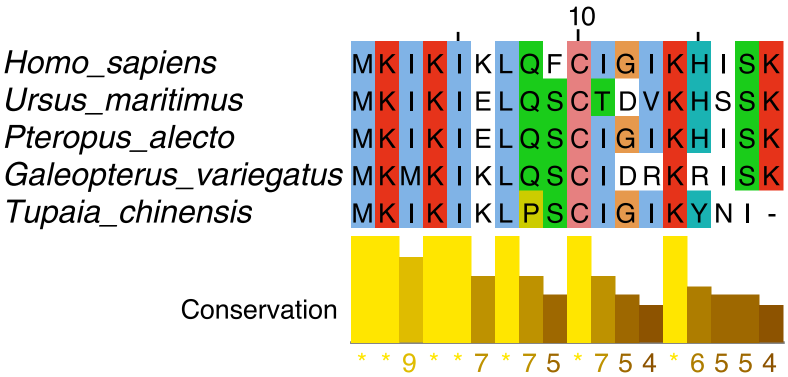

**HG1016**

ENSG00000254004 (ZNF260)

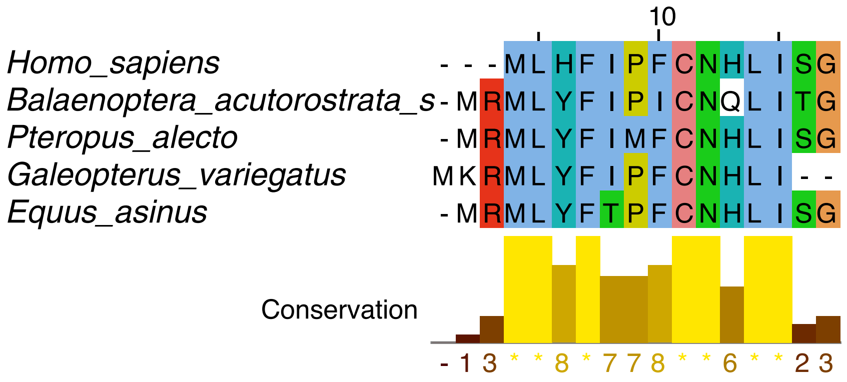

**HG1017**

ENSG00000255154 (HTD2)

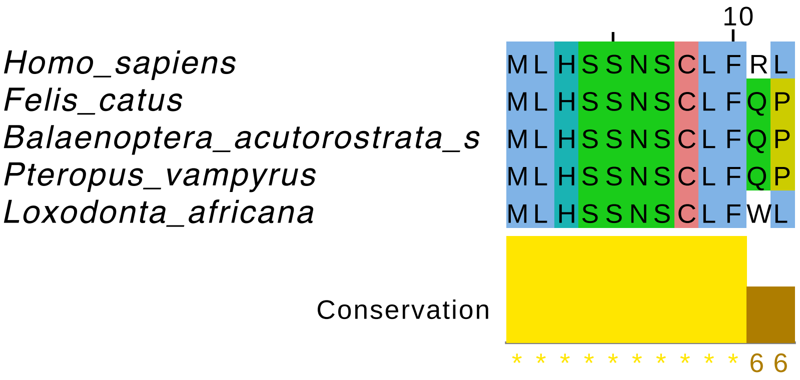

**HG1018**

ENSG00000259120 (SMIM6)

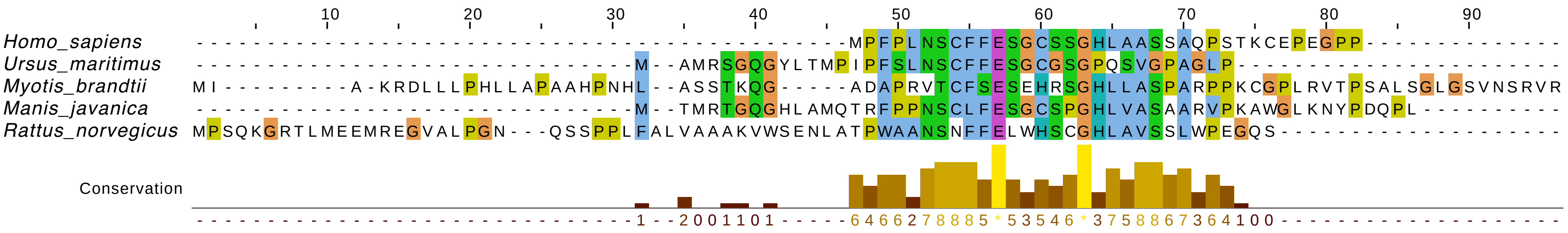

**HG1019**

ENSG00000019995 (ZRANB1)

```
                                       10        20        30        40
Homo_sapiens                  MASLKLNISISGFPLISSFLDQTSL-YLLPISFAFQSSVLLNPRV---
Odobenus_rosmarus_divergens   ----------------------MSL-YLLLISFAF-------------
Lipotes_vexillifer            MTWLRLNISVRGFLI---------------------------------
Orycteropus_afer_afer         MSWLKLNISIGGLPILTNPFPLKTFSTDLPLSLAYYLPCTAKYNFITS

Conservation
                              ------------------------------------------------
```

**HG1020**

ENSG00000023902 (PLEKHO1)

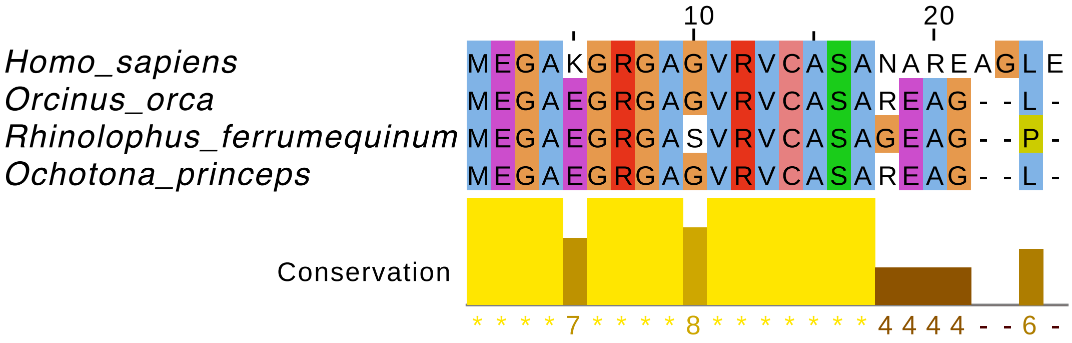

**HG1021**

ENSG00000064692 (SNCAIP)

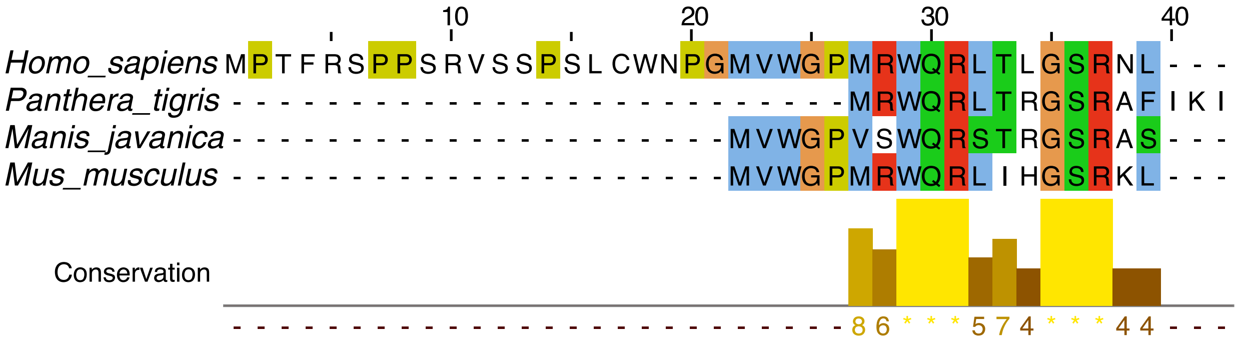

**HG1022**

ENSG00000065357 (DGKA)

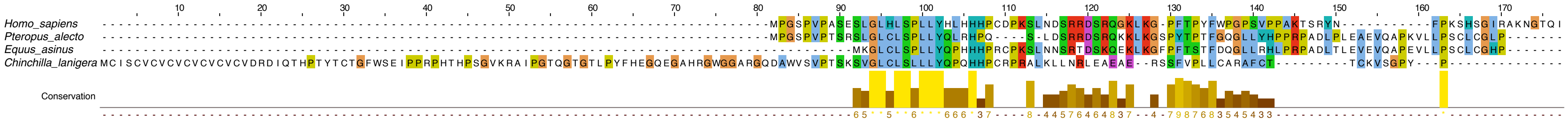

**HG1023**

ENSG00000068001 (HYAL2)

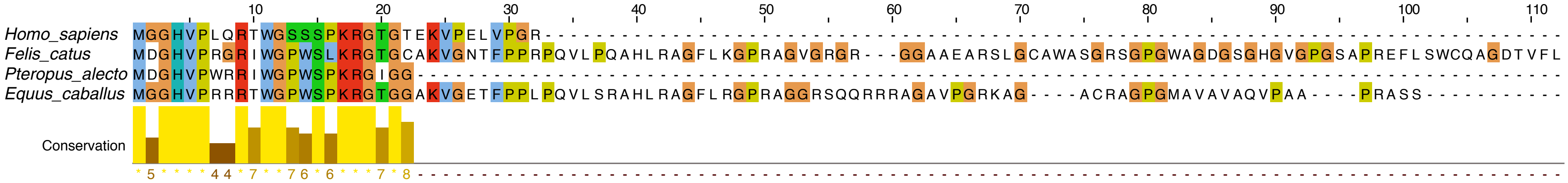

**HG1024**

ENSG00000073464 (CLCN4)

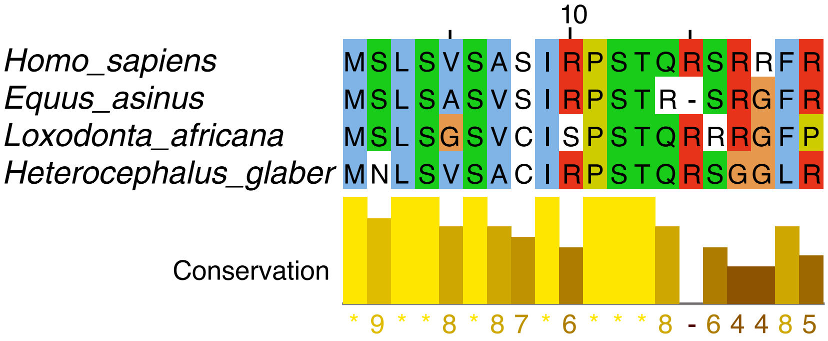

**HG1025**

ENSG00000074621 (SLC24A1)

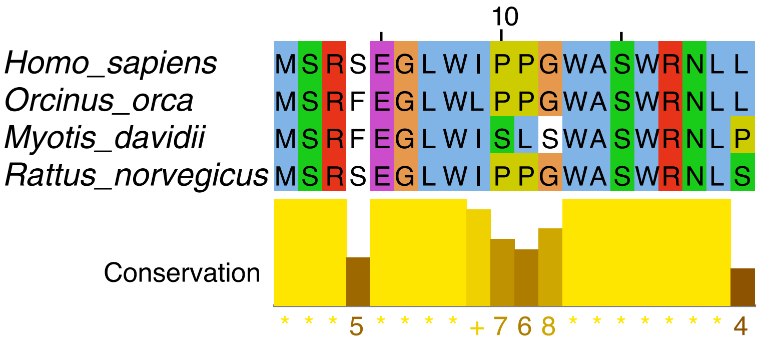

**HG1026.1**

ENSG00000075035 (WSCD2)

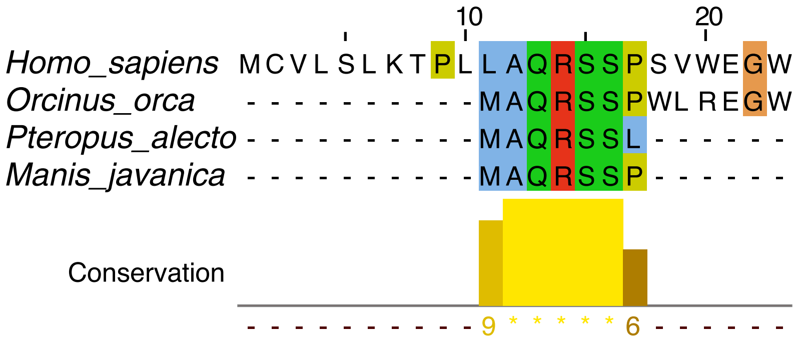

**HG1026.2**

ENSG00000179314 (WSCD1)

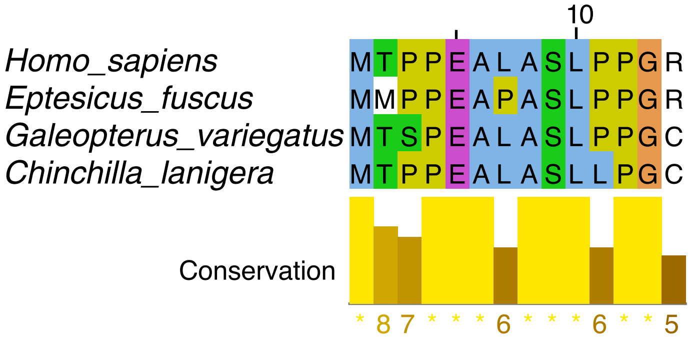

**HG1026.3**

ENSG00000179314 (WSCD1)

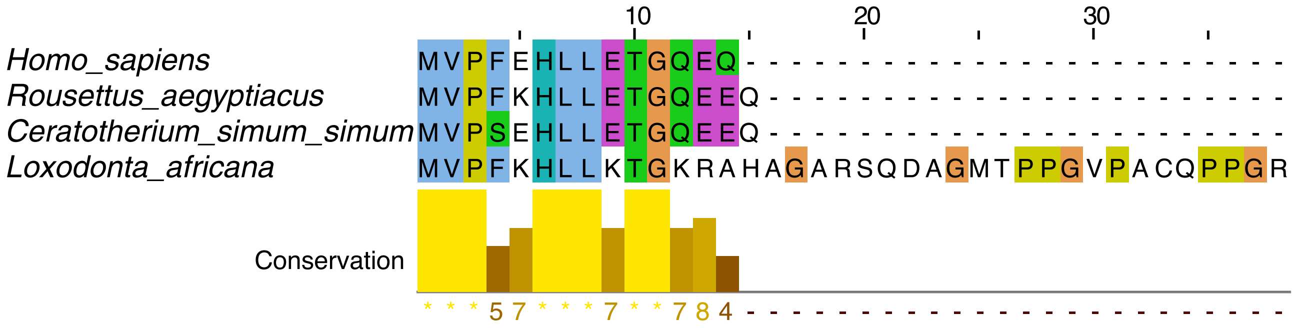

**HG1027**

ENSG00000075223 (SEMA3C)

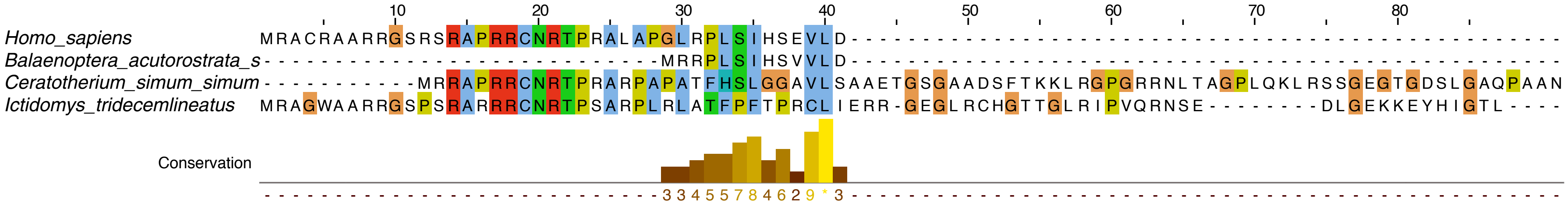

**HG1028**

ENSG00000076356 (PLXNA2)

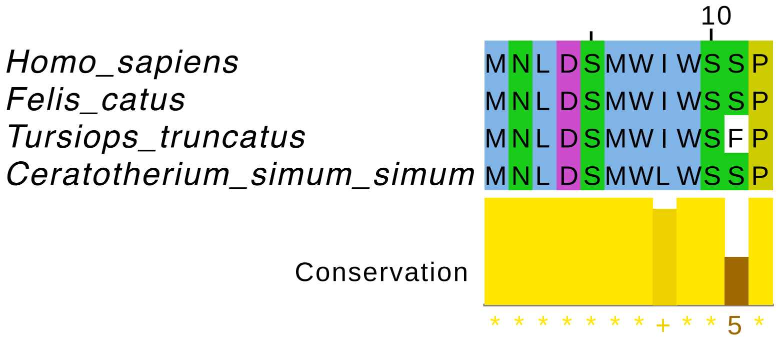

**HG1029**

ENSG00000079482 (OPHN1)

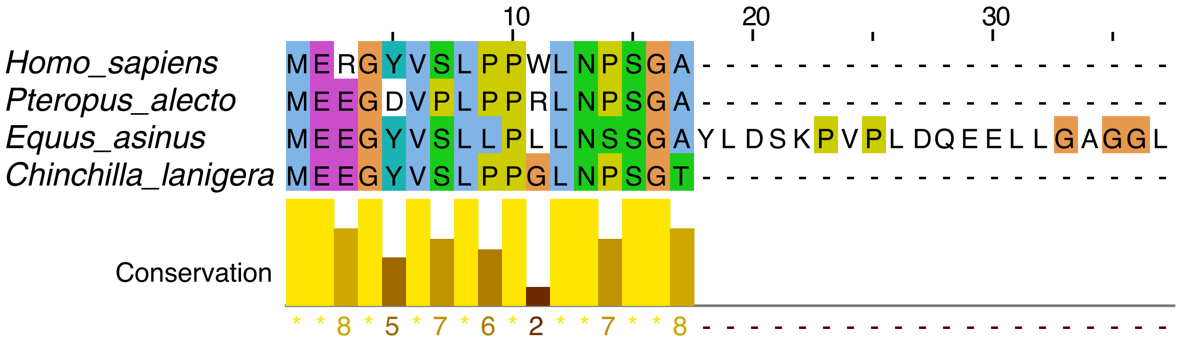

**HG1030**

ENSG00000082175 (PGR)

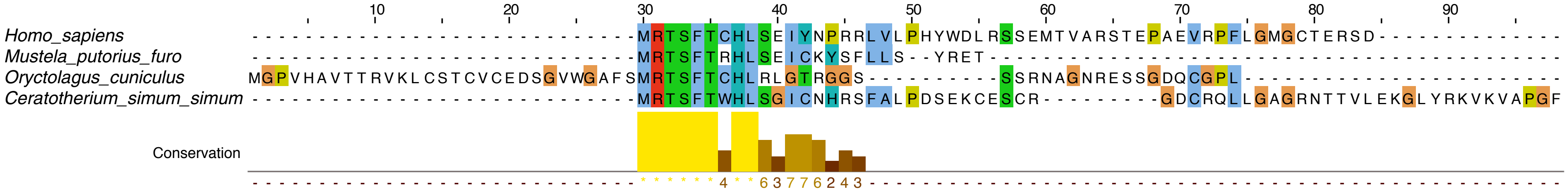

**HG1031**

ENSG00000082269 (FAM135A)

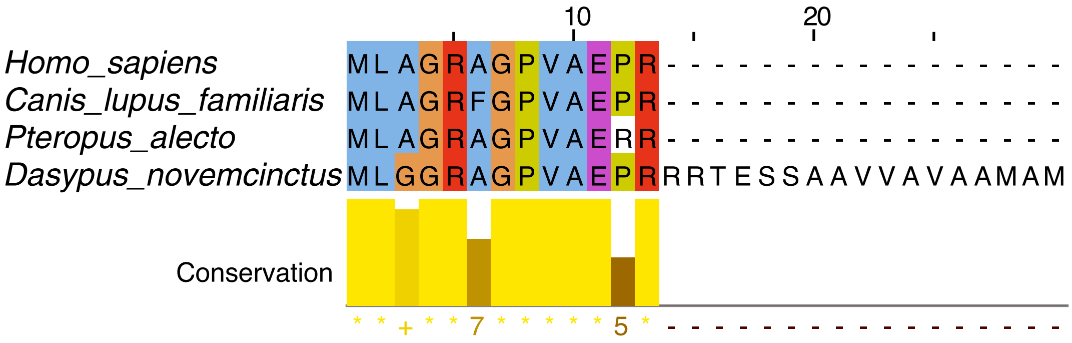

**HG1032**

ENSG00000084453 (SLCO1A2)

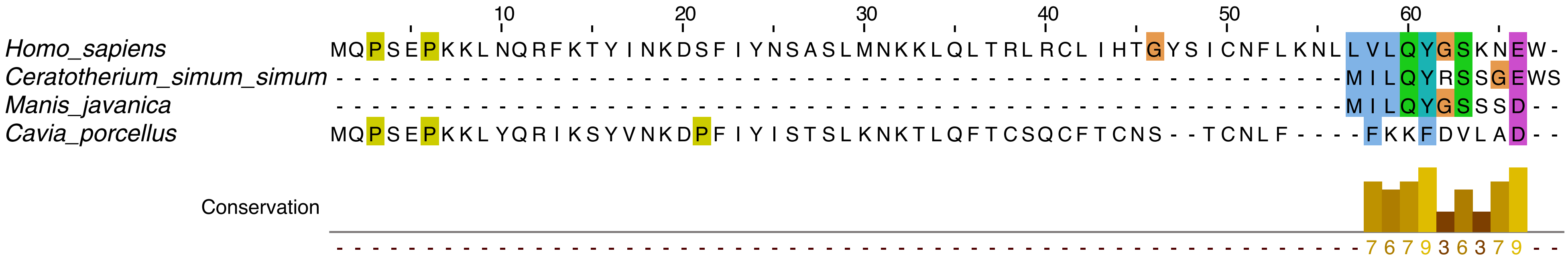

**HG1033**

ENSG00000089225 (TBX5)

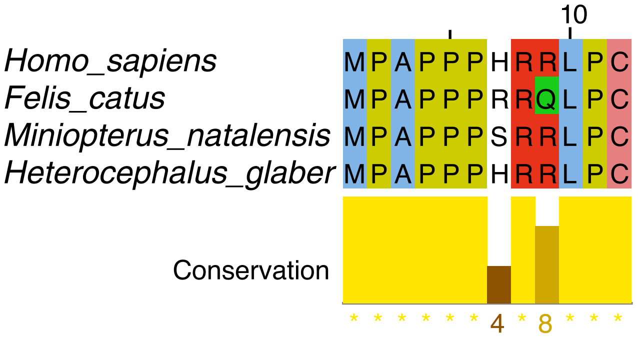

**HG1034**

ENSG00000090487 (SPG21)

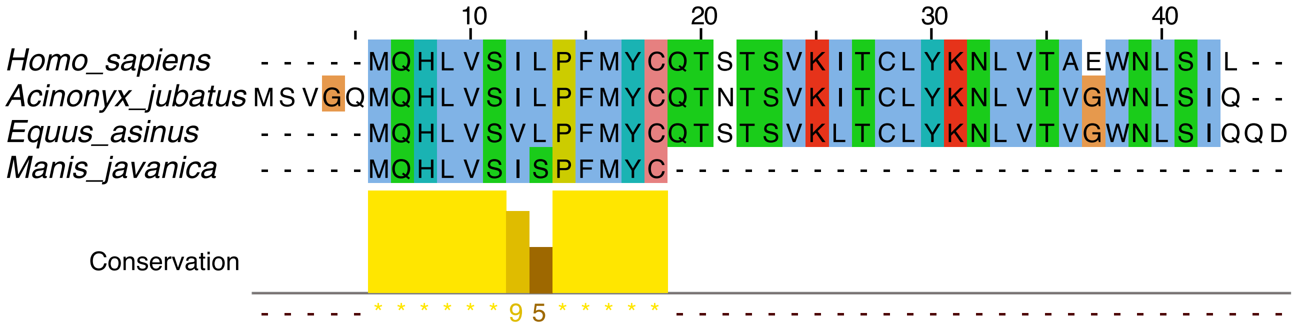

**HG1035**

ENSG00000093000 (NUP50)

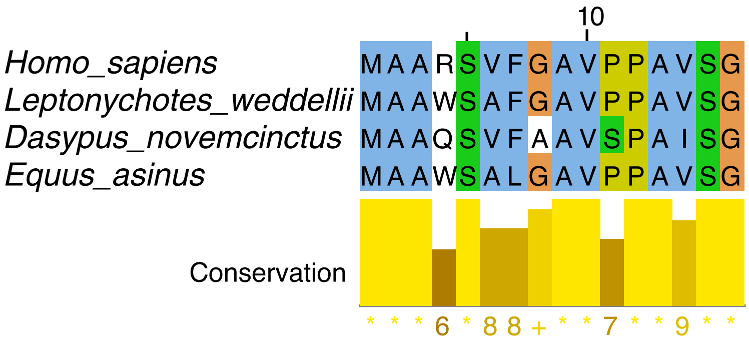

**HG1036**

ENSG00000100813 (ACIN1)

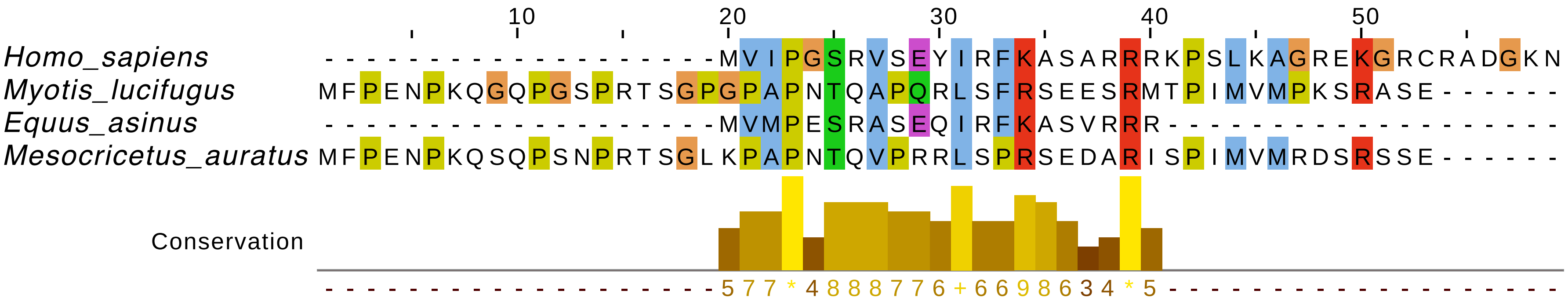

**HG1037**

ENSG00000102710 (SUPT20H)

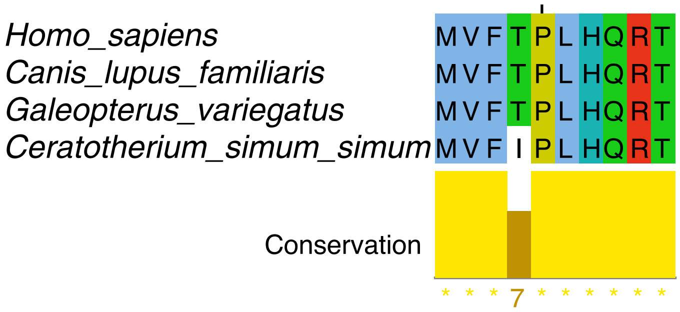

**HG1038**

ENSG00000104331 (IMPAD1)

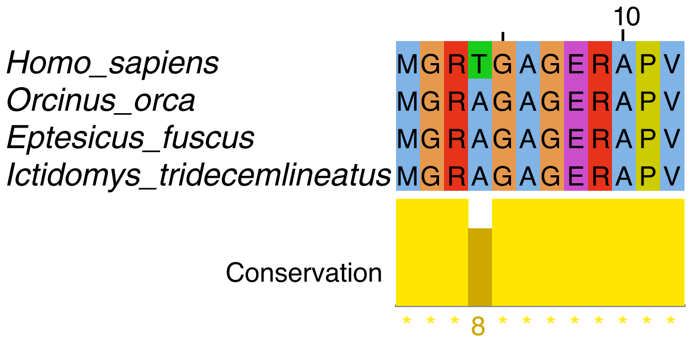

**HG1039**

ENSG00000104804 (TULP2)

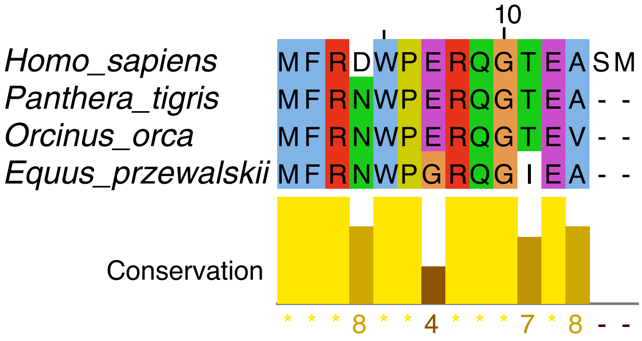

**HG1040**

ENSG00000105223 (PLD3)

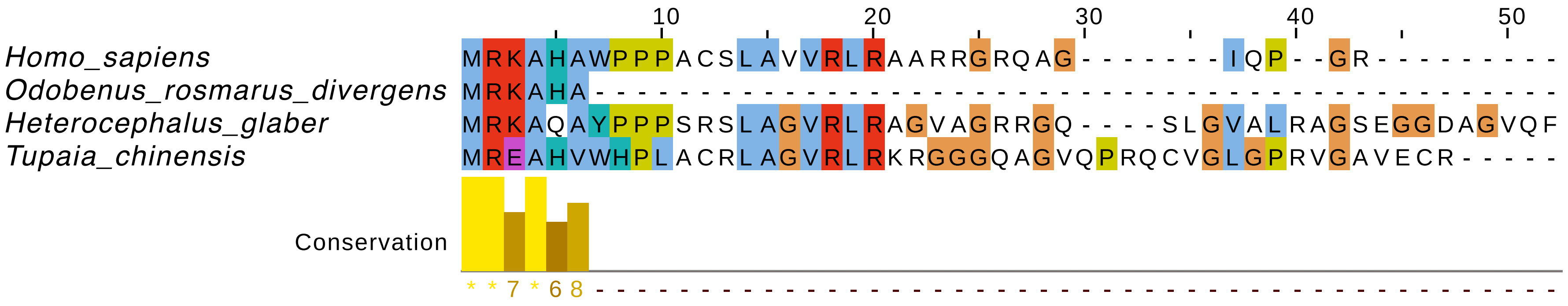

**HG1041**

ENSG00000106536 (POU6F2)

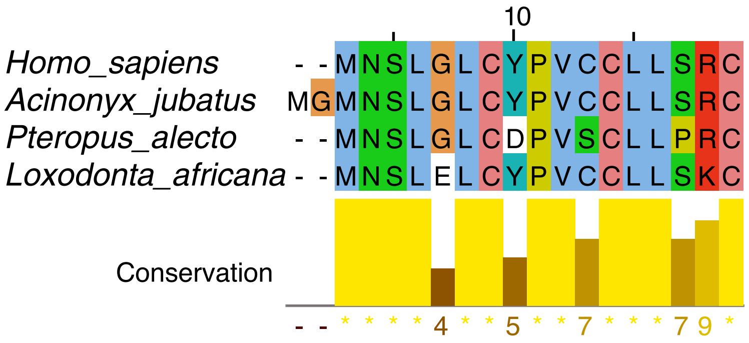

**HG1042**

ENSG00000107249 (GLIS3)

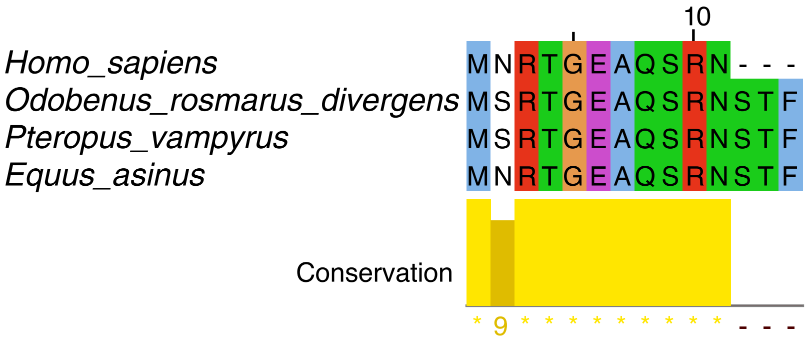

**HG1043**

ENSG00000107736 (CDH23)

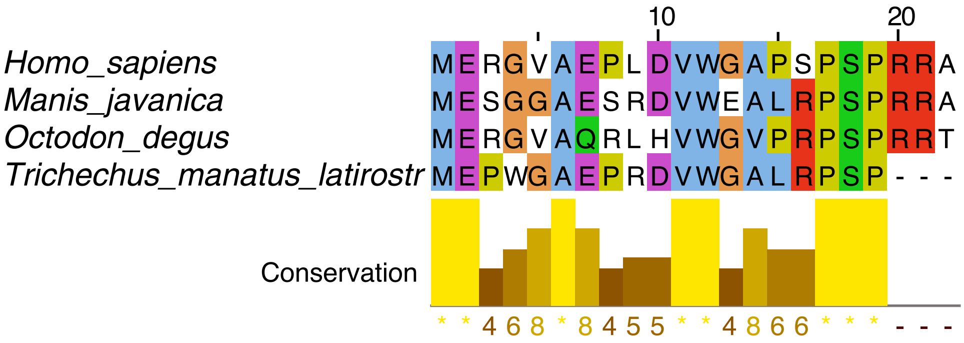

**HG1044**

ENSG00000109381 (ELF2)

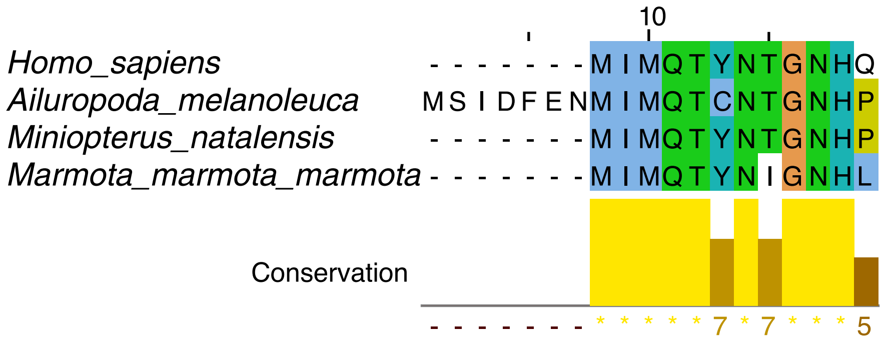

**HG1045**

ENSG00000110925 (CSRNP2)

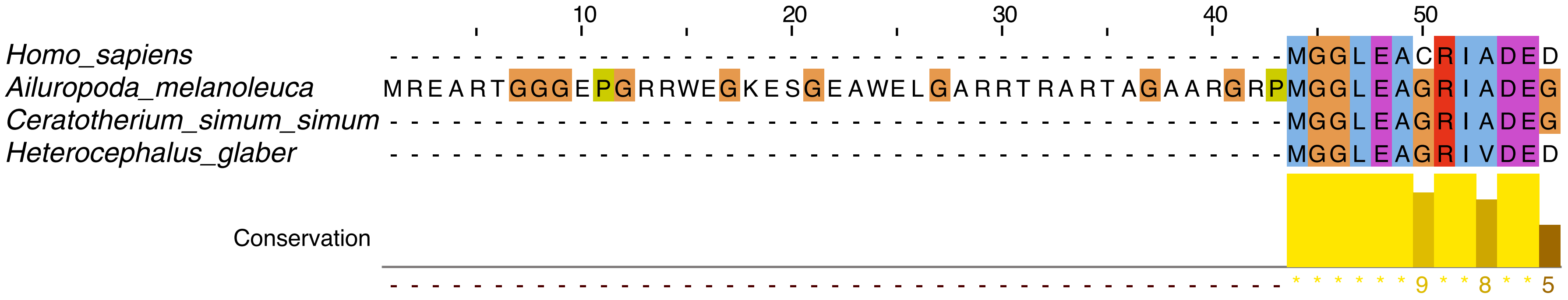

**HG1046**

ENSG00000112701 (SENP6)

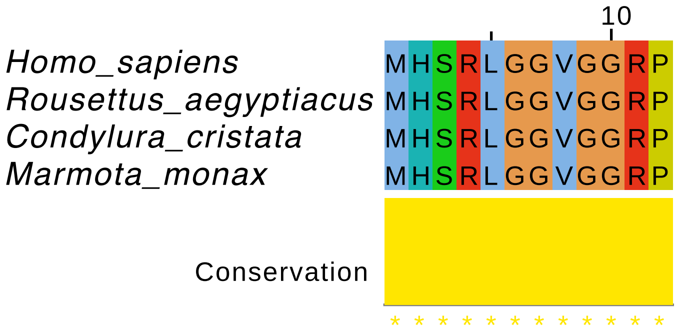

**HG1047**

ENSG00000112769 (LAMA4)

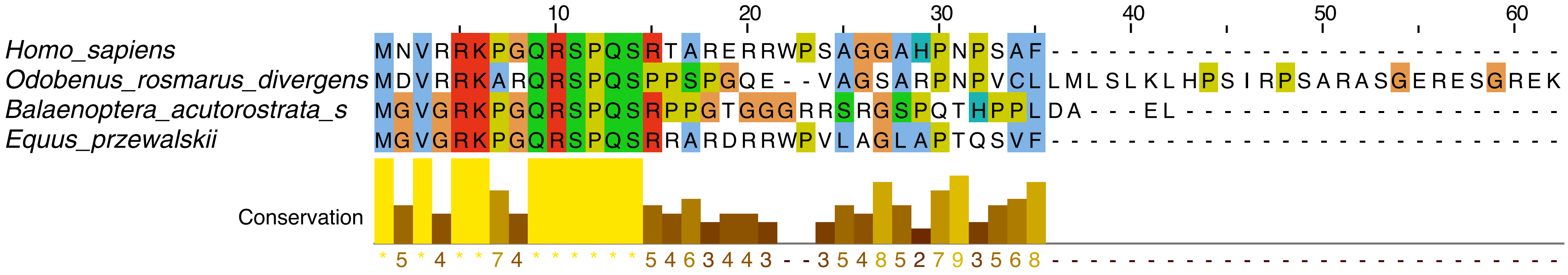

**HG1048**

ENSG00000115295 (CLIP4)

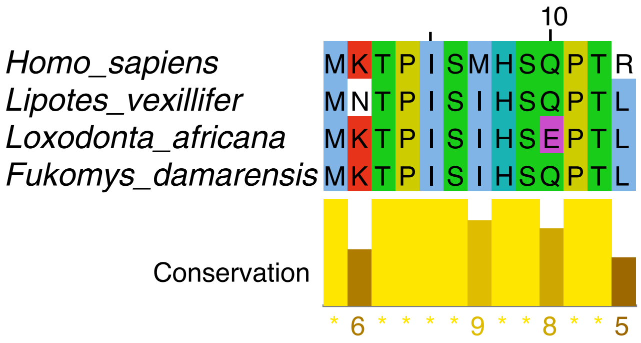

**HG1049**

ENSG00000116212

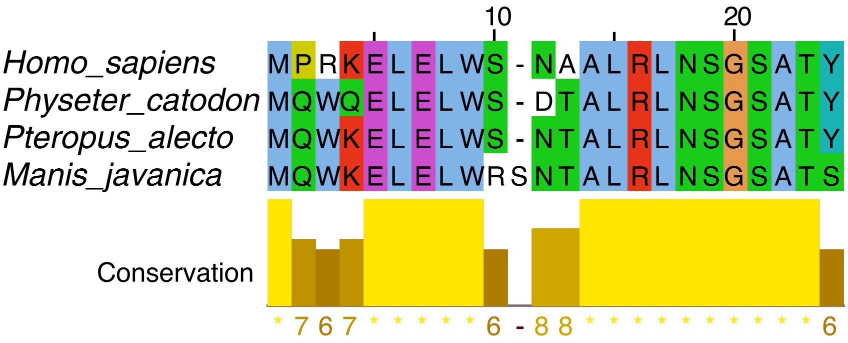

**HG1050**

ENSG00000116489 (CAPZA1)

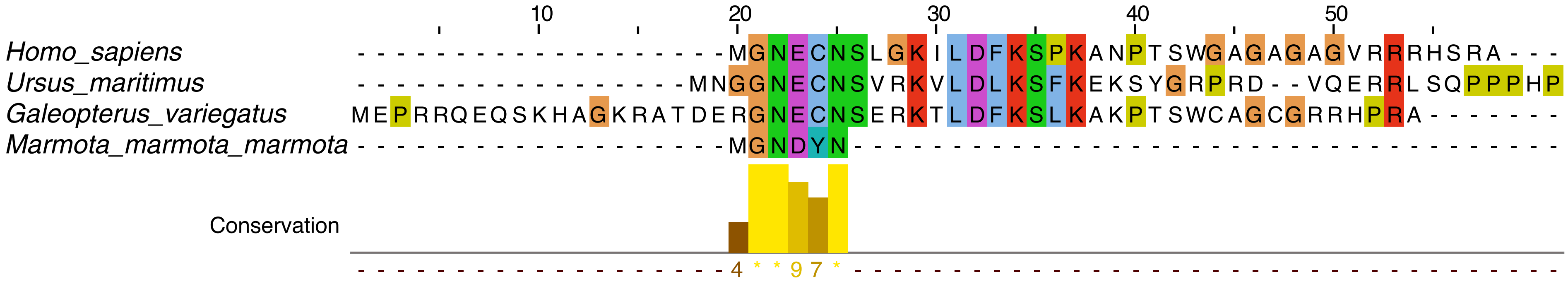

**HG1051**

ENSG00000116852 (KIF21B)

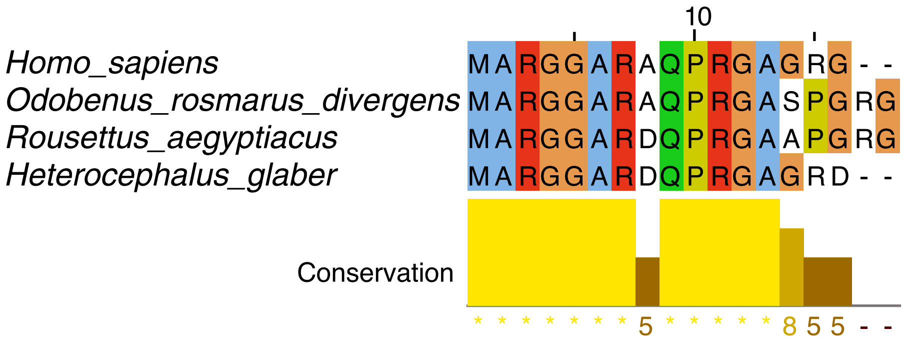

**HG1052**

ENSG00000118640 (VAMP8)

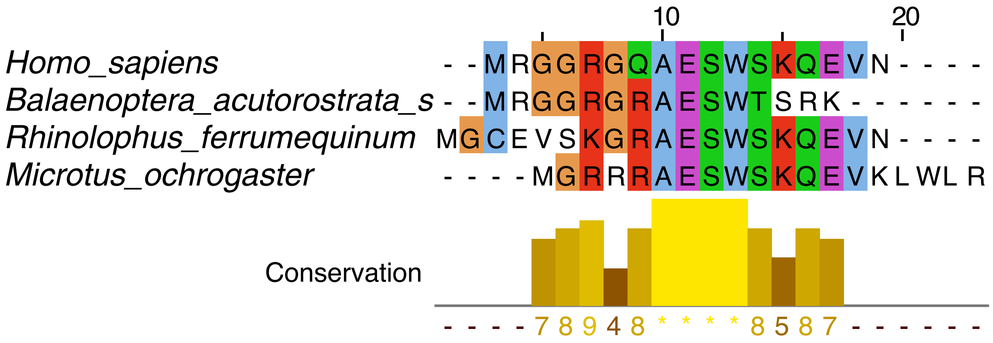

**HG1053**

ENSG00000119522 (DENND1A)

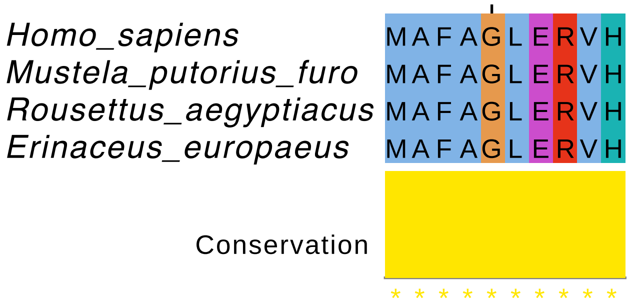

**HG1054**

ENSG00000120832 (MTERF2)

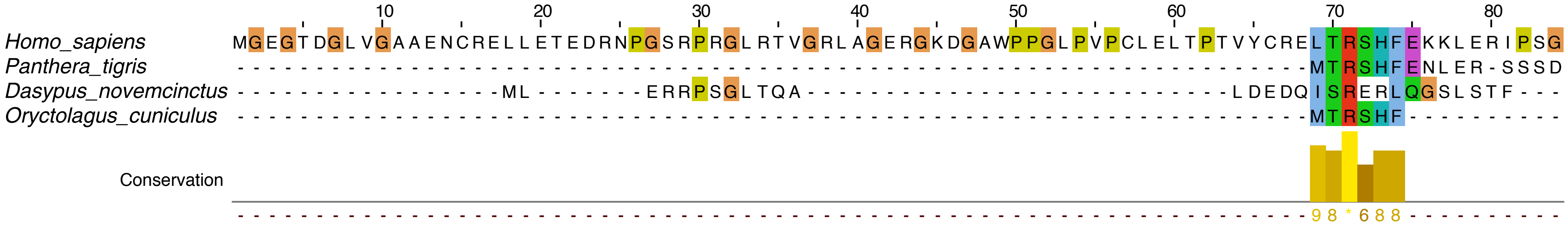

**HG1055**

ENSG00000122299 (ZC3H7A)

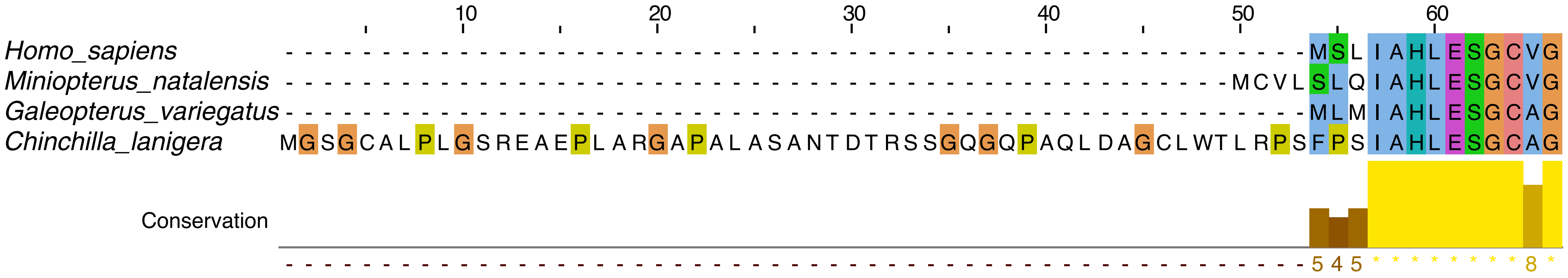

**HG1056**

ENSG00000128271 (ADORA2A)

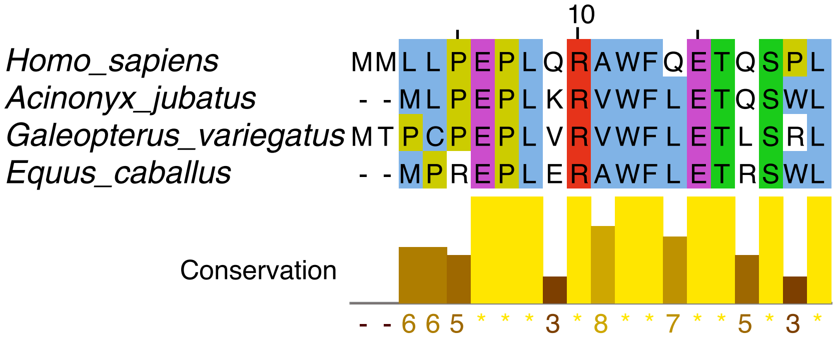

**HG1057**

ENSG00000131097 (HIGD1B)

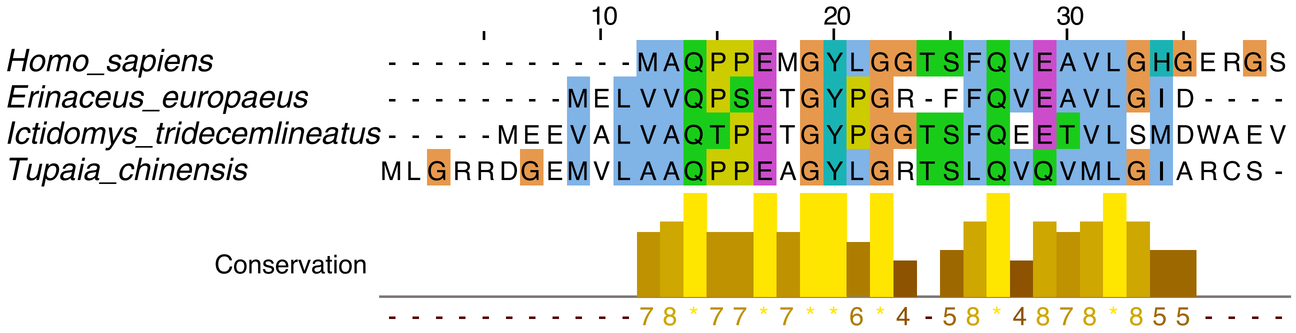

**HG1058**

ENSG00000131386 (GALNT15)

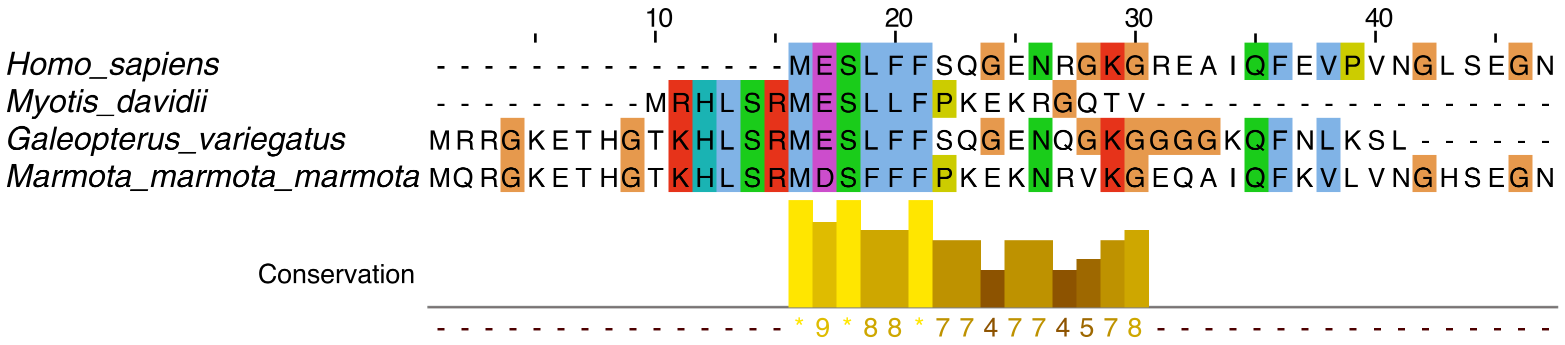

**HG1059.1**

ENSG00000132970 (WASF3)

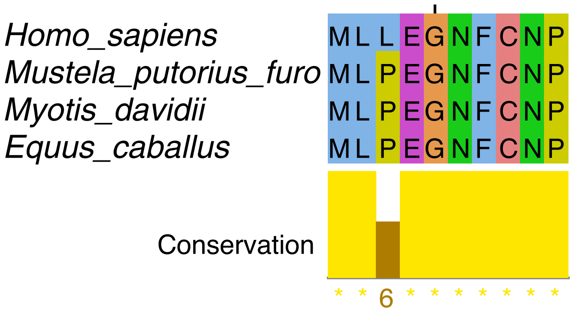

**HG1059.2**

ENSG00000112290 (WASF1)

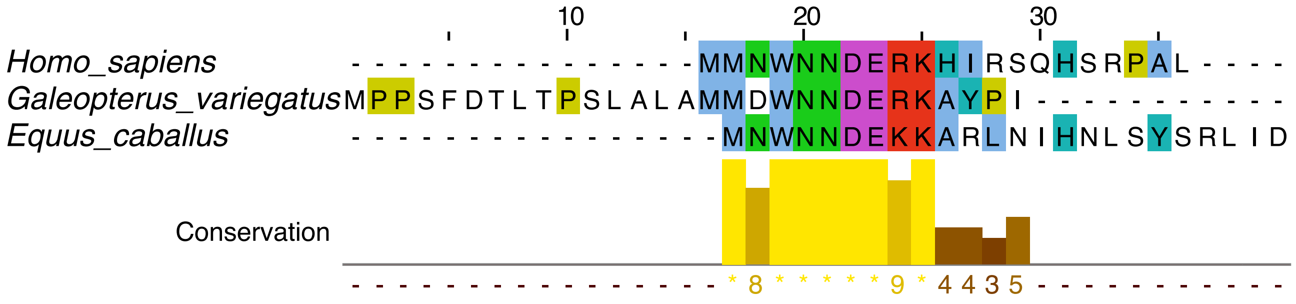

**HG1060**

ENSG00000134242 (PTPN22)

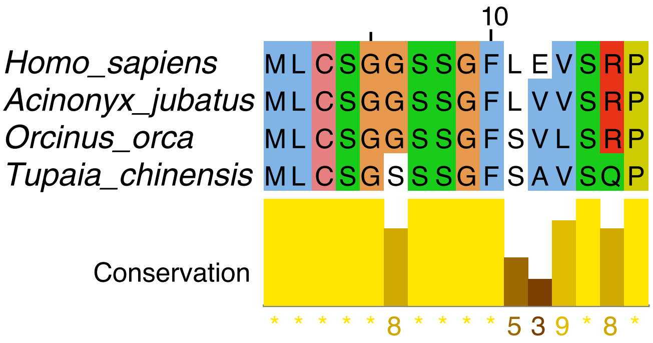

**HG1061**

ENSG00000134802 (SLC43A3)

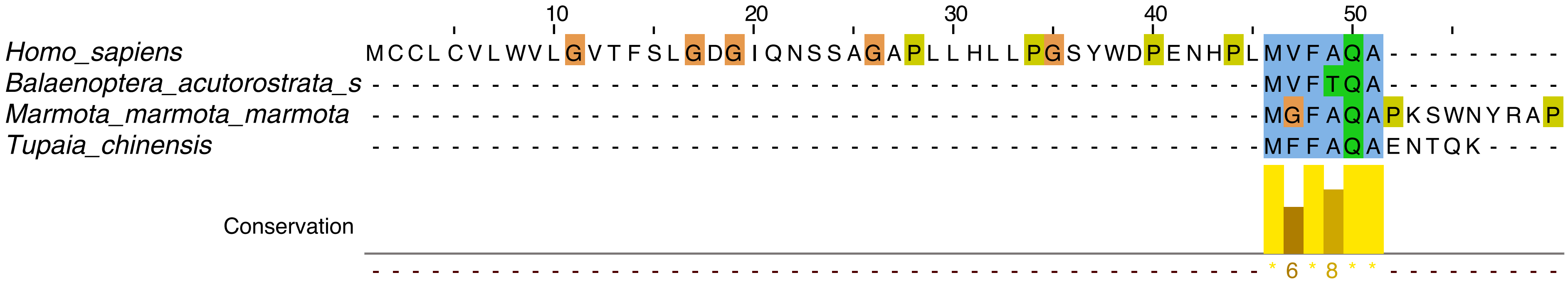

# HG1062

### ENSG00000137337 (MDC1)

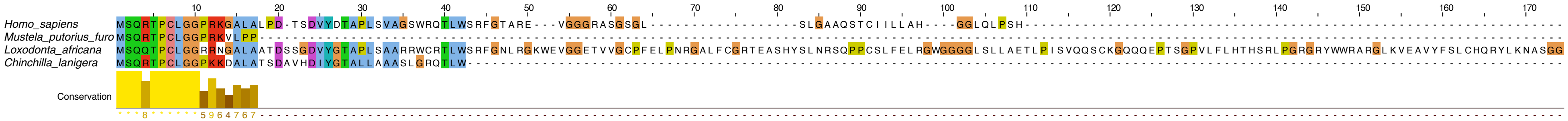

**HG1063**

ENSG00000138162 (TACC2)

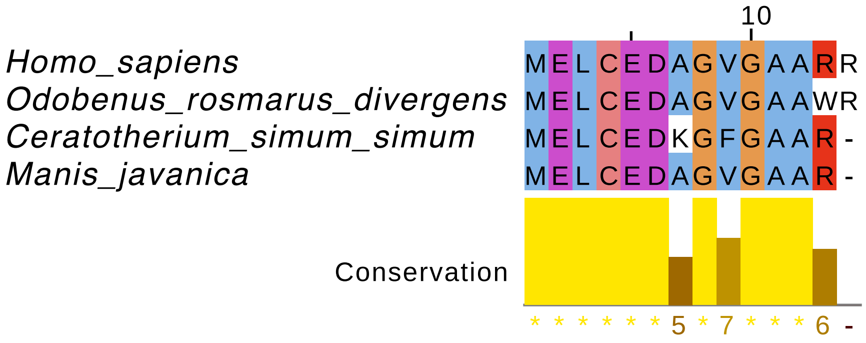

**HG1064**

ENSG00000139734 (DIAPH3)

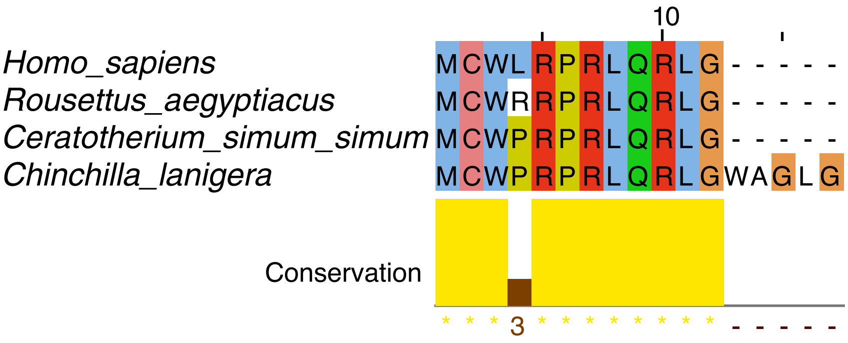

**HG1065**

ENSG00000140030 (GPR65)

```
                                 10        20        30        40        50        60        70        80        90       100       110
Homo_sapiens             MQAQIYQAPQSTNTSPEEIWKERNFKRKYQSLCKQSLVYSCLHQSTVRFMKKNKLRTTLYVHLQMPQLMLVGCLSAFCDNEHMDFCLLNSVDPFSQLPGAWIF-TSNC--------
Mustela_putorius_furo    ----------------------------------------------------------------------------------MELLKISQRLNARKCPAVWVSFSSRCKQIYPAPR
Galeopterus_variegatus   -----------------------------------------------------------MEAKELSQLILEGSLSVFCDNEHMDLCSLNLADHFS---------------------
Oryctolagus_cuniculus    -----------------------------------------------------------MNTKGISQLIIQSSSSVFCDHDHMDVCSLNSTDHFS---------------------
```

Conservation

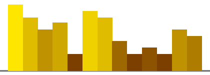

```
                         ----------------------------------------------------------------------------------*9783+9534376---------------------
```

# HG1066

### ENSG00000140553 (UNC45A)

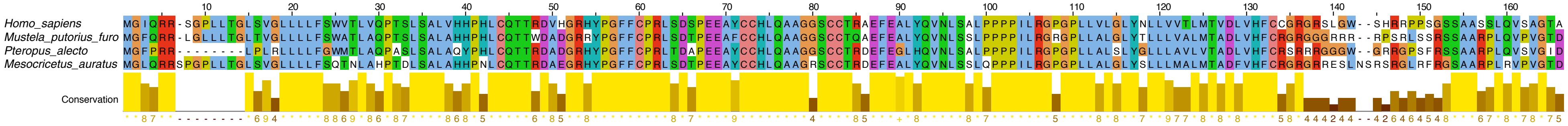

**HG1067**

ENSG00000143340 (FAM163A)

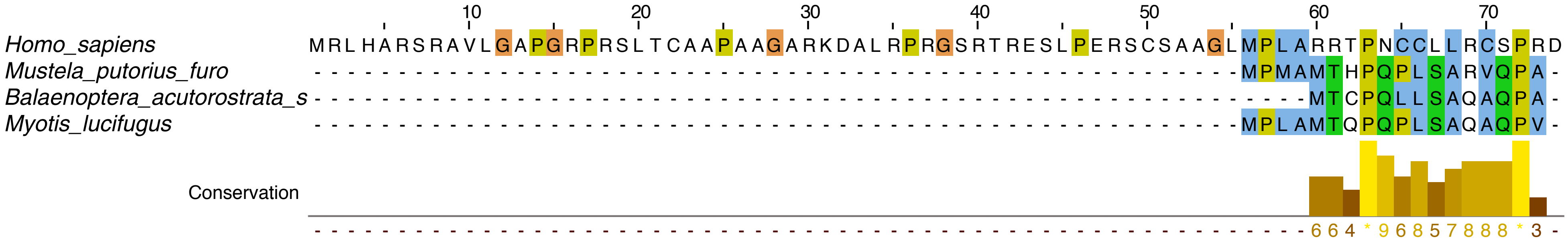

**HG1068**

ENSG00000144791 (LIMD1)

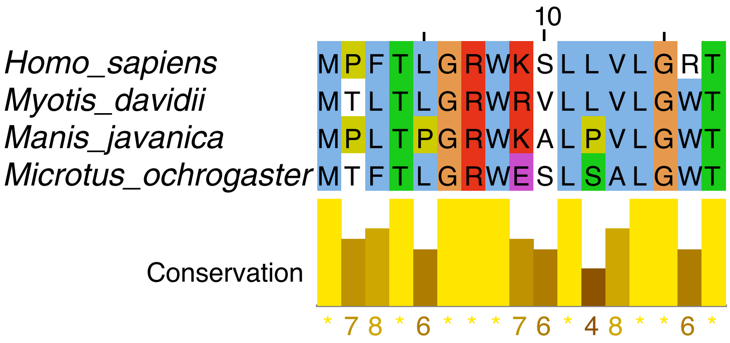

**HG1069**

ENSG00000146414 (SHPRH)

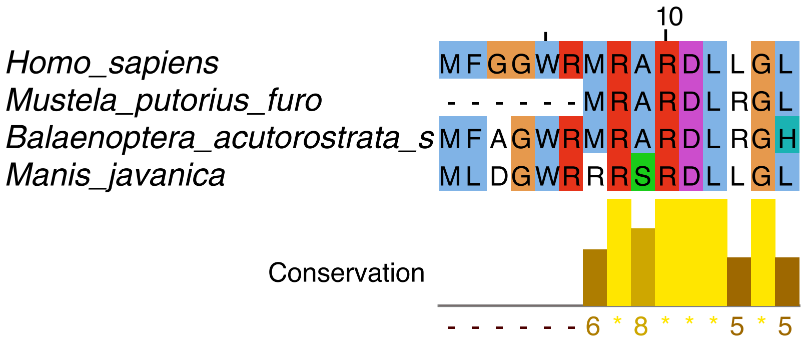

**HG1070**

ENSG00000147459 (DOCK5)

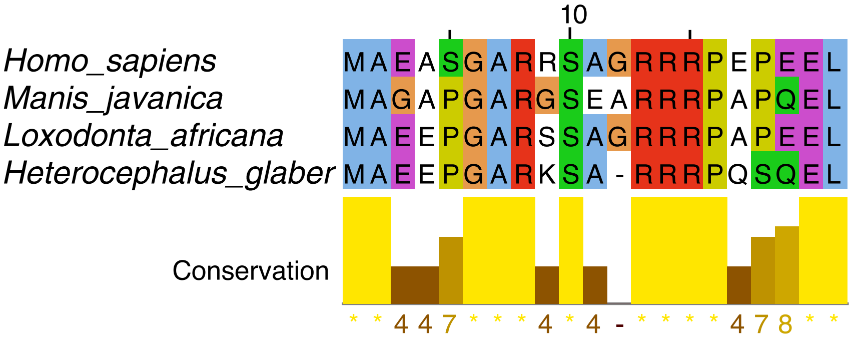

**HG1071**

ENSG00000147573 (TRIM55)

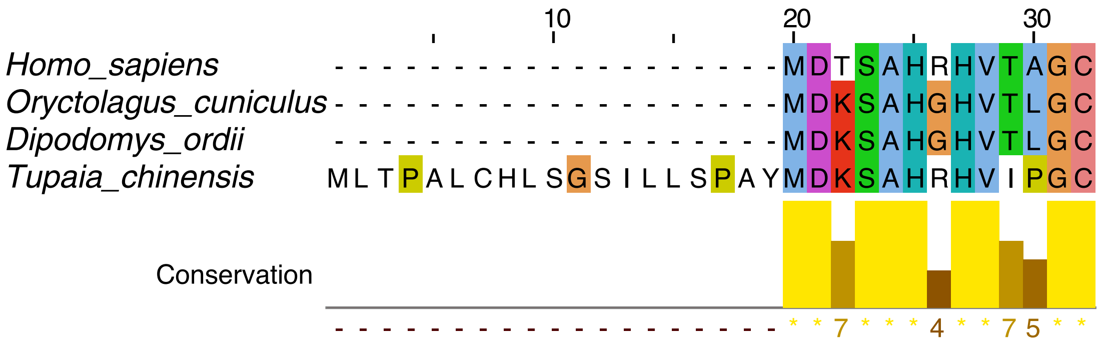

**HG1072**

ENSG00000147874 (HAUS6)

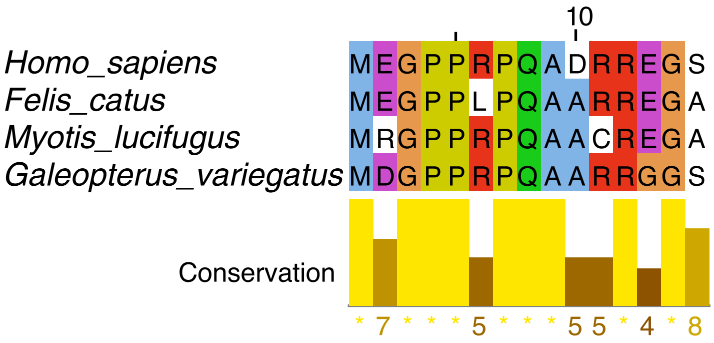

**HG1073**

ENSG00000149187 (CELF1)

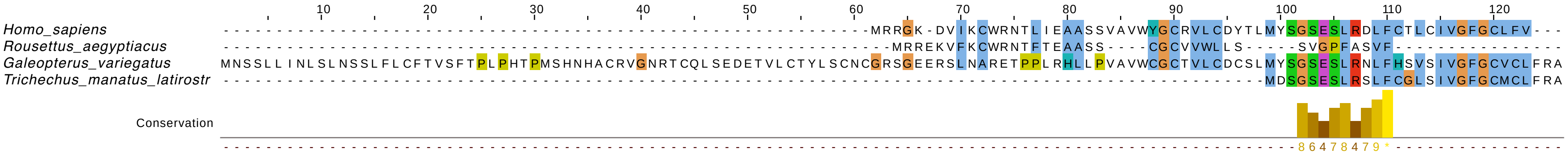

**HG1074**

ENSG00000150275 (PCDH15)

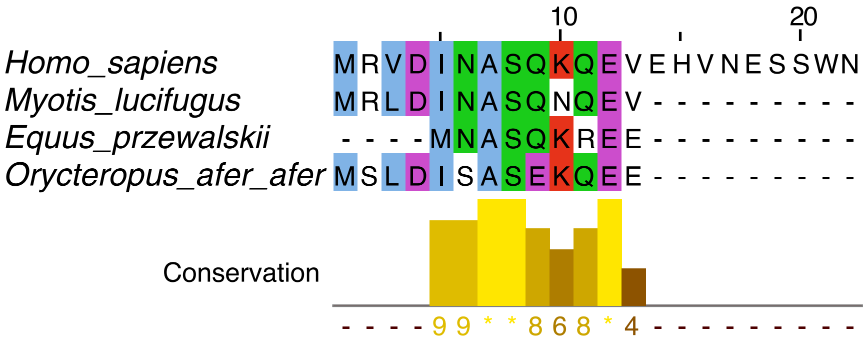

**HG1075**

ENSG00000150347 (ARID5B)

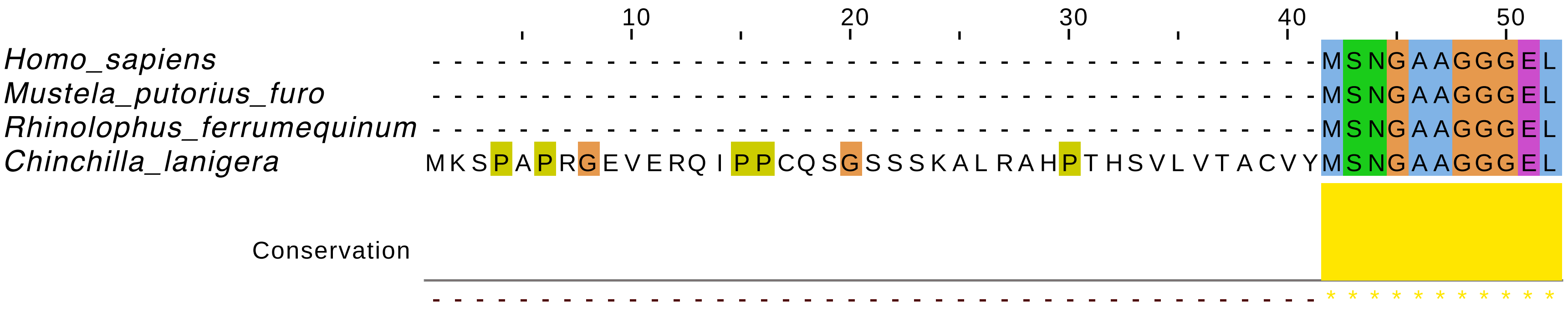

**HG1076**

ENSG00000150967 (ABCB9)

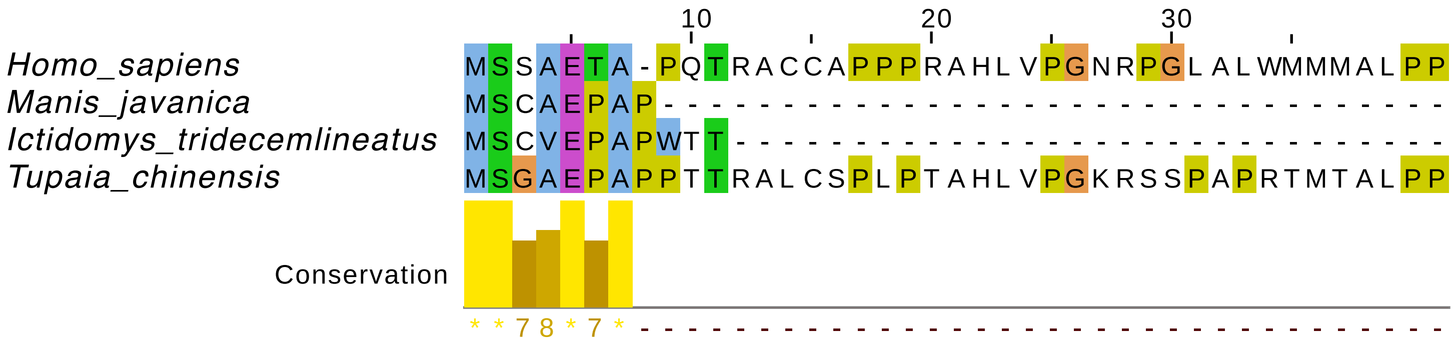

**HG1077**

ENSG00000151474 (FRMD4A)

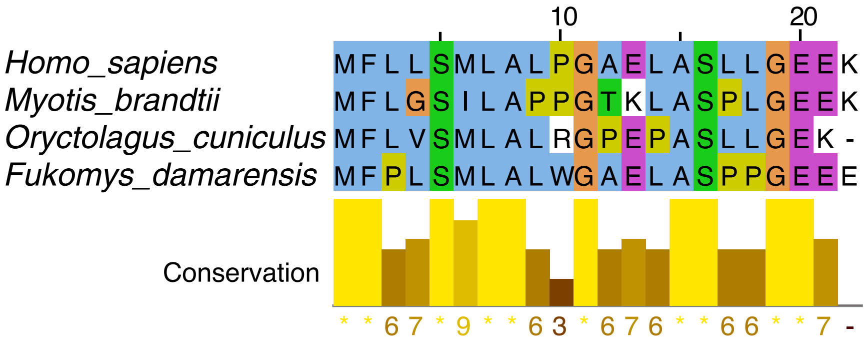

**HG1078**

ENSG00000162607 (USP1)

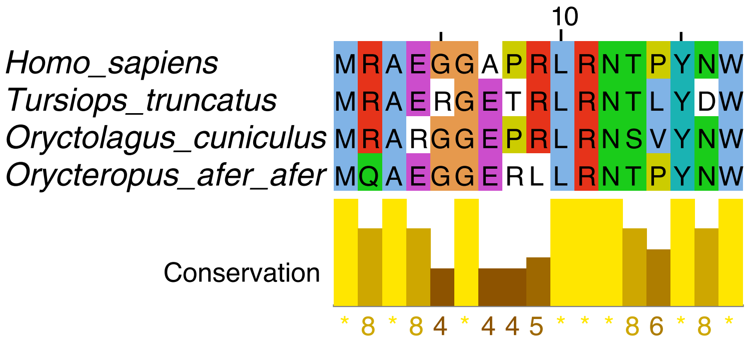

**HG1079**

ENSG00000162975 (KCNF1)

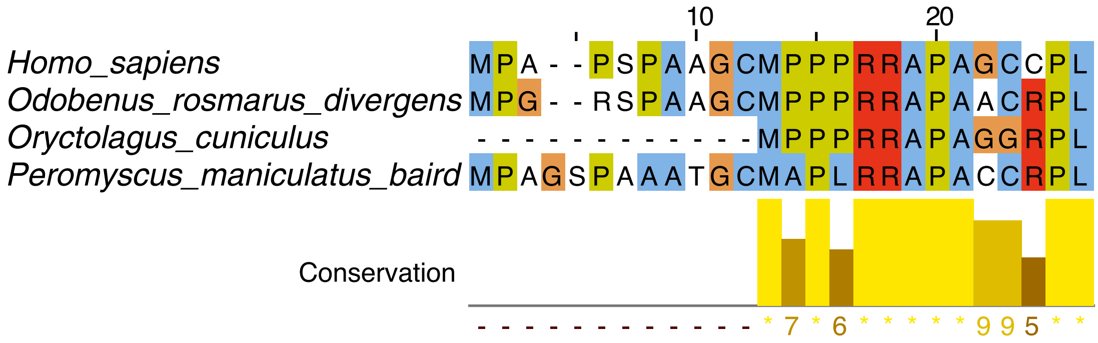

**HG1080**

ENSG00000164168 (TMEM184C)

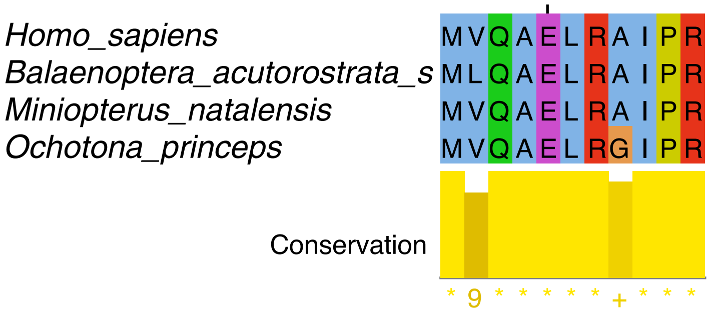

**HG1081**

ENSG00000165392 (WRN)

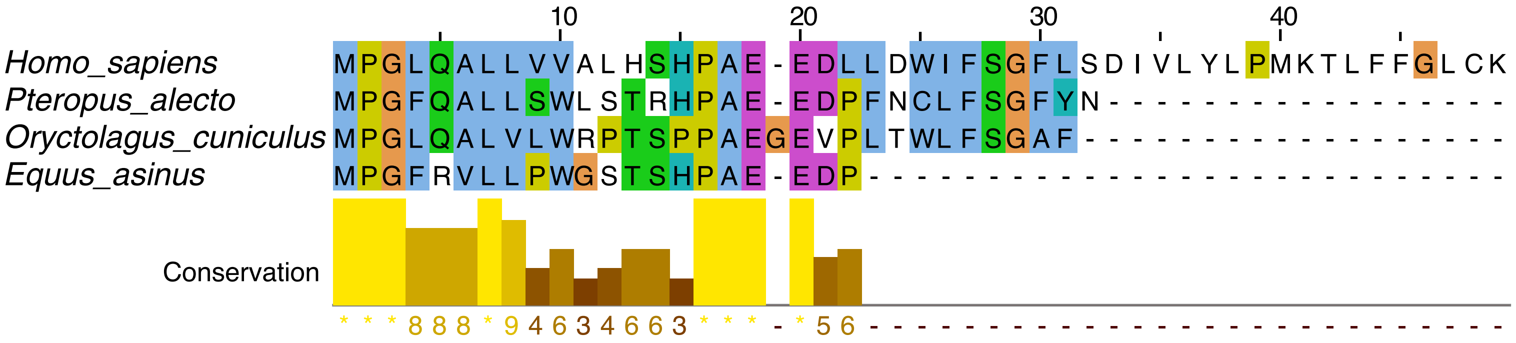

**HG1082**

ENSG00000165434 (PGM2L1)

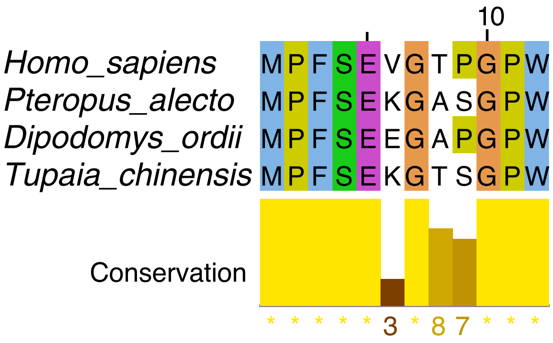

**HG1083**

ENSG00000165804 (ZNF219)

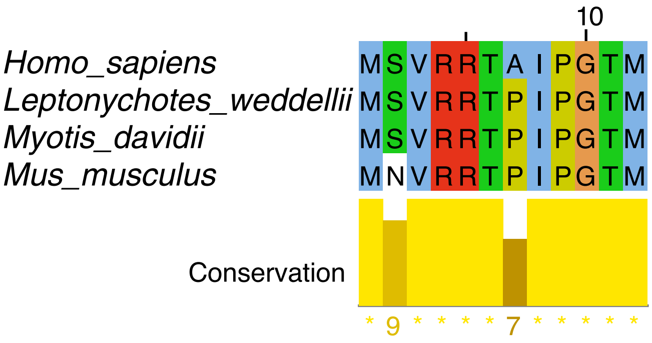

**HG1084**

ENSG00000165923 (AGBL2)

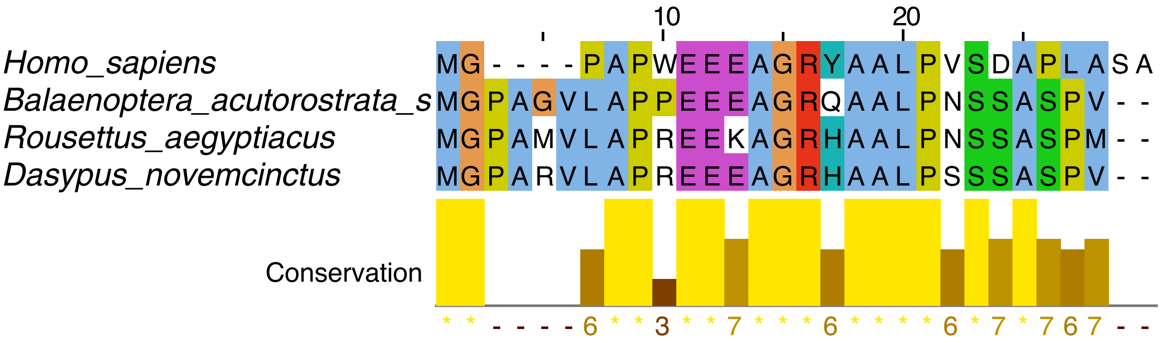

**HG1085**

ENSG00000166532 (RIMKLB)

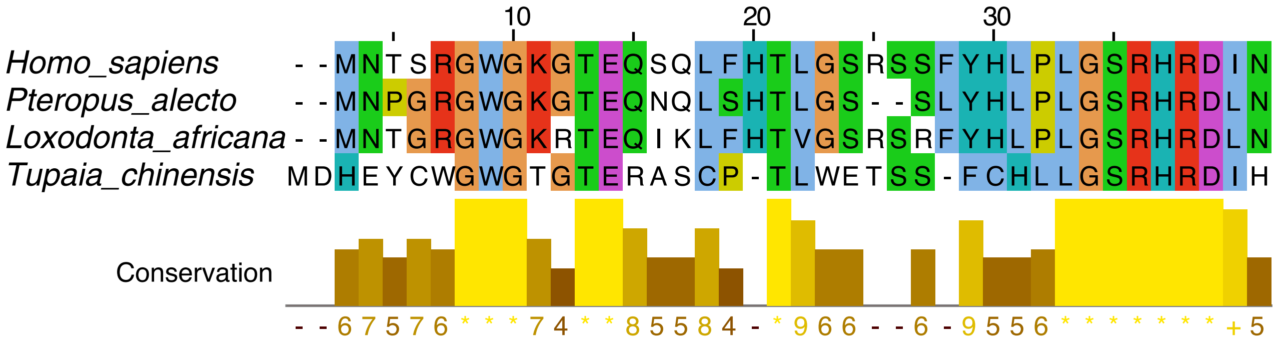

**HG1086**

ENSG00000167377 (ZNF23)

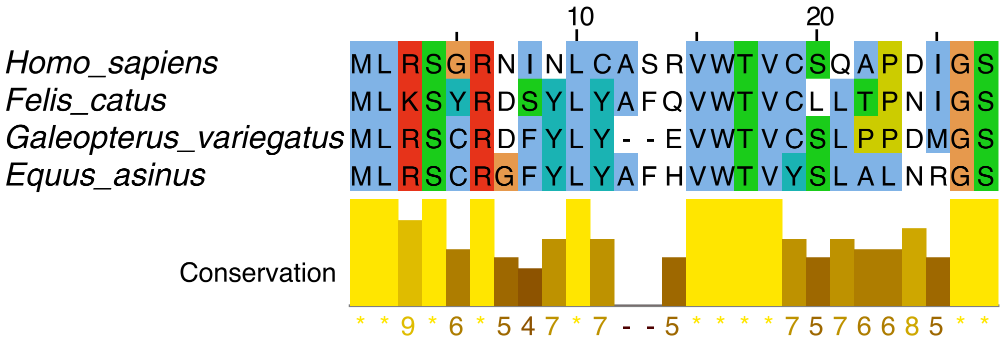

**HG1087**

ENSG00000168675 (LDLRAD4)

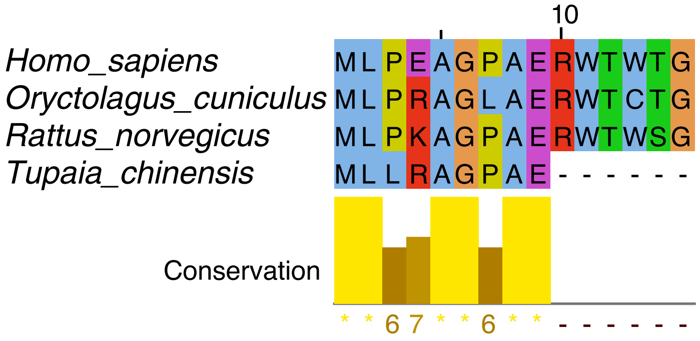

**HG1088**

ENSG00000169740 (ZNF32)

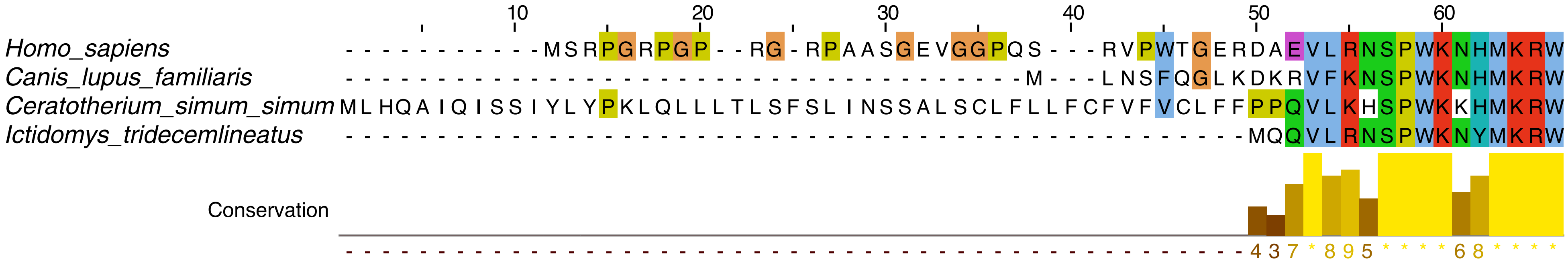

**HG1089**

ENSG00000169813 (HNRNPF)

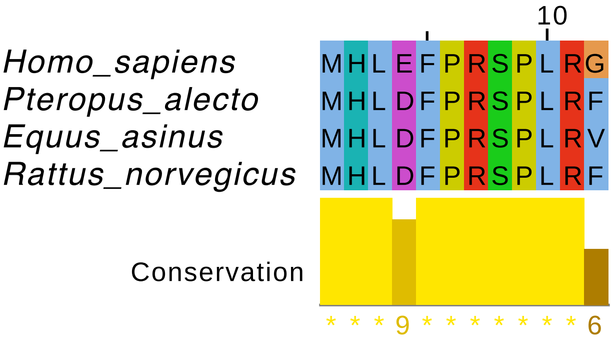

**HG1090.1**

ENSG00000170458 (CD14)

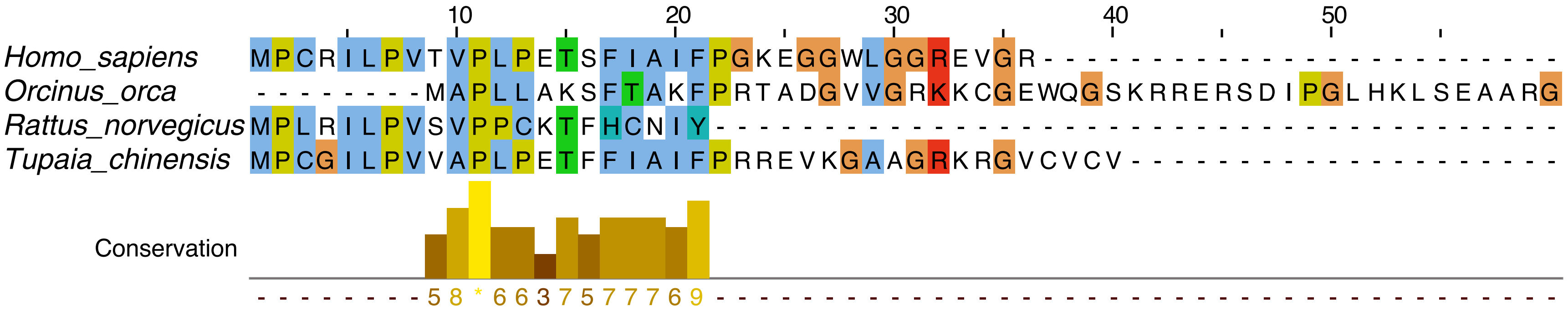

**HG1090.2**

ENSG00000170458 (CD14)

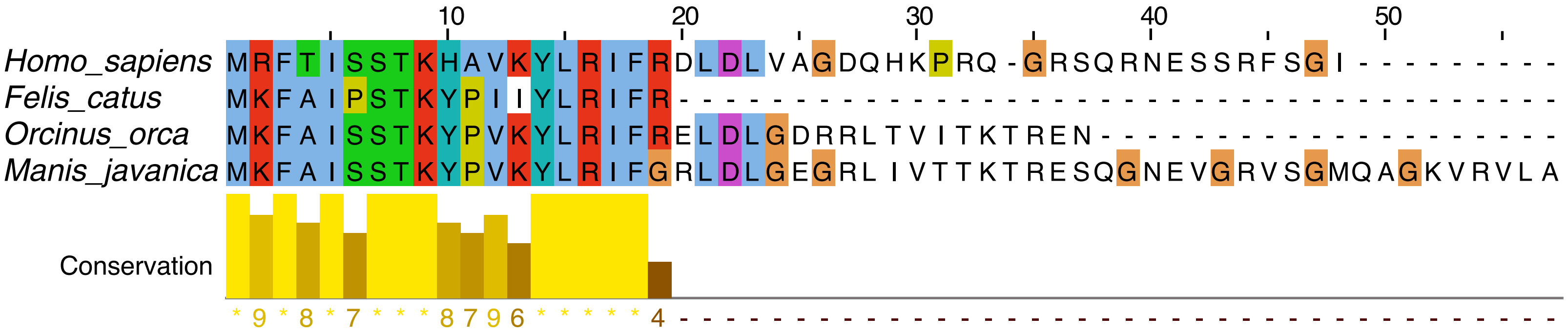

**HG1091**

ENSG00000172183 (ISG20)

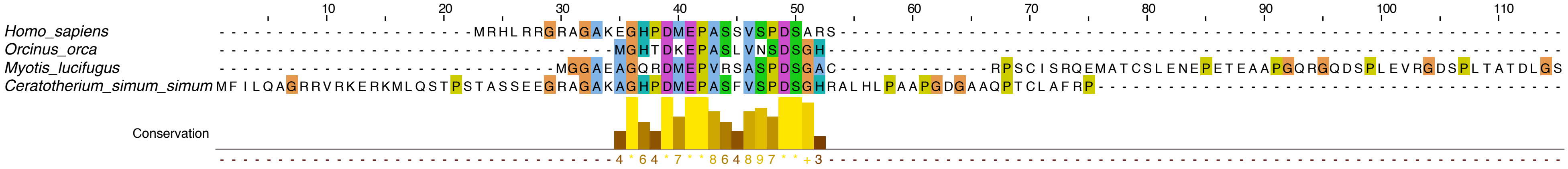

# HG1092

### ENSG00000173327 (MAP3K11)

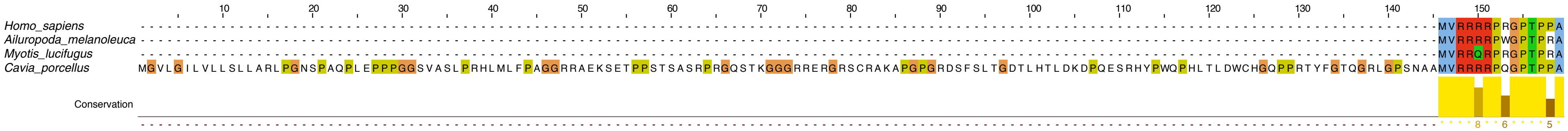

**HG1093**

ENSG00000174371 (EXO1)

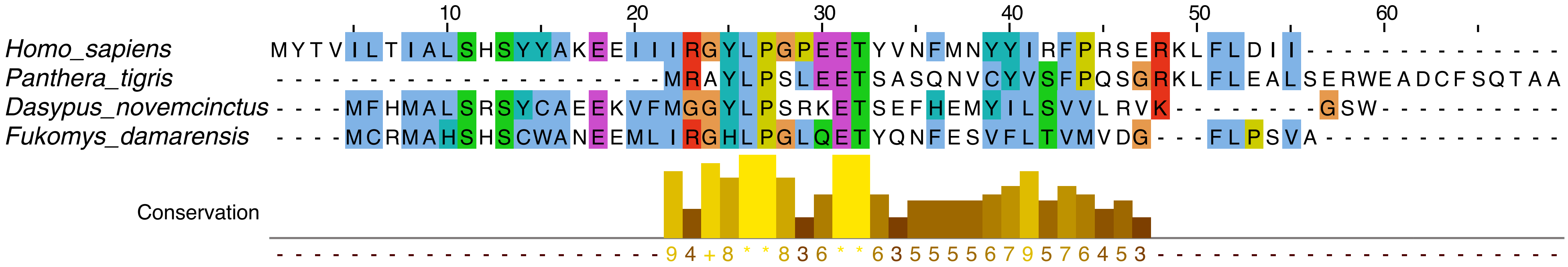

**HG1094**

ENSG00000177917 (ARL6IP6)

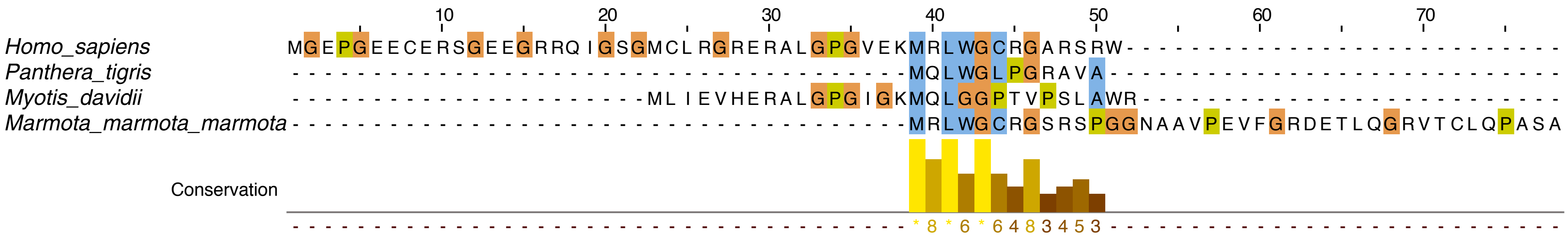

**HG1095**

ENSG00000179921 (GPBAR1)

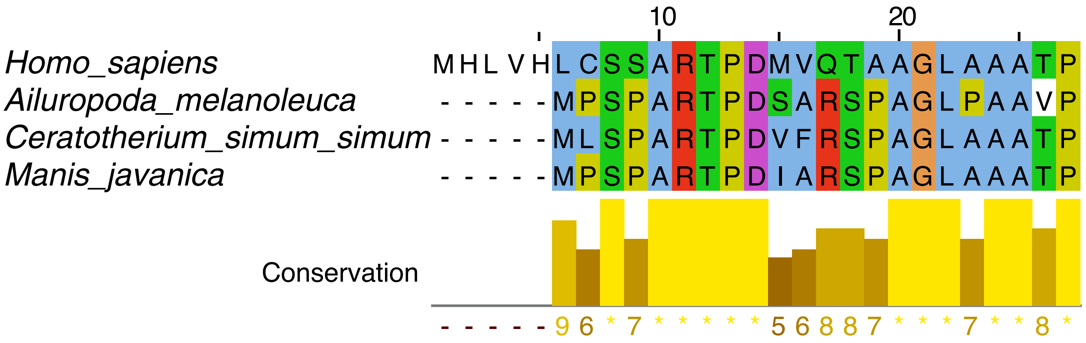

**HG1096**

ENSG00000181544 (FANCB)

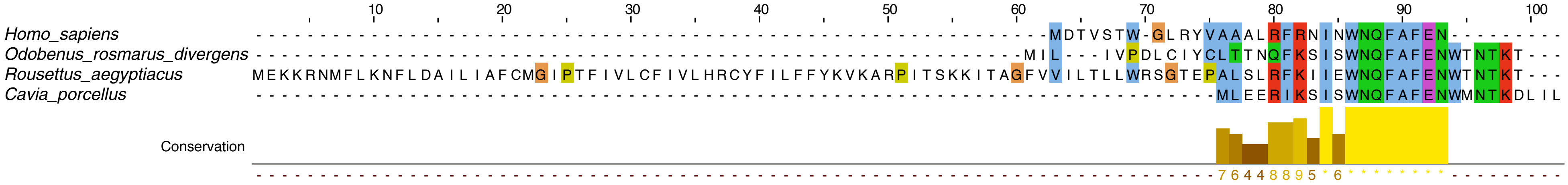

**HG1097**

ENSG00000182481 (KPNA2)

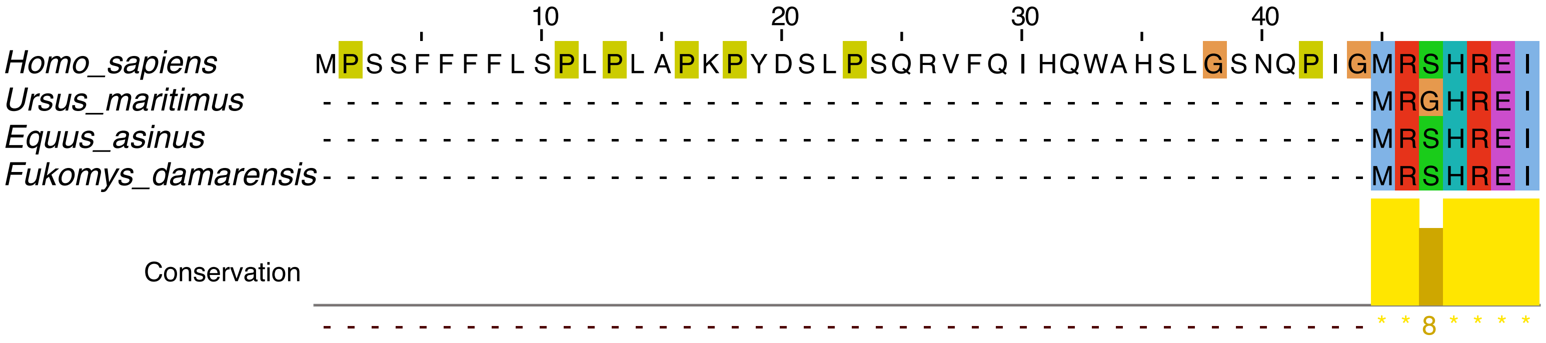

**HG1098**

ENSG00000183775 (KCTD16)

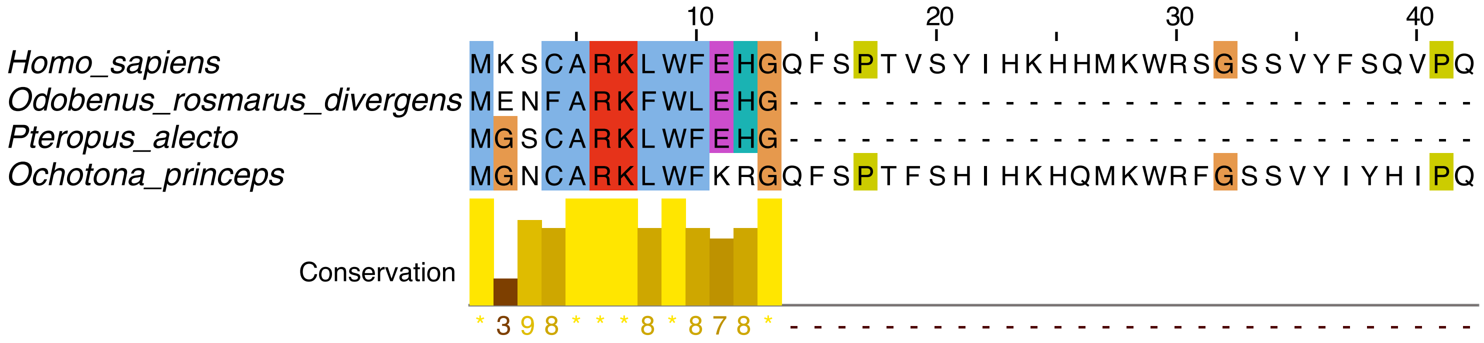

**HG1099**

ENSG00000186334 (SLC36A3)

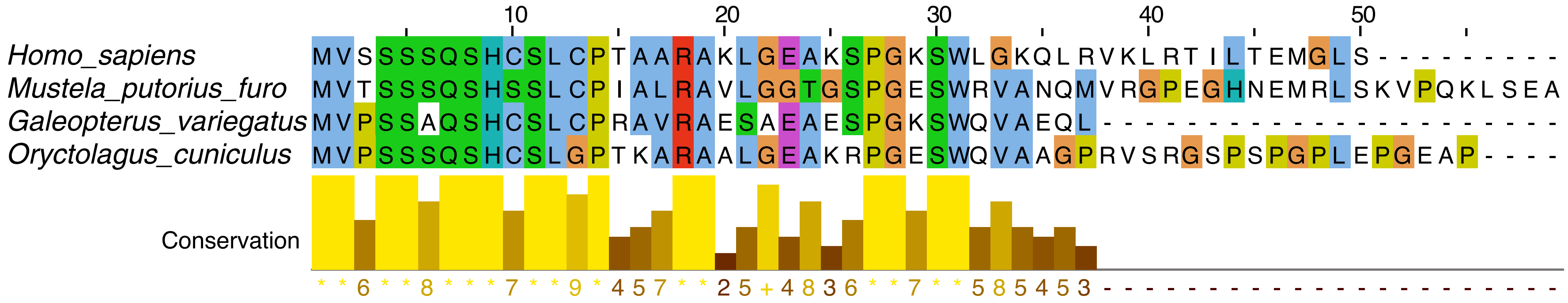

**HG1100**

ENSG00000187815 (ZFP69)

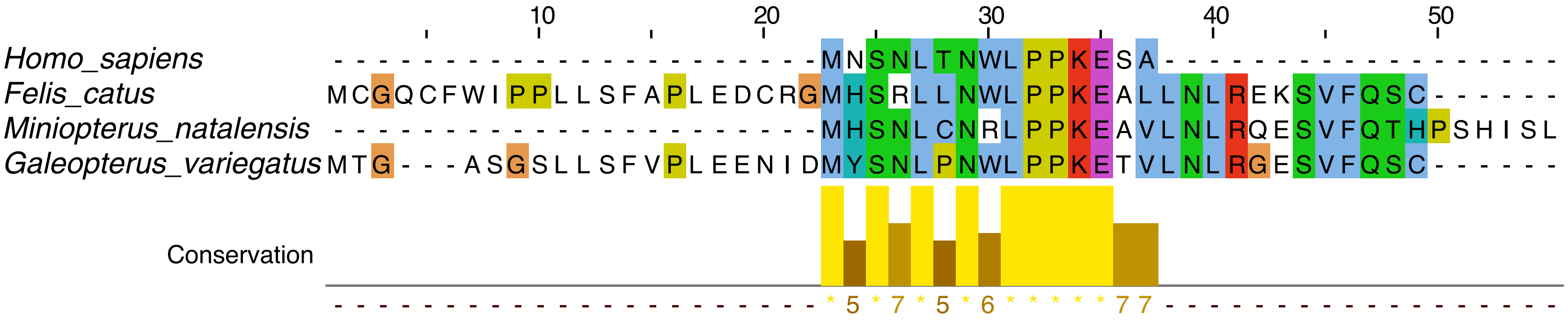

**HG1101**

ENSG00000189334 (S100A14)

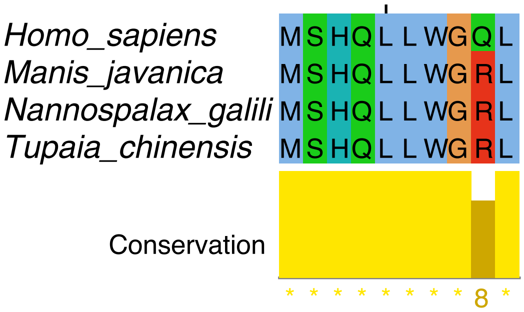

**HG1102**

ENSG00000196704 (AMZ2)

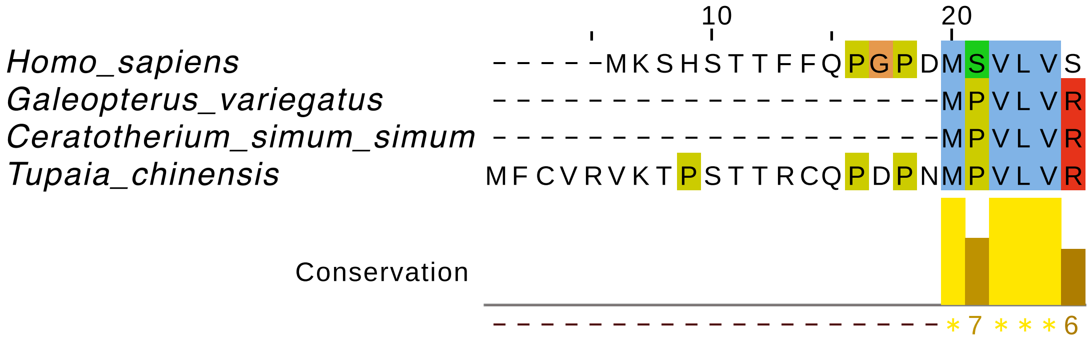

**HG1103**

ENSG00000197620 (CXorf40A)

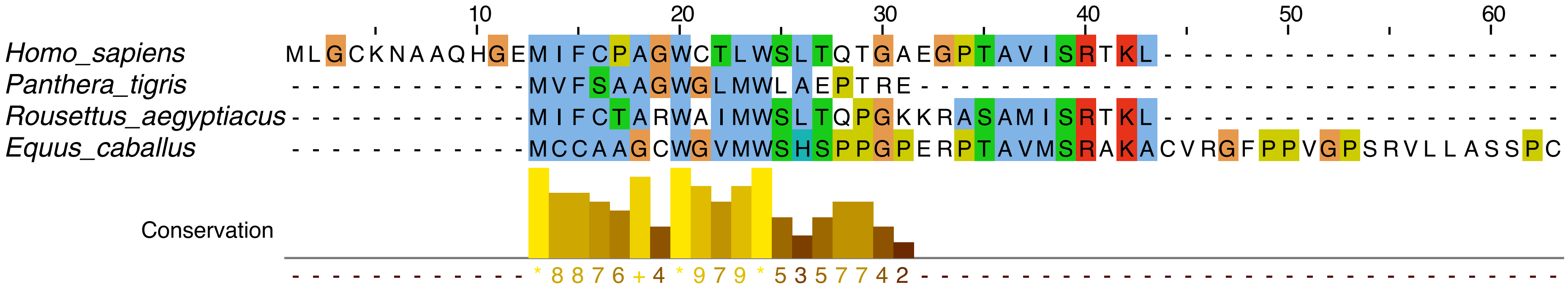

**HG1104**

ENSG00000205155 (PSENEN)

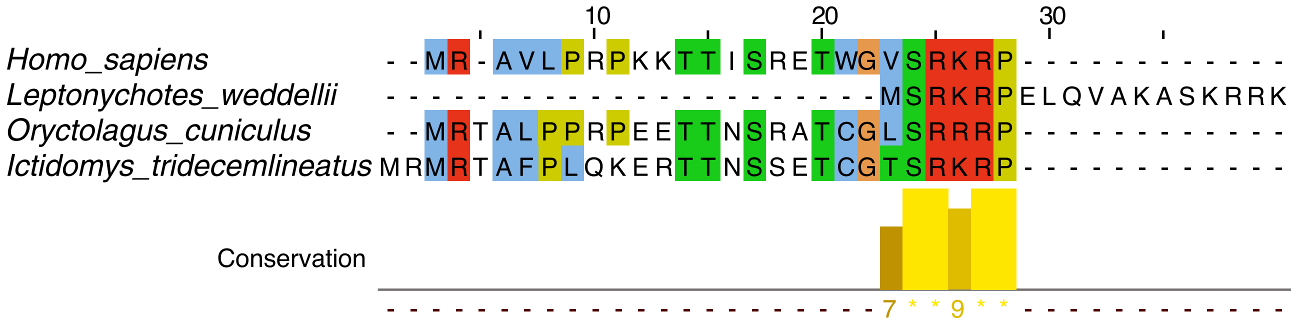

**HG1105**

ENSG00000213654 (GPSM3)

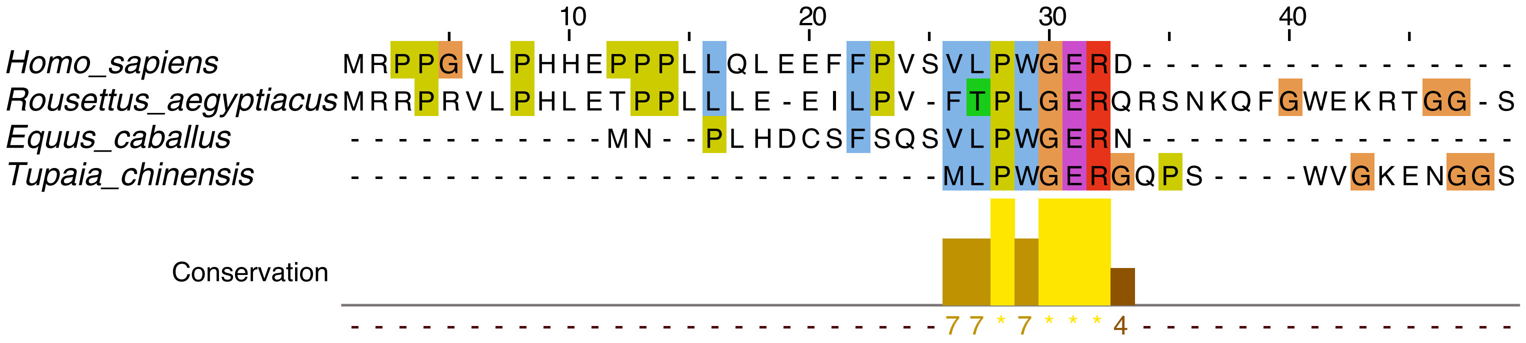

**HG1106**

ENSG00000215021 (PHB2)

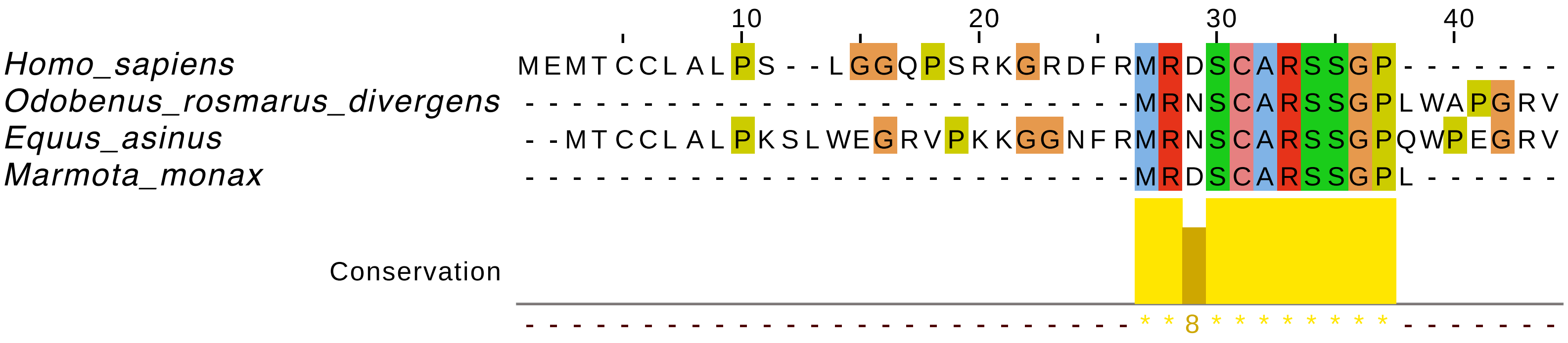

**HG1107**

ENSG00000217930 (PAM16)

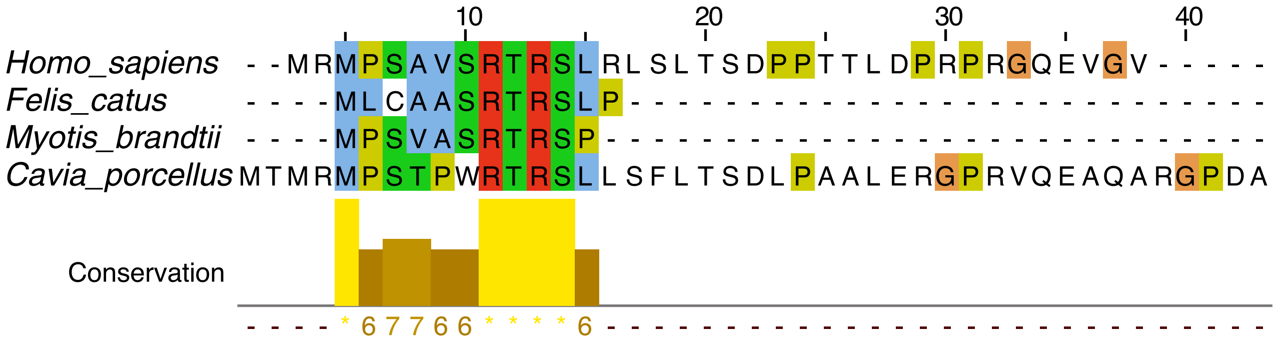

**HG1108**

ENSG00000251247 (ZNF345)

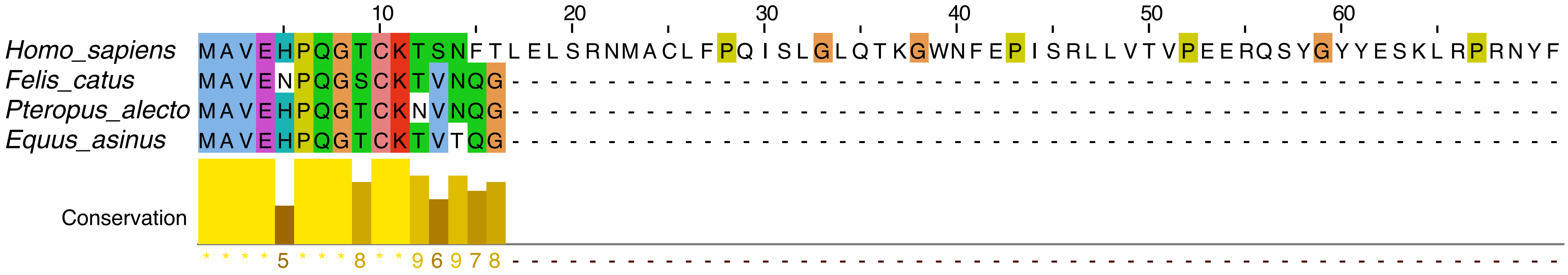

**HG1109**

ENSG00000278311 (GGNBP2)

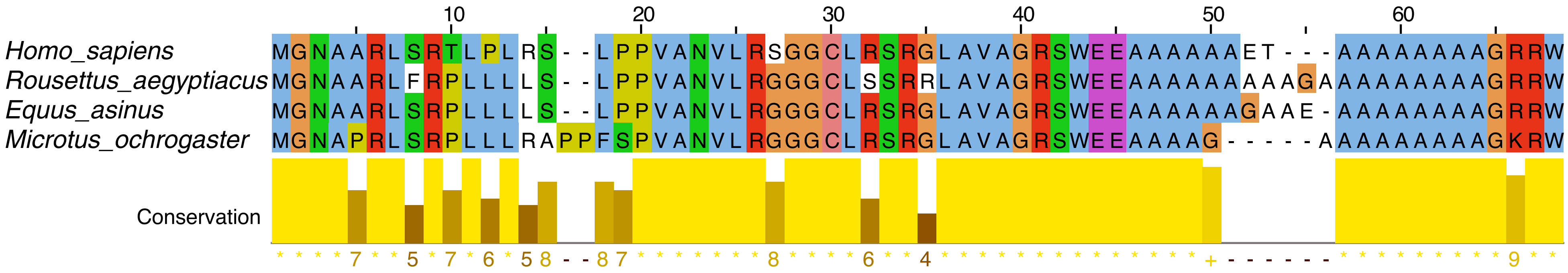

**HG1110**

ENSG00000034152 (MAP2K3)

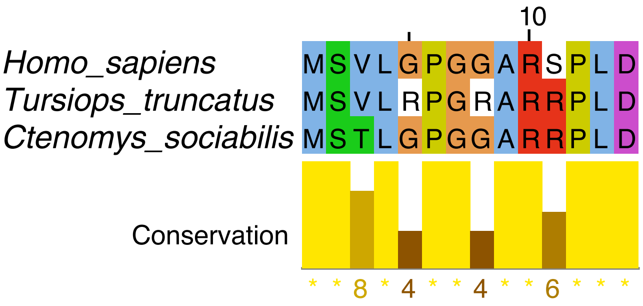

# HG1111

### ENSG00000040531 (CTNS)

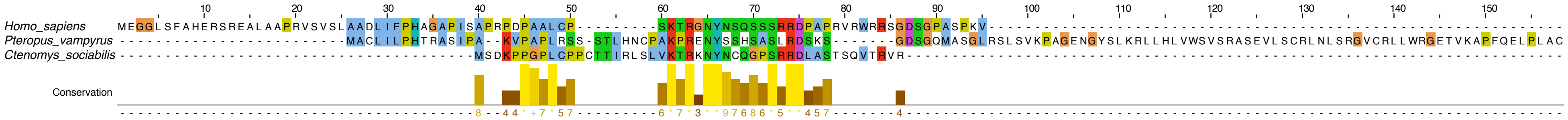

**HG1112**

ENSG00000068885 (IFT80)

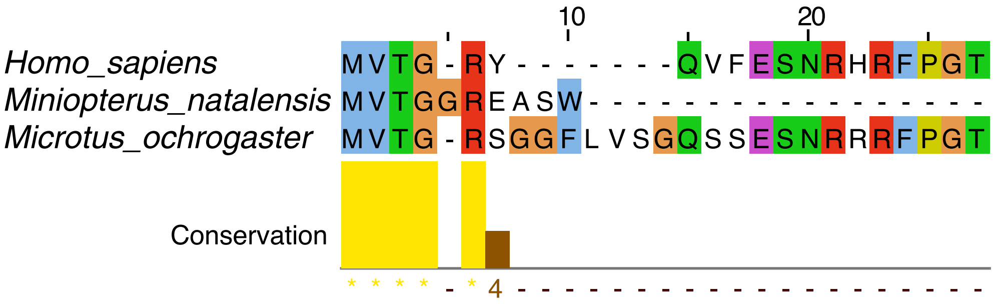

**HG1113**

ENSG00000072201 (LNX1)

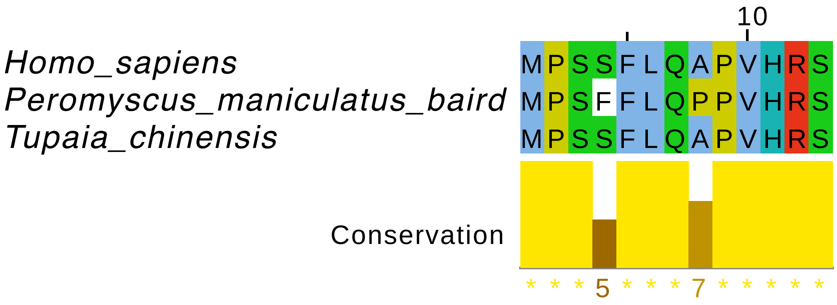

**HG1114**

ENSG00000073737 (DHRS9)

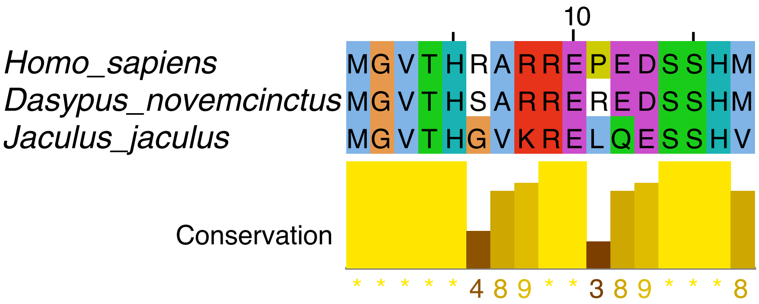

**HG1115**

ENSG00000074219 (TEAD2)

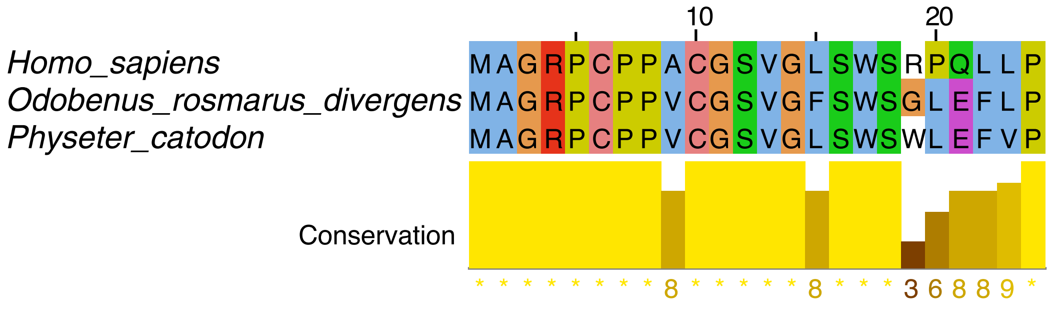

**HG1116**

ENSG00000084090 (STARD7)

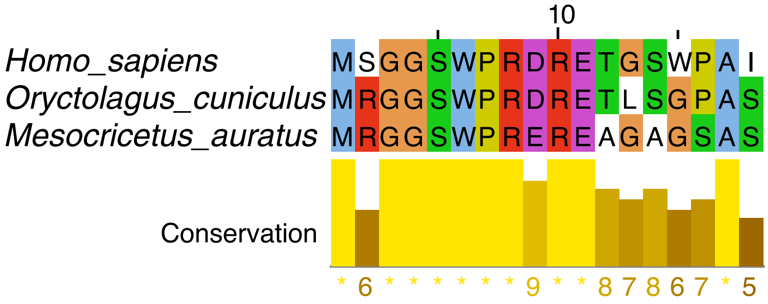

**HG1117**

ENSG00000089902 (RCOR1)

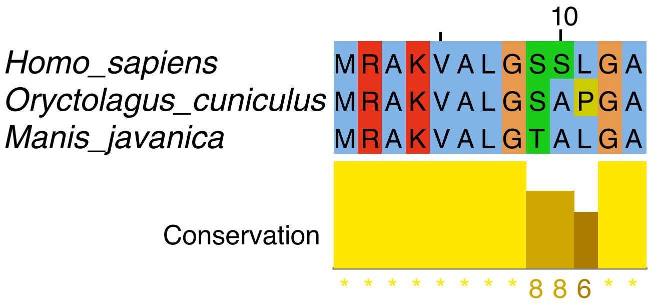

**HG1118**

ENSG00000090975 (PITPNM2)

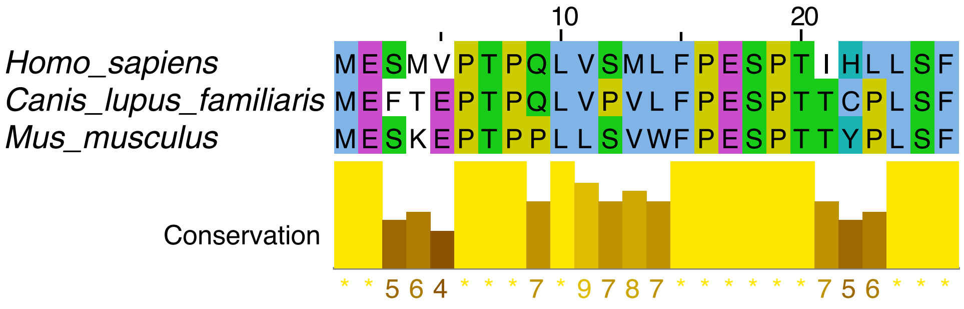

**HG1119**

ENSG00000092020 (PPP2R3C)

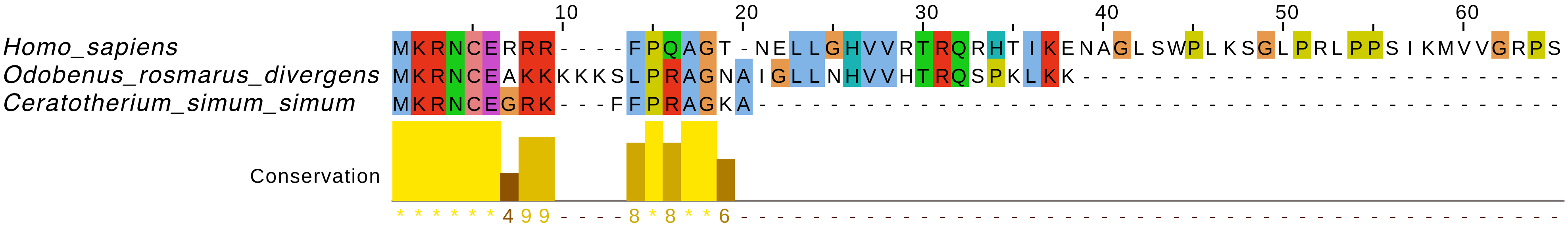

**HG1120**

ENSG00000095585 (BLNK)

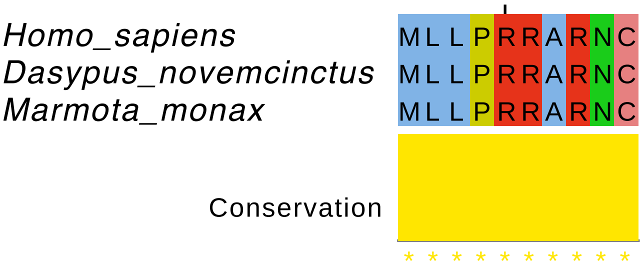

**HG1121**

ENSG00000100350 (FOXRED2)

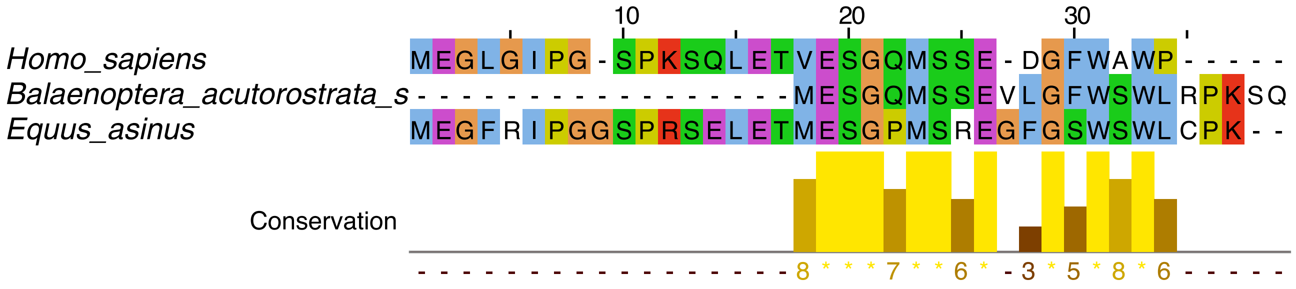

**HG1122**

ENSG00000100410 (PHF5A)

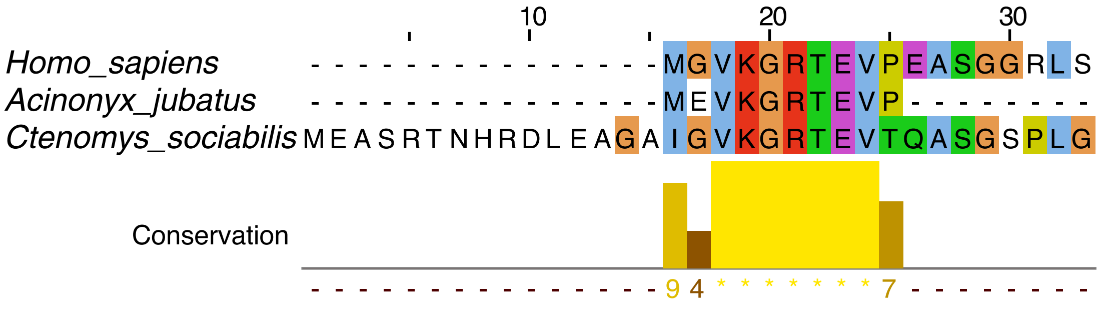

**HG1123**

ENSG00000100884 (CPNE6)

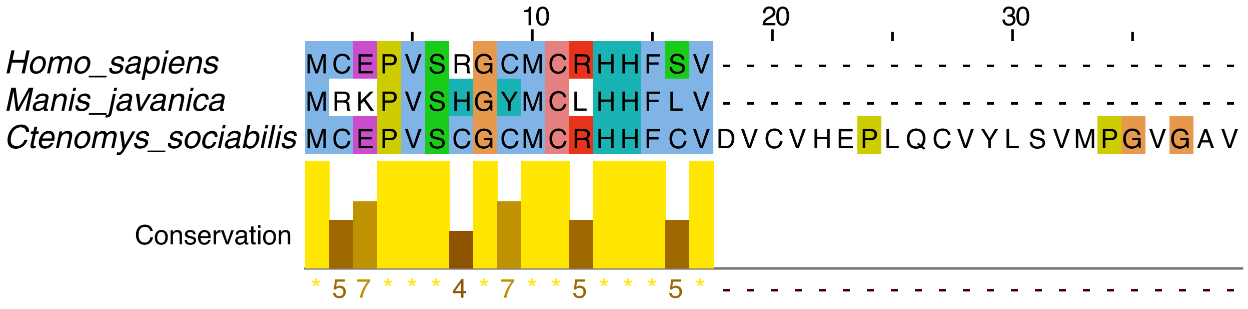

**HG1124**

ENSG00000100934 (SEC23A)

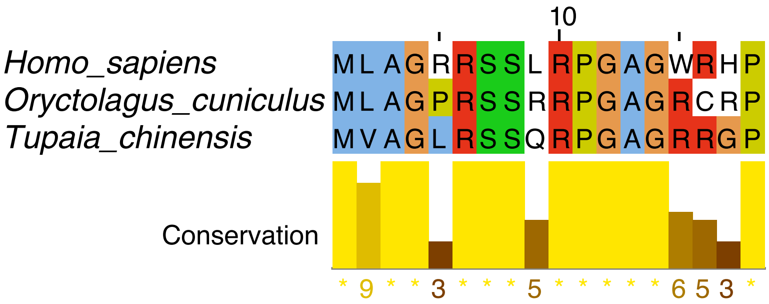

**HG1125**

ENSG00000101574 (METTL4)

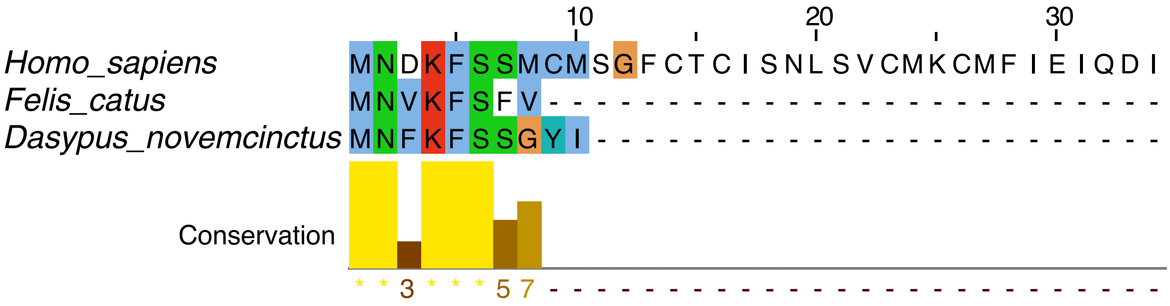

**HG1126**

ENSG00000101890 (GUCY2F)

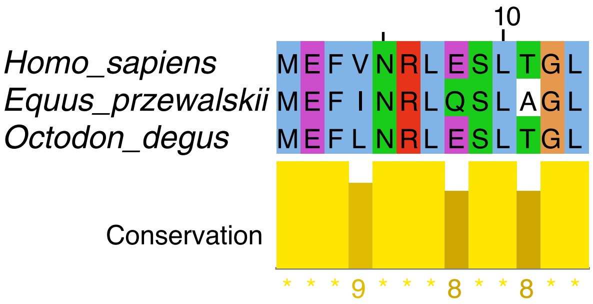

**HG1127**

ENSG00000103064 (SLC7A6)

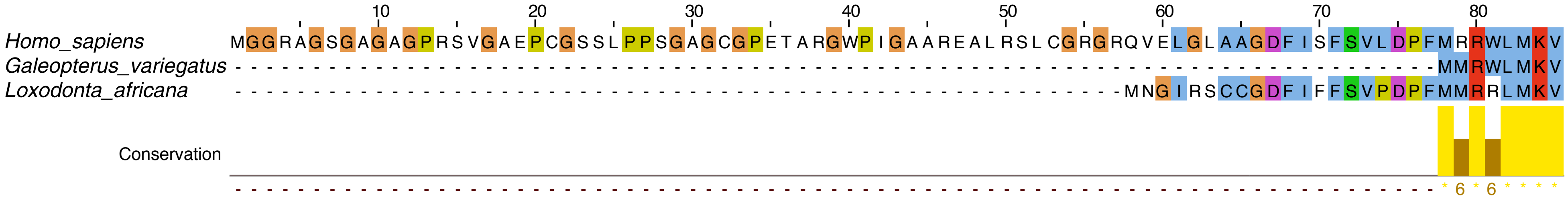

**HG1128**

ENSG00000103852 (TTC23)

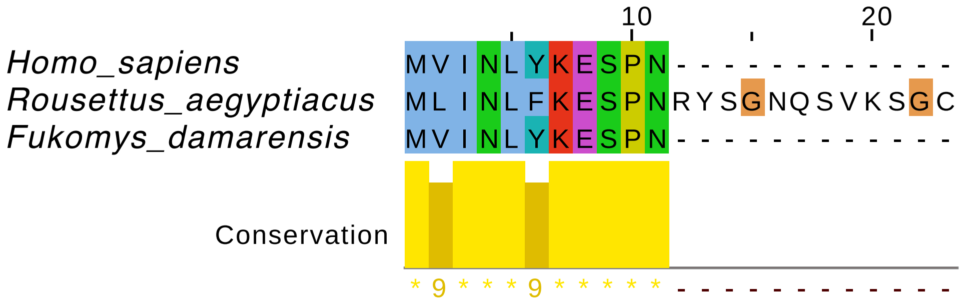

**HG1129**

ENSG00000104432 (IL7)

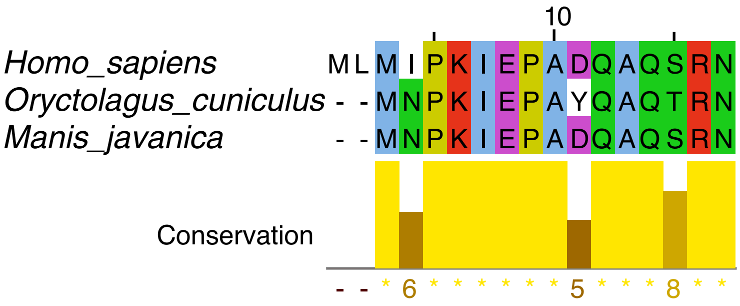

**HG1130**

ENSG00000105559 (PLEKHA4)

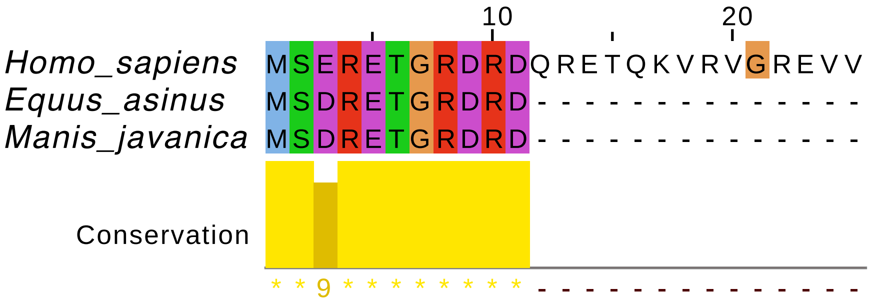

**HG1131**

ENSG00000106780 (MEGF9)

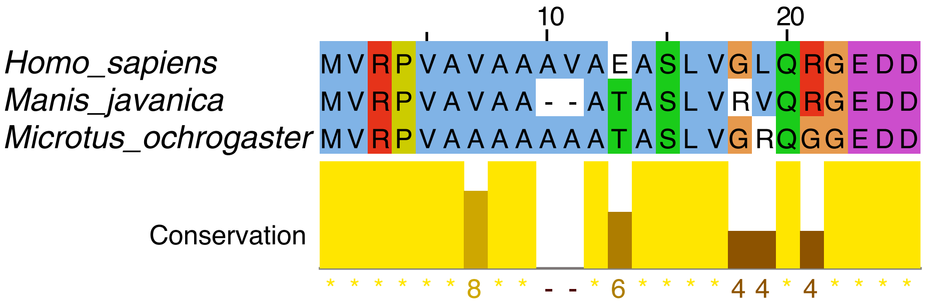

**HG1132**

ENSG00000109771 (LRP2BP)

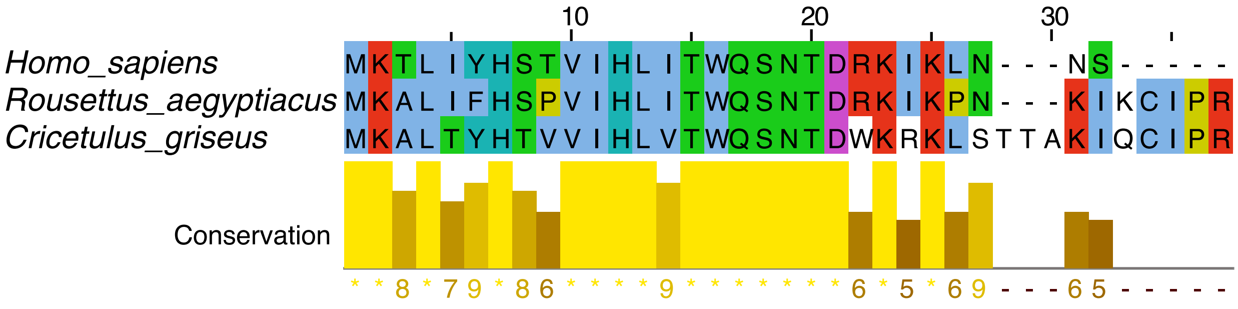

**HG1133**

ENSG00000111328 (CDK2AP1)

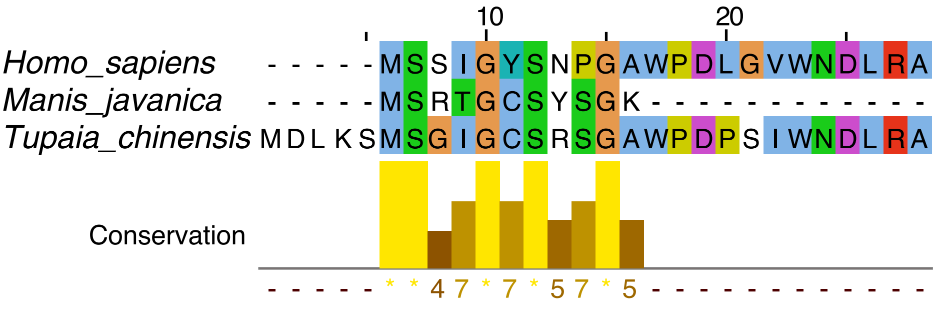

**HG1134**

ENSG00000112984 (KIF20A)

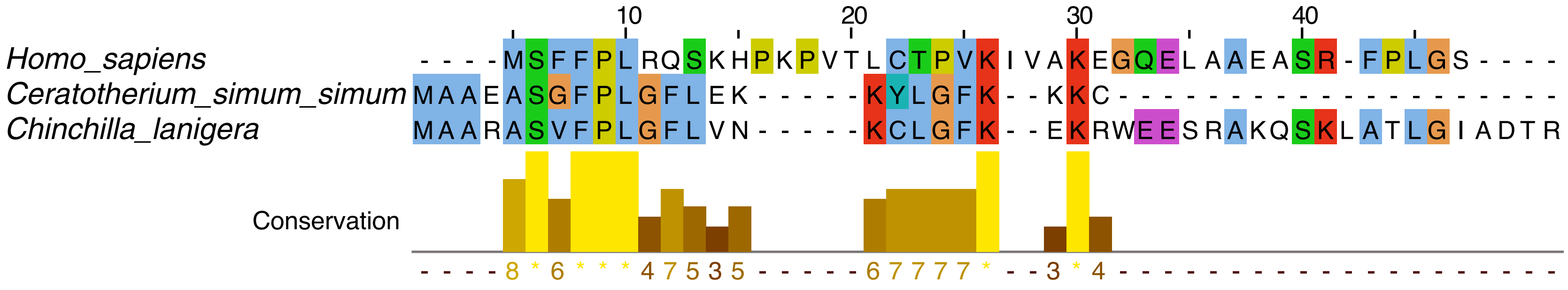

**HG1135**

ENSG00000113597 (TRAPPC13)

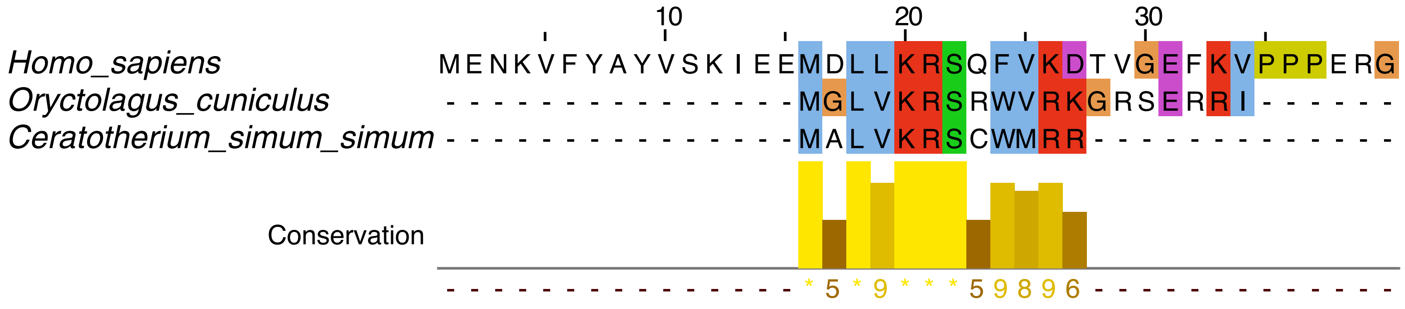

**HG1136**

ENSG00000113749 (HRH2)

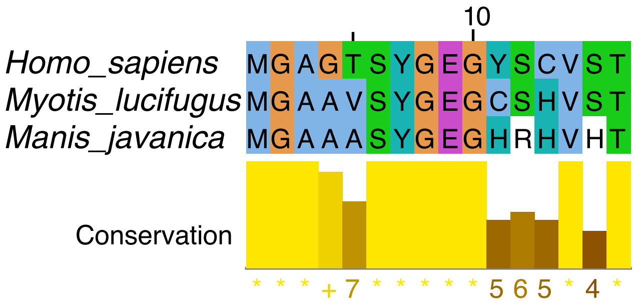

**HG1137**

ENSG00000115307 (AUP1)

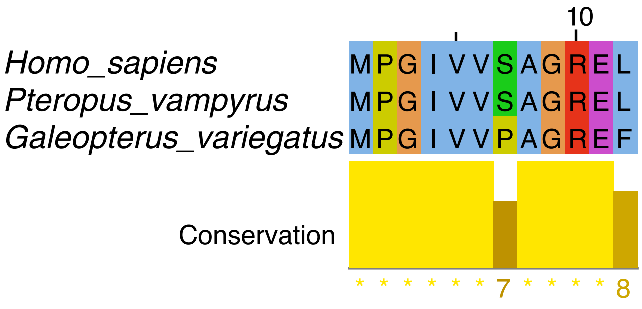

**HG1138**

ENSG00000116871 (MAP7D1)

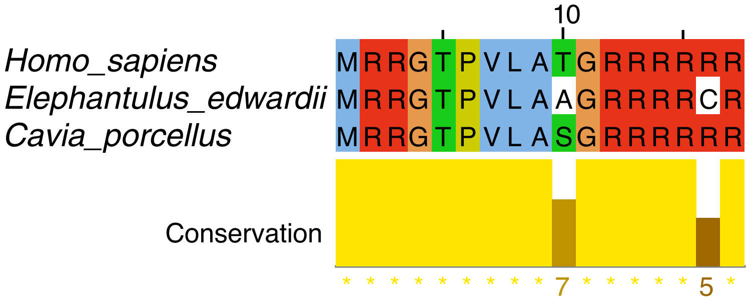

**HG1139.1**

ENSG00000118308 (LRMP)

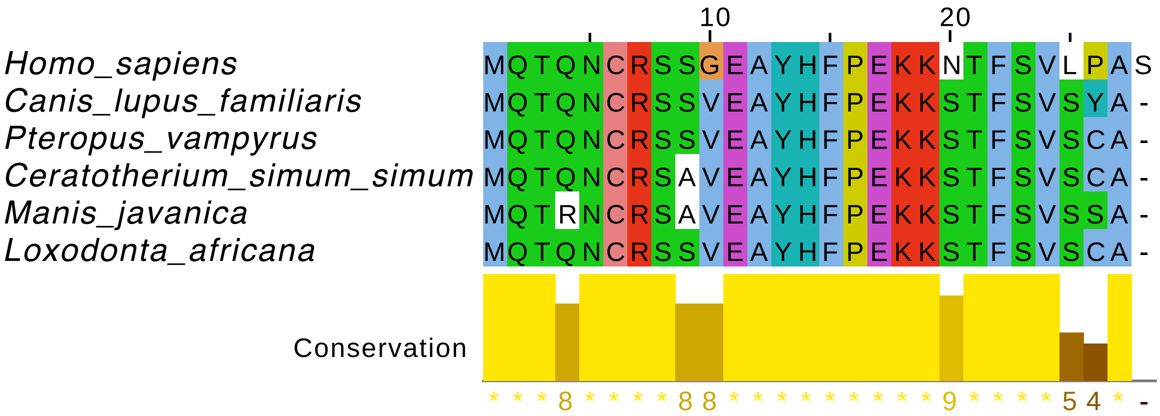

**HG1139.2**

ENSG00000118308 (LRMP)

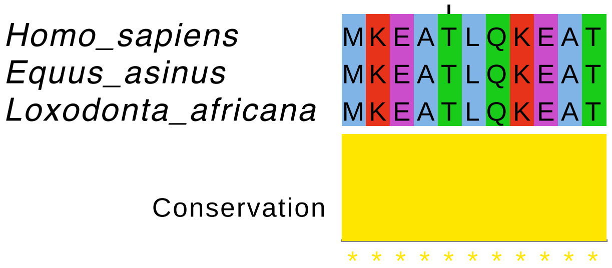

**HG1140**

ENSG00000118495 (PLAGL1)

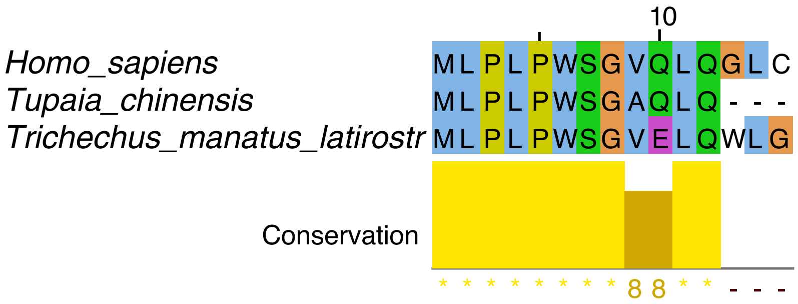

**HG1141**

ENSG00000121964 (GTDC1)

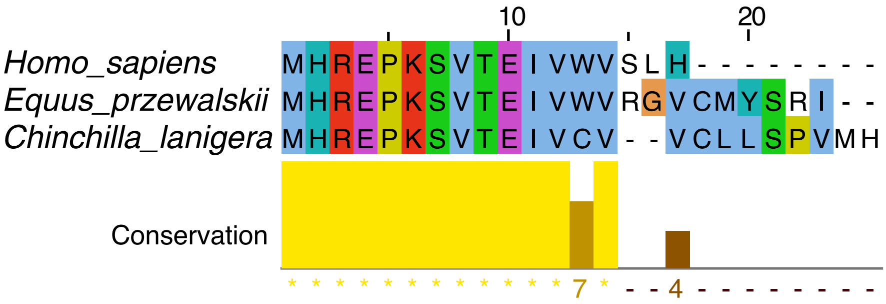

**HG1142**

ENSG00000124613 (ZNF391)

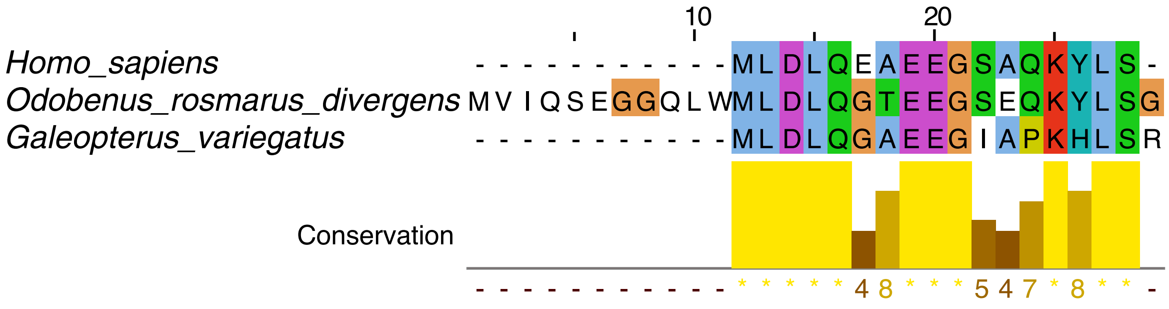

**HG1143**

ENSG00000125245 (GPR18)

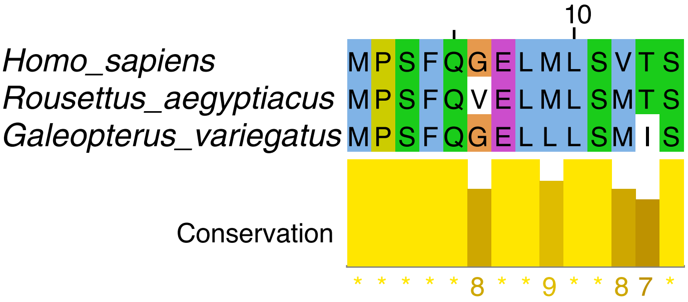

**HG1144**

ENSG00000128203 (ASPHD2)

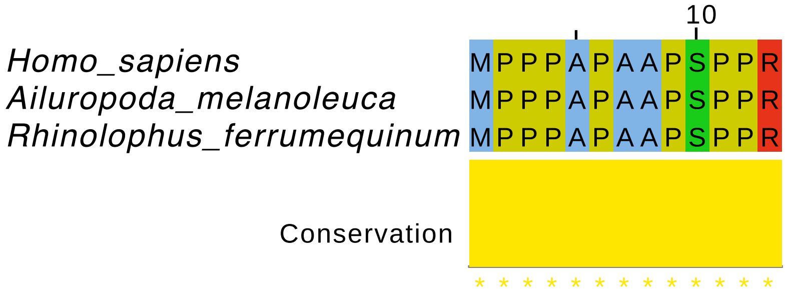

**HG1145**

ENSG00000128585 (MKLN1)

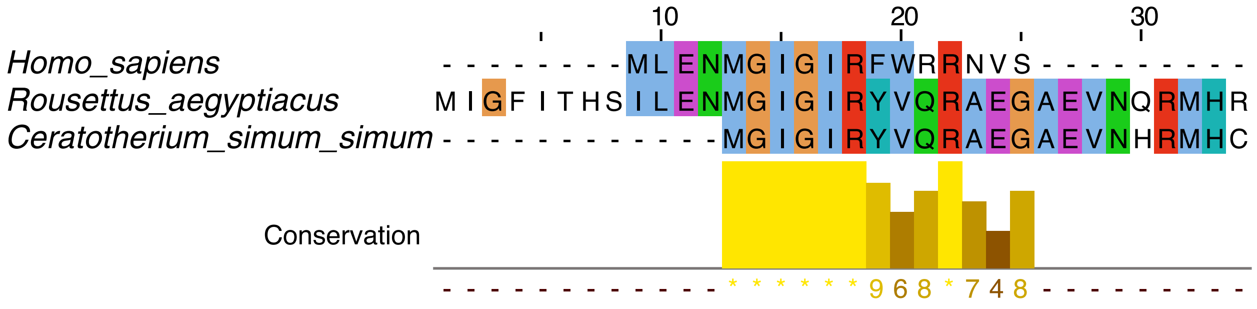

**HG1146**

ENSG00000130413 (STK33)

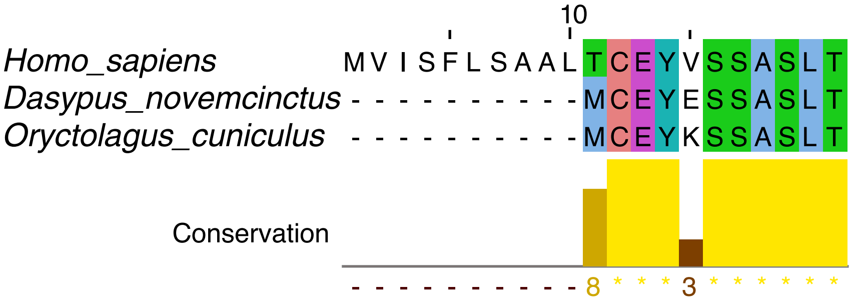

**HG1147**

ENSG00000131781 (FMO5)

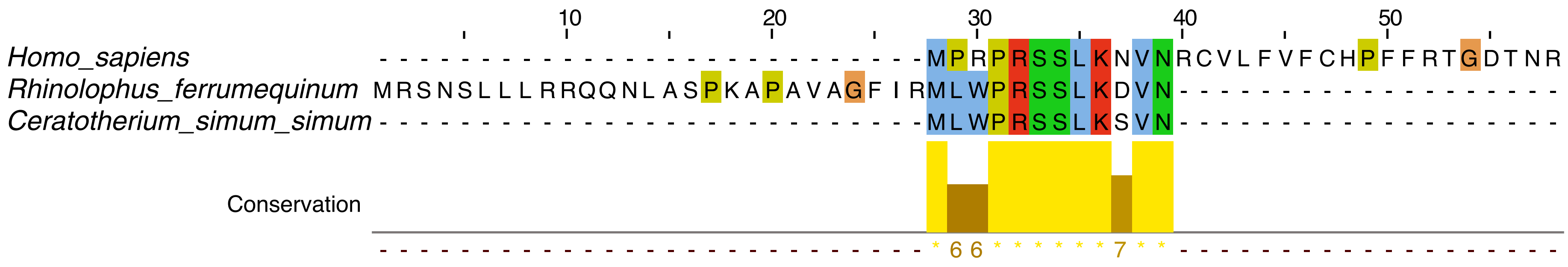

**HG1148**

ENSG00000132938 (MTUS2)

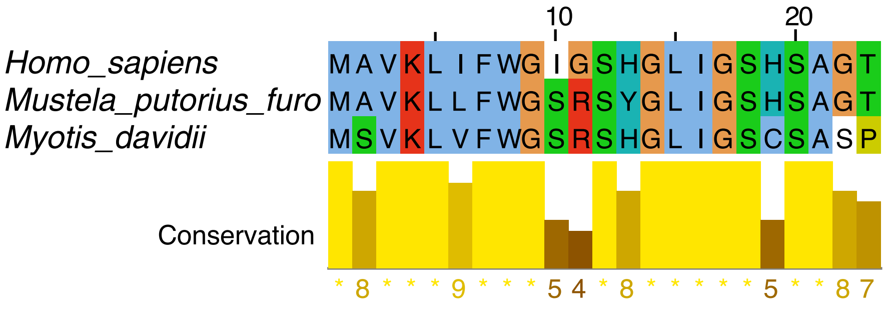

**HG1149**

ENSG00000132975 (GPR12)

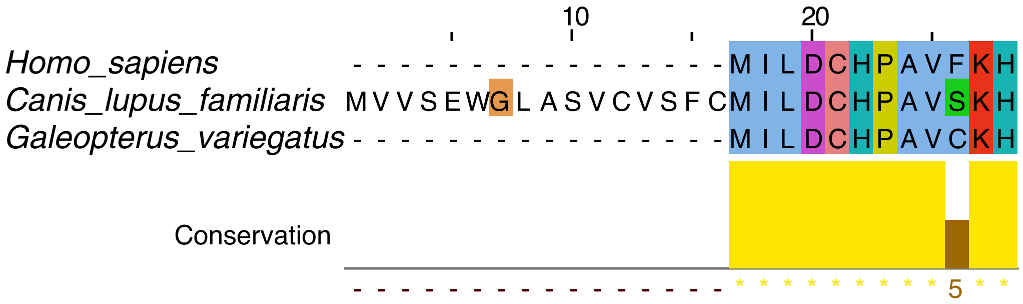

**HG1150**

ENSG00000135299 (ANKRD6)

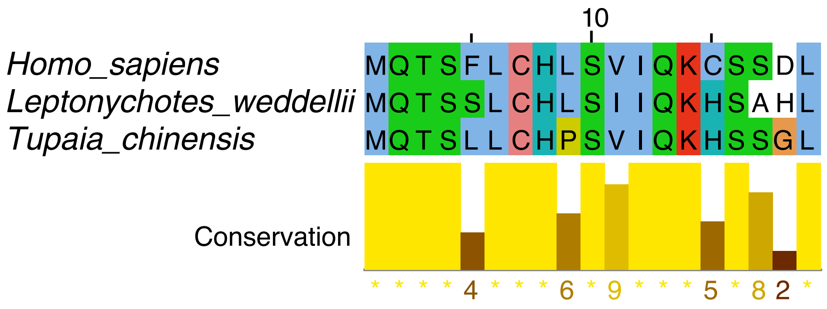

**HG1151**

ENSG00000137168 (PPIL1)

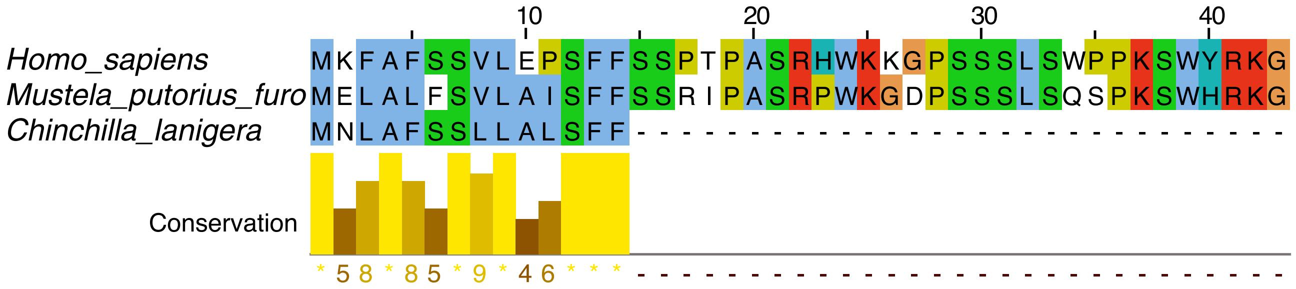

**HG1152**

ENSG00000138658 (ZGRF1)

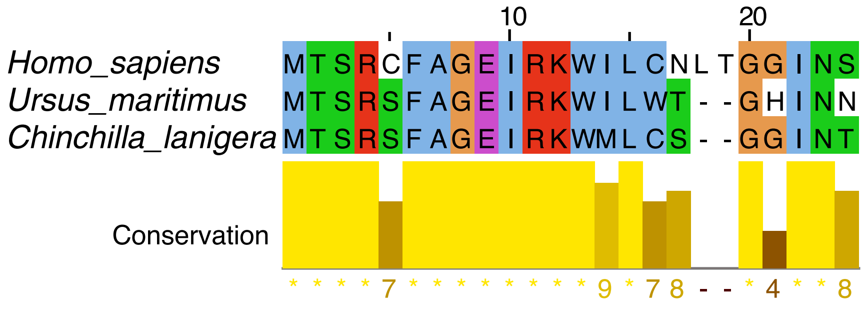

**HG1153**

ENSG00000138738 (PRDM5)

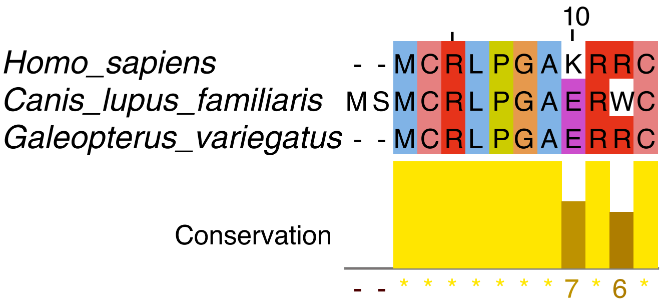

**HG1154**

ENSG00000139725 (RHOF)

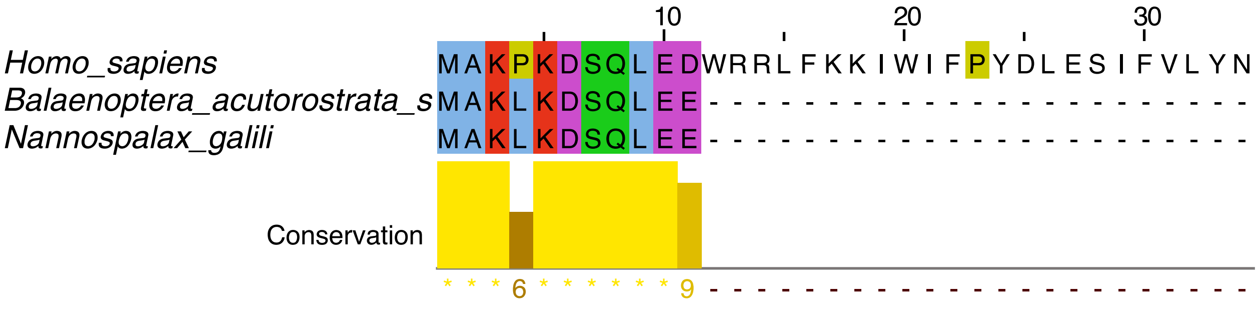

**HG1155**

ENSG00000141668 (CBLN2)

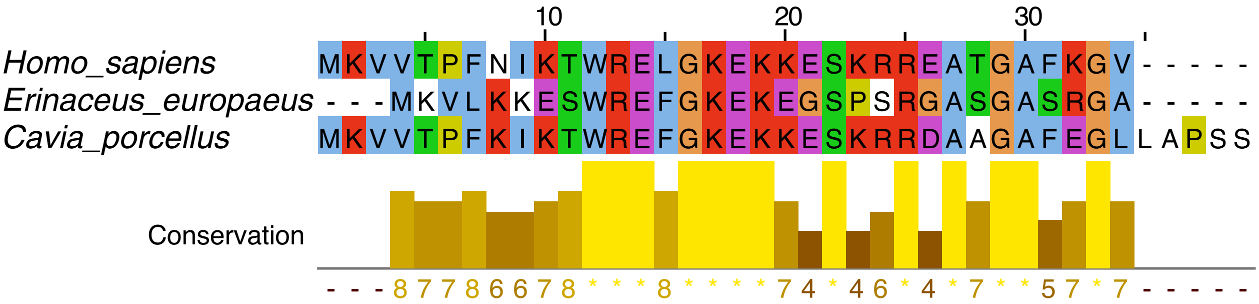

**HG1156**

ENSG00000144741 (SLC25A26)

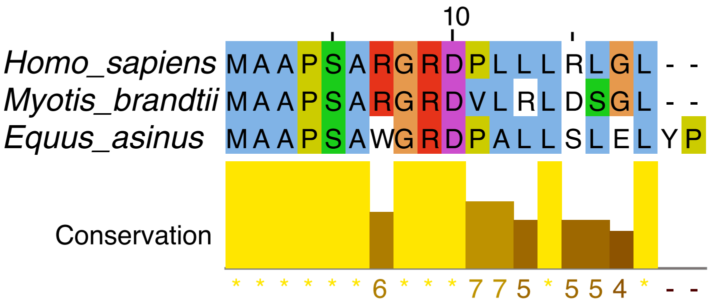

**HG1157**

ENSG00000148123 (PLPPR1)

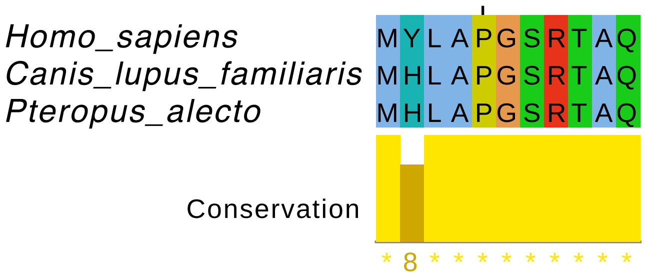

**HG1158**

ENSG00000149571 (KIRREL3)

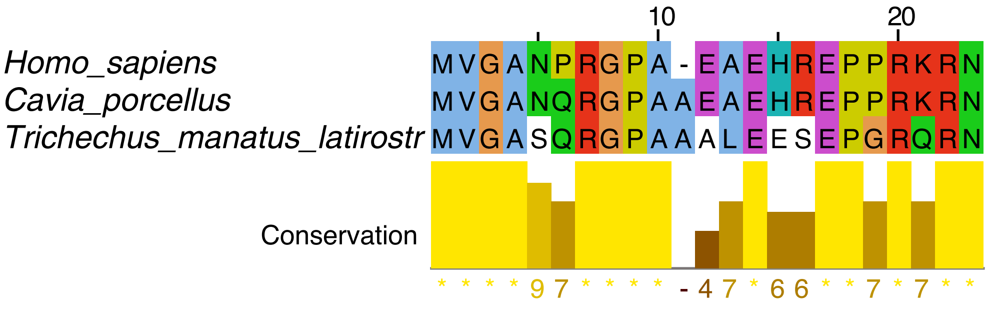

**HG1159**

ENSG00000152822 (GRM1)

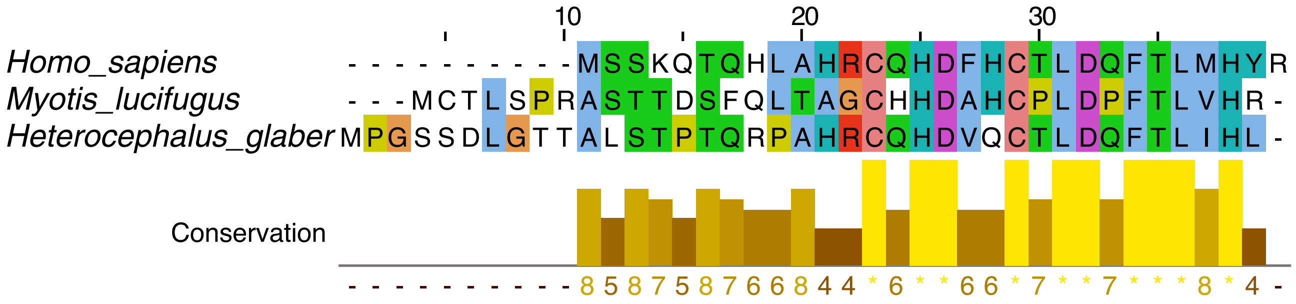

**HG1160**

ENSG00000154162 (CDH12)

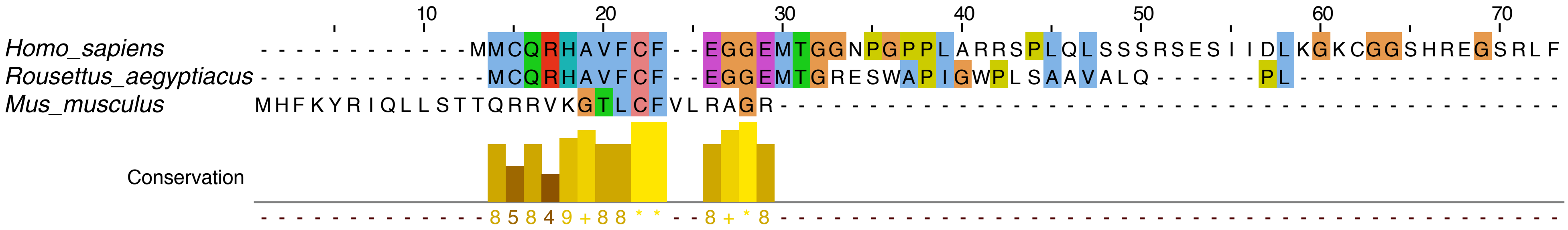

**HG1161**

ENSG00000154227 (CERS3)

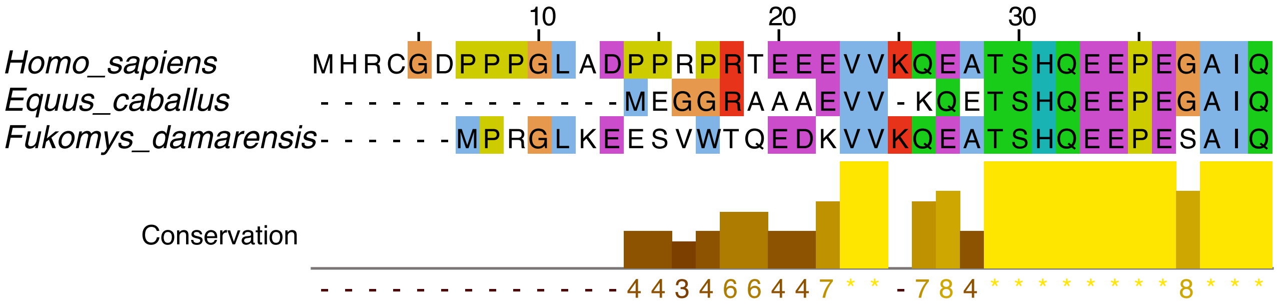

**HG1162**

ENSG00000155380 (SLC16A1)

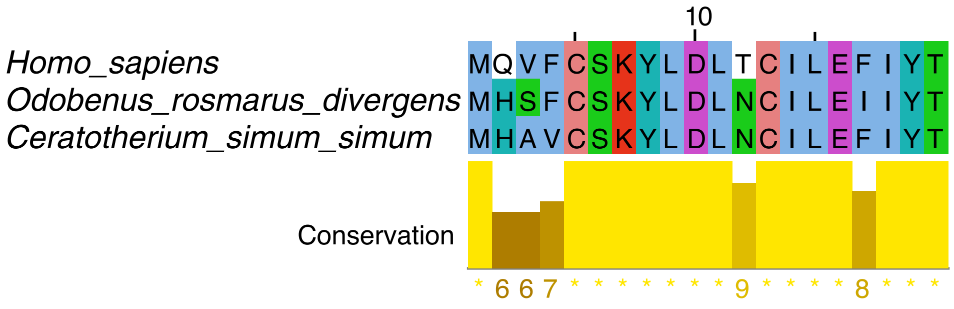

**HG1163**

ENSG00000157637 (SLC38A10)

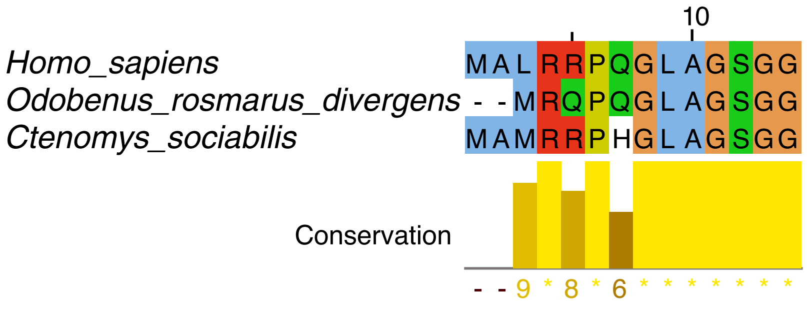

**HG1164**

ENSG00000162639 (HENMT1)

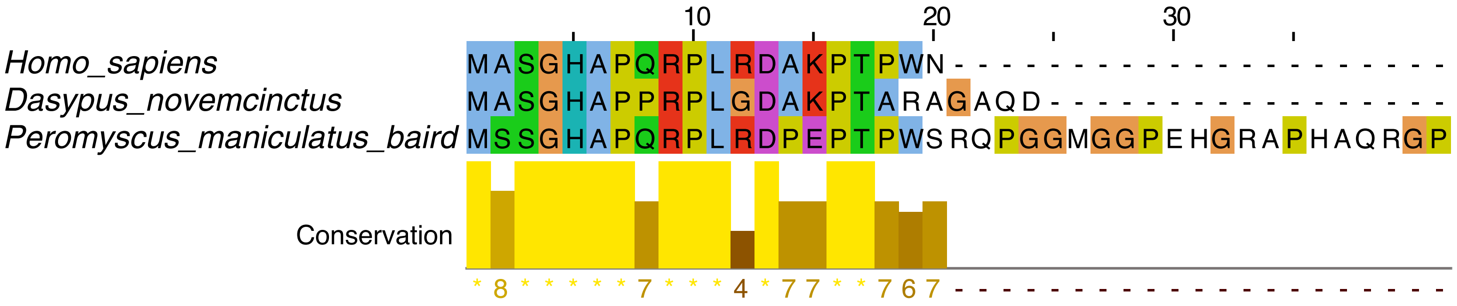

**HG1165**

ENSG00000162694 (EXTL2)

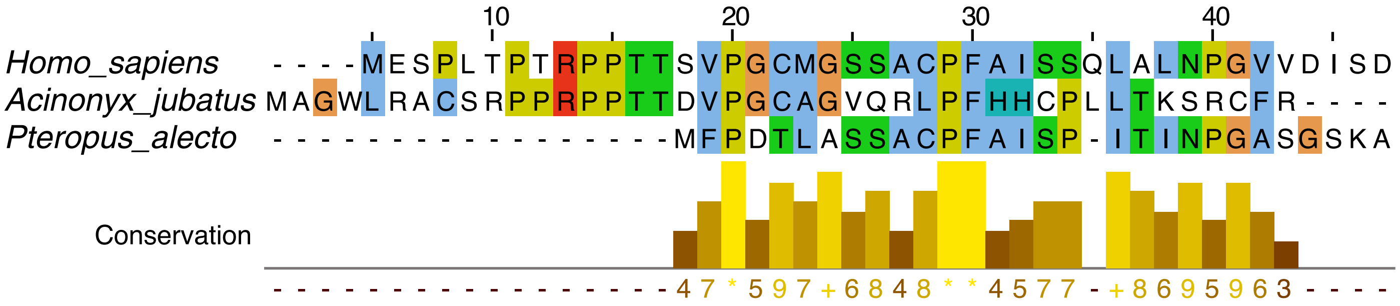

**HG1166**

ENSG00000165071 (TMEM71)

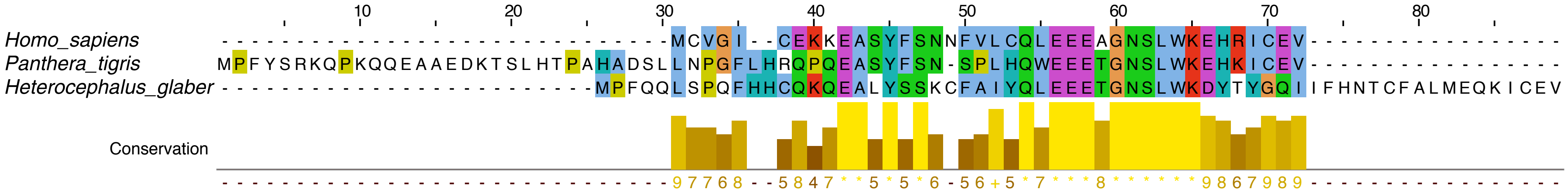

**HG1167**

ENSG00000165934 (CPSF2)

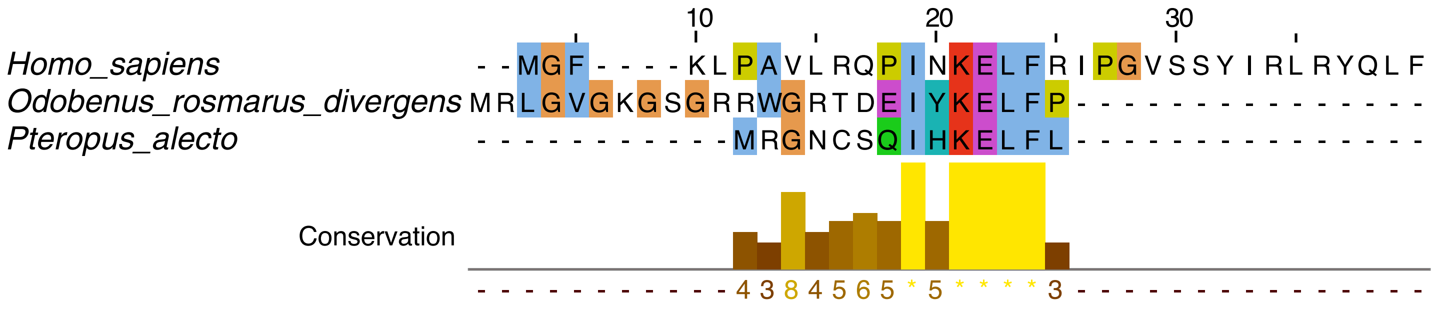

**HG1168**

ENSG00000166669 (ATF7IP2)

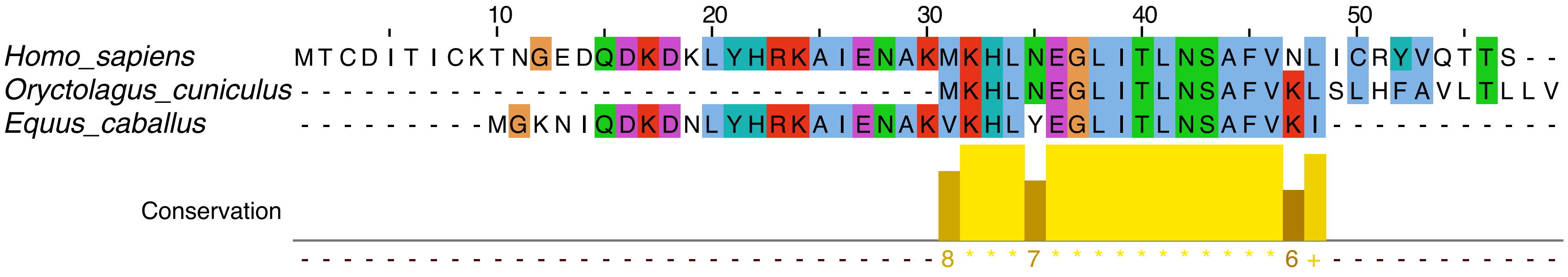

**HG1169**

ENSG00000167637 (ZNF283)

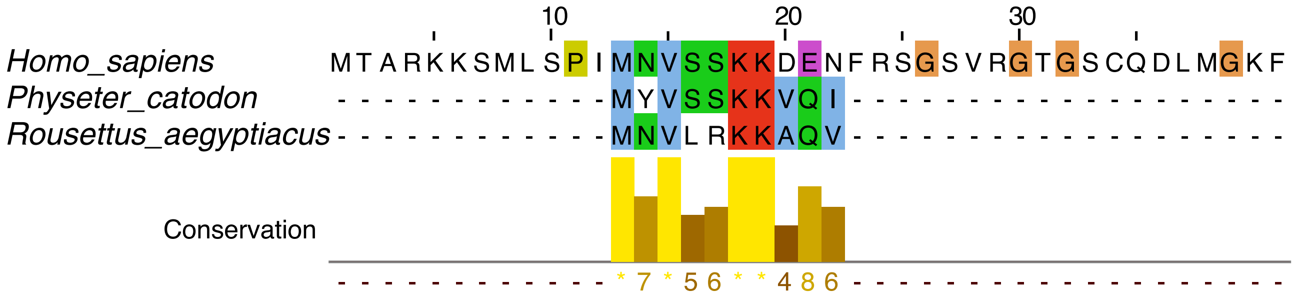

**HG1170**

ENSG00000167858 (TEKT1)

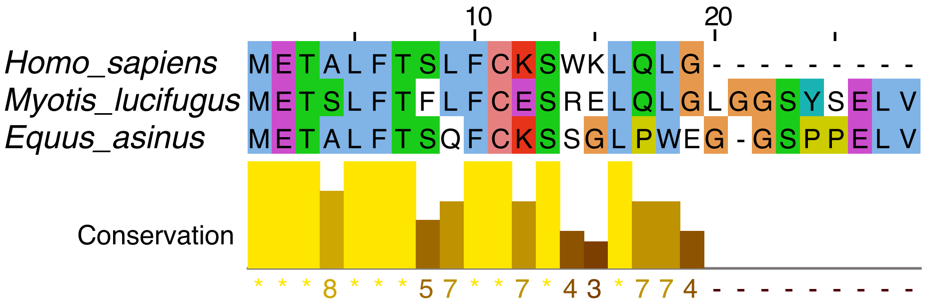

**HG1171**

ENSG00000167968 (DNASE1L2)

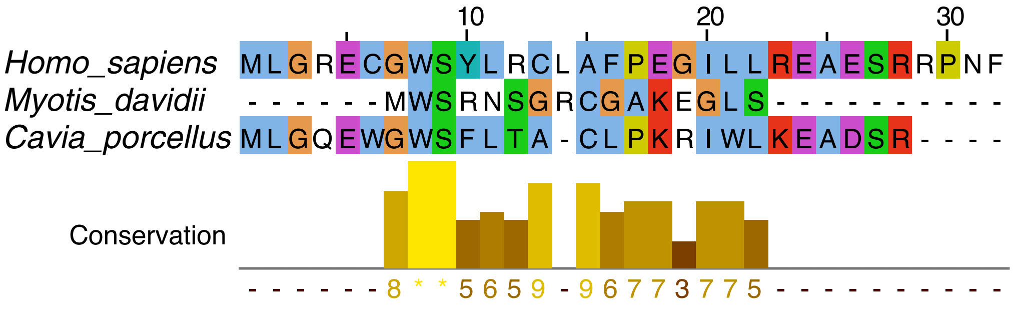

**HG1172**

ENSG00000170381 (SEMA3E)

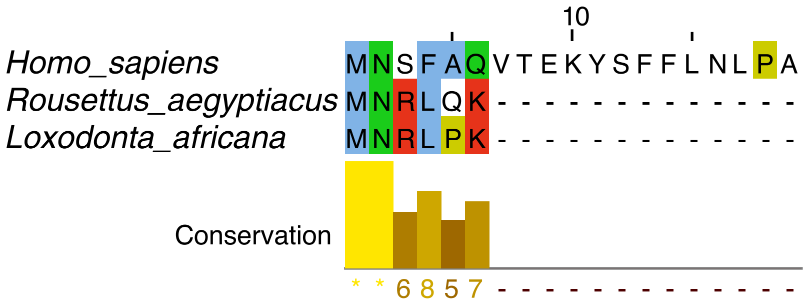

**HG1173**

ENSG00000170464 (DNAJC18)

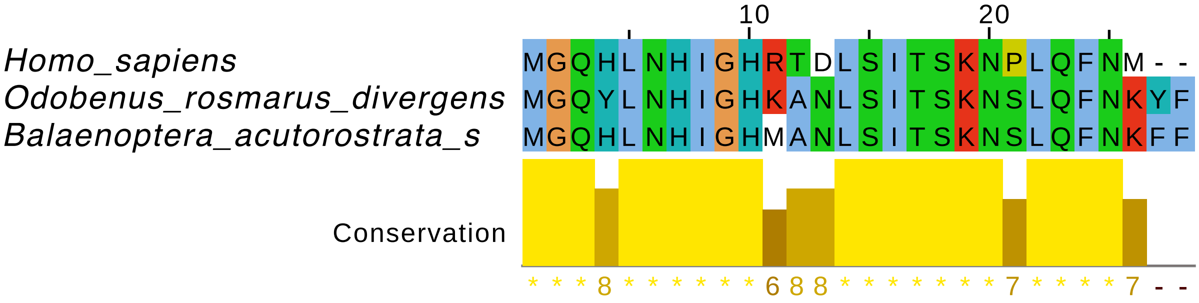

**HG1174**

ENSG00000173918 (C1QTNF1)

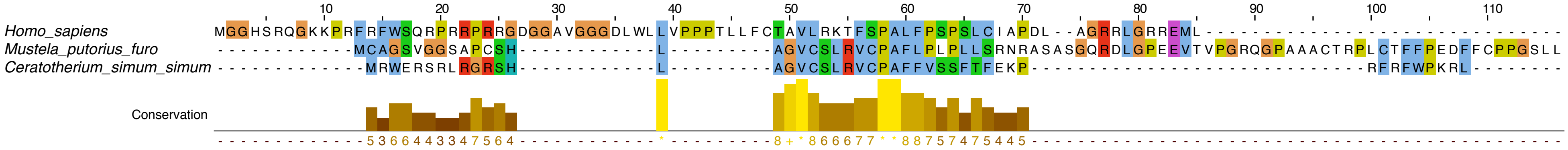

**HG1175**

ENSG00000174473 (GALNTL6)

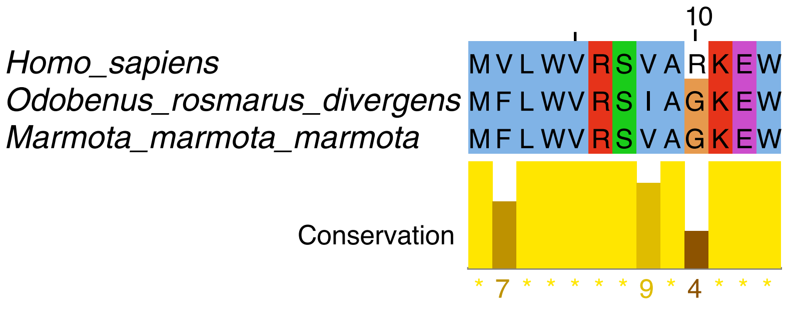

**HG1176**

ENSG00000176472 (ZNF575)

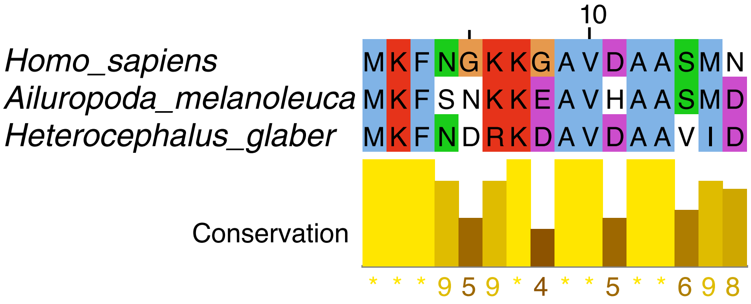

**HG1177**

ENSG00000178295 (GEN1)

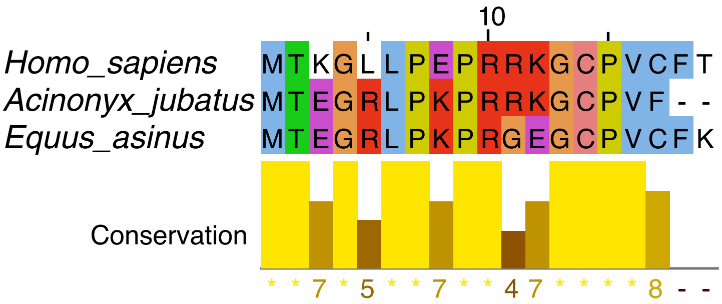

**HG1178**

ENSG00000182670 (TTC3)

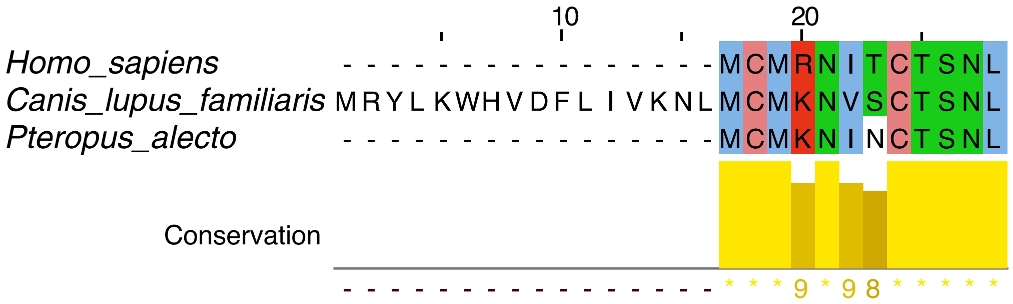

**HG1179**

ENSG00000185933 (CALHM1)

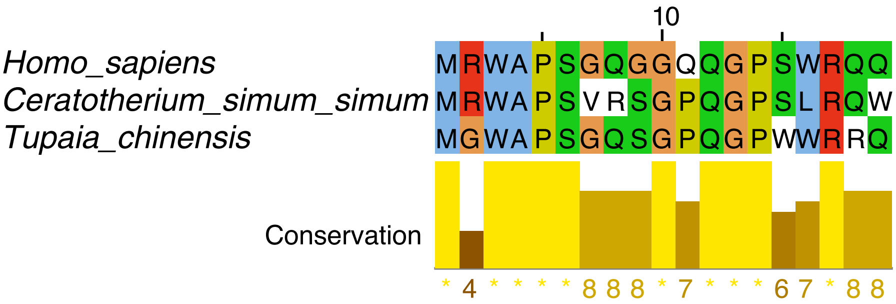

**HG1180**

ENSG00000188674

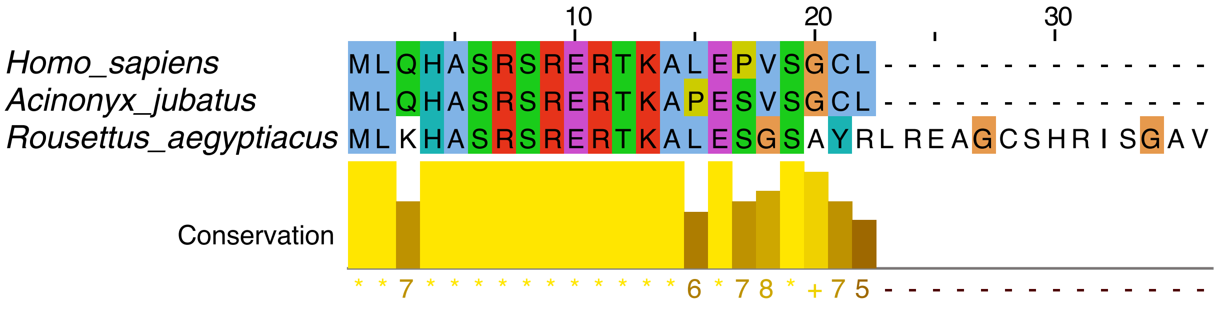

**HG1181**

ENSG00000189058 (APOD)

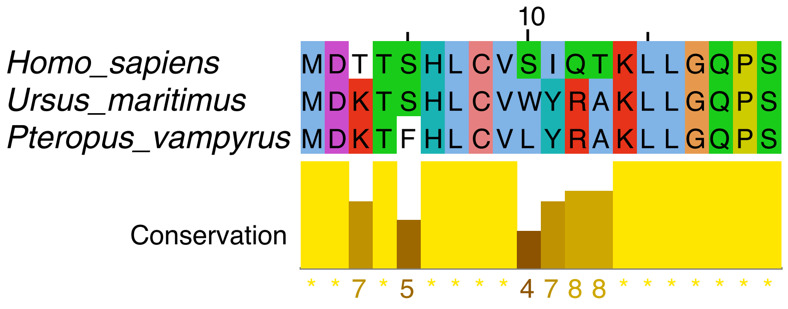

**HG1182**

ENSG00000196155 (PLEKHG4)

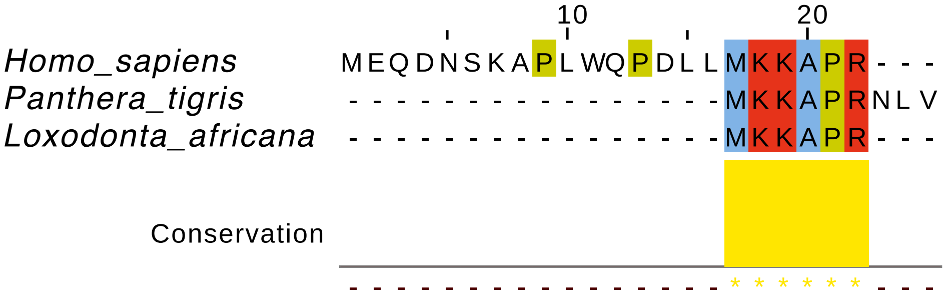

**HG1183**

ENSG00000198133 (TMEM229B)

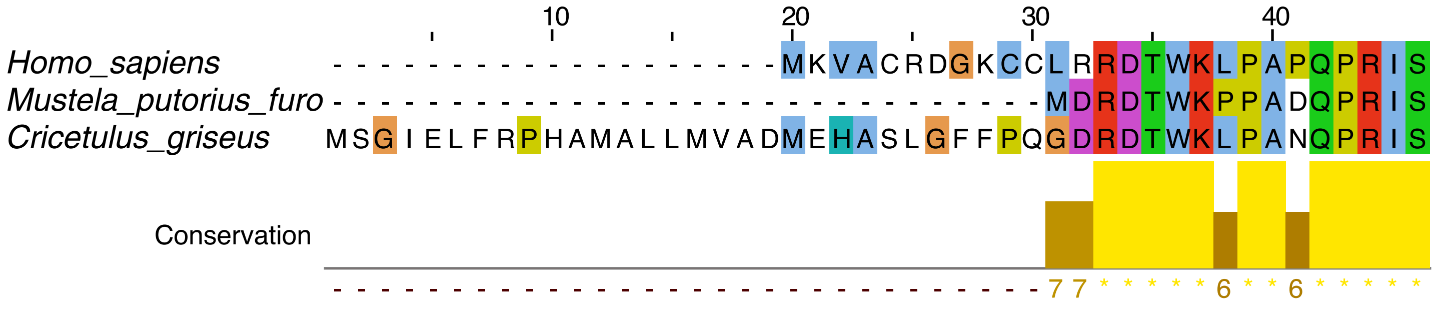

**HG1184**

ENSG00000198894 (CIPC)

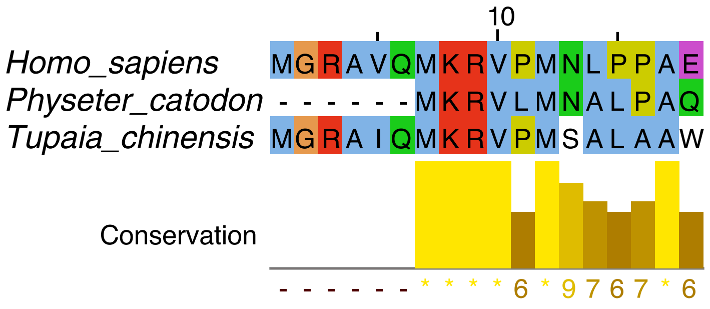

**HG1185**

ENSG00000198920 (KIAA0753)

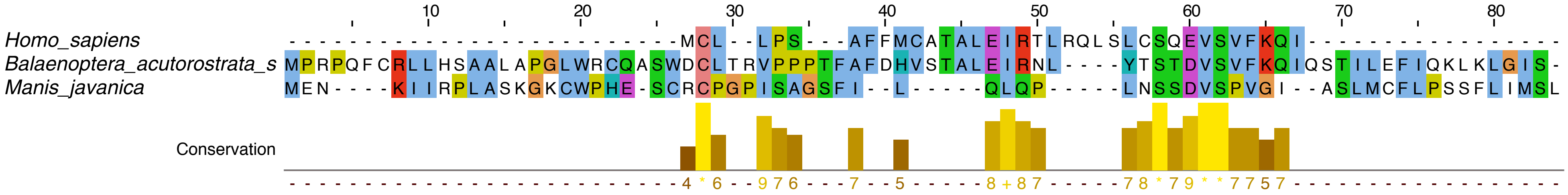

**HG1186**

ENSG00000204923

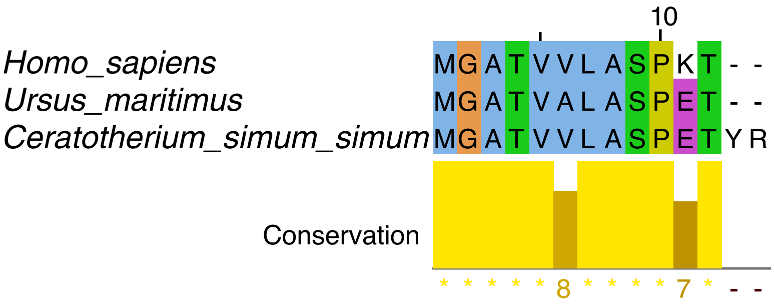

**HG1187**

ENSG00000213762 (ZNF134)

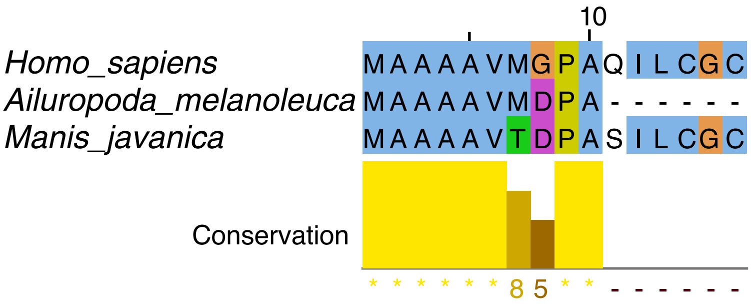

**HG1188**

ENSG00000214415 (GNAT3)

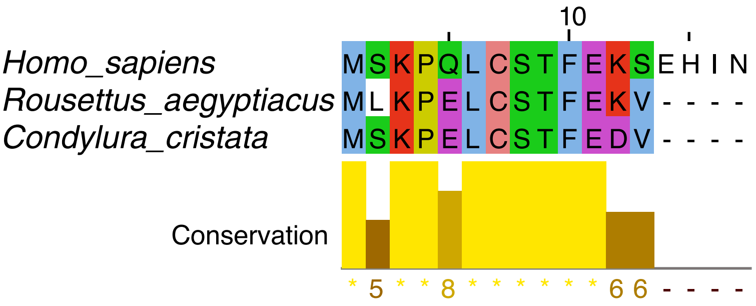

**HG1189**

ENSG00000279909

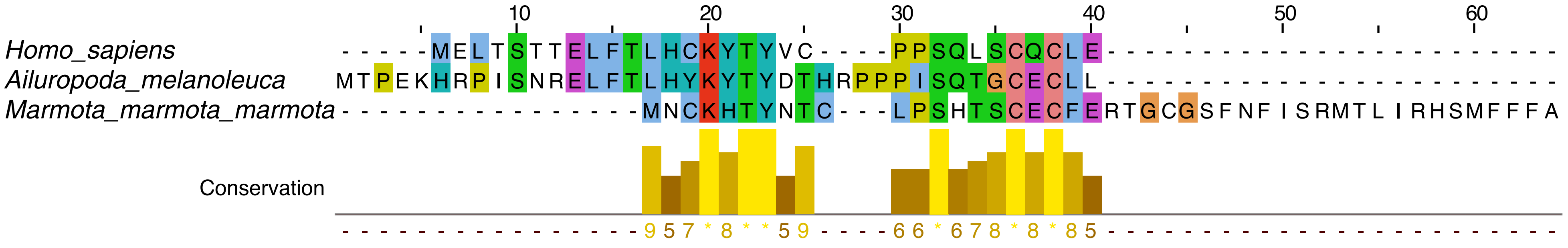

**HG1190**

ENSG00000123684 (LPGAT1)

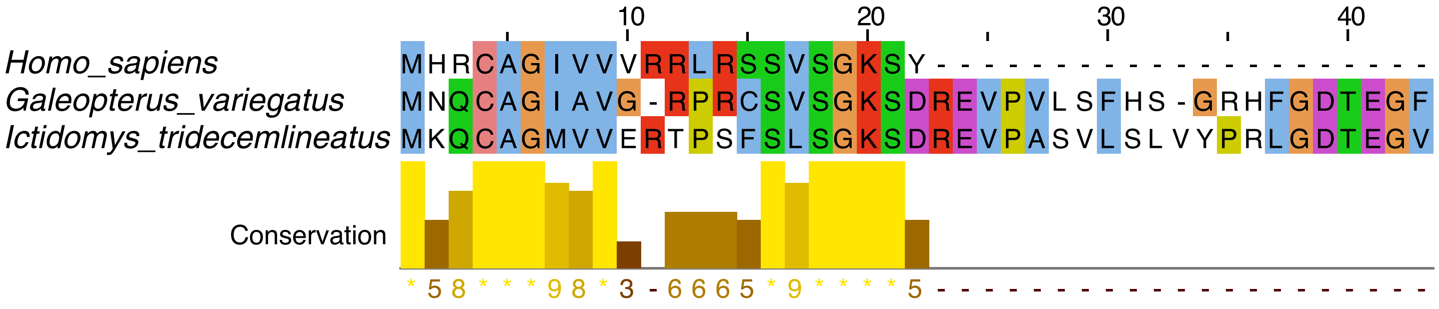

**HG1191**

ENSG00000164331 (ANKRA2)

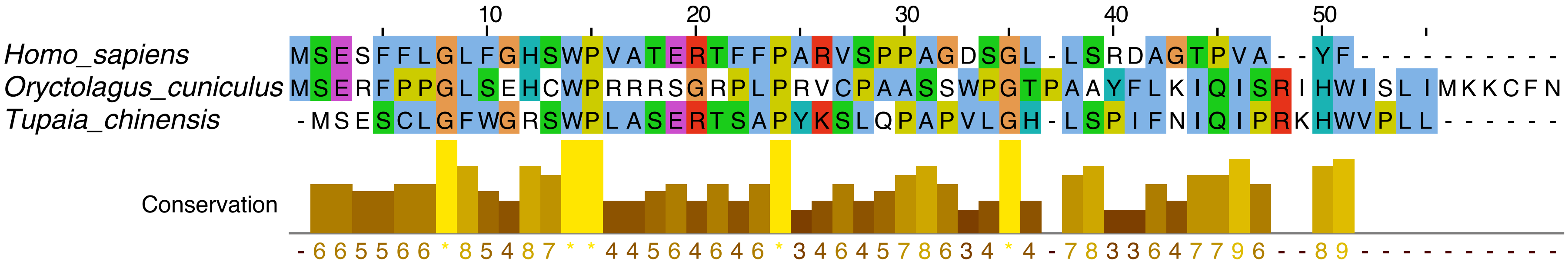

**HG1192**

ENSG00000168758 (SEMA4C)

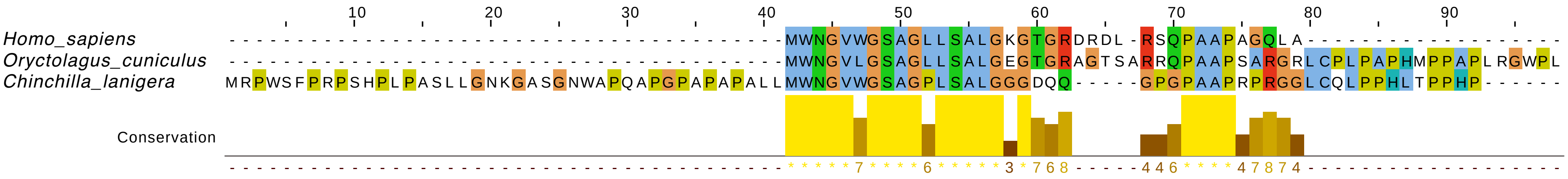

**HG1193**

ENSG00000177706 (FAM20C)

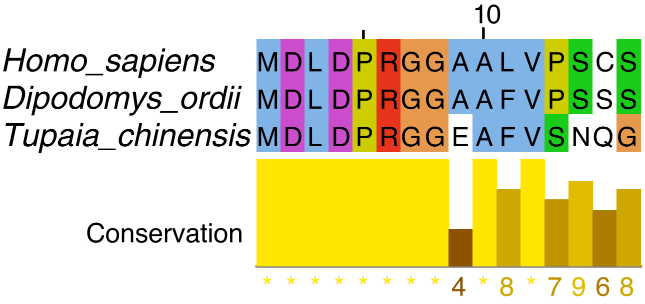

**HG1194**

ENSG00000214694 (ARHGEF33)

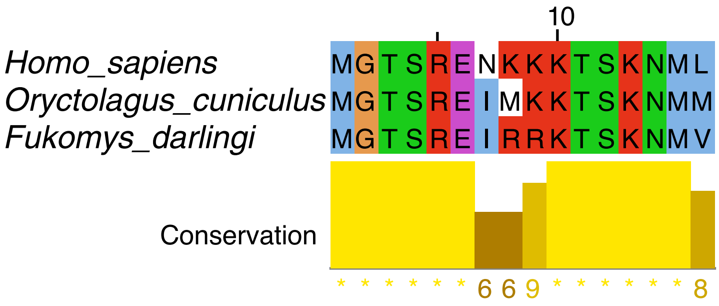


**Supplementary Figure S4.** Alignments of the extracted CPuORF sequences by ESUCA. The amino acid sequences of the extracted CPuORFs and their homologous putative uORFs were aligned using Clustal Ω ver. 1.2.2 and displayed using Jalview ver. 2.10.2. For each alignment, a putative uORF sequence was selected from each order in which uORF-tBLASTn and mORF-tBLASTn hits were found. In the alignments of the ENSG00000112245 (HG0001), ENSDARG00000003077 (HG0002), ENSGALG00000041706 (HG0004.1), ENSG00000031003 (HG0004.1), ENSGALG00000009013 (HG0010.1), ENSG00000189266 (HG0018), ENSDARG00000087059 (HG0026), ENSG00000172995 (HG0043), ENSDARG00000061687 (HG0044), ENSG00000139329 (HG0056), ENSG00000280987 (HG0057), ENSDARG00000018060 (HG0059.1), ENSDARG00000040008 (HG0076), ENSG00000164600 (HG0076), ENSG00000204406 (HG0079), ENSGALG00000016176 (HG0105), ENSG00000181690 (HG0125.2), ENSG00000017427 (HG0135), ENSG00000180530 (HG0170), ENSG00000177311 (HG0171.2), ENSG00000023445 (HG0262.1), ENSG00000204256 (HG0274.1), ENSG00000100239 (HG0275.1), ENSG00000175928 (HG0279.1), ENSGALG00000007717 (HG0403.1), ENSGALG00000026736 (HG0436), ENSG00000180008 (HG0460), ENSG00000166444 (HG0484), ENSG00000144460 (HG0504), ENSG00000177853 (HG0506.1), ENSG00000126860 (HG0513), ENSG00000273841 (HG0556), ENSG00000010818 (HG0579), ENSG00000164930 (HG0599), ENSG00000003147 (HG0604.2), ENSG00000074657 (HG0607), ENSG00000122203 (HG0614), ENSG00000259332 (HG0635), ENSG00000100359 (HG0641), ENSG00000166845 (HG0661), ENSG00000111837 (HG0683), ENSG00000085998 (HG0719), ENSG00000104783 (HG0724), ENSG00000115825 (HG0727), ENSG00000163794 (HG0816), ENSG00000127324 (HG0946.2), ENSG00000136235 (HG0956), ENSG00000154319 (HG0968), ENSG00000065357 (HG1022), ENSG00000082175 (HG1030), ENSG00000084453 (HG1032), ENSG00000140030 (HG1065), ENSG00000166532 (HG1085), ENSG00000174371 (HG1093), ENSG00000177917 (HG1094), ENSG00000092020 (HG1119), ENSG00000101574 (HG1125) CPuORFs, some putative uORF sequences were manually removed. After manual removal, $K_a/K_s$ analysis was conducted using the putative uORF sequences shown in the alignments.
