## Supplementary Figure S5 for "Exhaustive identification of conserved upstream open reading frames with potential translational regulatory functions from animal genomes"

```
HG0001 ENSDARG00000006242 ptp4a1

>Danio_rerio
-------------------------------------------------------------------MIKLQSSMSKHI-P--FCGVLGHTFMEF-LKGSGDYCQ--AQHGVYAEK-----
>Acipenser_sinensis
-------------------------------------------------------------------MIKLQSSMSKHIPQ--FCGVLGHTFMEF-LKGSGDYCQ--AQHEVYADK-----
>Actinia_tenebrosa
--------------------------------------------------------------------------MWT-TNI--CYDHCGHSLMEF-LIDSGDWCQ--AQHN----------
>Acanthogorgia_aspera
--------------------------------------------------------------------------MSR-AYCLFGFRSRFEKFMEYFLKDSGEFCQ--AQHE----------
>Amia_calva
-------------------------------------------------------------------MIKLQSSMSKHIPQ--FCGVLGHTFMEF-LKGSGDYCQ--AQHDVYADK-----
>Channa_argus
MS-------------------QQTEEAPP----QQQFQISDENSSC-HIEEEQKGDLTVVSALPPLSLRSHGISMFILLFG--FCVDLSRTYMAC-V--SGEFCQ--AQHLTSVEK-----
>Kaupichthys_hyoproroides
-------------------------------------------------------------------MIKLQSSMSKHIPQ--FCGVLGHTFMEF-LKDSGDYCQ--AQHEVYADK-----
>Anser_cygnoides
--------------------------------------------------------------------------------------------MEF-LKGSGDYCQ--AQHDLYADK-----
>Podocoryna_carnea
--------------------------------------------------------------------------------------------MEF-LRGSGDYCF--AQHRT---------
>Xenopus_tropicalis
MLFTGHGEEPQGRRSGPRQLSPDHRALRL---GDSSLL-YAPSPPAPLLFHRLFL--------GTGCMIKIQSSMSKHS-P--FCGVLGHTFMEF-LKGSGDYCQ--AQHELYADK-----
>Chaetura_pelagica
--------------------------------------------------------------------------------------------MEF-LKGSGDYCQ--AQHDLYADK-----
>Apteryx_australis_mantelli
--------------------------------------------------------------------------------------------MEF-LKGSGDYCQ--AQHDLYADK-----
>Stegodyphus_lineatus
---------------------------------------------------------------------MLDASMC-IDIA--VNQIQTFLLMEV-LVGSDEYCR--AQHK----------
>Meinertellus_cundinamarcensi
--------------------------------------------------------------------------MCR-DLV--FLGETGLTYMEF-QLDSGEYCQ--AQHK----------
>Scapharca_broughtonii
--------------------------------------------------------------------------MFR-KIL--FFGEFGHTFMEF-VTNSGDYCQ--AQHV----------
>Porichthys_notatus
-------------------------------------------------------------------MIKLQSSMSKHIPQ--FCGVLGHTFMEF-LKGRGDYCQ--AQHAADADE-----
>Oryzias_melastigma
-------------------------------------------------------------------MIKLQSSMSKHIPQ--FCGVLGHTFMEF-LKGSGDYCQ--AQHAAYADE-----
>Tripterygion_delaisi
--------------------------------------------------------------------------MFSLLFG--FCVDLSRTYMAC-V--SGEFCQ--AQHLTSVEK-----
>Buceros_rhinoceros_silvestri
--------------------------------------------------------------------------------------------MEF-LKGSGDYCQ--AQHDLYADK-----
>Caprimulgus_carolinensis
--------------------------------------------------------------------------------------------MEF-LKGSGDYCQ--AQHDLYADK-----
>Scyliorhinus_torazame
--------------------------------------------------------------------------------------------MEF-LKGSGDYCQ--AQHAVYADK-----
>Cariama_cristata
--------------------------------------------------------------------------------------------MEF-LKGSGDYCQ--AQHDLYADK-----
>Canis_lupus_familiaris
-------------------------------------------------------------------MIRPQSSMSKHI-P--FCGVLGHTFMEF-LKGSGDYCQ--AQHDLYADK-----
>Alatina_alata
---------------------------------------------------------------------MRRPCLC-T-------CFCGKTFMEF-LKDSGDWCQ--SQHK----------
>Hynobius_retardatus
MRAPELGRSEARAQCGGGSVAADTATLPPRSPAARPVW-YTARRACP--PATRAR--------GTGRMIQPRSSMSNHSPQ--FCGVLGHTFMEF-LKGSGDYCQ--AQHDLYADK-----
>Micropterus_floridanus
--------------------------------------------------------------------------------------------MEF-VRRSGDYCH--AQHI----------
>Balaenoptera_acutorostrata_s
--------------------------------------------------------------------------------------------MEF-LKGSGDYCQ--AQHDLYADK-----
>Astyanax_mexicanus
-------------------------------------------------------------------MVKLQSSMSKHIPQ--FCGVLGHTFMEF-LKGSGDYCQ--AQHGVYADK-----
>Calidris_pugnax
--------------------------------------------------------------------------------------------MEF-LKGSGDYCQ--AQHDLYADK-----
>Callorhinchus_milii
--------------------------------------------------------------------------------------------MEF-LKGSGDYCQ--AQHDVYADK-----
>Miniopterus_natalensis
-------------------------------------------------------------------MIRPQSSMSKHI-P--FCGVLGHTFMEF-LKGSGDYCQ--AQHDLYADK-----
>Haplochromis_burtoni
-------------------------------------------------------------------MIKLQSSMSKHIPQ--FCGVLGHTFMEF-LKGSGDYCQ--AQHAAYADE-----
>Pterocles_gutturalis
--------------------------------------------------------------------------------------------MEF-LKGSGDYCQ--AQHDLYADK-----
>Eucidaris_tribuloides
--------------------------------------------------------------------------MCK-HTE--VLAVINKPFMAL-VLDSGEFCQ--SQHK----------
>Dasypus_novemcinctus
-------------------------------------------------------------------MIRPQSSMSKHI-P--FCGVLGHTFMEF-LKGSGDYCQ--AQHDLYADK-----
>Alosa_alosa
-------------------------------------------------------------------MIKLQSSMSKHIPQ--FCGVLGHTFMEF-LKGSGDYCQ--AQHGVYADE-----
>Latimeria_menadoensis
-------------------------------------------------------------------MIKIQSSMSKHIPQ--FCGVLGHTFMEF-LKGSGDYCQ--AQHDLYADK-----
>Pogonus_chalceus
--------------------------------------------------------------------------MCM-ALV--FLGEKVFAFMEF-QLDSAEYCQ--AQHK----------
>Colius_striatus
--------------------------------------------------------------------------------------------MEF-LKGSGDYCQ--AQHDLYADK-----
>Columba_livia
--------------------------------------------------------------------------------------------MEF-LKGSGDYCQ--AQHDLYADK-----
>Leptosomus_discolor
--------------------------------------------------------------------------------------------MEF-LKGSGDYCQ--AQHDLYADK-----
>Ricordea_yuma
--------------------------------------------------------------------------MWR-GNP--VYGNRAHTFMEF-IIDSGDWCQ--AQHE----------
>Alligator_mississippiensis
-------------------------------------------------------------------MIRLQSSMSNRR-P--FCGVLGHTFMEF-LKGSGDYCQ--AQHDLYADK-----
>Cuculus_canorus
--------------------------------------------------------------------------------------------MEF-LKGSGDYCQ--AQHDLYADK-----
>Poecilia_formosa
-------------------------------------------------------------------MIKLQSSMSKHIP---FCGVLGRTFMEF-LKGSGDYCQ--AQHAAYADE-----
>Sarcophilus_harrisii
--------------------------------------------------------------------------------------------MEF-LKGSGDYCQ--AQHDLYADK-----
>Macrobrachium_nipponense
--------------------------------------------------------------------MKFKYTRS-NNAQ--LSSGRGYTLMEL-LKGSGEFCK--AQYHIK--------
>Forficula_auricularia
--------------------------------------------------------------------------------------------MEF-RLDSAEYCQ--AQHK----------
>Galeopterus_variegatus
--------------------------------------------------------------------------------------------MEF-LKGSGDYCQ--AQHDLYADK-----
>Monodelphis_domestica
-------------------------------------------------------------------MIRPQSSMSKHI-P--FCGVLGHTFMEF-LKGSGDYCQ--AQHDLYADK-----
>Occasjapyx_japonicus
--------------------------------------------------------------------------MCR-DLV--CLGKIGLTFMEF-FLDSGEYCQ--AQHK----------
>Macropus_eugenii
----------------------------------------------------------------------------MKAKQ--FCGVLGHTFMEF-LKGSGDYCQ--AQHDLYADK-----
>Arachnocampa_luminosa
--------------------------------------------------------------------------------------------MEF-HLDSAEYCQ--AQHK----------
>Eurylophella_sp._AD-2013
--------------------------------------------------------------------------MCR-DLV--FLGEIGLTYMEF-QLDSGEYCQ--AQHK----------
>Esox_lucius
-------------------------------------------------------------------MIKLQSSMSKHIP---FCGVLGHTFMEF-LKGSGDYCQ--AQHVAAYADE----
>Meganyctiphanes_norvegica
--------------------------------------------------------------------MKFKYCGN-NNTQ--LSCGRGYTLMEL-LKGSGEFCK--AQYHIK--------
>Haliaeetus_leucocephalus
-------------------------------------------------------------------MIRLQSSMSNHIPQ--FCGVLGHTFMEF-LKGSGDYCQ--AQHDLYADK-----
>Coturnix_japonica
-------------------------------------------------------------------MIRLQSSMSNHIP---FCGVLGHTFMEF-LKGSGDYCQ--AQHDLYADK-----
>Gavia_stellata
--------------------------------------------------------------------------------------------MEF-LKGSGDYCQ--AQHDLYADK-----
>Acanthogobius_hasta
-------------------------------------------------------------------MIKLQSSMSKHIPQ--FCGVLGHTFMEF-LKGSGDYCQ--AQHVVDADE-----
>Eurypyga_helias
--------------------------------------------------------------------------------------------MEF-LKGSGDYCQ--AQHDLYADK-----
>Apteronotus_leptorhynchus
--------------------------------------------------------------------------------------------MAF-LQRAGDYCH--AQHWCC--------
>Okanagana_villosa
--------------------------------------------------------------------------MCM-AIV--FLGETWLTFMEF-HLDSAEYCQ--AQHK----------
>Megachile_rotundata
--------------------------------------------------------------------------------------------MEF-QLDSAEYCQ--AQHK----------
>Erinaceus_europaeus
-------------------------------------------------------------------MIKPQSSMSKHIP---FCGVLGHTFMEF-LKGSGDYCQ--AQHDLYADK-----
>Oryctolagus_cuniculus
-------------------------------------------------------------------MIRPQSSMSKHI-P--FCGVLGHTFMEF-LKGSGDYCQ--AQHDLYADK-----
>Rhodinia_newara
--------------------------------------------------------------------------------------------MEF-HLDSAEYCQ--AQHK----------
>Protopterus_aethiopicus
-------------------------------------------------------------------MIKLQSSMSKHIPQ--FCGVIGHTFMEF-LKGSGDYCQ--AQHALYADK-----
>Lingula_anatina
-------------------------------------------------------------------------------MQ--FFGEFGHTFMEF-VTNSGEYCQ--AQHK----------
>Elephantulus_edwardii
-------------------------------------------------------------------MIRPQSSMSKHIPQ--FCGVLGHTFMEF-LKGSGDYCQ--AQHDLYADK-----
>Metallyticus_splendidus
--------------------------------------------------------------------------MCM-ALV--FLGETVLTYMEF-QLDSGEYCQ--AQHK----------
>Boreus_hyemalis
--------------------------------------------------------------------------------------------MEF-QLDSAEYCQ--AQHK----------
>Ornithorhynchus_anatinus
-------------------------------------------------------------------MIATHSSMSKHS-P--FCGVLGHTFMEF-LKDSGDYCQ--AQHDLYADK-----
>Cypridininae_sp._BMR-2011
-------------------------------------------------------------MEDSETSLLYKYRLCM-RPQ--VLAGVCCSFMEF-VLADGEYCQ--AQHK----------
>Perna_viridis
--------------------------------------------------------------------------MCR-NIL--FFGVLGHTFMEF-VTNSGDYCQ--AQHV----------
>Eptatretus_burgeri
--------------------------------------------------------------------------MNRHNAQ--FCGVLGRTFMEF-VKGRGEYCE--TQHIALADK-----
>Conwentzia_psociformis
---------------------------------------------------------------------------MV-NFN--SFSERVLTYMEF-QLDSAEYCQ--AQHK----------
>Triops_newberryi
--------------------------------------------------------------------------MCT-TGQ--RWVEWRHKFLAV-LLGSGDYCE--AQHK----------
>Hapalochlaena_maculosa
--------------------------------------------MC-RNILDLLGGI--VEKTDSPSTLAWPHHLCR-SIK--FFGEFGHTFMEF-VTNSGDFCQ--AQHV----------
>Epiophlebia_superstes
--------------------------------------------------------------------------------------------MEF-QLDSGEYCQ--AQHK----------
>Opisthocomus_hoazin
--------------------------------------------------------------------------------------------MEF-LKGSGDYCQ--AQHDLYADK-----
>Stenobothrus_lineatus
MV-------------------V------E----ASAVALADEGRTAGRHVMRLERAS--SPSSMVDSQAAGGGRLCA-HQQ--FRGEIVLTFMEF-QLDSGEYCQ--AQHK----------
>Plecoglossus_altivelis
--------------------------------------------------------------------------------------------MEF-LQGSGDYCH--AQHPSI--------
>Scleropages_formosus
--------------------------------------------------------------------------MCGLLFR--FCVDLGRPFMAF-VSVSGDFCQ--AQHLASADK-----
>Crassostrea_gigas
------------------------------------------MRDP-RGPVTVRGKF--VTGFV---LCSSLLNCWQ-HTE--FRSNFGHTYMEF-VTNSGDYCQ--AQHV----------
>Endeis_spinosa
--------------------------------------------------------------------------------------------MEF-LKESGDYCYCRAQHV----------
>Zonotrichia_albicollis
-------------------------------------------------------------------MIRLQSSMSNHI-P--FCGVLGHTFMEF-LKGSGDYCQ--AQHDLYADK-----
>Pecten_maximus
--------------------------------------------------------------------------------------------MEF-VTNSGDYCQ--AQHAL---------
>Pelecanus_crispus
--------------------------------------------------------------------------------------------MEF-LKGSGDYCQ--AQHDLYADK-----
>Sebastes_caurinus
-------------------------------------------------------------------MIKLQSSMSKHIP---FCGVLGHTFMEF-LKGSGDYCQ--AQHAAYADE-----
>Equus_przewalskii
-------------------------------------------------------------------MIRPQSSMSKHSPQ--FCGVLGHTFMEF-LKGSGDYCQ--AQHDLYADK-----
>Peruphasma_schultei
--------------------------------------------------------------------------------------------MEF-LPNSGEYCQ--AQHK----------
>Menopon_gallinae
--------------------------------------------------------------------MNFRFSGCM-ALK--NFHSTSSRLMEF-FLDSAEYCQ--AQHTK---------
>Glycera_dibranchiata
-------------------------------------------------------------ME------QCHPVLKT-HSK--FFGVLGHTFMEF-LCDSGEYCQ--AQHV----------
>Leuctra_sp._AD-2013
--------------------------------------------------------------------------MCM-ALV--LVGEIVLTYMEF-QLDSGEYCQ--AQHRK---------
>Cynoglossus_semilaevis
--------------------------------------------------------------------------------------------MEF-LKGSGDYCQ--AQHAAYADE-----
>Gorilla_gorilla
-------------------------------------------------------------------MIRPQSSMSKHIPQ--FCGVLGHTFMEF-LKGSGDYCQ--AQHDLYADK-----
>Loxodonta_africana
-------------------------------------------------------------------MIRPQSSMSKHI-P--FCGVLGHTFMEF-LKGSGDYCQ--AQHDLYADK-----
>Fulmarus_glacialis
--------------------------------------------------------------------------------------------MEF-LKGSGDYCQ--AQHDLYADK-----
>Nestor_notabilis
--------------------------------------------------------------------------------------------MEF-LKGSGDYCQ--AQHDLYADK-----
>Pinctada_martensii
--------------------------------------------------------------------------MCR-NIV--FFGEYGHTFMEF-VTNSGDYCQ--AQHK----------
>Leucoraja_erinacea
--------------------------------------------------------------------------MRKHIPQ--FCGVLGHTFMEF-LKGSGDYCQ--AQHDVYADK-----
>Inocellia_crassicornis
--------------------------------------------------------------------------------------------MEF-QLDSAEYCQ--AQHK----------
>Rhopilema_esculentum
-------------------------------------------------------------------------MNS-YHLL--LSKYFGKPFMEF-LKDSAEWCK--AQHVE---------
>Rattus_norvegicus
-------------------------------------------------------------------MIRPQSSMSRHI-P--FCGVLGHTFMEF-LKGSGDYCQ--AQHDLYADK-----
>Salmo_salar
-------------------------------------------------------------------MIKLQSSMSKHIPQ--FCGVLGHTFMEF-LKGSGDYCQ--AQHAAAYADE----
>Acropora_digitifera
--------------------------------------------------------------------------MWR-FSL--TFGSRAHTFMEF-IIDSGDWCQ--AQHE----------
>Tityus_serrulatus
--------------------------------------------------------------------------------------------MEY-LKTTGEYCR--AQHK----------
>Aurelia_aurita
-------------------------------------------------------------------------MTS-YHLQ--L-KCFGETFMEF-LKDSAEWCK--AQHVE---------
>Lepisosteus_oculatus
-------------------------------------------------------------------MIKLQSSMSKHIPQ--FCGVLGHTFMEF-LKGSGDYCQ--AQHGVYADK-----
>Sepia_pharaonis
--------------------------------------------MC-RNILDLLGGI--VKKTDSPSTLAWPHHLWR-LVK--FFGEFGHTFMEF-VTNSGDFCQ--AQHV----------
>Sepioloidea_lineolata
--------------------------------------------------------------------------------------------MEF-VTNSGDFCQ--AQHV----------
>Ictalurus_punctatus
-------------------------------------------------------------------MVKLQSSMSKHIPQ--FCGVLGHTFMEF-LKGSGDYCQ--AQHGVYADK-----
>Oropsylla_silantiewi
--------------------------------------------------------------------------------------------MEF-QLDSAEYCQ--AQHK----------
>Trichechus_manatus_latirostr
-------------------------------------------------------------------MIRPQSSMSKHIPQ--FCGVLGHTFMEF-LKGSGDYCQ--AQHDLYADK-----
>Aptenodytes_forsteri
--------------------------------------------------------------------------------------------MEF-LKGSGDYCQ--AQHDLYADK-----
>Streblospio_benedicti
--------------------------------------------------------------------------MCR-IIL--FFGQIGYTFMEF-LCNSGEYCQ--AQHVAIPSVLSRLH
>Squalus_acanthias
--------------------------------------------------------------------------MNRLVTQ--FSGISGCSFMEF-LKGSRDYCQ--AQHGLLADK-----
>Python_bivittatus
-------------------------------------------------------------------MIRLQSSMSNHI-P--FCGVLGHTFMEF-LKGSGDYCQ--AQHGLYADK-----
>Stylops_melittae
--------------------------------------------------------------------------MCI-GIL--NLGNKVVEYMEF-HFGSSEYCK--AQHK----------
>Tyto_alba
--------------------------------------------------------------------------------------------MEF-LKGSGDYCQ--AQHDLYADK-----
>Struthio_camelus_australis
--------------------------------------------------------------------------------------------MEF-LKGSGDYCQ--AQHDLYADK-----
>Deroceras_reticulatum
--------------------------------------------------------------------------MSS-FVL--FVGEFGHTFMEF-VTNSEDFCQ--AQHYRLVE------
>Pelodiscus_sinensis
-------------------------------------------------------------------MIRLQSSMSNHI-P--FCGVLGHTFMEF-LKGSGDYCQ--AQHDLYADK-----
>Takifugu_rubripes
-------------------------------------------------------------------MIKLQSSMSKHIPQ--FCGVLGHTFMEF-LKGSGDYCQ--AQHAVYADE-----
>Watasenia_scintillans
--------------------------------------------------------------------------------------------MEF-VTNSGDFCQ--AQHV----------
>Gynaikothrips_ficorum
----------------------------------------------MVHCAKFNGEN--SISMVIRRVIFDNKWPSV-GYQ--TCGEILPTFMEF-HLDSAEYCQ--AQHK----------
>Tinamus_guttatus
--------------------------------------------------------------------------------------------MEF-LKGSGDYCQ--AQHDLYADK-----
>Tetronarce_californica
--------------------------------------------------------------------------------------------MEF-VRNSSGYCP--AQHSIFADK-----
>Hydroptila_sp._AD-2013
--------------------------------------------------------------------------------------------MEF-RLDSGEYCQ--AQHK----------
>Calypte_anna
--------------------------------------------------------------------------------------------MEF-LKGSGDYCQ--AQHDLYADK-----
>Apaloderma_vittatum
--------------------------------------------------------------------------------------------MEF-LKGSGDYCQ--AQHDLYADK-----
>Orycteropus_afer_afer
-------------------------------------------------------------------MIRPQSSMSKHIPQ--FCGVLGHTFMEF-LKGSGDYCQ--AQHDLYADK-----
>Ruditapes_philippinarum
--------------------------------------------------------------------------MNR-IIL--FFGELGHTFMEF-LTNSGEYCL--AQHVHV--------
>Limulus_polyphemus
--------------------------------------------------------------------------------------------MEF-LKTAGEYCQ--SQHK----------
>Palythoa_caribaeorum
--------------------------------------------------------------------------MR--RDY--LAYDRGRTLMEF-LIDSGDWCQ--AQHE----------
>Thermobia_domestica
--------------------------------------------------------------------------------------------MEF-QLDSGEYCQ--AQHK----------
```

HG0001 ENSDARG00000035676 ptp4a2b

```
>Danio_rerio
---------------------------------------------------------------------MVGPFFCFRVDSDHTYMAFVTVCREFCQAQHLTFADK-
>Acipenser_sinensis
-------------------------------------------------------------M-IKLQSSMSKHIPQFCGVLGHTFMEFLKGSGDYCQAQHEVYADK-
>Amia_calva
-------------------------------------------------------------M-IKLQSSMSKHIPQFCGVLGHTFMEFLKGSGDYCQAQHDVYADK-
>Channa_argus
MS-QQTEEAPPQQ---QFQISDENSSCHIEEEQKGDLTV-VSALPPLS-------------L-RSHGISMFILLFGFCVDLSRTYMACV--SGEFCQAQHLTSVEK-
>Kaupichthys_hyoproroides
-------------------------------------------------------------M-IKLQSSMSKHIPQFCGVLGHTFMEFLKDSGDYCQAQHEVYADK-
>Anas_platyrhynchos
-------------------------------------------------------------------------------------MAILSVRADFCQAQHSILADK-
>Podocoryna_carnea
---------------------------------------------------------------------MNSLLSCFCVDLGRTFMAFLSVSGDFCQAQHLAFADK-
>Xenopus_tropicalis
MLFTGHGEEPQGRRSGPRQLSPDHRA-----LRLGDSSLLYAPSPPAPLLFHR-LFLGTGCM-IKIQSSMSKH-SPFCGVLGHTFMEFLKGSGDYCQAQHELYADK-
>Chaetura_pelagica
-------------------------------------------------------------------------------------MEFLKGSGDYCQAQHDLYADK-
>Apteryx_australis_mantelli
-------------------------------------------------------------------------------------MAILSVRADFCQAQHSTLADK-
>Scapharca_broughtonii
----------------------------------------------------------------------MFRKILFFGEFGHTFMEFVTNSGDYCQAQHV------
>Menidia_menidia
---------------------------------------------------------------------MFNLLFGFCVDVSRSYMACV--TGEFCQAQHLTSAEK-
>Porichthys_notatus
-------------------------------------------------------------M-IKLQSSMSKHIPQFCGVLGHTFMEFLKGRGDYCQAQHAADADE-
>Oryzias_latipes
---------------------------------------------------------------------MFSLFFGFCVDSSRTYMACV--TGEFCQAQHLTSAEK-
>Tripterygion_delaisi
-----------------------------------------------MVCLHPGA---------VTFSCIHSMNPSY--QHSYTCMAFARLGGEFCQAQHLLIIVDK
>Buceros_rhinoceros_silvestri
-------------------------------------------------------------------------------------MEFLKGSGDYCQAQHDLYADK-
>Caprimulgus_carolinensis
-------------------------------------------------------------------------------------MEFLKGSGDYCQAQHDLYADK-
>Trachinotus_blochii
-----------------------------------------------MVCLHPGA---------VTFSCVRSMNHLLSSHHSCTCMAFARLGGEFCQAQHLLLIVDK
>Scyliorhinus_torazame
-------------------------------------------------------------------------------------MEFLKGSGDYCQAQHAVYADK-
>Cariama_cristata
-------------------------------------------------------------------------------------MAILSVRADFCQAQHSILADK-
>Mustela_putorius_furo
-------------------------------------------------------------------------------------MAILSVRADFCQAQHSVFADK-
>Hynobius_chinensis
-------------------------------------------------------------M-IQPRSSMSNHSPQFCGVLGHTFMEFLKGSGDYCQAQHDLYADK-
>Micropterus_floridanus
-----------------------------------------------MVCLHPGA---------VTFSCVRSMNHLLSSHQSCTCMAFVKLGGEFCQAQHLLLIVDK
>Orcinus_orca
-------------------------------------------------------------------MPSPQLASVFRWNIRCTFMAILSVRADFCQAQHSVSADK-
>Astyanax_mexicanus
---------------------------------------------------------------------MFSLFFCFCVDLGRTFMAFV--SGDFCQAQHLSFENK-
>Calidris_pugnax
-------------------------------------------------------------------------------------MAILSVRADFCQAQHSILADK-
>Callorhinchus_milii
-------------------------------------------------------------------------------------MEFLKGSGDYCQAQHDVYADK-
>Rousettus_aegyptiacus
-------------------------------------------------------------------MPSPQLASVFRWNIRCTFMAILSVRADFCQAQHSVFADK-
>Haplochromis_burtoni
-------------------------------------------------------------------------------------MACV--SGEFCQAQHLTSVEK-
>Pterocles_gutturalis
-------------------------------------------------------------------------------------MEFLKGSGDYCQAQHDLYADK-
>Dasypus_novemcinctus
-------------------------------------------------------------------MPSPQLASVFRWNIRFTFMAILSVRADFCQAQHSVFADK-
>Clupea_harengus
---------------------------------------------------------------------MFNLLFCFCVELGRTYMAFVSVSGDFCQAQHLAFVDK-
>Latimeria_chalumnae
-------------------------------------------------------------M-IKIQSSMSKHIPQFCGVLGHTFMEFLKGSGDYCQAQHDLYADK-
>Colius_striatus
-------------------------------------------------------------------------------------MEFLKGSGDYCQAQHDLYADK-
>Columba_livia
-------------------------------------------------------------------------------------MAILSVRADFCQAQHSTLADK-
>Merops_nubicus
-------------------------------------------------------------------------------------MAILSVRADFCQAQHSPLADK-
>Corynactis_australis
----------------------------------------------------------------------MWRGNRVYGNRAHTFMEFIIDSGDWCQAQHE------
>Alligator_sinensis
-------------------------------------------------------------------------------------MAILSVRADFCQAQHSTLADK-
>Cuculus_canorus
-------------------------------------------------------------------------------------MEFLKGSGDYCQAQHDLYADK-
>Austrofundulus_limnaeus
---------------------------------------------------------------------MFSLLFGFCVDFNRTYMACV--TGEFCQAQHLTSVEK-
>Sarcophilus_harrisii
-------------------------------------------------------------------------------------MEFLKGSGDYCQAQHDLYADK-
>Galeopterus_variegatus
-------------------------------------------------------------------MPSPQLASVFRWNIRCTFMAILSVRADFCQAQHSVFADK-
>Monodelphis_domestica
-------------------------------------------------------------------------------------MAILSVRADFCQAQHSAFADK-
>Macropus_eugenii
-----------------------------------------------------------------------MKAKQFCGVLGHTFMEFLKGSGDYCQAQHDLYADK-
>Esox_lucius
-------------------------------------------------------------------------------------MAYVSVSGDFCQAQHLAFVDK-
>Haliaeetus_leucocephalus
------------------------------------------------------------------------MPRVFCWNIRRTFMAILSVRADFCQAQHSILADK-
>Asterias_amurensis
----------------------------------------------------------------------MSHLTGISSVPVSSYMAFVAISGEYCRSQHK------
>Coturnix_japonica
------------------------------------------------------------------------MPGVFWWNSRRTFMAILSVRAEFCQAQHSALADK-
>Gavia_stellata
-------------------------------------------------------------------------------------MAILSVRADFCQAQHSILADK-
>Acanthogobius_hasta
-------------------------------------------------------------M-IKLQSSMSKHIPQFCGVLGHTFMEFLKGSGDYCQAQHVVDADE-
>Chlamydotis_macqueenii
-------------------------------------------------------------------------------------MAILSVRADFCQAQHRILADK-
>Apteronotus_leptorhynchus
-------------------------------------------------------------------------------------MAFV--SGDFCQAQHLAFVNK-
>Glossoscolex_paulistus
----------------------------------------------------------------------MYTIASLSKASGHHLMALVYLCGEFCQAQHV------
>Cerapachys_biroi
-------------------------------------------------------------------------------------MEFQLDSAEYCQAQHK------
>Sorex_araneus
-------------------------------------------------------------------------------------MAILSVRADFCQAQHSVFADK-
>Ochotona_princeps
-------------------------------------------------------------------------------------MEFLKGSGDYCQAQHDLYADK-
>Protopterus_aethiopicus
-------------------------------------------------------------M-IKLQSSMSKHIPQFCGVIGHTFMEFLKGSGDYCQAQHALYADK-
>Lingula_anatina
-------------------------------------------------------------------------------------MEFVTNSGEYCQAQHK------
>Elephantulus_edwardii
-------------------------------------------------------------M-IRPQSSMSKHIPQFCGVLGHTFMEFLKGSGDYCQAQHDLYADK-
>Ornithorhynchus_anatinus
-------------------------------------------------------------M-IATHSSMSKHSPQFCGVLGHTFMEFLKDSGDYCQAQHDLYADK-
>Tauraco_erythrolophus
-------------------------------------------------------------------------------------MAILSVRADFCQAQHSILADK-
>Perna_viridis
----------------------------------------------------------------------MCRNILFFGVLGHTFMEFVTNSGDYCQAQHV------
>Eptatretus_burgeri
---------------------------------------------------------------------MNRHNAQFCGVLGRTFMEFVKGRGEYCETQHIALADK-
>Octopus_bimaculoides
-------------------------------------------------------------------------------------MEFVTNSGDFCQAQHV------
>Opisthocomus_hoazin
-------------------------------------------------------------------------------------MEFLKGSGDYCQAQHDLYADK-
>Scleropages_formosus
---------------------------------------------------------------------MCGLLFRFCVDLGRPFMAFVSVSGDFCQAQHLASADK-
>Crassostrea_gigas
-----------------------------------------MRDPRGPVTVRGKFVTGF---VLCSSLLNCWQHTEFRSNFGHTYMEFVTNSGDYCQAQHV------
>Corvus_brachyrhynchos
-------------------------------------------------------------------------------------MAILSVRADFCQAQHSTLAAK-
>Pecten_maximus
-------------------------------------------------------------------------------------MEFVTNSGDYCQAQHAL-----
>Egretta_garzetta
-------------------------------------------------------------------------------------MAILSVRADFCQAQHSILADK-
>Lateolabrax_maculatus
-----------------------------------------------MVCLHPGA---------VTSSCVRSMNHLLGSHRGRTCMAFARLGGEFCQAQHLLLIVDK
>Notothenia_coriiceps
-------------------------------------------------------------------------------------MACV--SGEFCQAQHLTSVEK-
>Ceratotherium_simum_simum
-------------------------------------------------------------------------------------MAILSVRADFCQAQHSVFADK-
>Perinereis_aibuhitensis
---------------------------------------------------------------MSSEAGVTTKLHQFFGELGHTFMEFLCNSGEYCQAQHV------
>Picoides_pubescens
-------------------------------------------------------------------------------------MAILSVRADFCQAQHSILADK-
>Platichthys_flesus
---------------------------------------------------------------------MFSLLFNFCVDLSHTYMACV--SGEFCQAQHLTSVEK-
>Homo_sapiens
-------------------------------------------------------------------MPSPQLASVFRWNIRCTFMAILSVRADFCQAQHSIFADK-
>Loxodonta_africana
-------------------------------------------------------------------------------------MAILSVRADFCQAQHSIFADK-
>Fulmarus_glacialis
-------------------------------------------------------------------------------------MAILSVRADFCQAQHSILADK-
>Nestor_notabilis
-------------------------------------------------------------------------------------MAILSVRAEFCQAQHSILADK-
>Leucoraja_erinacea
---------------------------------------------------------------------MRKHIPQFCGVLGHTFMEFLKGSGDYCQAQHDVYADK-
>Peromyscus_maniculatus_baird
-------------------------------------------------------------------MPSPQLASVFRWNIRCTFMAILSVRADFCQAQHSVFADK-
>Oncorhynchus_mykiss
---------------------------------------------------------------------MSSLLFCFCVDLDHTYMAYVSVSGDFCQAQHLAFVDK-
>Tupaia_chinensis
-------------------------------------------------------------------------------------MAILSVRADFCQAQHSVFADK-
>Ctenactis_echinata
----------------------------------------------------------------------MWRFNLVYGSRAHTFMEFIIDSGDWCQAQHE------
>Lepisosteus_oculatus
---------------------------------------------------------------------MSSLPFCFCVDLVRTFMAFLSVSGDFCQAQHLAFADK-
>Sepia_pharaonis
-------------------------------------------MCRNILDLLGGIVKKTDSPSTLAWPHHLWRLVKFFGEFGHTFMEFVTNSGDFCQAQHV------
>Sepioloidea_lineolata
-------------------------------------------------------------------------------------MEFVTNSGDFCQAQHV------
>Trichomycterus_areolatus
---------------------------------------------------------------------MFSLFFCFRADLSRTFMAFV--SGDFCHAQHLTIVNK-
>Trichechus_manatus_latirostr
-------------------------------------------------------------------MPSSQLASVFRWNIRCTFMAILSVRADFCQAQHSIFADK-
>Aptenodytes_forsteri
-------------------------------------------------------------------------------------MAILSVRADFCQAQHSILADK-
>Squalus_acanthias
---------------------------------------------------------------------MNRLVTQFSGISGCSFMEFLKGSRDYCQAQHGLLADK-
>Micrurus_fulvius
------------------------------------------------------------------------MLPVFHLNPSHTFMAILSVRADFCRAQHSTLAEK-
>Tyto_alba
-------------------------------------------------------------------------------------MEFLKGSGDYCQAQHDLYADK-
>Struthio_camelus_australis
-------------------------------------------------------------------------------------MAILSVRADFCQAQHSTLADK-
>Deroceras_reticulatum
----------------------------------------------------------------------MSSFVLFVGEFGHTFMEFVTNSEDFCQAQHYRLVE--
>Chrysemys_picta
-------------------------------------------------------------------------------------MAILSIRADFCQAQHSTLADK-
>Takifugu_rubripes
---------------------------------------------------------------------MFSLLFGFSVDLSHTFMACV--SGEFCQAQHLTSAEK-
>Watasenia_scintillans
-------------------------------------------------------------------------------------MEFVTNSGDFCQAQHV------
>Tinamus_guttatus
------------------------------------------------------------------------MPRVFCWNIRRTFMAILRVRADFCQAQHSTLADK-
>Tetronarce_californica
-------------------------------------------------------------------------------------MEFVRNSSGYCPAQHSIFADK-
>Calypte_anna
-------------------------------------------------------------------------------------MEFLKGSGDYCQAQHDLYADK-
>Apaloderma_vittatum
-------------------------------------------------------------------------------------MEFLKGSGDYCQAQHDLYADK-
>Orycteropus_afer_afer
-------------------------------------------------------------M-IRPQSSMSKHIPQFCGVLGHTFMEFLKGSGDYCQAQHDLYADK-
>Eurhomalea_rufa
-------------------------------------------------------------------------------------MEFLTNSGEYCLAQHVHV----
>Limulus_polyphemus
-------------------------------------------------------------MSSTNDSAKVPLLLPFEVRRHHTLMEFLKTAGEYCQSQHK------
```

HG0001 ENSDARG00000039997 ptp4a3

```
>Danio_rerio
-------------------------------------------------------------------------------------------MAFLQRTGDYCHAQHLHI--------
>Acipenser_sinensis
--------------------------------------------------------------------MIKLQSSMSKHIPQFCGVLGHTFMEFLKGSGDYCQAQHEVYADK-----
>Metridium_senile
----------------------------------------------------------------------------MWTGIAYRHSREYTLMEFLTNSGDWCQAQHH----------
>Amia_calva
--------------------------------------------------------------------MIKLQSSMSKHIPQFCGVLGHTFMEFLKGSGDYCQAQHDVYADK-----
>Kaupichthys_hyoproroides
--------------------------------------------------------------------MIKLQSSMSKHIPQFCGVLGHTFMEFLKDSGDYCQAQHEVYADK-----
>Anser_cygnoides
-------------------------------------------------------------------------------------------MEFLKGSGDYCQAQHDLYADK-----
>Podocoryna_carnea
-------------------------------------------------------------------------------------------MEFLRGSGDYCFAQHRT---------
>Xenopus_laevis
--------------------------------------------------------------------MIKIHSSMSKHS-PFCGVLGYTFMEFLKGSGDYCQAQHALYADK-----
>Chaetura_pelagica
-------------------------------------------------------------------------------------------MEFLKGSGDYCQAQHDLYADK-----
>Apteryx_australis_mantelli
-------------------------------------------------------------------------------------------MEFLKGSGDYCQAQHDLYADK-----
>Scapharca_broughtonii
----------------------------------------------------------------------------MFRKILFFGEFGHTFMEFVTNSGDYCQAQHV----------
>Porichthys_notatus
--------------------------------------------------------------------MIKLQSSMSKHIPQFCGVLGHTFMEFLKGRGDYCQAQHAADADE-----
>Oryzias_latipes
-------------------------------------------------------------------------------------------MEFVRQSGDYCHAQHI----------
>Buceros_rhinoceros_silvestri
-------------------------------------------------------------------------------------------MEFLKGSGDYCQAQHDLYADK-----
>Caprimulgus_carolinensis
-------------------------------------------------------------------------------------------MEFLKGSGDYCQAQHDLYADK-----
>Scyliorhinus_torazame
-------------------------------------------------------------------------------------------MEFLKGSGDYCQAQHAVYADK-----
>Cariama_cristata
-------------------------------------------------------------------------------------------MEFLKGSGDYCQAQHDLYADK-----
>Canis_lupus_familiaris
--------------------------------------------------------------------MIRPQSSMSKHI-PFCGVLGHTFMEFLKGSGDYCQAQHDLYADK-----
>Hynobius_retardatus
MRAPELGRSEARAQCGGGSVAADTATLPPRSPAARPVWYTARRACPPAT----------RARGT--GRMIQPRSSMSNHSPQFCGVLGHTFMEFLKGSGDYCQAQHDLYADK-----
>Micropterus_floridanus
-------------------------------------------------------------------------------------------MEFVRRSGDYCHAQHI----------
>Physeter_catodon
-------------------------------------------------------------------------------------------MEFLKGSGDYCQAQHDLYADK-----
>Pygocentrus_nattereri
-------------------------------------------------------------------------------------------MAFLQRAGDYCHAQHFQS--------
>Charadrius_vociferus
-------------------------------------------------------------------------------------------MEFLKGSGDYCQAQHDLYADK-----
>Callorhinchus_milii
-------------------------------------------------------------------------------------------MEFLKGSGDYCQAQHDVYADK-----
>Miniopterus_natalensis
--------------------------------------------------------------------MIRPQSSMSKHI-PFCGVLGHTFMEFLKGSGDYCQAQHDLYADK-----
>Neolamprologus_brichardi
-------------------------------------------------------------------------------------------MEFLKGSGDYCQAQHAAYADE-----
>Pterocles_gutturalis
-------------------------------------------------------------------------------------------MEFLKGSGDYCQAQHDLYADK-----
>Dasypus_novemcinctus
-------------------------------------------------------------------------------------------MEFLKGSGDYCQAQHDLYADK-----
>Clupea_harengus
-------------------------------------------------------------------------------------------MAFLQRAGDYCHAQHPRV--------
>Latimeria_menadoensis
--------------------------------------------------------------------MIKIQSSMSKHIPQFCGVLGHTFMEFLKGSGDYCQAQHDLYADK-----
>Colius_striatus
-------------------------------------------------------------------------------------------MEFLKGSGDYCQAQHDLYADK-----
>Columba_livia
-------------------------------------------------------------------------------------------MEFLKGSGDYCQAQHDLYADK-----
>Leptosomus_discolor
-------------------------------------------------------------------------------------------MEFLKGSGDYCQAQHDLYADK-----
>Alligator_sinensis
-------------------------------------------------------------------------------------------MEFLKGSGDYCQAQHDLYADK-----
>Cuculus_canorus
-------------------------------------------------------------------------------------------MEFLKGSGDYCQAQHDLYADK-----
>Fundulus_heteroclitus
--------------------------------------------------------------------MIKLQSSMSKHIP-FCGVLGRTFMEFLKGSGDYCQAQHAAYADE-----
>Sarcophilus_harrisii
-------------------------------------------------------------------------------------------MEFLKGSGDYCQAQHDLYADK-----
>Galeopterus_variegatus
-------------------------------------------------------------------------------------------MEFLKGSGDYCQAQHDLYADK-----
>Monodelphis_domestica
-------------------------------------------------------------------------------------------MEFLKGSGDYCQAQHDLYADK-----
>Macropus_eugenii
-----------------------------------------------------------------------------MKAKQFCGVLGHTFMEFLKGSGDYCQAQHDLYADK-----
>Esox_lucius
-------------------------------------------------------------------------------------------MAFLQRTGDYCHAQHPRA--------
>Haliaeetus_leucocephalus
--------------------------------------------------------------------MIRLQSSMSNHIPQFCGVLGHTFMEFLKGSGDYCQAQHDLYADK-----
>Leptasterias_sp._AR-2014
---------------------------------------------------------------------MSHS---------TGISSVPVSMAFVSISGEYCRSQHK----------
>Meleagris_gallopavo
-------------------------------------------------------------------------------------------MEFLKGSGDYCQAQHDLYADK-----
>Gavia_stellata
-------------------------------------------------------------------------------------------MEFLKGSGDYCQAQHDLYADK-----
>Acanthogobius_hasta
--------------------------------------------------------------------MIKLQSSMSKHIPQFCGVLGHTFMEFLKGSGDYCQAQHVVDADE-----
>Balearica_regulorum_gibberic
-------------------------------------------------------------------------------------------MEFLKGSGDYCQAQHDLYADK-----
>Apteronotus_leptorhynchus
-------------------------------------------------------------------------------------------MAFLQRAGDYCHAQHWCC--------
>Condylura_cristata
--------------------------------------------------------------------MIRPQSSMSKHI-PFCGVLGHTFMEFLKGSGDYCQAQHDLYADK-----
>Oryctolagus_cuniculus
--------------------------------------------------------------------MIRPQSSMSKHI-PFCGVLGHTFMEFLKGSGDYCQAQHDLYADK-----
>Protopterus_aethiopicus
--------------------------------------------------------------------MIKLQSSMSKHIPQFCGVIGHTFMEFLKGSGDYCQAQHALYADK-----
>Lingula_anatina
--------------------------------------------------------------------------------MQFFGEFGHTFMEFVTNSGEYCQAQHK----------
>Elephantulus_edwardii
--------------------------------------------------------------------MIRPQSSMSKHIPQFCGVLGHTFMEFLKGSGDYCQAQHDLYADK-----
>Ornithorhynchus_anatinus
--------------------------------------------------------------------MIATHSSMSKHS-PFCGVLGHTFMEFLKDSGDYCQAQHDLYADK-----
>Perna_viridis
----------------------------------------------------------------------------MCRNILFFGVLGHTFMEFVTNSGDYCQAQHV----------
>Opisthocomus_hoazin
-------------------------------------------------------------------------------------------MEFLKGSGDYCQAQHDLYADK-----
>Plecoglossus_altivelis
-------------------------------------------------------------------------------------------MEFLQGSGDYCHAQHPSI--------
>Scleropages_formosus
---------------------------------------------------------------------------MCGLLFRFCVDLGRPFMAFVSVSGDFCQAQHLASADK-----
>Crassostrea_gigas
-------------------------MRDPRGP------------------------VTVR-GKFVTGFVLCSSLLNCWQHTEFRSNFGHTYMEFVTNSGDYCQAQHV----------
>Endeis_spinosa
-------------------------------------------------------------------------------------------MEFLKESGDYCYCRAQHV--------
>Lepidothrix_coronata
--------------------------------------------------------------------MIRLQSSMSNHIPQFCGVLGHTFMEFLKGSGDYCQAQHDLYADK-----
>Pecten_maximus
-------------------------------------------------------------------------------------------MEFVTNSGDYCQAQHAL---------
>Egretta_garzetta
-------------------------------------------------------------------------------------------MEFLKGSGDYCQAQHDLYADK-----
>Sebastes_caurinus
--------------------------------------------------------------------MIKLQSSMSKHIP-FCGVLGHTFMEFLKGSGDYCQAQHAAYADE-----
>Equus_przewalskii
--------------------------------------------------------------------MIRPQSSMSKHSPQFCGVLGHTFMEFLKGSGDYCQAQHDLYADK-----
>Extatosoma_tiaratum
-----------------------------------------------------------------------------------------MHMEFLLSSGEYCQAQHK----------
>Perinereis_aibuhitensis
---------------------------------------------------------------------MSSEAGVTTKLHQFFGELGHTFMEFLCNSGEYCQAQHV----------
>Cynoglossus_semilaevis
-------------------------------------------------------------------------------------------MEFLKGSGDYCQAQHAAYADE-----
>Gorilla_gorilla
--------------------------------------------------------------------MIRPQSSMSKHIPQFCGVLGHTFMEFLKGSGDYCQAQHDLYADK-----
>Loxodonta_africana
--------------------------------------------------------------------MIRPQSSMSKHI-PFCGVLGHTFMEFLKGSGDYCQAQHDLYADK-----
>Fulmarus_glacialis
-------------------------------------------------------------------------------------------MEFLKGSGDYCQAQHDLYADK-----
>Melopsittacus_undulatus
-------------------------------------------------------------------------------------------MEFLKGSGDYCQAQHDLYADK-----
>Leucoraja_erinacea
---------------------------------------------------------------------------MRKHIPQFCGVLGHTFMEFLKGSGDYCQAQHDVYADK-----
>Rattus_norvegicus
--------------------------------------------------------------------MIRPQSSMSRHIP-FCGVLGHTFMEFLKGSGDYCQAQHDLYADK-----
>Salmo_salar
-------------------------------------------------------------------------------------------MAFLQRTGDYCHAQHPRA--------
>Tityus_serrulatus
-------------------------------------------------------------------------------------------MEYLKTTGEYCRAQHK----------
>Lepisosteus_oculatus
------------MACEG-----------------SFSWFRAETVISVSSSGSLLRVIPLRDTDF--IRSRKQHDTKVRAAKTSCTVTGGSLMEFLRGSGDYCHAQHHSV--------
>Sepia_pharaonis
-------------MCRN--------ILDLLGG------------------------IVKK-TD---SPSTLAWPHHLWRLVKFFGEFGHTFMEFVTNSGDFCQAQHV----------
>Ictalurus_punctatus
-----------------------------------------------------------------------------------------MLMEFLQGSGDYCHAQHRNA--------
>Trichechus_manatus_latirostr
--------------------------------------------------------------------MIRPQSSMSKHIPQFCGVLGHTFMEFLKGSGDYCQAQHDLYADK-----
>Aptenodytes_forsteri
-------------------------------------------------------------------------------------------MEFLKGSGDYCQAQHDLYADK-----
>Streblospio_benedicti
----------------------------------------------------------------------------MCRIILFFGQIGYTFMEFLCNSGEYCQAQHVAIPSVLSRLH
>Crotalus_horridus
--------------------------------------------------------------------MIRLQSSMSNHIPQFCGVLGHTFMEFLKGSGDYCQAQHGLYADK-----
>Tyto_alba
-------------------------------------------------------------------------------------------MEFLKGSGDYCQAQHDLYADK-----
>Struthio_camelus_australis
-------------------------------------------------------------------------------------------MEFLKGSGDYCQAQHDLYADK-----
>Chrysemys_picta
--------------------------------------------------------------------MIRLQSSMSNHLPQFCGVLGHTFMEFLKGSGDYCQAQHDLYADK-----
>Takifugu_rubripes
--------------------------------------------------------------------MIKLQSSMSKHIPQFCGVLGHTFMEFLKGSGDYCQAQHAVYADE-----
>Tinamus_guttatus
-------------------------------------------------------------------------------------------MEFLKGSGDYCQAQHDLYADK-----
>Calypte_anna
-------------------------------------------------------------------------------------------MEFLKGSGDYCQAQHDLYADK-----
>Apaloderma_vittatum
-------------------------------------------------------------------------------------------MEFLKGSGDYCQAQHDLYADK-----
>Orycteropus_afer_afer
--------------------------------------------------------------------MIRPQSSMSKHIPQFCGVLGHTFMEFLKGSGDYCQAQHDLYADK-----
>Ruditapes_philippinarum
----------------------------------------------------------------------------MNRIILFFGELGHTFMEFLTNSGEYCLAQHVHV--------
>Limulus_polyphemus
-------------------------------------------------------------------------------------------MEFLKTAGEYCQSQHK----------
>Palythoa_caribaeorum
-----------------------------------------------------------------------------MRRDYLAYDRGRTLMEFLIDSGDWCQAQHE----------
```

HG0001 ENSDARG00000054814 PTP4A3

>Danio_rerio
----------------------------------------------------------------------------------MEFWQGSGDYCHAQHHTA--------
>Acipenser_sinensis
-----------------------------------------------------------MIKLQSSMSKHIPQFCGVLGHTFMEFLKGSGDYCQAQHEVYADK-----
>Metridium_senile
---------------------------------------------------------MWTG----------IAYRHSREYTLMEFLTNSGDWCQAQHH----------
>Amia_calva
----------------------------------------------------------------------------------MESSRGRGDYCHAQHPTA--------
>Kaupichthys_hyoproroides
-----------------------------------------------------------MIKLQSSMSKHIPQFCGVLGHTFMEFLKDSGDYCQAQHEVYADK-----
>Anser_cygnoides
----------------------------------------------------------------------------------MEFLKGSGDYCQAQHDLYADK-----
>Podocoryna_carnea
----------------------------------------------------------------------------------MEFLRGSGDYCFAQHRT---------
>Xenopus_tropicalis
MLFTGHGEEPQGRRSGPRQLSPDHR-ALRL---GDSSLLYAPSPPAPLLFHRLFLGTGCMIKIQSSMSKHS-PFCGVLGHTFMEFLKGSGDYCQAQHELYADK-----
>Chaetura_pelagica
----------------------------------------------------------------------------------MEFLKGSGDYCQAQHDLYADK-----
>Apteryx_australis_mantelli
----------------------------------------------------------------------------------MEFLKGSGDYCQAQHDLYADK-----
>Stegodyphus_mimosarum
---------------------------------------------------M-LDASMCID----------IAVNQIQTFLLMEVLVGSDEYCRAQHK----------
>Machilis_hrabei
---------------------------------------------------------MCKDLVFLG----------ETALTYMEFQLDSGEYCQAQHK----------
>Scapharca_broughtonii
---------------------------------------------------------MFRK----------ILFFGEFGHTFMEFVTNSGDYCQAQHV----------
>Porichthys_notatus
-----------------------------------------------------------MIKLQSSMSKHIPQFCGVLGHTFMEFLKGRGDYCQAQHAADADE-----
>Oryzias_latipes
-----------------------------------------------------------MIKLQSSMSKHIPQFCGVLGHTFMEFLKGSGDYCQAQHAAYADE-----
>Buceros_rhinoceros_silvestri
----------------------------------------------------------------------------------MEFLKGSGDYCQAQHDLYADK-----
>Caprimulgus_carolinensis
----------------------------------------------------------------------------------MEFLKGSGDYCQAQHDLYADK-----
>Scyliorhinus_torazame
----------------------------------------------------------------------------------MEFLKGSGDYCQAQHAVYADK-----
>Cariama_cristata
----------------------------------------------------------------------------------MEFLKGSGDYCQAQHDLYADK-----
>Mustela_putorius_furo
-----------------------------------------------------------MIRPQSSMSKHIPQFCGVLGHTFMEFLKGSGDYCQAQHDLYADK-----
>Alatina_alata
---------------------------------------------------M-RRPCLCT---------------CFCGKTFMEFLKDSGDWCQSQHK----------
>Hynobius_retardatus
MRAPELGRSEARAQCGGGSVAADTA-TLPPRSPAARPVWYTARRACP--PATRARGTGRMIQPRSSMSNHSPQFCGVLGHTFMEFLKGSGDYCQAQHDLYADK-----
>Micropterus_salmoides
----------------------------------------------------------------------------------MEFVRRSGDYCHAQHI----------
>Orcinus_orca
----------------------------------------------------------------------------------MEFLKGSGDYCQAQHDLYADK-----
>Astyanax_mexicanus
----------------------------------------------------------------------------------MEFLQGSGDYCHAQHRNA--------
>Calidris_pugnax
----------------------------------------------------------------------------------MEFLKGSGDYCQAQHDLYADK-----
>Callorhinchus_milii
----------------------------------------------------------------------------------MEFLKGSGDYCQAQHDVYADK-----
>Eptesicus_fuscus
-----------------------------------------------------------MIRPQSSMSKHIPQFCGVLGHTFMEFLKGSGDYCQAQHDLYADK-----
>Neolamprologus_brichardi
----------------------------------------------------------------------------------MEFLKGSGDYCQAQHAAYADE-----
>Pterocles_gutturalis
----------------------------------------------------------------------------------MEFLKGSGDYCQAQHDLYADK-----
>Dasypus_novemcinctus
-----------------------------------------------------------MIRPQSSMSKHI-PFCGVLGHTFMEFLKGSGDYCQAQHDLYADK-----
>Alosa_alosa
-----------------------------------------------------------MIKLQSSMSKHIPQFCGVLGHTFMEFLKGSGDYCQAQHGVYADE-----
>Latimeria_chalumnae
-----------------------------------------------------------MIKIQSSMSKHIPQFCGVLGHTFMEFLKGSGDYCQAQHDLYADK-----
>Colius_striatus
----------------------------------------------------------------------------------MEFLKGSGDYCQAQHDLYADK-----
>Columba_livia
----------------------------------------------------------------------------------MEFLKGSGDYCQAQHDLYADK-----
>Leptosomus_discolor
----------------------------------------------------------------------------------MEFLKGSGDYCQAQHDLYADK-----
>Alligator_mississippiensis
-----------------------------------------------------------MIRLQSSMSNRR-PFCGVLGHTFMEFLKGSGDYCQAQHDLYADK-----
>Cuculus_canorus
----------------------------------------------------------------------------------MEFLKGSGDYCQAQHDLYADK-----
>Cyprinodon_variegatus
-----------------------------------------------------------MIKLQSSMSKHIP-FCGVLGRTFMEFLKGSGDYCQAQHAAYADE-----
>Sarcophilus_harrisii
----------------------------------------------------------------------------------MEFLKGSGDYCQAQHDLYADK-----
>Cancer_borealis
--------------------------------------------------MK-FKYCGDNN----------AQLSSGRGYTLMELLKGSGEFCKAQYHIK--------
>Galeopterus_variegatus
----------------------------------------------------------------------------------MEFLKGSGDYCQAQHDLYADK-----
>Monodelphis_domestica
-----------------------------------------------------------MIRPQSSMSKHI-PFCGVLGHTFMEFLKGSGDYCQAQHDLYADK-----
>Occasjapyx_japonicus
---------------------------------------------------------MCRDLVCLG----------KIGLTFMEFFLDSGEYCQAQHK----------
>Macropus_eugenii
--------------------------------------------------------------------MKAKQFCGVLGHTFMEFLKGSGDYCQAQHDLYADK-----
>Rhagoletis_zephyria
------------------------------------------------------------MVVTRKQISDRPKELARPVPTSMEFHLDSAEYCQAQHK----------
>Isonychia_bicolor
---------------------------------------------------------MCRDLVFLG----------EIGLTYMEFQLDSGEYCQAQHK----------
>Esox_lucius
------------MASRGSLNWLRAETAICVRDS-GRLRVALPNRAPGLILTGTLL---------------NIKLCVPAGRSLMEFFQGSGDYCHAQHPSA--------
>Meganyctiphanes_norvegica
--------------------------------------------------MK-FKYCGNNN----------TQLSCGRGYTLMELLKGSGEFCKAQYHIK--------
>Haliaeetus_leucocephalus
-----------------------------------------------------------MIRLQSSMSNHIPQFCGVLGHTFMEFLKGSGDYCQAQHDLYADK-----
>Meleagris_gallopavo
----------------------------------------------------------------------------------MEFLKGSGDYCQAQHDLYADK-----
>Gavia_stellata
----------------------------------------------------------------------------------MEFLKGSGDYCQAQHDLYADK-----
>Acanthogobius_hasta
-----------------------------------------------------------MIKLQSSMSKHIPQFCGVLGHTFMEFLKGSGDYCQAQHVVDADE-----
>Balearica_regulorum_gibberic
----------------------------------------------------------------------------------MEFLKGSGDYCQAQHDLYADK-----
>Apteronotus_leptorhynchus
----------------------------------------------------------------------------------MAFLQRAGDYCHAQHWCC--------
>Trialeurodes_vaporariorum
---------------------------------------------------------MCVVVVNVRE---------LCCLPLMEFQLDSSEYCQAQHK----------
>Dinoponera_quadriceps
----------------------------------------------------------------------------------MEFQLDSAEYCQAQHK----------
>Condylura_cristata
-----------------------------------------------------------MIRPQSSMSKHIP-FCGVLGHTFMEFLKGSGDYCQAQHDLYADK-----
>Oryctolagus_cuniculus
-----------------------------------------------------------MIRPQSSMSKHIP-FCGVLGHTFMEFLKGSGDYCQAQHDLYADK-----
>Rhodinia_newara
----------------------------------------------------------------------------------MEFHLDSAEYCQAQHK----------
>Protopterus_aethiopicus
-----------------------------------------------------------MIKLQSSMSKHIPQFCGVIGHTFMEFLKGSGDYCQAQHALYADK-----
>Lingula_anatina
----------------------------------------------------------------------------------MEFVTNSGEYCQAQHK----------
>Elephantulus_edwardii
-----------------------------------------------------------MIRPQSSMSKHIPQFCGVLGHTFMEFLKGSGDYCQAQHDLYADK-----
>Metallyticus_splendidus
---------------------------------------------------------MCMALVFLG----------ETVLTYMEFQLDSGEYCQAQHK----------
>Ornithorhynchus_anatinus
-----------------------------------------------------------MIATHSSMSKHS-PFCGVLGHTFMEFLKDSGDYCQAQHDLYADK-----
>Perna_viridis
---------------------------------------------------------MCRN----------ILFFGVLGHTFMEFVTNSGDYCQAQHV----------
>Eptatretus_burgeri
---------------------------------------------------------MNR---------HNAQFCGVLGRTFMEFVKGRGEYCETQHIALADK-----
>Hapalochlaena_maculosa
-------------MCRNILDLLGGI---VEK----------TDSP---STLA-WPHHLCRS----------IKFFGEFGHTFMEFVTNSGDFCQAQHV----------
>Epiophlebia_superstes
----------------------------------------------------------------------------------MEFQLDSGEYCQAQHK----------
>Opisthocomus_hoazin
----------------------------------------------------------------------------------MEFLKGSGDYCQAQHDLYADK-----
>Prosarthria_teretrirostris
---------------------------------------------------------MCMAIVSRG----------EIVLTYMEFQLDSGEYCQAQHK----------
>Plecoglossus_altivelis
----------------------------------------------------------------------------------MEFLQGSGDYCHAQHPSI--------
>Crassostrea_gigas
-------------MCNNVLLMRDPRGPVTVR----------GKFVTGFVLCS-SLLNCWQH----------TEFRSNFGHTYMEFVTNSGDYCQAQHV----------
>Pseudopodoces_humilis
-----------------------------------------------------------MIRLQSSMSNHIPQFCGVLGHTFMEFLKGSGDYCQAQHDLYADK-----
>Pecten_maximus
----------------------------------------------------------------------------------MEFVTNSGDYCQAQHAL---------
>Egretta_garzetta
----------------------------------------------------------------------------------MEFLKGSGDYCQAQHDLYADK-----
>Sebastes_caurinus
-----------------------------------------------------------MIKLQSSMSKHIP-FCGVLGHTFMEFLKGSGDYCQAQHAAYADE-----
>Equus_przewalskii
-----------------------------------------------------------MIRPQSSMSKHSPQFCGVLGHTFMEFLKGSGDYCQAQHDLYADK-----
>Extatosoma_tiaratum
--------------------------------------------------------------------------------MHMEFLLSSGEYCQAQHK----------
>Menopon_gallinae
--------------------------------------------------MN-FRFSGCMAL----------KNFHSTSSRLMEFFLDSAEYCQAQHTK---------
>Perinereis_aibuhitensis
-------------------------------------------------MSS-EAGVTTKL----------HQFFGELGHTFMEFLCNSGEYCQAQHV----------
>Leuctra_sp._AD-2013
---------------------------------------------------------MCMALVLVG----------EIVLTYMEFQLDSGEYCQAQHRK---------
>Cynoglossus_semilaevis
----------------------------------------------------------------------------------MEFLKGSGDYCQAQHAAYADE-----
>Homo_sapiens
----------------------------------------------------------------------------------MEFLKGSGDYCQAQHDLYADK-----
>Loxodonta_africana
-----------------------------------------------------------MIRPQSSMSKHI-PFCGVLGHTFMEFLKGSGDYCQAQHDLYADK-----
>Fulmarus_glacialis
----------------------------------------------------------------------------------MEFLKGSGDYCQAQHDLYADK-----
>Melopsittacus_undulatus
----------------------------------------------------------------------------------MEFLKGSGDYCQAQHDLYADK-----
>Ectopsocus_briggsi
----------------------------------------------------------------------------------MEFFLDSAEYCQAQHRK---------
>Leucoraja_erinacea
------------------------------------------------------------------MRKHIPQFCGVLGHTFMEFLKGSGDYCQAQHDVYADK-----
>Mus_musculus
-----------------------------------------------------------MIRPQSSMSRHIPQFCGVLGHTFMEFLKGSGDYCQAQHDLYADK-----
>Salmo_salar
----------------------------------------------------------------------------------MEFLQGSGDYCHAQHLSA--------
>Porites_astreoides
-------------------------------------------------------------------MWSSNLEFGSRAHTFMEFIIDSGDWCQAQHE----------
>Tityus_serrulatus
----------------------------------------------------------------------------------MEYLKTTGEYCRAQHK----------
>Lepisosteus_oculatus
------------MACEGSFSWFRAETVISVSSS-GSLLRVIPLRDTDFIRSRKQHDTKVR---------AAKTSCTVTGGSLMEFLRGSGDYCHAQHHSV--------
>Sepia_pharaonis
-------------MCRNILDLLGGI---VKK----------TDSP---STLA-WPHHLWRL----------VKFFGEFGHTFMEFVTNSGDFCQAQHV----------
>Sepioloidea_lineolata
----------------------------------------------------------------------------------MEFVTNSGDFCQAQHV----------
>Ictalurus_punctatus
--------------------------------------------------------------------------------MLMEFLQGSGDYCHAQHRNA--------
>Trichechus_manatus_latirostr
-----------------------------------------------------------MIRPQSSMSKHIPQFCGVLGHTFMEFLKGSGDYCQAQHDLYADK-----
>Aptenodytes_forsteri
----------------------------------------------------------------------------------MEFLKGSGDYCQAQHDLYADK-----
>Streblospio_benedicti
---------------------------------------------------------MCRI----------ILFFGQIGYTFMEFLCNSGEYCQAQHVAIPSVLSRLH
>Squalus_acanthias
---------------------------------------------------------MNR---------LVTQFSGISGCSFMEFLKGSRDYCQAQHGLLADK-----
>Crotalus_horridus
-----------------------------------------------------------MIRLQSSMSNHIPQFCGVLGHTFMEFLKGSGDYCQAQHGLYADK-----
>Tyto_alba
----------------------------------------------------------------------------------MEFLKGSGDYCQAQHDLYADK-----
>Struthio_camelus_australis
----------------------------------------------------------------------------------MEFLKGSGDYCQAQHDLYADK-----
>Chrysemys_picta
-----------------------------------------------------------MIRLQSSMSNHLPQFCGVLGHTFMEFLKGSGDYCQAQHDLYADK-----
>Takifugu_rubripes
-----------------------------------------------------------MIKLQSSMSKHIPQFCGVLGHTFMEFLKGSGDYCQAQHAVYADE-----
>Watasenia_scintillans
----------------------------------------------------------------------------------MEFVTNSGDFCQAQHV----------
>Tinamus_guttatus
----------------------------------------------------------------------------------MEFLKGSGDYCQAQHDLYADK-----
>Hydroptila_sp._AD-2013
----------------------------------------------------------------------------------MEFRLDSGEYCQAQHK----------
>Calypte_anna
----------------------------------------------------------------------------------MEFLKGSGDYCQAQHDLYADK-----
>Apaloderma_vittatum
----------------------------------------------------------------------------------MEFLKGSGDYCQAQHDLYADK-----
>Orycteropus_afer_afer
-----------------------------------------------------------MIRPQSSMSKHIPQFCGVLGHTFMEFLKGSGDYCQAQHDLYADK-----
>Ruditapes_philippinarum
---------------------------------------------------------MNRI----------ILFFGELGHTFMEFLTNSGEYCLAQHVHV--------
>Limulus_polyphemus
----------------------------------------------------------------------------------MEFLKTAGEYCQSQHK----------
>Palythoa_caribaeorum
--------------------------------------------------------------------MRRDYLAYDRGRTLMEFLIDSGDWCQAQHE----------
>Tricholepidion_gertschi
----------------------------------------------------------------------------------MEFQLDSGEYCQAQHK----------

HG0001 ENSDARG00000087443 ptp4a2a

```
>Danio_rerio
----------------------------------------------------------------------------MQSSQLVFC-S-LIEGCTFMAFVSVSGDFCQAQHLVIV-DK----
>Acipenser_sinensis
----------------------------------------------------------------------MIKLQS--SMSKHIPQFCGVLGHTFMEFLKGSGDYCQAQHEVYA-DK----
>Metridium_senile
-------------------------------------------------------------------------------MWTGIA-YRHSREYTLMEFLTNSGDWCQAQHH----------
>Amia_calva
----------------------------------------------------------------------MIKLQS--SMSKHIPQFCGVLGHTFMEFLKGSGDYCQAQHDVYA-DK----
>Hyalella_azteca
---------------------------------------------------------------------MKLKY-C--GGNCNQL-VLSGRGYSLMELLQASGEFCKAQYTVLLPHD----
>Channa_argus
-------------------MSQQTEEAPPQQQF---QISDENSSCHIEEEQKGDLTVVSAL------PPLSLRSHGISMFILLFG-FCVDLSRTYMACV--SGEFCQAQHLTSV-EK----
>Kaupichthys_hyoproroides
----------------------------------------------------------------------MIKLQS--SMSKHIPQFCGVLGHTFMEFLKDSGDYCQAQHEVYA-DK----
>Anser_cygnoides
-----------------------------------------------------------------------------------------------MAILSVRADFCQAQHSILA-DK----
>Podocoryna_carnea
------------------------------------------------------------------------------MNSLLSC-FCVDLGRTFMAFLSVSGDFCQAQHLAFA-DK----
>Xenopus_laevis
----------------------------------------------------------------------MIKIHS--SMSKHS-PFCGVLGYTFMEFLKGSGDYCQAQHALYA-DK----
>Chaetura_pelagica
-----------------------------------------------------------------------------------------------MEFLKGSGDYCQAQHDLYA-DK----
>Apteryx_australis_mantelli
-----------------------------------------------------------------------------------------------MEFLKGSGDYCQAQHDLYA-DK----
>Meinertellus_cundinamarcensi
-------------------------------------------------------------------------------MCRDLV-FLGETGLTYMEFQLDSGEYCQAQHK----------
>Scapharca_broughtonii
-------------------------------------------------------------------------------MFRK-ILFFGEFGHTFMEFVTNSGDYCQAQHV----------
>Menidia_menidia
------------------------------------------------------------------------------MFNLLFG-FCVDVSRSYMACV--TGEFCQAQHLTSA-EK----
>Porichthys_notatus
----------------------------------------------------------------------MIKLQS--SMSKHIPQFCGVLGHTFMEFLKGRGDYCQAQHAADA-DE----
>Oryzias_melastigma
----------------------------------------------------------------------MIKLQS--SMSKHIPQFCGVLGHTFMEFLKGSGDYCQAQHAAYA-DE----
>Tripterygion_delaisi
---------------------------------------------------------MVCL------HPGAVTFSC--IHSMNPS-Y--QHSYTCMAFARLGGEFCQAQHLLIIVDK----
>Buceros_rhinoceros_silvestri
-----------------------------------------------------------------------------------------------MEFLKGSGDYCQAQHDLYA-DK----
>Caprimulgus_carolinensis
-----------------------------------------------------------------------------------------------MEFLKGSGDYCQAQHDLYA-DK----
>Trachinotus_blochii
---------------------------------------------------------MVCL------HPGAVTFSC--VRSMNHL-LSSHHSCTCMAFARLGGEFCQAQHLLLIVDK----
>Scyliorhinus_torazame
-----------------------------------------------------------------------------------------------MEFLKGSGDYCQAQHAVYA-DK----
>Cariama_cristata
-----------------------------------------------------------------------------------------------MAILSVRADFCQAQHSILA-DK----
>Canis_lupus_familiaris
-----------------------------------------------------------------------------------------------MAILSVRADFCQAQHSVFA-DK----
>Alatina_alata
-------------------------------------------------------------------------------MRRPCL-CTCFCGKTFMEFLKDSGDWCQSQHK----------
>Hynobius_retardatus
MRAPELGRSEARAQCGGGSVAADTATLPPRSPAARPVWYTARRACPPATRARG--------------TGRMIQPRS--SMSNHSPQFCGVLGHTFMEFLKGSGDYCQAQHDLYA-DK----
>Micropterus_floridanus
---------------------------------------------------------MVCL------HPGAVTFSC--VRSMNHL-LSSHQSCTCMAFVKLGGEFCQAQHLLLIVDK----
>Orcinus_orca
----------------------------------------------------------------------------MPSPQLASV-FRWNIRCTFMAILSVRADFCQAQHSVSA-DK----
>Astyanax_mexicanus
------------------------------------------------------------------------------MRRSRLF-FTHTESCTFMAFV--SGDFCQAQHLVIV-DK----
>Calidris_pugnax
---------------------------------------------------------------------------------MPRV-SRWNIRRTFMAILSVRADFCQAQHSILA-DK----
>Callorhinchus_milii
-----------------------------------------------------------------------------------------------MEFLKGSRDYCQAQHGVLA-DK----
>Rousettus_aegyptiacus
----------------------------------------------------------------------------MPSPQLASV-FRWNIRCTFMAILSVRADFCQAQHSVFA-DK----
>Oreochromis_niloticus
---------------------------------------------------------MVCL------HPGAVTFSC--VRSMNHL-LSSHHGCTCMAFARLGGEFCQAQHLLLIVDK----
>Pterocles_gutturalis
-----------------------------------------------------------------------------------------------MEFLKGSGDYCQAQHDLYA-DK----
>Eucidaris_tribuloides
-------------------------------------------------------------------------------MCKHTE-VLAVINKPFMALVLDSGEFCQSQHK----------
>Dasypus_novemcinctus
-----------------------------------------------------------------------------------------------MEFLKGSGDYCQAQHDLYA-DK----
>Clupea_harengus
----------------------------------------------------------------------------MRNS-YRVF-FTPDESCTFMAFVSVSGDFCQAQHLVIV-DK----
>Latimeria_chalumnae
----------------------------------------------------------------------MIKIQS--SMSKHIPQFCGVLGHTFMEFLKGSGDYCQAQHDLYA-DK----
>Colius_striatus
-----------------------------------------------------------------------------------------------MEFLKGSGDYCQAQHDLYA-DK----
>Columba_livia
-----------------------------------------------------------------------------------------------MEFLKGSGDYCQAQHDLYA-DK----
>Leptosomus_discolor
-----------------------------------------------------------------------------------------------MEFLKGSGDYCQAQHDLYA-DK----
>Corynactis_australis
-------------------------------------------------------------------------------MWRGNR-VYGNRAHTFMEFIIDSGDWCQAQHE----------
>Alligator_mississippiensis
----------------------------------------------------------------------MIRLQS--SMSNRR-PFCGVLGHTFMEFLKGSGDYCQAQHDLYA-DK----
>Cuculus_canorus
-----------------------------------------------------------------------------------------------MEFLKGSGDYCQAQHDLYA-DK----
>Poeciliopsis_prolifica
-----------------------------------------------------------------------------------------------MAFARLGGAFCQAQHLLLIVDK----
>Sarcophilus_harrisii
-----------------------------------------------------------------------------------------------MEFLKGSGDYCQAQHDLYA-DK----
>Sclerodactyla_briareus
-----------------------------------------------MYDRRKPSISERFYSTSAVDSPNNITPMY--RFNTA---SQLHTSLSWMAFLGAFGEFCRSQHT----------
>Galeopterus_variegatus
----------------------------------------------------------------------------MPSPQLASV-FRWNIRCTFMAILSVRADFCQAQHSVFA-DK----
>Monodelphis_domestica
----------------------------------------------------------------------MIRPQS--SMSKHI-PFCGVLGHTFMEFLKGSGDYCQAQHDLYA-DK----
>Occasjapyx_japonicus
-------------------------------------------------------------------------------MCRDLV-CLGKIGLTFMEFFLDSGEYCQAQHK----------
>Macropus_eugenii
---------------------------------------------------------------------------------MKAKQFCGVLGHTFMEFLKGSGDYCQAQHDLYA-DK----
>Evechinus_chloroticus
-------------------------------------------------------------------------------MCKPAE-VSAISNLAFMASVLDSGEYCQSQHK----------
>Ephemera_danica
-----------------------------------------------------------------------------------------------MEFQLDSGEYCQAQHK----------
>Esox_lucius
-----------------------------------------------------------------------------------------------MAYVSVSGDFCQAQHLAFV-DK----
>Aquila_chrysaetos_canadensis
---------------------------------------------------------------------------------MPRV-FCWNIRRTFMAILSVRADFCQAQHSILA-DK----
>Asterias_amurensis
-------------------------------------------------------------------------------MSHLTG-ISSVPVSSYMAFVAISGEYCRSQHK----------
>Coturnix_japonica
----------------------------------------------------------------------MIRLQS--SMSNHI-PFCGVLGHTFMEFLKGSGDYCQAQHDLYA-DK----
>Gavia_stellata
-----------------------------------------------------------------------------------------------MAILSVRADFCQAQHSILA-DK----
>Acanthogobius_hasta
----------------------------------------------------------------------MIKLQS--SMSKHIPQFCGVLGHTFMEFLKGSGDYCQAQHVVDA-DE----
>Chlamydotis_macqueenii
-----------------------------------------------------------------------------------------------MAILSVRADFCQAQHRILA-DK----
>Apteronotus_leptorhynchus
-----------------------------------------------------------------------------------------------MAFV--SGDFCQAQHLAFV-NK----
>Glossoscolex_paulistus
-------------------------------------------------------------------------------MYTIAS-LSKASGHHLMALVYLCGEFCQAQHV----------
>Planococcus_citri
---------------------------------------------------------------------------------MDIV-TKSALCFTFMEFLLDSSEYCQAQHK----------
>Sorex_araneus
-----------------------------------------------------------------------------------------------MAILSVRADFCQAQHSVFA-DK----
>Oryctolagus_cuniculus
----------------------------------------------------------------------MIRPQS--SMSKHI-PFCGVLGHTFMEFLKGSGDYCQAQHDLYA-DK----
>Protopterus_aethiopicus
----------------------------------------------------------------------MIKLQS--SMSKHIPQFCGVIGHTFMEFLKGSGDYCQAQHALYA-DK----
>Lingula_anatina
-----------------------------------------------------------------------------------------------MEFVTNSGEYCQAQHK----------
>Elephantulus_edwardii
----------------------------------------------------------------------MIRPQS--SMSKHIPQFCGVLGHTFMEFLKGSGDYCQAQHDLYA-DK----
>Metallyticus_splendidus
-------------------------------------------------------------------------------MCMALV-FLGETVLTYMEFQLDSGEYCQAQHK----------
>Ornithorhynchus_anatinus
----------------------------------------------------------------------------MPSPRLASA-FRWNIRCTFMAILSVRADFCQAQHSVCA-DK----
>Tauraco_erythrolophus
-----------------------------------------------------------------------------------------------MAILSVRADFCQAQHSILA-DK----
>Cypridininae_sp._BMR-2011
----------------------------------------------------------------MEDSETSLLYKY--RLCMRPQ-VLAGVCCSFMEFVLADGEYCQAQHK----------
>Perna_viridis
-------------------------------------------------------------------------------MCRN-ILFFGVLGHTFMEFVTNSGDYCQAQHV----------
>Eptatretus_burgeri
------------------------------------------------------------------------------MNRHNAQ-FCGVLGRTFMEFVKGRGEYCETQHIALA-DK----
>Octopus_bimaculoides
-----------------------------------------------------------------------------------------------MEFVTNSGDFCQAQHV----------
>Epiophlebia_superstes
-----------------------------------------------------------------------------------------------MEFQLDSGEYCQAQHK----------
>Opisthocomus_hoazin
-----------------------------------------------------------------------------------------------MEFLKGSGDYCQAQHDLYA-DK----
>Prosarthria_teretrirostris
-------------------------------------------------------------------------------MCMAIV-SRGEIVLTYMEFQLDSGEYCQAQHK----------
>Plecoglossus_altivelis
-----------------------------------------------------------------------------------------------MEFLQGSGDYCHAQHPSI--------
>Scleropages_formosus
-----------------------------------------------------------------------------------------------MAFVSVSGDFCQAQHLASA-DK----
>Crassostrea_gigas
-----------------------------------------------MRDPRGPVTVRGKF------VTGFVLCSSLLNCWQHTE-FRSNFGHTYMEFVTNSGDYCQAQHV----------
>Parus_major
----------------------------------------------------------------------MIRLQS--SMSNHI-PFCGVLGHTFMEFLKGSGDYCQAQHDLYA-DK----
>Pecten_maximus
-----------------------------------------------------------------------------------------------MEFVTNSGDYCQAQHAL---------
>Egretta_garzetta
-----------------------------------------------------------------------------------------------MAILSVRADFCQAQHSILA-DK----
>Lateolabrax_maculatus
---------------------------------------------------------MVCL------HPGAVTSSC--VRSMNHL-LGSHRGRTCMAFARLGGEFCQAQHLLLIVDK----
>Sebastes_caurinus
----------------------------------------------------------------------MIKLQS--SMSKHIP-FCGVLGHTFMEFLKGSGDYCQAQHAAYA-DE----
>Ceratotherium_simum_simum
-----------------------------------------------------------------------------------------------MAILSVRADFCQAQHSVFA-DK----
>Perinereis_aibuhitensis
-----------------------------------------------------------------MSSE-----AG--VTTKLHQ-FFGELGHTFMEFLCNSGEYCQAQHV----------
>Picoides_pubescens
-----------------------------------------------------------------------------------------------MAILSVRADFCQAQHSILA-DK----
>Perla_marginata
-----------------------------------------------------------------------------------------------MEFQLDSGEYCQAQHK----------
>Cynoglossus_semilaevis
---------------------------------------------------------MVCP------HHRAVTTPC--VRSMNNL-LSSHDSCTCMAFARVGGEFCQAQHLLLIVDK----
>Cercocebus_atys
----------------------------------------------------------------------------MPSPQLASV-FRWNIRCTFMAILSVRADFCQAQHSVFA-DK----
>Loxodonta_africana
----------------------------------------------------------------------MIRPQS--SMSKHI-PFCGVLGHTFMEFLKGSGDYCQAQHDLYA-DK----
>Fulmarus_glacialis
-----------------------------------------------------------------------------------------------MAILSVRADFCQAQHSILA-DK----
>Nestor_notabilis
-----------------------------------------------------------------------------------------------MAILSVRAEFCQAQHSILA-DK----
>Leucoraja_erinacea
-------------------------------------------------------------------------------MRKHIPQFCGVLGHTFMEFLKGSGDYCQAQHDVYA-DK----
>Rhopilema_esculentum
------------------------------------------------------------------------------MNSYHLL-LSKYFGKPFMEFLKDSAEWCKAQHVE---------
>Heterocephalus_glaber
----------------------------------------------------------------------------MPSPQLASV-FRWNIRCTFMAILSVRADFCQAQHSVFA-DK----
>Salmo_salar
------------------------------------------------------------------------------MFNLLFC-FCVDLGHTYMAYVSVSGDFCQAQHLAFV-DK----
>Tupaia_chinensis
-----------------------------------------------------------------------------------------------MAILSVRADFCQAQHSVFA-DK----
>Porites_astreoides
-------------------------------------------------------------------------------MWSSNL-EFGSRAHTFMEFIIDSGDWCQAQHE----------
>Aurelia_aurita
------------------------------------------------------------------------------MTSYHLQ-L-KCFGETFMEFLKDSAEWCKAQHVE---------
>Lepisosteus_oculatus
------------------------------------------------------------------------------MSSLPFC-FCVDLVRTFMAFLSVSGDFCQAQHLAFA-DK----
>Sepia_pharaonis
-------------------------------------------MCRNILDLLGGIVKKTD-------SPSTLAWPH--HLWRLVK-FFGEFGHTFMEFVTNSGDFCQAQHV----------
>Sepioloidea_lineolata
-----------------------------------------------------------------------------------------------MEFVTNSGDFCQAQHV----------
>Ictalurus_punctatus
------------------------------------------------------------------------------MQSSRHF-FPCTESCTFMAFVSVSGDFCQAQHLVIV-DK----
>Trichechus_manatus_latirostr
----------------------------------------------------------------------------MPSSQLASV-FRWNIRCTFMAILSVRADFCQAQHSIFA-DK----
>Aptenodytes_forsteri
-----------------------------------------------------------------------------------------------MAILSVRADFCQAQHSILA-DK----
>Echinaster_spinulosus
-------------------------------------------------------------------------------MCQLTAILNTISVSTLMAFAAVSGEYCRSQHK----------
>Streblospio_benedicti
-------------------------------------------------------------------------------MCRIIL-FFGQIGYTFMEFLCNSGEYCQAQHVAIPSVLSRLH
>Squalus_acanthias
-------------------------------------------------------------------------------MNRLVTQFSGISGCSFMEFLKGSRDYCQAQHGLLA-DK----
>Python_bivittatus
----------------------------------------------------------------------MIRLQS--SMSNHI-PFCGVLGHTFMEFLKGSGDYCQAQHGLYA-DK----
>Tyto_alba
-----------------------------------------------------------------------------------------------MEFLKGSGDYCQAQHDLYA-DK----
>Struthio_camelus_australis
-----------------------------------------------------------------------------------------------MAILSVRADFCQAQHSTLA-DK----
>Deroceras_reticulatum
-------------------------------------------------------------------------------MSSFVL-FVGEFGHTFMEFVTNSEDFCQAQHYRLVE------
>Sphaerechinus_granularis
-------------------------------------------------------------------------------MCKPAE-DSAISNQAFMATVLDSGEYCQSQHK----------
>Pelodiscus_sinensis
----------------------------------------------------------------------MIRLQS--SMSNHI-PFCGVLGHTFMEFLKGSGDYCQAQHDLYA-DK----
>Takifugu_rubripes
----------------------------------------------------------------------MIKLQS--SMSKHIPQFCGVLGHTFMEFLKGSGDYCQAQHAVYA-DE----
>Watasenia_scintillans
-----------------------------------------------------------------------------------------------MEFVTNSGDFCQAQHV----------
>Tinamus_guttatus
---------------------------------------------------------------------------------MPRV-FCWNIRRTFMAILRVRADFCQAQHSTLA-DK----
>Tetronarce_californica
-----------------------------------------------------------------------------------------------MEFVRNSSGYCPAQHSIFA-DK----
>Hydroptila_sp._AD-2013
-----------------------------------------------------------------------------------------------MEFRLDSGEYCQAQHK----------
>Calypte_anna
-----------------------------------------------------------------------------------------------MEFLKGSGDYCQAQHDLYA-DK----
>Apaloderma_vittatum
-----------------------------------------------------------------------------------------------MEFLKGSGDYCQAQHDLYA-DK----
>Orycteropus_afer_afer
----------------------------------------------------------------------MIRPQS--SMSKHIPQFCGVLGHTFMEFLKGSGDYCQAQHDLYA-DK----
>Ruditapes_philippinarum
-------------------------------------------------------------------------------MNRI-ILFFGELGHTFMEFLTNSGEYCLAQHVHV--------
>Limulus_polyphemus
----------------------------------------------------------------MSSTNDS----A--KVPLLLP-FEVRRHHTLMEFLKTAGEYCQSQHK----------
>Protopalythoa_variabilis
--------------------------------------------------------------------------------MRRDY-LAYDRGRTLMEFLIDSGDWCQAQHE----------
>Thermobia_domestica
-----------------------------------------------------------------------------------------------MEFQLDSGEYCQAQHK----------
```

HG0001 ENSG00000112245 PTP4A1

>Homo_sapiens
-------------------------------------------------------------------MIRPQSSMSKHIPQF--CGVLGHTFMEF-LKGSGDYCQAQHDL-YADK-----
>Acipenser_sinensis
-------------------------------------------------------------------MIKLQSSMSKHIPQF--CGVLGHTFMEF-LKGSGDYCQAQHEV-YADK-----
>Actinia_tenebrosa
---------------------------------------------------------------------------MWTTNIC--YDHCGHSLMEF-LIDSGDWCQAQHN-----------
>Acanthogorgia_aspera
---------------------------------------------------------------------------MSRAYCLFGFRSRFEKFMEYFLKDSGEFCQAQHE-----------
>Amia_calva
-------------------------------------------------------------------MIKLQSSMSKHIPQF--CGVLGHTFMEF-LKGSGDYCQAQHDV-YADK-----
>Channa_argus
-------------------MSQQTEEAPPQQQF---QISDENSSC-HIEEEQKGDLTVVSALPPLSLRSH-GISMFILLFGF--CVDLSRTYMAC---VSGEFCQAQHLT-SVEK-----
>Kaupichthys_hyoproroides
-------------------------------------------------------------------MIKLQSSMSKHIPQF--CGVLGHTFMEF-LKDSGDYCQAQHEV-YADK-----
>Anser_cygnoides
--------------------------------------------------------------------------------------------MEF-LKGSGDYCQAQHDL-YADK-----
>Podocoryna_carnea
--------------------------------------------------------------------------------------------MEF-LRGSGDYCFAQHRT----------
>Megophrys_sangzhiensis
-------------------------------------------------------------------MIKPQSSMSKHSPQF--CGVLGHTFMEF-LKGSGDYCQAQHDL-YADK-----
>Chaetura_pelagica
--------------------------------------------------------------------------------------------MEF-LKGSGDYCQAQHDL-YADK-----
>Apteryx_australis_mantelli
--------------------------------------------------------------------------------------------MEF-LKGSGDYCQAQHDL-YADK-----
>Nephilengys_cruentata
-------------------------------------------------------------------MLD-ASMC--IDLAV--NQIHPFSLMEI-LVGSDEYCRAQHK-----------
>Lepismachilis_y-signata
---------------------------------------------------------------------------MCKDLVF--LGETALTYMEF-QLDSGEYCQAQHK-----------
>Scapharca_broughtonii
---------------------------------------------------------------------------MFRKILF--FGEFGHTFMEF-VTNSGDYCQAQHV-----------
>Porichthys_notatus
-------------------------------------------------------------------MIKLQSSMSKHIPQF--CGVLGHTFMEF-LKGRGDYCQAQHAA-DADE-----
>Oryzias_melastigma
-------------------------------------------------------------------MIKLQSSMSKHIPQF--CGVLGHTFMEF-LKGSGDYCQAQHAA-YADE-----
>Tripterygion_delaisi
--------------------------------------------------------------------------MFSLLFGF--CVDLSRTYMAC---VSGEFCQAQHLT-SVEK-----
>Buceros_rhinoceros_silvestri
--------------------------------------------------------------------------------------------MEF-LKGSGDYCQAQHDL-YADK-----
>Caprimulgus_carolinensis
--------------------------------------------------------------------------------------------MEF-LKGSGDYCQAQHDL-YADK-----
>Scyliorhinus_torazame
--------------------------------------------------------------------------------------------MEF-LKGSGDYCQAQHAV-YADK-----
>Cariama_cristata
--------------------------------------------------------------------------------------------MEF-LKGSGDYCQAQHDL-YADK-----
>Leptonychotes_weddellii
-------------------------------------------------------------------MIRPQSSMSKHIPQF--CGVLGHTFMEF-LKGSGDYCQAQHDL-YADK-----
>Alatina_alata
-------------------------------------------------------------------MRR-PCLC---------TCFCGKTFMEF-LKDSGDWCQSQHK-----------
>Hynobius_retardatus
MRAPELGRSEARAQCGGGSVAADTATLPPRSPAARPVWYTARRAC-PPATRARG----------TGRMIQPRSSMSNHSPQF--CGVLGHTFMEF-LKGSGDYCQAQHDL-YADK-----
>Micropterus_salmoides
--------------------------------------------------------------------------------------------MEF-VRRSGDYCHAQHI-----------
>Physeter_catodon
--------------------------------------------------------------------------------------------MEF-LKGSGDYCQAQHDL-YADK-----
>Astyanax_mexicanus
-------------------------------------------------------------------MVKLQSSMSKHIPQF--CGVLGHTFMEF-LKGSGDYCQAQHGV-YADK-----
>Calidris_pugnax
--------------------------------------------------------------------------------------------MEF-LKGSGDYCQAQHDL-YADK-----
>Callorhinchus_milii
--------------------------------------------------------------------------------------------MEF-LKGSGDYCQAQHDV-YADK-----
>Eptesicus_fuscus
-------------------------------------------------------------------MIRPQSSMSKHIPQF--CGVLGHTFMEF-LKGSGDYCQAQHDL-YADK-----
>Oreochromis_niloticus
-------------------------------------------------------------------MIKLQSSMSKHIPQF--CGVLGHTFMEF-LKGSGDYCQAQHAA-YADE-----
>Pterocles_gutturalis
--------------------------------------------------------------------------------------------MEF-LKGSGDYCQAQHDL-YADK-----
>Eucidaris_tribuloides
---------------------------------------------------------------------------MCKHTEV--LAVINKPFMAL-VLDSGEFCQSQHK-----------
>Dasypus_novemcinctus
-------------------------------------------------------------------MIRPQSSMSKHIPQF--CGVLGHTFMEF-LKGSGDYCQAQHDL-YADK-----
>Alosa_alosa
-------------------------------------------------------------------MIKLQSSMSKHIPQF--CGVLGHTFMEF-LKGSGDYCQAQHGV-YADE-----
>Latimeria_menadoensis
-------------------------------------------------------------------MIKIQSSMSKHIPQF--CGVLGHTFMEF-LKGSGDYCQAQHDL-YADK-----
>Gyrinus_marinus
---------------------------------------------------------------------------MCMAIVW--PGETVLTFMEF-QLDSAEYCQAQHK-----------
>Colius_striatus
--------------------------------------------------------------------------------------------MEF-LKGSGDYCQAQHDL-YADK-----
>Columba_livia
--------------------------------------------------------------------------------------------MEF-LKGSGDYCQAQHDL-YADK-----
>Leptosomus_discolor
--------------------------------------------------------------------------------------------MEF-LKGSGDYCQAQHDL-YADK-----
>Ricordea_yuma
---------------------------------------------------------------------------MWRGNPV--YGNRAHTFMEF-IIDSGDWCQAQHE-----------
>Alligator_mississippiensis
-------------------------------------------------------------------MIRLQSSMSNRRPQF--CGVLGHTFMEF-LKGSGDYCQAQHDL-YADK-----
>Cuculus_canorus
--------------------------------------------------------------------------------------------MEF-LKGSGDYCQAQHDL-YADK-----
>Cyprinus_carpio
-------------------------------------------------------------------MIKLQSSMSKHIPQF--CGVLGHTFMEF-LKGSGDYCQAQHGV-YAEK-----
>Fundulus_grandis
-------------------------------------------------------------------MIKLQSSMSKHIPQF--CGVLGRTFMEF-LKGSGDYCQAQHAA-YADE-----
>Sarcophilus_harrisii
--------------------------------------------------------------------------------------------MEF-LKGSGDYCQAQHDL-YADK-----
>Cancer_borealis
--------------------------------------------------------------------MK-FKYCGDNNAQL--SSGRGYTLMEL-LKGSGEFCKAQYHI-K--------
>Galeopterus_variegatus
--------------------------------------------------------------------------------------------MEF-LKGSGDYCQAQHDL-YADK-----
>Monodelphis_domestica
-------------------------------------------------------------------MIRPQSSMSKHIPQF--CGVLGHTFMEF-LKGSGDYCQAQHDL-YADK-----
>Occasjapyx_japonicus
---------------------------------------------------------------------------MCRDLVC--LGKIGLTFMEF-FLDSGEYCQAQHK-----------
>Macropus_eugenii
----------------------------------------------------------------------------MKAKQF--CGVLGHTFMEF-LKGSGDYCQAQHDL-YADK-----
>Chironomus_riparius
-----------------------------------------------------------------------------MDIVL--LRQLALTSMEF-KLDSAEYCQAQHK-----------
>Isonychia_bicolor
---------------------------------------------------------------------------MCRDLVF--LGEIGLTYMEF-QLDSGEYCQAQHK-----------
>Esox_lucius
-------------------------------------------------------------------MIKLQSSMSKHIPQF--CGVLGHTFMEF-LKGSGDYCQAQHVAAYADE-----
>Meganyctiphanes_norvegica
--------------------------------------------------------------------MK-FKYCGNNNTQL--SCGRGYTLMEL-LKGSGEFCKAQYHI-K--------
>Haliaeetus_leucocephalus
-------------------------------------------------------------------MIRLQSSMSNHIPQF--CGVLGHTFMEF-LKGSGDYCQAQHDL-YADK-----
>Marthasterias_glacialis
---------------------------------------------------------------------------MSHLTGI--CSVPVSSYMAF-VAISGEYCRSQHK-----------
>Coturnix_japonica
-------------------------------------------------------------------MIRLQSSMSNHIPQF--CGVLGHTFMEF-LKGSGDYCQAQHDL-YADK-----
>Gavia_stellata
--------------------------------------------------------------------------------------------MEF-LKGSGDYCQAQHDL-YADK-----
>Acanthogobius_hasta
-------------------------------------------------------------------MIKLQSSMSKHIPQF--CGVLGHTFMEF-LKGSGDYCQAQHVV-DADE-----
>Eurypyga_helias
--------------------------------------------------------------------------------------------MEF-LKGSGDYCQAQHDL-YADK-----
>Apteronotus_leptorhynchus
--------------------------------------------------------------------------------------------MAF-LQRAGDYCHAQHWC-C--------
>Planococcus_citri
-----------------------------------------------------------------------------MDIVT--KSALCFTFMEF-LLDSSEYCQAQHK-----------
>Orussus_abietinus
--------------------------------------------------------------------------------------------MEF-QLDSAEYCQAQHK-----------
>Condylura_cristata
-------------------------------------------------------------------MIRPQSSMSKHIPQF--CGVLGHTFMEF-LKGSGDYCQAQHDL-YADK-----
>Ornithodoros_turicata
--------------------------------------------------------------------------MKSDAAGT--NGEPSHSWMEVSLGPSGEYCLAQHK-----------
>Ochotona_princeps
-------------------------------------------------------------------MIRPQSSMSKHIPQF--CGVLGHTFMEF-LKGSGDYCQAQHDL-YADK-----
>Antheraea_yamamai
--------------------------------------------------------------------------------------------MEF-RLDSAEYCQAQHK-----------
>Protopterus_aethiopicus
-------------------------------------------------------------------MIKLQSSMSKHIPQF--CGVIGHTFMEF-LKGSGDYCQAQHAL-YADK-----
>Lingula_anatina
-------------------------------------------------------------------------------MQF--FGEFGHTFMEF-VTNSGEYCQAQHK-----------
>Elephantulus_edwardii
-------------------------------------------------------------------MIRPQSSMSKHIPQF--CGVLGHTFMEF-LKGSGDYCQAQHDL-YADK-----
>Metallyticus_splendidus
---------------------------------------------------------------------------MCMALVF--LGETVLTYMEF-QLDSGEYCQAQHK-----------
>Nannochorista_philpotti
---------------------------------------------------------------------------MCMAIVV--FPTVALRYMEF-HLDSAEYCQAQHK-----------
>Ornithorhynchus_anatinus
-------------------------------------------------------------------MIATHSSMSKHSPQF--CGVLGHTFMEF-LKDSGDYCQAQHDL-YADK-----
>Cypridininae_sp._BMR-2011
----------------------------------------------------------MED---SETSLL-YKYRLCMRPQV--LAGVCCSFMEF-VLADGEYCQAQHK-----------
>Perna_viridis
---------------------------------------------------------------------------MCRNILF--FGVLGHTFMEF-VTNSGDYCQAQHV-----------
>Eptatretus_burgeri
--------------------------------------------------------------------------MNRHNAQF--CGVLGRTFMEF-VKGRGEYCETQHIA-LADK-----
>Osmylus_fulvicephalus
---------------------------------------------------------------------------MCMARVF--LGETVLTYMEF-QLDSAEYCQAQHK-----------
>Triops_newberryi
---------------------------------------------------------------------------MCTTGQR--WVEWRHKFLAV-LLGSGDYCEAQHK-----------
>Hapalochlaena_maculosa
-------------------------------------------MC-RNILDLLGGIVEKTDSPST---LA-WPHHLCRSIKF--FGEFGHTFMEF-VTNSGDFCQAQHV-----------
>Epiophlebia_superstes
--------------------------------------------------------------------------------------------MEF-QLDSGEYCQAQHK-----------
>Opisthocomus_hoazin
--------------------------------------------------------------------------------------------MEF-LKGSGDYCQAQHDL-YADK-----
>Stenobothrus_lineatus
-------------------MV------VEASAV---ALADEGRTAGRHVMRLERASSPSSM---VDSQAA-GGGRLCAHQQF--RGEIVLTFMEF-QLDSGEYCQAQHK-----------
>Plecoglossus_altivelis
--------------------------------------------------------------------------------------------MEF-LQGSGDYCHAQHPSI---------
>Scleropages_formosus
--------------------------------------------------------------------------MCGLLFRF--CVDLGRPFMAF-VSVSGDFCQAQHLA-SADK-----
>Crassostrea_gigas
-----------------------------------------MRDP-RGPVTVRGKFVTGFV---L---CS-SLLNCWQHTEF--RSNFGHTYMEF-VTNSGDYCQAQHV-----------
>Parus_major
-------------------------------------------------------------------MIRLQSSMSNHIPQF--CGVLGHTFMEF-LKGSGDYCQAQHDL-YADK-----
>Pecten_maximus
--------------------------------------------------------------------------------------------MEF-VTNSGDYCQAQHAL----------
>Pelecanus_crispus
--------------------------------------------------------------------------------------------MEF-LKGSGDYCQAQHDL-YADK-----
>Sebastes_caurinus
-------------------------------------------------------------------MIKLQSSMSKHIP-F--CGVLGHTFMEF-LKGSGDYCQAQHAA-YADE-----
>Equus_przewalskii
-------------------------------------------------------------------MIRPQSSMSKHSPQF--CGVLGHTFMEF-LKGSGDYCQAQHDL-YADK-----
>Peruphasma_schultei
--------------------------------------------------------------------------------------------MEF-LPNSGEYCQAQHK-----------
>Menopon_gallinae
--------------------------------------------------------------------MN-FRFSGCMALKN--FHSTSSRLMEF-FLDSAEYCQAQHTK----------
>Perinereis_aibuhitensis
----------------------------------------------------------------M---SS-EAGVTTKLHQF--FGELGHTFMEF-LCNSGEYCQAQHV-----------
>Perla_marginata
--------------------------------------------------------------------------------------------MEF-QLDSGEYCQAQHK-----------
>Cynoglossus_semilaevis
--------------------------------------------------------------------------------------------MEF-LKGSGDYCQAQHAA-YADE-----
>Loxodonta_africana
-------------------------------------------------------------------MIRPQSSMSKHIPQF--CGVLGHTFMEF-LKGSGDYCQAQHDL-YADK-----
>Fulmarus_glacialis
--------------------------------------------------------------------------------------------MEF-LKGSGDYCQAQHDL-YADK-----
>Nestor_notabilis
--------------------------------------------------------------------------------------------MEF-LKGSGDYCQAQHDL-YADK-----
>Ectopsocus_briggsi
--------------------------------------------------------------------------------------------MEF-FLDSAEYCQAQHRK----------
>Leucoraja_erinacea
--------------------------------------------------------------------------MRKHIPQF--CGVLGHTFMEF-LKGSGDYCQAQHDV-YADK-----
>Inocellia_crassicornis
--------------------------------------------------------------------------------------------MEF-QLDSAEYCQAQHK-----------
>Rhopilema_esculentum
--------------------------------------------------------------------------MNSYHLLL--SKYFGKPFMEF-LKDSAEWCKAQHVE----------
>Mesocricetus_auratus
-------------------------------------------------------------------MIRPQSSMSRHIPQF--CGVLGHTFMEF-LKGSGDYCQAQHDL-YADK-----
>Salmo_salar
-------------------------------------------------------------------MIKLQSSMSKHIPQF--CGVLGHTFMEF-LKGSGDYCQAQHAAAYADE-----
>Acropora_digitifera
---------------------------------------------------------------------------MWRFSLT--FGSRAHTFMEF-IIDSGDWCQAQHE-----------
>Tityus_serrulatus
--------------------------------------------------------------------------------------------MEY-LKTTGEYCRAQHK-----------
>Aurelia_aurita
--------------------------------------------------------------------------MTSYHLQL---KCFGETFMEF-LKDSAEWCKAQHVE----------
>Lepisosteus_oculatus
-------------------------------------------------------------------MIKLQSSMSKHIPQF--CGVLGHTFMEF-LKGSGDYCQAQHGV-YADK-----
>Sepia_pharaonis
-------------------------------------------MC-RNILDLLGGIVKKTDSPST---LA-WPHHLWRLVKF--FGEFGHTFMEF-VTNSGDFCQAQHV-----------
>Sepioloidea_lineolata
--------------------------------------------------------------------------------------------MEF-VTNSGDFCQAQHV-----------
>Ictalurus_punctatus
-------------------------------------------------------------------MVKLQSSMSKHIPQF--CGVLGHTFMEF-LKGSGDYCQAQHGV-YADK-----
>Oropsylla_silantiewi
--------------------------------------------------------------------------------------------MEF-QLDSAEYCQAQHK-----------
>Trichechus_manatus_latirostr
-------------------------------------------------------------------MIRPQSSMSKHIPQF--CGVLGHTFMEF-LKGSGDYCQAQHDL-YADK-----
>Aptenodytes_forsteri
--------------------------------------------------------------------------------------------MEF-LKGSGDYCQAQHDL-YADK-----
>Streblospio_benedicti
---------------------------------------------------------------------------MCRIILF--FGQIGYTFMEF-LCNSGEYCQAQHVA-IPSVLSRLH
>Squalus_acanthias
--------------------------------------------------------------------------MNRLVTQF--SGISGCSFMEF-LKGSRDYCQAQHGL-LADK-----
>Crotalus_horridus
-------------------------------------------------------------------MIRLQSSMSNHIPQF--CGVLGHTFMEF-LKGSGDYCQAQHGL-YADK-----
>Stylops_melittae
---------------------------------------------------------------------------MCIGILN--LGNKVVEYMEF-HFGSSEYCKAQHK-----------
>Tyto_alba
--------------------------------------------------------------------------------------------MEF-LKGSGDYCQAQHDL-YADK-----
>Struthio_camelus_australis
--------------------------------------------------------------------------------------------MEF-LKGSGDYCQAQHDL-YADK-----
>Deroceras_reticulatum
---------------------------------------------------------------------------MSSFVLF--VGEFGHTFMEF-VTNSEDFCQAQHYR-LVE------
>Pelodiscus_sinensis
-------------------------------------------------------------------MIRLQSSMSNHIPQF--CGVLGHTFMEF-LKGSGDYCQAQHDL-YADK-----
>Takifugu_rubripes
-------------------------------------------------------------------MIKLQSSMSKHIPQF--CGVLGHTFMEF-LKGSGDYCQAQHAV-YADE-----
>Watasenia_scintillans
--------------------------------------------------------------------------------------------MEF-VTNSGDFCQAQHV-----------
>Gynaikothrips_ficorum
---------------------------------------------MVHCAKFNGENSISMV---IRRVIF-DNKWPSVGYQT--CGEILPTFMEF-HLDSAEYCQAQHK-----------
>Tinamus_guttatus
--------------------------------------------------------------------------------------------MEF-LKGSGDYCQAQHDL-YADK-----
>Tetronarce_californica
--------------------------------------------------------------------------------------------MEF-VRNSSGYCPAQHSI-FADK-----
>Hydroptila_sp._AD-2013
--------------------------------------------------------------------------------------------MEF-RLDSGEYCQAQHK-----------
>Calypte_anna
--------------------------------------------------------------------------------------------MEF-LKGSGDYCQAQHDL-YADK-----
>Apaloderma_vittatum
--------------------------------------------------------------------------------------------MEF-LKGSGDYCQAQHDL-YADK-----
>Orycteropus_afer_afer
-------------------------------------------------------------------MIRPQSSMSKHIPQF--CGVLGHTFMEF-LKGSGDYCQAQHDL-YADK-----
>Eurhomalea_rufa
--------------------------------------------------------------------------------------------MEF-LTNSGEYCLAQHVH-V--------
>Limulus_polyphemus
--------------------------------------------------------------------------------------------MEF-LKTAGEYCQSQHK-----------
>Palythoa_caribaeorum
---------------------------------------------------------------------------MRRDY-L--AYDRGRTLMEF-LIDSGDWCQAQHE-----------
>Thermobia_domestica
--------------------------------------------------------------------------------------------MEF-QLDSGEYCQAQHK-----------

HG0001 ENSG00000184007 PTP4A2

```
>Homo_sapiens
-----------------------MAILSVRADFCQAQHSIFADK
>Acipenser_sinensis
MIKLQSSMSKHIPQFCGVLGHTFMEFLKGSGDYCQAQHEVYADK
>Amia_calva
MIKLQSSMSKHIPQFCGVLGHTFMEFLKGSGDYCQAQHDVYADK
>Kaupichthys_hyoproroides
MIKLQSSMSKHIPQFCGVLGHTFMEFLKDSGDYCQAQHEVYADK
>Anser_cygnoides
-----------------------MAILSVRADFCQAQHSILADK
>Podocoryna_carnea
-------MNSLLSCFCVDLGRTFMAFLSVSGDFCQAQHLAFADK
>Leptobrachium_boringii
------MNSSPVTDSFWNIRLTFMAIRRVRTDYCQAQHSVGADK
>Chaetura_pelagica
-----------------------MEFLKGSGDYCQAQHDLYADK
>Apteryx_australis_mantelli
-----------------------MAILSVRADFCQAQHSTLADK
>Porichthys_notatus
MIKLQSSMSKHIPQFCGVLGHTFMEFLKGRGDYCQAQHAADADE
>Oryzias_latipes
MIKLQSSMSKHIPQFCGVLGHTFMEFLKGSGDYCQAQHAAYADE
>Buceros_rhinoceros_silvestri
-----------------------MEFLKGSGDYCQAQHDLYADK
>Caprimulgus_carolinensis
-----------------------MEFLKGSGDYCQAQHDLYADK
>Scyliorhinus_torazame
-----------------------MEFLKGSGDYCQAQHAVYADK
>Cariama_cristata
-----------------------MAILSVRADFCQAQHSILADK
>Acinonyx_jubatus
-----------------------MAILSVRADFCQAQHSVFADK
>Hynobius_retardatus
-----MTGPSRRTVSCRNIRLTFMAIVSVRADFCQAQHCPVAGK
>Orcinus_orca
-----MPSPQLASVFRWNIRCTFMAILSVRADFCQAQHSVSADK
>Astyanax_mexicanus
MVKLQSSMSKHIPQFCGVLGHTFMEFLKGSGDYCQAQHGVYADK
>Calidris_pugnax
----------MPRVSRWNIRRTFMAILSVRADFCQAQHSILADK
>Callorhinchus_milii
-----------------------MEFLKGSGDYCQAQHDVYADK
>Myotis_davidii
-----------------------MAILSVRADFCQAQHSVFADK
>Oreochromis_niloticus
MIKLQSSMSKHIPQFCGVLGHTFMEFLKGSGDYCQAQHAAYADE
>Pterocles_gutturalis
-----------------------MEFLKGSGDYCQAQHDLYADK
>Dasypus_novemcinctus
-----MPSPQLASVFRWNIRFTFMAILSVRADFCQAQHSVFADK
>Clupea_harengus
------MRNSYRVFFTPDESCTFMAFVSVSGDFCQAQHLVIVDK
>Latimeria_chalumnae
------------------------MAECVRADFCQAQHGILAEK
>Colius_striatus
-----------------------MEFLKGSGDYCQAQHDLYADK
>Columba_livia
-----------------------MAILSVRADFCQAQHSTLADK
>Merops_nubicus
-----------------------MAILSVRADFCQAQHSPLADK
>Alligator_sinensis
-----------------------MAILSVRADFCQAQHSTLADK
>Cuculus_canorus
-----------------------MEFLKGSGDYCQAQHDLYADK
>Sinocyclocheilus_anshuiensis
-----------------------MAFVSVSGDFCQAQHLVIVDK
>Kryptolebias_marmoratus
MIELHSSMSKHIPQFCGVLGHTFMEFLKGSGDYCQAQHAAYADE
>Sarcophilus_harrisii
-----------------------MEFLKGSGDYCQAQHDLYADK
>Galeopterus_variegatus
-----MPSPQLASVFRWNIRCTFMAILSVRADFCQAQHSVFADK
>Monodelphis_domestica
-----------------------MAILSVRADFCQAQHSAFADK
>Macropus_eugenii
---------MKAKQFCGVLGHTFMEFLKGSGDYCQAQHDLYADK
>Esox_lucius
-------MFSLLFYFCVELGHPYMAYVSVSGDFCQAQHLAFVDK
>Haliaeetus_leucocephalus
----------MPRVFCWNIRRTFMAILSVRADFCQAQHSILADK
>Numida_meleagris
----------MPGVFWWKIRRTFMAILSVRADFCQAQHSALADK
>Gavia_stellata
-----------------------MAILSVRADFCQAQHSILADK
>Acanthogobius_hasta
MIKLQSSMSKHIPQFCGVLGHTFMEFLKGSGDYCQAQHVVDADE
>Chlamydotis_macqueenii
-----------------------MAILSVRADFCQAQHRILADK
>Apteronotus_leptorhynchus
-------------------------MAFVSGDFCQAQHLAFVNK
>Condylura_cristata
-----MPSPQLASVFRWNIRCTFMAILSVRADFCQAQHSVFADK
>Ochotona_princeps
-----------------------MEFLKGSGDYCQAQHDLYADK
>Protopterus_aethiopicus
MIKLQSSMSKHIPQFCGVIGHTFMEFLKGSGDYCQAQHALYADK
>Elephantulus_edwardii
MIRPQSSMSKHIPQFCGVLGHTFMEFLKGSGDYCQAQHDLYADK
>Ornithorhynchus_anatinus
-----MPSPRLASAFRWNIRCTFMAILSVRADFCQAQHSVCADK
>Tauraco_erythrolophus
-----------------------MAILSVRADFCQAQHSILADK
>Opisthocomus_hoazin
-----------------------MEFLKGSGDYCQAQHDLYADK
>Scleropages_formosus
-----------------------MAFVSVSGDFCQAQHLASADK
>Corvus_cornix_cornix
-----------------------MAILSVRADFCQAQHSTLAAK
>Egretta_garzetta
-----------------------MAILSVRADFCQAQHSILADK
>Sebastes_caurinus
MIKLQSSMSKHIP-FCGVLGHTFMEFLKGSGDYCQAQHAAYADE
>Ceratotherium_simum_simum
-----MPSPQLASVFRWNIRCTFMAILSVRADFCQAQHSVFADK
>Picoides_pubescens
-----------------------MAILSVRADFCQAQHSILADK
>Cynoglossus_semilaevis
-----------------------MEFLKGSGDYCQAQHAAYADE
>Loxodonta_africana
-----------------------MAILSVRADFCQAQHSIFADK
>Fulmarus_glacialis
-----------------------MAILSVRADFCQAQHSILADK
>Nestor_notabilis
-----------------------MAILSVRAEFCQAQHSILADK
>Leucoraja_erinacea
-------MRKHIPQFCGVLGHTFMEFLKGSGDYCQAQHDVYADK
>Microtus_ochrogaster
---------MREKVFRWNIRCTFMAILSVRADFCQAQHSIFADK
>Coregonus_maraena
-------MFNLLFCFCVDLGHTYMAYVSVSGDFCQAQHLAFVDK
>Tupaia_chinensis
-----------------------MAILSVRADFCQAQHSVFADK
>Lepisosteus_oculatus
-------MSSLPFCFCVDLVRTFMAFLSVSGDFCQAQHLAFADK
>Trichomycterus_areolatus
-----------------------MAFVSVSGDFCQAQHLVIVDK
>Trichechus_manatus_latirostr
-----MPSSQLASVFRWNIRCTFMAILSVRADFCQAQHSIFADK
>Aptenodytes_forsteri
-----------------------MAILSVRADFCQAQHSILADK
>Squalus_acanthias
-------MNRLVTQFSGISGCSFMEFLKGSRDYCQAQHGLLADK
>Micrurus_fulvius
----------MLPVFHLNPSHTFMAILSVRADFCRAQHSTLAEK
>Tyto_alba
-----------------------MEFLKGSGDYCQAQHDLYADK
>Struthio_camelus_australis
-----------------------MAILSVRADFCQAQHSTLADK
>Chrysemys_picta
-----------------------MAILSIRADFCQAQHSTLADK
>Takifugu_rubripes
MIKLQSSMSKHIPQFCGVLGHTFMEFLKGSGDYCQAQHAVYADE
>Tinamus_guttatus
----------MPRVFCWNIRRTFMAILRVRADFCQAQHSTLADK
>Tetronarce_californica
-----------------------MEFVRNSSGYCPAQHSIFADK
>Calypte_anna
-----------------------MEFLKGSGDYCQAQHDLYADK
>Apaloderma_vittatum
-----------------------MAILSVRADFCQAQHSILADK
>Orycteropus_afer_afer
MIRPQSSMSKHIPQFCGVLGHTFMEFLKGSGDYCQAQHDLYADK
```

HG0001 ENSGALG00000003265 PTP4A2

>Gallus_gallus
-----------------------------------------------------MPGVFCWNLRRTFMAILSVRAEFCQAQHSALADK-----
>Acipenser_sinensis
-------------------------------------------MIKLQSSMSKHIPQFCGVLGHTFMEFLKGSGDYCQAQHEVYADK-----
>Amia_calva
-------------------------------------------MIKLQSSMSKHIPQFCGVLGHTFMEFLKGSGDYCQAQHDVYADK-----
>Channa_argus
MSQQTEEAPPQQQFQISDENSSCHIEEEQKGDLTVVSALPPLSLRSHGISMFILLFGFCVDLSRTYMACV--SGEFCQAQHLTSVEK-----
>Kaupichthys_hyoproroides
-------------------------------------------MIKLQSSMSKHIPQFCGVLGHTFMEFLKDSGDYCQAQHEVYADK-----
>Anser_cygnoides
------------------------------------------------------------------MAILSVRADFCQAQHSILADK-----
>Podocoryna_carnea
--------------------------------------------------MNSLLSCFCVDLGRTFMAFLSVSGDFCQAQHLAFADK-----
>Pelophylax_nigromaculatus
---------------------------------------------------MNSSPGSSWNIRRTFMAIRRVSTEYCQAQHSVCADK-----
>Chaetura_pelagica
------------------------------------------------------------------MEFLKGSGDYCQAQHDLYADK-----
>Apteryx_australis_mantelli
------------------------------------------------------------------MAILSVRADFCQAQHSTLADK-----
>Menidia_menidia
--------------------------------------------------MFNLLFGFCVDVSRSYMACV--TGEFCQAQHLTSAEK-----
>Porichthys_notatus
-------------------------------------------MIKLQSSMSKHIPQFCGVLGHTFMEFLKGRGDYCQAQHAADADE-----
>Oryzias_latipes
-------------------------------------------MIKLQSSMSKHIPQFCGVLGHTFMEFLKGSGDYCQAQHAAYADE-----
>Tripterygion_delaisi
--------------------------------------------------MFSLLFGFCVDLSRTYMACV--SGEFCQAQHLTSVEK-----
>Buceros_rhinoceros_silvestri
------------------------------------------------------------------MEFLKGSGDYCQAQHDLYADK-----
>Caprimulgus_carolinensis
------------------------------------------------------------------MEFLKGSGDYCQAQHDLYADK-----
>Scyliorhinus_torazame
------------------------------------------------------------------MEFLKGSGDYCQAQHAVYADK-----
>Cariama_cristata
------------------------------------------------------------------MAILSVRADFCQAQHSILADK-----
>Acinonyx_jubatus
------------------------------------------------------------------MAILSVRADFCQAQHSVFADK-----
>Hynobius_chinensis
------------------------------------------------MTGPSRRTVSCWNIRLTFMAIVSVRADFCQAQHCPVAGK-----
>Orcinus_orca
------------------------------------------------MPSPQLASVFRWNIRCTFMAILSVRADFCQAQHSVSADK-----
>Pygocentrus_nattereri
--------------------------------------------------MFSLFFCFCVDLGRTFMAFV--SGDFCQAQHLAFENK-----
>Calidris_pugnax
-----------------------------------------------------MPRVSRWNIRRTFMAILSVRADFCQAQHSILADK-----
>Callorhinchus_milii
------------------------------------------------------------------MEFLKGSGDYCQAQHDVYADK-----
>Miniopterus_natalensis
------------------------------------------------MPSPQLASVFRWNIRCTFMAILSVRADFCQAQHSVFADK-----
>Neolamprologus_brichardi
-------------------------------------------MIKLQSSMSKHIPQFCGVLGHTFMEFLKGSGDYCQAQHAAYADE-----
>Pterocles_gutturalis
------------------------------------------------------------------MEFLKGSGDYCQAQHDLYADK-----
>Eucidaris_tribuloides
---------------------------------------------------MCKHTEVLAVINKPFMALVLDSGEFCQSQHK----------
>Dasypus_novemcinctus
------------------------------------------------MPSPQLASVFRWNIRFTFMAILSVRADFCQAQHSVFADK-----
>Clupea_harengus
--------------------------------------------------MFNLLFCFCVELGRTYMAFVSVSGDFCQAQHLAFVDK-----
>Latimeria_chalumnae
-------------------------------------------MIKIQSSMSKHIPQFCGVLGHTFMEFLKGSGDYCQAQHDLYADK-----
>Colius_striatus
------------------------------------------------------------------MEFLKGSGDYCQAQHDLYADK-----
>Columba_livia
------------------------------------------------------------------MAILSVRADFCQAQHSTLADK-----
>Merops_nubicus
------------------------------------------------------------------MAILSVRADFCQAQHSPLADK-----
>Alligator_sinensis
------------------------------------------------------------------MAILSVRADFCQAQHSTLADK-----
>Cuculus_canorus
------------------------------------------------------------------MEFLKGSGDYCQAQHDLYADK-----
>Sinocyclocheilus_anshuiensis
--------------------------------------------------MFGPFFHFCVDLDHTYMAFVTICGEFCQAQHLTFVDK-----
>Fundulus_heteroclitus
-------------------------------------------MIKLQSSMSKHIP-FCGVLGRTFMEFLKGSGDYCQAQHAAYADE-----
>Sarcophilus_harrisii
------------------------------------------------------------------MEFLKGSGDYCQAQHDLYADK-----
>Galeopterus_variegatus
------------------------------------------------MPSPQLASVFRWNIRCTFMAILSVRADFCQAQHSVFADK-----
>Monodelphis_domestica
------------------------------------------------------------------MAILSVRADFCQAQHSAFADK-----
>Macropus_eugenii
----------------------------------------------------MKAKQFCGVLGHTFMEFLKGSGDYCQAQHDLYADK-----
>Esox_lucius
--------------------------------------------------MFSLLFYFCVELGHPYMAYVSVSGDFCQAQHLAFVDK-----
>Aquila_chrysaetos_canadensis
-----------------------------------------------------MPRVFCWNIRRTFMAILSVRADFCQAQHSILADK-----
>Gavia_stellata
------------------------------------------------------------------MAILSVRADFCQAQHSILADK-----
>Acanthogobius_hasta
-------------------------------------------MIKLQSSMSKHIPQFCGVLGHTFMEFLKGSGDYCQAQHVVDADE-----
>Chlamydotis_macqueenii
------------------------------------------------------------------MAILSVRADFCQAQHRILADK-----
>Apteronotus_leptorhynchus
------------------------------------------------------------------MAFV--SGDFCQAQHLAFVNK-----
>Xenophysella_greensladeae
---------------------------------------------------MCMATVFLGDACATFMEFHLDSAEYCQAQHK----------
>Sorex_araneus
------------------------------------------------------------------MAILSVRADFCQAQHSVFADK-----
>Ochotona_princeps
------------------------------------------------------------------MEFLKGSGDYCQAQHDLYADK-----
>Protopterus_aethiopicus
-------------------------------------------MIKLQSSMSKHIPQFCGVIGHTFMEFLKGSGDYCQAQHALYADK-----
>Elephantulus_edwardii
-------------------------------------------MIRPQSSMSKHIPQFCGVLGHTFMEFLKGSGDYCQAQHDLYADK-----
>Ornithorhynchus_anatinus
------------------------------------------------MPSPRLASAFRWNIRCTFMAILSVRADFCQAQHSVCADK-----
>Tauraco_erythrolophus
------------------------------------------------------------------MAILSVRADFCQAQHSILADK-----
>Eptatretus_burgeri
--------------------------------------------------MNRHNAQFCGVLGRTFMEFVKGRGEYCETQHIALADK-----
>Octopus_bimaculoides
------------------------------------------------------------------MEFVTNSGDFCQAQHV----------
>Opisthocomus_hoazin
------------------------------------------------------------------MEFLKGSGDYCQAQHDLYADK-----
>Scleropages_formosus
------------------------------------------------------------------MAFVSVSGDFCQAQHLASADK-----
>Zonotrichia_albicollis
-----------------------------------------------------MPTVFCWRILGTFMAILSVRAEFCQAQHSTLAAK-----
>Pecten_maximus
------------------------------------------------------------------MEFVTNSGDYCQAQHAL---------
>Egretta_garzetta
------------------------------------------------------------------MAILSVRADFCQAQHSILADK-----
>Lateolabrax_maculatus
-------------------------MVCL---------H--PGAVTSS-CVRSMNHLLGSHRGRTCMAFARLGGEFCQAQHLLLIVDK----
>Notothenia_coriiceps
------------------------------------------------------------------MACV--SGEFCQAQHLTSVEK-----
>Equus_asinus
------------------------------------------------MPSPQLASVFRWNIRCTFMAILSVRADFCQAQHSVFADK-----
>Perinereis_aibuhitensis
-------------------------------------------MSSEA-GVTTKLHQFFGELGHTFMEFLCNSGEYCQAQHV----------
>Picoides_pubescens
------------------------------------------------------------------MAILSVRADFCQAQHSILADK-----
>Cynoglossus_semilaevis
------------------------------------------------------------------MEFLKGSGDYCQAQHAAYADE-----
>Macaca_nemestrina
------------------------------------------------------------------MAILSVRADFCQAQHSVFADK-----
>Loxodonta_africana
------------------------------------------------------------------MAILSVRADFCQAQHSIFADK-----
>Fulmarus_glacialis
------------------------------------------------------------------MAILSVRADFCQAQHSILADK-----
>Nestor_notabilis
------------------------------------------------------------------MAILSVRAEFCQAQHSILADK-----
>Leucoraja_erinacea
--------------------------------------------------MRKHIPQFCGVLGHTFMEFLKGSGDYCQAQHDVYADK-----
>Fukomys_damarensis
------------------------------------------------------------------MAILSVRADFCQAQHSVFADK-----
>Salmo_salar
--------------------------------------------------MFSLLFCFCVDLDHTYMAYVSVSGDFCQAQHLAFVDK-----
>Tupaia_chinensis
------------------------------------------------------------------MAILSVRADFCQAQHSVFADK-----
>Lepisosteus_oculatus
--------------------------------------------------MSSLPFCFCVDLVRTFMAFLSVSGDFCQAQHLAFADK-----
>Sepia_pharaonis
---------------------MCRNILDLLGGIVKKTDS--PSTLAWP-HHLWRLVKFFGEFGHTFMEFVTNSGDFCQAQHV----------
>Ictalurus_punctatus
-------------------------------------------MVKLQSSMSKHIPQFCGVLGHTFMEFLKGSGDYCQAQHGVYADK-----
>Trichechus_manatus_latirostr
------------------------------------------------MPSSQLASVFRWNIRCTFMAILSVRADFCQAQHSIFADK-----
>Aptenodytes_forsteri
------------------------------------------------------------------MAILSVRADFCQAQHSILADK-----
>Streblospio_benedicti
---------------------------------------------------MCRIILFFGQIGYTFMEFLCNSGEYCQAQHVAIPSVLSRLH
>Squalus_acanthias
--------------------------------------------------MNRLVTQFSGISGCSFMEFLKGSRDYCQAQHGLLADK-----
>Gekko_japonicus
-------------------------------------------------MRKAACEVFLWNPRHTFMAILSIRADFCRAQHSTLADK-----
>Tyto_alba
------------------------------------------------------------------MEFLKGSGDYCQAQHDLYADK-----
>Struthio_camelus_australis
------------------------------------------------------------------MAILSVRADFCQAQHSTLADK-----
>Deroceras_reticulatum
---------------------------------------------------MSSFVLFVGEFGHTFMEFVTNSEDFCQAQHYRLVE------
>Chrysemys_picta
------------------------------------------------------------------MAILSIRADFCQAQHSTLADK-----
>Takifugu_rubripes
-------------------------------------------MIKLQSSMSKHIPQFCGVLGHTFMEFLKGSGDYCQAQHAVYADE-----
>Watasenia_scintillans
------------------------------------------------------------------MEFVTNSGDFCQAQHV----------
>Tinamus_guttatus
-----------------------------------------------------MPRVFCWNIRRTFMAILRVRADFCQAQHSTLADK-----
>Tetronarce_californica
------------------------------------------------------------------MEFVRNSSGYCPAQHSIFADK-----
>Calypte_anna
------------------------------------------------------------------MEFLKGSGDYCQAQHDLYADK-----
>Apaloderma_vittatum
------------------------------------------------------------------MAILSVRADFCQAQHSILADK-----
>Orycteropus_afer_afer
-------------------------------------------MIRPQSSMSKHIPQFCGVLGHTFMEFLKGSGDYCQAQHDLYADK-----

HG0001 ENSGALG00000016271 PTP4A1

```
>Gallus_gallus
--------------------------------------------------------------------------MIRLQSSMS--NHIPQFCGVLGHTFME-FLKGSGDYCQAQHDL-YADK----
>Acipenser_sinensis
--------------------------------------------------------------------------MIKLQSSMS--KHIPQFCGVLGHTFME-FLKGSGDYCQAQHEV-YADK----
>Actinia_tenebrosa
----------------------------------------------------------------------------MWTTNIC--------YDHCGHSLME-FLIDSGDWCQAQHN----------
>Acanthogorgia_aspera
----------------------------------------------------------------------MSRAYCLFG------------FRSRFEKFMEYFLKDSGEFCQAQHE----------
>Amia_calva
--------------------------------------------------------------------------MIKLQSSMS--KHIPQFCGVLGHTFME-FLKGSGDYCQAQHDV-YADK----
>Channa_argus
---------------------MSQQTEEAPPQQQFQISDENSSCHIEEE----------QKGDLTVVSALPPLSLRSHGISMF--ILLFGFCVDLSRTYMA-C--VSGEFCQAQHLTSVEK-----
>Kaupichthys_hyoproroides
--------------------------------------------------------------------------MIKLQSSMS--KHIPQFCGVLGHTFME-FLKDSGDYCQAQHEV-YADK----
>Anser_cygnoides
---------------------------------------------------------------------------------------------------ME-FLKGSGDYCQAQHDL-YADK----
>Podocoryna_carnea
---------------------------------------------------------------------------------------------------ME-FLRGSGDYCFAQHRT---------
>Pseudacris_regilla
MLFTGHG----GEVCERRSGPHPARH-RSPPHRALATGF--SRLLYAPSPPAPRPHHRFFLGT---G------CMIKLQSSMS--KHSPQFCGVLGHTFME-FLKGSGDYCQAQHDL-YADK----
>Chaetura_pelagica
---------------------------------------------------------------------------------------------------ME-FLKGSGDYCQAQHDL-YADK----
>Apteryx_australis_mantelli
---------------------------------------------------------------------------------------------------ME-FLKGSGDYCQAQHDL-YADK----
>Nephilengys_cruentata
----------------------------------------------------------------------------MLDASMC--IDLAV-NQIHPFSLME-ILVGSDEYCRAQHK----------
>Meinertellus_cundinamarcensi
---------------------------------------------------------------------------------MC--RDLVF-LGETGLTYME-FQLDSGEYCQAQHK----------
>Scapharca_broughtonii
---------------------------------------------------------------------------------MF--RKILF-FGEFGHTFME-FVTNSGDYCQAQHV----------
>Menidia_menidia
---------------------------------------------------------------------------------MF--NLLFGFCVDVSRSYMA-C--VTGEFCQAQHLTSAEK-----
>Porichthys_notatus
--------------------------------------------------------------------------MIKLQSSMS--KHIPQFCGVLGHTFME-FLKGRGDYCQAQHAA-DADE----
>Oryzias_melastigma
--------------------------------------------------------------------------MIKLQSSMS--KHIPQFCGVLGHTFME-FLKGSGDYCQAQHAA-YADE----
>Tripterygion_delaisi
---------------------------------------------------------------------------------MF--SLLFGFCVDLSRTYMA-C--VSGEFCQAQHLTSVEK-----
>Buceros_rhinoceros_silvestri
---------------------------------------------------------------------------------------------------ME-FLKGSGDYCQAQHDL-YADK----
>Caprimulgus_carolinensis
---------------------------------------------------------------------------------------------------ME-FLKGSGDYCQAQHDL-YADK----
>Scyliorhinus_torazame
---------------------------------------------------------------------------------------------------ME-FLKGSGDYCQAQHAV-YADK----
>Cariama_cristata
---------------------------------------------------------------------------------------------------ME-FLKGSGDYCQAQHDL-YADK----
>Mustela_putorius_furo
--------------------------------------------------------------------------MIRPQSSMS--KHIPQFCGVLGHTFME-FLKGSGDYCQAQHDL-YADK----
>Alatina_alata
------------------------------------------------------------------------------MRRPC--LC----TCFCGKTFME-FLKDSGDWCQSQHK----------
>Hynobius_retardatus
MRAPELGRSEARAQCGG--GSVAADTATLPPRSPAA-----RPVWYTARR--ACPPATRARGT---G------RMIQPRSSMS--NHSPQFCGVLGHTFME-FLKGSGDYCQAQHDL-YADK----
>Micropterus_floridanus
---------------------------------------------------------------------------------------------------ME-FVRRSGDYCHAQHI----------
>Orcinus_orca
---------------------------------------------------------------------------------------------------ME-FLKGSGDYCQAQHDL-YADK----
>Astyanax_mexicanus
--------------------------------------------------------------------------MVKLQSSMS--KHIPQFCGVLGHTFME-FLKGSGDYCQAQHGV-YADK----
>Calidris_pugnax
---------------------------------------------------------------------------------------------------ME-FLKGSGDYCQAQHDL-YADK----
>Callorhinchus_milii
---------------------------------------------------------------------------------------------------ME-FLKGSGDYCQAQHDV-YADK----
>Eptesicus_fuscus
--------------------------------------------------------------------------MIRPQSSMS--KHIPQFCGVLGHTFME-FLKGSGDYCQAQHDL-YADK----
>Maylandia_zebra
--------------------------------------------------------------------------MIKLQSSMS--KHIPQFCGVLGHTFME-FLKGSGDYCQAQHAA-YADE----
>Pterocles_gutturalis
---------------------------------------------------------------------------------------------------ME-FLKGSGDYCQAQHDL-YADK----
>Eucidaris_tribuloides
---------------------------------------------------------------------------------MC--KHTEV-LAVINKPFMA-LVLDSGEFCQSQHK----------
>Dasypus_novemcinctus
--------------------------------------------------------------------------MIRPQSSMS--KHIPQFCGVLGHTFME-FLKGSGDYCQAQHDL-YADK----
>Alosa_alosa
--------------------------------------------------------------------------MIKLQSSMS--KHIPQFCGVLGHTFME-FLKGSGDYCQAQHGV-YADE----
>Latimeria_chalumnae
--------------------------------------------------------------------------MIKIQSSMS--KHIPQFCGVLGHTFME-FLKGSGDYCQAQHDL-YADK----
>Gyrinus_marinus
---------------------------------------------------------------------------------MC--MAIVW-PGETVLTFME-FQLDSAEYCQAQHK----------
>Colius_striatus
---------------------------------------------------------------------------------------------------ME-FLKGSGDYCQAQHDL-YADK----
>Columba_livia
---------------------------------------------------------------------------------------------------ME-FLKGSGDYCQAQHDL-YADK----
>Leptosomus_discolor
---------------------------------------------------------------------------------------------------ME-FLKGSGDYCQAQHDL-YADK----
>Ricordea_yuma
----------------------------------------------------------------------------MWRGNPV--------YGNRAHTFME-FIIDSGDWCQAQHE----------
>Alligator_mississippiensis
--------------------------------------------------------------------------MIRLQSSMS--NRRPQFCGVLGHTFME-FLKGSGDYCQAQHDL-YADK----
>Cuculus_canorus
---------------------------------------------------------------------------------------------------ME-FLKGSGDYCQAQHDL-YADK----
>Cyprinus_carpio
--------------------------------------------------------------------------MIKLQSSMS--KHIPQFCGVLGHTFME-FLKGSGDYCQAQHGV-YAEK----
>Fundulus_heteroclitus
--------------------------------------------------------------------------MIKLQSSMS--KHIPQFCGVLGRTFME-FLKGSGDYCQAQHAA-YADE----
>Sarcophilus_harrisii
---------------------------------------------------------------------------------------------------ME-FLKGSGDYCQAQHDL-YADK----
>Carcinus_maenas
-----------------------------------------------------------------------------MKFKYCGNNNAQL-SSGRGYTLME-LLKGSGEFCKAQYHIK--------
>Galeopterus_variegatus
---------------------------------------------------------------------------------------------------ME-FLKGSGDYCQAQHDL-YADK----
>Monodelphis_domestica
--------------------------------------------------------------------------MIRPQSSMS--KHIPQFCGVLGHTFME-FLKGSGDYCQAQHDL-YADK----
>Occasjapyx_japonicus
---------------------------------------------------------------------------------MC--RDLVC-LGKIGLTFME-FFLDSGEYCQAQHK----------
>Macropus_eugenii
-------------------------------------------------------------------------------------MKAKQFCGVLGHTFME-FLKGSGDYCQAQHDL-YADK----
>Chironomus_riparius
-------------------------------------------------------------------------------------MDIVL-LRQLALTSME-FKLDSAEYCQAQHK----------
>Isonychia_bicolor
---------------------------------------------------------------------------------MC--RDLVF-LGEIGLTYME-FQLDSGEYCQAQHK----------
>Esox_lucius
--------------------------------------------------------------------------MIKLQSSMS--KHIPQFCGVLGHTFME-FLKGSGDYCQAQHVAAYADE----
>Meganyctiphanes_norvegica
-----------------------------------------------------------------------------MKFKYCGNNNTQL-SCGRGYTLME-LLKGSGEFCKAQYHIK--------
>Haliaeetus_leucocephalus
--------------------------------------------------------------------------MIRLQSSMS--NHIPQFCGVLGHTFME-FLKGSGDYCQAQHDL-YADK----
>Marthasterias_glacialis
---------------------------------------------------------------------------------MS---HLTGICSVPVSSYMA-FVAISGEYCRSQHK----------
>Gavia_stellata
---------------------------------------------------------------------------------------------------ME-FLKGSGDYCQAQHDL-YADK----
>Acanthogobius_hasta
--------------------------------------------------------------------------MIKLQSSMS--KHIPQFCGVLGHTFME-FLKGSGDYCQAQHVV-DADE----
>Eurypyga_helias
---------------------------------------------------------------------------------------------------ME-FLKGSGDYCQAQHDL-YADK----
>Apteronotus_leptorhynchus
---------------------------------------------------------------------------------------------------MA-FLQRAGDYCHAQHWCC--------
>Planococcus_citri
-------------------------------------------------------------------------------------MDIVT-KSALCFTFME-FLLDSSEYCQAQHK----------
>Lasius_neglectus
---------------------------------------------------------------------------------------------------ME-FQLDSAEYCQAQHK----------
>Condylura_cristata
--------------------------------------------------------------------------MIRPQSSMS--KHIPQFCGVLGHTFME-FLKGSGDYCQAQHDL-YADK----
>Ornithodoros_turicata
-----------------------------------------------------------------------------MKSDAAG-----T-NGEPSHSWMEVSLGPSGEYCLAQHK----------
>Ochotona_princeps
--------------------------------------------------------------------------MIRPQSSMS--KHIPQFCGVLGHTFME-FLKGSGDYCQAQHDL-YADK----
>Antheraea_yamamai
---------------------------------------------------------------------------------------------------ME-FRLDSAEYCQAQHK----------
>Protopterus_aethiopicus
--------------------------------------------------------------------------MIKLQSSMS--KHIPQFCGVIGHTFME-FLKGSGDYCQAQHAL-YADK----
>Lingula_anatina
---------------------------------------------------------------------------------------MQF-FGEFGHTFME-FVTNSGEYCQAQHK----------
>Elephantulus_edwardii
--------------------------------------------------------------------------MIRPQSSMS--KHIPQFCGVLGHTFME-FLKGSGDYCQAQHDL-YADK----
>Metallyticus_splendidus
---------------------------------------------------------------------------------MC--MALVF-LGETVLTYME-FQLDSGEYCQAQHK----------
>Nannochorista_philpotti
---------------------------------------------------------------------------------MC--MAIVV-FPTVALRYME-FHLDSAEYCQAQHK----------
>Ornithorhynchus_anatinus
--------------------------------------------------------------------------MIATHSSMS--KHSPQFCGVLGHTFME-FLKDSGDYCQAQHDL-YADK----
>Cypridininae_sp._BMR-2011
--------------------------------------------------------------------MEDSETSLLYKYRLC--MRPQV-LAGVCCSFME-FVLADGEYCQAQHK----------
>Perna_viridis
---------------------------------------------------------------------------------MC--RNILF-FGVLGHTFME-FVTNSGDYCQAQHV----------
>Eptatretus_burgeri
--------------------------------------------------------------------------------MNR--HNAQF-CGVLGRTFME-FVKGRGEYCETQHIALADK-----
>Osmylus_fulvicephalus
---------------------------------------------------------------------------------MC--MARVF-LGETVLTYME-FQLDSAEYCQAQHK----------
>Triops_newberryi
---------------------------------------------------------------------------------MC--TTGQR-WVEWRHKFLA-VLLGSGDYCEAQHK----------
>Hapalochlaena_maculosa
----------------------------------------------------MCRNILDLLGGIVEK--TDSPSTLAWPHHLC--RSIKF-FGEFGHTFME-FVTNSGDFCQAQHV----------
>Epiophlebia_superstes
---------------------------------------------------------------------------------------------------ME-FQLDSGEYCQAQHK----------
>Opisthocomus_hoazin
---------------------------------------------------------------------------------------------------ME-FLKGSGDYCQAQHDL-YADK----
>Stenobothrus_lineatus
------------------------------------MVVEASAVALADEGRTAGRHVMRLERA--SSPSSMVDSQAAGGGRLC--AHQQF-RGEIVLTFME-FQLDSGEYCQAQHK----------
>Plecoglossus_altivelis
---------------------------------------------------------------------------------------------------ME-FLQGSGDYCHAQHPSI--------
>Scleropages_formosus
---------------------------------------------------------------------------------MC--GLLFRFCVDLGRPFMA-FVSVSGDFCQAQHLASADK-----
>Crassostrea_gigas
---------------------------------------------MR-----DPRGPVTVRGKFVTGFVLCSSLLNCW-------QHTEF-RSNFGHTYME-FVTNSGDYCQAQHV----------
>Endeis_spinosa
---------------------------------------------------------------------------------------------------ME-FLKESGDYCYCRAQHV--------
>Lepidothrix_coronata
--------------------------------------------------------------------------MIRLQSSMS--NHIPQFCGVLGHTFME-FLKGSGDYCQAQHDL-YADK----
>Pecten_maximus
---------------------------------------------------------------------------------------------------ME-FVTNSGDYCQAQHAL---------
>Pelecanus_crispus
---------------------------------------------------------------------------------------------------ME-FLKGSGDYCQAQHDL-YADK----
>Sebastes_caurinus
--------------------------------------------------------------------------MIKLQSSMS--KHIP-FCGVLGHTFME-FLKGSGDYCQAQHAA-YADE----
>Equus_przewalskii
--------------------------------------------------------------------------MIRPQSSMS--KHSPQFCGVLGHTFME-FLKGSGDYCQAQHDL-YADK----
>Peruphasma_schultei
---------------------------------------------------------------------------------------------------ME-FLPNSGEYCQAQHK----------
>Menopon_gallinae
---------------------------------------------------------------------------MNFRFSGC--MALKN-FHSTSSRLME-FFLDSAEYCQAQHTK---------
>Perinereis_aibuhitensis
--------------------------------------------------------------------------MSSEAGVTT--KLHQF-FGELGHTFME-FLCNSGEYCQAQHV----------
>Perla_marginata
---------------------------------------------------------------------------------------------------ME-FQLDSGEYCQAQHK----------
>Cynoglossus_semilaevis
---------------------------------------------------------------------------------------------------ME-FLKGSGDYCQAQHAA-YADE----
>Gorilla_gorilla
--------------------------------------------------------------------------MIRPQSSMS--KHIPQFCGVLGHTFME-FLKGSGDYCQAQHDL-YADK----
>Loxodonta_africana
--------------------------------------------------------------------------MIRPQSSMS--KHIPQFCGVLGHTFME-FLKGSGDYCQAQHDL-YADK----
>Fulmarus_glacialis
---------------------------------------------------------------------------------------------------ME-FLKGSGDYCQAQHDL-YADK----
>Nestor_notabilis
---------------------------------------------------------------------------------------------------ME-FLKGSGDYCQAQHDL-YADK----
>Ectopsocus_briggsi
---------------------------------------------------------------------------------------------------ME-FFLDSAEYCQAQHRK---------
>Leucoraja_erinacea
---------------------------------------------------------------------------------MR--KHIPQFCGVLGHTFME-FLKGSGDYCQAQHDV-YADK----
>Inocellia_crassicornis
---------------------------------------------------------------------------------------------------ME-FQLDSAEYCQAQHK----------
>Rhopilema_esculentum
----------------------------------------------------------------------------------MNSYHLLL-SKYFGKPFME-FLKDSAEWCKAQHVE---------
>Heterocephalus_glaber
--------------------------------------------------------------------------MIRPQSSMS--RHIPQFCGVLGHTFME-FLKGSGDYCQAQHDL-YADK----
>Salmo_salar
--------------------------------------------------------------------------MIKLQSSMS--KHIPQFCGVLGHTFME-FLKGSGDYCQAQHAAAYADE----
>Acropora_digitifera
----------------------------------------------------------------------------MWRFSLT--------FGSRAHTFME-FIIDSGDWCQAQHE----------
>Tityus_serrulatus
---------------------------------------------------------------------------------------------------ME-YLKTTGEYCRAQHK----------
>Aurelia_aurita
----------------------------------------------------------------------------------MTSYHLQL--KCFGETFME-FLKDSAEWCKAQHVE---------
>Lepisosteus_oculatus
--------------------------------------------------------------------------MIKLQSSMS--KHIPQFCGVLGHTFME-FLKGSGDYCQAQHGV-YADK----
>Sepia_pharaonis
----------------------------------------------------MCRNILDLLGGIVKK--TDSPSTLAWPHHLW--RLVKF-FGEFGHTFME-FVTNSGDFCQAQHV----------
>Sepioloidea_lineolata
---------------------------------------------------------------------------------------------------ME-FVTNSGDFCQAQHV----------
>Ictalurus_punctatus
--------------------------------------------------------------------------MVKLQSSMS--KHIPQFCGVLGHTFME-FLKGSGDYCQAQHGV-YADK----
>Oropsylla_silantiewi
---------------------------------------------------------------------------------------------------ME-FQLDSAEYCQAQHK----------
>Trichechus_manatus_latirostr
--------------------------------------------------------------------------MIRPQSSMS--KHIPQFCGVLGHTFME-FLKGSGDYCQAQHDL-YADK----
>Aptenodytes_forsteri
---------------------------------------------------------------------------------------------------ME-FLKGSGDYCQAQHDL-YADK----
>Streblospio_benedicti
---------------------------------------------------------------------------------MC--RIILF-FGQIGYTFME-FLCNSGEYCQAQHVAIPSVLSRLH
>Squalus_acanthias
---------------------------------------------------------------------------------MN--RLVTQFSGISGCSFME-FLKGSRDYCQAQHGLLAD-K----
>Takydromus_sexlineatus
--------------------------------------------------------------------------MIRLQSSMS--NHIPQFCGVLGHTFME-FLKGSGDYCQAQHGL-YADK----
>Stylops_melittae
---------------------------------------------------------------------------------MC--IGILN-LGNKVVEYME-FHFGSSEYCKAQHK----------
>Tyto_alba
---------------------------------------------------------------------------------------------------ME-FLKGSGDYCQAQHDL-YADK----
>Struthio_camelus_australis
---------------------------------------------------------------------------------------------------ME-FLKGSGDYCQAQHDL-YADK----
>Deroceras_reticulatum
---------------------------------------------------------------------------------MS--SFVLF-VGEFGHTFME-FVTNSEDFCQAQHYRLVE------
>Pelodiscus_sinensis
--------------------------------------------------------------------------MIRLQSSMS--NHIPQFCGVLGHTFME-FLKGSGDYCQAQHDL-YADK----
>Takifugu_rubripes
--------------------------------------------------------------------------MIKLQSSMS--KHIPQFCGVLGHTFME-FLKGSGDYCQAQHAV-YADE----
>Watasenia_scintillans
---------------------------------------------------------------------------------------------------ME-FVTNSGDFCQAQHV----------
>Gynaikothrips_ficorum
-----------------------------------------------------MVHCAKFNGE--NSISMVIRRVIFDNKWPS--VGYQT-CGEILPTFME-FHLDSAEYCQAQHK----------
>Tinamus_guttatus
---------------------------------------------------------------------------------------------------ME-FLKGSGDYCQAQHDL-YADK----
>Tetronarce_californica
---------------------------------------------------------------------------------------------------ME-FVRNSSGYCPAQHSIFADK-----
>Annulipalpia_sp._AD-2013
---------------------------------------------------------------------------------MC--LAERLEACSLATTLME-FRLDSSEYCQAQHK----------
>Calypte_anna
---------------------------------------------------------------------------------------------------ME-FLKGSGDYCQAQHDL-YADK----
>Apaloderma_vittatum
---------------------------------------------------------------------------------------------------ME-FLKGSGDYCQAQHDL-YADK----
>Orycteropus_afer_afer
--------------------------------------------------------------------------MIRPQSSMS--KHIPQFCGVLGHTFME-FLKGSGDYCQAQHDL-YADK----
>Ruditapes_philippinarum
---------------------------------------------------------------------------------MN--RIILF-FGELGHTFME-FLTNSGEYCLAQHVHV--------
>Limulus_polyphemus
----------------------------------------------------------------------MSST--NDSAKVP--LLLPF-EVRRHHTLME-FLKTAGEYCQSQHK----------
>Palythoa_caribaeorum
-------------------------------------------------------------------------------------MRRDYLAYDRGRTLME-FLIDSGDWCQAQHE----------
>Thermobia_domestica
---------------------------------------------------------------------------------------------------ME-FQLDSGEYCQAQHK----------
```

HG0001 FBgn0024734 PRL-1

```
>Drosophila_melanogaster
-------------------------------------------MCM--------------------AIV--L-VRPVPTSME-FHLDSAEYCQAQHK------
>Acipenser_sinensis
--------------------------------------------------------MIKLQSSMSKHI-PQFCGVLGHTFME-FLKGSGDYCQAQHEVYADK-
>Exaiptasia_pallida
-----------------------------------------MW-----------------------RGL--ALGRRDYSLME-FLIDSGDWCQAQHYK-----
>Acanthogorgia_aspera
---------------------------------------------------------------MSRAYCLFGFRSRFEKFMEYFLKDSGEFCQAQHE------
>Amia_calva
--------------------------------------------------------MIKLQSSMSKHI-PQFCGVLGHTFME-FLKGSGDYCQAQHDVYADK-
>Anser_cygnoides
--------------------------------------------------------------------------------ME-FLKGSGDYCQAQHDLYADK-
>Xenopus_tropicalis
----------------------------------MYVRFDTRAQRPA------DNGMPTQGGVRDWAA-RQFCGVLGHTFME-FLKGSGDYCQAQHELYADK-
>Chaetura_pelagica
--------------------------------------------------------------------------------ME-FLKGSGDYCQAQHDLYADK-
>Apteryx_australis_mantelli
--------------------------------------------------------------------------------ME-FLKGSGDYCQAQHDLYADK-
>Nephilengys_cruentata
--------------------------------------MLDASMCID---------------L-----A--VNQIHPFSLME-ILVGSDEYCRAQHK------
>Meinertellus_cundinamarcensi
-------------------------------------------MCR--------------------DLV--FLGETGLTYME-FQLDSGEYCQAQHK------
>Scapharca_broughtonii
-----------------------------------------MFRKI-----------------------L-FFGEFGHTFME-FVTNSGDYCQAQHV------
>Buceros_rhinoceros_silvestri
--------------------------------------------------------------------------------ME-FLKGSGDYCQAQHDLYADK-
>Caprimulgus_carolinensis
--------------------------------------------------------------------------------ME-FLKGSGDYCQAQHDLYADK-
>Scyliorhinus_torazame
--------------------------------------------------------------------------------ME-FLKGSGDYCQAQHAVYADK-
>Cariama_cristata
--------------------------------------------------------------------------------ME-FLKGSGDYCQAQHDLYADK-
>Mustela_putorius_furo
--------------------------------------------------------MIRPQSSMSKHI-PQFCGVLGHTFME-FLKGSGDYCQAQHDLYADK-
>Alatina_alata
------------------------------------------MRRP-----------------------CLCTCFCGKTFME-FLKDSGDWCQSQHK------
>Hynobius_retardatus
MRAPELGRSEARAQCGGGSVAADTATLPPRSPAARPVWYTARRACPPATRARGTGRMIQPRSSMSNHS-PQFCGVLGHTFME-FLKGSGDYCQAQHDLYADK-
>Orcinus_orca
--------------------------------------------------------------------------------ME-FLKGSGDYCQAQHDLYADK-
>Astyanax_mexicanus
--------------------------------------------------------MVKLQSSMSKHI-PQFCGVLGHTFME-FLKGSGDYCQAQHGVYADK-
>Charadrius_vociferus
--------------------------------------------------------------------------------ME-FLKGSGDYCQAQHDLYADK-
>Callorhinchus_milii
--------------------------------------------------------------------------------ME-FLKGSGDYCQAQHDVYADK-
>Myotis_lucifugus
--------------------------------------------------------MIRPQSSMSNHI-P-FCGVLGHTFME-FLKGSGDYCQAQHDLYADK-
>Oreochromis_niloticus
--------------------------------------------------------MIKLQSSMSKHI-PQFCGVLGHTFME-FLKGSGDYCQAQHAAYADE-
>Pterocles_gutturalis
--------------------------------------------------------------------------------ME-FLKGSGDYCQAQHDLYADK-
>Eucidaris_tribuloides
-------------------------------------------MCKH---------------T-----E--VLAVINKPFMA-LVLDSGEFCQSQHK------
>Dasypus_novemcinctus
--------------------------------------------------------MIRPQSSMSKHI-P-FCGVLGHTFME-FLKGSGDYCQAQHDLYADK-
>Latimeria_chalumnae
--------------------------------------------------------MIKIQSSMSKHI-PQFCGVLGHTFME-FLKGSGDYCQAQHDLYADK-
>Dastarcus_helophoroides
-------------------------------------------MCM--------------------AIV--YFGELIPSFME-FHLDSAEYCQAQHK------
>Colius_striatus
--------------------------------------------------------------------------------ME-FLKGSGDYCQAQHDLYADK-
>Columba_livia
--------------------------------------------------------------------------------ME-FLKGSGDYCQAQHDLYADK-
>Leptosomus_discolor
--------------------------------------------------------------------------------ME-FLKGSGDYCQAQHDLYADK-
>Corynactis_australis
-----------------------------------------MWRGN-----------------------R-VYGNRAHTFME-FIIDSGDWCQAQHE------
>Alligator_mississippiensis
--------------------------------------------------------MIRLQSSMSNRR--PFCGVLGHTFME-FLKGSGDYCQAQHDLYADK-
>Cuculus_canorus
--------------------------------------------------------------------------------ME-FLKGSGDYCQAQHDLYADK-
>Sinocyclocheilus_grahami
--------------------------------------------------------MIKLQSSMSKHI-PQFCGVLGHTFME-FLKDSGDYCQAQHGVYAEK-
>Austrofundulus_limnaeus
--------------------------------------------------------------------------------MA-FARLGGEFCQAQHLLLIVDK
>Sarcophilus_harrisii
--------------------------------------------------------------------------------ME-FLKGSGDYCQAQHDLYADK-
>Forficula_auricularia
--------------------------------------------------------------------------------ME-FRLDSAEYCQAQHK------
>Galeopterus_variegatus
--------------------------------------------------------------------------------ME-FLKGSGDYCQAQHDLYADK-
>Monodelphis_domestica
--------------------------------------------------------MIRPQSSMSKHI--PFCGVLGHTFME-FLKGSGDYCQAQHDLYADK-
>Occasjapyx_japonicus
-------------------------------------------MCR--------------------DLV--CLGKIGLTFME-FFLDSGEYCQAQHK------
>Macropus_eugenii
-----------------------------------------------------------------MKA-KQFCGVLGHTFME-FLKGSGDYCQAQHDLYADK-
>Evechinus_chloroticus
-------------------------------------------MCKP---------------A-----E--VSAISNLAFMA-SVLDSGEYCQSQHK------
>Eurylophella_sp._AD-2013
-------------------------------------------MCR--------------------DLV--FLGEIGLTYME-FQLDSGEYCQAQHK------
>Esox_lucius
--------------------------------------------------------MIKLQSSMSKHI-P-FCGVLGHTFME-FLKGSGDYCQAQHVAAYADE
>Falco_cherrug
--------------------------------------------------------------------------------ME-FLKGSGDYCQAQHDLYADK-
>Leptasterias_sp._AR-2014
-------------------------------------------MSH-----------------------S-TGISSVPVSMA-FVSISGEYCRSQHK------
>Meleagris_gallopavo
--------------------------------------------------------------------------------ME-FLKGSGDYCQAQHDLYADK-
>Gavia_stellata
--------------------------------------------------------------------------------ME-FLKGSGDYCQAQHDLYADK-
>Acanthogobius_hasta
--------------------------------------------------------MIKLQSSMSKHI-PQFCGVLGHTFME-FLKGSGDYCQAQHVVDADE-
>Balearica_regulorum_gibberic
--------------------------------------------------------------------------------ME-FLKGSGDYCQAQHDLYADK-
>Okanagana_villosa
-------------------------------------------MCM--------------------AIV--FLGETWLTFME-FHLDSAEYCQAQHK------
>Leptopilina_boulardi
-------------------------------------------MCM--------------------AIL--FPDECILTYME-FQLDSAEYCQAQHK------
>Condylura_cristata
--------------------------------------------------------MIRPQSSMSKHI--PFCGVLGHTFME-FLKGSGDYCQAQHDLYADK-
>Ochotona_princeps
--------------------------------------------------------------------------------ME-FLKGSGDYCQAQHDLYADK-
>Azaleodes_micronipha
-------------------------------------------MCM--------------------AAV--LEVSSLATLME-FRLDSAEYCQAQHK------
>Lingula_anatina
--------------------------------------------------------------------------------ME-FVTNSGEYCQAQHK------
>Elephantulus_edwardii
--------------------------------------------------------MIRPQSSMSKHI-PQFCGVLGHTFME-FLKGSGDYCQAQHDLYADK-
>Metallyticus_splendidus
-------------------------------------------MCM--------------------ALV--FLGETVLTYME-FQLDSGEYCQAQHK------
>Nannochorista_philpotti
-------------------------------------------MCM--------------------AIV--VFPTVALRYME-FHLDSAEYCQAQHK------
>Ornithorhynchus_anatinus
--------------------------------------------------------MIATHSSMSKH--SPFCGVLGHTFME-FLKDSGDYCQAQHDLYADK-
>Cypridininae_sp._BMR-2011
---------------------MED----SETS---LLYKYRLCMRP-----------------------Q-VLAGVCCSFME-FVLADGEYCQAQHK------
>Perna_viridis
-----------------------------------------MCRNI-----------------------L-FFGVLGHTFME-FVTNSGDYCQAQHV------
>Osmylus_fulvicephalus
-------------------------------------------MCM--------------------ARV--FLGETVLTYME-FQLDSAEYCQAQHK------
>Epiophlebia_superstes
--------------------------------------------------------------------------------ME-FQLDSGEYCQAQHK------
>Opisthocomus_hoazin
--------------------------------------------------------------------------------ME-FLKGSGDYCQAQHDLYADK-
>Prosarthria_teretrirostris
-------------------------------------------MCM--------------------AIV--SRGEIVLTYME-FQLDSGEYCQAQHK------
>Crassostrea_gigas
MRDPR-----GPVTVRGKFVTGF-------VL---CSSLLNCWQHT-----------------------E-FRSNFGHTYME-FVTNSGDYCQAQHV------
>Sturnus_vulgaris
--------------------------------------------------------MIRLQSSMSNHI-PQFCGVLGHTFME-FLKGSGDYCQAQHDLYADK-
>Pecten_maximus
--------------------------------------------------------------------------------ME-FVTNSGDYCQAQHAL-----
>Egretta_garzetta
--------------------------------------------------------------------------------ME-FLKGSGDYCQAQHDLYADK-
>Ceratotherium_simum_simum
--------------------------------------------------------------------------------ME-FLKGSGDYCQAQHDLYADK-
>Medauroidea_extradentata
-------------------------------------------MCM--------------------ALV--FTAGVAVRHME-FLLDSGEYCQAQHK------
>Menopon_gallinae
-------------------------------------MNFRFSGCM--------------------ALK--NFHSTSSRLME-FFLDSAEYCQAQHTK-----
>Glycera_dibranchiata
-----------------------------------------MEQCHP---------------VLKTHSK--FFGVLGHTFME-FLCDSGEYCQAQHV------
>Leuctra_sp._AD-2013
-------------------------------------------MCM--------------------ALV--LVGEIVLTYME-FQLDSGEYCQAQHRK-----
>Homo_sapiens
--------------------------------------------------------------------------------ME-FLKGSGDYCQAQHDLYADK-
>Loxodonta_africana
--------------------------------------------------------MIRPQSSMSKHI-P-FCGVLGHTFME-FLKGSGDYCQAQHDLYADK-
>Fulmarus_glacialis
--------------------------------------------------------------------------------ME-FLKGSGDYCQAQHDLYADK-
>Melopsittacus_undulatus
--------------------------------------------------------------------------------ME-FLKGSGDYCQAQHDLYADK-
>Ectopsocus_briggsi
--------------------------------------------------------------------------------ME-FFLDSAEYCQAQHRK-----
>Leucoraja_erinacea
---------------------------------------------------------------MRKHI-PQFCGVLGHTFME-FLKGSGDYCQAQHDVYADK-
>Inocellia_crassicornis
--------------------------------------------------------------------------------ME-FQLDSAEYCQAQHK------
>Rhopilema_esculentum
----------------------------------------------------------------MNSYHLLLSKYFGKPFME-FLKDSAEWCKAQHVE-----
>Rattus_norvegicus
--------------------------------------------------------MIRPQSSMSRHI-P-FCGVLGHTFME-FLKGSGDYCQAQHDLYADK-
>Ctenactis_echinata
-----------------------------------------MWRFN-----------------------L-VYGSRAHTFME-FIIDSGDWCQAQHE------
>Aurelia_aurita
----------------------------------------------------------------MTSYHLQL-KCFGETFME-FLKDSAEWCKAQHVE-----
>Lepisosteus_oculatus
--------------------------------------------------------MIKLQSSMSKHI-PQFCGVLGHTFME-FLKGSGDYCQAQHGVYADK-
>Sepia_pharaonis
--MCR-----NILDLLGGIVKKTD----SPST---LAWPHHLWRLV-----------------------K-FFGEFGHTFME-FVTNSGDFCQAQHV------
>Sepioloidea_lineolata
--------------------------------------------------------------------------------ME-FVTNSGDFCQAQHV------
>Trichomycterus_areolatus
--------------------------------------------------------------------------------ME-FLQGSGDYCHAQHRDA----
>Oropsylla_silantiewi
--------------------------------------------------------------------------------ME-FQLDSAEYCQAQHK------
>Trichechus_manatus_latirostr
--------------------------------------------------------MIRPQSSMSKHI-PQFCGVLGHTFME-FLKGSGDYCQAQHDLYADK-
>Aptenodytes_forsteri
--------------------------------------------------------------------------------ME-FLKGSGDYCQAQHDLYADK-
>Henricia_sp._AR-2014
-----------------------------------------MCQLT-----------------------AILNTISVPTLMA-FAGISGEYCRSQHK------
>Crotalus_horridus
--------------------------------------------------------MIRLQSSMSNHI-PQFCGVLGHTFME-FLKGSGDYCQAQHGLYADK-
>Stylops_melittae
-------------------------------------------MCI--------------------GIL--NLGNKVVEYME-FHFGSSEYCKAQHK------
>Tyto_alba
--------------------------------------------------------------------------------ME-FLKGSGDYCQAQHDLYADK-
>Struthio_camelus_australis
--------------------------------------------------------------------------------ME-FLKGSGDYCQAQHDLYADK-
>Sphaerechinus_granularis
-------------------------------------------MCKP---------------A-----E--DSAISNQAFMA-TVLDSGEYCQSQHK------
>Chelonia_mydas
--------------------------------------------------------------------------------ME-FLKGSGDYCQAQHDLYADK-
>Watasenia_scintillans
--------------------------------------------------------------------------------ME-FVTNSGDFCQAQHV------
>Gynaikothrips_ficorum
MVHC-------------AKFNGEN----SISMVIRRVIFDNKWPSV--------------------GYQ--TCGEILPTFME-FHLDSAEYCQAQHK------
>Tinamus_guttatus
--------------------------------------------------------------------------------ME-FLKGSGDYCQAQHDLYADK-
>Hydroptila_sp._AD-2013
--------------------------------------------------------------------------------ME-FRLDSGEYCQAQHK------
>Calypte_anna
--------------------------------------------------------------------------------ME-FLKGSGDYCQAQHDLYADK-
>Apaloderma_vittatum
--------------------------------------------------------------------------------ME-FLKGSGDYCQAQHDLYADK-
>Orycteropus_afer_afer
--------------------------------------------------------MIRPQSSMSKHI-PQFCGVLGHTFME-FLKGSGDYCQAQHDLYADK-
>Eurhomalea_rufa
--------------------------------------------------------------------------------ME-FLTNSGEYCLAQHVHV----
>Limulus_polyphemus
--------------------------------------------------------------------------------ME-FLKTAGEYCQSQHK------
>Protopalythoa_variabilis
---------------------------------------------------------------MR-RDY--LAYDRGRTLME-FLIDSGDWCQAQHE------
>Tricholepidion_gertschi
--------------------------------------------------------------------------------ME-FQLDSGEYCQAQHK------
```

HG0002 ENSDARG00000003077

>Danio_rerio
---------------------------------------------------------------------------------------------MAANSVYDFTVETL----EE-----KSVSLDIFRGKVLLIMNVATF-----------------------------------------
>Acipenser_sinensis
---------------------------------------------------------------------------------------MSLKMAGGIKKFYDFSAKIL----NT----GEALNFSKFKGKVVLIENVASL-----------------------------------------
>Anemonia_viridis
------------------------------MLRTVFQFSS--------------------V----SR----PVS----RLIK--------SQIRSMASVHDFTVQ-N----LQ----GEPVSLSKYKGKVILIENVASL-----------------------------------------
>Talitrus_saltator
---------------------MK---PILALLATAGLANA--------------------------L----YNPDNATECC------------AVDGSLFDFKLKNL----TE----DGSYDLSKLRGKVVLVVNVATF-----------------------------------------
>Channa_striata
-------------------------------------------------------------------------------------------MAGNVTGFYEITAKLL----S-----GETFSFSTLKGKVVLLENVASL-----------------------------------------
>Anguilla_anguilla
-------------------------------------------------------------------------------------------MAGNIKKFYDFSLKLL----S-----GDVLNLSSLKGKVVLIENVASL-----------------------------------------
>Hydractinia_symbiolongicarpu
-----------------------------------------------------------------------------------------MNSSARMASFYDFSVKTL----K-----GETKSMSDYKGKVVLIENTATL-----------------------------------------
>Microhyla_fissipes
---------------------------------------------------------------------------------------MVMCRVMAVRSVFEFSARLL----S-----GEAVSLERYRGKVLLIENVASL-----------------------------------------
>Nephilengys_cruentata
------------------MGWLG---LAFATLACGVIL----------------------------G----SAAKVPSRSCF--------RPGQSDLTVYNFTLSDL----LQ----TRNISLSEYQDKILLIVNVATY-----------------------------------------
>Meinertellus_cundinamarcensi
--------------MGAWSPWMA---ALSVLVAY-------------------------------VG----ATDVVKSRPCY--------HNRRANTTIYEFTAQDM----MK----TKNISLSDYRGKVVLIVNVATY-----------------------------------------
>Scapharca_broughtonii
-------------------------------------------MAGRASQGALLLATV-LG----IV----DATLPRKAVCT--------QPKNATATIYDYSLKEI----YG----VKTLNMSMFQGQVVLFVNVATY-----------------------------------------
>Argulus_siamensis
---------------------------MNRRIADKSLLTT--------------LLLCLLA----GW----SRADLPSVG----------CRGNFPGTIHHYNVTAL----DG----QSLIDFKLFKNQVLLIVNVATY-----------------------------------------
>Menidia_menidia
---------------------------------------------------------------------------------------------MENKSVYDFSVETL----DG-----HQVPLRRFRGKVLLIVNVATF-----------------------------------------
>Opsanus_beta
-----------------------------------------------------------------------------------------MAAVRKVKRFYDLTAKLL----S-----GETLTFSALKGKVVLIENVASL-----------------------------------------
>Oryzias_latipes
---------------------------------------------------------------------------------------------MGEKSVYDFFATSL----DG-----QQIPLGTFRNKVLLIVNVATF-----------------------------------------
>Cryptocercus_wrighti
---MITIEKRPCTILGTNMHL-------------AGILAL-------------LLLTLHGS----HG----KVERVASVQCD--------NEDLIENSIYNYTLPDL----WE----VQNVSLSDFRGKVVLIVNVATY-----------------------------------------
>Tripterygion_delaisi
-----------------------------------------------------------------------------------------------MSTIYDFTSKLL----T-----GETFSFSSLKGKVVLIENVASL-----------------------------------------
>Calanus_helgolandicus
-----------------------------------------------------------------------------------------------MPAFYDIRAKKI----G-----GEVVSMADYKGKVVLIENTASL-----------------------------------------
>Arenicola_marina
---------------------------------------------------------------------------MAESKEK--------DWWKKAGSIYEFSAQDI----DG-----NDTSLEKYKGHVCLIVNVATK-----------------------------------------
>Trachinotus_blochii
-------------------------------------------------------------------------------------------MAGNLKRFYGLTAKLL----S-----GETFSFPALRGKVVLIENVASL-----------------------------------------
>Acinonyx_jubatus
-------------------------------------------------------------------------------MCAAPLAAA-ALAAAAPRSVYSFSARPL----AG----GEPMSLGSLRGKVLLIENVASL-----------------------------------------
>Hynobius_retardatus
-------------------------------------------------------------------------------------------MAYVAKSFYDLKATNL----E-----GETVDFNIFRGRVVLIENVASL-----------------------------------------
>Micropterus_floridanus
----------------------------------------------------------------------------------------------MATTLYNFTAKLL----T-----GETFNFSSLQGKVVLIENVASL-----------------------------------------
>Balaenoptera_acutorostrata_s
------------------------MTPVARLLKGGAS----------------RTPCSSSP-----R--LSPVSLGTHTMCAAQRSAA-ALAAAAPRSVYAFSARPL----AG----GEPVNLGSLRGKVLLIENVASL-----------------------------------------
>Pygocentrus_nattereri
--------------------------------------------------------------------------------------------MANVKSFYDITAKTL----T-----GEEFKFSTLQGKVVLIENVASL-----------------------------------------
>Callorhinchus_milii
--------------------------------------------------------------------------------------------MANSRKFYGFSTKLL----N-----GQTLNFSKFKGKVVLIENVASL-----------------------------------------
>Rhinolophus_ferrumequinum
--------MGRGRGGGAAGKVLSPHDPIARLLKGGAA----------------RDPFCRVP-----S--ATPS--ETSAMCAAQLVAA-ARAAAAPRSVYDFSARPL----AG----GDPVNLGSLRGKVLLIENVASL-----------------------------------------
>Haplochromis_burtoni
----------------------------------------------------------------------------------------------MATSFYDLTAKLL----T-----GETFNFSSLKGKVVLIENVASL-----------------------------------------
>Dasypus_novemcinctus
-------------------------------------------------------------------------------MCAAALSAA---ALAAQRSVHAFSARPL----AG----GEPVSLGSLRGKVLLIENVASL-----------------------------------------
>Clupea_harengus
--------------------------------------------------------------------------------------------MAAIKTLYDISAKLL----T-----GELLNFSSLKGKVILIENVASL-----------------------------------------
>Latimeria_chalumnae
-------------------------------------------------------------------------------------------MAVGMKKFYDFSAKIL----NT----GELLSFSKFKGKVVLLENVASL-----------------------------------------
>Rhodactis_indosinensis
-----------------MYLWYV---FVAFCLYWGNIGGT--------------------------E----RQK----SRSL--------CREEHKKSFYDFSMK-N----LD----GELVSMSDFRGHVVMAVNVATF-----------------------------------------
>Alligator_mississippiensis
------------MNWGRLAKRAL-----------------------LCG----------------AVAARGLG-RSMCAQVD--------EDWRSAKSIYYFHAKDI----DG-----NDVSLEKYQGQVCIITNVASK-----------------------------------------
>Echinococcus_multilocularis
-------------------------------------------------------------------------------MCASR------LSAAAQSTVFSFSARPL----AG----GEPVSLGSLRGKVLLIENVASL-----------------------------------------
>Cyprinodon_variegatus
---------------------------------------------------------------------------------------------MEAKSIYDFSVETL----E-----GENVPLSVYRGKVLLIVNVATF-----------------------------------------
>Sarcophilus_harrisii
-------------------------------------------------------------------------------MC---------AAAAVPRSVHAFSARLL----GS----GELVSLASLRGKVLLIENVASL-----------------------------------------
>Procambarus_clarkii
------------------MLW-------------AGLVSLAAVG----------V-----------V----AQDQVASRACF--------HHHSDGGNIYDFEEMDL----LE----TRNISLADFRGKVLMIVNVATY-----------------------------------------
>Sclerodactyla_briareus
-------------------------------------------------------------------------------------------MSSKARTIYSFSAKAL----EG----GGIVDMARYRGKVLLIENVASL-----------------------------------------
>Galeopterus_variegatus
-------------------------------------------------------------------------------MCATR------LAAVAHSSVHAFSARPL----AG----GEPVSLGSLQGKVLLIENVASL-----------------------------------------
>Occasjapyx_japonicus
--------------MRVDGRGVS---PTPRNP-------------LAGGVAALVLLGCMPWSS----WAASPPDQVPSRPCY--------HNLSLGGDVYDFSEEDL----DR----ASNVSLADFRGKVLLVVNVATY-----------------------------------------
>Evechinus_chloroticus
---------MVSRVLHRISRAVP---QFG--R-------------TACGVGAILSVGVYFYASRNSPFVTTMA-AGAGKNDP--------NWWKTAQNIYAFSANDI----DG-----NEVSLEKYKGHVCLIVNVASS-----------------------------------------
>Ciona_intestinalis
----------------------------------------------MHQKGAVPAVLLVLG----TL----AVA-TSIENS---------TCFHNTFSVYDHSFPTL----SG----E-TVNLGEFRGNVSMVINVATY-----------------------------------------
>Esox_lucius
----MYVRYC-YKMWR-INPVLV-----------------------VFG------------------TLASCG-LLAMSHPT--------EDWQKATSIYDFSAKDI----DG-----NVVSLEKYRGDVVIIVNVASK-----------------------------------------
>Gadus_morhua
----------------------------------------------------------------------------------------------MASKFFEITAKRL----T-----GEVLNFSSLKGKVVLIQNVASL-----------------------------------------
>Glomeris_pustulata
------------------------------------------------------------------------------------------MNTSCCTSFYEFSAKDI----Q-----GKDVDFSKYKGKVVLIENVASL-----------------------------------------
>Acanthogobius_hasta
---------------------------------------------------------------------------------------------MASAEFYKFSPKLL----S-----GDTLNFSSLQGKVVLIENVASL-----------------------------------------
>Sipunculus_nudus
----MLTGYHSGRLLTKLNEG----------V-------------LACA---------------LTGCVNRFS-AMAEAKDK--------DSWKKAQSIYEFTVKDI----DG-----NEVSLEKYKGHVCLIVNVASK-----------------------------------------
>Chanos_chanos
--------------------------------------------------------------------------------------------MAESKTLYDFSTKSL----T-----GEVFNFSALKGKVVLIENVASL-----------------------------------------
>Galloisiana_yuasai
--------MAPS------------------FLGLGGILAV-------------FLLALAGS----DA----LVKRVPKVPCQ--------DHREINGSIHDYTIQRF----WD----SCNISLSDYQGKVVLLANVATY-----------------------------------------
>Apteronotus_leptorhynchus
----------------------------------------------------------------------------------------------MAKSFYDIKAKTL----E-----GAEFAFSSLQGKVVLIENVASL-----------------------------------------
>Glossoscolex_paulistus
---------------------------------------------------------------------------MADARPK--------DWWKKAASIYDFTVKDI----DG-----NEVSLEKYRGHVCLIVNVATK-----------------------------------------
>Cimex_lectularius
---------------------------------------------MMKAQIFLAFATLFVG----SGLATSTVARVAKRKC---------LDAGDDANVYNFSFMDL----SG----KTNVSLGDFKGKMILVVNVATY-----------------------------------------
>Oscarella_carmela
--------------------------------------------------------------------------------------------MSAIKSFFQLAARSL----T-----GELVAFSRYKGKVLLVENVATA-----------------------------------------
>Lasius_turcicus
---------------------------------------------------------------------------------------------MAATSIYDFKANDI----DG-----HEVSLEKYKGHVVLICNVACK-----------------------------------------
>Erinaceus_europaeus
-------------------------------------------------------------------------------------------MCAAVRSVHAFTARPL----SG----GDALDLGSLRGKVLLIENVASL-----------------------------------------
>Armadillidium_vulgare
--------------------------MIRYLLLLGTL----------------------LG----PS----MGESYDSVNC------------TAEEDIYQFSAKLL----DG----SATVNFSDYRGKVFVIYNVATY-----------------------------------------
>Ixodes_persulcatus
----------------------------------------------------------------------------MATSAS--------DGWKTAASIYDFDAVDI----DG-----NKVSLSKYRGHVALIVNVATK-----------------------------------------
>Oryctolagus_cuniculus
-------------------------------------------------------------------------------MCAAR-------MAAAAQSVYSFSAHPL----AG----GEPVNLGSLRGKVLLIENVASL-----------------------------------------
>Sarsinebalia_urgorrii
------------------------------------------------------------------------------------------------MAFYSFTAKLL----S-----GEAVSMSRYKGKVILIENTASL-----------------------------------------
>Clytia_hemisphaerica
MAKSILFDLKVSTLLGIIVTVVA---MLA--I-------------YKA------FTGYLFPT--LS-AKETAN-TNLLHSNP--------QSPEKASSIYDFSALDI----DG-----NMVNLSKYNGLVTYIVNVASE-----------------------------------------
>Lingula_anatina
----------------------------------------------------------------------------------------MARSPVKIRNFFDFSAKLL----S-----GEDINMSRYKGKVILVENVASL-----------------------------------------
>Lithobius_forficatus
------------------------------------------------------------------------------------------------MSFYDLAAKNL----V-----GQQVSFSKYQGKVVLIENVASL-----------------------------------------
>Lutjanus_erythropterus
-------------------------------------------------------------------------------------------MARNVKRFYDLTAKLL----S-----GEMFSFSALKGKVVLIENVASL-----------------------------------------
>Macrostomum_lignano
--------------------------------------------------------------------------------M---------ASSAETSTVYDFEAVDI----DG----N-NVSMDKYKGRVLIVVNVACK-----------------------------------------
>Ornithorhynchus_anatinus
-------------------------------------------------------------------------------------------MCASPNSVYGFSGRLL----RS----GEPFSLAALRGKVLLIENVASL-----------------------------------------
>Mytilus_californianus
-----------------ME---------------GF-VDILTI----------VLATLWSV----SG----QDLSSRQVACY--------HHDDNLHSVYNFSMPDI----HG----EKTIDFSHYRGKVLLLVNVASF-----------------------------------------
>Triops_newberryi
------------------------------------MASRLLLP---------VLLF---V----AG----VRSDTHVGSTL--------CNSSTTGTIYPFSQKRI----LG----GEVVNFSDFEGKVVLIVNVATY-----------------------------------------
>Hapalochlaena_maculosa
MVEPLTTSMEPSKLMFLSHLLAS---FLSICLCEGFSITTGLTEEMTTRQNH-VHSHGHSH----GH----LHRSTNFLTCY--------QSDNHSTSLYDYKSYDI----HG----VEEIPLSVYRNKVLLIMNVASF-----------------------------------------
>Calopteryx_splendens
--------------MGEGKHWLA---PLLCLVSLGLG-GV-------------------------D----AGDRVASRMCS---NFSPNSVKSPRNGVYNFSIPDV----YG----RRNISLSEYQGKVLLIVNVATY-----------------------------------------
>Ophiocoma_echinata
---------MFASVLRVLSHTVS---KYP--P--------------------SLPLGF--A---GAAVVTKSL-CTMAEGGP--------SAPEKASSIYEFTVKDI----DG-----NDVSLEKYKGHVCLIVNVASK-----------------------------------------
>Clonorchis_sinensis
------------------------------------------------------------------------------MAS---------CQTKPEETIYNFTVKSI----DG----E-SICLSQYAGFVCLVVNVASK-----------------------------------------
>Tetrix_subulata
---------------------------MEALASFVFLVT--------------LLSPCALA----AS----KVERVSSLPCS--------TDYTDGDTIFNFTAEDI----HK----QRNISLSKFRGKVVLIANVATF-----------------------------------------
>Osmerus_mordax
-------------------------------------------------------------------------------------------MSGTIKTFYDFTAKLL----S-----GNILKLSSLQGKVVLVENVASL-----------------------------------------
>Scleropages_formosus
--------------------------------------------------------------------------------------------MAGEKTFYDFSAALV----T-----GELFSFSSLRGKVVLIENVASL-----------------------------------------
>Crassostrea_gigas
--------------------------MAGACCSLAFLAFLGV------------------------I----CRVSGTEVVCN--------QPPSSAESVYSFTAPDI----YQ----TKNVSLSKYRGKVLVITNLASF-----------------------------------------
>Endeis_spinosa
-----------------------------------------------------------------------------------------------------------------------------------MIVNTAAF-----------------------------------------
>Pecten_maximus
---------------------------------------------------------------------------MASVVNV--------RSPVKIQNFFDFHATTL----EG-----EKINFSTYRGKVILVMNVASL-----------------------------------------
>Lateolabrax_japonicus
----------------------------------------------------------------------------------------------MAASFYDFTSKLL----T-----GETFDFSALKGKVVLVENVASL-----------------------------------------
>Sebastes_rastrelliger
-----------------------------------------------------------------------------------------MTMAGNVKRFYDLTAKLL----S-----GKSLSFSALKGKVVLIENVASL-----------------------------------------
>Ceratotherium_simum_simum
-------------------------------------------------------------------------------MFAA---HL---ASVAPRSVYAFSARPL----AG----GELVSLGSLRGKVLLIENVASL-----------------------------------------
>Laternula_elliptica
---------------------------------------------MAVQRGQLLVAAL-VG----LL----APAHGARVQCT--------MPTNFTDTVYNYNAMNA----YE----NQTIDLSQYRGKVLAIMNVATY-----------------------------------------
>Manis_javanica
-------------------------------------------------------------------------------MC----------ASAALRSVYAFSARPL----AG----GEPVSLGSLRGKVLLIENVASL-----------------------------------------
>Spadella_cephaloptera
---------------------------------------------------------------------------------------MVRLTTSLLKSFYDLSAKTLACGGYT----ASNVTFDAFKGKAVLIENVASL-----------------------------------------
>Glycera_dibranchiata
------------------------------MIRDGGVLLL--------------PLLVASI----SL----AVDCLQMSTC------------SPEESIYDHYLN-D----LD----GRPVSLEKYAGNVLLIINTATF-----------------------------------------
>Paralichthys_olivaceus
-------------------------------------------------------------------------------------------MAGNAKRFYELTAKLL----S-----GESFSFAALKGKVVLVENVASL-----------------------------------------
>Rhinopithecus_bieti
-------------------------------------------------------------------------------MCAAG------LAAAAAQSVYAFSARPL----AG----GESVSLGSLRGKVLLIENVASL-----------------------------------------
>Loxodonta_africana
-------------------------------------------------------------------------------MCAAP---V---AAAALRSVHAFSARPL----AG----GEPVNLGSLRGKVLLIENVASL-----------------------------------------
>Pinctada_maxima
-------------------------------------------------------------------------------------MYSVCDERTDGKTLYSMDTKDL----D-----GGVKELSDYRGAVTMVINVATF-----------------------------------------
>Fukomys_darlingi
-------------------------------------------------------------------------------MCAAR------LAEAAQATVYSFSARPL----AG----EEPLSLGSLRGKVLLIENVASL-----------------------------------------
>Owenia_fusiformis
-------------------------------M-------------TMGLLTSALLLLCGFL-------APGLAHNEAKTACY--------HSRRNNASIYDYRVETL----SKYFPDKDIVNLSDYRGKVVLIVNVATY-----------------------------------------
>Oncorhynchus_kisutch
--------------------------------------------------------------------------------------------MACVKSIYDFSAETL----NG-----QSVPLSIYRGKVLLIVNLATF-----------------------------------------
>Tupaia_chinensis
-------------------------------------------------------------------------------MCAAQA--------AAVAQATVYXARPL----GA----EEPMSLGSLRGKVLLIENVASL-----------------------------------------
>Porites_astreoides
-------------------MWVV---NLAYLLATAFVASAMTIT---------QTLTTTSP----RK----FHKSNNLLTCY--------QSDNHTTSVYDFVMKDV----YG----EKTIPLSIYKGKVLMIVNVASF-----------------------------------------
>Thunnus_thynnus
---------------------------------------------------------------------------------------------MAVRRFYDLTVKLL----S-----GETFSLSALKGKVVLIENVASL-----------------------------------------
>Tityus_serrulatus
---------------------------------------------------------------------------------------------MAKRSFYEFSAKLL----N-----GETLNFSQYRGKVILIENVASL-----------------------------------------
>Lepisosteus_oculatus
-------------------------------------------------------------------------------------------MAGGMKKFYDFSMNLL----ST----GEAYNFSTLKGKVVLIENVASL-----------------------------------------
>Sepia_pharaonis
-------------------------------------------------------------------------------------------------------MQ-N----LD----GNVVSLKKFKGKYLMIVNVATF-----------------------------------------
>Euprymna_scolopes
-------------------------------------------------------------------------------------MSRSPATPTGRKSFFDFSAKTL----A-----GENIDFSRFKGKVVLVENVASL-----------------------------------------
>Ictalurus_furcatus
----------------------------------------------------------------------------------------------MAKSFYDLSVKTL----T-----GEELKLSCLQGKVVLIENVASL-----------------------------------------
>Caligus_rogercresseyi
----------------------------------------------------------------------------------------------MENSFYDFKVEKL-----N----GSTMHLNTLKGKIVLIENTASL-----------------------------------------
>Trichechus_manatus_latirostr
-------------------------------------------------------------------------------MCAALGADVAALAAAVPRSVHAFSARRL----AG----GEPVSLGSLRGKVLLIENVASL-----------------------------------------
>Pagrus_major
-----------------------------------------------------------------------------------------------MAKFYDLTAKLL----T-----GETFKFSSLRGKVVLIENVASL-----------------------------------------
>Henricia_sp._AR-2014
-------------MRGIGTA--------------ALAVAL-------------ILLPGY-------G----SALGLLESRC------------AGNTSIYQHTFTTL----NG----SQEITLSQYKGKILLVVNVASF-----------------------------------------
>Streblospio_benedicti
---------------------------------------------------------------------------------------------MAARSIFQFTAKLI----N-----GQEVNLNSFKGKVVLVENVASL-----------------------------------------
>Squalus_acanthias
-----------------------------------------------------------------------------MSAQP--------ADWQKATSIFEFDAKDI----DG-----NEVSLEKYKGFVSIITNVASK-----------------------------------------
>Protobothrops_mucrosquamatus
------------------------------------------------------------------------------------------MTTPIAKSFYDLSATDL----Q-----GEKVDFNHFRGRVVLIENVASF-----------------------------------------
>Schistosoma_mansoni
---------------------------------------------------------------------------------MS-------SSHKSWNSIYEFTVKDI----NG-----VDVSLEKYRGHVCLIVNVACK-----------------------------------------
>Nesiohelix_samarangae
--------MGPW---------------------------------STTGPGGIMIAFCGLF----AA----ASAGVIMLDCK--------QPVSDKTTLYDFTLMDV----HR----VKNISLSDYRGKVVLIVNVATY-----------------------------------------
>Symphylella_vulgaris
----------------------------MKSQ-------------MLVHL-AFIVLFCHLASA----L------KLDFTNC---------TAPEANQDIYKFVAEDL----NR----TKNVSLSEFRGKVLLIINVATFCDSTTQYKGLRTLQTEFGSENFSVLAFPCNHLACKTSSQCH
>Monopterus_albus
-------------------MNLI-----------------------QST------------------VLFSFL-LQAMSGLT--------EDWQRATSIYDFSATDI----DG-----NVVSLEKYRGNVVIITNVASK-----------------------------------------
>Alvinella_pompejana
---------------------------------------------------------------------MATA-SEEKSEKH--------DDWKKATSIYDFTCKDI----DG-----NEVSLEKYRGHVCLIVNVATN-----------------------------------------
>Chelonia_mydas
---------------------------------------------------------------------MGLQ-FTYSCAQA--------DDWRAAKSIYEFHAKDI----DG-----SDVSLEKYRGFVCIITNVASK-----------------------------------------
>Takifugu_rubripes
-------------------------------------------------------------------------------------------MAGSVKKFYELTAKLL----S-----GEVLSFSALRGKVVLIENVASL-----------------------------------------
>Watasenia_scintillans
----------------------------------------------------------------------------------------MARTPATAKTFFDFSAKTL----A-----GENVDFSRYKGKVILVENVASL-----------------------------------------
>Clio_pyramidata
--------MG-R---------------------------------TSVNSGGLLACLCGLV----VA----AAGSAVMLDCH--------QPANDPKSIYDFTLMDI----HH----QKNVSLSDYRGKVVLIVNVATY-----------------------------------------
>Phagocata_gracilis
-----------------------------------------------------------------------------------------------------------------------------------MVVNTACK-----------------------------------------
>Orycteropus_afer_afer
-------------------------------------------------------------------------------MCAAPLTAAAARAAAAPRSVYAFSARPL----AG----GDPVSLGSLRGKVLLIENVASL-----------------------------------------
>Villosa_lienosa
---------------------------------------------------------------------------------------MAVRSHVKPQNFFDFTAKTL----K-----GQDVNLSNYKGKVILVENVASL-----------------------------------------
>Mastigoproctus_giganteus
--------------------------------------------------------------------------------------------MFASKSFYDFAAKTL----Q-----GETIEFSLYKDRVVLIQNVASL-----------------------------------------
>Patiria_pectinifera
------------------------------MA-------------TAGLWMPASAVLMILWSV-----AAAAG-GTLESIC---------RADGAASTVHQYSLHPL----DE---TSPPVPLSDFAGKILLLINVATY-----------------------------------------
>Sinonovacula_constricta
---------------------------------------------MAVPAGQFVLLSMA-G----LL----ALASGTRVSC---------VPGTPAKTVYDFSAMSI----YE----NETISFDRYKGKVLAIINLATF-----------------------------------------
>Limulus_polyphemus
--------------------------------------------------------------------------------------------MSIAKSFYDLAAKSL----T-----GEIINFSKYEGKVVLIENVASL-----------------------------------------
>Palythoa_caribaeorum
-----MEAFEASRLKS------------LALILFGFVVFA--------------------------Y----ATDR-PKQVSD--------CQNRQKDSFHDFAALDL----H-----EEYIPFKRYEGKVVLVVNVATF-----------------------------------------
>Thermobia_domestica
-----------------------------------------------------------MG----NK----MP---WKSNK---------LLTDIPNSLYDLSAIDI----DG----Q-KVGLSKFAGKCVMVVNVASK-----------------------------------------

HG0003 ENSGALG00000014906 MOCS2

>Gallus_gallus
------------------------MSCQVTVLYFARSAELVGLRSES-V--SVPQRIT--SL-Q-----------LW---EEIVKIHPRLA--VIRD---QVVFAVRQEYVLLGD--QLLVLQTGDEVAIIPPISGG
>Acipenser_sinensis
------------------------MTTEVVVLYFAKSAELAEVRSEN-I--TVSHQVT--SL-Q-----------LW---QELVRKHPRLA--AIRD---QVVLAVRQEYVSLSD--ELLILHPGDEVAVIPPLSGG
>Adineta_ricciae
-----------------------MSTIQVAIIYFGRLRELVGISSEK-I--DFDSAIS--YTQQ----------SIL---DRIIEHRPILKSFLVTN---TFRLAVNQQYLLEGD---LIQFSKESEVALLPPFGGG
>Acanthogorgia_aspera
------------------------------------------------------------------------------------------------------MLALNEIYIDLE---ETVNLRSTDKIAVIPPLSGG
>Amia_calva
------------------------MNTEVVVLYFAKSAEIADVRSET-I--SVPQEIT--SL-Q-----------LW---QELVNRHPSLS--VLRD---QVVLAVRQEYVALGE--QLLSLQSGDEVAVIPPLSGG
>Talitrus_saltator
------------------MTQYGLEIVPVKIVFFASARELSGCSEVE-IDLSTPTT-----AKK-----------LF---SEVLNKHPELQ--KLEK---SVMVAVNQEYIDD-DT-NQVSLRAGDEVAFIPPISGG
>Podocoryna_carnea
--------------------MEESFKVPVSILFFAKSRELVGKTSDT-I--YLEQKTY---LKK-----------LL---EEIIHAYPSLK--NIGN---NFIIAYNQKYLTE-SE-DCITLKASDEIAIIPPISGG
>Megophrys_sangzhiensis
------------------------MTCEVVVLYFAKSSELTGVRSEI-I--NVPQEIT---SQN-----------LW---EEIVRFHPRLC--TIKD---QVVFAVRQEYVAI-GD-ELLSLRPGDEIAVIPPISGG
>Milnesium_tardigradum
------------------------MEISVKVLFFAKAKDLVGTGQGI-L--TLKEDIL--SLGS-----------LL---SNLYSQFPALL--PLQN---QVVLSLNQKYIDADDLQRPVSINIGDEIAVLPPISGG
>Apteryx_australis_mantelli
-------------------------MYDVLEVYWK--QNCVSLRSID-L--SCPRLCT--SL-LFKEISWLNANTLFRNLFVLLPIPLRLA--VIRD---QVVFAVQQEYVLLGD--QLLVLHPGDEVAIIPPISGG
>Machilis_hrabei
--------------------MAALCESEIRLLFFAKSRELAGKSEAS-I--CVPSLVH---VQD-----------LL---DLIVRE-FSLD--LIKN---NLILAHNQEYLDRE---QSVNIRTGDEIAVIPPLSGG
>Scapharca_broughtonii
--------------------MATPEEVQVTLLFFAKSRELSGLKTSQ-L--KTPTQVN---FQQ-----------LL---GLIVTAFPSLE--IISE---NIVLSLNEDYIDK--D-SSVNLNSGDEIAVIPPISGG
>Oryzias_latipes
------------------------MTAQVSVIYFGQSAELTGLKGEE-L--VSVPTPI--SSRE-----------LW---ELLQQRHPRLA--VLRG---HVVLAVAQQYIPIAE--QLVVLRDGDEVAVVPPLSGG
>Prorhinotermes_simplex
--------------------MADVSVVEIQVLFFAKARELCGQKQSC-L--LIP-SVT--SYNN-----------LL---DEIVTK-FSLE--SIRD---NLLLAINEEYTTSE---SEVQLKEGDEVAVIPPLSGG
>Eurytemora_affinis
-----------------------MADVDVKLILFAKARELVGASEVN-V--RFPQELS---SEH-----------LK---SKIIQSFPVLD--KLGG---TWLLALNEAYIDT----DRVTLNTGDTLAVIPPISGG
>Alatina_alata
---------------------MSNITVTVEVRFFAKARELAGIQKTH-V--RTRERIT---AHE-----------LI---SQIIDAFPKLD--VIRQ---CFLLAHNQEYCSL--D-QELNISDGDEIAVIPPISGG
>Hynobius_chinensis
------------------------MNSEVIVLYFAKSAELTGVRSET-L--SVPQQIT--PS-Q-----------LW---DEIVNRQPRLA--AIRD---NVVLAVRQEYVILGD--QLLTLQPGDEVAVIPPISGG
>Siniperca_chuatsi
------------------------MTAQVSVLYFAQSVELTGVKEEE-L--VAVPTPI--SSQD-----------LW---RLLLQRHPRLS--LLQD---QVVLAVRQQYVAIGD--QPVTLGDGDEVAVVPPLSGG
>Astyanax_mexicanus
------------------------MKSEVLVLYFAKSAELTGIKSET-I--SVQSELS--TL-Q-----------LW---QELEKLHPKLS--AVRD---QVVLAVRQEYVSLGD--QRVILRDGDEVAIIPPLSGG
>Callorhinchus_milii
------------------------MTAEVSVLYFARSAELSGVRSET-L--SVPQQLT--SM-Q-----------LW---EKIVNKHPSLI--AIRD---HVLLAVRQEYILLGD--QILFLQPGDEIAVIPPLSGG
>Oreochromis_niloticus
------------------------MTAQVCVLYFAKSVELTGVKEEK-T--VAVPTPI--SSHD-----------LW---KLLVQRHPRLC--TLQD---GVVLAVRQEYVPIGD--QVLTLGDGDEVAVVPPLSGG
>Alosa_alosa
------------MFGSA-VLQAPFMISEVSVLYFAKGAELTGVKSET-V--KLPSELT--TI-Q-----------LW---LELEKRHPKLC--VLRE---HVVLAVRQEYVDLCD--QTVSLHDGDEVAVIPPLSGG
>Latimeria_chalumnae
------------------------MTAQISVLYFAKSAELAGIRSEV-I--TVAQQIT--SQ-Q-----------LW---QEIVNRHPRLA--AIHD---HVILAVHQEYILLGD--QFLLLQPGDEVAIIPPISGG
>Priacma_serrata
------------------------MSVTVTILFFAKAREIVGKNSSS-L--EVPPALT--CT-D-----------LL---NTIVNE-YSLH--VIRN---NIILAVNEEFCAGN---DRLSLKTGDEIAVIPPLSGG
>Corynactis_australis
--------------MEH-INNDTTGKVEVTVLFFAKSRELVEQRSAE-V--LLSKRLS---GRE-----------LI---ASIVDNFPSLS--VIQD---NLVVAVNQSYVEK--E-SEIEFQGGEEVAIIPPISGG
>Cuculus_canorus
-------------------MTAELPLVQVSVLYFARSAELAGLRRET-L--RVPRQVT--AL-Q-----------LW---EEIVSLHPRLA--VIRD---QVVFAVRQEYVLLGD--QLLVLQPGDEVAIIPPISGG
>Ctenopharyngodon_idella
------------------------MNTEVSVLYFAKSAELTGLKAEN-I--TVPSNIT--SL-Q-----------LW---QDLENRHPRLS--VLRD---QVVLAVRQEYVPLGA--QPLTLRQGDEVAIIPPLSGG
>Fundulus_heteroclitus
------------------------MSAQVTVLYFAKSAELTGLKEEP-V--AVPTPIS--S-GD-----------LW---ALLLRKQPRLV--ALQG---HVILAVRQRYVAIGE--QQVTLEDGDEVAVVPPLSGG
>Petrolisthes_cinctipes
---------------------MTEREVSVSVLFFAGARDLAGVERSA-I--TLPVVTT--P-EQ-----------LL---ARVVRSFGELD--IIRD---SVIISLNQEYCVPG---QELVLSAEDEVAIIPPISGG
>Sclerodactyla_briareus
------------------------MPVDVTVQYYAISRDITGTREDR-L--TVPTHVT--L-KQ-----------LF---GCLTERHPGLK--QIET---NCLLAINGDWLTEES--GVVELKPGDEVAVIPPLSGG
>Forficula_auricularia
----------------M-NQDLNCDQVTVEVLFFAKARELSGVNKAI-I--QTPSLTN--S-EE-----------LL---DSLINR-FNLQ--QISS---CLVLAVDENWVESG----PLKLNSGSIIAVIPPLSGG
>Occasjapyx_japonicus
-----------------------MQTTELRVLFFASARELAKCNAAV-LRLDSAQ--Q--SGES-----------LL---EQVVLSFPDLQ--RVRG---SLMLALNQNYVQL--D-QRLTVSSGDELAVIPPISGG
>Sarcophaga_crassipalpis
-------------------MNSELPVINIHILFFAKARELSGLSESN-HQISTPKIIA---S-D-----------LL---EQIC-STYNLS--VIRD---NLIIAINESYCEN-LN-TELQLQEGDEIAIIPPISGG
>Echiniscus_testudo
--------------------------MQITVLLFARMRDLAGGRSEILI--DVPADCENGSV---------SSKRLL---QLIIAQCPRLK--EIK----HYQLAVNRQYVDLNSD-NLVQLGPTSEIALIPPVSGG
>Ciona_intestinalis
----------------------MGKNVQVNIKFFAKSRELVGIESEQ-L--ILPQDVT--SF-Q-----------LI---ETINSKYPRLS--DLQG---CFTLAVNGEYIKTDES-YSILLTDQSEIAVIPPLSGG
>Baetis_sp._AB-2009
----------------------MGQPIVVKVLLFASAREISGERESE-I--QVQQEET--Y-DT-----------LF---RKIVQR-FKLE--KLQN---SLILAINEDYPDAEN--TQLNLKPGDHIAVIPPLSGG
>Esox_lucius
------------------------MNTEVTVLYFAKSAELTGIKSET-I--RVPSELT--TL-D-----------LW---LDLVSRHPRLS--TVRE---SVVLAVRQEYVAVGE--LRVSLREGDEVAVIPPLSGG
>Leptasterias_sp._AR-2014
------------------MASICADVVEINVLFFAKSRELTGVKEGH-L--QVKTNLK--PE-E-----------LL---LQILSQYPQLS--IIQD---TVVLSVNQEYIAKGD--GTLNLQPGDEVAVIPPISGG
>Gadus_morhua
------------------------MTARVTVLYFAKSAELTGHRSEV-V--PLPAATS--SC-Q-----------LW---ALLLELHPSLA--ALQD---QVVLAVRQQYVALGD--QALLLADGDEVAVVPPLSGG
>Chlamydotis_macqueenii
--------------------------------------------------------------------------------------------------------------MLLGD--QLLVLQPGDEVAVIPPISGG
>Galloisiana_yuasai
--------------------MDLNLKISVNVLFFAKARELCGKKESK-L--QVVPSIS--Y-NL-----------LL---DHIIKE-FSLE--SIKG---NFILAINEEYTTTD---RELDLKEGDEIAIIPPLSGG
>Tigriopus_californicus
------------------------MSVSIKVLLFAKARELAGTHELT-L--NVVPNLS---ASD-----------LV---TQIEAQCPDLA--ILRG---SFVLALNQDYVDH--N-QVLDLKSGDELAVIPPISGG
>Acanthosoma_haemorrhoidale
----------------------MDCNVKIKILFFAKALELSGTKESF-L--SVSPNIS---TPL-----------LV---NKIV-DLFGLH--SISN---SILIAKNEEYVIKDGQ-EPLNLQEGDEIAIIPPISGG
>Myrmica_ruginodis
-------------------MDETATPIQATVLFFAKARELTGCKECK-L--CVPRKLS--SA-D-----------LL---DKIIHA-FELG--SIRN---QIILAVNEEFVVPD---SILVLSEKDKIAVIPPLSGG
>Rhipicephalus_appendiculatus
----------------------MRPEVPVRLLLFAKARELAGTSEAS-L--VVPAVLR--DVEA-----------LK---EVIFDAFPKLA--SLQR---SAVIAVNESYVDS--G-VEVTLRQGDEVAFIPPISGG
>Dyseriocrania_subpurpurella
--------------------MDDLQLVQVKVLFFAQSREVSGRREAS-I--NLPSKLS--CA-E-----------LL---DIFVDN-FSLS--AIKN---NILIAKNEELCCED---NLLEIKTGDEFAVIPPLSGG
>Lutjanus_erythropterus
------------------------MTAQVCVLYFAKSVELTGAKEEQ-I--VAVPTPI--SSQD-----------LW---KLLLQRHPRLS--VLQD---QVVLAVRQQYVAIGD--QVVTLGDGDEVAVVPPLSGG
>Mantis_religiosa
------------------MDVSSQQRVQIKVLFFAKARELSGIKESD-F--ISETPTS--Y-EL-----------LL---QQIVDK-YSLS--AIKS---SVVLAINEEYCSPS---EHFKLNEGDEVAVIPPLSGG
>Nannochorista_philpotti
----------------------MTEEVEVKLLFFANARDLAGINETL-L--LTRSKIL--CS-E-----------LL---RDITIN-YSLE--AIKN---NILLAVNEEYCESPN--TELNLKAGDEIAIIPPLSGG
>Dermanyssus_gallinae
-------------------------MVRVKLLFFAKARELAGTAERS-V--DVAERVF--SLDQ-----------LK---ELIAQLEPSVK--TLLD---TSILAVDLNYVVDSV----EIGTRTGEIAVIPPVSGG
>Ornithorhynchus_anatinus
-----------------------MVLSPVTVLYFAKSAELAGVRSET-I--SVPQEIK--SL-Q-----------LW---EEIESRHPRLA--DVRD---QVIFAVRQEYVQLGD--QVLLLQPGDEIAIIPPISGG
>Perna_viridis
---------------------MANNKVQVSLLFFAKSRELSGVSNSQ-I--KVSSQCS---FFQ-----------LF---ETIVQQYPSLS--VISE---NIILSLNEEYISR--D-GSITVQEGDEIAVIPPISGG
>Osmylus_fulvicephalus
------------------------MSVEVRILFFAKAKDFAQKSEST-L--NIPNTLT--F-KE-----------LL---TIIVDN-YNLQ--AIRN---TINISLNANFCDRD---ATLKFNTGDEIAVIPPLSGG
>Hapalochlaena_maculosa
--------------MDH-SFAGNKDPVKINVLFFAKSRELVGQNKSE-V--SITSRSV--TTAQ-----------LT---SFLLETFPRLK--PLKQ---SMIIALNEEYLSGD---ETIHLSNGDEIAVIPPLSGG
>Epiophlebia_superstes
--------------------MGDKDFVTVKVLFFAKARELSGIKFGQ-LKVNAAP--S---YTF-----------LL---EEIV-TKFSLE--KIRN---NILLAINEEYSGS--E-LQINLKDGDEIAVIPPISGG
>Opisthorchis_felineus
----------------------MKQAVRARVILFGEAKDLTGFQSAL-I--LVPVTCT--A-KE-----------LY---DLILGHFPKLK--LIEK---SLLLTLDFQYIDIEDNNTLLHISDSSELGVLPPLGGG
>Stenobothrus_lineatus
--------------------MDNQKKIEVKILFFAKARELAGCSESS-L--KVASHIS---YEE-----------LC---TLIV-ENFSLA--AIKK---NFILAVNEEYLSS--E-STLDFKEGDEIAVIPPISGG
>Osmerus_mordax
----------------------MTTKVSVLVLYFAKSADLTGLKSEV-I--HVQPELT--TL-A-----------LW---QELVNRHSRLS--AVRE---QIVLAVSQEYVTIGC--QLVSLKNGDEVAVIPPLSGG
>Scleropages_formosus
------------------------MNTEILVLYFAKSAELTGLRSEN-V--RVPPGLT--SL-Q-----------LW---EELVTRHPRLS--MLRT---QVVLAVREEYVPLGD--QPVRLQEGDEVAVIPPLSGG
>Crassostrea_gigas
----------------------MEGDVTINLLFFAKSRELVGSGSGE-L--VLTANTN---FDT-----------LK---QALLLKYPSLQ--VIAE---NIILSLNEEYIDD--N-QQLTLREGDEVAVIPPISGG
>Hypsibius_dujardini
--------------------------MDVNLKFLALAKDLAGTGDAT-I--RITAEESPFTVGS-----------FL---AKLIRTFPRLG--EIKD---QLILAHNLEYLDLQNPSASFELKEGDEIAVLPPVSGG
>Taeniopygia_guttata
------------------------MRCQVSVLYFARSAELAGLRCET-L--SVPQEIT--SL-Q-----------LW---EEIVKVHPRLA--LIRD---QVVFAVRQEYVLLGD--QLLVLQPGDEVAIIPPISGG
>Pecten_maximus
-----------------------MADVEVSVLFFAKSRELAGKKTAQ-IIVESPTN-----LQQ-----------LI---STIVSQFPSLQ--IIAD---NLVVSINEDYLQA--G-QEVHLNTGDEVAVIPPISGG
>Dissostichus_mawsoni
------------------------MTAQVTVLYFAKSAELSGLKEEE-L--VAVPTPI--SSQD-----------LW---SLLLQRHPRLC--VLQH---QVVLAVRQQYVAIGD--ELVALGDGDEVAVVPPLSGG
>Equus_asinus
----------------------MVPRCQVEVLYFAKSAEIAGIRSET-I--SVPQEIK--AL-Q-----------LW---NEIETRHPGMS----------------------------------------------
>Timema_cristinae
-------------------MMTNNLKVAVKVLFFAKARELSGWKESD-L--EVECSLT--Y-EC-----------LL---DTLVKE-FSLE--LVQN---HLILAVNQEYKSAG---QTIVLEAGDEIAVIPPLSGG
>Menopon_gallinae
--------------------MNDSSHVSVKLLFFAKARELTGKNEAI-V--DVDRNVK--G-AD-----------LL---DFLVSK-FELD--CIRK---NVILALNGDYVDLE---CDLELNSADELAIIPPLSGG
>Perinereis_aibuhitensis
------------------------MAVQVKILFFAKSKEIAQTNESE-L--ALPNSIS---LAQ-----------LK---EVLCQQYNGLS--AIIE---NCIISLNQNYVEE----GDLNLNSGDEIAVVPPISGG
>Fasciola_hepatica
----------------------MSTVISVSVLFFGEAREVVKLHRVC-L--SLPTTCT--P-AE-----------LK---REILKHFPQLE--LLSS---SLGLALDLQYLDCSSETDRITLSDLSEVSVFPPLSGG
>Colobus_angolensis_palliatus
----------------------MVPLCQAEVLYFAKSVEITGVRSET-I--SVPQEIK--AL-Q-----------LW---KEIETRHPGLA--DVRN---QILFAVHQEYVELGD--QLLLLQPGDEIAVIPH----
>Acerentomon_sp._AD-2013
----------------------MAYDVSVNVLFFAKAKELTNLSEVL-L--VLKTETT---GNE-----------IL---SALEKKFPVLS--RIKE---IIHLAVNEAYCED-LS-CSITLNSGDTVAVITPISGG
>Melopsittacus_undulatus
------------------------MRCQVSVLYFARSAELAGLRRET-L--SVPRQIT--AI-Q-----------LW---EEIVKVHPRLA--VIRD---QVVFAVRQEYVLLGD--QFLVLQPGDEVAIIPPISGG
>Liposcelis_bostrychophila
--------------MID-GFDEEATQVSIKILFFALVKDLVGKNEDR-I--YVPSTIV--G-SA-----------LL---DLIIKK-YNLE--TVKN---NVILAVNEDWVDLD---SLLILKDSDEVAVIPPLSGG
>Xanthostigma_xanthostigma
------------------------MSVEVKILFFASASTISNLSESI-L--NVSGQIA---YDD-----------LL---ELIV-EKYSLQ--NIKQ---NILLAHNESICEK--N-QTLNLSDGDEIAVVPPFSGG
>Rhopilema_esculentum
--------------------MDAVNEVSVRLLFFAKSRDIVGLHEAH-I--DISSVVT---GQE-----------LL---RAIVTKFPGLH--ILRE---NLLLSCNQEYLES--D-QVVELAGGEEVAVIPPISGG
>Marmota_marmota_marmota
----------------------MVPRCQVEILYFAKSAEITGVRSET-I--SVPQEIK--AL-Q-----------LW---HEIETRHPGLA--DVRN---QVIFAVRQEYVELGD--QLLLLHPGDEIAIIPPISGG
>Coregonus_clupeaformis
------------------------MNTEVSVFYFAKSAELTGLKSET-I--RVPSELT--TL-E-----------LW---QDLVSRHPRLS--AVRD---SVVLAVRQEYVALGD--QCVSLREGDEVAVIPPLSGG
>Favia_lizardensis
--------------MDH-FSNDTTEQVEVTVLFFAKSRELVEKRSAQ-I--SVGKQIT---GTD-----------LI---HYILNNFPSLA--VIQE---NLVIAVNQSYIEK--D-SEITFEGGEEVAVIPPISGG
>Sepioloidea_lineolata
---------------------------------------------------------------------------------------------------------------------------------MLPPLSGG
>Ictalurus_punctatus
------------------------MNTEVTVLYFAKSAELTGVKSEI-I--RVNSELT--SL-Q-----------LW---QELEHKHPKLS--AVRE---QVVLAVRREYVSVGE--QVVRLRHGDEVAVIPPLSGG
>Ceratophyllus_gallinae
---------------------MGNGEVSIKVIFFAKSRELSGVSETS-L--KVNSNIS--C-GE-----------LI---QLICAI-HGLQ--SIQN---NIIIALNEVYYQDLA--AILTLKTGDEVAIIPPLSGG
>Streblospio_benedicti
--------MLLFLIMLT-VLNILKLLYLVKVLFFAKSRELVGHSDSY-I--NLPLICL---ESQ-----------IR---TSIVENNPSLS--PISD---SFVLALNQEYIGE--G-EEVHISEGDEIAVIPPISGG
>Squalus_acanthias
------------------------MIAQVSVLYFARSAELAGVRSET-L--SVPQQLT--SL-Q-----------LW---QEIVKRHPRKQRENLREIQPRSVLNIFYHPVL-------------------------
>Takydromus_sexlineatus
------------------------MSCEVVVLYFARSAELAGVRTET-I--SVPQQIT--SL-Q-----------LW---EEIVRRHSRLT--AIQD---QVVFAVRQEYVLLGD--QLVVLNPGDEVAVIPPISGG
>Stylops_melittae
-----------------------MNTVTIKVLFFAKSRQLSKRKEGH-I--TLPAEIA---YDD-----------LL---TLIC-NQFQLE--EIKK---TIFLALNETYLQS--N-TLVHLHSDDIVAVIPPISSG
>Schistosoma_mansoni
-----------------------MTICNKKVLLFGEASELVGCKEIT-I--QFPLSCT--N-KT-----------LK---DIIIEQLVKLE--PLKD---SLTVAVNWKYTDINTSLNQISIEEIEELAILPPISGG
>Alvinella_pompejana
MKAYVICKPKYDHYDDKNFCNDTNCILKVTVFFFAKGRELVGKKEDQ-I--DIAPHNS--GT-A-----------LY---KLVVDRYPCLK--PIQE---NFALALNEEYIDKTD--EEIILSNGDQLAEIPPISGG
>Pelodiscus_sinensis
------------------------MSCEVVALYFARSAELAGVRSET-I--SVPQQIT--SL-Q-----------LW---EEIVKRHPRLA--AIQD---QVVFAVCQEYVLLGD--QLLVLQSGDEVAIIPPISGG
>Hydroptila_sp._AD-2013
---------------------MEVSKVVVKVLFFAASREIVGAEERE-L--EVPARSS--Y-SS-----------LL---HLLVHT-FKLD--IIKD---NFIVAKNEELCLDNG--HEIVFEKNDEIAIIPPLSGG
>Villosa_lienosa
-----------------------------------------------------------------------------------------------------------------------MQLHQGDEIAVIPPISGG
>Sinonovacula_constricta
----------------------MAAANVIQVLFFAKSRELTGIKQTN-L--LVNSSLT---AHE-----------LL---KKIISEFPGLQ--VLAD---TLILSLNEEYLALEDP-KILNLQEGDTVAVIPPISGG
>Limulus_polyphemus
----------------------MSSGVKVKLLFFAKARELVGVTETG-L--VVPNGLY--TQKK-----------LL---SIIVETFPVLC--SIQG---NFIVALNEECMDQS---QSIELKEGDQLAVIPPLSGG
>Palythoa_caribaeorum
--------------MEL-ANDAGCEKVQVTVLFFAKCRELVGTSSVQ-I--YLARRLS---GSK-----------LI---ETLLNLFPSLS--EIER---NLIVAVNQEYIEK--E-SNIELFGNDEIAVIPPISGG
>Thermobia_domestica
--------------------MAESEKVIVKLLFFAKARELSGHKDIT-I--NLPSITT---YSI-----------LV---SLIA-KEFSLE--PIQE---NLILALNQEYIES--D-RDISLKNGDEIAIIPPISGG

HG0003 FBgn0039280 Mocs2

>Drosophila_melanogaster
-------------------MNADGPVVNVHVLFFAKSRELANTPR-STVEVPTE---ITA--T-ELLDHLVSK-FGLTS--IR---DNLILAHNESYIDNLS--DRILFKEGDELAIIPPLSGG
>Acipenser_sinensis
------------------------MTTEVVVLYFAKSAELAEVRS-ENITVSHQ---VTS--L-QLWQELVRKHPRLAA--IR---DQVVLAVRQEYVSLSD--ELLILHPGDEVAVIPPLSGG
>Anemonia_viridis
----------------MDNKYNGTEDVPVILLFFAKSRELTECAS-KEVQLPRL---VTR--E-DLISSIIDIFPSLVQ--IE---KTIVVAVNQEYLDTGK---KVELRKGDTVAIIPPLSGG
>Adineta_ricciae
-----------------------MSTIQVAIIYFGRLRELVGISS-EKIDFDSA---ISYTQQ-SILDRIIEHRPILKSFLVT---NTFRLAVNQQYLLEGD---LIQFSKESEVALLPPFGGG
>Acanthogorgia_aspera
-----------------------------------------------------------------------------------------MLALNEIYIDLEE---TVNLRSTDKIAVIPPLSGG
>Amia_calva
------------------------MNTEVVVLYFAKSAEIADVRS-ETISVPQE---ITS--L-QLWQELVNRHPSLSV--LR---DQVVLAVRQEYVALGE--QLLSLQSGDEVAVIPPLSGG
>Talitrus_saltator
------------------MTQYGLEIVPVKIVFFASARELSGCSE-VEIDLSTP---TTA--K-KLFSEVLNKHPELQK--LE---KSVMVAVNQEYIDDD--TNQVSLRAGDEVAFIPPISGG
>Hydractinia_symbiolongicarpu
---------------------MEDCSVEVNILFFAKSKELTGKSV-DVVNLKKS---NCL--D-NLLKEIIHTYPALNI--LK---GRFIIAHNQAYLRDT--KEVVILETGDEIAIIPPISGG
>Xenopus_tropicalis
------------------------MTCEVSVLYFAKSSELAGVRS-ETLTVSQE---ITS--Q-QLWDQLVHSHPSLCA--IR---DQVVLAVRQEYVPIG--NELLTLNTGDEIAVIPPISGG
>Milnesium_tardigradum
------------------------MEISVKVLFFAKAKDLVGTGQ-GILTLKED---ILS-LG-SLLSNLYSQFPALLP--LQ---NQVVLSLNQKYIDADDLQRPVSINIGDEIAVLPPISGG
>Machilis_hrabei
--------------------MAALCESEIRLLFFAKSRELAGKSE-ASICVPSL---VHV--Q-DLLDLIVRE-FSLDL--IK---NNLILAHNQEYLDREQ---SVNIRTGDEIAVIPPLSGG
>Scapharca_broughtonii
--------------------MATPEEVQVTLLFFAKSRELSGLKT-SQLKTPTQ---VNF--Q-QLLGLIVTAFPSLEI--IS---ENIVLSLNEDYIDKD---SSVNLNSGDEIAVIPPISGG
>Argulus_siamensis
-----------------------MSLITVSVLFFAKARELVGAHQ-TTITLPQS---LIG--H-KLVALILDYYEALKP--IE---KNFILALNENYIELD---EEISLSEGDVIAVIPPISGG
>Oryzias_latipes
------------------------MTAQVSVIYFGQSAELTGLKGEELVSVPTP---ISS--R-ELWELLQQRHPRLAV--LR---GHVVLAVAQQYIPIAE--QLVVLRDGDEVAVVPPLSGG
>Prorhinotermes_simplex
--------------------MADVSVVEIQVLFFAKARELCGQKQ-SCLLIPSV----TS-YN-NLLDEIVTK-FSLES--IR---DNLLLAINEEYTTSES---EVQLKEGDEVAVIPPLSGG
>Tripterygion_delaisi
------------------------MSARVVVLYFAQSAELAGLKEEEIVAVPSP---ISS--L-DLWNLLVQRHPRLGV--LR---GHVVLAVRQQYVAIGD--EQVALGDGDEVAVVPPLSGG
>Eurytemora_affinis
-----------------------MADVDVKLILFAKARELVGASE-VNVRFPQE---LSS--E-HLKSKIIQSFPVLDK--LG---GTWLLALNEAYIDTD----RVTLNTGDTLAVIPPISGG
>Seriola_quinqueradiata
------------------------MTAQVCVLYFAKSVELTGVKQEDIVAVPTP---ISS--R-DLWSLLLQRHPRLSV--LQ---DQVVLAVRQHYVAIGD--QVVTLGDGDEVAVVPPLRGG
>Alatina_alata
---------------------MSNITVTVEVRFFAKARELAGIQK-THVRTRER---ITA--H-ELISQIIDAFPKLDV--IR---QCFLLAHNQEYCSLD---QELNISDGDEIAVIPPISGG
>Hynobius_retardatus
------------------------MNSEVIVLYFAKSAELTGVRS-ERLSVPQQ---ITS--S-QLWDEIVNRQPRLAA--IR---DNVVLAVRQEYVVLGD--QLLTLQPGDEVAVIPPISGG
>Siniperca_chuatsi
------------------------MTAQVSVLYFAQSVELTGVKEEELVAVPTP---ISS--Q-DLWRLLLQRHPRLSL--LQ---DQVVLAVRQQYVAIGD--QPVTLGDGDEVAVVPPLSGG
>Astyanax_mexicanus
------------------------MKSEVLVLYFAKSAELTGIKS-ETISVQSE---LST--L-QLWQELEKLHPKLSA--VR---DQVVLAVRQEYVSLGD--QRVILRDGDEVAIIPPLSGG
>Callorhinchus_milii
------------------------MTAEVSVLYFARSAELSGVRS-ETLSVPQQ---LTS--M-QLWEKIVNKHPSLIA--IR---DHVLLAVRQEYILLGD--QILFLQPGDEIAVIPPLSGG
>Oreochromis_niloticus
------------------------MTAQVCVLYFAKSVELTGVKEEKTVAVPTP---ISS--H-DLWKLLVQRHPRLCT--LQ---DGVVLAVRQEYVPIGD--QVLTLGDGDEVAVVPPLSGG
>Alosa_alosa
-------------MFGSAVLQAPFMISEVSVLYFAKGAELTGVKS-ETVKLPSE---LTT--I-QLWLELEKRHPKLCV--LR---EHVVLAVRQEYVDLCD--QTVSLHDGDEVAVIPPLSGG
>Latimeria_chalumnae
------------------------MTAQISVLYFAKSAELAGIRS-EVITVAQQ---ITS--Q-QLWQEIVNRHPRLAA--IH---DHVILAVHQEYILLGD--QFLLLQPGDEVAIIPPISGG
>Priacma_serrata
------------------------MSVTVTILFFAKAREIVGKNS-SSLEVPPA---LTC--T-DLLNTIVNE-YSLHV--IR---NNIILAVNEEFCAGND---RLSLKTGDEIAVIPPLSGG
>Ricordea_yuma
---------------MEYTNHVSTDKVEINVLFFAKSRELVEQRS-AKVLLAKR---LSG--R-ELIASLVDSFPSLTV--IR---DNLVVAVNQSYVEKET---EIEFEGGEEVAIIPPISGG
>Cuculus_canorus
-------------------MTAELPLVQVSVLYFARSAELAGLRR-ETLRVPRQ---VTA--L-QLWEEIVSLHPRLAV--IR---DQVVFAVRQEYVLLGD--QLLVLQPGDEVAIIPPISGG
>Ctenopharyngodon_idella
------------------------MNTEVSVLYFAKSAELTGLKA-ENITVPSN---ITS--L-QLWQDLENRHPRLSV--LR---DQVVLAVRQEYVPLGA--QPLTLRQGDEVAIIPPLSGG
>Fundulus_grandis
------------------------MSAQVTVLYFAKSSELTGLKE-EPVAVPTP---ISS--G-DLWALLLRKQPRLVA--LQ---GHVILAVRQRYVAIGE--QQVTLEDGDEVAVVPPLSGG
>Macrobrachium_nipponense
---------------------MTESEVSVNILFFASARDLAGVDR-SILTVPVN---TTP--E-ELLSQVVRSFGEISS--IK---DSVILSHNQQYCEPG---QGLTLSTGDEVAIIPPISGG
>Sclerodactyla_briareus
------------------------MPVDVTVQYYAISRDITGTRE-DRLTVPTH---VTL--K-QLFGCLTERHPGLKQ--IE---TNCLLAINGDWLTEES--GVVELKPGDEVAVIPPLSGG
>Forficula_auricularia
-----------------MNQDLNCDQVTVEVLFFAKARELSGVNK-AIIQTPSL---TNS--E-ELLDSLINR-FNLQQ--IS---SCLVLAVDENWVESGP----LKLNSGSIIAVIPPLSGG
>Occasjapyx_japonicus
-----------------------MQTTELRVLFFASARELAKCNA-AVLRLDSA---QQS--GESLLEQVVLSFPDLQR--VR---GSLMLALNQNYVQLD---QRLTVSSGDELAVIPPISGG
>Echiniscus_testudo
--------------------------MQITVLLFARMRDLAGGRSEILIDVPADCENGSVSSK-RLLQLIIAQCPRLKE--IK----HYQLAVNRQYVDLNS-DNLVQLGPTSEIALIPPVSGG
>Evechinus_chloroticus
------------------MADDLAGKVEVTVLFFAKSRDLAGTKE-ASLQIQSD---IKG--E-VLLNVISEKFPRLDA--IR---DSIVLCINQEYISSES---SYHLSPGDEVAVIPPLSGG
>Ciona_intestinalis
----------------------MGKNVQVNIKFFAKSRELVGIES-EQLILPQD---VTS--F-QLIETINSKYPRLSD--LQ---GCFTLAVNGEYIKTDE-SYSILLTDQSEIAVIPPLSGG
>Baetis_sp._AB-2009
----------------------MGQPIVVKVLLFASAREISGERE-SEIQVQQE---ETY--D-TLFRKIVQR-FKLEK--LQ---NSLILAINEDYPDAEN--TQLNLKPGDHIAVIPPLSGG
>Esox_lucius
------------------------MNTEVTVLYFAKSAELTGIKS-ETIRVPSE---LTT--L-DLWLDLVSRHPRLST--VR---ESVVLAVRQEYVAVGE--LRVSLREGDEVAVIPPLSGG
>Asterias_rubens
------------------MASICADVVEINLLFFAKSRELAGVKE-GHLQVQTN---LKP--E-ELLLTIVNHYPQLSV--IQ---DTVVLSLNQEYIAKG--DDTLNLRPGDEVAVIPPISGG
>Gadus_morhua
------------------------MTARVTVLYFAKSAELTGHRS-EVVPLPAA---TSS--C-QLWALLLELHPSLAA--LQ---DQVVLAVRQQYVALGD--QALLLADGDEVAVVPPLSGG
>Gallus_gallus
------------------------MSCQVTVLYFARSAELVGLRS-ESVSVPQR---ITS--L-QLWEEIVKIHPRLAV--IR---DQVVFAVRQEYVLLGD--QLLVLQTGDEVAIIPPISGG
>Chlamydotis_macqueenii
-------------------------------------------------------------------------------------------------MLLGD--QLLVLQPGDEVAVIPPISGG
>Galloisiana_yuasai
--------------------MDLNLKISVNVLFFAKARELCGKKE-SKLQVVPS---ISY--N-LLLDHIIKE-FSLES--IK---GNFILAINEEYTTTDR---ELDLKEGDEIAIIPPLSGG
>Tigriopus_californicus
------------------------MSVSIKVLLFAKARELAGTHE-LTLNVVPN---LSA--S-DLVTQIEAQCPDLAI--LR---GSFVLALNQDYVDHN---QVLDLKSGDELAVIPPISGG
>Trialeurodes_vaporariorum
------------------------MNVSVEVLLFGRAKELAGEEK-VRVQVPSS---SSR--S-HLLKNLVTR-FSLQA--IE---SILILAINEEYVLESD---EIKLQDGDRVAIIPPLSGG
>Diadromus_collaris
-----------------MTALMAPNPRNIRVVFFAKARELAGLSE-STIDVDGI---LSC--D-QLFARLIDK-FGLDC--IR---DNLILAINEQFVERGA---TLDLKEGDEIAVIPPLSGG
>Ornithodoros_turicata
----------------------MLSNVAIRVLFFAKARELLGKSE-VTLLIPNN---VQD--EKALKDFIFSSFPLLSV--LV---KCTILAVNELYLEPG---ASVDLKEGDEIAVIPPISGG
>Helicoverpa_armigera
--------------------MEIENDVTVKLLFFARSKELAGTKE-STIRLPSK---ISY--S-KLLNIICDN-YNLQT--IK---DNILLAKNEEVCDEDI---DIDIKEKDSIAVIPPISGG
>Lutjanus_erythropterus
------------------------MTAQVCVLYFAKSVELTGAKEEQIVAVPTP---ISS--Q-DLWKLLLQRHPRLSV--LQ---DQVVLAVRQQYVAIGD--QVVTLGDGDEVAVVPPLSGG
>Metallyticus_splendidus
------------------MDGNGQHKVRIKVLFFAKARELSGTRE-SSLISDSV---ISY--E-SLLQQIIET-YSLSS--IK---SSVVLAINEEYCSAAT---NVMLNEGDEVAVIPPLSGG
>Boreus_hyemalis
-----------------------MGTAKIKIIFFAKSRDLSGKTE-DTFSVPTN---ITC--S-ELLSLISSN-YGLDA--IK---DNVLLALNEEYCDDLN--AQLILKTGDEIAVIPPLSGG
>Dermanyssus_gallinae
-------------------------MVRVKLLFFAKARELAGTAE-RSVDVAER----VFSLD-QLKELIAQLEPSVKT--LL---DTSILAVDLNYVVDSV---EIG-TRTGEIAVIPPVSGG
>Ornithorhynchus_anatinus
-----------------------MVLSPVTVLYFAKSAELAGVRS-ETISVPQE---IKS--L-QLWEEIESRHPRLAD--VR---DQVIFAVRQEYVQLGD--QVLLLQPGDEIAIIPPISGG
>Perna_viridis
---------------------MANNKVQVSLLFFAKSRELSGVSN-SQIKVSSQ---CSF--F-QLFETIVQQYPSLSV--IS---ENIILSLNEEYISRD---GSITVQEGDEIAVIPPISGG
>Osmylus_fulvicephalus
------------------------MSVEVRILFFAKAKDFAQKSE-STLNIPNT---LTF--K-ELLTIIVDN-YNLQA--IR---NTINISLNANFCDRDA---TLKFNTGDEIAVIPPLSGG
>Hapalochlaena_maculosa
---------------MDHSFAGNKDPVKINVLFFAKSRELVGQNK-SEVSITSR---SVT-TA-QLTSFLLETFPRLKP--LK---QSMIIALNEEYLSGDE---TIHLSNGDEIAVIPPLSGG
>Epiophlebia_superstes
--------------------MGDKDFVTVKVLFFAKARELSGIKF-GQLKVNAA---PSY--T-FLLEEIVTKF-SLEK--IR---NNILLAINEEYSGSE---LQINLKDGDEIAVIPPISGG
>Opisthorchis_felineus
----------------------MKQAVRARVILFGEAKDLTGFQS-ALILVPVT---CTA--K-ELYDLILGHFPKLKL--IE---KSLLLTLDFQYIDIEDNNTLLHISDSSELGVLPPLGGG
>Ceuthophilus_sp._AD-2013
--------------------MADGHQVDVKILLFAKARELSGVKE-SNLEITSS---ISY--S-SLFDILVKT-FSLDA--IK---GNLILAINQEYSSADS---ILNLKAGDEIAVIPPLSGG
>Osmerus_mordax
----------------------MTTKVSVLVLYFAKSADLTGLKS-EVIHVQPE---LTT--L-ALWQELVNRHSRLSA--VR---EQIVLAVSQEYVTIGC--QLVSLKNGDEVAVIPPLSGG
>Crassostrea_gigas
----------------------MEGDVTINLLFFAKSRELVGSGS-GELVLTAN---TNF--D-TLKQALLLKYPSLQV--IA---ENIILSLNEEYIDDN---QQLTLREGDEVAVIPPISGG
>Hypsibius_dujardini
--------------------------MDVNLKFLALAKDLAGTGD-ATIRITAEESPFTV--G-SFLAKLIRTFPRLGE--IK---DQLILAHNLEYLDLQNPSASFELKEGDEIAVLPPVSGG
>Taeniopygia_guttata
------------------------MRCQVSVLYFARSAELAGLRC-ETLSVPQE---ITS--L-QLWEEIVKVHPRLAL--IR---DQVVFAVRQEYVLLGD--QLLVLQPGDEVAIIPPISGG
>Pecten_maximus
-----------------------MADVEVSVLFFAKSRELAGKKT-AQIIVESP---TNL--Q-QLISTIVSQFPSLQI--IA---DNLVVSINEDYLQAG---QEVHLNTGDEVAVIPPISGG
>Dissostichus_mawsoni
------------------------MTAQVTVLYFAKSAELSGLKEEELVAVPTP---ISS--Q-DLWSLLLQRHPRLCV--LQ---HQVVLAVRQQYVAIGD--ELVALGDGDEVAVVPPLSGG
>Equus_asinus
----------------------MVPRCQVEVLYFAKSAEIAGIRS-ETISVPQE---IKA--L-QLWNEIETRHPGMS-----------------------------------------------
>Sipyloidea_sipylus
--------------------MTEGNKVVIKVLFFAKARELSGKTE-STILIEPK---TTY--S-KVLDHIAKE-FSLEL--IK---NNVILAVNQSYASADQ---TLELKTGDELAVIPPLSGG
>Menopon_gallinae
--------------------MNDSSHVSVKLLFFAKARELTGKNE-AIVDVDRN---VKG--A-DLLDFLVSK-FELDC--IR---KNVILALNGDYVDLEC---DLELNSADELAIIPPLSGG
>Fasciola_hepatica
----------------------MSTVISVSVLFFGEAREVVKLHR-VCLSLPTT---CTP--A-ELKREILKHFPQLEL--LS---SSLGLALDLQYLDCSSETDRITLSDLSEVSVFPPLSGG
>Colobus_angolensis_palliatus
----------------------MVPLCQAEVLYFAKSVEITGVRS-ETISVPQE---IKA--L-QLWKEIETRHPGLAD--VR---NQILFAVHQEYVELGD--QLLLLQPGDEIAVIPH----
>Acerentomon_sp._AD-2013
----------------------MAYDVSVNVLFFAKAKELTNLSE-VLLVLKTE---TTG--N-EILSALEKKFPVLSR--IK---EIIHLAVNEAYCEDL--SCSITLNSGDTVAVITPISGG
>Melopsittacus_undulatus
------------------------MRCQVSVLYFARSAELAGLRR-ETLSVPRQ---ITA--I-QLWEEIVKVHPRLAV--IR---DQVVFAVRQEYVLLGD--QFLVLQPGDEVAIIPPISGG
>Liposcelis_bostrychophila
---------------MIDGFDEEATQVSIKILFFALVKDLVGKNE-DRIYVPST---IVG--S-ALLDLIIKK-YNLET--VK---NNVILAVNEDWVDLDS---LLILKDSDEVAVIPPLSGG
>Inocellia_crassicornis
------------------------MSVEVKLLFFASARTISNLNE-SVLKVPDE---ISY--K-DLLGLIVEKY-SLQK--IR---HSVLLAHNESICDKN---QILSLSEGDEVAVIPPFSGG
>Rhopilema_esculentum
--------------------MDAVNEVSVRLLFFAKSRDIVGLHE-AHIDISSV---VTG--Q-ELLRAIVTKFPGLHI--LR---ENLLLSCNQEYLESD---QVVELAGGEEVAVIPPISGG
>Marmota_marmota_marmota
----------------------MVPRCQVEILYFAKSAEITGVRS-ETISVPQE---IKA--L-QLWHEIETRHPGLAD--VR---NQVIFAVRQEYVELGD--QLLLLHPGDEIAIIPPISGG
>Coregonus_clupeaformis
------------------------MNTEVSVFYFAKSAELTGLKS-ETIRVPSE---LTT--L-ELWQDLVSRHPRLSA--VR---DSVVLAVRQEYVALGD--QCVSLREGDEVAVIPPLSGG
>Favia_lizardensis
---------------MDHFSNDTTEQVEVTVLFFAKSRELVEKRS-AQISVGKQ---ITG--T-DLIHYILNNFPSLAV--IQ---ENLVIAVNQSYIEKDS---EITFEGGEEVAVIPPISGG
>Sepioloidea_lineolata
--------------------------------------------------------------------------------------------------------------------MLPPLSGG
>Trichomycterus_areolatus
----------------------MNPEVQVLVLYFAKSSELSGVKS-ETISVQSE---LNS--L-KLWQQLELRHPKLSV--IR---EQVVLAVRREYVALGE--HRVHLQNGDEVAVIPPLSGG
>Oropsylla_silantiewi
---------------------MEVGQVSIKVKFFAKARELSGVSE-TLLKVNSN---FSC--G-ELVQLICIK-YGLQS--IQ---SNLIIALNETYCKDLA--AILDLKTGDEIAVIPPLSGG
>Streblospio_benedicti
---------MLLFLIMLTVLNILKLLYLVKVLFFAKSRELVGHSD-SYINLPLI---CLE--S-QIRTSIVENNPSLSP--IS---DSFVLALNQEYIGEG---EEVHISEGDEIAVIPPISGG
>Squalus_acanthias
------------------------MIAQVSVLYFARSAELAGVRS-ETLSVPQQ---LTS--L-QLWQEIVKRHPRKQRENLREIQPRSVLNIF--YH--------------------PVL---
>Atractaspis_aterrima
------------------------MSCEVTVLYFARSAELAGLRT-ETISVPQQ---ITS--L-QLWREIVKRHSSLAD--IE---DQVIFAICQEYVPLGD--QQFILHTGDEVAIIPPLSGG
>Stylops_melittae
-----------------------MNTVTIKVLFFAKSRQLSKRKE-GHITLPAE---IAY--D-DLLTLICNQF-QLEE--IK---KTIFLALNETYLQSN---TLVHLHSDDIVAVIPPISSG
>Trichobilharzia_regenti
-----------------------MSVCSKKVLLFGEASELVGCKE-IHLPFPKS---CTQ--K-ALKNIVIDQLAELEP--LK---DCITLAVNWCYVDTEDSINSISVEEIEEIAVLPPISGG
>Lytechinus_variegatus
------------MAEDAVSDSIAGEKVEVTVLFFAKSRDLTGTKE-ANIRVQSK---INT--G-SLLNVLSEKYPRLSA--IQ---DSIVLCINQEYISAEG---SFHLSPGDEIAVIPPLSGG
>Alvinella_pompejana
MKAYVICKPKYDHYDDKNFCNDTNCILKVTVFFFAKGRELVGKKE-DQIDIAPH---NSG--T-ALYKLVVDRYPCLKP--IQ---ENFALALNEEYIDKTD--EEIILSNGDQLAEIPPISGG
>Trachemys_scripta
------------------------MSCEVVALYFARSAELAGVRS-EIISVPQQ---LTS--L-QLWEEIVKRHPRLAA--IQ---DQVVFAVRQEYVLLGD--QLLVLQSGDEVAIIPPISGG
>Hydroptila_sp._AD-2013
---------------------MEVSKVVVKVLFFAASREIVGAEE-RELEVPAR---SSY--S-SLLHLLVHT-FKLDI--IK---DNFIVAKNEELCLDNG--HEIVFEKNDEIAIIPPLSGG
>Villosa_lienosa
----------------------------------------------------------------------------------------------------------MQLHQGDEIAVIPPISGG
>Meretrix_meretrix
---------------------MDDDKVEVTVLFFAKSRELAGTRE-TTLQTFKE---ITA--A-GLLNQIISQYNSLDI--LR---ESIVLSLNEEYLSLTD-SEQLTLRSGDTVAVIPPISGG
>Limulus_polyphemus
----------------------MSSGVKVKLLFFAKARELVGVTE-TGLVVPNG---LYT-QK-KLLSIIVETFPVLCS--IQ---GNFIVALNEECMDQSQ---SIELKEGDQLAVIPPLSGG
>Protopalythoa_variabilis
---------------MELANDAGCEKVQVTVLFFAKCRELVGTSS-VQIYLARR---LSG--S-KLIETLLNLFPSLSE--IE---RNLIVAVNQEYIEKE---SNIELFGNDEIAVIPPISGG
>Tricholepidion_gertschi
----------------MGVFLMELSLITVKVLFFAKARELSGKVE-DNFQIVCS---LSY--P-ELLSQIVKE-FCLEP--IE---RNIILAINEEYIGTEE---KLCLKEGDEIAVIPPLSGG

HG0004.1 ENSDARG00000076779 fam13b

```
>Danio_rerio
---------------------------------------------------------------MRNYRLIFCHNLKQP
>Acipenser_sinensis
---------------------------------------------------------------MRNYRLIFCHNLKQP
>Amia_calva
---------------------------------------------------------------MRNYRLIFCHNLKQP
>Channa_punctata
---------------------------------------------------------------MRNYRLIFCHNLKQP
>Anas_platyrhynchos
---------------------------------------------------------------MRNYRLIFCHNLKQP
>Xenopus_laevis
MVTSCSDCGPLAVNAFKCIHKILTERDCWGQRELKGLSKRKPQQNQS-YRHTAEELIELWISSMRNYRLIFCHNLKQP
>Chaetura_pelagica
---------------------------------------------------------------MRNYRLIFCHNLKQP
>Menidia_menidia
---------------------------------------------------------------MRNYRLIFCHNLKQP
>Porichthys_notatus
---------------------------------------------------------------MRNYRLIFCHNLKQP
>Oryzias_latipes
---------------------------------------------------------------MRNYRLIFCHNLKQP
>Tripterygion_delaisi
---------------------------------------------------------------MRNYRLIFCHNLKQP
>Buceros_rhinoceros_silvestri
---------------------------------------------------------------MRNYRLIFCHNLKQP
>Caprimulgus_carolinensis
---------------------------------------------------------------MRNYRLIFCHNLKQP
>Cariama_cristata
---------------------------------------------------------------MRNYRLIFCHNLKQP
>Canis_lupus_familiaris
---------------------------------------------------------------MRNYRLIFCHNLKQP
>Hynobius_retardatus
---------------------------------------------------------------MRNYRLIFCHNLKQP
>Micropterus_floridanus
---------------------------------------------------------------MRNYRLIFCHNLKQP
>Balaenoptera_acutorostrata_s
---------------------------------------------------------------MRNYRLIFCHNLKQP
>Pygocentrus_nattereri
---------------------------------------------------------------MRNYRLIFCHNLKQP
>Calidris_pugnax
---------------------------------------------------------------MRNYRLIFCHNLKQP
>Callorhinchus_milii
---------------------------------------------------------------MRNYRLIFCHNLKQP
>Pteropus_vampyrus
---------------------------------------------------------------MRNYRLIFCHNLKQP
>Maylandia_zebra
---------------------------------------------------------------MRNYRLIFCHNLKQP
>Pterocles_gutturalis
---------------------------------------------------------------MRNYRLIFCHNLKQP
>Clupea_harengus
--------------------------------------MRKPFQ-------LSEVLHGICGLSMRNYRLIFCHNLKQP
>Latimeria_chalumnae
---------------------------------------------------------------MRNYRLIFCHNLKQP
>Colius_striatus
---------------------------------------------------------------MRNYRLIFCHNLKQP
>Columba_livia
---------------------------------------------------------------MRNYRLIFCHNLKQP
>Merops_nubicus
---------------------------------------------------------------MRNYRLIFCHNLKQP
>Alligator_mississippiensis
---------------------------------------------------------------MRNYRLIFCHNLKQP
>Cuculus_canorus
---------------------------------------------------------------MRNYRLIFCHNLKQP
>Kryptolebias_marmoratus
---------------------------------------------------------------MRNYRLIFCHNLKQP
>Sarcophilus_harrisii
---------------------------------------------------------------MRNYRLIFCHNLKQP
>Galeopterus_variegatus
---------------------------------------------------------------MRNYRLIFCHNLKQP
>Monodelphis_domestica
---------------------------------------------------------------MRNYRLIFCHNLKQP
>Anopheles_funestus
---------------------------------------------------------------MLNYRLVSTHFLKQP
>Esox_lucius
---------------------------------------------------------------MRNYRLIFCHNLKQP
>Falco_cherrug
---------------------------------------------------------------MRNYRLIFCHNLKQP
>Gallus_gallus
---------------------------------------------------------------MRNYRLIFCHNLKQP
>Gavia_stellata
---------------------------------------------------------------MRNYRLIFCHNLKQP
>Balearica_regulorum_gibberic
---------------------------------------------------------------MRNYRLIFCHNLKQP
>Erinaceus_europaeus
---------------------------------------------------------------MRNYRLIFCHNLKQP
>Oryctolagus_cuniculus
---------------------------------------------------------------MRNYRLIFCHNLKQP
>Eudarcia_simulatricella
---------------------------------------------------------------MLNYRLVSTHFLKQP
>Tauraco_erythrolophus
---------------------------------------------------------------MRNYRLIFCHNLKQP
>Opisthocomus_hoazin
---------------------------------------------------------------MRNYRLIFCHNLKQP
>Plecoglossus_altivelis
----------------------------------------MLFAPVVFDLLTSEILNRICGLSMRNYRLIFCHNLKQP
>Scleropages_formosus
---------------------------------------------------------------MRNYRLIFCHNLKQP
>Lepidothrix_coronata
---------------------------------------------------------------MRNYRLIFCHNLKQP
>Nipponia_nippon
---------------------------------------------------------------MRNYRLIFCHNLKQP
>Trematomus_bernacchii
---------------------------------------------------------------MRNYRLIFCHNLKQP
>Ceratotherium_simum_simum
---------------------------------------------------------------MRNYRLIFCHNLKQP
>Manis_javanica
---------------------------------------------------------------MRNYRLIFCHNLKQP
>Glycera_dibranchiata
---------------------------------------------------------------MHGYRLILCRTLKQP
>Picoides_pubescens
---------------------------------------------------------------MRNYRLIFCHNLKQP
>Platichthys_stellatus
---------------------------------------------------------------MRNYRLIFCHNLKQP
>Homo_sapiens
---------------------------------------------------------------MRNYRLIFCHNLKQP
>Loxodonta_africana
---------------------------------------------------------------MRNYRLIFCHNLKQP
>Fulmarus_glacialis
---------------------------------------------------------------MRNYRLIFCHNLKQP
>Nestor_notabilis
---------------------------------------------------------------MRNYRLIFCHNLKQP
>Mus_musculus
---------------------------------------------------------------MRNYRLIFCHNLKQP
>Salmo_salar
---------------------------------------------------------------MRNYRLIFCHNLKQP
>Tupaia_chinensis
---------------------------------------------------------------MRNYRLIFCHNLKQP
>Lepisosteus_oculatus
---------------------------------------------------------------MRNYRLIFCHNLKQP
>Ameiurus_nebulosus
---------------------------------------------MTVFCLTTDVLYRICGPSMRNYRLIFCHNLKQP
>Trichechus_manatus_latirostr
---------------------------------------------------------------MRNYRLIFCHNLKQP
>Aptenodytes_forsteri
---------------------------------------------------------------MRNYRLIFCHNLKQP
>Crotalus_horridus
---------------------------------------------------------------MRNYRLIFCHNLKQP
>Tyto_alba
---------------------------------------------------------------MRNYRLIFCHNLKQP
>Struthio_camelus_australis
---------------------------------------------------------------MRNYRLIFCHNLKQP
>Chelonia_mydas
---------------------------------------------------------------MRNYRLIFCHNLKQP
>Takifugu_rubripes
---------------------------------------------------------------MRNYRLIFCHNLKQP
>Tinamus_guttatus
---------------------------------------------------------------MRNYRLIFCHNLKQP
>Calypte_anna
---------------------------------------------------------------MRNYRLIFCHNLKQP
>Apaloderma_vittatum
---------------------------------------------------------------MRNYRLIFCHNLKQP
>Orycteropus_afer_afer
---------------------------------------------------------------MRNYRLIFCHNLKQP
```

HG0004.1 ENSG00000031003 FAM13B

>Homo_sapiens
---------------------------------------------------------------MRNYRLIFCHNLKQP
>Acipenser_sinensis
---------------------------------------------------------------MRNYRLIFCHNLKQP
>Amia_calva
---------------------------------------------------------------MRNYRLIFCHNLKQP
>Channa_punctata
---------------------------------------------------------------MRNYRLIFCHNLKQP
>Anas_platyrhynchos
---------------------------------------------------------------MRNYRLIFCHNLKQP
>Xenopus_laevis
MVTSCSDCGPLAVNAFKCIHKILTERDCWGQRELKGLSKRKPQQNQS-YRHTAEELIELWISSMRNYRLIFCHNLKQP
>Chaetura_pelagica
---------------------------------------------------------------MRNYRLIFCHNLKQP
>Menidia_menidia
---------------------------------------------------------------MRNYRLIFCHNLKQP
>Porichthys_notatus
---------------------------------------------------------------MRNYRLIFCHNLKQP
>Oryzias_latipes
---------------------------------------------------------------MRNYRLIFCHNLKQP
>Tripterygion_delaisi
---------------------------------------------------------------MRNYRLIFCHNLKQP
>Buceros_rhinoceros_silvestri
---------------------------------------------------------------MRNYRLIFCHNLKQP
>Caprimulgus_carolinensis
---------------------------------------------------------------MRNYRLIFCHNLKQP
>Cariama_cristata
---------------------------------------------------------------MRNYRLIFCHNLKQP
>Canis_lupus_familiaris
---------------------------------------------------------------MRNYRLIFCHNLKQP
>Hynobius_retardatus
---------------------------------------------------------------MRNYRLIFCHNLKQP
>Micropterus_floridanus
---------------------------------------------------------------MRNYRLIFCHNLKQP
>Balaenoptera_acutorostrata_s
---------------------------------------------------------------MRNYRLIFCHNLKQP
>Pygocentrus_nattereri
---------------------------------------------------------------MRNYRLIFCHNLKQP
>Calidris_pugnax
---------------------------------------------------------------MRNYRLIFCHNLKQP
>Callorhinchus_milii
---------------------------------------------------------------MRNYRLIFCHNLKQP
>Pteropus_vampyrus
---------------------------------------------------------------MRNYRLIFCHNLKQP
>Maylandia_zebra
---------------------------------------------------------------MRNYRLIFCHNLKQP
>Pterocles_gutturalis
---------------------------------------------------------------MRNYRLIFCHNLKQP
>Clupea_harengus
--------------------------------------MRKPFQ-------LSEVLHGICGLSMRNYRLIFCHNLKQP
>Latimeria_chalumnae
---------------------------------------------------------------MRNYRLIFCHNLKQP
>Colius_striatus
---------------------------------------------------------------MRNYRLIFCHNLKQP
>Columba_livia
---------------------------------------------------------------MRNYRLIFCHNLKQP
>Merops_nubicus
---------------------------------------------------------------MRNYRLIFCHNLKQP
>Alligator_mississippiensis
---------------------------------------------------------------MRNYRLIFCHNLKQP
>Cuculus_canorus
---------------------------------------------------------------MRNYRLIFCHNLKQP
>Danio_rerio
---------------------------------------------------------------MRNYRLIFCHNLKQP
>Kryptolebias_marmoratus
---------------------------------------------------------------MRNYRLIFCHNLKQP
>Sarcophilus_harrisii
---------------------------------------------------------------MRNYRLIFCHNLKQP
>Galeopterus_variegatus
---------------------------------------------------------------MRNYRLIFCHNLKQP
>Monodelphis_domestica
---------------------------------------------------------------MRNYRLIFCHNLKQP
>Esox_lucius
---------------------------------------------------------------MRNYRLIFCHNLKQP
>Falco_cherrug
---------------------------------------------------------------MRNYRLIFCHNLKQP
>Gallus_gallus
---------------------------------------------------------------MRNYRLIFCHNLKQP
>Gavia_stellata
---------------------------------------------------------------MRNYRLIFCHNLKQP
>Balearica_regulorum_gibberic
---------------------------------------------------------------MRNYRLIFCHNLKQP
>Erinaceus_europaeus
---------------------------------------------------------------MRNYRLIFCHNLKQP
>Oryctolagus_cuniculus
---------------------------------------------------------------MRNYRLIFCHNLKQP
>Tauraco_erythrolophus
---------------------------------------------------------------MRNYRLIFCHNLKQP
>Opisthocomus_hoazin
---------------------------------------------------------------MRNYRLIFCHNLKQP
>Plecoglossus_altivelis
----------------------------------------MLFAPVVFDLLTSEILNRICGLSMRNYRLIFCHNLKQP
>Scleropages_formosus
---------------------------------------------------------------MRNYRLIFCHNLKQP
>Lepidothrix_coronata
---------------------------------------------------------------MRNYRLIFCHNLKQP
>Nipponia_nippon
---------------------------------------------------------------MRNYRLIFCHNLKQP
>Trematomus_bernacchii
---------------------------------------------------------------MRNYRLIFCHNLKQP
>Ceratotherium_simum_simum
---------------------------------------------------------------MRNYRLIFCHNLKQP
>Manis_javanica
---------------------------------------------------------------MRNYRLIFCHNLKQP
>Picoides_pubescens
---------------------------------------------------------------MRNYRLIFCHNLKQP
>Platichthys_stellatus
---------------------------------------------------------------MRNYRLIFCHNLKQP
>Loxodonta_africana
---------------------------------------------------------------MRNYRLIFCHNLKQP
>Fulmarus_glacialis
---------------------------------------------------------------MRNYRLIFCHNLKQP
>Nestor_notabilis
---------------------------------------------------------------MRNYRLIFCHNLKQP
>Mus_musculus
---------------------------------------------------------------MRNYRLIFCHNLKQP
>Salmo_salar
---------------------------------------------------------------MRNYRLIFCHNLKQP
>Tupaia_chinensis
---------------------------------------------------------------MRNYRLIFCHNLKQP
>Lepisosteus_oculatus
---------------------------------------------------------------MRNYRLIFCHNLKQP
>Ictalurus_punctatus
---------------------------------------------------------------MRNYRLIFCHNLKQP
>Trichechus_manatus_latirostr
---------------------------------------------------------------MRNYRLIFCHNLKQP
>Aptenodytes_forsteri
---------------------------------------------------------------MRNYRLIFCHNLKQP
>Crotalus_horridus
---------------------------------------------------------------MRNYRLIFCHNLKQP
>Tyto_alba
---------------------------------------------------------------MRNYRLIFCHNLKQP
>Struthio_camelus_australis
---------------------------------------------------------------MRNYRLIFCHNLKQP
>Chelonia_mydas
---------------------------------------------------------------MRNYRLIFCHNLKQP
>Takifugu_rubripes
---------------------------------------------------------------MRNYRLIFCHNLKQP
>Tinamus_guttatus
---------------------------------------------------------------MRNYRLIFCHNLKQP
>Calypte_anna
---------------------------------------------------------------MRNYRLIFCHNLKQP
>Apaloderma_vittatum
---------------------------------------------------------------MRNYRLIFCHNLKQP
>Orycteropus_afer_afer
---------------------------------------------------------------MRNYRLIFCHNLKQP

HG0004.1 ENSGALG00000041706

```
>Gallus_gallus
---------------------------------------------------------------MRNYRLIFCHNLKQP
>Acipenser_sinensis
---------------------------------------------------------------MRNYRLIFCHNLKQP
>Amia_calva
---------------------------------------------------------------MRNYRLIFCHNLKQP
>Channa_punctata
---------------------------------------------------------------MRNYRLIFCHNLKQP
>Anas_platyrhynchos
---------------------------------------------------------------MRNYRLIFCHNLKQP
>Xenopus_laevis
MVTSCSDCGPLAVNAFKCIHKILTERDCWGQRELKGLSKRKPQQNQS-YRHTAEELIELWISSMRNYRLIFCHNLKQP
>Chaetura_pelagica
---------------------------------------------------------------MRNYRLIFCHNLKQP
>Menidia_menidia
---------------------------------------------------------------MRNYRLIFCHNLKQP
>Porichthys_notatus
---------------------------------------------------------------MRNYRLIFCHNLKQP
>Oryzias_latipes
---------------------------------------------------------------MRNYRLIFCHNLKQP
>Tripterygion_delaisi
---------------------------------------------------------------MRNYRLIFCHNLKQP
>Buceros_rhinoceros_silvestri
---------------------------------------------------------------MRNYRLIFCHNLKQP
>Caprimulgus_carolinensis
---------------------------------------------------------------MRNYRLIFCHNLKQP
>Cariama_cristata
---------------------------------------------------------------MRNYRLIFCHNLKQP
>Canis_lupus_familiaris
---------------------------------------------------------------MRNYRLIFCHNLKQP
>Hynobius_retardatus
---------------------------------------------------------------MRNYRLIFCHNLKQP
>Micropterus_floridanus
---------------------------------------------------------------MRNYRLIFCHNLKQP
>Balaenoptera_acutorostrata_s
---------------------------------------------------------------MRNYRLIFCHNLKQP
>Pygocentrus_nattereri
---------------------------------------------------------------MRNYRLIFCHNLKQP
>Calidris_pugnax
---------------------------------------------------------------MRNYRLIFCHNLKQP
>Callorhinchus_milii
---------------------------------------------------------------MRNYRLIFCHNLKQP
>Pteropus_vampyrus
---------------------------------------------------------------MRNYRLIFCHNLKQP
>Maylandia_zebra
---------------------------------------------------------------MRNYRLIFCHNLKQP
>Pterocles_gutturalis
---------------------------------------------------------------MRNYRLIFCHNLKQP
>Clupea_harengus
--------------------------------------MRKPFQ-------LSEVLHGICGLSMRNYRLIFCHNLKQP
>Latimeria_chalumnae
---------------------------------------------------------------MRNYRLIFCHNLKQP
>Colius_striatus
---------------------------------------------------------------MRNYRLIFCHNLKQP
>Columba_livia
---------------------------------------------------------------MRNYRLIFCHNLKQP
>Merops_nubicus
---------------------------------------------------------------MRNYRLIFCHNLKQP
>Alligator_mississippiensis
---------------------------------------------------------------MRNYRLIFCHNLKQP
>Cuculus_canorus
---------------------------------------------------------------MRNYRLIFCHNLKQP
>Danio_rerio
---------------------------------------------------------------MRNYRLIFCHNLKQP
>Kryptolebias_marmoratus
---------------------------------------------------------------MRNYRLIFCHNLKQP
>Sarcophilus_harrisii
---------------------------------------------------------------MRNYRLIFCHNLKQP
>Galeopterus_variegatus
---------------------------------------------------------------MRNYRLIFCHNLKQP
>Monodelphis_domestica
---------------------------------------------------------------MRNYRLIFCHNLKQP
>Esox_lucius
---------------------------------------------------------------MRNYRLIFCHNLKQP
>Falco_cherrug
---------------------------------------------------------------MRNYRLIFCHNLKQP
>Gavia_stellata
---------------------------------------------------------------MRNYRLIFCHNLKQP
>Balearica_regulorum_gibberic
---------------------------------------------------------------MRNYRLIFCHNLKQP
>Erinaceus_europaeus
---------------------------------------------------------------MRNYRLIFCHNLKQP
>Oryctolagus_cuniculus
---------------------------------------------------------------MRNYRLIFCHNLKQP
>Tauraco_erythrolophus
---------------------------------------------------------------MRNYRLIFCHNLKQP
>Opisthocomus_hoazin
---------------------------------------------------------------MRNYRLIFCHNLKQP
>Plecoglossus_altivelis
----------------------------------------MLFAPVVFDLLTSEILNRICGLSMRNYRLIFCHNLKQP
>Scleropages_formosus
---------------------------------------------------------------MRNYRLIFCHNLKQP
>Lepidothrix_coronata
---------------------------------------------------------------MRNYRLIFCHNLKQP
>Nipponia_nippon
---------------------------------------------------------------MRNYRLIFCHNLKQP
>Trematomus_bernacchii
---------------------------------------------------------------MRNYRLIFCHNLKQP
>Ceratotherium_simum_simum
---------------------------------------------------------------MRNYRLIFCHNLKQP
>Manis_javanica
---------------------------------------------------------------MRNYRLIFCHNLKQP
>Picoides_pubescens
---------------------------------------------------------------MRNYRLIFCHNLKQP
>Platichthys_stellatus
---------------------------------------------------------------MRNYRLIFCHNLKQP
>Homo_sapiens
---------------------------------------------------------------MRNYRLIFCHNLKQP
>Loxodonta_africana
---------------------------------------------------------------MRNYRLIFCHNLKQP
>Fulmarus_glacialis
---------------------------------------------------------------MRNYRLIFCHNLKQP
>Nestor_notabilis
---------------------------------------------------------------MRNYRLIFCHNLKQP
>Mus_musculus
---------------------------------------------------------------MRNYRLIFCHNLKQP
>Salmo_salar
---------------------------------------------------------------MRNYRLIFCHNLKQP
>Tupaia_chinensis
---------------------------------------------------------------MRNYRLIFCHNLKQP
>Lepisosteus_oculatus
---------------------------------------------------------------MRNYRLIFCHNLKQP
>Ictalurus_punctatus
---------------------------------------------------------------MRNYRLIFCHNLKQP
>Trichechus_manatus_latirostr
---------------------------------------------------------------MRNYRLIFCHNLKQP
>Aptenodytes_forsteri
---------------------------------------------------------------MRNYRLIFCHNLKQP
>Crotalus_horridus
---------------------------------------------------------------MRNYRLIFCHNLKQP
>Tyto_alba
---------------------------------------------------------------MRNYRLIFCHNLKQP
>Struthio_camelus_australis
---------------------------------------------------------------MRNYRLIFCHNLKQP
>Chelonia_mydas
---------------------------------------------------------------MRNYRLIFCHNLKQP
>Takifugu_rubripes
---------------------------------------------------------------MRNYRLIFCHNLKQP
>Tinamus_guttatus
---------------------------------------------------------------MRNYRLIFCHNLKQP
>Calypte_anna
---------------------------------------------------------------MRNYRLIFCHNLKQP
>Apaloderma_vittatum
---------------------------------------------------------------MRNYRLIFCHNLKQP
>Orycteropus_afer_afer
---------------------------------------------------------------MRNYRLIFCHNLKQP
```

HG0004.1 FBgn0028494

```
>Drosophila_melanogaster
MLRYRLVSTHFLKQP
>Steatoda_grossa
MLSLRLVSCRFLKQP
>Prorhinotermes_simplex
MLTLRLVSTHFLKQP
>Anoplophora_glabripennis
MLKFRLVSTHFLKQP
>Eurylophella_sp._AD-2013
MLTLRLVSTHFLKQP
>Cimex_lectularius
MLTLRLVSTHFLKQP
>Leptopilina_clavipes
MLKLRLVSTHFLKQP
>Eudarcia_simulatricella
MLNYRLVSTHFLKQP
>Metallyticus_splendidus
MLTLRLVSTHFLKQP
>Corydalus_cornutus
MLTLRLVSTHFLKQP
>Conwentzia_psociformis
MLTLRLVSTHFLKQP
>Medauroidea_extradentata
MLTLRLVSTHFLKQP
>Tityus_serrulatus
MLSLRLVSCRFLKQP
>Oropsylla_silantiewi
MLTYRLVSTHFLKQP
>Annulipalpia_sp._AD-2013
MLTYRLVSTHFLKQP
>Limulus_polyphemus
MLSMRLVSCRFLKQP
>Zorotypus_caudelli
MLTLRLVSTHFLKQP
```

HG0004.2 ENSG00000138640 FAM13A

```
>Homo_sapiens
---------MLEAALRLPWGLCIGSCFSHQRICAEWNFSILLSF
>Mustela_putorius_furo
MRSKTSLPKMLGAALRLP--------------------------
>Physeter_catodon
---------MLGAALRSPWGLRIWSRVSHRSISTEWNSSVLLSF
>Eptesicus_fuscus
---------MLGAALRLRWGLRIGSCFSNQSIPTGWNFSLLLSF
>Loxodonta_africana
MRSTASLPKLLGTALHLPWGLWIGSRFLNQSISTEWNFSILLLF
```

HG0005 ENSDARG00000011055 fbxo9

>Danio_rerio
---------------------------------------------------------------------------------------------------------MAAVWRARKFA-RCVVRFS-DRDL----
>Acipenser_sinensis
---------------------------------------------------------------------------------------------------------MAAVWRARNFA-RCVVRFS-DRELN---
>Channa_argus
---------------------------------------------------------------------------------------------------------MAAFLRARKYV-CCVVRFS-DREL----
>Kaupichthys_hyoproroides
---------------------------------------------------------------------------------------------------------MAAVWRARNFS-RCVVRFS-DRELY---
>Podocoryna_carnea
---------------------------------------------------------------------------------------------------------MAAVRRARNFA-RCVVCFS-DRELN---
>Xenopus_laevis
---------------------------------------------------------------------------------------------------------MAAVRRSRSDS-RCVLRFS-DRELC---
>Apteryx_australis_mantelli
---------------------------------------------------------------------------------------------------------MAAVRRARSFS-RCVVRFS-DRELC---
>Stegodyphus_mimosarum
---------------------------------------------------------------------------------------------------------MAAVLIARRYV-RCVIRFNSDRELID--
>Menidia_menidia
---------------------------------------------------------------------------------------------------------MAAFLRARKYV-RCVVRFS-DREL----
>Opsanus_beta
---------------------------------------------------------------------------------------------------------MAAFLRARKYV-RCVVRFS-DREL----
>Oryzias_latipes
---------------------------------------------------------------------------------------------------------MAAFLRARKYV-RCVVRFS-DRDL----
>Prorhinotermes_simplex
---------------------------------------------------------------------------------------------------------MAAVIRAKKCF-RSVVRYS-DRESLP--
>Calanus_finmarchicus
---------------------------------------------------------------------------------------------------------MAAVWMAKCFT-RSVVRFS-DRDS----
>Trachinotus_blochii
---------------------------------------------------------------------------------------------------------MAAFLRARKYV-RCVVRFS-DREL----
>Panthera_tigris
---------------------------------------------------------------------------------------------------------MAAVRRARSYC-RCLVRFS-DRELC---
>Hynobius_retardatus
---------------------------------------------------------------------------------------------------------MAATRRARAHP-RGVIRFS-DRDLY---
>Micropterus_floridanus
---------------------------------------------------------------------------------------------------------MAAFLRARKYV-CCVVRFS-DREL----
>Pygocentrus_nattereri
---------------------------------------------------------------------------------------------------------MAAVWRARKFA-RCVVRFS-DRDL----
>Miniopterus_natalensis
---------------------------------------------------------------------------------------------------------MAAVRRARSYS-RCLVRFS-DRELC---
>Ophthalmotilapia_ventralis
---------------------------------------------------------------------------------------------------------MAAFLRARKYV-RCVVRFS-DREL----
>Clupea_harengus
---------------------------------------------------------------------------------------------------------MAAVWRARKFP-RSVVRFS-DREL----
>Latimeria_chalumnae
---------------------------------------------------------------------------------------------------------MAAVWRARNFI-GCVVRFS-DRELC---
>Alligator_mississippiensis
---------------------------------------------------------------------------------------------------------MAAVRRARSDS-RCVVRFS-DRELC---
>Fundulus_heteroclitus
---------------------------------------------------------------------------------------------------------MAAFLRARKYV-RCVVRFS-DREL----
>Sarcophilus_harrisii
---------------------------------------------------------------------------------------------------------MAAVRRARSHC-RCLVRFS-DRELC---
>Homarus_americanus
---------------------------------------------------------------------------------------------------------MAAVVRSRIYF-SCVVRFS-DRESIA--
>Sclerodactyla_briareus
---------------------------------------------------------------------------------------------------------MAAVARFNRYH-QNIVRYS-DRD-----
>Macropus_eugenii
---------------------------------------------------------------------------------------------------------MAAVRRARSHC-RCLVRFS-DRELC---
>Haploembia_palaui
---------------------------------------------------------------------------------------------------------MAAVLQAKGFF-ASVIKYS-DREFMP--
>Ephemera_danica
---------------------------------------------------------------------------------------------------------MAAVQRAAKLAGQLVVRFS-DRELN---
>Esox_lucius
---------------------------------------------------------------------------------------------------------MAAFLRYLQFP-RCVVRFS-DRDL----
>Meganyctiphanes_norvegica
---------------------------------------------------------------------------------------------------------MAAVARSYMYF-ASVVRFS-DRECIA--
>Falco_cherrug
---------------------------------------------------------------------------------------------------------MAAVRRARSYS-RCVVRFS-DRELC---
>Leptasterias_sp._AR-2014
---------------------------------------------------------------------------------------------------------MAAFLRFLEFS-NCIVRFS-DRDCKA--
>Coturnix_japonica
---------------------------------------------------------------------------------------------------------MAAVRRARSYS-RCVVRFS-DRELC---
>Gillichthys_mirabilis
---------------------------------------------------------------------------------------------------------MAAFLRARKYV-SCVVRFS-DREL----
>Chanos_chanos
---------------------------------------------------------------------------------------------------------MAAVWRSRKFA-RCVVRFS-DRDL----
>Euschistus_heros
-----------------------------------------------------------------------------------------------------MATNMAAVYVCLRLG-EIILRFS-DRDSKI--
>Apis_dorsata
---------------------------------------------------------------------------------------------------------MAAVLLARLYF-RCVVRFS-DREYKG--
>Erinaceus_europaeus
---------------------------------------------------------------------------------------------------------MAAVRRARSYC-RCLVRFS-DRELC---
>Rhipicephalus_appendiculatus
---------------------------------------------------------------------------------------------MASGPLVGNYKSMAAVVKARRYR-NCVLRYNSDRECIA--
>Ochotona_princeps
---------------------------------------------------------------------------------------------------------MAAVRRARSYC-RCLVRFS-DRELC---
>Lutjanus_erythropterus
---------------------------------------------------------------------------------------------------------MAAFLRARKYV-RCVVRFS-DREL----
>Calopteryx_splendens
---------------------------------------------------------------------------------------------------------MAAVLRAKELS-TCVVRYS-DRESSA--
>Ophiocoma_echinata
---------------------------------------------------------------------------------------------------------MAAVNKSLLYF-SCFVRFS-DRDWLA--
>Locusta_migratoria
---------------------------------------------------------------------------------------------------------MAAVFMARKYF-ESVIRFS-DRESVP--
>Plecoglossus_altivelis
---------------------------------------------------------------------------------------------------------MAAFQRSRKFV-RSVVRFS-DRDLVSLI
>Scleropages_formosus
---------------------------------------------------------------------------------------------------------MAAVWRARNFA-RCVVRFS-DRELN---
>Junco_hyemalis
---------------------------------------------------------------------------------------------------------MAAVRRARRDS-RCVVRFS-DRELC---
>Lateolabrax_japonicus
---------------------------------------------------------------------------------------------------------MAAFLRARKYV-RCVVRFS-DREL----
>Trematomus_bernacchii
---------------------------------------------------------------------------------------------------------MAAFLRARKYV-RCVIRFS-DRDL----
>Equus_asinus
---------------------------------------MLGALAQVPGQKGAAPQRRLGSPAGPILRCSHSLRPRLAAGAPPLRESPAA--APARPSLAPCPAKMAAVRRARSYC-RCLVRFS-DRELC---
>Medauroidea_extradentata
---------------------------------------------------------------------------------------------------------MAAALKARKFI-GSVIRYS-DRESLP--
>Perla_marginata
--------------------------------------------------------------------------------------------------------------MAKKYF-KSVIRFT-DRECLP--
>Cynoglossus_semilaevis
---------------------------------------------------------------------------------------------------------MAAFLRARKYV-RCVVRFS-DREL----
>Microcebus_murinus
MVAKKYVRILAQTPCGTAKGLVRKIPRPSPQHPPRRASNRPGSSPSSRRIGGAAPLRRPGSPIGPTRRCSHGPRPRLAAGVPLLRESPAAAAAPARPDLAPCPAKMAAVRRARSYC-RCLVRFS-DRELC---
>Acerentomon_sp._AD-2013
------------------------------------------------------------------------------------------------------------------MF-ECVIRFT-DREIYG--
>Melopsittacus_undulatus
-----------------------------------------------------------------------------------------MCLCCSRPACPSLAGKMAAVRRARSYS-RCVVRFS-DRELC---
>Ectopsocus_briggsi
---------------------------------------------------------------------------------------------------------MAAVLNAKFYF-GLVIRYS-DRELIR--
>Leucoraja_erinacea
---------------------------------------------------------------------------------------------------------MAAVWRARNFL-DYVVRFS-DRELQ---
>Cavia_porcellus
---------------------------------------------------------------------------------------------------------MAAVRRARSYC-RCLVRFS-DRELC---
>Salvelinus_alpinus
---------------------------------------------------------------------------------------------------------MAAFLRSLKFP-RCVVRFS-DRDL----
>Tupaia_chinensis
---------------------------------------------------------------------------------------------------------MAAVRRARSYC-RCLVRFS-DRELC---
>Tityus_serrulatus
---------------------------------------------------------------------------------------------------------MAAVIKGRRYF-GCVIRFNSDREYVA--
>Lepisosteus_oculatus
---------------------------------------------------------------------------------------------------------MAAMWRARNFA-RCVVRFS-DRELY---
>Trichomycterus_areolatus
----------------------------------------------------------------MYMRETPGTSGRLGMSVTLLYK--ADLQRPCRLDSTQTLSVMAAVWRARKFS-RCVVRFS-DRDL----
>Sparus_aurata
---------------------------------------------------------------------------------------------------------MAAFLRARKYV-RCVVRFS-DREL----
>Gekko_japonicus
-------------------------------------------------------MTQTN----------------------THTGSRAHDLALSHLLALSRVGKMAAVRRARSDS-RCVVRFS-DRELC---
>Pelodiscus_sinensis
---------------------------------------------------------------------------------------------------------MAAVRRARSYS-RCVVRFS-DRELC---
>Takifugu_rubripes
---------------------------------------------------------------------------------------------------------MAAFLRARKFV-RSVVRFS-DREL----
>Patiria_miniata
-------------------------------------------------------------------------------------------MYCFGRPSADRRRNMAAFLRFLEFS-KCITRFS-DRDCEA--
>Zorotypus_caudelli
---------------------------------------------------------------------------------------------------------MAAVWMARKYY-ESIIRFT-DREFFP--
>Thermobia_domestica
------------------------------------------------------------------------------------------------------MTKMAAVLRARKYC-GSVIRYS-DRESFS--

HG0005 ENSGALG00000016320

>Gallus_gallus
-------------------------------------------------------------------------------MAAVRRARSYS-RCVVRFSD-RELC---
>Acipenser_sinensis
-------------------------------------------------------------------------------MAAVWRARNFA-RCVVRFSD-RELN---
>Channa_argus
-------------------------------------------------------------------------------MAAFLRARKYV-CCVVRFSD-REL----
>Kaupichthys_hyoproroides
-------------------------------------------------------------------------------MAAVWRARNFS-RCVVRFSD-RELY---
>Podocoryna_carnea
-------------------------------------------------------------------------------MAAVRRARNFA-RCVVCFSD-RELN---
>Xenopus_tropicalis
-------------------------------------------------------------------------------MAAVRRSRSDS-RCVLRFSD-RELC---
>Apteryx_australis_mantelli
-------------------------------------------------------------------------------MAAVRRARSFS-RCVVRFSD-RELC---
>Stegodyphus_mimosarum
-------------------------------------------------------------------------------MAAVLIARRYV-RCVIRFNSDRELID--
>Apostichopus_japonicus
-------------------------------------------------------------------------------MAAVGIANRYP-GSVTRYSD-RELVD--
>Menidia_menidia
-------------------------------------------------------------------------------MAAFLRARKYV-RCVVRFSD-REL----
>Opsanus_beta
-------------------------------------------------------------------------------MAAFLRARKYV-RCVVRFSD-REL----
>Oryzias_melastigma
-------------------------------------------------------------------------------MAAFLRARKYV-RCVVRFSD-REL----
>Prorhinotermes_simplex
-------------------------------------------------------------------------------MAAVIRAKKCF-RSVVRYSD-RESLP--
>Trachinotus_blochii
-------------------------------------------------------------------------------MAAFLRARKYV-RCVVRFSD-REL----
>Panthera_tigris
-------------------------------------------------------------------------------MAAVRRARSYC-RCLVRFSD-RELC---
>Micropterus_floridanus
-------------------------------------------------------------------------------MAAFLRARKYV-CCVVRFSD-REL----
>Pygocentrus_nattereri
-------------------------------------------------------------------------------MAAVWRARKFA-RCVVRFSD-RDL----
>Miniopterus_natalensis
-------------------------------------------------------------------------------MAAVRRARSYS-RCLVRFSD-RELC---
>Ophthalmotilapia_ventralis
-------------------------------------------------------------------------------MAAFLRARKYV-RCVVRFSD-REL----
>Clupea_harengus
-------------------------------------------------------------------------------MAAVWRARKFP-RSVVRFSD-REL----
>Latimeria_chalumnae
-------------------------------------------------------------------------------MAAVWRARNFI-GCVVRFSD-RELC---
>Alligator_mississippiensis
-------------------------------------------------------------------------------MAAVRRARSDS-RCVVRFSD-RELC---
>Sinocyclocheilus_grahami
-------------------------------------------------------------------------------MAAVWRARKFA-RCVVRFSD-RDL----
>Fundulus_heteroclitus
-------------------------------------------------------------------------------MAAFLRARKYV-RCVVRFSD-REL----
>Sarcophilus_harrisii
-------------------------------------------------------------------------------MAAVRRARSHC-RCLVRFSD-RELC---
>Homarus_americanus
-------------------------------------------------------------------------------MAAVVRSRIYF-SCVVRFSD-RESIA--
>Sclerodactyla_briareus
-------------------------------------------------------------------------------MAAVARFNRYH-QNIVRYSD-RD-----
>Macropus_eugenii
-------------------------------------------------------------------------------MAAVRRARSHC-RCLVRFSD-RELC---
>Haploembia_palaui
-------------------------------------------------------------------------------MAAVLQAKGFF-ASVIKYSD-REFMP--
>Ephemera_danica
-------------------------------------------------------------------------------MAAVQRAAKLAGQLVVRFSD-RELN---
>Esox_lucius
-------------------------------------------------------------------------------MAAFLRYLQFP-RCVVRFSD-RDL----
>Meganyctiphanes_norvegica
-------------------------------------------------------------------------------MAAVARSYMYF-ASVVRFSD-RECIA--
>Falco_cherrug
-------------------------------------------------------------------------------MAAVRRARSYS-RCVVRFSD-RELC---
>Leptasterias_sp._AR-2014
-------------------------------------------------------------------------------MAAFLRFLEFS-NCIVRFSD-RDCKA--
>Gillichthys_mirabilis
-------------------------------------------------------------------------------MAAFLRARKYV-SCVVRFSD-REL----
>Chanos_chanos
-------------------------------------------------------------------------------MAAVWRSRKFA-RCVVRFSD-RDL----
>Grylloblatta_bifratrilecta
-------------------------------------------------------------------------------MAADLRCKKYS-TPVIRYSD-RELLP--
>Apis_cerana
-------------------------------------------------------------------------------MAAVLLARLYF-RCVVRFSD-REYKG--
>Erinaceus_europaeus
-------------------------------------------------------------------------------MAAVRRARSYC-RCLVRFSD-RELC---
>Ixodes_persulcatus
-------------------------------------------------------------------------------MAAAVKARRYR-NCVLRFNSDRECIA--
>Ochotona_princeps
-------------------------------------------------------------------------------MAAVRRARSYC-RCLVRFSD-RELC---
>Lutjanus_erythropterus
-------------------------------------------------------------------------------MAAFLRARKYV-RCVVRFSD-REL----
>Calopteryx_splendens
-------------------------------------------------------------------------------MAAVLRAKELS-TCVVRYSD-RESSA--
>Ophiocoma_echinata
-------------------------------------------------------------------------------MAAVNKSLLYF-SCFVRFSD-RDWLA--
>Locusta_migratoria
-------------------------------------------------------------------------------MAAVFMARKYF-ESVIRFSD-RESVP--
>Osmerus_mordax
-------------------------------------------------------------------------------MAAFQRYRKFV-RSVVRFSD-RDLVSVI
>Scleropages_formosus
-------------------------------------------------------------------------------MAAVWRARNFA-RCVVRFSD-RELN---
>Corvus_cornix_cornix
-------------------------------------------------------------------------------MAAVRRARRDS-RCVVRFSD-RELC---
>Lateolabrax_japonicus
-------------------------------------------------------------------------------MAAFLRARKYV-RCVVRFSD-REL----
>Sebastes_caurinus
-------------------------------------------------------------------------------MAAFLRARKYV-RCVVRFSD-REL----
>Ceratotherium_simum_simum
MRAAAAAAAAALRMLGASAQVPVAERAAPQPRPGSPAGPTLRCSHSPRPRLAAGAPSLRESPAA--APARPSLAPSPAKMAAVRRARSYC-RCLVRFSD-RELC---
>Ramulus_artemis
-------------------------------------------------------------------------------MAAILKARKYI-GSVIRYSD-RESLP--
>Menopon_gallinae
-------------------------------------------------------------------------------MAALLSAKRHF-GLVIRFSD-REFQG--
>Perla_marginata
-------------------------------------------------------------------------------------MAKKYF-KSVIRFTD-RECLP--
>Cynoglossus_semilaevis
-------------------------------------------------------------------------------MAAFLRARKYV-RCVVRFSD-REL----
>Papio_anubis
-------------MLGASAQVCAAEGAAPQRRPGSPVGLTLRCSHNSLLRLATGVPPLPESPAAAATPARPDLAPCPAKMAAVRRARSYC-RCLVRFSD-RELC---
>Acerentomon_sp._AD-2013
----------------------------------------------------------------------------------------MF-ECVIRFTD-REIYG--
>Melopsittacus_undulatus
---------------------------------------------------------------MCLCCSRPACPSLAGKMAAVRRARSYS-RCVVRFSD-RELC---
>Ectopsocus_briggsi
-------------------------------------------------------------------------------MAAVLNAKFYF-GLVIRYSD-RELIR--
>Leucoraja_erinacea
-------------------------------------------------------------------------------MAAVWRARNFL-DYVVRFSD-RELQ---
>Microtus_ochrogaster
-------------------------------------------------------------------------------MAAVRRARSYC-RCVVRFSD-RELC---
>Salvelinus_alpinus
-------------------------------------------------------------------------------MAAFLRSLKFP-RCVVRFSD-RDL----
>Tupaia_chinensis
-------------------------------------------------------------------------------MAAVRRARSYC-RCLVRFSD-RELC---
>Tityus_serrulatus
-------------------------------------------------------------------------------MAAVIKGRRYF-GCVIRFNSDREYVA--
>Lepisosteus_oculatus
-------------------------------------------------------------------------------MAAMWRARNFA-RCVVRFSD-RELY---
>Trichomycterus_areolatus
--------------------------MYMRETPGTS------------GRLGMSVTLLY--KADLQRPCRLDSTQTLSVMAAVWRARKFS-RCVVRFSD-RDL----
>Sparus_aurata
-------------------------------------------------------------------------------MAAFLRARKYV-RCVVRFSD-REL----
>Echinaster_spinulosus
-------------------------------------------------------------------------------MAAFLRFLEFA-NCIVRFSD-RDWKA--
>Streblospio_benedicti
-------------------------------------------------------------------------------MAAFLRYRMFS-SCVICFSW-RELTV--
>Gekko_japonicus
--------------------------------------MT-------------QTNTHTGSRAHDLALSHLLALSRVGKMAAVRRARSDS-RCVVRFSD-RELC---
>Chelonia_mydas
-------------------------------------------------------------------------------MAAVRRARSYS-RCVVRFSD-RELC---
>Takifugu_rubripes
-------------------------------------------------------------------------------MAAFLRARKFV-RSVVRFSD-REL----
>Thermobia_domestica
----------------------------------------------------------------------------MTKMAAVLRARKYC-GSVIRYSD-RESFS--

HG0006 FBgn0029971 CG18624

>Drosophila_melanogaster
----------------------------------MGSGIVVQLARKYGAVIFFPTVAVGSIYADWSHTRQWKRQQLQLAHRAQLKD--QR---------
>Echiuropus_macronychus
-------------------------------------MGVLEIVRRFRVVVIFPAFVTTCIWADYKHTAEWKKRNFLERDQLRDRES-LRN--------
>Machilis_hrabei
---------------------------------MALWSSFRQFWSRYCVVIIFPTACVTSIYADYSHTQKWKAQQLRIQEDKLKAE-NWS---------
>Blaberus_atropos
-------------------------------------MAVSQLFRRYWIVVLFPTVTAGSIFADLIHTKQWKEQKAQLQTAAELEE--TL---------
>Hynobius_chinensis
---------------------------------MGQLSNIGVFWRRYRVSILFPAFSIGTIYADYAHTQKYKKSLADFQRKQQLQ--------------
>Trypoxylus_dichotomus
-----------------------------------MSFKIGKLVNKYWVVTLLPTFVASTIYADYTHTQEWKKSLNKLN--------------------
>Litopenaeus_vannamei
--------------------------------MAPRLTALG-FLRKYKVVIFFPLVTFSAITADILHTRKWKAEKAAAALAESA---------------
>Apachyus_charteceus
------------------------MVLLSIPVISVYMSSIINFCRKYSLSILFPTLSLSIIIADFNHSRKWKKSLKNE---------------------
>Daphnia_pulex
-------------------------------------MSVLAFWRRYRVSIIFPVLSFSFIYADWSYTRRWKQENLKKLAEAVDPE--VIKA-------
>Campodea_augens
MADS----------RIVTSKLPLDAPPVDPTSFSARCKRANFLWRYYRASVLFPTVTVGLILLDYNHTQKWKREQKELSEKQAEK--------------
>Haploembia_palaui
----------------------MDK-------PPSVLRRTGYLIRKYFIVVALPTALGYAVIADYLHTQKWKREQAALQNKPSM---------------
>Isonychia_bicolor
-------------------------------------MAVGYYIRRYGISVLFPTFVALSIFADYNHTRRWKQQHKLLTSQESNLS-------------
>Onuphis_iridescens
---------------------------------MSGLAYYKYLIRRNFVSVIFPVGLGLVIYADWSRTQKYKASQK-----------------------
>Galloisiana_yuasai
-------------------------------MASGKLSTINKLWKTYRVVIFFPTFATAAIVADLVHTRNWKVQKASLEAKENN---------------
>Amynthas_koreanus
----------------------------------MQLSFAYAVFKRNFVSVIFPVCTVWAIYADWSRTRRYKAAKAALPSTHFQS--------------
>Tigriopus_californicus
-----------------------------------MFKQGLALFRKYAVSVIVPTVVFGSIAWDFNRTQRWKRNQIEFDESQKIANQ------------
>Adelphocoris_lineolatus
-----------------------------------MALPIRSFVRKYGPSIVFPVAAFSAIYADYSHTQNWKKEQLLKAKTSELL--------------
>Lineus_viridis
-----------------------MEKE-----VVSAPVKVLRFAKRNFVSVIFPVVAAWAIYSDYSRTQRFKAKKARELLVTGAVQKELIKDSF-----
>Tenthredo_koehleri
-------------------------------MAAPGKVSIGKLISRYAAVVFFPTFSATTIFLDWNHTRKWKIAESERVSALNSELQ------------
>Armadillidium_vulgare
--------------------------------MEGKTHPIIKFWKRAGRSIIFSTVSISLITADYLHTRKWKKELETKYENFYV---------------
>Dermacentor_variabilis
----------------------MAATGTA----LEKVRPLLKVCRRYAATVLFPTISGWLIYADYSRTQRWKKSKASAE--------------------
>Triodia_sylvina
-------------------------------------MAIGKFLRRYGVVIFFPTFAATSIYLDWSHTKKWKQQNKLFVNSKEIV--------------
>Mantis_religiosa
------------------------------MEV-SRLAQFRRFWKTYWVVITLPVVTAGSIFADLNHTRKWKKEKRELAKADILKR--TLSEEQ-----
>Tanzaniophasma_sp._AD-2013
-------------------------------MAVGILSSARNFWRRYCVVILFPTFTITTIAADLIHTRNWKAKTVAAEVLDKTVSEDTVYS-------
>Boreus_hyemalis
-------------------------------MALGKASAIRHFWKKYSVVIFFPTFTLATIYADYSHTQQWKAQQGSTK--------------------
>Sialis_lutaria
--------------------------------MAVRLSTIGSFWRKYRVVIIFPVITISSIAADYQHTQKWKASQTNQ---------------------
>Mytilus_galloprovincialis
----------------------MAGS--------RTISKILRIVRRHSLTVIFPVLCFGAIYLDYSRTQEYKRQRKLDH--------------------
>Xibalbanus_tulumensis
-------------------------------MDVTLVGRVFRFLRFYKFVIIFPVACGGAIFADYNHTRKWKEEKARLEALAYQAEN-PENIVQ-----
>Euroleon_nostras
------------------------------MSVGRKLKTVYGYWRYYRIVILFPTFTIGTISADLLHTKHWKEQQLALENKSKSEF--D----------
>Triops_newberryi
----------------------------------MALSALRAFWRRYNVVIVFPTLTAASIYADYSHTQQYKAAKALAASRFSELLSE-----------
>Epiophlebia_superstes
---------------------------------MAKGNAAVKFLKKYGISIFFPSATVAAIYADLSHTKKWKALQVQVAKVSQE---------------
>Phalangium_opilio
----------------------------------MNFKIALRTLKRYKVVILFPTITIWSIYADYTRTLRYKATKLQILQRQTVL--------------
>Hypochthonius_rufulus
----------------------MANESPSLNHNKLWYERLVKEFRRYRRTILFPTVLLGLVYLDFSHTQKWKKQKQLSEENQSQIVLN-----------
>Stenobothrus_lineatus
-----------------------------------MLKATGKFLRRYGVVIFFPTVVTAAIITDLSYTRQWKRLKAEKLRQQELLTD------------
>Phyllium_philippinicum
-------------------------------MEVSKLVAAKNLVKKYFWVVFFPTFAAVTISADWQHTRKWKLKKAALAEEQSKQA--VLAEDQSSNYQ
>Glycera_tridactyla
---------------------------------MSPAAGPLRLLKVNLVSITFPTVVAWAIYADWSRTRRFKAQKALERVKEHY---------------
>Cosmioperla_kuna
------------------------------MAAVKEVSAIGKFLRRYKVVIFFPTFTAATIFADLRHTRNWKAQETAQESTVKYTAEKLL---------
>Acerentomon_sp._AD-2013
MVHQTTTPHGSSSASKYGKQPTLVATQAEQDANLSFFRRALKTAWKYSLVVTFPSIAFAAIYADYSRTQKWKAEQARLKETE-----------------
>Ectopsocus_briggsi
------------------------------------MANVRYLLRRYSVVVIFPTLTAWSIYADLSHTRKWKAQEEARKAKEE----------------
>Inocellia_crassicornis
-------------------------------MAVQKVGFIRAFWRKYGVVITFPTLSFSAILADYTHTYKWKKQQAALLQELKDPE--LLKS-------
>Trichomycterus_areolatus
------------------------------MAVAKTPSALGLFWRKYRVSIIFPVLSVSAIFADYSRTSSWKKISKFER--------------------
>Oropsylla_silantiewi
---------------------------------MAYLASFRQLWKRYSIVVLFPSFALGSIYADWSNTQKWKQNTKKQ---------------------
>Gluvia_dorsalis
-------------------------------MASGSARGILYQLWRYKYVILFPTGTALSIYADYSHTKQWEALNKNKE--------------------
>Stylops_melittae
---------------------------------MIKIRFRLRSITRHISTALFPLVAAGLIWADYSATQNWKARKGLREAK------------------
>Alvinella_pompejana
----------------------MEGGT-----ALNRFQTLVAKLRRNFITIIFPVLTTWAIYADWSHTQKYKAAKAAALLQASEESQSIG---------
>Gynaikothrips_ficorum
------------------------------------MVKVANIVKRYFLTGFFPVLFVTPIYLDWSRTQRYKAEKLKESGGSS----------------
>Rhyacophila_fasciata
-------------------------------------MGLGKLVKRYSVVVLFPTVTVTSIVADLRHTYLWKKQKAELLKQ------------------
>Zorotypus_caudelli
---------------------------------MAVLKTARELFSRYFISVLFPTIAISSIVADYKHTQKWKAKQQAIIQSS-----------------
>Atelura_formicaria
----------------------------------MAAGGVGLFWRRYRVVIIFPTIVAACISADYAHTREWKAQQAAELKSRESVH-------------

HG0007 FBgn0034372 Gint3

>Drosophila_melanogaster
------------------------------------------MSFNFYVESFNS---T-------L--------QSIAR---QVEQFAGVALTAAVD------LRGNLE-------------SLTG---------DFRWEN-GPIAFTDLQATLLKVLIVLLLGTCVLAGYSWSIYGKVITEKFVRPST--LK-EI--EELKLSVAKLKLPKEHSPRI-
>Stegodyphus_lineatus
-----------------------------------------------------------------------MKSL-------------------DII--SNLITPSGVTE-------------YLYSVFPEDIPDA-AISLVTSFNTVDEFQLKILRVSLCILLINLFLIWCAWRKYGEKITERFMK-PA--SSKSL--EDLKREATELKLPKEHTPRI-
>Machilis_hrabei
MS---------S-----------------------VIDFE-------------------------SETSSLFE---ALHQLFQELTGFICETF-GLSLEVSQEE---------FKSYFWS---------VYHSII-DWLSFDDFQWKLLQAFLIILFTNIVLIYIAWRVYGERIADRFMYPES--VR-AI--EELKASVQRLKLPREHTPRI-
>Argulus_siamensis
---------------------------------------------------------------------------------------MFLD-----FD--------------------NLYLLLYDLY---IQ-LVEKLYE-ELYTKNDFRLRTIQCFSFIICVNIILIIIAWKIYGPRIAEKFMKPVP--TV--V--EQLSKDIASLKLPPEYSPRV-
>Cryptocercus_wrighti
------------------------------------------------------------------------------------------------------------------------------------------------------------------------------MSWKIYGKRISEKFMRPAT--CK-TI--EELKRSVSSLKLPKEHSPRV-
>Meloe_violaceus
MN---------A-----------------------FLNDFSTNSSLFD-----------VIYTYLTTINIYF-----------------IDTL-SLLRDHPKH---------------KDLLSVF------DNLP-ELLDFNEFQLTLLKTFLIVLFANLILIFIAWTIYGKRICDRFMKPAT--NK-AI--EELKASVSKLKLPKEHTPRI-
>Eucyclops_serrulatus
MNIWESLTNFTSSISTLILEDNIQPGRETDLLTDLFNFDSSIFSIESLYS-----------VANSFQNTLVGIFNSTCEISSVFGQVAGSIL----NEATTGNIS-----------SA-------VSLTFLESEKLF-DWFNFTEFQVKLLREFSIILLGNIVLVIFAWKIYGPRISSKFSVSPSRGSRQVI--EDLRLSMSELQLPNEHDYKLK
>Apachyus_charteceus
-------------------------------------------------------------------------------------------------------------------------------------------------------------------------------------------MKPAT--SK-TI--EDLKKSVSLLKLPKEHSPRV-
>Daphnia_magna
----------------M--------------------VDEKSFGDQQSTLWVEL-------FVEYFVNLIYVLVT---SVIHLTEVL----V-----------E-----YG----EHFLQSIFLLLISLSETIVAI-FDLAFSDFQIKLLQVFFCILAIHILAICVAWKIYSERITQRFLKPSL--YEISS--EQLKTAVSELKLPAEHTPRW-
>Occasjapyx_japonicus
-------------------------------------------------------------------------------------------------------------------------------------------------------------------------------------------MKPAS--LR-AV--EELRSSVAELRLPQEHSPRI-
>Aposthonia_japonica
MI---------P-----------------------LFGNYF-NSTVLD-----------ATTYYVFKLFALA-----------------NEVV-GISSSKTFDDS-------ESS--FTIQI---------FDLG-TWFAFDDFQITLLKVFLSFLCVNLICIYISWKIYGRRISEKFMKPAT--PK-AI--EELKKSVFLLKLPKEHSPRV-
>Isonychia_bicolor
MS---------S-----------------------LSGDVSANST-----------------VIDSIFQYL----------GKFCLLLQQAL-GLGPVVPLPEPLQDL----PHLPTTDILTDFIEW----------FAFDDFQFKLLKGFLVVLTFNIILIFIAWRFYGRNISEKFMRPAT--SK-AI--QDLKNSVSQLKLPSEHTPRI-
>Galloisiana_yuasai
MT---------S-----------------------INEDDSNSLSFIN-----------VITNYIYEISTVI-----------------YTAF-GFTSNIKVENN-------IPLHFIIECY---------YNII-NWLAFDEFQIKLVKGFTFILFANILFIYMSWKVYGKRISEKFMRPAT--SK-AI--EELKKSVSLLKLPKEHSPRI-
>Graphocephala_atropunctata
MT---------S-----------------------NVDEQLDSSSLWN-----------LLYFYCNQIYFAI-----------------FSNL-NITSNFTTSTLTSFIGNEEHSLNILSYL---------RYIA-SLLPYDEFQIELLKLFCAIILTNICLVAIAWKVYGKYICQRFMKPAT--PK-VI--EELKKSVSQLKLPKQHSPRI-
>Tenthredo_koehleri
MS---------S---SV------------------CNDVAPGSGSLLD-----------IINKFACDLIGLV-----------------FSTI-GITSNINSSNLDDDTSSEHHRHTIIQFY---------EEFS-AWLAFDEFQLQLFKVFAFLLVSNVALIYVAWHVYGARISNRFMKPAT--SA-AV--EELKKSVSRLKLPKEHSPRV-
>Rhipicephalus_annulatus
-----------------------------------------------------------------------MFWF---------------SASQK--EEEAESPLW-----WTSSLLTGPLVEAY---------EQLDTWCLGCDHFERSVAQALLVVFAINVVLILCVWRRCGDQIADQFLNLAT--SSSQL--EEFQKEASSLELPEEHSPRI-
>Enteucha_acetosae
MS---------A-----------------------FFETYIINSTLFE-----------VLSLTVNDSFGYF----------SEL---VIRCF-ESV----------------KVLEISNKLEKICENTFSKEFL-SSLEFDDFQFLLLKTLLIIFFATFLLICFVWKLYGKRISERFMKPAT--SA-AI--QELKDSVSKLKLPKEHTPRI-
>Empusa_pennata
-------------------------------------------------------------------------------------------------------------------------------------------------------------------------------------------MKPAT--SK-AI--EDLKKSVSLLKLPKEHSPRV-
>Panorpa_vulgaris
------------------------------------------------------------------------------------------------------------------------------------------------------------------------MNILFIYFAWNFYKDDIWEYFMRPTS--VD-ED--AELQKRIELLNLPREHTPRI-
>Sialis_lutaria
MT---------S-----------------------YITDNLFNTSYYT---D-------VILKYSEEVNNIYL---KVIQLFSEHETAIHP------L---IL-------------FYQS---------VVDFAF-SLVHFNEFQISILKYLLLVLIINLFLICIYWKLYGKAIGERFMKPAT--SR-AI--EELKASVDRLKLPKVHTPRI-
>Euroleon_nostras
-------------------------------------------------------------------------------------------------------------------------------------------------------------------------------------------MRPVT--SR-AI--EELKASVARLKLPKVHSPRI-
>Triops_newberryi
MK---------------------------------FWLPNFDGVSLWSWWESAVDSVSDLSENVKSSVFPWIP-----VSLFSLFWF-LVES-----VTS-----------------AWGSL-----TYNLSVVG-ESFSFDDFQTKFLQALFVVLFCNIGLTLLAWRVYGKKITERFMQTSS--DV-LK--EQLKTAVSELKLPNEHTPRL-
>Calopteryx_splendens
MT---------S-----------------------FNDALFSNSTLLD-----------VAHYFIHEWSNIL-----------------LELL-GIGSEHEIEKAD--------SNTLLHYC---------EDLI-QWFAFDEFQTTLLKGFAIILSANIVLICISWKVYGKRISERFMRPPS--TK-AI--EELRKSVSQLKLPKEHSPRV-
>Nothrus_palustris
--------------------------------------------------------------------------------------------------------------------------------------------------------------MIDTLLFWLLIILVSNLLLIVIFWNIYGKRIDVLITRKEE--KQQIL--DQLRREAFELKLPKEHSPRI-
>Ceuthophilus_sp._AD-2013
MT---------S-----------------------VAEDAI-NSSFVD-----------VIRVYVFELFTAV-----------------TKIF-GLASNIPLQDT-------TSTRLFFECY---------NSVI-SWLAFDDFQVKLLKGFVFILCANVVLIFISWKVYGKKISERFMKPPT--SK-AI--EELKKSVSLLKLPKEHSPRV-
>Crassostrea_gigas
-----------------------------------------MSSESLSS-----------A---TTQEAL------------------VIVDYISG--V----------------GKVIWDFL-------VDAFLL-LRHEQYDVQSKLLVVLIFWLFFNLFLIGIAWQYYGQNICEKLNRSKEK----------------------------
>Ramulus_artemis
MT---------S-----------------------VNGDIWFNFSFLE-----------PVLHCICELYKSL-----------------NEYL-ADSLNFNYSAP---------AYSRVETL---------VVLG-SWVTFDDFQINLLKGFILFFFINILLIFISWKYYGKKISEKFMGPVT--PK-AI--EELKKSVSLLKLPREHSPRV-
>Menopon_gallinae
MD---------------------------------QDSSTLGQTTFLD-----------NLYACFSKIFQEA-----------------YELL-SSVIQSEI----------LSYFEGVSFN---------LNVL-QYFAFDELQIKLLKGFLLIIVFNSVLIWITWRIYGKRISNRFMKTAS--SK-TI--EDLKKSASMLKLPKEHNPRV-
>Platynereis_dumerilii
MS---------------------------------------------------ERAP----I--D------------------SKIAEFNSTEAEGNTW-------------VDFG---YD---L-CKEFGKFWDDKLEFRVKLLVVLIICLAVNLNLIHCAWRKYGNKITERFMKSGS--SRASS--EN--KEVPEQRLHRE------
>Leuctra_sp._AD-2013
-------------------------------------------------------------------------------------------------------------------------------------------------------------------------------------------MRPPT--SK-SI--EELKKSVSLLKLPKEHSPRV-
>Acerentomon_sp._AD-2013
-------------------------------------------------------------------------------------------------------------------------------------------------------------------------------------------MRPAS--LK-VI--EELKRSIAELKLPREHSPKI-
>Liposcelis_bostrychophila
-------------------------------------------------------------------------------------------------------------------------------------------------------------------------------------------MRTAS--HK-AI--EDLKKSISMLKLPKEHSPRV-
>Xanthostigma_xanthostigma
MT---------S-----------------------FSYLDFVNASYND------------INYYAYEVYDYIL---QILNLFVFWKDCST------------------------------EH---------YLSLYQ-SFIVFNEFQVNILKSLLIIFLCINLLIFIYWKIYGHQICERFMRPAT--SR-AI--EELKASVAKLKLPKVHSPRI-
>Tityus_serrulatus
---------------------------------------------------------------------------------------MV--SHLF-----------------------Y--FT-DI--I-YEYLVIPFINIEEFEWKLLRSLLVFLLINLVFITVAWNKYGNKISERFMK-PV--ANKIL--EDLKKEATELKLPKEHTPRI-
>Trichomycterus_areolatus
-------------------------------------------------------------------------------------------------------------------------------------------------------------------------------------------MRPAT--SK-AI--QELKESVSRLKLPKEHTPRI-
>Archaeopsylla_erinacei
---------------------------------------------------------MSLSNYNFTYFNTVYDQL------EYLYTVLWENLIKTLYSTES----------------EVSFFDPI--------QETFE-YLINFDEYQLTLLKVLFVLICLNLLFIYVAWNYYKDVIWDCSIRPTT--LD-ED--VELIERNAKLNLPRVHSPRL-
>Streblospio_benedicti
---------------------------------------------------------MTEQEL----QNEK---------------VIHADTLQTL-------DLE-------------SLKET--------FCL-IVTDFWGKSDFQIKLFCILTVWLLINLVLIKIAWSTYGERLSEMFSGLTSRQTSRQTSEERVVEGVRAENIQRE------
>Stylops_melittae
------------------------------------------------------------------------------------------MV-NLQTNLTWVF------N-EFKQLFVSEI---------LYLE-SLTSLSNLQIRIVQVFTLLTILNLTLILIVWNIYGQRISNRFIKVAT--SK-EI--EELKLSVSKLKLPKEHSPRF-
>Frankliniella_cephalica
MS---------T-----------------------FTEESLFNTSAL------------LMNSLVSSISNTV-----------------LYAI-GLTSNVTEGC----------SHLSALPL---------FDWQ-ALFSYDDFQVVLLKVFLFILAFNLVSICIAWKIYGKNIYETFMKPVT--PK-II--EDLKKSVSRLNLPKEHSPRV-
>Annulipalpia_sp._AD-2013
-----------------------------------------------------------------------MSWW-------------RAFSEE-----G--LPLP----------LSLPSQESLSEFVQYFVPEIPL-GSVQFDDFHALLLKTFLVILFTTLFLIFVSWKLYGKKISERFMKPAT--SK-AI--QELKESVSRLKLPKEHTPRI-
>Limulus_polyphemus
MS---------------------------------------------------QAVDAA-NQSEL------------------EKTENSTNADPQNITW-------------LTGSLLGSAREGL-ELWISDNFSDIDEFQWNILKTLLTFLCINLTLICLAWKKYGGKITERFMK-PD--SSKAL--DDLRKEATELKLPKEHCPRI-
>Zorotypus_caudelli
-----------------------------------------------------------------------------------------------------------------------------------------------------------------------------MFICIAWKIYGEQISERFIKPAN--AK-FI--EDLKKSVSQLKLPKEHSPRV-
>Thermobia_domestica
-------------------------------------------------------------------------------------------------------------------------------------------------------------------------------------------MK-PA--STKSF--EDLAKGASELKLPKEHSPRV-

HG0008 ENSG00000175567 UCP2

>Homo_sapiens
--------------------------------------MTIRCFVSHPFSMENQGDR-AMIATGSFEERDTFREA-----
>Acipenser_sinensis
--------------------------------------MLVQRFVFGRLSMEQRGNL--IVTESNFSERDTFREEC----
>Podocoryna_carnea
--------------------------------------MLVQRFVFGRLSMEPRGNL--IITESNFSERDTFREEC----
>Megophrys_sangzhiensis
--------------------------------------MFIQHLVLHQFSMESKGYR-IFTASGKCDGRDAFREA-----
>Oryzias_latipes
--------------------------------------MILGLSTSEISSMEKQGQT--VKCAANFSQRDTFRES-----
>Acinonyx_jubatus
--------------------------------------MTIRCFVSQPFSMENQGDR-AMIATGSFEEWDAFREA-----
>Hynobius_retardatus
--------------------------------------MITRTAGSKPSPMEDQGYR-----ERKFDMRDTFRET-----
>Micropterus_floridanus
--------------------------------------MILRLITSEKSSMEKQGNQ--FIAGTKFSQRDTFRES-----
>Orcinus_orca
MCLEQMALSLLGCGPLRSISLSKLSSF---ALTCRTHSMTIRCFISQPFSMENQRDQ-AMIATGSFEEWDAFREA-----
>Astyanax_mexicanus
--------------------------------------MFFRCSFSERFSMEDRGNE--IIVESHFNQRDAFRES-----
>Miniopterus_natalensis
--------------------------------------MTFRCFVSRPFSMENQQDR-AMIATGSFEEWDAFREA-----
>Clupea_harengus
--------------------------------------MLLYCLVLERFSMEEQGNV--IIVESNFNQRDTFRES-----
>Latimeria_chalumnae
--------------------------------------MIIRYSDSAEFSMDQ-EKT--SIVKNNFNERDIFRES-----
>Alligator_sinensis
--------------------------------------MVIRCPACKRAPMERNGHR-MVVASSKCDGRDAFRES-----
>Sinocyclocheilus_grahami
--------------------------------------MFLICTGSNRFSMEERGNK--IQVEGHFNQRDTFRES-----
>Kryptolebias_marmoratus
---------MIKPDYLKKYYLLFFFLFLLKCLFSRSWSMILRLKVSERSSMEKQGNQ--FIGGTRFNQRDTFRES-----
>Sminthopsis_crassicaudata
--------------------------------------MIIRCSVSQQFSMENQGKC-TMVARSNFEERDTFREA-----
>Galeopterus_variegatus
--------------------------------------MTFGCFVSQPFSMENQGDR-AMIATGSFEERDAFREA-----
>Monodelphis_domestica
--------------------------------------MIIRCSVSQQFSMENQAKW-NMVAVSNFEERDTFREA-----
>Trichosurus_vulpecula
--------------------------------------MIIRCSVSQQLSMENQGKW-AMVAVSNFEQRDTFREA-----
>Esox_lucius
--------------------------------------MFLGHIVFERCSMEEQGNQ--IIVESNFDQRDAFRES-----
>Acanthogobius_hasta
--------------------------------------------------MENKRTQ--IIAGTKFSQRDAFREA-----
>Chanos_chanos
---------------------------------------MFMCPVFKHFSMEEQGNE--IIAESTFNQRDTFRES-----
>Condylura_cristata
--------------------------------------MTVRGFVSQPFFMENQRDQ-AMLTADSFEEWDAFREA-----
>Oryctolagus_cuniculus
--------------------------------------MNFRCFVSQPFSMENPGDR-GMIATGGFEERDAFREA-----
>Elephantulus_edwardii
--------------------------------------MTIRYSVSQPFSMENQKDQMIMIATGSFKEWDVFREA-----
>Scleropages_formosus
--------------------------------------MFLKCYTLKRVSMEDQVNE--IIAESQFSERDTFREA-----
>Lateolabrax_japonicus
--------------------------------------MILRLITSKRSSMEKQGYQ--FIAGTKFSQRDTFRES-----
>Sebastes_caurinus
--------------------------------------MILKLITSESCSMERQGNQ--CIAGTKFSQRDAFRES-----
>Equus_asinus
--------------------------------------MTIRCFVSQPFSMENQGDR-AMIATGSFEERDAFREA-----
>Manis_javanica
--------------------------------------MTIRCFVSQPFSMENQRDL-AMIATGSFEEWDAFKEA-----
>Loxodonta_africana
--------------------------------------MTIRCSVSQPFSMENQRDQ-IMIATGSFEEWDVFREA-----
>Leucoraja_ocellata
--------------------------------------MII-WRFISELSMEQRQSQ--TVVKRGYNERDTFRESSPARL
>Psammomys_obesus
--------------------------------------MIFKCFVSKPFSMENQGDR-AMIATGSFEERDTFREA-----
>Oncorhynchus_tshawytscha
--------------------------------------MFLRYIVFERFSMEEKGNQ--IIVESNFDQRDAFRES-----
>Tupaia_chinensis
--------------------------------------MTIRCSVSQPFSMENQGDW-AMIATGSFKEWDVFREA-----
>Lepisosteus_oculatus
--------------------------------------MLFTSLVLGIFSMEQQGTQ--IIVESNFNERDTFRES-----
>Ictalurus_punctatus
--------------------------------------MFLKCPFSELFSMEEQGNE--IIVDSHFNQRDAFREA-----
>Trichechus_manatus_latirostr
--------------------------------------MTIRCSVSQPFSMENQRDQ-IMIATGSFEEWDVFREA-----
>Squalus_acanthias
--------------------------------------MII-RRFISELSME-RQSQ--IIVKKSYNERDTFRES-----
>Gekko_japonicus
--------------------------------------MIIRCRLSAAFSMASKGSR-AAVALSNFDERDTFRES-----
>Chrysemys_picta
--------------------------------------MILKCLVSKRVSMENQGDQ-IVVALSKFDERDTFRES-----
>Orycteropus_afer_afer
--------------------------------------MTIRCFVSQPFSMENQRDHQIMIATGSFEEWDVFREA-----

HG0009 FBgn0050100 CG30100

>Drosophila_melanogaster
-----MKMVQPSRQQILRLYKHLIRYGNHLKLTDKNYFLGRVRHEFRDNRSLTNPVEVEFSFKRGETLLKKGRIL
>Talitrus_saltator
-----------------------------MELSDRRFFEKRIRREFTKQRNS--PEELDTAVQRAEAFLEKGRLL
>Latrodectus_geometricus
--------MTVSRQKVLSLYKELLKYSKNLKYTDKSYYLNRLKNEFLENRNLQNASEINFQIQRGKNFLEKKRLL
>Meinertellus_cundinamarcensi
-------MQIPNKQQVLCLYKSLLRYGETLRLTDKQYFRTRVRSEFQKNKSSDDSKEIVFQYEKGLKVLNSKRIL
>Blattella_germanica
-------MSNPTAREVLKLYRDILKYGRQLQFTDKNYFGQRIRNEFKAGKDLKNQEDINFHFQRGEALLKNKRVV
>Calanus_finmarchicus
------------MSQATLLYKSLLRYGRTLQMTDKDYFYRRVRFEFEKNSVLLEKQEVQFQLDKARQFLKNKRLV
>Photinus_pyralis
--------MHPNAKDILKLYKNLLRYGQELQYTDKNYFARRIKNEFRRNRDITDEADIQFNYKRGISLLEKRAVV
>Apachyus_charteceus
-----MASTLPNTRQILSLYKNLMRYGQKLKFTNQSYFYERVRREFENNKTIPED-KREFYYKKGLSLLEKKRVV
>Daphnia_pulex
MNTIFIIMNVPNKTQIFELFRSCIRYSGQLKLSDKRLVSKRIKSEFHKNKTLSNPEEICKAYEKGQSFLSKRRLL
>Occasjapyx_japonicus
---MAAYMGVPSPVQVLRLYRSLVRYGKELKYTDRDFYQRRIRNEFERHRLLTKDEDIRFQYEKGLAFISRKRII
>Haploembia_palaui
-------MSSPSASEVLRLYRELLRFGRQLQFTDQKYFLRRIKTQFQENKHLEDKEQIEFQFNKGQSLLKRKRVV
>Baetis_sp._AD-2013
--------MSVARSEVLNLYRSILKYGKDIKLTDKDYFRKRIRQEFRSNQKVTNAEEIQFHVERAKAFLANKRVV
>Glomeris_pustulata
--------MTIQRQSVLKLYRELLRYGQTLQYTDKKFYFRRIRKEFEQNRSLTAPADIQYYYEKGLTFLTNKRLI
>Galloisiana_yuasai
-------MTNIGGRQIIKLYRDLLRYGKRLEFTDINYFNRRIKTEFLKNKNLASLDEIQFQFERGISLLKNKRVV
>Tigriopus_californicus
---------MSQRSEALRLYKSLLRYGQSLRLTDSAYLHRRCTAEFRTPRST--PAQIERALAQARALLSQRRLL
>Cercopis_vulnerata
-------MNKPTVKQIVQLYKDLLRYGNNLKLTDKEYYIDRIHLEFRKNQSLDNPKDIQFCFDRGLALLKNKRVV
>Argochrysis_armilla
-----------MRSAILRLYKEILRYGETLKYTDKTYFRERIRTTFRNNKDLTNEKDIDFQFKKGQVFLEKRRVV
>Rhipicephalus_appendiculatus
------MSKPPSKQAVLSLYRSLVRYGHRLELSDKNYYFKRLREEFEARKTLTEPKELQFYFNKGKAFLERKRLI
>Papilio_machaon
--------MSITRSEILNLYKKLMQYSQSLNLTDVQYFRTRVSNEFRRNKTLTKPEDISFAYKKGEALLRKGSVL
>Mantis_religiosa
-------MSIPSPRQVLQLYKSILRYGQELKFTDKEYFYYRINQEFKNNKDIKRTEDIIFNFQRGQALLKNKRVV
>Panorpa_vulgaris
-------MSVPNSSHILRLYKDLLRYGQSLQLTDKLYFRSKIVNEFKKNKTIENHKDISFNYEKGKSLLKKGRVV
>Dermanyssus_gallinae
---MGPVSGAPSVQAMRSLYKQLARYGQQLVLTDRRWFERQLRNEFAKPRVT--AEEQNFAYRKGQRFLELKRLV
>Osmylus_fulvicephalus
-------MLIPNNGQVRNLYRHILRYGQTLRYTDKIYFYNRVRNEFKQNKSLTDPADIVFNFKKGEELLRKKTVC
>Triops_newberryi
-------MSGPSPYQVLKLYRSLLRYSSTLQYTDKSFYIKRIRSEFKKKKDLTNEEEIRKAFEKGLSFLSKKRIV
>Calopteryx_splendens
-------MCQPSPSQVLSLYRSLLRYGNQLKYTDKKYFIKRIRKEFESNRNLIP-KERGFQYERGLVLLHNKRVI
>Hypochthonius_rufulus
--------MSVSREHILRLYKNILRYSNSLVYTDKSYYLQRVKREFRENKQLTDDSDIRFYYEKGITFLNKKRLI
>Prosarthria_teretrirostris
-------MNAPTAHEVLRLYRKLLQYGKQLKLTDKKYFFDRIRKEFKTNKTLTNQEDVTFQFKRGLALLENKRVV
>Ramulus_artemis
-------MNTPARVQVLRLYRDLLRYGQNLQLTDKDYFCKRVREEFRNNRVLENKNDLLFHFNKGESLLKTKRVV
>Menopon_gallinae
-------MCVPTTREVLTLYKRLLRYGRQLKYTDQDYYSKRIKQEFQSNKDVKGN-EAIFHYEKGLQLLKLKRVV
>Perla_marginata
-------MSGPTAKQVIQLYRDLLRYSKHLKFTDKNFFTKRVRKEFKTNKRLSNEEDIQFHYQKGLILLKNKSVI
>Polydesmus_angustus
-----MSASVVGKSQILHLYRQLLRYGRTLTYTDKNYYYGRIRAEFEKHREAEL-KTQLFQYKKGKALLENGRLI
>Acerentomon_sp._AD-2013
--MCWQQMGIPTNGQILKLYRSLLRYGRTLQYTDKDFFKRKVQQEFHRNGAITAPNEIKFQYEKGISFLSRKRLV
>Liposcelis_bostrychophila
-------MSAPSRTQVLRLYKDILRYGEELTYTNKTYFKNRVKNEFLSAKSAKDNKEIDNLYQKGVALLMRRRVV
>Xanthostigma_xanthostigma
-------MEVPSSTQVLRLYRQLIRYGQKLQYTDKNYYYKRVIAEFKRNRNLEDPKEIKFQFEKGQSLLKRKSVI
>Trichomycterus_areolatus
---------MPSRSEILKLYKQLLRYGQQLRLTDKSYYQERINQEFKQHKNLASNKDITFNFEKGKKLLEKGLVM
>Ctenocephalides_felis
-------MNTPSAAQVLRLYKNLLRYGQNLQYTDKDYFCNRIRKEFRNNKTCSRTEDLIFNFEVCI---------
>Stylops_melittae
-----------MRLKVLNLYKSLMRYGKSLRLTDNEYFLERIRSEFRRNKDLYSNQEIDYHFKKGETLLKKKLLV
>Symphylella_vulgaris
--------MNPTRVQILSLYKTLLRYADNLKYSDTNYLKQRVRVEFSQHRSLTAKKDIEFYFKRGKSFLERKRLV
>Frankliniella_cephalica
------MSTLINSNQVLQLYRSLIRYGQSLKLTDQKYYLKRVRKEFRANKDLQDPEKIEFMYKKGVSLLDRKRLI
>Philopotamus_ludificatus
---------MPSKREILKLYKQLLRYGQQLQLTDKTYFQQRIVNEFKKNKDLSKSDDITFNFKKGQQLLNKGLVL
>Zorotypus_caudelli
--------MNPGKPGILILYKSLIRYGKNLKYTDKKYFIERIRQEFKNRKELENEEDIKFAFQRGVSVLNNKRII
>Atelura_formicaria
-------MVAPSSKQVLQLYRNLMRYGRQLKYTGKDYYSKRVREEFKRNKSLENDEKKIFHFEKGQALLSQKRIV

HG0010.1 ENSG00000125863 MKKS

>Homo_sapiens
-----------------------------------------------MSLRNLWRDYKVLVVMVPLVGLIHLGWYRIKSSPVFQIPK---N-DDIPEQDSLGLSNLQ-KSQIQG-----K--------
>Amia_calva
-----------------------------------------------MGLSSVWKNYKVLIVMGTSLGLIHWGWYSIKANPIFHKSE---T-NYFPEPGIVSFVSEP-QHQPKG-----K--------
>Micruropus_glaber
------------------------------------------MVFRWGRVIESLKTNVVLVVMVPAIVVIHLGWRKLQDNPNLVDPNMKRE-----IPPVEGFSYLKNKIVEELFTTTDK--------
>Anas_platyrhynchos
-----------------------------------------------MGLKTFWKSYKVLIVMGTGLGLVHWGWFHIKSSPIFQVKT---E-DFVPEPAIVAYVMQK-DNKNKE-----K--------
>Xenopus_laevis
-----------------------------------------------MGLGNLWKNYKVLIVMGTGLGLVHWGWYNIKSNPAFQPAK---EDSLEPEPAISAEKNTM----KKG-----K--------
>Apteryx_australis_mantelli
-----------------------------------------------MGLKTFWKNYKVLIVMGPSLGLVHWGWFHIKSSPIFQVKT---EENFVPEPGIVAYMMRN-EHKNKE-----KQQ------
>Mastotermes_darwiniensis
---------------------------------------------MSGKHGGFFNRNAVTVVMIPALVLMHWGWYKLQDVEVLVKKEEKKE-----LPVVVGARYLVHILEEKLQSFRGGSQSSGEK-
>Acinonyx_jubatus
-----------------------------------------------MSFQNFWRDYKVLVVMVPLVGLIHLGWHRIKSSPVFQMPN---K-DDISEPDSPVLGRPQ-KSHIQG-----K--------
>Hynobius_retardatus
-----------------------------------------------MGLKNLWKNYKVIIVMGTGLSLMHWGWYHVKTNPLFHPKR---D-EFVPEPSIVTYVLQG-NQQTKA-----K--------
>Physeter_catodon
-----------------------------------------------MSFQKFWRDYKVLVVMVPLVGLIHLGWHRIKSSPVFQIPN---K-DDLPEPDSLALTSPE-KSHIPG-----K--------
>Calidris_pugnax
-----------------------------------------------MGLKTIWKDYKVLIVMGTSLGLVHWGWFYVKSSPIFQVKT---E-DIVPEPGIVAYVMQG-DHKSKE-----K--------
>Miniopterus_natalensis
-----------------------------------------------MSFQSFWRDYKVLVVMVPLIGLIHLGWHRIKSSPVFQVPS---K-DNVPEPDSLALASLQ-KDHIQK-----K--------
>Echinarachnius_parma
-------------------------------------------MQPNPPKKNFFMRNRVLFVGLISLTFIHVGWFHIANDPVFNRDAVSGQASR--KLESVPAYIKS-KTGSASAE---KSS------
>Latimeria_chalumnae
-----------------------------------------------MGLKSIWSNYKVLIVVGTGLGLIHWGWYSIKSNPVFHKNR---E-DYAPEPGIATSIMQQ-DKQPKG-----KQQ------
>Columba_livia
-----------------------------------------------MGLKTIWKDYKVLIVMGTGLGLVHWGWFYMKSSPIFQVKT---E-KFVPEPGIVAYVMQS-DQKNKE-----K--------
>Alligator_sinensis
-----------------------------------------------MGLKTIWKNYKVLIVMGPSLVLIHWGWYNIQSNPIFKMKR---E-DLDQEPGIVTHVMQQ-DSKNQG-----K--------
>Callinectes_sapidus
------------------------------------------MGTGMGRAREFLRRNIVLVVMFPILVGVHFGWQKIQDVEMFVPKQQRID-----LPVIEGARYLREGVEKKLGLKKEE--------
>Sclerodactyla_briareus
-----------------------------------------------MTKPSFIRRYRVLIVAFAGLSLIHLGWYRLSKNPDFTGIAVGAE------PEIKPSYLKQ-KS-GKG-----TSA------
>Monodelphis_domestica
-----------------------------------------------MSLKTFWRDYKVLLVMIPSIGLIHIGWKSIKSNPIFQTAN--ANEDSIPEPGIVAHMIPQ-KNQNQV-----K--------
>Strongylocentrotus_purpuratu
---------------------------------------------MNPPKKNFFVRNKVLFVGLISLSLIHVGWFHIANDPVFNKGEK----KT--VSDLLPSYIKS-KALKAPAE---KSN------
>Meganyctiphanes_norvegica
--------------------------------------------MMWGRVSEFAKRNLVMIIMVPTLFGLHLGWLKIQDTEAFVPKHQKME-----LPAVTAAARLQEEIKRKVKGDNNDQ-------
>Aquila_chrysaetos_canadensis
-----------------------------------------------MGLKTIWKDYKVLIVMGTSLGLVHWGWFYIKSSPIFQVKT---E-DFVPEPGIVAYVMQS-DHKNKE-----K--------
>Marthasterias_glacialis
--------------------------------------------------MSFFARYRVLTVAAVGLTMVHFGWYYIASDPVFNPDK------K--VLGITPEYFKS-KQAQQKKA---AVEGVTKSQ
>Numida_meleagris
----------------------------------------------MMSLKTLWKDYKVLIVMGTSLGLVHWSWFHIKSSPLFQVKT---E-EIVPEPGIVAYVMQR-DRKNKE-----K--------
>Meccus_pallidipennis
---------------------------------------------MNNNKDKFFKKHLVMIVFIPLIGLIHYSWHLLQQNEILVKNRKHNE-----LPIITIFKRLTQGRESD---------------
>Oryctolagus_cuniculus
-----------------------------------------------MNLQKFWRDYKVLVVMVPLVGLIHLGWHRIKSSPIFQVPN---K-DNIPEPDSLALASLP-KGPPQG-----K--------
>Ornithorhynchus_anatinus
-----------------------------------------------MSLKTIWRDYKVLIVMVPSIGLIHLGWHSIKNSPIFQIPN---K-DTIPEPAIVTHVMQQ-EASKKG-----K--------
>Octopus_bimaculoides
---------------------------------------------MLKQLKAFAKNYTVLLVMIPSVVALHVGWYYMQFNPDLVEPEKQEI-------RILGKIFKK-T--PKE--------------
>Ceuthophilus_sp._AD-2013
-------------------------------------------MLDKGKMQGFINRNLVSIVMVPGLILLHWGWTRIQDIEVFVKKEEKKN-----LPVFLGVHYIQE----KFLKWKEGDGVHPDE-
>Scleropages_formosus
-----------------------------------------------MGLGNVWRQYRVLIVMSASLGLIHWGWFQLKGHPALHGKQQREDDEYIPEPAIVSYVASG-RQSG-G-----K--------
>Lepidothrix_coronata
-----------------------------------------------MGLKTIWKDYKVLIVMGTGLGLVHWGWFYIKSSPIFQVKT---E-DYVPEPGIVTYVMQS-DQKNKE-----K--------
>Ceratotherium_simum_simum
-----------------------------------------------MSFQNFWRDYKVLVVMVPLVGLIHLGWHRIKSSPVFQIPN---K-DKVPEPDNLALASSQ-KSHIQE-----K--------
>Manis_javanica
-----------------------------------------------MSFQTFWRDYKVLVVMVPLVGLIHLGWHRIKSSPVFQIPN---K-DDGSEPDSLVLASPQ-KKHIQG-----K--------
>Glycera_dibranchiata
----------------------------------------------MMSVAAFLRKNIVLVVCIPSLALIHYGWYNLQFKDNFIPQDQRHI-----RIA--GIDIDP---EKR-RTTTSQE-------
>Leuctra_sp._AD-2013
------------------------------------------MTGASSKFAGFINRNLVAIVMVPVLIGIHWGWNRIQNVEVFVTKEEKKD-----LPIILGARYLEQKIEEKVATWKDGDKAVTKE-
>Chinchilla_lanigera
-----------------------------------------------MSFQNFWRDYKVLLVMVPLIGLIHVGWHRIKSSPVFRVPT---K-DDIPEPHSLGLASPP-KSQIQG-----K--------
>Tupaia_chinensis
MRRVARGAKVPGNRGGSGRRFGLWGLLSVAAQGGAAAGGQVHQCCITMSFQNFWRDYKVLVVMVPLVGLIHLGWHRIKSSPVFQIPS---K-DDISEPDSLALASSQ-KSQIQG-----K--------
>Aptenodytes_forsteri
-----------------------------------------------MGLKTIWKDYKVLIVMGTSLGLVHWGWFYIKSSPIFQVKT---E-DFVPEPGIVAYVMRS-DHKNKE-----K--------
>Henricia_sp._AR-2014
--------------------------------------------------MSFFSRYRVLTIGVVGLAAIHAGWFYIAHNPTFNQEK------E--VVNITPEYYKS-KRSKLGAT---TASSDAKNI
>Takydromus_sexlineatus
-----------------------------------------------MGIKTIWKNYKVLIVMVPSLGLIHWGWYNIKSNPMFQPKR---E-DTVPVPGIVTYALQQ-DNKSKG-----K--------
>Pelodiscus_sinensis
-------------------------MILQYSHGCVC----RTGVDASMSLKTIWKNYKVLIVMGTSLGLIHWGWYKIKSNSIFQTKR---E-DYVPEPGIVTYVMQH-ESQNKG-----K--------
>Zorotypus_caudelli
-------------------------------------------MGQKGRLWEFVNRNIVTLVMVPGLVLIHWGWLRLQENDIFVPPQDRKE-----FVIFRSFRDMKTNLEERYRSIVDGNSKD----

HG0010.1 ENSGALG00000009013 MKKS

>Gallus_gallus
----------------------------------------------MMSLKTLWKDYKVLIVMGISLGLVHWSWFHIKSSPLFQVK---TE-EIVPEPGIVTYVMQSDHKNKEK----------
>Amia_calva
-----------------------------------------------MGLSSVWKNYKVLIVMGTSLGLIHWGWYSIKANPIFHKS---ET-NYFPEPGIVSFVSEPQHQPKGK----------
>Anas_platyrhynchos
-----------------------------------------------MGLKTFWKSYKVLIVMGTGLGLVHWGWFHIKSSPIFQVK---TE-DFVPEPAIVAYVMQKDNKNKEK----------
>Xenopus_tropicalis
-----------------------------------------------MGLGNLWKNYKVLIVMGTGLGLVHWGWYNIKSNPIFQPE---REDTLEPEPAITGYAVEKNTAKKAK----------
>Apteryx_australis_mantelli
---------------------------------------MAGRCQFTMGLKTFWKNYKVLIVMGPSLGLVHWGWFHIKSSPIFQVK---TEENFVPEPGIVAYMMRNEHKNKEKQQ--------
>Canis_lupus_familiaris
-----------------------------------------------MSFQNFWKDYKVLIVMVPLVGLIHMGWHRIKSSPVFQMPN----KDNVSEPENPELGCPPKNHIQGK----------
>Hynobius_retardatus
-----------------------------------------------MGLKNLWKNYKVIIVMGTGLSLMHWGWYHVKTNPLFHPK---RD-EFVPEPSIVTYVLQGNQQTKAK----------
>Orcinus_orca
-----------------------------------------------MSFQKFWRDYKVLIVMVPLVGLIHLGWHRIKSSPVFQIPN----KDDLPEPGRVAVTSPEKNHLPGK----------
>Calidris_pugnax
-----------------------------------------------MGLKTIWKDYKVLIVMGTSLGLVHWGWFYVKSSPIFQVK---TE-DIVPEPGIVAYVMQGDHKSKEK----------
>Miniopterus_natalensis
-----------------------------------------------MSFQSFWRDYKVLVVMVPLIGLIHLGWHRIKSSPVFQVPS----KDNVPEPDSLALASLQKDHIQKK----------
>Echinarachnius_parma
-------------------------------------------MQPNPPKKNFFMRNRVLFVGLISLTFIHVGWFHIANDPVFNRDAVSGQ--ASRKLESVPAYIKSKTGSASAEKSS------
>Latimeria_chalumnae
-----------------------------------------------MGLKSIWSNYKVLIVVGTGLGLIHWGWYSIKSNPVFHKN---RE-DYAPEPGIATSIMQQDKQPKGKQQ--------
>Columba_livia
-----------------------------------------------MGLKTIWKDYKVLIVMGTGLGLVHWGWFYMKSSPIFQVK---TE-KFVPEPGIVAYVMQSDQKNKEK----------
>Alligator_mississippiensis
-----------------------------------------------MGLKTIWKNYKVLIVMGPSLVLIHWGWYNIQSNPIFKMK---RE-DLDQEPGIVTHVMQQDSKNQGK----------
>Sclerodactyla_briareus
-----------------------------------------------MTKPSFIRRYRVLIVAFAGLSLIHLGWYRLSKNPDFTGIAV------GAEPEIKPSYLKQKSGKGTSA---------
>Monodelphis_domestica
-----------------------------------------------MSLKTFWRDYKVLLVMIPSIGLIHIGWKSIKSNPIFQTANA--NEDSIPEPGIVAHMIPQKNQNQVK----------
>Strongylocentrotus_purpuratu
---------------------------------------------MNPPKKNFFVRNKVLFVGLISLSLIHVGWFHIANDPVFNKGEK------KTVSDLLPSYIKSKALKAPAEKSN------
>Aquila_chrysaetos_canadensis
-----------------------------------------------MGLKTIWKDYKVLIVMGTSLGLVHWGWFYIKSSPIFQVK---TE-DFVPEPGIVAYVMQSDHKNKEK----------
>Marthasterias_glacialis
--------------------------------------------------MSFFARYRVLTVAAVGLTMVHFGWYYIASDPVFNPDK--------KVLGITPEYFKSKQAQQKKAAVEGVTKSQ
>Oryctolagus_cuniculus
-----------------------------------------------MNLQKFWRDYKVLVVMVPLVGLIHLGWHRIKSSPIFQVPN----KDNIPEPDSLALASLPKGPPQGK----------
>Ornithorhynchus_anatinus
-----------------------------------------------MSLKTIWRDYKVLIVMVPSIGLIHLGWHSIKNSPIFQIPN----KDTIPEPAIVTHVMQQEASKKGK----------
>Octopus_bimaculoides
---------------------------------------------MLKQLKAFAKNYTVLLVMIPSVVALHVGWYYMQFNPDLVEPEK-------QEIRILGKIF-KKTPKE------------
>Laupala_kohalensis
--------------------------------------------MISGKIRKFVNRNLVMIVMIPSLIFVHWGWLQIQQKDIFVKEEE---RKGLPIIGMAKIVMKKIMP-AEKEDSTPTQPE-
>Scleropages_formosus
-----------------------------------------------MGLGNVWRQYRVLIVMSASLGLIHWGWFQLKGHPALHGKQQREDDEYIPEPAIVSYVASGRQSGGK-----------
>Manacus_vitellinus
-----------------------------------------------MGLKTIWKDYKVLIVMGTGLGLVHWGWFYIKSSPLFQVK---TE-DYVPEPGIVTYVMQSDQKNKEK----------
>Ceratotherium_simum_simum
-----------------------------------------------MSFQNFWRDYKVLVVMVPLVGLIHLGWHRIKSSPVFQIPN----KDKVPEPDNLALASSQKSHIQEK----------
>Manis_javanica
-----------------------------------------------MSFQTFWRDYKVLVVMVPLVGLIHLGWHRIKSSPVFQIPN----KDDGSEPDSLVLASPQKKHIQGK----------
>Glycera_dibranchiata
----------------------------------------------MMSVAAFLRKNIVLVVCIPSLALIHYGWYNLQFKDNFIPQDQ-------RHIRIAGIDIDPEKRRTTTSQE-------
>Nomascus_leucogenys
-----------------------------------------------MSLRNLWKDYKVLVVMVPLVGLIHLGWYRIKSSPVFQIPR----NDDIPEQDSLGLSNLQKSQIQGK----------
>Fukomys_darlingi
-----------------------------------------------MSLQKFWRDYKVLVVMVPLIGLIHLGWHRIKSSPIFQIPT----KDNIPEPDSPALASPPKRQIQGK----------
>Tupaia_chinensis
MRRVARGAKVPGNRGGSGRRFGLWGLLSVAAQGGAAAGGQVHQCCITMSFQNFWRDYKVLVVMVPLVGLIHLGWHRIKSSPVFQIPS----KDDISEPDSLALASSQKSQIQGK----------
>Sepia_pharaonis
---------------------------------------------MASRLTAFLKEYKVLLIMLPAVAGIHYVWYKMQKSPVFMEGEK-------TKI--FGKDLVSDSPK-------------
>Aptenodytes_forsteri
-----------------------------------------------MGLKTIWKDYKVLIVMGTSLGLVHWGWFYIKSSPIFQVK---TE-DFVPEPGIVAYVMRSDHKNKEK----------
>Henricia_sp._AR-2014
--------------------------------------------------MSFFSRYRVLTIGVVGLAAIHAGWFYIAHNPTFNQEK--------EVVNITPEYYKSKRSKLGATTASSDAKNI
>Takydromus_sexlineatus
-----------------------------------------------MGIKTIWKNYKVLIVMVPSLGLIHWGWYNIKSNPMFQPK---RE-DTVPVPGIVTYALQQDNKSKGK----------
>Pelodiscus_sinensis
-----------------------------------------------MSLKTIWKNYKVLIVMGTSLGLIHWGWYKIKSNSIFQTK---RE-DYVPEPGIVTYVMQHESQNKGK----------

HG0010.2 ENSG00000125863 MKKS

```
>Homo_sapiens
--------------------------------------------------------MKNTSWIRKNWLLVAGISFIGVHLGTYFLQRSAKQSVKFQSQSKQKSIEE----
>Amia_calva
--------------------------------------------------------MRNRSWLRKNWLLIAGLSFLGIHLGTYVMQKTMKNSVRSDLQ--LKRKDASE--
>Anser_cygnoides
--------------------------------------------------------MRKASWSRKNLLLVAGLSFIGVHFGTMLVNFVAKKSVRSHSETKKDDRSE----
>Xenopus_tropicalis
-------------------------------------------------------MIKKQTWLRKNWLWVAGISFLGIHFGTYFIQKLAKQSAKSNLQLKESKNSE----
>Apteryx_australis_mantelli
---------------------------------------------------------MKTSWSRKNFLLVAGLSLIGVHVGSMLVNLVAKKSVQSHSEIKKDRRE-----
>Ursus_maritimus
---------------------------------------------------MLKEEMKNTSWIRKNWLLVAGVSFIGVHLGTYFIQRAAKQSVKSQSQSRGKQQNIEE--
>Ambystoma_mexicanum
--------------------------------------------------------MKSKSWLRKNWLLVAGISFVGLHIGTYIIQKVAKTSARSGSELKQQKDYDE---
>Physeter_catodon
--------------------------------------------------------MKNSNWIRKNWLLVAGISFTGIHLGTYFIQRAAKQSVKSQPEGKQKSLNNELK-
>Calidris_pugnax
--------------------------------------------------------MRKTSWSRKNFLLLAGLSVIAVHFGSMLVNFVAKKSARSHSEAKKEDRRE----
>Miniopterus_natalensis
--------------------------------------------------------MKNTSWIRKNWLLVAGVSFIGVHLGTYFIQRAAKQSVKSQSGDKQK--SIRE--
>Dasypus_novemcinctus
---------------------------------------------------------MKISWIKKNWLLVAGISFLGVHLGTYIIQRTAKQSVKSQSGVKPKHIEE----
>Latimeria_chalumnae
--------------------------------------------------------MKNHSWMRKNWLLVAGLSFLGIHLGTYIMQKSVKNSTKSTENTVLKAKDI----
>Columba_livia
--------------------------------------------------------MRKTTWNRKNLLLVAGLSVIGVHFGSMLVNFVAKKSARSHSEAKKEDHRE----
>Alligator_mississippiensis
--------------------------------------------------------MKKTSWLRKNWLLVAGLSFLGVHIGTYLIQRAAKNSVKSTVEIKHKNLDE----
>Sarcophilus_harrisii
--------------------------------------------------------MKQTSWIRRNWLLVAGLTFIGVHFSTYFLQIAAKHSAKSQIEYRQKNIKEIK--
>Galeopterus_variegatus
--------------------------------------------------------MKNTSWIRKNWLLVAGVSFIGVHLGTYFIQRTAKQSVKSQSGNK----------
>Monodelphis_domestica
--------------------------------------------------------MKQTSWIRKNWLLVAGLTFIGVHLGTYFIQVAAKHSAKSQSGSKEKNIKEMK--
>Aquila_chrysaetos_canadensis
--------------------------------------------------------MKKASWSRKNFLLVAGLSLIGVHFGSMLVNFVAKKSVRSHSEAKKGDRHE----
>Asterias_rubens
--------------------------------------------------------MKLPPWVARNKLLSVGIALLSIHLTIYIVQKTALATKSTKRLR-GMDMEQTERK
>Meleagris_gallopavo
--------------------------------------------------------MRKASWSKKNFLLVAGLSLIGVHLGTMLVNFVAKKSARSHSETKRDDHRE----
>Condylura_cristata
--------------------------------------------------------MKNTSWIRKNWLLVAGVSFIGVHLGTYFLQWSAKQSVKSQSIGKQKNIEE----
>Oryctolagus_cuniculus
--------------------------------------------------------MRNTSWIRKNWLLVAGVSFVGVHLGTYFIQRAAKQSVKSQSGSKQKSTEE----
>Ornithorhynchus_anatinus
--------------------------------------------------------MRKSSWIRRNWLLVAGLSFIGVHLGTYIIQITAKNSAKSVLETKKKSIQE----
>Scleropages_formosus
------------------------------------------------------MTMKNQSWLRKNWLWVAAASFLGIHVGTWLMQKAMKSSAHSQTRTKQKS--VDE--
>Ficedula_albicollis
--------------------------------------------------------MKKTTWNRKNFLLVAGLSLIGVHLGSMVVNYVAKKSAQSQSEAKRSS-------
>Ceratotherium_simum_simum
--------------------------------------------------------MKNISWIRKNWLLVAGVSFIGVHLGTYFIQRTAKQSVKYQSGGNNVLLNNEVK-
>Manis_javanica
--------------------------------------------------------MKNTSWIRKNWLLIAGVSFIGVHLGTYFMQRAAKQSVKSQSRNKQENIEE----
>Melopsittacus_undulatus
--------------------------------------------------------MRKANWNRKNFLLVVGLSVIGVHFGSMLVNFVAKKSAHSHSEAKKDRHE-----
>Marmota_marmota_marmota
--------------------------------------------------------MKSTSWIRKNWLLVAGVSFIGVHLGTYFIQRAAKQSIKSQSVGKQKSIEE----
>Tupaia_chinensis
MRRVARGAKVPGNRGGSGRRFGLWGLLSVAAQGGGAAAGGGQKEATLNLSLILKEDMRNTSWIRKNWLLVAGVSFIGVHLGTYIIQRTAKHSVKSQSGDKQKNIEE----
>Lepisosteus_oculatus
--------------------------------------------------------MSNRSWLQKNWLLVAGLSFLGIHLGTYLLQKTVKSSTRSDLA--LKQKDVK---
>Pygoscelis_adeliae
--------------------------------------------------------MRKASWSRKNFLLVAGLSLIGFHFGSMLVNFVAKKSVRSHSEAKKEDHHE----
>Echinaster_spinulosus
--------------------------------------------------------MKLPVWVANNKLLSIGIALATLHLTIYTIQKMAHSTSKSKRSLTGPPQHSRE--
>Takydromus_sexlineatus
--------------------------------------------------------MKKISWIRKNWLLVAGLSFVGVHLGTYFIQRAAKSSAKSGLNSKSKGLDE----
>Struthio_camelus_australis
---------------------------------------------------MLQKEEMKTSWSRKNFLLVAGLSLIGVHIGSTLVNLVAKKSVHSHSEIKKDRRE-----
>Pelodiscus_sinensis
--------------------------------------------------------MRKTSWIRKNWLLVAGFSFLGVHLGTYFIQRAAKNSAKSALEVKQKNLEE----
>Patiria_miniata
--------------------------------------------------------MKLPVWVAKHKLLSVGIALLTIHLTIYTVQKTALSTSKTRRSLTGPSPPADKED
```

HG0010.2 ENSGALG00000009013 MKKS

```
>Gallus_gallus
-----MRKASWSKKNFLLVAGLSLIGVHFGTMLVNFVAKKSARSHSETK--RDNHRE---
>Amia_calva
-----MRNRSWLRKNWLLIAGLSFLGIHLGTYVMQKTMKNSVRSDLQ--LKRKDASE---
>Anser_cygnoides
-----MRKASWSRKNLLLVAGLSFIGVHFGTMLVNFVAKKSVRSHSETK--KDDRSE---
>Xenopus_tropicalis
----MIKKQTWLRKNWLWVAGISFLGIHFGTYFIQKLAKQSAKSNLQLK--ESKNSE---
>Apteryx_australis_mantelli
------MKTSWSRKNFLLVAGLSLIGVHVGSMLVNLVAKKSVQSHSEIK--KDRRE----
>Mustela_putorius_furo
-----MKNISWIRKNWLLVAGVSFIGVHLGTYFIQRAAKQSVKSQSTGK--QESIKE---
>Ambystoma_mexicanum
-----MKSKSWLRKNWLLVAGISFVGLHIGTYIIQKVAKTSARSGSELK--QQKDYDE--
>Balaenoptera_acutorostrata_s
-----MKNSNWIRKNWLLVAGISFTGIHLGTYLIQRAAKQSVKSQSEGK--QKSLNNELK
>Calidris_pugnax
-----MRKTSWSRKNFLLLAGLSVIAVHFGSMLVNFVAKKSARSHSEAK--KEDRRE---
>Eptesicus_fuscus
-----MKTTSWIRKNWLLVAGVSFIGVHLGTYVIQWAAKQSVKSQSGNK--QKSIKE---
>Dasypus_novemcinctus
------MKISWIKKNWLLVAGISFLGVHLGTYIIQRTAKQSVKSQSGVK--PKHIEE---
>Latimeria_chalumnae
-----MKNHSWMRKNWLLVAGLSFLGIHLGTYIMQKSVKNSTKSTENTVLKAKDI-----
>Columba_livia
-----MRKTTWNRKNLLLVAGLSVIGVHFGSMLVNFVAKKSARSHSEAK--KEDHRE---
>Alligator_mississippiensis
-----MKKTSWLRKNWLLVAGLSFLGVHIGTYLIQRAAKNSVKSTVEIK--HKNLDE---
>Sarcophilus_harrisii
-----MKQTSWIRRNWLLVAGLTFIGVHFSTYFLQIAAKHSAKSQIEYR--QKNIKEIK-
>Galeopterus_variegatus
-----MKNTSWIRKNWLLVAGVSFIGVHLGTYFIQRTAKQSVKSQSGNK-----------
>Monodelphis_domestica
-----MKQTSWIRKNWLLVAGLTFIGVHLGTYFIQVAAKHSAKSQSGSK--EKNIKEMK-
>Aquila_chrysaetos_canadensis
-----MKKASWSRKNFLLVAGLSLIGVHFGSMLVNFVAKKSVRSHSEAK--KGDRHE---
>Condylura_cristata
-----MKNTSWIRKNWLLVAGVSFIGVHLGTYFLQWSAKQSVKSQSIGK--QKNIEE---
>Oryctolagus_cuniculus
-----MRNTSWIRKNWLLVAGVSFVGVHLGTYFIQRAAKQSVKSQSGSK--QKSTEE---
>Ornithorhynchus_anatinus
-----MRKSSWIRRNWLLVAGLSFIGVHLGTYIIQITAKNSAKSVLETK--KKSIQE---
>Scleropages_formosus
---MTMKNQSWLRKNWLWVAAASFLGIHVGTWLMQKAMKSSAHSQTRTKQKS--VDE---
>Corvus_brachyrhynchos
-----MKKTTWSRKNFLLVAGLSLIGVHLGSMLVNFVAKKSAQSHSEAK--RDRHE----
>Equus_asinus
-----MKNISWIRKNWLLVAGVSFLGVHLGTYFIQRAAKQSVKSQSGGN--KVLLNDEVK
>Manis_javanica
-----MKNTSWIRKNWLLIAGVSFIGVHLGTYFMQRAAKQSVKSQSRNK--QENIEE---
>Microcebus_murinus
-----MKNTSWVRKNWLLVAGLSLIGVHLGTYFIQRAAKQSVKSQSGAK--QKSIEE---
>Melopsittacus_undulatus
-----MRKANWNRKNFLLVVGLSVIGVHFGSMLVNFVAKKSAHSHSEAK--KDRHE----
>Dipodomys_ordii
-----MKNPSWIRKNWLLVAGVSFIGVHLGTYFMQRIAKQSVRSQSADK--QRNVEE---
>Tupaia_chinensis
-----MRNTSWIRKNWLLVAGVSFIGVHLGTYIIQRTAKHSVKSQSGDK--QKNIEE---
>Lepisosteus_oculatus
-----MSNRSWLQKNWLLVAGLSFLGIHLGTYLLQKTVKSSTRSDLA--LKQKDVK----
>Pygoscelis_adeliae
-----MRKASWSRKNFLLVAGLSLIGFHFGSMLVNFVAKKSVRSHSEAK--KEDHHE---
>Anolis_carolinensis
-----MKKISWIRKNWLLVAGLTFVGVHLGTYLVQRAAKSSAKSGLDIK--SKGLDE---
>Struthio_camelus_australis
MLQKEEMKTSWSRKNFLLVAGLSLIGVHIGSTLVNLVAKKSVHSHSEIK--KDRRE----
>Trachemys_scripta
-----MRKTSWIKKNWLLVAGLSFLGVHLGTYFIQRAAKNSAKSALEIK--QKDLGE---
```

HG0010.3 ENSG00000125863 MKKS

>Homo_sapiens
------------MLISFINIELFDFIATMLHIHTLIPKE-----------------
>Canis_lupus_familiaris
------MSLSHYMLISFINLELFSFIATLLHVQDFHS-------------------
>Miniopterus_natalensis
------MSLNHYMLISFINMKLFNFITTMLRIQTVIPEEYFFKMLNRKLWLTK---
>Dasypus_novemcinctus
-----------------MNMKLFNFIPTMLHVQTFILRVIFLNVRTHDNVIYAKCV
>Galeopterus_variegatus
------MSLNHFVLINFINIELFNFIDTMLHA------------------------
>Condylura_cristata
------MSLNLYILITFINMELFNFIATYLDFCS----------------------
>Oryctolagus_cuniculus
MWNHQHVIKSLSVSISFIIIEPFHFLASVLHIQTFTPKE-----------------
>Equus_caballus
------MSLNHYMLISFVNIELFNFMATMLHIQTFIPEE-----------------
>Manis_javanica
-------------------MEPFNFIAIVPHIPIFIP-------------------
>Fukomys_damarensis
------MLLSHYVLISFRNTESYNFIATTLNIPEEYFKKYFKTHGHK---------
>Tupaia_chinensis
------MSLNHYILISFTNIELFNFIAMMLHVALSVLQNNFFKC------------

HG0011 FBgn0027360 Tim10

```
>Drosophila_melanogaster
--------------------------------------------MRVRERMRKANQGDYDSSAASDRRFDY---------
>Euphrynichus_bacillifer
-------MVSKLVKINIAWAVIATIGISSFWLAKSSVDKNRYEAMKVRERMAKANVGEYEPSYRKFTS------------
>Stegodyphus_tentoriicola
------MALSKLTKINIIGATLAISGILSFFVVKKLVTIQRVEAMKARERMAKSNVGEYEVKRKFA--------------
>Machilis_hrabei
---MQLMITSKVAKIGISWLLITGAGVGGFVWTKRLVDKERYESMKIRERIRQSNFGEYKPSSRKFTG------------
>Prorhinotermes_simplex
------MPLGNVTKIAVGWAVVTAAGLYSFVLSKQSVDKHRYEAMKVRERIRRSNEGEYDSGPRKFTG------------
>Hynobius_chinensis
------MRLGKLGKIGVGWTILVVVGVSGFTYSKTSVDKKRYDNMRVRERMKKSNEGLYEATERFVGGETNKSQ------
>Dascillus_cervinus
------MLLNKTAKIGIGWGFVVSAGIYGFYLSKENITKKRYEHMKVRERMRQANFGEYEPSPRKFD-------------
>Scylla_paramamosain
------MLTSKWARIAVGWSAVIVIGVGGFVFTKQTIEQQRRESMKVRERMRNANTGDYESKRNFTG-------------
>Forficula_auricularia
-----MLILGTHAKVALGWSLIVATGIYSFVLAKKSIEKNRYENMKVRERMRNSNIGEYISTEKRKF-------------
>Aposthonia_japonica
---MSIMSLGHVQKIAIGWTLVISGSLYAIVWARDSINKSRYEVMKSRERMKRSNIGDYEVQDRKY--------------
>Isonychia_bicolor
------MPLTKVTKVGIGWAIITFAGISGFILSKKSIEKRRYEHMQVRERMRRSNYGEYAQSERKF--------------
>Meganyctiphanes_norvegica
------MFRTKTAKIVVGWAAAISCGVGGFILTKISVEDQRRDAMKVRERMRNANTGEYEAKRTFTG-------------
>Grylloblatta_bifratrilecta
MYNVSIMPLGNHAKIGIGWTIIIVSGLYSFVLSKRSIDKHRYDAMKVRERMRKSNAGDYGETARTFTG------------
>Meccus_pallidipennis
------MPMNVIHKICISWSLIVAAGIYSFVLSKRSIERNRYENMKVRERMRNANKGNYIASERKFTG------------
>Cephus_cinctus
------MRLNNHSKIAIGWLVVISTGLYGFVLSKQRVEVHRYESMKARERMRQAPQGDLSTRVQSST-------------
>Ixodes_persulcatus
------MRVGTVRRITIAWALVTVAGIGSFVYAKKLVDKQRYEAMKVRERMKNCNTGDYEPQRKFTF-------------
>Biston_suppressaria
------MPLGKFGKITLSWLAVTVTSLSLFVVAKDHINGQRLENMRVRERMRKSNQGEYEDVYRKL--------------
>Lithobius_forficatus
--------------------------------------------MKIRQRMRKSNSGEYEQSPRHSD-------------
>Mantis_religiosa
--------------------------------------------MKVRERMRKANYGEYETTRKFSG-------------
>Tanzaniophasma_sp._AD-2013
-----MMQPGSIKKVVIGWTVVTIVGLYGFVLSKKSVDKNRYEAMKVRERMRKANEGSYETERKFTG-------------
>Nannochorista_philpotti
-----MLQLTKATKIGICWSVITVSSLFGFVKSKEHIDKRRYENMKVRERMRKSNCGDYDKSTSRYI-------------
>Corydalinae_sp._KMRSPBM-2012
--------------------------------------------MKVRERMRKSNFGEYKESNRKFNG------------
>Metaseiulus_occidentalis
-----MLWKSPIFKICVGWGIAVVGGLGAFVVARDKIIAQRYEAMKSRERMRNANAGSYEQGNRVFRKDGTTEKIQNFKF
>Xibalbanus_tulumensis
------MALGNVAKVAVGWTIVISAGVGAFVLAKISIDKNRFASMKARERIRKANEGDYVPVRKYTT-------------
>Conwentzia_psociformis
------MKLGKVAKIGIGWTIITTASLYAFYLSKQSIDNQRYEHMKARERMRNANIGDYKSSSRNF--------------
>Triops_newberryi
--------MNRVTKVNIGWAILISVGIGSFVVAKNLVDTQRLEHMRSRERMRNANKGEYEPSERKFVD------------
>Epiophlebia_superstes
------MVLNNAGKIAIGWIVVITVGISSFVLSKQSIEQKRLENMRVRERMRMANSGDYPESYRKFNG------------
>Prosarthria_teretrirostris
------MPLNNVAKIGISWAFIVAASLYTFVLSKRYVDNHRYESMKARERMRKANSGDYEPSPRKFTG------------
>Phyllium_philippinicum
--------------------------------------------MKVRQRIRDANKGVYESSERKFD-------------
>Cosmioperla_kuna
---MSIMPLGKYAKIGIGWSFITAAGLYGFVLSKRSVDKRRYDSMKVRERMRNSNYGKYDTSARKFTG------------
>Ectopsocus_briggsi
----MPIKLGRDAKVAIGWAVVVVGGLYSFVLSKQSIDRKRYDNMKIRERMRNANVGEYEPSPRSFH-------------
>Raphidia_ariadne
------MVLGNSSKILIGWSVITFAGIYAFVLSKRSIEAKRYEHMKIRERMRKSNVGDYQITGRYSKSS-----------
>Tityus_serrulatus
------MPLNTLAKIGIGWVVLSIAGIGGFFVAKNSVNKHRYESMKVRERMRKANQGPYKPSPRFSR-------------
>Trichomycterus_areolatus
------MKLGNVGKVGVGWTIFVILGISGFAASKNSVDKKRYDSMKVRERMRKANEGEYQIPETRRFEA-----------
>Ceratophyllus_gallinae
-----MVAIGKFAKIGLSWAALTFASISAFVLSKQHIDAQRYENMKIRERIRNSNTGEYEKSYRHFD-------------
>Hydroptila_sp._AD-2013
------MKLSNFAKITISWAFLGVAGIGSFILSKNSVVHRRREDMVIRERMRNSNVGEYEASSRKF--------------
>Tricholepidion_gertschi
------MPVSKVAKICIGWAIASTAGVSGFIFCKHIIEKQRHESMKVRERMRDSNKGEYEPSYRKFTE------------
```

HG0012 FBgn0064116 CG33713

```
>Drosophila_melanogaster
MATAAAVAKVGKSVHRIFVGNLPWTVGHQELRGYFREFGRVVSA--NVIFDKRTGCSKGYGFVSFNSLTALEKIENEQKHILEGNYLNIQKS-------
>Boeckaxelia_carpenterii
--------MSMRSFSKIFTGNLPWSVGNRELGQYFSQFGRVLSA--QVHFDKKLGKSKSYGFVTFKDRQSAESALSHQNHILEGNILRVDRTENF----
>Machilis_hrabei
--------MAARNLQKIFVGNLPWTVGQRELRNYFSEFGHVVTA--NVVFDKNTGMSRGYGFVQFGTRDGYNSASNKQQHFLEGNYLNIQPASN-----
>Blaberus_atropos
--------MATKAIRKIFVGNLPWTVSHRELRNYFSEFGHVISA--NVIFDRNTGMSRGYGFVVFSTPGGFENASNKQSHILEGSKITIHPASN-----
>Gyrinus_marinus
------MAVTARSLYRLFVGNLPWTISNTELKLYFSKFGHVYNA--SVVFDKNTGLSRNYGFVTFSNQNGFDSATNVQTHKLEGNILKVQPAASS----
>Macrobrachium_rosenbergii
--------MAARSIHKLFIGNLPWTVGHRELRQHFSQFGYVITA--NVIFDRSKGMSKGFGFVQFGNRNAYEAALNKSPQFLEGNTLNVQPLHP-----
>Forficula_auricularia
------MANLIRKGPKIFVGNLPWTVGHFDLEKYFSKFGPVVHS--KVIFNKNTGLSKGYGFVEFDSKESIPKVLG-QKHILEGKLISTEININSN---
>Campodea_augens
---------MASSRLKLFVGNIPWTICHQELRQYFSEFGHVITS--NVMFDRNNGLSKGFGFITFSSQEAFNKAINKQTHFLEGNYLNIQPASTT----
>Aposthonia_japonica
--------MASRAIRKLFVSNLPWTVGHRELKTYFSEFGHVVSA--YVVFDRNTGLTRGYGFVVFGSPESYTNAVDYQNHFLEGNVLQVQNSAN-----
>Eurylophella_sp._AD-2013
--------MAARSIQKLFVGNLPWTVGNRELRQYFSEFGHVVSA--NVVFDKNTGMSRGYGFIMFGNQNGYSNATNKQHHVLEGNQITVQSSAN-----
>Meganyctiphanes_norvegica
--------MAARNLNKIFVNNIPWTIGHQELRQHFAQFGFVTSA--NVVFDRSTGMSRGYGFVQFANRQGVEAASAKSLQFLEGNMVNIRDANSK----
>Galloisiana_yuasai
--------MASRAIRKLFVGNLPWTISHRELKTYFSEFGHVVSA--NVIFDRNTGMSRGYGFIIFSTQGGYDSATNKQSHLLEGNQLTVQVASN-----
>Xenophysella_greensladeae
-----MASAAARSAQKLFVGNLPWTISSRELRQYFSEFGHVSSA--IVVFDRDTGLSRGYGFITFGTSGAYDSVARQQKHVLEGNDLTIQPAST-----
>Tenthredo_koehleri
-----MASKVGIIGNKLYVGNLPWTIANKQLKEYFSQFGHISRA--IVVFDKNSGMSKGFGFVTFSTGKSINTVQNKSVHMLEGHTLNIGSVDVSQQTE
>Metaphatus_ochraceus
------MASSGRALNRLFIGNLPWTVGHRQLREYFSQFGPVQSA--RVIFDKSTGLSKGYGFIEFATKSAAADATNKQVHTVEGHNILVQPQNN-----
>Mantis_religiosa
--------MAARALHKLFVGNLPWTIGHRELRTYFSEFGNVVSA--NVVFDKNTGMSRGYGFVIFGTKGGYESASKKQSHILEGHTLIVQPASN-----
>Tanzaniophasma_sp._AD-2013
----MAANATGRAIRKLFVSNLPWTVGHRELKTYFSEFGHVVSA--HVVFDRNTGMSRGYGFVIFSTQGGLDSATIKQNHLLEGNTLAVQVAA------
>Boreus_hyemalis
-----MASSVRNALHKLFVGNLPWTVGHRELRTYFSEFGPVASA--NVIYDRSSGVSRGYGFVVFSGRSGFENATNKQTHALEGHTLTIQPASN-----
>Corydalus_cornutus
--MATLPRQSINYVKRLFVGNLPWTVSHKELQHYFSKFGSVNTAGTRVVFDKKTGFSKKFGFVQFNNRNSIDDVI-KQTHTLEGNVLTVHETNQ-----
>Chrysopa_pallens
-MGIRKVATAAKALNRVFVGNLPWTISNRELKQYFSQFGLVSQV--NVIFDKKNGFSKGYGFVQFAKKESIQLLTETPNHNLEGNVLTIQPATF-----
>Triops_newberryi
--------MASRSAGRLFVSNLAWTVGSRELRQYFSQFGAVAQA--NVIFDKNTGMSRNYAFVVFATREGYNKALTHPTHTVEGQTVNIQPTGNTTE--
>Cordulegaster_boltonii
-------MAGRLVARKLFVGNLPWTVGHQELRQYFSEFGHVASA--NVVFNRETGISRGYGFVVFGNKSGIAAAMDRGNHQLEGNQITIQMASN-----
>Stenobothrus_lineatus
--------MAARAAYKLFVGNLPWTVGHRELRMYFSEFGPVASA--VVIFDKNTGLSRGYGFVLLGNKSGFSKVTNKQTHVLEGNQITVQAASH-----
>Ramulus_artemis
--------MASKVMKKLFVGNLPWTVGSRELKNYFSEFGHLVSA--NVVFDRNTGISRGYGFVIFSAQRGLESVLEKQSHILDGSTMSVQAATN-----
>Menopon_gallinae
-----MSLLNAPIRHKFLIKNLPWTVGDKQLTSYFVRFGGVVKS--CVEVDTKTGFSKGFGYVFFSRFL--SGDINSQPHILHNEVLEVIPVNL-----
>Leuctra_sp._AD-2013
-------MAARNAVNKLFVGNLPWTIGHEELFRYFSEFGSISSA--RVIFDNVTGMSKGYGFVMFNSKGALDNALGKQNHKLEGNKINIQRSTD-----
>Acerentomon_sp._AD-2013
---------MSSKITTVFVAYLSWSVSRLELGRRFSEFGKVLAA--DVVFDKETGFSKRYGFVRFADRHSAERALQTRNVFIDGKLAVVKPSGSGGSA-
>Liposcelis_bostrychophila
------MASRVVRSPKLVIFNLPWSVGPKELRQHFAQFGAVTYA--DVKFDRATGFSKGFGFIEFGAPEDTHKALEAPVHKLEGNLLNIRRSENE----
>Xanthostigma_xanthostigma
------MARFPRALNRLFVGNIPWTVGNKELRKYFSKFGLVNSV--NVMFDKQTGFSKNYGFVYYSMKDGFDNALNSKPHELEGSTLIVEPVSNPNGN-
>Trichomycterus_areolatus
-----MASTSVRALNRLFVGNLPWTVSHRELTQYFSSFGIVKSA--KVVFDRSSGLSRGYGFITFASRTGYDTAVNKPQHRLEGSVLTVNEASQ-----
>Ceratophyllus_gallinae
------MASTARSLNKLFVANLPWTIGHRELRKYFAEFGHVSTA--NVIFDKSTGISKGFGFVFFSAKGGLEKALNQQVHNLEGRVLNVKPSNN-----
>Gynaikothrips_ficorum
-----------MSSYLIKVSKLPWTVGAKELSAYCSSFGPVHSV--RVPVDYSTGFQYGFGWVYFANADSKSRFLE-KKHYLEGHQLIVDGSRSKIE--
>Rhyacophila_fasciata
-----MASQVGRSVSKLFVGNLPWTIGHKELGEYFSRFGRVQVA--KVVFDRSTGVSRGYGFVEFSSKTAYENAINQQMHSLEGKVLSVNPSSA-----
>Zorotypus_caudelli
--------MAARAVNKFFIGNLPWTVGPKELRNYFSEFGAVTSA--NVLFDKNTGFSKGFGFITFGSNENSEAIKNKQTFVLEGNNLVVEPAE------
>Atelura_formicaria
--------MAARAVNKLFVANLPWTVGHLELRQYFAQFGHVQSA--HVVFDKKTGMSRGYGFIQYATQDACNSATDKRNHVLEGNQLAVQISAAPK---
```

HG0013 FBgn0261381 mtTFB1

```
>Drosophila_melanogaster
------------------MSASEQGPKIKYGE--SAPKLDKAQLQFMKLIEEQNLDRVQKLKRIRRNNLLTAGALGVSVLAIYGYSIFSVQQEKFLDDFEEPKKVSS-----
>Artemia_franciscana
------------------MAGNDQMPKVDLAK--DRPKFSEAQLDHMRLVEKLNALRVTELQRLKRNNYITAGVIATCVLGIYGYSMFAVKQEKFLDELE-------------
>Meinertellus_cundinamarcensi
------------------------MQKVDLEK--DKPKLSRAQLDYMKLLEKQNAERVMKLQRIRRNNIITGSLLGVGVLSIYFYSIFAVKQEKFLDDFDAPKKISASSS--
>Argulus_siamensis
------------------MANDNVMPKLNLQN---DPQLKQIQLEYIKKIERENLIRVNRMKKLKMKNVYAGCGLLGFVLGVYTYTLYSVKQETFLDDLEDPSHV-------
>Mastotermes_darwiniensis
------------------MGKNEQMPKVDLVK--DRANISRTTLDYMKIVERENLRRVQALQKLRQKNVRTGLMLGAGVLGIYLYTIYSVRQEKFLDDFDEPATTSN-----
>Hynobius_chinensis
--------------------------------------------------------------------------------------MYAVKQEKFLDDFEEPAKEELTQKQL
>Dastarcus_helophoroides
-----------------MMSEQPKAP----KL--DYSKLKEAELKFVKLVEKQNFERVQKLKAIRKRNLITAGAVGAAVLGIYGYTLFAVHQESFLDDFDEPAKTVQ-----
>Forficula_auricularia
-------------------MSSNQMPKVDFVK--DKELLKNTAVDYIKHIEKLNIDRAKKYIRIRRNNIITGCLLGLGITSIFAFTIIGVKQEKFLDDFEEPAKISS-----
>Campodea_augens
-----------------MAESAEKMPKLDLQK--DKPQLHQAQLYYMQMIENKNLERVQKLQKLRKRNLATAGILGIAVVSIFSYSMLAVKQEKFLDEVFDETEAKATLKQ-
>Haploembia_palaui
------------------MADSEQMPKVDFAK--DKAELKHTAIDYVKLINKVNEERALKFQRIRKSNIITGCVLGFAALGIYSYTIYAVKQERFLDDFEEPERINIAN---
>Eurylophella_sp._AD-2013
------------------------MPKVDFVK--DKANLNPASLDYIRLIERQNRERVLKLQKLRKVNLWTAGILGVSVFSIYFYSMFAVKQETFLDDFNEPERGSNPN---
>Grylloblatta_bifratrilecta
------------------MGDSDQMPKVDFVK--DKDKLKFTAVKYVEILEKKNLERVQKLLRTRRNNNITGWLLGGGVFGIYLYSMFAVQQEKFLDDFNEPEKTED-----
>Anoplocnemis_curvipes
------------------MGDADQMPKLNLAK----DKVDKNVLDYIRLIEEQNRSRVEKLRRIRRNNLLTAFGLGIGVLSIYGYSIFAVRQETFLDDFERPAIKE------
>Neodiprion_sertifer
-------------------MADEQMPKIHEGE--NFRKLTLVEQEYMRLIEERNLERVTRLKRMQRNNRYTGLALGALVLGIYGYSMYAVRQENFLDDFEEPVKTTE-----
>Acanthopteroctetes_unifascia
------------------MGGNDVMPKINLDK-EKPEKLKKFELEYMKVINQKNLERVQKLERIRRNNILTGSILGVGVLGIYFYSMYAVKQESFLDDFNEPEKIIRNN---
>Empusa_pennata
-------------------MDKDQMQKVDLVK--DKANISRVVVDYMKLMEKENLERVSRLKRLRKRNNLTGLLLGGAVFGVYLYTILSVKQEKFLDDFEPPELVEKN----
>Tanzaniophasma_sp._AD-2013
--------------------------------------------------------------------------------------MFAVQQENFLDDFEEPAKNVE-----
>Boreus_hyemalis
------------------MSGKDVMPKLNLSD-KSQQKLQPTELDFMKIIEKQNLARVQKLKRIRRNNLITAGILATGVFGIYGYSIFAVQQEKFLDDFKEPEKLTTNN---
>Corydalus_cornutus
------------------MDNNEQMPKVNLKS----SNIRQSDIDFMKVIEKQNIERVRKLQRMKRNNLITAGLLGSGVFGIYLYSMLAVKQETFLDDFNEPAKVVE-----
>Conwentzia_psociformis
-----------------------------MLD-EEKKKLTPTQLRFMQEIEQKNLDRVQKLQRIRKNNILVGCSIGLSVLAIYGYSIYSVQQEKFLDDFNPPSKVQS-----
>Triops_cancriformis
-------------------MASDQMPKVDLVK--DATKLRQPQLDYMKLIEKQNLERVTKLKRIRKNNLLTGAALFASVFGIYVYSIAAVKQEKFLDDLETPSSKPSH----
>Cordulegaster_boltonii
---------------MAGKENSNEMPKVDFEK--DRGQLGRAELGYIRQIEEQTRTRAERLKKMRRGNLITGFALTAGVLGIYIYSIVAVKQEKFLDDFEPPKLEPKQ----
>Schistocerca_gregaria
-------------------MENKQDPK-------NKEFISKSMAEYMKLIEKQNLERVQKLQKTRRNNLITAGILGAAVTGIYFYSMYSVKQETFLDDFEEPKKISQ-----
>Timema_cristinae
------------------MADKESMPKVDFEK--DKAKFKNTAVDYIKLIEKLNLERVQKLQRIRRNNIITGCVLGGSVLGIYAYSMYSVKQETFLDDFDEPATTSN-----
>Menopon_gallinae
------------------MTSSEDMPRADLSD----EKLKN-AQAYAKYLEHKNYMRSLYGKRILRRKRFMAGFCCAFVVGVYCFTLFRVQQEAFLDDFEEPPRVNRDEKN-
>Perla_marginata
------------------MGDNERMPKVDFTK--DKENIKLSVIDYMKLVEKQNLERVEKLRRIQRRNLITGCILGVGVLGIYGYSMYAVKQEKFLDDFDEPAKTEAKLS--
>Acerentomon_sp._AD-2013
--------------MALTEKPGDVMPEVDLKK--DAADINQVHLDYMKLLEAKNTKRAHRYVRMIKKNKLLGSGLIFGVASIYVYSMYAVKQEKFLDDFEHPDANIPPPQ--
>Liposcelis_bostrychophila
------------------MARENEMPEVDFKD-IDKNKKLRQAYIYAKQLEERNVFRALIGARNLKWSRIIGITCGLFAAGVYSLTINRVQQETFLEDWEEPEIPGRS----
>Inocellia_crassicornis
------------------MSGSDQMPKVSFGT----SKINKTELDFMKFIEKQNLDRVEKLKILRRKNVITGCVLGAGILSIYAYSILSVKQEKFLDDFEEPKKVYAETQ--
>Ceratophyllus_gallinae
------------------MADNKLVQKPSISD--EPRKLSKVELDFMRIIEKENLARVEKLKRVRRNNLITGTLLTGSVFGIYLYTILSVKQEKFLDDFNEPAKISD-----
>Caligus_rogercresseyi
MSVNAGLLRALRAPSRNFVRGAKTMKQVNVDDSETRAKLGSIQMDLIHRAEKMNSRRATQFSKYRRHDVVVGGVCFSMAIAIYLYTIFAIKQEKFLDDFDMPDPMLPVQSKE
>Stylops_melittae
----------------MQVSG----AKVNPKD----KEFTPEQLELFKQFDNDKLRRLNRIRLRNSNNKKIGLALGFTVAAFYAYSMVAVQQEHFLDDFDKPEIVQD-----
>Gynaikothrips_ficorum
------------------MDKNEAMVKVDLSK----AQVDPTTLEHMKIMEKYNLARVQQMKRIKRKSNFMSLGLVSLAVGTYLYSMYAVRQEGFLDDFEEPKLKTK-----
>Platycentropus_radiatus
------------------MDSTEQKGKI--VS-TSKPAFSKGQIDYMKVVENLNLERVQKLKRIKRNNFITGSVLGGIVLGIYAYSIFSVKQEHFLDDFIEPTTVHKET---
>Atelura_formicaria
------------------MGDSERMPKVNLGK----DHMKQTELDYMKLVEKQNLERVQKLKTLRRRNVITGSLLGLAVVGIYGYSIYAVKQERFLDDFDEPQKTYEQK---
```

HG0014 FBgn0260392 CG42518

```
>Drosophila_melanogaster
MMDLSPNNQIEDRKPILTADGLVQTSNSPFEPTISQETQTSNG--IGGQCHLTVDQLDIEILPIIYDIVRCVEKDPLEN-AVKLRESQDCNHKIFELQKRFESAREQIRQLPGIDFNKEEQQQRLELLRNQLKLKQQLIRKYKDTEF--------------------------------------------------
>Echinogammarus_veneris
--------------------------------------------MSITGATVAAEDIDLDLLPLIYQYLRCLEKEQN--QTDLNRVALDATQKLTELNNKITLAREQVPKLAGVEYSPADQLQKLSALRAQLTLKKQLLAKYKSK-ASADG-------VTTRD----------------------------------
>Reticulitermes_flavipes
-----------------------------------MDSSEQNG--AALTCKLTVDDVDIEFLPLIYEIIRSVERDPH-DTSQKTRESQDTSQKVLELQKKLDQARNQIRRLPGVEHSKEEQLQKLETLRKQLQLKKELLLKYRN-MCTFDIPKS-------------------------------------------
>Agrilus_planipennis
----------------------------------MDSNNESN---SDLQKRMTVDDLNIEILPVFYEIIKSVERDHHDN-TAKTRESQDCSQKVLELQKRFDQARSQISLLPGIDYSKEQQLQHLEALKTQLHLKQELLYKYRF-MYPFQMQK--------------------------------------------
>Petrolisthes_cinctipes
----------------------------------------------MSSPVVAAEDVDVDFLPVIYQYLRCLEKEQ----TDLGRVSQESSQKVLELQRKIQTAREQVRKLPGVEYSREEQLRRLEALRKQLALKKRLLIKYKSK-SEATH----------------------------------------------
>Apachyus_charteceus
-----------------------------------MDISEST-S-QSLIQNLTVDDLDVEFLPIIYDIIRSVERDPL-DPSQKNRESQDSSQKVLELQKKLDQARNQVKQLQGIEYSKAEQLKKVEILRKQLQLKRDLLNKYRT-MCTFDIPK--------------------------------------------
>Daphnia_pulex
------------------------------------------------MATQLVEGGDFDLLSTIYDIIKSVEKEQQ-DNAQKQKDSQEAGQKVLELQRKLDHAREVVKKLPGIELSKDEQLLQIDLLRRQLTLKRDLLNKYRL-ITSFDREAQSEFQIHQH-----------------------------------
>Campodea_augens
-----------------------------------------------MAASNPIEQIDVEFLPTIYEIIRNIERDSG-DTTQKTRDLQEAGQKTLELQRKLDQAREQIQKFPGIEYNKEEQLRRLETLRKQLTLKRELLLKYKN-MCSFEVPKA-------------------------------------------
>Haploembia_palaui
----------------------------------MEEQQNLNGS-VAASTQLTVQDLDLEFLPLIYEILRSIEKDPH-DSSQKVRESQDISQKILELQKKLDTARSQVQQLPGVEYSKEEQLQKLETLRKQLQLKKDLLLKYRH-MCSFEVPK--------------------------------------------
>Baetis_sp._AD-2013
-----------------------------------MDSAEQP---TVTNGHLSAEEVDVELLPLIYELMRGIERESI-ETAQKPRESQDTSSKVLDLHKKLNLLREQIRKLPGIEYNREDQLKQLDHLRKQLRLKRELLLKYRN-MCSFEMPKA-------------------------------------------
>Tigriopus_californicus
-------------------------------------------------------------------------------------------------------MRTGIYGLSGIENSPKEQLKQLEILREQLRMKKKLIHKYKE-LNLKVFGLSGTY----------------------------------------
>Cuerna_arida
---------------------------MDVH---AINNDQQE-QTPTDYKPLTVENLDVDFLPLIYDIIRSVERDPH-DNAQKSSQSQDISQKVIELHNKLEQAKKQVRRLPGVEYSKEEQLEKLETLRKQLRLKRELLQKYRN-MYLLDSPKP-------------------------------------------
>Tenthredo_koehleri
----------------------------------MMEVVEIAE-ETPARSQFTVDDLDIEILPLVYEIIRSVEKDPH-DNTQKAKESQDTSHKILDLQKKLDSARSQIKRLPGIEYSKEEQLQKLETLRRQLRLKQELLLKYRN-MCTFDVPKV-------------------------------------------
>Armadillidium_vulgare
----------------------------------------------MSSHTIAAEEVDVDFLPLIYQYLRCLEKDQ----NDLNRISVESSQKILEIQKKIDIAREQVQRLPGVEYSKEEQLQRLEALRKQLSLKKKLLPKYKSK-AQPEQ-------SH-------------------------------------
>Plutella_xylostella
----------------------------------MENNTDHAE--QNSCAPLSVQDVDVDFLPIVYEIIRSVERDLHDN-SAKVRESQDCSQKVLELQKKFDAARAQILRLPGIDYNKQDQLKQFEILRTQLKLKRELLQKYRNNMCPF------------------------------------------------
>Empusa_pennata
-----------------------------MF---SEDQNGSAG--SALPAQLTVDDVDVEFLPLIYEIIKSVERDPH-DTSQKTRESQDTSQKVLELQKKLDQARNQIRRLPGIEYSKEEQIQKLETLRKQLQLKKELLLKYRH-MCTFDIPKST------------------------------------------
>Tanzaniophasma_sp._AD-2013
-----------------------------------MDVGEQNG--TSMPTQLTVDDVDVEFLPLVYEIIRSVERDPH-DTSQKTRESQDTSQKVLELQKKLDQARSQISRLPGVQYSKEEQLMKLETLHKQLQLKKELLLKYRH-MCSFDIPKS-------------------------------------------
>Panorpa_vulgaris
-----------------------------ME---PMVTDQSE---LKNNTTLTVDQVEIDILPTIYDIIRSVERDPHDS-AGKLRESQDCATKVLELQKKLLKARNQIKQLPGIDLNPEEQLRKVDTLRNQLKLKEELLHKYRN-MYSFDSSKI-------------------------------------------
>Corydalus_cornutus
-----------------------------ME---VSEQNDSN---NRSQEPLTVEEVDIEILPIIYDIIRSIERDPHDN-AAKARESQDCSQKVLELQKKLEFSRDQIRRLTGINCSREEQIAKLTSLRTQLKLKQDLLRKYRE-MVSLESTKS-------------------------------------------
>Pseudomallada_prasinus
-----------------------------ME---VSESNELN---NRNSTQLNIEELDFEILPVIYEIIRSVERDPHDN-AAKARESQDCSQKVLELQKKLDQARAQIRRLPGIEFSREQQLQRLEALKTQLRLKQELLQKYRH-MCSLDSIKA-------------------------------------------
>Triops_newberryi
-------------------------------------------------MTATSEEVDIEFLPLIYDIIKSFERDPV-DNTQKLKDSQETYQKVLELQKKLDHAREQIKKLPGVEFSKEEQLEQLDLLRQQLTLKRELLLKYRN-LGPFDLPK--------------------------------------------
>Epiophlebia_superstes
-----------------------------------MDSEEQKT--SLSPSQLSAEEVDVEILPLIYEIIRSIERDPH-DATQKPRESQDTAQKILELQKKLDQAREQVKRLPGVDLSREEQLKRLEFLRTQLRLKRELLLKYRT-TCTFEIPSNPTGNSNSFSSPSLSSSTQQQQQINTNSLPTSNSQPIVNNTGKA
>Locusta_migratoria
-----------------------------------MESIEQNGS-TATPSSLTVDDVDVDFLPLIYDIIRSVERDPH-DSTQKTRESQDISQKILELHKKFDEARSQIGRLPGVEYSKDEQLRKLETFRKQLQLKKDLLRKYRE-MCTFEIPK--------------------------------------------
>Phyllium_philippinicum
-------------------MEQKNATDSCEH---DMEGCEQNS--STHASELSVKDVDVEFLPLIYDIIRGVEKDPQ-DTSQKSRDSQDTSQKVLDLHKKLDQARNQIRRLTGIEYSKEEQLKKLETLRKQLQLKKDLLLKYRH-MCSFDVPK--------------------------------------------
>Cosmioperla_kuna
-----------------------------ME---ASDQSNQEG--SSIPTYLTVNDVDVEFLPLIYEIIRSVEKDPH-DISQKTRESQDTSHKVLELQKKLDQARSQIQRLPGVEYSKEEQLLKLETLRKQLHLKRELLQKYRN-MCSFEIPKV-------------------------------------------
>Acerentomon_franzi
------------------------------------------------MADVETDEIDTDFLPLIYDIIRSIEREPTGDLSAKPKEGPDANQKILDLNKKLEQAREQIRQFPGIECNKEEQLRRFDALKQQLTLKEQLLAKYRT-LCSFELQ---------------------------------------------
>Ectopsocus_briggsi
---------------MAEQTALSNAVKIEME---NLPNGNST-TDLVANNSTNNKEFDFEFLPLIYEVIKSVGPDQ--DPTGKGRESQDASIKILELERKLNEARQQILVLPGIEHSKEEQLRQLELCRKQVILKQKLIAKYRS-FCNFSLAD--------------------------------------------
>Xanthostigma_xanthostigma
-----------------------------ME---VTD--QSE---TLNCKQLSVEELDIDILPIVYEIIRSIERDHHDN-TAKTREAQDCSQKVIELQKKLEQARSQIRQLPGIEYSREQQLQKLEALRTQLKLKQELLQKYRH-MCTLDFNKS-------------------------------------------
>Trichomycterus_areolatus
-----------------------------ME------SVEEGE--GEGVGVVAESPLQLELLPALYDIVRSVEREVSEGGANKARESQDCSLKVLELHRQLEAARQQVRSLAGISQTKREQLRQFEILRTQLFLKRKLLKKYRA-MCLTESSFK-------------------------------------------
>Oropsylla_silantiewi
-----------------------------ME---SMNTELSEQ--QTNANPLTVDQVEIDILPTIYDIIRSVERDPHDN-AGKARESQDCSTKVLELQKKLDKIRTQVALLPGIDYNKEEQLQQLDNLRKQLKLKQELLHKYRT-MYSFDSMKI-------------------------------------------
>Mengenilla_moldrzyki
-----------------------------MD------ISN-----TEIEQGDSAEQITSEILPAVYEVIRSLEKDSYD----ISFKIHDCSQKILHLKDKFDRARQLIEDFPGIRYSIDDNKALMKSLRQQLKLNHELIQKYRN-IRDFGTTKP-------------------------------------------
>Platycentropus_radiatus
-----------------------------MD---VINNDLNEP--QNNFSPLTVQDVDVDILPIVYDIIRSVERDVHDN-SAKARESQDCSLKVLELQKKLDSARSQIKRLPGIDYNKQEQLRQFEILRRQLHLKRELLQKYRN-MCSFDAPFK-------------------------------------------
>Zorotypus_caudelli
-----------------------------------MDSNDQPAVPTVINNPATVDDIDVDFLPLIYEIIRSVERDPH-DTSQKSRESVDTSQKVLELQKKLDLARSQVRLLPGVELSKSEQLAHLDTLYKQLQLKRDLLHKYRN-MCSFDISKVQ------------------------------------------
>Atelura_formicaria
-----------------------------ME---SVEQNPVPTV-APSPSTITVDDADVEFLPVIYEILRSIERDPH-DSTQKSRESQDTSQKVLELQKKLDQAREQIVKLPGIEYSKEDQLKKLDTLRKQLRLKQELLLKYRN-MCTFELPKV-------------------------------------------
```

HG0015 ENSDARG00000007377 odc1

```
>Danio_rerio
MRFKNSCLREV----HTGPGLEAYLKGFNF-TGEPPWLSDIC
>Acipenser_sinensis
MLFKNAPFKKVYNGFSEGLGLALYLKEFNF-TGEPPW-----
>Amia_calva
MLFKNTRFSKAHNGFLEGPDFAFYLKVFNF-KGEPPWL----
>Channa_striata
MLFKNHHSKKSVTAYSSGPGLESYLKGFNF-TGEPPWLLCT-
>Podocoryna_carnea
MLFKNAPFKKVYNGFSVGLGLALYLKEFNF-TGEPPW-----
>Xenopus_laevis
MLFKNTNSRTAQSGSIESVGLVLYLPGFYF-TGEPPWE----
>Basilichthys_microlepidotus
MLFKNPHSRKSHSGCSTGPGLESYLKGFNF-TGEPPWLLCT-
>Porichthys_notatus
MLVKTHHCRTAHSDYSAGPGLESYLKGVNF-TGDPPWILCT-
>Oryzias_latipes
MLFQNLHSRKSYRGSSTGPSIESYLKGFNFFTGEPPWPQRLD
>Hynobius_chinensis
MLFKNSCSTTSYHGSSGGLRLSLYLPGFYF-TGEPPWQQS--
>Micropterus_floridanus
MLFKNHPLRKTHTGYSAGPGLESYLKGFNF-TGEPPWLLCT-
>Astyanax_mexicanus
MLFKNTRLRKAHTGY-AGPGLEAYLKGFNF-TGEPPWVL---
>Callorhinchus_milii
MFYKNNQIRKLYNGYSEGLRFLLYLKVFSF-VGEPPWRRS--
>Haplochromis_burtoni
MLFKNLHSRKSHTGYSAGPGLESYLKGFNF-TGEPPWLLCT-
>Alosa_alosa
MLLKNNHLRTLHT-YSEGNGLESYLKGFNF-TGEPPWLLYT-
>Latimeria_chalumnae
MLFKN-YISKLYNGCTEGSGLALYRKGFDF-AGAPPWEHG--
>Poecilia_reticulata
MLFKNLHSRKAHSSCSTGPGLELYLKGFNF-TGEPPWLLCT-
>Esox_lucius
MLFKNNRFRKAHTGYSAGPGLESYLKGFNF-TGEPPWLLCT-
>Gadus_morhua
MLFKINHFRKVHTGYSAGRGLESYLKGFNF-TGEPPWLPLT-
>Acanthogobius_hasta
MLFKNLHLRRS----ETGPGLEAYLKGFNF-TGEPPWLLCT-
>Apteronotus_leptorhynchus
MLFKTTCLRKAHTGYSAGPGLEVYLKGFNF-TGEPPWVI---
>Erinaceus_europaeus
MRFKGPRLRNCHTSSGC-------CTSFEV-TGDPP------
>Lutjanus_erythropterus
MLFKNHRSRQAHTGYSAGPGLESYLKGFNF-TGEPPWLLCT-
>Osmerus_mordax
MLFKTNCFRKAHTGYSAGPGLESYLKGFNF-TGEPPWLLCT-
>Scleropages_formosus
MLFKNSRLRKAHFRYSTGCGLESYLKGFNF-TGEPPWLLCT-
>Lateolabrax_japonicus
MLFKNNHSRKTHT--GAGPGLESYLKGFNF-TGEPPWLLCT-
>Perca_flavescens
MLFKNNHSRKTHT----GPGLECYLKGFNF-TGEPPWLLCT-
>Paralichthys_olivaceus
MLFKNFNSRRSHTGYSTGPGLESYLKGFNF-TGEPPWLLCT-
>Leucoraja_erinacea
MFFKNERVKRVFHGYTESPVLTLYLRGFNS-VGKPPWRTS--
>Salmo_salar
MLFKNNRFRKAHTGYSAGPGLESYLKGFNF-TGEPPWLLCT-
>Lepisosteus_oculatus
MLFKNAHFRKADIGFLEGPGLASYLKGFNF-TGEPPWLSG--
>Trichomycterus_areolatus
MLFKTSRLSKAH----TGPGLEVYLKGFNF-TGEPPWGL---
>Squalus_acanthias
MFFKNEQARKVYNGYVESPDLALYLRGFNS-VGKPPWRDS--
>Takifugu_rubripes
MLFQNPRFRKTNTGYSAGPGLESYLKGFNF-TGEPPWALYS-
```

HG0016 ENSGALG00000010399 SELENOT

>Gallus_gallus
---------------------------------------------------MAYATGPLLKFQICVS
>Acipenser_sinensis
--------------MAKIWLSFSVLFLGVFYLY-S-AVADNG--APGKKLKMQFATGPQLKFQICVS
>Kaupichthys_hyoproroides
--------------MGRRWLSFCLLFLGVF-FLSCA-SADNGG---VKKLKMQFATGPLLKFQICIS
>Podocoryna_carnea
--------------MAKRWLSFSLLFLGVFYLY-S-ALADNG--APGKKLKMQFSTGPRLKFQICVS
>Xenopus_laevis
---------------------------------------------------MQYTAGPLLKFQICVS
>Acinonyx_jubatus
--------------MRF--LLVLL--VAASAVIRSDASANLGG-VPSKRLKMQYATGPLLKFQICVS
>Ambystoma_mexicanum
---------------------------------------------------MHYTSGPLLRFQICVS
>Tursiops_truncatus
------------------MRLLLFLLVAASAVVRSDASANLGG-VPGKRLKMQYATGPLLKFQICVS
>Eptesicus_fuscus
------------------MRLLLFLLVAASALVRSDASANLGG-VPSKRLKMQYATGPLLKFQICVS
>Dasypus_novemcinctus
-------------------MRLLLLLLAASAVVRSEASANLGG-LPSKRLKMQYATGPLLKFQICVS
>Alligator_mississippiensis
---------------------------------------------------MAYATGPLLKFQICVS
>Misgurnus_anguillicaudatus
----------------MRWLPLSVLILWAC-SLYS-VSADNSG---LKKMKMQFATGPLLKFQICVS
>Galeopterus_variegatus
------------------MRLLLLLLVAASAMVRSEASANRGG-VPSKRLKMQYATGPLLKFQICVS
>Monodelphis_domestica
------------------MRFLLLL-LMAAAALQGEASADTGG-VPGKRLKMQYATGPLLKFQICVS
>Trichosurus_vulpecula
------------------MRFLLLL-LMAAAALRGEASADTGG-VPGKRLKMQYATGPLLKFQICVS
>Aegypius_monachus
--------------MRAAVLLLLLLALAAGPGGG---SAEQGG-LPAKKLRMAYATGPLLKFQICVS
>Gadus_morhua
-------------MASSRWLRFSLLFLGVF-SLYCATSADNSG---VKKMKMQFATGPLLKFQICIS
>Condylura_cristata
------------------MRLLLLLLVAASAVVGTDASANLGG-VPSKRLKMQYATGPLLKFQICVS
>Oryctolagus_cuniculus
------------------MRLLLLLLVAASAVARSEASANLGG-VPSKRLKMQYATGPLLKFQICVS
>Pseudopodoces_humilis
--------MRAAGLRRLLLLLLLLAGLAAAPGGGGAAAAEQGG-LPAKKLRMAYATGPLLKFQICVS
>Trematomus_bernacchii
----------------MKWLQFSLL-VGLS-SLLCASSGDNSG---VKKMKMQFANGPLLKFQICIS
>Ceratotherium_simum_simum
------------------MRLLVLLLVAASAVVRSDASANLGG-VPGKRLKMQYATGPLLKFQICVS
>Manis_javanica
------------------MRLLLLLLVAASAMIRSDASANLGG-MPSKRLKMQYATGPLLKFQICVS
>Platichthys_stellatus
--------------MEMRWPRFLLLALGAF-SLCCATSGDISG---VKKMKMQFATGPLLKFQICIS
>Aotus_nancymaae
------------------MRLLLLLLVAASAVVRSEASANLGG-VPSKRLKMQYATGPLLKFQICVS
>Loxodonta_africana
------------------MRLLLLLLMAASAVARSEASANLGG-MPSKRLKMQYATGPLLKFQICVS
>Melopsittacus_undulatus
----------------MRAVLLLLLALAARPAGG---SAEQGG-LPAKKLRMAYATGPLLKFQICVS
>Marmota_marmota_marmota
MKPNINFEFWVVNREITRKRLLLLLLVAASAVARSEASANLGG-VPSKRLKMQYATGPLLKFQICVS
>Oncorhynchus_mykiss
-------------------------------------------------MKMQFATGPLLKFQICIS
>Tupaia_chinensis
------------------MRLLLLFLVAASAVVRSEASANLGG-VPSKRLKMQYATGPLLKFQICVS
>Trichechus_manatus_latirostr
------------------MRFLLLLLMAVSAVVRSEASANLGG-VPSKRLKMQYATGPLLKFQICVS
>Gekko_japonicus
-------------MKRALALLL-LLSV-----GLDWASAQPGGLPPAKKLRMAYATGPLLKFQICVS
>Mauremys_reevesii
---------------------------------------------------MAYTTGPLLKFQICVS
>Orycteropus_afer_afer
------------------MRLLLLLLMAASAVVRSEASANLGG-VPNKRLKMQYASGPLLKFQICVS

HG0017 ENSDARG00000043154 ucp2

>Danio_rerio
------------------------------------------------------MFFICTSSQRFSMEERGNKI-QVESHFNQRDAFRES-----
>Acipenser_sinensis
------------------------------------------------------MLVQRFVFGRLSMEQRGNLI-VTESNFSERDTFREEC----
>Podocoryna_carnea
------------------------------------------------------MLVQRFVFGRLSMEPRGNLI-ITESNFSERDTFREEC----
>Oryzias_latipes
------------------------------------------------------MILGLSTSEISSMEKQGQTV-KCAANFSQRDTFRES-----
>Micropterus_floridanus
------------------------------------------------------MILRLITSEKSSMEKQGNQF-IAGTKFSQRDTFRES-----
>Astyanax_mexicanus
------------------------------------------------------MFFRCSFSERFSMEDRGNEI-IVESHFNQRDAFRES-----
>Callorhinchus_milii
------------------------------------------------------MII-QRFISGFSMEQPETQI-IVKKSFNERDIFWES-----
>Neolamprologus_brichardi
------------------------------------------------------MIIRLI-AVKKSMEKQEKLF-IAGTKFSQRDAFRES-----
>Alosa_alosa
------------------------------------------------------MLLYCLVLERFSMEDQGNVI-IVESNFNQRDTFRES-----
>Latimeria_menadoensis
------------------------------------------------------MIIRYSDSAEFSMDQE-KTS-IVKNNFNERDIFRES-----
>Kryptolebias_marmoratus
-----------------MIKPDYLKKYYLLFFFLFLLK--------CLFSRSWSMILRLKVSERSSMEKQGNQF-IGGTRFNQRDTFRES-----
>Sminthopsis_crassicaudata
------------------------------------------------------MIIRCSVSQQFSMENQGKCTMVARSNFEERDTFREA-----
>Galeopterus_variegatus
------------------------------------------------------MTFGCFVSQPFSMENQGDRAMIATGSFEERDAFREA-----
>Monodelphis_domestica
------------------------------------------------------MIIRCSVSQQFSMENQAKWNMVAVSNFEERDTFREA-----
>Trichosurus_vulpecula
------------------------------------------------------MIIRCSVSQQLSMENQGKWAMVAVSNFEQRDTFREA-----
>Esox_lucius
------------------------------------------------------MFLGHIVFERCSMEEQGNQI-IVESNFDQRDAFRES-----
>Acanthogobius_hasta
------------------------------------------------------------------MENKRTQI-IAGTKFSQRDAFREA-----
>Chanos_chanos
-------------------------------------------------------MFMCPVFKHFSMEEQGNEI-IAESTFNQRDTFRES-----
>Oryctolagus_cuniculus
------------------------------------------------------MNFRCFVSQPFSMENPGDRGMIATGGFEERDAFREA-----
>Osmerus_mordax
------------------------------------------------------MFFGYTVFEACSMEE-GKQI-IGESNFDQRDTFRES-----
>Scleropages_formosus
------------------------------------------------------MFLKCYTLKRVSMEDQVNEI-IAESQFSERDTFREA-----
>Lateolabrax_japonicus
------------------------------------------------------MILRLITSKRSSMEKQGYQF-IAGTKFSQRDTFRES-----
>Trematomus_bernacchii
------------------------------------------------------MILRLNTSEKSSMEKQGNLF-KAGTTFNQRDTFRES-----
>Equus_asinus
------------------------------------------------------MTIRCFVSQPFSMENQGDRAMIATGSFEERDAFREA-----
>Propithecus_coquereli
------------------------------------------------------MTIRCFVSQPFSMENQGNRAMIATGSFEEWDAFREA-----
>Leucoraja_ocellata
------------------------------------------------------MII-WRFISELSMEQRQSQT-VVKRGYNERDTFRESSPARL
>Ctenomys_sociabilis
MQAPPPALKSSASRADRIASPAAALFGHRHRWCPLTADCQRPLSRVLPAAGTRSMIFKCLVSQSFSMENQGDQAMIATGNFEERDTFREA-----
>Salmo_salar
------------------------------------------------------MFFGYVVFERFSMEEQGNQI-IIESNFDQRDAFRES-----
>Lepisosteus_oculatus
------------------------------------------------------MLFTSLVLGIFSMEQQGTQI-IVESNFNERDTFRES-----
>Ictalurus_furcatus
------------------------------------------------------MFLKCPFSELFSMEERGNEI-IVESHFNQRDAFREA-----
>Squalus_acanthias
------------------------------------------------------MII-RRFISELSME-RQSQI-IVKKSYNERDTFRES-----
>Gekko_japonicus
------------------------------------------------------MIIRCRLSAAFSMASKGSRAAVALSNFDERDTFRES-----
>Chrysemys_picta
------------------------------------------------------MILKCLVSKRVSMENQGDQIVVALSKFDERDTFRES-----

HG0018 ENSDARG00000053291 pnrc2

```
>Danio_rerio
---------------------------------------MFNSVPRQNN--AHWHMKTSDLA------GHTCAARSR-S-TTSRTWSPIGEEKLKSEKNHA--LNFFTTER-
>Acipenser_sinensis
---------------------------------------MYDSSTRFN--TNFRHMKTKTTA------GLTRSERSGEI-PTSPTWSQTGEKVSRKEEANK--SCYFTITKR
>Amia_calva
---------------------------------------MFGSAARPD--TAVRHMSAPSPA------GPTRATRSGFI-STSRTWSPVGEEKLKRPD-------FCTTKR-
>Channa_striata
---------------------------------------MFNSAPRENN--AHRHMKTSDFA------GHKRAARSRQK-TISRTWGPIGGERLKRD-LCF--VDFQTTKR-
>Anguilla_japonica
---------------------------------------MFGSASRQNC-SYFWHMKTPKLA------GLRRTERSGPT-PTSRTWSPIGEEKLRVEEVRT--RYFLTTKR-
>Podocoryna_carnea
---------------------------------------MYDSSTRFN--TNFRHMKTKTTA------GLRRSERSGEI-PTSPTWSQTGDKVSRKEEVNK--PCFFTITKR
>Basilichthys_microlepidotus
---------------------------------------MFSSASRENN--ALWHMRTSDLA------GHTSAARSRQK-ATSRTWGPIGGERLKRD-LCF--DNFLTTKR-
>Opsanus_beta
---------------------------------------MFSSAPREND--ANWHMKTSHCT----------HKGRQK-AASRTWGPTGGERLKRN-LCSV-----TTKR-
>Oryzias_melastigma
---------------------------------------MFSSVPRENI--AHWHMKTSDLA------GQKSAVRSRQK-ATSRTWGPTGGERLKKD-LCF--VNIPPTKR-
>Tripterygion_delaisi
---------------------------------------MFYSAPRDIN--AHCHMKTSDLA------GHNTSAARSRLRTTSRTWGPVGGERLKRD-LCSVFTT----KR-
>Trachinotus_blochii
---------------------------------------MFSSAPRENN--AHWHMKTSDLA------GHKSAARSRQK-TTSRTWGPIGGERLKID-LCF--INFLTTKR-
>Mustela_putorius_furo
---------------------------------------MLASAPRLNS--ADWPMKTSVLR------QRKGSSRKQH--LLSWAWQQGRGQVVEILQSE------KQTER-
>Micropterus_dolomieu
---------------------------------------MFSPAPRENN--ALWHMKTTDLA------GNKIAARSRQR-TTSRTWGPIGGERLKRD-HCF--FNFLTTKR-
>Pygocentrus_nattereri
---------------------------------------MFNSASRQNI--ARWHMKTSDTV------GLKRAVRSRA--ATSRTWSPIGEEKLRNKDIYAVIFN---TKR-
>Miniopterus_natalensis
---------------------------------------MLASAPRLNS--ADWPMKTSVLR------QRKGSVRKQH--LLSWAWQQGIGQVVEILQSE------KQTER-
>Haplochromis_burtoni
---------------------------------------MFSSASRENN--AYWHMKTSDPA------GHKCAARGRQK-TTSRTWGPIGGERLKRD-VYS--VNFLTTKR-
>Alosa_alosa
---------------------------------------MFSSVTRQYT--AHWHMKTSDLA------GHKCAARSRFS-TTSRTWSPVGEEKLRSEIIHV--LNFLTTKR-
>Poecilia_mexicana
---------------------------------------MFSSASRQNN--DHWHMKTSDLA------GHTSAARSRQR-TTSRTWGPIGGERLKQD-LIV--FSLTTKR--
>Esox_lucius
---------------------------------------MFSSASLENN--AHWHMKTSDLA------GLKSAARSRSK-TTLRTWSPIGEEKLRNH-LCT--FNILITKR-
>Acanthogobius_hasta
---------------------------------------MFSSVPRQNI--ALWHMRTSVLA------GHKSAVRSRR--TTSRTWGPIGGVRLKRA-HCNT---FLTTKR-
>Chanos_chanos
-------------------------------------------------------MKTSGLA------GRKCAARSR-T-TTSRTWSPIGEEKLRSKDIDE--LNLVTTKR-
>Plecoglossus_altivelis
MNTSFTRQPDQYLTKGFYHPSQKKSRFSSSRSTVFRGELMFSSGTRENI--ADRHMKTADLA------GHKSAARSRSK-TTLRTWSPIGEEKLRKKALCS--FHFLTTKR-
>Lateolabrax_japonicus
---------------------------------------MFSSAPRENN--APWHMKTSDLA------GHRSAARSRQR-TTSRTWGPIGGERLKRD-HCF--VDFLTTKR-
>Sebastes_caurinus
---------------------------------------MFSSVPRENNSTTHWHMKTSDLA------GHRSTARSRQKITTSRTWGPIGGERLKGD-LSFFYFNFLTTKR-
>Cynoglossus_semilaevis
---------------------------------------MFSSAARVNN--AHWHMKTSDPA------GHISAVRSRQK-TTSRTWGPIGGERLKED-SCS--TNFLTTKR-
>Microcebus_murinus
---------------------------------------MLASAPRLNS--ADWPMKTSVLT------QRKGSFRKQH--LLSWAWQQGRGQVVEILQSE------KQTER-
>Ictidomys_tridecemlineatus
---------------------------------------MLASAPRLNS--ADWPMKTSVLR------QRKGSVRKQH--LLSWAWQQGRGQVVEILQSE------KQTER-
>Oncorhynchus_tshawytscha
---------------------------------------MFYSASREND--AHWHMTTPDLA------GPKSAARSRPK-TTSRTWSPIGKEKLRRKYLCT--FNFLTTKR-
>Lepisosteus_oculatus
---------------------------------------MFGSASRYS--PDFRHMKTPNLA------GLTPSERSGLR-PTSRTWSPVGEEKLRGKESNR--LCFSTAKR-
>Ictalurus_punctatus
---------------------------------------MFSSASRQNA--AFWHMNTINTITTINIELNTCAVRSRR--TTSRTWRQISDETVRRRNLCT--GNFTTTTER
>Trichechus_manatus_latirostr
---------------------------------------MLASAPRLNS--ADWPMKTSVFR------QRKGSVRKQH--LLSWAWQQGRGQVLEILQSE------KQTER-
>Takifugu_rubripes
---------------------------------------MFSSVPRQNN--ALWHMKTLGLA------GHRSAVRSRSR-AAPRTRSPVEDPSKIGS------AHFATTKR-
```

HG0018 ENSG00000189266 PNRC2

```
>Homo_sapiens
---------------MLASAPRLNSADRPMKTSVLRQRKGSVRKQHLLSWAWQQGRGQVVEILQSEKQTER
>Anas_platyrhynchos
MQLDFFPWSFSAENPMLGSAPWLNTADHPMKTSIPSQRKESSERHHLLFQAWKQERGQELDGVESEKATER
>Chaetura_pelagica
---------------MLGSATWLDTADQPMKTSIPSQRKESSKRQHLLFQAWKQERGQELESVESEKTTER
>Apteryx_australis_mantelli
---------------MLGSATWLKAAYQPMKTSIPSQRKESSKRQHLRFQAWKQERGQELESVESEKTTER
>Buceros_rhinoceros_silvestri
---------------MLGSAAWLKSADQPMNTSVRSQRQESGKRQHLLFQAWKQERGHELESVGSEKRTER
>Caprimulgus_carolinensis
---------------MLGSATWLNTADQPMKTSIPSQRKESSKRQHLLFQAWKQERGQEWESVESEKTTER
>Cariama_cristata
---------------MLGSATWLNTAHQPMKTSIPSQRKESSKRQHLLFQAWKQERGQELESVESEKTTER
>Odobenus_rosmarus_divergens
---------------MLASAPRLNSADRPMKTSVLRQRKGSGRKQHLLSWAWQQGRGQVVEILQSEKQTER
>Orcinus_orca
---------------MLASAPRLNSADRPMKTSVLRQRKGSIRKQHLLSWAWQQGRGQVVEILQSEKQTER
>Charadrius_vociferus
---------------MLGSATWLNTADQPMKTSIPSQRKESSKRQHLLFQAWKQERGQELESVESEKTTER
>Eptesicus_fuscus
---------------MLASAPRLNSADWPMKTSVLRQRRGSVRKQHLLSWAWQQGRGQVVEILQSEKQTER
>Pterocles_gutturalis
---------------MLGSATWLNTADQPMKTSIPSQRKESSKRQHLLFQAWKQERGQELESVESEKTTER
>Dasypus_novemcinctus
---------------MLTSAPRLNSADQPMKTSVFRQRKGSVRKQHLLTWAWQQGRGQVVEILQSEKQTER
>Colius_striatus
---------------MLGSAPWLNTAEQPMKTSIPSQRKESSERQHLLFQAWKQERGQELESVESEKTTER
>Columba_livia
---------------MLGSAPWLNTAHQPMKTSIPGQRKESSERQHLLFQAWKQERGQELERAESEKTTER
>Merops_nubicus
---------------MLGSATWLNTADQPMKTSIPSQRKESSKRQHLLFQAWKQERGQELESVESEKTTER
>Alligator_mississippiensis
---------------MWGSASWLDTADRPMKTSIPSQRKRSGKRQHLLFQAWQQKRGREVAAVESEKKTER
>Cuculus_canorus
---------------MLGSAPWLNTADQPMKTYICSPRKESSERQHLLFQAWKQERGQELESVESEKTTER
>Sarcophilus_harrisii
---------------------MLNSADQPMKTSILRQRNRSGRKQHLLSWAWQQGRRQVVEILQSAKQTER
>Galeopterus_variegatus
---------------MLASAPRLNSTDWPMKTSVLRQRKGSTRRQHLLSWAWQQGRGQVVEILQSEKQTER
>Monodelphis_domestica
---------------------MLNSVDQPMKTSILRERNGSGRKQYLLSWAWQQGRRQVMEILQSAKRTER
>Macropus_eugenii
---------------------MLNSADQPMKTSILRQRNRSGRKQHLLSWAWQQGRRQVVEILQSAKQTER
>Falco_cherrug
---------------MLASATWLNTADQPMKTSIPSQRKESSKRQHLLFQAWKQERGQELESVESEKTTER
>Gallus_gallus
---------------MLGSAAWLNTADRPMKTSVPSQRKESSERQHLLFQAWKQERGQELESVESEKTTER
>Gavia_stellata
---------------MLGSATWLNTADQPMKTSIPSQRKESSKRQHLLFQAWKQERGQELESVESEKTTER
>Balearica_regulorum_gibberic
---------------MLGSATWLNTADQPMKTSIPSQRKESSKRQHLLFQAWKQERGQELESVESEKTTER
>Condylura_cristata
---------------MLASAPRLNSADRPMKTYVLRQRKGSVRKQHLLSWAWQQGRGQVVEILQSEKQTER
>Oryctolagus_cuniculus
---------------MLASAPRLNSADRPMNTSVLRQRRGSARKQHLGPWAWQRGRGQAVGTLHSGQQTER
>Ornithorhynchus_anatinus
---------------MLGSASWLNTADQPMKTSTRRQRERSGGKQHLLSQARQQGRGQAVEILWSEKQTER
>Tauraco_erythrolophus
---------------MLGSATWLNTADRPMKTSIPSQRKESSKRQHLLFQAWKQERGQELETVESEKTTER
>Opisthocomus_hoazin
---------------MLGSAPWLNTADQPMKTSIPSQRKESSKRQHLLFQAWKQERGQELESVQSEKTTER
>Sturnus_vulgaris
---------------MLGSAPWLKTADQPMKTSIPSQRKESSTRQHLFFQAWKQERGQELESVESEKTTER
>Phaethon_lepturus
---------------MLGSATWLNTADQPMKTSIPSQRKESSKRQHLLFQAWKQERGQALESVESEKTTER
>Ceratotherium_simum_simum
---------------MLASAPRLNSADRPMKTSVLRQRKGSVRKQHLLSWAWQQGRGQVVEILQSEKQTER
>Manis_javanica
---------------MLASAPRLNSADRPMKTSVLRQRKGSVRKQHLLSWAWQQGRGQVLEILRSAKQTER
>Picoides_pubescens
---------------MLGSAPWLNTADQPMKTSIPSQRKESSKRQHLLFQAWKQERGRELESVESEKTTER
>Loxodonta_africana
---------------MLASAPRLNSADRPMKTSVFRQRKGSVRKQHLLAWALQQGRGQVVEILRSEKQTER
>Fulmarus_glacialis
---------------MLGSATWLNTADQPMKTSIPSQRKESSKRQHLLFQAWKQERGQELESVESEKTTER
>Melopsittacus_undulatus
---------------MLGSAPRLNTADQPMNTSVLSQRKESSKRQHLRFQAWKQERGQGLESGESEKTTER
>Ictidomys_tridecemlineatus
---------------MLASAPRLNSADWPMKTSVLRQRKGSVRKQHLLSWAWQQGRGQVVEILQSEKQTER
>Tupaia_chinensis
---------------MLASAPRLNSADRPMKTYALRQRKGSVRKQLLLPWAWQPGRGQVVEILQSEKQTER
>Trichechus_manatus_latirostr
---------------MLASAPRLNSADWPMKTSVFRQRKGSVRKQHLLSWAWQQGRGQVLEILQSEKQTER
>Aptenodytes_forsteri
---------------MLGSATWLNTADQPMKTSIPSQRKESSKRQHLLFQAWKQERGQELESVESEKTSER
>Gekko_japonicus
---------------MLGSASWLKTADRPMRTTIPKQRKGSNRSQHLFFQVWQQERGQEVETVDSEKKTER
>Tyto_alba
---------------MWGSATWLNTANQPMKTSIPSQRKESSKRQHLLFQAWKQERGQELESVESEKTTER
>Struthio_camelus_australis
---------------MLGSATWLNAADQPMKTSIPSQRQESSKRQHLFFQAWTQERGQELESVESEKTTER
>Trachemys_scripta
---------------MLGSASWLNTADQPMKTSILSQRKGSNRRQHLLFQAWQQERGREVEAGESEKKTER
>Tinamus_guttatus
---------------MLGSATWLNAADQPMKTSIPSQRKESSKRQHLLFQAWKQERGQDLESVESEKTSER
>Calypte_anna
---------------MLGSAAWLNTADQPMKTSIPSQRKESSKRQHLLFQAWKQERGHELESVESEKTTER
>Apaloderma_vittatum
---------------MLGSAAWLKTADQPMKTSIPSQRKESSKRQHLLFQAWKQERGQELESVESEKTTER
>Orycteropus_afer_afer
---------------MLASAPRLNSADRPMKTSVLRQRRGLVRKQHLLSWAWQQGRRQVVEILQSEKQTER
```

HG0018 ENSGALG00000004122 PNRC2

```
>Gallus_gallus
----------------------------------------------------MLGSAAWLNTADRPMKTSVPSQRKESSERQHLLFQAWKQERGQELESVESEKTTER
>Anas_platyrhynchos
-------------------------------------MQLDFFPWSFSAENPMLGSAPWLNTADHPMKTSIPSQRKESSERHHLLFQAWKQERGQELDGVESEKATER
>Chaetura_pelagica
----------------------------------------------------MLGSATWLDTADQPMKTSIPSQRKESSKRQHLLFQAWKQERGQELESVESEKTTER
>Apteryx_australis_mantelli
MCFQKRVQERHGGACDARGWLDGEGRLRSAGVSRGPVIWLRLLPWSLCAENPMLGSATWLKAAYQPMKTSIPSQRKESSKRQHLRFQAWKQERGQELESVESEKTTER
>Buceros_rhinoceros_silvestri
----------------------------------------------------MLGSAAWLKSADQPMNTSVRSQRQESGKRQHLLFQAWKQERGHELESVGSEKRTER
>Caprimulgus_carolinensis
----------------------------------------------------MLGSATWLNTADQPMKTSIPSQRKESSKRQHLLFQAWKQERGQEWESVESEKTTER
>Cariama_cristata
----------------------------------------------------MLGSATWLNTAHQPMKTSIPSQRKESSKRQHLLFQAWKQERGQELESVESEKTTER
>Odobenus_rosmarus_divergens
----------------------------------------------------MLASAPRLNSADRPMKTSVLRQRKGSGRKQHLLSWAWQQGRGQVVEILQSEKQTER
>Balaenoptera_acutorostrata_s
----------------------------------------------------MLASAPRLNSADRPMKTSVLRQRKGSIRKQHLLSWAWQQGRGQVVEILQSEKQTER
>Charadrius_vociferus
----------------------------------------------------MLGSATWLNTADQPMKTSIPSQRKESSKRQHLLFQAWKQERGQELESVESEKTTER
>Eptesicus_fuscus
----------------------------------------------------MLASAPRLNSADWPMKTSVLRQRRGSVRKQHLLSWAWQQGRGQVVEILQSEKQTER
>Pterocles_gutturalis
----------------------------------------------------MLGSATWLNTADQPMKTSIPSQRKESSKRQHLLFQAWKQERGQELESVESEKTTER
>Dasypus_novemcinctus
----------------------------------------------------MLTSAPRLNSADQPMKTSVFRQRKGSVRKQHLLTWAWQQGRGQVVEILQSEKQTER
>Colius_striatus
----------------------------------------------------MLGSAPWLNTAEQPMKTSIPSQRKESSERQHLLFQAWKQERGQELESVESEKTTER
>Columba_livia
----------------------------------------------------MLGSAPWLNTAHQPMKTSIPGQRKESSERQHLLFQAWKQERGQELERAESEKTTER
>Merops_nubicus
----------------------------------------------------MLGSATWLNTADQPMKTSIPSQRKESSKRQHLLFQAWKQERGQELESVESEKTTER
>Alligator_mississippiensis
----------------------------------------------------MWGSASWLDTADRPMKTSIPSQRKRSGKRQHLLFQAWQQKRGREVAAVESEKKTER
>Cuculus_canorus
----------------------------------------------------MLGSAPWLNTADQPMKTYICSPRKESSERQHLLFQAWKQERGQELESVESEKTTER
>Sarcophilus_harrisii
----------------------------------------------------------MLNSADQPMKTSILRQRNRSGRKQHLLSWAWQQGRRQVVEILQSAKQTER
>Galeopterus_variegatus
----------------------------------------------------MLASAPRLNSTDWPMKTSVLRQRKGSTRRQHLLSWAWQQGRGQVVEILQSEKQTER
>Monodelphis_domestica
----------------------------------------------------------MLNSVDQPMKTSILRQRNGSGRKQYLLSWAWQQGRRQVVEILQSAKRTER
>Macropus_eugenii
----------------------------------------------------------MLNSADQPMKTSILRQRNRSGRKQHLLSWAWQQGRRQVVEILQSAKQTER
>Aquila_chrysaetos_canadensis
----------------------------------------------------MLGSATWLNTADQPMKTSIPSQRKESSKRQHLLFQAWKQERGQELESVESEKTTER
>Gavia_stellata
----------------------------------------------------MLGSATWLNTADQPMKTSIPSQRKESSKRQHLLFQAWKQERGQELESVESEKTTER
>Mesitornis_unicolor
----------------------------------------------------MLGSAAWLNTADQPMKTSVPSQRKESSERQHLLFQAWKQERGQGLESVESEKTTER
>Condylura_cristata
----------------------------------------------------MLASAPRLNSADRPMKTYVLRQRKGSVRKQHLLSWAWQQGRGQVVEILQSEKQTER
>Oryctolagus_cuniculus
----------------------------------------------------MLASAPRLNSADRPMNTSVLRQRRGSARKQHLGPWAWQRGRGQAVGTLHSGQQTER
>Ornithorhynchus_anatinus
----------------------------------------------------MLGSASWLNTADQPMKTSTRRQRERSGGKQHLLSQARQQGRGQAVEILWSEKQTER
>Tauraco_erythrolophus
----------------------------------------------------MLGSATWLNTADRPMKTSIPSQRKESSKRQHLLFQAWKQERGQELETVESEKTTER
>Opisthocomus_hoazin
----------------------------------------------------MLGSAPWLNTADQPMKTSIPSQRKESSKRQHLLFQAWKQERGQELESVQSEKTTER
>Serinus_canaria
----------------------------------------------------MLGSAPWLKTAHQPMKTSIPSQRKESSTRQHLFFQAWKQERGQELESVESEKTTER
>Phalacrocorax_carbo
----------------------------------------------------MLGSATWLNTADQPMKTSVPSQRKESSKRQHLLFQAWKQERGQELESVESEKTTER
>Ceratotherium_simum_simum
----------------------------------------------------MLASAPRLNSADRPMKTSVLRQRKGSVRKQHLLSWAWQQGRGQVVEILQSEKQTER
>Manis_javanica
----------------------------------------------------MLASAPRLNSADRPMKTSVLRQRKGSVRKQHLLSWAWQQGRGQVLEILRSAKQTER
>Picoides_pubescens
----------------------------------------------------MLGSAPWLNTADQPMKTSIPSQRKESSKRQHLLFQAWKQERGRELESVESEKTTER
>Nomascus_leucogenys
----------------------------------------------------MLASAPWLNSADRPMKTSVLRQRKGSVRKQHLLSWAWQQGRGQVVEILQSEKQTER
>Loxodonta_africana
----------------------------------------------------MLASAPRLNSADRPMKTSVFRQRKGSVRKQHLLAWALQQGRGQVVEILRSEKQTER
>Fulmarus_glacialis
----------------------------------------------------MLGSATWLNTADQPMKTSIPSQRKESSKRQHLLFQAWKQERGQELESVESEKTTER
>Nestor_notabilis
----------------------------------------------------MLGSAPWLNTADRPMNTSVPSQRKESSKRQHLRFQAWKQERGQDLESGESEKTTER
>Ictidomys_tridecemlineatus
----------------------------------------------------MLASAPRLNSADWPMKTSVLRQRKGSVRKQHLLSWAWQQGRGQVVEILQSEKQTER
>Tupaia_chinensis
----------------------------------------------------MLASAPRLNSADRPMKTYALRQRKGSVRKQLLLPWAWQPGRGQVVEILQSEKQTER
>Trichechus_manatus_latirostr
----------------------------------------------------MLASAPRLNSADWPMKTSVFRQRKGSVRKQHLLSWAWQQGRGQVLEILQSEKQTER
>Aptenodytes_forsteri
----------------------------------------------------MLGSATWLNTADQPMKTSIPSQRKESSKRQHLLFQAWKQERGQELESVESEKTSER
>Gekko_japonicus
----------------------------------------------------MLGSASWLKTADRPMRTTIPKQRKGSNRSQHLFFQVWQQERGQEVETVDSEKKTER
>Tyto_alba
----------------------------------------------------MWGSATWLNTANQPMKTSIPSQRKESSKRQHLLFQAWKQERGQELESVESEKTTER
>Struthio_camelus_australis
----------------------------------------------------MLGSATWLNAADQPMKTSIPSQRQESSKRQHLFFQAWTQERGQELESVESEKTTER
>Trachemys_scripta
----------------------------------------------------MLGSASWLNTADQPMKTSILSQRKGSNRRQHLLFQAWQQERGREVEAGESEKKTER
>Tinamus_guttatus
----------------------------------------------------MLGSATWLNAADQPMKTSIPSQRKESSKRQHLLFQAWKQERGQDLESVESEKTSER
>Calypte_anna
----------------------------------------------------MLGSAAWLNTADQPMKTSIPSQRKESSKRQHLLFQAWKQERGHELESVESEKTTER
>Apaloderma_vittatum
----------------------------------------------------MLGSAAWLKTADQPMKTSIPSQRKESSKRQHLLFQAWKQERGQELESVESEKTTER
>Orycteropus_afer_afer
----------------------------------------------------MLASAPRLNSADRPMKTSVLRQRRGLVRKQHLLSWAWQQGRRQVVEILQSEKQTER
```

HG0019 ENSG00000178397 FAM220A

>Homo_sapiens
-----MAA----ALSGLAVRLSRSAAARSYGVFCKGLTRTLLIFFDLAWRLRINFPYLYIVASMMLNVRLQVHIEIH--
>Canis_lupus_familiaris
-----MAA----ALSGLAVRLSRSAAARSYGVFCKGLTRTLLIFFDLAWRLRINFPYLYIVASMMLNVRLQVHIEIH--
>Hynobius_retardatus
------MA----ALAVLAARVSQSAAVRSYCVFSKGLSRTLLIFFDMAWRLRINFPYLYVIASMMLNVRLQVHIEIH--
>Balaenoptera_acutorostrata_s
---------------------------------------------------------------MMLNVRLQVHIEIH--
>Pteropus_vampyrus
-----MAA----ALSGLALRLSRSAAARSYGVFCKGLTRTLLIFFDLAWRLRINFPYLYIVASMMLNVRLQVHIEIH--
>Echinarachnius_parma
MAATYSSPWLLSQYQERRERRLKLAALQTVQP-FPLLTRTLNFFFNLAYQCRLRIPTIIFVGLLAFDVRLQLQINVVQI
>Latimeria_menadoensis
------MA----GLTALVVRFSGSAAARSYGVFSKNLTRTLLIFFDLAWRLRLRFPYLYIITSLMLNVRLQVHIEIH--
>Sarcophilus_harrisii
-----MAA----ALSALALRLSRSAAARSYGVFCKGLTRTLLIFFDLAWRLRINFPYLYIVASMMLNVRLQVHIEIH--
>Monodelphis_domestica
-----MAA----ALSALALRLSRSAAARSYGVFCKGLTRTLLIFFDLAWRLRINFPYLYIVASMMLNVRLQVHIEIH--
>Arachnocampa_luminosa
----------------MIFNYLAKSLLNIYPSVSGRLSPLMLFCFNFAWRAFTLFPYMFLATTMATGFQIRVEIEFE--
>Strongylocentrotus_purpuratu
-----MAPWMIYSRFQRHHERKLTAVIRTPPP-FPLLTRTLSFFFNLAYQCRMRIPTIVFVGLLAFDVRLQLQINVVQI
>Marthasterias_glacialis
---------MFKMLGMMRDPDSMIQVGASMPV-RPFLSRTLTFFFDMAFRCRFKLPSIVAFVLFAFDIQLQLQINVNH-
>Condylura_cristata
---------------------------------------------------------------MMLNVRLQVHIEIH--
>Oryctolagus_cuniculus
---------------------------------------------------------------MMLNVRLQVHIEIH--
>Lingula_anatina
------------MLE---NYIP-------QRMGPSTLSDVCMFFFKLAWQSRIRLPVILTVVFMILDTRLQLEINININ
>Crassostrea_gigas
--------------------MVLQTLNQSYASFARSLSRILLFFFRFAWQTKFSLPYLLNSTMLALDFRVRVVILL---
>Equus_asinus
-----MAA----ALSGLAVRLSRSAAARSYGVFCKGLTRTLLIFFDLAWRLRINFPYLYIVASMMLNVRLQVHIEIH--
>Manis_javanica
---------------------------------------------------------------MMLNVRLQVHIEIH--
>Brachionus_manjavacas
----------------MIFNYLAKSLLNIYPSVSGRLSPLMLFCFNFAWRAFTLFPYMFLATTMATGFQIRVEIEFE--
>Heterocephalus_glaber
-----MAA----VLSGLAVRLSRSAAARSYGVFCKGLTRTLLIFFDLAWRLRINFPYLYIVASMMLNVRLQVHIEIH--
>Tupaia_chinensis
---------------------------------------------------------------MMLNVRLQVHIEIH--
>Echinaster_spinulosus
-------------MLRISDPDSFIQVCTVLPV-RNVMSRTLTFFFDMAFRCRFKLPSIVAFVLFAFDIQLQLQINVNH-
>Takydromus_sexlineatus
MAVTAAAT----LLSALVARFSQSAAARSYGVFCKGLTRTLLIFFDLAWKLRINFPYLYIVASMMLNVRLQVHIEIH--
>Sphaerechinus_granularis
----MMAPWLMYLRYQRSQQRKVQ----TVSP-FPLLTRTLSFFFNLAYQCRMRIPTIVFVGLLAFDVRLQLQINVVQI
>Pelodiscus_sinensis
-----MAA----DLSALAARLSQSAAARSYGVFCKGLTRTLLIFFDLAWKLRINFPYLYIVASMMLNVRLQVHIEIH--
>Villosa_lienosa
------------------MLKLFSALWSTYSNFTRHLTPILMFFFRFAWLSRFELPGILTTVFLALNIRVTVEIHM---

HG0020 FBgn0035436 CG12016

```
>Drosophila_melanogaster
------------------------------------------------MASNAQLGKIILISAIAVFFYYFFWVAVLPFMLIDEGNPIRLFFPPLKYAFIVPTVFGVIFLGGIAAFSFYHIWSLRVKRD-----
>Meinertellus_cundinamarcensi
------------------------------------------------MISDAAAGRLAIVVGSSIFSYYTLWVIVLPFVDREHA--VRDFFPDVTYALVLPNILGVVFIGSLVYYACASLLKERCTS------
>Macrobrachium_nipponense
-----------------------------------------------MYGHDRVIGKCVLSVTGVALLYYLTWVVCLPFADDDSS--IRLLFPDPLVALIVPTIVFVFLITLTGFYIGHSLYVDYTTTSISS--
>Forficula_auricularia
-------------------------------------------------MSDPIYGKLIVISSVIVLIYFSCWVFLTPFADATDY--FFKVFPSLRYALIIPAVSGLIFVGIISLFILCSVI------------
>Daphnia_pulex
----------------------------------------------MAVGHDVLIGKILLFLCVIFVLYYTFWTICLPLLEPNFF--AHQYFPDSFYAIGVPLSVLSVMVTVLSVYALWLMKFQ----------
>Aposthonia_japonica
-------------------------------------------------MEDATLGKVILFCCGLFFIYYSMWVLVLPFVDENHV--IRRFFPNVFHAIFTPAVLGLSFISSLIFFTIYAMTFEQS--------
>Eurylophella_sp._AD-2013
------------------------------------------------MVHTRTVGQLILLISSLLLLYYTLWVVALPFADSDHF--LVQLFPSRRYAIIVPAISGLTFIGSLAVFTVWSMYSC----------
>Meganyctiphanes_norvegica
-----------------------------------------------MLGCDQLIGKCVLCVFLSLSLYYIVWVVGLPFTDEDSF--LRALFPDPIYAIVGPTLVFVALLLLNGVYMLKAYFKVQKLKAQ----
>Grylloblatta_bifratrilecta
-----------------------------------------------MVVTDATCGKAILILTLLPFCYYTVWVLILPFTDAEDV--LNAVFPSNRYALVVPALAGLCFVGGLVAFTLYSLAS-----------
>Xenophysella_greensladeae
----------------------------------------------MGALTDSTLGALLLTGASVFLLYYTLWVILSPFYEP-----ISSLFPPIRYALLVPAFTGLTFISSLIFFTVWALLSS----------
>Sceliphron_caementarium
-----------------------------------------------MFDSDREKGKLILIVTIILFLYYTIWIIGLPFIDDDRL---RSLFCSHDFALVVPAVSGLCFIGGLVLFTLYHVRPYLLSDRASKIQ
>Asellus_aquaticus
-----------------------------------------------MYGKDYMLGKVLLISSIVFVTYYLSWTILLPFVKDDSV--IRIFFPGPLLAILIPTSLFLFLLVTTFGYLLYTVAVNKKELL-----
>Pararge_aegeria
-----------------------------------------------------------------------------------------------------------------MSFTLYHIYPYL---------
>Metallyticus_splendidus
-------------------------------------------------MNDSALGKILLIASCLFFCYYTTWVIVLPFSDRDHI--INILFPPVQHALLVPACMGVAFIGSLIIYTVYNIKQ-----------
>Tanzaniophasma_sp._AD-2013
-------------------------------------------------MNDSTLGTFVVIFSVLFFMYYTFWVLVLPFVEQDHV--IHTIFTSVDFALMVPAVAGLFFIGGLVIFTLFALRH-----------
>Euroleon_nostras
-------------------------------------------------MKDVVLGRTLLIFSGIFVLYYTFWVVVLPLLPLADDNTFLLFFPDVTYALLLPAILGTLFIGSLSAFTLYHVLPHLKLKKK----
>Calopteryx_splendens
-----------------------------------------------MGISDAQLGKIILVLSASLFVYYTLWLWVPPFISPSSFNTFPTVFPPTRYALILPAISGLLFIGGLLVFTIAALQNA----------
>Gryllotalpa_sp._AD-2013
------------------------------------------------MLSDATLGKIILFSSCSFFIYYTAWVMFLPFADNIQV--IKS-FPPIWIALLVPAIMGLLFVGALMMYTVHAIRRK----------
>Ramulus_artemis
MILLCLLKKILRSLKKSILSVEHCLLHAFFLKYQQLFNFINCDTEYVIMVSDSTVGKLVLTATCIFFTYYSFWVLVLPFTDSEDK--INSFFPSVWYALVIPAVMGLIFIGSLVVFTLCAVETRGRS-------
>Cosmioperla_kuna
-------------------------------------------------MSDALIGKVILIASCTFFCYYTAWTLLLPFADGDST--INSLFPSIWYALFVPAYMGLAFIGSLMVFTYIALRT-----------
>Liposcelis_bostrychophila
-----------------------------------------------------MLAKLILALAVSFFSWYTFCVIALPFFEPTHL--IVSYFPGPRYTLIIPALFGLTFFGSLLIFTIAVLWKEKP--------
>Trichomycterus_areolatus
-----------------------------------------------MILPDRILGKSIVLITTFAFVYLVIWLTLLPFTDFEKDHWLYKGFLDREYALILCISWGFIFIGSLMLFTLYQIYPYL---------
>Thrips_palmi
-------------------------------------------------MNDKTVGSLLIALSASFSFYYSVWVLGLPFFDSDHF--IHSLFPAVEFALLVPALMGLTFIGSLIIFCIIELRKAGAASK-----
>Hydroptila_sp._AD-2013
-----------------------------------------------MIISDSNLGKLIVLAAVSIFTYLVIWLTFLPFTDFDEDHWLYKGFLPREYALIFCIGWGFLFIGSLMGFTLYHLYPFLK--------
>Thermobia_domestica
-----------------------------------------------MTIADETLGKLLLIGSLLFTLYYTFWIIILPFVDPDHI--LHDLFPPVTYGVMTPAVMGLCFIGSLFVFTVVCFKSEK---------
```

HG0021.1 ENSDARG00000068708 ifrd1

```
>Danio_rerio
MYRSRSKSNTGIAFNTFLFTVPLEYTQTKRHYSKNWSCIG
>Amia_calva
MCRFRSQTFTGIAFYSYLLTVPLDGFHLERQYSKNWSSIG
>Channa_punctata
MYRCSQQSFSRIAFDCFLSTVQREYFHFTRQYNKSWSCIG
>Anguilla_japonica
MYRFRPQIFTGTAFNSYLLTLLLKGFYLKRHCTRNWSSIG
>Podocoryna_carnea
MCRFCSPTFTRIAFYSSLLAVPFEDFQLMRHYSKNCLSVG
>Menidia_menidia
MYRLGQQTSFKIAFDFFLSTVPREHFHFTRQYNKSRSCIG
>Opsanus_beta
MYRLGLQLIAKITFTSIFCTVPREYSYFTRQYSKSWSCIG
>Oryzias_melastigma
MYRLGQQFISKIAFDYFLSTVPREHFTFTRQYNKSWSCIG
>Tripterygion_delaisi
MHRLGQQSFSRIAFNSFLPTVPREYFLFTRQYNKGWSCIG
>Astyanax_mexicanus
MYRFRSQALIGIAFNSLLFTVPLDDIQFKRRYSKNWSCIG
>Neolamprologus_brichardi
MYRLGPQSISKIAFDFFLSTVPREYFHFTRQYNKGWSCIG
>Clupea_harengus
MYRLRSKSFTGIALNTFFAAVPLEEIYFRRQYSKNWSSIG
>Fundulus_heteroclitus
MYRLGRQSISKIAFNFFLSTVPRERFHFTRQYNKSWSCIG
>Esox_lucius
MYRFRTQIFSRTAFNSFLTTVPREDCQFMRQFNKNWSCIG
>Gadus_morhua
----------------------------MRQYNKGWSCIG
>Acanthogobius_hasta
MYRLAQQVFSKIALISSLPAVPREYFHFTRQYNKGWSCIG
>Apteronotus_leptorhynchus
MYRFRLQAFIGIAFYSLPFTVALEDGKFKRHYSKNWSCIG
>Plecoglossus_altivelis
MYRLGSQSFSRIAFNSFLSTVPREDLHFIRQYNKNWSCIG
>Perca_flavescens
MYRLGQQCIPKIAFNSFLSAVPREYFHFTRHYNKSWSCIG
>Cynoglossus_semilaevis
MYRLGQQFISKIAFNSFLSTVPREYFHFTRRCNKNWSCIG
>Coregonus_clupeaformis
MYRLRPQSFSRIAFNSFLSTVPREDCQFMRQFNKNWSCIG
>Lepisosteus_oculatus
MCRFRLQTFTGIAFYSYLSTVPCGDFHLERQYSKNWSSIG
>Ictalurus_punctatus
MYRSRSQAFIGIAFNPLLFTVPFEDFQFKRHYSKNWSCIG
>Sparus_aurata
MYRLGQQTFSKIAFDSFLSTKPREYFHFTRQYSKSCSCIG
>Takifugu_rubripes
MYRLGQLTISKIAFNSFLSAVPREYFHFTRQYNKSWSCIG
```

HG0021.1 ENSG00000006652 IFRD1

```
>Homo_sapiens
------------------------------------------MYRFRSQLFTGISAAATAHSYP---RRFSTLLLAEDSPLSRPPHRR-TS--KKCSSIG
>Xenopus_laevis
------------------------------------------MYRLHSQAFTGVSV----TSTF---RHFYPIPIEDS---KLIPERR-FS--KKCSSIG
>Apteryx_australis_mantelli
------------------------------------------MHRFRSRAFTGISAFATDSS-FACFRQSPPLPRSGK----SASQRR-CSKKRSCSSIG
>Leptonychotes_weddellii
------------------------------------------MYRFRPQLFTGISAAATAHSYP---RRFSPLLLAEDSPLSRPPHRR-TS--KKCSSIG
>Ambystoma_mexicanum
------------------------------------------MYRFRSQALIGFPWSRRRCSPA---RRCSPLPLGDL---NLIPERR-FS--KKCSSIG
>Physeter_catodon
------------------------------------------MYRFRSQLFTGISAAATAHSYP---RRFSPLLLVENSPLSRPPHRR-TS--KKCSSIG
>Rhinolophus_ferrumequinum
------------------------------------------MYRFRSQLFTGISAAATARSHP---RRSSPLLLAEASPLSRPPHRP-TS--KKCSSIG
>Dasypus_novemcinctus
------------------------------------------MCRFRSQLFTGISATATTHSYP---RRFSPLLLVEDSPLSRPPHRR-TS--KKCSSIG
>Latimeria_menadoensis
------------------------------------------MIRSGSQTVTRIA------SCY---HHLLAVPF---GDLHLVPERR-YS--KKCSSIG
>Sarcophilus_harrisii
------------------------------------------MYRFRSQLFTGISAAAIAHSSP---RRFSPLLLAEDLSLSQPPHRR-TS--KNCSSIG
>Galeopterus_variegatus
MCRDWLVRVRVPPTCGDDRRRSCYLRRRGLDRPALAPALERNMYRFRSQLFTGISAAATAHSYP---RRFSPLLLAEDSPLSRPPHRR-TS--KKCSSIG
>Monodelphis_domestica
------------------------------------------MYRFRSQLFTGISAAAIAYSSP---RRFSPLLLAEDLSLSHPPHRR-TS--KKCSSIG
>Coturnix_japonica
------------------------------------------MHRLRSRAFTGISAIAAGSS-FACLRQSPPLPRPEK----SASQRL-WNKKRSCSSIG
>Sorex_araneus
------------------------------------------MYRFRSQLFTGISAAATAHSHP---RRFSPLLLAEDSPLCRPPHRR-TS--KKCSSIG
>Ochotona_princeps
------------------------------------------MYRFRSQLFTGISAAATAHSYP---RRFSPLLLAQDSPLSRPTHRR-TS--KKCSSIG
>Elephantulus_edwardii
------------------------------------------MHRFRSQLFTGISAAATAHSHP---RRFSPLLLAEDSPLSLQPHRR-TS--KKCSSIG
>Equus_przewalskii
------------------------------------------MYRFRLQLFTGISAAATAHSYP---RRFSPLLLAEDSPLSRLPHRR-TS--KKCSSIG
>Manis_javanica
------------------------------------------MCRFRSQLLTGISAAATTHFYP---RRCSPLLLAEDSPFSRPPHRR-TS--KKCSSIG
>Loxodonta_africana
------------------------------------------MYRFRSQLFTGISAAATAHSHP---RRFSPLLLVEYSPLSRLPHRR-TS--KKCSSIG
>Marmota_monax
------------------------------------------MYRFRSQLFTGISAAATAHFYP---RRFSPLLLAEDSPLSRPPHRR-TS--RKCSSIG
>Tupaia_chinensis
------------------------------------------MYRFRSQLFTGISAAATAHSYP---RRFSPLLLAEDSSLSRPPHRR-TS--KKCSSIG
>Trichechus_manatus_latirostr
------------------------------------------MYRFRSQLFTEISAAATAHSHP---RRFSPLLLVEDSLLNRLPHRR-TS--KKCSSIG
>Anolis_carolinensis
------------------------------------------MYRFRSRASTGISAAATALPIIARLPQSHPLPRSGK----PASQRRFYSKKRSCSSIG
>Struthio_camelus_australis
------------------------------------------MHRFRSRAFTGISAFATDSS-FACFRQSPPLPRSGK----SASQRR-CSKKRSCSSIG
>Chelonia_mydas
------------------------------------------MHRFCSRAFTGISTATTALS-FACLRQSPPLPRSGK----SAPQRR-CSKKRSCSSIG
```

HG0021.1 ENSGALG00000009448 IFRD1

```
>Gallus_gallus
-----------------------------------------------------MHRLRSRAFTGISAIAASSSFACLRQSPPLPRPEKS----ASQRRW-NKKRSCSSIG
>Xenopus_laevis
-----------------------------------------------------MYRLHSQAFTGVSVTS----T--FRHFYPIPIED---SKLIPERRF---SKKCSSIG
>Apteryx_australis_mantelli
-----------------------------------------------------MHRFRSRAFTGISAFATDSSFACFRQSPPLPRSGKS----ASQRRC-SKKRSCSSIG
>Canis_lupus_familiaris
-----------------------------------------------------MYRFRSQLFTGISAAATAHSY--PRRFSPLLLAEDSPLSRPPHRRT---SKKCSSIG
>Hynobius_chinensis
-----------------------------------------------------MHRFRSQALTGVSSRC-------PPHTAPLPLGD---LHLIPERRF---SKKCSSIG
>Lipotes_vexillifer
-----------------------------------------------------MYRFRSQLFTGISAAATAHSY--PRRFSPVLLVENSPLSRPPHRRT---SKKCSSIG
>Calidris_pugnax
-----------------------------------------------------MHRLRSRAFTGISAFAAGSSSACFRQSPPLPRPEKS----ASLTRW-NKKRSCSSIG
>Rousettus_aegyptiacus
-----------------------------------------------------MCRFRSQLFTGISAAAAARSH--PRRSSPLSLATASPLSRPPHRRA---SRKCSSIG
>Dasypus_novemcinctus
-----------------------------------------------------MCRFRSQLFTGISATATTHSY--PRRFSPLLLVEDSPLSRPPHRRT---SKKCSSIG
>Alligator_mississippiensis
-----------------------------------------------------MHRFRSRASTGISAAS-ASSIACRRQSPPLPRPTKP----AAHRRC-SKKRSCFSIG
>Cuculus_canorus
-----------------------------------------------------MHRLRSRASTGISAFAADSSSSCFRQSPPLPRPGKS----TSQKRW-NKKRSCSSIG
>Sarcophilus_harrisii
-----------------------------------------------------MYRFRSQLFTGISAAAIAHSS--PRRFSPLLLAEDLSLSQPPHRRT---SKNCSSIG
>Galeopterus_variegatus
-----------MCRDWLVRVRVPPTCGDDRRRSCYLRRRGLDRPALAPALERNMYRFRSQLFTGISAAATAHSY--PRRFSPLLLAEDSPLSRPPHRRT---SKKCSSIG
>Monodelphis_domestica
-----------------------------------------------------MYRFRSQLFTGISAAAIAYSS--PRRFSPLLLAEDLSLSHPPHRRT---SKKCSSIG
>Falco_peregrinus
-----------------------------------------------------MHRLRSRASTGISAFAAGSSSACFRQSPPLPRFGKS----ASQRRW-SKKRSCSSIG
>Sorex_araneus
-----------------------------------------------------MYRFRSQLFTGISAAATAHSH--PRRFSPLLLAEDSPLCRPPHRRT---SKKCSSIG
>Ochotona_princeps
-----------------------------------------------------MYRFRSQLFTGISAAATAHSY--PRRFSPLLLAQDSPLSRPTHRRT---SKKCSSIG
>Elephantulus_edwardii
-----------------------------------------------------MHRFRSQLFTGISAAATAHSH--PRRFSPLLLAEDSPLSLQPHRRT---SKKCSSIG
>Junco_hyemalis
-----------------------------------------------------MHRLRSRASTGISAFAAGSPSACSRQSPPLPRPEKS----ASQRRW-SKKRSCSSIG
>Ceratotherium_simum_simum
MSRAKWDSGGECYGYWLGRVRVPPTDGDDRRRSCYLRRRGLDRPALAPALKRNMYRFRLQLFTGISAAATARSY--PRRFSPLLLAEDSPLSRRPHRRT---SKKCSSIG
>Picoides_pubescens
-----------------------------------------------------MHRLRSRASTGISAFAACLSSVCFRQSPPLPRTGKS----TSQRRW-SKKPSCSSIG
>Otolemur_garnettii
-----------------------------------------------------MYRFRSQLFTGISATATAHSY--PRRFSPLLLTEDSPLSRPPHRRT---SKKCSSIG
>Melopsittacus_undulatus
-----------------------------------------------------MHRLRSRASTGISAFAASSSSACFRQSPPLPRTAKS----TSQRRW-SKKRSCSSIG
>Marmota_monax
-----------------------------------------------------MYRFRSQLFTGISAAATAHFY--PRRFSPLLLAEDSPLSRPPHRRT---SRKCSSIG
>Tupaia_chinensis
-----------------------------------------------------MYRFRSQLFTGISAAATAHSY--PRRFSPLLLAEDSSLSRPPHRRT---SKKCSSIG
>Takydromus_sexlineatus
-----------------------------------------------------MYRFRSRASTGISAAATALPIACFPQSHPLPRSGKP----ASQRRFYNKKRSCSSIG
>Struthio_camelus_australis
-----------------------------------------------------MHRFRSRAFTGISAFATDSSFACFRQSPPLPRSGKS----ASQRRC-SKKRSCSSIG
>Trachemys_scripta
-----------------------------------------------------MHRFCSRAFTGISTATTDSSFACLRQSPPLPRSGKS----APQRRC-SKKRSCSSIG
```

HG0021.2 ENSDARG00000036811 ifrd2

>Danio_rerio
MVESGAATCTILRPKHFQKHLPSQPGIRHRLPGDNHR
>Acipenser_sinensis
MVEFTRATCSDIQ--LLQTACLSQPRIRYILPGDNHR
>Menidia_menidia
MVDLATVACSELRPKHFEKYLPSQPGVRYRLPGDNHR
>Oryzias_latipes
MVDLAKGACSKLRHKHFETYLTSQLSVRYRLPGDNHR
>Trachinotus_blochii
MVDFATAACSKLRPKHFKKYLQSQPGSRYRLPGDNHR
>Pygocentrus_nattereri
MVESAAATCTILRPKHFEKSLPSQPGIRYRLPGDKHR
>Oreochromis_niloticus
MVDLATTACSKLRPKRFEKYLPSHPSIRYRLPGDNHR
>Alosa_alosa
MVESAAATCTIPRPKLSEKYLSSQPGIRYWLPGDNHR
>Cyprinodon_variegatus
MVDLATAACSKLRPKHFEKYLPSQPGVRYRLPGDNHR
>Esox_lucius
MVEFGTATCSILLPKHFEKYCISQPGIRYRLPGDNHR
>Gadus_morhua
MVDFATATCSKLRPNYLLISLSSQPGIKYRLPGYNQR
>Acanthogobius_hasta
MVISETAACSKLRPKHCPRTSLSQPGIRYRLPGDNHR
>Plecoglossus_altivelis
MVEFATATCSILRPKHFEKYLPSQPGIRYMLPGDNHR
>Scleropages_formosus
MVEFAAATCAILRPKHFENYIPSQPGIRYWLPGDNHR
>Lateolabrax_japonicus
MVDFATTSCSELRPKHFGKDLLSQPGIRFRLPGDTHR
>Trematomus_bernacchii
MVDLATTACSLLRPKHFGKTFISPPGIRYRLPGDNHR
>Cynoglossus_semilaevis
MVDFVTAACSKLRPKLSEKISLSQPGIRYRLPGDNHR
>Salmo_salar
MVEFETVMCSILCPKHFEKYFLSQPGIRYRLPGDNHR
>Lepisosteus_oculatus
MVDLGAAACTILRPKRSEKYLPSQPGIRYRLPGDNHR
>Ictalurus_punctatus
MVEAAAATCTNLRPKHFGKYLPSQPGIRYWLPGDKHR
>Takifugu_rubripes
MVDLATAACSKLRPKHFEKYLLSQPGSRYRLPGDNHR

HG0022 FBgn0061359 CG33671

```
>Drosophila_melanogaster
MSKYDSKYLEEKLKRELQTEYVSVTDESDGCGGKFSAVIVSPAFS-GKTLLQKHRLVNSTLAEELKEIHAFSQKSYTPEEWEKVKAQ---
>Talitrus_saltator
MSEYTAEYIKAKLEKELDSTHVSVVDESDGCGGKFSAIIVSSAFQ-GKPLLQRHRLVNSTLAEELKQIHAFSMKTYTPEQWEQQQSKEQ-
>Meinertellus_cundinamarcensi
-MVFTEQYIREKLVKELEAVHVEVIDESDGCGAKFNVLIVSFLFE-GKPLLQRHRLVNSILEEELKTIHAFSQKTLTPEQWEKQKSS---
>Prorhinotermes_simplex
-MSYTADYIKSKLLKELEAVYVEVQDESDGCGAKFSVVIVSDKFD-GKPLLQRHRLVNSILEEELKSIHAFSQKTLTVQQWEKQKSS---
>Leptinotarsa_decemlineata
-MTYTEEHLRNKLIKELEASHVEVIDESDGCGGKFSCVIVSEVFK-GKPLLQRHRLVNSVLSEELKSIHAFSQKTYTPEQWEQQK-----
>Farfantepenaeus_aztecus
MTTYTEDYIRDKLTKELDAKHVSVEDMSDGCGAKFNVVVVSPKFE-GKPLLQRHRLVNGALEEELKTIHAFSMKTLTPEQWEKQGQ----
>Meganyctiphanes_norvegica
MTSYTEEYIKEKLTRELEASHVAVEDMSDGCGAKFSVVIVSPKFS-GKPLLARHRLVNSTLEEELKEIHAFSMKTLTPEQWQQQQQ----
>Acyrthosiphon_pisum
-MTINEDLLKSKLTVALDASYVSVNDDSDGCGAKFSVVVVSDKFN-GKTLLQRHRMVNEVLAEELKEIHAFSQKTLTTEQWNNLNKV---
>Ganaspis_sp._G1
-MPYTAEHIRNKLITELKAEHVEVVDESDGCGGKFSAIIVSEQFK-GKPLLQRHRLVNAALAEELKDIHAFSQKTLTPEQWEKEQS----
>Triodia_sylvina
-MVYTEEYLKEKLVRGLDAVHVEVVDESDGCGGKFSVVVVSDKFQ-GKPLLARHRLVNSVLQEELKEIHAFTQKTYTIEQWSNQTNAVNE
>Mantis_religiosa
-MPLTEEYIKNKLMKELEATHVEVQDESDGCGAKYNVLIVSDKFE-GKPLLQRHRLVNAVLEEELKLVHAFSQRTLTPAQWEEQKAA---
>Boreus_hyemalis
-MKYTEEYIREKLIKELNAEHVEVTDDSDGCGGKFSAIVVASIFQ-GKALLARHKLVNAALAEELKTIHAFSQKTFTPQQWQEKNTS---
>Corydalus_cornutus
-MSVTEEQIKQKLLRELDATHVKVVDESDGCGAKFSVVIVSDKFV-GKSLLQRHRLVNTTLQEELKNIHAFSQKTFTIDQWEKENSSN--
>Conwentzia_psociformis
-MSYTEGHIKDKLTKELDAVHCEVIDESDGCGGKFSAIIVSDKFN-GKPLLQRHRLVNAVLQEELKSIHAFSQKTFTREQWEAAKKDSH-
>Cordulegaster_boltonii
MSVYTTNYIKEKLRTGVEATHVEVQDDSDGCGAKFSVLVVSPKFE-GLPLLQRHRLVNSALAEELPNIHAFSQKTFTPKQWDDQKNKDAN
>Teleogryllus_commodus
-MSYTEEYVKNKLLKELEATHVEVEDLSDGCGAKFNAIIVSSKFE-GKPLLQRHRLVNSILSEELKSIHAFSQKTMTPKEWEKQNSQ---
>Gasterosteus_aculeatus
------------------------------------MLGNTSKIKSRRQYKLITLQINSILEEELKTIHALTLKTLTPHQWETRQK----
>Menopon_gallinae
-MPYTKDYIQSKLEKELDPVYIKVEDESDGCGAKFSAIIVSPKFQ-GKPLLQRQRLVNSILSEELKDIHAFSQRTLTPEQWEKENS----
>Perla_marginata
-MPYTENDIKTKLIKELDATHVDVRDDSDGCGAKFSVVIVSEKFE-GKPLLARHRLVNSVLEEELKEIHAFSQKTLTPQQWDQQKSA---
>Liposcelis_bostrychophila
-MPYTEDYIQNKLQKELDAIYVKVEDQSDGCGAKFSAIIVSPKFE-GKPLLQRHRLVNSVLEEELKDIHAFSQRTLTPQQWEQQNSS---
>Ctenocephalides_felis
-MSYTKQYLEEKLKTQLQAEHVEVVDLSDGCGDKFSAVIVSSVFE-GKPLLARHKLVNSTLAEELKTIHAFSQKPLTPEQWKAQQ-----
>Thrips_palmi
-MSISVDHIKDKLSKELQPVHIEVVDESDGCGGKFNVIVVSDKFE-GKPLLQRHRLVNSILEEELKVIHAFSQKTLTPEQWAKQSSQ---
>Hydroptila_sp._AD-2013
--MVSEEEIRSKLLTQLEAAHVEVTDESDGCGGKFSAVIVSDKFI-GKPLLARHRLVNAALEEELKQIHAFSQKTFTVEQWEKSKAL---
>Thermobia_domestica
-MAYTEDYIKDKLIKELEAIHVEVEDISDGCGAKFNAVIVSPKFE-GKPLLQRHRLVNSVLEEELKTIHAFSQKTLTPEQWEKQKQT---
```

HG0023.1 ENSG00000175197 DDIT3

```
>Homo_sapiens
-MLKMSGW----QRQ-----------SQNQSWNLRRECSRRKCIFIHHHT
>Xenopus_laevis
-MINMSPKSLNVQPQTSS---RCRFIRQKRTDPTWRKCWRRKTIFIHHHT
>Ursus_americanus
-MLKMSGW----QPQ-----------SQNQSRNLRRECSRRKCIFIHHHT
>Hynobius_retardatus
-MVNMSLRSPSE-PSTAARPDPPRALQRPQAGCRRRKCWRTKTIFIHHHT
>Tursiops_truncatus
-MLKMSGW----QRQ-----------SQNQSRNLRRECSRRKCIFIHHHT
>Rousettus_aegyptiacus
-MLKMSGW----QRQ-----------SQNQSRKLRRECSRRKCIFIHHHT
>Dasypus_novemcinctus
MMLKMSGW----QRQ-----------SQSQSRNLRRECSRRKCIFIHHHT
>Alligator_mississippiensis
MQARMSR--------------GQRNRS-GDERWPRRKSSRRKCIFIHHHT
>Sarcophilus_harrisii
MMMKMTWW----QRQ-----------SQNQSRDPRRECSRRKSIFIHHHT
>Galeopterus_variegatus
-MLKMSGW----QRQ-----------SQNQSRNLRRECSRRKCIFIHHHT
>Haliaeetus_leucocephalus
MRSRMSG--------------GQRSGWSRGEPGPPRKASRRKCVLVDHHT
>Oryctolagus_cuniculus
---------------------------MANICVCSPQCSRRKCIFIHHHT
>Eogystia_hippophaecolus
-MLKMSGW----QRQ-----------SQNNSRNLRRECSRRKCIFIHHHT
>Parus_major
MRTRMNR--------------GQRPERGRGEPGPPRKASRRKCVLADHHT
>Ceratotherium_simum_simum
-MLKMSGW----QPQ-----------SQNQSRNLRRECSRRKCIFIHHHT
>Manis_javanica
-MLKMSGW----QRQ-----------SQNQSWNPRRECSRRKCIFIHHHT
>Loxodonta_africana
-MLKMSGW----QRQ-----------SQSQSRNLRRKCSRRKCIFIHHHT
>Chinchilla_lanigera
-MLKMSRW----QRQ-----------SQSQSWNLRRECSRRKCIFIHHHT
>Trichechus_manatus_latirostr
-MLKMNGW----QRQ-----------SQNQSRNLRRKCSRRKCIFIHHHT
>Trachemys_scripta
MRARMSR--------------RQRGSRGQGEQGPRRKSSRRKCVFIHHHT
```

HG0023.2 ENSDARG00000059836 ddit3

>Danio_rerio
MVNMSDQPSLQKHTQTLNQKQPRKRNNKKKRSYWDKISPTTHIQT
>Amia_calva
MVNMSYHQSTPENQNTANPRQSRKRNNKRKRICWGTPSPPTHRQT
>Kaupichthys_hyoproroides
MVNMSYHQSQQKYIHTPTPRQTRKRNNKRKRICWDKLSPTAHRQT
>Oryzias_melastigma
MVNMSYQQFLLKYLHS-SSRQSRRRANKRKRNHWDKLSPTHFQT-
>Pygocentrus_nattereri
MVNMSYQQPQQKYIHTPSPRETRKRHNKRKRNFWDKISPTTHNHT
>Haplochromis_burtoni
MVNMSYHQFLLKYLHS-NSRQSRRRANKRKRNYWDKLSPTNCQT-
>Alosa_alosa
MVNMCCHQSQLKYIHTPNPRQSRKRHNKRKRNCWDTLSPTTHRQT
>Austrofundulus_limnaeus
MVNMSDHQFLLKYLLS-ASRQSRRRANKRKRNYWDNISPTIFQT-
>Esox_lucius
MVNMSYHQSQLKYIHTPNPRQTRKRNNKRKRTCWDKLSPTPNCQT
>Apteronotus_leptorhynchus
MVNMSYHQSQQKYTHTPNPRQLRRRHNKRKRNFWDKISPTTHNHT
>Scleropages_formosus
MVNMSYQQSQLKYIHTPNPRQSKKRNNKRKRTFWDKLSPTTHQQT
>Notothenia_coriiceps
MVNMSYHQFLLKYPHS-SSRQSRRRTNKRKRDCWDKLSPTNWQT-
>Oncorhynchus_mykiss
MVNMSYHQSQLKYIHTPNPRQSRKRNNKRKRTCWDKLSPTTNCQT
>Lepisosteus_oculatus
MVNMSYHHSTPENKNSPNHRQSRKRNNKRKRIVSDTPTPPNHRQT
>Trichomycterus_areolatus
MVNMNYQQSQQKHIHSPSLRQSRKRHNKRKRNYWDTISPTTHNHT

HG0024.1 ENSDARG00000093406

```
>Danio_rerio
MGPPRCRSRRLLPERTEDFGRFVSLS
>Acipenser_sinensis
MGPPGSRNCCVTFERMDDFGWLVSLS
>Amia_calva
MGPPGSKPRCVSFERMEDFGWFVNLS
>Channa_argus
MGPPGDRTQHMSTERTEDSSWIVDLS
>Podocoryna_carnea
MGPPGSRNCCVTFERMEDFGWLVSLS
>Menidia_menidia
MGPPGDRTCHMTPERTEDSRWIVDLS
>Oryzias_latipes
MGSPGDKTGSMSTERTEDFSWIVDLS
>Astyanax_mexicanus
MGPPRGKSSRLLTERRENFSWFVHLS
>Haplochromis_burtoni
MGPPGDRKLHMSTERTEDLSWIVDLS
>Alosa_alosa
MGPPRSKSRRISPERTEDFCWFVTLS
>Kryptolebias_marmoratus
MGPPGDRTRHMSTKRTEDSRWIVDLS
>Esox_lucius
MGPPGDKPRRVSSERTEDFIWFVNLS
>Plecoglossus_altivelis
MGPPGDKPRRVSTERTEDFSWFVNLS
>Trematomus_bernacchii
MGPPGDRTGHMSTERTEDFCWIVDLS
>Cynoglossus_semilaevis
MGPPGDRRRHLLTERTEDFSMIVDLS
>Salmo_salar
MGPPGDNPRRVSSERTEDFSWFVNLS
>Lepisosteus_oculatus
MGPPGSKPRRVSFERMEDFGWFVNLS
>Ictalurus_punctatus
MGPPRSKSSRLLSERRENLNWFVSLS
>Takifugu_rubripes
MGPPADRTCYMSTERTEDFGWIVDLS
```

```
HG0024.2 ENSG00000112308 C6orf62

>Homo_sapiens
MDFRD-WSGVSCVLLHT
>Mustela_putorius_furo
MDFKD-WTGVSCVLLHT
>Tursiops_truncatus
MDFKD-WTDVSCVLLHT
>Pteropus_vampyrus
MDFKD-WTGVTCVLLHT
>Dasypus_novemcinctus
MDFKD-WTGVSCVLLHT
>Sarcophilus_harrisii
MDLKD-WAGVSCVLLYT
>Galeopterus_variegatus
MDFKD-WTGVSCILLHT
>Monodelphis_domestica
MDLKD-WAGVSCVLLYT
>Erinaceus_europaeus
MDFKD-WTDVSCVLLHT
>Oryctolagus_cuniculus
MDFED-WTGVVLSHT--
>Elephantulus_edwardii
MGFKDF-----------
>Phalacrocorax_carbo
---MD-WAGASCFLLYT
>Equus_asinus
MDSKD-WTGVSCVLLHT
>Manis_javanica
MDFKD-RTGVSFVLLHP
>Loxodonta_africana
MNFKDFWTGVSCVLLHT
>Dipodomys_ordii
MDFKD-WTGVSCVLLHT
>Orycteropus_afer_afer
MDFKDFWTGVSCVLLHT
```

HG0024.2 ENSGALG00000013628 C6orf62

```
>Gallus_gallus
MDWACASCFLLYT
>Anser_cygnoides
MDWACASCFLLYT
>Apteryx_australis_mantelli
MDWACASCFLLYT
>Buceros_rhinoceros_silvestri
MDWACASCFLLYT
>Caprimulgus_carolinensis
MDWACASCFLLYT
>Cariama_cristata
MDWACASCFLLYT
>Calidris_pugnax
MDWACASCFLLYT
>Pterocles_gutturalis
MDWACASCFLLYT
>Colius_striatus
MDWACASCFLLYT
>Columba_livia
MDWACASCFLLYT
>Leptosomus_discolor
MDWACASCFLLYT
>Alligator_sinensis
MDWACASCFLLYT
>Aquila_chrysaetos_canadensis
MDWACASCFLLYT
>Gavia_stellata
MDWACASCFLLYT
>Balearica_regulorum_gibberic
MDWACASCFLLYT
>Tauraco_erythrolophus
MDWACASCFLLYA
>Opisthocomus_hoazin
MDWVCASCSLSYT
>Taeniopygia_guttata
MDWACASCFLLYT
>Pelecanus_crispus
MDWACASCFLLYT
>Picoides_pubescens
MDWACASCFLLYT
>Fulmarus_glacialis
MDWACASCFLLYT
>Melopsittacus_undulatus
MDWACASCFLLYT
>Aptenodytes_forsteri
MDWACASCFLLYT
>Tyto_alba
MDWACASCFLLYT
>Struthio_camelus_australis
MDWACASCFLLYT
>Chrysemys_picta
MDWACASCFLLYT
>Tinamus_guttatus
MDWDCASCFLLFT
```

```
HG0024.3 ENSGALG00000013628 C6orf62

>Gallus_gallus
---------MAGQLLILFLDPSNTTFP-
>Anser_cygnoides
MVKPVCCPAMAGQLL-LFLDPSDTTFS-
>Apteryx_australis_mantelli
---------MVGQLLILFLDPSNKAFS-
>Buceros_rhinoceros_silvestri
---------MAGQLFSLFLDPSNTTFS-
>Caprimulgus_carolinensis
---------MAGQLLTLVFDPSDTTSS-
>Cariama_cristata
---------MAGQLLRLFLDPSDTTTFS
>Calidris_pugnax
---------MAGQLLTLFLDPSNTTFS-
>Pterocles_gutturalis
---------MAGQLLTLFRDPSDTTFS-
>Colius_striatus
---------MAGQLLILFRGPSDTTVS-
>Columba_livia
---------MAGQLLTLFLGPSDTTFS-
>Leptosomus_discolor
---------MTGQLLTLFLDPSDTTFS-
>Alligator_mississippiensis
---------MVGQRLILFLDPFNMTVS-
>Cuculus_canorus
---------MAGQLLTLFLDPSNTMFS-
>Falco_cherrug
---------MAGQLLTLFLDPSNTTFS-
>Gavia_stellata
---------MAGQLLSLFLDPSDTTFS-
>Chlamydotis_macqueenii
---------MAGQLLTLFLDPSDTTFS-
>Tauraco_erythrolophus
---------MAGQLLTLFLDPSDTTFS-
>Opisthocomus_hoazin
---------MAGRLLTLFLDPSNATFS-
>Corvus_cornix_cornix
---------MAGQLLSLFLDPSNTTFS-
>Pelecanus_crispus
---------MAGQLLTLFLDPSNTTFS-
>Picoides_pubescens
---------MAEQLLSWFLDPSDTTFS-
>Fulmarus_glacialis
---------MAGQLLTLFLDPSDTTFS-
>Nestor_notabilis
---------MAGQLLTLFLDPSERPFS-
>Aptenodytes_forsteri
---------MAGQLLTLFLDPSDTTLS-
>Anolis_carolinensis
---------MVGRLLILFYDPSNKTVS-
>Tyto_alba
---------MAGQLLTLFLDPSDTTFS-
>Struthio_camelus_australis
---------MVGQLLILFLDPSNTTFS-
>Chelonia_mydas
---------MVGQLHSLFLDPSNMIAS-
>Tinamus_guttatus
---------MVGQLLILFLDPSNTTFS-
```

HG0025 ENSDARG00000060054 epc1b

>Danio_rerio
-----------------------------------------------------------------------------------MRTLIGPAGNIRGSDRFMLHLEL
>Anguilla_anguilla
-----------------------------------------------------------------------------------MRTKTGKVVNFRGSDRFMLHLEL
>Podocoryna_carnea
-----------------------------------------------------------------------------------MITQAGTVLNFRGSDRFMLNLEL
>Oryzias_latipes
-----------------------------------------------------------------------------------MKAPLGTVVNIRRSDRFILQLEL
>Pygocentrus_nattereri
-----------------------------------------------------------------------------------MTALIGSVVNIRRSDRFMLHLEL
>Haplochromis_burtoni
-----------------------------------------------------------------------------------MKAPLGTVVNIRRSDRFILHLEL
>Clupea_harengus
-----------------------------------------------------------------------------------MRTKICTVVNRRGSDRYMLHLSL
>Kryptolebias_marmoratus
-----------------------------------------------------------------------------------MRTRSRPAVNLSRSDRFMLHLLL
>Esox_lucius
-----------------------------------------------------------------------------------MRTQISTVVHLRRSDRFMLHLVL
>Acanthogobius_hasta
MRPGLAAIGRGAVGAGPGAERSGSLLNVRHVVHGALKRRGAARSGASHKHSPPESDQSRPSPPRRRPTHFHSPTVDPDPPHTSMKAPLGSAGDLCTSDRFMSHLEL
>Apteronotus_leptorhynchus
-----------------------------------------------------------------------------------MTTLIGSVVNIRGSDRFMLHLEL
>Scleropages_formosus
-----------------------------------------------------------------------------------MRTLTGTVLNSRGSDRFILHLEL
>Notothenia_coriiceps
-----------------------------------------------------------------------------------MKAPLGTVVNIRRSDRFILHLEL
>Salvelinus_alpinus
-----------------------------------------------------------------------------------MKAQLETDVNLRRSDRFMLHLEF
>Lepisosteus_oculatus
-----------------------------------------------------------------------------------MRAQTGTVVNFRGSDRFMLHLEI
>Takifugu_rubripes
-----------------------------------------------------------------------------------MKALLGTVVNIRRSDRFILHLEL

HG0026 ENSDARG00000087059 fam213ab

```
>Danio_rerio
-----------MELV----TRAVGSVGLTV--IEA-LRSF-TELFLTQPVC-ATLTQLADTDLRTLD-----------GD-DRVFKARELWESSGAVIMAVRRPG
>Oxyacanthus_curtus
----------MSLVPEMLGSWGWLLTGGTLA-GIITYCNLPPRFWFGHVSKSNDIELLRDAPLQRLL-----------PAEGSVMTGQQLWKDTGAVVMAIRRPG
>Scapharca_broughtonii
----------MALIS----PTLLGIAGLTA--GAAILLNL-PASWLFSTKK-STLSYLESAKLQTLH-----------SSNNRNFSARELWKDKPAVIMAVRRPG
>Micropterus_dolomieu
MTFILEGELYNM--------WY-GALAFVATLMVVVALAN-TDLFLTKSAP-ASLEELGATDLQTTT-----------GD-GKTVKGQLLWERSGAVILAVRRPG
>Rhodactis_indosinensis
-----------MA-------WFLGAGGAVA--IAG-IVLA-HWDWYSNSAR-PDLAYLSSIKLKALN-----------GT-NKDVMASELWRESGAVIMVVRRPG
>Strongylocentrotus_purpuratu
------------------MALLWGAAAAAV--GG-LLVAN-MDFWLPRGGAATTEAYLAGAVLQLLGETKLQAQLSLVDEAVRTFKAKDLWKDTGAVVLAVRRPG
>Meganyctiphanes_norvegica
--MLLDGGLSIMALSTMLPNWPWLVTGALT--GFVAYCNL-PAYWLNAPQK-ATTEYLSGAPLMKLD-----------GS-GGQILSNELWQKNGALVMAVRRPG
>Perna_viridis
-----------MVFS----KALLGVAGFTA--GAAILINL-PTRFFMASEK-ATLSYLTTAKLQTID-----------EA-KKNFSASDLWKNNGIVMMAVRRPG
>Octopus_bimaculoides
-----------------------------------------------MAEK-ATLAYLQGATLKTLG-----------DD-PKEFGAGELWKNNGAIIMVVRRPG
>Saccostrea_glomerata
-----------MGIS----PVLVGFAGLTA--GIAVIGNL-PVTWFTTVEK-ATLKYLEKGKLQTIE-----------AS-PRIFSVKDLWKDHGVVIMAVRRPG
>Pecten_maximus
-----------MGVT----KALLGMAGVTA--MAATLCNV-PVSWFMSSER-ATLNYLENARVQTLD-----------SQ-PKNITASELWKKNGVVMMAVRRPG
>Anoplopoma_fimbria
--------------M----MEIIAAVGGFI--TDV-MNFF-TDKFLTPPLK-ATLKHLEEAELKTLK-----------GE-TRTLKGRSLWERSGAVIMAVRRPG
>Montastraea_faveolata
-----------MA-------WFLGAGGVAA--LTG-VIIA-NWDWYKNSPR-PDLAYLSSIKLKALD-----------GT-GKEVMASELWSKAGAVIMVVRRPG
>Sepia_officinalis
-----------------------------------------------MAEK-ATVGYLAGATLKTLG-----------DD-PKEFGAAELWRKSGAVIMVVRRPG
>Sepioloidea_lineolata
-----------------------------------------------MAEK-ATVGYLAGATLKTLG-----------DD-PKEFGAAELWKNTGAVIMVVRRPG
>Doryteuthis_pealeii
-----------------------------------------------MAEK-ATVGYLAGATLKTLG-----------DD-PKEFGAAELWKKTGAVIMVVRRPG
```

HG0027 ENSG00000077254 USP33

```
>Homo_sapiens
---------------------------MERNNRYQPLQKHAKL------------------
>Channa_argus
MLREFVTTRPALEELMEEALIWGGGVGRGRKNRHQPLQKHTKVERPLAL------------
>Millepora_alcicornis
---------------------------MERKNWYQQLQKHTKI------------------
>Latrodectus_hesperus
---------------------------MERNNWYQPLQKHTKL------------------
>Hynobius_retardatus
MLREFITTRPALQELLKEALN------MERKDHYQPLQKHTEVHRPVTL------------
>Chironomus_riparius
MLRDFVTTRPALQELLKEALN------MERNNWYQPLQKHTKCKDHRHYEETASTNGKITS
```

HG0028.1 ENSDARG00000032103 mapk6

>Danio_rerio
-----------------------------------------------------------------------MEDSFDCDCGELEPPDH
>Channa_argus
-----------------------------------------------------------------------MEDSFDCDCGELEPPDH
>Kaupichthys_hyoproroides
-----------------------------------------------------------------------MEDCFDCDCGELEPPDH
>Anas_platyrhynchos
-----------------------------------------------------------------------MDGSFDCDCGELEPPDH
>Microhyla_fissipes
-----------------------------------------------------------------------MDASFDCDCGELEPPDH
>Chaetura_pelagica
-----------------------------------------------------------------------MDGSFDCDCGELEPPDH
>Apteryx_australis_mantelli
-------------------------------------MYWDIDL----------------FSLLFRLSRNKMDGSFDCDCGELEPPDH
>Porichthys_notatus
MELYLADTDTWSRIRHRPDCNHTHRETDKELLDRTDPFLLSADSSTPSLMDSYSLHQDGSDGFWPRLYSSKMEDSFDCDCGELEPPDH
>Oryzias_latipes
-----------------------------------------------------------------------MEDSFDCDCGELEPPDH
>Buceros_rhinoceros_silvestri
-----------------------------------------------------------------------MDGSFDCDCGELEPPDH
>Caprimulgus_carolinensis
-----------------------------------------------------------------------MDGSFDCDCGELEPPDH
>Trachinotus_blochii
-----------------------------------------------------------------------MEDSFDCDCGELEPPDH
>Cariama_cristata
-----------------------------------------------------------------------MDGSFDCDCGELEPPDH
>Canis_lupus_familiaris
-----------------------------------------------------------------------MDGSFDCDCGELEPPDH
>Hynobius_retardatus
-----------------------------------------------------------------------MDSSFDCDCGELEPPDH
>Micropterus_floridanus
-----------------------------------------------------------------------MEDSFDCDCGELEPPDH
>Tursiops_truncatus
-----------------------------------------------------------------------MDGSFDCDCGELEPPDH
>Pygocentrus_nattereri
-----------------------------------------------------------------------MEDSFDCDCGELEPPDH
>Charadrius_vociferus
-----------------------------------------------------------------------MDGSFDCDCGELEPPDH
>Callorhinchus_milii
-----------------------------------------------------------------------MDDSFDCDCGELEPPDH
>Rousettus_aegyptiacus
---------------------MPLKVS-LD----INFFLWVQDDLLRE---KINSGQ-NGAQTASGLYRSKMDGSFDCDCGELEPPDH
>Haplochromis_burtoni
-----------------------------------------------------------------MLYPSKMEDSFDCDCGELEPPDH
>Pterocles_gutturalis
-----------------------------------------------------------------------MDGSFDCDCGELEPPDH
>Dasypus_novemcinctus
-----------------------------------------------------------------------MDGSFDCDCGELEPPDH
>Clupea_harengus
-----------------------------------------------------------------------MEDSFDCDCGELEPPDH
>Latimeria_chalumnae
-----------------------------------------------------------------------MDDSFDCDCGELEPPDH
>Colius_striatus
-----------------------------------------------------------------------MDGSFDCDCGELEPPDH
>Columba_livia
-----------------------------------------------------------------------MDGSFDCDCGELEPPDH
>Leptosomus_discolor
-----------------------------------------------------------------------MDGSFDCDCGELEPPDH
>Alligator_mississippiensis
-----------------------------------------------------------------------MDGSFDCDCGELEPPDH
>Cuculus_canorus
-----------------------------------------------------------------------MDGSFDCDCGELEPPDH
>Kryptolebias_marmoratus
-----------------------------------------------------------------------MEDSFDCDCGELEPPDH
>Sarcophilus_harrisii
-----------------------------------------------------------------------MDGSFDCDCGELEPPDH
>Galeopterus_variegatus
-----------------------------------------------------------------------MDGSFDCDCGELEPPDH
>Monodelphis_domestica
-----------------------------------------------------------------------MDGSFDCDCGELEPPDH
>Esox_lucius
-----------------------------------------------------------------------MEDSFDCDCGELEPPDC
>Haliaeetus_albicilla
-----------------------------------------------------------------------MDGSFDCDCGELEPPDH
>Gallus_gallus
-----------------------------------------------------------------------MDGSFDCDCGELEPPDH
>Gavia_stellata
-----------------------------------------------------------------------MDGSFDCDCGELEPPDH
>Acanthogobius_hasta
-----------------------------------------------------------------------MEDSFDCDCGELEPPDH
>Balearica_regulorum_gibberic
-----------------------------------------------------------------------MDGSFDCDCGELEPPDH
>Apteronotus_leptorhynchus
-----------------------------------------------------------------------MEDSFDCDCGELEPPDQ
>Condylura_cristata
-----------------------------------------------------------------------MDGSFDCDCGELEPPDH
>Oryctolagus_cuniculus
-----------------------------------------------------------------------MDGSFDCDCGELEPPDH
>Elephantulus_edwardii
-----------------------------------------------------------------------MDGSFDCDCGELEPPDH
>Ornithorhynchus_anatinus
-----------------------------------------------------------------------MDGSFDCDCGELEPPDQ
>Tauraco_erythrolophus
-----------------------------------------------------------------------MDGSFDCDCGELEPPDH
>Opisthocomus_hoazin
-----------------------------------------------------------------------MDGSFDCDCGELEPPDH
>Scleropages_formosus
-----------------------------------------------------------------------MEDSFDCDCGELEPPDH
>Lepidothrix_coronata
-----------------------------------------------------------------------MDGSFDCDCGELEPPDH
>Egretta_garzetta
-----------------------------------------------------------------------MDGSFDCDCGELEPPDH
>Notothenia_coriiceps
-----------------------------------------------------------------------MEDSFDCDCGELEPPDH
>Equus_asinus
----MAATAA-KREPRRRRCRQPSRRPEPT----LGAV----R--GPR---TPSMSR-RSPVSVCVLYQSKMDGSFDCDCGELEPPDH
>Manis_javanica
-----------------------------------------------------------------------MDGSFDCDCGELEPPDH
>Picoides_pubescens
-----------------------------------------------------------------------MDGSFDCDCGELEPPDH
>Cynoglossus_semilaevis
-----------------------------------------------------------------------MEDSFDCDCGELEPPDH
>Aotus_nancymaae
----MAATAA-KREPRRRRCRQHSRRPEPT----LGAVR------GLR---TLSMNR-RSPVSVCVLYQSKMDGSFDCDCGELEPPDH
>Loxodonta_africana
-----------------------------------------------------------------------MDGSFDCDCGEVEPPDH
>Fulmarus_glacialis
-----------------------------------------------------------------------MDGSFDCDCGELEPPDH
>Melopsittacus_undulatus
-----------------------------------------------------------------------MDGSFDCDCGELEPPDH
>Mus_musculus
----MAATAA-KREPRKRRCRQHSRRPEPT----LRAVR------GPC---TLSMSR-RSP--VSVLYQSKMDGSFDCDCGELEPPDH
>Oncorhynchus_kisutch
-----------------------------------------------------------------------MEDSFDCDCGELEPPDC
>Tupaia_chinensis
-----------------------------------------------------------------------MDGSFDCDCGELEPPDH
>Lepisosteus_oculatus
-----------------------------------------------------------------------MEDSFDCDCGELEPPDH
>Ictalurus_punctatus
-----------------------------------------------------------------------MEDSFDCDCGELEPPDH
>Trichechus_manatus_latirostr
-----------------------------------------------------------------------MDGSFDCDCGELEPPDH
>Pygoscelis_adeliae
-----------------------------------------------------------------------MDGSFDCDCGELEPPDH
>Squalus_acanthias
-----------------------------------------------------------------------MDDSFDCDCGELEPPDH
>Hypsiglena_sp._JMG-2014
-----------------------------------------------------------------------MDGSFDCDCGELEPPDH
>Tyto_alba
-----------------------------------------------------------------------MDGSFDCDCGELEPPDH
>Struthio_camelus_australis
-----------------------------------------------------------------------MDGSFDCDCGELEPPDH
>Pelodiscus_sinensis
----------------------------------------------------------MMAGTVLRLSRNKMDGSFDCDCGELEPPDH
>Takifugu_rubripes
-----------------------------------------------------------------------MEDSFDCDCGELEPPDH
>Tinamus_guttatus
-----------------------------------------------------------------------MDGSFDCDCGELEPPDH
>Calypte_anna
-----------------------------------------------------------------------MDGSFDCDCGELEPPDH
>Apaloderma_vittatum
-----------------------------------------------------------------------MDGSFDCDCGELEPPDH
>Orycteropus_afer_afer
-----------------------------------------------------------------------MDGSFDCDCGELEPPDH

HG0028.1 ENSG00000069956 MAPK6

>Homo_sapiens
-------------------------------------------------------------------------------------------------------MDGSFDCDCGELEPPDH
>Channa_argus
-------------------------------------------------------------------------------------------------------MEDSFDCDCGELEPPDH
>Kaupichthys_hyoproroides
-------------------------------------------------------------------------------------------------------MEDCFDCDCGELEPPDH
>Anas_platyrhynchos
-------------------------------------------------------------------------------------------------------MDGSFDCDCGELEPPDH
>Xenopus_laevis
-------------------------------------------------------------------------------------------------------MDGSFDCDCGELEPPDH
>Chaetura_pelagica
-------------------------------------------------------------------------------------------------------MDGSFDCDCGELEPPDH
>Apteryx_australis_mantelli
-------------------------------------------------------------------------------------MYWDIDLFSLLFRLSRNKMDGSFDCDCGELEPPDH
>Porichthys_notatus
---------------MELYLADTDTWSRIRHRPDCNHTHRETDKELL----DRTDPFL-----------LSADSSTPSLMDSYSLHQDGSD--GFWPRLYSSKMEDSFDCDCGELEPPDH
>Oryzias_latipes
-------------------------------------------------------------------------------------------------------MEDSFDCDCGELEPPDH
>Buceros_rhinoceros_silvestri
-------------------------------------------------------------------------------------------------------MDGSFDCDCGELEPPDH
>Caprimulgus_carolinensis
-------------------------------------------------------------------------------------------------------MDGSFDCDCGELEPPDH
>Trachinotus_blochii
-------------------------------------------------------------------------------------------------------MEDSFDCDCGELEPPDH
>Cariama_cristata
-------------------------------------------------------------------------------------------------------MDGSFDCDCGELEPPDH
>Ailuropoda_melanoleuca
-------------------------------------------------------------------------------------------------------MDGSFDCDCGELEPPDH
>Hynobius_retardatus
-------------------------------------------------------------------------------------------------------MDSSFDCDCGELEPPDH
>Micropterus_floridanus
-------------------------------------------------------------------------------------------------------MEDSFDCDCGELEPPDH
>Tursiops_truncatus
-------------------------------------------------------------------------------------------------------MDGSFDCDCGELEPPDH
>Pygocentrus_nattereri
-------------------------------------------------------------------------------------------------------MEDSFDCDCGELEPPDH
>Charadrius_vociferus
-------------------------------------------------------------------------------------------------------MDGSFDCDCGELEPPDH
>Callorhinchus_milii
-------------------------------------------------------------------------------------------------------MDDSFDCDCGELEPPDH
>Miniopterus_natalensis
-------------------------------------------------------------------------------------------------------MDGSFDCDCGELEPPDH
>Haplochromis_burtoni
-------------------------------------------------------------------------------------------------MLYPSKMEDSFDCDCGELEPPDH
>Pterocles_gutturalis
-------------------------------------------------------------------------------------------------------MDGSFDCDCGELEPPDH
>Dasypus_novemcinctus
-------------------------------------------------------------------------------------------------------MDGSFDCDCGELEPPDH
>Clupea_harengus
-------------------------------------------------------------------------------------------------------MEDSFDCDCGELEPPDH
>Latimeria_chalumnae
-------------------------------------------------------------------------------------------------------MDDSFDCDCGELEPPDH
>Colius_striatus
-------------------------------------------------------------------------------------------------------MDGSFDCDCGELEPPDH
>Columba_livia
-------------------------------------------------------------------------------------------------------MDGSFDCDCGELEPPDH
>Leptosomus_discolor
-------------------------------------------------------------------------------------------------------MDGSFDCDCGELEPPDH
>Alligator_mississippiensis
-------------------------------------------------------------------------------------------------------MDGSFDCDCGELEPPDH
>Cuculus_canorus
-------------------------------------------------------------------------------------------------------MDGSFDCDCGELEPPDH
>Sinocyclocheilus_rhinocerous
MTPVRRLTKWLSAGRPSFPSHTTATWLNH-PQPLCLQTQTGTRANPLHMCYEHCGVFYSLQEKQESMTSDSVDTSELGFMNNHTQETLPKR--AVMSVVSQCKMEDSFDCDCGELEPPDH
>Poecilia_formosa
--------------------------------------------------------------------------------MTKSKTSHGNS--TGYHRLYPSKMEDSFDCDCGELEPPDH
>Sarcophilus_harrisii
-------------------------------------------------------------------------------------------------------MDGSFDCDCGELEPPDH
>Galeopterus_variegatus
-------------------------------------------------------------------------------------------------------MDGSFDCDCGELEPPDH
>Monodelphis_domestica
-------------------------------------------------------------------------------------------------------MDGSFDCDCGELEPPDH
>Esox_lucius
-------------------------------------------------------------------------------------------------------MEDSFDCDCGELEPPDC
>Haliaeetus_albicilla
-------------------------------------------------------------------------------------------------------MDGSFDCDCGELEPPDH
>Coturnix_japonica
-------------------------------------------------------------------------------------------------------MDGSFDCDCGELEPPDH
>Gavia_stellata
-------------------------------------------------------------------------------------------------------MDGSFDCDCGELEPPDH
>Acanthogobius_hasta
-------------------------------------------------------------------------------------------------------MEDSFDCDCGELEPPDH
>Balearica_regulorum_gibberic
-------------------------------------------------------------------------------------------------------MDGSFDCDCGELEPPDH
>Apteronotus_leptorhynchus
-------------------------------------------------------------------------------------------------------MEDSFDCDCGELEPPDQ
>Condylura_cristata
-------------------------------------------------------------------------------------------------------MDGSFDCDCGELEPPDH
>Oryctolagus_cuniculus
-------------------------------------------------------------------------------------------------------MDGSFDCDCGELEPPDH
>Elephantulus_edwardii
-------------------------------------------------------------------------------------------------------MDGSFDCDCGELEPPDH
>Ornithorhynchus_anatinus
-------------------------------------------------------------------------------------------------------MDGSFDCDCGELEPPDQ
>Tauraco_erythrolophus
-------------------------------------------------------------------------------------------------------MDGSFDCDCGELEPPDH
>Opisthocomus_hoazin
-------------------------------------------------------------------------------------------------------MDGSFDCDCGELEPPDH
>Scleropages_formosus
-------------------------------------------------------------------------------------------------------MEDSFDCDCGELEPPDH
>Manacus_vitellinus
-------------------------------------------------------------------------------------------------------MDGSFDCDCGELEPPDH
>Egretta_garzetta
-------------------------------------------------------------------------------------------------------MDGSFDCDCGELEPPDH
>Notothenia_coriiceps
-------------------------------------------------------------------------------------------------------MEDSFDCDCGELEPPDH
>Equus_asinus
-----------------------------------MAATAAKREPRRRRCR----------------QPSRRPEPTLGAVRGPRTPSMSRRSPVSVCVLYQSKMDGSFDCDCGELEPPDH
>Manis_javanica
-------------------------------------------------------------------------------------------------------MDGSFDCDCGELEPPDH
>Picoides_pubescens
-------------------------------------------------------------------------------------------------------MDGSFDCDCGELEPPDH
>Platichthys_stellatus
-------------------------------------------------------------------------------------------------------MEDSFDCDCGELEPPDH
>Loxodonta_africana
-------------------------------------------------------------------------------------------------------MDGSFDCDCGEVEPPDH
>Fulmarus_glacialis
-------------------------------------------------------------------------------------------------------MDGSFDCDCGELEPPDH
>Melopsittacus_undulatus
-------------------------------------------------------------------------------------------------------MDGSFDCDCGELEPPDH
>Mus_musculus
-----------------------------------MAATAAKREPRKRRCR----------------QHSRRPEPTLRAVRGPCTLSMSRRSPVS--VLYQSKMDGSFDCDCGELEPPDH
>Oncorhynchus_kisutch
-------------------------------------------------------------------------------------------------------MEDSFDCDCGELEPPDC
>Tupaia_chinensis
-------------------------------------------------------------------------------------------------------MDGSFDCDCGELEPPDH
>Lepisosteus_oculatus
-------------------------------------------------------------------------------------------------------MEDSFDCDCGELEPPDH
>Ictalurus_punctatus
-------------------------------------------------------------------------------------------------------MEDSFDCDCGELEPPDH
>Trichechus_manatus_latirostr
-------------------------------------------------------------------------------------------------------MDGSFDCDCGELEPPDH
>Aptenodytes_forsteri
-------------------------------------------------------------------------------------------------------MDGSFDCDCGELEPPDH
>Squalus_acanthias
-------------------------------------------------------------------------------------------------------MDDSFDCDCGELEPPDH
>Python_bivittatus
-----------------------------------------------MVCF----------------QTLR-------SQHG-----IQATTSHLQEGFSQNKMDGSFDCDCGELEPPDH
>Tyto_alba
-------------------------------------------------------------------------------------------------------MDGSFDCDCGELEPPDH
>Struthio_camelus_australis
-------------------------------------------------------------------------------------------------------MDGSFDCDCGELEPPDH
>Chrysemys_picta
-------------------------------------------------------------------------------------------------------MDGSFDCDCGELEPPDH
>Takifugu_rubripes
-------------------------------------------------------------------------------------------------------MEDSFDCDCGELEPPDH
>Tinamus_guttatus
-------------------------------------------------------------------------------------------------------MDGSFDCDCGELEPPDH
>Calypte_anna
-------------------------------------------------------------------------------------------------------MDGSFDCDCGELEPPDH
>Apaloderma_vittatum
-------------------------------------------------------------------------------------------------------MDGSFDCDCGELEPPDH
>Orycteropus_afer_afer
-------------------------------------------------------------------------------------------------------MDGSFDCDCGELEPPDH

HG0028.1 ENSGALG00000031448 MAPK6

>Gallus_gallus
-------------------------------------------------------------------------------------------------------MDGSFDCDCGELEPPDH
>Channa_argus
-------------------------------------------------------------------------------------------------------MEDSFDCDCGELEPPDH
>Kaupichthys_hyoproroides
-------------------------------------------------------------------------------------------------------MEDCFDCDCGELEPPDH
>Anas_platyrhynchos
-------------------------------------------------------------------------------------------------------MDGSFDCDCGELEPPDH
>Xenopus_laevis
-------------------------------------------------------------------------------------------------------MDGSFDCDCGELEPPDH
>Chaetura_pelagica
-------------------------------------------------------------------------------------------------------MDGSFDCDCGELEPPDH
>Apteryx_australis_mantelli
-------------------------------------------------------------------------------------MYWDIDLFSLLFRLSRNKMDGSFDCDCGELEPPDH
>Porichthys_notatus
---------------MELYLADTDTWSRIRHRPDCNHTHRETDKELL----DRTDPFL-----------LSADSSTPSLMDSYSLHQDGS--DGFWPRLYSSKMEDSFDCDCGELEPPDH
>Oryzias_latipes
-------------------------------------------------------------------------------------------------------MEDSFDCDCGELEPPDH
>Buceros_rhinoceros_silvestri
-------------------------------------------------------------------------------------------------------MDGSFDCDCGELEPPDH
>Caprimulgus_carolinensis
-------------------------------------------------------------------------------------------------------MDGSFDCDCGELEPPDH
>Trachinotus_blochii
-------------------------------------------------------------------------------------------------------MEDSFDCDCGELEPPDH
>Cariama_cristata
-------------------------------------------------------------------------------------------------------MDGSFDCDCGELEPPDH
>Ailuropoda_melanoleuca
-------------------------------------------------------------------------------------------------------MDGSFDCDCGELEPPDH
>Hynobius_retardatus
-------------------------------------------------------------------------------------------------------MDSSFDCDCGELEPPDH
>Micropterus_floridanus
-------------------------------------------------------------------------------------------------------MEDSFDCDCGELEPPDH
>Tursiops_truncatus
-------------------------------------------------------------------------------------------------------MDGSFDCDCGELEPPDH
>Pygocentrus_nattereri
-------------------------------------------------------------------------------------------------------MEDSFDCDCGELEPPDH
>Charadrius_vociferus
-------------------------------------------------------------------------------------------------------MDGSFDCDCGELEPPDH
>Callorhinchus_milii
-------------------------------------------------------------------------------------------------------MDDSFDCDCGELEPPDH
>Miniopterus_natalensis
-------------------------------------------------------------------------------------------------------MDGSFDCDCGELEPPDH
>Haplochromis_burtoni
-------------------------------------------------------------------------------------------------MLYPSKMEDSFDCDCGELEPPDH
>Pterocles_gutturalis
-------------------------------------------------------------------------------------------------------MDGSFDCDCGELEPPDH
>Dasypus_novemcinctus
-------------------------------------------------------------------------------------------------------MDGSFDCDCGELEPPDH
>Clupea_harengus
-------------------------------------------------------------------------------------------------------MEDSFDCDCGELEPPDH
>Latimeria_chalumnae
-------------------------------------------------------------------------------------------------------MDDSFDCDCGELEPPDH
>Colius_striatus
-------------------------------------------------------------------------------------------------------MDGSFDCDCGELEPPDH
>Columba_livia
-------------------------------------------------------------------------------------------------------MDGSFDCDCGELEPPDH
>Leptosomus_discolor
-------------------------------------------------------------------------------------------------------MDGSFDCDCGELEPPDH
>Alligator_mississippiensis
-------------------------------------------------------------------------------------------------------MDGSFDCDCGELEPPDH
>Cuculus_canorus
-------------------------------------------------------------------------------------------------------MDGSFDCDCGELEPPDH
>Sinocyclocheilus_rhinocerous
MTPVRRLTKWLSAGRPSFPSHTTATWLNH-PQPLCLQTQTGTRANPLHMCYEHCGVFYSLQEKQESMTSDSVDTSELGFMNNHTQETLPK--RAVMSVVSQCKMEDSFDCDCGELEPPDH
>Poecilia_formosa
--------------------------------------------------------------------------------MTKSKTSHGN--STGYHRLYPSKMEDSFDCDCGELEPPDH
>Sarcophilus_harrisii
-------------------------------------------------------------------------------------------------------MDGSFDCDCGELEPPDH
>Galeopterus_variegatus
-------------------------------------------------------------------------------------------------------MDGSFDCDCGELEPPDH
>Monodelphis_domestica
-------------------------------------------------------------------------------------------------------MDGSFDCDCGELEPPDH
>Esox_lucius
-------------------------------------------------------------------------------------------------------MEDSFDCDCGELEPPDC
>Haliaeetus_albicilla
-------------------------------------------------------------------------------------------------------MDGSFDCDCGELEPPDH
>Gavia_stellata
-------------------------------------------------------------------------------------------------------MDGSFDCDCGELEPPDH
>Acanthogobius_hasta
-------------------------------------------------------------------------------------------------------MEDSFDCDCGELEPPDH
>Balearica_regulorum_gibberic
-------------------------------------------------------------------------------------------------------MDGSFDCDCGELEPPDH
>Apteronotus_leptorhynchus
-------------------------------------------------------------------------------------------------------MEDSFDCDCGELEPPDQ
>Condylura_cristata
-------------------------------------------------------------------------------------------------------MDGSFDCDCGELEPPDH
>Oryctolagus_cuniculus
-------------------------------------------------------------------------------------------------------MDGSFDCDCGELEPPDH
>Elephantulus_edwardii
-------------------------------------------------------------------------------------------------------MDGSFDCDCGELEPPDH
>Ornithorhynchus_anatinus
-------------------------------------------------------------------------------------------------------MDGSFDCDCGELEPPDQ
>Tauraco_erythrolophus
-------------------------------------------------------------------------------------------------------MDGSFDCDCGELEPPDH
>Opisthocomus_hoazin
-------------------------------------------------------------------------------------------------------MDGSFDCDCGELEPPDH
>Scleropages_formosus
-------------------------------------------------------------------------------------------------------MEDSFDCDCGELEPPDH
>Manacus_vitellinus
-------------------------------------------------------------------------------------------------------MDGSFDCDCGELEPPDH
>Egretta_garzetta
-------------------------------------------------------------------------------------------------------MDGSFDCDCGELEPPDH
>Notothenia_coriiceps
-------------------------------------------------------------------------------------------------------MEDSFDCDCGELEPPDH
>Equus_asinus
-----------------------------------MAATAAKREPRRRRCRQPS----------------RRPEPTLGAVRGPRTPSMSRRSPVSVCVLYQSKMDGSFDCDCGELEPPDH
>Manis_javanica
-------------------------------------------------------------------------------------------------------MDGSFDCDCGELEPPDH
>Picoides_pubescens
-------------------------------------------------------------------------------------------------------MDGSFDCDCGELEPPDH
>Platichthys_stellatus
-------------------------------------------------------------------------------------------------------MEDSFDCDCGELEPPDH
>Aotus_nancymaae
-----------------------------------MAATAAKREPRRRRCRQHS----------------RRPEPTLGAVRGLRTLSMNRRSPVSVCVLYQSKMDGSFDCDCGELEPPDH
>Loxodonta_africana
-------------------------------------------------------------------------------------------------------MDGSFDCDCGEVEPPDH
>Fulmarus_glacialis
-------------------------------------------------------------------------------------------------------MDGSFDCDCGELEPPDH
>Melopsittacus_undulatus
-------------------------------------------------------------------------------------------------------MDGSFDCDCGELEPPDH
>Mus_musculus
-----------------------------------MAATAAKREPRKRRCRQHS----------------RRPEPTLRAVRGPCTLSMSRRSPVS--VLYQSKMDGSFDCDCGELEPPDH
>Oncorhynchus_kisutch
-------------------------------------------------------------------------------------------------------MEDSFDCDCGELEPPDC
>Tupaia_chinensis
-------------------------------------------------------------------------------------------------------MDGSFDCDCGELEPPDH
>Lepisosteus_oculatus
-------------------------------------------------------------------------------------------------------MEDSFDCDCGELEPPDH
>Ictalurus_punctatus
-------------------------------------------------------------------------------------------------------MEDSFDCDCGELEPPDH
>Trichechus_manatus_latirostr
-------------------------------------------------------------------------------------------------------MDGSFDCDCGELEPPDH
>Aptenodytes_forsteri
-------------------------------------------------------------------------------------------------------MDGSFDCDCGELEPPDH
>Squalus_acanthias
-------------------------------------------------------------------------------------------------------MDDSFDCDCGELEPPDH
>Python_bivittatus
-----------------------------------------------MVCFQT-----------------------LRSQHGI-----QATTSHLQEGFSQNKMDGSFDCDCGELEPPDH
>Tyto_alba
-------------------------------------------------------------------------------------------------------MDGSFDCDCGELEPPDH
>Struthio_camelus_australis
-------------------------------------------------------------------------------------------------------MDGSFDCDCGELEPPDH
>Chrysemys_picta
-------------------------------------------------------------------------------------------------------MDGSFDCDCGELEPPDH
>Takifugu_rubripes
-------------------------------------------------------------------------------------------------------MEDSFDCDCGELEPPDH
>Tinamus_guttatus
-------------------------------------------------------------------------------------------------------MDGSFDCDCGELEPPDH
>Calypte_anna
-------------------------------------------------------------------------------------------------------MDGSFDCDCGELEPPDH
>Apaloderma_vittatum
-------------------------------------------------------------------------------------------------------MDGSFDCDCGELEPPDH
>Orycteropus_afer_afer
-------------------------------------------------------------------------------------------------------MDGSFDCDCGELEPPDH

HG0028.2 ENSG00000069956 MAPK6

```
>Homo_sapiens
MFIEACCVHIYSHFC--------
>Anser_cygnoides
MLIEAYYVHIYSHFC--------
>Rhacophorus_omeimontis
MMIEAYILHIYSHFCSMSTCFAH
>Chaetura_pelagica
MLIEAYYVHIYSHFC--------
>Apteryx_australis_mantelli
MLIEAYYVHIYSHFC--------
>Buceros_rhinoceros_silvestri
MLIEAYYVHIYSHFC--------
>Caprimulgus_carolinensis
MLIEAYYVHIYSHFC--------
>Cariama_cristata
MLIEAYYVHIYSHFC--------
>Acinonyx_jubatus
MFIEACCVHIYSHFC--------
>Balaenoptera_acutorostrata_s
MFIEACCVHIYSHFC--------
>Calidris_pugnax
MLIEAYYVHIYSHFC--------
>Eptesicus_fuscus
MFIEACCVHIYSHFC--------
>Pterocles_gutturalis
MLIEAYYVHIYSHFC--------
>Dasypus_novemcinctus
MFIEACCVHIYSHFC--------
>Colius_striatus
MLIEAYYVHIYSHFC--------
>Columba_livia
MLIEAYYVHIYSHFC--------
>Leptosomus_discolor
MLIEAYYVHIYSHFC--------
>Alligator_sinensis
MLIEAYYVHIYSYFC--------
>Cuculus_canorus
MLIEAYYVHIYSHFC--------
>Sarcophilus_harrisii
MFREAYCVHIYSLFC--------
>Galeopterus_variegatus
MFIEACCVHIYSHFC--------
>Aquila_chrysaetos_canadensis
MLIEAYYVHIYSHFC--------
>Coturnix_japonica
MLIEAYYVHIYSHFC--------
>Gavia_stellata
MLIEAYYVHIYSHFC--------
>Chlamydotis_macqueenii
MFIEAYYVHIYSHFC--------
>Condylura_cristata
MFIEACCVHIYSHFC--------
>Oryctolagus_cuniculus
MFIEACCVHIYSHFC--------
>Elephantulus_edwardii
MSIEACCVHIYSHFC--------
>Ornithorhynchus_anatinus
MLIEACFVHIYSHFC--------
>Tauraco_erythrolophus
MLKEAYYVHIYSHFC--------
>Opisthocomus_hoazin
MLIETYYVHIYSHFC--------
>Acanthisitta_chloris
MLIEAYYVHIYSHFC--------
>Egretta_garzetta
MLIEAYYVHIYSHFC--------
>Ceratotherium_simum_simum
MFIEACCVHIYSHFC--------
>Picoides_pubescens
MLIEAYYVHIYSHFC--------
>Loxodonta_africana
MFIEACCVHIYSHFC--------
>Fulmarus_glacialis
MLIEAYYVHIYSHFC--------
>Melopsittacus_undulatus
MLIEAYYVHIYSHFC--------
>Marmota_marmota_marmota
MFIEACCVHIYSHFC--------
>Tupaia_chinensis
MFLEACCVHIYSHFC--------
>Trichechus_manatus_latirostr
MFIEARCVHIYSRFC--------
>Pygoscelis_adeliae
MLIEAYYVHIYSHFC--------
>Takydromus_sexlineatus
MLIEAYYVHIYSHFC--------
>Tyto_alba
MLIEAYYVHIYSHFC--------
>Struthio_camelus_australis
MLIEAYYVHIYSHFC--------
>Pelodiscus_sinensis
MLLEAHYVHIYSHFC--------
>Tinamus_guttatus
MLIEAYYVHIYSHFC--------
>Calypte_anna
MLIEAYYVHIYSHFC--------
>Apaloderma_vittatum
MLIEAYYVHIYSHFC--------
>Orycteropus_afer_afer
MFIEACCVHIYSHFC--------
```

HG0029 ENSDARG00000007523 kmt2e

```
>Danio_rerio
----------------------------------------------------MHLDVCNECCG------------------
>Amia_calva
----------------------------------------------------MHLDVCNEHCGWHAPVFWIQMLRTQ----
>Channa_punctata
----------------------------------------------------MHLDVCNDCCGWHASVFWTPILWTQ----
>Xenopus_tropicalis
----------------------------------------------------MHLDVCNEHCGWHAPVFWIPMQRTP----
>Apteryx_australis_mantelli
------------------------------------------------MGLWMHLDVCNEHCGWHAPVFWIPMHRTP----
>Menidia_menidia
----------------------------------------------------MHLDVCNECCGWHASVFWTPFLWTQLDKN
>Porichthys_notatus
----------------------------------------------------MHLDVCNECCGWHASVFWTSILWTQ----
>Oryzias_melastigma
----------------------------------------------------MHLDVCNDCCGWHASVFWTPFLWTQ----
>Canis_lupus_familiaris
------------------------------------------------MGLWMHLDVCNEHCGWHAPVFWIPMHRTP----
>Hynobius_retardatus
----------------------------------------------------MHLDVCNEHCGCYASVLWPPMHRTP----
>Micropterus_dolomieu
----------------------------------------------------MHLDVCNDCCGWHASVFWTPILWTQ----
>Orcinus_orca
------------------------------------------------MGLWMHLDVCNEHCGWHAPVFWIPMHRTP----
>Pygocentrus_nattereri
----------------------------------------------------MHLDVCNECCGWHASVFWIV-LWTQ----
>Charadrius_vociferus
----------------------------------------------------MHLDVCNEHCGWHAPVFWIPMHRTP----
>Callorhinchus_milii
----------------------------------------------------MHLDVCNEHCGWHAPVFWMPMYRTP----
>Myotis_brandtii
------------------------------------------------MGLWMHLDVCNEHCGWHAPVFWIPMHRTP----
>Maylandia_zebra
----------------------------------------------------MHLDVCNECCGWHASVFWTPFLWTQ----
>Dasypus_novemcinctus
------------------------------------------------MGLWMHLDVCNEHCGWHAPVFWIPMHRTP----
>Alosa_alosa
----------------------------------------------------MHLDVCNECCGWHASVYWIPLLWTQ----
>Latimeria_chalumnae
----------------------------------------------------MHLDVCNEHCGWHAPVFWIPMHRTP----
>Columba_livia
----------------------------------------------------MHLDVCNEHCGWHAPVFWIPMHRTP----
>Cuculus_canorus
----------------------------------------------------MHLDVCNEHCGWHAPVFWIPMHRTP----
>Poecilia_formosa
----------------------------------------------------MHLDVCNDCCGWHASVFRTPFLWTQ----
>Esox_lucius
----------------------------------------------------MHLDVCNECCG------------------
>Falco_cherrug
----------------------------------------------------MHLDVCNEHCGWHAPVFWIPMHRTP----
>Gadus_morhua
----------------------------------------------------MHLDVCNECCG------------------
>Acanthogobius_hasta
----------------------------------------------------MHLDVCNDCCGWYASVFRTAILWTQ----
>Chlamydotis_macqueenii
----------------------------------------------------MHLDVCNEHCGWHAPVFWIPMHRTP----
>Apteronotus_leptorhynchus
----------------------------------------------------MHLDVCNESCGWHASVFWIAVLWTQ----
>Condylura_cristata
------------------------------------------------MDLWMHLDVCNEHCGWHAPVFWIPMHRTP----
>Ochotona_princeps
----------------------------------------------------MHLDVCNEHCGWHAPVFWIPMHRTP----
>Elephantulus_edwardii
------------------------------------------------MGLWMHLDVCNEHCGWHAPVFWIPMHRTP----
>Ornithorhynchus_anatinus
----------------------------------------------------MHLDVCNEHCGWHAPVFWIPMHRTP----
>Plecoglossus_altivelis
----------------------------------------------------MHLDVCNECCGWHASVFWTPMLWTQ----
>Scleropages_formosus
MAWQDGLGLYHGGPLSLAAFAHLRAHRGNFKEDPQYEAASRPVGSTQGQAKWMHLDVCNECCGWYASVFWIPTLWTL----
>Manacus_vitellinus
----------------------------------------------------MHLDVCNEHCGWHAPVFWIPMHRTP----
>Lateolabrax_japonicus
----------------------------------------------------MHLDVCNDCCGWHASVFWTPILWTQ----
>Sebastes_caurinus
----------------------------------------------------MHLDVCNDCCGWHASVFWTPILWTQ----
>Ceratotherium_simum_simum
------------------------------------------------MGLWMHLDVCNEHCGWHAPVFWIPMHRTP----
>Manis_javanica
------------------------------------------------MGLWMHLDVCNEHCGWHAPVFWIPMHRTP----
>Picoides_pubescens
----------------------------------------------------MHLDVCNEHCGWHAPVFWIPMHRTP----
>Platichthys_stellatus
----------------------------------------------------MHLDVCNDCCGWHASVFWTPILWTQ----
>Aotus_nancymaae
------------------------------------------------MGLWMHLDVCNEHCGWHAPVFWIPMHRTP----
>Loxodonta_africana
----------------------------------------------------MHLDVCNEHCGWHAPVFWIPMHRTP----
>Leucoraja_erinacea
----------------------------------------------------MHLDVCNEHCGWHAPVFWMPMCRTP----
>Mesocricetus_auratus
----------------------------------------------------MHLDVCNEHCGWHAPVFWIPMHRTP----
>Salmo_salar
----------------------------------------------------MHLDVCNECCGWHASVFWTPMLWTQ----
>Tupaia_chinensis
------------------------------------------------MGLWMHLDVCNEHCGWHAPVFWIPMHRTP----
>Lepisosteus_oculatus
----------------------------------------------------MHLDVCNEHCGWHAPVFWIPMLRTQ----
>Trichechus_manatus_latirostr
------------------------------------------------MDLWMHLDVCNEHCGWHAPVFWIPMHRTP----
>Aptenodytes_forsteri
----------------------------------------------------MHLDVCNEHCGWHAPVFWIPMHRTP----
>Squalus_acanthias
----------------------------------------------------MHLDVCNEHCGWHAPVFWMPMYRTP----
>Thamnophis_sirtalis
------------------------------------------------MSLWMHLDVCNEHCGWHAPVFWIPMHRNP----
>Struthio_camelus_australis
----------------------------------------------------MHLDVCNEHCGWHAPVFWIPMHRTP----
>Chelonia_mydas
----------------------------------------------------MHLDVCNEHCGWHAPVFWIPMHRTP----
>Takifugu_rubripes
----------------------------------------------------MHLDVCNDCCGWHASVFWTPILWTQ----
>Tinamus_guttatus
----------------------------------------------------MHLDVCNEHCGWHAPVFWISMHRTP----
>Calypte_anna
----------------------------------------------------MHLDVCNEHCGWHAPVFWIPMHRTP----
>Apaloderma_vittatum
----------------------------------------------------MHLDVCNEHCGWHAPVFWIPMHRTP----
>Orycteropus_afer_afer
------------------------------------------------MGFWMHLDVCNEHCGWHAPVFWIPMHRTP----
```

HG0029 ENSG00000005483 KMT2E

```
>Homo_sapiens
---------------------------MGLWMHLDVCNEHCGWHAPVFWIPMHRTP----
>Acipenser_sinensis
-------------------------------MHLDVCNEHFGWHASVFWIPMLRTP----
>Amia_calva
-------------------------------MHLDVCNEHCGWHAPVFWIQMLRTQ----
>Channa_punctata
-------------------------------MHLDVCNDCCGWHASVFWTPILWTQ----
>Anser_cygnoides
-------------------------------MHLDVCNEHCGWHAPVFWIPMHRTL----
>Xenopus_tropicalis
-------------------------------MHLDVCNEHCGWHAPVFWIPMQRTP----
>Chaetura_pelagica
-------------------------------MHLDVCNEHCGWHAPVFWIPMHRTP----
>Apteryx_australis_mantelli
---------------------------MGLWMHLDVCNEHCGWHAPVFWIPMHRTP----
>Menidia_menidia
-------------------------------MHLDVCNECCGWHASVFWTPFLWTQLDKN
>Porichthys_notatus
-------------------------------MHLDVCNECCGWHASVFWTSILWTQ----
>Oryzias_latipes
-------------------------------MHLDVCNDCCGWHASVFWTPFLWTQ----
>Buceros_rhinoceros_silvestri
-------------------------------MHLDVCNEHCGWHAPVFWIPMHRTP----
>Caprimulgus_carolinensis
-------------------------------MHLDVCNEHCGWHAPVFWIPMHRTP----
>Cariama_cristata
-------------------------------MHLDVCNEHCGWHAPVFWIPMHRTP----
>Mustela_putorius_furo
MLYRSILDSRFCEDIYSFLDLRFARHLMGLWMHLDVCNEHCGWHAPVFWIPMHRTP----
>Hynobius_retardatus
-------------------------------MHLDVCNEHCGCYASVLWPPMHRTP----
>Micropterus_dolomieu
-------------------------------MHLDVCNDCCGWHASVFWTPILWTQ----
>Orcinus_orca
---------------------------MGLWMHLDVCNEHCGWHAPVFWIPMHRTP----
>Pygocentrus_nattereri
-------------------------------MHLDVCNECCGWHASVFWIV-LWTQ----
>Calidris_pugnax
-------------------------------MHLDVCNEHCGWHAPVFWIPMHRTP----
>Callorhinchus_milii
-------------------------------MHLDVCNEHCGWHAPVFWMPMYRTP----
>Myotis_brandtii
---------------------------MGLWMHLDVCNEHCGWHAPVFWIPMHRTP----
>Haplochromis_burtoni
-------------------------------MHLDVCNECCGWHASVFWTPFLWTQ----
>Pterocles_gutturalis
-------------------------------MHLDVCNEHCGWHAPVFWIPMHRTP----
>Dasypus_novemcinctus
---------------------------MGLWMHLDVCNEHCGWHAPVFWIPMHRTP----
>Clupea_harengus
-------------------------------MHLDVCNECCGWHASVFWIPLLWTQ----
>Latimeria_chalumnae
-------------------------------MHLDVCNEHCGWHAPVFWIPMHRTP----
>Columba_livia
-------------------------------MHLDVCNEHCGWHAPVFWIPMHRTP----
>Merops_nubicus
-------------------------------MHLDVCNEHCGWHAPVFWIPMHRTP----
>Alligator_mississippiensis
-------------------------------MHLDVCNEHCGWHAPVFWIPMHRTP----
>Cuculus_canorus
-------------------------------MHLDVCNEHCGWHAPVFWIPMHRTP----
>Sinocyclocheilus_angustiporu
-------------------------------MHLDVCNECCG------------------
>Austrofundulus_limnaeus
-------------------------------MHLDVCNDCCGWHASVFWTPFLWTQ----
>Sarcophilus_harrisii
-------------------------------MHLDVCNEHCGWHAPVFWIPMHRTP----
>Galeopterus_variegatus
---------------------------MGLWMHLDVCNEHCGWHAPVFWIPMHRTP----
>Monodelphis_domestica
-------------------------------MHLDVCNEHCGWHAPVFWIPMHRTP----
>Esox_lucius
-------------------------------MHLDVCNECCG------------------
>Falco_cherrug
-------------------------------MHLDVCNEHCGWHAPVFWIPMHRTP----
>Gadus_morhua
-------------------------------MHLDVCNECCG------------------
>Coturnix_japonica
-------------------------------MHLDVCNEHCGWHAPVFWIPMHRTL----
>Acanthogobius_hasta
-------------------------------MHLDVCNDCCGWYASVFRTAILWTQ----
>Chlamydotis_macqueenii
-------------------------------MHLDVCNEHCGWHAPVFWIPMHRTP----
>Apteronotus_leptorhynchus
-------------------------------MHLDVCNESCGWHASVFWIAVLWTQ----
>Erinaceus_europaeus
---------------------------MGLWMHLDVCNEHCGWHAPVFWIPMHRTP----
>Ochotona_princeps
-------------------------------MHLDVCNEHCGWHAPVFWIPMHRTP----
>Elephantulus_edwardii
---------------------------MGLWMHLDVCNEHCGWHAPVFWIPMHRTP----
>Ornithorhynchus_anatinus
-------------------------------MHLDVCNEHCGWHAPVFWIPMHRTP----
>Plecoglossus_altivelis
-------------------------------MHLDVCNECCGWHASVFWTPMLWTQ----
>Scleropages_formosus
-------------------------------MHLDVCNECCGWYASVFWIPTLWTL----
>Parus_major
-----------------------MHLFMGLWMHLDVCNEHCGWHAPVFWIPMHRTP----
>Nipponia_nippon
-------------------------------MHLDVCNEHCGWHAPVFWIPMHRTP----
>Lateolabrax_japonicus
-------------------------------MHLDVCNDCCGWHASVFWTPILWTQ----
>Sebastes_caurinus
-------------------------------MHLDVCNDCCGWHASVFWTPILWTQ----
>Ceratotherium_simum_simum
---------------------------MGLWMHLDVCNEHCGWHAPVFWIPMHRTP----
>Manis_javanica
---------------------------MGLWMHLDVCNEHCGWHAPVFWIPMHRTP----
>Picoides_pubescens
-------------------------------MHLDVCNEHCGWHAPVFWIPMHRTP----
>Platichthys_stellatus
-------------------------------MHLDVCNDCCGWHASVFWTPILWTQ----
>Loxodonta_africana
---------------------------MGLWMHLDVCNEHCGWHAPVFWIPMHRTP----
>Leucoraja_erinacea
-------------------------------MHLDVCNEHCGWHAPVFWMPMCRTP----
>Mesocricetus_auratus
---------------------------MGLWMHLDVCNEHCGWHAPVFWIPMHRTP----
>Salmo_salar
-------------------------------MHLDVCNECCGWHASVFWTPMLWTQ----
>Tupaia_chinensis
---------------------------MGLWMHLDVCNEHCGWHAPVFWIPMHRTP----
>Lepisosteus_oculatus
-------------------------------MHLDVCNEHCGWHAPVFWIPMLRTQ----
>Trichomycterus_areolatus
-------------------------------MHLDVCHACSGWYASVFGIAQLWTH----
>Trichechus_manatus_latirostr
---------------------------MDLWMHLDVCNEHCGWHAPVFWIPMHRTP----
>Aptenodytes_forsteri
-------------------------------MHLDVCNEHCGWHAPVFWIPMHRTP----
>Squalus_acanthias
-------------------------------MHLDVCNEHCGWHAPVFWMPMYRTP----
>Thamnophis_sirtalis
---------------------------MSLWMHLDVCNEHCGWHAPVFWIPMHRNP----
>Tyto_alba
-------------------------------MHLDVCNEHCGWHAPVFWIPMHRTP----
>Struthio_camelus_australis
-------------------------------MHLDVCNEHCGWHAPVFWIPMHRTP----
>Chelonia_mydas
-------------------------------MHLDVCNEHCGWHAPVFWIPMHRTP----
>Takifugu_rubripes
-------------------------------MHLDVCNDCCGWHASVFWTPILWTQ----
>Tinamus_guttatus
-------------------------------MHLDVCNEHCGWHAPVFWISMHRTP----
>Calypte_anna
-------------------------------MHLDVCNEHCGWHAPVFWIPMHRTP----
>Apaloderma_vittatum
-------------------------------MHLDVCNEHCGWHAPVFWIPMHRTP----
>Orycteropus_afer_afer
---------------------------MGFWMHLDVCNEHCGWHAPVFWIPMHRTP----
```

HG0029 ENSG00000168137

>Homo_sapiens
---------------------------------------------------------------------------------MSITCWMRLCSSPPLSEWSVSINWTP
>Anas_platyrhynchos
---------------------------------------------------------------------------------MSVTCWPRLCGSSPPSERPVAALWTP
>Xenopus_tropicalis
---------------------------------------------------------------------------------MSVKCLSRLCGSPP-SVWSVAAIPTP
>Apteryx_australis_mantelli
---------------------------------------------------------------------------------MSVTCWPRLCGSSPPSERPVAALWTP
>Menidia_menidia
---------------------------------------------------------------------------------MTVSGAVRLCGAPP-LEWSAAGVWTL
>Buceros_rhinoceros_silvestri
---------------------------------------------------------------------------------MSVTCWPRLCGSSPPSERPVAALWTP
>Caprimulgus_carolinensis
---------------------------------------------------------------------------------MSVTCWPRLCGSSPPSERPVAALWTP
>Cariama_cristata
---------------------------------------------------------------------------------MSVTCWPRLCGSSPPSERPVAALWTP
>Canis_lupus_familiaris
---------------------------------------------------------------------------------MSITCWLRLCSSPPLSEWPVSINWTP
>Physeter_catodon
---------------------------------------------------------------------------------MSITCWLRLCSSPPLSEWPVSDNWTP
>Charadrius_vociferus
---------------------------------------------------------------------------------MSVTCWPRLCGSSPPSERPVAALWTP
>Pteropus_vampyrus
---------------------------------------------------------------------------------MSITCWLRLCSSPPLSEWPVSINWTP
>Pterocles_gutturalis
---------------------------------------------------------------------------------MSVTCWPRLCGSSPPSERPVAALWTP
>Dasypus_novemcinctus
---------------------------------------------------------------------------------MSITCCLRLCSAPPLSEWSVSLNWIP
>Latimeria_chalumnae
---------------------------------------------------------------------------------MSMKCWSRHCGSPP-LVWSVAAIWTP
>Colius_striatus
---------------------------------------------------------------------------------MSVTCWPRLCGSSPPSERPVAVLWTP
>Columba_livia
---------------------------------------------------------------------------------MSVTCWPRLCGSSPPSERPVAALWTP
>Leptosomus_discolor
---------------------------------------------------------------------------------MSVTCWPRLCGSSPPSERPVAALWTP
>Alligator_mississippiensis
---------------------------------------------------------------------------------MSVTCWPRLCGSSPPLERPVATLWTP
>Poecilia_mexicana
---------------------------------------------------------------------------------MTVSRGLRLDGAPL-LEWSVAISWTL
>Sarcophilus_harrisii
---------------------------------------------------------------------------------MSLTCWLRLCGSPPLSEWPVSINWTP
>Monodelphis_domestica
---------------------------------------------------------------------------------MSLTCWSRLCGSAPLSEWPVSINWIP
>Aquila_chrysaetos_canadensis
---------------------------------------------------------------------------------MSVTCWPRLCGSSPPSERPVAALWTP
>Gallus_gallus
---------------------------------------------------------------------------------MSVTCWPRLCGSSPPSERPVAALWTP
>Gavia_stellata
---------------------------------------------------------------------------------MSVTCWPRLCGASPPSGRPVAALWTP
>Mesitornis_unicolor
---------------------------------------------------------------------------------MSVTCWPRLCGSSPPSERPVAALWTP
>Condylura_cristata
---------------------------------------------------------------------------------MSITCWLRLCSSPPLSEWPVSINWTP
>Oryctolagus_cuniculus
---------------------------------------------------------------------------------MSVTCWPRLCSSPPLSEWSVSINWTP
>Ornithorhynchus_anatinus
------------------------MLTRSSESAAVM-------------------IF--RRRWQRGYW-DFTSTGQYKGFLMSVTCWLMLCGSPPLSEWSVSIKWTP
>Opisthocomus_hoazin
---------------------------------------------------------------------------------MSVTCWPRLCGSPPPPERPVAALWTP
>Plecoglossus_altivelis
---------------------------------------------------------------------------------MSVSRRLRLCGAPP-LEWSVAAVRTP
>Zonotrichia_albicollis
---------------------------------------------------------------------------------MSVTCWPRLCGSSPPSERPVAALWTP
>Egretta_garzetta
---------------------------------------------------------------------------------MSVTCWPRLCGSSPPSERPVAALWTP
>Equus_asinus
---------------------------------------------------------------------------------MSITCWLRLCSSPPLSEWPVSINWTP
>Manis_javanica
---------------------------------------------------------------------------------MSIICWLRLCSSPPLSEWPVSINWTP
>Picoides_pubescens
---------------------------------------------------------------------------------MSVTCWPRLCGSSPPSERPVAALWTP
>Loxodonta_africana
---------------------------------------------------------------------------------MSVTWWSRLCSSPPLSEWSVSINWTP
>Fulmarus_glacialis
---------------------------------------------------------------------------------MSVTCWPRLCGSSPPSERPVAALWTP
>Melopsittacus_undulatus
---------------------------------------------------------------------------------MSVTCWPRLCGSSPPSERPVAALWTP
>Cricetulus_griseus
----------------------------------------------------------------------------MAGFLMSITCWLRLCSSPPLSEWSVSINWTP
>Tupaia_chinensis
---------------------------------------------------------------------------------MSITCWLRLRSSPPLSEWSVSVNWTP
>Trichechus_manatus_latirostr
---------------------------------------------------------------------------------MSITCWLRLCSSPPLSEWSVSINWTP
>Aptenodytes_forsteri
---------------------------------------------------------------------------------MSVTCWPRLCGSSPPSERPVAALWTP
>Anolis_carolinensis
---------------------------------------------------------------------------------MSVTCWPRLCGCSPPLELPVATLWTP
>Struthio_camelus_australis
---------------------------------------------------------------------------------MSVTCWPRLCGSSPPSERPVAALWTP
>Trachemys_scripta
MGRGSGRVRIAPGASAGAPRALSAVSARPVEEAQPVVLRGRGRGCRRRRGSRSRLLPRKRRGAEPGLVARGARATGRPGFPMSVTCWPRLCGSSPPLERPVATFWTP
>Tinamus_guttatus
---------------------------------------------------------------------------------MSVTCWPRLCGSSPPSERPVAALWTP
>Calypte_anna
---------------------------------------------------------------------------------MSVTCWPRLCGSSPPSERPLAALWTP
>Apaloderma_vittatum
---------------------------------------------------------------------------------MSVTCWPGLCGSSPPSERPVAALWTP
>Orycteropus_afer_afer
---------------------------------------------------------------------------------MSISCWLRLCSLPPLPEWSISINWTP

HG0029 ENSGALG00000008167

```
>Gallus_gallus
-------------------------------MHLDVCNEHCGWHAPVFWIPMHRTL----
>Acipenser_sinensis
-------------------------------MHLDVCNEHFGWHASVFWIPMLRTP----
>Amia_calva
-------------------------------MHLDVCNEHCGWHAPVFWIQMLRTQ----
>Channa_punctata
-------------------------------MHLDVCNDCCGWHASVFWTPILWTQ----
>Anser_cygnoides
-------------------------------MHLDVCNEHCGWHAPVFWIPMHRTL----
>Xenopus_tropicalis
-------------------------------MHLDVCNEHCGWHAPVFWIPMQRTP----
>Chaetura_pelagica
-------------------------------MHLDVCNEHCGWHAPVFWIPMHRTP----
>Apteryx_australis_mantelli
---------------------------MGLWMHLDVCNEHCGWHAPVFWIPMHRTP----
>Menidia_menidia
-------------------------------MHLDVCNECCGWHASVFWTPFLWTQLDKN
>Porichthys_notatus
-------------------------------MHLDVCNECCGWHASVFWTSILWTQ----
>Oryzias_latipes
-------------------------------MHLDVCNDCCGWHASVFWTPFLWTQ----
>Buceros_rhinoceros_silvestri
-------------------------------MHLDVCNEHCGWHAPVFWIPMHRTP----
>Caprimulgus_carolinensis
-------------------------------MHLDVCNEHCGWHAPVFWIPMHRTP----
>Cariama_cristata
-------------------------------MHLDVCNEHCGWHAPVFWIPMHRTP----
>Mustela_putorius_furo
MLYRSILDSRFCEDIYSFLDLRFARHLMGLWMHLDVCNEHCGWHAPVFWIPMHRTP----
>Ambystoma_mexicanum
-------------------------------MHLDVCNEHCGWNASVFWIPMQRTP----
>Micropterus_dolomieu
-------------------------------MHLDVCNDCCGWHASVFWTPILWTQ----
>Orcinus_orca
---------------------------MGLWMHLDVCNEHCGWHAPVFWIPMHRTP----
>Pygocentrus_nattereri
-------------------------------MHLDVCNECCGWHASVFWIV-LWTQ----
>Calidris_pugnax
-------------------------------MHLDVCNEHCGWHAPVFWIPMHRTP----
>Callorhinchus_milii
-------------------------------MHLDVCNEHCGWHAPVFWMPMYRTP----
>Myotis_brandtii
---------------------------MGLWMHLDVCNEHCGWHAPVFWIPMHRTP----
>Haplochromis_burtoni
-------------------------------MHLDVCNECCGWHASVFWTPFLWTQ----
>Pterocles_gutturalis
-------------------------------MHLDVCNEHCGWHAPVFWIPMHRTP----
>Dasypus_novemcinctus
---------------------------MGLWMHLDVCNEHCGWHAPVFWIPMHRTP----
>Clupea_harengus
-------------------------------MHLDVCNECCGWHASVFWIPLLWTQ----
>Latimeria_chalumnae
-------------------------------MHLDVCNEHCGWHAPVFWIPMHRTP----
>Columba_livia
-------------------------------MHLDVCNEHCGWHAPVFWIPMHRTP----
>Merops_nubicus
-------------------------------MHLDVCNEHCGWHAPVFWIPMHRTP----
>Alligator_mississippiensis
-------------------------------MHLDVCNEHCGWHAPVFWIPMHRTP----
>Cuculus_canorus
-------------------------------MHLDVCNEHCGWHAPVFWIPMHRTP----
>Sinocyclocheilus_grahami
-------------------------------MHLDVCNECCG------------------
>Austrofundulus_limnaeus
-------------------------------MHLDVCNDCCGWHASVFWTPFLWTQ----
>Sarcophilus_harrisii
-------------------------------MHLDVCNEHCGWHAPVFWIPMHRTP----
>Galeopterus_variegatus
-------------------------------MHLDVCNEHCGWHAPVFWIPMHRTP----
>Monodelphis_domestica
-------------------------------MHLDVCNEHCGWHAPVFWIPMHRTP----
>Esox_lucius
-------------------------------MHLDVCNECCG------------------
>Falco_cherrug
-------------------------------MHLDVCNEHCGWHAPVFWIPMHRTP----
>Gadus_morhua
-------------------------------MHLDVCNECCG------------------
>Acanthogobius_hasta
-------------------------------MHLDVCNDCCGWYASVFRTAILWTQ----
>Chlamydotis_macqueenii
-------------------------------MHLDVCNEHCGWHAPVFWIPMHRTP----
>Apteronotus_leptorhynchus
-------------------------------MHLDVCNESCGWHASVFWIAVLWTQ----
>Condylura_cristata
---------------------------MDLWMHLDVCNEHCGWHAPVFWIPMHRTP----
>Ochotona_princeps
-------------------------------MHLDVCNEHCGWHAPVFWIPMHRTP----
>Elephantulus_edwardii
---------------------------MGLWMHLDVCNEHCGWHAPVFWIPMHRTP----
>Ornithorhynchus_anatinus
-------------------------------MHLDVCNEHCGWHAPVFWIPMHRTP----
>Plecoglossus_altivelis
-------------------------------MHLDVCNECCGWHASVFWTPMLWTQ----
>Scleropages_formosus
-------------------------------MHLDVCNECCGWYASVFWIPTLWTL----
>Manacus_vitellinus
-------------------------------MHLDVCNEHCGWHAPVFWIPMHRTP----
>Nipponia_nippon
-------------------------------MHLDVCNEHCGWHAPVFWIPMHRTP----
>Lateolabrax_japonicus
-------------------------------MHLDVCNDCCGWHASVFWTPILWTQ----
>Sebastes_caurinus
-------------------------------MHLDVCNDCCGWHASVFWTPILWTQ----
>Ceratotherium_simum_simum
---------------------------MGLWMHLDVCNEHCGWHAPVFWIPMHRTP----
>Manis_javanica
---------------------------MGLWMHLDVCNEHCGWHAPVFWIPMHRTP----
>Picoides_pubescens
-------------------------------MHLDVCNEHCGWHAPVFWIPMHRTP----
>Platichthys_stellatus
-------------------------------MHLDVCNDCCGWHASVFWTPILWTQ----
>Pan_paniscus
-------------------------------MHLDVCNEHCGWHAPVFWIPMHRTP----
>Loxodonta_africana
-------------------------------MHLDVCNEHCGWHAPVFWIPMHRTP----
>Leucoraja_erinacea
-------------------------------MHLDVCNEHCGWHAPVFWMPMCRTP----
>Mesocricetus_auratus
---------------------------MGLWMHLDVCNEHCGWHAPVFWIPMHRTP----
>Salmo_salar
-------------------------------MHLDVCNECCGWHASVFWTPMLWTQ----
>Tupaia_chinensis
---------------------------MGLWMHLDVCNEHCGWHAPVFWIPMHRTP----
>Lepisosteus_oculatus
-------------------------------MHLDVCNEHCGWHAPVFWIPMLRTQ----
>Ictalurus_punctatus
-------------------------------MHLDVCHACSGWYASVFGIARLCSQ----
>Trichechus_manatus_latirostr
---------------------------MDLWMHLDVCNEHCGWHAPVFWIPMHRTP----
>Aptenodytes_forsteri
-------------------------------MHLDVCNEHCGWHAPVFWIPMHRTP----
>Squalus_acanthias
-------------------------------MHLDVCNEHCGWHAPVFWMPMYRTP----
>Thamnophis_sirtalis
---------------------------MSLWMHLDVCNEHCGWHAPVFWIPMHRNP----
>Tyto_alba
-------------------------------MHLDVCNEHCGWHAPVFWIPMHRTP----
>Struthio_camelus_australis
-------------------------------MHLDVCNEHCGWHAPVFWIPMHRTP----
>Chelonia_mydas
-------------------------------MHLDVCNEHCGWHAPVFWIPMHRTP----
>Takifugu_rubripes
-------------------------------MHLDVCNDCCGWHASVFWTPILWTQ----
>Tinamus_guttatus
-------------------------------MHLDVCNEHCGWHAPVFWISMHRTP----
>Calypte_anna
-------------------------------MHLDVCNEHCGWHAPVFWIPMHRTP----
>Apaloderma_vittatum
-------------------------------MHLDVCNEHCGWHAPVFWIPMHRTP----
>Orycteropus_afer_afer
---------------------------MGFWMHLDVCNEHCGWHAPVFWIPMHRTP----
```

HG0030.1 ENSG00000100664 EIF5

>Homo_sapiens
-----------------------------------------------------------MSSLLLQTEDTKKLQSKISSSY-
>Acipenser_sinensis
MSLFYENGEYPDDRSDGWERAVA--LVPVWRKLGLQARSLQPQWRKLFHVRDRVHLQSFMSSALLQTEDTKKLQAKIFLSY-
>Kaupichthys_hyoproroides
-----------------------------------------------------------MSSTLLQTEDTKKLQSKIFLSC-
>Anser_cygnoides
-----------------------------------------------------------MSSRLLQTEDTKKLQSKISSSY-
>Megophrys_sangzhiensis
-----------------------------------------------------------MSSSLLQTEDTKKLQSKIFSSY-
>Chaetura_pelagica
-----------------------------------------------------------MSSRLLQTEDTKKLQSKISSSY-
>Apteryx_australis_mantelli
-----------------------------------------------------------MSSRLLQTEDTKKLQSKISSSY-
>Porichthys_notatus
-----------------------------------------------------------MSSLLFQTEDPEKSPTTILLSY-
>Oryzias_latipes
-----------------------------------------------------------MSSPFFQTEDPKKSPTKILLPY-
>Buceros_rhinoceros_silvestri
-----------------------------------------------------------MSSRLLQTEDTKKLQSKISSSY-
>Caprimulgus_carolinensis
-----------------------------------------------------------MSSRLLQTEDTKKLQSKISSSY-
>Seriola_quinqueradiata
-----------------------------------------------------------MSSPLFQTEDPEKSPTKILLSY-
>Cariama_cristata
-----------------------------------------------------------MSSRLLQTEDTKKLQSKISSSY-
>Mustela_putorius_furo
-----------------------------------------------------------MSSLLLQTEDTKKLQSKISSSY-
>Hynobius_chinensis
-----------------------------------------------------------MSSPLLQTEDTKKLLSQIFSSY-
>Siniperca_chuatsi
-----------------------------------------------------------MSSPLFQTEDPEKSPTKILLSY-
>Lipotes_vexillifer
-----------------------------------------------------------MSSLLLQTEDTKKLQSKISSSY-
>Astyanax_mexicanus
-----------------------------------------------------------MSSLLLQTEDTKKSPSKIFLSC-
>Calidris_pugnax
-----------------------------------------------------------MSSRLLQTEDTKKLQSKISSSY-
>Callorhinchus_milii
-----------------------------------------------------------MSSILLQTEDTKKFNSKIFSLYS
>Miniopterus_natalensis
-----------------------------------------------------------MSSLLLQTEDTKKLQSKISSSYR
>Haplochromis_burtoni
-----------------------------------------------------------MSSPLFQTEDPEKSPTKILLSY-
>Pterocles_gutturalis
-----------------------------------------------------------MSSRLLQTEDTKKLQSKISSSY-
>Dasypus_novemcinctus
-----------------------------------------------------------MSSLLLQTEDTKKLQSKISSSY-
>Latimeria_chalumnae
-----------------------------------------------------------MSSILLQAEDTKKLQSKIFSSY-
>Colius_striatus
-----------------------------------------------------------MSSRLLQTEDTKKLQSKISSSY-
>Columba_livia
-----------------------------------------------------------MSSRLLQTEDTKKLQSKISSSY-
>Merops_nubicus
-----------------------------------------------------------MSSRLLQTEDTKKLQSKISSSY-
>Alligator_mississippiensis
-----------------------------------------------------------MSSRLLQTEDTKKLQSKISSSY-
>Cuculus_canorus
-----------------------------------------------------------MSSRLLQTEDTKKLQSKISSSY-
>Ctenopharyngodon_idella
MSLFCENGEYLDDRSYGWESAIVQLTEPVWGELSQRARSLEPQWPYPFHVRDKVLLQSFMSSPLLQTEDTNNCQFKIFSSC-
>Fundulus_heteroclitus
-----------------------------------------------------------MSSPLFQTEDPEKSPTKILLSY-
>Sarcophilus_harrisii
-----------------------------------------------------------MSSLLLQTEDTKKLQSKISSSY-
>Galeopterus_variegatus
-----------------------------------------------------------MSSLLLQTEDTKKLQSKISSSY-
>Monodelphis_domestica
-----------------------------------------------------------MSSLLLQTEDTKKLQSKISSSY-
>Esox_lucius
-----------------------------------------------------------MSPPLCQTEDTKNLQSKIFLSY-
>Aquila_chrysaetos_canadensis
-----------------------------------------------------------MSSRLLQTEDTKKLQSKISSSY-
>Gallus_gallus
-----------------------------------------------------------MSSRLLQTEDTKKLQSKISSSY-
>Gavia_stellata
-----------------------------------------------------------MSSRLLQTEDTKKLQSKVSSSY-
>Chanos_chanos
-----------------------------------------------------------MSSSLLQTEDTKNRQSRIFSSC-
>Chlamydotis_macqueenii
-----------------------------------------------------------MSSRLLQTEDTKKLQSKISSSY-
>Apteronotus_leptorhynchus
-----------------------------------------------------------MSSLLLQTEDTPKSPSKIFLFC-
>Condylura_cristata
-----------------------------------------------------------MSSLLLQTEDTKKLQSKISSSY-
>Oryctolagus_cuniculus
-----------------------------------------------------------MSSLLLQTEDTKKLQSKISSSY-
>Elephantulus_edwardii
-----------------------------------------------------------MSSLLLQTEDTKKL---------
>Ornithorhynchus_anatinus
-----------------------------------------------------------MSSLLLQTEDTKKLQSKISSPY-
>Tauraco_erythrolophus
-----------------------------------------------------------MSSRLLQTEDTKKLQSKISSSY-
>Opisthocomus_hoazin
-----------------------------------------------------------MSSRLLQTEDTKKLQSKISSSY-
>Osmerus_mordax
MSLFYENGEYPDDRSDGWDRAVAQLLEPVWGKLSLQARSLQPQWQDLFHVRARLLLQLFMSSPLGQTEDTKNCQSKIFLSY-
>Scleropages_formosus
-MFGYLSLAYLATSPVFKLQAVAQLLEPVWDTRNLQARKPQPRGQNLFHVRDRVLLQPFMSSTLLQTEDTKKSPSKIFLPY-
>Acanthisitta_chloris
-----------------------------------------------------------MSSSLLQTEDTKKLQSKISSPY-
>Nipponia_nippon
-----------------------------------------------------------MSSRLLQTEDTKKLQSKISSSY-
>Notothenia_coriiceps
-----------------------------------------------------------MSSPLFQTEDPKKSPTKILLSY-
>Ceratotherium_simum_simum
-----------------------------------------------------------MSSLLLQTEDTKKLQSKISSSY-
>Manis_javanica
-----------------------------------------------------------MSSLLLQTEDTKKLQSKISSSY-
>Picoides_pubescens
-----------------------------------------------------------MSSRLLQTEDTKKLQSKISSSY-
>Loxodonta_africana
-----------------------------------------------------------MSSLLLQTEDTKKLQSKISSSY-
>Fulmarus_glacialis
-----------------------------------------------------------MSSRLLQTEDTKKLQSKISSSY-
>Melopsittacus_undulatus
-----------------------------------------------------------MSSRLLQTEDTKKLQSKISSSY-
>Leucoraja_erinacea
-----------------------------------------------------------MSSILLQTEDTKKFHSKIFSLHS
>Jaculus_jaculus
-----------------------------------------------------------MSSLLLQTEDTKKLQSKISSSY-
>Salmo_salar
MSLFCENGEYPDDRSDGWERAVAQLLEPVWGELNLQARSLQPHWQDLFHVRARTLFQSFMSSHLRQTEDTKNLQSKIFLSY-
>Tupaia_chinensis
-----------------------------------------------------------MSSLLLQTEDTKKLQSKISSSY-
>Lepisosteus_oculatus
MSLFYENGEYPDDRSDGWERAVAQLLEPVWGKLSLQARSLQPQWRKLFHVRDRVHLQSFMSSTLLQTEDTKKLQSKIFLSC-
>Ictalurus_furcatus
-----------------------------------------------------------MSSRLLQTEDTQKSPSRIFLFC-
>Trichechus_manatus_latirostr
-----------------------------------------------------------MSSLLLQTEDTKKLQSKISSSY-
>Aptenodytes_forsteri
-----------------------------------------------------------MSSRLLQTEDTKKLQSKISSSY-
>Gekko_japonicus
-----------------------------------------------------------MSSRLLQTEDTKKLQSKISASY-
>Tyto_alba
-----------------------------------------------------------MSSRLLQTEDTKKLQSKISSSY-
>Struthio_camelus_australis
-----------------------------------------------------------MSSRLLQTEDTKKLQSKISSSY-
>Chrysemys_picta
-----------------------------------------------------------MSSRLLQTEDTKKLQSKISSSY-
>Tinamus_guttatus
-----------------------------------------------------------MSSRLLQTEDTKKLQSKISSSY-
>Calypte_anna
-----------------------------------------------------------MSSRLLQTEDTKKLQSKISSSY-
>Apaloderma_vittatum
-----------------------------------------------------------MSSRLLQTEDTKKLQSKISSSY-
>Orycteropus_afer_afer
-----------------------------------------------------------MSSLLLQTEDTKNLQSKISSSY-

HG0030.1 ENSGALG00000011464 EIF5

```
>Gallus_gallus
-----------------------------------------------------------MSSRLLQTEDTKKLQSKISSSY-
>Acipenser_sinensis
MSLFYENGEYPDDRSDGWERAVA--LVPVWRKLGLQARSLQPQWRKLFHVRDRVHLQSFMSSALLQTEDTKKLQAKIFLSY-
>Kaupichthys_hyoproroides
-----------------------------------------------------------MSSTLLQTEDTKKLQSKIFLSC-
>Anas_platyrhynchos
-----------------------------------------------------------MSSRLLQTEDTKKLQSKISSSY-
>Leptobrachium_boringii
-----------------------------------------------------------MSSSLLQTEDTKKLQSKIFSSY-
>Chaetura_pelagica
-----------------------------------------------------------MSSRLLQTEDTKKLQSKISSSY-
>Apteryx_australis_mantelli
-----------------------------------------------------------MSSRLLQTEDTKKLQSKISSSY-
>Oryzias_latipes
-----------------------------------------------------------MSSPFFQTEDPKKSPTKILLPY-
>Buceros_rhinoceros_silvestri
-----------------------------------------------------------MSSRLLQTEDTKKLQSKISSSY-
>Caprimulgus_carolinensis
-----------------------------------------------------------MSSRLLQTEDTKKLQSKISSSY-
>Seriola_quinqueradiata
-----------------------------------------------------------MSSPLFQTEDPEKSPTKILLSY-
>Cariama_cristata
-----------------------------------------------------------MSSRLLQTEDTKKLQSKISSSY-
>Mustela_putorius_furo
-----------------------------------------------------------MSSLLLQTEDTKKLQSKISSSY-
>Hynobius_chinensis
-----------------------------------------------------------MSSPLLQTEDTKKLLSQIFSSY-
>Siniperca_chuatsi
-----------------------------------------------------------MSSPLFQTEDPEKSPTKILLSY-
>Lipotes_vexillifer
-----------------------------------------------------------MSSLLLQTEDTKKLQSKISSSY-
>Astyanax_mexicanus
-----------------------------------------------------------MSSLLLQTEDTKKSPSKIFLSC-
>Calidris_pugnax
-----------------------------------------------------------MSSRLLQTEDTKKLQSKISSSY-
>Callorhinchus_milii
-----------------------------------------------------------MSSILLQTEDTKKFNSKIFSLYS
>Miniopterus_natalensis
-----------------------------------------------------------MSSLLLQTEDTKKLQSKISSSYR
>Oreochromis_niloticus
-----------------------------------------------------------MSSPLFQTEDPEKSPTKILLSY-
>Pterocles_gutturalis
-----------------------------------------------------------MSSRLLQTEDTKKLQSKISSSY-
>Dasypus_novemcinctus
-----------------------------------------------------------MSSLLLQTEDTKKLQSKISSSY-
>Latimeria_chalumnae
-----------------------------------------------------------MSSILLQAEDTKKLQSKIFSSY-
>Colius_striatus
-----------------------------------------------------------MSSRLLQTEDTKKLQSKISSSY-
>Columba_livia
-----------------------------------------------------------MSSRLLQTEDTKKLQSKISSSY-
>Merops_nubicus
-----------------------------------------------------------MSSRLLQTEDTKKLQSKISSSY-
>Alligator_sinensis
-----------------------------------------------------------MSSRLLQTEDTKKLQSKISSSY-
>Cuculus_canorus
-----------------------------------------------------------MSSRLLQTEDTKKLQSKISSSY-
>Danio_rerio
MSLFCENGDYLDDRSYGWESAIVQLPEPVWGELSQRARSLEPQWPYPFHVRDKVFLQSFMSSPLLQTEDTKNCQFKIFSSC-
>Cyprinodon_variegatus
-----------------------------------------------------------MSSPLFQTEDPEKSPTKILLSY-
>Sarcophilus_harrisii
-----------------------------------------------------------MSSLLLQTEDTKKLQSKISSSY-
>Galeopterus_variegatus
-----------------------------------------------------------MSSLLLQTEDTKKLQSKISSSY-
>Monodelphis_domestica
-----------------------------------------------------------MSSLLLQTEDTKKLQSKISSSY-
>Esox_lucius
-----------------------------------------------------------MSPPLCQTEDTKNLQSKIFLSY-
>Aquila_chrysaetos_canadensis
-----------------------------------------------------------MSSRLLQTEDTKKLQSKISSSY-
>Gavia_stellata
-----------------------------------------------------------MSSRLLQTEDTKKLQSKVSSSY-
>Chanos_chanos
-----------------------------------------------------------MSSSLLQTEDTKNRQSRIFSSC-
>Chlamydotis_macqueenii
-----------------------------------------------------------MSSRLLQTEDTKKLQSKISSSY-
>Apteronotus_leptorhynchus
-----------------------------------------------------------MSSLLLQTEDTPKSPSKIFLFC-
>Condylura_cristata
-----------------------------------------------------------MSSLLLQTEDTKKLQSKISSSY-
>Oryctolagus_cuniculus
-----------------------------------------------------------MSSLLLQTEDTKKLQSKISSSY-
>Elephantulus_edwardii
-----------------------------------------------------------MSSLLLQTEDTKKL---------
>Ornithorhynchus_anatinus
-----------------------------------------------------------MSSLLLQTEDTKKLQSKISSPY-
>Tauraco_erythrolophus
-----------------------------------------------------------MSSRLLQTEDTKKLQSKISSSY-
>Opisthocomus_hoazin
-----------------------------------------------------------MSSRLLQTEDTKKLQSKISSSY-
>Plecoglossus_altivelis
-----------------------------------------------------------MSSPLGQTEDTKNCQSKIFLSY-
>Scleropages_formosus
-MFGYLSLAYLATSPVFKLQAVAQLLEPVWDTRNLQARKPQPRGQNLFHVRDRVLLQPFMSSTLLQTEDTKKSPSKIFLPY-
>Serinus_canaria
-----------------------------------------------------------MSSRLLQTEDTKKLQSKISSSY-
>Nipponia_nippon
-----------------------------------------------------------MSSRLLQTEDTKKLQSKISSSY-
>Notothenia_coriiceps
-----------------------------------------------------------MSSPLFQTEDPKKSPTKILLSY-
>Equus_asinus
-----------------------------------------------------------MSSLLLQTEDTKKLQSKISSSY-
>Manis_javanica
-----------------------------------------------------------MSSLLLQTEDTKKLQSKISSSY-
>Picoides_pubescens
-----------------------------------------------------------MSSRLLQTEDTKKLQSKISSSY-
>Macaca_nemestrina
-----------------------------------------------------------MSSLLLQTEDTKKLQSKISSFY-
>Loxodonta_africana
-----------------------------------------------------------MSSLLLQTEDTKKLQSKISSSY-
>Fulmarus_glacialis
-----------------------------------------------------------MSSRLLQTEDTKKLQSKISSSY-
>Melopsittacus_undulatus
-----------------------------------------------------------MSSRLLQTEDTKKLQSKISSSY-
>Leucoraja_erinacea
-----------------------------------------------------------MSSILLQTEDTKKFHSKIFSLHS
>Jaculus_jaculus
-----------------------------------------------------------MSSLLLQTEDTKKLQSKISSSY-
>Salmo_salar
MSLFCENGEYPDDRSDGWERAVAQLLEPVWGELNLQARSLQPHWQDLFHVRARTLFQSFMSSHLRQTEDTKNLQSKIFLSY-
>Tupaia_chinensis
-----------------------------------------------------------MSSLLLQTEDTKKLQSKISSSY-
>Lepisosteus_oculatus
MSLFYENGEYPDDRSDGWERAVAQLLEPVWGKLSLQARSLQPQWRKLFHVRDRVHLQSFMSSTLLQTEDTKKLQSKIFLSC-
>Ictalurus_punctatus
-----------------------------------------------------------MSSRLLQTEDTQKSPSRIFLFC-
>Trichechus_manatus_latirostr
-----------------------------------------------------------MSSLLLQTEDTKKLQSKISSSY-
>Aptenodytes_forsteri
-----------------------------------------------------------MSSRLLQTEDTKKLQSKISSSY-
>Gekko_japonicus
-----------------------------------------------------------MSSRLLQTEDTKKLQSKISASY-
>Tyto_alba
-----------------------------------------------------------MSSRLLQTEDTKKLQSKISSSY-
>Struthio_camelus_australis
-----------------------------------------------------------MSSRLLQTEDTKKLQSKISSSY-
>Chrysemys_picta
-----------------------------------------------------------MSSRLLQTEDTKKLQSKISSSY-
>Tinamus_guttatus
-----------------------------------------------------------MSSRLLQTEDTKKLQSKISSSY-
>Calypte_anna
-----------------------------------------------------------MSSRLLQTEDTKKLQSKISSSY-
>Apaloderma_vittatum
-----------------------------------------------------------MSSRLLQTEDTKKLQSKISSSY-
>Orycteropus_afer_afer
-----------------------------------------------------------MSSLLLQTEDTKNLQSKISSSY-
```

HG0030.2 FBgn0030719 eIF5

>Drosophila_melanogaster
--------------------------------------------------------------------------------------------------------------------------------------------------------------MNSKGRTKNHRTSSNRHRKLQPFRYT-
>Hyalella_azteca
-----------------------------------------------------------------------------------------------------------MIP-EGQ--FSRVV-----------------YDYPRACVSPTPLHRMCAHNNDSPINIRNRQFSSNRHRKLQPFRTS-
>Meinertellus_cundinamarcensi
-----------------------------------------MTVLSWKEP----------------------LLARGGS------GIDHTS-REHHPASQGPAVDRCMTS-PKR---------------------------------------QHPL------SYRTSKFSSNRHRKLQPFRET-
>Argulus_siamensis
-----------------------------------------MNLYSWNDL----------------------LLSPGNP------ASFRKQ-LVA---SCGPDPSGCMFA-AKK---------------------------------------YEPL------SYRKTKFSSNRHRKLQPFRS--
>Coptotermes_gestroi
-----------------------------------------------------------------------------------------------------------MIS-SKR---------------------------------------QEPI------SYRTSKFSSNRHRKLQPFRET-
>Agrilus_planipennis
------------------------------------------------------------------------------------------------------------MT-SQR---------------------------------------QEL------HCNRTTKSSSNRHRKLQPFRET-
>Callinectes_sapidus
-----------------------------------------------------------------------MILFSWHK--------------------GHISPGNCILSVEPFGGLLT-----------------------QS--------CMCACASDTPLDIRNRQFSSNRHRKLQPFRTS-
>Apachyus_charteceus
----------------------------------------MTLLSICRET----------------------TLSRRKS------GSFHI--ALEP---IRGLDSRCMSS-SKQ---------------------------------------QERI--NHRPVYRTSKFSTNRHRKLQPFRET-
>Occasjapyx_japonicus
-----------------------------------------------------------------------------------------------------------MIF-FQR---------------------------------------QEPI------SFRTSKFSSNRHRKLQPFRET-
>Haploembia_palaui
-----------------------------------------MTLNTCRET----------------------TLTRGKS------GSFHL--SLEP---FRGLDSTCMIS-SKR---------------------------------------QEPI------TYRTSKFSSNRHRKLQPFRET-
>Isonychia_bicolor
-----------------------------------------MTFHRCNEI----------------------SLSRGKS------GRLHI--ALEP---IFGLDPRCMSS-SKW---------------------------------------QNPI------SCRTSKFSSNRHRKLQPFRET-
>Meganyctiphanes_norvegica
-----------------------------------------------------------------------MILISWHK--------------------GHISPGHSILLVEPLGGLLT-----------------------KS--------CMCACASDTPLDIRNRQFSSNRHRKLQPFRTS-
>Galloisiana_yuasai
----------------------------------------MTLTSTCRET----------------------TLSRRKS------GSFHI--SLEP---IRGLDSRCMIS-SKR---------------------------------------QEPI------SYRTSKFSTNRHRKLQPFRET-
>Tupiocoris_notatus
-----------------------------------------------------------------------------------------------------------MIS-SKL---------------------------------------QWPV------SYRTSKFSSNRHRKSQPFRET-
>Copidosoma_floridanum
MTFLRQRETSRISSRGKGTVYNTGGGGGSTSSLRRTSRAV-DGLLSSRRTIASTTTSPSDVKITGKQRNNINVLSSGSKEECGQCQYRRCSPLFL--SSSCSAPAACMII-EGRGRLSSRSSSSSSSCCGCASHKQQLQDEDDHCQNAA---LRVHLNRLLLVDHRTAKFSSNRHRKLQPFRET-
>Eudarcia_simulatricella
-----------------------------------------------------------------------MTLA-RSTETSIQ----------------CGEKSCCIE------------------------RVQR----AADCARS-----CMTLYGPRTTGRRTSKFSSNRHRKLQPFRET-
>Metallyticus_splendidus
----------------------------------------MTLLPTCRET----------------------TLSRGKS------GSFHI--ALEP---VRGLDSRCMIS-SKR---------------------------------------QEPI------SYRTSKFSSNRHRKLQPFRET-
>Tanzaniophasma_sp._AD-2013
-----------------------------------------------------------------------MTLTLTR--------IQR----------FSRKFSGCFYI-SPKPSFG----------------------------LEI---SRMILSKWQELLVHRTKFSSNRHRKSQPFREC-
>Panorpa_vulgaris
-----------------------------------------------------------------------MTLERYRETS-VLRGVPR------SPTALSPRGTKCMI------------------------------------------------PPRRHLSYRTSKFSSNRHRKLQPFRET-
>Corydalus_cornutus
-----------------------------------------MTLAGCIET----------------------LLSRRKP------GITQYS-VLEP---VLGIDTRCMIS-SKF---------------------------------------QEPI------SYRISKFSTNRHRKLQPFRET-
>Pseudomallada_prasinus
-MRQRRAETFRISSRGKGSVCSKGSGGRTLSRHRRNNSTTIISYIRCNRSL---------------------RWCDGSR------LSTWCSEIIED---TRYGPAACMII-ENYPPLPTRSRHNSN--------------------QDT---TQHFLNHLVLPKHRTSKFSSNRHRKLQPFRET-
>Calopteryx_splendens
----------------------------------------MT-LLSSKGI----------------------SLSRGRS------GFFSI--ALEP---ILGLDSRCMIS-SKR---------------------------------------QEPI------SYRTSKFSSNRHRKLQPFRET-
>Gryllotalpa_sp._AD-2013
----------------------------------------MTLLPTCRET----------------------TLTRGKS------GSFHI--ALAP---TRGLDSRCMIS-SKR---------------------------------------QEPI------SYRTSKFSSNRHRKLQPFRET-
>Clitarchus_hookeri
-----------------------------------------MTLLTCRET----------------------ILSRGKS------GSFHI--AEEP---ARGQDSRCMIS-TKR---------------------------------------QEPI------CYRTSKFSSNRHRKLQPFRET-
>Menopon_gallinae
-----------------------------------------------------------------------MTLKHFKE---ASLTRRKCRPNLSP----GRGQF------------------------------------------------RCMISHKGQPSYRKSKLSTNRHRKLQPFRET-
>Leuctra_sp._AD-2013
----------------------------------------MTLLPTCRET----------------------ILTRGKS------GSFHR--ALEP---VRGLDSRCMIS-YKR---------------------------------------RDPI------SHRISKFSTNRHRKLQPFRET-
>Polydesmus_angustus
-----------------------------------------------------------------------MTLISLK-------------ELLL----LRGLNCCCKLT-D-----------------------PHWESVLSRC--------MFSTEKFKPVSHRTPKFQSNRHRKLQPFRS--
>Acerentomon_sp._AD-2013
-----------------------------------------------------------------------MTLINSKEEKRSKDGSGKLYIRLQ--TFCGPGFINCMNLYQPHS-------------------------------SASPLSALCCT----EPSKSHYSSHSNQHRKLLPFREES
>Liposcelis_bostrychophila
-----------------------------------------------------------------------MTLTHFKE---ASLTRRKCRPNLSP----GRGQF------------------------------------------------RCMISHKGQPSYRKSKLSTNRHRKLQPFRET-
>Inocellia_crassicornis
-----------------------------------------MTLVNHTET----------------------LLSRGKS------GSSRYS-ALEP---ALGIDTRCMSS-SKR---------------------------------------QGPI------SYRISKYSTSRHRKLQPFRET-
>Ctenocephalides_felis
-----------------------------------------------------------------------MTLERYRETS-VLRGVDR------SACAPSSHSAKCMT------------------------------------------------PTRRRHCYRQSKFSSNRHRKLQPFRET-
>Thrips_palmi
-----------------------------------------MTFRSCKETW---------------------TVSRGRS------GSSPI--AQEP---GRGPGGRCMIS-YQP---------------------------------------HGPI------SHRTTKFSSNRHRKSQPFRET-
>Zorotypus_caudelli
----------------------------------------MTLLPLCRE-----------------------TLTRRKS------GNFHI--ALVP---TGGLDSGCMIS-SKR---------------------------------------KQPI------CYRTSKFSSNRHRKLQPFRET-
>Tricholepidion_gertschi
-----------------------------------------MTLLSWKEI----------------------SLSRGKS------GSFHI--TLEP---ILGLDTRCMIS-SKR---------------------------------------QEPI------SYRTSKFSSNRHRKLQPFRET-

HG0031 ENSDARG00000000540 asnsd1

>Danio_rerio
---------------------------------------------MSSKNIQRT-E-----PGASEERAQKEELNQKIKEQKIVVDELSNLKKNRRVYTQQPNSNIFFLADKGETLSSCKQDLDNMRKEYQDI-------------
>Acipenser_sinensis
---------------------------------------------MST-K-SGQDN-----TSEV-NTKQKEKLNKKIKEQQVLVDELSNLKKNRKVYIQQPNSNIFFLGDKSETLSESKEMLDNMKKEYQDL-------------
>Amia_calva
---------------------------------------------MSNKS-QAKDD-----QING-KAKQKEELNKKMKEQKVVIDELSNLKKNRKVYVQQQNSNIFFLEDKSETLATCKKSLDHMKKEYQDL-------------
>Channa_striata
-----------------------------------------------------------------------------------------------------------------------------MKKELQDM-------------
>Anser_cygnoides
--------------------------------------MAARRGEEEEKKEEEEG------EGESAVLRPRDQLSRRIKEQKVIIDELSNLKKNRKVYKQQPNSNIFFLVDRTETLSQCKIVLDELKKAHQEMENSEKTKIKK---
>Polypedates_megacephalus
---------------------------------------------MSDVG-VVLGE-----DCRSWTDKQKEEFYAKFREQNVVVDELSNLKKDRKVYRQQPNSSIYFLADRAKTLSDCKKTLDDLKHTHQLLENSENIQDTD---
>Porichthys_notatus
---------------------------------------------MSSKL-HDIDA-----SFG--GTPSIEVLNRKISEQKVVLDELSNLKKNRKVYIQQQNSNIFFLADRKQTLSSCKKELDNLKRDLQDS-------------
>Oryzias_latipes
-------------------------------------------MKMSSKK-YNNSD-----NSQE-QTTYKEELSKKIKEQKVVVDELSNLKKNRKVYIQQRNSNIFFLADRSQTLSTSRKELDYLKKELQDI-------------
>Trachinotus_blochii
---------------------------------------------MSSKN-RDNDD-----NFEG-QSALKEELNKKIKEQKVVVDELSNLKKNRKVYIQQRNSNIFFLGDRSQTLGSCKKELDNMKKELQDM-------------
>Cariama_cristata
------------------------------------------------------------------------------------------------------------------MLSQCKNTLDELKKAHQEMENSENTKVKK---
>Phoca_largha
---------------------------------------------MPSRGSRPEDGAGLF--SADHSTLHKEDLSSKIKEQKIVVDELSNLKKNRKVYRQQQNSNIFFLADRTEMLSESKNILDELKKEYQEIENSEKTKIKK---
>Hynobius_retardatus
---------------------------------------------MSGTKPKN------KEDGSDSAAWTKEDLDTKIREQTVVVDELSNLKPNRKVYTQQPNSNIFFLADRTATLSESKKTLDDLKTAHQELEKSQKARVK----
>Micropterus_salmoides
------------------------------------------MNKMSSKI-RDNDD-----NCEG-RCALKEELNKKIKEQRVVVDELSNLKKNRKVYIQQRNSNIFFLADRSQTLASCKREMDNMKKDLQDM-------------
>Lipotes_vexillifer
---------------------------------------------MPSRGARPEDGSGLV--PTDNSTQHKEDLSSKIKEQKIVVDELSNLKKNRKVYRQQQNSDIFFLADRTEMLSESKNTLDELRKEYQEIENSEKTKIKK---
>Astyanax_mexicanus
---------------------------------------------MSIRSSEQPED-----PAQE-ERPTREELNTKIKEQKVLVDELSNLKKNRKVYLQQPNSNIFFLSDKGETLSSCKKELDKMKKDYQDM-------------
>Calidris_pugnax
---------------------------------------------MSGRAARQEEEEEKEKREESAKARHKEELSRRIKEQKVVVDELSNLKKNRKVYRQQPNSNIFFLVDRTETLSQCKNTLDELKKAHQEMENSEKTKVKK---
>Callorhinchus_milii
------------------------------------------MEAAETRAAPGGEE-----RETRDQATRRQELHNKIKEQNVAINELSNLKETRRVYKQQPNSNIFFLTDKTTALTECKRAFDELKKEHQDLENPDKMMK-----
>Myotis_lucifugus
-------------------------------------------MEMPAREARPEDGAGRA-------APGREDLSSRIKEQKIVVDELSNLKKNRTVYRQQQNSNIFFLADRTELLSESKNILDELKKEYQEIENSEKTKTKK---
>Oreochromis_niloticus
---------------------------------------------MSSKG-CSSDE-----QFEG-QAAVKEELNKKIKEQKVLVDELCNLKKNRKVYIQQRNSNIFFVGDRSQTLTSCKKELDYMKKELQDI-------------
>Clupea_harengus
---------------------------------------------MTTKRNLPAAN-----TVQ-DETPQKEELSKKIKEQKILVDELSNLKKNRRVYLQQPNSNIFFLTDRGQMLSTSKKELDNLKKDYQDM-------------
>Latimeria_chalumnae
---------------------------------------------MSGRV-QGEDN-----ESQ-LRTSHKEELHKKIKEQKVLVDELSNLKKNRKVYQQQPNSNIFFLADRSKTLSDGKKKLDDMKREYQEL-------------
>Columba_livia
---------------------------------------------MSGRPPRQEE------GGDSAKLRHKDELSRKIKEQKVVVDELSNLKKNRKVYRQQPNSNIFFLVDRTETLSQCKNTLDELKKAHQEMENSEKTKIKK---
>Alligator_sinensis
MRRERARVRNHPAAWVGGASCAEGVWLGGAEVRSQSKGAARLVSGMSGRAPRR--------DEEEARLRRKEELNGKIKEQKIVVDELSNLKKNRKVYRQQPNSNIFFLADRTETLSQCKNKLDELKKAHQEIENSEKSKIKK---
>Kryptolebias_marmoratus
---------------------------------------------MSLKS-PDSSD-----NSAE-QSALKEELNKKIKEQKIVLDELSNLKKNRRVFIQQRNSNIFFLADRSQTLSSCKNDLDVLKKELQDV-------------
>Sarcophilus_harrisii
---------------------------------------------MPGRGTRVEDGPEPA--ISAKLPLNKEELNSKIKEQKVVVDELSNLKKNRKVYKQQPNSNIFFLADRTEMLSESKNTLDELKKIHQEIENTEKNKSKK---
>Monodelphis_domestica
---------------------------------------------MPGKGTRREDATEPG--TFASLPLNKEELNSKIKEQKVVVDELSNLKKNRKVYKQQPNSNIFFLSDRTEMLSESKNTLDELKKVHQEIENTEKNKIKK---
>Trichosurus_vulpecula
------------------------------------------------------------------------------------------------------------------MLSESKNTLDELKKIHQEIENTEKNKIKK---
>Esox_lucius
---------------------------------------------MSV-----KNE-----IFDEEESALKVELNKKIKEQKVVVDELSNLKKSRRVYVQQPNSNIFFLADRGQTLSSCKRARQHEEGSPKYVIF-----------
>Aquila_chrysaetos_canadensis
---------------------------------------------MSGRAPRQEEG-------ESAKLRRKEELSRRIKEQKVVVDELSNLKKNRKVYRQQPNSNIFFLVDRTETLSQCKNTLDELKKAHQEMENSEKAKIKK---
>Gadus_morhua
---------------------------------------------MTSN--SREDE-----ISEA-HSAIKEELNKKIKEQKIVVDELSNLKKNRKVYVQQRNSNIFFLADKGQMLGTNKKELDILKKELQDM-------------
>Numida_meleagris
--------------------------------------------------------MSRREEEEGAGVCRKEELSRRIKEQKVVVDELSNLKKNRKVYKQQPNSNIFFLVDRTETLSQCKNTLDQLKKAHQELENSEKTKVKK---
>Acanthogobius_hasta
------------------------------------------------MS-RNTED-----NNEG-LSALKEEMNKKIREQKVVLDELSNLKKGRKVYTQQRNSNIFFLAERSQTLSACKKELDVMKKELQDI-------------
>Condylura_cristata
---------------------------------------------MPSQ--GPEEGARLV--SSDNSSLIKEDLSSKIKEQKIVVDELSNLKKNRKVYRQQQNSNVFFLADRTEMLSESKNILDELKKEYQEIENLDKTQIKK---
>Oryctolagus_cuniculus
---------------------------------------------MPGRGVHPEDSAVPV--PAGGRTLHKEDLCNKIKEQKVVVDELSNLKKNRRVYRQQQNSNIFFLADRTEMLSESKNILDELKKEYQEIENSEKSSIKK---
>Lutjanus_erythropterus
-----------------------------------------------------------------------------------------------------------------------------MKKELQDM-------------
>Ornithorhynchus_anatinus
------------------------------------------------------------------------------------------------------------------MLSESKNTLDELKKTHQEMDSLEKTKAKK---
>Plecoglossus_altivelis
---------------------------------------------MSSKCQLDLEN-----LFAGQESKRKEELNKKINEQKVVVDELNNLKKNRKVYSQQPNSNIFFLADRQQCLSSYKNKVDNMKKELLDI-------------
>Scleropages_formosus
---------------------------------------------MSVRE-QEKEG-----ASLG-DTSLKEELNKKIKEQKVVVDELSNLKKNRRVYVQQPNSNIFFLADKAETLGSCRRTLDNMKKEYQDL-------------
>Pseudopodoces_humilis
---------------------------------------------MSGRPPRQEDG-------ESAALRHKEELSRRIKEQKIVVDELSNLKKNRKVYKQQPNSNIFFLADRTEMLSQCKNTLDELKKAHQELENSEKAKIKK---
>Lateolabrax_japonicus
---------------------------------------------MSS-K-SDNSD-----NFDG-QSAAKEELNTKIKEQKVVVDELSNLKKNRKVYIQQRNSNIFFLADRGQTLGSCKKELDNMKKELLDM-------------
>Anoplopoma_fimbria
---------------------------------------------MSSKI-HSNED-----NCEG-TSALKEELNIKIKEQKVVVDELSNLKKNRKVYVQQRNSNIFFLADRGQTLGSCKKELDNMKKHLQDM-------------
>Ceratotherium_simum_simum
---------------------------------------------MPSRGTRPEDGAGLV--PTDDSAPHKEDLSNKIKEQKIVVDELSNLKKNRKVYRQQQNSNIFFLADRTEMLSESKNILDELKKEYQEIENSEKTKIKK---
>Manis_javanica
---------------------------------------------MPSGGSRPEDGAAPV--PTENSILHKEDLSSKIKEQKIVVDELSNLKKNRKVYRQQQNSNIFFLADRTEMLSESKNILDELKKEYQEIENSEKTKIKK---
>Hippoglossus_hippoglossus
---------------------------------------------MSSKG-RDNED-----SFEW-QSALKEELNKKIKEQKVVVDELSNLKKNRNVYIQQRNSNIFFLADRGQTLGSCKKELDNMKKELQDM-------------
>Pongo_abelii
---------------------------------------------MPSRGTRPEDSSVLI--PTDNSTPHKEDLSSKIKEQKIVVDELSNLKKNRKVYRQQQNSNIFFLADRTEMLSESKNILDELKKEYQEIENLDKTKIKK---
>Loxodonta_africana
-------------------------------------------MEMPSSRAHPEDSAELV--PSDNSTPHKEELNSKIKEQKIVVDELSNLKKNRKVYKQQQNSNIFFLADRTEMLSESKNILDELKKEYQEIENLEKTKIKK---
>Callospermophilus_lateralis
---------------------------------------MLRGVEMPIRGTRSDDSAVLF--STDYSTAQKEELSNKIKEQKIVVDELSNLKKNRKVYRQQQNSNIFFLSDRTEMLSENKNILDELKKEYQDIENSEKTKIKK---
>Oncorhynchus_mykiss
---------------------------------------------MSAKNQREN-E-----NIDEEESTLKEELNKKIKEQKIVFDELSNLKKNRRVYVQQPNSNIFFLADRGQTLASCKKELDNMKKDM----------------
>Tupaia_chinensis
-------------------------------------------MEMPSRGARLEDSAVRG--PTDDSTPHREDLHSKIKEQKIVVDELSNLKKNRRVYKQQQNSNIFFLADRAELLSESKNVLDELKKEYQEIENSEKTKIKK---
>Lepisosteus_oculatus
---------------------------------------------MSNKN-QVKDD-----VVQA-KAKQKEELNKKIREQKIVIDELSNLKKTRRVYVQQPNSHIFFLANKAETLSASKNSLENMKKEYQDV-------------
>Ictalurus_punctatus
----------------------------------------------------MSTN-----LVHE-ERATKEELNKQIKEQKVVIDELSNLKKNRKVFVQQPNSNIFFLADKGEMLSTCKKDLDKMKKEYQDM-------------
>Trichechus_manatus_latirostr
-------------------------------------------MEMPSSRAHPEDSTELV--PSDNSTPHREDLSSKIKEQKIVVDELSNLKKNRKVYKQQQNSNIFFLADRTEMLSESKNTLDELKKEYQEIENLEKTKIKKSSS
>Squalus_acanthias
--------------------------------------------------MEGTSD-----KVQSSEADRKEELNKQMKEQNVVIDELSNLKKSRRVYKQQPNSNIFFSADRTQTLSECKCALEEMKKEYQDLENSEKMKK-----
>Thamnophis_sirtalis
-------------MIISATGKEPGLLLT------------TQQEQMSS-------QALLREGGEGSKLPDKEELNRKIKEQKIVVDEISNLKKNRKVYRQQPNSNIFFLADKTQTLSECKITLDELKKAHQEIENPEKTKTKK---
>Tyto_alba
-------------------------------------------------------------------------------------------------------------------------MLDELKKAHQEMENSEKTKSKK---
>Trachemys_scripta
---------------------------------------------MSGREPRR--------EEEGAKPHQKEELNRKIKEQKVVVDELSNLKKNRKVYKQQPNSNIFFLVDRTETLSECKNTLDDLKRAHQEIENSEKTKIRK---
>Takifugu_rubripes
---------------------------------------------------MAFND-----HFDE-QSTLKEELNRKLKEQKVVVDELSNLKKNRKVYTQQRNSNIFFLADRAETLSSNKKKLDFMKKEVQNMKD-----------
>Orycteropus_afer_afer
------------------------------------------------------------------------------------------------------------------MLSESKNILDELKKEYQEIENSEKTKIKK---

HG0031 ENSG00000138381 ASNSD1

>Homo_sapiens
-----------------------------------------------MPSRGTRPE---DSSVLIPTDNSTPHKEDLSSKIKEQKIVVDELSNLKKNRTEQKCCLRARIYWMN------------------------------------
>Acipenser_sinensis
------------------------------------------------M----STK---SGQD-NTSEVNTKQKEKLNKKIKEQQVLVDELSNLKKNRKVYIQQPNSNIFFLGDKSETLSESKEMLDNMKKEYQDL-------------
>Amia_calva
-----------------------------------------------MS----NKS---QAKD-DQINGKAKQKEELNKKMKEQKVVIDELSNLKKNRKVYVQQQNSNIFFLEDKSETLATCKKSLDHMKKEYQDL-------------
>Channa_argus
-----------------------------------------------MT----SKT---QDNDINVEGQ-CGLKEELNRKIKEQKVAVDELSNLKKNRKVYIQQRNSNIFFLADRSQTLGSCKKELDNMKKELQDM-------------
>Anser_cygnoides
-----------------------------------------------MAARRGEEEEKKEEEEGEGESAVLRPRDQLSRRIKEQKVIIDELSNLKKNRKVYKQQPNSNIFFLVDRTETLSQCKIVLDELKKAHQEMENSEKTKIKK---
>Nanorana_parkeri
-----------------------------------------------MSD---------IVAGEDCRGWTDKQKEDFYTKFREQKVVIDELTNLKKDRKVYRQQPNSNIFFVADKVKTLSECKKTLADLEHTYQILENSEKLQDKK---
>Porichthys_notatus
-----------------------------------------------MS----SKL---HDID-ASFGG-TPSIEVLNRKISEQKVVLDELSNLKKNRKVYIQQQNSNIFFLADRKQTLSSCKKELDNLKRDLQDS-------------
>Oryzias_latipes
-------------------------------------------MK--MS----SKK---YNNSDNSQEQ-TTYKEELSKKIKEQKVVVDELSNLKKNRKVYIQQRNSNIFFLADRSQTLSTSRKELDYLKKELQDI-------------
>Trachinotus_blochii
-----------------------------------------------MS----SKN---RDNDDNFEGQ-SALKEELNKKIKEQKVVVDELSNLKKNRKVYIQQRNSNIFFLGDRSQTLGSCKKELDNMKKELQDM-------------
>Acinonyx_jubatus
-----------------------------------------------MPSRGSQPE---DGAALSSADNSTPHKEDLSSKIKEQKIVVDELSNLKKNRKVYRQQQNSNIFFLADRTEMLSESKNILDELKKEYQEIENSEKTKIKK---
>Hynobius_chinensis
-----------------------------------------------MSGTKPKNK-------EAGSDSAAWTKEDLDTKIREQTVVVDELSNLKPNRKVYTQQPNSNIFFLADRTATLSESKKTLDDLKTARQELEKSQEVRVK----
>Micropterus_salmoides
------------------------------------------MNK--MS----SKI---RDNDDNCEGR-CALKEELNKKIKEQRVVVDELSNLKKNRKVYIQQRNSNIFFLADRSQTLASCKREMDNMKKDLQDM-------------
>Orcinus_orca
-----------------------------------------------MPSRGARPE---DGSGLVPTDNSTQHKEDLSSKIKEQKIVVDELSNLKKNRI----------HWMS------------------------------------
>Astyanax_mexicanus
-----------------------------------------------MS----IRSS--EQPE-DPAQEERPTREELNTKIKEQKVLVDELSNLKKNRKVYLQQPNSNIFFLSDKGETLSSCKKELDKMKKDYQDM-------------
>Calidris_pugnax
-----------------------------------------------MSGRAA-RQEEEEEKEKREESAKARHKEELSRRIKEQKVVVDELSNLKKNRKVYRQQPNSNIFFLVDRTETLSQCKNTLDELKKAHQEMENSEKTKVKK---
>Callorhinchus_milii
-----------------------------------------------MEAAETRAAP--GGEE-RETRDQATRRQELHNKIKEQNVAINELSNLKETRRVYKQQPNSNIFFLTDKTTALTECKRAFDELKKEHQDLENPDKMMK-----
>Rousettus_aegyptiacus
---------------------------------------------MEMPFCETRPE---NGAELVPTDNSAPHKVDLSSKIKEQKVVVDELSNLKKNRI----------YWMS------------------------------------
>Neolamprologus_brichardi
-----------------------------------------------MS----SKS---YSNDEQFEGQ-AAVKEELNKKIKEQKVLVDELCNLKKNRKVYIQQRNSNIFFVGERSQTLTSCKKELDYMKKELQDI-------------
>Clupea_harengus
-----------------------------------------------MT----TKRN--LPAANTVQDE-TPQKEELSKKIKEQKILVDELSNLKKNRRVYLQQPNSNIFFLTDRGQMLSTSKKELDNLKKDYQDM-------------
>Latimeria_chalumnae
-----------------------------------------------MS----GRV---QGED-NESQLRTSHKEELHKKIKEQKVLVDELSNLKKNRKVYQQQPNSNIFFLADRSKTLSDGKKKLDDMKREYQEL-------------
>Columba_livia
-----------------------------------------------MSGRPP-RQEE------GGDSAKLRHKDELSRKIKEQKVVVDELSNLKKNRKVYRQQPNSNIFFLVDRTETLSQCKNTLDELKKAHQEMENSEKTKIKK---
>Alligator_sinensis
MRRERARVRNHPAAWVGGASCAEGVWLGGAEVRSQSKGAARLVSG--MSGRAP-RRD--------EEEARLRRKEELNGKIKEQKIVVDELSNLKKNRKVYRQQPNSNIFFLADRTETLSQCKNKLDELKKAHQEIENSEKSKIKK---
>Danio_rerio
-----------------------------------------------MS----SKNI--QRTE-PGASEERAQKEELNQKIKEQKIVVDELSNLKKNRRVYTQQPNSNIFFLADKGETLSSCKQDLDNMRKEYQDI-------------
>Kryptolebias_marmoratus
-----------------------------------------------MS----LKS---PDSSDNSAEQ-SALKEELNKKIKEQKIVLDELSNLKKNRRVFIQQRNSNIFFLADRSQTLSSCKNDLDVLKKELQDV-------------
>Sarcophilus_harrisii
-----------------------------------------------MPGRGTRVE---DGPEPAISAKLPLNKEELNSKIKEQKVVVDELSNLKKNRTEQKCSLKVKTH---------------------------------------
>Monodelphis_domestica
-----------------------------------------------MPGKGTRRE---DATEPGTFASLPLNKEELNSKIKEQKVVVDELSNLKKNRIEQKCSLKAKIH---------------------------------------
>Esox_lucius
-----------------------------------------------MS----------VKNEIFDEEE-SALKVELNKKIKEQKVVVDELSNLKKSRRVYVQQPNSNIFFLADRGQTLSSCKRARQHEEGSPKYVIF-----------
>Aquila_chrysaetos_canadensis
-----------------------------------------------MSGRAP-RQEEG-------ESAKLRRKEELSRRIKEQKVVVDELSNLKKNRKVYRQQPNSNIFFLVDRTETLSQCKNTLDELKKAHQEMENSEKAKIKK---
>Meleagris_gallopavo
-----------------------------------------------------------MSRREEEESAELGHKEELSRRIKEQKVVVDELSNLKKNRKVYRQQPNSNIFFLVDRTETLSQCKNTLDQLKKAHQELENSDKTKVKK---
>Acanthogobius_hasta
------------------------------------------------------MS---RNTEDNNEGL-SALKEEMNKKIREQKVVLDELSNLKKGRKVYTQQRNSNIFFLAERSQTLSACKKELDVMKKELQDI-------------
>Condylura_cristata
-----------------------------------------------MPSQ--GPE---EGARLVSSDNSSLIKEDLSSKIKEQKIVVDELSNLKKNRKVYRQQQNSNVFFLADRTEMLSESKNILDELKKEYQEIENLDKTQIKK---
>Oryctolagus_cuniculus
-----------------------------------------------MPGRGVHPE---DSAVPVPAGGRTLHKEDLCNKIKEQKVVVDELSNLKKNRRVYRQQQNSNIFFLADRTEMLSESKNILDELKKEYQEIENSEKSSIKK---
>Plecoglossus_altivelis
-----------------------------------------------MS----SKCQ--LDLENLFAGQESKRKEELNKKINEQKVVVDELNNLKKNRKVYSQQPNSNIFFLADRQQCLSSYKNKVDNMKKELLDI-------------
>Scleropages_formosus
-----------------------------------------------MS----VRE---QEKE-GASLGDTSLKEELNKKIKEQKVVVDELSNLKKNRRVYVQQPNSNIFFLADKAETLGSCRRTLDNMKKEYQDL-------------
>Zonotrichia_albicollis
-----------------------------------------------MSGRPP-RQE-------DGDSAALRHKEELSRRIKEQKVVVDELSNLKKNRKVYKQQPNSNIFFLADRTEMLSQSKNTLDELKKAHQELENSEKAKIKK---
>Lateolabrax_japonicus
------------------------------------------------M----SSK---SDNSDNFDGQ-SAAKEELNTKIKEQKVVVDELSNLKKNRKVYIQQRNSNIFFLADRGQTLGSCKKELDNMKKELLDM-------------
>Perca_flavescens
-----------------------------------------------MS----SKS---RDNDDNFEGQ-SAHKEEMNRKIKEQKIVVDELSNLKKNRKVYIQQRNSNIFFLADRSQTLGLCKKELDNMKKDLQDM-------------
>Ceratotherium_simum_simum
-----------------------------------------------MPSRGTRPE---DGAGLVPTDDSAPHKEDLSNKIKEQKIVVDELSNLKKNRKVYRQQQNSNIFFLADRTEMLSESKNILDELKKEYQEIENSEKTKIKK---
>Manis_javanica
-----------------------------------------------MPSGGSRPE---DGAAPVPTENSILHKEDLSSKIKEQKIVVDELSNLKKNRKVYRQQQNSNIFFLADRTEMLSESKILFPSLHRYIG---------------
>Hippoglossus_hippoglossus
-----------------------------------------------MS----SKG---RDNEDSFEWQ-SALKEELNKKIKEQKVVVDELSNLKKNRNVYIQQRNSNIFFLADRGQTLGSCKKELDNMKKELQDM-------------
>Loxodonta_africana
---------------------------------------------MEMPSSRAHPE---DSAELVPSDNSTPHKEELNSKIKEQKIVVDELSNLKKNRTEQKCFLKAKIYWMN------------------------------------
>Ictidomys_tridecemlineatus
-----------------------------------------MLRGVEMPIRGTRSD---DSAVSVSTDNSTAHKEELSNKIKEQKIVVDELSNLKKNRKVYRQQQNSNIFFLADRTEMLSENKNILDELKKEYQDIENSEKTKIKK---
>Oncorhynchus_mykiss
-----------------------------------------------MS----AKNQ--RENENIDEEE-STLKEELNKKIKEQKIVFDELSNLKKNRRVYVQQPNSNIFFLADRGQTLASCKKELDNMKKDM----------------
>Tupaia_chinensis
---------------------------------------------MEMPSRGARLE---DSAVRGPTDDSTPHREDLHSKIKEQKIVVDELSNLKKNRRVYKQQQNSNIFFLADRAELLSESKNVLDELKKEYQEIENSEKTKIKK---
>Lepisosteus_oculatus
-----------------------------------------------MS----NKN---QVKD-DVVQAKAKQKEELNKKIREQKIVIDELSNLKKTRRVYVQQPNSHIFFLANKAETLSASKNSLENMKKEYQDV-------------
>Ictalurus_punctatus
------------------------------------------------------------MST-NLVHEERATKEELNKQIKEQKVVIDELSNLKKNRKVFVQQPNSNIFFLADKGEMLSTCKKDLDKMKKEYQDM-------------
>Trichechus_manatus_latirostr
---------------------------------------------MEMPSSRAHPE---DSTELVPSDNSTPHREDLSSKIKEQKIVVDELSNLKKNRKVYKQQQNSNIFFLADRTEMLSESKNTLDELKKEYQEIENLEKTKIKKSSS
>Squalus_acanthias
-------------------------------------------------------ME--GTSD-KVQSSEADRKEELNKQMKEQNVVIDELSNLKKSRRVYKQQPNSNIFFSADRTQTLSECKCALEEMKKEYQDLENSEKMKK-----
>Protobothrops_mucrosquamatus
-----------------------------------------------MSSPA--------LLGEGGEESKLPDKEELNRKIKEQKIVVDELSNLKKNRKVYRQQPNSNIFFLADRTQTLSECRITLDELKKAHQEIENSEKTKTKK---
>Trachemys_scripta
-----------------------------------------------MSGREP-RRE--------EEGAKPHQKEELNRKIKEQKVVVDELSNLKKNRKVYKQQPNSNIFFLVDRTETLSECKNTLDDLKRAHQEIENSEKTKIRK---
>Takifugu_rubripes
-----------------------------------------------------------MAFNDHFDEQ-STLKEELNRKLKEQKVVVDELSNLKKNRKVYTQQRNSNIFFLADRAETLSSNKKKLDFMKKEVQNMKD-----------

HG0031 ENSGALG00000041666

>Gallus_gallus
----------------------------------------------------------MSRREEEETAELGYKGELSRRIKEQKVVVDELSNLKKNRKVYRQQPNSNIFFLVDRTEALSQCKNTLDQLKKAHQELENSEKTKVKK---
>Acipenser_sinensis
-----------------------------------------------MSTK-SGQD------NTS-EVNTKQKEKLNKKIKEQQVLVDELSNLKKNRKVYIQQPNSNIFFLGDKSETLSESKEMLDNMKKEYQDL-------------
>Amia_calva
-----------------------------------------------MSNKSQAKD------DQI-NGKAKQKEELNKKMKEQKVVIDELSNLKKNRKVYVQQQNSNIFFLEDKSETLATCKKSLDHMKKEYQDL-------------
>Channa_striata
--------------------------------------------------------------------------------------------------------------------------------MKKELQDM-------------
>Anas_platyrhynchos
--------------------------------------------------------------------------------------------------------------------------------------MENSEKTKIKK---
>Pelophylax_nigromaculatus
-----------------------------------------------MTDV------EVEKREDCRAWTDKQKEEFYTKFREQKVVVDELSNLKQDRKVYRQQPNSNIFFLADKAKTLSECKKTLDDLKKTSQILENSEKIQDQK---
>Porichthys_notatus
-----------------------------------------------MSSKLHDID--------ASFGGTPSIEVLNRKISEQKVVLDELSNLKKNRKVYIQQQNSNIFFLADRKQTLSSCKKELDNLKRDLQDS-------------
>Oryzias_latipes
---------------------------------------------MKMSSKKYNNS------DNS-QEQTTYKEELSKKIKEQKVVVDELSNLKKNRKVYIQQRNSNIFFLADRSQTLSTSRKELDYLKKELQDI-------------
>Caprimulgus_carolinensis
--------------------------------------------------------------------------------------------------------------------------------------MENSEKTKIKK---
>Trachinotus_blochii
-----------------------------------------------MSSKNRDND------DNF-EGQSALKEELNKKIKEQKVVVDELSNLKKNRKVYIQQRNSNIFFLGDRSQTLGSCKKELDNMKKELQDM-------------
>Cariama_cristata
--------------------------------------------------------------------------------------------------------------------MLSQCKNTLDELKKAHQEMENSENTKVKK---
>Acinonyx_jubatus
-----------------------------------------------MPSRGSQPED--GAALSSADNSTPHKEDLSSKIKEQKIVVDELSNLKKNRKVYRQQQNSNIFFLADRTEMLSESKNILDELKKEYQEIENSEKTKIKK---
>Hynobius_retardatus
-----------------------------------------------MSGTKPK------NKEDGSDSAAWTKEDLDTKIREQTVVVDELSNLKPNRKVYTQQPNSNIFFLADRTATLSESKKTLDDLKTAHQELEKSQKARVK----
>Micropterus_salmoides
--------------------------------------------MNKMSSKIRDND------DNC-EGRCALKEELNKKIKEQRVVVDELSNLKKNRKVYIQQRNSNIFFLADRSQTLASCKREMDNMKKDLQDM-------------
>Lipotes_vexillifer
-----------------------------------------------MPSRGARPED--GSGLVPTDNSTQHKEDLSSKIKEQKIVVDELSNLKKNRKVYRQQQNSDIFFLADRTEMLSESKNTLDELRKEYQEIENSEKTKIKK---
>Pygocentrus_nattereri
-----------------------------------------------MSVRSSEQT------DDLAQDERSAREELNRRIKEQKVVIDELSNLKKNRKVYIQQPNSNIFFLTDKGETLSSCKKELDKMKRHYQDM-------------
>Calidris_pugnax
-----------------------------------------------MSGRAARQEEEEEKEKREESAKARHKEELSRRIKEQKVVVDELSNLKKNRKVYRQQPNSNIFFLVDRTETLSQCKNTLDELKKAHQEMENSEKTKVKK---
>Callorhinchus_milii
--------------------------------------------MEAAETRAAPGG-----EERETRDQATRRQELHNKIKEQNVAINELSNLKETRRVYKQQPNSNIFFLTDKTTALTECKRAFDELKKEHQDLENPDKMMK-----
>Myotis_lucifugus
---------------------------------------------MEMPAREARPED--GAGRA-----APGREDLSSRIKEQKIVVDELSNLKKNRTVYRQQQNSNIFFLADRTELLSESKNILDELKKEYQEIENSEKTKTKK---
>Oreochromis_niloticus
-----------------------------------------------MSSKGCSSD------EQF-EGQAAVKEELNKKIKEQKVLVDELCNLKKNRKVYIQQRNSNIFFVGDRSQTLTSCKKELDYMKKELQDI-------------
>Pterocles_gutturalis
--------------------------------------------------------------------------------------------------------------------------------------MENSEKTKIKK---
>Alosa_alosa
--------------------------MCGIQVDNIS---YIVLFLLTMTTKRNLPA------ADTVQEETTQKEELSKQIKEQNILVDELSNLKKNRRVYLQQPNSNIFFLTDREQMLSTCKKDLDNLKKEYQAM-------------
>Latimeria_chalumnae
--------------------------------------------------------------------------------------------------------------------------------MKREYQEL-------------
>Columba_livia
-----------------------------------------------MSGRPPR------QEEGGDSAKLRHKDELSRKIKEQKVVVDELSNLKKNRKVYRQQPNSNIFFLVDRTETLSQCKNTLDELKKAHQEMENSEKTKIKK---
>Alligator_sinensis
MRRERARVRNHPAAWVGGASCAEGVWLGGAEVRSQSKGAARLVSG--MSGRAPR------RD--EEEARLRRKEELNGKIKEQKVIVVDELSNLKKNRKVYRQQPNSNIFFLADRTETLSQCKNKLDELKKAHQEIENSEKSKIKK---
>Cuculus_canorus
--------------------------------------------------------------------------------------------------------------------------------------MENSQKTEIKK---
>Pimephales_promelas
-----------------------------------------------MSSKNIHQP------EPGTQEELAQKEELNKKIKEQKILVDELSNLKKNRRVYTQQPNSNIFFLADKGESLSSCKKDLDNMRKEYQDI-------------
>Poecilia_formosa
-----------------------------------------------MSQTNSGSS------DNV-SKYSALKEELNRKVKEQKIIVDELSNLKKNRKVYIQQRNSNVFFLADRSQTLSSCKKQLDYMKKEQQDL-------------
>Sarcophilus_harrisii
-----------------------------------------------MPGRGTRVED--GPEPAISAKLPLNKEELNSKIKEQKVVVDELSNLKKNRKVYKQQPNSNIFFLADRTEMLSESKNTLDELKKIHQEIENTEKNKSKK---
>Monodelphis_domestica
-----------------------------------------------MPGKGTRRED--ATEPGTFASLPLNKEELNSKIKEQKVVVDELSNLKKNRKVYKQQPNSNIFFLSDRTEMLSESKNTLDELKKVHQEIENTEKNKIKK---
>Trichosurus_vulpecula
--------------------------------------------------------------------------------------------------------------------MLSESKNTLDELKKIHQEIENTEKNKIKK---
>Esox_lucius
-----------------------------------------------MSVKN----------EIFDEEESALKVELNKKIKEQKVVVDELSNLKKSRRVYVQQPNSNIFFLADRGQTLSSCKRARQHEEGSPKYVIF-----------
>Aquila_chrysaetos_canadensis
-----------------------------------------------MSGRAPR------QEE-GESAKLRRKEELSRRIKEQKVVVDELSNLKKNRKVYRQQPNSNIFFLVDRTETLSQCKNTLDELKKAHQEMENSEKAKIKK---
>Gadus_morhua
-----------------------------------------------MTSNS-RED------EIS-EAHSAIKEELNKKIKEQKIVVDELSNLKKNRKVYVQQRNSNIFFLADKGQMLGTNKKELDILKKELQDM-------------
>Gavia_stellata
--------------------------------------------------------------------------------------------------------------------------------------MENSEKTKIKK---
>Acanthogobius_hasta
--------------------------------------------------MSRNTE------DNN-EGLSALKEEMNKKIREQKVVLDELSNLKKGRKVYTQQRNSNIFFLAERSQTLSACKKELDVMKKELQDI-------------
>Balearica_regulorum_gibberic
--------------------------------------------------------------------------------------------------------------------------------------MENSEKTKIKK---
>Condylura_cristata
-----------------------------------------------MPSQ--GPEE--GARLVSSDNSSLIKEDLSSKIKEQKIVVDELSNLKKNRKVYRQQQNSNVFFLADRTEMLSESKNILDELKKEYQEIENLDKTQIKK---
>Oryctolagus_cuniculus
-----------------------------------------------MPGRGVHPED--SAVPVPAGGRTLHKEDLCNKIKEQKVVVDELSNLKKNRRVYRQQQNSNIFFLADRTEMLSESKNILDELKKEYQEIENSEKSSIKK---
>Lutjanus_erythropterus
--------------------------------------------------------------------------------------------------------------------------------MKKELQDM-------------
>Ornithorhynchus_anatinus
--------------------------------------------------------------------------------------------------------------------MLSESKNTLDELKKTHQEMDSLEKTKAKK---
>Plecoglossus_altivelis
-----------------------------------------------MSSKCQLDL-----ENLFAGQESKRKEELNKKINEQKVVVDELNNLKKNRKVYSQQPNSNIFFLADRQQCLSSYKNKVDNMKKELLDI-------------
>Scleropages_formosus
-----------------------------------------------MSVREQEK-------EGASLGDTSLKEELNKKIKEQKVVVDELSNLKKNRRVYVQQPNSNIFFLADKAETLGSCRRTLDNMKKEYQDL-------------
>Taeniopygia_guttata
------------------------------------------------MSGGPA------RQEDGESAALRHKEELSRRIKEQKVVVDELSNLKKNRKVYKQQPNSNIFFLADRTEMLSQCKNTLDELKKAHQELENSEKAKIKK---
>Nipponia_nippon
--------------------------------------------------------------------------------------------------------------------------------------MENSEKTRNKK---
>Lateolabrax_japonicus
-----------------------------------------------MSSK-SDNS------DNF-DGQSAAKEELNTKIKEQKVVVDELSNLKKNRKVYIQQRNSNIFFLADRGQTLGSCKKELDNMKKELLDM-------------
>Perca_flavescens
-----------------------------------------------MSSKSRDND------DNF-EGQSAHKEEMNRKIKEQKIVVDELSNLKKNRKVYIQQRNSNIFFLADRSQTLGLCKKELDNMKKDLQDM-------------
>Ceratotherium_simum_simum
-----------------------------------------------MPSRGTRPED--GAGLVPTDDSAPHKEDLSNKIKEQKIVVDELSNLKKNRKVYRQQQNSNIFFLADRTEMLSESKNILDELKKEYQEIENSEKTKIKK---
>Manis_javanica
-----------------------------------------------MPSGGSRPED--GAAPVPTENSILHKEDLSSKIKEQKIVVDELSNLKKNRKVYRQQQNSNIFFLADRTEMLSESKNILDELKKEYQEIENSEKTKIKK---
>Picoides_pubescens
--------------------------------------------------------------------------------------------------------------------------------------MENPEKTKIKK---
>Hippoglossus_hippoglossus
-----------------------------------------------MSSKGRDNE------DSF-EWQSALKEELNKKIKEQKVVVDELSNLKKNRNVYIQQRNSNIFFLADRGQTLGSCKKELDNMKKELQDM-------------
>Microcebus_murinus
-----------------------------------------------MPSRATHPED--SAVLVSTDESTPYKGDLSNKIKEQKIVVDELSNLKKNRKVYKQQQNSNIFFLADRTEMLSQSKNTLDELKKEYQEIENSEKTKIKK---
>Loxodonta_africana
---------------------------------------------MEMPSSRAHPED--SAELVPSDNSTPHKEELNSKIKEQKIVVDELSNLKKNRKVYKQQQNSNIFFLADRTEMLSESKNILDELKKEYQEIENLEKTKIKK---
>Fulmarus_glacialis
--------------------------------------------------------------------------------------------------------------------------------------MENSEKTKIKK---
>Melopsittacus_undulatus
--------------------------------------------------------------------------------------------------------------------------------------MENSEKTKIEK---
>Callospermophilus_lateralis
-----------------------------------------MLRGVEMPIRGTRSDD--SAVLFSTDYSTAQKEELSNKIKEQKIVVDELSNLKKNRKVYRQQQNSNIFFLSDRTEMLSENKNILDELKKEYQDIENSEKTKIKK---
>Oncorhynchus_mykiss
-----------------------------------------------MSAKNQREN------ENIDEEESTLKEELNKKIKEQKIVFDELSNLKKNRRVYVQQPNSNIFFLADRGQTLASCKKELDNMKKDM----------------
>Tupaia_chinensis
---------------------------------------------MEMPSRGARLED--SAVRGPTDDSTPHREDLHSKIKEQKIVVDELSNLKKNRRVYKQQQNSNIFFLADRAELLSESKNVLDELKKEYQEIENSEKTKIKK---
>Lepisosteus_oculatus
-----------------------------------------------MSNKNQVKD-------DVVQAKAKQKEELNKKIREQKIVIDELSNLKKTRRVYVQQPNSHIFFLANKAETLSASKNSLENMKKEYQDV-------------
>Ictalurus_punctatus
-----------------------------------------------------MST------NLV-HEERATKEELNKQIKEQKVVIDELSNLKKNRKVFVQQPNSNIFFLADKGEMLSTCKKDLDKMKKEYQDM-------------
>Trichechus_manatus_latirostr
---------------------------------------------MEMPSSRAHPED--STELVPSDNSTPHREDLSSKIKEQKIVVDELSNLKKNRKVYKQQQNSNIFFLADRTEMLSESKNTLDELKKEYQEIENLEKTKIKKSSS
>Aptenodytes_forsteri
--------------------------------------------------------------------------------------------------------------------------------------MENSEKTKIKK---
>Squalus_acanthias
-----------------------------------------------ME----G-T-----SDKVQSSEADRKEELNKQMKEQNVVIDELSNLKKSRRVYKQQPNSNIFFSADRTQTLSECKCALEEMKKEYQDLENSEKMKK-----
>Anolis_carolinensis
-----------------------------------------------MSSQMPQ------REEEGEGAKLGDKEELSRKIKEQKVIVDELSNLKKNRKVYRQQQNSNIFFLADRTQTLSECKNTLDELKKAHQEMENSEKTGIKK---
>Tyto_alba
---------------------------------------------------------------------------------------------------------------------------MLDELKKAHQEMENSEKTKSKK---
>Chelonia_mydas
-----------------------------------------------MSGREPR------REE--EGAKLHQKDELTRKIKEQKVVVDELSNLKKNRKVYKQQPNSNIFFLVDRTEMLSECKNTLDDLKRAHQEIENSEKTKIRK---
>Takifugu_rubripes
-----------------------------------------------------MAF-----NDHF-DEQSTLKEELNRKLKEQKVVVDELSNLKKNRKVYTQQRNSNIFFLADRAETLSSNKKKLDFMKKEVQNMKD-----------
>Calypte_anna
--------------------------------------------------------------------------------------------------------------------------------------MENSDKTKVKK---
>Orycteropus_afer_afer
--------------------------------------------------------------------------------------------------------------------MLSESKNILDELKKEYQEIENSEKTKIKK---

HG0032 ENSDARG00000104981 SLC45A4

>Danio_rerio
MGILRCFPSGFSRPSPLGLPS--TLY-----------STASIQIMKLVLTEWQLIWIFTVSSRQ
>Acipenser_sinensis
MGINRGAPTCVA-FPSLCCQP--TLF-----------STCSIAIMKLVLTEWQLIWIL-NCSRQ
>Amia_calva
MGIHRGAPTCVA-RPLLHCSP--SLF-----------STSSSEIMKLVLTEWQLIWIFTNCSRQ
>Channa_argus
MGIFRGAPPTAAACPPPPSH------CTPTHSIIQPSIQPFLYKMTLELTEWSSIWIFSACNRQ
>Pseudacris_regilla
MGINSASPVCVA-FPACSRKT--ENF-----------TTSSIEIMKLVLTEWQL-WIFNSNSQ-
>Chaetura_pelagica
--------------------------------------------MKLVLTEWQL-WIFNSNSQ-
>Oryzias_melastigma
MGIFRGAPPAVVTCPPPSRP-PCSLLPAPPQTVIEPSIQPPQHRMTLELTEWSSIRIFTACNRQ
>Mustela_putorius_furo
MGSNGGPPSCVS-FPPLCSRT--NNF-----------STASITMMELVLTEWQL-WIFNSNSQ-
>Hynobius_retardatus
MGINSGAPLCVA-FTTRCSQT--NHF-----------TTSSIEIMKLVLTEWQL-WIFNSSSQ-
>Pygocentrus_nattereri
MGIHRRAPTCVS-CPPACRLP--SLY-----------NTSSIEIMKLVLTEWQMIWIFTVSSRQ
>Calidris_pugnax
MGINSGAPLCVA-FPPLCSKN--NNF-----------STSSIIIMKLVLTEWQL-WIFNSNSQ-
>Callorhinchus_milii
MGSNSGAPPCVA-VPALYYKQ---NL-----------NTSSIKIMKLVLTEWQLIWIFNSNRQ-
>Myotis_brandtii
MGSNGGAPSRVS-APPLRSPR--NHA-----------CTPSVMMMKLVLTEWQL-WICNSHSQ-
>Haplochromis_burtoni
MGIFRGAPPTVAAFPPPPTPSPRDLLIAPTHSVIQPSIQPSIQKMTLELTEWSSIWIFTACNRQ
>Pterocles_gutturalis
--------------------------------------------MKLVLTEWQL-WIFNSNSQ-
>Dasypus_novemcinctus
MGIHSGPPFCVT-SPPLCSKK--NNF-----------STSSIIMMKLVLTEWQL-WIFNSNSQ-
>Alosa_alosa
MGIHRGAPTCAV-CPPPRRLP--SLY-----------RTSSIQIMKLVLTEWQSIWIFTNFSRQ
>Latimeria_chalumnae
MGIHGGPPLCVA-FPPLCSKK--NIF-----------NTSSIKIMKLILTEWQLLWIFINSSRQ
>Colius_striatus
--------------------------------------------MKLVLTEWQL-WIFNSNSQ-
>Columba_livia
MGINSGAPLCVA-FPPLCSKN--NNF-----------STSSIIIMKLVLTEWQL-WIFNSNSQ-
>Leptosomus_discolor
--------------------------------------------MKLVLTEWQL-WIFNSNSQ-
>Alligator_mississippiensis
MGINSGAPLCVA-FPPLCNKK---NF-----------STSSIIIMKLVLTEWQL-WIFNSNSQ-
>Cuculus_canorus
--------------------------------------------MKLVLTEWQL-WIFNSNSQ-
>Poecilia_formosa
MGIFAGAPPAIATRPSPPS--HCSLSPVPTHTI----IQPPRHKMTLELTEWSSIWIFTACNRQ
>Sarcophilus_harrisii
MGINSGPPFCDT-FSPLCSKK--NNF-----------STSSIIMMKLVLTEWQL-WIFNSNSQ-
>Monodelphis_domestica
MGINSGPPFSDT-FPLLCSKK--NNF-----------STSSIIMMKLVLTEWQL-WIFNSNSQ-
>Esox_lucius
MGIHRGALPCFASCPPPRCLP--SLF-----------CTSSTQIMKLVLTEWQLIWIFTDSSRQ
>Aquila_chrysaetos_canadensis
MGINSGAPLCVA-FPPLCSKN--NNF-----------STSSIIIMKLVLTEWQL-WIFNSNSQ-
>Coturnix_japonica
MGINSGAPLCVA-CPPLCSKN--NNF-----------STSSIIIMKLVLTEWQL-WIFNSNSQ-
>Eurypyga_helias
--------------------------------------------MKLVLTEWQL-WIFNSNSQ-
>Condylura_cristata
MGSNRGPPLCVA-LPPLRGTE--SSF-----------STASAIMMKLVLTEWQL-WIFNSDSQ-
>Oryctolagus_cuniculus
MGISRAPAVSVT-VPPLHSKK--RAC-----------STCSVMMMKLVLTEWQL-WIFNSNSQ-
>Opisthocomus_hoazin
--------------------------------------------MKLVLTEWQL-WIFNSNSQ-
>Scleropages_formosus
MGIYRGAPTCVA-SPPLHRLP--PLF-----------STSSTEIMKLVLTEWQLIWIFTDHSRQ
>Acanthisitta_chloris
MGINSGAPLCVA-FPPLCSTN--NNF-----------STSSIIIMKLVLTEWQL-WIFNTNSQ-
>Phaethon_lepturus
MGINSGAPLCVA-FPPLCSKN--NNF-----------STSSIIIMKLVLTEWQL-WIFNSNSQ-
>Equus_asinus
MGSNGGPPFCVT-FPPLCSKK--NNF-----------STSSIIMMKLVLTEWQL-WIFNSNSQ-
>Manis_javanica
MGSNGGPPFCVT-FPPLCSKK--SNF-----------STTSVIMMKLVLTEWQL-WIFNSNSQ-
>Picoides_pubescens
--------------------------------------------MKLVLTEWQL-WIFNSNSQ-
>Platichthys_stellatus
MGIFRGAPPTIAACPPPAPS-PR----APTLSV----IQPSIHKMTLELTEWSSIWIFTACNRQ
>Macaca_fascicularis
MGITSGTPVCVT-FPPLCSKK--IHF-----------STSSVIMMKLVLTEWQL-WIFNSNSQ-
>Loxodonta_africana
MGINSGLSFCVT-IPPLYSKK--NNF-----------STSSVIMMKLVLTEWQL-WIFNSNSQ-
>Melopsittacus_undulatus
--------------------------------------------MKLVLTEWQL-WIFNSNSQ-
>Cavia_porcellus
MGIHRGLPVFVT-FPPLSSKR--SDC-----------SASSVMMMKLVLTEWHL-WIFNSNLQ-
>Oncorhynchus_mykiss
MGIHRGALTCVASCPPPCRLP--SLF-----------CTSSTHIMKLVLTEWQLIWIFTDSSRQ
>Tupaia_chinensis
MGSVSAAAVGAA-SKPLCSKS--HNR-----------STSSVKMMKLVLTEWQL-WICNSHSQ-
>Lepisosteus_oculatus
MGIFRGAPTGVA-SPPLRRLP--SLF-----------RTSSTEIMKLVLTEWQLIWIFINCSRQ
>Ictalurus_punctatus
MGIHRRAPSCVS-CPAARCLP--SLF-----------NTSSTEIMKLVLTEWQMIWIFTVSNRQ
>Aptenodytes_forsteri
--------------------------------------------MKLVLTEWQL-WIFNSNSQ-
>Anolis_carolinensis
MGITSGAAVCVT-IPPLCSKQ--NNF-----------STSSIIIMKLVLTEWQL-WIFNSNSQ-
>Tyto_alba
MGINSGAPLCVA-FPPLCSKN--NNF-----------STSSIIIMKLVLTEWQL-WIFNSNSQ-
>Struthio_camelus_australis
MGINSGAPLCVA-FPPLCSKK--NNF-----------STSSIIIMKLVLTEWQL-WIFNSNSQ-
>Chrysemys_picta
MGINSGAPFCVA-FPPLCSKN--NNF-----------STLSIIIMKLVLTEWQL-WIFNSNSQ-
>Takifugu_rubripes
MGIIRGAPPNCTACPSPPSP-PSGLYFTPTHSDIQPTILPSIPKMTLELTEWSSIWIFTAYNRQ
>Tinamus_guttatus
MGINSGAPLCVA-FPPLCSKK--NNF-----------STSSIIIMKLVLTEWQL-WIFNSNSQ-
>Calypte_anna
--------------------------------------------MKLVLTEWQL-WIFNSNSQ-

HG0033 ENSG00000112144 ICK

>Homo_sapiens
----------------------------------------------MYLGDSHVLLNTLCWCCHRKIWLHSGEDCYHCRTEPLRP---------
>Anser_cygnoides
----------------------------------------------MIPATVHTVLSALYLCCHRKNWLHSEEK-HKKH---------------
>Apteryx_australis_mantelli
--------------------------MQLKLPEDGGLSDQRLSLENMIPATIHTVLSALYLCCHRKNWLHSEEK-HKNH---------------
>Menidia_menidia
----------------------------------------------MSPPA---LPTV-FCCCHRKIWLHSATGRERICFPPN-----------
>Oryzias_melastigma
----------------------------------------------MTPPA---LPPG-FCCCHRKIWLHSAAGEEHVGFEVLSKDHVDG----
>Buceros_rhinoceros_silvestri
----------------------------------------------MIPATVHTVLSALYLCCHRKNWLHLEEKQTKR----------------
>Caprimulgus_carolinensis
----------------------------------------------MIPATIHSVLSALYLCCHRKNWLHSEEK-HQKH---------------
>Cariama_cristata
----------------------------------------------MIPATVHTVLSALYLCCHRKNWLQSEEK-HKKH---------------
>Odobenus_rosmarus_divergens
----------------------------------------------MYLGDSHVLLNTLCWCCHRKIWLHSGEACRLCRTTPPRP---------
>Physeter_catodon
----------------------------------------------MYLGDSHVIWNTLCWCCHRKIWLHSGEVCYGCRTDPPRP---------
>Charadrius_vociferus
----------------------------------------------MIPATVHTVLSALYLCCHRKNWLHSEEK-HKKH---------------
>Pteropus_vampyrus
----------------------------------------------MYLGDSHVVLNTLCWCCHRKIWLHSGEACCHCRTEPPRP---------
>Pterocles_gutturalis
----------------------------------------------MIPATVHTVLSALYLCCHRKNWLHSEEK-YEKHKKR------------
>Colius_striatus
----------------------------------------------MIPATVHTVLSALYLCCHRKDWLHSEEK-H------------------
>Columba_livia
----------------------------------------------MIPATVHTVLSALYLCCHRKNWLHSEKK-H------------------
>Merops_nubicus
----------------------------------------------MIPATVHTVLSALYLCCHRKNWLHSEEK-HRKHQGLKCACS-------
>Alligator_mississippiensis
----------------------------------------------MILAAFHGVLSALYLCCHRKNWLHSEEEHREVYPKD------------
>Cuculus_canorus
----------------------------------------------MIPATVHTVLSALYLCCHRKNWLHSEEK-HKKHEGLKCACS-------
>Xiphophorus_maculatus
----------------------------------------------MNTPA---QPNV-FCCCHRKIWLHSAVEGEHKCFSS------------
>Sarcophilus_harrisii
----------------------------------------------MYLRDCHDISNTLCWCCHRKVWLHSEENCFHDRREQSQD---------
>Galeopterus_variegatus
----------------------------------------------MYLGDSHVLLNALCWCCHRKIWLHSGEACCHCRTEPPRP---------
>Monodelphis_domestica
----------------------------------------------MYLRHCPAIPNTLCWCWHRKVWLHSGEDCCSHERKGQSQD--------
>Esox_lucius
----------------------------------------------MSPPA---VPVV-HCCCHRKIWLHSTAARGLPFVFSSSP---------
>Aquila_chrysaetos_canadensis
----------------------------------------------MIPATVHTVLSALYLCCHRKNWLHSEEK-HKKH---------------
>Gallus_gallus
-------------------------------------MYQRLSLENMIPATVDTVLSALCLCCHRKNWLQSEKKH-------------------
>Gavia_stellata
----------------------------------------------MIPATVHTVLSALYLCCHRKNWLHSEEK-HKNH---------------
>Eurypyga_helias
----------------------------------------------MIPATVHTILSALYLCCHRKNWLHSEEE-HKKH---------------
>Erinaceus_europaeus
----------------------------------------------MYLGDSHVLLNTLCWCCHRKIWLHSGEACCHCRTEPPQP---------
>Oryctolagus_cuniculus
----------------------------------------------MYLGDSHVLLTTLCWCCHRKIWLHSGEACYHCRTEPPQP---------
>Ornithorhynchus_anatinus
----------------------------------------------MRLGDHHVVSDSLFLCCHRKIWLRSGERRVLGN---------------
>Tauraco_erythrolophus
----------------------------------------------MIPATVHTVLSALYLCCHRKNWLHSEEK-HKKH---------------
>Plecoglossus_altivelis
----------------------------------------------MNPST---LPAV-HCCCHRKIWLHSAAGGGHLSLFLLVPAHPHLQFAE
>Parus_major
MSRARPALGRSCSAGRRGRGAAIPGLSAGRAGAGRHVRDQRLSLENMIPATVHTVLSALYLCCHRKNWLHSEEKKHRKRQGLKCACS-------
>Egretta_garzetta
----------------------------------------------MIPATVHIVLSALYLCCHRKNWLHSEEK-H------------------
>Lateolabrax_japonicus
----------------------------------------------MSPPA---LPVVQLCCCHRKIWLHSAARGISRQ---------------
>Ceratotherium_simum_simum
----------------------------------------------MYLGDSHVLLNTLCWCCHRKIWLHSGEACCHCRTEPPRP---------
>Manis_javanica
----------------------------------------------MYLGDSGVVLNTLGWCCHRKIWLHSGEACFCCGTAPPRL---------
>Picoides_pubescens
----------------------------------------------MIPATVDTVLSALYLCCHRKNWLHSEKK-HRKH---------------
>Cynoglossus_semilaevis
----------------------------------------------MSPPA---WPSV-SCCCHRKIWLHSAAGHHHSDAFNS-----------
>Loxodonta_africana
---------------------------MEQTSLSTVFISRRLIFENMYLGDSCVVLNPLCWCCHRKVWLHAGEACYRCRTEPPQP---------
>Fulmarus_glacialis
----------------------------------------------MIPATVHTVLSALYLCCHRKNRLHSEEK-RKKH---------------
>Nestor_notabilis
----------------------------------------------MIPATVHTVLSALYLCCHRKNWLHSEEK-HKKH---------------
>Mus_musculus
----------------------------------------------MYLGDNHVLLNTLCWCCHRKIWLQSGEACYHCRTEPPRP---------
>Oncorhynchus_nerka
----------------------------------------------MSPTA---VPVV-HCCCHRKIWLHSTDAGGFKPATSVFLLAIVI----
>Tupaia_chinensis
----------------------------------------------MYLGDSRVLLNTLCWCCHRKIWLHSGEACDHGRTEPPRP---------
>Trichechus_manatus_latirostr
----------------------------------------------MYLRDSHVVLNTLCWCCHRKVWLHSGEACYHCRTEPPQP---------
>Pygoscelis_adeliae
----------------------------------------------MIPATVHTVLSALYLCCHRKNWLHSEEK-HKKH---------------
>Anolis_carolinensis
----------------------------------------------MSPENFHAALEILCLCCHRKNWLHSEEKCFSCS---------------
>Tyto_alba
----------------------------------------------MIPATVHTVLSALYLCCHRKNWLHSEKK-Y------------------
>Struthio_camelus_australis
--------------------MYMHVICNRDDLQLNSLSHQRLRLENMIPA-VHTVLSALYLCCHRKNWLHSEEKHKKH----------------
>Chrysemys_picta
----------------------------------------------MTLVAIHAVLSTLYLCCHRKNWLHSEEKHVSCREAFTKD---------
>Calypte_anna
----------------------------------------------MIPATVHTVLSALYLCCHRKNWLHSEEK-LKKH---------------
>Apaloderma_vittatum
----------------------------------------------MIPATVHTVLSALYLCCHRKNWLHSEEKPRNTKD--------------
>Orycteropus_afer_afer
----------------------------------------------MYLGDRHV-LNTLCWCCHRKIWLHSGGACYHCRTEPPQP---------

HG0033 ENSGALG00000016321 ICK

>Gallus_gallus
--------------------------MIPATVDTVLSALCLCCHRKNWLQSEKKH-----------
>Anas_platyrhynchos
----------MRVWSGQADQRLSLENMIPATVDTVLSALYLCCHRKNWLQSEEQHKNSGH------
>Apteryx_australis_mantelli
--MQL--KLPEDGGLSGNDQRLSLENMIPATIHTVLSALYLCCHRKNWLHSEEKHKNH--------
>Buceros_rhinoceros_silvestri
--------------------------MIPATVHTVLSALYLCCHRKNWLHLEEKQTKR--------
>Caprimulgus_carolinensis
--------------------------MIPATIHSVLSALYLCCHRKNWLHSEEKHQKH--------
>Cariama_cristata
--------------------------MIPATVHTVLSALYLCCHRKNWLQSEEKHKKH--------
>Ailuropoda_melanoleuca
--------------------------MYLGDSHVLLNTLCWCCHRKIWLHSGEACRLCRTAPPRP-
>Balaenoptera_acutorostrata_s
--------------------------MYLGDSHVLLSTLCWCCHRKIWLHSGEVCCGCRTDPPRP-
>Charadrius_vociferus
--------------------------MIPATVHTVLSALYLCCHRKNWLHSEEKHKKH--------
>Pteropus_alecto
--------------------------MYLGDSHVVLNTLCWCCHRKIWLHSGEACCHCRTEPPRP-
>Pterocles_gutturalis
--------------------------MIPATVHTVLSALYLCCHRKNWLHSEEKYEKHKKR-----
>Latimeria_menadoensis
--------------------------MSTVDLPVLLCYMYCCCHRKGWLHSKK-------------
>Colius_striatus
--------------------------MIPATVHTVLSALYLCCHRKDWLHSEEKH-----------
>Columba_livia
--------------------------MIPATVHTVLSALYLCCHRKNWLHSEKKH-----------
>Leptosomus_discolor
--------------------------MIPATVHTVLSALYLCCHRKNWLYSEKKH-----------
>Alligator_mississippiensis
--------------------------MILAAFHGVLSALYLCCHRKNWLHSEEEHREVYPKD----
>Cuculus_canorus
--------------------------MIPATVHTVLSALYLCCHRKNWLHSEEKHKKHEGLKCACS
>Sarcophilus_harrisii
--------------------------MYLRDCHDISNTLCWCCHRKVWLHSEENCFHDRREQSQD-
>Galeopterus_variegatus
--------------------------MYLGDSHVLLNALCWCCHRKIWLHSGEACCHCRTEPPRP-
>Haliaeetus_leucocephalus
--------------------------MIPATVHTVLSALYLCCHRKNWLHSEEKH-----------
>Gavia_stellata
--------------------------MIPATVHTVLSALYLCCHRKNWLHSEEKHKNH--------
>Balearica_regulorum_gibberic
--------------------------MIPATVHTVLSALYLCCHRKNWLHSEEKHKKH--------
>Erinaceus_europaeus
--------------------------MYLGDSHVLLNTLCWCCHRKIWLHSGEACCHCRTEPPQP-
>Oryctolagus_cuniculus
--------------------------MYLGDSHVLLTTLCWCCHRKIWLHSGEACYHCRTEPPQP-
>Tauraco_erythrolophus
--------------------------MIPATVHTVLSALYLCCHRKNWLHSEEKHKKH--------
>Opisthocomus_hoazin
--------------------------MIHATVLTVLSALYLCCHRKNWLHLEEKHRKHYGLKCACS
>Lepidothrix_coronata
--------------------------MIPATVHTVLSALYLCCHRKNWLQSKEKHKKH--------
>Nipponia_nippon
--------------------------MIPATVHTVLSALYLCCHRKNWLHSEEKHKKH--------
>Equus_asinus
--------------------------MYLGDSHVLLNTLCWCCHRKIWLHSGEACCRCRTGPPRP-
>Manis_javanica
--------------------------MYLGDSGVVLNTLGWCCHRKIWLHSGEACFCCGTAPPRL-
>Picoides_pubescens
--------------------------MIPATVDTVLSALYLCCHRKNWLHSEKKHRKH--------
>Homo_sapiens
--MG-YSFFPRSCGSLLPGRRLIFGNMYLGDSHVLLNTLCWCCHRKIWLHSGEDCYHCRTEPLRP-
>Loxodonta_africana
-------MEQTSLSTVFISRRLIFENMYLGDSCVVLNPLCWCCHRKVWLHAGEACYRCRTEPPQP-
>Fulmarus_glacialis
--------------------------MIPATVHTVLSALYLCCHRKNRLHSEEKRKKH--------
>Nestor_notabilis
--------------------------MIPATVHTVLSALYLCCHRKNWLHSEEKHKKH--------
>Cricetulus_griseus
--------------------------MYLGDNHVLLNTLCWCCHRKIWLQSGEACYCCRTDPPRP-
>Tupaia_chinensis
--------------------------MYLGDSRVLLNTLCWCCHRKIWLHSGEACDHGRTEPPRP-
>Trichechus_manatus_latirostr
--------------------------MYLRDSHVVLNTLCWCCHRKVWLHSGEACYHCRTEPPQP-
>Aptenodytes_forsteri
--------------------------MIPATVHTVLSALYLCCHRKNWLHSEEKHKKH--------
>Anolis_carolinensis
--------------------------MSPENFHAALEILCLCCHRKNWLHSEEKCFSCS-------
>Tyto_alba
--------------------------MIPATVHTVLSALYLCCHRKNWLHSEKKY-----------
>Struthio_camelus_australis
MYMHVICNRDDLQLNSLSHQRLRLENMIPA-VHTVLSALYLCCHRKNWLHSEEKHKKH--------
>Chrysemys_picta
--------------------------MTLVAIHAVLSTLYLCCHRKNWLHSEEKHVSCREAFTKD-
>Calypte_anna
--------------------------MIPATVHTVLSALYLCCHRKNWLHSEEKLKKH--------
>Apaloderma_vittatum
--------------------------MIPATVHTVLSALYLCCHRKNWLHSEEKPRNTKD------
>Orycteropus_afer_afer
--------------------------MYLGDRHV-LNTLCWCCHRKIWLHSGGACYHCRTEPPQP-

HG0034 ENSDARG00000060991 mief1

```
>Danio_rerio
------------------------------MGGWSRSAVLQLYRALLRAGQHLQYTDRDYYRRVVTREFRRCQT-LSAPAEREDALKRGQFFLNSGLGDLV
>Acipenser_sinensis
------------------------------MGGWSRSAVLQLYRALLRAGRPLQHTDRNYYRAALSREFRRCQG-LREEGARELALKRGLYFLHSGLGGLV
>Channa_punctata
MLHVQLYSLAVNRVPTSRCQGVPPLHVWSLMGGWSRCAVLELYRALLRAGRYLQYTDRNYYRRAVAREFRRCQD-LTIPQDKEEALKRGQFFLSSRLGGLM
>Anser_cygnoides
------------------------------MAAWSREAVLSLYRALLRRGRALRYTDRDFYRAAVRSEFRRQRG-LQRPEDRERQLEKGQAFLRSGLGGLI
>Rhacophorus_omeimontis
------------------------------MAGWSREAVLKLYRALLREGRKLQYTDRDYYSSYIRREFRKNQN-LANLEEKEKQLAKGQFFLQTKLGGLV
>Basilichthys_microlepidotus
------------------------------MGSWSRSAVLELYRALLRAGRHLQYTDRNYYRRAVAREFRRCQT-LTVPEDKEEALKRGQFFLNSRLGGLM
>Porichthys_notatus
------------------------------MGGWSRSAVLELYRALLRAGRRLQYTDRNYYYRVVALEFRRSQA-LTAPDQKEAALKRGQFFLSSKLGGLM
>Oryzias_melastigma
------------------------------MGSWSRVAVLELYRALLRAGRHLQYTDRNYYRQAVAREFRRCQA-LTVPEDKEEALKRGQFFLNSRLGGLM
>Tripterygion_delaisi
------------------------------MGSWSRGAVLELYRALLRAGRQLQHTERGYYRRAVAREFRRCQT-LTLPQDKEDALRRGQFFLSSRLGGLM
>Trachinotus_blochii
------------------------------MGGWSRSAVLELYRALLRAGRHLQYTDRNYYRRAVAREFRRCQA-LTVPADKEDALKRGQFFLNSRLGGLM
>Ailuropoda_melanoleuca
------------------------------MAPWSREAVLSLYRALLRQGRELRYTDRDFYLASIRREFRKNQK-LEDPEAREKQLEKGLVFLHSKLGGII
>Ambystoma_mexicanum
------------------------------MAAWSRTAVLNLYRTLLREGRHLKYTDRDFYFATVRREFRRNQD-LTEPDEKARQLEKGQYFLQTRLGGLV
>Micropterus_floridanus
------------------------------MGGWSRSAVLELYRALLRAGRHLQYTDRNYYRRAVAREFRRCQA-LTVPEDKEDALRRGQFFLRSRLGGLM
>Orcinus_orca
------------------------------MAPWSREAVLSLYRALLRQGRELRYTDRDFYFASIRREFRKNQK-LEDPEARERQLEKGLVFLHSKLGGII
>Astyanax_mexicanus
------------------------------MGGWSRTAVLELYRALLRAGRHLQYTDRNYYRRAVTREFRRCQA-LSTPGEKEDALRRGQFFLTSRLGGLV
>Calidris_pugnax
------------------------------MAAWSRQAVLTLYRALLRQGRGLRYTDQDFYLASIRREFRKNQG-LQQLEEKERQLEKGQAFLQSKLGGLV
>Callorhinchus_milii
------------------------------MATWSRTAVLNLYRALLRAGQHLRYTDREYYYSYVKREFRKNQH-LTEEQQRTQQLKKGIYFLNNKLGGLV
>Pteropus_vampyrus
------------------------------MAPWSREAVLSLYRALLRQGRELRYTDRDFYLASIRREFRKNQK-LEDPEARERQLEKGLVFLHSKLGGII
>Haplochromis_burtoni
------------------------------MGSWSRSAVLELYRALLRAGRHLQYTDRNYYRRAVAREFRRCQA-LTVPEDKEEALKRGRFFLSSRLGGLM
>Alosa_alosa
------------------------------MGGWSRSAVLELYRALLRAGRHLQYTDRNYYRRAVAREFRRCQA-LSAPQEKEDALKRGQFFLNSRLGGLV
>Latimeria_chalumnae
------------------------------MAGWSRTAVLNLYRALLREGRRLRYTDRDYYYNYVKREFRKNQG-LSQPEDREKQLEKGMFFLQNKLGGLI
>Alligator_mississippiensis
------------------------------MAAWSRQAVLSLYRTLLRQGRRLRFTDRDFYLATIRREFRKNKQ-LEQLEDKERQLEKGQAFLHSKLGGLV
>Cyprinodon_variegatus
------------------------------MGSWSRGAVLELYRALLRAGRHLQYTDRNYYRRAVTREFRRCQA-LTVPEDKEEALKRGQFFLSSRLGGLM
>Sarcophilus_harrisii
------------------------------MAEPGRGAVLGLYRTLLRRARELRLTDRDFYVETIRREFRRPLPPGAGPDAVQRQLEKGRAFLRGERGRLV
>Galeopterus_variegatus
------------------------------MAPWSREAVLSLYRALLRQGRELRYTDRDFYLAFIRREFRKNQK-LEDPEARERQLEKGLVFLHSKLGGII
>Monodelphis_domestica
------------------------------MAEPGRGAVLGLYRALLRRARGLRYTDRDFYVEAVRREFRRPLPPGVGPEAVQRQLDKGRAFLRGERGRLV
>Esox_lucius
------------------------------MGGWSRSAVLDLYRALLRAGRHLQYTDRDYYRRAVAHEFRRCQT-LTAPSDREDALKRGQFFLHSRLGGLM
>Numida_meleagris
------------------------------MAAWSRQAVLTLYRALLREGRGLRYTDRDFYLASIRREFRKNQE-LQRLEDKERQLEKGQAFLQSKLGGLV
>Acanthogobius_hasta
------------------------------MGGWSRSAVLELYRALLRAGRHLQYTDRNYYRRAVTREFRRCQA-LSGSQEKEDALKRGQFFLNSKLGGLM
>Apteronotus_leptorhynchus
------------------------------MGGWSRSAVLELYRALLRAGRHLQYTDRSYYQRAVTREFRRCQA-LNAPEEKEDALNRGRFFLSSRLGGLV
>Condylura_cristata
------------------------------MAPWSRSAVLSLYRALLRQGRELRYSDRDFYLASIRREFRKNQK-LEDPEAREKQLEKGLVFLHSKLGGLV
>Ochotona_princeps
------------------------------MAPWSRQAVLSLYRALLRQGRELRYTDRDFYLASIRREFRKNQK-LEEPEGRERQLEKGLVFLRSKLGGLV
>Plecoglossus_altivelis
------------------------------MGGWSRSAVLDLYRALLRAGRHLQHTDRNYYRRAVAREFRRCQA-LSAPSDKEDALKRGQFFLSSRLGGLV
>Scleropages_formosus
------------------------------MGGWSKTAVLELYRALLRAGRHLQYTDRNYYRHTLAREFRRCQA-LTNPQAREEALKRGQFFLRSRLGGLV
>Zonotrichia_albicollis
------------------------------MAAWSREAVLTLYRALLRRGRGLRYTDRDFYLASIRREFRRNQG-LQRLEDKERQLEKGQAFLRSSLGGLV
>Lateolabrax_japonicus
------------------------------MGGWSRSAVLELYRALLRAGRHLQYTDRNYYRRAVAREFRRCQA-LTIPEDKEDALKRGQFFLSSRLGGLM
>Anoplopoma_fimbria
------------------------------MGGWSRSAVLELYRALLRAGRHLQYTDRNYYRRAVAHEFRRCQA-LTIPEDKEEALRRGQFFLNSRLGGLM
>Ceratotherium_simum_simum
------------------------------MAPWSREAVLSLYRALLRQGRELRYTDRDFYLASIRREFRKNQK-LEDPEARERQLEKGLVFLHSKLGGLI
>Cynoglossus_semilaevis
------------------------------MGGWSRSAVLGLYKALLRAGRNLQYTDRNYYRRAVAREFRRCQA-LTIPEDKEEALKRGQFFLSSRLGGLM
>Pongo_abelii
------------------------------MAPWSREAVLSLYRALLRQGRQLRYTDRDFYFASIRREFRKNQK-LEDPVARERQLEKGLVFLNGKLGRII
>Leucoraja_erinacea
------------------------------MAAWSRVAVLNLYRSLLREGRRLRYTDREFYYTFIKREFRKNQH-LSGEEEKAQQLEKGLYFLESKIGGLV
>Jaculus_jaculus
------------------------------MAPWSREAVLSLYRALLRQGRELRYTDRDFYFASVRREFRKNQK-LQDPVAREKQLEKGLVFLHSKLGGLV
>Salvelinus_alpinus
------------------------------MGGWSRSAVLELYRALLRAGRHLQYTDRNYYRRAVAGEFRRCQA-LTVPSDKEDALKRGQFFLSSRLGGLM
>Lepisosteus_oculatus
------------------------------MGGWSRTAVLELYRALLRAGHHLQHTDRNYYRRAVAREFRRCQG-LTAPGDKERALQRGQFFLSSRLGGLV
>Ictalurus_punctatus
------------------------------MGGWSRGAVLELYRALLRAGRHLQYTDRNYYRRTIAREFRRCQS-LSEPREKEEALKRGQFFLNSRLGGLI
>Trichechus_manatus_latirostr
------------------------------MAPWSREAVLSLYRTLLRRGRELRYTDRDFYLASIRREFRKNQK-LEDPEARERQLEKGLFFLHSKLGGIV
>Squalus_acanthias
------------------------------MATWSRVAVLNLYRALLREGRNLQYTDREYYYTFIKREFRKNQR-LTGEEEKVKQLEKGIYFLKSKLGGLV
>Crotalus_adamanteus
------------------------------MATWSREAVLNLYRALLRQGRGLRYTDRDFYLSSIRKEFRKNQH-LTSPEDKERHLKKGQTFLHSKLGGLI
>Chrysemys_picta
------------------------------MAAWSREAVLNLYRALLRQGRGLRYTDRDFYLASIRREFRKNQQ-LERLEDRERQLEKGQAFLHSRLGGLV
>Takifugu_rubripes
------------------------------MGGWSRSAVLELYRALLRAGRHLQYTDRNYYRHAVVREFRRCQA-LTAPEEKEEALKRGQFFLKSRLGGLM
>Orycteropus_afer_afer
------------------------------MAPWSRAAVLSLYRTLLRQGRQLRYTDRDFYFASIRQEFRKNQK-LEDPAARERQLEKGLFFLHSKLGGII
```

HG0034 ENSG00000100335 MIEF1

```
>Homo_sapiens
------------------------------MAPWSREAVLSLYRALLRQGRQLRYTDRDFYFASIRREFRKNQK-LEDAEARERQLEKGLVFLNGKLGRII
>Acipenser_sinensis
------------------------------MGGWSRSAVLQLYRALLRAGRPLQHTDRNYYRAALSREFRRCQG-LREEGARELALKRGLYFLHSGLGGLV
>Channa_punctata
MLHVQLYSLAVNRVPTSRCQGVPPLHVWSLMGGWSRCAVLELYRALLRAGRYLQYTDRNYYRRAVAREFRRCQD-LTIPQDKEEALKRGQFFLSSRLGGLM
>Anser_cygnoides
------------------------------MAAWSREAVLSLYRALLRRGRALRYTDRDFYRAAVRSEFRRQRG-LQRPEDRERQLEKGQAFLRSGLGGLI
>Xenopus_tropicalis
------------------------------MAPWSREAVLSLYRALLRQGRQLRYTDRDFYFASIRREFRKNQK-LEDAEARERQLEKGLVFLNGKLGRII
>Basilichthys_microlepidotus
------------------------------MGSWSRSAVLELYRALLRAGRHLQYTDRNYYRRAVAREFRRCQT-LTVPEDKEEALKRGQFFLNSRLGGLM
>Porichthys_notatus
------------------------------MGGWSRSAVLELYRALLRAGRRLQYTDRNYYYRVVALEFRRSQA-LTAPDQKEAALKRGQFFLSSKLGGLM
>Oryzias_latipes
------------------------------MGSWSRVAVLELYRALLRAGRHLQYTDRNYYRQAVAREFRRCQA-LTVAEDKEEALKRGQFFLNSRLGGLM
>Tripterygion_delaisi
------------------------------MGSWSRGAVLELYRALLRAGRQLQHTERGYYRRAVAREFRRCQT-LTLPQDKEDALRRGQFFLSSRLGGLM
>Trachinotus_blochii
------------------------------MGGWSRSAVLELYRALLRAGRHLQYTDRNYYRRAVAREFRRCQA-LTVPADKEDALKRGQFFLNSRLGGLM
>Ursus_maritimus
------------------------------MAPWSREAVLSLYRALLRQGRELRYTDRDFYLASIRREFRKNQK-LEDPEAREKQLEKGLVFLHSKLGGII
>Hynobius_chinensis
------------------------------MAAWSRQAVLSLYRALLRGGRRLQYTDRDFYFASVRREFRRQQG-LTRPEDKERQLEKGLFFLRTQLGGLL
>Micropterus_floridanus
------------------------------MGGWSRSAVLELYRALLRAGRHLQYTDRNYYRRAVAREFRRCQA-LTVPEDKEDALRRGQFFLRSRLGGLM
>Orcinus_orca
------------------------------MAPWSREAVLSLYRALLRQGRELRYTDRDFYFASIRREFRKNQK-LEDPEARERQLEKGLVFLHSKLGGII
>Pygocentrus_nattereri
------------------------------MGGWSRSAVLELYRALLRAGRQLQYTDRNYYQRAVTQEFRRCQA-LSAPGEKEDALKRGQFFLSSRLGGLV
>Calidris_pugnax
------------------------------MAAWSRQAVLTLYRALLRQGRGLRYTDQDFYLASIRREFRKNQG-LQQLEEKERQLEKGQAFLQSKLGGLV
>Callorhinchus_milii
------------------------------MATWSRTAVLNLYRALLRAGQHLRYTDREYYYSYVKREFRKNQH-LTEEQQRTQQLKKGIYFLNNKLGGLV
>Pteropus_vampyrus
------------------------------MAPWSREAVLSLYRALLRQGRELRYTDRDFYLASIRREFRKNQK-LEDPEARERQLEKGLVFLHSKLGGII
>Haplochromis_burtoni
------------------------------MGSWSRSAVLELYRALLRAGRHLQYTDRNYYRRAVAREFRRCQA-LTVPEDKEEALKRGRFFLSSRLGGLM
>Alosa_alosa
------------------------------MGGWSRSAVLELYRALLRAGRHLQYTDRNYYRRAVAREFRRCQA-LSAPQEKEDALKRGQFFLNSRLGGLV
>Latimeria_chalumnae
------------------------------MAGWSRTAVLNLYRALLREGRRLRYTDRDYYYNYVKREFRKNQG-LSQPEDREKQLEKGMFFLQNKLGGLI
>Alligator_mississippiensis
------------------------------MAAWSRQAVLSLYRTLLRQGRRLRFTDRDFYLATIRREFRKNKQ-LEQLEDKERQLEKGQAFLHSKLGGLV
>Sinocyclocheilus_rhinocerous
------------------------------MGSWSRSAVLQLYRALLRAGQHLQYTDRDYYRRAVTREFRRCQT-LSAPAEREDALKKGQFFLNSGLGRLV
>Kryptolebias_marmoratus
------------------------------MGSWSRSAVLELYRALLRAGRRLQYTDRNYYRWAVAREFRRCQA-LTGAEDKEEALKRGRFFLRSGLGGLM
>Sarcophilus_harrisii
------------------------------MAEPGRGAVLGLYRTLLRRARELRLTDRDFYVETIRREFRRPLPPGAGPDAVQRQLEKGRAFLRGERGRLV
>Galeopterus_variegatus
------------------------------MAPWSREAVLSLYRALLRQGRELRYTDRDFYLAFIRREFRKNQK-LEDPEARERQLEKGLVFLHSKLGGII
>Monodelphis_domestica
------------------------------MAEPGRGAVLGLYRALLRRARGLRYTDRDFYVEAVRREFRRPLPPGVGPEAVQRQLDKGRAFLRGERGRLV
>Esox_lucius
------------------------------MGGWSRSAVLDLYRALLRAGRHLQYTDRDYYRRAVAHEFRRCQT-LTAPSDREDALKRGQFFLHSRLGGLM
>Numida_meleagris
------------------------------MAAWSRQAVLTLYRALLREGRGLRYTDRDFYLASIRREFRKNQE-LQRLEDKERQLEKGQAFLQSKLGGLV
>Acanthogobius_hasta
------------------------------MGGWSRSAVLELYRALLRAGRHLQYTDRNYYRRAVTREFRRCQA-LSGSQEKEDALKRGQFFLNSKLGGLM
>Apteronotus_leptorhynchus
------------------------------MGGWSRSAVLELYRALLRAGRHLQYTDRSYYQRAVTREFRRCQA-LNAPEEKEDALNRGRFFLSSRLGGLV
>Erinaceus_europaeus
------------------------------MAPWSREAVLSLYRALLRQGRELRYTDRDFYLASIRREFRKNQK-LEDPAAREKQLEKGLVFLHSKLGGII
>Ochotona_princeps
------------------------------MAPWSRQAVLSLYRALLRQGRELRYTDRDFYLASIRREFRKNQK-LEEPEGRERQLEKGLVFLRSKLGGLV
>Plecoglossus_altivelis
------------------------------MGGWSRSAVLDLYRALLRAGRHLQHTDRNYYRRAVAREFRRCQA-LSAPSDKEDALKRGQFFLSSRLGGLV
>Scleropages_formosus
------------------------------MGGWSKTAVLELYRALLRAGRHLQYTDRNYYRHTLAREFRRCQA-LTNPQAREEALKRGQFFLRSRLGGLV
>Sturnus_vulgaris
------------------------------MAAWSREAVLSLYRALLRRGRGLRYTDRDFYLASIRREFRRNQG-LQRLEDKERQLEKGQAFLQNRLGGLV
>Lateolabrax_japonicus
------------------------------MGGWSRSAVLELYRALLRAGRHLQYTDRNYYRRAVAREFRRCQA-LTIPEDKEDALKRGQFFLSSRLGGLM
>Trematomus_bernacchii
------------------------------MGGWSRGAVLELYRALLRAGRQLQYTDRNYYRRAVAHEFRRCQD-LKKAQDKEEALKRGQFFLDSRLGGLM
>Equus_asinus
------------------------------MAPWSREAVLSLYRALLRQGRELRYTDRDFYLASIRREFRKNQK-LEDPEARERQLEKGLVFLHSKLGGII
>Cynoglossus_semilaevis
------------------------------MGGWSRSAVLGLYKALLRAGRNLQYTDRNYYRRAVAREFRRCQA-LTIPEDKEEALKRGQFFLSSRLGGLM
>Leucoraja_erinacea
------------------------------MAAWSRVAVLNLYRSLLREGRRLRYTDREFYYTFIKREFRKNQH-LSGEEEKAQQLEKGLYFLESKIGGLV
>Psammomys_obesus
------------------------------MAPWSREAVLSLYRALLRQGRELRYTDRDFYFASIRREFRKNQK-LEDLEAREKQLEKGLVFLHSKLGGLI
>Salmo_salar
------------------------------MGGWSRSAVLELYRALLRAGRHLQYTDRNYYRRAVAGEFRRCQA-LTVSRDKEAALKRGQFFLSSRLGGLM
>Lepisosteus_oculatus
------------------------------MGGWSRTAVLELYRALLRAGHHLQHTDRNYYRRAVAREFRRCQG-LTAPGDKERALQRGQFFLSSRLGGLV
>Trichomycterus_areolatus
------------------------------MGGWSRGAVLQLYRALLRAGRNLQYTDRDYYRRAIAREFRRHQA-LTEPQEKEEALRRGQFFLSSRLGGLV
>Trichechus_manatus_latirostr
------------------------------MAPWSREAVLSLYRTLLRRGRELRYTDRDFYLASIRREFRKNQK-LEDPEARERQLEKGLFFLHSKLGGIV
>Squalus_acanthias
------------------------------MATWSRVAVLNLYRALLREGRNLQYTDREYYYTFIKREFRKNQR-LTGEEEKVKQLEKGIYFLKSKLGGLV
>Gekko_japonicus
------------------------------MAAWTREAVLNLYRSLLRQGRGLRYTDRDFYFTSIRKEFRKNQQ-LERLEDKERQLEKGQAFLRSKLGGLV
>Chrysemys_picta
------------------------------MAAWSREAVLNLYRALLRQGRGLRYTDRDFYLASIRREFRKNQQ-LERLEDRERQLEKGQAFLHSRLGGLV
>Takifugu_rubripes
------------------------------MGGWSRSAVLELYRALLRAGRHLQYTDRNYYRHAVVREFRRCQA-LTAPEEKEEALKRGQFFLKSRLGGLM
>Orycteropus_afer_afer
------------------------------MAPWSRAAVLSLYRTLLRQGRQLRYTDRDFYFASIRQEFRKNQK-LEDPAARERQLEKGLFFLHSKLGGII
```

```
HG0035 ENSG00000197147 LRRC8B

>Homo_sapiens
MKTNTEMVCLRNFKRCRPPD
>Anser_cygnoides
MKTNVKMACPRNFKRCRPPD
>Leptobrachium_boringii
MKSNAEMSRIKHFKRCRPPD
>Chaetura_pelagica
MKTNVKMACPRNFKRCRPPD
>Buceros_rhinoceros_silvestri
MKTNVKMACPRNFKRCRPPD
>Caprimulgus_carolinensis
MKTNVKMACPRNFKRCRPPD
>Cariama_cristata
MKTNVKMACPRNFKRCRPPD
>Acinonyx_jubatus
MKTNTEMVCLRNFKRCRPPD
>Ambystoma_mexicanum
------MQCTRKFKRCRPPD
>Physeter_catodon
MKTITEMVCLRNFKRCRPPD
>Charadrius_vociferus
MKTNVKMACPRNFKRCRPPD
>Myotis_ricketti
MKTNTEMVFLRNFKRCRPPD
>Pterocles_gutturalis
MKTNVKMACPRNFKRCRPPD
>Dasypus_novemcinctus
MKTNTEMVCLRNFKRCRPPD
>Colius_striatus
MKTNVKMACPRNFKRCRPPD
>Columba_livia
MKTNVKMACPRNFKRCRPPD
>Merops_nubicus
MKTNVKMACPRNFKRCRPPD
>Alligator_mississippiensis
MKANAKMACPRNFKRCRPPD
>Cuculus_canorus
MKANVKMACPRNFKRCRPPD
>Sarcophilus_harrisii
MKTKAEMVCLRNFKRCRPPD
>Galeopterus_variegatus
MKTNTEMVCLRNFKRCRPPD
>Monodelphis_domestica
MKTKAEMVCLRNFKRCRPPD
>Macropus_eugenii
MKTKAEMVCLRNFKRCRPPD
>Aquila_chrysaetos_canadensis
MKTNVKMACPRNFKRCRPPD
>Coturnix_japonica
MKTNVKMACPRNFKRCRPPD
>Gavia_stellata
MKTNVKMACPRNFKRCRPPD
>Balearica_regulorum_gibberic
MKTNVKMACPRNFKRCRPPD
>Condylura_cristata
MKTNTEMVCLRNFKRCRPPD
>Oryctolagus_cuniculus
MKKNTEMVCLRNFKRCRPPD
>Elephantulus_edwardii
MKTNTEMVCLRNFKRCRPPD
>Tauraco_erythrolophus
MKTNVKMACPRNFKRCRPPD
>Opisthocomus_hoazin
MKTNVKMACPRNFKKCRPPD
>Acanthis_flammea
MKTNVKMACPRNFKRCRPPD
>Egretta_garzetta
MKTNVKMACPRNFKRCRPPD
>Ceratotherium_simum_simum
MKTNTEMVCLRNFKRCRPPD
>Manis_javanica
MKTNTEMVCLRNFKRCRPPD
>Picoides_pubescens
MKTNVKMACPRNFKRCRPPD
>Loxodonta_africana
MKTNTEMVCLRNFKRCRPPD
>Melopsittacus_undulatus
MKTNVKMACPRNFKRCRPPD
>Nannospalax_galili
MKTNSEMVCLRNFKRCRPPD
>Tupaia_chinensis
MKTNTDMVCLRNFKRCRPPD
>Trichechus_manatus_latirostr
MKTNTEMVCLRNFKRCRPPD
>Aptenodytes_forsteri
MKTNVKMACPRNFKRCRPPD
>Gekko_japonicus
MKTVVKMACLRNFKRCRPPD
>Tyto_alba
MKTNVKMACPRNFKRCRPPD
>Struthio_camelus_australis
MKTNAKMACPRNFKRCRPPD
>Chrysemys_picta
MKINAKMACPRNFKRCRPPD
>Tinamus_guttatus
MKTNAKMACPRNFKRCRPPD
>Calypte_anna
MKTNVKMACPRNFKRCRPPD
>Apaloderma_vittatum
MKTNVKMACPRNFKRCRPPD
>Orycteropus_afer_afer
MKTNTEMVCLRNFKRCRPPD
```

HG0035 ENSGALG00000006167 LRRC8B

>Gallus_gallus
----------------------MKTNVKMACPRNFKRCRPPD
>Anser_cygnoides
----------------------MKTNVKMACPRNFKRCRPPD
>Leptobrachium_boringii
----------------------MKSNAEMSRIKHFKRCRPPD
>Chaetura_pelagica
----------------------MKTNVKMACPRNFKRCRPPD
>Buceros_rhinoceros_silvestri
----------------------MKTNVKMACPRNFKRCRPPD
>Caprimulgus_carolinensis
----------------------MKTNVKMACPRNFKRCRPPD
>Cariama_cristata
----------------------MKTNVKMACPRNFKRCRPPD
>Felis_catus
--------------MLAAHKCRMKTNTEMVCLRNFKRCRPPD
>Ambystoma_mexicanum
----------------------------MQCTRKFKRCRPPD
>Physeter_catodon
----------------------MKTITEMVCLRNFKRCRPPD
>Charadrius_vociferus
----------------------MKTNVKMACPRNFKRCRPPD
>Myotis_ricketti
----------------------MKTNTEMVFLRNFKRCRPPD
>Pterocles_gutturalis
----------------------MKTNVKMACPRNFKRCRPPD
>Dasypus_novemcinctus
----------------------MKTNTEMVCLRNFKRCRPPD
>Colius_striatus
----------------------MKTNVKMACPRNFKRCRPPD
>Columba_livia
----------------------MKTNVKMACPRNFKRCRPPD
>Merops_nubicus
----------------------MKTNVKMACPRNFKRCRPPD
>Alligator_mississippiensis
----------------------MKANAKMACPRNFKRCRPPD
>Cuculus_canorus
----------------------MKANVKMACPRNFKRCRPPD
>Sarcophilus_harrisii
----------------------MKTKAEMVCLRNFKRCRPPD
>Galeopterus_variegatus
----------------------MKTNTEMVCLRNFKRCRPPD
>Monodelphis_domestica
----------------------MKTKAEMVCLRNFKRCRPPD
>Macropus_eugenii
----------------------MKTKAEMVCLRNFKRCRPPD
>Aquila_chrysaetos_canadensis
----------------------MKTNVKMACPRNFKRCRPPD
>Gavia_stellata
----------------------MKTNVKMACPRNFKRCRPPD
>Balearica_regulorum_gibberic
----------------------MKTNVKMACPRNFKRCRPPD
>Condylura_cristata
----------------------MKTNTEMVCLRNFKRCRPPD
>Oryctolagus_cuniculus
----------------------MKKNTEMVCLRNFKRCRPPD
>Elephantulus_edwardii
----------------------MKTNTEMVCLRNFKRCRPPD
>Tauraco_erythrolophus
----------------------MKTNVKMACPRNFKRCRPPD
>Opisthocomus_hoazin
----------------------MKTNVKMACPRNFKKCRPPD
>Acanthis_flammea
----------------------MKTNVKMACPRNFKRCRPPD
>Egretta_garzetta
----------------------MKTNVKMACPRNFKRCRPPD
>Ceratotherium_simum_simum
----------------------MKTNTEMVCLRNFKRCRPPD
>Manis_javanica
----------------------MKTNTEMVCLRNFKRCRPPD
>Picoides_pubescens
----------------------MKTNVKMACPRNFKRCRPPD
>Homo_sapiens
----------------------MKTNTEMVCLRNFKRCRPPD
>Loxodonta_africana
----------------------MKTNTEMVCLRNFKRCRPPD
>Melopsittacus_undulatus
----------------------MKTNVKMACPRNFKRCRPPD
>Nannospalax_galili
----------------------MKTNSEMVCLRNFKRCRPPD
>Tupaia_chinensis
----------------------MKTNTDMVCLRNFKRCRPPD
>Trichechus_manatus_latirostr
----------------------MKTNTEMVCLRNFKRCRPPD
>Aptenodytes_forsteri
----------------------MKTNVKMACPRNFKRCRPPD
>Anolis_carolinensis
----------------------MKTLVRMACPRNFKRCRPPD
>Tyto_alba
----------------------MKTNVKMACPRNFKRCRPPD
>Struthio_camelus_australis
----------------------MKTNAKMACPRNFKRCRPPD
>Chelonia_mydas
MRVLFWKREPTELESHLKCLSRMKINAKMACPRNFKRCRPPD
>Tinamus_guttatus
----------------------MKTNAKMACPRNFKRCRPPD
>Calypte_anna
----------------------MKTNVKMACPRNFKRCRPPD
>Apaloderma_vittatum
----------------------MKTNVKMACPRNFKRCRPPD
>Orycteropus_afer_afer
----------------------MKTNTEMVCLRNFKRCRPPD

HG0036 ENSG00000155096 AZIN1

>Homo_sapiens
MIGGFHLPTCPFCCIF
>Anser_cygnoides
MIVGFHLLTCPFCCII
>Chaetura_pelagica
MIVGFHLLTCPFCCII
>Apteryx_australis_mantelli
MIVGFHLLTCPFCCII
>Buceros_rhinoceros_silvestri
MIVGFHLLTYPFCCII
>Caprimulgus_carolinensis
MIVGFHLLTCPFCCII
>Cariama_cristata
MIVGFHLLTCPFCCII
>Canis_lupus_familiaris
MIGGFHLLTCPFCCIF
>Ambystoma_mexicanum
MIVGFHLLTCSICCIF
>Lipotes_vexillifer
MIGGFHLLTCPFCCIF
>Calidris_pugnax
MIVGFHLLTCPFCCII
>Rousettus_aegyptiacus
MIGGFHLLTCPFCCIF
>Pterocles_gutturalis
MIVGFHLLTCPFCCII
>Dasypus_novemcinctus
MIGGFHLLTCPFCCIF
>Columba_livia
MIVGFHLLTCPFCCII
>Leptosomus_discolor
MIVGFHLLTCPFCCII
>Alligator_mississippiensis
MIVGFHLLTCPFCCII
>Cuculus_canorus
MIVGFHLLTCPFCCII
>Sarcophilus_harrisii
MIGGFHLLTCPFCCIF
>Galeopterus_variegatus
MIGGFHLLTCPFCCIF
>Monodelphis_domestica
MIGGFHLLTCPFCCIF
>Macropus_eugenii
MIGGFHLLTRPFCCIF
>Aquila_chrysaetos_canadensis
MIVGFHLLTCPFCCII
>Coturnix_japonica
MIVGFHLLTCPFCCII
>Mesitornis_unicolor
MIVGFHLLTCPFCCII
>Condylura_cristata
MIGGFHLLTCPFCCIF
>Elephantulus_edwardii
MIGGFHLLTCPFCCIF
>Tauraco_erythrolophus
MIVGFHLLTCPFCCII
>Opisthocomus_hoazin
MIVGFHLLTCPFCCII
>Ficedula_albicollis
MIVGFHLLTCPFCCII
>Phalacrocorax_carbo
MIVGFHLLTCPFCCII
>Equus_przewalskii
MIGGFHLLTCPFCCIF
>Manis_javanica
MIGGFHLLTCPFCCIF
>Picoides_pubescens
MIVGFHLLTCPFCCII
>Loxodonta_africana
MIGGFHLLTCPFCCIF
>Fulmarus_glacialis
MIVGFHLLTCPFCCII
>Melopsittacus_undulatus
MIVGFHLLTCPFCCII
>Cricetulus_griseus
MIGGFHLLTCPFCCIF
>Tupaia_chinensis
MIGGFHLLTCPFCCIF
>Trichechus_manatus_latirostr
MIGGFHLLTCPFCCIF
>Aptenodytes_forsteri
MIVGFHLLTCPFCCII
>Boiga_irregularis
MIVGFHLLTCPFCCII
>Tyto_alba
MIVGFHLLTCPFCCII
>Struthio_camelus_australis
MIVGFHLLTCPFCCII
>Chelonia_mydas
MIVGFHLLTCPFCCII
>Tinamus_guttatus
MIVGFHLLTCPFCCII
>Calypte_anna
MIVGFHLLTCPFCCII
>Apaloderma_vittatum
MIVGFHLLTCPFCCII
>Orycteropus_afer_afer
MIGGFHLLTCPFCCIF

HG0036 ENSGALG00000029295

>Gallus_gallus
MIVGFHLLTCPFCCII
>Anser_cygnoides
MIVGFHLLTCPFCCII
>Chaetura_pelagica
MIVGFHLLTCPFCCII
>Apteryx_australis_mantelli
MIVGFHLLTCPFCCII
>Buceros_rhinoceros_silvestri
MIVGFHLLTYPFCCII
>Caprimulgus_carolinensis
MIVGFHLLTCPFCCII
>Cariama_cristata
MIVGFHLLTCPFCCII
>Canis_lupus_familiaris
MIGGFHLLTCPFCCIF
>Hynobius_chinensis
MIVVFHLLTCPICCIL
>Lipotes_vexillifer
MIGGFHLLTCPFCCIF
>Calidris_pugnax
MIVGFHLLTCPFCCII
>Eptesicus_fuscus
MIGGFHLLTCPFCCIF
>Pterocles_gutturalis
MIVGFHLLTCPFCCII
>Dasypus_novemcinctus
MIGGFHLLTCPFCCIF
>Columba_livia
MIVGFHLLTCPFCCII
>Leptosomus_discolor
MIVGFHLLTCPFCCII
>Alligator_mississippiensis
MIVGFHLLTCPFCCII
>Cuculus_canorus
MIVGFHLLTCPFCCII
>Sarcophilus_harrisii
MIGGFHLLTCPFCCIF
>Galeopterus_variegatus
MIGGFHLLTCPFCCIF
>Monodelphis_domestica
MIGGFHLLTCPFCCIF
>Macropus_eugenii
MIGGFHLLTRPFCCIF
>Aquila_chrysaetos_canadensis
MIVGFHLLTCPFCCII
>Balearica_regulorum_gibberic
MIVGFHLLTCPFCCII
>Condylura_cristata
MIGGFHLLTCPFCCIF
>Elephantulus_edwardii
MIGGFHLLTCPFCCIF
>Tauraco_erythrolophus
MIVGFHLLTCPFCCII
>Opisthocomus_hoazin
MIVGFHLLTCPFCCII
>Ficedula_albicollis
MIVGFHLLTCPFCCII
>Phalacrocorax_carbo
MIVGFHLLTCPFCCII
>Ceratotherium_simum_simum
MIGGFHLLTCPFCCIF
>Manis_javanica
MIGGFHLLTCPFCCIF
>Picoides_pubescens
MIVGFHLLTCPFCCII
>Aotus_nancymaae
MIGGFHLPTCPFCCIF
>Loxodonta_africana
MIGGFHLLTCPFCCIF
>Fulmarus_glacialis
MIVGFHLLTCPFCCII
>Melopsittacus_undulatus
MIVGFHLLTCPFCCII
>Jaculus_jaculus
MIGGFHLLTCPFCCIF
>Tupaia_chinensis
MIGGFHLLTCPFCCIF
>Trichechus_manatus_latirostr
MIGGFHLLTCPFCCIF
>Aptenodytes_forsteri
MIVGFHLLTCPFCCII
>Boiga_irregularis
MIVGFHLLTCPFCCII
>Tyto_alba
MIVGFHLLTCPFCCII
>Struthio_camelus_australis
MIVGFHLLTCPFCCII
>Chelonia_mydas
MIVGFHLLTCPFCCII
>Tinamus_guttatus
MIVGFHLLTCPFCCII
>Calypte_anna
MIVGFHLLTCPFCCII
>Apaloderma_vittatum
MIVGFHLLTCPFCCII
>Orycteropus_afer_afer
MIGGFHLLTCPFCCIF

HG0037 ENSG00000140092 FBLN5

```
>Homo_sapiens
MKCSFLTFLLQLV
>Anser_cygnoides
MKCSFLTFLLQLV
>Xenopus_laevis
MKCSFLTFLQQLV
>Chaetura_pelagica
MKCSFLTFLLQLV
>Apteryx_australis_mantelli
MKCSFLTFLLQLV
>Buceros_rhinoceros_silvestri
MKCSFLTFLLQLV
>Caprimulgus_carolinensis
MKCSFLTFLLQLV
>Cariama_cristata
MKCSFLTFLLQLV
>Felis_catus
MKCSFLKFLLQLV
>Hynobius_retardatus
MKCSFLTFLLQLV
>Balaenoptera_acutorostrata_s
MKCSFLTFLLQLV
>Calidris_pugnax
MKCSFLTFLLQLV
>Myotis_brandtii
MKCSFLTFLLQLV
>Pterocles_gutturalis
MKCSFLTFLLQLV
>Dasypus_novemcinctus
MKCSFLAFLLQLV
>Colius_striatus
MKCSFLTFLLQLV
>Columba_livia
MKCSFLTFLLQLV
>Leptosomus_discolor
MKCSFLTFLLQLV
>Cuculus_canorus
MKCSFLTFLLQLV
>Monodelphis_domestica
MKCSFLTFLLQLV
>Trichosurus_vulpecula
MKCSFLTFLLQLV
>Aquila_chrysaetos_canadensis
MKCSFLTFLLQLV
>Coturnix_japonica
MKCSFLTFLLQLV
>Gavia_stellata
MKCSFLTFLLQLV
>Chlamydotis_macqueenii
MKCSFLTFLLQLV
>Condylura_cristata
MKCSFLTFLLQLV
>Oryctolagus_cuniculus
MKCSFLTFLLQLV
>Ornithorhynchus_anatinus
MKCSFLTFLLQLV
>Tauraco_erythrolophus
MKCSFLTFLLQLV
>Opisthocomus_hoazin
MKCSFLTFLLQLV
>Corvus_brachyrhynchos
MKCSFLTFLLQLV
>Egretta_garzetta
MKCSFLTFLLQLV
>Ceratotherium_simum_simum
MKCSFLTFLLQLV
>Picoides_pubescens
MKCSFLTFLLQLV
>Loxodonta_africana
MKCSFLTFLLQLV
>Fulmarus_glacialis
MKCSFLTFLLQLV
>Nestor_notabilis
MKCSFLTFLLQLV
>Chinchilla_lanigera
MKCSFLTFLLQLV
>Tupaia_chinensis
MKCSFLTFLLQLV
>Trichechus_manatus_latirostr
MKCSFLTFLLQLV
>Aptenodytes_forsteri
MKCSFLTFLLQLV
>Tyto_alba
MKCSFLTFLLQLV
>Struthio_camelus_australis
MKCSFLTFLLQLV
>Pelodiscus_sinensis
MKCSFLTFLLQLV
>Tinamus_guttatus
MKCSFLTFLLQLV
>Calypte_anna
MKCSFLTFLLQLV
>Apaloderma_vittatum
MKCSFLTFLLQLV
```

HG0037 ENSGALG00000010747 FBLN5

```
>Gallus_gallus
MKCSFLTFLLQLV
>Anser_cygnoides
MKCSFLTFLLQLV
>Xenopus_laevis
MKCSFLTFLQQLV
>Chaetura_pelagica
MKCSFLTFLLQLV
>Apteryx_australis_mantelli
MKCSFLTFLLQLV
>Buceros_rhinoceros_silvestri
MKCSFLTFLLQLV
>Caprimulgus_carolinensis
MKCSFLTFLLQLV
>Cariama_cristata
MKCSFLTFLLQLV
>Felis_catus
MKCSFLKFLLQLV
>Hynobius_retardatus
MKCSFLTFLLQLV
>Balaenoptera_acutorostrata_s
MKCSFLTFLLQLV
>Calidris_pugnax
MKCSFLTFLLQLV
>Myotis_brandtii
MKCSFLTFLLQLV
>Pterocles_gutturalis
MKCSFLTFLLQLV
>Dasypus_novemcinctus
MKCSFLAFLLQLV
>Colius_striatus
MKCSFLTFLLQLV
>Columba_livia
MKCSFLTFLLQLV
>Leptosomus_discolor
MKCSFLTFLLQLV
>Cuculus_canorus
MKCSFLTFLLQLV
>Monodelphis_domestica
MKCSFLTFLLQLV
>Trichosurus_vulpecula
MKCSFLTFLLQLV
>Aquila_chrysaetos_canadensis
MKCSFLTFLLQLV
>Gavia_stellata
MKCSFLTFLLQLV
>Chlamydotis_macqueenii
MKCSFLTFLLQLV
>Condylura_cristata
MKCSFLTFLLQLV
>Oryctolagus_cuniculus
MKCSFLTFLLQLV
>Ornithorhynchus_anatinus
MKCSFLTFLLQLV
>Tauraco_erythrolophus
MKCSFLTFLLQLV
>Opisthocomus_hoazin
MKCSFLTFLLQLV
>Corvus_brachyrhynchos
MKCSFLTFLLQLV
>Egretta_garzetta
MKCSFLTFLLQLV
>Ceratotherium_simum_simum
MKCSFLTFLLQLV
>Picoides_pubescens
MKCSFLTFLLQLV
>Aotus_nancymaae
MKCSFLTFLLQLV
>Loxodonta_africana
MKCSFLTFLLQLV
>Fulmarus_glacialis
MKCSFLTFLLQLV
>Nestor_notabilis
MKCSFLTFLLQLV
>Chinchilla_lanigera
MKCSFLTFLLQLV
>Tupaia_chinensis
MKCSFLTFLLQLV
>Trichechus_manatus_latirostr
MKCSFLTFLLQLV
>Aptenodytes_forsteri
MKCSFLTFLLQLV
>Tyto_alba
MKCSFLTFLLQLV
>Struthio_camelus_australis
MKCSFLTFLLQLV
>Pelodiscus_sinensis
MKCSFLTFLLQLV
>Tinamus_guttatus
MKCSFLTFLLQLV
>Calypte_anna
MKCSFLTFLLQLV
>Apaloderma_vittatum
MKCSFLTFLLQLV
```

HG0038 ENSGALG00000006571

>Gallus_gallus
-----------------------------------------------------------------------------------MSVTCWPRLCGSSPPSERPVAALWTP
>Anas_platyrhynchos
-----------------------------------------------------------------------------------MSVTCWPRLCGSSPPSERPVAALWTP
>Xenopus_tropicalis
-----------------------------------------------------------------------------------MSVKCLSRLCGSPP-SVWSVAAIPTP
>Apteryx_australis_mantelli
-----------------------------------------------------------------------------------MSVTCWPRLCGSSPPSERPVAALWTP
>Buceros_rhinoceros_silvestri
-----------------------------------------------------------------------------------MSVTCWPRLCGSSPPSERPVAALWTP
>Caprimulgus_carolinensis
-----------------------------------------------------------------------------------MSVTCWPRLCGSSPPSERPVAALWTP
>Cariama_cristata
-----------------------------------------------------------------------------------MSVTCWPRLCGSSPPSERPVAALWTP
>Ailuropoda_melanoleuca
-----------------------------------------------------------------------------------MSITCWLRLCSSPPLSEWPVSINWTP
>Orcinus_orca
-----------------------------------------------------------------------------------MSITCWLRLCSSPPLSEWPVSDNWTP
>Charadrius_vociferus
-----------------------------------------------------------------------------------MSVTCWPRLCGSSPPSERPVAALWTP
>Rousettus_aegyptiacus
-----------------------------------------------------------------------------------MSITCWLRLCGSPPLSEWPVSINWTP
>Pterocles_gutturalis
-----------------------------------------------------------------------------------MSVTCWPRLCGSSPPSERPVAALWTP
>Dasypus_novemcinctus
-----------------------------------------------------------------------------------MSITCCLRLCSAPPLSEWSVSLNWIP
>Latimeria_chalumnae
-----------------------------------------------------------------------------------MSMKCWSRHCGSPP-LVWSVAAIWTP
>Colius_striatus
-----------------------------------------------------------------------------------MSVTCWPRLCGSSPPSERPVAVLWTP
>Columba_livia
-----------------------------------------------------------------------------------MSVTCWPRLCGSSPPSERPVAALWTP
>Merops_nubicus
-----------------------------------------------------------------------------------MSVTCWPRLCGSSPPSERPVAALWTP
>Alligator_mississippiensis
-----------------------------------------------------------------------------------MSVTCWPRLCGSSPPLERPVATLWTP
>Sarcophilus_harrisii
-----------------------------------------------------------------------------------MSLTCWLRLCGSPPLSEWPVSINWTP
>Monodelphis_domestica
-----------------------------------------------------------------------------------MSLTCWSRLCGSAPLSEWPVSINWIP
>Aquila_chrysaetos_canadensis
-----------------------------------------------------------------------------------MSVTCWPRLCGSSPPSERPVAALWTP
>Gavia_stellata
-----------------------------------------------------------------------------------MSVTCWPRLCGASPPSGRPVAALWTP
>Chlamydotis_macqueenii
-----------------------------------------------------------------------------------MSVTCWPRLCGSSPPSERPVAALWTP
>Condylura_cristata
-----------------------------------------------------------------------------------MSITCWLRLCSSPPLSEWPVSINWTP
>Oryctolagus_cuniculus
-----------------------------------------------------------------------------------MSVTCWPRLCSSPPLSEWSVSINWTP
>Ornithorhynchus_anatinus
-----------------------------------------------MLTRSSESAAVMIFRRRW-QRGYWDFTSTGQYKGFLMSVTCWLMLCGSPPLSEWSVSIKWTP
>Opisthocomus_hoazin
-----------------------------------------------------------------------------------MSVTCWPRLCGSPPPPERPVAALWTP
>Ficedula_albicollis
-----------------------------------------------------------------------------------MSVTCWPRLCGSSPPSERPVAALWTP
>Nipponia_nippon
-----------------------------------------------------------------------------------MSVTCWPRLCGSSPPSERPVAALWTP
>Equus_asinus
-----------------------------------------------------------------------------------MSITCWLRLCSSPPLSEWPVSINWTP
>Manis_javanica
-----------------------------------------------------------------------------------MSIICWLRLCSSPPLSEWPVSINWTP
>Picoides_pubescens
-----------------------------------------------------------------------------------MSVTCWPRLCGSSPPSERPVAALWTP
>Callithrix_jacchus
-------------------------------MCSSMLFF-SILICVSTFSRSSELL-----ADNF-GGLHIQKSIKVTDARFLMSVTCWMRLCSSPPLSEWSVSIKWTP
>Loxodonta_africana
-----------------------------------------------------------------------------------MSVTWWSRLCSSPPLSEWSVSINWTP
>Fulmarus_glacialis
-----------------------------------------------------------------------------------MSVTCWPRLCGSSPPSERPVAALWTP
>Melopsittacus_undulatus
-----------------------------------------------------------------------------------MSVTCWPRLCGSSPPSERPVAALWTP
>Fukomys_darlingi
-----------------------------------------------------------------------------------MSVTCWLRICGSPPLSEWSVSISWTP
>Tupaia_chinensis
-----------------------------------------------------------------------------------MSITCWLRLRSSPPLSEWSVSVNWTP
>Trichechus_manatus_latirostr
-----------------------------------------------------------------------------------MSITCWLRLCSSPPLSEWSVSINWTP
>Aptenodytes_forsteri
-----------------------------------------------------------------------------------MSVTCWPRLCGSSPPSERPVAALWTP
>Anolis_carolinensis
-----------------------------------------------------------------------------------MSVTCWPRLCGCSPPLELPVATLWTP
>Struthio_camelus_australis
-----------------------------------------------------------------------------------MSVTCWPRLCGSSPPSERPVAALWTP
>Trachemys_scripta
MGRGSGRVRIAPGASAGAPRALSAVSARPVEEAQPVVLRGRGRGCRR--RRGSRSRLLPRKRRGAEPGLVARGARATGRPGFPMSVTCWPRLCGSSPPLERPVATFWTP
>Tinamus_guttatus
-----------------------------------------------------------------------------------MSVTCWPRLCGSSPPSERPVAALWTP
>Calypte_anna
-----------------------------------------------------------------------------------MSVTCWPRLCGSSPPSERPLAALWTP
>Apaloderma_vittatum
-----------------------------------------------------------------------------------MSVTCWPGLCGSSPPSERPVAALWTP
>Orycteropus_afer_afer
-----------------------------------------------------------------------------------MSISCWLRLCSLPPLPEWSISINWTP

HG0039.1 ENSG00000077092 RARB

>Homo_sapiens
MIIWINYRLLAGLSVIIHDSGLGKRPTAYVPKKGQSLMELGGLFY-AICLHT--
>Acipenser_sinensis
-------------------------------------MELGPLLH-AISLYT--
>Amia_calva
MIIWINYRLLAVLSVIIHDSSLGKRPRSSVPKRGQSLMELGPLLH-AISLHT--
>Anser_cygnoides
MIIWINYRLLAVLSVIIHDSGLGKRPTAYVPKKGQSLMELGVLLC-AIFLHT--
>Xenopus_tropicalis
MIIWINYRLLAVLSVIIHDSGLGKRPTAYVPKKGAEFDGVGCTFSIAICCAPKG
>Chaetura_pelagica
MIIWINYRLLAVLSVIIHDSGLGKRPTAYVPKKGQSLMELGVLSC-AIFLHT--
>Apteryx_australis_mantelli
MIIWINYRLLAVLSVIIHDSGLGKRPTAYVPKKGQSLMELGVLLC-AIFLLT--
>Buceros_rhinoceros_silvestri
MIIWINYRLLAVLSVIIHDSGLGKRPTAYVPKKGQSLMELGVLLC-AIFLHT--
>Ursus_maritimus
MIIWINYRLLAGLSVIIHDSGLGKRPTAYVPKKGQSLMELGGLFY-AICLHT--
>Orcinus_orca
MIIWINYRLLAGLSVIIHDSRLGKRPTAYVPKRGQSLMELGGLFY-AICLHT--
>Calidris_pugnax
MIIWINYRLLAVLSVIIHDSGLGKRPTAYVPKKGQSLMELGVLLC-AIFLHT--
>Callorhinchus_milii
MIIWINYRGF--------------------------------------------
>Eptesicus_fuscus
MIIWINYRLLAGLSVIIHDSGLGKRPTAYVPKKGQSLMELGGLFY-AICLHT--
>Dasypus_novemcinctus
MIIWINYRLLAGVSVIIHDSGPAGSPRACVPKKGQSLMELGGLVLDAICLRT--
>Latimeria_chalumnae
MIIWINYRLLAVLSVIIHDSGLGKRPRAYVPKKRGRVSWSWVYFYMPFSTTPRG
>Colius_striatus
-------------------------------------MELGVLLC-AIFLHT--
>Columba_livia
MIIWINYRLLAVLSVIIHDSGLGKRPTAYVPKKGQSLMELGVLLC-AIFLHT--
>Cuculus_canorus
-------------------------------------MELGVLLC-AIFLHT--
>Galeopterus_variegatus
MIIWINYRLLAGLSVIIHDSGLGKRPTAYVPKKGQSLMELGGLFY-AICLYT--
>Falco_cherrug
MIIWINYRLLAVLSVIIHDSGLGKRPTAYVPKKGQSLMELGVLLC-AIFLHT--
>Coturnix_japonica
MIIWINYRLLAVLSVIIHDSGLGKRPTAYVPKKGQSLMELGVLLC-AIFLHT--
>Gavia_stellata
MIIWINYRLLAVLSVIIHDSGLGKRPTAYVPKKGQSLMELGVLLC-AIFLHT--
>Mesitornis_unicolor
MIIWINYRLLAVLSVIIHDSGLGKRPTAYVPKKGQSLMELGVLLC-AIFLHT--
>Apteronotus_leptorhynchus
MITWIVSKLLATLSVIIRDSGLGKGARNMSNQR---------------------
>Condylura_cristata
MIIWINYRLLAGLSVIIHDSGLGKRPTAYVPKKGQSLMELGGLFY-AICLHT--
>Oryctolagus_cuniculus
MIIWINYRLLAGLSVIIHDSGLGKEPTAYVPKKGQSLMELGGLFY-AICLHT--
>Elephantulus_edwardii
MIIWINYRLLAGLSVIIHDSGLGKRPTAYVPKRGQSLMELGGLFP-AVCLRT--
>Opisthocomus_hoazin
-------------------------------------MELGLLLR-AIFLRA--
>Geospiza_fortis
MIIWINYRLLAVLSVIIHDSGLGKRPTAYVPKKGQSLMELGVLLC-AIFLHT--
>Nipponia_nippon
MIIWINYRLLAVLSVIIHDSGLGKRPTAYVPKKGQSLMELGVLLC-AIFLHT--
>Ceratotherium_simum_simum
MIIWINYRLLAGLSVIIHDSGLGKRPTAYVPKKGQSLMELGGLFY-AICLHT--
>Manis_javanica
MIIWINYRLLAGLSVIIHDSGLGKRPTAYVPKKGQSLMELGGLFY-AICLHT--
>Picoides_pubescens
MIIWINYRLLAVLSVIIHDSGLGKRPTAYVPKKGQSLMELGVLLC-AIFLHT--
>Loxodonta_africana
MTIWINYRLLAGLSVIIHDSGLGKRPTAYVPKKGQSLMELGGLFY-AICLRT--
>Melopsittacus_undulatus
MIIWINYRLLAVLSVIIHDSGLGKRPTAYVPKKGQSLMELGVLLC-AIFLYT--
>Callospermophilus_lateralis
MIIWINYRLLAGLSVIIHDSGLGKRPTAYVPKKGQSLMELGGLFY-AICLHT--
>Salmo_salar
-------------------------------------MELCALMF-AIIIHT--
>Tupaia_chinensis
MIIWINYRLLAGLSVIIHDSGLGKSPTAYVPKKGQSLMELGGLFY-AICLLT--
>Lepisosteus_oculatus
MIIWINYRLLVVLSVIIHDSSLGKRPRSYVPKKGQSLMELVPLLH-AISLHT--
>Trichechus_manatus_latirostr
MIIWINYRLLAGLSVIIHDSGLGKRPTAYVPKKGQSLMELGGLFY-AICLRT--
>Pygoscelis_adeliae
MIIWINYRLLAVLSVIIHDSGLGKRPTAYVPKKGQSLMELGVLLC-AIFLHT--
>Squalus_acanthias
MIIWINYRLLAVLSVIIHDSGLGKRPTAQRAKKRGEEKI---------------
>Struthio_camelus_australis
MIIWINYRLLAVLSVIIHDSGLGKRPTAYVPKKGQSLMELGVLLC-AIFLHT--
>Trachemys_scripta
MIIWINYRLLAVLSVIIHDSGLGKRRTAYVPKKGQSLMELGVLLY-AIFLHT--
>Tinamus_guttatus
MIIWINYRLLAVLSVIIHDSGLGKRPTAYVPKKGQSLMELGVLLC-AIFLHT--
>Calypte_anna
MIIWINYRLLAVLSVIIHDSGLGKRPTAYVPKKGQSLMELGVLLC-AIFLHT--
>Orycteropus_afer_afer
MIIWINYRLLAGLSVIIHDSGLGKRPTAYVPKKGQSLMELGGLFY-AICLRT--

HG0039.2 ENSG00000077092 RARB

>Homo_sapiens
MPRTRAIRAGFVWAPSG
>Odobenus_rosmarus_divergens
MPRTRASRAGFVWAPSG
>Orcinus_orca
MPRTRASRAGLVWAPSG
>Rousettus_aegyptiacus
MPRTRASRAGLVWAPSG
>Galeopterus_variegatus
MPRTRAIRAGFVWAPSG
>Condylura_cristata
MPRTRASRAGFVWAPSG
>Ochotona_princeps
MPRTRATQAGSLWAPSG
>Elephantulus_edwardii
MRRTRASRAGAVWAPSG
>Equus_przewalskii
MPRTRASRAGFVWAPSG
>Marmota_marmota_marmota
MPRTRASRAGFVWAPSG
>Tupaia_chinensis
MPRTRASRAGLVWAPSG
>Trichechus_manatus_latirostr
MPRTRAIRAGFVWAPSG
>Orycteropus_afer_afer
MPRTQTIRAGFVWAPSG

HG0040.1 ENSG00000164190 NIPBL

>Homo_sapiens
------------------------------------------------------------------------------------------------------------------------------------------------------------------------------MGSNDSWHLN
>Anser_cygnoides
------------------------------------------------------------------------------------------------------------------------------------------------------------------------------MGSNDSWHLN
>Xenopus_laevis
------------------------------------------------------------------------------------------------------------------------------------------------------------------------------MGSNDSWHFN
>Chaetura_pelagica
------------------------------------------------------------------------------------------------------------------------------------------------------------------------------MGSNDSWHLN
>Apteryx_australis_mantelli
------------------------------------------------------------------------------------------------------------------------------------------------------------------------------MGSNDSWHLN
>Buceros_rhinoceros_silvestri
------------------------------------------------------------------------------------------------------------------------------------------------------------------------------MGSNDSWHLN
>Caprimulgus_carolinensis
------------------------------------------------------------------------------------------------------------------------------------------------------------------------------MGSNDSWHLN
>Cariama_cristata
------------------------------------------------------------------------------------------------------------------------------------------------------------------------------MGSNDSWHLN
>Felis_catus
------------------------------------------------------------------------------------------------------------------------------------------------------------------------------MGSNDSWHLN
>Balaenoptera_acutorostrata_s
------------------------------------------------------------------------------------------------------------------------------------------------------------------------------MGSNDSWHLN
>Calidris_pugnax
------------------------------------------------------------------------------------------------------------------------------------------------------------------------------MGSNDSWHLN
>Miniopterus_natalensis
------------------------------------------------------------------------------------------------------------------------------------------------------------------------------MGSNDSWHLN
>Pterocles_gutturalis
------------------------------------------------------------------------------------------------------------------------------------------------------------------------------MGSNDSWHLN
>Dasypus_novemcinctus
----------------------------------------------------------------------------------------------------------------------------------------------------------------------MHKYSVWEMGSNDSWHLN
>Columba_livia
------------------------------------------------------------------------------------------------------------------------------------------------------------------------------MGSNDSWHLN
>Leptosomus_discolor
------------------------------------------------------------------------------------------------------------------------------------------------------------------------------MGSNDSWHLN
>Cuculus_canorus
------------------------------------------------------------------------------------------------------------------------------------------------------------------------------MGSNDSWHLN
>Sarcophilus_harrisii
MVDNWRNWCISRGKQIPRSISVSSAGASFGLLGSIIPPKQKVCIYTYRCIIISKLCQIFLKWHIHLVARRSPENPIESRGYPKSGHPPAAFPKSGEKPGHRSHPISFILHPSGPLRFSRERGRRLEGKPPNAFLWSPSREARRRPKLRSSHFPHSRLGLGLPTDSNADSPSVWEMGSNDSWHLS
>Galeopterus_variegatus
------------------------------------------------------------------------------------------------------------------------------------------------------------------------------MESNDSWHLN
>Monodelphis_domestica
------------------------------------------------------------------------------------------------------------------------------------------------------------------------------MGSNDSWHLS
>Haliaeetus_leucocephalus
------------------------------------------------------------------------------------------------------------------------------------------------------------------------------MGSNDSWHMN
>Coturnix_japonica
------------------------------------------------------------------------------------------------------------------------------------------------------------------------------MGSNDSWHLN
>Gavia_stellata
------------------------------------------------------------------------------------------------------------------------------------------------------------------------------MGSNDSWHLN
>Chlamydotis_macqueenii
------------------------------------------------------------------------------------------------------------------------------------------------------------------------------MGSNDSWHLN
>Condylura_cristata
------------------------------------------------------------------------------------------------------------------------------------------------------------------------------MGSNDSWHLN
>Oryctolagus_cuniculus
------------------------------------------------------------------------------------------------------------------------------------------------------------------------------MGSNDSWHLN
>Elephantulus_edwardii
------------------------------------------------------------------------------------------------------------------------------------------------------------------------------MGSNDSWHLN
>Ornithorhynchus_anatinus
------------------------------------------------------------------------------------------------------------------------------------------------------------------------------MGSNDSWHLN
>Tauraco_erythrolophus
------------------------------------------------------------------------------------------------------------------------------------------------------------------------------MGSNDSWHLN
>Opisthocomus_hoazin
------------------------------------------------------------------------------------------------------------------------------------------------------------------------------MGSNDSWHLN
>Lepidothrix_coronata
------------------------------------------------------------------------------------------------------------------------------------------------------------------------------MGSNDSWHLN
>Nipponia_nippon
------------------------------------------------------------------------------------------------------------------------------------------------------------------------------MGSNDSWHLN
>Equus_przewalskii
------------------------------------------------------------------------------------------------------------------------------------------------------------------------------MGSNDSWHLN
>Picoides_pubescens
------------------------------------------------------------------------------------------------------------------------------------------------------------------------------MGSNDSWHLN
>Loxodonta_africana
------------------------------------------------------------------------------------------------------------------------------------------------------------------------------MGSNDSWHLN
>Melopsittacus_undulatus
------------------------------------------------------------------------------------------------------------------------------------------------------------------------------MGSNDSWHLN
>Peromyscus_maniculatus_baird
------------------------------------------------------------------------------------------------------------------------------------------------------------------------------MGSNDSWHLN
>Tupaia_chinensis
------------------------------------------------------------------------------------------------------------------------------------------------------------------------------MGSNDSWHLN
>Aptenodytes_forsteri
------------------------------------------------------------------------------------------------------------------------------------------------------------------------------MGSNDSWHLN
>Gekko_japonicus
------------------------------------------------------------------------------------------------------------------------------------------------------------------------------MGSNDSWHLN
>Tyto_alba
------------------------------------------------------------------------------------------------------------------------------------------------------------------------------MGSNDSWHLN
>Pelodiscus_sinensis
------------------------------------------------------------------------------------------------------------------------------------------------------------------------------MGSNDSWHLN
>Tinamus_guttatus
------------------------------------------------------------------------------------------------------------------------------------------------------------------------------MGSNDSWHLN
>Calypte_anna
------------------------------------------------------------------------------------------------------------------------------------------------------------------------------MGSNDSWHLN
>Apaloderma_vittatum
------------------------------------------------------------------------------------------------------------------------------------------------------------------------------MGSNDSWHLN
>Orycteropus_afer_afer
------------------------------------------------------------------------------------------------------------------------------------------------------------------------------MGSNDSWHLN

HG0040.2 ENSG00000164190 NIPBL

```
>Homo_sapiens
MFPALRRRKRSRSHPPSRPGL--------------------------
>Apteryx_australis_mantelli
MFAALRR-RRKRSQPPSRPGLLLLYYLPSSSCSAAAAAAQQRRRLL-
>Mustela_putorius_furo
MFPALRRRKRSRSHPPSRPGL--------------------------
>Physeter_catodon
MFPALRRRKRSRSHPPSRPGL--------------------------
>Miniopterus_natalensis
MFPALRRRKRSRSHPPSRPGL--------------------------
>Alligator_mississippiensis
MFAALRR-RRKRSQPPSRPGLLLLYYLPSSSCSAAAAAAQQRRRLL-
>Galeopterus_variegatus
MFPALRRRKRSRSHPPSRPGL--------------------------
>Aquila_chrysaetos_canadensis
MFAALRR-RRKRSQPPSRPGLLLLYYLPSSSCSAAAAAAQRRRRRLL
>Coturnix_japonica
MFAALRR-RRKRSQPPSRPGLLLLYYLPSSSCSAAAAAAQRRRRLL-
>Condylura_cristata
MFPALRRRKRSRSHPPSRPGL--------------------------
>Oryctolagus_cuniculus
MFPALRRRKRSRSHPPSRPGL--------------------------
>Elephantulus_edwardii
MFPALRSRKRSRSHPPSRPGL--------------------------
>Ficedula_albicollis
MFAALRR-RRKRSQPPSRPGLLLLYYLPSSSCSAAAAAQRRRRLL--
>Equus_caballus
MFPALRRRKRSRSHPPSRPGL--------------------------
>Loxodonta_africana
MFPALRRRKRSRSHPPSRPGL--------------------------
>Melopsittacus_undulatus
MFAALRR-RRKRSQPPSRPGLLLLYYLPSSSCSAAAAVAQRRRRLL-
>Chinchilla_lanigera
MFPALRRRKRSRSHPPSRPGL--------------------------
>Trichechus_manatus_latirostr
MFPALRRRKRSRSHPPSWPGL--------------------------
>Anolis_carolinensis
MFAALRR-RRKRSQPPSRPGLLLLYYLPSSSSSAAQRRIL-------
>Pelodiscus_sinensis
MFAALRR-RRKRSQPPSRPGLLLLYYLPSSSCSAAAAAQQRRRLL--
>Orycteropus_afer_afer
MFPALRRRKRSRSHPPSRPGL--------------------------
```

HG0041 ENSDARG00000061352 bcl11aa

>Danio_rerio
----------------------------------------------------------------------------------------------------------MTAIPQSIFPAP-SLYYF
>Acipenser_sinensis
----------------------------------------------------------------------------------------------------------MTAIPQSIFPAP-SLYYF
>Anser_cygnoides
----------------------------------------------------------------------------------------------------------MTAIPQFIFPAP-SLYYF
>Pelophylax_nigromaculatus
----------------------------------------------------------------------------------------------------------MTAVPQFIFPAP-SLYYF
>Chaetura_pelagica
----------------------------------------------------------------------------------------------------------MTAIPQFIFPAP-SLYYF
>Apteryx_australis_mantelli
----------------------------------------------------------------------------------------------------------MTAIPQFIFPAP-SLYYF
>Oryzias_melastigma
------------------------MSGVRPQPDVQVRRERHVRALELAQLEGGGGEGEGKGGFCHFFP-FSLSIHLSLILSLSLFSL------------LSFLKKAMTAIPRSIFASAQSLYYF
>Tripterygion_delaisi
----------------------------------------------------------------------------------------------------------MTAIPRSIFASAPSLYYF
>Mustela_putorius_furo
----------------------------------------------------------------------------------------------------------MTALPQFIFPAP-SLYYF
>Lipotes_vexillifer
----------------------------------------------------------------------------------------------------------MTALPQFIFPAP-SLYYF
>Pygocentrus_nattereri
MSVHWFCYFHSLLPLPGFALPMFCMSVSIHLLTFKFAGTSRP-HL----------NLQLRGAFAIFS-SLS----------------------------LSLFKKAMTAIPQSIFPAP-SFYYF
>Calidris_pugnax
---------------------MFCMSLSIQAPDVQVRRDVTS-AL----------ELAAQGGFCHFFH-LSLSLLLSLPLSPS-PSLSLFFFPC-----TKKKKKAMTAIPQFIFPAP-SLYYF
>Rousettus_aegyptiacus
----------------------------------------------------------------------------------------------------------MTALPQFIFPAP-SLYYF
>Dasypus_novemcinctus
----------------------------------------------------------------------------------------------------------MTALPQFIFPAP-SLYYF
>Alosa_alosa
----------------------------------------------------------------------------------------------------------MTAIPQSIFPAP-SLYYF
>Latimeria_chalumnae
----------------------------------------------------------------------------------------------------------MTAIPQSIFPAP-SLYYF
>Alligator_mississippiensis
----------------------------------------------------------------------------------------------------------MTAIPQFIFPAP-SLYYF
>Sarcophilus_harrisii
----------------------------------------------------------------------------------------------------------MTALPQFIFPAP-SLYYF
>Galeopterus_variegatus
----------------------------------------------------------------------------------------------------------MTALPQFIFPAP-SLYYF
>Esox_lucius
----------------------------------------------------------------------------------------------------------MTAIPQSIFPAP-SLYYF
>Haliaeetus_leucocephalus
---------------------MFCMSLSIQAPDVQVRRDVTS-AL----------ELAAQGGFCHFFH-LSLSFSLSLSLPL--PLSLSFFFPL-----HKKKKKAMTAIPQFIFPAP-SLYYF
>Gallus_gallus
------------------------MSLSIQAPDVQVRGDVTS-AL----------ELAAQGGFCHFFH-LALFL-PSFPLSPS--LSPSLFFPR-----TQKKKKAMTAIPQFIFPAP-SLYYF
>Erinaceus_europaeus
----------------------------------------------------------------------------------------------------------MTALPQFIFPAP-SLYYF
>Ochotona_princeps
----------------------------------------------------------------------------------------------------------MTALPQFIFPAP-SLYYF
>Elephantulus_edwardii
----------------------------------------------------------------------------------------------------------MTALPQFIFPAP-SLYYF
>Ornithorhynchus_anatinus
----------------------------------------------------------------------------------------------------------MTALPQFIFPAP-SLYYF
>Scleropages_formosus
----------------------------------------------------------------------------------------------------------MSAVPQSIFSAP-SLHYP
>Geospiza_fortis
------------------------MSLSIQAPDVQVRRDVTS-AL----------ELAAQGGFCHFFH-LSLSPSLSPSLSLP-LPLSLFFLAQKKKKKKKKKKKAMTAIPQFIFPAP-SLYYF
>Egretta_garzetta
----------------------------------------------------------------------------------------------------------MTAIPQFIFPAP-SLYYF
>Ceratotherium_simum_simum
----------------------------------------------------------------------------------------------------------MTALPQFIFPAP-SLYYF
>Manis_javanica
----------------------------------------------------------------------------------------------------------MTALPQFIFPAP-SLYYF
>Callithrix_jacchus
MCVHWCELPLPSSPSLLLALPMFCVSLSIQTPDVQVRRDVTS-AL----------ELAAQGGFCHFFSSLSLSLSLSPPLSLFSLSLSFFFFFF-----LLKKKKAMTALPQFIFPAP-SLYYF
>Loxodonta_africana
----------------------------------------------------------------------------------------------------------MTALPQFIFPAP-SLYYF
>Melopsittacus_undulatus
------------------------MSLSIQAPDVQVRRDVTS-AL----------ELAAQGGFCHFFH-LSLSFSPSLSLSLS----SFFFFPFSL--HKKKKKKAMTAIPQFIFPAP-SLYYF
>Mus_musculus
----------------------------------------------------------------------------------------------------------MTALPQFIFPAP-SLYYF
>Salmo_salar
----------------------------------------------------------------------------------------------------------MTAIPQSIFPAP-SLYYF
>Lepisosteus_oculatus
----------------------------------------------------------------------------------------------------------MTAIPQSIFPAP-SLYYF
>Ictalurus_punctatus
----------------------------------------------------------------------------------------------------------MTAIPQSIFSAP-SLYYF
>Trichechus_manatus_latirostr
----------------------------------------------------------------------------------------------------------MTALPQFIFPAP-SLYYF
>Thamnophis_sirtalis
---------------------MFCMSLSIQPLDVQVRRDVTS-AL----------ELAAQGGFCHFFH-LSLSLSLLSPPTVS---------PSLAQKGKKEKEKAMTAIPQFIFPAP-SLYYF
>Pelodiscus_sinensis
----------------------------------------------------------------------------------------------------------MTAIPQFIFPAP-SLYYF
>Calypte_anna
----------------------------------------------------------------------------------------------------------MTAIPQFIFPAP-SLYYF
>Orycteropus_afer_afer
----------------------------------------------------------------------------------------------------------MTALPQFIFPAP-SLYYF

HG0041 ENSG00000119866 BCL11A

```
>Homo_sapiens
-------------------------------------------------------------------MTALPQFIFP-APSLYYF
>Acipenser_sinensis
-------------------------------------------------------------------MTAIPQSIFP-APSLYYF
>Anser_cygnoides
-------------------------------------------------------------------MTAIPQFIFP-APSLYYF
>Pelophylax_nigromaculatus
-------------------------------------------------------------------MTAVPQFIFP-APSLYYF
>Chaetura_pelagica
-------------------------------------------------------------------MTAIPQFIFP-APSLYYF
>Apteryx_australis_mantelli
-------------------------------------------------------------------MTAIPQFIFP-APSLYYF
>Tripterygion_delaisi
------------------------------------------------------------------MTAIPRSIFASAPSLYYF
>Canis_lupus_familiaris
-------------------------------------------------------------------MTALPQFIFP-APSLYYF
>Lipotes_vexillifer
-------------------------------------------------------------------MTALPQFIFP-APSLYYF
>Astyanax_mexicanus
-------------------------------------------------------------------MTAIPQFIFP-APSLYYF
>Calidris_pugnax
MFCMSLSIQAPDVQVRRDVTSALELAAQGGFCHFFHLSLSLLLSLPLSPSPSLSLFFFPCTKKKKKAMTAIPQFIFP-APSLYYF
>Rousettus_aegyptiacus
-------------------------------------------------------------------MTALPQFIFP-APSLYYF
>Dasypus_novemcinctus
-------------------------------------------------------------------MTALPQFIFP-APSLYYF
>Alosa_alosa
-------------------------------------------------------------------MTAIPQSIFP-APSLYYF
>Latimeria_chalumnae
-------------------------------------------------------------------MTAIPQSIFP-APSLYYF
>Alligator_mississippiensis
-------------------------------------------------------------------MTAIPQFIFP-APSLYYF
>Sinocyclocheilus_rhinocerous
-------------------------------------------------------------------MTAIPQSIFP-APSLYYF
>Sarcophilus_harrisii
-------------------------------------------------------------------MTALPQFIFP-APSLYYF
>Galeopterus_variegatus
-------------------------------------------------------------------MTALPQFIFP-APSLYYF
>Esox_lucius
-------------------------------------------------------------------MTAIPQSIFP-APSLYYF
>Haliaeetus_leucocephalus
MFCMSLSIQAPDVQVRRDVTSALELAAQGGFCHFFHLSLSFSLSLSLPL-PLSLSFFFPLHKKKKKAMTAIPQFIFP-APSLYYF
>Gallus_gallus
---MSLSIQAPDVQVRGDVTSALELAAQGGFCHFFHLALFL-PSFPLSPS-LSPSLFFPRTQKKKKAMTAIPQFIFP-APSLYYF
>Erinaceus_europaeus
-------------------------------------------------------------------MTALPQFIFP-APSLYYF
>Ochotona_princeps
-------------------------------------------------------------------MTALPQFIFP-APSLYYF
>Elephantulus_edwardii
-------------------------------------------------------------------MTALPQFIFP-APSLYYF
>Ornithorhynchus_anatinus
-------------------------------------------------------------------MTALPQFIFP-APSLYYF
>Scleropages_formosus
-------------------------------------------------------------------MSAVPQSIFS-APSLHYP
>Sturnus_vulgaris
-------------------------------------------------------------------MTAIPQFIFP-APSLYYF
>Egretta_garzetta
-------------------------------------------------------------------MTAIPQFIFP-APSLYYF
>Ceratotherium_simum_simum
-------------------------------------------------------------------MTALPQFIFP-APSLYYF
>Manis_javanica
-------------------------------------------------------------------MTALPQFIFP-APSLYYF
>Loxodonta_africana
-------------------------------------------------------------------MTALPQFIFP-APSLYYF
>Melopsittacus_undulatus
---MSLSIQAPDVQVRRDVTSALELAAQGGFCHFFHLSLSFSPSLSLSLSSFFFFPFSLHKKKKKKAMTAIPQFIFP-APSLYYF
>Dipodomys_ordii
-------------------------------------------------------------------MTALPQFIFP-APSLYYF
>Salmo_salar
-------------------------------------------------------------------MTAIPQSIFP-APSLYYF
>Lepisosteus_oculatus
-------------------------------------------------------------------MTAIPQSIFP-APSLYYF
>Ictalurus_punctatus
-------------------------------------------------------------------MTAIPQSIFS-APSLYYF
>Trichechus_manatus_latirostr
-------------------------------------------------------------------MTALPQFIFP-APSLYYF
>Thamnophis_sirtalis
MFCMSLSIQPLDVQVRRDVTSALELAAQGGFCHFFHLSLSLSLLSPPT---VSPSLAQKGKKEKEKAMTAIPQFIFP-APSLYYF
>Pelodiscus_sinensis
-------------------------------------------------------------------MTAIPQFIFP-APSLYYF
>Calypte_anna
-------------------------------------------------------------------MTAIPQFIFP-APSLYYF
>Orycteropus_afer_afer
-------------------------------------------------------------------MTALPQFIFP-APSLYYF
```

HG0041 ENSGALG00000034048

```
>Gallus_gallus
---------------------------------------------------------------------MTAIPQFIFP-APSLYYF
>Acipenser_sinensis
---------------------------------------------------------------------MTAIPQSIFP-APSLYYF
>Anser_cygnoides
---------------------------------------------------------------------MTAIPQFIFP-APSLYYF
>Pelophylax_nigromaculatus
---------------------------------------------------------------------MTAVPQFIFP-APSLYYF
>Chaetura_pelagica
---------------------------------------------------------------------MTAIPQFIFP-APSLYYF
>Apteryx_australis_mantelli
---------------------------------------------------------------------MTAIPQFIFP-APSLYYF
>Tripterygion_delaisi
---------------------------------------------------------------------MTAIPRSIFASAPSLYYF
>Mustela_putorius_furo
---------------------------------------------------------------------MTALPQFIFP-APSLYYF
>Lipotes_vexillifer
---------------------------------------------------------------------MTALPQFIFP-APSLYYF
>Astyanax_mexicanus
---------------------------------------------------------------------MTAIPQFIFP-APSLYYF
>Calidris_pugnax
MFCMSLSIQAPDVQVRRDVTSALELAAQGGFCHFFHL-SLSLLLSLPLSP-SPSLSLFFFPCTKKKKKAMTAIPQFIFP-APSLYYF
>Rousettus_aegyptiacus
---------------------------------------------------------------------MTALPQFIFP-APSLYYF
>Dasypus_novemcinctus
---------------------------------------------------------------------MTALPQFIFP-APSLYYF
>Alosa_alosa
---------------------------------------------------------------------MTAIPQSIFP-APSLYYF
>Latimeria_chalumnae
---------------------------------------------------------------------MTAIPQSIFP-APSLYYF
>Alligator_sinensis
---------------------------------------------------------------------MTAIPQFIFP-APSLYYF
>Catla_catla
---------------------------------------------------------------------MTAIPQSIFP-APSLYYF
>Sarcophilus_harrisii
---------------------------------------------------------------------MTALPQFIFP-APSLYYF
>Galeopterus_variegatus
---------------------------------------------------------------------MTALPQFIFP-APSLYYF
>Esox_lucius
---------------------------------------------------------------------MTAIPQSIFP-APSLYYF
>Haliaeetus_leucocephalus
MFCMSLSIQAPDVQVRRDVTSALELAAQGGFCHFFHL-SLSFSLSLSLPL--PLSLSFFFPLHKKKKKAMTAIPQFIFP-APSLYYF
>Erinaceus_europaeus
---------------------------------------------------------------------MTALPQFIFP-APSLYYF
>Ochotona_princeps
---------------------------------------------------------------------MTALPQFIFP-APSLYYF
>Elephantulus_edwardii
---------------------------------------------------------------------MTALPQFIFP-APSLYYF
>Ornithorhynchus_anatinus
---------------------------------------------------------------------MTALPQFIFP-APSLYYF
>Scleropages_formosus
---------------------------------------------------------------------MSAVPQSIFS-APSLHYP
>Sturnus_vulgaris
---------------------------------------------------------------------MTAIPQFIFP-APSLYYF
>Egretta_garzetta
---------------------------------------------------------------------MTAIPQFIFP-APSLYYF
>Ceratotherium_simum_simum
---------------------------------------------------------------------MTALPQFIFP-APSLYYF
>Manis_javanica
---------------------------------------------------------------------MTALPQFIFP-APSLYYF
>Callithrix_jacchus
MFCVSLSIQTPDVQVRRDVTSALELAAQGGFCHFFSSLSLSLSLSPPLSLFSLSLSFFFFFFLLKKKKAMTALPQFIFP-APSLYYF
>Loxodonta_africana
---------------------------------------------------------------------MTALPQFIFP-APSLYYF
>Melopsittacus_undulatus
---MSLSIQAPDVQVRRDVTSALELAAQGGFCHFFHL-SLSFSPSLSLSL-SSFFFFPFSLHKKKKKKAMTAIPQFIFP-APSLYYF
>Mus_musculus
---------------------------------------------------------------------MTALPQFIFP-APSLYYF
>Salmo_salar
---------------------------------------------------------------------MTAIPQSIFP-APSLYYF
>Lepisosteus_oculatus
---------------------------------------------------------------------MTAIPQSIFP-APSLYYF
>Ictalurus_punctatus
---------------------------------------------------------------------MTAIPQSIFS-APSLYYF
>Trichechus_manatus_latirostr
---------------------------------------------------------------------MTALPQFIFP-APSLYYF
>Gekko_japonicus
---------------------------------------------------------------------MTAIPQFIFP-APSLYYF
>Chrysemys_picta
---------------------------------------------------------------------MTAIPQFIFP-APSLYYF
>Calypte_anna
---------------------------------------------------------------------MTAIPQFIFP-APSLYYF
>Orycteropus_afer_afer
---------------------------------------------------------------------MTALPQFIFP-APSLYYF
```

HG0042.1 ENSG00000140937 CDH11

>Homo_sapiens
----MGG-VCYRNWQQVSNG
>Anas_platyrhynchos
----MCG-ICYVNWQQVSSG
>Chaetura_pelagica
----MCG-ICYENWQRVSSG
>Apteryx_australis_mantelli
----MCG-ICYVSWQQVSSG
>Buceros_rhinoceros_silvestri
----MCG-ICYINWQQVSSG
>Caprimulgus_carolinensis
----MCG-ICYINWQQVSSG
>Acinonyx_jubatus
----MGG-VCYRNWQQVSNG
>Balaenoptera_acutorostrata_s
MTNQMGGVLRYRNWQQVSNG
>Charadrius_vociferus
----MCG-ICYINWQQVSSG
>Eptesicus_fuscus
----MGG-VCYRNWQQVSNG
>Pterocles_gutturalis
----MCG-ICYINWQQVSSG
>Dasypus_novemcinctus
----MGE-LSYRNWQQVSNG
>Columba_livia
----MCG-ICYINWQQVSSG
>Leptosomus_discolor
----MCG-ICYINWQQVSSG
>Alligator_mississippiensis
----MGG-ICYINWQQVTNG
>Galeopterus_variegatus
----MGR-VCYRNWQQVSNG
>Monodelphis_domestica
----MGG-VCYRNWQQVSNG
>Falco_cherrug
----MCG-ICYINWQQVSSG
>Coturnix_japonica
----MCG-ICYINWQRVSSG
>Gavia_stellata
----MCG-ICYINWQQVSSG
>Chlamydotis_macqueenii
----MCG-ICYINWQQVSSG
>Sorex_araneus
----MGG-VCYRNWQQVSNG
>Oryctolagus_cuniculus
----MGG-VCYRNWQQVSNG
>Tauraco_erythrolophus
----MCG-ICYINWQQVSSG
>Opisthocomus_hoazin
----MCG-ICYINWQQVSSG
>Pseudopodoces_humilis
----MCG-ICYINWQQVSSG
>Phaethon_lepturus
----MCG-ICYINWQQVSNG
>Ceratotherium_simum_simum
----MGG-VCYRNWQQVSNG
>Manis_javanica
----MGG-VCYRNWQQVSNG
>Picoides_pubescens
----MCG-ICYINWQQVPSG
>Loxodonta_africana
----MGG-VYYRNWQQVSNG
>Fulmarus_glacialis
----MCG-ICYINWQQVSSG
>Melopsittacus_undulatus
----MCG-ICYINWQQVSSG
>Heterocephalus_glaber
----MGG-VCYRNWQQVSNG
>Tupaia_chinensis
----MGG-LCYRNWQQVSNG
>Aptenodytes_forsteri
----MSG-ICYITWQQVSSG
>Takydromus_sexlineatus
----MGG-ICYFNWQWVSSG
>Struthio_camelus_australis
----MCG-ICYRSWQQVSSG
>Chelonia_mydas
----MGG-ICYINWQQVSSG
>Tinamus_guttatus
----MCG-ICYISWQQVSSG
>Calypte_anna
----MCG-ICYINWQQVSSG
>Apaloderma_vittatum
----MCG-TCYINWQQVSSG
>Orycteropus_afer_afer
MTNQMGR-VCYRNWQQVSNG

HG0042.2 ENSGALG00000039985

>Gallus_gallus
MHSRCGGSVCAEHFI
>Anas_platyrhynchos
MHSMCGCNLCAEHFI
>Charadrius_vociferus
MCLMCGGILCAEHFI
>Columba_livia
MCLMCGGILCAEHFI
>Falco_cherrug
MCLMCGGILCAEHFI
>Balearica_regulorum_gibberic
MCLMCGGILCAEHFI
>Opisthocomus_hoazin
MCLMCGGLLCAEHFI
>Phalacrocorax_carbo
MCLMCGGILCAEHFI
>Fulmarus_glacialis
MCSVCGGILCAEHFI
>Melopsittacus_undulatus
MHLMCGGILCAQHFI
>Aptenodytes_forsteri
MCLMCGGILCAEHFI
>Struthio_camelus_australis
MHFMCGAILCAEHFI
>Tinamus_guttatus
MHFMCGGILCAERFI
>Apaloderma_vittatum
MCLMCGGILCAEHFI

HG0042.3 ENSG00000140937 CDH11

>Homo_sapiens
MPRGPLAAAADL
>Leptonychotes_weddellii
MPRGPLAAAADL
>Orcinus_orca
MPRGPLAAAADL
>Rousettus_aegyptiacus
MPRGPLAAAADL
>Manis_javanica
MPRGPLAAAADL
>Loxodonta_africana
MPRGPLAAAADL
>Octodon_degus
MPRGPLAAAADL
>Orycteropus_afer_afer
MPRGPLAAAADL

HG0042.4 ENSG00000140937 CDH11

>Homo_sapiens
MSSTSRDIPSQERSA
>Mustela_putorius_furo
MYSVSRDIPFQERSA
>Pteropus_vampyrus
MCSASRDIPFQERSA
>Condylura_cristata
MLSASRDIPLQDRSA
>Ceratotherium_simum_simum
MCSASRDIPLQERSA
>Octodon_degus
MCSESKDIPLQERSA

HG0042.5 ENSG00000140937 CDH11

```
>Homo_sapiens
------------------------------MQKSLCFQLSGPV-TFLRVVIC
>Leptonychotes_weddellii
MGPGLGSGLTPLRSSCARDRRPVARAHPPGPQKSLYFRLSGPV-TLVPVVIC
>Eptesicus_fuscus
--------------------------------------------MIIPVAVC
>Erinaceus_europaeus
MGPGLGPGVPPLRSP--RDRPPEARAHPLGPQKSFPFWLSGPE-AFSPVGIC
>Peromyscus_maniculatus_baird
------------------------------MTKSLCFWLCGPVKTFISVVIC
```

HG0043 ENSG00000172995 ARPP21

>Homo_sapiens
--------------------------------------------MLFNEGASASLSNKRSMREDRAVHGYGYWTRIFGKVD-DVSHCVRESHGDGHSELLMGTWDSSCL--------------
>Anas_platyrhynchos
MLIQHSHCAVQFNYCYFPAFIYRLSQRGEHTHIQGSFLCARCVLAPLLI------LLSLRFQVCAVLFQFIPLLLCHIQVDDDLSNSLKNQHGDWHAELLMGTWDSSCH--------------
>Chaetura_pelagica
----------------------------------------------------------------------------------------------------MGTWDSGCH--------------
>Buceros_rhinoceros_silvestri
----------------------------------------------------------------------------------------------------MGTWDSRCH--------------
>Caprimulgus_carolinensis
----------------------------------------------------------------------------------------------------MGTWDSRCH--------------
>Cariama_cristata
----------------------------------------------------------------------------------------------------MGTWDSRCR--------------
>Mustela_putorius_furo
-------------------------------MLHWGVLCCLFL-MLFNEGASASLRNKRSMMEDRAVHGFGYWRRIYEKVD-DVSNRVRESHGDGHSELLMGTWDSSCL--------------
>Hynobius_retardatus
----------------------------------------------------------------------------------------------------MGTWDSHCH--------------
>Lipotes_vexillifer
----------------------------------------------------------------------------------------------------MGTWDSRCL--------------
>Callorhinchus_milii
----------------------------------------------------------------------------------------------------MRTWDSLCYCSAVLISQVNSIR-
>Rousettus_aegyptiacus
-----------------------------------------------MKEQAPHLANKRSMMEDRAVRGYGYWRRIFGKVD-DVSNCVRESHGHGRSDLLMGTWDSSFL--------------
>Pterocles_gutturalis
----------------------------------------------------------------------------------------------------MGTWDSRRH--------------
>Dasypus_novemcinctus
----------------------------------------------------------------------------------------------------MGTWDSSCR--------------
>Latimeria_chalumnae
----------------------------------------------------------------------------------------------------MGTWDSRCH--------------
>Colius_striatus
----------------------------------------------------------------------------------------------------MGTWDSHCH--------------
>Columba_livia
----------------------------------------------------------------------------------------------------MGTWDSRCHLSLAWT--------
>Leptosomus_discolor
----------------------------------------------------------------------------------------------------MGTWDSHCH--------------
>Alligator_sinensis
----------------------------------------------------------------------------------------------------MGTWDAHCLS-------------
>Cuculus_canorus
----------------------------------------------------------------------------------------------------MGTWDSRCH--------------
>Galeopterus_variegatus
-------------------------------MLHRGVLCSLFL-MLFNEGASASLSNKRSMREDRAVRGYGYWRTIFGKVDDDVSKCVKESHGDGHSELLKGTWDSSCL--------------
>Aquila_chrysaetos_canadensis
-----------------------------------------------------------------MKHSLPSPPFLRRQVDYDPSNSLKDQHGDWHSELLMGTWDSRCH--------------
>Gallus_gallus
----------------------------------------------------------------------------------------------------MGTWDSSCR--------------
>Balearica_regulorum_gibberic
----------------------------------------------------------------------------------------------------MGTWDSRCH--------------
>Erinaceus_europaeus
----------------------------------------------------------------------------------------------------MGTWDSSCL--------------
>Oryctolagus_cuniculus
----------------------------------------------------------------------------------------------------MGTWDSSCL--------------
>Elephantulus_edwardii
----------------------------------------------------------------------------------------------------MGTWDSGHP--------------
>Tauraco_erythrolophus
----------------------------------------------------------------------------------------------------MRTWDSRCH--------------
>Opisthocomus_hoazin
----------------------------------------------------------------------------------------------------MGTWDSRCH--------------
>Parus_major
----------------------------------------------------------------------------------------------------MGTWDSRCR--------------
>Egretta_garzetta
----------------------------------------------------------------------------------------------------MGTWDSRCH--------------
>Equus_asinus
-------------------------------MLHWGVLCSLFL-MLFNEGASASLSNKRSMMEDRAVRGYGYWRRISGKVD-DVSNCVRESHGDGRSELLMGTWDSSCL--------------
>Manis_javanica
-------------------------------------------------------------------------------MVDDVSHCVRKPHGDEHSELLVGTWDPSCLCSLVWILLVHDSRR
>Picoides_pubescens
----------------------------------------------------------------------------------------------------MGTWDSRCH--------------
>Loxodonta_africana
----------------------------------------------------------------------------------------------------MGTWDSSCL--------------
>Fulmarus_glacialis
----------------------------------------------------------------------------------------------------MGTWDSRCH--------------
>Nestor_notabilis
----------------------------------------------------------------------------------------------------MGTWDSRCH--------------
>Heterocephalus_glaber
-------------------------------MLRWGVLCSLFL-MPFNERASTSLSNKRRMRKDSSVHGYGSWRRIFGKVDDDVSKCVREPHGDGHSALLMGTWDSSCF--------------
>Pygoscelis_adeliae
----------------------------------------------------------------------------------------------------MGTWDSRCH--------------
>Python_bivittatus
----------------------------------------------------------------------------------------------------MGSWDSHCH--------------
>Tyto_alba
----------------------------------------------------------------------------------------------------MGTWDSRCH--------------
>Tinamus_guttatus
----------------------------------------------------------------------------------------------------MGTWDSHCR--------------
>Calypte_anna
----------------------------------------------------------------------------------------------------MGTWDSRCH--------------
>Orycteropus_afer_afer
----------------------------------------------------------------------------------------------------MGTWDSGCL--------------

HG0044 ENSDARG00000061687 atxn1a

>Danio_rerio
----------------------------------------------MIPHELHGPLSRS-----RCDFNCL-----------RLKSVSIRP----
>Acipenser_sinensis
----------------------------------------------MIPHEVPGSLSDS-----RCDFQCI-----------RLRSVTSDTEASQ
>Amia_calva
----------------------------------------------MIPHEVHGSLSES-----RRDFPSL-----------RLRSMLVRH----
>Anser_cygnoides
----------------------------------------------MIPHEVTGSLAEI-----KCDFPFI-----------RLKSEPARQ----
>Rhacophorus_dennysi
----------------------------------------------MIPHEVVGVPSGI-----KCDFPFT-----------RLKSVSVRQ----
>Apteryx_australis_mantelli
----------------------------------------------MIPHEVTGSLSEI-----KCDFPFI-----------RLKSKPARQ----
>Oryzias_latipes
----------------------------------------------MIPHEVHGAHFRS-----WSNSIFS-----------CVKPGAGQQ----
>Canis_lupus_familiaris
----------------------------------------------MIPHEEPGSPTEI-----KCDFPCI-----------RLKPEPARQ----
>Hynobius_retardatus
----------------------------------------------MIPHEVSGSSSGI-----KCDFLSL-----------RLKPIPARQ----
>Balaenoptera_acutorostrata_s
----------------------------------------------MIPHEEPGSPTEI-----KCDFPFI-----------RLKAEPARE----
>Pygocentrus_nattereri
----------------------------------------------MIPHEPQAQLFCG-----RHVFSCF-----------RLRSASIQP----
>Calidris_pugnax
----------------------------------------------MIPHEVTGSLAEI-----KCDFPFI-----------RLKSEPARQ----
>Miniopterus_natalensis
----------------------------------------------MIPHEEPGSPTEI-----KCDFPFI-----------RLKSEPARQ----
>Dasypus_novemcinctus
----------------------------------------------MIPHEAPGAPAAV-----PCDFPCS-----------RLKSQPARQ----
>Latimeria_chalumnae
----------------------------------------------MIPHEVPGSPSEN-----QCDFTFTF----------RLKSEQKRH----
>Columba_livia
----------------------------------------------MIPHEVTGSLAEI-----KCDFPFI-----------RLKSEPARQ----
>Alligator_mississippiensis
----------------------------------------------MIPHEVTGSPSEI-----KCDFPFI-----------RLKSEPARQ----
>Sarcophilus_harrisii
----------------------------------------------MIPHEVSGSPSEI-----KCDFPFT-----------RLKPELARQ----
>Monodelphis_domestica
----------------------------------------------MIPHEVSGSPSEI-----KCDFPFT-----------RLKSELARQ----
>Aquila_chrysaetos_canadensis
----------------------------------------------MIPHEVTGSLAEI-----KCDFPFI-----------RLKSEPARQ----
>Coturnix_japonica
----------------------------------------------MIPHEVTGSLAEI-----KCDFPFI-----------RLKSEPARQ----
>Eurypyga_helias
----------------------------------------------MIPHEVTGSLAEI-----KCDFPFI-----------RLKSEPARQ----
>Condylura_cristata
----------------------------------------------MIPHEGPGSPTEI-----KCDFPFI-----------RLKSEPARQ----
>Oryctolagus_cuniculus
----------------------------------------------MIPHEEPGSPTEI-----KCDFPFI-----------RLKSEPASQ----
>Opisthocomus_hoazin
----------------------------------------------MIPHEVTGSLAEI-----KCDFPFI-----------RLKSEPARQ----
>Scleropages_formosus
----------------------------------------------MIPHEVCGPLTNS-----RRCLPFF-----------RLRPAPIRR----
>Lepidothrix_coronata
----------------------------------------------MIPHEVTGSLAEI-----KCDFPFI-----------RLKSEPDRQ----
>Egretta_garzetta
----------------------------------------------MIPHEVTGSLAEI-----KCDFPFI-----------RLKSEPARQ----
>Equus_asinus
----------------------------------------------MIPHEEPGSPTDI-----QCDFPFI-----------RLKSEPARQ----
>Manis_javanica
----------------------------------------------MIPHEGPGSPTEI-----KCDFPFI-----------RLKSEPARQ----
>Cercocebus_atys
----------------------------------------------MIPHEEPGSPTEI-----KCDFPFI-----------RLKSEPARQ----
>Loxodonta_africana
----------------------------------------------MIPHEEPGSPTEI-----KCDFPFI-----------RLKPEPASQ----
>Melopsittacus_undulatus
----------------------------------------------MIPHEVTGSLAEI-----KCDFPFI-----------RLKSEPARQ----
>Ictidomys_tridecemlineatus
----------------------------------------------MIPHEEPGSPTEI-----KCDFPFI-----------RLKSEPARQ----
>Salmo_salar
MQQEKA----TPEGYSRR---LLQ--KATPDTLVPAAIQEASSQREMIPHEVDGRLSSSFHGNGRCDSRCSHGNSRCAPLCFSLSQSQASDRR--
>Tupaia_chinensis
MLFPGADVSISLPSFPLSSLPFAPSLPSSSFSQKPELLSADCTHGEMVPHEEPGSPTEI-----KCDFSFI-----------RLKAESARQ----
>Lepisosteus_oculatus
----------------------------------------------MIPHEVHGSHTDS-----RRDFPCI-----------RLRSVPVRH----
>Ictalurus_punctatus
----------------------------------------------MIPHELWQVLSHS-----RHVFCCF-----------QLRSVSIQP----
>Aptenodytes_forsteri
----------------------------------------------MIPHEVTGSLAEI-----KCDFPFI-----------RLKSEPARQ----
>Anolis_carolinensis
-----------------------------------MRCLQRFVLTEMIPHEVSGSASQI-----KCDFPLS-----------RLKPEPARQ----
>Struthio_camelus_australis
MLF-------------SSLKPFVQWSSSTKMAPNPEELSKGCAHGEMIPHEVTGSLSEI-----KCDFPFI-----------RLKSEPARQ----
>Pelodiscus_sinensis
----------------------------------------------MIPHEVTGSLSEI-----KCDFPFI-----------RLKSEPARQ----
>Tinamus_guttatus
----------------------------------------------MIPHEVTGSLSDI-----KCDFPFI-----------RLKPEAARQ----

HG0044 ENSG00000124788 ATXN1

>Homo_sapiens
----------------------------------------------MIPHEEPGSPTEIKCDFPFI-RLKSEPSRQ----
>Acipenser_sinensis
----------------------------------------------MIPHEVPGSLSDSRCDFQCI-RLRSVTSDTEASQ
>Amia_calva
----------------------------------------------MIPHEVHGSLSESRRDFPSL-RLRSMLVRH----
>Anser_cygnoides
----------------------------------------------MIPHEVTGSLAEIKCDFPFI-RLKSEPARQ----
>Pelophylax_nigromaculatus
----------------------------------------------MIPHEVVGVPSGIKCDFPFT-RLKSVSVRQ----
>Apteryx_australis_mantelli
----------------------------------------------MIPHEVTGSLSEIKCDFPFI-RLKSKPARQ----
>Felis_catus
----------------------------------------------MIPHEEPGSPTEIKCDFPCI-RLKPEPARQ----
>Hynobius_retardatus
----------------------------------------------MIPHEVSGSSSGIKCDFLSL-RLKPIPARQ----
>Balaenoptera_acutorostrata_s
----------------------------------------------MIPHEEPGSPTEIKCDFPFI-RLKAEPARE----
>Calidris_pugnax
----------------------------------------------MIPHEVTGSLAEIKCDFPFI-RLKSEPARQ----
>Pteropus_vampyrus
----------------------------------------------MIPHEEPGSPTEIKCDFPFI-RLKSEPAKQ----
>Dasypus_novemcinctus
----------------------------------------------MIPHEAPGAPAAVPCDFPCS-RLKSQPARQ----
>Latimeria_chalumnae
----------------------------------------------MIPHEVPGSPSENQCDFTFTFRLKSEQKRH----
>Columba_livia
----------------------------------------------MIPHEVTGSLAEIKCDFPFI-RLKSEPARQ----
>Alligator_mississippiensis
--------------------MQDSQEAEQGEMENPEQLLEDCAHKEMIPHEVTGSPSEIKCDFPFI-RLKSEPARQ----
>Sinocyclocheilus_anshuiensis
-------------------------MAS-LSTKVLPAVLETHSQREMIPHELHGPLSHSRCDFHCL-RLKSVSIRP----
>Sarcophilus_harrisii
----------------------------------------------MIPHEVSGSPSEIKCDFPFT-RLKPELARQ----
>Monodelphis_domestica
----------------------------------------------MIPHEVSGSPSEIKCDFPFT-RLKSELARQ----
>Aquila_chrysaetos_canadensis
----------------------------------------------MIPHEVTGSLAEIKCDFPFI-RLKSEPARQ----
>Coturnix_japonica
----------------------------------------------MIPHEVTGSLAEIKCDFPFI-RLKSEPARQ----
>Eurypyga_helias
----------------------------------------------MIPHEVTGSLAEIKCDFPFI-RLKSEPARQ----
>Condylura_cristata
----------------------------------------------MIPHEGPGSPTEIKCDFPFI-RLKSEPARQ----
>Oryctolagus_cuniculus
----------------------------------------------MIPHEEPGSPTEIKCDFPFI-RLKSEPASQ----
>Opisthocomus_hoazin
----------------------------------------------MIPHEVTGSLAEIKCDFPFI-RLKSEPARQ----
>Scleropages_formosus
----------------------------------------------MIPHEVCGPLTNSRRCLPFF-RLRPAPIRR----
>Corvus_brachyrhynchos
----------------------------------------------MIPHEVTGSPAEIKCDFPFI-RLKSEPARQ----
>Egretta_garzetta
----------------------------------------------MIPHEVTGSLAEIKCDFPFI-RLKSEPARQ----
>Equus_asinus
----------------------------------------------MIPHEEPGSPTDIQCDFPFI-RLKSEPARQ----
>Manis_javanica
----------------------------------------------MIPHEGPGSPTEIKCDFPFI-RLKSEPARQ----
>Loxodonta_africana
----------------------------------------------MIPHEEPGSPTEIKCDFPFI-RLKPEPASQ----
>Melopsittacus_undulatus
----------------------------------------------MIPHEVTGSLAEIKCDFPFI-RLKSEPARQ----
>Marmota_monax
----------------------------------------------MIPHEEPGSPTEIKCDFPFI-RLKSEPARQ----
>Tupaia_chinensis
MLFPGADVSISLPSFPLSSLPFAPSLPSSSFSQKPELLSADCTHGEMVPHEEPGSPTEIKCDFSFI-RLKAESARQ----
>Lepisosteus_oculatus
----------------------------------------------MIPHEVHGSHTDSRRDFPCI-RLRSVPVRH----
>Aptenodytes_forsteri
----------------------------------------------MIPHEVTGSLAEIKCDFPFI-RLKSEPARQ----
>Thamnophis_sirtalis
------------------------------------MLVEDCGHKEMIPHEVSGSTQEIKCDFPLG-RLKSEPARQ----
>Struthio_camelus_australis
----------------------------------------------MIPHEVTGSLSEIKCDFPFI-RLKSEPARQ----
>Chrysemys_picta
-----------------------------MVLWNPEQLLEHCAHREMIPHEVTGSPSEIKCDFPFI-RLKSEPARQ----
>Tinamus_guttatus
----------------------------------------------MIPHEVTGSLSDIKCDFPFI-RLKPEAARQ----

HG0045.1 ENSG00000083168 KAT6A

>Homo_sapiens
MVNWFPASENLVFHGVKLTQHQDKG
>Anser_cygnoides
MVSCFPSSENLFSHRVKLVQHRDKG
>Apteryx_australis_mantelli
MVNCFPSSENLFSHGVKLVQHRDKG
>Buceros_rhinoceros_silvestri
MVSWFLSFENLFSHRVKLVQHRDKG
>Canis_lupus_familiaris
MVNWFPASENVVFHGVKLTQHQDKG
>Physeter_catodon
MVNWFPASGNVVFHGVKLTQHQDKG
>Charadrius_vociferus
MVSWFPSFENLFSHCVKLVQHRDKG
>Callorhinchus_milii
MVNCFLSRRNLFLLCVELVQHRDKG
>Eptesicus_fuscus
MVNWFPASENVVFHGVKLTQHQDKG
>Latimeria_chalumnae
MVNWFTSSENLFSHCVEFVQHQDKG
>Colius_striatus
MVSCFPSLENLFSHRVTLVQHRDKG
>Columba_livia
MVSWFLSFENLFSHRVKLVQHQDKG
>Leptosomus_discolor
MVSWFPSFENLFSHRVKLVQHRDKG
>Alligator_sinensis
MVSWLLSSENLFSHGVKLVQHRDKG
>Cuculus_canorus
MVSWFPSFENLFSHRVKLVQHRDKG
>Sarcophilus_harrisii
MVNWFPTSENLFFHGVKHIQHQDKG
>Galeopterus_variegatus
MVDWFPASENLIFRGVKLIQHQDKG
>Monodelphis_domestica
MVNWFPASENLVFHGVKLIQHQDKG
>Aquila_chrysaetos_canadensis
MVSWFPSFENLFSHCVKLVQHRDKG
>Coturnix_japonica
MVSCFPSSENLFSHCVKLVQHQDKG
>Balearica_regulorum_gibberic
MVSWFPSFENLFSHRVKLVQHRDKG
>Erinaceus_europaeus
MVNWFPVSENVVFHGVKLTQHQDKG
>Oryctolagus_cuniculus
MVSWFPASESRVSHGVGLTQHQDKG
>Elephantulus_edwardii
MVNWFPGSENLVFHGVKFTQHQDKG
>Tauraco_erythrolophus
MVSWFPSFENLFSHCVKLVQHRDKG
>Pseudopodoces_humilis
MVSWFPSFEDLLSHHLKLVQHQDKG
>Phaethon_lepturus
MVSWFPSFENLFSHCVKLVQHQDKG
>Equus_asinus
MVNWFPASEKVVFHGVKLTQHQDKG
>Manis_javanica
MVSWFPASENVVFHGVKLTEHQDKG
>Loxodonta_africana
MVNWFPASENLVFHGVKLTQHQDKG
>Fulmarus_glacialis
MVSWFPSFENLFSHRVKLVQHRDKG
>Melopsittacus_undulatus
MVSWFLSLENLFSHRVRLVQHRDKG
>Ctenomys_sociabilis
MVNWFPSSENLVFHGVKLTQHQDKG
>Tupaia_chinensis
MVNWFPASENLVFHGVKFTQHQDKG
>Trichechus_manatus_latirostr
MVNWFPASENLVFHGVKLSQHQDKG
>Aptenodytes_forsteri
MVSWFPSFENLFSHRVKLFQHRDKG
>Anolis_carolinensis
MVNWSPFSDNLFSYCVKLLQHRDKG
>Tyto_alba
MVSWFSSFENLFSHCVKLVQLRDKG
>Struthio_camelus_australis
MVNWFPSSENLFSHGVKLVQHRDKG
>Chelonia_mydas
MVNWFPSSKNLFSHGVKLIQHRDKG
>Tinamus_guttatus
MVNCFPSSENLFSHGVKLIQHRDKG

HG0045.2 ENSG00000083168 KAT6A

>Homo_sapiens
MAAVSVWGISAVRPGAPGSRLSFLPA---
>Mustela_putorius_furo
MAAVSVWGISAVRPGAPGSRFSSPPA---
>Balaenoptera_acutorostrata_s
MAAVSVWGISAVRPGAPLTRRSSPPA---
>Rousettus_aegyptiacus
MAAVSVWGISAVRPGAPGSRLSSPPA---
>Haliaeetus_leucocephalus
MAAVSVSGFAAVRPLPLGSRLSQASA---
>Gallus_gallus
MAAVSVSGFAAVRPLLLGSRLSQSSV---
>Ochotona_princeps
MAAVSVWGISAVRLGAPGSRLSFPPA---
>Elephantulus_edwardii
MAAVSVWGISAVRPGAPGSRLSSPPV---
>Pseudopodoces_humilis
MAAVSVSGFAAVRPSPPGSRRSRASA---
>Ceratotherium_simum_simum
MAAVSVWGISAVRPGAPGSRLCFPPA---
>Ctenomys_sociabilis
MAAVSVWGISAVRPGAPGSRLSPPPDSFY
>Gekko_japonicus
MAAVSVWVFAAVRPQAPGLRLS-------
>Chrysemys_picta
MAAVSVWGFAAVRPPARGLRLSQAPI---

HG0046 ENSDARG00000018817 bdnf

```
>Danio_rerio
MMTSNKTHYLPASVGVCGY
>Anser_cygnoides
MMTSNKTHYLPASVGETGF
>Nanorana_parkeri
MMTSNKRHYLPASVGETGF
>Chaetura_pelagica
MMTSNKTHYLPASVGETGF
>Canis_lupus_familiaris
MMTSNKTHYLPASVGETGF
>Lipotes_vexillifer
MMTSNKTHYLPASVGETGF
>Calidris_pugnax
MMTSNKTHYLPASVGETGF
>Miniopterus_natalensis
MMTSNKTHYLPASVGETGF
>Haplochromis_burtoni
MMTSNKTHYLPASVGVSGY
>Columba_livia
MMTSNKTHYLPASVGETGF
>Cuculus_canorus
MMTSNKTHYLPASVGETGF
>Austrofundulus_limnaeus
MMTSNKTHYLPASVGVSGY
>Galeopterus_variegatus
MMTSNKTHYLPASVGETRF
>Monodelphis_domestica
MMTSNKTHYLPASVGETGF
>Aquila_chrysaetos_canadensis
MMTSNKTHYLPASVGETGF
>Coturnix_japonica
MMTSNKTHYLPASVGETRF
>Condylura_cristata
MMTSNKTHYLPASVGETGF
>Ochotona_princeps
MMTSNKTHYLPASVGETRF
>Elephantulus_edwardii
MMTSNKTHYLPASVGETGF
>Opisthocomus_hoazin
MMTSNKTHYLPASVRETGF
>Corvus_brachyrhynchos
MMTSNKTHYLPASVGETGF
>Egretta_garzetta
MMTSNKTHYLPASVGETGF
>Notothenia_coriiceps
MMTSNKTHYLPASVGVSGY
>Ceratotherium_simum_simum
MMTSNKTHYLPASVGETGF
>Manis_javanica
MMTSNKTHYLPASVGETGF
>Cynoglossus_semilaevis
MMTSNKTHYLPASVGVSGY
>Aotus_nancymaae
MMTSNKTHYLPASVGETRF
>Loxodonta_africana
MMTSNKTHYLPASVGETGF
>Melopsittacus_undulatus
MMTSNKTHYLPASVGETGF
>Dipodomys_ordii
MMTSNKTHYLPASVGETGF
>Salmo_salar
MMTSNKTHYLPASVGVSGY
>Lepisosteus_oculatus
MMTSNKTHYLPASVGESGF
>Trichechus_manatus_latirostr
MMTSNKTHYLPASVGETGF
>Aptenodytes_forsteri
MMTSNKTHYLPASVGETGF
>Anolis_carolinensis
MMTSNETHSLPASGGETGL
>Struthio_camelus_australis
MMTSNKTHYLPASVGETGF
>Chelonia_mydas
MMTSNKTHYLPASVGETGF
>Tinamus_guttatus
MMTSNKTHYLPASVGETGF
>Calypte_anna
MMTSNKTHYLPASVGETGF
>Orycteropus_afer_afer
MMTSNKTHYLPASVGETRF
```

HG0046 ENSG00000176697 BDNF

>Homo_sapiens
MMTSNKTHYLPASVGETRF
>Anser_cygnoides
MMTSNKTHYLPASVGETGF
>Nanorana_parkeri
MMTSNKRHYLPASVGETGF
>Chaetura_pelagica
MMTSNKTHYLPASVGETGF
>Canis_lupus_familiaris
MMTSNKTHYLPASVGETGF
>Lipotes_vexillifer
MMTSNKTHYLPASVGETGF
>Calidris_pugnax
MMTSNKTHYLPASVGETGF
>Eptesicus_fuscus
MMTSNKTHYLPASVGETGF
>Haplochromis_burtoni
MMTSNKTHYLPASVGVSGY
>Columba_livia
MMTSNKTHYLPASVGETGF
>Cuculus_canorus
MMTSNKTHYLPASVGETGF
>Danio_rerio
MMTSNKTHYLPASVGVCGY
>Austrofundulus_limnaeus
MMTSNKTHYLPASVGVSGY
>Galeopterus_variegatus
MMTSNKTHYLPASVGETRF
>Monodelphis_domestica
MMTSNKTHYLPASVGETGF
>Aquila_chrysaetos_canadensis
MMTSNKTHYLPASVGETGF
>Gallus_gallus
MMTSNKTHYLPASVGETRF
>Condylura_cristata
MMTSNKTHYLPASVGETGF
>Ochotona_princeps
MMTSNKTHYLPASVGETRF
>Elephantulus_edwardii
MMTSNKTHYLPASVGETGF
>Opisthocomus_hoazin
MMTSNKTHYLPASVRETGF
>Acanthis_flammea
MMTSNKTHYLPASVGETGF
>Egretta_garzetta
MMTSNKTHYLPASVGETGF
>Notothenia_coriiceps
MMTSNKTHYLPASVGVSGY
>Ceratotherium_simum_simum
MMTSNKTHYLPASVGETGF
>Manis_javanica
MMTSNKTHYLPASVGETGF
>Cynoglossus_semilaevis
MMTSNKTHYLPASVGVSGY
>Loxodonta_africana
MMTSNKTHYLPASVGETGF
>Melopsittacus_undulatus
MMTSNKTHYLPASVGETGF
>Rattus_norvegicus
MMTSNKTHYLPASVGETRF
>Salmo_salar
MMTSNKTHYLPASVGVSGY
>Lepisosteus_oculatus
MMTSNKTHYLPASVGESGF
>Trichechus_manatus_latirostr
MMTSNKTHYLPASVGETGF
>Aptenodytes_forsteri
MMTSNKTHYLPASVGETGF
>Anolis_carolinensis
MMTSNETHSLPASGGETGL
>Struthio_camelus_australis
MMTSNKTHYLPASVGETGF
>Chelonia_mydas
MMTSNKTHYLPASVGETGF
>Tinamus_guttatus
MMTSNKTHYLPASVGETGF
>Calypte_anna
MMTSNKTHYLPASVGETGF
>Orycteropus_afer_afer
MMTSNKTHYLPASVGETRF

HG0047 ENSGALG00000015422 NRG1

>Gallus_gallus
MLLLLSLLP------LPLLL----QILLLLLLPHDNC-FPRLSRYQPQMLKVRVLPFALGYWFT----
>Anser_cygnoides
MLLLLSLLP------LPLLL----QILLLLLLPHDNC-FPRLSRYQPQMLKGDVLSLPQKSWNCLRGV
>Xenopus_laevis
MVLLLSLLP--------RLLLFPLLLAFLLLLLHDSY-FLP-NRYHLQMLQVREQQFTLGYWFT----
>Chaetura_pelagica
MLLLLSLLP------LP--L----QILLLLLLLHDNC-FPRLSRYQPQMLKVRVLPFALGYWFT----
>Buceros_rhinoceros_silvestri
------------------------------------------------MLKVRVMPFALGYWFT----
>Caprimulgus_carolinensis
MLLLLSLLP------LPLLL----QILLLLLLPLDNC-FPRLSRYQPQMLKVRVLPFALGYWFT----
>Mustela_putorius_furo
MLLLLSLLPLLPPPLLPPP-PPPLVLLLLLLLLHDSC-FLHLSRHQLQMLEVRNMPFSLGYWFT----
>Hynobius_retardatus
MRWLLSLLP------LRL---------PVLLSPHV-TSFLQRSGCPSPVLQVRSLHTALGYWFT----
>Physeter_catodon
------------------------------------------------MLEVRNMPFSLGYWFT----
>Calidris_pugnax
MLLLLSLLP------LPLLL----QILLLLLLPHDNC-FPRLSRYQPQMLKVRILPFALGYWFT----
>Eptesicus_fuscus
MLLLLSLLPLLPPPLLPPP-PPALVLLLLLLLLHDSC-FLHLSRYQLQMLEVRNMPFSLGYWFT----
>Pterocles_gutturalis
------------------------------------------------MLKVRILPFALGYWFT----
>Dasypus_novemcinctus
------MLGLL--SPPLPPPPPPLVLLLLAPLPHDSC-FRHLSGQQLQGLEGSPLPFSLGSWFT----
>Columba_livia
MLLLLSLLP------LPLLL----QILLLLLLPHDSC-FPRLSRYQPQTLKVRVLPFALGYWFT----
>Leptosomus_discolor
------------------------------------------------MLKVRVLAFALGYWFT----
>Cuculus_canorus
MLLLLSLLP------LPLLL----QILLLLLLPHDNC-FPHLSRYQPQMLKVRVLPFALGYWFT----
>Galeopterus_variegatus
------------------------------------------------MLEVRNMPFTLGYWFT----
>Haliaeetus_leucocephalus
MLLLLSLLP------LPLLL----QILLLLLLPHDNC-FPRLSRYQPQMLKVRVLPFALGYWFT----
>Gavia_stellata
MLLLLSLLP------LPLLL----QILLLLLLPHDNC-FPRLSRYQPQMLKVRVLPFALGYWFT----
>Balearica_regulorum_gibberic
------------------------------------------------MLKVRVLPFALGYWFT----
>Condylura_cristata
MLLLLSLLPLLPPPLLPPP-PPPLVLLLLLLLLHDSC-FLHLSRQQLQMLEVRNIPFSLGYWFT----
>Oryctolagus_cuniculus
MLLLLSLLPLLPPPLLP---PPPLLLFLLLLLLHDSC-FLHLSRHQLQMLGVRNLPFSLGYWFT----
>Elephantulus_edwardii
MLLLLSLLPLLPPPLLPPP-PPPLVLLWLLLLLHDSC-FLHLSRQQLQMLEVRNLPFSLGYWFT----
>Ornithorhynchus_anatinus
MLLLLSLLYLLLLL-LLLL-LLPLLLLLLLLLLRDSC-FLHLSRHQPQMLEVRNLPFTLGYWFT----
>Opisthocomus_hoazin
MLLLLSLLP------LPLLL----QILLLLLLPHDNC-FPRLSRYQPQTLKVRVLPFALGYWFT----
>Parus_major
MLLLLSLLP------LPLLL----EILLLLLLPHDNC-FPRLSRYQPQMLKVRVLPFALGYWFT----
>Egretta_garzetta
MLLLLSLLP------LPLML----QILLLLLLPHDNC-FPRLSRYQPQMLKVRVLPFALGYWFT----
>Ceratotherium_simum_simum
MLLLLSLLPLLPPPLLPPP-PPPLILLLLLLLLHDSC-FLHLSRHQLQMLEVRNVPFSLGYWFT----
>Manis_javanica
ML--------VLPAPLPLLPP-PPVLPLLLLLPHDSG-FLHLSRPQLQTLEVRNMPFRLGYWFT----
>Mandrillus_leucophaeus
MLLLLSLLPLLPPPLLPPP-PPALVLLLLLLLLHDSC-FLHLSRHQLQMLEVRNMPFSLGYWFT----
>Loxodonta_africana
MLLLLSLLPLLPPPLLPPP-PPPLVLLLLLLLLHDSC-FLHLSRQQLQMLEVRNLPFSLGYWFT----
>Melopsittacus_undulatus
MLLLLSLLP------LPPLL----QILLLLLLPHDNC-FPRLSRYQPQMLKVRVLPFALGYWFT----
>Chinchilla_lanigera
MLLLLSLLPLLPPLLLPPPPPPPLVLLLLLLLLHDSC-FLHLSRHQLQMLEVRNVPFSLGYWFT----
>Trichechus_manatus_latirostr
MLLLLSLLPLLPPPL---LPPPPLVLLLLLLLLHDSC-FLHLSRQQLQMLEVRNLPFSLGYWFT----
>Pygoscelis_adeliae
MLLLLSLLP------LPLLL----QILLLLLLPHDNC-FPRLSRYQPQMLKVRVLPFALGYWFT----
>Anolis_carolinensis
MLVLLSLLL------LLPLPPLPFLLLFLLLLLHDGCVFLHLSSHQPQMLKVRYQPFTLGCWFT----
>Chrysemys_picta
MLLLLSLLP------LPLLPPLLLFLILLLLLLHDNC-FLHLSRHQPQMLKVRSQPFALGYWFT----
>Calypte_anna
MLLLLSLLP------LPLLL----QILLLLLLPHDNC-FPRLSRYQPQMLKVRVLPFALGYWFT----
>Apaloderma_vittatum
------------------------------------------------MLKVRVLPFALGYWFT----
>Orycteropus_afer_afer
MLLLLSLLPLLPPPLLPP-PPPPLVLLLLLLLLHDSC-FLHLSRQQLQMLEVRNLPFSLGYWFT----

HG0048 ENSG00000177565 TBL1XR1

>Homo_sapiens
----------------------------------------------MGYILCCDLMV
>Anser_cygnoides
----------------------------------------------MGCVLCCDLMV
>Chaetura_pelagica
----------------------------------------------MGCVLCCDLMV
>Apteryx_australis_mantelli
----------------------------------------------MGCVLCCDLMV
>Cariama_cristata
----------------------------------------------MGCVLCCDLMV
>Acinonyx_jubatus
----------------------------------------------MGYILCCDLMV
>Lipotes_vexillifer
----------------------------------------------MGYILCCDLMV
>Charadrius_vociferus
----------------------------------------------MGCVLCCDLMV
>Myotis_brandtii
----------------------------------------------MGYILCCDLMV
>Colius_striatus
----------------------------------------------MGCVLCCDLMV
>Columba_livia
----------------------------------------------MGCVLCCDLMV
>Leptosomus_discolor
----------------------------------------------MGCVLCCDLMV
>Alligator_sinensis
----------------------------------------------MGYILRCDLMV
>Cuculus_canorus
----------------------------------------------MGCVLCCDLMV
>Sarcophilus_harrisii
----------------------------------------------MGYILCCDLMV
>Galeopterus_variegatus
----------------------------------------------MGYILCCDLMV
>Monodelphis_domestica
----------------------------------------------MGYILCCDLMV
>Aquila_chrysaetos_canadensis
----------------------------------------------MGCVLCCDLMV
>Coturnix_japonica
MTFSCSLFGSAWKGRQTCYLRMRHSASPLCGKRHLKGKLLHINALPMGCVLCCDLMV
>Balearica_regulorum_gibberic
----------------------------------------------MGCVLCCDLMV
>Condylura_cristata
----------------------------------------------MGYILCCDLMV
>Oryctolagus_cuniculus
---------------------MHQQQQLVIGGTGYSVAHSHINALPMGYILCCDLMV
>Tauraco_erythrolophus
----------------------------------------------MGCVLCCDLMV
>Opisthocomus_hoazin
----------------------------------------------MGCVLCCDLMV
>Corvus_cornix_cornix
----------------------------------------------MGCVLCCDLMV
>Phaethon_lepturus
----------------------------------------------MGCVLCCDLMV
>Ceratotherium_simum_simum
----------------------------------------------MGYILCCDLMV
>Picoides_pubescens
----------------------------------------------MGCVLCCDLMV
>Fulmarus_glacialis
----------------------------------------------MGCVLCCDLMV
>Melopsittacus_undulatus
----------------------------------------------MGCVLCCDLMV
>Cricetulus_griseus
----------------------------------------------MGYILCCDLMV
>Tupaia_chinensis
----------------------------------------------MGHVLCCGLMV
>Trichechus_manatus_latirostr
----------------------------------------------MGYILCCDLMV
>Pygoscelis_adeliae
----------------------------------------------MGCVLCCDLMV
>Anolis_carolinensis
----------------------------------------------MGYILCRDLMV
>Tyto_alba
----------------------------------------------MGCVLCCDLMV
>Struthio_camelus_australis
----------------------------------------------MGCVLCCDLMV
>Chelonia_mydas
----------------------------------------------MGYILCRDLMV
>Orycteropus_afer_afer
----------------------------------------------MGYILCCDLMV

HG0048 ENSGALG00000039182

>Gallus_gallus
-------------------------MGCVLCCDLMV
>Anser_cygnoides
-------------------------MGCVLCCDLMV
>Chaetura_pelagica
-------------------------MGCVLCCDLMV
>Apteryx_australis_mantelli
-------------------------MGCVLCCDLMV
>Cariama_cristata
-------------------------MGCVLCCDLMV
>Ailuropoda_melanoleuca
-------------------------MGYILCCDLMV
>Lipotes_vexillifer
-------------------------MGYILCCDLMV
>Calidris_pugnax
-------------------------MGCVLCCDLMV
>Rousettus_aegyptiacus
-------------------------MGYILCCDLMV
>Colius_striatus
-------------------------MGCVLCCDLMV
>Columba_livia
-------------------------MGCVLCCDLMV
>Leptosomus_discolor
-------------------------MGCVLCCDLMV
>Cuculus_canorus
-------------------------MGCVLCCDLMV
>Sarcophilus_harrisii
-------------------------MGYILCCDLMV
>Galeopterus_variegatus
-------------------------MGYILCCDLMV
>Monodelphis_domestica
-------------------------MGYILCCDLMV
>Aquila_chrysaetos_canadensis
-------------------------MGCVLCCDLMV
>Balearica_regulorum_gibberic
-------------------------MGCVLCCDLMV
>Condylura_cristata
-------------------------MGYILCCDLMV
>Oryctolagus_cuniculus
MHQQQQLVIGGTGYSVAHSHINALPMGYILCCDLMV
>Tauraco_erythrolophus
-------------------------MGCVLCCDLMV
>Opisthocomus_hoazin
-------------------------MGCVLCCDLMV
>Acanthisitta_chloris
-------------------------MGCVLCCDLMV
>Egretta_garzetta
-------------------------MGCVLCCDLMV
>Equus_asinus
-------------------------MGFILCCDLMV
>Picoides_pubescens
-------------------------MGCVLCCDLMV
>Cercocebus_atys
-------------------------MGYILCCDLMV
>Fulmarus_glacialis
-------------------------MGCVLCCDLMV
>Melopsittacus_undulatus
-------------------------MGCVLCCDLMV
>Chinchilla_lanigera
-------------------------MGYVLCCDLMV
>Tupaia_chinensis
-------------------------MGHVLCCGLMV
>Trichechus_manatus_latirostr
-------------------------MGYILCCDLMV
>Aptenodytes_forsteri
-------------------------MGCVLCCDLMV
>Gekko_japonicus
-------------------------MGCILRCDLMV
>Tyto_alba
-------------------------MGCVLCCDLMV
>Struthio_camelus_australis
-------------------------MGCVLCCDLMV
>Apaloderma_vittatum
-------------------------MDCVLCCDLMV
>Orycteropus_afer_afer
-------------------------MGYILCCDLMV

HG0049 ENSDARG00000044485 sall4

```
>Danio_rerio
---------------MINRNALLIIIITLTCAFLLEGLISPNLENF----------------
>Channa_argus
MCEPLDWSQSAVRGTVINRNALLIIIITLTCAFLLEGLIFRGEGFVEEAEEN----------
>Anser_cygnoides
---------------MINRNALLIIIITLTCAFRPMGVISQL--------------------
>Xenopus_laevis
------------MGGSI---------QQLLEFFFFFGGGERE--------------------
>Oryzias_latipes
MCERSGLEPISCQGLVINRNALLIIIITLTCAFPTRGANFPSRRICRRRAGKLSATLCS---
>Cyprinodon_variegatus
MCEPLDRSQSAVREPAINRNALLIIIITLTCAFLLEREGG----------------------
>Coturnix_japonica
---------------MINRNALLIIIITLTCAFRPMGVISQL--------------------
>Plecoglossus_altivelis
---------------MINRRALLII-ITGTCAVDGL--DFRNFELIFLV--IFYSFLADV--
>Ficedula_albicollis
---------------MINRNALLIIIITLTCAFRPKFPNSRNLL------------------
>Cebus_capucinus_imitator
-----------------------MSRIAGTCAFLLKGGNFRTPGIRCVK-------------
>Rattus_norvegicus
---------------MINRNALLIIIITGTCAFLPKGGKIS---------------------
>Lepisosteus_oculatus
---------------MINRNALLIIIITLTCAFLLEGLISQL--------------------
>Ictalurus_punctatus
---------------MINRNALLIIIITLTCAFLLEGLISADLENLLRRRSKACYYLPQMVV
>Chrysemys_picta
---------------MINRNALLIIIITLTCAFRPRGVISQL--------------------
```

HG0049 ENSGALG00000039238

```
>Gallus_gallus
MINRNALLIIIIIAGHMRGARRTGEAGGGRLDFRNFE-----KIFF--------
>Acipenser_sinensis
MINRNALLIIIIIEGHMRGGWARFP-----------------------------
>Anser_cygnoides
----------------MRGARRIGEVGGGRLDFRNFE-----KIFF--------
>Xenopus_tropicalis
MINRNALLIIIIMAGHMRAGWGSLPAIGSTFWE---------------------
>Chaetura_pelagica
MINRNALLIIIIIAGHMRGARRIGEAGGGRLDFRNFE-----KIFF--------
>Apteryx_australis_mantelli
MINRNALLIIIIIAGHMRGARRIGEVGGGRLDFRNFE-----KIFF--------
>Menidia_menidia
MINRRALLII--ITGHMRGGWARFT-----------------------------
>Oryzias_latipes
MINRRALLII--ITGHMRGGWARFT-----------------------------
>Charadrius_vociferus
MINRNALLIIIIIAGHMRGARRIGEAGGGRLDFRNFE-----KIFF--------
>Pteropus_vampyrus
MINRNALLIIIIIAGHMRAAPGAGEPGRGRAGRRAGGAAAAAAARFP-------
>Haplochromis_burtoni
MINRRALLII--ITGHMRGGCSRFT-----------------------------
>Alosa_alosa
MINRKALLII--ITGHMRGGWARFS-----------------------------
>Columba_livia
MINRNALLIIIIIAGHMRGARRIGEAGGGRLDFRNFE-----KIFF--------
>Cuculus_canorus
MINRNALLIIIIIAGHMRGARRIGEAGGGGSISVILR-----RFFFSNFLFCPS
>Ctenopharyngodon_idella
MINREALLII--ITGHMRGGWARFP-----------------------------
>Kryptolebias_marmoratus
MINRTALLII--ITGHMRGGWGSIY---EIFEEVL-------------------
>Monodelphis_domestica
MINRNALLIIIIIAGHMRAARRAGESGRGAGRREGGATA---AARFP-------
>Esox_lucius
MINRQALLII--ITGHMRGGWARFP-----------------------------
>Falco_cherrug
MINRNALLIIIIIAGHMRGARRIGEAGGGEARFP--------------------
>Chanos_chanos
MINRKALLII--ITGHMRGGWARFP-----------------------------
>Apteronotus_leptorhynchus
MINREALLII--ITGHMRGGWTRYP-----------------------------
>Plecoglossus_altivelis
MINRRALLII--ITGTCAVDGLDFRNFELIFLVIFYSF-------LADV-----
>Scleropages_formosus
----------------MRAGWAPLGRALS----------EKVCGNFP-------
>Corvus_brachyrhynchos
MINRNALLIIIIIAGHMRGARRIGEAGGGRLDFRNFE-----KIFF--------
>Egretta_garzetta
MINRNALLIIIIIAGHMRGARRIGEAGGGRLDFRNFE-----KIFF--------
>Trematomus_bernacchii
MINRRALLII--ITGHMRGGWARFT-----------------------------
>Picoides_pubescens
MINRNALLIIIIIAGHMRGARRIGEAGGGRLDFRNFE-----KIFF--------
>Cynoglossus_semilaevis
MINRTALLII--ITGHMRGGWARFF---T-------------------------
>Cebus_capucinus_imitator
MINRNALLIIIIIEGHMRAAPGAGEPGRGRAGRRAGGAAAAAAARFP-------
>Melopsittacus_undulatus
MINRNALLIIIIIAGHMRGARRTGEAGGGGSISVILR-----RFFFSNFLFCPS
>Peromyscus_maniculatus_baird
MINRNALLIIIIIAGHMRAVPGAGEPGRGRAGRRAGGAAAA-AARFP-------
>Salvelinus_alpinus
MINRKALLII--ITGHMRGGWTRFS-----------------------------
>Ictalurus_punctatus
MINRNALLIIIITLTCAFLLEGLISADLENLL----RRRSKACYYLPQMVV---
>Aptenodytes_forsteri
MINRNALLIIIIIAGHMRGARRIGEAGGGEARFP--------------------
>Struthio_camelus_australis
MINRNALLIIIIIAGHMRGARRIGEAGGGRLDFRNFE-----KIFF--------
>Chelonia_mydas
MINRNALLIIIIIAGHMRGAAWLGE--GGGSISVILR-----RFFFSNFLFCPS
>Tinamus_guttatus
MINRNALLIIIIIAGHMRGARRTGAVGGGEARFP--------------------
>Calypte_anna
MINRNALLIIIIIAGHMRGARRIGEAGGGRLDFRNFE-----KIFF--------
```

HG0050.1 ENSG00000074054 CLASP1

>Homo_sapiens
------------------------------MVHCCHSPDCIFETQALS--NLQRT--KRQPPRYVCWEGV----IVTAL------
>Amia_calva
------------------------------MLHCCHTPACISILQPLC--DLQRI--SGQPLG----------------------
>Anser_cygnoides
------------------------------MVHCCHSQNCIFETQTLS--NLHRT--KRRPPASVRGGICRTVCPLAFLLVLCF-
>Rhacophorus_dennysi
------------------------------MVNCCHSQDCIFDGQALR--KLFQTSDDH--EEVIC-------------------
>Apteryx_australis_mantelli
------------------------------MVHCCHSRNCIFETQTLS--NLHRT--KRRPPAYVCRRIFWTCCPLAFLLVLYF-
>Ailuropoda_melanoleuca
------------------------------MVHCCHSPDCIFETQALS--NFQRT--KRQPPGYVYWEGV----TLTAL------
>Hynobius_retardatus
------------------------------MLNCFHSPHCILGTQAHSIRNLHRT--KRRPRGYDC--------PLAFIVVVLDW
>Tursiops_truncatus
------------------------------MVHCCHSPDCISETQALS--NLQQT--KRPPPGDACWEGV----TPTAL------
>Pygocentrus_nattereri
------------------------------MVHCCHSQACIPKAQPLA--SLRCT------------------------------
>Calidris_pugnax
------------------------------MVHCCHSQNCIFETQTLS--NLHRT--KRRPPAYVYRGIFRTYCPLAFLLVLYF-
>Callorhinchus_milii
------------------------------MLHCCRSQDCICSPRPLD--NLQET--DRRPLG----------------------
>Rousettus_aegyptiacus
------------------------------MVHCCHSPDCIFETQALS--NLYRT--NRQPPGYVYWEGV----TLTAL------
>Dasypus_novemcinctus
------------------------------MVHCCHSPDCIFETQALS--NLWRARATRQPPGHVYQEGSS--------------
>Columba_livia
------------------------------MVHCCHSQNCIFETQTLS--NLHRT--KRRPPAYVYRGIFRTYCPLAFLLVLYF-
>Alligator_mississippiensis
------------------------------MVHCWHSQNCIFETQAFS--NLHRT--KRQPTGFICRGIFPTCCPLASFSLVVYL
>Sarcophilus_harrisii
------------------------------MVHCCHSPDCIFETKALS--NLYQT--QRQPPGSLPSGRFSLTAL----------
>Galeopterus_variegatus
------------------------------MVHCCHSPDCIFETQALS--NLQRT--KRQPPRYVYWEGV----ILTAL------
>Monodelphis_domestica
------------------------------MVHCCHSPDCISETKALS--NLCQT--KRQPPGSLPSKEIQPDCPLAFFGLCCI-
>Aquila_chrysaetos_canadensis
------------------------------MVHCCHSQNCIFETQTLS--NLHRT--KRQPPAYVYRGIFRTYCPLAFLLVLYF-
>Coturnix_japonica
------------------------------MVHCCHSQNCIFETQALS--NLH--------------------------------
>Condylura_cristata
------------------------------MVHCCHSPDCIFETQTLS--NLQRT--KRQPPGSFYLEGV----TLTAL------
>Oryctolagus_cuniculus
------------------------------MVHCCHSPDCIFETQALS--NLQQT--KRQLPRYGYWEGV----ILTAL------
>Elephantulus_edwardii
MYDILLPLTSSSSRYLNWTFNIQPAALQTHMVHCCHSPDCIFETQALG--NSQQT--KRQLLGSVYREGV----VLTAL------
>Taeniopygia_guttata
------------------------------MVHCCHSQNCIFETQALS--NLHRR--KRRPPAYICRGSFWPCCPLAFWLLLYF-
>Ceratotherium_simum_simum
------------------------------MVHCCHSPDCIFETQALS--NLQQT--KRQPPRYVYREGV----TLTAP------
>Manis_javanica
------------------------------MVHCCHSLDCIFEAQALS--NLQQT--KRQPPGHVYWEGV----TLTAL------
>Loxodonta_africana
------------------------------MVHCCHSLDCIFETQTLS--NLLQT--KR--------------------------
>Dipodomys_ordii
------------------------------MVHCCHSPDCIFETQALS--NLQRT--KRPPPGLGYREGV----ILTAL------
>Tupaia_chinensis
------------------------------MVHCCHSPDCIFETQALS--NLQRT--KRQPPGYIYREGV----ILTAL------
>Lepisosteus_oculatus
------------------------------MVHCFHSQACISVLQPLS--NLQHI------------------------------
>Trichechus_manatus_latirostr
------------------------------MVHCCHSPDCIFETQTLS--NLQQT--KRQPPGSLYQEGV----ILTAL------
>Aptenodytes_forsteri
------------------------------MVHCCHSQNCIFETQTLS--NLH--------------------------------
>Squalus_acanthias
------------------------------MVHCCRSPDRICSPELLS--NLQQA--DQRPLG----------------------
>Anolis_carolinensis
------------------------------MVHCCHSQNHIFEIQALG--NLHQT--KRRPRGSS-RRTAQTCRPLASLLPVLLF
>Struthio_camelus_australis
------------------------------MVHCCHSQNCIFETQTLS--NLHRT--KRQPPEYVCRGIFWTCCPLAFLLVLYF-
>Chelonia_mydas
------------------------------MVHCCHSQNCIFETQALS--NLHRT--KRRPTGYIYSGVSQTYCPLALFSPVLYF
>Tinamus_guttatus
------------------------------MVHCCHSQNCIFEIQTLS--NLHRT--KRRPPAYSCRGIF---------------
>Orycteropus_afer_afer
------------------------------MVHCCHSPDYIFETQALS--NLQQT--KRQPPGSVYQEGN----ILTAL------

HG0050.2 ENSG00000074054 CLASP1

>Homo_sapiens
MKYFRPSS-RTSLCHSLLYLLLVI
>Acinonyx_jubatus
MKNFRPKS-RTSLCHSLLYLLLVI
>Tursiops_truncatus
MKYSRPRR-RTSLCHSLPSPLPVI
>Myotis_brandtii
MKYFRPNT-RTSLCHSLLYRLLVI
>Dasypus_novemcinctus
MKYFRPNS-VASLCHSLLYRLLVI
>Galeopterus_variegatus
MKYFRPNS-RTSLCHSLLYLLLVI
>Condylura_cristata
MKYFRANS-RTSLCHSLLYLLLVI
>Oryctolagus_cuniculus
MKYFKPSSRRTSLCHSLLYLLLVI
>Elephantulus_edwardii
MKYFRPLC-MTSFCHSLHLLLDI-
>Ceratotherium_simum_simum
MKYFRPNS-RTSLCHSLLYLQLVI
>Fukomys_darlingi
MKYFRPSS-RTSLCHSLLYLLLVI
>Orycteropus_afer_afer
MKYFRHNS-IASLCHSLLYLLLDI

HG0051 ENSG00000116679 IVNS1ABP

>Homo_sapiens
------------MNVSISSEMNQIMMHHYHRRNSCL
>Anas_platyrhynchos
------------------------MMHDYHRRNSCL
>Apteryx_australis_mantelli
-------------------------MHDYHRRNSCL
>Ailuropoda_melanoleuca
------------MSFSISSEMNQIMMHDYHRRNSCL
>Hynobius_chinensis
------------MSLTVCAQRNQIMIHDYHKRNSCL
>Physeter_catodon
------------MSFSISSEMNQIMMHDYHRRSSCL
>Callorhinchus_milii
------------------------MMQHHHSRNSCL
>Rousettus_aegyptiacus
------------MSFSISSEMNQIMMHDYHRRNSCL
>Dasypus_novemcinctus
------------------------MMHDYHRRNSCL
>Latimeria_menadoensis
----------MRMNFIVYSTRNQIMMQDYHSRKQSL
>Alligator_sinensis
------------MSSLVRADRHQFMMHYYHRRNSCL
>Sarcophilus_harrisii
------------MNFFVVSQRNQIIMHDYHRRNSCL
>Galeopterus_variegatus
------------------------MMHHYHRRNSCL
>Macropus_eugenii
------------MNFFVLSQRNQIMMHDYHRRNSCL
>Aquila_chrysaetos_canadensis
------------MNSIVYAERHQIMMHDYHRRNSCL
>Numida_meleagris
------------MSCIVRAERQKIMMHDYHRRNSCL
>Erinaceus_europaeus
------------MSFPISSEMNQIMMHDYHRRNSCL
>Oryctolagus_cuniculus
MRSWRTANSDEKHGTQRSPGVQEIMMHHSHRRNSCL
>Elephantulus_edwardii
------------MNFSISTERNQIMMHDYHRRNSCL
>Ornithorhynchus_anatinus
------------------------MMHDYHRRNSCL
>Pseudopodoces_humilis
------------MFTAVRAQRHQVMMHDYHRRNSCL
>Ceratotherium_simum_simum
------------MSFSISSEMNQIMMHDYHRRNSCL
>Manis_javanica
------------MSVSISTEMNQIIMHDYHRRNSCL
>Loxodonta_africana
------------MNFSVSAERSQTMMQDYHRRNSCL
>Heterocephalus_glaber
------------MSFSISSEMNQIMMHRYHRRNSCL
>Tupaia_chinensis
------------MNFSISSEMNQIMMYHYHRRNWCL
>Trichechus_manatus_latirostr
------------MNFSISTERNQIMMHDYHRRNSCL
>Aptenodytes_forsteri
------------------------MMHDYHRRNSCL
>Anolis_carolinensis
------------MSAVVCGERDQSMMNDYHRRNSCL
>Struthio_camelus_australis
------------------------MMHDYHRRNSCL
>Chrysemys_picta
------------MNGVVCAERHQIMMHDYHRRNSCL
>Tinamus_guttatus
------------------------MMHDYHRRNSCL
>Calypte_anna
------------------------MMHDYHRRNSCL
>Orycteropus_afer_afer
------------MNFSTSTERNQIMMHDYHRRNSCL

HG0051 ENSGALG00000037162

>Gallus_gallus
------------MSCIIRAERQKIMMHDYHRRNSCL
>Acipenser_sinensis
------------MSTVVYSHRNQVIMRDYHRRTTFL
>Anas_platyrhynchos
------------------------MMHDYHRRNSCL
>Apteryx_australis_mantelli
-------------------------MHDYHRRNSCL
>Ailuropoda_melanoleuca
------------MSFSISSEMNQIMMHDYHRRNSCL
>Cynops_pyrrhogaster
------------MSLIVCAERNQIMIHDYHKRSSCL
>Physeter_catodon
------------MSFSISSEMNQIMMHDYHRRSSCL
>Rousettus_aegyptiacus
------------MSFSISSEMNQIMMHDYHRRNSCL
>Dasypus_novemcinctus
------------------------MMHDYHRRNSCL
>Latimeria_chalumnae
----------MRMNSIVYSTRNQIMMQDYHSRKQSL
>Alligator_sinensis
------------MSSLVRADRHQFMMHYYHRRNSCL
>Rutilus_rutilus
------------MSTIVNRQPDQVIMHDQHRKGLSL
>Sarcophilus_harrisii
------------MNFFVVSQRNQIIMHDYHRRNSCL
>Galeopterus_variegatus
------------------------MMHHYHRRNSCL
>Macropus_eugenii
------------MNFFVLSQRNQIMMHDYHRRNSCL
>Aquila_chrysaetos_canadensis
------------MNSIVYAERHQIMMHDYHRRNSCL
>Erinaceus_europaeus
------------MSFPISSEMNQIMMHDYHRRNSCL
>Oryctolagus_cuniculus
MRSWRTANSDEKHGTQRSPGVQEIMMHHSHRRNSCL
>Elephantulus_edwardii
------------MNFSISTERNQIMMHDYHRRNSCL
>Ornithorhynchus_anatinus
------------------------MMHDYHRRNSCL
>Sturnus_vulgaris
------------MFSTVRAQRHQVMMHDYHRRNSCL
>Ceratotherium_simum_simum
------------MSFSISSEMNQIMMHDYHRRNSCL
>Manis_javanica
------------MSVSISTEMNQIIMHDYHRRNSCL
>Pan_paniscus
------------MNVSISSEMNQIMMHHYHRRNSCL
>Loxodonta_africana
------------MNFSVSAERSQTMMQDYHRRNSCL
>Mus_musculus
------------MNFAISSEMTQIMMHDYHRRNSCL
>Salmo_salar
------------MSTTTRCQKPQVIMHDQHRKSLTL
>Tupaia_chinensis
------------MNFSISSEMNQIMMYHYHRRNWCL
>Trichechus_manatus_latirostr
------------MNFSISTERNQIMMHDYHRRNSCL
>Aptenodytes_forsteri
------------------------MMHDYHRRNSCL
>Squalus_acanthias
------------MIYIVRFVPRKDNMQDHHNRNSCL
>Anolis_carolinensis
------------MSAVVCGERDQSMMNDYHRRNSCL
>Struthio_camelus_australis
------------------------MMHDYHRRNSCL
>Pelodiscus_sinensis
------------MSGVVCAERHQIMMHDYHRRNSCL
>Tinamus_guttatus
------------------------MMHDYHRRNSCL
>Calypte_anna
------------------------MMHDYHRRNSCL
>Orycteropus_afer_afer
------------MNFSTSTERNQIMMHDYHRRNSCL

HG0052 ENSG00000178235 SLITRK1

>Homo_sapiens
MNLNCVCGEQDGRCYFVMRSGMNCSL
>Anser_cygnoides
-----------------MKSGMNCSV
>Apteryx_australis_mantelli
MNWNCFRCEEDGCCYFVMRSGMNRSV
>Buceros_rhinoceros_silvestri
MNWNCFPCEEDGCCYFVMRSGMNCSV
>Caprimulgus_carolinensis
MNWNSFPCEEDGCCYFVMRSGMNCSV
>Ailuropoda_melanoleuca
MNLNCVCGEQDGRCYFVMRSGMNCSL
>Lipotes_vexillifer
MNLNCVCGEQDGRCYFVMRSGMNCSL
>Calidris_pugnax
MNWNCFPCEEDGCCYFVMRSGMNCSV
>Miniopterus_natalensis
MNLNCVCGEQDGCCYFVMRSG-----
>Pterocles_gutturalis
MNWNCFPCEEDGCCYFVMRSGMNSSV
>Columba_livia
MNWNSFPWEEDGCCYFVMRSGMHCWV
>Leptosomus_discolor
MNWNCFPREEDGCCYFVMRSGMNCSV
>Galeopterus_variegatus
MNLNCVCGEQDGRCYFVMRSGMNCSL
>Monodelphis_domestica
MNLNCVCGKQDDCCYFVMKSGMNCSV
>Falco_cherrug
MNWNCFPCEKDGCCYFVMRSGMNCSV
>Coturnix_japonica
MNWNCFPREEDGCCYFVMRSGMNCSV
>Balearica_regulorum_gibberic
MNWNCFPFEEDGCCYFVMRSGMNCSV
>Erinaceus_europaeus
MNLNCVCGEQDGRCYFVMRSGMNCSL
>Oryctolagus_cuniculus
MNLNCVCGEQDGCCYFVMRSGMNCSL
>Elephantulus_edwardii
MNLDCVCGEQDGRCYFVMKSGMNCSL
>Tauraco_erythrolophus
-----------------MRSGMNCSV
>Lepidothrix_coronata
MNWNCFPCEEDGCCYFVMRSGMNCSV
>Egretta_garzetta
MNWNCFPCEEDGCCYFVMRSGMNCSV
>Ceratotherium_simum_simum
MNLNCVCGEQDGRCYFVMRSGMNCSL
>Manis_javanica
-----------------MRSGMNCSL
>Picoides_pubescens
-----------------MRSGMNCSV
>Loxodonta_africana
-----------------MRSGMNCSL
>Melopsittacus_undulatus
MNWNCFPCEEDGCCYFVTRSGMNCSV
>Jaculus_jaculus
MNLNCVCGEQDGGCYFVMRSGMNCSL
>Tupaia_chinensis
MNLNCVCGEQDGRCYFVMRSGMNCSL
>Trichechus_manatus_latirostr
-----------------MRSGMNCSL
>Aptenodytes_forsteri
MNWNCFPCEEDGCCYFVMRSGMNRSV
>Struthio_camelus_australis
MNWNCFRCEEDDCCYFVMRSGMNRSV
>Chrysemys_picta
MNWNCFRCEDDGCCYFVMTSGMNCSV
>Tinamus_guttatus
MDWNCFRYEEDGGCYFVMRSGMNCSV
>Apaloderma_vittatum
MNWNCFPCKEDGCCYFVMRSGMNCSV
>Orycteropus_afer_afer
MNLNCVCGEQDGRCYFVMRSGMNCSL

HG0053.1 ENSG00000184564 SLITRK6

>Homo_sapiens
---MVNYMQKSQHSCEVCREF
>Anser_cygnoides
-------MQKSQHSCEVCCEF
>Chaetura_pelagica
-------MQKSQHSSEVCCEF
>Buceros_rhinoceros_silvestri
MLQTVNYMQKSQHSCEVCCEF
>Caprimulgus_carolinensis
-------MQKSQHSCEVCCEF
>Cariama_cristata
-------MQKSQHSCEVCCEF
>Acinonyx_jubatus
---MVNYMQKSQHSCEVCREF
>Charadrius_vociferus
-------MQKKQRSCEVCCEF
>Eptesicus_fuscus
---MVNYMQKSQHSCEVCPEF
>Pterocles_gutturalis
-------MQKSQHSCEVCCEF
>Colius_striatus
-------MQKSQHSCEVCSEF
>Columba_livia
-------MQKTQHSCEVCCEF
>Merops_nubicus
-------MQKSQHSCEVCCEF
>Cuculus_canorus
-------MQKSQHCCEVCCEF
>Falco_cherrug
-------MQKSQHSCEVCCEF
>Gallus_gallus
-------MRKSQHSCEVCCEF
>Gavia_stellata
-------MQKSQHSCEVCCEF
>Mesitornis_unicolor
-------MQKSQRSCEVCREF
>Condylura_cristata
---MVNYMQKTQHSCEVCCEF
>Ornithorhynchus_anatinus
---MVNYMQKSQHSCEVCCEF
>Tauraco_erythrolophus
-------MQKSQHSYEVCCEC
>Opisthocomus_hoazin
-------MQKGQHSCEVCFEY
>Acanthisitta_chloris
-------MQKSQPSCEVCCEF
>Pelecanus_crispus
-------MQKSQHSCEVCCEF
>Equus_caballus
---MVNYMQKSQHSCEVCCEF
>Manis_javanica
---MVNYMQKSQHSCEVCCEF
>Picoides_pubescens
-------MQKSQHSCEVCCEF
>Loxodonta_africana
---MVNYMQKSQRSCEVCREF
>Fulmarus_glacialis
-------MQKSQHSCEVCCEF
>Nestor_notabilis
-------MQISQHSCEVCCEF
>Marmota_marmota_marmota
---MVNYMQKSQHSCEVCCEF
>Trichechus_manatus_latirostr
---MVNYMQKSQHSCEVCREF
>Aptenodytes_forsteri
-------MQKSQHSCEVCCEF
>Chelonia_mydas
---MVNYMQKSQHRWEVCCEF
>Calypte_anna
-------MQKSQHSCEVCCEF
>Apaloderma_vittatum
-------MQTTQHSCEVCCEF
>Orycteropus_afer_afer
---MVNYMQKSQHSCEVCREF

HG0053.2 ENSG00000184564 SLITRK6

>Homo_sapiens
----------MLSILSLTRGFNFMKLWDLTKSDI------------
>Canis_lupus_familiaris
-----------------------MKQWDLPKSDI------------
>Physeter_catodon
----------MLSILSLTRGFNFIKRWDLPKSDI------------
>Rousettus_aegyptiacus
----------MLSILGLTRGFHFIK---------------------
>Galeopterus_variegatus
----------MLSVLSLTRGFNFMKLWDLPKSDI------------
>Condylura_cristata
----------MLSILSLNRGFNFMKQRNCQKVIFQKKMHSGWCFLF
>Oryctolagus_cuniculus
----------MLSIFSWTRSLNFLMLGNTPQSDI------------
>Equus_asinus
MHSARNIFNEMLSILSLTKGFNFMKQWDLPKSDI------------
>Manis_javanica
----------MLSILSLTRGFNFMKQWDLPKSDII-----------
>Loxodonta_africana
MHSARNIFSEMLSILSLIGAVNFRKLWDLPKSDI------------
>Fukomys_darlingi
----------MLSILSLTRDFNFPKLWDLPKSDI------------
>Tupaia_chinensis
----------MLSIVSLTRGFNFMKLWDLPKSDI------------
>Orycteropus_afer_afer
----------MLSILSLTGVFNFKKLWDMPKSDI------------

HG0054 ENSG00000124479 NDP

>Homo_sapiens
-------MTTRKLQPDPALSLNGTGS------
>Anser_cygnoides
-------MTAKRLKPDFSLSWRGTGSHDLEFF
>Leptonychotes_weddellii
-------MTTRKLQPDPALCLKGTGS------
>Balaenoptera_acutorostrata_s
-------MTTRKLQPDPALSLKGTGA------
>Miniopterus_natalensis
-------MTTRKLQPDPALSLEGTGS------
>Dasypus_novemcinctus
-------MITRKLQSDPALSLRGTRS------
>Galeopterus_variegatus
-------MTTRKLQPDPALSLKGTGS------
>Meleagris_gallopavo
-------MTTKRLKPDFSLSWRGTGSQYLVFF
>Gavia_stellata
-------MTAKRLKPDFLLSWKGTGCQYLEFF
>Sorex_araneus
-------MTARKLQPDPALSLEGTGS------
>Oryctolagus_cuniculus
-------MTTRKLQPDPALSLKGSGA------
>Ceratotherium_simum_simum
-------MTTRKLQPDPALSLKGTGS------
>Loxodonta_africana
-------MTTRKLQPDPALSLKGTGS------
>Ctenomys_sociabilis
-------MTTTKLQPDPALALKGTGS------
>Tupaia_chinensis
-------MTTRKLQPDPTLSLKGTGS------
>Trichechus_manatus_latirostr
-------MTTRKLQPDPALSLKGTGS------
>Python_bivittatus
MSNPKPAMTTWRLKTDPLLSWQGTGPQSLEGF
>Orycteropus_afer_afer
-------MTTRKLQPDPTLSLKGTGS------

HG0054 ENSGALG00000016222 NDP

```
>Gallus_gallus
-----------MTAKRLKPDFSLSWRGTGSQYLEFF
>Anser_cygnoides
-----------MTAKRLKPDFSLSWRGTGSHDLEFF
>Apteryx_australis_mantelli
-----------MTAKRPKPDFSPSWKETGSQYLEFF
>Cariama_cristata
-----------MTAKRLKPDFSPSWKGTGCQYLEFF
>Felis_catus
-----------MTTRKLRPDRALCWEGTGS------
>Balaenoptera_acutorostrata_s
-----------MTTRKLQPDPALSLKGTGA------
>Calidris_pugnax
-----------MTAKRLKPDFSPSWKGTGCQYLEFF
>Pteropus_vampyrus
-----------MTTRKLQPDPALSLKGTGS------
>Pterocles_gutturalis
-----------MTAKRLKPDFSPSWKGTGCQYLEFF
>Colius_striatus
-----------MTSKRLKPVFSPSWEGTGGQYLEFC
>Leptosomus_discolor
-----------MTAKRLKPDFSPSWKGTGCQCLEFF
>Cuculus_canorus
-----------MPAKRLKPGFSPSWKGTDCQYLEFF
>Galeopterus_variegatus
-----------MTTRKLQPDPALSLKGTGS------
>Haliaeetus_albicilla
-----------MTAKRLKPDFSPSWRGTGCQYLEFF
>Gavia_stellata
-----------MTAKRLKPDFLLSWKGTGCQYLEFF
>Balearica_regulorum_gibberic
-----------MTAKRLKPDFSPSWKGTGCQYLEFF
>Sorex_araneus
-----------MTARKLQPDPALSLEGTGS------
>Oryctolagus_cuniculus
-----------MTTRKLQPDPALSLKGSGA------
>Tauraco_erythrolophus
-----------MTAKRLKPDFSPSWKGTGCQDLEFF
>Opisthocomus_hoazin
-----------MTAKRLKPDFSPSWKGTGCPYLEFF
>Nipponia_nippon
MISVNSNSRRRMTAKRLKPDFSPSWKGTGCQYLEFF
>Ceratotherium_simum_simum
-----------MTTRKLQPDPALSLKGTGS------
>Aotus_nancymaae
-----------MTTRKLQPDPALSLKGTGS------
>Loxodonta_africana
-----------MTTRKLQPDPALSLKGTGS------
>Fulmarus_glacialis
-----------MTAKRLKPDFSPSWKGTGCQYLEFF
>Nestor_notabilis
-----------MTAKALKPDFSPSWKGTGCHYLEFF
>Heterocephalus_glaber
-----------MTTKKLRPDPALTLKGTGS------
>Tupaia_chinensis
-----------MTTRKLQPDPTLSLKGTGS------
>Trichechus_manatus_latirostr
-----------MTTRKLQPDPALSLKGTGS------
>Pygoscelis_adeliae
-----------MTAKRLKPDFSSSWKGTGCQYLEFF
>Gekko_japonicus
----MSTPKLAMTTQRLKTDLSLSSKGTGPQSLECF
>Struthio_camelus_australis
-----------MTAKRLKPDFSPSWKGTGSRYLEFF
>Chrysemys_picta
-----------MTTKRLKPGFSSSLKGTESQYLEFF
>Tinamus_guttatus
-----------MTVKRTNPDFSLSWKGMGSQYLEFF
>Calypte_anna
-----------MTAKRLKPDSSPSWKGTDCQVLEFF
>Orycteropus_afer_afer
-----------MTTRKLQPDPTLSLKGTGS------
```

HG0055.1 ENSGALG00000010533 ELAVL4

```
>Gallus_gallus
MLHIVFQRVSCHDLIKRGGGCVCCSA-----------------------------------------
>Anser_cygnoides
MLHIVFQRVSCHDLIKRGGGLCVLQRLVYVDF-VASLGSRAVRRNVSFYITLSLLPTR---------
>Chaetura_pelagica
MLHIVFQRVSCHDLIKRGGGLYVLQRLVYVDF-VASLGSSAVRRNVSFYITLSLLFDSVRGCKGPSL
>Apteryx_australis_mantelli
MLHIVFQRVSCHDLIKRGGGLYVLQRLVYVDF-VASLGSGTVRRNVSFYITFFLLLTL---------
>Caprimulgus_carolinensis
MLHIVFQRVSCHDLIKRGGGLYVLQRLVYVDF-VASLGSSAVRRNVSFYITLFLLLTL---------
>Canis_lupus_familiaris
MLHLVFQRVSCHDLIKKRRRLVCVAVPGVLSCCFLPWAAAL--------------------------
>Hynobius_retardatus
MLHIVFQRVSCHDLIKRGGGLYVLQRIVYVDL-LHRWAAVL--------------------------
>Lipotes_vexillifer
MLHLVFQRVSCHDLIKKRRRLVCVAVPGVLSCCFLLWAAAL--------------------------
>Calidris_pugnax
MLHIVFQRVSCHDLIKRGGGLYVLQRLVYVDF-VASLGSSAVRRNVSFYITLFLLLTL---------
>Myotis_lucifugus
MLHLVFQRVSCHDLIKKRRRLVCVAVPGVLSCCFLLWAAAL--------------------------
>Dasypus_novemcinctus
MLHLVFQRVSCHDLIKKRRRLVCVAVPGVLSCCFLLWAAAL--------------------------
>Cuculus_canorus
MLHIVFQRVSCHDLIKRGGGVYVLQCLVYVDF-VASLGSSAVRRNVSFYITLFLLSTL---------
>Galeopterus_variegatus
MLHLVFQRVSCHDLIKKRRRLVCVAVPGVLSCCFLLWAAAL--------------------------
>Aquila_chrysaetos_canadensis
MLHIVFQRVSCHDLIKRGGGLYVLQRLVYVDF-VASLGSSAVRRNVSFYITLFLLLTL---------
>Balearica_regulorum_gibberic
MLHIVFQRVSCHDLIKRGGGLYVLQRLVYVDF-VASLGSSAVRRNVSFYITLFLLLTL---------
>Condylura_cristata
MLHLVFQRVSCHDLIKKRRRLVCVAVPGVLSCCFLLWAAAL--------------------------
>Oryctolagus_cuniculus
MLHLVFQRVSCHDLIKKRRRLVCVAVPGVLSCCFLLWAAAL--------------------------
>Elephantulus_edwardii
MLHLVFQRVSCHDLIKKRRRLVCVAVPGVLNCCFLLWAAAL--------------------------
>Opisthocomus_hoazin
MLHIVFQRVSCHDLIKRGGGLYVLQRLVYVDF-VASLGSSAVRRNVSFYITLFLLLTL---------
>Corvus_brachyrhynchos
MLHIVFQRVSCRDLIKRGGGLYVLQCLVYVDF-VASLGSSAVRRNVSFYITLFLLLTL---------
>Egretta_garzetta
MLHIVFQRVSCHDLIKRGGGLYVLQRLVYVDF-VASLGSSAVRRNVSFYITLFLLLTL---------
>Ceratotherium_simum_simum
MLHLVFQRVSCHDLIKKRRRLVCVAVPGVLSCCFLLWAAAL--------------------------
>Manis_javanica
MLHLVFQRVSCHDLIKKRRRLVCVAVPGVLSCCFLLWAAAL--------------------------
>Aotus_nancymaae
MLHLVFQRVSCHDLIKKRRRLVCVAVPGVLSCCFLLWAAAL--------------------------
>Loxodonta_africana
MLHLVFQRVSCHDLIKKRRRLVCVAVPGVLSCCFLLWAAAL--------------------------
>Fulmarus_glacialis
MLHIVFQRVSCHDLIKRGGGLYVLQRLVYVDF-VASLGSSAVRRNVSFYITLFLLLTL---------
>Melopsittacus_undulatus
MLHIVFQRVSCRDLIKRGGGLYVLQCLVYVDF-VASLGSSAVRRNVSFYITLFLLSTL---------
>Dipodomys_ordii
MLHLVFQRVSCHDLIKRGGGLYV--------------------------------------------
>Trichechus_manatus_latirostr
MLHLVFQRVSCHDLIKKRRRLVCVAVPGVLSCCFLLWAAAL--------------------------
>Aptenodytes_forsteri
MLHIVFQRVSCHDLIKRGGGLYVLQRLVYVDF-VASLGSSAVRRNVSFYITLFLLLTL---------
>Anolis_carolinensis
MLHIVFQRVSCHDLIKRGGGLYVYQRLVYADL-LHRWAAVL--------------------------
>Tyto_alba
MLHIVFQRVSCHDLIKRGGGLYVLQRLVYVDF-VASLGSSAVRRNVSFYITLFLLLTL---------
>Struthio_camelus_australis
MLHIVFQRVSCHDLIKRGGGLYVLQRLVYVDF-VASLGSSTVRRNVSFYITFFLLLTL---------
>Chelonia_mydas
MLHIVFQRVSCHDLIKRGGGLYVLQRLVYVDF-VASLGSSTVRRNVSFYITFFLLLTL---------
>Tinamus_guttatus
MLHIVFQRVSCHDLIKRGGGSCVLRRLVYVDF-VASLGSGTVRRNVSFYITLSLLPTL---------
>Orycteropus_afer_afer
MLHLVFQRVSCHDLIKKRRRLVCVAVPGVLSCCFLLWAAAL--------------------------
```

HG0055.2 ENSGALG00000010533 ELAVL4

>Gallus_gallus
-------MSILLHRWAAAL
>Anser_cygnoides
-------MSILLHRWAAAL
>Chaetura_pelagica
MCCSAWYMSILLHRWAAAL
>Apteryx_australis_mantelli
-------MSILLHRWAAVL
>Caprimulgus_carolinensis
-------MSILLHRWAAAL
>Calidris_pugnax
-------MSILLHRWAAAL
>Cuculus_canorus
-------MSILLHRWAAAL
>Aquila_chrysaetos_canadensis
-------MSILLHRWAAAL
>Balearica_regulorum_gibberic
-------MSILLHRWAAAL
>Opisthocomus_hoazin
-------MSILLHRWAAAL
>Corvus_brachyrhynchos
-------MSILLHRWAAAL
>Egretta_garzetta
-------MSILLHRWAAAL
>Fulmarus_glacialis
-------MSILLHRWAAAL
>Melopsittacus_undulatus
-------MSILLHRWAAAL
>Aptenodytes_forsteri
-------MSILLHRWAAAL
>Tyto_alba
-------MSILLHRWAAAL
>Struthio_camelus_australis
-------MSILLHRWAAAL
>Pelodiscus_sinensis
-------MSILLHRWAAVL
>Tinamus_guttatus
-------MSILLHRWAAVL

HG0056 ENSG00000139329 LUM

>Homo_sapiens
----------------MSWAVLSPHVHLHFVREQCSHAT--PQDPHNDITPFRDWRDWAG-SPHPPSALVSLRIWQPVPS----------------
>Ursus_maritimus
----------------MSGAVLAPHVHLHFVREQNAHMP--PCK----IPTMT-------------------------------------------
>Hynobius_chinensis
----------------MGWPFLS--IHLHFVREQYFHMP--HHK----FTMMT-------------------------------------------
>Orcinus_orca
-------------------------------------MP--HHK----IPTMT-------------------------------------------
>Calidris_pugnax
MSTSIKNNRWVVLVQNMVSTCT--------SLESCIFGHATPQDHHDDIAASENSVTRAFF--FFFFLFVTERIWQTFTS----------------
>Myotis_ricketti
----------------MYWAVLAHQSTCTSLESRAFTCH--TTRSHNDITPFRVWRDWAG-SPPFSSCITKNLAASSALTEFTAYIGGCLSILHLL
>Haliaeetus_leucocephalus
----------------MGCPCTKYGIHLHFIRESCIFVHATPQDHHDDIAASENSVTRAFF--FF-FSLVTQRIWQTFTS----------------
>Mesitornis_unicolor
----------------MVSTCT--------SLESYIFVHATPQDHHDDIAASQNSVTRAFF--F---SLVTQRIWQTLTS----------------
>Ficedula_albicollis
----------------MVSTCT--------SLESCIFVHATPQDHHDDIAASENSVTRAFF--FF-SPLVTQRIWQSFTS----------------
>Ceratotherium_simum_simum
----------------MSWADFAHHVHLHFVREQSVHMP--HHK----IPTMT-------------------------------------------
>Picoides_pubescens
----------------MGSTCT--------SLELHFFVHATPQDPHDDIAASHNSLTRAFLSFFFFPPAVTQRIWQIFTS----------------
>Loxodonta_africana
----------------MSRAVLSLHVHLHFVRELSFYTP--HHE----IPTMK-------------------------------------------
>Nestor_notabilis
----------------MVSTCT--------SLESCIFVHATPQDHHDDIAASENSVTRAFF--F---SLVTQRIWQTCTS----------------
>Marmota_marmota_marmota
----------------MSWAVVAQHVHLHFVREQSVNMP--HHK----SPTMT-------------------------------------------
>Tupaia_chinensis
----------------MSRAVLVHHVHLHFVREQCVHMP--HHK----LPTMT-------------------------------------------
>Pygoscelis_adeliae
----------------MGCPCTKYGIHLHFIRELHFCTCHTTRSP---------------------------------------------------
>Python_bivittatus
----------MGRIDGLSFTIFS--LHLHLIRQQQFHIP--HAR----IP----------------------------------------------
>Orycteropus_afer_afer
----------------MSWAVLALHVHLHFVREQSFHMP--HHK----IPTMK-------------------------------------------

HG0056 ENSGALG00000011271 LUM

```
>Gallus_gallus
MGCPCTECGIHLHFIKRAAFSYMPHHKITMMT
>Anser_cygnoides
----------------------MPHHKITMMT
>Apteryx_australis_mantelli
----------------------MPHHKITMMT
>Caprimulgus_carolinensis
----------------------MPHHKITMMT
>Cariama_cristata
----------------------MPHHKITMMT
>Hynobius_chinensis
MGWPF--LSIHLHFVRE-QYFHMPHHKFTMMT
>Charadrius_vociferus
----------------------MPHHKITMMT
>Myotis_ricketti
----------------------MPHHKIPQ--
>Pterocles_gutturalis
----------------------MPHHKITMMT
>Latimeria_chalumnae
---------MHLHLVKE-LCFFIPHHKLIMMT
>Colius_striatus
MDCPYTKYGIHLHFIRE-LHFCSCHTTISMMT
>Columba_livia
----------------------MPHHKITMMT
>Merops_nubicus
----------------------MPHHKITMMT
>Alligator_sinensis
----------------------MPHHKITMMT
>Haliaeetus_leucocephalus
MVS-------TCTSLERAAFLYMPHHRITMMT
>Gavia_stellata
----------------------MPHHKITMMT
>Chlamydotis_macqueenii
MGCPCTKCGIHLHFIRE-LHFRTCYTTNSP--
>Condylura_cristata
----------------------MPHHKIPTMT
>Ficedula_albicollis
MGCPCTKYGIHLHFIRE-LHFCTCHTTRSP--
>Nipponia_nippon
MGCPCTKYGIHLHFIRD-LHFSTCHTTRSP--
>Equus_przewalskii
----------------------MPHHKIPTMT
>Picoides_pubescens
MGCPYTKYGIHLHFIRA-AFFCPCHTTRSP--
>Propithecus_coquereli
MNWAVLAPHVHLHFVRE-QSVHMPHHKIPTMT
>Loxodonta_africana
MSRAVLSLHVHLHFVRE-LSFYTPHHEIPTMK
>Fulmarus_glacialis
----------------------MPHHKITMMT
>Nestor_notabilis
----------------------MPHHKITMMT
>Tupaia_chinensis
MSRAVLVHHVHLHFVRE-QCVHMPHHKLPTMT
>Pygoscelis_adeliae
----------------------MPHHKITMMT
>Gekko_japonicus
----------------------MPHHQNPMMT
>Tyto_alba
----------------------MPHHKITMMT
>Struthio_camelus_australis
----------------------MPHHKITMMT
>Pelodiscus_sinensis
MGCPCTVYTVHLHFIRE-QHLYIPHHKIIMMT
>Tinamus_guttatus
MGCPCTKYGIHLHFITE-LHFCTYHTTRSL--
>Apaloderma_vittatum
----------------------MPHHKITMMT
>Orycteropus_afer_afer
MSWAVLALHVHLHFVRE-QSFHMPHHKIPTMK
```

HG0057 ENSG00000280987 MATR3

```
>Homo_sapiens
---------------------------MLGAQWRRNQPSRAAESCLLVLSLKKILQSLLILFLGVTIFEAKLT
>Anas_platyrhynchos
--------------------------------MTK--TACFSTELSGVSQLEEDLQSLLILFLGVTIFEVKFA
>Apteryx_australis_mantelli
--------------------------------MTK--TACFSTELSGVSQLEEDLQSLLILFLGVTIFEVKFA
>Buceros_rhinoceros_silvestri
----------------MMHK--LNCICLKGL-MTK--TACFSTELSGVSQLEEDLQSLLILFLGVTLLEVKFA
>Caprimulgus_carolinensis
--------------------------------MTK--TACFSTELSGVSQLEEDLQSLLILFLGVTIFELKFA
>Cariama_cristata
--------------------------------MTK--PACFSTELSGVSQLEEDLQSLLILFLGVTIFEVKFA
>Odobenus_rosmarus_divergens
---------------------------MLGAQWRRNQPPRAAESCLLVLSLKKILQSLLILFLGVTIF-----
>Calidris_pugnax
--------------------------------MTK--TACFSTELSGVSQLEEDLQSLLILFLGVTVFEVKFA
>Miniopterus_natalensis
---------------------------MLGAQWRRNQPPRAAESCLLVLSLKKILQSLLILFLGVTIFEAQLT
>Pterocles_gutturalis
--------------------------------MTK--TACFSTELSGVSQLEEDLQSLLILFLGVTIFEVKFA
>Colius_striatus
--------------------------------MTK--TACFSTELSGVSQLEEDLQSLLILFLGVTVVEVKFA
>Columba_livia
--------------------------------MTK--TACFSTELSGVSQLEEDLQSLLILFLGVTIFEVKFA
>Merops_nubicus
--------------------------------MTK--PACFSTELSGVSQLEEDLQSLLILFLGVTIFEVKFA
>Cuculus_canorus
--------------------------------MTK--TACFSTELSGVSQLEEDLQSLLILFLGVTILK----
>Monodelphis_domestica
---------------------------MLGTQW----------------------------------------
>Gallus_gallus
MLAVQGRPPRCRFALLFFTPLPEHXIAPRGPGARS--HRRXRAELSGVSQLEEDLQSLLILFLGVTIFEVKFA
>Gavia_stellata
--------------------------------MTK--TACFSTELSGVSQLEEDLQSLLILFLGVTIFEVKFA
>Chlamydotis_macqueenii
--------------------------------MTK--TACFSTELSGVSQLEEDLQSLLILFLGVTIFEVKFA
>Condylura_cristata
---------------------------MLGAQWRRNQPPRAAESCLLVLSLKKILQSLLILFLGVTIF-----
>Oryctolagus_cuniculus
---------------------------MLGAQWRRRQSPRAAESCLLVLSLKKILQSLLILFLGVTIF-----
>Tauraco_erythrolophus
--------------------------------MTK--TACFSTELSGVSQLEEDLQSLLILFLGVTVLK----
>Opisthocomus_hoazin
--------------------------------MTK--TACFSTELSGVSQLEEDLQSLLILFLGVTVF-----
>Acanthisitta_chloris
--------------------------------MTK--TACFSTELSGVSQLEEDLQSLLILFLGVTVFEVKFA
>Egretta_garzetta
--------------------------------MTK--TACFSTELSGVSQLEEDLQSLLILFLGVTIFEVKFA
>Ceratotherium_simum_simum
---------------------------MLGAQWRRKQPLRAAESCLLFLSLKKILQSLLILFLGVTIF-----
>Picoides_pubescens
--------------------------------MTK--TACFSTELSGVSQLDEDLQSLLILFLGVTIFEVKFA
>Loxodonta_africana
---------------------------MLGAQGRRSQPPRAAESCLLVLSLKKILQSLLILFLGVTIF-----
>Fulmarus_glacialis
--------------------------------MTK--TACFSTELSGVSQLEEDLRSLLILFLGVTIFEVKFA
>Melopsittacus_undulatus
--------------------------------MTK--PACFSTELSGVSQLEEDLQSLLILFLGVTIFEVKFA
>Jaculus_jaculus
-----------------MRA-SRSAAAMLGAQWRRRQPRRAAESCLLVLSLKKILQSLLILFLGVTIF-----
>Trichechus_manatus_latirostr
---------------------------MLGAQWRRSQPPRAAESCLLVLSLKKILQSLLILFLGVTIF-----
>Aptenodytes_forsteri
--------------------------------MTK--TACFSTELSGVSQLEEDLQSLLILFLGVTIFEVKFA
>Tyto_alba
--------------------------------MTK--TACFSTELSGVSQLEEDLQSLLILFLGVTIFEVKFA
>Struthio_camelus_australis
--------------------------------MTK--TACFSTELSGVSQLEEDLQSLLILFLGVTIFEVKFA
>Tinamus_guttatus
--------------------------------MTK--TACFSTELSGVSQLEEDLQSLLILFLGVTIFEVKFA
```

HG0058 ENSG00000180332 KCTD4

>Homo_sapiens
--------------------------------MLPAFPFHHQAGAE------
>Anser_cygnoides
--------------------------------MLAAFPFQHRAGAEERCF--
>Xenopus_laevis
MQIVFFPAVLVISCCLHSSYATYCKSELRAHAMLAAFPFHHRGGAG------
>Apteryx_australis_mantelli
--------------------------------MPAAFPFQHRAGAE------
>Caprimulgus_carolinensis
--------------------------------MRAAFPFQHRAGAE------
>Cariama_cristata
--------------------------------MLAAFPFQHRAGAE------
>Ailuropoda_melanoleuca
--------------------------------MLPAFPFHHQAGAE------
>Charadrius_vociferus
--------------------------------MLAAFPFQHRAGAE------
>Miniopterus_natalensis
--------------------------------MLPAFPFHHQAGAE------
>Colius_striatus
--------------------------------MIAAFPFQHRAGAE------
>Columba_livia
--------------------------------MLAAFPFQHRAGAE------
>Leptosomus_discolor
--------------------------------MLAAFPFQHRAGAE------
>Galeopterus_variegatus
--------------------------------MLPAFPFHHQAGAE------
>Haliaeetus_albicilla
--------------------------------MLAAFPFQHRAGAE------
>Gavia_stellata
--------------------------------MLAAFPFQHRAGAE------
>Chlamydotis_macqueenii
--------------------------------MLAAFPFQHRAGAE------
>Erinaceus_europaeus
--------------------------------MLPAFPFHHQAGAE------
>Ochotona_princeps
--------------------------------MLPAFPFPRQAGTE------
>Tauraco_erythrolophus
--------------------------------MLAAFPF-------------
>Opisthocomus_hoazin
--------------------------------MLAAFPFQHRAGAE------
>Manacus_vitellinus
--------------------------------MLAAFPFQHRSGAE------
>Egretta_garzetta
--------------------------------MLAAFPFQHRAGAE------
>Ceratotherium_simum_simum
--------------------------------MLPAFPFHHQAGAE------
>Manis_javanica
--------------------------------MLPAFSFHHQAGLNKGVFVW
>Fulmarus_glacialis
--------------------------------MLAAFPFQHRAGAE------
>Nestor_notabilis
--------------------------------MLAAFPFQHRAGAE------
>Ictidomys_tridecemlineatus
--------------------------------MLPAFPFHRQAGTE------
>Tupaia_chinensis
--------------------------------MLPAFPFHHQAGAE------
>Trichechus_manatus_latirostr
--------------------------------MLPAFPFHHQAGTE------
>Pygoscelis_adeliae
--------------------------------MLAAFPFQHRAGAE------
>Tyto_alba
--------------------------------MLAAFPFQHRAGAE------
>Trachemys_scripta
--------------------------------MLAALPFHHRAGAE------
>Calypte_anna
--------------------------------MLSAFPFQHRAGAE------
>Apaloderma_vittatum
--------------------------------MLAAFPFQHRAGAE------
>Orycteropus_afer_afer
--------------------------------MLPVFPFHHQQAGAE------

HG0059.1 ENSDARG00000018060 pik3r2

>Danio_rerio
----------MLHSLCSIIWLNRGFKLRKVACTANTGQV---------------------------
>Acipenser_sinensis
--------MLVKLSLCSIVWLNRGFKRRGVLLTMGLGHH---------------------------
>Amia_calva
--------MLPTSSLCSIVWLNRGFKLRKVLSS-NTGQN---------------------------
>Channa_punctata
--------MLPTLSLCSIIWLNRGIKLRKVHRSTNTGQV---------------------------
>Anas_platyrhynchos
MPSRERWIMVVSFSLCSIIWLNRGFKLRKRGCT---------------------------------
>Xenopus_tropicalis
MIPEKSWIMFVSLFLCSIIWLNRGFKLRITGT----------------------------------
>Menidia_menidia
--------MLPTLSLCSIIWLNRGIKLRKVHRFTNTDQV---------------------------
>Tripterygion_delaisi
--------MLPTRSLCSIIWLNRGIKLRKVHQSANTGQV---------------------------
>Cariama_cristata
MPAHQCWIMFVSFSLCSIIWLNRGFKLRKRGCT---------------------------------
>Hynobius_chinensis
--------MIVAPSLCSIIWLNRGFKLRKGGT----------------------------------
>Micropterus_salmoides
--------MLPTLSLCSIIWLNRGIKLRKVHQFTNTGQV---------------------------
>Calidris_pugnax
MPARQCWIMFVSFSLCSIIWLNRGFKLRKRGCT---------------------------------
>Haplochromis_burtoni
--------MLPTISLCSIIWLNRGIKLRKVHQSTNTGQV---------------------------
>Clupea_harengus
--------MFPFLSLCSIIWLNRGFKLRKVLCVANTGQI---------------------------
>Latimeria_menadoensis
MVVVKCWSML-ALSLCSIIWLNRGLKLRKGVPLQDCALLPMWTEELDITSASKIELPVIWGRGPPI
>Columba_livia
MPAHKCWIMFVSFSLCSIIWLNRGFKLRKRGCT---------------------------------
>Poeciliopsis_prolifica
--------MLPTLSLCSIIWLNRGIKLRKVQQSSNTGQL---------------------------
>Monodelphis_domestica
---------MVRRALCSIVWLSRGFKLRKSG-----------------------------------
>Esox_lucius
--------MLPTLSLCSIVWLNRGIKLRKVLHCANTGQL---------------------------
>Aquila_chrysaetos_canadensis
MPAHKCWIMFVSFSLCSIIWLNRGFKLRKRGCT---------------------------------
>Coturnix_japonica
MPTRECWIMFVSFSLCSIIWLNRGFKLRKRGCT---------------------------------
>Lutjanus_erythropterus
--------MLPTLSLCSIIWLNRGIKLRKVHQATNTGQV---------------------------
>Scleropages_formosus
--------MLPTLSLCSIVWLNRGIKLRKVLYSANTGQF---------------------------
>Lepidothrix_coronata
MPAHQCWIMFVSFSLCSIIWLNRGFKLRKRGCT---------------------------------
>Lateolabrax_japonicus
--------MLPTLSLCSIIWLNRGIKLRKVHQSTNTGQV---------------------------
>Sebastes_caurinus
--------MLPTLSLCSIIWLNRGIKLRKVHQPTNTGQV---------------------------
>Cynoglossus_semilaevis
--------MLPSLSLCSIIWLNRGIKLRKLHLSTNTGQI---------------------------
>Salmo_salar
--------MLPTLSLCSIIWLNRGIKLRKVLHCANTGQL---------------------------
>Lepisosteus_oculatus
--------MLPTLSLCSIVWLNRGFKLRKALLS-NTGQH---------------------------
>Aptenodytes_forsteri
MPAHQCWIMFVSFSLCSIIWLNRGFKLRKRGCT---------------------------------
>Gekko_japonicus
--------MLLTFSLCSIIWLNRGFKLRERGCT---------------------------------
>Struthio_camelus_australis
MPARNCWIMFVSFSLCSIIWLNRGFKLRKRGCT---------------------------------
>Chrysemys_picta
MPSHECWIMFVAFSLCSIIWLNRGFKLRKRGCTSACGSLSVCGAILTRLR--STWLVQRW------
>Takifugu_rubripes
--------MLPTLSLCSIIWLNRGFKLRKVHQSSNTGQV---------------------------

HG0059.1 ENSGALG00000003428

```
>Gallus_gallus
MPTHECWIMFVSFSLCSIIWLNRGFKLRKR-GCT--------------------------------
>Acipenser_sinensis
--------MLVKLSLCSIVWLNRGFKRRGVLLTMGLGHH---------------------------
>Amia_calva
--------MLPTSSLCSIVWLNRGFKLRKVLSSNTG-QN---------------------------
>Channa_argus
--------MLPTLSLCSIIWLNRGIKLRKMYRSTNTGQV---------------------------
>Anas_platyrhynchos
MPSRERWIMVVSFSLCSIIWLNRGFKLRKR-GCT--------------------------------
>Megophrys_sangzhiensis
MVPVKSWIMFVSLFLCSIIWLNRGFKLRIR-GT---------------------------------
>Menidia_menidia
--------MLPTLSLCSIIWLNRGIKLRKVHRFTNTDQV---------------------------
>Ambystoma_mexicanum
MCGGKCWIMFVALSLCSIIWLNRGFKLRKG-GT---------------------------------
>Micropterus_salmoides
--------MLPTLSLCSIIWLNRGIKLRKVHQFTNTGQV---------------------------
>Haplochromis_burtoni
--------MLPTISLCSIIWLNRGIKLRKVHQSTNTGQV---------------------------
>Clupea_harengus
--------MFPFLSLCSIIWLNRGFKLRKVLCVANTGQI---------------------------
>Latimeria_chalumnae
MVVVKCWSML-ALSLCSIIWLNRGLKLRKGVPLQDCALLSMWTEELDITSASKIELPVIWGRGPPI
>Danio_rerio
----------MLHSLCSIIWLNRGFKLRKVACTANTGQV---------------------------
>Poecilia_mexicana
--------MLPMLSLCSIIWLNRGIKLRKVQQSTTLVSSEAAATTV-IT-AATL------------
>Monodelphis_domestica
---------MVRRALCSIVWLSRGFKLRKS-G----------------------------------
>Esox_lucius
--------MLPTLSLCSIVWLNRGIKLRKVLHCANTGQL---------------------------
>Falco_cherrug
MPTHKCWIMFVSFSLCSIIWLNRGFKLRKR-GCT--------------------------------
>Lutjanus_erythropterus
--------MLPTLSLCSIIWLNRGIKLRKVHQATNTGQV---------------------------
>Scleropages_formosus
--------MLPTLSLCSIVWLNRGIKLRKVLYSANTGQF---------------------------
>Lateolabrax_japonicus
--------MLPTLSLCSIIWLNRGIKLRKVHQSTNTGQV---------------------------
>Sebastes_caurinus
--------MLPTLSLCSIIWLNRGIKLRKVHQPTNTGQV---------------------------
>Platichthys_stellatus
--------MLPTISLCSIIWLNRGIKLRKSHQSTNTGQV---------------------------
>Oncorhynchus_mykiss
--------MLPTLSLCSIIWLNRGIKLRKVLHSANTGLL---------------------------
>Lepisosteus_oculatus
--------MLPTLSLCSIVWLNRGFKLRKALLSNTG-QH---------------------------
>Crotalus_adamanteus
--------MLVTFCLCFIIWLNRGFKLRERRGCT--------------------------------
>Takifugu_rubripes
--------MLPTLSLCSIIWLNRGFKLRKVHQSSNTGQV---------------------------
```

HG0059.2 ENSG00000145675 PIK3R1

>Homo_sapiens
--------------------------MDSVTKVSERIGPRCESQPGFKVLTSC
>Canis_lupus_familiaris
MDLQGGNHWEFGEARVRSDTVQIRRQMDSVTRVSERIGPRCENQPGGQVLTSC
>Balaenoptera_acutorostrata_s
--------------------------MDSVTRASERIGPRRESQPRVNVLTSC
>Myotis_brandtii
--------------------------MDSVTRVSERIGPRCESQPGVQVLTSC
>Dasypus_novemcinctus
--------------------------MDSVTRASERIGPRCESQPGGQVLTSR
>Erinaceus_europaeus
--------------------------MDSVTGVSERIGPRCENQPGVKMLTSC
>Oryctolagus_cuniculus
--------------------------MDSVARASERIGPGRERQPRVWVLTSC
>Equus_asinus
--------------------------MDSVTGVSERIGPRCESQPGVKVLASC
>Manis_javanica
--------------------------MDSVTRVSERIGPRCESQSEVQVLTSC
>Loxodonta_africana
--------------------------MDSVTRASERIGPRGESQPGVKC----
>Mus_musculus
--------------------------MDSVTGTSERIGPRCESQPGVKVLTSC
>Tupaia_chinensis
--------------------------MDSVTRASERIGPRCESRPGVRVLTSC

HG0059.3 ENSG00000105647 PIK3R2

```
>Homo_sapiens
MVAACTLGSLVWCSQQSRQPWAPMALRVRA-------
>Felis_catus
MVTACALGPLVWRSRKSRQPRAPMAPRVRE-------
>Balaenoptera_acutorostrata_s
MVTACALGPLAWRSWKSRQPREPMAPHVRE-------
>Rousettus_aegyptiacus
MVTACALGSLVWRSRKSRQPWAPMAPRVRE-------
>Condylura_cristata
MVTSSALGPLVWRSWKSRQSRAPMAPHVKE-------
>Equus_asinus
MVTACVLGPLIWRSWKRRQPWEPMAAHVRE-------
>Manis_javanica
MVTACALGPLVWRSWKSRQPRAPMAHHVRE-------
>Chinchilla_lanigera
MVAALTLCSLVWRSWKSWQPRVPMALYHRRMANPVAK
```

HG0059.4 ENSG00000145675 PIK3R1

>Homo_sapiens
MDSRMASHLSS
>Dasypus_novemcinctus
MDSRMASHLSS
>Equus_asinus
MDSRMANHLSS
>Loxodonta_africana
MDSRMASHLSS
>Nannospalax_galili
MDSRMASHLSS
>Tupaia_chinensis
MDSRMASHLSS

HG0059.5 ENSDARG00000038524 pik3r1

```
>Danio_rerio
-----------------------------------------------------MGRVCMKHLEGAPNHWTISFT
>Astyanax_mexicanus
-----------------------------------------------------MGKLCVKHREGALDHWRISFT
>Alosa_alosa
-----------------------------------------------------MGSLCLKHHAGAMEHWPINMT
>Apteronotus_leptorhynchus
MLNCIFKLTPLVSCDSYRKERFSRADFDGFLFNLAIYQSRGCFGQSTGADFSQMGTLCLKHREGAQEHWRIRLT
>Ictalurus_punctatus
-----------------------------------------------------METLCLKHHEAPLDHWRISFT
```

HG0060.1 ENSG00000049192 ADAMTS6

>Homo_sapiens
--------------------------------MNPDIEESAGCWIKVALKQWNIQ
>Anser_cygnoides
--------------------------------MTADLEESTGCWKRIPVKKWNVQ
>Chaetura_pelagica
--------------------------------MTADVEEGTGCWKRIPVKKWNI-
>Buceros_rhinoceros_silvestri
--------------------------------MTADIEESTGCWKRILVKKWNNQ
>Caprimulgus_carolinensis
--------------------------------MTADIEESTGCWKRIPVKKWNIQ
>Cariama_cristata
--------------------------------MTTDIEESTGYWKRIPVKKWNIQ
>Odobenus_rosmarus_divergens
-----------------MKILHHSFFPFGLKKMNPDIEESTGCWKKVPVKQWNI-
>Lipotes_vexillifer
-----------------MKILPSLSFSFWTEKMNPDTEESTGCWKKIPVKQWNI-
>Charadrius_vociferus
--------------------------------MTADIEESTGCWKRIPVKMWNIQ
>Pteropus_alecto
MLKNGHTLVTNQPNALLMKILPSLIFSFWIEKMNPDIEESTGCWKKVPVKQWTD-
>Dasypus_novemcinctus
----------------------------MKTKMNQDTEESMGCWTKVPVKQRNI-
>Columba_livia
--------------------------------MTADIEESTGCWKRIPVKRWNIQ
>Leptosomus_discolor
--------------------------------MTADIEEGTGCWKRIPVEKWNVQ
>Sarcophilus_harrisii
--------------------------------MNPERGESTGCWKRAPAKKWNI-
>Galeopterus_variegatus
--------------------------------MNPDLEESMGCWKKVLVKQWNT-
>Monodelphis_domestica
--------------------------------MNPERGESTGCWKRAPAKKWNI-
>Aquila_chrysaetos_canadensis
--------------------------------MTADIEESTGCWKRIPVKKWNIQ
>Gallus_gallus
--------------------------------MTADIEE--SCWKRVPVKKWNIQ
>Eurypyga_helias
--------------------------------MTADLEENTGCWKRIPVKKWNVQ
>Sorex_araneus
--------------------------------MNPVIEESMGCWKKVSIIQWNI-
>Oryctolagus_cuniculus
--------------------------------MNPDIEESTGCWKKLLVKQWNI-
>Tauraco_erythrolophus
--------------------------------MTADIEESTGCWKRIPVKKWNIQ
>Opisthocomus_hoazin
--------------------------------MTADIEEGTGFWKRIPVKKWNIQ
>Nipponia_nippon
--------------------------------MTADIEESTGCWKRIPFKKWNIQ
>Ceratotherium_simum_simum
-----------------MKILPSLFFSFWIEKMNPDIEESMGCWKKVPVKQ----
>Manis_javanica
--------------------------------MNPHLEESMGCWKKVPAKQWNI-
>Loxodonta_africana
--------------------------------MNPDTEESMGCWEKVPVKQWNIY
>Fulmarus_glacialis
--------------------------------MTADIEESTGCWKRIPVKKWNIQ
>Ictidomys_tridecemlineatus
--------------------------------MNPDIEESAGCWKKEPVKQWNI-
>Tupaia_chinensis
--------------------------------MNPDMDDSMGFWKKVPVKQWKM-
>Pygoscelis_adeliae
--------------------------------MTADIEESTGCWKRIPVKKWNLQ
>Struthio_camelus_australis
--------------------------------MTADIEESTGCRKRIPVKKWNIQ
>Chrysemys_picta
--------------------------------MTADIEESTGCWKRVPVKKWNIQ
>Orycteropus_afer_afer
--------------------------------MNPDTEESTEYWEKIPVKQWNI-

HG0060.1 ENSGALG00000014751

>Gallus_gallus
--------------------------------MTADIEE--SCWKRVPVKKWNIQ
>Anser_cygnoides
--------------------------------MTADLEESTGCWKRIPVKKWNVQ
>Chaetura_pelagica
--------------------------------MTADVEEGTGCWKRIPVKKWNI-
>Buceros_rhinoceros_silvestri
--------------------------------MTADIEESTGCWKRILVKKWNNQ
>Caprimulgus_carolinensis
--------------------------------MTADIEESTGCWKRIPVKKWNIQ
>Cariama_cristata
--------------------------------MTTDIEESTGYWKRIPVKKWNIQ
>Odobenus_rosmarus_divergens
-----------------MKILHHSFFPFGLKKMNPDIEESTGCWKKVPVKQWNI-
>Lipotes_vexillifer
-----------------MKILPSLSFSFWTEKMNPDTEESTGCWKKIPVKQWNI-
>Charadrius_vociferus
--------------------------------MTADIEESTGCWKRIPVKMWNIQ
>Pteropus_alecto
MLKNGHTLVTNQPNALLMKILPSLIFSFWIEKMNPDIEESTGCWKKVPVKQWTD-
>Dasypus_novemcinctus
----------------------------MKTKMNQDTEESMGCWTKVPVKQRNI-
>Columba_livia
--------------------------------MTADIEESTGCWKRIPVKRWNIQ
>Leptosomus_discolor
--------------------------------MTADIEEGTGCWKRIPVEKWNVQ
>Sarcophilus_harrisii
--------------------------------MNPERGESTGCWKRAPAKKWNI-
>Galeopterus_variegatus
--------------------------------MNPDLEESMGCWKKVLVKQWNT-
>Monodelphis_domestica
--------------------------------MNPERGESTGCWKRAPAKKWNI-
>Aquila_chrysaetos_canadensis
--------------------------------MTADIEESTGCWKRIPVKKWNIQ
>Eurypyga_helias
--------------------------------MTADLEENTGCWKRIPVKKWNVQ
>Oryctolagus_cuniculus
--------------------------------MNPDIEESTGCWKKLLVKQWNI-
>Tauraco_erythrolophus
--------------------------------MTADIEESTGCWKRIPVKKWNIQ
>Opisthocomus_hoazin
--------------------------------MTADIEEGTGFWKRIPVKKWNIQ
>Nipponia_nippon
--------------------------------MTADIEESTGCWKRIPFKKWNIQ
>Equus_asinus
-----------------MKILPSLFFSFWIEKMNPDIEESMRCWKKVPVKQWNI-
>Manis_javanica
--------------------------------MNPHLEESMGCWKKVPAKQWNI-
>Cercocebus_atys
-----------------MKTFTIIIFSFWIEKMNPDIEESAGCWIKVPLKQWNIQ
>Loxodonta_africana
--------------------------------MNPDTEESMGCWEKVPVKQWNIY
>Fulmarus_glacialis
--------------------------------MTADIEESTGCWKRIPVKKWNIQ
>Ictidomys_tridecemlineatus
--------------------------------MNPDIEESAGCWKKEPVKQWNI-
>Pygoscelis_adeliae
--------------------------------MTADIEESTGCWKRIPVKKWNLQ
>Struthio_camelus_australis
--------------------------------MTADIEESTGCRKRIPVKKWNIQ
>Chrysemys_picta
--------------------------------MTADIEESTGCWKRVPVKKWNIQ
>Orycteropus_afer_afer
--------------------------------MNPDTEESTEYWEKIPVKQWNI-

HG0060.2 ENSG00000049192 ADAMTS6

>Homo_sapiens
MLRFWYGIDSLAHR
>Anas_platyrhynchos
MVRFWSGVSSIAHR
>Chaetura_pelagica
MVRFWSGVNSVSHR
>Canis_lupus_familiaris
MLRFWYGIDSLAHR
>Physeter_catodon
MLRFWHGIDSLAHR
>Charadrius_vociferus
MVQFWSGVNSVAHR
>Dasypus_novemcinctus
MLRFWYGVDSLAHR
>Columba_livia
MVRYWSGANSIAHR
>Leptosomus_discolor
MVRFWSGVNFIAHR
>Galeopterus_variegatus
MLRLWYDIDSLTHR
>Aquila_chrysaetos_canadensis
MVRFWSGVNSVAHR
>Gallus_gallus
MVQFWSGVNPIAHR
>Eurypyga_helias
MVRFWSGVNSVAHR
>Sorex_araneus
MLRFWYDIDSLAHR
>Oryctolagus_cuniculus
MLRFWYGIDSLAHR
>Tauraco_erythrolophus
MVPFWSGVNSIAHR
>Opisthocomus_hoazin
MVRFWSGVNSIAHR
>Acanthisitta_chloris
MVRFWSGVNSIAHR
>Phaethon_lepturus
MVRFWSGVNSLAHR
>Ceratotherium_simum_simum
MLRFWYGIDSLAHR
>Manis_javanica
MLRFWYGIDSLAHR
>Loxodonta_africana
MLRFWYGIDSLAHR
>Fulmarus_glacialis
MVRFWSGVNSTAHR
>Nannospalax_galili
MLRFWYGIDSLAHR
>Tupaia_chinensis
MLRFGYDSDSLAHR
>Pygoscelis_adeliae
MVRFWSGVNSVAHR
>Thamnophis_sirtalis
MVQFWSGVKAIAHR
>Struthio_camelus_australis
MVRFWSGVKSIAHR
>Chrysemys_picta
MVRFWSGVNSIAHR
>Orycteropus_afer_afer
MLRFWHGVDSLAHR

HG0060.2 ENSGALG00000014751

>Gallus_gallus
MVQFWSGVNPIAHR
>Anser_cygnoides
MVRFWSGVSSIAHR
>Chaetura_pelagica
MVRFWSGVNSVSHR
>Buceros_rhinoceros_silvestri
MVLFWSGVNFIAHR
>Caprimulgus_carolinensis
MVRFWSGVNSIARR
>Cariama_cristata
MVRFWSGVNSVAHK
>Charadrius_vociferus
MVQFWSGVNSVAHR
>Dasypus_novemcinctus
MLRFWYGVDSLAHR
>Columba_livia
MVRYWSGANSIAHR
>Leptosomus_discolor
MVRFWSGVNFIAHR
>Aquila_chrysaetos_canadensis
MVRFWSGVNSVAHR
>Eurypyga_helias
MVRFWSGVNSVAHR
>Tauraco_erythrolophus
MVPFWSGVNSIAHR
>Opisthocomus_hoazin
MVRFWSGVNSIAHR
>Manacus_vitellinus
MVQFWSGVNSIAHK
>Nipponia_nippon
MVRFWSGVNSIAHR
>Microcebus_murinus
MLRFWYGIDSIAHR
>Loxodonta_africana
MLRFWYGIDSLAHR
>Fulmarus_glacialis
MVRFWSGVNSTAHR
>Chinchilla_lanigera
MLRFWYGINSLAHR
>Pygoscelis_adeliae
MVRFWSGVNSVAHR
>Thamnophis_sirtalis
MVQFWSGVKAIAHR
>Struthio_camelus_australis
MVRFWSGVKSIAHR
>Chrysemys_picta
MVRFWSGVNSIAHR
>Orycteropus_afer_afer
MLRFWHGVDSLAHR

HG0061.1 ENSG00000166398

```
>Homo_sapiens
----MVTCLMEMPFLKLRFAELHYFYT
>Anas_platyrhynchos
-----------MPFVKPMYTVLHCYCT
>Xenopus_laevis
-----------MPFTKPVFADFIYYLT
>Caprimulgus_carolinensis
-----------MPFVKLMYTVLHCYCT
>Felis_catus
----MVTCLMEMPFLKLRFAELHYFYT
>Hynobius_retardatus
-----------MPFTKPSYPVPHYYT-
>Physeter_catodon
----MVTCLMEMPFLKLRFAELHYFYT
>Cynopterus_sphinx
----MVTCLMEMPFLKLIFAELHYFYT
>Columba_livia
-----------MPFVKPMYAVLHCYCT
>Alligator_mississippiensis
-----------MPFMKPAYAVLHYYYT
>Sarcophilus_harrisii
-----------MPFMELLFAVLYCYYT
>Galeopterus_variegatus
----MATCLMEMPFLKLGLAELHYFYT
>Monodelphis_domestica
-----------MPFMELLFAVLHYYYT
>Condylura_cristata
----MVTCLMKMPFLKLRFAELHYFYT
>Oryctolagus_cuniculus
----MVICLLEMSFLKLTFAEVHYFYT
>Phaethon_lepturus
-----------MPFVKPMATVLHCYYT
>Ceratotherium_simum_simum
----MVICLMEMPFLKLRFAELHYFYT
>Manis_javanica
----MVTFLMEMPFLKLRFAELHYFYT
>Melopsittacus_undulatus
-----------MPFVKPTYTVLHFYCT
>Ctenomys_sociabilis
MFDHMVTCLMEMPFLKLQFAELHYFYT
>Tupaia_chinensis
---------MEMPFLKHRFAELQYFYT
>Tyto_alba
-----------MPFVKLMYMVLHCYCT
>Struthio_camelus_australis
-----------MPFVKPMYAVLHCYCT
>Pelodiscus_sinensis
-----------MPFLKRMYTVLHCYYT
>Calypte_anna
-----------MPFVKPMYTVLHCYCT
```

HG0061.1 ENSGALG00000004949

>Gallus_gallus
MPFVKPMYTVLHCYCT----
>Anser_cygnoides
MPFVKPMYTVLHCYCT----
>Leptobrachium_boringii
MPFTKPMYVALLCYLT----
>Chaetura_pelagica
MPFVKPMYTVLHCYCT----
>Apteryx_australis_mantelli
MPFVKPMYTVLHCYCT----
>Buceros_rhinoceros_silvestri
MPFVEPMYTVLHCYCT----
>Caprimulgus_carolinensis
MPFVKLMYTVLHCYCT----
>Cariama_cristata
MPFVKPMYTVLHCYCT----
>Charadrius_vociferus
MPFVKPMYTVLHCYCTKLYI
>Pterocles_gutturalis
MPFVKPMYTVLHCYCT----
>Colius_striatus
MPFVKPMYTVHHCYCT----
>Columba_livia
MPFVKPMYAVLHCYCT----
>Merops_nubicus
MPFVKPMYTVLHCYCT----
>Alligator_sinensis
MPFMKPAYAVLHYYYT----
>Cuculus_canorus
MPFVKPMYTVLHCYCT----
>Falco_cherrug
MPFVKPMYTVLHCYCT----
>Gavia_stellata
MPFVKPMYTVLHCYCT----
>Balearica_regulorum_gibberic
MPFVKPMYTVLHCYCT----
>Ornithorhynchus_anatinus
MPFVKPGFTVLHRYHT----
>Tauraco_erythrolophus
MPFVKPMYTVLHCYCT----
>Opisthocomus_hoazin
MPFVKRMCAVLDCYCTELYI
>Corvus_brachyrhynchos
MPFVKPMYTVLHCYCT----
>Egretta_garzetta
MPFVKPMYTVLHCYCT----
>Picoides_pubescens
MPFVTPMYTAPHRRCT----
>Fulmarus_glacialis
MPFVKPMYTVLHCYCT----
>Nestor_notabilis
MPFVKPVYTVLHCYCT----
>Aptenodytes_forsteri
MPFVKPMYTVLHCYCT----
>Tyto_alba
MPFVKLMYMVLHCYCT----
>Struthio_camelus_australis
MPFVKPMYAVLHCYCT----
>Pelodiscus_sinensis
MPFLKRMYTVLHCYYT----
>Tinamus_guttatus
MPFVKPMYTVLHCYCT----
>Calypte_anna
MPFVKPMYTVLHCYCT----
>Apaloderma_vittatum
MPFVKPMYTVLHCYCT----

HG0061.2 ENSGALG00000004949

>Gallus_gallus
--------------------------------------------------------MLQRKQLPQQTRKGEEGLQ----------------------------------
>Anas_platyrhynchos
--------------------------------------------------------MLQRKQLPQQTRKGEEGLQ----------------------------------
>Calidris_pugnax
--------------------------------------------------------MLQRKQLPQQRRKGGKGKKACNNK-HSNF------------------------
>Columba_livia
--------------------------------------------------------MLQRKQLPQQTRKGEEGLQ----------------------------------
>Sarcophilus_harrisii
--------------------------------------------------------MLQRKQLPQKTKGKEEAAEQLEITNTPNF------------------------
>Monodelphis_domestica
--------------------------------------------------------MLQRKQLPQKTEGKEEAPWQLEITNTPVFNNLTCEKRRHFIFFSCSIKKKHYR
>Aquila_chrysaetos_canadensis
--------------------------------------------------------MLQRKQLPQQTRKGEEGLQ----------------------------------
>Corvus_brachyrhynchos
--------------------------------------------------------MLQRKQLPQQTGKGEEGLQ----------------------------------
>Aptenodytes_forsteri
--------------------------------------------------------MLQRKQLPQQTRKGEEGLQ----------------------------------
>Struthio_camelus_australis
MNLLLLKYSKPVNSLESLLTEHRSCGNDLEALQIHLCIFVPSLRLRANAIWKHHFLTLQRKQLPQQTRKGEEGLQ----------------------------------

HG0061.3 ENSG00000166398

```
>Homo_sapiens
MLQGNRYLSKQKEWRRRLTTNA-I------------------------------
>Pteropus_alecto
MLQGNRYLSKQKERRRRLITNA-IF-----------------------------
>Galeopterus_variegatus
MLQGNCYLSKQKERRRRLIITNGIFKKTC--FRKTLEE-I--------------
>Condylura_cristata
MLQGNRYLSKQKERRRLIIYNMPFF-----------------------------
>Oryctolagus_cuniculus
MLQGNHYLSKQKERSRRLKITNDIFTKKQTCFQKTLEEIWTFLASGRTFVSKWI
>Ceratotherium_simum_simum
MLQGSHYLNKQKERRRRLIITNAIFKKKLG-FRKY-------------------
>Nannospalax_galili
MLQGNRYLSKEKERRRRRTVTDAM-FKENL--FSENPTGDL-------------
>Tupaia_chinensis
MLQGNRYLSKQKERRRRLTTTNAIFKKQTNSFPENIEE-I--------------
```

HG0062.1 ENSDARG00000075569 gpbp1l1

>Danio_rerio
MPPSTDGRHWT
>Pygocentrus_nattereri
MPRSTDGRHWT
>Clupea_harengus
MPRSTEGRHWT
>Poecilia_formosa
MPPSTDQKHWT
>Apteronotus_leptorhynchus
MPRSTDGRHWT
>Salmo_salar
MPRSTDGRHWT
>Trichomycterus_areolatus
MPRSTDGRHWT

HG0062.1 ENSGALG00000010276 GPBP1L1

>Gallus_gallus
MPHFIEGRHWT
>Anser_cygnoides
MPHFIEGRHWT
>Chaetura_pelagica
MPHFIEGRHWT
>Apteryx_australis_mantelli
MPHFIEGRHWT
>Buceros_rhinoceros_silvestri
MPHFIEGRHWT
>Caprimulgus_carolinensis
MPHFVEGRHWT
>Cariama_cristata
MPHFIEGRHWT
>Physeter_catodon
MPHLVEKRHWT
>Charadrius_vociferus
MPHFIEGRHWT
>Pterocles_gutturalis
MPHFIEGRHWT
>Colius_striatus
MPHFIEGRHWT
>Columba_livia
MPHFIEGRHWT
>Leptosomus_discolor
MPHFIEGRHWT
>Alligator_mississippiensis
MPHFIEGRHWT
>Cuculus_canorus
MPHFIEGRHWT
>Falco_cherrug
MPHFIEGRHWT
>Eurypyga_helias
MPHFIEGRHWT
>Tauraco_erythrolophus
MPHFIEGRHWT
>Opisthocomus_hoazin
MPHFVEGRHWT
>Geospiza_fortis
MPHFIEGRHWT
>Egretta_garzetta
MPHFIEGRHWT
>Picoides_pubescens
MPHFIEGRHWT
>Fulmarus_glacialis
MPHFIEGRHWT
>Melopsittacus_undulatus
MPHFIEGRHWT
>Oncorhynchus_mykiss
MPHSTDGRHWT
>Aptenodytes_forsteri
MPHFIEGRHWT
>Gekko_japonicus
MPRFIEGRHWT
>Tyto_alba
MPHFIEGRHWT
>Struthio_camelus_australis
MPHFIEGRHWT
>Pelodiscus_sinensis
MPHFIEGRHWT
>Tinamus_guttatus
MPHFIERRHWT
>Calypte_anna
MPHFVEGRHWT
>Apaloderma_vittatum
MPHFIEGRHWT

HG0062.2 ENSGALG00000010276 GPBP1L1

>Gallus_gallus
MSCPADLGVSVQY
>Anser_cygnoides
MNYPAELGVSVQY
>Caprimulgus_carolinensis
MNYPADLSVSVQY
>Charadrius_vociferus
MNYPADLSVSVQY
>Pterocles_gutturalis
MNYPADLSVSVQY
>Colius_striatus
MSYPADLGVSAQY
>Columba_livia
MNDPADLSVSVQY
>Cuculus_canorus
MNYPADLSVSVQY
>Haliaeetus_leucocephalus
MNYPADLSVSVQY
>Gavia_stellata
MNYPADLSVSVQY
>Chlamydotis_macqueenii
MNYPADLSVSVQY
>Opisthocomus_hoazin
MNYPADLSVSVQY
>Acanthisitta_chloris
MNCPAALSVSVQY
>Phaethon_lepturus
MNYPADLSVSVQY
>Fulmarus_glacialis
MNYPADLSVSVQY
>Melopsittacus_undulatus
MNYPADLGVSVQY
>Pygoscelis_adeliae
MNYPADLSVSVQY
>Tyto_alba
MNYPADLSVSVQY
>Struthio_camelus_australis
MNYPADLGASVQY
>Tinamus_guttatus
MNYPADLGASVQY
>Calypte_anna
MNYPADLSVSVQY

HG0062.3 ENSG00000159592 GPBP1L1

>Homo_sapiens
--MSYLIKQHHSAITQGS
>Leptonychotes_weddellii
MSMYYLFKQHHSAITQGS
>Balaenoptera_acutorostrata_s
MSMYYLFKQHHRAITQGS
>Rousettus_aegyptiacus
--MYYLFKQHHSAITQGS
>Dasypus_novemcinctus
MSMYYPFKQHHSTITQGS
>Galeopterus_variegatus
MSMYYLFKQHHSAITQGS
>Condylura_cristata
MSLNYLFKQHHSDITQGS
>Oryctolagus_cuniculus
MPMYYPFKQQHSAITQGS
>Ceratotherium_simum_simum
MSMYYLFKQHHSAITQGS
>Loxodonta_africana
MSMHYLFKQHHSAITQRS
>Marmota_marmota_marmota
MSMYYLFKQHHSAITQGS
>Trichechus_manatus_latirostr
MSMYYLFKQHHSTITQGS
>Orycteropus_afer_afer
MSMYYLFKQHHSAITQRS

HG0063.1 ENSG00000162630 B3GALT2

```
>Homo_sapiens
------------------MCACSIASFLLK-----------------------------------------
>Anas_platyrhynchos
------------------MCACSIASFLLK-----------------------------------------
>Chaetura_pelagica
------------------MCACSIASFLLK-----------------------------------------
>Buceros_rhinoceros_silvestri
------------------MCACSIASFLLK-----------------------------------------
>Caprimulgus_carolinensis
------------------MCACSRASFLLK-----------------------------------------
>Cariama_cristata
------------------MCACSIASFLLK-----------------------------------------
>Mustela_putorius_furo
------------------MCACSIASFLLK-----------------------------------------
>Charadrius_vociferus
------------------MCACSIASFLLK-----------------------------------------
>Pterocles_gutturalis
---MGYREQQPQEADGWGMCACSIASFLLK-----------------------------------------
>Dasypus_novemcinctus
------------------MCACSIASFWLK-----------------------------------------
>Colius_striatus
------------------MCACSIASFLLK-----------------------------------------
>Columba_livia
------------------MCACSIASFLLK-----------------------------------------
>Cuculus_canorus
------------------MCACSIASILLK-----------------------------------------
>Galeopterus_variegatus
------------------MCACSIASFLLK-----------------------------------------
>Monodelphis_domestica
------------------MCACSIASFLLK-----------------------------------------
>Macropus_eugenii
------------------MCACSIASFLLK-----------------------------------------
>Aquila_chrysaetos_canadensis
------------------MCACSIASFLLK-----------------------------------------
>Gallus_gallus
------------------MCACSIASFLLK-----------------------------------------
>Gavia_stellata
------------------MCACSIASFLLK-----------------------------------------
>Mesitornis_unicolor
------------------MCACSIASFLLK-----------------------------------------
>Condylura_cristata
------------------MCACSIASFLLK-----------------------------------------
>Ornithorhynchus_anatinus
MDGSDTGEEPSREAERWGMCACSMASF--------------------------------------------
>Tauraco_erythrolophus
------------------MCACSIASFLLK-----------------------------------------
>Opisthocomus_hoazin
------------------MCACSIASFLLK-----------------------------------------
>Phalacrocorax_carbo
MDRLDTVKKQSQEAGCWGMCACSIASFLLK-----------------------------------------
>Ceratotherium_simum_simum
------------------MCACSIASFLLK-----------------------------------------
>Manis_javanica
------------------MCACSIASLLLK-----------------------------------------
>Loxodonta_africana
------------------MCACSLTSFLLK-----------------------------------------
>Fulmarus_glacialis
------------------MCACSIASFLLK-----------------------------------------
>Nannospalax_galili
------------------MCACSIASFLLK-----------------------------------------
>Aptenodytes_forsteri
------------------MCACSIASFLLK-----------------------------------------
>Chelonia_mydas
------------------MCACSIASFLLK-----------------------------------------
>Apaloderma_vittatum
------------------MCACSIASFLPKRWRAKTIFWGYNPRTINGLTKEVSHLQTTTQHKQRKMNTKE
```

HG0063.1 ENSGALG00000002477

>Gallus_gallus
------------------MCACSIASFLLK-----------------------------------------
>Anas_platyrhynchos
------------------MCACSIASFLLK-----------------------------------------
>Chaetura_pelagica
------------------MCACSIASFLLK-----------------------------------------
>Buceros_rhinoceros_silvestri
------------------MCACSIASFLLK-----------------------------------------
>Caprimulgus_carolinensis
------------------MCACSRASFLLK-----------------------------------------
>Cariama_cristata
------------------MCACSIASFLLK-----------------------------------------
>Mustela_putorius_furo
------------------MCACSIASFLLK-----------------------------------------
>Charadrius_vociferus
------------------MCACSIASFLLK-----------------------------------------
>Pterocles_gutturalis
---MGYREQQPQEADGWGMCACSIASFLLK-----------------------------------------
>Dasypus_novemcinctus
------------------MCACSIASFWLK-----------------------------------------
>Colius_striatus
------------------MCACSIASFLLK-----------------------------------------
>Columba_livia
------------------MCACSIASFLLK-----------------------------------------
>Cuculus_canorus
------------------MCACSIASILLK-----------------------------------------
>Galeopterus_variegatus
------------------MCACSIASFLLK-----------------------------------------
>Monodelphis_domestica
------------------MCACSIASFLLK-----------------------------------------
>Macropus_eugenii
------------------MCACSIASFLLK-----------------------------------------
>Aquila_chrysaetos_canadensis
------------------MCACSIASFLLK-----------------------------------------
>Gavia_stellata
------------------MCACSIASFLLK-----------------------------------------
>Mesitornis_unicolor
------------------MCACSIASFLLK-----------------------------------------
>Condylura_cristata
------------------MCACSIASFLLK-----------------------------------------
>Ornithorhynchus_anatinus
MDGSDTGEEPSREAERWGMCACSMASF--------------------------------------------
>Tauraco_erythrolophus
------------------MCACSIASFLLK-----------------------------------------
>Opisthocomus_hoazin
------------------MCACSIASFLLK-----------------------------------------
>Phalacrocorax_carbo
MDRLDTVKKQSQEAGCWGMCACSIASFLLK-----------------------------------------
>Ceratotherium_simum_simum
------------------MCACSIASFLLK-----------------------------------------
>Manis_javanica
------------------MCACSIASLLLK-----------------------------------------
>Cercocebus_atys
------------------MCACSIASFLLK-----------------------------------------
>Loxodonta_africana
------------------MCACSLTSFLLK-----------------------------------------
>Fulmarus_glacialis
------------------MCACSIASFLLK-----------------------------------------
>Nannospalax_galili
------------------MCACSIASFLLK-----------------------------------------
>Aptenodytes_forsteri
------------------MCACSIASFLLK-----------------------------------------
>Chelonia_mydas
------------------MCACSIASFLLK-----------------------------------------
>Apaloderma_vittatum
------------------MCACSIASFLPKRWRAKTIFWGYNPRTINGLTKEVSHLQTTTQHKQRKMNTKE

HG0063.2 ENSG00000162630 B3GALT2

>Homo_sapiens
MD-TETRTQHCIVQPGK---
>Anser_cygnoides
MDTEEKRTQHCIVRQGKHQM
>Canis_lupus_familiaris
MD-TEKRIQHCIVRPGKQQM
>Balaenoptera_acutorostrata_s
MD-TEKRIQHCIVRPGKQQM
>Rousettus_aegyptiacus
MD-TEKRTQHCIVRPGKQQM
>Dasypus_novemcinctus
MD-TEKRTQHCIVQPGKQQM
>Galeopterus_variegatus
MD-TEKRTQHCIVQPGKQQM
>Monodelphis_domestica
MD-TEKRIQHCIVRQGKYKM
>Macropus_eugenii
MD-TEKRIQHCIVRQGKYEM
>Aquila_chrysaetos_canadensis
MDTGEKRTQHCIVRQGKHQM
>Gallus_gallus
MDTGERRTQHCIVRQGKHQM
>Erinaceus_europaeus
MD-TQKRIQHCIVRPGKQQM
>Oryctolagus_cuniculus
MD-TEKRTQHCIVQPGKQQM
>Lepidothrix_coronata
MDTGERRTQHCIVRHGKHQM
>Ceratotherium_simum_simum
MD-TEKRTQHCIVRPGKQQM
>Manis_javanica
MD-TEKRIQHCIVRPGKQQM
>Loxodonta_africana
MD-TEKRIKHCIVRPGKQQT
>Dipodomys_ordii
MD-TEKRTQHCIVQPGKQQM
>Tupaia_chinensis
MD-TGKRTQHRIVQPGKQQL
>Trichechus_manatus_latirostr
MD-TEKRIQHCIVQPGKQQM
>Gekko_japonicus
MD-TEKRTQHCIVQQGKHQL
>Chelonia_mydas
MD-TEKRIQHCIVRQGKCQM
>Orycteropus_afer_afer
MD-TEKRIQHCIVQPGKQQM

HG0064.1 ENSG00000102678 FGF9

```
>Homo_sapiens
MVILISNQMEKLRIFFPYY-GRMG-----
>Anas_platyrhynchos
MVILILNQMEKLRIFFFITVGWAEDLIAC
>Chaetura_pelagica
MVILILNQMEKLRIFFFITVGWAEDRIAC
>Canis_lupus_familiaris
MVILISNQMEKLRIFFPYY-GRMG-----
>Balaenoptera_acutorostrata_s
MVILISNQMEKLRIFFPYY-GRMG-----
>Charadrius_vociferus
MVILILNQMEKLRIFFFITVGWAEDRIAC
>Rousettus_aegyptiacus
MVIPIWNQMEKLRIFSLMTASWDEDLPVC
>Dasypus_novemcinctus
MVILISNPMEKLRIFFLITVGWDEDLTAC
>Leptosomus_discolor
MVILILNQMEKLRIFFFITVGWAEDRIAC
>Cuculus_canorus
MVILTLNQMEKLRIFFFITVGWAEDRIAC
>Galeopterus_variegatus
MVILISNQMEKLQIFFLITVGWDEDLPAY
>Macropus_eugenii
MVILISNQMEKLRIFFLITVGWDEDLTAC
>Aquila_chrysaetos_canadensis
MVILILNQMEKLRIFFFITVGWAEDRIAC
>Coturnix_japonica
MVILFLNQMEKLRIFFLYY-GWMG-----
>Eurypyga_helias
MVILILNQMEKLRIFFLITVGWAEDRIAC
>Condylura_cristata
MVILISNQMEKLRIFFPYY-GRMG-----
>Oryctolagus_cuniculus
MVILISKQMEKLRIFFPYY-GRMG-----
>Elephantulus_edwardii
MVILISNQMEKLRIFFLIMAGWDEDLTAC
>Opisthocomus_hoazin
MVILVLNQMEKLRIFFLDY-GWMG-----
>Lepidothrix_coronata
MIILILNQMEKLRIFFLYY-SWMG-----
>Egretta_garzetta
MVILILNQMEKLRIFFFITVGWAEDRIAC
>Equus_przewalskii
MVILISNQMEELRIFFPYY-GRMG-----
>Manis_javanica
MVILISNQMEKLRIFFLITVGWDEDLSAC
>Picoides_pubescens
MVILILNQMEKLRIFFLITVGWAEDRIAC
>Loxodonta_africana
MVILISSQMEKLRIFFLITVGWDEDLTAC
>Peromyscus_maniculatus_baird
MVILISKQMEKLRIFFPYY-GRMG-----
>Tupaia_chinensis
MVILISNPMEKLRIFFPYY-GRMG-----
>Aptenodytes_forsteri
MVILILNQMEKLRIFFFITVGWAEDRIAC
>Struthio_camelus_australis
MVILILNQMEKLRIFFFITVGWAEDLIAC
>Chelonia_mydas
MVILILNQMEKLRIFFLIAVGWAEDLTAC
>Tinamus_guttatus
MVFLILNQMEKLRIFFLYY-SWMG-----
>Calypte_anna
MVILILNQMEKLRIFFFITVGWAEDRIAC
>Orycteropus_afer_afer
MVILISNQMDKLQIFFLITVGWDEDFTAC
```

HG0064.2 ENSG00000113578 FGF1

>Homo_sapiens
MVCGFTQFFSWGFSELHSYQKGSRRERTSGSNSLRSRGIRASPPFLGGHSLLMVNGNSLPPAAACLQLSW---------------------------
>Panthera_tigris
----------------------------------------------------MENGNSLPPAAACLQHSR---------------------------
>Myotis_brandtii
----------------------------------------------------MANGNSLPAAAAGLPCSW---------------------------
>Galeopterus_variegatus
----------------------------------------------------MVNGNSLLPAAACLRCSWRGSVDTAPEKKLLRAKAKSSKSATTSS
>Condylura_cristata
----------------------------------------------------MENGNSLPPAAASPQRSW---------------------------
>Ceratotherium_simum_simum
----------------------------------------------------MENGNSLPPAAACLQRSC---------------------------
>Nannospalax_galili
----------------------------------------------------MVNGNSLPQSL--LAASW---------------------------
>Orycteropus_afer_afer
----------------------------------------------------MGNGNSLPPAPACLK------------------------------

HG0065 ENSGALG00000029927 STRBP

>Gallus_gallus
------------------------------------------------------------------------------------------------------------------------------------MKRFCQAKQLLSKTPRRREKEKERES-----
>Anser_cygnoides
------------------------------------------------------------------------------------------------------------------------------------MKRFCQAKQRLAKTPRRREKEKGREKLKH--
>Chaetura_pelagica
------------------------------------------------------------------------------------------------------------------------------------MKRFCQAKQCLAKTPRRREKEKGRERELKH-
>Apteryx_australis_mantelli
------------------------------------------------------------------------------------------------------------------------------------MKRFCQAKQCLAKTPRRREKEKGRGRELKH-
>Buceros_rhinoceros_silvestri
------------------------------------------------------------------------------------------------------------------------------------MKRFCQTKQCLAKTPRRREKEKGRERELKH-
>Caprimulgus_carolinensis
------------------------------------------------------------------------------------------------------------------------------------MKRFCQAKQCLAKTPRRREKEKGRERELKH-
>Cariama_cristata
------------------------------------------------------------------------------------------------------------------------------------MKRFCQAKQCLAKTPRRREKEKGRERELKH-
>Balaenoptera_acutorostrata_s
------------------------------------------------------------------------------------------------------------------------------------MRRYCHRKQCLVKTTRKREKDXGRRS-----
>Charadrius_vociferus
------------------------------------------------------------------------------------------------------------------------------------MKRFCQAKQCLAKTPRRREKEKGRERELKH-
>Colius_striatus
------------------------------------------------------------------------------------------------------------------------------------MKRFYQAKQCLAKTPRRREKEKGRERELKR-
>Columba_livia
------------------------------------------------------------------------------------------------------------------------------------MKRFCQTKQGLAKTPRRREKEKGRESELKH-
>Merops_nubicus
------------------------------------------------------------------------------------------------------------------------------------MKRFCQAKQCLAKTPRRREKEKGREKELKH-
>Alligator_mississippiensis
------------------------------------------------------------------------------------------------------------------------------------MRRFCQAKQCLAKTPRRREKEKGEGERS---
>Cuculus_canorus
------------------------------------------------------------------------------------------------------------------------------------MKRFCQAKQCLAKTPRRREKEKRRRRELKH-
>Monodelphis_domestica
------------------------------------------------------------------------------------------------------------------------------------MRRFCLTKQGPLKTTRKREKERGKS------
>Aquila_chrysaetos_canadensis
------------------------------------------------------------------------------------------------------------------------------------MKRFCQAKQCLAKTPRRREKEKGRERELKH-
>Gavia_stellata
------------------------------------------------------------------------------------------------------------------------------------MKRFCQAKQCLAKTPRRREKEKGRERELKH-
>Balearica_regulorum_gibberic
------------------------------------------------------------------------------------------------------------------------------------MKRFCQAKQCLAKTPRRREKEKGRERELKH-
>Condylura_cristata
------------------------------------------------------------------------------------------------------------------------------------MRRYCHTKQCLVKTPRKRGKDHRKKKLKTL-
>Tauraco_erythrolophus
------------------------------------------------------------------------------------------------------------------------------------MKRFCQAKQCLAKTPRRREKEKGRERELKH-
>Zonotrichia_albicollis
MEHVCHVGFPVFLIPEAGIARWLTNLPGISSNWHGKVVTCVWAHHKEVAFHSDICLLRYKWISDCPIWKLGCFDQWFASCAASSTSGINTPS--LLISSCFSLFAGELEKVTP--RAPWPYELLSWQGTERLMKRFCQAKQCLAKTPRRREKEKGRERELQH-
>Phaethon_lepturus
------------------------------------------------------------------------------------------------------------------------------------MKRFCQAKQCLAKTPRRREKEKGRERELKH-
>Picoides_pubescens
------------------------------------------------------------------------------------------------------------------------------------MKRFCQAKQCLAKTPRRREKEKGRERELKH-
>Otolemur_garnettii
------------------------------------------------------------------------------------------------------------------------------------MRRYCHTKQCLVKTTRKREKDREKS------
>Fulmarus_glacialis
------------------------------------------------------------------------------------------------------------------------------------MKRFCQAKQCLAKTPRRREKEKGRERELKH-
>Melopsittacus_undulatus
------------------------------------------------------------------------------------------------------------------------------------MKRFCQAKQCLAKTPRRREKEKGRERELKH-
>Aptenodytes_forsteri
------------------------------------------------------------------------------------------------------------------------------------MKRFCQAKQCLAKTPRRREKEKGREKELKH-
>Anolis_carolinensis
---------------------------------MGARGAVTWGPERRLA-------------------------------------AVELPERSERRDGEDAAAAGVLQSRIPGASVPLLRDLLSRQETERHMRSSWQAKQSLAKTPRRREKEKADRRRS---
>Struthio_camelus_australis
------------------------------------------------------------------------------------------------------------------------------------MKRFCQAKQCLAKTPRRREKEKGRGKELKH-
>Chrysemys_picta
------------------------------------------------------------------------------------------------------------------------------------MRRFCQAKTVPSKDTEEEGKGEGGGGAELRL
>Tinamus_guttatus
------------------------------------------------------------------------------------------------------------------------------------MKRFCQTKQCLAKTPRRREKEKGRGRDLKH-
>Calypte_anna
------------------------------------------------------------------------------------------------------------------------------------MKRLCQAKQCLAKTPRRREKEKGREKELKH-
>Apaloderma_vittatum
------------------------------------------------------------------------------------------------------------------------------------MKRFCQAKQCLAKTPRRREKEKGWERELKH-

```
HG0066 ENSG00000112182 BACH2

>Homo_sapiens
-----MDCQSLLISPFAQFPAS
>Anser_cygnoides
-----MDCCSLLISPLILFPAS
>Mustela_putorius_furo
-----MDCWSLLISPFAQFPAS
>Orcinus_orca
-----MDCRSLLFSPFAQFPAS
>Charadrius_vociferus
-----MDCCSLLISPLILFPAS
>Eptesicus_fuscus
-----MDCCSLLISPFAQFPAS
>Dasypus_novemcinctus
-----MDCCSLLISPFAQFSAS
>Columba_livia
-----MDCCSLLISPLILFPAS
>Alligator_mississippiensis
-----MDCCSLLISPFTQFPAS
>Cuculus_canorus
-----MDCCSLLISPLIPFPAS
>Sarcophilus_harrisii
-----MDCCNLLISPFIQFPAS
>Monodelphis_domestica
-----MDCCSLFISPFIQFPAS
>Macropus_eugenii
-----MDCCSLLISPFIQFPAS
>Falco_cherrug
-----MDCCSLLISPLILFPAS
>Gallus_gallus
-----MDCCSLLISPLILFPAS
>Mesitornis_unicolor
-----MDCCSLLISPVILFPAS
>Condylura_cristata
-----MDCCSFLISPFAQCPAS
>Oryctolagus_cuniculus
-----MDCCSLLISPFAQSPAS
>Elephantulus_edwardii
-----MDCCSLLISPFAQFPAS
>Opisthocomus_hoazin
-----MDCCSLLISPLILFPAS
>Pseudopodoces_humilis
-----MDCSSLLISPLILFPAS
>Egretta_garzetta
-----MDCCSLLISPLILFPAS
>Equus_asinus
-----MDCCSLLISPFAQFPAS
>Manis_javanica
-----MDCCSHLISPFAQFPAS
>Melopsittacus_undulatus
-----MDCCSLLLSPLILFPAS
>Heterocephalus_glaber
-----MDCCSLLISPFAQFPAS
>Trichechus_manatus_latirostr
-----MDCCSLLISPFAQFPAS
>Aptenodytes_forsteri
-----MDCCSLLISPLILFPAS
>Gekko_japonicus
-----MDCCKLLISPFVHFLVS
>Struthio_camelus_australis
-----MDCCSLLISPLILFPAS
>Chrysemys_picta
MVHILMDCCSLLISPFIQFPAS
>Tinamus_guttatus
-----MDCCSLFISPLIPFPAS
>Orycteropus_afer_afer
-----MDCCSILISPFAQFPAS
```

HG0067 ENSG00000166862 CACNG2

>Homo_sapiens
METGDQNFRKRD-------------------------------------------
>Channa_punctata
METGDQNFRKKENLNGYFDFFPVFIAFCLLSLPVWRKPANYLINSLSLDFLTPSG
>Anser_cygnoides
METGDQNFRKRD-------------------------------------------
>Microhyla_fissipes
METEDQNFRKRD-------------------------------------------
>Oryzias_latipes
METGDRNFRKKEKRSC---------------------------------------
>Canis_lupus_familiaris
METGDQNFRKRD-------------------------------------------
>Balaenoptera_acutorostrata_s
METGDQNFRKRD-------------------------------------------
>Astyanax_mexicanus
MESGDQNFRKKEN------------------------------------------
>Calidris_pugnax
METGDQNFRKRD-------------------------------------------
>Rousettus_aegyptiacus
METGDQNFRKRD-------------------------------------------
>Maylandia_zebra
METGDQNFRKKENLSG---------------------------------------
>Dasypus_novemcinctus
METGDQNFRKRD-------------------------------------------
>Alosa_alosa
METGDQNFRKKEN------------------------------------------
>Sinocyclocheilus_anshuiensis
METGDQNFRKKDI------------------------------------------
>Poecilia_mexicana
METGDQNFRKKENLSG---------------------------------------
>Galeopterus_variegatus
METGDQNFRKRD-------------------------------------------
>Monodelphis_domestica
METGDQNFRKRD-------------------------------------------
>Aquila_chrysaetos_canadensis
METGDQNFRKRD-------------------------------------------
>Meleagris_gallopavo
METGDQNFRKRD-------------------------------------------
>Apteronotus_leptorhynchus
METGDQNFRKKDIYQLFILFNFRFHFSVY--------------------------
>Zonotrichia_albicollis
METGDQNFRKRD-------------------------------------------
>Ceratotherium_simum_simum
METGDQNFRKRD-------------------------------------------
>Manis_javanica
METGDQNFRKRD-------------------------------------------
>Melopsittacus_undulatus
METGDQNFRKRD-------------------------------------------
>Nannospalax_galili
METGDQNFRKRD-------------------------------------------
>Salmo_salar
METGDQNFRKKEI------------------------------------------
>Tupaia_chinensis
METGDQNFRKRD-------------------------------------------
>Ictalurus_punctatus
METGDQNFRKKLN------------------------------------------
>Thamnophis_sirtalis
METGDQNFRKRD-------------------------------------------
>Struthio_camelus_australis
METGDQNFRKRD-------------------------------------------
>Chelonia_mydas
METGDQNFRKRD-------------------------------------------
>Takifugu_rubripes
METGDQNFRKKENPCVSFGFSF---SSCVPTRTI---------------------
>Tinamus_guttatus
METGDQNFRKRD-------------------------------------------

HG0068 ENSDARG00000071235

>Danio_rerio
--------------------------------------------------------MRFFRL-TFKCFVDCF
>Amia_calva
--------------------------------------------------------MRFFRL-TFKCFVDCF
>Xenopus_tropicalis
--------------------------------------------------------MRFFRL-TFKCFVDCF
>Menidia_menidia
--------------------------------------------------------MRFFRL-TFKCFVDCF
>Porichthys_notatus
--------------------------------------------------------MRFFRL-TFKCFVDCF
>Oryzias_melastigma
--------------------------------------------------------MRFFRL-TFKCFVDCF
>Hynobius_chinensis
--------------------------------------------------------MRFFRL-TFKCFVDGF
>Micropterus_floridanus
--------------------------------------------------------MRFFRL-TFKCFVDCF
>Astyanax_mexicanus
--------------------------------------------------------MRFFRL-TFKCFVDCF
>Rousettus_aegyptiacus
--------------------------------------------------------MRFFRL-TFKCFVDCF
>Haplochromis_burtoni
--------------------------------------------------------MRFFRL-TFKCFVDCF
>Clupea_harengus
--------------------------------------------------------MRFFRL-TFKCFVDCF
>Latimeria_chalumnae
MLLPYPSVQCRELSQTRSAAREILSYRDSKCSQEGISPAAWLPSFYPYFPLLLFAHMRYFRL-TFKCFVDCF
>Cyprinodon_variegatus
--------------------------------------------------------MRFFRL-TFKCFVDCF
>Monodelphis_domestica
--------------------------------------------------------MRFFRL-TFKCFVDCF
>Ciona_intestinalis
--------------------------------------------------------MRYFRLFTFKCFIDCF
>Esox_lucius
--------------------------------------------------------MRFFRL-TFKCFVDCF
>Gallus_gallus
--------------------------------------------------------MRFFRL-TFKCFVDCF
>Erinaceus_europaeus
--------------------------------------------------------MRFFRL-TFKCFVDCF
>Scleropages_formosus
--------------------------------------------------------MRFFRL-TFKCFVDCF
>Notothenia_coriiceps
--------------------------------------------------------MRFFRL-TFKCFVDCF
>Platichthys_stellatus
--------------------------------------------------------MRFFRL-SFKCFVDCF
>Papio_anubis
--------------------------------------------------------MRFFRL-TFKCFVDCF
>Leucoraja_erinacea
--------------------------------------------------------MRFFRL-TFKCFVDCF
>Psammomys_obesus
--------------------------------------------------------MRFFRL-TFKCFVDCF
>Salmo_salar
--------------------------------------------------------MRFFRL-SFKCFVDCF
>Lepisosteus_oculatus
--------------------------------------------------------MRFFRL-TFKCFVDCF
>Macropisthodon_rudis
--------------------------------------------------------MRFFRL-TFKCFVDCF
>Botryllus_schlosseri
--------------------------------------------------------MHYFKLTTFKCFIDCF
>Tetronarce_californica
--------------------------------------------------------MRFFRL-TFKCFVDCF

HG0068 ENSG00000130821 SLC6A8

```
>Homo_sapiens
--------------------------------------------------------MRFFRL-TFKCFVDCF
>Xenopus_tropicalis
--------------------------------------------------------MRFFRL-TFKCFVDCF
>Menidia_menidia
--------------------------------------------------------MRFFRL-TFKCFVDCF
>Porichthys_notatus
--------------------------------------------------------MRFFRL-TFKCFVDCF
>Oryzias_latipes
--------------------------------------------------------MRFFRL-TFKCFVDCF
>Tripterygion_delaisi
--------------------------------------------------------MRFFRL-TFKCFVDCF
>Hynobius_chinensis
--------------------------------------------------------MRFFRL-TFKCFVDGF
>Micropterus_floridanus
--------------------------------------------------------MRFFRL-TFKCFVDCF
>Pygocentrus_nattereri
--------------------------------------------------------MRFFRL-TFKCFVDCF
>Rousettus_aegyptiacus
--------------------------------------------------------MRFFRL-TFKCFVDCF
>Haplochromis_burtoni
--------------------------------------------------------MRFFRL-TFKCFVDCF
>Clupea_harengus
--------------------------------------------------------MRFFRL-TFKCFVDCF
>Latimeria_chalumnae
MLLPYPSVQCRELSQTRSAAREILSYRDSKCSQEGISPAAWLPSFYPYFPLLLFAHMRYFRL-TFKCFVDCF
>Danio_rerio
--------------------------------------------------------MRFFRL-TFKCFVDCF
>Cyprinodon_variegatus
--------------------------------------------------------MRFFRL-TFKCFVDCF
>Monodelphis_domestica
--------------------------------------------------------MRFFRL-TFKCFVDCF
>Ciona_intestinalis
--------------------------------------------------------MRYFRLFTFKCFIDCF
>Esox_lucius
--------------------------------------------------------MRFFRL-TFKCFVDCF
>Gallus_gallus
--------------------------------------------------------MRFFRL-TFKCFVDCF
>Erinaceus_europaeus
--------------------------------------------------------MRFFRL-TFKCFVDCF
>Notothenia_coriiceps
--------------------------------------------------------MRFFRL-TFKCFVDCF
>Platichthys_stellatus
--------------------------------------------------------MRFFRL-SFKCFVDCF
>Leucoraja_erinacea
--------------------------------------------------------MRFFRL-TFKCFVDCF
>Psammomys_obesus
--------------------------------------------------------MRFFRL-TFKCFVDCF
>Salmo_salar
--------------------------------------------------------MRFFRL-SFKCFVDCF
>Lepisosteus_oculatus
--------------------------------------------------------MRFFRL-TFKCFVDCF
>Macropisthodon_rudis
--------------------------------------------------------MRFFRL-TFKCFVDCF
>Botryllus_schlosseri
--------------------------------------------------------MHYFKLTTFKCFIDCF
>Tetronarce_californica
--------------------------------------------------------MRFFRL-TFKCFVDCF
```

HG0068 ENSGALG00000041419

>Gallus_gallus
--------------------------------------------------------MRFFRL-TFKCFVDCF
>Amia_calva
--------------------------------------------------------MRFFRL-TFKCFVDCF
>Xenopus_tropicalis
--------------------------------------------------------MRFFRL-TFKCFVDCF
>Menidia_menidia
--------------------------------------------------------MRFFRL-TFKCFVDCF
>Porichthys_notatus
--------------------------------------------------------MRFFRL-TFKCFVDCF
>Oryzias_latipes
--------------------------------------------------------MRFFRL-TFKCFVDCF
>Tripterygion_delaisi
--------------------------------------------------------MRFFRL-TFKCFVDCF
>Hynobius_chinensis
--------------------------------------------------------MRFFRL-TFKCFVDGF
>Micropterus_floridanus
--------------------------------------------------------MRFFRL-TFKCFVDCF
>Pygocentrus_nattereri
--------------------------------------------------------MRFFRL-TFKCFVDCF
>Rousettus_aegyptiacus
--------------------------------------------------------MRFFRL-TFKCFVDCF
>Haplochromis_burtoni
--------------------------------------------------------MRFFRL-TFKCFVDCF
>Clupea_harengus
--------------------------------------------------------MRFFRL-TFKCFVDCF
>Latimeria_chalumnae
MLLPYPSVQCRELSQTRSAAREILSYRDSKCSQEGISPAAWLPSFYPYFPLLLFAHMRYFRL-TFKCFVDCF
>Danio_rerio
--------------------------------------------------------MRFFRL-TFKCFVDCF
>Cyprinodon_variegatus
--------------------------------------------------------MRFFRL-TFKCFVDCF
>Monodelphis_domestica
--------------------------------------------------------MRFFRL-TFKCFVDCF
>Ciona_intestinalis
--------------------------------------------------------MRYFRLFTFKCFIDCF
>Esox_lucius
--------------------------------------------------------MRFFRL-TFKCFVDCF
>Erinaceus_europaeus
--------------------------------------------------------MRFFRL-TFKCFVDCF
>Scleropages_formosus
--------------------------------------------------------MRFFRL-TFKCFVDCF
>Taeniopygia_guttata
--------------------------------------------------------MRFFRL-TFKCFVDCF
>Notothenia_coriiceps
--------------------------------------------------------MRFFRL-TFKCFVDCF
>Platichthys_stellatus
--------------------------------------------------------MRFFRL-SFKCFVDCF
>Cebus_capucinus_imitator
--------------------------------------------------------MRFFRL-TFKCFVDCF
>Leucoraja_erinacea
--------------------------------------------------------MRFFRL-TFKCFVDCF
>Mus_musculus
--------------------------------------------------------MRFFRL-TFKCFVDCF
>Salmo_salar
--------------------------------------------------------MRFFRL-SFKCFVDCF
>Lepisosteus_oculatus
--------------------------------------------------------MRFFRL-TFKCFVDCF
>Macropisthodon_rudis
--------------------------------------------------------MRFFRL-TFKCFVDCF
>Botryllus_schlosseri
--------------------------------------------------------MHYFKLTTFKCFIDCF
>Tetronarce_californica
--------------------------------------------------------MRFFRL-TFKCFVDCF

HG0069 ENSG00000100697 DICER1

```
>Homo_sapiens
-MSKCSSDKSNTEISNIKT
>Apteryx_australis_mantelli
MMSKCSSGSINTEVAEIKT
>Cariama_cristata
MMSKCSSGSINTETAEIKT
>Ailuropoda_melanoleuca
-MSKCSSDNSNTEISNIKT
>Balaenoptera_acutorostrata_s
-MSKCSSDNSNTEISNIKT
>Charadrius_vociferus
MMSKCSSGSINTETAEIKT
>Pteropus_alecto
-MSKCSSDNSNTEISNIKT
>Pterocles_gutturalis
MMSKCSSGSINTETAEIKT
>Dasypus_novemcinctus
-MSKCSSDNSNTEISNIKT
>Columba_livia
MMSKCSSGSINTETAEIKT
>Alligator_sinensis
MMSKCSSENTNTKVSEIKT
>Galeopterus_variegatus
-MSKCSSDNSNTEISNIKT
>Aquila_chrysaetos_canadensis
MMSKCSSGSINTETAEIKT
>Gavia_stellata
MMSKCSSGSINTETAEIKT
>Mesitornis_unicolor
MMSKCSSGSVNTETAEIKT
>Erinaceus_europaeus
MMSKCSSDNSNTEIANIKT
>Elephantulus_edwardii
-MSKCGSDSSNTDIADIKT
>Ornithorhynchus_anatinus
MMSKCSSGNINTEVSEIKT
>Ficedula_albicollis
MMSKCSSGGINTETAEIKT
>Phaethon_lepturus
MMSKCSSGSINTEAAEIKT
>Ceratotherium_simum_simum
-MSKCSSDNSNTEISNIKT
>Loxodonta_africana
-MSKCSSDNSNAEISNIKT
>Melopsittacus_undulatus
MMSKCSSGSINTETAEIKT
>Ictidomys_tridecemlineatus
-MSKCSSDSSNTEISNIKT
>Trichechus_manatus_latirostr
-MSKCSSDNSNTEISNIKT
>Aptenodytes_forsteri
MMSKCSSGSINTETAEIKT
>Tyto_alba
MMSKCSSGSINTETAEIKT
>Struthio_camelus_australis
MMSKCNSGSINTEVAEIKT
>Chrysemys_picta
MMSKCSSKNTNTEISEIKT
>Tinamus_guttatus
MMSKCSSGNINTEVAEIKT
>Apaloderma_vittatum
MMSKCSSGSINTETAEIKT
>Orycteropus_afer_afer
-MSGCSSDGSNTEASHIKT
```

HG0069 ENSGALG00000010999 DICER1

>Gallus_gallus
MMSKCSLESISTEAAEIKT--------------------------
>Anser_cygnoides
MMSKCSLESISTEAAEIKT--------------------------
>Chaetura_pelagica
MMSKCSLGSIKTETAEIKTAVLEQKCNERNTG-------------
>Apteryx_australis_mantelli
MMSKCSSGSINTEVAEIKT--------------------------
>Buceros_rhinoceros_silvestri
MMSKCSSGSINTATAEIKT--------------------------
>Cariama_cristata
MMSKCSSGSINTETAEIKT--------------------------
>Calidris_pugnax
MMSKCSLGSINTETAEIKT--------------------------
>Pterocles_gutturalis
MMSKCSSGSINTETAEIKT--------------------------
>Colius_striatus
MMSKCSLGSINTETAEIKMAVLEQKCNERNTG-------------
>Columba_livia
MMSKCSSGSINTETAEIKT--------------------------
>Merops_nubicus
MMSKCSLGSINTETAEIKT--------------------------
>Alligator_sinensis
MMSKCSSENTNTKVSEIKT--------------------------
>Cuculus_canorus
MMSRCSLGSINSETAEIKTAVLKQKCK------------------
>Aquila_chrysaetos_canadensis
MMSKCSSGSINTETAEIKT--------------------------
>Gavia_stellata
MMSKCSSGSINTETAEIKT--------------------------
>Balearica_regulorum_gibberic
MMSKCSSGSINTETAEIKTTVLEQKCNERNTG-------------
>Erinaceus_europaeus
MMSKCSSDNSNTEIANIKT--------------------------
>Ornithorhynchus_anatinus
MMSKCSSGNINTEVSEIKT--------------------------
>Opisthocomus_hoazin
MMSKCSLGIINTETAEIKT--------------------------
>Acanthisitta_chloris
MMSKCSLGSINTETAEIKTAVLEQKCNERNTG-------------
>Phaethon_lepturus
MMSKCSSGSINTEAAEIKT--------------------------
>Picoides_pubescens
MMSKCSSGSINTETAKIKTAVLEQKCNERNTG-------------
>Fulmarus_glacialis
MMSKCSSGSINTETTEIKTAVLEQKCNERNTG-------------
>Melopsittacus_undulatus
MMSKCSSGSINTETAEIKT--------------------------
>Jaculus_jaculus
-MSKCSSDNSSTEVADIKT--------------------------
>Aptenodytes_forsteri
MMSKCSSGSINTETAEIKT--------------------------
>Tyto_alba
MMSKCSSGSINTETAEIKT--------------------------
>Struthio_camelus_australis
MMSKCNSGSINTEVAEIKT--------------------------
>Pelodiscus_sinensis
MMSKCSSKNTNTEVSEIKTQVRKYPSRTPAVNNTTWVIEEALQRQ
>Tinamus_guttatus
MMSKCSSGNINTEVAEIKT--------------------------
>Calypte_anna
MMSKCSLESINTETAEIKTAVLEQKCNERNTG-------------
>Apaloderma_vittatum
MMSKCSSGSINTETAEIKT--------------------------

HG0070 ENSDARG00000040926 nr2f2

>Danio_rerio
MDDCKCPWTWPLRARLLFGLLQF----------------------------
>Amia_calva
MADCRQPWTWPLSARLLFSLSSHQV--------------------------
>Oryzias_latipes
MADCKRPWTWPLSARQRLSSPTHQF--------------------------
>Tripterygion_delaisi
MADCKRPWTWPLRARQRLSSPLLHHF-------------------------
>Mustela_putorius_furo
MADCGLPWTWPPDSRLSSSSSSSSSSS-------YS------NSSAHQL--
>Orcinus_orca
MADCGLPWTWPPDSRLSSSTSSSSSSS-------SSS---SNSSAAHQL--
>Pygocentrus_nattereri
MADCRRPWTWPLRAGPRFGSPSPHF--------------------------
>Haplochromis_burtoni
MADCKRPWTWPLSARQRLSSPSHQF--------------------------
>Clupea_harengus
MDECKRPWTWPSRARLRFGSSSVLTIC--S--------------ATF----
>Latimeria_chalumnae
MADCELPWTWPLDTRLFFNSTVRSGLNIFE--------------AIFLLFA
>Austrofundulus_limnaeus
MADCKRPWTWPLSARQPLRSPSHQF--------------------------
>Esox_lucius
MADCKRPWTWPLSARLRFSSLSHPF--------------------------
>Acanthogobius_hasta
MADCTRPWTWPLSAGQRLGSPSHHF--------------------------
>Osmerus_mordax
MADCKRPWTWPLRERQRLSSPSHQF--------------------------
>Scleropages_formosus
MADCRRIWTWPLSHRTTFNT-------------------------------
>Lepidothrix_coronata
MADCELPWTWPPDTRLSSFNYSAADQL------------------------
>Notothenia_coriiceps
MADCKRPWTWPLSARQRLSSPSHQF--------------------------
>Platichthys_stellatus
MADCKRPWTWPLSARQRLSSPSHQF--------------------------
>Cercocebus_atys
MADCGLPWTWPPDSRLSSSSTSSFTTTSSSSSSSSSSSSSANSSAAHQL--
>Loxodonta_africana
MADCGLPWTWPPDSRLSSFSSSSSS-S-------SSF---SNSSAAHQL--
>Nannospalax_galili
MADCGLPWTWPPDSRLSSSSSSISSFS-------SSSSFSSSNSAAH----
>Salmo_salar
MADCKRPWTWPLSARLRFSSPSHQF--------------------------
>Lepisosteus_oculatus
MADCRQRWTWPLNAQLLSSSSTHQV--------------------------
>Ictalurus_punctatus
MDDCRRPWTWPLRAGLRFGSSCCQF--------------------------
>Trichechus_manatus_latirostr
MADCGLPWTWPPDSRLSSFSSSSFS-S-------S-----SNSSAAHQL--
>Takifugu_rubripes
MADCKRPWTWPLSARQRLGSPSHQF--------------------------

HG0070 ENSDARG00000052695 nr2f1a

```
>Danio_rerio
MADCNLPWNWPPDNCIS--------------------------------
>Acipenser_sinensis
MAGCILPWNWPPDTCIS--------------------------------
>Xenopus_laevis
MADCILPWIWPPDTYKP--------------------------------
>Menidia_menidia
MADCNLPWNWPPDTCIS--------------------------------
>Scyliorhinus_torazame
MADCVLPWIWPPDTCVP--------------------------------
>Mustela_putorius_furo
MADCGLPWTWPPDSRLSSSSSSSS----------SSSYS---NSSAHQL
>Ambystoma_mexicanum
MAACGLPWTWPPDTRRP--------------------------------
>Orcinus_orca
MADCGLPWTWPPDSRLSSSTSSSS----------SSSSSSSNSSAAHQL
>Pygocentrus_nattereri
MADCNLPWNWPPDSCIS--------------------------------
>Haplochromis_burtoni
MADCNLPWNWPPDTCIS--------------------------------
>Alosa_alosa
MADCNFPWNWPPDTCIS--------------------------------
>Latimeria_chalumnae
MADCVLPWTWPPDTCIS--------------------------------
>Cyprinodon_variegatus
MADCNLPWNWPPDTCIS--------------------------------
>Esox_lucius
MADCNLPWNWPPDACIS--------------------------------
>Gadus_morhua
MAVCNLPWNWPPDTCTS--------------------------------
>Acanthogobius_hasta
MAGCVLPWTWPPDTCVS--------------------------------
>Apteronotus_leptorhynchus
MADCNLPWNWSPDNCIS--------------------------------
>Osmerus_mordax
MADCNIPWNWPPDTCIS--------------------------------
>Lepidothrix_coronata
MADCELPWTWPPDTRLSSFNYSAADQL----------------------
>Perca_flavescens
MAGCNLPWNWPPDTCIS--------------------------------
>Manis_javanica
MADCGLRWTWPPDSRLSSSASSSA--------SSSNSSA-------PQV
>Cynoglossus_semilaevis
MADCNLPWNWPPDTCIS--------------------------------
>Cercocebus_atys
MADCGLPWTWPPDSRLSSSSTSSFTTTSSSSSSSSSSSSSANSSAAHQL
>Loxodonta_africana
MADCGLPWTWPPDSRLSSFSSSSS----------S-SSSFSNSSAAHQL
>Mesocricetus_auratus
MADCGLPWTWPPDSRLSFSSSSSS--------SSSSSFSSSNSSAAH--
>Oncorhynchus_mykiss
MANCNLPWNWPPDTCIS--------------------------------
>Lepisosteus_oculatus
MADCNLPWNWPPDTCIS--------------------------------
>Trichomycterus_areolatus
MADCELPWNWPPDNCIS--------------------------------
>Trichechus_manatus_latirostr
MADCGLPWTWPPDSRLSSFSSSSF----------S-SS--SNSSAAHQL
>Takydromus_sexlineatus
MADCGLLWIWPPDTSFSSCNSAWLPEREGGGDL----------------
>Takifugu_rubripes
MADCNLPWNWPPDTCIS--------------------------------
```

HG0070 ENSG00000175745 NR2F1

```
>Homo_sapiens
MAGCVRPWTWPPDTRAP-------------------------------------
>Acipenser_sinensis
MAGCILPWNWPPDTCIS-------------------------------------
>Xenopus_laevis
MADCILPWIWPPDTYKP-------------------------------------
>Menidia_menidia
MADCNLPWNWPPDTCIS-------------------------------------
>Oryzias_latipes
MADCKRPWTWPLSARQRLSSPTHQF-----------------------------
>Scyliorhinus_torazame
MADCVLPWIWPPDTCVP-------------------------------------
>Canis_lupus_familiaris
MADCGLPWTWPPDSRLSSSSSSSSSSSSSSSSSSSSSSSSSSSSSYSNSSAHQL
>Hynobius_retardatus
MAGCALPWTWPPDTSEP-------------------------------------
>Orcinus_orca
MADCGLPWTWPPDSRLSSSTSSSSSSSSSSSNS---------------SAAHQL
>Pygocentrus_nattereri
MADCNLPWNWPPDSCIS-------------------------------------
>Haplochromis_burtoni
MADCNLPWNWPPDTCIS-------------------------------------
>Clupea_harengus
MAGCNIPWNWPPDTCIS-------------------------------------
>Latimeria_chalumnae
MADCVLPWTWPPDTCIS-------------------------------------
>Catostomus_commersonii
MAGCNLPWNWPPDNCIS-------------------------------------
>Austrofundulus_limnaeus
MADCKRPWTWPLSARQPLRSPSHQF-----------------------------
>Esox_lucius
MADCKRPWTWPLSARLRFSSLSHPF-----------------------------
>Gadus_morhua
MAVCNLPWNWPPDTCTS-------------------------------------
>Acanthogobius_hasta
MAGCVLPWTWPPDTCVS-------------------------------------
>Osmerus_mordax
MADCNIPWNWPPDTCIS-------------------------------------
>Lepidothrix_coronata
MADCELPWTWPPDTRLSSFNYSAADQL---------------------------
>Perca_flavescens
MAGCNLPWNWPPDTCIS-------------------------------------
>Manis_javanica
MADCGLRWTWPPDSRLSSSASSSASSSNSSAPQV--------------------
>Platichthys_stellatus
MAGCNIPWNWPPDTCIS-------------------------------------
>Loxodonta_africana
MADCGLPWTWPPDSRLSSFSSSSSS-SSSFSNS---------------SAAHQL
>Nannospalax_galili
MADCGLPWTWPPDSRLSSSSSSISSFSSSSSFSSSN------------SAAH--
>Salmo_salar
MANCNLPWNWPPDTCIS-------------------------------------
>Lepisosteus_oculatus
MADCNLPWNWPPDTCIS-------------------------------------
>Trichomycterus_areolatus
MADCELPWNWPPDNCIS-------------------------------------
>Trichechus_manatus_latirostr
MADCGLPWTWPPDSRLSSFSSSSFS-SS--SNS---------------SAAHQL
>Takifugu_rubripes
MADCNLPWNWPPDTCIS-------------------------------------
```

HG0070 ENSG00000185551 NR2F2

>Homo_sapiens
MADCGLPWTWPPDSRLSSSSTSSFTTTSSSSSSSSSSSANSSAAHQL
>Menidia_menidia
MADCNLPWNWPPDTCIS------------------------------
>Mustela_putorius_furo
MADCGLPWTWPPDSRLSSSSSSSS--------SSSYS---NSSAHQL
>Ambystoma_mexicanum
MAACGLPWTWPPDTRRP------------------------------
>Orcinus_orca
MADCGLPWTWPPDSRLSSSTSSSS--------SSSSSSSNSSAAHQL
>Pygocentrus_nattereri
MADCNLPWNWPPDSCIS------------------------------
>Haplochromis_burtoni
MADCNLPWNWPPDTCIS------------------------------
>Latimeria_chalumnae
MADCVLPWTWPPDTCIS------------------------------
>Ctenopharyngodon_idella
MADCNLPWNWPPDNCIS------------------------------
>Fundulus_grandis
MADCNLPWNWPPDTCTS------------------------------
>Esox_lucius
MADCNLPWNWPPDACIS------------------------------
>Osmerus_mordax
MADCNIPWNWPPDTCIS------------------------------
>Lepidothrix_coronata
MADCELPWTWPPDTRLSSFNYSAADQL--------------------
>Manis_javanica
MADCGLRWTWPPDSRLSSSASSSASSSNSSAP-------------QV
>Cynoglossus_semilaevis
MADCNLPWNWPPDTCIS------------------------------
>Loxodonta_africana
MADCGLPWTWPPDSRLSSFSSSSS--------S-SSSFSNSSAAHQL
>Mesocricetus_auratus
MADCGLPWTWPPDSRLSFSSSSSSSSS------SSFSSSNSSAAH--
>Lepisosteus_oculatus
MADCNLPWNWPPDTCIS------------------------------
>Trichomycterus_areolatus
MADCELPWNWPPDNCIS------------------------------
>Trichechus_manatus_latirostr
MADCGLPWTWPPDSRLSSFSSSSF--------S-SS--SNSSAAHQL
>Squalus_acanthias
MADCVLPWSWPPDSRLFSADRV-------------------------
>Takydromus_sexlineatus
MADCGLLWIWPPDTSFSSCNSAWLPEREGGGDL--------------
>Takifugu_rubripes
MADCNLPWNWPPDTCIS------------------------------

HG0071.1 ENSDARG00000098240 meis2a

>Danio_rerio
MRHDSLVTCRLLMGRL
>Acipenser_sinensis
MRHDSLVTCGLLMGRL
>Oryzias_latipes
MRHDSLVTCRLLMGRL
>Odobenus_rosmarus_divergens
MKHDSSVTCGLLTGRS
>Lipotes_vexillifer
MKHDSSVTCGLLTGRS
>Pygocentrus_nattereri
MRHDSLVTCTLLMGRL
>Myotis_brandtii
MKHDSSVTCGLLTGRS
>Lipochromis_sp._matumbi_hunt
MRHDSLVTCRLLMGRL
>Clupea_harengus
MRHDSLVTCRLLMGRL
>Latimeria_chalumnae
MRHDSLVTCGLLTGRS
>Alligator_sinensis
MRHDSLVTCGLLTGRS
>Kryptolebias_marmoratus
MRHDSLVTCRLLMGRL
>Esox_lucius
MRHDSLVTCRLLMGRL
>Condylura_cristata
MKHDSSVTCGLLTGRS
>Ochotona_princeps
MKHDSSVTCGLLTGRS
>Elephantulus_edwardii
MKHDSSVTCGLLTGRS
>Zonotrichia_albicollis
MRHDSLVTCGLLTGRS
>Notothenia_coriiceps
MRHDSLVTCRLLMGRL
>Ceratotherium_simum_simum
MKHDSSVTCGLLTGRS
>Cynoglossus_semilaevis
MRHDSLVTCRLLMGRL
>Colobus_angolensis_palliatus
MKHDSSVTCGLLTGRS
>Loxodonta_africana
MKHDSSVTCGLLTGRS
>Jaculus_jaculus
MKHDSSVTCGLLTGRS
>Salmo_salar
MRHDSLVTCRLLMGRL
>Lepisosteus_oculatus
MRHDSLVTCGLLMGRL
>Ictalurus_punctatus
MRHDSLVTCRLLMGRL
>Python_bivittatus
MRHDSLVTCGLLTGRS
>Struthio_camelus_australis
MRHDSLVTCGLLTGRS
>Takifugu_rubripes
MRHDSLVTCRLLMGRL
>Orycteropus_afer_afer
MKHDSSVTCGLLTGRS

HG0071.1 ENSG00000134138 MEIS2

>Homo_sapiens
MKHDSSVTCGLLTGRS
>Acipenser_sinensis
MRHDSLVTCGLLMGRL
>Oryzias_latipes
MRHDSLVTCRLLMGRL
>Odobenus_rosmarus_divergens
MKHDSSVTCGLLTGRS
>Lipotes_vexillifer
MKHDSSVTCGLLTGRS
>Astyanax_mexicanus
MRHDSLVTCTLLMGRL
>Myotis_davidii
MKHDSSVTCGLLTGRS
>Haplochromis_burtoni
MRHDSLVTCRLLMGRL
>Clupea_harengus
MRHDSLVTCRLLMGRL
>Latimeria_chalumnae
MRHDSLVTCGLLTGRS
>Alligator_sinensis
MRHDSLVTCGLLTGRS
>Danio_rerio
MRHDSLVTCRLLMGRL
>Poecilia_reticulata
MRHDSLVTCRLLMGRL
>Esox_lucius
MRHDSLVTCRLLMGRL
>Erinaceus_europaeus
MKHDSSVTCGLLTGRS
>Ochotona_princeps
MKHDSSVTCGLLTGRS
>Elephantulus_edwardii
MKHDSSVTCGLLTGRS
>Zonotrichia_albicollis
MRHDSLVTCGLLTGRS
>Notothenia_coriiceps
MRHDSLVTCRLLMGRL
>Ceratotherium_simum_simum
MKHDSSVTCGLLTGRS
>Cynoglossus_semilaevis
MRHDSLVTCRLLMGRL
>Loxodonta_africana
MKHDSSVTCGLLTGRS
>Jaculus_jaculus
MKHDSSVTCGLLTGRS
>Salmo_salar
MRHDSLVTCRLLMGRL
>Lepisosteus_oculatus
MRHDSLVTCGLLMGRL
>Ictalurus_punctatus
MRHDSLVTCRLLMGRL
>Python_bivittatus
MRHDSLVTCGLLTGRS
>Struthio_camelus_australis
MRHDSLVTCGLLTGRS
>Takifugu_rubripes
MRHDSLVTCRLLMGRL
>Tetronarce_californica
MRDDSFVTSGLLTGRS
>Orycteropus_afer_afer
MKHDSSVTCGLLTGRS

HG0071.2 ENSG00000143995 MEIS1

>Homo_sapiens
MLSDSGALLQVP
>Panthera_tigris
MLSDSGALLQVP
>Balaenoptera_acutorostrata_s
MLSDSGALLQVP
>Myotis_davidii
MLSDSGALLQVP
>Galeopterus_variegatus
MLSDSGALLQVP
>Condylura_cristata
MLSDSGALLQVP
>Ochotona_princeps
MLSDSGGLLQVP
>Equus_caballus
MLSDSGALLQVP
>Manis_javanica
MLSDSGALLQVP
>Loxodonta_africana
MLSDSGALLQVP
>Mus_musculus
MLNDAGALLQVP
>Tupaia_chinensis
MLSDSGALLQVP

HG0072 ENSG00000104435 STMN2

>Homo_sapiens
---MGGADREGLISISNCILVLCTPAPLADQYLMLGDSDSAGVMSGDLGSQSA----
>Chaetura_pelagica
---------------------------------MPRDSGSAGVMSGDL---------
>Odobenus_rosmarus_divergens
---------------------------------MLGDSDSAGVMSGDLGSQSA----
>Hynobius_retardatus
-------------------------------------------MSGDLWSQSAGAEP
>Tursiops_truncatus
---------------------------------MLGDSDSAGVMSGDLGSQSA----
>Charadrius_vociferus
---------------------------------MPGDSGSAGVMSGDL---------
>Myotis_davidii
---------------------------------MLGGSDSARVMSGDLGSQSARASG
>Dasypus_novemcinctus
-------------------------------------------MSGDLGSQSA----
>Columba_livia
---------------------------------MPRDSGSAGVMSGDL---------
>Cuculus_canorus
---------------------------------MPRDSGSAGVMSGDL---------
>Galeopterus_variegatus
---MGGAAREGLISISNCILVLCTPAPLADQYLMLGDSDSAGVMSGDLGSQSA----
>Falco_cherrug
---------------------------------MPGDSGSAGVMSGDL---------
>Gallus_gallus
---------------------------------MPGDSGSAGVMSGDL---------
>Condylura_cristata
---------------------------------MLGDSDSAGVVSGDLGSQSA----
>Ochotona_princeps
---MGGAAREGLISISNCIVVLCTPAPLADQYLMLRDSDSAGVMSGDLGSQSA----
>Opisthocomus_hoazin
---------------------------------MPGGSGSAGVMSGDL---------
>Sturnus_vulgaris
---------------------------------MPGDSGSAGVMSGDLSSQSAWVEE
>Egretta_garzetta
---------------------------------MPGDSGSAGVMSGDL---------
>Ceratotherium_simum_simum
---------------------------------MLGDSDSAGVMSGDLGSQSA----
>Picoides_pubescens
---------------------------------MPGDSGSAGVMSGDL---------
>Loxodonta_africana
---MGGAAREGLISISNCILVLCTLAPLADQYLMLGDSDSAGVMSGDLGSQSA----
>Melopsittacus_undulatus
---------------------------------MPGDSGSAGVMSGDL---------
>Marmota_marmota_marmota
---MGGAAREGLISISNCILVLCTPAPLADQYSMLGDSDSAGVMSGDLGSQSA----
>Tupaia_chinensis
---------------------------------MLGDSDSAGVMSGDLGSQSA----
>Aptenodytes_forsteri
---------------------------------MPGDSGSAGVMSGDL---------
>Squalus_acanthias
MFHWKKRGGPILICISNCILALCTLASLVDQYLMPGDPENAPVMSRDLFSQSALLGR
>Anolis_carolinensis
---------------------------------MLADSGSAGVMSGELWSQSAWLEE
>Struthio_camelus_australis
---------------------------------MPGDSGSAGVMSGDL---------
>Chelonia_mydas
---------------------------------MLGDSSSAGVMSGDL---------
>Tinamus_guttatus
---------------------------------MPGDSGSAGVMSGDL---------
>Calypte_anna
---------------------------------MPGDSGSAGVMSGDL---------

HG0072 ENSGALG00000039690 STMN2

>Gallus_gallus
------------------------------MPGDSGSAGVMSGDL---------
>Chaetura_pelagica
------------------------------MPRDSGSAGVMSGDL---------
>Canis_lupus_familiaris
------------------------------MLGDSDSAGVMSGDLGSQSA----
>Lipotes_vexillifer
------------------------------MLGDSDSAGVMSGDLGSQSA----
>Charadrius_vociferus
------------------------------MPGDSGSAGVMSGDL---------
>Cynopterus_sphinx
------------------------------MPGDSDSAGVMSGDLGSQSAWVSE
>Columba_livia
------------------------------MPRDSGSAGVMSGDL---------
>Cuculus_canorus
------------------------------MPRDSGSAGVMSGDL---------
>Galeopterus_variegatus
MGGAAREGLISISNCILVLCTPAPLADQYLMLGDSDSAGVMSGDLGSQSA----
>Haliaeetus_leucocephalus
------------------------------MPGDSGSAGVMSGDL---------
>Opisthocomus_hoazin
------------------------------MPGGSGSAGVMSGDL---------
>Pseudopodoces_humilis
------------------------------MPGDSGSAGVMSGDLSSQSAWVEE
>Egretta_garzetta
------------------------------MPGDSGSAGVMSGDL---------
>Ceratotherium_simum_simum
------------------------------MLGDSDSAGVMSGDLGSQSA----
>Picoides_pubescens
------------------------------MPGDSGSAGVMSGDL---------
>Cebus_capucinus_imitator
MGGAAREGLITISNCILVLCTPALLADQYLMLGDSDSAGVMSGDLGSQSA----
>Loxodonta_africana
MGGAAREGLISISNCILVLCTLAPLADQYLMLGDSDSAGVMSGDLGSQSA----
>Melopsittacus_undulatus
------------------------------MPGDSGSAGVMSGDL---------
>Ictidomys_tridecemlineatus
------------------------------MLGDSDSAGVMSGDLGSQSA----
>Tupaia_chinensis
------------------------------MLGDSDSAGVMSGDLGSQSA----
>Aptenodytes_forsteri
------------------------------MPGDSGSAGVMSGDL---------
>Anolis_carolinensis
------------------------------MLADSGSAGVMSGELWSQSAWLEE
>Struthio_camelus_australis
------------------------------MPGDSGSAGVMSGDL---------
>Chelonia_mydas
------------------------------MLGDSSSAGVMSGDL---------
>Tinamus_guttatus
------------------------------MPGDSGSAGVMSGDL---------
>Calypte_anna
------------------------------MPGDSGSAGVMSGDL---------

HG0073 ENSDARG00000041708 csrp2bp

>Danio_rerio
------------------------------MSTASKVVLGLSIVLTISTVAGVHIKQNWDRQRLREGVVRDLERLERKRENLRALEEQIQLTRELVTERNRRQAEASETHDS--------------
>Amia_calva
------------------------------MSTTSKVVLGVSLALTLGTVAGVHIKQSLDRERLREGVYRDLERQARKKENLRQLEEQIALTRELEKERQRRAAGTQGS-----------------
>Channa_striata
------------------------------MSTTSKVVLGLSVSLTLTTVAGVHLKQALDRQRLHEGVVQDLERLERKKENLRLLQEQRVLTRHLEEERQRREAELHPQDHN--------------
>Xenopus_laevis
------------------------------MSTISKVVLGISVVLSVGTVVGVHVKQRLDRERLREGVLQDLERQSRKQENLRLLEEQIHLTKRLEAEREKILLTEGSSKF---------------
>Porichthys_notatus
------------------------------MSRVSKVVLGLSVVLTLGTVTGVHLLQARDRQRLHDGVLRDLERVQRKKDNLRRLQDQKVLSRRLEAERQRREAELQTTGHTHTD-----------
>Tripterygion_delaisi
------------------------------MSTTSKVVLGVSVVLSLSIVAGVHLKQDWDRQRLHEGVIRDLERLAKKKENLRLLEEQRILTIQLEEERQQREAQLSSQES---------------
>Odobenus_rosmarus_divergens
------------------------------MSRSSKAVLGLSVVLTAATVAGVHLKQRQDRQRLHDGVIRDIERQNRKKENIRLLGEQIILTEQLEAEREKMLLAKGSQKT---------------
>Hynobius_chinensis
------------------------------MSAASKVVLGVSVLASVATVAGVHLKQKLDRERLREGVFRDIERQHRKQENLRLLEEQIVLTRQLEVEREKMLMTKGSQKS---------------
>Pygocentrus_nattereri
------------------------------MSTASKVVLGVSVVLTVSTVVGVHVKQNWDRQRLREGVLRDLERLERKRENLRALEEQIQLTRELVAEQAYQQAEAARGS----------------
>Calidris_pugnax
-----------------------------------------------------------------------------------------------MEERDKALMAKGSQQP---------------
>Callorhinchus_milii
------------------------------MSAVSKVVLGVSVAVTAGTVAGVHLRQQLDREKLREGVFRDLERQEHKKKNLRLLEEQIILTNQLEADRVASESIQKSEKHKSDF-----------
>Rhinolophus_ferrumequinum
------------------------------MSRTSKVVLGLSVVLTVATVAGVHLKQRQDQQRLRDGVIRDIERQNRKKENIRLLGEQIILTEQLEAEREKTLLAKDLKKH---------------
>Maylandia_zebra
------------------------------MSAVSKVVLGVSVVLTVSTVAAVHLKQAWDRERLREGVVRDLERLERRKENLRMLEEQRLLTRQLEEERRRREARLHPQDT---------------
>Alosa_alosa
------------------------------MSTASKVVLGLSVVLTISTVAGVHIKQNWDRQRLHEGVLRDLERVERKRENLRLLEEQIQLTQELETERQRREASAASGATQSP------------
>Merops_nubicus
-----------------------------------------------------------------------------------------------MEERDKALMEKGSQQS---------------
>Cuculus_canorus
--------------------------------------------------------MSRKKQRLHDGVLRDLERQHQKKENIRLLEEQIALTKQLVEERDKGLMSKGSQES---------------
>Nothobranchius_furzeri
MNIADESSVVGIWLT---------EVRPNVMSAASKVVLGVSVLLTLGTVAAVHLNQAWDRERLREGVVQDLERVERKKENLRLLEEQRILTRHLEEERRRGEERLQSQDT---------------
>Numida_meleagris
------------------------------MSRSSRAVLVVTALLSAATIAAVHVQQRRERERLHSGVLRDLERQNQKKENIRLLEEQIALTKQLTEERDKMLMEKGSQQS---------------
>Acanthogobius_hasta
------------------------------MSRASRAVLGVSVVLTLSTVAGVHLKQAWDRERLHQGVVRDLERLQRKKENLQLLEEQRELTRILQQDRDQGRDQGQGRDQGQDQGRGQGQGQGRD
>Apteronotus_leptorhynchus
------------------------------MTVVSKVVLGISVVLTIGSVVGVHIKQNWDRERLREGVLRDLERLERKRKNLQDLEDQIQLTKVLVAERERRETQTANSAQGS-------------
>Plecoglossus_altivelis
------------------------------MSKTSKLVLGVSVVLTLSTVAGVHLKQTWDRERLHQGVLRDLERMERKKENLRVLEEQRTLTKELEAERDRRTTALQSPDQPHT------------
>Zonotrichia_albicollis
------------------------------MSRGSRLALAVSALLSAAIVAAVHIQQRRELERLRSGVVRDLERQKQKKENVRLLEEQIALTKRLIEERDKALMEKRSQQP---------------
>Gasterosteus_aculeatus
------------------------------MTTVSKVVLGVSVVLTLSTVAAVHLNQSWDRQRLHQGVVRDLERLERKKENLRLLEEQRILTSQLEEERLRREAERRPQDH---------------
>Ceratotherium_simum_simum
------------------------------MSRSSKVVLGLSVVLTAATVAGVHLKQRQDQQRLRDGVIRDIERQNRKKENIRLLGEQIILTEQLEAEREKMVLAKGSQKT---------------
>Papio_hamadryas
------------------------------MSRTSKVVLGLSVLLTAATLAGVHVKQQWDRQRLRDGVIRDIERQIRKKENIRLLGEQIILTEQLEAEREKTLLAKGSQKS---------------
>Mus_musculus
------------------------------MSRSSKAVLGLSVLLTAATVAGVHLKQRQDRQRLRDGVIRDIERQNQKKENIRLLGEQIILTKQLEAEREKMLEKGSQN-T---------------
>Salmo_salar
------------------------------MSTTSKVVLGVSVILTVSTVAGVHLKQNWDQERLHAGVLRDIERLERKRENLRVLEQQRSLTKELKAERDRRESGQQGSDQPQE------------
>Ictalurus_punctatus
------------------------------MSTASKVVLGLSVVLTVGTVAGVHIKQNWDRQRLREGVLRDLERVERKLENQRALEEQIRLTTELVAHREQQEAGETAGSN---------------
>Takydromus_sexlineatus
------------------------------MSAASKAVLAASVAVSGAVVAAVHLQQSRVRERLREGVLRDFERQNRKQENLRLLEQQIALTEQLEMERSKMLMGKESPH----------------
>Trachemys_scripta
----------MHWPLELCGSRSANGPVGEGMSTASKVALGISVLLSAGTVAAVHIQQRRVKERLHEGVIRDLERQSRKLENIRLLQEQIILTTQLEAERDKMLMEKGSQQS---------------
>Takifugu_rubripes
------MLSVSSWLA---------KTNRNVMSRASKVVLGVSVVLTLSTVTAVHLKQTWDRQRLHDGVLRDLERLEKKKENLRLLEEQRVLTRQLEYERQGRET----------------------

HG0074 ENSDARG00000078624 arhgef9b

>Danio_rerio
-------------------------------------------------------------------------------MTLDVQTVAVFTVIAILLLVNVILIFFMGSS
>Maylandia_zebra
-------------------------------------------------------------------------------MALDVQTCAVFTVIAVLVLVNVLLMFILGTR
>Fundulus_heteroclitus
-------------------------------------------------------------------------------MALDVQTCAVFTVIAVLVLVNVLLMFILGTR
>Esox_lucius
-------------------------------MVDAIWTVKFKQPRSRAEQTSNANLPSSRQSTHRR-GSGVLLALQHCTMALDVQTFVVFTVIAVLVLVNVLLMFILGTR
>Cynoglossus_semilaevis
-------------------------------------------------------------------------------MALDVQTCVVFTVIAFLVLVNVLLMFILGTR
>Salmo_salar
MSCNSACRVKTRSVAWEAVFPAVFFFSICSQMVDAVWTVKFKQSRSRPEQSSSARLPSSGQAAHRHRSSRIYLALQHYTMALDVQTFVVFTVIAVLVLVNVLLMFILGTR

HG0074 ENSG00000131089 ARHGEF9

```
>Homo_sapiens
------MDSLTEQ----RLTS-PNLPAPH--------LEHYSVLHCTMTLDVQTVVVFAVIVVLLLVNVILMFFLGTR
>Anser_cygnoides
------MDSLTEQ----RLTS-PNLPAPH--------LEHYSVLHCTMTLDVQTVVVFAVIVVLLLVNVILMFFLGTR
>Nanorana_parkeri
------MDSFAEQ----RLTP-QNLPAPR--------LDHYNVLHCTMTLDVQTVVVFAVIVVLLLVNVILMFFLGTR
>Apteryx_australis_mantelli
------MDSLTEQ----RLTS-PNLPAPH--------LEHYSVLHCTMTLDVQTVVVFAVIVVLLLVNVILMFFLGTR
>Oryzias_latipes
-------------------MARGGGGAGR--WSRQRMGFLNTVLRCNLPPETQKVVVFFIMMLLIIINVVLMFLLAFQ
>Panthera_tigris
------MDSLTEQ----RLTS-PNLPAPH--------LEHYSVLHCTMTLDVQTVVVFAVIVVLLLVNVILMFFLGTR
>Hynobius_retardatus
------MDSFTEQ----RLTT-QNLQAPH--------LEQYSVLHCTMTLDVQTVVVFAVIVVLLLVNVILMFFLGTR
>Orcinus_orca
------MDSLTEQ----RLTS-PNLPAPH--------LEHYSVLHCTMTLDVQTVVVFAVIVVLLLVNVILMFFLGTR
>Calidris_pugnax
------MDSLTEQ----RLTS-PNLPAPH--------LEHYSVLHCTMTLDVQTVVVFAVIVVLLLVNVILMFFLGTR
>Myotis_brandtii
------MDSLTEQ----RLTS-PNLPAPH--------LEHYSVLHCTMTLDVQTVVVFAVIVVLLLVNVILMFFLGTR
>Neolamprologus_brichardi
--------------------MARGGGSGR--WSRQRMGFFNTMLRCNLPPETQKVVVFLIMMLLIIINVVLMFLLAFQ
>Danio_rerio
-----------------------------------------------MTLDVQTVAVFTVIAILLLVNVILIFFMGSS
>Fundulus_heteroclitus
--------------------MARGNGSGR--WSRQKMGFFNTLLRCNLPPDTQKVVVFVIMMLLIIINVVLMFLLAFQ
>Monodelphis_domestica
--------------------MA---KGPD--CSRPQWSILHTFLRCNMNPETQRVVVFVILLFLIIINVVLMFLLAFR
>Macropus_eugenii
-----------------------------------------------MTLDVQTIVVFAVIVVLLLVNVILMFFLGTR
>Esox_lucius
MVDAIWTVKFKQPRSRAEQTSNANLPSSRQSTHRRGSGVLLALQHCTMALDVQTFVVFTVIAVLVLVNVLLMFILGTR
>Aquila_chrysaetos_canadensis
------MDSLTEQ----RLTS-PNLPAPH--------LEHYSVLHCTMTLDVQTVVVFAVIVVLLLVNVILMFFLGTR
>Meleagris_gallopavo
------MDSLTEQ----RLTS-PNLPAPH--------LEHYSVLHCTMTLDVQTVVVFAVIVVLLLVNVILMFFLGTR
>Condylura_cristata
------MDSLTEQ----RLTS-PNLPAPH--------LEHYSVLHCTMTLDVQTVVVFAVIVVLLLVNVILMFFLGTR
>Ochotona_princeps
------MDSLTEQ----RLTS-PNLPAPH--------LEHYSVLHCTMTLDVQTVVVFAVIVVLLLVNVILMFFLGTR
>Elephantulus_edwardii
------MDSLTEQ----RLTS-PNLPAPH--------LEHYSVLHCTMTLDVQTVVVFAVIVVLLLVNVILMFFLGTR
>Pseudopodoces_humilis
------MDSLTEQ----RLTS-PNLPAPH--------LEHYSVLHCTMTLDVQTVVVFAVIVVLLLVNVILMFFLGTR
>Ceratotherium_simum_simum
------MDSLTEQ----RLTS-PNLPAPH--------LEHYSVLHCTMTLDVQTVVVFAVIVVLLLVNVILMFFLGTR
>Cynoglossus_semilaevis
-----------------------------------------------MALDVQTCVVFTVIAFLVLVNVLLMFILGTR
>Cricetulus_griseus
------MDSLTEQ----RLTS-PNLPAPH--------LEHYSVLHCTMTLDVQTVVVFAVIVVLLLVNVILMFFLGTR
>Salmo_salar
--------------------MARGG-GAH--RARQRLGYLNTLLRCNLHPETQKVVVFMIMMLLVIINVVLMFLLAF-
>Lepisosteus_oculatus
------MDSYTEP----RLTSENNLPAQH--------LEYYSLLHCTMTLDVQTVVVFTVIVVLVLVNVILMFFLGSR
>Trichechus_manatus_latirostr
------MDSLTEQ----RLTS-PNLPAPH--------LEHYSVLHCTMTLDVQTVVVFAVIVVLLLVNVILMFFLGTR
>Struthio_camelus_australis
------MDSLTEQ----RLTS-PNLPAPH--------LEHYSVLHCTMTLDVQTVVVFAVIVVLLLVNVILMFFLGTR
>Pelodiscus_sinensis
------MDSLTEQ----RLTS-PNLPAPH--------LEHYSVLHCTMTLDVQTVVVFAVIVVLLLVNVILMFFLGTR
```

HG0075 ENSG00000165699 TSC1

>Homo_sapiens
---------------------------------------------MKDTRLTALETEVPVVARTVW-----------
>Anas_platyrhynchos
---------------------------------------------MKGVWLTALQTEAQRIHRTVW-----------
>Apteryx_australis_mantelli
---------------------------------------------MKGVWLIAVETEARLIRRTAW-----------
>Leptonychotes_weddellii
---------------------------------------------MKDTRLTALETEVPIVPRTVW-----------
>Balaenoptera_acutorostrata_s
MATCQGCAVGKRWKPANPELPLLCLGSHFIVRLVLTDACFFSSVTMKDTRLTALETEVPIVPRTVW-----------
>Calidris_pugnax
---------------------------------------------MKGVWLTALQTEARLVHRTVW-----------
>Miniopterus_natalensis
---------------------------------------------MKDTRLTALETELPIVPRTVW-----------
>Dasypus_novemcinctus
---------------------------------------------MKDTKLTALESEVSFVPRTVW-----------
>Columba_livia
---------------------------------------------MKGVWLTALQTEAQLIHRTVW-----------
>Alligator_mississippiensis
---------------------------------------------MKGARLTALKTEALLVHRTVW-----------
>Sarcophilus_harrisii
---------------------------------------------MKDTRLRALETEVLIVHRTVW-----------
>Galeopterus_variegatus
---------------------------------------------MKDSRLTALETEVPIIPRTAW-----------
>Monodelphis_domestica
---------------------------------------------MKDTRLTALETEVLIVHRTVW-----------
>Haliaeetus_leucocephalus
---------------------------------------------MKGVWLTALQTEAQLVHRTVW-----------
>Coturnix_japonica
---------------------------------------------MKGVWLTALQTEAQGIHRTVW-----------
>Condylura_cristata
---------------------------------------------MRDTRLTALETDVPVVPRTGWSRR--------
>Oryctolagus_cuniculus
---------------------------------------------MRDNKLAALGTEAPVVHRAVYSWPQ-------
>Elephantulus_edwardii
---------------------------------------------MKDTRLTALETEVSVVPRTV------------
>Ornithorhynchus_anatinus
---------------------------------------------MKVSRLTTLETEGPIIYRTVW-----------
>Parus_major
---------------------------------------------MKGVWLTALQTEAQVIHRTVW-----------
>Ceratotherium_simum_simum
---------------------------------------------MKDTRLTALETEVPVVPRTVW-----------
>Manis_javanica
---------------------------------------------MKDTRLTALETEVPVVPRTV------------
>Loxodonta_africana
---------------------------------------------MKDTRLTALETEISIVPRAVL-----------
>Mesocricetus_auratus
---------------------------------------------MKDTRLTALETDVPTVPRTVW-----------
>Tupaia_chinensis
---------------------------------------------MRDTGLTALETEISIVPRTVW-----------
>Trichechus_manatus_latirostr
---------------------------------------------MKDTRLTALETEVSIVPRAVW-----------
>Aptenodytes_forsteri
---------------------------------------------MKGVWLTALQTEARLVHRTVW-----------
>Struthio_camelus_australis
----------------------------------MDVQTRMKFLIMKGVWLIALETEAQLIHRTVW-----------
>Chrysemys_picta
---------------------------------------------MKGARLTALETEVLLVHRTVW-----------
>Orycteropus_afer_afer
---------------------------------------------MKETRLTALETEVSIVPQSSLVVAPMKSLWSL

HG0076 ENSDARG00000040008 neurod6a

```
>Danio_rerio
MGTSSGSWHQCHFS---------------------------------------
>Anser_cygnoides
MGTSSGSWHQCHFLEINEIVQIY--TDLDSRRRCDTQTEKIWKAKNEN-----
>Microhyla_fissipes
MGTSSGSWHQCHFF---------------------------------------
>Canis_lupus_familiaris
MGTSSGSWHQCHFLEINEIVQIY--TDLETPGDDATLSLKRFGRYK--MKTDY
>Hynobius_retardatus
MGTSSGSWHQCHFLEINEIVQLH--TDPDSWRRRGIQTEQTWKPKKWKLIID-
>Lipotes_vexillifer
MGTSSGSWHQCHFLEINEIVQIH--TDLETSGDDATLSLKRCGRSK--MKTDY
>Calidris_pugnax
MGTSSGSWHQCHFLEINEIVQIH--TDLDSRRRCDIQTEKIWKAKNEN-----
>Pteropus_alecto
MGTSSGSWHQCHFLEINEIVQIY--TDLETPGDDATLSLKRFGRSK--MKTDY
>Sarcophilus_harrisii
MGTSSGSWHQCHFLEINEIVQRSIPRSRDSRRRCDIRTEKIWKTKK-------
>Galeopterus_variegatus
MGTSSGSWHQCHFLEINEIVQIY--TDLETPGDDATLSLKRFGRPK--MKTDY
>Macropus_eugenii
MGTSSGSWHQCHFLEINEIVQRSIPRSRDSRRRCDIRTEKIWKTKK-------
>Aquila_chrysaetos_canadensis
MGTSSGSWHQCHFLEINEIVQIY--TDLDSRRRCDIQTEKIWKAKNEN-----
>Gallus_gallus
MGTSSGSWHQCHFLEINEIVQIY--TDLGSRRRCDTQTEKIWKAKNEN-----
>Condylura_cristata
MGTSSGSWHQCHFLEINEIVQIY--TDLETPGDDATLSLKRFGRSK--MKTDY
>Ochotona_princeps
MGTSSGSWHQCHFLEINEIVQIY--TDLETPGDDATFSLKRFGRPK--MKTDY
>Elephantulus_edwardii
MGTSSGSWHQCHFLEINEIVQIY--TDLETRGDDATLSLKRFGRPK--MRTDY
>Ficedula_albicollis
MGTSSGSWHQCHFLEINEIVQIY--TDLDSSRRCDIQTEKIWKAKNEN-----
>Ceratotherium_simum_simum
MGTSSGSWHQCHFLEINEIVQIY--TDLETPGDDATLSLKRFGRSK--MKTDY
>Pongo_abelii
MGTSSGSWHQCHFLEINEIVQIY--TDLETPGDDATLSLKRFGRPK--MKTDY
>Loxodonta_africana
MGTSSGSWHQCHFLEINEIVQIH--TDLETPGDDATLSLKRFGRPK--MRTDY
>Peromyscus_maniculatus_baird
MGTSSGSWHQCHFLEINEIVQIH--TDLEAPGDDATLSLKRFGRSK--MKTDY
>Tupaia_chinensis
MGTSSGSWHQCHFLEINEIVQIH--TDLETPGDDATLSLKRFGRPK--MKTDY
>Trichechus_manatus_latirostr
MGTSSGSWHQCHFLEINEIVQIY--TDLETPGDDATLSLKRFGRPK--MRTDY
>Anolis_carolinensis
MGTSSGSWHQCHFLEINEIVQIY--TDLDSRRRCDIQIEKIWKAKK-------
>Struthio_camelus_australis
MGTSSGSWHQCHFLEINEIVQIH--TDLDSRRRCDIQTEKIWKAKNEN-----
>Chrysemys_picta
MGTSSGSWHQCHFLEINEIVQRY--TDLDSRRRCDIQTEKIWKAKNEN-----
>Tinamus_guttatus
MGTSSGSWHQCHFLEINEIVQIH--TDLGSRRRCDIQTEKIWKAKNEN-----
>Calypte_anna
MGTSSGSWHQCHFLEINEIVQIY--TDLDSRRRCDIQTEKIWKAKNEN-----
>Orycteropus_afer_afer
MGTSSGSWHQCHFLEINEIVQIY--TDLETSGDDATLSLKRFGRPK--MRTDY
```

HG0076 ENSG00000164600 NEUROD6

```
>Homo_sapiens
---------------MGTSSGSWHQCHFLEINEIVQIY--TDLETPGDDATLSLKRFG--RPKMKTDY
>Anser_cygnoides
---------------MGTSSGSWHQCHFLEINEIVQI--YTDLDSRRRCDTQTEKIWKAKNEN-----
>Microhyla_fissipes
MHHVVLCPVTSQADGMGTSSGSWHQCHFF---------------------------------------
>Panthera_tigris
---------------MGTSSGSWHQCHFLEINEIVQIY--TDLETPGDDATLSLKRFG--RYKMKTDY
>Hynobius_retardatus
---------------MGTSSGSWHQCHFLEINEIVQL--HTDPDSWRRRGIQTEQTWKPKKWKLIID-
>Orcinus_orca
---------------MGTSSGSWHQCHFLEINEIVQIH--TDLETPGDDATLSLKRCG--RSKMKTDY
>Calidris_pugnax
---------------MGTSSGSWHQCHFLEINEIVQI--HTDLDSRRRCDIQTEKIWKAKNEN-----
>Pteropus_alecto
---------------MGTSSGSWHQCHFLEINEIVQIY--TDLETPGDDATLSLKRFG--RSKMKTDY
>Sarcophilus_harrisii
---------------MGTSSGSWHQCHFLEINEIVQRSIPRSRDSRRRCDIRTEKIWKTKK-------
>Galeopterus_variegatus
---------------MGTSSGSWHQCHFLEINEIVQIY--TDLETPGDDATLSLKRFG--RPKMKTDY
>Macropus_eugenii
---------------MGTSSGSWHQCHFLEINEIVQRSIPRSRDSRRRCDIRTEKIWKTKK-------
>Aquila_chrysaetos_canadensis
---------------MGTSSGSWHQCHFLEINEIVQI--YTDLDSRRRCDIQTEKIWKAKNEN-----
>Gallus_gallus
---------------MGTSSGSWHQCHFLEINEIVQI--YTDLGSRRRCDTQTEKIWKAKNEN-----
>Condylura_cristata
---------------MGTSSGSWHQCHFLEINEIVQIY--TDLETPGDDATLSLKRFG--RSKMKTDY
>Ochotona_princeps
---------------MGTSSGSWHQCHFLEINEIVQIY--TDLETPGDDATFSLKRFG--RPKMKTDY
>Elephantulus_edwardii
---------------MGTSSGSWHQCHFLEINEIVQIY--TDLETRGDDATLSLKRFG--RPKMRTDY
>Scleropages_formosus
---------------MGTSSGSWHQCHCSKREEQGETERK--------DGPGTQRVYGT---------
>Lepidothrix_coronata
---------------MGTSSGSWHQCHFLEINEIVQI--YTDLDSRRRCDIQTEKIWKAKNEN-----
>Ceratotherium_simum_simum
---------------MGTSSGSWHQCHFLEINEIVQIY--TDLETPGDDATLSLKRFG--RSKMKTDY
>Loxodonta_africana
---------------MGTSSGSWHQCHFLEINEIVQIH--TDLETPGDDATLSLKRFG--RPKMRTDY
>Ictidomys_tridecemlineatus
---------------MGTSSGSWHQCHFLEINEIVQIY--TDLEAPGDDATLSLKRFG--RPKMKTDY
>Tupaia_chinensis
---------------MGTSSGSWHQCHFLEINEIVQIH--TDLETPGDDATLSLKRFG--RPKMKTDY
>Trichechus_manatus_latirostr
---------------MGTSSGSWHQCHFLEINEIVQIY--TDLETPGDDATLSLKRFG--RPKMRTDY
>Anolis_carolinensis
---------------MGTSSGSWHQCHFLEINEIVQI--YTDLDSRRRCDIQIEKIWKAKK-------
>Struthio_camelus_australis
---------------MGTSSGSWHQCHFLEINEIVQI--HTDLDSRRRCDIQTEKIWKAKNEN-----
>Chrysemys_picta
---------------MGTSSGSWHQCHFLEINEIVQR--YTDLDSRRRCDIQTEKIWKAKNEN-----
>Tinamus_guttatus
---------------MGTSSGSWHQCHFLEINEIVQI--HTDLGSRRRCDIQTEKIWKAKNEN-----
>Calypte_anna
---------------MGTSSGSWHQCHFLEINEIVQI--YTDLDSRRRCDIQTEKIWKAKNEN-----
>Orycteropus_afer_afer
---------------MGTSSGSWHQCHFLEINEIVQIY--TDLETSGDDATLSLKRFG--RPKMRTDY
```

HG0077.1 ENSGALG00000005074

```
>Gallus_gallus
------------------MIP-QPDPTTKEKKIVHLRLGRHCN
>Chaetura_pelagica
MLQRFDLTSSSSPEAAVAMIP-QPDPTTKDKRIVRLKLGRRCN
>Apteryx_australis_mantelli
------------------MIP-QPDPTTKEKRIVRLKLGRHCN
>Buceros_rhinoceros_silvestri
------------------MIP-QPDPTTKEKRIVRLKLGRRCN
>Caprimulgus_carolinensis
------------------MIP-QPDPTTKEKRIVCLKLGRRCN
>Cariama_cristata
------------------MIP-QPDPTTKEKRIVCLKLGRRCN
>Charadrius_vociferus
------------------MIP-QPDPTTKEKRIVHLKLGRRCN
>Pterocles_gutturalis
------------------MIP-QPDPTTKEKRIVCLKLGRHCN
>Colius_striatus
------------------MTP-QPDPTTKEKRIVRLKLGHRCN
>Columba_livia
------------------MIP-QPDPTTKEKRIVRLKLGRHCN
>Leptosomus_discolor
------------------MIP-QPDPTTKEKRVVRLKLGRRCN
>Alligator_mississippiensis
------------------MIP-QPDPTTSERRASYFEAEQSLN
>Cuculus_canorus
------------------MIP-QPDPTTKEKRIVRLKLGRRCN
>Falco_cherrug
------------------MIP-QPDPTTKEKRIVRLKLGRRCN
>Gavia_stellata
------------------MIP-QPDPTTKEKRIVRLKLGRRCN
>Chlamydotis_macqueenii
------------------MIP-QPDPTTKEKRIVCLKLGRRCN
>Opisthocomus_hoazin
------------------MIP-QPDPTTKEKRIVCLKLGRRCN
>Manacus_vitellinus
------------------MIP-QPDPTTKEKRIVRLKLGRHCN
>Phaethon_lepturus
------------------MIP-QPDPTTKEKRIVHLKLGRRCN
>Picoides_pubescens
------------------MIP-QPDPTTKEKRVVCLKLGRCCN
>Fulmarus_glacialis
------------------MIP-QPDPTTKEKRIVRLKLGRRCN
>Melopsittacus_undulatus
------------------MIP-QPDPTTKEKRIVCLKLGRRCN
>Pygoscelis_adeliae
------------------MIP-QPDPTTKEKRIVHLKLGRRCN
>Tyto_alba
------------------MIP-QPDPTTKEKRIVHLKLGRHCN
>Struthio_camelus_australis
------------------MIP-QPDPTTKEKRIVHLKLGRHCN
>Chrysemys_picta
MFQRFTLSSSSSPEACQGNDSTARSDHQREKRIVHLKLGRHCN
>Tinamus_guttatus
------------------MTP-QPDPSTKEKRIVRLKQGRHCN
>Calypte_anna
------------------MIP-QPDPTTKEKRIVHLKLGHPCN
>Apaloderma_vittatum
------------------MIP-QPDPTTKEKRIVCLKLGHRCN
```

HG0077.2 ENSGALG00000005074

>Gallus_gallus
MPSGGVLLKLGHCNL
>Caprimulgus_carolinensis
MLSGGVVLKTEHCSL
>Calidris_pugnax
MLSGGVVLKTEHCNL
>Pterocles_gutturalis
MLSGGVVLKTEHCSL
>Columba_livia
MLSGGVVLKTEHCNL
>Aquila_chrysaetos_canadensis
MLSGGVVLKTEHCSL
>Manacus_vitellinus
MLSGGVVLKTEHCSL
>Nipponia_nippon
MLSGGVVLKTEHCSL
>Nestor_notabilis
MLSGGVVLKTEHCSL

HG0078.1 ENSGALG00000016633 HMBOX1

>Gallus_gallus
MVNNTEDPQEAL
>Anas_platyrhynchos
MVNNTEDPQEAL
>Chaetura_pelagica
MVNNTEDPQEAL
>Apteryx_australis_mantelli
MVNNTEDPQEAL
>Buceros_rhinoceros_silvestri
MVNNTEDPQEAL
>Cariama_cristata
MVNNTEDPQEAL
>Charadrius_vociferus
MVNNTEDPQEAL
>Pterocles_gutturalis
MVNNTEDPREAL
>Colius_striatus
MVNNTEDLQEAL
>Leptosomus_discolor
MVNNTEDPQEAL
>Alligator_mississippiensis
MVNNTEDPHEAF
>Cuculus_canorus
MVNNTEDPQEAL
>Falco_cherrug
MVNNTEDPQEAL
>Gavia_stellata
MVNNTEDPQEAL
>Chlamydotis_macqueenii
MVNNTEDPQEAL
>Tauraco_erythrolophus
MVNNTEDPQEAL
>Opisthocomus_hoazin
MVNNTEDPQEAL
>Zonotrichia_albicollis
MVNNTEDPQEAL
>Egretta_garzetta
MVNNTEDPQEAL
>Picoides_pubescens
MRNNTEDPQEAL
>Fulmarus_glacialis
MVNNTEDPREAL
>Melopsittacus_undulatus
MVNNTEDPQEAL
>Aptenodytes_forsteri
MVNNTEDPQEAL
>Tyto_alba
MVNNTEDPQEAL
>Struthio_camelus_australis
MVNNTEDPQEAL
>Pelodiscus_sinensis
MVNNAEDPQEAF
>Tinamus_guttatus
MVNNTEDPYEAL
>Calypte_anna
MVNNTEDPQEAL
>Apaloderma_vittatum
MVNNTEDPQEAL

HG0078.2 ENSG00000147421 HMBOX1

>Homo_sapiens
MVDNADHLWKGY
>Felis_catus
MINNADHL----
>Pteropus_alecto
MVNNADHL----
>Dasypus_novemcinctus
MINYANHLWKGY
>Erinaceus_europaeus
MVDNTDHLWKGY
>Oryctolagus_cuniculus
MINHIDHLWKGY
>Equus_asinus
MVNNADHL----
>Loxodonta_africana
MVNIADHLWKGY
>Heterocephalus_glaber
MVNNADHLWKGY
>Trichechus_manatus_latirostr
MVNNADHLWKGY
>Orycteropus_afer_afer
MINNADHLWKGY

HG0079 ENSG00000204406 MBD5

>Homo_sapiens
MSTIFWREKRYSLIGTRKDIEHLEINCLHYTAESHTTLVMFSSLLSRRLHYWNFEVMKTSDEPTCNTLDSDWKIKRKFKECGL---
>Anas_platyrhynchos
--------------------------------------------------------MKETQE---ESLDNERKIKRKQKECGI---
>Xenopus_laevis
---------------------------------------------------------MKSPGKHCLNLSTDWKIRRKQEECDK---
>Odobenus_rosmarus_divergens
--------------------------------------------------------MKTSDEPTYNILDSDWKMKKKFKECGL---
>Ambystoma_mexicanum
------------MV-SSPEAGSSDLPSL-YTEE--PSSSEFVSLETERLCNQDLGIMRIIQEKASEVFVMDWKIKRKLKESCF---
>Orcinus_orca
------------------------------------MLCFY-QFLCRRLHYWNFEDMKTSDEPTCNILDSDWKMKRKFKECGL---
>Calidris_pugnax
--------------------------------------------------------MKETQE---DNLDNERKIKRKQKECGI---
>Eptesicus_fuscus
MSTVFWREKRYSHIGIHKDIEQLEICCLHYAAGSHSTCYVF-KFLSRRLQYWNFEDMNTSDEPTCNILDSDWKMKRKFKECGL---
>Dasypus_novemcinctus
--------------------------------------------------------MKTSNEPTCNALDSDWKIKRKYKECDL---
>Columba_livia
--------------------------------------------------------MKETQE---DNLDHEWKIKRKQKECGI---
>Alligator_sinensis
--------------------------------------------------------MKESQE---ETLDNEWKIKRKQKECGI---
>Galeopterus_variegatus
--------------------------------------------------------MKTSDEPTCNTLDSDWKIKRKFKECGL---
>Monodelphis_domestica
--------------------------------------------------------MNKSEERIYNTLNIEKKI--KLKECGI---
>Falco_cherrug
--------------------------------------------------------MKETQE---DNLDNERKIKRKQKECGI---
>Gallus_gallus
--------------------------------------------------------MKGTQE---ENLDNERKIKRKQKECGI---
>Condylura_cristata
--------------------------------------------------------MKTSEEPTCNILNSVWKMKRKFKECGL---
>Oryctolagus_cuniculus
--------------------------------------------------------MKTSDEPTCNTLDSDWKIKRKFKECGL---
>Manacus_vitellinus
--------------------------------------------------------MKESQE---DNLDSEQKIKSKQKECGMQRV
>Equus_przewalskii
--------------------------------------------------------MKTSDEPACNILNSDWKMKRKLKECGL---
>Manis_javanica
--------------------------------------------------------MKTSDEPTCSILNSDWKMKRKFKECGL---
>Loxodonta_africana
--------------------------------------------------------MKTSEEPACNTLNSDWKIERKFKECGQ---
>Tupaia_chinensis
--------------------------------------------------------MKTSDEPTCNTLDSDWKIK--FKGCGL---
>Trichechus_manatus_latirostr
--------------------------------------------------------MKTSDEPTCNTLDSDWKIERKFKECGQ---
>Aptenodytes_forsteri
--------------------------------------------------------MKETQE---DNLDNERKIKRKQKECGI---
>Python_bivittatus
--------------------------------------------------------------------------MRKQKECGI---
>Struthio_camelus_australis
--------------------------------------------------------MKESQE---ENLDNEWKIKRKQKECGI---
>Chrysemys_picta
--------------------------------------------------------MKESQEDTLSTLDNEWKIKRKQKECGI---
>Tinamus_guttatus
--------------------------------------------------------MKESQE---ENLDNEWKIKRKQKECGI---
>Orycteropus_afer_afer
--------------------------------------------------------MKTSDEPTCNTLDSDWKIERKFKECGQ---

HG0080.1 ENSG00000179603 GRM8

>Homo_sapiens
---------------------------MGPDGSLQKVQPQVVPFLLWQE
>Anser_cygnoides
---------------------MVLYYLLGTDGSLQKLQFQMVNLIL---
>Canis_lupus_familiaris
---------------------------MGPDGSLQKVQPLVVPLLLWQE
>Orcinus_orca
---------------------------MGPDGSLQKVQPQVVPLLLWQE
>Calidris_pugnax
---------------------MILYYLLGTDGSLQKVQLQMVNLIL---
>Myotis_lucifugus
MEIKLNLRGLTVTFQSRKLAEAHRAFLRGPDGSLQKVRPQVVPLLLWQE
>Dasypus_novemcinctus
---------------------------MGPDGSLQKVQPQVVPLLLWRE
>Galeopterus_variegatus
---------------------------MGPDGSLQKEQPQVVPLLLWRE
>Monodelphis_domestica
---------------------------MGPDGSLQKV------------
>Aquila_chrysaetos_canadensis
---------------------MVLYYLLGTDGSLQKVQRQMVNLIL---
>Coturnix_japonica
---------------------MVLYHLLGTDGSLQKVQFQMVNLSL---
>Sorex_araneus
---------------------------MGPDGRLQTVEPQVVLFLLWQE
>Oryctolagus_cuniculus
---------------------------MGPDGSLQKVQPQAVPLLLWQE
>Elephantulus_edwardii
---------------------------MGPNGGLQQTQPQVVLLLLWQE
>Ornithorhynchus_anatinus
---------------------------MGPDDTRWKVRLQTVPLLLQQE
>Lepidothrix_coronata
---------------------MVLYHLLGTDGSLQKVQLQMVSLIL---
>Equus_asinus
---------------------------MGPDGSLQKVQPQVVPLLLWQE
>Manis_javanica
---------------------------MGPDGSLQKVQALVVPLLLRQK
>Loxodonta_africana
---------------------------MGPDGSLQKVQPQVVPLFLWQE
>Mus_musculus
---------------------------MGPDGSLQKVQPQVVPLLLWQE
>Tupaia_chinensis
---------------------------MGPDGSLQKVQAQVVPFLLWQE
>Trichechus_manatus_latirostr
---------------------------MGPDGSLRKVQPQVVPLFLWQE
>Anolis_carolinensis
--------------------MVLYSASLGTDGSLQKGKLQIVHFLL---
>Struthio_camelus_australis
---------------------MVLYYLLGTDGSLQKVQLQMVNLIL---
>Trachemys_scripta
----------------------------------------MVHLLF---
>Tinamus_guttatus
---------------------MVLYYLLGTDGSLQKVQLQMVNFTL---
>Calypte_anna
----------------------------------------MVNLIL---
>Orycteropus_afer_afer
---------------------------MGPDGSLQKVQPQVVPILLWQE

HG0080.1 ENSGALG00000039403

```
>Gallus_gallus
--------------MVLY-HLLGTDGSLQKVQFQMVNLNL---
>Anser_cygnoides
--------------MVLY-YLLGTDGSLQKLQFQMVNLIL---
>Leptonychotes_weddellii
---------------------MGPDGSLQKVQPLVVPLLLWQE
>Orcinus_orca
---------------------MGPDGSLQKVQPQVVPLLLWQE
>Calidris_pugnax
--------------MILY-YLLGTDGSLQKVQLQMVNLIL---
>Dasypus_novemcinctus
---------------------MGPDGSLQKVQPQVVPLLLWRE
>Galeopterus_variegatus
---------------------MGPDGSLQKEQPQVVPLLLWRE
>Aquila_chrysaetos_canadensis
--------------MVLY-YLLGTDGSLQKVQRQMVNLIL---
>Oryctolagus_cuniculus
---------------------MGPDGSLQKVQPQAVPLLLWQE
>Lepidothrix_coronata
--------------MVLY-HLLGTDGSLQKVQLQMVSLIL---
>Equus_asinus
---------------------MGPDGSLQKVQPQVVPLLLWQE
>Aotus_nancymaae
MGIHCDVQIKKTCCCPQSLFPMGPDGSLQKVQPQVVPLLLWQE
>Loxodonta_africana
---------------------MGPDGSLQKVQPQVVPLFLWQE
>Fukomys_damarensis
---------------------MGPDGSLQKVQPQVVPLPLCQQ
>Tupaia_chinensis
---------------------MGPDGSLQKVQAQVVPFLLWQE
>Trichechus_manatus_latirostr
---------------------MGPDGSLRKVQPQVVPLFLWQE
>Anolis_carolinensis
--------------MVLYSASLGTDGSLQKGKLQIVHFLL---
>Struthio_camelus_australis
--------------MVLY-YLLGTDGSLQKVQLQMVNLIL---
>Tinamus_guttatus
--------------MVLY-YLLGTDGSLQKVQLQMVNFTL---
>Calypte_anna
----------------------------------MVNLIL---
>Orycteropus_afer_afer
---------------------MGPDGSLQKVQPQVVPILLWQE
```

HG0080.2 ENSG00000179603 GRM8

```
>Homo_sapiens
----------------------------------------------------MTRNNSATRLISRT---------------
>Odobenus_rosmarus_divergens
----------------------------------------------------MTRNNSATGLTSRT---------------
>Myotis_brandtii
----------------------------------------------------MPRNNSAAGLISRTWIVDLIPTSEPRRLF
>Condylura_cristata
----------------------------------------------------MTRNNSATGLTSRT---------------
>Oryctolagus_cuniculus
----------------------------------------------------MTRNNSATGLTSRT---------------
>Chinchilla_lanigera
----------------------------------------------------MTRNNSATGLTSRT---------------
>Tupaia_chinensis
----------------------------------------------------MTRNNSATGLTSRT---------------
>Trichechus_manatus_latirostr
MGDPRVTGRGQARLGCSDGADRGRILEHSFSDCLPQVTSWTCPGGTRTCPAEMTRNNSATGLTSRT---------------
```

HG0080.3 ENSG00000179603 GRM8

>Homo_sapiens
-------------------------------------------------MEETWRLLGQLSPLSPEISHKCGRCARLTRRRRRRRRR--------------------------------
>Mustela_putorius_furo
-------------------------------------------------MEDTWRRPGQLSPLSPETSHKCGRGACLARRRRRR-----------------------------------
>Erinaceus_europaeus
-------------------------------------------------MEATWGWPAQLCPLGPQISHKCGRGAGQARRRRRRHGSAGFL--TAPSAGSRE-----------------
>Oryctolagus_cuniculus
-------------------------------------------------MEETWRRLGQLSPLNPESSHKCGRGARLARGGGGGGGGADMELRAPQRASSPRPLRGAGNNSATGLTSRT
>Mus_musculus
MVGAKRLVIPGAIPAAAPRAAWGEWPRAEGREAPGVDQSRPPVGVFVIHMEETRRWLGRLSPLSPETSHKCGRGARLSRWPRRRQR---------------------------------

HG0081.1 ENSDARG00000028228 zbtb18

>Danio_rerio
MMSEQLTGCHPPVFSSNAQTGS-------------------------------------------------------------------------------------------------------------
>Anser_cygnoides
MMSEQLTACHPSAFYSNTQTGS-------------------------------------------------------------------------------------------------------------
>Xenopus_laevis
MMSEQLTACHPSAFYSNTQTGS-------------------------------------------------------------------------------------------------------------
>Apteryx_australis_mantelli
MMSEQLTACHPSAFYSNTQTGS-------------------------------------------------------------------------------------------------------------
>Canis_lupus_familiaris
MMSEQLTACHPSAFYSNTQTGS-------------------------------------------------------------------------------------------------------------
>Balaenoptera_acutorostrata_s
MMSEQLTACHPSAFYSNTQTGS-------------------------------------------------------------------------------------------------------------
>Astyanax_mexicanus
MMSEQLTGCHPPVFSSTAQTGS-------------------------------------------------------------------------------------------------------------
>Calidris_pugnax
MMSEQLTACHPSAFYSNTQTGS-------------------------------------------------------------------------------------------------------------
>Miniopterus_natalensis
MMSEQLTACHPSAFYSNTQTGS-------------------------------------------------------------------------------------------------------------
>Dasypus_novemcinctus
MMSEQLTACHPSAFYSNTQTGS----------------SCVR-SCAGGGI--------------------SFLSNIIS-----------------------TLHLSS------------------------
>Columba_livia
MMSEQLTACHPSAFYSNTQTGS-------------------------------------------------------------------------------------------------------------
>Kryptolebias_marmoratus
MMSEQLTGCHPPHLLLHRLHPSPVLYCESQRQPGRVSALCVCVCVCVRVFAAADQRQLEGGGGRRGGGGRWYLVPPPTSPPLCICPSQFEQRGFFLFVCLFSLPLSLSFFFFFAYLPSSSALLFYPLIDSE
>Monodelphis_domestica
MMSEQLTACHPSAFYSNTQTGS-------------------------------------------------------------------------------------------------------------
>Esox_lucius
MMSEQLTGCHPPVFYSNTQTGS-------------------------------------------------------------------------------------------------------------
>Coturnix_japonica
MMSEQLTACHPSAFYSNTQTGS-------------------------------------------------------------------------------------------------------------
>Erinaceus_europaeus
MMSEQLTACHPSAFYSNTQTGS-------------------------------------------------------------------------------------------------------------
>Oryctolagus_cuniculus
MMSEQLTACHPSAFYSNTQTGS-------------------------------------------------------------------------------------------------------------
>Corvus_brachyrhynchos
MMSEQLTACHPSAFYSNTQTGS-------------------------------------------------------------------------------------------------------------
>Ceratotherium_simum_simum
MMSEQLTACHPSAFYSNTQTGS-------------------------------------------------------------------------------------------------------------
>Manis_javanica
MMSEQLTACHPSAFYSNTQTGS-------------------------------------------------------------------------------------------------------------
>Cercocebus_atys
MMSEQLTACHPSAFYSNTQTGS-------------------------------------------------------------------------------------------------------------
>Cricetulus_griseus
MMSEQLTACHPSAFYSNTQTGS-------------------------------------------------------------------------------------------------------------
>Salmo_salar
MMSEQLTGCHPPVFYSNAQTGS-------------------------------------------------------------------------------------------------------------
>Tupaia_chinensis
MMSEQLTACHPSAFYSNTQTGS-------------------------------------------------------------------------------------------------------------
>Ictalurus_punctatus
MMSEQLTGCHPPVFSSAAQTAS-------------------------------------------------------------------------------------------------------------
>Anolis_carolinensis
MMSEQLTACHPSAFYSNTQTGS----------------ECVCRSVCKEGI--------------------SFLSNIIS-----------------------TLHLSLS-----------------------
>Chelonia_mydas
MMSEQLTACHPSAFYSNTQTGS-------------------------------------------------------------------------------------------------------------
>Tinamus_guttatus
MMSEQLTACHPSAFYSNTQTGS-------------------------------------------------------------------------------------------------------------

HG0081.2 ENSDARG00000028228 zbtb18

```
>Danio_rerio
MIVPIVSYFFFPPPLSVVQRVSKMNDGNGAASEP
>Astyanax_mexicanus
-----------------------MNDSNGAASEP
>Esox_lucius
-----------------------MNDGNGEGSEP
```

HG0082 ENSDARG00000062379 slc35a4

>Danio_rerio
--------------------------------MAEDKDPVTKLKDLVHLKDQLEEIQKKVENEVQAGVPQGGSLLASPFLKGFLAGYVVSRLRSSAVLGVALGTLSGIFAAQNYQVPNIEATLRDILNSLKKRPGE
>Acipenser_sinensis
--------------------------------MADDKDALSKLKDLAHLKNQLEDIQKRVEEEVQAGIPQGGSLLASPFLKGFLAGYVVAKLRSSAVMGVLIGTCTGIYAAQSFDVPNIEKTVKDYFNSIKKGPN-
>Amia_calva
--------------------------------MAGDKDPLSKLKDLAQLKDQLEDIQKRVENEVQAGIPQGGSLLASPFLKGFLAGYVVARLRSSAVLGVVVGTCTGIYAAQNFDVPNIEKSVKDYYNSMKKRPN-
>Odorrana_margaretae
MVAISTCSRPQSVSCSPAVWLSVR--LPWLDNMA-DDKGTPKLKDLAFLKDQLEGLQRKLENEVQAGVSQGGSFLSSPFLKGFLAGYIISKLRASAVLGFAVGAFTGIYAAQAYAVPNVEKTIKDYLGSLRKGPD-
>Porichthys_notatus
--------------------------------MANDKDPLKQLKDLTQLRNQLQEIQRQVENELSVGIPQGSTVWGSPFLKGFLAGYVVAKLRSSAVIGVLLGTVTGMYVAQNYQVPIIDKTFKEYMNHVKNAPK-
>Oryzias_melastigma
------------------------------------------------------------------------------------------------------------------------MTIREYMNSLKKGPK-
>Trachinotus_blochii
--------------------------------MADDKDPLKQLKDLTQLRNQLEEIQRRVENEVSVGIPQGGSVLGSPFLKGFLAGYVVAKLRSSAILGVLLGTITGMYAAQNYQVPNIERTVKEYLNNMKKGPK-
>Ursus_maritimus
--------------------------------MADDKDSLPKLKDLAFLKNQLERLQQRVEDEVSSGVSQDGSLMSSPFLKGFLAGYVVAKLRASAVLGFAVGTCTGIYAAQAYAVPNVEKTLRDYLRSLRKGPD-
>Micropterus_floridanus
---------------------------------------------------------------------------------------MVAKLRSSAIAGVLLGTITGMYVAQSYQVPNIERTVRDYLNNMKKGPK-
>Pteropus_vampyrus
-----------MAGALDIRWEGCACATSSTGKMADDKDSLPKLKDLAFLKNQLEHLQQRVEDEVNSGVGQDGSLLSSPFLKGFLAGYVVAKLRASAVLGFAVGTCTGIYAAQTYAVPNVEKTLRDYLRSLRKGPD-
>Maylandia_zebra
--------------------------------MADDKDPLKQLRDLTQLKNQLDEIQRRVEHEVSAGIPQGGSVLGSPFLKGFLAGYVVARLRSSAFLGVLLGTVTGMYAAQNYQVPNIERTFKEYMNNLKKGPK-
>Dasypus_novemcinctus
--------------------------------MADDKDSLPKLKDLAFLKNQLERLQRRVEDEVNSGVGQDGSLLSSPFLKGFLAGYVVAKLRASAVMGFVVGTCTGIYIAQAYAVPNVEKTLRDYLRSLRKGPD-
>Latimeria_chalumnae
--------------------------------MADDKDTLRKLKDLAQLKDQLEDIQKRVEEEVNAGIPQGGSLLASPFLKGFLTGYVVARLRSSALLGVVVGTCTGIYAAQNYKVPNIENTVKDYISSLKKGSN-
>Fundulus_grandis
--------------------------------MADNKDPLRQLKDLTQLKNQLEAIQRRVENEVSSGIPQGSSVLGSPFLKGFLAGYVVAKLRSSAFLGVLLGTVTGIYAAQNYQVPNVERTVKDYMSTLRKGPK-
>Sarcophilus_harrisii
-----------------------------------------------------------------MADDKEGSFLSSPFLKGFLAGYVVAKLRASAVMGFAVGLCTGIYAAQSYAVPNVEKTLQDYLRSLRKGPD-
>Esox_lucius
--------------------------------MADDKDPFKQMKDLNQLKNQLESIQKRVESEFATGIPQGGSVLGSPFLKGFLAGYVVAKLRSSAIIGVVLGTFTGIYAAQNYQVPNIERTLKDYMSLFKKGP--
>Gallus_gallus
--------------------------------MADDKDSLPKLKDLALLKGQLESLQRRVEDEVHAGVGQGGSLLASPFLKGFLAGYLVAKLRFSAVLGFAVGTCTGIYAAQNYAVPDVEKTVRDYLSALRKGGE-
>Acanthis_flammea
--------------------------------MADDKDSLPKLKDLAFLKGQLESLQRRVEDEVQAGVGQDGSLLASPLLKGFLAGYLVAKLRFSAVLGFVAGTCTGIYAAQNYAVPNVEKTIRDYVNSLKKGRD-
>Lateolabrax_japonicus
-----------------------------------------------------------------------------------------------------------MYAAQNYQVPNIERTMKDCMNNLRKGPK-
>Trematomus_bernacchii
--------------------------------MAGDKDPLKQLKDLTKLKSQLEEIQRRVENEVTLGIPQGGSVLGSPFLKGFLAGYAVAKLRSSAFLGLLVGTATGIYAAQNYPVPNIERTVKDYINSLKKGPK-
>Equus_asinus
--------------------------------MADDKDSLPKLKDLAFLKDQLESLQRRVEDEVNSGVGQDGSLLSSPFLKGFLAGYVVAKLRASAVLGFAVGTCTGIYAAQAYAVPNVEKTLRDYLRSLRKGPD-
>Manis_javanica
--------------------------------MADDKDSLPKLKDLAFLKNQLERLQQRVEDEVNSGVSQDGSLLSSPFLKGFLAGYVVAKLRASAVLGFAVGTCTGIYAAQAYAVPNVEKTLRDYLRSLRKGPD-
>Platichthys_stellatus
--------------------------------MADDKDPLKQLRDLTHLKSQLEEIQRRVESEASLGIPQGGSVLGSPLLKGFLAGYVVAKLRSSAILGVLLGTLTGIYAAQSYQVPNIERTVKDYMNTFKKGPK-
>Nomascus_leucogenys
--------------------------------MADDKDSLPKLKDLAFLKNQLERLQRRVEDEVSSGVDQDGSLLSSPFLKGFLAGYVVAKLRASAVLGFAVGTCTGIYAAQAYAVPNVEKTLRDYLQSLRKGPD-
>Marmota_monax
--------------------------------MADDKDSLPKLKDLAFLKNQLERLQQRVEDEVSSGVGQDGSLLSSPFLKGFLAGYVVAKLRASAVLGFAVGTFTGIYAAQAYAVPNVEKTLRDYFRSLRKGPD-
>Salmo_salar
--------------------------------MAGDKDPFKQMKDLNQLKNQLEDIQKRVETEFAAGIPQGGSVLASPFLKGFLAGYVVAKLRSSAIIGVLLGTFTGIYAAQNYQVPNIESTLKDYMSLFRKGPK-
>Trichomycterus_areolatus
--------------------------------MANDKDPLSQVKDLALLKDQLADIQKRVENEVQAGIPQGGSLLASPFLKGFLAGYVVARLRASAVLGVALGTLTGIYAAQNYEVPNIERTIKDYLNSLKKGPGN
>Gekko_japonicus
--------------------------------MADDKDSLPKLKDLAFLKDQLANLQQRVENEVQTGVGQEGSLLASPFLKGFLAGYVVAKLRSSAVLGFLVGTCSGVYVAQTYAIPNVEKTIKEFFSSFRKGPD-

HG0082 ENSG00000176087 SLC35A4

>Homo_sapiens
-------------------MADDKDSLPKLKDLAFLKNQLESLQRRVEDEVNSGVGQDGSLLSSPFLKGFLAGYVVAKLRASAVLGFAVGTCTGIYAAQAYAVPNVEKTLRDYLQLLRKGPD-
>Acipenser_sinensis
-------------------MADDKDALSKLKDLAHLKNQLEDIQKRVEEEVQAGIPQGGSLLASPFLKGFLAGYVVAKLRSSAVMGVLIGTCTGIYAAQSFDVPNIEKTVKDYFNSIKKGPN-
>Amia_calva
-------------------MAGDKDPLSKLKDLAQLKDQLEDIQKRVENEVQAGIPQGGSLLASPFLKGFLAGYVVARLRSSAVLGVVVGTCTGIYAAQNFDVPNIEKSVKDYYNSMKKRPN-
>Pelophylax_nigromaculatus
-------------------MADD-KGSPKLKDLAFLKNQLEGLQRKLENEVQAGVSQEGSFLSSPFLKGFLAGYIVSKLRASAVLGFAVGAFTGIYAAQAYAVPNVEKTIKDYLGSLRKGPD-
>Porichthys_notatus
-------------------MANDKDPLKQLKDLTQLRNQLQEIQRQVENELSVGIPQGSTVWGSPFLKGFLAGYVVAKLRSSAVIGVLLGTVTGMYVAQNYQVPIIDKTFKEYMNHVKNAPK-
>Oryzias_melastigma
-----------------------------------------------------------------------------------------------------------MTIREYMNSLKKGPK-
>Trachinotus_blochii
-------------------MADDKDPLKQLKDLTQLRNQLEEIQRRVENEVSVGIPQGGSVLGSPFLKGFLAGYVVAKLRSSAILGVLLGTITGMYAAQNYQVPNIERTVKEYLNNMKKGPK-
>Ursus_maritimus
-------------------MADDKDSLPKLKDLAFLKNQLERLQQRVEDEVSSGVSQDGSLMSSPFLKGFLAGYVVAKLRASAVLGFAVGTCTGIYAAQAYAVPNVEKTLRDYLRSLRKGPD-
>Micropterus_floridanus
--------------------------------------------------------------------------MVAKLRSSAIAGVLLGTITGMYVAQSYQVPNIERTVRDYLNNMKKGPK-
>Rhinolophus_ferrumequinum
-------------------MADDKDSLPKLKDLAFLKNQLERLQQRVEDEVNSGMGQDGSLLSSPFLKGFLAGYVVAKLRASAVLGFAVGTCTGIYAAQAYAVPNVEKTLRDYLRSLRKGPD-
>Maylandia_zebra
-------------------MADDKDPLKQLRDLTQLKNQLDEIQRRVEHEVSAGIPQGGSVLGSPFLKGFLAGYVVARLRSSAFLGVLLGTVTGMYAAQNYQVPNIERTFKEYMNNLKKGPK-
>Dasypus_novemcinctus
-------------------MADDKDSLPKLKDLAFLKNQLERLQRRVEDEVNSGVGQDGSLLSSPFLKGFLAGYVVAKLRASAVMGFVVGTCTGIYIAQAYAVPNVEKTLRDYLRSLRKGPD-
>Latimeria_chalumnae
-------------------MADDKDTLRKLKDLAQLKDQLEDIQKRVEEEVNAGIPQGGSLLASPFLKGFLTGYVVARLRSSALLGVVVGTCTGIYAAQNYKVPNIENTVKDYISSLKKGSN-
>Pimephales_promelas
------------------------------------------------------------------------------MRSSAILGVALGTLTGIYAVQNYQVPNIEQTVRDFLNSLKKGPRD
>Fundulus_grandis
-------------------MADNKDPLRQLKDLTQLKNQLEAIQRRVENEVSSGIPQGSSVLGSPFLKGFLAGYVVAKLRSSAFLGVLLGTVTGIYAAQNYQVPNVERTVKDYMSTLRKGPK-
>Sarcophilus_harrisii
----------------------------------------------------MADDKEGSFLSSPFLKGFLAGYVVAKLRASAVMGFAVGLCTGIYAAQSYAVPNVEKTLQDYLRSLRKGPD-
>Esox_lucius
-------------------MADDKDPFKQMKDLNQLKNQLESIQKRVESEFATGIPQGGSVLGSPFLKGFLAGYVVAKLRSSAIIGVVLGTFTGIYAAQNYQVPNIERTLKDYMSLFKKGP--
>Gallus_gallus
-------------------MADDKDSLPKLKDLALLKGQLESLQRRVEDEVHAGVGQGGSLLASPFLKGFLAGYLVAKLRFSAVLGFAVGTCTGIYAAQNYAVPDVEKTVRDYLSALRKGGE-
>Acanthis_flammea
-------------------MADDKDSLPKLKDLAFLKGQLESLQRRVEDEVQAGVGQDGSLLASPLLKGFLAGYLVAKLRFSAVLGFVAGTCTGIYAAQNYAVPNVEKTIRDYVNSLKKGRD-
>Lateolabrax_japonicus
----------------------------------------------------------------------------------------------MYAAQNYQVPNIERTMKDCMNNLRKGPK-
>Trematomus_bernacchii
-------------------MAGDKDPLKQLKDLTKLKSQLEEIQRRVENEVTLGIPQGGSVLGSPFLKGFLAGYAVAKLRSSAFLGLLVGTATGIYAAQNYPVPNIERTVKDYINSLKKGPK-
>Equus_asinus
-------------------MADDKDSLPKLKDLAFLKDQLESLQRRVEDEVNSGVGQDGSLLSSPFLKGFLAGYVVAKLRASAVLGFAVGTCTGIYAAQAYAVPNVEKTLRDYLRSLRKGPD-
>Manis_javanica
-------------------MADDKDSLPKLKDLAFLKNQLERLQQRVEDEVNSGVSQDGSLLSSPFLKGFLAGYVVAKLRASAVLGFAVGTCTGIYAAQAYAVPNVEKTLRDYLRSLRKGPD-
>Cynoglossus_semilaevis
--------MSRIPSTPPSKMANDKDPLKQLKDLTQLKNQLEEIQRRVENEISAGIPQGGSVLGSPFLKGFLAGYVVAKLRSSAILGLVIGTCTGIYAAQNYQVPNIEQTVKEYMRNIKKGPK-
>Heterocephalus_glaber
-------------------MADDKDSLPKIKDLAFLKNQLERLQQRVENEVNSGVGQDGSLLSSPFLKGFLAGYVVAKLRASAVLGFAVGTCTGIYAAQAYALPNVEKTLRDYFQSLRKGPD-
>Salmo_salar
MLLSTPGIRVDHRAFNRAKMADDKDPFKQMKNLNQLKNQLEEIQKRVESEFAEGIPQGGSVLASPFLKGFLAGYMVAKLRSSAVIGVVLGTFTGIYAAQNYQVPNIESTLKDYMSLFRKGPK-
>Ictalurus_punctatus
-------------------MANGKDPLTQLKDLAQLKDQLEDIQKRVENEVQIGIPQGGSLLASPFLKGFLAGYVVAKLRASAVLGVALGTLTGIYTAQTYQVPNIETTIKDYLNSLKKGPNN
>Gekko_japonicus
-------------------MADDKDSLPKLKDLAFLKDQLANLQQRVENEVQTGVGQEGSLLASPFLKGFLAGYVVAKLRSSAVLGFLVGTCSGVYVAQTYAIPNVEKTIKEFFSSFRKGPD-

HG0083.1 ENSGALG00000008039 MFSD13A

```
>Gallus_gallus
MDWPRQRCNRCCEKTSHLKWR
>Anser_cygnoides
MDWPQQRCSRCWGKTSHLKWR
>Chaetura_pelagica
MDWLQQRHSRCWEKTSHLKWR
>Apteryx_australis_mantelli
MDWPQQRSSRCWEKTSHLKWR
>Buceros_rhinoceros_silvestri
MDWPRQRCSRCWEKTSHPKWR
>Cariama_cristata
MDWPQQRCSRCWEKTSHLKWR
>Charadrius_vociferus
MDWPQQRCSRCWEKTSHLKWR
>Pterocles_gutturalis
MDWLQQGCSRCWEKTSHLKWR
>Colius_striatus
MDWPQQRCRRCWEKTSHVKWR
>Columba_livia
MDWPQQRCSRCWDETSHLKWR
>Merops_nubicus
MDWPQQRCSRCWEKTSHLKWR
>Cuculus_canorus
MDLPQQRCSRCWEKTSHLKWR
>Falco_cherrug
MDWPQQRCSRCWEKTSHLTWR
>Gavia_stellata
MDWPQKICSRCWEKTSHLKWR
>Eurypyga_helias
MDWPQQRCSRCWEKTSHLKWR
>Tauraco_erythrolophus
MDWSQQRCSRCWEKTFHLKWR
>Opisthocomus_hoazin
MDWPQQRCGRCWEKASHLKWR
>Sturnus_vulgaris
MDWPQQRCSRCWEKTSHLKWR
>Egretta_garzetta
MDWPQQICSRCWEKTSHLKWR
>Picoides_pubescens
MDQLQQRCSWCWEKTSHVKWR
>Fulmarus_glacialis
MDWPQQISSRCWEKTSHVKWR
>Nestor_notabilis
MDWPQKRCRRCWDKTSHLKWR
>Aptenodytes_forsteri
MDWPQQICSRCWEKTSHLKWR
>Tyto_alba
MDWPQQRCSRCWEKTSHLKWR
>Struthio_camelus_australis
MDWPQQRSSRCWEKTSHLKWR
>Chelonia_mydas
MDWPPTKVNRCCGKVSHLKWR
>Calypte_anna
MDWPQQRCSRCWEKTSHLKWR
>Apaloderma_vittatum
MDWPQQRCNRCWEKISHLKWR
```

HG0083.2 ENSG00000138111 MFSD13A

>Homo_sapiens
MGRPPVAMSPPRCSWG
>Ailuropoda_melanoleuca
MARLLLDMSAPRCSWG
>Balaenoptera_acutorostrata_s
MARLPVAMSAPQYSWG
>Dasypus_novemcinctus
MARLPVAMSPPQCSCG
>Ceratotherium_simum_simum
MARLLVAMSAPRCFWG
>Cavia_porcellus
MARPPVISSPPRCSWG

HG0084.1 ENSGALG00000009207 TRIM2

```
>Gallus_gallus
MMFYSFGLLRNCSSIPARILSSALH-
>Anser_cygnoides
MMFCSFGLLRNCSSIPARIWSSALH-
>Chaetura_pelagica
MMVCSFGLLRNCSSIPARILSSALH-
>Apteryx_australis_mantelli
MMFYSFGLLRNCSSVPARILSSALH-
>Buceros_rhinoceros_silvestri
MMVCSFGLLRNCSSSLQGFYPQHCID
>Caprimulgus_carolinensis
MMVCSFGFLRNCSSVPARILSSALH-
>Cariama_cristata
MMVCSFGLLRNCSSVPARILSSALH-
>Calidris_pugnax
MMVYSFGLLRNCSSVPARILSSALH-
>Pterocles_gutturalis
MMVCSFGFLRNYSSVPARILSSALH-
>Colius_striatus
MMVCSFGLLRNCSSVPARILSSALH-
>Merops_nubicus
MMVCSFGLLRNCSSVPARILSSALH-
>Cuculus_canorus
MMVCSFGLLRNCSSVPARILSSARH-
>Aquila_chrysaetos_canadensis
MMVCSFGLLRNCSSVPARILSSALH-
>Gavia_stellata
MMVCSFGLLRNCSSVPARILSSALH-
>Balearica_regulorum_gibberic
MMVCSFGLLRNCSSVPARILSSALH-
>Tauraco_erythrolophus
MMVCSLGLLRNCSSVPARILSSALH-
>Opisthocomus_hoazin
MMVCSFGLLRNCSSVPARILSSALH-
>Lepidothrix_coronata
MMVCSFGLLRNCSSVPARILSSALH-
>Pelecanus_crispus
MMVCSFGLLRNCSSVPARILSSALH-
>Picoides_pubescens
MMVCSFGLLRNCSSVPARILSSALH-
>Fulmarus_glacialis
MMVCSFGLLRNCSSVPARILSSALH-
>Melopsittacus_undulatus
MMVCSFGLLRNCSSVPARILSSALH-
>Tyto_alba
MMVCSFGLLRNCSSVPARILSSALH-
>Struthio_camelus_australis
MMICSFGLLRNCSSVPARILSSALH-
>Chelonia_mydas
-MFCSFGLLRNCSSIPERILYAALL-
>Tinamus_guttatus
MMFCSPGLLRNCSSVPARILSSALY-
>Calypte_anna
MMVCSLGLLWNCSSVPARILSSALH-
>Apaloderma_vittatum
MMVCSFGLLRNCSSVSARILSSALH-
```

HG0084.2 ENSG00000154447 SH3RF1

>Homo_sapiens
MSHFKGCTQL
>Canis_lupus_familiaris
MSHFKGCTEL
>Equus_asinus
MNHFKGCTQL
>Nannospalax_galili
MNHFKGCTQL
>Orycteropus_afer_afer
MSHYKGCTQL

HG0085 ENSG00000107779 BMPR1A

>Homo_sapiens
MHQFNTVLEFMRWKHRSKLFGENQKYSFI------------------------------
>Polypedates_megacephalus
MHQFDTVLELMNWKYG-------------------------------------------
>Chaetura_pelagica
MHFHNITFELMKWKYR-------------------------------------------
>Odobenus_rosmarus_divergens
MHQFNTVLEFMRWKHRSKLFGENWKYSFI------------------------------
>Orcinus_orca
MHQFNTVLEFMRWKHRSKLFGENWKYSFI------------------------------
>Calidris_pugnax
MHFHNTTFELMKWKYR-------------------------------------------
>Rousettus_aegyptiacus
MHQFNTVLEFMRWKHRSKLFGENWKSSFI------------------------------
>Dasypus_novemcinctus
MHQFNTVLEFMRWKHRSKLFGENWKYNFI------------------------------
>Alligator_mississippiensis
MHCYNIILELMKWKYR-------------------------------------------
>Monodelphis_domestica
MCHFNIALELMKWKYRSKQFGENRKSSFIQQTLLRKQETAVEVGIIIKLLQAEFH----
>Falco_cherrug
MHFHNITFELMKWKYR-------------------------------------------
>Gallus_gallus
MRFHNTTFELMKWKYR-------------------------------------------
>Gavia_stellata
MHFHNITFELMKWKYR-------------------------------------------
>Chlamydotis_macqueenii
MHCHSITFELMKWKYR-------------------------------------------
>Erinaceus_europaeus
MHQFNTVLEFMRWKHRSKVFGENWKYSFI------------------------------
>Oryctolagus_cuniculus
MHPFNTVLEFMRWKHRSKPFRENWKCSSVWAHL--------------------------
>Ornithorhynchus_anatinus
MHFFNTVLELMKWKYRSKQFEEIWKYSLARPFSKESGNCSWSSS--------NY-----
>Opisthocomus_hoazin
MHFHNITFELMKWKYR-------------------------------------------
>Parus_major
MHFHNTTFELMKWKYR-------------------------------------------
>Equus_asinus
MHQFNTVLEFMRWKHRSKLFGENWKYSFI------------------------------
>Manis_javanica
MHQFNTVLEFMRWKHRSKLFGENWKHSFI------------------------------
>Picoides_pubescens
MHFHNTTFELMKWKYR-------------------------------------------
>Loxodonta_africana
MHQFNTVLEFMRWKHRSKLFGENWKHSFI------------------------------
>Melopsittacus_undulatus
MHFHNITFELMKWKYR-------------------------------------------
>Marmota_monax
MHQLNTVLEFMRWKHRSKLFGENWKYSFIATSL-RSEEKAIRGGVIVKWLWPSTRNLIE
>Tupaia_chinensis
MHPFNTVLEFMRWKHRSKPRGEDLKYGFI------------------------------
>Trichechus_manatus_latirostr
MHQFNTVLEFMRWKHRSKLFGENWKYSFI------------------------------
>Tinamus_guttatus
MHFHNTTFELMKWKYR-------------------------------------------

HG0085 ENSGALG00000002003 BMPR1A

```
>Gallus_gallus
------MRFHNTTFELMKWKYR---------------------------------------
>Anas_platyrhynchos
MEIDRNMHFHSITFELMKWKYR---------------------------------------
>Chaetura_pelagica
------MHFHNITFELMKWKYR---------------------------------------
>Calidris_pugnax
------MHFHNTTFELMKWKYR---------------------------------------
>Alligator_mississippiensis
------MHCYNIILELMKWKYR---------------------------------------
>Monodelphis_domestica
------MCHFNIALELMKWKYRSKQFGENRKSSFIQQTLLRKQETAVEVGIIIKLLQAEFH
>Falco_cherrug
------MHFHNITFELMKWKYR---------------------------------------
>Gavia_stellata
------MHFHNITFELMKWKYR---------------------------------------
>Balearica_regulorum_gibberic
------MHFHNITFELMKWKYR---------------------------------------
>Ornithorhynchus_anatinus
------MHFFNTVLELMKWKYRSKQFEEIWKYSLARPFSKESGNCSWSSS--------NY-
>Opisthocomus_hoazin
------MHFHNITFELMKWKYR---------------------------------------
>Parus_major
------MHFHNTTFELMKWKYR---------------------------------------
>Picoides_pubescens
------MHFHNTTFELMKWKYR---------------------------------------
>Melopsittacus_undulatus
------MHFHNITFELMKWKYR---------------------------------------
>Struthio_camelus_australis
------MQFHNITFELMKWKYR---------------------------------------
>Chrysemys_picta
------MHCYNIILELMKWKYR---------------------------------------
>Tinamus_guttatus
------MHFHNTTFELMKWKYR---------------------------------------
```

```
HG0086 ENSGALG00000009835 XPOT

>Gallus_gallus
MLVATSELWGGSRHRRDGGVTGKMIVISTLEEVGKPVSRSRKRRV-
>Anser_cygnoides
-----------------------MIVGSPLEEVGKPVSRSKKPIV-
>Rana_clamitans
-----------------------MLVDSAVPERGKPIVRNRELEV-
>Chaetura_pelagica
-----------------------MIVASPLEEVGKPVRSRKQRI--
>Apteryx_australis_mantelli
-----------------------MIVVSPLEEVGKPVRNRKQRV--
>Buceros_rhinoceros_silvestri
-----------------------MIVVSPLEEVGKPVRSRKQRV--
>Caprimulgus_carolinensis
-----------------------MIVVSPLKEVGKPVESRKRRVQQ
>Cariama_cristata
-----------------------MIVVSPLEEVGKPVRSRKQRV--
>Hynobius_chinensis
-----------------------MFVVYAEGDGGEPSKENRKLEL-
>Charadrius_vociferus
-----------------------MIVVSLLEEVGKPVRSRKQRV--
>Colius_striatus
-----------------------MIVVSPVEEVGKPVRSRKQKV--
>Columba_livia
-----------------------MIVGSPLEEVGKPVRSRKQRV--
>Merops_nubicus
-----------------------MIVVSPLEEVGKPVRSRKQRA--
>Cuculus_canorus
-----------------------MIVVSLLEEVGKPVRSRKRSV--
>Haliaeetus_leucocephalus
-----------------------MIVVSSLEEVGKPVRSRKQRV--
>Gavia_stellata
-----------------------MIVVFLLEEVGKPVRSRKQRV--
>Mesitornis_unicolor
-----------------------MIVVSPLEEVGKPVRNRKQRV--
>Tauraco_erythrolophus
-----------------------MVVVSPLEEVGKPVGSRKQRV--
>Opisthocomus_hoazin
-----------------------MIVASPLEEVGKPVTCRKQRVQQ
>Manacus_vitellinus
-----------------------MIVVSPLEEVGKPVRSRKQRV--
>Phaethon_lepturus
-----------------------MVVVSPLEEVGKPVRSRKQRV--
>Fulmarus_glacialis
-----------------------MIVVSPLEEVGKPVRSRKQRVYQ
>Melopsittacus_undulatus
-----------------------MMVVPPLEEVGRPVRSRKQRV--
>Pygoscelis_adeliae
-----------------------MIVVSPLEEVGKPVRSRKQRV--
>Tyto_alba
-----------------------MFVVSPLEEVGKPVRSRKQRV--
>Struthio_camelus_australis
-----------------------MIVVSPLEEVGKAVRNRKQRV--
>Chrysemys_picta
-----------------------MFVVSTIGEGGKPVKNRKQKV--
>Apaloderma_vittatum
-----------------------MIVVSPLEEVGKPVRSRRQKAEQ
```

HG0087 ENSG00000164548 TRA2A

>Homo_sapiens
-------------------------MWKINQDKRRLMKSIKDQKIKIEWGKKLNRRLVR---------
>Xenopus_tropicalis
-----MSDVEGRVNVREEMEEFFPRILKIKQDKRRLMKSFKDQKKSIEWGNFLNRRLGR---------
>Canis_lupus_familiaris
MSDVEENNFEGRVNVREEIEEFFPRMWKINQDKRRLMKSIKDQKIKIEWGKKLNRRLVR---------
>Orcinus_orca
MSDVEENNFEGRVNVREEIEEFFPRMWKINQDKRRLMKSIKDQKIKIEWGKKLNRRLVR---------
>Calidris_pugnax
MSDVEENNFEGRVNVREEIEEFFPRMWKINQDKRRLMKSIKDQKIKIEWGKKLNRRLVR---------
>Myotis_brandtii
MSDVEENNFEGRVNVREEIEEFFPRMWKINQDKRRLMKSIKDQKIKIEWGKKLNRRLVR---------
>Haplochromis_burtoni
------------------------------------MKTYTEQNIELEREEVYRKNGVR---------
>Dasypus_novemcinctus
MSDVEENNFEGRVNVREEIEEFFPRMWKINQDKRRLMKSIKDQKIKIEWGKKLNRRLVR---------
>Latimeria_chalumnae
MSDVEENTFEGRVNVREEIEEFYPRMWKNNQDKRRLMKNVKDQKIKIEWGKKLNRRNVR---------
>Columba_livia
MSDVEENNFEGRVNVREEIEEFFPRMWKINQDKRRLMKSIKDQKIKIEWGKKLNRRLVR---------
>Sarcophilus_harrisii
-------------------------MWKINQDKRRLMKSIKDQKIKIEWGKKLNRRLVR---------
>Aquila_chrysaetos_canadensis
MSDVEENNFEGRVNVREEIEEFFPRMWKINQDKRRLMKSIKDQKIKIEWGKKLNRRLVR---------
>Coturnix_japonica
MSDVEENNFEGRVNVREEIEEFFPRMWKINQDKRRLMKSIKDQKIKIEWGKKLNRRLVR---------
>Condylura_cristata
MSDIEENNFEGRVNVREEIEEFFPRMWKINQDKRRLMKSIKDQKIKIEWGKKLNRRLVR---------
>Ochotona_princeps
MSDVEENNFEGRVNVREEIEEYFPRMWKINQDKRRLMKSIKDQKIKIEWGKKLNRRLVR---------
>Scleropages_formosus
MSDIEDGNFGGRVIGRDGFEEFYPRMWKNIQDKRRLMKTCTDQKI--RWEGKNERKNGQIEALALCHV
>Parus_major
MSDVEENNFEGRVNVREEIEEFFPRMWKINQDKRRLMKSIKDQKIKIEWGKKLNRRLVR---------
>Lateolabrax_japonicus
------------------------------------MKTCTEQNIEVEREEM----------------
>Anoplopoma_fimbria
------------------------------------MKTCTEQNIEVEREEM----------------
>Equus_asinus
MSDVEENNFEGRVNVREEIEEFFPRMWKINQDKRRLMKSIKDQKIKIEWGKKLNRRLVR---------
>Manis_javanica
MSDIEENNFEGRVNVREEIEEFFPRMWKINQDKRRLMKSIKDQKIKIEWGKKLNRRLVR---------
>Cynoglossus_semilaevis
------------------------------------MKTCTEQYIEVQRVENVTEEWCQIEALVV---
>Marmota_marmota_marmota
MSDVEENNFEGRVNVREEIEEFFPRMWKINQDKRRLMKSIKDQKIKIEWGKKLNRRLVR---------
>Trichechus_manatus_latirostr
-------------------------MWKINQDKRRLMKSIKDQKIKIEWGKKLNRRLVR---------
>Pygoscelis_adeliae
MSDVEENNFEGRVNVREEIEEFFPRMWKINQDKRRLMKSIKDQKIKIEWGKKLNRRLVR---------
>Struthio_camelus_australis
-------------------------MWKINQDKRRLMKSIKDQKIKIEWGKKLNRRLVR---------
>Pelodiscus_sinensis
-------------------------MWKINQDKRRLMKSIKDQKIKIEWGKKLNRRLVR---------
>Tinamus_guttatus
-------------------------MWKINQDKRRLMKSIKDQKIKIEWGKKLNRRLVR---------

HG0088 ENSG00000273079 GRIN2B

>Homo_sapiens
---------------------------------------------------------------------MQARRRDWTF----PTCSLP
>Anas_platyrhynchos
---------------------------------------------------------------------MQARRGDWTF----PTCSLP
>Chaetura_pelagica
---------------------------------------------------------------------MQARRGDWTF----PTCSLP
>Apteryx_australis_mantelli
---------------------------------------------------------------------MQARRGDWTF----PTCSLP
>Canis_lupus_familiaris
---------------------------------------------------------------------MPARRGDWTF----PTCSLP
>Orcinus_orca
-------------------------------------------------------MLFLLFILRLSYVDMQARRHDWIY----PTCSLP
>Charadrius_vociferus
---------------------------------------------------------------------MQARRGDWTF----PTCSLP
>Eptesicus_fuscus
---------------------------------------------------------------------MQARRGAWTF----PTCSLP
>Columba_livia
---------------------------------------------------------------------MQARRGDWTF----PTCSLP
>Cuculus_canorus
---------------------------------------------------------------------MQARRGDWTF----PTCSLP
>Haliaeetus_leucocephalus
---------------------------------------------------------------------MQARRGDWTF----PTCSLP
>Coturnix_japonica
---------------------------------------------------------------------MQARRGAWTF----PTCSLP
>Condylura_cristata
---------------------------------------------------------------------MQARREDWTF----STCLLP
>Ochotona_princeps
-------------------------------------------------------MILRSFILCLPYVDMQARRGDCTFPTSFPTCSLP
>Elephantulus_edwardii
-------------------------------------------------------MIFLSFTLRLSYVDMQARRGDWTS----PTCSLP
>Opisthocomus_hoazin
---------------------------------------------------------------------MQARRGDWTF----PTCSLP
>Manacus_vitellinus
---------------------------------------------------------------------MQARRGDWTF----PTCSLP
>Nipponia_nippon
---------------------------------------------------------------------MQARRGDWTF----PTCSLP
>Manis_javanica
---------------------------------------------------------------------MQARRGDWTP----PPCSLP
>Ctenomys_sociabilis
---------------------------------------------------------------------MQARRGDWTF----PTCSLP
>Tupaia_chinensis
-------------------------------------------------------MISRSFILRLSYVDMKARRGDWTS----PTCPLP
>Trichechus_manatus_latirostr
-------------------------------------------------------MIFLSFILRLSYVDMQARRGDWTF----PTCSLP
>Aptenodytes_forsteri
---------------------------------------------------------------------MQARRGDWTF----PTCSLP
>Thamnophis_sirtalis
MISLDPAYFDFPLYWIRFILEGDILLCLCTTSRRKIASVEASCKKAVSMCIWHRPFPICSPNLILLYVDMQARRGDWIF----PTCSLP
>Struthio_camelus_australis
---------------------------------------------------------------------MQARRGDWTF----PTCSLP
>Chrysemys_picta
---------------------------------------------------------------------MQARREVWIV----PTCSLL
>Tinamus_guttatus
---------------------------------------------------------------------MQARRGDWTF----PTCSLP
>Orycteropus_afer_afer
-------------------------------------------------------MIFLSFTLRLSYVDMQARRGDWTF----PTCSLP

HG0089.1 ENSG00000092421 SEMA6A

```
>Homo_sapiens
MTKACEGESAA--RGPERSPR----
>Chaetura_pelagica
MTKDCKGET--GSSFPERSP-----
>Canis_lupus_familiaris
MTKACRGESAA--RGPERSPR----
>Hynobius_retardatus
MTKDCQGESAAGSRFPERSP-----
>Balaenoptera_acutorostrata_s
MTKACEGESAA--RGPERSPR----
>Calidris_pugnax
MTKDCKGES--GSSFPERSP-----
>Miniopterus_natalensis
MTKACEGESAA--RGPERSPR----
>Columba_livia
MTKDCKGES--GSSFPERSP-----
>Cuculus_canorus
MTKDCQGES--GSSFPERSP-----
>Galeopterus_variegatus
MTKACEGENTA--RGPKRAPR----
>Monodelphis_domestica
MTKDCEGESGP--RSPERSPI----
>Falco_cherrug
MTKDCKGES--GSSFPERSP-----
>Coturnix_japonica
MTKDCKGES--GSSFPERSP-----
>Sorex_araneus
MTKACKGESAA--RGPERSPSIMDY
>Oryctolagus_cuniculus
MTKACGGESAA--RGPERAPR----
>Lepidothrix_coronata
MTKDCKGES--GSSFPERSPKM-DY
>Egretta_garzetta
MTKDCKGES--GSSFPERSP-----
>Ceratotherium_simum_simum
MTKACEGESAA--RGPERSPR----
>Manis_javanica
MTKACEAESAA--RGPESSPR----
>Picoides_pubescens
MTKDCKGES--GSSFPERSP-----
>Loxodonta_africana
MTKACEGESAA--RSPERSPSIMDY
>Heterocephalus_glaber
MTKACKAESAA--RGPEKSPR----
>Trichechus_manatus_latirostr
MTKACEGESAA--RGPERSPR----
>Python_bivittatus
MTKDCKGES--GSSFAERSP-----
>Struthio_camelus_australis
MTKDCKGES--GSSFPERSP-----
>Chrysemys_picta
MTKDCKGES--GSSFPERSP-----
>Orycteropus_afer_afer
MTKACKGESEA--RCPEKSPR----
```

HG0089.1 ENSGALG00000002294 SEMA6A

```
>Gallus_gallus
MTKDCKGES--GSSFPERSP-------------------
>Nanorana_parkeri
MTKDCQRVP-TGSAFPERSLSV---DY------------
>Chaetura_pelagica
MTKDCKGET--GSSFPERSP-------------------
>Odobenus_rosmarus_divergens
MTKACKGESAA--CGPERSPR------------------
>Hynobius_retardatus
MTKDCQGESAAGSRFPERSP-------------------
>Lipotes_vexillifer
MTKACEGESAA--RGPERSPR------------------
>Calidris_pugnax
MTKDCKGES--GSSFPERSP-------------------
>Miniopterus_natalensis
MTKACEGESAA--RGPERSPR------------------
>Columba_livia
MTKDCKGES--GSSFPERSP-------------------
>Alligator_mississippiensis
MTKDCTGERGCGV--PRAVPGHGLLNGTLPQLGLSSGCR
>Cuculus_canorus
MTKDCQGES--GSSFPERSP-------------------
>Monodelphis_domestica
MTKDCEGESGP--RSPERSPI------------------
>Falco_cherrug
MTKDCKGES--GSSFPERSP-------------------
>Sorex_araneus
MTKACKGESAA--RGPERSPSI--MDY------------
>Oryctolagus_cuniculus
MTKACGGESAA--RGPERAPR------------------
>Ficedula_albicollis
MTKDCKGES--GSSFPERSPKM---DY------------
>Egretta_garzetta
MTKDCKGES--GSSFPERSP-------------------
>Equus_caballus
MTKACKGESTA--RGPERSPR------------------
>Picoides_pubescens
MTKDCKGES--GSSFPERSP-------------------
>Cebus_capucinus_imitator
MTKACEGESAA--RGPERSPR------------------
>Loxodonta_africana
MTKACEGESAA--RSPERSPSI--MDY------------
>Heterocephalus_glaber
MTKACKAESAA--RGPEKSPR------------------
>Trichechus_manatus_latirostr
MTKACEGESAA--RGPERSPR------------------
>Takydromus_sexlineatus
MTKDCKGES--GSSFAERSP-------------------
>Struthio_camelus_australis
MTKDCKGES--GSSFPERSP-------------------
>Chrysemys_picta
MTKDCKGES--GSSFPERSP-------------------
>Orycteropus_afer_afer
MTKACKGESEA--RCPEKSPR------------------
```

HG0089.2 ENSG00000092421 SEMA6A

>Homo_sapiens
MNTSRLHPRK
>Balaenoptera_acutorostrata_s
MNTARLHPRK
>Rhinolophus_ferrumequinum
MNTERLHPRK
>Galeopterus_variegatus
MNTARLHPRK
>Sorex_araneus
MNTARLHPRK
>Oryctolagus_cuniculus
MNTARLHPRK
>Equus_asinus
MNTARLHPRK
>Manis_javanica
MNTARLHPRK
>Jaculus_jaculus
MNTARLHPRK
>Trichechus_manatus_latirostr
MNTARLHPRK

HG0089.3 ENSGALG00000002294 SEMA6A

>Gallus_gallus
MNKLHLQFRK
>Chaetura_pelagica
MNKLHLQFRK
>Calidris_pugnax
MNKLHLQFRK
>Columba_livia
MNKLHLQFRK
>Falco_cherrug
MNKLHLQFRK
>Egretta_garzetta
MNKLHLQFRK
>Picoides_pubescens
MNKLHLQFRK
>Struthio_camelus_australis
MNKLHLQFRK

HG0090.1 ENSG00000172765 TMCC1

>Homo_sapiens
MKIGMKPSSHSCMRSHC
>Mustela_putorius_furo
----MKPPSHSCMCSYC
>Balaenoptera_acutorostrata_s
----MKPSSQSCMCSYC
>Miniopterus_natalensis
----MKPSSHSCMCSYC
>Dasypus_novemcinctus
----MKPPSYSCMCSHC
>Sorex_araneus
----MKPSSHSCMCSCW
>Ceratotherium_simum_simum
----MKPPSHSCMCSYC
>Loxodonta_africana
MKIGMKPPSHSSMCSHC
>Chinchilla_lanigera
----MKPPSHSCMHSHC
>Tupaia_chinensis
MKIGMKPSSHCCMCSHY
>Trichechus_manatus_latirostr
----MKPPSHSCMCSNC

HG0090.1 ENSGALG00000008487 TMCC1

>Gallus_gallus
-MEISMRSPWHSCMCSHS
>Anser_cygnoides
-MEISMRSPWHSCMCSHS
>Apteryx_australis_mantelli
-MEISMRSPWHNCMCSH-
>Buceros_rhinoceros_silvestri
-MEISMRSPWHSCMCSHS
>Mustela_putorius_furo
-----MKPPSHSCMCSYC
>Calidris_pugnax
-MEISMRSPWHSCMCSHS
>Eptesicus_fuscus
-----MKPPSHSCMCSYY
>Merops_nubicus
-MEISMRSPWHSCMCSHS
>Alligator_mississippiensis
-----MRSPWHCCMRSHS
>Cuculus_canorus
-----MRSPWHSCMCSHS
>Sarcophilus_harrisii
MEEISMRPSSHSCMCRHS
>Aquila_chrysaetos_canadensis
-MEISMRSPWHSCMCSHS
>Mesitornis_unicolor
-MEISMRSPWHSCMCSRS
>Condylura_cristata
-----MKPPSHSCMCSYW
>Tauraco_erythrolophus
-----MRSPWHSCMCSHS
>Opisthocomus_hoazin
-MEISMRSPWHSCMCSHS
>Geospiza_fortis
MMEISMRSPWHSCMCSHS
>Phaethon_lepturus
-MEISMRSPWHSCMCSHS
>Ceratotherium_simum_simum
-----MKPPSHSCMCSYC
>Picoides_pubescens
-MEISMRSPWHSCMCSHS
>Nestor_notabilis
-----MRSPWHSCMYSHS
>Trichechus_manatus_latirostr
-----MKPPSHSCMCSNC
>Aptenodytes_forsteri
-----MRSPWHSCMCSHC
>Takydromus_sexlineatus
-----MRPPWHSCMCIHS
>Tyto_alba
-MEISMRSPWHSCMCSHS
>Struthio_camelus_australis
-MEISMRSPWHSCMCSHS
>Chrysemys_picta
-MEISMRPPWHSCMCSRS

HG0090.2 ENSDARG00000060954 tmcc1b

>Danio_rerio
--------------------------MTSMTLRQTTDACTLSEST----------
>Channa_argus
-----------------------------MTLSQVTDACPL--------------
>Basilichthys_microlepidotus
--------------------------MTSMTLSQATDACPLKRPN----------
>Oryzias_melastigma
--------------------------MTSMTLSQATDACPLKGSD----------
>Trachinotus_blochii
--------------------------MTSMTLSQATDACPLKRPN----------
>Micropterus_floridanus
--------------------------MTSMTLSQATDACALKRPN----------
>Pygocentrus_nattereri
---------MDRAIPELRARGGQAGLMTSMTLSQSADACTLAEST----------
>Maylandia_zebra
--------------------------MTSMTLSQATDACPLKRPN----------
>Austrofundulus_limnaeus
--------------------------MTSMTLSQATDACPLKRPN----------
>Esox_lucius
--------------------------MTSMTLSQSADACTGSK------------
>Gadus_morhua
--------------------------MTSMTLRQTTDACTLV-------------
>Apteronotus_leptorhynchus
---------------------------MSMTLSQSADACTLTEST----------
>Plecoglossus_altivelis
--------------------------MTSMTLSQSSDACTLAVPI----------
>Lateolabrax_japonicus
--------------------------MTSMTLSQATDACPLKRPN----------
>Notothenia_coriiceps
--------------------------MTSMTLSQATDACPLKRPN----------
>Ictalurus_punctatus
MNGHSAQSSQGSPIPELRARGGRAGPMTSMTLSQSTDACTLTESTRVPSPLLPGF
>Takifugu_rubripes
--------------------------MTSMTLSQATDACPHKGH-----------

HG0090.3 ENSGALG00000008487 TMCC1

```
>Gallus_gallus
----------------MLSFVQWRDCCMSAMLKGK
>Anas_platyrhynchos
---------------------------MSAMLKGK
>Apteryx_australis_mantelli
---------------------------MSAMLKDN
>Charadrius_vociferus
-------MQRDELASAILSFVQWRDCCMSAMLKGK
>Merops_nubicus
---------------------------MSAMLKGK
>Cuculus_canorus
---------------------------MSTMLKGK
>Aquila_chrysaetos_canadensis
---------------------------MSAMLKGK
>Tauraco_erythrolophus
---------------------------MSAMLKGK
>Opisthocomus_hoazin
---------------------------MSAMLKGK
>Ficedula_albicollis
MSCARCSCVRVLTAFTPVSSVQWRDCCESAMLKGK
>Nipponia_nippon
---------------------------MSAMLKGK
>Nestor_notabilis
---------------------------MSAMLKGK
>Tyto_alba
---------------------------MSAMLKGK
>Struthio_camelus_australis
---------------------------MPAMLKDA
```

HG0091.1 ENSG00000172209 GPR22

>Homo_sapiens
-----------------------------------MAHYRHIFSLRSFSKRCL---------------------------------------------
>Nanorana_parkeri
-----------------------------------MAQHKHIFSLRSFSGEICSTCFPVGFAYK----------------------------------
>Chaetura_pelagica
-----------------------------------MAQYRHKFSLRSFFEDMLITLFPFYSANK----------------------------------
>Caprimulgus_carolinensis
-----------------------------------MAQYRHKFSLRSFFEDMLITLFLSLLLC-----------------------------------
>Odobenus_rosmarus_divergens
-----------------------------------MAQYRHIFSLRSFLKRCL---------------------------------------------
>Hynobius_retardatus
-----------------------------------MAQHKHIFSLRSFSEEMLMALFPWYSAYK----------------------------------
>Orcinus_orca
MYFQMLIDSTVGRVRDCSSLQAALHVPYISSTAETMAQYRHTFSLRSFSKRCL---------------------------------------------
>Calidris_pugnax
-----------------------------------MAHYRHKFSLRSFLEDMLITLFPFYSANK----------------------------------
>Rousettus_aegyptiacus
-----------------------------------MAQYRHIFSLRSFQRDVYEPLPQVL--------------------------------------
>Pterocles_gutturalis
-----------------------------------MAHYRHKFSLRSFLEDMLITLFPFYSANK----------------------------------
>Colius_striatus
-----------------------------------MAQYRHKFSLRSFFEDMLITIFPFYSANK----------------------------------
>Merops_nubicus
-----------------------------------MAQYRHKFSLRSFLEDMFITLFPFYSANK----------------------------------
>Galeopterus_variegatus
-----------------------------------MAQYRHIFSLRSFLKRCL---------------------------------------------
>Aquila_chrysaetos_canadensis
-----------------------------------MAQYRHKFSLRSFFEDMLITLFPFYSANK----------------------------------
>Coturnix_japonica
-----------------------------------MAHYRHKFSLRSFFEDVLFTLFPFFSANK----------------------------------
>Erinaceus_europaeus
-----------------------------------MAQYRHIFSLRSFRRDVYDPLPPILY-------------------------------------
>Oryctolagus_cuniculus
-----------------------------------MAQHRHIFSLRSFLKRCL---------------------------------------------
>Tauraco_erythrolophus
-----------------------------------MAQYRHKFSLRSFFEDMLITLFKNSANK-----------------------------------
>Zonotrichia_albicollis
-----------------------------------MAQYRHKFSLRSFFEDMLITLFPFYSANKVNKMNPSNLELLHLHREFTKSLVEGMQQLIQFYY
>Phaethon_lepturus
-----------------------------------MAQYRHKFSLRSFFEEMLITLFSFYSANK----------------------------------
>Ceratotherium_simum_simum
-----------------------------------MAQYRHIFSLRSFSKRCL---------------------------------------------
>Manis_javanica
-----------------------------------MAQYRHIFSLRSFPKRCL---------------------------------------------
>Melopsittacus_undulatus
-----------------------------------MAQYRHKFSLRSFFEDMLITLFPFYSANK----------------------------------
>Ictidomys_tridecemlineatus
-----------------------------------MAQYRHIFSLRSFLKRCL---------------------------------------------
>Aptenodytes_forsteri
-----------------------------------MAQYRHKFSLRSFFEDMLITLFPFYSANK----------------------------------
>Calypte_anna
-----------------------------------MAHYRHKFSLRSFFEDMLITLLPFYSANK----------------------------------
>Apaloderma_vittatum
-----------------------------------MAQYRHKFSLRSFFGDTLITLFPFYSANK----------------------------------

HG0091.2 ENSG00000172209 GPR22

```
>Homo_sapiens
--------------MSKYILEATNVMFPFLFSHILENKEILSCTPSKGNIIPII----------------------------
>Canis_lupus_familiaris
--------------MSKYILEARNRMFPFLFSHVLENKETLSYIPSEGNIIPIIEEVSLGRDFLRQNKHKYFVLVFRNPSAN
>Orcinus_orca
--------------MSKYILEARNLMF-LLFSHVLENKETSSHILSKGNNSYHLRKFLSVVTF-------------------
>Pteropus_alecto
--------------MSKYILGARNLMFPFLFSHVLENKKTL-----------------------------------------
>Dasypus_novemcinctus
--------------MAKYIFETRNLMYPFLFSRILENKETL-----------------------------------------
>Galeopterus_variegatus
--------------MSKYILEARNLMFPFLFSHILENKETLSYIPSKGNIISII----------------------------
>Monodelphis_domestica
-------MFSCNKGMFKYISKARNQMSQFLFSLILENKNTQPHSPQKGNIINVT----------------------------
>Oryctolagus_cuniculus
--------------MSKYILEARKLVFPFLFSHILENRDNTIYSLKGKSNSYHLRKFLLVTTF-------------------
>Ornithorhynchus_anatinus
--------------MLKYISEARDRLSGFLFPAILENKNTRPHQARG-NRINIS----------------------------
>Ceratotherium_simum_simum
MRTAIETDRFVTKGCLNIFWKLETLCFLFCYHYVLENKETPSYKPSKGNIILII----------------------------
>Manis_javanica
--------------MSKYILEARNLMFPFLFSNVLGNRHFHIFPQREM----------------------------------
>Heterocephalus_glaber
--------------MSKYILEVRNLMFPFLFSHILENKETLS----------------------------------------
>Chrysemys_picta
--------------MPKYIFEARNWMSLFLFSFILENKNMLP----------------------------------------
```

HG0091.3 ENSG00000172209 GPR22

```
>Homo_sapiens
-------------MYPLLMKTNKIKQQMVFIKRTWPGFII-----
>Odobenus_rosmarus_divergens
-----------MTMYPLLMKTNQIKEQMVFIKRTWTGFII-----
>Lipotes_vexillifer
-------------MYPLLMKTNQIK--------------------
>Pteropus_alecto
-------------MYPLLMKTNQIK--------------------
>Galeopterus_variegatus
MTNGLFHLSFITNLYLLLMKTNQIKEQMVFIKKNMDWIYNIKQVR
>Monodelphis_domestica
--MDIFSLNIMICYVSFLMKISQIK--------------------
>Erinaceus_europaeus
-------------MYLLLMKTSQIKQQMAFIKRT-----------
>Oryctolagus_cuniculus
-------------MYPLLMKTNQIK--------------------
>Ornithorhynchus_anatinus
-------------MYPPLMKINQIKWQMVSLKSTRTGFYSLMQLM
>Ceratotherium_simum_simum
-------------MYPVLMRTNQIKYQMAFIKRTWTGFII-----
>Manis_javanica
-------------MYLLLMKTNQIK--------------------
>Heterocephalus_glaber
-------------MYPLLMKTNQIK--------------------
```

HG0092.1 ENSGALG00000009966 FRS2

>Gallus_gallus
--------------------------------------MISSVTRQHDTLKYDRSPDAATEAGNINGQI
>Anser_cygnoides
--------------MVIKQLLSCSAAVC----LHVLGQMISSVTRQHDTLKYDRTPDAATEAGNINGQI
>Xenopus_laevis
--------------------------------------------MRHVILV------------------
>Apteryx_australis_mantelli
------------MLHGAKCLMLMTLGWK----VTFLDQMITSVTRQHDTLKYDRSPDAAAEAGNINGQI
>Buceros_rhinoceros_silvestri
MI-SSVTRQHDTLSMTEALMLQQRLGILMAESRDHMVF-------------------------------
>Charadrius_vociferus
MI-SSVTRQHDALSMTEALMLQQRLGILMAKSRDHMVF-------------------------------
>Pteropus_alecto
MRGIWFKLQYREFRLDIKNFQKIKYQILKR-------FVMSSNKKKHGILKYD----------------
>Dasypus_novemcinctus
--------------MQLKLFKQIGTKLHLFFSEESSTRLSQQT--KHVILKYD----------------
>Latimeria_menadoensis
-------------------------------------------------MKYERGPDVARGIRNLNGQI
>Colius_striatus
MI-SSMTRQHDTLSMTEALMLQQRLGILMAKSRDLMVF-------------------------------
>Columba_livia
--------------------------------------MISSVTRHHDTLKYDRSPDAAAEPGNINGQI
>Leptosomus_discolor
MI-SSVTRQHDTLSMTEALMLQQRLGILMAKSRDHMVF-------------------------------
>Alligator_mississippiensis
------------MSPFDDC---SDMGACI---ECILDPVVAHVKRQHDILKYDRSPDAATEARNINGQI
>Cuculus_canorus
MI-SSVTRQHDTSSMTEALMLQQRLGILMAKSRDHMVF-------------------------------
>Sarcophilus_harrisii
--------------MQLLLFSHIRRKLHLIFSKEKFARLCKQA--KYGILKYD----------------
>Aquila_chrysaetos_canadensis
MI-SSVTRQHDTLSMTEALMLQQRLGILMAKSRDHMVF-------------------------------
>Gavia_stellata
MI-SSVTRQHDPLSMTEALMLQQRLGILMAKSRDHMVF-------------------------------
>Balearica_regulorum_gibberic
--------------------------------------MISSVTRQHDTLKYDRSPDAAAEAGNINGQI
>Condylura_cristata
--------------MQLWLFNQIRAKLYLFFSEDNSTRISHHTEKSHGILKYD----------------
>Tauraco_erythrolophus
MI-SSVTRQHDTLSMTEALMLQQRLGILMAKCRDHMVF-------------------------------
>Opisthocomus_hoazin
--------------------------------------MISSVTRQHDTLKYDRSPDAAAEAGNINGQI
>Zonotrichia_albicollis
MI-SSVTRQHDTLSMTEALMLQQRLGILMAKSRDQMVF-------------------------------
>Phaethon_lepturus
--------------------------------------MISSVTRQHDTLKYDRSPDAAAEAGNINGQI
>Fulmarus_glacialis
MI-SSVTRQYDTLSMTEALMLQQRLGILMARSRDHMVF-------------------------------
>Aptenodytes_forsteri
MI-SSVTRQHDTLSMTEALMLQQRLGILMAKSRDHMVF-------------------------------
>Pelodiscus_sinensis
--------------------------------------MFVYVRRQHAILKYDRSPDAATEARNINGQI
>Calypte_anna
MI-SSVARQHDTLSMTEALMLQQRLGILMAKSRDPMVF-------------------------------

HG0092.2 ENSG00000166225 FRS2

>Homo_sapiens
------MKQSICHKVRCSCGMSTSLEQNCICFSQKRIPQG--------------------------------
>Panthera_tigris
------MKQSVCHKVRCSCGFSTRLEQNCICFSQKRIPQG--------------------------------
>Orcinus_orca
------MKQSICHKVRCSCGFSTRLGQNCIYFSQKRIPQEKTKMRGIWFKLQYKEFRLDIKSFQKIKYQILK
>Rousettus_aegyptiacus
------MKQSICHKVRCSCGFSTRLEQNCICFSQKRIPQGARQVYCIKQRFEDS------------------
>Dasypus_novemcinctus
------MKQSICHKVRCS------------------------------------------------------
>Monodelphis_domestica
------MKLFICDEVKCSRGFLVISEGNCI------------------------------------------
>Condylura_cristata
------MKQSICHKLRCNCGFSTRLEQNCISFSQKIIPQG--------------------------------
>Oryctolagus_cuniculus
------MKQPICHKVRCSSGFSTRLEQNCICFSQKRTPQG--------------------------------
>Ceratotherium_simum_simum
------MKQSICPKVRCSCGFSTRLEQNCIYFSQKRIPQG--------------------------------
>Manis_javanica
----MQSDETICHKVRCSCGFSTRLDQNCVSFSQRRILQG--------------------------------
>Loxodonta_africana
------MKQTICHKVKCSCGFSTRLEQNCICFSQKRIPQG--------------------------------
>Fukomys_damarensis
------MKQSICHKIRCSCGFSTRLEQNYICFSQKRIPQG--------------------------------
>Tupaia_chinensis
MPGYQVMKQSIWYKVRCSCGYSTRLEQNYICFSQKRILQG--------------------------------

HG0093 ENSG00000169554 ZEB2

```
>Homo_sapiens
MLFLLTISGQNYKQDFAAPCLN-------------
>Anas_platyrhynchos
MLLLLTISGQNYNQDFAESACC-------------
>Chaetura_pelagica
MLFLLTISGQNYKQDLAEPACC-------------
>Apteryx_australis_mantelli
MLFLLTISGQNYKQDFAEPACC-------------
>Odobenus_rosmarus_divergens
MLFLLTISGQNYKQDFADRACVLPKQGAESMRTAI
>Lipotes_vexillifer
MLFLLTISGQNYKQDFADRACVLPKQGAESMRTAI
>Myotis_brandtii
MLFLLTISGQNYKQDFADRACVLPEQGAESMRTAI
>Dasypus_novemcinctus
MLFLLTISGQNYKQDFADRACVLPKQGAESMRTAI
>Columba_livia
MLFLLTISGQNYKQDFAEPACC-------------
>Alligator_sinensis
MLFLLPISGQNYNQDFAESACC-------------
>Galeopterus_variegatus
MLFLLTISGQNYKQDFTDRACVLPKQGAESMRTAI
>Falco_peregrinus
MLFLLTISGQNYKQDFAEPACC-------------
>Coturnix_japonica
MLFLLTISGQNYKQDFAESACC-------------
>Erinaceus_europaeus
MLFLLTISGQNYKQDFTDRACVLPEPGAESMRAAI
>Oryctolagus_cuniculus
MLFLLTISGQNYKQDFADRACVLPKQGAESMRTAI
>Parus_major
MLFLLTISGQNYKQDFAEPACCCCCRRRCC-----
>Egretta_garzetta
MLFLLTISGQNYKQDFAEPACC-------------
>Ceratotherium_simum_simum
MLFLLTISGQNYKQDFADRACVLPKQGAESMRTAI
>Loxodonta_africana
MLFLLTISGQNYKQDFADRACVLPKQGAESMRTAI
>Peromyscus_maniculatus_baird
MLFLLTISGQNYKQDFAARCLN-------------
>Trichechus_manatus_latirostr
MLFLLTISGQNYKQDFADRACVLPKQGAESMRTAI
>Pygoscelis_adeliae
MLFLLTISGQNYKQDFAESACC-------------
>Thamnophis_sirtalis
MLLLLFTAGQNYKQDFAESACC-------------
>Struthio_camelus_australis
MLFLLTISGQNYRQDFAEPACC-------------
>Chelonia_mydas
MLFLLTISGQNYKQHFAGRVCVLLNQGVESMRTAI
>Orycteropus_afer_afer
MLFLLTISGQNYKQDFADRACVLPKQGAESMRTAI
```

HG0093 ENSGALG00000040465 Sip1

```
>Gallus_gallus
MLFLLTISGQNYKQDFAESACC-------------
>Anas_platyrhynchos
MLLLLTISGQNYNQDFAESACC-------------
>Chaetura_pelagica
MLFLLTISGQNYKQDLAEPACC-------------
>Apteryx_australis_mantelli
MLFLLTISGQNYKQDFAEPACC-------------
>Odobenus_rosmarus_divergens
MLFLLTISGQNYKQDFADRACVLPKQGAESMRTAI
>Lipotes_vexillifer
MLFLLTISGQNYKQDFADRACVLPKQGAESMRTAI
>Myotis_brandtii
MLFLLTISGQNYKQDFADRACVLPEQGAESMRTAI
>Dasypus_novemcinctus
MLFLLTISGQNYKQDFADRACVLPKQGAESMRTAI
>Latimeria_chalumnae
MSLLLVIASPN------------------------
>Columba_livia
MLFLLTISGQNYKQDFAEPACC-------------
>Alligator_mississippiensis
MLFLLPISGQNYNQDFAESACC-------------
>Galeopterus_variegatus
MLFLLTISGQNYKQDFTDRACVLPKQGAESMRTAI
>Falco_cherrug
MLFLLTISGQNYKQDFAEPACC-------------
>Condylura_cristata
MLFLLTISGQSYKQDFADRACVLPKQGAESMRTAI
>Oryctolagus_cuniculus
MLFLLTISGQNYKQDFADRACVLPKQGAESMRTAI
>Parus_major
MLFLLTISGQNYKQDFAEPACCCCCRRRCC-----
>Egretta_garzetta
MLFLLTISGQNYKQDFAEPACC-------------
>Ceratotherium_simum_simum
MLFLLTISGQNYKQDFADRACVLPKQGAESMRTAI
>Otolemur_garnettii
MLFLLTISGQNYKQDFADRACVLPKQGAESMRTAI
>Loxodonta_africana
MLFLLTISGQNYKQDFADRACVLPKQGAESMRTAI
>Peromyscus_maniculatus_baird
MLFLLTISGQNYKQDFADRACVLPKQGAESMRTAI
>Trichechus_manatus_latirostr
MLFLLTISGQNYKQDFADRACVLPKQGAESMRTAI
>Pygoscelis_adeliae
MLFLLTISGQNYKQDFAESACC-------------
>Python_bivittatus
MLLLLFTAGQNYKQDFAESACC-------------
>Struthio_camelus_australis
MLFLLTISGQNYRQDFAEPACC-------------
>Chelonia_mydas
MLFLLTISGQNYKQHFAESACC-------------
>Orycteropus_afer_afer
MLFLLTISGQNYKQDFADRACVLPKQGAESMRTAI
```

HG0094.1 ENSG00000112175 BMP5

>Homo_sapiens
---------------------MQQKLLTTVDNWKSEFQLS
>Anser_cygnoides
-----MPRSADK--ILVFCSINAEKTITTVENWKSEFQLS
>Apteryx_australis_mantelli
-----MSRSADK--ILVFCSISAEKTITTVENWKSEFQLS
>Mustela_putorius_furo
---------------------MQQKLLTTVDNWKSKFQLS
>Balaenoptera_acutorostrata_s
---------------------MQQKLLTTVDNWKSEFQLS
>Calidris_pugnax
-----MPRSADK--ILVFCSINAEKTITTVENWKSEFQLS
>Pteropus_alecto
---------------------MQQKLLTTVDNWKSEFQPS
>Columba_livia
-----MPRSADK--ILVFCSINAEKTITTVENWKSEFQLS
>Alligator_mississippiensis
-----MPRSADK--VLFFCSINAEKTITTVENWKSEFQLS
>Monodelphis_domestica
-----MHRSFDQIILAVFCYINAVQTITTVDNWKSEFQLS
>Aquila_chrysaetos_canadensis
-----MPRSADK--ILVFCSINAEKTITTVENWKSEFQLS
>Coturnix_japonica
-----MPRSADK--ILVFCSINAEKTITTVENWKSEFQLS
>Sorex_araneus
---------------------MQQKLLTTVDNWKSEFQLS
>Ochotona_princeps
---------------------MKQKLLTTVDNWRSEFQLS
>Elephantulus_edwardii
---------------------MQQKLLTTVDNWKFEFQLS
>Corvus_brachyrhynchos
-----MPRSADK--ILVFCSINAEKTITTVENWKSEFQLS
>Ceratotherium_simum_simum
---------------------MQQKLLTTVDNWKSEFQLS
>Manis_javanica
---------------------MQQKLLTTVDNWKSEFQLS
>Loxodonta_africana
---------------------MQQKLLTTVDNWKSKFQLS
>Marmota_marmota_marmota
---------------------MQQKLLATVDNWKSEFQLS
>Trichechus_manatus_latirostr
---------------------MQQKLLTTVDNWKSKFQLS
>Aptenodytes_forsteri
-----MPRSADK--ILVFCSINAEKTITTVENWKSEFQLS
>Anolis_carolinensis
MLCTGMPASTNQ--VLVFCSIHVEKTVMTVENWKSEFQLS
>Struthio_camelus_australis
-----MPRSADK--ILVFCSINAEKTITTVENWKSEFQLS
>Pelodiscus_sinensis
-----MPRSADK--ILVFCSINAEKAITTVENWKSEFQLS
>Tinamus_guttatus
-----MSRSADK--ILVFCSINAEKTITTVENWKSEFQLS
>Orycteropus_afer_afer
---------------------MQQKLLTTVDNWKYEFQLS

HG0094.2 ENSG00000112175 BMP5

>Homo_sapiens
MDLQEGFQVNSGKHIYLNSTT----------------------------------------------------------------------------------------
>Ailuropoda_melanoleuca
MDLQEGFQVNSGKHIYLNSTTLSLILH----------------------------------------------------------------------------------
>Pteropus_vampyrus
MDLQEGFQVNTGKHIYLNSTTLSLILH----------------------------------------------------------------------------------
>Dasypus_novemcinctus
MDLQEGFQVNSGKHIYLNSTTLSLILH----------------------------------------------------------------------------------
>Galeopterus_variegatus
MDLQEGFQVNSGKHIYLNSTTLSLILH----------------------------------------------------------------------------------
>Monodelphis_domestica
MDFREGLQVNSGKHIYFTSTILSFILH----------------------------------------------------------------------------------
>Sorex_araneus
MDLKEGFQVNSGKHIYLNSRTLNLVLH----------------------------------------------------------------------------------
>Oryctolagus_cuniculus
MDLQEGFQVNSGEHIYLNSTTLSVILH----------------------------------------------------------------------------------
>Ceratotherium_simum_simum
MDLQEGFQVNSGKHIYLNSTTLSLILH----------------------------------------------------------------------------------
>Manis_javanica
MDLQEGFQGNSGKHIYLNSTTLSLILHQDDTKDVKVITKLPDKYISQHQGADEHRSVIQKSGSVLERTQGLRRTQEQVKITLEIQFIRRSTLLNSNTKGLIIINSYRNA
>Loxodonta_africana
MDLQEGFQVNSGKHIYLNSRTLSLILH----------------------------------------------------------------------------------
>Ictidomys_tridecemlineatus
MDLQEGFQVNSGKHIYLNSTTLSLILH----------------------------------------------------------------------------------
>Trichechus_manatus_latirostr
MDLQEGFQINSGKHIYLNSRTLSLISH----------------------------------------------------------------------------------
>Chelonia_mydas
MDLQKFFQVNSGKHMN---------------------------------------------------------------------------------------------
>Orycteropus_afer_afer
MDLQEGFQVNSGKHIYLNSTTLSLTLH----------------------------------------------------------------------------------

HG0095 ENSDARG00000013708 usp9

```
>Danio_rerio
MEERSSLILGYAMVSATYPVPQVVLSLSEVCRFSVA-----PEE----KLISSGVPAT
>Amia_calva
------------MVSAKHPVLQVVLSYTDVCWFSVI-----VEE----AIISSLTTTT
>Channa_argus
------------MVSAAHPDLQVLVSLSCVFG--FSSRCSTAEVA----VSSLLACAP
>Microhyla_fissipes
------------MVSAKRSVFEVELCKADICRLAF------LEG----TIISSLVTAT
>Menidia_menidia
------------MVSAAHPDLQVLVSLSRAFG--LSSLCSTAEVA----VISLLVSAP
>Porichthys_notatus
------------MVSAAHPDLQVLVSLSRVLG--LDSLCSTAELA----VISLLASAP
>Oryzias_melastigma
------------MVSATHPDLQVLVSLSCVFG--LSSLCSTAEVA----VISLLASAP
>Ambystoma_mexicanum
------------MVSAKCGVHEVVLCQPHTCWYYVY-----IKG----SPISSLTLAT
>Pygocentrus_nattereri
------------MVSAAHPVLQVVLSLSDVCRFSIS-----PEE----AVIFSCTPAT
>Callorhinchus_milii
------------MVSARQPELEVVLLCTDLNWFSIF-----TEE----AIHSSLATTT
>Haplochromis_burtoni
------------MVSAAHPDLQVLVSLSRVFR--LSSLCSTAEVA----VISLLASAP
>Dasypus_novemcinctus
------------MVSARWFILDLELFKTDICWYFVF-----YKW----TIISFFKTTT
>Alosa_alosa
------------MVSAIPPVLQVVLSLSRACRLSVS-----LEE----AVISPRAPAT
>Latimeria_chalumnae
------------MLSAKQSVLEVVLLYTDTCWFAIF-----TEG----VIISSLKTTT
>Poeciliopsis_prolifica
------------MVSAVHPDLQVLATLSRVFR--LSFLRCTAEVA----VISLLASAP
>Sarcophilus_harrisii
------------MVSARWFVLNLELFKTDICWYFIF-----YKR----TLISPLTTTT
>Esox_lucius
------------MVSAARPDLQVVLSLSLAPRPQCSSLRSVSEVAEAPPVISLLASAP
>Gadus_morhua
------------MVSAAHPDLQVLVSLSCVFG--LVYLCSTAEAA----VIPLVASAP
>Apteronotus_leptorhynchus
------------MVTATQPKPQVLLSCLLALWIRPA-----ALQ----AVISRLAPTT
>Ornithorhynchus_anatinus
------------MVAATWLALELELFKTDTCRYCIV-----SKR----TIIPSLPTTT
>Scleropages_formosus
------------MVAASRPALQLVLSHTDVCWFSVT-----TDE----ALISSPAIAT
>Trematomus_bernacchii
------------MVSAAHPDLQVLVSLSRAFG--LSSLCSTAEVA----VISLLASAP
>Platichthys_stellatus
------------MVSAAHPDLQVLVSLSRVFG--LSSLCSTAEVA----VISLLASAP
>Macaca_nemestrina
------------MVSARWFILELDLFKTDKCWYFIF-----YKW----TIISFLKTTT
>Oncorhynchus_mykiss
------------MVSAARPDLQVVLSLSLVPRPLLSSLCSSSEVAEAPPVISLVVSAA
>Lepisosteus_oculatus
------------MVSAKHPVLQVVLSSTDACWFSII-----VEE----AIISSLTTTT
>Ictalurus_punctatus
------------MVS------ASALALSAVCYFSIS-----PEE----AVISSRAPVT
```

HG0096 ENSGALG00000014186 MPPED1

```
>Gallus_gallus
-------------------------------MVFSCRCRKIADMIA
>Anas_platyrhynchos
-------------------------------MVFFHQCRKIADMIA
>Chaetura_pelagica
-------------------------------MVFFRGCRKIADMIA
>Apteryx_australis_mantelli
-------------------------------MVFFCRCRKIADMIA
>Buceros_rhinoceros_silvestri
-------------------------------MVFFHRSRKIAEMIA
>Caprimulgus_carolinensis
-------------------------------MVFFCRCRKIADMIA
>Cariama_cristata
-------------------------------MVFFHRCRKIADMIA
>Calidris_pugnax
MLVVSQPLMQEENPRLKGFSEGSVLIRSGDVMVFFRRCRKIADMIA
>Pterocles_gutturalis
-------------------------------MVFFHRCRKIAAMIA
>Colius_striatus
-------------------------------MVFFHRCRKIADMIA
>Columba_livia
-------------------------------MVFFHRCRKIADMIA
>Leptosomus_discolor
-------------------------------MVFFRRCRKIADMIA
>Cuculus_canorus
-------------------------------MVVFRRCRKIADMIA
>Haliaeetus_albicilla
-------------------------------MVFSRRCRKIADMIA
>Gavia_stellata
-------------------------------MVFFHRCRKIADMIA
>Mesitornis_unicolor
-------------------------------MVFFHRCRKIADMIA
>Tauraco_erythrolophus
-------------------------------MVFSRRCRKIADMIA
>Opisthocomus_hoazin
-------------------------------MVFFRRCRKIADMIA
>Acanthisitta_chloris
-------------------------------MVFFHRCRKIADMIA
>Nipponia_nippon
-------------------------------MVFFRRCRKIADMIA
>Fulmarus_glacialis
-------------------------------MVFFHRCRKIADMIA
>Nestor_notabilis
-------------------------------MVFFRRCRKIADMIA
>Aptenodytes_forsteri
-------------------------------MVFFHRCRKIADMIA
>Tyto_alba
-------------------------------MVFFRRCRKIADMIA
>Struthio_camelus_australis
-------------------------------MVFFRRCRKIADMIA
>Chrysemys_picta
-------------------------------MVFFCRCRKIAQMII
>Tinamus_guttatus
-------------------------------MVFFCRYRKIADMIA
```

HG0097 ENSGALG00000037496

>Gallus_gallus
MELQPRAGAADTPSAFLRSSHHERGRFHLVKMAVWDRCNL
>Ailuropoda_melanoleuca
-------------------------------MAVRDRCNL
>Orcinus_orca
-------------------------------MAVRDRCNL
>Calidris_pugnax
-------------------------------MAVWDRCNL
>Rousettus_aegyptiacus
-------------------------------MAVRDRCNL
>Dasypus_novemcinctus
-------------------------------MAVRGRCHL
>Columba_livia
-------------------------------MAVWDRCNL
>Alligator_mississippiensis
-------------------------------MAVGDRCNL
>Sarcophilus_harrisii
-------------------------------MAVRDHCNL
>Galeopterus_variegatus
-------------------------------MAVRDRCNL
>Monodelphis_domestica
-------------------------------MAVRDHCNL
>Falco_peregrinus
-------------------------------MAVWDRCNL
>Erinaceus_europaeus
-------------------------------MAVRDRCNL
>Elephantulus_edwardii
-------------------------------MAVRDRCNL
>Lepidothrix_coronata
-------------------------------MAVWDRCNL
>Nipponia_nippon
-------------------------------MAVWDRCNL
>Ceratotherium_simum_simum
-------------------------------MAVRDRCNL
>Manis_javanica
-------------------------------MAVRDRCNL
>Pongo_abelii
-------------------------------MAVRDRCNL
>Loxodonta_africana
-------------------------------MAVRDRCNL
>Melopsittacus_undulatus
-------------------------------MAVRDRCNL
>Microtus_ochrogaster
-------------------------------MAVCDRCNL
>Trichechus_manatus_latirostr
-------------------------------MAVRDRCNL
>Aptenodytes_forsteri
-------------------------------MAVWDRCNL
>Gekko_japonicus
-------------------------------MAVWDRCNL
>Pelodiscus_sinensis
-------------------------------MAVWDHCNL
>Calypte_anna
-------------------------------MAVWDRYNL

HG0098 ENSG00000198739 LRRTM3

>Homo_sapiens
MQKRNIHEDPIT
>Anser_cygnoides
MQKRNIHEDPIT
>Panthera_tigris
MQKRNIHEDPIT
>Tursiops_truncatus
MQKRNIHEDPIT
>Calidris_pugnax
MQKRNIHEDPIT
>Pteropus_alecto
MQKRNIHEDPIT
>Dasypus_novemcinctus
MQKRNIHEDPIT
>Galeopterus_variegatus
MQKRNIHEDPIT
>Monodelphis_domestica
MQKRNIHEDPIT
>Aquila_chrysaetos_canadensis
MQKHNIHEDPIT
>Coturnix_japonica
MQKRNIHEDPIT
>Condylura_cristata
MQKRNIHEDPIT
>Oryctolagus_cuniculus
MQKRNIHEDPIT
>Elephantulus_edwardii
MQKRNIHEDPIT
>Ornithorhynchus_anatinus
MQKRNIHEDPIT
>Lepidothrix_coronata
MQKRNIHEDPIT
>Ceratotherium_simum_simum
MQKRNIHEDPIT
>Manis_javanica
MQKRNIHEDPIT
>Loxodonta_africana
MQKRNIHEDPIT
>Marmota_marmota_marmota
MQKRNIHEDPIT
>Tupaia_chinensis
MQKRNIHEDPIT
>Trichechus_manatus_latirostr
MQKRNIHEDPIT
>Anolis_carolinensis
MQKHIIHEDPIT
>Struthio_camelus_australis
MQKHNIHEDPIT
>Chrysemys_picta
MQNRNIHEDPIT
>Tinamus_guttatus
MQKRNIHEDPIT
>Orycteropus_afer_afer
MQKRNIHEDPIT

HG0099.1 ENSG00000168575 SLC20A2

>Homo_sapiens
--------------------------------------------MRKT---NGSEKVYRIIGGDITAFPDREEKYGMCFTAD---------
>Anas_platyrhynchos
----------------------------------------------------------------------------MCFTADREQANEQYL
>Mustela_putorius_furo
MFLTAPLLTQLLLCIIFELLSSTAIAGSPLMNRVLFHVDGELVATRRN---GKGYRNQLQLGGDITAFPDWEEKYGMCFTAD---------
>Balaenoptera_acutorostrata_s
----------------------------------------------------------------------------MCFTAD---------
>Miniopterus_natalensis
--------------------------------------------MRENPSFSMSSVEIKEPGGDITAFPDREEKYGMCFTAD---------
>Dasypus_novemcinctus
----------------------------------------------------------------------------MCFTAD---------
>Cuculus_canorus
----------------------------------------------------------------------------MCFTADRRQPNEQYL
>Sarcophilus_harrisii
----------------------------------------------------------------------------MCFTAD---------
>Galeopterus_variegatus
----------------------------------------------------------------------------MCFNAD---------
>Monodelphis_domestica
------------MCQK------------KLIIEQV--QSLDFLQPRKKDACKRGY----------HSVSSLREKYGMCFTAD---------
>Haliaeetus_albicilla
----------------------------------------------------------------------------MCFTADRRQPNEQYL
>Condylura_cristata
----------------------------------------------------------------------------MCFTAD---------
>Oryctolagus_cuniculus
----------------------------------------------------------------------------MHFPAD---------
>Elephantulus_edwardii
----------------------------------------------------------------------------MCFTTDCTQPNELD-
>Ornithorhynchus_anatinus
----------------------------------------------------------------------------MCFTAD---------
>Sturnus_vulgaris
----------------------------------------------------------------------------MCFTADRRQPNEQYL
>Equus_asinus
----------------------------------------------------------------------------MCFTAD---------
>Manis_javanica
----------------------------------------------------------------------------MCFTPD---------
>Loxodonta_africana
----------------------------------------------------------------------------MCFTAD---------
>Cavia_porcellus
----------------------------------------------------------------------------MCFTAD---------
>Tupaia_chinensis
-----------MGCSILELIRPLTG-----------TDSCRHVHPQAP---PSSDVGSCMAGGDITAFPGWEEKYGMCFPAD---------
>Trichechus_manatus_latirostr
----------------------------------------------------------------------------MCFTAD---------
>Struthio_camelus_australis
----------------------------------------------MPSCAGGSFIPGLLVTG-HHSCTSLGAKYGMCFTADRTQPNEQYL
>Chrysemys_picta
-------------------------------------------------------------MTGHCSCTSFEAKYGMCFTAD---------
>Tinamus_guttatus
----------------------------------------------------------------------------MCFTADRMQPNEQYL
>Orycteropus_afer_afer
----------------------------------------------------------------------------MCFTADRTQPNELY-

HG0099.2 ENSG00000144136 SLC20A1

>Homo_sapiens
MNLRPFSSPPWNSAPCF
>Leptonychotes_weddellii
MNLRPFSSPPWNS-PCF
>Tursiops_truncatus
MNLRPFSSPPWNSAPCF
>Condylura_cristata
MNLRPFSSPPWNSAPC-
>Oryctolagus_cuniculus
MNLRPFSSPPWNFAPCF
>Equus_przewalskii
MNLRPFSSQPWNSAPCF
>Manis_javanica
MNRRPCSSKPWNSAPLL
>Rattus_norvegicus
MNLRPFSNPPWNSAPCF
>Tupaia_chinensis
MNLRPFSSPPWNSAPCF
>Trichechus_manatus_latirostr
MNLRPCSSLPWNSAPCF
>Orycteropus_afer_afer
MNLRPCFSRPWNSAPCF

HG0099.3 ENSG00000168575 SLC20A2

>Homo_sapiens
-----------------------------------MMLCIVHQGTPLGRISH-------
>Leptonychotes_weddellii
-----------------------------------MMLCIVQQSTPLGQISH-------
>Eptesicus_fuscus
-----------------------------------MMLCIVHQSTPLGQISH-------
>Galeopterus_variegatus
MPVAKKSLLFLFVSTCKNHLKEDRVIGPNSIISIIVMLCIVH-----------------
>Erinaceus_europaeus
MPVVKKLALFSLLVLAKTTYKKLVVLGLNSVICIIMMLCIVHQNTPLSQISH-------
>Elephantulus_edwardii
MPVVKGAYYSIFVSTCKSNLKEVSAVGLTSIISIIVMLCIVHQSTPLCQISH-------
>Ceratotherium_simum_simum
MPVVKKSLLFLFVSTCKNNLKEVSIIGLKSIISIIMMLCIVHRARPLARFPIKRTPTLS
>Heterocephalus_glaber
-------------------------------MGIIMMLCIVHQGTPLGHISH-------
>Tupaia_chinensis
MPVVKELTL--SVSTCKINLKEVSIIGLHSVISIIVMLCIVHQGTPLGQISH-------
>Orycteropus_afer_afer
-----------------------------------MMLCIVHQSTPLCQISH-------

HG0099.4 ENSDARG00000010641 slc20a1b

>Danio_rerio
MDVRCICLIVQAKTNRNRSSLFRG
>Trachinotus_blochii
MAVHSLCLIAQDPKKSNRSSIITG
>Astyanax_mexicanus
MDVRSVCLITQTKTHCSSTSLFRG
>Clupea_harengus
----MGCLIHQSQTNCSKSSLLRG
>Apteronotus_leptorhynchus
MDVCSVCLIAQTKTHCSRYSLFRG
>Lutjanus_erythropterus
MAVCSLCLIAQHQSNSNRSSIITG
>Sebastes_caurinus
MAVCSICLISEHQRNRNRSSLITG
>Platichthys_stellatus
MAVMSLCLIAQRSKNSNRSSLILG
>Trichomycterus_areolatus
MDVRSVCLIGQTKTHCSSTSLFRG

HG0100.1 ENSDARG00000104148 crebbpb

>Danio_rerio
------------------------------------------------------MRVLGVGLSWGEWRWRSGALLSRV----
>Acipenser_sinensis
------------------------------------------------------MWVLGVELSWGEWRWRGAVATFLRFFVA
>Channa_punctata
------------------------------------------------------MWVLGVGLSWGEWRWRSGALLHLF----
>Menidia_menidia
---------------------------------------------------MIIMWVLGVGLSWGEWRWRSGALLHFF----
>Porichthys_notatus
------------------------------------------------------MWVLGVGLSWVDLRRRSVALLPFF----
>Oryzias_latipes
MKTFDVTQNGYVVEHLAIARLPKCCSVVPDFLNGRFETRPVHGDRCSWGYLMTIMWVLGVGLTWGEWRWRSGALLHLF----
>Trachinotus_blochii
------------------------------------------------------MWVLGVGLSWGEWRWRSGALLHLF----
>Micropterus_floridanus
----------------------------------------------MLGHLTIIMWVLGVGLSWGEWRWRSGALLHLF----
>Pygocentrus_nattereri
------------------------------------------------------MWVLGVGLSWGEWRWRRAALLLLF----
>Maylandia_zebra
------------------------------------------------------MWVLGVGLSWGEWRWRSGALLHLF----
>Alosa_alosa
----------------------------------------------------MIMWVLGVGLSWGEWRWRSGALLLLF----
>Nothobranchius_furzeri
MKKFDVTQNGYFEH-LELRLYPRNAEIHLNIPDRHLKARFLHGDPFISGYLMIIMWVLGVGLSWGEWRWRSGALLHLF----
>Esox_lucius
------------------------------------------------------MWVLGVGLSWGDCQWRCVALLPFF----
>Acanthogobius_hasta
----------------------------------------------------MIMWVLGVGLSWGEWRWRSGALLHLF----
>Apteronotus_leptorhynchus
------------------------------------------------------MMVFGVGLSWGEWQWRSGALLLVL----
>Lutjanus_erythropterus
------------------------------------------------------MWVLGVGLSWGEWRWRSGALLHLF----
>Plecoglossus_altivelis
------------------------------------------------------MWVLGVGLSWGEWRWRSGALLHLF----
>Scleropages_formosus
------------------------------------------------------MWVLGVGLSWGERRWRSGALLLLF----
>Lateolabrax_japonicus
------------------------------------------------------MWVLGVGLSWGEWRWRSGALLHLF----
>Trematomus_bernacchii
------------------------------------------------------MWVLGVGLSWGEWRWRSGALLHLF----
>Cynoglossus_semilaevis
-----------------------------MLSNCRAEVGSLHGEYFIVRYLTIIMWVLGVGLSWGEWRWRSGALLHLF----
>Salvelinus_alpinus
------------------------------------------------------MWVLVVGLSWVEWRWRSGALLHLF----
>Lepisosteus_oculatus
------------------------------------------------------MWVLGVGLSWGEWFWRSGAL--------
>Trichomycterus_areolatus
------------------------------------------------------MSVIGVDLSWVEWQWRSGALLLVF----
>Takifugu_rubripes
MNTFDVTQNDFFTH-LDITFFLGNAAVHLTSLDCHPEVRFLHGEYFILGYLTMIMWVLGVGLSWGEWRWRSGALLHLF----

HG0100.1 ENSDARG00000104609 crebbpa

>Danio_rerio
----------------------------------------------------MIMWVLGVGLSWGEWWWRRGALLLLF----
>Acipenser_sinensis
------------------------------------------------------MWVLGVELSWGEWRWRGAVATFLRFFVA
>Channa_punctata
------------------------------------------------------MWVLGVGLSWGEWRWRSGALLHLF----
>Menidia_menidia
---------------------------------------------------MIIMWVLGVGLSWGEWRWRSGALLHFF----
>Porichthys_notatus
------------------------------------------------------MWVLGVGLSWVDLRRRSVALLPFF----
>Oryzias_latipes
MKTFDVTQNGYVVEHLAIARLPKCCSVVPDFLNGRFETRPVHGDRCSWGYLMTIMWVLGVGLTWGEWRWRSGALLHLF----
>Trachinotus_blochii
------------------------------------------------------MWVLGVGLSWGEWRWRSGALLHLF----
>Micropterus_floridanus
----------------------------------------------MLGHLTIIMWVLGVGLSWGEWRWRSGALLHLF----
>Pygocentrus_nattereri
------------------------------------------------------MWVLGVGLSWGEWRWRRAALLLLF----
>Maylandia_zebra
------------------------------------------------------MWVLGVGLSWGEWRWRSGALLHLF----
>Clupea_harengus
----------------------------------------------------MIMWVLGVGLSWGEWRWRSGALLLLF----
>Latimeria_chalumnae
-----------------------------------------MKCVLTVGFLSAIMWVLGVVLSGGDRWWRRRGALLLLH---
>Nothobranchius_furzeri
MKKFDVTQNGYFEH-LELRLYPRNAEIHLNIPDRHLKARFLHGDPFISGYLMIIMWVLGVGLSWGEWRWRSGALLHLF----
>Esox_lucius
------------------------------------------------------MWVLGVGLSWGDCQWRCVALLPFF----
>Acanthogobius_hasta
----------------------------------------------------MIMWVLGVGLSWGEWRWRSGALLHLF----
>Apteronotus_leptorhynchus
------------------------------------------------------MMVFGVGLSWGEWQWRSGALLLVL----
>Lutjanus_erythropterus
------------------------------------------------------MWVLGVGLSWGEWRWRSGALLHLF----
>Plecoglossus_altivelis
------------------------------------------------------MWVLGVGLSWGEWRWRSGALLHLF----
>Scleropages_formosus
------------------------------------------------------MWVLGVGLSWGERRWRSGALLLLF----
>Lateolabrax_japonicus
------------------------------------------------------MWVLGVGLSWGEWRWRSGALLHLF----
>Sebastes_caurinus
------------------------------------------------------MWVLGVGLSWGEWRWRSGALLHLF----
>Cynoglossus_semilaevis
-----------------------------MLSNCRAEVGSLHGEYFIVRYLTIIMWVLGVGLSWGEWRWRSGALLHLF----
>Oncorhynchus_kisutch
------------------------------------------------------MWVLVVGLSWVEWRWRSGALLHLF----
>Lepisosteus_oculatus
------------------------------------------------------MWVLGVGLSWGEWFWRSGAL--------
>Trichomycterus_areolatus
------------------------------------------------------MSVIGVDLSWVEWQWRSGALLLVF----
>Takifugu_rubripes
MNTFDVTQNDFFTH-LDITFFLGNAAVHLTSLDCHPEVRFLHGEYFILGYLTMIMWVLGVGLSWGEWRWRSGALLHLF----

HG0100.2 ENSDARG00000061108 ep300b

```
>Danio_rerio
---------------------------------------------MAEKTDASSEDKSRENCS--VAEEAT
>Channa_punctata
---------------------------------------------MADRTECSIEDGSWDNFC--VAEEAT
>Micropterus_floridanus
---------------------------------------------MADRTECSTEDGSWDSFC--VAEEAT
>Pygocentrus_nattereri
---------------------------------------------MAVKMDPALEERSRHQVS--VAHEAT
>Pundamilia_nyererei
---------------------------------------------MPGRTEYSPEDRSWDGFC--VAEEAT
>Clupea_harengus
MRRTPERRYRGAFGLHSLSCFPSSLTRYDALAQRNRTELLISPPTMAEKTDTAMEDKSWDNLP--VAEEAT
>Esox_lucius
---------------------------------------------MAEKTDAAMEDRSWDNVS--VTEEAK
>Apteronotus_leptorhynchus
---------------------------------------------MAENTDTAIEDKSRDNVF--VADEAT
>Plecoglossus_altivelis
---------------------------------------------MAEKTDATMEDSSWDNFS--VAEEAT
>Scleropages_formosus
---------------------------------------------MAEKTDAPAEDRSWDNLNFTVAEEAT
>Notothenia_coriiceps
---------------------------------------------MAERTESSTEDGSWDSFC--VAEEAT
>Salmo_salar
---------------------------------------------MAEKTDAAMEDRSWDNFS--AAKEAK
```

HG0101 ENSG00000099250 NRP1

>Homo_sapiens
MARAVAPGRGTSVGKGRGSRS
>Apteryx_australis_mantelli
MVGAVAPAEGTSVGKGRGSLS
>Odobenus_rosmarus_divergens
MARAVAPGRGTSVGKERGNRS
>Orcinus_orca
MARAVAPGRGTSVGKERGNRS
>Calidris_pugnax
MAGAAAPGEGTSVGKGRGSRS
>Rousettus_aegyptiacus
MARAVAPGRGTSVGKERGNRS
>Dasypus_novemcinctus
MARAVAPGRGTSVGKGRGNRS
>Monodelphis_domestica
MARAAALGSGTSVGKERGNRS
>Aquila_chrysaetos_canadensis
MAGAAAPGEGTSVGKGRGSRS
>Coturnix_japonica
MAGAAALGAGTSVGKGRGSRS
>Condylura_cristata
MARAVAPGRGTSVGKERGNRS
>Oryctolagus_cuniculus
MARAVAPGRGTSVGKGRGSRS
>Elephantulus_edwardii
MARAVAPGRGTSVGKGRGNRS
>Ornithorhynchus_anatinus
MARAASSGRGTSVGREIGNRS
>Geospiza_fortis
MARAAAPGEGTSVGKGRGSRS
>Ceratotherium_simum_simum
MARAVAPGRGTSVGKERGNRS
>Manis_javanica
MARAVAP--------------
>Loxodonta_africana
MARAVALGRGTSVGKERGNRS
>Melopsittacus_undulatus
MAGAAAPGEETSVGKGKGNRS
>Microtus_ochrogaster
MARAVAPGRGTSVGKGRGNRS
>Tupaia_chinensis
MARAVAPGRGTSVGKGRGNRS
>Lepisosteus_oculatus
-MATPAPGRGTSLGKERGNQS
>Aptenodytes_forsteri
--MAGAPGEGTSVGKGRGSRS
>Anolis_carolinensis
MAKAVTPGKGTSVGKGRGNWS
>Struthio_camelus_australis
MVGAVAPGEGTSVGKGRGSRS
>Pelodiscus_sinensis
MAGAATPGKGATVGKGRGNQS

HG0102 ENSG00000182263 FIGN

```
>Homo_sapiens
-----------------------------MISNHRHLHLTLASIHQQT----HRIDATLTYRVLSPRVQVK
>Acipenser_sinensis
-----------------------------MISNRRHPHLTRAPIHQQT----HRIDVTLTSEAALPHTQA-
>Anser_cygnoides
MLAPCKAAVLGLKFQHGRGCSQSSLQKLGMISNHRHLHLTRAPIHQQT----HRIDATLTYRVLVPNLQVK
>Apteryx_australis_mantelli
-----------------------------MISNHRHLHLTRAPIHQQT----RRIDATLTYRVLVPNLQVK
>Acinonyx_jubatus
-----------------------------MISNHRHLHLTLASIHQQT----HRIDATLTYRVFAPRVQVK
>Physeter_catodon
-----------------------------MISNHRHLHLTLASIHQQT----HRIDATLTYRVFAPRVQVK
>Calidris_pugnax
-----------------------------MISNHRHLHLTRAPIHQQT----HRIDATLTYRVLVPNLQVK
>Rousettus_aegyptiacus
-----------------------------MISNHRHLHLTLASIHQQT----HRIDATLTYRLFAPRVQVK
>Dasypus_novemcinctus
-----------------------------MISNHRHLHLTRASIHQQT----HRIDATLTYRVLARRVQVK
>Alligator_mississippiensis
MLAPCKAAVLGLKFQHGRGCSQSSLQKLGMISNHRHLHLTRAPIHQQT----HRIDAILTYRVLIPNLQVK
>Galeopterus_variegatus
-----------------------------MISNHRHLHLTLASIHQQT----HRIDATLTYRVLAPRVQVK
>Gallus_gallus
MLAPCKAAVLGLKFQHGRGCSQSSLQKLGMISNHRHLHLTRAPIHQQT----HRIDAALTYRVLEPNLQVK
>Apteronotus_leptorhynchus
-----------------------------MVSNRMHLRLTRTTIHQRT----NRIDVALTSEVILSYTQV-
>Ochotona_princeps
-----------------------------MISNHKHLRLTLASIHQQT----QRIDATLTYRVLAPRVQVK
>Ornithorhynchus_anatinus
-----------------------------MISNHRHLHLTRAPIHQQT----HRIDATLTYRVPGPRIQVK
>Scleropages_formosus
-----------------------------MVSNGKHPRLTRAAVHQKKKIQHSGIDVTLTCEAALSHTQV-
>Manacus_vitellinus
-----------------------------MISNHRHLHLTRAPIHQQT----HRIDATLTYRVLVPNLQVK
>Equus_asinus
-----------------------------MISNHRHLHLTLASIHQQT----HRIDATLTYRVFAPRVQVK
>Manis_javanica
-----------------------------MISNHRHLHLTRASIHQQT----HRIDATLTYRVFAPRVQVK
>Ictidomys_tridecemlineatus
-----------------------------MISNHRHLHLTLASIHQQT----HRIDATLTYRVLAPRVQVK
>Tupaia_chinensis
-----------------------------MISNHRHLHLTLASIHQQT----HRIDATLTYRVLAPRVQVK
>Lepisosteus_oculatus
-----------------------------MISNRRHPRLTLAPIHQQT----YRIDVTLTSEAALPHIQA-
>Trichechus_manatus_latirostr
-----------------------------MISNHRHLHLTRASIHQQT----HRIDATLTYRVLPPRVQVK
>Struthio_camelus_australis
-----------------------------MISNHRHLHLTRAPIHQQT----PRIDATLTYRVLVPNLQVK
>Chrysemys_picta
MLAPCKAAVLGLKFQHGRGCSQSSLQKLGMISNHRHLHLTRAPIHQQT----QRIDATLTYRVLVPNLQVK
>Tinamus_guttatus
-----------------------------MISNHRHLHLTRAPVHQQT----RRIDATLTYRVLLPNLQVK
```

HG0103 ENSG00000165197 VEGFD

```
>Homo_sapiens
-----------------------------------------------------MPFETIFSSCLMSTA-
>Canis_lupus_familiaris
-----------------------------------------------------MPFETVFSSCLMSTAW
>Orcinus_orca
-----------------------------------------------------MPFEPVCSSCLRSTAW
>Rousettus_aegyptiacus
-----------------------------------------------------MPFETVFSSCLMSTSW
>Dasypus_novemcinctus
-----------------------------------------------------MPFDTIFSSCLTSTA-
>Sorex_araneus
-----------------------------------------------------MPFETVFSSCLMSTAW
>Oryctolagus_cuniculus
-----------------------------------------------------MPFETIFSACLMSTA-
>Equus_asinus
-----------------------------------------------------MPFETVFSSCLMSSAW
>Manis_javanica
-----------------------------------------------------MPFETVSSSCLMSTVW
>Loxodonta_africana
-----------------------------------------------------MPFEAIFSICLMSAAC
>Octodon_degus
-----------------------------------------------------MPFETVFSSCLMSTA-
>Tupaia_chinensis
-----------------------------------------------------MPFATIFSSCLMSTV-
>Trichechus_manatus_latirostr
MRTFYIFFFSLSPPPTRLCKNVPCLIEVISLDFDHNYLFFHVRSWGFQLSGEEMPFEAIFSICLMSTAC
```

HG0103 ENSGALG00000016558 VEGFD

```
>Gallus_gallus
----------------MHFKRDASSCMTTTA
>Anser_cygnoides
MTEKLESFLVCIGDREMHFKRDASSCVTTTA
>Chaetura_pelagica
----------------MHFETDTSSCMTTTA
>Apteryx_australis_mantelli
----------------MHFKTATSSCMTTTV
>Buceros_rhinoceros_silvestri
----------------MHFETDASICMTTTA
>Cariama_cristata
----------------MHFETDASSCMTTTA
>Charadrius_vociferus
----------------MHFETDASSCMTTTA
>Pterocles_gutturalis
----------------MHFETDASSCMTTTA
>Colius_striatus
----------------MHFKTDVSSCMTTTA
>Columba_livia
----------------MHFETDTSSYMTTTA
>Merops_nubicus
----------------MHFETDASSCMTTTE
>Alligator_mississippiensis
----------------MHFKTDSSSCTISTA
>Cuculus_canorus
----------------MHFETDASSCVMTTA
>Aquila_chrysaetos_canadensis
----------------MHFETDASSCMTTTA
>Mesitornis_unicolor
----------------MHFKTDASSCMTTTA
>Ornithorhynchus_anatinus
----------------MHFERDSSSCTISTA
>Tauraco_erythrolophus
----------------MHFKTDACSCMTTTA
>Opisthocomus_hoazin
----------------MHFKTDASSCMMTTA
>Lepidothrix_coronata
----------------MHFETDASSCMTTTA
>Egretta_garzetta
----------------MHFETDASSCMTTTA
>Melopsittacus_undulatus
----------------MHFETDASSCMTTSA
>Aptenodytes_forsteri
----------------MHFETDASSCMTTTA
>Struthio_camelus_australis
----------------MHFETDASSCMTITA
>Chelonia_mydas
----------------MHFETEPSSCMISTA
>Apaloderma_vittatum
----------------MHFKTDASSCMTLTA
```

HG0104.1 ENSGALG00000038097 EI24

>Gallus_gallus
-----------------------------------------------------MMREAWVRLFPSPCS
>Anas_platyrhynchos
-----------------------------------------------------MMREAWVRLFPSPCS
>Chaetura_pelagica
------------------------------------------------------MREVWDRLFPSPCS
>Cariama_cristata
------------------------------------------------------MREVWVRLFPSPCS
>Charadrius_vociferus
------------------------------------------------------MREVWVRLFPSPCS
>Colius_striatus
------------------------------------------------------MREVWVLLFPSPCS
>Columba_livia
------------------------------------------------------MREVRVRLFPSPRS
>Merops_nubicus
------------------------------------------------------MREVWGRLFPSPCS
>Sarcophilus_harrisii
------------------------------------------------------MRDFGVLLFPSPCS
>Monodelphis_domestica
--------------------------MGNPR----------PREGKLCVSRLFMMRDFGILLFPSPCS
>Trichosurus_vulpecula
------------------------------------------------------MRDFGILLFPSPCS
>Haliaeetus_leucocephalus
------------------------------------------------------MREVWVRLFPSPCS
>Gavia_stellata
------------------------------------------------------MREVWVRLFPSPCS
>Balearica_regulorum_gibberic
------------------------------------------------------MREVWVRLFPSPCS
>Tauraco_erythrolophus
------------------------------------------------------MREVWAQLFPSPRS
>Sturnus_vulgaris
MVVPSGKPVRAGNPVGNHQGVTELVIPGQPTVSSSAMALCVLSPQAPSVSHLSRMREAWVRLSPSPCS
>Phalacrocorax_carbo
------------------------------------------------------MREVWVRLFPSPCS
>Fulmarus_glacialis
------------------------------------------------------MREVWVRLFPSPCS
>Melopsittacus_undulatus
------------------------------------------------------MREVWVRLFPSPCS
>Aptenodytes_forsteri
---------------------------------MPAAVRCFLSPQAPSVSHLSRMREVWVRLFPSPCS
>Anolis_carolinensis
------------------------------------------------------MRQGWVHFFPSPCS
>Tyto_alba
------------------------------------------------------MREVWVRFFPSPCS
>Chelonia_mydas
-----------------------------------------------------MMTEVWALLFPSPCS
>Calypte_anna
------------------------------------------------------MREVWVGLSPSPCS
>Apaloderma_vittatum
------------------------------------------------------MREVWVRLFPSPCS

HG0104.2 ENSG00000149547 EI24

```
>Homo_sapiens
---------MLVLK-------------QKSSLQSLSKNKSRPRSFMMRDLGTLLSPVCS-
>Leptonychotes_weddellii
---------------------------------------------MMKDFETLLPPLCSW
>Lipotes_vexillifer
---------------------------------------------MMRHLGTLLSPLCS-
>Miniopterus_natalensis
---------------------------------------------MMKEFGMLLYPLCSW
>Condylura_cristata
---------------------------------------------MMKVLGALLSPLCNW
>Oryctolagus_cuniculus
MPSAGSVAPALGAPRRSERRARRSRAAGGGRGTGPSRGGRGAGPFMMRDLGAFLSPLCSR
>Ceratotherium_simum_simum
---------------------------------------------MMKDFGTLLSPPCSW
>Manis_javanica
---------------------------------------------MMKDFGMLLPLLCSW
```

HG0105 ENSG00000135298 ADGRB3

```
>Homo_sapiens
------------------------------------------------MGNGDWDRRGVSPSLLLGRVPVLVS--
>Anser_cygnoides
MWLDLVDPEPKLLFSLGGVCVCRAGVFSPRFLSSAAFFNLGFEPEPGKMGNGASDRRGISPSLLLGRVPVVLVF-
>Chaetura_pelagica
------------------------------------------------MGNGASDRRGISPSLLLGRVPVVLVF-
>Odobenus_rosmarus_divergens
------------------------------------------------MGNGDWDRRGVSPSLLLGRVPVLVS--
>Lipotes_vexillifer
------------------------------------------------MGNGTWDRRGVSPSLLLGRVPVLVS--
>Pteropus_alecto
------------------------------------------------MGNGDWDRRGVSPSLLLGRVPVLVS--
>Dasypus_novemcinctus
------------------------------------------------MGNGDWDRRGVSPSLLLGRVPVLVS--
>Galeopterus_variegatus
------------------------------------------------MGNGDWDRRGVSPSLLLGRVPVLVS--
>Monodelphis_domestica
------------------------------------------------MGNGDSARRGVSPSLLLGRVPVLGS--
>Aquila_chrysaetos_canadensis
------------------------------------------------MGNGASDRRGISPSLLLGRVPVVLVF-
>Coturnix_japonica
------------------------------------------------MGNGASDRRGISPSLVSGRVPVVLEV-
>Condylura_cristata
------------------------------------------------MGNGDWDRRGVSPSLLLGRVPVLVS--
>Oryctolagus_cuniculus
------------------------------------------------MGNGDWERRGVSPSLLLGRVPVLVS--
>Parus_major
------------------------------------------------MGNGASVRRGISPSLLLGRVPVVLVF-
>Nipponia_nippon
------------------------------------------------MGNGASDRRGISPSLLLGRVPVVLVF-
>Equus_caballus
------------------------------------------------MGNGDWDRRGVSPSLLLGRVPVLVS--
>Manis_javanica
------------------------------------------------MGNGAWDRRGVSPSLLPGRVPALVS--
>Loxodonta_africana
------------------------------------------------MGNGDGARRGVSPSLLLGRVAVLVS--
>Ictidomys_tridecemlineatus
------------------------------------------------MGNGDWDRRGVSPSLLLGRVPVLVS--
>Trichechus_manatus_latirostr
------------------------------------------------MGNGDWDRRGVSPSLLLGRVPVLVS--
>Pygoscelis_adeliae
------------------------------------------------MGNGASDRRGISPSLLLGRVPVVLVF-
>Thamnophis_sirtalis
------------------------------------------------MGNDASARRGMSPCLFLGHVLVVLGSS
>Struthio_camelus_australis
MWLDLVDPEPKLLFFLGGVYVWRAGVFSLRFSSSAAFFNLGFEPEPGKMGNGASDRRGISPSLLSGRVPVVLVF-
>Chrysemys_picta
------------------------------------------------MGNGASDRRGISPSLLLGRVPVVLVF-
>Calypte_anna
MWLDLVDPESKLLFFLCGVCVWRAGVFSLRFLSSTAFLNLGFEPEPGKMGNGASDRRGISPSLLLGRVPVVLVF-
```

HG0105 ENSGALG00000016176

```
>Gallus_gallus
MWLDLADPEPELLFFRGAVGVREACGGYLAAFFSLLLPFFFFFFFFLVGFEPEPGKMGNGASDRRGISPSLVSGRVPVVLEV-
>Anser_cygnoides
MWLDLVDPEPKLLFSLGGVCVCRAG------VFS--PRFLSSAAFFNLGFEPEPGKMGNGASDRRGISPSLLLGRVPVVLVF-
>Chaetura_pelagica
--------------------------------------------------------MGNGASDRRGISPSLLLGRVPVVLVF-
>Leptonychotes_weddellii
--------------------------------------------------------MGNGDWDRRGVSPSLLLGRVPVLVS--
>Orcinus_orca
--------------------------------------------------------MGNGAWDRRGVSPSLLLGRVPVLVS--
>Dasypus_novemcinctus
--------------------------------------------------------MGNGDWDRRGVSPSLLLGRVPVLVS--
>Galeopterus_variegatus
--------------------------------------------------------MGNGDWDRRGVSPSLLLGRVPVLVS--
>Monodelphis_domestica
--------------------------------------------------------MGNGDSARRGVSPSLLLGRVPVLGS--
>Aquila_chrysaetos_canadensis
--------------------------------------------------------MGNGASDRRGISPSLLLGRVPVVLVF-
>Condylura_cristata
--------------------------------------------------------MGNGDWDRRGVSPSLLLGRVPVLVS--
>Ochotona_princeps
--------------------------------------------------------MGNGGWERRGISRSLLLGRVPVLVS--
>Lepidothrix_coronata
MWLDLVDPEPKLLFFLGGVCVWRAG------IFS--LRFLSSAAFFNLGFEPEPGKMGNGASVRRGISPSLLLGRVPVVLVF-
>Nipponia_nippon
--------------------------------------------------------MGNGASDRRGISPSLLLGRVPVVLVF-
>Manis_javanica
--------------------------------------------------------MGNGAWDRRGVSPSLLPGRVPALVS--
>Microcebus_murinus
--------------------------------------------------------MGNGDWDRRGVSPSLLLGRVPVLVS--
>Loxodonta_africana
--------------------------------------------------------MGNGDGARRGVSPSLLLGRVAVLVS--
>Ictidomys_tridecemlineatus
--------------------------------------------------------MGNGDWDRRGVSPSLLLGRVPVLVS--
>Trichechus_manatus_latirostr
--------------------------------------------------------MGNGDWDRRGVSPSLLLGRVPVLVS--
>Pygoscelis_adeliae
--------------------------------------------------------MGNGASDRRGISPSLLLGRVPVVLVF-
>Thamnophis_sirtalis
--------------------------------------------------------MGNDASARRGMSPCLFLGHVLVVLGSS
>Struthio_camelus_australis
MWLDLVDPEPKLLFFLGGVYVWRAG------VFS--LRFSSSAAFFNLGFEPEPGKMGNGASDRRGISPSLLSGRVPVVLVF-
>Pelodiscus_sinensis
--------------------------------------------------------MGNGASDRRGISPSLVLGRVPVVLVF-
>Calypte_anna
MWLDLVDPESKLLFFLCGVCVWRAG------VFS--LRFLSSTAFLNLGFEPEPGKMGNGASDRRGISPSLLLGRVPVVLVF-
```

HG0106 ENSG00000147548 NSD3

>Homo_sapiens
----------------------------------------------MEQTEGPMRERES
>Pseudacris_regilla
----------------------------------------------MEQTEGPMRERES
>Canis_lupus_familiaris
----------------------------------------------MEQTEGPMRERES
>Hynobius_retardatus
----------------------------------------------MDQTEGPMRERES
>Balaenoptera_acutorostrata_s
----------------------------------------------MEQSEGPMRERES
>Calidris_pugnax
----------------------------------------------MEQTEGPMRRRES
>Miniopterus_natalensis
----------------------------------------------MEQTEGPMRERES
>Latimeria_chalumnae
----------------------------------------------MEQTEGPMRKRES
>Sarcophilus_harrisii
----------------------------------------------MEQTEGPMRERES
>Galeopterus_variegatus
----------------------------------------------MEQTEGPMRERES
>Aegypius_monachus
MRRGALRAPLGARRKGSHLPARRGRRRPRRRPLSPLGWVDAAGAVRMEQTEGPMRKRES
>Gallus_gallus
----------------------------------------------MEQTEGPMRKRES
>Condylura_cristata
----------------------------------------------MEQTEGPMRERES
>Oryctolagus_cuniculus
----------------------------------------------MEQTEGPMRERES
>Elephantulus_edwardii
----------------------------------------------MEQTEVPMREREN
>Ornithorhynchus_anatinus
MRLRPAPAPS------PSAPGSSCGPTLLRAPVSLLGSHRHGRGVRMEQTEGPMRERES
>Lepidothrix_coronata
----------------------------------------------MEVTEGPMRKRES
>Ceratotherium_simum_simum
----------------------------------------------MEQTEGPMRERES
>Manis_javanica
----------------------------------------------MEQTEGPMRERES
>Ictidomys_tridecemlineatus
----------------------------------------------MEQTEGPMRERES
>Lepisosteus_oculatus
----------------------------------------------MEHAEGPVRERES
>Trichechus_manatus_latirostr
----------------------------------------------MEQTEGPMREREN
>Takydromus_sexlineatus
----------------------------------------------MEQTDGPMRKRES
>Chrysemys_picta
----------------------------------------------MEQTDGPMRERES
>Orycteropus_afer_afer
----------------------------------------------MEQTEGPMREREN

HG0107.1 ENSG00000058668 ATP2B4

>Homo_sapiens
MHILFTPSSGQVGLS
>Chaetura_pelagica
MHILFTPLSGEVGLG
>Ailuropoda_melanoleuca
MHILFTPPSGQVGLS
>Balaenoptera_acutorostrata_s
MHILFTPPSGQVGLS
>Charadrius_vociferus
MHILFTPLSGEVGLG
>Callorhinchus_milii
MHFLTTPSSGEVGLS
>Rhinolophus_ferrumequinum
MHILFTPPSGQVGLS
>Dasypus_novemcinctus
MHILFILQSGQVGLS
>Latimeria_chalumnae
MHFLFTPLSGEVGLG
>Columba_livia
MHILFTPLSGEVGLG
>Alligator_mississippiensis
MHILFTPLSGEVGLG
>Galeopterus_variegatus
MHILFTPSSGQVGLS
>Aquila_chrysaetos_canadensis
MHILFTPLSGEVGLG
>Gallus_gallus
MHILFTPSSGGLGLG
>Condylura_cristata
MHILFTPPSGQVGLS
>Ornithorhynchus_anatinus
MHILFTPRSREVGLG
>Lepidothrix_coronata
MHILFTPLSGEVGLG
>Equus_asinus
MHILFTPPSGQVGLS
>Nestor_notabilis
MHILFTPLSGEVGLG
>Ictidomys_tridecemlineatus
MHILFTPSSGQVSLN
>Tupaia_chinensis
MFILFIPPSGQVGLS
>Pygoscelis_adeliae
MHILFTPLSGEVGLG
>Gekko_japonicus
MCILFTPSSEEVGLG
>Chrysemys_picta
MHILFTPLSGEVGLS

HG0107.1 ENSGALG00000003601 ATP2B4

```
>Gallus_gallus
MHILFTPSSGGLGLG
>Anser_cygnoides
MRILFTPASGGEGLG
>Chaetura_pelagica
MHILFTPLSGEVGLG
>Ailuropoda_melanoleuca
MHILFTPPSGQVGLS
>Balaenoptera_acutorostrata_s
MHILFTPPSGQVGLS
>Calidris_pugnax
MHILFTPLSGEVGLG
>Rhinolophus_ferrumequinum
MHILFTPPSGQVGLS
>Latimeria_chalumnae
MHFLFTPLSGEVGLG
>Columba_livia
MHILFTPLSGEVGLG
>Alligator_sinensis
MHILFTPLSGEVGLG
>Galeopterus_variegatus
MHILFTPSSGQVGLS
>Aquila_chrysaetos_canadensis
MHILFTPLSGEVGLG
>Condylura_cristata
MHILFTPPSGQVGLS
>Ornithorhynchus_anatinus
MHILFTPRSREVGLG
>Ficedula_albicollis
MHILFTPSSPQLGLG
>Equus_asinus
MHILFTPPSGQVGLS
>Cebus_capucinus_imitator
MHILFTPSSGQVGLS
>Nestor_notabilis
MHILFTPLSGEVGLG
>Ictidomys_tridecemlineatus
MHILFTPSSGQVSLN
>Pygoscelis_adeliae
MHILFTPLSGEVGLG
>Gekko_japonicus
MCILFTPSSEEVGLG
>Chrysemys_picta
MHILFTPLSGEVGLS
```

HG0107.2 ENSG00000070961 ATP2B1

>Homo_sapiens
-------------------------------------------MIDMEKLVMGQRSRYFSGKCCC
>Odobenus_rosmarus_divergens
-------------------------------------------MIDMKKLVMGQRSRYFSGKCC-
>Lipotes_vexillifer
-------------------------------------------MTDRKKLVMGQRSRYFSGKCC-
>Pteropus_vampyrus
-------------------------------------------MIDMRKLIMGQRSRYFSGKCC-
>Dasypus_novemcinctus
MRREPSPRSRLLVSVTGPRPFFSFLFLSPSENGSCYRARYVHLMIDMKKLVMGQRSRHFLGKSCC
>Galeopterus_variegatus
---------------------------------------------------MGQRSRYFSGKCCY
>Monodelphis_domestica
---------------------------------------------------MGQSSRDLHGKCC-
>Condylura_cristata
-------------------------------------------MIDVKKLVMGQRSRYFSGKCC-
>Oryctolagus_cuniculus
-------------------------------------------MIEMKKQVMGQRSRYFSGKCCC
>Elephantulus_edwardii
-------------------------------------------MIDRKKLVMGQRSRYFSGKCCC
>Ceratotherium_simum_simum
-------------------------------------------MIDRKKLVMGQRSRYFSGKCC-
>Manis_javanica
-------------------------------------------MIDTKKLVMGQRSRYFSGKCC-
>Loxodonta_africana
----------------------------------------MHLMIDMKKLVMGQRSRYFSGKCCC
>Cavia_porcellus
-------------------------------------------MIDMKKQVMGQRSRYFSGKCCC
>Tupaia_chinensis
-------------------------------------------MIDMKKLVMGQKSRYFSEKCCY
>Orycteropus_afer_afer
----------------------------------------MHLMIDMKKLVMGHMSRYFSGKCCC

HG0107.2 ENSGALG00000030550

```
>Gallus_gallus
----------MGRPQRYLSKMGLLLG
>Anas_platyrhynchos
----------MGRPQRYLSRRGLLLG
>Caprimulgus_carolinensis
----------MGQAERYLSRWGLLLR
>Calidris_pugnax
----------MGQPQRYLSRCGLLLG
>Colius_striatus
----------MGQPQRYLSRRGLLLE
>Columba_livia
----------MGQPQRYLSRRGLLLG
>Merops_nubicus
----------MGPPQRYLSRRGLLLG
>Cuculus_canorus
----------MGQPQRYLSRRGLLLG
>Aquila_chrysaetos_canadensis
----------MGQPQRYLSRRGLLLG
>Eurypyga_helias
----------MGQPQRYLSRRDLLLG
>Tauraco_erythrolophus
----------MGQPQRYLSRRDLLLG
>Opisthocomus_hoazin
----------MGQPQRYLSRRGLLLG
>Corvus_brachyrhynchos
----------MGQPQRYLSRRGLLLG
>Nipponia_nippon
----------MGQPQRYLSRRGLLLG
>Fulmarus_glacialis
----------MGQPQRYLSRRGLLLG
>Nestor_notabilis
----------MGQPQRFLSRRGFLLG
>Aptenodytes_forsteri
----------MGQPQRYLSRQGLLLG
>Tyto_alba
----------MGQPQRYLSRRGLLLG
>Struthio_camelus_australis
MPFGLGTELTMGQHQRYISRWGLLLG
>Tinamus_guttatus
MPFGLGTELAMGQHQRYLSRRGLLLG
>Calypte_anna
----------MGQPERFLSRRGLLLG
>Apaloderma_vittatum
----------MGQPQRYLSRRGLLLG
```

HG0107.3 ENSDARG00000063433 atp2b2

```
>Danio_rerio
MGNHVWSDAGLQPW
>Acipenser_sinensis
MGNCIWSEAGLQP-
>Channa_argus
MGSHTWSRSGLQPW
>Menidia_menidia
MGSHTWSGSGLQPW
>Porichthys_notatus
MGSHTWSMSGLQPW
>Oryzias_latipes
MGSHTWSGSGRQPW
>Micropterus_salmoides
MGNHTWSGSGLQPW
>Pygocentrus_nattereri
MGNHVWSDAGLQPW
>Haplochromis_burtoni
MGSHTWSGSGLQPW
>Austrofundulus_limnaeus
MGSHTWSGSGLQPW
>Lateolabrax_japonicus
MGSHTWSGSGLQPW
>Platichthys_stellatus
MGSHTWSGSGLQPW
>Lepisosteus_oculatus
MGNYIWSGAGLQP-
>Ictalurus_punctatus
MGNAVWSDAALQPR
```

HG0107.3 ENSG00000157087 ATP2B2

```
>Homo_sapiens
-----------MPVPGSSMGPTEWRLLGLQP
>Canis_lupus_familiaris
-----------MPAPGSSMGPTEWRLLGLQP
>Lipotes_vexillifer
-----------MPAPGSSMGPAEWRLLGLQP
>Myotis_davidii
-----------MPAPGSSMGPTEWRLLGLDP
>Dasypus_novemcinctus
-----------MSAPGSSMGPAEWRLLGLQP
>Condylura_cristata
------------------MGPTEWRLLGLQP
>Oryctolagus_cuniculus
-----------MPALGSSMGPTEWRLLGLQP
>Equus_asinus
-----------MQAPGSSMGPTEWRLLGLQP
>Manis_javanica
MILGATTPDHVDASSGSSMGPTEWRLLGLQP
>Nannospalax_galili
-----------MPAPGSSMGPTEWRLLGLQP
>Tupaia_chinensis
------------------MGPTGWRLLGLQP
>Trichechus_manatus_latirostr
-----------MPALGSSMGATEWRLLGLQP
>Orycteropus_afer_afer
-----------MPALGNSMGATEWRLLGLQP
```

HG0107.4 ENSG00000070961 ATP2B1

>Homo_sapiens
MSVWFKKDFHT
>Mustela_putorius_furo
MSVWFKKDFHI
>Balaenoptera_acutorostrata_s
MSVWFKKDFHI
>Rousettus_aegyptiacus
MSVWFKKDFHI
>Galeopterus_variegatus
MSVWFKKDFHI
>Condylura_cristata
MSVWLKKDFHI
>Ceratotherium_simum_simum
MSVWFKKDFHI
>Loxodonta_africana
MSVWFKEDFHI
>Mus_musculus
MSVWFKEDFHI
>Tupaia_chinensis
MSVWFKKDFHI
>Trichechus_manatus_latirostr
MSVWFKEDFHI

HG0108.1 ENSGALG00000007396 TAOK3

```
>Gallus_gallus
-------------------MKEETDWDLKAPTSS
>Anas_platyrhynchos
-------------------MKEEIDWELKARASS
>Apteryx_australis_mantelli
-------------------MKEEIDWESKARTSS
>Buceros_rhinoceros_silvestri
-------------------MKEETDWKSKAQTGS
>Cariama_cristata
-------------------MKEEIVWKLKARTSS
>Calidris_pugnax
-------------------MKEEIDWELKARTSS
>Colius_striatus
-------------------MKEEIDWKLKAQTSS
>Columba_livia
-------------------MKEETDWKLKARTGS
>Merops_nubicus
-------------------MKEEIDWELKAQTS-
>Alligator_mississippiensis
-------------------MKAETDWEFKARTRS
>Cuculus_canorus
-------------------MKEESDWELKARTSS
>Falco_cherrug
-------------------MKEEIDWELKARTSS
>Gavia_stellata
-------------------MKEEIDWELKVRTSS
>Eurypyga_helias
-------------------MKEETDWELKARASS
>Tauraco_erythrolophus
-------------------MKEEIDWKLKARTSS
>Opisthocomus_hoazin
-------------------MKEEVDWELKGRTSS
>Corvus_cornix_cornix
-------------------MKEEIGWDLRAQTSS
>Phaethon_lepturus
-------------------MKEEIDWELKARTSS
>Nestor_notabilis
-------------------MKEETDWELKAQTSL
>Aptenodytes_forsteri
-------------------MKEEIDWELKARTSS
>Takydromus_sexlineatus
MLIAIVTQILNCYLQTPLSMREETDWELEAGTSS
>Tyto_alba
-------------------MKEEIGWKLKARTSS
>Pelodiscus_sinensis
-------------------MREETDWEFKARTRS
>Tinamus_guttatus
-------------------MKEEVDWESKARTSS
```

```
HG0108.2 ENSG00000135090 TAOK3

>Homo_sapiens
----------------MNLSNMEYFVPHTKRY
>Acinonyx_jubatus
MQKTEVLKEDKGFAAKMNLSNTVYFVPFTKRY
>Balaenoptera_acutorostrata_s
----------------MSLSNTVYFVPYTKRY
>Eptesicus_fuscus
----------------MNLSNAVYFVPYTKRY
>Condylura_cristata
----------------MNLSNTVHFVPYTKRC
>Oryctolagus_cuniculus
----------------MNLSNPVHFLPHTKRY
>Equus_asinus
----------------MNLSNTVYFVPYTKRY
>Manis_javanica
----------------MNLRNTVYFVPYTKRY
>Marmota_marmota_marmota
----------------MNLSNTAHFVPHTKRY
>Tupaia_chinensis
----------------MNLSNT-NFVPHTKRY
>Trichechus_manatus_latirostr
----------------MNLSNTVYFVPCTKRC
```

HG0108.3 ENSDARG00000098304 taok1b

>Danio_rerio
MSNQRGLTCRSRTAR
>Pygocentrus_nattereri
MSHQRGLARRSRTAR
>Ictalurus_punctatus
MSHHRGLACRSRTAR

HG0109.1 ENSG00000048540 LMO3

```
>Homo_sapiens
MNGSHRQTSDCSHLQAFLKGEGCVLFPGYSGEALQSSHCAY
>Anser_cygnoides
----------------------MLLFDLSQGEELQSSHCAY
>Chaetura_pelagica
----------------------MLLFDLSQGEELQSSHCAY
>Ursus_maritimus
----------------------MLLFDLSQGEALQSSHCAF
>Charadrius_vociferus
----------------------MLLFDLFQGEELQSSHCAY
>Myotis_lucifugus
----------------------MLLFDLSQGEALQSSHCAF
>Columba_livia
----------------------MLLFDLSQGEELQSSHCAY
>Alligator_sinensis
----------------------MLLFDLSQGEELQSSHCAY
>Cuculus_canorus
----------------------MLLFDLSQGEELQSSHCAY
>Galeopterus_variegatus
----------------------MLLFDLSQGEALQSSHCAY
>Aquila_chrysaetos_canadensis
----------------------MLLFDLFQGEELQSSHCAY
>Coturnix_japonica
----------------------MLLFDLSQGEELQSSHCAY
>Condylura_cristata
----------------------MLLFDLSQGEALQSSHCAY
>Lepidothrix_coronata
----------------------MLLFDLFQGEELQSSHCAY
>Nipponia_nippon
----------------------MLLFDLFQGEELQSSHCAY
>Equus_asinus
----------------------MLLFDLSQGEALQSSHCAF
>Picoides_pubescens
----------------------MLLFDLSQGEELQSSHCAY
>Melopsittacus_undulatus
----------------------MLLFDLFQGEELQSSHCAY
>Marmota_marmota_marmota
----------------------MLLFDLSQGEALQSSHCAY
>Struthio_camelus_australis
----------------------MLLFDLSQGEELQSSHCAY
>Chelonia_mydas
----------------------MLLFDLSQGEELQSSHCAY
>Tinamus_guttatus
----------------------MLLFDLSQGEELQSSHCAY
>Calypte_anna
----------------------MLLFDLSQGEELQSSHCAY
>Orycteropus_afer_afer
----------------------MLLFDLSQGEALQSSHCAY
```

HG0109.2 ENSG00000048540 LMO3

```
>Homo_sapiens
MQHIFYG----DYSDCCYEIQVRFMIFFILNMICYANAACDSLIKSICYQEAKRGRWDFRITGRVREACGFVIRHH--FAHCEGL----RV-CALSCLAVLIPSPTPCRHS------DPLPKWLHNSRSSAV----------
>Mustela_putorius_furo
------------------------MIFFILNMICYANAACDSLIKSICYHEAKRGRWDFRITGRVREACGFVIRHH--FAHCHTARGSVCALCLLWPPSSLVPSPTPCRHS------DPLPKWLHNSRSSAV----------
>Hynobius_retardatus
-------------------------------MIWYANAACDSLIKSICYQPAKRDHWDYRITGRVREACGFVIRHH--LHTAATCEGLAACLCLCVCVPE------------------------------------------
>Tursiops_truncatus
------------------------MIFFILNMICYANAACDSLIKSICYHDAKRGRWDFRITGRVREACGFVIRHH--FAHCHTARGSVCAPCLVWPPSSFPPPPAPPLRP------A--PKVASQ----------------
>Myotis_brandtii
MQHIFIGWGEKNFPQSCYEIQLRFMIFFILNMICYANAACDSLIKSICYREAQRGRWDFRIMGRVREACGFVIRHR--FAHRHPARGL--SVCAASRLAALVPSPTPCRHS------DPLPKWLHNSRSSSGVGCAPGPPSS
>Galeopterus_variegatus
------------------------MIFFILNMICYANAACDSLIKSICYQEAKRGLWDFRITGRVREACGFVIRHY--FAHCHTARGSVCV-CAVSCLAVLLPSPPPCRHS------DPLPKWLHNTRSSAV----------
>Oryctolagus_cuniculus
------------------------MIFFILNMICYANAACDSLIKSICYQEAKGGRWDFRITGRVREACGFVIRHR--FAHRHTARGSVCAPCLVWQPSPFPH-PLPPLRP------A--PKVASQ----------------
>Loxodonta_africana
----------------------------------------------------------------------MWLCHSPSFCTLPHCEGL--RVCAVSRLAVLIPSPTLRRHS------DPLPKWLHNSRSSAV----------
>Ctenomys_sociabilis
------------------------MTFFILNMICYANAACDSLIKSICYQEAKKGRWDFRIAGRVREACGFVIRHH--FAHCHAARGSVCALCLVRPPSSLPPPPASTPTLSQSGFTVVDPQWWCRLRTRSTLEPN------
>Trichechus_manatus_latirostr
----------------------------------------------------------------------MWLCRSPSFCTLPPCEGL--RVCAVSLLAVLIPSPTLRRHS------DPLPKWLHNSRSSAV----------
>Orycteropus_afer_afer
----------------------------------------------------------------------MWLCHSPSFCTLPHCEGL--LVRALSRLAVLFPSPTPRRH--------------------------------
```

HG0109.3 ENSG00000048540 LMO3

```
>Homo_sapiens
-----------MYACLDGPEKPNLRNLPMYRTSQLRGSH---------------------------------
>Canis_lupus_familiaris
-----------MCACLDGPEKPNLRNLPMYRTNQLKGFLLIRRL----------------------------
>Orcinus_orca
-----------MCACLDRPEKPSLRELPMYRTNPLKGFLLICRL----------------------------
>Rousettus_aegyptiacus
-----------MCACLDGPEKPNLRNLPMYRTNQLKGFLLICRL----------------------------
>Dasypus_novemcinctus
-----------MCACSDGPAKPNLRKLPMYRTNQLKGVLLICRL----------------------------
>Galeopterus_variegatus
------MG-EVRVRLLRGPEQPNLRNLPMHRPNQLKGFPLICRLSVNFVIFQEKRYKAATVLTERPPPISLN
>Monodelphis_domestica
MSKDLFIQMRCMCACISIRERQKLRNLLIYRSNQLKGVVLTC------------------------------
>Oryctolagus_cuniculus
-----------MCACIDGPGKPNLRNLPMHRTSQRKGFSLICRL----------------------------
>Elephantulus_edwardii
----------------------------MYRTNQLKGVSLICRL----------------------------
>Equus_asinus
-----------MCSCLDGPEKPNLRNLPMYRTNQLKGSY---------------------------------
>Manis_javanica
------MGEVHMCLLR-PARKAKHRKLPMYRTNQLKGFLLICRL----------------------------
>Cavia_porcellus
----------------------------MFRTNQLKGFSLICRL----------------------------
>Tupaia_chinensis
--------MRCMCACLDGPEKPNLRISAYVQNKSAE-GVLINLPSVNFVLFQEKRYKAASVLTEQPPPVSLN
>Trichechus_manatus_latirostr
-----------MCACLNRPAKPNLRNLPMYRTNQLKGVLLFCRL----------------------------
>Orycteropus_afer_afer
----------------------------MYRTNQLKGVLLICRL----------------------------
```

HG0110.1 ENSG00000091656 ZFHX4

>Homo_sapiens
MPPTFCSSVFIFTQ
>Anas_platyrhynchos
MPPTSCSIIFIFTQ
>Chaetura_pelagica
MPPTSCSIIFIFTQ
>Canis_lupus_familiaris
MPPTFCFSVFIFTQ
>Orcinus_orca
MPPTFCSSVFIFTQ
>Myotis_brandtii
MPPTFCSSVFIFTQ
>Columba_livia
MPPTSCSIIFIFTQ
>Cuculus_canorus
MPPTSCSIIFIFTQ
>Falco_cherrug
MPPTSCSIIFIFTQ
>Gallus_gallus
MPPTSCSIIFIFTQ
>Condylura_cristata
MPPTFCSSVFIFTQ
>Ochotona_princeps
MPPTFCSSVFIFTQ
>Opisthocomus_hoazin
MPPTSCSVIFIFTQ
>Corvus_cornix_cornix
MPPTSCSIIFIFTQ
>Egretta_garzetta
MPPTSCSIIFIFTQ
>Ceratotherium_simum_simum
MPPTFCSSVFIFTQ
>Loxodonta_africana
MPPTFCSSVFIFTQ
>Melopsittacus_undulatus
MPPTSCSIIFIFTQ
>Ictidomys_tridecemlineatus
MPPTFCSSVFIFTQ
>Tupaia_chinensis
MPPTFCSSVFIFTQ
>Aptenodytes_forsteri
MPPTSCSIIFIFTQ
>Struthio_camelus_australis
MPPTSCSIIFIFTQ
>Calypte_anna
MPPTSCSIIFIFTQ

HG0110.2 ENSDARG00000073944

>Danio_rerio
---MRSVLIPAI-FCTKVISRNKDRTDFFSLSPPSSN---------RSQGASVVCSTT----------------------KKQTLL--------NALRNTEIPRLTSAA---------------------------------------------------------------------------------------------
>Oryzias_latipes
MKHMRTELITAI-FCRKLISRNKDPKSFFFSITRSHFHLFLAR------------S--------------DTRLIGCFASKLFLYC-------L------------------------------------------------------------------------------------------------------------
>Odobenus_rosmarus_divergens
MKHMRTELITAI-FCRKLISRIKIQRLFF-LFT-HQLPLPARPLRPLQPP-----PPPPPPPPPGRS--RRRHRLLCIPITA------------------------------------------------------------------------------------------------------------------------
>Ambystoma_mexicanum
MKHMRTALITAI-FCRKLISRIKIQRVSWLRCSGSVQQQQQPPLSGQRRGLGAACTGDRGDPAPGAASAKGRGPGGCVQRASQGAGD------PRAAEDPEQPV----PGTGAQPWPRTRSPRRASGVQRTGHTQGTAAPYPGTLRPRRTPPAAGGCRGRAGEEGGRAGAAAAAAAMERGRRRRCCRAGLPAAPLLSPPSAA
>Astyanax_mexicanus
MKHMRTELITAIFFCRKLISRNKDPKSFFSLFN-HQISLSPLPLSL------------------------------------------------------------------------------------------------------------------------------------------------------------
>Neolamprologus_brichardi
---MRTELITAI-FCRKLISRNKDPKSFFFSIIRSHFHLFLAP------------S--------------NTRLIACFASKLFLYC-------L------------------------------------------------------------------------------------------------------------
>Latimeria_chalumnae
MKHMRTELITAI-FCRKLISRIKIQRLFF-LFN-HQISLSPLPLSLQ------------------------HRHIACFASKLFVTFSLLLFFANSLPNNNALPILMQTDPKELPSWDL------------------------------------------------------------------------------------
>Nothobranchius_furzeri
MKHMRTELITAI-FCRKLISRNKDPKSFFFSIIRSHFHLFLAP------------S--------------NTRLIACFASKLFLYC-------L------------------------------------------------------------------------------------------------------------
>Notothenia_coriiceps
MKHMRTELITAI-FCRKLISRNKDPKSFFFSIIRSHFHLFLAP------------S--------------NTRLIACFASKLFLYC-------L------------------------------------------------------------------------------------------------------------
>Salmo_salar
MKHMRTELITAI-FCRKLISRNKDPKSFF-LFN-HQISLSPLPRSL------------------------------------------------------------------------------------------------------------------------------------------------------------
>Lepisosteus_oculatus
MKHMRTELITAI-FCRKLISRNKDPKSFFSFQS-PDLTFTSSSLPLTQDT-----SPALHP------------NYFYIPFTTNTS----FFYTPQQQ---------------------------------------------------------------------------------------------------------
>Takifugu_rubripes
MKHMRTELITAI-FCRKLISRNKDPKSFFFSITRSHFHLFLAR------------S--------------NTRLIACFASKLFLYR-------L------------------------------------------------------------------------------------------------------------

HG0111 ENSGALG00000034944

>Gallus_gallus
MEVISWGGPRALWARVLLSDHWLPPE
>Anser_cygnoides
MEVISWGGPRAPWARVLLSDHWLPPA
>Caprimulgus_carolinensis
MEVISWGGPRALWARVLLSDHWLPPA
>Calidris_pugnax
MEVISWGGPRALWARVLLSDHWLPPA
>Pterocles_gutturalis
MEVISWGGPRALWARVLLSDHWLPPA
>Leptosomus_discolor
MEVISWGGPRALWARVLLSDHWLPPA
>Alligator_sinensis
MEVIIQGGPRALWAQVLLSDHWLPPA
>Cuculus_canorus
MEVISWGGPRALWAGVLLSDHWLPTA
>Aquila_chrysaetos_canadensis
MEVISWGGPRALWARVLLSDHWLPPA
>Gavia_stellata
MEVISWGGPRALWARVLLSDHWLPPA
>Balearica_regulorum_gibberic
MEVISWGGPRALWARVLLSDHWLPPA
>Tauraco_erythrolophus
MEVISWGGPRALWARVLLSDHWLPPA
>Opisthocomus_hoazin
MEVISWGGPRALWARVLLSDHWLPPA
>Lepidothrix_coronata
MEVISWGGPRALWARVLLSDHWLPPA
>Egretta_garzetta
MEVISWGGPRALWARVLLSDHWLPPA
>Melopsittacus_undulatus
MEVISWGGPRALWARVLLGDHWLPPA
>Aptenodytes_forsteri
MEVISWGGPRALWARVLLSDHWLPPA
>Protobothrops_mucrosquamatus
-----MGGLRVLWVRVLLNNHWLIAT
>Struthio_camelus_australis
MEVISWGGPRALWARVLLSDHWLPPA
>Chrysemys_picta
MEVISWGGPRALWARVLLSYHWLPPA
>Tinamus_guttatus
MEGMSWGGPRALWARVLLSDHWLPPA
>Calypte_anna
MEVISWGGPRALWAGVLLSDHWLPPA
>Apaloderma_vittatum
MEVISWGGPRALWARVLLGDHWLPPA

HG0112 ENSG00000105997 HOXA3

>Homo_sapiens
----------------------------------------------------------------------MKRSARGGHWRRSVT
>Anser_cygnoides
MC--KDT-ASWLSAKKMYMLLVENEERTAKYPMADFRRLR---------------------------------SARRGHWRKSAT
>Xenopus_laevis
---------------------------------------------------MEMN-AGFLVPGVISDPSCFNRSDQGGHWRRNAT
>Ailuropoda_melanoleuca
---------------------------------------------------M----PRVGDN-HWIS-LHLSRSARGGHWRRSVT
>Calidris_pugnax
------------------------------------------------------------------M-DFILRSARRGHWRKSAT
>Eptesicus_fuscus
----------------MPRLLVEIECRAAQPHMADCHQLRAPGHRKRRFPER----SGVGDNWGFRL-KYLSRSARGSHWRRSVT
>Columba_livia
------------------------------------------------------------------M-DFILRSARRGHWRKSAT
>Poecilia_reticulata
------M-GIHVLTSRVEQPLFGHLTFPFRPPLTDRRPSRPPGRTASPCVTE-IRPSNFR--PSLGNTLSCLRFTRGGHWRRGVT
>Esox_lucius
--------------------------------MEDKQVVKTAVYETLVAQRHEIYQNKSLPYYC----VHETRFTQGGHWRRGVT
>Coturnix_japonica
------------------------------------------------------------------M-DFILRSARSGHWRSSAT
>Erinaceus_europaeus
---------------------------------------------------M----CRR------GS-CGNPRSARGSHWRRSIT
>Scleropages_formosus
----------------MRKLHISHE--------LCGMELEHPKHVKKRLAGQDFSLNKVITIYDLGEKFPPLRSTRGGHWRRGVT
>Sturnus_vulgaris
------------------------------------------------------------------M-DFILRSARRGHWRKSAT
>Notothenia_coriiceps
-----------------------MEKKQDLDVMADGE-------------EHEIQPSKFRASHALQDTLNCLRFTRGGHWRRGVT
>Equus_asinus
------------------------------------------------------------------M-EFILRSARGGHWRRSVT
>Cynoglossus_semilaevis
----------------------------------------------------MRKYCSV-----LCVRRQRKRFTRGGHWRRGVT
>Chinchilla_lanigera
------------------------------------------------------------------M-EFILRSARGGHWRRSVT
>Salmo_salar
--------------------------------MA----------VDEILKRYEFCLNKSHPSYSLHEILPCHRFTRGGHWRRGVT
>Pygoscelis_adeliae
------------------------------------------------------------------M-DFILRSARRGHWRKSAT
>Anolis_carolinensis
MCSCRKTQSLGCALMKMYTFWEENPARREKIHMAGCHHPSSPVHWKIRFKDS-LSLEII-----LPGVGNARRSTRRGHWRRNVT
>Struthio_camelus_australis
MC--KDT-ASWLSAKKMYTLSVEIEPRTAKYPMADYHHLREPVHRKTPICLADRSLEQFS----PTL-SDFKRSARRGHWRKSAT
>Takifugu_rubripes
------------------------MKREADLGMA-------------ECTMHEIQPSKFTPSLGARKTINCLRFTRGGHWRRGVT
>Tinamus_guttatus
------------------------------------------------------------------M-DFILRSARRGHWRKSAT

HG0113 ENSDARG00000099727 ssbp2

>Danio_rerio
-------------------MGGIAPAVAMVTGLCPGRL
>Acipenser_sinensis
-------------------MGSDAPCVAMVTGLCPGRL
>Polypedates_megacephalus
----------------------------MVTGLFPGRL
>Porichthys_notatus
-------MERGRSRTAGHPMRVAAACVAMVTGLCPGRL
>Oryzias_latipes
-------------------MEYDAACVTMVTGLCPGRL
>Canis_lupus_familiaris
-------------------MVGVAAWVAMGTGLFPGRL
>Orcinus_orca
----------------------------MGTGLFPGRL
>Haplochromis_burtoni
-------------------MGDDAACVTMVTGLCPGRL
>Dasypus_novemcinctus
----------------------------MGTGLFPGRL
>Alosa_alosa
MSVRGVTKQNGRFFCARCPMGAIGPAVTMVTGLCPGRL
>Latimeria_menadoensis
----------------------------MVTGLFPGRL
>Austrofundulus_limnaeus
-------------------MEDAAACVTMVTGLCPGRL
>Falco_peregrinus
-------------------MGCVCGSVSMVTGLFPGRL
>Sorex_araneus
----------------------------MGTGLFPGRL
>Ornithorhynchus_anatinus
-------------------MAGVAAWVAMGTGLFPGRL
>Trematomus_bernacchii
----------------------------MVTGLCPRRL
>Ceratotherium_simum_simum
----------------------------MGTGLFPGRL
>Platichthys_stellatus
-------MASGRSRPAGHPMGGDAACVAMVTGPCPGRL
>Callithrix_jacchus
----------------------------MGTGLFPGRL
>Microtus_ochrogaster
-------------------MVGVAAWVAMGTGLFPGRL
>Tupaia_chinensis
----------------------------MGTGLFPGRL
>Pelodiscus_sinensis
-------------------MGCVCGSVSMVTGLFPGRL

HG0113 ENSGALG00000023089 SSBP2

>Gallus_gallus
------------------------MGCVCGGVAMATGLFPGRL
>Odorrana_margaretae
---------------------------------MVTGLFPGRL
>Canis_lupus_familiaris
------------------------MVGVAAWVAMGTGLFPGRL
>Hynobius_retardatus
-------------MGGPPACAPSNRRRRCMRVAMATGLFQGRL
>Orcinus_orca
---------------------------------MGTGLFPGRL
>Pteropus_vampyrus
---------------------------------MGTGLFPGRL
>Dasypus_novemcinctus
---------------------------------MGTGLFPGRL
>Latimeria_chalumnae
---------------------------------MVTGLFPGRL
>Monodelphis_domestica
---------------------------------MGTGLFPGRL
>Falco_peregrinus
------------------------MGCVCGSVSMVTGLFPGRL
>Erinaceus_europaeus
MEGEACPSRALPGRRGRGLRAGRPIVGVAAWVAMGTGLFPGRL
>Ochotona_princeps
---------------------------------MGTGLFPGRL
>Ornithorhynchus_anatinus
------------------------MAGVAAWVAMGTGLFPGRL
>Ceratotherium_simum_simum
---------------------------------MGTGLFPGRL
>Cercocebus_atys
---------------------------------MGTGLFPGRL
>Marmota_marmota_marmota
---------------------------------MGTGLFPGRL
>Tupaia_chinensis
---------------------------------MGTGLFPGRL
>Anolis_carolinensis
------------------------MARVCGGVSMATGLFPGRL
>Pelodiscus_sinensis
------------------------MGCVCGSVSMVTGLFPGRL

HG0114.1 ENSGALG00000003701 SGMS1

```
>Gallus_gallus
--------------------------------------------------------MKPRRSLLM-LWSRSDCCPL
>Anser_cygnoides
--------------------------------------------------------MKPRKSFLM-HWSHSDCCPI
>Apteryx_australis_mantelli
------------------------------MLSPACGCSRQGLIVGQT---GSSDKMKKPRRSLLLPWCHSGCCPV
>Cariama_cristata
--------------------------------------------------------MKPRRSFPT-RWNRSDCCPI
>Calidris_pugnax
MNRVLWVEKHLQEEECQHSYEVTSFLCLPQHLDARIRQLRQGLAYCQT---ASLDKMKPRRSFLM-RWSRSDCCPI
>Pterocles_gutturalis
--------------------------------------------------------MKPRRSFLM-RWSRSDCCPI
>Columba_livia
--------------------------------------------------------MKPRRSFAT-CQSRADRCPI
>Merops_nubicus
--------------------------------------------------------MKPRRSFPTRRWSRSDCCPI
>Cuculus_canorus
--------------------------------------------------------MKPRRSFLT-CWSHLDCCPI
>Haliaeetus_albicilla
--------------------------------------------------------MKPRRSFPT-RWSRSDCCPV
>Gavia_stellata
--------------------------------------------------------MKPRRSFPT-RWSRSDCCPI
>Balearica_regulorum_gibberic
--------------------------------------------------------MKPRRSFPT-RWSRSDCCPI
>Opisthocomus_hoazin
--------------------------------------------------------MKPRRSFLT-HWSHSDCCPV
>Pseudopodoces_humilis
--------------------------------------------------------MKPRRSFPP-RWSRSDCCPL
>Nipponia_nippon
--------------------------------------------------------MKPRRSFPM-RWSRSDCCPI
>Fulmarus_glacialis
--------------------------------------------------------MKPRRSFLT-RWSRSDCCPI
>Nestor_notabilis
--------------------------------------------------------MKPRRSFPT-HWSRSDCCPI
>Aptenodytes_forsteri
--------------------------------------------------------MKPRRSFPT-RWSCSDCCPI
>Tyto_alba
--------------------------------------------------------MKPRRSFPM-CWSRSDCCPI
>Chrysemys_picta
------------------MYKVFRNL---------------YLSTCQTGFCCTSDRMNKPKRGFLLYWKNSDCCLV
>Calypte_anna
--------------------------------------------------------MKPRKSFPV-HWSRSDCCPI
>Apaloderma_vittatum
--------------------------------------------------------MKPRRSFVT-RQNRSDCCPV
```

HG0114.2 ENSG00000198964 SGMS1

>Homo_sapiens
MTRMFLNNPRKNPAQK
>Felis_catus
MTRIFLNNQRKTPGAP
>Myotis_davidii
MTRIFLSNPRKNLVEK
>Galeopterus_variegatus
MTRIFLNNPRKNPVQK
>Condylura_cristata
MTRIFLNNQRKNLVQK
>Equus_asinus
MTRIFLNNQRKNLAQK
>Fukomys_damarensis
MTRIFLHNLRKNPVQK

HG0115.1 ENSGALG00000010976 CCDC126

>Gallus_gallus
MESRPAK-PRRCVQDDPARRLLTL------------
>Anas_platyrhynchos
MESRPTK-LRRCVQDDPARRLFTL------------
>Apteryx_australis_mantelli
MESRPTKTWKRCIQDDLAGGLFTL------------
>Buceros_rhinoceros_silvestri
--------MRRCIQDDPARRFFIS------------
>Caprimulgus_carolinensis
--------MRRCIQDDSARRLFNS------------
>Cariama_cristata
--------MRRCIQDDPARRFFTS------------
>Charadrius_vociferus
--------MRRCIQDDPARRFFTS------------
>Colius_striatus
--------MRRCIQDDPDRRLFTSQKFGFV------
>Columba_livia
--------MRRCIQDDPSRRLFTS------------
>Leptosomus_discolor
--------MRSCIQDDPARRFFTS------------
>Alligator_mississippiensis
MGSRLSIIWRECVQDGPSRRFVTL------------
>Aquila_chrysaetos_canadensis
--------MRRCIQDDPARRFFTS------------
>Gavia_stellata
--------MRRCIQDDPARRFFTSY-----------
>Balearica_regulorum_gibberic
--------MRRCIQDDPARRFFTSQKSYFCLICNII
>Tauraco_erythrolophus
--------MRRCIQDDPATRFFTSQKS---------
>Opisthocomus_hoazin
--------MRRCIQDDPARR----------------
>Lepidothrix_coronata
--------MRRCIQDDPTRRFFTS------------
>Phaethon_lepturus
--------MRRCIQDDPARRFFTS------------
>Melopsittacus_undulatus
--------MRRCIQDD--------------------
>Tyto_alba
---------MRCNQDDPARRFFTSQKSYFCLICNII
>Struthio_camelus_australis
MESRPTKTWKRCIQDDQARGFFTL------------
>Apaloderma_vittatum
--------MRRYIQDDPARRFFTS------------

HG0115.2 ENSG00000169193 CCDC126

>Homo_sapiens
MSANRVNRVDTTLLKMKNIQY
>Ursus_maritimus
---------------MKNIQY
>Pteropus_alecto
---------------MKNIQY
>Galeopterus_variegatus
---------------MENIKY
>Oryctolagus_cuniculus
---------------MKNIQY
>Marmota_marmota_marmota
---------------MKIIPY
>Tupaia_chinensis
---------------MKDIQY

HG0116 ENSG00000010270 STARD3NL

>Homo_sapiens
-------------MCLLFRDGEVGKRLL
>Ursus_maritimus
MRSPTSFQCLGWDAGTGFRDGEVGERLL
>Myotis_ricketti
------------MRPLLSRDGEVGKRLL
>Condylura_cristata
------------MRPFLSRDGEIGKRLL
>Ceratotherium_simum_simum
------------MRSLLSRDGEVGKRLL
>Marmota_marmota_marmota
------------MRCLLSRDGEVGKRFL

HG0116 ENSGALG00000033038

>Gallus_gallus
--------------------------------------------------------------------------------------------MDQHSSDGETGKGLL
>Apteryx_australis_mantelli
--------------------------------------------------------------------------------------------MDQPSSDGETGRRLL
>Buceros_rhinoceros_silvestri
--------------------------------------------------------------------------------------------MDQPSSDGETGRPPL
>Caprimulgus_carolinensis
--------------------------------------------------------------------------------------------MDQSSSDGETGRRPL
>Cariama_cristata
--------------------------------------------------------------------------------------------MDQPSSDGETGRRPL
>Calidris_pugnax
--------------------------------------------------------------------------------------------MDQPSSDGETGRRPL
>Pterocles_gutturalis
--------------------------------------------------------------------------------------------MDQPSSDGETGRRPL
>Colius_striatus
--------------------------------------------------------------------------------------------MDQPSSDGETGRRPL
>Columba_livia
--------------------------------------------------------------------------------------------MDQPSSDGETGRRPL
>Leptosomus_discolor
--------------------------------------------------------------------------------------------MDQPSSDGETGRRPL
>Alligator_mississippiensis
--------------------------------------------------------------------------------------------MDQPSSDGEAGRRLL
>Cuculus_canorus
--------------------------------------------------------------------------------------------MDQPSRDGETARRLL
>Aquila_chrysaetos_canadensis
--------------------------------------------------------------------------------------------MDQPSSDGETGRRPL
>Gavia_stellata
--------------------------------------------------------------------------------------------MDQPSSDGETGRRLL
>Balearica_regulorum_gibberic
--------------------------------------------------------------------------------------------MDQPSSDGETGRRPL
>Opisthocomus_hoazin
--------------------------------------------------------------------------------------------MDQPSSDGETGRRQL
>Lepidothrix_coronata
--------------------------------------------------------------------------------------------MDQPSSDGETGRRPL
>Egretta_garzetta
--------------------------------------------------------------------------------------------MDQPSSDGETGRRPL
>Tyto_alba
--------------------------------------------------------------------------------------------MDQHSSDGETGRLPL
>Struthio_camelus_australis
MGKDDIAVPSDVPLTQKIDILLLTSFFFLLLLACKLSPALLSTFFQYFQKNIVEHVCESCVTVQLLLLGCHESIAIWGKVWLKETRKRSSSCMDQPSSDGETGRRLP
>Chrysemys_picta
--------------------------------------------------------------------------------------------MDQPASDGEAGKRLL
>Tinamus_guttatus
--------------------------------------------------------------------------------------------MDQPSSDGETGRRLL

HG0117.1 ENSG00000140945 CDH13

>Homo_sapiens
MLSAAACMNENAAGRF
>Anser_cygnoides
MLSAAECMNEKASGTF
>Ursus_maritimus
MLSAAKCMNENAAGRF
>Lipotes_vexillifer
MLSAAACMNENAARRF
>Calidris_pugnax
MLSAAECMNEKASGTF
>Rhinolophus_ferrumequinum
MLSAAACMNENAAGRF
>Dasypus_novemcinctus
MLSAAACMNENAAGRF
>Columba_livia
MLSAAECMNEKASGTF
>Numida_meleagris
MLSAAECMNEKASGTF
>Condylura_cristata
MLSAAACMNENAAGRF
>Oryctolagus_cuniculus
MLSAAACMNENAAGRC
>Sturnus_vulgaris
MLSAAECMNEKASGTF
>Ceratotherium_simum_simum
MLSAAACMNENAAGRF
>Manis_javanica
MLSAAKCMNENAAGRV
>Loxodonta_africana
MLSAAACVNENAAGRF
>Ctenomys_sociabilis
MLSAAACMNENAAGRF
>Tupaia_chinensis
MLSAAACMNENAAGRF
>Trichechus_manatus_latirostr
MLSAAASVNENAAGLF
>Aptenodytes_forsteri
MLSAAECMNEKASGTF
>Anolis_carolinensis
MLSAAECMKEKASGVF
>Struthio_camelus_australis
MLSAAECMNEKASGTF
>Pelodiscus_sinensis
MLSAAECMNEKASGTF

HG0117.2 ENSG00000140945 CDH13

```
>Homo_sapiens
MPPPRGSLLARCSPRICHAKRGSVRKESVL
>Dasypus_novemcinctus
------------------------MKESVL
>Condylura_cristata
----------MYSPHICHAERGSFMKESVL
>Oryctolagus_cuniculus
------------------------MKESVL
>Mus_musculus
------------------------MKESVL
>Tupaia_chinensis
------------------------MKESVL
```

HG0118.1 ENSG00000183662 FAM19A1

```
>Homo_sapiens
--------------------------------------------MTRFPPLKPWLLF
>Anas_platyrhynchos
-----------------------------------MLGDDLKVKMRRFPPLKPWPLF
>Mustela_putorius_furo
--------------------------------------------MTRFLPLKPWLLF
>Hynobius_retardatus
--------------------------------------------MTRFPPLESWLLF
>Lipotes_vexillifer
--------------------------------------------MTRFPALKPWFLF
>Pteropus_vampyrus
--------------------------------------------MTRFPPLKPWLLF
>Monodelphis_domestica
--------------------------------------------MTRFPPLKPWLLF
>Aquila_chrysaetos_canadensis
-----------------------------------MLGDDLKVKMRRFPPLKPWPLF
>Erinaceus_europaeus
--------------------------------------------MTRFPPLKPWLLF
>Oryctolagus_cuniculus
--------------------------------------------MTRFPPLKPWLLF
>Elephantulus_edwardii
--------------------------------------------MTRFPPLKPWLLF
>Parus_major
-----------------------------------MLGNDLKVKMRRFPPLKPWPLF
>Equus_asinus
--------------------------------------------MTRFPPLKPWLLF
>Manis_javanica
--------------------------------------------MTRFPPIKPWLLL
>Loxodonta_africana
MHWSGDRSVTDYKLNLVRKLHIISPSLQRSRQAEVTPGDDLKVKMTRFPPLKAWLLF
>Fulmarus_glacialis
-----------------------------------MLGDDLKVKMRRFPPLKPWPLF
>Nannospalax_galili
--------------------------------------------MTRFPPLKPWLLF
>Tupaia_chinensis
--------------------------------------------MTRFPPLKPWLLF
>Trichechus_manatus_latirostr
--------------------------------------------MTRFPPLKPWLLF
>Struthio_camelus_australis
--------------------------------------------MRRFPPLKPWPLF
>Chrysemys_picta
--------------------------------------------MTRFPPLKPWPLF
>Orycteropus_afer_afer
--------------------------------------------MTRFPPLKPWLLF
```

HG0118.2 ENSG00000183662 FAM19A1

>Homo_sapiens
MHWSGDGPSATIN-------------------------------------------
>Lipotes_vexillifer
MHWSGDGPSLTIN-------------------------------------------
>Pteropus_alecto
MHWNGDGPSLTIN-------------------------------------------
>Condylura_cristata
MHWRGDGPSLTIN-------------------------------------------
>Ochotona_princeps
MHWSGDGPAVAQLIPIWRLHILSPPLQKSLQAEVTPGDYLKVKMTRFLLLKPWLLF
>Elephantulus_edwardii
MHWSGGAPSLTIN-------------------------------------------
>Tupaia_chinensis
MHWSGNGPSVTIN-------------------------------------------
>Orycteropus_afer_afer
MHWSGDGPSLTIN-------------------------------------------

HG0119 ENSGALG00000043570 SGK3

>Gallus_gallus
------------------------MMEFTFSQEELRILDQYR
>Anser_cygnoides
------------------------MMEFRFSQEERRISDQHR
>Chaetura_pelagica
------------------------MMEFTFSEEKLRVVDQYR
>Buceros_rhinoceros_silvestri
------------------------MMEFTFSREKLRLLGQHR
>Caprimulgus_carolinensis
------------------------MMEFTFSQEKLRILDQYR
>Cariama_cristata
------------------------MMGFTFSPEKLCILDQYR
>Calidris_pugnax
------------------------MMEFTFSQEKLRILDQDR
>Pterocles_gutturalis
------------------------MMEFTFPREKLRILDQYH
>Leptosomus_discolor
------------------------MMEFTFSEEKLHISDQYR
>Alligator_mississippiensis
------------------------MMESRFSQDKLHLLDQYR
>Aquila_chrysaetos_canadensis
MDNLTVLLNTKYVKYGGRGGREDYMMEFTFSQEKLRILDQYR
>Gavia_stellata
-------------------------MEFTFFQEKLRILDQYR
>Chlamydotis_macqueenii
------------------------MMEFTFSQEKLRILDQYR
>Tauraco_erythrolophus
------------------------MMEFTFSQEKLRILDQYR
>Ficedula_albicollis
------------------------MMEFTFSQEKLRILDQYR
>Pelecanus_crispus
------------------------MMEFTFSQEKLRILDQYH
>Fulmarus_glacialis
------------------------MMEFTFSQEKLRILDQYR
>Pygoscelis_adeliae
---------------MLSCRDSDYMMEFTFSQEKLRILDQYR
>Takydromus_sexlineatus
------------------------MMEFKFTQERLHVLDQYC
>Struthio_camelus_australis
------------------------MMELTFFQEKLHILDQYR
>Chrysemys_picta
------------------------MMEFKFSQEKLHILDQYR
>Apaloderma_vittatum
------------------------MMEFTFSQEKLRIKDQYH

HG0120 ENSG00000022355 GABRA1

>Homo_sapiens
MKSPPISHPVCPGTYYYCLGKEKSFVV------------------------------
>Mustela_putorius_furo
MKSPPISHPVCPGTYYYCLGKEKSFVV------------------------------
>Calidris_pugnax
MKSQPISHPVYPGTYYYCLGKEKAFVV------------------------------
>Rhinolophus_ferrumequinum
MKSPPISHPVCPGTYYYCLGKEKSFVV------------------------------
>Dasypus_novemcinctus
MKSPPISHPVCPGTYYYCLGKEKSFVVSLQIGSWEANLGVKSSAEEHAESMMAQTE-
>Galeopterus_variegatus
MKSPPISHPVCPGTYYYCLGREKSFVV------------------------------
>Monodelphis_domestica
MKSLPISHPVYPGTYYYCLGKERSFVVYLQTEYWEEIWV------------------
>Macropus_eugenii
MKSLPISHPVYPGTYYYCLGKEKSFVVYLQTEYWEEIWV------------------
>Esox_lucius
MKSQPISHPVYPGTYYYCPGKVKLHLAAICVFRV--DFLVTNLLGT-EKSLTPHTNE
>Condylura_cristata
MKSPPISHPVCPGTYYYCLGKEKSFVV------------------------------
>Ochotona_princeps
MKSPPISHPVCPGTYYYCLGKEKSFVV------------------------------
>Scleropages_formosus
MKSQPIPHPVYSGTHYYCLG-------------------------------------
>Zonotrichia_albicollis
MKSQPISHPVYPGTYYYCLGKEKAFVV------------------------------
>Ceratotherium_simum_simum
MKSPPISHPVCPGTYYYCLGKEKSFVV------------------------------
>Manis_javanica
MKSPPISHPVCPGTYYYCLGKEKSFVV------------------------------
>Cavia_porcellus
MKSPPISHPVCPGTYYYCLGKEKSFVV------------------------------
>Salmo_salar
MKSQPISHPVYPGTYYYCPGKVKLYLAAISVFTV--DFVVR----------------
>Tupaia_chinensis
MKSPPISHPVCPGTYYYCLGKEKSFVV------------------------------
>Lepisosteus_oculatus
MKSQPISHPVYPGTYYYCLGKENLYLAAISVFRV--DFAVRLPIYR-DR--------
>Trichechus_manatus_latirostr
MKSPPISHPVCPGTYYYCLGKEISFVV------------------------------
>Thamnophis_sirtalis
MKSQPISHPVYPGTYYYCLGKVQSFLL------------------------------
>Chrysemys_picta
MKSQPISHPVYPGTYYYCLGKEKAFVV------------------------------

HG0121 ENSG00000122786 CALD1

```
>Homo_sapiens
--------MCSLSMDFWEGPRKGGLEFY---
>Anas_platyrhynchos
--------MYSLSIDFWEGPRKGCSFIE---
>Apteryx_australis_mantelli
--------MYSLSIDFWEGPRKGCSFIE---
>Canis_lupus_familiaris
--------MCSLSTDFWEGPRKGGLEFY---
>Balaenoptera_acutorostrata_s
MPTVNIGDMCSLSMDFWEGLRKGGLEFY---
>Calidris_pugnax
--------MYSLSIDFWEGPRKSCSFIE---
>Dasypus_novemcinctus
--------MCSLSMDFWEGLRKGGLQFY---
>Columba_livia
--------MYSLSIDFWEGPRKSCSFIE---
>Galeopterus_variegatus
--------MCSLSMDFWEGPRKGGLEFY---
>Haliaeetus_leucocephalus
--------MYSLSIDFWEGPRKSCSFIE---
>Coturnix_japonica
--------MYSLSIDFWEGPKQGCSFIE---
>Condylura_cristata
--------MCSLSMGFWEELRKGRAVWSLVE
>Oryctolagus_cuniculus
--------MCSLSMDFWEGPRKGGLQFY---
>Corvus_brachyrhynchos
--------MYSLSIDFWEGPRKSCTFIE---
>Nipponia_nippon
--------MYSLSIDCWEGPRKSCSFIE---
>Equus_asinus
--------MCSLSMDFWEGLRKGGLEFY---
>Manis_javanica
-------------MDFWEGLREGGLEFY---
>Loxodonta_africana
--------MCSLSVDCWERLRKGGLQFIE--
>Nannospalax_galili
--------MCSLSMDFWEGPRKGGLQFY---
>Tupaia_chinensis
--------MCSLSLDFWEGPRKGGLEFY---
>Aptenodytes_forsteri
--------MYSLSIDFWEGPRKSCSFIE---
>Struthio_camelus_australis
--------MYSLSIDFWEGPRKGCSFIE---
```

HG0122 ENSG00000169925 BRD3

>Homo_sapiens
-----------------------------MLSCDRPSSCLEEAGLQPGSL---------PLCPD-HPHIPKAVAWLGPEMGRRAFSGSVEAARASPSRCCDRK-
>Felis_catus
-------------------------------------------------------------------------------MGRRASERSTVEARASPSRCCDGQ-
>Balaenoptera_acutorostrata_s
-------------------------------------------------------------------------------MGCRTSAGSTAEARASPSCCCRDGQ
>Calidris_pugnax
-------------------------------------------------------------------------------MGCQSRSGSIKEERKSPSRCCRDSE
>Pteropus_alecto
-------------------------------------------------------------------------------MGCHASAGSTEEARAPPSRCCDGQ-
>Cuculus_canorus
-------------------------------------------------------------------------------MGCQSRSGNIKEERKSPSRCCCDSE
>Sarcophilus_harrisii
-------------------------------------------------------------------------------MGCQSYSGSIKKERESPSCCCDSK-
>Galeopterus_variegatus
MQMPRRQTCRAGTTASCLPDWLRASPCHSLLPSQTPEL-LLLVMIEPRSVPCSRAPAPHAVCGSAEVWGAAGVAQLGPEMGCHTSPGSIGAARASRSRCCDRK-
>Monodelphis_domestica
-------------------------------------------------------------------------------MGCQSYSGSIKKERESPSCCCDSK-
>Macropus_eugenii
-------------------------------------------------------------------------------MGCQSYSGSIKKERESPSCCCDSK-
>Balearica_regulorum_gibberic
-------------------------------------------------------------------------------MGCQSHSGSIKEERKSPSCCCCDRE
>Erinaceus_europaeus
-------------------------------------------------------------------------------MGCHAPKGSTEHISTSLSRCCDN--
>Oryctolagus_cuniculus
-------------------------------------------------------------------------------MGRRASLGSVDAARAPARRRCCGRK
>Ornithorhynchus_anatinus
-------------------------------------------------------------------------------MGCQSYSGSIKQVKESPSCCCDDSE
>Equus_caballus
-------------------------------------------------------------------------------MGRRASRGSTEELRASPSRCCDGQ-
>Manis_javanica
-------------------------------------------------------------------------------MGCHASAGSTEETRASLSRCCDGQ-
>Loxodonta_africana
-------------------------------------------------------------------------------MGCRAASGGVEGERAPAGCGCDRK-
>Ictidomys_tridecemlineatus
-------------------------------------------------------------------------------MGCQASSGSTEGAGAFWSCSCDGK-
>Tupaia_chinensis
-------------------------------------------------------------------------------MGCRGPSGSIAAARAPASPCRDGK-
>Trichechus_manatus_latirostr
-------------------------------------------------------------------------------MGCRAASGSLEGARASAGRGCDGK-
>Chelonia_mydas
-------------------------------------------------------------------------------MGCQSHSGSIKEDRESPSCCCDSK-
>Orycteropus_afer_afer
-------------------------------------------------------------------------------MGCRASSGIVEGTWASLGRGCDE--

HG0123 ENSG00000173276 ZBTB21

```
>Homo_sapiens
MTSSLVWDDVKFKMDGITAAAAAAARASRV-
>Amia_calva
-------------MDGITTATRSTEF-----
>Channa_argus
MTSSLVWDDVKFKMDGITTSAQKSTQH----
>Xenopus_laevis
MTSSVVWDDVKFKMDGITAAAARQRLAE---
>Menidia_menidia
-------------MDGITISSQKFTQH----
>Oryzias_latipes
-------------MDGITTKTQKSCTTVNKV
>Hynobius_retardatus
-------------MDGITAAARAGQF-----
>Myotis_lucifugus
MTSSLVWDDVKFKMDGITAAAAAAARASRV-
>Oreochromis_niloticus
MTSSLVWDDVKFKMDGITTSAEKSTQH----
>Pimephales_promelas
MTSSLEWDDVKYKMDGITTASCSTQL-----
>Fundulus_heteroclitus
MTSSLVWDDVKFKMDGITTSRLKATQH----
>Galeopterus_variegatus
MTSSLVWDDVKFKMDGITAAAAAVARASRV-
>Esox_lucius
-------------MDGITTTTHSTEF-----
>Coturnix_japonica
MTSSLVWDDVKFKMDGITAAAAARAAEP---
>Lateolabrax_japonicus
MTSSLVWDDVKFKMDGITTSAQKSTQH----
>Trematomus_bernacchii
----MVWDDVKFKMDGITTSAQKSTQQH---
>Platichthys_stellatus
MTSSLVWDDVKFKMDGITTSAQKAPQQ----
>Heterocephalus_glaber
MTSSLVWDDVKFKMDGITAAAAAAARASRV-
>Salmo_salar
MTSSLE-------------------------
>Thamnophis_sirtalis
MTSSLVWDDVKFKMDGITAVAAARAGES---
>Chrysemys_picta
MTSSLVWDDVKFKMDGITAVAAARAGES---
>Takifugu_rubripes
MTSSLVWDDVKFKMDGITTAA----------
```

HG0124.1 ENSDARG00000031763 smad6b

```
>Danio_rerio
------------------MQYSIVPSKLMLWLCHRYTVLGCARAARHSANI-NVARAQK-----
>Acipenser_sinensis
MHFDFNVAVCNCVTLAIYLARYCICPNLMLQSGCKYTAFGCAIVARHLQKL-HVLRRPWN----
>Channa_argus
----------------------------MLRLCHQCLSVGCARIARHFTDI-NVLQRP------
>Porichthys_notatus
--MQQNSLFFILLLFLFPPSLPLFSRSLMVRLGHRCSSLGCARVARRSTHI-NVLQRP------
>Eptesicus_fuscus
----------------------------MLQPCGAPSTTGCARSARRPAAVLHVGLRPAQPAPR
>Oreochromis_niloticus
----------------------------MLRFCHQCSSLGCACTARHFSVI-NVLQRPEKCL--
>Dasypus_novemcinctus
----------------------------MLQPCGAPSTTGCARSARHPTAELHVGLRPAQPAPR
>Alosa_alosa
------------------MQHYIVASNLMLCFSHQYSAFGCACVARQSTNI-NVVQRA------
>Esox_lucius
------------------MQYTTVNYILMLRVGHKSTTQGCACSARRSQNL-DVVRRP------
>Apteronotus_leptorhynchus
------------------MLHNIVPSK-LMWFYHQHRVLGCACIAQQSANI-NVRSQKLF----
>Oryctolagus_cuniculus
----------------------------MLQPCGAPSTTGCARSARRSAAELHVGLRPAPR---
>Lateolabrax_japonicus
----------------------------MLRLCHQCSSVGCACIARHFTDI-NVLQRP------
>Notothenia_coriiceps
----------------------------MLRFCHQCSSVGCAGIARHFTDI-NVLQRPEKYL--
>Equus_asinus
----------------------------MLQPCGAPSTTGCARSARRPAAELHVGLRPA-----
>Cynoglossus_semilaevis
----------------------------MLRLCHQCSSLGCACAARRFTYI-NVLQRR------
>Callithrix_jacchus
----------------------------MLQPCGAPSTTGCARSARRSAAVLHVGLRPAQPAPR
>Microtus_ochrogaster
----------------------------MLQPCGAPSTTGCARSARRPAAELHVGLRPAQPAPR
>Oncorhynchus_kisutch
------------------MQHTTVNYILMLRVSHKSTTPGCACSARRLQNI-NVVRRP------
>Lepisosteus_oculatus
----------------------------MLQFCCKYTALSCAFIARHLRKM-NVWRRP------
>Ictalurus_punctatus
MHFVFSVALHFIDALYILMLPNNVPFNPMLWFCYRPTVLCCARAAQRSANL-NVAQAQKQL---
>Takifugu_rubripes
----------------------------MLRLCHQCSSVSCAWSARHFTHI-NVLQRP------
```

HG0124.2 ENSDARG00000053209 smad6a

>Danio_rerio
MMGINQWLHCSPQG----PRVCVSEEERE---------------------------------------------------------------------
>Pygocentrus_nattereri
------MRECVRECDCVYVWDCVRISRRRKRGRETAQGNNRRSPLILSAARNERIRRVAGARAFTLPVEGSFVRACFFAPLHEASFCAVRADWNVNSS
>Latimeria_chalumnae
MMGGSQ--------------------------------------------------------------------------------------------
>Scleropages_formosus
MMGGSQWLLSYPRG----PRVCVSGGGKKRREKR-----GKANPRVLSEQI-----------------------------------------------
>Salmo_salar
MMGINQWLQSFPGA----P-VCVRASVSEKNSRK----------------------------------------------------------------

HG0125.1 ENSDARG00000051926 plag1

>Danio_rerio
MRKKLLKIREIVESQEKYRRKDVNAPAGLMSLLC
>Channa_argus
MRKKLLKIRKIVESQEKLNRKDVKASGALESGRC
>Xenopus_laevis
MGKKLLKNRKGI----------------------
>Menidia_menidia
MRKKLLKIRKIVESPEKFNRKDVEASGALKYRQC
>Oryzias_melastigma
MRKKFVKIRKTVESPEKHSRKDVKATRAFESRQC
>Mustela_putorius_furo
MRKKLLEIPRAV----------------------
>Pygocentrus_nattereri
MRKKLLKIRRTVESPEKRPRKDVKAPTGLVSLWC
>Myotis_lucifugus
MRKKLLEIPRAV----------------------
>Maylandia_zebra
MRKKLLKIRKIVESEEKLNRKDDKASGTPVSRQC
>Clupea_harengus
MRKKLLKIRKIVETQEKHRRKDVNASADVGSLRC
>Poecilia_reticulata
MRKKLQKIRKIVESQEKVHRKDVEAPRAP-----
>Esox_lucius
MRKKLLKIRKIVESQEKLHRKDVKAPVAPESWRC
>Lateolabrax_japonicus
MRKKLLKIRKIVESQEKLNRKNVKAPGAPESGQC
>Notothenia_coriiceps
MRKKLLKIRKIVESEEKPNRKNVKATGAPESGRC
>Manis_javanica
MRKKLLEIPRAV----------------------
>Platichthys_stellatus
MRKKLLKIRKIVESEEKLNRKDVKASGAPESGRC
>Rhinopithecus_bieti
MRKKLLEIPRAV----------------------
>Rattus_norvegicus
MRKKLLEIPRAV----------------------
>Lepisosteus_oculatus
MRKKLLKIRKILQS--------------------
>Trichomycterus_areolatus
MRKNLLKIRKIVESQEKGSRKDAKAPKCLVSLWC
>Takifugu_rubripes
MRKKLIKILKLVESQEKLNRKDAEAPRAPGSSWC

HG0125.2 ENSG00000181690 PLAG1

>Homo_sapiens
MLKPRDWPKWEGLDSTLFHEESMGLAKIKV
>Mustela_putorius_furo
MLKPRDWPKWEGLDSALFHEESMGLAKIKV
>Lipotes_vexillifer
MLKPRDWPKWEGLDSALFHEESMELAKIKV
>Myotis_davidii
MLKPRDWPKWEGLDSALFHEESVGLAKIKV
>Dasypus_novemcinctus
----------------------MGLAKIKV
>Galeopterus_variegatus
MLKPRDWPKWEGLDSTLFHEESMELAKIKV
>Erinaceus_europaeus
MLKPRDWPKWEGLDSTLLHEESMELAKIKV
>Ochotona_princeps
MLKPRDWPKWEGLDSTLFHEELMGLAKIKI
>Elephantulus_edwardii
MVKPRDWPKWEGLDSTLFHKD---------
>Equus_caballus
MLKPRDWPKWEGLNSALFHEESMGLAKIKV
>Manis_javanica
MLKPRDWSKWERLDSALFYEESMGLAKIKV
>Loxodonta_africana
MVKPRDWPKWEGLDSALFHKESVGLAKIKV
>Jaculus_jaculus
MLKPRDWPKWEGLDSTLFHEELMGLAKIRV
>Tupaia_chinensis
MLKPRDWPKWEGLDSTLFHEESMGLAKI--
>Trichechus_manatus_latirostr
MVKLCDWPKWEGLDSALFHKESMGLAKIKV
>Anolis_carolinensis
--------------------MSVELVKIKT
>Orycteropus_afer_afer
MVKPRDWPKWEELDSALFHKESMGLAKIKV

HG0126.1 ENSGALG00000010899 G6PC2

```
>Gallus_gallus
MNCEKIQANMFQLFIRRIA---------------------------
>Anser_cygnoides
MNCEKIQANMFQLFIERIA---------------------------
>Chaetura_pelagica
---------MFQLFIQRIS---------------------------
>Apteryx_australis_mantelli
MNCEKIQANMFQLFIQRIA---------------------------
>Buceros_rhinoceros_silvestri
MDCEKIRANMFQLFTQRIA---------------------------
>Caprimulgus_carolinensis
---------MFQLFIQRIA---------------------------
>Merops_nubicus
---------MFQLFIQRTA---------------------------
>Cuculus_canorus
MNCEKNQANMFQLFIQRIS---------------------------
>Haliaeetus_albicilla
MNCEKIQANMFQLFIQRIA---------------------------
>Gavia_stellata
MNCEKIQANMFQLFIQRIA---------------------------
>Chlamydotis_macqueenii
MNCEKIQANMFQLFIQRIA---------------------------
>Opisthocomus_hoazin
MNCEKIQANMFQLFIQRIAPVTHCASTKLSPQFAAAQHVLHLVSSN
>Acanthisitta_chloris
MNCDKIQANMFQLFIQSIA---------------------------
>Phalacrocorax_carbo
MNCEKIQANMFQLFIQRIA---------------------------
>Picoides_pubescens
MNCEKIQANMFQLFIQRIA---------------------------
>Fulmarus_glacialis
MNCEKIQANMFQLCIQRIA---------------------------
>Nestor_notabilis
MNCEKIQANMFQLFIERIA---------------------------
>Aptenodytes_forsteri
MNCEKIQANMFQLFIQRIA---------------------------
>Tyto_alba
MNCEKIQANMFQLFIQRIA---------------------------
>Struthio_camelus_australis
MNCEKIQANMFQLFIQRIA---------------------------
>Chrysemys_picta
MNCVKIQANIFQLFIQRVA---------------------------
```

HG0126.2 ENSGALG00000010899 G6PC2

```
>Gallus_gallus
MCFPRDSEPRVPGMAGYATLIQYKAQPSGWY----------------
>Anas_platyrhynchos
MCFPGDLEPRVPGMAGYTPPIRYKGNVNHLLSLFNSSLPWLSFSLET
>Apteryx_australis_mantelli
MCFLGDLESHVPGMAGYTTLIQYKAQSSGWY----------------
>Buceros_rhinoceros_silvestri
MCFPGDLEPRVPGMAG-------------------------------
>Caprimulgus_carolinensis
-------------MAGYTTPIQYKAQPSDWY----------------
>Cuculus_canorus
MCFPGDLEPRVPGMSGYTMAIQYKAQPSGWY----------------
>Haliaeetus_albicilla
-------------MAGYTTPIQYKAQPSGWY----------------
>Gavia_stellata
MCFPGDLEPRVPGMAGYTTPIQYKAQPSGWY----------------
>Opisthocomus_hoazin
MCFPGDLETRVPGMAGYTTAIQYKAQPSGWY----------------
>Corvus_cornix_cornix
MCFPRDLEPRVSGMAGYTTPI--------------------------
>Phalacrocorax_carbo
MCFPGDLEPRVPGMAGCTTLIQYKAQPSGWY----------------
>Fulmarus_glacialis
MCFPGDLEPRVPGMAGYTTPIQYKAQPSGWY----------------
>Aptenodytes_forsteri
-------------MAGYTTLIQYKAQPSGWY----------------
>Tyto_alba
-------------MAGCTTPIQYKAQPSGWY----------------
>Struthio_camelus_australis
MCFLRDLESRVPGMAGTPL----------------------------
>Tinamus_guttatus
MCFPGDLESHVPGMAGYTTLIQYKGNVNHFLSFFNSSLPWFSFSLI-
```

HG0127.1 ENSGALG00000029709

```
>Gallus_gallus
MQNQSTH-EEKSPLRGNVRGIAVGDNLLHSPTP-----
>Anas_platyrhynchos
MQNPSAH-EEKSPLHGNVRGIALGDNLLHSAAP-----
>Buceros_rhinoceros_silvestri
MQNSSTH-EEKSPL--VVMSEE----------------
>Caprimulgus_carolinensis
MQNPST--------------------------------
>Cariama_cristata
MQNPSTH-EEPSHG--NVRGIAIGDNLLHSATL-----
>Calidris_pugnax
MQNPSTH-EEKSPL--IVMLEE----------------
>Colius_striatus
MQNPSTH-EEKSSL--VVMLEE----------------
>Columba_livia
-----------MISRGNVRGIAAGDNPLHSAAL-----
>Falco_cherrug
-----------MISRGNVRGIAVGDNPLHSATVQDKHR
>Gavia_stellata
MQNPSTH-EENSPL--VVMLEE----------------
>Eurypyga_helias
MQNPSTH-EEKSPL--VVMLEE----------------
>Corvus_brachyrhynchos
-----ME-DFEGERHGNVRGIAVGDNLLHSATL-----
>Pelecanus_crispus
MQNPSTH-EEKSPL--VVMLEE----------------
>Picoides_pubescens
MQNPSSH-EEKSPLL--VTLEE----------------
>Fulmarus_glacialis
MQNPSTH-EEESPL--VVMLEE----------------
>Pygoscelis_adeliae
MQNPSTH-EEKSPL--VVMLEE----------------
>Tyto_alba
MQNPSTH-EEKSPL--VVMLEE----------------
>Struthio_camelus_australis
MQNLFTQ-EERS-----CLGIATGDNLLHSDPLQEKHH
>Trachemys_scripta
MQNHTVSQEENSPHYA----------------------
>Tinamus_guttatus
MQNPSTQ-EEKSPL------------------------
>Apaloderma_vittatum
MQNPSTH-EEKSPL--VIMLKE----------------
```

HG0127.2 ENSG00000109452 INPP4B

>Homo_sapiens
-------------------------------MVLPGKRSSVFYEKLDKGFTGGIVLEICS
>Canis_lupus_familiaris
-------------------------------MVLPGKRSVVFYEKSDQGFQRGI------
>Myotis_brandtii
-------------------------------MVLPGKRSALSYEKSDQEFQSGISGL---
>Dasypus_novemcinctus
-------------------------------MVLPGKKSAVFYEKSSQEL----------
>Condylura_cristata
--------------------------MEPKEMVLPGRRFAVFYEKTDQEL----------
>Oryctolagus_cuniculus
-------------------------------MVLPGKRSVL-YEKSDQEP----------
>Fukomys_damarensis
MKYNLTRNCTIVTLRIFHLLKVEIRHQEPKEMVLPGKRSAVFYEKSDQES----------
>Tupaia_chinensis
-------------------------------MVLPGKRAAVFSEKSDQEP----------

HG0128 ENSDARG00000091029 phox2bb

>Danio_rerio
------------------------------------------------MRTICSRDLGLGGSLGGFS
>Anas_platyrhynchos
------------------------------------------------MRTICSRDLRQGIS-----
>Apteryx_australis_mantelli
------------------------------------------------MRTICSRDLRQGIS-----
>Oryzias_latipes
------------------------------------------------MRTICSRDLGLGGS-----
>Mustela_putorius_furo
------------------------------------------------MRTICSRDLRQGIS-----
>Rousettus_aegyptiacus
MQESMIGRLDSSVPQCQPNRRMSYCHVKSASNKTNRFAIVQVERAKFIMRTICSRDLRQGIS-----
>Haplochromis_burtoni
------------------------------------------------MRTICSRDLGLGGS-----
>Austrofundulus_limnaeus
------------------------------------------------MRTICSRDLGLGGS-----
>Monodelphis_domestica
------------------------------------------------MRTICSRDLRQGFS-----
>Gallus_gallus
MQESAIGRLDSSVPQCQPNRLMSYCHVKSASNKTNRFAIVQVERAKFIMRTICSRDLRQGIS-----
>Oryctolagus_cuniculus
MQESMIGRLDSSVPQCQPNRRMSYCHVKSASNKTNRFAIVQVERAKFIMRTICSRDLRQGIS-----
>Corvus_brachyrhynchos
MQESAIGRLDSSVPQCQPNRLMSYCHVKSASNKTNRFAIVQVERAKFIMRTICSRDLRQGIS-----
>Ceratotherium_simum_simum
------------------------------------------------MRTICSRDLRQGIS-----
>Cercocebus_atys
MQESMIGRLDSSVPQCQPNRRMSYCHVKSASNKTNRFAIVQVERAKFIMRTICSRDLRQGIS-----
>Microtus_ochrogaster
MQESMIGRLDSSVPQCQPNRRMSYCHVKSASNKTNRFAIVQVERAKFIMRTICSRDLRQGISQEAFS
>Salmo_salar
------------------------------------------------MRTICSRDLRLGGS-----
>Lepisosteus_oculatus
------------------------------------------------MRTICSRDLRQGCS-----
>Pelodiscus_sinensis
---------------------MSYCHVKSASNKTNRFAIVQVERAKFIMRTICSRDLRQGIS-----

HG0128 ENSG00000109132 PHOX2B

>Homo_sapiens
MQESMIGRLDSSVPQCQPNRRMSYCHVKSASNKTNRFAIVQVERAKFIMRTICSRDLRQGIS-----
>Anas_platyrhynchos
------------------------------------------------MRTICSRDLRQGIS-----
>Xenopus_tropicalis
---------------------MSYCHVKSTSNKTNRFAIVQVERAKFIMRTLCSGDPRRGSS-----
>Apteryx_australis_mantelli
------------------------------------------------MRTICSRDLRQGIS-----
>Oryzias_latipes
------------------------------------------------MRTICSRDLGLGGS-----
>Mustela_putorius_furo
------------------------------------------------MRTICSRDLRQGIS-----
>Hynobius_retardatus
---------------------MSYCHVKSASPETNRPAIVQVQRAQCTSGAAML-------------
>Rousettus_aegyptiacus
MQESMIGRLDSSVPQCQPNRRMSYCHVKSASNKTNRFAIVQVERAKFIMRTICSRDLRQGIS-----
>Haplochromis_burtoni
------------------------------------------------MRTICSRDLGLGGS-----
>Rutilus_rutilus
------------------------------------------------MRTICSRDLGLGGSLGGFS
>Poecilia_formosa
------------------------------------------------MRTICSRDLGLGGS-----
>Monodelphis_domestica
------------------------------------------------MRTICSRDLRQGFS-----
>Meleagris_gallopavo
MQESAIGRLDSSVPQCQPNRLMSYCHVKSASNKTNRFAIVQVERAKFIMRTICSRDLRQGIS-----
>Apteronotus_leptorhynchus
------------------------------------------------MRTICSRDLGLGGSLGGF-
>Oryctolagus_cuniculus
MQESMIGRLDSSVPQCQPNRRMSYCHVKSASNKTNRFAIVQVERAKFIMRTICSRDLRQGIS-----
>Corvus_brachyrhynchos
MQESAIGRLDSSVPQCQPNRLMSYCHVKSASNKTNRFAIVQVERAKFIMRTICSRDLRQGIS-----
>Ceratotherium_simum_simum
------------------------------------------------MRTICSRDLRQGIS-----
>Microtus_ochrogaster
MQESMIGRLDSSVPQCQPNRRMSYCHVKSASNKTNRFAIVQVERAKFIMRTICSRDLRQGISQEAFS
>Salmo_salar
------------------------------------------------MRTICSRDLRLGGS-----
>Lepisosteus_oculatus
------------------------------------------------MRTICSRDLRQGCS-----
>Pelodiscus_sinensis
---------------------MSYCHVKSASNKTNRFAIVQVERAKFIMRTICSRDLRQGIS-----

HG0129 ENSDARG00000099437

>Danio_rerio
--------------------------------------------------------------MPAVFLLLRSLVVRLFGSRLAASGVQLLRRILTTATGHLGTVLRNIWERISSQQSKEAILGCVLCLLNMHKKVDN------
>Amia_calva
----------------------------------------------------------------------------------------------------------------------------------MHKKVDN------
>Channa_argus
--------------------------------------------------------------MPAVFILLRSLVIRLLSSRLAGSVAQFLRRSLSTGIAHLGTALRHVWDCVRSQESKEAILGCVLCILNMHKKVEN------
>Xenopus_laevis
--------------------------------------------------------------MPVPFLLIRTLLVRAAGSRLAVSGARQLTKGARWARARLLILLQRLWDRLTSEETRQALLSCVLCLLNLKHKSNT------
>Menidia_menidia
--------------------------------------------------------------MPAVFILLRSLVIRLLSSRLAGSLAQFLRRSLSTGAAHLGTALRHIWDRVRSQESKEAILGCVLCILNMHKKVEN------
>Porichthys_notatus
----------------------------------------------------------------------------------------------------------------------------------MHKKVEN------
>Oryzias_melastigma
----------------------------------------------------------------------------------------------------------------------------------MHKKVEN------
>Ambystoma_mexicanum
--------------------------------------------------------------MPLPFLLLRTLLIRLAGSRLAASGVQFLRKRLAWAGARLAPLLRRVWRRATSEESRQAVLGCVLCVLNLRKRVDGE-----
>Astyanax_mexicanus
----------------------------------------------------------------------------------------------------------------------------------MHKKVDN------
>Oreochromis_niloticus
--------------------------------------------------------------MPAVFILLRSLVIRLLSSRLAGSVAQYLRRILSTGATHLGTALRHIWDRVRSQESKEAILGCVLCILNMHKKVEN------
>Alosa_alosa
--------------------------------------------------------------MPAVFLLLRSLVIRLLSSKLAGSAAQFLRRSLSAAVSHLGTALRHIWDRVRSQESKEAILGCVLCILNMHKKVDN------
>Latimeria_chalumnae
--------------------------------------------------------------MPALFLLLRTLFIRLLGSQLAVSAGLFLRRSLAAAGGRLAALLRRVWDRIRSEESRQVALSCVLCILNLHKKVDE------
>Nothobranchius_furzeri
--------------------------------------------------------------MPAVFILLRSLVIRLLSSRLAGSLAQFLRRSLTTGAAHLGTALRHIWERVRSQESKEAVLGCVLCILNMHKKVEN------
>Esox_lucius
--------------------------------------------------------------MPAVLVLLRA----LLSSRLTGSLVQFLRRTLSSTFGHVGAALRYVWDRISSQESKEAILGCVLCLLNMHKKVDEPEAESR
>Apteronotus_leptorhynchus
----------------------------------------------------------------------------------------------------------------------------------MQKKVDN------
>Lutjanus_erythropterus
----------------------------------------------------------------------------------------------------------------------------------MHKKVEN------
>Sebastes_caurinus
--------------------------------------------------------------MPAVFILLRSLVIRLLSSRLAGSVAQFLRRTLSTGAAHLGTALRHVWDRIRSQESKEAVLGCVLCILNMHKKVET------
>Platichthys_stellatus
---------------------------------------------------------------MPVFILLRSLVIRLLSSKLAGSLAQYLRRGLSAGAAHLGTALRYVWDRIRSQESKEAVLGCVLCVLGMAKKVEK------
>Salmo_salar
MQERENPCKRERIPVREREREKSKTRSRSQRGGDFIPVLFLSFLSVFLLLISLCVSGCGSSIMPAVFLFLL---RTLLSSRMIGSLLQFLRRTLSSTFSHVGAALRYVWDRISSQESKEAILGCVLCLLNMHKKVDQPAAESH
>Lepisosteus_oculatus
--------------------------------------------------------------MPAFFLLLRSLLIRLLSSKLAGSAAQFLRRSLSATASHIGSLLRHIWDRIRSQESREAILSCVLCILNMHKKVDN------
>Trachemys_scripta
--------------------------------------------------------------MPAPLLLLPLLR-------------TLLARLLLLARKRLLPLLRRLGARLGSRESREAVLTCLLCILNLRKKADD------

HG0130 ENSGALG00000002993

>Gallus_gallus
MSCWEIRRIFFSLVEEEPCPGG-WLH-GL-------F-----WRAKEAKGSCLSRTQR----ELGHVLSAHL-GNTVCKSRRKIAGIIQTAILIFVIFFRRGMHLA-------------------------------
>Anas_platyrhynchos
MSCWEIWWIFYSLVEEEPCAGG-RLR-AL-------F-----WRAKEGKGSRLSRTER----ELGHVLSAHL-GNTGCKSERKIAGIVQTAILIFVIFFRRGMHLA-------------------------------
>Panthera_tigris
MSSWGARRIGVSFLQEEPGAGGFRWLPPPLL---GLFPPLVRGGEETGDRGDLRGS-------EASDMCQHIWGAHPSSPRGDLPEQFKLRY---------------------------------------------
>Orcinus_orca
MTSWGSRRIGVSFLQEEPRAGGFRWLPPPLL---VLFPPLV-QGVETGEQGDFPGS-------EASDMCQHIWGTHPSSPRGDLLEQFKLRY---------------------------------------------
>Calidris_pugnax
MSCWEIWWIFFSLVEEEPGAGS-WLR-GL-------F-----WRAKEGKGSHLSRGER----ELGHVLSAHL-GNTVCKSERKFAGIVQTAILIFLIFFRRGMHLA-------------------------------
>Pteropus_alecto
MSSWGARRIGVSFLQKEPGAGGFRWLPPPLL---VLFPPLRRG-AETGERGHLRGS-------EASDMCQHIWGAHPSSPRGDLPEQFKLRY---------------------------------------------
>Columba_livia
MSCWEIWWIFFSLVEEEPGAGS-WLR-GL-------F-----WRAKEGKGYRLSHAES----SDMC--CQHIWGIQSASPKGNLLGSFK-------VRY--------------------------------------
>Aquila_chrysaetos_canadensis
MSCWEIWWIFFSLVEEEPGAGG-WLR-GL-------F-----WRVKEGKGSRLSRAER----ELGHVLSAHL-GNTVCKSERKFAGIVQTAILIFVIFFRRGMHLA-------------------------------
>Chlamydotis_macqueenii
MSRWEIWWIFFLLVEEEPGAGS-WLR-GV-------F-----WRAKEGKGSRLSRAER----ELGHVLSAHL-GNTVCKSERKFAGIVQTAILIFVIFFRCGMHLA-------------------------------
>Condylura_cristata
MSSWGARRIGVSFLQEEPRAGGFRWLLPPLL---VLFPPLVLG-AETGERGDLRSS-------EASDMCQHIWGAHQSSPRGDLPEQFKLRY---------------------------------------------
>Oryctolagus_cuniculus
MSSWGARRIGVSFLQERAQRGWLQVAGAAAA--------AARADSAAGPRSRDAGAGRPRRQRGIGHVSAHL-GRTSVEPEGSFAGTLQTAILILGVTVPGRLSGGSASVRPLESARACVCVCRVGPPLPPPCFPVE
>Elephantulus_edwardii
MSSWGARRIGVSLVPEEPSAGGFRWLPPPPL---LVLFPALGRRAETGERGDLQGG-------EASDMCQHIWGAQPSSPRGDLPEQFKLRY---------------------------------------------
>Lepidothrix_coronata
MSCWEIWWIFFSLVEEEPGAGS-WLR-GL-------F-----WRAKEGKGSRLSRAER----EPGHVLSAHL-GNTVCKSERKFAGIVQTAILIFVIFFRRGMHLA-------------------------------
>Ceratotherium_simum_simum
MSSWGARRIGVSFLPEEPSAGGFRWLPPPLL---VLFPPLVRG-GETGERGDLRGS-------EASDMCQHIWGAHPSSPRGDLPEQFKLRY---------------------------------------------
>Carlito_syrichta
MSSWGARRIGVSFLQEEPSAGGFRWLPPPSL---LVLSPPRVRGAETRERGDLRGS-------EASDMCQHIWGAHPSSPRGDLPEQFKLRY---------------------------------------------
>Loxodonta_africana
MSSWGARRIGVSLVQEEPSAGGFRWLPPPLL---VLFPPHV-RRAETGEWGDLLGS-------EASDMCQHIWGAHLSSPRGDLLEQFKLRY---------------------------------------------
>Microtus_ochrogaster
MSSWGARRIGVSVLQEEPRAGGFRWLTPPPPRSPQVLFPPLVRGAKTQEQGDLRGK-------EASDMCQHIWGAHPKSPRGDLPKQFKLRY---------------------------------------------
>Trichechus_manatus_latirostr
MSSWGDRQIGVSLVQEEPIAGGFRWLLPPLL---VLFPPLV-RRAETGERGDVLGS-------EASDMCQHTWGAHPSSPRGDLPEQFKLQY---------------------------------------------
>Chrysemys_picta
MSCWEIWWIFVSLLEEEPRAGGCRLL-GL-------L-----WRAKEGKGSRLSRAER----ELGHVLSAHL-GNTVCKSERRFAGIVQSAILIFVISFRRGMHLA-------------------------------
>Calypte_anna
MSCWEIWWIFFSLVEEEPGAGS-WLR-GL-------F-----WRLKEGKGSRLSRAGR----ELGHVLSAHL-GNTVCKSQRKFAGIVQTAILIFVIFFRRGMHLV-------------------------------
>Orycteropus_afer_afer
MSSWGARRIVVSLVKEEASTGSFRWLLPPPPL--LVLCSPLVRRAETGERGDLLGS-------EASDMCQHICGARPSSPRGDLPEQFELRY---------------------------------------------

HG0131 ENSGALG00000013135 GALNT1

>Gallus_gallus
------------------------------------MAVNLQRRYR-TWSLKLD---
>Anas_platyrhynchos
------------------------------------MAVNLQRRYR-TWSLKLD---
>Apteryx_australis_mantelli
------------------MLYSNAVGSHHQCLWTIQMAVNIQRRYR-TWSLKLD---
>Buceros_rhinoceros_silvestri
------------------------------------MAVNLQRRYR-TWSLKLD---
>Charadrius_vociferus
------------------------------------MAVNLQRRYR-TVCLKLD---
>Colius_striatus
------------------------------------MAVNLQRRYR-TWSVKLD---
>Columba_livia
--MSEWPRPVCYSMVPTTGALQQCIGILPPVLVDLQMAVNLQRRYRRTWSLKLD---
>Leptosomus_discolor
MFMSEWPRPVCS-MVPTTGALQQCIGILPPVLVDHPDAVNLQRRYRT-WSLKLD---
>Alligator_mississippiensis
------------------------------------MAVNLQKRYRTWSQTGLEQSC
>Cuculus_canorus
------------------------------------MAVNLQRRYR-TWCLKLD---
>Aquila_chrysaetos_canadensis
------------------------------------MAVNLQRRYR-TWSLKLD---
>Mesitornis_unicolor
------------------------------------MAVNLQRRYR-T---------
>Tauraco_erythrolophus
------------------------------------MAVNLQRRYR-TWCLKLD---
>Opisthocomus_hoazin
------------------------------------MAVNLQRRYR-TWSLKLD---
>Acanthisitta_chloris
------------------------------------MAVNLQRRYR-TWSLKLD---
>Phaethon_lepturus
-------------MVPTTGAPRQCIGILPPVLVDHPDAVNLQRRYRT-WSLKLD---
>Melopsittacus_undulatus
------------------------------------MAINLQKRYRT-WCLKLD---
>Pygoscelis_adeliae
------------------------------------MAVNLQRRYR-TWSLKLD---
>Struthio_camelus_australis
--MSEWPRPIWILPQ---VLYSNAVGSRHQCWWTIQMAVNIQRRYR-TWSLKLD---
>Chelonia_mydas
------------------------------------MAVNLQKRYRT-WSRKLD---
>Tinamus_guttatus
--------------------------------------MAIQRRYRM-WCLKLD---

HG0132 ENSG00000151067 CACNA1C

```
>Homo_sapiens
----MRYSFTRGGSSG-KEQFLGFDAIMGIR
>Anas_platyrhynchos
----MHCNFTKGGSSG-KDQFLVFDAIMGIR
>Xenopus_laevis
----MQSFSAKGGSRG-TDQTRGFDAIMGIR
>Panthera_tigris
----MHYNFTKGGSSG-KEQFLVFDAIMGIR
>Hynobius_retardatus
----MHCYPTRGDCSGEKEQFLGFDSIMGIR
>Tursiops_truncatus
----MHYNFTEGGSSG-KDQFLVFDAIMGIR
>Calidris_pugnax
----MHCNFTKGGSSG-KDQFLVFDAIMGIR
>Pteropus_vampyrus
----MHYNFTQGGSSG-KDQFLVFDAIMGIR
>Dasypus_novemcinctus
----MHYNFTQGGSSG-KDQFSVFDAIMGIR
>Galeopterus_variegatus
----MHDSLSEGGGSG-RQQFAVFDAIMGIR
>Gallus_gallus
----MHCNFTKGGSSG-KDQFLVFDAIMGIR
>Condylura_cristata
----MHCNLTKGGGRG-KAQFLVFDAIMGIR
>Oryctolagus_cuniculus
----MHCKLTRGGSSG-KEQFLVFDAIMGIR
>Scleropages_formosus
MYRGMRCSVLLGGSSE--CLFLFFNSIMGIR
>Lepidothrix_coronata
----MHCNFTKGGSSG-KDQFLVFDAIMGIR
>Ceratotherium_simum_simum
----MHYNSTQGGSSG-KDQFLVFDAIMGIR
>Manis_javanica
----MYYNFTKGGSSG-KEQFWVFDAIMGIR
>Chinchilla_lanigera
----MHYNLTKGGSSG-KEQFLVFDAIMGIR
>Anolis_carolinensis
----MLCNCTKGGSSG-KDQILGFDAIMGIR
>Struthio_camelus_australis
----MHCNFTKGGSSG-KDQFLVFDAIMGIR
>Tinamus_guttatus
----MHCNLTKGGSSG-KEQFLVFDAIMGIR
```

HG0133.1 ENSDARG00000051886 ankrd11

>Danio_rerio
------MESTVDDDAMEA
>Acipenser_sinensis
------MESTVDDDAMEA
>Channa_striata
------MESTVDDDAMAA
>Menidia_menidia
------MESTVDDDAMAA
>Porichthys_notatus
------MESTVDDDAMAA
>Oryzias_melastigma
------MESTVDDDAMAA
>Astyanax_mexicanus
------MESTVDDDAMEA
>Haplochromis_burtoni
------MESTVDDDAMAA
>Alosa_alosa
------MESTVDDDAMAA
>Kryptolebias_marmoratus
------MESTVDDDAMAA
>Esox_lucius
------MESTVDDDAIEA
>Acanthogobius_hasta
------MESTVDDDAMAA
>Scleropages_formosus
------MESTVDDDAMEA
>Anoplopoma_fimbria
------MESTVDDDAMAA
>Platichthys_stellatus
------MESTVDDDAMAA
>Salmo_salar
------MESTVDDDAMEA
>Lepisosteus_oculatus
------MESTVDDDAMEP
>Ictalurus_punctatus
------MESTVDDDAMEA
>Squalus_acanthias
MPWWRAMESTVDDDAMDA
>Takifugu_rubripes
------MESTVDDDAMAA

HG0133.1 ENSG00000167522 ANKRD11

>Homo_sapiens
MESQTVDDEAPAV
>Canis_lupus_familiaris
MESQTVDDEAMAV
>Miniopterus_natalensis
MESQTVDDEAMAV
>Ceratotherium_simum_simum
MESQTVDDEAMAV
>Loxodonta_africana
MESQTVDDEAMAV
>Chinchilla_lanigera
MESQTVDDEAMAV

HG0133.2 ENSDARG00000051886 ankrd11

>Danio_rerio
MGKTSIGREYQV
>Pygocentrus_nattereri
MGKSSIGREYQA
>Esox_lucius
MGKTSIGQEYQG

HG0133.3 ENSG00000167522 ANKRD11

>Homo_sapiens
-MGSARA--------PPLSRRGAAQPSEHEYIQPCSGTHLHWI------
>Rousettus_aegyptiacus
MAGPNGEGQIVNRTEVEIIYLTKLYSLVSEYTQHCSGHTLYWI------
>Psammomys_obesus
-MGSARA--------PPPFRRGAASPSEPEHTV-----LLEFIPSPEQH
>Pelodiscus_sinensis
--------------------------MEEEQIHSCFGTHLYWI------

HG0134.1 ENSGALG00000041267 BMI1

>Gallus_gallus
MAAWLTFIFC
>Odobenus_rosmarus_divergens
MAAWLTFIFC
>Ambystoma_mexicanum
MAAWLTFIFC
>Lipotes_vexillifer
MAAWLTFIFC
>Astyanax_mexicanus
MAAWVAFIFC
>Calidris_pugnax
MAAWLTFIFC
>Latimeria_chalumnae
MAAWLTFIFC
>Alligator_mississippiensis
MAAWLTFIFC
>Danio_rerio
MAAWLVFIFC
>Sarcophilus_harrisii
MAAWLTFIFC
>Ochotona_princeps
MAAWLAFIFC
>Corvus_brachyrhynchos
MAAWLTFIFC
>Ceratotherium_simum_simum
MAAWLTFIFC
>Manis_javanica
MAAWLTFIFC
>Loxodonta_africana
MAAWLTFIFC
>Chinchilla_lanigera
MAAWLTFIFC
>Pygoscelis_adeliae
MAAWLTFIFC
>Anolis_carolinensis
MAAWLTFIFC
>Chelonia_mydas
MAAWLTFIFC
>Orycteropus_afer_afer
MAAWLTFIFC

HG0134.2 ENSG00000168283 BMI1

```
>Homo_sapiens
MRGRDRGETMGMWAREPRSGLAAPPSPAE
>Odobenus_rosmarus_divergens
---------MGMLAREPRSGLETPPSPAE
>Orcinus_orca
---------MGMLAREPRSGLETPPSPAE
>Pteropus_vampyrus
---------MGMLAREPRSGLETPPSPAE
>Ochotona_princeps
---------MGMLARAPRSGLETPPGPAE
>Ceratotherium_simum_simum
---------MGLLAREPRSGLETPPSPAE
>Loxodonta_africana
---------MGRLAREPRSGLETPPSPAE
>Dipodomys_ordii
---------MGMWAREPRSGLETPPRPAE
>Orycteropus_afer_afer
---------MGLLAREPQSGLETPPSPAE
```

HG0135 ENSDARG00000094132 igf1

>Danio_rerio
MLPQLFPVENVSVM
>Acipenser_sinensis
MFPQLFPVENVCVM
>Amia_calva
MFLQLFPVENVCVM
>Xenopus_laevis
MLPQLFPVYSVCVM
>Menidia_menidia
MLPQLFPVENVCVM
>Oryzias_melastigma
MLPQLFPVENVCVM
>Austrofundulus_limnaeus
MLPQLFPVENVCVM
>Meleagris_gallopavo
MLPQLFPVYSVCVM
>Parus_major
MLPQLFPVYSVCVM
>Mesocricetus_auratus
MLPQLFPVYSVCVL
>Oncorhynchus_kisutch
MLPQLFPVENVCVM
>Lepisosteus_oculatus
MFPQLFPVENVCVM

HG0135 ENSG00000017427 IGF1

>Homo_sapiens
MFPQLFPVYSVCVL
>Acipenser_sinensis
MFPQLFPVENVCVM
>Amia_calva
MFLQLFPVENVCVM
>Xenopus_laevis
MLPQLFPVYSVCVM
>Menidia_menidia
MLPQLFPVENVCVM
>Oryzias_melastigma
MLPQLFPVENVCVM
>Ailuropoda_melanoleuca
MFPQLFPVYSVCVL
>Myotis_brandtii
MFPQLFPVYSVCVL
>Dasypus_novemcinctus
MFPQLFPVYSVCVL
>Austrofundulus_limnaeus
MLPQLFPVENVCVM
>Monodelphis_domestica
MFPQLFPVYSVCVL
>Meleagris_gallopavo
MLPQLFPVYSVCVM
>Erinaceus_europaeus
MFPQLFPVYSVCVL
>Oryctolagus_cuniculus
MFPQLFPVYSVCVL
>Parus_major
MLPQLFPVYSVCVM
>Equus_caballus
MFPQLFPVYSVCVL
>Nannospalax_galili
MFPQLFPVYSVCVL
>Oncorhynchus_kisutch
MLPQLFPVENVCVM
>Tupaia_chinensis
MFPQLFPVYSVCVL
>Lepisosteus_oculatus
MFPQLFPVENVCVM

HG0136.1 ENSG00000125845 BMP2

```
>Homo_sapiens
------------------------------------------------------MWTLFQWTCPRV---LLRRTAVS
>Felis_catus
------------------------------------------------------MWRLFQWTCPRV---LLRRSAVS
>Physeter_catodon
------------------------------------------------------MWRLFQWTCPRV---LLRRSAVS
>Eptesicus_fuscus
------------------------------------------------------MWRLFQWTCPRV---LLRRSAVS
>Dasypus_novemcinctus
------------------------------------------------------MRRRSRWTCPRV---LPRRSAAS
>Latimeria_chalumnae
------------------------------------------------------MWILFQWTCSSV---LLSQSAAS
>Galeopterus_variegatus
------------------------------------------------------MWRLFQWTCPRV---LLRRTAVS
>Monodelphis_domestica
------------------------------------------------------MWRLFQWTCPRV---LLSQSAVS
>Condylura_cristata
------------------------------------------------------MWRFFQWTCPRV---LLRRSAVS
>Oryctolagus_cuniculus
------------------------------------------------------MWRLFQWTCPRG---LLRRTAVS
>Ornithorhynchus_anatinus
MRKKTHREQRRLPQTPAPPPSPSSSRSPSRSPWSPPRPAPDPPAAPVSSARDFSMWRLFRWTCPRV---LLRQSAVS
>Ceratotherium_simum_simum
------------------------------------------------------MWRLFQWTCPRV---LLRRSAVS
>Manis_javanica
------------------------------------------------------MWRLFQWTCPRV---LLRRSAVS
>Loxodonta_africana
------------------------------------------------------MWRLFQWTCPRV---LLRRSAVS
>Dipodomys_ordii
------------------------------------------------------MWRLFQWTCPRV---LLRRTAVS
>Tupaia_chinensis
------------------------------------------------------MWRLFQWTCHRV---LLRRTAVS
>Lepisosteus_oculatus
------------------------------------------------------MWKLFQWTCQRV---LLSRSPLS
>Trichechus_manatus_latirostr
------------------------------------------------------MWRLFQWTCLRV---LLRRSAVS
>Trachemys_scripta
------------------------------------------------------MWSPFQWTCPSAPPLLLRRSAGS
>Orycteropus_afer_afer
------------------------------------------------------MWRLFQWTCPRV---LLRRSAVS
```

HG0136.2 ENSG00000125845 BMP2

>Homo_sapiens
MRSTYCRRSGAWDALAEV
>Oryctolagus_cuniculus
MPSTSCRGSGARDVPAEV
>Ceratotherium_simum_simum
MPITSCRSSGAWDAPAEV
>Marmota_monax
MPSTSCRGSGAWDVMVEV

HG0137.1 ENSG00000181722 ZBTB20

>Homo_sapiens
---------------MNICTRPKPWSNLPEE
>Amia_calva
---------------MKICCRPHTWTDLPEE
>Anser_cygnoides
---------------MNVCTRSKAWSNLPEE
>Panthera_tigris
---------------MNICTRPKPWSNLPEE
>Balaenoptera_acutorostrata_s
---------------MNICTRPKPWSDLPEE
>Rousettus_aegyptiacus
---------------MNICTRPKPWSDLPEE
>Dasypus_novemcinctus
---------------MNIFTRPKRWSDLPEE
>Latimeria_menadoensis
---------------MKIRVRPKRWTDLPEK
>Gallus_gallus
---------------MNVCTKSKAWSNLPEE
>Erinaceus_europaeus
---------------MNISTRPKSWSDLPEE
>Oryctolagus_cuniculus
---------------MNICTRPQPWSDLPEE
>Elephantulus_edwardii
---------------MNICTRPKPWSDLPEE
>Lepidothrix_coronata
---------------MNVCTRSKAWSNLPEE
>Equus_asinus
---------------MNICTRPKPWSDLPEE
>Manis_javanica
---------------MNICTRPQPWSDLPEE
>Cavia_porcellus
MSLFLLCRSSLFPAPMNICTRPKPWSDLPEE
>Tupaia_chinensis
---------------MNICTRPKPWSDLPEE
>Trichechus_manatus_latirostr
---------------MNICTRPKPWSDLPEE
>Pelodiscus_sinensis
---------------MNICTRAKAWSDLPEE
>Orycteropus_afer_afer
---------------MNICTRPKPWSNLPEE

HG0137.2 ENSG00000181722 ZBTB20

>Homo_sapiens
----------------MSWKCRNLSLPSRKNL-------------------------
>Mustela_putorius_furo
MRSLNSFHLLLTVQSSMSWKCRNLSLPSRKTVAGTFPPRFERKRIGWNKTLPSIWST
>Lipotes_vexillifer
----------------MSWKCRNLSLPSRKIP-------------------------
>Rousettus_aegyptiacus
MRSLNSFHLLLTVQSSMSWKCRNLSLPSRKTVTGTFPPRFERKRIGWNKTLLSIWST
>Erinaceus_europaeus
----------------MSWKCRNLSLPSRKIP-------------------------
>Elephantulus_edwardii
----------------MSWKCRNLSLPSRQIL-------------------------
>Equus_przewalskii
MRSLNSFHLLLTVQSSMSWKCRNLSLPSRKTVAGTFPPRFERKRIGWNKTLLSIWST
>Cricetulus_griseus
MSSQNSFHLLLTVQSSMSWKCRNLSLPSRKTLWNTVNSTN-----------------
>Tupaia_chinensis
MRSLSSFHLLLTVQSSMSWKCRNLSLPSRKIL-------------------------

HG0138.1 ENSG00000185070 FLRT2

>Homo_sapiens
MPRDPRCCRASVFPFHLPDPVGFG
>Anser_cygnoides
MP--EPSCSATFSPFNLPDPVGFG
>Felis_catus
MPREPRCCRASFFPFHLPDPVGFG
>Lipotes_vexillifer
MPREPRCCRASFLPFHLPDPVGFG
>Calidris_pugnax
MP--EPSCSATFSPFNLPDPVGFG
>Myotis_davidii
MPREPRCCRASFFPFHLPDPVGFG
>Dasypus_novemcinctus
MPREPRCCRASFFPFHLPDPVGFG
>Monodelphis_domestica
MQQEPSYYCTSFFSFHLPDPVGFG
>Oryctolagus_cuniculus
MPREPRCYRAGFFPFHLPDPVGFG
>Corvus_brachyrhynchos
MP--EPSCSATFSPFNLPDPVGFG
>Equus_asinus
MPREPGCCRASFFPFHLPDPVGFG
>Loxodonta_africana
MPREPRYCHASFFPFHLPDPVGFG
>Chinchilla_lanigera
MPREPHCCRASFFPFHLPDPVGFG
>Tupaia_chinensis
MPREPRSCRASFFPFHLPDPVGFG
>Trichechus_manatus_latirostr
MPRKPRYCLASFFPFHLPDPVGFG
>Crotalus_adamanteus
------MPGLSCFALHLPDPVGFG
>Struthio_camelus_australis
MP--EPSCSATFSPFNLPDPVGFG
>Pelodiscus_sinensis
MQ--EASASASVSPFHLPDPAGFG
>Tinamus_guttatus
MP--EPSSSAAFSPLNLPDPVGFG
>Orycteropus_afer_afer
MPREPRCCRASFFPFHLPDPVGFG

HG0138.2 ENSG00000185070 FLRT2

>Homo_sapiens
-------------------------------------------------------------------------------MGRRDLGGDP
>Odobenus_rosmarus_divergens
MFETCRCPLRVSHLPLSPQRNERTEITSVATISAPRRCAASARGASLALGSGCLAAARRQHCACRRVHPAPGSGWRFSTMGRRDLGGDP
>Orcinus_orca
-------------------------------------------------------------------------------MGRRDLGGDP
>Pteropus_alecto
-------------------------------------------------------------------------------MGRRDLGGDP
>Dasypus_novemcinctus
-------------------------------------------------------------------------------MGRRDLGGDP
>Oryctolagus_cuniculus
-------------------------------------------------------------------------------MGRRDLGGDP
>Equus_asinus
-------------------------------------------------------------------------------MGRRDLGGDP
>Loxodonta_africana
-------------------------------------------------------------------------------MGRRDLGGDP
>Cricetulus_griseus
-------------------------------------------------------------------------------MGRRDLGGDP

HG0139 ENSGALG00000006379 SHH

>Gallus_gallus
MCSVISDLKAAIPAPVLLGIGG-----------
>Anas_platyrhynchos
MCSVISDLKPAIPAPVLLGIGG-----------
>Apteryx_australis_mantelli
MCSVISDLKAAIPAPVLIGIGG-----------
>Charadrius_vociferus
MCSVISDLKPAIPAPVLLGIGG-----------
>Myotis_brandtii
MCSVISDQQPAAPAPVWVGIGRESPALVAGRVL
>Latimeria_chalumnae
MCSVINDLKPVISAPTLIGIAG-----------
>Columba_livia
MCSVISDLTPAIPAPVLLGIGG-----------
>Falco_cherrug
MCSVISDLKPAIPAPVLLGIGG-----------
>Oryctolagus_cuniculus
MCSVISDRQPAIRAPVWVGIGGESPAAAAGKVI
>Opisthocomus_hoazin
MCSVISDLKPAIPAPVLLGIGG-----------
>Parus_major
MCSVISDLKPAIPAPVLLGIGG-----------
>Nipponia_nippon
MCSVISDLKPAIPAPVLLGIGG-----------
>Ceratotherium_simum_simum
MCSVISDQQPAVPAPVWVGISGESPAAAAGKVI
>Manis_javanica
MCSVISAQQPAEPAPVWVGIAGQSPAAAAGKVV
>Chlorocebus_sabaeus
MCSVISGRQPAIPAPVWVGSGGECPAAAAGKVI
>Fukomys_damarensis
MCSAISDRQPAIAAPVWAEIGGEYTAAAAGKVI
>Lepisosteus_oculatus
MYSVINEQKADISAPASVGIGG-----------
>Pygoscelis_adeliae
MCSVISDLKPAIPAPVLLGIGG-----------
>Chrysemys_picta
MCSVISDLKPAIPTPVLLGIGG-----------
>Tinamus_guttatus
MCSVISDLKAAIPAPVLLGIGG-----------

HG0140 ENSG00000009307 CSDE1

>Homo_sapiens
--MKCNRTDFTT
>Apteryx_australis_mantelli
--MKCNRTEFTT
>Canis_lupus_familiaris
--MKCNRTDFTT
>Physeter_catodon
--MKCNRTDFTT
>Myotis_davidii
--MKCNRTDFTT
>Dasypus_novemcinctus
--MKCNRTDFTT
>Alligator_mississippiensis
--MKCNRTEFTT
>Sarcophilus_harrisii
--MKCNRTDFTT
>Galeopterus_variegatus
--MKCNRTDFTT
>Monodelphis_domestica
--MKCNRTDFTT
>Sorex_araneus
--MKCNRTDFTT
>Oryctolagus_cuniculus
--MKCNRTDFTT
>Ceratotherium_simum_simum
--MKCNRTDFTT
>Manis_javanica
MSMKCNRTDFTT
>Loxodonta_africana
--MKCNRTDFTT
>Jaculus_jaculus
--MKCNRTDFTT
>Trichechus_manatus_latirostr
--MKCNRTDFTT
>Python_bivittatus
--MKCNRTEFTT
>Chrysemys_picta
--MKCNRTEFTT
>Orycteropus_afer_afer
--MKCNRTDFTT

```
HG0141.1 ENSGALG00000010435 RASGRP3

>Gallus_gallus
MFVLNTSAEPSTIFLEA-QLRMQPTIK-----------
>Anser_cygnoides
MFVLNTSAEPSTIFLEERKLRMQPTIK-----------
>Charadrius_vociferus
MFVLNTSAEPSTIFLEV-ELRMQPTIK-----------
>Pterocles_gutturalis
MFVLNTSAEPSTIFLE-VQLRMQPTIK-----------
>Columba_livia
MFVLNTSAEPSTIFLEG-RLRMQPTIK-----------
>Alligator_mississippiensis
---------------------MQPAIKR----------
>Cuculus_canorus
MFVLNTSAEPSTIFLE-VQLRMQPTIK-----------
>Aquila_chrysaetos_canadensis
MFVLNTSAEPSTIFP-----------------------
>Eurypyga_helias
MFVLNTSAEPSTIFLE-VKLRMQPKIK-----------
>Tauraco_erythrolophus
MFVLNTSAEPSTIFLE-VQLRMQPTIK-----------
>Opisthocomus_hoazin
---------------------MQPTIK-----------
>Zonotrichia_albicollis
MFVLNTSAEPSTIFIE-VQLRMQPTK------------
>Egretta_garzetta
MFVLNTSAEPSTIFLE-VQLRMQPTIK-----------
>Picoides_pubescens
MFVLNTSAEPSTISPE-VQLRMQPTIK-----------
>Python_bivittatus
MFVLNTSAKLSTFFEASQR--MQLTVIEWGPSCFSLLW
>Struthio_camelus_australis
MFVLNTSAEPSTLFCRPACL------------------
>Chrysemys_picta
MFVLNTSAEPSTIFHEA---------------------
>Tinamus_guttatus
MFVLNTSAEPSTIFPE-GQLRMQPTIK-----------
>Apaloderma_vittatum
---------------------MQPTIK-----------
```

HG0141.2 ENSG00000152689 RASGRP3

>Homo_sapiens
--MIMEMVYLMLKMSLVF
>Canis_lupus_familiaris
----MEMVYLMLKMCLVF
>Balaenoptera_acutorostrata_s
--MIMEMVYMMLKMPLVF
>Rousettus_aegyptiacus
--MIMEIVCLMLKMSLVF
>Dasypus_novemcinctus
----MKMVYLMLKMSLVF
>Erinaceus_europaeus
----MEMVCLMWKMSLLF
>Ochotona_princeps
MYMIMEMVYLMLKMSLVF
>Equus_przewalskii
--MIIEMVCLMLKMSLVF
>Loxodonta_africana
--MIMEMVYLMLKMSIVF
>Tupaia_chinensis
----MEMAYLMLKMSLVF
>Orycteropus_afer_afer
--MIMEMVYLMPKMSTVF

HG0141.3 ENSG00000152689 RASGRP3

>Homo_sapiens
MLMRWLSQLRGYEVVCE---
>Mustela_putorius_furo
--MKWLSQLRGYEA-CELGE
>Equus_asinus
MLTKWLSQLRGYEKVCKLRE
>Manis_javanica
MLMKWLNQLRG---------
>Tupaia_chinensis
MLMRWLSQLRG---------

HG0142.1 ENSG00000204217 BMPR2

>Homo_sapiens
MFSPRPGYFFDIVKLRGK---
>Apteryx_australis_mantelli
MFPPKPVCFFDIVKLRGK---
>Odobenus_rosmarus_divergens
MFSPRPGYFFDIVKLRGK---
>Hynobius_retardatus
MFHPKPGCFFDIVKLRGK---
>Lipotes_vexillifer
MFSPRPGYFFDIVKLRGK---
>Pteropus_alecto
MFSPRPGYFFDIVKLRGK---
>Monodelphis_domestica
MFPSKSGFFFDIVKLRGKSSW
>Aquila_chrysaetos_canadensis
MFPSQPGSFFDIVKLRGK---
>Condylura_cristata
MFSPRPGYFFDIVKLRGK---
>Oryctolagus_cuniculus
MFSPRPGYFFDIVKLRGK---
>Elephantulus_edwardii
MFSPRPGSFFDIVKLRGE---
>Acanthis_flammea
MFPSQPGAFFDVVKLRGK---
>Loxodonta_africana
MFSLRPGSFFDIVKLRGE---
>Jaculus_jaculus
MFSPRPGYFFDIVKLRGK---
>Trichechus_manatus_latirostr
MFSPRPGYFFDIVKLRGE---
>Pygoscelis_adeliae
MFPSPPGCFFDIVKPRRK---
>Struthio_camelus_australis
MFAPTPGCFFDIVKLRGK---
>Chelonia_mydas
MFHAKPGCFFDIVKLRGK---
>Orycteropus_afer_afer
MFSPRPGYFFHIVKLRGE---

HG0142.2 ENSG00000204217 BMPR2

>Homo_sapiens
--------MKALQLGPL-ISHLSFQTVL
>Acinonyx_jubatus
--------MKALQLGPL-ICHLSFQTAL
>Lipotes_vexillifer
--------MKALQLGPL-ICHLSFQTAL
>Miniopterus_natalensis
--------MKALQLGPL-ICHLSFQTAL
>Monodelphis_domestica
MKGIFCSGVRALQTGPLSICHLSFQTAL
>Condylura_cristata
--------MNALQLGPR-ICHLSFQTAL
>Oryctolagus_cuniculus
--------MKALPLRPL-PCHLSFQTAL
>Elephantulus_edwardii
--------MNALQLGLL-ICHLSFQTAL
>Ictidomys_tridecemlineatus
--------MKALQLGPL-ICHLSFQTAL
>Trichechus_manatus_latirostr
--------MKALQLGPL-ICHLSFQTAL
>Orycteropus_afer_afer
--------MKALQLGPL-ICHLSFQTAL

HG0143 ENSG00000101958 GLRA2

>Homo_sapiens
MMPRTGTFSFFSA-NCTKP-NLFLIFKETRFLPNFESGQ
>Mustela_putorius_furo
MMPRTGTFCFFSA-NCTNP-NLFLIFKETRFLPDFESGQ
>Orcinus_orca
MMPSTGTFCFFSA-NCTSP-NLFLIFKETRFLPDFESGQ
>Rousettus_aegyptiacus
-MPRTGTFCFFSA-NCTNP-NLFLIFKETRFLPDFEFGQ
>Dasypus_novemcinctus
MMPRTGTFCFFSA-NCTNP-NLFLIFKEIRFLPDFQSGQ
>Galeopterus_variegatus
MMPRTGTFCFFSV-NCTNP-NLFLIFKETRFLPDFESGR
>Monodelphis_domestica
MIPRTGTFYFFSP-NYTNP-HLFLIFKAIRLLTDFESGQ
>Sorex_araneus
MMPRTGTFCFFSA-NCTNP-NLFLIFKETRFLPNFEPGQ
>Oryctolagus_cuniculus
MMPRTGTFCFFSA-NCSIP-NLFLIFKETRFLPDFESGQ
>Elephantulus_edwardii
MMPRSGTFCFFLV-NCTNP-NLFLVFKEIRFLPDFESGQ
>Equus_caballus
MMPRTGTFCFFSA-NCANP-NLFLIFKETRLLSDFESGQ
>Manis_javanica
MMPRTGTFCFFSA-NCTNR-NLFLIFKETRFLTDFESGQ
>Loxodonta_africana
MMPRTGTFCFFSV-NCTNP-NLFLVFKEIRFLPDFESGQ
>Peromyscus_maniculatus_baird
MMPRTGTFCFFSA-NCTTP-NLFLIFKETRFLPDFESGQ
>Tupaia_chinensis
MMPKTGTFCFFSA-N------------------------
>Trichechus_manatus_latirostr
MMPRTGTFCFFSV-NCTNP-NLFLVFKEIRFLPDFESGQ
>Anolis_carolinensis
MIPRTGTFDFLPSPRHADEGLLFLILKNFKLLLDLGI--
>Chelonia_mydas
MIPRTGTSYFLPP-ICRHPDLLFLIFKKIKLLLDLGI--
>Orycteropus_afer_afer
MMPKTGTFCFFLV-NCTNP-NLFLVFKEIRFLPDFGSGQ

HG0144 ENSG00000117114 ADGRL2

>Homo_sapiens
MMQTDGFDEAGHL
>Acinonyx_jubatus
MMQTAGFDEAGHL
>Ambystoma_mexicanum
-MQTDGFDEVGNA
>Balaenoptera_acutorostrata_s
MMQTAGFDEAGHL
>Pteropus_alecto
MMQIAGFDEAGHI
>Alligator_mississippiensis
-MQTTGFDDAGHL
>Galeopterus_variegatus
MMQTAGFDEAGHL
>Monodelphis_domestica
MMYTAGFNEAGHL
>Sorex_araneus
MMQTAVFDEAGHL
>Oryctolagus_cuniculus
MMQTAGFDEAGHL
>Elephantulus_edwardii
MMQTAGFDEAGHI
>Equus_caballus
MMQTPGFDEAGHL
>Manis_javanica
MMQTAGFDEAGHL
>Loxodonta_africana
MMQTAGFDEAGHL
>Ictidomys_tridecemlineatus
MMQTAGFDEAGHL
>Tupaia_chinensis
MMQTAGFDEAGHL
>Trichechus_manatus_latirostr
MMQTAGFDEAGHL
>Chrysemys_picta
MIQTTGFNDAGHL
>Orycteropus_afer_afer
MMQTAGFNEAGHL

HG0145 ENSG00000162599 NFIA

>Homo_sapiens
MIPYGMDRTTCCPTCHCKCFLN
>Canis_lupus_familiaris
MIPYGMDRTTCCPTCHCKCFLN
>Pygocentrus_nattereri
MIPYGMERATCCPTCQCKCFLN
>Charadrius_vociferus
MIPYGMDRATCCPTCQCKCFLN
>Alosa_alosa
MIPYGMERATCCPTCQCKCFLN
>Cuculus_canorus
MIPYGMDRATCCPTCQCKCFLN
>Sinocyclocheilus_anshuiensis
MIPYGMEWATCCPTCQCKCFLN
>Monodelphis_domestica
MIPYGMDRTTCCPTCHRKCFLN
>Haliaeetus_leucocephalus
MIPYGMDRATCCPTCQCKCFLN
>Apteronotus_leptorhynchus
MIPYGMERATCCPTCQCKCFLN
>Sturnus_vulgaris
MIPYGMDRATCCPTCQCKCFLN
>Nipponia_nippon
MIPYGMDRATCCPTCQCKCFLN
>Lateolabrax_maculatus
MIPYGME-ATCCPTCQCKCFLN
>Manis_javanica
MIPYGMDRTTCCPTCHCKCFLN
>Melopsittacus_undulatus
MIPYGMDRATCCPTCQCKCFLN
>Mus_musculus
MIPYGMDRTTCCPTCHCKCFLN
>Salmo_salar
MIPYGMERATCCPTCQCKCFLN
>Lepisosteus_oculatus
MIPYGMDWATCCPTCQCKCFLN
>Ameiurus_nebulosus
-----MERATCCPTCQCKCFLN

HG0146.1 ENSG00000163681 SLMAP

```
>Homo_sapiens
MLQ---CCNVSATKGEKRPRSQTKHKNWRVTHLLGYLQSPNCVP-GVFLAEAARCLSLFFPEKKMD
>Anas_platyrhynchos
-----------------------------MTHLLGFETFYPSFKEPLLLFEGIRCLFLFFQRRMD-
>Apteryx_australis_mantelli
-----------------------------MTHLVGFKTSYPSFKEPLLLFEGIRCLLLFFQRRMD-
>Panthera_tigris
MLQ---CCNISATEGEKRPRSQTKHKNWCVTHLLGYLQSLNCLP-GVFLAGADRCLSLFFPGKRMD
>Calidris_pugnax
-----------------------------MTHLLGFKTFYPSFKELLLLFEGIRCLFLFFQRRMD-
>Pteropus_alecto
MLQ---CCNISATEGEKRPRSQTKHKNWCVTHLLGYLQSSNCLP-GVFLAEAVQCLSLFFPRKKMD
>Galeopterus_variegatus
MLQ---CCNVSATKGEKRPRSQTKHKNWCVTHLLGYLQSPNCLP-GVFLAEAA-------------
>Haliaeetus_albicilla
-----------------------------MTHLLGFKTFYPSFKEPLLLFEGIRCLFLFFQRRMD-
>Meleagris_gallopavo
-----------------------------MTHLPGFKTFSPSVEEPFLLSEEIRCLFLFFQRRMD-
>Condylura_cristata
MLQ---CCNISATKGEKRPRSQTKRKNWRVTHLLGYLQSPNCLP-GLFLAEAVRCLNLFFLGKKMD
>Oryctolagus_cuniculus
MLQ---CCNVSASEGAPRPRSHTEPRNRRVTHLLGDLQPQNCLP-GVFLAEAARCLSLSFPGKMD-
>Pseudopodoces_humilis
-----------------------------MTHLLGFKTLYPSFPGPLLLSEGIRCLFLFFQRRMD-
>Equus_asinus
MLQ---CCNISATEGERRPRSQTKHKNWFVTHLLGYLQSPNCLP-GVFLAEAVRCLSLFFPGKRMD
>Manis_javanica
MLQ---GCNISATEGERRPRSQTKPANCCVTHLCGYLQSPNCLP-GVL-AETVRCLSLFFPGKKMD
>Marmota_marmota_marmota
MLQ---CCNVSATKGEKRPRSQTKHKNWCVTHLLGYLQSPNCLP-GVFLAEAARCLSLFFPGKMD-
>Tupaia_chinensis
MLQ---CCNVSATEGEKRPRSQTKHKNWCVTHLLGYLQSPNCLP-GVFLAEAARCSSLFFPGKKMD
>Pygoscelis_adeliae
-----------------------------MTHLLGFKTFCPSFKEPLLLFEGIRCLFLFFRRRMD-
>Struthio_camelus_australis
MLHRSGFWSVLFTEGGKKKAVEIPNKSICMTHLVGFKTS---------------------------
```

HG0146.2 ENSG00000163681 SLMAP

>Homo_sapiens
MLRQCAVCVNLKFWVG
>Felis_catus
MLRQCAVCFNLKFWGG
>Pteropus_alecto
MLRQCAFCFNLKFWGG
>Galeopterus_variegatus
MLRQCVVCFNLEFWGG
>Condylura_cristata
MLRQCAVCFDLKFWGG
>Ceratotherium_simum_simum
MLRQCTVCFNLKFWGG
>Manis_javanica
MLRQCAVCFNLKFGGG
>Marmota_marmota_marmota
MLRQCAVCFNLKFWGG
>Tupaia_chinensis
MLRQCAVCVNLQFWGG

HG0146.3 ENSG00000163681 SLMAP

>Homo_sapiens
MAAGAGKVGKFPASSGAA
>Felis_catus
MAAGAGKVGKFPALSGTA
>Rousettus_aegyptiacus
MAAGAGKVGKFSASSGTA
>Condylura_cristata
MAAGAGKVGKFPASSDMV
>Oryctolagus_cuniculus
MAAGAGKVGKFPASSGTA
>Mus_musculus
MAANAGKVGKFPASSATA

HG0147 ENSG00000108984 MAP2K6

>Homo_sapiens
MKIARLQLASLLQN
>Acinonyx_jubatus
MKIARLQLASLLQN
>Orcinus_orca
MKIARLQLASLLQN
>Eptesicus_fuscus
MKIARLQLASLLQN
>Dasypus_novemcinctus
MKIARLQLASLLQN
>Macropus_eugenii
MKTARLQLESLLQN
>Meleagris_gallopavo
MKIERLQFASLLQN
>Erinaceus_europaeus
MKIARLQLASLLQN
>Oryctolagus_cuniculus
MKIARLQLASLLQN
>Elephantulus_edwardii
MKIARLQLASLLQN
>Equus_asinus
MKIARLQLASLLQN
>Manis_javanica
MKIARLQLASLLQN
>Loxodonta_africana
MKIARLQLASLLQN
>Nannospalax_galili
MKIARLQLASLLQN
>Trichechus_manatus_latirostr
MKIARLQLASLLQN
>Gekko_japonicus
MKIERLQFASLLQN
>Struthio_camelus_australis
MKIERLQFASLLQN
>Pelodiscus_sinensis
MKIGRLQFASLLQN

HG0147 ENSGALG00000004370 MAP2K6

>Gallus_gallus
MKIERLQFASLLQN
>Acinonyx_jubatus
MKIARLQLASLLQN
>Orcinus_orca
MKIARLQLASLLQN
>Eptesicus_fuscus
MKIARLQLASLLQN
>Dasypus_novemcinctus
MKIARLQLASLLQN
>Latimeria_chalumnae
MKITSLHFASLLQN
>Erinaceus_europaeus
MKIARLQLASLLQN
>Ochotona_princeps
MKIARLQLASLLQN
>Elephantulus_edwardii
MKIARLQLASLLQN
>Equus_asinus
MKIARLQLASLLQN
>Manis_javanica
MKIARLQLASLLQN
>Otolemur_garnettii
MKIARLQLASLLQN
>Loxodonta_africana
MKIARLQLASLLQN
>Dipodomys_ordii
MKIARLQFASLLQN
>Trichechus_manatus_latirostr
MKIARLQLASLLQN
>Gekko_japonicus
MKIERLQFASLLQN
>Struthio_camelus_australis
MKIERLQFASLLQN
>Trachemys_scripta
MKIERLQFASLLQN

HG0148.1 ENSGALG00000011855 RHOJ

>Gallus_gallus
MLEIRWKMEWI
>Apteryx_australis_mantelli
MLEMKWKMEWI
>Caprimulgus_carolinensis
MLEMKWKMEWI
>Charadrius_vociferus
MLEMKWKMEWI
>Columba_livia
MLEMKWKMEWI
>Merops_nubicus
MLEMRWKMEWI
>Cuculus_canorus
MLEMKWKMEWI
>Falco_cherrug
MLEMKWKMEWI
>Gavia_stellata
MLEMKWKMEWI
>Eurypyga_helias
MLEMKWKMEWI
>Opisthocomus_hoazin
MLEMKWKMEWI
>Corvus_brachyrhynchos
MLEMKWKMEWI
>Nipponia_nippon
MLEMKWKMEWI
>Picoides_pubescens
MLEMKWKMEWI
>Melopsittacus_undulatus
MLEMKWKMEWI
>Tyto_alba
MLEMKWKMEWI
>Pelodiscus_sinensis
MLEMKWKMEWI
>Calypte_anna
MLEMKWKMEWI

HG0148.2 ENSGALG00000011855 RHOJ

```
>Gallus_gallus
-----------MILWTALVIVRSSCDSLETLV
>Anas_platyrhynchos
-----------MILLTALVIVRSGCDSLETLV
>Apteryx_australis_mantelli
--MYIFNLFTSMILLTALVIVRSGCDSLETLV
>Caprimulgus_carolinensis
-----------MILLTDLVIVRSGCDSLETLV
>Cariama_cristata
-----------MILLTDLVIVRSGCDSLETLV
>Calidris_pugnax
MIFLSVNLFTSMILLTDLVIVKSGCDSLETLV
>Columba_livia
-----------MILLTDLAIVRSGCDSLETLV
>Merops_nubicus
-----------MLLLTDLVIVRSGCDSLETLV
>Cuculus_canorus
-----------MILLTDLVIVRSGCDSLETLV
>Aquila_chrysaetos_canadensis
-----------MILLTDLVIVRSGCDSLETLV
>Gavia_stellata
-----------MILLTDLVIVRSGCDSLETLV
>Mesitornis_unicolor
-----------MILSTDLVIVRSGCDSLETLV
>Opisthocomus_hoazin
-----------MILLTDLVIVRSGCDSLETLV
>Manacus_vitellinus
-----------MILLTDLVIVRSGCDSLETLL
>Nipponia_nippon
-----------MILLTDLVIVRSGCDSLETLV
>Picoides_pubescens
-----------MILLTDLVIVRSGCDSLETLV
>Melopsittacus_undulatus
-----------MILFTDLVIVRSGCDSLETLV
>Aptenodytes_forsteri
-----------MILLTDLVIVRSGFDSLETLV
>Tyto_alba
-----------MILLTDLVIVRSGCDSLETLV
>Struthio_camelus_australis
-----------MILLTALVIVRSGCDSLETLV
>Calypte_anna
-----------MILWTDLVIVRSGCDSLEALV
```

HG0149.1 ENSGALG00000014645 MEF2C

>Gallus_gallus
MILTIKRVYVQERMQGTVN
>Anas_platyrhynchos
MILTIKRVYVQERMQGTVN
>Calidris_pugnax
------------MQGTVN
>Columba_livia
MILTIKRVYV-ERMQGTVN
>Alligator_mississippiensis
MILRIKKEYVQERMQGTVN
>Cuculus_canorus
------------MQGTVN
>Monodelphis_domestica
----MKFLFLQERMQGSVN
>Aquila_chrysaetos_canadensis
MILTIKRVYVQERMQGTVN
>Balearica_regulorum_gibberic
------------MQGTVN
>Ornithorhynchus_anatinus
-------MYVQERMQGSVN
>Lepidothrix_coronata
MILTIKRVYVQERMQRTVN
>Phaethon_lepturus
------------MQGTVN
>Otolemur_garnettii
------------MQGFGN
>Fulmarus_glacialis
------------MQGTVN
>Aptenodytes_forsteri
MILTIKRVYVQEQMQGTVN
>Anolis_carolinensis
MILNKRKGYVQERMPGSVN
>Chrysemys_picta
MILTIKRVYVQERMQGSVN
>Tinamus_guttatus
------------MQGTVN

HG0149.2 ENSG00000081189 MEF2C

>Homo_sapiens
MWNILKDNIKVNECRNLGTELCKC
>Ursus_maritimus
----------MNECRNLGTEQCKC
>Orcinus_orca
----------MNECRNLGTEQCKC
>Rousettus_aegyptiacus
----------MNECRNLGTEQCKC
>Dasypus_novemcinctus
----------MNECRNLRTELCKC
>Condylura_cristata
----------MNECRNLGTEQCKC
>Ochotona_princeps
----------MNECRNLGTELCKC
>Ceratotherium_simum_simum
----------MNECRNLGTELCKC
>Manis_javanica
----------MNECRNSGTEQGKC
>Loxodonta_africana
----------MNECRNLGTELCKC
>Heterocephalus_glaber
----------MNECRNLGTELCKC
>Tupaia_chinensis
----------MNECRNLGTELCKC
>Trichechus_manatus_latirostr
----------MNECGNLGTELCKC
>Orycteropus_afer_afer
----------MNECRNLGTELCKC

HG0150.1 ENSGALG00000032289 IKZF5

>Gallus_gallus
---------MAAAAEGGGRGAGRRRRHQQMEHKLT
>Anser_cygnoides
MGRESFGSFVCGALRFNAFLLEKMRRHQQMEHKLT
>Cariama_cristata
-----------------------------MEHKLT
>Charadrius_vociferus
-----------------------------MEHKLT
>Pterocles_gutturalis
-----------------------------MEHKLT
>Merops_nubicus
-----------------------------MEHKLT
>Alligator_mississippiensis
-----------------------------MEHKLT
>Haliaeetus_leucocephalus
-----------------------------MEYKLT
>Gavia_stellata
-----------------------------MEHKLT
>Mesitornis_unicolor
-----------------------------MEHKLT
>Tauraco_erythrolophus
-----------------------------MEHKLT
>Opisthocomus_hoazin
-----------------------------MEHKLT
>Parus_major
-----------------------------MEHKLT
>Egretta_garzetta
-----------------------------MEHKLT
>Fulmarus_glacialis
-----------------------------MEHKLT
>Nestor_notabilis
-----------------------------MEHKLT
>Aptenodytes_forsteri
-----------------------------MEHKLT
>Chelonia_mydas
-----------------------------MEHKLT

HG0150.2 ENSG00000095574 IKZF5

```
>Homo_sapiens
MAFWSPLLYLRSKFQLLYISSMLEWQEKGGTDLG-------------
>Mustela_putorius_furo
MAFWSPLVCLQSKFRLLYISSMLEWQEKGGTDLG-------------
>Orcinus_orca
MAFWSPLVYLQSKFRLLYISSMLEWQEKGGTDLG-------------
>Pteropus_alecto
MAFWSPLVCLQSKFHLLYISSMLEWQEKGGTDLG-------------
>Dasypus_novemcinctus
MAFWSPMVCLQSKFRLLYISSMLEWQEKGGADLGEKLFFLGITNCKL
>Sarcophilus_harrisii
---------------------MLEWKEKGGIDLG-------------
>Galeopterus_variegatus
MAFWSPLACHQSEFRLLHISSMLQWQEKGT-DLG-------------
>Monodelphis_domestica
MAFWSQLMYFQRGFQLLYISSMLEWKEKSGIDLG-------------
>Condylura_cristata
MAFWSPWVCLQSKFR--------------------------------
>Oryctolagus_cuniculus
---------------------MLEWQEKGT-DLG-------------
>Ornithorhynchus_anatinus
MACWSPLAHRQSGFLILGYSSMLEREERSGTDLG-------------
>Equus_asinus
MAFWSPLVCLQSKFLLLYISSMLEWQDKGGTDLG-------------
>Manis_javanica
---------------------MLEWQEKGS-DLG-------------
>Loxodonta_africana
MAFWSPWLCLQSKFPLLFISSMVEWQEKGGTDLG-------------
>Marmota_marmota_marmota
MAFWSPLVCLQNKFQILYISSMLEWQEKGGTDLG-------------
>Tupaia_chinensis
MAFWSPVVYLQNKFQLLYISSMLEWQEKGITDLG-------------
```

HG0151 ENSG00000006468 ETV1

>Homo_sapiens
MFLLNLEGSFCSLSPSVAFSPVRCKA--SLVD----R----HC-------
>Acinonyx_jubatus
MFLLNLEGIFCSRSPSVAFSPVRCKA--SLVD----R----HC-------
>Balaenoptera_acutorostrata_s
MFLLKLEGIFCSRSPSVAFSPVRCKA--SLVD----R----HC-------
>Miniopterus_natalensis
MFLLNLEGIFCSSSPSVAFSLKAILFPSRLIEVQIPRKSPAKCSKYKTLS
>Dasypus_novemcinctus
MVLLNLEGIFCSRSPSVAFSPVRCKA--SLVD----R----HC-------
>Alligator_mississippiensis
MFLLNLEGIFCSRSPCVTFRL---------LAVQILRKAADQMPRK----
>Galeopterus_variegatus
MFLLNLEGIFCSCSPSVAFSPVRCKA--SLVD----R----HC-------
>Haliaeetus_leucocephalus
MFLLNLEGIFCSRPPRVAFRL---------IGVQIPRKSPDQMPSK----
>Condylura_cristata
MFLLNLEGFFCSRSPSVAFRL---------IDVQIPRKSPA---------
>Pseudopodoces_humilis
MFLLNLEGIFCSRSPRVAFRL---------IGVQIPRKSPDQMPSK----
>Ceratotherium_simum_simum
MFLLNLEGIFCSRSPSVAFSPVRCKA--SLVD----R----HC-------
>Manis_javanica
MFLLNLKGFFCSSSPSVAFSPVRCKA--SLVD----R----HC-------
>Loxodonta_africana
MFLLNLEGIFCSRSPSVAFSPVRCKA--SLVD----R----HC-------
>Chinchilla_lanigera
MFLLNLEGIFCSRSPSVAFSPVRCKA--SLVD----R----HC-------
>Tupaia_chinensis
MFLLNLEGIVCSRSPSVAFSPVRCKA--SLAD----R----HC-------
>Struthio_camelus_australis
MFLLNLEGIFCSRSPGVAFRL---------IGVQIPKKSPDQMPSK----
>Chelonia_mydas
MFLLNLEGIFCSRSPCVTFRL---------IGEQIPRKSSGQMPRK----
>Orycteropus_afer_afer
MFLPNLEGIFCSSSPTVAF---------RLIEVQIPRKSPAKCSKYKILS

HG0151 ENSGALG00000035504 ER81

```
>Gallus_gallus
MFLLNLEGNFLQPIAARGLQADWSPDPEEISRSNAH-
>Anas_platyrhynchos
MFLLNLEGIFSQPTAARGLQADWRPDPEEISRSNAQ-
>Rousettus_aegyptiacus
MFLLNLEGIFLQQFA----------------------
>Mesitornis_unicolor
MFLLNLEGIFLQPIAAPGLQADWSPDPEEISRSNAQ-
>Manacus_vitellinus
MFLLNLEGIFLQPIAARGLQADWSPDPEEISRSNAQ-
>Nomascus_leucogenys
MFLLNLEGSFCSLSPSVAF----SPVRCKASLVDRHC
>Struthio_camelus_australis
MIA----WSW-LAKTSLWEKADWSPDPEEISRSNAQ-
```

HG0152 ENSDARG00000015536 sox6

```
>Danio_rerio
--------MLHDGKCFEYLRHGRLH--------------------------------------------------------
>Megophrys_sangzhiensis
--------MLHDGKCLEYLQPWLLTFTAKCDSPPAPPNQVTAQSPKAPLALQTYANDLDSGDLPAAHPYLLWGSLGFLIHE
>Menidia_menidia
--------MLHDGKCFEYLRHGCAH--------------------------------------------------------
>Felis_catus
--------MLHDGKCFEYLRHGRSH--------------------------------------------------------
>Pygocentrus_nattereri
--------MLRDGKCFEYLRHGRSH--------------------------------------------------------
>Miniopterus_natalensis
--------MLHDGKCFEYLRHGRSH--------------------------------------------------------
>Haplochromis_burtoni
--------MLHDGKCFEYLRHGCAH--------------------------------------------------------
>Latimeria_chalumnae
--------MLHDGKCFEYLRHGRSH--------------------------------------------------------
>Fundulus_heteroclitus
--------MLHDGKCFEYLRHGCAH--------------------------------------------------------
>Equus_asinus
--------MLHDGKCFEYLRHGRSH--------------------------------------------------------
>Platichthys_stellatus
--------MLHDGKCFEYLRHGCAH--------------------------------------------------------
>Cebus_capucinus_imitator
--------MLHDGKCFEYLRHGRSH--------------------------------------------------------
>Peromyscus_maniculatus_baird
--------MLHDGKCFEYLRHGRSH--------------------------------------------------------
>Salmo_salar
--------MLHDGKCFEYLRHGRSH--------------------------------------------------------
>Lepisosteus_oculatus
-------MMLHDGKCFEYLRHGR----------------------------------------------------------
>Ictalurus_punctatus
MPLATHFLMLHDGKCFECLRHGRAH--------------------------------------------------------
>Pelodiscus_sinensis
--------MLHDGKCFEYLRHGRSH--------------------------------------------------------
>Takifugu_rubripes
--------MLHDGKCFEYLRHGCAH--------------------------------------------------------
```

HG0153 ENSGALG00000003136 IKZF2

```
>Gallus_gallus
MEEVPAGLPLATAPLPQFP-----------
>Odobenus_rosmarus_divergens
MEEVKTGLPLATDPLP--------------
>Ambystoma_mexicanum
MEEVRAGLPLATAAALP-------------
>Orcinus_orca
MEEVKTGLPLATDPLP--------------
>Rousettus_aegyptiacus
MEEVKTGLPLATDPLP--------------
>Galeopterus_variegatus
MEEVKTGLPLATDPLP--------------
>Erinaceus_europaeus
MEEVKTGLPLATDPLP--------------
>Oryctolagus_cuniculus
MEEVKTGLPLATDPLP--------------
>Elephantulus_edwardii
MEEVKTGLPLATDPLP--------------
>Ceratotherium_simum_simum
MEEVKTGLPLATDPLP--------------
>Manis_javanica
MEEVKTGLPLATEPLP--------------
>Macaca_mulatta
MEEVKTGLPLATDPLP--------------
>Loxodonta_africana
MEEVKTGLPLATEPLP--------------
>Mus_musculus
MEEVKTGLPLATDPLP--------------
>Tupaia_chinensis
MEEVKTGLPLATDPLP--------------
>Trichechus_manatus_latirostr
MEEVKTGLPLATDPLP--------------
>Chrysemys_picta
MEEVKTGLPLATDPLLTVSITNCSAQTHKH
>Orycteropus_afer_afer
MEEVKTGLPLATDPLP--------------
```

HG0154 ENSG00000128573 FOXP2

```
>Homo_sapiens
MPSVWDVIGQRCTHSSVNTAVNSCLMVALTVS--
>Leptobrachium_boringii
MPSVWDLISQRCIHSSVNTAVNSYWMVTVTAS--
>Mustela_putorius_furo
MPSVWDVIGQRCTHSSVNTAVNSCLMVALTVS--
>Hynobius_retardatus
MPSVWDVIGQRCIHSSVNTAVNSCLKVALTVS--
>Calidris_pugnax
-------------------------MVALTVS--
>Pteropus_alecto
MPSVWDVIGQRCTHSSVNTAVNSCLMVALTVS--
>Kryptolebias_marmoratus
--------------------------MSLTVSAS
>Galeopterus_variegatus
MPSVWDVIGQRCTHSSVNTAVNSCLMVALTVS--
>Monodelphis_domestica
MPSVWDVIGQRCTHSSVNTAVNSCLMVALTVS--
>Condylura_cristata
MPSVWDVIGQRCTHSSVNTAVNSCLMVALTVS--
>Oryctolagus_cuniculus
MPSVWDVIGQRCTHSSVNTAVNSCLMVALTVS--
>Lepidothrix_coronata
MPSVWDVIGQRCTHSSVNTAVNSCLMVALTVS--
>Mus_musculus
MPSVWDVIGQRCTHSSVNTAVNSCLMVALTVS--
>Salmo_salar
-MGLGREQAEVLSHSSVNTAGNTFLMMSLTVSVS
>Lepisosteus_oculatus
MPSVWDVIGQRCTHSSVNTAVNSCLIMALTVS--
>Ictalurus_punctatus
-------------------------MMALTVSVS
>Anolis_carolinensis
MPSVWDVIGQRCSRRSVNTAVNSCLMVALTVS--
>Pelodiscus_sinensis
MPSVWDVIGQRCTHSSVNTAVNSCLMVALTVS--
```

HG0155 ENSG00000136535 TBR1

>Homo_sapiens
MAGAVSSAAQAARSKVSLF
>Anser_cygnoides
MAEAVSSSPQAARSKVSLS
>Canis_lupus_familiaris
MAGAVSSAAQAARSKVSLF
>Orcinus_orca
MAGAVSSAAQAARSKVSLF
>Pteropus_vampyrus
MAGAVSSAAQAARSKVSLF
>Sarcophilus_harrisii
MAGAVSSAAQAARSKVSLF
>Monodelphis_domestica
MAEAVSSAAQAARSKVSLF
>Aquila_chrysaetos_canadensis
MAEAVSSSPQAARSKVSLS
>Gallus_gallus
MAGAVSSSPQAARSKVSLS
>Condylura_cristata
MAGAVSSAAQAARSKVSLF
>Ochotona_princeps
MAGAVSSAAQAVRSKVSLF
>Elephantulus_edwardii
MAGAVSSAAQAARSKVSLF
>Ceratotherium_simum_simum
MAGAVSSAAQAARSKVSLF
>Ictidomys_tridecemlineatus
MAGAVSSAAQAARSKVSLF
>Trichechus_manatus_latirostr
MAGAVSSAAQAARSKVSLF
>Struthio_camelus_australis
MAEAVSSSPQAARSKVSLS
>Tinamus_guttatus
MAGAVSSSPQAARSKVSLS
>Orycteropus_afer_afer
MARAVSSAAQAARSKVSLF

HG0156 ENSG00000156113 KCNMA1

>Homo_sapiens
MAVDGCLGWRWRREELPRAPAPAARC
>Apteryx_australis_mantelli
MAGDGFFGGRWRREELSEAPARPARC
>Odobenus_rosmarus_divergens
MAVDGCSGWRWRREELPRAPAPAARC
>Orcinus_orca
MAVDGCSGWRWRREELPRAPAPAARC
>Pteropus_vampyrus
MAVDGCSGWRWRREELPRAPAPAARC
>Macropus_eugenii
MAVDGCLGWRWRREELPGAPAPPAHC
>Aquila_chrysaetos_canadensis
MAGDGFFGGRWRREELSEAPARPARC
>Gallus_gallus
MAGDGLFGGRWRREELSAAPARPARC
>Condylura_cristata
MAVDGCSGWRWRREELPRAPAPAARC
>Zonotrichia_albicollis
MAGDGLFGGRWRREELSEAPARPARC
>Ceratotherium_simum_simum
MAVDGCSGWRWRREELPRAPAPAARC
>Manis_javanica
MAVDGCSGWRWRREELPRAPAPAARC
>Loxodonta_africana
MAVDGCSGWRWRREELPRAPAPAARC
>Melopsittacus_undulatus
MAGDGVFGGRWRREELSEAPARPARC
>Microtus_ochrogaster
MAVDGCSGWRWRREELPRAPAPAARC
>Trichechus_manatus_latirostr
MAVDGCSGWRWRREELPRAPAPAARC
>Chrysemys_picta
MAGEGCFGGRWRWGELSGAPARPARC
>Orycteropus_afer_afer
MAVDGCSGWRWRREELPRAPAPAARC

HG0157 ENSG00000173926 MARCH3

>Homo_sapiens
MCPGPLGREQPPTGGNLALRTAAGTSVCYRDSARTMEILYECGSH
>Phoca_largha
-----------------------------------MEILYECGSH
>Balaenoptera_acutorostrata_s
-----------------------------------MEILYECGSH
>Pteropus_alecto
-----------------------------------MEILYECGSH
>Dasypus_novemcinctus
-----------------------------------MEILYECGSH
>Galeopterus_variegatus
-----------------------------------MEILYECGSH
>Condylura_cristata
MCPGALGPEQPPVGATWDSVPAPGASVCYLDSARRMEIL------
>Oryctolagus_cuniculus
-----------------------------------MEILHERGSH
>Elephantulus_edwardii
-----------------------------------MEILYECGSH
>Ceratotherium_simum_simum
-----------------------------------MEILYECGSH
>Manis_javanica
-----------------------------------MEILYECGSH
>Loxodonta_africana
-----------------------------------MAILYECGSH
>Heterocephalus_glaber
MCPASLGREQPPVRDNLGPRTCGRASVGDPESAGRMEILYQCGSH
>Tupaia_chinensis
MCPGPLGSEQPPIGATWDSGPAAGASVCDRDTARKMEILYECGSH
>Trichechus_manatus_latirostr
-----------------------------------MEILYECGSH
>Gekko_japonicus
-----------------------------------MEILYECGSH
>Pelodiscus_sinensis
-----------------------------------MEILYECGSH
>Orycteropus_afer_afer
-----------------------------------MEILYECGSH

HG0158 ENSG00000198939 ZFP2

>Homo_sapiens
---------------------------------------------------------------------------------------MTVYGEVMLENYRNPVSLGNHLSKPNGTSQLERE-LSLMQKEPPE--GGFRD-------------------------------------------------------------------------
-------------------------------------------------------------------------------------------------------------------------------------FLGYCCQI----------------------------------------------------------------NLTFSPL
IILSQSRGESID----------------------------------------------------------------
--------------------------------------------------------------------------------------------------------------------------------------------------------------------------------------------------------------------
-----------------------------------------------------------------------------------------------------

>Acinonyx_jubatus
--------------------------------------------------MAARPLPLGTQFQESVTFKDVSADFTWEEWLRLDSAQGNLYGKVMLENYRDLVSLGHRLSKPNVSSQLELEELSQVTKELPAC--LSPG-------------------------------------------------------------------------
-------------------------------------------------------------------------------------------------------------------------------------LLF---------------------------------------------------------------SRFSSM-LL----
-----PD---------------------------------------------------------------------
--------------------------------------------------------------------------------------------------------------------------------------------------------------------------------------------------------------------
-----------------------------------------------------------------------------------------------------

>Lipotes_vexillifer
----------------------------------------------------------------------------------------------MLGNYSNLVSLGHQLSKPNVISQLD---------------------------------------------------------------------------------------------
--------------------------------------------------------------------------------------------------------------------------------------------------------------------------------------------------------------------
----------------------------------------------------------------------------
--------------------------------------------------------------------------------------------------------------------------------------------------------------------------------------------------------------------
-----------------------------------------------------------------------------------------------------

>Rousettus_aegyptiacus
----------------------------------------------------MAPGLLTTRDQALMAFQDIAVAFSEKEWKLLSPAQRTLYRDVMLENYSHLVSLGIAFSKPQLITQLEQEDEPWKDKSECLL-DLCPG-------------------------------------------------------------------------
-------------------------------------------------------------------------------------------------------------------------------------ENIQCNDE----------FKW----------------------------------------------------------
----------------------------------------------------------------------------
--------------------------------------------------------------------------------------------------------------------------------------------------------------------------------------------------------------------
-----------------------------------------------------------------------------------------------------

>Dasypus_novemcinctus
----------------------------------------------------------MTKAQGLLSFEDVAVVFTWQEWQLLDPAQKDLYKDVMLENYNNLLSVGYQATKPDSLFKLEQGEPPWMVEGKIQSWPYPAHSMAGT-L-AFIQNSAVTN-------------------------------------------------------
-------------------------------------------------------------------------------HSTALGKKSYGCNEDAVKA---------------------------CKPLVTVPERTHAGKKPHECCDYKRAFASKSLLIPHQT-PIEERLYGCSQCQKAFITKSALIYHQKTYHREGESYGCNKCGKAFPWE
SKLIL--HQRIHSGERPFECKVCGKAFMVRTHLTVHERTHTGEKPYECSDCKNAFSKKAQLLIHQRIHTGERPYGC
SECQQAFIQKSDLTNHKKIHLVGRKSHGCSECGKVFPWKSKLLVHQRTHSGERPYRCNACDKAFTSKAHLIVHQRLHTGEKPYECCDCRKAFSTKAHLIIHQRTHTGERPYECKECQQAFIQKSGLTNHLQTCHPRVKLYGCHECGKVLSCKSTLIIHQRTHTGEKPF----KCNVCDKAFTAKSYLSVHQRIHTGEKPYECSECKKAFTTL
STLIGHQRTHTGERPYECNECQKAFFRKSALTNHQQTQHTGGNVGI-------------------------------------------------------
>Sarcophilus_harrisii
----------------------------------------------------------------------------------------------MLENYRNLFFLGLPVSKPEVISQLEQGEVPWIAEKEATF-RIHTG-------------------------------------------------------------------------
-------------------------------------------------------------------------------------------------------------------------------------EKP---------------------------------------------------------------YACNECTKAFCDG
SSLTK--HQRIHTGEKPYKCSDCGKSFSDSSSLA------------------------------------------
--------------------------------------------------------------------------------------------------------------------------------------------------------------------------------------------------------------------
-----------------------------------------------------------------------------------------------------

>Galeopterus_variegatus
----------------------------------------------------------------------------------------------MLENYTNLVSLGYSASKPDVITLLEQGKEPCMVVKDVTG-GWCPG-------------------------------------------------------------------------
-------------------------------------------------------------------------------------------------------------------------------------LFSRHKTK----------TLSPEKDIYEISLSK----------------------------------------------
----------------------------------------------------------------------------
--------------------------------------------------------------------------------------------------------------------------------------------------------------------------------------------------------------------
-----------------------------------------------------------------------------------------------------

>Monodelphis_domestica
--------------------------------------------------MASVPLPT-GPCQESVTFQDVAVEFTWEEWRYLNPSQKKLYKDVMLENYQSLISLGFEVSKPDVIHQLERKETSWMPEADVSQ-SSYPG-------------------------------------------------------------------------
-------------------------------------------------------------------------------------------------------------------------------------MKEFTGEKPYECSECGNVFSSRTQLTMHQRIHTRKFLYECKECGKACHCNSELIRHQRIHT-GEKPYECTVCGKAFSMN
GNLIV--HQRIHTGEKPFECSECGKSFRQRTHLYVHQTVHTQEISHECSECGKAFMRKTQLAVHQRIHTGGRPYGC
SVCGKDFFCSSDLIKHQKIHI-IEKSYECSECGKVFRSKIQLTVHQRIHPGKSLYECKECGKEFHCNSNLIKHQRIHTEEKPYQCNDCGKAFRRKAHLTLHQRIHTGDILYGCQECEKTFLYISDLTRHQR-IHTEEKPYECAVCVKAFRSKTQLSVHQRIHTGEKPY----ECTECGKFFRQRSHLYIHQRIHTVDILYECKECGKAFPYN
SDLIQHQKVHTGEKPHECTVCTKAFRSKTQLTVHQRIH-TGEKPYECTECGKFFRQRTHLYIHQRIHTGEILYECKDYEKAFPYKLELIQH----------
>Trichosurus_vulpecula
-------------------------------------------------------------MLESVTFNDVAVDFTQEEWKQPDPAQRSLYRDVLLENYRNFISLGQQVSKPHMISQLEQGEEPWMVESIIQK-GICPE-------------------------------------------------------------------------
-------------------------------------------------------------------------------------------------------------------------------------FSFPLPME---------------QKTFAFSIPFRL-------------------------------------------G
G---------------------------------------------------------------------------
--------------------------------------------------------------------------------------------------------------------------------------------------------------------------------------------------------------------
-----------------------------------------------------------------------------------------------------

>Sorex_araneus
-----------------------------------------------------MTHPASRSAPGTVAFEDVAVYFTRLEWSCLFPEQRTPYRDVMLENYRHLASLGFLASKPALVSLLEQGREPGALILQ----------------------------------------------------------------------------------
--------------------------------------------------------------------------------------------------------------------------------------------------------------------------------------------------------------------
----------------------------------------------------------------------------
--------------------------------------------------------------------------------------------------------------------------------------------------------------------------------------------------------------------
-----------------------------------------------------------------------------------------------------

>Ochotona_princeps
----------------------------------------------------------------------------------------------MLENYRNVASVGCQLFKPGLISWLEEELESMIGERGVLQ-GVTTH-------------------------------------------------------------------------
-------------------------------------------------------------------------------------------------------------------------------------VQSNAGEKLYECKECGKAFITSSRLSEHLRSHTGEKPYGCYECGKAFASSSYLTAHLRTHT-GEKPFVCTTCGKAFTRS
CYLRV--HMRTHTGEKPYECQECGKAFTGRSWLTIHLRTHTGEKPYECKECEKAFTSFALLNEHIKTHTGEKPFVC
NICTKSFRNSS-----------------------------CLKTHFRIHTGIKPYQCKDCGKAFSGRAGFTKHVLTHTGEKPYECKECGKTFRTSSGLTEHVRIHTGEKPFECYQCGKAMAHSSSLVAHLR-THTGEKPFECNMCEKAFTSSSYLRVHMRSHTGEKPY----VCKDCG----------------------------KAFIAH
SGLSQHLRTHTGEKPNECKQ---------------------------------------------------------------------------------
>Equus_asinus
----------------------------------------------------------------------------------------------MLENYRNLVSLGFPFSKPGIIS------------------------------------------------------------------------------------------------
--------------------------------------------------------------------------------------------------------------------------------------------------------------------------------------------------------------------
----------------------------------------------------------------------------
--------------------------------------------------------------------------------------------------------------------------------------------------------------------------------------------------------------------
-----------------------------------------------------------------------------------------------------

>Manis_javanica
----------------------------------------------------MHNEFLQTVSKGSVMFRDVSINFCQEEWECLDSGQMNLYKEVMLENFSNLVSVGLSNSKPAVISLLEQGKEPWMVERKLTR-SLCSD-------------------------------------------------------------------------
-------------------------------------------------------------------------------------------------------------------------------------LVSMCETK----------ILSLKKRHFSQVISTHE--------DMPTFIQPFLTPHQKAIN-EEKVCECKICGKAFNQN
SQLTQ--YQRIHSAEKNYECKECGKAFI------------------------------------------------
--------------------------------------------------------------------------------------------------------------------------------------------------------------------------------------------------------------------
-----------------------------------------------------------------------------------------------------

>Loxodonta_africana
----------------------------------------------------------MIEAQESLAFEDVAVKFTWEEWQLLDPAQKNLYWDVMLENFSNLVSMGYQVTTPDALSRLEREELPWNILDEINSQNISEIRKVVDHLPEHLRNESRVNRLERCQEHNTWENIFHQRRSHFSLRQNHNIFDLDEESLKSNLTLIDQSRSYEVKN
SAEFNRDEKTFLHASHEQFHTEIKFPQSQKSNSTKSKIIKHQNT-------HKIEKPHVCSECGKAFIRKSWLISHQIIHSGERPHRGYSCGEAFSRKFKLTEHHQRKHGGEEFLECMGRGKAFLKKSQLNTHRKIHTDEKWYKCSECGKGFTWKRNLISHQRIHTEEKPYMCSECGKGFIRKGNLIIHQRIHT-GEKPYVCGQCGKGFMWK
GNLVL--HQRIHTGEKSYICIECGKGFIKKGDFIIHQRTHTGEKPYACSECGKGFSQKGNLIIHQRIHSGEKPYIC
AECGKGFIQKRDFIIHQRIHT-GEKPYACGECGKDFIRKQYLIIHQRIHSGEKPYVCTDCGKRFIQKRDFVIHQRTHTGEKPYVCGQCGKGFIRKGHLIIHQRIHSGEKPYICTKCGKGFIQKRKLIVHQR-SHSGEKPYICTECGKGFIQKRDYDIHQRIHTGEKLY----ICRECGKAFTRKGNLTIHQRIHTGEKPYICSECGKGFIQK
GNLIIHQRIHSGEKPYICTECGKGFNQKGSLITHQRIH-TGEKPYVCGKCGKGFRKKSHLITHQRFHTXKSHLACSICGQSYSQESSFIEHQKIHTVEKPF
>Marmota_marmota_marmota
--------------------------------------------------------------------------------MHLDSAQRNLYGKVMLENYRNLVSLGHQLSKPLVISQLEQEELNLKNKELPGC--ASPD-------------------------------------------------------------------------
-------------------------------------------------------------------------------------------------------------------------------------FLG---------------------------------------------------------------YYCQTN-LLFSPL
SILSKPRRERID----------------------------------------------------------------
--------------------------------------------------------------------------------------------------------------------------------------------------------------------------------------------------------------------
-----------------------------------------------------------------------------------------------------

>Tupaia_chinensis
MKSLPLVTCRSRPPHRMMGTALPPSFCACAAPPTPPSYSGNSSSCGFSCACPADLGMGSVFLKEFVSFEDVAVGFTWEEWQDLDHAQRTLYRDVMLETYSNLVSLGHCITKPELILKMEQGAEPWMVEEPPNQ-SLPDVQKVDD----------LNETSEKSQDKTLWQLVITNNNIST--KERIELEKTFSLSLNHVLKLITNNENYIGMI
PEEFSVCQNMLLSSNFDELHAQEE-----PDLNPTGKFLRHPEHFSQLPKIPVGQQPFEYSGQEKAFHREAVLLTQKRIHM------YGACKYAYAKAF--------------------------DNSALIAQERTQEGRKPFEYDVYGNIFYKNCDLTKHPIIHTGEKPYRFSEFEKSFIKKSYLTKHQRTQT-REKPYGCNKCRKFFHEK
SDLIV--HQKTHIGEKPHECNECKKSFYRKSDLTVHQRTHTGEKPYECNECRKSFNQKSNLSRHQRIHTGEKPYEC
NRCKKSFCQKSDLTFHQRTHT-GERPHECKECRKSFYKKSDLTVHKRTHTGEKPYECNVCRKTFYQKSKLTVHQRTHTGEKPFECNECRKSFNRKADLTVHQRTHTGEKPYECNECRKSFYQKSVLTVHQR-THTGEKPYECNNCRKAFCQKSHLSRHQKTHTGGKPYRKTFSCNVCGKMFYEKSKFTKHQKIHTGEKPYKCKECEKSFISK
SKVIRHERTHTGEKPYRCNKCEKSFRQKSNLTVHQRIH-TGGKSYKCFECGKTFGQKSTLIVHHRIHTGRNPMNVKNVGNLST------------------
>Trichechus_manatus_latirostr
----------------------------------------------------------------------------------------------MLENYRILVSLGLCFSKPSVILLLEQGKEPWMVKRELTE-GLCSG-------------------------------------------------------------------------
-------------------------------------------------------------------------------------------------------------------------------------E------------------------------------------------------------------------------
----------------------------------------------------------------------------
--------------------------------------------------------------------------------------------------------------------------------------------------------------------------------------------------------------------
-----------------------------------------------------------------------------------------------------

>Gekko_japonicus
-------------------------------------------------------------MLSPVTFEDVAVYFTEGQVALLDPNQRALYREVMMENYENMASLGFLFIKPALISQLEQGEELWVPESLDWD-AAETL-------------------------------------------------------------------------
-------------------------------------------------------------------------------------------------------------------------------------DTSGEGSR-----------------------------------------------------------------------
----------------------------------------------------------------------------
--------------------------------------------------------------------------------------------------------------------------------------------------------------------------------------------------------------------
-----------------------------------------------------------------------------------------------------

HG0159.1 ENSG00000182197 EXT1

>Homo_sapiens
MQENYSPLLARPGCEIFSPALSRALCNPKMKDRRGES
>Acinonyx_jubatus
MQENYSPLLARPG-----------------------
>Balaenoptera_acutorostrata_s
MQENYSPLLARPGCEIFSPALSRLLCNPKMKDRRGES
>Pteropus_vampyrus
MQENYSPLLARPGCEIFSPALSRPLCNPKMKDRRGES
>Dasypus_novemcinctus
MQENSSPLLARPGCEVFSPALSRPLCNPKMKDRRGES
>Sarcophilus_harrisii
----------------------------MKDRRGES
>Galeopterus_variegatus
----------------------------MKDRRGES
>Monodelphis_domestica
----------------------------MKDRRGES
>Condylura_cristata
MQENYSLLVARPGCEIFCPALSRPLCNPKMKDRRGES
>Oryctolagus_cuniculus
----------------------------MKDRRGES
>Ceratotherium_simum_simum
MQENYSPLLARPGCEIFSLALSRPLCNPKMKDRRGES
>Manis_javanica
MRGNYSPLLARPGCKIFSPALSRPLCNPKMKDRRGES
>Loxodonta_africana
MQENYCPLLARPGCEIFRPALSRPLCNPKMKDRRGES
>Marmota_marmota_marmota
MQENYSPLLARPGCEIFSPGLSRALCNPKMKDRRGES
>Trichechus_manatus_latirostr
MQENYRQLLARPGCEIFRPALSRPLCNPKMKDRRGES
>Takydromus_sexlineatus
----------------------------MKDRRGES
>Chelonia_mydas
----------------------------MKDRRGES

HG0159.2 ENSG00000182197 EXT1

>Homo_sapiens
MQGFVRLRRPSCKMH
>Ailuropoda_melanoleuca
MQGFVRLRRPSCKMH
>Balaenoptera_acutorostrata_s
MQGFVRLRRPSCKMH
>Pteropus_alecto
MQGFVRLRRPSCKMH
>Dasypus_novemcinctus
MQGFVRLRRSSCRMH
>Galeopterus_variegatus
MQGFVRLRRPSCKMH
>Monodelphis_domestica
MQGFVRLRRSSWKM-
>Sorex_araneus
MQGFVRLRRPSCKMH
>Oryctolagus_cuniculus
MQGFVRLRRPSCRMH
>Ceratotherium_simum_simum
MQGFVRLRRPSCKMH
>Manis_javanica
MQGFVRLRRPGCKMH
>Loxodonta_africana
MQGFVRLPRSGCEMH
>Cricetulus_griseus
MQGFVRLRRPSCKMH
>Trichechus_manatus_latirostr
MQGFVRLRRSGCKMH
>Takydromus_sexlineatus
MQGFVRLRRSGWKM-
>Pelodiscus_sinensis
MQGFVRLRRESWKM-

HG0160 ENSDARG00000060089 btaf1

>Danio_rerio
MAPHIKPAAWLLFCDLQNYFSPFVRLKYRFRGSA--EG
>Amia_calva
MAPHIKPASWVLSSEQQKYPLFLIRMKYLSARSS--GG
>Channa_punctata
MAPHIKTVFWVAFNVQQNNSLHLIRLKYLSARTS--EG
>Megophrys_sangzhiensis
MAPHIKPGVWLLRCHRQISSL--IRFKRLLGSCRGPDG
>Menidia_menidia
MAPHIKTVFWVGLVAQQNNSLILIRLKYLSASSS--EG
>Oryzias_melastigma
MAPHIKTVFWIALYVEQKRALILIRLKYLSAKTS--EG
>Micropterus_salmoides
MAPHIKTVFWVALNVLQNNSLHLIRLKYLSSKSS--EG
>Astyanax_mexicanus
MAPRIKPASWLLFCDQQNYFSLLIRLKYLFGRSA--EG
>Haplochromis_burtoni
MAPHIKTVFWVVLNVQQNNSLNLIRLKYLSAKSS--EG
>Alosa_alosa
MAPHIKPASWLLFCDQQNYFFLLIRLKYLSGRSS--EG
>Latimeria_chalumnae
MAPHIKSGFWVLSSNHQIFSLNLLNDQSMFRCCP--EG
>Kryptolebias_marmoratus
MAPHIKTVFWVSPNEQRNKSLFLVRLKYLSADSS--EG
>Esox_lucius
MAPHIKSASWVPLGEQQNYSLYLIRLKYLSGSSS--EG
>Notothenia_coriiceps
MAPHIKTVFWVALGLLQNNSLHLIRLRYLSSKSS--EG
>Salmo_salar
MAPHIKSASWVPFSEHKNYSLNLIRLKYLSGSSS--EG
>Lepisosteus_oculatus
MAPHIKPDSWVLFSEQQNYCLFLIRIKYLSGCSS--AG
>Ictalurus_punctatus
MAPHIKSASRLLFCEQQNYFSLLIRVKYLFGTSA--EG

HG0161 ENSGALG00000017191

>Gallus_gallus
MCWLFFVPYTFPNS
>Anas_platyrhynchos
MCWLLFVLYTFPNS
>Caprimulgus_carolinensis
MCWLLFVP------
>Cariama_cristata
MCWLLFVQ------
>Charadrius_vociferus
MCWLLFVL------
>Pterocles_gutturalis
MSWLLFVLQTFPNS
>Columba_livia
MCWLLFVL------
>Leptosomus_discolor
MCWLLFVL------
>Haliaeetus_leucocephalus
MCWLLFVL------
>Gavia_stellata
MCWLLFVL------
>Chlamydotis_macqueenii
MCWLLFVL------
>Opisthocomus_hoazin
MCWLLFALQTFPNS
>Zonotrichia_albicollis
MCWLPFVLQTFPNS
>Fulmarus_glacialis
MCWLLFVL------
>Aptenodytes_forsteri
MCWLLFVL------
>Tyto_alba
MCWLLFVL------
>Chrysemys_picta
MCWLRFVLYTFPNT

HG0162 ENSG00000138347 MYPN

>Homo_sapiens
----MFHRHMVYTAQNNLGPVS------------------------------------------------------------
>Anas_platyrhynchos
----MLRRHMVYAAQNNLGPAN------------------------------------------------------------
>Xenopus_laevis
----MLYRHMVHRAQNNTQPASDKCCCCFSGDRLTQLHTIIFPKFQDNPCSLSSAFTKIGLFHFHPHPWTISIEETAVRAGH
>Canis_lupus_familiaris
MPGGMFHRHMVYTAQNNLGPVS------------------------------------------------------------
>Lipotes_vexillifer
----MFHRHMVYTAQNNLGPVS------------------------------------------------------------
>Eptesicus_fuscus
----MFDRHMVYTAQNNLGPVS------------------------------------------------------------
>Gallus_gallus
----MLCRHMVYAAQNNLGPAN------------------------------------------------------------
>Erinaceus_europaeus
----MFHRHMVYTAQNNLGPVS------------------------------------------------------------
>Oryctolagus_cuniculus
----MFHRHMVYTAQNNLGPAS------------------------------------------------------------
>Parus_major
----MLRGHMVYTAQNNLGPAN------------------------------------------------------------
>Equus_asinus
----MFHRHMVYTAQNNLGPAS------------------------------------------------------------
>Manis_javanica
----MFHRHMVYTAQNNLGPVS------------------------------------------------------------
>Loxodonta_africana
----MFHRHMVYTAQNNLGPVS------------------------------------------------------------
>Ictidomys_tridecemlineatus
----MFHRHMVYTAQNNLGPVS------------------------------------------------------------
>Pygoscelis_adeliae
---------MVYAAQNNLGPAN------------------------------------------------------------
>Anolis_carolinensis
----MFHKHMVYVSQNNLGPAN------------------------------------------------------------
>Orycteropus_afer_afer
----MFHRHMVYTAQNNLGPVS------------------------------------------------------------

HG0163 ENSG00000146285 SCML4

>Homo_sapiens
MFAAGAQGCWRLTVSY-----------------------------------------
>Xenopus_tropicalis
MFAAGERSCWRLSVC------------------------------------------
>Ailuropoda_melanoleuca
MFAAGEQGCWRLTVSY-----------------------------------------
>Hynobius_retardatus
MFAAGESGCWRLTVC------------------------------------------
>Orcinus_orca
MFAAGEQGCWRLTVSY-----------------------------------------
>Rousettus_aegyptiacus
MFAAGEQGCWRLTVSY-----------------------------------------
>Dasypus_novemcinctus
MFAVGEQGCWRQTVSS-----------------------------------------
>Gallus_gallus
MFAAGERSCWRLARC------------------------------------------
>Sorex_araneus
MFATGELGCWRLTVSSSLPSGRLASAGLGLHVLKVFSLLKQHSWLGGEPARALVNKL
>Elephantulus_edwardii
MFAAGEQGCWRLTVNDSHTSGKQVLQFSVCMCRKSSRFRVNIPDLGKNLSRALVNKP
>Corvus_brachyrhynchos
MFAAGELSCWRLTLC------------------------------------------
>Ceratotherium_simum_simum
MFAAGEQGCWRLTVSS-----------------------------------------
>Manis_javanica
MFAAGEQGCCRLTVSC-----------------------------------------
>Loxodonta_africana
MFAAGEQGCWRLTVSDSHTSGRQVLQYSVCMCRKSSVF-------------------
>Ictidomys_tridecemlineatus
MFAAGEQGCWRLTVSY-----------------------------------------
>Anolis_carolinensis
MFAAGEQSCWRCTVR------------------------------------------
>Chrysemys_picta
MFAAGELSCWRLTVC------------------------------------------

HG0164 ENSG00000151292 CSNK1G3

```
>Homo_sapiens
MLTLPIWYSYLVMYLFSQYPVSVCLS
>Xenopus_laevis
------------MFPFSQHPI-----
>Felis_catus
------------MYLFSQYPISVCLS
>Lipotes_vexillifer
MLTLPTWYSYLVMYLFSQYPISVWLS
>Eptesicus_fuscus
MLTLPAWYSYLVMYLFSQYPISVCLS
>Dasypus_novemcinctus
MLTLPTWYSYLVMYLFSQHPISVCLS
>Alligator_sinensis
------------MFPFSHYPI-----
>Galeopterus_variegatus
MLTLPTWCSYLVMYLFSQYPISVCLS
>Erinaceus_europaeus
MLTLPTWYSYLVMYLFSQYPISVCLS
>Oryctolagus_cuniculus
MLTLPTWCSTLVMYLLSQYPISVCWS
>Ceratotherium_simum_simum
MLTLPTWYSYLVMYLFSQYPISVCLS
>Manis_javanica
------------MYLFSQYPISVCLS
>Loxodonta_africana
MLTLRTWYSYLVMYLFSEYPISACLS
>Octodon_degus
MLTLPGWYSYLVMYLFSQYPVSVCLS
>Tupaia_chinensis
MLTLPSWHSDLVMYLFPQYLISVCLS
>Trichechus_manatus_latirostr
MLTLSTWYSYLVMYLFLQYPISACLS
>Orycteropus_afer_afer
MLTLSTWYSYLVMYLFSQYPISVCLS
```

HG0165 ENSG00000177508 IRX3

>Homo_sapiens
MSIRARSPGNLRQSVRT
>Canis_lupus_familiaris
MSIRARSPGNLRQSVRT
>Orcinus_orca
MSIRARSPGNLRQSVRT
>Pygocentrus_nattereri
MSIRARSPNNLRQSVRT
>Callorhinchus_milii
MSIRPSTPNNLHESVRT
>Latimeria_chalumnae
MSIRAVSPNNLKQSVRT
>Catostomus_commersonii
MSITKRSSNNLKQSVRT
>Esox_lucius
MSIRARSPNNLRQSVRT
>Gallus_gallus
MSIRARSPNNLRQSVRT
>Erinaceus_europaeus
MSIRAQSPGNRGQSVRT
>Scleropages_formosus
MSIRAKSPNNLRQSLRT
>Parus_major
MSIRAQSPNNLRQSVRT
>Ceratotherium_simum_simum
MSIRARSPGNLRQSVRT
>Heterocephalus_glaber
MSIRARSPGNLRQSVRT
>Lepisosteus_oculatus
MSIRAKSPNNLRQSVRT
>Trichechus_manatus_latirostr
MSIRARSPGNLRQSVRT
>Trachemys_scripta
MSIRTQSPNNLRQSVRT

HG0166.1 ENSG00000148948 LRRC4C

>Homo_sapiens
---------------------------MKLQRRWREGDRI
>Anas_platyrhynchos
---------------------------MKMQRSWREGKRI
>Felis_catus
---------------------------MKLQRRWREGDRI
>Rousettus_aegyptiacus
---------------------------MKLQRRWREGDRI
>Dasypus_novemcinctus
---------------------------MKLQRRWRKGDRI
>Aquila_chrysaetos_canadensis
---------------------------MKMQRSWREGKRI
>Gallus_gallus
---------------------------MKMQRSWREGKRI
>Condylura_cristata
---------------------------MKLQRR-------
>Ochotona_princeps
---------------------------MKLQRRRREGDRI
>Corvus_brachyrhynchos
---------------------------MKLQRSWREGKRI
>Equus_przewalskii
---------------------------MKLQRRCGEGDRI
>Microtus_ochrogaster
---------------------------MKSQRRWREGDRI
>Trichechus_manatus_latirostr
---------------------------MKLQRRWREGDRI
>Struthio_camelus_australis
MDLFLTAGGNLEFFPLPPSLYSIYCEIMKMQRSWREGKRI
>Pelodiscus_sinensis
---------------------------MKMQRSWREGKRI
>Orycteropus_afer_afer
---------------------------MKLQRRWREGDRI

HG0166.2 ENSG00000148948 LRRC4C

>Homo_sapiens
MQDAAFSWNRTQWIN------------
>Leptonychotes_weddellii
MRDAALSWNRTQWIN------------
>Balaenoptera_acutorostrata_s
MRDAALSWNRTQWIN------------
>Rousettus_aegyptiacus
MRDAALSWNRTQWIN------------
>Dasypus_novemcinctus
MRDAALSWNRTQWIN------------
>Sorex_araneus
MQDAAPSGERTQWIN------------
>Elephantulus_edwardii
MRDAALSQKRTQWRN------------
>Equus_przewalskii
MQDAALSWNRTQWIN------------
>Microtus_ochrogaster
MRDAALSWSRTQRIN------------
>Trichechus_manatus_latirostr
MRDAALSWNRTQWINCAREKEERSFFL
>Orycteropus_afer_afer
MRDAAFSWNRTQWINCAREKEGRSFFF

HG0166.3 ENSG00000148948 LRRC4C

```
>Homo_sapiens
MIGNEEPRIHKEKSMFIFLYKGESEPRRYFWNEKFGAFLVK------------------------------------------
>Acinonyx_jubatus
--------------MFVFLYKGESKPRRYFWN---------------------------------------------------
>Balaenoptera_acutorostrata_s
--------------MFIFFCKGESKPRRYFWNEKFGAFFSKVKT---------------------------------------
>Miniopterus_natalensis
MISNGEPTSHKEKVCSLFFYTGESQPRIYFWNEKFGAFFFFFFFRRVKDLI--------------------------------
>Dasypus_novemcinctus
--------------MFVFLYKGESKPRRLFYFFIFGTKSLGLY----------------------------------------
>Condylura_cristata
MIGNGERRIHKEKVCS-FFSKGESKPRR-------------------------------------------------------
>Oryctolagus_cuniculus
--------------MFVFLCKGESKPRRYFWNEKFGAFFRKVKDLRRWCFLFSFEFPTRGEEINNTSAKKFQRRKVDRGRMRH
>Equus_przewalskii
--------------MFIFLSKGESEPRRYFCNEKFGAFLGK------------------------------------------
>Nannospalax_galili
--------------MFLFLYKEESTPRRYFWNEKFGAFFRKVKT---------------------------------------
```

HG0166.4 ENSG00000148948 LRRC4C

>Homo_sapiens
MEGDWGFCRAGSREPGRWR
>Leptonychotes_weddellii
MEGDWGFCRAGNREPGRWR
>Dasypus_novemcinctus
MEGEKGFRSAASREPGRWR
>Condylura_cristata
MEGDPGCCRAGSPEPGRWR
>Ochotona_princeps
MEGDWGFCCVASREPGRWR
>Peromyscus_maniculatus_baird
MEGDLGFRGTESREPGLWH
>Trichechus_manatus_latirostr
MEGDWGFCRAGSREPGRWR
>Orycteropus_afer_afer
MEGDQCFCRAGSREPGRWR

HG0167 ENSDARG00000104005 slc4a3

>Danio_rerio
----------------------MPPHRLTGNPAVKEKG
>Menidia_menidia
----------------------MPPQRLPGTPAVKEKG
>Porichthys_notatus
----------------------MPPQRLPGTPAVKEKG
>Oryzias_latipes
----------------------MPSQRLPGSPAEKEKG
>Micropterus_floridanus
----------------------MPPKRLPGSPTVKEKG
>Pygocentrus_nattereri
----------------------MPPHRLTGNPTVKEKG
>Haplochromis_burtoni
----------------------MPPQRLPGSPAVKEKG
>Alosa_alosa
----------------------MPPHRLTGSPTVKEKG
>Latimeria_chalumnae
----------------------MPPHRILGRATVKEKG
>Fundulus_heteroclitus
----------------------MPPQRLPGSPVVKEKG
>Apteronotus_leptorhynchus
----------------------MPPHRLTGNPTVKEKG
>Scleropages_formosus
MQRESRSAGCSELRAQRACAAVMPPHRLTGTPTVKGNG
>Platichthys_stellatus
----------------------MPPQGLRGPPAVKEKG
>Salvelinus_alpinus
----------------------MPPHRLPGTPTVKEKG
>Lepisosteus_oculatus
----------------------MPPHSFTGIPTVKEKG
>Ictalurus_punctatus
-------------------MQVMPPHRLTGNPTVKEKG

HG0168 ENSGALG00000006997 BRINP1

>Gallus_gallus
MPAIWIKEGDFLSEHEKDC
>Anas_platyrhynchos
MPAIWIKEGDFLSEHEKDC
>Hynobius_chinensis
MPEIWSRDWAFLSEHEKDC
>Calidris_pugnax
MPAIWIKEGDFLSEHEKDC
>Columba_livia
MPAIWIKEGDFLSEHEKDC
>Alligator_mississippiensis
MPASRIQGRDFLSEHEKDC
>Cuculus_canorus
MPAIWIKEEDFLSEHEKDC
>Monodelphis_domestica
MPAIWITEGDLKAEYEKDC
>Haliaeetus_leucocephalus
MPAIWIKEGDFLSEHEKDC
>Ornithorhynchus_anatinus
MPAIWIREGDLTAEHEKGC
>Lepidothrix_coronata
MPAIWIKEGDFLSEHEKDC
>Picoides_pubescens
MPAICIKEGDFLSEHEKDC
>Melopsittacus_undulatus
MPAIWIKEGDFLSEHEKDC
>Aptenodytes_forsteri
MPAIWIKEGDFLSEHEKDC
>Pelodiscus_sinensis
MPAIWIKERDFLSEHEKDC
>Calypte_anna
MPAIWIKEGYFLSEHEKDC

HG0169 ENSG00000114933 INO80D

>Homo_sapiens
MVGKLSFRRAEEQLSGVMSSQCL
>Anas_platyrhynchos
MVGKLLFWRAEEPLSDVMSSRCL
>Chaetura_pelagica
MVGKLLFWRAEEPLSDVMSSQCL
>Hynobius_retardatus
MVGKLLFWRAEELLSGVMSSQCL
>Physeter_catodon
MVGKLSFRRAEELLSGVMSSQCL
>Calidris_pugnax
MVGKLLFWRAEEPLSDVMSSQCL
>Pteropus_vampyrus
MVGKLSFRRAEERLSGVMSSQCL
>Columba_livia
MVGKLLFWRAEEPLSDVMYSQCL
>Monodelphis_domestica
MVGKLSFRRAEERLSGVMSSQCR
>Aquila_chrysaetos_canadensis
MVGKLLFWRAEEPLSDVMSSQCL
>Gallus_gallus
MVGKLLFWRAEEPLCDVMSSRCL
>Serinus_canaria
MVGKLLFWRAEEPLSDVMSSQCL
>Egretta_garzetta
MVGKLLFWRAEEPLSDVMSSQCL
>Picoides_pubescens
MVGKLLFWRAEDPLSDVMSSQCL
>Melopsittacus_undulatus
MVGKLLFWRAEEPLSDVMSSQCL
>Aptenodytes_forsteri
MVGKLLFWRAEEPLSDVMSSQCL

HG0170 ENSG00000180530 NRIP1

>Homo_sapiens
MKALRSAEQIDYSGVPV----LGTTAVRERTLIFAFNGEQKMKK--------EKEYI---H------
>Anas_platyrhynchos
MKALLSAEQIDCYSVPV----LGTTAVKERNSMCLMEMEKDEKQTGFMEKGEEGKYLMELHYLRSVF
>Canis_lupus_familiaris
MKALLSAEQIDCSGVPV----LGTTAVKERTPLLAFNWKQKMKK--------EKQCI---H------
>Balaenoptera_acutorostrata_s
--------------------------------MFAFIVEQKVKK--------EKQYI---R------
>Rhinolophus_ferrumequinum
-----------------------------------------MKK--------EKQDI---H------
>Dasypus_novemcinctus
-----------------------------------------MKK--------EKQYI---H------
>Galeopterus_variegatus
-----------------------------------------MKE--------EKHYI---H------
>Oryctolagus_cuniculus
-----------------------------------------MKT--------AKQYI---H------
>Equus_asinus
----MTSERLFNSPLNFLSQKSETQILEIEELMFAFNGQQKMKK--------EKRYI---H------
>Manis_javanica
-----------------------------------------MKK--------EKQYI---H------
>Loxodonta_africana
-----------------------------------------MKK--------EKQYI---H------
>Ictidomys_tridecemlineatus
-----------------------------------------MKM--------EKQYI---H------
>Tupaia_chinensis
-----------------------------------------MKK--------ENQCI---H------
>Trichechus_manatus_latirostr
-----------------------------------------MKK--------EKQYI---H------
>Anolis_carolinensis
MKALLSAEQIDCYSVPV----LGTTAVKERLLALTLHINHLEGE--------EKKHI---NCVQSSV
>Orycteropus_afer_afer
-----------------------------------------MKK--------VKQYI---H------

HG0171.1 ENSG00000177311 ZBTB38

>Homo_sapiens
-----------------------------------------MYS--------KEKRTLAMVQLNQDSEADMQCHLQH
>Acinonyx_jubatus
----------------------------------------------------------------------MQCHLQH
>Balaenoptera_acutorostrata_s
----------------------------------------------------------------------MQCHLQH
>Myotis_davidii
----------------------------------------------------------------------MQCHFQH
>Dasypus_novemcinctus
----------------------------------------------------------------------MQCHLQH
>Columba_livia
--------MM---------------SECHCATVAKKSLRGERRERKKTRSFASSNNMRCAYTLNQDSVADTQCHLQH
>Galeopterus_variegatus
----------------------------------------------------------------------MQCHLQH
>Monodelphis_domestica
----------------------------------------------------MYSSLLEFTVLNQDSVADMQCHFQH
>Condylura_cristata
----------------------------------------------------------------------MQCHLQH
>Corvus_cornix_cornix
----------------------------------------------------------------------MQCHLQH
>Equus_asinus
----------------------------------------------------------------------MQCHLQH
>Manis_javanica
MPTRAHRAMMWGWGRWFHHWVRLGATVSVRATFISGVLVGERCA--------LKRRGLVVVQLNQDSEADMQCHLQH
>Heterocephalus_glaber
----------------------------------------------------------------------MQCHLQH
>Struthio_camelus_australis
----------------------------------------------------------------------MQCHLQH
>Tinamus_guttatus
----------------------------------------------------------------------MQCHLQH

HG0171.2 ENSG00000177311 ZBTB38

>Homo_sapiens
MRNFFFQSCFPVDKFSSVITVHSSCSAYDFCVYVTIASKTCEQEDREISWSPSLYIKMAECLVMLALH-----
>Felis_catus
---------------------------------------------------------MAECLVMLALH-----
>Pteropus_vampyrus
---------------------------------------------------------MAECLVMLALH-----
>Galeopterus_variegatus
---------------------------------------------------------MAECLVMLALH-----
>Monodelphis_domestica
---------------------------------------------------------MAECAAMLVRH-----
>Erinaceus_europaeus
---------------------------------------------------------MAECPVMLALH-----
>Sturnus_vulgaris
---------------------------------------------------------MAGCIAMLTLL-----
>Equus_asinus
MRNFFFQSCFPVDKFSSRDHSTLKLFCVRFCVCVTIASKTCEQEDGEISWSPSLYIKMAECLVMLALH-----
>Manis_javanica
----------------------------MICVCVTIASKTCEQEDGEISWSPSLYIKMAECHASTALSDTYRF
>Trichechus_manatus_latirostr
---------------------------------------------------------MAECLVMLALH-----
>Tinamus_guttatus
---------------------------------------------------------MAGCIATLTLH-----

HG0171.3 ENSG00000177311 ZBTB38

>Homo_sapiens
MLAVLQKAFRAADLDVEEMEDSEPRISK
>Acinonyx_jubatus
------------------MEDSEPRISK
>Physeter_catodon
------------------MEDSEPRISK
>Myotis_davidii
------------------MEDSEPRISK
>Galeopterus_variegatus
------------------MEDKEPRISK
>Erinaceus_europaeus
------------------MEDSEPRISM
>Equus_asinus
------------------MEDSEPRISK
>Manis_javanica
------------------MEDSEPRISK
>Heterocephalus_glaber
------------------MEDSEPRISK
>Trichechus_manatus_latirostr
------------------MEDSESRISK

HG0171.4 ENSG00000177311 ZBTB38

```
>Homo_sapiens
MGDIIRKICGADHKK---------------------------------
>Acinonyx_jubatus
MGDIIRKICGADLNKK--------------------------------
>Balaenoptera_acutorostrata_s
MGDIIRKICGADLDKK--------------------------------
>Rousettus_aegyptiacus
MGDIIRKICGADFNKK--------------------------------
>Condylura_cristata
MRDVIRKICGADLDEK--------------------------------
>Equus_asinus
MGDIIRKICGAALNK---------------------------------
>Manis_javanica
MGDIIRKICGADLNKK--------------------------------
>Heterocephalus_glaber
MSDIIRKICGADLNRVKNPEQESVCDSWAQGVGSVWASEEQLIRIRLS
```

HG0172.1 ENSG00000196782 MAML3

>Homo_sapiens
MQLSSQSTLRTRGGINTKTFLWKENAISK---LAVFRCGVE--------------------------------------
>Mustela_putorius_furo
MQLSSQSTLRTRGGINTKTFLWKENAISK---LAVFRCGVE--------------------------------------
>Balaenoptera_acutorostrata_s
MQLSSQSTLRTRGGINTKTFFVEGKCNKQACRFPM-RCGVSVCRACPHWRHMLVCVHGVCFSVRREKMLANHRKSPGIY
>Myotis_brandtii
MQLSSQSTLRTRGGINTKTFLWKENAISK---LAVFRCGVE--------------------------------------
>Dasypus_novemcinctus
MQLNSQSTLRTRGGINTKTFLWKENAISK---LAVF-------------------------------------------
>Monodelphis_domestica
MQPRSQSTLRTRGGINTKTFLWKENAIRK---LAVF-------------------------------------------
>Gallus_gallus
MQPGSASTLRARRGTSAQTFL----------------------------------------------------------
>Condylura_cristata
MQLSSQSTLRTRGGINTKTFLWKENAISR---LAVFRCGVE--------------------------------------
>Oryctolagus_cuniculus
MQLSSQSTLRTRGGINTKTFLWKENAISK---LAVLRCAVE--------------------------------------
>Ficedula_albicollis
MQLSSQSTLRTRCGINTKTFFQKENTISK---RLCLSFDAAECE-----------------------------------
>Ceratotherium_simum_simum
MQLSSQSTLRTRGGINTKTFLWKENAISK---LAVFRCGVE--------------------------------------
>Manis_javanica
MQLSSQSTLRTRGGINTKTFLWKENAISK---LAVFRCGVE--------------------------------------
>Mus_musculus
MQLSSQSTLRTRGGINTKTFLWKENAISK---LAVFRCGVE--------------------------------------
>Tupaia_chinensis
MQLSSQSTLRTRGGINTKTFLWKENAISK---LAVFRCGVE--------------------------------------
>Pelodiscus_sinensis
MQLSSQSTLSTRCGINTKTFFLEGKYNKKAALPVFWCCGV---------------------------------------

HG0172.2 ENSG00000196782 MAML3

>Homo_sapiens
MDYKKKASKERISP------------------
>Mustela_putorius_furo
MDYKKKAGEERISP------------------
>Balaenoptera_acutorostrata_s
MDYKKKAGEERIFP------------------
>Myotis_brandtii
MDYKKKAGEERISP------------------
>Dasypus_novemcinctus
MDYKKKAGEERSTP------------------
>Condylura_cristata
MDYKKKAGDERIAP------------------
>Oryctolagus_cuniculus
MDYKKKAGEERRSP------------------
>Ceratotherium_simum_simum
MDYKKKAGKERISP------------------
>Nannospalax_galili
MDYKKKAGEERIAP------------------
>Tupaia_chinensis
MDYKKKAGEERLSPGVAIRFAAGSVCCDSFC

HG0173 ENSG00000101746 NOL4

>Homo_sapiens
---------------------------------------MIELACHRHWYLS
>Anser_cygnoides
---------------------------------------MIELACHRHWYLS
>Lipotes_vexillifer
MQERLALTVQRREFRGSQTNPWVHLERKVGPRLRDAVSAMIELACHRHWYLS
>Calidris_pugnax
---------------------------------------MIELACHRHWYLS
>Pteropus_alecto
---------------------------------------MIELAFHRHWYLS
>Aquila_chrysaetos_canadensis
---------------------------------------MIELACHRHWYLS
>Sorex_araneus
---------------------------------------MIELACHRYWYLS
>Oryctolagus_cuniculus
---------------------------------------MIELVRHRHWYLS
>Elephantulus_edwardii
---------------------------------------MIELACHRHWYLS
>Corvus_cornix_cornix
---------------------------------------MIELACHRHWYLS
>Loxodonta_africana
---------------------------------------MIELACHRHWYLS
>Fukomys_damarensis
---------------------------------------MIELACHRHWYLS
>Tupaia_chinensis
---------------------------------------MIELACHRHWYLS
>Trichechus_manatus_latirostr
---------------------------------------MIELACHRHWYLS
>Chrysemys_picta
---------------------------------------MIELACHRHWYLS

HG0174 ENSG00000144355 DLX1

>Homo_sapiens
--------MRLTRGRAASPALVQRSPSASDQSPVIMQDSGPISSASS
>Anas_platyrhynchos
-----------------------------------MQDGGPISSAVS
>Mustela_putorius_furo
-----------------------------------MQDSGPISSASS
>Rousettus_aegyptiacus
-----------------------------------MQGSGPISSASS
>Galeopterus_variegatus
-----------------------------------MQDSGPISSASS
>Monodelphis_domestica
MITRRGSGTHLSQVLLASLPLVQPSQSTSDQSQAIMQDGGPIRSASS
>Coturnix_japonica
-----------------------------------MQDGEPISSAGS
>Ochotona_princeps
-----------------------------------MQDGGPISSASS
>Ornithorhynchus_anatinus
-----------------------------------MQDGGPIRSASS
>Zonotrichia_albicollis
-----------------------------------MQDGGPISSAVS
>Equus_przewalskii
---------------------------------MIMQDSGPISSASS
>Loxodonta_africana
-----------------------------------MQDSGPISSANL
>Octodon_degus
-----------------------------------MQDGGPISSASS
>Tupaia_chinensis
-----------------------------------MQDSGPISSASS
>Chrysemys_picta
-----------------------------------MQDGGPISSASS

HG0175 ENSG00000151320 AKAP6

>Homo_sapiens
MCCKHQKKTVGSGDAVLERSEV---
>Anas_platyrhynchos
MCCKHQKRMI---------------
>Felis_catus
MCCKHQKETVGAGDAVLERSKV---
>Balaenoptera_acutorostrata_s
MCCKHQKKAVGAGDAVLERSKA---
>Calidris_pugnax
MCRKHQKRMI---------------
>Rousettus_aegyptiacus
MCCKHQKKTVGAGDAVLERSKV---
>Dasypus_novemcinctus
MCRKHQKRTVGAGDAVLERSKV---
>Monodelphis_domestica
MCRKHQKRMVRAGDALLERSMV---
>Condylura_cristata
MCCKHQKRTVRAGDAVLERSKI---
>Parus_major
MCRKHQKRMI---------------
>Equus_caballus
MCCKHQKKTVGPGDAVLERSKV---
>Manis_javanica
MCCKHQKQTVGAGDELPEEGLGHLR
>Fukomys_darlingi
MCCKHQKKTVGAGDAVLERSKV---
>Struthio_camelus_australis
MCRKHQKGTI---------------
>Pelodiscus_sinensis
MCRKHQKRMV---------------

HG0176 ENSG00000170962 PDGFD

>Homo_sapiens
---------------------------------------------MQEVWGELGD
>Apteryx_australis_mantelli
---------------------------------------------MQEVWGKLGE
>Physeter_catodon
---------------------------------------------MQEVWGELGD
>Pteropus_alecto
---------------------------------------------MQEVWGELGD
>Galeopterus_variegatus
---------------------------------------------MQEVWGELGE
>Monodelphis_domestica
---------------------------------------------MQEVWGELGD
>Macropus_eugenii
MLGLDQDRHRHIWILLQLPLLATGGAVGRKSTCCSSSPTGLTNFSMQEVWGELGD
>Haliaeetus_leucocephalus
---------------------------------------------MQEVWGQLGE
>Oryctolagus_cuniculus
---------------------------------------------MQEVWGELGD
>Ceratotherium_simum_simum
---------------------------------------------MQEVWGELSD
>Loxodonta_africana
---------------------------------------------MQEVWGELGD
>Heterocephalus_glaber
---------------------------------------------MQEVWGKLSD
>Tupaia_chinensis
---------------------------------------------MQEVWGELGD
>Trichechus_manatus_latirostr
---------------------------------------------MQEVWGELGD
>Orycteropus_afer_afer
---------------------------------------------MQEVWGELGD

HG0177 ENSG00000171843 MLLT3

>Homo_sapiens
MLRNHLLYPGAA--AAAAAFAFGAEFNKRASEQASAGGKRQRMSAIYPPLLGALSFIAASS
>Felis_catus
------------------------------------------MSAIYPPLLGALSFIAASS
>Orcinus_orca
------------------------------------------MSAIYPPLLGALSFIAASS
>Myotis_brandtii
------------------------------------------MSAIYPLLLGALSFIAASS
>Dasypus_novemcinctus
------------------------------------------MSAIYPPLLGALSFIAASS
>Alligator_mississippiensis
MCRAHALTAALQALELELELLLGAQFNKSERK-RQGEKERKRMSAISPPLLGALPVIAASS
>Galeopterus_variegatus
------------------------------------------MSAIYPPLLGALSFIAASS
>Monodelphis_domestica
------------------------------------------MSAIYPLLLGALSFISASS
>Condylura_cristata
------------------------------------------MSAIYPPLLGALSFIAASS
>Oryctolagus_cuniculus
------------------------------------------MSAIYPPLPGALSFIAASS
>Elephantulus_edwardii
------------------------------------------MSAIYPPLPGALSFIAASS
>Equus_caballus
MLRNLILWPWSCCCCCCCFCFWG--------------------------------------
>Ictidomys_tridecemlineatus
------------------------------------------MTAIYPPFLGALSFIAASS
>Pelodiscus_sinensis
------------------------------------------MSAIYPPLPCAPSVTSASS
>Orycteropus_afer_afer
------------------------------------------MSAIYPLLLGALSFIAASP

HG0178.1 ENSG00000131634 TMEM204

>Homo_sapiens
------------------------------MGPE-ASSFSTGLALLQVQEGG
>Odobenus_rosmarus_divergens
--------------------------------MGREAGCSSGLALLQVQEGG
>Balaenoptera_acutorostrata_s
--------------------------------MGSSEGFSSGLALLQVQEGG
>Pteropus_vampyrus
--------------------------------MGQAAAFSSGLALFQVQEGG
>Galeopterus_variegatus
------------------------------MGLTAATSFNTGLALLQVQEGG
>Monodelphis_domestica
------------------------------MGLS-AAWFCTGLALLKVQAGG
>Trichosurus_vulpecula
------------------------------MGLS-AAWFSTGLALLKVQAGG
>Oryctolagus_cuniculus
------------------------------MGLE-AAGFSTGLALLQVQEGG
>Ceratotherium_simum_simum
--------------------------------MGRAGGFSSGLALLQVQEGG
>Ctenomys_sociabilis
MFVASIASGSQKDGILTTLDSCWCPGFFPNMGLA-ATGFSTGLALLQVQEGG
>Tupaia_chinensis
------------------------------MGFT-AAAFSTGLALLQVQEAG

HG0178.1 ENSGALG00000029047 TMEM204

>Gallus_gallus
------------------------------MDLHVFQFITGLALLIKVQEGG
>Anser_cygnoides
------------------------------MEVPIFQFITGLALLIKVQEGG
>Apteryx_australis_mantelli
------------------------------MDLPIFQFITGLALLIKVQEGG
>Calidris_pugnax
------------------------------MDLPILQFSTGLALLIKVQEGG
>Callorhinchus_milii
------------------------------MDLCAHQLLTAASPVKKAQEGG
>Columba_livia
------------------------------MDLPVIQFITGLALLIKVQEGG
>Alligator_mississippiensis
------------------------------MDLPVLQFITGLALLIKVQEGG
>Aquila_chrysaetos_canadensis
------------------------------MDLPIVQFITGLALLIKVQEGG
>Corvus_brachyrhynchos
------------------------------MDLPVIQFTTGLALLIKVQEGG
>Egretta_garzetta
------------------------------MDLPVVQFITGLALLIKVQEGG
>Fukomys_damarensis
MFVASLASGSQKDGILRTLDSRRGPCFFPEMDLAATGFSTGLALLQV-QQGG
>Aptenodytes_forsteri
------------------------------MDLPIVQFITGLALLIKVQEGG
>Struthio_camelus_australis
------------------------------MDLPVFQFITGLALLIKVQEGG
>Chelonia_mydas
------------------------------MDLPVLQFITRLALLIQVQEGG

HG0178.2 ENSGALG00000029047 TMEM204

>Gallus_gallus
----------------------------------------------------------------------------MFDCRGSGSVCCSRLRPEAHQNSHKNSHFL----------------------------------------------------------------
---------------------------------------------------------------------
>Anser_cygnoides
----------------------------------------------------------------------------MFDCCGSGSVCCSRLRPEAHQKQPQKQPFPLSWGK---GGGRQT-----FSSQQ----------------------PVLWS-------------
LPLAVRRSETDTA-EQ-----IPQD--T------PNFH--SSLG-------------------------
>Apteryx_australis_mantelli
----------------------------------------------------------------------------MFDCRGSGSVCCSRLRPEAHQKQPQKQQFPLSGETEREGGKKTL-----FLTAV----------------------SALQS-------------
PPLAALMFREGHSGAN-----SSRDSKLSEFLWDENIR--VEFGLALR---MTILSVCGIF--------
>Calidris_pugnax
----------------------------------------------------------------------------MFDCRGSGSVCCSRLRPEAHQNQPQKQPFPLRGEVG---GPFPR-----SSC------------------------------------------
----SSIFREGHGTAN-----SSRYFKLAELLWDASIH--IQLGLALR---E-----------------
>Aquila_chrysaetos_canadensis
----------------------------------------------------------------------------MFDCCGSGSVCCSRLRPEAHQKQPQKQPFPLRGEVG---GAFSS---Q-QWLFSGPCTRSSD-----------LQRGTRQS-------------
KLLEILQTCRAPLG-------------------------------------------------------
>Lepidothrix_coronata
----------------------------------------------------------------------------MFDCRGSGSVCCSRLRPEAHQKQPQKQPFPLGGEVG---GPFPH-----CSGCSPPGTHSSD-----------LRRGTRQS-------------
SPLEILQAC----------------------------R--APLG-------------------------
>Microcebus_murinus
MSTASLGPVSQHRSLPVSTAMGETAGLPCAWDPWPLLWDRKVGRGQQRGWPGAGPQ---KSP---QAFPIRKHILLVSGCGGRAAICCSWLRPPAHQNSLKNNHFLSAERRGELELEHTHTSLQRALCFPGPAIRCGAPGKGPGSRGAAAKPALLRARPGAGLPRCRLPP
ELLSVFCRCRGRGTRSLLGPLLPRTTA-RSRLWIRAGAPVSSPGWASQQLAAAQARPCSRCRKEDKAGP
>Mus_musculus
--------------------MSVLAGIPD----------YSYGTG---RWGGAASGGLEQASEVSPGLSHQEAHPVSVWLCGRAAICCSWLRPPAHQKQPQKQPFPLC--------------------------------------------------------------
---------------------------------------------------------------------
>Struthio_camelus_australis
----------------------------------------------------------------------------MFDCRGSGSVCCSRLRPEAHQKQPQKQPFPLS--------------------------------------------------------------
---------------------------------------------------------------------

HG0179 ENSDARG00000036549 agpat3

```
>Danio_rerio
-------MLSHIERLP-CYNRLR----DTAVAQYE---REKLEKVAGFPPTGSVVLSEGPCKRGL
>Opsanus_beta
--------------------MLN----QSAVMIFTGICGALVVMAMAYYIYWSVVLAEGPCKRGL
>Oryzias_latipes
--------------------MLN----QSAVMIFTGICGAMVLMAMAYYVYWSVILADGPCKRGL
>Trachinotus_blochii
-------------------------------------------------MCMSVVLAEGPCKRGL
>Pygocentrus_nattereri
------------------------------------------MTAGALQHNGSVVLSEGPCKRGL
>Neolamprologus_brichardi
--------------------MLN----QSAVMIFTGICGAMVFMAMAYYVYWSVVLAEGPCKRGL
>Clupea_harengus
--------------------MHT----Q-SLLHWSKSVS-QIFFKPKKGPVRSVVLSEGPCKRGL
>Fundulus_heteroclitus
--------------------MLN----QSAVMIFTGICGAMVFMAMAYYVYWSVVLAEGPCKRGL
>Esox_lucius
--------------------MLS----QSTVMIFSGICGAFVVLAMGYYVYWTIVLSEGPCRRGL
>Scleropages_formosus
-------MLC------GCYQLRCGVFRGIVAVRWTGS-GLVPGGQVSWKRTRSVILSEGPCKRGL
>Ficedula_albicollis
---------------------------MEDCLLWEG----------LHPGTGTIILSDGPCSRF-
>Salmo_salar
MLKNKYVYGSGVQRHQCCFSVELK---TNALDLFGNV------NRRQDPEIRSVVLADGPCKRGL
>Lepisosteus_oculatus
-----------------------MLANQTSVLIFTGICAALLILAVAYYTYWSVVLSEGPCKRY-
>Pelodiscus_sinensis
-----------------------MRANGKDMETTQ-----NSKEHLGSPAVMTVILSEGPCKAY-
```

HG0179 ENSG00000160216 AGPAT3

>Homo_sapiens
-------MLRWVSVASPEFLLNEVCRVIWTAVLSEGPCTRS-
>Opsanus_beta
-MLNQSAVMIFTGICGALVVMAMAYYIYWSVVLAEGPCKRGL
>Canis_lupus_familiaris
------------------MSCFSARGRIRTAVLSEGPCTRS-
>Astyanax_mexicanus
-MLNQGAVMIFTGICGALVVMAVAYYAYWSVVLSEGPCKRGL
>Myotis_lucifugus
-MLASYSTCLFSGVCAALVAVALTYYLYWTAVLSEGPCSRY-
>Haplochromis_burtoni
-MLNQSAVMIFTGICGAMVFMAMAYYVYWSVVLAEGPCKRGL
>Danio_rerio
-MLNQGAVMIFTGVCGALVVMAVAYYAYWSVVLSEGPCKRGL
>Fundulus_heteroclitus
-MLNQSAVMIFTGICGAMVFMAMAYYVYWSVVLAEGPCKRGL
>Monodelphis_domestica
-MLDSYSTSLFSAVCATLFVMALAYYLYWTSVLSVGPCTKS-
>Macropus_eugenii
-MLANYSVCLFSAICANLFVMALAYYLYWTSVLSEGPCTHY-
>Esox_lucius
-MLSQSTVMIFSGICGAFVVLAMGYYVYWTIVLSEGPCRRGL
>Equus_asinus
----------MVGSTSP-------ELWPRTAVLSEGPCTRY-
>Rattus_norvegicus
-MLASSSTCLFSGVCVALLAVALAYYFYWTAVLSEGPCGRP-
>Lepisosteus_oculatus
MLANQTSVLIFTGICAALLILAVAYYTYWSVVLSEGPCKRY-

HG0180.1 ENSG00000112246 SIM1

```
>Homo_sapiens
--MSIFRGPGFAAGSRRKG-----WLLRGPTGI
>Anas_platyrhynchos
--MTLFRGAMLSATSRRKGQKYLIWLLRGAASF
>Odobenus_rosmarus_divergens
--MSIFWGPGFASPNRRKG-----WLLRGPAGI
>Orcinus_orca
--MSISREPGFAARNRRKG--------------
>Myotis_brandtii
--MSLFQGPGFAARNRRKG-----WLLRGPAGI
>Monodelphis_domestica
--MSVFRVLVFAAWNRREA-----WFLDGPIDI
>Sorex_araneus
--MSVFRAPGCAAWNRKKG-----WPRRAPAGL
>Oryctolagus_cuniculus
--MSTFRAPRFAAQNGRKG-----WLLRGPAGI
>Equus_asinus
MIMSIFRGPGFAARNRRKG-----WLLGDLAGI
>Manis_javanica
-MMSTFREPGYEAPNRRKG-----WLLRSPAGI
>Loxodonta_africana
--MSIFRGPGFAARNRRKG-----WLLWGASGI
>Heterocephalus_glaber
--MSIFQGTGFAAENRRKR-----WLLRGSASI
>Tupaia_chinensis
--MSIFRGPGCASPNRKKN-----WLPWGLAGI
>Trichechus_manatus_latirostr
--MSIFRGPGFAARNRRKV-----WLLRGPAGI
```

HG0180.2 ENSG00000112246 SIM1

>Homo_sapiens
---MRTVKRWAPPKPLPLVGTEGP
>Mustela_putorius_furo
MVGMSTIKRWAPLHPLPPAGTEVA
>Pteropus_alecto
---MNTIKRRASPHPLPPAGTEGR
>Loxodonta_africana
---MSAIKRWAPPHPLPPVGTEAA
>Tupaia_chinensis
---MIAIRRWAPAHPLPPAGTEVA

HG0181.1 ENSG00000143776 CDC42BPA

```
>Homo_sapiens
----------MHSVFFLPF--------L-FFFFPGSELLMQIGLYSLMMQPDSFQDYVARVEF
>Ailuropoda_melanoleuca
---------------------------------------MRIGPDSLMMQPDSLRDCVARVEF
>Pteropus_vampyrus
---------------------------------------MQIGLNSLMMQPDSFQDYVARIEF
>Galeopterus_variegatus
---------------------------------------MQIGLYSFMMQPDSFQD-------
>Monodelphis_domestica
MHYSSVGGGNLHLGFRLPLVTYFVFLLLFSIHPSKSELLMPSGLYSLMMQRDSFQDCVA----
>Coturnix_japonica
-------------------------------------MRPEWGFYTLMMQWDSFQGCVA----
>Condylura_cristata
---------------------------------------MHLGLNSLVMQPDSFQDYVARVEF
>Ochotona_princeps
---------------------------------------MQMGLYSLMMQPDSFQDYVARVEF
>Equus_asinus
---------------------------------------MQIGLNSLMMQPDSFQDYVARVEF
>Loxodonta_africana
---------------------------------------MRIGLYSLMMQPASFQDYVA----
>Mesocricetus_auratus
---------------------------------------MPRGLYSLMMQPDSFQDYVARIEF
>Tupaia_chinensis
---------------------------------------MQIRLYSLMMQPDSFQDYVARVEF
>Thamnophis_sirtalis
---------------------------------------MEWGLYTLMMQKDSFRDWVS----
>Chrysemys_picta
-------------------------------------MQTEWGFYTLMMQWDSFQDYVA----
```

HG0181.2 ENSG00000143776 CDC42BPA

>Homo_sapiens
MYMGEIEDASVK
>Balaenoptera_acutorostrata_s
MYMCEIEDASVK
>Rhinolophus_ferrumequinum
MYMCEIEDASVK
>Erinaceus_europaeus
MYMCEIEDASVK
>Ochotona_princeps
MYMGEIEDASVK
>Ceratotherium_simum_simum
MYMCEIEDASVK
>Cricetulus_griseus
MFMGEIEDASVK
>Tupaia_chinensis
MFMGEIEDASVK
>Trichechus_manatus_latirostr
MYMCEIEDASVK

HG0182 ENSGALG00000031991

```
>Gallus_gallus
MDVNQPKQEHLLALKGPVSH---------------------
>Ailuropoda_melanoleuca
MEVNPPKQEHLLALKGPVSL---------------------
>Lipotes_vexillifer
MEVNPPKQEHLLALKGPVSL---------------------
>Dasypus_novemcinctus
MEVNPPKQEHLLALKGPVFL---------------------
>Sarcophilus_harrisii
MEVNPPKQEHLLALKGPVSSFSSVQNVERRTRKSNKF----
>Trichosurus_vulpecula
MEVNPPKQEHLLALKGTVSSFSSVQNVERKTRKSYNYEFKF
>Oryctolagus_cuniculus
MEVNPPKQEHLLALKGPVSL---------------------
>Sturnus_vulgaris
MDVNQPKQEHLLALKGPVFH---------------------
>Manis_javanica
MEVNPPKQEHLLALKGPVSL---------------------
>Papio_anubis
MEVNPPKQEHLLALKGPVSL---------------------
>Cricetulus_griseus
MEVNPPKQEHLLALKGPVSSDLARETVL-------------
>Tupaia_chinensis
MEVNPPKQEHLLALKGPVSL---------------------
>Aptenodytes_forsteri
MDVNQPKQEHLLALKGPVIH---------------------
>Protobothrops_mucrosquamatus
MDVTQSKQEHLFALKCPVLSYTAHRIRFF------------
```

HG0183 ENSG00000011465 DCN

>Homo_sapiens
-----------------------MW---VSAGLCSLRETLFFNCAME------------------------------------
>Leptonychotes_weddellii
MGRKKADKLSGCNTRDISLPYDLMW---VSAGLCSLRETLFFNCAME------------------------------------
>Lipotes_vexillifer
-----------------------MW---VSAGLCSLRETLFFNCAME------------------------------------
>Rousettus_aegyptiacus
-----------------------MW---VSAG---------------------------------------------------
>Dasypus_novemcinctus
-----------------------MW---VSAGLCSLRETFIF-----------------------------------------
>Galeopterus_variegatus
-----------------------MW---VSAGLCSLRETLFFNCAME------------------------------------
>Erinaceus_europaeus
-----------------------MW---VSAGLSSLRETVFFLTAMQ------------------------------------
>Elephantulus_edwardii
-----------------------MW---VSAGLCSLRETFFFFSVCYGVEAGGFQPSDSHRAAPNPLLLSTPANSFAWAEYLV
>Manis_javanica
-----------------------MW---VSAGLCSLRETLFFNCAME------------------------------------
>Loxodonta_africana
-----------------------MW---VSAGLCSLRETLFFNCAME------------------------------------
>Dipodomys_ordii
-----------------------MW---VSAGLCSLRETLFFNCAME------------------------------------
>Protobothrops_mucrosquamatus
----------------------MMYCGCQQPGVFSLRKTLLFNCAME------------------------------------
>Pelodiscus_sinensis
-----------------------MW---VSAWIVFIEETLFFNCAVE------------------------------------
>Orycteropus_afer_afer
-----------------------MW---VSAGLCSLRETLFFNCAME------------------------------------

HG0184 ENSG00000136542 GALNT5

>Homo_sapiens
-----------------------------------------------------------------------------MLCYEMPRRQRLTSGRN-
>Odobenus_rosmarus_divergens
----------------------------------------------------------------------------------MPRRQRLTSSRS-
>Orcinus_orca
-----------------------------------------------------------------------------MLCCEMPRRQRLTSGRS-
>Pteropus_alecto
-----------------------------------------------------------------------------MLCCEMPRRQRLTSGRS-
>Dasypus_novemcinctus
-----------------------------------------------------------------------------MLCCEMPRRQRLTSSRS-
>Monodelphis_domestica
-----------------------------------------------------------------------------MVCCEMPRRQRLTSSSRS
>Condylura_cristata
-----------------------------------------------------------------------------MLCYEMPRRQTDKG----
>Oryctolagus_cuniculus
MRSGVELQRADRERGGEKKHQPQPAPPFLGLVGRWTRVHRENTSDRLNYLVAGIACSWNDLRAVFTRVQPGPAEQAPMLYCEMPRRQRLTSSRS-
>Equus_caballus
-----------------------------------------------------------------------------MLCYEMPRRQRLTSGRS-
>Manis_javanica
-----------------------------------------------------------------------------MLCCEMPRRQRLTSGRS-
>Loxodonta_africana
-----------------------------------------------------------------------------MLCCEMPRRQRLTSSRS-
>Ictidomys_tridecemlineatus
-----------------------------------------------------------------------------MRCCEMPRRQRLTGGRS-
>Pelodiscus_sinensis
-----------------------------------------------------------------------------MPCCKMPRRQRLTGSRI-
>Orycteropus_afer_afer
-----------------------------------------------------------------------------MLCCEMPRRQRLTSSRS-

HG0185 ENSG00000143033 MTF2

>Homo_sapiens
MHRQSAGNQNGEGLY---------------------------------------
>Canis_lupus_familiaris
MHRQSAGNQNGEGL----------------------------------------
>Lipotes_vexillifer
MHRQSAGNQNGEGL----------------------------------------
>Miniopterus_natalensis
MHRQSAGNQNGEGL----------------------------------------
>Galeopterus_variegatus
MHRQSAGNQNGEGL----------------------------------------
>Erinaceus_europaeus
MHRQSAGNQNGEGL----------------------------------------
>Ochotona_princeps
MHRQSAGNQNGEGL----------------------------------------
>Equus_asinus
MHRQSAGNQNGEGL----------------------------------------
>Loxodonta_africana
MHRQSAGNQNGEGLY---------------------------------------
>Heterocephalus_glaber
MHRQSAGNQNGEGL----------------------------------------
>Tupaia_chinensis
MHRQSAGNQNGEGL----------------------------------------
>Trichomycterus_areolatus
MHRDSAGNQNGETKSRKIFV------GSNWNNS---RLLTCDSFF---------
>Trichechus_manatus_latirostr
MHRQSAGNQNGEGLYGSGLCLRLVYERPLYPNPSPQDFGLCVRLCGVPGGAGAQ
>Orycteropus_afer_afer
MHRQSAGNQNGDGLY---------------------------------------

HG0186 ENSG00000167552 TUBA1A

>Homo_sapiens
-----------------------------------------------MASLGRLGVWTNRKRPRPIPMVTELYIGSRIRLCAAGSLTSTA----
>Mustela_putorius_furo
-------------------------------------------------------------------MVTELYIGSRIRPCAAGSLTSTA----
>Orcinus_orca
-------------------------------------------------------------------MVTELYIGSRIRPCAAGSLTSTA----
>Rousettus_aegyptiacus
-------------------------------------------------------------------MVTELYIGSRIRPCAAGSLTSTA----
>Sarcophilus_harrisii
-------------------------------------------------------------------MVTELYIGSCIGPCAVSSLTSTS----
>Sorex_araneus
-------------------------------------------------------------------MVTESYLGSRIRPCAAGSLTSTA----
>Ochotona_princeps
MDARAAGFCSLSLSEGCRKVYGYWAGGGGGTHYLIPCRLHQSAPSLHGSPGADAGVWTNRNRPRPIPMATGLYIGSLRRPHRRFSYIDRLRVAL
>Equus_asinus
------------------------------------MSLAPITTLLSWLLWADRGVWTNRKRPQPIPMVTELYIGSRIRPCAAGSLTSTA----
>Manis_javanica
-------------------------------------------------------------------MVTELYIGSCILPCTAGFLTSTA----
>Loxodonta_africana
-------------------------------------------------------------------MVTELYIGSCIRPCAAGSLTSTA----
>Marmota_marmota_marmota
-------------------------------------------------------------------MVTELYIGSRIRLCAAGSLTSTA----
>Trichechus_manatus_latirostr
------------------------------------MSLAPITTLLSWLLWADRGVWTNRKRPRPIPMVTELYIGSRIRPCAAGSLTSTA----
>Python_bivittatus
-------------------------------------------------------------------MVTGLYIGSRVILCATGSLTSTS----
>Orycteropus_afer_afer
-------------------------------------------------------------------MVTELYIGSRIRSSAAGSLTSTA----

```
HG0187.1 ENSGALG00000004058 GPR146

>Gallus_gallus
MDKMFLPAST-------GQQGTAYQKSW
>Anser_cygnoides
MGKMLLPAIK-------KLQGGAYQESW
>Caprimulgus_carolinensis
MGKMFLPAIK-------EQQGCAYQKSW
>Pterocles_gutturalis
---MFLPAIT-------EHQGSAYQKSW
>Alligator_mississippiensis
MSKMLLQAIKKH---SFKQQGSAYQKSW
>Aquila_chrysaetos_canadensis
MGKMFLPTIK-------GQQGTAYQKSW
>Gavia_stellata
MGKMFLPAIE-------EQ---------
>Lepidothrix_coronata
---MFLPAVK-------GQQGRAYQKSW
>Fulmarus_glacialis
MGKMFLPAIK-------EQQGSAYQKSW
>Aptenodytes_forsteri
MGKMFLPAIK-------EQQGSAYQKSW
>Takydromus_sexlineatus
---MFLPAVN-------LRKGSAYQESW
>Struthio_camelus_australis
MGKMFLPAVRKH---SLKQQGSAYQKSW
>Chrysemys_picta
---MFLPAIKKPICYSLKQQGRAYRKSW
```

HG0187.2 ENSDARG00000059610 gpr146

>Danio_rerio
MSMEIQQNPMPP
>Pygocentrus_nattereri
MSVEIQQNPMPP
>Esox_lucius
MSVEVQQNPMPP
>Chanos_chanos
MSMEVQQNPMPP
>Apteronotus_leptorhynchus
MSMEIKQNPMPP
>Oncorhynchus_kisutch
MSVEVQQNPMPP
>Trichomycterus_areolatus
MSVEIKQDPMPP

HG0187.3 ENSGALG00000004058 GPR146

>Gallus_gallus
MCMYVQGKSHVNTDSVP-------------
>Anas_platyrhynchos
MCMYVQGKSHVNTDSVPRTGLVQPSAGVF-
>Caprimulgus_carolinensis
MCMYVQGKSHVYTEFSATTEGSAAASAVFF
>Pterocles_gutturalis
MRMYVQGKSHVYTEFGATTQGDAAVSAVFS
>Haliaeetus_leucocephalus
MCIYVQGKSRVYTKFGAATEGSAAASAVFF
>Fulmarus_glacialis
MCMYVQGKSHVYTEFGATTEGSAATSAVLF
>Pygoscelis_adeliae
MCMYVQGKSRVYTEFGATTEASAAASAVFF

HG0188 ENSG00000157625 TAB3

>Homo_sapiens
------------MTVVLEDHYFWKAG
>Felis_catus
MTLSVLKRTDQKMTAVLEDHYYWKAG
>Lipotes_vexillifer
------------MTAVLEDHYYWKTD
>Pteropus_vampyrus
------------MTVVLEDYYYWKAG
>Dasypus_novemcinctus
------------MTVVLKDRYYWKAG
>Sarcophilus_harrisii
------------MTETLEHHFYWKAN
>Galeopterus_variegatus
MTLTVLRSTDQKMTVVLEDHYFWKAH
>Erinaceus_europaeus
------------MTVVLEDHYYWKAG
>Oryctolagus_cuniculus
------------MTVVLEVHYFWKAG
>Ceratotherium_simum_simum
------------MTAVLEDHYYWKAG
>Manis_javanica
------------MTAVLEDHYYWKAG
>Loxodonta_africana
------------MTAVFEDHCYWKAS
>Fukomys_darlingi
------------MTVVLEDHYFWKAS
>Tupaia_chinensis
MTWTVLRSTDQKMTMVLEDHYFWKAG
>Trichechus_manatus_latirostr
------------MTAVLEDHCYWKAS

HG0188 ENSGALG00000016284 TAB3

>Gallus_gallus
--------------------MPEASEDHNHWKVC
>Anas_platyrhynchos
--------------------MPEASEDHNHWKAC
>Pelophylax_nigromaculatus
--------------------MQEGSEDHNHWRAL
>Apteryx_australis_mantelli
-----------------MQKMPEASEDHNHWKAY
>Merops_nubicus
--------------------MPEASEDHNHWKAC
>Alligator_mississippiensis
MVLPPEP------SP-PRTKMPEASEDHNHWKAC
>Aquila_chrysaetos_canadensis
--------------------MPEASEDHNHWKAC
>Gavia_stellata
--------------------MPEASEDHNHWKAC
>Parus_major
-MQPPEPQPQPEPSPQRSTKMPEASEDHNHWKAC
>Gekko_japonicus
-MLLLE------PV-PPRATMPEASEDHNHWEAC
>Struthio_camelus_australis
--------------------MPEASEDHNHWKAY
>Chrysemys_picta
--------------------MPEASEDHIHWKTY
>Tinamus_guttatus
--------------------MPEASEDHNHWKAY

HG0189 ENSDARG00000001807 tnfrsf21

```
>Danio_rerio
--------MGIIAFVFQNQALIPYYLKPVHSGAQTIKRREGTEFVH---------
>Amia_calva
--------MGIIPC-LQNQALIPYYLKPVHFATLTLKNCRWKKDQC---------
>Trachinotus_blochii
--------MGITTAFFLNQALIPYYLKPVHLETQTIKHCS---------------
>Micropterus_salmoides
--------MGITTAFFLNQALIPYYLKPVHLETRTIKRCS---------------
>Oreochromis_niloticus
-MPPSVVAMGINTASSVKQTLTLHYLKPVHLKTQSIKHCSGDCHCSEPGSASLLR
>Latimeria_chalumnae
--------MGYRPTGFRNQALLPYYLKPVHLVKVIFTNWAESYF-----------
>Esox_lucius
--------MGIKPF-FQNQALIPYYLKPVHFETQTIKNCRLKEHQQ---------
>Lutjanus_erythropterus
--------MGISTAFLLKPALIPYYLKPVHLETRTIKHCS---------------
>Trematomus_bernacchii
--------MGITTAFFLNQALIPYYLKPVHLETRTIKHCS---------------
>Leucoraja_ocellata
--------MGINAADIDDQSLLPYYLKPVDLVKAAARCLQRWSFLYQ--------
>Salmo_salar
--------MGIIPF-FQNQALIPYYLKPVHFETQTIKTAG---------------
>Lepisosteus_oculatus
MNSHSGIAMGIIPF-FQNRALIPYYLKPVHFDTMTLKDWRWNKYQY---------
>Squalus_acanthias
--------MGINAADIDDQSLLPYYLKPVDLVKVAARFLQWWNRWCQ--------
```

HG0190 ENSGALG00000006949 CHD2

>Gallus_gallus
MSFFPRTFLY
>Anser_cygnoides
MSFFPRTFLY
>Apteryx_australis_mantelli
MSFFPRTFLY
>Cariama_cristata
MSFFPRTFLY
>Cuculus_canorus
MSFFPRTFLY
>Monodelphis_domestica
MSFFPRTFLY
>Aquila_chrysaetos_canadensis
MSFFPRTFLY
>Gavia_stellata
MSFFPRTFLY
>Chlamydotis_macqueenii
MSFFPRTFLY
>Manacus_vitellinus
MSFFPRTFLY
>Egretta_garzetta
MSFFPRTFLY
>Pygoscelis_adeliae
MSFFPRTFLY
>Struthio_camelus_australis
MSFFPRTFLY

HG0191 ENSGALG00000036097

>Gallus_gallus
MNKRKNPFWTVFLNQASPSISRTF
>Anas_platyrhynchos
MNKQKNPFWTVFLNEASPSISGTY
>Apteryx_australis_mantelli
MNKQKKTFWTVFLNQASPSILGTY
>Calidris_pugnax
MNKQKNPFWTVFLNQASPSVSGTY
>Alligator_mississippiensis
MDRLKKKFWTVFINQASPSLLETY
>Monodelphis_domestica
MDKLKNSFWIVFPNETSPSSFSAQ
>Aquila_chrysaetos_canadensis
MSKQKNPFWTVFLNQASPSVLGTY
>Ornithorhynchus_anatinus
MDKLKKPLWTVFLPEASLSLLGCH
>Lepidothrix_coronata
MNKRKTPIWTVFLNQASPSVSGTY
>Anolis_carolinensis
MDRIKKPFWTVFLNQFFPSPMEAC
>Struthio_camelus_australis
MNKQKKTFWTVFLNQASPSILGTY
>Pelodiscus_sinensis
MDILKKTFWIVFLSQASPSFLGTY
>Tinamus_guttatus
MNKQKKTFWTVFLNQASPSILGTY

HG0192 ENSG00000125107 CNOT1

>Homo_sapiens
MQKLPRSCQAISQLSIQKETYHSKDLRSCMPGNGMKTKCAMFQNRVRPLVKKN
>Felis_catus
----------------------------------------MFQNRVIPLVKKN
>Lipotes_vexillifer
----------------------------------------MFQNRVRPLVKN-
>Myotis_brandtii
----------------------------------------MFQNRVRPLVKKN
>Dasypus_novemcinctus
----------------------------------------MFQNRVRLLVKKN
>Sarcophilus_harrisii
----------------------------------------MFQNRIRSLVKKN
>Monodelphis_domestica
----------------------------------------MFQNRIRSLVKKN
>Condylura_cristata
----------------------------------------MFQNRVRPLVKKN
>Ceratotherium_simum_simum
----------------------------------------MFQNRLRPLVKKN
>Loxodonta_africana
----------------------------------------MFQNRVRPLVKKN
>Ictidomys_tridecemlineatus
----------------------------------------MFQNRVRPLVKKN
>Trichechus_manatus_latirostr
----------------------------------------MFQNRVRPLVKKN
>Pelodiscus_sinensis
----------------------------------------MFQNRIRPLGKKS

HG0193 ENSG00000134853 PDGFRA

>Homo_sapiens
MISFRRMDYNIESITKRGF--------
>Anser_cygnoides
MISFGRMDYNIESITGRGL--------
>Ursus_maritimus
MISFRRMDYNIESITKRGF--------
>Ambystoma_mexicanum
MIRLGRMGYNIESITRRAFLGRLQSGN
>Lipotes_vexillifer
MISFRRMDYNIESITKRGF--------
>Myotis_ricketti
MISFRRMDYNIESITKRGF--------
>Dasypus_novemcinctus
MISFRRMDYNIESITKRGF--------
>Monodelphis_domestica
MISF-RIDFNIESITKRGV--------
>Oryctolagus_cuniculus
MISFRRMDYNIESITKRGF--------
>Lepidothrix_coronata
MISFGRMDYNIESITERGL--------
>Equus_asinus
MISFRRMDYNIESITKSGF--------
>Fukomys_damarensis
MISFRRMDYNIESITKSGFCAHYS---
>Pelodiscus_sinensis
MISFGRMDSNIESITRLGL--------

HG0194 ENSG00000143507 DUSP10

```
>Homo_sapiens
--------MCESIAEEERQMVEEYTYL---
>Ailuropoda_melanoleuca
--------MGESLAEAERQMVEGYTYL---
>Balaenoptera_acutorostrata_s
--------MCESTAEAERQMVEGYTYL---
>Sarcophilus_harrisii
MKLSEWGLDAPSIATVKRQMVEGYTYL---
>Monodelphis_domestica
MKLSEWGFNVQSIATVKRQMVEGYTYL---
>Macropus_eugenii
--------MYKSIATVKRQMVEGYTYL---
>Sorex_araneus
------------------MVEGYTY----
>Elephantulus_edwardii
------------------MVKGYYFIKLF
>Equus_asinus
--------MRESIAEAERQMVEGYTYL---
>Marmota_marmota_marmota
--------MCQSIAEAERQMVEGYTYL---
>Tupaia_chinensis
--------MYETVAQEERQMVEGSTYL---
>Trichechus_manatus_latirostr
--------MCESIAEEERQMVEGYTHL---
>Chrysemys_picta
------------------MLKGNTYL---
```

HG0195 ENSG00000165966 PDZRN4

>Homo_sapiens
MKNSSQVWRVIPFQFTCLSVLGKLEDLLSLSAS
>Odobenus_rosmarus_divergens
MKNSSQVWRVIPFQFTCLSVLGKLEDLLSLSAS
>Lipotes_vexillifer
MKNSSQVWRVIPFQFTCLSVLGKLEDLLSLSAC
>Pteropus_alecto
MKNSSQVWRVIPFQFTCLSVLGKLEDLLSLSAS
>Latimeria_chalumnae
MKNWSQEWREIPCQCHYLSVLGKLESFALVC--
>Gallus_gallus
MKNSSQVWRVIPCQCICLSVLGK----------
>Erinaceus_europaeus
MKNSSQVWRVIPFQFTCLSVLGELEDLLSLSAA
>Oryctolagus_cuniculus
MKNSSRDWRVIHFQLPCRSVLGTLEDQLPRSAS
>Elephantulus_edwardii
MKNSSQVWRVIPFQFTCLSVLGKLEDLLSLSAS
>Manis_javanica
MKTGGQVWRVLPFRLACLPVPGKLEDSRARSAP
>Loxodonta_africana
MKNSSQVWRVIPFQFTCLSVLGKLEDLLSLSAS
>Ictidomys_tridecemlineatus
MKNSSQVWRVIPFQFTCLSVLGKLEDLLSLSAS
>Trichechus_manatus_latirostr
MKNSSQVWRVIPFQFTCLSVLGKLEDLLSLSAS

HG0196 ENSG00000189403 HMGB1

```
>Homo_sapiens
MLQSGESEE-AASGSRSHSHCSTLSSIETAPGQVRAGRALGDSVPR----
>Megophrys_sangzhiensis
MLQIGERVKRLRIDPASHSHCSTLSSIDTALETS----------------
>Odobenus_rosmarus_divergens
MLQSGESEE-AASGSRSHSHCSTLSSIETAPGQVRARRALGDSVPR----
>Pteropus_vampyrus
MLQSGESEE-AASGSRSHSHCSTLSSIETAPGQVRAGRALGDSVLAEEK-
>Dasypus_novemcinctus
MLQSGESEE-AASGSRSHSHCSTLSSIETAPGKCEPGGRWATRSLADEK-
>Galeopterus_variegatus
MLQSGESEE-AASGSRSHSHCSTLSSIETAPGQVRAGRALGDSVPR----
>Macropus_eugenii
MLQNGESEE-AASGSRSHSHCSTLSSIETAPGQVRAGRALGYSVRR----
>Condylura_cristata
MLQSGESEE-AASGSRSHSHCSTLSSIETAPGQVRAGRALGDSVPR----
>Ochotona_princeps
MLQSGESEE-AASGSRSHSHCSTLSSIETAPGQARAGRALGDSVPRGGK-
>Equus_przewalskii
MLQSGESEE-AASGSRSHSHCSTLSSIETAPGQVRAGRALGDSVPR----
>Manis_javanica
MLQSGESEE-AASGSRSHSHCSTLSSIETAPGQVRGRRAPGDSVPR----
>Ictidomys_tridecemlineatus
MLQSGESEE-AASGSRSHSHCSTLSSIETAPGQVRAGRALGDSVPR----
>Trichechus_manatus_latirostr
MLQSGESEE-AASGSRSHSHCSTLSSIETAPGQAEPDGTGRLCASRTKNY
```

HG0197 ENSG00000196132 MYT1

>Homo_sapiens
MIAFRDVILKEDRQ
>Odobenus_rosmarus_divergens
MIAFRDVILKEDRQ
>Balaenoptera_acutorostrata_s
MIAFRDVILKEDRQ
>Myotis_davidii
MIAFRDVILKEDRQ
>Galeopterus_variegatus
MIAFRDVILKEDRQ
>Monodelphis_domestica
MIAFRDVILKGERQ
>Coturnix_japonica
MIGFRDVILQRGRQ
>Oryctolagus_cuniculus
MIAFRDVILKEDRQ
>Equus_asinus
MIAFRDVILKEDRQ
>Manis_javanica
MIAFRDVILKEDRQ
>Chinchilla_lanigera
MIAFRDVILKEDRQ
>Tupaia_chinensis
MIAFRDVILKEDRQ
>Pelodiscus_sinensis
MIAFRDVMLKRARQ

HG0198.1 ENSDARG00000018688 elk3

```
>Danio_rerio
------------------------MCVCVRIRVCAR-VPECRMVARGGKACLHRRAERLGNRARREEVSRLAVCLLQLPAHRERERKSE-----------------------------
>Calidris_pugnax
---------------------MLQPC-------C---ASVARARGEGRKSLFTQTANRLGNNA-GEEVSRLAV---------------------------------------------
>Latimeria_menadoensis
-------------------MCVLYYC-------N---VEHQWAGREGRKSLFTQTAKCLGNNA-GKEVSRLSV---------------------------------------------
>Fundulus_heteroclitus
MCGGFEANEASLKASQRSRRCEFKDCRSFRQHVCRQRVGPLLAAEEGRKSVFTQTASCLGNRR-GEEVSRLCVCLLQLPALRKKKPRIRFQRSTTVQTRSQHGKTGERTGFDVKSFAE
>Monodelphis_domestica
------------------------MCVCVCVCVCVC-VCVC----------------------VVTVTILLCVCLILQPC--------------------------------------
>Coturnix_japonica
---------------------MLQPC-------R---TSVARARGEGRKSLFTQTANRLGNNA-GKEVSRLSV---------------------------------------------
>Ornithorhynchus_anatinus
---------------------MLQPC-------C---TSAARERGAGRKSLFTQTANRLGNNA-VKEVSRLSV---------------------------------------------
>Manacus_vitellinus
---------------------MLQPC-------C---TSTARARGAGRKRLFTQTANRPGNSA-GEEVSRAAV---------------------------------------------
>Nannospalax_galili
------------------------MRASEGSRGT---QPRPRRTARGGKACLHRLHTAWGIMQ-------------------------------------------------------
>Lepisosteus_oculatus
-------------------MCVIIER-------C---WTSVGRGRRGEESLFTQTANRLGNNA-GKEVSRLSV---------------------------------------------
>Ictalurus_punctatus
----------------MGTDSGYSITVCVRVCVG---PRWAGREERGGKACLHRRPSAWGTE--------------------------------------------------------
>Python_bivittatus
------------------------MCLMLQQCCL---HISGQRGEKGGKACLHRLQTAWGIMQERK----------------------------------------------------
```

HG0198.2 ENSG00000111145 ELK3

>Homo_sapiens
MESRGLSPGSS
>Mustela_putorius_furo
MESRGLSPGSS
>Lipotes_vexillifer
MESRGLSPGSS
>Rousettus_aegyptiacus
MESRGLSPGSS
>Dasypus_novemcinctus
MESRGVSPGSS
>Galeopterus_variegatus
MESRGLSPGSS
>Ochotona_princeps
MESRGLSPGSS
>Elephantulus_edwardii
MESRGVSPGSS
>Ceratotherium_simum_simum
MESRGLSPGSS
>Manis_javanica
MESRGLSPGSS
>Microtus_ochrogaster
MESRGLSPGSS
>Tupaia_chinensis
MESRGLSPGSS
>Trichechus_manatus_latirostr
MESRGVSPGSS

HG0199 ENSGALG00000007989 ADNP

>Gallus_gallus
MLRQKGNEQDMTLLLCPGFGLTQ
>Anas_platyrhynchos
----------MMLLLCPGFGLTQ
>Apteryx_australis_mantelli
MLRQKGKEQDMILLLCPGFGLTQ
>Calidris_pugnax
MLRQKGKEQDMILLLCPGFGLTQ
>Columba_livia
-----MLNQDMILLLCPGFGLTQ
>Alligator_mississippiensis
----------MSLLLCPGFGLTQ
>Aquila_chrysaetos_canadensis
MLRQKGKEQDMILLLCPGFGLTQ
>Manacus_vitellinus
MLRQKGKEQDMILLLCPGWGLPQ
>Aptenodytes_forsteri
MLRQKGKEQDMILLLCPGFGLTQ
>Struthio_camelus_australis
----------MILLLCPGFGLTQ
>Chrysemys_picta
----------MSLLLCPGFGLTQ
>Tinamus_guttatus
----------MILLLCPGFGLTQ

HG0200 ENSGALG00000012015 NDST4

>Gallus_gallus
MHCNGHMNVSGEESAPLRLIVIMAL
>Anser_cygnoides
MYCNGHMNISGGESAPLHLLIIMAL
>Apteryx_australis_mantelli
MHCNGHMNNSVGESAPLRLLVTMAL
>Mustela_putorius_furo
MVCNGYMNLWGVIFAPSHTLFTMAL
>Hynobius_retardatus
MHCNGHMNCWGAAPAQSCTLFTMSL
>Calidris_pugnax
MHCNGHMNVLGGESAPPHLLIPMAL
>Lepidothrix_coronata
MLCNGHMNVSEGESAPLQLLIPMAL
>Nannospalax_galili
MVCNGYMNLWEVESAPSHALATMAL
>Thamnophis_sirtalis
MNCNGHMNIWRVVFIPSRGLFMMAL
>Struthio_camelus_australis
MYCNGHMNNSVGESAPLRLLLTMAL
>Chrysemys_picta
MHCNGHMNILGVVPAPFFVLSTMAL
>Tinamus_guttatus
MHCNGHMNSSGPIST--HLLVTMAL

HG0201 ENSGALG00000015256 PAM

```
>Gallus_gallus
--------------------MVCQKLEDACRAHNNVVMN
>Anser_cygnoides
--------------------MVCQKLEDACRAHNNEVVN
>Calidris_pugnax
--------------------MVCQKLEDARRAHNNEVVN
>Columba_livia
--------------------MVCQKLEDACRAHNNEVVN
>Alligator_mississippiensis
--------------------MVCQKL-------------
>Sarcophilus_harrisii
MIRDNIASSTFLIYRASKTDGLSSAVGDACRAHDNEVVN
>Falco_cherrug
--------------------MVCQKLEDACGAHNNEVVN
>Pseudopodoces_humilis
--------------------MVCQKLEDACRAHNNEVVK
>Aptenodytes_forsteri
--------------------MVCQKLEDAHRAHNNEAVN
>Takydromus_sexlineatus
--------------------MVCQKL-------------
>Struthio_camelus_australis
--------------------MVCQKL-------------
>Trachemys_scripta
--------------------MVCQKL-------------
```

HG0202 ENSG00000124766 SOX4

>Homo_sapiens
----------------------------------------------------------------------------MRSDLELLIARGDYYCIGFQANGKPGGGVWHEEALVTAADWLQPRL-----------------------------------------
>Channa_punctata
----------------------------------------------------------------------------MRSDTRLLIARADYY------------------------------------------------------------------------
>Oryzias_melastigma
----------------------------------------------------------------------------MRSDTRLLIAHADYYCVSFSTNE----------------------------------------------------------------
>Mustela_putorius_furo
----------------------------------------------------------------------------MRSDLELLIARGDYYCIGFQANGKAGGGAWHEEALVTAADWPRPRL-----------------------------------------
>Ambystoma_mexicanum
MLLAGSHPCTVCTPPLPPLHPGSGYTAHSPNPLEHIRTPSQINGSLPAPPKRFTLNPKISASPRVPFFEKFALWLGMRSALEPFIARGDYYCIGCRANGGGRGRDRACGNVGY-SD-----------------------------------------------
>Pteropus_vampyrus
----------------------------------------------------------------------------MRSDLELLIARGDYYCIGFQANGKPRGGAWHEEALVTAADWPWPRL-----------------------------------------
>Pimephales_promelas
----------------------------------------------------------------------------MRSDPKLLIARADYY------------------------------------------------------------------------
>Austrofundulus_limnaeus
----------------------------------------------------------------------------MRSDTRLLIARADYY------------------------------------------------------------------------
>Scleropages_formosus
----------------------------------------------------------------------------MRSDITLLIARAHDY------------------------------------------------------------------------
>Ceratotherium_simum_simum
----------------------------------------------------------------------------MRSDMELLIARGDYYCIGFQANGKPGGGAWHEEALVTAADWPRPRL-----------------------------------------
>Cavia_porcellus
----------------------------------------------------------------------------MRSDLELLIARGDYYCIGFQANRKGGGGAWHEEALVTAADWLRPKDKEARGGIGVCSKLQQEKQCVRERGLSHSQVSSSCTLFKSLH
>Chrysemys_picta
----------------------------------------------------------------------------MRSDLELLIARGDYYCIGFQANSSRRGGVWHEEALDTASDWPGRGL-----------------------------------------

HG0203 ENSG00000128606 LRRC17

>Homo_sapiens
------------MSL----WHP-FPKFSLGTPRTPAGSHCSRTALVKITQTWISKEYFHCSVCNTK
>Xenopus_laevis
------------MSL----WHP-FPEPRARIPSTPAAHTEYL----IITG----------------
>Ursus_maritimus
------------MSL----WHP-FPKFRD-SHRTSAGSHHS-------------------------
>Physeter_catodon
------------MRLEREAETR-RPRVSKGLHRTSAGSHCS-------------------------
>Rousettus_aegyptiacus
------------MSL----WHP-FPKFRDSTTPQLDLQALHLH-----------------------
>Dasypus_novemcinctus
------------MSL----WHP-FPKFSDS--TDPEGSQCSLTAQVTIAQTWISKKYIHCFICHIK
>Condylura_cristata
MVSA-----GMAPCEACEGWIRGRPHSLVWTTCTSAGSHCS-------------------------
>Oryctolagus_cuniculus
------------MSL----WHP-FPKLRTSTEPSWVPLFL----------NYVS------------
>Equus_caballus
------------MSL----WHP-FPKFRE-LHGTSAGSHCSSTALVKLTHTRISKEYFHCSICHTK
>Manis_javanica
MLPRLIGQDQFLLITWPQSWLR--KIPEMGLHCTSAVSHCS-------------------------
>Marmota_monax
------------MSL----WHP-FPKFRNSESPSWVSLFL----------NCI-------------
>Tupaia_chinensis
------------MSL----WHP-FPKFRDS--TCWVALFL----------SCIS------------

HG0204 ENSG00000162599 NFIA

>Homo_sapiens
MPLTPAYSRHV
>Felis_catus
MPLTPAYSRHV
>Myotis_lucifugus
MPLTPAYSRHV
>Galeopterus_variegatus
MPLTPAYSRHV
>Condylura_cristata
MPLTPAYSRHV
>Ochotona_princeps
MPLTPAYSRHV
>Elephantulus_edwardii
MPLTPAYSRHV
>Loxodonta_africana
MPLTPAYSRHV
>Nannospalax_galili
MPLTPAYSRHV
>Thamnophis_sirtalis
MPLTPGYSRHV
>Chrysemys_picta
MPLTPGYSRHV
>Orycteropus_afer_afer
MPLTPAYSRHV

HG0205 ENSG00000184611 KCNH7

>Homo_sapiens
MGREFWFQKRMLCHKERLARWPGL
>Leptonychotes_weddellii
MGRDFWFQKRMLCHKERLARWPGL
>Physeter_catodon
MGRDFWFQKRMLCHKERLARWPGL
>Rousettus_aegyptiacus
MGRDFWFQKRMLCYKERLARWPGL
>Monodelphis_domestica
----------MICHKEKLAQWSGL
>Erinaceus_europaeus
MGRDFWFQKKMLCHKERLARWPGL
>Ochotona_princeps
MGRAFWFQMRMLCQPERLARWPGL
>Equus_caballus
MGRDFWFQKRMLCHKERLARWPGL
>Loxodonta_africana
MGRDFWFPKRMLCHKERLARWPGL
>Marmota_marmota_marmota
MGRDFWFQKRMLCHKERLARWPGL
>Trichechus_manatus_latirostr
MGRDFWFQKRMLCHKERLARWPGL
>Pelodiscus_sinensis
MGRDAVIQRRMICHKG--------

HG0206.1 ENSGALG00000002069

```
>Gallus_gallus
MVGAIPRQPRRFWGVPGWQVSSASCANDRPLWDGEIDPEWSGILLQREPHTGGLTWTS
>Anser_cygnoides
MVGAIPRQPRRFWGVPGWQVSSASCANDSPRRDGEVDPEHGGVLLQRDAALQHIP---
>Calidris_pugnax
MVGAIPRQPRRFWGVPRWQVSSANCANDRPLRDKEVNPEQSRVLLQRDAALQCTP---
>Columba_livia
MVGAIPRQPRRFWGVPDWQVSSASCANDRPLWDRETDPEQSRVLLQRDDALQSTL---
>Alligator_mississippiensis
MVGAIPRQPRRFWGVPGWQVSPASCANDRVLRGHQGYLARRGGRLRSDAVLQRAP---
>Falco_cherrug
MVGAIPRQPRRFWGVPGWQVSSASCANDRLLRDREVNPEQSGVLLQRDAVLHRTL---
>Manacus_vitellinus
MVGAIPRQPRRFWGVPDWQVSSAGCANDRLLWDREVNPEISRVLLQRDAVLQTTS---
>Pygoscelis_adeliae
MVGAIPRQPRRFWGVPGWQVSSASCANGRPLRDPEVNPEESGVLLQRDAALQCTP---
>Boiga_irregularis
MVGAIPRQPRRFWGVPGWQVSPASFTSHRRLLQQEKTPARGRDPVQWDVAVQPTS---
>Struthio_camelus_australis
MVGAIPRQPRRFWGVPGWQVSSASCANDYPLQDGEPNLEPCGVLLQRDAALQRTP---
>Chrysemys_picta
MVGAIPRQPRRFWGVPGWQVSPASCANDRLLLGQESCSARGG----------------
```

HG0206.2 ENSG00000005238 FAM214B

>Homo_sapiens
--------------------------------MGLVLLSSGAARNGGSHLASAAGMDQTAS
>Canis_lupus_familiaris
-------------------------------------------------------MDQTAS
>Lipotes_vexillifer
-------------------------------------------------------MDQTAS
>Myotis_davidii
-------------------------------------------------------MDQTTS
>Sarcophilus_harrisii
-------------------------------------------------------MGQTTS
>Galeopterus_variegatus
-------------------------------------------------------MDQTAS
>Monodelphis_domestica
MEPPAREGED-------TFSTKPYSPRGVLSSHMESHLSQGAARDGGSDRASGVGMGQTTS
>Condylura_cristata
-------------------------------------------------------MDQTAS
>Ochotona_princeps
-------------------------------------------------------MDQTAS
>Elephantulus_edwardii
-------------------------------------------------------MDQTAN
>Ceratotherium_simum_simum
MEGVVWGGRGGRGGRWGCWGAGRGGRPHQLVGNGLVQLSPGATRNGGSHLASAAGMDQTAS
>Manis_javanica
-------------------------------------------------------MDQTAS
>Marmota_marmota_marmota
-------------------------------------------------------MDQTAS
>Tupaia_chinensis
-------------------------------------------------------MDQTAS

HG0207.1 ENSGALG00000007121 TNFSF8

```
>Gallus_gallus
-----------------------------------MSQDDGENPARNRDGLRPEPDQGEIQTSTLTAI
>Anser_cygnoides
--------------------------------------MKGENPARNRDGLQPEPDQGEIQSSTLRAI
>Canis_lupus_familiaris
--------------------------------------MQGENPSWNRDGLRTESDEER---------
>Pteropus_vampyrus
MKNEPLPHPCCCLSPHAPREEFLRKAAREGFFPSHMIQDAGENPSWNRDGPRTESDEER---------
>Sarcophilus_harrisii
--------------------------------------MEGENPTWNRDGL-----------------
>Monodelphis_domestica
--------------------------------------MKGENPAWNRDGL-----------------
>Trichosurus_vulpecula
--------------------------------------MKGENPAWNRDGL-----------------
>Parus_major
--------------------------------------MKGENPARSRDGLQLEPDRGEPQSSAWRAI
>Ceratotherium_simum_simum
-----------------------------------MIQDAGENPSWNRDGPRTESDEER---------
>Pan_paniscus
MKNEPLPHPCCRLSPHAPREEFLRQAAGEGFFPSHMIQDSGENPSWNRDGPRTESDEER---------
>Chrysemys_picta
--------------------------------------MKGENPARSGDGLRMRI-------------
```

HG0207.2 ENSG00000106952 TNFSF8

>Homo_sapiens
----------MGPELNQMKRDKV------
>Ailuropoda_melanoleuca
----------MGPEVNQMRRDKVCCGKAI
>Physeter_catodon
----------MGPEPNQMRRDKV------
>Rousettus_aegyptiacus
MQGRILLGTEMGPEPNQMRRDKV------
>Dasypus_novemcinctus
MQGENPSWNRDGPEPTQMRRDKV------
>Galeopterus_variegatus
----------MGPELNQMKTDKV------
>Sorex_araneus
----------MGPEPNQMRRDKV------
>Oryctolagus_cuniculus
----------MSPEPSQMKRDKV------
>Ceratotherium_simum_simum
MQGRILLGTEMGPEPNQMRRDKV------
>Manis_javanica
----------MGPEPNQMRRDKV------
>Loxodonta_africana
----------MGPEPTQMRRDKV------
>Marmota_marmota_marmota
----------MGPEPDQMRRDKV------

HG0208 ENSGALG00000013272 PRMT8

>Gallus_gallus
MPSWSLHAQVQFSWCFSPV
>Anser_cygnoides
MPSWSLHAQVRFSWCFSPV
>Chaetura_pelagica
MPSWSLHAQVQFSWCFSPI
>Charadrius_vociferus
MPGWSLHAQVQFSWCFSPV
>Aquila_chrysaetos_canadensis
MPSWSLHAQVQFSWCFSPV
>Mesitornis_unicolor
MPSWSLHAQVQFSWCSSPV
>Zonotrichia_albicollis
MPSWSLHAQVQFSWCFSPV
>Nipponia_nippon
MPSWSLHAQVQISWCFSPV
>Struthio_camelus_australis
MPSWSLYAQVQFSSCFSPI
>Chrysemys_picta
MPRWSLHAQVQFSWGFFAF
>Calypte_anna
MPSWSLYAQVQFSWCFSPV

HG0209 ENSG00000101040 ZMYND8

```
>Homo_sapiens
MHPQSPKSASLFSLPNCGKMRRLLEHLLAWLKRK-----------
>Canis_lupus_familiaris
-------------------MRRLLEHLLAWPKRK-----------
>Rousettus_aegyptiacus
-------------------MRRLLEHVG-----------------
>Dasypus_novemcinctus
MHPQSPKSASLFSLPNCGKMRRLLEHLLAWLKRK-----------
>Oryctolagus_cuniculus
-------------------MRRLLEHLLAWLKRK-----------
>Elephantulus_edwardii
MHPQSPHSASLFSLPNSGKMRRLLEHLVAWLKRK-----------
>Ceratotherium_simum_simum
-------------------MRRLLEHLLAWLKRK-----------
>Loxodonta_africana
-------------------MRRLLEHLLAWLKRK-----------
>Chinchilla_lanigera
MHPQSPKSASLFSLPNCGKMRRLLEHWLAWLKRK-----------
>Trichechus_manatus_latirostr
MHPQSPKSASLFSLPNCGKMRRLLEHLLAAFLPKVFSFSRLLGAK
>Chrysemys_picta
-------------------MRWLLEQLLG----------------
```

HG0210 ENSG00000155849 ELMO1

```
>Homo_sapiens
--------------MPCRCKMCLWPAGGRKRK
>Felis_catus
--------------MPCRCKMCLWPAGGRKRK
>Balaenoptera_acutorostrata_s
--------------MPCRCKMCLWPAGGRKRK
>Rhinolophus_ferrumequinum
--------------MPRSCKTCLWPAGGRKRK
>Dasypus_novemcinctus
--------------MPCRCKMCLWPAGGRKRK
>Condylura_cristata
--------------------MCLWPAGGRKRK
>Oryctolagus_cuniculus
--------------MPCRCKMCLWPAGGRKRK
>Equus_asinus
--------------MPCRCKMCLWPAGGRKRK
>Manis_javanica
--------------------MCLWPAGGRKRK
>Nannospalax_galili
--------------MPCRCKMSLRPAGGRKRK
>Anolis_carolinensis
MAEARLLSSVPRREPPCRCKSRLQPAGGRKRK
```

HG0211 ENSG00000176204 LRRTM4

>Homo_sapiens
MDKPSLHSF
>Mustela_putorius_furo
MDKPSLHSF
>Rousettus_aegyptiacus
MDKPSLHSF
>Condylura_cristata
MDKPSLHSF
>Oryctolagus_cuniculus
MDKPSLHSF
>Lepidothrix_coronata
MDKPSLHSF
>Manis_javanica
MDKPSLHSF
>Loxodonta_africana
MDKPSLHSF
>Heterocephalus_glaber
MDKPSLHSF
>Tupaia_chinensis
MDKPSLHSF
>Trichechus_manatus_latirostr
MDKPSLHSF

HG0212 ENSDARG00000100244 ebf3a

>Danio_rerio
MLPSILQRRAKGPSSSSPVSGNTSTVIIYAFICLLEPEVVDSAVSRSLF---------
>Hynobius_retardatus
------------------------MVIIFSFICLLEQESLARSARFRSVLKLF-----
>Alosa_alosa
MLPSILQ-DAKGPSSTSPVSGNTSMVIIFSFICLLEPERLARSGRFSSVLKLF-----
>Latimeria_chalumnae
------------------------MVIIFSFICLLEQESLARSGRFSSVLKLF-----
>Esox_lucius
MLLSILQ-DAKGPSSTSPVSGNTSMVIIFFVYLSCGAGAPCQKWSIQQCPEAFLML---
>Apteronotus_leptorhynchus
MLPSILQ-DAKGPSSTSPVSGNTSMVIIFSFICLLEPERLARSGRFSSVLKLF-----
>Lepisosteus_oculatus
MLPSILQ-DAKGPSSTSPVSGNTSMVIIFSFICLLEQERLARSGRFSSVLKVFDAVVG
>Anolis_carolinensis
------------------------MVIIFSFICLLEQSRPALPEVVDSAVS-------
>Pelodiscus_sinensis
------------------------MVIIFSFICLLEQESLARSGRFSSVLKLF-----
>Tetronarce_californica
------------------------MVIIFRLFVFWSRRTLSE--VVDSAVS-------

HG0213.1 ENSGALG00000001946 RSPO1

>Gallus_gallus
MGLDPSSCAPYGDEGRTMLPAT
>Anser_cygnoides
MGLDPSSCAPYGDEGRTMLPAT
>Xenopus_laevis
MGLDPNSCGPYKDESRTMLPVT
>Apteryx_australis_mantelli
MGLDPRSCAPYGVDGRIMLPAT
>Hynobius_retardatus
MGLHLSTFDPYGDENRAMLPAT
>Calidris_pugnax
MGLEPSPCAPYGDEGRAMLPAT
>Lepidothrix_coronata
MGPRPSSCAPHRDEGRAMLPAT
>Struthio_camelus_australis
MGLDPHSCAPYGVDGRIMLPAT
>Pelodiscus_sinensis
MGLDPSSSAPYGVEGRLMLPAT

HG0213.2 ENSGALG00000001946 RSPO1

```
>Gallus_gallus
--------------------------MGMESTALRPGFINVQIALI
>Anser_cygnoides
--------------------------MGMESTALRPGFINVQIALI
>Calidris_pugnax
--------------------------MGMESTALRPGFINVQIALI
>Lepidothrix_coronata
MQEFALDAFTKLDTLVIIIIKSPDPVDGDGVYGLRPGFINVQIALI
>Struthio_camelus_australis
--------------------------MGMESTALKPGFINVQIALI
>Chelonia_mydas
--------------------------MGMESAALWAGVINVPIALI
```

HG0214.1 ENSGALG00000012917 CDH6

>Gallus_gallus
METNQIESPLGSGCLHAGVKKCCRSSGSPSERRGG
>Anas_platyrhynchos
METNQIESPFGFSSLAA------------------
>Ambystoma_mexicanum
METNQIESPLGFACLPASVKSDVGRASAS------
>Calidris_pugnax
METNQTESPFGFSSLPA------------------
>Alligator_sinensis
METNQIESPLGFSSLHARVKSDAGREGAPLRCQ--
>Monodelphis_domestica
METNQIESPFGFSFPACLS----------------
>Acanthis_flammea
METNQTESPFGSCSLPA------------------
>Struthio_camelus_australis
METNQIESPFGCSSLPA------------------
>Chrysemys_picta
METNQIESPLGFGSLHARVKSDAGRERAPLRCE--

HG0214.2 ENSG00000040731 CDH10

>Homo_sapiens
-----------------------------------------------------MNKTLMKCWKPVETQ--
>Mustela_putorius_furo
-----------------------------------------------------MSKTLMRCWKPVETQ--
>Miniopterus_natalensis
MLRESWSCIEVEAGGENNAGNTQKRDVVTLSPGLSGLAPTSFIPRLRKKAFGNMSKTLMKCWKPVETQ--
>Galeopterus_variegatus
-----------------------------------------------------MYKTLMRCWKPVEMQ--
>Monodelphis_domestica
---------------------------------------MLLPSPEGESKIGNMNKILINCWKPIGYPTY
>Oryctolagus_cuniculus
-----------------------------------------------------MNKPSMKCWKPVETQ--
>Equus_caballus
-----------------------------------------------------MSEALMKCWKPVETQ--
>Cavia_porcellus
-----------------------------------------------------MNKALMKCWKPAETQ--
>Tupaia_chinensis
-----------------------------------------------------MYKTLRNCWKPVETQ--

HG0215 ENSDARG00000079280 dhrsx

>Danio_rerio
MQSLISPVTKAILVALFIFAILLILYVILWYICRDVDCDHGI
>Channa_argus
MQSLISPVTKAILVALFIFAILLILYVILWYICRDVDCDHGI
>Pygocentrus_nattereri
MQSLISPVTKAVLVALFIFAILLILYVILWYICRDVDCDHGI
>Maylandia_zebra
MQSFISPVTKAILVALFIFAILLILYVILWYICRDVDCDHGI
>Poecilia_latipinna
MQSLISPVTKAILVALFIFAILLILYVILWYICRDVDCDHGI
>Esox_lucius
MQSFLSPFNKAILVALFIFAILLILYVILWYICRDVDCDHGI
>Platichthys_stellatus
MQSIISPVTKAILVALFIFAILLILYVILWYICRDVDCDHGI

HG0215 ENSGALG00000016681 DHRSX

>Gallus_gallus
----MQSLISPVTKAILVALFIFAILLILYVILWYICRDVDCDHGI
>Astyanax_mexicanus
----MQSLISPVTKAVLVALFIFAILLILYVILWYICRDVDCDHGI
>Maylandia_zebra
----MQSFISPVTKAILVALFIFAILLILYVILWYICRDVDCDHGI
>Sinocyclocheilus_rhinocerous
----MQSLISPVTKAILVALFIFAILLILYVILWYICRDVDCDHGI
>Austrofundulus_limnaeus
----MQSLISPVTKAILVALFIFAILLILYVILWYICRDVDCDHGI
>Esox_lucius
MSSVMQSFISPVTKAILVALFIFAILLILYVILWYICRDVDCDHGI
>Zonotrichia_albicollis
----MQSLISPVTKAILVALFIFAILLILYVILWYICRDVDCDHGI
>Platichthys_stellatus
----MQSIISPVTKAILVALFIFAILLILYVILWYICRDVDCDHGI
>Salmo_salar
MSSVMQSFLSPVNKAILVALFIFAILLILYVILWYICRDVDCDHGI

HG0216.1 ENSGALG00000020922

>Gallus_gallus
-------------------------------------MICGSVQLELSE
>Anas_platyrhynchos
-------------------------------------MICGSVQLELSE
>Calidris_pugnax
-------------------------------------MICGWVQLELSE
>Columba_livia
-------------------------------------MICGWVQLELSE
>Alligator_mississippiensis
-------------------------------------MICGSVQLELTE
>Zonotrichia_albicollis
MPGAEILGWFCLTKVEGTREPRREVDSSCQDERASLEMICGSVQLELSE
>Aptenodytes_forsteri
-------------------------------------MICGSVQLELSE
>Takydromus_sexlineatus
-------------------------------------MVCGSVKLELTK
>Chrysemys_picta
-------------------------------------MICGSVQLELTE

HG0216.2 ENSG00000116667 C1orf21

```
>Homo_sapiens
------------------------------MICSSARLKLTE
>Canis_lupus_familiaris
------------------------------MICSSARLKLTE
>Orcinus_orca
------------------------------MLCSSARLKLTE
>Eptesicus_fuscus
------------------------------MICSSARLKLSE
>Dasypus_novemcinctus
------------------------------MICCSARLKLTE
>Sarcophilus_harrisii
MSFYLMFSLILVDFFFLFFKKQKYVPVSVEMICSSARLELTE
>Monodelphis_domestica
------------------------------MICSSARLELTE
>Macropus_eugenii
------------------------------MICSSPRLELTE
>Condylura_cristata
------------------------------MICSSARLKLTE
>Elephantulus_edwardii
------------------------------MICSSARLKLTE
>Ceratotherium_simum_simum
------------------------------MICSSARLKLTE
>Manis_javanica
------------------------------MICSSARLKLTE
>Loxodonta_africana
------------------------------MICSSARLKLTE
>Ictidomys_tridecemlineatus
------------------------------MICSSARLKLTE
>Tupaia_chinensis
------------------------------MICSSARLKLTE
>Trichechus_manatus_latirostr
------------------------------MICSSARLKLTE
```

HG0217.1 ENSG00000150394 CDH8

>Homo_sapiens
-------------------------------------------------------------------------------MNFQVVDLPLRSAQ
>Odobenus_rosmarus_divergens
-------------------------------------------------------------------------------MNFQVVDLPLRSAQ
>Orcinus_orca
-------------------------------------------------------------------------------MNFQVVDLPLRSAQ
>Galeopterus_variegatus
MFLSSASALMLELGPPRAARTARKGASSPGGSDGGGALGVLGSEERKVVDPLGRGKGSHRASATAKGEGGRGRGGGTAYMNFQVIDLPLRSAQ
>Gallus_gallus
-------------------------------------------------------------------------------MNFQALDLPPRSAQ
>Loxodonta_africana
-------------------------------------------------------------------------------MNFQVVDLPLRSAQ
>Ctenomys_sociabilis
-------------------------------------------------------------------------------MNFQVVDLPLRSAQ
>Trichechus_manatus_latirostr
-------------------------------------------------------------------------------MNFQVVDLPLRSAQ
>Orycteropus_afer_afer
-------------------------------------------------------------------------------MNFQVVDLPLRSAQ

HG0217.2 ENSGALG00000005319 CDH8

>Gallus_gallus
MRRKSGESCLFTGTGEWLFYLTLSIRV
>Chaetura_pelagica
MKRKSGGSCLFTGTG------------
>Apteryx_australis_mantelli
MKRKSGESCLFTGTG------------
>Buceros_rhinoceros_silvestri
MKRKSGKACLFTGTG------------
>Calidris_pugnax
MRRKSGESCLFTGTG------------
>Pterocles_gutturalis
MTRRSGESCLFAGTG------------
>Columba_livia
MKRKSAKSCLFPGTG------------
>Falco_cherrug
MKRKSGESCLFTGTG------------
>Gavia_stellata
MKRQSGESCLFTGTG------------
>Chlamydotis_macqueenii
MKRKSGESCLFAGTG------------
>Sturnus_vulgaris
MRRKSGAPCLFPATG------------
>Egretta_garzetta
MKRKSGESCLFTGTG------------
>Picoides_pubescens
MKRKSGKSCLFTGTG------------
>Nestor_notabilis
MKRKPGESCVFTGTG------------
>Pygoscelis_adeliae
MKRKSGESCLFAGSG------------
>Struthio_camelus_australis
MKRKSGESCLFTGTG------------
>Tinamus_guttatus
MKRKAGESCLFTGTG------------
>Calypte_anna
MKRKSGESCLFRGTG------------

HG0218 ENSDARG00000100149 axin2

>Danio_rerio
--------------------------------------MDERSSPADLRERGDYS--
>Channa_argus
---------------------------------MRGDGQPKRSSPADIRE-------
>Xenopus_laevis
MQL-------QKWGALAESSGCTEW-----STLHTHEGRGRRGSPADLRGRADYRGI
>Haplochromis_burtoni
---------------------------------MREAGQPERSPPADIRE-------
>Cyprinodon_variegatus
---------------------------------MREDGQPERSPPADIRE-------
>Gadus_morhua
MRIRALDTPTVDYGMFLCGYQTGVFFEELTHGSMRENGQTERSSPADIRE-------
>Lutjanus_erythropterus
---------------------------------MREDGQPERSSPADIRE-------
>Notothenia_coriiceps
---------------------------------MREDGQPERSSPADIRE-------
>Cynoglossus_semilaevis
---------------------------------MGEDGQPERSSPADI---------

HG0219 ENSDARG00000104082 trps1

>Danio_rerio
MAHAKAPPSFRLQRLLQFN-QQ----------------------
>Pygocentrus_nattereri
---------------MQFN-QQVLL-------------------
>Clupea_harengus
MAYAKAPPSFRLQRLLQFN-QQ----------------------
>Sarcophilus_harrisii
MALATAPPSFGLLRLLLPINQQIYRSALKMSSVSSVHLLRQLH-
>Monodelphis_domestica
MALATAPPSFGLLRLLLPINQQIYRSALKMSSVSSVHLLRQLH-
>Esox_lucius
MAYAKAPPSFRLQRLLQFN-QQRAVAGR--KTS-----------
>Apteronotus_leptorhynchus
MVHAKAPPSFRLQRQLQFN-QQPGLAV-----------------
>Salmo_salar
MAYAKAPPSFRLQRLLQFN-QQREKHCD--SCCGGTQDCYYL--
>Trichomycterus_areolatus
---------------MLFN-QQAGLSSDCAKQCDGPWTLIQNHR

HG0220 ENSGALG00000008418 LARGE2

>Gallus_gallus
-------------MLGPAGLGLPT
>Anser_cygnoides
-------------MLGPAGLGLPT
>Calidris_pugnax
-------------MLGPVGLGLPT
>Aquila_chrysaetos_canadensis
MPSGGVAPVPGSRMLGPAGLGLPT
>Acanthis_flammea
-------------MPGPAGLGLPT
>Aptenodytes_forsteri
-------------MLGPAGLGLPT
>Struthio_camelus_australis
-------------MLGPAGLGLPT
>Pelodiscus_sinensis
-------------MMGPAGLDLPT
>Tinamus_guttatus
-------------MLGPAGLGLPT

HG0221 ENSGALG00000010859 EPGN

>Gallus_gallus
------------------------------------------------------------MPTGNRKLGFQEKTRSQC
>Canis_lupus_familiaris
MFDSYPGMLLPGLQISELSFVGLLGRWLATLELVQVCPGNGAFPEFRFFFVVLSTFTYTIMPTGNRKPVFQKKT----
>Calidris_pugnax
------------------------------------------------------------MPTGNRKPGFQKKI----
>Galeopterus_variegatus
----------------------------------------------------------MIMPTGNRKPVFPKKT----
>Erinaceus_europaeus
----------------------------------------------------------MIMPTGNRKPGFQKKA----
>Oryctolagus_cuniculus
------------------------------------------------------------MPTGNRKPVFQKKT----
>Lepidothrix_coronata
------------------------------------------------------------MPTGNRKPGFQKKT----
>Homo_sapiens
----------------------------------------------------------MIMPTGNRKPVFQKKT----
>Struthio_camelus_australis
------------------------------------------------------------MPTGNRKLEFQKKS----

HG0222 ENSGALG00000011564 RUFY3

>Gallus_gallus
MQKPGFLWLLRINSL---
>Anser_cygnoides
MRKSGFLWLLRINSLQHI
>Alligator_sinensis
MQRARFLWMLRIHCL---
>Aquila_chrysaetos_canadensis
MQKPGFLWLL--------
>Corvus_brachyrhynchos
--------MLRISSL---
>Pygoscelis_adeliae
MQKPGFLWSLRINSL---
>Struthio_camelus_australis
MQKPGFL-----------
>Pelodiscus_sinensis
MQWPGFL-----------
>Calypte_anna
MQKPGFLWWLRIGSLQHS

HG0223 ENSG00000091656 ZFHX4

>Homo_sapiens
--------------------------------------------MDGRVRERLRLPLPFRVLLRLRLRLGCDWDATVTRASIASIESNAKTQPGSLTGWMK
>Anas_platyrhynchos
--------------------------------------------MDGRGSEKPL--QPCLLLVRLRQRLGCDWDATVTWASIASIESNAKTQPGSLTGWMK
>Monodelphis_domestica
--------------------------------------------MDGRGRERLQMPLPFRLLVRLRQRLGCDWDTTVTWASIASIESNAKTQPGSLTGWMK
>Coturnix_japonica
MVYFFFFFFFQLCFSWIFSGSILGSGAGVFKKIICSIIFLVWWWMDGRGSEKPL--QPCLLLVRLRQRLGCDWDATVTWASIASIESNAKTQPGSLTGWMK
>Oryctolagus_cuniculus
--------------------------------------------MDGRVRERLRLPLPFRVLLRLRLRLGCDWDATVTRASIASIESNAKTQPGSLTGWMK
>Ficedula_albicollis
--------------------------------------------MDGRGSEKPL--QPCLLLVRLRQRLGCDWDATVTWASIASIESNAKTQPGSLTGWMK
>Ceratotherium_simum_simum
--------------------------------------------MDGRVRERLRLPLPFLVLLRLRLRLGCDWDTTVTRASIASIESNAKTQPGSLTGWMK
>Rattus_norvegicus
---------------------------------------------MDGVRERLRLPLPFRLVLRLRLRLGCDWDATVTQASIASIESNAKTQPGSLTGWMK
>Tinamus_guttatus
--------------------------------------------MDGRGSEKPL--QPCLLLVRLRQRLGCDWDATVTWASIASIESNAKTQPGSLTGWMK

HG0224 ENSG00000118007 STAG1

>Homo_sapiens
MPPDGGLGLQLR
>Ailuropoda_melanoleuca
MPPDGGLGLQLR
>Lipotes_vexillifer
MPPDGGLGLQLR
>Rousettus_aegyptiacus
MPPDGGLGLQLR
>Gallus_gallus
MPPDGGLGLRLR
>Condylura_cristata
MPPDGGLGLQLR
>Ochotona_princeps
MPPDGGLGLQLR
>Equus_asinus
MPPDGGLGLQLR
>Mus_musculus
MPPDGGLGLQLR

HG0225 ENSG00000150051 MKX

```
>Homo_sapiens
MFAGWDRQSGNMRRPRYSCSRAARAGLGK-----------
>Amia_calva
MFAGRAA---------------------------------
>Mustela_putorius_furo
MFAGWAGQSRNMRRPRYSCSRAARARLGK-----------
>Pteropus_alecto
MFAGWARQSGNMRRPRYSCSRAARARLGK-----------
>Oreochromis_niloticus
MFAVSGCVLGNMSRVRYSESRAARRRPLDKAEEETAVRAV
>Poecilia_latipinna
MFAGSASVQGNMSRRRYSESRAAG----------------
>Manis_javanica
MFAGWAGQSGNMRRPQHSCSRAARAPPGK-----------
>Ictidomys_tridecemlineatus
MFAGWARQSGNMRRPRYSCSRAARARLGK-----------
>Salmo_salar
MFAGWGSVLGNMRRVTYSVSRAENRRPLNKAERRPRFVWF
```

HG0226.1 ENSG00000180592

>Homo_sapiens
MERFIYRLYYLLWEGRWQYKGMLISGRFWGEGVEY-QFLLHHLSGVSNQRWL--------------------
>Mustela_putorius_furo
MERFIYRLYYLLWEGRWQYKGMLISGRFWGEGVEY-QFLLHHLSGVSNQRWL--------------------
>Miniopterus_natalensis
MERFIYRLYYLLWEGRWQYKGMLISGRFWGEGVEY-QFLLHHLSGVSNQRWL--------------------
>Coturnix_japonica
---------------------MLISGRFLGGRGGISIFIASPVRSVQSALALGELIDEGVSGRAHFTLSFYL
>Ceratotherium_simum_simum
MERFIYRLYYLLWEGRWQYKGMLISGRFWGEGVEY-QFLLHHLSGVSNQRWL--------------------
>Manis_javanica
MERFIYRLYYLLWEGRWQYKGMLISGRFWGEGVEY-QFLLHHLSGVSNQRWL--------------------
>Cavia_porcellus
MERFIYRLYYLLWEGRWQYKGMLISGRFWGEGVEY-QFLLHHLSGVSNQRWL--------------------
>Salmo_salar
--------------------MYLTNGGGGAQGVEY-PFLLHHLSVASNRIRV--------------------

HG0226.2 ENSG00000180592

```
>Homo_sapiens
MAEQPIESADMVNKF-IKETPWL-------------LKNSTLYIWICTSGKAQKYPFILLITRMCLFLEEQGNT
>Mustela_putorius_furo
---------------------------------------------------------------MCLFLEEQGNI
>Manis_javanica
---------------------------------------------------------------MCLFLEEQGNM
>Chinchilla_lanigera
----------MSESFFFGRGEYLLVRPEQGKVELLNLKKEIIVHFFFL--EGSEISVYSLTTRMCLFLEEEGNI
```

HG0226.3 ENSG00000180592

```
>Homo_sapiens
----------------------------------MRLGGPLLSFGNILDILENDTKICLNLTKKRLNHIDFHPK
>Mustela_putorius_furo
MPGLHLCCSCKHRTVKKILPNFAKEPTSHLGILIMQLGRPLLSFGNILDILGNDTKICFNLTKKRLNNIDIHPK
>Chinchilla_lanigera
MPGLHLCCSCKHRTVKKVLPNSAKQPTKRLGTLVIQVGRLLLSFGNILDSLENDSKLCFNLTKKRLNHTDIHPK
```

HG0227.1 ENSGALG00000031929

>Gallus_gallus
-MTALNNWWHFGGYGWSSVY-
>Anser_cygnoides
-MTALNNWWHFGGYHWIPVY-
>Ambystoma_mexicanum
MMTTFNNSWQFGGFHWIRAY-
>Calidris_pugnax
-MTALNNWWQFGGFGWISVYS
>Aquila_chrysaetos_canadensis
-MTALNNRWRFGGFRWISVC-
>Lepidothrix_coronata
-MTALNNWWQFGGFRWISV--
>Struthio_camelus_australis
-MTALNNWWHFGRFHWIPVY-
>Chrysemys_picta
MMTAFNNWWQFGGFHWISVC-

HG0227.2 ENSG00000275163

>Homo_sapiens
--------------MGKVSEEE-PFLP--FVAISTGLQGTSFLLGFYVCFTHCLSFLCLRHFHE----
>Anser_cygnoides
--------------MEKVSNWECSLLPCLSVAISAGLLGVAFLLGFCVCFTYCFPLPFGQEPFYDQAR
>Coturnix_japonica
MRSLQQNCFVTDVSMEKVSNWECSLFPCLSVAISAGL-------------------------------
>Lepidothrix_coronata
--------------MEKVSNWECSLLPSLAVAISAGLQGVAFLLGFCVCFTYCFPLPFGQELFYDLAK

HG0228 ENSDARG00000087887 phf21b

```
>Danio_rerio
MSLTASKEARTCCAARVV-LAIVIVAFQNTSLTPQITRIASEVEEKCS
>Astyanax_mexicanus
MSLVASKEARTCSAA---------------------------------
>Alosa_alosa
MSPVPSKEARTCFAA---------------------------------
>Latimeria_chalumnae
MSLVPSKEARTCCAAGVVVLSIVIVGFF--------------------
>Apteronotus_leptorhynchus
MSLVPSKEARTCFAA---------------------------------
>Scleropages_formosus
MSPIGSKETRTCFGD---------------------------------
>Salmo_salar
MSPLHSKEARTCFAD---------------------------------
>Lepisosteus_oculatus
MSPIPSKEARTCFAD---------------------------------
```

HG0229 ENSGALG00000005474

>Gallus_gallus
MCIHFWWTLPFLGV---------------
>Apteryx_australis_mantelli
MCMHFGWTLPFLAV---------------
>Calidris_pugnax
MCIHFGWTLPLLCI---------------
>Haliaeetus_leucocephalus
MCIHFEWTLPFLGI---------------
>Lepidothrix_coronata
MCIRFGWTLPFLGI---------------
>Phalacrocorax_carbo
MCIHFGWTLPLLLI---------------
>Pygoscelis_adeliae
MCIHFGWTLPLLCI---------------
>Pelodiscus_sinensis
MCAHFRWTLPCLGFSDLVLNSPEADIY

HG0230 ENSGALG00000011007 OSBPL3

>Gallus_gallus
----------------------------------------------MQIMLLECIWVRDEFCITLRSLPIHLLDYLLMTF
>Anser_cygnoides
----------------------------------------------MQIMPLECI-------------------------
>Calidris_pugnax
MVAVDLPFLPSSSCWLWGAAAGGHGGGARWAVARPAAGERLRNVHGIQIVPLDCIWVKDESRVSLRSLPTHFLNYLLMVF
>Aquila_chrysaetos_canadensis
----------------------------------------------MQIMPLDCIWIKDEFCISFRSLPTHFLNCFLMVF
>Sturnus_vulgaris
-----------------------MGEECSIAVKRQQFSPPLGSFHGIQIMPLDCIWVKDEFSISLRSLPTHFPDYLLMVF
>Pygoscelis_adeliae
----------------------------------------------MQIMPLDCIWVKDEFCVSLRSLPTHFLNYLFMVF
>Anolis_carolinensis
-------------------------------------------------MLLECIWEHNDFCLPDPRLPVPLMVLQSD--
>Chrysemys_picta
-------------------------------------------------MPLECIWVNNDSCLSRPCLPIHFLNSLWMVF

HG0231 ENSG00000150636

```
>Homo_sapiens
MQEILLFTHVAATETL-EQPSKPRGS-----
>Anas_platyrhynchos
MQEILLFTRLCNTEKQ-EEPRKPRG------
>Canis_lupus_familiaris
MQEILLFTHVSVTLSR---------------
>Dasypus_novemcinctus
MQEILLFTHVSVTLRR---------------
>Monodelphis_domestica
MQEIQLFTHVSVTLRSKSSPASPEGALGRCL
>Pseudopodoces_humilis
MQEILLFTHVSVTLRSKRSPAN---------
>Struthio_camelus_australis
MQEILLFTHVFVTLRSKSSPASPEGVKLD--
>Pelodiscus_sinensis
MQE-ILFTHVSAALSSKNSPASPKGVKSH--
```

HG0232 ENSG00000180875 GREM2

>Homo_sapiens
-------------------------------MGVCCSLAAVELKAAAGDQT
>Alligator_mississippiensis
-------------------------------MGICSSLAKEELKAAAGYI-
>Monodelphis_domestica
-------------------------------MGICCSPAKAELNPAEGDQT
>Oryctolagus_cuniculus
-------------------------------MGVCWSPPAVELKAAAGDQT
>Equus_caballus
MQRREESELSPGLLPHLQGCQPGLHTRPALLMGVCWRPAAVQLSAAAGDQT
>Ictidomys_tridecemlineatus
-------------------------------MGFCWSLAAVELKAAAGDQT
>Tupaia_chinensis
-------------------------------MGIGWSPAAIELQAAAGDQT
>Pelodiscus_sinensis
-------------------------------MGICSNLVEAELKAAVGDHT

HG0233 ENSG00000206432 TMEM200C

>Homo_sapiens
MSGGEEGSEAGCGMRDAEDLTLLGNSWKWLPPQRRRGLR-AWGGRSPPRGC------------------------
>Mustela_putorius_furo
-------------MRAAEDLTLLGNSWKWLPPQPRRGLR-TWGGRSPPLGC------------------------
>Balaenoptera_acutorostrata_s
-------------MQDAEDLTLLGNSWKWLPPRHRRGLR-IPGGRSPPLGC------------------------
>Rousettus_aegyptiacus
-------------MQDAEDLTLLGNSWKWLPPQHRRGLR-AWGGRSPPRGC------------------------
>Monodelphis_domestica
-------------MKDGEGITLLGNSSPWLLPQAWRRGSENITGATSPAGSPGILSRGEMRDAGRLHLRATVHSG
>Oryctolagus_cuniculus
-------------MQDAADLTLLGNSWKWLPPLRRRGLR-ARGGRSPPRGC------------------------
>Equus_asinus
-------------MQDAEDLTLLGNSRKWLPPQHQRGLR-TRGGRSPPRGC------------------------
>Chrysemys_picta
-------------MKDGEGTTLLGNSLEWLCPGAPRGLTS--SGRSLPLSLLGL---------------------

HG0234.1 ENSGALG00000011250 ASIC4

>Gallus_gallus
MLTCLELLSIPWILLAHDA
>Calidris_pugnax
MLTCLELLSIPWILLAHDA
>Columba_livia
MLTCLELLSIPWILLAHDA
>Aquila_chrysaetos_canadensis
MLTCLELLSIPWILLAHDA
>Lepidothrix_coronata
MLTCLELLSIPWILLAHDA
>Thamnophis_sirtalis
MLTCLELLSIPWILLAHDA
>Tinamus_guttatus
MLTCLELLSIPWILLAHDA

HG0234.2 ENSG00000108684 ASIC2

```
>Homo_sapiens
---MVNMGHYALVNRLRAAQRLVLSIFSLALA-FNLVSSGVCRGEDFISIG
>Mustela_putorius_furo
MPGMLHAGHYARVNRLRAAERLV----------------------------
>Monodelphis_domestica
---MLNTGHYALVNRLGAAERLVLSILSCLLLILSLVVCVCVQGKDFISIG
>Condylura_cristata
---MLNTGHYALVNRLRAAERLVLSIFSLALA-FNLVSSGVCRGEDFISIG
>Rattus_norvegicus
---MLNTGHYALVNRLRAAQRLVLSIFSLALA-FNLVSSGVCRGEDFISIG
```

HG0235.1 ENSGALG00000035927

>Gallus_gallus
MLATSESLGNFTVGALRAVLEL-----TAKEARRGQTMGCPVLGLLHEQTPRAPLGCPAKRA
>Anser_cygnoides
MFPTSESPGNFTVEASRCAQSWSSHPKGARKGQQGRTMGYVVLGLLREKPANVPLGFPAKRA
>Calidris_pugnax
-------------------------------------MGYLVLGLLRDKPASVPLRFPAKRA
>Aquila_chrysaetos_canadensis
MLPTSESPGNFTMEASNCAQSWS---------------------------------------
>Sturnus_vulgaris
MLPTSESPGNFTMEASHCSQSWS---------------------------------------
>Struthio_camelus_australis
MLPTSESPGNFKLEVSHCAQSWS---------------------------------------
>Pelodiscus_sinensis
MFPTSDAPGNFTVETSDCAQFFS---------------------------------------

HG0235.2 ENSG00000101638 ST8SIA5

>Homo_sapiens
MPPAPAAPTRNFAPRSPPPA--------------------
>Mustela_putorius_furo
MPPAPSPPGGNFAPPSPPPG--------------------
>Myotis_lucifugus
---MPATPAGNFAPRSPPPPDCLVGLVPP-PPPPPTGPRA
>Ceratotherium_simum_simum
MLPAPAGPATNFARRSPPPD--------------------
>Ictidomys_tridecemlineatus
MPPARAAPAGNFALRSPPPAGPRCETPVRPRAAGVSEPRA

HG0236 ENSGALG00000004437 LHFPL2

>Gallus_gallus
MVAAGAGQGEPGGARAGGGGGSGAEPQRTGGAGERALLAGGIARLLKWLTKAKNRGHWCVMSSSGQSPRPSPPVRSVMCQYEQ
>Anas_platyrhynchos
------------------------------------------------------------MSNSGQSLQPALPARSVMCQYEQ
>Calidris_pugnax
------------------------------------------------------------MSNSGQSLRPPPPACSVMCQYEY
>Alligator_mississippiensis
------------------------------------------------------------MSRSGQSLGPPPHYCNVMCQYEY
>Aquila_chrysaetos_canadensis
------------------------------------------------------------MSKSGQSLRPPPPACSIMCQYEY
>Manacus_vitellinus
------------------------------------MKNTSIARHLKWLTKPKDLEQWCVMSNSAQSFQPPPPACSVICQYEY
>Struthio_camelus_australis
-----------------------------------------------------------------------------MCQYEH

HG0237 ENSGALG00000012123 PAX6

```
>Gallus_gallus
---------MCGVISGF-EKR-APARPHFPLWLRRREEVFCWKMMTEVRLC--
>Cyprinodon_variegatus
MEKTTPYFLSTQFFLSFSSSL-LLPWLCFPLWLWHRGEVFIWGMMTEVGSR--
>Esox_lucius
---------MCGLISVFWRPVSPLSPHRFPLWLWRREKSLRGADDDRGRLMGK
>Callithrix_jacchus
-------------------------------------------MMTEVRLR--
>Salmo_salar
---------MCRVISRF-EKL-ALALPRFPLCTAE------------------
>Lepisosteus_oculatus
-------------------------------------------MMTEVRLC--
>Chrysemys_picta
---------MCGVISGF-EKR-ALAQPHFPLWLRRREEVFCWKMMTEVRLC--
```

HG0238 ENSG00000049618 ARID1B

```
>Homo_sapiens
------------------------------------------MLGPIGSASRVFTHIRAR-EEK-R---KVSFKPELKTFTMKAHSRSRPRA--
>Channa_argus
--------------------------------MPCWATRAP------GDAAAYYREPGWRRGGGG-KEKKVSFKPGRETFNMKLH---------
>Latimeria_chalumnae
MKKLWSMSLQGLHASTPFHIFQALCSNKREVDNSCWSSLSHAMLGQRGSGRHVFTHIRVR-EEKKK---KVSFKPEQKTFNMKSHSRSRHRHRQ
>Kryptolebias_marmoratus
----------------------------------------------------------MEKRRRG-ERKKVSFKPGRETFNMKPH---------
>Esox_lucius
--------------------------------------------------------MHLPISGLE-ERKKVSFKPGRQTSNMKPH---------
>Parus_major
------------------------------------------MLGQRGSGRRVFTHIRVR-EERRRKKKKFHLNLNKKTFNMKSHSGNGHGP--
>Lepisosteus_oculatus
--------------------------------------------------------MHLPISGLERKGRKVSFKTWTKTFNMKSHRGSRQSQ--
```

HG0239 ENSG00000149256 TENM4

>Homo_sapiens
MEPGRLKLPCVQIKIAYFKDYLKDYS
>Mustela_putorius_furo
MEPGRLKLPCVQIKIAYLKNYSKDYS
>Physeter_catodon
MEPGRLKLPCVQIKIAYFKDYLKDYS
>Pteropus_alecto
MEPGRLKLPCVQIKIAYFKDYLKDYS
>Condylura_cristata
MEPGRLKLPCVQIKIAYFKDYLKDYS
>Chinchilla_lanigera
MEPGRLKLPCVQIKIAYFKDYLKDYS
>Anolis_carolinensis
MESGKLQSAYDQVKTAYFKAHLMDYS

HG0240 ENSG00000166342 NETO1

>Homo_sapiens
MTDPFFLPSMPQRKIPKGWSEERCRWTSSRGGCSDLLRGASETGD
>Calidris_pugnax
MTDPLIPSSMHQ-KKPKGWKSLGS---ESCRKCFHLLLVLYETGD
>Myotis_lucifugus
MTDPFLLPPMPQRKIPQGWTEEGCRSSSRRARLPQPAAR---GG-
>Falco_peregrinus
MTDPLIPSSMHQ-KKPKGWKSLGS---ESCRKCFDLLLVLYETGD
>Pseudopodoces_humilis
MTDPLIPSSMHQ-KKPKGWKSLGS---ESCRKCFDLLLVLYETGD
>Mesocricetus_auratus
MTDPCLLPSMPQQKTPKGWREKSCGWTSSRRRCSDLVLLVDETGD
>Chrysemys_picta
MTDPLILPSMHQ-KNPKVWETLGL---KSYRKCFDRLLVLYETGD

HG0241 ENSG00000198561 CTNND1

>Homo_sapiens
------------------------------------------------------------------ME-----VSGWLCSLLPASSLLWWLGCFFHDFLNLDWAVLCVKPISCDLLLTV-----------------
>Balaenoptera_acutorostrata_s
-------------------------------------------------------------MSAP-LK-----VSQICSSLLPASSLLWWLGCFSHDILNLDWAVLC-----------------------------
>Coturnix_japonica
---------MGAVTQP-----------TACPRAEAV-RSEG-RTGGRPTAAAGPSRRYPACRSHPKYELTACLFALCGESSLPPASSLWWLGYFSHDSLRLDLAVPCLCQSHPSFLL-------------------
>Oryctolagus_cuniculus
-----------------------------------------------------------------------------------------------MIVLNLDWAVLCVKSISYDILTVV-----------------
>Lepidothrix_coronata
---------MS-----------------------A---------VGSPGLAAQP------------APMGQRLAPKSPSALFPPASSLWWLGCFSHDSLRLDLAVPVPLPIKPELPPLTVKRGAPAPSGSLFLPSP
>Ictidomys_tridecemlineatus
-----------------------------------------------------------------------------------------------MIFLNLDWAVLCVKTINYDPSLNSVK---------------
>Python_bivittatus
MIRKDVAEQLGPWHSPKEKVLFFLSVALFPPRPDSFLNLQGSEANLLPEAAAKGTVRFPERSARG----------TKAQAFFLLLPCCWWLGCFSHDFLHLDQAVP------------------------------

HG0242.1 ENSGALG00000007112 LINS1

```
>Gallus_gallus
---------MSRGNLEEECLL--ACLFLLLDKNNFKTDLQQSEIIGEDLQRSVIY
>Apteryx_australis_mantelli
-----------------------------MDKNNFETNLQ---------------
>Columba_livia
-----------------------MLLFLLLDKKNFEADLH---------------
>Haliaeetus_albicilla
----MVEEVRSLGNIEEKCLI--VLLFLLLDKNNFKADLQRP-------------
>Sturnus_vulgaris
-----MRSLHLPRNLEEKCPV--VLLFLLLDESNWEAALQ---------------
>Pelodiscus_sinensis
MSLSSRGFCIIRAILQKECLYRATNPFLLLDRSNFKSDP----------------
```

HG0242.2 ENSG00000140471 LINS1

>Homo_sapiens
-----MREEVPACTFCELRWPRAGICVPS---NGGVKRVRLSQ---------------IPGFTEETS
>Mustela_putorius_furo
-----MRRKRSACAFPELRWPGG-WCCSF---NGGVKAGSPNPDLRQKLQERRPACVIKQSFPEETS
>Manis_javanica
MWPLRMREEGPTCTFRELRLPGGGSCALEWRFEGRVRFAR---SLAGK-------------------

HG0243.1 ENSG00000163697 APBB2

>Homo_sapiens
---MGSLLSRQLIFQIITYFLHHC
>Physeter_catodon
------MALSQDTFQIITYFLHHC
>Equus_asinus
MVPRGSVSSRQLIFQVITYFLHLC
>Manis_javanica
---------MQLVFQIITYFLHHC
>Chinchilla_lanigera
MVPGGSLFSRQLILQIITYFRHHC
>Struthio_camelus_australis
----------MKRFQIIIYFLVHC

HG0243.2 ENSG00000163697 APBB2

>Homo_sapiens
MLWSDLPSRGTETFAGWKIHKRKLL
>Felis_catus
MLWSDLPSRGTETFAGWKIHKRKLL
>Balaenoptera_acutorostrata_s
MLWSALPSRGTETFAGWKIHKRKLL
>Rousettus_aegyptiacus
MLWSDLPSRGTETFAGWKIHKRKLL
>Dasypus_novemcinctus
MLWLDLPSRGTEIFAGWKIHKRKLL
>Galeopterus_variegatus
MLWSDLPSRGTETFAGWKIHKRKLL
>Condylura_cristata
MLGSDLPSCGTEAFAGWKIHKRKLL
>Ceratotherium_simum_simum
MLWSDLPSRGTETFAGWKIHKRKLL
>Manis_javanica
MLWSDLPSRGTETFAGWKIHKRKLL
>Loxodonta_africana
MLWSDLPSRGRETFAGWKIHKRKLL
>Fukomys_damarensis
MLWSDLPSRGTETFAGWKIHKRKLL
>Tupaia_chinensis
MLWSDLPSCCTETFAGWKIHKRKFL

HG0244 ENSGALG00000010837 ASB5

```
>Gallus_gallus
MLTGTLRSLKAVVMKQPKYQHFSRIDPSEQLMLMVVF-
>Xenopus_tropicalis
MLTGTIRSLNAVVIKHPKITFRKKPYSMSDI-------
>Calidris_pugnax
MRTGTLRSLKAVVMKQPKCWCFSRIDPSQQLMIRAVF-
>Columba_livia
-------------------------------MIRVVF-
>Tauraco_erythrolophus
------------------------------MMIRVVF-
>Saimiri_boliviensis
-------------MKQPKDQLFRSTTDESQAHLRLGSN
```

HG0245 ENSG00000091831 ESR1

>Homo_sapiens
----------------------------------------------------------------MQPHTPFYLPYLGYSVV--LPRVILCTHYVFLANEEGESNRKRDKQRYIGVWHGAHKAAH-------------------
>Mustela_putorius_furo
MRKL---RAG-----------TILLLAKHSSSYRYTLTPPPPFSPDVSSQSPQLLNIAVCRWQHGQPHSPFYLPYLSYRVA--LPRVILCTHYILPANEESESNRKRDKQR--------------------------------
>Pteropus_alecto
-----------------------------------------------------------------------------------------MYTLHGPANEKSASSSERHKQR--------------------------------
>Monodelphis_domestica
-----------------------------MSSC---LF-PSLQRRLLSSPAVAKYSLAVCGWLHRQPHTQFYLPYLGYRID--LPRVIFCTHYIFLENMKSESNRGKR-----------------------------------
>Rattus_norvegicus
MAVLEGTPWGVFFFFFPGEETEGFLDSKCSSSYTRSL--SARLALDASYLRPLMLNIAVCGWLHMQSYTPSYLPYLSYRYSYCPSRVILCTHHAVPAN-KTESNRKGDKQRQIGAWHRYTGNTSGKKERKETSLWLSLHLLQT
>Thamnophis_sirtalis
--------------------------------------------------------------------------------------------------MKMNKTRSLDKRR--------------------------------

HG0246 ENSG00000162676 GFI1

>Homo_sapiens
MSKPLGLGVKVSSGLG-L--------------------SKP---SE---------------------------------------------------------
>Mustela_putorius_furo
MSKPLGLGVKVFSALLGVFIPKCFHNKLFRGSGAGFALRTP---RPLAGLQAGGTT--R-R------------------------------------------
>Rousettus_aegyptiacus
MSKPLGLGVKVSA------------------------------------------------------------------------------------------
>Ochotona_princeps
MSKPLGLGVKISAGPG-L--------------------RKL---WP---------------------------------------------------------
>Parus_major
MSKPLGLGLKVFSAPLGVFIPEEFSTTSYF-------------------------------------------------------------------------
>Rattus_norvegicus
MSKPLGLGVKVFSASLGVFILKCFHNKLFRGPEAQVSLCGRGGQWPGSSPAAPGFGDSRLRSGSDACPSRRPGCVAHGQRVQELCELPARSARQLPSICVLNL

HG0247.1 ENSDARG00000014746 rbfox1

>Danio_rerio
MLALGRCCRDWD----
>Apteryx_australis_mantelli
MLALGRCCMAWD----
>Calidris_pugnax
MLALGRCCMAWD----
>Corvus_brachyrhynchos
MLALGRCCMAWDQSSC
>Chrysemys_picta
MLALGRCCMAWD----

HG0247.2 ENSG00000100320 RBFOX2

>Homo_sapiens
MGGSWRKKEGERLPERKRKERKKH
>Canis_lupus_familiaris
MGGSWREKEGERLPERKRKERKKR
>Orcinus_orca
MGGSWRKKEGERLPERKRKERKKH
>Myotis_brandtii
MGGSWRTKEGERLPERKRKERKKH
>Dasypus_novemcinctus
MGGSWRKKEGERLPERKRKERKKH
>Galeopterus_variegatus
MGGSWRK-EGERLPERKRKERKKH
>Macropus_eugenii
MGGSWRKKEGERLPERKRKERKKH
>Condylura_cristata
MGGSWRKKEGERLPERKRKERKKH
>Ceratotherium_simum_simum
MGGSWRRKEGERLPERKRKERKKH
>Manis_javanica
MGGSWRKKEGERLPERKRKERKKH
>Cricetulus_griseus
MGGSWRKKEGERLPERKRKERKKH
>Tupaia_chinensis
MGGSWRKKEGERLPERKRKERKKH

HG0248.1 ENSGALG00000004570 NCOA3

>Gallus_gallus
MVSDSGDPVKSILCLNAC
>Anser_cygnoides
MTSDSEGPMKNMLCLNVC
>Aquila_chrysaetos_canadensis
MTSDSEGPLKNMLCLNVC
>Manacus_vitellinus
MTSDSEGPLKNRLCLNVC
>Chrysemys_picta
MITGPDGPLKSLLCLNVC

HG0248.2 ENSG00000124151 NCOA3

```
>Homo_sapiens
---------MAAGGSKYLLDGGLRDQ
>Leptonychotes_weddellii
---------MAAGGSKYLLNSGVTDQ
>Orcinus_orca
---------MAAGGSKYLLNSGIRDQ
>Myotis_lucifugus
---------MAAGGSKYLLNSGIRDQ
>Galeopterus_variegatus
MASKSEHADISTAGSKYLMNSGLRDQ
>Condylura_cristata
---------MAAGGSKYLLNSGVRDQ
>Oryctolagus_cuniculus
---------MAAGGSKYLLNSEHRDQ
>Equus_asinus
---------MAAGGSKYLLNSGVRDQ
>Manis_javanica
---------MAAGGSEYLLNSGVRDQ
>Loxodonta_africana
---------MAAVGSKHLLNSGPRDQ
>Nannospalax_galili
---------MAAGGSKYLLNSGLRDN
```

HG0249 ENSDARG00000062550 rc3h1a

>Danio_rerio
--------MLCFSHYVLNAHNARKYLRTVIFGRALSVPVEPLGLTAVSFYAKRT
>Pygocentrus_nattereri
---------------MLRVHNA--------------------------------
>Apteronotus_leptorhynchus
MNRFR-HILLCIQHCVLNVHSA--------------------------------
>Scleropages_formosus
----------------MNVYNVRIY-----------------------------
>Gekko_japonicus
MNTWSSRDKGYISHYVLTVCTYRRYLKTIFFPKNLI-PVESVL-----------

HG0250 ENSGALG00000014861

>Gallus_gallus
-------------MNWKHLDLCRALNQILV-----------------
>Anser_cygnoides
-------------MNWKHLGLFRAVNQLFV-----------------
>Haliaeetus_leucocephalus
MKENYALKSQLFKMNWKHPDLFRVVNQLFV-----------------
>Manis_javanica
-------------MNWKHIDLCRRMPCECSPRRRSLAQKAAGRHSLG
>Aptenodytes_forsteri
-------------MNWKRLDLFRAVNQLFV-----------------

HG0251 ENSG00000185053 SGCZ

>Homo_sapiens
MMLTREAAFFLGMQQFAGF
>Mustela_putorius_furo
MMLSREAAFFLGMQQFAGF
>Rousettus_aegyptiacus
MMLSREAAFFLGMQQFAGF
>Chinchilla_lanigera
MMLSREAAFFLGMQQFAGF
>Chrysemys_picta
-MLSSEAVFFLGMQQFAGF

HG0252.1 ENSGALG00000009645 ESRRG

>Gallus_gallus
MSLHGVVDFPSKETGFGFFKIPDDSPD
>Alligator_mississippiensis
MSLHGVVDFPSQETGFGFLQIPDDSPD
>Sturnus_vulgaris
MSLHGVVDFPSKETGFGFFKIPDDSPD
>Microcebus_murinus
MFLHGGVL-------------------

HG0252.2 ENSG00000119715 ESRRB

>Homo_sapiens
---------------------------MVVRLFFFFFFFCQLPVISITESGSIYRLG--VLCP-----------------------------
>Myotis_davidii
------------------------------MVVGIFFFFCQLPVISITESDLFIGWEYCVL--SH--KETQTGDCQLVLEELSSESSATGST
>Corvus_brachyrhynchos
MPTGDCGGLLPGHSAELKAQLLLPCLRPGYFFFFFFFFFCQLPVISITELGSIYRLGLGVLCPSHKERQRHRGDCRLVQAEPKDI-------
>Mus_musculus
------------------------------MVVGFFFLFCQLPVISITESGSIYSLR--VLCP-----------------------------

HG0252.3 ENSG00000196482 ESRRG

>Homo_sapiens
MCNVPAWGFCRLS-----------------------
>Mustela_putorius_furo
MCNVPAWGFCRLS-----------------------
>Orcinus_orca
MCNVPAWGFCRLSQPKRPASASLKSPTTHLLNLLQF
>Monodelphis_domestica
MCKVPAWGFGGVGCRLS-------------------
>Manis_javanica
MCNVPAWGFCRLS-----------------------
>Mus_musculus
MCDVPAWGVCRLS-----------------------

HG0253.1 ENSGALG00000015184

```
>Gallus_gallus
MYQNVNDEALLNLVKGVTSAERAVGAARSGGAGRRAGQGRVGQGGPGRAGHGAPSSAAAASRSLDRQRRARKPNTNKAARTVPSSSSLSLFLIF
>Lipotes_vexillifer
MYQNVND---------------------------------------------------------------------------------------
>Pteropus_vampyrus
MYQNVND---------------------------------------------------------------------------------------
>Anolis_carolinensis
MYQNVNDPPLLNLVKGVTSAESVVRAVPSGGAVPGLPLLLLLPRG--RD---------------------------------------------
```

HG0253.2 ENSG00000140332 TLE3

>Homo_sapiens
MSAQSQALTRSTQPREGGRPLPAEP
>Odobenus_rosmarus_divergens
MSAQPRALTRSTQWGEGGRPQPAEP
>Orcinus_orca
MSAQPPALTRSTQRGEGGRPQPAEP
>Rousettus_aegyptiacus
MSAQSPALTCSTQPREGGRLQPAEP
>Oryctolagus_cuniculus
MSAQPPALTRSTQPREGGRPKPAEP
>Elephantulus_edwardii
MSSQSPALIRQTQPREGYRPQPADP
>Equus_caballus
MSAQPPALTRSTQPGEGGRPQPAEP
>Manis_javanica
MSARSQALTRTTQPGEGGRPQPTEP
>Jaculus_jaculus
MSAQPRALTLSTQPWEGGRPQPAEP
>Trichechus_manatus_latirostr
MSAQSPALTRSTQPGEGGRPQPAEP
>Orycteropus_afer_afer
MSAQSPALTRSTQPEEGGRLQLAEP

HG0253.3 ENSG00000140332 TLE3

>Homo_sapiens
MPGRLICTY
>Odobenus_rosmarus_divergens
MPGRLICTY
>Equus_caballus
MPGRLICTY
>Manis_javanica
MPGRLICTY

HG0254.1 ENSGALG00000035919

```
>Gallus_gallus
MLDGACGCQTQPEAAADGR-R--------DPSCAPPVPPRS--------------------------
>Aquila_chrysaetos_canadensis
MLDGACGCQTQPEAAAAAGYRDIPLRVSRGPFPAPPAPSASTFAATDWAWVVSVLRYMKHYCCGEVY
>Pseudopodoces_humilis
MLDGACGCQAQPAAAGHRDTGTPR---------PPRARSDEDFFSQI--------------C-----
>Trachemys_scripta
MLAGACGCQSQPEAAAAAGQPDPR-----APLPAPRRRRQQVMVSCVLHLFL---EERK--------
```

HG0254.2 ENSG00000072134 EPN2

>Homo_sapiens
MTIQGSYSSLITETEPS
>Galeopterus_variegatus
MTVQGSCSKSITETEPS
>Loxodonta_africana
MIVQGCSSRLITETDPS
>Nannospalax_galili
MTIQGSCSIWITETEPS
>Orycteropus_afer_afer
MIVQGSSSWLITETDPS

HG0255 ENSGALG00000012139 ARL14EP

```
>Gallus_gallus
-------------MSATGSAAVGGAARKPGAGVPAQRRRKVHGKGDLHRNH
>Alligator_mississippiensis
-------------MAASSTAAVVGGTRESGSGAAGQGRKKAYGKGDPHRNY
>Haliaeetus_leucocephalus
MVSRRVGGGEGRLMAATSSAAVAEGTRKSGAGVPAQRRKKVHGKGDLHRNY
>Trachemys_scripta
-------------MAATSTTALVGGTRESGSGVAAQGRKKAYGKGDPHRNY
```

HG0256 ENSGALG00000039901

```
>Gallus_gallus
MAATTTISLRHSQPCNRFPRDVPELLEFKPPKGPRSTQKQCP
>Calidris_pugnax
MAATTTISLRNSQPCNKFPTNVPELLEFKPP-----------
>Alligator_mississippiensis
MAATTTISLRNSHPCNTFPRMCLNSWSSNRLKGPTTAQTQYS
>Haliaeetus_leucocephalus
MAATTTISLRNSQPCNKFPTNVPELLEFKPP-----------
```

HG0257 ENSG00000198791 CNOT7

>Homo_sapiens
------------------------------------------MLSPVMRRPLVSLRGFWDCSREREGFEISLPKNQQE---LCL-----------------------------
>Physeter_catodon
------------------------------------------MLPPVMRGPLMPLRGFWNCSRERWRKAVFN-YRVFP---LCVCVFSEEGYKGRGPGSFHCVCVCVCVCVCV
>Galeopterus_variegatus
--------------------------MPLCCPLVSLGACSTEMLPPVMRRLWVSFRGFWNCSKERGGGKHC------------------------------------------
>Corvus_brachyrhynchos
MGQVRRDHSGSPGPTSLLEQGHPGAYSTRLCPDSSGMSSLRQTPKPFWAVRSRSLHGFWSCSSSCSGFPLKLSVHQFLPIISCPIVWHHQDEPGTRPLGIHTH----------

HG0258.1 ENSGALG00000016797 FHL2

>Gallus_gallus
MSNCLITKYIVWVIQ-------
>Columba_livia
MPDCLTTKHIVWAIQ-------
>Pelodiscus_sinensis
MPDCLITKYIVWVIQCSNKPLY

HG0258.2 ENSGALG00000016797 FHL2

>Gallus_gallus
MDVTLCPWSRR
>Chaetura_pelagica
MDVTPCPWSRR
>Apteryx_australis_mantelli
MDVTPCPWSRR
>Cariama_cristata
MDVTPCPWSRR
>Calidris_pugnax
MDVTPCPWSRR
>Colius_striatus
MDVTPCPWSRR
>Columba_livia
MDVTPCPWSRR
>Cuculus_canorus
MDVTPCPWSRR
>Mesitornis_unicolor
MDVTPCPWSRR
>Opisthocomus_hoazin
MDVTPCPWSRR
>Acanthisitta_chloris
MDVTPCPWSRR
>Nipponia_nippon
MDVTPCPWSRR
>Aptenodytes_forsteri
MDVTPCPWSRR
>Struthio_camelus_australis
MDVTPCPWSRR

HG0259 ENSGALG00000009936 AP4S1

>Gallus_gallus
MLKSKMAAPGKGAQFRP-------
>Gekko_japonicus
MLESKMAAPGKAVQPREPRGGCAE
>Pelodiscus_sinensis
-----MAAPGKGAQFGP-------

HG0260.1 ENSGALG00000016890 MBNL2

>Gallus_gallus
MLLSMIFFSQKSTSGT
>Anas_platyrhynchos
MLLSMVVFSQKSTSGT
>Apteryx_australis_mantelli
MLLSMVFFSQKSTSGT
>Buceros_rhinoceros_silvestri
MQLSMVFPSQKSTTGT
>Calidris_pugnax
MRLSMVFPSQKSTSGT
>Columba_livia
MRLSMVFPSQKSTSGT
>Merops_nubicus
MRLSMVFPSQKSTSGT
>Cuculus_canorus
MQLSMFFPSQKSTSGT
>Aquila_chrysaetos_canadensis
MRLSMFFPSQKSTSGT
>Mesitornis_unicolor
MRLSMVFPSQKSTSGT
>Tauraco_erythrolophus
MQLSMVFPSQKSASGT
>Opisthocomus_hoazin
MRLSMVFRSQKSTSGT
>Taeniopygia_guttata
MRLSMCFPSQKSTSGT
>Egretta_garzetta
MQLSMVFPSQKSTSGT
>Fulmarus_glacialis
MQLSMVFPSQKSTSGT
>Nestor_notabilis
MRLSMVFPSQKSTSGT
>Aptenodytes_forsteri
MRLSMVFPSQKSTSGT
>Struthio_camelus_australis
MLLSMVFFSRKSTSGT
>Tinamus_guttatus
MLLSMFVFSQKSTSGT
>Apaloderma_vittatum
MRLSMVFPSQKSTSGT

HG0260.2 ENSG00000139793 MBNL2

```
>Homo_sapiens
MSDCLSIHSLIILFLGLDFTKQFITHLQTYMWEFSQQ
>Mustela_putorius_furo
------------------------------MWGFSQQ
>Lipotes_vexillifer
MSDYLPINSLLTLFPGLGFFKQFVTYLQTYMWDFSQQ
>Pteropus_alecto
MSDCLPIII-SSLLLGLDFIKQFITYLQTYMWDFSQ-
>Dasypus_novemcinctus
MSDCLPINSLI-LFCGLDFIKQFIIYLKTYMWDFSQQ
>Galeopterus_variegatus
MSDCLPINSLITLLPGLDFIKRFITYLQTYMWEFSQQ
>Condylura_cristata
MSDRLLINSLLTLSPGLDFIKQFITYLQTYMWDFPPQ
>Oryctolagus_cuniculus
------------------------------MWEFSRQ
>Elephantulus_edwardii
------------------------------MWDLSQH
>Equus_asinus
MSDCPPINSLLTLFPGLDFIKQFITYLQTYMWDFSQ-
>Manis_javanica
MSDCLPINSLFTLLPRLNFIKTFIPYLQTYMWDFSQQ
>Loxodonta_africana
------------------------------MWDFSQQ
>Chinchilla_lanigera
MSDCLPIHSLSTLSPGLDFIKQLITYLQTYMWEFSQQ
>Tupaia_chinensis
MSDCPLINSLITLFPRLDFIKQFITYLQTYMWEFSQQ
>Trichechus_manatus_latirostr
------------------------------MWDCSQQ
```

HG0260.3 ENSG00000152601 MBNL1

>Homo_sapiens
MTPTMPWALGRQWGRL
>Odobenus_rosmarus_divergens
MTPTMPWALGRQSGCL
>Lipotes_vexillifer
MTPTMPWALARQSGCL
>Pteropus_alecto
----MPWALGRQSGCL
>Condylura_cristata
----MPWAPGRQSGCL
>Manis_javanica
----MPWALGRQSGRL
>Heterocephalus_glaber
MTPTMPWALGRQSGCL
>Tupaia_chinensis
MTPTMPWALGRLSGCL
>Orycteropus_afer_afer
MTPTMPWVLCLQSGCL

HG0260.4 ENSG00000152601 MBNL1

>Homo_sapiens
MSCGAHNAPMTRS
>Odobenus_rosmarus_divergens
MSCGAHNAPVTGN
>Lipotes_vexillifer
MSCGAHNAPVTGN
>Pteropus_alecto
MSCGAHNAPVTGN
>Dasypus_novemcinctus
MSCGAHNAPVTRN
>Condylura_cristata
MSSGAHNAPVTRN
>Tupaia_chinensis
MSCGAHNAAVTRN
>Orycteropus_afer_afer
MSCGAHNAPVTRN

HG0260.5 ENSG00000152601 MBNL1

```
>Homo_sapiens
MFFPPVSPPGSKTVWCILRRQSHQIAGF-------------------------
>Odobenus_rosmarus_divergens
MFFPPVSHPGSKTVWCILKWQPHQILTVFCRSDSHLQPNLAGFLTKSPIRRPL
>Pteropus_vampyrus
MFFPPASHPGSKTVWCICKEAAPSDLTVLS-----------------------
>Orycteropus_afer_afer
MFFPPVLPPGSKTMWCILRRQPHQIPGFYSNLI--LTPS--------------
```

HG0260.6 ENSG00000152601 MBNL1

```
>Homo_sapiens
--MGLRFICSGDNMAYLPWNIPVFQK----------------------------------------------------
>Odobenus_rosmarus_divergens
MIWGCGVFCSNNNMT---------------------------------------------------------------
>Orcinus_orca
-------------MTYLPWTITVFKKMNILC-----------------------------------------------
>Pteropus_vampyrus
-------------MTYLPWNSPVFQKMVFYFLKRGPLLSINHIMQSWGERGFYISNIIYKLDGKLVCECLCLCDNQFY
```

HG0261.1 ENSGALG00000005867 MARCH8

>Gallus_gallus
MSAAFKPPSCK-
>Anas_platyrhynchos
MSAAFKPPSCK-
>Buceros_rhinoceros_silvestri
MRAAFKPPSCKS
>Charadrius_vociferus
MSAAFKPPSCK-
>Columba_livia
MSAAFKPPSCK-
>Haliaeetus_leucocephalus
MSTAFKPPSCK-
>Eurypyga_helias
MSTAFKPPSCK-
>Corvus_cornix_cornix
MSTAFKPPSCK-
>Nipponia_nippon
MSAAFKPPSCK-
>Picoides_pubescens
MSTAFKPPSCKQ
>Pygoscelis_adeliae
MSAAFKPPSCK-
>Apaloderma_vittatum
MSIAFKPPSCK-

HG0261.2 ENSGALG00000009520 MARCH1

```
>Gallus_gallus
-----------------MEYLTQRQEMKPGSNVKGHVKLLHSRNLLKDCFT
>Anser_cygnoides
MGFRRCRYFHCRKRNMRLPQDENCMEMNPGSNVKGHVKLCHSRNLLKDCFA
>Calidris_pugnax
--------------------------MNPGRNVKGHVKLCHSRSLLKDCFT
>Zonotrichia_albicollis
--------------------------MNPGSNVKGHVKLCHSRNFLKDCFT
>Struthio_camelus_australis
--------------------------MNPRSRLKGHVKLCHSRNLLKDCFT
```

HG0261.3 ENSG00000145416 MARCH1

```
>Homo_sapiens
MPAPRDAGRTQPGGRLSYFTLRR-------------
>Pteropus_alecto
MPAPRDAGYAQPGGCLSDFALRR-------------
>Monodelphis_domestica
MPEPRDMGTVQTCGRLSDFTLRWKCTQG--------
>Nannospalax_galili
MPAPRDAGCAQPGGRLNHFILPRKCTQEEF------
>Tupaia_chinensis
MPASGAAGCAQPGGRLSHFTPRGFQQRLSESVKSTA
```

HG0261.4 ENSG00000165406 MARCH8

>Homo_sapiens
MLEAGQGGSAASSEPT
>Ailuropoda_melanoleuca
MLRAGWGGPAASSAPT
>Pteropus_vampyrus
MLQAGPGGLAASSAPT

HG0262.1 ENSG00000023445 BIRC3

```
>Homo_sapiens
MFLLPPGNKALKSAVEKPNVCEIQETMLANHW-----------------------
>Canis_lupus_familiaris
---MPPGNKALKVWSRGPSVGGPDMD-----PREGTHHDLFVGRC----------
>Balaenoptera_acutorostrata_s
MFLLPPGNKALKSAVESPNICGTQEACLQNTSRKKNHGDLFIGYY----------
>Dasypus_novemcinctus
MFLLPPGNKVLESAVENPNICASQQTMLAKSLVEKHA-VYWLSLVLLKYLFTNAR
>Oryctolagus_cuniculus
MFLPPPGHRVLKSAVRKPQHL----------------------------------
>Equus_asinus
MFLLPPGNK-LKSAVESANLCGTQETMLANHCLENQD-DLFVGHC----------
>Cavia_porcellus
MFLLLPGNQVLKSAPRLQILVEKAL------------------------------
```

HG0262.2 ENSG00000023445 BIRC3

>Homo_sapiens
-------------------MSLILNLICGNKGKVILAGAYC
>Ailuropoda_melanoleuca
MNRGRVTKSSHRSCNCKVTITLNLNVTEGHEGEVILAGTYC
>Balaenoptera_acutorostrata_s
---------------------------------MIVAGTYC
>Dasypus_novemcinctus
---------------------------------MTVAGAYC
>Cavia_porcellus
-------------------MILILNLICGSKE-VITMEAYC

```
HG0262.3 ENSGALG00000017186

>Gallus_gallus
MHSTRKAHNVTVAQEKVLFDIPVFVNG--VDQLLH------
>Haliaeetus_albicilla
-----------MIQGEVLFDMLSSIVRKGRGSFCTESSCVS
>Fulmarus_glacialis
-----------MIQGKVLFDMLSSIVKMGWGSFCTESSCVN
```

HG0262.4 ENSG00000023445 BIRC3

>Homo_sapiens
MNTKKIFITMCKIFGQGKEY
>Rousettus_aegyptiacus
---------MCKLFGQ----
>Equus_asinus
MNTKKVFITMCKVFEQ----

HG0263.1 ENSG00000169826 CSGALNACT2

>Homo_sapiens
MEILILFLIFDNFLLKV
>Leptonychotes_weddellii
MEILILFLIFDNLLLKV
>Physeter_catodon
MEILILFLLFYNLSQKV
>Pteropus_alecto
MEILVLFLIFDNLLLKV
>Equus_asinus
MEILILFLIFDNLLLKV
>Manis_javanica
MEILILFSVFDNLLLKV
>Microtus_ochrogaster
MEILIFFLIFDNFLLKL

HG0263.2 ENSGALG00000002543 CSGALNACT2

>Gallus_gallus
--------------------------------MANVGTCDLWLCNDT
>Calidris_pugnax
------------------------------MVMDDIGTSDLSLCNDT
>Sturnus_vulgaris
------------------------------MVMDDIGTSDLHLCNDT
>Struthio_camelus_australis
MSPFPFNGFLFSLGWDILNKSFGVNHAYKLMVMDDVGTSNLYLCNDT

HG0263.3 ENSGALG00000002543 CSGALNACT2

>Gallus_gallus
MIPGWCLPFCSPSAVFFVWDKKS
>Calidris_pugnax
MTPWWCLPSSSSSAGFFVWGRKS
>Corvus_brachyrhynchos
MIPCWCLPSSSPSAGFSCLG---
>Pygoscelis_adeliae
MTPWWCLPPSSPSAGFFVWGRKS

HG0263.4 ENSG00000147408 CSGALNACT1

>Homo_sapiens
MDASSSGGPKRMLPVV
>Balaenoptera_acutorostrata_s
MDESSSGTPGRVLPVI
>Cynopterus_sphinx
MDESSSGGPKRMLPVV
>Ceratotherium_simum_simum
MDESSSRGFRRMLPVV

HG0264.1 ENSG00000151458 ANKRD50

>Homo_sapiens
MELCDSLKFRKSVAELWNQAFLPQFNN--
>Acinonyx_jubatus
MELCDPSRFRKSIATLWNQAIFHQFNN--
>Lipotes_vexillifer
MEFCDPSRYRKSVGQGLEPGSFPSVQQFI
>Rousettus_aegyptiacus
MELCDPSRYRKTVAELWNRAIFYQFNP--
>Dasypus_novemcinctus
MALCDLPKHRKSVTKF-------------
>Sarcophilus_harrisii
MELYDLLSHMKSIIEFWDQTSFHQINN--
>Galeopterus_variegatus
MQLCDSSRCRKSVAELWNQAFFPSVQQLN
>Oryctolagus_cuniculus
MELCDLSSYWKGVTKLWS-----------
>Equus_asinus
MELCDPSRYRKRVAKLWNQATFHQFNN--
>Manis_javanica
MELCDPSRYRKSVAKFWNQAIFYQFNS--
>Loxodonta_africana
MELCDPSRHRKSVTEFWN-----------
>Fukomys_darlingi
MELYGPSRYSKSVAELWHQSCFHQFNN--
>Trichechus_manatus_latirostr
MELCDPSRHRKSVTELWN-----------
>Orycteropus_afer_afer
MKLCDPSKHKKSVTELWNKAFFPSVKQ--

HG0264.2 ENSG00000151458 ANKRD50

>Homo_sapiens
MYCLISVLETSTIT----------
>Mustela_putorius_furo
MYCLICTLETSTIT----------
>Orcinus_orca
MYCLICILETSTIT----------
>Myotis_lucifugus
MYCLICILEGSTIT----------
>Sarcophilus_harrisii
MYCLISILETSTISINSRCSHGVI
>Galeopterus_variegatus
MYCLIYILEISTIT----------
>Ceratotherium_simum_simum
MYCLICIFETSTVT----------
>Manis_javanica
MYCLIHILETSTVT----------
>Cavia_porcellus
MYCLIYVLETSTIT----------
>Tupaia_chinensis
MYCLICILETSTIT----------
>Orycteropus_afer_afer
MYCLICVLETFTIT----------

HG0264.3 ENSGALG00000011831 ANKRD50

>Gallus_gallus
MDSFFWIYIRRD
>Anser_cygnoides
MDSFFWIYIQRD
>Calidris_pugnax
MDSFFWMYIQRD
>Zonotrichia_albicollis
MDSFFCVYIQRD
>Aptenodytes_forsteri
MDSFFWMYIQRD

HG0264.4 ENSG00000151458 ANKRD50

>Homo_sapiens
MERPVLMYPDSSK---
>Physeter_catodon
------MYPDSSKC--
>Manis_javanica
------MYPDSSKC--
>Trichechus_manatus_latirostr
------MYPDSSKCRR

HG0265.1 ENSG00000144285 SCN1A

>Homo_sapiens
MELFLQINTSGVMEAGGS
>Mustela_putorius_furo
MELFLQINTSGTMEAGGS
>Rhinolophus_ferrumequinum
MELFLQINTSGAMEAGGS
>Condylura_cristata
MQLFLQINTSGAMEAGRS
>Oryctolagus_cuniculus
MGLFLLINTSGAMEAEGS
>Equus_asinus
MELLLQINTSGAMEAGGS
>Manis_javanica
MELFLQINTSGAMEAGGS
>Heterocephalus_glaber
MELFLQINTSGAMEAGGS

HG0265.2 ENSG00000196876 SCN8A

>Homo_sapiens
MAPTAVPPAASSAPLQSGRASRAPR
>Odobenus_rosmarus_divergens
MAPTAVPPAAASAPLQSGRASRAPR
>Orcinus_orca
MAPTAVPPAASSAPQQSGRASRAPR
>Oryctolagus_cuniculus
MAPTAVPPAAASAPLQSGRASRAPR
>Elephantulus_edwardii
MAPTAVPPAAASVPLQSGRASRAPR
>Equus_asinus
MAPTAVPPAAASAPLQSGRASRAPR
>Mus_musculus
MAPTAVPPAAASAPLQSGRASRAPR

HG0265.3 ENSDARG00000018032 scn8ab

>Danio_rerio
MATSAVSPGSFKHP----------
>Astyanax_mexicanus
MATSAVSPGSLRHP----------
>Austrofundulus_limnaeus
MATAAVTPDSFLHPQFTDSCRDAC
>Apteronotus_leptorhynchus
MATSAVSPGSFRHP----------
>Salmo_salar
MATTAVSPESFRHLQSNNAFLGSL
>Lepisosteus_oculatus
MATTAVSPESFRHLQSSDAVLDFC

HG0266.1 ENSG00000184185 KCNJ12

>Homo_sapiens
--------------------------------------------MAATTNSALKTVPCPLSTIT-
>Odobenus_rosmarus_divergens
---------------------MELICAPSQGLLATDSSPAGVWEMAAAADTALKTATCPLSTVT-
>Orcinus_orca
--------------------------------------------MAASASAALKTVTCLVSTVT-
>Rhinolophus_ferrumequinum
MQPARRRPWPGTAARSSEDAERAAPGGLTEGLLATDLSPIGAWEMAAATNAALKTVTCPLSTVT-
>Galeopterus_variegatus
--------------------------------------------MAAAANSALKTVTCPLSTIT-
>Oryctolagus_cuniculus
---------------------------------MIGSSPADAWEMAVTAHSALKTVKCSLSTIT-
>Equus_przewalskii
--------------------------------------------MAATANAALKTATCPLSLLRR
>Fukomys_damarensis
--------------------------------------------MAAAANSALKTVTCPLSTIT-
>Tupaia_chinensis
--------------------------------------------MAAAANSALKTVTCLLSTVT-

HG0266.1 ENSG00000260458 KCNJ18

>Homo_sapiens
-------------------------------------------------------------------------------------------------------------MAATTNSALKTVPCLLSTIT
>Canis_lupus_familiaris
MWQPRAPTPPLPGRTVPRAPGRHLVIFRGLNSHHGLLWPSRGGAGGSIGRGVQRPGFLCPSCHSLQCSFVTLGKSLLCSEPLFPCRVELILPTSQGLLVTDSSPAGVQEMEAAADTALKTATCLLSTVT
>Orcinus_orca
-------------------------------------------------------------------------------------------------------------MAASASAALKTVTCLVSTVT
>Rhinolophus_ferrumequinum
-----------------------------MQPARRRPWPGTAARSS---------------------------------E---DAERAAPGGLTEGLLATDLSPIGAWEMAAATNAALKTVTCPLSTVT
>Galeopterus_variegatus
-------------------------------------------------------------------------------------------------------------MAAAANSALKTVTCPLSTIT
>Condylura_cristata
-----------------------------MRPARRRPGPGTAAR------------------------------------SSEDAECAAPGALTEGLPVTSSFAAGTWEMAASASAALKTAACLLSTVT
>Oryctolagus_cuniculus
--------------------------------------------------------------------------------------------------MIGSSPADAWEMAVTAHSALKTVKCSLSTIT
>Fukomys_damarensis
-------------------------------------------------------------------------------------------------------------MAAAANSALKTVTCPLSTIT
>Tupaia_chinensis
-------------------------------------------------------------------------------------------------------------MAAAANSALKTVTCLLSTVT

HG0266.2 ENSGALG00000001181 KCNJ5

>Gallus_gallus
MFCHAGQREGRRRRPRDQKTTTACW
>Calidris_pugnax
MFSPNRQREEGRGQPRDPKTSPALW
>Aquila_chrysaetos_canadensis
MFSNNRQREEGRRQPRDQKTTPAIW
>Parus_major
MFAASRQREGGRRQPGAPRTIPAPW
>Struthio_camelus_australis
MFCNNRQREEGEREPRDKKTTPALW

HG0267 ENSDARG00000028173 slc4a2a

>Danio_rerio
MLPESSQLSLSKREKG
>Pygocentrus_nattereri
MLPESSQLSLSEREKG
>Apteronotus_leptorhynchus
MLPKNCQLSLSEKEKG
>Scleropages_formosus
MLPKSTPLSLSEKEKG
>Lepisosteus_oculatus
MLPKSSPLSLSEKEKG
>Trichomycterus_areolatus
MLPRTTQLSLSKREKG

HG0267 ENSDARG00000052330 slc4a2b

>Danio_rerio
MLPVNSCLSLSEKEKG
>Trachinotus_blochii
MLP-KSCLSPSEKEKG
>Pygocentrus_nattereri
MLPESSQLSLSEREKG
>Pundamilia_nyererei
MLP-KSCLSPSEKEKG
>Alosa_alosa
MLPISSCLSPSEKEKG
>Kryptolebias_marmoratus
MLP-KSCLSPSEKEKG
>Apteronotus_leptorhynchus
MLPKNCQLSLSEKEKG
>Plecoglossus_altivelis
MLPKRSCLSPSEKEKG
>Scleropages_formosus
MLPKSTPLSLSEKEKG
>Sebastes_caurinus
MLP-KSCLSPSEKEKG
>Platichthys_stellatus
MLP-KSCLSPSEKEKG
>Lepisosteus_oculatus
MLPKSSPLSLSEKEKG

HG0267 ENSG00000164889 SLC4A2

>Homo_sapiens
------------------------MPPPRCPESRSDSEKG
>Panthera_tigris
------------------------MPPPCCPESRSDSEKG
>Balaenoptera_acutorostrata_s
------------------------MPPPCCPESRSYSEKG
>Myotis_brandtii
MVSRSQRPRGHLCTGSHFSPGHLVMPPPCCPESRSDSEKG
>Galeopterus_variegatus
------------------------MPPPCCPESRSDSEKG
>Oryctolagus_cuniculus
------------------------MPPPCCSESRSDSEKG
>Equus_asinus
------------------------MPPPCCPESRSDSEKG
>Manis_javanica
------------------------MPPPCCLESRSDSEKG
>Loxodonta_africana
------------------------MPPPCCPESRSDSEKG
>Mus_musculus
------------------------MPPPCCPESRSDSEKG
>Trichechus_manatus_latirostr
------------------------MPPPCCPESRSDSEKG
>Orycteropus_afer_afer
------------------------MPPPCCPESLSDSEKG

HG0268.1 ENSDARG00000011703 clocka

>Danio_rerio
MSTSRLKVLSLQLRP
>Pygocentrus_nattereri
MSTSRLKLLSLPLQP
>Clupea_harengus
MSTSRLKVVSLPLRP
>Lepisosteus_oculatus
MSTSRLKCDSLRLRP
>Trichomycterus_areolatus
MSTSRLKLLSVPLQP
>Takifugu_rubripes
MSTSRLKVLSPPLRP

HG0268.2 ENSG00000134852 CLOCK

>Homo_sapiens
----------------------MVGHIEEYLEPIVS
>Panthera_tigris
----------------------MVSHIEEYIEPIVF
>Myotis_lucifugus
-------MP-PYVTLSDRILRLMVSHIEEYIEPIVF
>Galeopterus_variegatus
----------------------MIGHTEEYLEPIVF
>Equus_asinus
MNPGASFMPPNSHHILRRILRLMVSHIEEYIEPVVF
>Heterocephalus_glaber
----------------------MVGHIEEALEPIVF

HG0268.3 ENSG00000134852 CLOCK

>Homo_sapiens
------------------------------MITRRQLVLRRSTNVYYKTKT
>Physeter_catodon
--------------------------------------MRRSTNVYYKTKT
>Pteropus_alecto
-------------------------------MLIPQIKKRRSTNVYHKTKT
>Loxodonta_africana
------------MKETFDV-RDKDQTKRENTKCGKWRPYRRSTNVYHKTKT
>Heterocephalus_glaber
MTRTAEFVSGKPAAASFSRKINSMQSSPILPLCRPATILRRSTNVYHKTKT

HG0269.1 ENSGALG00000028685 SEMA5A

>Gallus_gallus
MVTERCFSPMSARRVSS
>Anser_cygnoides
MVTKRCFSHMSARRVAG
>Columba_livia
MVTKRCFSHMSARRVAS
>Corvus_brachyrhynchos
MVTKRCFSHMSARRVDS
>Aptenodytes_forsteri
MVTKRCFSHMSARRVVS
>Struthio_camelus_australis
MVTKRCFSHMSARRTAS

HG0269.2 ENSDARG00000061471 sema5ba

>Danio_rerio
MLVMPLCIPEAHLGS
>Salmo_salar
--MTSCCIPEAHLGS
>Ictalurus_punctatus
-MSLWKCIPEAHLGS

HG0269.3 ENSGALG00000011686 SEMA5B

>Gallus_gallus
MPLTQTEITKDFPWCLTSNMTLQ
>Lepidothrix_coronata
---------------MTLNMTLQ
>Aptenodytes_forsteri
---------------MTSNMTLQ

```
HG0270.1 ENSGALG00000015147 ALDH1A1

>Gallus_gallus
MVNVTVFKMFHKCRSRITIFPSCFLFFFFSFLFFFFFIS--SLARMSE----LD--------
>Calidris_pugnax
MVNLTGFKMFCKCRSRITIFPCF-LFFFF----FFFSIS--SLARMSE----LD--------
>Pterocles_gutturalis
MVNLTGFKMFRKYRSRITIFPCF-LFLFF-----FFLIS--SLARMSE----LD--------
>Columba_livia
MVNLTGFKTFHKCRSRITIFSCFLFFFF---------LFVFSFHHWQESVNWIDMEVIEEEK
>Aquila_chrysaetos_canadensis
MVNLTGFKMFCKCRSRITIFPHF-LFFFLS-FFFFFFIS--SLARMSE----LD--------
>Gavia_stellata
MVNLTGFKMFRKCRSRITIFPCF-LFFFF--FFFFFLIS--SLGRMSE----LD--------
>Chlamydotis_macqueenii
MVNLTGFKMFRKSRSRITIFPCFLFFFF---------FFFFSFHHWQE--------------
>Opisthocomus_hoazin
MINLTGFKMPHKCRSRITNFPCFLFFLF-SFFHFIIGKN--------E--------------
>Manacus_vitellinus
MVNLTGFKMYRKCRSGITIFPCFLFFPF-FFFHFIIGKN--------E--------------
>Phaethon_lepturus
MVNLTGFKMFRKCRSRITIFPCF-LFFFF-----FFFIS--SLARMSE----LD--------
>Melopsittacus_undulatus
MVIITGFKMFCKCRSRITILPSLFFLFFFLFCLW----------------------------
>Aptenodytes_forsteri
MVNLTGFKMFRKCRFKITIFLCFLFFFF------FIFFSFFSFHHWQE--------------
>Tyto_alba
MVNLTGFKMFHKCRSRITIFPCFLFFFF-SFLFFFFFFFFFFFFPHWQE--------------
>Apaloderma_vittatum
MVNLTGFKMFCKRRSRITIFPFS-F------FFFFFFIS--SLARMSE----LD--------
```

HG0270.2 ENSGALG00000015147 ALDH1A1

>Gallus_gallus
MISFLFLALPPPSLSLISLT
>Calidris_pugnax
MISFLFLALPPPSLSH----
>Pterocles_gutturalis
MISFLFLALPPPSLSH----
>Columba_livia
MISFLSLALPPPSLSH----
>Falco_cherrug
MISFLFLALPPPSLSLISLT
>Gavia_stellata
MISFLFLALPPPSLSH----
>Chlamydotis_macqueenii
MISFLFLALLPPSLSH----
>Opisthocomus_hoazin
MISFLFLALPPP-LSLISLT
>Lepidothrix_coronata
MISFLFLALPPPSLSLV---
>Nipponia_nippon
MISFLFLALPPPSLSH----
>Fulmarus_glacialis
MISFLFLALPPPSLSH----
>Aptenodytes_forsteri
MISFLFLAVPPPSLSLISLT
>Tyto_alba
MMSFLFLALPPPSLSH----

HG0270.3 ENSG00000128918 ALDH1A2

>Homo_sapiens
MEAAPGVLGRMDQPPAPRPSRPIRRAPA
>Leptonychotes_weddellii
----------MDQPPAPRPSRPIRKAPA
>Lipotes_vexillifer
----------MDQPPAPRPSRPIRRAPA
>Eptesicus_fuscus
----------MDQPPRPGPHVQ------
>Dasypus_novemcinctus
----------MDQPPAPRPSRPIRRAPA
>Monodelphis_domestica
----------MDQPPLPLPSCPIRRDPA
>Macropus_eugenii
----------MDQPPLPLPSCPIRRDPA
>Condylura_cristata
----------MDQPPAPRASRPIRRAPA
>Elephantulus_edwardii
----------MDQPPTPRPSRPIRRAPA
>Ceratotherium_simum_simum
----------MDQPPAPRPSRPIRRAPA
>Ictidomys_tridecemlineatus
----------MDQPPAPRPSRPIRRVPA

HG0271.1 ENSGALG00000004064 GPER1

>Gallus_gallus
MELELCHLPTCTKENC-
>Caprimulgus_carolinensis
MELKFCYLPACMKKLC-
>Cariama_cristata
MELKFYYLPTCTKKHC-
>Calidris_pugnax
MELKFCRLPTCPEKRCR
>Pterocles_gutturalis
MELKFYYLPTCMKKHC-
>Merops_nubicus
MELKFYYLPPCTKKHC-
>Haliaeetus_albicilla
MELKFYHLPTCVKKHC-
>Gavia_stellata
MELKFYYLPTCMKKHC-
>Eurypyga_helias
MELKFYYLPTCMKKHC-
>Pelecanus_crispus
MELKFYYLPTCMKKHC-
>Fulmarus_glacialis
MELKFYYLPTCMKKHC-
>Apaloderma_vittatum
MEIKFYYLPTCMKKHC-

HG0271.2 ENSG00000164850 GPER1

>Homo_sapiens
MPGSLQGESRFFLKMDSPFKTELWEPFGKS
>Orcinus_orca
MPGSLPRASPFLLKRDFLLKNGFWEPCGKS
>Equus_asinus
MSGSLQRAAPFFLKMDLLFKTDFCGPWGKS
>Mesocricetus_auratus
MPGSLPRESPFFVKMQLLLKTDFCDPFSKS
>Tupaia_chinensis
MLDSLQRESPFFLTMDSWFKTDFCDFFGNS

HG0271.3 ENSDARG00000074661 gper1

>Danio_rerio
MLMDWRKQDCAGLNTCEKQSVQKKKKKQDP
>Pygocentrus_nattereri
MPVDWRRQGSAGIKTCEEQSVQKNQERHNP
>Salmo_salar
MHMDWMRQASTGIRTCEKQTAQKKRKKQSP
>Ictalurus_punctatus
MLLDWRRQSSTGIKMCKEQTVQKKHSL---

HG0272.1 FBgn0034649 CG9865

```
>Drosophila_melanogaster
------MSDTRKDPCKPNACRIQACLTKNHYQEDKCLDVLEAMRQCCLKWHKES--LCCSGIDLEKTYLPDASKEKNATGKSNK--------------------------
>Boeckaxelia_potanini
MDREYNSVLKPRDPCKPKACAIQRCLERTHYNVVLCRPFYEQLTECCRQFAARS--DACGSEEQPPASALDREPVA----------------------------------
>Parasteatoda_tepidariorum
--------MQKKEPCQKYACELQKCLKENNFQQEHCLSVIKSLLQCCEKWNFKS--KCCEGFIESSDMVPDL--------------------------------------
>Meinertellus_cundinamarcensi
---------MKKDPCKIFACEIQRCLQENKFQESACQAAINALKKCCKKWREDS--LVCSGIKIDDELNKDIPQK-----------------------------------
>Mastotermes_darwiniensis
---------MKKDPCKPHACSIQKCLKENNYQEASCQEFIELLKECCRKWSSAS--ISCSGIDTQEAPSQTPEDKKQ---------------------------------
>Carabus_granulatus
-------MPVTKDPCKATACKIQKCLKENNYQESACADVLEEMKECCRKWAKVS--LCCSGIKIEDKAGSSSFNQSVK--------------------------------
>Carcinus_maenas
---------MKKDPCQRQACDIQDCLQRNKYQEAACEIQINALQECCRKWHEES--GCCAGVKLKGTVTSDAALKTKPSSQKSNVK------------------------
>Forficula_auricularia
-----MPRIDPKNPCMHYSCDYNKCLTETNYKEDKCKYYLDQLRNCCKKFAHLNS-LSCEGIKFDPPKENDHTETVTK--------------------------------
>Occasjapyx_japonicus
------MPAKPNDPCKPLACEIQRCLVANDFLESKCEEEVKRMLACCDLWGETSGGRSCSGFWINRPGEAPRKRQEAKK-------------------------------
>Aposthonia_japonica
--------MPKKDPCEPHACKIQKCLTENNFQEQKCKAVIDDMINCCKKWGGES--LCCSGFLSKGNKTDSSKEGSSAVTNEGKPSSET---------------------
>Eurylophella_sp._AD-2013
---------MKGDPCQLQACAIQKCLQVNNYKEAACPKEFEEMRKCCEKWGQLS--GCCSGFKDPELISSENRQGDKVASK-----------------------------
>Galloisiana_yuasai
----------MKDPCKPYACKIQKCLKEHNYQENACKDVIEEMVICCRKWGDIS--LCCSGIDTGKTNGSSGSKTVSGKT------------------------------
>Pachypsylla_venusta
-------MSKKKDPCKVHACAIQKCLKENDFQESKCLQYFEYMRQCCVKFHADS--ICCSGIDINKSYLKDNEKSSPSKTDHIERSTLVAKISSYYSSRSNNNLFERFFQ
>Brachycistis_timberlakei
--------MTTTDPCKPYACKLQKCLKDNVYQPSRCEEVIEEIRQCCIKHSAKS--IVCEGIDTSKQYEHNTVDYKKATK------------------------------
>Ixodes_persulcatus
--------MKAVDPCQKEACAIQKCLQEHNYQEDACVKFIKEMLKCCEKWREMS--GCCSGFLKDEASKKAKGELKSKT-------------------------------
>Antheraea_yamamai
---------MSKDPCKRFACDIQRCLVENNYQEVRCASVFEAMRRCCLKFKPSS--LVCEGYDLEPRKFAPVTDRAKK--------------------------------
>Mantis_religiosa
---------MKKDPCKPYACEIQKCLKEHNYQIEPCQKFVEYLRDCCRKWGDVS--ISCSGIDTRTSPNKASDKE-----------------------------------
>Panorpa_vulgaris
------MSNVPKDPCKPSACKIQQCLQEHNFQEQSCIDVIEQMRLCCVKWEGKS--FVCNGINTAEPADLAISPKHTKHS----RY------------------------
>Dermanyssus_gallinae
--------MPKGDPCQKQACDIQKCLQANNYQEALCVEAFRAMDLCCEKWGLQS--GCCTGFIKEQQLKKKIANESTKTPISFN--------------------------
>Pseudomallada_prasinus
------MPKPYIDPCKEFACRIQKCLRENNFQEHRCKQVIEELRQCCVKWKSES--ISCSGIDINSPKEEQ---------------------------------------
>Triops_newberryi
------------------------------------------MKECCSKWQERS--QSCAGFLVPEIEKKKNEPLS----------------------------------
>Calopteryx_splendens
------------------------------------------MQKCCDKWFDES--LCCSGFKKPSES------------------------------------------
>Stenobothrus_lineatus
---------MRKDPCKPIACKIQKCLTENRFQEPACQRVLEEMKECCRKWAHIS--LCCSGIDTSDKPKD----------------------------------------
>Extatosoma_tiaratum
----------MKDPCKPQACKIQKCLTENNYQEAACKDVIKEMVECCRKWADVS--LCCSGINTKETTANSPEEI-----------------------------------
>Menopon_gallinae
--------MTKKDPCKSLACALQKCLKENDFQEERCKKVLDDIRTCCKKFKEIS--VVCSGIDVSEKIK-----------------------------------------
>Cosmioperla_kuna
--------MKKRDPCKAHACRIQKCLAENNFQESACQAVLEDMRSCCRKFKNVS--LCCSGIDIGDTPTLETASEPKK--------------------------------
>Ectopsocus_briggsi
-------MPKKVEPCKPLACALQKCLTENNYQEAACKEAISALKDCCAKWGETSP-VVCSGFNK----------------------------------------------
>Xanthostigma_xanthostigma
----------MKDPCKLTACKIQKCLQENHFQEDACADVLEEMRQCCLKWYKES--LCCSGIRLDEPKIKEKKSINQSKTN-----------------------------
>Ctenocephalides_felis
--------MRSKDPCKANACMIQKCLSENNFQESACTDVIELMRLCCVKWGTKS--FVCNGIKTEKETSSTSSTVENN----KIPYKLDHKS------------------
>Thrips_palmi
------MSKRSKDPCKPFACAIQDCLNAKNFQESACVDVIENLNECCRHWKHKS--LVCSGMLIGPAEVEKSDVKSDGTSGVEVRKA-----------------------
>Hydroptila_sp._AD-2013
--------MKQKDPCKATACLIQKCLRERGFQEERCADVLEEMRQCCLKWHKVS--LCCSGIKLDPPKIEKETK------------------------------------
>Zorotypus_caudelli
-------MKKKKDPCELYACMLQKCLTENNFRESCCIAVLEQLRECCRIWGNAS--ICCSGFDNSPKAEKEPENKEPESSAKK---------------------------
>Thermobia_domestica
--------MKGGDPCQKQACDIQKCLKIHNYQVSACKDFFEAMKKCCEKWGDLS--GCCSGFKKERELEEAQKTK-----------------------------------
```

HG0272.2 FBgn0034649 CG9865

```
>Drosophila_melanogaster
---MESVRKANQRIRNYPVLLSKCADKATAYAVCVSRD--------LNVQHKICDTEFKEFLSCIRKTALEMKTKL
>Oxyacanthus_curtus
---MESVRKARERMKRFPMLLKECHDPATSYARCVALK--------ENVLKDDCSQEFQMFKNCLASAAKKMGTKM
>Machilis_hrabei
---MESVRKAKVRLQMYPKLLAQCGREGSAYAQCVITI--------ENLKKGDCSKEFQDFRVCIQKAACKLNTRL
>Argulus_siamensis
---MDSVKNAQLKLSKYTQSIVKCNQEASMYAQCVLVQ--------DDLKKHACSEEFKKLVICINKVIGKK----
>Periplaneta_americana
---MESVLKAKQRLRRYPELLVQCTGPATAYAMCVLAK--------ESVRHAECDREFQEFQLCLQKAARHLGTRL
>Euwallacea_fornicatus
---MESVSKAKQRFRQYPVIFAKCSREASVYATCVLKQ--------DNIKKNDCAEEFSGFRTCLRKAAVSLKTKI
>Farfantepenaeus_aztecus
---MESVKKAQNRLRNYPKLVANCAPVATEYARCVALK--------GNVLKGDCAQEFEAFKRCILESAKKMKTKI
>Occasjapyx_japonicus
---MESVKRARSRFQMYPKLLAECGGPAAAYAKCVATH--------EDLPKGACEKEFQGLKECLQAAAKKMGTRL
>Aposthonia_japonica
---MESVRKAKVRMRRYPELLKTCSQPAAAYAKCVAKS--------GNIRRHECEELFIKFMSCMKTAAKNLGSRL
>Eurylophella_sp._AD-2013
---MESVSRARARLRMFPRFLAECAAEGTSYAQCVAQQ--------DNPQRGACAKEFEQFSRCVRGAAQKHGTRI
>Meganyctiphanes_norvegica
---MESVKRARNRLKAYPKLAAQCTGVATEYARCVALK--------ENVMKGDCNAEFEAFKKCLMDSAKRMNTRL
>Grylloblatta_bifratrilecta
---MDSVNKGRQRLQRYPGLLVECNQPAITYATCVVSK--------DNVKKAECEKEFKLLLECLHKAAKKRGTRL
>Kerria_lacca
---MDSVSKAKNRIHKYPVLFIECRNEGKLYAACIASK--------QDIQFNACVNEFRNFKNCLMKSAIKLGTKL
>Pseudomasaris_vespoides
---MEAVKNAKDRLRKYPLLLAECHDAATKYAVCVLAK--------KNLQKNDCEAEFHEFKICLTKAAAKHRTKL
>Metaphatus_ochraceus
---MESVQKARDRLAKYPLIFGKCGKQATLYARCVCLK--------DNVKKDDCAPEFKEFMSCLKTTAKDMKVRF
>Mantis_religiosa
---MESVRKAKQRLRHYPKLLAECHGPATAYASCVLAK--------ENVKRGDCEQEFQALRKCLHQAAKHIGSRL
>Boreus_hyemalis
MQVMDSVRSARQRLSQYPKFLAECTQSATQYAMCVCRD--------TNVQKDQCINEFNAFKQCFQKAALKAKVKL
>Chrysopa_pallens
---MESVKKAHKRFRNYPVLLTECRNEASEYAACVLNS--------GGIKYKECAKEFTNFKNCIQKAAVKHKTKL
>Triops_newberryi
---MEPLRKARTRLHNYPLVIAECGPQATSYGRCIALKL-------DDLKKDTCVKEFEAFKGCLQQASKKLKTKL
>Epiophlebia_superstes
---MEAVNRAKGRMRRFPRLLSLCTAEGNAYAVCILAAHESGGPRGTGLTKGQCDAEFQALKLCVSEAARKAGTRI
>Stenobothrus_lineatus
---MEAVQAARSRLRKYPLLLSKCSTEASLYATCVLSR--------DNIKHSECEQEFRKLTTCLQKAAQKAGTRL
>Medauroidea_extradentata
---MDSVRKAHSRLQRYPELLRRCSKEATAYAGCVIGK--------ENVRKWDCDEEFKQLMSCLRSAAYKAKIRL
>Menopon_gallinae
MGKVEVLKMAKTRLRKYPDLFAECRTEGAAYAACVVKQ--------ENVTRNSCAAEFQNFKNCAMSAAKRLSTRL
>Cosmioperla_kuna
---MESVRRARARLRRYPELFAVCHAPATKYAACVVNK--------DNIKHNECGKEFLEFKQCLEQAAHKCGTRL
>Ectopsocus_briggsi
---MEAVQRSRARLARYPFHLTACAIEGGAYATCVLEK--------ENLTKDACVKEFRALVKCMKVSAAKSGSRL
>Xanthostigma_xanthostigma
---MESVKLARTRFRKYPLLLTECHNFAAAYASCVLVK--------ENIQQNDCAKEFEAFKNCLKKTALKHKTRL
>Oropsylla_silantiewi
---MDSVRKARARFSQYPSLLGECATQATTYASCVCRD--------LNVEKDRCLKEFTELQKCVKKAASKAKVRI
>Gynaikothrips_ficorum
---MESVKRAKLRLSRYPALLAECREPAKQYAACVSRQ--------DNVVKGSCQTEFVRFKGCLLKASAKMATRM
>Annulipalpia_sp._AD-2013
---MESVKKAHTRLRQYPVLLAKCGESASAYAKCVIRL--------DTVDKNDCSKEFNSFKSCLMKAAKDMKTKM
>Zorotypus_caudelli
---MDALTKARNRLKKYPQHLKECTEPAAAYASCIVRK--------DNLKKGDCEKEFKELKKCLLLVAEKRKTKL
>Atelura_formicaria
---MESVKKARGRIRKYPQLLAGCAKEGTKYAKCVMDN--------ANVQKGCCTKEFEELKTCLIKAAKKQGTRI
```

HG0272.3 ENSG00000143315 PIGM

```
>Homo_sapiens
MPDVIVCSRFGWQPGGWEMRRPPAAKNRREG
>Rhinolophus_ferrumequinum
------------------MRRQPEGRSRS--
>Ceratotherium_simum_simum
------------------MRRPPEARNRRED
>Manis_javanica
------------------MRGPPEARNAS--
```

HG0273.1 ENSGALG00000003293 MBTPS1

```
>Gallus_gallus
MSLFIRLYLKRQ
>Anas_platyrhynchos
MSLFIRLYLKRP
>Chaetura_pelagica
MSLFIRLYLKRP
>Apteryx_australis_mantelli
MSLFIRLYLKRP
>Cariama_cristata
MSLFIRLYLKRP
>Calidris_pugnax
MSLFIRLYLKRP
>Merops_nubicus
MSLFIRLYLKRP
>Cuculus_canorus
MSLFIRLYLKRP
>Aquila_chrysaetos_canadensis
MSLFIRLYLKRP
>Gavia_stellata
MSLFIRLYLKRP
>Eurypyga_helias
MSLFIRLYLKRP
>Opisthocomus_hoazin
MSLFIRLYLKRP
>Nipponia_nippon
MSLFTRLYLKRQ
>Fulmarus_glacialis
MSLFIRLYLKRP
>Aptenodytes_forsteri
MSLFIRLYLKRP
>Struthio_camelus_australis
MSLFIRLYLKRP
>Tinamus_guttatus
MSLFIRLYLKRP
```

HG0273.2 ENSG00000140943 MBTPS1

>Homo_sapiens
MECPGVFAAEAPLSRARRP--
>Felis_catus
METRGVFAAEAPLSRARGTRA
>Lipotes_vexillifer
MEILGVFAAEAPLSRAGRTRA
>Rhinolophus_ferrumequinum
METRGVFAAEAPLSRAHRTRA
>Condylura_cristata
MESRGVFAAETPLS-------
>Equus_asinus
MESRGVFAAEAPLSRARRTRA
>Manis_javanica
MEARGVLAAEAPLSPARRN--
>Psammomys_obesus
METRGVFAAEAPFSRARQLRA
>Trichechus_manatus_latirostr
METRGVLSVEARLSRARRTGA

HG0273.3 ENSG00000140943 MBTPS1

>Homo_sapiens
MLQLLAETKIHF
>Chinchilla_lanigera
MLQLLAETKIHF
>Tupaia_chinensis
MLQLLAETKIHF

HG0274.1 ENSG00000204256 BRD2

>Homo_sapiens
MTGPSGQYRPPRSPWSSPFSGPAQSSESVHPLYSALKRISKMEAAAP
>Ailuropoda_melanoleuca
MTGPGGQYRPHRSPRRSPFSGPAQSSESVHPLYSALKRISKMEAAAP
>Orcinus_orca
MTGPGGQYRPHRSPRSSPFSSPAQSSESVHPLYSALKRISKMEAAAP
>Myotis_davidii
MTVPGGQYRPHRSPRSSPFSGPAQSSESVHPLYSALKRISKMEAAAP
>Dasypus_novemcinctus
MTGPGGQY------------------------------------------
>Galeopterus_variegatus
MTGPCGQCRPHRSPRSSPFSGPAQSSESVHPLYSALKRISKMEAAAP
>Condylura_cristata
--------------------------------------------MEAAAP
>Oryctolagus_cuniculus
MTGPGGQYRPYRSPRSSPFSGPAQSSESVHPLYSALKRISKMEAASP
>Elephantulus_edwardii
MTRPGGQYRPHRSPRSSPFPGPAQSSESIHPLYSALKRISKMEAAAP
>Equus_caballus
MTGPGGQYRPHRSPRSSPFSGPAQSSESIHPLYSALKRISKMEAAAP
>Manis_javanica
MIGPGGQYRPHRSPRSFPFSGPAQSSESVHPLYSALERISKMVAAAP
>Loxodonta_africana
MTDTGGQYRPHRSPRSSPFPGPAQSSESIHPLYSALKRISKMEAAAP
>Trichechus_manatus_latirostr
MTGPGGQYRPHRSPRSSPFPGPAQSSESVRPLYSALKRISKMEAAAP
>Orycteropus_afer_afer
MTGPGGQYRPHRSPRSSPFLGPAQSSESIHPLYSALKRISKMEAAAP

HG0274.2 ENSDARG00000100129

>Danio_rerio
----------------------------MLLRNKSYPKICKDNATGAARSESNKVQHIHLRLAHSDRSSRSV
>Astyanax_mexicanus
---------------------------MLLRCNKSDPKICEENAAGAARTESNKTC----------------
>Ictalurus_punctatus
MGGQTHTHTHTHTHTHTQTTESSQPCIMPLRCNKKLPENLHGNATGAAWSESNKVVLTL-RY----------

HG0275.1 ENSG00000100239 PPP6R2

>Homo_sapiens
--------------------------------------------MKKIFLKLHILQEKNHKCFSVLPEYMI-------------------------
>Canis_lupus_familiaris
--------------------------------------------MKKSFLKLHIVQEKNPKCFHILFELRIKKFP---------------------
>Balaenoptera_acutorostrata_s
--------------------------------------------MKKFFLKLHILQEKIPNVFIFCLNTKFKQGVSTEVRRLRRKCASPCSGLLTG
>Rousettus_aegyptiacus
MCVRSAAGRPGFASASALAATRASQEPPRRPARGAGEEIINLILNEKSFLKVAYTPRKNPKCFHILPEYKI-------------------------
>Ceratotherium_simum_simum
--------------------------------------------MKKFFLKLHVLQEKNRSFLNILPEFKI-------------------------
>Manis_javanica
MCVRPTPGRPGFASASVLASTRASEEPPQRPARGAGEEIINLLLNEKIILKVTYSPRKNPKCFYILPENKI-------------------------
>Cavia_porcellus
--------------------------------------------MKKRILKVGYTPRKKPKCFSVLPDYKI-------------------------

HG0275.2 ENSDARG00000045540 ppp6r2a

>Danio_rerio
MLGHSVPSAFCHSNACRPVEEV
>Pygocentrus_nattereri
MPGRSVLSAFCHSNACLLVEEV
>Ictalurus_punctatus
-----MPSAVCHSNACLLVEKV

HG0276.1 ENSGALG00000008912

```
>Gallus_gallus
MDSSSSITVTWNIQHHM
>Anas_platyrhynchos
MDLSLSVTVTWNIQHCL
>Calidris_pugnax
MDSSLHVTATWSIQHHL
>Manacus_vitellinus
MDSSLSVTVTWSIQQHL
>Pygoscelis_adeliae
MDSSLSVTVTWSTQHHL
```

HG0276.2 ENSG00000073734 ABCB11

>Homo_sapiens
MMKTEVGKGCETF
>Equus_asinus
-MKTEVGKGCKTL
>Trichechus_manatus_latirostr
MMKTEVGKGCKTL

HG0277.1 ENSG00000102531 FNDC3A

>Homo_sapiens
MAAAAFVVPRVKQKAGATRRGSEERG
>Felis_catus
MAAAAFVVPRVKQKAGATRRGSEERG
>Orcinus_orca
MAAAAFVVPRVKQKAGATRRGSEERG
>Pteropus_vampyrus
MAAAAFVVPRVKQKAGATRRGSEERG
>Monodelphis_domestica
MAAVGFVISRVKQKEGATGRLGEERG
>Condylura_cristata
MAAAAFVVPRVKQKAGAARRGSEERG
>Ochotona_princeps
MAAAAFVVPRVKQKAGATRRGSEERG
>Elephantulus_edwardii
MAAAAFVVPRVKQKAGATRRGSGERG
>Ceratotherium_simum_simum
MAAAAFVVPRVKQKAGATRRGSEERG
>Octodon_degus
MAAAAFVVPRVKQKAGATRRGSEERG
>Trichechus_manatus_latirostr
MAAAAFVVPRVKQKAGATRRGSEERG
>Orycteropus_afer_afer
MAAAAFVVPRVKQKAGATRRGSEERG

HG0277.2 ENSGALG00000004169

>Gallus_gallus
MAERSVGGRQRRG
>Calidris_pugnax
MAERSVGGRQRRG
>Haliaeetus_leucocephalus
MAERSVGGRQRRG

HG0278.1 ENSG00000119946 CNNM1

>Homo_sapiens
-----------------------------------------------------------MPGHLQPPAGTSCPGGV----------------
>Myotis_lucifugus
MGNFPGQWGRRGAGRELLPGRSSLRCCLLLKQEALLQAKLSLGCSSDIPECVRVMHRAVMRGHFQPPAGTSCPDRVRTPTASSPRAQGACTW
>Dasypus_novemcinctus
------------------------------------------------------MLCAVMRGHLQPSSGTSCPG------------------
>Condylura_cristata
-----------------------------------------------------------MRGHLWPPAGTTCPGRV----------------
>Oryctolagus_cuniculus
-----------------------------------------------------------MRGHLQPPAPTPCPGRVCAPTASSPRAQGARTW
>Equus_caballus
----------------------------------MLQGKLSPDCNSDTTKCVWVMQSAVMRGHLQPPAKTSCPGRV----------------

HG0278.2 ENSDARG00000078733 cnnm2b

>Danio_rerio
-------------------------------------------------------------------------------MHKSRRRAAEMKP--DWL
>Channa_argus
MYIKKSRVESCRSFYMRCTCFGISFVEEAADAPPTSQRRFPVGRRGRSSRGHQGRHELARKPALASRQCRTRSGNEQVPMHKSRQPSAEMKADSDWL
>Alosa_alosa
-------------------------------------------------------------------------------MHKLKRRTAEMKADSDWL
>Lateolabrax_japonicus
-------------------------------------------------------------------------------MHKSRQPSAEMKADSDWL

HG0279.1 ENSG00000175928 LRRN1

>Homo_sapiens
MFTFSARCPTYHNIVFTFC
>Leptonychotes_weddellii
MFTFSACCPTYHYLVFNFC
>Balaenoptera_acutorostrata_s
MFTFSARCPSYHYTVFTFC
>Rhinolophus_ferrumequinum
MFTFSARCPT---------
>Galeopterus_variegatus
MFTFSACCPTCHRTVVTFC
>Ceratotherium_simum_simum
MFTFSACCATYHYIVFTCC
>Manis_javanica
MFTFLVCCPTYHQTVSAFC
>Nannospalax_galili
MFTFSACCST---------
>Orycteropus_afer_afer
MLAFPACCPTHHYIGFTFC

HG0279.2 ENSGALG00000035099 Lrrn1

```
>Gallus_gallus
MSAPPLRGGEERLRG-QRPPGCRNVFRHGFVEDPGTQRSS
>Anas_platyrhynchos
---------MTRLRTKANGTIYRNIFRHRFAEDPGTQRSG
>Calidris_pugnax
-----MLSGTP--NSRLCSRAVRNLFRHRFAEDSGTQRNS
```

HG0280.1 ENSG00000156299 TIAM1

```
>Homo_sapiens
----------------MAHLEILTKPLLFPMGFLAHSHRLR
>Acinonyx_jubatus
------------------------------MGFLAPPHRLR
>Oryctolagus_cuniculus
MLCILVPQLGPEQPAWMAPLEILTKLLLFCMGFSAHYPR--
>Equus_caballus
------------------------------MGFLAHYHRFR
>Ictidomys_tridecemlineatus
----------------MAPLEILTKPLLFGMGFLAHSSRLR
>Tupaia_chinensis
----------------MAYLKILTQLLLSEMGILAHSHRLR
```

```
HG0280.2 FBgn0085447 sif

>Drosophila_melanogaster
MGAKESVEAAAKMNADGWGHVERSSRYHNHRH-NQRSQSLNRGQALSQ------ADVSGMW
>Rhynchophorus_ferrugineus
MGAKGSVEAA-KINAQNWSNFEHNHSKQFVGH-HQRSQSLNRGVREQWSGRGADLGGSDMW
>Trichoplusia_ni
MGAKESVEAA-KFSAQNWNNFEHAAVKHPYRHGTQRSQSLNRGHSLSR------TD--GTW
>Chrysopa_pallens
MGAKGSVEAA-KYNAQQWNSMEGSRHRH-----SQRSHSLNRGHSLSR------AP--DMW
>Oropsylla_silantiewi
MGAKGSVEAA----KENWEHIERGR--HHHRL-HQRSQSLNRGHSLSR------ADTAGMW
```

HG0281.1 ENSGALG00000006766 DEPDC5

>Gallus_gallus
MCRMSSTVFTRNS
>Anas_platyrhynchos
---MSSTVFTQSS
>Apteryx_australis_mantelli
---MSSTVFAQSS
>Buceros_rhinoceros_silvestri
---MTSTVFTQSS
>Calidris_pugnax
MCTMSSTVFTQSS
>Colius_striatus
---MSSTVFAQSS
>Aquila_chrysaetos_canadensis
---MSSTVFTQSS
>Chlamydotis_macqueenii
MCKMSSTVFTQGS
>Pelecanus_crispus
---MSSTVFTQSS
>Picoides_pubescens
---MSSNVFTQSS
>Fulmarus_glacialis
---MSSTVFTQSS
>Melopsittacus_undulatus
---MSSTVFTQSS
>Calypte_anna
---MSSTIFTQSS

HG0281.2 ENSG00000100150 DEPDC5

>Homo_sapiens
MTSLPQAWNS
>Leptonychotes_weddellii
MTSLPQAWNS
>Balaenoptera_acutorostrata_s
MISLPQAWNS
>Miniopterus_natalensis
MTSLPQAWNS
>Dasypus_novemcinctus
MTSLPLAWNS
>Galeopterus_variegatus
MTSLPQAWNS
>Condylura_cristata
MTSLPQAWNS
>Oryctolagus_cuniculus
MTSLPQAWNS
>Equus_przewalskii
MTSLPQAWNS
>Manis_javanica
MTSHPQAWNS

HG0282.1 ENSDARG00000061328 cdon

>Danio_rerio
----------------------------------MEGPVRICHRENVFRC
>Channa_argus
---------------------MFFPSARTSSPCLMEGPVRICHQKNAFCC
>Trachinotus_blochii
----------------------------------MEGPVRICHQKNASCC
>Micropterus_floridanus
----------------------------------MEGPVRICLQKNAFCC
>Pygocentrus_nattereri
----------------------------------MEGPVRICCQKNAFCC
>Neolamprologus_brichardi
---------------------MSLKPSRTLSPFLMEGPVRICQQKNAFCC
>Esox_lucius
----------------------------------MEGPVRICHQKNELFF
>Lateolabrax_maculatus
MWVEWNSSLTPPPPPPPPAVFMFLPSSRTLSPCLMEGPVRICLQKNAFCC
>Sebastes_caurinus
---------------------MLLPSSRKHSPCLMEGPVRICLQKNAFCC
>Cynoglossus_semilaevis
---------------------MSRPLSYTFCPCLMEGPVRICHQKNASCC
>Oncorhynchus_kisutch
----------------------------------MEGPVRICHQKNELFC
>Trichomycterus_areolatus
----------------------------------MEGPVRICCYRNAFCC

HG0282.2 ENSG00000144857 BOC

>Homo_sapiens
MKSCRHLYRLRVAARK
>Condylura_cristata
MKSCQHSHRLRVAAVK
>Loxodonta_africana
MKSCQHSHRRRVAATK

HG0283.1 ENSGALG00000010270 AREL1

>Gallus_gallus
MQVPGVPFSVTAFSMASLC
>Anser_cygnoides
MQIPGVSFSVIIFSMASSC
>Apteryx_australis_mantelli
MQVPGVSLSVMNFSLASSC
>Charadrius_vociferus
MQVPGVSLSVIIFSMDSSC
>Colius_striatus
MQVPGN-------------
>Leptosomus_discolor
MHVPGISLSVIIFSMAASC
>Haliaeetus_leucocephalus
MQVPGVSPSVIIFSMASSC
>Mesitornis_unicolor
MQVPRVSLSVIIFSMASSC
>Opisthocomus_hoazin
MQAPGVSPSVIIFSMASSC
>Phaethon_lepturus
MQVPGVSLSVIIFSMASSC
>Picoides_pubescens
MQVPGVSLSVIIFSMTSSC
>Fulmarus_glacialis
MQVPGVSLSVTIFSMASSC
>Struthio_camelus_australis
MRVPGVSLSVMNFSMASSC
>Apaloderma_vittatum
MQIPGVSLSVIIFSMASSC

HG0283.2 ENSG00000119682 AREL1

>Homo_sapiens
MDRPFLLTLWRFSFMWKLEDKVKGAKSYLLW-
>Panthera_tigris
MDRHFPLTLQQFSFLWKLEDKVNGARSYLLW-
>Balaenoptera_acutorostrata_s
MDRHFLLTLQLFSFMWKLEDKVKVAKSYLLW-
>Pteropus_alecto
MDRHFLLTLQLFSLMWKLKTR-----------
>Dasypus_novemcinctus
MDRPILLTLWLFSSMWKLEDKVKGGKSYLFW-
>Galeopterus_variegatus
MDRPFLLTLRLFSFMWKLEDKAKGAKSYLLW-
>Condylura_cristata
MDRHFLLTLQLFSFMWNLEDKVTATKSTCSGE
>Oryctolagus_cuniculus
MDNDLPMTLWLFSFMWKLEDKVKEARDYLLW-
>Equus_asinus
MDRHFLLTLWLFSFLWKLEDNVKGAKNYPLW-
>Manis_javanica
MDRCFPSTLQLFSSMWKLEDKVKGAPEYLLC-
>Dipodomys_ordii
MDRPFLVTLWLFFFMWKLEDKVKGAKSYLLR-
>Tupaia_chinensis
MDRGFLLTLQLFSFMWQTRWKTR---------
>Trichechus_manatus_latirostr
MDRPFLLTLQLFSPMWKLEDRVKGAEVICFGE
>Orycteropus_afer_afer
MDRPFLLTLQLFSSMWKLEDKVEGARSDLLW-

HG0284.1 ENSG00000007168 PAFAH1B1

>Homo_sapiens
----------------------------------------------------------MGVKDGRGPAEAAVQRSGGMNLTC
>Leptonychotes_weddellii
----------------------------------------------------------MGVKDGRGPAEAAVQRSGGMNPIC
>Physeter_catodon
----------------------------------------------------------MGVKDGRGPAEAAVQRSGGMNLTC
>Miniopterus_natalensis
----------------------------------------------------------MGVKDGRGPAEAAVQRSGGMNLTC
>Galeopterus_variegatus
----------------------------------------------------------MGVKDGRGPAEAAVQRSGGMNLTC
>Erinaceus_europaeus
----------------------------------------------------------MGVKDGRGPAEAAVQRSGGMNLAC
>Ochotona_princeps
----------------------------------------------------------MGVKDGRGPAEAAVQRSGGMNLTC
>Ceratotherium_simum_simum
----------------------------------------------------------MGVKDGRGPAESAVQRSGGMNLTC
>Mesocricetus_auratus
MPCFLSLLSGNSDAAWLFVFVVMVTFTLSLSLSLTPSLPPSLPPSFLPSLSLPLSRRWMGVKDGRGPAEAAVQRSGGMNLTC
>Tupaia_chinensis
----------------------------------------------------------MGVKDGRGPAEAAGQRSGGMNLTC
>Trichechus_manatus_latirostr
----------------------------------------------------------MGVKDGRGPAEAAVQRSGGMNLTC
>Orycteropus_afer_afer
----------------------------------------------------------MGVKDGRGPAEAAVQRSGGMNLTC

HG0284.2 ENSGALG00000005834 PAFAH1B1

>Gallus_gallus
MPPGFSVGFPIWRNICALSHCGKLNCGS
>Anser_cygnoides
MPLGFPIGFHIWRNISDLSHSVAD----
>Haliaeetus_albicilla
MPPDFLLDFTCGKIL-------------
>Melopsittacus_undulatus
MPPDFLSGFTSGKLL-------------

HG0285.1 ENSGALG00000012156 DPP10

>Gallus_gallus
MLRGRPRAAGQEEHGGGADTGLGFPPAGAVAPRLPPAFLFLYFSSSSAAAPLRGEVGVMRLAPREGCPGPQRQ
>Calidris_pugnax
----------------------------------------------------------MRLAPREGCPGPQRQ
>Acanthis_flammea
----------------------------------------------------------MRLAPREGCPGPQQQ

HG0285.2 ENSG00000175497 DPP10

>Homo_sapiens
-----------------MNQVNALFRAK
>Loxodonta_africana
MELSLAFCYISPRALRMMNQVNALFRAK
>Ctenomys_sociabilis
-----------------MNQVNALFRAK

HG0286.1 ENSG00000116698 SMG7

>Homo_sapiens
MSLQSAQYLRF--------------
>Leptonychotes_weddellii
MSLQSAQYLRF--------------
>Lipotes_vexillifer
MSLQSAQYLRF--------------
>Eptesicus_fuscus
MSLQSAQYLRF--------------
>Erinaceus_europaeus
MSLQSAQYLRF--------------
>Elephantulus_edwardii
MSLQSAQYLRF--------------
>Heterocephalus_glaber
MSLQSAQYLRFMDSKWKAMGFYMLD
>Trichechus_manatus_latirostr
MSLQSAQYLRF--------------
>Orycteropus_afer_afer
MSLQSAQYLRF--------------

HG0286.2 ENSDARG00000060767 smg7

>Danio_rerio
-MDNMAPGVDLACESVVV
>Pygocentrus_nattereri
MMDNMAPGVDLASGSVVV
>Apteronotus_leptorhynchus
----MAPGVDLASGPVVV
>Trichomycterus_areolatus
-MDNMAPGIDLAPEPAVV

```
HG0287.1 ENSG00000170325 PRDM10

>Homo_sapiens
MLQTDLSVLHEHTCGRQMTVLYPVLL
>Acinonyx_jubatus
-----------------MTVPHPVLF
>Balaenoptera_acutorostrata_s
-----------------MTVLHPELL
>Rousettus_aegyptiacus
-----------------MTVLHPMLL
>Oryctolagus_cuniculus
-----------------MTALSPVLL
>Equus_asinus
-----------------MTVLHPLLP
>Loxodonta_africana
-----------------MSVVCP---
>Heterocephalus_glaber
MLQTDPSVFHAHACAGQMTVPHPVLL
>Tupaia_chinensis
-MLRSPSVLHEHVCGGQMTALLPVLL
```

HG0287.2 ENSDARG00000104251 prdm10

>Danio_rerio
MQGSLLCSYHSEGRPTSTFS----------------------------
>Amia_calva
MQCGHLCSYHSEVRPTWPFPLKTQISPPISEPKPKSRRRGT-------
>Channa_punctata
MQCGHLCSYHSEVRTTCTFSRKCKKTRLCFVIF---FVRQLLLVLLLI
>Oryzias_latipes
MQCSRLCSYHSEVRTACTFFAKCKITQNSRATSSTTLVQTIIFS----
>Pygocentrus_nattereri
MQCGHLCSYHSEVRPTSTILLK--------------------------
>Plecoglossus_altivelis
MQRSLLCSYHSEELCDNKVTGGEESS----------------------
>Cynoglossus_semilaevis
MQRDHLCSYHSEVRTTYRESAGDSCY----------------------

HG0288.1 ENSGALG00000006655 MAP3K13

>Gallus_gallus
MIRYWLPASRLNCPVL--
>Apteryx_australis_mantelli
MIRYWVHMLHLNCVVLIL
>Calidris_pugnax
MIRYWVRVSHLNCVVLIL
>Falco_cherrug
MIRYWLHVSHLNCVVLIL
>Aptenodytes_forsteri
MIRYWVCVSHLNCVVLIL

HG0288.2 ENSG00000073803 MAP3K13

>Homo_sapiens
-----------------------MYCGIQI--LALWERFWSPLLSQNSVPKIF
>Mustela_putorius_furo
MERAKDRAELVGLPVFLVSTYQLLRCAVAVRGRTHTHRFWSPLLHPNSVPLIL
>Rhinolophus_ferrumequinum
------------------------------MCRNSSLLFWSPLLRQNSIRVIL
>Mus_musculus
------------------------------------MPFWSPLLHQTSVAAIF
>Tupaia_chinensis
-------------------------------------------MHQNSLPATL

HG0289 ENSG00000072415 MPP5

>Homo_sapiens
MDFILQNQENWLIGKIDL
>Odobenus_rosmarus_divergens
MDFILQNQEIGFIGEIDL
>Balaenoptera_acutorostrata_s
MDFILQNQEIGFIGKIEL
>Miniopterus_natalensis
MDFILQNQEIGFIGKIDL
>Dasypus_novemcinctus
MDFILQNQENWLYRKF--
>Galeopterus_variegatus
MDSILQNQENWLYRKN--
>Condylura_cristata
MDFILQNQEIGFIGKFEL
>Ceratotherium_simum_simum
MDFILQNQEIGFIGKIDL
>Manis_javanica
MDFMLQNQEIGFLGKIDL
>Loxodonta_africana
MDFILQNQENWLYRKN--
>Nannospalax_galili
MDFILQNQENWLYRKI--
>Tupaia_chinensis
MDFILQNQEHWLYRKN--
>Trichechus_manatus_latirostr
MDFILQNQENWLYRKN--
>Orycteropus_afer_afer
MDFILQNQENWLYRKIDL

HG0289 ENSGALG00000009581 MPP5

```
>Gallus_gallus
--------MDFTQQRNENWPNISISI---
>Anser_cygnoides
--------MDFTQHHNKNWPNSISV----
>Chaetura_pelagica
--------MDFTQHHNENWPHSSISV---
>Apteryx_australis_mantelli
--------MDFIPHHNKNWPYSSICV---
>Buceros_rhinoceros_silvestri
--------MDFTQHHNENWPNSSISVCGE
>Caprimulgus_carolinensis
--------MDFTQHHNENWPNRSISV---
>Cariama_cristata
--------MDFTQHRNKNWPNSSISV---
>Calidris_pugnax
--------MDFTQHHNENWPNISISV---
>Pterocles_gutturalis
--------MDFTQHHNENWPSSSISV---
>Colius_striatus
--------MDSTQHHNKNCPNSSISV---
>Columba_livia
--------MDFTQHRNKNWPSSSISV---
>Leptosomus_discolor
--------MDFMQRHNENWPGSSISV---
>Cuculus_canorus
--------MDFTQHHNENWPNSSISV---
>Aquila_chrysaetos_canadensis
--------MDFTQHHNENWPNSSISV---
>Gavia_stellata
--------MDFTQHHNKNWPSSSISV---
>Mesitornis_unicolor
--------MDFMQHHNENWPNSSISV---
>Tauraco_erythrolophus
--------MDFTQHHNENCPNSSISV---
>Opisthocomus_hoazin
--------MDSTQHHNKNWPNSTISV---
>Lepidothrix_coronata
-------MMDFTQHHNKNWPNSSICV---
>Phalacrocorax_carbo
--------MDFTQHRNENWPNSSISV---
>Picoides_pubescens
--------MDFTHHHNENWLNSSIFA---
>Fulmarus_glacialis
--------MDSTQQHNKNWPNSSISV---
>Melopsittacus_undulatus
--------MDFTQCHSENWPNNSISI---
>Aptenodytes_forsteri
--------MDFTQHHNKNWPNSSISV---
>Tyto_alba
--------MDFTQHHNENWPNSSISD---
>Struthio_camelus_australis
--------MDFMPHHNENWPYSSICV---
>Tinamus_guttatus
--------MDFIPHHNENWPFSTICV---
>Calypte_anna
MGNIFQVVMDFRQHQNESWPSSSISA---
>Apaloderma_vittatum
--------MDFTQHPNENWPNSSFSV---
```

HG0290.1 ENSG00000116539 ASH1L

>Homo_sapiens
MLVSSFGDPCFLWNTVL
>Acinonyx_jubatus
MLVASFGDPCFLWNTVL
>Balaenoptera_acutorostrata_s
MLVASFGDPCFLWNTVL
>Rhinolophus_ferrumequinum
MLVAHFGDPCFLWNTVL
>Dasypus_novemcinctus
MLVASFGDPCFLWNTVL
>Sarcophilus_harrisii
MLVASFGEQPFLFNTTF
>Monodelphis_domestica
MLVASFGEQPFLFKTVL
>Sorex_araneus
MLVASFGDPCFLWNRVL
>Ochotona_princeps
MLVPSFGDPCFLWNTIL
>Ceratotherium_simum_simum
MLVASFGRPCFLWNTVL
>Manis_javanica
MLVASFGDPCFLWNTVL
>Loxodonta_africana
MLVASLGDLCFLWNIVL
>Dipodomys_ordii
MLVASFGKSCFLWNTVL
>Tupaia_chinensis
MLVSSFGDPCFLWNTVL
>Trichechus_manatus_latirostr
MLVASLGGLCFLWNVVL
>Orycteropus_afer_afer
MLVASLGDLCFLWNIVL

HG0290.2 ENSDARG00000070981 ash1l

>Danio_rerio
MEARLTREARSDEPELH
>Astyanax_mexicanus
MEARLTPQARSDSPELD
>Haplochromis_burtoni
MEARLTREARADTSRPS
>Austrofundulus_limnaeus
MEARLTREARADTSRPS
>Notothenia_coriiceps
MEARLTREARADTSRPS
>Cynoglossus_semilaevis
MEARLTREARADTSRPS

HG0291.1 ENSGALG00000009173 GFRA1

```
>Gallus_gallus
---------------MDRNLESLHLGGFGISTC
>Anser_cygnoides
---------------MDVNLESLDLGGFGISTR
>Apteryx_australis_mantelli
---------------MDLNLESLDLGGCGISTC
>Buceros_rhinoceros_silvestri
---------------MDVNLESLDLGGFGISTC
>Calidris_pugnax
---------------MDLNSESLDLGGFGTSTC
>Pterocles_gutturalis
---------------MDLNLESLDLGGFGISTC
>Colius_striatus
MFFPFSFFFFSPVNSMDLNLESLDLGGFGISTC
>Leptosomus_discolor
---------------MDLNLESLDLGGFGISTC
>Cuculus_canorus
---------------MDLNLESLDLGGFGISTC
>Balearica_regulorum_gibberic
---------------MDLNLESLDLGGFGISTC
>Opisthocomus_hoazin
---------------MDLNLEPSDLGGFGISTC
>Lepidothrix_coronata
---------------MDVNLEPLDLGGFGISTC
>Pelecanus_crispus
---------------MDLNLESLDLGGFGISTC
>Calypte_anna
---------------MDRNLESSNVGGSRISTC
>Apaloderma_vittatum
---------------MDVNLESLDLDSFGISTC
```

HG0291.2 ENSG00000151892 GFRA1

```
>Homo_sapiens
MELNFGRPEQHSCPGIAAR
>Mustela_putorius_furo
MELNFGRPEQRGCPGIAVC
>Lipotes_vexillifer
MELNFGRPEQRGRPGIAAR
>Galeopterus_variegatus
MELNFGRPEERSRPGTAAC
>Condylura_cristata
MELNFGRPEQRGCPGIAAC
>Ochotona_princeps
MELNFGRPEEHDLPGIAAC
>Equus_przewalskii
MELNFGRPEQRGSPGIAAC
>Rattus_norvegicus
MELNFEWPEERSRPGIAAR
>Trichechus_manatus_latirostr
MELNFGRPEQRGRPSFAAC
>Orycteropus_afer_afer
MELNFGRREQRGRPSIAAC
```

HG0292.1 ENSGALG00000006502 PCGF5

>Gallus_gallus
MPGKVPLRRAAPQLS
>Chaetura_pelagica
MPGKVELHRAAPQLL
>Buceros_rhinoceros_silvestri
MPGEVHLHRAAPQLL
>Caprimulgus_carolinensis
MPGEVQLHRAAPQLL
>Cariama_cristata
MPGEVQLHRAAPQL-
>Calidris_pugnax
MPGEVQPRRAAPQLS
>Leptosomus_discolor
MPGEVQLPRAAPQLS
>Falco_cherrug
MPGEVQLHRAAPQLP
>Balearica_regulorum_gibberic
MPGEVQLHRAAPQLL
>Ficedula_albicollis
MPGEVQLHRAAPQLL
>Nipponia_nippon
MPGEVQLHRAAPQLL
>Aptenodytes_forsteri
MPGEVQLRRAAPQLL

HG0292.2 ENSG00000180628 PCGF5

```
>Homo_sapiens
MALSPLRRRWDRLGEDAAEETNSQDMGKRNHQKE
>Felis_catus
---------------------MVSDMGKQNHQKE
>Lipotes_vexillifer
-------------------------MGKQNHQKE
>Pteropus_vampyrus
-------------------------MGKRNHQKE
>Dasypus_novemcinctus
-------------------------MGKQNHQKE
```

HG0293 ENSG00000152127 MGAT5

>Homo_sapiens
----MEVRKGNQLTQEPE
>Acinonyx_jubatus
----MEMRKGNQLTQEPE
>Orcinus_orca
----METRKGNRLTQEPE
>Pteropus_alecto
----MEARKGNWLTQEPE
>Sarcophilus_harrisii
----MELKEGNRLTQEPE
>Galeopterus_variegatus
----MGVKKGNRLTQEPE
>Monodelphis_domestica
----MEMKEGNWLTQEPE
>Condylura_cristata
----MEARKGNRLTQEPE
>Ochotona_princeps
----MEVRKGSLLTQEPE
>Ceratotherium_simum_simum
----MEVSKGNRLTQEPE
>Loxodonta_africana
----MELRKGHRLTQEPE
>Chinchilla_lanigera
MKQRMEARKGNQLTQEPE
>Tupaia_chinensis
----MDVRKGNWLTQEPE
>Orycteropus_afer_afer
----MELRKGHQLTQEPE

HG0293 ENSGALG00000012187 MGAT5

>Gallus_gallus
MLRKAHWPTNRMELKYGYMLTKESE
>Anser_cygnoides
-----------MELRYRYTLTKESE
>Chaetura_pelagica
-----------MELKYGYTLTKELE
>Buceros_rhinoceros_silvestri
-----------MELKYGYTLTKELE
>Caprimulgus_carolinensis
-----------MELKYRYVLTKELE
>Cariama_cristata
-----------MELKYGYTLTKELE
>Calidris_pugnax
-----------MELKYGHTLTKELE
>Haliaeetus_albicilla
-----------MELKYGYTLTKELE
>Gavia_stellata
-----------MELKYGYTLTKELE
>Eurypyga_helias
-----------MELKYGYMLTKELE
>Sturnus_vulgaris
-----------MELKYGYVLTKEWE
>Pelecanus_crispus
-----------MELKYGYMLTKELE
>Fulmarus_glacialis
-----------MDLKYGYTLTKELE
>Nestor_notabilis
-----------MELKHGYMLTKELE
>Tinamus_guttatus
-----------MEVKYNYMLTQESD
>Calypte_anna
-----------MELKHGYTLTKELE
>Apaloderma_vittatum
-----------MELKYGYTLTEELE

HG0294.1 ENSGALG00000017036 LHFP

```
>Gallus_gallus
---------MMGRNILLAFRIRASPSVRICQG
>Anser_cygnoides
---------MMGGNIPLAVRIRASPSARICQG
>Apteryx_australis_mantelli
---------MMGENIIMAVKIQASPRARIYQG
>Calidris_pugnax
---------MMGGNIPPAVRIQASPSARICQG
>Columba_livia
---------MMGGNIPPAVRIRASPSVRICQG
>Aquila_chrysaetos_canadensis
MGQKRIFDFMMGGNIPLAVRIRASPSARICQG
>Zonotrichia_albicollis
---------MMGGNIPLALRIRASPSVRICQG
>Struthio_camelus_australis
---------MMGENIFMAVRIQASPGARIYQG
>Tinamus_guttatus
---------MMGENISMAVKIRVSPGARIYQG
```

HG0294.2 ENSG00000183722 LHFP

>Homo_sapiens
MFPPEKDSKLCVRPGRGREGGRRPPAMDHHL----------
>Myotis_ricketti
MFRPGKTASCVCAPARG-EGGRRPPAMDHHPYYGGPEVEAQ
>Galeopterus_variegatus
MFPPARESKLCVRPGPGREGGRRPPAMDHRSYYRGWEVDAQ
>Oryctolagus_cuniculus
-----MFRQVVCAPGTHREGGRRPPTMDHRSYHREWEVDAQ
>Dipodomys_ordii
MCVSGKDLKLCARPGPGRKGGCRPPAPRPHPTTLSHG----
>Tupaia_chinensis
MFPPGNDSKLCARSGPGREGGRRPPAMDHHPYYREWEVDTQ

HG0295.1 ENSG00000166888 STAT6

```
>Homo_sapiens
MYVCALSYFSFWWWWWKGGGASRASLELAGQSYRPMGPGSAR
>Ailuropoda_melanoleuca
-----------------------------------MGPGSAR
>Balaenoptera_acutorostrata_s
-----------------------------------MGPGSAR
>Pteropus_alecto
-----------------------------------MGPRSAR
>Sarcophilus_harrisii
-----------------------------------MGPGSVH
>Galeopterus_variegatus
---------------------MQVSPELAGQSYRPMGPGSAH
>Monodelphis_domestica
-----------------------------------MGPGSVH
>Erinaceus_europaeus
-----------------------------------MGPGSAH
>Ceratotherium_simum_simum
-----------------------------------MGPGSAR
>Manis_javanica
-----------------------------------MGPGSAC
>Loxodonta_africana
-----------------------------------MGPGSAR
>Nannospalax_galili
-----------------------------------MGPGSAR
>Tupaia_chinensis
-----------------------------------MGPGSAR
>Trichechus_manatus_latirostr
-----------------------------------MGPGSAR
>Orycteropus_afer_afer
-----------------------------------MGPGSAC
```

HG0295.2 ENSGALG00000003282 STAT5B

>Gallus_gallus
-----------------------------------------------------------MRAGVPSLCP-----------------------QLRCAGGGRKQPEHSAARLRRQLTFWKEIAALSSFL
>Odobenus_rosmarus_divergens
------------------------------------------------------------------------------------------------------------MVYRRQLTFEEEIAVLCSFL
>Orcinus_orca
------------------------------------------------------------------------------------------------------------MVYQRQLTFEEEIAALGSFL
>Pteropus_alecto
------------------------------------------------------------------------------------------------------------MVYQRQLTFEEEIAALCSFL
>Elephantulus_edwardii
------------------------------------------------------------------------------------------------------------MVYRRQLTFEEEIAALCSFL
>Ceratotherium_simum_simum
MRKLRPWFGISRARDFPALPPPSSGHSCA--S-----GPPQSRAAGRWDSRGSPPSGQSPRPAPPSLATPGPAAAAAAHSPGSQRPRAEAGRERQRLGGVLAKARALCMVYRRQLTFEEEIAALCSFL
>Chlorocebus_sabaeus
MRKLRPRFGISRARDFPAPPTNHIPPAIL--TRPGSGTLPRSRSRGPAGTPGGPRRRGSPHARRRPALPPAGTAAAVAHSQGSRRPRAEAGRERAGLGGVLARARALCMVYRRQLTFEEEIAALRSFL
>Cricetulus_griseus
------------------------------------------------------------------------------------------------------------MVYRRQLTFEEEIAALCSFL
>Trichechus_manatus_latirostr
MRKLRPWFGISRARGFPTLPLSPQRPLQRLLPATISGRFRGAAGKGWLGLPGDPAVGAAPRAAPPRLAARGPATAAAGHSPGSQQPRAEAGRELGSREGELGRARALCMVYRRQLTFEEEIAALRSFL

HG0296 ENSGALG00000006912 CPEB3

>Gallus_gallus
MQIARGRRWRRRRVLGKTGTRISFSLQGESSEFRCKRKETWRNCCISLPYKHISHLD
>Mustela_putorius_furo
MQITG-GGW----CLGKNRNQNQFLIAG-----------------------------
>Rousettus_aegyptiacus
MQINR-GGW----CLGKNRNQNQFLIAG-----------------------------
>Monodelphis_domestica
MQITG-GGW----CLEKTGTRISFSLQGERSEFR-----------------------
>Papio_anubis
MQITG-GGW----CLGKNRNQNQFLIAGRTL--------------------------
>Mesocricetus_auratus
MQITG-GGW----CLGKNRNQNQFLIAG-----------------------------

HG0297 ENSGALG00000031684

>Gallus_gallus
MCFCFVLQRFFWRLIFTF
>Lipotes_vexillifer
MCFCFVL-----------
>Rousettus_aegyptiacus
MCFCFVL-----------
>Oryctolagus_cuniculus
MCFCFVL-----------
>Elephantulus_edwardii
MCFCFVL-----------
>Cebus_capucinus_imitator
MCFCFVL-----------
>Jaculus_jaculus
MCFCFVL-----------

HG0298 ENSGALG00000042308

>Gallus_gallus
MESIVWECG
>Acinonyx_jubatus
MESIVWECG
>Myotis_davidii
MESVVWECG
>Cercocebus_atys
MESVVWECG

HG0299 ENSGALG00000040465 Sip1

```
>Gallus_gallus
-------------------------------------MLGKEKEKNVIRRGVTRQSVPRFVFPGVAERDQS---
>Notothenia_coriiceps
MLTFGSP-----QLSPPVKIGGVRFR--GLLLCVTRQRKGGGKKKNVIRRGVTRQSLPRFVSWSGCKTSVVNLP
>Platichthys_stellatus
MLTFGSP-----QLSPPVKIGGVRFR--GLLLCVTWQRKGGGKKKNVIRRGVTRQSLPRFVSWSGCKTSVVNLP
>Takifugu_rubripes
MFNPEDAHVWLSSIIPSREDRWRAFQGASPPLCST-AKKRRWKEKNVIRRGVTRQSLPRFVSWSGCKTSVVNLP
```

HG0300 FBgn0031668 CG31917

>Drosophila_melanogaster
------------------------MVNVMKGVLVECDPAMKQFLLHLDEKLALGRKFIIQDLDENHLFISTDIVEVLQARVDDLMDRISFPLHDKDA---------
>Eulimnogammarus_cruentus
------------------------MVNVIKGILIKCDPVMKQLLLHLDETLALGSKFVLANLDDCHLFVAQDVYEQLKVRVDELYEKVYNNYSASNPKSFE-----
>Latrodectus_tredecimguttatus
------------------------MVYVMKGILVECDPAMKQFLLHLDETNAFKKKFIIEDLDETHLFIAPDMLEILQTRIDDLMDQISFPLAEK-----------
>Machilis_hrabei
------------------------MVNVKKGILVECDQAMKQFLLHLDETLGLGQKFIVQDLDETHVFIAPEILETLQSRIDDLMDQISFPLADKGM---------
>Argulus_siamensis
------------------------MVNFLKGILVECDPAMKQFLLHLDETFALGKKFIIQDLDDNHLFISSDIIETLKARIDDLMDQISFPLAEKGA---------
>Mastotermes_darwiniensis
---------MKYIQYL-RRKSTEIMVNVMKGVLIECDQAMKQFLLHLDETMALGRKFIIQDLDETHLFISADILETLQSRVDDLMDQISFPLADRGL---------
>Sitophilus_oryzae
------------------------MVNVMKGVLVRCDNAMKQFLLHLDETLKLGRKFIIQDLDENHLFISADILDTLQAKIDDLMDQISFPLADKGN---------
>Eriocheir_sinensis
------------------------MVNVMKGILIKCDPVMKQFLLHLDETMKLGSKFVLVDLDECHLFVQPEVYNSLKNLIDEQYEKVYYNQKSSLE---------
>Forficula_auricularia
------------------------MVNVMKGILIECDHAMKQFLLHLDETSALGKRFIIQDLDETHLFISSDILETLQARVDDLMDKISFPLADRGL---------
>Daphnia_magna
---------------------------------------MKQLLLHLDETHALGNKFIVQDLDETHLFIAPEYLEVLQTRIDDLMDRISFPLAERETD--------
>Occasjapyx_japonicus
------------------------MVNVMKGILVECDPAMKQFLLHLDETSALGRKFIIQDLDETHIFISADIIEVLQTQIDDLMDQISFPLADKNN---------
>Haploembia_palaui
------------------------MVHVTKGILIECDHAMKQFLLHLDETLALGKKFILQDLDETHLFISADILTTLQARIDDLMDQISFPLADKGF---------
>Eurylophella_sp._AD-2013
------------------------MVNVTKGSLITCDPAMKQFLLHLDDTLALGKKFIIQDLDETHLFVSSDIVEILRNRVDDLMDQISFTDKQQQ----------
>Meganyctiphanes_norvegica
------------------------MVNVMKGLLIKCDPVMKQLILHLDETLGLGSKFVLVDLDECHLFVQSEVYKTLKTRIDELYEKVYYNKSTSDSSWQASRYYE
>Grylloblatta_bifratrilecta
------------------------MVNVMKGILIECDQAMKQFLLHLDETTALGRKFIIQDLDETHLFISADILETLQARVDDLMDQISFPLADKGL---------
>Cuerna_arida
------------------------MVNVMKGILVECDHAMKQFLLHLDETVALGRKFIIQDLDETHLFISADILDVLQARVDDLMDQISFPLAEKGL---------
>Copidosoma_floridanum
------------------------MVNVMKGVLVECDPAMKQFLLHLDETLALGRKFILQDLDETHLFISADIVETLQGRVDDLMDQLSIPVHEKGL---------
>Asellus_aquaticus
------------------------MVQVQKGILIKCDPVMKQFLLHLDEKQTLGQRFVLVDLDDCHLFVQSDIYNSLKVRIDELYEKIYYNISGSNQQKNFE----
>Antheraea_assama
------------------------MVNVMKGVLVECDPAMKQFLLHLDETLALGRKFILQDLDETHLFISADIVETLQARVDDLMDQLSIPVHDKGL---------
>Mantis_religiosa
------------------------MVNVMKGVLVECDQAMKQFLLHLDDKLALGRKFIIQDLDDTHLFISADILDALKARVDDLMDQISFPLADKGF---------
>Tanzaniophasma_sp._AD-2013
---------------------------------------MKQFLLHLDDTTALGRKFILQDLDETHLFIAADILETLQARVDDLMDQISFPLADKGL---------
>Boreus_hyemalis
------------------------MVNVMKGVLVECDQAMKQFLLHLDETLALGRKFIIQDLDETHVFISSDIIETLQARVDDLMDQISFPLHDKGL---------
>Corydalinae_sp._KMRSPBM-2012
------------------------MVNVMKGILVQCDQPMKQFLLHLDETLKLGRRFIIQDLDETHLFISADVIEILQAKIDDLMDQISFPLADKGN---------
>Cypridininae_sp._BMR-2011
------------------------MVNALKGILVECDPAMKQFLLHLDEKEILGRRFIIQDLDETKLFISSDILDTLQAKIDDLMDKISFPITET-----------
>Osmylus_fulvicephalus
------------------------MVNVMKGVLVQCDQAMKQFLLHLDETLKLGRRFIIHDLDETHLFISADVIETLQAKIDDLMDQISFPLSEKGY---------
>Triops_newberryi
------------------------MVNVVKGILVECDPAMKQFLLHLDETNALGRKFILQDLDDTHVFISTEILEVLQARIDDLMDQISFPLHEKDP---------
>Epiophlebia_superstes
------------------------MVNVMKGILVECDHAMKQFLLHLDDTVALGRKFIIQDLDETHLFISADILGVLRARIDDLMDQISFPLADKGY---------
>Platynothrus_peltifer
---------------------------------------MKQFLLHLDETNALGTRFIIQDLDANHLFISSDILDKLKNSIDDLMDQISFVDVN------------
>Prosarthria_teretrirostris
------------------------MVNVMKGVLVECDQAMKQFLLHLDETLALGKKFIIQDLDETHLFISADIVETLQARVDDLMDQISFPLADRGL---------
>Timema_cristinae
------------------------MVNVMKGILVECDQAMKQFLLHLDETIALGRKFIIQDLDETHLFISADIVETLQARVDDLMDQISFPLSDKGL---------
>Menopon_gallinae
----------------------------MKGILVECDQAMKQFLLHLDETLALGRKFIIQDLDETHLFVSADVLNILQTKVDDLIDEISAPITEK-----------
>Leuctra_sp._AD-2013
------------------------MVNVLKGVLIGCDQAMKQFLLHLDETNALGRKFIIQDLDETHLFISADILGTLQAKIDDLMDQISFPLADKGL---------
>Polydesmus_angustus
------------------------MVNVMKGVLIECDPAMKQFLVYLDDTNALKRKFIIRDLDDTHLFVSVDIIPILQTKIDDLMDQISFPLAEK-----------
>Acerentomon_sp._AD-2013
------------------------MVNVVKGVLVECDPAMKQFLLQLDETNALNRKFIIQDLDETHLFISSDILETLQEKIDDLMDQISFPLSEK-----------
>Liposcelis_bostrychophila
------------------------MVNVTKGILVECDPAMKQFLLHLDETSALGGKFIIQDLDDTHLFIAAEYLSILRSKVDDLMDQISAPITDK-----------
>Inocellia_crassicornis
------------------------MVNVMKGVLIECDQAMKQFLLHLDESLKLGRRFIIQDLDDTHLFVSADVIEILQTKIDDLMDQISFPIAEKAQ---------
>Ceratophyllus_gallinae
------------------------MVNLKKGILIECDQAMKQFLLHLDETQALGRKFIIEDLDETHVFISADILETLQARIDDLMDQISFPLHEKGL---------
>Archispirostreptus_gigas
------------------------MVNVMKGVLIECDPAMKQFLLHLDETNALGKHFIIRDLDETHLFITADSLETLQGKIDDLMDQISFPLSEKGI---------
>Stylops_melittae
MSLKLVKYRQKFFDNIRPHRNILLYVNVVKGVLIKCDPAIKQFLIVLDETLKLGRRFIVQDLDETHLFVSSDIVDTLQTKLDDILDQINFTVVEKS----------
>Thrips_palmi
------------------------MVNVVKGVLVECDQAMKQFLLHLDETMALGCRFIIQNLDDTHLFISADILTTLQSRIDDLMDQVSFPLVEKDA---------
>Platycentropus_radiatus
------------------------MVNVMKGILVECDQAMKQFLLHLDETLALGRKFILQDLDETHVFISTDIVETLQARIDDLMDQINFPLHEKGQ---------
>Zorotypus_gurneyi
------------------------MVNVMKGILVECDHAMKQFLLHLDETIALGRKFIIQDLDETHLFISADILETLQARVDDLMDQISFPLAEK-----------
>Atelura_formicaria
------------------------MVNVTKGVLVSCDRAMKQFLLHLDETLALGRKFIVHDLDETHLFISADILETLQARVDDLMDQISFPLADKY----------

HG0301 FBgn0036856 CG9666

>Drosophila_melanogaster
----------------------------------MAFP---TTSAQQA-ETNRKILEEIQTKKQLLAGGIINLGLSPPNQMPA--PQLLGQPTTVNPDFQAGV--------GIATNATSTARSAFNPTSSTTLGFFIPQDSYFGNSFIPVLPRLEPLPSPATTPTTPNAP
PSHSISK
>Machilis_hrabei
----------------------------MTLDFDMAFQ--PANNKQEM-Q-NRKILQELQLKKQLLLKEGAAPTSPL---STP--PLS---PN-----PQGFPLLP-----SEAHMLPSSQRTALQCAHSQSPGYFITQDSSFGNIILPVLPRFEQNSK-----------
-------
>Periplaneta_americana
----------------------------------MAFQ--PPSARQEL-A-NRKILEEIQLKKQQILKQGVAPSLNSPTL-TP--SLSAVGGG-----VNLPVSMS-----PEAHMMSSSQRAALQHAHSQSFGYFVTQDSSFGNLILPVLPRFDIK-------------
-------
>Pogonus_chalceus
----------------------------------MAFT--QTNAQREI-T-SRKILEELHLKKQMLLKQGAVTSLNNTPAP--------ILPLVSPQTGTIPFTGAPAAALLGEAQTTSAQRTALHSAHSQSFGFFITQDSSFGNLILPVIPRFDNK-------------
-------
>Procambarus_clarkii
----------------------------------------------------------------MILKQGVSQALNAPATSQQ--PT-G------------------WSSGTDGHIAGNSQRSNLLQANSSSFGYFIPQDSSFGNFILPVLPRLEPK-------------
-------
>Apachyus_charteceus
----------------------------------MAFQ--PPSARQEI-A-NRKILEELQLKKQILLKQGATSTLNTQLTPNV--PVSGTTPSFVSCGLNRGPLVSASSPMSSEPYMTAAQRTA----LQQSYGYFVTQDSLYGNSILPVLPRFESK-------------
-------
>Campodea_augens
----------------------------------MAFQ--PVNNRQDL-Q-NRKILEELQMKKQMLKQGAAPVSLNSTTSPSG--M------GTVTAANSAFSHL-----PPEMQMLQHNQRNALQLANSQSFGHFVTQDSAFGNTLLPVLPRLETPKP-----------
-------
>Haploembia_palaui
----------------------------------MAYQ--PPTARQDV-ATSRKILEELQLKKQLLLKQGVQSTLTSTSLSSL--SMPAT------------DTAN-----ITANMMSSAQRAALNHAHSTSFGYFISQDSSFGNIILPVLPRHESK-------------
-------
>Isonychia_bicolor
----------------------------------MAFQ--PPSNRQEL-Q-NKKILEELHLKKQLLLKQGAVTALNSSNFQS----------S-----PSVPTYGS-----SDTHMMSNNQRTQLQIANTQSYGYFVPQESSFGNFILPVLPRFDTPPSK----------
-------
>Meganyctiphanes_norvegica
----------------------------------MASQ-APGRGAADS-NTQRKILEDLQLKKQMILKQGASQALNAPAPQQT--AT-G------------------WSSGTDGHIAGNSQRQV-QQTTNQSFGYFVTQDSSFGNFILPVLPRLEPTK------------
-------
>Galloisiana_yuasai
----------------------------------MALQ--PTNVRQEL-A-NRKILEELQLKKQLLLKQGAAPSLINTPTITT--SLSMT--A-----GTGPLSMS-----PEAHMLSSNQRAALQHAHAQSFGFFVSQDSSFGNLIIPVLPRFDVAK------------
-------
>Tigriopus_californicus
MAANPSHHTPNSTSGGASGPAVSAGTAAAATGGSLAFG---NPSSGKI-DQQRKILEELERQKKQLAKGNLGPSAANASPAPNHTPNSGG--------PASGTLSKAMEQGDVPSILTQAQKAAMELANKSSFGYFIPQDSAFGNSILPVVPRLQPK-------------
-------
>Clastoptera_arizonana
----------------------------------MDFR--QSQSAREL-A-NRKILEELALKKQMLLKQGAQPLGST---------P--ISPMPV-----TPATNIPANIAPDVHVMSSTTRAALAHANSQSFGFFVPQDSSFGNLILPVLPRLEK--------------
-------
>Leptopilina_clavipes
----------------------------------MAFS--QTN-SREL-Q-NRKILEELQLKKQMLLKQGVTTNLGSSLTSTAGS-----------------ATTVPATQTSDGVTISASQRAALHNAHSNSYGFFVTQDSSFGNLILPVLPRFDNK-------------
-------
>Bombyx_mori
----------------------------------MAFP--QPNAQREL-T-NRKILEDLQLKKQMLLKQGAVAPLTAA---LSQPNP--VSQLPMCSIPPV-SATAGFPQLPEANVINTSHRAALQHANATSCGYFVSQDSSFGNQILPVLPRFENK-------------
-------
>Mantis_religiosa
----------------------------------MAFQ--PPSARQEI-A-NRKILEEIQLKKQLLKQGVVTQNLS-----TP--IAPLVGAG-----VNVPMTVS-----PEAHMLSSSQRAAFQHALLQSFGYFIPQDSSFGNLILPVLPRFDK--------------
-------
>Tanzaniophasma_sp._AD-2013
----------------------------------MAFQ--P-ASRQEI-A-NRKILEELQLKKQLLLKQGVTPTLNTPAL-SQ--SIPGVSPS-----PILPTSLP-----SDAHMMTSSQRVALQHAHQMSFGYFVTQDSSFGNLILPVLPRFDTK-------------
-------
>Nannochorista_philpotti
----------------------------------MAFP--Q-TNTQD----HRKILEELQAKKQQLLKGGAATLTTTPQLPN-------------------ILTTPVANQTIDTHAINSTQRNALNNANSTSFGYFVPTDSSFGNPILPVLPRFDPKV------------
-------
>Corydalus_cornutus
----------------------------------MAFP--QPNAQRDL-A-ARKILEELQLKKQLLLKQGVQATTTANSVTT----N--LSTLPTTTGAPNSTGGP---LSTDSHSMSSAQRAALHAAHTQSFGYFVTQDSSFGNLILPVLPRYDTK-------------
-------
>Euroleon_nostras
----------------------------------MAFP--QPN-QREL-A-NRKILEELQLKKQILLKQGVQATANTNNLTT----P--LGPSVITSGTVINQTGP---GTPQEHNLTSAQRAALHSAHTQSFGYFIPQDSAFGNLILPVLPRFDVK-------------
-------
>Triops_newberryi
----------------------------------MSFQ--PAPGKQEL-A-SRKILEEIQMQKKLLKQGGPLPTPTTVNP-VP--LASQ-----------TGQVSAPTANPELQSLNPTQRAALQHAHAHSFGYYVTQESAYGNLILPVIPRLELVAPSK----------
-------
>Calopteryx_splendens
----------------------------------MAFQ--PPSSRQEI-Q-NRKILEELQLKKQLLLKQGAASALNTTPIPPP--PMPV---G-----NTMPL-PT-----GEQHLVSSTQRSALQFAHTHSFGFFVTQDSLFGNLILPVLPRFDMK-------------
-------
>Ceuthophilus_sp._AD-2013
----------------------------------MAFQ--PPSARQEL-Q-NRKILEELQLKKQILLKQGVAPTLSI----SP--SLSV---G-----GSNAVAMT-----PEAHMMSSSQRAALQHAHTQSFGFFVSQDSLFGNLILPVLPRFDK--------------
-------
>Peruphasma_schultei
----------------------------------MAYQ--PPSVRQEM-A-TRKILEELQLKKQMLLKQGVAPTLSSQNLSA----------------IPMSGGLT-----PDAHMMSSAQRTALQNANSQSCGYFITQDSSFGNLILPVLPRFDSK-------------
-------
>Menopon_gallinae
----------------------------------MAFQ---PSTRPDIQNPQKKILEELNLRKQLLKIG-TAQTLNPSALNTP---I--------------GVPLINSNPVSTDNQMSTTARAALQHANSTSFGYFVPQESSFGNNIIAVLPRFNESK------------
-------
>Perla_marginata
----------------------------------MAFQ--APNLRQEL-A-NRKILEELQLKKQMLLKQGVTQATLNTSIPSS--IPIG---M-----PASGLGVP-----AELHLVSPAQRQALQNANSQSFGVFVCQDSSFGNLILPVLPRFEVKPSL----------
-------
>Acerentomon_sp._AD-2013
----------------------------------MAFQQTTPNSRPDL-QQSKKILEELQMKKQQLLKGVNSTNQSSCP------PA-----------STAPISG-----TPDVHMMSPSQRSALQNANATSYGFFIAQDSSFGNLILPVLPRYDNK-------------
-------
>Ectopsocus_briggsi
----------------------------------MAFQ---SNSAGDFPNPQRKILQELNLKKQLLKSG-TAQTLNPSALNAP---I--------------GLPSLNP-PVAPENQMSNIARSALNSANTTSFGYFVAQESSFGNNILAVLPRFPEAK------------
-------
>Xanthostigma_xanthostigma
----------------------------------MAFP--QPNAQREL-A-NRKILEELQLKKQLLLKQGVQASTTTVASNIS---P--IGASTLPAPGPTVQSGA---PSMDTHNLSSAQRAALHSAHTQSFGFFIAQDSSFGNLILPVLPRFDTK-------------
-------
>Ceratophyllus_gallinae
----------------------------------MALP--PTNAQQD----HRKILEELQQKKQQLLKGGAPTLNTTLPTSN-------------------IPNPQGIIPGNDMHMINSTQRIALRNANSTSFGYFVPTDSSFGNQILPVLPRIDAE-------------
-------
>Gynaikothrips_ficorum
----------------------------------MSFQ--PTN-SRKD-P-NRKILEELQQKKQMLLKSGAGVVSPPSVP-----------------------------NLASEPQANKVQCMVMGAAIAQSFAYYIPQESNFGNPILPVIPRFDKN-------------
-------
>Annulipalpia_sp._AD-2013
----------------------------------MAFN--QPNAQREL-T-NRKILEELQLKKQILLKQGPAPNLTGSSVSLTSAPT--PAQLPVSTIPPI-SATTILPQISDTHLINTSHRAALNNANATSCGFFVPQDSSFGNQILPVLPRFDSK-------------
-------
>Zorotypus_gurneyi
----------------------------------MAFQ--PPSSRQEQ-A-NRKLLEELQMKKQLLLKQGVSQTLTVPSITP-------LSVS-----TPVISGLT-----PEAHMMSPAQRVALQHAHSQSFGYFITQDSSFGNQILPVLPRFDSK-------------
-------
>Thermobia_domestica
MFRLLNHLRIERW-NG--------KENLGIRVCMMAFQ--APNARQEI-Q-NRKILEEIQMKKQLLLKQGAAPNLNSPPNLTP--TN--IPVT-----GPMPMYMS-----PEAHMMSSSQRTALQHAQCQSFGYFITQDSSFGNLILPVLPRFDNK-------------
-------

HG0302 FBgn0031454 CG9960

>Drosophila_melanogaster
-------MDSDSTVTSLEENTENFC----TNPTRDILAEGITNLFKPTIERLDERVASTIQLQAELRGQLDALAAQLRDIEKAQ-SQIPEFADKVKELLNVKHKVTVISNVLVTSQERLTGLHKLIEKEQRRRQALLDSALSTNIS----------------------
>Oxyacanthus_flavus
------MADSDSVSTSVGDQTEDLV---SP--TRDALVDGIMGMLKPCLDQLDDRVRQTRASQSELQQQLEGLQQQLTSVQQKT-QCPLHLDQYIKKLQNAKRRVVVVNNILQTTQERLNKLNNQMIKENTRRQALLDPSMTAS------------------------
>Machilis_hrabei
-----MEVDTESTNTSLDDKTEDFG----ENPTRDTLAEGIMCVIKPTIDQLDERVKVTRISQVELKQQIEALAEELKKIADAQ-QCPLDLDLYVKKLLNAKRRVTVVSNVLQNAQDRLNKIHQLVAREGSRRRALLEPSPVTTPGSGP-GT---------SRQVNPI
>Cryptocercus_wrighti
-----MDLDTESTTTSIDDKTEDFC----ENPTRDTLAEGLMCLLKPTIDQLDERVRATRISQVELKQQIEGLAEELKKISESQ-QCPLDLDLYVRKLINAKLKVTVVSNILQTAQDRLNKVHQQIAREQSRRRALLEPSPITTPLEAT-NA---------SIQ----
>Pogonus_chalceus
-----MDVDSESTATSIDENTENFC----DNPTRDTLAEGLMGLIKPTVDQLDERVRATRISQLELKQCIESLSEELKKVSETQ-QTVVDLDAYVKKLINAKHRVTVLSNVLQNAQDRLNKVHQSIERETIKRKAFLESTPGEATTSTTSSVI---------------
>Astacus_astacus
------MADTESSCTSVDDRTDDLV---SP--TRDALVDGIVGLLKPCLDQLDDRVRATRVSQSELHQQLEGLAAELSRVQEQT-RCPLQLDAYIKKLQNAKRRVVVVNNILQNTQDRLNKLHSQVNKENARRRALLETTSPLPITPSTSLHA-P-------------
>Occasjapyx_japonicus
-----MEVDTASTTTSLEDKTEDFGFGDGDNPTRDSLAEGLVALLKPTLDQLDERVKNTKISQVELRQQIEALTEELRGIAGEQ-QCPIDLDHYVKKLINARRRVTVVSTVLQTTQERLNKLHQLVQRETGRRRPLLEPSPGAPPSASALPFQ-P-------------
>Baetis_sp._AD-2013
-----------------------------------------MCLLKPTIDSLDERVRATRISQVELRQQIDGLADEIKKISDEQ-QCPLDLESYVKKLLNAKRRVTVVSNILQVAQERLNKVHQLVERESARRKALTEPSPVISPSAHP-IE---------TNPSIQ-
>Meganyctiphanes_norvegica
MIQSFNMADTESTCTSVDDRTEDLV---SP--TRDALVDGIVSLLKPCLDQLDERVRTTRLSQSELHQQLEGLAVELSRVKDQV-ECPLQLDSYIKKLQNAKRRVVVVNNILQNTQDRLNKLHSQVNKETTRRRALLETTST-ASTPSTSLHA-P-------------
>Galloisiana_yuasai
-----MDLDTESTTTSIDDKTEDFG----ENPTRDTLAEGLICLLKPTIDQLDERVRATRISQVELKQKIEGLSDELKKISGEQ-QCPLDLDLYVKKLANAKQKVTVVSNVLQAVQERLNKVLIQIDKDEARRRALLEPSPVISPLDVE-ST---------STQ----
>Clastoptera_arizonana
-----MDVETGSTNTSLDDKTEDFC----ENPTRDTLAEGLMCMLKPTIDQLDERVKVTRISQLELKQQIEGLSEELKKISAEQ-SSPLDWEAYANKLTNAKHKITVVSNILQAAQERLYKVSQQINKETARRQALLKQSSPTSSTDSKPLS---------SVQ----
>Ganaspis_sp._G1
-----MDLDTASDNTSLDDKTEDFC----DNPTRDALTEGLMSLLKPTVDQLDERIRSTRISQIELKQQIDSLNEELVKISESL-QQPLELDSYLKKLVNAKQKVTIVSNILQTTQDRLNKIHQAIERESAKKRALLDHSSSYNDDETQKSAK-ERETGHSSVDVQQ-
>Asellus_aquaticus
------MADSDSTCTSVNDRTEDLI---SPKPTRDALVDGIIGLLKPCLDQLDEKVRSTRLSQSELQQQLETLSTQLNRVQDQV-QCPLQLDSYIKKLQNAKRRVVIVNNVLQSTQDRLNKLHGQVGKETSRRKALLESQPSSSTALI--------------------
>Triodia_sylvina
-----MDADTESTTTSAEENTGNLC----DNPTRDTLAEGILGLLKPTVDQLDERVRATRISQVELKQQIDTLTEELKKINQAL-SNHPDIESYVKKLITCKHKVTVVLNVLQTSQERLNEIRRMISLEEGVRRAVLPALEAQSEPEPSTSATLISPEGS--IN----
>Mantis_religiosa
-----------------------------------------MCLLKPTIDQLDERVRATRISQVELRQQIEGLAEELKKISEAQ-QCPLDLDVYVRKLINAKLKVTVVSNILQTAQERLNKIHQQIARETSRRKALMEPSPITTPLEAG-NP---------SIQ----
>Tanzaniophasma_sp._AD-2013
-----------------------------------------MCLLKPTIDQLDERVRATRISQVELKQQIEGLSEELRKISDGQ-QCPLDLDVYVKKLLNAKQKVTVVSNILQTAQERLNKVHIQIAKETARRRALLEPSPISTPLEVS-NT---------TVQ----
>Nannochorista_philpotti
-----MDADSESTATSLDENTENFC----DNPTRDTLSEGLMGILKPTIDQLDERVRATRVSQLELKQQIDRVSEDVRKILDNK-KPPAELDDYVKKLVNTKLKVTVITNILQSVQERLNKIHSQIDRETARREAQLSSQTEPYDSEVASGSN-Q-------LQNNLN
>Corydalus_cornutus
------MDDTESTATSIDENTENFC----ENPTRDTLAEGLMGLLKPTIDQLDERVRAMRISQSELKQQIEILHEELRKIAELQ-QYDLDLDVYVKKLVTAKHKVTIVLNILQCAQDRLNKIHAGIEKETIRRKALLEKLPTPVQSPPPVPQTSSDDLGH--FGG---
>Chrysopa_pallens
------MEDTESTATSIDENTENSC----DNPIRDTLAEGLMGLLKPTIVQLNERANAVRISQSELKQQIETLQEEIKKISELQ-QSDVDIDSYVKKLMTAKHKVVAVSTVLQNAQDRLNKIHSGIEKETNRRKIVLEKSNTTLQEPSSSSIS---------------
>Triops_newberryi
-----MDSEVISNDTSLDDKIEDFG----DYPTRDILAEGIVNLLRPTVEGLDQSIRDTKQSQAELRTQIEALTSALNVVAENH-SCPIDLEPYIKKLLDSKRRITVISSVLQTSQDRLVKVHHLAQKDIARRKSSLDTQASVPAP----------------------
>Calopteryx_splendens
-----MEQDNDNKVSASVSKTNSFG----ENPTGDALAEGLICLFKPTIDLLDERITATRNSQLELKDKIEILAEELRKISELH-KCPLDLELYVKKLAGAKRKVTVVSNVLQNTQERLSKVQVQVAREAQRRRALLEPSPAMSPVSVESNA---------SLQ----
>Prosarthria_teretrirostris
-----MELDTESTTTSIDDKTEDFC----ENPTRDTLAEGLMCLLKPTVDQLDERVRATRITQVELKQQIEGLAEELRKLSATQ-QCPLDLDLYVKKLMNAKQKVTVVSNILQTAQDRLNKVHLLIEKESSRRRTLLEPSPSTTPLEAT-NP---------SIQ----
>Medauroidea_extradentata
-----------------------------------------MCLLKPTIDQLDDRVRAVRISQVELKQQIDGLSEELKKIAESQ-SCPLELDAYVRKLMNAKLKVTVVSNILQTAQERLNKVHANIAKEAAKRKALLEPSQKTTPLEVS-QP---------SIQ----
>Menopon_gallinae
-----MEVDTESTNTSVEDKTEDFC----DNPTRDALTEGLMRLFKPTIDQLDEKVRTIRISQVELKQQIESLSDELKKINESSTDKEIDLDVYVKKLMTVKHKVTVVSNILQNSEERLLKLQQNISKDSTSRKALLESTSVTQVSVTEPDD--------SSVQ----
>Leuctra_sp._AD-2013
-----MDLDDASTTTSIDDKTEDFC----ENPIRDTLTEGLMCLLKPTIDQLDERVRATRISQVELKQQIEGLSDELRKISDAQ-QCPLNLDLYVRKLTNAKLKVTVVSNILQTAQERLNKVHQMIAREAAKRRSLLEPSPISSPLETV-NP---------SIQ----
>Acerentomon_sp._AD-2013
-----MEIDTDSRNTSLDDKTEDYD----DNPTRDALVDGMISLLKPAIDQIDERVKATRLSQLELGEQIGSLAEEIKRVSEEQ-NCPLQLDQYVQKLQNAKKRITVVSNVLQSVQDRLNQLHNIATQELTRQQSLTEVEAPGQSSEDTQ------------------
>Liposcelis_bostrychophila
-----------------------------------------MRLFKPTIDQLDDKVRTIRISQVELKQQIESLSEELKRISESNVDKELDLDVYVKKLMNVKHKVTVVSNILQNSEERLLKLQQNISKDSTSHKALLETTETSQATQAEPDS--------SAIQ----
>Inocellia_crassicornis
------MDDTESTATSIDENTENFC----DNPTRDTLAEGLMGLLKPTVDQLDERVRATRISQCELKQQIEGLSEELKKISEAQ-QTNLELDAYVKKLINSKQKVILVSNILQAAQDRLNKIHLAIDKESAKRKALLEKTPAGSTSTLETSQTNR-------------
>Oropsylla_silantiewi
-----MEPDSESTNTSLDENTENFC----DYPTRDTLSEGLMGILKPTVDQLEERVQATKASQIALKSQIDEVSNEISKILSNR-KPPLELDEYLKKLVNTKNKITVVTNILQATQERLNRIHISIEKETQRRESLLATNISLPS---TSGSQ-----------INL-
>Gynaikothrips_ficorum
-----MEGDTASTNTSNDDKTDDYC----DNPTRDTLADGLMCLLKPTVDQLDERVRATRISQNEVKQHLDNLSEELKKISDAQ-QCPIDLDLYVQKLNNAKQKITIVSTFLNSAQDRLSKLSQQISREQSRRKALIEPSQSTLFDASS-GS---------VI-----
>Platycentropus_radiatus
-----MDGDTESTATSVEENTENFC----DNPTRDTLVEGLMGLLKPTVDKLDESVRATRISQVELKQQIDGLADELKKISDSQ-SNFLDLEEYVKKLMNSKRKITAISTILQSTQERLDKVHHLVSREAARKQALLESSEQSASTSGFNDV--SQ-EGN--IN----
>Atelura_formicaria
-----MDVDTESTNTSIDDKTEDFG----ENPTRDTLAEGFMCLLKPTIDQLDERVRATRISQVELRQQIDGLAEELRKISEAQ-HCPLDLDVYVKKLLDAKRRVTVVSNILQNAQDRLNKVHQLVARESVRRRALMEPSPVTTPSTAL-PE---------AKSSQQ-

HG0303 FBgn0260464

```
>Drosophila_melanogaster
MHTDL-SSHLHTPACNKLIEELQACHEN-NAFAKFVGVCNSIDDKVVKCLKGERIARSAANRAKARERQAKYKEKLLQQESN---------
>Hyalellopsis_costata
MHPDL-SPHLHTPECNEIIQKLFNCHKE-HSLLKFLGTCNDLDHQVWTCLKAERVERRALNRFKAEQRYQQLKEHMKSS------------
>Machilis_hrabei
MHPDL-STHLHTDECNQLIQLLKDCHDR-NIFNKFLGVCNGEDNAVVKCLRKEREENRRKNLEKSREKQRKISQARAAQQ-----------
>Mastotermes_darwiniensis
MHTDL-STHLHTEECNKLIELLKDCHKE-HSFGKFLGYCNDADREVIKCLRKEREQNRKANQQKAQLMQDKVRTSYPE-------------
>Gyrinus_marinus
MHTDL-SPHLHSKECNELIKQLQTCHTE-FPIGKFFGYCNSVDAAMVRCLKAERIERSKLNREKARERQEQIKRIL---------------
>Astacus_leptodactylus
MHPDL-SPHLHSDECNELISKLHACHKE-NKFLKFVGICNDIDSAVWKCMKAERLQRRAKNRALGEEQRKKVFAKLDEGYSWKKM------
>Forficula_auricularia
MHPDL-SPHLHGETCLKIIEAYKTCLIE-HPFGKFINYCGKLDAELNRCLKEERETRRRQNFEKSSYYKDMLKKGDQEK------------
>Daphnia_pulex
MHSNL-SPHLHTPECNQLIQLLLACHKE-NPVGRFVGFCNKQDTLMVACLRKEREANRSVNAEKSKIKQEEIRRRLKEQESI---------
>Occasjapyx_japonicus
MHPDL-SAHLHTPECNVLIEALKACHSE-HKFAKFVGYCNEADSAVLRCLRDERENRRKANFEKSAEKRRRLRNAADQARSE---------
>Isonychia_bicolor
MHPDL-SSHLHTEKCNLLITDLKACHSQ-NKFGRFLGFCNDLDKELVKCLREEREQNRKKNFDKSRAKQKAVLERL---------------
>Meganyctiphanes_norvegica
MHPDL-SPHLHLEECNILIAKLKDCHNE-NKFAKFMGACNDIDKAVNKCLKKEREIKRNANRAAGKEKREATLKKIEEGHSWKEL------
>Galloisiana_yuasai
MHPDL-SAHLHTTECNILITKLKECHEE-NKFRKFLGICNDIDTKLAACLRQERADNRKANQEKARQTQE----KLKRVALKAKSQNL---
>Cercopis_vulnerata
MHPDL-SSHLHTPECNELITQLLECHKN-HNFLRYFGHCTHLDNVMLKCLKRERLTRRAANAAKAVEKNKQVRERMRELYN----------
>Fopius_arisanus
MSDTYSRLPLHQGACNELYYQVQECEKSHHALSKIFGKCSDVHVALTKCLKQNKLNLAAANREQSRIRQENARKKLKAEKNSNKQ------
>Biston_suppressaria
MHTDL-SAHLHTDECNVLIKKLQDCHSE-HSFLKFVGYCNDFDYEMRKCLKGERLRRQKANLDESRKRHAEIRARIVAQEQAKH-------
>Mantis_religiosa
MHPDL-SPHLHTEECQNLINKFHECHKN-YPYKKFLGVCNDLDKAVNKCLKNERIERRRLNQLKARELQEKAKKSKPSA------------
>Tanzaniophasma_sp._AD-2013
MHPDL-SPNLHSAECNILVTKLRECHQE-HKFGRFLGLCNDADLELVRCLRKEREANRLNNPNWSRERQKNICESKASN------------
>Nannochorista_philpotti
MHPDL-SPHLHTEECNIIVQQLKECHNE-HQFRKFVGFCNSIDSKLIKCFKAERENNRRRNKELADTRRETTYRLMEAAKE----------
>Corydalus_cornutus
MHTDL-SPHLHTEECNYIINLLKACHTD-NPFKRFLGVCNDIDRDLIKCLRKEREQNRKLNFEKSREKQERLKQTQKLT------------
>Euroleon_nostras
MHPDL-SPHLHTPECNELIQQLHECHRD-HPLRKFVGYCNDIDRVLLKCLKRERQERSRLNRQKAFEKQERIKRLLSEQEQKS--------
>Triops_newberryi
MHSDL-SAHLHSPECNALIELLLKCHKE-NPVRKFTGYCNSQDTAVTVCLKKEREQNRKLNADKSAQRQKAIQERLRQAQQEKAQS-----
>Cordulegaster_boltonii
MHPDL-SPHLHTEECNILITKLKTCHSE-NRFAKFMGICNDIDRELLDCLRKERNDRRKANFEKSREKRQIVQEKLKQEKVK---------
>Ceuthophilus_sp._AD-2013
MHPDL-SPHLHTDECNVLIRKLHECHRV-NNIGKFFGYCNDVDNELVKCLRKERDIRRKSNLENSRARQEKYKDILKVQKEGQS-------
>Peruphasma_schultei
MSKNV-LKHNYSGNCTGLLEKLTECRED-NKYTKFFGVCNDAFKELDKCLQKEYEDRRTENHAKARALQERKLGKQSQQQT----------
>Perla_marginata
MHPDL-SPHLHTEECNVLIKKLKDCHEQ-NTFGKFLGFCNSADNELLKCLRREREIRRKANFDKAKEFQAKIRQPIPQV------------
>Acerentomon_sp._AD-2013
MHTNL-APHLHTDECNELIQKLFQCRLD-HPVLRFVGYCNDLDILMTKCLKRERLERRRISIEQSTKSRLERLELRRQERQKETESANQQA
>Inocellia_crassicornis
MHSDL-SPHLHSEDCNKVIAELHKCHAD-HPFKKFIGHCNDIDRALVKCLKKERLERTRINYEKSIIRQKEIRERMKNMS-----------
>Ceratophyllus_gallinae
MHSDL-STHLHSEECNAIVQRLTACRQE-NSFKKFLGACNDIDRELVKCFRKDREARRKKNKEASDLRYNKKLNI----------------
>Rhyacophila_fasciata
MHTDL-SPQLHTEECNKIITALKTCHEQ-HAFRKFIGFCNSIDDQMIKCLKEERLRRRAANIKASAEKHAAILKRIEQNTNA---------
>Zorotypus_gurneyi
MHTDL-SRHLHTEECNILIQKLQECHTG-HPFAKFIGFCNDFDSEMRRCLKKERLERKRINLEQAVARRERMNKAIHADVNKN--------
>Tricholepidion_gertschi
MHPDL-SAHLHTDECNILISKLKECHKE-YKFRRFVGACNDIDHELTKCLRAEREANRKVNLERARRQQEVVRQKMREV------------
```

>Drosophila_melanogaster
---------------MKIMISQVGRELYKVPLRILDHRVFVNGEIEAFLENFEVRRNDSEVEKLFQVTETVGSLKYDLSRCSATGRGSGAE-NLAQLDTEVSHLLDGVNALLAKA-KV-ERTASTQLQEARLAREQRRAEFLTNLEHGYRRIENSFEEKEEEIAELYSDL
QLKLNIAK-------
>Hyalella_azteca
--------------------MAITSEVEEVFSRLFDHRPFLKGEISFFKREFEEKRGDREVEELFRALELTTEIKQAQVEK---VVEASDA-NLPRTIADIQVALRMCHTSLDSD--SRTSRLSSEIERQREDRQQRLAVAKAEVEAKLASINAAYDLKEKELREKFAKL
DSSNTCDS-------
>Machilis_hrabei
----------------------------------------------------------------------------------------------------------MCGRIAEKE--D-SRRSVDTLEENRVSREHEWETFVTDMAEKCSKIDQTFGNKGDEMKEYYSEL
EKKLHLDQ-------
>Mastotermes_darwiniensis
-------------------MAAVVKDVGEIWTRLFDHRPFLSGEIKFFLKEFEEKRSDREVERLFQILERATEIKETQLDR---VKLASDV-HLPTLNANLEVAVGMCNRILEKE--E-EHQSDTALTARREIRKAEWESFIEDMTQKCSKVDATFQEKEEELKEFYSDL
EKKLHIGK-------
>Calanus_finmarchicus
---------------MSANNRSGFEDITEVHSRLFNHRPFLQGEVKHFVREFEEKRGDREVENIFSCLERVSELRDLGLPG---LLSSAGSNSSTSLVANLEVAESMLGRILEQG--E-TGEVEKTLVASREAREKEWTNFLKEVQIRCKKVDQSFKEQEDSIRKKYQDL
ESGS-----------
>Pogonus_chalceus
-------------------MSQLSKDIGEIWNRLFDHRAFLNGEINFVLKEFEQKRLDKEVEYLFRLLEQITDIKDTQIVR---LRQVGNT-SLPSINKNLDFALNACKRIEDIE--L-KYKEDTSLALAEEERKLRWEKFVDDMSHKCSRVDNTFDEKEEELHEFYTDL
ERKLQIAK-------
>Eucyclops_serrulatus
---------------------MVSKDVFGVYSRLFNHRPFLQGEARHFVKEFEEKRNDREVENIFNTLETVTELKDCQIDK---VKGQCEL-NLTTLSANLLVANSMCSKILDQE--R-TSDFDQALESSRQVREKEWSQFLSDIQNKCEKIDDAYKQKEKDLRLHYQKL
ENVFLQD--------
>Farfantepenaeus_aztecus
-------------------MASAASDVNEVFSRLFDHRPFLKGEIEYFKKEFEEKRGDREVEQLFRSLELITEIKEGQIEK---IVNSSDD-NLPRTIADIQVALHMLEDTIDTE--KKFNS-EELLAKKRAERKSKLTATQQEVQEKLNFLENNYLEKEQALRAQFESL
EKSAGF---------
>Forficula_auricularia
----------------------------------------------------------------------------------------------------------MCNRIVEKE--Q-LTRKAITLQKHRESRRAEWEKFIGDMTTTCSKVDQTFLEKEDELRHFYSDL
EDKLHIAN-------
>Occasjapyx_japonicus
----------------------------------------------------------------------------------------------------------MATRISACD--T-QFDSESILCERRVAREQEWERFLQDTSEKSVRVDNAFRDKELQLRGYYSLL
EDKMRP---------
>Baetis_sp._AD-2013
-------------------MSSIPKDVGEIWSRLFDHKPFVQGEVKFFLKEFEEKREDREVEKVFQLLEQVADIKDVQIEK---TIKSCGN-QLSILNSNLEVATGMCDSILSRE--E-ENSVDVALMVKRESRKAEWEAFIRDMGEKCARVDKTFKEREEEIREDYQEL
ERKLHLDKTPE----
>Meganyctiphanes_norvegica
-------------------MAAAVGDVNEVFSRLFDHRPFLKGEIEYFRKEFEEKRGDREVEQLFRCLELTTEIKEAQIDK---VVGSSDD-NLPRTIADIQVALHMCEDTIQVE--KKFAT-EELLSKKRTERQIRLQSAQADVSEKLALLQNTFQDKELELRRQFESL
EREAGYS--------
>Galloisiana_yuasai
----------------------------------------------------------------------------------------------------------MCNRILEKE--E-VYRSDTAIFAKREIRKTEWQNFIEGMTHKCVFVDQTFQEKEDQIAEFYSDL
EQKLHLEK-------
>Graphocephala_atropunctata
-------------------MNNIIRDCGEIWNRLFDHRPFLNGEILYFQEEFEVKRSDREITKLFEVLEHVTEIRGSQIEK---VKSLCET-NLPDYQTKLSALLSSCDAILDDE--N-KYDIDTNLEVNREIRKSDWERFQSEMQVKYSNVDETFNDKEKELLEFYSDL
EKKLHINNTTISRKE
>Solenopsis_invicta
-------------------MASITKDTGEIWSRLFDHRPFVQGEITFFLREFQEKRADREVERLFKILEYSTELKESQLDR---TEQLGDC-HLPSLKANVDVALSMCNLVLQKE--E-NFDSDSVLNENRLARRTEWEKFINDMSDKCQRVDQTFQEKENEIQEFYVDL
EKKLHITP-------
>Eogystia_hippophaecolus
-------------------MTELSREIAEVWSRLFDHRPFLNGEIKFMLKEFEEKRNDREVENLFVILEKLTDIKDTQVDR---IKRTSE--SLPVLSEKLEQALQLSEKVEKDFLEN-HKTIEEKRLANQSKRQKEWDQFIDDMNFKCQRIDNAFEEKEEELRDLYADL
KHKLNIAK-------
>Tanzaniophasma_sp._AD-2013
------------------------------------------------------------------------------------------------------------------------------------------------MTEKCAKVDATFEEKEQELREFYTDL
ERKLHIGK-------
>Panorpa_vulgaris
-------------------MTEIIKDVGEIWSRLFDHRPFLNGEIKFFLREFEEKRSDHEVENLFSTLEIATDIKDTDISR---TTLAADS-HLPVLIENIQAAVTLCDKIVDLG--H-SEDLTKQLEENREKRKAEWVLFVDDMTHKCSRIDNTFEEKEEELKDFYSDL
ERKLHIHK-------
>Corydalus_cornutus
------------------MSAEYLKDIGVIWTRLFDHRPFLNGEINFALKEFEEKRSDKEVEQLFTLLEHAADINDSQIKR---FKTAVDT-EIPQLKNQIESILKVTDNILEQA--E-RYKQDETLEINNRARKKEWEEFVEDMTYKCRQVDNTFEEKEEELQEFYADL
EKKLIRQ--------
>Osmylus_fulvicephalus
-------------------MSDIFKDIGNLWTRLFDHRSFLHGEKQFFLREFEEKRTDTEVKQLFNILETITDIKDTEIDR---MKLIANN-HLPSVEINLLLAFDLVDGILQNE--S-NDAQKEKLIQNRTLRDVSLKEFLKDIDFKKERVDRTFEEKEVEILELYLNL
ERKLRLSN-------
>Triops_newberryi
----------------------------------------------------------------------------------------------------------------------------------------------------MSALDNLIKEKELSYRQYYASL
EEQLKAK--------
>Epiophlebia_superstes
----------------------------------------------------------------------------------------------------------MCNRILEKE--E-EYCPDRALVNNRERRKAEWEAFVEDMTQKCSKVEITFGEKEKEMREFYEQL
GQKLHID--------
>Stenobothrus_lineatus
-------------------MAAIIKDTGELWTRLFDHRPFLNGEIKFVLREFEEKRSDGEVEKLFKILECVSEIKDTQIER---VKLASDI-HLPNLNGNLEVAVSMCNRILEKE--Q-QHRLDTTLEAKREIRKAEWENFIEDMAQKCTKVDTTFQEKEQELKEFYLDL
EKKLHITK-------
>Ramulus_artemis
----------------------------------------------------------------------------------------------------------MSCRILDKE--Q-QHLADNSLEARRELRKAEWHDFITDMTDKCFKVDRTFEEKERELSDFYVDL
ERKLHLDQ-------
>Cosmioperla_kuna
----------------------------------------------------------------------------------------------------------MCNRILEND--E-RHRSDTTLVAKRELRKVEWEKFIEDMTQKCSRVDKTFEEKEEELKEFYADL
EKKLHLNN-------
>Inocellia_crassicornis
--------------MSEVVKDTIICDIAGIWTRLFDHRPFLHGEIQFMLKEFQEKRSDSEIENLFKILECGTDIKDTQTDR---IKHRINN-NLPELKTKIDEALNICNDINQQE--D-LYKSDKTLAVNRITRKKEWELFIEDMTYKCSKVDNTFEEKEGELQEFYTDL
ERKLHITK-------
>Ctenocephalides_felis
-------------------MTEIFKDINETWMRLFDHRPFLNGEINYMQQEFEGKRKDIEIEHLFSTLQVSTEIVETHIPR---AKICSES-YLPSIKSSVDKALEVSESVFSED--F-EKSVKEALLEKRAKREAGWESFLEDVNQRYQSIDLTFEEKEQELNEFYSDL
NRKLHLHT-------
>Lepeophtheirus_salmonis
MPGNSAITAGGILEYSGKRTMSVAKDIFDIHDRLFNHRPFLQGEIRHFIKEFEEKREDREVHNLFNVLEKVTELRESEIDK---IITQCDG-IFPNLSANLLVAQSMSNKILDQG--R-TREIEQALENSKDTREKEWTSFMSEIQDTCAKIDDAYKRKEEDITTQYSKL
EESIEKKN-------
>Frankliniella_cephalica
----------------------------------------------------------------------------------------------------------MCNRMIKSE--E-EHLADNSLEAKRALRKADWENFISDMAGRCSRVDSTFQEKEEELREFYEDL
EAKLHVVK-------
>Rhyacophila_fasciata
-------------------MSDLTKDCGKIWMRLFDHRPFLNGEIKFMLREFEEKRSDREVTNLFSALENATEIKDNQVER---MKESAAL-QLPELATHLVKALSAYEDIEGYK--T-ELSEDTTLEKRREIRKKEWARFTEEMSDTYTRIDNAFEQKEEELKDFYADL
EHKLNITTK------
>Atelura_formicaria
----------------------------------------------------------------------------------------------------------MCSRIIEKE--E-EHCIDTALLDNRLSRKAEWELFIGDMTDRCSKVDVTFEDKEGELREYYADL
EKKLRTEQ-------

HG0305 FBgn0050290 Ppcdc

>Drosophila_melanogaster
MTSQIYSQKKFVPTPPEKGSFPLDHEGLCKKQFLLYASCLRKNAQDTSQCRQDAQNYLACRMDNNLMEKTEWSKLGFHDQSTKTDQKEPEVQKQ
>Asprogammarus_rhodophthalmus
MGSMTLGTRSFQPTPPAKGSFPLDHEGECKTFMIKYMECMHQTGGNNSECREKARDYLSCRMEKGLMAREEWSKLGYADLASSAARDGSE----
>Meinertellus_cundinamarcensi
MTTMTFGQKSFKPTPPDKGSFPLDHEGECKRTMLKYMLCLRENNNDNSQCRVQAKDYLQCRMDNNLMAKEDWKTLGLGDIVPEK----------
>Prorhinotermes_simplex
MSSVTFSQKKFTPIPPDKGSFPLDHEGQCKKAMVKYMICLKTNKNDSSPCREDAMAYLDCRMSNHLMAKEEWSKLGFGDVKTKTSEK-------
>Dastarcus_helophoroides
MSAATFSQQRFVPTPPEKGSFPLDHEGLCKKFMVKYMGCLRSNKNNNSACREEARAYLACRIDNNLMTKEEWSKLGFSDSEPKEQH--------
>Astacus_leptodactylus
MASMTFSQKSFTPTPPQKGSFPLDHEGECKKFMIKYMLCLRENQNENNKCREESREYLGCRMEKGLMAKEEWSKLGYSDLTTSAHHTS------
>Forficula_auricularia
MSAMTFSQKQFIPTAPDKGSFPLDHEGICKKVMLKYMICLNENDNKNSLCRQEAKDYLSCRMNNNLMAKEEWETLGYKDEKN------------
>Daphnia_pulex
MASNTFGQKNFTPTGPLKGSFPLDHDGECKINMLRYMICLRENSNQSSECRQLSKEYLDCRMQKQLMVREEWTKLGFGDISEENKQ--------
>Occasjapyx_japonicus
MSSTTFGQKTFTPTPPEKGSFPLDHEGLCRRFMLRYLLCIERQSHDNSACRGEARDYLECRMDRQLMAREDWGKLGFSDSSAAPPAEKP-----
>Aposthonia_japonica
MSSMTFGQKTFTPTPPDKGSFPLDHEGLCKRQMLKYMICLREYDNDGAKCRAESKDYLSCRMENNLMAKEDWSKLGFDDNVKMKNL--------
>Meganyctiphanes_norvegica
MSSMTFGQKSFTPTPPQKGSFPLDHEGECKKFMIKYMLCLRESKNENSLCRQQSKDYLECRMEKGLMAKEEWTKLGLADVGSAEPSNSEQ----
>Grylloblatta_bifratrilecta
MSSVTFGQKSFTPTPPDKGSFPLDHQGQCKKEMLKYMICLRRNRNDNSACREESMRYLGCRMDNNLMAREEWSKLGFSDLELKPEKNN------
>Halyomorpha_halys
MSSLTFGQKKFTPTPPEKGSFPLDHDGQCKRTMLKYMICLRNTNSDSSKCRQEAKEYLACRMDNNLMAKEDWSKLGFKEEEVKQEN--------
>Psyttalia_concolor
MSSYTFSQKIFTPIPPQKGSFPLDHEGYCKLPMIKYMRCLHTNKNDNTKCRDDAREYLGCRMENNLMSREEWSKLGFHDDDKIEEGEKPKNT--
>Armadillidium_vulgare
MSSMTFGQKLFKPIPPEEGSFPLDHEGLCKKSMINYMQCLSKYNNNNSECRLEAKEYLDCRMRNNLMASEEWSKLGFSDKDKQQSEVTEVKDN-
>Metallyticus_splendidus
MSTMTFGQKQFAPVPPQKGSFPLDHEGQCKKAMLKYMICLNANSNDNSSCRAEAKVYLQCRMDNNLMAKEEWSKLGLSDVEAKLQKE-------
>Boreus_hyemalis
MSSMTFGQKTFIPTAPDKGSFPLDHEGLCKKLMLKYMHCLRMSKNDNSLCRQEAKEYLGCRMDNELMAKEDWSKLGFKESDVVK----------
>Chrysopa_pallens
MSTMTFGQKSFTPTPPDKGSFPLDHEGICKETMYNYMRCLRLNKNNNSACRIEAKDYLACRMDNNLMSKEEWSKLGFQNQNKSTINK-------
>Gryllotalpa_sp._AD-2013
MSSMTFGQKTFTPVPPEKGSFPLDHEGVCRTLMLKYMLCLRRNDNDNSQCREEAKSYLGCRMDNHLMAKEEWSKLGFKDLDMKDDSKEN-----
>Medauroidea_extradentata
MSSVTFGQKSFTPTPPDKGSFPLDHEGQCKREMLKYLLCLRANNNDNSACRREAKTYLGCRMDNNLMAKEEWSKLGFSDLNLDSKGSDKSL---
>Menopon_gallinae
---MAYQGKVFKPVPPMKGSFPLDHDRSCRPEMLKYMKCLMENKYDSEFCRTEIKTYLECRMDNNLMAKESWEALGLKDIAESNSEYSS-----
>Cosmioperla_kuna
MSSMTFGQKSFKPTAPDKGSFPLDHEGQCKHEMIKYMLCLSTNNNNNSACREEAKSYLGCRMDNQLMAKEDWSKLGFDDLRSDSNRVEH-----
>Acerentomon_sp._AD-2013
MGTRSFQQKPLQSIPPEKGSFPLDHENECKRSMLEYMICLQQNKNDNSACRMKAKGYLKCRMDNGLMAIEDWPSLGFKDGNVEASVSAKL----
>Ectopsocus_briggsi
MSSMTFNQKKFTPTAPDKGSFPLDHENLCKQQMMKYMQCLRVSKADNSKCRLESKDYLACRMEHQLMTKEPWSKLGLEDLEDQKTVQ-------
>Inocellia_crassicornis
MSAMNF-QRTFIPTPPDKGSFPLDHEGLCKDQMIRYMTCLRQNRNDNSCCRIEAKDYLGCRMDNNLMLKEDWSKLGYSDNKNNVKKD-------
>Ceratophyllus_gallinae
MSSMTFSQKSFTPTAPQKGSFPLDHEGQCKKFMIKYMSCLRQHKNDNSSCRNEAQAYLGCRMDNELMAKEDFSKLGFNSNEINK----------
>Caligus_rogercresseyi
MASNVFGAKNFKPVAPEKGSFPLDYEGKCKKEYLKFMICLKDNKYVVENCREESKNYLVCRMDHGLMAKETLPNLGFKEEENKKADK-------
>Mengenilla_moldrzyki
MSNPLLRQKVSKPIPPQKGSFPLDHNAICKKEMLKYIICLRNNENGGSNCRELAKSYLKCRMDNNLMASEEWSHLGF-----------------
>Frankliniella_cephalica
--MAQFAKSRFVPTPPEKGSFPLDHDGVCKDLMRKYMTCLSSNKQDNTKCRKEIKTYLDCRMNNNLMAREEWKLLGLEDVGAKSEVVQK-----
>Zorotypus_gurneyi
MSSATFGKKIFTPTPPDKGSFPLDHEGQCKRVMLQYMSCLTKNNNDNSFCRLEAKNYLACRMENNLMAKEEWSKLGFQD---------------

HG0306 FBgn0259726

>Drosophila_melanogaster
--------MSIPQ--------------YRPPK------E--------IQDLLDQ---KEQQ----------QANQRRRTAVLPK---RTWIQRNPRLFQISFITGSLLIFFSKPLFDCFIADPLPPPENKVPPHKR----------------------------------
----------------------
>Micruropus_glaber
-----------------------MSNAKTTGRRSSVPP---LHGQ--KYDWLYQKPALKVRTQEHFDAEEYLKRRIEVPKTVSNPWWLKFYNRNPRRFFITFCVVTCGTFYSRFLYDAFVVPTLYIVSSLKEIVFQPPVPAIEPLQLASVTTNSTCDSVQPSRTEPTKRV
SESASATAQTAKPEAVASETSK
>Machilis_hrabei
---------------------------------------------------------MAIK----YGSG-------RVSPHLPK---RTLMQKYPRIISGLLITTGLLVFFSKPLYDAFCGDTVTVETLRARKAAKSQSQ------------------------------
----------------------
>Reticulitermes_flavipes
-----------------------------------------------------------MK----------QQPPFVRQKVLKD---RNWIQRNPRTFVCLFTASSLLVLFSRPLYDIFVREYPPVPVDPQLPK------------------------------------
----------------------
>Calanus_finmarchicus
MADRGWHSPGHPS--------------YKKDRVGQTDPS--------RGTSLDE---IKRRSKQKNADKRFNSPFAFHEPKYNK---VNLINRYPRLFGVAFTTVCVLLLYSKPIYDATLREPTQYELDRANFLKRRME--VGGWWDNPFWKR---SSDKPE-----K--
----------------------
>Carabus_granulatus
---------------------------------------------------------MNDNR----------------RTVIRK---RSWLQRHPKIFFWGTVSSSLLIFFSRPIYDIFIRTDFIPPPTPKEKSKQ----------------------------------
----------------------
>Farfantepenaeus_aztecus
--------MSKPVKSPYPRAKIDLPGSYRSGPTGSQRPGRETEGSNSRSRLLQQQDPSHYR----YNPESVKS---NKPEVSRF---RRWIYRNPRRFQIIGLGIGLTIFFSRPLYDIFLREHIEGPLPVKQK-------------------------------------
----------------------
>Occasjapyx_japonicus
-----------------------------------------------------MPGP---------RPTPVVR---RGTYNLPK---RSWRQRNPRAFTGVFVGTCLLVLFSRPLYDAFLRDEGSPSAPRPN--------------------------------------
----------------------
>Eurylophella_sp._AD-2013
--------------------------------------------------MNRPKPP---------KK----P---HYLNQIGK---RSWVQRNPRAFVIIFTTTCLTIFYSRPLYDIFFREYPEGHIPKNYIFQKKEE-------------------------------
----------------------
>Grylloblatta_bifratrilecta
-----------------------------------------------------------MK----------QQPIVRRKLILKD---SPWYQRNPRLFVGLFTTGCLLVLFSRPLYDVFLRPAPTGPAPHPSKR------------------------------------
----------------------
>Meccus_pallidipennis
---------------------------------------------------------MWKE----------KVETPKFRKTSKD---LPWVNRNPRIFIILTTAFALSIFFSKPAYDLMYAERVPHPSLSRSLDAGKGQ-------------------------------
----------------------
>Chyphotes_mellipes
--------------------------------MTALDPCKK----------------FACKLQQCLKDNVYQP--SRCEGVIEE-------LRQCCIKNSTHSVVCEGIDISKPYDHKTVDYKTAIR-------------------------------------------
----------------------
>Palaephatus_luteolus
------------------------------------------------------------------------MDQRRRSKVLGK---RSWMYRYPKTFQITFTTLGLGIFFSKPIYDIFIRKYEDFDYTEPPTTLSQRRN------------------------------
----------------------
>Mantis_religiosa
-----------------------------------------------------------MN----------PKP-FIRRKVLKD---RSWIYRYPRAFVGITTAASMLIFFSRPLYDAFVRDLPEAPPEPRVPRLKPSHF------------------------------
----------------------
>Tanzaniophasma_sp._AD-2013
-----------------------------------------------------------MK----------EQPIVRRK-VLKD---SPWYIRNPRTFVGIFTTSCLLVLFSRPLYDAFLRDYTSSPPPAAPRSLK----------------------------------
----------------------
>Boreus_hyemalis
-------------------------------------------------------------------------MSVKRGPTMIH---RTWIQRNPRLFSGVFITGSLLILFSRPIYDMFFRTDFAPPRQSLYKK------------------------------------
----------------------
>Corydalus_cornutus
---------------------------------------------------------MSNRK----------------NPVMVK---HSWARRNPKLFSAIFVTTSLLIFWSRPIYDIFFRPVNEMAPPRKVKFD-----------------------------------
----------------------
>Chrysopa_pallens
-----------------------------------------------------------MKE----------------RTVLKK---RSWAQRNPRAFFAIFATTSLLILYSRPIYDIFIREDYDTYKKRMEKERLSRQ--AAE--------------------------
----------------------
>Triops_newberryi
---------------------------------------------------------MDPSN---------RKVVWSRFGEKVK---LPWYRRKPILFQKIFISVSLLIFFSRPLYDIFLRSYIEGPNPIRDPYSINER--LGSRVVLK--------PKTPE-----EKS
E---------------------
>Cordulegaster_boltonii
--------------------------------------------------MNEFKPW---------KEKKFPP---PSRKLLKE---HSWIRRNPGIFSSVFTLGCLMILFSRPLYDAFLRDSVEGRQVLPSNSK-----------------------------------
----------------------
>Schistocerca_gregaria
---------------------------------------------------------MKES----------K-ESPVRRKVLKD---QPWIKRNPRAFAGLLTASFLLAFFSRPLYDIFLRKDNVVPLESPRIAK-----------------------------------
----------------------
>Timema_cristinae
-----------------------------------------------------------------------MKEKLVRRQVLKD---RPFYQRNPRLFGGVFTTVCLLIYFSRPIYDIFFRPDFAPAPIDQIRAIRKKVSD-----------------------------
----------------------
>Leuctra_sp._AD-2013
---------------------------------------------------------MDRK----------PS-PVVRRQVLKD---LPWTRRNPRLFAGIFLTGSMLLLFSRPIYDIFIRDNPEQTPLSAN--------------------------------------
----------------------
>Raphidia_ariadne
--------------------------------------------------------------------------MQKRNTILHK---PTFLQRNPRIIGFSLITLSMLMLFSKPIYDTFFAPNYKFPPKPKN--------------------------------------
----------------------
>Oropsylla_silantiewi
-----------------------------------------------------------------------MNQGPRRRAIRKD---LPWAYRNPRKAANIIVIGSILTIFSKPIYDIFLQEPIDYTVKWVQKIKANIESKKGSE-------------------------
----------------------
>Annulipalpia_sp._AD-2013
--------MSYPY--------------GKPPK------P--------VDPPI--------K----------RDPLKQRRPVLKN---QSFVYRYPGTFTGVMTFLGLAIFFSRPIYDVFFSEMFPDLADPILCDHP-TRGVKGKRYVAP--------EQQ---------Q
----------------------
>Zorotypus_caudelli
-----------------------------------------------------------MN----------SEGPKIRKAVLKD---RSWIYRHPQLFIYSCATIGFLIIFHRPIYEMFNPLPPRIKTQ-LDEAKSKKNSVKISE-------------------------
----------------------
>Atelura_formicaria
-----------------------------------------------------------------------MQ---PRQKPLKD---RSWVQRNPRTFTVTLTSTCLLILFSRPLYDAFLRPAEPPSVPMKK--------------------------------------
----------------------

HG0307 FBgn0037822 CG14683

>Drosophila_melanogaster
-MSAQYQRFLKVLEKWPAEKSKV--GRDLGEQIRKQVTKLTSLEG---GATDKELDRQINSLERLSNNVYAKKYPRTFE-STATGLTAAQCSQVLSSEFLQYLNEDSKKKK--------------------
>Hyalella_azteca
MSGSSYHRFVELLKLWPVDKTKI-GGRDLGEYIQHRISKTFVQGVAT-SVDEEKCSAIVSSLTRLAKDTHKNKFPRTRE-SSALGLTKEECSGLLSNEFLALLHDREK-PFFNRFSNFPNKKT---------
>Machilis_hrabei
MASIYYKKFIEILGKWPLDSTKK--GRDFGEVIRKWVSGTFSHGEAS-SVDVKECEKVYVTLNSLANNKYRDTYKRTHS-TSATGLTRKECQDIVSTEFLQKLQEEENRGILRSVWSSMKSK----------
>Argulus_siamensis
------------------------------------------------------MLKQSDALSKMLNNSYKKMYPRKFK-STALGLDYETCKVANSNEFLKSMSK---------------------------
>Cryptocercus_wrighti
-MARSYRNFLQLLELWPVDSTKV--GRDLGQHIREQVKVAFKTGEVF-PGDETECNRIYASLKRLADNRYGQMYPREST-TCASGLLADECQAILSHQFLEELRESER-GFFSKLFHRKS------------
>Carabus_granulatus
-MAASYKRMLKLLEKWPVDKNKR--GQDLGEHLRKQVQAAFAARQF--DTNQEECDRKYLSLKRIAEDHHLKTHPRTLK-SSATGLTAEQCSQILSPEFLEALEREDQ-SVFRRIFSF-------GSKK---
>Litopenaeus_vannamei
MARNSYRNFLRLLEKWPLDESKK-GGRDLGEHLRIRVSEAFRLGDAT-PTDEVECQRVCASLNRLANNTYATRYPRKLT-STSTGVTIQECHQIASTEFLKAMEDSEK-GLLSGMFKQG-------------
>Forficula_auricularia
-MAHSYKIALQILEKWPICSSKK--GRDLGEFIRNQFNIGFKDQQIL-FGSEVECNRNLESIKRISENYHANKYKRTLN-SSSTGLTAEECEKSLSNEFLKHLNK---------------------------
>Occasjapyx_japonicus
MSGATYRNYLKLLEKWPVDANKK--GRDLGEHLRTRISEAFHRGAAT-QVDEADCASKLSSLTRLTVDHHAKQYPRRYE-SAATGLSADKCRQIVSTEFINQLLEEEK-GLVEKLLATKLEKKSKD------
>Eurylophella_sp._AD-2013
-MAQHYKNFLRILEKWPLDKSKG--ERDLGLIVRQRVKQAFPSGEST-PVNIDECDKTYGILKRLVDNHHLNLHKRKLE-SSSTGLSAEECRQVLSTEFLKELEKQDR-SLWSRLFNRNKE-----------
>Meganyctiphanes_norvegica
MSQKSYRTFLRLLENWPMDTTKT-GGRDLGEHIRIRVSEAFHKGETT-PVNETECMRIYSSLNRIASNTYFNQYPRSRN-STSTGLSGDICRQVISTEGLASMRVKEP-PTFEKKDN---------------
>Galloisiana_yuasai
-MAKTYRQFLILLEKWPIDSSKR--GRDLGEHLREEVKKAFTAGEVT-PVDEAKCSRYQASLQRLVNNHYGQLYRRSSS-SSASGLTAQECSDLVSSEFLDYLQKEEK-GFFARYFKRK-------------
>Clastoptera_arizonana
-MANHYRNFLRLLEKWPVDTSKS--SRDLGFFIREQLKVNFPQGEIARNIDEKKCNEAYNSLNRLADNYYFDKYQRKFH-SSATGLDAEQCKTVLSDEFLEYLKEENNQTIVSKVFGNKNKEQ---------
>Argochrysis_armilla
-MAGNYKNYLKLLELWPLDKSKA--GRDLGQHIRDQVKVAFSKGDVGNDINREQCNRYYQSLQRIVGNHYGKTYPRTLS-SSASGLTKEQCNEILSPEFNEYLKYYDQ-SFFEKVRLSLALIFTRSNKAKKS
>Danaus_plexippus
-MTSQYKKFTKLIAKWPIDNNKA--ERDLGKFIRDKVKAAFEGGNSK-NLDSELCTRQLYSLNKIADNHYRNKYKRIHD-SSATGLSSEECNLVLSSEVLQYLKEENK-GFFKNIFKKD-------------
>Boreus_hyemalis
-MAASYKKYLQLLEKWPIDKTKN--ARDLGLYLRQQVKLTFGGGVVS-RAEEEKCERQLHALNRLADNTYGKLFSRTST-STATGLSADQCRQILSTDFLELLQEEDK-GMMSKIMSKAKPKT---------
>Euroleon_nostras
-MSNSYRRFLNLLKDWPLDTTKQ--NRDLGQHLRDRIKVAFATGEQF-QGDQQKCDLEYLALRRLADNYYAKKYPRTLQ-SSAIGLSLKECKDIISNEFLESVKEDEK-SVLQKLFSKKSRVVAN-------
>Calopteryx_splendens
-MAGRYKLFLRLLESWPVDNSKK--GRDLGYHIRKKLLEAFPMGETS-SFNAEECEKVYLSLKKINDNYYGDKYKRHTN-VTSTGLTYEQCAGVTSTEFLTEVSAGRR------LFKLSQKR----------
>Stenobothrus_lineatus
-MANNYRKFVKLLESWPVDPSKT--ERDIGFYLRERVKKAFTAGELF-EGDQEKCAREHASLKRLADNVYGKLYKRTTS-SSATGLTAEECNSVLASEFLTLLQKENR-SIFSRLFRGK-------------
>Clitarchus_hookeri
-MANHYRNFLKLLEGWPIDASKK--GRDLGEHLRDQVKTAFNAGEVS-RIDKAKCEKIYMSLKKLADNHYGNLYHRTSS-SSAMGLTAEQCGSVMSTEFLKSLEEEEK-GFFKRIFKRD-------------
>Leuctra_sp._AD-2013
-MAGIYRDFLRILEKWPVDTTKK--GRDIGEYIRQQVKVAFPAGEVF-KGDEEHCRKSAASLKRLSENHYGELYKRTSQ-SSATGLTAEQCNGVLSSEFMEMVEESNK-SFLSKLFGRNKDS----------
>Acerentomon_sp._AD-2013
MSAPLYHKYMKLLERWPVDLSKK--GRDLGEFLRTKIFSEFKYGETS-NVDVLNCERKFEALNRLSADHYMTKYPRLYT-STSSGISVEQCKLVLSTETLKQLEYEDT-GVLKRTFWNLTSYKNKVDDSKK-
>Xanthostigma_xanthostigma
-MASNYRRFLELLKKWPLDKTKE--GRDLGQHIRDKVKVAFAEGDIS-QTNQEECDKQYNSLKRLSESYYANTYPRKVS-SSSTGLKSDQCTQILSNEFLELLEQEDR-SFVSKLFNRSKK-----------
>Oropsylla_silantiewi
-MASSYKTFMKILKQWPIDKTKK--DRDLGRYLRDSIKTVFTKGEVS-TVDEVKCSKQLEALDRIANNTNFVKYKREFA-SSATGLTSEQCNTVLSTDFLEYLEYENK-GIVTKLVTKPPAVES--------
>Caligus_rogercresseyi
-MSASSSRWLKLLQSWPSEPSKK--SRDLSQILLKRKSPSS--------------EAEVRALETLLSNKLALKYSRVYSEATATALGREDLKTLTSTDFLESSTAYANLSFWERIKLKFFSK----------
>Thrips_palmi
-MAQLYERFLKVLQKWPLNADYPPPLRSLPAHLRDQGKAAFSARGEF-SGDTLQCEKRLSSLERLTTNQIARAYPRTQQ-TGALGLTSDQCVFALSDSYQKSLETSATTSLFQRIVSSFRQGSGKEKA----
>Platycentropus_radiatus
-MTSQYRNFLKLLEKWPLDPSKK--GRDLGDRIRVCIKAAFNTEQGF-TGDLKECQQKYESLNRLISNQHGNLFKRTKA-HSSTGLTAQECNLVLSSDFLEYLQQEER-SIFANILPQKD------------

HG0308 FBgn0264743 CG44001

```
>Drosophila_melanogaster
MPYIIIRGNLASYS--HKYPWRVLVSGLKADDIEQLNKFSCGG---YSDESTIVYLVHPCRILSALEILGFRVVASSSTAVK------QDYNEYMWTMRKEFDEPEPLEAESVVRENLSNIGREAASLGNYHKVDSPE--------
>Meinertellus_cundinamarcensi
MPYAIVTGNLKQSEGAGRPPWQVSVAGLKDKEIAQLNRFQCLT----KDKYTVVFYQHPCIILTALEVIGYRVVSSCPAGEG----NENRSDEYVWTVRKDFSEPEPSDAEEENGI------------------------------
>Coptotermes_gestroi
MPYILVRGNLSSYG--KDYPWRVLVSGLKACDIEQLKRFASGG---YCDESTIVYLQHPCVVLTALEVLGYQVVASSSTSIK------QDYNEYMWTMRKEFSEPEPMVTKPTLKQ------------------------------
>Lepicerus_sp._AD-2013
MPYILVRGNLASYG--HKYPFRVLVSGLKAGDIDQLNRFASGG---YSDDYTIVYLQHPCVILTALEVLGYRVVASSSTSVK------QDYNEHMWTMRKEFAEPEPDPDADTETDV-TTLC-E----------------------
>Apachyus_charteceus
MPYILVRGNLASYG--HKHPWRVLVSGLKASDIEQLKRFASGG---YTDESTIVYLQHPCVILTALEVLDYRVVASSSTSVK------QDYNEYMWTMRKEFSEPAPLLTCKPTN-------------------------------
>Aposthonia_japonica
----------------------------------------------------------------------------------------------MWTMRREFTEPDPTSKPGF---------------------------------
>Ephemera_danica
MPYILVRGTVTQHP-------CAHVSGLKASDIEQLKHFGSARKAVPADDLTVSYALHPCVVLTALEVLGYSVVSCTAGPTG-----PGGETQYLWTLHKDFPEPEPDSDLYNHH-------------------------------
>Grylloblatta_bifratrilecta
MPYIIVRGNLASYG--HRNPWRVLVSGLKAGDLEQLKGFSSGG---YSDDSTIVYCHHPCVVLTALEVLGYKVIASSSTSVK------QDYNEYMWTMRKEFPEPEPDSQTKQPVLT----CN-----------------------
>Lygus_lineolaris
MPYIIVRGNLASYD--NRYPWRVLVSGLKADDLEQLQRFNCGG---YCDNSTIVYLVHPTVILSALEVLGYQVVASSSTAVK------QDYNEYMWTMRKEFAEPEPTKEKEKTNSA-----------------------------
>Bombus_terrestris
MPYILVRGNLASYG--HRYPWRVLVSGLKASDIEQLSRFASGG---YCDDSTIVYLQHPCVILTALEVLGYKVVASSSTSVK------QDYNEYMWTMRKDFSEPEPEPVIKTGKLP-RR--------------------------
>Zygaena_fausta
MPYILIRGNLASYG--HKNPWRVLVSGLKAADIEQLKRFACGG---YSDDSTIVYLQHPCVILSALEVLGYKVVASSSTAVK------QDYNEYMWTMRKEFSEPEPSIIVEQDNIN-NYSK-EAMHLGNYQKDDEV---------
>Mantis_religiosa
----------------------------------------------------------------------------------------------MWTMRKDFASPEPQLAKPTLKQ------------------------------
>Tanzaniophasma_sp._AD-2013
MPYILVRGNLASYG--DRNPWRVLVSGLKAAEIEQLKGFASGG---YCDNSTIVYLQHPCVVLTALEVLDYKVVASSSTSIK------QDYNEYMWTMRKEFSEPNPQTKL-----------------------------------
>Boreus_hyemalis
MPYILIRGNLASYS--HKYPWRVLVSGLKATDIEQLKRFACGG---YCDDSTIVYLQHPCVILTALEVLGYKVVASSSTSVK------QDYNEYMWTMRKEVFEPETCIAPSTVTTI-SPSVDERDNIANISKEALTTITPNKIDE
>Corydalus_cornutus
MPYILVRGHLTSYG--HRYTPRVLVSGLKANDIEQLARFPSGG---YSDDLTIVYLSHPCVILTALEVLGYKVIASSSTSVK------QDYNEYMWTMRKDFTEPEPVNVKEVAHDA-----------------------------
>Chrysopa_pallens
MPYILVRGHLTSYG--HKYTPRVLVSGLKAPDIEQLSRFPSGG---YADDLTIVYHTHPCVILTALEVLGYKVIASSSTSVK------QDYNEYMWTMRKDFAEPEPDPLSK----------------------------------
>Triops_newberryi
MPYVIVSGNLTPYK--SGTPWKVGISGLKGIELQELDRFILEG---QADPHTVVFKHHPIVLLNALEVLGYRVISSYSTDKPTINISQEPVPISVWTMRREFPEPEPSDDDSDSNHPTDASL------------------------
>Epiophlebia_superstes
MPYIIIRGNLASYS--NRNPWRVLVSGLKADDITQLKRFPSGG---YCDDSTIVYMQHPCVILTALEVIGYRVVASSSTSVK------QDYNEYMWTMRKEFTEPEPSVTSFDSGIT-EPLP------------------------
>Ceuthophilus_sp._AD-2013
MPYILVRGNLASYG--HRHPWRVLVSGLKATDIEQLKRFASGG---YSDDSTIVYLQHPCVILTALEVLGYKVVASSSTAVK------QDYNEYMWTMRGEFAEPEPLLSKPTSPTS-----------------------------
>Peruphasma_schultei
MPYILVRGNLASYG--HKHPWRVLVSGLKASDIDQLKGFASGG---YCDESTIVYLQHPCVILTALEVLDYRVVASSSTSVK------QDYNEYMWTMRKEFSEPEPHAAKPGY--------------------------------
>Leuctra_sp._AD-2013
MPYILVRGNLASYG--HKNPWRVLVSGLKAADIEQLKRFASGG---YCDNSTIVYLQHPCVILTALEVLDYKVVASSSTVIK------QDYNEYMWTMKKDFAEPEPQTVSNPE--------------------------------
>Ectopsocus_briggsi
MPYIIVRGNLASYG--HRDPFRVIVSGLKAQDIEQLKQYPCGG---YTDEVTIVYLAHPCIILNVLEVLGYRVIASSTTGVK------QDYNEYMWTMRKEFHEPEPLERN-----------------------------------
>Ctenocephalides_felis
----------------------------------------------------------------------------------------------MWTMTKEFSEPDPTKCGESKHE------------------------------
>Thrips_palmi
MPYIIVRGNLASYD--NRYPWRVLVSGLKAGDIEQLSRFASGG---YCDDSTIVYLQHPCVILSALEVLGYHVVASSSTSVK------QDYNEYMWTMRKDFSEPEPVPAPITAKPS-A---------------------------
>Annulipalpia_sp._AD-2013
MPYILIRGNLASYG--HKNPWRVLVSGLQAADIEQLKKFACGG---FCDTSTIVYLQHPCVILSALEVLGFRVVASSSTAVK------QDYNEYMWTMRREFEEPEPSVFMERDNIT-NIT-KEAMNIADIQNIKANDL-------
>Thermobia_domestica
MPYVIVRGNLTALGNNHSQPWRVLVSGLKADDILNLKRFQCGD---YPDDFTIAFLQHPCVILTALEVLGYRVVSSSSSS----------NNEYMWTMRKEFSEPEPIPQNSESFD------------------------------
```

HG0309 FBgn0037689 CG8135

```
>Drosophila_melanogaster
------MSSVEAEVLQKIDLIEP-TTGKLFFE--HKTELLFCRPHLMPLKTQALERLEEMHRDTARQLQKKR----QQKKKSTEPTGSL-------------------------------
>Talitrus_saltator
--------------MSGLKEVLP-RKGLLYQE--ESYQLALCRPKLMPLKSITLEKLEAMQQQLQERSKEQL----QQQELSTQQQ----------------------------------
>Argulus_siamensis
---------MSEEDLKLLKEVIP-YQGVVFQE--DVPNFILCKPKLLPMKSVTLEKLEKMQKESQERVKNQQ----NEVQDANIDSS---------------------------------
>Cryptocercus_wrighti
--------MSEERKIPALKEYIP-YKGLLHHE--EIPEFLLCKPKLMPLKSVTLEKLENMQKEAEEKLKKQQ----MNEQRS--------------------------------------
>Oxysternon_conspicillatum
------------MSEGEIKPIVP-KTGQLLIE--NPEELILCKPRLIPLKSYTLEKLEKMQEEAIKTLRESR----EKTEG---------------------------------------
>Eucyclops_serrulatus
--MPKG-EVKVTVQSEPFREVLP-KRGLIFQE--SSPQLVLCKPKLLPLKSFTLEKLDKMQKEATEKAREQM----IEQQKQEDEDERLFGSVSENAETHAPESVSFE------------
>Farfantepenaeus_aztecus
-------MPEGSEAGQVLKEVLP-RKGLLYQE--DSSPLALCRPKLMPLKSVTLEKLEAMQKAAQEKGRQQQ----EEQ-LERQATAPA------NGSRI------F-------------
>Occasjapyx_japonicus
-MSSEGSSKTTAAPQQLFKEILP-RKGLLHQE--DIPTYALCKPKLMPLKSFTLEKLERVQREAQEKVLDRS----NQPEQN--------------------------------------
>Eurylophella_sp._AD-2013
----------MGDSKDSVKEILP-KKGLLFSE--DIASLILCKPKLMPLKTLTLEKLEKMQKEAEVRGREEH----EKEANESF-----------------------------------
>Tigriopus_californicus
--------------------------------------MVLCKPKLLPLKSFTLQKLEAMQSEAKIKAQAQL----RPNEDLEHNDEDE-QRLAADGAAFISESVSFE------------
>Nilaparvata_lugens
-----M-ANLEGSSDSNIQEYLP-TTGLVHNE--EIKELIFCKPKLMPLKSVTLEKLERMQKEADQKLKEQQ----RLPTAAPLPTFQS-------------------------------
>Argochrysis_armilla
-----M-SQAADNISEEIDIVLP-KQGLLYKE--ESLDYILCKPKLMPLKSVTLEKLEKMQKDAEQKLKDTE----VTTNGYKENK----------------------------------
>Metaphatus_ochraceus
------------------------------------------------------MRFERMQQEASRQWKEKL----KSAKKN--------------------------------------
>Boreus_hyemalis
MSINDLNKNITLDSKTVPQYVLP-PYGEICLK--YESKLKLSHPHIMPLKSVTLEKLEKMQQEAFKQVQENA----KLKEDY--------------------------------------
>Corydalus_cornutus
--------MSNTLNEKNVKYVIPSNTGQLFFE--DTRQLVLCKPRLMALKSLTLEKLEKMQIDAEEKMKLHK----SNYS----------------------------------------
>Osmylus_fulvicephalus
--------MSSGSSTTPMRPVLPINTGQLFFE--DTKELILCKPRLMPLKSLTLEKLEEMQKLAEAEVRAHR----NSTVEEHK------------------------------------
>Triops_newberryi
-------MEESKNTPEVPQRILP-EHGLIHLE-CNNSSWSLCKPKLMPLKSVTLDKLELMQKAAQEKMQQMK----DSSSQS--------------------------------------
>Cordulegaster_boltonii
--------MSDGFKIDR--ENFSVPRGLIQYQ--DFPHFVLCKPMLMPLKPVNLRKLEEMQREILKKVKSQE----NAQHE---------------------------------------
>Teleogryllus_commodus
---------MSKDQTPVLKDFIP-NSGMIYHE--DLPQYILCKPKLMPLKSVTLEKLEKMQKDAEDKLKQQR----LEQINEGNVNNVP-------------------------------
>Sipyloidea_sipylus
---------MSSHNQPALKEFIP-RKGFVVPE--DETQYILCKPKLMPLKSVTLMKLEKMQRDANEKLKKQK----AEPQNDTC-----------------------------------
>Menopon_gallinae
-------MSSSHDNDTDLKYTLP-TTGYIFEE--DVNRYVLCKPKLMPMKSITLEKLEKMQKEAEDKFKSHM----NSKCEE--------------------------------------
>Liposcelis_bostrychophila
---------MSTVNNENLKFTLP-TTGFIYEE--DISQYILCKPKLMPLKSITLEKLEKMQKEAEEKLKSQQ----AYTKKE--------------------------------------
>Caligus_rogercresseyi
-----------------------------------------------------------MQEESQARAALVLREEGQHLLRSTGPQGIP--------PQHDAEILSMTTQTHRGNKDEEE
>Frankliniella_cephalica
-----------MSEPANVKHVIL-STGVIQQQEKDIFQPTLCKPKLMPLKSVTLQRLEDMQKLAEEKLRQQQ----INSSTGESTQL---------------------------------
```

HG0310 FBgn0039339 CG5116

>Drosophila_melanogaster
-MNDIKRLKDQQREKHPGFDGYMDCMTRSLFTGLATFCLSFSGTYFAQKIVQ-SKIR--YPIKYNILISSLVATGVSYQITSTRTKACQAAWMAFEDKHSVLKEKTF----------
>Poekilogammarus_pictoides
MSDRKSRMVNQFGEEHPAIGSYLECVSRAYLTGLAGFSLSFACTYSVQRLTA-HRLP--YTANGQLLLAAIVASAVAYKVTANHVQHCHNFWMAAEDKHTALNPIQDKGDTQQL---
>Machilis_hrabei
-MNDLNKLKEEYGDKYPAFGSYTECMSRALLTGLSSFVISFSGVYIFQTLMK-KHLP--YGSKANVLISSGVASIVSYYITAKRTQACQSAWMAAEDKHTYLNPISERGNKKDT---
>Argulus_siamensis
-------------------------MKTNYFYGFGYFVIGLGCAEVIQKVVG-AEKY--YGKSIKIGTSLMAAVLVAFKFTLNGAKECSAILKDRSH--------------------
>Prorhinotermes_simplex
-MENINELKKQQRDIHPGFGSYLECMTRALFTGVAAFTLAFSGSFIGQSLLK-KHIP--YGKNTYILLSALLATGISYQVTTQRTKACQAAWMAAEDKHTYFKRGSQAEE-------
>Eucyclops_serrulatus
MLLAQEASSETKKDS--IPASQVECQTRSVMYGVATGTISFVGVYFAQRFLTRHYKG--ASVSLFLATSLFFGGFSGYMVTDAKLKDCAKASKKFKEFSSN----PNK---------
>Farfantepenaeus_aztecus
-MNNNTELINQYKDAFPAFGGYMECMSRTLLTGLSSFCLAFGGTYITQHLTN-KYLP--YGRKNHIIVSSLIACAVAYKVTSIRAEACQAGWMAAEDKHTYLNPISEREKTVDVEEE
>Occasjapyx_japonicus
-MQNMNEVVKKTEEKYPSFGSYTECMSRALLSGMSAFALAFSGTYLLQTALS-RHLP--YGGKGFVLVSSFVAVGASYRTTTTRTRACQRAWVRADGIKDDF---------------
>Isonychia_bicolor
-MNNVKQLKEQYSGEHRGFGIYLECMSRALLMGLSSFCIAFATSYIPQSLLK-KQLPTSYGVGGRILTSSVLAAGVSYYITSQRTKTCQAAWMAVEEKYTYLTPISVIENIESK---
>Meganyctiphanes_norvegica
-MQNTSDTIKQYKGAFPAFGSYMECMSRTFLTGLSAFSLGFGGTYVAQHLTA-KHLP--YDRKLHILFSSIIAVAVSYKVTSVRTNACQAAWMAAEDKHTFLNPISERPKTVDEEE-
>Cuerna_arida
MSRGKERLLELYGDDHPGFGSYLECMTRANFTGLATFTIAFSSTYIAQSILK-NKLP--YSSKFFILSSTAIAAYFSYKVSKARSQVCQAAWLAAEDKSTYFTELEKTSPTPVS---
>Tenthredo_koehleri
-MNSNDNSDTIISAKFPGYGSYLECMTRSLFTGLSTFAIVFSGAFFGQRFMH-KSTP--YSKKSTILISTSLAALASYYITSSYTVACKKSWARANGN-------------------
>Mantis_religiosa
-MENINELKRQQSEKHPGFGSYLECMTRTLFTGLASFTLAFSGSFIGQSLLK-NKLP--YGKNGSILISSLLAVGVSYQVTSQRTKACQAAWMASEDKHTYLKDIPAST--------
>Boreus_hyemalis
-MNDIKKLKLQQKDAHPGFGSYLECMTRALFTGLATFSITFSGIYFIQKSQS-KRLP--YTSNGCILVSAGFAVLASYYITASKTKICQAAWMAAEDKHTYLKELDKVS--------
>Triops_newberryi
-MNNIGELKERYTQIYPAFGSYTECMSRALLTGLSASIFGFSASYFAQYLLK-NRIP--YGLKGQVLTSSAIAILVSYKVTSDRSKACQKAWMAAEDKHTYLQDVEKPRSS------
>Calopteryx_splendens
-MNNINALKEQYKEVHPGFGSYLECMSRTLLTGLASFTLAFAGTYIGQSLLK-KSLP--YKKKSFVIISSLVAVAVSYRVTSVRAKACQAGWLAAEEKHTFFSENPTSDK-------
>Ceuthophilus_sp._AD-2013
--------------------------------------MVFASTYLSQKLLQ-KKLP--YGQNMFVLVSSLLAVGVSYQVTAHRTRNCQSGWMAAEDKCTYFTQNMPDE--------
>Extatosoma_tiaratum
-MNDINQLKREQSEKHPGFGSYLECSSRGLLTGLASFTLAFSATYISQTLLQ-NRLP--YSRNSIILCSACAAIAVSYHVTSRRMQACQAAWLAAEDKHTYLSGIGEAD--------
>Leuctra_sp._AD-2013
-MNDIKKLKEQHSEAHPGFGSYLECMTRSLFTGLAAFTLAFSGSFLAQSLLK-KHLP--YRMNGYILVSSGLAVATSYHITAKRTSACQAAWMAVEDKHTYLTDQPEQKN-------
>Inocellia_crassicornis
-MNDIKILKEQQRESHPAFGSYLECMTRALFTGLASFTIVFSSTFFTQKLLVNKIFP--YRLPRAILISSACAVLSSYQITATRTKSCQAAWMAAEDKYTYLTETK-----------
>Thermobia_domestica
-MSS-SNQRKGLSQNHHAHGSYLECMTRAALTGLSSFTIAFSAAYLSQYTFR-HRLPF-SSRRGPIICSLVVASAFTYYTTSRTLKECELASVPSEEKTTSFRIIQEKQ--------

HG0311 FBgn0263251 vnc

```
>Drosophila_melanogaster
------------------------MSLQTGLAFIFTILASVSIVAAIILLGWFLIWKAFLSKFRLVRELLGQEEPEDQQPLAWDHQNQQPHHHGRARKARRD---
>Cryptocercus_wrighti
--------------------------MKELLLAVFSIISSLFLVLILVCLGWYLVWRFFLSRFRFVRELLGGMTESTSVS-------DLKNGRSRIKKTRRE---
>Aleochara_curtula
--------------------------MKDIILAITTVISSLAIVLFLILLGWYIVWKLFLSRFRFVRELLGGGSPAGETP-------E---NKNRSKRSRRD---
>Forficula_auricularia
MS------IDLVPNEGNSVFTPDDMSIQKILAIIAYSIVSIIIVLAIVAFVWYLLWKFFLSKFRFVREFVGAIYETPTEN-------E-NTKSTKSRKSRTRRLD
>Aposthonia_japonica
--------------------------MKEFLLAVFSVISSLLLVLGLVGFGWFLVWKLFLSRFRFVRELLGGSSESSSVN-------DLKNGRSRIKKTRKD---
>Graminella_nigrifrons
---------------------------MEFVLAIFSFLSSFVVVIAIVCLGWYLVWKVFLSNFRFVRELLGGMSDTPPVE-------G-ETRKARIKKARRE---
>Pseudomasaris_vespoides
--------------------------MQEIVLAAFSVISSLLIVFGLVLLGWYLVWKFFLSRFRFVRELLGGMSESSSVN-------DLKNGRSRIKKVRRD---
>Tischeria_quercitella
--------------------------MKQTFLAIVALISSVALVCLFVLICWFILWKCCLSKFRFVRELLGSMSESPPAS-------EVRQ-PTRAKKLRKE---
>Mantis_religiosa
--------------------------MKEFFLAIFSIISSVFLVLVFVCLGWYFVWKLFLSRFRFVRELLGGMSESNSAS-------DLKDGRSRMKKMRRD---
>Bittacus_pilicornis
------------------------------------------------------------------------MTESSSVN-------DLKNGRNRIKKVRLD---
>Sialis_lutaria
------------------------------------------------------------------------MSETSSQT-------NLKNEKGRVKKVRRE---
>Osmylus_fulvicephalus
-------------------MSKNIMNVENMILMVLTIFVTFSTVIVFIFLGWFIVWKCCLSRFRFVKELLSEKQEEEVPV----------RKSSRSRRVHRE---
>Laupala_kohalensis
--------------------------MMELFLAVFSIISSVLLVLVLVFLGWYLVWKLFLSRFRFVRELLGAMSESASVN-------DLKNGRSRMKKTRRD---
>Medauroidea_extradentata
--------------------------MKDFLLAFVSTVVSVAIVVTLVIFGWYLVWKLFLSRFRFVRELLGGANDTPPGS-------E----GKRAKKVRRD---
>Leuctra_sp._AD-2013
---------------------------MEILLTIFSVLFSLALVVSLAGLGWYLVWKFFLSRFRFVRELLGGMSESQSVA-------DLKSGRSRIKKTRLD---
>Ectopsocus_briggsi
--------------------------MISAIFSIFSFLIAFVVVIGLVTFGWFLLWKYLLSRFRFVRDLFGDSRDSTSDS-------P-ESTRPRTRPNRKARID
>Xanthostigma_xanthostigma
------------------------------------------------------------------------MNGNEHPP-------NQRKSSRSSKKVRRD---
>Ctenocephalides_felis
--------------------------MKDFLLACVSIISSVLIVLILVSAGWYIVWKLFLSRFRFVRELLGGMSESSTVV-------DLKNSRNKIKKVRRD---
>Frankliniella_cephalica
--------------------------MKEILLAVFSLFSSLTLVLALVALGWFLVWKFFLSRFRFVRDLLGSRSESRTVA-------ELKTGRSRAKKSRRE---
>Annulipalpia_sp._AD-2013
MCFTHSQKLGKCRNHFRPMFE--GSSTSSLVWAAALSAGLPILLVLLIATGWFLVWKCFLSKFRFVRELLGGNADQSPSA-------EPKDNRNRSKKVRRE---
>Zorotypus_caudelli
--------------------------MNEFLLAAFSIISSFILVLILVSVGWYLVWILFLSRFRFVRELLGNMSENQVSS-------K----KSRLE--------
```

HG0312 FBgn0038641 CG7708

>Drosophila_melanogaster
----------MCGLHFRTRNSPSQKTSLRWGDCCLA-------------------------------------------------
>Hyalella_azteca
----------MCGLGFRLKNTPSKKTSLRWGDCCGCY------------------------------------------------
>Parasteatoda_tepidariorum
----------MVSLRTRLRNFPSKKTSLRWGDSCVRFFP-FSLTFQ---------------------------------------
>Argulus_siamensis
---------MTCNLGFRTKNTPCQKTSLRWGDYC---------------------------------------------------
>Prorhinotermes_simplex
----------MCGLAFRTRNSPSQKTSLRWGDCCTNYSFSHNYPLH-HRQLHSLQSVQKLKTSAMRVFTAFALHQRFHHSVSLKI
>Calanus_finmarchicus
--------MAVCSLGYQTENTPSKKTSLRWGDDCRVYAVFSLHACK-QDR-----------------------------------
>Nicrophorus_vespilloides
----------MCALGFRTRNSPSQKTSLRWGDCCLAS------------------------------------------------
>Homarus_americanus
----------MCGLGFRLKNTPSKKTSLRWGDCCGRGVVKRSCSSRR---QEGI-------------------------------
>Tigriopus_californicus
--------MTTATLGFTTENSPSKRTSLRWGDDCLQIKQTHLDRSC---------------------------------------
>Aphis_glycines
----------MCSLGFKTRNSPSQKTSLRWGDCCPLF------------------------------------------------
>Polistes_dominula
----------MGGLAFRTRNSPSQKTSLRWGDCCPPNPNPPSSSALWKNRQRDMPPVPV--------------------------
>Ornithodoros_turicata
----------MVSPRYRLRNFPSKRTSLRWGDSCTRGLHRHLLSAQ---------------------------------------
>Tischeria_quercitella
----------MCGLRFRTRNSPSQKTSLRWGDCC---------------------------------------------------
>Varroa_jacobsoni
MPYMSPGQKSNGNNLVRHRNFPSKRTSLRWGDSCPSAYPQLPPLHCAASL---LGPF--------------------H-------
>Triops_newberryi
----------MCGLGFRTRNSPSQKTSLRWGDCCFSRNSTFVQSF----------------------------------------
>Nothrus_palustris
---------MFRLIRLRLRNFPSKKTSLRWGDSCGQNVQLKNY------------------------------------------
>Locusta_migratoria
--------MSSCGPAFRTRNSPSQKTSLRWGDSCAAAAAHL--------------------------------------------
>Liposcelis_bostrychophila
----------MCGLGFKTRNSPSQKTSLRWGDCCFNYNY----------------------------------------------
>Oropsylla_silantiewi
----------MCGLGFRTRNSPSQKTSLRWGDCC---------------------------------------------------
>Caligus_rogercresseyi
-----------MSVGYGTENSPSKRTSLRWGDDCCLLAINSED------------------------------------------

HG0313 FBgn0259725

```
>Drosophila_melanogaster
------MSDTRKDPCKPNACRIQACLTSKHRR-----------------------------------------------------
>Gmelinoides_fasciatus
MDREYNSVLKPKDPCKPKACAIQRCLERTRYNVVLCQPYYEKLTECCRQFAARSDA--CGSEEQPPAS-A---------------
>Meinertellus_cundinamarcensi
---------MKKDPCKIFACEIQRCLQENKFQESACQAAINALKKCCKKWREDSLV--CSGIKIDDELNKDIPQK----------
>Reticulitermes_flavipes
---------MKKDPCKHHACNIQKCLKENSYQEQSCQQFIELLRECCCKWGNASVS--CSGIDTRKTP-SQTVRSLACLA-----
>Lepicerus_sp._AD-2013
--------MSSRDPCKPFACNIQKCLKGKEIKRIICFITLF--IVYLRKKLPRAGL--SKGNNTNDKM-L---------------
>Occasjapyx_japonicus
------MPAKPNDPCKPLACEIQRCLVANDFLESKCEEEVKRMLACCDLWGETSGGRSCSGFWINRPGEAPRKRQEAKK------
>Clastoptera_arizonana
---------MKKDPCKKFACEIQKCLKENEFQEDKCKQYIEYLRNCCKTFGPQSTS--CSGIKTSSPPKVD--------------
>Lasius_neglectus
--------MTTVDPCKQFACKLQKCLTDNVYQPSRCQEIIEELRKCCIKHSSISTV--CDGINTSKRYEHNTVDYRKVLK-----
>Papilio_xuthus
----MEFLYMPADPCKKFACEIQRCLSENKFQEVRCARVFESMRQCCLKFKHISLV--CEGYRLEPREFAPVTDRPDKN------
>Tanzaniophasma_sp._AD-2013
----------MKDPCKPFACRIQKCLFVYTCHC----------------------------------------------------
>Nannochorista_philpotti
------MPLPSKDPCKLNACKIQKCLKEHNYQEDACLDIIEQMRLCCVKWGNKSYV--CNGIKTESKQPKYETSNESVKI-----
>Cordulegaster_boltonii
------MKRTKKDPCKVHACRIQQCLKENKFQEDNCKDIINEMQKCCDKWFYESLC--CSGFKKPLQS-----------------
>Tetrix_subulata
--------MKKKDPCKPIACKIQKCLVV---------------------------------------------------------
>Ramulus_artemis
---------MKKDPCKVQACKIQKCLAENNYQESACKEVIKEMVECCRKWSDVSLC--CSGINTKVST-ASSAEV----------
>Cosmioperla_kuna
--------MKKRDPCKAHACRIQKCLAENNFQESACQAVLEDMRSCCRKFKNVSLC--CSGIDIGDTP-TLETASEPKK------
>Inocellia_crassicornis
----------MKDPCKLTACKIQKCLKENRYQEDACAEVLEEMRQCCLKWHQVSLC--CSGIRLDEPKIKD-KKLSSKDDNN---
>Oropsylla_silantiewi
--------MRPKDPCKANACMIQKCLNENNFQESACGEVLELMRLCCVKWGDKSFV--CNGINLEKAENSTDNTKSTKLKPQQKS
>Rhyacophila_fasciata
----------MKDPCKASACKIQTCLKENGYQESRCADVLEAMRQCCLKWHKESLC--CSGIRLEPRPVLQTRQAETHSENTDKN
```

HG0314 FBgn0260468 CG7950

```
>Drosophila_melanogaster
-----------------------------------MSGYQYLDVKIKLRDPDSVTLTPVFFCGCIHSSLASIFGEIGGQTILEIVKFSSSQKRSILRVPENVLDRVRVAIALIGYYQEVPC-HFQVLSTSRKPLDFEESPEEFVAFN-
>Blattella_germanica
-----------------------------------MSGYYYLDVSLEFEGGSRPELNAVYLKNHILQALNTLFGEVGASTRVDILKYNPNTQRAVLRCPSASYVKLRSSLSLCGNYNAVAC-VYRIHNTSSCLLSLLGTSRTYKH---
>Anoplophora_glabripennis
-----------------------------------MSKYYYLDVSLQLAKTSENTITATYFKNNIVNAVNQLFGEVATAVNIDILKYNSTNQRAILRVPKTHYIKLRSSLTLCGTYEGEPC-AYRIHKASPLLLALQGDSRDYQH---
>Aposthonia_japonica
-----------------------------------------------------------------MLSMKTLFGETGTCIDVDIIKYNEKSRRAVIRCPSSSYIKLRSALTLSGNYSDIPC-VYRVHNASSSPLSLLGNSRNYKH---
>Xenophysella_greensladeae
-----------------------------------------------------------------MQALKSLFGEAGQSTCVDILKYDPSHKRAILRCPSYAYVKLRSALTLCGNCSGEEC-AFRIHCCSPSLLALTANSRTYSHR--
>Psyttalia_concolor
--------------------------------------MQYLDVSLDLPSNPSREISPVFLKSHVTQSLKQLFGEKGADCSIDILKYNSTTRRFILRCSSQDYVRLRASLTLASTYEEDNC-IYTVNRSSASPFGLLANGRDYNH---
>Nemophora_degeerella
-----------------------------------MTDYFYLDVSLELDTSDNVEISALYLKKHIIDSLNTLFGEVGASIPVDILKYEPTTKRSILRVPEEFYVKLYSSLTLCGKYQNILC-AYRVHKTTPSLLSLLGDSRTHKF---
>Mantis_religiosa
-----------------------------------MSDYYYLDVSLEIAGKEKQEVTAVYFKNHVMQSLRTLFGEVGGCIIVDVLKYNPEEQRAVLRVPASSYVKLHSSLTLCGEYENTVC-AYRVHNTSPCLLSFLGNSRTYHH---
>Panorpa_vulgaris
-----------------------------------MSNYYYLDVSLELSAA-KSKVSAVLFKKHVIESLQSLFGSISSATQIDILRYNREQIRAIIRVPEKYYVKLRSALTLSGGYEKFQC-AYTIHSSSPCLLSILADSRSYEH---
>Chrysopa_pallens
-----------------------------------MSNYHYLDLTLELPKGLKREISAIFFKKHVMQALNALFGEVGASTTVDLLKYESIPHRAIIRVPSVYYVKLRSSLTLAGDYEGIPC-AYRIHKSSTYLLNLLGDSRNYNHE--
>Prosarthria_teretrirostris
MGRLAKCPCRVLQIQRNVCVAVRRFDNSTNGF--IMSDYYYMDISLICSSPTEEKVSAVYFKDHVMQSLRTLFGEVGSSINVDVLKYDIEKQRAVLRCPSASYVKLHSSLTLSGDYCGSPC-VYKVHQTSAYLLSLLGNSRTYKHI--
>Menopon_gallinae
-----------------------------------MKGYYYLDVLIKFEQG-KISLDENKLKTIIVYSLNTLFGECGSAIEVDVLKYTQTDQRAILRCLADKYVKVRASLSLLNEYAGVQL-QLQVVKASASLLSLQSNSRTYKFA--
>Liposcelis_bostrychophila
-----------------------------------MSSYYYLDVQTKFHQGETIQLTPSEAKETIRKNLKELFGECGTLQSVEILKSYLKEGRFILRCSCSFYVKLRSSLIVCSSLG---N-NFKIIKTSPSLLSLLSKPWETDF---
>Raphidia_ariadne
---------------------MNIFCNYFDIFVILLNVYK----YKEFEKGINKEVTVAFFKKHVLSALNTLHGEVGASTTVDILKYDSTKRRAVLRIPISNYIQLRNALTLVGDYEGIPC-AYRIHNCSAYLLNLIGDSSLYIH---
>Stylops_melittae
-----------------------------------MSNYFYFDISLELEANSTFKPSINNINSHIQQSIKLLFGELGLPHEFHILKYNKAENRFIIRIPKDDFIKFHTALTLSDTFNHVPIITYRIHKITPFLL-FLSNSRTFEY---
>Megalurothrips_sjostedti
-----------------------------------MSKFFYLDVSLEFEDKQKSQVDAVAFKNYIWTAIRTLFGDFGASINVDLLKYNEKDTRGIIRCPAREYVKVRGSLTFCGKIQGTTC-RFLVHQSSSLLLSLCGNSRTYKHC--
>Hydroptila_sp._AD-2013
------------------------------------MTYHYLDVTLDVNIQSAFNITPSSFMKNVTNSVEELFGEIGGAIDIKLLKYNQEDKRAILRVQSNSFEKLYAALTLKGVYESFEC-AYRIHKTSQSLLALTSSSLTYTH---
>Zorotypus_gurneyi
-----------------------------------MNDYSYFDVSLEVADE-NISISKNSFKRHVLSSLNTLFGEVGESVDLKILKYNVQLHRGIIKCPAESAVKLHSALTLSGKFDKYYS-VYKVHKSSTSLLSLAANSRTYNHVVK
```

HG0315 FBgn0260234 CG42508

>Drosophila_melanogaster
MKPKKSAFANSSTGYGRNKHGKGHSGKDKVEGGGKGKSR--DEKD-KHKHEQKVDEKDDKDLNSGFGDYLRTPEAFEMMKLFVFANTIMLIVTMAWPHIKEQFYMVNQWLESFREEHQQ
>Leptinotarsa_decemlineata
MI-----------------------KSRQLKRTKNS-----LNKPIKDREKKSEADRIDSDDEQGFSKWIRSEEGLDNLKLFVLGNTLIMFIAISWPQIKEVLDSAYYLYVEFTERR--
>Aposthonia_japonica
MK---------------------------------------KLMRSNSAMPAATNENRRDNNEEGFTAWLRSPEGIKFMHTFVLANTFVVILTVAWPHMTEVVDIIKSYIFD-------
>Isonychia_bicolor
------------------------------------------------------------------------------MKLFVIANSIIMLTTVGWPHITKTWEILQELLSGEL-----
>Acanthocasuarina_muelleriana
MKNKAAKSGSTAYRGG-------TAYKKLAPSAESKKET--N--PRPKVKKAPAPLTDHINENEGFGGWLSSEEGIGTMKMFVMANSVMMLITVGWPYLYQLLNYLHDSADNTPVRLY-
>Polistes_canadensis
MKKRINNTDNKNTKN------------RSNEERGKQKND--DEKDNEKNTIERLEQNEDEYSNQGFGDWLRSNDGAEMMRLFVIANSLLLFVTMAWPNMKESFYFIRDYFMGEEDDY--
>Micropterix_calthella
MKPRN-----KKSQN------------KANFNTGAALHK--T--ESKLLAKENTEKFVDEDEEKGFGGWLRSTEGIEIMRLFVIANSVVAFLTLAWPQMQKAFGIFQELVAENF-----
>Empusa_pennata
MKKRKSPNTV------------------AATE-----------DKVKPKI---KVDDVSDEENEGFGGWLKSSEGVAYMRLFVLVNSLVVFMTMTWPHIVECYDIVYSYFTE-------
>Euroleon_nostras
MKKHGRKQKKV-TI--------NEKQKQLQEKEKAKEKE--NKKPSDTVDKKN--DIDSDDENQGFSVWLRSSEGIELIQLFVLSNSIVLFVTVAWPQMQEAINIIISFFKETFST---
>Triops_newberryi
MKCKTGNLKK--SK--------------SIEA-----------KDIA-TQPSDTASGGPEEAAGGWTGWLASPEGVGWMQFFVITNAIIILTTTGLPHISAVWDIVKSWF---------
>Prosarthria_teretrirostris
------------------------MKKRKQENNKAHTTN--KDTSSSELVDNP--DISDTETNQGFGNWLRSNEGMEYMRLFVLANSLVVFLTMAWPQMSQAFNTIGSYFTSD------
>Leuctra_sp._AD-2013
MKKKKAGKLT------------------SPPATEV--EK--NISKINTNIVAKQELNEDDEENQGFSDWLKSSEGVSYMRLFVIANSLMVFMTMTWPHLSSVFEIISSFFTDD------
>Ectopsocus_briggsi
MQ----------------------HQHRKQQKEGAAKRKVAKSVVNAAESRKPDVDPDPYDEDAGFGGWLQSGDGVTYLRLFVISNSLMLFLTLGWPHIKESIEIITSFFYD-------
>Hydroptila_sp._AD-2013
------------------------------------------------------------------------------MRLFVIGNSIVVFLTMTWPHMQSTAKIVREMIFGEEE----
>Zorotypus_gurneyi
------------------------------------------------------------------------------MKLFVISNSVVMVLTMLWPQISSAVEVIGSILNDQN-----

HG0316 FBgn0041164 armi

>Drosophila_melanogaster
-----------------------------------------------------------------------------------------MSS---EASGSITIK--E-----EAS-----------TSGTATNIPVESSETEAKKPEIGADKAESQSETTEGGKAAETT
ENHLKRLQNLRKELSYLSETDWMYESLDKKAAQ----
>Homarus_americanus
MASGNSSGLGASLFRGALHGNTSNTNNLSGNSSTSNQQHHHHHHTAGHHSQQGGVTGSPHIHHPQQPQP-------------------PPPQ---PTLGPVTFSTEETSRNRGTP-----------GSPDHMSTV------------MAKLWF---DHNVESSLSSLKK
EEHDKRTRDLRKQLDYITETNWKYSPVEKYIGQQ---
>Meganyctiphanes_norvegica
-MASNSSQFGTSIFRGGGSNNNSQNSNNSINNNNNNNNNNNNNTTPQNQQLSGALQQA---SHPFQSQQTTPIQQQEQQQQLQQAQPSGSSS---QVQYQPGIDNTPNDYADNRS-----------QSPDFMATL------------MAKLWF---DHNVESSLSSLRK
EENEKRMRDLRKQLDYINDTNWKYAPVEKFLGQQ---
>Graphocephala_atropunctata
-----------------------------------------------------MVRVSPGDHGIMGPMN-------------------TIPK---TILSSPSQS--E-AQGSLNS-----------NLNNTSSAV------------LSRLKF---EEAIEKNSVHIEK
EAHLKRVKNLRKELDYLQNTAWKYPSIDHYIGQ----
>Argochrysis_armilla
------------------------------------------------------------------------------------MNSSTLSP---STLGHTSSQRTSESQDSEIT-----------TSGTQTSKI------------LNRMKS---ELQEQKQQKRLLK
ELHLKRVQSLRKELDYLKATEWRYQPIDRYLDQSNRN
>Phymatopus_californicus
-----------------------------------------------------------------------------------MSSNQSISQ---TLLTPPGKI--E-SQST---------------DGPVAH---------DIDQALSHIRL---EHNIERSLRNCHK
EIHLKRLQSLRKELEFLSSTDWQFEDRDKGTSQY---
>Teleogryllus_commodus
-----------------------------------------------------------------MSST-------------------GLPK---TVLSSPTQS--E-SQASELS-----------T--TSLNSE------------LVHLKF---EHSFDQCLLQMQR
DAHNKRMKLLRKELDYIQVTAWKYPVIDKYIGQ----
>Ramulus_artemis
-----------------------------------------------------------------MSTS-------------------RLSK---TVLAPPGQG--E-GQSSELS-----------SSSTLTPNQ------------SNLLKF---EYVMDQVMSASPR
EQHEKRMKVLRKEVEYLQNTAWKYQPIDKFIGP----
>Leuctra_sp._AD-2013
-----------------------------------------------------------------MSSS-------------------GLSK---AVLSSPGQR--E-TQNTEQN-----------NRPTSAGDP------------LGLLKV---DCSVDQSLAQLQR
DDHKKRMKVLRKELNNLQSTAWQYPPIDKFVGQ----
>Thrips_palmi
-----------------------------------------------------------------MSSP-------------------NINR---SVLSTTRAPRKE-TGSSELM--------------NNQLSL------------LARYKV---EHAVDQARSQVLK
EAHQKRLKVLRKELDYLKNSNWMYQPVDKFIGQ----
>Hydroptila_sp._AD-2013
------------------------------------------------------------------------------------MSSSPSTSPVPSVSTAPSPSPAP-SAHSESSAVAHVHPHAAGSSSTVTNIG-----RSFHEDILKQVKL---DSKFNLLAKQAIT
AEHKTRMKSLRKETEYLNTTAWQFESVDKF-------
>Atelura_formicaria
------------------------------------------------------------------------------------------------------------------------------------MMA------------FSRFLF---EHTVDHSHAFVQR
DEHQKRLKALRKELDYLQNTAWKYQPIEKYIGQ----

HG0317 FBgn0050011 gem

```
>Drosophila_melanogaster
MEK--SEIRLQRMSNEYQS----------QS-SYMYL-----RTKML---LKIENTL--LRSHRQRETTGIKKLYNSFFVLF
>Nicrophorus_vespilloides
-----------------------------MD-ILCNH-----LQVKM---IKIDNTF--LQNHRQRETTGIKKLYNSFFVMF
>Trachymyrmex_zeteki
------------------MRCSQGVAMIFEDAFVVFS-----RFLSL---KSRGCTL--LETHRQRETTGIKKLYNSFFVMF
>Papilio_polytes
MEAIPSHQRTYLIGNRFTSKPFPGLVPHIQY-HLCRS-----NTNML---LKIENEL--LQSHRQRETTGIKKLYNSFFVLF
>Osmylus_fulvicephalus
------------------------MDSGI-R-CHYCL-----VQIKM---NRIENSL--LQSHRQRETTGLKKLYNSFFVLF
>Xanthostigma_xanthostigma
------------------------MDTR--Y-FAFYF-----IQDKM---NTIENVL--LQSHRQRETTGIKKLYNSFFVLF
>Ceratophyllus_gallinae
-------MELKLLQFDF-------------S-STRIF-----VNKML---LKIENSL--LQSHRQRETTGIKKLYNSFFVLF
>Stylops_melittae
-------MENKLFG---------KMLSFY-AQFLSYKIFIRNRQRLMQPLLYHQVLLITKHSNQHQKMIDIKCLYNSFFVMF
```

HG0318 FBgn0035293 CG5687

>Drosophila_melanogaster
MKYRN-NRGTILQIQFRTFLRYSAANLIPFES-
>Tribolium_castaneum
MKYRSRSNRQIIQIQFRSFLR------------
>Copidosoma_floridanum
MRYAK-KTSKIIQLQFRTFLRNSSNTGYY----
>Boreus_hyemalis
MKYRN-KSGQILQIQFRTFLIVESGKRS-----
>Corydalus_cornutus
MKYNC-SNRQIIQIQFRTFLLDKRIIDNRILIR
>Osmylus_fulvicephalus
MKYRNRSNRQIIQIQFRTFLSTDKHSR------
>Oropsylla_silantiewi
MKYRN-SCGLILQIQFRTFLLDTRIYDNRG---

HG0319 FBgn0026778 Rad1

>Drosophila_melanogaster
MA---SESENTTSYFVPHLVDQVFELYRSKAESNSLTKEERGQFFYDLSILLGQELVLRSLHLLDDHN-FSFFHAKNNRSVCVVEISK-GTEYFRLIPGVSFCKCEFFQCHVLQLPRGILYQDLPSGEPGILEDWSEESRVSYTCQHILALRLHQFLKHTGGKTTEKILK
KEDIKELRTDVFRD------
>Gyrinus_marinus
-----------------------------------------------MHSLYGQV-FVRALELLEKYN-FTIFQT-TDGLRKVLKVIGT-TEEYALFHNINFCPCRAFQYQVLS------------------------SKSSYTCKHVLAAKLAEI----SGKFKVETYT
SEQITEVLNNQIPDG-----
>Polyommatus_icarus
-------------------------------MRNSTTKQLSDQDLITLHSVFGPV-LQRALEILEKQPLLEVYTT-ANKSRTLLEVKGEKDLRYRIFPMVNFCPCQAFKHQVLE------------------------KRIQLTCKHILAGKIALL----LGKTKEHEVT
QEQYQMLLHSMFEQQQ----
>Panorpa_vulgaris
-------MSKEKSTFLPHT---LVELIAKVKKSYSANKSFSEEDFLSLHSIFGTV-LEKALEILDRYN-ITVYRT-SS-SRCLIQVASH-DTYYKLFQDVNYCSCQAFKFQVLG------------------------TQKQYTCKHVLAAILADI----TGKIITEQIS
DEQFNVLIESMVSL------
>Annulipalpia_sp._AD-2013
MSYDSRTKKLSTCKILPRI---AFDLLKKVQNESSESKKISDDNLLTLHSVFGNV-LERALDVNSKRNSVTVYTL-QNGSRSLIEVRGSNGQTYRLLPQVNYCQCPAFQNYALQ------------------------LKTELTCKHVLAAYVADV----TQSAVENIVN
REQFDLLFDSFVASDEFELL

HG0320 FBgn0040993

>Drosophila_melanogaster
-MKT---PDTKAKLLNNISLSKHLSSKKGGSRTSNCGNCLEFSVLTRLSSVP
>Cosmioperla_kuna
MNSRAFVYQETKRFLPTVRLSKHLNTRKG-SRRGNAGNRLELQVLTQLTHVP
>Frankliniella_cephalica
MNNR---ADTKAKLLNNLSLSKHLSSKQGASRTTNCGNRLEFALLTLRQNIP

HG0321 ENSDARG00000070426 chac1

```
>Danio_rerio
--MHQAKKSVLRDCTITTAKQLGSDTVRGIAVSQPIPLLLDRRKLTLAT---
>Channa_punctata
MAMHRARQNQLRDSIIATAKQLGSNTVRGIPASQPVPLLLDRRKLTLAS---
>Anguilla_japonica
--MHQAKQKIRRYSIAKTAKQLGT--TRGITASQPIPLLLDREK--------
>Basilichthys_microlepidotus
MAMHSARQNQPRNSIFATAKQLGSNTVRGIPASQPVPLLLDRRKLTLAT---
>Oryzias_melastigma
MAMHSARQTRPRASTRATAKQLGSGTVRGIPASQPVPLLLYRRRPQLAAAAS
>Tripterygion_delaisi
MAMHRARQTQPRPSISAAARQSASSTVRGIPANQPVPLLLDRRKITIAG---
>Seriola_quinqueradiata
--MHRARQNQLRNSIFATAKQLGSNTVRGIPASQPVPLLLDRRKLTLAS---
>Micropterus_salmoides
--MHSARQNQLRNSIFATAKQLGSNTVRGIPASQPVPLLLDRRKLTLAS---
>Pygocentrus_nattereri
--MHQAKQKHPRTSILTTAKQLGSSAVRGIAVSQPIPLLLDRRKLVLAS---
>Haplochromis_sp._redtail_she
MAMHSARQNKQRNSIFATAKQLGSNTVRGIPASQPVPLLLDRRKLTLAS---
>Clupea_harengus
--MHSARQN-VNLSIFTTAKQLGSNTVRGIAVSQPIPLLLDRRKQILAS---
>Xiphophorus_maculatus_x_Xiph
MAMHSARQNQPRNSIFATAKQLGSNTGRGIPASQPVPLLLDRRKLTLTTS--
>Esox_lucius
--MHRARKNRLRNSIFTTAKQLGSNTVRGITVSQPIPLLLDRRKLTLAS---
>Apteronotus_leptorhynchus
--MHQAKKNGLRNSIFTTAKQLGSNTVRGIAVSQPIPLLLDRRKLTLGS---
>Lutjanus_erythropterus
MAMHSARRNQLRNSIFATAKQLGSSTVRGIPASQPVPLLLDRRKLTLAS---
>Plecoglossus_altivelis
--MHLARQNRQRNSIFTTAKQLGSNTVRGISVSQPIPLLLDRRKLILAS---
>Scleropages_formosus
--MHRTRQ----TCITKTAKQFGSHT-----ASRPIALLLDRRKYSTP----
>Lateolabrax_maculatus
--MHRARQHRARDSISAAAKQLGSSAVRGIPASQPVPLLLDRRKLTLAS---
>Gasterosteus_aculeatus
--MHSARQNPLRNSIFATAKQLGSNTVRGIPASQPVPLLLDRRKLTLAS---
>Solea_solea
MAMHQAKKYQLRDSILATAKQLGSNTVRGIPVSQPVPLLLDRRKLTLAS---
>Oncorhynchus_nerka
--MHRAKQNGLRYSIFTTAKQLGSNSVRGITVSQPIPLLLDRRKLTLPS---
>Ictalurus_punctatus
--MHQAKQTDLRNSIFTTAKQLGSSTVRGIAVSQPIPLLLDRRKLALAG---
>Pagrus_major
MAMHSARQNKLRNSIFAPAKQSGSNTVRGIPASQPVPLLLDRRKLTLAS---
>Takifugu_rubripes
MAMHSARQH-HRISILATAKQLGSNTVRGIPASQPVPLLLDRRKLTLAS---
```

HG0322 ENSDARG00000070917 kitlga

>Danio_rerio
MMILDMYLWDEENPFPARRSKIK
>Channa_punctata
-----MCTQDDDSPFPPRRSKIK
>Anguilla_anguilla
-----MCARNYENPFPPVRSRIK
>Menidia_menidia
-----MCTQDDDNPFPHRRSKIK
>Opsanus_beta
-----MCTQDDENPFPPRRSKIK
>Oryzias_latipes
-----MCTQDDDNPFPPRRSKIK
>Tripterygion_delaisi
-----MCTQDEDNPFPTRRSKIK
>Micropterus_salmoides
-----MCTQDDDNPFPPRRSKIK
>Pygocentrus_nattereri
-MSLDMRKRDEENPFPARRSKIK
>Haplochromis_burtoni
-----MCTQDDDNPFPPRRSKIK
>Austrofundulus_limnaeus
-----MCLQDDENPFPLRRSKIK
>Esox_lucius
-----MCTRDDENPFPPRRSKIK
>Gadus_morhua
-----MCTRDDESLFPHRRSKIK
>Apteronotus_leptorhynchus
-MILDMCRRHEESSFPARRSKIK
>Scleropages_formosus
-----MCTRNDESPFPPQRSKIK
>Lateolabrax_japonicus
-----MCTQDDDNPFPPRRSKIK
>Sebastes_caurinus
-----MCTQDDDNPFPPRRSKIK
>Cynoglossus_semilaevis
-----MCTQDDDSPFPPRRSKIK
>Salmo_salar
-----MCTRDDENPFPPRRSKIK
>Ictalurus_punctatus
-MVLDMHERDEEKSFPARRSKIK
>Takifugu_rubripes
-----MCTQDDDNPFPPRRSKIK

HG0323 ENSDARG00000071197 usp40

```
>Danio_rerio
---MFIPVKSSYARV-SPLIVNRELGKTKMR
>Channa_punctata
---MFSPVKSSYFRV-SPAIVNRELGKTKMR
>Menidia_menidia
---MFSPVKSSYFRV-SPAIVNRELGKTKMR
>Porichthys_notatus
---MFSPVKSSYFRV-SPAIVNRELGKTKMR
>Oryzias_latipes
---MFSPVNSSYFRV-SPAIVNRELGKTKMR
>Trachinotus_blochii
---MFSPVKSSYFRV-SPAIVNRELGKTKMR
>Micropterus_floridanus
---MFSPVKSSYFRV-SPAIVNRELGKTKMR
>Haplochromis_burtoni
---MFSPVKSSYFRV-SPAIVNRELGKTKMR
>Clupea_harengus
MIAMVSPVKSVHSGV-NPMVVNRELGKTKMR
>Poecilia_mexicana
---MFSPVKSSYFRV-SPTIINRELGKTKMR
>Esox_lucius
---MFTPVKSSYFRV-SPTTINRELGKTKMR
>Acanthogobius_hasta
---MFSPIKSSYFRV-SPAIINTELGKTKMR
>Apteronotus_leptorhynchus
---MFSPVKSAYLRV-SPTTVNRELGKTKMR
>Lutjanus_erythropterus
---MFSPVKSSYFRV-SPAIVNRELGKTKMR
>Plecoglossus_altivelis
---MFSPVKSSYFRV-SPTTVNRELGKTKMR
>Scleropages_formosus
---MFSAVQSEYPRLTLPRAVNRELGKTKMR
>Lateolabrax_japonicus
---MFSPVKSSYFRV-SPAIVNRELGKTKMR
>Notothenia_coriiceps
---MFSPVKSSYFRV-SPAIVNRELGKTKMR
>Hippoglossus_hippoglossus
---MFSPVKSSYFRV-SPATVNRELGKTKMR
>Salmo_salar
---MFTPVKSSYFRV-SPTTINRELGKTKMR
>Takifugu_rubripes
---MFSPVKSSYFRV-SPTIVNRELGKTKMR
```

HG0324 ENSDARG00000094965 nfil3-5

>Danio_rerio
MTVVFVEFHLFLIFLTSLDT
>Channa_punctata
MSEVFVEFHLFLIFLANLDT
>Menidia_menidia
MSEVFVEFHLFLIFLANLDT
>Porichthys_notatus
MSEIIVEVRLFLIFLANVDT
>Oryzias_latipes
MSEVFVEFHLFLIFLANLDT
>Micropterus_floridanus
MSEVFVEFHLFLIFLANMDT
>Astyanax_mexicanus
MRDIFIEFHLFLIFLANLDT
>Oreochromis_niloticus
MSEVFVEFHLFLIFLANLDT
>Clupea_harengus
MKEVYVEFHLFLIFLSNFDT
>Austrofundulus_limnaeus
MSEVFVEFHLFLIFLANLDT
>Acanthogobius_hasta
MYKVILEFHLFLIFLANLDT
>Lutjanus_erythropterus
MRVANAELQLFFIFLANLDT
>Plecoglossus_altivelis
MCEVFVEFHLFLIFLANFDT
>Scleropages_formosus
MRKLYVEFHLLLIFLANLDT
>Lateolabrax_japonicus
MSEVFVEFHLFLIFLAYLDT
>Sebastes_caurinus
MSEVFVEFHLFLIFLANMDT
>Cynoglossus_semilaevis
MSKIFVEFHLFLIFLANLDT
>Oncorhynchus_mykiss
MRGINAEFHCFLIFLASLDT
>Lepisosteus_oculatus
MWEIFVESHLFLIFLANLDT
>Takifugu_rubripes
MSEVFVEFHLFLIFLANMDT

HG0325 ENSDARG00000079738 znf219

```
>Danio_rerio
----------------MYLIQVKRNHDVCCH
>Channa_argus
----------------MYLIQVKRNHDVSCH
>Menidia_menidia
-MLVYNPDFHFFSQLVMYLIQVKRNHDVSCH
>Porichthys_notatus
----------------MYLIPVKRNHDVSCH
>Oryzias_melastigma
----------------MYLIQVKRNHDVWCH
>Micropterus_salmoides
----------------MYLIQVKRNHDVSCH
>Pygocentrus_nattereri
----------------MYLIQVKRNHDVCCH
>Haplochromis_burtoni
----------------MYLIQVKRNHDVSCH
>Alosa_alosa
----------------MYLIQVKRNHDVSCH
>Austrofundulus_limnaeus
----------------MYLIQVKRNHDVSCH
>Esox_lucius
----------------MYLIQVKRNHDVSCH
>Apteronotus_leptorhynchus
----------------MYLIQVKRNHDVSCH
>Scleropages_formosus
----------------MYLIQVKRNHDVSCH
>Lateolabrax_japonicus
----------------MYLIQVKRNHDVSCH
>Sebastes_caurinus
----------------MYLIQVKRNHDVSCH
>Cynoglossus_semilaevis
-MLVYKPDFHFFPQLVMYLIQVKRNHDVSCH
>Salmo_salar
MLVYNNPDFHFFPQLVMYLIQVKRNHDVSCH
>Ictalurus_punctatus
----------------MYLIQAKRNHDVSCH
>Takifugu_rubripes
----------------MYLIQVKRNHDVSCH
```

HG0326 ENSDARG00000012848 arih2

>Danio_rerio
---------MSDGECDFFCFFRCC
>Channa_argus
---------MSDGECDFFCFFRCC
>Menidia_menidia
---------MSDGECDFFCFFRCC
>Oryzias_latipes
---------MSDGECDFFCFFRCC
>Tripterygion_delaisi
---------MSDGECDFFCFFRCC
>Micropterus_salmoides
---------MSDGECDFFCFFRCC
>Pygocentrus_nattereri
---------MSDGECDFFCFFRCC
>Oreochromis_niloticus
---------MSDGECDFFCFFRCC
>Austrofundulus_limnaeus
---------MSDGECDFFCFFRCC
>Acanthogobius_hasta
---------MSDGECDFFCFFRCC
>Apteronotus_leptorhynchus
---------MSDGECDFFCFFRCC
>Plecoglossus_altivelis
---------MSDGECDFFCFFRCC
>Scleropages_formosus
---------MSDGECDFFCFFRCC
>Lateolabrax_japonicus
---------MSDGECDFFCFFRCC
>Anoplopoma_fimbria
---------MSDGECDFFCFFRCC
>Cynoglossus_semilaevis
---------MSDGECDFFCFFRCC
>Ictalurus_punctatus
---------MSDGECDFFCFFRCC
>Takifugu_rubripes
MNIQSSAVQMSDGECDFFCFFRCC

HG0327 ENSDARG00000022203 gtf2ird1

```
>Danio_rerio
-------------MTFNFTGLQNKE-RGKGTHSSVKENREKTGQTHQIN-HTECFSTD---YRPFEIMTFSKVAVAFL-----------------------------------
>Amia_calva
-------------MGFNFTGLQNKE-RGKGTHSSAKEKRKKGKNR-AII-HRRTVHKAPTISKCFGLQCFGLRWPLYRDRGTN----RAVSIGLNGEEGCVAGLTADGI----
>Channa_punctata
MLLLSWAGISRRRMSFNFTGLQNKE-RGKGHPALLREKRNK---LRQNT-QIIDY----KPRERSQQCDFSDYGASAGGTATLWMGIVSVSDGLKRGINLFRD----------
>Menidia_menidia
MLVLSWAGISRRRMSFNFTGLQNKE-RGKGHPAL-------------------------------------------------------------------------------
>Porichthys_notatus
MLLLSGAGISRRRMSFNFTGLQNKE-RGKGHPAQLKQRNKKKEKNLKGK-RDRL-----------------------------------------------------------
>Oryzias_latipes
-------------MSFNFTGLQNKE-KEKGTPCSAKTGKTGINWEKSKE--SETAVQ----GAMIRIRVFF------NGREL-------------------------------
>Tripterygion_delaisi
MLLLSWAGISRRRMSFNFTGLQNKE-RGKGHPALVKGAKKQ---E--------------------------------------------------------------------
>Pygocentrus_nattereri
-------------MSFNFTGLQNKE-RGKGTLCSVKEKQKEKRTGKQD-----------------------------------------------------------------
>Oreochromis_niloticus
MLLLSWAGISRRRMSFNFTGLQNKE-RGKGHPALLRGKNRHKLRKKKNT-QIADN----GTWARSRRGDDPDYDVLADGTATLWMGIVSVSDGLKRGNKPIPGLTAATRLFNR
>Fundulus_heteroclitus
MLLLRWAGISRRRMSFNFTGLQNKEKRGKGHPALLKGINKKQNSENKTG-ETPGL----QES-------KPSPGQ--------------------------------------
>Esox_lucius
-------------MSFNFTGLQNKE-RGKGTHSTVKENSKKRGTFK-------------------------------------------------------------------
>Apteronotus_leptorhynchus
-------------MSFNFTGLQNKE-RGKGTLSSVIENRKKRTKIDKII-SPKIFKQATKKRNMLQIMRFHKLAIAFL-----------------------------------
>Scleropages_formosus
-------------MAFNFTGLQNKE-RGERGPCAALKKTERNGHKH-------------------------------------------------------------------
>Cynoglossus_semilaevis
MLLLSWAGINRRRMSFNFTGLQNKE-RGKGHPALLKG----------------------------------------------------------------------------
>Salmo_salar
-------------MSFNFTGLATEQ--RNGTHSTVKENRKNGGNRKIISVIT-----------------FC------------------------------------------
>Lepisosteus_oculatus
-------------MSFNFTGLQNKD-GKGDTLLR-------------------------------------------------------------------------------
>Ictalurus_punctatus
-------------MSFNFTGLQNKE-KGKGDTVPALKTRKKDGGRKKKRSKSRCNLTRKGRRTRSRLCGFCEVSVALF-----------------------------------
>Takifugu_rubripes
MLLLSWAGISRRRMSFNFTGLQNKE-RGKGTPDRVKGEKQE------------------------------------------------------------------------
```

HG0328 ENSDARG00000099999 tcf3b

>Danio_rerio
----------MWMEVVKMSTRRMNSPQL
>Menidia_menidia
----------MWTEVVKMSTRRMKSSQL
>Oryzias_latipes
----------MWTEVVKMSTRRMKSSQL
>Tripterygion_delaisi
----------MWTEVVKMSTRRMKSSQL
>Astyanax_mexicanus
----------MWMEVVKMSTRRMNSSQL
>Neolamprologus_brichardi
----------MWTEVVKMSTRRMKSSQL
>Alosa_alosa
MRKDTLLLPTMWMEVVKMSTRRMKSLQL
>Kryptolebias_marmoratus
----------MWTEVVKMSTRRMKSSQL
>Esox_lucius
MRRD-TLLLTMWTEVVKMSTRRMKSSQL
>Gadus_morhua
----------MWTEVVKMSARRMKSSQL
>Apteronotus_leptorhynchus
MRRNTLLLLTMWMVVAKMSTRRMNSSQL
>Plecoglossus_altivelis
----------MCSEVVKMSSRRMKSAQL
>Scleropages_formosus
----------MWMEVVKMSSRRMK----
>Lateolabrax_japonicus
----------MWTEVVKMSTRRMKSSQL
>Notothenia_coriiceps
----------MWTEVVKMSTRRMKSSQL
>Cynoglossus_semilaevis
----------MWTEVVKMSTRRMKSSQL
>Oncorhynchus_mykiss
MRRDTLLLLTMWTEVVKMSTRRMKSS--
>Takifugu_rubripes
----------MWTEVVKMSTRRMKSSQL

HG0329 ENSDARG00000036442 atp11c

```
>Danio_rerio
MRMVREDSIQLIWKRCSFTAALHIARGNR-LESR--------------------------
>Channa_argus
-----MNRNQTKWKRCSVAAVFHIARTNARVETR--------------------------
>Menidia_menidia
MRMVRGDRNQTKWRRCSVAAAFHIARTNVRVESR--------------------------
>Oryzias_melastigma
MRMVRRYRSQTKWKRCSVAAAFHIARTNDRLESR--------------------------
>Tripterygion_delaisi
MRMVRRSRNQTKWKPCSGAAASPIARTRVGTR----------------------------
>Micropterus_floridanus
MRMVRRYRNQTKWKRCSVAAAFHIARTNVRVENR--------------------------
>Pygocentrus_nattereri
MQMVREERSQPEWKRCSFTAALHIARRNR-VENR--------------------------
>Maylandia_zebra
MRMVRRYRNQTKWKRCSVAAAFHIARTNVRVENR--------------------------
>Alosa_alosa
MRMVRGERLPRKWKQCSNTATFHVARTNWIE-NR--------------------------
>Austrofundulus_limnaeus
MRMVRRYRNQTKWKRCSVAPAFHIARXNVRVENR--------------------------
>Esox_lucius
MRMVRRDRIQTKWKQCSIAAAFHIARWTSRVENR--------------------------
>Gadus_morhua
MRMVRRARNQTKWKRCSLDAVPHIARRHVRVETR--------------------------
>Apteronotus_leptorhynchus
MRMIREQRSETKWKQCSFTAAFHIDWKHR-LGNR--------------------------
>Plecoglossus_altivelis
MRMVRRERIQTKWKQCSIAAAFHIARTRV--GNR--------------------------
>Notothenia_coriiceps
MRMVRRYRNQTKWKRCSDAAAFHIARTNVRVENR--------------------------
>Salmo_salar
MRMVRRDRIQTKWKQCSIAAAFHIAISTTRVENRELVNKYKGNVTYLSEKLKQSIVNWFK
>Takifugu_rubripes
MRMVPRSRNRAQWEACSVAVAFHIARYSARVESR--------------------------
```

HG0330 ENSDARG00000039392 wnk1b

>Danio_rerio
MDEVDLTVFVCH--HQCLPLSNCVLFFYVYKRTPPLLGSGPL-HCKSNC----
>Acipenser_sinensis
MDGVVWTVLVSQ--CLRLLFCDCILFFSNYKRTPPVQGPDPL-HCNCIHCPY-
>Amia_calva
MDGVDLMVLVCQ--HPCLIFCDSVLFFSIYKRTPPLQGPDPV-RCKILNFLC-
>Menidia_menidia
MDGVDLKRLEYQ--QQHPAFSCSVLSLSIFKRTPPILHHQSPNYICKDIL---
>Pygocentrus_nattereri
MDGVDLTVLVCH--HQRLLLCDCVLFFYIYKRTPPLLSSDHK-YCNSNCL---
>Clupea_harengus
MDGVDVTVLQCP--LQCLPVCDCVLFFSIYKRTRPIQGFGPQ-CCLYSRNCSC
>Kryptolebias_marmoratus
MDGVNLKILLYR--HQCSAFCCSVRSFSTYKRTPPIIHPQSLRYIYKDSL---
>Esox_lucius
MDGVDLTVLVCH--HQRLLLCDCFLFFSIYKRTPPLLGPTPPERCKSNRDCCC
>Acanthogobius_hasta
MEENYLK---SP--PRHCGLYNSDQSFSNYKRTPPILHIVNL-HCKSRNYKQ-
>Apteronotus_leptorhynchus
MDGVDLTVLVCR--HQCLLLCDCILFFYVYKRTPPLLGLCPT-YCISNCL---
>Plecoglossus_altivelis
MDEVDLKSLVCL--HPCPPFCDCVL-FSIYKRTPPVLHLHRE-PCLHSNWL--
>Scleropages_formosus
MGCVDLMVLVCQQQHQCLLFCDGVLFFSTYKRTPPLVGPDPV-WCKSIRYFC-
>Lateolabrax_japonicus
MDGVDLKRLVYQ--QQHSEFCCGVQSFTIYKRTPPLLHLQSLHYIYKDRL---
>Trematomus_bernacchii
MDGVDLKRLVYQ--HQHSVFFCGVQSFTIYKRTPPILYPQSLHYIYKDRQ---
>Oncorhynchus_mykiss
MDGVDLTVLVCH--HQRLLLCDCFLFFSIYKRTPPLLGPTTPERCKSYSNYCC
>Lepisosteus_oculatus
MDCVNLMVIVCL--HQHLIFCDIILFFFIYKRTPPLQGPDPL-HCKILSFHC-
>Trichomycterus_areolatus
MDGVDLTLLIYH--RQRLCLSDCILFYYAYKRTPPLLG--------------

HG0331 ENSDARG00000044899 tmem183a

>Danio_rerio
MYATLSVSSIMAEDPGGAFFNFTLILTDFSLPLL--ERWQG----
>Channa_punctata
----------MAEDPGGTFFNFSSIFLFFSLPQLAIETWQDRKPR
>Oryzias_melastigma
---------MMAEDSGGTFFNFYLIFLPFFVPKLVIETWLDL---
>Micropterus_floridanus
---------MMAEDPGGTFFNFSLIFFSFSLPPLAIETWQDSQAR
>Pygocentrus_nattereri
---------MMAEDPGGAYRIFSFIFLRFSPPLL--QTWQGAQSR
>Neolamprologus_brichardi
---------MMAEDPGGTFFNFSVIFLSFSLPQLAIETWQDSQSR
>Cyprinodon_variegatus
---------MMAEDPGGTFFNLSVIFLSFSLTRLAIEPWQDSQAR
>Esox_lucius
---------MMAEDPGGTLFNFSPTFFSFSLLLLVLETWQDTQP-
>Gadus_morhua
---------MMAEDPGGTFLKFTQFFISFYLGLLVLETWQHTQSR
>Scleropages_formosus
---------MMAEDPGGPYFNLSFIFLSFSLSLLVVPTWQEPQLR
>Lateolabrax_maculatus
---------MMAEDPGGTLFKFSLIFLSFSLPPLP----------
>Sebastes_caurinus
----------MAEDPGGTFSNLSFIFLSFSLPPLAIETWQDPQAR
>Scophthalmus_maximus
---------MMAEDPGGALFNFSIIFLSFSLPRLAIEPWQDSHPR
>Salmo_salar
---------MMAEDPGGTIFNFSPTFLFFSLPLLVIETWQDTQPR
>Lepisosteus_oculatus
----------MAEDPGGVDFNFSFISFRFSLSLLVVATWQEPQSR
>Ictalurus_punctatus
----------MAEDPGGAYFNFSLNFLCFLLPLL--EIWQGTQSR
>Takifugu_rubripes
---------MMAEDPGGTISNFSLFFLFFALPPLAIATWQDSQVC

HG0332 ENSDARG00000102893 atg7

>Danio_rerio
MKRNSVYRFLSVFSCLLVCVFLF-FWGFAFYSMVLAP--
>Amia_calva
MKRNGVYQFFSVFSCLVVFLFLFLFWFFVFYTMVMTP--
>Channa_argus
MKRNGAYRFLSVFSCFVVCAFLFLFWFFIFYSLVLAP--
>Menidia_menidia
MKRNGAYRFLSVFSCFIVCVFLFLFWFFVFYSMMHAP--
>Porichthys_notatus
MKRNGAYRFLSVFSCFVVCVFLFLFWFFVFYSLVLAP--
>Oryzias_melastigma
MKRNGAYRFLSVFSCFIVCAFLFLFWSFVFYSMLRAP--
>Pygocentrus_nattereri
MKRTGAYRFLSVFSCLLVCVFLFLFWFFVFYSMVLAP--
>Haplochromis_burtoni
MKRNGAYRFLSVFSCFIVCAFLFMFWFFVFYSMVLAP--
>Poecilia_formosa
MKRNGAYRFLSVFSCFVVCVFLFLFWFFVFYSMVLAP--
>Esox_lucius
-MRSGAYKFLTVFSCLIVCGFMFMFWFFVFYSFAP----
>Apteronotus_leptorhynchus
MKRDGAYRLLSVFSCLVIFVFLLFFWVFVFYSMVLAP--
>Plecoglossus_altivelis
-MKRGAYKFLSVFSCFVVCAFLFLFWLFIFYSMMLAP--
>Scleropages_formosus
MKKICAYRLLSVLSCVGICLFLFLFWFFVFYTLVMAP--
>Lateolabrax_japonicus
MKRNGAYKFLSVFSCFIVCAFLLLFWFFIFYSMVLAP--
>Platichthys_stellatus
MKRSGAYRFLSVLSFFAVCASLLLFWFIIFYTMMHVT--
>Trichomycterus_areolatus
MKRNGAYRFLSVFSCFAVCAFIFIFWFFVFYSMFLAPEN
>Takifugu_rubripes
MKRNGAYKFLSVFSCFIVCAFLFIFWFFIFYSMVHAP--

HG0333 ENSDARG00000042518 hipk2

```
>Danio_rerio
MQEIAAIL----SAAQDGRFLLSCEQCCRFIFCHE
>Porichthys_notatus
MRE--ATADTRSSLLQDGRFLLSFVQKARFLLLP-
>Astyanax_mexicanus
MQETAAIL----SAAQDGRFLLSSELKYLFIFCHE
>Oreochromis_niloticus
MQETAS---TLSSAFQDGRFLLSFEQKYRFIFCHE
>Kryptolebias_marmoratus
MQETAATLSSPSSLFQDGRFLLSFKQKARFIFCHE
>Esox_lucius
MQETATL-----SVAQDGRFLLSSEQKSRFIFCNE
>Acanthogobius_hasta
MQEIAATLSSP--VSQDGRFLLSFERKSRCIPLP-
>Plecoglossus_altivelis
MQETATL-----TVAQDGRFLLSSEISLYFLP---
>Scleropages_formosus
MQEEPLL-----SADQDGRFLLSSELRSPPIYLP-
>Notothenia_coriiceps
MQETAA---TLSSLFQDGRFSLSFEQKSRFIVLP-
>Cynoglossus_semilaevis
MQENTATL-----SSQDGRFLLSFEQNRRFIFCHV
>Oncorhynchus_kisutch
MQETATL-----SGAQDGRFLLSSEQISRFIFCHE
>Lepisosteus_oculatus
MQETATL-----TVAQDGRFLLSSKLKSRFIFLP-
>Trichomycterus_areolatus
MR----DQDRIVSAAQDGRFLLSSGLKDLLTVLQ-
>Takifugu_rubripes
-MNWASNAATLSSALQDGRFSLSSEKKPLFIFLP-
```

HG0334 ENSDARG00000075349

```
>Danio_rerio
-------------------------MTSLCGYHRNQGSTADG----------------------
>Anguilla_japonica
-------------------------MTSLHGYHRNQGPSANG----------------------
>Trachinotus_blochii
-------------------------MTSLLGFNKSQSSTADG----------------------
>Micropterus_floridanus
MRVKLFRQEKKQHGLLCFGSFLLCDMTSVLGYNRSQSSTAAGGRHLVRCVFLQQCKALPRQSPQ
>Astyanax_mexicanus
-------------------------MTSPSGYHRNRRSTADS----------------------
>Maylandia_zebra
-------------------------MTSRLGYKRSSSSTADG----------------------
>Alosa_alosa
-------------------------MTSPCGCHRNQGCPADG----------------------
>Esox_lucius
-------------------------MRSLHGFYRNQGSTVDG----------------------
>Acanthogobius_hasta
-------------------------MTSLIEYTRNQSSSSDG----------------------
>Apteronotus_leptorhynchus
-------------------------MTSPSSCHRNQGSTADS----------------------
>Plecoglossus_altivelis
-------------------------MTSLHGYHRNQGSTAHG----------------------
>Lateolabrax_japonicus
-------------------------MTSLLGSNKSQSSTADG----------------------
>Anoplopoma_fimbria
-------------------------MTSLFGYNKSQSSTADG----------------------
>Oncorhynchus_mykiss
-------------------------MTSLSGYHRNQGSTAEG----------------------
>Ictalurus_furcatus
-------------------------MTSSSGCHRNQGSTADS----------------------
```

HG0335 ENSDARG00000034700 vegfab

```
>Danio_rerio
MWIVIALEEKFDLLLNNVRGPASWILIFLDILKIIGNS
>Channa_striata
MWIATPLQDDKSLLRNNVRRCASWILVFLDILKIVGNS
>Porichthys_notatus
MWIAIALKENKDLLRNNVTRRTSWILVFLDILKVVGNS
>Oryzias_latipes
MWIATP-QESQDLPRNDVRRCASWILVFLDILKIIGNS
>Pygocentrus_nattereri
MWIVIPFEEILDLLQNNVRGFASWVLTFLDILKTVSNG
>Ophthalmotilapia_ventralis
MWIATPLQENKDLLRNNVRRCASWILVFLDILKIIGNS
>Poecilia_latipinna
-----MWRENKDLLRNDVRRCASWILAFLDILKIVGNS
>Gadus_morhua
MWIAIPLQENDSLLQNNVKRYASWILVFLDILKNIGNS
>Apteronotus_leptorhynchus
MWIVILFEENISLPENNVRGFATWVLTFLDILKIIGNS
>Notothenia_coriiceps
MWIAIPLQEDKGLLRNNVRRFASRILVFLDILKIIGNS
>Scophthalmus_maximus
MWIATALQENTGPLRNNVRRYASWIFVFLDILKSIGNS
>Oncorhynchus_mykiss
MWIAISFEENKGLLQNNVKRYASWILMFLDILKIIGNS
>Ictalurus_punctatus
MWIVRAFEQKLSLRRDHVSGFASWLLTFLDILKIIGNS
>Takifugu_rubripes
MRILISLQENKGRLRKNVTSCASWILVFLDILKIIGNS
```

HG0336 ENSDARG00000028676 nuak1b

>Danio_rerio
-------------------------------------------MRVIYVQNAGVVAAFSPEIFCGRLK----QRNNGEIMTERATGKRTRHNGVGYVIME
>Channa_argus
---------------------------------MRSCTWAVDKMRVIFIQNAGFVAAFSPRRCESVMLAAFRTTDNVNEAAELTTARSCQS------SEE
>Menidia_menidia
---------------------------------MRSRTWAADKMRVIFIQNAGFVAAFSPFS--------------------------------------
>Oryzias_latipes
---------------------------------MRCSTWAVDKMRVIFIQNAGFVAAFSPFS--------------------------------------
>Pundamilia_nyererei
---------------------------------MRSFTWAVDKMRVIFIQNAGFVAAFSPFS--------------------------------------
>Clupea_harengus
-------------------------------------------MRVIYIQNAGFVAAFSGGNDG------------------------------------
>Kryptolebias_marmoratus
-------------------------------------------MRVLFTQNAGFVAAFSPLL--------------------------------------
>Esox_lucius
-------------------------------------------MRVIFIQNAGFVAAFSGGNDAEGFAC-------------------------------
>Apteronotus_leptorhynchus
-------------------------------------------MRVIYIQNAGCVAAFSGSNDG------------------------------------
>Notothenia_coriiceps
---------------------------------MRSCTWAVDKMRVIFIQNAGFVAAFSPFRCESVMLAAFRTTDKVNEP-ELTTARSCKS------TEE
>Cynoglossus_semilaevis
---------------------------------MRVYTWAKDKMRVIFIQNAGFVAAFSPRRCESVTLAAFRVTNTA-EEPELNTNKSCQS------AE-
>Salmo_salar
MAIFKTSELDISDHSASLKSTWAWSTCAIHRVQQLNDVAPEDKMRVIFIQNAGFVAAFSGGNDAEGFAC-------------------------------
>Takifugu_rubripes
----------------------------------MNGQPHLDKMRVIFIQNAGFVAAFSPFRRESVMSAAFRTTDKVTDLQ-SASAGSCQA------IKE

HG0337 ENSDARG00000054906

```
>Danio_rerio
------------------MFLHRADLSLRPRTKYRDSD-------------------
>Channa_argus
------------------MFLHRADLNLSPRTKRFPLVRLRVGFLLPERSWRQTGGV
>Micropterus_salmoides
------------------MFLHRADLNLSPRTKRFPLR-------------------
>Oreochromis_niloticus
------------------MCLHRAHLNLSPRTKRAPLR-------------------
>Alosa_alosa
------------------MCLHRLDSNLRPRTKYRGLH-------------------
>Esox_lucius
------------------MFYHRADLSLRPRTKYRDFH-------------------
>Lutjanus_erythropterus
------------------MCLHRADLNLSPPTKRFPLR-------------------
>Plecoglossus_altivelis
------------------MFLHRTDLSLRPRTKYRDFH-------------------
>Lateolabrax_maculatus
------------------MFLHRADLNLSLRTKRAPLI-------------------
>Salmo_salar
------------------MFHYRADLSLRPRTKYRDFH-------------------
>Ictalurus_punctatus
MIAGARSLGGSGQGRSCSMFHHRANLSLRPRTKYRAFD-------------------
>Sparus_aurata
------------------MFLHRADLSFSPRTKRVPLK-------------------
```

HG0338 ENSDARG00000000151 thraa

```
>Danio_rerio
MLTCRGSSPPQSAVIGCWMESR
>Channa_argus
-------------MIGCWMELR
>Pygocentrus_nattereri
-------------MIGCWMELR
>Neolamprologus_brichardi
-------------MIGCWMELR
>Alosa_alosa
-------------MIGCWMELR
>Poecilia_reticulata
-------------MIGCWMELR
>Esox_lucius
-------------MIGCWMELR
>Lateolabrax_japonicus
-------------MIGCWMELR
>Trematomus_bernacchii
-------------MIGCWMELR
>Salmo_salar
-------------MIGCWMELR
>Ictalurus_punctatus
-------------MIGCWMELR
```

HG0339 ENSDARG00000063144 shisa7b

```
>Danio_rerio
MNPDLFMPVQWAQPCCPVRTGPKPPFVLFCWVELNPEQRHSRKRVQHT---
>Menidia_menidia
MNPDIFMSVLRAEPRCPARTGPKPSFVLFCWAKLNSEQRHSREGIVRTQTR
>Oryzias_latipes
MNPDIFMSALWAEPRCPARTGPKPSFVLFCWAKLNSEQRHSRDGVST----
>Pygocentrus_nattereri
MNPTSFMSVQWAQSRCPARTGPKPPFVLFCWAELNPEQRHSRKRVQHT---
>Oreochromis_niloticus
MNPDSFMSVLWAEPRCPARTGPKPSFVLFCWAKLNSEQRHSREGIKRT---
>Alosa_alosa
MNLSQFMGFLWTEPCSPARDGPKPTFVLFCWAELSPEQRHLCKGATHT---
>Kryptolebias_marmoratus
MNPDIFMSLLWAEPRCPARTGPKPSFVLFCWAKLNSEQRHSREGIERT---
>Esox_lucius
MNPDIFMSVLWAEPRCPARTGPKPTIVLFCWAELNSEQRHSRKGIKHT---
>Lateolabrax_japonicus
MNPDIFMSVLWAEPRCPARTGPKPSFVLFCWAKLNSEQRHSHKGIERT---
>Platichthys_stellatus
MNPDIFMSVLWAEPRCPARTGPKPSFVLFCWAKLNSEQRHSHKGIERTFL-
>Salmo_salar
MNPDIFMSVLWAEPRCPARTGPKPTFVLFCWAELNSEQRHSRKGIKRT---
```

HG0340 ENSDARG00000017591 fat1a

```
>Danio_rerio
--MQRLGTYKLHEKPPNKGR--
>Channa_punctata
--MPRSGLHKLHAKPPNLRVYQ
>Menidia_menidia
--MPRFGLHKLHAKPPNIRD--
>Neolamprologus_brichardi
--MPRFGIHKLHAKPPNLRI--
>Poecilia_reticulata
--MRRFGLQKLHAKPPNLRDY-
>Esox_lucius
--MQTFGPYKLHEKPPNPRR--
>Anoplopoma_fimbria
--MPRFGLHKLHAKPPNIRD--
>Oncorhynchus_masou
--MQTFVPYKLHEKPPNPRR--
>Ictalurus_furcatus
MYMYKLGTYKLHEKPPNHGR--
```

HG0341 ENSDARG00000039430 msl2b

>Danio_rerio
----------------MMTRIWSYFVPWKRGKR
>Amia_calva
-----------------MMFIWAYFVPWKRGKR
>Anguilla_japonica
MSLLFFNYIQVKDVHLMMMLVWAYFVPWKRGKR
>Alosa_alosa
-----------------MMILWAHFVPWKRGKR
>Esox_lucius
-----------------MMIIWAYFVPWKRGKR
>Apteronotus_leptorhynchus
------------MDFYPMMIVYAYFLPWKREKR
>Plecoglossus_altivelis
-----------------MMLIWAYFVPWKRGKR
>Salmo_salar
-----------------MMIIWAYFVPWKRGKR
>Lepisosteus_oculatus
------------------MLIWAYFVPWKRGKR

HG0342 ENSDARG00000045801 fbxl14b

```
>Danio_rerio
------------------------------------MYDRLDPRVERRLLSKRQTGSSAA
>Amia_calva
MQSETSLLHDCGRKRRSDRPGLFLLYITGLVEVFLKMYDRLDPRGELEGLSQDKVGSAV-
>Alosa_alosa
------------------------------------MYDRLDPRVDRGRLSEHKMGSVA-
>Esox_lucius
------------------------------------MYDRLDPRRERRHVSKGIIGTVV-
>Apteronotus_leptorhynchus
------------------------------------MYDRLDPRVERRRLSRHKMGSAV-
>Scleropages_formosus
------------------------------------MYDRLDPRQEAEGSSEHTSGCAAV
>Salmo_salar
------------------------------------MYDRLDPRREPRRVSKGIIGSVV-
>Lepisosteus_oculatus
------------------------------------MYDRLDPRGEREGLSQDKVGSAV-
>Trichomycterus_areolatus
------------------------------------MYDRLDPRVERRRLSEHIMGSNAV
```

HG0343 ENSDARG00000057062

```
>Danio_rerio
---------------MDYSRTDANLGFPTELLP---
>Menidia_menidia
------------MPVMDYGRTSANLGWHS-------
>Porichthys_notatus
------------MSVMDYSRTDANLSRHSQSSIRLH
>Trachinotus_blochii
------------MPVMDYSRTDANLSWHSQNSIPPY
>Micropterus_floridanus
------------MQVMDYSRTDANLSWHSQNSIPPY
>Pygocentrus_nattereri
---------------MDYSRTYANLGFPAKLSP---
>Alosa_alosa
MKRVGYCKVPVQAQAMDYSRTDANLGFPAKLSS---
>Notothenia_coriiceps
------------MLVMDYSRTDANLSWHSQNSIPPN
>Ictalurus_punctatus
------------MPVMDYRRTDANLAFSAEVSP---
```

HG0344 ENSDARG00000032072 cdk16

>Danio_rerio
MAEDYWERAAALGYYYCC
>Acipenser_sinensis
MAEDYWESTAALGKH---
>Amia_calva
MAEDYWEQTAAPGHY---
>Astyanax_mexicanus
MAEDYWEHTAALGHC---
>Clupea_harengus
MAEDYWESTVALGYYC--
>Esox_lucius
MAEDYWESTVALGYC---
>Salmo_salar
MAEDYWESTLALGYFC--
>Lepisosteus_oculatus
MAEDYWELPAARGHY---

HG0345 ENSDARG00000035910 znf281b

```
>Danio_rerio
MEEANSCKGTLFWL
>Channa_punctata
MEEPSSCKRTMFWL
>Trachinotus_blochii
MEEPSSCGKTMFWL
>Micropterus_salmoides
MEEPSSCGRTMFWL
>Astyanax_mexicanus
MEEGNSCERTMFWH
>Maylandia_zebra
MEEPSSCGRTMFWL
>Notothenia_coriiceps
MEEPSSCGRTMFWL
>Ictalurus_punctatus
MEEGNSCRRTMFWQ
```

HG0346 ENSDARG00000041982 adamts6

```
>Danio_rerio
------------------------------MFVCTVAASASAGKQMDSSAQVPNIDLLLWNNMKGVVMDSCPAHG
>Channa_punctata
-------------------------------------------MQMDRSRQVPNIELPVWSYT------------
>Oryzias_melastigma
---------------------------------------------MDRSGQVSSIGLPVWSST------------
>Pygocentrus_nattereri
---------------------------------------------MDSSRQPPNIDLLLWNYLK-----------
>Apteronotus_leptorhynchus
---------------------------------------------MDRSRQAPHIGLLLWN--------------
>Lateolabrax_maculatus
-------------------------------------------MQMDRSRQVSNIELPVWSYT------------
>Notothenia_coriiceps
-------------------------------------------MQMDRSRQVSNIELPVWSYT------------
>Lepisosteus_oculatus
MRKILKRKGEEGSELTAELQCRRVLLSVSSSVGNASDNEDQNGTQMDSSRQTSSIDLLFWNYLKSFN--------
```

HG0347 ENSDARG00000063194

```
>Danio_rerio
----------------------MADLSKSKPGLK---
>Pygocentrus_nattereri
----------------------MADLSKSNPGLK---
>Oreochromis_niloticus
----------------------MADLSKSNPGLK---
>Poecilia_formosa
MGEEYEMDSVSQRANRVDDEVKMADLSKSNPGLK---
>Apteronotus_leptorhynchus
----------------------MADLSKSNPGLK---
>Cynoglossus_semilaevis
----------------------MADLSKSNPGLK---
>Salmo_salar
----------------------MADLSKSNPGLKIET
>Lepisosteus_oculatus
----------------------MADLSKSNPGLK---
```

HG0348 ENSDARG00000036139 lctla

>Danio_rerio
MSFLLAALDKLAYLLSLVAFCVSLSYSWLK
>Opsanus_beta
MSLILTVLDKLAYFLSLVAFCVSLSSSWMK
>Pygocentrus_nattereri
MSFLLAALDKLTYFLSLIAFCVSVSYSWLK
>Oreochromis_niloticus
MSLILTVLDKLAYFLSLIAFCVSVSYSWLK
>Fundulus_heteroclitus
MSLILTVLDKLSYFLSLVAFCVSASHGWLR
>Esox_lucius
MSLLLTVLDKLTYLLSLIAICVSASNSWLR
>Salmo_salar
MSLILTVLDKLTYFLSLIAFCVSVSYGWLK

HG0349 ENSDARG00000028699 crkl

```
>Danio_rerio
------------------------------MHQSWLTLVAACLVKRQVAVQ
>Channa_punctata
-MVVVSFFVFIF-LYFGREHLLTAQRRNHIMHQSWIKLVAACL--------
>Micropterus_dolomieu
------------------------------MLQSWIKPVAACL--------
>Alosa_alosa
------------------------------MHQSWLMLVPACL--------
>Apteronotus_leptorhynchus
------------------------------MQESWFMPVAACL--------
>Notothenia_coriiceps
MFVCRKLFCFSILAAQTITQPLTAEHRNHIMFQSWIKPVAACL--------
```

HG0350 ENSDARG00000036194 pitx2

```
>Danio_rerio
---------------------------------------------------------------------------------MQVDIISHTCPCHSGSPCSRLKPRYLLPSRNGKKIFP
>Lipochromis_sp._matumbi_hunt
------------------------------------------MGGRNMVTWRSGERRSWIDSAQATAEMNSEWCS----VCMPLDLISPTCPCHSDSQAR------------------
>Poecilia_reticulata
MLLFGRGKISASPGINTLADISRFCELRLVGHIHMDAPVRENRGGGKAVTWRCGERRSWMGRAQATAEMSRERWS----VCKPLDLISPTCPCHSDSQPR------------------
>Esox_lucius
---------------------------------------------------------------------------------MALDLISPTCPCHSDTPRS------------------
>Apteronotus_leptorhynchus
----------------------------------MSSLMTCQKEGGFCVTSGCRERLSWTAKEQGTAEMKSERCALCENVCMQVDIISPTCPCHSGFIAARGMKLR------------
>Platichthys_stellatus
--------------------------------------------------------------------MNTERCG----VCMPLDLISPTCPCHSDSQQR------------------
```

HG0351 ENSDARG00000068096 atf5a

>Danio_rerio
MRTKPCFLWLNSQTSLSCTLSSGI--
>Pygocentrus_nattereri
MLTQPCFLWLNSQTTISCTINSAV--
>Esox_lucius
MLTQPCSLWLNSQTPPTHIVPSDYSD
>Apteronotus_leptorhynchus
MLTQPFLLWLNSQTTRSCTVNSAV--
>Salmo_salar
MLTQPCSLWLNSQTPPTHILRSD---
>Ictalurus_punctatus
MHTQPCFLWLNSLTTLTCITSSDS--

HG0352 ENSDARG00000070214 slc41a1

>Danio_rerio
MGNIIQVLPKFRQDLTQAWRRGTHLPLFLDL---------------
>Astyanax_mexicanus
MGNIICMLPKFRQDLKQARRGVMHLPLLLNL---------------
>Alosa_alosa
MGSIICMLPKYRQDLSRAWSTGMHLPFLSDL---------------
>Esox_lucius
MENILELLPKHRQGLTCAWGRAATHTCLRGRTCSLVAPSNLFFITT
>Salmo_salar
MENILELLPKHRQEGTHAWGRAATHTCLCRRKSSLVAPLNRFFITT
>Ictalurus_punctatus
MGNIICMLPKFRQDLMQSLERAKAPASFTQPVEREL----------

```
HG0353 ENSDARG00000078864 lats2

>Danio_rerio
-----------------------------------MANMHTRLGWKKHADVNPHMSLEEGECEP
>Astyanax_mexicanus
--------------------------------------MQTMPEWNQPTSVTPFMSWEEGECDP
>Alosa_alosa
-----------------------------------MAKMCTRLAWSKAVLLTPHKTWDEGERDP
>Esox_lucius
-----------------------------------MAQMGARMVWGKGTLVTPHMTWNKEECDP
>Salmo_salar
-----------------------------------MAQMGARTVWGKGTLVTPHMTWNKEECDP
>Trichomycterus_areolatus
MSFFKTTRVKQSINELSDGKVSSVTLLTTVRNISLMAMMQTRLEWNQPTCMNPFMSAKEGKCDP
```

HG0354 ENSDARG00000019897 gpat4

>Danio_rerio
MIIFPYMPMKSMFQRRLQKLRHSRGVQQSVFR
>Astyanax_mexicanus
--------MKALFQRRLQKLRL----------
>Esox_lucius
--------MKALFQRRLQKLRLYMD-------
>Apteronotus_leptorhynchus
--------MKTLFQRRLQKLRL----------
>Ictalurus_punctatus
--------MKALFQRRLQKLRL----------

HG0355 ENSDARG00000019930 tal1

>Danio_rerio
MDDPPQNGSMHTLTSTARRR---
>Acipenser_sinensis
MDGPTQNGSTQFLTSKYKR----
>Amia_calva
MDGPPQNGSTHGIISTHKR----
>Osmerus_mordax
MDDHRQNGSTHTPSTTFKQRLRT
>Salmo_salar
MDDPRQNGSTHTLSTTLRLW---

HG0356 ENSDARG00000020475 znrf1

>Danio_rerio
MIRPGKWAVASARHSCFSQT
>Alosa_alosa
MFRPGERAVASHRHSWRSQT
>Esox_lucius
MFRSGKWAVASVRHSCFSQT
>Salmo_salar
MFLSGKWAVASVIHSCFSQT
>Ictalurus_punctatus
MIRPRKRALASPGHSCFSQT

HG0357 ENSDARG00000037285 mipa

>Danio_rerio
MSAGHAHHLSIRVPLLNLS-----------------------------
>Oreochromis_niloticus
MSAGHAYRLSIHVLLLTTSGVGL-------------------------
>Cyprinodon_variegatus
MSAGHT------------------------------------------
>Esox_lucius
MSAGHAQPLSIHVPLLTTIRVSILMQRDRYSAPRRQHPGLYDKAKKRL
>Salmo_salar
MSAGHAQPLSINVPLLTTSRVGI-------------------------

HG0358 ENSDARG00000062506 zc3h18

```
>Danio_rerio
MAALTVSCRVYAFFR------------------
>Pygocentrus_nattereri
MAALTVSCGVYAFLR------------------
>Alosa_alosa
MAALTLSCRVYYFLR------------------
>Plecoglossus_altivelis
MAALSVSCRFFVFFNIADKSEIHSFTLESKKLK
>Ictalurus_punctatus
MAALTVPCRACAFLR------------------
```

HG0359 ENSDARG00000070348 hoxc10a

>Danio_rerio
----------------------MSASYCPYPLRHKGDCFESKEKQDK
>Astyanax_mexicanus
----------------------MSASYCPNPLRRKRDCFKT------
>Alosa_alosa
---------MLLIF---LPPQKMSASYCPYPSRKKRNCCKTKE----
>Apteronotus_leptorhynchus
----------------------MSASYCPYPLRHKRDCLKT------
>Ictalurus_punctatus
MTGRLESIIMLPIFSLPPTSQKMSASYCPYPFRHKRACFKT------

HG0360 ENSDARG00000070801 opa1

>Danio_rerio
MCPIWKDELNFILQRDPKLIYICH-
>Pygocentrus_nattereri
MPPIWKDELQFVLK-----------
>Clupea_harengus
MSTIWKDDLNCVFTFKLKKLYICQL
>Cyprinodon_variegatus
MCPVWKDELNCTRR-----------
>Anoplopoma_fimbria
MCTIWKDELNRA-------------

HG0361 ENSDARG00000077581 zzz3

>Danio_rerio
MCSKRFAEVRCPHTAAEERAPR---
>Oryzias_latipes
--MFIIGQTRCPHSAAEERAPG---
>Haplochromis_burtoni
----MSPAGRCPHLAAEERAPR---
>Apteronotus_leptorhynchus
MCSRRFAEVRWPHSAAEESTTMRSQ
>Ictalurus_punctatus
--------MRCPYSAEEESTRMRSQ

```
HG0362 ENSDARG00000078355 zc3h4

>Danio_rerio
MSFISHRNCRT-ENLRNVFNLFLF--------------------
>Esox_lucius
MSFILQRNCRT-ENHRTQFQPISLTTVSGELTTMN---------
>Apteronotus_leptorhynchus
MSFISHRNCKDGENFRNVSACFSFKLLMGINCYELKLFLKPINK
>Scleropages_formosus
MSFILQRSCRT-ENLRTHFNLSLFKQLMGN--------------
>Salmo_salar
MSFILQRNCRT-ENLRTQFQPISLTTVSGELTTMN---------
```

HG0363 ENSDARG00000095896 pou3f3b

>Danio_rerio
MRYNLPALKKFLQKGGGDQQTYS
>Alosa_alosa
MRHNLPALKKFLRKGGGLHQTYS
>Esox_lucius
MRYNLPARKK-------------
>Notothenia_coriiceps
MRYNLPARKKRL-----------
>Cynoglossus_semilaevis
MRYNLPARNR-------------

HG0364 ENSDARG00000004988 ppp3ca

>Danio_rerio
MPLHQPHLVTANTRRE
>Alosa_alosa
MPLHQPHLAPAKTYRE
>Esox_lucius
MPLHQQHLATAKTHRE
>Salmo_salar
MPLHQPHLATAKTHRE

HG0365 ENSDARG00000016132 keap1a

```
>Danio_rerio
-----MPTWLETESCYSGLLTRQKKHGSLSLNDNQLCLYFFMARGWV
>Pygocentrus_nattereri
MYHSVMPLWSEMDHCHDGLRTRQKKHGSLSPSDHQLCLYFFMTRGWV
>Apteronotus_leptorhynchus
-----MPIRSETVPCHHGLQTRQNKRSSLSPSDHQLCLYFLMTRGWV
>Trichomycterus_areolatus
-----MPIWSAQDPSHDGLRLRLKKPGSLSPSDHQLCLYYFMARGWI
```

HG0366 ENSDARG00000017803 gsk3b

>Danio_rerio
MELSGSERP-RISVLKLGNWKILIRIAG
>Pygocentrus_nattereri
MELAGAERP-RVSVSKPGNWKTLLRILG
>Apteronotus_leptorhynchus
MELRGTERP-WISVFKLGNWKILIRIVG
>Ictalurus_punctatus
MELRGAEVQWWISVFKSGDW--------

HG0367 ENSDARG00000034056 csnk1g2b

```
>Danio_rerio
-----MFTVLPRTDFKETMRSTRGRSLVGFH---------------------------------------------
>Astyanax_mexicanus
MKNVCLFAPFPKTDLEETMRSLRGRFAVAFWIQ-------------------------------------------
>Esox_lucius
-----MSAVLPKTDQKEAMRGSGGSCQAVLWIQLLGF---------------------------------------
>Ictalurus_punctatus
------------------MRSPTGRSALADQIQYQEACVCACICERVSECACMRAHAITSCLMHAVMCSTPQKEEP
```

HG0368 ENSDARG00000058606 sik1

>Danio_rerio
MTCKCKCKNHLCCGRRWKLFAS
>Clupea_harengus
MTCKSAVVNLFDCGRRWNLFVC
>Chanos_chanos
MTCKFVSKSDFSCGRRWKLFVY
>Apteronotus_leptorhynchus
MTYKYASENHFGCERRWNLLVL

HG0369 ENSDARG00000059276 hoxd4a

>Danio_rerio
------MKSLHLDWPSWSHGRLTLFS
>Pygocentrus_nattereri
MASSALIGFLHLDWPSWSHGCPTLFS
>Lateolabrax_japonicus
------MDQHHLYWPIRSHGSLTLFS
>Sebastes_caurinus
------MNQHHLHWPIWSHGSLTLFS

HG0370 ENSDARG00000062693 nrxn3b

>Danio_rerio
-----MWQNNRIRSNMENLCFNPERRLKGTSSVKHF--
>Pygocentrus_nattereri
MLSFTMQRYCRVKSNMEDLCYKPGSRFKRMSSGKQIGH
>Esox_lucius
-----MGHGCGSEAKMATVCFNPCSRLKGNSSNGSLYH
>Salmo_salar
---------------MVAVCFNPCSRLKGKSSDGPLYH

HG0371 ENSDARG00000062765 whsc1l1

>Danio_rerio
-------------------MEHAEAGPVRKRET
>Pygocentrus_nattereri
-------------------MEH-AEGPVRKRET
>Alosa_alosa
MIVHLCEQKQGSNIWHLFRMEH-AEGPVRKRET
>Scleropages_formosus
-------------------MEH-AEGPVRERET

HG0372 ENSDARG00000069467 igsf9bb

>Danio_rerio
MQGMSKRVSAADSASNDQGLG-
>Apteronotus_leptorhynchus
---MSKRISADDSASNVLGARL
>Lepisosteus_oculatus
---MSKRVLAGDCANTVQGARR
>Ameiurus_nebulosus
---MSERIPAAASASSIPGARL

HG0373 ENSDARG00000074611 wdr37

>Danio_rerio
MTVALRSAEAVINALGFLFEVFLFFITLSSRLRMCLLCLSFTALHGCFR---------
>Pygocentrus_nattereri
---------------------------------MRLLCLSFTALHCPVPDRKCAVGRH
>Apteronotus_leptorhynchus
----------------------------MIALKMCLLCLSFTAPHCPVPGGTSAEGRH
>Trichomycterus_areolatus
---------------------------------MCLLCFSCTALHCPVAGRKCAVGRH

HG0374 ENSDARG00000075147 lrrc38a

```
>Danio_rerio
MISWKIELRVFVAAFRCTSLCRLQSRRQGKMPSRQT------------
>Pygocentrus_nattereri
MISWKTAFRVFVAVVRSRSFFSL-------------------------
>Esox_lucius
MISWKTLFRVSLA-ARWPGLCRLHTKRKKDASARTLTPFFLLLPPPLP
>Apteronotus_leptorhynchus
MISWKVASHISVA-VRFWSFCRL-------------------------
```

HG0375 ENSDARG00000076171 znf827

>Danio_rerio
MGKDLFSSYP------
>Porichthys_notatus
MGKNLFSSYPQTRTRM
>Plecoglossus_altivelis
MGKNLFSSYP------
>Oncorhynchus_kisutch
MGKNLFSSYP------

HG0376 ENSDARG00000077229 ano8b

>Danio_rerio
MTHNNLHFTLTFI
>Alosa_alosa
MSHNNLHFTFTFI
>Apteronotus_leptorhynchus
MTHNNLHFTFTFI
>Lepisosteus_oculatus
MSHGNLHFTFTFI

HG0377 ENSDARG00000079549 cdc42ep1a

```
>Danio_rerio
MQPRFSGIGVMHIFSSHLSSPNLSGYEQRPMAKTGHS----------------------------------
>Astyanax_mexicanus
----------MHIFNSHPSQLEHFGHGQEPATKELCVFLMEKAFLRNISLSCSFFSFALRSPEPGIVLHEW
>Alosa_alosa
----------MHIFNSHFS----------------------------------------------------
>Ictalurus_punctatus
----------MSIFISHLSQWDHFGYGQRPVAKNLCVHTYGKFSLRNISLSCTFFSLTFLEKP--------
```

HG0378 ENSDARG00000080009 bahcc1b

>Danio_rerio
MIGTYFFGRQSQNCAECKT--------
>Apteronotus_leptorhynchus
MTGTYFFGRQIKNCAECETWAKLVTGL
>Scleropages_formosus
MIGSYFSGWGSQQYRECNT--------
>Trichomycterus_areolatus
MTGTYFFGRHTQNGAECKT--------

HG0379 ENSDARG00000104372 gnb1b

>Danio_rerio
MRGFDSHTIVFFGQCHINKGHLRLRTDQNVSRQ------
>Esox_lucius
MRGLEGYTVTFSSQCQKTQSLRQLRKGSSVPAAGERQKY
>Apteronotus_leptorhynchus
MRGFDSYTIVFFGQCHIKQGQLRLSTDQCVSRQQE----
>Ameiurus_nebulosus
MRGFDSNTVFSFGQCHIKHGQLRLNTDPCVSRQ------

HG0380 ENSDARG00000005350 dip2ba

```
>Danio_rerio
-------------MAEGVVLRMDCYSPFNSRFMP
>Plecoglossus_altivelis
-------------MAEGPIVRMDCLPDLNACSMS
>Trichomycterus_areolatus
MGSRDQRSASSHTMAEGAAVRMDCSSHFNTRFES
```

HG0381 ENSDARG00000006124 kdm5c

```
>Danio_rerio
---------------MACSSDRMNGG
>Astyanax_mexicanus
MGLEGLCLGCDEKPKMACSLDRMNGG
>Clupea_harengus
---------------MACSTDRMNGG
```

HG0382 ENSDARG00000017242

>Danio_rerio
MTEPFFTADACGGGLTCRC
>Alosa_alosa
MQEPFFRTNECGGG-----
>Apteronotus_leptorhynchus
MKQPLFTADECGGGLTCLC

HG0383 ENSDARG00000036501 rab39bb

```
>Danio_rerio
-------MMLLLSLPFHLHKTSKHFSIFQTWIKT-------
>Alosa_alosa
MRDESGYMMLLMLLPFHQ-QNSKHFFIFKTWSENNTSQYHN
>Platichthys_stellatus
MRDESGDMMLLLALPFHQ-QTSKHTQPAPT-----------
```

HG0384 ENSDARG00000054748 cuedc1b

>Danio_rerio
MQLNKQKKRRKGFLSLQWTRFFIV
>Pygocentrus_nattereri
MQLNKQKKRKKAFSFLSGLT----
>Apteronotus_leptorhynchus
MQLKKQKRRRKGLLLPPWTPTTLH

HG0385 ENSDARG00000059549 akap10

>Danio_rerio
MNEGNRESSGCCL
>Salmo_salar
MNEGTRESTGCCH
>Ictalurus_punctatus
MNEGTRESTRCCH

HG0386 ENSDARG00000062084 clcn1a

>Danio_rerio
MCEEETERLSVCVCMCPCHPRVGMLRQTGSLFLGQ
>Alosa_alosa
-----------------------MLRQTSGLFLGQ
>Salmo_salar
-----------------------MLRQTAGLFLGQ

HG0387 ENSDARG00000071150 cica

>Danio_rerio
MQERNRVEDIDRLGLTGNQED
>Astyanax_mexicanus
MREGNRVEDIDRLRLTENLED
>Apteronotus_leptorhynchus
MRVGNRVEVLDRLGHTGNLED

HG0388 ENSDARG00000077228 ntrk3a

```
>Danio_rerio
MQPVEEDSLGALIFEPFLH
>Alosa_alosa
MQPGDEDSIGVSIFQPFHL
>Salmo_salar
MQPREEDKPGVVIFEPFLI
```

HG0389 ENSDARG00000077361 bptf

>Danio_rerio
MAAASDLKRGAGRSF
>Alosa_alosa
MAAMSDPKRDAGRSF
>Apteronotus_leptorhynchus
MAAASDSKRDAGHSF

HG0390 ENSDARG00000090035 rtn4r

>Danio_rerio
MTERGT-TGRRCSPSRARTATVG
>Esox_lucius
MTERGS-TRRRCSPTRFSRKTSG
>Apteronotus_leptorhynchus
MTERGRINGRLCSPSRAFKATVG

HG0391 ENSDARG00000099634 elna

>Danio_rerio
MIHRTKGFALS
>Pygocentrus_nattereri
MIHRTEGFALS
>Trichomycterus_areolatus
MIHRTEGFAFS

HG0392 ENSDARG00000100003 glulb

```
>Danio_rerio
MRMRDKQPGLPYKTLWKYSHSYSDYLYFLIGLI
>Astyanax_mexicanus
--MREKQARKSSRSSWTHSQSGIDYLFLLKGFL
>Ictalurus_punctatus
--MREKEFVLSY---------------------
```

HG0393 ENSGALG00000002090 GPSM2

>Gallus_gallus
MELKGPQMKELERVSDSLRCV--
>Anser_cygnoides
MELKGPQMQELEHVSESVRCVWQ
>Chaetura_pelagica
-------MKELEHVSEGLRCV--
>Apteryx_australis_mantelli
MELKGPQMQELERVSESLRCV--
>Buceros_rhinoceros_silvestri
MELKGPQMKEREHVSEGLRCVWQ
>Caprimulgus_carolinensis
MELKGPQMKEREHISEGLRCV--
>Cariama_cristata
MELKGPQMKELEHVSEGLRCV--
>Calidris_pugnax
MELKGPQMKELEHVSEGLRCV--
>Pterocles_gutturalis
MELKGPQMKELEHVSEALR----
>Colius_striatus
MELKGPQMKELENVSEGLRCV--
>Columba_livia
-------MKELEHVSEGFRGV--
>Leptosomus_discolor
MELKGPQMKELEHVSEGLRRV--
>Cuculus_canorus
MELKGPQMKELEHVSEGFRCV--
>Aquila_chrysaetos_canadensis
MELKGPQMKELEHVSEGLRCV--
>Gavia_stellata
MELKGPQMKELEHISEGLRCV--
>Mesitornis_unicolor
MEPKGPQMKELERVTEGLRCV--
>Tauraco_erythrolophus
MELKGPQMKELEHVSEGLRCV--
>Opisthocomus_hoazin
MELKGPEMRKPEHVSEGV-----
>Ficedula_albicollis
MELKGPQMKERTRTSLTV-----
>Phaethon_lepturus
MELKGPQMKELEHVSEGLRCV--
>Picoides_pubescens
MDLKGPQMKELEHVAEGLRCV--
>Fulmarus_glacialis
MELKGPQMKELEHVSEGLRCV--
>Nestor_notabilis
MELKGPQMKELEHVSEGLRCV--
>Aptenodytes_forsteri
MELKGPQMKELGTPF--------
>Tyto_alba
MELKGPQMKELEHVTEGLRCV--
>Struthio_camelus_australis
MELKGPQMQELEHVSESLRCV--
>Tinamus_guttatus
MELKGPQMQELEHGAECLRCI--
>Calypte_anna
MEIRGPEMKEQEHISEDLRCV--
>Apaloderma_vittatum
-------MKELEHVSEGL-----

HG0394 ENSGALG00000002479 MAT1A

```
>Gallus_gallus
MRSPCMHTQIQAHADFTD----
>Anser_cygnoides
MHTPCVHAQIQAHAEFTR----
>Chaetura_pelagica
MHSPCMCTQMQAHADFTH----
>Apteryx_australis_mantelli
MHTPCTHTQIQAHADFTH----
>Caprimulgus_carolinensis
MHTPCMRTQMQAHADFTH----
>Cariama_cristata
MHAPCMRTQMQAHAEFTH----
>Charadrius_vociferus
MHTPCMRTQMQARADFTY----
>Pterocles_gutturalis
MHTPCMRTQMQAHADFTH----
>Colius_striatus
MHTPCMHAQMQAHAGFTH----
>Columba_livia
MHTPCMCTQMQAHADFTH----
>Merops_nubicus
---------MQAHAEFTHSNLS
>Aquila_chrysaetos_canadensis
MHAPCMHTQMQAHADFTH----
>Gavia_stellata
MHTACMRTQMQAHADFTH----
>Chlamydotis_macqueenii
MHAPCMRTQMQAHADFTR----
>Tauraco_erythrolophus
MHTPCMCTRMQAHADFTH----
>Manacus_vitellinus
MRSPCVHAQMQAYAAFTH----
>Egretta_garzetta
MHTPCMCTQMQAHADFTH----
>Fulmarus_glacialis
MHTPCMRTQMQAHADFTH----
>Nestor_notabilis
MHTPCMCTQMQAHAVFTH----
>Pygoscelis_adeliae
MHTPCMRTQMQAHADFTH----
>Tyto_alba
MHTPCMRTQMQAHADFTH----
>Struthio_camelus_australis
MHAPCTHT--------------
>Tinamus_guttatus
MHTACTHSQLQARADFTH----
>Calypte_anna
MHAPCMCTQMKAHADFIH----
>Apaloderma_vittatum
MHTPCVCTQMQTHADFTH----
```

HG0395 ENSGALG00000014936 ESCO1

>Gallus_gallus
MLLAHRDKLLWPKHRLPLVKT
>Anser_cygnoides
MLLAHLDKSLWPEHRLPLVKT
>Apteryx_australis_mantelli
MLLAHLDKLLWPKHRLPLVKT
>Buceros_rhinoceros_silvestri
MLLAHLDKLLWPQHPLHLVKT
>Caprimulgus_carolinensis
MLLAHLDELLWPKHRLPLVKT
>Cariama_cristata
MLLAHLDKLLWPKHRLPLVKT
>Calidris_pugnax
MLLAHLDKLLWPKHRLPLVKT
>Colius_striatus
MLLAHLAKLLWPKHRLPFVKT
>Columba_livia
MLLAHLDKLLWPKHRLPLVKT
>Merops_nubicus
MLLAHLDKLLWPKHRFPLVKT
>Aegypius_monachus
MLLAHLDKLLWPKHRLPLVKT
>Gavia_stellata
MLLAHLNKLLWPKHRLPLVKT
>Chlamydotis_macqueenii
MLLAHLDKLRWPKHRLPLVKT
>Tauraco_erythrolophus
MLLAHLDKLLWPKHRLPLVKT
>Opisthocomus_hoazin
MPLAHLDKLLWPKHRLPLVKT
>Geospiza_fortis
MLLSHVDKLLWPKHRLPLVKT
>Egretta_garzetta
MLLAHLDKLLWPKHRLPLVKT
>Fulmarus_glacialis
MLLAHLDKLLWPKHRLPLVKT
>Nestor_notabilis
MLLAHLDKLPWPKHRLPLVET
>Pygoscelis_adeliae
MLLAHLDKLLWPKHRLPLVKT
>Tyto_alba
MLLARLDKLLWPKHRLPLVKS
>Struthio_camelus_australis
MLLAHLDKLLWPKHRLPLVKT
>Tinamus_guttatus
MLLAHLDKLLWPKHRLPLVET

HG0396.1 ENSGALG00000034253

>Gallus_gallus
------------------MKDHCKEPWTFVVSYNTVIQRV
>Chaetura_pelagica
MQIKQRKKECQEWPSLISAKDHCRVPWILAVSYNVLIQRV
>Caprimulgus_carolinensis
------------------MKDHCREPWILVYKCT------
>Cariama_cristata
------------------MKDHYREPWILVVSYNILIQRV
>Charadrius_vociferus
------------------MKDHFREPWILVVSYNILIQRV
>Pterocles_gutturalis
------------------MKDHSREPWILGVSYNILIQRV
>Colius_striatus
------------------MKDHCREPWILVVSYNILIQRV
>Merops_nubicus
------------------MKDHCKEPWLLVVSYNVLIQRV
>Gavia_stellata
------------------MKDHCREPWILVVSYNILIQRV
>Balearica_regulorum_gibberic
------------------MKDHCREPWMLVVSCNILIQRV
>Tauraco_erythrolophus
------------------MKDHCREPWILVVSFNILIQRI
>Opisthocomus_hoazin
------------------MKDHCREPWILVYKCT------
>Pelecanus_crispus
------------------MKDHCREPWILVVSYNILIQRV
>Picoides_pubescens
------------------MKGHCREPWILVVS--------
>Fulmarus_glacialis
------------------MKDHCREPWILVVSYNILIQRV
>Nestor_notabilis
------------------MKDHCREPWILLVSCNILIQKV
>Aptenodytes_forsteri
------------------MKDHCREPWILVVSYNILIQRV
>Tyto_alba
------------------MKDQCRKSWILAVSYNILIQRV
>Struthio_camelus_australis
------------------MKDHCEEPWTFVYKCT------
>Tinamus_guttatus
---------------MVSMKDHCEELWTFVYKCT------
>Calypte_anna
------------------MKDCYREPWILVVSYNILIQSV
>Apaloderma_vittatum
------------------MKDHCREPWILVVSYNVLIQRV

HG0396.2 ENSGALG00000034253

>Gallus_gallus

---------------------------------------------------------------------MQKKTNKGRKDVKHDLP

>Anas_platyrhynchos

MKTNLSWLLQLHQLSATSSERPAQTPQVVDQSSFSSSSHLGAQSFYVVLGLNTRLLKLFVLMRKWISQLSTNKTNKGKKDVKHGLP

>Calidris_pugnax

-------------------------------------------------------------MWKWISQLNANQTNKGKKDVKRGLP

>Pterocles_gutturalis

-------------------------------------------------------------MWKWISQLNANQTNKGKKDVEHGLP

>Columba_livia

-------------------------------------------------------------MWKRILQLNANQTNKGKKDVKRGLP

>Gavia_stellata

-------------------------------------------------------------MWKWISQLNANQTNKGKKDVKRGLP

>Nipponia_nippon

-------------------------------------------------------------MWKWISQLNANQTNKGKKDVKRGLP

>Fulmarus_glacialis

-------------------------------------------------------------MWKWISQLNANQTNKGKKDVKCGLP

>Pygoscelis_adeliae

MKTNPSWLLQLHQLSAT----------------------------------------RFVLMWKWISQLNANQTNKGKKDVKRGLP

HG0397 ENSGALG00000009438 SLC39A9

>Gallus_gallus
MAESPSWDLGVCTG
>Apteryx_australis_mantelli
MAEGLSWDLGVCTG
>Buceros_rhinoceros_silvestri
MAESPSWELGVCTG
>Cariama_cristata
MAESLSWELGVCTG
>Colius_striatus
MAESLCWELGVCTG
>Leptosomus_discolor
MAESLSWELGVCTG
>Cuculus_canorus
MAESLCWELGVCTG
>Falco_cherrug
MAESLSWELGVCTG
>Gavia_stellata
MAESLSWELGVCTG
>Eurypyga_helias
MAESLSWELGVCTG
>Tauraco_erythrolophus
MAESLSWELGVCTG
>Opisthocomus_hoazin
MAESLSWELGVCTS
>Corvus_cornix_cornix
MAESLSWELGVCTG
>Egretta_garzetta
MAESLSWELGVCTG
>Fulmarus_glacialis
MAESLSWELGVCTG
>Aptenodytes_forsteri
MAESLSWELGVCTG
>Tyto_alba
MAERLSWELGVCTG
>Struthio_camelus_australis
MAGGLSWDLGVCTG
>Apaloderma_vittatum
MAESRSWELGVCTG

HG0398 ENSGALG00000005369 BTBD10

>Gallus_gallus
------------MMEEKVRLARAALNY
>Chaetura_pelagica
------------MMEEKVRLADGAVNY
>Buceros_rhinoceros_silvestri
------------MMEEKVRLADGAVNY
>Caprimulgus_carolinensis
------------MMEEKVRLADGAVNY
>Cariama_cristata
------------MMEEKVRLADGAVNY
>Charadrius_vociferus
------------MMEEKVRLADGALNY
>Pterocles_gutturalis
------------MMEEKVRLADGGMNY
>Colius_striatus
------------MMEEKVRLADGAVNY
>Leptosomus_discolor
------------MMEEKVRLADGAVNY
>Aquila_chrysaetos_canadensis
------------MMEEKVRLADGAVNY
>Chlamydotis_macqueenii
------------MMEEKVRLADGAVNY
>Tauraco_erythrolophus
------------MMEEKVRLAGGAVNY
>Opisthocomus_hoazin
------------MMEEKVRLADGALNY
>Corvus_brachyrhynchos
------------MMEEKVRLADGAVKY
>Egretta_garzetta
------------MMEEKVRLADGAVNY
>Nestor_notabilis
------------MMEEKVRLADGAVNY
>Struthio_camelus_australis
MYSYNPLSLEGKMMEEKVRLADVAVNY
>Calypte_anna
------------MMEEKVRLADGAVNY

HG0399 ENSGALG00000032659

>Gallus_gallus
-------MTNHFCVCTTRIQNS------
>Anser_cygnoides
-------MTNHICLCTTRIQNS------
>Chaetura_pelagica
-------MTNNICLCTTRIQEFLNRRLV
>Caprimulgus_carolinensis
MIFKCIFMTNHICLCTTRIQNS------
>Cariama_cristata
-------MTNHTCLCTTQIRNS------
>Charadrius_vociferus
-------MTNHIC---------------
>Pterocles_gutturalis
----MHFMTNHICLCTTQIQNS------
>Columba_livia
-------MTDHICLCTTREQNS------
>Leptosomus_discolor
MIFKCIFMTNHICLCTTRIQNS------
>Falco_cherrug
-------MTNHICLCTTQIQN-------
>Gavia_stellata
-------MTNHICLCTTRIQNS------
>Mesitornis_unicolor
-------MTNHICLCTTRIQNS------
>Tauraco_erythrolophus
-------MPNHICLCTTRIQNS------
>Taeniopygia_guttata
-------MTNYICLCTTRIQNS------
>Pelecanus_crispus
-------MTNHICLCTTRIQNS------
>Fulmarus_glacialis
-------MTNHICLCTTRIQNS------
>Aptenodytes_forsteri
-------MTNHICLCTTRIQNS------
>Apaloderma_vittatum
-------MTNRICLCTTRIQNS------

HG0400 ENSGALG00000004261 AQP9

>Gallus_gallus
MISLAVQHEISYFSV
>Anser_cygnoides
MISLAVQDEISYFNV
>Apteryx_australis_mantelli
MISLAVQDEI-----
>Caprimulgus_carolinensis
MTSLAVQDEISYFNV
>Calidris_pugnax
MISLAVQDEISYFNV
>Columba_livia
MISLAVQDEISYFNV
>Cuculus_canorus
MISLAVQDEISYFNV
>Aquila_chrysaetos_canadensis
MISLAVQDEISYFNV
>Balearica_regulorum_gibberic
MISLAVQDEISYFNV
>Geospiza_fortis
MISLAVQDEISYFNV
>Egretta_garzetta
MISLAVQDEISYFNV
>Picoides_pubescens
MISLAAQDEISYFNV
>Fulmarus_glacialis
MISLAVQDQISYFNV
>Aptenodytes_forsteri
MISLAVQDEISYFNV
>Tyto_alba
MISLAVQDEISYFNV
>Struthio_camelus_australis
MISLAVQDEISYFNV
>Tinamus_guttatus
MISLALRDEISYFNV

HG0401 ENSGALG00000009057 TASP1

>Gallus_gallus
--MILLQNLFLNGSMLFASRRC
>Anas_platyrhynchos
MQMIPLQNLFLNGSMLFSNRRW
>Apteryx_australis_mantelli
--MIPLQSLFLNGSMLFSSRRW
>Charadrius_vociferus
--MIPLQGLFLNGRMLFSSRRW
>Leptosomus_discolor
--MIPLQSLFLNGSMPFSSRRW
>Falco_cherrug
--MILLQSLFLNGKVLFSSRRW
>Gavia_stellata
--MIPLQSLFLDGRMLFSSRRW
>Balearica_regulorum_gibberic
--MIPLQSLFLNGRMLFSSRRW
>Sturnus_vulgaris
--MISLQSLFLNGSVLLPSRRW
>Egretta_garzetta
--MIPLQSLFLNGRMLFSSRRW
>Fulmarus_glacialis
--MIPLQSLFLNGRMLFSSRRW
>Nestor_notabilis
--MILLQSLFLNGRVLFSSRRW
>Pygoscelis_adeliae
--MIPLQSLFLNGRMLFSSRRW
>Tyto_alba
--MIPLLSLFLNGRMLFSSRRW
>Struthio_camelus_australis
--MIPLQSLLLNGSMIFSSRRW
>Tinamus_guttatus
--MIPLQRLFLNGSMFFSSRRW
>Apaloderma_vittatum
--MIPLQSLFLNGRMLFSSRRW

HG0402 ENSGALG00000035584

>Gallus_gallus
MARCWKGREQPMLLI
>Anas_platyrhynchos
MARCWKGRERPMLLL
>Chaetura_pelagica
MAQCWKGREQPMLLF
>Calidris_pugnax
MARCWKGREQPMLLF
>Pterocles_gutturalis
MARCWKGREQPMLLF
>Columba_livia
MARCWKGREQPMLLF
>Leptosomus_discolor
MALCWKGREQPMLLF
>Falco_cherrug
MARCWKGREQPMLLF
>Chlamydotis_macqueenii
MARCWKGREQPMLLF
>Tauraco_erythrolophus
MARCWKGREQPMLLF
>Taeniopygia_guttata
MARCWKGREQPMLLF
>Pelecanus_crispus
MARCWKGREQPMLLF
>Fulmarus_glacialis
MARCWKGREQPMLLF
>Aptenodytes_forsteri
MARCWKGREQPMLLF
>Tyto_alba
MAQCWKGREQPMLLF
>Struthio_camelus_australis
MARCWKGREQPMLLF
>Calypte_anna
MARCWKGREQPMLLF

HG0403.1 ENSGALG00000007717 GCNT7

>Gallus_gallus
MNPPPNTNPFSRMLSGKEAFIQHEIQSIPITCMPN-------------------------
>Anser_cygnoides
------------MLSAEEAFFQHVIQCIPVRIMR--------------------------
>Chaetura_pelagica
------------MLSGKEEFFQHEIQCLPTI-----------------------------
>Cariama_cristata
MNLPPNIQLLSGMLSGKEAFFQHEIQCIPII-----------------------------
>Charadrius_vociferus
------------MLSGKEAFFQHEIQCIPII-----------------------------
>Pterocles_gutturalis
------------MLSGKEAFFHPEIQCIQII-----------------------------
>Columba_livia
------------MLSGKEAFFQHGIQCIPII-----------------------------
>Merops_nubicus
------------MLSGKEEFFQHEIQCIPII-----------------------------
>Aquila_chrysaetos_canadensis
------------MLSGKEAFFQREIQCLPII-----------------------------
>Gavia_stellata
------------MLSGKEAFFQHEIQCIPII-----------------------------
>Balearica_regulorum_gibberic
------------MLSGKEAFFQHEIQCIPII-----------------------------
>Tauraco_erythrolophus
------------MLSGKKAFFQHEIQCIPIL-----------------------------
>Lepidothrix_coronata
------------MLSGKEAFFQHEIQCIPIISMFN-------------------------
>Fulmarus_glacialis
------------MLSGKEAFFQHEIQCIPVI-----------------------------
>Tyto_alba
------------MLSGKEAFFQNEIQCIPII-----------------------------
>Tinamus_guttatus
------------MLSGKKAFFQHQIQCIPEFCLGSSTSKASRKRPHAVATGFRTSVLSCF

HG0403.2 ENSGALG00000007717 GCNT7

>Gallus_gallus

MLSIAGN-------QVMLMFNPFQLYEISTQATE

>Anas_platyrhynchos

MLSIAGSQMKP-----------------------

>Apteryx_australis_mantelli

MLSIAGSQMKP-----------------------

>Calidris_pugnax

MLSIAGSQMNSQMKTVMLMFNPFHLRQISPQATE

>Struthio_camelus_australis

MLSIAGSQMKP-----------------------

HG0404 ENSGALG00000008700 PDCD4

>Gallus_gallus
MEHSTRKDKTVLTR---
>Apteryx_australis_mantelli
MEHSTGKNKTVLTW---
>Buceros_rhinoceros_silvestri
MEHSARKNKSVLTG---
>Caprimulgus_carolinensis
MEHSTRKNKTVLT----
>Charadrius_vociferus
MEHSTRKNKTVLTR---
>Colius_striatus
MEHSTRKNKTVLTR---
>Leptosomus_discolor
MEHSTRKNKIVLTR---
>Haliaeetus_leucocephalus
MEHSIRKNKTVLTR---
>Gavia_stellata
MEHSPRKDKTVLTR---
>Chlamydotis_macqueenii
MEHSTRKNKTVLTRQFA
>Tauraco_erythrolophus
MEHSTRRSKTVLTR---
>Ficedula_albicollis
MEHSTRKDKTVLTR---
>Fulmarus_glacialis
MEHSARKNKTVLTR---
>Nestor_notabilis
MEHSTTKTKTVLTR---
>Aptenodytes_forsteri
MEHSTRKNNTVLTR---
>Tyto_alba
MEHSTRKNKTVITR---

HG0405.1 ENSGALG00000009185

>Gallus_gallus
------------------------------------------------------------------------MLLSSLWAFLDLRFFS
>Anser_cygnoides
------------MW------NTLL-AYLRVRE--AVQEVLCRLYKYSVRQ-----AVISLSLLQISAACLGHMLLSRLRAFLDLRFFS
>Cariama_cristata
------------------------------------------------------------------------MLLISLRAFLDLRFFS
>Charadrius_vociferus
MKERLARVGDLKSFSPSPLTHSQTEELHRHQEITLHQDARSR-YFLPCTQQVTSFEILFSPLKQISAACLGYMLLISLRAFLDLRFFS
>Pterocles_gutturalis
------------------------------------------------------------------------MLLSSLQAFLDLRCFS
>Colius_striatus
------------------------------------------------------------------------MLLIGLRAFLDLRFFS
>Leptosomus_discolor
------------------------------------------------------------------------MLLISLRAFLDLRFFS
>Aquila_chrysaetos_canadensis
----------------------------------------------------------MDNHGKISTACLGYMLLISLRAFLDLRFFS
>Gavia_stellata
----------------------------------------------------------------MSTACLGYMLLISLRAFLDLRFFS
>Chlamydotis_macqueenii
------------------------------------------------------------------------MLLISLQAFLDLGFFS
>Opisthocomus_hoazin
----------------------------------------------------------------MSTACLGCMLLISLRAFLDLRFSS
>Acanthisitta_chloris
------------------------------------------------------------------------MLLISLLASLDLRFFS
>Phalacrocorax_carbo
----------------------------------------------------------------MSTACLGYMLLISLQAFLDLRFSS
>Struthio_camelus_australis
------------------------------------------------------------------------MLLSSLRAFLDLRFFS

HG0405.2 ENSGALG00000009185

>Gallus_gallus
-----------------------------------------MPHPDIFSPELSKLPVL------
>Charadrius_vociferus
-----------------------------------------MPDPGIFSPVLSKLPVSRSCSVL
>Pterocles_gutturalis
-----------------------------------------MPHPGILCPVLSKLQVLRSCSVL
>Colius_striatus
-----------------------------------------MPNPGIFFPVLSKLAVLRSCPGL
>Leptosomus_discolor
-----------------------------------------MPDPGIFSPVLSKLPFLRSCSVL
>Haliaeetus_albicilla
MKERLARAGELKSFSPSPLVSQCKDRRTTETSEISTLAFLRMPDPGIFSPALSKLPVSRLCSVL
>Chlamydotis_macqueenii
-----------------------------------------MPDPDIFYPVLSKLPVLRSCSVL
>Phalacrocorax_carbo
-----------------------------------------MPDPGIFSPILSKLPVLRSGSVL
>Picoides_pubescens
-----------------------------------------MLDPGIFSPVLSKLPVLRSCSVL
>Fulmarus_glacialis
-----------------------------------------MPDPGIFSPVLSKLPVLRSCSVL
>Nestor_notabilis
-----------------------------------------MPDPGIFSTVLSKLPVSRSCSVL
>Pygoscelis_adeliae
-----------------------------------------MPDPGIFSPVLSKLPVLRSCSVL

HG0406 ENSGALG00000004935 LSM14A

>Gallus_gallus

MKAAAMLGASGRFWCGVC

>Chaetura_pelagica

-----MLGASGRFRCGVC

>Apteryx_australis_mantelli

MKAAAMLGASGCFWCCLC

>Calidris_pugnax

MKAAAMLGASGRFWCGLC

>Columba_livia

MKAAAMLGASGRFWCGVC

>Leptosomus_discolor

MKAAAMLGASGRFWCGVC

>Aquila_chrysaetos_canadensis

MKAAAMLGASGCFWCGVC

>Tauraco_erythrolophus

MKPSSMWRASGRFRCGVC

>Pseudopodoces_humilis

MKAAAMLGAAGRGPC---

>Nipponia_nippon

MKAAAMFGASGRFWCGVC

>Picoides_pubescens

MKAAAMLGGSGRFWCGVC

>Melopsittacus_undulatus

MTAAAMLGASGSFWCGVC

>Pygoscelis_adeliae

MKAAAMFGASGWFWCGVC

>Calypte_anna

MKAAAMLGASGRFRCGVC

HG0407 ENSGALG00000015027 JAK2

>Gallus_gallus
MERARVLRGSRQLALLRRAPAGYIMFYHCLCACFI
>Anas_platyrhynchos
------------------------MFYHSLCACFV
>Chaetura_pelagica
------------------------MFHHCLCTCFL
>Buceros_rhinoceros_silvestri
------------------------MFHHCLCACFV
>Cariama_cristata
------------------------MFQHCLCACFV
>Calidris_pugnax
------------------------MFHHCLCACFA
>Columba_livia
------------------------MFHHCLCACFV
>Cuculus_canorus
------------------------MFHHCLCACIV
>Aquila_chrysaetos_canadensis
-----------------------MMFHHCLCACFV
>Eurypyga_helias
------------------------MFHHCLCACFA
>Tauraco_erythrolophus
------------------------MFHHCVCACFV
>Lepidothrix_coronata
MGRERA--DAACAPGCTASAAGYIMSHHCLCACFV
>Egretta_garzetta
------------------------MFYHCLCACFV
>Nestor_notabilis
------------------------MFHHCSCACFV

HG0408 ENSGALG00000036255 WNT7B

>Gallus_gallus

MTGRSPEAAAAARTA-AAAHTLPREQH

>Anas_platyrhynchos

MTGRSPEAAAAARAA-AAAHTLHREQH

>Chaetura_pelagica

MTGRSPEAAAAARAA-AAAHTLHREQH

>Apteryx_australis_mantelli

MTGRSPEAAAAARAA-AAAHTLHREQH

>Charadrius_vociferus

MTGRSPEAAAAARAA-AAAHTLHREQH

>Columba_livia

MTGRSPEAAAAARTA-AAAHTLPREQH

>Cuculus_canorus

MTGRSPEAAAAARAAAAAAHTLHREQH

>Aquila_chrysaetos_canadensis

MTGRSPEAAAAARAA-AAAHTLHREQH

>Lepidothrix_coronata

MTGRSPEAAAAARAA-APAHTLHREQH

>Nipponia_nippon

MTGRSPEAAAAARAA-AAAHTLHREQH

>Melopsittacus_undulatus

MTGRSPEAAAAARAA-AAAHTLHREQH

>Aptenodytes_forsteri

MTGRSPEAAAAARAA-AAAHTLHREQH

>Struthio_camelus_australis

MTGRSPEAAAAARAA-AAAHTLHREQH

>Calypte_anna

MTGRSPEAAAAARAA-AAAHTLHREQH

HG0409 ENSGALG00000002315

>Gallus_gallus
MVSTPCEPPLQK---------
>Anser_cygnoides
MDSTSCEPPLQK---------
>Apteryx_australis_mantelli
MNSTSCEPPLQKATLAQKWNL
>Cariama_cristata
MDSTSCEPPLQK---------
>Pterocles_gutturalis
MNSTSCEPPLQK---------
>Columba_livia
MDSTSCEPPLQK---------
>Chlamydotis_macqueenii
MDSTSCEPPLQK---------
>Opisthocomus_hoazin
MDSTSCEPPLQK---------
>Zonotrichia_albicollis
MDSTSCEPPLQK---------
>Phalacrocorax_carbo
MDSTSCEPPLQK---------
>Fulmarus_glacialis
MDSTSCEPPLQK---------
>Melopsittacus_undulatus
MDSTSCEPPLQK---------
>Pygoscelis_adeliae
MDSTSCEPPLQK---------

HG0410 ENSGALG00000008148 MYBPC3

>Gallus_gallus
MGASTEWFFWGSQKPCSLICV
>Anas_platyrhynchos
MGASTKSFFRGSQKPWRLIPV
>Cariama_cristata
MGASTKCFFRGSEKPCSLIPK
>Calidris_pugnax
MGASTECFFRGSQKPCRLIPV
>Columba_livia
MEASTKCFFRGFQKPCSLIPV
>Leptosomus_discolor
MGASTKCFFRGSQKPCSFIPV
>Aquila_chrysaetos_canadensis
MGASTKCFFRGSQKPCSLIPV
>Eurypyga_helias
MGSSTECFFRGSQKPHILIPD
>Parus_major
MGASIERFFRGSQKPCSHIPV
>Phaethon_lepturus
MGASTGCFFRGSQKPCRLIPV
>Fulmarus_glacialis
MGASTECFFRGSQKPCSLISV
>Melopsittacus_undulatus
MGASTKCFFRGSQKPCTLIPV
>Aptenodytes_forsteri
MGASTECFFRGSQKPCSLIPV

HG0411 ENSGALG00000012166 SLC35F5

>Gallus_gallus

--------MAIPTLTLLKWRLSSL

>Anser_cygnoides

MAEKWFYKMTTPTLTLLKWRLSSL

>Cariama_cristata

--------MTAPTLTLLKWRLSSL

>Calidris_pugnax

--------MTAPNLTQLKWRLSSL

>Aquila_chrysaetos_canadensis

--------MTAPTLTLLKWRLSSL

>Gavia_stellata

--------MTAPTLTLLKWRLSSL

>Eurypyga_helias

--------MTAPTLTLLRWRLSSL

>Tauraco_erythrolophus

--------MTASTLTLLKWRLSSL

>Sturnus_vulgaris

--------MTVPTLRLLEWRLSSL

>Nipponia_nippon

--------MTPPTLRLLKWRLSSL

>Picoides_pubescens

--------MTAPTVILLKWRLSSL

>Fulmarus_glacialis

--------MTAPTLTSLKWRLSSL

>Aptenodytes_forsteri

--------MTAPTLTLLKWRLSSL

HG0412 ENSGALG00000003446 PRLR

>Gallus_gallus
-----------------------------MNERNKIFLCTHESFREPLM
>Chaetura_pelagica
-----------------------------MKERNKNLPCTHESFSEHLI
>Caprimulgus_carolinensis
-----------------------------MKERNKNLPCTHESFSEPLI
>Cariama_cristata
-----------------------------MKERNRNLPCTHQSFCKPLI
>Calidris_pugnax
-----------------------------MKERNKNLHRTHESFSEPLI
>Pterocles_gutturalis
-----------------------------MKERNKNLPRTHESFSEPLI
>Colius_striatus
-----------------------------MKERNKNLPCTHESFSEPLI
>Columba_livia
-----------------------------MKERNKNLPCTHETFSEPLI
>Haliaeetus_leucocephalus
-----------------------------MKEKNKNLPCTHESFSEPLI
>Chlamydotis_macqueenii
MTENLQGVSPMAAWDEDGGYIRSSTDLTFMKKRNKNLHNTHESFREPLI
>Lepidothrix_coronata
-----------------------------MKERNKKLPYIHESFSEPLI
>Nipponia_nippon
-----------------------------MKEKNKNLPHTHESFSEPLI

HG0413 ENSGALG00000005475

>Gallus_gallus

MRSCCWLLLDFVSG

>Anser_cygnoides

MQSCCWLLLDFVSG

>Charadrius_vociferus

MQSCCQLLLEFVSG

>Colius_striatus

MQSCCQLLLGFVSG

>Falco_cherrug

MQSCCQLLMDFVSG

>Gavia_stellata

MQSCCQLLLDFVSG

>Balearica_regulorum_gibberic

MQSCCQLLLDFVSG

>Opisthocomus_hoazin

MQSCCQCLLDFVFG

>Egretta_garzetta

MQSCCQLLLDFVSG

>Melopsittacus_undulatus

MQSCCQLLLDSVSG

>Struthio_camelus_australis

MQSCCQFLLDFFSG

>Tinamus_guttatus

MQSCCQFLLVFLSG

HG0414 ENSGALG00000032599

>Gallus_gallus
MMKALQWERAVVQG
>Anser_cygnoides
MMKAPQWEHAVVQG
>Chaetura_pelagica
MMKALQWDRAVVQG
>Buceros_rhinoceros_silvestri
MMKALQWERAVVQG
>Charadrius_vociferus
MMKALQWERAVVQG
>Columba_livia
MMKALQWERAVVQG
>Cuculus_canorus
MMKAPQWERAVVQG
>Aquila_chrysaetos_canadensis
MMKALQWERAVVQG
>Sturnus_vulgaris
MMKALQWERAVVQG
>Phaethon_lepturus
MMKALQWEHAVVQG
>Pygoscelis_adeliae
MMKALQWERAVVQG
>Calypte_anna
MMKALQWDRAVVQG

HG0415 ENSGALG00000034970

>Gallus_gallus
MLSRTVMTREASLAKD
>Anas_platyrhynchos
MLSATVMTREASPAKD
>Chaetura_pelagica
MLSTTVMKREAPLARD
>Cariama_cristata
MLSATVMKKESPLARD
>Charadrius_vociferus
MLSATVMKKEAPLARD
>Pterocles_gutturalis
MLSTTVMKREAPLARD
>Leptosomus_discolor
MLSTTVMKREAPLARD
>Cuculus_canorus
MLSRTVMKIEALLAKD
>Gavia_stellata
MLSTTVMKREATLDRD
>Balearica_regulorum_gibberic
MLSTTVMKREAALARD
>Pelecanus_crispus
MLSTTVMKREASLARD
>Fulmarus_glacialis
MLSTTVMKREAPLARD

HG0416 ENSGALG00000015874 TTK

>Gallus_gallus

MKDYISCRCWILGII

>Anas_platyrhynchos

MESYISCLCWVLGII

>Charadrius_vociferus

MESYISCLCWILEII

>Leptosomus_discolor

MESYVSCLCWILGII

>Gavia_stellata

MESYISCLCWILGII

>Eurypyga_helias

MENYLSCLCWILRII

>Egretta_garzetta

MENYISCLCWILGII

>Fulmarus_glacialis

MESYISCLCWILGII

>Tyto_alba

MESYVSCLCWILGII

>Struthio_camelus_australis

MENYVSCLCWISGNI

HG0417 ENSGALG00000001522 PHF20

>Gallus_gallus

----------------MDFSALTIPDGGKHVVGGELGE

>Anas_platyrhynchos

----------------MDFSGLTIPDGGMHIVGGELGE

>Apteryx_australis_mantelli

MAVVIHLQGLPIVVGTMDFSGVTIPDGGVHIVGGELGE

>Calidris_pugnax

----------------MDLSGLSIPDGGVQIVGGELGE

>Columba_livia

MAVVIHLRGLPGVVGTMDFSGLTIPAGGVHVVGGELGE

>Haliaeetus_albicilla

MAVFIHLQGLPIVVGTMDFSGLTIPDGGVHIVGGELGE

>Parus_major

MAVVIHLQGVPIAVGTMDFSGLTIPDGGVHIVGGELGE

>Pygoscelis_adeliae

MAVVIHLQGLLIVVGTMDFSGLTIPDGGMHIVGGELGE

>Struthio_camelus_australis

MAVVIHLQGLPIVVGTMDFSGLTIPDGGVHIVGGELGE

HG0418 ENSGALG00000010337 GPR149

>Gallus_gallus
-------------------MKNCIFSSTFCTSLLPDFSRA-----------
>Anser_cygnoides
-------------------MKNCIFSSTFGTSLLPDFSRACVVLVIGDA--
>Caprimulgus_carolinensis
-------------------MKNCIFFSTFSTSLLPDFS-------------
>Calidris_pugnax
-------------------MKNCIFFSTFDASLLPDFSRA-----------
>Haliaeetus_albicilla
-------------------MKNCIFFSTFDTSLLPEFSRA-----------
>Eurypyga_helias
MACYYLNSLNYPSFSAERIMKNCFLFSTFATSLLPDFSRARVVLVIKGTFT
>Tauraco_erythrolophus
-------------------MRNCIFSATFDTSLLPDFSRA-----------
>Phaethon_lepturus
-------------------MKNCIFFSIFDTSLLPDFSRA-----------
>Tyto_alba
-------------------MKNCIFFSTFGTSLLPDFLRA-----------

HG0419 ENSGALG00000012542 RASD2

>Gallus_gallus

MGGRPLEPCSPSQSC

>Anas_platyrhynchos

MGGRPLEPCGPSHSC

>Apteryx_australis_mantelli

MGDRPLQSCSPSQSC

>Calidris_pugnax

MGEHPLKPCSPSQSS

>Columba_livia

MGECPLEPCGPSQSC

>Haliaeetus_albicilla

MGERPLEPCGPSQSC

>Corvus_brachyrhynchos

MGERPLEPCGPSQSC

>Aptenodytes_forsteri

MGERPLEPCGPSQSC

>Tinamus_guttatus

MGDSPLHICSPSQSC

HG0420 ENSGALG00000000336 PPP1R12B

>Gallus_gallus

MPKIIHLNRQGLITRAANSLISLF

>Anser_cygnoides

MPKIIHPKQQGLITRAVNSLISLF

>Apteryx_australis_mantelli

MPKIIHPKQRGLITRAGNSLIGLF

>Calidris_pugnax

MPKIIHPKQQGLITRAVNSLISLF

>Aquila_chrysaetos_canadensis

MPKIIHPKQQGLITRAVNSLISLF

>Corvus_brachyrhynchos

MPKIIHLKQQGLITRAVNSLISLF

>Nipponia_nippon

MPKITHPKQQGLITRAVNSLISLF

>Calypte_anna

MPKIIHPKQQGLITRAVNSLISLF

HG0421 ENSGALG00000003492

>Gallus_gallus
-------------------------------------------------------------MEANLKRAKKGDLKVSVHRMEIERIRYVLSSYLRCRLAKIEKFFP
HVLEKEKSRAKGEPSILSPEEFAFAKEYMANTETYLKNVALKHMPPNLQKVALLKSVPKPNLDSFVFLRVLKRQENILVEPETDEQREYTIDLEEGSQHLIRYKTI
APLVASGAVQLI
>Anser_cygnoides
-------------------------------------------------------------MESNLKRAKAGDLKVSVHRMEIERIRYVLSSYLRCRLVKIEKFFP
HVLEKEKSRAEGEPSILSPEEFAFAKEYMANTETYLKNVALKHMPPNLQKVSLLKSVPKPNLDSFVFLRVLERQENILVEPETDEQREYTIDLEEGSQHLIRYKTI
APLVASGAVQLI
>Apteryx_australis_mantelli
--------------------------------------------------------------------------------MEIERIRYVLSSYLRCRLVKIEKFFP
HVLEKEKSRAEGEPSILSPEEFAFAKEYMANTETYLRNVALKHMPPNLQKVSLLKSVPKPNLDSFVFLRVLERQENILVEPETDEQREYTIDLEEGSQHLIRYKTI
APLVASGAVQLI
>Charadrius_vociferus
-------------------------------------------------------------MEANLKRAKRGDLKVSVHRMEIERIRYVLSSYLRCRLVKIEKFFP
HVLEKEKSRAEGEPSILSPEEFAFAKEYMANTETYLKNVALKHMPPNLQKVSLLKSVPKPNLDSFVFLRVLERQENILVEPETDEQREYTIDLEEGSQHLIRYKTI
APLVASGAVQLI
>Falco_cherrug
-------------------------------------------------------------MEANLKRAKRGDLKVSIHRMEIERIRYVLSSYLRCRLVKIEKFFP
HVLEKEKSRTEGEPSILSPEEFAFAKEYMANTETYLKNVALKHMPPNLQKVSLLKSVPKPNLDSFVFLRVLERQENILVEPETDEQREYAIDLEEGSQHLIRYRTI
APLVASGAVQLI
>Geospiza_fortis
MAGAMAAAAGGDSDGESEELVLTPAQLIHSLEQAWLNEKFAPELLESKPEIIECVVEQLDHMEANLKRAKKGDLKVSVHHMEIERIRFVLSSYLRCRLVKIEKFFP
HILEKEKSRAEGEPSILSPEEFAFAKEYMANTETYLKNVALKHMPPNLQKVSLLKSVPKPNLDSFVFLRVLERQENILVEPETDEQREYTIDLEEGSQHLIRYRTV
APLVASGAVQLI
>Nipponia_nippon
MAAASGEGTAADSDGDSEELVLTPAQLIRSLEQAWLNEKFAPELLESKPEIIECVVEQLDHMEANLKRAKRGDLKVSVHRMEIERIRYVLSSYLRCRLVKIEKFFP
HVLEKEKSRAEGEPSILSPEEFAFAKEYMANTETYLKNVALKHMPPNLQKVSLLKSVPKPNLDSFVFLRVLERQENILVEPETDEQREYTIDLEEGSQHLIRYKTI
APLVASGAVQLI
>Struthio_camelus_australis
------------------------------------------------------------MNEANLKRAKKGDLKVSIHHMEIERIRYVLSSYLRCRLVKIEKFFP
HVLEKEKSRAEGEPSILSPEEFAFAKEYMANTETYLKNVALKHMPPNLQKVSLLKSVPKPNLDSFVFLRVLERQENILVEPETDEQREYTIDLEEGSQHLIRYKTI
APLVASGAVQLI

HG0422 ENSGALG00000005772 BOK

>Gallus_gallus
--------------------------MKGKMKNRLWESSSWEL
>Anas_platyrhynchos
------------------------------MKDRLFENSCWKL
>Leptosomus_discolor
MLIDAASPKKHESNPKKKSLRGGKCNMKGKMKNWPFENSCWEM
>Aquila_chrysaetos_canadensis
--------------------------MKGKMKNWPFENSCWEL
>Gavia_stellata
--------------------------MRGKMKNWPFENSCWEL
>Balearica_regulorum_gibberic
--------------------------MRGKMKNWPFESSCWEL
>Fulmarus_glacialis
--------------------------MRGKMKNWSFESSCWDL
>Aptenodytes_forsteri
--------------------------MRGKMKNWPFENSCWEL

HG0423 ENSGALG00000006329

>Gallus_gallus
MLSKLVGKKGGHLVSVL
>Leptosomus_discolor
MLSKLVGKNSGHLVSVL
>Aquila_chrysaetos_canadensis
MLSKLVGKNSGHLVSVL
>Gavia_stellata
MLSKLVGKNSGHLVSVL
>Balearica_regulorum_gibberic
MLSKLVGKNSGHLVSVL
>Phaethon_lepturus
MLSKLVGKNSGHLVSVL
>Tyto_alba
MLSKLVGKNSGHLVSVL
>Tinamus_guttatus
MLSKLVGRNSGHLVSVL

HG0424 ENSGALG00000039553

>Gallus_gallus
MTIYLSCIRQRCCLQ
>Anser_cygnoides
MTIYLSCIRKRFCLQ
>Calidris_pugnax
MTIFLSCIRKRFCL-
>Cuculus_canorus
MTIFLSCIRKRFC--
>Haliaeetus_leucocephalus
MTIFLSCIRKRFCL-
>Acanthisitta_chloris
MTIFLSCIRKRFCL-
>Phalacrocorax_carbo
MTIFLSCIRRRFCL-
>Aptenodytes_forsteri
MTIFLSCIRKRVCL-

HG0425 ENSGALG00000043044

>Gallus_gallus
MIQKTVRSCDCKMSSFYLMLKGGKLKLQEL
>Caprimulgus_carolinensis
------------MSSFHLMLKGGKLELQE-
>Falco_cherrug
------------MSSFHSLLKGGKLKLQE-
>Mesitornis_unicolor
------------MSSFHLMLKGGKLKHQE-
>Tauraco_erythrolophus
------------------MLKEGKRKLQE-
>Opisthocomus_hoazin
------------MSSFQLMPKGGKLKLQE-
>Ficedula_albicollis
--------MDCEMNSFHLILKEGKLKLQK-
>Tyto_alba
------------MSSFHLMLKGGKLKLQE-

HG0426 ENSGALG00000002192 PTPRC

>Gallus_gallus
-----------------------------------------------------------------MLLRKDTPDFQK
>Anser_cygnoides
-----------------------------------------------------------------MLLHKDTPDFQK
>Calidris_pugnax
-----------------------------------------------------------------MLFHKDTPDFQK
>Columba_livia
-----------------------------------------------------------------MLFHKDTPDFQK
>Falco_cherrug
-----------------------------------------------------------------MLFHKDTPDFQK
>Ficedula_albicollis
MPVELRAGHSSLTSGWRWRKKVLAAVAAGLVKRSLVPLMYRMKAEISGCPQRTNPFNQTNQRERHMLFHKDTPDFQK
>Aptenodytes_forsteri
-----------------------------------------------------------------MLFHKDTPDFQK

HG0427 ENSGALG00000005550 ST6GAL1

>Gallus_gallus
MKVFLELFSASRGCCVWGYPALNPGQKSANSQK
>Anser_cygnoides
MKVFLELFRVSHNRCVLVYPALTPGRKSANSQK
>Calidris_pugnax
MKVFLECFGASQNCCVLGYSA------------
>Columba_livia
MKLFLELFSASQNCCVLRCPA------------
>Aquila_chrysaetos_canadensis
MKVFLELFSASQNCCVLRYPA------------
>Ficedula_albicollis
MKVFLELFNTSRSCCVLGYPA------------
>Aptenodytes_forsteri
MKVFLELFSASQNCCVLGYPA------------

HG0428 ENSGALG00000005976 TCEA2

>Gallus_gallus
MSHPVAGRFAAPLLK
>Charadrius_vociferus
MSVPVDGRFSTPLLK
>Leptosomus_discolor
MSVPVDGRFSAPLLK
>Haliaeetus_albicilla
MSVPVDGRFSTPLLK
>Gavia_stellata
MSVPVDGRFSTPLLK
>Pelecanus_crispus
MSVPVDGRFSTPLLK
>Fulmarus_glacialis
MSVPVDGRFSTPLLK

HG0429 ENSGALG00000006626 PDE6C

>Gallus_gallus

----MPLYSMYKVPFSY

>Anser_cygnoides

---------MYKVPFSY

>Columba_livia

MYYACELYNMYKVPFSY

>Gavia_stellata

MYYACELYNMYKVPFSY

>Opisthocomus_hoazin

MYYACELYNMYKVPFSY

>Pelecanus_crispus

MYYACELYNMYKVPFSY

>Tinamus_guttatus

---------MYKVPFSY

HG0430 ENSGALG00000015490

>Gallus_gallus

-------------------------------------------MLSMGTSASDALTLLKTYFVAWLKRR

>Anser_cygnoides

-------------------------------------------MLSTGTSASDARTLLKTYFVVWLKGR

>Apteryx_australis_mantelli

MKVYVLKELTCLRRSQWYQGNGAQQKEGSPKTTNLAVLGHRCKMLSMGTFASDAR-ILKIYFFAWLTGR

>Calidris_pugnax

-----------MGIAPYF-----------FLVNAVNRLIQRSKMLSVGTSASDARTLQKAYFVAWLKGR

>Columba_livia

-------------------------------------------MLSLETSAYDAGTLLKTYFVAQLKGR

>Haliaeetus_albicilla

---------------------------------MKIFSLIRSKMLSMETSASDARTLLKTYFVAWLKGR

>Zonotrichia_albicollis

-------------------------------------------MLSMGTSASDARTLLKTSFVAWFKGR

HG0431 ENSGALG00000026542 FAM217B

>Gallus_gallus
MPAEAESKDVVMFAFGTLCCVDRKEWSRWKIM
>Aquila_chrysaetos_canadensis
----------MMFAFDTLCLANPKDWSRLKIM
>Manacus_vitellinus
-----------MSAFDTLCLANPKDWSRLKIM
>Melopsittacus_undulatus
------MQEVVMFAFDTLCLTNPKDWS-----
>Aptenodytes_forsteri
-----------MFAFDTLCLTNPKDWSRLKIM
>Struthio_camelus_australis
-----------MPAFDALCLINPKHWSPLKVM
>Tinamus_guttatus
---------MVMFAFDTLCLVNPKHWSPLKIM

HG0432 ENSGALG00000040546

>Gallus_gallus
MLENFTSVLIVISEHPPKN----
>Anser_cygnoides
MLENFTSVLIVISEHPPKN----
>Calidris_pugnax
MLEYFTSVLIVISEHPQRTETLH
>Columba_livia
MLENFTSVLIVISEHPQRTETLH
>Haliaeetus_albicilla
MLENFTSVLIVISEHPQRTETLH
>Corvus_cornix_cornix
MLENSTSVLTVIREHPQRSAAFH
>Aptenodytes_forsteri
MLENFTSVLIVISEHPQRTETLH

HG0433 ENSGALG00000002952 NRG4

>Gallus_gallus

MQISGGVDSADCPVRV

>Calidris_pugnax

MQMRDYVDSADCPVRV

>Columba_livia

MQIRDYVDSADCPVHV

>Aquila_chrysaetos_canadensis

MQIRDYVDSADCPVHV

>Zonotrichia_albicollis

MRIRDYVDSADCPVHV

>Aptenodytes_forsteri

MQIRDYVDSADCPVHV

HG0434 ENSGALG00000008357 TPCN1

>Gallus_gallus
MGARTFAFWWLEDGHDGRFPGKRF
>Calidris_pugnax
MGTRPCIFLCLEDTPDGRLPGESF
>Columba_livia
MGTRTFIFLCLEDTPDGRLSRDTF
>Aquila_chrysaetos_canadensis
MGTHTFIFLCLEDTPDGRLSGDSF
>Pygoscelis_adeliae
MSTHTFIFLCLEDTPDGRLSGESF
>Struthio_camelus_australis
MGTRVFVRFCLEDTPDGRLPGGRF

HG0435 ENSGALG00000010778 CHRM3

>Gallus_gallus
--MAKSFSSVWYIQLQAG
>Anser_cygnoides
--MAKSFSSAWYDGLQVG
>Pterocles_gutturalis
--MAKSFISVWYNGLPAG
>Aquila_chrysaetos_canadensis
MVMAKSFISIWYNGLPVG
>Egretta_garzetta
MVMAKSFISVWYNGLPVG
>Aptenodytes_forsteri
MVMAKSFISVWYNGLPVG

HG0436 ENSGALG00000026736 OGN

>Gallus_gallus
MLKLSTAPCSRILPLTLGTECWQSIFYPMESTFKTVNLGKLSHCSNLDVIHCWIVHNFTLLSVCMGLTYA
>Anser_cygnoides
----------------------------MESIFKTVNLGRSIHCSICIS---------------------
>Apteryx_australis_mantelli
----------------------------MESTFKTVNL--------------------------------
>Calidris_pugnax
----------------------------MESTFKTVKL--------------------------------
>Columba_livia
----------------------------MESTFKTVKLRMSIHFNNS-----------------------
>Pygoscelis_adeliae
----------------------------MESTFKTVKL--------------------------------

HG0437 ENSGALG00000001728 NACC2

>Gallus_gallus
MRGAAPARRPGPPFSPGPRRPASAAAARRKTMGRTMCPTLDTPALNA
>Haliaeetus_leucocephalus
-----------------------------------MCPTLDIPALNV
>Parus_major
-----------------------------------MCPTLDIPALNV
>Melopsittacus_undulatus
-----------------------------------MCPTLDIPALNV
>Aptenodytes_forsteri
-----------------------------------MCPTLDIPALNV

HG0438 ENSGALG00000008545 SENP5

>Gallus_gallus

MWYTARCCVTCKERQFCT

>Anser_cygnoides

MWYTAWC-----------

>Falco_cherrug

MWYAGWCCLTCKERWFCL

>Corvus_brachyrhynchos

MWFAGRCCLTCKERWFCV

>Aptenodytes_forsteri

MWYAGWCCLTCKERWFCV

HG0439 ENSGALG00000009172 OSBPL6

>Gallus_gallus
MQKSIPCIFVHE
>Calidris_pugnax
MQKTIPCLFVHE
>Aquila_chrysaetos_canadensis
MQKTIPCLFVHE
>Taeniopygia_guttata
MQKTIPCIFVHE
>Pygoscelis_adeliae
MQKAIPCLFVHE

HG0440 ENSGALG00000010090

>Gallus_gallus

MMLGHLLLTTGFSKEIQHGLMC

>Anas_platyrhynchos

-MLGHLLLSTGFSKEIQHGLDG

>Calidris_pugnax

MMLEHL----------------

>Corvus_cornix_cornix

MILQHL----------------

>Aptenodytes_forsteri

MMLEHL----------------

HG0441 ENSGALG00000015069 B4GALT4

>Gallus_gallus
MSHRTTRSAAALPTCSFPSLL
>Apteryx_australis_mantelli
MNCKTRGSVAAIPTSGFPSLL
>Corvus_brachyrhynchos
MNFKTAGSVAALPTSSFPSLL
>Struthio_camelus_australis
MNCKTRGSVAAVPTSGFPSLL
>Tinamus_guttatus
MNCKTRGSVAAVPTSAFPSLL

HG0442 ENSGALG00000030250

>Gallus_gallus

-----------MTFTSMMN-QEKPVKSS

>Calidris_pugnax

---MKYFLVLDMTFSITMNPQEKPVKSS

>Aquila_chrysaetos_canadensis

MAVMKYFLVLDMTFTTTMN-QEKPIKSS

>Corvus_brachyrhynchos

-----------MTFSTTMN-QEKPVKSS

>Struthio_camelus_australis

-----------MTFTIRMN-QEKPAKSS

HG0443 ENSGALG00000039454

>Gallus_gallus

MGWVSNHVSQDSVLL------------------------------------

>Anas_platyrhynchos

MGWVSNHVSQDSLFCNDGLRRCFPGEVRAGGWEAKRDVLVGFVVVGMEVNN

>Calidris_pugnax

MGWVSNHVS------------------------------------------

>Struthio_camelus_australis

MGWVSNHVS------------------------------------------

>Tinamus_guttatus

MGWVSNHVS------------------------------------------

HG0444 ENSGALG00000000558 SLC1A6

>Gallus_gallus

--------MLYLPKEKILHPAFELRESRNKTTQNNPPR------RKRPTVLLPDH

>Aquila_chrysaetos_canadensis

--------MLYLSKEKILHPAFELKELKNETTQNNPPR------RKRPAALPA--

>Lepidothrix_coronata

MLQSTMRATVYKTRENILLPAFELKELKNETTQNNPPR------RKRPAALPA--

>Struthio_camelus_australis

--------MLYLSKEKILQPAFDLKELQNKPTQNNPQEIRRVFRPKRPAVLPA--

HG0445 ENSGALG00000004017

>Gallus_gallus

MVALPGVGLGQPPQQGGDESLYKQGRK

>Anas_platyrhynchos

MVVLLGVGLGQPPHQQGDKSLHKQRRK

>Acanthis_flammea

MVAVRRAGLGQPPEQQGETAPYERRRK

>Pygoscelis_adeliae

--MVSVRGLGQPLEQQGDKSPYDQRRK

HG0446 ENSGALG00000006807 PLSCR5

>Gallus_gallus

MLPNTSLCSPARSAWALPFCSHASYLSAAHMCV-ISMATWALEVEI

>Anser_cygnoides

------------------------------------MATWALEVEI

>Calidris_pugnax

-MPHSSL--PGSVHRLCHFCSHASYLSADRMCVMISMATWALEAEI

>Parus_major

---------------------------------MISTATWAFKAGV

HG0447 ENSGALG00000008784

>Gallus_gallus
MKLNFKECFNQKEN
>Anser_cygnoides
MKLILK--------
>Calidris_pugnax
MKLILKECFTQKEN
>Corvus_cornix_cornix
MKLALRECFNQKEN

HG0448 ENSGALG00000011762 UBA6

>Gallus_gallus
MRPGLPAVPAC
>Aquila_chrysaetos_canadensis
MNPGLPAIPAC
>Lepidothrix_coronata
MSPGLPAIPAC
>Aptenodytes_forsteri
MNPGLPAIPAC

HG0449 ENSGALG00000014106 PPFIBP1

>Gallus_gallus
MDQWNWLKRNGGDQH
>Apteryx_australis_mantelli
MGWWYWLKSDGGDQH
>Aquila_chrysaetos_canadensis
MDQWYLLKSDSGDQH
>Pseudopodoces_humilis
MDQWYMLKSNSGDQH

HG0450 ENSGALG00000015528 JAKMIP1

>Gallus_gallus

---------MRDKTHMADGRDFCERFG

>Anas_platyrhynchos

MGMNGKRKDMRNS--THNGRDLCERFY

>Calidris_pugnax

---------MRDR--SHDGRDLCERFC

>Aptenodytes_forsteri

---------MRDC--SHDGRDLCERFC

HG0451 ENSGALG00000019489 CHST9

>Gallus_gallus

MGL----------------CTD---FR----LYRSESGQVTEEDAGTVRTSEK

>Apteryx_australis_mantelli

----------------------------------------------MLKTCEK

>Aquila_chrysaetos_canadensis

MGLLAGFQQRFRQMLSVSCCMNNTSFVWGKTEEVRVSGEDTEEDAGTLKTSEK

>Lepidothrix_coronata

-------------MLPMSCYLNNTAFVYSETEEVGVSGEDTEEDAGMLKTSEK

HG0452 ENSGALG00000004227 FAM20B

>Gallus_gallus
MLPWQSVFYSASYE
>Chaetura_pelagica
MLQWQYLFHSASYE
>Apteryx_australis_mantelli
MLQWQYLFYSASYE

HG0453 ENSGALG00000005995 SLC13A5

>Gallus_gallus

MQVLADSPLSARLCAGVKFPLC

>Calidris_pugnax

MRVLADSPLSARL---------

>Manacus_vitellinus

MQVLADSPLSARLCTGVKFPLC

HG0454 ENSGALG00000007186 TMEM268

>Gallus_gallus
MLPCQWRAESQGSRQWESE
>Anas_platyrhynchos
MPPCSWKAESRGSLQWECE
>Falco_cherrug
MSPCSRKAGSPGSPQWECE

HG0455 ENSGALG00000010718 THSD7A

>Gallus_gallus

MTAAASQRKITCSVSWSPLRTAVSLLADGKLLLTAEWKSVEQQPY

>Anas_platyrhynchos

MTAAASQRKITCSVSWSPLRTAVSLLADGKLQLTAEWKSVE----

>Zonotrichia_albicollis

MTAAASQSKITCSVSWSPLRTAVSLLADGKLL-------------

HG0456 ENSGALG00000033256

>Gallus_gallus
MHCSKLFLVFSSG
>Columba_livia
MHYSKLFLVSSSG
>Pseudopodoces_humilis
MHYSKLFLVSSSG

HG0457 ENSGALG00000039538 CLDND1

>Gallus_gallus
MEQLGPAGSWRCVLLAYCRTPVARCL
>Apteryx_australis_mantelli
MEQLGQIGSLHFMLLAYYRTQIYL--
>Gavia_stellata
MEQLGQIGSLHFMLSAYHGTQTTLCL

HG0458 ENSG00000060339 CCAR1

>Homo_sapiens
MWRRGAAWRKRGKLAHAPKADGFEMASMLAGTRLRSML-----
>Acinonyx_jubatus
MRRRGEAWRKRGKLAHAPKADGFEMASVLAGTRFRSLL-----
>Balaenoptera_acutorostrata_s
------------------------MASVLAGTGFRSLLLKTNK
>Miniopterus_natalensis
------------------------MASVLAGTRFRSLL-----
>Dasypus_novemcinctus
MGRKGEAWRKREKLAHAPKADGFEMASVLAGSRLRLLF-----
>Sarcophilus_harrisii
------------------------MASALAATRLRSPL-----
>Galeopterus_variegatus
------------------------MASVLTGTRLR--------
>Monodelphis_domestica
------------------------MASALAATRLRSFL-----
>Trichosurus_vulpecula
------------------------MASALAATRLRSLL-----
>Sorex_araneus
------------------------MASLLAGTRFRSLP-----
>Oryctolagus_cuniculus
------------------------MASVLAGTRLRTLL-----
>Elephantulus_edwardii
------------------------MASVLTGFRLRSLL-----
>Equus_caballus
------------------------MASVLAGTRFRSLL-----
>Manis_javanica
------------------------MASVLAGTRFRSLL-----
>Loxodonta_africana
------------------------MASLLTRSRLRSLL-----
>Fukomys_damarensis
------------------------MASVLAGTRFRSLL-----
>Tupaia_chinensis
------------------------MASALAGTRLR--------
>Trichechus_manatus_latirostr
------------------------MASVLTGIRFRSLL-----
>Orycteropus_afer_afer
------------------------MASVLAGTRLRSLL-----

HG0459.1 ENSG00000163848 ZNF148

>Homo_sapiens
MKGRNSTMISFSTLQHSFP
>Canis_lupus_familiaris
MKGRNSTMISFSTLQHSLP
>Lipotes_vexillifer
MKGRNSTMISFSTLQHSLP
>Eptesicus_fuscus
MKGRNSTMISFSTVQNSFP
>Dasypus_novemcinctus
MKGGNSTMISFSILQHFLS
>Sarcophilus_harrisii
-------MISFSTLQHPLP
>Galeopterus_variegatus
-------MISFSTLQRSLP
>Monodelphis_domestica
-------MISFSTLQHPLP
>Condylura_cristata
-------MISFSTLQHSLP
>Oryctolagus_cuniculus
MKGRNSTMISVSTLQHSLP
>Elephantulus_edwardii
-------MISFSTLQHSLP
>Ceratotherium_simum_simum
MKGRNSTMISFSTLQHSLP
>Manis_javanica
-------MISFSTVQHCLP
>Loxodonta_africana
-------MISFSTLQHFLP
>Marmota_marmota_marmota
-------MISFSTLQHSLP
>Tupaia_chinensis
MKGRNSAMISFSTLQHSLP
>Trichechus_manatus_latirostr
-------MISFSTLQHSLP
>Orycteropus_afer_afer
-------MISFSTLQHSLP

HG0459.2 ENSG00000163848 ZNF148

>Homo_sapiens

MGVCLTYLMEESDGISD----

>Acinonyx_jubatus

MGICLTYYIEDLDGISD----

>Orcinus_orca

MGICLTYFVEELDGIND----

>Myotis_brandtii

MGICLTYFIEESDGISGLRWY

>Dasypus_novemcinctus

MGICLTYFIEELECISD----

>Galeopterus_variegatus

MGICLTYFIEELNGISD----

>Erinaceus_europaeus

MGICLTYLTEELDGIND----

>Oryctolagus_cuniculus

MGICLVDLIEELDGVSD----

>Ceratotherium_simum_simum

MGICLTYFIEELDGVSD----

>Jaculus_jaculus

MGICLTYFIEEFDGISD----

>Tupaia_chinensis

MGICSTYFIEELDGISD----

>Trichechus_manatus_latirostr

MGICLLYFIEELDDISD----

>Orycteropus_afer_afer

MGICLTYFIEELDGISD----

HG0459.3 ENSG00000163848 ZNF148

>Homo_sapiens
MSASLLLWEEEEKEEE-EEEEEGEEEEKGAGVGAVAEAATAKVLPPPPFPSSQGP----------------------------------------------
>Lipotes_vexillifer
MSASLLLWEEEEKEEE--EEEEGEEEEKGAGVGAVAEAATAKVLPPPPFPSS-------------------------------------------------
>Rousettus_aegyptiacus
MSASLLPWEEEEKEEEEEEEEEGEEEEKGAGVGAVAEAATAKVLPPPPFPSPQGP----------------------------------------------
>Galeopterus_variegatus
MSASLLLWEEEEKEEE-EEEEEGEEEEKGAGVGAVAEAATAKVLLPPPFPSSQGP----------------------------------------------
>Trichechus_manatus_latirostr
MSASLLLWEEEEKEE---EEEEGEEEEKGAGVGAVAEAATAKVLPPPPFPSSPGPSEGWSCRRLQSVTARLGARPRIKGGICSLEIQPSDLAGWAFACYTL
>Orycteropus_afer_afer
MSASLLLWEEEEKEE---EEEEGEEEEKGAGVGAVAEAATAKVLPPPPFPSSQGP----------------------------------------------

HG0460 ENSG00000180008 SOCS4

>Homo_sapiens
MAMV----MAAVTRTTPVKLRSGRYIQKVPRRNFLLEKMKKQYL
>Ursus_maritimus
MAMV----MAAVTRTAPVKLRSERYIQRVPRRNFLLEKLKKQYL
>Balaenoptera_acutorostrata_s
MAVV----MAAVTRTAPVKLRSERYIQKVPRRNILPEKLKEQYL
>Eptesicus_fuscus
MAMV----MAAVTRTAPIKLRSERYIQKVPRRNFLPEKLKKQYL
>Dasypus_novemcinctus
-MAM----MAPVTRTASVKLCSGRYIQKVPRRNFLLEKLKKQYL
>Sarcophilus_harrisii
MFLLPGRASHIQRWQLISPLGYVRYFLKRPRRNFLLESLKNQHI
>Galeopterus_variegatus
MAMV----MAAVIWTAPAKLRSGRYIQKVSRRNFLQEKLKKPYL
>Monodelphis_domestica
--------MEAAPWTARVKLRCGRYFHKGLRRNFLLKSLKNQHL
>Macropus_eugenii
--------MEAAPWTARVRRRCGRYFHKGPRRNFLLESLKNQHM
>Condylura_cristata
--------MAAVTPTAPVKLRSGRYIQKVPRRTFLLEKLKKQYL
>Ornithorhynchus_anatinus
MA----ASGAPTPRAARVKRRSGRYFQKVPGRSSPPEKRKGGRF
>Ceratotherium_simum_simum
MAMV----MAAVPRAAPVKLRFGRYIQKVPRRNFLPEKLKKQYS
>Manis_javanica
MAMT----MAAVTRTAPIKLRSEKYIQNVPRRNFLPEKLKKQYL
>Loxodonta_africana
----------------------MRYMQKVPRRNFLPEKLKKQYL
>Nannospalax_galili
MEAV----MAAAPRTAPVKLHSGRFIQKVPRRNFLSEKLKKQYL
>Tupaia_chinensis
MAMV----MAAVTRTAPVKLRSGRYIQKVPRRNFLPEKLEKQYL
>Trichechus_manatus_latirostr
MAVV----MAAVTRTAIVELRSKRYIQKVPRRNFLPEKLKKQYL
>Orycteropus_afer_afer
MAMV----MAAVTQRAPVKLRSGRYIQKVPGRNYLPEKLKKQYL

HG0461 ENSG00000064419 TNPO3

>Homo_sapiens
MAPVRERPWPRQ
>Ursus_maritimus
MAPVRERPWPTQ
>Lipotes_vexillifer
MAPVRERPWPTK
>Miniopterus_natalensis
MAPVGERPWPTQ
>Dasypus_novemcinctus
MAPVRERPWPRQ
>Sarcophilus_harrisii
MAPARERPWPRQ
>Galeopterus_variegatus
MAPVRERPWPRQ
>Monodelphis_domestica
MAPARERPWPRQ
>Erinaceus_europaeus
MAPVRERPWPTQ
>Oryctolagus_cuniculus
MAPVRERPWPRQ
>Elephantulus_edwardii
MAPARERLWPRQ
>Equus_asinus
MAPVRERPWPTQ
>Manis_javanica
MAPVWERPWPTQ
>Loxodonta_africana
MAPVRERSWPRQ
>Dipodomys_ordii
MAPVRERPWPSQ
>Tupaia_chinensis
MAPVRERPWPRQ
>Trichechus_manatus_latirostr
MAPVRERPWPRQ

HG0462 ENSG00000125686 MED1

>Homo_sapiens
MAAASSTLLFLPPGTSD
>Acinonyx_jubatus
MAAAGSTLLFLPPGTSD
>Orcinus_orca
MAAAASTLLFLPPGTSD
>Pteropus_alecto
MAAAASTLLFLPPGTSD
>Dasypus_novemcinctus
MAAAASSLIFFPPGSSD
>Sarcophilus_harrisii
MAAAAGTLQLLPPAASD
>Monodelphis_domestica
MAAAAGTLQLLPPGASD
>Sorex_araneus
MAAAGSTLLFLPPGTTD
>Oryctolagus_cuniculus
MAAAAGTLLFLPPGTSD
>Elephantulus_edwardii
MAAAESTLLFLPPGTSD
>Ceratotherium_simum_simum
MAAAASTLLFLPPGTSD
>Manis_javanica
MAAAASTLLFLPPGTSD
>Loxodonta_africana
MAAAASTLHFLPPGTSD
>Marmota_marmota_marmota
MAAAASTLLFLPPGTSD
>Tupaia_chinensis
MAAAASTLLFLPPGTSD
>Trichechus_manatus_latirostr
MAAAASTLLFLPPGTSD
>Orycteropus_afer_afer
MAAVASTLPFLPPGTSE

HG0463 ENSG00000129473 BCL2L2

>Homo_sapiens
--------------MKGPSWGLLATSAVS
>Ailuropoda_melanoleuca
--------------MKGPSWGLLATSAGS
>Balaenoptera_acutorostrata_s
--------------MKGPSWGLLATSAGS
>Eptesicus_fuscus
--------------MKGPSWGLLATSAGS
>Dasypus_novemcinctus
--------------MKGPSWGLLATGAGS
>Sarcophilus_harrisii
--------------MKGLSWGLPATSAVS
>Galeopterus_variegatus
--------------MKGPSWGLLATSAGS
>Monodelphis_domestica
--------------MKGLSWGLPATSAVS
>Erinaceus_europaeus
-------------MMKGPSWGLLATSAGS
>Oryctolagus_cuniculus
--------------MKGPSWGLLATGAGS
>Elephantulus_edwardii
--------------MKGPSWGLLATSAGS
>Ceratotherium_simum_simum
--------------MKGPSWGLLATSAGS
>Manis_javanica
--------------MKGPSWGLFATSAGS
>Loxodonta_africana
MDSATQLGSTLAGRMKGPSWGLLATGAGS
>Dipodomys_ordii
--------------MKGPSWGLLATSAGS
>Trichechus_manatus_latirostr
--------------MKGPSWRLFATSAGS
>Orycteropus_afer_afer
--------------MKGPSWGLLATSAGS

HG0464 ENSG00000188215 DCUN1D3

>Homo_sapiens
---------------------------------------------------------------------------------------MLRYMGQRQRGHPIQRGPL
>Odobenus_rosmarus_divergens
-----------------------------------------MVGGLDPPGRRRFQKGFD------------WRNLWSSCWLAPLADGMLRHMGQWQRGHLIQRGPL
>Physeter_catodon
---------------------------------------------MRSPLRGHFWSMPPHLAVFLPLHCPPTMLHSAEGWLAPLADGMLRHVGQWQQGHPIQRGPL
>Pteropus_alecto
-----------------------------------------MVGGLDPPGRRRFQKGFD------------WRNLWSSCRLAPLADGMLRFKGQWQRGHPVQRGPL
>Dasypus_novemcinctus
---------------------------------------------------------------------------------------MLRHMGQGQQGHPVQRGPL
>Sarcophilus_harrisii
---------------------------------------------------------------------------------------MLRHAGQHQRGCPIRRGPL
>Galeopterus_variegatus
---------------------------------------------------------------------------------------MLRYMGQWQRGHPIQRGPL
>Monodelphis_domestica
---------------------------------------------------------------------------------------MLRHAGHHQRGCPIRRGPL
>Macropus_eugenii
---------------------------------------------------------------------------------------MLRHAGQHQRGCPIRRRPL
>Condylura_cristata
-----------------------------------------MVGDLDPPGRRRFQSGFD------------WRNLWSSCWLTPLADGMLRHMGQRQKGRPIQGGPL
>Oryctolagus_cuniculus
-----------------------------------------MVGGLDPPGRRRFQKGFD------------WRNLWSSCWLVPLADGMLKYMGQRQRGHPFQRGPL
>Ceratotherium_simum_simum
MEGAGLRGGRRRWAADEEEAAAVVEEEAEAAGVGSLETRAGMVGGLDPPGRRRFQKGFD------------WRNLWSSCWLAPLADGMLRHMGQWQRGHPVQRGPL
>Manis_javanica
-----------------------------------MCVQGGRQKQLELRLKEESWRLIS-----VQRKLSRATVFDLSGWLAPLADGMLRHKGQWQRGHPIQRGPL
>Loxodonta_africana
-----------------------------------------MVGGLDPPSRRRFQKGSD------------WRNLWSSCGLAPLADGMLRHTGQRQQGHLIPRGPL
>Chinchilla_lanigera
---------------------------------------------------------------------------------------MLRYMGQWQRGHPIQRGPL
>Tupaia_chinensis
-------------------------------------------------------------------------MSHEGGWLAPLADGMLTYMGQWQQGHLIQRGPL
>Trichechus_manatus_latirostr
-----------------------------------------MVGGLDPPSRRRFQKGFD------------WRNLWSSCWLAPLADGMLRHMGQRQRGHLIPRGPL

HG0465 ENSG00000132153 DHX30

>Homo_sapiens
----------------------------------------------------------------------------------------------------------
--------------------------------------------MRNSKERIQLQFKSLQNLRLSLLL
>Mustela_putorius_furo
MLRRWEGAPCAALTSAVQTRSLRCPRGARAPTHSAPLPEVPLAPAAAAPHWARRRGLAAGWPPSGVSPRRPLRAPGGGPAGDPLRRGVGAGGGAGSAGAAGGTSAS
GPGRGRAAAGGSCSPAVVGGARGSCAVHRGLFHICNNRRRPSLVMRNSKERIQLQFKNLQNLRLSLLV
>Orcinus_orca
----------------------------------------------------------------------------------------------------------
--------------------------------------------MRNSKERIQLQFKSPQNLRLSLLL
>Rousettus_aegyptiacus
----------------------------------------------------------------------------------------------------------
--------------------------------------------MRNSKERIQLQFKSLQNLRLSLLL
>Dasypus_novemcinctus
----------------------------------------------------------------------------------------------------------
--------------------------------------------MRNSKERIQLQLKAYKI-------
>Sarcophilus_harrisii
----------------------------------------------------------------------------------------------------------
--------------------------------------------MKNRTERIYFPFKSRRNLRFSLLL
>Galeopterus_variegatus
----------------------------------------------------------------------------------------------------------
--------------------------------------------MRNSKDRIQLQFKSLQNLRLSLLS
>Condylura_cristata
----------------------------------------------------------------------------------------------------------
--------------------------------------------MRNSKETIQLQFKSLQNLRPSLLL
>Elephantulus_edwardii
----------------------------------------------------------------------------------------------------------
--------------------------------------------MRNSKERIQLQFKSLQNLRLSLLL
>Equus_caballus
----------------------------------------------------------------------------------------------------------
--------------------------------------------MRNSKERIQLQFRSLQDLRLSLLS
>Manis_javanica
----------------------------------------------------------------------------------------------------------
--------------------------------------------MRRSKERIQLQLKSLHDLGVSLLL
>Loxodonta_africana
----------------------------------------------------------------------------------------------------------
--------------------------------------------MRNSKERIQLQFRSLQNLKLSLLL
>Rattus_norvegicus
----------------------------------------------------------------------------------------------------------
--------------------------------------------MRNSKERIQLQFESLQNLSPSLLL
>Tupaia_chinensis
----------------------------------------------------------------------------------------------------------
--------------------------------------------MRNSKERTQLQFKSLHNLRLSLHL
>Trichechus_manatus_latirostr
----------------------------------------------------------------------------------------------------------
-------------------------------------------MRNNSKERIQLQFKSLQNLRSSLLL
>Orycteropus_afer_afer
----------------------------------------------------------------------------------------------------------
--------------------------------------------MRNSKERIQLQFKSLQNLRLSLLL

HG0466 ENSG00000139679 LPAR6

>Homo_sapiens

MFLLGISEKKEIFIKPSKDPDNLQINWRL

>Mustela_putorius_furo

MFLTEISKKKEIFIKPSKDTKNLHILWRL

>Lipotes_vexillifer

---MSLLEEKEIFIKSRKDPKDLQIHWRL

>Miniopterus_natalensis

MFLLEILEEKEIFIKPPKDPENLQIHWGF

>Dasypus_novemcinctus

MFLSEILEKKEILIKPPKDPKNLHISWRL

>Sarcophilus_harrisii

MFLLEISKGKEILMKPSKDPKHLQINKRS

>Galeopterus_variegatus

MFLLEISEKKEIFIKPPRDPNNLQINWRL

>Monodelphis_domestica

-------------MKPTKDHLHLQVNRRL

>Macropus_eugenii

MFLLEISKGKEILMKPSKDPKHLQIN---

>Erinaceus_europaeus

MFLLETSEKKEILIKPSKDPKNLQLHWRL

>Oryctolagus_cuniculus

MFHLEISEKTEIFLKPRLDPNNLQIN---

>Equus_przewalskii

MFLLEISEKEEIFIKPPKDPKNLQIRWRL

>Manis_javanica

MFLFEISEKKEIFIKSPKDPRNLHILWRL

>Loxodonta_africana

MFLLEIPEKKEIFIKPPKEPKNLQVNWRL

>Marmota_marmota_marmota

MFLSYILEKKEIFIKPPKDPNNLRINWML

>Orycteropus_afer_afer

MLLSEISEKKEILIKPPQELKNL------

HG0467.1 ENSG00000163320 CGGBP1

>Homo_sapiens
-----------------MTSWHHRENRVQAQKLFPHHHLLKN
>Mustela_putorius_furo
-----------------MTSRHRRENRVQAQKLFPHHHLLKN
>Tursiops_truncatus
-----------------MTSQHLRENRVQAQKLFPHHHLLKN
>Pteropus_vampyrus
-----------------MTSRHRRENRVQAQKLFPHHHLLKN
>Dasypus_novemcinctus
-----------------MTFRHRRENRVQAQKLFPHHHLLKN
>Sarcophilus_harrisii
MPNSTFPEYSGLGTGKGMTTRYASENRVQAQKPFLYHHLLRN
>Galeopterus_variegatus
-----------------MTSRHRRENRVQAQKPFPHHHFLKN
>Monodelphis_domestica
-----------------MTARYASENRVQAQKPFLYHHLLRN
>Condylura_cristata
-----------------MISRHRRENRVQAQKLFPHHHLLKN
>Oryctolagus_cuniculus
-----------------MTSWHLRENRVQAQKLFPHHHLLKN
>Equus_asinus
-----------------MTSWHRRENRVQAQKLFPHHHLLKN
>Manis_javanica
-----------------MTSWHRRENRVQAQKLFLHHHLLKN
>Loxodonta_africana
-----------------MTFRYRRENRVQAQKLFPHHHLLKN
>Nannospalax_galili
-----------------MTSRHRRENRVQAQKLFPHHHLLKN
>Tupaia_chinensis
-----------------MISRHRRENRVQAQKLFPHHHLLKN
>Trichechus_manatus_latirostr
-----------------MTFRYRRENRVQAQKLFPHHHLLKN

HG0467.2 ENSG00000163320 CGGBP1

>Homo_sapiens
MRLWFRSGDHSLLSIKTKKTGN
>Orcinus_orca
MRLWFCSGDPSLLGIKTKKTGN
>Miniopterus_natalensis
MRLWFCAGGPLLLSIKTKKTGN
>Oryctolagus_cuniculus
MRLWLCAGDHSLLSIKTKKTGN
>Ceratotherium_simum_simum
MRLWICAGDPSLLSIKKKKTGN
>Marmota_marmota_marmota
MCLRFCAGDPSLLSIKKKKTGN
>Tupaia_chinensis
-MLWFCAGDPSLLSIQTKKTGI

HG0468 ENSG00000187778 MCRS1

>Homo_sapiens
MLLPRVLETDLDRSPTLGVHSWENEGRRVRISSQPLSSPGQNPWMSLWDRSS------------
>Acinonyx_jubatus
--------------------------------------------MSLWDPSS------------
>Orcinus_orca
--------------------------------------------MNLWDSS-F-----------
>Myotis_brandtii
--------------------------------------------MSLWDPSS------------
>Dasypus_novemcinctus
--------------------------------------------MSPWDVSF------------
>Galeopterus_variegatus
--------------------------------------------MSLWDPSS------------
>Macropus_eugenii
--------------------------------------------MNLWDSS-YPFGLALRDW--
>Erinaceus_europaeus
--------------------------------------------MSLWDPSS------------
>Oryctolagus_cuniculus
--------------------------------------------MSLWDPSSWPRGLKPVALGL
>Equus_asinus
--------------------------------------------MSLWDPSS------------
>Manis_javanica
--------------------------------------------MSLWEPSS------------
>Loxodonta_africana
--------------------------------------------MSLWDPSSWPCGLEPVALGL
>Ictidomys_tridecemlineatus
--------------------------------------------MRLWDPSS------------
>Tupaia_chinensis
--------------------------------------------MSLWDPSF------------
>Trichechus_manatus_latirostr
--------------------------------------------MSLWDPSSWPCGLEPVALGL
>Orycteropus_afer_afer
--------------------------------------------MSLCDPSF------------

HG0469 ENSG00000198369 SPRED2

>Homo_sapiens
MKGAKLPAAGQASR
>Ailuropoda_melanoleuca
MKGAKLPAAGQASR
>Physeter_catodon
MKGAKLPAAGQASR
>Eptesicus_fuscus
MKGAKLPAAGQASR
>Dasypus_novemcinctus
MKGAKLPAAGQASR
>Sarcophilus_harrisii
MKGTELPAAGHSSR
>Monodelphis_domestica
MKGTELPAAGHSSR
>Condylura_cristata
MKGAKLPAAGQASR
>Oryctolagus_cuniculus
MKGAKLPAAGQASR
>Ornithorhynchus_anatinus
MKGTKLPAAGHSSR
>Ceratotherium_simum_simum
MKGAKLPAAGQASR
>Manis_javanica
MKGAKLPAAGQASR
>Loxodonta_africana
MKGAKLPAAGQASR
>Cavia_porcellus
MKGAKLPAAGQASR
>Tupaia_chinensis
MKGAKLLAAGQASR
>Trichechus_manatus_latirostr
MKGAKLPAAGQASR

HG0470 ENSG00000224470 ATXN1L

>Homo_sapiens

----MWAPQPEAERGTGKLQRSPF

>Panthera_tigris

----MWAPQPAAERGAGEPQRSPF

>Balaenoptera_acutorostrata_s

----MWAPQPAAERGAGEPQRSPF

>Rousettus_aegyptiacus

----MWAPQPSAERGAGEPQRSPF

>Dasypus_novemcinctus

----MWPPQPAAERGAGKPQRSCF

>Sarcophilus_harrisii

MEAQMWVPQSAVEGGTRKSQRSLF

>Galeopterus_variegatus

----MWAPQPAAERGAGKPQRSPF

>Condylura_cristata

----MWAPQPAAERGAGEPQRSPF

>Ochotona_princeps

----MWAPQPAAERGAGKPQRSPF

>Elephantulus_edwardii

----MWAPRPTAARGAGKPQRSPF

>Equus_przewalskii

----MWAPQPAAERGAGEPQRSPF

>Manis_javanica

----MWAPQPAAERGAGEPQRSPF

>Loxodonta_africana

----MWAPQPAAERGAGKPQRSPF

>Heterocephalus_glaber

----MWAPQPTAERGAGKPQRSPF

>Trichechus_manatus_latirostr

----MWAPQPATERGAGKPQRSPF

>Orycteropus_afer_afer

----MWAPQPAAERVAGKPQRSPF

HG0471 ENSG00000023041 ZDHHC6

>Homo_sapiens
-------------------------------------------------------------------MTIKEHTWTCGTWKCAHRKRKSIILLK
>Ursus_americanus
-------------------------------------------------------------------MTIKEHTWTCGTWKCVHRKRKSIIF--
>Balaenoptera_acutorostrata_s
-------------------------------------------------------------------MTIKEHTWTCGTWKCVHRKRKSVVL--
>Pteropus_alecto
-------------------------------------------------------------------MSIKEHTWTCGTWKCVHRKRKAVIL--
>Dasypus_novemcinctus
-------------------------------------------------------------------MTVKGDTWTHGTWKCVHRQRKSIIL--
>Galeopterus_variegatus
-------------------------------------------------------------------MTIKEHTWTCGTWKCMHRKRKSIVL--
>Macropus_eugenii
-------------------------------------------------------------------MSINEDTWTFDTWKCVYKKGTLQGF--
>Condylura_cristata
MLVFGEQPQPGDLDHVWYSRLHWRLSLEPCKIEFHTKRLDSPSYHKELCVRNAENGIICINIRQLEGMTAKEHTWTWGTWKYVHRKRKSIIL--
>Oryctolagus_cuniculus
-------------------------------------------------------------------MTVKEHTWTCGTWKCVHRKRKSVIL--
>Elephantulus_edwardii
-------------------------------------------------MLNSEDDILFV--NILESMTVEEDMWTCGMWKCMHRKRKSVIL--
>Ceratotherium_simum_simum
-------------------------------------------------------------------MTVKEHTWTCGTWKCVHRKRKSIIL--
>Loxodonta_africana
------------------------------------------------------------------MTIKEEDTWTCATWKCVHRKRTSINL--
>Chinchilla_lanigera
-------------------------------------------------------------------MTIKEHTWTCGTWKCVHRKRKSIIL--
>Trichechus_manatus_latirostr
-------------------------------------------------------------------MTVKEDTWTCGTWKCVHRKRKSISL--
>Orycteropus_afer_afer
-------------------------------------------------------------------MTVQEDKWTWGTWKCVPRKRKPVIL--

HG0472 ENSG00000105991 HOXA1

>Homo_sapiens
--------MEEVRKLAQSRRASQDQVTQ
>Leptonychotes_weddellii
--------MEEVRKLARSPRASQDPVPQ
>Tursiops_truncatus
--------MEEVRKLARSPRASQDPVTQ
>Rousettus_aegyptiacus
--------MEEVRKLARSPRASQDQVTQ
>Monodelphis_domestica
--------MEEVRKLARSPRARQDPVTV
>Condylura_cristata
--------MEEVRKLARSSRASQDQVTQ
>Oryctolagus_cuniculus
--------MEEVRKLARSPRASQDPVTQ
>Elephantulus_edwardii
--------MEEVRKLARSSRASQQPATQ
>Equus_przewalskii
--------MEEVRKLARSPRASQDPVTQ
>Manis_javanica
MSFFFSGPMEEVRKLARSPRASQNPVTQ
>Loxodonta_africana
--------MEEVRKLARSPRASQDPVTQ
>Peromyscus_maniculatus_baird
--------MEEVRKLARSPRASQDPVTQ
>Tupaia_chinensis
--------MEEVRKLAQSPRASQDPVTQ
>Trichechus_manatus_latirostr
--------MEEVRKLARSRRASQDPVTQ
>Orycteropus_afer_afer
--------MEEVRKLARSPRASQDPVTQ

HG0473.1 ENSG00000138650 PCDH10

>Homo_sapiens

MRKGDACHPSCAKI---------------

>Odobenus_rosmarus_divergens

MRKGDACHPSCAKIEK-------------

>Lipotes_vexillifer

MRKGDACHPSCAKTEKL--G---------

>Pteropus_alecto

MRKGDACHPSCAKIEK-------------

>Dasypus_novemcinctus

MRKRDVCHPSCTKTEK-------------

>Galeopterus_variegatus

MRKWDACHPSCAKIEK-------------

>Monodelphis_domestica

MRSGDVCLPSCAKREKGEAGWWGVEP---

>Condylura_cristata

MRRGDACRPACAKVEERG-----------

>Ochotona_princeps

MRIGDAGHPSCAEIEK-------------

>Equus_asinus

MRKGDACHPSCAKIEK-------------

>Manis_javanica

MRKGDACHPSCAKIEK-------------

>Loxodonta_africana

MRKGDACHPSCAKIEK-------------

>Marmota_marmota_marmota

MRKGDVCHLSCAKIEKL--DWIEDSFKMK

>Tupaia_chinensis

MRKGDACHPSCAKIEK-------------

>Trichechus_manatus_latirostr

MRKGDACHPSCAKIEK-------------

HG0473.2 ENSG00000240184 PCDHGC3

>Homo_sapiens
MSDSAPSAQALTR
>Leptonychotes_weddellii
MSDSAPSAQALTR
>Orcinus_orca
MSDSEPSARALTR
>Myotis_davidii
MSDSAPSAQALTR
>Dasypus_novemcinctus
MSDSAPSARALTR
>Erinaceus_europaeus
MSDSAPSAQALNR
>Oryctolagus_cuniculus
MSDSAPSAQALSR
>Equus_przewalskii
MSDSAPSAQALTR
>Manis_javanica
MSDSAPSAPALTR
>Loxodonta_africana
MSDSAPSAQALSH
>Mus_musculus
MSDSAPSAQALTR
>Trichechus_manatus_latirostr
MSDSAPSAQALTR

HG0474 ENSG00000139083 ETV6

>Homo_sapiens
MTASGWP-WSLSGLGRGKESGKNLRTS-----
>Ailuropoda_melanoleuca
MTASGWP-WSASGLGRGKESGKKLRTS-----
>Balaenoptera_acutorostrata_s
MTASGRP-WSVSGLGRGKESGKKLRTS-----
>Pteropus_alecto
MTASGWP-WSVSGLGRGKESGKKLRTS-----
>Sarcophilus_harrisii
MIASGCLFGGFLGWGRGKESGKKLRTS-----
>Monodelphis_domestica
MIASGWLFGGFLGWGRGKESGKKLRTS-----
>Condylura_cristata
MTASGRP-WSASGLGRGKESGKKLRTS-----
>Oryctolagus_cuniculus
MTASGWP-RSRSGLGRGKESGKTLRTCCSPPL
>Elephantulus_edwardii
MTASGWP-WSISGLGRGKESGKNQRTS-----
>Equus_przewalskii
MTASGWP-WSVSGLGRGKESGKKLRTS-----
>Loxodonta_africana
MTASGWP-WSISGLGRGKESGKKLRTS-----
>Octodon_degus
MTASGCP-WSVSGLGRGKESGKKLRTS-----
>Tupaia_chinensis
MTASGWP-WSFSGLGRGKESGKKLRTS-----
>Trichechus_manatus_latirostr
MTASGWP-WSLSGLGRGKESEKN---------
>Orycteropus_afer_afer
MTASGWP-WSISGLGRGKESGKKLRTS-----

HG0475 ENSG00000109118 PHF12

>Homo_sapiens
MAAPASLPGRTGCRQCRL
>Canis_lupus_familiaris
MAAPASLPGRTSGRQRRL
>Orcinus_orca
MAAPASFPGRTSGRQHRL
>Rousettus_aegyptiacus
MAAPASLPSRTSSRQRRL
>Dasypus_novemcinctus
MAAPASLPGRTSGHQRRL
>Sarcophilus_harrisii
MAAPASLPGRAGGRQRRL
>Galeopterus_variegatus
MAAPANLPGPTSGRQRRL
>Erinaceus_europaeus
MAAPANLPGRTSGRQRRL
>Oryctolagus_cuniculus
MAAPASLPGRTSGRQRRL
>Elephantulus_edwardii
MAAPDSLPGRTSGRQRRL
>Ceratotherium_simum_simum
MAAPASLPGRTSGRQRRL
>Heterocephalus_glaber
MAAPASLPGRTSGRQHRL
>Trichechus_manatus_latirostr
MAAPASLPGRTSSRQRRL
>Orycteropus_afer_afer
MAAPASLPGRTSGLQRRL

HG0476 ENSG00000130939 UBE4B

>Homo_sapiens
MNNTWWGEGER-VGVEG-------------------------------
>Mustela_putorius_furo
MNNTWWGEGVR-VGVEG-------------------------------
>Orcinus_orca
MNNTWWGEGVR-VGVEG-------------------------------
>Pteropus_alecto
MNNTWWGEGVR-VGVEG-------------------------------
>Dasypus_novemcinctus
MNNTWWGEGVR-VGVEG-------------------------------
>Galeopterus_variegatus
MNNTWWGEGVR-VGVEGRRIYSSSESIAVLPLPPARAPAWRRGVAGLF
>Monodelphis_domestica
MNNPWWGQGGQRRAEG--------------------------------
>Condylura_cristata
MNNTWWGEGVR-VGVEG-------------------------------
>Ochotona_princeps
MNNTWWGEGVR-VGVEG-------------------------------
>Equus_przewalskii
MNNTWWGEGVR-LGVEG-------------------------------
>Ictidomys_tridecemlineatus
MNNTWWGEGVR-VGVEG-------------------------------
>Tupaia_chinensis
MNNTCWGEGVRRVGVEG-------------------------------
>Trichechus_manatus_latirostr
MNNTWWGEGVR-VEVEG-------------------------------
>Orycteropus_afer_afer
MNNTWWGEGVK-VRVER-------------------------------

HG0477 ENSG00000166925 TSC22D4

>Homo_sapiens
MSDPDVPRGSDVPAMWPPCSRGA
>Phoca_largha
MSDPDVPRGPDAPAMWPPCSRGA
>Balaenoptera_acutorostrata_s
MSDPDVPRGPDVPTMWPPCSRGA
>Rhinolophus_ferrumequinum
MSDSDVPRGPDVPAMWPPCSRGA
>Dasypus_novemcinctus
MSDPDVPRGPDAPAMWPPCSRGA
>Sarcophilus_harrisii
MSDPDVARGPDPTAIWPPSARGA
>Galeopterus_variegatus
MSDPDVARGPDAPTMWPPCSRGA
>Monodelphis_domestica
MLDPDVPRGPDPTAIWPPSARGA
>Condylura_cristata
MSDPDVPRGPDAPAMWPPCSRGA
>Oryctolagus_cuniculus
MSDPDVPRGPDAPAMWPPCSRGA
>Equus_caballus
MSDPDVPRGPDVPAMWPPCSRGA
>Manis_javanica
MSDPDIPRGPDAPTMWPPCSRGA
>Loxodonta_africana
MSDPDVPRGSDVPAMWPPCSRGA
>Cricetulus_griseus
MSDPDVPRGPDVPAMWPPCSRGA

HG0478 ENSG00000168214 RBPJ

>Homo_sapiens
MISKRKENDTDTLAEVF
>Acinonyx_jubatus
MISKGKENDTDTLAEVF
>Lipotes_vexillifer
MISKGKENDTDTLAEVF
>Myotis_brandtii
MISKGKENDTDTLAEVF
>Dasypus_novemcinctus
MISKGKENDTDTLAEVF
>Sarcophilus_harrisii
MISRGKEND--TLAEVF
>Galeopterus_variegatus
MISKGKENETDTLAEVF
>Monodelphis_domestica
MISKGKEND--TLAEVF
>Oryctolagus_cuniculus
MISKGKENDTDTLAEVF
>Ornithorhynchus_anatinus
MISKGKEND--TLAEVF
>Ceratotherium_simum_simum
MISKGKENDTDTLAEVF
>Manis_javanica
MISKGKENDTDTLAEVF
>Dipodomys_ordii
MISKEKENDTDILAEVF
>Orycteropus_afer_afer
MISKGKENDTDTLAEVF

HG0479.1 ENSG00000168453 HR

>Homo_sapiens
----------------------------------------------------------------------------------------------------------
------------------------------------------MAQPTASAQKLVRPIRAVCRILQIPESDPSNLRP
>Leptonychotes_weddellii
----------------------------------------------------------------------------------------------------------
------------------------------------------MAQPTASAQKLVRPIRAVCRILQIPESDPSNLRP
>Lipotes_vexillifer
----------------------------------------------------------------------------------------------------------
------------------------------------------MAQPTASAQKLVRPIRAVCRILQIPEPDPSNLRP
>Myotis_lucifugus
MRSRWHRRGVGGSPRGTGGGRPRRGRRITGSCFPQGAGGGNRRSSFLTLSCASGDGDQRSCCAPRCALPPLAALPRFAPSLVLSSGFQDRRPLPGTTPSPGPSAFP
PLGRLRPGARVPPARLPLSLRGHRSLAPARPSPRRSTESPRPMAQPTASAQKLVRPIRAVCRILQIPESDPSNLRP
>Sarcophilus_harrisii
----------------------------------------------------------------------------------------------------------
------------------------------------------MAQPTTSAQKLVRPIRAVCRILQIPESDPSNLRP
>Monodelphis_domestica
----------------------------------------------------------------------------------------------------------
------------------------------------------MAQPTTSAQKLVRPIRAVCRILQIPESDPSNLRP
>Condylura_cristata
----------------------------------------------------------------------------------------------------------
------------------------------------------MAQPTASAQKLVRPIRAVCRILQIPESDPSNLRP
>Oryctolagus_cuniculus
----------------------------------------------------------------------------------------------------------
------------------------------------------MAQPTASAQKLVRPIRAVCRILQIPESDPSNLRP
>Ceratotherium_simum_simum
----------------------------------------------------------------------------------------------------------
------------------------------------------MAQPTASAQKLVRPIRAVCRILQIPESDPSNLRS
>Manis_javanica
----------------------------------------------------------------------------------------------------------
------------------------------------------MAQPTASAQKLVRPIRAVCRILQIPESDPSNLRP
>Ictidomys_tridecemlineatus
----------------------------------------------------------------------------------------------------------
------------------------------------------MAQPTASAQKLVRPIRAVCRILQIPESDPSNLRP
>Tupaia_chinensis
----------------------------------------------------------------------------------------------------------
------------------------------------------MAQPTASAQKLVRPIRAVCRILQIPESDPSNLRP
>Trichechus_manatus_latirostr
----------------------------------------------------------------------------------------------------------
------------------------------------------MAQPTASAQKLVRPIRAVCRILQIPESDPSNLRS
>Orycteropus_afer_afer
----------------------------------------------------------------------------------------------------------
------------------------------------------MAQPTASAQKLVRPIRAVCRILQIPESDPSNLRP

HG0479.2 ENSG00000168453 HR

>Homo_sapiens
MAIRGPAALSAALYLH----
>Odobenus_rosmarus_divergens
MAMRGPAALSAALSLH----
>Lipotes_vexillifer
MAVRGPAALSAGHSLH----
>Myotis_lucifugus
MAIRGPAALPAALSLH----
>Monodelphis_domestica
MAIGGPAALSVPLGLH----
>Condylura_cristata
MAIRGPAALSAALSLH----
>Oryctolagus_cuniculus
MAIGGPEALSAALSLH----
>Ceratotherium_simum_simum
MAIRGPAALSAALSLH----
>Manis_javanica
MAIGGPAALSAALSLH----
>Ictidomys_tridecemlineatus
MAIRGPAALSAALSLH----
>Trichechus_manatus_latirostr
MAIRGPAALSAALFAELSLH
>Orycteropus_afer_afer
MAIRSPAALSAALSLH----

HG0480 ENSG00000198018 ENTPD7

>Homo_sapiens
MPGTRRPPSQGKRKRR
>Canis_lupus_familiaris
MPETRRPVSQGKKKRR
>Balaenoptera_acutorostrata_s
MTGTRRPPSQGKRKRR
>Myotis_davidii
MPGTRRPPSQGKRKRR
>Galeopterus_variegatus
MPGTRRPPSQGKRKRR
>Monodelphis_domestica
MPGTGKPPCQGKRKRR
>Condylura_cristata
MPGTRRPSAQGKRKRR
>Oryctolagus_cuniculus
MPGTRRPPSQGKRKRR
>Equus_asinus
MPGTRRPPSQGKRKRR
>Manis_javanica
MSGTRRPPSQGKRKMR
>Loxodonta_africana
MPGTRRPHSQGKRKRR
>Dipodomys_ordii
MPGTRRPPSQGRRKRR
>Tupaia_chinensis
MPGTRRPPSQGKRKRR
>Trichechus_manatus_latirostr
MPGTRRPHSQGKRKRR

HG0481 ENSG00000198963 RORB

>Homo_sapiens
MVFSAEQLFADHLLHSC
>Ursus_maritimus
MVFSAEQLFADHLLHSC
>Balaenoptera_acutorostrata_s
MVFSAEQLFADYLPHSC
>Rousettus_aegyptiacus
MVFSAEQLFADHLLHSC
>Monodelphis_domestica
MVFSAEQIFGCSL-HSC
>Condylura_cristata
MVFSAKQLFAFHLLHSC
>Elephantulus_edwardii
MVFSAEQLFTDHLLPSC
>Ceratotherium_simum_simum
MIFSAEQLFADHLLHSC
>Manis_javanica
MVLSVEQLFADHLLQCC
>Loxodonta_africana
MVFSAEQLFTDHLLHLC
>Peromyscus_maniculatus_baird
MLLSAEQLFADRLLPSC
>Tupaia_chinensis
MVFSAEQLFTDYLLHSC
>Trichechus_manatus_latirostr
MVSSAEQLFTNHLLHSC
>Orycteropus_afer_afer
MVFSAEQPFVDHLVHSC

HG0482 ENSG00000067167 TRAM1

>Homo_sapiens
-----------------------------MGSEAPASGCSGAVRSSQREAAASR---------------
>Odobenus_rosmarus_divergens
MLTPAVGSPSFFFFFFFFGRPRRLRTGCGMGSEAPASGSGGAVRSSQGEAAAAASR-------------
>Physeter_catodon
-----------------------------MGSEAPASGSGGAVRSSQGEAAAASR--------------
>Pteropus_alecto
-----------------------------MGSKAPASGSGGAVRSSQGEAAAASR--------------
>Dasypus_novemcinctus
-----------------------------MGSGAPASGCGWAVSSSQGEAGAAASQ-------------
>Galeopterus_variegatus
-----------------------------MGSEAPASGCGGAVRSSQGAAAAASR--------------
>Monodelphis_domestica
-----------------------------MGSEVPESGCGWAENSSQGEETAGAATAEATAAAAAASCP
>Condylura_cristata
-----------------------------MGSEAPANGSSGAVRSSQGEAAAAFSR-------------
>Oryctolagus_cuniculus
-----------------------------MGSEAPASGCGGAVRSGQGEAAAAAVSR------------
>Equus_asinus
-----------------------------MGSEAPASGGGGAVRSGRGEEAEAAARR------------
>Manis_javanica
-----------------------------MGSEAPASCSDGAVRSSQGEAAAVACR-------------
>Rattus_norvegicus
-----------------------------MGSEAPASGCGGAVRSTQREAAVAALR-------------
>Trichechus_manatus_latirostr
-----------------------------MGSEALASGCGGAVRSSQGEVAAAACR-------------

HG0483 ENSG00000164463 CREBRF

>Homo_sapiens
MERGRGPCSGTGPSP
>Ailuropoda_melanoleuca
MERGRGPCSGTGPSP
>Orcinus_orca
MERGRGPCSGTGPSP
>Rousettus_aegyptiacus
MERGRGPCSGTGPSP
>Dasypus_novemcinctus
MERGRGPCSGTGPSS
>Sarcophilus_harrisii
MERGRGPRTRTGPSP
>Galeopterus_variegatus
MERGRGPCSGTGPSP
>Macropus_eugenii
MERGRGPRSRTGPSP
>Oryctolagus_cuniculus
MERGRGPCSGTGPSP
>Equus_przewalskii
MERGRGPCSGTGPRP
>Manis_javanica
MERGRGPCSGTGPRP
>Ictidomys_tridecemlineatus
MERGRGPCSGTGPIP
>Trichechus_manatus_latirostr
MERGRGPCSGTAPSP

HG0484 ENSG00000166444 ST5

>Homo_sapiens
MNQLASISLTNHRKMRDCLGATTSTMSPCTG-
>Canis_lupus_familiaris
--------------MRDCLGTSNSTMSPCTG-
>Eptesicus_fuscus
--------------MRDCLGTSTSTMRPCTG-
>Galeopterus_variegatus
--------------MRDCLGTSTSTMGPCTG-
>Monodelphis_domestica
--------------MRDCPGSASSTMSTACTD
>Elephantulus_edwardii
--------------MRDYLSANTSTMSPCTG-
>Ceratotherium_simum_simum
--------------MRDCLSTSPSTMSPCTG-
>Manis_javanica
--------------MRDCLGTSTSTRNPCTG-
>Loxodonta_africana
--------------MRDCLGASTSTMSPCTG-
>Chinchilla_lanigera
--------------MRDCLGSSSSTMSPCTG-
>Tupaia_chinensis
--------------MQDCFGASSSAMSPCTG-
>Trichechus_manatus_latirostr
--------------MRDCLGASTSTMSPCTG-
>Orycteropus_afer_afer
--------------MRDCLGASTSIMSPCTG-

HG0485.1 ENSG00000182667 NTM

>Homo_sapiens
MSGDNYGEVILSHTLGFLVVSFSKTVDLNLLAQA
>Mustela_putorius_furo
MSRDNY----------------------------
>Orcinus_orca
MRRDNY----------------------------
>Pteropus_alecto
MSRDNY----------------------------
>Monodelphis_domestica
MSRDNY----------------------------
>Macropus_eugenii
MSRDNY----------------------------
>Erinaceus_europaeus
MSRDNY----------------------------
>Ochotona_princeps
MSGDNYGEVILSHTLGFLVVSFSETVDLNLRA--
>Equus_caballus
MSRDNY----------------------------
>Manis_javanica
MSRDNY----------------------------
>Microtus_ochrogaster
MSGDNYGEVRLSHALGFLVVSFSKTVDLNLLGRA
>Tupaia_chinensis
MSGDNYGEVALSHTLGFL-VSFSETVDLNLLA--
>Trichechus_manatus_latirostr
MSGDNY----------------------------

HG0485.2 ENSG00000183715 OPCML

>Homo_sapiens
MQEGASGSARLES
>Mustela_putorius_furo
MQEGASGSARLES
>Orcinus_orca
MQEGASGSARLES
>Ochotona_princeps
MQEGASGSARLES
>Equus_asinus
MQEGASGSARLES
>Mesocricetus_auratus
MQEGASGSARLES

HG0486 ENSG00000067900 ROCK1

>Homo_sapiens
MRLWASERLWVAVAPGPGSGE-RRSPQ
>Canis_lupus_familiaris
MRLWASERLWVAVAPGPGSGE-RRSPQ
>Lipotes_vexillifer
MRLWASERLWVAVAPGPGSGE-RRSPQ
>Myotis_brandtii
MRLWASERLWVALAPGPGSGE-RRSPQ
>Dasypus_novemcinctus
MRLWASERLWVAVAPSPGSGE-WRSPQ
>Sarcophilus_harrisii
MRLWDSVRLRVAVAPRARPGERRAAQ-
>Sorex_araneus
MRLWASERLWVAVAPGPGSPAERRSPQ
>Ochotona_princeps
MRLWACERLWVAVVPGPGSGEPRC-PQ
>Equus_asinus
MRLWASERLWVAVAPGPGSGE-RRSPQ
>Dipodomys_ordii
MRLWASGRLWVAVAPGPGSGERRWSPQ
>Tupaia_chinensis
MRLWASERLWVAVAPGPGSGE-RRSPQ
>Orycteropus_afer_afer
MRLWASERLWVAVAPGPGSGE-RRSPQ

HG0487 ENSG00000091831 ESR1

>Homo_sapiens
----------------------------------MEHFWKDVLDPAGWPAGF
>Mustela_putorius_furo
----------------------------------MEHFWKDALDPAGWPLGC
>Lipotes_vexillifer
----------------------------------MEHFWKDALDPAGWPVGC
>Rousettus_aegyptiacus
----------------------------------MEPFWKDALDPAGWPVGC
>Monodelphis_domestica
MNPCNFAHAYRATCTYTHILSFILTLSAFEFCISMEHFRKDALCPVGWPTGC
>Condylura_cristata
----------------------------------MEHFWKDALDPAGWPVCC
>Oryctolagus_cuniculus
----------------------------------MEHFWKDTLDPAGWPVGC
>Ceratotherium_simum_simum
----------------------------------MEHFWKDALDPAGWPAGC
>Manis_javanica
----------------------------------MDHFWKDALDPAGWRVGC
>Ictidomys_tridecemlineatus
----------------------------------MEHFWKDTLAPAGWPTGC
>Tupaia_chinensis
----------------------------------MEHFWKDDLDPAGWPVGC
>Trichechus_manatus_latirostr
----------------------------------MEHLWKDAFDPAGWPVGC

HG0488.1 ENSG00000116194 ANGPTL1

>Homo_sapiens
----------------------------------MWQTEQSHNAFSAGLKTAHRPELPLYYLKLHRWLVKFN----
>Canis_lupus_familiaris
----------------------------------MWQTEQSHNAFSAGLKTAHRPELPLYYLKLHRWLVKFNRLVL
>Balaenoptera_acutorostrata_s
----------------------------------MWQTEQSHNAFSAGLKTARRPGTSTILLKIT-----------
>Myotis_lucifugus
----------------------------------MWQTEQSHNAFSAGLKTAHRPELPLHYLRLHRWLVKFD----
>Dasypus_novemcinctus
----------------------------------MWQTEQSHNAFSAGLKTALRPELPLYYLKLHRWLVKFN----
>Sarcophilus_harrisii
----------------------------------MKQTEERHNTFSAGKKDISHMEICAVLSKII-----------
>Condylura_cristata
----------------------------------MWLTEQSHNAFSAGLKTTHRVDRRLCYLKLHRWLVKFN----
>Oryctolagus_cuniculus
----------------------------------MWQTEQSHNAFSAGLRTAHRPELLLYDLKLHRWLVKLN----
>Equus_przewalskii
MWAELESTEKERESLFKGTVYTIWVSCSWLLHFSMWQTEQNHNAFSAGLKTAHRPELPLYYLKLHRWLVKFN----
>Manis_javanica
----------------------------------MWQTEQSHSAFSAGLQIAHRPELPLYYLKLHRWLVKFN----
>Callospermophilus_lateralis
----------------------------------MWQTEQSHKAFSEGLDTAHRPELPLHYLKLHRWLVKFN----
>Tupaia_chinensis
MWAALESTEKERESLFKGTVYTGRVSCSSLLHFSMWQTAQGHSAFSAGLKTAHRPELHHTT---------------

HG0488.2 ENSG00000116194 ANGPTL1

>Homo_sapiens
MQLPLKSQMNKLLH
>Felis_catus
MQLPLKSQMNKLLH
>Balaenoptera_acutorostrata_s
MQLPLKSQTNKLLH
>Rousettus_aegyptiacus
MQLPLKSQMNKLLH
>Galeopterus_variegatus
--------MNKLLH
>Oryctolagus_cuniculus
MQLPLKSQMNKLLN
>Equus_caballus
MQLPLKSQTSKLLH
>Loxodonta_africana
MQLPLESQMNKLHF
>Marmota_marmota_marmota
MQLPLKSQMNKLLH
>Tupaia_chinensis
MQLPLKSQMKKLLH

HG0489 ENSG00000143847 PPFIA4

>Homo_sapiens
------------------------------------------MLWEKPLPTCPLAWHCPL
>Odobenus_rosmarus_divergens
------------------------------------------MPWEKPLPTCPLAWRCPL
>Balaenoptera_acutorostrata_s
------------------------------------------MPWEKPLPICSLTWHCLL
>Myotis_brandtii
--------------------------------------------MGGAPPTCLLAWHCPL
>Dasypus_novemcinctus
--------------------------------------------MGEPLPSCPPTWHCPL
>Galeopterus_variegatus
------------------------------------------MPWEKPLPTCPLAWHCLF
>Monodelphis_domestica
----------------------------------------MASAWESPLPAYPLAWHCPL
>Condylura_cristata
------------------------------------------MPWEKPLPIRPLAWHCPL
>Equus_asinus
------------------------------------------MPWEKPLPTCPLAWHCLL
>Loxodonta_africana
------------------------------------------MPWEKLLSTHPPTWHCPL
>Heterocephalus_glaber
MLSAGLGTQGTQPQPREVKPGFLAPFHRGTAGPRLLSEQEQLMAWEKPLPTCPLSWHCPL
>Tupaia_chinensis
------------------------------------------MPWEKPLLICPLAWHCPL

HG0490.1 ENSG00000152413 HOMER1

>Homo_sapiens

MAPFILFGRMNRSCLAFSSSSPPLLCSES

>Odobenus_rosmarus_divergens

MAPFILLGRMNRSCPAFSSSSPPPLCSES

>Myotis_lucifugus

MAPFI-LGRMNRSCPAFSASSPPLLGSES

>Dasypus_novemcinctus

MAPFILLGRMNRSCPAFSSSSPPPLCS--

>Galeopterus_variegatus

MAPFILLERINRSCPAFSSSSTPPLCFES

>Monodelphis_domestica

MASFILFWRMNRSSPAFSSCSPPPLSSDS

>Sorex_araneus

MAPFILLGRMNRSCPVFSSSSPPPLCSES

>Manis_javanica

MVPFILLGRMNRSCPAFSSSSPPPLCSES

>Loxodonta_africana

MAPFILLGRVNRSCPVFSSFSLPPLCSES

>Dipodomys_ordii

MAPFLLLGRMNRSCLAFSSTSPPLR-SES

>Tupaia_chinensis

MAPFI-LGRMNRSCPAFSSSSSPPLSSES

>Orycteropus_afer_afer

MAPFILLGSMSRSCPVFSSSSLPPLCSES

HG0490.2 ENSG00000152413 HOMER1

>Homo_sapiens

MPPLFERFGIYKREIILPKRTN

>Acinonyx_jubatus

MPPLLERFGIYEREILLPKGTN

>Balaenoptera_acutorostrata_s

MPPLFEHFGISQREIILPKGTN

>Pteropus_vampyrus

MPPLFEHFGIYGREIILPKGTN

>Galeopterus_variegatus

MPPLFERFGVYQREIILPKRTN

>Condylura_cristata

MPPLFERFGVYEREIILPKGTN

>Ceratotherium_simum_simum

MPPLFERFGIYEGEIILPKGTN

>Manis_javanica

MSPLFERFDIYKREIILPKGTN

>Dipodomys_ordii

MPPLFERFGVYERELILPKRNN

>Tupaia_chinensis

MPPLFERFGIYEREIILPKRTN

HG0491 ENSG00000154114 TBCEL

>Homo_sapiens
-------MQDTGLRRPEWGTR-----------
>Mustela_putorius_furo
-------MQDTGLRRPEWGTRMPDIIRYRQLA
>Orcinus_orca
-------MQDTGLRRPEWGTR-----------
>Rousettus_aegyptiacus
MKAASGSMQDTGLLRAELGTR-----------
>Monodelphis_domestica
-------MQDTGLRRPELGTR-----------
>Condylura_cristata
-------MEDTGLRRPEWGTR-----------
>Ochotona_princeps
-------MQDTGLRRLDWGTR-----------
>Elephantulus_edwardii
-------MQDTWLRRSEWGPR-----------
>Ceratotherium_simum_simum
-------MQDTGLRRPEWGPR-----------
>Loxodonta_africana
-------MQDTGLGRPEWGTR-----------
>Ictidomys_tridecemlineatus
-------MQDTGLRRPEWGTR-----------
>Trichechus_manatus_latirostr
-------MQGTGLRPPELGTR-----------

HG0492 ENSG00000162670 BRINP3

>Homo_sapiens
-------------------------MEYLFESWMYFSHSLLAS------
>Mustela_putorius_furo
-------------------------MEYLFESWIYFSHSHLAS------
>Orcinus_orca
-------------------------MEYLLASWMYFSHSHLAS------
>Pteropus_alecto
-------------------------MEYLFECWMYFSHCHLAS------
>Galeopterus_variegatus
-------------------------MEYLFESWMYFPHGHLAS------
>Monodelphis_domestica
-------------------------MECLFDFWIHYSHSHLAS------
>Erinaceus_europaeus
-------------------------MDYLFESWMYFSHSHRAS------
>Equus_asinus
MPPGTADYFKVPPPCSYHCTLIWKIMEYLCESWMYISHSHLASWNFLWC
>Loxodonta_africana
------------------------MMEYLFESWTYFSHSHLAS------
>Heterocephalus_glaber
-------------------------MEYLFESWMYFSHGHLAS------
>Trichechus_manatus_latirostr
-------------------------MEYLFESWTYFSHSHLAS------
>Orycteropus_afer_afer
---------------------------------MYFSHSHLAY------

HG0493 ENSG00000164707 SLC13A4

>Homo_sapiens
MPPFIILSQFLFAGHKTMLINPT
>Canis_lupus_familiaris
MPPSITLPRCLSAGPRTMLSNPT
>Balaenoptera_acutorostrata_s
MPPFIILSRFLFAGHKTMLTNPT
>Rhinolophus_ferrumequinum
MSRFTILCRFLSAGHKTMLTNPT
>Dasypus_novemcinctus
MPPFTILSQYLFAGHKTMLINPT
>Galeopterus_variegatus
MPPFIILSQFLSAGHKTMLINPT
>Monodelphis_domestica
MPQFVILGQFPFPAQKTLLINPT
>Oryctolagus_cuniculus
MPPHRTLSQFLFAGHKTMLINPT
>Ceratotherium_simum_simum
MPPFIIFSHFLFAGHKTMLANPT
>Manis_javanica
-----------------MLTDPS
>Peromyscus_maniculatus_baird
-----------------MLLNPT
>Tupaia_chinensis
MPPFIILSQFLFAGHKTMLINPT

>Homo_sapiens
-----------------------------------------MHAAVENIPELKKCNYVLCVLNAQKN---P---DPVGKRDEMWRE
>Mustela_putorius_furo
-----------------------------------------MLDAVENTPELKKCNGMLC--VCVKYQKSP---DPVGNRDEMWRE
>Orcinus_orca
-----------------------------------------MFDAVENTPELKKCNWMLCVR------------------------
>Rousettus_aegyptiacus
-----------------------------------------MLDPVENIPELKKCNWMLCVLNTRRAQTSWVIETKCGEND-----
>Dasypus_novemcinctus
MRNFAGNCHLGEQPESPHHPNSDVSYQKLETSAVSISDGNRLLDAVGNTPALKKCSWMLCVC------------------------
>Monodelphis_domestica
-----------------------------------------MQSTVENIPELKKCNWMLCVCSIQGAPIS------CG--------
>Condylura_cristata
-----------------------------------------MLDAVENTPELKKCNWMVCVCVCVKYQKSP---DPMGNKDEMWRE
>Ceratotherium_simum_simum
-----------------------------------------MLDAVENTPELKKCNWMLC--VC----------------------
>Loxodonta_africana
-----------------------------------------MLDAVENTPELKKCNWMVCVCEIPE---EP---RPSG--------
>Nannospalax_galili
-----------------------------------------MLDAVENTPELKKCNWMWCLLNTRS----P---DPVGNRDEMWRE
>Trichechus_manatus_latirostr
-----------------------------------------MLDAVENSLELKKCNWMLCVC------------------------
>Orycteropus_afer_afer
-----------------------------------------MLGAVENTPEVKKCNWVLCVC------------------------

HG0495.1 ENSG00000196730 DAPK1

>Homo_sapiens

----------------------MHEGATEAQERW

>Felis_catus

----------------------MHEGPTEAQERW

>Balaenoptera_acutorostrata_s

----------------------MHEGPTEAQERW

>Rousettus_aegyptiacus

----------------------MHEGPTEAQERW

>Galeopterus_variegatus

----------------------MHEGPTEAQERW

>Erinaceus_europaeus

----------------------MYEGPTEAQKRW

>Oryctolagus_cuniculus

MVGLRQRSGEIRGVLGARDWRPMHEGSTEAREPW

>Equus_asinus

----------------------MHEGPTEAQERW

>Loxodonta_africana

----------------------MHEGPTTAQERW

>Ictidomys_tridecemlineatus

MIGLRQRPERPGELRDAGDWRLMHEGPTSVQERW

HG0495.2 ENSG00000196730 DAPK1

>Homo_sapiens
-----------------------------------MVWEAELKCPGLW
>Mustela_putorius_furo
-----------------------------------MVWEAELKCAGLW
>Orcinus_orca
-----------------------------------MVWEAELKCAGLP
>Eptesicus_fuscus
-----------------------------------MVWEAELKCAGLW
>Dasypus_novemcinctus
MVGLRQRSGGTRGSPGRPGLETDARGARGGAGAAVMVWEAELKCAGLR
>Galeopterus_variegatus
-----------------------------------MVWEVEPKCPGLR
>Monodelphis_domestica
---------------------------------MVMVWEVELKSVGLW
>Condylura_cristata
-----------------------------------MVWDAELKCAGLW
>Equus_przewalskii
-----------------------------------MVWEAELKCAGLW
>Loxodonta_africana
-----------------------------------MVWEAELKCAGLW
>Nannospalax_galili
----------------------MHEGPAEAQEAVVMVWEAELKCPGLW
>Tupaia_chinensis
-----------------------------------MVWEEELKCARLW

HG0496 ENSG00000135913 USP37

>Homo_sapiens
MSSVPRRVLCLWHKDQNKGELLLHRVFR
>Canis_lupus_familiaris
MSSSPRRVHCLWHKDQNKGELLLHRVFQ
>Physeter_catodon
MSSSPRRAHCLWHKDQNKGELLLHRVFQ
>Pteropus_alecto
MSSSPRRVHCLWHKHQNKGELLLHRVFQ
>Monodelphis_domestica
MGSSPRRVLYLWHKHQNKDYWK------
>Oryctolagus_cuniculus
MSSFPRRVHCLWHKDQHKGELLLHRVFQ
>Equus_asinus
MSSFPRRVHCLWHKDQNKGELLLHRVFQ
>Manis_javanica
MSSSPRRVHCLWHKDQNKESFR------
>Loxodonta_africana
MSSSPRRVLCLWHKDQNKGELLLHRVFQ
>Octodon_degus
MSSFPRRVPCLWHKDQNKGELLLHRVFQ
>Trichechus_manatus_latirostr
MSSSPRRVLCLWHKDQNKGELLLHRVFQ

HG0497 ENSG00000147571 CRH

>Homo_sapiens
----------------------MIYAGAEAARNRAVKRASAY
>Mustela_putorius_furo
MGPGIKYTQAGQLALSPTDHGPMIYAGAEAARNRAVKRASAY
>Pteropus_alecto
----------------------MIYAGAEAARNRAVKRASAY
>Galeopterus_variegatus
----------------------MIYAGAEAARNRAVKRASAY
>Monodelphis_domestica
----------------------MIYSGADRARNLAVKKASAY
>Condylura_cristata
----------------------MIYAGAEAARNRAVKRASAY
>Oryctolagus_cuniculus
----------------------MIYAGAEAARNRAVKRASAY
>Equus_asinus
----------------------MIYAGAEAARNRAVKRASAY
>Loxodonta_africana
----------------------MIYAGAEAARNRAVKRASAY
>Peromyscus_maniculatus_baird
----------------------MIYAGAEAARNRAVKRASAY
>Tupaia_chinensis
----------------------MIYAGAEAARNRAVKRASAY

HG0498 ENSG00000005379 TSPOAP1

>Homo_sapiens
--------------------------------------MRQAAHRPRPP
>Mustela_putorius_furo
--------------------------------------MRQAAHRPRPP
>Balaenoptera_acutorostrata_s
--------------------------------------MRQAAHRPRPP
>Rousettus_aegyptiacus
--------------------------------------MRQAAHRPRPP
>Dasypus_novemcinctus
--------------------------------------MRQAAHRPRPP
>Monodelphis_domestica
MHTLMLGPSPIPRTLGHRELLGNPGFPRAEPLTSKGETMRQAAHRPRPP
>Equus_przewalskii
--------------------------------------MRQAAHRPRPP
>Manis_javanica
--------------------------------------MRQAAHRPRPP
>Mus_musculus
--------------------------------------MRQAAHRPRPP
>Tupaia_chinensis
--------------------------------------MRQAAHRPRPP

HG0499 ENSG00000011258 MBTD1

>Homo_sapiens
MLMLLISWGHPS
>Mustela_putorius_furo
MLMLLISWGHPS
>Orcinus_orca
MLMLLISWGHPS
>Myotis_lucifugus
MLMLLISWGHPS
>Dasypus_novemcinctus
MLMLLISWGHQS
>Sarcophilus_harrisii
MLMLLISWGHPN
>Oryctolagus_cuniculus
MLMLLISWGHPS
>Equus_caballus
MLMLLISWGHPS
>Manis_javanica
MLMLLISWGHPS
>Mus_musculus
MLMLLISWGHPS

HG0500 ENSG00000070778 PTPN21

>Homo_sapiens

MLPSSGHWRLRCFLMAAAAAG

>Ailuropoda_melanoleuca

MLPSPGLWGLSCFLMAAASAG

>Pteropus_vampyrus

MLPSPGLWRLSCILMEAAAAG

>Dasypus_novemcinctus

MLPSPGLWKRNCFLMAAAAAG

>Sarcophilus_harrisii

MIPSLRPWRIRCFLMAADAAG

>Monodelphis_domestica

MIPSPGPWRIRCFLMAADTAG

>Ceratotherium_simum_simum

MLPSPGLWRLRCFLMAAAAAG

>Manis_javanica

MLPSPGLWRLSCFLMAAAAAG

>Loxodonta_africana

MLPSPGLWRLDCFLMAAAAPG

>Rattus_norvegicus

MLPSLGRWRLGCFLMAAAAVG

HG0501 ENSG00000110400 NECTIN1

>Homo_sapiens

MTRATCGGHLPSL

>Mustela_putorius_furo

MTRATCGGHPPSL

>Pteropus_vampyrus

MTRATCGGHLPSF

>Sminthopsis_crassicaudata

MTRLTCGGHLPRL

>Monodelphis_domestica

MTRLTCGGHLPRL

>Condylura_cristata

MTRATCGGHLPSL

>Ochotona_princeps

MTRATCGGHLPSL

>Ceratotherium_simum_simum

MTRATCGGHLPSL

>Manis_javanica

MTRATCGGHLPSL

>Marmota_marmota_marmota

MTRATCGGHLPSL

HG0502 ENSG00000126882

>Homo_sapiens
MSEPHNYSLNYKLFLTQRISVIFVHCEEKKREPRAHEMQQNPYL
>Canis_lupus_familiaris
MSEQHDYPLTSRLSFF----------------------------
>Rhinolophus_ferrumequinum
-------------------------------------MQQNPYL
>Galeopterus_variegatus
-------------------------------------MQQSPRL
>Monodelphis_domestica
-------------------------------------MQQNPYL
>Erinaceus_europaeus
MSEQYDYLLTSKLFFF----------------------------
>Oryctolagus_cuniculus
-------------------------------------MQQNPYL
>Equus_caballus
-------------------------------------MQQNPYL
>Microtus_ochrogaster
MSEQHAYPLNYKLFFF----------------------------
>Orycteropus_afer_afer
-------------------------------------MQQNPYL

HG0503 ENSG00000141564 RPTOR

>Homo_sapiens

MGRSWLWLQQLRRVRDPQG

>Felis_catus

MGRSWLWLQQLRRVRDPQG

>Orcinus_orca

MGPSWLWLQQLRRVRDPQG

>Rhinolophus_ferrumequinum

MGRSWLWLQQLRRVRDPQG

>Galeopterus_variegatus

MGRSWLWLQQLRRVRDPQG

>Monodelphis_domestica

MGRSWLQLQQLGRMWGPQG

>Ceratotherium_simum_simum

MGRSWLWLQQLRRLRDPQG

>Manis_javanica

MGRSWLPLQQLRRVRDPQG

>Loxodonta_africana

MGRSWLCLQQLRRVRDPQG

>Ictidomys_tridecemlineatus

MGRSWLLPQQLRRVRDPQG

HG0504 ENSG00000144460 NYAP2

>Homo_sapiens

MQTDCRNREERLLLISEEYNQMRYISQLGDCFWSCNHLCSKVGKIIQFLPCWTWKERGCVFIPTH

>Mustela_putorius_furo

MQTDCRNREERLLLISEEYNQMRHLSQPG------------------------------------

>Myotis_lucifugus

MQTDCRNREERLLLISEEYNQMRHLSQPG------------------------------------

>Galeopterus_variegatus

---------------------MKYLSQPGDCFGSCNHLCSKVGKIIQFLPCWTWKERGCVFIPTH

>Condylura_cristata

MQTDCRNREERLLLISEEYNQMRHLSQPG------------------------------------

>Oryctolagus_cuniculus

---------------------MKYLSQPGDCFGSCNHLCSKVGKIIQFLLCWTWKERGCVFIPTH

>Manis_javanica

MQTDCTHREERLLLISEEYNQMRHLSQPG------------------------------------

>Peromyscus_maniculatus_baird

MQADCRSRGERLLLISEEHDQMRYLSQLGDCFGSCNHLCSKVGKIIQFFPCWTWKERGCALIPTH

>Tupaia_chinensis

---------------------MRHLSQPGDCFGSCNHLCSKVGKIILFLPCWTWKEQGCVFIPTH

HG0505 ENSG00000157470

>Homo_sapiens

MGSRGAPTPRADVNY

>Odobenus_rosmarus_divergens

MGSRGAPAPRADVNY

>Lipotes_vexillifer

MGSRGAPAPRADVNY

>Rhinolophus_ferrumequinum

MGSRGAPAPREDVNY

>Dasypus_novemcinctus

MGCRGAPAPRADVNY

>Monodelphis_domestica

MGSRIPPAPSADVNY

>Condylura_cristata

MGPRGAPDSRADVNY

>Oryctolagus_cuniculus

MGSRGAPGPRADVNY

>Manis_javanica

MGSRRAPAPCADVNY

>Rattus_norvegicus

MGSRGAPDPRADVNY

HG0506.1 ENSG00000177853 ZNF518A

>Homo_sapiens

MAAIRDSQAMGTTTDIISSFDHCS

>Ursus_maritimus

MAAIRDSQAMGTTTDIVSSFDHCS

>Physeter_catodon

MAAIRDSQAMGTTTDIISSFDHCS

>Miniopterus_natalensis

MAAIRDSQAMGTTTDIISSFDHCS

>Galeopterus_variegatus

MAAIRDSQAMGSTTDIISSFDHCS

>Trichosurus_vulpecula

MAAIGDSPAMELTPDFVSSFDQCS

>Oryctolagus_cuniculus

MAAIRDSQAMGTTTDI-ISFDHCS

>Equus_asinus

MAAIRDSQAMGTTTDI-ISFDHCS

>Manis_javanica

MAAIRDSQAMGTTTDTISSFDHCS

>Mesocricetus_auratus

MAAIWDSQAMGTTTDV-ISFDHCA

HG0506.2 ENSG00000177853 ZNF518A

>Homo_sapiens
MSLHSIVSCLNYR
>Mustela_putorius_furo
MSWHSIVSCLNYK
>Balaenoptera_acutorostrata_s
MSRHSIVSCSNYK
>Miniopterus_natalensis
MSQHNIVSCLNYK
>Dasypus_novemcinctus
MCQHSIVSCLNYE
>Galeopterus_variegatus
MSWHSIVSCLNYK
>Equus_asinus
MSRHSIVSCLNYK
>Manis_javanica
MSQHSIVSCLNYK
>Loxodonta_africana
MSQHNIVSCLNFK
>Marmota_marmota_marmota
MSRHSIISCLNYR

HG0507 ENSG00000060237 WNK1

>Homo_sapiens

MTAAPLLPPPARPPLAAGWMRTVRR

>Odobenus_rosmarus_divergens

MTAAPLLLQPSRPPLAAGWMRTVRR

>Rousettus_aegyptiacus

MTAAPLLLQPSRPPLAAGWMWTARR

>Monodelphis_domestica

MTAVPQLLVPARPPPAAGWMRSVRR

>Oryctolagus_cuniculus

MTAAPLLPQPSRPPLAAGWMRTVQR

>Loxodonta_africana

MTAAPLLLQLSWPPLAAGWMRTVRR

>Rattus_sp.

MTAALLLPQPYRPPLAAGWMRTVRR

>Trichechus_manatus_latirostr

MTAALLLLQPSWPTVAAGWMRTVRR

>Orycteropus_afer_afer

MTAAPLLLQSSWLPLAAGWMRTVRR

HG0508 ENSG00000061936 SFSWAP

>Homo_sapiens

MAAVLRLGTGCGVFD

>Acinonyx_jubatus

MAAVSSLGPGCGAFD

>Pteropus_alecto

MAAVSRLGPGCGACD

>Dasypus_novemcinctus

MAAVLRLVLGCGTFD

>Monodelphis_domestica

MAAVLRLRPGCGASD

>Oryctolagus_cuniculus

MAAVLRLGPGCGAFD

>Equus_asinus

MAAVLRLGPGCGAFD

>Manis_javanica

MAAMPRLGLGCGASD

>Marmota_monax

MAAVLRLRTGCGAFD

HG0509 ENSG00000068024 HDAC4

>Homo_sapiens
---------------------------MCWWEFGARWSYRFRGNFEPFRIT
>Canis_lupus_familiaris
MPRPAPRGRQDWGFPRLRANTPCRCTCMCPWNLGALWSCRLRGSCEPL---
>Balaenoptera_acutorostrata_s
---------------------------MCRWDSGALWSSRSCGSFEPF---
>Pteropus_alecto
---------------------------MCPWDFGALWSYCLRGNFEPS---
>Dasypus_novemcinctus
---------------------------MCWWNFGALWSYRLCGSIKPF---
>Galeopterus_variegatus
---------------------------MCWWNFGALCSYRLLGNFEPF---
>Ornithorhynchus_anatinus
---------------------------MYWWSIGALWSCGLCGNFDPF---
>Ceratotherium_simum_simum
---------------------------MCRWNF------------------
>Peromyscus_maniculatus_baird
---------------------------MCWWKFGAGWSYCLRGHFEPF---

HG0510 ENSG00000134769 DTNA

>Homo_sapiens
MVLKSSCCPFIEF
>Acinonyx_jubatus
MVLESSCCFFIEF
>Balaenoptera_acutorostrata_s
MVLKSSCCFFIEF
>Pteropus_alecto
MVLESSCCFFIEF
>Dasypus_novemcinctus
MVLESSCCCFIEF
>Monodelphis_domestica
MVLESSCCAFIEL
>Ceratotherium_simum_simum
MVLESSCCFFIEF
>Tupaia_chinensis
MVLESSCCFFIEF
>Orycteropus_afer_afer
MVLESSCCSFIEF

HG0511 ENSG00000143337 TOR1AIP1

>Homo_sapiens
MAETSSSPFSSSLVPPARRRERSTAADPAQALPLSQPSRHTMGPEAGLLHSSD----DAGGGPSDSQLPP
>Felis_catus
-----------------------------------------MGPEAGLQHSSD----DAGGGPGDSQLPP
>Balaenoptera_acutorostrata_s
-----------------------------------------MGPEAGLQHSSD----DAGGSPGDSQLPP
>Rhinolophus_ferrumequinum
-----------------------------------------MGPEAGLQHSSD----DAGGSPGDSQLPP
>Dasypus_novemcinctus
-----------------------------------------MGPEAGLQYSSD----DAGGSPGDSQLPP
>Sarcophilus_harrisii
-----------------------------------------MGPEAAGQRGSGSGRSGGPTGPGDSQLPP
>Oryctolagus_cuniculus
-----------------------------------------MGPGAASRHSSD----DAGGGPGDSQLPP
>Equus_asinus
-----------------------------------------MGPEAGLRHSSD----DAGGGPGDSQLPP
>Marmota_marmota_marmota
-----------------------------------------MGPEAGLRHSSD----DAGGGPGDSQLPP

HG0512.1 ENSG00000112319 EYA4

>Homo_sapiens

MSCFHQRHSAKGKLRHWKERE---

>Canis_lupus_familiaris

MSCLQQRHNAKGKLRRWKERE---

>Balaenoptera_acutorostrata_s

MSSFHQRHSVKGKLQHWKARE---

>Miniopterus_natalensis

MSCFHQRHSAKGKLRHWKERQ---

>Monodelphis_domestica

MSCFYQKNSREGKLRHQEGRRINK

>Equus_przewalskii

MSCFHQRHSAKGKLRHWKERE---

>Manis_javanica

MSCFRQRHSAEGRLRHWKERE---

>Mesocricetus_auratus

MSCFHQRHSAKGKLRHWKERE---

HG0512.2 ENSG00000104313 EYA1

>Homo_sapiens

------------MLSAAAVWWGRSAGRKAVTKQNGGS

>Leptonychotes_weddellii

------------MLSAAAVWWGKSAGLKAVTKQNGGS

>Physeter_catodon

--MVLFLCLGFSDAISCCCLEGKSAGLKAVTKQNGGS

>Pteropus_alecto

------------MLSAAAVWWGRSAGLKAVSPQHGGS

>Galeopterus_variegatus

------------MLSAAAVWWGRSAGRKAVAKQNGGS

>Oryctolagus_cuniculus

------------MLSAAAVRWGESAGRKAVTARSGGS

>Elephantulus_edwardii

MRLCCFFVSGSQMLSAAAVQWGKSAGLQAVTTQNGGS

>Equus_przewalskii

------------MLSAAAVEWGQSAGLKAVAKQNGGS

>Loxodonta_africana

------------MLSAAAV------------------

>Cavia_porcellus

------------MLSAAAVRWGKRAGRRAVPKQNSGS

HG0512.3 ENSG00000104313 EYA1

>Homo_sapiens
MPFTAAVSSFFSFG
>Panthera_tigris
MPFTAAVSSFFSFG
>Lipotes_vexillifer
MPFTAAVSSFFSFG
>Myotis_brandtii
MPFTAAVSSFFSFG
>Sorex_araneus
MPFTAAVFSFFSFG
>Elephantulus_edwardii
MPFTAAVSSFFSFG
>Ceratotherium_simum_simum
MPFTAAVSSFFSFG
>Loxodonta_africana
MPFTAAVSSFFSFG
>Ictidomys_tridecemlineatus
MPFTAAVSSFFSFG

HG0513 ENSG00000126860 EVI2A

>Homo_sapiens

MCFSCSGKEMCHLWFGFKSGKLAAHILFYCRFTLRLIFSKSILL-------------

>Odobenus_rosmarus_divergens

---------MCHLWFGFKSGKLAVHILFFF---------------------------

>Physeter_catodon

---------MCHLWLVES---------------------------------------

>Miniopterus_natalensis

MCFSCSEKEMCHLWFGLKSGKLAVHVLFCSRFTLTFIFLQVCSTWKLCLKTRKEVAY

>Manis_javanica

---------MCHLWFGFKTGKLAVHILFYCRFTFKVHDL------------------

>Loxodonta_africana

MCFSCSRKEMCHPSVVWL---------------------------------------

>Chinchilla_lanigera

MCLSCSGKEMCHLWFGFKSGKLAVRTLFCCRCTLHFVFSKSILL-------------

HG0514 ENSG00000147570 DNAJC5B

>Homo_sapiens

MKKRRDLKRDNQLAGKVTNEK

>Panthera_tigris

---MPPVQRDNQLADEVTNEK

>Physeter_catodon

---MTPVQRDNQLADRVTNQK

>Myotis_lucifugus

---MGPLQRDNQLADRVTNEK

>Ornithorhynchus_anatinus

MSSFRVFQRDNQLAARVTNDK

>Ceratotherium_simum_simum

---MPPVQRDNQLADRVTNEK

>Microtus_ochrogaster

---MPPVQRDNQLADKVTNEK

>Tupaia_chinensis

---MAHMQRDNQLAGRVTNEK

HG0515 ENSG00000170471 RALGAPB

>Homo_sapiens

MRAPPVASGAIWIVL

>Leptonychotes_weddellii

MRAPPVASGAIWIVL

>Balaenoptera_acutorostrata_s

MRAPPVASGAIWIVL

>Myotis_brandtii

MRAPPVASGAIWIVL

>Monodelphis_domestica

MRAGPLGAGTIWIVL

>Erinaceus_europaeus

MRAPPVASGAIWIVL

>Mesocricetus_auratus

MRAPPVASGAIWIVL

>Tupaia_chinensis

MRAPPVASGAIWIVL

HG0516 ENSG00000041515 MYO16

>Homo_sapiens
MRMSLK-LFAGSWNRASMQ-----
>Panthera_tigris
MRMSLK-LSAGSWNRASVR-----
>Rousettus_aegyptiacus
MRMSLK-LFA--------------
>Monodelphis_domestica
MRMSLKLLFARSWTRSCIQ-----
>Oryctolagus_cuniculus
MRMSGK-LLAGSWSGASVQ-----
>Manis_javanica
MRMSLK-LFAPSWNRASVQGCVLR
>Chinchilla_lanigera
MRMSLE-LFA--------------

HG0517 ENSG00000086717 PPEF1

>Homo_sapiens

MIDCVLSPQVRLKESTTSKVKI--

>Canis_lupus_familiaris

-MDCVLSLQVRLKDSATNKVKI--

>Monodelphis_domestica

-MDCVLSLLVRPKDSTMVKVEIWE

>Oryctolagus_cuniculus

-MDCVLSQQIRLRVSAANKVKI--

>Ceratotherium_simum_simum

-MDCVLSLQIRLKDTAASKMKI--

>Mus_musculus

-MDCALSLQVRMNDSATSEMKI--

>Tupaia_chinensis

MMDSVLNLQVQLKYSATSKAKI--

HG0518 ENSG00000111254 AKAP3

>Homo_sapiens

-----------MEGHRKKQDLELSKNIPASSLTL-------KVYSG----VFRGSLFQYYRDSLYYMAGGG------------------------------

>Canis_lupus_familiaris

MSLFSPNKGRYMEGWRRKQDLELSKNIPASSLTF-------KV----------------------------------------------------------

>Pteropus_vampyrus

-----------MEGWVRKQDLELSKNIPASSLTL-------KLYSG----VFCGHYPS-TTEIPYITWQEGWVNLGIVKTKNN------------------

>Sarcophilus_harrisii

-----------MEGSGRKQDLELNKNITASLIDF-------KKCILESFVVHCSSAA----EVPD-----NYMAGGVGKLKNCEDYKTIQDPSLYLRICTQ

>Oryctolagus_cuniculus

-----------MEGSGRKQDLELSKDIPASSLTL-------KIYSG----VSCGSLLKCYRDS--------------------------------------

>Equus_asinus

-----------MEGWGRKQDLELSTNIFIDFQSIFWSLLWFTIPVLQRFHILHGRRG------------------G-------------------------

>Chinchilla_lanigera

-----------MEGWGRKQDLELSKNIPASSLTL-------KVYCG----IFCGSLVQSYRDSLYITWQEV------------------------------

HG0519 ENSG00000127074 RGS13

>Homo_sapiens
MVDCSTGYICQWNVSTYYTTNMRKK---
>Acinonyx_jubatus
MVDRSTGYICQWNFSTYHTTNMRKK---
>Pteropus_vampyrus
MVDCSSGYICQWNFSTCHTTNMRKK---
>Trichosurus_vulpecula
-MFCATRHICQWNVSTYHISQYEGKIYC
>Loxodonta_africana
MVDCCTGYICQRNVSTYTT-NMRKK---
>Cavia_porcellus
MVDSSTRYIC------------------
>Orycteropus_afer_afer
MVDCSSGYIC------------------

HG0520 ENSG00000197415 VEPH1

>Homo_sapiens

----MNNFRFTELSGFHSL-CPRS-LPESPAQSQTMQLSNCKGAFEE

>Panthera_tigris

----MNNFRFTELRGFHLL-CPRSPLPESRAQAQNL-----------

>Lipotes_vexillifer

----MNNFRFTELSRFHSL-CPWSPLPESLTQSQNRPLSSSKCAFEE

>Rousettus_aegyptiacus

MGSAMNNFRFTKLSGFCPLSVPRSPLPESPAQSQNIQLSNNKPAFEE

>Monodelphis_domestica

----MNNWRFPGLPDFS------------------------------

>Manis_javanica

----MNNFRFTELSGFHSL-CPRSQLPKSPTQSQHRQLSNSRHAFEE

>Tupaia_chinensis

----MNHSKFTELSGFHSL-CPRNLLPEFLTQSQNIPLSNRMGALEE

HG0521 ENSG00000038274 MAT2B

>Homo_sapiens

-----------------------MWRTARDSGIGIKMLI

>Myotis_lucifugus

-----------------------MLKTARVSGIGIKMLI

>Sarcophilus_harrisii

-----------------------MWETARDPETGI----

>Ceratotherium_simum_simum

-----------------------MWKTARDSGIGIKMLI

>Ictidomys_tridecemlineatus

MKHETSHWVFWKLPRGISCQLTALWKTARDSGIGIKMLI

HG0522 ENSG00000140396 NCOA2

>Homo_sapiens

MAAAASATASAAKVSADGSRHLTA

>Rousettus_aegyptiacus

MAAAASAAASAAKVSADGSRHLTA

>Monodelphis_domestica

MAAT--ASAPAGKVSAVGSRHLID

>Ochotona_princeps

MAAAASATATAAKVSADGSRHLTA

>Dipodomys_ordii

MAATVSAPAWAAKVSADGSRHLTA

HG0523 ENSG00000075213 SEMA3A

>Homo_sapiens

MGCSWYSYKVNMLLAAHIYP

>Monodelphis_domestica

MGCSLYSYKVNKLLTAYIYD

>Trichosurus_vulpecula

MGCSLYSYKVNKLLTAYIYD

>Mus_musculus

-----------MLLAAHVCA

HG0524 ENSG00000107929 LARP4B

>Homo_sapiens
MRTGRSVGDLESRG
>Macropus_eugenii
MRTGRSVGDLESRG
>Manis_javanica
MRTGRSVGATESRG
>Microtus_ochrogaster
MRTGRSVGDLESRG

HG0525 ENSG00000136167 LCP1

>Homo_sapiens
MTSPEIAVVQRKQSRGIELLQNARVSNIVSAAASYLTKVEEILYFQLRKLKIPPASDDQAPLDTEHCAFLLQVTSCLVTTHPGLTKLFCRSERRGSWSYTSTTKDS
FFPARSVT
>Canis_lupus_familiaris
-----------------------------------------------------------------MICHSSHFLPC--DRTAGVDSCSADPRGGGFWSHASTTEDS
FSTRSVA-
>Ornithorhynchus_anatinus
------------------------------------------------------------------------MTSCLVTT--------------------------
--------
>Mus_musculus
------------------------------------------MLLERRKWK----------VASRFHCDLLSVTSCLVTTQPCLTRLLCRSEGLGFLTTQQCC---
--------

HG0526 ENSG00000143569 UBAP2L

>Homo_sapiens
MRLEPERGERERARASEAWALPEYSTL
>Felis_catus
MRLEPERGERGRARASAAWAPPEYSTL
>Monodelphis_domestica
MRLGP-GGWREREGASAAWAWPEHSTL
>Chinchilla_lanigera
MRLQPEGGERERARASAAWALPEYSTL

HG0527 ENSG00000198382 UVRAG

>Homo_sapiens

MS----APGLSGVEGLHCGNMALP

>Monodelphis_domestica

MSALGLGSGRPGREVLHCGNMALP

>Oryctolagus_cuniculus

MS----GPGLCRVEGLHCGNMALP

>Chinchilla_lanigera

MS----APGLGPVEGLHCGNMALP

HG0528 ENSG00000204767

>Homo_sapiens

MGMFPWEHHVSSNKQTSRSFISPPVYLLWKEDAFKFPAVSPWKLMLWETKSCMF-------------------------------------------

>Rousettus_aegyptiacus

MGMSPSGHHVSSKKPTRRSLISLPVYMLWKEDAFKFPVVSHWKSMLRERKPSVPEVAGFFWNLHFGSLALSSKWYFCCLFVQDSKRFPGSELPASLP

>Ornithorhynchus_anatinus

--MLLVDHHVSSEKQTYQSLIMLIIYT----------------------------------------------------------------------

HG0529 ENSG00000112851 ERBIN

>Homo_sapiens
--------MLAQLRNIREINSTQCV
>Canis_lupus_familiaris
--------MLAQLRNIREINSTQCV
>Lipotes_vexillifer
--------MLAQLRNIREINSTQCV
>Pteropus_alecto
--------MLAQLRNSREINSAQCV
>Dasypus_novemcinctus
--------MLAQLRNIRQIDLLQCV
>Galeopterus_variegatus
--------MLAQLRNIREINSTQCV
>Sorex_araneus
--------MLAQLRNIRETNST---
>Oryctolagus_cuniculus
--------MLAQLRNTREINSTQCV
>Elephantulus_edwardii
--------MLAQLRNTSKINSALCV
>Equus_asinus
MREPLIHRMLAQLRNIREINSTQCV
>Manis_javanica
--------MLAQLRNIREINSTKSV
>Loxodonta_africana
--------MLAQLRNIREINSTQCV
>Cricetulus_griseus
--------MLAQLRNIREINSTQCF
>Tupaia_chinensis
--------MLAQLRSIREINSTKCV
>Trichechus_manatus_latirostr
--------MLAQLRNIREINSTQCV
>Orycteropus_afer_afer
--------MLAQLRNIIEINSTQSV

HG0530 ENSG00000128594 LRRC4

>Homo_sapiens
MREQSAWLPPATPRARREEETKSEWERGAQLPWRKLKDTTPVLSCLPSEMERGAPSSLKPG
>Felis_catus
-------------------------------------------------MERGAPSSFKPG
>Lipotes_vexillifer
-------------------------------------------------MERGAPSSLKPG
>Myotis_brandtii
-------------------------------------------------MERGAPSSLKPG
>Dasypus_novemcinctus
-------------------------------------------------MERGALSSLKPG
>Galeopterus_variegatus
-------------------------------------------------MERRAPSSLKPG
>Condylura_cristata
-------------------------------------------------MERGAPSSLKPG
>Oryctolagus_cuniculus
-------------------------------------------------MERGAPSSLKPG
>Elephantulus_edwardii
-------------------------------------------------MERGAPSSLKPG
>Ceratotherium_simum_simum
-------------------------------------------------MERGAPSSLKPG
>Manis_javanica
-------------------------------------------------MERGAPSSLKPG
>Loxodonta_africana
-------------------------------------------------MERGAPSSLKPG
>Dipodomys_ordii
-------------------------------------------------MERGAPSSLKPG
>Tupaia_chinensis
-------------------------------------------------MERGAPSSLKPG
>Trichechus_manatus_latirostr
-------------------------------------------------MERGAPSSLKPG
>Orycteropus_afer_afer
-------------------------------------------------MERGAPSLLKPG

HG0531 ENSG00000146834 MEPCE

>Homo_sapiens
MRAQPGWWRSAHAWFPGEGNARSPCALAAGW
>Felis_catus
MRAQPGWWRSAHAWFRGKRNAL---------
>Balaenoptera_acutorostrata_s
MRAQPGWWRSAHA------------------
>Pteropus_alecto
MRARPGWWRSAHA------------------
>Dasypus_novemcinctus
MRAQPGW------------------------
>Galeopterus_variegatus
MSAQPGWWRSAHAWFGGERNALQQCALAAKC
>Condylura_cristata
MRAQPGWWRNAHA------------------
>Oryctolagus_cuniculus
MRAQPGWWRSAHAWPRG--------------
>Elephantulus_edwardii
MRAQPGCWRSAHASSWGQRNAL---------
>Equus_caballus
MRAQPGWWRSAHA------------------
>Manis_javanica
MRAQPGWWRSAHA------------------
>Loxodonta_africana
MRAQPGWWRCAHA------------------
>Nannospalax_galili
MRAQPGWWRSAHAWFRGERNARPPCALSAKW
>Tupaia_chinensis
MRAQPGWWQSAHA------------------
>Trichechus_manatus_latirostr
MRAQLGWWRSAHA------------------
>Orycteropus_afer_afer
MRAQPGWGRSAHA------------------

HG0532 ENSG00000105221 AKT2

>Homo_sapiens

MVPGTVLCPVHGASCMSCCPELSRAR

>Mustela_putorius_furo

---------------MSCCPELSQAR

>Physeter_catodon

---------------MSCCPELSRAR

>Pteropus_vampyrus

---------------MSCCPELSRAR

>Dasypus_novemcinctus

---------------MSCCPELPQAR

>Galeopterus_variegatus

---------------MSCCPEVSQAR

>Erinaceus_europaeus

---------------MSCCPELSLAR

>Oryctolagus_cuniculus

---------------MSCCPELPQAR

>Elephantulus_edwardii

---------------MSCCHILT---

>Equus_asinus

---------------MSCCPELSRAR

>Manis_javanica

---------------MSCCPELSQAR

>Loxodonta_africana

---------------MSCCLKVP---

>Cricetulus_griseus

---------------MSCCPELSQAR

>Trichechus_manatus_latirostr

---------------MSCCPELP---

>Orycteropus_afer_afer

---------------MSCCLELS---

HG0533 ENSG00000114209 PDCD10

>Homo_sapiens
MKIPLHFSTAFSTVVGF
>Acinonyx_jubatus
MKIPLHFSTAFSTVVGF
>Lipotes_vexillifer
MKIPLHFSTAFSTVVGF
>Eptesicus_fuscus
MKIPLHFSTAFSTVVGF
>Dasypus_novemcinctus
MKIPLHFSTAFSRVVGF
>Galeopterus_variegatus
MKIPLHFSAAFSAVVGF
>Condylura_cristata
MKIPLNFSTAFSTVVGL
>Oryctolagus_cuniculus
MKIPLHFSTAFSTVVGF
>Ceratotherium_simum_simum
MKIPLHFSTAFSTVVGF
>Manis_javanica
MKIPLHFSTAFLL----
>Loxodonta_africana
MKIPLHFSTAFSPVVGF
>Ictidomys_tridecemlineatus
MKIPLHFSIAFSTVVGF
>Tupaia_chinensis
MKITLHFSTAFSTVVGF
>Trichechus_manatus_latirostr
MKIPLHFSTAFSPVVGF
>Orycteropus_afer_afer
MKIPLHFSTAFSPVVGF

HG0534 ENSG00000117408 IPO13

>Homo_sapiens

MPALGPPSPHHSLGTQAGV

>Felis_catus

MPALGPLSPHRSLGTQAGV

>Physeter_catodon

MLALGPLSPHHSLGTEAGV

>Myotis_davidii

MPTLGPLSPHHSLGPQAGV

>Dasypus_novemcinctus

MPALGPPSPHLSLGTQAGV

>Erinaceus_europaeus

MPALGSLSHHQSLGTQAGV

>Ochotona_princeps

MPALGAPSPHCSLDTQAGV

>Elephantulus_edwardii

MSALGAPSPHHSLGTQAGV

>Ceratotherium_simum_simum

MPALGPLSPRHSLGTRAGV

>Manis_javanica

MPAQGPLSPHHSLGTQAGV

>Loxodonta_africana

MPALGPPSPHYSLGTQAGV

>Cavia_porcellus

MPALGTPSPQHSLGTQAGV

>Tupaia_chinensis

MPTLGSPSPHHSLGTQAGV

>Trichechus_manatus_latirostr

MPALGPPSLHHSVGTQAGV

>Orycteropus_afer_afer

MPALGAPSPHHSLGTQAGV

HG0535 ENSG00000118407 FILIP1

>Homo_sapiens
MTSAIGWSWERKWEKLFPLYSFGLDSF-------------------
>Acinonyx_jubatus
MTSAIGQS--------------------------------------
>Orcinus_orca
MTSAIG----------------------------------------
>Eptesicus_fuscus
MTSAIGQSWESKWEKLFPLYSFGLDSFPGLQSLSWAVGQATGFFQG
>Dasypus_novemcinctus
MTSAIG----------------------------------------
>Galeopterus_variegatus
MTSAIG----------------------------------------
>Condylura_cristata
MTSAIGYS--------------------------------------
>Oryctolagus_cuniculus
MTSAIGWS--------------------------------------
>Elephantulus_edwardii
MTSAIGWS--------------------------------------
>Ceratotherium_simum_simum
MTSAIGQS--------------------------------------
>Manis_javanica
MTSAIGWS--------------------------------------
>Loxodonta_africana
MTSAIG----------------------------------------
>Cavia_porcellus
MTSAIGWS--------------------------------------
>Tupaia_chinensis
MTSAIG----------------------------------------
>Orycteropus_afer_afer
MTSAIG----------------------------------------

HG0536 ENSG00000181751 C5orf30

>Homo_sapiens
--------------------------------------MSVFPWDSLEIVQ
>Phoca_largha
--------------------------------------MPVFPWDSLDIVQ
>Balaenoptera_acutorostrata_s
--------------------------------------MPVFPWDSLEIVQ
>Myotis_davidii
--------------------------------------MPVFPWDSLEIVQ
>Dasypus_novemcinctus
--------------------------------------MPVFPWDSVEIVH
>Galeopterus_variegatus
--------------------------------------MTVCPWDSLEIAQ
>Condylura_cristata
--------------------------------------MPVFLWDSLEIVQ
>Oryctolagus_cuniculus
--------------------------------------MPAFPGDSLEIVQ
>Ceratotherium_simum_simum
--------------------------------------MPVFSWDSLEIVQ
>Manis_javanica
--------------------------------------MPVFPWDSLEIVQ
>Loxodonta_africana
--------------------------------------MPVFPWDSLEIVH
>Nannospalax_galili
MTASPLPLASVVQPAVGTPSFSAHATLHRYLEYRRKIIMPVLPWDSLEIVQ
>Tupaia_chinensis
--------------------------------------MPVFPWDSLEIVQ
>Trichechus_manatus_latirostr
--------------------------------------MPVFPWDSLEIVH
>Orycteropus_afer_afer
--------------------------------------MPVFPWDSLEIVH

HG0537 ENSG00000184481 FOXO4

>Homo_sapiens

MCLLALDASPLRGGIVRDCGRLH

>Felis_catus

MCLLALGASPLGGGIVRDCGRLL

>Balaenoptera_acutorostrata_s

MCLLALGASPLGGGIARDCGRLL

>Miniopterus_natalensis

MCLLALGASPLGGGIVRDCGRLL

>Dasypus_novemcinctus

MCPLALDASPLGGGIVRDCGRLH

>Galeopterus_variegatus

MCLLALGASPLGGGIVRDCGRLH

>Sorex_araneus

MCLLALGASPLGGGIVRDCGRLL

>Oryctolagus_cuniculus

MCLLALSASPLGGGIVRDCGRLQ

>Ceratotherium_simum_simum

MCLLALSASPLGGGIVRDCGRLL

>Manis_javanica

MCLLALGASPLGGGIVRDCGRLL

>Loxodonta_africana

MCPLALGASPLRGGIVRDCSRLH

>Peromyscus_maniculatus_baird

MCLLALVASPLRGGMVRDCGRLH

>Tupaia_chinensis

MCLLALGASPLGGGIARDCGRLH

>Trichechus_manatus_latirostr

MCPLALGASPLRGGIVKDCSRLQ

>Orycteropus_afer_afer

MCPLALGASPLGGGIVRDCSWLH

HG0538 ENSG00000006116 CACNG3

>Homo_sapiens
MRTNPPEPAPFRGP
>Canis_lupus_familiaris
MWTNPLEPAPFQGP
>Physeter_catodon
MWTNPLEPAPFRGP
>Miniopterus_natalensis
MRTNPLEPAPFRGP
>Dasypus_novemcinctus
MWTNPLEPLPFRGP
>Galeopterus_variegatus
MWTSPLEPAPFRGP
>Condylura_cristata
MWTNPLEPAPFRGP
>Ochotona_princeps
MWTNPLEPAPFRGP
>Elephantulus_edwardii
MWTNPLESAPFRGP
>Equus_asinus
MWTNPLEPAPFRGP
>Loxodonta_africana
MWTNPLKPAPFRGP
>Chinchilla_lanigera
MRTNPLEPAPSRGP
>Tupaia_chinensis
MWTNPLEPAPFRGP
>Trichechus_manatus_latirostr
MWTNPLEAAPFRGP

HG0539 ENSG00000115966 ATF2

>Homo_sapiens
---------------MAHQLESILAMRKRM
>Acinonyx_jubatus
---------------MAHQLESILAMKKRT
>Lipotes_vexillifer
---------------MAHQLESILAMKKRT
>Myotis_brandtii
---------------MAHQLESILAMKKRT
>Dasypus_novemcinctus
---------------MAHQLESIFAMKKRT
>Galeopterus_variegatus
---------------MAHQLESILAVRKRT
>Erinaceus_europaeus
---------------MAHQLESIFVMKKRT
>Elephantulus_edwardii
---------------MAHQLESILAMEKRM
>Equus_asinus
---------------MAHQLESILAMKKRT
>Manis_javanica
---------------MAHQLESILAMGKRT
>Loxodonta_africana
---------------MAHQLESILAMIKRM
>Nannospalax_galili
---------------MAHQLQSILAMKKRT
>Tupaia_chinensis
---------------MAHQLESILAMKKRT
>Orycteropus_afer_afer
MIRAEGFLAAEGIKLMAHLLESILAMKKRM

HG0540 ENSG00000120519 SLC10A7

>Homo_sapiens

MSGGTKESVETLQGLGDRSLKRKKAKQINKTHPP

>Panthera_tigris

MSGGTVESVETPRGLSDRSLKRKKAKQINKTHPP

>Tursiops_truncatus

MSGGTVESVETPRGLSDSSLKRKKAKQINKTHPP

>Miniopterus_natalensis

MSRGTVESVETPRGLSDRSLKRKKAKQINKTHPP

>Dasypus_novemcinctus

MSGGTEETVETPRGLSDRSLKRKKAKQINKTHPP

>Condylura_cristata

MSGGAVESVETSRGLSDRSLKRKKAKQINKTHPP

>Oryctolagus_cuniculus

MSGGTEESVETPRRLRDSSSKRKKAKQTNKTHPP

>Elephantulus_edwardii

MSGGIEQSVESPREIRDRSLKRKKAKQINKTHPP

>Equus_przewalskii

MSGGTVESVETPRGLSDRSLKRKKAKQINKTHPP

>Manis_javanica

MSGGTVESVETPRGLSDSSLKRKKAKQINKTHPP

>Loxodonta_africana

MSGGNEESVETPRVLSDRSLKRKKAKQINKTHPP

>Marmota_marmota_marmota

MSGGIQESVETPRGFRDRSLKRKKAKQINKTHPP

>Tupaia_chinensis

MSGGTAESVETPRGLSARSLEREKTKQINKTHPP

>Trichechus_manatus_latirostr

MSGGTEESVETPRGLSDRSLKRKKAKQINKTHPP

HG0541.1 ENSG00000122584 NXPH1

>Homo_sapiens
METERLGGHVRGAGDARPE
>Mustela_putorius_furo
MEMKRQGGHVRGAGDARPE
>Orcinus_orca
MEMKRQGGHVRDAGDARPE
>Pteropus_vampyrus
METKRQGGHVRGAGDARPE
>Dasypus_novemcinctus
METQRQGRRVRGAGDARPE
>Sorex_araneus
METKRQGGHVRSAGDARPE
>Ochotona_princeps
METERQGGHVWGAGDARPQ
>Elephantulus_edwardii
METKRQGGHVRGAGDARPQ
>Ceratotherium_simum_simum
METKRQGGHVRGAGDARP-
>Loxodonta_africana
METKRQRGHVRGAGDARPE
>Marmota_marmota_marmota
METELQGGHVRGAGDARPE
>Tupaia_chinensis
METERQGGHVRGAGDARPE
>Trichechus_manatus_latirostr
METNRQGGHVRGAGEARPE
>Orycteropus_afer_afer
METKRQGGHVRGAGDARPE

HG0541.2 ENSG00000122584 NXPH1

>Homo_sapiens
--------------MEPAQSGPFSQDSPQVLCGC
>Leptonychotes_weddellii
MSGAPETPGPSSSSMEPAQSGPCSPVSPLVLCGC
>Orcinus_orca
MFGTPETPGPSSSSMEPSQSGPCLLDSPPVLCGC
>Eptesicus_fuscus
MSGAPETPGPSSSSMEPAQSGPCSPDSPPVLCGC
>Dasypus_novemcinctus
--------------MDPGQNWTCSPDSTPFLCGC
>Condylura_cristata
MSGAPETPGPSSSSMEPAQSGPCSPDSPLILCGC
>Elephantulus_edwardii
--------------MEPIQSGPCSQDLPPVLCGC
>Equus_asinus
MFGAPETPGPSSSSMEPAQSHPCSPDSPPVLCGC
>Marmota_marmota_marmota
--------------MEPAQSGQCSPDSPPVLCGC
>Tupaia_chinensis
--------------MEPVQSGPCSPDSPPVQCGC
>Trichechus_manatus_latirostr
MSGAPEKPGPSSSSMEPAQSGPCSPDSPSVLCGC
>Orycteropus_afer_afer
--------------MEPAQSDSCSPASPPVYSGC

HG0542 ENSG00000126581 BECN1

>Homo_sapiens
-MVVLEASLRGRPEATVPPR
>Felis_catus
-MAVLEASLRGRPDATVPPR
>Eptesicus_fuscus
MMAVLEASLQGSPEAVVPPR
>Dasypus_novemcinctus
-MGVLEAFPRGRPQATVPPR
>Galeopterus_variegatus
-MAVLEALLPGRPEAAVPPR
>Condylura_cristata
-MAILEAWVRVRPAATVPPR
>Oryctolagus_cuniculus
-MAILEASFRGRPEATVPPR
>Ceratotherium_simum_simum
-MAVLEASLRDRPEAAVPPR
>Manis_javanica
-MAVLEVPLRGRPEATVPLR
>Loxodonta_africana
-MAVLGALFRGRPEATVPPR
>Mus_musculus
-MGTLEVSLRGGPEATVPPR
>Tupaia_chinensis
-MAVLDASLWGRPAATVPPR
>Trichechus_manatus_latirostr
-MAVLGALFRCRPEAAVPPR
>Orycteropus_afer_afer
-MAVLEALLRGRPEATVPPR

HG0543 ENSG00000131931 THAP1

>Homo_sapiens
-----------------------------------MGDGRWLWSFC
>Ailuropoda_melanoleuca
-----------------------------------MGDGRWLWSFC
>Lipotes_vexillifer
-----------------------------------MGDGRWLWSFC
>Rousettus_aegyptiacus
-----------------------------------MGDGRWLWSFC
>Dasypus_novemcinctus
MACFPRNVTSDLGTERQWDELLARGEVEAASAILEMGDGRWLWFYC
>Galeopterus_variegatus
-----------------------------------MGDRRWLWSFC
>Condylura_cristata
-----------------------------------MGDGRRLWSFC
>Ochotona_princeps
-----------------------------------MGEGRWLWPFC
>Ceratotherium_simum_simum
-----------------------------------MGDGRWLCSFC
>Manis_javanica
-----------------------------------MGDGRWLWSFC
>Loxodonta_africana
-----------------------------------MGDGRWLWSFC
>Marmota_marmota_marmota
-----------------------------------MGDGRWLWSFC
>Trichechus_manatus_latirostr
-----------------------------------MGDGRGLWSFC
>Orycteropus_afer_afer
-----------------------------------MGDGRWLWSFC

HG0544 ENSG00000136802 LRRC8A

>Homo_sapiens
---------------------------------------------MEQKECQDPAQADAGQPQHRQPTRR-
>Panthera_tigris
---------------------------------------------MEREECQDPAQTDVGQPQHRQPARR-
>Orcinus_orca
---------------------------------------------MEQKECQDPAQTDVGQPQHRQPTRR-
>Rousettus_aegyptiacus
---------------------------------------------MERKECQDPAQADVGQPRHRQPTRR-
>Dasypus_novemcinctus
---------------------------------------------MEQRECQDPAQTDVGQPQHRHLTRR-
>Galeopterus_variegatus
---------------------------------------------MEQKECQDPVQTDVSQPQHRQPTCR-
>Erinaceus_europaeus
---------------------------------------------MEHKECQDPAQTDVGQPQHQQPTRR-
>Ochotona_princeps
MLSGAVRDKQKEAPGERGGGRRDGAAGARAACWAERCGKPKQAVSMEQQECQDPAQTDVGQPQPRQPTRR-
>Ceratotherium_simum_simum
---------------------------------------------MEREECQDPGQTDVGQPQRRQQTRR-
>Manis_javanica
---------------------------------------------MERKECQDPAQTDMGQPQHQEPAHR-
>Loxodonta_africana
---------------------------------------------MDQRECEDPAQTDVGHPQHRQLTPRR
>Microtus_ochrogaster
---------------------------------------------MEQKECQDPVHPDRDQPQHQQPTRR-
>Tupaia_chinensis
---------------------------------------------MEQKECQDPAQADVGQPRHRQPTRR-
>Orycteropus_afer_afer
---------------------------------------------MERGECQDLAQTDVGLPQHHQPTCR-

HG0545 ENSG00000142784 WDTC1

>Homo_sapiens

MCILWTWAWLECSGVLKILL

>Acinonyx_jubatus

MCVLWTWAWLECSRVLKILL

>Balaenoptera_acutorostrata_s

MCVLWTWAWLECSRVLKILL

>Pteropus_alecto

MCVVWTWAWLECSRVLKTLL

>Dasypus_novemcinctus

MCVLWTWALLECSRVLKILL

>Galeopterus_variegatus

MCILWTWAWLECSRVLKILL

>Erinaceus_europaeus

MCVLWTWAWLECSRVLKILL

>Oryctolagus_cuniculus

MCVLWTWAWLECSRVLKILL

>Ceratotherium_simum_simum

MCVLWTWAWLECSRVLKILL

>Manis_javanica

MCVLWTWAWLECSRVLKIPL

>Loxodonta_africana

MYFLWTWAWLECTRVLKILL

>Chinchilla_lanigera

MCILWTWAWLECASVLKILL

>Tupaia_chinensis

MCALWTCTWLECSRVLKILR

>Trichechus_manatus_latirostr

MCVLWTWAWLECTKVLKILL

HG0546 ENSG00000148143 ZNF462

>Homo_sapiens
MRLPNNFHNNNPSRKRRKRKRIRRRWGWRTRSTSRRRDASSVPNVRG
>Mustela_putorius_furo
MRLPNNFHNNNPSRKRRKRKRIRRRWGWRTRNTSRRRDASSVPNVRR
>Physeter_catodon
MRLPNNFHNNNPSRKRRKRKRIRRRWGWRTRNTSRRRDASSVPNVRE
>Pteropus_alecto
MRLPNNFHNNNPSRKRRKRKRIRRRWGWRTRNTSRRRDASSVPNVRG
>Dasypus_novemcinctus
MRLPNNFHNNNPSRKRRKRKRIRRRRSRRTRNTSRRRDASSVPNVRG
>Condylura_cristata
MRLPNNFHNNNPSRKRRKRKRIRRRWGWRTRNTSRRRDASSVPDVRG
>Oryctolagus_cuniculus
MRLPNNFHNNNPSRKRRKRKRIRRRWGWRTRSTFRRRDASSVPNVRG
>Elephantulus_edwardii
MRLQNNFHNNNPSRKRRKRKRIRRRWGWGSRDTSRRRDASSVPNVKG
>Equus_asinus
MRLPNNFHNNNPSRKRRKRKRIRRRWGWRTRNTSRRRDASSVPNVRG
>Manis_javanica
MRLPNNFHNNNPSRKRRKRKRIRRRWDRRTRNTSRRRDASSVPNVRE
>Loxodonta_africana
MRLPNNFHNNNPSRKRRKRKRIRRRWGWRTRNTSRRRDA-SVPNVRG
>Octodon_degus
MRLPNNFHNNNPSRKRRKRKRIRRRWGWRTRSTSRRRDASSVPNVRG
>Tupaia_chinensis
MRLPNNFHNNNPSRKRRKRKRIRRRWGWRTRSTSRRRDAPSVPNVRG
>Trichechus_manatus_latirostr
MRLLNNFHNNNPSRKRRKRKRIRRRWGWRTRKHLPAPGRFFCS----

HG0547 ENSG00000149596 JPH2

>Homo_sapiens
MALTSRPGADLAPGLFQPLL-ASCFPPPGVQW------
>Panthera_tigris
MALTSRPSADLAPGLIQSLV-ASCFPPPGAQW------
>Orcinus_orca
MALTSRPSADLAPGFLQPRV-ASCLPPPGAQ-------
>Pteropus_alecto
MALTSRPSADLAPGLNQPLV-ASCFPPAGARWW-----
>Dasypus_novemcinctus
MALSSRPGAGLA-GPSRPLL-AARSPPPGARW------
>Galeopterus_variegatus
MALTSHPSADLAPGLFPPLL-APCFPPPGVRW------
>Oryctolagus_cuniculus
MALPSPPGTDLAAGLLQPLL-ASRSPCCGARW------
>Equus_przewalskii
MALTSRPSADLAPGLLQPLL-ASRVPPPGALWQW----
>Manis_javanica
MALTGPSGADLARGLLPL-R-ASCCPVLGHGGELGAQP
>Loxodonta_africana
MALTSWPSADLAPGLSQPLVAAG-CPPPGAPW------
>Cavia_porcellus
MALTSTPNADLAPGLFQPLL-ASCFPPPGARW------
>Tupaia_chinensis
MALTSPPSAHLAPGLFQPLR-AACFPPPGARW------
>Trichechus_manatus_latirostr
MALTSRPSADLAPGLSQPLVAARCCPPPGAPW------
>Orycteropus_afer_afer
MALTSWPSADLAPGFSQPLLVAR-CPPPRARW------

HG0548 ENSG00000155744 FAM126B

>Homo_sapiens
--------------------------------------------------------------------------MNSPLELCRIPVPPPPLEN
>Canis_lupus_familiaris
----------------------------------MLPLEE------TGPRV--PGTSWHIFWELLSWKIFKSQQMNSTLELCRIPVPPPPLEN
>Orcinus_orca
--------------------------------------------------------------------------MNSTLELCRIPVPPPPLEH
>Myotis_lucifugus
--------------------------------------------------------------------------MNSTLELCRIPVPPPPLEN
>Galeopterus_variegatus
--------------------------------------------------------------------------MNSILELCCIPVPPPPLEN
>Erinaceus_europaeus
--------------------------------------------------------------------------MNSTLDFCCIPVPPPPLEN
>Ochotona_princeps
--------------------------------------------------------------------------MNSTLELCLIPVPPPPLEN
>Elephantulus_edwardii
----------------------------------------------------------------------MSQQMNSTREICHIPVPPPPLEN
>Ceratotherium_simum_simum
--------------------------------------------------------------------------MNSTLELCRIPVPPPPLEN
>Manis_javanica
--------------------------------------------------------------------------MNSTLELCCIPVPPPPLEN
>Loxodonta_africana
--------------------------------------------------------------------------MNSTLEICHIPVPPPPLEN
>Mus_musculus
--------------------------------------------------------------------------MNSTLELCRIPVPPPPLEN
>Tupaia_chinensis
MFSLHGDRNPKTLHRGGRAAAVFQPMAEAVVESVDLPNERVACYAAVGRLLREAVAVGSRRQRGDSWKIFNPQQMNSALELCCIPVPPPPLEN
>Trichechus_manatus_latirostr
--------------------------------------------------------------------------MNSTLETCRIPVPPPPLEN

HG0549 ENSG00000156256 USP16

>Homo_sapiens

-MQLWALSPWIVILCQ-

>Acinonyx_jubatus

-MQLWALSPWIVILCQ-

>Balaenoptera_acutorostrata_s

-MQLWALSPWIVILCQ-

>Pteropus_alecto

-MQLWVLSPWIVILGQ-

>Dasypus_novemcinctus

-MQLWALSPWIIILCQ-

>Condylura_cristata

-MQLWALSPWIVILCQ-

>Oryctolagus_cuniculus

-MQLWVLSPWIVFCVSK

>Ceratotherium_simum_simum

-MQLWALSLWMVILCQ-

>Manis_javanica

-MQLWVLSPWIVILCQ-

>Loxodonta_africana

-MQLWALSPWIVILCH-

>Heterocephalus_glaber

MMQLWALSPWIVILCQ-

>Tupaia_chinensis

-MQLWALPPWIVILCQ-

>Trichechus_manatus_latirostr

-MQLWALSPWIVILCQ-

>Orycteropus_afer_afer

-MQLWSLLPWIVTLCH-

HG0550.1 ENSG00000156650 KAT6B

>Homo_sapiens
--------------------MTFKCASSVKYKENLYG
>Canis_lupus_familiaris
--------------------MTFKCSSSVKYKENLYG
>Lipotes_vexillifer
--------------------MTFKCASSVKYKENLYG
>Myotis_brandtii
MFVRCCLICFGVLCPFVELLMTFKCASSVKYKENLYG
>Galeopterus_variegatus
--------------------MTFKCASSVKYKENLYG
>Sorex_araneus
--------------------MTFKCASSVKYKENLYG
>Oryctolagus_cuniculus
--------------------MTFKCASPVKYKENLYG
>Elephantulus_edwardii
--------------------MTFTCVSSVKYKENLYG
>Equus_asinus
--------------------MTFKCASSVKYKENLYG
>Manis_javanica
--------------------MTFKCSSSVKYKENLYG
>Loxodonta_africana
--------------------MTFKCASSVKYKENLYG
>Nannospalax_galili
--------------------MTFKCASSDKYKENLYG
>Tupaia_chinensis
--------------------MTFNCASSVKNKENLYG
>Orycteropus_afer_afer
--------------------MTFKCATSVKYKENLYG

HG0550.2 ENSG00000156650 KAT6B

>Homo_sapiens

MFFTRMQSFLLVK

>Odobenus_rosmarus_divergens

MFFARMQSFLLVK

>Orcinus_orca

MFFTRMQFFLLVK

>Myotis_lucifugus

MFFARMQSFLLVK

>Sorex_araneus

MFFTRMQSFLLVK

>Ceratotherium_simum_simum

MVFTRMQSFVLVR

>Loxodonta_africana

MFFARMQSFLLVK

>Trichechus_manatus_latirostr

MFFARTQSFLLVK

>Orycteropus_afer_afer

MFFARMQSFLLVK

HG0551 ENSG00000162951 LRRTM1

>Homo_sapiens
MSEPALCSCPGREWQAVSAE-----------
>Odobenus_rosmarus_divergens
MSEPALRSRPGREWQAVSAE-----------
>Orcinus_orca
MSEPALRSRPGREWQAVSAEQKVAPVSGPFQ
>Rousettus_aegyptiacus
MSEPALRSRPGREWQVVSAD-----------
>Galeopterus_variegatus
MSEPAPRSCPGREWRAVSAEQKVAPVRGRFQ
>Condylura_cristata
MSEPALRSRPEREWQAVSEK-----------
>Ochotona_princeps
MSEPVLCSCPGREWPAVSAE-----------
>Equus_asinus
MSEPALRSRPGREWQAVSAE-----------
>Manis_javanica
MSEPALRSLPGREWQAVSAE-----------
>Loxodonta_africana
MSEPAPCSRPGREWQAVSAE-----------
>Fukomys_darlingi
MSEPALRSCPGREWQAVSAE-----------
>Tupaia_chinensis
MSEPALRSWPRREWQAVSAESKVARSSGRFQ
>Trichechus_manatus_latirostr
MSEPAPGSRPGREWQAVSAE-----------
>Orycteropus_afer_afer
MSELAPRSRPGREWQAVSVEQKVETVSGLFQ

>Homo_sapiens
--------------------------------------------------------------------------------MMVSSCQRRLWLPRER
>Acinonyx_jubatus
----------------------------------------------------------------------------MTKLIMVSSCQRRLWMPRER
>Physeter_catodon
---------------------------------------------------------------------------------MVSSCQRRLWMPRER
>Rousettus_aegyptiacus
MRLSSASSCHAAMGGVLHYVDISSLSAIKNQWCPASLLRFIHSTNLGTRITYGNSFYTWELKYRFYICCLTCFLLSFCRLIMVSSCQRRLWMPRER
>Dasypus_novemcinctus
---------------------------------------------------------------------------------MVSSCQRRLWVPRKM
>Galeopterus_variegatus
---------------------------------------------------------------------------------MVSSCQRRLWMPRER
>Condylura_cristata
--------------------------------------------------------------------------------MMVSSCQRRLWIPRDR
>Elephantulus_edwardii
---------------------------------------------------------------------------------MVSSCQRKLWMPRER
>Ceratotherium_simum_simum
---------------------------------------------------------------------------------MVSSCQRRLWMPGER
>Manis_javanica
---------------------------------------------------------------------------------MVSSCQRRLWMPRER
>Loxodonta_africana
---------------------------------------------------------------------------------MISSCRRRLWMPRER
>Nannospalax_galili
---------------------------------------------------------------------------------MVSSCQRRLWIPRER
>Tupaia_chinensis
---------------------------------------------------------------------------------MVSSCQRRLWMPRER
>Orycteropus_afer_afer
---------------------------------------------------------------------------------MVSSCRRRLWMPRER

HG0553 ENSG00000183475 ASB7

>Homo_sapiens
---MVTRQSVHQEKGHSSIRLIPVT-----
>Leptonychotes_weddellii
MCTMVIRQSTYQEKGHSSIRLFPVT-----
>Physeter_catodon
MCTMVFRQSTYQEKGHSSRKADPCDLMNPL
>Rhinolophus_ferrumequinum
MCTMVIRQSTCQEKGHSSVRLIPVT-----
>Dasypus_novemcinctus
MYTMVIRQSIYQEKGHSSVRLIPVT-----
>Galeopterus_variegatus
MYTMVIRQSIYQEKGHPPIRLIPVT-----
>Condylura_cristata
MCTMVIRQSTCQDEEHLSRRLIPVT-----
>Oryctolagus_cuniculus
MCTMVTRQSTYQERGHSSVRLIPVT-----
>Equus_asinus
MCTMIIRQSIYQEKGHSSVRLIPVT-----
>Manis_javanica
MCTMVIRQSAYQEKEHFSVRLIPVT-----
>Loxodonta_africana
MCTMVIRQHLYQDKGHSIIRLIPVT-----
>Ictidomys_tridecemlineatus
---MVIRQSIHQEKGHSSVRLIPVT-----
>Tupaia_chinensis
---MVVRQSTCHEKGHAIVRPFPVT-----
>Orycteropus_afer_afer
---MVIRQSLYQEKGHSKV--EPCDLMNPW

HG0554 ENSG00000196914 ARHGEF12

>Homo_sapiens
---MTEWSFELDFCVPDGVGPDPRALGVGRRCYCKMQVG
>Felis_catus
---MTEWSFELDFCVPYGVGPDPRALGVGWRCYCKMQVG
>Physeter_catodon
---MTEWSFELDSCVPYGVGPDPGALGVGWRCYCKMQVG
>Rousettus_aegyptiacus
---MTEWSFELNFSSLSESGLIPELWGVGWRCYCKMQVG
>Galeopterus_variegatus
---MTEWSFELDFCVPDEVGPDLSALGVGRRCYCKMQVG
>Condylura_cristata
---MTEWSFELDFCVPYGVGPDPRARGVGWRCSCKMQVG
>Oryctolagus_cuniculus
---MTAWSVELDFCVPDGVGPDPRALGVGWRCYCKMQVG
>Elephantulus_edwardii
---MTEWSFELDFCVLYGIGPDPRALGVGWRCYWKMQVG
>Ceratotherium_simum_simum
---MTAWSFELDFCVPYGVGPDPRALGVGWRCSCKMQVG
>Manis_javanica
---MTEWSFEVDFCVPYGVGLDPRALGVGWGCYCKMQVG
>Loxodonta_africana
---MTEWSFELDFCVPYGVGPDPRALGVGWRCYCKMQVG
>Chinchilla_lanigera
---MTEWSFELDFCVPAGIGPDPRALGVGWRCFCKMQVG
>Tupaia_chinensis
MFWMTEWSFELDFCVPDGVGPDPRALGVGWRCSCKMQVG
>Trichechus_manatus_latirostr
---MTEWSFELDFCV----GPDPSALGVGWRCYCKMQVG

HG0555 ENSG00000204310 AGPAT1

>Homo_sapiens
MGPTSHSHLPSIGRG
>Ursus_maritimus
MGPASHSHLPSIGRG
>Orcinus_orca
MGPASHSHLPSIGRG
>Myotis_lucifugus
MGPASHSHLPSIGRG
>Dasypus_novemcinctus
MGPASHSHLPSSGRG
>Galeopterus_variegatus
MGPTSHSHLPSIGRG
>Erinaceus_europaeus
MGPASHSHLPSIGRG
>Oryctolagus_cuniculus
MGPASHAHLPSIGRG
>Equus_caballus
MGPASHSHLPSIGRG
>Manis_javanica
MGPASQSHLPYIGRG
>Loxodonta_africana
MGPASHSHLPSIGRG
>Marmota_marmota_marmota
MGPTSHSHLPSIGRG
>Trichechus_manatus_latirostr
MGPASHSHLPSIGRG
>Orycteropus_afer_afer
MGPASHSHLLSTGRG

HG0556 ENSG00000273841 TAF9

>Homo_sapiens
MLLPNILLTGTPGVGKTTLGKELASKSGLKYINVGDLAREV
>Felis_catus
MLLPNILLTGTPGVGKTTLGKELASRSGLKYINVGDLAREV
>Orcinus_orca
MLLPNILLTGTPGVGKTTLGKELASRSGLKYVNVGDLAREV
>Pteropus_vampyrus
MLLPNILLTGTPGVGKTTLGKELASRSGLKYINVGDLAREV
>Dasypus_novemcinctus
MLQPNILLTGTPGVGKTTLGKELASRSGLKYVNVGDLAKEV
>Galeopterus_variegatus
MLLSNILLTGTPGVGKTTLGKELTARSGLKYFNVGDLAQEV
>Sorex_araneus
MLLPNILLTGTPGVGKTTLGKELASRSGLKYVNVGDLAREV
>Ochotona_princeps
MLRPNILLTGTPGVGKTTLGKELASRSGLKYVNVGDLAREV
>Elephantulus_edwardii
MLLPNILLTGTPGVGKTTLGKELASRSGLKYINVGDLAKEV
>Equus_przewalskii
MLLPNILLTGTPGVGKTTLGKELASRSGLKYINVGDLAREV
>Loxodonta_africana
MLQPNILLTGTPGVGKTTLGKELASRSGLKYVNVGDLAKEV
>Marmota_marmota_marmota
MLLPNILLTGTPGVGKTTLGKELASRSGLKYINVGDLAQEV
>Trichechus_manatus_latirostr
MLQPNILLTGTPGVGKTTLGKELASRSGLKYINVGDLAKEV
>Orycteropus_afer_afer
MLRPNILLTGTPGVGKTTLGKELASRSGLKYINVGDLAKEV

HG0557 ENSG00000012232 EXTL3

>Homo_sapiens

MHICWSAKPRNKSYGI

>Ailuropoda_melanoleuca

MHICWSAKPRNKSYGS

>Balaenoptera_acutorostrata_s

MHMCWSAKPRNESYGS

>Eptesicus_fuscus

MHICWSAKPRNKSYGC

>Galeopterus_variegatus

MHTCWSAEPRNKSYGR

>Condylura_cristata

MHVCWPAKRRSKSHGR

>Equus_asinus

MHICWSAKPRNKSYGS

>Manis_javanica

MHTCLSAKPRNKSYGS

>Loxodonta_africana

MHNCWPAKPRNKSYGT

>Ictidomys_tridecemlineatus

MHICWSAKPRNKSYGL

>Tupaia_chinensis

MNICWSAKPRNKSYGS

>Trichechus_manatus_latirostr

MHICWPAKPRNKSYGT

>Orycteropus_afer_afer

MHLCWPAKPRNKSDGS

HG0558 ENSG00000029363 BCLAF1

>Homo_sapiens
MVFLAFLSREMAAVWLQRR
>Canis_lupus_familiaris
MVFLPFLSREMAAV-----
>Balaenoptera_acutorostrata_s
MVFLPCLSREMAAV-----
>Miniopterus_natalensis
MVFLPFLSRVMAAV-----
>Dasypus_novemcinctus
MVFLPFLSREMAAV-----
>Galeopterus_variegatus
MVFLSFLSREMAAVWLQRR
>Condylura_cristata
MVFLPFLSREMAAV-----
>Ochotona_princeps
MVFLSFSSREMAAV-----
>Ceratotherium_simum_simum
MVFLPFSSREMAAV-----
>Manis_javanica
MVFLPFLSREMAAV-----
>Loxodonta_africana
MVFLSVLSREMAAV-----
>Chinchilla_lanigera
MVFLSFLSREMAAV-----
>Tupaia_chinensis
MVFLSFLSREMAAV-----

HG0559 ENSG00000078747 ITCH

>Homo_sapiens
MHFTVALWRQRLNPRK
>Panthera_tigris
MHLTVAFWRQRLNPKK
>Balaenoptera_acutorostrata_s
MHLTVAFWRQRLNPKK
>Miniopterus_natalensis
MHLTVAFWRQRLNPKK
>Dasypus_novemcinctus
MHLTVAFWRQRFNPKK
>Galeopterus_variegatus
MHFTVAFWRQRLNPKK
>Condylura_cristata
MHLTVAFWRQRLNPKK
>Oryctolagus_cuniculus
MPFTVAFWRQRLNQKK
>Equus_asinus
MHLTVAFWRQRLNPKK
>Loxodonta_africana
MHVTVAFWRQRLNPKK
>Dipodomys_ordii
MHFTVAFWRQRLNPKK
>Tupaia_chinensis
MHFTVAFWRQRLNPKK
>Trichechus_manatus_latirostr
MHLTVAFWRQHLNPKK

HG0560.1 ENSG00000092051 JPH4

>Homo_sapiens

MDAVPGRQPEPQPTCHSG

>Leptonychotes_weddellii

MDAVPGRQPEPQPTCHSG

>Balaenoptera_acutorostrata_s

MDAVPGRQPEPQSTCHSG

>Rousettus_aegyptiacus

MDAVPGRQPEPQPTCHSG

>Dasypus_novemcinctus

MDAVPGKQPEPQPTCHSG

>Galeopterus_variegatus

MDAVPERQPEPQTTCHSG

>Condylura_cristata

MDAVPGRQPEPQPTCHSG

>Ochotona_princeps

MDVEPGRRPAPQPTCHSG

>Elephantulus_edwardii

MDAVPGRQPEPQPTRHS-

>Ceratotherium_simum_simum

MDAVPGRQPEPQPTCHSG

>Manis_javanica

MDDLPGKRLEPQPTCHSG

>Dipodomys_ordii

MDAVPGRQPEPQPTCHSG

>Orycteropus_afer_afer

MDVILRRQPEPQPTCHSG

>Homo_sapiens
----------------------------------------------------------------------------------------------------------
---------------------------------------------------MEQTLICAGMESEF
>Mustela_putorius_furo
MGLPPPSFSSGISPSLALTPAPLSLYSSRPFLASPASLLPRLASASPSLPIFSSVSSLSLCSPPLSPPPSVSPHSLPAYQDFFPPWGSLCFSFSPLPASSLTSDLA
GTHLFQPGNPPPQLLTSPLYLLPLLLSPPSPLLSSPCPFKSSWPSLPRDAGMEQTLICAGMESEF
>Orcinus_orca
----------------------------------------------------------------------------------------------------------
---------------------------------------------------MEQTLICAGMESEF
>Myotis_lucifugus
----------------------------------------------------------------------------------------------------------
---------------------------------------------------MEQTLICAGMESEF
>Dasypus_novemcinctus
----------------------------------------------------------------------------------------------------------
---------------------------------------------------MEQTLICAGMESEF
>Galeopterus_variegatus
----------------------------------------------------------------------------------------------------------
---------------------------------------------------MEQTLICAGMESEF
>Erinaceus_europaeus
----------------------------------------------------------------------------------------------------------
---------------------------------------------------MEQTLICAGMESEF
>Oryctolagus_cuniculus
----------------------------------------------------------------------------------------------------------
---------------------------------------------------MEQTLICARMESEF
>Elephantulus_edwardii
----------------------------------------------------------------------------------------------------------
---------------------------------------------------MEQTLICAGMESEF
>Ceratotherium_simum_simum
----------------------------------------------------------------------------------------------------------
---------------------------------------------------MEQTLICAGMESEF
>Manis_javanica
----------------------------------------------------------------------------------------------------------
---------------------------------------------------MEQTLICAGMESEF
>Dipodomys_ordii
------------------------------------------------MLVF--LSTLGVSVPLFLLPTCLIPH----------------------PG--VS----
-------KSPSSSTWTSPLHPHHRHRLSSPLLLAPLCPFKSAWPSLPRDAGMEQTLICAGMESEF

HG0561 ENSG00000100393 EP300

>Homo_sapiens
MRRRRTAPRRKRLMAAAELRETSAGQGPAVAGRGLRL
>Mustela_putorius_furo
MRRRRTAPRRKRLMAAAELRETSAGQGPAVAGRGLRL
>Physeter_catodon
MRRRRTAPRRKKLMAAAELRETSAGQGPAVAGRGLRL
>Rousettus_aegyptiacus
MRRRRTAPRRKRLMAAAELRETSAGQGPAVADRGLRL
>Dasypus_novemcinctus
MRRRRTAPRRKRLMAAAELRETSAGQGPAVAGRGLRL
>Erinaceus_europaeus
MRRRRTAPRRKRLMAAAELRETSAGQGPAVAGRGLRL
>Oryctolagus_cuniculus
MRRRRTAPRRKRLMAAAELRETSAGQGPAAAGRGLRL
>Equus_asinus
MRRRRTAPRRKRLMAAAELRETSAGQGPAVADRGLRL
>Manis_javanica
MRRRRTAPRRKRLMAAAELRETSAGQGPAVAGRGLRL
>Loxodonta_africana
MRRRRTAPRRKRLMAA-ALRETSAGQGPAVAGRGLRL
>Octodon_degus
-------------MAAAELRETSAGQGPAVAGRGQRF
>Trichechus_manatus_latirostr
MRRRRTAPRRKRLMAA-ALRETSAGQGPAVAGRGLRL
>Orycteropus_afer_afer
MRRRRTAPRRKRLMAA-ALRETSAGQEPAVAGRGLHL

HG0562 ENSG00000105821 DNAJC2

>Homo_sapiens
-----------------------MRRFLGVLEPRREARSER-------------
>Leptonychotes_weddellii
-----------------------MRRFLGVVEPRREARTER-------------
>Physeter_catodon
-----------------------MRRFLDIVEPRREARTGR-------------
>Rousettus_aegyptiacus
-----------------------MRRFLGVVDPRREARTGR-------------
>Galeopterus_variegatus
-----------------------MRRFLGVVERRREARTGRQSRRGR-------
>Oryctolagus_cuniculus
-----------------------MRRFVGVVEPRREARTGR-------------
>Elephantulus_edwardii
-----------------------MRRFLGVE-QRREARSGG-------------
>Ceratotherium_simum_simum
-----------------------MRRFLGVLEPRREARTGR-------------
>Manis_javanica
-----------------------MRRFLGAVESRREARTGR-------------
>Loxodonta_africana
-----------------------MRRFLGVVEPRREARTEGTAGGNAKHPPPGP
>Ictidomys_tridecemlineatus
MSGGGIGVHTTKDPSRKCLLLVSTRRFLGVVEPRREARTGR-------------
>Tupaia_chinensis
-----------------------MRRVLGDVEPRREAQKER-------------
>Orycteropus_afer_afer
-----------------------MRRFLGVVEPRREARTGR-------------

HG0563 ENSG00000107249 GLIS3

>Homo_sapiens
---------------------MDDSIRLITHTMDNLLDFA
>Leptonychotes_weddellii
--------------------------------MDNLLDFA
>Lipotes_vexillifer
---------------------MDDSIRLINHTMDNLLDFA
>Rousettus_aegyptiacus
--------------------------------MDNLLDFA
>Dasypus_novemcinctus
--------------------------------MDNLLDFA
>Galeopterus_variegatus
MLHRLKTHCTLFLFLLLKMPAFIFLNRLRTYTMDNLLDFA
>Sorex_araneus
--------------------------------MDNLLDFA
>Ochotona_princeps
--------------------------------MDNLLDLA
>Ceratotherium_simum_simum
--------------------------------MDNLLDFA
>Loxodonta_africana
--------------------------------MDNVLDLA
>Chinchilla_lanigera
--------------------------------MDNLLDFA
>Tupaia_chinensis
--------------------------------MDNLLDFA
>Trichechus_manatus_latirostr
--------------------------------MDNVLDLA

HG0564 ENSG00000109171 SLAIN2

>Homo_sapiens
----------------------------------------------MAAGAGYWRRLPQSRSRRR
>Odobenus_rosmarus_divergens
----------------------------------------------MAAKAGYWRRLPQSWSRRR
>Balaenoptera_acutorostrata_s
----------------------------------------------MAVEAGYWRRLPQSWSRRR
>Pteropus_alecto
----------------------------------------------MAAEAGYWRRLPQSRSRRR
>Dasypus_novemcinctus
----------------------------------------------MAAEAGYWRRLPQSGSRRR
>Galeopterus_variegatus
----------------------------------------------MAAEAGCWRRLPQSRSRRR
>Condylura_cristata
----------------------------------------------MAAEAGYWRRLPQSWSRRR
>Oryctolagus_cuniculus
----------------------------------------------MAAEAGYWRRLPQSWSRRR
>Elephantulus_edwardii
----------------------------------------------MAAEAGYWHRLPKSGSRRR
>Ceratotherium_simum_simum
----------------------------------------------MAAEAGYWRRLPQSWSRRR
>Ictidomys_tridecemlineatus
----------------------------------------------MAAEAGYWRRLPQSRSRRR
>Trichechus_manatus_latirostr
----------------------------------------------MAAEAGYWRRLPQSWSRRR
>Orycteropus_afer_afer
MTPPTAPPGSELSARFSQCHPGAVSTGDALRSPTGPNSRGRRDGGKMAAEAGYWRRLPQSGSRRR

HG0565 ENSG00000120533 ENY2

>Homo_sapiens
MRVLAFCVLRCPSY
>Leptonychotes_weddellii
MRVLAFCVLRCPSY
>Lipotes_vexillifer
MRVLAFCVLRCPSY
>Rousettus_aegyptiacus
MRVLAFCVLRCPSY
>Dasypus_novemcinctus
MRVLAFCVLRCPSY
>Condylura_cristata
MRVLAFCVLRCPSY
>Ochotona_princeps
MRVLAFCVLRCPSY
>Elephantulus_edwardii
MRVLALYVLRCPSY
>Equus_asinus
MRVLAFCVLRCPSY
>Fukomys_damarensis
MRVLAFCVLRCPSY
>Tupaia_chinensis
MRVLAFCVLRCPSY
>Trichechus_manatus_latirostr
MRVLAFCVLRCPSY
>Orycteropus_afer_afer
MRVLAFCVLRCPSY

HG0566 ENSG00000124496 TRERF1

>Homo_sapiens
MHLLQKQPRPWLWRMLKEEDAEAGRP-------------------------
>Canis_lupus_familiaris
MHLLQKQPRPWLWRMLKEEDMEAGRP-------------------------
>Balaenoptera_acutorostrata_s
MHLLQKQPRPRLWRMLKEEDTEAGRP-------------------------
>Pteropus_alecto
MHLLQKQPRPWLWRMLKEEDMEAGRP-------------------------
>Dasypus_novemcinctus
MHLLQRQPRPWLWRMLEEEDTEAVGPERLSSRS------------------
>Condylura_cristata
MHLLQKQPRPWLSRMLKEEDTEARRP-------------------------
>Oryctolagus_cuniculus
MHLLQKQPRPWLRRPLKEEDEEAGRL-------------------------
>Ceratotherium_simum_simum
MHLLQKQPRPWLWRMLKEEDTEAARP-------------------------
>Manis_javanica
MHLLQKQPRPRLWRTLKEEDTEAGRP-------------------------
>Loxodonta_africana
MHLLLTQPRPWFWRMLKRKTRKQDSPEILSFLTLKQNQVSSSHLSSSMRAA
>Marmota_marmota_marmota
MHLLQKQPRPWLWRMLKEEDAEAGRP-------------------------
>Tupaia_chinensis
MHLLQKQPRPWLWRMLKEEDTEAGQP-------------------------
>Trichechus_manatus_latirostr
MHLLLRQPRRCLWRMLKEEDPEAGRS-------------------------

HG0567 ENSG00000132155 RAF1

>Homo_sapiens

-MRGLLGAPSPGLGTRRM

>Canis_lupus_familiaris

-MRGLPGAPSPGLGTRRM

>Physeter_catodon

-MRGLPGAPSPGLGTRRM

>Rousettus_aegyptiacus

-MRGLPGAPSPGLGTRRM

>Dasypus_novemcinctus

-MRGLPGVPSPGLGTRRM

>Galeopterus_variegatus

MMRGLPGAPSPGLGTRRM

>Condylura_cristata

-MRGLPGAPSPGLGTRRM

>Ochotona_princeps

-MRGLPGAPSPGLGTRRM

>Ceratotherium_simum_simum

-MRGLPGAPSPGLGTRRM

>Manis_javanica

-MLGLPGAPSPGLGTRRM

>Nannospalax_galili

-MRGLLGAPSPGLGTRRM

>Trichechus_manatus_latirostr

-MRGLPGAPSPGLGTRRM

>Orycteropus_afer_afer

-MRWLLGAPSPGLGTRRM

HG0568.1 ENSG00000138271 GPR87

>Homo_sapiens

MKEIKP--GITYAEPTPQSSPSVS

>Canis_lupus_familiaris

MKGLKP--GITYAKPFPQPSPSVS

>Physeter_catodon

MKEIKP--GITYAEPLPQSSPSVS

>Pteropus_alecto

MKEIKP--GITYAEPLPQPSPSVS

>Galeopterus_variegatus

MKEIKP--GIAYAELMPQPSPSVS

>Condylura_cristata

MKESKPTNKTTYAEPMRQSSPSVS

>Oryctolagus_cuniculus

MKEIKP--GIPYAEPMPPSSPSVS

>Equus_asinus

MKEIKP--GTTYAGPLPQSSPSVS

>Manis_javanica

MKEIKP--GITYAEPLPQSSPSVS

>Heterocephalus_glaber

MKEIKP--GLTHAEPMPQSSPSVS

>Tupaia_chinensis

MKEIKP--GITYAKPMPELSPSVS

>Trichechus_manatus_latirostr

MKEIKQ--GITYAEPKLQLSPSVS

>Orycteropus_afer_afer

MKEIKQ--GITYAEPKLQLSPSVS

HG0568.2 ENSG00000138271 GPR87

>Homo_sapiens
MFRFHALPENPLPCRP
>Canis_lupus_familiaris
MLIFHAVPENPLLCQP
>Miniopterus_natalensis
MFIFHALPENPHPCQP
>Ceratotherium_simum_simum
MFLFHALPESPLLCQP
>Manis_javanica
MLIFHAFPENPLPCQP
>Tupaia_chinensis
MILFSALPENPLSCQP

HG0569 ENSG00000139651 ZNF740

>Homo_sapiens
------------------------------------------------------MATGLMGHEGVATVIW
>Acinonyx_jubatus
------------------------------------------------------MAAGLKGHEGVVTVIW
>Orcinus_orca
------------------------------------------------------MAAGLKGHEGAVTVIW
>Myotis_brandtii
------------------------------------------------------MAAGLKGHEGVVTVIW
>Galeopterus_variegatus
------------------------------------------------------MAAGLKGHEGVITVTW
>Erinaceus_europaeus
MKQTSWTSVREFGQKTKTGRLFLLTIDGSQAEPKDSCGIYLVFFFFLTQRKFNMMAAGLKGHAGVVTVIW
>Elephantulus_edwardii
------------------------------------------------------MAAGLKGHKGVVTVIW
>Ceratotherium_simum_simum
------------------------------------------------------MAAGLRGHEGVVTVIW
>Manis_javanica
------------------------------------------------------MAAELKGHKEVVTVIW
>Loxodonta_africana
------------------------------------------------------MAAGLKGHKGVVTVIW
>Marmota_marmota_marmota
------------------------------------------------------MAAGLRGHEGAITVIW
>Trichechus_manatus_latirostr
------------------------------------------------------MAAGLKGHRGVITVIW
>Orycteropus_afer_afer
------------------------------------------------------MAAGLKGHRGVVTVIW

HG0570 ENSG00000152214 RIT2

>Homo_sapiens
MRPARAEDQQPARR---------------
>Ursus_maritimus
MRPARAEDQQPARR---------------
>Rousettus_aegyptiacus
MRPARAQDQQPARR---------------
>Dasypus_novemcinctus
MRPAPAEDQQPARR---------------
>Condylura_cristata
MRPAGAEDQQPARR---------------
>Ochotona_princeps
MRPAGAEDQQPAGH---------------
>Elephantulus_edwardii
MRPAQAEDQQPAWELMLLPACQTFLLPQF
>Equus_asinus
MRPAGAEDQQPARR---------------
>Loxodonta_africana
MRPAGAEDQQPARR---------------
>Octodon_degus
MRPAPAQDQQPARR---------------
>Tupaia_chinensis
MRPARAEDQQPARC---------------
>Trichechus_manatus_latirostr
MRPAGAEDQQLARR---------------
>Orycteropus_afer_afer
MRSAGAEDQQPAWR---------------

HG0571 ENSG00000153266 FEZF2

>Homo_sapiens

----------------------------MARNASPWWWGKET

>Canis_lupus_familiaris

----------------------------MARNASPWRWGKET

>Orcinus_orca

----------------------------MARNASPWRWGKET

>Pteropus_alecto

----------------------------MARNASPWRWGKET

>Galeopterus_variegatus

MSAECKRPVISRGGEDVCHQNVTEEIRCMARNASPWWWGKET

>Erinaceus_europaeus

----------------------------MARNASPWRWGKET

>Ochotona_princeps

----------------------------MARNASPWQWGKET

>Equus_asinus

----------------------------MARNASPWRWGKET

>Manis_javanica

----------------------------MARNASPWRWGKET

>Loxodonta_africana

----------------------------MARNASPWRWGKET

>Mesocricetus_auratus

----------------------------MARNASPWRWGKET

>Trichechus_manatus_latirostr

----------------------------MARNASPWRWGKET

>Orycteropus_afer_afer

----------------------------MARNASPWRWGKET

HG0572 ENSG00000158161 EYA3

>Homo_sapiens
MRIRVTLTPRFPPLDWFYCGSGLISMSCCGAFTAFGL--
>Canis_lupus_familiaris
MRIRVTLTPRFPPSAWFSCESGPVPMSCFGASAAVGQ--
>Physeter_catodon
MRIRVTLTPPFPAASWFSCVFGPVPMSCCGASAAFGL--
>Myotis_brandtii
MRFRVTLTPRFPPLAWFYCGSGPVPMSCSGAPAAFGL--
>Dasypus_novemcinctus
-------------------------MSCCGTSAAFGL--
>Galeopterus_variegatus
-------------------------MPYCGASAAFGL--
>Condylura_cristata
MRIRVTLTPRFPPPAWFYCGSGPVPMSCCGASASFGL--
>Oryctolagus_cuniculus
MRFRVTLTPRFPPLARSYRGSGPVPMSCCGSSVALGL--
>Ceratotherium_simum_simum
MCIRVTLTPRFPPPAWFCCGSGPVPMSCCGASGAFGL--
>Manis_javanica
-------------------------MSCCVAFAGFGRAR
>Marmota_marmota_marmota
MRIRVTLTPRFPPLCWFYRGSGPVPMSCCGASAAFGL--
>Tupaia_chinensis
-------------------------MSRCGAPAAFGL--
>Trichechus_manatus_latirostr
MRIRVMLTPPFPPLAWFCCGSGPVLMSCCGASAAFGL--

HG0573.1 ENSG00000162526 TSSK3

>Homo_sapiens

-------------------------MKSLGRKGAGQGRWWSRAASQRQHELRG

>Leptonychotes_weddellii

MALSLPTTRGCCVQTENALALGRLRMKSLGGKGAGQGRWWSRAASQRQHELRG

>Eptesicus_fuscus

-------------------------MKSLGRKGPGQGLWWSRAASQRQHELRG

>Galeopterus_variegatus

-------------------------MKSLGGKEAGQG----------------

>Condylura_cristata

-------------------------MKSLGGKGAGQGRWWSRAASRRQHELRG

>Oryctolagus_cuniculus

MSDPLLPPTRGCVQTENALALGRLWMKSLGGRGAGRGRWCSRAASRRQHALSG

>Elephantulus_edwardii

-------------------------MKSLGGKEAGQGRWWSRAASLRQHELRG

>Ceratotherium_simum_simum

-------------------------MKSLGGKVAGQGQWWSRAASWRQHELRG

>Manis_javanica

-------------------------MKSLGGKSAGQGRWWSRAASQRQHELGV

>Loxodonta_africana

-------------------------MKSLGGKEAGQGRWWSRAASRRQHELRG

>Ictidomys_tridecemlineatus

-------------------------MKSLGGKGAGQGRWWSRAASQRQHELRG

>Trichechus_manatus_latirostr

-------------------------MKSLGGKGAGQGRWWSRAASRRQHELRG

>Orycteropus_afer_afer

-------------------------MKSLGGKGAGQGRWWSRAASWRQHELRG

HG0573.2 ENSG00000206203 TSSK2

>Homo_sapiens
MRTMPAGPHDGGM
>Canis_lupus_familiaris
MRTMPAGPRGEGM
>Miniopterus_natalensis
MRTMPAGPRGKGM
>Dasypus_novemcinctus
MRTMPAGPRGEGM
>Galeopterus_variegatus
MRTMPAGPRGEGM
>Condylura_cristata
MRTMPTGPRGEGM
>Equus_asinus
MRTMPAGPRREGM
>Cricetulus_griseus
MRTMPAGPCGEGM
>Tupaia_chinensis
MRTMPAGPRGEGM

HG0574 ENSG00000164663 USP49

>Homo_sapiens

MEEDGKNLFNGLRKWRLL

>Panthera_tigris

MEEDGKNLFNGLKDWSLL

>Pteropus_vampyrus

---MEKNLFNGLKEWRLL

>Dasypus_novemcinctus

MDEDGKKLFNGLTEWRLL

>Galeopterus_variegatus

MEEDGKNLFNGLRERQLL

>Oryctolagus_cuniculus

MEEDGKNLFNGLRAWQLL

>Elephantulus_edwardii

MEEDGKSLLNGLKEWRLL

>Ceratotherium_simum_simum

MEEDGKNLFNGLKEWRLP

>Manis_javanica

MEEDRKNLFNGLKEWRLL

>Psammomys_obesus

MEEDGNNLFNGLREWRLH

>Tupaia_chinensis

MEEDGKKLLNGVSEWRLL

>Trichechus_manatus_latirostr

---MEKTLFNGLKEWRLL

>Orycteropus_afer_afer

MEEEGKNLFNGLKEWRLL

HG0575 ENSG00000164663 USP49

>Homo_sapiens
--------------------------------------------------------------------MMAILWMGRDLERTSSLSCNSSQYPT------------
-
>Felis_catus
MRWGVREENRVKGSWRSPAGGRTGLACTGVGGSQMCMLWGDATEGLHGSGHRCCSEGPSVGSLWCRRGGTMILWMGRDLERTSSLSCNFSQYPT------------
-
>Pteropus_vampyrus
--------------------------------------------------------------------MMAILWMGRNLERTSSLSCNFSQYPT------------
-
>Dasypus_novemcinctus
--------------------------------------------------------------------MMAILWMGRDLERTSSLSATSVTIPHKSSQVTEKWKSN
Q
>Galeopterus_variegatus
--------------------------------------------------------------------------MGRDLERTSSLSCNFSHYPT------------
-
>Oryctolagus_cuniculus
--------------------------------------------------------------------------MERDLDRTSSLSCNFSQYLT------------
-
>Elephantulus_edwardii
--------------------------------------------------------------------MMAILWMGRDLERTSTLSCNVSQYPT------------
-
>Ceratotherium_simum_simum
--------------------------------------------------------------------------MGRDLERTSSLSCNFSQYPT------------
-
>Manis_javanica
--------------------------------------------------------------------------MGRDLERTSSLSCNFSQYPT------------
-
>Peromyscus_maniculatus_baird
--------------------------------------------------------------------------MGRDLERTLGLSCNFSQYPT------------
-
>Tupaia_chinensis
----------------------------------------------------------------------MILWMERGLGRMCSLPCSFSQYPT------------
-
>Trichechus_manatus_latirostr
--------------------------------------------------------------------MMAILWMGRDLERT----CSFSQYPT------------
-
>Orycteropus_afer_afer
--------------------------------------------------------------------MMANLWMGRDLERTSSLSCNFSHYPT------------
-

HG0576 ENSG00000175029 CTBP2

>Homo_sapiens
-------------------MKVYSLKKTDSPSCEL
>Ailuropoda_melanoleuca
-------------------MKLESLKKTDSPSCEL
>Lipotes_vexillifer
MPGSLALSGGFIALNGLEPMKLDSLKKTDSPSCEL
>Rhinolophus_ferrumequinum
-------------------MKLNSLQKTDNPSCDL
>Dasypus_novemcinctus
---MYSKINGFIALNGFEPMKLYSLKKTDSTSCEL
>Oryctolagus_cuniculus
-------------------MKLCSLRETDSPSCEL
>Elephantulus_edwardii
-------------------MTLYTLKQTDSSSCEL
>Ceratotherium_simum_simum
-------------------MKLDSLKKTDSPSCEL
>Loxodonta_africana
-------------------MKLHSLKKTDSPSCEL
>Chinchilla_lanigera
-------------------MKLYPLKKTDSPSCEP
>Tupaia_chinensis
-------------------MKLYSLKKTDSRLCEL
>Trichechus_manatus_latirostr
-------------------MKLRSLKKTDSPSSEL
>Orycteropus_afer_afer
-------------------MKLYSLKKTDSPSLEL

HG0577 ENSG00000250510 GPR162

>Homo_sapiens

MLSTGVSLGAACLLTGRGLWGSWECL

>Ursus_maritimus

MLSSRVGLGAACFTIGQGLYASWVCV

>Lipotes_vexillifer

MLSTRVGLGAACLTPGQGLWASWGCV

>Myotis_brandtii

MLSIRVGRGAACLTTGQGLWASWRCV

>Dasypus_novemcinctus

MLSSGAGQGAAHLTPGPGLWAAWGCA

>Galeopterus_variegatus

MLSSRVGLGAAHLTTGQGLWAPWGRV

>Oryctolagus_cuniculus

MLSSRAGLGPAGLSTGQAPWASWGCA

>Elephantulus_edwardii

MLSSRVGLEAACLTTGQGWWAPWGCV

>Equus_asinus

MLSTRVGLGAACLTTGQGLWASWGCV

>Manis_javanica

MLSSRIGLGAVCLTTGQGLWASWRCV

>Mus_musculus

MLRSKQDLGSACLATRQGLWASWGCV

>Trichechus_manatus_latirostr

MLSSGVGLGAACLTTGQGLWASWGCV

>Orycteropus_afer_afer

MLSSEGGLGAACLTTGQELWASWGCV

HG0578 ENSG00000004897 CDC27

>Homo_sapiens

MSRGWPGRSRYRGGLRHCRKWA

>Ailuropoda_melanoleuca

MSRGWPGRSRYGGGLRHCRKWA

>Orcinus_orca

MSRGWPGRSRYGGGLRHCRKWA

>Rousettus_aegyptiacus

MSRGWPGRSRYGGGLRHCRKWA

>Dasypus_novemcinctus

MSRDWPGRSRYGGGLRHCRKWA

>Condylura_cristata

MSRGWPGRSRYGGGLRHCRKWA

>Oryctolagus_cuniculus

MSRGWPGRSRYGGGA-------

>Equus_asinus

MSRGWPGRSRYGGGLRHCRKWA

>Ictidomys_tridecemlineatus

MSRGWPGRSRYGGGLRHCRKWA

>Tupaia_chinensis

MSRGWPGRSRYGGCLRHCRKWA

>Trichechus_manatus_latirostr

MSRGWPGWSRYGGGLRHCRKWA

>Orycteropus_afer_afer

MSRGWPSRSRYGGGA-------

HG0579 ENSG00000010818 HIVEP2

>Homo_sapiens

MPIDCGVRVEGSKEKEQSASLQTPWNMEATKL

>Felis_catus

--------------------------MEANKL

>Balaenoptera_acutorostrata_s

MPIDYGVGVEGSKEKEHSASLQTPWNMEASKL

>Myotis_lucifugus

--------------------------MEASKL

>Galeopterus_variegatus

MTIDYGVGVEGSREKEQSASLQTPWNMEATKL

>Ochotona_princeps

--------------------------MEAAKL

>Ceratotherium_simum_simum

--------------------------MEASKL

>Manis_javanica

MPRDYRVDVEGSKEQEHSASLQTPWTMETSRL

>Loxodonta_africana

--------------------------MEATKL

>Nannospalax_galili

MPIDYRVGVEGSKEKEQSANSQTSWNMEATQL

>Tupaia_chinensis

MPIDCGEGVGGSKEKEQSAGLQTPWNMEATKL

>Orycteropus_afer_afer

MPVDCGEGVEGSKEKEKS-SLQTSGNMEATKL

HG0580 ENSG00000073711 PPP2R3A

>Homo_sapiens
-------------------MCSSYCILIVTKPQI--------------------------
>Acinonyx_jubatus
-------------------MCSSYCILTVTKPQI--------------------------
>Balaenoptera_acutorostrata_s
-------------------MCNGYCILTVTKPQILMISLCGKGHYIYKKPLNIVIKYIFS
>Rhinolophus_ferrumequinum
-------------------MCNSYCILTVTKPQI--------------------------
>Galeopterus_variegatus
-------------------MCSIYCILTVTKPQI--------------------------
>Erinaceus_europaeus
MIKLKRSVVKLLYLLTQADLCSSCCALTVSKPQI--------------------------
>Equus_caballus
-------------------MCNSYCILTVNKPQI--------------------------
>Manis_javanica
-------------------MCNSYCILTVTKPQI--------------------------
>Mesocricetus_auratus
-------------------MCSSYCILIVTKPQI--------------------------
>Tupaia_chinensis
-------------------MCSNYCILIVTKPQI--------------------------
>Trichechus_manatus_latirostr
-------------------MCNSYCILTVTHPQI--------------------------
>Orycteropus_afer_afer
-------------------MCNSYCIMTVTQPQI--------------------------

HG0581 ENSG00000087074 PPP1R15A

>Homo_sapiens
----------------------------------------------------MNALASLTVR--TCDRFWQTEPALLPPG
>Canis_lupus_familiaris
----------------------------------------------------MNALASRAVL--SCDRWWQPEPALLPPG
>Orcinus_orca
----------------------------------------------------MNALASRAVR--TCDRLWQPQPALLPPG
>Rhinolophus_ferrumequinum
----------------------------------------------------MKALASRAVR--TCDRLWQPEPALLPPG
>Dasypus_novemcinctus
MDYKSLAVCVLAQRLFSSASLFFSVAPPDPARVCRHRACGSWRSPSPTPEREMKAPASRAAR--TCDRSWQPAPALLPPG
>Galeopterus_variegatus
----------------------------------------------------MNALASRAVR--TCDRFWQPEPALLPPG
>Erinaceus_europaeus
----------------------------------------------------MTAPAPRTVRSPTRDRLWQPEPALLPPG
>Oryctolagus_cuniculus
----------------------------------------------------MNALASRAVR--TYDRFWQPEPALLPPG
>Equus_caballus
----------------------------------------------------MNALASRAVR--TCDRLWKTEPALLPPG
>Loxodonta_africana
----------------------------------------------------MNAPDSRTAR--TYDRFWLREPALLSPG
>Ctenomys_sociabilis
----------------------------------------------------MNALASRTVR--TCDRFWQPEPALLPPG
>Tupaia_chinensis
----------------------------------------------------MNTLDSRAVR--TCDRFWKPEPALLSPG

HG0582 ENSG00000087338 GMCL1

>Homo_sapiens
METV-ASAATVGAAREKVAV
>Felis_catus
MEAV-ASAATVGAAREKVAV
>Physeter_catodon
MEAV-ASNTTVGAAREKVAV
>Rousettus_aegyptiacus
MEVV-ASAATVGAAREKVAV
>Dasypus_novemcinctus
MEAV-SWAATVGAAREKVAV
>Galeopterus_variegatus
MEAV-ASAETVGAAREKVAV
>Condylura_cristata
---MEVVAATVGAAREKVAA
>Oryctolagus_cuniculus
MEVV-ASAATVGAAREKVAV
>Ceratotherium_simum_simum
MEVV-ACAATVGAGREKVAV
>Loxodonta_africana
MEAV-SSAATVGAARETVAV
>Ictidomys_tridecemlineatus
MEAV-ASSSTVGAAREKVAV
>Orycteropus_afer_afer
MEAVSSFAATVGAARERVAV

HG0583 ENSG00000087502 ERGIC2

>Homo_sapiens

MAVGWDHNTSMTI

>Acinonyx_jubatus

MAVGWYHNTSMTI

>Orcinus_orca

MAVGWDRSTSMTI

>Myotis_brandtii

MAVGWDRHTSMTI

>Galeopterus_variegatus

MAVGWERNTSMTI

>Oryctolagus_cuniculus

MAVGWDRNTSMTI

>Elephantulus_edwardii

MAVGWDHNTSITI

>Ceratotherium_simum_simum

MAVGWDRNTSMTI

>Loxodonta_africana

MAVGWDSNRSITI

>Cricetulus_griseus

MAVGWDRSTSMTI

>Tupaia_chinensis

MAVGWDRSTSMTI

>Orycteropus_afer_afer

MAVGWDRNTSITI

HG0584 ENSG00000092051 JPH4

>Homo_sapiens
MDAVPGRQPEPQPT----
>Leptonychotes_weddellii
MDAVPGRQPEPQPTCHSG
>Balaenoptera_acutorostrata_s
MDAVPGRQPEPQSTCHSG
>Rousettus_aegyptiacus
MDAVPGRQPEPQPTCHSG
>Dasypus_novemcinctus
MDAVPGKQPEPQPTCHSG
>Galeopterus_variegatus
MDAVPERQPEPQTTCHSG
>Condylura_cristata
MDAVPGRQPEPQPTCHSG
>Oryctolagus_cuniculus
MDAEPGRRPEPQPTGHSG
>Elephantulus_edwardii
MDAVPGRQPEPQPTRHS-
>Ceratotherium_simum_simum
MDAVPGRQPEPQPTCHSG
>Dipodomys_ordii
MDAVPGRQPEPQPTCHSG
>Orycteropus_afer_afer
MDVILRRQPEPQPTCHSG

HG0585 ENSG00000106038 EVX1

>Homo_sapiens
MGLPPRGGAAQRSARFTR
>Odobenus_rosmarus_divergens
MGLPPRGGAAQRSARITR
>Lipotes_vexillifer
MGLPPRGGAAQRSARITR
>Miniopterus_natalensis
MGLPPRGGAAQRSGRITR
>Dasypus_novemcinctus
MGLPPRGGAAQRNARITR
>Galeopterus_variegatus
MGLPPRGGAAQRSARITR
>Sorex_araneus
MGLPPRGGAAQRSARITR
>Oryctolagus_cuniculus
MGLPPRGGAAQRRARITR
>Elephantulus_edwardii
MGLPPRGGAAQSSTHITR
>Ceratotherium_simum_simum
MGLPPRGGAAQRSARITR
>Mus_musculus
MGLPPRGGAAQRSARITR
>Orycteropus_afer_afer
MGLPPRGGAAQRSARITR

HG0586.1 ENSG00000108375 RNF43

>Homo_sapiens
----------------------MGENDIPAADVG
>Acinonyx_jubatus
----------------------MGENDIPAAEVG
>Balaenoptera_acutorostrata_s
----------------------MGENDIPEAEVG
>Myotis_lucifugus
----------------------MGENDIPAAEVG
>Erinaceus_europaeus
----------------------MGENDIPAAEVG
>Oryctolagus_cuniculus
----------------------MGENDIPAAEVG
>Elephantulus_edwardii
----------------------MGENNIPAAEVG
>Ceratotherium_simum_simum
----------------------MGENDIPAAEVG
>Manis_javanica
----------------------MGENDIPAAEVG
>Loxodonta_africana
----------------------MGENDIPATEVG
>Heterocephalus_glaber
MFNLKQSWKFQREVVLLLVRVKMGENDIPAAEVG
>Tupaia_chinensis
----------------------MGENDIPVAEVG

HG0586.2 ENSG00000108375 RNF43

>Homo_sapiens
MLFGEKKNFC
>Canis_lupus_familiaris
MLFGEKKNFC
>Miniopterus_natalensis
MLFGEKKNFC
>Condylura_cristata
MLFGEKKNFC
>Equus_asinus
MLFGEKKNFC
>Tupaia_chinensis
MLFGEKKNFC

HG0587 ENSG00000119138 KLF9

>Homo_sapiens

MPERCGPAGAGVQKPSVDSGKSVS--

>Odobenus_rosmarus_divergens

MPERCGPAKAGAQKASVDWGKSVS--

>Orcinus_orca

MPERCGPARAGAQKPSVDWGKSVS--

>Rousettus_aegyptiacus

MPERCGPARAGAQKPSGDWGKSVS--

>Dasypus_novemcinctus

MPERCGPAGAGAQKPSMDWGQSVS--

>Condylura_cristata

MPERCGPPGAGAQKLSVDWGESVY--

>Equus_asinus

MPERCGPAREGAPKPSADWGKSIS--

>Manis_javanica

MPERCGPARAGAQKPSTDWGKSAA--

>Loxodonta_africana

MPERCGPAGAGAQKPSVDWGKSVSSN

>Marmota_marmota_marmota

MPERCGPARAGAQKPSVDWGKSVS--

>Trichechus_manatus_latirostr

MPERCGPAGAGAQKPSVDWGKSVS--

>Orycteropus_afer_afer

MPERCGPADTGAQKPSVD--------

HG0588 ENSG00000120162 MOB3B

>Homo_sapiens
-------------------------------MRFPWKSFKRKMEGAV
>Felis_catus
-------------------------------MRFSWKLFKRKMEGAV
>Lipotes_vexillifer
-------------------------------MRFPWKSFKRKMEGAV
>Myotis_davidii
-------------------------------MRFPWKSLERKMEGAI
>Dasypus_novemcinctus
-------------------------------MRFPWKSLKRKMEGAV
>Galeopterus_variegatus
-------------------------------MRFPWKSFKRKMEGAI
>Condylura_cristata
-------------------------------MRFSWKLFKRKMEGAA
>Ochotona_princeps
-------------------------------MRLPWKSFKRKMEGAV
>Ceratotherium_simum_simum
-------------------------------MRFPWKLLKRKMEGAL
>Manis_javanica
-------------------------------MRSPWKLFKRKMEGAI
>Mesocricetus_auratus
MRNSRFREVKRRVRRTPHHSLAATYKQMGRNMRFPWRSFKRKMEGAI
>Tupaia_chinensis
-------------------------------MRFPWKSFKRKMEGTI

HG0589.1 ENSG00000121871 SLITRK3

>Homo_sapiens
------MMDTIQLRREFAPPAG---------------------------------
>Mustela_putorius_furo
------MLDTIQPPREFAPPAGQPLGADLGDLERSAPIVL---------------
>Balaenoptera_acutorostrata_s
------MMDTIQPPREFAPLAGKPLGVNLGDREPSARIVL---------------
>Pteropus_alecto
------MMDTIEPSREFAAPAGKTLGVNLGDRECSARIVL---------------
>Galeopterus_variegatus
MLQGYSMMDTIQPRREFAPPAGKPLGANLGDRERSAPIVL---------------
>Condylura_cristata
-MQAYDTMDTIQPPREFAHPVGKPLGANLGDRDHSAP------------------
>Oryctolagus_cuniculus
------MVDTIQPPREFAPPAGKPLAANLGDGERSAPLVL---------------
>Equus_caballus
-------MDTIQPPREFAPPAGKPLGANLGDRERSAPIVL---------------
>Manis_javanica
------MMDTIHSLRVFAAPAGKPFGANLGDLERSARIAV---------------
>Nannospalax_galili
------MTDTIQPRGEFAPPAGKPLGANLGDRERSAPIVL---------------
>Tupaia_chinensis
-------MDTIQPPREFAPPAGKPLGANLGDCERSAPIGTRRHRFLLRLIEISVF
>Orycteropus_afer_afer
------MMDTIQPLREFIPPAGKPLGANLGDRERSEAIVL---------------

HG0589.2 ENSG00000121871 SLITRK3

>Homo_sapiens
MQRPFPCKSQRWKHLNGLQA------
>Mustela_putorius_furo
MQRPFPFKSQRWKHLKGLQA------
>Lipotes_vexillifer
MQRPFPCKSQRWKHLKGLQA------
>Rousettus_aegyptiacus
MQRPFLCKSQRWKHLEGLQA------
>Galeopterus_variegatus
MQRPFPRKSQRCKYLKGLQA------
>Condylura_cristata
MRRPFLCKSQRWETLKRITGEIWWPF
>Equus_asinus
MQRPFACKSQRWKHLKGLQA------
>Manis_javanica
MQRPFRCKSRRWKHLKGLQA------
>Ictidomys_tridecemlineatus
MQRPFPCKSQRWKYLKGFRL------
>Tupaia_chinensis
MQRPFPCKSPRWKH------------
>Orycteropus_afer_afer
MQLPFPCKSPRWKHLKGLQA------

HG0589.3 ENSG00000121871 SLITRK3

>Homo_sapiens
----------------------------------------------------MDLHSGSSPAFPILTPYTRPGAPV-------
>Rhinolophus_ferrumequinum
----------------------------------------------------MDLHSGSSPAFSMLPPNTRLGVPSVGADVVL
>Galeopterus_variegatus
MRGDRTAAADLHSWRGGGAGGAEGTSWRRLWRWADLGAPERRSLGAPIWGWLMDLHSGSSPAFPILSANTRPGAPSLGADVVL
>Condylura_cristata
----------------------------------------------------MDLHSDSCLAFRILPPSTQPALPV-------
>Equus_asinus
----------------------------------------------------MALHSGSSPAFPILPPSTRPGAPCVRADVVL
>Manis_javanica
----------------------------------------------------MDLHSGSSPAFRILPPNSRPGAPSLGADVAV
>Loxodonta_africana
----------------------------------------------------MDL-LGSSPAFGYWLPNTRPALPV-------
>Marmota_marmota_marmota
----------------------------------------------------MDLHLCSSPAFPILPPNTRPGAPSLGADVVL
>Tupaia_chinensis
----------------------------------------------------MDLHSGSSPALPTSPPNTRPGSRSFGADVVL

HG0590 ENSG00000122557 HERPUD2

>Homo_sapiens
----------------------------------------------------------------------------------------------------------
---------------------------------------------------------------MGFVSSSALVTDLDTEY
>Felis_catus
----------------------------------------------------------------------------------------------------------
---------------------------------------------------------------MAFVSPDALVTDLDTEY
>Balaenoptera_acutorostrata_s
----------------------------------------------------------------------------------------------------------
---------------------------------------------------------------MAFVSPDALVTDLDKEY
>Rhinolophus_ferrumequinum
----------------------------------------------------------------------------------------------------------
---------------------------------------------------------------MAFVSSDALVTDLDTEY
>Condylura_cristata
----------------------------------------------------------------------------------------------------------
---------------------------------------------------------------MAFLSPDALVTDLDTE-
>Oryctolagus_cuniculus
----------------------------------------------------------------------------------------------------------
---------------------------------------------------------------MAFVSPDALVTDLNTEY
>Elephantulus_edwardii
----------------------------------------------------------------------------------------------------------
---------------------------------------------------------------MAFVSSDALVTDLDSRF
>Equus_asinus
MTHFRPALPGLKGTQTFRTTTTKPARSRPAPRLWSPPSCHLPPPGPRPACAPLPCPRGQAESGPHPSANFSRSQRSASRDPGPGCGRHRAAGRERPPRKTFGRRPC
PGVKGGGGGPHGLLSVRSPAEPGPGGENCTHGRAPRPRRRRRRRHDHHHHHHYRHCPSSLVPAMAFVSPDALVTDVDTEY
>Loxodonta_africana
----------------------------------------------------------------------------------------------------------
---------------------------------------------------------------MAFVSPDALVTDLDTKY
>Chinchilla_lanigera
----------------------------------------------------------------------------------------------------------
---------------------------------------------------------------MAFVSPDALVTDLDTEY
>Trichechus_manatus_latirostr
----------------------------------------------------------------------------------------------------------
---------------------------------------------------------------MAFVSPDALVTDLDTKY
>Orycteropus_afer_afer
----------------------------------------------------------------------------------------------------------
---------------------------------------------------------------MAFVSPEALVIVLDTKY

HG0591.1 ENSG00000127124 HIVEP3

>Homo_sapiens
MNAGFQREQRFSFGHRKWCLQHRRRA------
>Leptonychotes_weddellii
MKAGFQREQCFSFGHRKWCLQHRRRA------
>Physeter_catodon
MNADFQREQRFSFGHRKWCLQHRRA-------
>Myotis_davidii
MNAGFQREQRFSFGHRKWCLQHRSA-------
>Galeopterus_variegatus
MNAGFQREQRLSFGHRKWCLQHRRA-------
>Condylura_cristata
MNAGFQREQRFSFGHRKWCLQHRRA-------
>Oryctolagus_cuniculus
MNAGFQREQRFSFGHRKWCLQHRRVSLLIQDC
>Equus_asinus
MNAGFQREQRFSFGHRKWCLQHRRA-------
>Manis_javanica
MNAGFPREQRFSFGHRKWCLQRRRA-------
>Ictidomys_tridecemlineatus
MNAGFQREQRFSFGHRKWCLQHRRRV------
>Tupaia_chinensis
MSAGFQREQRFSFGHRKWCLQHRRV-------
>Orycteropus_afer_afer
MNAGFQREERFSFGHRKWCLQHRRRA------

HG0591.2 ENSG00000127124 HIVEP3

>Homo_sapiens
MWPKLDTSLGHWDYS
>Acinonyx_jubatus
MWPKPDASLGHLDYS
>Balaenoptera_acutorostrata_s
MWPKPDASLGRLDYS
>Pteropus_alecto
MWPKPDSSLGHLDYS
>Galeopterus_variegatus
MWPKLDTSLGHLDYS
>Condylura_cristata
MWPQPDASLGHLDYS
>Equus_asinus
MWPKADASLGRLDYS
>Manis_javanica
MWPKPEASLGRLDYS
>Fukomys_darlingi
MWPKLDNSLGHEDYS
>Tupaia_chinensis
MWPKLDTSSGHLDYS
>Orycteropus_afer_afer
MWPKPDTSLGHSDSP

HG0591.3 ENSG00000127124 HIVEP3

>Homo_sapiens

MEEPRGVLAKQARPSGGKSGA---------------------

>Canis_lupus_familiaris

MAELQEILARQALPSGGKSRA---------------------

>Pteropus_alecto

MEEPQEILARQALPSKGKSRE---------------------

>Oryctolagus_cuniculus

MEEPPELPARQAVPSRGKSGVEEDGGCSRSGHERWFPERTAF

>Ceratotherium_simum_simum

MEEPREILARQALPSRGKSRA---------------------

>Ictidomys_tridecemlineatus

MEQPQETLARQALPSRGQSGA---------------------

>Orycteropus_afer_afer

MEDPQEILAGQALPSRGKSCV---------------------

HG0591.4 ENSG00000127124 HIVEP3

>Homo_sapiens
MDTDVSNLQPFA
>Acinonyx_jubatus
MHTDVSSLQPFA
>Galeopterus_variegatus
MDTDASNLQPFA
>Cavia_porcellus
MDTEVSNFQPFA

HG0592 ENSG00000133401 PDZD2

>Homo_sapiens
MNTGKADDGQGPQGTWGPVGSGPQEQDL-
>Mustela_putorius_furo
MNTGKADDSQGSQGTWGLPGSSPKEQVLS
>Balaenoptera_acutorostrata_s
MNTGKAEDSPRSRGTWGPAGPSPQEQVLS
>Myotis_davidii
MNMNTADDSQGPQGTWGPAGSGPQEQVLS
>Dasypus_novemcinctus
MSTGQADGSQGPPGTWGP-----QERVLS
>Erinaceus_europaeus
MNLGHVDDSQGPQGMWGLVGPSPPKQALS
>Oryctolagus_cuniculus
MNTCKAHDNQASQGTWGPAGPGPQERVLS
>Ceratotherium_simum_simum
MNTGKADDSQGSQGTWGPLGSGPQEQVVS
>Manis_javanica
MHTDGASGSQGSHGPWGPLGSSSQEQPVP
>Fukomys_darlingi
MNTGRADDHQDSQGTWGPAGSGPQEQDLP
>Tupaia_chinensis
MNTGKADDSQCSQGSRDPVGSGPQKQDLS
>Trichechus_manatus_latirostr
MNIGQADDSQGSQGTWGPMGPGPQEQALS

HG0593 ENSG00000135048 TMEM2

>Homo_sapiens
MERRSGPEDARDSSPSPWTAV
>Felis_catus
MERRSGPGDARSSAPSLWTAV
>Balaenoptera_acutorostrata_s
MEGRSRPGDARSSALSLWTAV
>Myotis_brandtii
MERRAGPGDAQNSAPSLWTSV
>Dasypus_novemcinctus
MERRSELGDARSSAPSLGNAV
>Condylura_cristata
MERRSGPADARSSAPAFWIAV
>Elephantulus_edwardii
MERGSGPRVARNPALSLWTSV
>Ceratotherium_simum_simum
MEGRSGPGDARSSAPSLWTAV
>Manis_javanica
MERGSGPGDTRSSAWSLWTPV
>Loxodonta_africana
MERGSGPGDARSSALSLWISV
>Marmota_marmota_marmota
MERRSGPGDARSSTQSLWIIA
>Trichechus_manatus_latirostr
MERGFGPGDARSSALSLWISV

HG0594 ENSG00000136169 SETDB2

>Homo_sapiens
-------------------------------------------------------MFHPFVDQKMELVFIFKKIMLMI----
>Canis_lupus_familiaris
MGSWPSSGIGGRPAVPRRIGVSREPEAVKGAAGRGLRRNNPEAQLGGVAADPCLRMFHASMDQKMELVFIFKKILSMI----
>Physeter_catodon
-------------------------------------------------------MFHAFMDRKMELVFIFKMILSMI----
>Pteropus_alecto
-------------------------------------------------------MFHAFMDQKMELVFIFKKILSVI----
>Galeopterus_variegatus
----------------------------------------------------------------MELVFIF-KDTVNDLIQL
>Condylura_cristata
------------------------------------------------------------MDQKMALVFIF-KDSVSHLVQL
>Oryctolagus_cuniculus
----------------------------------------------------------------MELVYIFKKKLSVL----
>Equus_asinus
------------------------------------------------------------MDQKMKLVFIFKKILSMN----
>Loxodonta_africana
-------------------------------------------------------MFYVFVDQKMKSFFYFKKDTVNDLIEL
>Ictidomys_tridecemlineatus
---------------------------------------------------------MYFEDQKMELVFIFKKILSMI----
>Tupaia_chinensis
-------------------------------------------------------MFHAFVDQKMELVFIFKKLLSVI----
>Trichechus_manatus_latirostr
-------------------------------------------------------MFYAFVDQKMELFSFLKKILSMI----

HG0595 ENSG00000141098 GFOD2

>Homo_sapiens
MPACQEPCLTRTFGRSYPCQRLL
>Felis_catus
MPACQEPCLTRTVGRSYSCWRLL
>Lipotes_vexillifer
MPACQEPCVTRTVGWSYSCWRLL
>Myotis_lucifugus
MPACQEPCLTRTVGWSSSCWRLL
>Dasypus_novemcinctus
------------MGRSYPCWRLL
>Erinaceus_europaeus
MPECQEPCLTRTVGTSYSCWKPL
>Oryctolagus_cuniculus
MPACQEPCLTMPAGISDPCWRHL
>Ceratotherium_simum_simum
MPACQEPCLTRTVGRSYSCWRLL
>Manis_javanica
------------MGRSYSCWRLL
>Loxodonta_africana
MPACQEPCLTRTLGRWDPHQRLL
>Heterocephalus_glaber
MPACQEPCLTRTVGRSAPCWRPL
>Tupaia_chinensis
MPACQEPCLTRTMGNPFPCWRCS

HG0596 ENSG00000157483 MYO1E

>Homo_sapiens
MESPLGSHWLL
>Felis_catus
MESPQGSHWLL
>Orcinus_orca
MESPQGSHWFL
>Rhinolophus_ferrumequinum
MECPQGSHWLL
>Dasypus_novemcinctus
MESRLGSHWLL
>Galeopterus_variegatus
MESPLGSHWLL
>Condylura_cristata
MECPLGSHWLL
>Oryctolagus_cuniculus
MQSPLGSHWLL
>Equus_asinus
MESPQGSHWLL
>Ictidomys_tridecemlineatus
MESPLGSHWLL
>Tupaia_chinensis
MESPLGSHWLL
>Orycteropus_afer_afer
MESPLGSHWLL

HG0597 ENSG00000163032 VSNL1

>Homo_sapiens
MGFPICSRDLPERVAAAAGLARRDPRGI
>Mustela_putorius_furo
MGFPICSRDLPELVAAAAGLARRDPRGI
>Balaenoptera_acutorostrata_s
MGFPICSRDLPERAAATAGLARRDPRGI
>Pteropus_alecto
MGFPICSRDLPERVAAAAGLKRRDPKGI
>Condylura_cristata
MGFPIRSRDLPERVAAAAGLARRDPRGI
>Oryctolagus_cuniculus
MGFPICSRDLPERVAATAGLARRDPTGI
>Equus_caballus
MGFPICSRDLPERVAAAAGLARRDPGGI
>Manis_javanica
MGFPICSRDLPEIVAAVAGLARRDPKGI
>Loxodonta_africana
MGFPICSRDLSERVAAAAVVARRDPRGI
>Psammomys_obesus
MGFPICSRDSPERVAAAAGLARRDPRGI
>Tupaia_chinensis
MGFPICSRDLPERVAAATGLARRDPRGI
>Trichechus_manatus_latirostr
MGFPICGRDLPERVAAAAGLARRDPRGI

HG0598 ENSG00000163788 SNRK

>Homo_sapiens
--------------------------------------------------------------------MTTLKMNFLYSPDILI
>Odobenus_rosmarus_divergens
--------------------------------------------------------------------MATLKMNSLFSPDILI
>Lipotes_vexillifer
------------------------------------------------------------------MSMATLKMNSLFSPDI--
>Miniopterus_natalensis
------------------------------------------------------------------MPMATLKMNSLFSPDILV
>Dasypus_novemcinctus
------------------------------------------------------------------MSMATLKMNSLFSPDSLI
>Galeopterus_variegatus
------------------------------------------------------------------MSMASLKMNSLFSPDILV
>Condylura_cristata
MISTGCGESAAGRESSAPRPLRTDDRLAAENLLRFCFGTSSTWPLLRRKTTSKTVITRTIQHAVCRMSMATLKMNSLSLPDILI
>Equus_caballus
------------------------------------------------------------------MSMVTLKMNSLFSPDILI
>Loxodonta_africana
------------------------------------------------------------------MSMATLKMNSLFSPDILI
>Ictidomys_tridecemlineatus
------------------------------------------------------------------MSVATLKMNSLFSADILI
>Trichechus_manatus_latirostr
------------------------------------------------------------------MSMVALKMNSLFSPDTLI
>Orycteropus_afer_afer
------------------------------------------------------------------MSMATLKMNSLFSPDILI

HG0599 ENSG00000164930 FZD6

>Homo_sapiens
MRNLNILSLDGDLLRMQRVIHPSHVVKSGI
>Odobenus_rosmarus_divergens
---------------MKRVINPSHVVKPGI
>Balaenoptera_acutorostrata_s
---------------MKRAINPSHVVKLGI
>Rousettus_aegyptiacus
MRNSNILSLDGN-LRMKRVINSSHVVKTGI
>Condylura_cristata
---------------MKRAINSSHVVKAGI
>Oryctolagus_cuniculus
MRTLNILPLAGNPLRMKRAVNLRHVVISGI
>Equus_asinus
MRNLNILSLDGN-LRMKSAITPSHVVKSGI
>Loxodonta_africana
MRNLNISYLDGNLLRMRRLIQAMW------
>Dipodomys_ordii
---------------MNGVVNPSHVVKPEI
>Tupaia_chinensis
MRNLNVLFLDGNLLRMKRATNPSHVVKSGI
>Trichechus_manatus_latirostr
MRNLNILYLDGSLLRT--------------
>Orycteropus_afer_afer
---------------MKRAINLGHVVKSGI

HG0600 ENSG00000173065 FAM222B

>Homo_sapiens

MRAGPSHRPVSHCPMLC

>Felis_catus

MRAGPSHRHVSDCLMLC

>Rousettus_aegyptiacus

MRAGPSHRHVSDCPMLC

>Dasypus_novemcinctus

MRAGPSHRHVSDCLMLC

>Condylura_cristata

MRAGPSHRHVSGCPMLC

>Ochotona_princeps

MRAGPSQRHVSHCPMLC

>Elephantulus_edwardii

MRAGPSHRHVSDCLMLC

>Equus_asinus

MRAGPSHRHVSDCPMLC

>Manis_javanica

MRAGPSHRHVNGCPMLC

>Nannospalax_galili

MRAGPSHRHVSHCPMLC

>Tupaia_chinensis

MRAGPSHRHVSHCPMLC

>Trichechus_manatus_latirostr

MRAGPSYRHVSDCPMLC

HG0601 ENSG00000175893 ZDHHC21

>Homo_sapiens

MNQKKISKCRFQKNYSTSQ

>Canis_lupus_familiaris

MNQDKTSKCKYQKNYSTSW

>Physeter_catodon

MNQNKSSKCKYQKNYNTSQ

>Eptesicus_fuscus

MNQDKTSKCKYQKNYNTSQ

>Dasypus_novemcinctus

MNQDKPSKCKYQKNYNTSQ

>Galeopterus_variegatus

MNQEKTSKCKYQKNYNTSQ

>Condylura_cristata

MNQDTTSKCKYQRNCHTSQ

>Oryctolagus_cuniculus

MNQDKTFKCKYQKNYNTFP

>Ceratotherium_simum_simum

MNQDKTSKCKYQKNYNTSQ

>Manis_javanica

MNQDKTSKCKYQKNYNTSQ

>Marmota_marmota_marmota

MNQDKTSRCKYQKNYNTSQ

>Tupaia_chinensis

MNQGKTSKCKYQKKYSTSQ

HG0602 ENSG00000179195 ZNF664

>Homo_sapiens
MQEETFRRKTGREKLRLTLNLPNRASLR
>Panthera_tigris
MQEETFHRKTGREKLRPTIALPNRASLR
>Lipotes_vexillifer
MQGGTFHRKTGRGELRPASNLPSRASLR
>Miniopterus_natalensis
MQEEAFHRKTGRAKLRLTLSLPSRVSLR
>Dasypus_novemcinctus
MQEETFHRKTGREKLRPTLNLPNRASLR
>Erinaceus_europaeus
MQEETFHRKTGREKLRLTFNLPNRASLR
>Oryctolagus_cuniculus
MQEEAFRRNAGRETLRLAARPPSGASPR
>Ceratotherium_simum_simum
MQEETFHRKTGREKLRLTLNLRIRASLR
>Manis_javanica
MQEEIFHRKSGREKLRLTVKLPNRVSLR
>Loxodonta_africana
MQEEAFHRKIGREKLRPTLNLLTRASLR
>Heterocephalus_glaber
MQEEAFHRKTGREKLRLTLNLPHRASLR
>Tupaia_chinensis
MQEETFHRKTGREKLRLSLNLPNRASLR

HG0603 ENSG00000197329 PELI1

>Homo_sapiens
MSVTQRRCEAEGPSGGAAAAASTSSSGSSPSPAVRSNWSPALRERTTKQPQRLSLRPRLLSARLRGQLPLAGLGWRARRAAAE-----RRAVC----SRSPRTTLHP
LASGLGGRSGIPATSNCPEIMQKVSQGS--
>Mustela_putorius_furo
-----------------------------------------------------------------------------------------------------------
--------------------MQKVSQGS--
>Lipotes_vexillifer
-----------------------------------------------------------------------------------------------------------
--------------------MQKVSQGS--
>Rousettus_aegyptiacus
-----------------------------------------------------------------------------------------------------------
--------------------MQKVSKAPDQ
>Dasypus_novemcinctus
-----------------------------------------------------------------------------------------------------------
--------------------MQKVSPSS--
>Condylura_cristata
-----------------------------------------------------------------------------------------------------------
--------------------MQKVSQGS--
>Ochotona_princeps
MSVTQRQCEAEGPSGGAAAAASTSS--SSSGAARAELSRALRERSTKRAPRLALGPRLLSAGLLGQLPPQRA-WLLPGEAGGGGGGGTPGCLLARPPPHPRAHSLHP
LAAGLGSRAGILATSDCPEIMQKVSQGS--
>Ceratotherium_simum_simum
-----------------------------------------------------------------------------------------------------------
--------------------MQKVSQGS--
>Loxodonta_africana
-----------------------------------------------------------------------------------------------------------
--------------------MQKVSQAS--
>Ictidomys_tridecemlineatus
-----------------------------------------------------------------------------------------------------------
--------------------MQKVSQGS--
>Trichechus_manatus_latirostr
-----------------------------------------------------------------------------------------------------------
--------------------MQKVSQAS--
>Orycteropus_afer_afer
-----------------------------------------------------------------------------------------------------------
--------------------MQKVSQAS--

HG0604.1 ENSG00000003147 ICA1

>Homo_sapiens
-----------------------------------MLFSCPFSPNHQQ
>Acinonyx_jubatus
-----------------------------------MLFPCPFSPNHQQ
>Tursiops_truncatus
-----------------------------------MLFPCPFSPNHQQ
>Myotis_davidii
-----------------------------------MLFPCPFSPNHQQ
>Galeopterus_variegatus
MGVEKRPQGVERRRLTLPTT--TRLKQGYNITYPLMLLLYPFSPNHQQ
>Condylura_cristata
-----------------------------------MLFPCPFSPNHQQ
>Elephantulus_edwardii
-----------------------------------MLFLCPFFPNHQL
>Equus_asinus
-----------------------------------MLFPCPFSPNHQQ
>Dipodomys_ordii
-----------------------------------MCFPCPFSPNHQQ
>Tupaia_chinensis
-----------------------------------MLFLCPFSPNHQQ
>Trichechus_manatus_latirostr
MGVKERPQGVERGRLIPPTPVPPHPKEGCNITYLLMLFPCPFSPNHQQ

HG0604.2 ENSG00000003147 ICA1

>Homo_sapiens
MSDPEGRSDRGREAPGGRGSEVII
>Odobenus_rosmarus_divergens
MKELEGRSDRGQEAPGGRGSEVII
>Tursiops_truncatus
MKELEGRSDRGREAPRGRGSEVII
>Rousettus_aegyptiacus
MRELEGRSDRGREAPGGRGSEVII
>Condylura_cristata
MKELEGRSDRGREAPGGRGSEVII
>Ceratotherium_simum_simum
MRELEGRSDRGREAPGGRGSEVII
>Ictidomys_tridecemlineatus
MWDLEGHSDRGREAPGGQGSEV--

HG0604.3 ENSG00000003147 ICA1

>Homo_sapiens

MATWNTCRRAGAGSLWAADPSGGYPRRWGSRRDPGE

>Galeopterus_variegatus

MDTWNTCGRAGAGSPRAADPSKGYPKRWGSRKDPRE

>Ceratotherium_simum_simum

MDTWNTCGRAGRGHPGRQMRAGGYPRRWGSRKDPRE

>Mus_musculus

MDTWNTC-----------------------------

>Orycteropus_afer_afer

MDTWNTCRKAGAGSPWTAGPRGGYPRRWGSRKDPRE

HG0605 ENSG00000053254 FOXN3

>Homo_sapiens
MKKNQPLEDQHSRDYGGPSVFSVTGAMLTSVSWTHSDSGAFT
>Felis_catus
--------------------------MLTSVSWTHSDSGAFT
>Physeter_catodon
--------------------------MLTSVSWTHSDSGAFT
>Miniopterus_natalensis
--------------------------MLTSVSWTHSDSGAFT
>Condylura_cristata
--------------------------MLTSVSWTHSDSGAFT
>Oryctolagus_cuniculus
--------------------------MRTSVSWTHCDSGAFT
>Ceratotherium_simum_simum
--------------------------MLTSVSWTHSDSGAFT
>Manis_javanica
--------------------------MLTSVSWTHSDSGAFT
>Loxodonta_africana
--------------------------MLTSVSWTHSDSGAFT
>Nannospalax_galili
--------------------------MLTSISWTHSDSGAFT
>Tupaia_chinensis
--------------------------MLTSVSWTHSNSGAFT

HG0606 ENSG00000055130 CUL1

>Homo_sapiens

MRFHSLHLKDILSELL

>Odobenus_rosmarus_divergens

MRFHSLHLKDILSELL

>Balaenoptera_acutorostrata_s

MRFHSLPLKDVLSELL

>Cynopterus_sphinx

MKVHSLQLKDILSELL

>Dasypus_novemcinctus

MRSHSLHLKDILSELL

>Galeopterus_variegatus

MRFHFLHLKDILF---

>Ceratotherium_simum_simum

MRFHSLHLKDILSELL

>Manis_javanica

MRFHSLHLKDILSALL

>Heterocephalus_glaber

MRFHSLHLKDILSELL

>Tupaia_chinensis

MRFASLRLKDILSELL

>Trichechus_manatus_latirostr

MRAPSLHLKDTASELL

HG0607 ENSG00000074657 ZNF532

>Homo_sapiens
MRFQRPPCPLSHWPCLRPPPPPQTAPVFFPRGEGIWTSPPRCLKEEKGMFCLVLFLLEATKGCWGCF---------
>Odobenus_rosmarus_divergens
------------------------------------------------MFCFVLFLLEATKGCWGYF---------
>Orcinus_orca
------------------------------------------------MFCFVLFLLEATKGFWGCF---------
>Miniopterus_natalensis
------------------------------------------------MFCFVLFLLEATKGCWGYF---------
>Dasypus_novemcinctus
------------------------------------------------MFCFVLFLLEATKGCWGYV---------
>Galeopterus_variegatus
------------------------------------MAAKSRGACRKAMFCFVLFLLEATKGCWGYF---------
>Condylura_cristata
------------------------------------------------MFCFVLFLVEATKGVGDTSDCYEGRKAC
>Oryctolagus_cuniculus
---------MNWWVCDGS---------LFVAGECYFACLSSL---AGCVLFCAFFLLEATKGCWGYF---------
>Ceratotherium_simum_simum
------------------------------------------------MFCFVLFLLEATKGCWGYF---------
>Ictidomys_tridecemlineatus
------------------------------------------------MFCFVLFLLEATKGCWGYF---------
>Orycteropus_afer_afer
------------------------------------------------MFYFVFFLLEAAKGCWGDF---------

HG0608 ENSG00000084731 KIF3C

>Homo_sapiens

MNDPPAWGEDSR

>Leptonychotes_weddellii

MNDPPARGEDSR

>Orcinus_orca

MNDPPAGGEDSR

>Rousettus_aegyptiacus

MNDPPAPGEDSR

>Galeopterus_variegatus

MNDPPSVGEDSR

>Condylura_cristata

MNDPPAGGADSR

>Elephantulus_edwardii

MNDPPAGGDDSR

>Loxodonta_africana

MNDPPAEGEDSR

>Peromyscus_maniculatus_baird

MNDPPAGGEDSR

>Trichechus_manatus_latirostr

MNDPPASGEDSR

>Orycteropus_afer_afer

MNDPPAWGEDSR

HG0609 ENSG00000095539 SEMA4G

>Homo_sapiens

----MATQPCDSMSPRFQDPPWPHDSLTPMTL

>Ailuropoda_melanoleuca

----MATQPCDSMSPRFQDPPWPHDP------

>Physeter_catodon

----MATQPCDSMSPQFQDPPWPHDS------

>Pteropus_alecto

----MATQSCDSMSPQFQDPPWPHDA------

>Galeopterus_variegatus

----MATQPCDSMSPQFQDPPWPHDSLAPMAL

>Oryctolagus_cuniculus

----MATQPCDSMSPPFQDPPWPHDSLTPMAS

>Ceratotherium_simum_simum

----MATQPCDSMSPQFQDPPWPHDP------

>Manis_javanica

MWDSMATQPCDSMSPRFQDPPWPHDL------

>Loxodonta_africana

----MATQPCDPMSPRFQDPPWPHDSVTPMPS

>Rattus_norvegicus

----MATQPCGSMSPPFQDPPWPHDYLTPMTS

>Tupaia_chinensis

----MAIQPCDSMSPRFQDPPWPHDSLTPTAL

HG0610 ENSG00000106069 CHN2

>Homo_sapiens
MGWGPRRLRAAGRGQRRRRPHRG
>Odobenus_rosmarus_divergens
MGWGPRRLRAVARGQLRRRPHRG
>Orcinus_orca
MGWGPRRLRAAGSGQRRRRPHRG
>Myotis_lucifugus
MGWGPRQPGAAGRGQRRLRPHRG
>Dasypus_novemcinctus
MGWGPRRLRAAGRGQRRRRPHRG
>Sorex_araneus
MGWGPRRLRAAGPGQRRRRPLRG
>Ochotona_princeps
MGWGPRRLRAAGPGQRRQRLQRG
>Elephantulus_edwardii
MGWGPRRLRAAGPGQRRPA-HRG
>Equus_asinus
MGWGPRRLRAAGRGQRRRRPHRG
>Loxodonta_africana
MGWGPRRLRAAGRGQLRRRPHWG
>Ictidomys_tridecemlineatus
MGWGPRRLRAAGQGQRQWRPHRG

HG0611 ENSG00000108309 RUNDC3A

>Homo_sapiens

MAMEGLRGPVGQRAQP

>Odobenus_rosmarus_divergens

MAVQASRGPGGQRAQP

>Orcinus_orca

MALEGSRGPGGQRAQP

>Eptesicus_fuscus

MAVEGSRGPDGQKAQP

>Galeopterus_variegatus

MAAEGSRRPGGQRAQP

>Sorex_araneus

MAAEGSRGPGGQRARP

>Ochotona_princeps

MALEGSRGSGGQSAQP

>Elephantulus_edwardii

MAAEGSRGPGGRRAQP

>Equus_asinus

MAVEGSRGPGGQRAQP

>Manis_javanica

MALEGSRGPGGQRAQR

>Ictidomys_tridecemlineatus

MAMERSMGPGGQRAQP

HG0612.1 ENSG00000110422 HIPK3

>Homo_sapiens
-----------------MAASRALWPRPQRRGAAGLQGVNSGDVRACV------------------------------------------------------------
-----
>Mustela_putorius_furo
-----------------MAASLALWPSHQRRRAAGLEGVNSGDVSACDCWGGEM----------------------------------------------------
-----
>Lipotes_vexillifer
-----------------MAASLALWPSPQRRRAAGLEGVNSGDVSACDCWGGEM----------------------------------------------------
-----
>Rousettus_aegyptiacus
-----------------MAASLALWPSHQRRRAAGLEGANSGDVSACDCWGGEM----------------------------------------------------
-----
>Condylura_cristata
-----------------MAASPAWWPSHRRRRAAGLQSVHSGDVSACDCWGGEM----------------------------------------------------
-----
>Ochotona_princeps
-----------------MASSRALGPDRQRREAAESEGVNPRDVRAWDCRGGEM----------------------------------------------------
-----
>Elephantulus_edwardii
-----------------MAASLAWWPRRWRRSAAWLQGVSSPGPECV-------------------------------------------------------------
-----
>Manis_javanica
-----------------MAASVSLRPSRQRRRAAGLEGGNSGDWSACDCWGGEMRSARSASPAAAGGRRGGRAAGLLPASLVTLLFAALGELAERLRRRMCVFLSG
GRRLF
>Rattus_norvegicus
MGANTNVHTRRRSAQPRGCAAEAWWPCHRRRGAAGLEGVNSGDVRACDCWGGEM----------------------------------------------------
-----
>Trichechus_manatus_latirostr
-----------------MAASPALGPRHQRRRAAGLEGVNSGDVSACDCWAGEM----------------------------------------------------
-----
>Orycteropus_afer_afer
-----------------MAASPALWPRHQRRRAAGLEGVDSGDLSACDCWAGEM----------------------------------------------------
-----

HG0612.2 ENSG00000110422 HIPK3

>Homo_sapiens
MLRAAAARRGESAGL
>Odobenus_rosmarus_divergens
MLRAAAARRGESSGL
>Eptesicus_fuscus
MLRAAAARRGESAGL
>Dasypus_novemcinctus
MLRAAAARRGESAGL
>Galeopterus_variegatus
MLRAAAARRGESAGL
>Condylura_cristata
MLRVAAARRGESAGL
>Oryctolagus_cuniculus
MLRAAAARRGESAGL
>Manis_javanica
MLRAAAARRGESAGL
>Loxodonta_africana
MLRVAAAGRGESSGL
>Ictidomys_tridecemlineatus
MLRAAAARRGESAGL

HG0612.3 ENSG00000110422 HIPK3

>Homo_sapiens

MKMSVSLSGGRRLL

>Ursus_maritimus

MGMSVFLSGGRRLF

>Lipotes_vexillifer

--MSVFLSGGRRLF

>Eptesicus_fuscus

--MSVSPSGGRRLL

>Mesocricetus_auratus

--MSVFLSGGRRLL

HG0613 ENSG00000114861 FOXP1

>Homo_sapiens
MVRASIPSLICCQP--------
>Canis_lupus_familiaris
MVRASIPSLICCQP--------
>Balaenoptera_acutorostrata_s
MVRASIPCLICCQP--------
>Pteropus_alecto
MVRASIPSLICCQP--------
>Galeopterus_variegatus
MVRALIPSLICFQPCTAGCDAC
>Ochotona_princeps
MVRASIPSLICCQP--------
>Ceratotherium_simum_simum
MVRASIPSLICCQP--------
>Loxodonta_africana
MVRASIPSRFCCQP--------
>Jaculus_jaculus
MVRAALPSLICCRP--------
>Tupaia_chinensis
MVRASILSLICCQP--------
>Orycteropus_afer_afer
MVRASVRSPICCQP--------

HG0614 ENSG00000122203 KIAA1191

>Homo_sapiens
MGRGRVSRSWTTTPSMPCAPLGQVSREIPRSPEALSAVLPARWVVLLPSGSWGPSKSPPLSQWLGSRLRHSLSGDVGPRNLFSAGLRAAGSPAAAGSQNLWMDSSW
EALLLCSSWTMFLINICLSA
>Odobenus_rosmarus_divergens
-----------------------------------------------------------------------------------------------------MDSSQ
ADLLICSSWTMFLINTCLSA
>Galeopterus_variegatus
-----------------------------------------------------------------------------------------------------MDSSW
EALLISSSWTMFLINICLSA
>Erinaceus_europaeus
----------------------------------------------------------------------------------------------------MDTSSW
TASLLHSSWTMDD-HLCLSA
>Oryctolagus_cuniculus
---------------------------------------------MFPGQCWGLMFLLLPLPWFESGQCHSLSGYLGLWSLCSAGVQAAGGFLSVGEQNLWMD-SW
KASLAGSSWKKFLISIC---
>Elephantulus_edwardii
-----------------------------------------------------------------------------------------------------MDFSW
DALLTCSSWTMFLINICLSA
>Ceratotherium_simum_simum
----------------------------------------------------------------------------------------------------------
----MCSSWTMFLINNCLSA
>Chinchilla_lanigera
-----------------------------------------------------------------------------------------------------MDSSG
KALLICSSWTMFLINICLSA
>Tupaia_chinensis
----------------------------------------------------------------------------------------------------------
----------MLLITICLSA
>Trichechus_manatus_latirostr
-----------------------------------------------------------------------------------------------------MDSSW
EALLTCSSWTMFHI-ICLSA
>Orycteropus_afer_afer
-----------------------------------------------------------------------------------------------------MDSSW
EVLLTCSSWTMFLINICLSA

HG0615 ENSG00000128191 DGCR8

>Homo_sapiens

MKTDSLSRQSLKLSAL

>Acinonyx_jubatus

MKTDSLSRQSFKLSAL

>Pteropus_alecto

MKTDSLSRQSFKLSAL

>Galeopterus_variegatus

MKTDSLSRQSFKLSAL

>Erinaceus_europaeus

MKTDSLSRQSFKLSAL

>Ochotona_princeps

MKTESLSRQSFKLSAL

>Equus_asinus

MKTDSLSRQSFKLSAL

>Manis_javanica

MKTDSLSRQSFKLSAL

>Loxodonta_africana

MKTDSLSRQSFKLSAL

>Marmota_marmota_marmota

MKTDSLSRQSFKLSAL

>Orycteropus_afer_afer

MKTDSLSRQSFKLSAL

HG0616 ENSG00000128739 SNRPN

>Homo_sapiens
-------------------------------------------------MERARDRLHLRRT--TEQHVPEVEVQVKRRRTASLSNQECQLYPRRSQQ-QQVPVVD
FQAELRQAFLAETPRGG
>Leptonychotes_weddellii
-------------------------------------------------MERARDRLHLRRT--TEQHVPEVEVQVKRRRTASLSNQECQLYPRRSQQ-QQIPVVD
FQAELRQAFLAETPRGG
>Tursiops_truncatus
-------------------------------------------------MERARDRLHLRRT--TEQHVPEVEVQVKRRRTASLSNQECHLYPRRSQQQQQVPVVD
FQAELRQAFLAETPRGG
>Miniopterus_natalensis
-------------------------MGAC--------SRRPASRRPSSTMSRGACAGPLRRIRFKGQGLCFLAL-RPRGPPDACPPSRCQQFPRRSPQ-PRARAAD
FAAELRPAFVAETPRGG
>Galeopterus_variegatus
-------------------------------------------------MERARDRLHLRRT--TEQHVPEVEVQVKRRRTASLSNQECHLYPRRSQQ-QPVPVVD
FQVELRQAFLAETPRGG
>Erinaceus_europaeus
-------------------------------------------------MERARDRLNLRRT--TEQQVPEVEVQVKRRRTASLSNQECHLYPRRSQQ-QQVPVVD
FQAELRQAFLAETPRGG
>Ochotona_princeps
-------------------------------------------------MERARDRLHLRRT--TEQHVPEVEVQVKRRRTASLSNQECQLYPRRSQQ-QQVPVLD
FQAELRQAFLAETPRGG
>Equus_przewalskii
-------------------------------------------------MERPRDRLHLRRT--TEQHVPEVEVQVKRRRTASLSNQECQLYPRRSQQ-QQIPLLD
FQAELRQAFLAETPRGG
>Ictidomys_tridecemlineatus
-------------------------------------------------MERARDRLHLRRT--TEQHVPEVEVQVKRRRTASLSNQECHLYPRRSQQ-QQVPVVD
FQAELRQAFLAETPRGG
>Trichechus_manatus_latirostr
-------------------------------------------------MERVRDRLHLRRT--TEQHVPEVEVQVKRRRTASLNNQECQLYPRRSQQ-QQVPVVD
FQAELRQAFLAETPRGG
>Orycteropus_afer_afer
MTDNFSTGLVQGEFLPHNVAVMWEYTGLRSLSMPDNIGSSPPP-RSAAVKWGGKDRLHLRRT--TEQHVPEIEVQVKRRRTASLSNQECQLYPRRSQQ-QQVPVLD
FQAELRQAFLTETPRGG

HG0617 ENSG00000135164 DMTF1

>Homo_sapiens
MRATWVKKLPVNSHHFCPSLGNTFRECCGDRNMGEKQSG
>Mustela_putorius_furo
--------------------------------MGEKQSG
>Physeter_catodon
--------------------------------MEEKQSG
>Miniopterus_natalensis
--------------------------------MEEKQSG
>Galeopterus_variegatus
--------------------------------MGEKQSG
>Oryctolagus_cuniculus
--------------------------------MGEKQLG
>Equus_asinus
--------------------------------MGEKQSG
>Loxodonta_africana
--------------------------------MGKKQSG
>Dipodomys_ordii
--------------------------------MGETQSG
>Tupaia_chinensis
--------------------------------MGEIQSG
>Orycteropus_afer_afer
--------------------------------MGEKQSG

HG0618 ENSG00000137817 PARP6

>Homo_sapiens
MPISLALVRLVVWRGAA--
>Leptonychotes_weddellii
MPIPLACVQLVVWRGAA--
>Balaenoptera_acutorostrata_s
MPISLAWVQLVVWRGAA--
>Eptesicus_fuscus
MPISLAWVQLVVWRGAA--
>Dasypus_novemcinctus
MPISLALVQLVVWRGAA--
>Condylura_cristata
MPISLARVQLVVWRGAA--
>Oryctolagus_cuniculus
MPILLAWLRLVAWRGAAEH
>Ceratotherium_simum_simum
MPISLAWVQLVVWRGAA--
>Manis_javanica
MPISLAWVQLVVWRGAA--
>Loxodonta_africana
MPISLAWVQLVVWRGAA--
>Heterocephalus_glaber
MPISLAWVQLVVWRGAA--

HG0619 ENSG00000138593 SECISBP2L

>Homo_sapiens
MEKAARGAVYTD
>Felis_catus
MEKAARGALHTD
>Lipotes_vexillifer
MEKAARGALDTD
>Miniopterus_natalensis
MEKAARGALYTD
>Galeopterus_variegatus
MEKAARGALHTN
>Condylura_cristata
MEKAARGALHTD
>Elephantulus_edwardii
MEKAARGALHTD
>Equus_asinus
MEKAARGALHVD
>Manis_javanica
MEKAARGALHTN
>Chinchilla_lanigera
MEKAARGALHTD
>Trichechus_manatus_latirostr
MEKAARIALYTD

HG0620 ENSG00000145555 MYO10

>Homo_sapiens
MCHQLFSACERCAQ-
>Odobenus_rosmarus_divergens
MCHQLFSARERCAQK
>Physeter_catodon
MCHQLFSARERGAQE
>Rousettus_aegyptiacus
MCHQLFSARERCAPK
>Galeopterus_variegatus
MCHQLFSARERFAQE
>Oryctolagus_cuniculus
MCHQLFPARERSVQK
>Ceratotherium_simum_simum
MCHQLFSARERGAQK
>Manis_javanica
MCHQLFSARQRCAQE
>Rattus_norvegicus
MCHQLFSAREHFAQK
>Tupaia_chinensis
MCHQLFTAREHLAQK
>Trichechus_manatus_latirostr
MCHQLFSARERCALK

HG0621.1 ENSG00000146352 CLVS2

>Homo_sapiens
MPLSPPPPLLKPRLMDARECRIA
>Odobenus_rosmarus_divergens
MLLSPPP--PKPGLMDARECRIA
>Pteropus_alecto
-MLSSPPPLLKPRLMDAGECRIA
>Dasypus_novemcinctus
--------------MDTRECRIA
>Condylura_cristata
--------------MDARECRIA
>Oryctolagus_cuniculus
---------MKPGLMDARECRIA
>Equus_caballus
--MLLPPTPLKPRLMDARECRIA
>Loxodonta_africana
---MLLSPLLKSQLMDAGECRIA
>Rattus_norvegicus
MVLSPPRPLRKPGLMDAGECRIA
>Tupaia_chinensis
--------------MDARECRIA
>Orycteropus_afer_afer
MLLS---PPPKPRLMDARECRIA

HG0621.2 ENSG00000146352 CLVS2

>Homo_sapiens
MAGPGENLGQRWP
>Odobenus_rosmarus_divergens
MAGPGEDLGRRWP
>Orcinus_orca
MAGPGENLGQRWP
>Rousettus_aegyptiacus
MAGPGEDPGQPWP
>Galeopterus_variegatus
MAGPGEHLGQRWP
>Oryctolagus_cuniculus
MAGPGEDLGQQWS
>Ceratotherium_simum_simum
MAGPGEDLGQRWP
>Loxodonta_africana
MAGPSEDLEQRWL
>Ictidomys_tridecemlineatus
MAGPGGDLGQRWP

HG0621.3 ENSG00000146352 CLVS2

>Homo_sapiens

MPPKLPSYP------------------------NEAGKGAERARELSECLRRSEICAGIIREELG

>Odobenus_rosmarus_divergens

MPRKLPSYPAES--GREERGACAS-----------------------------------------

>Galeopterus_variegatus

MPPKLPSQPNKA--GREELSACAS-----------------------------------------

>Condylura_cristata

MPPEPPAEASGPEAGGTRLRGCLNATDVAILLKRVGERAAECVRELSECLRRREIEAGIMREELG

>Marmota_marmota_marmota

MPQKLPSFPKRQGEKS-------------------------------------------------

HG0622 ENSG00000148677 ANKRD1

>Homo_sapiens

MKIHLPLSWLLMGV-GVLLRFPGWKIISPGPSYIS---------

>Leptonychotes_weddellii

MKIHLPLSWPLVGA-GVLLGFPGWEITSPGPSYISRPVWRAPQG

>Orcinus_orca

MKLHLPLSWPRVGA-GVLLGFPGWKISSPGPSYIS---------

>Pteropus_alecto

MKIHLPLSWPVVGA-GMLLGFPGWKIISPSPSYIS---------

>Dasypus_novemcinctus

MRIHLPLSWLLVRV-GVLLGFPGWKIISPSPSYIS---------

>Galeopterus_variegatus

MKIHLLLSWPLVGA-GVLLGFPGWKIISPSPSYIS---------

>Condylura_cristata

MKIHLPLSWPLVGG-GVLLGFPGWKISSPVLAI-----------

>Oryctolagus_cuniculus

MKIHLPLSWLLAGGQGCYLGSQVGRLSHPAQTI-----------

>Ceratotherium_simum_simum

MKIHLPLSWPLVGA-GVLLGFPGWKIISPSPVYIS---------

>Loxodonta_africana

MKIHLPLSWLLR-GRGCYLGSQVGRLSHPAQAI-----------

>Jaculus_jaculus

MKIHLPLSWPLVGA-GALLGFPGWKIISPSPSYITG--------

HG0623 ENSG00000151422 FER

>Homo_sapiens

MTIFARLHCTFSQVSSLSLMCD-TSSWTRYFS-------

>Felis_catus

MVIPVRLHCTFSQVCSSSLMCN-PSTWTCYFS-------

>Orcinus_orca

MTIPVRLHCTFSQVSSSSLMCN-PSTWTCYFS-------

>Pteropus_vampyrus

MIIPVRLH-------------------------------

>Dasypus_novemcinctus

---------MFSQVSSSSLMCS--STWTCYFS-------

>Condylura_cristata

MIITVSLHCTFSNVSSLLLMCS-PSIWTFYFS-------

>Oryctolagus_cuniculus

----MSLHSTFFQVFSSSLMSNHTSTWTCYFS-------

>Equus_asinus

MIIPVRLHCTFSQVSSSSLMCN-PSTWTCYFS-------

>Loxodonta_africana

-------------------MCN-TTTWTCYFS-------

>Dipodomys_ordii

-------------------MCN-TSTWTCYFR-------

>Tupaia_chinensis

MIPPVRLHCTFSQVSS-SRMCG-TSPWTCCFSLDVASNE

HG0624 ENSG00000156853 ZNF689

>Homo_sapiens

MALRSIKSIAGSCLCSRQRRCGSSAAIFPEGIFRCLSPKFGQEFPE

>Acinonyx_jubatus

MALRSIKSIAGSCLCSRLRRCGSSAAIFPEGIFRCLSPKFGQEFPE

>Tursiops_truncatus

MALRSIKSIAGSCLCFRQRRCSSGAAIFPEGIFRCLSPKFGQEFPE

>Eptesicus_fuscus

MALWSIKSIAGSCLCSRQRRCSSSAAIFAEGIFRCLSPKFGQEFPE

>Dasypus_novemcinctus

MALWSIKSIAGSCRCSREKRCSSSAAIFPEGILQCLSPKFGQEFPE

>Galeopterus_variegatus

MALRSIKSIAGSCLCSRQRRCNSSAAIFPEGIFRCLSPKFGQEFPE

>Oryctolagus_cuniculus

MALRSIKSIAGSRLCSRQRRCSCGAAIFPKGIFRCLSPKLGQEFPE

>Ceratotherium_simum_simum

MALRSIKSIAGSCLCSRHRRCSSSAAIFPEGIFRCLSPKFGQEFPE

>Manis_javanica

MALWSIKSIAGSCLCSRQRHWSSGAAIFPEGIFQCLSPKFGQEFPE

>Marmota_marmota_marmota

MALRSIKSIAGSCLCYRQRRCSSSAAIFPEGIFRCLSPKFGQEFPE

>Tupaia_chinensis

MALRSIKSIAGSCLCSRQRRCSSAAAIFPEGIFRCLSPKFGQEFPE

HG0625 ENSG00000164050 PLXNB1

>Homo_sapiens

MLLSPDPAGPAR

>Mustela_putorius_furo

MLLSPDPAGPAR

>Lipotes_vexillifer

MLLSPDPAGLAR

>Pteropus_vampyrus

MLLSPDPAGPAR

>Oryctolagus_cuniculus

MLLSPDPAGPAR

>Elephantulus_edwardii

MLLSPDPAGPAR

>Ceratotherium_simum_simum

MLLSPDPAGPAR

>Loxodonta_africana

MLLSPDPADPAR

>Dipodomys_ordii

MLLSPNPAGPAR

>Tupaia_chinensis

MQLSPDPAGPAR

>Orycteropus_afer_afer

MLLSPDPASPAR

HG0626 ENSG00000168939 SPRY3

>Homo_sapiens
-------MDKPFPQGFSHVP
>Acinonyx_jubatus
-------MDKPFPQGFSDVP
>Physeter_catodon
-------MDKPFLQGFSDVP
>Miniopterus_natalensis
-------MDRSFPQGFSDVP
>Galeopterus_variegatus
MTSAFPSMDKPFPQGFSDVP
>Erinaceus_europaeus
-------MDKPFPQGFSDVP
>Oryctolagus_cuniculus
-------MDKPFSQGFSDVP
>Equus_asinus
-------MDKPFPQGFSDVP
>Manis_javanica
-------MDKPFSQGFSDVP
>Heterocephalus_glaber
-------MDKPFPQGFSDVP
>Trichechus_manatus_latirostr
-------MDKPFPQGFSDVP

HG0627 ENSG00000169760 NLGN1

>Homo_sapiens
MRGLPLWIYLAEGYRPAANWI
>Odobenus_rosmarus_divergens
MRGFPLWIYLTEGCRPAANWI
>Balaenoptera_acutorostrata_s
MRGLLLWIYLTEGYRPAANWI
>Rhinolophus_ferrumequinum
MRGLPLWIFLTEGYRPAPNWI
>Sorex_araneus
MRGLPLWICLTEGYRPAANWI
>Equus_asinus
MKGLRLWI-------------
>Manis_javanica
MRVLPLWIYLTKGYRTAANWI
>Loxodonta_africana
MKGLPPWIPLTQGYRPAAKWI
>Marmota_marmota_marmota
MRGLPLWICLAEGYRPAANWI
>Tupaia_chinensis
MRGLAPRTRLPEGYRPTVNWI
>Orycteropus_afer_afer
MRGLPPWIYLTEGYRPATNWI

HG0628 ENSG00000170677 SOCS6

>Homo_sapiens

MKQEKYIDAALQPLQMFGDNIPDRNIDPLD

>Felis_catus

MKLEKYIDAALQPLQMFEDNIPERN-----

>Lipotes_vexillifer

------MQHCVQPLQMFEDNIPERN-----

>Dasypus_novemcinctus

---------------MFGDNIPQRS-----

>Galeopterus_variegatus

MKLEKYIDAALQPLQMFRDNIPERN-----

>Oryctolagus_cuniculus

MKLEKCIDAALQPLQMFGNNIPERN-----

>Ceratotherium_simum_simum

MKLEKYIDAALQPLQMFEDNTPERN-----

>Loxodonta_africana

MKLEKYIDAALQPLQMFGDNIPERN-----

>Heterocephalus_glaber

MKLEKHTDAALKPLQMFGDNIPERKLIPVD

>Tupaia_chinensis

MKLEKHTDAALQPLQMFGDTIPERR-----

>Trichechus_manatus_latirostr

MKLEKYVDAALQPLQMFGDNIPERN-----

HG0629 ENSG00000170961 HAS2

>Homo_sapiens
MLFLNFLFFLAGRLKFI
>Odobenus_rosmarus_divergens
MLFLNFLFFLAGRLKFI
>Lipotes_vexillifer
MLFLNFLFFLAGRLKFI
>Myotis_brandtii
MLFLNFLFFLAGRLKFI
>Dasypus_novemcinctus
MLFLNFLFFLAGRLKFI
>Galeopterus_variegatus
MLFLNFLFFLAGRLKFI
>Condylura_cristata
MLFLNFLFFLAGRLKFI
>Ceratotherium_simum_simum
MLFLNFLFFLAGRLKFI
>Mus_musculus
MLFLNFLFFLAGRLKFI
>Tupaia_chinensis
MLFLNFLFFLAGRLKFI
>Orycteropus_afer_afer
MLFLNFLFFLAGRLKFI

HG0630 ENSG00000171552 BCL2L1

>Homo_sapiens
MIPPAGAG--FFASHREAG
>Canis_lupus_familiaris
MIPPAGAP--FFASRREAG
>Lipotes_vexillifer
MIPPAGAG--FFASRREAG
>Pteropus_vampyrus
MIPPAGAG--LFASRREAG
>Dasypus_novemcinctus
MIPSAGAA--FFANRREAG
>Galeopterus_variegatus
MIPPAGAV--FFASRREAG
>Oryctolagus_cuniculus
MIPPAGAV--CFASRREAG
>Equus_caballus
MILPAA---AFFASHREAG
>Heterocephalus_glaber
MIPPAGAG--FFASRREAG
>Tupaia_chinensis
MIPPAGAA--YFASRREAG
>Trichechus_manatus_latirostr
MIPPAGAVLLFFASRREAG

HG0631 ENSG00000173548 SNX33

>Homo_sapiens

--MSPPCSQAYARRPDRAVSGPWTLL

>Physeter_catodon

MDMSSPCSQACAGRP---VSGPWTLL

>Cynopterus_sphinx

MDMSLSCLQACAGKPDRAVSGPWTQL

>Dasypus_novemcinctus

--MSPPCTQACAQRPERAFSGPWTLL

>Oryctolagus_cuniculus

MAGSPPSLEAHGQIPDWAAAGPWTLL

>Equus_asinus

MDMSPPCSHVCARRPERAVSGPWTLF

>Manis_javanica

MDVSPPCSQECARRSDRAVPGPWTLF

>Loxodonta_africana

--MSPPCSRSCAWRLDRAVSEPWTLL

>Ictidomys_tridecemlineatus

--MSPPCSQAYAQRLNRVVSGPWTLF

>Trichechus_manatus_latirostr

--MSPPCSQLCAWRPDRAVSGPWTLL

>Orycteropus_afer_afer

--MSPPYSQTFSWRLDRAVSGTWALL

HG0632.1 ENSG00000181315 ZNF322

>Homo_sapiens

MTVRATMGQEDQRKTP

>Ursus_maritimus

------MGQEDQWKTP

>Orcinus_orca

MTVKKMMGQEDQWKTP

>Dasypus_novemcinctus

MTSKTMMGQEDQWKTP

>Galeopterus_variegatus

MTMKAMMGQEGRWKTP

>Oryctolagus_cuniculus

MTVKVMMGQEDQWKTP

>Equus_caballus

MTVKKMMGQEDQWKTP

>Loxodonta_africana

MTAKTMMGQEDQWKTP

>Dipodomys_ordii

MTMKAKMGQEDQWKSP

>Trichechus_manatus_latirostr

------MGQEDQWKTP

>Orycteropus_afer_afer

MTVKTMMGQEDQWKTP

HG0632.2 ENSG00000181315 ZNF322

>Homo_sapiens

MYVFFHLDSRVGNTK-----------

>Canis_lupus_familiaris

MLLSFQLDSRVGNTK-----------

>Rhinolophus_ferrumequinum

MLLFIHLDSRVGNTK-----------

>Condylura_cristata

MLLIFHLDCRAGNTKQGISVYDGETL

HG0633 ENSG00000186174 BCL9L

>Homo_sapiens

MLRGWEAGTQAPSDPRIASPGPGWY

>Mustela_putorius_furo

MLGGWEAGTQAPSDPRIPSPGPGWY

>Orcinus_orca

MLRGWKAGTQAPSDPRIPTPGPGWY

>Pteropus_vampyrus

MLRGWKAGTQAPSNPRIPSPGPGWY

>Dasypus_novemcinctus

MLGDWEAGTQAPSDSRRPSLGPLWY

>Condylura_cristata

MFRGWEAGTQAPSDPRIPSPGPGWY

>Oryctolagus_cuniculus

MLRSWEAGTQAPRDSRTPSPGPGWD

>Equus_asinus

MLRGWEAGTQAPSDPRRP-PGPGWY

>Manis_javanica

MLRGWEAGTQAPSDPRIPSPGPGGY

>Marmota_marmota_marmota

MLRRREAGTQAPSDPRIPSPGPGWY

>Tupaia_chinensis

MLRGWETGTQAPSDPRIPSPGPGWC

HG0634 ENSG00000198026 ZNF335

>Homo_sapiens
-----------------MPESERNVATKASE
>Ursus_maritimus
------------MATSSMPESERSAATKASE
>Physeter_catodon
------------MATSSMPESERNAATKASE
>Rousettus_aegyptiacus
------------MATSSMPESERNAATKASE
>Dasypus_novemcinctus
MPIDSSAAGGKRMATSSMPESEKNADTKASE
>Galeopterus_variegatus
------------MATSSMPESEKNAATEASE
>Condylura_cristata
------------MATSSMPESERNAAAKTSE
>Ceratotherium_simum_simum
------------MATSSMPESERNAATKASE
>Manis_javanica
------------MATSSMPESERNAATQASE
>Cavia_porcellus
------------MATSSMPESERNAAKKASE
>Tupaia_chinensis
------------MATSSMPVSERNVATKASE

HG0635 ENSG00000259332 ST20-MTHFS

>Homo_sapiens
MAAETGYWRCLPESRTCAEEAVHLWPAVLRDPRWGNPGDGAWPCYMRGPRLE
>Leptonychotes_weddellii
---------------------------------------------MPAPRLE
>Rousettus_aegyptiacus
---------------------------------------------MPAPRLE
>Dasypus_novemcinctus
---------------------------------------------MPAPRLE
>Condylura_cristata
---------------------------------------------MPAPRLE
>Oryctolagus_cuniculus
---------------------------------------------MPAPRLE
>Elephantulus_edwardii
---------------------------------------------MPAPRLE
>Manis_javanica
---------------------------------------------MPAPRLE
>Fukomys_darlingi
---------------------------------------------MTAPRLE
>Trichechus_manatus_latirostr
---------------------------------------------MPAPRLE
>Orycteropus_afer_afer
---------------------------------------------MPAPQLE

HG0636 ENSG00000002587 HS3ST1

>Homo_sapiens

---------------------------------------------------------------------------------------------------MAQPRSL

FRAPDGLDRGAAQ

>Felis_catus

---------------------------------------------------------------------------------------------------MAQPESR

LRAPDRLDRGAAQ

>Physeter_catodon

---------------------------------------------------------------------------------------------------MAKPESR

LRAPDRLEGGAAQ

>Rhinolophus_ferrumequinum

---------------------------------------------------------------------------------------------------MAQPESR

LRTPDPLDRGAAQ

>Erinaceus_europaeus

---------------------------------------------------------------------------------------------------MAQPQSR

LGAPDWLDQRAAQ

>Oryctolagus_cuniculus

MPIPRARKPGQSEDPGPGGHPGAAPVSEDRRLPLLPGRHTEHRRLRTEHQNARLQGLRRAGNREPPPLPGLRGCLRPFHRALACLDIRSTCRGCSAVLPMAQPESR

LRAPDGLDGGVAQ

>Ceratotherium_simum_simum

---------------------------------------------------------------------------------------------------MAQPESR

LRAPDRLDRGAAQ

>Manis_javanica

---------------------------------------------------------------------------------------------------MAQPESR

LVAPDRLDRGAAQ

>Loxodonta_africana

---------------------------------------------------------------------------------------------------MAQPQSR

PRAPDGLDGGAAQ

>Cavia_porcellus

---------------------------------------------------------------------------------------------------MAQPESR

FRAPDGRDRRAAQ

HG0637 ENSG00000005102 MEOX1

>Homo_sapiens

MFSVIGDKYTYRGSQ

>Odobenus_rosmarus_divergens

MFSVIGDKYTSRGPQ

>Lipotes_vexillifer

MFSVIGDKYTYRGPH

>Pteropus_vampyrus

MFSVIGDKYTYRGPQ

>Dasypus_novemcinctus

MYSVIGDKYTHKGP-

>Galeopterus_variegatus

MFSVISDKYTYKGPQ

>Equus_asinus

MFSVIGDKYTYRGPQ

>Marmota_marmota_marmota

MFSAIGNKYMYKGPQ

>Tupaia_chinensis

---------MYKGPQ

>Trichechus_manatus_latirostr

MFSVISDKYTYKGPQ

HG0638 ENSG00000013561 RNF14

>Homo_sapiens

MSSTSWPLTQLQQSQLHQLENKWDLHELHSELKYRLRC

>Mustela_putorius_furo

MSSRSWPFIQLQQSHLHLLEDKWDLHKFQILN------

>Physeter_catodon

MSSTSWPLIQLQQSHLHHLENK----------------

>Myotis_davidii

MSSTSWPLIQLQQSHLHHLENKWDICMNSRL-------

>Dasypus_novemcinctus

MSPTSWPLIQHQQSHLHWPESKWGLHEFQTP-------

>Galeopterus_variegatus

MSPTSQPPIKLQQSQLHWLKNKWDRTNSQLN-------

>Condylura_cristata

MSSTSWPLIELLQSHLHRLENK----------------

>Ceratotherium_simum_simum

MSSTSWPLIHLQQSHLHRLENKWDFHEFQILN------

>Manis_javanica

MSSTSWPLTQLQ-FHLHCLENKWDLREFQTLN------

>Marmota_marmota_marmota

MSPTSWPLTQLQQSQLHWLENKWDLHKFQTELKFRLQC

HG0639 ENSG00000018236 CNTN1

>Homo_sapiens
MGAAVPACEVTA
>Canis_lupus_familiaris
MGAAVPASEVTA
>Orcinus_orca
MGAAVPASEVTA
>Myotis_brandtii
MGAAVPASEVTA
>Dasypus_novemcinctus
MGAAVPASEVTA
>Galeopterus_variegatus
MGAAVPASEVTA
>Equus_przewalskii
MGAAVPASEVTA
>Loxodonta_africana
MGAAVPASEVTA
>Rattus_norvegicus
MGAAVPASEVTA
>Tupaia_chinensis
MGAAVPASEVTA

HG0640 ENSG00000062194 GPBP1

>Homo_sapiens
------MLFRCEIFFLFLLFGS
>Canis_lupus_familiaris
------MLFRCEIFFLFLLFGS
>Balaenoptera_acutorostrata_s
------MLFRCEIFFLFLLFGS
>Miniopterus_natalensis
------MLFRCEIFFLFLLFGS
>Galeopterus_variegatus
------MLFRCEIFFLFLLFGS
>Oryctolagus_cuniculus
------MLFRCEIFFLFLLFGS
>Elephantulus_edwardii
MCSCLGMLFRCEIFF-------
>Ceratotherium_simum_simum
------MLFRCEIFFLFLLFGS
>Loxodonta_africana
------MLFRCEILFL--LFGS
>Orycteropus_afer_afer
------MFFRCEIFFLFLIFGS

HG0641 ENSG00000100359 SGSM3

>Homo_sapiens
MILDQMKPEEPSSSKIAG-
>Leptonychotes_weddellii
MFLDQMKPEEPSSCKIIG-
>Physeter_catodon
MFLDQMKLEEPSSCKILG-
>Miniopterus_natalensis
MFLDQMKPEKPSSCKILG-
>Dasypus_novemcinctus
MFLDQMKPEEPSSSKILGE
>Galeopterus_variegatus
MFLGQMKPEEPSSSKIG--
>Ceratotherium_simum_simum
MFLDQMKPEEPSNCKILV-
>Manis_javanica
MFLDQLKPEEPSGHRILD-
>Loxodonta_africana
-----MTSEEPSSSKILGD
>Tupaia_chinensis
MFLDQMKTEEPSSSHTAG-

HG0642 ENSG00000100815 TRIP11

>Homo_sapiens

MAAGVELAGVTHGTEESH

>Ursus_maritimus

MAAGVELAGVALGTEESH

>Orcinus_orca

MAAGVELAGVANGTGESH

>Myotis_lucifugus

MAAGVELAGVAHRTGASH

>Condylura_cristata

MAAGVELAGVAHGAGESH

>Ceratotherium_simum_simum

MAAGVELAGVAHGTGESH

>Manis_javanica

MAAGVELAGIAHGIGESH

>Loxodonta_africana

MAAGVELAGVAHGTGESH

>Heterocephalus_glaber

MAASVELAGTAFGTEESH

>Trichechus_manatus_latirostr

MAAGVELAGVARGIGESH

HG0643 ENSG00000103653 CSK

>Homo_sapiens
--------------------------------------MLFPTALMVPSDRLALL-------
>Acinonyx_jubatus
---------------------------------------------MVPSDRLAFL-------
>Physeter_catodon
--------------MLPAA--------------RWPGPQCPERALMVPSDRLAFL-------
>Myotis_davidii
---------------------------------------------MVPRDRLAFL-------
>Dasypus_novemcinctus
---------------------------------------------MVPSDRLAFL-------
>Condylura_cristata
--------------MLPAA--------------RWPGPQCPERALMVPSDRLAFL-------
>Oryctolagus_cuniculus
---------------------------------------------MVPGDRPALLTRGHQLS
>Equus_asinus
---------------------------------------------MVPSNRLAFL-------
>Manis_javanica
MMALPPSTSSGSLWHLFTLLWGRSGSSVGSHLTPLLHVPLLPVALMVPGDKWAFL-------
>Fukomys_damarensis
---------------------------------------------MVPSDRSAFL-------

HG0644 ENSG00000104299 INTS9

>Homo_sapiens

----------------------------------------------MLSAGRWRRLHRKTLPGFEEFSDCY

>Mustela_putorius_furo

----------------------------------------------MLSAGRWR-----------------

>Balaenoptera_acutorostrata_s

----------------------------------------------MLSAGRWRRLNRKTLSGSEEWSDWY

>Pteropus_alecto

----------------------------------------------MLSASEWRSLNRKTVSGFEESIDWC

>Oryctolagus_cuniculus

----------------------------------------------MLSAGQRRSLNRKTHAGVEAS----

>Equus_asinus

----------------------------------------------MLSAGRWQRLNRKTLSGYEGVE---

>Loxodonta_africana

----------------------------------------------MLCAGWWRRLNRKTYSGFEESSDCC

>Cricetulus_griseus

------------------------------MLR----KVREKGKLQLPRCSLWRRLNRKTLPCFEK-----

>Tupaia_chinensis

----------------------------------------------MLSAGQWRRLNRKTLPGVKESGGWY

>Trichechus_manatus_latirostr

MTEGGVRRGKGEKENITTVPMMHPGMTTFPMFPGGGAGREETACPLMLSAGWWRRLNRKTPFGFEGSSDCC

HG0645 ENSG00000107282 APBA1

>Homo_sapiens
MVALEAAAAGAELYSQAWHQRCH
>Canis_lupus_familiaris
MVALEVGAAGAERHSQAWPPRCH
>Rousettus_aegyptiacus
MVALEVGAAGAELYSQAWHQRCH
>Dasypus_novemcinctus
MVALEVGAAGAELHSQAWHHRCH
>Condylura_cristata
MVALEVGAAGAELSSQAGHQRCH
>Elephantulus_edwardii
MVALEVGPAGAELDSQARHRGCH
>Ceratotherium_simum_simum
MVALEVGAAGAELYSQAWHQRCH
>Manis_javanica
MVALEVGAARAELDSQARHQRCR
>Loxodonta_africana
MVALEVGAAGTELYSQAWHQRCH
>Heterocephalus_glaber
MVALEVGAAGAELYSQVWHRRCH

HG0646 ENSG00000108312 UBTF

>Homo_sapiens
MEQWEREAGRFPCKSPSGLTPSRRPRQGSEDPGEQMEEWSSL
>Balaenoptera_acutorostrata_s
---MGPGSRRFPRKSPSVPTSSRRTRQGSEEPGEQMEEWSSL
>Rousettus_aegyptiacus
-----------------------------------MEEWSSL
>Condylura_cristata
-----------------------------------MEEWSSL
>Ochotona_princeps
-----------------------------------MEEWSSL
>Elephantulus_edwardii
------------------------------------MEWSSL
>Ceratotherium_simum_simum
-----------------------------------MEEWSSL
>Dipodomys_ordii
---MGTGSGQVPSQIPLRLKPPRPHRQGSEEPGEQMEEWSSL
>Trichechus_manatus_latirostr
-----------------------------------MEEWSSL
>Orycteropus_afer_afer
-----------------------------------MEEWSSL

HG0647 ENSG00000108395 TRIM37

>Homo_sapiens

MGVGGAGPEEMGRAGGAVVRAP

>Felis_catus

----------MGRAGGAVVGAP

>Balaenoptera_acutorostrata_s

----------MGRAGGAVVGDP

>Pteropus_alecto

----------MGRADGAVVGAP

>Oryctolagus_cuniculus

----------MGRAGGAVVGVL

>Equus_asinus

----------MGRAGGAVVGAP

>Manis_javanica

----------MGRAGGAVVGAL

>Dipodomys_ordii

MGVAGLELEAMGRAGGAVVGAP

>Tupaia_chinensis

MGVAGAEPEAMGRAGGAVVGAP

>Trichechus_manatus_latirostr

----------MGRAGGAVVGAS

HG0648 ENSG00000114331 ACAP2

>Homo_sapiens
MTSGRGALRSSAQR-
>Odobenus_rosmarus_divergens
MTSGRGALRSSAQR-
>Orcinus_orca
MTSGRGALRSSAQR-
>Rousettus_aegyptiacus
MTSGRGVLPSSAQL-
>Dasypus_novemcinctus
MTGGRGALRSSVQR-
>Condylura_cristata
MTSGRGALRSSAQR-
>Ochotona_princeps
MTSGRGALRSPAQRS
>Elephantulus_edwardii
MTSGLGALRSSAQH-
>Ceratotherium_simum_simum
MTSGRGALRSSARP-
>Ictidomys_tridecemlineatus
MTSGRGALRSSAQC-

HG0649.1 ENSG00000114439 BBX

>Homo_sapiens
MWRIHWDIEKYQRRIVPL
>Panthera_tigris
MWRIHWDIEKYQRRVVLP
>Balaenoptera_acutorostrata_s
MWRSHWDIEKYQRRIVPL
>Rousettus_aegyptiacus
MWRIHWDIEKYQRRIVPL
>Dasypus_novemcinctus
MWRIHWDIEKYQRRIVPL
>Condylura_cristata
MWRSHWDIETYQRRFVPL
>Oryctolagus_cuniculus
MWRSHWDIEKYQRRIVPL
>Ceratotherium_simum_simum
MWRIHWDIEKYQRRIVPL
>Manis_javanica
MWRIHWDIEKYQRRIVPL
>Cavia_porcellus
MWRIHWDIEKYQRRIVPL

HG0649.2 ENSG00000114439 BBX

>Homo_sapiens
----MESLDLTGTLRKTPSL
>Canis_lupus_familiaris
----MESLDLTGTLRKTPSL
>Balaenoptera_acutorostrata_s
----MESLDLTGTLRKTPSL
>Myotis_brandtii
----MESLDLTGTLRKTPSL
>Dasypus_novemcinctus
MEITMESLDLTGNLRKTPAL
>Condylura_cristata
----MESLDLTGTLRKIPSL
>Equus_asinus
----MESLDLTGTLRKTPRL
>Manis_javanica
----MESLDLTGTLRKTPSL
>Microtus_ochrogaster
----MESLDLTGTLRKTPSL
>Tupaia_chinensis
----MESLDLTGTLGKTPSL

HG0650 ENSG00000115226 FNDC4

>Homo_sapiens

MGRPLGTEQPPEA

>Canis_lupus_familiaris

MGRPLGTEQPPEA

>Orcinus_orca

MGRPLGTEQPPEA

>Miniopterus_natalensis

MGRPLGTEQPPEA

>Dasypus_novemcinctus

MGRPLGTEQPPEA

>Sorex_araneus

MGRPLGTEQPPEA

>Equus_asinus

MGRPLGTEQPPEA

>Peromyscus_maniculatus_baird

MGRPLGTEQPPEA

>Tupaia_chinensis

MGRPLGTEQPPEA

>Orycteropus_afer_afer

MGRPLGTEQPPEA

HG0651 ENSG00000115935 WIPF1

```
>Homo_sapiens
MIPFSGTVFSLGSCPAGPLSACVPDHGSLQYPLTIYQQDC---
>Felis_catus
MMPFSGTVFSLGSHLAC-LSTCVPDHAFLQYPLTIHRQGC---
>Physeter_catodon
MTPFSGTVFSLISCLL---PVCLCP------------------
>Miniopterus_natalensis
MMPFSGTVFSLGSCFACSVSACVPGHAFLQYPLTIYQQDC---
>Oryctolagus_cuniculus
MMDFSGTVCSPGSGPSCSLSAGGPDHAFHGAP-----------
>Ceratotherium_simum_simum
MMPFSETVFSLGSYLACSLSACVPDHAFLQYPLTIYQDC----
>Manis_javanica
MMPFSGTVVTLGSYLAYSLSACVPDHALLHYPLKTHQQDC---
>Loxodonta_africana
MIPFSGMVFCLGSCLTCSLSACVPDHACPAVSFHDFSAGLLKV
>Cavia_porcellus
------MIFSLGSALACALSACIPDHASLQFLLTIYQQDC---
>Orycteropus_afer_afer
MMPFSGTIFSLGSCLVCSLSACVPDHACPAVSFNDLSARLLKV
```

HG0652 ENSG00000118515 SGK1

>Homo_sapiens

MMEGEKPLSVATESGERLLQ--------

>Odobenus_rosmarus_divergens

-MGSAKPLPVATESGERLLQ--------

>Balaenoptera_acutorostrata_s

MMEREEPLPVTTESGERLLQ--------

>Myotis_lucifugus

MMEREKPLPVAIESGERLLQSPLP----

>Galeopterus_variegatus

MMEREKPLPVATESGERLLQ--------

>Erinaceus_europaeus

-MEREKPLPVATESGERLLQ--------

>Oryctolagus_cuniculus

-MERERSLPVATERGERLRQSQALQNV-

>Loxodonta_africana

MMERESPPPVATEGGESLLQ--------

>Mus_musculus

MMEREKPLPVASESGASYCSRCRSRMSE

>Orycteropus_afer_afer

-MARENPPPVATESGEGLLQWLVLQNV-

HG0653 ENSG00000120334 CENPL

>Homo_sapiens

MIRESGMGDSVITAFY

>Ursus_maritimus

------MGDSVITAFY

>Lipotes_vexillifer

------MVDSVITAFY

>Pteropus_alecto

MIREAGLGDSVITAFH

>Dasypus_novemcinctus

MIREAGMGDSVISAFC

>Oryctolagus_cuniculus

MIAESGMGDSVDTAFY

>Ceratotherium_simum_simum

------MGDSVITAFY

>Manis_javanica

MIGAAGMGDSVITAFY

>Loxodonta_africana

MICEAGMRDSVVAAFY

>Fukomys_damarensis

------MGDRVTTAFY

HG0654 ENSG00000133794 ARNTL

>Homo_sapiens

----------MWNPGPSLVFGRS

>Acinonyx_jubatus

----------MWNPRPSLVSGRS

>Galeopterus_variegatus

MLLDPSFTFQMWNPRPLLVFGRS

>Elephantulus_edwardii

----------MWNPGPLLVFRRS

>Manis_javanica

----------MWNPRPSLVSERS

>Loxodonta_africana

MVLDSSLTFQMWNPRPLLVFERS

>Ictidomys_tridecemlineatus

MLLDPTSTTQMWNPRPSLVFGRS

>Tupaia_chinensis

MLLDPRLTFQMWNPRPSLVVGRS

>Trichechus_manatus_latirostr

MLLDPSLTFQMWNPRPLLIFGRS

>Orycteropus_afer_afer

----------MWNPRPFLVFGRS

HG0655 ENSG00000143061 IGSF3

>Homo_sapiens
---------------------------------------------------MWQEPGATSELLWTLPVA---
>Canis_lupus_familiaris
---------------------------------------------------MWEKPRGPSELFWTLPLA---
>Physeter_catodon
---------------------------------------------------MWQEPPAPSELFWTLPLA---
>Rhinolophus_ferrumequinum
---------------------------------------------------MWQKPRLSAELFWTLPLLRGL
>Condylura_cristata
MREITDTFTAKPRTFLYNQWMKGFSQTAFSLFLED-YPPRSTIPVETARHTMWPKPRAPSELIWTLPLA---
>Ochotona_princeps
---------------------------------------------------MWQEPGAPSE-FWTLPLA---
>Ceratotherium_simum_simum
MREITDTFTGKPRTFLYNQWMKGFSQTLFSPFSLRIKPLQSSIPVQTARHTMWQKPRAPSELFWTLPLA---
>Loxodonta_africana
---------------------------------------------------MWQKPGAPSELVWTLLLP---
>Peromyscus_maniculatus_baird
---------------------------------------------------MWQKPRAPSELLWTLPLA---
>Trichechus_manatus_latirostr
---------------------------------------------------MWQKPGAPSELVWTLHLP---

HG0656 ENSG00000150776 C11orf57

>Homo_sapiens

MGIFPLSSTWKPQPPLQAP

>Panthera_tigris

MGVFSLTSLWKLHPPLRAL

>Lipotes_vexillifer

MGVFSLSSTWKPQPPLQPL

>Rousettus_aegyptiacus

MGVFSLSSFWKPRSPLQAL

>Dasypus_novemcinctus

MGVFSLSSAWRPQPPL---

>Galeopterus_variegatus

MGVFSLSSTWKPQHPLQAP

>Oryctolagus_cuniculus

MGVFSLSSTWKPQRPRQAP

>Equus_caballus

MGVFFLSSTWKPQLPLQAR

>Marmota_marmota_marmota

MGVFSLFSTWKPQPPFQAP

>Tupaia_chinensis

MGVFPLFSTWKLQTSLQAP

HG0657 ENSG00000151276 MAGI1

>Homo_sapiens

MQTKLAGACAGLAGKPRKFVL

>Ursus_maritimus

MQTKLAGACAGLAGKPRKFVL

>Balaenoptera_acutorostrata_s

MQTKLAGACAGLAGKPRKFVL

>Pteropus_vampyrus

MQTKLAGACAGLAGKPRKFVL

>Condylura_cristata

MQTKLAGAFAELAGKPRKFVL

>Ochotona_princeps

MQTKLAGARAGLAGKPGKFVG

>Equus_asinus

MQTKLAGACAGLAGKPRKFVL

>Manis_javanica

MQTKLAGACARLAGKPRKFVL

>Ictidomys_tridecemlineatus

MQTKLAGACAGLAEEPRKFVL

>Tupaia_chinensis

MQTKLAGACAGLAGKPRKVVL

HG0658 ENSG00000153786 ZDHHC7

>Homo_sapiens
--------------------------------------MHLKILKKRHKMFK
>Canis_lupus_familiaris
--------------------------------------MHLKILKKRRKMFK
>Balaenoptera_acutorostrata_s
--------------------------------------MHLKILKKRHKMFK
>Rhinolophus_ferrumequinum
--------------------------------------MHLKILKKRRKMFK
>Condylura_cristata
--------------------------------------MHLKILKKRHKMFQ
>Equus_asinus
--------------------------------------MHLRVLKKRRKTFK
>Manis_javanica
MMRPCLSRSSRRRRRRRPGGAAEPGDPARASDPVTQDSMHIKILKKRRKMSK
>Loxodonta_africana
--------------------------------------MHLKTLKKRRKMFK
>Dipodomys_ordii
--------------------------------------MHLKILKKRHKMFA
>Tupaia_chinensis
--------------------------------------MHLKILKKRHKAFR

HG0659.1 ENSG00000158856 DMTN

>Homo_sapiens
----MEVYVTEEAAALLPHALGP
>Odobenus_rosmarus_divergens
MRRGMEVYVTEEAAVLLPHAPGP
>Balaenoptera_acutorostrata_s
----MEVYVTEEAA--EPHAPGP
>Rousettus_aegyptiacus
MRRGMEVYVTEEAAVLLPHAPGP
>Erinaceus_europaeus
----MEVYLTEEEAVLLPHAPGP
>Equus_asinus
MRRGMEVYVTEEAAVLLPHAPGP
>Manis_javanica
MRRGMEVYVTEEAAVLLPHAPGP
>Fukomys_darlingi
MRLEMEVYVTTEAAALLPHAPGP
>Trichechus_manatus_latirostr
MRRGMEVYVTEEAAVLLPHSPGP
>Orycteropus_afer_afer
MRRGMEVYVTEEAAELLPHAPGP

HG0659.2 ENSG00000158856 DMTN

>Homo_sapiens

MRTRQPGGTRQLLSRPQARSAQQPRRA

>Mustela_putorius_furo

MRTPQPGGTHSLLSWPGARSAQEPGRA

>Balaenoptera_acutorostrata_s

MRTPQPGGAPFLLSRPRARSAQEPDRA

>Pteropus_alecto

MRTLQPSGTPFLHSRPRARSAQKPGRA

>Condylura_cristata

MRTLQRSGPPSLLSRPRARSAPEPGRA

>Equus_asinus

MRTPRPGGTPFLLSRPRARSAQEPRSA

>Manis_javanica

MRPPQPAGTPFLLSPHRVRSAQEPGRA

>Trichechus_manatus_latirostr

MRTLQLGGTPFLLSRPRTRSAQERSRA

HG0660 ENSG00000165861 ZFYVE1

>Homo_sapiens

-----MEDETIKPNQRCC

>Canis_lupus_familiaris

-----MEDKTIKPNQRCC

>Lipotes_vexillifer

-----MEDETIKPSQRCC

>Rhinolophus_ferrumequinum

-----MEDETIKPNQRCC

>Dasypus_novemcinctus

-----MEDETIKPNQYCC

>Ceratotherium_simum_simum

-----MENKTIKPNQRCC

>Loxodonta_africana

-----MEDETIKPSQCCC

>Fukomys_darlingi

MSLHFTEDEPIKPNQHCC

>Tupaia_chinensis

-----MEDETITPNQHCC

>Trichechus_manatus_latirostr

-----MEDETVRPNQSCC

HG0661 ENSG00000166845 C18orf54

>Homo_sapiens

-MDFLLEIRTEACTEEMLFLREMNRNNPL

>Acinonyx_jubatus

MMDFLLEIGTEVFTEEM-FLREMNRNNPF

>Physeter_catodon

MMDFLLEIHLEVVTEEMLFLREMSRNNPF

>Rousettus_aegyptiacus

MMYFFWR--IRTVTEEMLFLREMNRNKPF

>Galeopterus_variegatus

MMNFLLEIRTQAFTEEMLFLREMNRNNPF

>Oryctolagus_cuniculus

-MGLLLGSRTGARMEEMLFLRGMNRNNPF

>Equus_asinus

MMDFSLKIRAEAFREKRLFLREMNRNNLF

>Manis_javanica

MMDFLLE--TEVFTEEMLFLREINRNNPF

>Loxodonta_africana

----------------------MNRNNPF

>Tupaia_chinensis

MMDLLSEIRSEAFVEEMLFLREMNRNNSF

HG0662 ENSG00000166897 ELFN2

>Homo_sapiens

MVAAPGRTVLPAPASGR

>Canis_lupus_familiaris

MVAAPGRTVPPAPASRC

>Orcinus_orca

MVAAPGRTVLPAPASGR

>Miniopterus_natalensis

MVAATGRTVLPPPASSC

>Galeopterus_variegatus

MVATPGRTVLPAPASGC

>Erinaceus_europaeus

MVATPGRTVLPAAASGL

>Oryctolagus_cuniculus

MVAAPGRTVLPAPTSGR

>Manis_javanica

MVAAPGRTVLPAPASGH

>Ictidomys_tridecemlineatus

MVAAPGRTVPPAPASGR

>Tupaia_chinensis

MVAAPGRTVLPTPASGR

HG0663 ENSG00000168615 ADAM9

>Homo_sapiens

MSGASSDRGLSCVSAARVLVGRACSSGARASRPDLGPRQGWKMMEEAEVEATEC

>Odobenus_rosmarus_divergens

MSGASSDRGRV------ASVPSACSSRARASRPALGPRQGWEMMEQPEAEATEC

>Orcinus_orca

------------------------------------------MVERAEAGATEC

>Rousettus_aegyptiacus

------------------------------------------MMEPADAEAIEC

>Dasypus_novemcinctus

-------------------------------------------MEEAEAGATEC

>Condylura_cristata

------------------------------------------MMEQAEAEVSEC

>Oryctolagus_cuniculus

MSGASLDRGPSCVSAARVLVPGRARP-----GQPWGPREGWKMMGEAE--ATEC

>Equus_asinus

MSGASSDWSRV------ASVPSACSPAGARTRPALGPRQGWEMMEPAEAQATDC

>Manis_javanica

MSGASS-DSRV------ASVPSACSPRARASLPALGPRQGWEVMERAEAEATVS

>Chinchilla_lanigera

---------MGCVSAARMPASGR------AHPGPHGARQGSEMMDKAETAR---

HG0664 ENSG00000169085

>Homo_sapiens
----------------------MIPEPDWIRMPRD
>Felis_catus
----------------------MIPEPDWIRKPRD
>Balaenoptera_acutorostrata_s
----------------------MIPEPDWIRKPRD
>Myotis_davidii
MLQRRLKLDELTAPACLLWTCTMIPEPDWIRKPCN
>Galeopterus_variegatus
----------------------MIPEPDWIRKPHD
>Oryctolagus_cuniculus
----------------------MIPEPDWISDARD
>Equus_caballus
----------------------MIPEPEWIRKPRD
>Manis_javanica
----------------------MIPEPDWIRKPRD
>Chinchilla_lanigera
----------------------MIAEPDWIRRPRD
>Tupaia_chinensis
--------MSSQPPACLLWTCTMIPEPDWIRKARD

HG0665 ENSG00000169515 CCDC8

>Homo_sapiens
MSAKMWIWTSSWKP
>Ailuropoda_melanoleuca
MSAKMWIWTSSWKP
>Balaenoptera_acutorostrata_s
----MWIWTSSWKP
>Rhinolophus_ferrumequinum
----MWIWTSSWKP
>Galeopterus_variegatus
MSAKMWIWTSSWKP
>Equus_caballus
MSAKMWIWTSSWRP
>Manis_javanica
MSAKMWIWTSSWKP
>Loxodonta_africana
----MWTWMSSWKP
>Marmota_monax
MSAKMWIWTSSWKP
>Tupaia_chinensis
----MWIWTSWWKP

HG0666 ENSG00000171943 SRGAP2C

>Homo_sapiens
MSDPRPASGRCSSPG
>Ailuropoda_melanoleuca
MSDPRPASGRCSSPG
>Physeter_catodon
MSDPRPASGRCSSPG
>Rousettus_aegyptiacus
MSDPRPASGRCSSPG
>Condylura_cristata
MSDPRPASGRCSSPG
>Oryctolagus_cuniculus
MSDPRPASGRCSSPG
>Ceratotherium_simum_simum
MSDPRPASGRCSSPG
>Manis_javanica
MSDPRPASGRCSSPG
>Loxodonta_africana
MSDPRPASGRCSSPG
>Cricetulus_griseus
MSDPRPASGRCSSPG

HG0667.1 ENSG00000182700 IGIP

>Homo_sapiens
MCQYTVRISLICTVWGPHLRMLNFAN
>Acinonyx_jubatus
MCQLYSRISLICTVWGPH--------
>Physeter_catodon
MCQLYSRISLICTVWGPH--------
>Myotis_brandtii
MCQLHSRISLICTVWGPH--------
>Galeopterus_variegatus
--------------------MLNFAN
>Condylura_cristata
MCQLYSRISLICTVWGPH--------
>Ceratotherium_simum_simum
MCQLYSRISLICTVWGPH--------
>Manis_javanica
MCQLYSRISLICTVWGPH--------
>Rattus_sp.
--------------------MLNFAN
>Tupaia_chinensis
--------------------MLNFAN

HG0667.2 ENSG00000182700 IGIP

>Homo_sapiens
--------------------------------MLRKSVGFSAVREGRSS-ALIWVTEANMSKTLVN
>Ailuropoda_melanoleuca
MGPFSISWGICTGQRLPAMRYELKPISLFSYSMLRKSVGFSAVREGRSSAALIWVTEAGWGKTLVN
>Lipotes_vexillifer
--------------------------------MLRKSVGFLAVREGRSSAALIWVTEASLSKSLVN
>Rousettus_aegyptiacus
--------------------------------MLRKSVGFSVVREGRSSAALIWVTEAGLSKTLVN
>Galeopterus_variegatus
--------------------------------MLRKSVGFSAVREGRSS-ALIWVTEAGLSKTSVN
>Sorex_araneus
---------MCTGRRLPAMRCELKPISLFSYSMLRKSVGFSAVREGRSSAALIWVTEAGLSKTLVN
>Ceratotherium_simum_simum
--------------------------------MLRKSVGFSAVREGRSSAALIWVTEAGLSKTLVN
>Manis_javanica
--------------------------------MLRKSVGFSAVREGRSSAALIWVTEAGLSKTLVN
>Chinchilla_lanigera
--------------------------------MMRKSVGFSAVREGRSSAALIWVTEAGLSKTLVN

HG0668 ENSG00000182944 EWSR1

>Homo_sapiens
MVAPGVARFAPAQCGA
>Tursiops_truncatus
MVAPEVAGFAPAQCGA
>Eptesicus_fuscus
MVAPEVAGFAPAQCGA
>Condylura_cristata
MVAPEVADFAPAQCGA
>Ochotona_princeps
MVAPEVASFAPAQCGA
>Ceratotherium_simum_simum
MVAPEVAGFAPAQCGA
>Loxodonta_africana
MVAPEVAGLAPAQCGA
>Marmota_marmota_marmota
MVAPEVASFAPAQCGA
>Trichechus_manatus_latirostr
MVAPEVAGFAPAHCGA
>Orycteropus_afer_afer
MVTPEVAGFAPAQCGA

HG0669 ENSG00000183876 ARSI

>Homo_sapiens

MVESTFWKLPRRVSGWPG

>Mustela_putorius_furo

MVESTFWKQPRRVSGWPG

>Myotis_lucifugus

MVESTFWKSPRRVPGWPG

>Dasypus_novemcinctus

MVESTFWKPPSRVSGWPG

>Condylura_cristata

MVQSTFGELPRRVSGWPG

>Oryctolagus_cuniculus

MVESAFWEPPRRVSGWPG

>Ceratotherium_simum_simum

MVESTFWKPPRRVSGWPG

>Loxodonta_africana

MVESTFWKPPSRVSGWPG

>Cavia_porcellus

MVESTFWRPPRYVPGWPG

>Tupaia_chinensis

MVESTFCKSPRRVSGWPD

HG0670 ENSG00000187164 SHTN1

>Homo_sapiens
---------------------------------------------------------------------------------------------------MISLARS
APGGGGAGADPTSG
>Leptonychotes_weddellii
---------------------------------------------------------------------------------------------------MISLARS
APGGGGAGADPTSG
>Lipotes_vexillifer
---------------------------------------------------------------------------------------------------MISPARS
APGGGGAGANPTSG
>Eptesicus_fuscus
---------------------------------------------------------------------------------------------------MISPVRS
APGGGGAGTNPTSG
>Dasypus_novemcinctus
---------------------------------------------------------------------------------------------------MISLARF
APGGGGAGANPTSG
>Oryctolagus_cuniculus
---------------------------------------------------------------------------------------------------MISPARS
APGGGGAGANPTSG
>Equus_caballus
MRVLMSRGWRRRKGSKSEPETRQTGLHALLPAHPAAPASVHKAEAQRPPQAKARAAVPVPPQVSPSTPSPVPGLLVRPRPPPRGLPAGEAPRPLSRCVRMISLVRS
APGGGGAGANPTSG
>Loxodonta_africana
---------------------------------------------------------------------------------------------------MISLARS
APGGGGAGANPTSG
>Marmota_marmota_marmota
---------------------------------------------------------------------------------------------------MISLARS
APGGGGAGADPTSG
>Tupaia_chinensis
---------------------------------------------------------------------------------------------------MISLARS
APGGGGAGADPTSG

HG0671 ENSG00000189184 PCDH18

>Homo_sapiens

------MPKVSGSRDTVCIKQKFASC

>Odobenus_rosmarus_divergens

MQGSDSTPIISGSRDTVCIKQKFASC

>Orcinus_orca

MQGLDSMPIISGSRDTVCIKQKFASC

>Pteropus_vampyrus

MRGLHGVPMLSGSRHTVCIKQKFASR

>Dasypus_novemcinctus

MQGLDSLPVISASRDTVCMKQKFANC

>Galeopterus_variegatus

MQGLDSMPTISGSRNTVCIKQKFASG

>Ceratotherium_simum_simum

MQGLDSMPVISGSRDTVCIKQKFASC

>Manis_javanica

MQGSDSMPIVSGSRDTVCIKQKFASC

>Cavia_porcellus

MQGI-GLHADSGNRDTVCMKQKFASC

>Trichechus_manatus_latirostr

MQGLDSMPIISGNRDTVCIKQKFASC

HG0672 ENSG00000196369 SRGAP2B

>Homo_sapiens
MSDPRPASGRCSSPG
>Ailuropoda_melanoleuca
MSDPRPASGRCSSPG
>Physeter_catodon
MSDPRPASGRCSSPG
>Rousettus_aegyptiacus
MSDPRPASGRCSSPG
>Condylura_cristata
MSDPRPASGRCSSPG
>Oryctolagus_cuniculus
MSDPRPASGRCSSPG
>Ceratotherium_simum_simum
MSDPRPASGRCSSPG
>Manis_javanica
MSDPRPASGRCSSPG
>Loxodonta_africana
MSDPRPASGRCSSPG
>Cricetulus_griseus
MSDPRPASGRCSSPG

HG0673 ENSG00000266028 SRGAP2

>Homo_sapiens

MSDPRPASGRCSSPG

>Odobenus_rosmarus_divergens

MSDPRPASGRCSSPG

>Physeter_catodon

MSDPRPASGRCSSPG

>Rousettus_aegyptiacus

MSDPRPASGRCSSPG

>Condylura_cristata

MSDPRPASGRCSSPG

>Oryctolagus_cuniculus

MSDPRPASGRCSSPG

>Ceratotherium_simum_simum

MSDPRPASGRCSSPG

>Manis_javanica

MSDPRPASGRCSSPG

>Loxodonta_africana

MSDPRPASGRCSSPG

>Cricetulus_griseus

MSDPRPASGRCSSPG

HG0674.1 ENSG00000006576 PHTF2

>Homo_sapiens
---------------MFFSLAQPKMTNV
>Ailuropoda_melanoleuca
---------------MFFSLAQPKMTNV
>Lipotes_vexillifer
MLFSLLLEGGSQVLCMFFSLAQPKMTNV
>Myotis_davidii
---------------MFFSLEQLKMTNV
>Erinaceus_europaeus
MLICLLLERGSKVVCVFFSLAQPKMTNV
>Ochotona_princeps
---------------MFFSLTQPKMTTV
>Elephantulus_edwardii
---------------MLFSLAQPKMTNV
>Equus_asinus
---------------MFFSLAQPKMTNV
>Orycteropus_afer_afer
---------------MFFFLAQPKMTNV

HG0674.2 ENSG00000116793 PHTF1

>Homo_sapiens

MRPAWPAASPPASRWDPGH

>Lipotes_vexillifer

MRPAWPGASPPASWPGRGH

>Nannospalax_galili

MRPTRPAASPPASRPGPSH

>Tupaia_chinensis

MRPAGPVASPRASRTGRGH

HG0675 ENSG00000048991 R3HDM1

>Homo_sapiens
---------------MQSISLQVLWKEHLLNSV--------
>Canis_lupus_familiaris
---------------MQSISLQILWKVHLHSLCMMKRTPVQ
>Orcinus_orca
---------------MQSISLQILWKVHLHSLCMMKQIPIR
>Myotis_brandtii
MPNQPAVCVVSSGGAMQSISLQILWKVHFHLLCVRKQIPIQ
>Galeopterus_variegatus
---------------MQSISLQILWKVHLH-----------
>Oryctolagus_cuniculus
---------------MQNISLQILWKVHLH-----------
>Ceratotherium_simum_simum
---------------MQSISLQILWKVHLHSLCMMKQIPIQ
>Fukomys_damarensis
--------MELSRRAMQSISLQILWKVHLH-----------
>Orycteropus_afer_afer
---------------MQSISLQILWKVHLH-----------

HG0676 ENSG00000058673 ZC3H11A

>Homo_sapiens
MAIMLLCLLQLAAPLCSYSITIRFYLFWLNTP
>Felis_catus
MAIMLLCLLQLAAPLCSYSITIRFYLFWLNTP
>Eptesicus_fuscus
MAVMLLCLLQLAAPLCSYSITIRFYLFWLNTP
>Oryctolagus_cuniculus
MALPPLCLPQLAALLCSYSVTLRFYLFWLNTP
>Equus_caballus
MAIMLLCLLQLAAPLCSYSVTIRFYLFWLNTP
>Loxodonta_africana
MAITLLCLLQLAAPLRSYSITVRFYLFWLNTP
>Cricetulus_griseus
MAIMLLCLLQLAAPLCSYSVTIRFYLFWLNTP
>Trichechus_manatus_latirostr
MAIMLLCLLQLAAPLCSYSVTVRFYLFWLNTP
>Orycteropus_afer_afer
MAIMLLCLLQLAAPLYSYSITIRFYLFWLNTP

HG0677 ENSG00000064989 CALCRL

>Homo_sapiens

-MIHQPLVLPSVKTLDGPTFRS

>Ailuropoda_melanoleuca

-MIHQPLVLPPVKTLDGPTFRS

>Balaenoptera_acutorostrata_s

MMIHQPLVLPPVKTMDGPTFRS

>Rousettus_aegyptiacus

-MIHQPLVLPPVKTLDGPTFRS

>Dasypus_novemcinctus

-MTHQPLVLPPVKTLDVPTFRS

>Galeopterus_variegatus

MMIYQPLVLSPVKTLDGPTFRS

>Ochotona_princeps

-MIHPPLILPPVKTVDDPTFRS

>Equus_asinus

-MMHQPLVLPPVKTLDGPTFRS

>Nannospalax_galili

-MIHQPLVLPAVKTSDGPTFRS

HG0678.1 ENSG00000090776 EFNB1

>Homo_sapiens
----------------------------------------------MGLARSAASPRSSKYTEQGPR--
>Canis_lupus_familiaris
MRAGQQWEVCEARTGSHWTFGRNRWGAGTRRRGCGATRPRLQKHKLMGLALLVASSRSSKYGKRGPR--
>Orcinus_orca
----------------------------------------------MGLARSAVSPRSSKYTERAPR--
>Eptesicus_fuscus
----------------------------------------------MGLARSAASPLSFKSTERGPR--
>Galeopterus_variegatus
----------------------------------------------MGLARSAASLR-SKCIERGPPLK
>Condylura_cristata
----------------------------------------------MGLACSAASPLSSKYTEWGPC--
>Ochotona_princeps
----------------------------------------------MGLARSPASPRSSKYRERGPALK
>Ictidomys_tridecemlineatus
----------------------------------------------MGLARSAASPRSFKYTERGPR--
>Tupaia_chinensis
----------------------------------------------MGLAHSAASPRSSKYTERGPC--

HG0678.2 ENSG00000108947 EFNB3

>Homo_sapiens
MPPPPGPRLPQSPT
>Mustela_putorius_furo
MPPPPGPRLPQSPT
>Physeter_catodon
MPPPPGPRLPQSPT
>Myotis_davidii
MPPPPGAWLPQSPT
>Oryctolagus_cuniculus
MPPPPGPRLPQSPT
>Mesocricetus_auratus
MLPPPGPRLPMSPT

HG0679 ENSG00000100281 HMGXB4

>Homo_sapiens
MAAIGGVEAGSGRAE
>Odobenus_rosmarus_divergens
MAAVGGAEAGSGRAE
>Rousettus_aegyptiacus
MAVVGGAEAGSGRAK
>Dasypus_novemcinctus
MAAVGGAEAGSRRAE
>Condylura_cristata
MAAVGGAEAGSGPAE
>Ochotona_princeps
MAAVGGAEAGSGRA-
>Ceratotherium_simum_simum
MAAVGGAEAGSGRAE
>Manis_javanica
MAAVGGAEAGSGRAE
>Ictidomys_tridecemlineatus
MAAVGGAEAGSGRAE

HG0680 ENSG00000100678 SLC8A3

>Homo_sapiens

MPRTRLRPAGSPRGR

>Odobenus_rosmarus_divergens

MPRTRLRPAGSPRGR

>Eptesicus_fuscus

MPRTRLRPAGSPRGR

>Erinaceus_europaeus

MPGTRLRPGGCPRGR

>Oryctolagus_cuniculus

MPRTRLRPAGSPRGR

>Equus_przewalskii

MPRTRLRPAGSPRGR

>Marmota_marmota_marmota

MPRTRLRPAGSPRGR

>Trichechus_manatus_latirostr

MPRTRLRPAGSPRGR

>Orycteropus_afer_afer

MPRTRLRPAGSPRGR

HG0681 ENSG00000101349 PAK5

>Homo_sapiens
---------------------------------------------------MLPPLAKMNTEEK---
>Mustela_putorius_furo
---------------------------------------------------MLPPLAKMNTEEK---
>Balaenoptera_acutorostrata_s
---------------------------------MRHRRCADSQRPEIHCGEMLPPLAKMNTEEK---
>Cynopterus_sphinx
-------------------------------------MRRRRQPPEVHCGKMLPPLAKMNTEEK---
>Dasypus_novemcinctus
MPSCKLQLLWSASLSTLFPELRGSAALHLISPSLNCSLEIFSRQQEIHCGEMFPPLAKMNAEEKEEN
>Erinaceus_europaeus
---------------------------------MRCRRVADNHPPVIHSGKMLPPLAKMNTEEK---
>Manis_javanica
---------------------------------------------------MLPPLAKMNTEEK---
>Ictidomys_tridecemlineatus
---------------------------------MSHCRFADSQPPEIHCGEMLLPLAKMNTEEK---
>Orycteropus_afer_afer
---------------------------------------------------MFPPLAKMNTEENEGN

HG0682 ENSG00000105829 BET1

>Homo_sapiens

MSWRFRGRSWCFAGPWY

>Ailuropoda_melanoleuca

MSWRFGERSWCFSGPWY

>Lipotes_vexillifer

MSWRFRGRSWCFAGPRY

>Myotis_davidii

MSWRYRGRSW-------

>Dasypus_novemcinctus

MPWHFRGRSWSSGGPWD

>Oryctolagus_cuniculus

MSWRLG-----------

>Ceratotherium_simum_simum

MSWRFRGRSWCFAGPWS

>Ictidomys_tridecemlineatus

MSWRFGARSWCCSGPWD

>Tupaia_chinensis

MSWRSWGRSWCSE----

HG0683 ENSG00000111837 MAK

>Homo_sapiens

MKHTRRRSFTRERTRVFIRMEVLFPCPIPSCYIS

>Canis_lupus_familiaris

-------------------MEALFPCQIPYWYIS

>Myotis_brandtii

MKHTRRRSLTPEQKRVFIRMDALFPCHIPYCCSS

>Erinaceus_europaeus

-------------------MEALFSCHTPYCSIS

>Oryctolagus_cuniculus

-------------------MESLFPCQIPYYCTA

>Equus_caballus

MKHTRRPSFTRQRRRVFIRMEALFPCQIAYCCIS

>Loxodonta_africana

-------------------MEALFPCQIHYYIS-

>Tupaia_chinensis

-------------------MEALFPCQIPYCCIS

>Orycteropus_afer_afer

-------------------MEALFP---------

HG0684 ENSG00000113916 BCL6

>Homo_sapiens

MRGLCHSPHCLLH-----------------------------------------------------------------

>Felis_catus

MRGLCHSPHCPLP-----------------------------------------------------------------

>Pteropus_vampyrus

MRGLCHSPHCPLP-----------------------------------------------------------------

>Erinaceus_europaeus

MRGLCHSPHCPLP-----------------------------------------------------------------

>Oryctolagus_cuniculus

MRGLCHSPHCPLP-----------------------------------------------------------------

>Ceratotherium_simum_simum

MRGLCHSPHCPLP-----------------------------------------------------------------

>Manis_javanica

MRGLCHSPHCPCLDGVTGKKRRIQCWLPRCTKRFI-------------------------------------------

>Ictidomys_tridecemlineatus

MRGLCHSPHCPLPWRSNRKKKKKAYTM-LATENLFECNEIKGRRSHTIRPPPRLRGLRRMGSGASVSPGAGRSLEDSF

>Tupaia_chinensis

MRGLCHSPHCPLP-----------------------------------------------------------------

HG0685 ENSG00000117262 GPR89A

>Homo_sapiens

MASGWRAAVPAAAPGRRQTV------------------

>Odobenus_rosmarus_divergens

MASGWRAAVPAVAPGRSPPL------------------

>Physeter_catodon

MASGWRAEVPAVARGRQLPV------------------

>Pteropus_vampyrus

MASGWRAAAPVAVLGQRPPV------------------

>Galeopterus_variegatus

MASDWSAALRAAAPGRRPPA------------------

>Erinaceus_europaeus

MASGSGARLSAAAPGRRPQV------------------

>Equus_asinus

MASGWRSAVPAVAPGRRLPV------------------

>Marmota_monax

MASGRSAAVPAAASGRRPAALVGRRPERSCPGEWEVET

>Tupaia_chinensis

MASGWLAAVPAVAPGRRLPV------------------

HG0686 ENSG00000118985 ELL2

>Homo_sapiens

----------------------------MTVRQQRDGPSSGGGCRSPSSRGRSGG

>Leptonychotes_weddellii

----------------------------MTVRQQRDGRSCGGGCRSPSSRGRSGG

>Balaenoptera_acutorostrata_s

----------------------------MTVRQQRDGRSCGGGCRSPSSRGRSGG

>Miniopterus_natalensis

----------------------------MTVRQQRDGRSCGGGCRSPSSRGHRGD

>Condylura_cristata

MNEPGAGRGPEGGAGVGGGSPRSGCAQYMTVRQQRDGRSCGGGCRSPSSRGRSGG

>Oryctolagus_cuniculus

----------------------------MTVRQQRDGRSCGGGCRSPSSRGRSGG

>Ceratotherium_simum_simum

----------------------------MTVRQQRDGRSCGGGCRSPSSRGRSGG

>Manis_javanica

----------------------------MTVRQQRDGRSCGGGCRSPSSRGRSGA

>Octodon_degus

----------------------------MTVRQQRDGRSCGGGCRSPSSRGRSGG

HG0687.1 ENSG00000120549 KIAA1217

>Homo_sapiens

------------------------------------------------------------------MQFSRARETLHRSGK

>Acinonyx_jubatus

------------------------------------------------------------------MQFSPARETLHRSGE

>Balaenoptera_acutorostrata_s

------------------------------------------------------------------MQFSPARETLYRSGE

>Myotis_brandtii

------------------------------------------------------------------MHFSPARETLHGSGE

>Condylura_cristata

------------------------------------------------------------------MQFSWARATLHRSGE

>Ceratotherium_simum_simum

------------------------------------------------------------------MRFSRASETLHRSGE

>Manis_javanica

------------------------------------------------------------------MQSSRASATLHRSGA

>Mus_musculus

MGPVLDCIQLRVPGNPAGCVLGQQSRHLLGGRGGDSHRLQDTAPSPGDSRRVADAPGPGPRRPGAGMRSSRARETLHRSGD

>Tupaia_chinensis

------------------------------------------------------------------MQFSRAKETLHRSGE

HG0687.2 ENSG00000120549 KIAA1217

>Homo_sapiens

MVFIGCEHSS-----------------------------------------------------

>Chinchilla_lanigera

MVFIGCEHSSWGTFQDLCVLCD--DFRL-----------------------------------

>Trichechus_manatus_latirostr

MVFIGCEHSSWERFRISLSFLIFSSLRRKLSKTSKCGTEDLDEASKQRNRKIGSLSQCSFSRR

HG0688 ENSG00000122420 PTGFR

>Homo_sapiens
----------------------MSGLQSCTVLRGR
>Ailuropoda_melanoleuca
--MPDEQL--SSAHDSSVFMLQMSGLQACTVLRGR
>Myotis_lucifugus
-------------------MPKMPGLQACTLLRGR
>Dasypus_novemcinctus
MEHPRARSSAPASAPPAARDAKMSGLQGCALLRGR
>Oryctolagus_cuniculus
-------------------MLQMSGLQACTVLRGR
>Equus_caballus
----------------------MSGLQACTVLRGR
>Manis_javanica
MALGAARFSAPPGAPSAAGDAKMSGLQTCTVLRER
>Loxodonta_africana
MAHRGARSSALFRASPAAGDTKMSGLQACTDLRGR
>Trichechus_manatus_latirostr
----------------------MSGLQACTDLRGR

HG0689 ENSG00000123352 SPATS2

>Homo_sapiens

MNVTTLVTSYNLPGKGDMNVCNDTS-----------

>Leptonychotes_weddellii

-----------------MNVCNDIS-----------

>Balaenoptera_acutorostrata_s

-----------------MNVCNDIS-----------

>Pteropus_alecto

-----------------MNVCNDIS-----------

>Oryctolagus_cuniculus

-----------------MNVCNAIS-----------

>Ceratotherium_simum_simum

-----------------MNVCNDIS-----------

>Fukomys_darlingi

-----------------MTVCNDSS-----------

>Tupaia_chinensis

-----------------MNVCHDTS-----------

>Trichechus_manatus_latirostr

-----------------MNVCNDIPDKKLIYKKERK

HG0690 ENSG00000125931 CITED1

>Homo_sapiens
MCPAAKLQKGHR
>Leptonychotes_weddellii
MFPAAQLQKGHR
>Orcinus_orca
MFPAAQLQKGHR
>Rhinolophus_ferrumequinum
MFPAAQLQKGHR
>Galeopterus_variegatus
MFPAAQLQKGHR
>Oryctolagus_cuniculus
MLPAAQLQKGHR
>Equus_caballus
MFPAAQLQKGHR
>Manis_javanica
MLPAAELQKGHR
>Ictidomys_tridecemlineatus
MFPAAQLQKGHR

HG0691 ENSG00000129535 NRL

>Homo_sapiens

MPFGLCAHSPLELSRGTRPCSMEPSVSWEAVPVWLWH

>Myotis_davidii

---------------------MVPPVSGEAVLAWLGH

>Galeopterus_variegatus

---------------------MGPLVSGEAVPVWL--

>Oryctolagus_cuniculus

---------------------MEPPVSGKTVAAWL--

>Elephantulus_edwardii

---------------------MGPPVLEEAVPVWF--

>Ceratotherium_simum_simum

---------------------MGPPVSGEAVPTWL--

>Loxodonta_africana

---------------------MGPPVSGEAVPAWLGH

>Ictidomys_tridecemlineatus

---------------------MGPPISGEAVPVWL--

>Trichechus_manatus_latirostr

---------------------MGPPVSGEAVPAWL--

HG0692.1 ENSG00000130338 TULP4

>Homo_sapiens

MALRGAPAGGGAVRWSPAFPGDT------GPSFEVESFWF

>Ailuropoda_melanoleuca

MALKSAPVCGGPVRWSPALCQDTATCLLGSQA---GP---

>Pteropus_alecto

MALRAAWACGGAVRWSPALPGDTATCQETGLGSEVGCFWF

>Dasypus_novemcinctus

MALRSGLAGGGAVGRGSVFPGDVNTGWLGAH----GLMWF

>Galeopterus_variegatus

MALRGNLSSGGVVQWSLMLPGDT------GPSFEVGNFWF

>Equus_asinus

MAPRGAWACGVAVRWSPALHGDTAACREAGPGSECGSFWF

>Manis_javanica

MALRSVLACGGAVRWSPALPRDTTTCWPTGPGSEVGSVWF

>Ictidomys_tridecemlineatus

MALRGAPAGGGVFLGSPAFLRDT------GPSFAVGSFWF

>Tupaia_chinensis

MALWGGPAGGGAGQWGPALPGDT------GPSFETGSFWF

HG0692.2 ENSG00000130338 TULP4

>Homo_sapiens

MDLILRLAFFIKMEKDL------------

>Felis_catus

MDLILRLVFSFKMEKYVIFRSRKGRGK--

>Dasypus_novemcinctus

MDLILRLVFSIKMERYVTFRSRRGGKCLG

>Galeopterus_variegatus

MDLILRLVFFSMMEKYVIFRNRKGEK---

HG0693 ENSG00000131653 TRAF7

>Homo_sapiens

MKRAGGAAAPGRPRAGGASQGP----

>Phoca_largha

MKREGGAAAPGRPRAGHAAQGP----

>Physeter_catodon

MKREGGAAVPERPRAGDAARGP----

>Pteropus_alecto

MKLEGGAAAPGRPRAGDAARGP----

>Dasypus_novemcinctus

MSREGGAAAPGRPRAGAAARGP----

>Ceratotherium_simum_simum

MKRVGGAAAPGRPRAGDATRGP----

>Manis_javanica

MKREGGGAAPGRPRAGDAAQGPWMPL

>Ictidomys_tridecemlineatus

MKRADGAAAPGQPRAGNAAQGPWMPL

>Tupaia_chinensis

MKREGGVAAPGRPRAGDAARRP----

HG0694 ENSG00000133265 HSPBP1

>Homo_sapiens
MAAPSRVRCGRLCL-
>Felis_catus
MAAPSRVRCGRLCL-
>Physeter_catodon
MAAPSIVRCGRVCL-
>Rousettus_aegyptiacus
MAAPTRGRCGQLCL-
>Dasypus_novemcinctus
MAAPSRVRCGPLCF-
>Condylura_cristata
MAAPSRVRCGRLCLY
>Oryctolagus_cuniculus
MAAPSRVRCGRL---
>Equus_przewalskii
MAAPTRVRCGRLCL-
>Marmota_marmota_marmota
MAAPSRVRCGRLCL-

HG0695 ENSG00000133704 IPO8

>Homo_sapiens
MESELWGGKREDRGRGVRVAQQVAAGGDGGGGWGVR
>Mustela_putorius_furo
MESELWGVEREDRGRGVRVAQQVAAGGDGGGGWGVR
>Physeter_catodon
MESELLGVKREDRGRGVRVAQQVAAGGDGGGGWGVR
>Galeopterus_variegatus
MESELWGGKREDRGRGVRVAQRVAAGGDGGGGWGVR
>Oryctolagus_cuniculus
MECELWGGKREARGHGVRVAQQVAAGGGGRGVR---
>Elephantulus_edwardii
MESELWGGKREEGGRGVRVAQQVAAGGEGGGEWGVR
>Ceratotherium_simum_simum
MESELWGVEREDRGRGVRVAQQVAAGGDGGGGWGVR
>Marmota_marmota_marmota
MESELWGGKREDRGRGVRVAQQVAAGGDRGGGWGVR
>Tupaia_chinensis
MESELWGGEREDRGRGVRVAQQVAAGGDGGGGWGVR

HG0696 ENSG00000138380 CARF

>Homo_sapiens
--------------MLEAWNLNLSHYLKLLNNRRKWN----------------------------------
>Physeter_catodon
--------------------------MKILNNRRKWN----------------------------------
>Myotis_brandtii
MNFSGQEGILRGSRMLEAWNLNLSYFLKLLNNRRKWN----------------------------------
>Condylura_cristata
--------------MLETWDLNLSYFLKVTHTKNTSEVAANTIKNTSAVSWKMKGVCICVCVREREKITTY
>Oryctolagus_cuniculus
--------------MLKAGHLDLSHCLKLLN-RRKWN----------------------------------
>Ceratotherium_simum_simum
--------------MLEAWNLNLSYCLKLLNNRRQWN----------------------------------
>Loxodonta_africana
--------------MPEAWNLNLNYCLQLLTNRRKWN----------------------------------
>Ictidomys_tridecemlineatus
-----MKEHSGGGRMLEDWNLNLSHCLKLLNSRRKWN----------------------------------
>Trichechus_manatus_latirostr
--------------MLEAWNLNLNHCLKLLTNRRKWD----------------------------------

HG0697 ENSG00000143344 RGL1

>Homo_sapiens

MDWLPDRVPCSW------

>Acinonyx_jubatus

MDWLPDQVPCSWQEYGAE

>Lipotes_vexillifer

MDWLPDQVPCSWQEYEAE

>Eptesicus_fuscus

MDWLSDQVPCSWQEYGAQ

>Dasypus_novemcinctus

MDWLPDRVPCSW------

>Condylura_cristata

MDWLPYPIPCSWQECEAE

>Equus_caballus

MDWLPDQVPCSWQEYGAE

>Chinchilla_lanigera

MDWLRDRVPCSWQEYGEE

>Tupaia_chinensis

MDWLPDRVPCIRQDCEE-

HG0698 ENSG00000157554 ERG

>Homo_sapiens

MTHREKRWQNQGQLKPSGSEQLVDGLA----Y------------

>Mustela_putorius_furo

MTRREKRWWGQGQLKPSSPRSGEKSMEPFWETDHKHADLEMVSH

>Balaenoptera_acutorostrata_s

MTHREKRWGSQGQLKLPQVL---NRW------------------

>Miniopterus_natalensis

MTHREKRWWSQGQLELSPLL---RGR------------------

>Ceratotherium_simum_simum

MTHREKRWWSRGQLKLPQVL---NGW------------------

>Manis_javanica

MTHREKRWCSQGQLKLSHAV---NGR------------------

>Loxodonta_africana

MTHREKRWWNQGQLKLPQVL---NGW------------------

>Peromyscus_maniculatus_baird

MTPREKRWRNQGQLKPVWVL---HGW------------------

>Trichechus_manatus_latirostr

MTHREKRWWNQGQLKLPQAL---NGW------------------

HG0699 ENSG00000160785 SLC25A44

>Homo_sapiens

MEEDA-TGRRHSLGTTDRLGAAA

>Tursiops_truncatus

MEEDA-TGGRQSPETEDRVGAAA

>Myotis_davidii

MEEDV-HRSWQSPGTADRLGAAA

>Galeopterus_variegatus

MEEDA-AGGQQSLGTEDRVGAAA

>Condylura_cristata

MEEDA-TRGPESPGTEDRLGAAA

>Oryctolagus_cuniculus

MEEDAPGGGRRSLGTEDRVGAAA

>Equus_asinus

MEEDA-TRSQKSRGTQDRVGAAA

>Fukomys_darlingi

MEEDA-TGGSQSLGTEGRAGAAA

>Tupaia_chinensis

MEEDA-IGGRQSLGVEDRVGAAA

HG0700 ENSG00000165156 ZHX1

>Homo_sapiens

MQDG-GGVALRGAAAPAGSGASRLRDQRARRLWAAGGNWSAESAAGTGKEKTDCCFGPVGNDGLS

>Odobenus_rosmarus_divergens

--------------------------------------------------MADRCFRPVGNDGLS

>Lipotes_vexillifer

MQDG-GGGALRGTAAPAGDGR--------------------------------------------

>Myotis_ricketti

--------------------------------------------------MADQCFGPAGNDGLS

>Erinaceus_europaeus

MQDG-GGRALRGAAAPAGHGR--------------------------------------------

>Ceratotherium_simum_simum

--------------------------------------------------MADRYFGPVGNDGLS

>Manis_javanica

--------------------------------------------------MADRYFGPVGNDGLS

>Ictidomys_tridecemlineatus

MQDG-GGGVLRGAAAPAGED---------------------------------------------

>Trichechus_manatus_latirostr

MQDGDGGGALRGAADPAGESR--------------------------------------------

HG0701 ENSG00000166292 TMEM100

>Homo_sapiens
-MDC-LS-PSFHPGETVQI
>Canis_lupus_familiaris
-MDC-LS-LSFHPGETPQV
>Balaenoptera_acutorostrata_s
-MDC-FS-LSFHPGETAQV
>Rhinolophus_ferrumequinum
-MDCFFS-LSFHPGEASQV
>Dasypus_novemcinctus
MMDC-LS-LSFCPGGTLQI
>Galeopterus_variegatus
-MDC-LS-LSSPPGETLQI
>Equus_asinus
-MDCFSPRDAFHPGETP--
>Manis_javanica
-MEC-FS-PSFHLGKAPQV
>Peromyscus_maniculatus_baird
-MDC-LS-VSFHPGQAFQI

HG0702 ENSG00000167377 ZNF23

>Homo_sapiens
MELALGQQEPIVSEVGDL
>Miniopterus_natalensis
MELALGQQEPVVSEVGDL
>Dasypus_novemcinctus
MELALGQQEPVVSEVGDL
>Galeopterus_variegatus
MELALGQQEPVVSEVGDL
>Manis_javanica
MELALGQQEPVVSEVGDL
>Loxodonta_africana
MELALGQQEPVVSEVGDL
>Cavia_porcellus
MELALGQQEPVVSEVGDL
>Tupaia_chinensis
MELALGQQEPVVSEVGDL
>Trichechus_manatus_latirostr
MELALGQQEPVVSEVGDF

HG0703 ENSG00000171862 PTEN

>Homo_sapiens

------MRCGRIRARRWDATALSSLLSEAAAMMEV

>Odobenus_rosmarus_divergens

MWRDSLCTAAGYAPGRRAAAALGSLLSEAAAMMEV

>Lipotes_vexillifer

MWRDSLCTAAGYTFGRWDAAALSSLLSEAAAMMEV

>Rousettus_aegyptiacus

MWRNSLCTAAGYMIGRWDAAALSSLLSEAAAMMEV

>Condylura_cristata

MWRDSLCTAAGYVLGRWDAAALSSLLSEAAAMMEV

>Ceratotherium_simum_simum

MWRDSLCTAAGYAFGRRDAAALSSLLSEAAAMMEV

>Ictidomys_tridecemlineatus

MWRDSLCTAAGYAFGRWDAAALSSLLSEAAAMMEV

>Trichechus_manatus_latirostr

MWRNSLCTAAGYAIGRRDTAALSSLLSEAAAMMEV

>Orycteropus_afer_afer

MWRNSLCTAAGYALGRRDTVALSSLLSEAAAMMEV

HG0704.1 ENSG00000173598 NUDT4

>Homo_sapiens

MESGGQLGRGGAFISAPWFQ

>Leptonychotes_weddellii

MESGGRPGRGGVCSPAPGLQ

>Lipotes_vexillifer

MESGGRPGRGGASSPAPGFQ

>Pteropus_vampyrus

MESGGRLGKGGASSPAPGLQ

>Erinaceus_europaeus

MESGGRPGRGGASSSAAGLQ

>Oryctolagus_cuniculus

MESGRRPGRGGASLPAPGLQ

>Mus_musculus

MESGGRRSRGGASTAAPGLQ

>Trichechus_manatus_latirostr

MESSGRPGRGGASTAAPGLQ

>Orycteropus_afer_afer

MENGGRPGRGGASTSAPGLQ

HG0704.2 ENSG00000272325 NUDT3

>Homo_sapiens

MRRPRRASGGAV

>Physeter_catodon

MRRPRRASGGAV

>Pteropus_alecto

MRRPRRASGGAV

>Oryctolagus_cuniculus

MRRPRRASGGAV

>Elephantulus_edwardii

MRRPRRASGGAV

>Heterocephalus_glaber

MKRPRRASGGVV

>Orycteropus_afer_afer

MRRPRRASGGAV

HG0705.1 ENSG00000175224 ATG13

>Homo_sapiens

-MAAPGSDRLLGLRRE

>Acinonyx_jubatus

-MAAPGGDRLLGLRQE

>Balaenoptera_acutorostrata_s

-MAAPGSDRLLG----

>Miniopterus_natalensis

-MAAPGSDRLLGLRRE

>Galeopterus_variegatus

-MAAPGQYRLLGLRQE

>Erinaceus_europaeus

-MAAPGSDRLRGLRQK

>Ceratotherium_simum_simum

MMAAPGSDRLLGLQQE

>Loxodonta_africana

-MAAPGADGLLGLRLK

>Ictidomys_tridecemlineatus

-MAAPGSDRLLG----

HG0705.2 ENSG00000175224 ATG13

>Homo_sapiens

MRARNHSLCRSFAVERPVETCDPFSLNKSLVLLCCPGWSVVV--

>Canis_lupus_familiaris

MRAPSHSLCRSFAVERPVESCDPFSFNKSLGFFW----PL----

>Orcinus_orca

MRAPSHSLCRSFTVE-----------------------------

>Rousettus_aegyptiacus

MRAPSHCLCRSFAVERPVETCDPFSLSKSLVMSPICPWLQEKRL

>Condylura_cristata

MRVPSHSLCRSFTGEGDC--------------------------

>Ceratotherium_simum_simum

MRAPSHSLCRSFAGKGDC--------------------------

>Loxodonta_africana

MRAPSHSLCRSVAGEGDC--------------------------

>Peromyscus_maniculatus_baird

MRARSHSLCRSFPGEGDC--------------------------

HG0706 ENSG00000178498 DTX3

>Homo_sapiens
-----------------------------------------------MGFPGHTTYK
>Leptonychotes_weddellii
-----------------------------------------------MGFPGHTTYK
>Physeter_catodon
-----------MVTFVILTQSPPL--PLPQSSSAECGSRGFLLLSSSMGFPGHTTYK
>Rousettus_aegyptiacus
-----------------------------------------------MGFPGHTTYK
>Galeopterus_variegatus
-----------------------------------------------MGFPGHTAYK
>Sorex_araneus
-----------------------------------------------MGLPGHTTYK
>Ochotona_princeps
-----------------------------------------------MGFPGHTTYK
>Equus_caballus
MAALTGGAAATLFAFQTDFRRRDWVRSQGSPGQLSVESEGLLLLSSSMGFPGHTTYK
>Dipodomys_ordii
-----------------------------------------------MGFPGHTTYK

HG0707 ENSG00000179796 LRRC3B

>Homo_sapiens
-------MGAQARKEIMKRHMC
>Felis_catus
----------------MKRHMC
>Myotis_davidii
----------------MKRHMC
>Galeopterus_variegatus
----------------MKKHMC
>Erinaceus_europaeus
----------------MKRRMC
>Equus_asinus
----------------MKSHMC
>Manis_javanica
----------------MKRPLC
>Fukomys_damarensis
MSAKPHSRGSQARTAIMKRCMC
>Tupaia_chinensis
----------------MKKHMC

HG0708 ENSG00000184613 NELL2

>Homo_sapiens

MMCALEGLSSSRAWGGLPFLS

>Ursus_americanus

MMCALEGLSSSRAWGVLPFFS

>Balaenoptera_acutorostrata_s

MMCALEGLSSARAWGVLPFFS

>Pteropus_alecto

MLCTLEGPSSSRAWGVLPFFC

>Elephantulus_edwardii

MMCALEGLSSSRAWGILAFSS

>Loxodonta_africana

MMCALEGLSSSLAWAVLAFSS

>Ctenomys_sociabilis

-MSAAQAPSSARAWGGRPFLS

>Trichechus_manatus_latirostr

MMCALEGLSSSRAWGVLVFSS

>Orycteropus_afer_afer

MMCALEDLSSSRA--------

HG0709 ENSG00000186439 TRDN

>Homo_sapiens
-----MRLFLFEADTSS------------------------------------
>Acinonyx_jubatus
-----MRLFLFEADTSA------------------------------------
>Tursiops_truncatus
-----MKLFLFEADTSA------------------------------------
>Eptesicus_fuscus
-----MRLFLFEADTSA------------------------------------
>Dasypus_novemcinctus
-----MRLFLYEADTSA------------------------------------
>Galeopterus_variegatus
MSPGEMRLFLFEADTSS------------------------------------
>Condylura_cristata
-----MRLFLFEADTSAEPQVLLGKGSGCFQGENSVEKRQSSTGQQRSSRNNQ
>Oryctolagus_cuniculus
-----MRLFLFEADTSS------------------------------------
>Dipodomys_ordii
-----MRLFLFEADTSS------------------------------------

HG0710 ENSG00000196505 GDAP2

>Homo_sapiens
MGPPECSKVSSEY
>Mustela_putorius_furo
MGPPECSKVCSEY
>Orcinus_orca
MGPPECSKVYSEY
>Oryctolagus_cuniculus
MGPPECSEVSSEY
>Ceratotherium_simum_simum
MGPPECSEVSSAY
>Loxodonta_africana
MGTPECSDISSEY
>Chinchilla_lanigera
MGPPECSKLSRDY
>Tupaia_chinensis
MGPPECTKVTSDY
>Orycteropus_afer_afer
MGTPKCSNVSSEY

HG0711 ENSG00000198300 PEG3

>Homo_sapiens
MSTLGWWRRRAPGAMRVR
>Canis_lupus_familiaris
MSTLGWWRRRAPGALWVC
>Balaenoptera_acutorostrata_s
MSTVGWWRRRVPGAVWGH
>Pteropus_alecto
MSTLGWWCRQARGAV---
>Galeopterus_variegatus
MSTLGWWRRRAPGAMWAH
>Equus_asinus
MSTLGWWRRRAPGAMWAR
>Manis_javanica
MSTLGWWRRRSP------
>Loxodonta_africana
MSTLDWWRHRAPGAGWPC
>Nannospalax_galili
MSTLGWWRRRAPGAVWVH

HG0712 ENSG00000213949 ITGA1

>Homo_sapiens
MKGRCPFKVCFYSPWTLA-------------------------------------------
>Canis_lupus_familiaris
MKGRCPFKVCFAAQPVDFSLNTNPRSWLYLSMSRTEPGRLPVGFQRPSAGRGGRWGSESGI
>Orcinus_orca
MKGRCPFKVCFAARGL---------------------------------------------
>Pteropus_alecto
MKGRCPFKVCLAARGL---------------------------------------------
>Oryctolagus_cuniculus
MKGRCPFKVCCAVRGL---------------------------------------------
>Ceratotherium_simum_simum
MKGRCPFKVCFAARGL---------------------------------------------
>Manis_javanica
MKGRCPFKVCFAARGL---------------------------------------------
>Loxodonta_africana
MKGRCPFKVCFAAGGL---------------------------------------------
>Nannospalax_galili
MKGRCPFKVCFAARGL---------------------------------------------

HG0713 ENSG00000008513 ST3GAL1

>Homo_sapiens

MPDSFDTLQRQRALDRREEEEEEERREKE-E--AM------------------------------------------

>Panthera_tigris

MPDSFDTLQRQRPEGGGRGGQGGRGGGGRGGRRDVSREGGQCSGQPPL-----------------------------

>Balaenoptera_acutorostrata_s

MPDSFDTLQRQRPEGGGRGGRRGRGGGEEKGAM--------------------------------------------

>Miniopterus_natalensis

MPDSFDTLQRQRPEGGGSGGERG-------GRGDVS-----------------------------------------

>Oryctolagus_cuniculus

MPDSFDTLQRQRDLEQRQPPEREQREAM-------------------------------------------------

>Manis_javanica

MPDSFDTLQRQRAERGGRGGRGARRGGGGDASFA---AGSARNTGPPLTLRPLSSPQRRELPRASSRDSAFWTWKCQ

>Loxodonta_africana

MPDSFDTLQRQRAGEQREEEEEDGEEEEE-EEEAL------------------------------------------

>Rattus_norvegicus

MPDTFDTLQRQRALDRREKRGGDRGGP--------------------------------------------------

HG0714 ENSG00000051825 MPHOSPH9

>Homo_sapiens

-----MTRIHFSQTAQEPLFSNKRI

>Acinonyx_jubatus

MKHLKMTRIQFLQTLKEPTFSNKRV

>Pteropus_alecto

MKHLKMTRIQFLQTVKEPTFSNKRA

>Oryctolagus_cuniculus

MKRLRKTRIQFPQTVKEPIFSNKRV

>Equus_asinus

MKHLKMTRTQFLQTVKKPTFSNKRV

>Loxodonta_africana

MKHLKMTRIQFLQTVKEPIFSTKRA

>Fukomys_damarensis

-----MTRIQFPQTVKEPLFSNKRL

>Tupaia_chinensis

MKHLKMTRIRFLQTVKQPLFSNKRD

HG0715 ENSG00000071246 VASH1

>Homo_sapiens
MCYLCSRCVGGLL
>Felis_catus
MCYLCSWCVGGLL
>Lipotes_vexillifer
MCYLCSWCVGGLL
>Myotis_brandtii
MCYLCSRCVGGLL
>Galeopterus_variegatus
MCYRCSWCVGGLL
>Condylura_cristata
MCYRCSWCVGGLL
>Ceratotherium_simum_simum
MCYLCSWCVGGLL
>Nannospalax_galili
MCYLCSWCMGGLL

HG0716 ENSG00000075413 MARK3

>Homo_sapiens
MSPGTRRGTARARPGI
>Odobenus_rosmarus_divergens
MSPGTRRGTARARPGI
>Orcinus_orca
MSPATRRGTARARPGI
>Rousettus_aegyptiacus
MSPGARRGTARARPGI
>Galeopterus_variegatus
MSPGTRRGTARARPGI
>Oryctolagus_cuniculus
MSPGSRRGTARARPGI
>Marmota_monax
MSPGARGGTARARPRI
>Tupaia_chinensis
MSPGTLRGTARARPGI

HG0717 ENSG00000083857 FAT1

>Homo_sapiens

MLKIPGRQSWYFKPGS

>Mustela_putorius_furo

MLKIGGRQSWYFRYGF

>Lipotes_vexillifer

MLKIRGKQSWYFIHGL

>Miniopterus_natalensis

MLKIPGRQSWYFRRGF

>Galeopterus_variegatus

MLKIGGRQSWHFRHG-

>Condylura_cristata

MLKIAGRQSWYFRRGF

>Ceratotherium_simum_simum

MLKIRGRQSWYFRPGF

>Rattus_norvegicus

MLKIQGRQSWYFEPGS

HG0718 ENSG00000085274 MYNN

>Homo_sapiens

-------------------------------MMGEDKQLQAPARSAQLASG-ERPAVLCCRASAQSPGRA--------PDADCRHQNARDEEDRRSAAS-------

-----------

>Acinonyx_jubatus

-------------------------------MMGETKKLQAPACSAQSGSR-KED--FR----AVLPDLGPEREMLTIGRPECLE---------------------

-----------

>Rousettus_aegyptiacus

MISLNSKIPPRPRAPVHKAAVPTRGAPKKGRMLGEDKQLQAPACSAQFGSS-KE--ALRCSSSRPRPRMRAERETPTVGRPECRE---------------------

-----------

>Galeopterus_variegatus

--------------------------------MGEDANYK-PQRSARKPAPGRRPAAHRHPASAQNPGLAPMLS---------AFQNAQNEEDRRSAVSQPPGLVS

MAPAIAGTVAC

>Ceratotherium_simum_simum

MVES------R----RTQSSCSHSWGPEEGEGDGRRQATTSPSMQCASWAPGKRSAAQRCAASASNGGRARDADFQ------------------------PSKMLR

MRK-TGGAL--

>Manis_javanica

-------------------------------MMGKEERLQAPACSAQCGGWKERPAAFRCPASAQKAGAA------------------------------------

-----------

>Ictidomys_tridecemlineatus

-------------------------------MTGEDKQLQAPACSAQRGL----PRVWRCRLPGLSPESG------------------------------------

-----------

>Trichechus_manatus_latirostr

-------------------------------MMGEDEQLQAPA---------------------------------------------------------------

-----------

HG0719 ENSG00000085998 POMGNT1

>Homo_sapiens
MTGQRPPLLQQPAHCLDALIQGHQLLQDLHIDGSQIPEVMGWPGHW
>Odobenus_rosmarus_divergens
---------------------------------------MGWPGQW
>Rousettus_aegyptiacus
---------------------------------------MGWPGQW
>Sorex_araneus
---------------------------------------MGWPGRW
>Equus_asinus
-----------------------------------MPEGMGWPGRW
>Manis_javanica
---------------------------------------MGWSGQW
>Ictidomys_tridecemlineatus
---------------------------------------MDWPGQW
>Orycteropus_afer_afer
---------------------------------------MGWPGQW

HG0720 ENSG00000087053 MTMR2

>Homo_sapiens
MAASGLPPGPALA
>Mustela_putorius_furo
MAASGLPPGPALA
>Orcinus_orca
MAASGLPPGPARA
>Pteropus_alecto
MAASGLPPGPALA
>Oryctolagus_cuniculus
MAASGLPPGPALA
>Loxodonta_africana
MAASGLPPGPALA
>Marmota_marmota_marmota
MAASGLPPGPALA
>Trichechus_manatus_latirostr
MAASGLPPRPALA

HG0721 ENSG00000095951 HIVEP1

>Homo_sapiens

MYVEFMELPFGGLLYLQFLRRKRRP

>Ailuropoda_melanoleuca

-----MELPFCDLLFLQFLRRKRGP

>Balaenoptera_acutorostrata_s

MDIGFMELPFCDLLFLQFLRRKRGP

>Cynopterus_sphinx

MDIKFMELPFCDLLFLQFLRRKRGP

>Dasypus_novemcinctus

-----MELPFCDLLFLQFLRGKSCP

>Ceratotherium_simum_simum

MDLEFMELPFCDLLFLQFLRRKRGP

>Ictidomys_tridecemlineatus

MYIEFMEPPFCDLLFLQFLRRRRSS

>Tupaia_chinensis

MSAEFMELLFCDLLFLQFLRRKRSL

HG0722 ENSG00000101445 PPP1R16B

>Homo_sapiens
MRPQPHQRPRAALAPGAPC
>Ailuropoda_melanoleuca
MRPQPRRSPRAALAPGAPR
>Balaenoptera_acutorostrata_s
MRPQPRRSPHAAPAPGALC
>Rousettus_aegyptiacus
MRPQPRRSPRAPRAVGAPC
>Dasypus_novemcinctus
MRPQAPRSPRAALAPGALC
>Condylura_cristata
MRPQPHRSPRAPLASGAPC
>Oryctolagus_cuniculus
MRPQPHGSPHAPLDPGALC
>Chinchilla_lanigera
MRPQPRGSPRAPLAPGAPC

HG0723 ENSG00000102974 CTCF

>Homo_sapiens
---------------------------------------------------------------------------------------------------------M
ITDLKPKNKMR
>Ailuropoda_melanoleuca
---------------------------------------------------------------------------------------------------------M
ITDLEPKNKMH
>Balaenoptera_acutorostrata_s
---------------------------------------------------------------------------------------------------------M
ITDLGPKNKMQ
>Eptesicus_fuscus
MLPCANPSPHLSSPALHSSPGQPPHPSLPLSAPAAPAAPTQAHCTTLPPFCVRACGAIKPCAELLLWRQRRRGGRCGCAELARDARWPERWSGGGAGAAGGVARRM
VTDLGPKNKMR
>Ceratotherium_simum_simum
---------------------------------------------------------------------------------------------------------M
ITDLGPKNKMH
>Manis_javanica
---------------------------------------------------------------------------------------------------------M
ITDLGPKNMMR
>Loxodonta_africana
---------------------------------------------------------------------------------------------------------M
ITNLGPKNKMR
>Marmota_marmota_marmota
---------------------------------------------------------------------------------------------------------M
ITDLEPKNKMH

HG0724 ENSG00000104783 KCNN4

>Homo_sapiens
-------------MDV------SFLLCPKTHPSPSLFSKGGAGEPPPTLGPPLPLPALPGRHHSPEGTGL-CCAS--EQKSQSQQSRLTTCKPQWLPCACCEVGDP
GQEAG
>Lipotes_vexillifer
-----------MWTHA------SSLLCPKTHPAPP-----------------------------------------------------------------------
-----
>Rousettus_aegyptiacus
MLCVRVSRGVWVWLRVGVYYRVQTVVCDHCQ----SVYPRGCMSL-PCSA-PRHTPAPP-----------------------------------------------
-----
>Galeopterus_variegatus
------------MDGC------LFLALPQDTPQPLLSLGRGVGEPPPTQGPPRSLPALPSRHHSPEGTGL-CCASEQKPQNKQSSQPATTCKPQWLLFACCEVGDL
GQEAG
>Condylura_cristata
-------------MHA------SSSL--------------------------------------------------------------------------------
-----
>Oryctolagus_cuniculus
-------------MDA------SSLLCPKTHPSPSLVWEGGGWGASPYPGASPPSPPCPAVTT-PQRAQS--CAE--PPSTAPE----PAEPRGWRPA--CHSGCP
LRAAG
>Ceratotherium_simum_simum
-------------MHA------SSLLCPKTHPSPSLV---------------------------------------------------------------------
-----
>Trichechus_manatus_latirostr
-------------MDV------SSLLCPETHPGPSLVWEGGLGSH-PCPGASLAPPRPAGQSPPPRGHSVPQCLRVKAPEGAQQADDL-----QATVAALCGL---
-----

HG0725 ENSG00000112218 GPR63

>Homo_sapiens

-----------MYFQAHQWDLILLFCVEMLEELLHLCLHLPSY

>Mustela_putorius_furo

-----------MCFQTHQ-------------------------

>Myotis_lucifugus

---MELCLFSRTCFQACQGNVKLLLGVEMLEEHLPSC------

>Oryctolagus_cuniculus

-----------MCFEACQWNLTLPFCFEMLEEPPHLCVYLPS-

>Ceratotherium_simum_simum

-----------MCFEAHQ-------------------------

>Manis_javanica

MEEGTLCPFSRMRFQAHQCNMILLLCVEMLEEHPLLCWHLPSY

>Marmota_marmota_marmota

-----------MCFQAHQWNLILLFCVEMLEEHTHLCLHPPSS

>Tupaia_chinensis

-----------MCFQAHQWNLMLLFCVEMLKK-LHLCLHLPPF

HG0726.1 ENSG00000115504 EHBP1

>Homo_sapiens
MWLRRGWRTRWRRR-LLLRRRRRSGYEKGTGPDSARQRD
>Leptonychotes_weddellii
MWLRRGWRTRWRRRRLLLRRRRRSGAEEGTGPDSARQRD
>Lipotes_vexillifer
MWLRRGWRTRWRRRRLLLRRRRRSGAEEGTGPDSARQRD
>Rousettus_aegyptiacus
MWLRRGWRTRWRRRRLLLRRRRRSGAEEGTGPDSARQRD
>Condylura_cristata
MWLRRGWRTRWRRRRLLLRRRRRSGAEEGTGPDSARQRD
>Oryctolagus_cuniculus
MWLRQGWRTRWRRRRLLLRRRRRSGSEEGTGPDSARQRD
>Equus_asinus
MWLRRGWRTRWRRRRLLLRRRRRSGAEEGTGPDSARQRD
>Heterocephalus_glaber
MWLPRGWRTRWRRRRLLLRWRWRSGSEEGTGPDSALQRD

HG0726.2 ENSG00000115504 EHBP1

>Homo_sapiens
MQSSFALDPLHY
>Oryctolagus_cuniculus
MQSSFALDPLLY
>Loxodonta_africana
MQSSFALDPLLY
>Fukomys_darlingi
MQSSFALDPLPY

HG0727 ENSG00000115825 PRKD3

>Homo_sapiens
MKYACTILKSFVFRHRKVHPPRMKMRTFEKVI
>Felis_catus
--------------MEKFTPPKMKMKTFEKVI
>Balaenoptera_acutorostrata_s
----------------------MKMRTFEKVI
>Condylura_cristata
----------------------MKMWVIEGI-
>Equus_przewalskii
----------------------MKMRTFENVI
>Marmota_marmota_marmota
MKYACTIFKSFLHHHRKVYPPQMKMRTFERVI
>Trichechus_manatus_latirostr
----------------------MKMRTFEKVI
>Orycteropus_afer_afer
----------------------MKMRTFEKVI

HG0728 ENSG00000116991 SIPA1L2

>Homo_sapiens

MLLKRTEAYNIITTTEKQVGFEETVF--------------------------

>Felis_catus

MLLKRTEAYKISSTSEKQAGFRKTIL--------------------------

>Balaenoptera_acutorostrata_s

MLLKRTEVFKIITATEKQVGFGETIL--------------------------

>Rhinolophus_ferrumequinum

MVLKRTEVYRIIT-TEKQVGFGESML--------------------------

>Dasypus_novemcinctus

MLLKRTEEYKIITITEKQVGFGKNCILTMLFAFG------------------

>Oryctolagus_cuniculus

MLLKRTEVYKFMT-TVEQAGFGDTMF--------------------------

>Equus_asinus

MLLKRTEVYKIITTTEKQVGFGETIL--------------------------

>Manis_javanica

MLLKRTEVYKIISTTEEQVGFKPGCGDVVCFWLKIEITTIEKDFISAFNITL

HG0729 ENSG00000119383 PTPA

>Homo_sapiens

---------------------MAAVFAVVTLTLGFRL

>Felis_catus

---------------------MAVVFTGVTLILGFRL

>Lipotes_vexillifer

---------------------MAGVLAGVTLILGFRL

>Pteropus_alecto

---------------------MAVVFAGVTLILGFRL

>Dasypus_novemcinctus

---------------------MAAVSAGVTLILGFRL

>Erinaceus_europaeus

---------------------MAGVFVGVTLILGFQL

>Manis_javanica

MAAASREPAVAGHARWGPRAGMAGVFAGVTLILGFRL

>Peromyscus_maniculatus_baird

---------------------MAAAFTRVTLTLGFRS

HG0730 ENSG00000125170 DOK4

>Homo_sapiens
MQARGEGAARTMLRRVR------
>Mustela_putorius_furo
-----------MLRRTR------
>Physeter_catodon
-----------MLRQPRPSISNR
>Rhinolophus_ferrumequinum
-----------MLRRVR------
>Galeopterus_variegatus
-----------MLRLPR------
>Condylura_cristata
-----------MLRRER------
>Ochotona_princeps
-----------MLRRVR------
>Nannospalax_galili
MQAPGKGAARTMLRRVR------

HG0731 ENSG00000129255 MPDU1

>Homo_sapiens

MAPSRRSGERQGCKSGFQTGSVNKGVGGLTAQACSFGKLYN--SNSASKTWSLTETYFTPLE---------------

>Panthera_tigris

MAPRRRR--REEAKCGLRIGSVNKGAKDGTAQAHSLSTPQHSNGEGASKTWPLDETYFTLQKVGRSKKKRGTGGGTA

>Rousettus_aegyptiacus

MAPSRTSGENQEDKSRI-----NKGAKDRIAQAYSFGTL--------------------------------------

>Dasypus_novemcinctus

MAPSRRSEKRHGNKSCFRTGTVNECAKALTAQA--------------------------------------------

>Oryctolagus_cuniculus

MAPSGRSGERQGTKCCFRSASANKSAKDLTAQACTSDTL--------------------------------------

>Equus_caballus

MAPSRSS----------------------------------------------------------------------

>Loxodonta_africana

MAPSRRN----------------------------------------------------------------------

>Heterocephalus_glaber

MAPSSRREESPKGNFSFQTATVNKGAENLTAQAYCFGTL--------------------------------------

HG0732 ENSG00000135914 HTR2B

>Homo_sapiens

----------------------MDRCVYALILENLN-FF-----F-------------------------------

>Ailuropoda_melanoleuca

----------------------MDRSVYTLIFENLKIFFSKEGVNFGFECLA------------------------

>Orcinus_orca

----------------------MDKCMYALIL--------------------------------------------

>Miniopterus_natalensis

----------------------MDQCMYALILENLE-GI-----NFGFKYLAWVQYLKKADDQFSH----------

>Ceratotherium_simum_simum

----------------------MDRHMYALILENLK-CF-----FKGRSQ-LWLQGFGMGTMP-------------

>Manis_javanica

----------------------MDRCMKALILPNLK-FF-------------------------------------

>Loxodonta_africana

----------------------MDRCMYALIFEGLK-FFSKEGVNFGFQCLAWVQCLKKADN--------------

>Rattus_norvegicus

MRVITLGQQRDHTWTLFRRPANMDRYVHSPILENLNLFRRKKATLAGSV--WRIPCFAKADDLLATDHADHCLELD

HG0733 ENSG00000136247 ZDHHC4

>Homo_sapiens

MLGRGMPELPAENYPTKRN---------VGWVP----------------

>Leptonychotes_weddellii

MLGGGMPELPVENYPNEKY---------VGWGP----------------

>Orcinus_orca

MFGGGMPELPAENYPSKRN---------M--------------------

>Myotis_brandtii

MLAGGMPELPAENHPDKKY---------VG-------------------

>Galeopterus_variegatus

MLGGGMPELPAEYYPTRRN---------M--------------------

>Manis_javanica

-----MPELPAESYPYEKH---------VGWGP----------------

>Chinchilla_lanigera

MLGRGMPELSAEIVSNQEKRCKLDTLNYLGLQEKDTPKAQHSIHLEGKY

>Tupaia_chinensis

MLGKGMPELPAENYPAKIN---------VQEMPPNRPQH----------

HG0734 ENSG00000139318 DUSP6

>Homo_sapiens
MCPAPPPLPPPLLSPSSRRARCCSLFALGLIRSGNSFPFL
>Ursus_maritimus
MCPAPPPPPPPLLSPSSRRARCCSLFALGLIRSGNSFPFL
>Rousettus_aegyptiacus
MCPAPPPPPPPLLSPSSRRARCCSLFALGLIRSGNSFPFL
>Sorex_araneus
MCPAPPPPPPPLLSPSSRGARCCSLFALGLIRSGNSFPFL
>Equus_asinus
MCPAPPPPPPPLLSPSSRRARCCSLFALGLIRSGNSFPFL
>Loxodonta_africana
MCPAPPPPPPPFLSPSSCGARCCSLFALGLIRSGNSFPFL
>Ctenomys_sociabilis
MCLLLLHRRLLFFLLPSRRARCCSLFALGLIRSGNSFPFL
>Trichechus_manatus_latirostr
MCPAPPPPPPPLLSLSSRGARCCSLFALGLIRSGNSFPFL

HG0735 ENSG00000139579 NABP2

>Homo_sapiens

MDRVPACRDEGSGRSGQ

>Ursus_maritimus

MDRVLGCPDEGSGRFGE

>Orcinus_orca

MHRVPGCPDEGSGGFGE

>Miniopterus_natalensis

MDQVPSCPDGGSGRFRN

>Dasypus_novemcinctus

MERVSGCRKEGSGMFGE

>Oryctolagus_cuniculus

MGRIPGCPDEGSGRVGE

>Equus_asinus

MDRVPGCPDEGSGRFGE

>Mus_musculus

MDRVLGCGDAGSGRSAE

HG0736 ENSG00000139826 ABHD13

>Homo_sapiens

MWEPAGAFAGDTRAGAGSGATARSRSSGLDRGAGSGPWAGGG-------------------------------------

>Felis_catus

MWEPAGTFAGDTERGPEVEEPR-----AAPQELRARPWRRRRPLGWRRTMRSDGSGAGGRCRAATGSVSAAHCAACEQV

>Physeter_catodon

MWEPAGVFAGYTERGPEVGEPR-----AAPQELRATLWRRRRPLGWRRTMRSDGSGAGGRFCAAPGSESAADCAACEQV

>Pteropus_alecto

MWEPAVAFAGDTEFGQEVEESR-----PAPQDLRARLWQQRWLFCWKRTMRSDGSGTAGRCCAALGCVFATDCAACEQI

>Dasypus_novemcinctus

MWDPAGVFAGDTRA-----------------------------------------------------------------

>Galeopterus_variegatus

MCEPAGAFAGDTQAGAGSGAADG-RAAGAPG------------------------------------------------

>Equus_asinus

MWESAGVFAGDTKRGPEVEEQT-----AAPQELRVRPWRRRRPLGWRRTMRNDGSGAGGSCCAAPGSVSAADCAACEQV

>Marmota_marmota_marmota

MWEPAGTFAGDTPALAGSRAAKS-RAAGSPG------------------------------------------------

HG0737 ENSG00000141639 MAPK4

>Homo_sapiens

MLPTSKGSRGSWPGS

>Balaenoptera_acutorostrata_s

MLPTSEASRGSWLGS

>Rousettus_aegyptiacus

MLSPSEASRGSWPGS

>Dasypus_novemcinctus

MLPTSETAKESWPGS

>Equus_asinus

MLPPSEASGGSWPGS

>Manis_javanica

MLPPSEASRGSWPGS

>Cavia_porcellus

MLPTSKCSGGSWPGS

>Tupaia_chinensis

MLPTSKACGGSWPGS

HG0738 ENSG00000143393 PI4KB

>Homo_sapiens

MEPGSGRCC

>Odobenus_rosmarus_divergens

MEPGSGRCC

>Galeopterus_variegatus

MEPGSGRCC

>Erinaceus_europaeus

MEPGSGRCC

>Ochotona_princeps

MEPGSGRCC

>Ceratotherium_simum_simum

MEPGSGRCC

>Loxodonta_africana

MEPGSGRCC

>Mesocricetus_auratus

MEPGSGRCC

HG0739 ENSG00000144331 ZNF385B

>Homo_sapiens

MICSEALPAGCFLAPHFTGAPG--------

>Panthera_tigris

MICSDALPVGCFLAPHFTEAPG--------

>Rousettus_aegyptiacus

MICSDALPVCCFLAPHFTGAPG--------

>Dasypus_novemcinctus

MICSGALPVGCFLAPHFTGAPG--------

>Ochotona_princeps

MICSDALPVGSFLALHFTEAPG--------

>Elephantulus_edwardii

MICSDPLPVGCFLAPHFTGAPG--------

>Ictidomys_tridecemlineatus

MICSDTLPVGCFLAPHFTGAPG--------

>Orycteropus_afer_afer

MICSDALPVGCFLAPHFTEAPASNDGIIYL

HG0740 ENSG00000144331 ZNF385B

>Homo_sapiens

MFYISTPSNIKLMFMG

>Mustela_putorius_furo

MFCISTSSKIKLIFMG

>Pteropus_alecto

MFCISISSKIKLMFMG

>Oryctolagus_cuniculus

MFCISTSSKIKLVFMG

>Equus_asinus

MFCIATSSKIKLMFMG

>Mus_musculus

MLYISGSSEIKLIFMG

>Trichechus_manatus_latirostr

MFCVSTSSEIKLMFMG

>Orycteropus_afer_afer

MFCVSTSSEIKLIFMG

HG0741 ENSG00000144834 TAGLN3

>Homo_sapiens
MRKSNQCAILPGARQQRLQFNMKGGFLPAC
>Myotis_brandtii
--------------MQHRQLTVKAAVLSAC
>Condylura_cristata
--------------MQPQLLSVKAAFFPAC
>Elephantulus_edwardii
-------------MMQHQPLHVRAALLFAC
>Equus_caballus
--------------MQHQQLNVKAAFLPAC
>Ictidomys_tridecemlineatus
--------------MQHPQL----------
>Trichechus_manatus_latirostr
--------------MQHQQLNLKAALLSAC
>Orycteropus_afer_afer
-------------MMQHQQLNVKAAVLSAC

HG0742 ENSG00000154556 SORBS2

>Homo_sapiens

---------------------------------------MFCKFP-----GGHEAPDTWIGLNLKPLEFKTF

>Odobenus_rosmarus_divergens

---------------------------------------MLCEYT-----GGHKAPDTWIGFNLKPLEFKTF

>Lipotes_vexillifer

---------------------------------------MLCKAA-----GGHEAFDTWVGLNLKPLEFKTF

>Miniopterus_natalensis

---------------------------------------------------MFALADTWIGLNLKSLEFKTF

>Dasypus_novemcinctus

MGWRVAGTAFSSRRTQLAAVTAENSANAPRQKQLPAAGILFPS--DQPSLLPFFPSDTWIGLNRKPLEFKTF

>Elephantulus_edwardii

-----------------------------------------------------------MGLNLKPLEFKTF

>Equus_caballus

-----------------------------MLAQRASACTLRPRLERRSLCIHLTDRDTWIGLNLKPLEVKTF

>Nannospalax_galili

--------------------------------------MLKLRWRTGIGAFHLSDRDTWIGLNLKPLEFKTF

HG0743 ENSG00000154814 OXNAD1

>Homo_sapiens

MEKLSMFQLLYIQKSQCNSRTKIFCQS--ADHILNDS------

>Felis_catus

METPSLFQLVYIQRSQYNTRTKVLHVSQSASHILNDF------

>Rousettus_aegyptiacus

METLSVFQLVCI-------------------------------

>Galeopterus_variegatus

-----MFQLVYPKVSV---------------------------

>Manis_javanica

METLFVFQPVYIQRSQFNTRTKILHVSQPASHSFNDS------

>Loxodonta_africana

MEKLPLFQLVCIKRSQYNTRTNILHKS--ASHTLNDSQNLMDI

>Ictidomys_tridecemlineatus

-----MFQPVYIQRCQCNTRSRL-CVSEPAIP-----------

>Orycteropus_afer_afer

MEKLSMFQLVYIKRSQYK-------------------------

HG0744 ENSG00000163053 SLC16A14

>Homo_sapiens

MIGDIATNLYTQQLRGLQAEQQVRGSGRGAE

>Odobenus_rosmarus_divergens

MLGDIATALYAQPLRGLRGKQQVRGSGPGAE

>Balaenoptera_acutorostrata_s

MIGDIATNLSAQPLRGLRGKQQVRGSGRGAQ

>Rousettus_aegyptiacus

MIGDIATNLYAQQLRGLRGKQQVRGSGPGAE

>Galeopterus_variegatus

MTSDIATNHYAQPPRGLGAEQQVRGSGPGAE

>Oryctolagus_cuniculus

MRGDPAARLCTRPLWGLGARQPARGSGRGAE

>Ceratotherium_simum_simum

MTGDIATNLYTQPLRGLRGKQQVRGSGPGGE

>Ictidomys_tridecemlineatus

MIGDIATHLYTQLLRGLRVEQQVRGSVPGAE

HG0745 ENSG00000163145 C1QTNF7

>Homo_sapiens

---------MLRRALYFSSSYFLHCFGVASNNKHIFLL------

>Canis_lupus_familiaris

---------MLRRALYFSSRYFLYCFEAASNKTHIFLL------

>Lipotes_vexillifer

MNSKLQKHSHAQTGTLFFIKFFLCCFEAALNDKHIFLL------

>Pteropus_alecto

---------MLRRDLYFSSSYFLHCFAAALNNKHIFLL------

>Galeopterus_variegatus

---------MLRQALYFSSSYFLHCFGAALNNRHIFLLYIF---

>Manis_javanica

---------MLRRALYFSSSYFLYCFEAALNNKHVFLLYIF---

>Loxodonta_africana

---------MLRRALYFSSSYFLHCFEAALNNKHLFLLYIF---

>Tupaia_chinensis

---------MIRRAFSFSSSYFLHWFGAALNNKHIFLLHIFEGF

HG0746 ENSG00000164741 DLC1

>Homo_sapiens

MISTTKWELRFWRN---------

>Leptonychotes_weddellii

MISTTKCELIFWRNSRGRTSFLA

>Lipotes_vexillifer

MISTTKCELIFWRN---------

>Myotis_brandtii

MISTTKCELIFWRN---------

>Ochotona_princeps

MISTTKCELRFWRNSLGDGLLR-

>Equus_przewalskii

MISTTKCELIFWRN---------

>Manis_javanica

MISTTKCELIFWRN---------

>Trichechus_manatus_latirostr

MISTTKCELIFWRN---------

HG0747 ENSG00000168036 CTNNB1

>Homo_sapiens
MIFSMNLGFCRSIHLQSQNPPGI--------
>Odobenus_rosmarus_divergens
----MNLGFCRSIHLQSQNPPGI--------
>Tursiops_truncatus
----MNLGFCRSIHLQSQNPPGL--------
>Erinaceus_europaeus
----MNLGFCRSIHLQSQNPPGI--------
>Ochotona_princeps
----MNLGFCRSVHLQSQNPPGILTFTKQLF
>Jaculus_jaculus
----MNLGFCRSIHLQSQNPPGI--------
>Trichechus_manatus_latirostr
----MNLGFCRSIHLQSQNPPGI--------
>Orycteropus_afer_afer
----MSLGFCRSLHLQSQNPPGI--------

HG0748 ENSG00000168830 HTR1E

>Homo_sapiens
MLWPFPLPTENGTQETT--------------------------------
>Ursus_maritimus
MLWCFLLPIQNGT------------------------------------
>Tursiops_truncatus
MLWCFPLPTRNGTQETAAEEKITASLQMRDFQVYIVSNCRKITISFSTV
>Eptesicus_fuscus
MLWRFPLPTQNGTQETT--------------------------------
>Galeopterus_variegatus
MLWSFPLPTESGTQETT--------------------------------
>Manis_javanica
MLWHFPLPIQKRTQETT--------------------------------
>Heterocephalus_glaber
MLWPFPLPTENETQETM--------------------------------
>Tupaia_chinensis
MLCPFPLLRENRTHETT--------------------------------

HG0749 ENSG00000169760 NLGN1

>Homo_sapiens

MVGFTWRWLWVPRILDRGCIWREKRINRHVLEVSVGE-----------

>Mustela_putorius_furo

MVGFTWRWLWVPRILDRGFIWREKRINRHVLEVSVGE-----------

>Physeter_catodon

MVGFTWRWL---------------------------------------

>Myotis_brandtii

MVGSPWRWLWVPRILDPGCSWREKRINRHVLEVSVGE-----------

>Condylura_cristata

MIGFTWRWLWVSRIQDRGCIWREKRINRHVLEVSVGE-----------

>Oryctolagus_cuniculus

MVGFTWRCLWVPRNLDRGCIWKEKRINRHVLEVSVGELKKKKKRRGYN

>Equus_caballus

MVGFIWRWLWVPRILDPGCIWREERINRHVLEVSVGE-----------

>Chinchilla_lanigera

MVGSTWRWLWVPRIADRGCIWKEKRINRDVWEVSVGE-----------

HG0750 ENSG00000170881 RNF139

>Homo_sapiens
---------------------------------------------------------MVAPRAGEAVEGGGVRP
>Mustela_putorius_furo
---------------------------------------------------------MVAPRAGEVLEGGGARP
>Lipotes_vexillifer
---------------------------------------------------------MVAPRAGEAVEGGGARP
>Myotis_lucifugus
MVTLTDVGYAVRDDDVGAPLASACDATAARSPRSRVTWGTASASQQPQRQRPAGEVKMVAPRAGEAVEGGGARP
>Dasypus_novemcinctus
---------------------------------------------------------MVAPRAGEAVEGGGARP
>Oryctolagus_cuniculus
---------------------------------------------------------MVAPRAGEAVEGGGARP
>Equus_caballus
---------------------------------------------------------MVAPRAGEAVEGGGARP
>Chinchilla_lanigera
---------------------------------------------------------MVAPRAGEAVEGGGVRS

HG0751 ENSG00000171791 BCL2

>Homo_sapiens

MGSLPYAFVLVLQKGNLTEDHAVLKKY----------------K-----------------------

>Acinonyx_jubatus

MESLPYAFVLFSQKRKLDRGSCCP-------------------------------------------

>Myotis_brandtii

MGSLPYAFVLSSQKGKLDRRSCRP-------------------------------------------

>Galeopterus_variegatus

-----MHLFWFCKKGNLTEDHAVLKKY----------------NIPEE---VDGYERYFVIIMCLMK

>Oryctolagus_cuniculus

MGSLPYAFVLVLQRRKLDRGSCCPLKTQHHVGKRLAFTTLMNDNVPRETEIRKGFE-----------

>Ceratotherium_simum_simum

------MHFYFHKKGNLTEDHAVLKKY----------------NITEE---VD--------------

>Manis_javanica

-----MHLFCFHKKGNLTEDHAVLKKY----------------NITEE---AD--------------

>Fukomys_darlingi

-----MHLFWFYKKGNLTEDHAVLKKY----------------K-----------------------

HG0752 ENSG00000172269 DPAGT1

>Homo_sapiens

MKFGITLLFHFHPHPLMPILKMAAAAFRSNGCSANKIRAETVDRRVQRGRT----------------------------------------------------

>Ailuropoda_melanoleuca

---------------------MAAAAFHDAGCSANKIREETVGMQVQWGHSRGCAIEARGSPEGGSSQSCWVSLRLLWEDPAFGERPLLPAGGSGLPPSLSIS

>Lipotes_vexillifer

----------------MPILKMAA-AFHNDSCSAN--------------------------------------------------------------------

>Rousettus_aegyptiacus

---------------------MAAAAFRNDSCFANKICAETVGRKVQWG------------------------------------------------------

>Oryctolagus_cuniculus

---------------------MAATAFRSKLLC------EWDPGGNRIGRCRKAALRLP--------------------------------------------

>Elephantulus_edwardii

----------------MPILKMSAIAFPSNSGSAEKIRAETVGR-----------------------------------------------------------

>Ceratotherium_simum_simum

---------------------MAASAFHNDSCSANKIRAETVRRQTQWGRARVDL------------------------------------------------

>Marmota_marmota_marmota

-------------------MKNGVASFRVDSSFAIKIRAETLDGQVQPGRAEVAL------------------------------------------------

HG0753 ENSG00000173406 DAB1

>Homo_sapiens

MGSRFLGGCGSGGGCSGLGEHSKRVNEMCSSECHF

>Mustela_putorius_furo

MGSRFLRGCDSGGGCCPRAERSEGVNEMCSSEHCF

>Balaenoptera_acutorostrata_s

MGNCFLGGCGSGGGWSRRA----------------

>Rousettus_aegyptiacus

---------MIEGALLHNTKHSKGVNEMCSSGCHF

>Condylura_cristata

---------------------------MHSSGCHF

>Ochotona_princeps

---------------------------MCGSGCCF

>Heterocephalus_glaber

---------------------------MSSSGCRF

>Tupaia_chinensis

---------------------------MCSLECCF

HG0754 ENSG00000175182 FAM131A

>Homo_sapiens
---------------------------------MRQECSPYVTRHRFQMGRGLAQHPEPKQE
>Felis_catus
---------------------------------MKQECSPSVTRHRFQVGRGLAQHPEPK--
>Rousettus_aegyptiacus
---------------------------------MRQECSPYVTRHRFQVGRGLAQHPEPK--
>Erinaceus_europaeus
---------------------------------MRQECSPYVTRHGFQVGRGLAQHPEPK--
>Oryctolagus_cuniculus
MYIACWHCLGDGPQFAKPQRALLGPWHSHLLKAMRRECSPYVTRHKFQVGRGLAQHPEPKQE
>Equus_caballus
---------------------------------MRQECSPYVTRHRFQVGRGLAQHPEPK--
>Manis_javanica
---------------------------------MRQECSPYVTRHRFQVGRGLAQHPEPK--
>Mus_musculus
---------------------------------MRQECSPYVTRHKFQVGRGLAQHPEPKQE

HG0755 ENSG00000176597 B3GNT5

>Homo_sapiens
MPVWKNFDVPVSSFYRVFHSSAISEKWD--------
>Ursus_maritimus
MPVWKHCDLPVFSSYWAFHSFAISEKMV--------
>Physeter_catodon
MSEWKHSDVPVSSFCSVFRLLPFPKKMVLKYFVK--
>Miniopterus_natalensis
MPAGEHLDVPVSSFYSVFHSFATSAKNAIKRKYFVK
>Oryctolagus_cuniculus
MPVWTNADVPVSSFQSVSILWPSQRKWC--------
>Equus_caballus
MPVWTHVDVPVSSFYSVFHSFAISEKMALKENIL--
>Loxodonta_africana
MPAQKSFDVPASSFCTVVHILFNKK-LLFHF-----
>Heterocephalus_glaber
MLLWKNFDIPVSSFYSMFHSFATLEKMVLKANIM--

HG0756 ENSG00000177051 FBXO46

>Homo_sapiens

MQRAAMVACGASHHHLGPEMR

>Mustela_putorius_furo

-----MVACGASPDHPGPEMQ

>Orcinus_orca

-----MVACGAPHRHSGPEMW

>Rhinolophus_ferrumequinum

-----MVACGIPYHHLGSEMQ

>Galeopterus_variegatus

MQREEMVVCGVPHHHSGPEMQ

>Equus_asinus

-----MVACGAPHHPPGPEMW

>Loxodonta_africana

-----MVARGAPHHHSAPKMR

>Marmota_marmota_marmota

-----MVACGASHHHPGPEIR

HG0757.1 ENSG00000177301 KCNA2

>Homo_sapiens

MLLCFLGPVAAP----------

>Ailuropoda_melanoleuca

MLLCFLGPQAAP----------

>Balaenoptera_acutorostrata_s

MLLCFLGP--------------

>Rousettus_aegyptiacus

MLLCFLGPSAAP----------

>Equus_asinus

MLLCFLGP--------------

>Loxodonta_africana

MLLCFLGPLAAP----------

>Mus_musculus

MLLCFLGPLSAPSSLLQAAGRH

>Tupaia_chinensis

MLLCFLGPLAAP----------

HG0757.2 ENSG00000182255 KCNA4

>Homo_sapiens
MNLKGEPGESS
>Equus_asinus
MNLRGEPGESA
>Trichechus_manatus_latirostr
MNLRGEPGESS

HG0758 ENSG00000179299 NSUN7

>Homo_sapiens

MLQMRGKPFPGTSEF

>Ailuropoda_melanoleuca

MLKVQGRPFPGTSEL

>Orcinus_orca

MLKVQGKPFPGTSEL

>Rousettus_aegyptiacus

MLTVQGKPFPGTSEL

>Oryctolagus_cuniculus

MLKVGGKPFPGTSEL

>Ceratotherium_simum_simum

MLKVQGKPFPGTSGL

>Loxodonta_africana

MLKVQGKPLPGTSEL

>Cavia_porcellus

MHKIRGKPFPGTSEV

HG0759 ENSG00000180929 GPR62

>Homo_sapiens
---------------------------------------------------------------------------------------------------MAAWASE
PPPGQPMR-------
>Panthera_tigris
---------------------------------------------------------------------------------------------------MAAWAME
PAPGHPVR-------
>Orcinus_orca
---------------------------------------------------------------------------------------------------MAAWVSE
PAPGQPVSTG-----
>Rousettus_aegyptiacus
---------------------------------------------------------------------------------------------------MVAWAAE
PGPGQPVR-------
>Oryctolagus_cuniculus
MQDRVWLPPKPTLDHLPSPPSAVTAQLLASGWQLLGPRGLAPGPAPPLPALCISRGHHQSDSQCHLRQRPGCWSWAGAGARASRLSLAPLHGAGQTSQKMAAWATE
PPPPQPMSSLHPSTG
>Ceratotherium_simum_simum
---------------------------------------------------------------------------------------------------MAAWASE
PAPGQPMR-------
>Ctenomys_sociabilis
---------------------------------------------------------------------------------------------------MAAWASE
PPSGQPMR-------
>Tupaia_chinensis
---------------------------------------------------------------------------------------------------MAAWASE
PPPGQPMRTG-----

HG0760 ENSG00000182463 TSHZ2

>Homo_sapiens
MFNKEIKCLNSHQKKKKPQKQNQKNS-KSK------NKKERGKKIQNKQTNKQGRTNL-Y---FKAAGTSHP--CLPPREGGLWPVVEELQGGSSGGQRPSDVLGA
TGQEAEETQRGQRDSGPQRAARGWGARSGTGAK
>Felis_catus
MFNKEIKCLNSHQKKKKKKKKKTQKPKISKKAKTKRERERERGKQNPKQTNKQTRQNQPL---LQTSQRKPP--VSAARRGGSLARGGGVAGGIARGTEAE-----
---------------------------------
>Pteropus_alecto
MCNKEIKCLNSHQKKKKKTKQTNKKIPKSK------NK--RERKKNPKQTNKQTRHSQPL---LQSSRHEPP--ESAIQRGGLRPLVEELQGGSSGGQRPRDALGA
TGQEAEDTQRGERDRGPPRAARGRGARSGTGAK
>Condylura_cristata
MFNKEIKCLNSHQKKKKNQKNSKKQKQ-----------KEREKGKKKSKTNKQTNQAEPTSPSNQPAQAT-R--VCHPERGVSGPWWRSCRGDRQGGRG-------
--RVTP---------------------------
>Oryctolagus_cuniculus
MFNKEIKCLNSHQKKKKQKQTKKNSKS---------KNKKERG--KKAKTNKQTRQNQPL---LPTSRHQPA--VSATQRGGSPARAAGVAGGIVRGTEAE-----
---------------------------------
>Equus_asinus
MFNKEIKCLNSHQKKKKQNKISKKQK------------QKEREEKKSKQTNKQGRTNL-Y---FKPAGASHP--CLPPREGGLWSVVEELQGGSSGGQRPSDALGA
TGQEAEETQRGERDSGPPRAARGWGARSGTGAK
>Cavia_porcellus
MFNKEIKCLTSHHKKKQKQQQQNIYIYTSKSKNK---NRER----KKKPQNKQTNKAEPS---SAASQPAPATVSAAAGEGGLRPEVAEWQGGSSGGQRPSDVLGA
SGRDTEETPRGERDSGPRARLGAGAPEVGLERK
>Tupaia_chinensis
MFNKEIKCFNSHQKKKQKQKKNSKSKN---------KKRKREREEKKSKTNKQTRQSQPL---LQPSRHQPP--CLPPERGVSGPWWRSCRGDRQGDRG-------
--RVTS---------------------------

HG0761 ENSG00000185985 SLITRK2

>Homo_sapiens

------------------MTWRLAATPREIRCKHVCSRGACVGGLGMGE---------------------------------------------------------

--------

>Leptonychotes_weddellii

------------------MTPRLAATRREIRCKHVCL---------------------------------------------------------------------

--------

>Lipotes_vexillifer

------------------MTPRLAATRREIRCKHVCL---------------------------------------------------------------------

--------

>Eptesicus_fuscus

MPQGRRRWVGFSGITRLLITPRLAETHREIRC--------------------------------------------------------------------------

--------

>Ochotona_princeps

------------------MTPRLAATRREIRCKLVCPVGT------------------------------------------------------------------

--------

>Ceratotherium_simum_simum

------------------MTPRLAATRREIRCKHVCSGTCVWVGWGGGGEERRNRFEKRSDFILPCG-PLLAFSIPPTPPPCFAILIGLPCFFLPRILLSPASPPP

RFACRYLS

>Manis_javanica

------------------MTPRLAATRREIRCRRVCSRTRVWVGGGGETGTALNDA-----VISICRAAPSPHSLPPTPPLALSF---------------------

--------

>Fukomys_damarensis

------------------MTPRLAATRREIRCKHVCSGTRVWVGWGSGKGVKKPL---------------------------------------------------

--------

HG0762 ENSG00000187147 RNF220

>Homo_sapiens
MHHTLRLLPSVS
>Canis_lupus_familiaris
MHHTHRLLPSVS
>Lipotes_vexillifer
MHHTHRLLPSVS
>Myotis_lucifugus
MHHTHRLLPSVS
>Equus_asinus
MHHTHRLLPSVS
>Loxodonta_africana
MHHTPRLLPSVS
>Marmota_marmota_marmota
MHHTLRLLPSVS
>Orycteropus_afer_afer
MHHTPRLLPSVS

HG0763 ENSG00000188050 RNF133

>Homo_sapiens

MLEDSGSCLYPEDFFLFVILN

>Acinonyx_jubatus

MPEDRGTCFHPEDFSHFIIFD

>Rousettus_aegyptiacus

MLEDGRTCFHPEDFFHVLTFN

>Ochotona_princeps

MLEDSGSCLYSEGFFLFFVLN

>Ceratotherium_simum_simum

MLEISGTYFHPEDFFHVIILN

>Peromyscus_maniculatus_baird

MLEDGGSCFHPEDIFSFLILN

>Tupaia_chinensis

MLEDSGSCFHPDDFFFFLTLN

>Trichechus_manatus_latirostr

MLEGSGTCFHLEHVFHFIILN

HG0764 ENSG00000188725 SMIM15

>Homo_sapiens

-----------------------MCTVEL-KKTDYQGSSEVVIGSLITPGQS-----------------------------

>Canis_lupus_familiaris

-------------------------MVEP-KKTSSQGSTEIVIGSLIISGQSWRSLLKHKT--------------------

>Tursiops_truncatus

-------------------------MVEL-KKTANQGSSEIVIG-------------------------------------

>Myotis_brandtii

-----------------------MCVVELK-KTTNQGSSI-----------------------------------------

>Oryctolagus_cuniculus

----------------------------------------------MIVGQS-----------------------------

>Ceratotherium_simum_simum

-------------------------MVELKKKTTNQGSSEIVIGSLITPGQS-----------------------------

>Manis_javanica

-------------------------MVEL-KKTTDQGWPKTVTGSLIIPGQDHCSVIR-----------------------

>Ictidomys_tridecemlineatus

MAASVLVSGCRAEVAAAQEFNALRSILWNQRRQNHQGSSEIVLGSLIIPGQNQRSLLKHKNFDSSYSLFPVVHPQLTLQFS

HG0765 ENSG00000196123

>Homo_sapiens
------------------------------MVKSLLQEYIAKAFLAEGPET-------
>Odobenus_rosmarus_divergens
------------------------------MVRSLLQGDIARAFLAKEPET-------
>Lipotes_vexillifer
MVKKPPGHWVPLRRWEGFPEEVRPKPSPEEWLRSLPRGDIGRAFLAEGPET-------
>Dasypus_novemcinctus
------------------------------MVRSLLQGDTGRTFLAEGRDLKEAGVCW
>Equus_przewalskii
------------------------------MVRCLLQGDIGRTFLAERPET-------
>Manis_javanica
------------------------------MVRSLLQGDIGRAFLAERPEI-------
>Loxodonta_africana
------------MCWEDFPKEVRPEPSPE--------GCIGRAFLAEGPET-------
>Nannospalax_galili
------------------------------MVEFAYREALARAFLVEGPER-------

HG0766 ENSG00000196189 SEMA4A

>Homo_sapiens
------------------------------------------------------------------------------------MMGTWRAALPFQPG
>Panthera_tigris
-------------------------------------------------------------------------------------MGTWRAALPAQPG
>Dasypus_novemcinctus
-------------------------------------------------------------------------------------MGTWRAVLPAQPG
>Oryctolagus_cuniculus
MSLVLSKRMHSESGWDRTEGPGFLDRLGRGEGGRKRCAVCWEAGEPRFVFPSQSTDFRPSQDLSQLERWKFWFPWPRAPSAGSGAMGTWRAALPFQPG
>Equus_przewalskii
-------------------------------------------------------------------------------------MGTWRAALPAQPG
>Manis_javanica
-------------------------------------------------------------------------------------MGTWRAALLAQPG
>Chinchilla_lanigera
------------------------------------------------------------------------------------MMGAWRAALPFQPG
>Orycteropus_afer_afer
-------------------------------------------------------------------------------------MGTWRAPLPAQPG

HG0767.1 ENSG00000197555 SIPA1L1

>Homo_sapiens
-----------------------------MLDSFCNILVSRKHGIMATTESQ---------
>Canis_lupus_familiaris
MFQPLNSSEASKMMLRTARGIKAAASEAGGRLARCLIIVERKHGIVAVTESP---------
>Balaenoptera_acutorostrata_s
---------------------------------------------MAITESQRYKLHSVFF
>Rousettus_aegyptiacus
---------------------------------------------MAVTTSQ---------
>Equus_caballus
---------------------------------------------MTIAESQ---------
>Loxodonta_africana
---------------------------------------------MAIAESQ---------
>Dipodomys_ordii
---------------------------------------------MAITESQ---------
>Orycteropus_afer_afer
---------------------------------------------MAVTESQ---------

HG0767.2 ENSG00000197555 SIPA1L1

>Homo_sapiens

-----------MERATQSCVLFLPSS

>Galeopterus_variegatus

MITVLCNLLGDMERATQSSILILPSS

>Jaculus_jaculus

MLTVLCDLVGDMERAARSSILFLPSS

HG0768 ENSG00000213023 SYT3

>Homo_sapiens

MGCRPP-PPRLPPRTRQR

>Balaenoptera_acutorostrata_s

MGYRQP-PPRLPSRTQQG

>Rhinolophus_ferrumequinum

MGCRQP--PRLPPRTRQG

>Galeopterus_variegatus

MGCRPPPPPRLPPRTRQR

>Condylura_cristata

MGCRQP-PPRLPSRTRQG

>Ochotona_princeps

MGCRQP-PPQLLPRTRQG

>Equus_caballus

MGCRQP-PPRLPSRTRQG

>Mus_musculus

MGYRPP-PTRLTPRTRQG

HG0769.1 ENSG00000280165 PCDH20

>Homo_sapiens
MNREGVSSAALPSRAL
>Mustela_putorius_furo
MNREGVSSAALPTGAR
>Pteropus_alecto
MNGEGVSSGALPSGAR
>Condylura_cristata
MNREGVSSVALPTGAR
>Oryctolagus_cuniculus
MNGEGVSSAALPAGAR
>Equus_asinus
MNREGVSWAALPSGAR
>Mus_musculus
MNREGVSSATLPSGAL
>Tupaia_chinensis
MNGEGVSSAALPSGAR

HG0769.2 ENSG00000197991

>Homo_sapiens

-----------------------------------MNVPFCPLAFLPLFEFPSLIEEPVTIG

>Lipotes_vexillifer

-----------------------------------MTVPRYHLTFLPPLELLSLTDEPVTIG

>Myotis_lucifugus

-----------------------------------MTVPFCPSPFCTPS-APVLIGEPVTIG

>Nannospalax_galili

MRSDLVPSHLRLQAQVSHSGSSGGEWTCEHKGVDVINVPFCPYAFLTS--------------

HG0770 ENSG00000010165 METTL13

>Homo_sapiens

MWLFFRGKNSHCSVPEPACGGQAPQMVSHRE---

>Orcinus_orca

MWLFVRGKNSRCGVPEPARG-ETPRMVPHRE---

>Pteropus_alecto

MWLLVGRKNSHRSVSEPAG-GEAPPLVPHRE---

>Ceratotherium_simum_simum

MWLFVRGKNSHCSVPEPACG-EAPPMVPRRVGES

>Manis_javanica

-------------------------MVPHRE---

>Loxodonta_africana

MGLFACGKNSHCSVPTPACG--------------

>Orycteropus_afer_afer

MWIFACGRNSHCSVPEPACG--------------

HG0771 ENSG00000013306 SLC25A39

>Homo_sapiens

MGPPDPGRRAARAPA----

>Felis_catus

MGPLGPGRRAARAPAWSQN

>Myotis_brandtii

MGPPDPGLRAARVPS----

>Dasypus_novemcinctus

MGPSGSGRRAARAPAWSQD

>Galeopterus_variegatus

MGPPGPGRRTVRVPAWSQD

>Manis_javanica

MGPPGPGTRTARAPA----

>Tupaia_chinensis

MGPPGPGRRAARAPAQSQD

HG0772 ENSG00000064309 CDON

>Homo_sapiens
--------------------MTWGWLSEPAAAQEVWRLSQPRGE
>Mustela_putorius_furo
MEGPVRIAAAAAVAAASARGMTWGWLSEPAAAQEVRGLRLPRRE
>Dasypus_novemcinctus
MEGPVRIAAA-----ARARGMSWGRPSEPAAAQEVWGLSRPRRE
>Oryctolagus_cuniculus
--------------------MTWGWLSEPAAPQEGLGA------
>Ceratotherium_simum_simum
MEGPVRIAAAATVAAARARGMTWGWLSEPAAAQEVWGLSLPR--
>Manis_javanica
MEGPVRIPAAAAVVAASARGMTWGWLSEPAAAQEVWGLSLPR--
>Peromyscus_maniculatus_baird
--------------------MTWGWLSEPAAAQEVWGLSQPRGE

HG0773 ENSG00000065548 ZC3H15

>Homo_sapiens

-----------------------------------------MLCDIDLSQRPRLRGQ

>Leptonychotes_weddellii

MSDITSDAKSFSHVPTLPPPGRPPQKRTRSSASGRRSSLPTMLCDTDLSQRPRFGGE

>Pteropus_alecto

-----------------------------------------MLCDIDLSQRPRFGGE

>Condylura_cristata

-----------------------------------------MLCGRDLSQRPHFGGE

>Ceratotherium_simum_simum

-----------------------------------------MLCGSDLSQRPRLGGE

>Chinchilla_lanigera

-----------------------------------------MLCDSDLSQTPRFGGE

>Tupaia_chinensis

-----------------------------------------MLCGTDLSQRPRFGGE

HG0774 ENSG00000066933 MYO9A

>Homo_sapiens

MCLLRPGGVSLTGRGMRRGRSPGP

>Mustela_putorius_furo

MCLPRPGGVSLTGRGMRGGSSLGP

>Orcinus_orca

MCLPRPGGVSLTGRGMRGGRSPGP

>Myotis_lucifugus

---------------MRGGRSPGP

>Condylura_cristata

MCLPRPGGVSLTGRGMRGGRWPGP

>Ceratotherium_simum_simum

MCLPRPGGVSLTGRGMRGGRSPGP

>Rattus_norvegicus

MCLPRPGGVSLTGRGMRGRRSLGP

HG0775 ENSG00000068001 HYAL2

>Homo_sapiens

----------MGFIKSSASCPTGSSWRK--------------------------

>Mustela_putorius_furo

----------MGFIKSPAGCPAGSSWCE--------------------------

>Lipotes_vexillifer

----------MGFIKLPAGCPEGSSWCE--------------------------

>Myotis_brandtii

----------MGFIKSPAGCPLGSSWRESLDWRGSNLRSGGLPYSS--------

>Dasypus_novemcinctus

----------MGFIKSPADCPKGSSWREWPGGRG--------------------

>Galeopterus_variegatus

MMGGGSGWEGMEFIKSPAGCPTGSSWRKQPGWRSTPQEGGRPPTSWGDRLPHSS

>Mus_musculus

----------MGFIKSPADCPEGSSWRKEPGWRDSTQKI---------------

HG0776 ENSG00000082014 SMARCD3

>Homo_sapiens

MALGAAGAVGSGPRHRGGTG

>Felis_catus

MALGAAGAVGSGPGHRGGTG

>Rousettus_aegyptiacus

MALGAAGALGSGPGHRGGTG

>Dasypus_novemcinctus

MALGAAGVTGSGPGHRGGTG

>Ceratotherium_simum_simum

MALGAAGAVGSGPRHRGGTG

>Manis_javanica

MALGAAGTVGSGPGHQGGTG

>Ictidomys_tridecemlineatus

MALGAAGAVGSGPGHRGGTG

HG0777 ENSG00000085644 ZNF213

>Homo_sapiens

----MPAQLPQGIPLGD

>Canis_lupus_familiaris

MRVQMPTRLPQGIPLGD

>Balaenoptera_acutorostrata_s

----MPTKLPLGIPLGD

>Myotis_davidii

----MPAQLPQGIPLGD

>Ceratotherium_simum_simum

----MPAQLPQEIPLGD

>Manis_javanica

----MLTQLPQGIPLGD

>Ictidomys_tridecemlineatus

----MPAQPPQGIPLGD

HG0778 ENSG00000089116 LHX5

>Homo_sapiens
MRELRGDQTKMRIRTA
>Leptonychotes_weddellii
MRELGGGQTKMRIGTA
>Rousettus_aegyptiacus
MRELGGGQTKMRIGTA
>Galeopterus_variegatus
MRELGGGQTKMRIGTA
>Condylura_cristata
MRELGGGQTKMRIGTA
>Ochotona_princeps
MRELGGGQTKMRIRTA
>Octodon_degus
MRELGGGQTKMRIRTA

HG0779 ENSG00000091140 DLD

>Homo_sapiens

MRREGRPWRSGGGAQRR

>Balaenoptera_acutorostrata_s

MRRAGRPWRGGGDA---

>Galeopterus_variegatus

MRRAGRPWRSGGGA---

>Erinaceus_europaeus

MRRAGRPWRSGGVV---

>Ochotona_princeps

MRRAGRPWESGGRAGPR

>Fukomys_damarensis

MRRAGRPWKSGGGAQRR

>Orycteropus_afer_afer

MRRAGRPWRGGDGALQR

HG0780 ENSG00000092199 HNRNPC

>Homo_sapiens
----------------------------------MPSLEQQQPG
>Canis_lupus_familiaris
----------------------------------MPSLEQQQPG
>Dasypus_novemcinctus
MAFSQKAIFLLFPLNPILLFLYTTPHKPQAAATWMPSLEQQQPG
>Condylura_cristata
----------------------------------MPSLEQQQPG
>Ceratotherium_simum_simum
----------------------------------MPSLEQQQPG
>Manis_javanica
----------------------------------MPSLEQQQPG
>Rattus_norvegicus
----------------------------------MPSLEQQQPG

HG0781 ENSG00000100099 HPS4

>Homo_sapiens
MHLPLLAFLERS-LKRWFCRQPQ---
>Odobenus_rosmarus_divergens
MHLPDWAFLARSFL-KKFCREPEC--
>Myotis_lucifugus
MHLPYCTFLARS-LKGSFCREPES--
>Dasypus_novemcinctus
MHVPCGAFLERSFLKSLSCRQPY---
>Ceratotherium_simum_simum
MHLPWWAFLARSSLKRSFCREPES--
>Nannospalax_galili
MHLLSGAFPERN-WKRSLCRQPCRCS
>Tupaia_chinensis
MFLSCRAFLERD-FKRPFCKQFH---

HG0782 ENSG00000100368 CSF2RB

>Homo_sapiens
MGLAWLPAGQEHRTSGH------
>Panthera_tigris
MGLAWPPAEQAYRTPGH------
>Myotis_davidii
MGLTWPPTGEGHRVPGQ------
>Oryctolagus_cuniculus
MGLAWPPPGREHKLPGQ------
>Ceratotherium_simum_simum
MGLAWLPAREGHKIPGQ------
>Manis_javanica
MGRAWPPAGEEEETPGPVHTWTM
>Tupaia_chinensis
MGLAWPPAGQEHKIPGH------

HG0783 ENSG00000105053 VRK3

>Homo_sapiens

----------------------------------MRAQRSPPSGSFRWATEKVAVKMEFPTQ--

>Felis_catus

MGVGDEEIIPPGHCPRAVRRPAAPARALSFLGSRMRAQRFPPPGSLRGTTKKVSSKMEFPAQGI

>Physeter_catodon

--------------------------------------------------------MEFPAQ--

>Pteropus_vampyrus

--------------------------------------------------------MEFPDQ--

>Condylura_cristata

--------------------------------------------------------MQLQAQ--

>Ceratotherium_simum_simum

----------------------------------MRAQRLPSSGSFRGATEKDC----------

>Cricetulus_griseus

--------------------------------------------------------MEFPTQ--

HG0784 ENSG00000105851 PIK3CG

>Homo_sapiens
MKGAPEKRKNLDAHWVA-----------
>Odobenus_rosmarus_divergens
MQGDPEKRKSLEAHWVDSNLFCFQKLNK
>Orcinus_orca
MKGDPEKRKSLEAHWVGLNLFCFQKLNK
>Galeopterus_variegatus
MKGAPEKRKSVEAYWVA-----------
>Oryctolagus_cuniculus
MKGAPEKRKSLEAHWAGLNLFCFQKLNK
>Ceratotherium_simum_simum
MKGDPEKRKNLEAHWVV-----------
>Dipodomys_ordii
MKGAPEKRKN------------------

HG0785 ENSG00000107242 PIP5K1B

>Homo_sapiens
----------------------------MSRPPITTNYR
>Rousettus_aegyptiacus
MIILMMVLKLQSVCVPETLNLGTKTVSTMNRPPITTNYR
>Galeopterus_variegatus
----------------------------MNRPPISTNYR
>Oryctolagus_cuniculus
----------------------------MNRPPITTSYR
>Equus_przewalskii
----------------------------MNRPPVVTNYR
>Peromyscus_maniculatus_baird
----------------------------MSRPPITTNYR
>Tupaia_chinensis
----------------------------MNRPPITTNYR

HG0786 ENSG00000107862 GBF1

>Homo_sapiens

MSSGCRSWD

>Physeter_catodon

MSSGCRSWD

>Pteropus_alecto

MSSGCRSWD

>Oryctolagus_cuniculus

MSSGCRSWD

>Equus_caballus

MSSGCRSWD

>Marmota_marmota_marmota

MSSGCRSWD

>Orycteropus_afer_afer

MSSGCRSWD

HG0787 ENSG00000108592 FTSJ3

>Homo_sapiens

----------------MAENLEPGTQGTQAEA--EVVSCILVRFQVWRKRLWV-----------------------

>Dasypus_novemcinctus

---------MLMATLKMAENLEPGT---------------------------------------------------

>Galeopterus_variegatus

----------------MAENLEPGTQGAQPEA--EVHLLPTCAGPSVVKTVLVRGERVEQGAECDCRAKWETENRI

>Oryctolagus_cuniculus

-MTGISSREKLTATPEMAENLEPGTQGRKRKRKWKCACCLHVRLQVWRRRRDRAGR-FS---QGDCRGTGS-----

>Equus_asinus

MWEQRIYTGTCVATLKMAENLESGTQATQAEA--EVRLLLHVPLQLGEKTALGRD---------------------

>Manis_javanica

----------------MAENLEPGTQTAEAEV--E-----------------------------------------

>Peromyscus_maniculatus_baird

----------------MAENLKPGTQETRAEA--EVR---------------------------------------

HG0788 ENSG00000113739 STC2

>Homo_sapiens
MLSGELREGAG
>Myotis_lucifugus
MLSGELREGAG
>Erinaceus_europaeus
MLSGELREGAG
>Ochotona_princeps
MLSGELREGAG
>Equus_przewalskii
MLSGELREGAG
>Loxodonta_africana
MLSGELREGAG
>Rattus_norvegicus
MLSGELREGAG

HG0789 ENSG00000115355 CCDC88A

>Homo_sapiens

MVSARGEGASPACWY

>Ailuropoda_melanoleuca

MVSARGEGASPACRW

>Pteropus_alecto

MVSVRGEGASPAYRW

>Sorex_araneus

MVSAQGEGASPACGW

>Oryctolagus_cuniculus

MVSARGEGASPACRW

>Ceratotherium_simum_simum

MMSARREGASPACRW

>Ictidomys_tridecemlineatus

MVSARGEGANPACRW

HG0790.1 ENSG00000115568 ZNF142

```
>Homo_sapiens
--MTMLHPKPSRLQTAFPVRWLL---------------------
>Ailuropoda_melanoleuca
--MTMLHPKLCRLQNGFPICWLL---------------------
>Physeter_catodon
--MTMLHPKPCRLQNVFPICWHL---------------------
>Rousettus_aegyptiacus
--MTMLHPKPCRLQSAFPFVGSCDSRSRSMMWTLSSPPLYR---
>Equus_asinus
MIMTMLYLKPCRLQNAFPIRRIF---------------------
>Manis_javanica
----MLHPKP----------------------------------
>Marmota_marmota_marmota
--MVVLHPKPCRLQNIFPVLWLLCASSRPTVWMLSSPPLFREES
```

HG0790.2 ENSG00000115568 ZNF142

>Homo_sapiens
-------------------------------MWTLSSPPFYR
>Mustela_putorius_furo
-------------------------------MWTLSSPPLYR
>Physeter_catodon
-------------------------------MWTLSSPPLHR
>Rhinolophus_ferrumequinum
MRLLPVLGRKLCRLQIGFPIRPVCDSASCPAMWTLSSPPLYR
>Oryctolagus_cuniculus
-------------------------------MWTLSSPPFHR
>Tupaia_chinensis
-------------------------------MWTLLSPPFYR

HG0791 ENSG00000131323 TRAF3

>Homo_sapiens

MRKMRPKEVMPLG

>Balaenoptera_acutorostrata_s

MRKLRPREVMPLG

>Oryctolagus_cuniculus

MRKLRPKEVTPLG

>Equus_asinus

MMKLRPKEVMPLG

>Chinchilla_lanigera

MRKLRPKEVIPFG

>Tupaia_chinensis

MRKLRPKEVTPLG

>Trichechus_manatus_latirostr

MRKLRSKEVMPLG

HG0792 ENSG00000132424 PNISR

>Homo_sapiens

MLRSCSLNQPKF

>Ursus_maritimus

MLPSCSLNQPKF

>Miniopterus_natalensis

MLPSCSLNQPKF

>Ochotona_princeps

MLPSCDLNQPKF

>Ceratotherium_simum_simum

MLPSCSLNQPEF

>Manis_javanica

MLPSCSLNQPKF

>Cavia_porcellus

MLSSCSLNQPKF

HG0793 ENSG00000132698 RAB25

>Homo_sapiens

MRGGAGPDLLRAEG

>Erinaceus_europaeus

MRGGVGPGLLRAEG

>Oryctolagus_cuniculus

MRGGAGPGLPRAEG

>Equus_caballus

MRGGADPGLLRTEG

>Loxodonta_africana

MRGGEGPSLLRAED

>Trichechus_manatus_latirostr

MRGGAGPSLLRAEG

>Orycteropus_afer_afer

MRGGAGPSLLKAGG

HG0794.1 ENSG00000135218 CD36

>Homo_sapiens

------------MLEHLIEKSFLAILKIAFQ---

>Acinonyx_jubatus

------------MFEHLIEKSFLAFLKIAYK---

>Balaenoptera_acutorostrata_s

------------MLEHLIEKSFLAFLKIAYN---

>Myotis_brandtii

------------MLEHLIEKSFLAFLKIADNA--

>Oryctolagus_cuniculus

------------MLEHLIEKSFLALLKLTNDWKN

>Equus_asinus

------------MLELLIEKFFLAFLQIAYK---

>Heterocephalus_glaber

MSFGPVKNCLFKVLEHLIENSFLMHIKAAYK---

HG0794.2 ENSG00000135218 CD36

>Homo_sapiens

---MSMAFRCQDNLKDR---------------------------------

>Canis_lupus_familiaris

-----MAFRCQDNLKDR---------------------------------

>Lipotes_vexillifer

---MSMAFTCQDNLKDRGVVKTLSLISEFSAGVLWKSCVYFLHLLLQQSL

>Erinaceus_europaeus

---MSMAFRRPDKLKDR---------------------------------

>Mus_musculus

MLRMSMAVRRQDNLEDR---------------------------------

HG0795 ENSG00000137601 NEK1

>Homo_sapiens
--------------------------------------------------------------------------------------------------MSFSKCGM
VPLLY
>Acinonyx_jubatus
--------------------------------------------------------------------------------------------------MSFSKCGM
MPLPH
>Myotis_lucifugus
--------------------------------------------------------------------------------------------------MSFSKCGM
MPLPL
>Dasypus_novemcinctus
MPPVPRPFKAARNYLFEVGEAVTQALAPLPPGVSRGLRGQSVVFIWVVWACLTGRCPRCDVIQPEASAKSWCQAWEARHHLSGWTVSCWVDRLDFSFRILFSKCGM
VPLPY
>Equus_asinus
--------------------------------------------------------------------------------------------------MSFSKCGM
VPLSL
>Loxodonta_africana
--------------------------------------------------------------------------------------------------MSFSKCGM
IPLP-
>Fukomys_damarensis
--------------------------------------------------------------------------------------------------MSFAKCRM
VPLLC

HG0796 ENSG00000138074 SLC5A6

>Homo_sapiens
MRPSPLSGILADPLFLFPFSEGLPRRRAASRSRLQTPSARCSPRPDAEVAASQKLGPLHPPSEKGAMLISGSTKGPS
>Odobenus_rosmarus_divergens
------------------------------------------------------------------MLISGQTTGPS
>Rhinolophus_ferrumequinum
------------------------------------------------------------------MLISGRTTGPS
>Galeopterus_variegatus
------------------------------------------------------------------MLISRRVKGPS
>Condylura_cristata
------------------------------------------------------------------MLISGRTTEPS
>Manis_javanica
------------------------------------------------------------------MLLSGRTKGPS
>Fukomys_darlingi
------------------------------------------------------------------MLISGSKKRSS

HG0797 ENSG00000140526 ABHD2

>Homo_sapiens
------------------------------------------MSAAGSRGAAGTRLGAS-------------
>Felis_catus
------------------------------------------MSAAGSRGASGTRLGANQDAVVNHAQLLG-
>Physeter_catodon
------------------------------------------MSAGGSR-----------------------
>Dasypus_novemcinctus
------------------------------------------MSAAGSRGAVGARLGASWTVVVTHAQLLG-
>Galeopterus_variegatus
------------------------------------------MSAAGSRGASGTGLGASWAAVGNQAQLLG-
>Ceratotherium_simum_simum
MNGRVPAVRGVAEAVVRGVAAVAWAGPVAAGAALARGSGCISMSAAGSCGAAGTGLGASWTAVVNHAQLLG-
>Nannospalax_galili
------------------------------------------MSAAGSRGAAGTRLGASWTARVHRAAFRLI

HG0798.1 ENSG00000142949 PTPRF

>Homo_sapiens

MRTDGPWIGGRSGGPGARPLVLSIQQE

>Orcinus_orca

MRTDGPWIGGRSRGPSALPSVLVVQE-

>Rhinolophus_ferrumequinum

MRTDGPWIGGRSGGPGALPSVLVVQE-

>Dasypus_novemcinctus

MRTDGPWIGGRNGGPSALPLMVIVQQV

>Oryctolagus_cuniculus

MRTDGPWIGG-----------------

>Manis_javanica

MRTDGPWMSGRSRGPCALASELGVQE-

>Chinchilla_lanigera

MRTDGPWIGGRSSGLGALPAVLIVQQE

HG0798.2 ENSG00000153707 PTPRD

>Homo_sapiens

--MLAPREAMGFGSRFLS

>Canis_lupus_familiaris

MLAPALKGVMGFGSRFLS

>Condylura_cristata

MLAPARREAMDFSSRFLS

>Mesocricetus_auratus

MLAPVLRDIMGFGSRFLS

HG0799 ENSG00000143398 PIP5K1A

>Homo_sapiens

MPGEWCGLELVQEPARRV

>Acinonyx_jubatus

MPAKWCGLELFQDPARRV

>Balaenoptera_acutorostrata_s

MPAEWCGPELVPEPARRI

>Rousettus_aegyptiacus

MPAEWCGSELVQEPARRV

>Oryctolagus_cuniculus

MLGEWCGPELVQELAGRV

>Equus_asinus

MLAEWCGPEVVLEPARRV

>Mesocricetus_auratus

MPGKWCGPELVPEPERRQ

HG0800 ENSG00000143612 C1orf43

>Homo_sapiens
----------------------------------------------MFSGRVWLGLPP
>Miniopterus_natalensis
----------------------------------------------MFSGRVSLGLSP
>Oryctolagus_cuniculus
---------------------------------MRRSPFLTTPPSAMFSGRVWLGLPP
>Loxodonta_africana
MHGRPPLPLTGVVPRLGRRRSRVPLQPESRAGGEAGRRRSPPPPAAMFLGRVFLGLPP
>Marmota_marmota_marmota
----------------------------------------------MFSGRVWLGLPP
>Trichechus_manatus_latirostr
----------------------------------------------MFSGQVSLGLPP
>Orycteropus_afer_afer
----------------------------------------------MFLGRVSLGLPP

HG0801 ENSG00000144278 GALNT13

>Homo_sapiens

MFANHFEIAKCHIQFNWI------------------------------

>Mustela_putorius_furo

MFANHFEIAKCHIHGMHPAFI---LEFTPVAWSYHC----SICLRSVC

>Balaenoptera_acutorostrata_s

MFANHFEIAKCHIHGMHPAFI---LEFTPVAWICHS----CICLRSLC

>Rhinolophus_ferrumequinum

MFANHFEIAKHHIQLSSLGNG---------------------------

>Condylura_cristata

MFANHFEIAKHHIHGMYPAFS---LEFTRWLGLITVAPVFDLCVN---

>Equus_asinus

MFANHFEIAKRHIHGMHLAFI---LEITSVAWFYHS----SVCLQSMY

>Cricetulus_griseus

MFSNHFEIAKHHIQCQRKMLVAILYHFLPYSTR----QGLSLNLHFIG

HG0802 ENSG00000144283 PKP4

>Homo_sapiens
MSLIPGATTAAA
>Pteropus_vampyrus
MSLIPGATTAAA
>Oryctolagus_cuniculus
MSLIPGATTAAA
>Equus_asinus
MSLIPGATTAAA
>Loxodonta_africana
MSLVPGATTAAA
>Ctenomys_sociabilis
MSLIPGATTAAA
>Trichechus_manatus_latirostr
MSLIPGATTAAA

HG0803 ENSG00000144802 NFKBIZ

>Homo_sapiens
MLSRTWVIIADYSWKQLICSYVFV--------------
>Pteropus_vampyrus
MLSRTWVIIADYSWKRFAFTYLHDLTLQTQRLTGRSGG
>Erinaceus_europaeus
MLSRTWVLIVDYSWKQLICLYVFV--------------
>Equus_asinus
MLSRTWAIIADHNWKQLICLYVFV--------------
>Nannospalax_galili
MLSRTWVIIAGYSWKQLIRSYVFV--------------
>Trichechus_manatus_latirostr
MLSRTWVIIADYSWKQFICFYVFV--------------
>Orycteropus_afer_afer
MLSRPWVIIADYSWK-----------------------

HG0804 ENSG00000144815 NXPE3

>Homo_sapiens

-------------------------MCFLNQLQSLLCIRALKGTSRIEANETESSSAAQKSKDMEF

>Panthera_tigris

-------------------------MCFLNQPQPLLCTKN---SS---------------------

>Myotis_davidii

-----------------------------MNLSSSFIPRTLKDASREEANETESSPAAQKNKDMEF

>Erinaceus_europaeus

-------------------------MCFLNQANVSSMQITLKDNNRKEAN----------------

>Ceratotherium_simum_simum

MILTANSLRTVSTKEQLGLVTERRKMCFLNQPQPLLSTKN---TSRKEAD----------------

>Dipodomys_ordii

---------------------------MLLNWLQSLLHITLKTASRIEAN----------------

>Tupaia_chinensis

-------------------------MCFWNQLQSLFYRKSS-------------------------

HG0805.1 ENSG00000145936 KCNMB1

>Homo_sapiens
-------------------------------------------------------------------MFGHGGLPQMGTKSCFPGDGRGGQTSC------------
--------------------

>Ailuropoda_melanoleuca
-------------------------------------------------------------------MFGHGGLSQMGTKGRFPRDGLAAQSSCCGSGSLFWGGGW
GPGPKS-LP-AEDSRPPDTK

>Physeter_catodon
-------------------------------------------------------------------MFGHGGSSQMGTKGRFPRDGLGAQTSCWSSGGLFWGGGW
GPGRKV-SA-EDSQGTGETK

>Myotis_lucifugus
-------------------------------------------------------------------MFGHGG---------------------------------
--------------------

>Galeopterus_variegatus
---------------------------MQVRGLGS-------SGCLRGW-VPALLSLCPEAVCSDLTMFGHGGLSQRGTKGCFPGDGLGGQTSCWSSASSFRVGAG
GQAERNLLPAEDPQGTRETK

>Oryctolagus_cuniculus
MKKKSFQIYQGGLFWFLSAVAVCCSTSVSPETLGNAGAGARAAAAVSGLGPRHLLSLFPRAVCSRLTMFGHGGSSQMGTQGCFPGDGLGGQTSC------------
--------------------

>Trichechus_manatus_latirostr
-------------------------------------------------------------------MFGHGGSLQMRTKGRFPGDGPGGRASCWSSGASFRVGAR
QKETVCQLKLTGDPGD----

HG0805.2 ENSG00000145936 KCNMB1

>Homo_sapiens
-------------------------------MEVQGLGNSCLPQ---GWVPALR-VCVSRLCAQT-----------------------------------------
-------------------
>Orcinus_orca
MSPNIPRWSFLTSQFCCL-FCEICFPLDSGNCRCRGLGNSCP---PLGWVPALPSLCPGAVCSDLTMFGHGGSSQMGTKGHFPRDGLGAQTSCWSSGGLFWGGGWG
--PGRKVSAEDSQGTRETK
>Myotis_davidii
-------------------------------MQVRGLGSSCHPPSSCGLVPALS-VCP-GLCAQT-----------------------------------------
-------------------
>Galeopterus_variegatus
-------------------------------MQVRGLGSSGC---LRGWVPALLSLCPEAVCSDLTMFGHGGLSQRGTKGCFPGDGLGGQTSCWSSASSFRVGAGG
QAERNLLPAEDPQGTRETK
>Oryctolagus_cuniculus
-------------------------------MQVQGPGQQ--LP-CPGWVPAISSVCFLGLCARA-----------------------------------------
-------------------
>Ceratotherium_simum_simum
------MVHFPASQFCCMLLFETCFPRDSGKCRCRGLVAAAILR-TPGWVPALPSFCPRLCAQT------------------------------------------
-------------------

HG0806 ENSG00000147481 SNTG1

>Homo_sapiens

MGRAVGVEMLKGPHFLK

>Mustela_putorius_furo

MGSAEGVEMLKGPHFMK

>Myotis_brandtii

MGSAVRIEVLKGPHFLE

>Condylura_cristata

MGAAAGVEMLKGPHFLK

>Manis_javanica

MGSAEGVNMLKAPYFLK

>Loxodonta_africana

MGSAVRVEMPRGPHFLK

>Mus_musculus

MGVAVGIEILKGPEFLK

HG0807.1 ENSG00000147488 ST18

>Homo_sapiens

----------MFLTECDLRGILEKIIKFLNGAAWRNPKRNSIRAFWNTSRKR

>Canis_lupus_familiaris

MEPFCREKGRMFLTECDLTGILEKAITFLNGAVWRNPKRNSIRAFWNISRKR

>Rousettus_aegyptiacus

MEPFCREKGRMFLTECDLTGVLEKVIKFLNGAVWRNPKRNSIRAFWNISRKR

>Dasypus_novemcinctus

----------MFLTECDLTGRLENIITFLNGAVWRNPKRNSIRAFWNISRKR

>Manis_javanica

MEPFCGEKGRMFLTECDLTGVLEKAIKFLNGAVWRSPKRNSIRTFWNISRKR

>Chinchilla_lanigera

MEPLCREKGRMFVTECDPTGVLEKIIKYLNGAAWRNPRRNSIRSFWNISRKR

>Tupaia_chinensis

----------MFLTECDLAGMLETVIKFLNGAAWRNPKRNSIRTFWNISRKR

HG0807.2 ENSG00000147488 ST18

>Homo_sapiens

MSGQILQETQKSMIKKLSTLIYCKG

>Mustela_putorius_furo

MSGQSLQAIQKSMVKKPRTSIYCKG

>Balaenoptera_acutorostrata_s

MSGQSFQVIQKPMVKKPGTPTYCKG

>Rousettus_aegyptiacus

MSGQSLQVIQRSMLKKLCTLIYCKG

>Dasypus_novemcinctus

MSGQSLQLIQISMVKKPGASIYCKG

>Oryctolagus_cuniculus

MSGQSLPVTQKSMVKKPGASLHCKG

HG0807.3 ENSG00000186487 MYT1L

>Homo_sapiens
MEKSRHREIK
>Oryctolagus_cuniculus
MEKSRHREIK
>Equus_przewalskii
MEKSRHREIK
>Manis_javanica
MEKSRHREIK

HG0808 ENSG00000149269 PAK1

>Homo_sapiens

MPGLAFPTSSCCWW-

>Ailuropoda_melanoleuca

MPGPAPPTSSCCWW-

>Miniopterus_natalensis

MPGPALPTSSCWWW-

>Equus_asinus

MPGPARPTSSCWWW-

>Loxodonta_africana

MPGSAPPTSSCWWW-

>Marmota_marmota_marmota

MPGLALRTSSCCWWW

>Trichechus_manatus_latirostr

MPGPAPPTSSCWWW-

HG0809 ENSG00000151287 TEX30

>Homo_sapiens

MCSFIVFVSYPCLPTHLSQQNLLFREMSFASQM

>Ursus_maritimus

MCSFIVLR-------------------------

>Dasypus_novemcinctus

--------------------------MSFASQM

>Ochotona_princeps

--------------------------MSFAPQV

>Equus_caballus

MCSFIVLVSYPCLPTHLSQQNLLFREMSFASQM

>Manis_javanica

MYSFIVLVSYPCLPTHLSQQNLLFRAMSFAPQM

>Ictidomys_tridecemlineatus

-------MSYPCLTTHLSQQNLLFREMSFASQM

HG0810 ENSG00000156504 FAM122B

>Homo_sapiens
MGARTRKNWPSGGSRF--------------------------------------------------------------
>Canis_lupus_familiaris
MGARTRKTWPRGGSRC--------------------------------------------------------------
>Pteropus_vampyrus
MGARTRKTWSRGGSRF--------------------------------------------------------------
>Oryctolagus_cuniculus
MGARTRKHWPC-------------------------------------------------------------------
>Ceratotherium_simum_simum
MGARTRKTWPRAGSRF--------------------------------------------------------------
>Manis_javanica
MGARTRKIWPRGGSPLLKRGRRKRRQVVDFALGPKGFPWAIRTQERKQRHRVGGRGAEPPTCSPGELGVGGEKYRTQG
>Heterocephalus_glaber
MGARTRKNWPHGGSGFRSEAWEV----I----EVAGYRY---------------------------------------

HG0811 ENSG00000157350 ST3GAL2

>Homo_sapiens

MGSRRARPTAERISISAAEFSHTHRASCFCPLRVGVWLPSSGRGFGPGGGTTLAWNPSPLKPLCSLL

>Ailuropoda_melanoleuca

MGTRRAEPSARRILVSAAEFSDTHRASWVSSLRTGALFPPLAEGLGQEGQFQWGRLSDGIQAT----

>Balaenoptera_acutorostrata_s

MGSRRAEPNARRILVS-EEFSGTHRASWVSSLWIGAWFPFLPWQRGWALRAN---------------

>Miniopterus_natalensis

--------------------------------------------------MPLAWNQGCSKPTCSLL

>Galeopterus_variegatus

--------------------------------------------------MTLAWNLGPLKPTCSPL

>Equus_asinus

MGRGTAEPNTRRILVSAAEFSDTHRASCVSSLRIGAWFPPLAEGLGQEGQQRWGRLLHGIQAA----

>Rattus_norvegicus

-------------------------------------------------MRTRAWDAGLLKPVCSLL

HG0812 ENSG00000157368 IL34

>Homo_sapiens

MGLAATAPRLSSTLPGR

>Felis_catus

MGLAATAPQLPSTLPGR

>Balaenoptera_acutorostrata_s

MGLTATAPRLPSALPGR

>Galeopterus_variegatus

MGLTATAPQLSSALPGR

>Equus_asinus

MGLTASAPRLPSALPGR

>Ctenomys_sociabilis

MGLGATAPRLSSALPGR

>Tupaia_chinensis

MGLAATTPRLSSTLPGR

HG0813 ENSG00000160199 PKNOX1

>Homo_sapiens

MKIGTQTPFAFHPR

>Odobenus_rosmarus_divergens

MKIGTQTPFAFDSR

>Orcinus_orca

MKIGTQTPFAFDSR

>Eptesicus_fuscus

MKIGTQIPFAFDSR

>Elephantulus_edwardii

MKIGTQTTFAFDSR

>Rattus_norvegicus

MKIGTQTPFASHSR

>Orycteropus_afer_afer

MKIGTQTTFAFDSR

HG0814 ENSG00000163071 SPATA18

>Homo_sapiens

MRRGGCGPGHLPSGFPNPARSDPRGRMETPAAL

>Acinonyx_jubatus

--------------------------MEEPAAL

>Physeter_catodon

--------------------------METPAAL

>Myotis_lucifugus

MRRGGCGPGHLPPASPNPARSDPRGRMETSAEL

>Equus_caballus

MRWGGCGPGHLPPGFPNPARSHPRGRMETPAAL

>Manis_javanica

MRLGGCGSGHLPPGFPNPARCDPRGTMEKPAAL

>Peromyscus_maniculatus_baird

--------------------------MDTPAAF

HG0815 ENSG00000163072 NOSTRIN

>Homo_sapiens

---MKVQGWDFLLGMFDKD

>Ursus_maritimus

---MNVLGWDFLLRVFDKD

>Rousettus_aegyptiacus

MNRMKVQGWDILLRVFDKG

>Equus_asinus

---MKVQGWDFLLGVFDKD

>Manis_javanica

---MKVQGWDFLLGVFDKA

>Nannospalax_galili

---MKVQDWDFLLGVFDKD

>Tupaia_chinensis

---MKVQGWDFLLGMFDKD

HG0816 ENSG00000163794 UCN

>Homo_sapiens

----------------------------------------------------------------------------------------------------------

------------------------------------------MFFADVSRSCPGTI

>Mustela_putorius_furo

----------------------------------------------------------------------------------------------------------

------------------------------------------MFLADVSRSCPGAI

>Orcinus_orca

----------------------------------------------------------------------------------------------------------

------------------------------------------MFLADVSRSCPGAI

>Rousettus_aegyptiacus

----------------------------------------------------------------------------------------------------------

------------------------------------------MFPADVSRSCPGAI

>Dasypus_novemcinctus

MVVTGCHQCPPDRLRLGKRWRGLETVRAAPRPPDASSPSGGGGHGQHHGQPQPPLAGGHHGPRLVWCGDGPEASLAVSSGRKDSAAFQSHRSFMPQPACRSLESGA

TFRAASSAPREMRSRVSAGAPPGGLNP-RDSGRGRHCFYNQCMLFADVSRSCPGAI

>Manis_javanica

----------------------------------------------------------------------------------------------------------

-------------------------------MRSCHCFYNQCMFLADVSRSCPGAI

>Peromyscus_maniculatus_baird

---------------------------------------------------------------------------------------------MPQPASPSSTARV

I----PKLAPGTRNLLSGCAPLRGLEAAGLEAAAVIAFYNQFMFFADVSRSCPGAI

HG0817 ENSG00000163888 CAMK2N2

>Homo_sapiens

MTSSFRRLPC

>Canis_lupus_familiaris

MTSSFRRLPC

>Orcinus_orca

MTSSFRRLPC

>Oryctolagus_cuniculus

MTSSFRRLPC

>Ceratotherium_simum_simum

MTSSFRRLPC

>Manis_javanica

MTSSFRRLPC

>Rattus_norvegicus

MTSSFRRLPC

HG0818 ENSG00000164125 FAM198B

>Homo_sapiens

------------------------------MVRNKPHFSLGRLII-

>Mustela_putorius_furo

MFYLHASAFKSSLHY-SQGLHSSWSQARGQLCKNKPHFSLGKLIGV

>Miniopterus_natalensis

MKYVLLTCLSLSKFFALESEIALFLEPSMGPAVNKPHFSLGRLIGV

>Galeopterus_variegatus

------------------------------MLRNKPHFSLGRLIAV

>Erinaceus_europaeus

------------------------------MVRTTPHFSLGKRSGV

>Ceratotherium_simum_simum

------------------------------MVRNKPHFSLGRLIGV

>Ictidomys_tridecemlineatus

---------------------------MARMVRINSHFSLGRLIEV

HG0819 ENSG00000164188 RANBP3L

>Homo_sapiens

-----MKELSCVFNQMNWHLYI

>Canis_lupus_familiaris

----MKEELSCVFIQTNWHLYM

>Balaenoptera_acutorostrata_s

MCFEMKEELSCVFIQMNWHLYM

>Pteropus_vampyrus

MCFEMKEELSCVFIQMNWHL--

>Condylura_cristata

MHFEMKEELSWVFIQMNWHLSL

>Equus_asinus

----MKEELSCVFIQMNWHLYM

>Marmota_marmota_marmota

---------------MNWHHFM

HG0820 ENSG00000164946 FREM1

>Homo_sapiens

MLRSRGTLSGPLTLPLGL--------------

>Acinonyx_jubatus

MLRSWGTLSGPLALPLGL--------------

>Miniopterus_natalensis

MLRSWGTLSGPLALPLGL--------------

>Equus_asinus

MLRSWGTLSGPFALPLGL--------------

>Manis_javanica

MLRSWGTLLEPLALPLGLGADKAGGSTEALLC

>Mus_musculus

MLRSWGTLSGPLVLPPGPQADTWGRSAGSLLC

>Tupaia_chinensis

MLRSQGTLSGPLALPLRL--------------

HG0821 ENSG00000165300 SLITRK5

>Homo_sapiens
MMTGPWRCETGR
>Mustela_putorius_furo
MMSGPWRCETGR
>Lipotes_vexillifer
MMSGPWRCETGR
>Pteropus_vampyrus
-MSGPWRCETGR
>Dasypus_novemcinctus
-MSGPWRCETGR
>Equus_caballus
MMSGPRRCETGR
>Ictidomys_tridecemlineatus
MMSGPWRCETGR

HG0822 ENSG00000166170 BAG5

>Homo_sapiens

MQLGAQLRCRTP

>Leptonychotes_weddellii

MQLGAQLRCRTP

>Balaenoptera_acutorostrata_s

MQLGAQLRCRTP

>Rousettus_aegyptiacus

MQLGAQLRHRTP

>Condylura_cristata

MTLGAQLRCRTP

>Equus_asinus

MLLGAQLRCRTP

>Tupaia_chinensis

MPLGAQLRCRTP

HG0823 ENSG00000166268 MYRFL

>Homo_sapiens
----------------------MAEDMLHSNKKKMKIFQEHS--
>Mustela_putorius_furo
----------------------------------MKIFREHS--
>Balaenoptera_acutorostrata_s
--------------------------MLHSNKKKMNIFQEHA--
>Myotis_lucifugus
MSFWFYNHLENLTISLGLWAQLDAEDTLPSSEKKMKIFREHS--
>Oryctolagus_cuniculus
----------------------MAEDMPHSPAKKMKVFRSTRRL
>Mus_musculus
----------------------MAEKPLHVNKKTMETFQEFS--
>Trichechus_manatus_latirostr
----------------------MSEDRLHSSKKKMKFFQEHS--

HG0824 ENSG00000167100

>Homo_sapiens
MERLEPARSPPRPG
>Felis_catus
MERLEPARSPPRPG
>Balaenoptera_acutorostrata_s
MERTEPARSPPRPG
>Condylura_cristata
MEQIEPARSPPHPG
>Ochotona_princeps
MERLEPARSPPRPG
>Equus_przewalskii
MKRIEPAWSPPRPG
>Peromyscus_maniculatus_baird
MERLEPARSPPRPG

HG0825 ENSG00000169180 XPO6

>Homo_sapiens
MGGGLPESAGTMT
>Odobenus_rosmarus_divergens
MGGGLPESAGTMT
>Pteropus_alecto
MGGGLPEPAGTMT
>Galeopterus_variegatus
MGGGLPEPAGTMT
>Ceratotherium_simum_simum
MGGGLPEPAGTMT
>Loxodonta_africana
MGRGLPEPAGTMT
>Mus_musculus
MGGGLPEPAGTMT

HG0826 ENSG00000169851 PCDH7

>Homo_sapiens

MTLRRVVKKKEKPALSGVRRD-----------------------------------------------------

>Pteropus_alecto

MTLRRVVKKKREKRALGGEKGLTLWAPLKMARAAFSRAGETRTRTAL---------------------------

>Dasypus_novemcinctus

MTLRRVVKKEK-PELSGVEKGLTLWASRKMARAALRRAGETRPRTAL---------------------------

>Equus_asinus

MTLRRVVKKKEKPELSGVRRD-----------------------------------------------------

>Manis_javanica

MTLRRVVKKKEKPELLGVRKD-----------------------------------------------------

>Loxodonta_africana

MTLRRVVKKKEKPELPAVRKG-----------------------------------------------------

>Ictidomys_tridecemlineatus

MTLRRVVKKKEKPELSEGEKRLTPGVSEDGSGCSLARRGNPDADRAVTRVVFTAPCPKRPAGDFFILLTGFSAA

HG0827 ENSG00000170365 SMAD1

>Homo_sapiens
------------------------------------------MCIREFAVEQLFLYSAPCL---------------------------------
>Odobenus_rosmarus_divergens
------------------------------------------MCICEFAVEQLFIYSALCLC--------------------------------
>Orcinus_orca
------------------------------------------MCICEFAVQQLFIYSAPGLCADFGLL-LKTLGMVISTFQDLSTKRLTRILCK
>Rousettus_aegyptiacus
MYSKVWRRRVIFFPSPLFRLHPPAPPSVVSFLSLDRIRLPHCMCICEFAVEQLFIYCAPWLCEGFSGLFLEEVGVVGVRETGRD----VGLFF-
>Equus_caballus
------------------------------------------MCICEFAAEQLFLCSAPCLCADLGLL-FETLGMVVSTLQDLKTKELRRTLRK
>Trichechus_manatus_latirostr
------------------------------------------MCICEFAVEQLFILPAPCLC--------------------------------
>Orycteropus_afer_afer
------------------------------------------MCICEFTVEQLFIYPAPCLC--------------------------------

HG0828 ENSG00000170390 DCLK2

>Homo_sapiens
MTWAARRCGHF
>Acinonyx_jubatus
MSWAARRCGHF
>Orcinus_orca
MSWAARRCGHF
>Rousettus_aegyptiacus
MSRAARRCGHF
>Ceratotherium_simum_simum
MSWAARRCGHF
>Trichechus_manatus_latirostr
MSWAAQRCDHF
>Orycteropus_afer_afer
MSWAAGRCGHF

HG0829 ENSG00000171502 COL24A1

>Homo_sapiens
MDSTCTLSSSELASVFR
>Odobenus_rosmarus_divergens
MASTCTLSSSELPSVFR
>Physeter_catodon
MDSTCTLSSSELASVFR
>Miniopterus_natalensis
MDSTCTLSRTELASVFR
>Dasypus_novemcinctus
MDSTGTLGTSELAFVFR
>Manis_javanica
MDSTCTLSSSELASVFR
>Ictidomys_tridecemlineatus
MDSTCTLSSSELASVFR

HG0830 ENSG00000171631 P2RY6

>Homo_sapiens

-----------MALEGGVQAEEMGAVLSEPLPP-------

>Ailuropoda_melanoleuca

-----------MALEGGVQAEEVGTALCQLAVCGRSIYVD

>Balaenoptera_acutorostrata_s

---------MTMALEGGVQAEEVGTVLSSEVWTFCSC---

>Rhinolophus_ferrumequinum

MDKKTAAKRRTMALEGGIQAEEVGAVLGQLTM--------

>Dasypus_novemcinctus

-----------MALEGGIQTKEVGAVLSQHTCLRM-----

>Equus_przewalskii

-----------MALEGGVQAEEPLWT--------------

>Chinchilla_lanigera

-----------MSLEGGVQAAEVGAIFSELMPLQM-----

HG0831.1 ENSG00000174482 LINGO2

>Homo_sapiens

MQLSRFVSSSWLVGL

>Canis_lupus_familiaris

MQLSRFTSSNWLVGL

>Condylura_cristata

MQLPRFISSTWLVGR

>Ochotona_princeps

MQLSRFLSSNWLVEL

>Elephantulus_edwardii

MQLSRFVSNSWRVGR

>Mus_musculus

MQCSRHVSSSWLVGL

>Tupaia_chinensis

MQLSRFISNSWLVGL

HG0831.2 ENSG00000174482 LINGO2

>Homo_sapiens

---MRRSSEQSTESGHYQHEEPNNSNW

>Canis_lupus_familiaris

---MQRGSEKNTESGHHQDEEPNKSDW

>Ochotona_princeps

---MKRGSEKNLKRGYHQHVQPNTPAW

>Fukomys_damarensis

---MRRRSEKNTASGHHHHEDPKNSNW

>Tupaia_chinensis

MRVMRRDSEQSTASGHHQHEEPNNPNW

HG0832 ENSG00000175048 ZDHHC14

>Homo_sapiens
MTVGLPGGLTWVSSAKLSPSREPV-
>Canis_lupus_familiaris
MTAGLPRGLTWVSSAKLSPSRQPA-
>Orcinus_orca
MTAGLPKGLTWVSSAKLSPSREPA-
>Ochotona_princeps
MPEGLPGGLSWASPAKWAASREPAW
>Equus_przewalskii
MTAGLPKGLTWGSSAKLSPTQEPA-
>Mesocricetus_auratus
MTAGLPKGLTWVSSAKLSPSREPV-
>Tupaia_chinensis
MTVGLPRGLTWVSSAKLPPSREPV-

HG0833 ENSG00000176641 RNF152

>Homo_sapiens

MSSRRRGERDSVTDTEMNTPVSTGCL

>Felis_catus

----------------MNTPEHALCL

>Balaenoptera_acutorostrata_s

----------------MNTPEYARCL

>Myotis_ricketti

----------------MNTPEHAPCL

>Ceratotherium_simum_simum

----------------MNTPEYARCL

>Manis_javanica

----------------MNTPEGASCP

>Heterocephalus_glaber

MSSCRRGERDSMTDSDMNTPANAKYL

HG0834 ENSG00000182489 XKRX

>Homo_sapiens

MLAFVLFKFHSKYVDDWNRGQSRVKFKVVLLGFPSQTPSR--I--------

>Ailuropoda_melanoleuca

MVAFALSRFHSTYVDDWNPGPMKSQVQSGTPWFSIPYSNRCNLKL------

>Myotis_davidii

------------MLTIGILSQSRIKFKVVLLGSPSQTSTS--LI-------

>Oryctolagus_cuniculus

MLAFSLFQFHSTYVDDWTLEPVKSQLQSGTPGLCIPDSEWNRSPEERDLLL

>Ceratotherium_simum_simum

MIAFALSSFHSTYVDDGILGQSRVKFKVVLLGSPSQTPTS--LI-------

>Manis_javanica

MLALPCPSFTALMLTIGILGQSRAKFKVVLLDSPSQAPTS--LI-------

>Marmota_marmota_marmota

MLAFALSKFLSTYIDDWILGPIKNQVQSGPLGLFIPDSTSNWV--------

HG0835 ENSG00000182901 RGS7

>Homo_sapiens

MKLPCLAAPLAAS

>Rousettus_aegyptiacus

MKLPCRAAPLAAS

>Erinaceus_europaeus

MKLPCLAAPLAAS

>Oryctolagus_cuniculus

MKLPCLAAPLAAS

>Loxodonta_africana

MKLPCRAPPLAAS

>Mesocricetus_auratus

MRLPCLAAPLAAS

>Tupaia_chinensis

MKLPCLAAPLAAS

HG0836 ENSG00000197461 PDGFA

>Homo_sapiens
MSANMQNYRPGRS
>Mustela_putorius_furo
----MQNYRPGRS
>Lipotes_vexillifer
----MQNYRPGRS
>Rousettus_aegyptiacus
----MQNYRPGRS
>Condylura_cristata
MPANMQNYRPGRS
>Ceratotherium_simum_simum
MAANMQNYRPGRS
>Dipodomys_ordii
MPTNMRNYRPGRS

HG0837 ENSG00000198690 FAN1

>Homo_sapiens

---------------------------MRSYRIPLLVISSQESKSKPLLSFTLNILCFIAQNIQFF

>Panthera_tigris

------------------------MKNMKPNRIPLCIVSRQGSKSKPLLSFSSDILGFTTGNTQFF

>Balaenoptera_acutorostrata_s

------------------------MNNMKSNRIPLFIISIQGSKSKPLLSFPGDFLCFTT------

>Miniopterus_natalensis

MVFICTCFT-------LQVEGIDYRGNMKSDRSPLFLISSQGSESKPLPSFPRDSLCFIT------

>Erinaceus_europaeus

MRFCWKLYKAQHSTAVFMEEETVQRNSMKPKRIRLFISR---------------------------

>Ceratotherium_simum_simum

---------------------------MKSNRIPLFIISSQGSKSKPLLSSLGDTLCFIT------

>Loxodonta_africana

---------------------------MKSNRTPLSVSSGPESPSKPLPSFT--------------

HG0838 ENSG00000198707 CEP290

>Homo_sapiens

-----------------MVSPSSRCRFGLLGTIWLDPESAWNRDRDLSGPAAGSSLVVAVVRGLRWLPGLV---------

>Felis_catus

-----------------MVSPSSRCRS-----------------------------------------------------

>Myotis_lucifugus

-----------------MVSPSSRRRYCLLGTSWRDPVSAWNRAGERSGIAAGPAWWMR---------------------

>Condylura_cristata

-----------------MVSPSSHRR------------------------------------------------------

>Oryctolagus_cuniculus

-----------------MVSPSSRRRLCLLGASWRD--------------------------------------------

>Ceratotherium_simum_simum

-----------------MVSPSSRRRYSLLRTSWREPVSGWNCAGERSGVAAGPGLGSVSA--CRAPPRLAPLDVRVPWW

>Heterocephalus_glaber

MWPFLASGPYSALAGWGMVSPSSCRRFY----------------------------------------------------

HG0839 ENSG00000198791 CNOT7

>Homo_sapiens

MGHPLPPPLPPPPPPPPPSAVSMARRRRRSASSATQVHK

>Leptonychotes_weddellii

MGHPLLPPLPPPPP--PLVAVSMARRKRRSAGSATQVQK

>Erinaceus_europaeus

MGHPLPPPPLPP-PPQPPGAFSMARRKRRSANSSTRVHK

>Oryctolagus_cuniculus

MGHPLPPPPL-PPPPLPPSAVSMARRKRRSASSATQVHK

>Elephantulus_edwardii

MGHPLPPPLPPP----PPGAVSMARRKRRSASSAIQEYK

>Microtus_ochrogaster

----------------------MARRKRRSASSETQVHK

>Tupaia_chinensis

----------------------MARRRMRSASSATQVHK

HG0840 ENSG00000278195 SSTR3

>Homo_sapiens

MITSLAARREALTPWAGK---------------------------------------------------------

>Canis_lupus_familiaris

MITSLAASRGALAPWAGQ---------------------------------------------------------

>Balaenoptera_acutorostrata_s

MITGLAARRGALTPWAGQ---------------------------------------------------------

>Myotis_brandtii

MITSLAARRGSLTPRTGQ-------CRSWKQPDASCCVASSAHAPAFLSPLSVLGAVLCTLQCNRSHPGRPIADC

>Dasypus_novemcinctus

MRTDLAARRAAFTPWAGP---------------------------------------------------------

>Equus_asinus

MITGLAARRGALTPWAESWNHRTAEYRSWKKPDASRRWPPRL-----T----------CQLA-------------

>Manis_javanica

MITGLAARREAVTQWAGQ---------------------------------------------------------

HG0841 ENSG00000278259 MYO19

>Homo_sapiens

MTRWIVLLVPPATQNPG

>Mustela_putorius_furo

MTRWIALLVP-VTQNPG

>Balaenoptera_acutorostrata_s

MTRWIALLVP-ATQNPG

>Myotis_brandtii

MTRWIALLVP-VTQNSG

>Galeopterus_variegatus

MTRWIALLVP-VTQNPG

>Ceratotherium_simum_simum

MTRWIALLVP-VTQNPG

>Chinchilla_lanigera

MTKWIALLGP-GTQNPG

HG0842 ENSG00000002746 HECW1

>Homo_sapiens

-------------------------------MKCAALPRIRRCCWSRNTVRL

>Odobenus_rosmarus_divergens

-------------------------------MKCAALPRTRRRCGSQNTVRL

>Rousettus_aegyptiacus

-------------------------------MKCAALQCIRRCCWSCNTVRL

>Oryctolagus_cuniculus

MRASPARGWFGSVVPTEEQARREGCRREDKAMKCAALPLVRRRGGSRNPVRL

>Ictidomys_tridecemlineatus

-------------------------------MKCAALPRIRRCCWNCNTVRL

>Orycteropus_afer_afer

-----------------------MRAREDKARKCAALPRIRRCVGSRNTVRL

HG0843 ENSG00000043093 DCUN1D1

>Homo_sapiens

MRVEELRSTDPHVP

>Canis_lupus_familiaris

MSVEQLKSTDPHVP

>Oryctolagus_cuniculus

MSVEQLKSTDPHVP

>Equus_asinus

MNVEQLKSTDPHVP

>Peromyscus_maniculatus_baird

MRVKQLKSADPHGP

>Tupaia_chinensis

MSVEQLKSTDPHVP

HG0844 ENSG00000091009 RBM27

>Homo_sapiens

MPGGLGHREL

>Canis_lupus_familiaris

MPGGLGHREL

>Lipotes_vexillifer

MPGGLGHREL

>Dasypus_novemcinctus

MPGGLGHREL

>Galeopterus_variegatus

MPGGLGHREL

>Oryctolagus_cuniculus

MPGGLGHREL

HG0845 ENSG00000099260 PALMD

>Homo_sapiens

---------------------------------------------MVALDLLPRLTVLDF

>Panthera_tigris

MCLSGLICTSAQRRSCRAGDFCLPPLVQRASPLSPLSSPQGGSCFMVALDFLRRLPALDF

>Galeopterus_variegatus

---------------------------------------------MVALDFLHRLPVLDF

>Equus_asinus

---------------------------------------------MVALDCLPRLPVLDL

>Fukomys_darlingi

---------------------------------------------MVALDFLQRLTVLDF

>Orycteropus_afer_afer

---------------------------------------------MVVLDFLPRLPILDF

HG0846 ENSG00000100852 ARHGAP5

>Homo_sapiens

---MGKREKEDDPYDLEDVSAQK------------------------

>Canis_lupus_familiaris

---MGKREKEDDPYDLEDLSAQK------------------------

>Pteropus_vampyrus

MVVGVMAVKEDDPHYLEDLSAQK------------------------

>Condylura_cristata

---MGKREKEDDLYHVEDLSTQK------------------------

>Ceratotherium_simum_simum

---MGKREKEDDPYDLEDLSAQK------------------------

>Microtus_ochrogaster

---MGSREKEDDPYDLQDLSLAMKELQRTKYNSEIQDLYFEMVLFSE

HG0847 ENSG00000106086 PLEKHA8

>Homo_sapiens
MALLLVLLASWGLGQ
>Odobenus_rosmarus_divergens
MALLPVLLASWGLGQ
>Lipotes_vexillifer
MALLPVLLASWGLGQ
>Condylura_cristata
MALLPVLLASWGLGQ
>Equus_przewalskii
MALLPVLLAAWGLGQ
>Trichechus_manatus_latirostr
MALLPVLLAYWGLGQ

HG0848 ENSG00000111817 DSE

>Homo_sapiens

MARRGRGGLGSEGAGSPGALE

>Orcinus_orca

MARRAR---------------

>Pteropus_vampyrus

MARRAR---------------

>Oryctolagus_cuniculus

MARRAR---------------

>Ceratotherium_simum_simum

MARRAR---------------

>Loxodonta_africana

MARRVR---------------

HG0849 ENSG00000111846 GCNT2

>Homo_sapiens

MTCRMVTSPTCNFMVGTGGPPGIRTSLRTSVFV------------------------------

>Panthera_tigris

----MVTSPFGDFNVGTGGHIRIRIFLRQLVFVW-----------------------------

>Oryctolagus_cuniculus

MTCRWVTPPAGDFPVGTGGALGIGTSQGRFVCV------------------------------

>Ceratotherium_simum_simum

----MVTSVLGDLTVGVGGHTGTRTSLSQLVFV------------------------------

>Ctenomys_sociabilis

----MVTSAVGDFAVGRRGRPGPWEWFCAAIKIQGVEGTNRARHMQRSYLRNVSEKSENATEL

>Tupaia_chinensis

MTCRMILSPLSGLAVRTGGPTGIRPLLRHSLFM------------------------------

HG0850 ENSG00000113916 BCL6

>Homo_sapiens

MCLQLSFAKEKILEVW--------------------

>Canis_lupus_familiaris

MCLQHSFSTEIIIELWLGPIWCVFLCPS--------

>Pteropus_vampyrus

MCLQHSFSKETITELWLMCQFSESVLPILS------

>Ceratotherium_simum_simum

MCLQHSFTEEIIIELWLGPI----------------

>Fukomys_damarensis

MCLQHSFSKEMTKELWLGPLFKQDFAL---------

>Tupaia_chinensis

MCLQHSFSKEMIVELWLGLSVSLLTHLELEWGEEER

HG0851 ENSG00000114279 FGF12

>Homo_sapiens
MHSVDLTLLGRTS
>Leptonychotes_weddellii
MHSVDLTQPGRTS
>Galeopterus_variegatus
MHSVDLAQPGRTS
>Elephantulus_edwardii
MHSVDLT------
>Equus_przewalskii
MHSVDLTQPGRTS
>Nannospalax_galili
MHSVDLTPPGRTS

HG0852 ENSG00000114942 EEF1B2

>Homo_sapiens

MPILLIYFQTQRK---VHGQFVGHLIQ---GPQFVRGEVGMQKYFDL

>Lipotes_vexillifer

MPILLIDLQTQRK---LGG----------------------------

>Pteropus_vampyrus

MPLLLMYFQTQRK---ATDDLWAI-----------------------

>Condylura_cristata

MPPLLKYFQRQRK---VRL----------------------------

>Loxodonta_africana

MMVLLIYFQTERK---KRG----------------------------

>Marmota_marmota_marmota

MPLLLIYFQKKKKHRNVRGPLMGHLIQNFRTPRFLRGRM--------

HG0853 ENSG00000121807 CCR2

>Homo_sapiens

----MYRVCLTHTQGCISKISKLILFWRPQPNVQCS-----------------------------------------------------

>Felis_catus

----MYRACLTHTQGCISKISKLILL---------------------------------------------------------------

>Oryctolagus_cuniculus

----MCTDCLTHTQGCTSKISK-------------------------------------------------------------------

>Ceratotherium_simum_simum

----MYRVCLTHTQGCISKISKLMLLWRA------------------------------------------------------------

>Manis_javanica

----MCRVCLTDAGLHKQKISKCILPWRPQPSV-QRSWLEKKNCIILFFFFSYLYHYLPLYLRLLFLQETLFPTSAHQVSTSRSVWFSY

>Marmota_marmota_marmota

MSEQMYRDSLTHNQGCTSKISKSILLWRPQPSSVQRSYLEKKNCIFLMLFF--------------------------------------

HG0854 ENSG00000127249 ATP13A4

>Homo_sapiens

MGAASFETAALRECNKQGAASNSS---------------------------

>Canis_lupus_familiaris

MGAACFGTAALRQCSKQG--RNSSEQRVTRGRS-----------SVC----

>Pteropus_alecto

MGAASFGTAALREVGKQGASRKSSEQRVTRGRS-----------SIC----

>Ceratotherium_simum_simum

MGAASFGTAALRECSKQGAGRNSSEQRVTRGRS-----------SVC----

>Manis_javanica

MGAASFGTAALGEHSRQGAGRHSSKQRVTCGRS-----------SVC----

>Mus_musculus

MGAASSETAALRECRSKEQPEIPPGTKCPGGGVHFAKYCHLLKEGVCSRSF

HG0855 ENSG00000127804 METTL16

>Homo_sapiens

MAARWERIICVSRSFAMRLL

>Acinonyx_jubatus

MAAHLERNICVSRRFALRVV

>Pteropus_vampyrus

MAARWERNICVSRSFAVRLA

>Equus_caballus

MAARWERNICVSRSFALRVA

>Manis_javanica

MAARWERIICVSRSFALRVA

>Cavia_porcellus

MAARGDRSICVSRSFALRVT

HG0856 ENSG00000130055 GDPD2

>Homo_sapiens

MGKPAQRNSSPQAP

>Orcinus_orca

MGKAAQRNSSPQAP

>Dasypus_novemcinctus

MGKAAQRNSSPQAP

>Equus_asinus

MGKAAQRNSSPRAP

>Ictidomys_tridecemlineatus

MGKAAQRNSSPQAP

>Trichechus_manatus_latirostr

MGKAAQRNSSPQAR

HG0857 ENSG00000130347 RTN4IP1

>Homo_sapiens

MDRILTGTLIRAFHMRRAE

>Leptonychotes_weddellii

MDRILTGTLIGAFQMGRAE

>Balaenoptera_acutorostrata_s

MDRILTGTLIGAFQM----

>Myotis_lucifugus

MDRILTGTLIGAFPKRRAI

>Galeopterus_variegatus

MDRILTGTLTGAF------

>Equus_asinus

MDKILTGKLIGALQMRIAE

HG0858 ENSG00000132702 HAPLN2

>Homo_sapiens

MRTQAPEGRGAQVPGF---------------------------------------

>Lipotes_vexillifer

MRTQAPEGRGAQVLGLLDAWTSQGPRQLLHLEPLPPAGQKAASHSFLHLCEEAHF

>Galeopterus_variegatus

MRTQALEGRGAQVLGL---------------------------------------

>Oryctolagus_cuniculus

MPTQAPGGGGAQVPGL---------------------------------------

>Loxodonta_africana

MCTQAPEGRGLQVLGL---------------------------------------

>Heterocephalus_glaber

MRTQGPGSRGAQVPGL---------------------------------------

HG0859 ENSG00000132842 AP3B1

>Homo_sapiens

MRRGWTARSAQGAVTSPEAPARAPPYEN

>Odobenus_rosmarus_divergens

MRRGWAARSA------------------

>Orcinus_orca

MRRGWIARSALGALTSPEAPPGAPPNET

>Rousettus_aegyptiacus

MRRGWTARSA------------------

>Heterocephalus_glaber

MRRTR-APGRRVAVMSPEAAARAPPREN

>Orycteropus_afer_afer

MRRGWTARSA------------------

HG0860 ENSG00000133056 PIK3C2B

>Homo_sapiens
MASPLSPAAAAIVSPGVA
>Oryctolagus_cuniculus
MASPRSPAARAIVSPGVA
>Ceratotherium_simum_simum
MASPRSPAAPAIVSPGVA
>Manis_javanica
MASRLSPAAPAIVSPGVA
>Mesocricetus_auratus
MASPRSPAARAIVSPGAA
>Tupaia_chinensis
MASPLSPAARAIVSPGVA

HG0861 ENSG00000134001 EIF2S1

>Homo_sapiens

MVNDEEGPLFVQRLR------MRGGSACAVE----------------------

>Leptonychotes_weddellii

MVSEEEGPLFSWRLRDARWFRMRGGVEPSARRGGSAAGAGGLSGLCD------

>Myotis_lucifugus

MVSEEEGPRFCWRLRAARWCRMRGGTERGAR----------------------

>Galeopterus_variegatus

MISGEEAPLLSAL--------ARGGSACAVGLSEAHAEEDRWLVRLDLV----

>Condylura_cristata

---------------------MRGGSACAVGSNEAHAAEARRLLRVDHI----

>Mus_musculus

MVSGNEGSFSV------GACAVRGGSACALGWAKRTLRSFGVRGWCSESPRCH

HG0862 ENSG00000134245 WNT2B

>Homo_sapiens
----------------------------------------------------------------------------MAPQGGEVGAA
>Leptonychotes_weddellii
----------------------------------------------------------------------------MAPQGGEVGAA
>Orcinus_orca
----------------------------------------------------------------------------MAPQGGEVGAA
>Oryctolagus_cuniculus
-----------------------------------------------------------------MTLGSQPPATSMAPQGGEVGAA
>Ceratotherium_simum_simum
MRARGQRGCPHTLPVAPLRAANPLTPHPAPGHRAPDPGLGAPSSASGHRSLRPSPGPCAPVPGLQVPLGPQPPASPMAPQGGEVGAA
>Manis_javanica
----------------------------------------------------------------------------MAPQGGEVGAA

HG0863 ENSG00000134313 KIDINS220

>Homo_sapiens

MAAGCGEGDALAVAVSCFPVL

>Ailuropoda_melanoleuca

MAAGFGEDEALEAAVSPFPVF

>Physeter_catodon

MAAGFGEGDSLKAAVSPFLVF

>Ceratotherium_simum_simum

MAAGCGEGDALKAGVSPLPVL

>Marmota_monax

MAAGCGEGDASKAAVFFSFVL

>Tupaia_chinensis

MAAGCGEGSPLKAAVSRFRVL

HG0864 ENSG00000134533 RERG

>Homo_sapiens

MVNPKTGTKKIYSVL-----

>Ailuropoda_melanoleuca

MVNPKAGTKKIFSALQCARQ

>Myotis_davidii

MVNPKAGTKTIFSAL-----

>Dasypus_novemcinctus

MVNPKAGTRKICSALWF---

>Oryctolagus_cuniculus

MVNPKTGTQKICSAL-----

>Ceratotherium_simum_simum

MVNPKTGTRKIFSAL-----

HG0865 ENSG00000135540 NHSL1

>Homo_sapiens

MPRRGVGLQFGMLGTRQRVGVPWLCVLQSPDKYS

>Mustela_putorius_furo

MLGTGAGSQEGMLGTHQRVGVPWFCSSPSPDKRS

>Miniopterus_natalensis

MPGRGVGLQVGMLGAHQRVGVPWFCSLQSPDKCS

>Oryctolagus_cuniculus

MPGRGAGLQFGVLGTPWSIGVPWFCSLQSTDKCA

>Manis_javanica

MPGRGVRSQCGVPGAPHRVRVPWFCSLQSPGKCS

>Cavia_porcellus

-----------MAGMHQRTPVPCFCSSQSPDKRA

HG0866 ENSG00000136011 STAB2

>Homo_sapiens
MRKNSLGVYQTKLK
>Balaenoptera_acutorostrata_s
MRKHSLGVHQTKLM
>Miniopterus_natalensis
MRENSLGVHQTKLK
>Oryctolagus_cuniculus
MRRNSLGVQQTMIK
>Equus_asinus
MRKNSLGVHQTKLK
>Peromyscus_maniculatus_baird
MRKNSPGTHQSKLK

HG0867 ENSG00000137203 TFAP2A

>Homo_sapiens
MDPSRAPALRGASAP
>Pteropus_vampyrus
MDQSRAPALGGASAR
>Oryctolagus_cuniculus
MDPSEAPALRGASAP
>Elephantulus_edwardii
MEPSGAPALGGASAP
>Tupaia_chinensis
MDPSEVPALQGASAW
>Trichechus_manatus_latirostr
MEPSRAPALGRASAP

HG0868 ENSG00000137843 PAK6

>Homo_sapiens

MPEDLMEKSQSEPGRAVEEGRAAAGA-AGP------------------------

>Ursus_maritimus

-----MEKSQSEPGCVVEEGRAAAGK-EAGARAPCSLPGSPGPAGAPRPRPRAL

>Rousettus_aegyptiacus

-----MEKSQSEPGCGVHCGGGARRDQEGG------------------------

>Equus_caballus

-----MEKSQSEPRGAVEEGRAAAE----PVARRCRLPGSPGPAGALWPRCL--

>Manis_javanica

-----MEKSLSEPGCAVEEEPSAASTGKAARARQCRLPGSPGPAGAPGPLYL--

>Microtus_ochrogaster

-----MEKSQSELGCTVEEGRAAAGA-TGP------------------------

HG0869 ENSG00000139910 NOVA1

>Homo_sapiens
MSFNGTDIASSFALESS
>Ailuropoda_melanoleuca
MSYNGTGIASSFALESS
>Physeter_catodon
MSFNGTGIASSFALESS
>Rousettus_aegyptiacus
MSYNGTGIASSFALESS
>Manis_javanica
MSYNGTGIASSFALESS
>Cavia_porcellus
MSYNGTGIASSFALESS

HG0870 ENSG00000140993 TIGD7

>Homo_sapiens

MALGSTRRTRKEYLRA-------------------

>Balaenoptera_acutorostrata_s

MALGGTRRT--------------------------

>Pteropus_vampyrus

MELGSTRRT--------------------------

>Galeopterus_variegatus

MALGSTRKTWKKYMRAQNYRRVFQQKKFMRNN---

>Ceratotherium_simum_simum

MALGSTRRTCKEYLRAQSYKGCQQKNLWETIDQKH

>Marmota_marmota_marmota

MALGSTRSS--------------------------

HG0871 ENSG00000141127 PRPSAP2

>Homo_sapiens

MPAQKGREPSRGTAEEESILEEFCCQRIKGGCP

>Ursus_maritimus

MSL-----LMALPFFCVLIQEEFCCQRIKGGCP

>Rousettus_aegyptiacus

MSL-----LMALPFFCVLIQEEFCCQRLKGGCP

>Oryctolagus_cuniculus

---------MALPFFCVLIQEEFCCQRLKGGCP

>Equus_caballus

MSL-----LMALPFFCVSIQEEFCCQSIKGVCP

>Mesocricetus_auratus

MFVMLKTPTARSPGRARVRLEEFCCQRITGGCP

HG0872 ENSG00000142583 SLC2A5

>Homo_sapiens

MLLAVIWHGQSAPRMSSLSIQCTRYFG

>Ursus_maritimus

--------------MSSLSIQRVCYFG

>Physeter_catodon

MLLAVTWLGQSTARMSSLAIQCICYFG

>Cynopterus_sphinx

MRLVVTWHGQSAVGMSSLSIQCMCYFG

>Dasypus_novemcinctus

--------------MSSLSI-------

>Ceratotherium_simum_simum

--------------MSSLSIQCICYFG

HG0873 ENSG00000142875 PRKACB

>Homo_sapiens

MLGSYALLLGASLFKTKESSFRFLCAACSSVCVYTIGSSPSRD

>Felis_catus

MLGSYALLLGAFLF-----------------------------

>Rousettus_aegyptiacus

MLGSYALLLGDFL------------------------------

>Galeopterus_variegatus

MLGSYALLLGAFLFKTKESSFHFLCAACSSVRVYTIGSSTSRD

>Manis_javanica

MLGSYALLLGAFLFKTKETAFHFLCAACSSVCVYTISSTSRD-

>Ictidomys_tridecemlineatus

MLGSYALLLGVFLFKTKESSSHFLCAACSSVCVYTIGSSTSRD

HG0874 ENSG00000143515 ATP8B2

>Homo_sapiens

MGSTGPPGRAPSAER

>Lipotes_vexillifer

MGRTWPPGRAPSAER

>Condylura_cristata

MGWTGPPGRAPGAER

>Oryctolagus_cuniculus

MGRTGPPGRAPSAER

>Ceratotherium_simum_simum

MGRTGPPGRAPSAER

>Mus_musculus

MGRTGPPGRAPRAER

HG0875 ENSG00000145284 SCD5

>Homo_sapiens

MGDTPATDAGRQDLHP

>Rhinolophus_ferrumequinum

MGNTPATDAGLPDLHP

>Condylura_cristata

MGGTPATDAGRQDLHP

>Ceratotherium_simum_simum

MGDTPATDAGRQDLHP

>Loxodonta_africana

MGGTPATDAGRLDLHL

>Orycteropus_afer_afer

MSDTPATDAGRPDLHP

HG0876 ENSG00000145908 ZNF300

>Homo_sapiens

-MAAPCGNSSREQPCCFLNKASEDAERAKPPSTERRS-------------LGSPVWCSRTGSRRVCQ-

>Ursus_maritimus

MLGSPVLCS----------PRQEAGE----------------FFNPR---------------------

>Dasypus_novemcinctus

-MAAPIRNSR----------------------------------------------------------

>Equus_caballus

--------------MLLLNKAAEDAERAKPPGAEPRRLGAPRFAHPGRKRESPPVWMRRPSQVKP---

>Manis_javanica

-MAAPRRTLRPGAPSCLLDKAAEDAEWAKPPGGTSAGSRASWFAHPDRKRERWPVWM-----------

>Chinchilla_lanigera

-MAAPLRIVAWEQPYCLLNKAPLTEEYRLQPSAALSHRK----------RERAHRY----G-CDSCHL

HG0877 ENSG00000146426 TIAM2

>Homo_sapiens

-------------MKKRLTSWTHFACLLNVLCCSQDHVKPTDH

>Felis_catus

-----------------------------MLPCSQDPVKPSDY

>Physeter_catodon

-----------------------------MLPCSQDPVKSTDH

>Pteropus_vampyrus

-----------------------------MLPCSQDHVKPAEH

>Ceratotherium_simum_simum

------------MSWQSIISRTHFLHVLTMLPCSQDHVKPTEH

>Cavia_porcellus

MITLSVLYSHFPDWTASRNTWIHFPCFLNMLHCSQDHVRSTDI

HG0878 ENSG00000146802 TMEM168

>Homo_sapiens

MAVAAGPVTEKVYADTGLY--------------------------

>Felis_catus

MAAAARPVTEKVSIETGLY--------------------------

>Lipotes_vexillifer

MAAVAGPVVEKVSDETGLC--------------------------

>Dasypus_novemcinctus

MAAVAGPVTEKVSAETGLWLAPRQTPRVGKGLVAAAKIIINKNFT

>Equus_asinus

MAAAAGLVTEKVSNDTGLC--------------------------

>Fukomys_darlingi

MAAAAGPVTEKVSTETGLY--------------------------

HG0879 ENSG00000147145 LPAR4

>Homo_sapiens

MDKNMHFQRASCPFTCLLANYLQASAHSLI

>Galeopterus_variegatus

----MHFQSASCPFTCLLANCLQA------

>Ochotona_princeps

MDKNMHFQSASCPFTCLLANCLQA------

>Loxodonta_africana

MDKNMHFQSASCPFTCLLANCLQA------

>Cricetulus_griseus

MDKNMRFQSASCPFTCLLANYLQA------

>Trichechus_manatus_latirostr

MDKNMHFQSANCPFTCLLANSLQA------

HG0880 ENSG00000151789 ZNF385D

>Homo_sapiens

MGSAAGSPSALSSRRGS

>Canis_lupus_familiaris

MGSAAGSPSARSSRGS-

>Myotis_lucifugus

MGSAAGSPGARSSRRGS

>Equus_asinus

MGSAAGSPSARSCRRGS

>Ictidomys_tridecemlineatus

MGSAAGSPSARSSRRGP

>Orycteropus_afer_afer

MGSAAGSPSAHSSRRGL

HG0881 ENSG00000153822 KCNJ16

>Homo_sapiens
---------------------------------------------------------------------MAVGADSFRRVLTGPREKGPEMLSYGALTR------
>Balaenoptera_acutorostrata_s
---------------------------------------------------------------------MAVGADSLRESLEEQRERSPGRLVAVLLYDAKDTGV
>Rhinolophus_ferrumequinum
---------------------------------------------------------------------MAVGADSLRESTEEQREKNSEVSSSELKRT------
>Oryctolagus_cuniculus
---------------------------------------------------------------------MVVGADSRRQSRKRAKRKSPEMPECGAPTR------
>Equus_caballus
MGREEGDFVFILVFCKPPLARLPTCRHGNQLKKVLIHWSEAENSVAFVCGECVRKSKEEQDSPVEFLISMAVGADSFRESTKEQKEKNPEMSSYSALRR------
>Manis_javanica
---------------------------------------------------------------MGFLISMAVGADSLRENMKGQRGKSPRCLAAVLL--------

HG0882.1 ENSG00000154832 CXXC1

>Homo_sapiens
--------------------------------------------------------------------------------MFPANIWIRPPYGYDKQDGGA-----
------------------------
>Odobenus_rosmarus_divergens
MAKMPAPSWKPRLGRCHVLPVNVRTRPSYGISFQDDGICVEVAHRRGHISCNVESVQAVTVRTQPTRWRCPRESSHIGAAMFPANLWTRPPYGLSSQDGGA-----
------------------------
>Oryctolagus_cuniculus
----------------------------------------------------------------------------MGAAMFPTGEWTLAPYVLDSQDGGT-----
------------------------
>Manis_javanica
----------------------------------MTASAYKSRGRRSHVSCNVEAVQVVTVRTQPTKWRRPQDSSETGAAMFPGTVCIRPPYGLGGQDGGTWESRS
SGIYVVAVKPTHWACAVGVSGKRD
>Loxodonta_africana
--------------------------------------------------------------------------------MFHGNLRTRSPYGLDSQDGGS-----
------------------------
>Fukomys_damarensis
--------------------------------------------------------------------------------MFPANI--------------------
------------------------

HG0882.2 ENSG00000154832 CXXC1

>Homo_sapiens

MWKGKETEWRNEYNLIQVVLRKRSLYL----------------------------------------------

>Mustela_putorius_furo

MWKGKVTEWKSECNLIQVVFREEPLFL----------------------------------------------

>Manis_javanica

-----MILWKKERNLIQAVLRKRPIFIPTSLKVPAYGISPQDGGTYAEAGSRTLPCVACERTDSAILRNHIPR

HG0883 ENSG00000156011 PSD3

>Homo_sapiens

MLGEAAWCFLS

>Rousettus_aegyptiacus

MLGEAAWCSLS

>Oryctolagus_cuniculus

MLGEAAWCFLS

>Equus_asinus

MLGEAAWCFLS

>Microtus_ochrogaster

MLGEAAWCFLS

>Tupaia_chinensis

MLGEAAWCFLS

HG0884 ENSG00000156463 SH3RF2

>Homo_sapiens

MWTLLLQVGTGVLK

>Pteropus_alecto

MWTLPLQVRTGVLK

>Oryctolagus_cuniculus

MWTLLLHVGTGVWK

>Ceratotherium_simum_simum

MWTLPLQVRTGVLK

>Manis_javanica

MWTLPLQVRTGVLK

>Mus_musculus

MWILLLQVGTGVLK

HG0885.1 ENSG00000157502 MUM1L1

>Homo_sapiens

MPTKPWKDSAT-TEVPSWVKALLDKQLE

>Mustela_putorius_furo

MPIEPQKGSVT-SEVLSREKALLDKQLE

>Balaenoptera_acutorostrata_s

MPVEPQKVSAT-TKVPSREKALLDKQLE

>Ceratotherium_simum_simum

MPVEPQKGSAT-TEALSREKALLDKQLE

>Nannospalax_galili

-----------------MKAPLGKLLE

>Tupaia_chinensis

MPVKPQKGSSTPTAVPSRVKAVLDKQS-

HG0885.2 ENSG00000157502 MUM1L1

>Homo_sapiens
MAVVNASYFQRVLEDLE
>Canis_lupus_familiaris
MAVVKASYFRRFLETFE
>Equus_asinus
MAVVKASYFRRFLETLK
>Chinchilla_lanigera
-----MSYFQRLLEDLV

HG0886 ENSG00000158850 B4GALT3

>Homo_sapiens
MAAAMPARLLGWR
>Leptonychotes_weddellii
MAVTMPARLSGWR
>Pteropus_alecto
MAAAVPARLSGWR
>Galeopterus_variegatus
MAATMPARLSGWR
>Ochotona_princeps
MAAAKPARLLGWR
>Mus_musculus
MAAAIPARLSGWR

HG0887 ENSG00000160959

>Homo_sapiens

--MIVEICPPAELGRGLVSGTFQEPVP

>Leptonychotes_weddellii

MNRPREICLQAEQGRGLAFGASQEPVP

>Balaenoptera_acutorostrata_s

MNRPREICPQAERGRGLTSRASQEPVP

>Pteropus_alecto

MNRPREICPQAERGRGLASGASQEPVP

>Ceratotherium_simum_simum

MNRPREICPQAERGRGLASGVSQEPVP

>Heterocephalus_glaber

---------MAEPGRGWVSGVSQEPVP

HG0888 ENSG00000164023 SGMS2

>Homo_sapiens
------MEETVIGWLPSTFCIRKQSPL
>Canis_lupus_familiaris
------MEGTVIRWLPSTFCARKQNQL
>Orcinus_orca
----------------MFCTRKQNQL
>Oryctolagus_cuniculus
MYNRELNKNNGIRWLPSMFCVKKQNQV
>Equus_caballus
------MAGTIIG--------------
>Marmota_marmota_marmota
------MAGTIIRWLPSAFCIRKQNQL

HG0889 ENSG00000164091 WDR82

>Homo_sapiens

MADTMSRRRAGCP

>Canis_lupus_familiaris

MADTMSRRRAGWP

>Rousettus_aegyptiacus

MADTMSRRRAGCP

>Galeopterus_variegatus

MADTMSRRLAGWP

>Oryctolagus_cuniculus

MADTMSRCRAGCP

>Ictidomys_tridecemlineatus

MADTMSRRRAGWP

HG0890 ENSG00000165813

>Homo_sapiens

MEKLAPEPELQQPC

>Mustela_putorius_furo

MEKLGPEPELQRPR

>Equus_caballus

MEKLSPEPELQRPR

>Callospermophilus_lateralis

MERLAPEPELQRPR

>Trichechus_manatus_latirostr

MEKLAPEPELQRPR

>Orycteropus_afer_afer

MKKLVPEPELQ---

HG0891 ENSG00000166211 SPIC

>Homo_sapiens
---------MKHLYKGLKCLPGLSTYFILFSSSNNC
>Ailuropoda_melanoleuca
----MILVLMKHFYKGLKCLPELSVY----------
>Lipotes_vexillifer
---------MKQLGKGLKCLPELSIY----------
>Oryctolagus_cuniculus
MQLCQCLMLMKHLCTGLKCVPGLPVY----------
>Equus_przewalskii
---------MKHLCKGLKCLPELSIY----------
>Tupaia_chinensis
---------MKQLCKGLKCLPESSIY----------

HG0892 ENSG00000168418 KCNG4

>Homo_sapiens

MGSPCLAEKQLETGSLRQEDW----------------------------

>Balaenoptera_acutorostrata_s

MGSPCPAEKQLERGSLRQEGCSSFSGRTL--------------------

>Rousettus_aegyptiacus

MGSPCLAKKQLETGSLRQEGCSSSSGHTL--------------------

>Equus_przewalskii

MGSPCPAEKQLETGSLRQEGCSSSSGRAL--------------------

>Loxodonta_africana

MGSPCLAKEQLESGCLRQEEG----------------------------

>Mesocricetus_auratus

MGSPCLAENWLGTRFLRQSAALLPLAVCCEPSNACEIRVHCSASVKLGP

HG0893 ENSG00000168769 TET2

>Homo_sapiens

MQVTSAPSARPPRDAGPAE-----------

>Odobenus_rosmarus_divergens

MRVTSAPSARPPRDAGPAE-----------

>Orcinus_orca

MRVTSAPSPRPPRDAGPAE-----------

>Pteropus_vampyrus

MRVTSAPSARPPREAGPAE-----------

>Oryctolagus_cuniculus

MRVTSAPSARPPRDASPAE-----------

>Ceratotherium_simum_simum

MRVTSAPSARPPWDAGPPAMRTDVKVPYEY

HG0894.1 ENSG00000169282 KCNAB1

>Homo_sapiens

MLRAPVEHPACGPGDAGGASCGKPALI

>Acinonyx_jubatus

MLRVPVEHPA-----------------

>Physeter_catodon

MLRAPVEHPACGPEGAGSASCGMPAWI

>Rousettus_aegyptiacus

MLRVPVEHPACGPEGAGGASCGMPAWI

>Equus_asinus

MLHAPVEHPACGPEGAGGASCGMPAWI

>Heterocephalus_glaber

MLRAPAAS-CLRSKDAGGASCGMPELI

HG0894.2 ENSG00000069424 KCNAB2

>Homo_sapiens

MALSPVVPVQEQNFPTVKNRASLDK

>Ursus_maritimus

-----MLLSQEQNLPSVKKRAGQDQ

>Rousettus_aegyptiacus

MVLSLAVLTQEQNLPSVKNRAGGGT

>Equus_caballus

MLSSLAVLIQEQNLPSVKKRAGRDQ

>Cricetulus_griseus

-MVSSTGPVPQQDFPSVKRRAGLDK

HG0895 ENSG00000170035 UBE2E3

>Homo_sapiens

MNLRCWPWKQLEQLFSKR

>Odobenus_rosmarus_divergens

MNLRCWPWKRLEQLFSKK

>Lipotes_vexillifer

MNLLCWP-----------

>Miniopterus_natalensis

MNLRCWPWKRLEQLFSKS

>Manis_javanica

MNLRCWPCKRLEQLVS--

>Trichechus_manatus_latirostr

MNLRCWPWKRLEQLFSKK

HG0896 ENSG00000170365 SMAD1

>Homo_sapiens

----------MCFRRGGGTNACQISQLQ-------------

>Panthera_tigris

----------MYFRKGGGTNACRILQLQ-------------

>Tursiops_truncatus

MKASRESVFLMYFGKGGGTNACRILQPQ-------------

>Myotis_davidii

----------MSLGREGGTDACRISQLQ-------------

>Fukomys_damarensis

----------MIFGKGGGTNPCQISHLQ-------------

>Trichechus_manatus_latirostr

----------MCFRKGGGTNACRILQLSSCGSCLVGGTWNF

HG0897 ENSG00000171435 KSR2

>Homo_sapiens

MPCPCNWEGGKPDFF------------

>Mustela_putorius_furo

MPCHCNWEGGKADFF------------

>Myotis_lucifugus

MPCHCNWEGGKPDFFFFLHKSQPAKNT

>Equus_caballus

MPCHCNWEGGKPDFF------------

>Manis_javanica

MPCHCHREGGKPDFF------------

>Tupaia_chinensis

MPRRCSGEGGKPDFF------------

HG0898 ENSG00000171729 TMEM51

>Homo_sapiens

MKSCLLACPESPQITTGPSTAAIRTGGLAIARNHPAGVQLIFSSVGREILWSAFKGFLL

>Ailuropoda_melanoleuca

MESCFQDCPEHPHVTTDPSATAVHTGCLESA----------------------------

>Balaenoptera_acutorostrata_s

MKSCCQACLEAP-----------------------------------------------

>Manis_javanica

MKSCFQACPVPLQVSTDLS--AVCMGRLTLARSRPWRGQLLFCECGTRGFVQRV-----

>Loxodonta_africana

MKSCFQACPESPQVTTDFSAAAVHTGCLTYTWNHPSRVWLIFF----------------

>Cavia_porcellus

MKSCVQVCWEPPPVTTELSAAAVCTRHPKSVRSHPLRVQRMTF----------------

HG0899 ENSG00000173320 STOX2

>Homo_sapiens

MCRGHMRVGAPTQPSTRAHAQRPEPRQG------ED--------------------------------------------------------------------

>Orcinus_orca

-----MRVGARAHTRTRTHARTPP--------------------------------------------------------------------------------

>Pteropus_vampyrus

MCRGHMRVGAHTHARAHTHTRAGTHTTLSLGRSESDETLQPRGLLPLFLDSAEPARGSCCL-------------------------------------------

>Elephantulus_edwardii

MCGGHMRVGTHTPTPSRASAEKEAGLTRRCRG-ERDR---GASTLPGLG--GTDCR-SRCLQSRPR-SRCTRDP-HSSLCDEDCLPRTAGEAEAACAGDCGPRD

>Equus_asinus

-----MRVGAHTRAHTRA------RTTLSLGRRKIDEALQSRGRLGLFLDSAGAGLPQ-LLSAEPARIPVHTRPRFEPL-------------------------

>Trichechus_manatus_latirostr

MCGGHMRVGTHTRTPPRASARKKEN-------------------------------------------------------------------------------

HG0900 ENSG00000173517 PEAK1

>Homo_sapiens

MVVAAAACAPLRYLPKLELML--------

>Mustela_putorius_furo

MVVTTAACAPFRYLPELELML--------

>Pteropus_vampyrus

MVVAAAACSPLRYLPKLELML--------

>Dasypus_novemcinctus

MVVPAAACAPHRCLSKLGLMLLTYLGWFI

>Loxodonta_africana

MVVPAAACAPFRYLPQLDLMLKTFLCGFV

>Ictidomys_tridecemlineatus

MVVTAAACATLRYLPKLELML--------

HG0901 ENSG00000175697 GPR156

>Homo_sapiens

MRNQSEGTRTVTAIWWLTRIITLCSLMVTKMSTL----DSHGMHSQQGTS----------

>Canis_lupus_familiaris

--------------------------MVTKMPTL----DSHDMHCQQGTRGAWRVSDEGD

>Myotis_davidii

------------------------------MPTP----DSHGMHWQQGTSRAWWVTEEGA

>Oryctolagus_cuniculus

--------------------------MVAKWSSL----DSPAAGDPLDNV----------

>Equus_przewalskii

--------------------------MVTKRPTL----DSHGMH----------------

>Manis_javanica

--------------------------MIAKMPPRLTWCDSVAEHK-----ESQRVSDEEA

HG0902 ENSG00000177182 CLVS1

>Homo_sapiens

MASVLLLFWMNTTT

>Rousettus_aegyptiacus

MASVLLLLWMSTAT

>Dasypus_novemcinctus

MASILLPFWMATTT

>Condylura_cristata

MASVLLLFWMATTK

>Equus_caballus

MALTLLLFWMATTT

>Jaculus_jaculus

MASVLLLFWMATTT

HG0903 ENSG00000180616 SSTR2

>Homo_sapiens

MSHWILGLQGPLR

>Miniopterus_natalensis

MSQWILGLQDPLR

>Oryctolagus_cuniculus

MSQWILGLQGPLR

>Ceratotherium_simum_simum

MSQWILCLQDPLR

>Nannospalax_galili

MSQWILGLQGPLR

>Tupaia_chinensis

MSQWILGLQGPLR

HG0904 ENSG00000182329

>Homo_sapiens

MYLIKGPVFVVFLGLL---------------------

>Canis_lupus_familiaris

MYLIKGPVFVVFLGSLLSSGNLEV-------------

>Rousettus_aegyptiacus

MYLIKGPVFVVSLGSLLSSGNLEV-------------

>Ceratotherium_simum_simum

MYLIKGPVFVVFLGSLLSSGNLEV-------------

>Manis_javanica

MYLIKGPVFVAFLGSLLSSGNLEV-------------

>Chinchilla_lanigera

MYLIKGPVFVVFLGFCGASEILRCNQLLAFKTKMNCP

HG0905 ENSG00000183337 BCOR

>Homo_sapiens
MLESVEDGAGRPED
>Canis_lupus_familiaris
MPESVEAGAGRPED
>Balaenoptera_acutorostrata_s
MPESVEAGAGRPED
>Rousettus_aegyptiacus
MPESVEAGAGRPED
>Manis_javanica
MPESVEAGAGRPED
>Fukomys_darlingi
MPESVEAGAGRPED

HG0906 ENSG00000183773 AIFM3

>Homo_sapiens

MAAPASLRPAGGPAPWGAP

>Eptesicus_fuscus

MAASAFLRAVGGPAPRGAL

>Galeopterus_variegatus

MAAPAFLRAAGCPAPWGAP

>Equus_przewalskii

MVAPASLRAVGGPAPLGAL

>Marmota_marmota_marmota

MATPASLRAAGDPAPWGAP

>Tupaia_chinensis

MADPASPRAAGRPALWGAP

HG0907 ENSG00000186862 PDZD7

>Homo_sapiens

MELEARRAGLSRASRFLLDL

>Felis_catus

MEPGARGALCSRASRFFLDL

>Orcinus_orca

MEPGARGALLSRASRFPLDL

>Myotis_lucifugus

MEPGARRALLPRASRFLLDL

>Ceratotherium_simum_simum

MELGARRAFLLRASRFLLDL

>Ictidomys_tridecemlineatus

MELRALGAGLWRVGRFLLDL

HG0908.1 ENSG00000188554 NBR1

>Homo_sapiens
MHCGTKGLGHTVPWKL
>Felis_catus
MHCGSKGLRHSVPWKL
>Pteropus_alecto
MHCGSKGLRHCVPWKL
>Oryctolagus_cuniculus
MHCVTKGLRHSVPWKL
>Ceratotherium_simum_simum
MHCGSKGLRHSVPWKL
>Microtus_ochrogaster
MHCGTKGLHHSIPWKL

HG0908.2 ENSG00000188554 NBR1

>Homo_sapiens

MERNIQYQSGHNSHGNPVDRLETSAGLYMSTVMDWSVVMFSYLESRD

>Felis_catus

--------------------------------MDWSV-RFSHLESGE

>Rousettus_aegyptiacus

--------------------------------MDWSL-MFSYLESGE

>Oryctolagus_cuniculus

MEKSSRLWKRARSSRHVVERLEPSARLCTSTAMDWSAVRCLYLKSGR

>Ceratotherium_simum_simum

--------------------------------MDWSV-MFSHLESGE

HG0909 ENSG00000189079 ARID2

>Homo_sapiens

MTEPRRRRPRNGLRALVGSAGSGGRF-------

>Rousettus_aegyptiacus

MTEPRRRRPRNGLRALVGSAGSGGRF-------

>Erinaceus_europaeus

MTEPRRRRPRNGLRALVGSAGSGGRL-------

>Ceratotherium_simum_simum

MTEPRRRRPRNGLRALVGSAGSGGRF-------

>Manis_javanica

MTEPRRRRPRNGLRALVGGAGSGGRFWTAIGAF

>Heterocephalus_glaber

MTEPRRRRPRNGLRALVGSAGRGGRF-------

HG0910 ENSG00000197935 ZNF311

>Homo_sapiens

---MNPKSRAVFDTLSQTVAALWVKKGPLAGTTKTCVLQRRLASTGATSQLV

>Ailuropoda_melanoleuca

MLLMNPKSRALCDT--------------------------------------

>Physeter_catodon

---MNPKSRAVFDTSSQKAAALWVRRGPLAGTTKA-----------------

>Oryctolagus_cuniculus

MLLIVPKSRAVFDTPKHRAAVFWVGKGPLTGLSETCVPQRRLLSTGTT----

>Equus_asinus

MLLMNPKSRAVFDTSRQVAAAL------------------------------

>Marmota_marmota_marmota

--MMNLKSRAMFKTSSQIAAAL------------------------------

HG0911 ENSG00000198673 FAM19A2

>Homo_sapiens

MPRRAAGGLRGALPAALRGGIEGLKLARSCC

>Felis_catus

MPRGAAAGLRGALPAALRGGIEGLQLARSCC

>Rousettus_aegyptiacus

MPRGSAVGPRGALPAALLGGIEGLQLARSCC

>Oryctolagus_cuniculus

MPRGVTGELRGALPAALGGSSDWLRLAGSCC

>Equus_przewalskii

MPRGAAVGLRGAAPAALGGGISGLQLDRSCC

>Rattus_norvegicus

MPRVAAGGLREAGPAALEGEIKGLQLARSCC

HG0912 ENSG00000198822 GRM3

>Homo_sapiens

-----------------------------MPGGPCFCQESQLDATASAWSR

>Mustela_putorius_furo

-------------MREG--GKEKENEKERMPEGPRLCRESQLDATV-----

>Tursiops_truncatus

-----------------------------MPEGPCLCRESQLDAKNSARPR

>Rousettus_aegyptiacus

MKGGRKEEGEREKENEGGRKEEKENEKERMPEGPCLC--------------

>Manis_javanica

-----------------------------MPEGPCLC--------------

>Nannospalax_galili

-----------------------------MPEGPCLCWESQLDATVSA---

HG0913.1 ENSG00000204599 TRIM39

>Homo_sapiens

MRLSLKFALQLRHKE

>Acinonyx_jubatus

MRTSLKFALLLRHKE

>Physeter_catodon

MRTALKFALLLRHEE

>Equus_przewalskii

MRTSLKFALLLRHKE

>Loxodonta_africana

MRTSLEFALPLKHEE

>Ictidomys_tridecemlineatus

MRTSLKFVLLLRHEE

HG0913.2 ENSG00000204599 TRIM39

>Homo_sapiens

MPQPQIWRKKMENGGGTELWTTSQERVARWDPEQW------------------------

>Oryctolagus_cuniculus

MPLPEIWREKMETRGGRSFGRRSGE-GRQVGPGTAMDAKSGRP----------------

>Equus_caballus

----------METFGGTELWTTLRR-GSQSGT-----PDSGRNKMRTSLKFALLLRLKE

>Manis_javanica

----------METGVGLSCGARSGE-GRRWDPGQW------------------------

>Loxodonta_africana

----------METGVGQSCGQRSGE-GCKMGPWTVVETQ--------------------

HG0914 ENSG00000227877 MRLN

>Homo_sapiens

MVFFIKERDHSLDFGLRLLNPWTESRLLRDHLC

>Panthera_tigris

---MIHSRDTI----------------------

>Orcinus_orca

MVFVTKERDHLLDFGLRLSNP------------

>Myotis_davidii

MVFFTKESDHLLNFGLRFLTPEQIQGC------

>Ceratotherium_simum_simum

MVFFTEERDHLLDFGLRFLNA------------

>Nannospalax_galili

----MGKRDHSLDFGLGLLNP------------

HG0915 ENSG00000254469 AP002495.2

```
>Homo_sapiens
MYFPGCSTCHLASVGLIFKNSSFLKPQNAFLCYGRRKSSSRGQKAKVRRPNWRENRFGGPRSEA-TVAPRGPRSRE----------
>Felis_catus
MHFTGYITCLLASVGIIFQKQRLLEAPGQIPGIPEAESSATNQKSEVRGPNWREKGFCGPRSEA-AMALPGVA-------------
>Balaenoptera_acutorostrata_s
MYFPGNSTYLLTSVGVIFRKQRLLEAPKRIHGLREAEEWRFGSEGG--GVT-----------------------------------
>Equus_caballus
-----------------------MRLQNEFPGFTRRKSGAPGQKAEVRGLNWRENGFVVRDLRQRWLFKVHCPG------------
>Manis_javanica
-----MPSCL---GGGHFQKHRFLRPRNEFPGFGRRKSDAPGQKAEVRRLNWRENGLLVRELRQPAVALGGPVSWGYHKDSFSSGS
>Peromyscus_maniculatus_baird
-----MSNCCLASVGAIFQKSGFFKPQNASLSFCRRGGARARRR-------RSPDRFGGRIRLE----PRS---------------
```

HG0916 ENSG00000020633 RUNX3

>Homo_sapiens

MIHSAYASVENLWLDIPVGPGIQILGS----------------------------------------------------

>Odobenus_rosmarus_divergens

MIHSAYASVENLWLDIPVGLGIQILGS----------------------------------------------------

>Loxodonta_africana

MIHSAYASVENLWVDIPVGPGIQILGS----------------------------------------------------

>Octodon_degus

MIHSAYAAVENLWLDIPVGPGIRILGYKQKLTFLGDFFYSLTHTLSRSFLFFFFVAVGGERTKQKTNKNQHPHPFTSNF

>Orycteropus_afer_afer

MIHSVYASVENLWVDIPVGPGI---------------------------------------------------------

HG0917 ENSG00000042980 ADAM28

>Homo_sapiens

MNQGVSFCLTGEEAGTDPAAPT

>Ailuropoda_melanoleuca

MKPGVSFCLT------------

>Orcinus_orca

MNPEVSFCLT------------

>Equus_asinus

MNLGVPFCLT------------

>Mus_musculus

MKQGVSFCLP------------

HG0918 ENSG00000047648 ARHGAP6

>Homo_sapiens

MDLLLLPPPGHLERTGGA

>Pteropus_vampyrus

MDLLLLSSPGHLECAGGA

>Ceratotherium_simum_simum

MDLLLLPPRGYLERAGGA

>Manis_javanica

MGLLPLSPLGHLERAGGA

>Cavia_porcellus

MDLLLLPPQGHLERVGGA

HG0919 ENSG00000051009 FAM160A2

>Homo_sapiens
MGSYWLLQTG
>Ursus_maritimus
MASYWLLQTG
>Sorex_araneus
MASYWLLQTG
>Microtus_ochrogaster
MASYWLLQTG
>Orycteropus_afer_afer
MASYWLLQTG

HG0920 ENSG00000051382 PIK3CB

>Homo_sapiens

---MGATILRAPWL

>Canis_lupus_familiaris

---MGAVILRAPWL

>Pteropus_alecto

MTEMGATILRAPWL

>Condylura_cristata

---MGAIILRAPWL

>Marmota_marmota_marmota

---MGTIILRAPWL

HG0921 ENSG00000058799 YIPF1

>Homo_sapiens

MRKLRPRGAR-IQQRSSRIGNQTQLCLTPEPVPLTTG--

>Leptonychotes_weddellii

MRKLRPREAHSPTARRSRIGNQTQVCWTPEPVPLTTG--

>Lipotes_vexillifer

MRKLRPPGAHSLTARSSRIGNQTQVCSTPEPVPLTTG--

>Loxodonta_africana

MRKLRPRGAHSHAARSSRIGSQTQFLQSFFLIIGRRPTG

>Microtus_ochrogaster

MRKLRPRGAHT-QQRSNRIGTQTQTPENSLPHPWERASG

HG0922 ENSG00000079841 RIMS1

>Homo_sapiens

MQTTMKDWVLALPGSAAAAAAAAAAAPPPA--AAARAPL

>Odobenus_rosmarus_divergens

MQTTMKDWVLSLPGSAAAAA---AAPPPPA--AARRAPL

>Rousettus_aegyptiacus

----MKDWVRSLPGSAAAAA---AAPPPAA--AARRAPL

>Condylura_cristata

MQTTMKDWVLCLPGSAAAA----ASAPPPA--AAGRAPL

>Nannospalax_galili

----MRDWVLALPGSAAAAATA---PPPAAAAATGRAPL

HG0923 ENSG00000081138 CDH7

>Homo_sapiens
---------------------------------MTLPSCLRSSTD----HGFHDRLRYLLHHSLAKAIQQW
>Odobenus_rosmarus_divergens
---------------------------------MTLPSCLRSSID----HGFYDRLGYLLHHSLAKVIQQW
>Myotis_brandtii
MDGLRPLLLSPSSSGFSCEGLHLSGGRWRRGARGCARVCARLAAEQLPFRQPSLGLRNLLHHSLAKAIQRW
>Dasypus_novemcinctus
----------------------------------------------------MSEFRCLLQHSLAKAIQQW
>Peromyscus_maniculatus_baird
---------------------------------ML--AQLRQPV-PPASAEVLAGRRYLLHHSLAKGTQHW

HG0924 ENSG00000081307 UBA5

>Homo_sapiens

MSATHRKRLRGRPVGVSETCLSVRRWVHVPRALG------------------

>Odobenus_rosmarus_divergens

MTATYRKRYPGRPAGAAATCCLCGAGGTSPGFWARKAAATCLLTYPWRGLQA

>Galeopterus_variegatus

MTAAHRKRCREMRVGAAETCL-------------------------------

>Ceratotherium_simum_simum

MTATYRKRYRVRPAGVAETWLAVRG---------------------------

>Orycteropus_afer_afer

MTAAHRKQHCGRSAGAEETCLSVRCWRYPGLS--------------------

HG0925 ENSG00000088766 CRLS1

>Homo_sapiens

-----------------------------------------------MEAAVAQCLSGCRVSMEKRLASVPGC

>Pteropus_vampyrus

MNYKRLPLPPGAQGSPAATVSQKRREDVRPPLAAPDHLYKRTCWGLKTSMEVGEVLGGFGYSREKRLFSAPGC

>Erinaceus_europaeus

-------------------------------------------------MEEAELPPDCRNSMEQRVACVPDC

>Manis_javanica

------------------MVSQKRRADVRPWR-LQATCISGVCRGLKTGMEAAEVLRACGKSMERRSAGVPGC

>Marmota_marmota_marmota

-----------------------------------------------MEASAAVLPRHFGVSMEKRLAGDPSC

HG0926 ENSG00000095397 WHRN

>Homo_sapiens

MPRTPCRTPAPLHRLPLPGVLWD

>Balaenoptera_acutorostrata_s

MPRTPCRTPVRLHQLRFGG-LWD

>Pteropus_vampyrus

MPRTPCRTAARLHRLQFGG-VWD

>Condylura_cristata

MPRTPWRTPARLHRLRFGG-L--

>Equus_asinus

MPRTPCRTPARLHQLRFGG-LWD

HG0927 ENSG00000095951 HIVEP1

>Homo_sapiens

MAACGRPLALLGALAARE--------PSSTWIN-

>Mustela_putorius_furo

MAACGRHLPLLGALAARERPELRSCEASST----

>Rousettus_aegyptiacus

MAACGRHLPLLGALAARERPELRSCELSSKWIN-

>Marmota_monax

MAACGSHLPLLGALTAREQPELRSCDPSSTWIN-

>Tupaia_chinensis

MAACGRHLPLLGALAARERPELRSCEQSSTWSSN

HG0928 ENSG00000102057 KCND1

>Homo_sapiens
MPQSPLHHCKGIPGSSAFPETLESAENSPRVSWPL
>Mustela_putorius_furo
--------MRKLQFPNLFPRSLEGTDDPPQLSWPL
>Oryctolagus_cuniculus
MPQSPRQPCQGVPGPFSIPKNSGECDSRSRVCWPL
>Manis_javanica
MSQFPLQPC--------------------------
>Heterocephalus_glaber
MPQSPLQPY--------------------------

HG0929 ENSG00000102239 BRS3

>Homo_sapiens

MGKHLAKHAGRHTELKQRSIWMSWIFFPFCSVLFS--

>Galeopterus_variegatus

---------------------MSWIFLPFCCILFS--

>Condylura_cristata

---------------------MSWICLPFCSVLPSRY

>Oryctolagus_cuniculus

-------MLGDTEPLKERSIRMSWIFFPFSSVLFS--

>Ictidomys_tridecemlineatus

---------------------MSWIFFPFCSVLFS--

HG0930 ENSG00000102524 TNFSF13B

>Homo_sapiens

MQKGRKEKIQDNSPEG--------------------

>Balaenoptera_acutorostrata_s

MQPGREKKIQENSPEGERRAGPCSQCRTSTNTHNRK

>Pteropus_vampyrus

MQQGREKKIQENSPEGKSRAGRWSQCRTSTNR----

>Oryctolagus_cuniculus

MQRGREKRIEENSPEGQSRVLRCSQCRAHTDNRK--

>Equus_asinus

MQRGREKKIQENSPEGDSQAGRCSQCRTSTNTDNRK

HG0931 ENSG00000104951 IL4I1

>Homo_sapiens

MCFLPSPDKWPFLKFLINSYEDNTFGD

>Canis_lupus_familiaris

MRFFASPDKRPLLKFLFIK--------

>Pteropus_alecto

MCFLASPDKWPPLTFLSVN--------

>Equus_przewalskii

MRLPARPDTWPLLNFLLIN--------

>Tupaia_chinensis

MHFLASPDTWPLLKFLFIN--------

HG0932 ENSG00000105997 HOXA3

>Homo_sapiens

MLRVRAHPAPRGPGRAAKPGRGPEPVSEPEPEPALVRS-----

>Felis_catus

MLRVGAHPAPRGPGRAAKPGRGPEPEPEPKPEPEPELAALIRS

>Pteropus_vampyrus

MLRVGAHPALRGPGRAAEPGRGPEPEP--------ALAALIRS

>Ceratotherium_simum_simum

MPRVGAHPVPRGPGRAAKPGRGLEPEPEPA------LAALFRS

>Cavia_porcellus

MLRVHPHPAPRGPGRAAQPGRGPEPEPEPA------LAAIVRS

HG0933 ENSG00000106330 MOSPD3

>Homo_sapiens

MTEAVRRKYRKGLSGTWQH---------------

>Odobenus_rosmarus_divergens

MTEAVRRKCRNGLSGRWQPRSGQKLMGRCSDQKL

>Rousettus_aegyptiacus

MTAVIRRKCRNGLSGRWQPRSGHKLIGRCCDQKL

>Galeopterus_variegatus

MTDVVRRKCRKGPSMRWHPWPGHRLIGRCFDHKP

>Fukomys_damarensis

MGEAVRRKYRKGLSGRWRH---------------

HG0934 ENSG00000107443 CCNJ

>Homo_sapiens
MSGAGVRRTTAGLGL
>Tursiops_truncatus
MSGAGVPRTLAGLGL
>Rousettus_aegyptiacus
MSGAGVPRTLAGLGL
>Ceratotherium_simum_simum
MSGAGVPRTTAGLGL
>Ictidomys_tridecemlineatus
MSGAGVPRTPAGLGL

HG0935 ENSG00000108518 PFN1

>Homo_sapiens

--------------------------MGSRPRPYRPLQIKI-----------------

>Eptesicus_fuscus

--------------------------MGSRPRPYRPLQIKI-----------------

>Condylura_cristata

MGGGIFKKLHLQSSVSGQAGDGGREGMGNKPRPYRPLQIKI-----------------

>Oryctolagus_cuniculus

--------------------------MGSRPRPYRPLQIKI-----------------

>Ictidomys_tridecemlineatus

--------------------------MGSRPRPYRPLQIKIQRGGEEQEVAVRGLLHS

HG0936 ENSG00000108799 EZH1

>Homo_sapiens

MEAGHLFCCCVLPFS

>Canis_lupus_familiaris

MEAEHLFCLCVLPFL

>Pteropus_vampyrus

MEAEHLFCLYVFPFL

>Galeopterus_variegatus

MEAEHFFCLSVLPFL

>Ceratotherium_simum_simum

MEAEHLFCLCVLPFL

HG0937 ENSG00000109846 CRYAB

>Homo_sapiens

MTSHWPAQTCLCLSFLSSVSTGYVSHCQIPDHKSP

>Odobenus_rosmarus_divergens

-------------MGSSSVSTGYVSHCQIPDHKSP

>Ochotona_princeps

MTSHWLAQTCLCFSFLSSVSTGYVSRCQIPDHKSP

>Equus_asinus

--------MCLCFSFLSSVSPGYVSHCQIPNHKSP

>Marmota_monax

MTSHWPAQTCLCFSFLSSVSTGYVSHCQIPDHKSP

HG0938 ENSG00000111665 CDCA3

>Homo_sapiens

--------------MGTTSAQPIGSGQGAWREV

>Canis_lupus_familiaris

----------------------MESRQGAWREV

>Myotis_lucifugus

MASLPVGGSPIADLSEPVSAQPIGSRQGAWREV

>Galeopterus_variegatus

MASCAGGGSHTAGLWESTSAQPIGSGQRAWREV

>Mesocricetus_auratus

----------------------MGSGQGAWREV

HG0939 ENSG00000111674 ENO2

>Homo_sapiens

--------------------------MG----A---LEERPRQPILFLAPRRNASLAAIPL

>Lipotes_vexillifer

--------------------------------------------MLFWAPRRNANLAAGPP

>Miniopterus_natalensis

--------------------------MR----AVDEGLQLCRLPTLFLAPRRRASLAAIPL

>Condylura_cristata

--------------------------MG----LLEERPHLPRQPI-FLAPT----------

>Trichechus_manatus_latirostr

MHVWERDWDLGEWVESGQAGVLSGGRMGGSHPRSRREAGQARQPLLFLGPRRNKSLAAILL

HG0940 ENSG00000115556 PLCD4

>Homo_sapiens

--------MGWTGESSHLSPSYCFPPAITCQSQQPAAVEEGRKQASARGAKEKRSQAGLPDPTASQSSGGTAHRHRKHRTAILLSCPR

>Balaenoptera_acutorostrata_s

--------MGWTEERAHTSPHPTLSIWL------------------------------------------------------------

>Rousettus_aegyptiacus

--------MGWTEERAHTSPHPLLPSGCNCQSQQQTAVEEGRKPEGAKGEK---ELAGFPHPTEESV--------TEHTDTGLLSCPR

>Equus_asinus

MSWAFKNWRGWTEESSHLSPPYSFPLAVTASHSSKLR---------------------------------------------------

>Chinchilla_lanigera

--------MGWTKERAYTSPHPAPSLCL------------------------------------------------------------

HG0941 ENSG00000116285 ERRFI1

>Homo_sapiens

MRGRAPRGSRRSSGGMKATG-----

>Mustela_putorius_furo

MRGRAPWGSRGSAGGMQATG-----

>Rhinolophus_ferrumequinum

MRGRAPWGSRGSAGGTKATSRPTNP

>Equus_asinus

MRGRAPWGSRGSAGGMKATG-----

>Ctenomys_sociabilis

MRGRAPPGSRRFPGGMKATG-----

HG0942 ENSG00000117477

>Homo_sapiens

MEDAEARCCHGSVVLAAAEAGAGVSFASM----------------

>Felis_catus

MEDAETRRCYGTWVLAAVGAGAGVSLASLSVRA------------

>Tursiops_truncatus

MEDAETRRCHGSRVLAAAA-RAGVSLALLSELRKRRKGELPWPTL

>Myotis_brandtii

MEDAETRRCHGSRVPAA----------------------------

>Oryctolagus_cuniculus

MEDSAPRRCHGSPVLAAAGVAAGVFHASLSELRKR-RGTLPRPSL

HG0943 ENSG00000120690 ELF1

>Homo_sapiens

----------------------------------------------MLVRPRSLREEKFQKNCLNLTRISMNQEN-----

>Balaenoptera_acutorostrata_s

----------------------------------------------MPLESRSSREEKFQKNCLNLSKIPVNQENLKHLP

>Rousettus_aegyptiacus

---------------MTWSSAVSFLLQPGLEIRIHFSTLKQKPLKTGSTWPRRSREEKFQKNCLRLNRISVNQEHSKFLP

>Oryctolagus_cuniculus

---------------------------------------------MCYPRPRRLREQNFQQHCLSLSRISTNQENLKHLS

>Heterocephalus_glaber

MNEGEAEEEAAAAAGAAWSSV------------------AEVAAAFMLVRSKSLREEKV---------------------

HG0944 ENSG00000120833 SOCS2

>Homo_sapiens
MVQERR-GGGVPGIFHVYFPTPTPAAGVQFGLTQPPAFSL
>Ursus_maritimus
MVQEHR-RGGVPGIFHVSVSPPTPQPQGSSLD--------
>Miniopterus_natalensis
MVQEHR-RAGVPGIFHVAVSPPTPRPGV------------
>Equus_przewalskii
MVREHR-CGGVPGIFHVCVFPPTPQP--------------
>Chinchilla_lanigera
MVQEHRLDGGVPGIFHFSVSPARPQGSSWD----------

HG0945 ENSG00000124782 RREB1

>Homo_sapiens

MIATAGESFIVYQWVRKWR----------------

>Odobenus_rosmarus_divergens

MIAIAGESLIVYQLGQKMEVIGSRGWVLLSTFWVL

>Myotis_lucifugus

MIAIARENLIVYQLGQKWSFMAEAS----------

>Equus_przewalskii

MIAIAGENLIVYQLGQKMEFYGRGFLEA-------

>Manis_javanica

MIAIAGESLVVYQLGQKMEFHGRGFLEA-------

HG0946.1 ENSG00000127324 TSPAN8

>Homo_sapiens
MTSKGTYLPGDSLCDI
>Mustela_putorius_furo
MTSKRTYSPSDSPCDI
>Pteropus_alecto
MTSKRTYLPKDSLCDI
>Dasypus_novemcinctus
MTSKRTYPPRDSPCDI
>Heterocephalus_glaber
MTSKRTYMPGDSLCDI

HG0946.2 ENSG00000127324 TSPAN8

>Homo_sapiens

MCGLIFLSSLITFYSSLYFFAFSFCSEAVDTEISAGKLLQSILQDKPVTNS

>Rousettus_aegyptiacus

MCGLIFYPL------------------------------------------

>Galeopterus_variegatus

MCGLIFYPL------------------------------------------

>Ceratotherium_simum_simum

MRGLIFYPL------------------------------------------

>Manis_javanica

MCWLIFYLL------------------------------------------

HG0947 ENSG00000129226 CD68

>Homo_sapiens

-MTKREVARAEAPESGCGWDHLQYRK-------------------

>Physeter_catodon

-MTKTLVAGAEVPESGWGLGCLSYRKSKIYFLFFQMNEGRGYRVD

>Equus_asinus

-MTKRLVAGGEVRESGWGWDY---LKSKTFISSFP-KGGE-----

>Loxodonta_africana

-MTKRLVAGAEVSESGLGVGIFSVTGSKTFISSFP-KEGRGNRVD

>Nannospalax_galili

MTKEGPVAGAEVSESGWGWDYLSYGK-------------------

HG0948 ENSG00000129595 EPB41L4A

>Homo_sapiens

MRGDAPPPSAPSSAGASPAAPGR

>Odobenus_rosmarus_divergens

MRGDAPPPSALSSAGASPAAPGR

>Miniopterus_natalensis

MRGDAPPPSAPSSAGASPAAPGR

>Equus_asinus

MRGDAPPPSAPSSAGASPAAPGR

>Marmota_marmota_marmota

MHGDA-PPSAPSSAGASPAAPGR

HG0949 ENSG00000131381 RBSN

>Homo_sapiens

------------------------MKPLPHHLCAA---

>Canis_lupus_familiaris

------------------------MIPLPHHLCAAWLS

>Physeter_catodon

MLWTWGLAWWFVDSYSREDTSSRAVMPLPHHLCAAWLS

>Myotis_lucifugus

------------------------MMPLPHHLCAAWLS

>Equus_asinus

------------------------MMPLPHHLCAAWLS

HG0950 ENSG00000132964 CDK8

>Homo_sapiens

MSLALSRGLLLLLPHQSGWCCGRRA

>Felis_catus

MSPALLRGLLLLLPHQSGWCCGRRA

>Oryctolagus_cuniculus

MSPALSRGLLLLLPHPSRWCCGRRA

>Ceratotherium_simum_simum

MSPALSRGLLLRLPHQSGWCCGRRV

>Mus_musculus

MSPAPARGLL-PSPREAGWCCGRRV

HG0951 ENSG00000134061 CD180

>Homo_sapiens

MLSSQQHFLFQDHPSEYLSGCQIARAFTV

>Balaenoptera_acutorostrata_s

MLSSQQHFLFSDYPSEHSLAARLSRPS--

>Galeopterus_variegatus

MLSRQQYFLFQDHPLSTSLAAKLQGPL--

>Ceratotherium_simum_simum

MLSHQQHFLF-------------------

>Trichechus_manatus_latirostr

MRNSQQHFLF-------------------

HG0952 ENSG00000134453 RBM17

>Homo_sapiens

MGRVGRAGRH

>Canis_lupus_familiaris

MGRVGRAGRH

>Dasypus_novemcinctus

MGKVGRAGRH

>Condylura_cristata

MGRVGRAGRH

>Equus_caballus

MGRVGRAGRH

HG0953 ENSG00000134815 DHX34

>Homo_sapiens
MNLCPSQRSLQMS
>Orcinus_orca
MNLCPSQRSPQES
>Cynopterus_sphinx
MNLCPSQRSLQDS
>Ceratotherium_simum_simum
MNLCPSQRSLQES
>Heterocephalus_glaber
MNLCPSLRSLQES

HG0954 ENSG00000134874 DZIP1

>Homo_sapiens

MSGPSRNSPVDWGMSGFSSAY

>Odobenus_rosmarus_divergens

MVVPSRDSPVD----------

>Physeter_catodon

MVGPSRDSPVD----------

>Myotis_brandtii

MVGPSQGSLVD----------

>Ceratotherium_simum_simum

MVGPSRDSPVD----------

HG0955 ENSG00000134982 APC

>Homo_sapiens

MLFPGTVVGCW

>Acinonyx_jubatus

MLFPGSVVGYW

>Ceratotherium_simum_simum

MLFPGSLVGYW

>Ictidomys_tridecemlineatus

MLFPGSAVGCR

>Orycteropus_afer_afer

MLFPGNAVGCW

HG0956 ENSG00000136235 GPNMB

>Homo_sapiens

MILPEA-LYLIKIEREAPQMPEEHCCSWWTGPEEFRVKP

>Canis_lupus_familiaris

MICPKAPLYLIKIRSGAQQRPEEHCCS------------

>Dasypus_novemcinctus

MTCPEA---------------------------------

>Equus_asinus

MICPEA-QSFIKIRSGAQQRPEEHG--------------

>Manis_javanica

MICPKALN-------------------------------

HG0957 ENSG00000136874 STX17

>Homo_sapiens

MTRRSMVVRLGGLGARGRKPLLAKPRGGWQKVPHLGSLLLANRGSERRLYFDFSLQIRFFYMSGEDSCYQGGHTTFF

>Myotis_lucifugus

-------------------------------------------------------------MSREDSCCK-------

>Equus_asinus

-------------------------------------------------------------MSGEDSCYK-------

>Manis_javanica

-------------------------------------------------------------MSGEDSCFKER-----

>Ictidomys_tridecemlineatus

-------------------------------------------------------------MSGEDRCYK-------

HG0958 ENSG00000137497 NUMA1

>Homo_sapiens

MNNHSCPFAPSLGGN-------------

>Felis_catus

MNKHSCPFAPSLGRG-------------

>Equus_asinus

MNKHSCPFAPSLGRG-------------

>Cavia_porcellus

MNKHSCPFVPSSLGWKLIRIQPIRLSVS

>Tupaia_chinensis

MNKHSCPFASSLLKA-HSKARPLRIALP

HG0959 ENSG00000137558 PI15

>Homo_sapiens

-------------------------MLPNDVFVNSYPLALFFF----IDPNSTFLL

>Leptonychotes_weddellii

-------------------------MLPNDDFVNSYPLALFLFFSFLIDPNSTFLL

>Lipotes_vexillifer

MYRYHYRLSSGLEF--SICLDFLKKVKSSNDFVNSYPLALFFFSFHRP--------

>Ochotona_princeps

MGRFHYNCLLGLSSIFWVWRKSDQAMLPNVDFVNSSPLALFPPYILTLPF------

>Trichechus_manatus_latirostr

-------------------------MLPKDNFINSHLLSLSPFFFSFIDNNSTFLL

HG0960 ENSG00000140297 GCNT3

>Homo_sapiens

MISLSLLSYLRAGSQKESDGATCCPLLCRY----------------------------------------------------------

>Myotis_lucifugus

MTYLSLLSYLRAGN--------------------------------------------------------------------------

>Equus_przewalskii

MLYLSLLFYLRTGS--------------------------------------------------------------------------

>Manis_javanica

MVGLSLLSYVRAGNQAG-DRATCCSLFSEIKEEPEA---VSWTSCEYQKIEFGAVKRRPGDMVLTGASLELNQPSPGNKYVMASGPPQ

>Mus_musculus

MVCLSSLPPLRTGSQKVSDRATCCSLLCIGLTGAWVSSLDGW----------------------------------------------

HG0961 ENSG00000140853 NLRC5

>Homo_sapiens

MARLEIAEGRREGEKWIPVLRSGLEGEVCKESALWKQPVNPAQGT-HSPDTHETLSVGTLEHLS

>Canis_lupus_familiaris

--------------------MASVSLYPHEEPVLWKQPANPAQSAGHGSGTHETISADTLEYL-

>Myotis_brandtii

----------------------MLSEADHWRPVLWKQPANPAQGTGLGPGTHETVSTDTLEHL-

>Oryctolagus_cuniculus

--------------------------------------------------------MGTLEHLP

>Nannospalax_galili

----------------------------------------------------------MLKHLS

HG0962.1 ENSG00000143319 ISG20L2

>Homo_sapiens
--------------------MVSRSLRGRRTWVRCMRRLPPIPAWSQGKGMPGFVSLLVVHAADAWVAQR---LSTPYFSLFLSIPRCSFPRRSIDRTCSSLLDSE
GSSSISPSTPFILQMRLKPRAR----------
>Physeter_catodon
------------------------------------MHVEISPTWRRGKWVPEFVSVLGTRARHAWVIQR---STMPNL---------------------------
--------------------------------
>Myotis_brandtii
-------------------------------------MWRWPPPGNGGKGYLELHLFVGPALGMPGKPR-------------------------------------
--------------------------------
>Ceratotherium_simum_simum
-------------------------------------MWRFSPPGGEGKGCLDCVSVCGTHAGAAWGI--------------------------------------
--------------------------------
>Ictidomys_tridecemlineatus
MCLLRWATSIVNPLPPGTAVVWCQRSPPGRTWVRCA--WRLLPPSRSERGVPGFVSLFVGSRLRCHLCRTPVSHSSPEH-----LRHC-LSGRITALACSGFP---
-------------DLRLRTRRAHWVLLFLWCW

HG0962.2 ENSG00000143319 ISG20L2

>Homo_sapiens

----------------MLDSRIQQRMSPTSSFHP

>Dasypus_novemcinctus

----------------MLVLGIQQRMIPTSSFHP

>Manis_javanica

----------------MLVRGIQQRMSPLSSFHR

>Dipodomys_ordii

MQVPESLWTPRFSPDHMLDNGIQQRMSSSSFFHT

>Tupaia_chinensis

----------------MLDAEIQQRMSPPASIHP

HG0963 ENSG00000143867 OSR1

>Homo_sapiens

MSESELGLSPARSQLLLVQTQRGSRERGSPSPACRIRG---C----------------------------------------------------------------

--

>Myotis_lucifugus

MSESELGLSPARSQPLPVQTQPGSREPRAASGTHHPRR-------PELRGR-------------------------------------------------------

--

>Manis_javanica

MSESELRLYPTRSQLLRVQTQRGSRERASPSRGRGTPG---QQSAPACRAPSVP---------AAGEGPGL-----------------------------------

--

>Marmota_marmota_marmota

MSESELGLSPARSQQLRFRTSEGTRSEAHDPSAGRTPRLL---SAPACGALGVP---------AAGEGSGLQNIRRVCGLQLLWLPGSRAASPRDALSSRTTAERD

RD

>Tupaia_chinensis

MSESKLGLYPARSQAFPVQTQRGSREQGSPTPGGLDSRAVGCSRVSDSRNPRNWREIWTLALQGYPRTPALLVSPVSC-----------SESRGDALNPGTAE---

--

HG0964 ENSG00000145246 ATP10D

>Homo_sapiens

MAGVEAQPAFTLRICA--------------

>Mustela_putorius_furo

MAGVEA------------------------

>Balaenoptera_acutorostrata_s

MAGIEA------------------------

>Myotis_brandtii

MAGVEAPPVSALRICAWLLSCFLGLGIVVL

>Ceratotherium_simum_simum

MAGVEAQAASTRRICAWFFSCTLGLAIVML

HG0965 ENSG00000146006 LRRTM2

>Homo_sapiens

MKPMLSRGIS

>Leptonychotes_weddellii

MKPMLSRGIS

>Balaenoptera_acutorostrata_s

MKPMLSRGIS

>Erinaceus_europaeus

MKPMLSRGIS

>Orycteropus_afer_afer

MKPMLSRGIS

HG0966 ENSG00000148356 LRSAM1

>Homo_sapiens

MFVVGRRLRLTAGKQGTAVR-------------------------

>Canis_lupus_familiaris

MLLVGRRLMLTAGEPCSAVLRFSVSDFVILV--CMTF--------

>Rhinolophus_ferrumequinum

MLLVGRRQRLTGGEPGTALLPLEETSLWIQGSSCQVVCDWRP---

>Oryctolagus_cuniculus

MFLVGRLLRLTAGEPGASR--------------------------

>Fukomys_damarensis

MFLMGRCLKLAAGKSGTARVTPE----------CQAPKPRAPPIL

HG0967 ENSG00000149925 ALDOA

>Homo_sapiens

MSGASGFPKYRSLPEDPWREKGRGH----------------

>Lipotes_vexillifer

MSGASGFPKYRSPARGSVARKGQGSPEKVGDIGWG------

>Ceratotherium_simum_simum

MSGASGFPKYRSLPGDLWRGKGRGHWRLLGTLAGGRAGAAL

>Manis_javanica

MSGASSFPKYRSLPGDLP-----------------------

>Heterocephalus_glaber

MSGASGFPKYRSLLGDPRREKGRG-----------------

HG0968 ENSG00000154319 FAM167A

>Homo_sapiens

MRFSSLPQGLQPPALPEKRELWAFFLPVWSSSAFRTDRPWHRLPRDSCDAVCSCLLWSMAPTGFQDGS

>Rousettus_aegyptiacus

----------------------------------------------------------MVPTGLGEES

>Equus_asinus

----------------------------------------------------------MAPTGLWEES

>Fukomys_damarensis

----------------------------------------------------------MAPTGFQD--

>Tupaia_chinensis

----------------------------------------------------------MAPTGFQDES

HG0969 ENSG00000154645 CHODL

>Homo_sapiens

MRAGGRAAGLGRRE

>Physeter_catodon

MRSGGRAAGFGRPE

>Myotis_lucifugus

MRAGDRAAGLGQRE

>Dasypus_novemcinctus

MRAGGRAAGLARQE

>Chinchilla_lanigera

MRAGARATGLGRRE

HG0970 ENSG00000154727 GABPA

>Homo_sapiens

--MGRRFSRSTLTGQEASRRQSATGRV

>Miniopterus_natalensis

--MGRRFSRSALTGQEASRRQSATGRL

>Oryctolagus_cuniculus

--MGRRFSRSALTGQEASRRQSATGRL

>Manis_javanica

--MGHRFNRSTFTEQEVSRRRSATRRL

>Trichechus_manatus_latirostr

MHYGPRCTRSSLTGQEASRRRSATGRL

HG0971 ENSG00000155592 ZKSCAN2

>Homo_sapiens

MHSGKPHCAVTSCLEKRPGFCILE--YTFTFSRAWHRFD-----------------------

>Oryctolagus_cuniculus

MHPGKPSGRDTSCLGKRLGFFLDLSVPGHLFRGLG---------------------------

>Ceratotherium_simum_simum

MHPGKPHRGDTSCLEKRPSLGTLV--HRHLFKGLG---------------------------

>Loxodonta_africana

MHPGKPHCGDTSCLEKRSGFWTLG--YTVLFSRA----------------------------

>Cavia_porcellus

MHSGKPHCQVTSCLEKRPGFWIWG--THSPFQGPTIDLINQPEVLFLLKLLNCGFGVRLDSF

HG0972 ENSG00000155926 SLA

>Homo_sapiens

MKNFNRKLLLQKIRSFAANGEHLRQP-------------------------------------------

>Canis_lupus_familiaris

-MNFNRKLLLQKIRSFSESGGFPGISGSRDAPRREACSEWLLKTGTYPSASPAHLPHF-----------

>Balaenoptera_acutorostrata_s

MKNFNRKLLLQKIRLFSANRAFPGISGSADVPRREVCSVPLLKTGTCLLQAF-LVSSRGSCNWFSNYAS

>Oryctolagus_cuniculus

MKDFNSERLLQELRSVSANAAFPRRAGCPAAPAFGRCHLRSRGS-------------------------

>Heterocephalus_glaber

MKNFNRKLLLQKIRSFPKNGALLGISSSPDALWREVCSEWLVETGTRPLPAK-RLPRF-----------

HG0973 ENSG00000156535 CD109

>Homo_sapiens

---------------------MFATAARSAIVMGMGGVEPPSPVSI

>Mustela_putorius_furo

---------------------MPAGAARSAIVMPVGGVEPPSPVSI

>Rousettus_aegyptiacus

---------------------MSARATRSAIVMRVGGVEPPSPVSI

>Marmota_marmota_marmota

--------------------------------MRVGRVEPPSPVSI

>Tupaia_chinensis

MLEPGWADRVERPLSNPGGFAIFAGAARSAIVIRVGGVEPPSPVSI

HG0974 ENSG00000157578 LCA5L

>Homo_sapiens
MQTNDHPYADHFEYLQPCFCHCLCRKATCFKKKSSWSAY--ERVGCFLQ-----------LIFPKQTIPPAISGFLKYILSLKRDQNLSKKYYRGKLINTTV
>Mustela_putorius_furo
-----------------------------------------------------------------------MMVFLKYILSLKRDQNMSKNTALETS-----
>Miniopterus_natalensis
---------------------------------------------------------------------------------------MSKNTTLGTS-----
>Equus_przewalskii
----------------------------------MWEFRLFVPQSFFFGFLERHQKRQVEMETDGHPCADHFASFLKYILSVKRDQNMSKNTTLETS-----
>Dipodomys_ordii
-------------------------------------MVVQFPFLCYF---------------------GALRCFLKYIF----------------------

HG0975.1 ENSG00000182158 CREB3L2

>Homo_sapiens

MRAGLLPSFWMHIPL

>Mustela_putorius_furo

MRAALLPSFWMRALL

>Equus_caballus

MRAYLLPSFWIHTLL

>Dipodomys_ordii

MRADLLPSLWMHIPR

>Tupaia_chinensis

MRADLLPLFWMLIPL

HG0975.2 ENSG00000157613 CREB3L1

>Homo_sapiens
MEASCAPGGAGGGGVG
>Odobenus_rosmarus_divergens
MEASCAPGGAGGGGVG
>Lipotes_vexillifer
MEASCAPGGAGGGGVG
>Ceratotherium_simum_simum
MEASCAPGGAGGGGVG
>Loxodonta_africana
MEASCAPGGAGGGGVG

HG0976 ENSG00000158008 EXTL1

>Homo_sapiens

-------------------------------------------------------------------------MGLSLPICTMTAQD

>Canis_lupus_familiaris

-------------------------------------------------------------------------MGLSLPICTMKALN

>Oryctolagus_cuniculus

MWPGGQRDTALWSGQGLHQPEQCPVAGALLLGLGCPEAEEAESGEQKGKSGKLGASPCLAVTCCSTLHRQPVSAGLSLPICTMRALD

>Loxodonta_africana

------------------------------------------------------------------------MPGLSLPICTMRALD

>Chinchilla_lanigera

------------------------------------------------------------MASDFLYGPRLVLMGLSLPICTMTVLD

HG0977 ENSG00000158321 AUTS2

>Homo_sapiens

MLPS-LGGLAPLGSLRSPGLPWLHFLKIWEPGSEFSPRRV

>Odobenus_rosmarus_divergens

MLPS-LGGLVPLRSLGSLGLPWMHFLKILEPGSEFSLRRV

>Rousettus_aegyptiacus

MLPS-LGGLVPLRSLGSPGLPWMHFLKIWEPGSEFSLRCV

>Galeopterus_variegatus

MLPS-LGGLAPRRSLWSPGLPWVHFLKILEPGSEFSPRRV

>Chinchilla_lanigera

MLPSQGGGLAPLRSLPSPGLPWMQFLKNFAAWE-------

HG0978 ENSG00000159023 EPB41

>Homo_sapiens

MCHCGDGSMASPGLAA

>Balaenoptera_acutorostrata_s

MCHCGDDSTASPLTAA

>Rousettus_aegyptiacus

MCHRGDGFMAPPGLAV

>Equus_przewalskii

MCHCGDGSMAFLLLAA

>Peromyscus_maniculatus_baird

MCHSDDGSTASP----

HG0979 ENSG00000159202 UBE2Z

>Homo_sapiens

MWEEVRVSGKWLFQADI

>Acinonyx_jubatus

---------MWLFQGDI

>Oryctolagus_cuniculus

MWEEVRVNGMWLFQGDI

>Equus_caballus

MWEEVRVNGMWLFRGDI

>Manis_javanica

MWEEVRVNGMWLFHGAI

HG0980 ENSG00000162670 BRINP3

>Homo_sapiens

MIQILGT-F-FCVLHRFVCHHAVKQVLRES

>Odobenus_rosmarus_divergens

MIQN-FE-I-FCMLFRFVCHHAVKQVLRES

>Equus_asinus

MIQIWGLFL-LRAI----------------

>Chinchilla_lanigera

MIRFWGLFFSFCLLCRFVCHHAVSRC----

>Tupaia_chinensis

MIQMLGTFF-FCVLFRFVCHPCS-------

HG0981 ENSG00000163629 PTPN13

>Homo_sapiens

MLTAPGDRRAGEPC-SVAPRSAS---AAVRVAYRS

>Pteropus_vampyrus

MITGPRDRRAGKQH-TLAPRSASATSAALEARSRS

>Oryctolagus_cuniculus

MITAPRDRRAGALH-SQAPGSAS---AALGVGCRS

>Ceratotherium_simum_simum

MVTVSGDRRVGEQH-SLAPRPASASRAALGAGCRS

>Chinchilla_lanigera

MVSGPGDRRAGEQQRPRAPRSASDAPAALGAGCRR

HG0982 ENSG00000164116 GUCY1A3

>Homo_sapiens
MCGFARRALELLEIRKHSPEVCEATKHPSYQCP
>Mustela_putorius_furo
MWGFAGRARELPEIRKRSRRVWEANEHPSHQYP
>Dasypus_novemcinctus
MCGFARRALELLEIGKRSREVWEAIEHSSHQCP
>Oryctolagus_cuniculus
MCGFARRAPQLLRIPQRSPEVWAVHRAPQL---
>Ictidomys_tridecemlineatus
MCGFARRAPELQQIWKHGPQVWETAKHTGQQCP

HG0983 ENSG00000164128 NPY1R

>Homo_sapiens

MDSNMGNKNKLNS

>Galeopterus_variegatus

MDSNMGNKNKLNS

>Erinaceus_europaeus

MDSNMGNKNKLNI

>Ochotona_princeps

MDSNMGYKNKLNS

>Ceratotherium_simum_simum

MDSNMGNKNKLNI

HG0984 ENSG00000164185 ZNF474

>Homo_sapiens

----------------------MLKGKIKRSWRHLSFNEDFPTFQTAVLQ--

>Ailuropoda_melanoleuca

-----------------------MTTNITRSWRHLRFNEDFPAFQTAALQ--

>Orcinus_orca

----------------------MIKAKIER----------------------

>Mus_musculus

---------------------------MSRSWRHLRFNGGFPTFRTAALQ--

>Tupaia_chinensis

MALCPHTVDSPRTAGTTRPGPTQRERGEAAGWRLLRLDGDFPTFQTCPVMRL

HG0985 ENSG00000164604 GPR85

>Homo_sapiens

MKACLLVL----GRRTGKLLLKILYHLHKQNSSEMVK

>Pteropus_alecto

MKARLLIF----WR-----------------------

>Elephantulus_edwardii

MSARIWGGFPWQGGGGGELSFKILNHPHKQNSSEMVK

>Loxodonta_africana

MSAWIFFLARWEE----EGEFKILYHPHKQNSSEMVK

>Ictidomys_tridecemlineatus

MSACF--W----GVGSGGWLLKILYHPHKKNSSEIVK

HG0986 ENSG00000164663 USP49

>Homo_sapiens

MVRGAGRDGPQNLKDEMGSKGREGAKRSWGSPDGCRTGVDCRGVGIECACYGELLQKVCKGQGRNAVVKDQV

>Acinonyx_jubatus

--------------MRWGVREENRVKGSWRSPAGGRTGLACTGVGVRCACCGEMLRKVCMGQGTGAVVKDQV

>Dasypus_novemcinctus

---------------------------------------------MKCACCGEILQKVWKGQGTKP------

>Oryctolagus_cuniculus

----------------------------------MSNSSRARRAGIERACGRERLQKACEGQGRKAGVRDPV

>Cricetulus_griseus

----------------MGSKRREAGRALMG------------------------VEPWWAAQGWGLHMYLP-

HG0987 ENSG00000164932 CTHRC1

>Homo_sapiens
MQPAAASERGGARR
>Physeter_catodon
MQPAVASERGGAGL
>Oryctolagus_cuniculus
MQPAAVSQRGGARS
>Equus_asinus
MQPAAASERGGARL
>Dipodomys_ordii
MQPAAASERGAARH

HG0988 ENSG00000166130 IKBIP

>Homo_sapiens

--------MELAAERLSEATRMPGRGKGLGSGRCQVCEGARSPPRVPTNAKAATSCSPW

>Physeter_catodon

---------------------MAGRGRGLVSGRCQV-----------------------

>Rousettus_aegyptiacus

MGSHVSQLEEFGEGGCCEAAGMVGRKRGLVYGRR-------------------------

>Equus_asinus

--------------MRCEAARMAGRGRGLVSGRRQIREGARSPPRLPAELRRPRPDPPG

>Ctenomys_sociabilis

---------------------MAGRGKGLVSGRGRVCEEARPPRDAPVATSRESLK---

HG0989 ENSG00000166603 MC4R

>Homo_sapiens
----------------------MAASRKIILNRLECIRLKLKQK
>Balaenoptera_acutorostrata_s
------------------------------------MRLKLKWK
>Myotis_lucifugus
MLVGASERQLMWACAHTLSPNGTAAPRRIILKRPECLRL-----
>Equus_caballus
------------------------------------MRPKLK--
>Trichechus_manatus_latirostr
------------------------------------MRLKLKQK

HG0990 ENSG00000166866 MYO1A

>Homo_sapiens

MKLGHLYTCEGAWAGRGSKRENQKSGHWQEEGSLFPSFLAPWTAPIKPPLHP-GARTLMPHSCQIEISSTILQGQ-----

>Canis_lupus_familiaris

----------------------------------------------------------MPPSCQIGISFTILQGTP----

>Dasypus_novemcinctus

----------------------------------------------------------MPPSCQIAISSTILQGGTEVTP

>Manis_javanica

--------------------------------------MTPWTAPINQGTSQPWGLGFLCPFVTLEFYPPFSKGHSEGTP

>Chinchilla_lanigera

----------------------------------------------------------MPPSCQTGISSTLLQGTLK---

HG0991 ENSG00000167654 ATCAY

>Homo_sapiens

MPSCTRAGLAFVCQ

>Odobenus_rosmarus_divergens

MSGCIRAGLAFVCA

>Myotis_brandtii

MPGCSRAGLAFVCA

>Equus_asinus

MLGCIRAGLAFVCA

>Manis_javanica

MPGCNRAGPAFVCA

HG0992 ENSG00000168000 BSCL2

>Homo_sapiens
MRHRYQRLDRHFTF
>Rousettus_aegyptiacus
MSYRYQLLDQHFVF
>Galeopterus_variegatus
MRYRYQLLDHHFTF
>Equus_asinus
MRHRYQLLDHHFMF
>Manis_javanica
MSYRYQLLDHHFMF

HG0993 ENSG00000169306 IL1RAPL1

>Homo_sapiens

------------MYSEDLQATLQFLS

>Ochotona_princeps

MLLGEQEDLKTYMYPEDLQARLQFLS

>Loxodonta_africana

------------MYSEDLQARLRFLS

>Mesocricetus_auratus

------------MYSEDLQARLRFLS

>Orycteropus_afer_afer

------------MYSEDLQARLRFLS

HG0994 ENSG00000170852 KBTBD2

>Homo_sapiens

MLQITASRFS--FISEVFT

>Acinonyx_jubatus

MLQITASRVSPFFISEAFT

>Tursiops_truncatus

MLQITASRVSPFFISEAFT

>Rousettus_aegyptiacus

MLQITASRVSPFFISEAFT

>Ceratotherium_simum_simum

MLQITASRVSPFFITEAFT

HG0995 ENSG00000171307 ZDHHC16

>Homo_sapiens

MITADTYLVLPHWQECPCWYL--

>Galeopterus_variegatus

MIAADTYLVLSNWQKCPC-----

>Oryctolagus_cuniculus

MVAADTYLVLLHWQECLCRCL--

>Equus_asinus

MIAADTYLVLSHWQECPCYLWDR

>Peromyscus_maniculatus_baird

-MRAANTLVLSHWQECPCWLL--

HG0996 ENSG00000173890 GPR160

>Homo_sapiens

----------------MKCRCIELTHIAYWYIKMKCKEPKIT

>Dasypus_novemcinctus

---------------------------------MKCKETKVR

>Galeopterus_variegatus

--------------------------MAFWYSKMKCKNQK--

>Ochotona_princeps

---------------------------------MKGKEPKIT

>Fukomys_damarensis

MQVDAGSAFSTQRALSIHQAHRYRTYIAFWHSKMKCKEPTVT

HG0997 ENSG00000177728 TMEM94

>Homo_sapiens

MAAAPGGGDVK

>Felis_catus

MAAAPGGGDVK

>Oryctolagus_cuniculus

MAAAPGGGDVK

>Nannospalax_galili

MAAAPGGGDVK

>Trichechus_manatus_latirostr

MAAAPGGGDVK

HG0998 ENSG00000178172 SPINK6

>Homo_sapiens

MYWLLGKCRLPVENDILY------------------------

>Eptesicus_fuscus

MYWLLGKFSLLVENDILY------------------------

>Galeopterus_variegatus

MYWLLGKCRLLVENNILLIGSKCFLGSVLTENLSWAKHPYSE

>Condylura_cristata

MYWLLGKCRLLVGNDRSTNKI---------------------

>Loxodonta_africana

MYWLLGKCRLPVENNILY------------------------

HG0999 ENSG00000178401 DNAJC22

>Homo_sapiens

MEVGAGRDGPFSLCGQRK

>Canis_lupus_familiaris

MEVGAE------------

>Pteropus_alecto

MEVEAE------------

>Equus_asinus

MAVGAE------------

>Marmota_monax

MEVGAE------------

HG1000 ENSG00000179083 FAM133A

>Homo_sapiens

---------------MLPIFFSATEQLPWYNKKDEYLLFIERMELA--------------------

>Myotis_davidii

MSGKPGLYPSARFANPCCRFFSATEQLPWYNPKDQCLLFIERT-----------------------

>Oryctolagus_cuniculus

---------------MLPIL-FCYRTTPLVNPKDQCLLFTERKIPRSGMAGSHEDPEKRTGKQLER

>Manis_javanica

---------MKILQMASLGFFSATEHLPWYNQKDQCLLFMKRRT----------------------

>Chinchilla_lanigera

MFGKLGLYPSTRFANPCCRFFSATELFSWYNTKDLCLLFIERIEPAWLTSR---------------

HG1001 ENSG00000179761 PIPOX

>Homo_sapiens
MTIIEPRIESHRDARVRLAAELG
>Ailuropoda_melanoleuca
MTIIEPRIES-------------
>Rousettus_aegyptiacus
MTIIEPRIES-------------
>Oryctolagus_cuniculus
MTIIEPRIES-------------
>Chinchilla_lanigera
MTIIEPRIES-------------

HG1002 ENSG00000181027 FKRP

>Homo_sapiens
--------------------------------------------------------------------------MAAAAAAGAQLSWAGTALLELPQPTT------
-
>Canis_lupus_familiaris
--------------------------------------------------------------------------MAAAAAAAMQLSWAGTTHPELPQQPQRMPWKP
S
>Pteropus_alecto
-------------------------------MPARPLSPLAGPKSPR-------------RPLSHQEPIGLHLQPAPGRDGAVQLSWAGTTPLELPQPGM------
-
>Condylura_cristata
------------------------------------------------------------MTLCSIMQNWLIWDFLVFLPAALPLSWAGTAPPELPQPAT------
-
>Heterocephalus_glaber
MPQVVPKFIGLQEKGFPEGGDLPHWFGCLLRKRKDPIRSLVEKKKEKGFPDLVGVTFLGTMKLGSIVPNQLTCGFLVFLPSVEQLSWTGTALLKLPQPTA------
-

HG1003 ENSG00000184867 ARMCX2

>Homo_sapiens

------------------------------------------------MLIHPPGFHCLTSTVYISALQSVLRMKL-

>Miniopterus_natalensis

-----------------------------------------------------------MSAVCISALQSVLKMKL-

>Oryctolagus_cuniculus

-----------------------------------------------------------MLAIYISALQSVLRMKLN

>Ceratotherium_simum_simum

MRSSRDQELTWAKVLKALTAVLRPPLGTNSNALRTAVRLTQARRRRRKFPQIRAGSQSPTSAVYISALQSVLRMKP-

>Peromyscus_maniculatus_baird

------------------------------------------------MMICPPGS---------------------

HG1004 ENSG00000185875 THNSL1

>Homo_sapiens

MFTSQWKLSTTHIN----

>Canis_lupus_familiaris

MFTSQWELSASHTC----

>Pteropus_vampyrus

MFTSQWELSTAHTS----

>Galeopterus_variegatus

MFTCQWELSTAHTS----

>Loxodonta_africana

MFTSQWELSTAHTSRKAK

HG1005 ENSG00000186472 PCLO

>Homo_sapiens

MEEAPAAGVTCERD

>Mustela_putorius_furo

MEEAPAAGVTCERD

>Myotis_lucifugus

MEEAPAAGVPCERD

>Galeopterus_variegatus

MEEAPAAGVTCERD

>Equus_asinus

MEEAPAAGVTCELD

HG1006.1 ENSG00000188227 ZNF793

>Homo_sapiens

---------------------------------------MDVTLPVVRKQIY-

>Mustela_putorius_furo

-----------------------------------MAARMDVTLPVVRKQRIY

>Balaenoptera_acutorostrata_s

MRGKWERFFLENVMTCPSVTASQESAWFLEIYSDSTAARMDVTLPVVRKQRVY

>Loxodonta_africana

-----------------------------------MAVGMDVTLPVVRKQRVY

>Cavia_porcellus

-----------------------------------MATKMDVTLPMVRKQKVY

HG1006.2 ENSG00000188227 ZNF793

>Homo_sapiens

---------------------------MGLFLAKVMTYASVPASQE

>Felis_catus

-----------------------------------MMYPSVTASQE

>Equus_asinus

MGRACPLARRRGTAIDLGLWARIQEKWERFFLAKEMTCPSVTASQE

>Cavia_porcellus

----------------------------------MMTYLSVTSFQE

HG1007 ENSG00000188910 GJB3

>Homo_sapiens
----------MATDAPISRLRKRRLRRATSPSGGWPGS--AWQRRV
>Mustela_putorius_furo
----------MAADAPISHMKKPRLRAATGPSEGGPGSGCPWH---
>Balaenoptera_acutorostrata_s
-----------------MKKKPRLRRASSPGEGCRGNE-------
>Equus_asinus
MRFICSLALPMATDALISHLKTRRLRRATSPSTSGPGN--PWLRGA
>Chinchilla_lanigera
----------MAASAPTPCRSKPRLGRARIPSGAGPGA--TWLRRD

HG1008 ENSG00000198252 STYX

>Homo_sapiens

MAAGCKTPDPPLPCLRRR--RCWSHWDPVVCVC--

>Felis_catus

MAAGYEPPNLPLPRCRRR--RRRSRWDPLVCVC--

>Physeter_catodon

MAAGHEPPSPPLPGCRRRHRRRRSRWDPLVCVC--

>Eptesicus_fuscus

MAAGYEPPNPPLPLCRRR--RRRSRRDPLVCVCQL

>Sorex_araneus

MAAGHEPPNPPLPHCRRR--RRRSRWDPSVCEC--

HG1009 ENSG00000198909 MAP3K3

>Homo_sapiens

MPQRPAAGGAGAGAWGGGGGPERSPRPPRGARPAAVPAARAPGMQPRLRR

>Odobenus_rosmarus_divergens

MPQRPAAGGAGAGAWGGGGGPERSPRPPRGARPAAVPAARAPGMQLRLRR

>Dasypus_novemcinctus

MPQRPAAGGAGAGAWGGGGGPERSPRPPRGARPATVPAARAPGMQLRLRR

>Oryctolagus_cuniculus

MPQSPVAAGAGAGAWAGGGGPERSPRPPRGARPAAVPAARAPGMQLRLRR

>Rattus_norvegicus

MPQRPAR--AGAGAWGGGGGPERSPRPPRGARPAAVPAACSPGMQPRLRR

HG1010 ENSG00000203740 METTL11B

>Homo_sapiens

-------MTVSSSFRKRRLIQRNSKASFLSL--

>Galeopterus_variegatus

-------MTVSSSFKKRSLIQRKSQANFLSL--

>Ochotona_princeps

-------MTVSSSFRKRRLIHRSSQASFLSQ--

>Equus_asinus

-------MTVSSSCRKRRLIQRNSQASFLSL--

>Chinchilla_lanigera

MGQDWFKMTVSSSFRKRRLIQRNSHTSFFSLQE

HG1011 ENSG00000204439 C6orf47

>Homo_sapiens

----------------------MTIQRREEPLLNSE----------

>Lipotes_vexillifer

----------------------MTIQGSGEPLLNSE----------

>Rousettus_aegyptiacus

MCLFSLPQKLKRKQAACPTLGCMTIQGRGEPLQNSE----------

>Manis_javanica

----------------------MTIQGNGEPLLNSEQDSPSKYLFF

>Ictidomys_tridecemlineatus

----------------------MTIQGSGEPLLNSE----------

HG1012 ENSG00000213694 S1PR3

>Homo_sapiens
MNVPGAPSRGFWS
>Balaenoptera_acutorostrata_s
MDVPRAPSGGFWS
>Condylura_cristata
MDVPGAPSAGIWN
>Equus_przewalskii
MGVPRAPSAGFWS
>Microtus_ochrogaster
MNVPGAPSRGLLR

HG1013 ENSG00000214022 REPIN1

>Homo_sapiens
MDGRRVPAQPAAEVRLASLQSEV
>Felis_catus
MDARRASAHPAGEVRLASLRSEV
>Rousettus_aegyptiacus
MDARRASAHPAGEVRLASLQSGA
>Dasypus_novemcinctus
MDGRRASAHPAGEVRLASQQSEV
>Heterocephalus_glaber
MDGRRAPARPAGEVRLASLRSEG

HG1014.1 ENSG00000214595 EML6

>Homo_sapiens

MQIPDVKGGVAPPRASPGGRSPQVQ

>Mustela_putorius_furo

MQMPDLKGGVTLPRTSPGGRSPQVQ

>Pteropus_vampyrus

MQISDLKGGATPARTSPRGRSPQVQ

>Manis_javanica

MQIPDLKGGVTPPRTSPGGRSPQVQ

>Mesocricetus_auratus

MQISDFKGGVTPGRTSPAGRSPQVQ

HG1014.2 ENSG00000214595 EML6

>Homo_sapiens

--MAELTRVACQIKAIAE

>Mustela_putorius_furo

MVMAKLTRVLCQIKAVTE

>Myotis_lucifugus

MVMAKLSRVPCQIKAVTE

>Manis_javanica

--MAKLTPASCQIKAVTE

>Chinchilla_lanigera

--MAKLTRAACQIETVTE

HG1015 ENSG00000240720 LRRD1

>Homo_sapiens

MKIKIKLQFCIGIKHISK

>Ursus_maritimus

MKIKIELQSCTDVKHSSK

>Pteropus_alecto

MKIKIELQSCIGIKHISK

>Galeopterus_variegatus

MKMKIKLQSCIDRKRISK

>Tupaia_chinensis

MKIKIKLPSCIGIKYNI-

HG1016 ENSG00000254004 ZNF260

>Homo_sapiens

---MLHFIPFCNHLISG

>Balaenoptera_acutorostrata_s

-MRMLYFIPICNQLITG

>Pteropus_alecto

-MRMLYFIMFCNHLISG

>Galeopterus_variegatus

MKRMLYFIPFCNHLI--

>Equus_asinus

-MRMLYFTPFCNHLISG

HG1017 ENSG00000255154 HTD2

>Homo_sapiens
MLHSSNSCLFRL
>Felis_catus
MLHSSNSCLFQP
>Balaenoptera_acutorostrata_s
MLHSSNSCLFQP
>Pteropus_vampyrus
MLHSSNSCLFQP
>Loxodonta_africana
MLHSSNSCLFWL

HG1018 ENSG00000259120 SMIM6

>Homo_sapiens

----------------------------------------------MPFPLNSCFFESGCSSGHLAASSAQPSTKCEPEGPP--------------

>Ursus_maritimus

-------------------------------M--AMRSGQGYLTMPIPFSLNSCFFESGCGSGPQSVGPAGLP-----------------------

>Myotis_brandtii

MI---------A-KRDLLLPHLLAPAAHPNHL--ASSTKQG-----ADAPRVTCFSESEHRSGHLLASPARPPKCGPLRVTPSALSGLGSVNSRVR

>Manis_javanica

-------------------------------M--TMRTGQGHLAMQTRFPPNSCLFESGCSPGHLVASAARVPKAWGLKNYPDQPL----------

>Rattus_norvegicus

MPSQKGRTLMEEMREGVALPGN---QSSPPLFALVAAAKVWSENLATPWAANSNFFELWHSCGHLAVSSLWPEGQS--------------------

HG1019 ENSG00000019995 ZRANB1

>Homo_sapiens

MASLKLNISISGFPLISSFLDQTSL-YLLPISFAFQSSVLLNPRV---

>Odobenus_rosmarus_divergens

----------------------MSL-YLLLISFAF-------------

>Lipotes_vexillifer

MTWLRLNISVRGFLI---------------------------------

>Orycteropus_afer_afer

MSWLKLNISIGGLPILTNPFPLKTFSTDLPLSLAYYLPCTAKYNFITS

HG1020 ENSG00000023902 PLEKHO1

>Homo_sapiens

MEGAKGRGAGVRVCASANAREAGLE

>Orcinus_orca

MEGAEGRGAGVRVCASAREAG--L-

>Rhinolophus_ferrumequinum

MEGAEGRGASVRVCASAGEAG--P-

>Ochotona_princeps

MEGAEGRGAGVRVCASAREAG--L-

HG1021 ENSG00000064692 SNCAIP

>Homo_sapiens

MPTFRSPPSRVSSPSLCWNPGMVWGPMRWQRLTLGSRNL---

>Panthera_tigris

--------------------------MRWQRLTRGSRAFIKI

>Manis_javanica

---------------------MVWGPVSWQRSTRGSRAS---

>Mus_musculus

---------------------MVWGPMRWQRLIHGSRKL---

HG1022 ENSG00000065357 DGKA

>Homo_sapiens
---------------------------------------------------------------------------------MPGSPVPASESLGLHLSPLLYHLHH
HPCDPKSLNDSRRDSRQGKLKG-PFTPYFWPGPSVPPAKTSRYN-----------FPKSHSGIRAKNGTQI
>Pteropus_alecto
---------------------------------------------------------------------------------MPGSPVPTSRSLGLCLSPLLYQLRH
PQ----S-LDSRRDSRQKKLKGSPYTPTFGQGLLYHPPRPADLPLEAEVQAPKVLLPSCLCGLP-------
>Equus_asinus
-------------------------------------------------------------------------------------------MKGLCLSPLLYQPHH
HPRCPKSLNNSRTDSKQEKLKGFPFTSTFDQGLLRHLPRPADLTLEVEVQAPEVLLPSCLCGHP-------
>Chinchilla_lanigera
MCISCVCVCVCVCVCVCVDRDIQTHPTYTCTGFWSEIPPRPHTHPSGVKRAIPGTQGTGTLPYFHEGQEGAHRGWGGARGQDAWVSVPTSKSVGLCLSLLLYQPQH
HPCRPRALKLLNRLEAEAE--RSSFVPLLCARAFCT---------TCKVSGPY---P--------------

HG1023 ENSG00000068001 HYAL2

>Homo_sapiens
MGGHVPLQRTWGSSSPKRGTGTEKVPELVPGR-------------------------------------------------------------------------
------
>Felis_catus
MDGHVPRGRIWGPWSLKRGTGCAKVGNTFPPRPQVLPQAHLRAGFLKGPRAGVGRGR---GGAAEARSLGCAWASGRSGPGWAGDGSGHVGPGSAPREFLSWCQA
GDTVFL
>Pteropus_alecto
MDGHVPWRRIWGPWSPKRGIGG-----------------------------------------------------------------------------------
------
>Equus_caballus
MGGHVPRRRTWGPWSPKRGTGGAKVGETFPPLPQVLSRAHLRAGFLRGPRAGGRSQQRRRAGAVPGRKAG----ACRAGPGMAVAVAQVPAA----PRASS----
------

HG1024 ENSG00000073464 CLCN4

>Homo_sapiens

MSLSVSASIRPSTQRSRRFR

>Equus_asinus

MSLSASVSIRPSTR-SRGFR

>Loxodonta_africana

MSLSGSVCISPSTQRRRGFP

>Heterocephalus_glaber

MNLSVSACIRPSTQRSGGLR

HG1025 ENSG00000074621 SLC24A1

>Homo_sapiens

MSRSEGLWIPPGWASWRNLL

>Orcinus_orca

MSRFEGLWLPPGWASWRNLL

>Myotis_davidii

MSRFEGLWISLSWASWRNLP

>Rattus_norvegicus

MSRSEGLWIPPGWASWRNLS

HG1026.1 ENSG00000075035 WSCD2

>Homo_sapiens

MCVLSLKTPLLAQRSSPSVWEGW

>Orcinus_orca

----------MAQRSSPWLREGW

>Pteropus_alecto

----------MAQRSSL------

>Manis_javanica

----------MAQRSSP------

HG1026.2 ENSG00000179314 WSCD1

>Homo_sapiens

MTPPEALASLPPGR

>Eptesicus_fuscus

MMPPEAPASLPPGR

>Galeopterus_variegatus

MTSPEALASLPPGC

>Chinchilla_lanigera

MTPPEALASLLPGC

HG1026.3 ENSG00000179314 WSCD1

>Homo_sapiens

MVPFEHLLETGQEQ------------------------

>Rousettus_aegyptiacus

MVPFKHLLETGQEEQ-----------------------

>Ceratotherium_simum_simum

MVPSEHLLETGQEEQ-----------------------

>Loxodonta_africana

MVPFKHLLKTGKRAHAGARSQDAGMTPPGVPACQPPGR

HG1027 ENSG00000075223 SEMA3C

>Homo_sapiens

MRACRAARRGSRSRAPRRCNRTPRALAPGLRPLSIHSEVLD--------------------------------------------------

>Balaenoptera_acutorostrata_s

----------------------------MRRPLSIHSVVLD--------------------------------------------------

>Ceratotherium_simum_simum

-----------MRRAPRRCNRTPRARPAPATFHSLGGAVLSAAETGSGAADSFTKKLRGPGRRNLTAGPLQKLRSSGEGTGDSLGAQPAAN

>Ictidomys_tridecemlineatus

MRAGWAARRGSPSRARRRCNRTPSARPLRLATFPFTPRCLIERR-GEGLRCHGTTGLRIPVQRNSE--------DLGEKKEYHIGTL----

HG1028 ENSG00000076356 PLXNA2

>Homo_sapiens

MNLDSMWIWSSP

>Felis_catus

MNLDSMWIWSSP

>Tursiops_truncatus

MNLDSMWIWSFP

>Ceratotherium_simum_simum

MNLDSMWLWSSP

HG1029 ENSG00000079482 OPHN1

>Homo_sapiens
MERGYVSLPPWLNPSGA--------------------
>Pteropus_alecto
MEEGDVPLPPRLNPSGA--------------------
>Equus_asinus
MEEGYVSLLPLLNSSGAYLDSKPVPLDQEELLGAGGL
>Chinchilla_lanigera
MEEGYVSLPPGLNPSGT--------------------

HG1030 ENSG00000082175 PGR

>Homo_sapiens

-----------------------------MRTSFTCHLSEIYNPRRLVLPHYWDLRSSEMTVARSTEPAEVRPFLGMGCTERSD--------------

>Mustela_putorius_furo

-----------------------------MRTSFTRHLSEICKYSFLLS--YRET-------------------------------------------

>Oryctolagus_cuniculus

MGPVHAVTTRVKLCSTCVCEDSGVWGAFSMRTSFTCHLRLGTRGGS----------SSRNAGNRESSGDQCGPL------------------------

>Ceratotherium_simum_simum

-----------------------------MRTSFTWHLSGICNHRSFALPDSEKCESCR---------GDCRQLLGAGRNTTVLEKGLYRKVKVAPGF

HG1031 ENSG00000082269 FAM135A

>Homo_sapiens

MLAGRAGPVAEPR-----------------

>Canis_lupus_familiaris

MLAGRFGPVAEPR-----------------

>Pteropus_alecto

MLAGRAGPVAERR-----------------

>Dasypus_novemcinctus

MLGGRAGPVAEPRRRTESSAAVVAVAAMAM

HG1032 ENSG00000084453 SLCO1A2

>Homo_sapiens

MQPSEPKKLNQRFKTYINKDSFIYNSASLMNKKLQLTRLRCLIHTGYSICNFLKNLLVLQYGSKNEW-

>Ceratotherium_simum_simum

--------------------------------------------------------MILQYRSSGEWS

>Manis_javanica

--------------------------------------------------------MILQYGSSSD--

>Cavia_porcellus

MQPSEPKKLYQRIKSYVNKDPFIYISTSLKNKTLQFTCSQCFTCNS--TCNLF----FKKFDVLAD--

HG1033 ENSG00000089225 TBX5

>Homo_sapiens
MPAPPPHRRLPC
>Felis_catus
MPAPPPRRQLPC
>Miniopterus_natalensis
MPAPPPSRRLPC
>Heterocephalus_glaber
MPAPPPHRRLPC

HG1034 ENSG00000090487 SPG21

>Homo_sapiens

-----MQHLVSILPFMYCQTSTSVKITCLYKNLVTAEWNLSIL--

>Acinonyx_jubatus

MSVGQMQHLVSILPFMYCQTNTSVKITCLYKNLVTVGWNLSIQ--

>Equus_asinus

-----MQHLVSVLPFMYCQTSTSVKLTCLYKNLVTVGWNLSIQQD

>Manis_javanica

-----MQHLVSISPFMYC---------------------------

HG1035 ENSG00000093000 NUP50

>Homo_sapiens

MAARSVFGAVPPAVSG

>Leptonychotes_weddellii

MAAWSAFGAVPPAVSG

>Dasypus_novemcinctus

MAAQSVFAAVSPAISG

>Equus_asinus

MAAWSALGAVPPAVSG

HG1036 ENSG00000100813 ACIN1

>Homo_sapiens

-------------------MVIPGSRVSEYIRFKASARRRKPSLKAGREKGRCRADGKN

>Myotis_lucifugus

MFPENPKQGQPGSPRTSGPGPAPNTQAPQRLSFRSEESRMTPIMVMPKSRASE------

>Equus_asinus

-------------------MVMPESRASEQIRFKASVRRR-------------------

>Mesocricetus_auratus

MFPENPKQSQPSNPRTSGLKPAPNTQVPRRLSPRSEDARISPIMVMRDSRSSE------

HG1037 ENSG00000102710 SUPT20H

>Homo_sapiens
MVFTPLHQRT
>Canis_lupus_familiaris
MVFTPLHQRT
>Galeopterus_variegatus
MVFTPLHQRT
>Ceratotherium_simum_simum
MVFIPLHQRT

HG1038 ENSG00000104331 IMPAD1

>Homo_sapiens

MGRTGAGERAPV

>Orcinus_orca

MGRAGAGERAPV

>Eptesicus_fuscus

MGRAGAGERAPV

>Ictidomys_tridecemlineatus

MGRAGAGERAPV

HG1039 ENSG00000104804 TULP2

>Homo_sapiens
MFRDWPERQGTEASM
>Panthera_tigris
MFRNWPERQGTEA--
>Orcinus_orca
MFRNWPERQGTEV--
>Equus_przewalskii
MFRNWPGRQGIEA--

HG1040 ENSG00000105223 PLD3

>Homo_sapiens

MRKAHAWPPPACSLAVVRLRAARRGRQAG-------IQP--GR---------

>Odobenus_rosmarus_divergens

MRKAHA----------------------------------------------

>Heterocephalus_glaber

MRKAQAYPPPSRSLAGVRLRAGVAGRRGQ----SLGVALRAGSEGGDAGVQF

>Tupaia_chinensis

MREAHVWHPLACRLAGVRLRKRGGGQAGVQPRQCVGLGPRVGAVECR-----

HG1041 ENSG00000106536 POU6F2

>Homo_sapiens

--MNSLGLCYPVCCLLSRC

>Acinonyx_jubatus

MGMNSLGLCYPVCCLLSRC

>Pteropus_alecto

--MNSLGLCDPVSCLLPRC

>Loxodonta_africana

--MNSLELCYPVCCLLSKC

HG1042 ENSG00000107249 GLIS3

>Homo_sapiens

MNRTGEAQSRN---

>Odobenus_rosmarus_divergens

MSRTGEAQSRNSTF

>Pteropus_vampyrus

MSRTGEAQSRNSTF

>Equus_asinus

MNRTGEAQSRNSTF

HG1043 ENSG00000107736 CDH23

>Homo_sapiens

MERGVAEPLDVWGAPSPSPRRA

>Manis_javanica

MESGGAESRDVWEALRPSPRRA

>Octodon_degus

MERGVAQRLHVWGVPRPSPRRT

>Trichechus_manatus_latirostr

MEPWGAEPRDVWGALRPSP---

HG1044 ENSG00000109381 ELF2

>Homo_sapiens
-------MIMQTYNTGNHQ
>Ailuropoda_melanoleuca
MSIDFENMIMQTCNTGNHP
>Miniopterus_natalensis
-------MIMQTYNTGNHP
>Marmota_marmota_marmota
-------MIMQTYNIGNHL

HG1045 ENSG00000110925 CSRNP2

>Homo_sapiens

-------------------------------------------MGGLEACRIADED

>Ailuropoda_melanoleuca

MREARTGGGEPGRRWEGKESGEAWELGARRTRARTAGAARGRPMGGLEAGRIADEG

>Ceratotherium_simum_simum

-------------------------------------------MGGLEAGRIADEG

>Heterocephalus_glaber

-------------------------------------------MGGLEAGRIVDED

HG1046 ENSG00000112701 SENP6

>Homo_sapiens
MHSRLGGVGGRP
>Rousettus_aegyptiacus
MHSRLGGVGGRP
>Condylura_cristata
MHSRLGGVGGRP
>Marmota_monax
MHSRLGGVGGRP

HG1047 ENSG00000112769 LAMA4

>Homo_sapiens

MNVRRKPGQRSPQSRTARERRWPSAGGAHPNPSAF---------------------------

>Odobenus_rosmarus_divergens

MDVRRKARQRSPQSPPSPGQE--VAGSARPNPVCLLMLSLKLHPSIRPSARASGERESGREK

>Balaenoptera_acutorostrata_s

MGVGRKPGQRSPQSRPPGTGGGRRSRGSPQTHPPLDA---EL--------------------

>Equus_przewalskii

MGVGRKPGQRSPQSRRARDRRWPVLAGLAPTQSVF---------------------------

HG1048 ENSG00000115295 CLIP4

>Homo_sapiens
MKTPISMHSQPTR
>Lipotes_vexillifer
MNTPISIHSQPTL
>Loxodonta_africana
MKTPISIHSEPTL
>Fukomys_damarensis
MKTPISIHSQPTL

HG1049 ENSG00000116212

>Homo_sapiens
MPRKELELWS-NAALRLNSGSATY
>Physeter_catodon
MQWQELELWS-DTALRLNSGSATY
>Pteropus_alecto
MQWKELELWS-NTALRLNSGSATY
>Manis_javanica
MQWKELELWRSNTALRLNSGSATS

HG1050 ENSG00000116489 CAPZA1

>Homo_sapiens

-------------------MGNECNSLGKILDFKSPKANPTSWGAGAGAGVRRRHSRA---

>Ursus_maritimus

-----------------MNGGNECNSVRKVLDLKSFKEKSYGRPRD--VQERRLSQPPPHP

>Galeopterus_variegatus

MEPRRQEQSKHAGKRATDERGNECNSERKTLDFKSLKAKPTSWCAGCGRRHPRA-------

>Marmota_marmota_marmota

-------------------MGNDYN------------------------------------

HG1051 ENSG00000116852 KIF21B

>Homo_sapiens

MARGGARAQPRGAGRG--

>Odobenus_rosmarus_divergens

MARGGARAQPRGASPGRG

>Rousettus_aegyptiacus

MARGGARDQPRGAAPGRG

>Heterocephalus_glaber

MARGGARDQPRGAGRD--

HG1052 ENSG00000118640 VAMP8

>Homo_sapiens

--MRGGRGQAESWSKQEVN----

>Balaenoptera_acutorostrata_s

--MRGGRGRAESWTSRK------

>Rhinolophus_ferrumequinum

MGCEVSKGRAESWSKQEVN----

>Microtus_ochrogaster

----MGRRRAESWSKQEVKLWLR

HG1053 ENSG00000119522 DENND1A

>Homo_sapiens
MAFAGLERVH
>Mustela_putorius_furo
MAFAGLERVH
>Rousettus_aegyptiacus
MAFAGLERVH
>Erinaceus_europaeus
MAFAGLERVH

HG1054 ENSG00000120832 MTERF2

>Homo_sapiens

MGEGTDGLVGAAENCRELLETEDRNPGSRPRGLRTVGRLAGERGKDGAWPPGLPVPCLELTPTVYCRELTRSHFEKKLERIPSG

>Panthera_tigris

--------------------------------------------------------------------MTRSHFENLER-SSSD

>Dasypus_novemcinctus

-----------------ML-------ERRPSGLTQA---------------------------LDEDQISRERLQGSLSTF---

>Oryctolagus_cuniculus

--------------------------------------------------------------------MTRSHF----------

HG1055 ENSG00000122299 ZC3H7A

>Homo_sapiens

-----------------------------------------------------MSLIAHLESGCVG

>Miniopterus_natalensis

-------------------------------------------------MCVLSLQIAHLESGCVG

>Galeopterus_variegatus

-----------------------------------------------------MLMIAHLESGCAG

>Chinchilla_lanigera

MGSGCALPLGSREAEPLARGAPALASANTDTRSSGQGQPAQLDAGCLWTLRPSFPSIAHLESGCAG

HG1056 ENSG00000128271 ADORA2A

>Homo_sapiens

MMLLPEPLQRAWFQETQSPL

>Acinonyx_jubatus

--MLPEPLKRVWFLETQSWL

>Galeopterus_variegatus

MTPCPEPLVRVWFLETLSRL

>Equus_caballus

--MPREPLERAWFLETRSWL

HG1057 ENSG00000131097 HIGD1B

>Homo_sapiens

-----------MAQPPEMGYLGGTSFQVEAVLGHGERGS

>Erinaceus_europaeus

--------MELVVQPSETGYPGR-FFQVEAVLGID----

>Ictidomys_tridecemlineatus

-----MEEVALVAQTPETGYPGGTSFQEETVLSMDWAEV

>Tupaia_chinensis

MLGRRDGEMVLAAQPPEAGYLGRTSLQVQVMLGIARCS-

HG1058 ENSG00000131386 GALNT15

>Homo_sapiens

---------------MESLFFSQGENRGKGREAIQFEVPVNGLSEGN

>Myotis_davidii

---------MRHLSRMESLLFPKEKRGQTV-----------------

>Galeopterus_variegatus

MRRGKETHGTKHLSRMESLFFSQGENQGKGGGGKQFNLKSL------

>Marmota_marmota_marmota

MQRGKETHGTKHLSRMDSFFFPKEKNRVKGEQAIQFKVLVNGHSEGN

HG1059.1 ENSG00000132970 WASF3

>Homo_sapiens

MLLEGNFCNP

>Mustela_putorius_furo

MLPEGNFCNP

>Myotis_davidii

MLPEGNFCNP

>Equus_caballus

MLPEGNFCNP

HG1059.2 ENSG00000112290 WASF1

>Homo_sapiens

---------------MMNWNNDERKHIRSQHSRPAL----

>Galeopterus_variegatus

MPPSFDTLTPSLALAMMDWNNDERKAYPI-----------

>Equus_caballus

----------------MNWNNDEKKARLNIHNLSYSRLID

HG1060 ENSG00000134242 PTPN22

>Homo_sapiens

MLCSGGSSGFLEVSRP

>Acinonyx_jubatus

MLCSGGSSGFLVVSRP

>Orcinus_orca

MLCSGGSSGFSVLSRP

>Tupaia_chinensis

MLCSGSSSGFSAVSQP

HG1061 ENSG00000134802 SLC43A3

>Homo_sapiens

MCCLCVLWVLGVTFSLGDGIQNSSAGAPLLHLLPGSYWDPENHPLMVFAQA---------

>Balaenoptera_acutorostrata_s

---------------------------------------------MVFTQA---------

>Marmota_marmota_marmota

---------------------------------------------MGFAQAPKSWNYRAP

>Tupaia_chinensis

---------------------------------------------MFFAQAENTQK----

HG1062 ENSG00000137337 MDC1

>Homo_sapiens

MSQRTPCLGGPRKGALALPD-TSDVYDTAPLSVAGSWRQTLWSRFGTARE---VGGGRASGSGL------------------SLGAAQSTCIILLAH----GGLQL
PSH-----------------------------------------------------------------

>Mustela_putorius_furo

MSQRTPCLGGPRKVLPP-----------------------------------------------------------------------------------------
--------------------------------------------------------------------

>Loxodonta_africana

MSQQTPCLGGRRNGALAATDSSGDVYGTAPLSAARRWCRTLWSRFGNLRGKWEVGGETVVGCPFELPNRGALFCGRTEASHYSLNRSQPPCSLFELRGWGGGGLSL
LAETLPISVQQSCKGQQQEPTSGPVLFLHTHSRLPGRGRYWWRARGLKVEAVYFSLCHQRYLKNASGG

>Chinchilla_lanigera

MSQRTPCLGGPKKDALATSDAVHDIYGTALLAAASLGRQTLW----------------------------------------------------------------
--------------------------------------------------------------------

HG1063 ENSG00000138162 TACC2

>Homo_sapiens
MELCEDAGVGAARR
>Odobenus_rosmarus_divergens
MELCEDAGVGAAWR
>Ceratotherium_simum_simum
MELCEDKGFGAAR-
>Manis_javanica
MELCEDAGVGAAR-

HG1064 ENSG00000139734 DIAPH3

>Homo_sapiens
MCWLRPRLQRLG-----
>Rousettus_aegyptiacus
MCWRRPRLQRLG-----
>Ceratotherium_simum_simum
MCWPRPRLQRLG-----
>Chinchilla_lanigera
MCWPRPRLQRLGWAGLG

HG1065 ENSG00000140030 GPR65

>Homo_sapiens

MQAQIYQAPQSTNTSPEEIWKERNFKRKYQSLCKQSLVYSCLHQSTVRFMKKNKLRTTLYVHLQMPQLMLVGCLSAFCDNEHMDFCLLNSVDPFSQLPGAWIF-TS

NC--------

>Mustela_putorius_furo

----------------------------------------------------------------------------------MELLKISQRLNARKCPAVWVSFSS

RCKQIYPAPR

>Galeopterus_variegatus

-----------------------------------------------------------MEAKELSQLILEGSLSVFCDNEHMDLCSLNLADHFS-----------

----------

>Oryctolagus_cuniculus

-----------------------------------------------------------MNTKGISQLIIQSSSSVFCDHDHMDVCSLNSTDHFS-----------

----------

HG1066 ENSG00000140553 UNC45A

>Homo_sapiens

MGIQRR-SGPLLTGLSVGLLLLFSWVTLVQPTSLSALVHHPHLCQTTRDVHGRHYPGFFCPRLSDSPEEAYCCHLQAAGGSCCTRAEFEALYQVNLSALPPPPILR

GPGPLLVLGLYNLLVVTLMTVDLVHFCCGRGRSLGW--SHRRPPSGSSAASSLQVSAGTA

>Mustela_putorius_furo

MGFQRR-LGLLLTGLTVGLLLLFSWATLAQPTSLSALAHHPHLCQTTWDADGRRYPGFFCPRLSDTPEEAFCCHLQAAGGSCCTQAEFEALYQVNLSSLPPPPILR

GRGPLLALGLYTLLLVALMTADLVHFCRGRGGGRRR--RPSRLSSRSSAARPLQVPVGTD

>Pteropus_alecto

MGFPRR--------LPLRLLLLFGWMTLAQPASLSALAQYPHLCQTTRDADGRHYPGFFCPRLTDAPEEAYCCHLQAAGGSCCTRDEFEGLHQVNLSALPPPPILR

GPGPLLALSLYGLLLAVLVTADLVHFCRSRRRGGGW--GRRGPSFRSSAARPLQVSVGID

>Mesocricetus_auratus

MGLQRRSPGPLLTGLSVGLLLLFSQTNLAHPTDLSALAHHPNLCQTTRDAEGRHYPGFFCPRLSDTPEEAYCCHLQAAGRSCCTRDEFEALYQVNLSSLQPPPILR

GPGPLLALGLYSLLLMALMTADFVHFCRGRGRRESLNSRSRGLRFRGSAARPLRVPVGTD

HG1067 ENSG00000143340 FAM163A

>Homo_sapiens
MRLHARSRAVLGAPGRPRSLTCAAPAAGARKDALRPRGSRTRESLPERSCSAAGLMPLARRTPNCCLLRCSPRD
>Mustela_putorius_furo
-------------------------------------------------------MPMAMTHPQPLSARVQPA-
>Balaenoptera_acutorostrata_s
-----------------------------------------------------------MTCPQLLSAQAQPA-
>Myotis_lucifugus
-------------------------------------------------------MPLAMTQPQPLSAQAQPV-

HG1068 ENSG00000144791 LIMD1

>Homo_sapiens
MPFTLGRWKSLLVLGRT
>Myotis_davidii
MTLTLGRWRVLLVLGWT
>Manis_javanica
MPLTPGRWKALPVLGWT
>Microtus_ochrogaster
MTFTLGRWESLSALGWT

HG1069 ENSG00000146414 SHPRH

>Homo_sapiens

MFGGWRMRARDLLGL

>Mustela_putorius_furo

------MRARDLRGL

>Balaenoptera_acutorostrata_s

MFAGWRMRARDLRGH

>Manis_javanica

MLDGWRRRSRDLLGL

HG1070 ENSG00000147459 DOCK5

>Homo_sapiens
MAEASGARRSAGRRRPEPEEL
>Manis_javanica
MAGAPGARGSEARRRPAPQEL
>Loxodonta_africana
MAEEPGARSSAGRRRPAPEEL
>Heterocephalus_glaber
MAEEPGARKSA-RRRPQSQEL

HG1071 ENSG00000147573 TRIM55

>Homo_sapiens

-------------------MDTSAHRHVTAGC

>Oryctolagus_cuniculus

-------------------MDKSAHGHVTLGC

>Dipodomys_ordii

-------------------MDKSAHGHVTLGC

>Tupaia_chinensis

MLTPALCHLSGSILLSPAYMDKSAHRHVIPGC

HG1072 ENSG00000147874 HAUS6

>Homo_sapiens

MEGPPRPQADRREGS

>Felis_catus

MEGPPLPQAARREGA

>Myotis_lucifugus

MRGPPRPQAACREGA

>Galeopterus_variegatus

MDGPPRPQAARRGGS

HG1073 ENSG00000149187 CELF1

>Homo_sapiens
-------------------------------------------------------------MRRGK-DVIKCWRNTLIEAASSVAVWYGCRVLCDYTLMYSGSESL
RDLFCTLCIVGFGCLFV---
>Rousettus_aegyptiacus
---------------------------------------------------------------MRREKVFKCWRNTFTEAASS----CGCVVWLLS-----SVGPF
ASVF----------------
>Galeopterus_variegatus
MNSSLLINLSLNSSLFLCFTVSFTPLPHTPMSHNHACRVGNRTCQLSEDETVLCTYLSCNCGRSGEERSLNARETPPLRHLLPVAVWCGCTVLCDCSLMYSGSESL
RNLFHSVSIVGFGCVCLFRA
>Trichechus_manatus_latirostr
--------------------------------------------------------------------------------------------------MDSGSESL
RSLFCGLSIVGFGCMCLFRA

HG1074 ENSG00000150275 PCDH15

>Homo_sapiens
MRVDINASQKQEVEHVNESSWN
>Myotis_lucifugus
MRLDINASQNQEV---------
>Equus_przewalskii
----MNASQKREE---------
>Orycteropus_afer_afer
MSLDISASEKQEE---------

HG1075 ENSG00000150347 ARID5B

>Homo_sapiens

-----------------------------------------MSNGAAGGGEL

>Mustela_putorius_furo

-----------------------------------------MSNGAAGGGEL

>Rhinolophus_ferrumequinum

-----------------------------------------MSNGAAGGGEL

>Chinchilla_lanigera

MKSPAPRGEVERQIPPCQSGSSSKALRAHPTHSVLVTACVYMSNGAAGGGEL

HG1076 ENSG00000150967 ABCB9

>Homo_sapiens

MSSAETA-PQTRACCAPPPRAHLVPGNRPGLALWMMMALPP

>Manis_javanica

MSCAEPAP---------------------------------

>Ictidomys_tridecemlineatus

MSCVEPAPWTT------------------------------

>Tupaia_chinensis

MSGAEPAPPTTRALCSPLPTAHLVPGKRSSPAPRTMTALPP

HG1077 ENSG00000151474 FRMD4A

>Homo_sapiens

MFLLSMLALPGAELASLLGEEK

>Myotis_brandtii

MFLGSILAPPGTKLASPLGEEK

>Oryctolagus_cuniculus

MFLVSMLALRGPEPASLLGEK-

>Fukomys_damarensis

MFPLSMLALWGAELASPPGEEE

HG1078 ENSG00000162607 USP1

>Homo_sapiens

MRAEGGAPRLRNTPYNW

>Tursiops_truncatus

MRAERGETRLRNTLYDW

>Oryctolagus_cuniculus

MRARGGEPRLRNSVYNW

>Orycteropus_afer_afer

MQAEGGERLLRNTPYNW

HG1079 ENSG00000162975 KCNF1

>Homo_sapiens

MPA--PSPAAGCMPPPRRAPAGCCPL

>Odobenus_rosmarus_divergens

MPG--RSPAAGCMPPPRRAPAACRPL

>Oryctolagus_cuniculus

------------MPPPRRAPAGGRPL

>Peromyscus_maniculatus_baird

MPAGSPAAATGCMAPLRRAPACCRPL

HG1080 ENSG00000164168 TMEM184C

>Homo_sapiens
MVQAELRAIPR
>Balaenoptera_acutorostrata_s
MLQAELRAIPR
>Miniopterus_natalensis
MVQAELRAIPR
>Ochotona_princeps
MVQAELRGIPR

HG1081 ENSG00000165392 WRN

>Homo_sapiens

MPGLQALLVVALHSHPAE-EDLLDWIFSGFLSDIVLYLPMKTLFFGLCK

>Pteropus_alecto

MPGFQALLSWLSTRHPAE-EDPFNCLFSGFYN-----------------

>Oryctolagus_cuniculus

MPGLQALVLWRPTSPPAEGEVPLTWLFSGAF------------------

>Equus_asinus

MPGFRVLLPWGSTSHPAE-EDP---------------------------

HG1082 ENSG00000165434 PGM2L1

>Homo_sapiens
MPFSEVGTPGPW
>Pteropus_alecto
MPFSEKGASGPW
>Dipodomys_ordii
MPFSEEGAPGPW
>Tupaia_chinensis
MPFSEKGTSGPW

HG1083 ENSG00000165804 ZNF219

>Homo_sapiens

MSVRRTAIPGTM

>Leptonychotes_weddellii

MSVRRTPIPGTM

>Myotis_davidii

MSVRRTPIPGTM

>Mus_musculus

MNVRRTPIPGTM

HG1084 ENSG00000165923 AGBL2

>Homo_sapiens

MG----PAPWEEEAGRYAALPVSDAPLASA

>Balaenoptera_acutorostrata_s

MGPAGVLAPPEEEAGRQAALPNSSASPV--

>Rousettus_aegyptiacus

MGPAMVLAPREEKAGRHAALPNSSASPM--

>Dasypus_novemcinctus

MGPARVLAPREEEAGRHAALPSSSASPV--

HG1085 ENSG00000166532 RIMKLB

>Homo_sapiens

--MNTSRGWGKGTEQSQLFHTLGSRSSFYHLPLGSRHRDIN

>Pteropus_alecto

--MNPGRGWGKGTEQNQLSHTLGS--SLYHLPLGSRHRDLN

>Loxodonta_africana

--MNTGRGWGKRTEQIKLFHTVGSRSRFYHLPLGSRHRDLN

>Tupaia_chinensis

MDHEYCWGWGTGTERASCP-TLWETSS-FCHLLGSRHRDIH

HG1086 ENSG00000167377 ZNF23

>Homo_sapiens

MLRSGRNINLCASRVWTVCSQAPDIGS

>Felis_catus

MLKSYRDSYLYAFQVWTVCLLTPNIGS

>Galeopterus_variegatus

MLRSCRDFYLY--EVWTVCSLPPDMGS

>Equus_asinus

MLRSCRGFYLYAFHVWTVYSLALNRGS

HG1087 ENSG00000168675 LDLRAD4

>Homo_sapiens

MLPEAGPAERWTWTG

>Oryctolagus_cuniculus

MLPRAGLAERWTCTG

>Rattus_norvegicus

MLPKAGPAERWTWSG

>Tupaia_chinensis

MLLRAGPAE------

HG1088 ENSG00000169740 ZNF32

>Homo_sapiens

-----------MSRPGRPGP--RG-RPAASGEVGGPQS---RVPWTGERDAEVLRNSPWKNHMKRW

>Canis_lupus_familiaris

-------------------------------------M---LNSFQGLKDKRVFKNSPWKNHMKRW

>Ceratotherium_simum_simum

MLHQAIQISSIYLYPKLQLLLTLSFSLINSSALSCLFLLFCFVFVCLFFPPQVLKHSPWKKHMKRW

>Ictidomys_tridecemlineatus

-------------------------------------------------MQQVLRNSPWKNYMKRW

HG1089 ENSG00000169813 HNRNPF

>Homo_sapiens
MHLEFPRSPLRG
>Pteropus_alecto
MHLDFPRSPLRF
>Equus_asinus
MHLDFPRSPLRV
>Rattus_norvegicus
MHLDFPRSPLRF

HG1090.1 ENSG00000170458 CD14

>Homo_sapiens

MPCRILPVTVPLPETSFIAIFPGKEGGWLGGREVGR------------------------

>Orcinus_orca

--------MAPLLAKSFTAKFPRTADGVVGRKKCGEWQGSKRRERSDIPGLHKLSEAARG

>Rattus_norvegicus

MPLRILPVSVPPCKTFHCNIY---------------------------------------

>Tupaia_chinensis

MPCGILPVVAPLPETFFIAIFPRREVKGAAGRKRGVCVCV--------------------

HG1090.2 ENSG00000170458 CD14

>Homo_sapiens

MRFTISSTKHAVKYLRIFRDLDLVAGDQHKPRQ-GRSQRNESSRFSGI---------

>Felis_catus

MKFAIPSTKYPIIYLRIFR--------------------------------------

>Orcinus_orca

MKFAISSTKYPVKYLRIFRELDLGDRRLTVITKTREN--------------------

>Manis_javanica

MKFAISSTKYPVKYLRIFGRLDLGEGRLIVTTKTRESQGNEVGRVSGMQAGKVRVLA

HG1091 ENSG00000172183 ISG20

>Homo_sapiens

----------------------MRHLRRGRAGAKEGHPDMEPASSVSPDSARS-----------------------------------------------------

---------

>Orcinus_orca

----------------------------------MGHTDKEPASLVNSDSGH------------------------------------------------------

---------

>Myotis_lucifugus

-----------------------------MGGAEAGQRDMEPVRSASPDSGAC-------------RPSCISRQEMATCSLENEPETEAAPGQRGQDSPLEVRGDS

PLTATDLGS

>Ceratotherium_simum_simum

MFILQAGRRVRKERKMLQSTPSTASSEEGRAGAKAGHPDMEPASFVSPDSGHRALHLPAAPGDGAAQPTCLAFRP-------------------------------

---------

HG1092 ENSG00000173327 MAP3K11

>Homo_sapiens
----------------------------------------------------------------------------------------------------------
---------------------------------------MVRRRRPRGPTPPA
>Ailuropoda_melanoleuca
----------------------------------------------------------------------------------------------------------
---------------------------------------MVRRRRPWGPTPRA
>Myotis_lucifugus
----------------------------------------------------------------------------------------------------------
---------------------------------------MVRRQRPRGPTPRA
>Cavia_porcellus
MGVLGILVLLSLLARLPGNSPAQPLEPPPGGSVASLPRHLMLFPAGGRRAEKSETPPSTSASRPRGQSTKGGGRRERGRSCRAKAPGPGRDSFSLTGDTLHTLDKD
PQESRHYPWQPHLTLDWCHGQPPRTYFGTQGRLGPSNAAMVRRRRPQGPTPPA

HG1093 ENSG00000174371 EXO1

>Homo_sapiens

MYTVILTIALSHSYYAKEEIIIRGYLPGPEETYVNFMNYYIRFPRSERKLFLDII--------------

>Panthera_tigris

---------------------MRAYLPSLEETSASQNVCYVSFPQSGRKLFLEALSERWEADCFSQTAA

>Dasypus_novemcinctus

----MFHMALSRSYCAEEKVFMGGYLPSRKETSEFHEMYILSVVLRVK--------GSW----------

>Fukomys_damarensis

----MCRMAHSHSCWANEEMLIRGHLPGLQETYQNFESVFLTVMVDG---FLPSVA-------------

HG1094 ENSG00000177917 ARL6IP6

>Homo_sapiens

MGEPGEECERSGEEGRRQIGSGMCLRGRERALGPGVEKMRLWGCRGARSRW---------------------------

>Panthera_tigris

--------------------------------------MQLWGLPGRAVA----------------------------

>Myotis_davidii

----------------------MLIEVHERALGPGIGKMQLGGPTVPSLAWR--------------------------

>Marmota_marmota_marmota

--------------------------------------MRLWGCRGSRSPGGNAAVPEVFGRDETLQGRVTCLQPASA

HG1095 ENSG00000179921 GPBAR1

>Homo_sapiens

MHLVHLCSSARTPDMVQTAAGLAAATP

>Ailuropoda_melanoleuca

-----MPSPARTPDSARSPAGLPAAVP

>Ceratotherium_simum_simum

-----MLSPARTPDVFRSPAGLAAATP

>Manis_javanica

-----MPSPARTPDIARSPAGLAAATP

HG1096 ENSG00000181544 FANCB

>Homo_sapiens

--------------------------------------------------------------MDTVSTW-GLRYVAAALRFRNINWNQFAFEN---------

>Odobenus_rosmarus_divergens

------------------------------------------------------------MIL---IVPDLCIYCLTTNQFKSISWNQFAFENWTNTKT---

>Rousettus_aegyptiacus

MEKKRNMFLKNFLDAILIAFCMGIPTFIVLCFIVLHRCYFILFFYKVKARPITSKKITAGFVVILTLLWRSGTEPALSLRFKIIEWNQFAFENWTNTKT---

>Cavia_porcellus

---------------------------------------------------------------------------MLEERIKSISWNQFAFENWMNTKDLIL

HG1097 ENSG00000182481 KPNA2

>Homo_sapiens

MPSSFFFFLSPLPLAPKPYDSLPSQRVFQIHQWAHSLGSNQPIGMRSHREI

>Ursus_maritimus

--------------------------------------------MRGHREI

>Equus_asinus

--------------------------------------------MRSHREI

>Fukomys_damarensis

--------------------------------------------MRSHREI

HG1098 ENSG00000183775 KCTD16

>Homo_sapiens

MKSCARKLWFEHGQFSPTVSYIHKHHMKWRSGSSVYFSQVPQ

>Odobenus_rosmarus_divergens

MENFARKFWLEHG-----------------------------

>Pteropus_alecto

MGSCARKLWFEHG-----------------------------

>Ochotona_princeps

MGNCARKLWFKRGQFSPTFSHIHKHQMKWRFGSSVYIYHIPQ

HG1099 ENSG00000186334 SLC36A3

>Homo_sapiens

MVSSSSQSHCSLCPTAARAKLGEAKSPGKSWLGKQLRVKLRTILTEMGLS---------

>Mustela_putorius_furo

MVTSSSQSHSSLCPIALRAVLGGTGSPGESWRVANQMVRGPEGHNEMRLSKVPQKLSEA

>Galeopterus_variegatus

MVPSSAQSHCSLCPRAVRAESAEAESPGKSWQVAEQL----------------------

>Oryctolagus_cuniculus

MVPSSSQSHCSLGPTKARAALGEAKRPGESWQVAAGPRVSRGSPSPGPLEPGEAP----

HG1100 ENSG00000187815 ZFP69

>Homo_sapiens

----------------------MNSNLTNWLPPKESA------------------

>Felis_catus

MCGQCFWIPPLLSFAPLEDCRGMHSRLLNWLPPKEALLNLREKSVFQSC------

>Miniopterus_natalensis

----------------------MHSNLCNRLPPKEAVLNLRQESVFQTHPSHISL

>Galeopterus_variegatus

MTG---ASGSLLSFVPLEENIDMYSNLPNWLPPKETVLNLRGESVFQSC------

HG1101 ENSG00000189334 S100A14

>Homo_sapiens
MSHQLLWGQL
>Manis_javanica
MSHQLLWGRL
>Nannospalax_galili
MSHQLLWGRL
>Tupaia_chinensis
MSHQLLWGRL

HG1102 ENSG00000196704 AMZ2

>Homo_sapiens
-----MKSHSTTFFQPGPDMSVLVS
>Galeopterus_variegatus
-------------------MPVLVR
>Ceratotherium_simum_simum
-------------------MPVLVR
>Tupaia_chinensis
MFCVRVKTPSTTRCQPDPNMPVLVR

HG1103 ENSG00000197620 CXorf40A

>Homo_sapiens

MLGCKNAAQHGEMIFCPAGWCTLWSLTQTGAEGPTAVISRTKL--------------------

>Panthera_tigris

------------MVFSAAGWGLMWLAEPTRE--------------------------------

>Rousettus_aegyptiacus

------------MIFCTARWAIMWSLTQPGKKRASAMISRTKL--------------------

>Equus_caballus

------------MCCAAGCWGVMWSHSPPGPERPTAVMSRAKACVRGFPPVGPSRVLLASSPC

HG1104 ENSG00000205155 PSENEN

>Homo_sapiens

--MR-AVLPRPKKTTISRETWGVSRKRP------------

>Leptonychotes_weddellii

----------------------MSRKRPELQVAKASKRRK

>Oryctolagus_cuniculus

--MRTALPPRPEETTNSRATCGLSRRRP------------

>Ictidomys_tridecemlineatus

MRMRTAFPLQKERTTNSSETCGTSRKRP------------

HG1105 ENSG00000213654 GPSM3

>Homo_sapiens

MRPPGVLPHHEPPPLLQLEEFFPVSVLPWGERD----------------

>Rousettus_aegyptiacus

MRRPRVLPHLETPPLLLE-EILPV-FTPLGERQRSNKQFGWEKRTGG-S

>Equus_caballus

-----------MN--PLHDCSFSQSVLPWGERN----------------

>Tupaia_chinensis

-------------------------MLPWGERGQPS----WVGKENGGS

HG1106 ENSG00000215021 PHB2

>Homo_sapiens

MEMTCCLALPS--LGGQPSRKGRDFRMRDSCARSSGP-------

>Odobenus_rosmarus_divergens

--------------------------MRNSCARSSGPLWAPGRV

>Equus_asinus

--MTCCLALPKSLWEGRVPKKGGNFRMRNSCARSSGPQWPEGRV

>Marmota_monax

--------------------------MRDSCARSSGPL------

HG1107 ENSG00000217930 PAM16

>Homo_sapiens

--MRMPSAVSRTRSLRLSLTSDPPTTLDPRPRGQEVGV-----

>Felis_catus

----MLCAASRTRSLP---------------------------

>Myotis_brandtii

----MPSVASRTRSP----------------------------

>Cavia_porcellus

MTMRMPSTPWRTRSLLSFLTSDLPAALERGPRVQEAQARGPDA

HG1108 ENSG00000251247 ZNF345

>Homo_sapiens

MAVEHPQGTCKTSNFTLELSRNMACLFPQISLGLQTKGWNFEPISRLLVTVPEERQSYGYYESKLRPRNYF

>Felis_catus

MAVENPQGSCKTVNQG-------------------------------------------------------

>Pteropus_alecto

MAVEHPQGTCKNVNQG-------------------------------------------------------

>Equus_asinus

MAVEHPQGTCKTVTQG-------------------------------------------------------

HG1109 ENSG00000278311 GGNBP2

>Homo_sapiens

MGNAARLSRTLPLRS--LPPVANVLRSGGCLRSRGLAVAGRSWEEAAAAAAET---AAAAAAAAGRRW

>Rousettus_aegyptiacus

MGNAARLFRPLLLLS--LPPVANVLRGGGCLSSRRLAVAGRSWEEAAAAAAAAGAAAAAAAAAAGRRW

>Equus_asinus

MGNAARLSRPLLLLS--LPPVANVLRGGGCLRSRGLAVAGRSWEEAAAAAAGAAE-AAAAAAAAGRRW

>Microtus_ochrogaster

MGNAPRLSRPLLLRAPPFSPVANVLRGGGCLRSRGLAVAGRSWEEAAAAG-----AAAAAAAAAGKRW

HG1110 ENSG00000034152 MAP2K3

>Homo_sapiens
MSVLGPGGARSPLD
>Tursiops_truncatus
MSVLRPGRARRPLD
>Ctenomys_sociabilis
MSTLGPGGARRPLD

HG1111 ENSG00000040531 CTNS

>Homo_sapiens

MEGGLSFAHERSREALAAPRVSVSLAADLIFPHAGAPISAPRPDPAALCP---------SKTRGNYNSQSSSRRDPAPRVRWRRSGDSGPASPKV-----------
----------------------------------------------------

>Pteropus_vampyrus

-------------------------MACLILPHTRASIPA--KVPAPLRSS-STLHNCPAKPRENYSSHSASLRDSKS-------GDSGQMASGLRSLSVKPAGEN
GYSLKRLLHLVWSVSRASEVLSCRLNLSRGVCRLLWRGETVKAPFQELPLAC

>Ctenomys_sociabilis

---------------------------------------MSDKPPGPLCPPCTTIRLSLVKTRKNYNCQGPSRRDLASTSQVTRVR--------------------
----------------------------------------------------

HG1112 ENSG00000068885 IFT80

>Homo_sapiens

MVTG-RY-------QVFESNRHRFPGT

>Miniopterus_natalensis

MVTGGREASW-----------------

>Microtus_ochrogaster

MVTG-RSGGFLVSGQSSESNRRRFPGT

HG1113 ENSG00000072201 LNX1

>Homo_sapiens
MPSSFLQAPVHRS
>Peromyscus_maniculatus_baird
MPSFFLQPPVHRS
>Tupaia_chinensis
MPSSFLQAPVHRS

HG1114 ENSG00000073737 DHRS9

>Homo_sapiens

MGVTHRARREPEDSSHM

>Dasypus_novemcinctus

MGVTHSARRERED SSHM

>Jaculus_jaculus

MGVTHGVKRELQESSHV

HG1115 ENSG00000074219 TEAD2

>Homo_sapiens

MAGRPCPPACGSVGLSWSRPQLLP

>Odobenus_rosmarus_divergens

MAGRPCPPVCGSVGFSWSGLEFLP

>Physeter_catodon

MAGRPCPPVCGSVGLSWSWLEFVP

HG1116 ENSG00000084090 STARD7

>Homo_sapiens

MSGGSWPRDRETGSWPAI

>Oryctolagus_cuniculus

MRGGSWPRDRETLSGPAS

>Mesocricetus_auratus

MRGGSWPREREAGAGSAS

HG1117 ENSG00000089902 RCOR1

>Homo_sapiens
MRAKVALGSSLGA
>Oryctolagus_cuniculus
MRAKVALGSAPGA
>Manis_javanica
MRAKVALGTALGA

HG1118 ENSG00000090975 PITPNM2

>Homo_sapiens

MESMVPTPQLVSMLFPESPTIHLLSF

>Canis_lupus_familiaris

MEFTEPTPQLVPVLFPESPTTCPLSF

>Mus_musculus

MESKEPTPPLLSVWFPESPTTYPLSF

HG1119 ENSG00000092020 PPP2R3C

>Homo_sapiens

MKRNCERRR----FPQAGT-NELLGHVVRTRQRHTIKENAGLSWPLKSGLPRLPPSIKMVVGRPS

>Odobenus_rosmarus_divergens

MKRNCEAKKKKKSLPRAGNAIGLLNHVVHTRQSPKLKK---------------------------

>Ceratotherium_simum_simum

MKRNCEGRK---FFPRAGKA---------------------------------------------

HG1120 ENSG00000095585 BLNK

>Homo_sapiens
MLLPRRARNC
>Dasypus_novemcinctus
MLLPRRARNC
>Marmota_monax
MLLPRRARNC

HG1121 ENSG00000100350 FOXRED2

>Homo_sapiens

MEGLGIPG-SPKSQLETVESGQMSSE-DGFWAWP-----

>Balaenoptera_acutorostrata_s

-----------------MESGQMSSEVLGFWSWLRPKSQ

>Equus_asinus

MEGFRIPGGSPRSELETMESGPMSREGFGSWSWLCPK--

HG1122 ENSG00000100410 PHF5A

>Homo_sapiens

---------------MGVKGRTEVPEASGGRLS

>Acinonyx_jubatus

---------------MEVKGRTEVP--------

>Ctenomys_sociabilis

MEASRTNHRDLEAGAIGVKGRTEVTQASGSPLG

HG1123 ENSG00000100884 CPNE6

>Homo_sapiens

MCEPVSRGCMCRHHFSV----------------------

>Manis_javanica

MRKPVSHGYMCLHHFLV----------------------

>Ctenomys_sociabilis

MCEPVSCGCMCRHHFCVDVCVHEPLQCVYLSVMPGVGAV

HG1124 ENSG00000100934 SEC23A

>Homo_sapiens
MLAGRRSSLRPGAGWRHP
>Oryctolagus_cuniculus
MLAGPRSSRRPGAGRCRP
>Tupaia_chinensis
MVAGLRSSQRPGAGRRGP

HG1125 ENSG00000101574 METTL4

>Homo_sapiens

MNDKFSSMCMSGFCTCISNLSVCMKCMFIEIQDI

>Felis_catus

MNVKFSFV--------------------------

>Dasypus_novemcinctus

MNFKFSSGYI------------------------

HG1126 ENSG00000101890 GUCY2F

>Homo_sapiens
MEFVNRLESLTGL
>Equus_przewalskii
MEFINRLQSLAGL
>Octodon_degus
MEFLNRLESLTGL

HG1127 ENSG00000103064 SLC7A6

>Homo_sapiens
MGGRAGSGAGAGPRSVGAEPCGSSLPPSGAGCGPETARGWPIGAAREALRSLCGRGRQVELGLAAGDFISFSVLDPFMRRWLMKV
>Galeopterus_variegatus
-----------------------------------------------------------------------------MMRWLMKV
>Loxodonta_africana
---------------------------------------------------------MNGIRSCCGDFIFFSVPDPFMMRRLMKV

HG1128 ENSG00000103852 TTC23

>Homo_sapiens

MVINLYKESPN------------

>Rousettus_aegyptiacus

MLINLFKESPNRYSGNQSVKSGC

>Fukomys_damarensis

MVINLYKESPN------------

HG1129 ENSG00000104432 IL7

>Homo_sapiens
MLMIPKIEPADQAQSRN
>Oryctolagus_cuniculus
--MNPKIEPAYQAQTRN
>Manis_javanica
--MNPKIEPADQAQSRN

HG1130 ENSG00000105559 PLEKHA4

>Homo_sapiens

MSERETGRDRDQRETQKVRVGREVV

>Equus_asinus

MSDRETGRDRD--------------

>Manis_javanica

MSDRETGRDRD--------------

HG1131 ENSG00000106780 MEGF9

>Homo_sapiens
MVRPVAVAAAVAEASLVGLQRGEDD
>Manis_javanica
MVRPVAVAA--ATASLVRVQRGEDD
>Microtus_ochrogaster
MVRPVAAAAAAAATASLVGRQGGEDD

HG1132 ENSG00000109771 LRP2BP

>Homo_sapiens

MKTLIYHSTVIHLITWQSNTDRKIKLN---NS-----

>Rousettus_aegyptiacus

MKALIFHSPVIHLITWQSNTDRKIKPN---KIKCIPR

>Cricetulus_griseus

MKALTYHTVVIHLVTWQSNTDWKRKLSTTAKIQCIPR

HG1133 ENSG00000111328 CDK2AP1

>Homo_sapiens

-----MSSIGYSNPGAWPDLGVWNDLRA

>Manis_javanica

-----MSRTGCSYSGK------------

>Tupaia_chinensis

MDLKSMSGIGCSRSGAWPDPSIWNDLRA

HG1134 ENSG00000112984 KIF20A

>Homo_sapiens

----MSFFPLRQSKHPKPVTLCTPVKIVAKEGQELAAEASR-FPLGS----

>Ceratotherium_simum_simum

MAAEASGFPLGFLEK-----KYLGFK--KKC--------------------

>Chinchilla_lanigera

MAARASVFPLGFLVN-----KCLGFK--EKRWEESRAKQSKLATLGIADTR

HG1135 ENSG00000113597 TRAPPC13

>Homo_sapiens

MENKVFYAYVSKIEEMDLLKRSQFVKDTVGEFKVPPPERG

>Oryctolagus_cuniculus

---------------MGLVKRSRWVRKGRSERRI------

>Ceratotherium_simum_simum

---------------MALVKRSCWMRR-------------

HG1136 ENSG00000113749 HRH2

>Homo_sapiens

MGAGTSYGEGYSCVST

>Myotis_lucifugus

MGAAVSYGEGCSHVST

>Manis_javanica

MGAAASYGEGHRHVHT

HG1137 ENSG00000115307 AUP1

>Homo_sapiens
MPGIVVSAGREL
>Pteropus_vampyrus
MPGIVVSAGREL
>Galeopterus_variegatus
MPGIVVPAGREF

HG1138 ENSG00000116871 MAP7D1

>Homo_sapiens

MRRGTPVLATGRRRRRR

>Elephantulus_edwardii

MRRGTPVLAAGRRRRCR

>Cavia_porcellus

MRRGTPVLASGRRRRRR

HG1139.1 ENSG00000118308 LRMP

>Homo_sapiens
MQTQNCRSSGEAYHFPEKKNTFSVLPAS
>Canis_lupus_familiaris
MQTQNCRSSVEAYHFPEKKSTFSVSYA-
>Pteropus_vampyrus
MQTQNCRSSVEAYHFPEKKSTFSVSCA-
>Ceratotherium_simum_simum
MQTQNCRSAVEAYHFPEKKSTFSVSCA-
>Manis_javanica
MQTRNCRSAVEAYHFPEKKSTFSVSSA-
>Loxodonta_africana
MQTQNCRSSVEAYHFPEKKSTFSVSCA-

HG1139.2 ENSG00000118308 LRMP

>Homo_sapiens
MKEATLQKEAT
>Equus_asinus
MKEATLQKEAT
>Loxodonta_africana
MKEATLQKEAT

HG1140 ENSG00000118495 PLAGL1

>Homo_sapiens
MLPLPWSGVQLQGLC
>Tupaia_chinensis
MLPLPWSGAQLQ---
>Trichechus_manatus_latirostr
MLPLPWSGVELQWLG

HG1141 ENSG00000121964 GTDC1

>Homo_sapiens
MHREPKSVTEIVWVSLH--------
>Equus_przewalskii
MHREPKSVTEIVWVRGVCMYSRI--
>Chinchilla_lanigera
MHREPKSVTEIVCV--VCLLSPVMH

HG1142 ENSG00000124613 ZNF391

>Homo_sapiens

-----------MLDLQEAEEGSAQKYLS-

>Odobenus_rosmarus_divergens

MVIQSEGGQLWMLDLQGTEEGSEQKYLSG

>Galeopterus_variegatus

-----------MLDLQGAEEGIAPKHLSR

HG1143 ENSG00000125245 GPR18

>Homo_sapiens

MPSFQGELMLSVTS

>Rousettus_aegyptiacus

MPSFQVELMLSMTS

>Galeopterus_variegatus

MPSFQGELLLSMIS

HG1144 ENSG00000128203 ASPHD2

>Homo_sapiens

MPPPAPAAPSPPR

>Ailuropoda_melanoleuca

MPPPAPAAPSPPR

>Rhinolophus_ferrumequinum

MPPPAPAAPSPPR

HG1145 ENSG00000128585 MKLN1

>Homo_sapiens

--------MLENMGIGIRFWRRNVS---------

>Rousettus_aegyptiacus

MIGFITHSILENMGIGIRYVQRAEGAEVNQRMHR

>Ceratotherium_simum_simum

------------MGIGIRYVQRAEGAEVNHRMHC

HG1146 ENSG00000130413 STK33

>Homo_sapiens

MVISFLSAALTCEYVSSASLT

>Dasypus_novemcinctus

----------MCEYESSASLT

>Oryctolagus_cuniculus

----------MCEYKSSASLT

HG1147 ENSG00000131781 FMO5

>Homo_sapiens

---------------------------MPRPRSSLKNVNRCVLFVFCHPFFRTGDTNR

>Rhinolophus_ferrumequinum

MRSNSLLLRRQQNLASPKAPAVAGFIRMLWPRSSLKDVN-------------------

>Ceratotherium_simum_simum

---------------------------MLWPRSSLKSVN-------------------

HG1148 ENSG00000132938 MTUS2

>Homo_sapiens

MAVKLIFWGIGSHGLIGSHSAGT

>Mustela_putorius_furo

MAVKLLFWGSRSYGLIGSHSAGT

>Myotis_davidii

MSVKLVFWGSRSHGLIGSCSASP

HG1149 ENSG00000132975 GPR12

>Homo_sapiens
----------------MILDCHPAVFKH
>Canis_lupus_familiaris
MVVSEWGLASVCVSFCMILDCHPAVSKH
>Galeopterus_variegatus
----------------MILDCHPAVCKH

HG1150 ENSG00000135299 ANKRD6

>Homo_sapiens

MQTSFLCHLSVIQKCSSDL

>Leptonychotes_weddellii

MQTSSLCHLSIIQKHSAHL

>Tupaia_chinensis

MQTSLLCHPSVIQKHSSGL

HG1151 ENSG00000137168 PPIL1

>Homo_sapiens

MKFAFSSVLEPSFFSSPTPASRHWKKGPSSSLSWPPKSWYRKG

>Mustela_putorius_furo

MELALFSVLAISFFSSRIPASRPWKGDPSSSLSQSPKSWHRKG

>Chinchilla_lanigera

MNLAFSSLLALSFF-----------------------------

HG1152 ENSG00000138658 ZGRF1

>Homo_sapiens
MTSRCFAGEIRKWILCNLTGGINS
>Ursus_maritimus
MTSRSFAGEIRKWILWT--GHINN
>Chinchilla_lanigera
MTSRSFAGEIRKWMLCS--GGINT

HG1153 ENSG00000138738 PRDM5

>Homo_sapiens
--MCRLPGAKRRC
>Canis_lupus_familiaris
MSMCRLPGAERWC
>Galeopterus_variegatus
--MCRLPGAERRC

HG1154 ENSG00000139725 RHOF

>Homo_sapiens

MAKPKDSQLEDWRRLFKKIWIFPYDLESIFVLYN

>Balaenoptera_acutorostrata_s

MAKLKDSQLEE-----------------------

>Nannospalax_galili

MAKLKDSQLEE-----------------------

HG1155 ENSG00000141668 CBLN2

>Homo_sapiens

MKVVTPFNIKTWRELGKEKKESKRREATGAFKGV-----

>Erinaceus_europaeus

---MKVLKKESWREFGKEKEGSPSRGASGASRGA-----

>Cavia_porcellus

MKVVTPFKIKTWREFGKEKKESKRRDAAGAFEGLLAPSS

HG1156 ENSG00000144741 SLC25A26

>Homo_sapiens

MAAPSARGRDPLLLRLGL--

>Myotis_brandtii

MAAPSARGRDVLRLDSGL--

>Equus_asinus

MAAPSAWGRDPALLSLELYP

HG1157 ENSG00000148123 PLPPR1

>Homo_sapiens
MYLAPGSRTAQ
>Canis_lupus_familiaris
MHLAPGSRTAQ
>Pteropus_alecto
MHLAPGSRTAQ

HG1158 ENSG00000149571 KIRREL3

>Homo_sapiens

MVGANPRGPA-EAEHREPPRKRN

>Cavia_porcellus

MVGANQRGPAAEAEHREPPRKRN

>Trichechus_manatus_latirostr

MVGASQRGPAAALEESEPGRQRN

HG1159 ENSG00000152822 GRM1

>Homo_sapiens

----------MSSKQTQHLAHRCQHDFHCTLDQFTLMHYR

>Myotis_lucifugus

---MCTLSPRASTTDSFQLTAGCHHDAHCPLDPFTLVHR-

>Heterocephalus_glaber

MPGSSDLGTTALSTPTQRPAHRCQHDVQCTLDQFTLIHL-

HG1160 ENSG00000154162 CDH12

>Homo_sapiens

------------MMCQRHAVFCF--EGGEMTGGNPGPPLARRSPLQLSSSRSESIIDLKGKCGGSHREGSRLF

>Rousettus_aegyptiacus

-------------MCQRHAVFCF--EGGEMTGRESWAPIGWPLSAAVALQ------PL---------------

>Mus_musculus

MHFKYRIQLLSTTQRRVKGTLCFVLRAGR--------------------------------------------

HG1161 ENSG00000154227 CERS3

>Homo_sapiens

MHRCGDPPPGLADPPRPRTEEEVVKQEATSHQEEPEGAIQ

>Equus_caballus

-------------MEGGRAAAEVV-KQETSHQEEPEGAIQ

>Fukomys_damarensis

------MPRGLKEESVWTQEDKVVKQEATSHQEEPESAIQ

HG1162 ENSG00000155380 SLC16A1

>Homo_sapiens
MQVFCSKYLDLTCILEFIYT
>Odobenus_rosmarus_divergens
MHSFCSKYLDLNCILEIIYT
>Ceratotherium_simum_simum
MHAVCSKYLDLNCILEFIYT

HG1163 ENSG00000157637 SLC38A10

>Homo_sapiens

MALRRPQGLAGSGG

>Odobenus_rosmarus_divergens

--MRQPQGLAGSGG

>Ctenomys_sociabilis

MAMRRPHGLAGSGG

HG1164 ENSG00000162639 HENMT1

>Homo_sapiens

MASGHAPQRPLRDAKPTPWN---------------------

>Dasypus_novemcinctus

MASGHAPPRPLGDAKPTARAGAQD-----------------

>Peromyscus_maniculatus_baird

MSSGHAPQRPLRDPEPTPWSRQPGGMGGPEHGRAPHAQRGP

HG1165 ENSG00000162694 EXTL2

>Homo_sapiens

----MESPLTPTRPPTTSVPGCMGSSACPFAISSQLALNPGVVDISD

>Acinonyx_jubatus

MAGWLRACSRPPRPPTTDVPGCAGVQRLPFHHCPLLTKSRCFR----

>Pteropus_alecto

-----------------MFPDTLASSACPFAISP-ITINPGASGSKA

HG1166 ENSG00000165071 TMEM71

>Homo_sapiens

------------------------------MCVGI--CEKKEASYFSNNFVLCQLEEEAGNSLWKEHRICEV-----------------

>Panthera_tigris

MPFYSRKQPKQQEAAEDKTSLHTPAHADSLLNPGFLHRQPQEASYFSN-SPLHQWEEETGNSLWKEHKICEV-----------------

>Heterocephalus_glaber

-------------------------MPFQQLSPQFHHCQKQEALYSSKCFAIYQLEEETGNSLWKDYTYGQIIFHNTCFALMEQKICEV

HG1167 ENSG00000165934 CPSF2

>Homo_sapiens

--MGF----KLPAVLRQPINKELFRIPGVSSYIRLRYQLF

>Odobenus_rosmarus_divergens

MRLGVGKGSGRRWGRTDEIYKELFP---------------

>Pteropus_alecto

-----------MRGNCSQIHKELFL---------------

HG1168 ENSG00000166669 ATF7IP2

>Homo_sapiens

MTCDITICKTNGEDQDKDKLYHRKAIENAKMKHLNEGLITLNSAFVNLICRYVQTTS--

>Oryctolagus_cuniculus

------------------------------MKHLNEGLITLNSAFVKLSLHFAVLTLLV

>Equus_caballus

---------MGKNIQDKDNLYHRKAIENAKVKHLYEGLITLNSAFVKI-----------

HG1169 ENSG00000167637 ZNF283

>Homo_sapiens

MTARKKSMLSPIMNVSSKKDENFRSGSVRGTGSCQDLMGKF

>Physeter_catodon

------------MYVSSKKVQI-------------------

>Rousettus_aegyptiacus

------------MNVLRKKAQV-------------------

HG1170 ENSG00000167858 TEKT1

>Homo_sapiens

METALFTSLFCKSWKLQLG---------

>Myotis_lucifugus

METSLFTFLFCESRELQLGLGGSYSELV

>Equus_asinus

METALFTSQFCKSSGLPWEG-GSPPELV

HG1171 ENSG00000167968 DNASE1L2

>Homo_sapiens

MLGRECGWSYLRCLAFPEGILLREAESRRPNF

>Myotis_davidii

------MWSRNSGRCGAKEGLS----------

>Cavia_porcellus

MLGQEWGWSFLTA-CLPKRIWLKEADSR----

HG1172 ENSG00000170381 SEMA3E

>Homo_sapiens

MNSFAQVTEKYSFFLNLPA

>Rousettus_aegyptiacus

MNRLQK-------------

>Loxodonta_africana

MNRLPK-------------

HG1173 ENSG00000170464 DNAJC18

>Homo_sapiens
MGQHLNHIGHRTDLSITSKNPLQFNM--
>Odobenus_rosmarus_divergens
MGQYLNHIGHKANLSITSKNSLQFNKYF
>Balaenoptera_acutorostrata_s
MGQHLNHIGHMANLSITSKNSLQFNKFF

HG1174 ENSG00000173918 C1QTNF1

>Homo_sapiens

MGGHSRQGKKPRFRFWSQRPRRPRRGDGGAVGGGDLWLLVPPPTLLFCTAVLRKTFSPALFPSPSLCIAPDL--AGRRLGRREML---------------------

----------

>Mustela_putorius_furo

------------MCAGSVGGSAPCSH------------L---------AGVCSLRVCPAFLPLPLLSRNRASASGQRDLGPEEVTVPGRQGPAAACTRPLCTFFPE

DFFCPPGSLL

>Ceratotherium_simum_simum

-------------MRWERSRLRGRSH------------L---------AGVCSLRVCPAFFVSSFTFEKP-----------------------------RFRFWPK

RL--------

HG1175 ENSG00000174473 GALNTL6

>Homo_sapiens
MVLWVRSVARKEW
>Odobenus_rosmarus_divergens
MFLWVRSIAGKEW
>Marmota_marmota_marmota
MFLWVRSVAGKEW

HG1176 ENSG00000176472 ZNF575

>Homo_sapiens

MKFNGKKGAVDAASMN

>Ailuropoda_melanoleuca

MKFSNKKEAVHAASMD

>Heterocephalus_glaber

MKFNDRKDAVDAAVID

HG1177 ENSG00000178295 GEN1

>Homo_sapiens

MTKGLLPEPRRKGCPVCFT

>Acinonyx_jubatus

MTEGRLPKPRRKGCPVF--

>Equus_asinus

MTEGRLPKPRGEGCPVCFK

HG1178 ENSG00000182670 TTC3

>Homo_sapiens
----------------MCMRNITCTSNL
>Canis_lupus_familiaris
MRYLKWHVDFLIVKNLMCMKNVSCTSNL
>Pteropus_alecto
----------------MCMKNINCTSNL

HG1179 ENSG00000185933 CALHM1

>Homo_sapiens

MRWAPSGQGGQQGPSWRQQ

>Ceratotherium_simum_simum

MRWAPSVRSGPQGPSLRQW

>Tupaia_chinensis

MGWAPSGQSGPQGPWWRRQ

HG1180 ENSG00000188674

>Homo_sapiens

MLQHASRSRERTKALEPVSGCL--------------

>Acinonyx_jubatus

MLQHASRSRERTKAPESVSGCL--------------

>Rousettus_aegyptiacus

MLKHASRSRERTKALESGSAYRLREAGCSHRISGAV

HG1181 ENSG00000189058 APOD

>Homo_sapiens

MDTTSHLCVSIQTKLLGQPS

>Ursus_maritimus

MDKTSHLCVWYRAKLLGQPS

>Pteropus_vampyrus

MDKTFHLCVLYRAKLLGQPS

HG1182 ENSG00000196155 PLEKHG4

>Homo_sapiens

MEQDNSKAPLWQPDLLMKKAPR---

>Panthera_tigris

----------------MKKAPRNLV

>Loxodonta_africana

----------------MKKAPR---

HG1183 ENSG00000198133 TMEM229B

>Homo_sapiens

-------------------MKVACRDGKCCLRRDTWKLPAPQPRIS

>Mustela_putorius_furo

------------------------------MDRDTWKPPADQPRIS

>Cricetulus_griseus

MSGIELFRPHAMALLMVADMEHASLGFFPQGDRDTWKLPANQPRIS

HG1184 ENSG00000198894 CIPC

>Homo_sapiens

MGRAVQMKRVPMNLPPAE

>Physeter_catodon

------MKRVLMNALPAQ

>Tupaia_chinensis

MGRAIQMKRVPMSALAAW

HG1185 ENSG00000198920 KIAA0753

>Homo_sapiens

--------------------------MCL--LPS---AFFMCATALEIRTLRQLSLCSQEVSVFKQI-----------------

>Balaenoptera_acutorostrata_s

MPRPQFCRLLHSAALAPGLWRCQASWDCLTRVPPPTFAFDHVSTALEIRNL----YTSTDVSVFKQIQSTILEFIQKLKLGIS-

>Manis_javanica

MEN----KIIRPLASKGKCWPHE-SCRCPGPISAGSFI--L-----QLQP-----LNSSDVSPVGI--ASLMCFLPSSFLIMSL

HG1186 ENSG00000204923

>Homo_sapiens

MGATVVLASPKT--

>Ursus_maritimus

MGATVALASPET--

>Ceratotherium_simum_simum

MGATVVLASPETYR

HG1187 ENSG00000213762 ZNF134

>Homo_sapiens
MAAAAVMGPAQILCGC
>Ailuropoda_melanoleuca
MAAAAVMDPA------
>Manis_javanica
MAAAAVTDPASILCGC

HG1188 ENSG00000214415 GNAT3

>Homo_sapiens

MSKPQLCSTFEKSEHIN

>Rousettus_aegyptiacus

MLKPELCSTFEKV----

>Condylura_cristata

MSKPELCSTFEDV----

HG1189 ENSG00000279909

>Homo_sapiens

-----MELTSTTELFTLHCKYTYVC----PPSQLSCQCLE------------------------

>Ailuropoda_melanoleuca

MTPEKHRPISNRELFTLHYKYTYDTHRPPPISQTGCECLL------------------------

>Marmota_marmota_marmota

----------------MNCKHTYNTC---LPSHTSCECFERTGCGSFNFISRMTLIRHSMFFFA

HG1190 ENSG00000123684 LPGAT1

>Homo_sapiens

MHRCAGIVVVRRLRSSVSGKSY---------------------

>Galeopterus_variegatus

MNQCAGIAVG-RPRCSVSGKSDREVPVLSFHS-GRHFGDTEGF

>Ictidomys_tridecemlineatus

MKQCAGMVVERTPSFSLSGKSDREVPASVLSLVYPRLGDTEGV

HG1191 ENSG00000164331 ANKRA2

>Homo_sapiens

MSESFFLGLFGHSWPVATERTFFPARVSPPAGDSGL-LSRDAGTPVA--YF----------

>Oryctolagus_cuniculus

MSERFPPGLSEHCWPRRRSGRPLPRVCPAASSWPGTPAAYFLKIQISRIHWISLIMKKCFN

>Tupaia_chinensis

-MSESCLGFWGRSWPLASERTSAPYKSLQPAPVLGH-LSPIFNIQIPRKHWVPLL------

HG1192 ENSG00000168758 SEMA4C

>Homo_sapiens

-----------------------------------------MWNGVWGSAGLLSALGKGTGRDRDL-RSQPAAPAGQLA-------------------

>Oryctolagus_cuniculus

-----------------------------------------MWNGVLGSAGLLSALGEGTGRAGTSARRQPAAPSARGRLCPLPAPHMPPAPLRGWPL

>Chinchilla_lanigera

MRPWSFPRPSHPLPASLLGNKGASGNWAPQAPGPAPAPALLMWNGVWGSAGPLSALGGGDQQ-----GPGPAAPRPRGGLCQLPPHLTPPHP------

HG1193 ENSG00000177706 FAM20C

>Homo_sapiens

MDLDPRGGAALVPSCS

>Dipodomys_ordii

MDLDPRGGAAFVPSSS

>Tupaia_chinensis

MDLDPRGGEAFVSNQG

HG1194 ENSG00000214694 ARHGEF33

>Homo_sapiens
MGTSRENKKKTSKNML
>Oryctolagus_cuniculus
MGTSREIMKKTSKNMM
>Fukomys_darlingi
MGTSREIRRKTSKNMV

**Supplementary Figure S5**. FASTA-formatted CPuORF peptide sequences with gaps used in Supplementary Figure S4.
