## Supplementary Figure S6 for "Exhaustive identification of conserved upstream open reading frames with potential translational regulatory functions from animal genomes"

### (a) *PTP4A1*

```
  1 GCTAGCCTGTGTCGCCGCCGCCTCGGGACCGGCTGTATGATTAGGCCACAATCTTCTGTG
                                          M  I  R  P  Q  S  S  V
                                          M  I  R  P  Q  S  S  V
 61 TCGCCGCCGCCTCGGGACCGGCTGTCAATGAGTAAACATATTCCTCAATTCTGTGGTGTT
    S  P  P  P  R  D  R  L  S  M  S  K  H  I  P  Q  F  C  G  V
    S  P  P  P  R  D  R  L  S  M  S  K  H  I  P  Q  F  C  G  V
                                   C
                                   ⬇
121 CTTGGTCACACATTTATGGAGTTTCTGAAGGGCAGTGGAGATTACTGCCAGGCACAGCAC
    L  G  H  T  F  M  E  F  L  K  G  S  G  D  Y  C  Q  A  Q  H
    L  G  H  T  F  M  E  F S  E  G  Q  W  R  L  L  P  G  T  A  R
           C
           ⬆
181 GACCTCTATGCAGACAAGTGAACTGTAGAAACTGATTACTGCTCCACCAAGAAGCCCCCA
    D  L  Y  A  D  K  *
      P  L  R  R  Q  *
241 TAAGAGTGGTTATCCTGGACACAGAAGTGTTGAATTGAAATCCACAGAGCATTTTACAAG
301 AGTTCTGACCTGGATGGGGTAAACCTCAGTGCACTTCTTTTCTGTTGGCCTCAGTATTAC
361 TGGATTGAAGAATTGCTGCTTCTTGTTAGGAGGTTCATTTCACTTATCATTACTTACAAC
421 TTCATACTCAAAGCACTGAGAATTTCAAGTGGAGTATATTGAAGTAGACTTCAGTTTCTT
481 TGCATCATTTCTGTATTCAATTTTTTTAATTATTTCATAACCCTATTGAGTGTTTTTTAA
541 CTAAATTAACATGGCTCAAGCTT
```

**(b)** ***MKKS***

A ⬇

1 GCTAGCTTGAATCTGAGCTTCATATCGAAAGAAGAG**ATGAAAAATACCAGTTGGATTAGA**

**M K N T S W I R**

**M K N T S W I R K**

61 **AAGAACTGGCTTCTTGTAGCTGGGATATCTTTCATAGGTGTCCATCTTGGAACATACTTT**

**K N W L L V A G I S F I G V H L G T Y F**

**E L A S C S W D I F H R C P S W N I L F**

121 **TTGCAGAGGTCTGCAAAGCAGTCTGTAAAATTTCAGTCTCAAAGCAAACAAAAGAGTATT**

**L Q R S A K Q S V K F Q S Q S K Q K S I**

**A E V C K A V C K I S V S K Q T K E Y**

181 **GAAGAGTGA**AGTAAAATAAATATTTGGAATTACTAATTTGTCATTAAATCATTCTATGCT

**E E ***

**E E ***

241 GATTAGCTTCATAAACATTGAACTTTTTGATTTTATAGCCACAATGCTGCATATTCATAC

301 TTTAATTCCTAAAGAATAATTTTTAATGTTAAAACGTGATAATGCAATAAATAGAAAAAT

361 GTGGTTTACAAAATAAAAACGGTCTTCACTAGTTACCACCTGAAGTAAGATGTCTCGTTA

421 AGCTT

**(c)** ***SLC6A8***

```
                                                    A
                                                    ↓
  1 GCTAGCCGCCCGCGCGCCCCCGGGCCCCCGACACACATGAGATTCTTCAGGCTCACTTTC
                                        M  R  F  F  R  L  T  F
                                        M  R I  L  Q  A  H  F  Q
 61 AAGTGCTTCGTGGACTGCTTCTGACTGCGCCGCCCGCGCCCCGCACCCCGCCGCCCGCCC
    K  C  F  V  D  C  F  *
      V  L  R  G  L  L   *
121 GCCGCCCCGTCCCCCGGCCCGGCCGCCCCCCGGCCCCCGGCCGGCCCGCGCCCTCGGGGC
181 CCTCCCCGGTGCCGCCGGTGCCCCCCGCCTGACCGCCGCCCCCCGTGAGGCGCCGCGACC
241 CCGGCCCGGCCGTGCGGCCCGCCGAGGCCATGGAAGCTT
```

**(d)** ***FAM13B***

A (↓)

1 GCTAGCGAAGTACTGATCGAGTTCTGCATTTCTTCA**ATGAGGAACTACAGGCTGATCTTC**

**M R N Y R L I F**

**M R K L Q A D L L**

A (↑)

61 **TGCCATAATCTCAAACAGCCATAA**ATGACAAAAGAATGCTTGCTCAGTGAGGGTAGCTGG

**C H N L K Q P ***

**P K S Q T A ***

121 TGCAGAAGCCATTTTTTAAACTTAGGTATTTAAGTACTGAAAGAAAAGACAGCTTTGATT

181 TCTGGCTGCAAAAAAGATATGAAAGCTT

**(e)** ***MIEF1***

```
  1 GCTAGCGGTGTGACACCCAGCCCCTGCCAGTCCCCCATGGCCCCGTGGAGCCGAGAGGCG
                                        M  A  P  W  S  R  E  A
                                        M  A  P  W  S  R  E  A
 61 GTGCTGAGTCTCTATCGGGCTCTGTTGCGCCAGGGCCGACAGCTTCGCTACACTGATCGA
    V  L  S  L  Y  R  A  L  L  R  Q  G  R  Q  L  R  Y  T  D  R
    V  L  S  L  Y  R  A  L  L  R  Q  G  R  Q  L  R  Y  T  D  R
121 GACTTCTACTTTGCCTCCATCCGCCGTGAATTCCGAAAAAATCAGAAGCTAGAGGACGCT
    D  F  Y  F  A  S  I  R  R  E  F  R  K  N  Q  K  L  E  D  A
    D  F  Y  F  A  S  I  R  R  E  F  R  K  N  Q  K  L  E  D  A
          T
          ⬇
181 GAGGCCCGGGAGAGGCAGCTGGAGAAGGGCCTGGTCTTTCTCAACGGCAAATTGGGGAGG
    E  A  R  E  R  Q  L  E  K  G  L  V  F  L  N  G  K  L  G  R
    E  A S  G  E  A  A  G  E  G  P  G  L  S  Q  R  Q  I  G  E  N

241 ATCATTTAGGATCCTCCAAGGGAAAGAGGACAAAGGTGCCTTCTGTAGACACTCCTGCTC
    I  I  *
       I  *
301 TCTTCCATCCCCATCTTACAGATGTATTAAGAAGCCTCAGATGAGCAATGGAAGCTT
```

**(f)** ***EIF5***

```
                                             C                                      A
 61 CCTCAGTGGCGAAAATTTTTTCATGTCAGAGACCGAGAACTCTTGCAGTCGTTTATGTCA
                                                                  M  S
                                                                  M  S  I
121 TCCCTTCTTCTCCAGACAGAAGATACCAAAAAGTTGCAATCAAAGATCTCTTCATCTTAT
    S  L  L  L  Q  T  E  D  T  K  K  L  Q  S  K  I  S  S  S  Y
      P  S  S  P  D  R  R  Y  Q  K  V  A  I  K  D  L  F  I  L
181 TGATAAAGCCACTAATAAGCCAAAATGTCCTCGAG
    *
    *
```

Line 1: 1 GCTAGCCAGCCATTGGTACCTGTATTGGGGAAACATAGCATACAAGCAAGAAGCTTACAG

### (g) *MAPK6*

1 GCTAGCGTCGGAGAAGTCCCGTTGTATCAGAGTAAG**ATGGACGGTAGCTTTGATTGTGAT**

**M D G S F D C D**

**M D V A L I V I**

A

61 **TGTGGTGAGCTGGAGCCACCTGATCACTAA**CAAAAGACATCTTCTGTTAACCAACAGCCG

**C G E L E P P D H ***

**V V S W S H L I T ***

121 CCAGGGCTTCCTGTTGAAATAAATATATAGCAACAAAGGAAAAAAAGAAGCAAAACGGAA

181 ATAGTGCTTACCAGCACCTTAGAATGATGCTGCTCAGGACCAGTCCAACACTGAATGTAT

241 CTGCACTGTGAGGAGAATGTTCATAGAAGCCTGTTGTGTGCATATTTATTCACATTTTTG

301 TTAAATGTTAAATCGTTTAGCACGGTAATCTGAGTGCACAGTATGTCATTTCATTCCGTT

361 TGAGTTTCTTGTTTTCGTTAAATGTCTGCAGAGTTGCTGCCCCTTTCTTGAACTATGAGT

421 ACTGCAATCTTTTTAATTCTCAATATGAATAGAGCTTTTTGAGCTTTAAATCTAAGGGGA

481 ACTCGACAGGCCTGTTTGGCATATGCAATGAACATCAAGAAACCATCTTGCTGTGGAAGC

541 ATAATTATTTTTCTTCTCCCTTTTTGAAAGATCTTTCCTTTTGATGCCAGTTTTCTTCCT

601 TGTTTACACAAGTTCAATTTGAAAGGAAAAGGCAATAGTAAGGGTTTCAAAATGGAAGCT

661 T

**(h)** ***MEIS2***

A C

1 GCTAGCTTTGATTGACAGCTGGAGTGGCAAAAAGCC**ATGAAACACGACAGTTCGGTTACA**

**M K H D S S V T**

**M K T R Q F G Y L**

61 **TGTGGGCTGCTGACGGGCCGCTCGTAA**CCTTCAGTTCGGGGGCTTGACAATTTTTTTCTT

**C G L L T G R S ***

**W A A D G P L ***

121 CTTTTTCTTCTTTTTCTTTTCTTTCTTTTTTTTTCCAACTGAGGGGAAGAGAAGAGAAAG

181 AGGGGGAAAGGAGGACCGAAGAGGAGGAGGAGGGGAGGGGGGAGGAGGAGGAGGTGGAGG

241 AGGAGGAGGAAGATCAGGAGGAGGAGGAAGAAGAGGAAAAAAGAGAAAAAGAAGAAATAT

301 CACAGAAAAAAAAATTCTTCGTTGTCTAGACTGGGCTTTTTTTCCCCCCTAAAAAATAGC

361 ATATTGGAGAATTGGGAGAAGTCTCTTTGGTTTGGAAAAAAAAAAAAGGAATCTTCAGCC

421 TAGATCACTTTCTTATCCGGACTGGGATATTAAATATACGACACATCCAGGAGTTTATTG

481 GAGCGCAGACTGATGGCTCGAG

### (i) *KAT6A*

```
                                                               A
                                                               ↓
  1 GCTAGCCAACAGGTTGTTTTGGTTTCTATAGTACAATTGGGGTGGCATTCTGTTTTGTGA

 61 AAGGAGGAAGGACTTAGGCCAGAAAACTCATATGCTATGGTTAACTGGTTCCCAGCCTCC
                                        M  V  N  W  F  P  A  S
                                        M  V K  L  V  P  S  L  R
                      C
                      ↑
121 GAGAATCTTGTTTTCCATGGTGTAAAACTTACTCAGCATCAGGATAAGGGATAACGACTC
    E  N  L  V  F  H  G  V  K  L  T  Q  H  Q  D  K  G  *
      E  S  C  F  P  R  C  K  T  Y  S  A  S  G   K  G  *
181 TATGGATATACAGAATCCTTCACCATGGTAAAACCTCGAG
```

**(j)** ***SLC35A4***

C

1 GAGCTCTGCGCCTGCGCAACAAGTTCGGCGGGGAAG**ATGGCGGATGACAAGGATTCTCTG**

**M A D D K D S L**

**M A T T R I L C**

A

61 **CCTAAGCTTAAGGACCTGGCATTTCTCAAGAACCAGCTGGAAAGCCTGCAGCGGCGTGTA**

**P K L K D L A F L K N Q L E S L Q R R V**

**L S L R T W H F S R T S W K A C S G V I R**

A C

121 **GAAGACGAAGTCAACAGTGGAGTGGGCCAGGATGGCTCGCTGTTGTCCTCCCCGTTCCTC**

**E D E V N S G V G Q D G S L L S S P F L**

**R R S H S G V G Q D G S L L S S P F L**

C

181 **AAGGGATTCCTGGCTGGCTATGTGGTGGCCAAACTGAGGGCATCAGCAGTATTGGGCTTT**

**K G F L A G Y V V A K L R A S A V L G F**

**K G F L A G Y V V A K L R A S A V L G F**

C C

241 **GCTGTGGGCACCTGCACTGGCATCTATGCGGCTCAGGCATATGCTGTGCCCAACGTGGAG**

**A V G T C T G I Y A A Q A Y A V P N V E**

**A V G T C T G I Y A A Q A Y A V P N V E**

301 **AAGACATTAAGGGACTATTTGCAGTTGCTACGCAAGGGGCCCGACTAG**CTCTAGGTGCCA

**K T L R D Y L Q L L R K G P D ***

**K T L R D Y L Q L L R K G P D ***

361 TGGAAGAGGCAGGATGAGCAGCTCAGCCTTCAGGTGGAGACACTTTATCTGGATTCCCCA

421 GCTGTCATCCATTTGCTATCTCCAACTTTCCTGCCACCTTCATCCTTGCCTCCCTTCCTG

481 CAGATTGTGGACAGTAGTTCCTCAGCCTGCACCCTGGATTCCTTCTTCCCCTTCCTAGCT

541 CCATGGGACTCGCCCCAAGACTGTGGCTTCAAGGACCACCAGCCCCTTACTCTTCAAGCC

601 CTGACTGTGGAGTTGGTAGATGCCTCTGATCCTCAGTATTCTCTCTGGCAATGTTCCACG

661 GCTTCTCCTTCCTGGGAGCTGGCTCCATAACTTGATTTTCCCCAAACGTGTTGCAATCCC

721 TGCTGCCCCTTAGCCACCCAGGGTCTTGTGTGGGTATGAGCTCGAG

**(k)** ***LRRC8B***

```
  1 GCTAGCTCAGAAGGTGATCTCTTTAATGCTTTCTTTTTAAGAATTTTTCAAATTGAGACT
 61 AATTGCAGAGGTTCCAGTTGACCAGCATTCATAGGAATGAAGACAAACACAGAGATGGTG
                                        M  K  T  N  T  E  M  V
                                        M  K  T  N  T  E  M  V
                                      A
                                      ⬇
121 TGTCTAAGAAACTTCAAAAGGTGTAGACCTCCTGACTGAAGCATATTGGATTTATTTAAT
    C  L  R  N  F  K  R  C  R  P  P  D  *
    C  L   K  T  S  K  G  V  D  L  L  N *
181 TTTTTTCACTGTATTTCTGTCCTCCTACAAGGGAAAGTCATGATTACACAAGCTT
```

**(I)** ***CDH11***

```
                                                        A
                                                        ↓
  1 GCTAGCCGTGTTGTCATTTGTTGAGTGACCAATCAGATGGGTGGAGTGTGTTACAGAAAT
                                        M  G  G  V  C  Y  R  N
                                        M  GR  S  V  L  Q  K  L
                         C
                         ↑
 61 TGGCAGCAAGTATCCAATGGGTGAAGAAGAAGCTAACTGGGGACGTGGGCAGCCCTGACG
    W  Q  Q  V  S  N  G  *
      A  A  S  I  Q  R  *
121 TGATGAGCTCAACCAGCAGAGACATTCCATCCCAAGAGAGGTCTGCGTGACGCGTCCGGG
181 AGGCCACCCTCAGCAAGACCACCGTACAGTTGGTGGAAGGGGTGACAGCTGCATTCTCCT
241 GTGCCTACCACGTAACCAAAAATGAAAGCTT
```

**(m)** ***PNRC2***

1 GCTAGCCTCTCAAACTTGTGTGCTGAGGAGACTCAG**ATGTTGGCCTCAGCTCCTAGGCTG**

**M L A S A P R L**

**M L A S A P R L**

A

61 **AACTCAGCAGATCGGCCCATGAAAACTTCTGTATTGAGACAAAGGAAGGGATCTGTCAGA**

**N S A D R P M K T S V L R Q R K G S V R**

**N S A D R P M K N F C I E T K E G I C Q K**

121 **AAGCAACACTTGTTATCTTGGGCTTGGCAGCAAGGAAGAGGACAGGTAGTGGAGATCCTG**

**K Q H L L S W A W Q Q G R G Q V V E I L**

**A T L V I L G L A A R K R T G S G D P A**

T T

181 **CAATCTGAAAAGCAGACTGAAAGGTGA**CAAAGAAGCTGAAGATGGAAGCTT

**Q S E K Q T E R ***

**I L K A D L K ***

**(n)** ***BACH2***

C T

1 GCTAGCGCCCCACAAAACTTTGGGGTCCATGTCTGA**ATGGATTGCCAGAGCCTTCTCATC**

**M D C Q S L L I**

**M D C P E P S H L**

61 **TCTCCCTTCGCCCAGTTCCCTGCATCCTAA**GACTCGAAGGCAGCACAGGACCTGGAAAAA

**S P F A Q F P A S ***

**S L R P V P C I ***

121 TTACATGGTGTGAACGGCATGTAAGCTT

**(o)** ***FGF9***

```
  1 GCTAGCCTATAATAACGCCTAGGCATTTAAGTTGCTATGGTCATTCTGATCTCAAACCAA
                                        M  V  I  L  I  S  N  Q
                                        M  V  I  L  I  S  N  Q
          A                                               T
 61 ATGGAGAAACTACGGATTTTTTTTCCTTATTACGGTCGGATGGGATGAAGACCTTCCTGC
    M  E  K  L  R  I  F  F  P  Y  Y  G  R  M  G  *
    M  E  KT T  D  F  F  S  L  L  R  S  M  G  *
121 CTGCTAAGAGCTGGGGATCTATCTATAGAGATACATAGATATGTTTATCAATATGTCAGT
181 GTGTGAGTATAAAGTGGTGGTTTCTTAGACTATCAGTGGTTTGACCTTGAACCTGTGCCA
241 GTGAAACAGCAGATTACTTTTATTTATGCATTTAATGGATTGAAGAAAAGAACCTTTTTT
301 TTCTCTCTCTCTCTGCAACTGCAGTAAGGGAGGGGAGTTGGATATACCTCGCCTAATATC
361 TCCTGGGTTGACACCATCATTATTGTTTATTCTTGTGCTCCAAAAGCCGAGTCCTCTGAT
421 GGAAGCTT
```

**(p)** ***PNISR***

```
  1 GCTAGCCTAGAGTGGGGCATACATAATCTTGCTGCTATGCTTCGAAGCTGTAGTCTGAAT
                                        M  L  R  S  C  S  L  N
                                        M  L  R  S  C  S  L  I

             A
             ⬇
 61 CAACCTAAGTTTTAAACAGAAGGTGAACCTCTGAGAAAATCAAGTATATTTTAAAAGAAG
    Q  P  K  F  *
     N  L  S  L *

121 GGATGTGGGAAGCTT
```

### (q) ***TMEM184C***

```
  1 GCTAGCTTCCTGGTCTGTGCTGCTCTCCTGGAAGCCATGGTACAGGCAGAGCTCAGGGCG
                                          M  V  Q  A  E  L  R  A
                                          M  V  Q   Q  S  S  G  R

              A
              ⬇
 61 ATCCCCAGGTGAGGGCAGCGGCTCTGCCTGGGATTCCACCGCAGTACAACCGGGTAGATG
    I  P  R  *
     S  P  G *
121 CGGGGTGGAGAAGAAAGGATGTTGCCTGCACTGCTCGCCAATAGCACCCTGAGAGGCTAC
181 ATTTGCAGAAGCAGCAGCAGCAGAAGACACAGCGCCGGTCCAGGAGGCGGCTCGAGCTGT
241 TCGTAAAGTCGCCCGACAGCTTTTTCTCCGTAGTATGCGAGTTGACAAAACAGCCAGAGA
301 ACAGGGCTCCCCATTACAATCTTTTCGAGATCTTTTCCCTTGCTAACCGGATCTGATTTG
361 TGCGAAAACATGCCTTGCAAAGCTT
```

**Supplementary Figure S6.** Nucleotide sequences of the 5′-UTRs and amino acid sequences of the CPuORFs analyzed in this study. **(a-q)** The 5′-UTRs of *PTP4A1* **(a)**, *MKKS* **(b)**, *SLC6A8* **(c)**, *FAM13B* **(d)**, *MIEF1* **(e)**, *EIF5* **(f)**, *MAPK6* **(g)**, *MEIS2* **(h)**, *KAT6A* **(i)**, *SLC35A4* **(j)**, *LRRC8B* **(k)**, *CDH11* **(l)**, *PNRC2* **(m)**, *BACH2* **(n)**, *FGF9* **(o)**, *PNISR* **(p)**, and *TMEM184C* **(q)**. The nucleotide sequences of the CPuORFs are shown in bold. The deduced amino acid sequences of the wild-type (WT-aa) and frameshift (fs) CPuORFs are indicated. The nucleotide sequences of other uORFs are underlined with a bold line. A dotted underline indicates the nucleotide sequences of other uORFs overlapping the CPuORFs. The initiation codons of these overlapping uORFs were changed to other codons, as indicated, while keeping the CPuORF amino acid sequence unaltered. The replaced nucleotides are shown as white letters in a black background. The nucleotides that were deleted and inserted in the frameshift mutants are shaded. The main coding sequences that were contained in the reporter constructs are boxed. The shaded boxes indicate the nucleotides changed to avoid the appearance of in-frame termination codons. Cloning sites were added at either end of the nucleotide sequences in controls to be subcloned into plasmid pGL4.10 with an SV40 promoter (pSV40:5'UTR::luc2) are underlined (Fig.2b).
