## Supplementary Table S1 for "Exhaustive identification of conserved upstream open reading frames with potential translational regulatory functions from animal genomes"

Supplementary Table S1. CPuORFs extracted from *D. melanogaster*, *D. rerio*, *G. gallus*, and *H. sapiens*.

| HG number | Species | Gene ID | Gene symbol | Gene description† | uORF-mORF fusion ratio | $K_a/K_s$ analysis before manual validation | | | $K_a/K_s$ analysis after manual validation | | Previous report* | CPuORF amino acid sequence |
|---|---|---|---|---|---|---|---|---|---|---|---|---|
| | | | | | | Median pairwise $K_a/K_s$ ratio | $U$-test $p$ value | $q$ value | Median pairwise $K_a/K_s$ ratio | $U$-test $p$ value | | |
| HG0001 | *D. melanogaster* | FBgn0024734 | PRL-1 | PRL-1 | 0.00 | 0.06 | 0.0E+00 | 0.0E+00 | 0.06 | 0.0E+00 | Crowe | MCMAIVLVRPVPTSMEFHLDSAEYCQAQHK* |
| | *D. rerio* | ENSDARG00000006242 | ptp4a1 | protein tyrosine phosphatase type IVA, member 1 | 0.04 | 0.10 | 0.0E+00 | 0.0E+00 | 0.10 | 0.0E+00 | Crowe | MIKLQSSMSKHIPFCGVLGHTFMEFLKGSGDYCQAQHGVYAEK* |
| | | ENSDARG00000035676 | ptp4a2b | protein tyrosine phosphatase type IVA, member 2b | 0.00 | 0.22 | 0.0E+00 | 0.0E+00 | 0.22 | 0.0E+00 | Crowe | MVGPFFCFRVDSDHTYMAFVTVCREFCQAQHLTFADK* |
| | | ENSDARG00000039997 | ptp4a3 | protein tyrosine phosphatase type IVA, member 3 | 0.00 | 0.07 | 0.0E+00 | 0.0E+00 | 0.07 | 0.0E+00 | Crowe | MAFLQRTGDYCHAQHLHI* |
| | | ENSDARG00000054814 | PTP4A3 | protein tyrosine phosphatase 4A3b | 0.00 | 0.08 | 0.0E+00 | 0.0E+00 | 0.08 | 0.0E+00 | Crowe | MEFWQGSGDYCHAQHHTA* |
| | | ENSDARG00000087443 | ptp4a2a | protein tyrosine phosphatase type IVA, member 2a | 0.00 | 0.14 | 0.0E+00 | 0.0E+00 | 0.14 | 0.0E+00 | Crowe | MQSSQLVFCSLIEGCTFMAFVSVSGDFCQAQHLVIVDK* |
| | *G. gallus* | ENSGALG00000003265 | PTP4A2 | protein tyrosine phosphatase type IVA, member 2 | 0.00 | 0.24 | 0.0E+00 | 0.0E+00 | 0.24 | 0.0E+00 | Crowe | MPGVFCWNLRRTFMAILSVRAEFCQAQHSALADK* |
| | | ENSGALG00000016271 | PTP4A1 | protein tyrosine phosphatase type IVA, member 1 | 0.00 | 0.08 | 0.0E+00 | 0.0E+00 | 0.08 | 0.0E+00 | Crowe | MIRLQSSMSNHIPQFCGVLGHTFMEFLKGSGDYCQAQHDLYADK* |
| | *H. sapiens* | ENSG00000112245 | PTP4A1 | protein tyrosine phosphatase type IVA, member 1 | 0.00 | 0.09 | 0.0E+00 | 0.0E+00 | 0.09 | 0.0E+00 | Crowe | MIRPQSSMSKHIPQFCGVLGHTFMEFLKGSGDYCQAQHDLYADK* |
| | | ENSG00000184007 | PTP4A2 | protein tyrosine phosphatase type IVA, member 2 | 0.00 | 0.22 | 0.0E+00 | 0.0E+00 | 0.22 | 0.0E+00 | Crowe | MAILSVRADFCQAQHSIFADK* |
| HG0002 | *D. rerio* | ENSDARG00000003077 | gpx9 | glutathione peroxidase 9 | 0.04 | 0.07 | 0.0E+00 | 0.0E+00 | 0.07 | 0.0E+00 | | MAANSVYDFTVETLEEKSVSLDIFRGKVLLIMNVATF* |
| HG0003 | *D. melanogaster* | FBgn0039280 | Mocs2 | Molybdenum cofactor synthesis 2 | 0.14 | 0.07 | 0.0E+00 | 0.0E+00 | 0.07 | 0.0E+00 | Hayden | MNADGPVVNVHVLFFAKSRELANTPRSTVEVPTEITATELLDHLVSKFGLTSIRDNLILAHNESYIDNLSDRILFKEGDELAIIPPLSGG* |
| | *G. gallus* | ENSGALG00000014906 | MOCS2 | molybdenum cofactor synthesis 2 | 0.02 | 0.07 | 0.0E+00 | 0.0E+00 | 0.07 | 0.0E+00 | Hayden | MSCQVTVLYFARSAELVGLRSESVSVPQRITSLQLWEEIVKIHPRLAVIRDQVVFAVRQEYVLLGDQLLVLQTGDEVAIIPPISGG* |
| HG0004.1 | *D. melanogaster* | FBgn0028494 | | | 0.00 | 0.01 | 9.6E-86 | 1.2E-85 | 0.01 | 9.6E-86 | Mackowiak | MLRYRLVSTHFLKQP* |
| | *D. rerio* | ENSDARG00000076779 | fam13b | family with sequence similarity 13, member B | 0.00 | 0.00 | 0.0E+00 | 0.0E+00 | 0.00 | 0.0E+00 | Mackowiak | MRNYRLIFCHNLKQP* |
| | *G. gallus* | ENSGALG00000041706 | | | 0.00 | 0.00 | 0.0E+00 | 0.0E+00 | 0.00 | 0.0E+00 | Mackowiak | MRNYRLIFCHNLKQP* |
| | *H. sapiens* | ENSG00000031003 | FAM13B | family with sequence similarity 13 member B | 0.00 | 0.00 | 0.0E+00 | 0.0E+00 | 0.00 | 0.0E+00 | Mackowiak | MRNYRLIFCHNLKQP* |
| HG0004.2 | *H. sapiens* | ENSG00000138640 | FAM13A | family with sequence similarity 13 member A | 0.15 | 0.34 | 2.4E-05 | 1.3E-04 | 0.34 | 2.4E-05 | | MLEAALRLPWGLCIGSCFSHQRICAEWNFSILLSF* |
| HG0005 | *D. rerio* | ENSDARG00000011055 | fbxo9 | F-box protein 9 | 0.16 | 0.04 | 0.0E+00 | 0.0E+00 | 0.04 | 0.0E+00 | Crowe, Mackowiak | MAAVWRARKFARCVVRFSDRDL* |
| | *G. gallus* | ENSGALG00000016320 | | | 0.22 | 0.04 | 0.0E+00 | 0.0E+00 | 0.04 | 0.0E+00 | Crowe, Mackowiak | MAAVRRARSYSRCVVRFSDRELC* |
| HG0006 | *D. melanogaster* | FBgn0029971 | CG18624 | GEO13364p1 | 0.09 | 0.08 | 0.0E+00 | 0.0E+00 | 0.08 | 0.0E+00 | Hayden, Mackowiak | MGSGIVVQLARKYGAVIFFPTVAVGSIYADWSHTRQWKRQQLQLAHRAQLKDQR* |
| HG0007 | *D. melanogaster* | FBgn0034372 | Gint3 | GDI interacting protein 3 | 0.25 | 0.06 | 0.0E+00 | 0.0E+00 | 0.06 | 0.0E+00 | Hayden | MSFNFYVESFNSTLQSIARQVEQFAGVALTAAVDLRGNLESLTGDFRWENGPIAFTDLQATLLKVLIVLLLGTCVLAGYSWSIYGKVITEKFVRPSTLKEIEELKLSVAKLKLPKEHSPRI* |
| HG0008 | *H. sapiens* | ENSG00000175567 | UCP2 | uncoupling protein 2 | 0.00 | 0.30 | 1.3E-225 | 2.6E-223 | 0.30 | 1.3E-225 | Crowe, Samandi | MTIRCFVSHPFSMENQGDRAMIATGSFEERDTFREA* |
| HG0009 | *D. melanogaster* | FBgn0050100 | CG30100 | AT22563p2 | 0.07 | 0.05 | 0.0E+00 | 0.0E+00 | 0.05 | 0.0E+00 | Hayden | MKMVQPSRQQILRLYKHLIRYGNHLKLTDKNYFLGRVRHEFRDNRSLTNPVEVEFSFKRGETLLKKGRIL* |
| HG0010.1 | *G. gallus* | ENSGALG00000009013 | MKKS | McKusick-Kaufman syndrome | 0.00 | 0.11 | 1.1E-270 | 7.2E-269 | 0.11 | 5.4E-265 | Akimoto | MMSLKTLWKDYKVLIVMGISLGLVHWSWFHIKSSPLFQVKTEEIVPEPGIVTYVMQSDHKNKEK* |
| | *H. sapiens* | ENSG00000125863 | MKKS | McKusick-Kaufman syndrome | 0.00 | 0.10 | 0.0E+00 | 0.0E+00 | 0.10 | 0.0E+00 | Akimoto | MSLRNLWRDYKVLVVMVPLVGLIHLGWYRIKSSPVFQIPKNDDIPEQDSLGLSNLQKSQIQGK* |
| HG0010.2 | *G. gallus* | ENSGALG00000009013 | MKKS | McKusick-Kaufman syndrome | 0.00 | 0.27 | 5.0E-187 | 2.3E-185 | 0.27 | 5.0E-187 | Crowe, Akimoto, Mackowiak | MRKASWSKKNFLLVAGLSLIGVHFGTMLVNFVAKKSARSHSETKRDNHRE* |
| | *H. sapiens* | ENSG00000125863 | MKKS | McKusick-Kaufman syndrome | 0.00 | 0.23 | 3.0E-245 | 7.0E-243 | 0.23 | 3.0E-245 | Crowe, Akimoto, Mackowiak | MKNTSWIRKNWLLVAGISFIGVHLGTYFLQRSAKQSVKFQSQSKQKSIEE* |
| HG0010.3 | *H. sapiens* | ENSG00000125863 | MKKS | McKusick-Kaufman syndrome | 0.00 | 0.44 | 7.0E-15 | 1.3E-13 | 0.44 | 7.0E-15 | | MLISFINIELFDFIATMLHIHTLIPKE* |
| HG0011 | *D. melanogaster* | FBgn0027360 | Tim10 | Translocase of inner membrane 10 | 0.03 | 0.05 | 3.5E-290 | 9.6E-290 | 0.05 | 3.5E-290 | Hayden | MRVRERMRKANQGDYDSSAASDRRFDY* |
| HG0012 | *D. melanogaster* | FBgn0064116 | CG33713 | GM05135p | 0.07 | 0.05 | 5.0E-301 | 1.4E-300 | 0.05 | 5.0E-301 | Hayden | MATAAAVAKVGKSVHRIFVGNLPWTVGHQELRGYFREFGRVVSANVIFDKRTGCSKGYGFVSFNSLTALEKIENEQKHILEGNYLNIQKS* |
| HG0013 | *D. melanogaster* | FBgn0261381 | mtTFB1 | Mitochondrial Transcription Factor B1 | 0.21 | 0.05 | 4.3E-307 | 1.4E-306 | 0.05 | 4.3E-307 | | MSASEQGPKIKYGESAPKLDKAQLQFMKLIEEQNLDRVQKLKRIRRNNLLTAGALGVSVLAIYGYSIFSVQQEKFLDDFEEPKKVSS* |
| HG0014 | *D. melanogaster* | FBgn0260392 | CG42518 | Uncharacterized protein, isoform A | 0.27 | 0.04 | 0.0E+00 | 0.0E+00 | 0.04 | 0.0E+00 | | MMDLSPNNQIEDRKPILTADGLVQTSNSPFEPTISQETQTSNGIGGQCHLTVDQLDIEILPIIYDIVRCVEKDPLENAVKLRESQDCNHKIFELQKRFESAREQIRQLPGIDFNKEEQQQRLELLRNQLKLKQQLIRKYKDTEF* |
| HG0015 | *D. rerio* | ENSDARG00000007377 | odc1 | ornithine decarboxylase 1 | 0.00 | 0.24 | 2.5E-211 | 4.1E-210 | 0.24 | 2.5E-211 | | MRFKNSCLREVHTGPGLEAYLKGFNFTGEPPWLSDIC* |
| HG0016 | *G. gallus* | ENSGALG00000010399 | SELENOT | selenoprotein T | 0.00 | 0.14 | 4.1E-254 | 2.4E-252 | 0.14 | 4.1E-254 | | MAYATGPLLKFQICVS* |
| HG0017 | *D. rerio* | ENSDARG00000043154 | ucp2 | uncoupling protein 2 | 0.04 | 0.30 | 2.3E-104 | 3.0E-103 | 0.30 | 2.3E-104 | Crowe | MFFICTSSQRFSMEERGNKIQVESHFNQRDAFRES* |
| HG0018 | *D. rerio* | ENSDARG00000053291 | pnrc2 | proline-rich nuclear receptor coactivator 2 | 0.00 | 0.47 | 9.8E-87 | 1.1E-85 | 0.47 | 9.8E-87 | Crowe, Samandi | MFNSVPRQNNAHWHMKTSDLAGHTCAARSRSTTSRTWSPIGEEKLKSEKNHALNFFTTER* |
| | *G. gallus* | ENSGALG00000004122 | PNRC2 | proline rich nuclear receptor coactivator 2 | 0.00 | 0.34 | 0.0E+00 | 0.0E+00 | 0.34 | 0.0E+00 | Crowe, Samandi | MLGSAAWLNTADRPMKTSVPSQRKESSERQHLLFQAWKQERGQELESVESEKTTER* |
| | *H. sapiens* | ENSG00000189266 | PNRC2 | proline rich nuclear receptor coactivator 2 | 0.00 | 0.32 | 0.0E+00 | 0.0E+00 | 0.34 | 0.0E+00 | Crowe, Samandi | MLASAPRLNSADRPMKTSVLRQRKGSVRKQHLLSWAWQQGRGQVVEILQSEKQTER* |
| HG0019 | *H. sapiens* | ENSG00000178397 | FAM220A | family with sequence similarity 220 member A | 0.06 | 0.07 | 8.8E-203 | 1.5E-200 | 0.07 | 8.8E-203 | | MAAALSGLAVRLSRSAAARSYGVFCKGLTRTLLIFFDLAWRLRINFPYLYIVASMMLNVRLQVHIEIH* |
| HG0020 | *D. melanogaster* | FBgn0035436 | CG12016 | SD05789p2 | 0.07 | 0.07 | 1.1E-163 | 1.7E-163 | 0.07 | 1.1E-163 | Hayden | MASNAQLGKIILISAIAVFFYYFFWVAVLPFMLIDEGNPIRLFFPPLKYAFIVPTVFGVIFLGGIAAFSFYHIWSLRVKRD* |
| HG0021.1 | *D. rerio* | ENSDARG00000068708 | ifrd1 | interferon-related developmental regulator 1 | 0.05 | 0.49 | 3.6E-60 | 3.5E-59 | 0.49 | 3.6E-60 | Zhao, Samandi | MYRSRSKSNTGIAFNTFLFTVPLEYTQTKRHYSKNWSCIG* |
| | *G. gallus* | ENSGALG00000009448 | IFRD1 | interferon related developmental regulator 1 | 0.02 | 0.45 | 2.8E-99 | 8.7E-98 | 0.45 | 2.8E-99 | Zhao, Samandi | MHRLRSRAFTGISAIAASSSFACLRQSPPLPRPEKSASQRRWNKKRSCSSIG* |
| | *H. sapiens* | ENSG00000006652 | IFRD1 | interferon related developmental regulator 1 | 0.00 | 0.38 | 3.8E-79 | 2.9E-77 | 0.38 | 3.8E-79 | Zhao, Samandi | MYRFRSQLFTGISAAATAHSYPRRFSTLLLAEDSPLSRPPHRRTSKKCSSIG* |
| HG0021.2 | *D. rerio* | ENSDARG00000036811 | ifrd2 | interferon-related developmental regulator 2 | 0.00 | 0.27 | 8.4E-109 | 1.1E-107 | 0.27 | 8.4E-109 | | MVESGAATCTILRPKHFQKHLPSQPGIRHRLPGDNHR* |
| HG0022 | *D. melanogaster* | FBgn0061359 | CG33671 | Mevalonate kinase | 0.00 | 0.03 | 2.7E-164 | 4.2E-164 | 0.03 | 2.7E-164 | Hayden | MSKYDSKYLEEKLKRELQTEYVSVTDESDGCGGKFSAVIVSPAFSGKTLLQKHRLVNSTLAEELKEIHAFSQKSYTPEEWEKVKAQ* |
| HG0023.1 | *H. sapiens* | ENSG00000175197 | DDIT3 | DNA damage inducible transcript 3 | 0.22 | 0.27 | 1.3E-37 | 5.8E-36 | 0.27 | 1.3E-37 | Crowe, Jousse, Mackowiak, Samandi | MLKMSGWQRQSQNQSWNLRRECSRRKCIFIHHHT* |
| HG0023.2 | *D. rerio* | ENSDARG00000059836 | ddit3 | DNA-damage-inducible transcript 3 | 0.06 | 0.34 | 8.9E-49 | 7.8E-48 | 0.34 | 8.9E-49 | Crowe, Jousse | MVNMSDQPSLQKHTQTLNQKQPRKRNNKKKRSYWDKISPTTHIQT* |
| HG0024.1 | *D. rerio* | ENSDARG00000093406 | C19H6orf62 | | 0.00 | 0.48 | 2.9E-39 | 2.4E-38 | 0.48 | 2.9E-39 | | MGPPRCRSRRLLPERTEDFGRFVSLS* |
| HG0024.2 | *G. gallus* | ENSGALG00000013628 | C6orf62 | chromosome 6 open reading frame 62 | 0.00 | 0.00 | 3.7E-13 | 2.2E-12 | 0.00 | 3.7E-13 | | MDWACASCFLLYT* |
| | *H. sapiens* | ENSG00000112308 | C6orf62 | chromosome 6 open reading frame 62 | 0.00 | 0.39 | 7.5E-12 | 1.0E-10 | 0.39 | 7.5E-12 | | MDFRDWSGVSCVLLHT* |
| HG0024.3 | *G. gallus* | ENSGALG00000013628 | C6orf62 | chromosome 6 open reading frame 62 | 0.00 | 0.48 | 5.2E-47 | 7.4E-46 | 0.48 | 5.2E-47 | | MAGQLLILFLDPSNTTFP* |
| HG0025 | *D. rerio* | ENSDARG00000060054 | epc1b | enhancer of polycomb homolog 1 (Drosophila) b | 0.10 | 0.41 | 8.5E-29 | 5.5E-28 | 0.41 | 8.5E-29 | | MRTLIGPAGNIRGSDRFMLHLEL* |
| HG0026 | *D. rerio* | ENSDARG00000087059 | fam213ab | family with sequence similarity 213, member Ab | 0.22 | 0.08 | 8.8E-53 | 7.9E-52 | 0.07 | 1.6E-51 | | MELVTRAVGSVGLTVIEALRSFTELFLTQPVCATLTQLADTDLRTLDGDDRVFKARELWESSGAVIMAVRRPG* |
| HG0027 | *H. sapiens* | ENSG00000077254 | USP33 | ubiquitin specific peptidase 33 | 0.00 | 0.47 | 1.0E-02 | 2.6E-02 | 0.47 | 1.0E-02 | | MERNNRYQPLQKHAKL* |
| HG0028.1 | *D. rerio* | ENSDARG00000032103 | mapk6 | mitogen-activated protein kinase 6 | 0.00 | 0.07 | 0.0E+00 | 0.0E+00 | 0.07 | 0.0E+00 | | MEDSFDCDCGELEPPDH* |
| | *G. gallus* | ENSGALG00000031448 | MAPK6 | Mitogen-activated protein kinase 6 | 0.00 | 0.07 | 0.0E+00 | 0.0E+00 | 0.07 | 0.0E+00 | | MDGSFDCDCGELEPPDH* |
| | *H. sapiens* | ENSG00000069956 | MAPK6 | mitogen-activated protein kinase 6 | 0.00 | 0.07 | 0.0E+00 | 0.0E+00 | 0.07 | 0.0E+00 | | MDGSFDCDCGELEPPDH* |
| HG0028.2 | *H. sapiens* | ENSG00000069956 | MAPK6 | mitogen-activated protein kinase 6 | 0.00 | 0.35 | 9.1E-15 | 1.6E-13 | 0.35 | 9.1E-15 | | MFIEACCVHIYSHFC* |
| HG0029 | *D. rerio* | ENSDARG00000007523 | kmt2e | lysine (K)-specific methyltransferase 2E | 0.00 | 0.37 | 4.7E-64 | 4.7E-63 | 0.37 | 4.7E-64 | | MHLDVCNECCG* |
| | *G. gallus* | ENSGALG00000008167 | | | 0.00 | 0.37 | 1.5E-134 | 5.6E-133 | 0.37 | 1.5E-134 | | MHLDVCNEHCGWHAPVFWIPMHRTL* |
| | *H. sapiens* | ENSG00000005483 | KMT2E | lysine methyltransferase 2E | 0.00 | 0.36 | 7.2E-199 | 1.2E-196 | 0.36 | 7.2E-199 | | MGLWMHLDVCNEHCGWHAPVFWIPMHRTP* |
| | | ENSG00000168137 | | | 0.00 | 0.30 | 3.0E-274 | 7.7E-272 | 0.30 | 3.0E-274 | | MSITCWMRLCSSPPLSEWSVSINWTP* |
| HG0030.1 | *G. gallus* | ENSGALG00000011464 | EIF5 | eukaryotic translation initiation factor 5 | 0.00 | 0.36 | 4.2E-136 | 1.6E-134 | 0.36 | 4.2E-136 | | MSSRLLQTEDTKKLQSKISSSY* |
| | *H. sapiens* | ENSG00000100664 | EIF5 | eukaryotic translation initiation factor 5 | 0.00 | 0.42 | 1.0E-91 | 9.7E-90 | 0.42 | 1.0E-91 | | MSSLLLQTEDTKKLQSKISSSY* |
| HG0030.2 | *D. melanogaster* | FBgn0030719 | eIF5 | eIF5 | 0.00 | 0.10 | 3.0E-116 | 4.0E-116 | 0.10 | 3.0E-116 | Mackowiak | MNSKGRTKNHRTSSNRHRKLQPFRYT* |
| HG0031 | *D. rerio* | ENSDARG00000000540 | asnsd1 | asparagine synthetase domain containing 1 | 0.00 | 0.09 | 0.0E+00 | 0.0E+00 | 0.09 | 0.0E+00 | Mackowiak | MSSKNIQRTEPGASEERAQKEELNQKIKEQKIVVDELSNLKKNRRVYTQQPNSNIFFLADKGETLSSCKQDLDNMRKEYQDI* |
| | *G. gallus* | ENSGALG00000041666 | | | 0.00 | 0.04 | 0.0E+00 | 0.0E+00 | 0.04 | 0.0E+00 | Mackowiak | MSRREEEETAELGYKGELSRRIKEQKVVVDELSNLKKNRKVYRQQPNSNIFFLVDRTEALSQCKNTLDQLKKAHQELENSEKTKVKK* |
| | *H. sapiens* | ENSG00000138381 | ASNSD1 | asparagine synthetase domain containing 1 | 0.00 | 0.07 | 0.0E+00 | 0.0E+00 | 0.07 | 0.0E+00 | Mackowiak | MPSRGTRPEDSSVLIPTDNSTPHKEDLSSKIKEQKIVVDELSNLKKNRTEQKCCLRARIYWMN* |
| HG0032 | *D. rerio* | ENSDARG00000104981 | SLC45A4 | | 0.00 | 0.36 | 0.0E+00 | 0.0E+00 | 0.36 | 0.0E+00 | | MGILRCFPSGFSRPSPLGLPSTLYSTASIQIMKLVLTEWQLIWIFTVSSRQ* |
| HG0033 | *G. gallus* | ENSGALG00000016321 | ICK | intestinal cell kinase | 0.00 | 0.47 | 3.1E-52 | 4.9E-51 | 0.47 | 3.1E-52 | Crowe | MIPATVDTVLSALCLCCHRKNWLQSEKKH* |
| | *H. sapiens* | ENSG00000112144 | ICK | intestinal cell kinase | 0.01 | 0.33 | 2.8E-117 | 3.2E-115 | 0.33 | 2.8E-117 | Crowe | MYLGDSHVLLNTLCWCCHRKIWLHSGEDCYHCRTEPLRP* |
| HG0034 | *D. rerio* | ENSDARG00000060991 | mief1 | mitochondrial elongation factor 1 | 0.00 | 0.12 | 0.0E+00 | 0.0E+00 | 0.12 | 0.0E+00 | Andreev, Mackowiak, Samandi | MGGWSRSAVLQLYRALLRAGQHLQYTDRDYYRRVVTREFRRCQTLSAPAEREDALKRGQFFLNSGLGDLV* |
| | *H. sapiens* | ENSG00000100335 | MIEF1 | mitochondrial elongation factor 1 | 0.00 | 0.14 | 0.0E+00 | 0.0E+00 | 0.14 | 0.0E+00 | Andreev, Mackowiak, Samandi | MAPWSREAVLSLYRALLRQGRQLRYTDRDFYFASIRREFRKNQKLEDAEARERQLEKGLVFLNGKLGRII* |
| HG0035 | *G. gallus* | ENSGALG00000006167 | LRRC8B | leucine rich repeat containing 8 family member B | 0.00 | 0.21 | 5.7E-305 | 4.7E-303 | 0.21 | 5.7E-305 | Mackowiak | MKTNVKMACPRNFKRCRPPD* |
| | *H. sapiens* | ENSG00000197147 | LRRC8B | leucine rich repeat containing 8 family member B | 0.00 | 0.20 | 0.0E+00 | 0.0E+00 | 0.20 | 0.0E+00 | Mackowiak | MKTNTEMVCLRNFKRCRPPD* |
| HG0036 | *G. gallus* | ENSGALG00000029295 | | | 0.00 | 0.34 | 2.2E-39 | 2.7E-38 | 0.34 | 2.2E-39 | | MIVGFHLLTCPFCCII* |
| | *H. sapiens* | ENSG00000155096 | AZIN1 | antizyme inhibitor 1 | 0.00 | 0.34 | 1.6E-44 | 7.9E-43 | 0.34 | 1.6E-44 | | MIGGFHLPTCPFCCIF* |
| HG0037 | *G. gallus* | ENSGALG00000010747 | FBLN5 | fibulin 5 | 0.00 | 0.06 | 2.4E-06 | 9.5E-06 | 0.06 | 2.4E-06 | | MKCSFLTFLLQLV* |
| | *H. sapiens* | ENSG00000140092 | FBLN5 | fibulin 5 | 0.00 | 0.06 | 1.9E-04 | 8.1E-04 | 0.06 | 1.9E-04 | | MKCSFLTFLLQLV* |
| HG0038 | *G. gallus* | ENSGALG00000006571 | | | 0.04 | 0.26 | 1.9E-221 | 9.8E-220 | 0.26 | 1.9E-221 | | MSVTCWPRLCGSSPPSERPVAALWTP* |
| HG0039.1 | *H. sapiens* | ENSG00000077092 | RARB | retinoic acid receptor beta | 0.02 | 0.39 | 1.2E-126 | 1.5E-124 | 0.39 | 1.2E-126 | Reynolds | MIIWINYRLLAGLSVIIHDSGLGKRPTAYVPKKGQSLMELGGLFYAICLHT* |
| HG0039.2 | *H. sapiens* | ENSG00000077092 | RARB | retinoic acid receptor beta | 0.04 | 0.29 | 2.2E-22 | 5.9E-21 | 0.29 | 2.2E-22 | Reynolds | MPRTRAIRAGFVWAPSG* |
| HG0040.1 | *H. sapiens* | ENSG00000164190 | NIPBL | NIPBL, cohesin loading factor | 0.00 | 0.00 | 1.8E-20 | 4.5E-19 | 0.00 | 1.8E-20 | | MGSNDSWHLN* |
| HG0040.2 | *H. sapiens* | ENSG00000164190 | NIPBL | NIPBL, cohesin loading factor | 0.00 | 0.40 | 3.3E-36 | 1.4E-34 | 0.40 | 3.3E-36 | Crowe | MFPALRRRKRSRSHPPSRPGL* |
| HG0041 | *D. rerio* | ENSDARG00000061352 | bcl11aa | B-cell CLL/lymphoma 11Aa | 0.00 | 0.14 | 2.1E-59 | 2.0E-58 | 0.14 | 2.1E-59 | | MTAIPQSIFPAPSLYYF* |
| | *G. gallus* | ENSGALG00000034048 | | | 0.00 | 0.14 | 4.0E-46 | 5.5E-45 | 0.14 | 4.0E-46 | | MTAIPQFIFPAPSLYYF* |
| | *H. sapiens* | ENSG00000119866 | BCL11A | B-cell CLL/lymphoma 11A | 0.00 | 0.14 | 3.0E-90 | 2.7E-88 | 0.14 | 3.0E-90 | | MTALPQFIFPAPSLYYF* |
| HG0042.1 | *H. sapiens* | ENSG00000140937 | CDH11 | cadherin 11 | 0.02 | 0.20 | 2.7E-227 | 5.4E-225 | 0.20 | 2.7E-227 | | MGGVCYRNWQQVSNG* |
| HG0042.2 | *G. gallus* | ENSGALG00000039985 | | | 0.02 | 0.37 | 1.4E-11 | 7.9E-11 | 0.37 | 1.4E-11 | | MHSRCGGSVCAEHFI* |
| HG0042.3 | *H. sapiens* | ENSG00000140937 | CDH11 | cadherin 11 | 0.00 | 0.00 | 1.3E-02 | 3.2E-02 | 0.00 | 1.3E-02 | | MPRGPLAAAADL* |
| HG0042.4 | *H. sapiens* | ENSG00000140937 | CDH11 | cadherin 11 | 0.00 | 0.29 | 2.1E-04 | 8.9E-04 | 0.29 | 2.1E-04 | | MSSTSRDIPSQERSA* |
| HG0042.5 | *H. sapiens* | ENSG00000140937 | CDH11 | cadherin 11 | 0.00 | 0.47 | 1.4E-03 | 4.9E-03 | 0.47 | 1.4E-03 | | MQKSLCFQLSGPVTFLRVVIC* |
| HG0043 | *H. sapiens* | ENSG00000172995 | ARPP21 | cAMP regulated phosphoprotein 21 | 0.00 | 0.17 | 3.1E-276 | 8.4E-274 | 0.17 | 2.4E-265 | | MLFNEGASASLSNKRSMREDRAVHGYGYWTRIFGKVDDVSHCVRESHGDGHSELLMGTWDSSCL* |
| HG0044 | *D. rerio* | ENSDARG00000061687 | atxn1a | ataxin 1a | 0.00 | 0.33 | 9.6E-100 | 1.2E-98 | 0.33 | 9.6E-100 | Crowe | MIPHELHGPLSRSRCDFNCLRLKSVSIRP* |
| | *H. sapiens* | ENSG00000124788 | ATXN1 | ataxin 1 | 0.00 | 0.36 | 2.1E-157 | 3.2E-155 | 0.36 | 2.1E-157 | Crowe | MIPHEEPGSPTEIKCDFPFIRLKSEPSRQ* |
| HG0045.1 | *H. sapiens* | ENSG00000083168 | KAT6A | lysine acetyltransferase 6A | 0.00 | 0.45 | 2.7E-102 | 2.8E-100 | 0.45 | 2.7E-102 | | MVNWFPASENLVFHGVKLTQHQDKG* |

| | | | | | | | | | | | | |
|---|---|---|---|---|---|---|---|---|---|---|---|---|
| HG0045.2 | *H. sapiens* | ENSG00000083168 | KAT6A | lysine acetyltransferase 6A | 0.04 | 0.37 | 7.0E-26 | 2.1E-24 | 0.37 | 7.0E-26 | | MAAVSVWGISAVRPGAPGSRLSFLPA* |
| HG0046 | *D. rerio* | ENSDARG00000018817 | bdnf | brain-derived neurotrophic factor | 0.01 | 0.18 | 4.1E-213 | 7.1E-212 | 0.18 | 4.1E-213 | | MMTSNKTHYLPASVGVCGY* |
| | *H. sapiens* | ENSG00000176697 | BDNF | brain derived neurotrophic factor | 0.00 | 0.18 | 2.0E-109 | 2.2E-107 | 0.18 | 2.0E-109 | | MMTSNKTHYLPASVGETRF* |
| HG0047 | *G. gallus* | ENSGALG00000015422 | NRG1 | neuregulin 1 | 0.00 | 0.34 | 1.3E-173 | 5.5E-172 | 0.34 | 1.3E-173 | | MLLLLSLLPLPLLLQILLLLLLPHDNCFPRLSRYQPQMLKVRVLPFALGYWFT* |
| HG0048 | *G. gallus* | ENSGALG00000039182 | | | 0.00 | 0.30 | 2.2E-39 | 2.7E-38 | 0.30 | 2.2E-39 | | MGCVLCCDLMV* |
| | *H. sapiens* | ENSG00000177565 | TBL1XR1 | transducin beta like 1 X-linked receptor 1 | 0.00 | 0.30 | 8.0E-29 | 2.6E-27 | 0.30 | 8.0E-29 | | MGYILCCDLMV* |
| HG0049 | *D. rerio* | ENSDARG00000044485 | sall4 | spalt-like transcription factor 4 | 0.05 | 0.31 | 2.6E-32 | 1.8E-31 | 0.31 | 2.6E-32 | Mackowiak | MINRNALLIIIITLTCAFLLEGLISPNLENF* |
| | *G. gallus* | ENSGALG00000039238 | | | 0.00 | 0.46 | 1.7E-100 | 5.3E-99 | 0.46 | 1.7E-100 | Mackowiak | MINRNALLIIIIIAGHMRGARRTGEAGGGRLDFRNFEKIFF* |
| HG0050.1 | *H. sapiens* | ENSG00000074054 | CLASP1 | cytoplasmic linker associated protein 1 | 0.00 | 0.45 | 2.7E-143 | 3.7E-141 | 0.45 | 2.7E-143 | | MVHCCHSPDCIFETQALSNLQRTKRQPPRYVCWEGVIVTAL* |
| HG0050.2 | *H. sapiens* | ENSG00000074054 | CLASP1 | cytoplasmic linker associated protein 1 | 0.00 | 0.45 | 1.9E-06 | 1.2E-05 | 0.45 | 1.9E-06 | | MKYFRPSSRTSLCHSLLYLLLVI* |
| HG0051 | *G. gallus* | ENSGALG00000037162 | | | 0.01 | 0.38 | 4.3E-59 | 7.8E-58 | 0.38 | 4.3E-59 | Crowe | MSCIIRAERQKIMMHDYHRRNSCL* |
| | *H. sapiens* | ENSG00000116679 | IVNS1ABP | influenza virus NS1A binding protein | 0.00 | 0.37 | 4.9E-63 | 3.3E-61 | 0.37 | 4.9E-63 | Crowe | MNVSISSEMNQIMMHHYHRRNSCL* |
| HG0052 | *H. sapiens* | ENSG00000178235 | SLITRK1 | SLIT and NTRK like family member 1 | 0.17 | 0.35 | 1.6E-124 | 2.0E-122 | 0.35 | 1.6E-124 | | MNLNCVCGEQDGRCYFVMRSGMNCSL* |
| HG0053.1 | *H. sapiens* | ENSG00000184564 | SLITRK6 | SLIT and NTRK like family member 6 | 0.00 | 0.41 | 3.9E-06 | 2.3E-05 | 0.41 | 3.9E-06 | | MVNYMQKSQHSCEVCREF* |
| HG0053.2 | *H. sapiens* | ENSG00000184564 | SLITRK6 | SLIT and NTRK like family member 6 | 0.00 | 0.31 | 8.6E-19 | 2.0E-17 | 0.31 | 8.6E-19 | | MLSILSLTRGFNFMKLWDLTKSDI* |
| HG0054 | *G. gallus* | ENSGALG00000016222 | NDP | NDP, norrin cystine knot growth factor | 0.19 | 0.23 | 1.2E-198 | 5.8E-197 | 0.23 | 1.2E-198 | | MTAKRLKPDFSLSWRGTGSQYLEFF* |
| | *H. sapiens* | ENSG00000124479 | NDP | NDP, norrin cystine knot growth factor | 0.00 | 0.29 | 2.2E-38 | 9.9E-37 | 0.29 | 2.2E-38 | | MTTRKLQPDPALSLNGTGS* |
| HG0055.1 | *G. gallus* | ENSGALG00000010533 | ELAVL4 | ELAV like RNA binding protein 4 | 0.00 | 0.06 | 3.0E-138 | 1.2E-136 | 0.06 | 3.0E-138 | | MLHIVFQRVSCHDLIKRGGGCVCCSA* |
| HG0055.2 | *G. gallus* | ENSGALG00000010533 | ELAVL4 | ELAV like RNA binding protein 4 | 0.00 | 0.00 | 1.3E-33 | 1.4E-32 | 0.00 | 1.3E-33 | | MSILLHRWAAAL* |
| HG0056 | *G. gallus* | ENSGALG00000011271 | LUM | lumican | 0.01 | 0.32 | 1.6E-51 | 2.5E-50 | 0.32 | 1.6E-51 | | MGCPCTECGIHLHFIKRAAFSYMPHHKITMMT* |
| | *H. sapiens* | ENSG00000139329 | LUM | lumican | 0.01 | 0.33 | 2.2E-15 | 4.0E-14 | 0.36 | 2.5E-09 | | MSWAVLSPHVHLHFVREQCSHATPQDPHNDITPFRDWRDWAGSPHPPSALVSLRIWQPVPS* |
| HG0057 | *H. sapiens* | ENSG00000280987 | MATR3 | matrin 3 | 0.00 | 0.42 | 5.6E-41 | 2.7E-39 | 0.42 | 1.1E-33 | | MLGAQWRRNQPSRAAESCLLVLSLKKILQSLLILFLGVTIFEAKLT* |
| HG0058 | *H. sapiens* | ENSG00000180332 | KCTD4 | potassium channel tetramerization domain containing 4 | 0.01 | 0.21 | 1.3E-146 | 1.8E-144 | 0.21 | 1.3E-146 | | MLPAFPFHHQAGAE* |
| HG0059.1 | *D. rerio* | ENSDARG00000018060 | pik3r2 | phosphoinositide-3-kinase, regulatory subunit 2 (beta) | 0.01 | 0.17 | 1.1E-275 | 2.3E-274 | 0.17 | 3.9E-270 | | MLHSLCSIIWLNRGFKLRKVACTANTGQV* |
| | *G. gallus* | ENSGALG00000003428 | | | 0.01 | 0.17 | 1.4E-168 | 5.6E-167 | 0.17 | 1.4E-168 | | MPTHECWIMFVSFSLCSIIWLNRGFKLRKRGCT* |
| HG0059.2 | *H. sapiens* | ENSG00000145675 | PIK3R1 | phosphoinositide-3-kinase regulatory subunit 1 | 0.00 | 0.48 | 1.3E-11 | 1.7E-10 | 0.48 | 1.3E-11 | | MDSVTKVSERIGPRCESQPGFKVLTSC* |
| HG0059.3 | *H. sapiens* | ENSG00000105647 | PIK3R2 | phosphoinositide-3-kinase regulatory subunit 2 | 0.13 | 0.50 | 2.5E-07 | 1.9E-06 | 0.50 | 2.5E-07 | | MVAACTLGSLVWCSQQSRQPWAPMALRVRA* |
| HG0059.4 | *H. sapiens* | ENSG00000145675 | PIK3R1 | phosphoinositide-3-kinase regulatory subunit 1 | 0.00 | 0.00 | 2.0E-04 | 8.6E-04 | 0.00 | 2.0E-04 | | MDSRMASHLSS* |
| HG0059.5 | *D. rerio* | ENSDARG00000038524 | pik3r1 | phosphoinositide-3-kinase, regulatory subunit 1 (alpha) | 0.18 | 0.26 | 6.7E-06 | 2.3E-05 | 0.26 | 6.7E-06 | | MGRVCMKHLEGAPNHWTISFT* |
| HG0060.1 | *G. gallus* | ENSGALG00000014751 | | | 0.00 | 0.48 | 1.1E-19 | 8.7E-19 | 0.48 | 1.1E-19 | | MTADIEESCWKRVPVKKWNIQ* |
| | *H. sapiens* | ENSG00000049192 | ADAMTS6 | ADAM metallopeptidase with thrombospondin type 1 motif [illegible] | 0.00 | 0.47 | 9.0E-55 | 5.3E-53 | 0.47 | 9.0E-55 | | MNPDIEESAGCWIKVALKQWNIQ* |
| HG0060.2 | *G. gallus* | ENSGALG00000014751 | | | 0.00 | 0.35 | 1.6E-16 | 1.1E-15 | 0.35 | 1.6E-16 | | MVQFWSGVNPIAHR* |
| | *H. sapiens* | ENSG00000049192 | ADAMTS6 | ADAM metallopeptidase with thrombospondin type 1 motif [illegible] | 0.00 | 0.36 | 4.0E-29 | 1.3E-27 | 0.36 | 4.0E-29 | | MLRFWYGIDSLAHR* |
| HG0061.1 | *G. gallus* | ENSGALG00000004949 | | | 0.00 | 0.27 | 4.3E-12 | 2.5E-11 | 0.27 | 4.3E-12 | | MPFVKPMYTVLHCYCT* |
| | *H. sapiens* | ENSG00000166398 | | | 0.02 | 0.27 | 2.2E-51 | 1.3E-49 | 0.27 | 2.2E-51 | | MVTCLMEMPFLKLRFAELHYFYT* |
| HG0061.2 | *G. gallus* | ENSGALG00000004949 | | | 0.00 | 0.34 | 3.2E-06 | 1.2E-05 | 0.34 | 3.2E-06 | | MLQRKQLPQQTRKGEEGLQ* |
| HG0061.3 | *H. sapiens* | ENSG00000166398 | | | 0.03 | 0.30 | 9.8E-10 | 1.0E-08 | 0.30 | 9.8E-10 | | MLQGNRYLSKQKEWRRRLTTNAI* |
| HG0062.1 | *D. rerio* | ENSDARG00000075569 | gpbp1l1 | GC-rich promoter binding protein 1-like 1 | 0.00 | 0.13 | 9.7E-04 | 2.6E-03 | 0.13 | 9.7E-04 | | MPPSTDGRHWT* |
| | *G. gallus* | ENSGALG00000010276 | GPBP1L1 | GC-rich promoter binding protein 1 like 1 | 0.00 | 0.12 | 1.8E-02 | 3.7E-02 | 0.12 | 1.8E-02 | | MPHFIEGRHWT* |
| HG0062.2 | *G. gallus* | ENSGALG00000010276 | GPBP1L1 | GC-rich promoter binding protein 1 like 1 | 0.00 | 0.38 | 5.3E-08 | 2.3E-07 | 0.38 | 5.3E-08 | | MSCPADLGVSVQY* |
| HG0062.3 | *H. sapiens* | ENSG00000159592 | GPBP1L1 | GC-rich promoter binding protein 1 like 1 | 0.00 | 0.29 | 3.0E-14 | 5.1E-13 | 0.29 | 3.0E-14 | | MSYLIKQHHSAITQGS* |
| HG0063.1 | *G. gallus* | ENSGALG00000002477 | | | 0.00 | 0.12 | 5.1E-09 | 2.4E-08 | 0.12 | 5.1E-09 | | MCACSIASFLLK* |
| | *H. sapiens* | ENSG00000162630 | B3GALT2 | beta-1,3-galactosyltransferase 2 | 0.00 | 0.12 | 5.9E-09 | 5.5E-08 | 0.12 | 5.9E-09 | | MCACSIASFLLK* |
| HG0063.2 | *H. sapiens* | ENSG00000162630 | B3GALT2 | beta-1,3-galactosyltransferase 2 | 0.00 | 0.41 | 1.9E-11 | 2.4E-10 | 0.41 | 1.9E-11 | | MDTETRTQHCIVQPGK* |
| HG0064.1 | *H. sapiens* | ENSG00000102678 | FGF9 | fibroblast growth factor 9 | 0.00 | 0.33 | 2.1E-133 | 2.7E-131 | 0.33 | 2.1E-133 | | MVILISNQMEKLRIFFPYYGRMG* |
| HG0064.2 | *H. sapiens* | ENSG00000113578 | FGF1 | fibroblast growth factor 1 | 0.01 | 0.38 | 1.2E-14 | 2.1E-13 | 0.38 | 1.2E-14 | | MVCGFTQFFSWGFSELHSYQKGSRRERTSGSNSLRSRGIRASPPFLGGHSLLMVNGNSLPPAAACLQLSW* |
| HG0065 | *G. gallus* | ENSGALG00000029927 | STRBP | Spermatid perinuclear RNA-binding protein | 0.00 | 0.36 | 1.6E-85 | 4.3E-84 | 0.36 | 1.6E-85 | | MKRFCQAKQLLSKTPRRREKEKERES* |
| HG0066 | *H. sapiens* | ENSG00000112182 | BACH2 | BTB domain and CNC homolog 2 | 0.22 | 0.18 | 5.0E-206 | 8.9E-204 | 0.18 | 5.0E-206 | | MDCQSLLISPFAQFPAS* |
| HG0067 | *H. sapiens* | ENSG00000166862 | CACNG2 | calcium voltage-gated channel auxiliary subunit gamma 2 | 0.00 | 0.01 | 2.6E-80 | 2.1E-78 | 0.01 | 2.6E-80 | | METGDQNFRKRD* |
| HG0068 | *D. rerio* | ENSDARG00000071235 | | | 0.02 | 0.00 | 3.2E-256 | 6.1E-255 | 0.00 | 3.2E-256 | Mackowiak | MRFFRLTFKCFVDCF* |
| | *G. gallus* | ENSGALG00000041419 | | | 0.03 | 0.00 | 1.7E-293 | 1.3E-291 | 0.00 | 1.7E-293 | Mackowiak | MRFFRLTFKCFVDCF* |
| | *H. sapiens* | ENSG00000130821 | SLC6A8 | solute carrier family 6 member 8 | 0.02 | 0.00 | 2.0E-232 | 4.3E-230 | 0.00 | 2.0E-232 | Mackowiak | MRFFRLTFKCFVDCF* |
| HG0069 | *G. gallus* | ENSGALG00000010999 | DICER1 | dicer 1, ribonuclease III | 0.00 | 0.42 | 2.9E-40 | 3.7E-39 | 0.42 | 2.9E-40 | | MMSKCSLESISTEAAEIKT* |
| | *H. sapiens* | ENSG00000100697 | DICER1 | dicer 1, ribonuclease III | 0.03 | 0.27 | 1.1E-78 | 8.0E-77 | 0.27 | 1.1E-78 | | MSKCSSDKSNTEISNIKT* |
| HG0070 | *D. rerio* | ENSDARG00000040926 | nr2f2 | nuclear receptor subfamily 2, group F, member 2 | 0.00 | 0.36 | 4.4E-41 | 3.7E-40 | 0.36 | 4.4E-41 | | MDDCKCPWTWPLRARLLFGLLQF* |
| | | ENSDARG00000052695 | nr2f1a | nuclear receptor subfamily 2, group F, member 1a | 0.03 | 0.13 | 1.7E-126 | 2.3E-125 | 0.13 | 1.7E-126 | | MADCNLPWNWPPDNCIS* |
| | *H. sapiens* | ENSG00000175745 | NR2F1 | nuclear receptor subfamily 2 group F member 1 | 0.03 | 0.12 | 6.3E-162 | 9.8E-160 | 0.12 | 6.3E-162 | | MAGCVRPWTWPPDTRAP* |
| | | ENSG00000185551 | NR2F2 | nuclear receptor subfamily 2 group F member 2 | 0.20 | 0.07 | 1.5E-119 | 1.8E-117 | 0.07 | 1.5E-119 | | MADCGLPWTWPPDSRLSSSSTSSFTTTSSSSSSSSSSSANSSAAHQL* |
| HG0071.1 | *D. rerio* | ENSDARG00000098240 | meis2a | Meis homeobox 2a | 0.00 | 0.39 | 2.0E-42 | 1.7E-41 | 0.39 | 2.0E-42 | | MRHDSLVTCRLLMGRL* |
| | *H. sapiens* | ENSG00000134138 | MEIS2 | Meis homeobox 2 | 0.00 | 0.39 | 1.1E-68 | 8.0E-67 | 0.39 | 1.1E-68 | | MKHDSSVTCGLLTGRS* |
| HG0071.2 | *H. sapiens* | ENSG00000143995 | MEIS1 | Meis homeobox 1 | 0.00 | 0.00 | 1.2E-08 | 1.1E-07 | 0.00 | 1.2E-08 | | MLSDSGALLQVP* |
| HG0072 | *G. gallus* | ENSGALG00000039690 | STMN2 | Stathmin-2 | 0.11 | 0.34 | 2.7E-35 | 3.0E-34 | 0.34 | 2.7E-35 | | MPGDSGSAGVMSGDL* |
| | *H. sapiens* | ENSG00000104435 | STMN2 | stathmin 2 | 0.12 | 0.40 | 1.7E-101 | 1.7E-99 | 0.40 | 1.7E-101 | | MGGADREGLISISNCILVLCTPAPLADQYLMLGDSDSAGVMSGDLGSQSA* |
| HG0073 | *D. rerio* | ENSDARG00000041708 | csrp2bp | csrp2 binding protein | 0.02 | 0.13 | 2.3E-203 | 3.7E-202 | 0.13 | 2.3E-203 | | MSTASKVVLGLSIVLTISTVAGVHIKQNWDRQRLREGVVRDLERLERKRENLRALEEQIQLTRELVTERNRRQAEASETHDS* |
| HG0074 | *D. rerio* | ENSDARG00000078624 | arhgef9b | Cdc42 guanine nucleotide exchange factor (GEF) 9b | 0.06 | 0.06 | 1.9E-10 | 8.0E-10 | 0.06 | 1.9E-10 | Mackowiak | MTLDVQTVAVFTVIAILLLVNVILIFFMGSS* |
| | *H. sapiens* | ENSG00000131089 | ARHGEF9 | Cdc42 guanine nucleotide exchange factor 9 | 0.01 | 0.04 | 8.1E-218 | 1.5E-215 | 0.04 | 8.1E-218 | Mackowiak | MDSLTEQRLTSPNLPAPHLEHYSVLHCTMTLDVQTVVVFAVIVVLLLVNVILMFFLGTR* |
| HG0075 | *H. sapiens* | ENSG00000165699 | TSC1 | tuberous sclerosis 1 | 0.00 | 0.45 | 1.6E-49 | 8.8E-48 | 0.45 | 1.6E-49 | Crowe | MKDTRLTALETEVPVVARTVW* |
| HG0076 | *D. rerio* | ENSDARG00000040008 | neurod6a | neuronal differentiation 6a | 0.01 | 0.30 | 5.9E-120 | 8.1E-119 | 0.50 | 7.1E-15 | | MKTTWTHCLLTGFDGRTTVKNTKANSTPGAVMMDAPLWTGHDGQQAALNRHAPGRVVRSSRHMPDDQGRNTGPLRASHVTSPCHVTSAADGMGTSSGSWHQCHFS* |
| | *H. sapiens* | ENSG00000164600 | NEUROD6 | neuronal differentiation 6 | 0.00 | 0.47 | 5.9E-31 | 2.1E-29 | 0.47 | 5.7E-09 | | MGTSSGSWHQCHFLEINEIVQIYTDLETPGDDATLSLKRFGRPKMKTDY* |
| HG0077.1 | *G. gallus* | ENSGALG00000005074 | | | 0.00 | 0.48 | 7.0E-33 | 7.1E-32 | 0.48 | 7.0E-33 | | MIPQPDPTTKEKKIVHLRLGRHCN* |
| HG0077.2 | *G. gallus* | ENSGALG00000005074 | | | 0.00 | 0.08 | 1.3E-10 | 6.8E-10 | 0.08 | 1.3E-10 | | MPSGGVLLKLGHCNL* |
| HG0078.1 | *G. gallus* | ENSGALG00000016633 | HMBOX1 | homeobox containing 1 | 0.00 | 0.01 | 6.7E-05 | 2.2E-04 | 0.01 | 6.7E-05 | | MVNNTEDPQEAL* |
| HG0078.2 | *H. sapiens* | ENSG00000147421 | HMBOX1 | homeobox containing 1 | 0.00 | 0.37 | 1.4E-02 | 3.4E-02 | 0.37 | 1.4E-02 | | MVDNADHLWKGY* |
| HG0079 | *H. sapiens* | ENSG00000204406 | MBD5 | methyl-CpG binding domain protein 5 | 0.00 | 0.30 | 4.9E-94 | 4.8E-92 | 0.31 | 2.0E-89 | | MSTIFWREKRYSLIGTRKDIEHLEINCLHYTAESHTTLVMFSSLLSRRLHYWNFEVMKTSDEPTCNTLDSDWKIKRKFKECGL* |
| HG0080.1 | *G. gallus* | ENSGALG00000039403 | | | 0.01 | 0.29 | 4.2E-68 | 9.5E-67 | 0.29 | 4.2E-68 | | MVLYHLLGTDGSLQKVQFQMVNLNL* |
| | *H. sapiens* | ENSG00000179603 | GRM8 | glutamate metabotropic receptor 8 | 0.02 | 0.29 | 1.0E-86 | 8.8E-85 | 0.29 | 1.0E-86 | | MGPDGSLQKVQPQVVPFLLWQE* |
| HG0080.2 | *H. sapiens* | ENSG00000179603 | GRM8 | glutamate metabotropic receptor 8 | 0.00 | 0.09 | 8.7E-03 | 2.3E-02 | 0.09 | 8.7E-03 | | MTRNNSATRLISRT* |
| HG0080.3 | *H. sapiens* | ENSG00000179603 | GRM8 | glutamate metabotropic receptor 8 | 0.09 | 0.34 | 1.0E-06 | 6.9E-06 | 0.34 | 1.0E-06 | | MEETWRLLGQLSPLSPEISHKCGRCARLTRRRRRRRRR* |
| HG0081.1 | *D. rerio* | ENSDARG00000028228 | zbtb18 | zinc finger and BTB domain containing 18 | 0.06 | 0.41 | 7.1E-19 | 3.9E-18 | 0.41 | 7.1E-19 | | MMSEQLTGCHPPVFSSNAQTGS* |
| HG0081.2 | *D. rerio* | ENSDARG00000028228 | zbtb18 | zinc finger and BTB domain containing 18 | 0.08 | 0.35 | 1.2E-02 | 2.2E-02 | 0.35 | 1.2E-02 | | MIVPIVSYFFFPPPLSVVQRVSKMNDGNGAASEP* |
| HG0082 | *D. rerio* | ENSDARG00000062379 | slc35a4 | solute carrier family 35, member A4 | 0.01 | 0.07 | 5.5E-233 | 9.9E-232 | 0.07 | 5.5E-233 | Andreev | MAEDKDPVTKLKDLVHLKDQLEEIQKKVENEVQAGVPQGGSLLASPFLKGFLAGYVVSRLRSSAVLGVALGTLSGIFAAQNYQVPNIEATLRDILNSLKKRPGE* |
| | *H. sapiens* | ENSG00000176087 | SLC35A4 | solute carrier family 35 member A4 | 0.01 | 0.07 | 2.7E-235 | 6.1E-233 | 0.07 | 2.7E-235 | Andreev | MADDKDSLPKLKDLAFLKNQLESLQRRVEDEVNSGVGQDGSLLSSPFLKGFLAGYVVAKLRASAVLGFAVGTCTGIYAAQAYAVPNVEKTLRDYLQLLRKGPD* |
| HG0083.1 | *G. gallus* | ENSGALG00000008039 | MFSD13A | major facilitator superfamily domain containing 13A | 0.00 | 0.31 | 1.3E-53 | 2.2E-52 | 0.31 | 1.3E-53 | | MDWPRQRCNRCCEKTSHLKWR* |
| HG0083.2 | *H. sapiens* | ENSG00000138111 | MFSD13A | major facilitator superfamily domain containing 13A | 0.07 | 0.49 | 1.1E-03 | 3.9E-03 | 0.49 | 1.1E-03 | | MGRPPVAMSPPRCSWG* |
| HG0084.1 | *G. gallus* | ENSGALG00000009207 | TRIM2 | tripartite motif containing 2 | 0.00 | 0.50 | 3.2E-35 | 3.6E-34 | 0.50 | 3.2E-35 | | MMFYSFGLLRNCSSIPARILSSALH* |
| HG0084.2 | *H. sapiens* | ENSG00000154447 | SH3RF1 | SH3 domain containing ring finger 1 | 0.00 | 0.12 | 4.4E-03 | 1.3E-02 | 0.12 | 4.4E-03 | | MSHFKGCTQL* |
| HG0085 | *G. gallus* | ENSGALG00000002003 | BMPR1A | bone morphogenetic protein receptor type 1A | 0.00 | 0.10 | 5.8E-17 | 4.2E-16 | 0.10 | 5.8E-17 | | MRFHNTTFELMKWKYR* |
| | *H. sapiens* | ENSG00000107779 | BMPR1A | bone morphogenetic protein receptor type 1A | 0.00 | 0.28 | 4.3E-74 | 3.1E-72 | 0.28 | 4.3E-74 | | MHQFNTVLEFMRWKHRSKLFGENQKYSFI* |
| HG0086 | *G. gallus* | ENSGALG00000009835 | XPOT | exportin for tRNA | 0.00 | 0.41 | 1.7E-90 | 4.8E-89 | 0.41 | 1.7E-90 | | MLVATSELWGGSRHRRDGGVTGKMIVISTLEEVGKPVSRSRKRRV* |
| HG0087 | *H. sapiens* | ENSG00000164548 | TRA2A | transformer 2 alpha homolog | 0.00 | 0.06 | 5.2E-81 | 4.2E-79 | 0.06 | 5.2E-81 | | MWKINQDKRRLMKSIKDQKIKIEWGKKLNRRLVR* |
| HG0088 | *H. sapiens* | ENSG00000273079 | GRIN2B | glutamate ionotropic receptor NMDA type subunit 2B | 0.02 | 0.35 | 1.5E-30 | 5.2E-29 | 0.35 | 1.5E-30 | | MQARRRDWTFPTCSLP* |
| HG0089.1 | *G. gallus* | ENSGALG00000002294 | SEMA6A | semaphorin 6A | 0.00 | 0.36 | 2.0E-69 | 4.8E-68 | 0.36 | 2.0E-69 | | MTKDCKGESGSSFPERSP* |
| | *H. sapiens* | ENSG00000092421 | SEMA6A | semaphorin 6A | 0.00 | 0.40 | 3.1E-68 | 2.1E-66 | 0.40 | 3.1E-68 | | MTKACEGESAARGPERSPR* |
| HG0089.2 | *H. sapiens* | ENSG00000092421 | SEMA6A | semaphorin 6A | 0.00 | 0.00 | 6.4E-18 | 1.4E-16 | 0.00 | 6.4E-18 | | MNTSRLHPRK* |
| HG0089.3 | *G. gallus* | ENSGALG00000002294 | SEMA6A | semaphorin 6A | 0.00 | 0.00 | 1.1E-14 | 7.5E-14 | 0.00 | 1.1E-14 | | MNKLHLQFRK* |
| HG0090.1 | *G. gallus* | ENSGALG00000008487 | TMCC1 | transmembrane and coiled-coil domain family 1 | 0.00 | 0.02 | 1.3E-53 | 2.3E-52 | 0.02 | 1.3E-53 | | MEISMRSPWHSCMCSHS* |
| | *H. sapiens* | ENSG00000172765 | TMCC1 | transmembrane and coiled-coil domain family 1 | 0.00 | 0.35 | 1.1E-02 | 2.9E-02 | 0.35 | 1.1E-02 | | MKIGMKPSSHSCMRSHC* |
| HG0090.2 | *D. rerio* | ENSDARG00000060954 | tmcc1b | transmembrane and coiled-coil domain family 1b | 0.00 | 0.34 | 1.2E-37 | 9.5E-37 | 0.34 | 1.2E-37 | | MTSMTLRQTTDACTLSEST* |
| HG0090.3 | *G. gallus* | ENSGALG00000008487 | TMCC1 | transmembrane and coiled-coil domain family 1 | 0.09 | 0.33 | 2.0E-08 | 9.2E-08 | 0.33 | 2.0E-08 | | MLSFVQWRDCCMSAMLKGK* |
| HG0091.1 | *H. sapiens* | ENSG00000172209 | GPR22 | G protein-coupled receptor 22 | 0.00 | 0.37 | 3.4E-42 | 1.7E-40 | 0.37 | 3.4E-42 | | MAHYRHIFSLRSFSKRCL* |
| HG0091.2 | *H. sapiens* | ENSG00000172209 | GPR22 | G protein-coupled receptor 22 | 0.00 | 0.47 | 2.3E-12 | 3.3E-11 | 0.47 | 2.3E-12 | | MSKYILEATNVMFPFLFSHILENKEILSCTPSKGNIIPII* |
| HG0091.3 | *H. sapiens* | ENSG00000172209 | GPR22 | G protein-coupled receptor 22 | 0.00 | 0.35 | 4.9E-11 | 6.0E-10 | 0.35 | 4.9E-11 | | MYPLLMKTNKIKQQMVFIKRTWPGFII* |
| HG0092.1 | *G. gallus* | ENSGALG00000009966 | FRS2 | fibroblast growth factor receptor substrate 2 | 0.00 | 0.11 | 8.3E-65 | 1.8E-63 | 0.11 | 8.3E-65 | | MISSVTRQHDTLKYDRSPDAATEAGNINGQI* |

| | | | | | | | | | | | | |
|---|---|---|---|---|---|---|---|---|---|---|---|---|
| HG0092.2 | *H. sapiens* | ENSG00000166225 | FRS2 | fibroblast growth factor receptor substrate 2 | 0.01 | 0.36 | 6.4E-07 | 4.4E-06 | 0.36 | 6.4E-07 | | MKQSICHKVRCSCGMSTSLEQNCICFSQKRIPQG* |
| HG0093 | *G. gallus* | ENSGALG00000040465 | Sip1 | zinc finger E-box binding homeobox 2 | 0.13 | 0.26 | 1.7E-35 | 2.0E-34 | 0.26 | 1.7E-35 | Crowe, Samandi | MLFLLTISGQNYKQDFAESACC* |
| | *H. sapiens* | ENSG00000169554 | ZEB2 | zinc finger E-box binding homeobox 2 | 0.22 | 0.44 | 2.7E-40 | 1.3E-38 | 0.44 | 2.7E-40 | Crowe, Samandi | MLFLLTISGQNYKQDFAAPCLN* |
| HG0094.1 | *H. sapiens* | ENSG00000112175 | BMP5 | bone morphogenetic protein 5 | 0.00 | 0.31 | 2.7E-60 | 1.7E-58 | 0.31 | 2.7E-60 | | MQQKLLTTVDNWKSEFQLS* |
| HG0094.2 | *H. sapiens* | ENSG00000112175 | BMP5 | bone morphogenetic protein 5 | 0.00 | 0.15 | 8.4E-26 | 2.6E-24 | 0.15 | 8.4E-26 | | MDLQEGFQVNSGKHIYLNSTT* |
| HG0095 | *D. rerio* | ENSDARG00000013708 | usp9 | ubiquitin specific peptidase 9 | 0.00 | 0.46 | 9.7E-57 | 8.9E-56 | 0.46 | 9.7E-57 | | MEERSSLILGYAMVSATYPVPQVVLSLSEVCRFSVAPEEKLISSGVPAT* |
| HG0096 | *G. gallus* | ENSGALG00000014186 | MPPED1 | metallophosphoesterase domain containing 1 | 0.01 | 0.42 | 6.1E-03 | 1.4E-02 | 0.42 | 6.1E-03 | | MVFSCRCRKIADMIA* |
| HG0097 | *G. gallus* | ENSGALG00000037496 | | | 0.01 | 0.25 | 1.6E-37 | 1.8E-36 | 0.25 | 1.6E-37 | | MELQPRAGAADTPSAFLRSSHHERGRFHLVKMAVWDRCNL* |
| HG0098 | *H. sapiens* | ENSG00000198739 | LRRTM3 | leucine rich repeat transmembrane neuronal 3 | 0.00 | 0.00 | 1.2E-06 | 8.1E-06 | 0.00 | 1.2E-06 | | MQKRNIHEDPIT* |
| HG0099.1 | *H. sapiens* | ENSG00000168575 | SLC20A2 | solute carrier family 20 member 2 | 0.00 | 0.01 | 5.3E-105 | 5.6E-103 | 0.01 | 5.3E-105 | | MRKTNGSEKVYRIIGGDITAFPDREEKYGMCFTAD* |
| HG0099.2 | *H. sapiens* | ENSG00000144136 | SLC20A1 | solute carrier family 20 member 1 | 0.00 | 0.28 | 3.1E-03 | 9.5E-03 | 0.28 | 3.1E-03 | | MNLRPFSSPPWNSAPCF* |
| HG0099.3 | *H. sapiens* | ENSG00000168575 | SLC20A2 | solute carrier family 20 member 2 | 0.00 | 0.45 | 1.0E-02 | 2.6E-02 | 0.45 | 1.0E-02 | | MMLCIVHQGTPLGRISH* |
| HG0099.4 | *D. rerio* | ENSDARG00000010641 | slc20a1b | solute carrier family 20 (phosphate transporter), member 1b | 0.00 | 0.36 | 5.5E-07 | 2.1E-06 | 0.36 | 5.5E-07 | | MDVRCICLIVQAKTNRNRSSLFRG* |
| HG0100.1 | *D. rerio* | ENSDARG00000104148 | crebbpb | CREB binding protein b | 0.00 | 0.48 | 5.8E-40 | 4.7E-39 | 0.48 | 5.8E-40 | | MRVLGVGLSWGEWRWRSGALLSRV* |
| | | ENSDARG00000104609 | crebbpa | CREB binding protein a | 0.00 | 0.50 | 2.6E-33 | 1.8E-32 | 0.50 | 2.6E-33 | | MIMWVLGVGLSWGEWWWRRGALLLLF* |
| HG0100.2 | *D. rerio* | ENSDARG00000061108 | ep300b | E1A binding protein p300 b | 0.00 | 0.47 | 2.3E-13 | 1.0E-12 | 0.47 | 2.3E-13 | | MAEKTDASSEDKSRENCSVAEEAT* |
| HG0101 | *H. sapiens* | ENSG00000099250 | NRP1 | neuropilin 1 | 0.00 | 0.45 | 2.0E-19 | 4.7E-18 | 0.45 | 2.0E-19 | | MARAVAPGRGTSVGKGRGSRS* |
| HG0102 | *H. sapiens* | ENSG00000182263 | FIGN | fidgetin, microtubule severing factor | 0.00 | 0.45 | 6.5E-32 | 2.4E-30 | 0.45 | 6.5E-32 | | MISNHRHLHLTLASIHQQTHRIDATLTYRVLSPRVQVK* |
| HG0103 | *G. gallus* | ENSGALG00000016558 | VEGFD | vascular endothelial growth factor D | 0.00 | 0.21 | 2.2E-93 | 6.6E-92 | 0.21 | 2.2E-93 | | MHFKRDASSCMTTTA* |
| | *H. sapiens* | ENSG00000165197 | VEGFD | vascular endothelial growth factor D | 0.00 | 0.48 | 1.7E-02 | 4.1E-02 | 0.48 | 1.7E-02 | | MPFETIFSSCLMSTA* |
| HG0104.1 | *G. gallus* | ENSGALG00000038097 | EI24 | EI24, autophagy associated transmembrane protein | 0.00 | 0.32 | 3.0E-25 | 2.6E-24 | 0.32 | 3.0E-25 | | MMREAWVRLFPSPCS* |
| HG0104.2 | *H. sapiens* | ENSG00000149547 | EI24 | EI24, autophagy associated transmembrane protein | 0.02 | 0.47 | 5.5E-05 | 2.7E-04 | 0.47 | 5.5E-05 | | MLVLKQKSSLQSLSKNKSRPRSFMMRDLGTLLSPVCS* |
| HG0105 | *G. gallus* | ENSGALG00000016176 | | | 0.00 | 0.33 | 6.9E-110 | 2.4E-108 | 0.35 | 2.0E-107 | | MWLDLADPEPELLFFRGAVGVREACGGYLAAFFSLLLPFFFFFFFFLVGFEPEPGKMGNGASDRRGISPSLVSGRVPVVLEV* |
| | *H. sapiens* | ENSG00000135298 | ADGRB3 | adhesion G protein-coupled receptor B3 | 0.00 | 0.33 | 1.1E-87 | 9.3E-86 | 0.33 | 1.1E-87 | | MGNGDWDRRGVSPSLLLGRVPVLVS* |
| HG0106 | *H. sapiens* | ENSG00000147548 | NSD3 | nuclear receptor binding SET domain protein 3 | 0.00 | 0.11 | 1.8E-29 | 6.2E-28 | 0.11 | 1.8E-29 | | MEQTEGPMRERES* |
| HG0107.1 | *G. gallus* | ENSGALG00000003601 | ATP2B4 | ATPase plasma membrane Ca2+ transporting 4 | 0.00 | 0.22 | 4.5E-81 | 1.1E-79 | 0.22 | 4.5E-81 | | MHILFTPSSGGLGLG* |
| | *H. sapiens* | ENSG00000058668 | ATP2B4 | ATPase plasma membrane Ca2+ transporting 4 | 0.00 | 0.20 | 1.7E-80 | 1.4E-78 | 0.20 | 1.7E-80 | | MHILFTPSSGQVGLS* |
| HG0107.2 | *G. gallus* | ENSGALG00000030550 | | | 0.00 | 0.30 | 6.0E-44 | 7.8E-43 | 0.30 | 6.0E-44 | | MGRPQRYLSKMGLLLG* |
| | *H. sapiens* | ENSG00000070961 | ATP2B1 | ATPase plasma membrane Ca2+ transporting 1 | 0.00 | 0.31 | 1.2E-12 | 1.7E-11 | 0.31 | 1.2E-12 | | MIDMEKLVMGQRSRYFSGKCCC* |
| HG0107.3 | *D. rerio* | ENSDARG00000063433 | atp2b2 | ATPase, Ca++ transporting, plasma membrane 2 | 0.01 | 0.28 | 3.5E-23 | 2.1E-22 | 0.28 | 3.5E-23 | Crowe | MGNHVWSDAGLQPW* |
| | *H. sapiens* | ENSG00000157087 | ATP2B2 | ATPase plasma membrane Ca2+ transporting 2 | 0.00 | 0.41 | 9.9E-07 | 6.6E-06 | 0.41 | 9.9E-07 | Crowe | MPVPGSSMGPTEWRLLGLQP* |
| HG0107.4 | *H. sapiens* | ENSG00000070961 | ATP2B1 | ATPase plasma membrane Ca2+ transporting 1 | 0.00 | 0.00 | 6.5E-03 | 1.8E-02 | 0.00 | 6.5E-03 | | MSVWFKKDFHT* |
| HG0108.1 | *G. gallus* | ENSGALG00000007396 | TAOK3 | TAO kinase 3 | 0.00 | 0.29 | 5.8E-53 | 9.5E-52 | 0.29 | 5.8E-53 | | MKEETDWDLKAPTSS* |
| HG0108.2 | *H. sapiens* | ENSG00000135090 | TAOK3 | TAO kinase 3 | 0.01 | 0.32 | 8.1E-10 | 8.7E-09 | 0.32 | 8.1E-10 | | MNLSNMEYFVPHTKRY* |
| HG0108.3 | *D. rerio* | ENSDARG00000098304 | taok1b | TAO kinase 1b | 0.07 | 0.23 | 1.6E-02 | 2.9E-02 | 0.23 | 1.6E-02 | | MSNQRGLTCRSRTAR* |
| HG0109.1 | *H. sapiens* | ENSG00000048540 | LMO3 | LIM domain only 3 | 0.00 | 0.34 | 1.3E-34 | 5.1E-33 | 0.34 | 1.3E-34 | | MNGSHRQTSDCSHLQAFLKGEGCVLFPGYSGEALQSSHCAY* |
| HG0109.2 | *H. sapiens* | ENSG00000048540 | LMO3 | LIM domain only 3 | 0.17 | 0.41 | 1.9E-23 | 5.3E-22 | 0.41 | 1.9E-23 | | MQHIFYGDYSDCCYEIQVRFMIFFILNMICYANAACDSLIKSICYQEAKRGRWDFRITGRVREACGFVIRHHFAHCEGLRVCALSCLAVLIPSPTPCRHSDPLPKWLHNSRSSAV* |
| HG0109.3 | *H. sapiens* | ENSG00000048540 | LMO3 | LIM domain only 3 | 0.08 | 0.43 | 1.7E-15 | 3.1E-14 | 0.43 | 1.7E-15 | | MYACLDGPEKPNLRNLPMYRTSQLRGSH* |
| HG0110.1 | *H. sapiens* | ENSG00000091656 | ZFHX4 | zinc finger homeobox 4 | 0.00 | 0.34 | 5.2E-28 | 1.6E-26 | 0.34 | 5.2E-28 | | MPPTFCSSVFIFTQ* |
| HG0110.2 | *D. rerio* | ENSDARG00000073944 | ZFHX3 (1 of many) | si:ch73-386h18.1 | 0.19 | 0.47 | 8.0E-08 | 3.1E-07 | 0.47 | 8.0E-08 | | MRSVLIPAIFCTKVISRNKDRTDFFSLSPPSSNRSQGASVVCSTTKKQTLLNALRNTEIPRLTSAA* |
| HG0111 | *G. gallus* | ENSGALG00000034944 | | | 0.04 | 0.25 | 5.3E-63 | 1.1E-61 | 0.25 | 5.3E-63 | | MEVISWGGPRALWARVLLSDHWLPPE* |
| HG0112 | *H. sapiens* | ENSG00000105997 | HOXA3 | homeobox A3 | 0.00 | 0.32 | 2.6E-50 | 1.4E-48 | 0.32 | 2.6E-50 | | MKRSARGGHWRRSVT* |
| HG0113 | *D. rerio* | ENSDARG00000099727 | ssbp2 | single-stranded DNA binding protein 2 | 0.00 | 0.40 | 4.7E-35 | 3.4E-34 | 0.40 | 4.7E-35 | | MGGIAPAVAMVTGLCPGRL* |
| | *G. gallus* | ENSGALG00000023089 | SSBP2 | single stranded DNA binding protein 2 | 0.00 | 0.38 | 2.5E-05 | 8.8E-05 | 0.38 | 2.5E-05 | | MGCVCGGVAMATGLFPGRL* |
| HG0114.1 | *G. gallus* | ENSGALG00000003701 | SGMS1 | sphingomyelin synthase 1 | 0.01 | 0.46 | 1.3E-14 | 8.5E-14 | 0.46 | 1.3E-14 | | MKPRRSLLMLWSRSDCCPL* |
| HG0114.2 | *H. sapiens* | ENSG00000198964 | SGMS1 | sphingomyelin synthase 1 | 0.00 | 0.36 | 3.6E-04 | 1.5E-03 | 0.36 | 3.6E-04 | | MTRMFLNNPRKNPAQK* |
| HG0115.1 | *G. gallus* | ENSGALG00000010976 | CCDC126 | coiled-coil domain containing 126 | 0.00 | 0.41 | 5.0E-11 | 2.6E-10 | 0.41 | 5.0E-11 | | MESRPAKPRRCVQDDPARRLLTL* |
| HG0115.2 | *H. sapiens* | ENSG00000169193 | CCDC126 | coiled-coil domain containing 126 | 0.02 | 0.01 | 6.5E-10 | 7.1E-09 | 0.01 | 6.5E-10 | | MSANRVNRVDTTLLKMKNIQY* |
| HG0116 | *G. gallus* | ENSGALG00000033038 | | | 0.01 | 0.30 | 4.0E-36 | 4.6E-35 | 0.30 | 4.0E-36 | | MDQHSSDGETGKGLL* |
| | *H. sapiens* | ENSG00000010270 | STARD3NL | STARD3 N-terminal like | 0.00 | 0.35 | 1.5E-05 | 8.0E-05 | 0.35 | 1.5E-05 | | MCLLFRDGEVGKRLL* |
| HG0117.1 | *H. sapiens* | ENSG00000140945 | CDH13 | cadherin 13 | 0.07 | 0.49 | 2.4E-19 | 5.6E-18 | 0.49 | 2.4E-19 | | MLSAAACMNENAAGRF* |
| HG0117.2 | *H. sapiens* | ENSG00000140945 | CDH13 | cadherin 13 | 0.00 | 0.00 | 1.1E-03 | 4.0E-03 | 0.00 | 1.1E-03 | | MPPPRGSLLARCSPRICHAKRGSVRKESVL* |
| HG0118.1 | *H. sapiens* | ENSG00000183662 | FAM19A1 | family with sequence similarity 19 member A1, C-C motif ch | 0.00 | 0.22 | 2.2E-18 | 4.8E-17 | 0.22 | 2.2E-18 | | MTRFPPLKPWLLF* |
| HG0118.2 | *H. sapiens* | ENSG00000183662 | FAM19A1 | family with sequence similarity 19 member A1, C-C motif ch | 0.00 | 0.38 | 1.3E-03 | 4.4E-03 | 0.38 | 1.3E-03 | | MHWSGDGPSATIN* |
| HG0119 | *G. gallus* | ENSGALG00000043570 | SGK3 | | 0.04 | 0.20 | 3.5E-51 | 5.4E-50 | 0.20 | 3.5E-51 | | MMEFTFSQEELRILDQYR* |
| HG0120 | *H. sapiens* | ENSG00000022355 | GABRA1 | gamma-aminobutyric acid type A receptor alpha1 subunit | 0.08 | 0.46 | 1.5E-03 | 5.1E-03 | 0.46 | 1.5E-03 | | MKSPPISHPVCPGTYYYCLGKEKSFVV* |
| HG0121 | *H. sapiens* | ENSG00000122786 | CALD1 | caldesmon 1 | 0.00 | 0.50 | 6.8E-12 | 9.1E-11 | 0.50 | 6.8E-12 | | MCSLSMDFWEGPRKGGLEFY* |
| HG0122 | *H. sapiens* | ENSG00000169925 | BRD3 | bromodomain containing 3 | 0.00 | 0.40 | 4.5E-75 | 3.3E-73 | 0.40 | 4.5E-75 | | MLSCDRPSSCLEEAGLQPGSLPLCPDHPHIPKAVAWLGPEMGRRAFSGSVEAARASPSRCCDRK* |
| HG0123 | *H. sapiens* | ENSG00000173276 | ZBTB21 | zinc finger and BTB domain containing 21 | 0.03 | 0.41 | 3.1E-36 | 1.4E-34 | 0.41 | 3.1E-36 | | MTSSLVWDDVKFKMDGITAAAAAAARASRV* |
| HG0124.1 | *D. rerio* | ENSDARG00000031763 | smad6b | SMAD family member 6b | 0.00 | 0.49 | 2.3E-13 | 1.0E-12 | 0.49 | 2.3E-13 | | MQYSIVPSKLMLWLCHRYTVLGCARAARHSANINVARAQK* |
| HG0124.2 | *D. rerio* | ENSDARG00000053209 | smad6a | SMAD family member 6a | 0.09 | 0.14 | 3.5E-04 | 9.9E-04 | 0.14 | 3.5E-04 | | MMGINQWLHCSPQGPRVCVSEEERE* |
| HG0125.1 | *D. rerio* | ENSDARG00000051926 | plag1 | pleiomorphic adenoma gene 1 | 0.00 | 0.33 | 2.7E-38 | 2.1E-37 | 0.33 | 2.7E-38 | | MRKKLLKIREIVESQEKYRRKDVNAPAGLMSLLC* |
| HG0125.2 | *H. sapiens* | ENSG00000181690 | PLAG1 | PLAG1 zinc finger | 0.00 | 0.48 | 3.3E-09 | 3.2E-08 | 0.50 | 3.5E-04 | | MLKPRDWPKWEGLDSTLFHEESMGLAKIKV* |
| HG0126.1 | *G. gallus* | ENSGALG00000010899 | G6PC2 | glucose-6-phosphatase catalytic subunit 2 | 0.00 | 0.30 | 2.5E-10 | 1.2E-09 | 0.30 | 2.5E-10 | | MNCEKIQANMFQLFIRRIA* |
| HG0126.2 | *G. gallus* | ENSGALG00000010899 | G6PC2 | glucose-6-phosphatase catalytic subunit 2 | 0.07 | 0.46 | 1.2E-15 | 8.2E-15 | 0.46 | 1.2E-15 | | MCFPRDSEPRVPGMAGYATLIQYKAQPSGWY* |
| HG0127.1 | *G. gallus* | ENSGALG00000029709 | | | 0.00 | 0.33 | 3.4E-34 | 3.7E-33 | 0.33 | 3.4E-34 | | MQNQSTHEEKSPLRGNVRGIAVGDNLLHSPTP* |
| HG0127.2 | *H. sapiens* | ENSG00000109452 | INPP4B | inositol polyphosphate-4-phosphatase type II B | 0.00 | 0.40 | 6.2E-09 | 5.8E-08 | 0.40 | 6.2E-09 | | MVLPGKRSSVFYEKLDKGFTGGIVLEICS* |
| HG0128 | *D. rerio* | ENSDARG00000091029 | phox2bb | paired-like homeobox 2bb | 0.00 | 0.25 | 4.1E-45 | 3.6E-44 | 0.25 | 4.1E-45 | | MRTICSRDLGLGGSLGGFS* |
| | *H. sapiens* | ENSG00000109132 | PHOX2B | paired like homeobox 2b | 0.12 | 0.39 | 5.7E-33 | 2.2E-31 | 0.39 | 5.7E-33 | | MQESMIGRLDSSVPQCQPNRRMSYCHVKSASNKTNRFAIVQVERAKFIMRTICSRDLRQGIS* |
| HG0129 | *D. rerio* | ENSDARG00000099437 | | | 0.00 | 0.14 | 2.5E-88 | 3.0E-87 | 0.14 | 2.5E-88 | | MPAVFLLLRSLVVRLFGSRLAASGVQLLRRILTTATGHLGTVLRNIWERISSQQSKEAILGCVLCLLNMHKKVDN* |
| HG0130 | *G. gallus* | ENSGALG00000002993 | | | 0.00 | 0.31 | 1.7E-69 | 4.1E-68 | 0.31 | 1.7E-69 | | MSCWEIRRIFFSLVEEEPCPGGWLHGLFWRAKEAKGSCLSRTQRELGHVLSAHLGNTVCKSRRKIAGIIQTAILIFVIFFRRGMHLA* |
| HG0131 | *G. gallus* | ENSGALG00000013135 | GALNT1 | polypeptide N-acetylgalactosaminyltransferase 1 | 0.00 | 0.35 | 1.1E-14 | 7.4E-14 | 0.35 | 1.1E-14 | | MAVNLQRRYRTWSLKLD* |
| HG0132 | *H. sapiens* | ENSG00000151067 | CACNA1C | calcium voltage-gated channel subunit alpha1 C | 0.00 | 0.32 | 1.0E-41 | 5.2E-40 | 0.32 | 1.0E-41 | Crowe | MRYSFTRGGSSGKEQFLGFDAIMGIR* |
| HG0133.1 | *D. rerio* | ENSDARG00000051886 | ankrd11 | ankyrin repeat domain 11 | 0.08 | 0.00 | 4.1E-82 | 4.3E-81 | 0.00 | 4.1E-82 | Mackowiak | MESTVDDDAMEA* |
| | *H. sapiens* | ENSG00000167522 | ANKRD11 | ankyrin repeat domain 11 | 0.13 | 0.05 | 2.0E-09 | 2.0E-08 | 0.05 | 2.0E-09 | Mackowiak | MESQTVDDEAPAV* |
| HG0133.2 | *D. rerio* | ENSDARG00000051886 | ankrd11 | ankyrin repeat domain 11 | 0.00 | 0.17 | 7.2E-03 | 1.5E-02 | 0.17 | 7.2E-03 | | MGKTSIGREYQV* |
| HG0133.3 | *H. sapiens* | ENSG00000167522 | ANKRD11 | ankyrin repeat domain 11 | 0.00 | 0.36 | 3.2E-04 | 1.3E-03 | 0.36 | 3.2E-04 | | MGSARAPPLSRRGAAQPSEHEYIQPCSGTHLHWI* |
| HG0134.1 | *G. gallus* | ENSGALG00000041267 | BMI1 | Polycomb complex protein BMI-1 | 0.09 | 0.24 | 1.8E-02 | 3.7E-02 | 0.24 | 1.8E-02 | | MAAWLTFIFC* |
| HG0134.2 | *H. sapiens* | ENSG00000168283 | BMI1 | BMI1 proto-oncogene, polycomb ring finger | 0.08 | 0.46 | 7.1E-07 | 4.9E-06 | 0.46 | 7.1E-07 | | MRGRDRGETMGMWAREPRSGLAAPPSPAE* |
| HG0135 | *D. rerio* | ENSDARG00000094132 | igf1 | insulin-like growth factor 1 | 0.00 | 0.31 | 6.6E-08 | 2.6E-07 | 0.31 | 6.6E-08 | | MLPQLFPVENVSVM* |
| | *H. sapiens* | ENSG00000017427 | IGF1 | insulin like growth factor 1 | 0.00 | 0.38 | 1.6E-13 | 2.6E-12 | 0.38 | 2.0E-17 | | MFPQLFPVYSVCVL* |
| HG0136.1 | *H. sapiens* | ENSG00000125845 | BMP2 | bone morphogenetic protein 2 | 0.03 | 0.15 | 2.0E-30 | 7.1E-29 | 0.15 | 2.0E-30 | Crowe | MWTLFQWTCPRVLLRRTAVS* |
| HG0136.2 | *H. sapiens* | ENSG00000125845 | BMP2 | bone morphogenetic protein 2 | 0.15 | 0.30 | 6.3E-04 | 2.4E-03 | 0.30 | 6.3E-04 | | MRSTYCRRSGAWDALAEV* |
| HG0137.1 | *H. sapiens* | ENSG00000181722 | ZBTB20 | zinc finger and BTB domain containing 20 | 0.00 | 0.32 | 1.1E-04 | 5.1E-04 | 0.32 | 1.1E-04 | | MNICTRPKPWSNLPEE* |
| HG0137.2 | *H. sapiens* | ENSG00000181722 | ZBTB20 | zinc finger and BTB domain containing 20 | 0.00 | 0.20 | 6.2E-03 | 1.8E-02 | 0.20 | 6.2E-03 | | MSWKCRNLSLPSRKNL* |
| HG0138.1 | *H. sapiens* | ENSG00000185070 | FLRT2 | fibronectin leucine rich transmembrane protein 2 | 0.00 | 0.13 | 5.3E-92 | 5.1E-90 | 0.13 | 5.3E-92 | Mackowiak | MPRDPRCCRASVFPFHLPDPVGFG* |
| HG0138.2 | *H. sapiens* | ENSG00000185070 | FLRT2 | fibronectin leucine rich transmembrane protein 2 | 0.00 | 0.00 | 2.6E-06 | 1.6E-05 | 0.00 | 2.6E-06 | Mackowiak | MGRRDLGGDP* |
| HG0139 | *G. gallus* | ENSGALG00000006379 | SHH | sonic hedgehog | 0.14 | 0.50 | 1.4E-02 | 3.0E-02 | 0.50 | 1.4E-02 | | MCSVISDLKAAIPAPVLLGIGG* |
| HG0140 | *H. sapiens* | ENSG00000009307 | CSDE1 | cold shock domain containing E1 | 0.00 | 0.00 | 1.3E-05 | 7.3E-05 | 0.00 | 1.3E-05 | | MKCNRTDFTT* |
| HG0141.1 | *G. gallus* | ENSGALG00000010435 | RASGRP3 | RAS guanyl releasing protein 3 | 0.02 | 0.43 | 2.5E-10 | 1.3E-09 | 0.43 | 2.5E-10 | | MFVLNTSAEPSTIFLEAQLRMQPTIK* |
| HG0141.2 | *H. sapiens* | ENSG00000152689 | RASGRP3 | RAS guanyl releasing protein 3 | 0.00 | 0.21 | 3.4E-13 | 5.3E-12 | 0.21 | 3.4E-13 | | MIMEMVYLMLKMSLVF* |
| HG0141.3 | *H. sapiens* | ENSG00000152689 | RASGRP3 | RAS guanyl releasing protein 3 | 0.00 | 0.28 | 3.0E-05 | 1.5E-04 | 0.28 | 3.0E-05 | | MLMRWLSQLRGYEVVCE* |
| HG0142.1 | *H. sapiens* | ENSG00000204217 | BMPR2 | bone morphogenetic protein receptor type 2 | 0.00 | 0.10 | 2.5E-68 | 1.8E-66 | 0.10 | 2.5E-68 | Mackowiak | MFSPRPGYFFDIVKLRGK* |
| HG0142.2 | *H. sapiens* | ENSG00000204217 | BMPR2 | bone morphogenetic protein receptor type 2 | 0.00 | 0.32 | 1.5E-13 | 2.5E-12 | 0.32 | 1.5E-13 | | MKALQLGPLISHLSFQTVL* |
| HG0143 | *H. sapiens* | ENSG00000101958 | GLRA2 | glycine receptor alpha 2 | 0.00 | 0.44 | 1.5E-32 | 5.6E-31 | 0.44 | 1.5E-32 | | MMPRTGTFSFFSANCTKPNLFLIFKETRFLPNFESGQ* |
| HG0144 | *H. sapiens* | ENSG00000117114 | ADGRL2 | adhesion G protein-coupled receptor L2 | 0.00 | 0.09 | 1.4E-37 | 6.2E-36 | 0.09 | 1.4E-37 | | MMQTDGFDEAGHL* |
| HG0145 | *H. sapiens* | ENSG00000162599 | NFIA | nuclear factor I A | 0.00 | 0.39 | 1.2E-06 | 7.9E-06 | 0.39 | 1.2E-06 | | MIPYGMDRTTCCPTCHCKCFLN* |
| HG0146.1 | *H. sapiens* | ENSG00000163681 | SLMAP | sarcolemma associated protein | 0.00 | 0.47 | 1.2E-06 | 8.0E-06 | 0.47 | 1.2E-06 | | MLQCCNVSATKGEKRPRSQTKHKNWRVTHLLGYLQSPNCVPGVFLAEAARCLSLFFPEKKMD* |
| HG0146.2 | *H. sapiens* | ENSG00000163681 | SLMAP | sarcolemma associated protein | 0.00 | 0.32 | 4.8E-04 | 1.9E-03 | 0.32 | 4.8E-04 | | MLRQCAVCVNLKFWVG* |
| HG0146.3 | *H. sapiens* | ENSG00000163681 | SLMAP | sarcolemma associated protein | 0.00 | 0.40 | 4.6E-05 | 2.3E-04 | 0.40 | 4.6E-05 | | MAAGAGKVGKFPASSGAA* |
| HG0147 | *G. gallus* | ENSGALG00000004370 | MAP2K6 | mitogen-activated protein kinase kinase 6 | 0.01 | 0.48 | 6.0E-03 | 1.4E-02 | 0.48 | 6.0E-03 | | MKIERLQFASLLQN* |
| | *H. sapiens* | ENSG00000108984 | MAP2K6 | mitogen-activated protein kinase kinase 6 | 0.01 | 0.48 | 1.5E-11 | 1.9E-10 | 0.48 | 1.5E-11 | | MKIARLQLASLLQN* |
| HG0148.1 | *G. gallus* | ENSGALG00000011855 | RHOJ | ras homolog family member J | 0.00 | 0.00 | 2.6E-60 | 5.0E-59 | 0.00 | 2.6E-60 | | MLEIRWKMEWI* |

| | | | | | | | | | | | | |
|---|---|---|---|---|---|---|---|---|---|---|---|---|
| HG0148.2 | *G. gallus* | ENSGALG00000011855 | RHOJ | ras homolog family member J | 0.00 | 0.41 | 2.3E-08 | 1.1E-07 | 0.41 | 2.3E-08 | | MILWTALVIVRSSCDSLETLV* |
| HG0149.1 | *G. gallus* | ENSGALG00000014645 | MEF2C | myocyte enhancer factor 2C | 0.00 | 0.10 | 3.0E-10 | 1.5E-09 | 0.10 | 3.0E-10 | | MILTIKRVYVQERMQGTVN* |
| HG0149.2 | *H. sapiens* | ENSG00000081189 | MEF2C | myocyte enhancer factor 2C | 0.00 | 0.34 | 9.3E-08 | 7.3E-07 | 0.34 | 9.3E-08 | | MWNILKDNIKVNECRNLGTELCKC* |
| HG0150.1 | *G. gallus* | ENSGALG00000032289 | IKZF5 | Zinc finger protein Pegasus | 0.25 | 0.00 | 1.5E-48 | 2.2E-47 | 0.00 | 1.5E-48 | | MAAAAEGGGRGAGRRRRHQQMEHKLT* |
| HG0150.2 | *H. sapiens* | ENSG00000095574 | IKZF5 | IKAROS family zinc finger 5 | 0.00 | 0.26 | 2.0E-48 | 1.1E-46 | 0.26 | 2.0E-48 | | MAFWSPLLYLRSKFQLLYISSMLEWQEKGGTDLG* |
| HG0151 | *G. gallus* | ENSGALG00000035504 | ER81 | ETS variant 1 | 0.00 | 0.32 | 1.1E-04 | 3.5E-04 | 0.32 | 1.1E-04 | | MFLLNLEGNFLQPIAARGLQADWSPDPEEISRSNAH* |
| | *H. sapiens* | ENSG00000006468 | ETV1 | ETS variant 1 | 0.00 | 0.21 | 4.0E-35 | 1.6E-33 | 0.21 | 4.0E-35 | | MFLLNLEGSFCSLSPSVAFSPVRCKASLVDRHC* |
| HG0152 | *D. rerio* | ENSDARG00000015536 | sox6 | SRY (sex determining region Y)-box 6 | 0.00 | 0.26 | 4.8E-14 | 2.3E-13 | 0.26 | 4.8E-14 | | MLHDGKCFEYLRHGRLH* |
| HG0153 | *G. gallus* | ENSGALG00000003136 | IKZF2 | IKAROS family zinc finger 2 | 0.00 | 0.18 | 2.4E-31 | 2.4E-30 | 0.18 | 2.4E-31 | | MEEVPAGLPLATAPLPQFP* |
| HG0154 | *H. sapiens* | ENSG00000128573 | FOXP2 | forkhead box P2 | 0.02 | 0.46 | 1.7E-04 | 7.4E-04 | 0.46 | 1.7E-04 | | MPSVWDVIGQRCTHSSVNTAVNSCLMVALTVS* |
| HG0155 | *H. sapiens* | ENSG00000136535 | TBR1 | T-box, brain 1 | 0.00 | 0.44 | 3.9E-09 | 3.8E-08 | 0.44 | 3.9E-09 | | MAGAVSSAAQAARSKVSLF* |
| HG0156 | *H. sapiens* | ENSG00000156113 | KCNMA1 | potassium calcium-activated channel subfamily M alpha 1 | 0.06 | 0.45 | 1.5E-11 | 1.9E-10 | 0.45 | 1.5E-11 | | MAVDGCLGWRWRREELPRAPAPAARC* |
| HG0157 | *H. sapiens* | ENSG00000173926 | MARCH3 | membrane associated ring-CH-type finger 3 | 0.00 | 0.00 | 1.1E-30 | 3.9E-29 | 0.00 | 1.1E-30 | | MCPGPLGREQPPTGGNLALRTAAGTSVCYRDSARTMEILYECGSH* |
| HG0158 | *H. sapiens* | ENSG00000198939 | ZFP2 | ZFP2 zinc finger protein | 0.26 | 0.32 | 2.6E-22 | 7.1E-21 | 0.32 | 2.6E-22 | | MTVYGEVMLENYRNPVSLGNHLSKPNGTSQLERELSLMQKEPPEGGFRDFLGYCCQINLTFSPLIILSQSRGESID* |
| HG0159.1 | *H. sapiens* | ENSG00000182197 | EXT1 | exostosin glycosyltransferase 1 | 0.01 | 0.08 | 7.8E-35 | 3.1E-33 | 0.08 | 7.8E-35 | Crowe | MQENYSPLLARPGCEIFSPALSRALCNPKMKDRRGES* |
| HG0159.2 | *H. sapiens* | ENSG00000182197 | EXT1 | exostosin glycosyltransferase 1 | 0.00 | 0.19 | 7.2E-08 | 5.7E-07 | 0.19 | 7.2E-08 | | MQGFVRLRRPSCKMH* |
| HG0160 | *D. rerio* | ENSDARG00000060089 | btaf1 | BTAF1 RNA polymerase II, B-TFIID transcription factor-ass | 0.00 | 0.33 | 9.3E-35 | 6.7E-34 | 0.33 | 9.3E-35 | | MAPHIKPAAWLLFCDLQNYFSPFVRLKYRFRGSAEG* |
| HG0161 | *G. gallus* | ENSGALG00000017191 | | | 0.02 | 0.25 | 2.7E-20 | 2.2E-19 | 0.25 | 2.7E-20 | | MCWLFFVPYTFPNS* |
| HG0162 | *H. sapiens* | ENSG00000138347 | MYPN | myopalladin | 0.01 | 0.37 | 1.9E-13 | 3.0E-12 | 0.37 | 1.9E-13 | | MFHRHMVYTAQNNLGPVS* |
| HG0163 | *H. sapiens* | ENSG00000146285 | SCML4 | sex comb on midleg like 4 (Drosophila) | 0.02 | 0.29 | 1.2E-25 | 3.6E-24 | 0.29 | 1.2E-25 | | MFAAGAQGCWRLTVSY* |
| HG0164 | *H. sapiens* | ENSG00000151292 | CSNK1G3 | casein kinase 1 gamma 3 | 0.00 | 0.30 | 4.8E-39 | 2.3E-37 | 0.30 | 4.8E-39 | | MLTLPIWYSYLVMYLFSQYPVSVCLS* |
| HG0165 | *H. sapiens* | ENSG00000177508 | IRX3 | iroquois homeobox 3 | 0.17 | 0.17 | 1.1E-58 | 6.7E-57 | 0.17 | 1.1E-58 | | MSIRARSPGNLRQSVRT* |
| HG0166.1 | *H. sapiens* | ENSG00000148948 | LRRC4C | leucine rich repeat containing 4C | 0.00 | 0.01 | 2.6E-28 | 8.3E-27 | 0.01 | 2.6E-28 | | MKLQRRWREGDRI* |
| HG0166.2 | *H. sapiens* | ENSG00000148948 | LRRC4C | leucine rich repeat containing 4C | 0.00 | 0.35 | 4.8E-06 | 2.8E-05 | 0.35 | 4.8E-06 | | MQDAAFSWNRTQWIN* |
| HG0166.3 | *H. sapiens* | ENSG00000148948 | LRRC4C | leucine rich repeat containing 4C | 0.00 | 0.50 | 4.7E-04 | 1.8E-03 | 0.50 | 4.7E-04 | | MIGNEEPRIHKEKSMFIFLYKGESEPRRYFWNEKFGAFLVK* |
| HG0166.4 | *H. sapiens* | ENSG00000148948 | LRRC4C | leucine rich repeat containing 4C | 0.01 | 0.31 | 5.8E-10 | 6.3E-09 | 0.31 | 5.8E-10 | | MEGDWGFCRAGSREPGRWR* |
| HG0167 | *D. rerio* | ENSDARG00000104005 | slc4a3 | | 0.04 | 0.31 | 1.8E-33 | 1.3E-32 | 0.31 | 1.8E-33 | | MPPHRLTGNPAVKEKG* |
| HG0168 | *G. gallus* | ENSGALG00000006997 | BRINP1 | BMP/retinoic acid inducible neural specific 1 | 0.00 | 0.15 | 3.1E-10 | 1.5E-09 | 0.15 | 3.1E-10 | | MPAIWIKEGDFLSEHEKDC* |
| HG0169 | *H. sapiens* | ENSG00000114933 | INO80D | INO80 complex subunit D | 0.00 | 0.47 | 2.0E-08 | 1.7E-07 | 0.47 | 2.0E-08 | | MVGKLSFRRAEEQLSGVMSSQCL* |
| HG0170 | *H. sapiens* | ENSG00000180530 | NRIP1 | nuclear receptor interacting protein 1 | 0.00 | 0.07 | 3.1E-51 | 1.7E-49 | 0.04 | 2.7E-48 | | MKALRSAEQIDYSGVPVLGTTAVRERTLIFAFNGEQKMKKEKEYIH* |
| HG0171.1 | *H. sapiens* | ENSG00000177311 | ZBTB38 | zinc finger and BTB domain containing 38 | 0.00 | 0.00 | 2.8E-59 | 1.7E-57 | 0.00 | 2.8E-59 | | MYSKEKRTLAMVQLNQDSEADMQCHLQH* |
| HG0171.2 | *H. sapiens* | ENSG00000177311 | ZBTB38 | zinc finger and BTB domain containing 38 | 0.00 | 0.10 | 5.2E-30 | 1.8E-28 | 0.47 | 2.6E-11 | | MRNFFFQSCFPVDKFSSVITVHSSCSAYDFCVYVTIASKTCEQEDREISWSPSLYIKMAECLVMLALH* |
| HG0171.3 | *H. sapiens* | ENSG00000177311 | ZBTB38 | zinc finger and BTB domain containing 38 | 0.00 | 0.01 | 4.1E-18 | 8.9E-17 | 0.01 | 4.1E-18 | | MLAVLQKAFRAADLDVEEMEDSEPRISK* |
| HG0171.4 | *H. sapiens* | ENSG00000177311 | ZBTB38 | zinc finger and BTB domain containing 38 | 0.00 | 0.31 | 9.9E-03 | 2.6E-02 | 0.31 | 9.9E-03 | | MGDIIRKICGADHKK* |
| HG0172.1 | *H. sapiens* | ENSG00000196782 | MAML3 | mastermind like transcriptional coactivator 3 | 0.07 | 0.48 | 2.2E-12 | 3.1E-11 | 0.48 | 2.2E-12 | | MQLSSQSTLRTRGGINTKTFLWKENAISKLAVFRCGVE* |
| HG0172.2 | *H. sapiens* | ENSG00000196782 | MAML3 | mastermind like transcriptional coactivator 3 | 0.01 | 0.21 | 3.2E-06 | 1.9E-05 | 0.21 | 3.2E-06 | | MDYKKKASKERISP* |
| HG0173 | *H. sapiens* | ENSG00000101746 | NOL4 | nucleolar protein 4 | 0.16 | 0.00 | 1.4E-02 | 3.4E-02 | 0.00 | 1.4E-02 | | MIELACHRHWYLS* |
| HG0174 | *H. sapiens* | ENSG00000144355 | DLX1 | distal-less homeobox 1 | 0.01 | 0.39 | 2.1E-06 | 1.3E-05 | 0.39 | 2.1E-06 | | MRLTRGRAASPALVQRSPSASDQSPVIMQDSGPISSASS* |
| HG0175 | *H. sapiens* | ENSG00000151320 | AKAP6 | A-kinase anchoring protein 6 | 0.00 | 0.39 | 3.9E-12 | 5.4E-11 | 0.39 | 3.9E-12 | | MCCKHQKKTVGSGDAVLERSEV* |
| HG0176 | *H. sapiens* | ENSG00000170962 | PDGFD | platelet derived growth factor D | 0.20 | 0.01 | 1.3E-23 | 3.6E-22 | 0.01 | 1.3E-23 | | MQEVWGELGD* |
| HG0177 | *H. sapiens* | ENSG00000171843 | MLLT3 | MLLT3, super elongation complex subunit | 0.09 | 0.46 | 5.3E-08 | 4.3E-07 | 0.46 | 5.3E-08 | | MLRNHLLYPGAAAAAAAFAFGAEFNKRASEQASAGGKRQRMSAIYPPLLGALSFIAASS* |
| HG0178.1 | *G. gallus* | ENSGALG00000029047 | TMEM204 | transmembrane protein 204 | 0.00 | 0.15 | 4.4E-45 | 5.8E-44 | 0.15 | 4.4E-45 | Crowe, Mackowiak | MDLHVFQFITGLALLIKVQEGG* |
| | *H. sapiens* | ENSG00000131634 | TMEM204 | transmembrane protein 204 | 0.05 | 0.24 | 6.4E-27 | 2.0E-25 | 0.24 | 6.4E-27 | Crowe, Mackowiak | MGPEASSFSTGLALLQVQEGG* |
| HG0178.2 | *G. gallus* | ENSGALG00000029047 | TMEM204 | transmembrane protein 204 | 0.02 | 0.49 | 7.3E-09 | 3.4E-08 | 0.49 | 7.3E-09 | | MFDCRGSGSVCCSRLRPEAHQNSHKNSHFL* |
| HG0179 | *D. rerio* | ENSDARG00000036549 | agpat3 | 1-acylglycerol-3-phosphate O-acyltransferase 3 | 0.00 | 0.09 | 8.0E-29 | 5.2E-28 | 0.09 | 8.0E-29 | Mackowiak | MLSHIERLPCYNRLRDTAVAQYEREKLEKVAGFPPTGSVVLSEGPCKRGL* |
| | *H. sapiens* | ENSG00000160216 | AGPAT3 | 1-acylglycerol-3-phosphate O-acyltransferase 3 | 0.12 | 0.08 | 8.3E-54 | 4.7E-52 | 0.08 | 8.3E-54 | Mackowiak | MLRWVSVASPEFLLNEVCRVIWTAVLSEGPCTRS* |
| HG0180.1 | *H. sapiens* | ENSG00000112246 | SIM1 | single-minded family bHLH transcription factor 1 | 0.01 | 0.33 | 7.1E-35 | 2.8E-33 | 0.33 | 7.1E-35 | | MSIFRGPGFAAGSRRKGWLLRGPTGI* |
| HG0180.2 | *H. sapiens* | ENSG00000112246 | SIM1 | single-minded family bHLH transcription factor 1 | 0.03 | 0.34 | 1.2E-06 | 7.9E-06 | 0.34 | 1.2E-06 | | MRTVKRWAPPKPLPLVGTEGP* |
| HG0181.1 | *H. sapiens* | ENSG00000143776 | CDC42BPA | CDC42 binding protein kinase alpha | 0.00 | 0.48 | 3.7E-15 | 6.6E-14 | 0.48 | 3.7E-15 | | MHSVFFLPFLFFFFPGSELLMQIGLYSLMMQPDSFQDYVARVEF* |
| HG0181.2 | *H. sapiens* | ENSG00000143776 | CDC42BPA | CDC42 binding protein kinase alpha | 0.00 | 0.04 | 4.5E-06 | 2.7E-05 | 0.04 | 4.5E-06 | | MYMGEIEDASVK* |
| HG0182 | *G. gallus* | ENSGALG00000031991 | | | 0.06 | 0.05 | 1.1E-56 | 1.9E-55 | 0.05 | 1.1E-56 | | MDVNQPKQEHLLALKGPVSH* |
| HG0183 | *H. sapiens* | ENSG00000011465 | DCN | decorin | 0.02 | 0.41 | 1.8E-12 | 2.6E-11 | 0.41 | 1.8E-12 | | MWVSAGLCSLRETLFFNCAME* |
| HG0184 | *H. sapiens* | ENSG00000136542 | GALNT5 | polypeptide N-acetylgalactosaminyltransferase 5 | 0.00 | 0.28 | 9.3E-07 | 6.2E-06 | 0.28 | 9.3E-07 | | MLCYEMPRRQRLTSGRN* |
| HG0185 | *H. sapiens* | ENSG00000143033 | MTF2 | metal response element binding transcription factor 2 | 0.00 | 0.18 | 3.0E-04 | 1.2E-03 | 0.18 | 3.0E-04 | | MHRQSAGNQNGEGLY* |
| HG0186 | *H. sapiens* | ENSG00000167552 | TUBA1A | tubulin alpha 1a | 0.08 | 0.41 | 3.3E-14 | 5.7E-13 | 0.41 | 3.3E-14 | | MASLGRLGVWTNRKRPRPIPMVTELYIGSRIRLCAAGSLTSTA* |
| HG0187.1 | *G. gallus* | ENSGALG00000004058 | GPR146 | G protein-coupled receptor 146 | 0.25 | 0.40 | 5.2E-14 | 3.3E-13 | 0.40 | 5.2E-14 | | MDKMFLPASTGQQGTAYQKSW* |
| HG0187.2 | *D. rerio* | ENSDARG00000059610 | gpr146 | G protein-coupled receptor 146 | 0.00 | 0.25 | 5.3E-04 | 1.5E-03 | 0.25 | 5.3E-04 | | MSMEIQQNPMPP* |
| HG0187.3 | *G. gallus* | ENSGALG00000004058 | GPR146 | G protein-coupled receptor 146 | 0.08 | 0.36 | 4.5E-06 | 1.7E-05 | 0.36 | 4.5E-06 | | MCMYVQGKSHVNTDSVP* |
| HG0188 | *G. gallus* | ENSGALG00000016284 | TAB3 | TGF-beta activated kinase 1 and MAP3K7 binding protein 3 | 0.03 | 0.02 | 2.4E-33 | 2.5E-32 | 0.02 | 2.4E-33 | | MPEASEDHNHWKVC* |
| | *H. sapiens* | ENSG00000157625 | TAB3 | TGF-beta activated kinase 1 and MAP3K7 binding protein 3 | 0.00 | 0.27 | 4.9E-28 | 1.5E-26 | 0.27 | 4.9E-28 | | MTVVLEDHYFWKAG* |
| HG0189 | *D. rerio* | ENSDARG00000001807 | tnfrsf21 | tumor necrosis factor receptor superfamily, member 21 | 0.00 | 0.25 | 7.0E-24 | 4.3E-23 | 0.25 | 7.0E-24 | | MGIIAFVFQNQALIPYYLKPVHSGAQTIKRREGTEFVH* |
| HG0190 | *G. gallus* | ENSGALG00000006949 | CHD2 | chromodomain helicase DNA binding protein 2 | 0.00 | 0.00 | 4.3E-08 | 1.9E-07 | 0.00 | 4.3E-08 | | MSFFPRTFLY* |
| HG0191 | *G. gallus* | ENSGALG00000036097 | | | 0.00 | 0.42 | 8.6E-16 | 6.0E-15 | 0.42 | 8.6E-16 | | MNKRKNPFWTVFLNQASPSISRTF* |
| HG0192 | *H. sapiens* | ENSG00000125107 | CNOT1 | CCR4-NOT transcription complex subunit 1 | 0.00 | 0.23 | 1.0E-18 | 2.3E-17 | 0.23 | 1.0E-18 | | MQKLPRSCQAISQLSIQKETYHSKDLRSCMPGNGMKTKCAMFQNRVRPLVKKN* |
| HG0193 | *H. sapiens* | ENSG00000134853 | PDGFRA | platelet derived growth factor receptor alpha | 0.03 | 0.22 | 8.1E-06 | 4.6E-05 | 0.22 | 8.1E-06 | | MISFRRMDYNIESITKRGF* |
| HG0194 | *H. sapiens* | ENSG00000143507 | DUSP10 | dual specificity phosphatase 10 | 0.01 | 0.37 | 5.3E-11 | 6.4E-10 | 0.37 | 5.3E-11 | | MCESIAEEERQMVEEYTYL* |
| HG0195 | *H. sapiens* | ENSG00000165966 | PDZRN4 | PDZ domain containing ring finger 4 | 0.01 | 0.44 | 9.5E-05 | 4.3E-04 | 0.44 | 9.5E-05 | | MKNSSQVWRVIPFQFTCLSVLGKLEDLLSLSAS* |
| HG0196 | *H. sapiens* | ENSG00000189403 | HMGB1 | high mobility group box 1 | 0.01 | 0.36 | 4.2E-09 | 4.0E-08 | 0.36 | 4.2E-09 | | MLQSGESEEAASGSRSHSHCSTLSSIETAPGQVRAGRALGDSVPR* |
| HG0197 | *H. sapiens* | ENSG00000196132 | MYT1 | myelin transcription factor 1 | 0.00 | 0.00 | 3.4E-04 | 1.4E-03 | 0.00 | 3.4E-04 | | MIAFRDVILKEDRQ* |
| HG0198.1 | *D. rerio* | ENSDARG00000018688 | elk3 | ELK3, ETS-domain protein | 0.06 | 0.30 | 2.3E-03 | 5.6E-03 | 0.30 | 2.3E-03 | | MCVCVRIRVCARVPECRMVARGGKACLHRRAERLGNRARREEVSRLAVCLLQLPAHRERERKSE* |
| HG0198.2 | *H. sapiens* | ENSG00000111145 | ELK3 | ELK3, ETS transcription factor | 0.12 | 0.00 | 7.2E-05 | 3.4E-04 | 0.00 | 7.2E-05 | | MESRGLSPGSS* |
| HG0199 | *G. gallus* | ENSGALG00000007989 | ADNP | activity dependent neuroprotector homeobox | 0.00 | 0.37 | 9.0E-03 | 2.0E-02 | 0.37 | 9.0E-03 | | MLRQKGNEQDMTLLLCPGFGLTQ* |
| HG0200 | *G. gallus* | ENSGALG00000012015 | NDST4 | N-deacetylase and N-sulfotransferase 4 | 0.02 | 0.40 | 1.1E-17 | 8.0E-17 | 0.40 | 1.1E-17 | | MHCNGHMNVSGEESAPLRLIVIMAL* |
| HG0201 | *G. gallus* | ENSGALG00000015256 | PAM | peptidylglycine alpha-amidating monooxygenase | 0.00 | 0.30 | 8.1E-14 | 5.0E-13 | 0.30 | 8.1E-14 | | MVCQKLEDACRAHNNVVMN* |
| HG0202 | *H. sapiens* | ENSG00000124766 | SOX4 | SRY-box 4 | 0.09 | 0.46 | 6.4E-15 | 1.1E-13 | 0.46 | 6.4E-15 | | MRSDLELLIARGDYYCIGFQANGKPGGGVWHEEALVTAADWLQPRL* |
| HG0203 | *H. sapiens* | ENSG00000128606 | LRRC17 | leucine rich repeat containing 17 | 0.09 | 0.21 | 3.0E-14 | 5.1E-13 | 0.21 | 3.0E-14 | | MSLWHPFPKFSLGTPRTPAGSHCSRTALVKITQTWISKEYFHCSVCNTK* |
| HG0204 | *H. sapiens* | ENSG00000162599 | NFIA | nuclear factor I A | 0.00 | 0.00 | 2.7E-09 | 2.7E-08 | 0.00 | 2.7E-09 | | MPLTPAYSRHV* |
| HG0205 | *H. sapiens* | ENSG00000184611 | KCNH7 | potassium voltage-gated channel subfamily H member 7 | 0.01 | 0.35 | 6.7E-04 | 2.5E-03 | 0.35 | 6.7E-04 | | MGREFWFQKRMLCHKERLARWPGL* |
| HG0206.1 | *G. gallus* | ENSGALG00000002069 | | | 0.00 | 0.48 | 3.4E-17 | 2.5E-16 | 0.48 | 3.4E-17 | | MVGAIPRQPRRFWGVPGWQVSSASCANDRPLWDGEIDPEWSGILLQREPHTGGLTWTS* |
| HG0206.2 | *H. sapiens* | ENSG00000005238 | FAM214B | family with sequence similarity 214 member B | 0.00 | 0.16 | 6.0E-30 | 2.1E-28 | 0.16 | 6.0E-30 | | MGLVLLSSGAARNGGSHLASAAGMDQTAS* |
| HG0207.1 | *G. gallus* | ENSGALG00000007121 | TNFSF8 | TNF superfamily member 8 | 0.04 | 0.40 | 1.5E-13 | 9.0E-13 | 0.40 | 1.5E-13 | | MSQDDGENPARNRDGLRPEPDQGEIQTSTLTAI* |
| HG0207.2 | *H. sapiens* | ENSG00000106952 | TNFSF8 | TNF superfamily member 8 | 0.02 | 0.24 | 1.3E-02 | 3.2E-02 | 0.24 | 1.3E-02 | | MGPELNQMKRDKV* |
| HG0208 | *G. gallus* | ENSGALG00000013272 | PRMT8 | protein arginine methyltransferase 8 | 0.00 | 0.41 | 4.5E-03 | 1.1E-02 | 0.41 | 4.5E-03 | | MPSWSLHAQVQFSWCFSPV* |
| HG0209 | *H. sapiens* | ENSG00000101040 | ZMYND8 | zinc finger MYND-type containing 8 | 0.22 | 0.31 | 1.9E-13 | 2.9E-12 | 0.31 | 1.9E-13 | | MHPQSPKSASLFSLPNCGKMRRLLEHLLAWLKRK* |
| HG0210 | *H. sapiens* | ENSG00000155849 | ELMO1 | engulfment and cell motility 1 | 0.03 | 0.36 | 2.4E-03 | 7.6E-03 | 0.36 | 2.4E-03 | | MPCRCKMCLWPAGGRKRK* |
| HG0211 | *H. sapiens* | ENSG00000176204 | LRRTM4 | leucine rich repeat transmembrane neuronal 4 | 0.00 | 0.00 | 1.1E-06 | 7.2E-06 | 0.00 | 1.1E-06 | | MDKPSLHSF* |
| HG0212 | *D. rerio* | ENSDARG00000100244 | ebf3a | early B-cell factor 3a | 0.00 | 0.32 | 2.9E-07 | 1.1E-06 | 0.32 | 2.9E-07 | | MLPSILQRRAKGPSSSSPVSGNTSTVIIYAFICLLEPEVVDSAVSRSLF* |
| HG0213.1 | *G. gallus* | ENSGALG00000001946 | RSPO1 | R-spondin 1 | 0.23 | 0.47 | 4.4E-05 | 1.5E-04 | 0.47 | 4.4E-05 | | MGLDPSSCAPYGDEGRTMLPAT* |
| HG0213.2 | *G. gallus* | ENSGALG00000001946 | RSPO1 | R-spondin 1 | 0.07 | 0.29 | 3.4E-03 | 8.6E-03 | 0.29 | 3.4E-03 | | MGMESTALRPGFINVQIALI* |
| HG0214.1 | *G. gallus* | ENSGALG00000012917 | CDH6 | cadherin 6 | 0.20 | 0.47 | 6.3E-08 | 2.7E-07 | 0.47 | 6.3E-08 | | METNQIESPLGSGCLHAGVKKCCRSSGSPSERRGG* |
| HG0214.2 | *H. sapiens* | ENSG00000040731 | CDH10 | cadherin 10 | 0.01 | 0.38 | 1.0E-02 | 2.7E-02 | 0.38 | 1.0E-02 | | MNKTLMKCWKPVETQ* |
| HG0215 | *D. rerio* | ENSDARG00000079280 | dhrsx | dehydrogenase/reductase (SDR family) X-linked | 0.12 | 0.01 | 1.2E-15 | 6.1E-15 | 0.01 | 1.2E-15 | | MQSLISPVTKAILVALFIFAILLILYVILWYICRDVDCDHGI* |
| | *G. gallus* | ENSGALG00000016681 | DHRSX | dehydrogenase/reductase X-linked | 0.03 | 0.01 | 1.7E-25 | 1.5E-24 | 0.01 | 1.7E-25 | | MQSLISPVTKAILVALFIFAILLILYVILWYICRDVDCDHGI* |
| HG0216.1 | *G. gallus* | ENSGALG00000020922 | | | 0.00 | 0.18 | 2.6E-04 | 8.0E-04 | 0.18 | 2.6E-04 | | MICGSVQLELSE* |
| HG0216.2 | *H. sapiens* | ENSG00000116667 | C1orf21 | chromosome 1 open reading frame 21 | 0.07 | 0.14 | 2.6E-09 | 2.6E-08 | 0.14 | 2.6E-09 | | MICSSARLKLTE* |
| HG0217.1 | *H. sapiens* | ENSG00000150394 | CDH8 | cadherin 8 | 0.00 | 0.04 | 1.0E-02 | 2.7E-02 | 0.04 | 1.0E-02 | | MNFQVVDLPLRSAQ* |
| HG0217.2 | *G. gallus* | ENSGALG00000005319 | CDH8 | cadherin 8 | 0.11 | 0.50 | 1.3E-09 | 6.5E-09 | 0.50 | 1.3E-09 | | MRRKSGESCLFTGTGEWLFYLTLSIRV* |
| HG0218 | *D. rerio* | ENSDARG00000100149 | axin2 | axin 2 (conductin, axil) | 0.02 | 0.45 | 9.2E-04 | 2.4E-03 | 0.45 | 9.2E-04 | | MDERSSPADLRERGDYS* |
| HG0219 | *D. rerio* | ENSDARG00000104082 | trps1 | trichorhinophalangeal syndrome I | 0.00 | 0.29 | 1.8E-04 | 5.3E-04 | 0.29 | 1.8E-04 | | MAHAKAPPSFRLQRLLQFNQQ* |
| HG0220 | *G. gallus* | ENSGALG00000008418 | LARGE2 | LARGE xylosyl- and glucuronyltransferase 2 | 0.06 | 0.01 | 2.1E-19 | 1.6E-18 | 0.01 | 2.1E-19 | | MLGPAGLGLPT* |
| HG0221 | *G. gallus* | ENSGALG00000010859 | EPGN | epithelial mitogen | 0.00 | 0.33 | 6.7E-14 | 4.2E-13 | 0.33 | 6.7E-14 | | MPTGNRKLGFQEKTRSQC* |
| HG0222 | *G. gallus* | ENSGALG00000011564 | RUFY3 | RUN and FYVE domain containing 3 | 0.00 | 0.39 | 7.0E-04 | 2.0E-03 | 0.39 | 7.0E-04 | | MQKPGFLWLLRINSL* |
| HG0223 | *H. sapiens* | ENSG00000091656 | ZFHX4 | zinc finger homeobox 4 | 0.00 | 0.50 | 1.9E-05 | 9.9E-05 | 0.50 | 1.9E-05 | | MDGRVRERLRLPLPFRVLLRLRLRLGCDWDATVTRASIASIESNAKTQPGSLTGWMK* |

| | | | | | | | | | | | | |
|---|---|---|---|---|---|---|---|---|---|---|---|---|
| HG0224 | *H. sapiens* | ENSG00000118007 | STAG1 | stromal antigen 1 | 0.00 | 0.09 | 9.2E-07 | 6.1E-06 | 0.09 | 9.2E-07 | | MPPDGGLGLQLR* |
| HG0225 | *H. sapiens* | ENSG00000150051 | MKX | mohawk homeobox | 0.21 | 0.30 | 5.2E-10 | 5.7E-09 | 0.30 | 5.2E-10 | | MFAGWDRQSGNMRRPRYSCSRAARAGLGK* |
| HG0226.1 | *H. sapiens* | ENSG00000180592 | | | 0.07 | 0.06 | 1.7E-05 | 9.1E-05 | 0.06 | 1.7E-05 | | MERFIYRLYYLLWEGRWQYKGMLISGRFWGEGVEYQFLLHHLSGVSNQRWL* |
| HG0226.2 | *H. sapiens* | ENSG00000180592 | | | 0.09 | 0.44 | 1.9E-02 | 4.5E-02 | 0.44 | 1.9E-02 | | MAEQPIESADMVNKFIKETPWLLKNSTLYIWICTSGKAQKYPFILLITRMCLFLEEQGNT* |
| HG0226.3 | *H. sapiens* | ENSG00000180592 | | | 0.26 | 0.40 | 8.1E-03 | 2.2E-02 | 0.40 | 8.1E-03 | | MRLGGPLLSFGNILDILENDTKICLNLTKKRLNHIDFHPK* |
| HG0227.1 | *G. gallus* | ENSGALG00000031929 | | | 0.05 | 0.48 | 5.6E-04 | 1.6E-03 | 0.48 | 5.6E-04 | | MTALNNWWHFGGYGWSSVY* |
| HG0227.2 | *H. sapiens* | ENSG00000275163 | | | 0.00 | 0.20 | 2.6E-04 | 1.1E-03 | 0.20 | 2.6E-04 | | MGKVSEEEPFLPFVAISTGLQGTSFLLGFYVCFTHCLSFLCLRHFHE* |
| HG0228 | *D. rerio* | ENSDARG00000087887 | phf21b | PHD finger protein 21B | 0.05 | 0.39 | 2.1E-13 | 9.4E-13 | 0.39 | 2.1E-13 | | MSLTASKEARTCCAARVVLAIVIVAFQNTSLTPQITRIASEVEEKCS* |
| HG0229 | *G. gallus* | ENSGALG00000005474 | | | 0.00 | 0.40 | 3.0E-03 | 7.7E-03 | 0.40 | 3.0E-03 | | MCIHFWWTLPFLGV* |
| HG0230 | *G. gallus* | ENSGALG00000011007 | OSBPL3 | oxysterol binding protein like 3 | 0.00 | 0.48 | 2.0E-05 | 7.2E-05 | 0.48 | 2.0E-05 | | MQIMLLECIWVRDEFCITLRSLPIHLLDYLLMTF* |
| HG0231 | *H. sapiens* | ENSG00000150636 | | | 0.13 | 0.48 | 1.3E-04 | 5.9E-04 | 0.48 | 1.3E-04 | | MQEILLFTHVAATETLEQPSKPRGS* |
| HG0232 | *H. sapiens* | ENSG00000180875 | GREM2 | gremlin 2, DAN family BMP antagonist | 0.20 | 0.34 | 9.4E-11 | 1.1E-09 | 0.34 | 9.4E-11 | | MGVCCSLAAVELKAAAGDQT* |
| HG0233 | *H. sapiens* | ENSG00000206432 | TMEM200C | transmembrane protein 200C | 0.17 | 0.45 | 1.2E-08 | 1.1E-07 | 0.45 | 1.2E-08 | | MSGGEEGSEAGCGMRDAEDLTLLGNSWKWLPPQRRRGLRAWGGRSPPRGC* |
| HG0234.1 | *G. gallus* | ENSGALG00000011250 | ASIC4 | acid sensing ion channel subunit family member 4 | 0.10 | 0.00 | 5.9E-03 | 1.4E-02 | 0.00 | 5.9E-03 | | MLTCLELLSIPWILLAHDA* |
| HG0234.2 | *H. sapiens* | ENSG00000108684 | ASIC2 | acid sensing ion channel subunit 2 | 0.03 | 0.43 | 1.5E-04 | 6.5E-04 | 0.43 | 1.5E-04 | | MVNMGHYALVNRLRAAQRLVLSIFSLALAFNLVSSGVCRGEDFISIG* |
| HG0235.1 | *G. gallus* | ENSGALG00000035927 | | | 0.00 | 0.48 | 5.7E-08 | 2.5E-07 | 0.48 | 5.7E-08 | | MLATSESLGNFTVGALRAVLELTAKEARRGQTMGCPVLGLLHEQTPRAPLGCPAKRA* |
| HG0235.2 | *H. sapiens* | ENSG00000101638 | ST8SIA5 | ST8 alpha-N-acetyl-neuraminide alpha-2,8-sialyltransferase | 0.03 | 0.37 | 9.2E-05 | 4.2E-04 | 0.37 | 9.2E-05 | | MPPAPAAPTRNFAPRSPPPA* |
| HG0236 | *G. gallus* | ENSGALG00000004437 | LHFPL2 | lipoma HMGIC fusion partner-like 2 | 0.02 | 0.47 | 1.0E-07 | 4.6E-07 | 0.47 | 1.0E-07 | | MVAAGAGQGEPGGARAGGGGGSGAEPQRTGGAGERALLAGGIARLLKWLTKAKNRGHWCVMSSSGQSPRPSPPVRSVMCQYEQ* |
| HG0237 | *G. gallus* | ENSGALG00000012123 | PAX6 | paired box 6 | 0.00 | 0.35 | 1.3E-04 | 4.1E-04 | 0.35 | 1.3E-04 | | MCGVISGFEKRAPARPHFPLWLRRREEVFCWKMMTEVRLC* |
| HG0238 | *H. sapiens* | ENSG00000049618 | ARID1B | AT-rich interaction domain 1B | 0.16 | 0.43 | 1.9E-05 | 1.0E-04 | 0.43 | 1.9E-05 | | MLGPIGSASRVFTHIRAREEKRKVSFKPELKTFTMKAHSRSRPRA* |
| HG0239 | *H. sapiens* | ENSG00000149256 | TENM4 | teneurin transmembrane protein 4 | 0.00 | 0.33 | 1.4E-02 | 3.5E-02 | 0.33 | 1.4E-02 | | MEPGRLKLPCVQIKIAYFKDYLKDYS* |
| HG0240 | *H. sapiens* | ENSG00000166342 | NETO1 | neuropilin and tolloid like 1 | 0.06 | 0.42 | 1.0E-08 | 9.2E-08 | 0.42 | 1.0E-08 | | MTDPFFLPSMPQRKIPKGWSEERCRWTSSRGGCSDLLRGASETGD* |
| HG0241 | *H. sapiens* | ENSG00000198561 | CTNND1 | catenin delta 1 | 0.06 | 0.49 | 1.4E-03 | 4.8E-03 | 0.49 | 1.4E-03 | | MEVSGWLCSLLPASSLLWWLGCFFHDFLNLDWAVLCVKPISCDLLLTV* |
| HG0242.1 | *G. gallus* | ENSGALG00000007112 | LINS1 | lines homolog 1 | 0.00 | 0.21 | 3.5E-08 | 1.6E-07 | 0.21 | 3.5E-08 | | MSRGNLEEECLLACLFLLLDKNNFKTDLQQSEIIGEDLQRSVIY* |
| HG0242.2 | *H. sapiens* | ENSG00000140471 | LINS1 | lines homolog 1 | 0.08 | 0.45 | 1.1E-02 | 2.7E-02 | 0.45 | 1.1E-02 | | MREEVPACTFCELRWPRAGICVPSNGGVKRVRLSQIPGFTEETS* |
| HG0243.1 | *H. sapiens* | ENSG00000163697 | APBB2 | amyloid beta precursor protein binding family B member 2 | 0.04 | 0.43 | 3.8E-04 | 1.5E-03 | 0.43 | 3.8E-04 | | MGSLLSRQLIFQIITYFLHHC* |
| HG0243.2 | *H. sapiens* | ENSG00000163697 | APBB2 | amyloid beta precursor protein binding family B member 2 | 0.00 | 0.13 | 2.9E-17 | 5.9E-16 | 0.13 | 2.9E-17 | Crowe | MLWSDLPSRGTETFAGWKIHKRKLL* |
| HG0244 | *G. gallus* | ENSGALG00000010837 | ASB5 | ankyrin repeat and SOCS box containing 5 | 0.00 | 0.22 | 9.7E-05 | 3.2E-04 | 0.22 | 9.7E-05 | | MLTGTLRSLKAVVMKQPKYQHFSRIDPSEQLMLMVVF* |
| HG0245 | *H. sapiens* | ENSG00000091831 | ESR1 | estrogen receptor 1 | 0.24 | 0.37 | 5.6E-05 | 2.7E-04 | 0.37 | 5.6E-05 | | MQPHTPFYLPYLGYSVVLPRVILCTHYVFLANEEGESNRKRDKQRYIGVWHGAHKAAH* |
| HG0246 | *H. sapiens* | ENSG00000162676 | GFI1 | growth factor independent 1 transcriptional repressor | 0.12 | 0.30 | 6.3E-08 | 5.1E-07 | 0.30 | 6.3E-08 | | MSKPLGLGVKVSSGLGLSKPSE* |
| HG0247.1 | *D. rerio* | ENSDARG00000014746 | rbfox1 | RNA binding protein, fox-1 homolog (C. elegans) 1 | 0.05 | 0.00 | 8.4E-07 | 3.1E-06 | 0.00 | 8.4E-07 | | MLALGRCCRDWD* |
| HG0247.2 | *H. sapiens* | ENSG00000100320 | RBFOX2 | RNA binding protein, fox-1 homolog 2 | 0.05 | 0.00 | 1.2E-05 | 6.8E-05 | 0.00 | 1.2E-05 | | MGGSWRKKEGERLPERKRKERKKH* |
| HG0248.1 | *G. gallus* | ENSGALG00000004570 | NCOA3 | nuclear receptor coactivator 3 | 0.00 | 0.40 | 2.1E-02 | 4.3E-02 | 0.40 | 2.1E-02 | | MVSDSGDPVKSILCLNAC* |
| HG0248.2 | *H. sapiens* | ENSG00000124151 | NCOA3 | nuclear receptor coactivator 3 | 0.00 | 0.41 | 1.8E-09 | 1.8E-08 | 0.41 | 1.8E-09 | | MAAGGSKYLLDGGLRDQ* |
| HG0249 | *D. rerio* | ENSDARG00000062550 | rc3h1a | ring finger and CCCH-type domains 1a | 0.00 | 0.08 | 7.9E-04 | 2.2E-03 | 0.08 | 7.9E-04 | | MLCFSHYVLNAHNARKYLRTVIFGRALSVPVEPLGLTAVSFYAKRT* |
| HG0250 | *G. gallus* | ENSGALG00000014861 | | | 0.02 | 0.30 | 3.4E-03 | 8.7E-03 | 0.30 | 3.4E-03 | | MNWKHLDLCRALNQILV* |
| HG0251 | *H. sapiens* | ENSG00000185053 | SGCZ | sarcoglycan zeta | 0.00 | 0.09 | 7.6E-05 | 3.6E-04 | 0.09 | 7.6E-05 | | MMLTREAAFFLGMQQFAGF* |
| HG0252.1 | *G. gallus* | ENSGALG00000009645 | ESRRG | estrogen related receptor gamma | 0.00 | 0.03 | 7.2E-04 | 2.1E-03 | 0.03 | 7.2E-04 | | MSLHGVVDFPSKETGFGFFKIPDDSPD* |
| HG0252.2 | *H. sapiens* | ENSG00000119715 | ESRRB | estrogen related receptor beta | 0.00 | 0.49 | 1.4E-03 | 4.9E-03 | 0.49 | 1.4E-03 | | MVVRLFFFFFFFCQLPVISITESGSIYRLGVLCP* |
| HG0252.3 | *H. sapiens* | ENSG00000196482 | ESRRG | estrogen related receptor gamma | 0.00 | 0.22 | 1.1E-04 | 5.1E-04 | 0.22 | 1.1E-04 | | MCNVPAWGFCRLS* |
| HG0253.1 | *G. gallus* | ENSGALG00000015184 | | | 0.02 | 0.28 | 6.1E-05 | 2.0E-04 | 0.28 | 6.1E-05 | | MYQNVNDEALLNLVKGVTSAERAVGAARSGGAGRRAGQGRVGQGGPGRAGHGAPSSAAAASRSLDRQRRARKPNTNKAARTVPSSSSLSLFLIF* |
| HG0253.2 | *H. sapiens* | ENSG00000140332 | TLE3 | transducin like enhancer of split 3 | 0.03 | 0.41 | 4.8E-16 | 9.1E-15 | 0.41 | 4.8E-16 | | MSAQSQALTRSTQPREGGRPLPAEP* |
| HG0253.3 | *H. sapiens* | ENSG00000140332 | TLE3 | transducin like enhancer of split 3 | 0.00 | 0.00 | 2.0E-03 | 6.5E-03 | 0.00 | 2.0E-03 | | MPGRLICTY* |
| HG0254.1 | *G. gallus* | ENSGALG00000035919 | | | 0.05 | 0.46 | 3.4E-04 | 1.0E-03 | 0.46 | 3.4E-04 | | MLDGACGCQTQPEAAADGRRDPSCAPPVPPRS* |
| HG0254.2 | *H. sapiens* | ENSG00000072134 | EPN2 | epsin 2 | 0.00 | 0.45 | 5.6E-03 | 1.6E-02 | 0.45 | 5.6E-03 | | MTIQGSYSSLITETEPS* |
| HG0255 | *G. gallus* | ENSGALG00000012139 | ARL14EP | ADP ribosylation factor like GTPase 14 effector protein | 0.00 | 0.20 | 1.6E-05 | 5.7E-05 | 0.20 | 1.6E-05 | | MSATGSAAVGGAARKPGAGVPAQRRRKVHGKGDLHRNH* |
| HG0256 | *G. gallus* | ENSGALG00000039901 | | | 0.23 | 0.35 | 7.7E-05 | 2.5E-04 | 0.35 | 7.7E-05 | | MAATTTISLRHSQPCNRFPRDVPELLEFKPPKGPRSTQKQCP* |
| HG0257 | *H. sapiens* | ENSG00000198791 | CNOT7 | CCR4-NOT transcription complex subunit 7 | 0.00 | 0.41 | 6.7E-04 | 2.5E-03 | 0.41 | 6.7E-04 | | MLSPVMRRPLVSLRGFWDCSREREGFEISLPKNQQELCL* |
| HG0258.1 | *G. gallus* | ENSGALG00000016797 | FHL2 | four and a half LIM domains 2 | 0.00 | 0.19 | 2.0E-02 | 4.1E-02 | 0.19 | 2.0E-02 | | MSNCLITKYIVWVIQ* |
| HG0258.2 | *G. gallus* | ENSGALG00000016797 | FHL2 | four and a half LIM domains 2 | 0.18 | 0.00 | 1.7E-14 | 1.1E-13 | 0.00 | 1.7E-14 | | MDVTLCPWSRR* |
| HG0259 | *G. gallus* | ENSGALG00000009936 | AP4S1 | adaptor related protein complex 4 sigma 1 subunit | 0.00 | 0.15 | 1.5E-02 | 3.1E-02 | 0.15 | 1.5E-02 | | MLKSKMAAPGKGAQFRP* |
| HG0260.1 | *G. gallus* | ENSGALG00000016890 | MBNL2 | muscleblind-like 2 isoform 7 | 0.00 | 0.36 | 3.6E-27 | 3.3E-26 | 0.36 | 3.6E-27 | | MLLSMIFFSQKSTSGT* |
| HG0260.2 | *H. sapiens* | ENSG00000139793 | MBNL2 | muscleblind like splicing regulator 2 | 0.00 | 0.47 | 2.1E-16 | 4.0E-15 | 0.47 | 2.1E-16 | | MSDCLSIHSLIILFLGLDFTKQFITHLQTYMWEFSQQ* |
| HG0260.3 | *H. sapiens* | ENSG00000152601 | MBNL1 | muscleblind like splicing regulator 1 | 0.00 | 0.35 | 3.3E-09 | 3.2E-08 | 0.35 | 3.3E-09 | | MTPTMPWALGRQWGRL* |
| HG0260.4 | *H. sapiens* | ENSG00000152601 | MBNL1 | muscleblind like splicing regulator 1 | 0.00 | 0.30 | 7.3E-08 | 5.8E-07 | 0.30 | 7.3E-08 | | MSCGAHNAPMTRS* |
| HG0260.5 | *H. sapiens* | ENSG00000152601 | MBNL1 | muscleblind like splicing regulator 1 | 0.10 | 0.38 | 7.5E-03 | 2.1E-02 | 0.38 | 7.5E-03 | | MFFPPVSPPGSKTVWCILRRQSHQIAGF* |
| HG0260.6 | *H. sapiens* | ENSG00000152601 | MBNL1 | muscleblind like splicing regulator 1 | 0.03 | 0.06 | 4.1E-03 | 1.2E-02 | 0.06 | 4.1E-03 | | MGLRFICSGDNMAYLPWNIPVFQK* |
| HG0261.1 | *G. gallus* | ENSGALG00000005867 | MARCH8 | membrane associated ring-CH-type finger 8 | 0.00 | 0.24 | 8.2E-03 | 1.9E-02 | 0.24 | 8.2E-03 | | MSAAFKPPSCK* |
| HG0261.2 | *G. gallus* | ENSGALG00000009520 | MARCH1 | membrane associated ring-CH-type finger 1 | 0.00 | 0.26 | 1.3E-04 | 4.1E-04 | 0.26 | 1.3E-04 | | MEYLTQRQEMKPGSNVKGHVKLLHSRNLLKDCFT* |
| HG0261.3 | *H. sapiens* | ENSG00000145416 | MARCH1 | membrane associated ring-CH-type finger 1 | 0.00 | 0.49 | 8.0E-05 | 3.7E-04 | 0.49 | 8.0E-05 | | MPAPRDAGRTQPGGRLSYFTLRR* |
| HG0261.4 | *H. sapiens* | ENSG00000165406 | MARCH8 | membrane associated ring-CH-type finger 8 | 0.03 | 0.48 | 2.1E-02 | 4.8E-02 | 0.48 | 2.1E-02 | | MLEAGQGGSAASSEPT* |
| HG0262.1 | *H. sapiens* | ENSG00000023445 | BIRC3 | baculoviral IAP repeat containing 3 | 0.00 | 0.37 | 1.3E-05 | 7.0E-05 | 0.35 | 3.2E-05 | | MFLLPPGNKALKSAVEKPNVCEIQETMLANHW* |
| HG0262.2 | *H. sapiens* | ENSG00000023445 | BIRC3 | baculoviral IAP repeat containing 3 | 0.00 | 0.39 | 5.6E-03 | 1.6E-02 | 0.39 | 5.6E-03 | | MSLILNLICGNKGKVILAGAYC* |
| HG0262.3 | *G. gallus* | ENSGALG00000017186 | | | 0.09 | 0.10 | 6.8E-03 | 1.6E-02 | 0.10 | 6.8E-03 | | MHSTRKAHNVTVAQEKVLFDIPVFVNGVDQLLH* |
| HG0262.4 | *H. sapiens* | ENSG00000023445 | BIRC3 | baculoviral IAP repeat containing 3 | 0.00 | 0.14 | 3.9E-03 | 1.2E-02 | 0.14 | 3.9E-03 | | MNTKKIFITMCKIFGQGKEY* |
| HG0263.1 | *H. sapiens* | ENSG00000169826 | CSGALNACT2 | chondroitin sulfate N-acetylgalactosaminyltransferase 2 | 0.14 | 0.48 | 7.1E-03 | 2.0E-02 | 0.48 | 7.1E-03 | | MEILILFLIFDNFLLKV* |
| HG0263.2 | *G. gallus* | ENSGALG00000002543 | CSGALNACT2 | chondroitin sulfate N-acetylgalactosaminyltransferase 2 | 0.00 | 0.20 | 1.5E-04 | 4.8E-04 | 0.20 | 1.5E-04 | | MANVGTCDLWLCNDT* |
| HG0263.3 | *G. gallus* | ENSGALG00000002543 | CSGALNACT2 | chondroitin sulfate N-acetylgalactosaminyltransferase 2 | 0.00 | 0.32 | 1.1E-04 | 3.6E-04 | 0.32 | 1.1E-04 | | MIPGWCLPFCSPSAVFFVWDKKS* |
| HG0263.4 | *H. sapiens* | ENSG00000147408 | CSGALNACT1 | chondroitin sulfate N-acetylgalactosaminyltransferase 1 | 0.00 | 0.34 | 1.5E-02 | 3.7E-02 | 0.34 | 1.5E-02 | | MDASSSGGPKRMLPVV* |
| HG0264.1 | *H. sapiens* | ENSG00000151458 | ANKRD50 | ankyrin repeat domain 50 | 0.00 | 0.45 | 2.0E-10 | 2.3E-09 | 0.45 | 2.0E-10 | | MELCDSLKFRKSVAELWNQAFLPQFNN* |
| HG0264.2 | *H. sapiens* | ENSG00000151458 | ANKRD50 | ankyrin repeat domain 50 | 0.00 | 0.31 | 1.4E-16 | 2.8E-15 | 0.31 | 1.4E-16 | | MYCLISVLETSTIT* |
| HG0264.3 | *G. gallus* | ENSGALG00000011831 | ANKRD50 | ankyrin repeat domain 50 | 0.00 | 0.01 | 3.4E-03 | 8.7E-03 | 0.01 | 3.4E-03 | | MDSFFWIYIRRD* |
| HG0264.4 | *H. sapiens* | ENSG00000151458 | ANKRD50 | ankyrin repeat domain 50 | 0.18 | 0.15 | 3.6E-03 | 1.1E-02 | 0.15 | 3.6E-03 | | MERPVLMYPDSSK* |
| HG0265.1 | *H. sapiens* | ENSG00000144285 | SCN1A | sodium voltage-gated channel alpha subunit 1 | 0.00 | 0.30 | 2.0E-05 | 1.0E-04 | 0.30 | 2.0E-05 | | MELFLQINTSGVMEAGGS* |
| HG0265.2 | *H. sapiens* | ENSG00000196876 | SCN8A | sodium voltage-gated channel alpha subunit 8 | 0.16 | 0.21 | 1.2E-06 | 7.9E-06 | 0.21 | 1.2E-06 | | MAPTAVPPAASSAPLQSGRASRAPR* |
| HG0265.3 | *D. rerio* | ENSDARG00000018032 | scn8ab | sodium channel, voltage gated, type VIII, alpha subunit b | 0.00 | 0.36 | 8.9E-04 | 2.4E-03 | 0.36 | 8.9E-04 | | MATSAVSPGSFKHP* |
| HG0266.1 | *H. sapiens* | ENSG00000184185 | KCNJ12 | potassium voltage-gated channel subfamily J member 12 | 0.03 | 0.48 | 4.0E-10 | 4.5E-09 | 0.48 | 4.0E-10 | | MAATTNSALKTVPCPLSTIT* |
| | | ENSG00000260458 | KCNJ18 | potassium voltage-gated channel subfamily J member 18 | 0.08 | 0.47 | 3.4E-09 | 3.4E-08 | 0.47 | 3.4E-09 | | MAATTNSALKTVPCLLSTIT* |
| HG0266.2 | *G. gallus* | ENSGALG00000001181 | KCNJ5 | potassium voltage-gated channel subfamily J member 5 | 0.28 | 0.39 | 2.4E-04 | 7.3E-04 | 0.39 | 2.4E-04 | | MFCHAGQREGRRRRPRDQKTTTACW* |
| HG0267 | *D. rerio* | ENSDARG00000028173 | slc4a2a | solute carrier family 4 (anion exchanger), member 2a | 0.00 | 0.33 | 7.2E-05 | 2.2E-04 | 0.33 | 7.2E-05 | Mackowiak | MLPESSQLSLSKREKG* |
| | | ENSDARG00000052330 | slc4a2b | solute carrier family 4 (anion exchanger), member 2b | 0.00 | 0.37 | 3.4E-14 | 1.6E-13 | 0.37 | 3.4E-14 | Mackowiak | MLPVNSCLSLSEKEKG* |
| | *H. sapiens* | ENSG00000164889 | SLC4A2 | solute carrier family 4 member 2 | 0.00 | 0.06 | 3.1E-31 | 1.1E-29 | 0.06 | 3.1E-31 | Mackowiak | MPPPRCPESRSDSEKG* |
| HG0268.1 | *D. rerio* | ENSDARG00000011703 | clocka | clock circadian regulator a | 0.00 | 0.34 | 7.6E-07 | 2.8E-06 | 0.34 | 7.6E-07 | | MSTSRLKVLSLQLRP* |
| HG0268.2 | *H. sapiens* | ENSG00000134852 | CLOCK | clock circadian regulator | 0.00 | 0.34 | 3.1E-04 | 1.3E-03 | 0.34 | 3.1E-04 | | MVGHIEEYLEPIVS* |
| HG0268.3 | *H. sapiens* | ENSG00000134852 | CLOCK | clock circadian regulator | 0.01 | 0.40 | 1.5E-02 | 3.8E-02 | 0.40 | 1.5E-02 | | MITRRQLVLRRSTNVYYKTKT* |
| HG0269.1 | *G. gallus* | ENSGALG00000028685 | SEMA5A | semaphorin 5A | 0.00 | 0.32 | 2.5E-06 | 9.7E-06 | 0.32 | 2.5E-06 | | MVTERCFSPMSARRVSS* |
| HG0269.2 | *D. rerio* | ENSDARG00000061471 | sema5ba | sema domain, seven thrombospondin repeats (type 1 and t | 0.29 | 0.04 | 5.2E-03 | 1.1E-02 | 0.04 | 5.2E-03 | | MLVMPLCIPEAHLGS* |
| HG0269.3 | *G. gallus* | ENSGALG00000011686 | SEMA5B | semaphorin 5B | 0.02 | 0.11 | 8.6E-03 | 2.0E-02 | 0.11 | 8.6E-03 | | MPLTQTEITKDFPWCLTSNMTLQ* |
| HG0270.1 | *G. gallus* | ENSGALG00000015147 | ALDH1A1 | aldehyde dehydrogenase 1 family member A1 | 0.00 | 0.46 | 2.0E-15 | 1.4E-14 | 0.46 | 2.0E-15 | | MVNVTVFKMFHKCRSRITIFPSCFLFFFFSFLFFFFFISSLARMSELD* |
| HG0270.2 | *G. gallus* | ENSGALG00000015147 | ALDH1A1 | aldehyde dehydrogenase 1 family member A1 | 0.00 | 0.32 | 2.3E-17 | 1.7E-16 | 0.32 | 2.3E-17 | | MISFLFLALPPPSLSLISLT* |
| HG0270.3 | *H. sapiens* | ENSG00000128918 | ALDH1A2 | aldehyde dehydrogenase 1 family member A2 | 0.01 | 0.05 | 3.5E-05 | 1.8E-04 | 0.05 | 3.5E-05 | | MEAAPGVLGRMDQPPAPRPSRPIRRAPA* |
| HG0271.1 | *G. gallus* | ENSGALG00000004064 | GPER1 | G protein-coupled estrogen receptor 1 | 0.00 | 0.29 | 7.8E-11 | 4.1E-10 | 0.29 | 7.8E-11 | | MELELCHLPTCTKENC* |
| HG0271.2 | *H. sapiens* | ENSG00000164850 | GPER1 | G protein-coupled estrogen receptor 1 | 0.10 | 0.38 | 1.3E-05 | 7.3E-05 | 0.38 | 1.3E-05 | | MPGSLQGESRFFLKMDSPFKTELWEPFGKS* |
| HG0271.3 | *D. rerio* | ENSDARG00000074661 | gper1 | G protein-coupled estrogen receptor 1 | 0.00 | 0.37 | 7.1E-04 | 1.9E-03 | 0.37 | 7.1E-04 | | MLMDWRKQDCAGLNTCEKQSVQKKKKKQDP* |
| HG0272.1 | *D. melanogaster* | FBgn0034649 | CG9865 | CG9865, isoform A | 0.09 | 0.06 | 5.2E-266 | 1.3E-265 | 0.06 | 5.2E-266 | Hayden | MSDTRKDPCKPNACRIQACLTKNHYQEDKCLDVLEAMRQCCLKWHKESLCCSGIDLEKTYLPDASKEKNATGKSNK* |
| HG0272.2 | *D. melanogaster* | FBgn0034649 | CG9865 | CG9865, isoform A | 0.00 | 0.05 | 2.2E-230 | 4.4E-230 | 0.05 | 2.2E-230 | Hayden | MESVRKANQRIRNYPVLLSKCADKATAYAVCVSRDLNVQHKICDTEFKEFLSCIRKTALEMKTKL* |
| HG0272.3 | *H. sapiens* | ENSG00000143315 | PIGM | phosphatidylinositol glycan anchor biosynthesis class M | 0.07 | 0.26 | 1.6E-04 | 6.8E-04 | 0.26 | 1.6E-04 | | MPDVIVCSRFGWQPGGWEMRRPPAAKNRREG* |
| HG0273.1 | *G. gallus* | ENSGALG00000003293 | MBTPS1 | membrane bound transcription factor peptidase, site 1 | 0.00 | 0.00 | 2.5E-25 | 2.2E-24 | 0.00 | 2.5E-25 | | MSLFIRLYLKRQ* |
| HG0273.2 | *H. sapiens* | ENSG00000140943 | MBTPS1 | membrane bound transcription factor peptidase, site 1 | 0.00 | 0.49 | 1.3E-06 | 8.4E-06 | 0.49 | 1.3E-06 | | MECPGVFAAEAPLSRARRP* |
| HG0273.3 | *H. sapiens* | ENSG00000140943 | MBTPS1 | membrane bound transcription factor peptidase, site 1 | 0.00 | 0.00 | 1.4E-03 | 4.7E-03 | 0.00 | 1.4E-03 | | MLQLLAETKIHF* |
| HG0274.1 | *H. sapiens* | ENSG00000204256 | BRD2 | bromodomain containing 2 | 0.00 | 0.41 | 1.2E-28 | 4.0E-27 | 0.44 | 1.5E-19 | | MTGPSGQYRPPRSPWSSPFSGPAQSSESVHPLYSALKRISKMEAAAP* |
| HG0274.2 | *D. rerio* | ENSDARG00000100129 | | | 0.13 | 0.43 | 1.2E-02 | 2.3E-02 | 0.43 | 1.2E-02 | | MLLRNKSYPKICKDNATGAARSESNKVQHIHLRLAHSDRSSRSV* |
| HG0275.1 | *H. sapiens* | ENSG00000100239 | PPP6R2 | protein phosphatase 6 regulatory subunit 2 | 0.00 | 0.35 | 2.4E-04 | 1.0E-03 | 0.37 | 1.9E-03 | | MKKIFLKLHILQEKNHKCFSVLPEYMI* |

| | | | | | | | | | | | | |
|---|---|---|---|---|---|---|---|---|---|---|---|---|
| HG0275.2 | *D. rerio* | ENSDARG00000045540 | ppp6r2a | protein phosphatase 6, regulatory subunit 2a | 0.03 | 0.29 | 2.8E-03 | 6.7E-03 | 0.29 | 2.8E-03 | | MLGHSVPSAFCHSNACRPVEEV* |
| HG0276.1 | *G. gallus* | ENSGALG00000008912 | | | 0.13 | 0.28 | 7.7E-04 | 2.2E-03 | 0.28 | 7.7E-04 | | MDSSSSITVTWNIQHHM* |
| HG0276.2 | *H. sapiens* | ENSG00000073734 | ABCB11 | ATP binding cassette subfamily B member 11 | 0.12 | 0.30 | 6.9E-03 | 1.9E-02 | 0.30 | 6.9E-03 | | MMKTEVGKGCETF* |
| HG0277.1 | *H. sapiens* | ENSG00000102531 | FNDC3A | fibronectin type III domain containing 3A | 0.10 | 0.02 | 5.4E-16 | 1.0E-14 | 0.02 | 5.4E-16 | | MAAAAFVVPRVKQKAGATRRGSEERG* |
| HG0277.2 | *G. gallus* | ENSGALG00000004169 | | | 0.06 | 0.00 | 7.2E-03 | 1.7E-02 | 0.00 | 7.2E-03 | | MAERSVGGRQRRG* |
| HG0278.1 | *H. sapiens* | ENSG00000119946 | CNNM1 | cyclin and CBS domain divalent metal cation transport med | 0.20 | 0.22 | 2.0E-09 | 2.0E-08 | 0.22 | 2.0E-09 | | MPGHLQPPAGTSCPGGV* |
| HG0278.2 | *D. rerio* | ENSDARG00000078733 | cnnm2b | cyclin and CBS domain divalent metal cation transport med | 0.00 | 0.19 | 2.7E-03 | 6.5E-03 | 0.19 | 2.7E-03 | | MHKSRRRAAEMKPDWL* |
| HG0279.1 | *H. sapiens* | ENSG00000175928 | LRRN1 | leucine rich repeat neuronal 1 | 0.00 | 0.35 | 1.0E-13 | 1.6E-12 | 0.38 | 1.3E-10 | | MFTFSARCPTYHNIVFTFC* |
| HG0279.2 | *G. gallus* | ENSGALG00000035099 | Lrrn1 | leucine rich repeat neuronal 1 | 0.28 | 0.47 | 2.3E-02 | 4.6E-02 | 0.47 | 2.3E-02 | | MSAPPLRGGEERLRGQRPPGCRNVFRHGFVEDPGTQRSS* |
| HG0280.1 | *H. sapiens* | ENSG00000156299 | TIAM1 | T-cell lymphoma invasion and metastasis 1 | 0.00 | 0.26 | 2.2E-05 | 1.2E-04 | 0.26 | 2.2E-05 | Crowe | MAHLEILTKPLLFPMGFLAHSHRLR* |
| HG0280.2 | *D. melanogaster* | FBgn0085447 | sif | still life | 0.00 | 0.04 | 9.4E-08 | 9.7E-08 | 0.04 | 9.4E-08 | | MGAKESVEAAAKMNADGWGHVERSSRYHNHRHNQRSQSLNRGQALSQADVSGMW* |
| HG0281.1 | *G. gallus* | ENSGALG00000006766 | DEPDC5 | DEP domain containing 5 | 0.01 | 0.12 | 4.8E-28 | 4.5E-27 | 0.12 | 4.8E-28 | | MCRMSSTVFTRNS* |
| HG0281.2 | *H. sapiens* | ENSG00000100150 | DEPDC5 | DEP domain containing 5 | 0.04 | 0.11 | 1.0E-09 | 1.1E-08 | 0.11 | 1.0E-09 | | MTSLPQAWNS* |
| HG0282.1 | *D. rerio* | ENSDARG00000061328 | cdon | cell adhesion associated, oncogene regulated | 0.07 | 0.29 | 2.2E-19 | 1.2E-18 | 0.29 | 2.2E-19 | | MEGPVRICHRENVFRC* |
| HG0282.2 | *H. sapiens* | ENSG00000144857 | BOC | BOC cell adhesion associated, oncogene regulated | 0.05 | 0.27 | 1.8E-02 | 4.2E-02 | 0.27 | 1.8E-02 | | MKSCRHLYRLRVAARK* |
| HG0283.1 | *G. gallus* | ENSGALG00000010270 | AREL1 | apoptosis resistant E3 ubiquitin protein ligase 1 | 0.10 | 0.29 | 3.9E-31 | 3.9E-30 | 0.29 | 3.9E-31 | | MQVPGVPFSVTAFSMASLC* |
| HG0283.2 | *H. sapiens* | ENSG00000119682 | AREL1 | apoptosis resistant E3 ubiquitin protein ligase 1 | 0.00 | 0.42 | 7.2E-24 | 2.1E-22 | 0.42 | 7.2E-24 | | MDRPFLLTLWRFSFMWKLEDKVKGAKSYLLW* |
| HG0284.1 | *H. sapiens* | ENSG00000007168 | PAFAH1B1 | platelet activating factor acetylhydrolase 1b regulatory subu | 0.13 | 0.10 | 4.4E-05 | 2.2E-04 | 0.10 | 4.4E-05 | Crowe | MGVKDGRGPAEAAVQRSGGMNLTC* |
| HG0284.2 | *G. gallus* | ENSGALG00000005834 | PAFAH1B1 | platelet activating factor acetylhydrolase 1b regulatory subu | 0.03 | 0.18 | 1.5E-02 | 3.3E-02 | 0.18 | 1.5E-02 | | MPPGFSVGFPIWRNICALSHCGKLNCGS* |
| HG0285.1 | *G. gallus* | ENSGALG00000012156 | DPP10 | dipeptidyl peptidase like 10 | 0.20 | 0.13 | 9.2E-03 | 2.1E-02 | 0.13 | 9.2E-03 | | MLRGRPRAAGQEEHGGGADTGLGFPPAGAVAPRLPPAFLFLYFSSSSAAAPLRGEVGVMRLAPREGCPGPQRQ* |
| HG0285.2 | *H. sapiens* | ENSG00000175497 | DPP10 | dipeptidyl peptidase like 10 | 0.00 | 0.00 | 9.9E-03 | 2.6E-02 | 0.00 | 9.9E-03 | | MNQVNALFRAK* |
| HG0286.1 | *H. sapiens* | ENSG00000116698 | SMG7 | SMG7, nonsense mediated mRNA decay factor | 0.00 | 0.00 | 5.8E-04 | 2.2E-03 | 0.00 | 5.8E-04 | | MSLQSAQYLRF* |
| HG0286.2 | *D. rerio* | ENSDARG00000060767 | smg7 | SMG7 nonsense mediated mRNA decay factor | 0.08 | 0.49 | 9.9E-03 | 2.0E-02 | 0.49 | 9.9E-03 | | MDNMAPGVDLACESVVV* |
| HG0287.1 | *H. sapiens* | ENSG00000170325 | PRDM10 | PR/SET domain 10 | 0.00 | 0.33 | 1.7E-12 | 2.4E-11 | 0.33 | 1.7E-12 | | MLQTDLSVLHEHTCGRQMTVLYPVLL* |
| HG0287.2 | *D. rerio* | ENSDARG00000104251 | prdm10 | PR domain containing 10 | 0.00 | 0.40 | 5.4E-07 | 2.0E-06 | 0.40 | 5.4E-07 | | MQGSLLCSYHSEGRPTSTFS* |
| HG0288.1 | *G. gallus* | ENSGALG00000006655 | MAP3K13 | mitogen-activated protein kinase kinase kinase 13 | 0.20 | 0.44 | 4.8E-03 | 1.2E-02 | 0.44 | 4.8E-03 | | MIRYWLPASRLNCPVL* |
| HG0288.2 | *H. sapiens* | ENSG00000073803 | MAP3K13 | mitogen-activated protein kinase kinase kinase 13 | 0.02 | 0.41 | 6.9E-04 | 2.6E-03 | 0.41 | 6.9E-04 | | MYCGIQILALWERFWSPLLSQNSVPKIF* |
| HG0289 | *G. gallus* | ENSGALG00000009581 | MPP5 | membrane palmitoylated protein 5 | 0.00 | 0.45 | 1.1E-32 | 1.1E-31 | 0.45 | 1.1E-32 | | MDFTQQRNENWPNISISI* |
| | *H. sapiens* | ENSG00000072415 | MPP5 | membrane palmitoylated protein 5 | 0.00 | 0.30 | 1.5E-16 | 2.9E-15 | 0.30 | 1.5E-16 | | MDFILQNQENWLIGKIDL* |
| HG0290.1 | *H. sapiens* | ENSG00000116539 | ASH1L | ASH1 like histone lysine methyltransferase | 0.00 | 0.48 | 1.9E-09 | 1.9E-08 | 0.48 | 1.9E-09 | | MLVSSFGDPCFLWNTVL* |
| HG0290.2 | *D. rerio* | ENSDARG00000070981 | ash1l | ash1 (absent, small, or homeotic)-like (Drosophila) | 0.04 | 0.28 | 6.6E-06 | 2.3E-05 | 0.28 | 6.6E-06 | | MEARLTREARSDEPELH* |
| HG0291.1 | *G. gallus* | ENSGALG00000009173 | GFRA1 | GDNF family receptor alpha 1 | 0.24 | 0.41 | 2.2E-03 | 5.9E-03 | 0.41 | 2.2E-03 | | MDRNLESLHLGGFGISTC* |
| HG0291.2 | *H. sapiens* | ENSG00000151892 | GFRA1 | GDNF family receptor alpha 1 | 0.14 | 0.44 | 3.0E-06 | 1.8E-05 | 0.44 | 3.0E-06 | | MELNFGRPEQHSCPGIAAR* |
| HG0292.1 | *G. gallus* | ENSGALG00000006502 | PCGF5 | polycomb group ring finger 5 | 0.17 | 0.24 | 1.0E-28 | 9.6E-28 | 0.24 | 1.0E-28 | | MPGKVPLRRAAPQLS* |
| HG0292.2 | *H. sapiens* | ENSG00000180628 | PCGF5 | polycomb group ring finger 5 | 0.00 | 0.11 | 5.2E-06 | 3.0E-05 | 0.11 | 5.2E-06 | | MALSPLRRRWDRLGEDAAEETNSQDMGKRNHQKE* |
| HG0293 | *G. gallus* | ENSGALG00000012187 | MGAT5 | mannosyl (alpha-1,6-)-glycoprotein beta-1,6-N-acetyl-gluco | 0.00 | 0.33 | 4.8E-27 | 4.4E-26 | 0.33 | 4.8E-27 | Mackowiak | MLRKAHWPTNRMELKYGYMLTKESE* |
| | *H. sapiens* | ENSG00000152127 | MGAT5 | mannosyl (alpha-1,6-)-glycoprotein beta-1,6-N-acetyl-gluco | 0.00 | 0.28 | 1.0E-23 | 2.8E-22 | 0.28 | 1.0E-23 | Mackowiak | MEVRKGNQLTQEPE* |
| HG0294.1 | *G. gallus* | ENSGALG00000017036 | LHFP | lipoma HMGIC fusion partner | 0.05 | 0.41 | 7.5E-14 | 4.7E-13 | 0.41 | 7.5E-14 | | MMGRNILLAFRIRASPSVRICQG* |
| HG0294.2 | *H. sapiens* | ENSG00000183722 | LHFP | lipoma HMGIC fusion partner | 0.12 | 0.43 | 3.7E-04 | 1.5E-03 | 0.43 | 3.7E-04 | | MFPPEKDSKLCVRPGRGREGGRRPPAMDHHL* |
| HG0295.1 | *H. sapiens* | ENSG00000166888 | STAT6 | signal transducer and activator of transcription 6 | 0.15 | 0.06 | 9.5E-36 | 3.9E-34 | 0.06 | 9.5E-36 | | MYVCALSYFSFWWWWWKGGGASRASLELAGQSYRPMGPGSAR* |
| HG0295.2 | *G. gallus* | ENSGALG00000003282 | STAT5B | signal transducer and activator of transcription 5B | 0.17 | 0.50 | 8.8E-05 | 2.9E-04 | 0.50 | 8.8E-05 | | MRAGVPSLCPQLRCAGGGRKQPEHSAARLRRQLTFWKEIAALSSFL* |
| HG0296 | *G. gallus* | ENSGALG00000006912 | CPEB3 | cytoplasmic polyadenylation element binding protein 3 | 0.23 | 0.17 | 2.1E-03 | 5.6E-03 | 0.17 | 2.1E-03 | | MQIARGRRWRRRRVLGKTGTRISFSLQGESSEFRCKRKETWRNCCISLPYKHISHLD* |
| HG0297 | *G. gallus* | ENSGALG00000031684 | | | 0.20 | 0.00 | 1.1E-05 | 4.0E-05 | 0.00 | 1.1E-05 | | MCFCFVLQRFFWRLIFTF* |
| HG0298 | *G. gallus* | ENSGALG00000042308 | | | 0.00 | 0.01 | 2.0E-04 | 6.2E-04 | 0.01 | 2.0E-04 | | MESIVWECG* |
| HG0299 | *G. gallus* | ENSGALG00000040465 | Sip1 | zinc finger E-box binding homeobox 2 | 0.00 | 0.39 | 1.2E-02 | 2.7E-02 | 0.39 | 1.2E-02 | | MLGKEKEKNVIRRGVTRQSVPRFVFPGVAERDQS* |
| HG0300 | *D. melanogaster* | FBgn0031668 | CG31917 | RE52596p2 | 0.00 | 0.02 | 0.0E+00 | 0.0E+00 | 0.02 | 0.0E+00 | Hayden | MVNVMKGVLVECDPAMKQFLLHLDEKLALGRKFIIQDLDENHLFISTDIVEVLQARVDDLMDRISFPLHDKDA* |
| HG0301 | *D. melanogaster* | FBgn0036856 | CG9666 | CG9666, isoform A | 0.15 | 0.05 | 0.0E+00 | 0.0E+00 | 0.05 | 0.0E+00 | Hayden | MAFPTTSAQQAETNRKILEEIQTKKQLLAGGIINLGLSPPNQMPAPQLLGQPTTVNPDFQAGVGIATNATSTARSAFNPTSSTTLGFFIPQDSYFGNSFIPVLPRLEPLPSPATTPTTPNAPPSHSISK* |
| HG0302 | *D. melanogaster* | FBgn0031454 | CG9960 | CG9960 | 0.20 | 0.04 | 1.2E-269 | 3.2E-269 | 0.04 | 1.2E-269 | Hayden | MDSDSTVTSLEENTENFCTNPTRDILAEGITNLFKPTIERLDERVASTIQLQAELRGQLDALAAQLRDIEKAQSQIPEFADKVKELLNVKHKVTVISNVLVTSQERLTGLHKLIEKEQRRRQALLDSALSTNIS* |
| HG0303 | *D. melanogaster* | FBgn0260464 | | | 0.00 | 0.05 | 1.4E-232 | 2.9E-232 | 0.05 | 1.4E-232 | Hayden | MHTDLSSHLHTPACNKLIEELQACHENNAFAKFVGVCNSIDDKVVKCLKGERIARSAANRAKARERQAKYKEKLLQQESN* |
| HG0304 | *D. melanogaster* | FBgn0260467 | CG7071 | CG7071, isoform A | 0.03 | 0.06 | 0.0E+00 | 0.0E+00 | 0.06 | 0.0E+00 | Hayden | MKIMISQVGRELYKVPLRILDHRVFVNGEIEAFLENFEVRRNDSEVEKLFQVTETVGSLKYDLSRCSATGRGSGAENLAQLDTEVSHLLDGVNALLAKAKVERTASTQLQEARLAREQRRAEFLTNLEHGYRRIENSFEEKEEEIAELYSDLQLKLNIAK* |
| HG0305 | *D. melanogaster* | FBgn0050290 | Ppcdc | Phosphopantothenoylcysteine decarboxylase | 0.29 | 0.04 | 4.7E-246 | 1.1E-245 | 0.04 | 4.7E-246 | Hayden | MTSQIYSQKKFVPTPPEKGSFPLDHEGLCKKQFLLYASCLRKNAQDTSQCRQDAQNYLACRMDNNLMEKTEWSKLGFHDQSTKTDQKEPEVQKQ* |
| HG0306 | *D. melanogaster* | FBgn0259726 | | | 0.06 | 0.08 | 0.0E+00 | 0.0E+00 | 0.08 | 0.0E+00 | Hayden | MSIPQYRPPKEIQDLLDQKEQQQANQRRRTAVLPKRTWIQRNPRLFQISFITGSLLIFFSKPLFDCFIADPLPPPENKVPPHKR* |
| HG0307 | *D. melanogaster* | FBgn0037822 | CG14683 | Probable methyltransferase-like protein 15 homolog | 0.12 | 0.06 | 1.1E-202 | 2.0E-202 | 0.06 | 1.1E-202 | | MSAQYQRFLKVLEKWPAEKSKVGRDLGEQIRKQVTKLTSLEGGATDKELDRQINSLERLSNNVYAKKYPRTFESTATGLTAAQCSQVLSSEFLQYLNEDSKKKK* |
| HG0308 | *D. melanogaster* | FBgn0264743 | CG44001 | LP22840p | 0.00 | 0.04 | 2.4E-197 | 4.3E-197 | 0.04 | 2.4E-197 | Hayden | MPYIIIRGNLASYSHKYPWRVLVSGLKADDIEQLNKFSCGGYSDESTIVYLVHPCRILSALEILGFRVVASSSTAVKQDYNEYMWTMRKEFDEPEPLEAESVVRENLSNIGREAASLGNYHKVDSPE* |
| HG0309 | *D. melanogaster* | FBgn0037689 | CG8135 | LMBR1 domain-containing protein 2 homolog | 0.23 | 0.06 | 2.3E-243 | 5.2E-243 | 0.06 | 2.3E-243 | | MSSVEAEVLQKIDLIEPTTGKLFFEHKTELLFCRPHLMPLKTQALERLEEMHRDTARQLQKKRQQKKKSTEPTGSL* |
| HG0310 | *D. melanogaster* | FBgn0039339 | CG5116 | CG5116, isoform A | 0.08 | 0.05 | 6.5E-119 | 8.7E-119 | 0.05 | 6.5E-119 | Hayden | MNDIKRLKDQQREKHPGFDGYMDCMTRSLFTGLATFCLSFSGTYFAQKIVQSKIRYPIKYNILISSLVATGVSYQITSTRTKACQAAWMAFEDKHSVLKEKTF* |
| HG0311 | *D. melanogaster* | FBgn0263251 | vnc | variable nurse cells | 0.01 | 0.06 | 5.0E-128 | 6.8E-128 | 0.06 | 5.0E-128 | Hayden | MSLQTGLAFIFTILASVSIVAAIILLGWFLIWKAFLSKFRLVRELLGQEEPEDQQPLAWDHQNQQPHHHGRARKARRD* |
| HG0312 | *D. melanogaster* | FBgn0038641 | CG7708 | High-affinity choline transporter 1 | 0.07 | 0.05 | 8.1E-98 | 1.0E-97 | 0.05 | 8.1E-98 | Mackowiak | MCGLHFRTRNSPSQKTSLRWGDCCLA* |
| HG0313 | *D. melanogaster* | FBgn0259725 | | | 0.12 | 0.06 | 9.8E-78 | 1.2E-77 | 0.06 | 9.8E-78 | | MSDTRKDPCKPNACRIQACLTSKHRR* |
| HG0314 | *D. melanogaster* | FBgn0260468 | CG7950 | CG7950, isoform A | 0.15 | 0.06 | 0.0E+00 | 0.0E+00 | 0.06 | 0.0E+00 | Hayden | MSGYQYLDVKIKLRDPDSVTLTPVFFCGCIHSSLASIFGEIGGQTILEIVKFSSSQKRSILRVPENVLDRVRVAIALIGYYQEVPCHFQVLSTSRKPLDFEESPEEFVAFN* |
| HG0315 | *D. melanogaster* | FBgn0260234 | CG42508 | GH25417p | 0.00 | 0.07 | 3.8E-68 | 4.5E-68 | 0.07 | 3.8E-68 | | MKPKKSAFANSSTGYGRNKHGKGHSGKDKVEGGGKGKSRDEKDKHKHEQKVDEKDDKDLNSGFGDYLRTPEAFEMMKLFVFANTIMLIVTMAWPHIKEQFYMVNQWLESFREEHQQ* |
| HG0316 | *D. melanogaster* | FBgn0041164 | armi | armitage | 0.10 | 0.07 | 6.3E-165 | 1.0E-164 | 0.07 | 6.3E-165 | | MSSEASGSITIKEEASTSGTATNIPVESSETEAKKPEIGADKAESQSETTEGGKAAETTENHLKRLQNLRKELSYLSETDWMYESLDKKAAQ* |
| HG0317 | *D. melanogaster* | FBgn0050011 | gem | gemini | 0.00 | 0.05 | 4.5E-17 | 5.0E-17 | 0.05 | 4.5E-17 | Mackowiak | MEKSEIRLQRMSNEYQSQSSYMYLRTKMLLKIENTLLRSHRQRETTGIKKLYNSFFVLF* |
| HG0318 | *D. melanogaster* | FBgn0035293 | CG5687 | CG5687, isoform A | 0.00 | 0.04 | 1.7E-13 | 1.8E-13 | 0.04 | 1.7E-13 | Mackowiak | MKYRNNRGTILQIQFRTFLRYSAANLIPFES* |
| HG0319 | *D. melanogaster* | FBgn0026778 | Rad1 | Radiation insensitive 1 | 0.00 | 0.07 | 5.4E-12 | 5.7E-12 | 0.07 | 5.4E-12 | Hayden | MASESENTTSYFVPHLVDQVFELYRSKAESNSLTKEERGQFFYDLSILLGQELVLRSLHLLDDHNFSFFHAKNNRSVCVVEISKGTEYFRLIPGVSFCKCEFFQCHVLQLPRGILYQDLPSGEPGILEDWSEESRVSYTCQHILALRLHQFLKHTGGKTTEKILKKEDIKELRTDVFRD* |
| HG0320 | *D. melanogaster* | FBgn0040993 | | | 0.00 | 0.06 | 1.0E-02 | 1.0E-02 | 0.06 | 1.0E-02 | Hayden | MKTPDTKAKLLNNISLSKHLSSKKGGSRTSNCGNCLEFSVLTRLSSVP* |
| HG0321 | *D. rerio* | ENSDARG00000070426 | chac1 | ChaC, cation transport regulator homolog 1 (E. coli) | 0.03 | 0.25 | 6.2E-160 | 9.3E-159 | 0.25 | 6.2E-160 | | MHQAKKSVLRDCTITTAKQLGSDTVRGIAVSQPIPLLLDRRKLTLAT* |
| HG0322 | *D. rerio* | ENSDARG00000070917 | kitlga | kit ligand a | 0.00 | 0.34 | 5.5E-27 | 3.4E-26 | 0.34 | 5.5E-27 | | MMILDMYLWDEENPFPARRSKIK* |
| HG0323 | *D. rerio* | ENSDARG00000071197 | usp40 | ubiquitin specific peptidase 40 | 0.02 | 0.03 | 7.8E-132 | 1.1E-130 | 0.03 | 7.8E-132 | Mackowiak | MFIPVKSSYARVSPLIVNRELGKTKMR* |
| HG0324 | *D. rerio* | ENSDARG00000094965 | nfil3-5 | nuclear factor, interleukin 3 regulated, member 5 | 0.00 | 0.10 | 6.6E-97 | 8.1E-96 | 0.10 | 6.6E-97 | | MTVVFVEFHLFLIFLTSLDT* |
| HG0325 | *D. rerio* | ENSDARG00000079738 | znf219 | zinc finger protein 219 | 0.13 | 0.07 | 2.2E-12 | 9.8E-12 | 0.07 | 2.2E-12 | | MYLIQVKRNHDVCCH* |
| HG0326 | *D. rerio* | ENSDARG00000012848 | arih2 | ariadne homolog 2 (Drosophila) | 0.00 | 0.00 | 5.5E-89 | 6.6E-88 | 0.00 | 5.5E-89 | | MSDGECDFFCFFRCC* |
| HG0327 | *D. rerio* | ENSDARG00000022203 | gtf2ird1 | GTF2I repeat domain containing 1 | 0.00 | 0.39 | 3.7E-20 | 2.2E-19 | 0.39 | 3.7E-20 | | MTFNFTGLQNKERGKGTHSSVKENREKTGQTHQINHTECFSTDYRPFEIMTFSKVAVAFL* |
| HG0328 | *D. rerio* | ENSDARG00000099999 | tcf3b | transcription factor 3b | 0.00 | 0.08 | 2.0E-63 | 2.0E-62 | 0.08 | 2.0E-63 | | MWMEVVKMSTRRMNSPQL* |
| HG0329 | *D. rerio* | ENSDARG00000036442 | atp11c | ATPase, Class VI, type 11C | 0.00 | 0.43 | 2.2E-20 | 1.3E-19 | 0.43 | 2.2E-20 | | MRMVREDSIQLIWKRCSFTAALHIARGNRLESR* |
| HG0330 | *D. rerio* | ENSDARG00000039392 | wnk1b | WNK lysine deficient protein kinase 1b | 0.00 | 0.32 | 3.7E-37 | 2.8E-36 | 0.32 | 3.7E-37 | | MDEVDLTVFVCHHQCLPLSNCVLFFYVYKRTPPLLGSGPLHCKSNC* |
| HG0331 | *D. rerio* | ENSDARG00000044899 | tmem183a | transmembrane protein 183A | 0.02 | 0.34 | 6.3E-58 | 5.9E-57 | 0.34 | 6.3E-58 | | MYATLSVSSIMAEDPGGAFFNFTLILTDFSLPLLERWQG* |
| HG0332 | *D. rerio* | ENSDARG00000102893 | atg7 | ATG7 autophagy related 7 homolog (S. cerevisiae) | 0.00 | 0.03 | 3.8E-84 | 4.2E-83 | 0.03 | 3.8E-84 | | MKRNSVYRFLSVFSCLLVCVFLFFWGFAFYSMVLAP* |
| HG0333 | *D. rerio* | ENSDARG00000042518 | hipk2 | homeodomain interacting protein kinase 2 | 0.00 | 0.47 | 5.0E-15 | 2.5E-14 | 0.47 | 5.0E-15 | | MQEIAAILSAAQDGRFLLSCEQCCRFIFCHE* |
| HG0334 | *D. rerio* | ENSDARG00000075349 | si:ch73-138n13.1 | si:ch73-138n13.1 | 0.05 | 0.46 | 3.0E-14 | 1.5E-13 | 0.46 | 3.0E-14 | | MTSLCGYHRNQGSTADG* |
| HG0335 | *D. rerio* | ENSDARG00000034700 | vegfab | vascular endothelial growth factor Ab | 0.00 | 0.32 | 2.3E-31 | 1.5E-30 | 0.32 | 2.3E-31 | | MWIVIALEEKFDLLLNNVRGPASWILIFLDILKIIGNS* |
| HG0336 | *D. rerio* | ENSDARG00000028676 | nuak1b | NUAK family, SNF1-like kinase, 1b | 0.03 | 0.34 | 1.1E-13 | 5.4E-13 | 0.34 | 1.1E-13 | | MRVIYVQNAGVVAAFSPEIFCGRLKQRNNGEIMTERATGKRTRHNGVGYVIME* |
| HG0337 | *D. rerio* | ENSDARG00000054906 | | | 0.06 | 0.47 | 8.6E-13 | 3.8E-12 | 0.47 | 8.6E-13 | | MFLHRADLSLRPRTKYRDSD* |
| HG0338 | *D. rerio* | ENSDARG00000000151 | thraa | thyroid hormone receptor alpha a | 0.04 | 0.00 | 1.3E-19 | 7.3E-19 | 0.00 | 1.3E-19 | | MLTCRGSSPPQSAVIGCWMESR* |
| HG0339 | *D. rerio* | ENSDARG00000063144 | shisa7b | shisa family member 7 | 0.11 | 0.21 | 2.3E-35 | 1.7E-34 | 0.21 | 2.3E-35 | Mackowiak | MNPDLFMPVQWAQPCCPVRTGPKPPFVLFCWVELNPEQRHSRKRVQHT* |
| HG0340 | *D. rerio* | ENSDARG00000017591 | fat1a | FAT atypical cadherin 1a | 0.06 | 0.15 | 2.6E-15 | 1.3E-14 | 0.15 | 2.6E-15 | | MQRLGTYKLHEKPPNKGR* |
| HG0341 | *D. rerio* | ENSDARG00000039430 | msl2b | male-specific lethal 2 homolog b (Drosophila) | 0.00 | 0.08 | 5.0E-20 | 2.9E-19 | 0.08 | 5.0E-20 | | MMTRIWSYFVPWKRGKR* |
| HG0342 | *D. rerio* | ENSDARG00000045801 | fbxl14b | F-box and leucine-rich repeat protein 14b | 0.08 | 0.30 | 2.2E-16 | 1.2E-15 | 0.30 | 2.2E-16 | | MYDRLDPRVERRLLSKRQTGSSAA* |
| HG0343 | *D. rerio* | ENSDARG00000057062 | si:dkey-157g16.6 | si:dkey-157g16.6 | 0.05 | 0.30 | 1.3E-09 | 5.1E-09 | 0.30 | 1.3E-09 | | MDYSRTDANLGFPTELLP* |
| HG0344 | *D. rerio* | ENSDARG00000032072 | cdk16 | cyclin-dependent kinase 16 | 0.19 | 0.16 | 1.8E-11 | 7.6E-11 | 0.16 | 1.8E-11 | | MAEDYWERAAALGYYYCC* |
| HG0345 | *D. rerio* | ENSDARG00000035910 | znf281b | zinc finger protein 281b | 0.05 | 0.39 | 4.4E-05 | 1.4E-04 | 0.39 | 4.4E-05 | | MEEANSCKGTLFWL* |
| HG0346 | *D. rerio* | ENSDARG00000041982 | adamts6 | ADAM metallopeptidase with thrombospondin type 1 motif, | 0.22 | 0.25 | 1.9E-13 | 8.9E-13 | 0.25 | 1.9E-13 | | MFVCTVAASASAGKQMDSSAQVPNIDLLLWNNMKGVVMDSCPAHG* |
| HG0347 | *D. rerio* | ENSDARG00000063194 | | | 0.00 | 0.00 | 2.2E-02 | 3.9E-02 | 0.00 | 2.2E-02 | | MADLSKSKPGLK* |
| HG0348 | *D. rerio* | ENSDARG00000036139 | lctla | lactase-like a | 0.08 | 0.11 | 7.7E-15 | 3.9E-14 | 0.11 | 7.7E-15 | | MSFLLAALDKLAYLLSLVAFCVSLSYSWLK* |
| HG0349 | *D. rerio* | ENSDARG00000028699 | crkl | v-crk avian sarcoma virus CT10 oncogene homolog-like | 0.00 | 0.36 | 1.4E-05 | 4.8E-05 | 0.36 | 1.4E-05 | | MHQSWLTLVAACLVKRQVAVQ* |
| HG0350 | *D. rerio* | ENSDARG00000036194 | pitx2 | paired-like homeodomain 2 | 0.12 | 0.46 | 1.5E-03 | 3.9E-03 | 0.46 | 1.5E-03 | | MQVDIISHTCPCHSGSPCSRLKPRYLLPSRNGKKIFP* |
| HG0351 | *D. rerio* | ENSDARG00000068096 | atf5a | activating transcription factor 5a | 0.00 | 0.42 | 5.5E-04 | 1.5E-03 | 0.42 | 5.5E-04 | | MRTKPCFLWLNSQTSLSCTLSSGI* |
| HG0352 | *D. rerio* | ENSDARG00000070214 | slc41a1 | solute carrier family 41 (magnesium transporter), member 1 | 0.05 | 0.33 | 3.5E-05 | 1.1E-04 | 0.33 | 3.5E-05 | | MGNIIQVLPKFRQDLTQAWRRGTHLPLFLDL* |
| HG0353 | *D. rerio* | ENSDARG00000078864 | lats2 | large tumor suppressor kinase 2 | 0.23 | 0.40 | 2.9E-06 | 1.0E-05 | 0.40 | 2.9E-06 | | MANMHTRLGWKKHADVNPHMSLEEGECEP* |
| HG0354 | *D. rerio* | ENSDARG00000019897 | gpat4 | glycerol-3-phosphate acyltransferase 4 | 0.05 | 0.18 | 1.6E-06 | 5.6E-06 | 0.18 | 1.6E-06 | | MIIFPYMPMKSMFQRRLQKLRHSRGVQQSVFR* |
| HG0355 | *D. rerio* | ENSDARG00000019930 | tal1 | T-cell acute lymphocytic leukemia 1 | 0.27 | 0.50 | 9.6E-04 | 2.5E-03 | 0.50 | 9.6E-04 | | MDDPPQNGSMHTLTSTARRR* |
| HG0356 | *D. rerio* | ENSDARG00000020475 | znrf1 | zinc and ring finger 1 | 0.08 | 0.31 | 1.2E-02 | 2.3E-02 | 0.31 | 1.2E-02 | | MIRPGKWAVASARHSCFSQT* |
| HG0357 | *D. rerio* | ENSDARG00000037285 | mipa | major intrinsic protein of lens fiber a | 0.05 | 0.40 | 2.3E-03 | 5.7E-03 | 0.40 | 2.3E-03 | | MSAGHAHHLSIRVPLLNLS* |

| | | | | | | | | | | | | |
|---|---|---|---|---|---|---|---|---|---|---|---|---|
| HG0358 | *D. rerio* | ENSDARG00000062506 | zc3h18 | zinc finger CCCH-type containing 18 | 0.15 | 0.28 | 4.7E-06 | 1.6E-05 | 0.28 | 4.7E-06 | | MAALTVSCRVYAFFR* |
| HG0359 | *D. rerio* | ENSDARG00000070348 | hoxc10a | homeobox C10a | 0.07 | 0.30 | 3.9E-07 | 1.5E-06 | 0.30 | 3.9E-07 | | MSASYCPYPLRHKGDCFESKEKQDK* |
| HG0360 | *D. rerio* | ENSDARG00000070801 | opa1 | optic atrophy 1 (autosomal dominant) | 0.04 | 0.10 | 7.6E-06 | 2.6E-05 | 0.10 | 7.6E-06 | | MCPIWKDELNFILQRDPKLIYICH* |
| HG0361 | *D. rerio* | ENSDARG00000077581 | zzz3 | zinc finger, ZZ-type containing 3 | 0.24 | 0.37 | 1.5E-03 | 3.9E-03 | 0.37 | 1.5E-03 | | MCSKRFAEVRCPHTAAEERAPR* |
| HG0362 | *D. rerio* | ENSDARG00000078355 | zc3h4 | zinc finger CCCH-type containing 4 | 0.01 | 0.42 | 7.7E-04 | 2.1E-03 | 0.42 | 7.7E-04 | | MSFISHRNCRTENLRNVFNLFLF* |
| HG0363 | *D. rerio* | ENSDARG00000095896 | pou3f3b | POU class 3 homeobox 3b | 0.22 | 0.31 | 2.3E-04 | 6.6E-04 | 0.31 | 2.3E-04 | | MRYNLPALKKFLQKGGGDQQTYS* |
| HG0364 | *D. rerio* | ENSDARG00000004988 | ppp3ca | protein phosphatase 3, catalytic subunit, alpha isozyme | 0.00 | 0.42 | 9.3E-03 | 1.9E-02 | 0.42 | 9.3E-03 | | MPLHQPHLVTANTRRE* |
| HG0365 | *D. rerio* | ENSDARG00000016132 | keap1a | kelch-like ECH-associated protein 1a | 0.00 | 0.17 | 3.3E-05 | 1.1E-04 | 0.17 | 3.3E-05 | | MPTWLETESCYSGLLTRQKKHGSLSLNDNQLCLYFFMARGWV* |
| HG0366 | *D. rerio* | ENSDARG00000017803 | gsk3b | glycogen synthase kinase 3 beta | 0.00 | 0.20 | 8.8E-05 | 2.7E-04 | 0.20 | 8.8E-05 | | MELSGSERPRISVLKLGNWKILIRIAG* |
| HG0367 | *D. rerio* | ENSDARG00000034056 | csnk1g2b | casein kinase 1, gamma 2b | 0.03 | 0.22 | 8.1E-04 | 2.2E-03 | 0.22 | 8.1E-04 | | MFTVLPRTDFKETMRSTRGRSLVGFH* |
| HG0368 | *D. rerio* | ENSDARG00000058606 | sik1 | salt-inducible kinase 1 | 0.23 | 0.28 | 2.1E-03 | 5.2E-03 | 0.28 | 2.1E-03 | | MTCKCKCKNHLCCGRRWKLFAS* |
| HG0369 | *D. rerio* | ENSDARG00000059276 | hoxd4a | homeobox D4a | 0.04 | 0.48 | 2.1E-02 | 3.8E-02 | 0.48 | 2.1E-02 | | MKSLHLDWPSWSHGRLTLFS* |
| HG0370 | *D. rerio* | ENSDARG00000062693 | nrxn3b | neurexin 3b | 0.02 | 0.32 | 2.2E-02 | 3.8E-02 | 0.32 | 2.2E-02 | | MWQNNRIRSNMENLCFNPERRLKGTSSVKHF* |
| HG0371 | *D. rerio* | ENSDARG00000062765 | whsc1l1 | Wolf-Hirschhorn syndrome candidate 1-like 1 | 0.00 | 0.24 | 9.7E-03 | 1.9E-02 | 0.24 | 9.7E-03 | | MEHAEAGPVRKRET* |
| HG0372 | *D. rerio* | ENSDARG00000069467 | igsf9bb | immunoglobulin superfamily, member 9Bb | 0.26 | 0.50 | 3.0E-02 | 5.0E-02 | 0.50 | 3.0E-02 | | MQGMSKRVSAADSASNDQGLG* |
| HG0373 | *D. rerio* | ENSDARG00000074611 | wdr37 | WD repeat domain 37 | 0.06 | 0.46 | 4.2E-03 | 9.4E-03 | 0.46 | 4.2E-03 | | MTVALRSAEAVINALGFLFEVFLFFITLSSRLRMCLLCLSFTALHGCFR* |
| HG0374 | *D. rerio* | ENSDARG00000075147 | lrrc38a | leucine rich repeat containing 38a | 0.18 | 0.17 | 2.7E-03 | 6.5E-03 | 0.17 | 2.7E-03 | | MISWKIELRVFVAAFRCTSLCRLQSRRQGKMPSRQT* |
| HG0375 | *D. rerio* | ENSDARG00000076171 | znf827 | zinc finger protein 827 | 0.00 | 0.18 | 1.6E-02 | 3.0E-02 | 0.18 | 1.6E-02 | | MGKDLFSSYP* |
| HG0376 | *D. rerio* | ENSDARG00000077229 | ano8b | anoctamin 8b | 0.07 | 0.01 | 9.4E-05 | 2.9E-04 | 0.01 | 9.4E-05 | | MTHNNLHFTLTFI* |
| HG0377 | *D. rerio* | ENSDARG00000079549 | cdc42ep1a | CDC42 effector protein (Rho GTPase binding) 1a | 0.00 | 0.28 | 2.5E-02 | 4.2E-02 | 0.28 | 2.5E-02 | | MQPRFSGIGVMHIFSSHLSSPNLSGYEQRPMAKTGHS* |
| HG0378 | *D. rerio* | ENSDARG00000080009 | bahcc1b | BAH domain and coiled-coil containing 1b | 0.21 | 0.16 | 6.4E-05 | 2.0E-04 | 0.16 | 6.4E-05 | | MIGTYFFGRQSQNCAECKT* |
| HG0379 | *D. rerio* | ENSDARG00000104372 | gnb1b | guanine nucleotide binding protein (G protein), beta polype | 0.00 | 0.47 | 1.7E-02 | 3.0E-02 | 0.47 | 1.7E-02 | | MRGFDSHTIVFFGQCHINKGHLRLRTDQNVSRQ* |
| HG0380 | *D. rerio* | ENSDARG00000005350 | dip2ba | disco-interacting protein 2 homolog Ba | 0.00 | 0.41 | 9.4E-03 | 1.9E-02 | 0.41 | 9.4E-03 | | MAEGVVLRMDCYSPFNSRFMP* |
| HG0381 | *D. rerio* | ENSDARG00000006124 | kdm5c | lysine (K)-specific demethylase 5C | 0.00 | 0.04 | 3.0E-03 | 7.1E-03 | 0.04 | 3.0E-03 | | MACSSDRMNGG* |
| HG0382 | *D. rerio* | ENSDARG00000017242 | | | 0.00 | 0.29 | 4.4E-03 | 1.0E-02 | 0.29 | 4.4E-03 | | MTEPFFTADACGGGLTCRC* |
| HG0383 | *D. rerio* | ENSDARG00000036501 | rab39bb | RAB39B, member RAS oncogene family b | 0.00 | 0.27 | 1.7E-02 | 3.1E-02 | 0.27 | 1.7E-02 | | MMLLLSLPFHLHKTSKHFSIFQTWIKT* |
| HG0384 | *D. rerio* | ENSDARG00000054748 | cuedc1b | CUE domain containing 1b | 0.06 | 0.04 | 1.2E-02 | 2.2E-02 | 0.04 | 1.2E-02 | | MQLNKQKKRRKGFLSLQWTRFFIV* |
| HG0385 | *D. rerio* | ENSDARG00000059549 | akap10 | A kinase (PRKA) anchor protein 10 | 0.00 | 0.15 | 3.2E-03 | 7.6E-03 | 0.15 | 3.2E-03 | | MNEGNRESSGCCL* |
| HG0386 | *D. rerio* | ENSDARG00000062084 | clcn1a | chloride channel, voltage-sensitive 1a | 0.00 | 0.25 | 5.9E-03 | 1.3E-02 | 0.25 | 5.9E-03 | | MCEEETERLSVCVCMCPCHPRVGMLRQTGSLFLGQ* |
| HG0387 | *D. rerio* | ENSDARG00000071150 | cica | capicua transcriptional repressor a | 0.16 | 0.28 | 1.2E-02 | 2.2E-02 | 0.28 | 1.2E-02 | | MQERNRVEDIDRLGLTGNQED* |
| HG0388 | *D. rerio* | ENSDARG00000077228 | ntrk3a | neurotrophic tyrosine kinase, receptor, type 3a | 0.00 | 0.35 | 1.4E-02 | 2.6E-02 | 0.35 | 1.4E-02 | | MQPVEEDSLGALIFEPFLH* |
| HG0389 | *D. rerio* | ENSDARG00000077361 | bptf | bromodomain PHD finger transcription factor | 0.08 | 0.36 | 1.5E-02 | 2.7E-02 | 0.36 | 1.5E-02 | | MAAASDLKRGAGRSF* |
| HG0390 | *D. rerio* | ENSDARG00000090035 | rtn4r | reticulon 4 receptor | 0.26 | 0.36 | 6.3E-03 | 1.3E-02 | 0.36 | 6.3E-03 | | MTERGTTGRRCSPSRARTATVG* |
| HG0391 | *D. rerio* | ENSDARG00000099634 | elna | elastin a | 0.00 | 0.15 | 2.5E-02 | 4.4E-02 | 0.15 | 2.5E-02 | | MIHRTKGFALS* |
| HG0392 | *D. rerio* | ENSDARG00000100003 | glulb | glutamate-ammonia ligase (glutamine synthase) b | 0.00 | 0.04 | 4.1E-03 | 9.4E-03 | 0.04 | 4.1E-03 | | MRMRDKQPGLPYKTLWKYSHSYSDYLYFLIGLI* |
| HG0393 | *G. gallus* | ENSGALG00000002090 | GPSM2 | G protein signaling modulator 2 | 0.00 | 0.43 | 5.0E-36 | 5.8E-35 | 0.43 | 5.0E-36 | | MELKGPQMKELERVSDSLRCV* |
| HG0394 | *G. gallus* | ENSGALG00000002479 | MAT1A | methionine adenosyltransferase 1A | 0.00 | 0.39 | 1.6E-46 | 2.2E-45 | 0.39 | 1.6E-46 | | MRSPCMHTQIQAHADFTD* |
| HG0395 | *G. gallus* | ENSGALG00000014936 | ESCO1 | establishment of sister chromatid cohesion N-acetyltransfer | 0.00 | 0.40 | 5.4E-03 | 1.3E-02 | 0.40 | 5.4E-03 | | MLLAHRDKLLWPKHRLPLVKT* |
| HG0396.1 | *G. gallus* | ENSGALG00000034253 | | | 0.00 | 0.43 | 2.9E-22 | 2.4E-21 | 0.43 | 2.9E-22 | | MKDHCKEPWTFVVSYNTVIQRV* |
| HG0396.2 | *G. gallus* | ENSGALG00000034253 | | | 0.00 | 0.37 | 1.0E-04 | 3.3E-04 | 0.37 | 1.0E-04 | | MQKKTNKGRKDVKHDLP* |
| HG0397 | *G. gallus* | ENSGALG00000009438 | SLC39A9 | solute carrier family 39 member 9 | 0.00 | 0.17 | 3.1E-46 | 4.3E-45 | 0.17 | 3.1E-46 | | MAESPSWDLGVCTG* |
| HG0398 | *G. gallus* | ENSGALG00000005369 | BTBD10 | BTB domain containing 10 | 0.00 | 0.13 | 3.0E-52 | 4.9E-51 | 0.13 | 3.0E-52 | | MMEEKVRLARAALNY* |
| HG0399 | *G. gallus* | ENSGALG00000032659 | | | 0.00 | 0.43 | 4.8E-07 | 2.0E-06 | 0.43 | 4.8E-07 | | MTNHFCVCTTRIQNS* |
| HG0400 | *G. gallus* | ENSGALG00000004261 | AQP9 | aquaporin 9 | 0.00 | 0.19 | 1.3E-13 | 7.7E-13 | 0.19 | 1.3E-13 | | MISLAVQHEISYFSV* |
| HG0401 | *G. gallus* | ENSGALG00000009057 | TASP1 | taspase 1 | 0.10 | 0.38 | 1.3E-16 | 9.5E-16 | 0.38 | 1.3E-16 | | MILLQNLFLNGSMLFASRRC* |
| HG0402 | *G. gallus* | ENSGALG00000035584 | | | 0.24 | 0.34 | 3.8E-03 | 9.5E-03 | 0.34 | 3.8E-03 | | MARCWKGREQPMLLI* |
| HG0403.1 | *G. gallus* | ENSGALG00000007717 | GCNT7 | glucosaminyl (N-acetyl) transferase family member 7 | 0.00 | 0.12 | 3.9E-50 | 5.9E-49 | 0.33 | 5.1E-19 | | MNPPPNTNPFSRMLSGKEAFIQHEIQSIPITCMPN* |
| HG0403.2 | *G. gallus* | ENSGALG00000007717 | GCNT7 | glucosaminyl (N-acetyl) transferase family member 7 | 0.00 | 0.33 | 1.1E-02 | 2.4E-02 | 0.33 | 1.1E-02 | | MLSIAGNQVMLMFNPFQLYEISTQATE* |
| HG0404 | *G. gallus* | ENSGALG00000008700 | PDCD4 | programmed cell death 4 | 0.00 | 0.32 | 1.4E-17 | 1.0E-16 | 0.32 | 1.4E-17 | | MEHSTRKDKTVLTR* |
| HG0405.1 | *G. gallus* | ENSGALG00000009185 | | | 0.00 | 0.49 | 5.3E-08 | 2.4E-07 | 0.49 | 5.3E-08 | | MLLSSLWAFLDLRFFS* |
| HG0405.2 | *G. gallus* | ENSGALG00000009185 | | | 0.00 | 0.40 | 2.9E-10 | 1.4E-09 | 0.40 | 2.9E-10 | | MPHPDIFSPELSKLPVL* |
| HG0406 | *G. gallus* | ENSGALG00000004935 | LSM14A | LSM14A, mRNA processing body assembly factor | 0.23 | 0.34 | 1.4E-04 | 4.3E-04 | 0.34 | 1.4E-04 | | MKAAAMLGASGRFWCGVC* |
| HG0407 | *G. gallus* | ENSGALG00000015027 | JAK2 | Janus kinase 2 | 0.05 | 0.28 | 2.5E-25 | 2.1E-24 | 0.28 | 2.5E-25 | | MERARVLRGSRQLALLRRAPAGYIMFYHCLCACFI* |
| HG0408 | *G. gallus* | ENSGALG00000036255 | WNT7B | Protein Wnt-7b | 0.06 | 0.21 | 1.5E-10 | 7.7E-10 | 0.21 | 1.5E-10 | | MTGRSPEAAAAARTAAAAHTLPREQH* |
| HG0409 | *G. gallus* | ENSGALG00000002315 | | | 0.00 | 0.32 | 1.1E-07 | 4.6E-07 | 0.32 | 1.1E-07 | | MVSTPCEPPLQK* |
| HG0410 | *G. gallus* | ENSGALG00000008148 | MYBPC3 | myosin-binding protein C, cardiac-type | 0.20 | 0.46 | 4.4E-11 | 2.4E-10 | 0.46 | 4.4E-11 | | MGASTEWFFWGSQKPCSLICV* |
| HG0411 | *G. gallus* | ENSGALG00000012166 | SLC35F5 | solute carrier family 35 member F5 | 0.00 | 0.27 | 7.8E-27 | 7.1E-26 | 0.27 | 7.8E-27 | | MAIPTLTLLKWRLSSL* |
| HG0412 | *G. gallus* | ENSGALG00000003446 | PRLR | prolactin receptor | 0.02 | 0.43 | 1.5E-05 | 5.4E-05 | 0.43 | 1.5E-05 | | MNERNKIFLCTHESFREPLM* |
| HG0413 | *G. gallus* | ENSGALG00000005475 | | | 0.19 | 0.36 | 5.6E-13 | 3.2E-12 | 0.36 | 5.6E-13 | | MRSCCWLLLDFVSG* |
| HG0414 | *G. gallus* | ENSGALG00000032599 | | | 0.18 | 0.17 | 2.7E-03 | 7.1E-03 | 0.17 | 2.7E-03 | | MMKALQWERAVVQG* |
| HG0415 | *G. gallus* | ENSGALG00000034970 | | | 0.04 | 0.46 | 4.8E-07 | 2.0E-06 | 0.46 | 4.8E-07 | | MLSRTVMTREASLAKD* |
| HG0416 | *G. gallus* | ENSGALG00000015874 | TTK | TTK protein kinase | 0.13 | 0.31 | 7.0E-06 | 2.6E-05 | 0.31 | 7.0E-06 | | MKDYISCRCWILGII* |
| HG0417 | *G. gallus* | ENSGALG00000001522 | PHF20 | PHD finger protein 20 | 0.00 | 0.48 | 8.0E-06 | 2.9E-05 | 0.48 | 8.0E-06 | | MDFSALTIPDGGKHVVGGELGE* |
| HG0418 | *G. gallus* | ENSGALG00000010337 | GPR149 | G protein-coupled receptor 149 | 0.14 | 0.36 | 1.3E-06 | 5.1E-06 | 0.36 | 1.3E-06 | | MKNCIFSSTFCTSLLPDFSRA* |
| HG0419 | *G. gallus* | ENSGALG00000012542 | RASD2 | RASD family member 2 | 0.03 | 0.48 | 1.7E-03 | 4.6E-03 | 0.48 | 1.7E-03 | | MGGRPLEPCSPSQSC* |
| HG0420 | *G. gallus* | ENSGALG00000000336 | PPP1R12B | | 0.00 | 0.38 | 4.4E-04 | 1.3E-03 | 0.38 | 4.4E-04 | | MPKIIHLNRQGLITRAANSLISLF* |
| HG0421 | *G. gallus* | ENSGALG00000003492 | | | 0.21 | 0.05 | 2.8E-20 | 2.2E-19 | 0.05 | 2.8E-20 | | MEANLKRAKKGDLKVSVHRMEIERIRYVLSSYLRCRLAKIEKFFPHVLEKEKSRAKGEPSILSPEEFAFAKEYMANTETYLKNVALKHMPPNLQKVALLKSVPKPNLDSFVFLRVLKRQENILVEPETDEQREYTIDLEEGSQHLIRYKTIAPLVASGAVQLI* |
| HG0422 | *G. gallus* | ENSGALG00000005772 | BOK | BOK, BCL2 family apoptosis regulator | 0.00 | 0.44 | 7.2E-03 | 1.7E-02 | 0.44 | 7.2E-03 | | MKGKMKNRLWESSSWEL* |
| HG0423 | *G. gallus* | ENSGALG00000006329 | | | 0.00 | 0.28 | 2.6E-03 | 6.9E-03 | 0.28 | 2.6E-03 | | MLSKLVGKKGGHLVSVL* |
| HG0424 | *G. gallus* | ENSGALG00000039553 | | | 0.00 | 0.27 | 2.8E-04 | 8.4E-04 | 0.27 | 2.8E-04 | | MTIYLSCIRQRCCLQ* |
| HG0425 | *G. gallus* | ENSGALG00000043044 | | | 0.00 | 0.26 | 6.2E-07 | 2.5E-06 | 0.26 | 6.2E-07 | | MIQKTVRSCDCKMSSFYLMLKGGKLKLQEL* |
| HG0426 | *G. gallus* | ENSGALG00000002192 | PTPRC | protein tyrosine phosphatase, receptor type C | 0.00 | 0.17 | 1.6E-03 | 4.4E-03 | 0.17 | 1.6E-03 | | MLLRKDTPDFQK* |
| HG0427 | *G. gallus* | ENSGALG00000005550 | ST6GAL1 | ST6 beta-galactoside alpha-2,6-sialyltransferase 1 | 0.01 | 0.45 | 1.3E-03 | 3.6E-03 | 0.45 | 1.3E-03 | | MKVFLELFSASRGCCVWGYPALNPGQKSANSQK* |
| HG0428 | *G. gallus* | ENSGALG00000005976 | TCEA2 | transcription elongation factor A2 | 0.00 | 0.04 | 1.8E-09 | 8.8E-09 | 0.04 | 1.8E-09 | | MSHPVAGRFAAPLLK* |
| HG0429 | *G. gallus* | ENSGALG00000006626 | PDE6C | phosphodiesterase 6C | 0.00 | 0.00 | 3.1E-12 | 1.8E-11 | 0.00 | 3.1E-12 | | MPLYSMYKVPFSY* |
| HG0430 | *G. gallus* | ENSGALG00000015490 | | | 0.00 | 0.39 | 2.1E-07 | 9.0E-07 | 0.39 | 2.1E-07 | | MLSMGTSASDALTLLKTYFVAWLKRR* |
| HG0431 | *G. gallus* | ENSGALG00000026542 | FAM217B | family with sequence similarity 217 member B | 0.00 | 0.33 | 6.8E-09 | 3.2E-08 | 0.33 | 6.8E-09 | | MPAEAESKDVVMFAFGTLCCVDRKEWSRWKIM* |
| HG0432 | *G. gallus* | ENSGALG00000040546 | | | 0.05 | 0.44 | 6.9E-03 | 1.6E-02 | 0.44 | 6.9E-03 | | MLENFTSVLIVISEHPPKN* |
| HG0433 | *G. gallus* | ENSGALG00000002952 | NRG4 | neuregulin 4 | 0.00 | 0.43 | 9.6E-05 | 3.1E-04 | 0.43 | 9.6E-05 | | MQISGGVDSADCPVRV* |
| HG0434 | *G. gallus* | ENSGALG00000008357 | TPCN1 | two pore segment channel 1 | 0.24 | 0.34 | 5.9E-05 | 2.0E-04 | 0.34 | 5.9E-05 | | MGARTFAFWWLEDGHDGRFPGKRF* |
| HG0435 | *G. gallus* | ENSGALG00000010778 | CHRM3 | cholinergic receptor muscarinic 3 | 0.00 | 0.14 | 2.6E-07 | 1.1E-06 | 0.14 | 2.6E-07 | | MAKSFSSVWYIQLQAG* |
| HG0436 | *G. gallus* | ENSGALG00000026736 | OGN | osteoglycin | 0.10 | 0.16 | 8.1E-07 | 3.3E-06 | 0.26 | 2.5E-03 | | MLKLSTAPCSRILPLTLGTECWQSIFYPMESTFKTVNLGKLSHCSNLDVIHCWIVHNFTLLSVCMGLTYA* |
| HG0437 | *G. gallus* | ENSGALG00000001728 | NACC2 | NACC family member 2 | 0.28 | 0.40 | 7.4E-04 | 2.1E-03 | 0.40 | 7.4E-04 | | MRGAAPARRPGPPFSPGPRRPASAAAARRKTMGRTMCPTLDTPALNA* |
| HG0438 | *G. gallus* | ENSGALG00000008545 | SENP5 | SUMO1/sentrin specific peptidase 5 | 0.12 | 0.16 | 3.0E-05 | 1.0E-04 | 0.16 | 3.0E-05 | | MWYTARCCVTCKERQFCT* |
| HG0439 | *G. gallus* | ENSGALG00000009172 | OSBPL6 | oxysterol binding protein like 6 | 0.14 | 0.04 | 1.6E-02 | 3.4E-02 | 0.04 | 1.6E-02 | | MQKSIPCIFVHE* |
| HG0440 | *G. gallus* | ENSGALG00000010090 | | | 0.00 | 0.28 | 5.3E-05 | 1.8E-04 | 0.28 | 5.3E-05 | | MMLGHLLLTTGFSKEIQHGLMC* |
| HG0441 | *G. gallus* | ENSGALG00000015069 | B4GALT4 | beta-1,4-galactosyltransferase 4 | 0.00 | 0.40 | 2.4E-04 | 7.5E-04 | 0.40 | 2.4E-04 | | MSHRTTRSAAALPTCSFPSLL* |
| HG0442 | *G. gallus* | ENSGALG00000030250 | | | 0.00 | 0.40 | 7.3E-03 | 1.7E-02 | 0.40 | 7.3E-03 | | MTFTSMMNQEKPVKSS* |
| HG0443 | *G. gallus* | ENSGALG00000039454 | | | 0.00 | 0.00 | 7.9E-04 | 2.3E-03 | 0.00 | 7.9E-04 | | MGWVSNHVSQDSVLL* |
| HG0444 | *G. gallus* | ENSGALG00000000558 | SLC1A6 | solute carrier family 1 member 6 | 0.13 | 0.39 | 3.0E-03 | 7.7E-03 | 0.39 | 3.0E-03 | | MLYLPKEKILHPAFELRESRNKTTQNNPPRRKRPTVLLPDH* |
| HG0445 | *G. gallus* | ENSGALG00000004017 | | | 0.03 | 0.41 | 1.3E-02 | 2.8E-02 | 0.41 | 1.3E-02 | | MVALPGVGLGQPPQQGGDESLYKQGRK* |
| HG0446 | *G. gallus* | ENSGALG00000006807 | PLSCR5 | phospholipid scramblase family member 5 | 0.00 | 0.32 | 5.4E-03 | 1.3E-02 | 0.32 | 5.4E-03 | | MLPNTSLCSPARSAWALPFCSHASYLSAAHMCVISMATWALEVEI* |
| HG0447 | *G. gallus* | ENSGALG00000008784 | | | 0.00 | 0.16 | 8.3E-03 | 1.9E-02 | 0.16 | 8.3E-03 | | MKLNFKECFNQKEN* |
| HG0448 | *G. gallus* | ENSGALG00000011762 | UBA6 | ubiquitin like modifier activating enzyme 6 | 0.00 | 0.09 | 5.9E-05 | 2.0E-04 | 0.09 | 5.9E-05 | | MRPGLPAVPAC* |
| HG0449 | *G. gallus* | ENSGALG00000014106 | PPFIBP1 | PPFIA binding protein 1 | 0.00 | 0.30 | 5.0E-03 | 1.2E-02 | 0.30 | 5.0E-03 | | MDQWNWLKRNGGDQH* |
| HG0450 | *G. gallus* | ENSGALG00000015528 | JAKMIP1 | janus kinase and microtubule interacting protein 1 | 0.00 | 0.09 | 8.9E-04 | 2.5E-03 | 0.09 | 8.9E-04 | | MRDKTHMADGRDFCERFG* |
| HG0451 | *G. gallus* | ENSGALG00000019489 | CHST9 | carbohydrate sulfotransferase 9 | 0.00 | 0.35 | 1.2E-03 | 3.3E-03 | 0.35 | 1.2E-03 | | MGLCTDFRLYRSESGQVTEEDAGTVRTSEK* |
| HG0452 | *G. gallus* | ENSGALG00000004227 | FAM20B | FAM20B, glycosaminoglycan xylosylkinase | 0.00 | 0.14 | 6.2E-03 | 1.5E-02 | 0.14 | 6.2E-03 | | MLPWQSVFYSASYE* |
| HG0453 | *G. gallus* | ENSGALG00000005995 | SLC13A5 | solute carrier family 13 member 5 | 0.07 | 0.08 | 1.4E-03 | 3.8E-03 | 0.08 | 1.4E-03 | | MQVLADSPLSARLCAGVKFPLC* |
| HG0454 | *G. gallus* | ENSGALG00000007186 | TMEM268 | transmembrane protein 268 | 0.08 | 0.36 | 1.8E-02 | 3.7E-02 | 0.36 | 1.8E-02 | | MLPCQWRAESQGSRQWESE* |
| HG0455 | *G. gallus* | ENSGALG00000010718 | THSD7A | thrombospondin type 1 domain containing 7A | 0.01 | 0.17 | 2.2E-02 | 4.5E-02 | 0.17 | 2.2E-02 | | MTAAASQRKITCSVSWSPLRTAVSLLADGKLLLTAEWKSVEQQPY* |
| HG0456 | *G. gallus* | ENSGALG00000033256 | | | 0.00 | 0.20 | 3.1E-03 | 8.0E-03 | 0.20 | 3.1E-03 | | MHCSKLFLVFSSG* |
| HG0457 | *G. gallus* | ENSGALG00000039538 | CLDND1 | claudin domain containing 1 | 0.00 | 0.28 | 2.3E-02 | 4.6E-02 | 0.28 | 2.3E-02 | | MEQLGPAGSWRCVLLAYCRTPVARCL* |

| | | | | | | | | | | | | |
|---|---|---|---|---|---|---|---|---|---|---|---|---|
| HG0458 | *H. sapiens* | ENSG00000060339 | CCAR1 | cell division cycle and apoptosis regulator 1 | 0.05 | 0.35 | 6.3E-39 | 2.9E-37 | 0.35 | 6.3E-39 | | MWRRGAAWRKRGKLAHAPKADGFEMASMLAGTRLRSML* |
| HG0459.1 | *H. sapiens* | ENSG00000163848 | ZNF148 | zinc finger protein 148 | 0.00 | 0.30 | 5.3E-10 | 5.8E-09 | 0.30 | 5.3E-10 | | MKGRNSTMISFSTLQHSFP* |
| HG0459.2 | *H. sapiens* | ENSG00000163848 | ZNF148 | zinc finger protein 148 | 0.03 | 0.47 | 1.3E-11 | 1.7E-10 | 0.47 | 1.3E-11 | | MGVCLTYLMEESDGISD* |
| HG0459.3 | *H. sapiens* | ENSG00000163848 | ZNF148 | zinc finger protein 148 | 0.00 | 0.21 | 1.3E-05 | 7.3E-05 | 0.21 | 1.3E-05 | | MSASLLLWEEEEKEEEEEEEEGEEEEKGAGVGAVAEAATAKVLPPPPFPSSQGP* |
| HG0460 | *H. sapiens* | ENSG00000180008 | SOCS4 | suppressor of cytokine signaling 4 | 0.01 | 0.34 | 7.7E-46 | 4.0E-44 | 0.40 | 1.5E-38 | Crowe, Samandi | MAMVMAAVTRTTPVKLRSGRYIQKVPRRNFLLEKMKKQYL* |
| HG0461 | *H. sapiens* | ENSG00000064419 | TNPO3 | transportin 3 | 0.02 | 0.35 | 8.0E-03 | 2.2E-02 | 0.35 | 8.0E-03 | | MAPVRERPWPRQ* |
| HG0462 | *H. sapiens* | ENSG00000125686 | MED1 | mediator complex subunit 1 | 0.08 | 0.32 | 6.4E-18 | 1.4E-16 | 0.32 | 6.4E-18 | | MAAASSTLLFLPPGTSD* |
| HG0463 | *H. sapiens* | ENSG00000129473 | BCL2L2 | BCL2 like 2 | 0.01 | 0.18 | 2.8E-30 | 9.8E-29 | 0.18 | 2.8E-30 | | MKGPSWGLLATSAVS* |
| HG0464 | *H. sapiens* | ENSG00000188215 | DCUN1D3 | defective in cullin neddylation 1 domain containing 3 | 0.01 | 0.48 | 1.3E-10 | 1.5E-09 | 0.48 | 1.3E-10 | | MLRYMGQRQRGHPIQRGPL* |
| HG0465 | *H. sapiens* | ENSG00000132153 | DHX30 | DExH-box helicase 30 | 0.00 | 0.35 | 2.8E-23 | 7.9E-22 | 0.35 | 2.8E-23 | Crowe | MRNSKERIQLQFKSLQNLRLSLLL* |
| HG0466 | *H. sapiens* | ENSG00000139679 | LPAR6 | lysophosphatidic acid receptor 6 | 0.04 | 0.49 | 1.5E-12 | 2.2E-11 | 0.49 | 1.5E-12 | | MFLLGISEKKEIFIKPSKDPDNLQINWRL* |
| HG0467.1 | *H. sapiens* | ENSG00000163320 | CGGBP1 | CGG triplet repeat binding protein 1 | 0.00 | 0.35 | 7.1E-15 | 1.3E-13 | 0.35 | 7.1E-15 | | MTSWHHRENRVQAQKLFPHHHLLKN* |
| HG0467.2 | *H. sapiens* | ENSG00000163320 | CGGBP1 | CGG triplet repeat binding protein 1 | 0.01 | 0.32 | 3.0E-07 | 2.2E-06 | 0.32 | 3.0E-07 | | MRLWFRSGDHSLLSIKTKKTGN* |
| HG0468 | *H. sapiens* | ENSG00000187778 | MCRS1 | microspherule protein 1 | 0.00 | 0.09 | 7.2E-27 | 2.2E-25 | 0.09 | 7.2E-27 | Samandi | MLLPRVLETDLDRSPTLGVHSWENEGRRVRISSQPLSSPGQNPWMSLWDRSS* |
| HG0469 | *H. sapiens* | ENSG00000198369 | SPRED2 | sprouty related EVH1 domain containing 2 | 0.02 | 0.01 | 2.1E-19 | 5.1E-18 | 0.01 | 2.1E-19 | | MKGAKLPAAGQASR* |
| HG0470 | *H. sapiens* | ENSG00000224470 | ATXN1L | ataxin 1 like | 0.00 | 0.39 | 1.4E-20 | 3.4E-19 | 0.39 | 1.4E-20 | | MWAPQPEAERGTGKLQRSPF* |
| HG0471 | *H. sapiens* | ENSG00000023041 | ZDHHC6 | zinc finger DHHC-type containing 6 | 0.01 | 0.43 | 2.4E-19 | 5.6E-18 | 0.43 | 2.4E-19 | | MTIKEHTWTCGTWKCAHRKRKSIILLK* |
| HG0472 | *H. sapiens* | ENSG00000105991 | HOXA1 | homeobox A1 | 0.06 | 0.23 | 1.2E-25 | 3.5E-24 | 0.23 | 1.2E-25 | | MEEVRKLAQSRRASQDQVTQ* |
| HG0473.1 | *H. sapiens* | ENSG00000138650 | PCDH10 | protocadherin 10 | 0.00 | 0.42 | 1.3E-02 | 3.2E-02 | 0.42 | 1.3E-02 | | MRKGDACHPSCAKI* |
| HG0473.2 | *H. sapiens* | ENSG00000240184 | PCDHGC3 | protocadherin gamma subfamily C, 3 | 0.02 | 0.18 | 6.6E-10 | 7.1E-09 | 0.18 | 6.6E-10 | | MSDSAPSAQALTR* |
| HG0474 | *H. sapiens* | ENSG00000139083 | ETV6 | ETS variant 6 | 0.00 | 0.39 | 1.3E-08 | 1.1E-07 | 0.39 | 1.3E-08 | | MTASGWPWSLSGLGRGKESGKNLRTS* |
| HG0475 | *H. sapiens* | ENSG00000109118 | PHF12 | PHD finger protein 12 | 0.02 | 0.42 | 6.9E-09 | 6.3E-08 | 0.42 | 6.9E-09 | | MAAPASLPGRTGCRQCRL* |
| HG0476 | *H. sapiens* | ENSG00000130939 | UBE4B | ubiquitination factor E4B | 0.00 | 0.15 | 1.8E-05 | 9.6E-05 | 0.15 | 1.8E-05 | | MNNTWWGEGERVGVEG* |
| HG0477 | *H. sapiens* | ENSG00000166925 | TSC22D4 | TSC22 domain family member 4 | 0.01 | 0.17 | 8.4E-52 | 4.8E-50 | 0.17 | 8.4E-52 | Mackowiak | MSDPDVPRGSDVPAMWPPCSRGA* |
| HG0478 | *H. sapiens* | ENSG00000168214 | RBPJ | recombination signal binding protein for immunoglobulin ka | 0.00 | 0.10 | 1.3E-05 | 6.9E-05 | 0.10 | 1.3E-05 | | MISKRKENDTDTLAEVF* |
| HG0479.1 | *H. sapiens* | ENSG00000168453 | HR | HR, lysine demethylase and nuclear receptor corepressor | 0.00 | 0.07 | 1.1E-28 | 3.6E-27 | 0.07 | 1.1E-28 | Crowe, Mackowiak | MAQPTASAQKLVRPIRAVCRILQIPESDPSNLRP* |
| HG0479.2 | *H. sapiens* | ENSG00000168453 | HR | HR, lysine demethylase and nuclear receptor corepressor | 0.00 | 0.38 | 5.9E-05 | 2.9E-04 | 0.38 | 5.9E-05 | | MAIRGPAALSAALYLH* |
| HG0480 | *H. sapiens* | ENSG00000198018 | ENTPD7 | ectonucleoside triphosphate diphosphohydrolase 7 | 0.00 | 0.20 | 1.2E-18 | 2.8E-17 | 0.20 | 1.2E-18 | | MPGTRRPPSQGKRKRR* |
| HG0481 | *H. sapiens* | ENSG00000198963 | RORB | RAR related orphan receptor B | 0.01 | 0.38 | 1.0E-06 | 6.7E-06 | 0.38 | 1.0E-06 | | MVFSAEQLFADHLLHSC* |
| HG0482 | *H. sapiens* | ENSG00000067167 | TRAM1 | translocation associated membrane protein 1 | 0.11 | 0.50 | 1.4E-17 | 2.9E-16 | 0.50 | 1.4E-17 | Crowe | MGSEAPASGCSGAVRSSQREAAASR* |
| HG0483 | *H. sapiens* | ENSG00000164463 | CREBRF | CREB3 regulatory factor | 0.07 | 0.22 | 3.8E-17 | 7.7E-16 | 0.22 | 3.8E-17 | Mackowiak | MERGRGPCSGTGPSP* |
| HG0484 | *H. sapiens* | ENSG00000166444 | ST5 | suppression of tumorigenicity 5 | 0.00 | 0.41 | 9.3E-17 | 1.9E-15 | 0.44 | 2.7E-11 | | MNQLASISLTNHRKMRDCLGATTSTMSPCTG* |
| HG0485.1 | *H. sapiens* | ENSG00000182667 | NTM | neurotrimin | 0.00 | 0.00 | 4.5E-16 | 8.6E-15 | 0.00 | 4.5E-16 | | MSGDNYGEVILSHTLGFLVVSFSKTVDLNLLAQA* |
| HG0485.2 | *H. sapiens* | ENSG00000183715 | OPCML | opioid binding protein/cell adhesion molecule like | 0.03 | 0.00 | 2.4E-03 | 7.6E-03 | 0.00 | 2.4E-03 | | MQEGASGSARLES* |
| HG0486 | *H. sapiens* | ENSG00000067900 | ROCK1 | Rho associated coiled-coil containing protein kinase 1 | 0.00 | 0.16 | 2.0E-25 | 5.9E-24 | 0.16 | 2.0E-25 | | MRLWASERLWVAVAPGPGSGERRSPQ* |
| HG0487 | *H. sapiens* | ENSG00000091831 | ESR1 | estrogen receptor 1 | 0.07 | 0.41 | 1.3E-11 | 1.6E-10 | 0.41 | 1.3E-11 | | MEHFWKDVLDPAGWPAGF* |
| HG0488.1 | *H. sapiens* | ENSG00000116194 | ANGPTL1 | angiopoietin like 1 | 0.00 | 0.48 | 1.5E-09 | 1.5E-08 | 0.48 | 1.5E-09 | | MWQTEQSHNAFSAGLKTAHRPELPLYYLKLHRWLVKFN* |
| HG0488.2 | *H. sapiens* | ENSG00000116194 | ANGPTL1 | angiopoietin like 1 | 0.03 | 0.24 | 1.1E-04 | 4.9E-04 | 0.24 | 1.1E-04 | | MQLPLKSQMNKLLH* |
| HG0489 | *H. sapiens* | ENSG00000143847 | PPFIA4 | PTPRF interacting protein alpha 4 | 0.00 | 0.48 | 6.5E-06 | 3.7E-05 | 0.48 | 6.5E-06 | | MLWEKPLPTCPLAWHCPL* |
| HG0490.1 | *H. sapiens* | ENSG00000152413 | HOMER1 | homer scaffolding protein 1 | 0.00 | 0.48 | 6.0E-07 | 4.2E-06 | 0.48 | 6.0E-07 | | MAPFILFGRMNRSCLAFSSSSPPLLCSES* |
| HG0490.2 | *H. sapiens* | ENSG00000152413 | HOMER1 | homer scaffolding protein 1 | 0.00 | 0.45 | 6.3E-04 | 2.4E-03 | 0.45 | 6.3E-04 | | MPPLFERFGIYKREIILPKRTN* |
| HG0491 | *H. sapiens* | ENSG00000154114 | TBCEL | tubulin folding cofactor E like | 0.03 | 0.26 | 3.4E-12 | 4.8E-11 | 0.26 | 3.4E-12 | Mackowiak | MQDTGLRRPEWGTR* |
| HG0492 | *H. sapiens* | ENSG00000162670 | BRINP3 | BMP/retinoic acid inducible neural specific 3 | 0.00 | 0.40 | 1.2E-06 | 7.6E-06 | 0.40 | 1.2E-06 | | MEYLFESWMYFSHSLLAS* |
| HG0493 | *H. sapiens* | ENSG00000164707 | SLC13A4 | solute carrier family 13 member 4 | 0.12 | 0.34 | 3.1E-17 | 6.3E-16 | 0.34 | 3.1E-17 | | MPPFIILSQFLFAGHKTMLINPT* |
| HG0494 | *H. sapiens* | ENSG00000184226 | PCDH9 | protocadherin 9 | 0.00 | 0.44 | 1.7E-14 | 2.9E-13 | 0.44 | 1.7E-14 | | MHAAVENIPELKKCNYVLCVLNAQKNPDPVGKRDEMWRE* |
| HG0495.1 | *H. sapiens* | ENSG00000196730 | DAPK1 | death associated protein kinase 1 | 0.00 | 0.13 | 2.1E-08 | 1.8E-07 | 0.13 | 2.1E-08 | | MHEGATEAQERW* |
| HG0495.2 | *H. sapiens* | ENSG00000196730 | DAPK1 | death associated protein kinase 1 | 0.00 | 0.26 | 1.1E-05 | 6.2E-05 | 0.26 | 1.1E-05 | | MVWEAELKCPGLW* |
| HG0496 | *H. sapiens* | ENSG00000135913 | USP37 | ubiquitin specific peptidase 37 | 0.01 | 0.42 | 1.1E-02 | 2.8E-02 | 0.42 | 1.1E-02 | | MSSVPRRVLCLWHKDQNKGELLLHRVFR* |
| HG0497 | *H. sapiens* | ENSG00000147571 | CRH | corticotropin releasing hormone | 0.09 | 0.00 | 3.9E-04 | 1.5E-03 | 0.00 | 3.9E-04 | | MIYAGAEAARNRAVKRASAY* |
| HG0498 | *H. sapiens* | ENSG00000005379 | TSPOAP1 | TSPO associated protein 1 | 0.00 | 0.00 | 4.8E-16 | 9.1E-15 | 0.00 | 4.8E-16 | | MRQAAHRPRPP* |
| HG0499 | *H. sapiens* | ENSG00000011258 | MBTD1 | mbt domain containing 1 | 0.02 | 0.13 | 2.0E-03 | 6.6E-03 | 0.13 | 2.0E-03 | | MLMLLISWGHPS* |
| HG0500 | *H. sapiens* | ENSG00000070778 | PTPN21 | protein tyrosine phosphatase, non-receptor type 21 | 0.00 | 0.24 | 6.4E-22 | 1.7E-20 | 0.24 | 6.4E-22 | | MLPSSGHWRLRCFLMAAAAAG* |
| HG0501 | *H. sapiens* | ENSG00000110400 | NECTIN1 | nectin cell adhesion molecule 1 | 0.11 | 0.19 | 5.4E-07 | 3.8E-06 | 0.19 | 5.4E-07 | | MTRATCGGHLPSL* |
| HG0502 | *H. sapiens* | ENSG00000126882 | | | 0.00 | 0.00 | 1.7E-22 | 4.6E-21 | 0.00 | 1.7E-22 | | MSEPHNYSLNYKLFLTQRISVIFVHCEEKKREPRAHEMQQNPYL* |
| HG0503 | *H. sapiens* | ENSG00000141564 | RPTOR | regulatory associated protein of MTOR complex 1 | 0.00 | 0.19 | 3.7E-18 | 8.2E-17 | 0.19 | 3.7E-18 | | MGRSWLWLQQLRRVRDPQG* |
| HG0504 | *H. sapiens* | ENSG00000144460 | NYAP2 | neuronal tyrosine-phosphorylated phosphoinositide-3-kinas | 0.00 | 0.28 | 2.6E-14 | 4.5E-13 | 0.36 | 3.7E-09 | | MQTDCRNREERLLLISEEYNQMRYISQLGDCFWSCNHLCSKVGKIIQFLPCWTWKERGCVFIPTH* |
| HG0505 | *H. sapiens* | ENSG00000157470 | | | 0.05 | 0.31 | 6.7E-07 | 4.6E-06 | 0.31 | 6.7E-07 | | MGSRGAPTPRADVNY* |
| HG0506.1 | *H. sapiens* | ENSG00000177853 | ZNF518A | zinc finger protein 518A | 0.00 | 0.18 | 4.0E-08 | 3.3E-07 | 0.33 | 1.1E-04 | | MAAIRDSQAMGTTTDIISSFDHCS* |
| HG0506.2 | *H. sapiens* | ENSG00000177853 | ZNF518A | zinc finger protein 518A | 0.00 | 0.49 | 2.0E-02 | 4.7E-02 | 0.49 | 2.0E-02 | | MSLHSIVSCLNYR* |
| HG0507 | *H. sapiens* | ENSG00000060237 | WNK1 | WNK lysine deficient protein kinase 1 | 0.01 | 0.44 | 1.9E-08 | 1.7E-07 | 0.44 | 1.9E-08 | | MTAAPLLPPPARPPLAAGWMRTVRR* |
| HG0508 | *H. sapiens* | ENSG00000061936 | SFSWAP | splicing factor SWAP homolog | 0.16 | 0.38 | 5.4E-10 | 5.9E-09 | 0.38 | 5.4E-10 | | MAAVLRLGTGCGVFD* |
| HG0509 | *H. sapiens* | ENSG00000068024 | HDAC4 | histone deacetylase 4 | 0.10 | 0.30 | 2.0E-08 | 1.8E-07 | 0.30 | 2.0E-08 | | MCWWEFGARWSYRFRGNFEPFRIT* |
| HG0510 | *H. sapiens* | ENSG00000134769 | DTNA | dystrobrevin alpha | 0.04 | 0.19 | 7.2E-05 | 3.4E-04 | 0.19 | 7.2E-05 | | MVLKSSCCPFIEF* |
| HG0511 | *H. sapiens* | ENSG00000143337 | TOR1AIP1 | torsin 1A interacting protein 1 | 0.25 | 0.45 | 8.1E-15 | 1.4E-13 | 0.45 | 8.1E-15 | | MAETSSSPFSSSLVPPARRRERSTAADPAQALPLSQPSRHTMGPEAGLLHSSDDAGGGPSDSQLPP* |
| HG0512.1 | *H. sapiens* | ENSG00000112319 | EYA4 | EYA transcriptional coactivator and phosphatase 4 | 0.00 | 0.34 | 6.1E-09 | 5.7E-08 | 0.34 | 6.1E-09 | | MSCFHQRHSAKGKLRHWKERE* |
| HG0512.2 | *H. sapiens* | ENSG00000104313 | EYA1 | EYA transcriptional coactivator and phosphatase 1 | 0.02 | 0.43 | 3.6E-13 | 5.6E-12 | 0.43 | 3.6E-13 | | MLSAAAVWWGRSAGRKAVTKQNGGS* |
| HG0512.3 | *H. sapiens* | ENSG00000104313 | EYA1 | EYA transcriptional coactivator and phosphatase 1 | 0.00 | 0.00 | 3.4E-03 | 1.0E-02 | 0.00 | 3.4E-03 | | MPFTAAVSSFFSFG* |
| HG0513 | *H. sapiens* | ENSG00000126860 | EVI2A | ecotropic viral integration site 2A | 0.02 | 0.20 | 1.5E-11 | 2.0E-10 | 0.23 | 1.6E-06 | | MCFSCSGKEMCHLWFGFKSGKLAAHILFYCRFTLRLIFSKSILL* |
| HG0514 | *H. sapiens* | ENSG00000147570 | DNAJC5B | DnaJ heat shock protein family (Hsp40) member C5 beta | 0.00 | 0.39 | 6.0E-07 | 4.2E-06 | 0.39 | 6.0E-07 | | MKKRRDLKRDNQLAGKVTNEK* |
| HG0515 | *H. sapiens* | ENSG00000170471 | RALGAPB | Ral GTPase activating protein non-catalytic beta subunit | 0.06 | 0.00 | 4.1E-03 | 1.2E-02 | 0.00 | 4.1E-03 | | MRAPPVASGAIWIVL* |
| HG0516 | *H. sapiens* | ENSG00000041515 | MYO16 | myosin XVI | 0.16 | 0.20 | 4.1E-09 | 3.9E-08 | 0.20 | 4.1E-09 | | MRMSLKLFAGSWNRASMQ* |
| HG0517 | *H. sapiens* | ENSG00000086717 | PPEF1 | protein phosphatase with EF-hand domain 1 | 0.00 | 0.42 | 1.5E-04 | 6.4E-04 | 0.42 | 1.5E-04 | | MIDCVLSPQVRLKESTTSKVKI* |
| HG0518 | *H. sapiens* | ENSG00000111254 | AKAP3 | A-kinase anchoring protein 3 | 0.06 | 0.44 | 2.0E-04 | 8.6E-04 | 0.44 | 2.0E-04 | | MEGHRKKQDLELSKNIPASSLTLKVYSGVFRGSLFQYYRDSLYYMAGGG* |
| HG0519 | *H. sapiens* | ENSG00000127074 | RGS13 | regulator of G protein signaling 13 | 0.01 | 0.23 | 2.9E-08 | 2.4E-07 | 0.23 | 2.9E-08 | | MVDCSTGYICQWNVSTYYTTNMRKK* |
| HG0520 | *H. sapiens* | ENSG00000197415 | VEPH1 | ventricular zone expressed PH domain containing 1 | 0.00 | 0.49 | 1.3E-04 | 5.9E-04 | 0.49 | 1.3E-04 | | MNNFRFTELSGFHSLCPRSLPESPAQSQTMQLSNCKGAFEE* |
| HG0521 | *H. sapiens* | ENSG00000038274 | MAT2B | methionine adenosyltransferase 2B | 0.08 | 0.21 | 2.7E-04 | 1.1E-03 | 0.21 | 2.7E-04 | | MWRTARDSGIGIKMLI* |
| HG0522 | *H. sapiens* | ENSG00000140396 | NCOA2 | nuclear receptor coactivator 2 | 0.00 | 0.25 | 1.4E-05 | 7.8E-05 | 0.25 | 1.4E-05 | | MAAAASATASAAKVSADGSRHLTA* |
| HG0523 | *H. sapiens* | ENSG00000075213 | SEMA3A | semaphorin 3A | 0.10 | 0.04 | 1.1E-03 | 3.9E-03 | 0.04 | 1.1E-03 | | MGCSWYSYKVNMLLAAHIYP* |
| HG0524 | *H. sapiens* | ENSG00000107929 | LARP4B | La ribonucleoprotein domain family member 4B | 0.13 | 0.13 | 1.1E-03 | 4.1E-03 | 0.13 | 1.1E-03 | | MRTGRSVGDLESRG* |
| HG0525 | *H. sapiens* | ENSG00000136167 | LCP1 | lymphocyte cytosolic protein 1 | 0.07 | 0.35 | 1.7E-02 | 4.0E-02 | 0.35 | 1.7E-02 | | MTSPEIAVVQRKQSRGIELLQNARVSNIVSAAASYLTKVEEILYFQLRKLKIPPASDDQAPLDTEHCAFLLQVTSCLVTTHPGLTKLFCRSERRGSWSYTSTTKDSFFPARSVT* |
| HG0526 | *H. sapiens* | ENSG00000143569 | UBAP2L | ubiquitin associated protein 2 like | 0.06 | 0.40 | 8.3E-04 | 3.0E-03 | 0.40 | 8.3E-04 | | MRLEPERGERERARASEAWALPEYSTL* |
| HG0527 | *H. sapiens* | ENSG00000198382 | UVRAG | UV radiation resistance associated | 0.08 | 0.08 | 1.5E-04 | 6.6E-04 | 0.08 | 1.5E-04 | | MSAPGLSGVEGLHCGNMALP* |
| HG0528 | *H. sapiens* | ENSG00000204767 | | | 0.00 | 0.32 | 7.8E-03 | 2.1E-02 | 0.32 | 7.8E-03 | | MGMFPWEHHVSSNKQTSRSFISPPVYLLWKEDAFKFPAVSPWKLMLWETKSCMF* |
| HG0529 | *H. sapiens* | ENSG00000112851 | ERBIN | erbb2 interacting protein | 0.10 | 0.50 | 6.4E-03 | 1.8E-02 | 0.50 | 6.4E-03 | | MLAQLRNIREINSTQCV* |
| HG0530 | *H. sapiens* | ENSG00000128594 | LRRC4 | leucine rich repeat containing 4 | 0.00 | 0.00 | 3.0E-11 | 3.7E-10 | 0.00 | 3.0E-11 | | MREQSAWLPPATPRARREEETKSEWERGAQLPWRKLKDTTPVLSCLPSEMERGAPSSLKPG* |
| HG0531 | *H. sapiens* | ENSG00000146834 | MEPCE | methylphosphate capping enzyme | 0.09 | 0.47 | 4.0E-13 | 6.2E-12 | 0.47 | 4.0E-13 | | MRAQPGWWRSAHAWFPGEGNARSPCALAAGW* |
| HG0532 | *H. sapiens* | ENSG00000105221 | AKT2 | AKT serine/threonine kinase 2 | 0.00 | 0.32 | 4.0E-19 | 9.2E-18 | 0.32 | 4.0E-19 | | MVPGTVLCPVHGASCMSCCPELSRAR* |
| HG0533 | *H. sapiens* | ENSG00000114209 | PDCD10 | programmed cell death 10 | 0.00 | 0.28 | 1.1E-05 | 6.0E-05 | 0.28 | 1.1E-05 | | MKIPLHFSTAFSTVVGF* |
| HG0534 | *H. sapiens* | ENSG00000117408 | IPO13 | importin 13 | 0.02 | 0.29 | 2.5E-36 | 1.1E-34 | 0.29 | 2.5E-36 | Mackowiak | MPALGPPSPHHSLGTQAGV* |
| HG0535 | *H. sapiens* | ENSG00000118407 | FILIP1 | filamin A interacting protein 1 | 0.00 | 0.21 | 1.6E-11 | 2.0E-10 | 0.21 | 1.6E-11 | | MTSAIGWSWERKWEKLFPLYSFGLDSF* |
| HG0536 | *H. sapiens* | ENSG00000181751 | C5orf30 | chromosome 5 open reading frame 30 | 0.04 | 0.20 | 2.7E-05 | 1.4E-04 | 0.20 | 2.7E-05 | | MSVFPWDSLEIVQ* |
| HG0537 | *H. sapiens* | ENSG00000184481 | FOXO4 | forkhead box O4 | 0.00 | 0.40 | 2.5E-16 | 4.8E-15 | 0.40 | 2.5E-16 | | MCLLALDASPLRGGIVRDCGRLH* |
| HG0538 | *H. sapiens* | ENSG00000006116 | CACNG3 | calcium voltage-gated channel auxiliary subunit gamma 3 | 0.04 | 0.46 | 5.6E-03 | 1.6E-02 | 0.46 | 5.6E-03 | | MRTNPPEPAPFRGP* |
| HG0539 | *H. sapiens* | ENSG00000115966 | ATF2 | activating transcription factor 2 | 0.00 | 0.21 | 1.3E-20 | 3.4E-19 | 0.21 | 1.3E-20 | | MAHQLESILAMRKRM* |
| HG0540 | *H. sapiens* | ENSG00000120519 | SLC10A7 | solute carrier family 10 member 7 | 0.04 | 0.32 | 2.8E-37 | 1.2E-35 | 0.32 | 2.8E-37 | Samandi | MSGGTKESVETLQGLGDRSLKRKKAKQINKTHPP* |
| HG0541.1 | *H. sapiens* | ENSG00000122584 | NXPH1 | neurexophilin 1 | 0.00 | 0.28 | 7.3E-24 | 2.1E-22 | 0.28 | 7.3E-24 | | METERLGGHVRGAGDARPE* |
| HG0541.2 | *H. sapiens* | ENSG00000122584 | NXPH1 | neurexophilin 1 | 0.00 | 0.41 | 7.8E-13 | 1.2E-11 | 0.41 | 7.8E-13 | | MEPAQSGPFSQDSPQVLCGC* |
| HG0542 | *H. sapiens* | ENSG00000126581 | BECN1 | beclin 1 | 0.03 | 0.34 | 1.1E-29 | 3.7E-28 | 0.34 | 1.1E-29 | | MVVLEASLRGRPEATVPPR* |
| HG0543 | *H. sapiens* | ENSG00000131931 | THAP1 | THAP domain containing 1 | 0.01 | 0.17 | 1.1E-04 | 4.8E-04 | 0.17 | 1.1E-04 | | MGDGRWLWSFC* |
| HG0544 | *H. sapiens* | ENSG00000136802 | LRRC8A | leucine rich repeat containing 8 family member A | 0.01 | 0.45 | 8.6E-17 | 1.7E-15 | 0.45 | 8.6E-17 | | MEQKECQDPAQADAGQPQHRQPTRR* |
| HG0545 | *H. sapiens* | ENSG00000142784 | WDTC1 | WD and tetratricopeptide repeats 1 | 0.01 | 0.43 | 3.7E-09 | 3.6E-08 | 0.43 | 3.7E-09 | Crowe | MCILWTWAWLECSGVLKILL* |
| HG0546 | *H. sapiens* | ENSG00000148143 | ZNF462 | zinc finger protein 462 | 0.00 | 0.42 | 6.5E-07 | 4.5E-06 | 0.42 | 6.5E-07 | | MRLPNNFHNNNPSRKRRKRKRIRRRWGWRTRSTSRRRDASSVPNVRG* |
| HG0547 | *H. sapiens* | ENSG00000149596 | JPH2 | junctophilin 2 | 0.05 | 0.48 | 6.4E-25 | 1.9E-23 | 0.48 | 6.4E-25 | Crowe | MALTSRPGADLAPGLFQPLLASCFPPPGVQW* |

| | | | | | | | | | | | | |
|---|---|---|---|---|---|---|---|---|---|---|---|---|
| HG0548 | *H. sapiens* | ENSG00000155744 | FAM126B | family with sequence similarity 126 member B | 0.00 | 0.30 | 1.5E-12 | 2.1E-11 | 0.30 | 1.5E-12 | | MNSPLELCRIPVPPPPLEN* |
| HG0549 | *H. sapiens* | ENSG00000156256 | USP16 | ubiquitin specific peptidase 16 | 0.16 | 0.20 | 2.0E-04 | 8.5E-04 | 0.20 | 2.0E-04 | | MQLWALSPWIVILCQ* |
| HG0550.1 | *H. sapiens* | ENSG00000156650 | KAT6B | lysine acetyltransferase 6B | 0.00 | 0.14 | 8.7E-24 | 2.5E-22 | 0.14 | 8.7E-24 | | MTFKCASSVKYKENLYG* |
| HG0550.2 | *H. sapiens* | ENSG00000156650 | KAT6B | lysine acetyltransferase 6B | 0.00 | 0.15 | 1.1E-06 | 7.2E-06 | 0.15 | 1.1E-06 | | MFFTRMQSFLLVK* |
| HG0551 | *H. sapiens* | ENSG00000162951 | LRRTM1 | leucine rich repeat transmembrane neuronal 1 | 0.03 | 0.48 | 1.9E-15 | 3.4E-14 | 0.48 | 1.9E-15 | | MSEPALCSCPGREWQAVSAE* |
| HG0552 | *H. sapiens* | ENSG00000169641 | LUZP1 | leucine zipper protein 1 | 0.00 | 0.25 | 6.3E-16 | 1.2E-14 | 0.25 | 6.3E-16 | | MMVSSCQRRLWLPRER* |
| HG0553 | *H. sapiens* | ENSG00000183475 | ASB7 | ankyrin repeat and SOCS box containing 7 | 0.00 | 0.49 | 6.0E-16 | 1.1E-14 | 0.49 | 6.0E-16 | | MVTRQSVHQEKGHSSIRLIPVT* |
| HG0554 | *H. sapiens* | ENSG00000196914 | ARHGEF12 | Rho guanine nucleotide exchange factor 12 | 0.00 | 0.47 | 2.2E-09 | 2.2E-08 | 0.47 | 2.2E-09 | Samandi | MTEWSFELDFCVPDGVGPDPRALGVGRRCYCKMQVG* |
| HG0555 | *H. sapiens* | ENSG00000204310 | AGPAT1 | 1-acylglycerol-3-phosphate O-acyltransferase 1 | 0.15 | 0.18 | 1.8E-16 | 3.5E-15 | 0.18 | 1.8E-16 | | MGPTSHSHLPSIGRG* |
| HG0556 | *H. sapiens* | ENSG00000273841 | TAF9 | TATA-box binding protein associated factor 9 | 0.00 | 0.12 | 1.5E-59 | 9.5E-58 | 0.11 | 2.2E-51 | | MLLPNILLTGTPGVGKTTLGKELASKSGLKYINVGDLAREV* |
| HG0557 | *H. sapiens* | ENSG00000012232 | EXTL3 | exostosin like glycosyltransferase 3 | 0.00 | 0.30 | 1.6E-03 | 5.5E-03 | 0.30 | 1.6E-03 | | MHICWSAKPRNKSYGI* |
| HG0558 | *H. sapiens* | ENSG00000029363 | BCLAF1 | BCL2 associated transcription factor 1 | 0.01 | 0.35 | 2.9E-12 | 4.1E-11 | 0.35 | 2.9E-12 | | MVFLAFLSREMAAVVWLQRR* |
| HG0559 | *H. sapiens* | ENSG00000078747 | ITCH | itchy E3 ubiquitin protein ligase | 0.00 | 0.13 | 3.2E-20 | 7.8E-19 | 0.13 | 3.2E-20 | | MHFTVALWRQRLNPRK* |
| HG0560.1 | *H. sapiens* | ENSG00000092051 | JPH4 | junctophilin 4 | 0.11 | 0.42 | 3.1E-04 | 1.2E-03 | 0.42 | 3.1E-04 | | MDAVPGRQPEPQPTCHSG* |
| HG0560.2 | *H. sapiens* | ENSG00000092051 | JPH4 | junctophilin 4 | 0.00 | 0.00 | 5.0E-03 | 1.4E-02 | 0.00 | 5.0E-03 | | MEQTLICAGMESEF* |
| HG0561 | *H. sapiens* | ENSG00000100393 | EP300 | E1A binding protein p300 | 0.04 | 0.34 | 1.9E-04 | 8.1E-04 | 0.34 | 1.9E-04 | | MRRRRTAPRRKRLMAAAELRETSAGQGPAVAGRGLRL* |
| HG0562 | *H. sapiens* | ENSG00000105821 | DNAJC2 | DnaJ heat shock protein family (Hsp40) member C2 | 0.07 | 0.45 | 8.4E-12 | 1.1E-10 | 0.45 | 8.4E-12 | | MRRFLGVLEPRREARSER* |
| HG0563 | *H. sapiens* | ENSG00000107249 | GLIS3 | GLIS family zinc finger 3 | 0.00 | 0.17 | 2.1E-08 | 1.8E-07 | 0.17 | 2.1E-08 | | MDDSIRLITHTMDNLLDFA* |
| HG0564 | *H. sapiens* | ENSG00000109171 | SLAIN2 | SLAIN motif family member 2 | 0.05 | 0.37 | 4.6E-05 | 2.3E-04 | 0.37 | 4.6E-05 | | MAAGAGYWRRLPQSRSRRR* |
| HG0565 | *H. sapiens* | ENSG00000120533 | ENY2 | ENY2, transcription and export complex 2 subunit | 0.00 | 0.00 | 1.8E-05 | 9.5E-05 | 0.00 | 1.8E-05 | | MRVLAFCVLRCPSY* |
| HG0566 | *H. sapiens* | ENSG00000124496 | TRERF1 | transcriptional regulating factor 1 | 0.00 | 0.31 | 4.9E-17 | 9.9E-16 | 0.31 | 4.9E-17 | | MHLLQKQPRPWLWRMLKEEDAEAGRP* |
| HG0567 | *H. sapiens* | ENSG00000132155 | RAF1 | Raf-1 proto-oncogene, serine/threonine kinase | 0.01 | 0.08 | 1.1E-35 | 4.6E-34 | 0.08 | 1.1E-35 | | MRGLLGAPSPGLGTRRM* |
| HG0568.1 | *H. sapiens* | ENSG00000138271 | GPR87 | G protein-coupled receptor 87 | 0.00 | 0.32 | 1.7E-22 | 4.7E-21 | 0.32 | 1.7E-22 | | MKEIKPGITYAEPTPQSSPSVS* |
| HG0568.2 | *H. sapiens* | ENSG00000138271 | GPR87 | G protein-coupled receptor 87 | 0.00 | 0.38 | 2.0E-04 | 8.4E-04 | 0.38 | 2.0E-04 | | MFRFHALPENPLPCRP* |
| HG0569 | *H. sapiens* | ENSG00000139651 | ZNF740 | zinc finger protein 740 | 0.00 | 0.30 | 1.8E-18 | 4.0E-17 | 0.30 | 1.8E-18 | | MATGLMGHEGVATVIW* |
| HG0570 | *H. sapiens* | ENSG00000152214 | RIT2 | Ras like without CAAX 2 | 0.02 | 0.38 | 7.9E-05 | 3.7E-04 | 0.38 | 7.9E-05 | | MRPARAEDQQPARR* |
| HG0571 | *H. sapiens* | ENSG00000153266 | FEZF2 | FEZ family zinc finger 2 | 0.13 | 0.00 | 3.0E-05 | 1.5E-04 | 0.00 | 3.0E-05 | | MARNASPWWWGKET* |
| HG0572 | *H. sapiens* | ENSG00000158161 | EYA3 | EYA transcriptional coactivator and phosphatase 3 | 0.00 | 0.45 | 1.1E-22 | 3.1E-21 | 0.45 | 1.1E-22 | | MRIRVTLTPRFPPLDWFYCGSGLISMSCCGAFTAFGL* |
| HG0573.1 | *H. sapiens* | ENSG00000162526 | TSSK3 | testis specific serine kinase 3 | 0.01 | 0.39 | 2.8E-11 | 3.4E-10 | 0.39 | 2.8E-11 | | MKSLGRKGAGQGRWWSRAASQRQHELRG* |
| HG0573.2 | *H. sapiens* | ENSG00000206203 | TSSK2 | testis specific serine kinase 2 | 0.00 | 0.27 | 1.1E-10 | 1.3E-09 | 0.27 | 1.1E-10 | | MRTMPAGPHDGGM* |
| HG0574 | *H. sapiens* | ENSG00000164663 | USP49 | ubiquitin specific peptidase 49 | 0.00 | 0.28 | 1.7E-17 | 3.5E-16 | 0.28 | 1.7E-17 | | MEEDGKNLFNGLRKWRLL* |
| HG0575 | *H. sapiens* | ENSG00000164663 | USP49 | ubiquitin specific peptidase 49 | 0.00 | 0.45 | 1.7E-02 | 4.0E-02 | 0.45 | 1.7E-02 | | MMAILWMGRDLERTSSLSCNSSQYPT* |
| HG0576 | *H. sapiens* | ENSG00000175029 | CTBP2 | C-terminal binding protein 2 | 0.00 | 0.37 | 3.8E-09 | 3.7E-08 | 0.37 | 3.8E-09 | | MKVYSLKKTDSPSCEL* |
| HG0577 | *H. sapiens* | ENSG00000250510 | GPR162 | G protein-coupled receptor 162 | 0.00 | 0.49 | 1.9E-13 | 3.0E-12 | 0.49 | 1.9E-13 | Crowe | MLSTGVSLGAACLLTGRGLWGSWECL* |
| HG0578 | *H. sapiens* | ENSG00000004897 | CDC27 | cell division cycle 27 | 0.05 | 0.22 | 6.1E-11 | 7.3E-10 | 0.22 | 6.1E-11 | | MSRGWPGRSRYRGGLRHCRKWA* |
| HG0579 | *H. sapiens* | ENSG00000010818 | HIVEP2 | human immunodeficiency virus type I enhancer binding pro | 0.08 | 0.25 | 1.5E-17 | 3.2E-16 | 0.28 | 9.6E-14 | | MPIDCGVRVEGSKEKEQSASLQTPWNMEATKL* |
| HG0580 | *H. sapiens* | ENSG00000073711 | PPP2R3A | protein phosphatase 2 regulatory subunit B"alpha | 0.00 | 0.40 | 1.8E-04 | 7.9E-04 | 0.40 | 1.8E-04 | | MCSSYCILIVTKPQI* |
| HG0581 | *H. sapiens* | ENSG00000087074 | PPP1R15A | protein phosphatase 1 regulatory subunit 15A | 0.14 | 0.39 | 3.3E-20 | 8.2E-19 | 0.39 | 3.3E-20 | Crowe, Mackowiak | MNALASLTVRTCDRFWQTEPALLPPG* |
| HG0582 | *H. sapiens* | ENSG00000087338 | GMCL1 | germ cell-less, spermatogenesis associated 1 | 0.01 | 0.39 | 1.2E-17 | 2.5E-16 | 0.39 | 1.2E-17 | | METVASAATVGAAREKVAV* |
| HG0583 | *H. sapiens* | ENSG00000087502 | ERGIC2 | ERGIC and golgi 2 | 0.04 | 0.18 | 4.1E-12 | 5.6E-11 | 0.18 | 4.1E-12 | | MAVGWDHNTSMTI* |
| HG0584 | *H. sapiens* | ENSG00000092051 | JPH4 | junctophilin 4 | 0.07 | 0.25 | 2.6E-05 | 1.4E-04 | 0.25 | 2.6E-05 | | MDAVPGRQPEPQPT* |
| HG0585 | *H. sapiens* | ENSG00000106038 | EVX1 | even-skipped homeobox 1 | 0.05 | 0.16 | 8.2E-03 | 2.2E-02 | 0.16 | 8.2E-03 | | MGLPPRGGAAQRSARFTR* |
| HG0586.1 | *H. sapiens* | ENSG00000108375 | RNF43 | ring finger protein 43 | 0.00 | 0.08 | 7.3E-05 | 3.4E-04 | 0.08 | 7.3E-05 | | MGENDIPAADVG* |
| HG0586.2 | *H. sapiens* | ENSG00000108375 | RNF43 | ring finger protein 43 | 0.00 | 0.00 | 1.4E-09 | 1.4E-08 | 0.00 | 1.4E-09 | | MLFGEKKNFC* |
| HG0587 | *H. sapiens* | ENSG00000119138 | KLF9 | Kruppel like factor 9 | 0.22 | 0.42 | 4.0E-19 | 9.2E-18 | 0.42 | 4.0E-19 | | MPERCGPAGAGVQKPSVDSGKSVS* |
| HG0588 | *H. sapiens* | ENSG00000120162 | MOB3B | MOB kinase activator 3B | 0.00 | 0.07 | 6.6E-29 | 2.2E-27 | 0.07 | 6.6E-29 | | MRFPWKSFKRKMEGAV* |
| HG0589.1 | *H. sapiens* | ENSG00000121871 | SLITRK3 | SLIT and NTRK like family member 3 | 0.01 | 0.45 | 6.5E-10 | 7.1E-09 | 0.45 | 6.5E-10 | | MMDTIQLRREFAPPAG* |
| HG0589.2 | *H. sapiens* | ENSG00000121871 | SLITRK3 | SLIT and NTRK like family member 3 | 0.04 | 0.44 | 2.8E-06 | 1.7E-05 | 0.44 | 2.8E-06 | | MQRPFPCKSQRWKHLNGLQA* |
| HG0589.3 | *H. sapiens* | ENSG00000121871 | SLITRK3 | SLIT and NTRK like family member 3 | 0.20 | 0.44 | 2.3E-12 | 3.3E-11 | 0.44 | 2.3E-12 | | MDLHSGSSPAFPILTPYTRPGAPV* |
| HG0590 | *H. sapiens* | ENSG00000122557 | HERPUD2 | HERPUD family member 2 | 0.00 | 0.25 | 6.0E-23 | 1.7E-21 | 0.25 | 6.0E-23 | | MGFVSSSALVTDLDTEY* |
| HG0591.1 | *H. sapiens* | ENSG00000127124 | HIVEP3 | human immunodeficiency virus type I enhancer binding pro | 0.00 | 0.12 | 1.8E-37 | 7.9E-36 | 0.12 | 1.8E-37 | Mackowiak | MNAGFQREQRFSFGHRKWCLQHRRRA* |
| HG0591.2 | *H. sapiens* | ENSG00000127124 | HIVEP3 | human immunodeficiency virus type I enhancer binding pro | 0.01 | 0.35 | 3.3E-10 | 3.7E-09 | 0.35 | 3.3E-10 | | MWPKLDTSLGHWDYS* |
| HG0591.3 | *H. sapiens* | ENSG00000127124 | HIVEP3 | human immunodeficiency virus type I enhancer binding pro | 0.00 | 0.49 | 1.4E-05 | 7.4E-05 | 0.49 | 1.4E-05 | | MEEPRGVLAKQARPSGGKSGA* |
| HG0591.4 | *H. sapiens* | ENSG00000127124 | HIVEP3 | human immunodeficiency virus type I enhancer binding pro | 0.06 | 0.19 | 1.3E-03 | 4.5E-03 | 0.19 | 1.3E-03 | | MDTDVSNLQPFA* |
| HG0592 | *H. sapiens* | ENSG00000133401 | PDZD2 | PDZ domain containing 2 | 0.00 | 0.46 | 1.7E-18 | 3.7E-17 | 0.46 | 1.7E-18 | | MNTGKADDGQGPQGTWGPVGSGPQEQDL* |
| HG0593 | *H. sapiens* | ENSG00000135048 | TMEM2 | transmembrane protein 2 | 0.01 | 0.44 | 1.4E-12 | 2.0E-11 | 0.44 | 1.4E-12 | | MERRSGPEDARDSSPSPWTAV* |
| HG0594 | *H. sapiens* | ENSG00000136169 | SETDB2 | SET domain bifurcated 2 | 0.00 | 0.26 | 6.0E-11 | 7.3E-10 | 0.26 | 6.0E-11 | | MFHPFVDQKMELVFIFKKIMLMI* |
| HG0595 | *H. sapiens* | ENSG00000141098 | GFOD2 | glucose-fructose oxidoreductase domain containing 2 | 0.00 | 0.40 | 6.8E-13 | 1.0E-11 | 0.40 | 6.8E-13 | | MPACQEPCLTRTFGRSYPCQRLL* |
| HG0596 | *H. sapiens* | ENSG00000157483 | MYO1E | myosin IE | 0.11 | 0.24 | 1.9E-04 | 8.3E-04 | 0.24 | 1.9E-04 | | MESPLGSHWLL* |
| HG0597 | *H. sapiens* | ENSG00000163032 | VSNL1 | visinin like 1 | 0.07 | 0.26 | 5.9E-24 | 1.7E-22 | 0.26 | 5.9E-24 | | MGFPICSRDLPERVAAAAGLARRDPRGI* |
| HG0598 | *H. sapiens* | ENSG00000163788 | SNRK | SNF related kinase | 0.03 | 0.35 | 7.8E-10 | 8.4E-09 | 0.35 | 7.8E-10 | | MTTLKMNFLYSPDILI* |
| HG0599 | *H. sapiens* | ENSG00000164930 | FZD6 | frizzled class receptor 6 | 0.00 | 0.36 | 4.8E-18 | 1.0E-16 | 0.36 | 2.2E-17 | | MRNLNILSLDGDLLRMQRVIHPSHVVKSGI* |
| HG0600 | *H. sapiens* | ENSG00000173065 | FAM222B | family with sequence similarity 222 member B | 0.06 | 0.29 | 5.9E-05 | 2.8E-04 | 0.29 | 5.9E-05 | | MRAGPSHRPVSHCPMLC* |
| HG0601 | *H. sapiens* | ENSG00000175893 | ZDHHC21 | zinc finger DHHC-type containing 21 | 0.00 | 0.42 | 6.9E-05 | 3.3E-04 | 0.42 | 6.9E-05 | | MNQKKISKCRFQKNYSTSQ* |
| HG0602 | *H. sapiens* | ENSG00000179195 | ZNF664 | zinc finger protein 664 | 0.00 | 0.32 | 1.7E-20 | 4.3E-19 | 0.32 | 1.7E-20 | | MQEETFRRKTGREKLRLTLNLPNRASLR* |
| HG0603 | *H. sapiens* | ENSG00000197329 | PELI1 | pellino E3 ubiquitin protein ligase 1 | 0.05 | 0.02 | 5.1E-21 | 1.3E-19 | 0.02 | 5.1E-21 | | MSVTQRRCEAEGPSGGAAAASTSSSGSSPSPAVRSNWSPALRERTTKQPQRLSLRPRLLSARLRGQLPLAGLGWRARRAAAERRAVCSRSPRTTLHPLASGLGGRSGIPATSNCPEIMQKVSQGS* |
| HG0604.1 | *H. sapiens* | ENSG00000003147 | ICA1 | islet cell autoantigen 1 | 0.03 | 0.33 | 5.9E-04 | 2.2E-03 | 0.33 | 5.9E-04 | | MLFSCPFSPNHQQ* |
| HG0604.2 | *H. sapiens* | ENSG00000003147 | ICA1 | islet cell autoantigen 1 | 0.13 | 0.48 | 1.8E-03 | 5.9E-03 | 0.42 | 7.4E-03 | | MSDPEGRSDRGREAPGGRGSEVII* |
| HG0604.3 | *H. sapiens* | ENSG00000003147 | ICA1 | islet cell autoantigen 1 | 0.22 | 0.35 | 6.2E-05 | 3.0E-04 | 0.35 | 6.2E-05 | | MATWNTCRRAGAGSLWAADPSGGYPRRWGSRRDPGE* |
| HG0605 | *H. sapiens* | ENSG00000053254 | FOXN3 | forkhead box N3 | 0.00 | 0.09 | 1.4E-20 | 3.4E-19 | 0.09 | 1.4E-20 | | MKKNQPLEDQHSRDYGGPSVFSVTGAMLTSVSWTHSDSGAFT* |
| HG0606 | *H. sapiens* | ENSG00000055130 | CUL1 | cullin 1 | 0.00 | 0.48 | 4.4E-05 | 2.2E-04 | 0.48 | 4.4E-05 | | MRFHSLHLKDILSELL* |
| HG0607 | *H. sapiens* | ENSG00000074657 | ZNF532 | zinc finger protein 532 | 0.00 | 0.44 | 4.5E-11 | 5.5E-10 | 0.49 | 8.0E-09 | | MRFQRPPCPLSHWPCLRPPPPPQTAPVFFPRGEGIWTSPPRCLKEEKGMFCLVLFLLEATKGCWGCF* |
| HG0608 | *H. sapiens* | ENSG00000084731 | KIF3C | kinesin family member 3C | 0.01 | 0.30 | 8.5E-04 | 3.1E-03 | 0.30 | 8.5E-04 | | MNDPPAWGEDSR* |
| HG0609 | *H. sapiens* | ENSG00000095539 | SEMA4G | semaphorin 4G | 0.00 | 0.16 | 7.5E-31 | 2.7E-29 | 0.16 | 7.5E-31 | Crowe | MATQPCDSMSPRFQDPPWPHDSLTPMTL* |
| HG0610 | *H. sapiens* | ENSG00000106069 | CHN2 | chimerin 2 | 0.22 | 0.38 | 1.2E-12 | 1.8E-11 | 0.38 | 1.2E-12 | | MGWGPRRLRAAGRGQRRRRPHRG* |
| HG0611 | *H. sapiens* | ENSG00000108309 | RUNDC3A | RUN domain containing 3A | 0.01 | 0.29 | 2.7E-18 | 6.0E-17 | 0.29 | 2.7E-18 | | MAMEGLRGPVGQRAQP* |
| HG0612.1 | *H. sapiens* | ENSG00000110422 | HIPK3 | homeodomain interacting protein kinase 3 | 0.00 | 0.50 | 5.0E-14 | 8.4E-13 | 0.50 | 5.0E-14 | | MAASRALWPRPQRRGAAGLQGVNSGDVRACV* |
| HG0612.2 | *H. sapiens* | ENSG00000110422 | HIPK3 | homeodomain interacting protein kinase 3 | 0.00 | 0.14 | 6.8E-07 | 4.6E-06 | 0.14 | 6.8E-07 | | MLRAAAARRGESAGL* |
| HG0612.3 | *H. sapiens* | ENSG00000110422 | HIPK3 | homeodomain interacting protein kinase 3 | 0.06 | 0.20 | 9.2E-03 | 2.4E-02 | 0.20 | 9.2E-03 | | MKMSVSLSGGRRLL* |
| HG0613 | *H. sapiens* | ENSG00000114861 | FOXP1 | forkhead box P1 | 0.00 | 0.17 | 2.7E-08 | 2.3E-07 | 0.17 | 2.7E-08 | | MVRASIPSLICCQP* |
| HG0614 | *H. sapiens* | ENSG00000122203 | KIAA1191 | KIAA1191 | 0.03 | 0.30 | 7.8E-29 | 2.6E-27 | 0.36 | 2.0E-21 | Crowe | MGRGRVSRSWTTTPSMPCAPLGQVSREIPRSPEALSAVLPARWVVLLPSGSWGPSKSPPLSQWLGSRLRHSLSGDVGPRNLFSAGLRAAGSPAAAGSQNLWMDSSWEALLLCSSWTMFLINICLSA* |
| HG0615 | *H. sapiens* | ENSG00000128191 | DGCR8 | DGCR8, microprocessor complex subunit | 0.00 | 0.07 | 9.9E-11 | 1.2E-09 | 0.07 | 9.9E-11 | | MKTDSLSRQSLKLSAL* |
| HG0616 | *H. sapiens* | ENSG00000128739 | SNRPN | small nuclear ribonucleoprotein polypeptide N | 0.00 | 0.06 | 4.5E-36 | 1.9E-34 | 0.06 | 4.5E-36 | | MERARDRLHLRRTTEQHVPEVEVQVKRRRTASLSNQECQLYPRRSQQQQVPVVDFQAELRQAFLAETPRGG* |
| HG0617 | *H. sapiens* | ENSG00000135164 | DMTF1 | cyclin D binding myb like transcription factor 1 | 0.00 | 0.01 | 3.5E-34 | 1.4E-32 | 0.01 | 3.5E-34 | | MRATWVKKLPVNSHHFCPSLGNTFRECCGDRNMGEKQSG* |
| HG0618 | *H. sapiens* | ENSG00000137817 | PARP6 | poly(ADP-ribose) polymerase family member 6 | 0.01 | 0.26 | 4.5E-12 | 6.2E-11 | 0.26 | 4.5E-12 | | MPISLALVRLVVWRGAA* |
| HG0619 | *H. sapiens* | ENSG00000138593 | SECISBP2L | SECIS binding protein 2 like | 0.04 | 0.38 | 9.7E-12 | 1.3E-10 | 0.38 | 9.7E-12 | | MEKAARGAVYTD* |
| HG0620 | *H. sapiens* | ENSG00000145555 | MYO10 | myosin X | 0.05 | 0.47 | 8.6E-04 | 3.1E-03 | 0.47 | 8.6E-04 | | MCHQLFSACERCAQ* |
| HG0621.1 | *H. sapiens* | ENSG00000146352 | CLVS2 | clavesin 2 | 0.03 | 0.40 | 1.8E-03 | 6.0E-03 | 0.40 | 1.8E-03 | | MPLSPPPPLLKPRLMDARECRIA* |
| HG0621.2 | *H. sapiens* | ENSG00000146352 | CLVS2 | clavesin 2 | 0.02 | 0.37 | 4.3E-03 | 1.3E-02 | 0.37 | 4.3E-03 | | MAGPGENLGQRWP* |
| HG0621.3 | *H. sapiens* | ENSG00000146352 | CLVS2 | clavesin 2 | 0.03 | 0.09 | 4.6E-06 | 2.7E-05 | 0.09 | 4.6E-06 | | MPPKLPSYPNEAGKGAERARELSECLRRSEICAGIIREELG* |
| HG0622 | *H. sapiens* | ENSG00000148677 | ANKRD1 | ankyrin repeat domain 1 | 0.13 | 0.43 | 1.4E-13 | 2.3E-12 | 0.43 | 1.4E-13 | | MKIHLPLSWLLMGVGVLLRFPGWKIISPGPSYIS* |
| HG0623 | *H. sapiens* | ENSG00000151422 | FER | FER tyrosine kinase | 0.00 | 0.45 | 5.5E-17 | 1.1E-15 | 0.45 | 5.5E-17 | | MTIFARLHCTFSQVSSLSLMCDTSSWTRYFS* |
| HG0624 | *H. sapiens* | ENSG00000156853 | ZNF689 | zinc finger protein 689 | 0.07 | 0.39 | 9.0E-10 | 9.6E-09 | 0.39 | 9.0E-10 | Crowe, Samandi | MALRSIKSIAGSCLCSRQRRCGSSAAIFPEGIFRCLSPKFGQEFPE* |
| HG0625 | *H. sapiens* | ENSG00000164050 | PLXNB1 | plexin B1 | 0.01 | 0.19 | 1.7E-04 | 7.2E-04 | 0.19 | 1.7E-04 | | MLLSPDPAGPAR* |
| HG0626 | *H. sapiens* | ENSG00000168939 | SPRY3 | sprouty RTK signaling antagonist 3 | 0.01 | 0.36 | 1.5E-05 | 7.9E-05 | 0.36 | 1.5E-05 | | MDKPFPQGFSHVP* |
| HG0627 | *H. sapiens* | ENSG00000169760 | NLGN1 | neuroligin 1 | 0.00 | 0.39 | 4.2E-08 | 3.4E-07 | 0.39 | 4.2E-08 | | MRGLPLWIYLAEGYRPAANWI* |
| HG0628 | *H. sapiens* | ENSG00000170677 | SOCS6 | suppressor of cytokine signaling 6 | 0.00 | 0.34 | 1.5E-14 | 2.6E-13 | 0.34 | 1.5E-14 | | MKQEKYIDAALQPLQMFGDNIPDRNIDPLD* |
| HG0629 | *H. sapiens* | ENSG00000170961 | HAS2 | hyaluronan synthase 2 | 0.00 | 0.00 | 2.7E-06 | 1.7E-05 | 0.00 | 2.7E-06 | | MLFLNFLFFLAGRLKFI* |
| HG0630 | *H. sapiens* | ENSG00000171552 | BCL2L1 | BCL2 like 1 | 0.00 | 0.25 | 3.2E-13 | 5.0E-12 | 0.25 | 3.2E-13 | | MIPPAGAGFFASHREAG* |
| HG0631 | *H. sapiens* | ENSG00000173548 | SNX33 | sorting nexin 33 | 0.00 | 0.49 | 7.7E-13 | 1.2E-11 | 0.49 | 7.7E-13 | | MSPPCSQAYARRPDRAVSGPWTLL* |
| HG0632.1 | *H. sapiens* | ENSG00000181315 | ZNF322 | zinc finger protein 322 | 0.01 | 0.27 | 1.1E-09 | 1.1E-08 | 0.27 | 1.1E-09 | | MTVRATMGQEDQRKTP* |
| HG0632.2 | *H. sapiens* | ENSG00000181315 | ZNF322 | zinc finger protein 322 | 0.00 | 0.40 | 9.9E-03 | 2.6E-02 | 0.40 | 9.9E-03 | | MYVFFHLDSRVGNTK* |
| HG0633 | *H. sapiens* | ENSG00000186174 | BCL9L | B-cell CLL/lymphoma 9 like | 0.02 | 0.49 | 2.0E-06 | 1.2E-05 | 0.49 | 2.0E-06 | | MLRGWEAGTQAPSDPRIASPGPGWY* |

| | | | | | | | | | | | | |
|---|---|---|---|---|---|---|---|---|---|---|---|---|
| HG0634 | *H. sapiens* | ENSG00000198026 | ZNF335 | zinc finger protein 335 | 0.04 | 0.23 | 1.0E-20 | 2.6E-19 | 0.23 | 1.0E-20 | | MPESERNVATKASE* |
| HG0635 | *H. sapiens* | ENSG00000259332 | ST20-MTHFS | ST20-MTHFS readthrough | 0.05 | 0.48 | 4.1E-10 | 4.6E-09 | 0.32 | 1.9E-18 | | MAAETGYWRCLPESRTCAEEAVHLWPAVLRDPRWGNPGDGAWPCYMRGPRLE* |
| HG0636 | *H. sapiens* | ENSG00000002587 | HS3ST1 | heparan sulfate-glucosamine 3-sulfotransferase 1 | 0.08 | 0.32 | 1.8E-19 | 4.4E-18 | 0.32 | 1.8E-19 | | MAQPRSLFRAPDGLDRGAAQ* |
| HG0637 | *H. sapiens* | ENSG00000005102 | MEOX1 | mesenchyme homeobox 1 | 0.03 | 0.42 | 1.7E-02 | 4.1E-02 | 0.42 | 1.7E-02 | | MFSVIGDKYTYRGSQ* |
| HG0638 | *H. sapiens* | ENSG00000013561 | RNF14 | ring finger protein 14 | 0.00 | 0.45 | 3.3E-12 | 4.7E-11 | 0.45 | 3.3E-12 | | MSSTSWPLTQLQQSQLHQLENKWDLHELHSELKYRLRC* |
| HG0639 | *H. sapiens* | ENSG00000018236 | CNTN1 | contactin 1 | 0.00 | 0.00 | 5.2E-11 | 6.3E-10 | 0.00 | 5.2E-11 | | MGAAVPACEVTA* |
| HG0640 | *H. sapiens* | ENSG00000062194 | GPBP1 | GC-rich promoter binding protein 1 | 0.00 | 0.00 | 5.5E-06 | 3.2E-05 | 0.00 | 5.5E-06 | | MLFRCEIFFLFLLFGS* |
| HG0641 | *H. sapiens* | ENSG00000100359 | SGSM3 | small G protein signaling modulator 3 | 0.00 | 0.41 | 1.9E-09 | 1.9E-08 | 0.40 | 2.2E-07 | | MILDQMKPEEPSSSKIAG* |
| HG0642 | *H. sapiens* | ENSG00000100815 | TRIP11 | thyroid hormone receptor interactor 11 | 0.00 | 0.44 | 1.1E-09 | 1.2E-08 | 0.44 | 1.1E-09 | | MAAGVELAGVTHGTEESH* |
| HG0643 | *H. sapiens* | ENSG00000103653 | CSK | CSK, non-receptor tyrosine kinase | 0.00 | 0.48 | 5.0E-08 | 4.1E-07 | 0.48 | 5.0E-08 | | MLFPTALMVPSDRLALL* |
| HG0644 | *H. sapiens* | ENSG00000104299 | INTS9 | integrator complex subunit 9 | 0.00 | 0.48 | 3.3E-10 | 3.7E-09 | 0.48 | 3.3E-10 | | MLSAGRWRRLHRKTLPGFEEFSDCY* |
| HG0645 | *H. sapiens* | ENSG00000107282 | APBA1 | amyloid beta precursor protein binding family A member 1 | 0.03 | 0.42 | 4.7E-09 | 4.4E-08 | 0.42 | 4.7E-09 | | MVALEAAAAGAELYSQAWHQRCH* |
| HG0646 | *H. sapiens* | ENSG00000108312 | UBTF | upstream binding transcription factor, RNA polymerase I | 0.28 | 0.00 | 2.4E-07 | 1.8E-06 | 0.00 | 2.4E-07 | | MEQWEREAGRFPCKSPSGLTPSRRPRQGSEDPGEQMEEWSSL* |
| HG0647 | *H. sapiens* | ENSG00000108395 | TRIM37 | tripartite motif containing 37 | 0.00 | 0.34 | 1.6E-08 | 1.4E-07 | 0.34 | 1.6E-08 | | MGVGGAGPEEMGRAGGAVVRAP* |
| HG0648 | *H. sapiens* | ENSG00000114331 | ACAP2 | ArfGAP with coiled-coil, ankyrin repeat and PH domains 2 | 0.02 | 0.35 | 2.9E-11 | 3.6E-10 | 0.35 | 2.9E-11 | Mackowiak | MTSGRGALRSSAQR* |
| HG0649.1 | *H. sapiens* | ENSG00000114439 | BBX | BBX, HMG-box containing | 0.00 | 0.16 | 5.9E-03 | 1.7E-02 | 0.16 | 5.9E-03 | | MWRIHWDIEKYQRRIVPL* |
| HG0649.2 | *H. sapiens* | ENSG00000114439 | BBX | BBX, HMG-box containing | 0.00 | 0.12 | 4.1E-04 | 1.6E-03 | 0.12 | 4.1E-04 | | MESLDLTGTLRKTPSL* |
| HG0650 | *H. sapiens* | ENSG00000115226 | FNDC4 | fibronectin type III domain containing 4 | 0.29 | 0.00 | 1.2E-10 | 1.4E-09 | 0.00 | 1.2E-10 | Mackowiak | MGRPLGTEQPPEA* |
| HG0651 | *H. sapiens* | ENSG00000115935 | WIPF1 | WAS/WASL interacting protein family member 1 | 0.01 | 0.45 | 2.0E-16 | 3.9E-15 | 0.45 | 2.0E-16 | | MIPFSGTVFSLGSCPAGPLSACVPDHGSLQYPLTIYQQDC* |
| HG0652 | *H. sapiens* | ENSG00000118515 | SGK1 | serum/glucocorticoid regulated kinase 1 | 0.18 | 0.37 | 9.6E-14 | 1.6E-12 | 0.37 | 9.6E-14 | | MMEGEKPLSVATESGERLLQ* |
| HG0653 | *H. sapiens* | ENSG00000120334 | CENPL | centromere protein L | 0.01 | 0.32 | 2.2E-04 | 9.4E-04 | 0.32 | 2.2E-04 | | MIRESGMGDSVITAFY* |
| HG0654 | *H. sapiens* | ENSG00000133794 | ARNTL | aryl hydrocarbon receptor nuclear translocator like | 0.12 | 0.41 | 3.0E-03 | 9.5E-03 | 0.41 | 3.0E-03 | | MWNPGPSLVFGRS* |
| HG0655 | *H. sapiens* | ENSG00000143061 | IGSF3 | immunoglobulin superfamily member 3 | 0.00 | 0.22 | 8.6E-23 | 2.4E-21 | 0.22 | 8.6E-23 | | MWQEPGATSELLWTLPVA* |
| HG0656 | *H. sapiens* | ENSG00000150776 | C11orf57 | chromosome 11 open reading frame 57 | 0.00 | 0.36 | 2.8E-13 | 4.3E-12 | 0.36 | 2.8E-13 | | MGIFPLSSTWKPQPPLQAP* |
| HG0657 | *H. sapiens* | ENSG00000151276 | MAGI1 | membrane associated guanylate kinase, WW and PDZ dor | 0.00 | 0.28 | 1.7E-04 | 7.3E-04 | 0.28 | 1.7E-04 | Crowe | MQTKLAGACAGLAGKPRKFVL* |
| HG0658 | *H. sapiens* | ENSG00000153786 | ZDHHC7 | zinc finger DHHC-type containing 7 | 0.00 | 0.39 | 1.3E-02 | 3.3E-02 | 0.39 | 1.3E-02 | | MHLKILKKRHKMFK* |
| HG0659.1 | *H. sapiens* | ENSG00000158856 | DMTN | dematin actin binding protein | 0.00 | 0.24 | 1.2E-19 | 2.9E-18 | 0.24 | 1.2E-19 | | MEVYVTEEAAALLPHALGP* |
| HG0659.2 | *H. sapiens* | ENSG00000158856 | DMTN | dematin actin binding protein | 0.01 | 0.46 | 5.3E-09 | 5.0E-08 | 0.46 | 5.3E-09 | | MRTRQPGGTRQLLSRPQARSAQQPRRA* |
| HG0660 | *H. sapiens* | ENSG00000165861 | ZFYVE1 | zinc finger FYVE-type containing 1 | 0.00 | 0.30 | 1.4E-03 | 4.8E-03 | 0.30 | 1.4E-03 | | MEDETIKPNQRCC* |
| HG0661 | *H. sapiens* | ENSG00000166845 | C18orf54 | chromosome 18 open reading frame 54 | 0.00 | 0.33 | 1.6E-17 | 3.4E-16 | 0.29 | 8.9E-16 | | MDFLLEIRTEACTEEMLFLREMNRNNPL* |
| HG0662 | *H. sapiens* | ENSG00000166897 | ELFN2 | extracellular leucine rich repeat and fibronectin type III dom | 0.00 | 0.40 | 5.3E-08 | 4.3E-07 | 0.40 | 5.3E-08 | | MVAAPGRTVLPAPASGR* |
| HG0663 | *H. sapiens* | ENSG00000168615 | ADAM9 | ADAM metallopeptidase domain 9 | 0.09 | 0.42 | 8.9E-11 | 1.1E-09 | 0.42 | 8.9E-11 | | MSGASSDRGLSCVSAARVLVGRACSSGARASRPDLGPRQGWKMMEEAEVEATEC* |
| HG0664 | *H. sapiens* | ENSG00000169085 | | | 0.01 | 0.25 | 1.9E-07 | 1.4E-06 | 0.25 | 1.9E-07 | | MIPEPDWIRMPRD* |
| HG0665 | *H. sapiens* | ENSG00000169515 | CCDC8 | coiled-coil domain containing 8 | 0.07 | 0.19 | 1.1E-05 | 6.0E-05 | 0.19 | 1.1E-05 | | MSAKMWIWTSSWKP* |
| HG0666 | *H. sapiens* | ENSG00000171943 | SRGAP2C | SLIT-ROBO Rho GTPase activating protein 2C | 0.00 | 0.00 | 1.8E-05 | 9.7E-05 | 0.00 | 1.8E-05 | | MSDPRPASGRCSSPG* |
| HG0667.1 | *H. sapiens* | ENSG00000182700 | IGIP | IgA inducing protein | 0.00 | 0.00 | 1.3E-11 | 1.7E-10 | 0.00 | 1.3E-11 | | MCQYTVRISLICTVWGPHLRMLNFAN* |
| HG0667.2 | *H. sapiens* | ENSG00000182700 | IGIP | IgA inducing protein | 0.17 | 0.45 | 5.1E-04 | 2.0E-03 | 0.45 | 5.1E-04 | | MLRKSVGFSAVREGRSSALIWVTEANMSKTLVN* |
| HG0668 | *H. sapiens* | ENSG00000182944 | EWSR1 | EWS RNA binding protein 1 | 0.03 | 0.21 | 3.1E-20 | 7.7E-19 | 0.21 | 3.1E-20 | | MVAPGVARFAPAQCGA* |
| HG0669 | *H. sapiens* | ENSG00000183876 | ARSI | arylsulfatase family member I | 0.16 | 0.41 | 8.2E-06 | 4.6E-05 | 0.41 | 8.2E-06 | | MVESTFWKLPRRVSGWPG* |
| HG0670 | *H. sapiens* | ENSG00000187164 | SHTN1 | shootin 1 | 0.04 | 0.27 | 9.0E-07 | 6.1E-06 | 0.27 | 9.0E-07 | | MISLARSAPGGGGAGADPTSG* |
| HG0671 | *H. sapiens* | ENSG00000189184 | PCDH18 | protocadherin 18 | 0.01 | 0.36 | 2.5E-10 | 2.8E-09 | 0.36 | 2.5E-10 | | MPKVSGSRDTVCIKQKFASC* |
| HG0672 | *H. sapiens* | ENSG00000196369 | SRGAP2B | SLIT-ROBO Rho GTPase activating protein 2B | 0.00 | 0.00 | 1.8E-05 | 9.7E-05 | 0.00 | 1.8E-05 | | MSDPRPASGRCSSPG* |
| HG0673 | *H. sapiens* | ENSG00000266028 | SRGAP2 | SLIT-ROBO Rho GTPase activating protein 2 | 0.00 | 0.00 | 1.8E-05 | 9.7E-05 | 0.00 | 1.8E-05 | | MSDPRPASGRCSSPG* |
| HG0674.1 | *H. sapiens* | ENSG00000006576 | PHTF2 | putative homeodomain transcription factor 2 | 0.01 | 0.19 | 3.3E-04 | 1.3E-03 | 0.19 | 3.3E-04 | | MFFSLAQPKMTNV* |
| HG0674.2 | *H. sapiens* | ENSG00000116793 | PHTF1 | putative homeodomain transcription factor 1 | 0.00 | 0.43 | 1.5E-03 | 5.2E-03 | 0.43 | 1.5E-03 | | MRPAWPAASPPASRWDPGH* |
| HG0675 | *H. sapiens* | ENSG00000048991 | R3HDM1 | R3H domain containing 1 | 0.01 | 0.46 | 6.2E-03 | 1.8E-02 | 0.46 | 6.2E-03 | | MQSISLQVLWKEHLLNSV* |
| HG0676 | *H. sapiens* | ENSG00000058673 | ZC3H11A | zinc finger CCCH-type containing 11A | 0.00 | 0.23 | 2.0E-16 | 3.9E-15 | 0.23 | 2.0E-16 | Mackowiak | MAIMLLCLLQLAAPLCSYSITIRFYLFWLNTP* |
| HG0677 | *H. sapiens* | ENSG00000064989 | CALCRL | calcitonin receptor like receptor | 0.06 | 0.34 | 3.3E-09 | 3.2E-08 | 0.34 | 3.3E-09 | | MIHQPLVLPSVKTLDGPTFRS* |
| HG0678.1 | *H. sapiens* | ENSG00000090776 | EFNB1 | ephrin B1 | 0.07 | 0.40 | 7.1E-14 | 1.2E-12 | 0.40 | 7.1E-14 | | MGLARSAASPRSSKYTEQGPR* |
| HG0678.2 | *H. sapiens* | ENSG00000108947 | EFNB3 | ephrin B3 | 0.08 | 0.06 | 1.2E-06 | 7.7E-06 | 0.06 | 1.2E-06 | | MPPPPGPRLPQSPT* |
| HG0679 | *H. sapiens* | ENSG00000100281 | HMGXB4 | HMG-box containing 4 | 0.20 | 0.38 | 6.5E-06 | 3.7E-05 | 0.38 | 6.5E-06 | | MAAIGGVEAGSGRAE* |
| HG0680 | *H. sapiens* | ENSG00000100678 | SLC8A3 | solute carrier family 8 member A3 | 0.01 | 0.00 | 1.2E-02 | 3.1E-02 | 0.00 | 1.2E-02 | | MPRTRLRPAGSPRGR* |
| HG0681 | *H. sapiens* | ENSG00000101349 | PAK5 | p21 (RAC1) activated kinase 5 | 0.00 | 0.21 | 7.2E-03 | 2.0E-02 | 0.21 | 7.2E-03 | | MLPPLAKMNTEEK* |
| HG0682 | *H. sapiens* | ENSG00000105829 | BET1 | Bet1 golgi vesicular membrane trafficking protein | 0.06 | 0.40 | 9.6E-04 | 3.5E-03 | 0.40 | 9.6E-04 | | MSWRFRGRSWCFAGPWY* |
| HG0683 | *H. sapiens* | ENSG00000111837 | MAK | male germ cell associated kinase | 0.02 | 0.39 | 1.2E-11 | 1.5E-10 | 0.38 | 2.5E-10 | | MKHTRRRSFTRERTRVFIRMEVLFPCPIPSCYIS* |
| HG0684 | *H. sapiens* | ENSG00000113916 | BCL6 | B-cell CLL/lymphoma 6 | 0.00 | 0.21 | 2.8E-10 | 3.2E-09 | 0.21 | 2.8E-10 | | MRGLCHSPHCLLH* |
| HG0685 | *H. sapiens* | ENSG00000117262 | GPR89A | G protein-coupled receptor 89A | 0.08 | 0.38 | 1.1E-11 | 1.4E-10 | 0.38 | 1.1E-11 | | MASGWRAAVPAAAPGRRQTV* |
| HG0686 | *H. sapiens* | ENSG00000118985 | ELL2 | elongation factor for RNA polymerase II 2 | 0.08 | 0.22 | 1.8E-07 | 1.4E-06 | 0.22 | 1.8E-07 | | MTVRQQRDGPSSGGGCRSPSSRGRSGG* |
| HG0687.1 | *H. sapiens* | ENSG00000120549 | KIAA1217 | KIAA1217 | 0.05 | 0.45 | 4.1E-05 | 2.1E-04 | 0.45 | 4.1E-05 | | MQFSRARETLHRSGK* |
| HG0687.2 | *H. sapiens* | ENSG00000120549 | KIAA1217 | KIAA1217 | 0.00 | 0.05 | 5.2E-03 | 1.5E-02 | 0.05 | 5.2E-03 | | MVFIGCEHSS* |
| HG0688 | *H. sapiens* | ENSG00000122420 | PTGFR | prostaglandin F receptor | 0.00 | 0.33 | 5.8E-09 | 5.5E-08 | 0.33 | 5.8E-09 | | MSGLQSCTVLRGR* |
| HG0689 | *H. sapiens* | ENSG00000123352 | SPATS2 | spermatogenesis associated serine rich 2 | 0.00 | 0.25 | 2.3E-08 | 2.0E-07 | 0.25 | 2.3E-08 | | MNVTTLVTSYNLPGKGDMNVCNDTS* |
| HG0690 | *H. sapiens* | ENSG00000125931 | CITED1 | Cbp/p300 interacting transactivator with Glu/Asp rich carbo | 0.25 | 0.18 | 6.0E-08 | 4.8E-07 | 0.18 | 6.0E-08 | | MCPAAKLQKGHR* |
| HG0691 | *H. sapiens* | ENSG00000129535 | NRL | neural retina leucine zipper | 0.22 | 0.48 | 5.6E-07 | 3.9E-06 | 0.48 | 5.6E-07 | | MPFGLCAHSPLELSRGTRPCSMEPSVSWEAVPVWLWH* |
| HG0692.1 | *H. sapiens* | ENSG00000130338 | TULP4 | tubby like protein 4 | 0.00 | 0.41 | 1.5E-14 | 2.7E-13 | 0.41 | 1.5E-14 | | MALRGAPAGGGAVRWSPAFPGDTGPSFEVESFWF* |
| HG0692.2 | *H. sapiens* | ENSG00000130338 | TULP4 | tubby like protein 4 | 0.00 | 0.37 | 6.5E-03 | 1.8E-02 | 0.37 | 6.5E-03 | | MDLILRLAFFIKMEKDL* |
| HG0693 | *H. sapiens* | ENSG00000131653 | TRAF7 | TNF receptor associated factor 7 | 0.05 | 0.37 | 4.1E-16 | 7.8E-15 | 0.37 | 4.1E-16 | | MKRAGGAAAPGRPRAGGASQGP* |
| HG0694 | *H. sapiens* | ENSG00000133265 | HSPBP1 | HSPA (Hsp70) binding protein 1 | 0.06 | 0.23 | 1.2E-09 | 1.2E-08 | 0.23 | 1.2E-09 | | MAAPSRVRCGRLCL* |
| HG0695 | *H. sapiens* | ENSG00000133704 | IPO8 | importin 8 | 0.05 | 0.45 | 1.5E-07 | 1.1E-06 | 0.45 | 1.5E-07 | | MESELWGGKREDRGRGVRVAQQVAAGGDGGGGWGVR* |
| HG0696 | *H. sapiens* | ENSG00000138380 | CARF | calcium responsive transcription factor | 0.11 | 0.32 | 1.6E-05 | 8.8E-05 | 0.32 | 1.6E-05 | | MLEAWNLNLSHYLKLLNNRRKWN* |
| HG0697 | *H. sapiens* | ENSG00000143344 | RGL1 | ral guanine nucleotide dissociation stimulator like 1 | 0.00 | 0.32 | 1.4E-07 | 1.0E-06 | 0.32 | 1.4E-07 | | MDWLPDRVPCSW* |
| HG0698 | *H. sapiens* | ENSG00000157554 | ERG | ERG, ETS transcription factor | 0.00 | 0.49 | 1.6E-07 | 1.2E-06 | 0.49 | 1.6E-07 | | MTHREKRWQNQGQLKPSGSEQLVDGLAY* |
| HG0699 | *H. sapiens* | ENSG00000160785 | SLC25A44 | solute carrier family 25 member 44 | 0.09 | 0.43 | 7.7E-12 | 1.0E-10 | 0.43 | 7.7E-12 | | MEEDATGRRHSLGTTDRLGAAA* |
| HG0700 | *H. sapiens* | ENSG00000165156 | ZHX1 | zinc fingers and homeoboxes 1 | 0.00 | 0.46 | 8.3E-06 | 4.7E-05 | 0.46 | 8.3E-06 | | MQDGGGVALRGAAAPAGSGASRLRDQRARRLWAAGGNWSAESAAGTGKEKTDCCFGPVGNDGLS* |
| HG0701 | *H. sapiens* | ENSG00000166292 | TMEM100 | transmembrane protein 100 | 0.05 | 0.46 | 7.2E-05 | 3.4E-04 | 0.46 | 7.2E-05 | | MDCLSPSFHPGETVQI* |
| HG0702 | *H. sapiens* | ENSG00000167377 | ZNF23 | zinc finger protein 23 | 0.02 | 0.00 | 6.8E-05 | 3.2E-04 | 0.00 | 6.8E-05 | | MELALGQQEPIVSEVGDL* |
| HG0703 | *H. sapiens* | ENSG00000171862 | PTEN | phosphatase and tensin homolog | 0.19 | 0.42 | 4.1E-10 | 4.6E-09 | 0.42 | 4.1E-10 | | MRCGRIRARRWDATALSSLLSEAAAMMEV* |
| HG0704.1 | *H. sapiens* | ENSG00000173598 | NUDT4 | nudix hydrolase 4 | 0.05 | 0.46 | 2.5E-07 | 1.9E-06 | 0.46 | 2.5E-07 | | MESGGQLGRGGAFISAPWFQ* |
| HG0704.2 | *H. sapiens* | ENSG00000272325 | NUDT3 | nudix hydrolase 3 | 0.00 | 0.00 | 7.5E-05 | 3.5E-04 | 0.00 | 7.5E-05 | | MRRPRRASGGAV* |
| HG0705.1 | *H. sapiens* | ENSG00000175224 | ATG13 | autophagy related 13 | 0.00 | 0.29 | 7.7E-09 | 7.1E-08 | 0.29 | 7.7E-09 | | MAAPGSDRLLGLRRE* |
| HG0705.2 | *H. sapiens* | ENSG00000175224 | ATG13 | autophagy related 13 | 0.00 | 0.29 | 1.3E-12 | 1.9E-11 | 0.29 | 1.3E-12 | Samandi | MRARNHSLCRSFAVERPVETCDPFSLNKSLVLLCCPGWSVVV* |
| HG0706 | *H. sapiens* | ENSG00000178498 | DTX3 | deltex E3 ubiquitin ligase 3 | 0.05 | 0.00 | 3.8E-06 | 2.3E-05 | 0.00 | 3.8E-06 | | MGFPGHTTYK* |
| HG0707 | *H. sapiens* | ENSG00000179796 | LRRC3B | leucine rich repeat containing 3B | 0.00 | 0.01 | 2.8E-12 | 3.9E-11 | 0.01 | 2.8E-12 | | MGAQARKEIMKRHMC* |
| HG0708 | *H. sapiens* | ENSG00000184613 | NELL2 | neural EGFL like 2 | 0.15 | 0.41 | 1.1E-06 | 7.4E-06 | 0.41 | 1.1E-06 | | MMCALEGLSSSRAWGGLPFLS* |
| HG0709 | *H. sapiens* | ENSG00000186439 | TRDN | triadin | 0.00 | 0.39 | 8.4E-03 | 2.3E-02 | 0.39 | 8.4E-03 | | MRLFLFEADTSS* |
| HG0710 | *H. sapiens* | ENSG00000196505 | GDAP2 | ganglioside induced differentiation associated protein 2 | 0.00 | 0.36 | 1.2E-04 | 5.2E-04 | 0.36 | 1.2E-04 | | MGPPECSKVSSEY* |
| HG0711 | *H. sapiens* | ENSG00000198300 | PEG3 | paternally expressed 3 | 0.01 | 0.35 | 4.2E-11 | 5.1E-10 | 0.35 | 4.2E-11 | Mackowiak | MSTLGWWRRRAPGAMRVR* |
| HG0712 | *H. sapiens* | ENSG00000213949 | ITGA1 | integrin subunit alpha 1 | 0.04 | 0.32 | 2.3E-12 | 3.2E-11 | 0.32 | 2.3E-12 | | MKGRCPFKVCFYSPWTLA* |
| HG0713 | *H. sapiens* | ENSG00000008513 | ST3GAL1 | ST3 beta-galactoside alpha-2,3-sialyltransferase 1 | 0.00 | 0.46 | 2.3E-04 | 9.5E-04 | 0.46 | 2.3E-04 | | MPDSFDTLQRQRALDRREEEEEEERREKEEAM* |
| HG0714 | *H. sapiens* | ENSG00000051825 | MPHOSPH9 | M-phase phosphoprotein 9 | 0.00 | 0.33 | 6.9E-08 | 5.5E-07 | 0.33 | 6.9E-08 | | MTRIHFSQTAQEPLFSNKRI* |
| HG0715 | *H. sapiens* | ENSG00000071246 | VASH1 | vasohibin 1 | 0.03 | 0.30 | 6.3E-03 | 1.8E-02 | 0.30 | 6.3E-03 | | MCYLCSRCVGGLL* |
| HG0716 | *H. sapiens* | ENSG00000075413 | MARK3 | microtubule affinity regulating kinase 3 | 0.03 | 0.29 | 1.5E-03 | 5.2E-03 | 0.29 | 1.5E-03 | | MSPGTRRGTARARPGI* |
| HG0717 | *H. sapiens* | ENSG00000083857 | FAT1 | FAT atypical cadherin 1 | 0.00 | 0.50 | 1.4E-02 | 3.4E-02 | 0.50 | 1.4E-02 | | MLKIPGRQSWYFKPGS* |
| HG0718 | *H. sapiens* | ENSG00000085274 | MYNN | myoneurin | 0.02 | 0.42 | 2.1E-07 | 1.5E-06 | 0.42 | 2.1E-07 | | MMGEDKQLQAPARSAQLASGERPAVLCCRASAQSPGRAPDADCRHQNARDEEDRRSAAS* |
| HG0719 | *H. sapiens* | ENSG00000085998 | POMGNT1 | protein O-linked mannose N-acetylglucosaminyltransferase | 0.03 | 0.29 | 6.6E-07 | 4.5E-06 | 0.16 | 1.7E-09 | | MTGQRPPLLQQPAHCLDALIQGHQLLQDLHIDGSQIPEVMGWPGHW* |
| HG0720 | *H. sapiens* | ENSG00000087053 | MTMR2 | myotubularin related protein 2 | 0.19 | 0.18 | 2.9E-05 | 1.5E-04 | 0.18 | 2.9E-05 | | MAASGLPPGPALA* |
| HG0721 | *H. sapiens* | ENSG00000095951 | HIVEP1 | human immunodeficiency virus type I enhancer binding pro | 0.00 | 0.27 | 8.7E-05 | 4.0E-04 | 0.27 | 8.7E-05 | | MYVEFMELPFGGLLYLQFLRRKRRP* |
| HG0722 | *H. sapiens* | ENSG00000101445 | PPP1R16B | protein phosphatase 1 regulatory subunit 16B | 0.16 | 0.42 | 2.1E-09 | 2.1E-08 | 0.42 | 2.1E-09 | | MRPQPHQRPRAALAPGAPC* |
| HG0723 | *H. sapiens* | ENSG00000102974 | CTCF | CCCTC-binding factor | 0.00 | 0.32 | 1.5E-03 | 5.1E-03 | 0.32 | 1.5E-03 | | MITDLKPKNKMR* |
| HG0724 | *H. sapiens* | ENSG00000104783 | KCNN4 | potassium calcium-activated channel subfamily N member | 0.30 | 0.33 | 5.7E-12 | 7.7E-11 | 0.49 | 7.6E-05 | | MDVSFLLCPKTHPSPSLFSKGGAGEPPPTLGPPLPLPALPGRHHSPEGTGLCCASEQKSQSQQSRLTTCKPQWLPCACCEVGDPGQEAG* |
| HG0725 | *H. sapiens* | ENSG00000112218 | GPR63 | G protein-coupled receptor 63 | 0.14 | 0.49 | 2.9E-08 | 2.5E-07 | 0.49 | 2.9E-08 | | MYFQAHQWDLILLFCVEMLEELLHLCLHLPSY* |
| HG0726.1 | *H. sapiens* | ENSG00000115504 | EHBP1 | EH domain binding protein 1 | 0.00 | 0.37 | 4.6E-08 | 3.7E-07 | 0.37 | 4.6E-08 | | MWLRRGWRTRWRRRLLLRRRRRSGYEKGTGPDSARQRD* |
| HG0726.2 | *H. sapiens* | ENSG00000115504 | EHBP1 | EH domain binding protein 1 | 0.00 | 0.13 | 1.5E-04 | 6.6E-04 | 0.13 | 1.5E-04 | | MQSSFALDPLHY* |

| | | | | | | | | | | | | |
|---|---|---|---|---|---|---|---|---|---|---|---|---|
| HG0727 | *H. sapiens* | ENSG00000115825 | PRKD3 | protein kinase D3 | 0.00 | 0.05 | 7.4E-17 | 1.5E-15 | 0.05 | 3.0E-11 | | MKYACTILKSFVFRHRKVHPPRMKMRTFEKVI* |
| HG0728 | *H. sapiens* | ENSG00000116991 | SIPA1L2 | signal induced proliferation associated 1 like 2 | 0.00 | 0.28 | 1.2E-10 | 1.4E-09 | 0.28 | 1.2E-10 | | MLLKRTEAYNIITTTEKQVGFEETVF* |
| HG0729 | *H. sapiens* | ENSG00000119383 | PTPA | protein phosphatase 2 phosphatase activator | 0.00 | 0.32 | 1.3E-08 | 1.1E-07 | 0.32 | 1.3E-08 | | MAAVFAVVTLTLGFRL* |
| HG0730 | *H. sapiens* | ENSG00000125170 | DOK4 | docking protein 4 | 0.00 | 0.30 | 2.9E-08 | 2.4E-07 | 0.30 | 2.9E-08 | | MQARGEGAARTMLRRVR* |
| HG0731 | *H. sapiens* | ENSG00000129255 | MPDU1 | mannose-P-dolichol utilization defect 1 | 0.06 | 0.47 | 1.0E-06 | 6.7E-06 | 0.47 | 1.0E-06 | | MAPSRRSGERQGCKSGFQTGSVNKGVGGLTAQACSFGKLYNSNSASKTWSLTETYFTPLE* |
| HG0732 | *H. sapiens* | ENSG00000135914 | HTR2B | 5-hydroxytryptamine receptor 2B | 0.21 | 0.28 | 4.1E-07 | 2.9E-06 | 0.28 | 4.1E-07 | | MDRCVYALILENLNFFF* |
| HG0733 | *H. sapiens* | ENSG00000136247 | ZDHHC4 | zinc finger DHHC-type containing 4 | 0.18 | 0.49 | 1.6E-06 | 1.0E-05 | 0.49 | 1.6E-06 | | MLGRGMPELPAENYPTKRNVGWVP* |
| HG0734 | *H. sapiens* | ENSG00000139318 | DUSP6 | dual specificity phosphatase 6 | 0.04 | 0.48 | 1.6E-04 | 6.8E-04 | 0.48 | 1.6E-04 | | MCPAPPPLPPPLLSPSSRRARCCSLFALGLIRSGNSFPFL* |
| HG0735 | *H. sapiens* | ENSG00000139579 | NABP2 | nucleic acid binding protein 2 | 0.22 | 0.49 | 1.2E-06 | 7.8E-06 | 0.49 | 1.2E-06 | | MDRVPACRDEGSGRSGQ* |
| HG0736 | *H. sapiens* | ENSG00000139826 | ABHD13 | abhydrolase domain containing 13 | 0.20 | 0.47 | 1.6E-02 | 3.9E-02 | 0.47 | 1.6E-02 | | MWEPAGAFAGDTRAGAGSGATARSRSSGLDRGAGSGPWAGGG* |
| HG0737 | *H. sapiens* | ENSG00000141639 | MAPK4 | mitogen-activated protein kinase 4 | 0.00 | 0.40 | 3.2E-06 | 2.0E-05 | 0.40 | 3.2E-06 | | MLPTSKGSRGSWPGS* |
| HG0738 | *H. sapiens* | ENSG00000143393 | PI4KB | phosphatidylinositol 4-kinase beta | 0.03 | 0.00 | 1.2E-08 | 1.1E-07 | 0.00 | 1.2E-08 | | MEPGSGRCC* |
| HG0739 | *H. sapiens* | ENSG00000144331 | ZNF385B | zinc finger protein 385B | 0.19 | 0.42 | 1.6E-06 | 1.0E-05 | 0.42 | 1.6E-06 | | MICSEALPAGCFLAPHFTGAPG* |
| HG0740 | *H. sapiens* | ENSG00000144331 | ZNF385B | zinc finger protein 385B | 0.00 | 0.34 | 1.0E-03 | 3.7E-03 | 0.34 | 1.0E-03 | | MFYISTPSNIKLMFMG* |
| HG0741 | *H. sapiens* | ENSG00000144834 | TAGLN3 | transgelin 3 | 0.04 | 0.35 | 9.4E-06 | 5.2E-05 | 0.35 | 9.4E-06 | | MRKSNQCAILPGARQQRLQFNMKGGFLPAC* |
| HG0742 | *H. sapiens* | ENSG00000154556 | SORBS2 | sorbin and SH3 domain containing 2 | 0.00 | 0.50 | 1.3E-04 | 5.8E-04 | 0.50 | 1.3E-04 | | MFCKFPGGHEAPDTWIGLNLKPLEFKTF* |
| HG0743 | *H. sapiens* | ENSG00000154814 | OXNAD1 | oxidoreductase NAD binding domain containing 1 | 0.24 | 0.35 | 1.8E-07 | 1.4E-06 | 0.35 | 1.8E-07 | | MEKLSMFQLLYIQKSQCNSRTKIFCQSADHILNDS* |
| HG0744 | *H. sapiens* | ENSG00000163053 | SLC16A14 | solute carrier family 16 member 14 | 0.04 | 0.35 | 4.7E-14 | 8.0E-13 | 0.35 | 4.7E-14 | | MIGDIATNLYTQQLRGLQAEQQVRGSGRGAE* |
| HG0745 | *H. sapiens* | ENSG00000163145 | C1QTNF7 | C1q and TNF related 7 | 0.00 | 0.31 | 4.7E-05 | 2.3E-04 | 0.31 | 4.7E-05 | | MLRRALYFSSSYFLHCFGVASNNKHIFLL* |
| HG0746 | *H. sapiens* | ENSG00000164741 | DLC1 | DLC1 Rho GTPase activating protein | 0.03 | 0.08 | 5.4E-04 | 2.1E-03 | 0.08 | 5.4E-04 | | MISTTKWELRFWRN* |
| HG0747 | *H. sapiens* | ENSG00000168036 | CTNNB1 | catenin beta 1 | 0.00 | 0.16 | 2.0E-04 | 8.5E-04 | 0.16 | 2.0E-04 | | MIFSMNLGFCRSIHLQSQNPPGI* |
| HG0748 | *H. sapiens* | ENSG00000168830 | HTR1E | 5-hydroxytryptamine receptor 1E | 0.03 | 0.46 | 1.7E-04 | 7.2E-04 | 0.46 | 1.7E-04 | | MLWPFPLPTENGTQETT* |
| HG0749 | *H. sapiens* | ENSG00000169760 | NLGN1 | neuroligin 1 | 0.00 | 0.47 | 6.8E-04 | 2.5E-03 | 0.47 | 6.8E-04 | | MVGFTWRWLWVPRILDRGCIWREKRINRHVLEVSVGE* |
| HG0750 | *H. sapiens* | ENSG00000170881 | RNF139 | ring finger protein 139 | 0.28 | 0.20 | 3.0E-06 | 1.8E-05 | 0.20 | 3.0E-06 | | MVAPRAGEAVEGGGVRP* |
| HG0751 | *H. sapiens* | ENSG00000171791 | BCL2 | BCL2, apoptosis regulator | 0.01 | 0.36 | 4.1E-06 | 2.4E-05 | 0.36 | 4.1E-06 | | MGSLPYAFVLVLQKGNLTEDHAVLKKYK* |
| HG0752 | *H. sapiens* | ENSG00000172269 | DPAGT1 | dolichyl-phosphate N-acetylglucosaminephosphotransferas | 0.05 | 0.44 | 1.6E-07 | 1.2E-06 | 0.44 | 1.6E-07 | | MKFGITLLFHFHPHPLMPILKMAAAAFRSNGCSANKIRAETVDRRVQRGRT* |
| HG0753 | *H. sapiens* | ENSG00000173406 | DAB1 | DAB1, reelin adaptor protein | 0.00 | 0.33 | 1.5E-04 | 6.7E-04 | 0.33 | 1.5E-04 | | MGSRFLGGCGSGGGCSGLGEHSKRVNEMCSSECHF* |
| HG0754 | *H. sapiens* | ENSG00000175182 | FAM131A | family with sequence similarity 131 member A | 0.05 | 0.28 | 2.5E-05 | 1.3E-04 | 0.28 | 2.5E-05 | | MRQECSPYVTRHRFQMGRGLAQHPEPKQE* |
| HG0755 | *H. sapiens* | ENSG00000176597 | B3GNT5 | UDP-GlcNAc:betaGal beta-1,3-N-acetylglucosaminyltransfe | 0.00 | 0.49 | 1.2E-06 | 7.7E-06 | 0.49 | 1.2E-06 | | MPVWKNFDVPVSSFYRVFHSSAISEKWD* |
| HG0756 | *H. sapiens* | ENSG00000177051 | FBXO46 | F-box protein 46 | 0.06 | 0.40 | 3.4E-05 | 1.7E-04 | 0.40 | 3.4E-05 | | MQRAAMVACGASHHHLGPEMR* |
| HG0757.1 | *H. sapiens* | ENSG00000177301 | KCNA2 | potassium voltage-gated channel subfamily A member 2 | 0.00 | 0.11 | 3.3E-15 | 5.9E-14 | 0.11 | 3.3E-15 | | MLLCFLGPVAAP* |
| HG0757.2 | *H. sapiens* | ENSG00000182255 | KCNA4 | potassium voltage-gated channel subfamily A member 4 | 0.00 | 0.16 | 5.5E-03 | 1.6E-02 | 0.16 | 5.5E-03 | | MNLKGEPGESS* |
| HG0758 | *H. sapiens* | ENSG00000179299 | NSUN7 | NOP2/Sun RNA methyltransferase family member 7 | 0.05 | 0.24 | 2.8E-12 | 3.9E-11 | 0.24 | 2.8E-12 | | MLQMRGKPFPGTSEF* |
| HG0759 | *H. sapiens* | ENSG00000180929 | GPR62 | G protein-coupled receptor 62 | 0.23 | 0.23 | 2.1E-10 | 2.3E-09 | 0.23 | 2.1E-10 | | MAAWASEPPPGQPMR* |
| HG0760 | *H. sapiens* | ENSG00000182463 | TSHZ2 | teashirt zinc finger homeobox 2 | 0.22 | 0.36 | 3.9E-08 | 3.2E-07 | 0.36 | 3.9E-08 | | MFNKEIKCLNSHQKKKKPQKQNQKNSKSKNKKERGKKIQNKQTNKQGRTNLYFKAAGTSHPCLPPREGGLWPVVEELQGGSSGGQRPSDVLGATGQEAEETQRGQRDSGPQRAARGWGARSGTGAK* |
| HG0761 | *H. sapiens* | ENSG00000185985 | SLITRK2 | SLIT and NTRK like family member 2 | 0.00 | 0.43 | 1.3E-10 | 1.5E-09 | 0.43 | 1.3E-10 | | MTWRLAATPREIRCKHVCSRGACVGGLGMGE* |
| HG0762 | *H. sapiens* | ENSG00000187147 | RNF220 | ring finger protein 220 | 0.00 | 0.14 | 1.5E-08 | 1.3E-07 | 0.14 | 1.5E-08 | | MHHTLRLLPSVS* |
| HG0763 | *H. sapiens* | ENSG00000188050 | RNF133 | ring finger protein 133 | 0.00 | 0.41 | 2.0E-07 | 1.5E-06 | 0.41 | 2.0E-07 | Crowe | MLEDSGSCLYPEDFFLFVILN* |
| HG0764 | *H. sapiens* | ENSG00000188725 | SMIM15 | small integral membrane protein 15 | 0.11 | 0.37 | 2.2E-09 | 2.2E-08 | 0.37 | 2.2E-09 | | MCTVELKKTDYQGSSEVVIGSLITPGQS* |
| HG0765 | *H. sapiens* | ENSG00000196123 | | | 0.08 | 0.44 | 1.1E-06 | 7.0E-06 | 0.44 | 1.1E-06 | | MVKSLLQEYIAKAFLAEGPET* |
| HG0766 | *H. sapiens* | ENSG00000196189 | SEMA4A | semaphorin 4A | 0.15 | 0.20 | 6.0E-13 | 9.1E-12 | 0.20 | 6.0E-13 | | MMGTWRAALPFQPG* |
| HG0767.1 | *H. sapiens* | ENSG00000197555 | SIPA1L1 | signal induced proliferation associated 1 like 1 | 0.00 | 0.28 | 5.5E-05 | 2.7E-04 | 0.28 | 5.5E-05 | | MLDSFCNILVSRKHGIMATTESQ* |
| HG0767.2 | *H. sapiens* | ENSG00000197555 | SIPA1L1 | signal induced proliferation associated 1 like 1 | 0.00 | 0.26 | 4.9E-03 | 1.4E-02 | 0.26 | 4.9E-03 | | MERATQSCVLFLPSS* |
| HG0768 | *H. sapiens* | ENSG00000213023 | SYT3 | synaptotagmin 3 | 0.13 | 0.30 | 2.0E-06 | 1.3E-05 | 0.30 | 2.0E-06 | | MGCRPPPPRLPPRTRQR* |
| HG0769.1 | *H. sapiens* | ENSG00000280165 | PCDH20 | protocadherin 20 | 0.11 | 0.34 | 3.7E-07 | 2.7E-06 | 0.34 | 3.7E-07 | | MNREGVSSAALPSRAL* |
| HG0769.2 | *H. sapiens* | ENSG00000197991 | | | 0.00 | 0.46 | 2.3E-03 | 7.4E-03 | 0.46 | 2.3E-03 | | MNVPFCPLAFLPLFEFPSLIEEPVTIG* |
| HG0770 | *H. sapiens* | ENSG00000010165 | METTL13 | methyltransferase like 13 | 0.00 | 0.45 | 4.3E-07 | 3.1E-06 | 0.45 | 4.3E-07 | | MWLFFRGKNSHCSVPEPACGGQAPQMVSHRE* |
| HG0771 | *H. sapiens* | ENSG00000013306 | SLC25A39 | solute carrier family 25 member 39 | 0.00 | 0.37 | 8.4E-07 | 5.7E-06 | 0.37 | 8.4E-07 | | MGPPDPGRRAARAPA* |
| HG0772 | *H. sapiens* | ENSG00000064309 | CDON | cell adhesion associated, oncogene regulated | 0.03 | 0.22 | 1.5E-11 | 1.9E-10 | 0.22 | 1.5E-11 | | MTWGWLSEPAAAQEVWRLSQPRGE* |
| HG0773 | *H. sapiens* | ENSG00000065548 | ZC3H15 | zinc finger CCCH-type containing 15 | 0.04 | 0.47 | 8.6E-05 | 4.0E-04 | 0.47 | 8.6E-05 | | MLCDIDLSQRPRLRGQ* |
| HG0774 | *H. sapiens* | ENSG00000066933 | MYO9A | myosin IXA | 0.00 | 0.42 | 1.9E-03 | 6.2E-03 | 0.42 | 1.9E-03 | | MCLLRPGGVSLTGRGMRRGRSPGP* |
| HG0775 | *H. sapiens* | ENSG00000068001 | HYAL2 | hyaluronoglucosaminidase 2 | 0.02 | 0.48 | 1.3E-03 | 4.5E-03 | 0.48 | 1.3E-03 | | MGFIKSSASCPTGSSWRK* |
| HG0776 | *H. sapiens* | ENSG00000082014 | SMARCD3 | SWI/SNF related, matrix associated, actin dependent regul | 0.16 | 0.33 | 2.2E-07 | 1.7E-06 | 0.33 | 2.2E-07 | | MALGAAGAVGSGPRHRGGTG* |
| HG0777 | *H. sapiens* | ENSG00000085644 | ZNF213 | zinc finger protein 213 | 0.20 | 0.25 | 5.9E-07 | 4.2E-06 | 0.25 | 5.9E-07 | | MPAQLPQGIPLGD* |
| HG0778 | *H. sapiens* | ENSG00000089116 | LHX5 | LIM homeobox 5 | 0.02 | 0.17 | 1.7E-02 | 4.1E-02 | 0.17 | 1.7E-02 | | MRELRGDQTKMRIRTA* |
| HG0779 | *H. sapiens* | ENSG00000091140 | DLD | dihydrolipoamide dehydrogenase | 0.09 | 0.45 | 2.8E-05 | 1.4E-04 | 0.45 | 2.8E-05 | | MRREGRPWRSGGGAQRR* |
| HG0780 | *H. sapiens* | ENSG00000092199 | HNRNPC | heterogeneous nuclear ribonucleoprotein C (C1/C2) | 0.02 | 0.00 | 1.8E-04 | 7.8E-04 | 0.00 | 1.8E-04 | | MPSLEQQQPG* |
| HG0781 | *H. sapiens* | ENSG00000100099 | HPS4 | HPS4, biogenesis of lysosomal organelles complex 3 subu | 0.00 | 0.50 | 9.0E-05 | 4.2E-04 | 0.50 | 9.0E-05 | | MHLPLLAFLERSLKRWFCRQPQ* |
| HG0782 | *H. sapiens* | ENSG00000100368 | CSF2RB | colony stimulating factor 2 receptor beta common subunit | 0.08 | 0.35 | 2.8E-06 | 1.7E-05 | 0.35 | 2.8E-06 | | MGLAWLPAGQEHRTSGH* |
| HG0783 | *H. sapiens* | ENSG00000105053 | VRK3 | vaccinia related kinase 3 | 0.17 | 0.04 | 8.6E-07 | 5.8E-06 | 0.04 | 8.6E-07 | | MRAQRSPPSGSFRWATEKVAVKMEFPTQ* |
| HG0784 | *H. sapiens* | ENSG00000105851 | PIK3CG | phosphatidylinositol-4,5-bisphosphate 3-kinase catalytic su | 0.11 | 0.43 | 7.3E-03 | 2.0E-02 | 0.43 | 7.3E-03 | | MKGAPEKRKNLDAHWVA* |
| HG0785 | *H. sapiens* | ENSG00000107242 | PIP5K1B | phosphatidylinositol-4-phosphate 5-kinase type 1 beta | 0.00 | 0.38 | 1.1E-03 | 4.1E-03 | 0.38 | 1.1E-03 | | MSRPPITTNYR* |
| HG0786 | *H. sapiens* | ENSG00000107862 | GBF1 | golgi brefeldin A resistant guanine nucleotide exchange fac | 0.00 | 0.00 | 4.4E-05 | 2.2E-04 | 0.00 | 4.4E-05 | | MSSGCRSWD* |
| HG0787 | *H. sapiens* | ENSG00000108592 | FTSJ3 | FtsJ homolog 3 | 0.18 | 0.23 | 9.8E-11 | 1.2E-09 | 0.23 | 9.8E-11 | | MAENLEPGTQGTQAEAEVVSCILVRFQVWRKRLWV* |
| HG0788 | *H. sapiens* | ENSG00000113739 | STC2 | stanniocalcin 2 | 0.10 | 0.00 | 2.5E-08 | 2.1E-07 | 0.00 | 2.5E-08 | | MLSGELREGAG* |
| HG0789 | *H. sapiens* | ENSG00000115355 | CCDC88A | coiled-coil domain containing 88A | 0.00 | 0.25 | 3.2E-08 | 2.6E-07 | 0.25 | 3.2E-08 | | MVSARGEGASPACWY* |
| HG0790.1 | *H. sapiens* | ENSG00000115568 | ZNF142 | zinc finger protein 142 | 0.00 | 0.47 | 2.9E-05 | 1.5E-04 | 0.47 | 2.9E-05 | | MTMLHPKPSRLQTAFPVRWLL* |
| HG0790.2 | *H. sapiens* | ENSG00000115568 | ZNF142 | zinc finger protein 142 | 0.00 | 0.20 | 6.9E-05 | 3.3E-04 | 0.20 | 6.9E-05 | | MWTLSSPPFYR* |
| HG0791 | *H. sapiens* | ENSG00000131323 | TRAF3 | TNF receptor associated factor 3 | 0.00 | 0.32 | 6.1E-04 | 2.3E-03 | 0.32 | 6.1E-04 | | MRKMRPKEVMPLG* |
| HG0792 | *H. sapiens* | ENSG00000132424 | PNISR | PNN interacting serine and arginine rich protein | 0.00 | 0.17 | 1.2E-05 | 6.7E-05 | 0.17 | 1.2E-05 | | MLRSCSLNQPKF* |
| HG0793 | *H. sapiens* | ENSG00000132698 | RAB25 | RAB25, member RAS oncogene family | 0.00 | 0.26 | 4.4E-07 | 3.1E-06 | 0.26 | 4.4E-07 | | MRGGAGPDLLRAEG* |
| HG0794.1 | *H. sapiens* | ENSG00000135218 | CD36 | CD36 molecule | 0.00 | 0.41 | 4.6E-04 | 1.8E-03 | 0.41 | 4.6E-04 | | MLEHLIEKSFLAILKIAFQ* |
| HG0794.2 | *H. sapiens* | ENSG00000135218 | CD36 | CD36 molecule | 0.00 | 0.17 | 6.0E-04 | 2.3E-03 | 0.17 | 6.0E-04 | | MSMAFRCQDNLKDR* |
| HG0795 | *H. sapiens* | ENSG00000137601 | NEK1 | NIMA related kinase 1 | 0.00 | 0.35 | 1.9E-05 | 9.8E-05 | 0.35 | 1.9E-05 | | MSFSKCGMVPLLY* |
| HG0796 | *H. sapiens* | ENSG00000138074 | SLC5A6 | solute carrier family 5 member 6 | 0.03 | 0.33 | 2.7E-11 | 3.3E-10 | 0.33 | 2.7E-11 | | MRPSPLSGILADPLFLFPFSEGLPRRRAASRSRLQTPSARCSPRPDAEVAASQKLGPLHPPSEKGAMLISGSTKGPS* |
| HG0797 | *H. sapiens* | ENSG00000140526 | ABHD2 | abhydrolase domain containing 2 | 0.01 | 0.34 | 5.0E-09 | 4.7E-08 | 0.34 | 5.0E-09 | | MSAAGSRGAAGTRLGAS* |
| HG0798.1 | *H. sapiens* | ENSG00000142949 | PTPRF | protein tyrosine phosphatase, receptor type F | 0.06 | 0.42 | 5.2E-05 | 2.5E-04 | 0.42 | 5.2E-05 | | MRTDGPWIGGRSGGPGARPLVLSIQQE* |
| HG0798.2 | *H. sapiens* | ENSG00000153707 | PTPRD | protein tyrosine phosphatase, receptor type D | 0.00 | 0.47 | 1.6E-02 | 3.8E-02 | 0.47 | 1.6E-02 | | MLAPREAMGFGSRFLS* |
| HG0799 | *H. sapiens* | ENSG00000143398 | PIP5K1A | phosphatidylinositol-4-phosphate 5-kinase type 1 alpha | 0.01 | 0.35 | 1.6E-06 | 1.0E-05 | 0.35 | 1.6E-06 | | MPGEWCGLELVQEPARRV* |
| HG0800 | *H. sapiens* | ENSG00000143612 | C1orf43 | chromosome 1 open reading frame 43 | 0.03 | 0.37 | 1.6E-02 | 3.9E-02 | 0.37 | 1.6E-02 | | MFSGRVWLGLPP* |
| HG0801 | *H. sapiens* | ENSG00000144278 | GALNT13 | polypeptide N-acetylgalactosaminyltransferase 13 | 0.00 | 0.39 | 6.0E-04 | 2.3E-03 | 0.39 | 6.0E-04 | | MFANHFEIAKCHIQFNWI* |
| HG0802 | *H. sapiens* | ENSG00000144283 | PKP4 | plakophilin 4 | 0.12 | 0.00 | 7.2E-05 | 3.4E-04 | 0.00 | 7.2E-05 | | MSLIPGATTAAA* |
| HG0803 | *H. sapiens* | ENSG00000144802 | NFKBIZ | NFKB inhibitor zeta | 0.05 | 0.40 | 1.8E-05 | 9.5E-05 | 0.40 | 1.8E-05 | | MLSRTWVIIADYSWKQLICSYVFV* |
| HG0804 | *H. sapiens* | ENSG00000144815 | NXPE3 | neurexophilin and PC-esterase domain family member 3 | 0.00 | 0.46 | 2.7E-03 | 8.4E-03 | 0.46 | 2.7E-03 | | MCFLNQLQSLLCIRALKGTSRIEANETESSSAAQKSKDMEF* |
| HG0805.1 | *H. sapiens* | ENSG00000145936 | KCNMB1 | potassium calcium-activated channel subfamily M regulator | 0.04 | 0.49 | 8.8E-05 | 4.1E-04 | 0.49 | 8.8E-05 | | MFGHGGLPQMGTKSCFPGDGRGGQTSC* |
| HG0805.2 | *H. sapiens* | ENSG00000145936 | KCNMB1 | potassium calcium-activated channel subfamily M regulator | 0.06 | 0.49 | 6.4E-03 | 1.8E-02 | 0.49 | 6.4E-03 | | MEVQGLGNSCLPQGWVPALRVCVSRLCAQT* |
| HG0806 | *H. sapiens* | ENSG00000147481 | SNTG1 | syntrophin gamma 1 | 0.00 | 0.42 | 7.5E-05 | 3.5E-04 | 0.42 | 7.5E-05 | | MGRAVGVEMLKGPHFLK* |
| HG0807.1 | *H. sapiens* | ENSG00000147488 | ST18 | ST18, C2H2C-type zinc finger | 0.00 | 0.31 | 6.6E-10 | 7.1E-09 | 0.31 | 6.6E-10 | Mackowiak | MFLTECDLRGILEKIIKFLNGAAWRNPKRNSIRAFWNTSRKR* |
| HG0807.2 | *H. sapiens* | ENSG00000147488 | ST18 | ST18, C2H2C-type zinc finger | 0.00 | 0.37 | 3.8E-07 | 2.8E-06 | 0.37 | 3.8E-07 | | MSGQILQETQKSMIKKLSTLIYCKG* |
| HG0807.3 | *H. sapiens* | ENSG00000186487 | MYT1L | myelin transcription factor 1 like | 0.00 | 0.00 | 1.5E-05 | 7.8E-05 | 0.00 | 1.5E-05 | | MEKSRHREIK* |
| HG0808 | *H. sapiens* | ENSG00000149269 | PAK1 | p21 (RAC1) activated kinase 1 | 0.15 | 0.42 | 7.2E-03 | 2.0E-02 | 0.42 | 7.2E-03 | | MPGLAFPTSSCCWW* |
| HG0809 | *H. sapiens* | ENSG00000151287 | TEX30 | testis expressed 30 | 0.04 | 0.38 | 5.2E-03 | 1.5E-02 | 0.38 | 5.2E-03 | | MCSFIVFVSYPCLPTHLSQQNLLFREMSFASQM* |
| HG0810 | *H. sapiens* | ENSG00000156504 | FAM122B | family with sequence similarity 122B | 0.00 | 0.44 | 6.2E-03 | 1.8E-02 | 0.44 | 6.2E-03 | | MGARTRKNWPSGGSRF* |
| HG0811 | *H. sapiens* | ENSG00000157350 | ST3GAL2 | ST3 beta-galactoside alpha-2,3-sialyltransferase 2 | 0.00 | 0.40 | 1.3E-08 | 1.2E-07 | 0.40 | 1.3E-08 | | MGSRRARPTAERISISAAEFSHTHRASCFCPLRVGVWLPSSGRGFGPGGGTTLAWNPSPLKPLCSLL* |
| HG0812 | *H. sapiens* | ENSG00000157368 | IL34 | interleukin 34 | 0.02 | 0.31 | 1.4E-08 | 1.2E-07 | 0.31 | 1.4E-08 | | MGLAATAPRLSSTLPGR* |
| HG0813 | *H. sapiens* | ENSG00000160199 | PKNOX1 | PBX/knotted 1 homeobox 1 | 0.00 | 0.32 | 3.4E-04 | 1.4E-03 | 0.32 | 3.4E-04 | | MKIGTQTPFAFHPR* |
| HG0814 | *H. sapiens* | ENSG00000163071 | SPATA18 | spermatogenesis associated 18 | 0.18 | 0.40 | 8.8E-05 | 4.1E-04 | 0.40 | 8.8E-05 | | MRRGGCGPGHLPSGFPNPARSDPRGRMETPAAL* |
| HG0815 | *H. sapiens* | ENSG00000163072 | NOSTRIN | nitric oxide synthase trafficking | 0.00 | 0.26 | 2.3E-03 | 7.5E-03 | 0.26 | 2.3E-03 | | MKVQGWDFLLGMFDKD* |
| HG0816 | *H. sapiens* | ENSG00000163794 | UCN | urocortin | 0.20 | 0.15 | 2.3E-15 | 4.3E-14 | 0.39 | 1.9E-02 | | MFFADVSRSCPGTI* |
| HG0817 | *H. sapiens* | ENSG00000163888 | CAMK2N2 | calcium/calmodulin dependent protein kinase II inhibitor 2 | 0.00 | 0.00 | 4.4E-04 | 1.7E-03 | 0.00 | 4.4E-04 | | MTSSFRRLPC* |
| HG0818 | *H. sapiens* | ENSG00000164125 | FAM198B | family with sequence similarity 198 member B | 0.00 | 0.49 | 4.2E-03 | 1.3E-02 | 0.49 | 4.2E-03 | | MVRNKPHFSLGRLII* |
| HG0819 | *H. sapiens* | ENSG00000164188 | RANBP3L | RAN binding protein 3 like | 0.00 | 0.36 | 3.5E-04 | 1.4E-03 | 0.36 | 3.5E-04 | | MKELSCVFNQMNWHLYI* |
| HG0820 | *H. sapiens* | ENSG00000164946 | FREM1 | FRAS1 related extracellular matrix 1 | 0.00 | 0.37 | 8.9E-06 | 5.0E-05 | 0.37 | 8.9E-06 | | MLRSRGTLSGPLTLPLGL* |

| | | | | | | | | | | | | |
|---|---|---|---|---|---|---|---|---|---|---|---|---|
| HG0821 | *H. sapiens* | ENSG00000165300 | SLITRK5 | SLIT and NTRK like family member 5 | 0.04 | 0.07 | 2.5E-08 | 2.1E-07 | 0.07 | 2.5E-08 | | MMTGPWRCETGR* |
| HG0822 | *H. sapiens* | ENSG00000166170 | BAG5 | BCL2 associated athanogene 5 | 0.18 | 0.26 | 7.4E-03 | 2.0E-02 | 0.26 | 7.4E-03 | | MQLGAQLRCRTP* |
| HG0823 | *H. sapiens* | ENSG00000166268 | MYRFL | myelin regulatory factor-like | 0.00 | 0.27 | 7.7E-07 | 5.2E-06 | 0.27 | 7.7E-07 | | MAEDMLHSNKKKMKIFQEHS* |
| HG0824 | *H. sapiens* | ENSG00000167100 | | | 0.04 | 0.26 | 1.7E-04 | 7.3E-04 | 0.26 | 1.7E-04 | | MERLEPARSPPRPG* |
| HG0825 | *H. sapiens* | ENSG00000169180 | XPO6 | exportin 6 | 0.00 | 0.00 | 3.6E-05 | 1.8E-04 | 0.00 | 3.6E-05 | | MGGGLPESAGTMT* |
| HG0826 | *H. sapiens* | ENSG00000169851 | PCDH7 | protocadherin 7 | 0.00 | 0.45 | 6.2E-04 | 2.3E-03 | 0.45 | 6.2E-04 | | MTLRRVVKKKEKPALSGVRRD* |
| HG0827 | *H. sapiens* | ENSG00000170365 | SMAD1 | SMAD family member 1 | 0.00 | 0.35 | 1.0E-08 | 9.2E-08 | 0.35 | 1.0E-08 | | MCIREFAVEQLFLYSAPCL* |
| HG0828 | *H. sapiens* | ENSG00000170390 | DCLK2 | doublecortin like kinase 2 | 0.00 | 0.17 | 6.8E-08 | 5.5E-07 | 0.17 | 6.8E-08 | | MTWAARRCGHF* |
| HG0829 | *H. sapiens* | ENSG00000171502 | COL24A1 | collagen type XXIV alpha 1 chain | 0.12 | 0.29 | 3.4E-04 | 1.4E-03 | 0.29 | 3.4E-04 | | MDSTCTLSSSELASVFR* |
| HG0830 | *H. sapiens* | ENSG00000171631 | P2RY6 | pyrimidinergic receptor P2Y6 | 0.25 | 0.41 | 6.6E-08 | 5.3E-07 | 0.41 | 6.6E-08 | | MALEGGVQAEEMGAVLSEPLPP* |
| HG0831.1 | *H. sapiens* | ENSG00000174482 | LINGO2 | leucine rich repeat and Ig domain containing 2 | 0.00 | 0.32 | 2.5E-03 | 8.0E-03 | 0.32 | 2.5E-03 | | MQLSRFVSSSWLVGL* |
| HG0831.2 | *H. sapiens* | ENSG00000174482 | LINGO2 | leucine rich repeat and Ig domain containing 2 | 0.21 | 0.44 | 1.3E-02 | 3.3E-02 | 0.44 | 1.3E-02 | | MRRSSEQSTESGHYQHEEPNNSNW* |
| HG0832 | *H. sapiens* | ENSG00000175048 | ZDHHC14 | zinc finger DHHC-type containing 14 | 0.13 | 0.37 | 1.6E-03 | 5.4E-03 | 0.37 | 1.6E-03 | Crowe | MTVGLPGGLTWSSAKLSPSREPV* |
| HG0833 | *H. sapiens* | ENSG00000176641 | RNF152 | ring finger protein 152 | 0.00 | 0.37 | 6.7E-06 | 3.8E-05 | 0.37 | 6.7E-06 | | MSSRRRGERDSVTDTEMNTPVSTGCL* |
| HG0834 | *H. sapiens* | ENSG00000182489 | XKRX | XK related, X-linked | 0.00 | 0.33 | 1.7E-02 | 4.2E-02 | 0.33 | 1.7E-02 | | MLAFVLFKFHSKYVDDWNRGQSRVKFKVVLLGFPSQTPSRI* |
| HG0835 | *H. sapiens* | ENSG00000182901 | RGS7 | regulator of G protein signaling 7 | 0.06 | 0.23 | 4.0E-04 | 1.6E-03 | 0.23 | 4.0E-04 | | MKLPCLAAPLAAS* |
| HG0836 | *H. sapiens* | ENSG00000197461 | PDGFA | platelet derived growth factor subunit A | 0.27 | 0.12 | 2.3E-05 | 1.2E-04 | 0.12 | 2.3E-05 | | MSANMQNYRPGRS* |
| HG0837 | *H. sapiens* | ENSG00000198690 | FAN1 | FANCD2 and FANCI associated nuclease 1 | 0.02 | 0.40 | 1.0E-06 | 6.8E-06 | 0.40 | 1.0E-06 | | MRSYRIPLLVISSQESKSKPLLSFTLNILCFIAQNIQFF* |
| HG0838 | *H. sapiens* | ENSG00000198707 | CEP290 | centrosomal protein 290 | 0.01 | 0.41 | 9.8E-03 | 2.6E-02 | 0.41 | 9.8E-03 | | MVSPSSRCRFGLLGTIWLDPESAWNRDRDLSGPAAGSSLVVAVVRGLRWLPGLV* |
| HG0839 | *H. sapiens* | ENSG00000198791 | CNOT7 | CCR4-NOT transcription complex subunit 7 | 0.04 | 0.42 | 5.2E-07 | 3.7E-06 | 0.42 | 5.2E-07 | | MGHPLPPPLPPPPPPPPPSAVSMARRRRRSASSATQVHK* |
| HG0840 | *H. sapiens* | ENSG00000278195 | SSTR3 | somatostatin receptor 3 | 0.16 | 0.43 | 1.8E-06 | 1.1E-05 | 0.43 | 1.8E-06 | | MITSLAARREALTPWAGK* |
| HG0841 | *H. sapiens* | ENSG00000278259 | MYO19 | myosin XIX | 0.03 | 0.16 | 6.9E-09 | 6.3E-08 | 0.16 | 6.9E-09 | | MTRWIVLLVPPATQNPG* |
| HG0842 | *H. sapiens* | ENSG00000002746 | HECW1 | HECT, C2 and WW domain containing E3 ubiquitin protein | 0.03 | 0.37 | 1.1E-03 | 4.0E-03 | 0.37 | 1.1E-03 | | MKCAALPRIRRCCWSRNTVRL* |
| HG0843 | *H. sapiens* | ENSG00000043093 | DCUN1D1 | defective in cullin neddylation 1 domain containing 1 | 0.12 | 0.44 | 1.0E-02 | 2.6E-02 | 0.44 | 1.0E-02 | | MRVEELRSTDPHVP* |
| HG0844 | *H. sapiens* | ENSG00000091009 | RBM27 | RNA binding motif protein 27 | 0.00 | 0.00 | 1.4E-03 | 4.8E-03 | 0.00 | 1.4E-03 | | MPGGLGHREL* |
| HG0845 | *H. sapiens* | ENSG00000099260 | PALMD | palmdelphin | 0.02 | 0.44 | 7.0E-04 | 2.6E-03 | 0.44 | 7.0E-04 | | MVALDLLPRLTVLDF* |
| HG0846 | *H. sapiens* | ENSG00000100852 | ARHGAP5 | Rho GTPase activating protein 5 | 0.00 | 0.33 | 3.8E-05 | 1.9E-04 | 0.33 | 3.8E-05 | Mackowiak | MGKREKEDDPYDLEDVSAQK* |
| HG0847 | *H. sapiens* | ENSG00000106086 | PLEKHA8 | pleckstrin homology domain containing A8 | 0.28 | 0.13 | 1.8E-08 | 1.6E-07 | 0.13 | 1.8E-08 | | MALLLVLLASWGLGQ* |
| HG0848 | *H. sapiens* | ENSG00000111817 | DSE | dermatan sulfate epimerase | 0.29 | 0.15 | 3.8E-04 | 1.5E-03 | 0.15 | 3.8E-04 | | MARRGRGGLGSEGAGSPGALE* |
| HG0849 | *H. sapiens* | ENSG00000111846 | GCNT2 | glucosaminyl (N-acetyl) transferase 2, I-branching enzyme | 0.01 | 0.49 | 3.2E-03 | 9.9E-03 | 0.49 | 3.2E-03 | | MTCRMVTSPTCNFMVGTGGPPGIRTSLRTSVFV* |
| HG0850 | *H. sapiens* | ENSG00000113916 | BCL6 | B-cell CLL/lymphoma 6 | 0.00 | 0.41 | 5.7E-03 | 1.6E-02 | 0.41 | 5.7E-03 | | MCLQLSFAKEKILEVW* |
| HG0851 | *H. sapiens* | ENSG00000114279 | FGF12 | fibroblast growth factor 12 | 0.00 | 0.19 | 3.9E-06 | 2.4E-05 | 0.19 | 3.9E-06 | | MHSVDLTLLGRTS* |
| HG0852 | *H. sapiens* | ENSG00000114942 | EEF1B2 | eukaryotic translation elongation factor 1 beta 2 | 0.05 | 0.25 | 1.8E-07 | 1.4E-06 | 0.25 | 1.8E-07 | | MPILLIYFQTQRKVHGQFVGHLIQGPQFVRGEVGMQKYFDL* |
| HG0853 | *H. sapiens* | ENSG00000121807 | CCR2 | C-C motif chemokine receptor 2 | 0.07 | 0.41 | 1.1E-04 | 4.9E-04 | 0.41 | 1.1E-04 | | MYRVCLTHTQGCISKISKLILFWRPQPNVQCS* |
| HG0854 | *H. sapiens* | ENSG00000127249 | ATP13A4 | ATPase 13A4 | 0.00 | 0.42 | 5.6E-04 | 2.1E-03 | 0.42 | 5.6E-04 | | MGAASFETAALRECNKQGAASNSS* |
| HG0855 | *H. sapiens* | ENSG00000127804 | METTL16 | methyltransferase like 16 | 0.10 | 0.49 | 1.6E-02 | 3.9E-02 | 0.49 | 1.6E-02 | | MAARWERIICVSRSFAMRLL* |
| HG0856 | *H. sapiens* | ENSG00000130055 | GDPD2 | glycerophosphodiester phosphodiesterase domain containi | 0.04 | 0.33 | 1.8E-02 | 4.3E-02 | 0.33 | 1.8E-02 | | MGKPAQRNSSPQAP* |
| HG0857 | *H. sapiens* | ENSG00000130347 | RTN4IP1 | reticulon 4 interacting protein 1 | 0.00 | 0.29 | 3.7E-07 | 2.6E-06 | 0.29 | 3.7E-07 | | MDRILTGTLIRAFHMRRAE* |
| HG0858 | *H. sapiens* | ENSG00000132702 | HAPLN2 | hyaluronan and proteoglycan link protein 2 | 0.04 | 0.23 | 1.8E-08 | 1.6E-07 | 0.23 | 1.8E-08 | | MRTQAPEGRGAQVPGF* |
| HG0859 | *H. sapiens* | ENSG00000132842 | AP3B1 | adaptor related protein complex 3 beta 1 subunit | 0.12 | 0.05 | 2.6E-05 | 1.4E-04 | 0.05 | 2.6E-05 | | MRRGWTARSAQGAVTSPEAPARAPPYEN* |
| HG0860 | *H. sapiens* | ENSG00000133056 | PIK3C2B | phosphatidylinositol-4-phosphate 3-kinase catalytic subunit | 0.00 | 0.24 | 5.9E-07 | 4.1E-06 | 0.24 | 5.9E-07 | | MASPLSPAAAAIVSPGVA* |
| HG0861 | *H. sapiens* | ENSG00000134001 | EIF2S1 | eukaryotic translation initiation factor 2 subunit alpha | 0.00 | 0.13 | 1.2E-07 | 9.3E-07 | 0.13 | 1.2E-07 | | MVNDEEGPLFVQRLRMRGGSACAVE* |
| HG0862 | *H. sapiens* | ENSG00000134245 | WNT2B | Wnt family member 2B | 0.02 | 0.00 | 3.0E-08 | 2.5E-07 | 0.00 | 3.0E-08 | | MAPQGGEVGAA* |
| HG0863 | *H. sapiens* | ENSG00000134313 | KIDINS220 | kinase D interacting substrate 220 | 0.00 | 0.45 | 1.2E-03 | 4.3E-03 | 0.45 | 1.2E-03 | | MAAGCGEGDALAVAVSCFPVL* |
| HG0864 | *H. sapiens* | ENSG00000134533 | RERG | RAS like estrogen regulated growth inhibitor | 0.00 | 0.46 | 1.3E-03 | 4.7E-03 | 0.46 | 1.3E-03 | | MVNPKTGTKKIYSVL* |
| HG0865 | *H. sapiens* | ENSG00000135540 | NHSL1 | NHS like 1 | 0.00 | 0.39 | 1.3E-06 | 8.4E-06 | 0.39 | 1.3E-06 | | MPRRGVGLQFGMLGTRQRVGVPWLCVLQSPDKYS* |
| HG0866 | *H. sapiens* | ENSG00000136011 | STAB2 | stabilin 2 | 0.09 | 0.18 | 4.8E-05 | 2.3E-04 | 0.18 | 4.8E-05 | | MRKNSLGVYQTKLK* |
| HG0867 | *H. sapiens* | ENSG00000137203 | TFAP2A | transcription factor AP-2 alpha | 0.09 | 0.41 | 9.6E-03 | 2.5E-02 | 0.41 | 9.6E-03 | | MDPSRAPALRGASAP* |
| HG0868 | *H. sapiens* | ENSG00000137843 | PAK6 | p21 (RAC1) activated kinase 6 | 0.09 | 0.41 | 3.5E-08 | 2.9E-07 | 0.41 | 3.5E-08 | | MPEDLMEKSQSEPGRAVEEGRAAAGAAGP* |
| HG0869 | *H. sapiens* | ENSG00000139910 | NOVA1 | NOVA alternative splicing regulator 1 | 0.04 | 0.35 | 1.1E-03 | 4.1E-03 | 0.35 | 1.1E-03 | | MSFNGTDIASSFALESS* |
| HG0870 | *H. sapiens* | ENSG00000140993 | TIGD7 | tigger transposable element derived 7 | 0.01 | 0.30 | 2.5E-05 | 1.3E-04 | 0.30 | 2.5E-05 | | MALGSTRRTRKEYLRA* |
| HG0871 | *H. sapiens* | ENSG00000141127 | PRPSAP2 | phosphoribosyl pyrophosphate synthetase associated prote | 0.04 | 0.37 | 6.8E-06 | 3.9E-05 | 0.37 | 6.8E-06 | | MPAQKGREPSRGTAEEESILEEFCCQRIKGGCP* |
| HG0872 | *H. sapiens* | ENSG00000142583 | SLC2A5 | solute carrier family 2 member 5 | 0.16 | 0.30 | 1.2E-05 | 6.6E-05 | 0.30 | 1.2E-05 | | MLLAVIWHGQSAPRMSSLSIQCTRYFG* |
| HG0873 | *H. sapiens* | ENSG00000142875 | PRKACB | protein kinase cAMP-activated catalytic subunit beta | 0.00 | 0.48 | 1.2E-04 | 5.5E-04 | 0.48 | 1.2E-04 | | MLGSYALLLGASLFKTKESSFRFLCAACSSVCVYTIGSSPSRD* |
| HG0874 | *H. sapiens* | ENSG00000143515 | ATP8B2 | ATPase phospholipid transporting 8B2 | 0.22 | 0.24 | 9.2E-05 | 4.2E-04 | 0.24 | 9.2E-05 | | MGSTGPPGRAPSAER* |
| HG0875 | *H. sapiens* | ENSG00000145284 | SCD5 | stearoyl-CoA desaturase 5 | 0.09 | 0.26 | 1.3E-05 | 7.0E-05 | 0.26 | 1.3E-05 | | MGDTPATDAGRQDLHP* |
| HG0876 | *H. sapiens* | ENSG00000145908 | ZNF300 | zinc finger protein 300 | 0.05 | 0.36 | 1.2E-03 | 4.3E-03 | 0.36 | 1.2E-03 | | MAAPCGNSSREQPCCFLNKASEDAERAKPPSTERRSLGSPVWCSRTGSRRVCQ* |
| HG0877 | *H. sapiens* | ENSG00000146426 | TIAM2 | T-cell lymphoma invasion and metastasis 2 | 0.00 | 0.33 | 3.3E-07 | 2.4E-06 | 0.33 | 3.3E-07 | | MKKRLTSWTHFACLLNVLCCSQDHVKPTDH* |
| HG0878 | *H. sapiens* | ENSG00000146802 | TMEM168 | transmembrane protein 168 | 0.01 | 0.37 | 1.8E-04 | 7.8E-04 | 0.37 | 1.8E-04 | | MAVAAGPVTEKVYADTGLY* |
| HG0879 | *H. sapiens* | ENSG00000147145 | LPAR4 | lysophosphatidic acid receptor 4 | 0.03 | 0.28 | 2.0E-07 | 1.5E-06 | 0.28 | 2.0E-07 | | MDKNMHFQRASCPFTCLLANYLQASAHSLI* |
| HG0880 | *H. sapiens* | ENSG00000151789 | ZNF385D | zinc finger protein 385D | 0.01 | 0.33 | 5.7E-05 | 2.8E-04 | 0.33 | 5.7E-05 | | MGSAAGSPSALSSRRGS* |
| HG0881 | *H. sapiens* | ENSG00000153822 | KCNJ16 | potassium voltage-gated channel subfamily J member 16 | 0.00 | 0.45 | 1.6E-02 | 3.9E-02 | 0.45 | 1.6E-02 | | MAVGADSFRRVLTGPREKGPEMLSYGALTR* |
| HG0882.1 | *H. sapiens* | ENSG00000154832 | CXXC1 | CXXC finger protein 1 | 0.01 | 0.30 | 4.5E-06 | 2.6E-05 | 0.30 | 4.5E-06 | | MFPANIWIRPPYGYDKQDGGA* |
| HG0882.2 | *H. sapiens* | ENSG00000154832 | CXXC1 | CXXC finger protein 1 | 0.01 | 0.26 | 7.0E-03 | 1.9E-02 | 0.26 | 7.0E-03 | | MWKGKETEWRNEYNLIQVVLRKRSLYL* |
| HG0883 | *H. sapiens* | ENSG00000156011 | PSD3 | pleckstrin and Sec7 domain containing 3 | 0.00 | 0.00 | 1.5E-05 | 7.9E-05 | 0.00 | 1.5E-05 | | MLGEAAWCFLS* |
| HG0884 | *H. sapiens* | ENSG00000156463 | SH3RF2 | SH3 domain containing ring finger 2 | 0.00 | 0.16 | 2.9E-08 | 2.4E-07 | 0.16 | 2.9E-08 | | MWTLLLQVGTGVLK* |
| HG0885.1 | *H. sapiens* | ENSG00000157502 | MUM1L1 | MUM1 like 1 | 0.00 | 0.37 | 2.7E-07 | 2.0E-06 | 0.37 | 2.7E-07 | | MPTKPWKDSATTEVPSWVKALLDKQLE* |
| HG0885.2 | *H. sapiens* | ENSG00000157502 | MUM1L1 | MUM1 like 1 | 0.00 | 0.43 | 3.6E-03 | 1.1E-02 | 0.43 | 3.6E-03 | | MAVVNASYFQRVLEDLE* |
| HG0886 | *H. sapiens* | ENSG00000158850 | B4GALT3 | beta-1,4-galactosyltransferase 3 | 0.05 | 0.33 | 4.4E-05 | 2.2E-04 | 0.33 | 4.4E-05 | | MAAAMPARLLGWR* |
| HG0887 | *H. sapiens* | ENSG00000160959 | | | 0.00 | 0.43 | 1.5E-06 | 9.7E-06 | 0.43 | 1.5E-06 | | MIVEICPPAELGRGLVSGTFQEPVP* |
| HG0888 | *H. sapiens* | ENSG00000164023 | SGMS2 | sphingomyelin synthase 2 | 0.00 | 0.27 | 1.5E-05 | 8.2E-05 | 0.27 | 1.5E-05 | | MEETVIGWLPSTFCIRKQSPL* |
| HG0889 | *H. sapiens* | ENSG00000164091 | WDR82 | WD repeat domain 82 | 0.08 | 0.24 | 1.4E-04 | 6.2E-04 | 0.24 | 1.4E-04 | | MADTMSRRRAGCP* |
| HG0890 | *H. sapiens* | ENSG00000165813 | | | 0.11 | 0.18 | 2.5E-07 | 1.8E-06 | 0.18 | 2.5E-07 | | MEKLAPEPELQQPC* |
| HG0891 | *H. sapiens* | ENSG00000166211 | SPIC | Spi-C transcription factor | 0.00 | 0.40 | 2.2E-05 | 1.1E-04 | 0.40 | 2.2E-05 | | MKHLYKGLKCLPGLSTYFILFSSSNNC* |
| HG0892 | *H. sapiens* | ENSG00000168418 | KCNG4 | potassium voltage-gated channel modifier subfamily G mer | 0.28 | 0.43 | 1.3E-06 | 8.4E-06 | 0.43 | 1.3E-06 | | MGSPCLAEKQLETGSLRQEDW* |
| HG0893 | *H. sapiens* | ENSG00000168769 | TET2 | tet methylcytosine dioxygenase 2 | 0.26 | 0.38 | 1.0E-02 | 2.7E-02 | 0.38 | 1.0E-02 | | MQVTSAPSARPPRDAGPAE* |
| HG0894.1 | *H. sapiens* | ENSG00000169282 | KCNAB1 | potassium voltage-gated channel subfamily A member regu | 0.00 | 0.50 | 4.8E-04 | 1.9E-03 | 0.50 | 4.8E-04 | | MLRAPVEHPACGPGDAGGASCGKPALI* |
| HG0894.2 | *H. sapiens* | ENSG00000069424 | KCNAB2 | potassium voltage-gated channel subfamily A regulatory be | 0.00 | 0.29 | 2.2E-04 | 9.2E-04 | 0.29 | 2.2E-04 | | MALSPVVPVQEQNFPTVKNRASLDK* |
| HG0895 | *H. sapiens* | ENSG00000170035 | UBE2E3 | ubiquitin conjugating enzyme E2 E3 | 0.00 | 0.45 | 1.2E-02 | 3.0E-02 | 0.45 | 1.2E-02 | | MNLRCWPWKQLEQLFSKR* |
| HG0896 | *H. sapiens* | ENSG00000170365 | SMAD1 | SMAD family member 1 | 0.00 | 0.40 | 2.1E-03 | 6.9E-03 | 0.40 | 2.1E-03 | | MCFRRGGGTNACQISQLQ* |
| HG0897 | *H. sapiens* | ENSG00000171435 | KSR2 | kinase suppressor of ras 2 | 0.10 | 0.20 | 2.9E-05 | 1.5E-04 | 0.20 | 2.9E-05 | | MPCPCNWEGGKPDFF* |
| HG0898 | *H. sapiens* | ENSG00000171729 | TMEM51 | transmembrane protein 51 | 0.02 | 0.34 | 1.2E-07 | 9.4E-07 | 0.34 | 1.2E-07 | | MKSCLLACPESPQITTGPSTAAIRTGGLAIARNHPAGVQLIFSSVGREILWSAFKGFLL* |
| HG0899 | *H. sapiens* | ENSG00000173320 | STOX2 | storkhead box 2 | 0.03 | 0.36 | 7.9E-05 | 3.7E-04 | 0.36 | 7.9E-05 | | MCRGHMRVGAPTQPSTRAHAQRPEPRQGED* |
| HG0900 | *H. sapiens* | ENSG00000173517 | PEAK1 | pseudopodium enriched atypical kinase 1 | 0.00 | 0.40 | 7.4E-05 | 3.5E-04 | 0.40 | 7.4E-05 | | MVVAAAACAPLRYLPKLELML* |
| HG0901 | *H. sapiens* | ENSG00000175697 | GPR156 | G protein-coupled receptor 156 | 0.00 | 0.49 | 5.2E-03 | 1.5E-02 | 0.49 | 5.2E-03 | | MRNQSEGTRTVTAIWWLTRIITLCSLMVTKMSTLDSHGMHSQQGTS* |
| HG0902 | *H. sapiens* | ENSG00000177182 | CLVS1 | clavesin 1 | 0.00 | 0.20 | 4.2E-08 | 3.5E-07 | 0.20 | 4.2E-08 | | MASVLLLFWMNTTT* |
| HG0903 | *H. sapiens* | ENSG00000180616 | SSTR2 | somatostatin receptor 2 | 0.05 | 0.09 | 7.0E-08 | 5.6E-07 | 0.09 | 7.0E-08 | | MSHWILGLQGPLR* |
| HG0904 | *H. sapiens* | ENSG00000182329 | | | 0.00 | 0.29 | 3.7E-04 | 1.5E-03 | 0.29 | 3.7E-04 | | MYLIKGPVFVVFLGLL* |
| HG0905 | *H. sapiens* | ENSG00000183337 | BCOR | BCL6 corepressor | 0.00 | 0.00 | 1.2E-05 | 6.4E-05 | 0.00 | 1.2E-05 | | MLESVEDGAGRPED* |
| HG0906 | *H. sapiens* | ENSG00000183773 | AIFM3 | apoptosis inducing factor, mitochondria associated 3 | 0.11 | 0.19 | 4.4E-09 | 4.2E-08 | 0.19 | 4.4E-09 | | MAAPASLRPAGGPAPWGAP* |
| HG0907 | *H. sapiens* | ENSG00000186862 | PDZD7 | PDZ domain containing 7 | 0.05 | 0.45 | 3.5E-05 | 1.8E-04 | 0.45 | 3.5E-05 | | MELEARRAGLSRASRFLLDL* |
| HG0908.1 | *H. sapiens* | ENSG00000188554 | NBR1 | NBR1, autophagy cargo receptor | 0.00 | 0.39 | 7.4E-04 | 2.7E-03 | 0.39 | 7.4E-04 | | MHCGTKGLGHTVPWKL* |
| HG0908.2 | *H. sapiens* | ENSG00000188554 | NBR1 | NBR1, autophagy cargo receptor | 0.00 | 0.25 | 4.4E-06 | 2.6E-05 | 0.25 | 4.4E-06 | | MERNIQYQSGHNSHGNPVDRLETSAGLYMSTVMDWSVVMFSYLESRD* |
| HG0909 | *H. sapiens* | ENSG00000189079 | ARID2 | AT-rich interaction domain 2 | 0.27 | 0.18 | 8.0E-04 | 2.9E-03 | 0.18 | 8.0E-04 | | MTEPRRRRPRNGLRALVGSAGSGGRF* |
| HG0910 | *H. sapiens* | ENSG00000197935 | ZNF311 | zinc finger protein 311 | 0.00 | 0.41 | 5.5E-06 | 3.2E-05 | 0.41 | 5.5E-06 | | MNPKSRAVFDTLSQTVAALWVKKGPLAGTTKTCVLQRRLASTGATSQLV* |
| HG0911 | *H. sapiens* | ENSG00000198673 | FAM19A2 | family with sequence similarity 19 member A2, C-C motif cl | 0.00 | 0.41 | 8.9E-04 | 3.2E-03 | 0.41 | 8.9E-04 | | MPRRAAGGLRGALPAALRGGIEGLKLARSCC* |
| HG0912 | *H. sapiens* | ENSG00000198822 | GRM3 | glutamate metabotropic receptor 3 | 0.11 | 0.35 | 2.4E-04 | 1.0E-03 | 0.35 | 2.4E-04 | | MPGGPCFCQESQLDATASAWSR* |
| HG0913.1 | *H. sapiens* | ENSG00000204599 | TRIM39 | tripartite motif containing 39 | 0.00 | 0.45 | 4.7E-04 | 1.8E-03 | 0.45 | 4.7E-04 | | MRLSLKFALQLRHKE* |
| HG0913.2 | *H. sapiens* | ENSG00000204599 | TRIM39 | tripartite motif containing 39 | 0.01 | 0.10 | 2.7E-04 | 1.1E-03 | 0.10 | 2.7E-04 | | MPQPQIWRKKMENGGGTELWTTSQERVARWDPEQW* |
| HG0914 | *H. sapiens* | ENSG00000227877 | MRLN | myoregulin | 0.01 | 0.33 | 8.5E-04 | 3.1E-03 | 0.33 | 8.5E-04 | | MVFFIKERDHSLDFGLRLLNPWTESRLLRDHLC* |
| HG0915 | *H. sapiens* | ENSG00000254469 | AP002495.2 | Putative short transient receptor potential channel 2-like pro | 0.10 | 0.47 | 9.2E-04 | 3.3E-03 | 0.47 | 9.2E-04 | | MYFPGCSTCHLASVGLIFKNSSFLKPQNAFLCYGRRKSSSRGQKAKVRRPNWRENRFGGPRSEATVAPRGPRSRE* |
| HG0916 | *H. sapiens* | ENSG00000020633 | RUNX3 | runt related transcription factor 3 | 0.17 | 0.26 | 6.3E-04 | 2.4E-03 | 0.26 | 6.3E-04 | | MIHSAYASVENLWLDIPVGPGIQILGS* |
| HG0917 | *H. sapiens* | ENSG00000042980 | ADAM28 | ADAM metallopeptidase domain 28 | 0.01 | 0.41 | 8.1E-03 | 2.2E-02 | 0.41 | 8.1E-03 | | MNQGVSFCLTGEEAGTDPAAPT* |

| | | | | | | | | | | | | |
|---|---|---|---|---|---|---|---|---|---|---|---|---|
| HG0918 | *H. sapiens* | ENSG00000047648 | ARHGAP6 | Rho GTPase activating protein 6 | 0.11 | 0.43 | 2.6E-03 | 8.3E-03 | 0.43 | 2.6E-03 | | MDLLLLPPPGHLERTGGA* |
| HG0919 | *H. sapiens* | ENSG00000051009 | FAM160A2 | family with sequence similarity 160 member A2 | 0.00 | 0.00 | 4.6E-05 | 2.3E-04 | 0.00 | 4.6E-05 | | MGSYWLLQTG* |
| HG0920 | *H. sapiens* | ENSG00000051382 | PIK3CB | phosphatidylinositol-4,5-bisphosphate 3-kinase catalytic su | 0.00 | 0.12 | 9.4E-06 | 5.2E-05 | 0.12 | 9.4E-06 | | MGATILRAPWL* |
| HG0921 | *H. sapiens* | ENSG00000058799 | YIPF1 | Yip1 domain family member 1 | 0.03 | 0.46 | 1.1E-04 | 4.9E-04 | 0.46 | 1.1E-04 | | MRKLRPRGARIQQRSSRIGNQTQLCLTPEPVPLTTG* |
| HG0922 | *H. sapiens* | ENSG00000079841 | RIMS1 | regulating synaptic membrane exocytosis 1 | 0.03 | 0.34 | 2.2E-07 | 1.6E-06 | 0.34 | 2.2E-07 | | MQTTMKDWVLALPGSAAAAAAAAAAAPPPAAAARAPL* |
| HG0923 | *H. sapiens* | ENSG00000081138 | CDH7 | cadherin 7 | 0.14 | 0.30 | 1.7E-04 | 7.5E-04 | 0.30 | 1.7E-04 | | MTLPSCLRSSTDHGFHDRLRYLLHHSLAKAIQQW* |
| HG0924 | *H. sapiens* | ENSG00000081307 | UBA5 | ubiquitin like modifier activating enzyme 5 | 0.21 | 0.50 | 2.8E-04 | 1.1E-03 | 0.50 | 2.8E-04 | | MSATHRKRLRGRPVGVSETCLSVRRWVHVPRALG* |
| HG0925 | *H. sapiens* | ENSG00000088766 | CRLS1 | cardiolipin synthase 1 | 0.30 | 0.49 | 3.3E-03 | 1.0E-02 | 0.49 | 3.3E-03 | | MEAAVAQCLSGCRVSMEKRLASVPGC* |
| HG0926 | *H. sapiens* | ENSG00000095397 | WHRN | whirlin | 0.00 | 0.34 | 5.8E-07 | 4.1E-06 | 0.34 | 5.8E-07 | | MPRTPCRTPAPLHRLPLPGVLWD* |
| HG0927 | *H. sapiens* | ENSG00000095951 | HIVEP1 | human immunodeficiency virus type I enhancer binding pro | 0.00 | 0.30 | 1.1E-05 | 6.1E-05 | 0.30 | 1.1E-05 | | MAACGRPLALLGALAAREPSSTWIN* |
| HG0928 | *H. sapiens* | ENSG00000102057 | KCND1 | potassium voltage-gated channel subfamily D member 1 | 0.05 | 0.29 | 1.7E-03 | 5.7E-03 | 0.29 | 1.7E-03 | | MPQSPLHHCKGIPGSSAFPETLESAENSPRVSWPL* |
| HG0929 | *H. sapiens* | ENSG00000102239 | BRS3 | bombesin receptor subtype 3 | 0.25 | 0.20 | 1.5E-03 | 5.1E-03 | 0.20 | 1.5E-03 | | MGKHLAKHAGRHTELKQRSIWMSWIFFPFCSVLFS* |
| HG0930 | *H. sapiens* | ENSG00000102524 | TNFSF13B | TNF superfamily member 13b | 0.12 | 0.31 | 2.9E-04 | 1.2E-03 | 0.31 | 2.9E-04 | | MQKGRKEKIQDNSPEG* |
| HG0931 | *H. sapiens* | ENSG00000104951 | IL4I1 | interleukin 4 induced 1 | 0.12 | 0.45 | 4.2E-03 | 1.3E-02 | 0.45 | 4.2E-03 | | MCFLPSPDKWPFLKFLINSYEDNTFGD* |
| HG0932 | *H. sapiens* | ENSG00000105997 | HOXA3 | homeobox A3 | 0.11 | 0.48 | 9.3E-05 | 4.3E-04 | 0.48 | 9.3E-05 | | MLRVRAHPAPRGPGRAAKPGRGPEPVSEPEPEPALVRS* |
| HG0933 | *H. sapiens* | ENSG00000106330 | MOSPD3 | motile sperm domain containing 3 | 0.05 | 0.34 | 7.1E-04 | 2.7E-03 | 0.34 | 7.1E-04 | | MTEAVRRKYRKGLSGTWQH* |
| HG0934 | *H. sapiens* | ENSG00000107443 | CCNJ | cyclin J | 0.00 | 0.19 | 4.4E-05 | 2.2E-04 | 0.19 | 4.4E-05 | | MSGAGVRRTTAGLGL* |
| HG0935 | *H. sapiens* | ENSG00000108518 | PFN1 | profilin 1 | 0.06 | 0.14 | 1.6E-03 | 5.5E-03 | 0.14 | 1.6E-03 | | MGSRPRPYRPLQIKI* |
| HG0936 | *H. sapiens* | ENSG00000108799 | EZH1 | enhancer of zeste 1 polycomb repressive complex 2 subun | 0.00 | 0.46 | 6.0E-03 | 1.7E-02 | 0.46 | 6.0E-03 | | MEAGHLFCCCVLPFS* |
| HG0937 | *H. sapiens* | ENSG00000109846 | CRYAB | crystallin alpha B | 0.03 | 0.44 | 1.1E-04 | 5.1E-04 | 0.44 | 1.1E-04 | | MTSHWPAQTCLCLSFLSSVSTGYVSHCQIPDHKSP* |
| HG0938 | *H. sapiens* | ENSG00000111665 | CDCA3 | cell division cycle associated 3 | 0.13 | 0.33 | 9.1E-04 | 3.3E-03 | 0.33 | 9.1E-04 | | MGTTSAQPIGSGQGAWREV* |
| HG0939 | *H. sapiens* | ENSG00000111674 | ENO2 | enolase 2 | 0.22 | 0.49 | 4.9E-03 | 1.4E-02 | 0.49 | 4.9E-03 | | MGALEERPRQPILFLAPRRNASLAAIPL* |
| HG0940 | *H. sapiens* | ENSG00000115556 | PLCD4 | phospholipase C delta 4 | 0.01 | 0.20 | 9.7E-03 | 2.5E-02 | 0.20 | 9.7E-03 | | MGWTGESSHLSPSYCFPPAITCQSQQPAAVEEGRKQASARGAKEKRSQAGLPDPTASQSSGGTAHRHRKHRTAILLSCPR* |
| HG0941 | *H. sapiens* | ENSG00000116285 | ERRFI1 | ERBB receptor feedback inhibitor 1 | 0.11 | 0.42 | 8.6E-04 | 3.1E-03 | 0.42 | 8.6E-04 | | MRGRAPRGSRRSSGGMKATG* |
| HG0942 | *H. sapiens* | ENSG00000117477 | | | 0.04 | 0.47 | 6.5E-05 | 3.1E-04 | 0.47 | 6.5E-05 | | MEDAEARCCHGSVVLAAAEAGAGVSFASM* |
| HG0943 | *H. sapiens* | ENSG00000120690 | ELF1 | E74 like ETS transcription factor 1 | 0.00 | 0.38 | 9.3E-04 | 3.4E-03 | 0.38 | 9.3E-04 | | MLVRPRSLREEKFQKNCLNLTRISMNQEN* |
| HG0944 | *H. sapiens* | ENSG00000120833 | SOCS2 | suppressor of cytokine signaling 2 | 0.00 | 0.50 | 4.6E-04 | 1.8E-03 | 0.50 | 4.6E-04 | | MVQERRGGGVPGIFHVYFPTPTPAAGVQFGLTQPPAFSL* |
| HG0945 | *H. sapiens* | ENSG00000124782 | RREB1 | ras responsive element binding protein 1 | 0.00 | 0.43 | 1.4E-03 | 4.9E-03 | 0.43 | 1.4E-03 | | MIATAGESFIVYQWVRKWR* |
| HG0946.1 | *H. sapiens* | ENSG00000127324 | TSPAN8 | tetraspanin 8 | 0.00 | 0.43 | 5.8E-03 | 1.7E-02 | 0.43 | 5.8E-03 | | MTSKGTYLPGDSLCDI* |
| HG0946.2 | *H. sapiens* | ENSG00000127324 | TSPAN8 | tetraspanin 8 | 0.00 | 0.08 | 3.4E-06 | 2.0E-05 | 0.27 | 7.4E-04 | | MCGLIFLSSLITFYSSLYFFAFSFCSEAVDTEISAGKLLQSILQDKPVTNS* |
| HG0947 | *H. sapiens* | ENSG00000129226 | CD68 | CD68 molecule | 0.14 | 0.49 | 1.3E-02 | 3.3E-02 | 0.49 | 1.3E-02 | | MTKREVARAEAPESGCGWDHLQYRK* |
| HG0948 | *H. sapiens* | ENSG00000129595 | EPB41L4A | erythrocyte membrane protein band 4.1 like 4A | 0.01 | 0.12 | 1.8E-02 | 4.3E-02 | 0.12 | 1.8E-02 | | MRGDAPPPSAPSSAGASPAAPGR* |
| HG0949 | *H. sapiens* | ENSG00000131381 | RBSN | rabenosyn, RAB effector | 0.00 | 0.16 | 5.9E-05 | 2.9E-04 | 0.16 | 5.9E-05 | | MKPLPHHLCAA* |
| HG0950 | *H. sapiens* | ENSG00000132964 | CDK8 | cyclin dependent kinase 8 | 0.13 | 0.44 | 6.1E-04 | 2.3E-03 | 0.44 | 6.1E-04 | | MSLALSRGLLLLLPHQSGWCCGRRA* |
| HG0951 | *H. sapiens* | ENSG00000134061 | CD180 | CD180 molecule | 0.00 | 0.29 | 1.1E-02 | 2.8E-02 | 0.29 | 1.1E-02 | | MLSSQQHFLFQDHPSEYLSGCQIARAFTV* |
| HG0952 | *H. sapiens* | ENSG00000134453 | RBM17 | RNA binding motif protein 17 | 0.05 | 0.00 | 7.1E-03 | 2.0E-02 | 0.00 | 7.1E-03 | | MGRVGRAGRH* |
| HG0953 | *H. sapiens* | ENSG00000134815 | DHX34 | DExH-box helicase 34 | 0.00 | 0.10 | 1.0E-04 | 4.7E-04 | 0.10 | 1.0E-04 | | MNLCPSQRSLQMS* |
| HG0954 | *H. sapiens* | ENSG00000134874 | DZIP1 | DAZ interacting zinc finger protein 1 | 0.00 | 0.27 | 9.6E-05 | 4.4E-04 | 0.27 | 9.6E-05 | | MSGPSRNSPVDWGMSGFSSAY* |
| HG0955 | *H. sapiens* | ENSG00000134982 | APC | APC, WNT signaling pathway regulator | 0.00 | 0.17 | 1.2E-03 | 4.1E-03 | 0.17 | 1.2E-03 | | MLFPGTVVGCW* |
| HG0956 | *H. sapiens* | ENSG00000136235 | GPNMB | glycoprotein nmb | 0.09 | 0.14 | 6.4E-05 | 3.1E-04 | 0.36 | 1.1E-02 | | MILPEALYLIKIEREAPQMPEEHCCSWWTGPEEFRVKP* |
| HG0957 | *H. sapiens* | ENSG00000136874 | STX17 | syntaxin 17 | 0.04 | 0.48 | 8.3E-03 | 2.2E-02 | 0.48 | 8.3E-03 | | MTRRSMVVRLGGLGARGRKPLLAKPRGGWQKVPHLGSLLLANRGSERRLYFDFSLQIRFFYMSGEDSCYQGGHTTFF* |
| HG0958 | *H. sapiens* | ENSG00000137497 | NUMA1 | nuclear mitotic apparatus protein 1 | 0.00 | 0.10 | 2.2E-05 | 1.2E-04 | 0.10 | 2.2E-05 | | MNNHSCPFAPSLGGN* |
| HG0959 | *H. sapiens* | ENSG00000137558 | PI15 | peptidase inhibitor 15 | 0.00 | 0.47 | 1.4E-03 | 4.9E-03 | 0.47 | 1.4E-03 | | MLPNDVFVNSYPLALFFFIDPNSTFLL* |
| HG0960 | *H. sapiens* | ENSG00000140297 | GCNT3 | glucosaminyl (N-acetyl) transferase 3, mucin type | 0.00 | 0.46 | 6.6E-04 | 2.5E-03 | 0.46 | 6.6E-04 | | MISLSLLSYLRAGSQKESDGATCCPLLCRY* |
| HG0961 | *H. sapiens* | ENSG00000140853 | NLRC5 | NLR family CARD domain containing 5 | 0.09 | 0.04 | 4.2E-07 | 3.0E-06 | 0.04 | 4.2E-07 | | MARLEIAEGRREGEKWIPVLRSGLEGEVCKESALWKQPVNPAQGTHSPDTHETLSVGTLEHLS* |
| HG0962.1 | *H. sapiens* | ENSG00000143319 | ISG20L2 | interferon stimulated exonuclease gene 20 like 2 | 0.11 | 0.11 | 4.8E-04 | 1.9E-03 | 0.11 | 4.8E-04 | | MVSRSLRGRRTWVRCMRRLPPIPAWSQGKGMPGFVSLLVVHAADAWVAQRLSTPYFSLFLSIPRCSFPRRSIDRTCSSLLDSEGSSSISPSTPFILQMRLKPRAR* |
| HG0962.2 | *H. sapiens* | ENSG00000143319 | ISG20L2 | interferon stimulated exonuclease gene 20 like 2 | 0.00 | 0.36 | 5.6E-04 | 2.2E-03 | 0.36 | 5.6E-04 | | MLDSRIQQRMSPTSSFHP* |
| HG0963 | *H. sapiens* | ENSG00000143867 | OSR1 | odd-skipped related transciption factor 1 | 0.13 | 0.49 | 6.2E-05 | 3.0E-04 | 0.49 | 6.2E-05 | | MSESELGLSPARSQLLLVQTQRGSRERGSPSPACRIRGC* |
| HG0964 | *H. sapiens* | ENSG00000145246 | ATP10D | ATPase phospholipid transporting 10D (putative) | 0.04 | 0.23 | 2.5E-04 | 1.0E-03 | 0.23 | 2.5E-04 | | MAGVEAQPAFTLRICA* |
| HG0965 | *H. sapiens* | ENSG00000146006 | LRRTM2 | leucine rich repeat transmembrane neuronal 2 | 0.00 | 0.00 | 6.2E-07 | 4.3E-06 | 0.00 | 6.2E-07 | | MKPMLSRGIS* |
| HG0966 | *H. sapiens* | ENSG00000148356 | LRSAM1 | leucine rich repeat and sterile alpha motif containing 1 | 0.05 | 0.43 | 5.4E-05 | 2.6E-04 | 0.43 | 5.4E-05 | | MFVVGRRLRLTAGKQGTAVR* |
| HG0967 | *H. sapiens* | ENSG00000149925 | ALDOA | aldolase, fructose-bisphosphate A | 0.11 | 0.49 | 8.3E-04 | 3.1E-03 | 0.49 | 8.3E-04 | | MSGASGFPKYRSLPEDPWREKGRGH* |
| HG0968 | *H. sapiens* | ENSG00000154319 | FAM167A | family with sequence similarity 167 member A | 0.02 | 0.27 | 3.6E-06 | 2.2E-05 | 0.32 | 4.8E-04 | | MRFSSLPQGLQPPALPEKRELWAFFLPVWSSSAFRTDRPWHRLPRDSCDAVCSCLLWSMAPTGFQDGS* |
| HG0969 | *H. sapiens* | ENSG00000154645 | CHODL | chondrolectin | 0.15 | 0.30 | 4.2E-05 | 2.1E-04 | 0.30 | 4.2E-05 | | MRAGGRAAGLGRRE* |
| HG0970 | *H. sapiens* | ENSG00000154727 | GABPA | GA binding protein transcription factor alpha subunit | 0.06 | 0.37 | 2.6E-03 | 8.2E-03 | 0.37 | 2.6E-03 | | MGRRFSRSTLTGQEASRRQSATGRV* |
| HG0971 | *H. sapiens* | ENSG00000155592 | ZKSCAN2 | zinc finger with KRAB and SCAN domains 2 | 0.15 | 0.47 | 1.9E-03 | 6.2E-03 | 0.47 | 1.9E-03 | | MHSGKPHCAVTSCLEKRPGFCILEYTFTFSRAWHRFD* |
| HG0972 | *H. sapiens* | ENSG00000155926 | SLA | Src like adaptor | 0.27 | 0.42 | 9.4E-03 | 2.5E-02 | 0.42 | 9.4E-03 | | MKNFNRKLLLQKIRSFAANGEHLRQP* |
| HG0973 | *H. sapiens* | ENSG00000156535 | CD109 | CD109 molecule | 0.16 | 0.24 | 6.9E-06 | 4.0E-05 | 0.24 | 6.9E-06 | | MFATAARSAIVMGMGGVEPPSPVSI* |
| HG0974 | *H. sapiens* | ENSG00000157578 | LCA5L | LCA5L, lebercilin like | 0.03 | 0.23 | 1.3E-02 | 3.3E-02 | 0.23 | 1.3E-02 | | MQTNDHPYADHFEYLQPCFCHCLCRKATCFKKKSSWSAYERVGCFLQLIFPKQTIPPAISGFLKYILSLKRDQNLSKKYYRGKLINTTV* |
| HG0975.1 | *H. sapiens* | ENSG00000182158 | CREB3L2 | cAMP responsive element binding protein 3 like 2 | 0.02 | 0.38 | 1.4E-04 | 6.2E-04 | 0.38 | 1.4E-04 | | MRAGLLPSFWMHIPL* |
| HG0975.2 | *H. sapiens* | ENSG00000157613 | CREB3L1 | cAMP responsive element binding protein 3 like 1 | 0.19 | 0.00 | 7.0E-08 | 5.6E-07 | 0.00 | 7.0E-08 | | MEASCAPGGAGGGGVG* |
| HG0976 | *H. sapiens* | ENSG00000158008 | EXTL1 | exostosin like glycosyltransferase 1 | 0.03 | 0.41 | 7.0E-03 | 1.9E-02 | 0.41 | 7.0E-03 | | MGLSLPICTMTAQD* |
| HG0977 | *H. sapiens* | ENSG00000158321 | AUTS2 | AUTS2, activator of transcription and developmental regula | 0.17 | 0.47 | 3.0E-04 | 1.2E-03 | 0.47 | 3.0E-04 | | MLPSLGGLAPLGSLRSPGLPWLHFLKIWEPGSEFSPRRV* |
| HG0978 | *H. sapiens* | ENSG00000159023 | EPB41 | erythrocyte membrane protein band 4.1 | 0.01 | 0.40 | 2.7E-03 | 8.6E-03 | 0.40 | 2.7E-03 | | MCHCGDGSMASPGLAA* |
| HG0979 | *H. sapiens* | ENSG00000159202 | UBE2Z | ubiquitin conjugating enzyme E2 Z | 0.00 | 0.37 | 1.2E-02 | 3.0E-02 | 0.37 | 1.2E-02 | | MWEEVRVSGKWLFQADI* |
| HG0980 | *H. sapiens* | ENSG00000162670 | BRINP3 | BMP/retinoic acid inducible neural specific 3 | 0.00 | 0.16 | 3.8E-06 | 2.3E-05 | 0.16 | 3.8E-06 | | MIQILGTFFCVLHRFVCHHAVKQVLRES* |
| HG0981 | *H. sapiens* | ENSG00000163629 | PTPN13 | protein tyrosine phosphatase, non-receptor type 13 | 0.00 | 0.48 | 5.5E-03 | 1.6E-02 | 0.48 | 5.5E-03 | | MLTAPGDRRAGEPCSVAPRSASAAVRVAYRS* |
| HG0982 | *H. sapiens* | ENSG00000164116 | GUCY1A3 | guanylate cyclase 1 soluble subunit alpha | 0.00 | 0.47 | 3.0E-04 | 1.2E-03 | 0.47 | 3.0E-04 | | MCGFARRALELLEIRKHSPEVCEATKHPSYQCP* |
| HG0983 | *H. sapiens* | ENSG00000164128 | NPY1R | neuropeptide Y receptor Y1 | 0.00 | 0.12 | 8.7E-04 | 3.2E-03 | 0.12 | 8.7E-04 | | MDSNMGNKNKLNS* |
| HG0984 | *H. sapiens* | ENSG00000164185 | ZNF474 | zinc finger protein 474 | 0.00 | 0.35 | 1.8E-03 | 5.9E-03 | 0.35 | 1.8E-03 | | MLKGKIKRSWRHLSFNEDFPTFQTAVLQ* |
| HG0985 | *H. sapiens* | ENSG00000164604 | GPR85 | G protein-coupled receptor 85 | 0.05 | 0.20 | 8.8E-06 | 5.0E-05 | 0.20 | 8.8E-06 | | MKACLLVLGRRTGKLLLKILYHLHKQNSSEMVK* |
| HG0986 | *H. sapiens* | ENSG00000164663 | USP49 | ubiquitin specific peptidase 49 | 0.00 | 0.39 | 1.0E-04 | 4.7E-04 | 0.39 | 1.0E-04 | | MVRGAGRDGPQNLKDEMGSKGREGAKRSWGSPDGCRTGVDCRGVGIECACYGELLQKVCKGQGRNAVVKDQV* |
| HG0987 | *H. sapiens* | ENSG00000164932 | CTHRC1 | collagen triple helix repeat containing 1 | 0.21 | 0.34 | 6.8E-03 | 1.9E-02 | 0.34 | 6.8E-03 | | MQPAAASERGGARR* |
| HG0988 | *H. sapiens* | ENSG00000166130 | IKBIP | IKBKB interacting protein | 0.07 | 0.46 | 6.6E-04 | 2.5E-03 | 0.46 | 6.6E-04 | | MELAAERLSEATRMPGRGKGLGSGRCQVCEGARSPPRVPTNAKAATSCSPW* |
| HG0989 | *H. sapiens* | ENSG00000166603 | MC4R | melanocortin 4 receptor | 0.00 | 0.34 | 1.5E-02 | 3.7E-02 | 0.34 | 1.5E-02 | | MAASRKIILNRLECIRLKLKQK* |
| HG0990 | *H. sapiens* | ENSG00000166866 | MYO1A | myosin IA | 0.23 | 0.41 | 5.1E-03 | 1.5E-02 | 0.41 | 5.1E-03 | | MKLGHLYTCEGAWAGRGSKRENQKSGHWQEEGSLFPSFLAPWTAPIKPPLHPGARTLMPHSCQIEISSTILQGQ* |
| HG0991 | *H. sapiens* | ENSG00000167654 | ATCAY | ATCAY, caytaxin | 0.16 | 0.37 | 4.4E-04 | 1.7E-03 | 0.37 | 4.4E-04 | | MPSCTRAGLAFVCQ* |
| HG0992 | *H. sapiens* | ENSG00000168000 | BSCL2 | BSCL2, seipin lipid droplet biogenesis associated | 0.11 | 0.21 | 9.2E-05 | 4.2E-04 | 0.21 | 9.2E-05 | | MRHRYQRLDRHFTF* |
| HG0993 | *H. sapiens* | ENSG00000169306 | IL1RAPL1 | interleukin 1 receptor accessory protein like 1 | 0.00 | 0.22 | 1.9E-03 | 6.3E-03 | 0.22 | 1.9E-03 | | MYSEDLQATLQFLS* |
| HG0994 | *H. sapiens* | ENSG00000170852 | KBTBD2 | kelch repeat and BTB domain containing 2 | 0.00 | 0.12 | 1.9E-07 | 1.4E-06 | 0.12 | 1.9E-07 | | MLQITASRFSFISEVFT* |
| HG0995 | *H. sapiens* | ENSG00000171307 | ZDHHC16 | zinc finger DHHC-type containing 16 | 0.10 | 0.26 | 1.6E-03 | 5.4E-03 | 0.26 | 1.6E-03 | | MITADTYLVLPHWQECPCWYL* |
| HG0996 | *H. sapiens* | ENSG00000173890 | GPR160 | G protein-coupled receptor 160 | 0.00 | 0.46 | 9.1E-03 | 2.4E-02 | 0.46 | 9.1E-03 | | MKCRCIELTHIAYWYIKMKCKEPKIT* |
| HG0997 | *H. sapiens* | ENSG00000177728 | TMEM94 | transmembrane protein 94 | 0.07 | 0.00 | 2.6E-04 | 1.1E-03 | 0.00 | 2.6E-04 | | MAAAPGGGDVK* |
| HG0998 | *H. sapiens* | ENSG00000178172 | SPINK6 | serine peptidase inhibitor, Kazal type 6 | 0.00 | 0.07 | 3.4E-06 | 2.1E-05 | 0.07 | 3.4E-06 | | MYWLLGKCRLPVENDILY* |
| HG0999 | *H. sapiens* | ENSG00000178401 | DNAJC22 | DnaJ heat shock protein family (Hsp40) member C22 | 0.06 | 0.15 | 5.1E-06 | 3.0E-05 | 0.15 | 5.1E-06 | | MEVGAGRDGPFSLCGQRK* |
| HG1000 | *H. sapiens* | ENSG00000179083 | FAM133A | family with sequence similarity 133 member A | 0.00 | 0.49 | 1.6E-03 | 5.6E-03 | 0.49 | 1.6E-03 | | MLPIFFSATEQLPWYNKKDEYLLFIERMELA* |
| HG1001 | *H. sapiens* | ENSG00000179761 | PIPOX | pipecolic acid and sarcosine oxidase | 0.00 | 0.00 | 1.2E-03 | 4.3E-03 | 0.00 | 1.2E-03 | | MTIIEPRIESHRDARVRLAAELG* |
| HG1002 | *H. sapiens* | ENSG00000181027 | FKRP | fukutin related protein | 0.02 | 0.44 | 8.7E-05 | 4.0E-04 | 0.44 | 8.7E-05 | | MAAAAAAGAQLSWAGTALLELPQPTT* |
| HG1003 | *H. sapiens* | ENSG00000184867 | ARMCX2 | armadillo repeat containing, X-linked 2 | 0.00 | 0.48 | 2.6E-04 | 1.1E-03 | 0.48 | 2.6E-04 | | MLIHPPGFHCLTSTVYISALQSVLRMKL* |
| HG1004 | *H. sapiens* | ENSG00000185875 | THNSL1 | threonine synthase like 1 | 0.00 | 0.27 | 6.7E-03 | 1.9E-02 | 0.27 | 6.7E-03 | | MFTSQWKLSTTHIN* |
| HG1005 | *H. sapiens* | ENSG00000186472 | PCLO | piccolo presynaptic cytomatrix protein | 0.04 | 0.07 | 7.8E-06 | 4.4E-05 | 0.07 | 7.8E-06 | | MEEAPAAGVTCERD* |
| HG1006.1 | *H. sapiens* | ENSG00000188227 | ZNF793 | zinc finger protein 793 | 0.00 | 0.39 | 7.0E-04 | 2.6E-03 | 0.39 | 7.0E-04 | | MDVTLPVVRKQIY* |
| HG1006.2 | *H. sapiens* | ENSG00000188227 | ZNF793 | zinc finger protein 793 | 0.00 | 0.44 | 2.0E-03 | 6.7E-03 | 0.44 | 2.0E-03 | | MGLFLAKVMTYASVPASQE* |
| HG1007 | *H. sapiens* | ENSG00000188910 | GJB3 | gap junction protein beta 3 | 0.22 | 0.32 | 4.2E-05 | 2.1E-04 | 0.32 | 4.2E-05 | | MATDAPISRLRKRRLRRATSPSGGWPGSAWQRRV* |
| HG1008 | *H. sapiens* | ENSG00000198252 | STYX | serine/threonine/tyrosine interacting protein | 0.16 | 0.31 | 5.7E-03 | 1.6E-02 | 0.31 | 5.7E-03 | | MAAGCKTPDPPLPCLRRRRCWSHWDPVVCVC* |
| HG1009 | *H. sapiens* | ENSG00000198909 | MAP3K3 | mitogen-activated protein kinase kinase kinase 3 | 0.12 | 0.41 | 3.2E-03 | 9.8E-03 | 0.41 | 3.2E-03 | | MPQRPAAGGAGAGAWGGGGGPERSPRPPRGARPAAVPAARAPGMQPRLRR* |
| HG1010 | *H. sapiens* | ENSG00000203740 | METTL11B | methyltransferase like 11B | 0.05 | 0.35 | 3.3E-04 | 1.3E-03 | 0.35 | 3.3E-04 | | MTVSSSFRKRRLIQRNSKASFLSL* |
| HG1011 | *H. sapiens* | ENSG00000204439 | C6orf47 | chromosome 6 open reading frame 47 | 0.00 | 0.27 | 4.1E-03 | 1.2E-02 | 0.27 | 4.1E-03 | | MTIQRREEPLLNSE* |
| HG1012 | *H. sapiens* | ENSG00000213694 | S1PR3 | sphingosine-1-phosphate receptor 3 | 0.04 | 0.30 | 1.4E-03 | 4.8E-03 | 0.30 | 1.4E-03 | | MNVPGAPSRGFWS* |
| HG1013 | *H. sapiens* | ENSG00000214022 | REPIN1 | replication initiator 1 | 0.13 | 0.29 | 8.5E-06 | 4.8E-05 | 0.29 | 8.5E-06 | | MDGRRVPAQPAAEVRLASLQSEV* |
| HG1014.1 | *H. sapiens* | ENSG00000214595 | EML6 | echinoderm microtubule associated protein like 6 | 0.10 | 0.44 | 1.4E-04 | 6.2E-04 | 0.44 | 1.4E-04 | | MQIPDVKGGVAPPRASPGGRSPQVQ* |
| HG1014.2 | *H. sapiens* | ENSG00000214595 | EML6 | echinoderm microtubule associated protein like 6 | 0.23 | 0.30 | 2.4E-03 | 7.8E-03 | 0.30 | 2.4E-03 | | MAELTRVACQIKAIAE* |
| HG1015 | *H. sapiens* | ENSG00000240720 | LRRD1 | leucine rich repeats and death domain containing 1 | 0.04 | 0.39 | 1.1E-02 | 2.8E-02 | 0.39 | 1.1E-02 | | MKIKIKLQFCIGIKHISK* |

| | | | | | | | | | | | | |
|---|---|---|---|---|---|---|---|---|---|---|---|---|
| HG1016 | *H. sapiens* | ENSG00000254004 | ZNF260 | zinc finger protein 260 | 0.00 | 0.06 | 6.6E-07 | 4.5E-06 | 0.06 | 6.6E-07 | | MLHFIPFCNHLISG* |
| HG1017 | *H. sapiens* | ENSG00000255154 | HTD2 | hydroxyacyl-thioester dehydratase type 2 | 0.00 | 0.21 | 2.6E-03 | 8.4E-03 | 0.21 | 2.6E-03 | | MLHSSNSCLFRL* |
| HG1018 | *H. sapiens* | ENSG00000259120 | SMIM6 | small integral membrane protein 6 | 0.12 | 0.47 | 1.6E-02 | 3.9E-02 | 0.47 | 1.6E-02 | | MPFPLNSCFFESGCSSGHLAASSAQPSTKCEPEGPP* |
| HG1019 | *H. sapiens* | ENSG00000019995 | ZRANB1 | zinc finger RANBP2-type containing 1 | 0.00 | 0.22 | 1.1E-03 | 4.0E-03 | 0.22 | 1.1E-03 | | MASLKLNISISGFPLISSFLDQTSLYLLPISFAFQSSVLLNPRV* |
| HG1020 | *H. sapiens* | ENSG00000023902 | PLEKHO1 | pleckstrin homology domain containing O1 | 0.12 | 0.43 | 9.2E-03 | 2.4E-02 | 0.43 | 9.2E-03 | | MEGAKGRGAGVRVCASANAREAGLE* |
| HG1021 | *H. sapiens* | ENSG00000064692 | SNCAIP | synuclein alpha interacting protein | 0.03 | 0.43 | 4.1E-04 | 1.6E-03 | 0.43 | 4.1E-04 | | MPTFRSPPSRVSSPSLCWNPGMVWGPMRWQRLTLGSRNL* |
| HG1022 | *H. sapiens* | ENSG00000065357 | DGKA | diacylglycerol kinase alpha | 0.06 | 0.47 | 3.2E-04 | 1.3E-03 | 0.47 | 7.0E-03 | | MPGSPVPASESLGLHLSPLLYHLHHHPCDPKSLNDSRRDSRQGKLKGPFTPYFWPGPSVPPAKTSRYNFPKSHSGIRAKNGTQI* |
| HG1023 | *H. sapiens* | ENSG00000068001 | HYAL2 | hyaluronoglucosaminidase 2 | 0.01 | 0.45 | 1.3E-02 | 3.3E-02 | 0.45 | 1.3E-02 | | MGGHVPLQRTWGSSSPKRGTGTEKVPELVPGR* |
| HG1024 | *H. sapiens* | ENSG00000073464 | CLCN4 | chloride voltage-gated channel 4 | 0.01 | 0.43 | 3.9E-03 | 1.2E-02 | 0.43 | 3.9E-03 | | MSLSVSASIRPSTQRSRRFR* |
| HG1025 | *H. sapiens* | ENSG00000074621 | SLC24A1 | solute carrier family 24 member 1 | 0.00 | 0.36 | 1.5E-02 | 3.7E-02 | 0.36 | 1.5E-02 | Crowe | MSRSEGLWIPPGWASWRNLL* |
| HG1026.1 | *H. sapiens* | ENSG00000075035 | WSCD2 | WSC domain containing 2 | 0.00 | 0.12 | 2.6E-03 | 8.2E-03 | 0.12 | 2.6E-03 | | MCVLSLKTPLLAQRSSPSVWEGW* |
| HG1026.2 | *H. sapiens* | ENSG00000179314 | WSCD1 | WSC domain containing 1 | 0.00 | 0.32 | 1.5E-02 | 3.6E-02 | 0.32 | 1.5E-02 | | MTPPEALASLPPGR* |
| HG1026.3 | *H. sapiens* | ENSG00000179314 | WSCD1 | WSC domain containing 1 | 0.00 | 0.11 | 2.2E-04 | 9.4E-04 | 0.11 | 2.2E-04 | | MVPFEHLLETGQEQ* |
| HG1027 | *H. sapiens* | ENSG00000075223 | SEMA3C | semaphorin 3C | 0.04 | 0.16 | 2.1E-03 | 6.7E-03 | 0.16 | 2.1E-03 | | MRACRAARRGSRSRAPRRCNRTPRALAPGLRPLSIHSEVLD* |
| HG1028 | *H. sapiens* | ENSG00000076356 | PLXNA2 | plexin A2 | 0.05 | 0.06 | 1.0E-02 | 2.7E-02 | 0.06 | 1.0E-02 | | MNLDSMWIWSSP* |
| HG1029 | *H. sapiens* | ENSG00000079482 | OPHN1 | oligophrenin 1 | 0.06 | 0.46 | 1.0E-02 | 2.7E-02 | 0.46 | 1.0E-02 | | MERGYVSLPPWLNPSGA* |
| HG1030 | *H. sapiens* | ENSG00000082175 | PGR | progesterone receptor | 0.09 | 0.10 | 2.3E-04 | 9.8E-04 | 0.13 | 3.3E-03 | | MRTSFTCHLSEIYNPRRLVLPHYWDLRSSEMTVARSTEPAEVRPFLGMGCTERSD* |
| HG1031 | *H. sapiens* | ENSG00000082269 | FAM135A | family with sequence similarity 135 member A | 0.00 | 0.33 | 2.6E-03 | 8.3E-03 | 0.33 | 2.6E-03 | | MLAGRAGPVAEPR* |
| HG1032 | *H. sapiens* | ENSG00000084453 | SLCO1A2 | solute carrier organic anion transporter family member 1A2 | 0.00 | 0.13 | 1.4E-02 | 3.4E-02 | 0.13 | 4.1E-02 | | MQPSEPKKLNQRFKTYINKDSFIYNSASLMNKKLQLTRLRCLIHTGYSICNFLKNLLVLQYGSKNEW* |
| HG1033 | *H. sapiens* | ENSG00000089225 | TBX5 | T-box 5 | 0.04 | 0.21 | 1.2E-03 | 4.1E-03 | 0.21 | 1.2E-03 | | MPAPPPHRRLPC* |
| HG1034 | *H. sapiens* | ENSG00000090487 | SPG21 | SPG21, maspardin | 0.00 | 0.19 | 3.3E-05 | 1.7E-04 | 0.19 | 3.3E-05 | | MQHLVSILPFMYCQTSTSVKITCLYKNLVTAEWNLSIL* |
| HG1035 | *H. sapiens* | ENSG00000093000 | NUP50 | nucleoporin 50 | 0.11 | 0.30 | 1.8E-03 | 6.1E-03 | 0.30 | 1.8E-03 | | MAARSVFGAVPPAVSG* |
| HG1036 | *H. sapiens* | ENSG00000100813 | ACIN1 | apoptotic chromatin condensation inducer 1 | 0.02 | 0.44 | 8.7E-03 | 2.3E-02 | 0.44 | 8.7E-03 | | MVIPGSRVSEYIRFKASARRRKPSLKAGREKGRCRADGKN* |
| HG1037 | *H. sapiens* | ENSG00000102710 | SUPT20H | SPT20 homolog, SAGA complex component | 0.00 | 0.00 | 1.2E-04 | 5.6E-04 | 0.00 | 1.2E-04 | | MVFTPLHQRT* |
| HG1038 | *H. sapiens* | ENSG00000104331 | IMPAD1 | inositol monophosphatase domain containing 1 | 0.21 | 0.10 | 9.8E-05 | 4.5E-04 | 0.10 | 9.8E-05 | Mackowiak | MGRTGAGERAPV* |
| HG1039 | *H. sapiens* | ENSG00000104804 | TULP2 | tubby like protein 2 | 0.00 | 0.01 | 4.6E-05 | 2.3E-04 | 0.01 | 4.6E-05 | | MFRDWPERQGTEASM* |
| HG1040 | *H. sapiens* | ENSG00000105223 | PLD3 | phospholipase D family member 3 | 0.00 | 0.24 | 5.1E-04 | 2.0E-03 | 0.24 | 5.1E-04 | | MRKAHAWPPPACSLAVVRLRAARRGRQAGIQPGR* |
| HG1041 | *H. sapiens* | ENSG00000106536 | POU6F2 | POU class 6 homeobox 2 | 0.16 | 0.20 | 6.5E-03 | 1.8E-02 | 0.20 | 6.5E-03 | | MNSLGLCYPVCCLLSRC* |
| HG1042 | *H. sapiens* | ENSG00000107249 | GLIS3 | GLIS family zinc finger 3 | 0.00 | 0.13 | 2.3E-03 | 7.3E-03 | 0.13 | 2.3E-03 | | MNRTGEAQSRN* |
| HG1043 | *H. sapiens* | ENSG00000107736 | CDH23 | cadherin related 23 | 0.24 | 0.42 | 5.2E-03 | 1.5E-02 | 0.42 | 5.2E-03 | | MERGVAEPLDVWGAPSPSPRRA* |
| HG1044 | *H. sapiens* | ENSG00000109381 | ELF2 | E74 like ETS transcription factor 2 | 0.00 | 0.11 | 2.4E-03 | 7.7E-03 | 0.11 | 2.4E-03 | | MIMQTYNTGNHQ* |
| HG1045 | *H. sapiens* | ENSG00000110925 | CSRNP2 | cysteine and serine rich nuclear protein 2 | 0.00 | 0.27 | 3.1E-03 | 9.6E-03 | 0.27 | 3.1E-03 | | MGGLEACRIADED* |
| HG1046 | *H. sapiens* | ENSG00000112701 | SENP6 | SUMO1/sentrin specific peptidase 6 | 0.15 | 0.00 | 5.5E-03 | 1.6E-02 | 0.00 | 5.5E-03 | | MHSRLGGVGGRP* |
| HG1047 | *H. sapiens* | ENSG00000112769 | LAMA4 | laminin subunit alpha 4 | 0.11 | 0.46 | 2.0E-02 | 4.6E-02 | 0.46 | 2.0E-02 | | MNVRRKPGQRSPQSRTARERRWPSAGGAHPNPSAF* |
| HG1048 | *H. sapiens* | ENSG00000115295 | CLIP4 | CAP-Gly domain containing linker protein family member 4 | 0.00 | 0.30 | 3.4E-03 | 1.0E-02 | 0.30 | 3.4E-03 | | MKTPISMHSQPTR* |
| HG1049 | *H. sapiens* | ENSG00000116212 | | | 0.00 | 0.39 | 2.7E-03 | 8.4E-03 | 0.39 | 2.7E-03 | | MPRKELELWSNAALRLNSGSATY* |
| HG1050 | *H. sapiens* | ENSG00000116489 | CAPZA1 | capping actin protein of muscle Z-line alpha subunit 1 | 0.00 | 0.13 | 9.0E-04 | 3.3E-03 | 0.13 | 9.0E-04 | | MGNECNSLGKILDFKSPKANPTSWGAGAGAGVRRRHSRA* |
| HG1051 | *H. sapiens* | ENSG00000116852 | KIF21B | kinesin family member 21B | 0.28 | 0.22 | 6.3E-05 | 3.0E-04 | 0.22 | 6.3E-05 | | MARGGARAQPRGAGRG* |
| HG1052 | *H. sapiens* | ENSG00000118640 | VAMP8 | vesicle associated membrane protein 8 | 0.09 | 0.41 | 3.5E-03 | 1.1E-02 | 0.41 | 3.5E-03 | | MRGGRGQAESWSKQEVN* |
| HG1053 | *H. sapiens* | ENSG00000119522 | DENND1A | DENN domain containing 1A | 0.02 | 0.00 | 4.4E-03 | 1.3E-02 | 0.00 | 4.4E-03 | | MAFAGLERVH* |
| HG1054 | *H. sapiens* | ENSG00000120832 | MTERF2 | mitochondrial transcription termination factor 2 | 0.02 | 0.19 | 1.7E-03 | 5.7E-03 | 0.19 | 1.7E-03 | | MGEGTDGLVGAAENCRELLETEDRNPGSRPRGLRTVGRLAGERGKDGAWPPGLPVPCLELTPTVYCRELTRSHFEKKLERIPSG* |
| HG1055 | *H. sapiens* | ENSG00000122299 | ZC3H7A | zinc finger CCCH-type containing 7A | 0.01 | 0.14 | 5.9E-04 | 2.2E-03 | 0.14 | 5.9E-04 | | MSLIAHLESGCVG* |
| HG1056 | *H. sapiens* | ENSG00000128271 | ADORA2A | adenosine A2a receptor | 0.00 | 0.38 | 4.7E-04 | 1.8E-03 | 0.38 | 4.7E-04 | | MMLLPEPLQRAWFQETQSPL* |
| HG1057 | *H. sapiens* | ENSG00000131097 | HIGD1B | HIG1 hypoxia inducible domain family member 1B | 0.11 | 0.40 | 2.2E-03 | 7.2E-03 | 0.40 | 2.2E-03 | | MAQPPEMGYLGGTSFQVEAVLGHGERGS* |
| HG1058 | *H. sapiens* | ENSG00000131386 | GALNT15 | polypeptide N-acetylgalactosaminyltransferase 15 | 0.00 | 0.43 | 1.2E-03 | 4.3E-03 | 0.43 | 1.2E-03 | | MESLFFSQGENRGKGREAIQFEVPVNGLSEGN* |
| HG1059.1 | *H. sapiens* | ENSG00000132970 | WASF3 | WAS protein family member 3 | 0.00 | 0.00 | 3.3E-05 | 1.7E-04 | 0.00 | 3.3E-05 | | MLLEGNFCNP* |
| HG1059.2 | *H. sapiens* | ENSG00000112290 | WASF1 | WAS protein family member 1 | 0.20 | 0.05 | 1.5E-02 | 3.7E-02 | 0.05 | 1.5E-02 | | MMNWNNDERKHIRSQHSRPAL* |
| HG1060 | *H. sapiens* | ENSG00000134242 | PTPN22 | protein tyrosine phosphatase, non-receptor type 22 | 0.00 | 0.37 | 1.7E-03 | 5.6E-03 | 0.37 | 1.7E-03 | | MLCSGGSSGFLEVSRP* |
| HG1061 | *H. sapiens* | ENSG00000134802 | SLC43A3 | solute carrier family 43 member 3 | 0.00 | 0.02 | 1.1E-05 | 6.2E-05 | 0.02 | 1.1E-05 | | MCCLCVLWVLGVTFSLGDGIQNSSAGAPLLHLLPGSYWDPENHPLMVFAQA* |
| HG1062 | *H. sapiens* | ENSG00000137337 | MDC1 | mediator of DNA damage checkpoint 1 | 0.26 | 0.41 | 7.9E-04 | 2.9E-03 | 0.41 | 7.9E-04 | | MSQRTPCLGGPRKGALALPDTSDVYDTAPLSVAGSWRQTLWSRFGTAREVGGGRASGSGLSLGAAQSTCIILLAHGGLQLPSH* |
| HG1063 | *H. sapiens* | ENSG00000138162 | TACC2 | transforming acidic coiled-coil containing protein 2 | 0.02 | 0.15 | 1.3E-02 | 3.3E-02 | 0.15 | 1.3E-02 | | MELCEDAGVGAARR* |
| HG1064 | *H. sapiens* | ENSG00000139734 | DIAPH3 | diaphanous related formin 3 | 0.14 | 0.21 | 7.1E-05 | 3.4E-04 | 0.21 | 7.1E-05 | | MCWLRPRLQRLG* |
| HG1065 | *H. sapiens* | ENSG00000140030 | GPR65 | G protein-coupled receptor 65 | 0.00 | 0.10 | 5.1E-04 | 2.0E-03 | 0.16 | 2.7E-03 | | MQAQIYQAPQSTNTSPEEIWKERNFKRKYQSLCKQSLVYSCLHQSTVRFMKKNKLRTTLYVHLQMPQLMLVGCLSAFCDNEHMDFCLLNSVDPFSQLPGAWIFTSNC* |
| HG1066 | *H. sapiens* | ENSG00000140553 | UNC45A | unc-45 myosin chaperone A | 0.26 | 0.32 | 1.2E-05 | 6.5E-05 | 0.32 | 1.2E-05 | | MGIQRRSGPLLTGLSVGLLLLFSWVTLVQPTSLSALVHHPHLCQTTRDVHGRHYPGFFCPRLSDSPEEAYCCHLQAAGGSCCTRAEFEALYQVNLSALPPPPILRGPGPLLVLGLYNLLVVTLMTVDLVHFCCGRGRSLGWSHRRPPSGSSAASSLQVSAGTA* |
| HG1067 | *H. sapiens* | ENSG00000143340 | FAM163A | family with sequence similarity 163 member A | 0.04 | 0.35 | 1.4E-12 | 2.0E-11 | 0.35 | 1.4E-12 | | MRLHARSRAVLGAPGRPRSLTCAAPAAGARKDALRPRGSRTRESLPERSCSAAGLMPLARRTPNCCLLRCSPRD* |
| HG1068 | *H. sapiens* | ENSG00000144791 | LIMD1 | LIM domains containing 1 | 0.00 | 0.33 | 9.4E-04 | 3.4E-03 | 0.33 | 9.4E-04 | | MPFTLGRWKSLLVLGRT* |
| HG1069 | *H. sapiens* | ENSG00000146414 | SHPRH | SNF2 histone linker PHD RING helicase | 0.21 | 0.19 | 2.7E-03 | 8.6E-03 | 0.19 | 2.7E-03 | | MFGGWRMRARDLLGL* |
| HG1070 | *H. sapiens* | ENSG00000147459 | DOCK5 | dedicator of cytokinesis 5 | 0.25 | 0.42 | 7.3E-03 | 2.0E-02 | 0.42 | 7.3E-03 | | MAEASGARRSAGRRRPEPEEL* |
| HG1071 | *H. sapiens* | ENSG00000147573 | TRIM55 | tripartite motif containing 55 | 0.00 | 0.22 | 4.4E-04 | 1.7E-03 | 0.22 | 4.4E-04 | | MDTSAHRHVTAGC* |
| HG1072 | *H. sapiens* | ENSG00000147874 | HAUS6 | HAUS augmin like complex subunit 6 | 0.15 | 0.25 | 1.4E-04 | 6.0E-04 | 0.25 | 1.4E-04 | | MEGPPRPQADRREGS* |
| HG1073 | *H. sapiens* | ENSG00000149187 | CELF1 | CUGBP Elav-like family member 1 | 0.09 | 0.33 | 1.7E-02 | 4.1E-02 | 0.33 | 1.7E-02 | | MRRGKDVIKCWRNTLIEAASSVAVWYGCRVLCDYTLMYSGSESLRDLFCTLCIVGFGCLFV* |
| HG1074 | *H. sapiens* | ENSG00000150275 | PCDH15 | protocadherin related 15 | 0.00 | 0.41 | 8.8E-03 | 2.3E-02 | 0.41 | 8.8E-03 | | MRVDINASQKQEVEHVNESSWN* |
| HG1075 | *H. sapiens* | ENSG00000150347 | ARID5B | AT-rich interaction domain 5B | 0.00 | 0.00 | 1.0E-02 | 2.6E-02 | 0.00 | 1.0E-02 | | MSNGAAGGGEL* |
| HG1076 | *H. sapiens* | ENSG00000150967 | ABCB9 | ATP binding cassette subfamily B member 9 | 0.05 | 0.17 | 9.1E-04 | 3.3E-03 | 0.17 | 9.1E-04 | | MSSAETAPQTRACCAPPPRAHLVPGNRPGLALWMMMALPP* |
| HG1077 | *H. sapiens* | ENSG00000151474 | FRMD4A | FERM domain containing 4A | 0.28 | 0.48 | 2.8E-03 | 8.9E-03 | 0.48 | 2.8E-03 | | MFLLSMLALPGAELASLLGEEK* |
| HG1078 | *H. sapiens* | ENSG00000162607 | USP1 | ubiquitin specific peptidase 1 | 0.00 | 0.38 | 2.0E-03 | 6.7E-03 | 0.38 | 2.0E-03 | | MRAEGGAPRLRNTPYNW* |
| HG1079 | *H. sapiens* | ENSG00000162975 | KCNF1 | potassium voltage-gated channel modifier subfamily F mem | 0.29 | 0.37 | 2.6E-04 | 1.1E-03 | 0.37 | 2.6E-04 | Crowe | MPAPSPAAGCMPPPRRAPAGCCPL* |
| HG1080 | *H. sapiens* | ENSG00000164168 | TMEM184C | transmembrane protein 184C | 0.03 | 0.04 | 3.8E-05 | 1.9E-04 | 0.04 | 3.8E-05 | Mackowiak | MVQAELRAIPR* |
| HG1081 | *H. sapiens* | ENSG00000165392 | WRN | Werner syndrome RecQ like helicase | 0.01 | 0.45 | 4.2E-03 | 1.3E-02 | 0.45 | 4.2E-03 | | MPGLQALLVVALHSHPAEEDLLDWIFSGFLSDIVLYLPMKTLFFGLCK* |
| HG1082 | *H. sapiens* | ENSG00000165434 | PGM2L1 | phosphoglucomutase 2 like 1 | 0.03 | 0.27 | 1.1E-02 | 2.8E-02 | 0.27 | 1.1E-02 | | MPFSEVGTPGPW* |
| HG1083 | *H. sapiens* | ENSG00000165804 | ZNF219 | zinc finger protein 219 | 0.00 | 0.14 | 2.6E-03 | 8.4E-03 | 0.14 | 2.6E-03 | | MSVRRTAIPGTM* |
| HG1084 | *H. sapiens* | ENSG00000165923 | AGBL2 | ATP/GTP binding protein like 2 | 0.13 | 0.08 | 1.2E-04 | 5.4E-04 | 0.08 | 1.2E-04 | | MGPAPWEEEAGRYAALPVSDAPLASA* |
| HG1085 | *H. sapiens* | ENSG00000166532 | RIMKLB | ribosomal modification protein rimK like family member B | 0.03 | 0.33 | 3.2E-04 | 1.3E-03 | 0.43 | 6.7E-03 | | MNTSRGWGKGTEQSQLFHTLGSRSSFYHLPLGSRHRDIN* |
| HG1086 | *H. sapiens* | ENSG00000167377 | ZNF23 | zinc finger protein 23 | 0.10 | 0.45 | 1.9E-02 | 4.4E-02 | 0.45 | 1.9E-02 | | MLRSGRNINLCASRVWTVCSQAPDIGS* |
| HG1087 | *H. sapiens* | ENSG00000168675 | LDLRAD4 | low density lipoprotein receptor class A domain containing 4 | 0.00 | 0.27 | 3.9E-04 | 1.5E-03 | 0.27 | 3.9E-04 | | MLPEAGPAERWTWTG* |
| HG1088 | *H. sapiens* | ENSG00000169740 | ZNF32 | zinc finger protein 32 | 0.02 | 0.37 | 5.7E-05 | 2.8E-04 | 0.37 | 5.7E-05 | | MSRPGRPGPRGRPAASGEVGGPQSRVPWTGERDAEVLRNSPWKNHMKRW* |
| HG1089 | *H. sapiens* | ENSG00000169813 | HNRNPF | heterogeneous nuclear ribonucleoprotein F | 0.00 | 0.12 | 1.2E-02 | 3.1E-02 | 0.12 | 1.2E-02 | | MHLEFPRSPLRG* |
| HG1090.1 | *H. sapiens* | ENSG00000170458 | CD14 | CD14 molecule | 0.09 | 0.10 | 1.9E-03 | 6.3E-03 | 0.10 | 1.9E-03 | | MPCRILPVTVPLPETSFIAIFPGKEGGWLGGREVGR* |
| HG1090.2 | *H. sapiens* | ENSG00000170458 | CD14 | CD14 molecule | 0.05 | 0.37 | 1.1E-02 | 2.7E-02 | 0.37 | 1.1E-02 | | MRFTISSTKHAVKYLRIFRDLDLVAGDQHKPRQGRSQRNESSRFSGI* |
| HG1091 | *H. sapiens* | ENSG00000172183 | ISG20 | interferon stimulated exonuclease gene 20 | 0.06 | 0.32 | 7.7E-05 | 3.6E-04 | 0.32 | 7.7E-05 | | MRHLRRGRAGAKEGHPDMEPASSVSPDSARS* |
| HG1092 | *H. sapiens* | ENSG00000173327 | MAP3K11 | mitogen-activated protein kinase kinase kinase 11 | 0.00 | 0.21 | 4.6E-05 | 2.3E-04 | 0.21 | 4.6E-05 | | MVRRRRPRGPTPPA* |
| HG1093 | *H. sapiens* | ENSG00000174371 | EXO1 | exonuclease 1 | 0.00 | 0.13 | 4.9E-05 | 2.4E-04 | 0.40 | 2.7E-02 | | MYTVILTIALSHSYYAKEEIIIRGYLPGPEETYVNFMNYYIRFPRSERKLFLDII* |
| HG1094 | *H. sapiens* | ENSG00000177917 | ARL6IP6 | ADP ribosylation factor like GTPase 6 interacting protein 6 | 0.11 | 0.20 | 4.4E-05 | 2.2E-04 | 0.21 | 3.8E-04 | | MGEPGEECERSGEEGRRQIGSGMCLRGRERALGPGVEKMRLWGCRGARSRW* |
| HG1095 | *H. sapiens* | ENSG00000179921 | GPBAR1 | G protein-coupled bile acid receptor 1 | 0.29 | 0.46 | 1.2E-02 | 3.1E-02 | 0.46 | 1.2E-02 | | MHLVHLCSSARTPDMVQTAAGLAAATP* |
| HG1096 | *H. sapiens* | ENSG00000181544 | FANCB | Fanconi anemia complementation group B | 0.04 | 0.20 | 1.0E-02 | 2.6E-02 | 0.20 | 1.0E-02 | | MDTVSTWGLRYVAAALRFRNINWNQFAFEN* |
| HG1097 | *H. sapiens* | ENSG00000182481 | KPNA2 | karyopherin subunit alpha 2 | 0.11 | 0.15 | 5.8E-05 | 2.8E-04 | 0.15 | 5.8E-05 | | MPSSFFFFLSPLPLAPKPYDSLPSQRVFQIHQWAHSLGSNQPIGMRSHREI* |
| HG1098 | *H. sapiens* | ENSG00000183775 | KCTD16 | potassium channel tetramerization domain containing 16 | 0.00 | 0.49 | 1.4E-02 | 3.4E-02 | 0.49 | 1.4E-02 | | MKSCARKLWFEHGQFSPTVSYIHKHHMKWRSGSSVYFSQVPQ* |
| HG1099 | *H. sapiens* | ENSG00000186334 | SLC36A3 | solute carrier family 36 member 3 | 0.08 | 0.42 | 3.9E-04 | 1.5E-03 | 0.42 | 3.9E-04 | | MVSSSSQSHCSLCPTAARAKLGEAKSPGKSWLGKQLRVKLRTILTEMGLS* |
| HG1100 | *H. sapiens* | ENSG00000187815 | ZFP69 | ZFP69 zinc finger protein | 0.00 | 0.38 | 1.1E-02 | 2.9E-02 | 0.38 | 1.1E-02 | | MNSNLTNWLPPKESA* |
| HG1101 | *H. sapiens* | ENSG00000189334 | S100A14 | S100 calcium binding protein A14 | 0.04 | 0.04 | 7.7E-05 | 3.6E-04 | 0.04 | 7.7E-05 | | MSHQLLWGQL* |
| HG1102 | *H. sapiens* | ENSG00000196704 | AMZ2 | archaelysin family metallopeptidase 2 | 0.04 | 0.00 | 3.5E-04 | 1.4E-03 | 0.00 | 3.5E-04 | | MKSHSTTFFQPGPDMSVLVS* |
| HG1103 | *H. sapiens* | ENSG00000197620 | CXorf40A | chromosome X open reading frame 40A | 0.12 | 0.39 | 1.0E-03 | 3.6E-03 | 0.39 | 1.0E-03 | | MLGCKNAAQHGEMIFCPAGWCTLWSLTQTGAEGPTAVISRTKL* |
| HG1104 | *H. sapiens* | ENSG00000205155 | PSENEN | presenilin enhancer gamma-secretase subunit | 0.12 | 0.29 | 2.8E-03 | 8.9E-03 | 0.29 | 2.8E-03 | | MRAVLPRPKKTTISRETWGVSRKRP* |
| HG1105 | *H. sapiens* | ENSG00000213654 | GPSM3 | G protein signaling modulator 3 | 0.23 | 0.15 | 3.3E-04 | 1.3E-03 | 0.15 | 3.3E-04 | | MRPPGVLPHHEPPPLLQLEEFFPVSVLPWGERD* |
| HG1106 | *H. sapiens* | ENSG00000215021 | PHB2 | prohibitin 2 | 0.19 | 0.46 | 7.2E-03 | 2.0E-02 | 0.46 | 7.2E-03 | | MEMTCCLALPSLGGQPSRKGRDFRMRDSCARSSGP* |
| HG1107 | *H. sapiens* | ENSG00000217930 | PAM16 | presequence translocase associated motor 16 homolog | 0.19 | 0.26 | 1.1E-03 | 3.8E-03 | 0.26 | 1.1E-03 | | MRMPSAVSRTRSLRLSLTSDPPTTLDPRPRGQEVGV* |
| HG1108 | *H. sapiens* | ENSG00000251247 | ZNF345 | zinc finger protein 345 | 0.07 | 0.41 | 2.4E-03 | 7.8E-03 | 0.41 | 2.4E-03 | | MAVEHPQGTCKTSNFTLELSRNMACLFPQISLGLQTKGWNFEPISRLLVTVPEERQSYGYYESKLRPRNYF* |
| HG1109 | *H. sapiens* | ENSG00000278311 | GGNBP2 | gametogenetin binding protein 2 | 0.14 | 0.36 | 1.7E-04 | 7.3E-04 | 0.36 | 1.7E-04 | | MGNAARLSRTLPLRSLPPVANVLRSGGCLRSRGLAVAGRSWEEAAAAAAETAAAAAAAAGRRW* |
| HG1110 | *H. sapiens* | ENSG00000034152 | MAP2K3 | mitogen-activated protein kinase kinase 3 | 0.13 | 0.04 | 8.8E-03 | 2.4E-02 | 0.04 | 8.8E-03 | | MSVLGPGGARSPLD* |
| HG1111 | *H. sapiens* | ENSG00000040531 | CTNS | cystinosin, lysosomal cystine transporter | 0.05 | 0.05 | 1.4E-03 | 4.7E-03 | 0.05 | 1.4E-03 | | MEGGLSFAHERSREALAAPRVSVSLAADLIFPHAGAPISAPRPDPAALCPSKTRGNYNSQSSSRRDPAPRVRWRRSGDSGPASPKV* |
| HG1112 | *H. sapiens* | ENSG00000068885 | IFT80 | intraflagellar transport 80 | 0.00 | 0.05 | 2.3E-03 | 7.5E-03 | 0.05 | 2.3E-03 | | MVTGRYQVFESNRHRFPGT* |
| HG1113 | *H. sapiens* | ENSG00000072201 | LNX1 | ligand of numb-protein X 1 | 0.00 | 0.09 | 1.5E-02 | 3.7E-02 | 0.09 | 1.5E-02 | | MPSSFLQAPVHRS* |
| HG1114 | *H. sapiens* | ENSG00000073737 | DHRS9 | dehydrogenase/reductase 9 | 0.00 | 0.24 | 5.5E-03 | 1.6E-02 | 0.24 | 5.5E-03 | | MGVTHRARREPEDSSHM* |

| | | | | | | | | | | | | |
|---|---|---|---|---|---|---|---|---|---|---|---|---|
| HG1115 | *H. sapiens* | ENSG00000074219 | TEAD2 | TEA domain transcription factor 2 | 0.26 | 0.24 | 2.9E-03 | 9.2E-03 | 0.24 | 2.9E-03 | | MAGRPCPPACGSVGLSWSRPQLLP* |
| HG1116 | *H. sapiens* | ENSG00000084090 | STARD7 | StAR related lipid transfer domain containing 7 | 0.18 | 0.37 | 5.9E-03 | 1.7E-02 | 0.37 | 5.9E-03 | | MSGGSWPRDRETGSWPAI* |
| HG1117 | *H. sapiens* | ENSG00000089902 | RCOR1 | REST corepressor 1 | 0.05 | 0.29 | 8.8E-03 | 2.3E-02 | 0.29 | 8.8E-03 | | MRAKVALGSSLGA* |
| HG1118 | *H. sapiens* | ENSG00000090975 | PITPNM2 | phosphatidylinositol transfer protein membrane associated | 0.24 | 0.38 | 3.4E-03 | 1.1E-02 | 0.38 | 3.4E-03 | | MESMVPTPQLVSMLFPESPTIHLLSF* |
| HG1119 | *H. sapiens* | ENSG00000092020 | PPP2R3C | protein phosphatase 2 regulatory subunit B"gamma | 0.00 | 0.26 | 2.3E-04 | 9.9E-04 | 0.39 | 2.8E-02 | | MKRNCERRRFPQAGTNELLGHVVRTRQRHTIKENAGLSWPLKSGLPRLPPSIKMVVGRPS* |
| HG1120 | *H. sapiens* | ENSG00000095585 | BLNK | B-cell linker | 0.02 | 0.00 | 7.3E-03 | 2.0E-02 | 0.00 | 7.3E-03 | | MLLPRRARNC* |
| HG1121 | *H. sapiens* | ENSG00000100350 | FOXRED2 | FAD dependent oxidoreductase domain containing 2 | 0.00 | 0.35 | 2.4E-03 | 7.6E-03 | 0.35 | 2.4E-03 | | MEGLGIPGSPKSQLETVESGQMSSEDGFWAWP* |
| HG1122 | *H. sapiens* | ENSG00000100410 | PHF5A | PHD finger protein 5A | 0.04 | 0.21 | 5.8E-03 | 1.6E-02 | 0.21 | 5.8E-03 | | MGVKGRTEVPEASGGRLS* |
| HG1123 | *H. sapiens* | ENSG00000100884 | CPNE6 | copine 6 | 0.00 | 0.25 | 1.3E-02 | 3.2E-02 | 0.25 | 1.3E-02 | | MCEPVSRGCMCRHHFSV* |
| HG1124 | *H. sapiens* | ENSG00000100934 | SEC23A | Sec23 homolog A, coat complex II component | 0.00 | 0.38 | 1.5E-02 | 3.6E-02 | 0.38 | 1.5E-02 | | MLAGRRSSLRPGAGWRHP* |
| HG1125 | *H. sapiens* | ENSG00000101574 | METTL4 | methyltransferase like 4 | 0.01 | 0.19 | 5.4E-05 | 2.6E-04 | 0.21 | 2.5E-03 | | MNDKFSSMCMSGFCTCISNLSVCMKCMFIEIQDI* |
| HG1126 | *H. sapiens* | ENSG00000101890 | GUCY2F | guanylate cyclase 2F, retinal | 0.00 | 0.22 | 1.0E-02 | 2.7E-02 | 0.22 | 1.0E-02 | | MEFVNRLESLTGL* |
| HG1127 | *H. sapiens* | ENSG00000103064 | SLC7A6 | solute carrier family 7 member 6 | 0.04 | 0.15 | 2.1E-03 | 6.8E-03 | 0.15 | 2.1E-03 | | MGGRAGSGAGAGPRSVGAEPCGSSLPPSGAGCGPETARGWPIGAAREALRSLCGRGRQVELGLAAGDFISFSVLDPFMRRWLMKV* |
| HG1128 | *H. sapiens* | ENSG00000103852 | TTC23 | tetratricopeptide repeat domain 23 | 0.00 | 0.05 | 1.9E-02 | 4.6E-02 | 0.05 | 1.9E-02 | | MVINLYKESPN* |
| HG1129 | *H. sapiens* | ENSG00000104432 | IL7 | interleukin 7 | 0.00 | 0.00 | 7.3E-03 | 2.0E-02 | 0.00 | 7.3E-03 | | MLMIPKIEPADQAQSRN* |
| HG1130 | *H. sapiens* | ENSG00000105559 | PLEKHA4 | pleckstrin homology domain containing A4 | 0.02 | 0.17 | 6.5E-03 | 1.8E-02 | 0.17 | 6.5E-03 | | MSERETGRDRDQRETQKVRVGREVV* |
| HG1131 | *H. sapiens* | ENSG00000106780 | MEGF9 | multiple EGF like domains 9 | 0.05 | 0.50 | 1.8E-02 | 4.3E-02 | 0.50 | 1.8E-02 | | MVRPVAVAAAVAEASLVGLQRGEDD* |
| HG1132 | *H. sapiens* | ENSG00000109771 | LRP2BP | LRP2 binding protein | 0.00 | 0.25 | 3.7E-03 | 1.1E-02 | 0.25 | 3.7E-03 | | MKTLIYHSTVIHLITWQSNTDRKIKLNNS* |
| HG1133 | *H. sapiens* | ENSG00000111328 | CDK2AP1 | cyclin dependent kinase 2 associated protein 1 | 0.09 | 0.37 | 1.8E-02 | 4.2E-02 | 0.37 | 1.8E-02 | | MSSIGYSNPGAWPDLGVWNDLRA* |
| HG1134 | *H. sapiens* | ENSG00000112984 | KIF20A | kinesin family member 20A | 0.00 | 0.10 | 6.2E-54 | 3.6E-52 | 0.10 | 6.2E-54 | | MSFFPLRQSKHPKPVTLCTPVKIVAKEGQELAAEASRFPLGS* |
| HG1135 | *H. sapiens* | ENSG00000113597 | TRAPPC13 | trafficking protein particle complex 13 | 0.00 | 0.06 | 1.4E-03 | 4.7E-03 | 0.06 | 1.4E-03 | | MENKVFYAYVSKIEEMDLLKRSQFVKDTVGEFKVPPPERG* |
| HG1136 | *H. sapiens* | ENSG00000113749 | HRH2 | histamine receptor H2 | 0.00 | 0.32 | 1.1E-02 | 2.8E-02 | 0.32 | 1.1E-02 | | MGAGTSYGEGYSCVST* |
| HG1137 | *H. sapiens* | ENSG00000115307 | AUP1 | ancient ubiquitous protein 1 | 0.22 | 0.31 | 9.6E-03 | 2.5E-02 | 0.31 | 9.6E-03 | | MPGIVVSAGREL* |
| HG1138 | *H. sapiens* | ENSG00000116871 | MAP7D1 | MAP7 domain containing 1 | 0.16 | 0.17 | 2.8E-03 | 8.8E-03 | 0.17 | 2.8E-03 | | MRRGTPVLATGRRRRRR* |
| HG1139.1 | *H. sapiens* | ENSG00000118308 | LRMP | lymphoid restricted membrane protein | 0.19 | 0.44 | 1.2E-02 | 3.0E-02 | 0.44 | 1.2E-02 | | MQTQNCRSSGEAYHFPEKKNTFSVLPAS* |
| HG1139.2 | *H. sapiens* | ENSG00000118308 | LRMP | lymphoid restricted membrane protein | 0.00 | 0.00 | 1.8E-03 | 6.1E-03 | 0.00 | 1.8E-03 | | MKEATLQKEAT* |
| HG1140 | *H. sapiens* | ENSG00000118495 | PLAGL1 | PLAG1 like zinc finger 1 | 0.00 | 0.16 | 2.8E-03 | 8.8E-03 | 0.16 | 2.8E-03 | | MLPLPWSGVQLQGLC* |
| HG1141 | *H. sapiens* | ENSG00000121964 | GTDC1 | glycosyltransferase like domain containing 1 | 0.00 | 0.23 | 3.3E-03 | 1.0E-02 | 0.23 | 3.3E-03 | | MHREPKSVTEIVWVSLH* |
| HG1142 | *H. sapiens* | ENSG00000124613 | ZNF391 | zinc finger protein 391 | 0.18 | 0.37 | 1.2E-02 | 2.9E-02 | 0.37 | 1.2E-02 | | MLDLQEAEEGSAQKYLS* |
| HG1143 | *H. sapiens* | ENSG00000125245 | GPR18 | G protein-coupled receptor 18 | 0.00 | 0.24 | 1.2E-02 | 3.0E-02 | 0.24 | 1.2E-02 | | MPSFQGELMLSVTS* |
| HG1144 | *H. sapiens* | ENSG00000128203 | ASPHD2 | aspartate beta-hydroxylase domain containing 2 | 0.27 | 0.00 | 2.3E-03 | 7.4E-03 | 0.00 | 2.3E-03 | | MPPPAPAAPSPPR* |
| HG1145 | *H. sapiens* | ENSG00000128585 | MKLN1 | muskelin 1 | 0.27 | 0.04 | 7.1E-03 | 2.0E-02 | 0.04 | 7.1E-03 | | MLENMGIGIRFWRRNVS* |
| HG1146 | *H. sapiens* | ENSG00000130413 | STK33 | serine/threonine kinase 33 | 0.02 | 0.15 | 4.5E-03 | 1.3E-02 | 0.15 | 4.5E-03 | | MVISFLSAALTCEYVSSASLT* |
| HG1147 | *H. sapiens* | ENSG00000131781 | FMO5 | flavin containing monooxygenase 5 | 0.00 | 0.33 | 1.7E-02 | 4.1E-02 | 0.33 | 1.7E-02 | | MPRPRSSLKNVNRCVLFVFCHPFFRTGDTNR* |
| HG1148 | *H. sapiens* | ENSG00000132938 | MTUS2 | microtubule associated scaffold protein 2 | 0.00 | 0.41 | 1.8E-02 | 4.3E-02 | 0.41 | 1.8E-02 | | MAVKLIFWGIGSHGLIGSHSAGT* |
| HG1149 | *H. sapiens* | ENSG00000132975 | GPR12 | G protein-coupled receptor 12 | 0.00 | 0.00 | 2.9E-03 | 9.1E-03 | 0.00 | 2.9E-03 | | MILDCHPAVFKH* |
| HG1150 | *H. sapiens* | ENSG00000135299 | ANKRD6 | ankyrin repeat domain 6 | 0.01 | 0.37 | 1.6E-02 | 3.8E-02 | 0.37 | 1.6E-02 | | MQTSFLCHLSVIQKCSSDL* |
| HG1151 | *H. sapiens* | ENSG00000137168 | PPIL1 | peptidylprolyl isomerase like 1 | 0.24 | 0.46 | 4.1E-03 | 1.2E-02 | 0.46 | 4.1E-03 | | MKFAFSSVLEPSFFSSPTPASRHWKKGPSSSLSWPPKSWYRKG* |
| HG1152 | *H. sapiens* | ENSG00000138658 | ZGRF1 | zinc finger GRF-type containing 1 | 0.01 | 0.41 | 1.6E-02 | 3.8E-02 | 0.41 | 1.6E-02 | | MTSRCFAGEIRKWILCNLTGGINS* |
| HG1153 | *H. sapiens* | ENSG00000138738 | PRDM5 | PR/SET domain 5 | 0.16 | 0.05 | 2.3E-03 | 7.4E-03 | 0.05 | 2.3E-03 | | MCRLPGAKRRC* |
| HG1154 | *H. sapiens* | ENSG00000139725 | RHOF | ras homolog family member F, filopodia associated | 0.00 | 0.21 | 8.9E-03 | 2.4E-02 | 0.21 | 8.9E-03 | | MAKPKDSQLEDWRRLFKKIWIFPYDLESIFVLYN* |
| HG1155 | *H. sapiens* | ENSG00000141668 | CBLN2 | cerebellin 2 precursor | 0.08 | 0.27 | 7.4E-03 | 2.0E-02 | 0.27 | 7.4E-03 | | MKVVTPFNIKTWRELGKEKKESKRREATGAFKGV* |
| HG1156 | *H. sapiens* | ENSG00000144741 | SLC25A26 | solute carrier family 25 member 26 | 0.18 | 0.18 | 8.8E-03 | 2.3E-02 | 0.18 | 8.8E-03 | | MAAPSARGRDPLLLRLGL* |
| HG1157 | *H. sapiens* | ENSG00000148123 | PLPPR1 | phospholipid phosphatase related 1 | 0.03 | 0.15 | 1.4E-02 | 3.4E-02 | 0.15 | 1.4E-02 | | MYLAPGSRTAQ* |
| HG1158 | *H. sapiens* | ENSG00000149571 | KIRREL3 | kin of IRRE like 3 (Drosophila) | 0.00 | 0.24 | 3.1E-03 | 9.6E-03 | 0.24 | 3.1E-03 | | MVGANPRGPAEAEHREPPRKRN* |
| HG1159 | *H. sapiens* | ENSG00000152822 | GRM1 | glutamate metabotropic receptor 1 | 0.00 | 0.26 | 6.2E-03 | 1.8E-02 | 0.26 | 6.2E-03 | | MSSKQTQHLAHRCQHDFHCTLDQFTLMHYR* |
| HG1160 | *H. sapiens* | ENSG00000154162 | CDH12 | cadherin 12 | 0.29 | 0.15 | 1.6E-02 | 3.9E-02 | 0.15 | 1.6E-02 | | MMCQRHAVFCFEGGEMTGGNPGPPLARRSPLQLSSSRSESIIDLKGKCGGSHREGSRLF* |
| HG1161 | *H. sapiens* | ENSG00000154227 | CERS3 | ceramide synthase 3 | 0.13 | 0.35 | 1.4E-02 | 3.5E-02 | 0.35 | 1.4E-02 | | MHRCGDPPPGLADPPRPRTEEEVVKQEATSHQEEPEGAIQ* |
| HG1162 | *H. sapiens* | ENSG00000155380 | SLC16A1 | solute carrier family 16 member 1 | 0.00 | 0.22 | 7.6E-03 | 2.1E-02 | 0.22 | 7.6E-03 | | MQVFCSKYLDLTCILEFIYT* |
| HG1163 | *H. sapiens* | ENSG00000157637 | SLC38A10 | solute carrier family 38 member 10 | 0.01 | 0.08 | 2.3E-03 | 7.6E-03 | 0.08 | 2.3E-03 | | MALRRPQGLAGSGG* |
| HG1164 | *H. sapiens* | ENSG00000162639 | HENMT1 | HEN1 methyltransferase homolog 1 | 0.20 | 0.29 | 8.7E-03 | 2.3E-02 | 0.29 | 8.7E-03 | | MASGHAPQRPLRDAKPTPWN* |
| HG1165 | *H. sapiens* | ENSG00000162694 | EXTL2 | exostosin like glycosyltransferase 2 | 0.05 | 0.08 | 2.2E-03 | 7.2E-03 | 0.08 | 2.2E-03 | | MESPLTPTRPPTTSVPGCMGSSACPFAISSQLALNPGVVDISD* |
| HG1166 | *H. sapiens* | ENSG00000165071 | TMEM71 | transmembrane protein 71 | 0.02 | 0.29 | 8.4E-03 | 2.3E-02 | 0.29 | 8.4E-03 | | MCVGICEKKEASYFSNNFVLCQLEEEAGNSLWKEHRICEV* |
| HG1167 | *H. sapiens* | ENSG00000165934 | CPSF2 | cleavage and polyadenylation specific factor 2 | 0.00 | 0.08 | 8.6E-03 | 2.3E-02 | 0.08 | 8.6E-03 | | MGFKLPAVLRQPINKELFRIPGVSSYIRLRYQLF* |
| HG1168 | *H. sapiens* | ENSG00000166669 | ATF7IP2 | activating transcription factor 7 interacting protein 2 | 0.00 | 0.36 | 1.5E-02 | 3.8E-02 | 0.36 | 1.5E-02 | | MTCDITICKTNGEDQDKDKLYHRKAIENAKMKHLNEGLITLNSAFVNLICRYVQTTS* |
| HG1169 | *H. sapiens* | ENSG00000167637 | ZNF283 | zinc finger protein 283 | 0.08 | 0.33 | 3.5E-03 | 1.1E-02 | 0.33 | 3.5E-03 | | MTARKKSMLSPIMNVSSKKDENFRSGSVRGTGSCQDLMGKF* |
| HG1170 | *H. sapiens* | ENSG00000167858 | TEKT1 | tektin 1 | 0.19 | 0.37 | 1.4E-02 | 3.4E-02 | 0.37 | 1.4E-02 | | METALFTSLFCKSWKLQLG* |
| HG1171 | *H. sapiens* | ENSG00000167968 | DNASE1L2 | deoxyribonuclease 1 like 2 | 0.06 | 0.15 | 1.5E-03 | 5.0E-03 | 0.15 | 1.5E-03 | | MLGRECGWSYLRCLAFPEGILLREAESRRPNF* |
| HG1172 | *H. sapiens* | ENSG00000170381 | SEMA3E | semaphorin 3E | 0.00 | 0.04 | 7.2E-03 | 2.0E-02 | 0.04 | 7.2E-03 | | MNSFAQVTEKYSFFLNLPA* |
| HG1173 | *H. sapiens* | ENSG00000170464 | DNAJC18 | DnaJ heat shock protein family (Hsp40) member C18 | 0.24 | 0.37 | 1.0E-02 | 2.6E-02 | 0.37 | 1.0E-02 | | MGQHLNHIGHRTDLSITSKNPLQFNM* |
| HG1174 | *H. sapiens* | ENSG00000173918 | C1QTNF1 | C1q and TNF related 1 | 0.30 | 0.43 | 2.3E-147 | 3.4E-145 | 0.43 | 2.3E-147 | | MGGHSRQGKKPRFRFWSQRPRRPRRGDGGAVGGGDLWLLVPPPTLLFCTAVLRKTFSPALFPSPSLCIAPDLAGRRLGRREML* |
| HG1175 | *H. sapiens* | ENSG00000174473 | GALNTL6 | polypeptide N-acetylgalactosaminyltransferase-like 6 | 0.00 | 0.12 | 2.2E-03 | 7.0E-03 | 0.12 | 2.2E-03 | | MVLWVRSVARKEW* |
| HG1176 | *H. sapiens* | ENSG00000176472 | ZNF575 | zinc finger protein 575 | 0.00 | 0.02 | 5.4E-03 | 1.6E-02 | 0.02 | 5.4E-03 | | MKFNGKKGAVDAASMN* |
| HG1177 | *H. sapiens* | ENSG00000178295 | GEN1 | GEN1, Holliday junction 5' flap endonuclease | 0.03 | 0.38 | 9.7E-03 | 2.5E-02 | 0.38 | 9.7E-03 | | MTKGLLPEPRRKGCPVCFT* |
| HG1178 | *H. sapiens* | ENSG00000182670 | TTC3 | tetratricopeptide repeat domain 3 | 0.00 | 0.01 | 7.6E-03 | 2.1E-02 | 0.01 | 7.6E-03 | | MCMRNITCTSNL* |
| HG1179 | *H. sapiens* | ENSG00000185933 | CALHM1 | calcium homeostasis modulator 1 | 0.26 | 0.36 | 1.6E-02 | 3.9E-02 | 0.36 | 1.6E-02 | | MRWAPSGQGGQQGPSWRQQ* |
| HG1180 | *H. sapiens* | ENSG00000188674 | | | 0.19 | 0.28 | 4.2E-03 | 1.3E-02 | 0.28 | 4.2E-03 | Mackowiak | MLQHASRSRERTKALEPVSGCL* |
| HG1181 | *H. sapiens* | ENSG00000189058 | APOD | apolipoprotein D | 0.00 | 0.43 | 1.2E-02 | 3.1E-02 | 0.43 | 1.2E-02 | | MDTTSHLCVSIQTKLLGQPS* |
| HG1182 | *H. sapiens* | ENSG00000196155 | PLEKHG4 | pleckstrin homology and RhoGEF domain containing G4 | 0.00 | 0.00 | 7.2E-03 | 2.0E-02 | 0.00 | 7.2E-03 | | MEQDNSKAPLWQPDLLMKKAPR* |
| HG1183 | *H. sapiens* | ENSG00000198133 | TMEM229B | transmembrane protein 229B | 0.00 | 0.16 | 1.5E-02 | 3.7E-02 | 0.16 | 1.5E-02 | | MKVACRDGKCCLRRDTWKLPAPQPRIS* |
| HG1184 | *H. sapiens* | ENSG00000198894 | CIPC | CLOCK interacting pacemaker | 0.01 | 0.16 | 1.4E-02 | 3.5E-02 | 0.16 | 1.4E-02 | | MGRAVQMKRVPMNLPPAE* |
| HG1185 | *H. sapiens* | ENSG00000198920 | KIAA0753 | KIAA0753 | 0.00 | 0.11 | 2.1E-03 | 6.8E-03 | 0.11 | 2.1E-03 | | MCLLPSAFFMCATALEIRTLRQLSLCSQEVSVFKQI* |
| HG1186 | *H. sapiens* | ENSG00000204923 | | | 0.00 | 0.21 | 6.5E-03 | 1.8E-02 | 0.21 | 6.5E-03 | | MGATVVLASPKT* |
| HG1187 | *H. sapiens* | ENSG00000213762 | ZNF134 | zinc finger protein 134 | 0.29 | 0.01 | 3.7E-03 | 1.1E-02 | 0.01 | 3.7E-03 | | MAAAAVMGPAQILCGC* |
| HG1188 | *H. sapiens* | ENSG00000214415 | GNAT3 | G protein subunit alpha transducin 3 | 0.04 | 0.22 | 4.5E-03 | 1.3E-02 | 0.22 | 4.5E-03 | | MSKPQLCSTFEKSEHIN* |
| HG1189 | *H. sapiens* | ENSG00000279909 | | | 0.00 | 0.34 | 7.0E-03 | 1.9E-02 | 0.34 | 7.0E-03 | | MELTSTTELFTLHCKYTYVCPPSQLSCQCLE* |
| HG1190 | *H. sapiens* | ENSG00000123684 | LPGAT1 | lysophosphatidylglycerol acyltransferase 1 | 0.11 | 0.41 | 1.3E-02 | 3.2E-02 | 0.41 | 1.3E-02 | | MHRCAGIVVVRRLRSSVSGKSY* |
| HG1191 | *H. sapiens* | ENSG00000164331 | ANKRA2 | ankyrin repeat family A member 2 | 0.05 | 0.11 | 5.2E-03 | 1.5E-02 | 0.11 | 5.2E-03 | | MSESFFLGLFGHSWPVATERTFFPARVSPPAGDSGLLSRDAGTPVAYF* |
| HG1192 | *H. sapiens* | ENSG00000168758 | SEMA4C | semaphorin 4C | 0.29 | 0.49 | 1.4E-02 | 3.4E-02 | 0.49 | 1.4E-02 | | MWNGVWGSAGLLSALGKGTGRDRDLRSQPAAPAGQLA* |
| HG1193 | *H. sapiens* | ENSG00000177706 | FAM20C | FAM20C, golgi associated secretory pathway kinase | 0.29 | 0.47 | 2.1E-02 | 4.9E-02 | 0.47 | 2.1E-02 | | MDLDPRGGAALVPSCS* |
| HG1194 | *H. sapiens* | ENSG00000214694 | ARHGEF33 | Rho guanine nucleotide exchange factor 33 | 0.00 | 0.19 | 6.5E-03 | 1.8E-02 | 0.19 | 6.5E-03 | | MGTSRENKKKTSKNML* |

[†] The gene description is based on information contained in Ensembl (https://www.ensembl.org/index.html) and Ensembl Metazoa (https://metazoa.ensembl.org/index.html) .

The fusion ratio (FR) was calculated as shown in Supplementary Fig. S2

[*] Akimoto: Akimoto et al., Biochim.Biophys. Acta (2013); Andreev: Andreev et al., eLife (2015); Crowe: Crowe et al., BMC Genomics (2006); Hayden: Hayden et al., BMC Genomics (2008); Jousse: Jousse et al., Nucleic Acids Res. (2001); Mackowiak: Mackowiak et al., Genome Biology (2015); Reynolds: Reynolds et al., J. Cell. Biol. (1996); Samandi: Samandiet al., eLife (2017); Zhao: Zhao et al., J Biol. Chem. (2010)
