## Supplementary Table S2 for "Exhaustive identification of conserved upstream open reading frames with potential translational regulatory functions from animal genomes"

Supplementary Table S2. Taxonomic range of sequence conservation of the CPuORFs.

| HG number | Species | Gene ID | Taxonomic category ** | | | | | | | | | | | | | | | | | | |
|---|---|---|---|---|---|---|---|---|---|---|---|---|---|---|---|---|---|---|---|---|---|
| | | | Metazoa | | | | | | | | | | | | | | | | | | |
| | | | | Eumetazoa*** | | | | | | | | | | | | | | | | | |
| | | | | | Bilateria | | | | | | | | | | | | | | | | |
| | | | | | | Protostomia | | | | Deuterostomia | | | | | | | | | | | |
| | | | | | | | Ecdysozoa | | | | Chordata | | | | | | | | | | |
| | | | | | | | | Arthropoda | | | | Vertebrata | | | | | | | | | |
| | | | | | | | | | | | | | Euteleostomi | | | | | | | | |
| | | | | | | | | | | | | | Actinopterygii | | Sarcopterygii | | | | | | |
| | | | | | | | | | | | | | | | | Tetrapoda | | | | | |
| | | | | | | | | | | | | | | | | | Amniota | | | | |
| | | | | | | | | | | | | | | | | | Sauropsida | | Mammalia | | |
| | | | | | | | | | | | | | | | | | | | | Eutheria | |
| | | | Metazoa* | Cnidaria | Bilateria* | Protostomia* | Ecdysozoa* | Arthropoda* | Insecta | Deuterostomia* | Chordata* | Vertebrata* | Actinopterygii* | Ostarioclupeomorpha | Sarcopterygii* | Amphibia | Sauropsida* | Aves | Mammalia* | Eutheria* | Euarchontoglires |
| HG0001 | D. melanogaster | FBgn0024734 | 0 | 8 | 0 | 10 | 0 | 4 | 22 | 5 | 0 | 3 | 7 | 3 | 1 | 2 | 3 | 27 | 4 | 10 | 4 |
| | D. rerio | ENSDARG00000006242 | 0 | 9 | 0 | 14 | 0 | 9 | 22 | 1 | 0 | 6 | 20 | 3 | 2 | 2 | 3 | 27 | 4 | 10 | 4 |
| | | ENSDARG00000035676 | 0 | 3 | 0 | 13 | 0 | 1 | 1 | 1 | 0 | 6 | 22 | 3 | 2 | 2 | 3 | 29 | 4 | 10 | 5 |
| | | ENSDARG00000039997 | 0 | 3 | 0 | 9 | 0 | 3 | 1 | 1 | 0 | 3 | 18 | 3 | 2 | 2 | 3 | 27 | 4 | 10 | 4 |
| | | ENSDARG00000054814 | 0 | 5 | 0 | 12 | 0 | 6 | 15 | 0 | 0 | 5 | 17 | 3 | 2 | 2 | 3 | 27 | 4 | 10 | 4 |
| | | ENSDARG00000087443 | 0 | 8 | 0 | 14 | 0 | 4 | 9 | 6 | 0 | 6 | 23 | 3 | 2 | 2 | 3 | 29 | 4 | 10 | 5 |
| | G. gallus | ENSGALG00000003265 | 0 | 1 | 0 | 7 | 0 | 0 | 1 | 1 | 0 | 6 | 20 | 4 | 2 | 2 | 3 | 28 | 4 | 10 | 5 |
| | | ENSGALG00000016271 | 0 | 9 | 0 | 13 | 0 | 10 | 22 | 2 | 0 | 6 | 21 | 4 | 2 | 2 | 3 | 26 | 4 | 10 | 4 |
| | H. sapiens | ENSG00000112245 | 0 | 9 | 0 | 13 | 0 | 9 | 22 | 2 | 0 | 6 | 20 | 4 | 2 | 2 | 3 | 27 | 4 | 10 | 3 |
| | | ENSG00000184007 | 0 | 1 | 0 | 0 | 0 | 0 | 0 | 0 | 0 | 5 | 16 | 4 | 2 | 2 | 3 | 29 | 4 | 10 | 4 |
| HG0002 | D. rerio | ENSDARG00000003077 | 1 | 6 | 5 | 22 | 0 | 18 | 8 | 6 | 1 | 2 | 27 | 4 | 1 | 2 | 3 | 0 | 2 | 10 | 5 |
| HG0003 | D. melanogaster | FBgn0039280 | 0 | 8 | 3 | 11 | 3 | 10 | 22 | 4 | 1 | 2 | 15 | 3 | 1 | 2 | 2 | 5 | 1 | 1 | 2 |
| | G. gallus | ENSGALG00000014906 | 0 | 7 | 3 | 12 | 3 | 9 | 23 | 2 | 1 | 2 | 14 | 3 | 1 | 2 | 2 | 5 | 1 | 1 | 2 |
| HG0004.1 | D. melanogaster | FBgn0028494 | 0 | 0 | 0 | 0 | 0 | 3 | 13 | 0 | 0 | 0 | 0 | 0 | 0 | 0 | 0 | 0 | 0 | 0 | 0 |
| | D. rerio | ENSDARG00000076779 | 0 | 0 | 0 | 1 | 0 | 0 | 2 | 0 | 0 | 1 | 19 | 2 | 1 | 2 | 3 | 28 | 2 | 9 | 5 |
| | G. gallus | ENSGALG00000041706 | 0 | 0 | 0 | 0 | 0 | 0 | 0 | 0 | 0 | 1 | 19 | 3 | 1 | 2 | 3 | 27 | 2 | 9 | 5 |
| | H. sapiens | ENSG00000031003 | 0 | 0 | 0 | 0 | 0 | 0 | 0 | 0 | 0 | 1 | 19 | 3 | 1 | 2 | 3 | 28 | 2 | 9 | 4 |
| HG0004.2 | H. sapiens | ENSG00000138640 | 0 | 0 | 0 | 0 | 0 | 0 | 0 | 0 | 0 | 0 | 0 | 0 | 0 | 0 | 0 | 0 | 0 | 4 | 0 |
| HG0005 | D. rerio | ENSDARG00000011055 | 0 | 1 | 0 | 0 | 0 | 7 | 12 | 4 | 0 | 1 | 23 | 3 | 1 | 2 | 3 | 5 | 2 | 4 | 4 |
| | G. gallus | ENSGALG00000016320 | 0 | 1 | 0 | 1 | 0 | 6 | 12 | 5 | 0 | 1 | 23 | 4 | 1 | 1 | 3 | 4 | 2 | 4 | 4 |
| HG0006 | D. melanogaster | FBgn0029971 | 0 | 0 | 0 | 6 | 0 | 13 | 27 | 0 | 0 | 0 | 0 | 1 | 0 | 1 | 0 | 0 | 0 | 0 | 0 |
| HG0007 | D. melanogaster | FBgn0034372 | 0 | 0 | 0 | 3 | 0 | 11 | 27 | 0 | 0 | 0 | 0 | 1 | 0 | 0 | 0 | 0 | 0 | 0 | 0 |
| HG0008 | H. sapiens | ENSG00000175567 | 0 | 1 | 0 | 0 | 0 | 0 | 0 | 0 | 0 | 2 | 12 | 4 | 1 | 2 | 3 | 0 | 3 | 10 | 4 |
| HG0009 | D. melanogaster | FBgn0050100 | 0 | 0 | 0 | 0 | 0 | 14 | 26 | 0 | 0 | 0 | 0 | 1 | 0 | 0 | 0 | 0 | 0 | 0 | 0 |
| HG0010.1 | G. gallus | ENSGALG00000009013 | 0 | 0 | 0 | 3 | 0 | 0 | 1 | 5 | 0 | 0 | 2 | 0 | 1 | 2 | 3 | 7 | 3 | 5 | 4 |
| | H. sapiens | ENSG00000125863 | 0 | 0 | 0 | 2 | 0 | 3 | 5 | 5 | 0 | 0 | 2 | 0 | 1 | 2 | 3 | 8 | 2 | 5 | 3 |
| HG0010.2 | G. gallus | ENSGALG00000009013 | 0 | 0 | 0 | 0 | 0 | 0 | 0 | 0 | 0 | 0 | 3 | 0 | 1 | 2 | 3 | 9 | 3 | 7 | 5 |
| | H. sapiens | ENSG00000125863 | 0 | 0 | 0 | 0 | 0 | 0 | 0 | 3 | 0 | 0 | 3 | 0 | 1 | 2 | 3 | 10 | 3 | 7 | 4 |
| HG0010.3 | H. sapiens | ENSG00000125863 | 0 | 0 | 0 | 0 | 0 | 0 | 0 | 0 | 0 | 0 | 0 | 0 | 0 | 0 | 0 | 0 | 0 | 6 | 4 |
| HG0011 | D. melanogaster | FBgn0027360 | 0 | 0 | 0 | 0 | 0 | 10 | 24 | 0 | 0 | 0 | 0 | 1 | 0 | 1 | 0 | 0 | 0 | 0 | 0 |
| HG0012 | D. melanogaster | FBgn0064116 | 0 | 0 | 0 | 0 | 0 | 6 | 27 | 0 | 0 | 0 | 0 | 1 | 0 | 0 | 0 | 0 | 0 | 0 | 0 |
| HG0013 | D. melanogaster | FBgn0261381 | 0 | 0 | 0 | 0 | 0 | 6 | 27 | 0 | 0 | 0 | 0 | 0 | 0 | 1 | 0 | 0 | 0 | 0 | 0 |
| HG0014 | D. melanogaster | FBgn0260392 | 0 | 0 | 0 | 0 | 0 | 8 | 24 | 0 | 0 | 0 | 0 | 1 | 0 | 0 | 0 | 0 | 0 | 0 | 0 |
| HG0015 | D. rerio | ENSDARG00000007377 | 0 | 1 | 0 | 0 | 0 | 0 | 0 | 0 | 0 | 3 | 22 | 3 | 1 | 2 | 0 | 0 | 0 | 1 | 0 |
| HG0016 | G. gallus | ENSGALG00000010399 | 0 | 1 | 0 | 0 | 0 | 0 | 0 | 0 | 0 | 0 | 6 | 1 | 0 | 2 | 3 | 3 | 2 | 10 | 5 |
| HG0017 | D. rerio | ENSDARG00000043154 | 0 | 1 | 0 | 0 | 0 | 0 | 0 | 0 | 0 | 3 | 14 | 3 | 1 | 0 | 2 | 0 | 3 | 1 | 4 |
| HG0018 | D. rerio | ENSDARG00000053291 | 0 | 1 | 0 | 0 | 0 | 0 | 0 | 0 | 0 | 0 | 22 | 3 | 0 | 0 | 0 | 0 | 0 | 3 | 2 |
| | G. gallus | ENSGALG00000004122 | 0 | 0 | 0 | 0 | 0 | 0 | 0 | 0 | 0 | 0 | 0 | 0 | 0 | 0 | 3 | 28 | 4 | 10 | 5 |
| | H. sapiens | ENSG00000189266 | 0 | 0 | 0 | 0 | 0 | 0 | 0 | 0 | 0 | 0 | 0 | 0 | 0 | 0 | 3 | 29 | 4 | 10 | 4 |
| HG0019 | H. sapiens | ENSG00000178397 | 0 | 0 | 0 | 4 | 0 | 0 | 1 | 5 | 0 | 0 | 0 | 0 | 1 | 1 | 2 | 0 | 2 | 6 | 3 |
| HG0020 | D. melanogaster | FBgn0035436 | 0 | 0 | 0 | 0 | 0 | 4 | 19 | 0 | 0 | 0 | 0 | 1 | 0 | 0 | 0 | 0 | 0 | 0 | 0 |
| HG0021.1 | D. rerio | ENSDARG00000068708 | 0 | 1 | 0 | 0 | 0 | 0 | 0 | 0 | 0 | 0 | 20 | 3 | 0 | 0 | 0 | 0 | 0 | 0 | 0 |
| | G. gallus | ENSGALG00000009448 | 0 | 0 | 0 | 0 | 0 | 0 | 0 | 0 | 0 | 0 | 0 | 0 | 0 | 2 | 3 | 8 | 2 | 7 | 5 |
| | H. sapiens | ENSG00000006652 | 0 | 0 | 0 | 0 | 0 | 0 | 0 | 0 | 0 | 0 | 0 | 0 | 1 | 2 | 2 | 3 | 2 | 10 | 4 |
| HG0021.2 | D. rerio | ENSDARG00000036811 | 0 | 0 | 0 | 0 | 0 | 0 | 0 | 0 | 0 | 0 | 18 | 2 | 0 | 0 | 0 | 0 | 0 | 0 | 0 |
| HG0022 | D. melanogaster | FBgn0061359 | 0 | 0 | 0 | 0 | 0 | 3 | 19 | 0 | 0 | 0 | 1 | 0 | 0 | 0 | 0 | 0 | 0 | 0 | 0 |
| HG0023.1 | H. sapiens | ENSG00000175197 | 0 | 0 | 0 | 0 | 0 | 0 | 1 | 0 | 0 | 0 | 0 | 0 | 0 | 2 | 2 | 2 | 1 | 8 | 3 |
| HG0023.2 | D. rerio | ENSDARG00000059836 | 0 | 0 | 0 | 0 | 0 | 0 | 0 | 0 | 0 | 0 | 11 | 3 | 0 | 0 | 0 | 0 | 0 | 0 | 0 |
| HG0024.1 | D. rerio | ENSDARG00000093406 | 0 | 1 | 0 | 0 | 0 | 0 | 0 | 0 | 0 | 0 | 15 | 2 | 0 | 0 | 0 | 0 | 0 | 0 | 0 |
| HG0024.2 | G. gallus | ENSGALG00000013628 | 0 | 0 | 0 | 0 | 0 | 0 | 0 | 0 | 0 | 0 | 0 | 0 | 0 | 0 | 2 | 24 | 0 | 0 | 0 |
| | H. sapiens | ENSG00000112308 | 0 | 0 | 0 | 0 | 0 | 0 | 0 | 0 | 0 | 0 | 0 | 0 | 0 | 0 | 0 | 1 | 2 | 10 | 3 |
| HG0024.3 | G. gallus | ENSGALG00000013628 | 0 | 0 | 0 | 0 | 0 | 0 | 0 | 0 | 0 | 0 | 0 | 0 | 0 | 0 | 3 | 25 | 0 | 0 | 0 |
| HG0025 | D. rerio | ENSDARG00000060054 | 0 | 1 | 0 | 0 | 0 | 0 | 0 | 0 | 0 | 0 | 12 | 2 | 0 | 0 | 0 | 0 | 0 | 0 | 0 |
| HG0026 | D. rerio | ENSDARG00000087059 | 0 | 2 | 0 | 8 | 0 | 2 | 0 | 1 | 0 | 0 | 2 | 0 | 0 | 0 | 0 | 0 | 0 | 0 | 0 |
| HG0027 | H. sapiens | ENSG00000077254 | 0 | 1 | 0 | 0 | 0 | 1 | 1 | 0 | 0 | 0 | 1 | 0 | 0 | 1 | 0 | 0 | 0 | 0 | 0 |
| HG0028.1 | D. rerio | ENSDARG00000032103 | 0 | 0 | 0 | 0 | 0 | 0 | 0 | 0 | 0 | 2 | 17 | 3 | 1 | 2 | 3 | 29 | 3 | 11 | 5 |
| | G. gallus | ENSGALG00000031448 | 0 | 0 | 0 | 0 | 0 | 0 | 0 | 0 | 0 | 2 | 17 | 4 | 1 | 2 | 3 | 28 | 3 | 11 | 5 |
| | H. sapiens | ENSG00000069956 | 0 | 0 | 0 | 0 | 0 | 0 | 0 | 0 | 0 | 2 | 17 | 4 | 1 | 2 | 3 | 29 | 3 | 11 | 4 |
| HG0028.2 | H. sapiens | ENSG00000069956 | 0 | 0 | 0 | 0 | 0 | 0 | 0 | 0 | 0 | 0 | 0 | 0 | 0 | 1 | 3 | 29 | 2 | 10 | 4 |

| | | | | | | | | | | | | | | | | | | | | | |
|---|---|---|---|---|---|---|---|---|---|---|---|---|---|---|---|---|---|---|---|---|---|
| HG0029 | *D. rerio* | ENSDARG00000007523 | 0 | 0 | 0 | 0 | 0 | 0 | 0 | 0 | 0 | 3 | 20 | 2 | 1 | 2 | 2 | 13 | 1 | 11 | 4 |
| | *G. gallus* | ENSGALG00000008167 | 0 | 0 | 0 | 0 | 0 | 0 | 0 | 0 | 0 | 3 | 21 | 4 | 1 | 2 | 3 | 22 | 3 | 11 | 5 |
| | *H. sapiens* | ENSG00000005483 | 0 | 0 | 0 | 0 | 0 | 0 | 0 | 0 | 0 | 3 | 21 | 4 | 1 | 2 | 3 | 23 | 3 | 11 | 4 |
| | | ENSG00000168137 | 0 | 0 | 0 | 0 | 0 | 0 | 0 | 0 | 0 | 0 | 3 | 0 | 1 | 1 | 3 | 25 | 3 | 10 | 3 |
| HG0030.1 | *G. gallus* | ENSGALG00000011464 | 0 | 0 | 0 | 0 | 0 | 0 | 0 | 0 | 0 | 2 | 13 | 5 | 1 | 2 | 3 | 28 | 3 | 11 | 5 |
| | *H. sapiens* | ENSG00000100664 | 0 | 0 | 0 | 0 | 0 | 0 | 0 | 0 | 0 | 2 | 14 | 5 | 1 | 2 | 3 | 29 | 3 | 11 | 4 |
| HG0030.2 | *D. melanogaster* | FBgn0030719 | 0 | 0 | 0 | 0 | 0 | 7 | 26 | 0 | 0 | 0 | 0 | 0 | 0 | 0 | 0 | 0 | 0 | 0 | 0 |
| HG0031 | *D. rerio* | ENSDARG00000000540 | 0 | 0 | 0 | 0 | 0 | 0 | 0 | 0 | 0 | 2 | 22 | 2 | 1 | 2 | 3 | 8 | 4 | 9 | 4 |
| | *G. gallus* | ENSGALG00000041666 | 0 | 0 | 0 | 0 | 0 | 0 | 0 | 0 | 0 | 2 | 22 | 3 | 1 | 2 | 3 | 18 | 4 | 9 | 4 |
| | *H. sapiens* | ENSG00000138381 | 0 | 0 | 0 | 0 | 0 | 0 | 0 | 0 | 0 | 2 | 20 | 3 | 1 | 2 | 3 | 6 | 2 | 8 | 3 |
| HG0032 | *D. rerio* | ENSDARG00000104981 | 0 | 0 | 0 | 0 | 0 | 0 | 0 | 0 | 0 | 1 | 13 | 2 | 1 | 2 | 3 | 20 | 2 | 7 | 4 |
| HG0033 | *G. gallus* | ENSGALG00000016321 | 0 | 0 | 0 | 0 | 0 | 0 | 0 | 0 | 0 | 0 | 0 | 0 | 1 | 0 | 3 | 26 | 1 | 9 | 5 |
| | *H. sapiens* | ENSG00000112144 | 0 | 0 | 0 | 0 | 0 | 0 | 0 | 0 | 0 | 0 | 8 | 0 | 0 | 0 | 3 | 26 | 3 | 9 | 4 |
| HG0034 | *D. rerio* | ENSDARG00000060991 | 0 | 0 | 0 | 0 | 0 | 0 | 0 | 0 | 0 | 3 | 21 | 3 | 1 | 2 | 3 | 4 | 2 | 7 | 4 |
| | *H. sapiens* | ENSG00000100335 | 0 | 0 | 0 | 0 | 0 | 0 | 0 | 0 | 0 | 3 | 21 | 4 | 1 | 2 | 3 | 4 | 2 | 7 | 3 |
| HG0035 | *G. gallus* | ENSGALG00000006167 | 0 | 0 | 0 | 0 | 0 | 0 | 0 | 0 | 0 | 0 | 0 | 0 | 0 | 2 | 3 | 26 | 3 | 11 | 5 |
| | *H. sapiens* | ENSG00000197147 | 0 | 0 | 0 | 0 | 0 | 0 | 0 | 0 | 0 | 0 | 0 | 0 | 0 | 2 | 3 | 27 | 3 | 11 | 4 |
| HG0036 | *G. gallus* | ENSGALG00000029295 | 0 | 0 | 0 | 0 | 0 | 0 | 0 | 0 | 0 | 0 | 0 | 0 | 0 | 1 | 3 | 26 | 3 | 11 | 4 |
| | *H. sapiens* | ENSG00000155096 | 0 | 0 | 0 | 0 | 0 | 0 | 0 | 0 | 0 | 0 | 0 | 0 | 0 | 1 | 3 | 27 | 3 | 11 | 3 |
| HG0037 | *G. gallus* | ENSGALG00000010747 | 0 | 0 | 0 | 0 | 0 | 0 | 0 | 0 | 0 | 0 | 0 | 0 | 0 | 2 | 1 | 28 | 3 | 8 | 4 |
| | *H. sapiens* | ENSG00000140092 | 0 | 0 | 0 | 0 | 0 | 0 | 0 | 0 | 0 | 0 | 0 | 0 | 0 | 2 | 1 | 29 | 3 | 8 | 3 |
| HG0038 | *G. gallus* | ENSGALG00000006571 | 0 | 0 | 0 | 0 | 0 | 0 | 0 | 0 | 0 | 0 | 0 | 0 | 1 | 1 | 3 | 24 | 3 | 10 | 4 |
| HG0039.1 | *H. sapiens* | ENSG00000077092 | 0 | 0 | 0 | 0 | 0 | 0 | 0 | 0 | 0 | 2 | 4 | 1 | 1 | 1 | 1 | 21 | 0 | 11 | 4 |
| HG0039.2 | *H. sapiens* | ENSG00000077092 | 0 | 0 | 0 | 0 | 0 | 0 | 0 | 0 | 0 | 0 | 0 | 0 | 0 | 0 | 0 | 0 | 0 | 8 | 4 |
| HG0040.1 | *H. sapiens* | ENSG00000164190 | 0 | 0 | 0 | 0 | 0 | 0 | 0 | 0 | 0 | 0 | 0 | 0 | 0 | 1 | 2 | 26 | 3 | 9 | 4 |
| HG0040.2 | *H. sapiens* | ENSG00000164190 | 0 | 0 | 0 | 0 | 0 | 0 | 0 | 0 | 0 | 0 | 0 | 0 | 0 | 0 | 3 | 5 | 0 | 9 | 3 |
| HG0041 | *D. rerio* | ENSDARG00000061352 | 0 | 0 | 0 | 0 | 0 | 0 | 0 | 0 | 0 | 0 | 8 | 2 | 1 | 1 | 3 | 10 | 2 | 11 | 4 |
| | *G. gallus* | ENSGALG00000034048 | 0 | 0 | 0 | 0 | 0 | 0 | 0 | 0 | 0 | 0 | 7 | 3 | 1 | 1 | 3 | 9 | 2 | 11 | 4 |
| | *H. sapiens* | ENSG00000119866 | 0 | 0 | 0 | 0 | 0 | 0 | 0 | 0 | 0 | 0 | 7 | 3 | 1 | 1 | 3 | 10 | 2 | 11 | 3 |
| HG0042.1 | *H. sapiens* | ENSG00000140937 | 0 | 0 | 0 | 0 | 0 | 0 | 0 | 0 | 0 | 0 | 0 | 0 | 0 | 0 | 3 | 25 | 1 | 9 | 4 |
| HG0042.2 | *G. gallus* | ENSGALG00000039985 | 0 | 0 | 0 | 0 | 0 | 0 | 0 | 0 | 0 | 0 | 0 | 0 | 0 | 0 | 0 | 13 | 0 | 0 | 0 |
| HG0042.3 | *H. sapiens* | ENSG00000140937 | 0 | 0 | 0 | 0 | 0 | 0 | 0 | 0 | 0 | 0 | 0 | 0 | 0 | 0 | 0 | 0 | 0 | 6 | 1 |
| HG0042.4 | *H. sapiens* | ENSG00000140937 | 0 | 0 | 0 | 0 | 0 | 0 | 0 | 0 | 0 | 0 | 0 | 0 | 0 | 0 | 0 | 0 | 0 | 4 | 1 |
| HG0042.5 | *H. sapiens* | ENSG00000140937 | 0 | 0 | 0 | 0 | 0 | 0 | 0 | 0 | 0 | 0 | 0 | 0 | 0 | 0 | 0 | 0 | 0 | 3 | 1 |
| HG0043 | *H. sapiens* | ENSG00000172995 | 0 | 0 | 0 | 0 | 0 | 0 | 0 | 0 | 0 | 1 | 0 | 0 | 1 | 1 | 2 | 24 | 0 | 10 | 3 |
| HG0044 | *D. rerio* | ENSDARG00000061687 | 0 | 0 | 0 | 0 | 0 | 0 | 0 | 0 | 0 | 0 | 5 | 2 | 1 | 2 | 3 | 14 | 2 | 8 | 4 |
| | *H. sapiens* | ENSG00000124788 | 0 | 0 | 0 | 0 | 0 | 0 | 0 | 0 | 0 | 0 | 4 | 1 | 1 | 2 | 3 | 14 | 2 | 8 | 3 |
| HG0045.1 | *H. sapiens* | ENSG00000083168 | 0 | 0 | 0 | 0 | 0 | 0 | 0 | 0 | 0 | 1 | 0 | 0 | 1 | 0 | 3 | 20 | 2 | 9 | 4 |
| HG0045.2 | *H. sapiens* | ENSG00000083168 | 0 | 0 | 0 | 0 | 0 | 0 | 0 | 0 | 0 | 0 | 0 | 0 | 0 | 0 | 2 | 3 | 0 | 5 | 2 |
| HG0046 | *D. rerio* | ENSDARG00000018817 | 0 | 0 | 0 | 0 | 0 | 0 | 0 | 0 | 0 | 0 | 6 | 0 | 0 | 1 | 2 | 15 | 1 | 10 | 4 |
| | *H. sapiens* | ENSG00000176697 | 0 | 0 | 0 | 0 | 0 | 0 | 0 | 0 | 0 | 0 | 6 | 1 | 0 | 1 | 2 | 15 | 1 | 10 | 3 |
| HG0047 | *G. gallus* | ENSGALG00000015422 | 0 | 0 | 0 | 0 | 0 | 0 | 0 | 0 | 0 | 0 | 0 | 0 | 0 | 2 | 2 | 19 | 1 | 11 | 4 |
| HG0048 | *G. gallus* | ENSGALG00000039182 | 0 | 0 | 0 | 0 | 0 | 0 | 0 | 0 | 0 | 0 | 0 | 0 | 0 | 0 | 1 | 22 | 2 | 7 | 5 |
| | *H. sapiens* | ENSG00000177565 | 0 | 0 | 0 | 0 | 0 | 0 | 0 | 0 | 0 | 0 | 0 | 0 | 0 | 0 | 3 | 22 | 2 | 7 | 4 |
| HG0049 | *D. rerio* | ENSDARG00000044485 | 0 | 0 | 0 | 0 | 0 | 0 | 0 | 0 | 0 | 0 | 5 | 1 | 0 | 1 | 1 | 3 | 0 | 0 | 2 |
| | *G. gallus* | ENSGALG00000039238 | 0 | 0 | 0 | 0 | 0 | 0 | 0 | 0 | 0 | 0 | 12 | 4 | 0 | 1 | 1 | 15 | 1 | 1 | 2 |
| HG0050.1 | *H. sapiens* | ENSG00000074054 | 0 | 0 | 0 | 0 | 0 | 0 | 0 | 0 | 0 | 2 | 2 | 1 | 0 | 2 | 3 | 10 | 2 | 11 | 4 |
| HG0050.2 | *H. sapiens* | ENSG00000074054 | 0 | 0 | 0 | 0 | 0 | 0 | 0 | 0 | 0 | 0 | 0 | 0 | 0 | 0 | 0 | 0 | 0 | 8 | 3 |
| HG0051 | *G. gallus* | ENSGALG00000037162 | 0 | 0 | 0 | 0 | 0 | 0 | 0 | 0 | 0 | 1 | 2 | 1 | 1 | 1 | 3 | 8 | 3 | 11 | 5 |
| | *H. sapiens* | ENSG00000116679 | 0 | 0 | 0 | 0 | 0 | 0 | 0 | 0 | 0 | 1 | 0 | 0 | 1 | 1 | 3 | 9 | 3 | 11 | 4 |
| HG0052 | *H. sapiens* | ENSG00000178235 | 0 | 0 | 0 | 0 | 0 | 0 | 0 | 0 | 0 | 0 | 0 | 0 | 0 | 0 | 1 | 20 | 1 | 10 | 4 |
| HG0053.1 | *H. sapiens* | ENSG00000184564 | 0 | 0 | 0 | 0 | 0 | 0 | 0 | 0 | 0 | 0 | 0 | 0 | 0 | 0 | 1 | 25 | 1 | 8 | 1 |
| HG0053.2 | *H. sapiens* | ENSG00000184564 | 0 | 0 | 0 | 0 | 0 | 0 | 0 | 0 | 0 | 0 | 0 | 0 | 0 | 0 | 0 | 0 | 0 | 8 | 4 |
| HG0054 | *G. gallus* | ENSGALG00000016222 | 0 | 0 | 0 | 0 | 0 | 0 | 0 | 0 | 0 | 0 | 0 | 0 | 0 | 0 | 2 | 20 | 0 | 8 | 5 |
| | *H. sapiens* | ENSG00000124479 | 0 | 0 | 0 | 0 | 0 | 0 | 0 | 0 | 0 | 0 | 0 | 0 | 0 | 0 | 1 | 3 | 0 | 9 | 4 |
| HG0055.1 | *G. gallus* | ENSGALG00000010533 | 0 | 0 | 0 | 0 | 0 | 0 | 0 | 0 | 0 | 0 | 0 | 0 | 0 | 1 | 2 | 17 | 0 | 11 | 4 |
| HG0055.2 | *G. gallus* | ENSGALG00000010533 | 0 | 0 | 0 | 0 | 0 | 0 | 0 | 0 | 0 | 0 | 0 | 0 | 0 | 0 | 1 | 17 | 0 | 0 | 0 |
| HG0056 | *G. gallus* | ENSGALG00000011271 | 0 | 0 | 0 | 0 | 0 | 0 | 0 | 0 | 0 | 0 | 0 | 0 | 1 | 1 | 3 | 22 | 0 | 5 | 2 |
| | *H. sapiens* | ENSG00000139329 | 0 | 0 | 0 | 0 | 0 | 0 | 0 | 0 | 0 | 0 | 0 | 0 | 0 | 1 | 1 | 7 | 0 | 6 | 2 |
| HG0057 | *H. sapiens* | ENSG00000280987 | 0 | 0 | 0 | 0 | 0 | 0 | 0 | 0 | 0 | 0 | 0 | 0 | 0 | 0 | 0 | 25 | 1 | 6 | 2 |
| HG0058 | *H. sapiens* | ENSG00000180332 | 0 | 0 | 0 | 0 | 0 | 0 | 0 | 0 | 0 | 0 | 0 | 0 | 0 | 1 | 1 | 21 | 0 | 7 | 4 |
| HG0059.1 | *D. rerio* | ENSDARG00000018060 | 0 | 0 | 0 | 0 | 0 | 0 | 0 | 0 | 0 | 0 | 18 | 0 | 1 | 2 | 2 | 9 | 1 | 0 | 0 |
| | *G. gallus* | ENSGALG00000003428 | 0 | 0 | 0 | 0 | 0 | 0 | 0 | 0 | 0 | 0 | 17 | 1 | 1 | 2 | 1 | 2 | 1 | 0 | 0 |
| HG0059.2 | *H. sapiens* | ENSG00000145675 | 0 | 0 | 0 | 0 | 0 | 0 | 0 | 0 | 0 | 0 | 0 | 0 | 0 | 0 | 0 | 0 | 0 | 8 | 3 |
| HG0059.3 | *H. sapiens* | ENSG00000105647 | 0 | 0 | 0 | 0 | 0 | 0 | 0 | 0 | 0 | 0 | 0 | 0 | 0 | 0 | 0 | 0 | 0 | 6 | 1 |
| HG0059.4 | *H. sapiens* | ENSG00000145675 | 0 | 0 | 0 | 0 | 0 | 0 | 0 | 0 | 0 | 0 | 0 | 0 | 0 | 0 | 0 | 0 | 0 | 3 | 2 |
| HG0059.5 | *D. rerio* | ENSDARG00000038524 | 0 | 0 | 0 | 0 | 0 | 0 | 0 | 0 | 0 | 0 | 1 | 3 | 0 | 0 | 0 | 0 | 0 | 0 | 0 |
| HG0060.1 | *G. gallus* | ENSGALG00000014751 | 0 | 0 | 0 | 0 | 0 | 0 | 0 | 0 | 0 | 0 | 0 | 0 | 0 | 0 | 1 | 16 | 2 | 8 | 4 |
| | *H. sapiens* | ENSG00000049192 | 0 | 0 | 0 | 0 | 0 | 0 | 0 | 0 | 0 | 0 | 0 | 0 | 0 | 0 | 1 | 17 | 2 | 9 | 4 |
| HG0060.2 | *G. gallus* | ENSGALG00000014751 | 0 | 0 | 0 | 0 | 0 | 0 | 0 | 0 | 0 | 0 | 0 | 0 | 0 | 0 | 2 | 17 | 0 | 3 | 2 |
| | *H. sapiens* | ENSG00000049192 | 0 | 0 | 0 | 0 | 0 | 0 | 0 | 0 | 0 | 0 | 0 | 0 | 0 | 0 | 2 | 15 | 0 | 8 | 4 |
| HG0061.1 | *G. gallus* | ENSGALG00000004949 | 0 | 0 | 0 | 0 | 0 | 0 | 0 | 0 | 0 | 0 | 0 | 0 | 0 | 1 | 2 | 28 | 1 | 0 | 0 |
| | *H. sapiens* | ENSG00000166398 | 0 | 0 | 0 | 0 | 0 | 0 | 0 | 0 | 0 | 0 | 0 | 0 | 0 | 2 | 2 | 8 | 2 | 6 | 4 |
| HG0061.2 | *G. gallus* | ENSGALG00000004949 | 0 | 0 | 0 | 0 | 0 | 0 | 0 | 0 | 0 | 0 | 0 | 0 | 0 | 0 | 0 | 7 | 2 | 0 | 0 |
| HG0061.3 | *H. sapiens* | ENSG00000166398 | 0 | 0 | 0 | 0 | 0 | 0 | 0 | 0 | 0 | 0 | 0 | 0 | 0 | 0 | 0 | 0 | 0 | 3 | 4 |
| HG0062.1 | *D. rerio* | ENSDARG00000075569 | 0 | 0 | 0 | 0 | 0 | 0 | 0 | 0 | 0 | 0 | 3 | 3 | 0 | 0 | 0 | 0 | 0 | 0 | 0 |
| | *G. gallus* | ENSGALG00000010276 | 0 | 0 | 0 | 0 | 0 | 0 | 0 | 0 | 0 | 0 | 1 | 0 | 0 | 0 | 3 | 27 | 0 | 1 | 0 |
| HG0062.2 | *G. gallus* | ENSGALG00000010276 | 0 | 0 | 0 | 0 | 0 | 0 | 0 | 0 | 0 | 0 | 0 | 0 | 0 | 0 | 0 | 20 | 0 | 0 | 0 |
| HG0062.3 | *H. sapiens* | ENSG00000159592 | 0 | 0 | 0 | 0 | 0 | 0 | 0 | 0 | 0 | 0 | 0 | 0 | 0 | 0 | 0 | 0 | 0 | 9 | 3 |

| | | | | | | | | | | | | | | | | | | | | | |
|---|---|---|---|---|---|---|---|---|---|---|---|---|---|---|---|---|---|---|---|---|---|
| HG0063.1 | *G. gallus* | ENSGALG00000002477 | 0 | 0 | 0 | 0 | 0 | 0 | 0 | 0 | 0 | 0 | 0 | 0 | 0 | 0 | 1 | 19 | 3 | 6 | 3 |
| | *H. sapiens* | ENSG00000162630 | 0 | 0 | 0 | 0 | 0 | 0 | 0 | 0 | 0 | 0 | 0 | 0 | 0 | 0 | 1 | 20 | 3 | 6 | 2 |
| HG0063.2 | *H. sapiens* | ENSG00000162630 | 0 | 0 | 0 | 0 | 0 | 0 | 0 | 0 | 0 | 0 | 0 | 0 | 0 | 0 | 2 | 4 | 2 | 10 | 4 |
| HG0064.1 | *H. sapiens* | ENSG00000102678 | 0 | 0 | 0 | 0 | 0 | 0 | 0 | 0 | 0 | 0 | 0 | 0 | 0 | 0 | 1 | 16 | 1 | 10 | 4 |
| HG0064.2 | *H. sapiens* | ENSG00000113578 | 0 | 0 | 0 | 0 | 0 | 0 | 0 | 0 | 0 | 0 | 0 | 0 | 0 | 0 | 0 | 0 | 0 | 5 | 2 |
| HG0065 | *G. gallus* | ENSGALG00000029927 | 0 | 0 | 0 | 0 | 0 | 0 | 0 | 0 | 0 | 0 | 0 | 0 | 0 | 0 | 3 | 25 | 1 | 2 | 1 |
| HG0066 | *H. sapiens* | ENSG00000112182 | 0 | 0 | 0 | 0 | 0 | 0 | 0 | 0 | 0 | 0 | 0 | 0 | 0 | 0 | 3 | 14 | 3 | 10 | 2 |
| HG0067 | *H. sapiens* | ENSG00000166862 | 0 | 0 | 0 | 0 | 0 | 0 | 0 | 0 | 0 | 0 | 7 | 4 | 0 | 1 | 2 | 8 | 1 | 6 | 3 |
| HG0068 | *D. rerio* | ENSDARG00000071235 | 0 | 0 | 0 | 0 | 0 | 0 | 0 | 0 | 2 | 2 | 14 | 1 | 1 | 2 | 1 | 1 | 1 | 2 | 2 |
| | *G. gallus* | ENSGALG00000041419 | 0 | 0 | 0 | 0 | 0 | 0 | 0 | 0 | 2 | 2 | 15 | 2 | 1 | 2 | 1 | 1 | 1 | 2 | 2 |
| | *H. sapiens* | ENSG00000130821 | 0 | 0 | 0 | 0 | 0 | 0 | 0 | 0 | 2 | 2 | 13 | 2 | 1 | 2 | 1 | 1 | 1 | 2 | 1 |
| HG0069 | *G. gallus* | ENSGALG00000010999 | 0 | 0 | 0 | 0 | 0 | 0 | 0 | 0 | 0 | 0 | 0 | 0 | 0 | 0 | 2 | 26 | 1 | 1 | 1 |
| | *H. sapiens* | ENSG00000100697 | 0 | 0 | 0 | 0 | 0 | 0 | 0 | 0 | 0 | 0 | 0 | 0 | 0 | 0 | 2 | 16 | 1 | 10 | 2 |
| HG0070 | *D. rerio* | ENSDARG00000040926 | 0 | 0 | 0 | 0 | 0 | 0 | 0 | 0 | 0 | 0 | 15 | 2 | 1 | 0 | 0 | 1 | 0 | 4 | 2 |
| | | ENSDARG00000052695 | 0 | 0 | 0 | 0 | 0 | 0 | 0 | 0 | 0 | 1 | 14 | 3 | 1 | 2 | 1 | 1 | 0 | 5 | 2 |
| | *H. sapiens* | ENSG00000175745 | 0 | 0 | 0 | 0 | 0 | 0 | 0 | 0 | 0 | 1 | 15 | 3 | 1 | 2 | 0 | 1 | 0 | 5 | 1 |
| | | ENSG00000185551 | 0 | 0 | 0 | 0 | 0 | 0 | 0 | 0 | 0 | 1 | 8 | 3 | 1 | 1 | 1 | 1 | 0 | 5 | 1 |
| HG0071.1 | *D. rerio* | ENSDARG00000098240 | 0 | 0 | 0 | 0 | 0 | 0 | 0 | 0 | 0 | 0 | 11 | 2 | 1 | 0 | 2 | 2 | 0 | 8 | 3 |
| | *H. sapiens* | ENSG00000134138 | 0 | 0 | 0 | 0 | 0 | 0 | 0 | 0 | 0 | 1 | 11 | 3 | 1 | 0 | 2 | 2 | 0 | 8 | 2 |
| HG0071.2 | *H. sapiens* | ENSG00000143995 | 0 | 0 | 0 | 0 | 0 | 0 | 0 | 0 | 0 | 0 | 0 | 0 | 0 | 0 | 0 | 0 | 0 | 7 | 4 |
| HG0072 | *G. gallus* | ENSGALG00000039690 | 0 | 0 | 0 | 0 | 0 | 0 | 0 | 0 | 0 | 0 | 0 | 0 | 0 | 0 | 2 | 14 | 0 | 5 | 4 |
| | *H. sapiens* | ENSG00000104435 | 0 | 0 | 0 | 0 | 0 | 0 | 0 | 0 | 0 | 1 | 0 | 0 | 0 | 1 | 2 | 15 | 0 | 7 | 4 |
| HG0073 | *D. rerio* | ENSDARG00000041708 | 0 | 0 | 0 | 0 | 0 | 0 | 0 | 0 | 0 | 1 | 12 | 3 | 0 | 2 | 2 | 5 | 0 | 3 | 2 |
| HG0074 | *D. rerio* | ENSDARG00000078624 | 0 | 0 | 0 | 0 | 0 | 0 | 0 | 0 | 0 | 0 | 5 | 0 | 0 | 0 | 0 | 0 | 0 | 0 | 0 |
| | *H. sapiens* | ENSG00000131089 | 0 | 0 | 0 | 0 | 0 | 0 | 0 | 0 | 0 | 0 | 7 | 1 | 0 | 2 | 1 | 7 | 2 | 7 | 2 |
| HG0075 | *H. sapiens* | ENSG00000165699 | 0 | 0 | 0 | 0 | 0 | 0 | 0 | 0 | 0 | 0 | 0 | 0 | 0 | 0 | 2 | 9 | 3 | 11 | 4 |
| HG0076 | *D. rerio* | ENSDARG00000040008 | 0 | 0 | 0 | 0 | 0 | 0 | 0 | 0 | 0 | 0 | 0 | 0 | 0 | 2 | 2 | 8 | 2 | 9 | 5 |
| | *H. sapiens* | ENSG00000164600 | 0 | 0 | 0 | 0 | 0 | 0 | 0 | 0 | 0 | 0 | 1 | 0 | 0 | 2 | 2 | 8 | 2 | 9 | 4 |
| HG0077.1 | *G. gallus* | ENSGALG00000005074 | 0 | 0 | 0 | 0 | 0 | 0 | 0 | 0 | 0 | 0 | 0 | 0 | 0 | 0 | 2 | 26 | 0 | 0 | 0 |
| HG0077.2 | *G. gallus* | ENSGALG00000005074 | 0 | 0 | 0 | 0 | 0 | 0 | 0 | 0 | 0 | 0 | 0 | 0 | 0 | 0 | 0 | 8 | 0 | 0 | 0 |
| HG0078.1 | *G. gallus* | ENSGALG00000016633 | 0 | 0 | 0 | 0 | 0 | 0 | 0 | 0 | 0 | 0 | 0 | 0 | 0 | 0 | 2 | 26 | 0 | 0 | 0 |
| HG0078.2 | *H. sapiens* | ENSG00000147421 | 0 | 0 | 0 | 0 | 0 | 0 | 0 | 0 | 0 | 0 | 0 | 0 | 0 | 0 | 0 | 0 | 0 | 8 | 2 |
| HG0079 | *H. sapiens* | ENSG00000204406 | 0 | 0 | 0 | 0 | 0 | 0 | 0 | 0 | 0 | 0 | 0 | 0 | 0 | 2 | 3 | 9 | 1 | 10 | 3 |
| HG0080.1 | *G. gallus* | ENSGALG00000039403 | 0 | 0 | 0 | 0 | 0 | 0 | 0 | 0 | 0 | 0 | 0 | 0 | 0 | 0 | 1 | 7 | 0 | 7 | 5 |
| | *H. sapiens* | ENSG00000179603 | 0 | 0 | 0 | 0 | 0 | 0 | 0 | 0 | 0 | 0 | 0 | 0 | 0 | 0 | 2 | 8 | 2 | 11 | 4 |
| HG0080.2 | *H. sapiens* | ENSG00000179603 | 0 | 0 | 0 | 0 | 0 | 0 | 0 | 0 | 0 | 0 | 0 | 0 | 0 | 0 | 0 | 0 | 0 | 4 | 3 |
| HG0080.3 | *H. sapiens* | ENSG00000179603 | 0 | 0 | 0 | 0 | 0 | 0 | 0 | 0 | 0 | 0 | 0 | 0 | 0 | 0 | 0 | 0 | 0 | 2 | 2 |
| HG0081.1 | *D. rerio* | ENSDARG00000028228 | 0 | 0 | 0 | 0 | 0 | 0 | 0 | 0 | 0 | 0 | 3 | 2 | 0 | 1 | 2 | 7 | 1 | 7 | 4 |
| HG0081.2 | *D. rerio* | ENSDARG00000028228 | 0 | 0 | 0 | 0 | 0 | 0 | 0 | 0 | 0 | 0 | 1 | 1 | 0 | 0 | 0 | 0 | 0 | 0 | 0 |
| HG0082 | *D. rerio* | ENSDARG00000062379 | 0 | 0 | 0 | 0 | 0 | 0 | 0 | 0 | 0 | 0 | 13 | 1 | 1 | 1 | 1 | 2 | 1 | 5 | 2 |
| | *H. sapiens* | ENSG00000176087 | 0 | 0 | 0 | 0 | 0 | 0 | 0 | 0 | 0 | 0 | 13 | 2 | 1 | 1 | 1 | 2 | 1 | 5 | 1 |
| HG0083.1 | *G. gallus* | ENSGALG00000008039 | 0 | 0 | 0 | 0 | 0 | 0 | 0 | 0 | 0 | 0 | 0 | 0 | 0 | 0 | 1 | 26 | 0 | 0 | 0 |
| HG0083.2 | *H. sapiens* | ENSG00000138111 | 0 | 0 | 0 | 0 | 0 | 0 | 0 | 0 | 0 | 0 | 0 | 0 | 0 | 0 | 0 | 0 | 0 | 4 | 1 |
| HG0084.1 | *G. gallus* | ENSGALG00000009207 | 0 | 0 | 0 | 0 | 0 | 0 | 0 | 0 | 0 | 0 | 0 | 0 | 0 | 0 | 1 | 26 | 0 | 0 | 0 |
| HG0084.2 | *H. sapiens* | ENSG00000154447 | 0 | 0 | 0 | 0 | 0 | 0 | 0 | 0 | 0 | 0 | 0 | 0 | 0 | 0 | 0 | 0 | 0 | 3 | 1 |
| HG0085 | *G. gallus* | ENSGALG00000002003 | 0 | 0 | 0 | 0 | 0 | 0 | 0 | 0 | 0 | 0 | 0 | 0 | 0 | 0 | 2 | 12 | 2 | 0 | 0 |
| | *H. sapiens* | ENSG00000107779 | 0 | 0 | 0 | 0 | 0 | 0 | 0 | 0 | 0 | 0 | 0 | 0 | 0 | 1 | 1 | 11 | 2 | 9 | 3 |
| HG0086 | *G. gallus* | ENSGALG00000009835 | 0 | 0 | 0 | 0 | 0 | 0 | 0 | 0 | 0 | 0 | 0 | 0 | 0 | 2 | 1 | 24 | 0 | 0 | 0 |
| HG0087 | *H. sapiens* | ENSG00000164548 | 0 | 0 | 0 | 0 | 0 | 0 | 0 | 0 | 0 | 0 | 5 | 0 | 1 | 1 | 1 | 8 | 1 | 8 | 2 |
| HG0088 | *H. sapiens* | ENSG00000273079 | 0 | 0 | 0 | 0 | 0 | 0 | 0 | 0 | 0 | 0 | 0 | 0 | 0 | 0 | 2 | 14 | 0 | 8 | 3 |
| HG0089.1 | *G. gallus* | ENSGALG00000002294 | 0 | 0 | 0 | 0 | 0 | 0 | 0 | 0 | 0 | 0 | 0 | 0 | 0 | 2 | 3 | 9 | 1 | 8 | 3 |
| | *H. sapiens* | ENSG00000092421 | 0 | 0 | 0 | 0 | 0 | 0 | 0 | 0 | 0 | 0 | 0 | 0 | 0 | 1 | 2 | 10 | 1 | 9 | 3 |
| HG0089.2 | *H. sapiens* | ENSG00000092421 | 0 | 0 | 0 | 0 | 0 | 0 | 0 | 0 | 0 | 0 | 0 | 0 | 0 | 0 | 0 | 0 | 0 | 6 | 3 |
| HG0089.3 | *G. gallus* | ENSGALG00000002294 | 0 | 0 | 0 | 0 | 0 | 0 | 0 | 0 | 0 | 0 | 0 | 0 | 0 | 0 | 0 | 7 | 0 | 0 | 0 |
| HG0090.1 | *G. gallus* | ENSGALG00000008487 | 0 | 0 | 0 | 0 | 0 | 0 | 0 | 0 | 0 | 0 | 0 | 0 | 0 | 0 | 3 | 17 | 1 | 5 | 0 |
| | *H. sapiens* | ENSG00000172765 | 0 | 0 | 0 | 0 | 0 | 0 | 0 | 0 | 0 | 0 | 0 | 0 | 0 | 0 | 0 | 0 | 0 | 8 | 2 |
| HG0090.2 | *D. rerio* | ENSDARG00000060954 | 0 | 0 | 0 | 0 | 0 | 0 | 0 | 0 | 0 | 0 | 13 | 3 | 0 | 0 | 0 | 0 | 0 | 0 | 0 |
| HG0090.3 | *G. gallus* | ENSGALG00000008487 | 0 | 0 | 0 | 0 | 0 | 0 | 0 | 0 | 0 | 0 | 0 | 0 | 0 | 0 | 0 | 13 | 0 | 0 | 0 |
| HG0091.1 | *H. sapiens* | ENSG00000172209 | 0 | 0 | 0 | 0 | 0 | 0 | 0 | 0 | 0 | 0 | 0 | 0 | 0 | 2 | 0 | 15 | 0 | 6 | 3 |
| HG0091.2 | *H. sapiens* | ENSG00000172209 | 0 | 0 | 0 | 0 | 0 | 0 | 0 | 0 | 0 | 0 | 0 | 0 | 0 | 0 | 1 | 0 | 2 | 6 | 3 |
| HG0091.3 | *H. sapiens* | ENSG00000172209 | 0 | 0 | 0 | 0 | 0 | 0 | 0 | 0 | 0 | 0 | 0 | 0 | 0 | 0 | 0 | 0 | 2 | 6 | 3 |
| HG0092.1 | *G. gallus* | ENSGALG00000009966 | 0 | 0 | 0 | 0 | 0 | 0 | 0 | 0 | 0 | 0 | 0 | 0 | 1 | 1 | 2 | 18 | 1 | 3 | 0 |
| HG0092.2 | *H. sapiens* | ENSG00000166225 | 0 | 0 | 0 | 0 | 0 | 0 | 0 | 0 | 0 | 0 | 0 | 0 | 0 | 0 | 0 | 0 | 1 | 8 | 3 |
| HG0093 | *G. gallus* | ENSGALG00000040465 | 0 | 0 | 0 | 0 | 0 | 0 | 0 | 0 | 0 | 0 | 0 | 0 | 1 | 0 | 3 | 9 | 0 | 9 | 4 |
| | *H. sapiens* | ENSG00000169554 | 0 | 0 | 0 | 0 | 0 | 0 | 0 | 0 | 0 | 0 | 0 | 0 | 0 | 0 | 3 | 10 | 0 | 9 | 3 |
| HG0094.1 | *H. sapiens* | ENSG00000112175 | 0 | 0 | 0 | 0 | 0 | 0 | 0 | 0 | 0 | 0 | 0 | 0 | 0 | 0 | 3 | 10 | 1 | 10 | 2 |
| HG0094.2 | *H. sapiens* | ENSG00000112175 | 0 | 0 | 0 | 0 | 0 | 0 | 0 | 0 | 0 | 0 | 0 | 0 | 0 | 0 | 1 | 0 | 1 | 9 | 3 |
| HG0095 | *D. rerio* | ENSDARG00000013708 | 0 | 0 | 0 | 0 | 0 | 0 | 0 | 0 | 0 | 1 | 15 | 3 | 1 | 2 | 0 | 0 | 2 | 1 | 1 |
| HG0096 | *G. gallus* | ENSGALG00000014186 | 0 | 0 | 0 | 0 | 0 | 0 | 0 | 0 | 0 | 0 | 0 | 0 | 0 | 0 | 1 | 25 | 0 | 0 | 0 |
| HG0097 | *G. gallus* | ENSGALG00000037496 | 0 | 0 | 0 | 0 | 0 | 0 | 0 | 0 | 0 | 0 | 0 | 0 | 0 | 0 | 3 | 8 | 2 | 10 | 3 |
| HG0098 | *H. sapiens* | ENSG00000198739 | 0 | 0 | 0 | 0 | 0 | 0 | 0 | 0 | 0 | 0 | 0 | 0 | 0 | 0 | 2 | 7 | 2 | 11 | 4 |
| HG0099.1 | *H. sapiens* | ENSG00000168575 | 0 | 0 | 0 | 0 | 0 | 0 | 0 | 0 | 0 | 0 | 0 | 0 | 0 | 0 | 1 | 6 | 3 | 11 | 4 |
| HG0099.2 | *H. sapiens* | ENSG00000144136 | 0 | 0 | 0 | 0 | 0 | 0 | 0 | 0 | 0 | 0 | 0 | 0 | 0 | 0 | 0 | 0 | 0 | 7 | 3 |
| HG0099.3 | *H. sapiens* | ENSG00000168575 | 0 | 0 | 0 | 0 | 0 | 0 | 0 | 0 | 0 | 0 | 0 | 0 | 0 | 0 | 0 | 0 | 0 | 6 | 3 |
| HG0099.4 | *D. rerio* | ENSDARG00000010641 | 0 | 0 | 0 | 0 | 0 | 0 | 0 | 0 | 0 | 0 | 5 | 3 | 0 | 0 | 0 | 0 | 0 | 0 | 0 |
| HG0100.1 | *D. rerio* | ENSDARG00000104148 | 0 | 0 | 0 | 0 | 0 | 0 | 0 | 0 | 0 | 0 | 21 | 3 | 0 | 0 | 0 | 0 | 0 | 0 | 0 |
| | | ENSDARG00000104609 | 0 | 0 | 0 | 0 | 0 | 0 | 0 | 0 | 0 | 0 | 21 | 3 | 1 | 0 | 0 | 0 | 0 | 0 | 0 |
| HG0100.2 | *D. rerio* | ENSDARG00000061108 | 0 | 0 | 0 | 0 | 0 | 0 | 0 | 0 | 0 | 0 | 9 | 2 | 0 | 0 | 0 | 0 | 0 | 0 | 0 |
| HG0101 | *H. sapiens* | ENSG00000099250 | 0 | 0 | 0 | 0 | 0 | 0 | 0 | 0 | 0 | 0 | 1 | 0 | 0 | 0 | 2 | 8 | 2 | 9 | 3 |

| | | | | | | | | | | | | | | | | | | | | | |
|---|---|---|---|---|---|---|---|---|---|---|---|---|---|---|---|---|---|---|---|---|---|
| HG0102 | *H. sapiens* | ENSG00000182263 | 0 | 0 | 0 | 0 | 0 | 0 | 0 | 0 | 0 | 0 | 3 | 1 | 0 | 0 | 2 | 7 | 1 | 7 | 4 |
| HG0103 | *G. gallus* | ENSGALG00000016558 | 0 | 0 | 0 | 0 | 0 | 0 | 0 | 0 | 0 | 0 | 0 | 0 | 0 | 0 | 2 | 21 | 1 | 0 | 0 |
| | *H. sapiens* | ENSG00000165197 | 0 | 0 | 0 | 0 | 0 | 0 | 0 | 0 | 0 | 0 | 0 | 0 | 0 | 0 | 0 | 0 | 0 | 9 | 3 |
| HG0104.1 | *G. gallus* | ENSGALG00000038097 | 0 | 0 | 0 | 0 | 0 | 0 | 0 | 0 | 0 | 0 | 0 | 0 | 0 | 0 | 2 | 19 | 3 | 0 | 0 |
| HG0104.2 | *H. sapiens* | ENSG00000149547 | 0 | 0 | 0 | 0 | 0 | 0 | 0 | 0 | 0 | 0 | 0 | 0 | 0 | 0 | 0 | 0 | 0 | 6 | 1 |
| HG0105 | *G. gallus* | ENSGALG00000016176 | 0 | 0 | 0 | 0 | 0 | 0 | 0 | 0 | 0 | 0 | 0 | 0 | 0 | 0 | 2 | 8 | 1 | 7 | 4 |
| | *H. sapiens* | ENSG00000135298 | 0 | 0 | 0 | 0 | 0 | 0 | 0 | 0 | 0 | 0 | 0 | 0 | 0 | 0 | 2 | 9 | 1 | 9 | 3 |
| HG0106 | *H. sapiens* | ENSG00000147548 | 0 | 0 | 0 | 0 | 0 | 0 | 0 | 0 | 0 | 0 | 1 | 0 | 1 | 2 | 2 | 4 | 2 | 9 | 3 |
| HG0107.1 | *G. gallus* | ENSGALG00000003601 | 0 | 0 | 0 | 0 | 0 | 0 | 0 | 0 | 0 | 0 | 0 | 0 | 1 | 0 | 3 | 8 | 1 | 5 | 3 |
| | *H. sapiens* | ENSG00000058668 | 0 | 0 | 0 | 0 | 0 | 0 | 0 | 0 | 0 | 1 | 0 | 0 | 1 | 0 | 3 | 8 | 1 | 6 | 3 |
| HG0107.2 | *G. gallus* | ENSGALG00000030550 | 0 | 0 | 0 | 0 | 0 | 0 | 0 | 0 | 0 | 0 | 0 | 0 | 0 | 0 | 0 | 21 | 0 | 0 | 0 |
| | *H. sapiens* | ENSG00000070961 | 0 | 0 | 0 | 0 | 0 | 0 | 0 | 0 | 0 | 0 | 0 | 0 | 0 | 0 | 0 | 0 | 1 | 10 | 4 |
| HG0107.3 | *D. rerio* | ENSDARG00000063433 | 0 | 0 | 0 | 0 | 0 | 0 | 0 | 0 | 0 | 0 | 11 | 2 | 0 | 0 | 0 | 0 | 0 | 0 | 0 |
| | *H. sapiens* | ENSG00000157087 | 0 | 0 | 0 | 0 | 0 | 0 | 0 | 0 | 0 | 0 | 0 | 0 | 0 | 0 | 0 | 0 | 0 | 9 | 3 |
| HG0107.4 | *H. sapiens* | ENSG00000070961 | 0 | 0 | 0 | 0 | 0 | 0 | 0 | 0 | 0 | 0 | 0 | 0 | 0 | 0 | 0 | 0 | 0 | 7 | 3 |
| HG0108.1 | *G. gallus* | ENSGALG00000007396 | 0 | 0 | 0 | 0 | 0 | 0 | 0 | 0 | 0 | 0 | 0 | 0 | 0 | 0 | 3 | 20 | 0 | 0 | 0 |
| HG0108.2 | *H. sapiens* | ENSG00000135090 | 0 | 0 | 0 | 0 | 0 | 0 | 0 | 0 | 0 | 0 | 0 | 0 | 0 | 0 | 0 | 0 | 0 | 7 | 3 |
| HG0108.3 | *D. rerio* | ENSDARG00000098304 | 0 | 0 | 0 | 0 | 0 | 0 | 0 | 0 | 0 | 0 | 0 | 2 | 0 | 0 | 0 | 0 | 0 | 0 | 0 |
| HG0109.1 | *H. sapiens* | ENSG00000048540 | 0 | 0 | 0 | 0 | 0 | 0 | 0 | 0 | 0 | 0 | 0 | 0 | 0 | 0 | 2 | 14 | 0 | 5 | 2 |
| HG0109.2 | *H. sapiens* | ENSG00000048540 | 0 | 0 | 0 | 0 | 0 | 0 | 0 | 0 | 0 | 0 | 0 | 0 | 0 | 1 | 0 | 0 | 0 | 6 | 3 |
| HG0109.3 | *H. sapiens* | ENSG00000048540 | 0 | 0 | 0 | 0 | 0 | 0 | 0 | 0 | 0 | 0 | 0 | 0 | 0 | 0 | 0 | 0 | 1 | 9 | 4 |
| HG0110.1 | *H. sapiens* | ENSG00000091656 | 0 | 0 | 0 | 0 | 0 | 0 | 0 | 0 | 0 | 0 | 0 | 0 | 0 | 0 | 0 | 13 | 0 | 6 | 3 |
| HG0110.2 | *D. rerio* | ENSDARG00000073944 | 0 | 0 | 0 | 0 | 0 | 0 | 0 | 0 | 0 | 0 | 7 | 1 | 1 | 1 | 0 | 0 | 0 | 1 | 0 |
| HG0111 | *G. gallus* | ENSGALG00000034944 | 0 | 0 | 0 | 0 | 0 | 0 | 0 | 0 | 0 | 0 | 0 | 0 | 0 | 0 | 3 | 19 | 0 | 0 | 0 |
| HG0112 | *H. sapiens* | ENSG00000105997 | 0 | 0 | 0 | 0 | 0 | 0 | 0 | 0 | 0 | 0 | 7 | 0 | 0 | 1 | 1 | 8 | 0 | 4 | 1 |
| HG0113 | *D. rerio* | ENSDARG00000099727 | 0 | 0 | 0 | 0 | 0 | 0 | 0 | 0 | 0 | 0 | 8 | 0 | 1 | 1 | 1 | 1 | 1 | 5 | 3 |
| | *G. gallus* | ENSGALG00000023089 | 0 | 0 | 0 | 0 | 0 | 0 | 0 | 0 | 0 | 0 | 0 | 0 | 1 | 2 | 2 | 1 | 2 | 6 | 4 |
| HG0114.1 | *G. gallus* | ENSGALG00000003701 | 0 | 0 | 0 | 0 | 0 | 0 | 0 | 0 | 0 | 0 | 0 | 0 | 0 | 0 | 1 | 20 | 0 | 0 | 0 |
| HG0114.2 | *H. sapiens* | ENSG00000198964 | 0 | 0 | 0 | 0 | 0 | 0 | 0 | 0 | 0 | 0 | 0 | 0 | 0 | 0 | 0 | 0 | 0 | 4 | 2 |
| HG0115.1 | *G. gallus* | ENSGALG00000010976 | 0 | 0 | 0 | 0 | 0 | 0 | 0 | 0 | 0 | 0 | 0 | 0 | 0 | 0 | 1 | 20 | 0 | 0 | 0 |
| HG0115.2 | *H. sapiens* | ENSG00000169193 | 0 | 0 | 0 | 0 | 0 | 0 | 0 | 0 | 0 | 0 | 0 | 0 | 0 | 0 | 0 | 0 | 0 | 2 | 4 |
| HG0116 | *G. gallus* | ENSGALG00000033038 | 0 | 0 | 0 | 0 | 0 | 0 | 0 | 0 | 0 | 0 | 0 | 0 | 0 | 0 | 2 | 19 | 0 | 0 | 0 |
| | *H. sapiens* | ENSG00000010270 | 0 | 0 | 0 | 0 | 0 | 0 | 0 | 0 | 0 | 0 | 0 | 0 | 0 | 0 | 0 | 0 | 0 | 4 | 1 |
| HG0117.1 | *H. sapiens* | ENSG00000140945 | 0 | 0 | 0 | 0 | 0 | 0 | 0 | 0 | 0 | 0 | 0 | 0 | 0 | 0 | 2 | 7 | 0 | 9 | 3 |
| HG0117.2 | *H. sapiens* | ENSG00000140945 | 0 | 0 | 0 | 0 | 0 | 0 | 0 | 0 | 0 | 0 | 0 | 0 | 0 | 0 | 0 | 0 | 0 | 2 | 3 |
| HG0118.1 | *H. sapiens* | ENSG00000183662 | 0 | 0 | 0 | 0 | 0 | 0 | 0 | 0 | 0 | 0 | 0 | 0 | 0 | 1 | 1 | 5 | 1 | 10 | 3 |
| HG0118.2 | *H. sapiens* | ENSG00000183662 | 0 | 0 | 0 | 0 | 0 | 0 | 0 | 0 | 0 | 0 | 0 | 0 | 0 | 0 | 0 | 0 | 0 | 5 | 2 |
| HG0119 | *G. gallus* | ENSGALG00000043570 | 0 | 0 | 0 | 0 | 0 | 0 | 0 | 0 | 0 | 0 | 0 | 0 | 0 | 0 | 3 | 18 | 0 | 0 | 0 |
| HG0120 | *H. sapiens* | ENSG00000022355 | 0 | 0 | 0 | 0 | 0 | 0 | 0 | 0 | 0 | 0 | 4 | 0 | 0 | 0 | 2 | 2 | 2 | 7 | 4 |
| HG0121 | *H. sapiens* | ENSG00000122786 | 0 | 0 | 0 | 0 | 0 | 0 | 0 | 0 | 0 | 0 | 0 | 0 | 0 | 0 | 0 | 10 | 0 | 7 | 4 |
| HG0122 | *H. sapiens* | ENSG00000169925 | 0 | 0 | 0 | 0 | 0 | 0 | 0 | 0 | 0 | 0 | 0 | 0 | 0 | 0 | 1 | 3 | 4 | 9 | 4 |
| HG0123 | *H. sapiens* | ENSG00000173276 | 0 | 0 | 0 | 0 | 0 | 0 | 0 | 0 | 0 | 0 | 12 | 1 | 0 | 2 | 2 | 1 | 0 | 1 | 2 |
| HG0124.1 | *D. rerio* | ENSDARG00000031763 | 0 | 0 | 0 | 0 | 0 | 0 | 0 | 0 | 0 | 0 | 12 | 2 | 0 | 0 | 0 | 0 | 0 | 3 | 3 |
| HG0124.2 | *D. rerio* | ENSDARG00000053209 | 0 | 0 | 0 | 0 | 0 | 0 | 0 | 0 | 0 | 0 | 2 | 1 | 1 | 0 | 0 | 0 | 0 | 0 | 0 |
| HG0125.1 | *D. rerio* | ENSDARG00000051926 | 0 | 0 | 0 | 0 | 0 | 0 | 0 | 0 | 0 | 0 | 12 | 2 | 0 | 1 | 0 | 0 | 0 | 3 | 2 |
| HG0125.2 | *H. sapiens* | ENSG00000181690 | 0 | 0 | 0 | 0 | 0 | 0 | 0 | 0 | 0 | 0 | 0 | 0 | 0 | 0 | 1 | 0 | 0 | 11 | 4 |
| HG0126.1 | *G. gallus* | ENSGALG00000010899 | 0 | 0 | 0 | 0 | 0 | 0 | 0 | 0 | 0 | 0 | 0 | 0 | 0 | 0 | 1 | 19 | 0 | 0 | 0 |
| HG0126.2 | *G. gallus* | ENSGALG00000010899 | 0 | 0 | 0 | 0 | 0 | 0 | 0 | 0 | 0 | 0 | 0 | 0 | 0 | 0 | 0 | 15 | 0 | 0 | 0 |
| HG0127.1 | *G. gallus* | ENSGALG00000029709 | 0 | 0 | 0 | 0 | 0 | 0 | 0 | 0 | 0 | 0 | 0 | 0 | 0 | 0 | 1 | 19 | 0 | 0 | 0 |
| HG0127.2 | *H. sapiens* | ENSG00000109452 | 0 | 0 | 0 | 0 | 0 | 0 | 0 | 0 | 0 | 0 | 0 | 0 | 0 | 0 | 0 | 0 | 0 | 4 | 3 |
| HG0128 | *D. rerio* | ENSDARG00000091029 | 0 | 0 | 0 | 0 | 0 | 0 | 0 | 0 | 0 | 0 | 5 | 0 | 0 | 0 | 1 | 4 | 1 | 3 | 3 |
| | *H. sapiens* | ENSG00000109132 | 0 | 0 | 0 | 0 | 0 | 0 | 0 | 0 | 0 | 0 | 5 | 2 | 0 | 2 | 1 | 4 | 1 | 3 | 2 |
| HG0129 | *D. rerio* | ENSDARG00000099437 | 0 | 0 | 0 | 0 | 0 | 0 | 0 | 0 | 0 | 0 | 14 | 2 | 1 | 2 | 1 | 0 | 0 | 0 | 0 |
| HG0130 | *G. gallus* | ENSGALG00000002993 | 0 | 0 | 0 | 0 | 0 | 0 | 0 | 0 | 0 | 0 | 0 | 0 | 0 | 0 | 1 | 7 | 0 | 9 | 3 |
| HG0131 | *G. gallus* | ENSGALG00000013135 | 0 | 0 | 0 | 0 | 0 | 0 | 0 | 0 | 0 | 0 | 0 | 0 | 0 | 0 | 2 | 18 | 0 | 0 | 0 |
| HG0132 | *H. sapiens* | ENSG00000151067 | 0 | 0 | 0 | 0 | 0 | 0 | 0 | 0 | 0 | 0 | 1 | 0 | 0 | 2 | 1 | 6 | 0 | 7 | 3 |
| HG0133.1 | *D. rerio* | ENSDARG00000051886 | 0 | 0 | 0 | 0 | 0 | 0 | 0 | 0 | 0 | 1 | 16 | 2 | 0 | 0 | 0 | 0 | 0 | 0 | 0 |
| | *H. sapiens* | ENSG00000167522 | 0 | 0 | 0 | 0 | 0 | 0 | 0 | 0 | 0 | 0 | 0 | 0 | 0 | 0 | 0 | 0 | 0 | 4 | 1 |
| HG0133.2 | *D. rerio* | ENSDARG00000051886 | 0 | 0 | 0 | 0 | 0 | 0 | 0 | 0 | 0 | 0 | 1 | 1 | 0 | 0 | 0 | 0 | 0 | 0 | 0 |
| HG0133.3 | *H. sapiens* | ENSG00000167522 | 0 | 0 | 0 | 0 | 0 | 0 | 0 | 0 | 0 | 0 | 0 | 0 | 0 | 0 | 1 | 0 | 0 | 1 | 1 |
| HG0134.1 | *G. gallus* | ENSGALG00000041267 | 0 | 0 | 0 | 0 | 0 | 0 | 0 | 0 | 0 | 0 | 0 | 2 | 1 | 1 | 3 | 3 | 1 | 6 | 2 |
| HG0134.2 | *H. sapiens* | ENSG00000168283 | 0 | 0 | 0 | 0 | 0 | 0 | 0 | 0 | 0 | 0 | 0 | 0 | 0 | 0 | 0 | 0 | 0 | 6 | 2 |
| HG0135 | *D. rerio* | ENSDARG00000094132 | 0 | 0 | 0 | 0 | 0 | 0 | 0 | 0 | 0 | 0 | 7 | 0 | 0 | 1 | 0 | 2 | 0 | 0 | 1 |
| | *H. sapiens* | ENSG00000017427 | 0 | 0 | 0 | 0 | 0 | 0 | 0 | 0 | 0 | 0 | 7 | 0 | 0 | 1 | 0 | 2 | 1 | 5 | 3 |
| HG0136.1 | *H. sapiens* | ENSG00000125845 | 0 | 0 | 0 | 0 | 0 | 0 | 0 | 0 | 0 | 0 | 1 | 0 | 1 | 0 | 1 | 0 | 2 | 10 | 4 |
| HG0136.2 | *H. sapiens* | ENSG00000125845 | 0 | 0 | 0 | 0 | 0 | 0 | 0 | 0 | 0 | 0 | 0 | 0 | 0 | 0 | 0 | 0 | 0 | 1 | 2 |
| HG0137.1 | *H. sapiens* | ENSG00000181722 | 0 | 0 | 0 | 0 | 0 | 0 | 0 | 0 | 0 | 0 | 1 | 0 | 1 | 0 | 1 | 3 | 0 | 10 | 3 |
| HG0137.2 | *H. sapiens* | ENSG00000181722 | 0 | 0 | 0 | 0 | 0 | 0 | 0 | 0 | 0 | 0 | 0 | 0 | 0 | 0 | 0 | 0 | 0 | 6 | 2 |
| HG0138.1 | *H. sapiens* | ENSG00000185070 | 0 | 0 | 0 | 0 | 0 | 0 | 0 | 0 | 0 | 0 | 0 | 0 | 0 | 0 | 2 | 5 | 1 | 8 | 3 |
| HG0138.2 | *H. sapiens* | ENSG00000185070 | 0 | 0 | 0 | 0 | 0 | 0 | 0 | 0 | 0 | 0 | 0 | 0 | 0 | 0 | 0 | 0 | 0 | 6 | 2 |
| HG0139 | *G. gallus* | ENSGALG00000006379 | 0 | 0 | 0 | 0 | 0 | 0 | 0 | 0 | 0 | 0 | 1 | 0 | 1 | 0 | 1 | 10 | 0 | 3 | 3 |
| HG0140 | *H. sapiens* | ENSG00000009307 | 0 | 0 | 0 | 0 | 0 | 0 | 0 | 0 | 0 | 0 | 0 | 0 | 0 | 0 | 3 | 1 | 2 | 10 | 3 |
| HG0141.1 | *G. gallus* | ENSGALG00000010435 | 0 | 0 | 0 | 0 | 0 | 0 | 0 | 0 | 0 | 0 | 0 | 0 | 0 | 0 | 3 | 15 | 0 | 0 | 0 |
| HG0141.2 | *H. sapiens* | ENSG00000152689 | 0 | 0 | 0 | 0 | 0 | 0 | 0 | 0 | 0 | 0 | 0 | 0 | 0 | 0 | 0 | 0 | 0 | 8 | 2 |
| HG0141.3 | *H. sapiens* | ENSG00000152689 | 0 | 0 | 0 | 0 | 0 | 0 | 0 | 0 | 0 | 0 | 0 | 0 | 0 | 0 | 0 | 0 | 0 | 3 | 1 |
| HG0142.1 | *H. sapiens* | ENSG00000204217 | 0 | 0 | 0 | 0 | 0 | 0 | 0 | 0 | 0 | 0 | 0 | 0 | 0 | 1 | 1 | 5 | 1 | 8 | 2 |
| HG0142.2 | *H. sapiens* | ENSG00000204217 | 0 | 0 | 0 | 0 | 0 | 0 | 0 | 0 | 0 | 0 | 0 | 0 | 0 | 0 | 0 | 0 | 1 | 7 | 2 |
| HG0143 | *H. sapiens* | ENSG00000101958 | 0 | 0 | 0 | 0 | 0 | 0 | 0 | 0 | 0 | 0 | 0 | 0 | 0 | 0 | 2 | 0 | 1 | 11 | 4 |
| HG0144 | *H. sapiens* | ENSG00000117114 | 0 | 0 | 0 | 0 | 0 | 0 | 0 | 0 | 0 | 0 | 0 | 0 | 0 | 1 | 2 | 0 | 1 | 10 | 4 |

| | | | | | | | | | | | | | | | | | | | | | |
|---|---|---|---|---|---|---|---|---|---|---|---|---|---|---|---|---|---|---|---|---|---|
| HG0145 | *H. sapiens* | ENSG00000162599 | 0 | 0 | 0 | 0 | 0 | 0 | 0 | 0 | 0 | 0 | 4 | 4 | 0 | 0 | 0 | 6 | 1 | 2 | 1 |
| HG0146.1 | *H. sapiens* | ENSG00000163681 | 0 | 0 | 0 | 0 | 0 | 0 | 0 | 0 | 0 | 0 | 0 | 0 | 0 | 0 | 0 | 8 | 0 | 5 | 4 |
| HG0146.2 | *H. sapiens* | ENSG00000163681 | 0 | 0 | 0 | 0 | 0 | 0 | 0 | 0 | 0 | 0 | 0 | 0 | 0 | 0 | 0 | 0 | 0 | 5 | 3 |
| HG0146.3 | *H. sapiens* | ENSG00000163681 | 0 | 0 | 0 | 0 | 0 | 0 | 0 | 0 | 0 | 0 | 0 | 0 | 0 | 0 | 0 | 0 | 0 | 3 | 2 |
| HG0147 | *G. gallus* | ENSGALG00000004370 | 0 | 0 | 0 | 0 | 0 | 0 | 0 | 0 | 0 | 0 | 0 | 0 | 1 | 0 | 2 | 1 | 0 | 10 | 3 |
| | *H. sapiens* | ENSG00000108984 | 0 | 0 | 0 | 0 | 0 | 0 | 0 | 0 | 0 | 0 | 0 | 0 | 0 | 0 | 2 | 2 | 1 | 10 | 2 |
| HG0148.1 | *G. gallus* | ENSGALG00000011855 | 0 | 0 | 0 | 0 | 0 | 0 | 0 | 0 | 0 | 0 | 0 | 0 | 0 | 0 | 1 | 16 | 0 | 0 | 0 |
| HG0148.2 | *G. gallus* | ENSGALG00000011855 | 0 | 0 | 0 | 0 | 0 | 0 | 0 | 0 | 0 | 0 | 0 | 0 | 0 | 0 | 0 | 20 | 0 | 0 | 0 |
| HG0149.1 | *G. gallus* | ENSGALG00000014645 | 0 | 0 | 0 | 0 | 0 | 0 | 0 | 0 | 0 | 0 | 0 | 0 | 0 | 0 | 3 | 11 | 2 | 0 | 1 |
| HG0149.2 | *H. sapiens* | ENSG00000081189 | 0 | 0 | 0 | 0 | 0 | 0 | 0 | 0 | 0 | 0 | 0 | 0 | 0 | 0 | 0 | 0 | 0 | 10 | 3 |
| HG0150.1 | *G. gallus* | ENSGALG00000032289 | 0 | 0 | 0 | 0 | 0 | 0 | 0 | 0 | 0 | 0 | 0 | 0 | 0 | 0 | 2 | 15 | 0 | 0 | 0 |
| HG0150.2 | *H. sapiens* | ENSG00000095574 | 0 | 0 | 0 | 0 | 0 | 0 | 0 | 0 | 0 | 0 | 0 | 0 | 0 | 0 | 0 | 0 | 3 | 8 | 4 |
| HG0151 | *G. gallus* | ENSGALG00000035504 | 0 | 0 | 0 | 0 | 0 | 0 | 0 | 0 | 0 | 0 | 0 | 0 | 0 | 0 | 0 | 4 | 0 | 1 | 1 |
| | *H. sapiens* | ENSG00000006468 | 0 | 0 | 0 | 0 | 0 | 0 | 0 | 0 | 0 | 0 | 0 | 0 | 0 | 0 | 2 | 3 | 0 | 9 | 3 |
| HG0152 | *D. rerio* | ENSDARG00000015536 | 0 | 0 | 0 | 0 | 0 | 0 | 0 | 0 | 0 | 0 | 7 | 2 | 1 | 1 | 1 | 0 | 0 | 3 | 2 |
| HG0153 | *G. gallus* | ENSGALG00000003136 | 0 | 0 | 0 | 0 | 0 | 0 | 0 | 0 | 0 | 0 | 0 | 0 | 0 | 1 | 1 | 0 | 0 | 10 | 5 |
| HG0154 | *H. sapiens* | ENSG00000128573 | 0 | 0 | 0 | 0 | 0 | 0 | 0 | 0 | 0 | 0 | 3 | 1 | 0 | 2 | 2 | 2 | 1 | 3 | 3 |
| HG0155 | *H. sapiens* | ENSG00000136535 | 0 | 0 | 0 | 0 | 0 | 0 | 0 | 0 | 0 | 0 | 0 | 0 | 0 | 0 | 0 | 5 | 2 | 8 | 2 |
| HG0156 | *H. sapiens* | ENSG00000156113 | 0 | 0 | 0 | 0 | 0 | 0 | 0 | 0 | 0 | 0 | 0 | 0 | 0 | 0 | 1 | 5 | 1 | 9 | 1 |
| HG0157 | *H. sapiens* | ENSG00000173926 | 0 | 0 | 0 | 0 | 0 | 0 | 0 | 0 | 0 | 0 | 0 | 0 | 0 | 0 | 2 | 0 | 0 | 11 | 4 |
| HG0158 | *H. sapiens* | ENSG00000198939 | 0 | 0 | 0 | 0 | 0 | 0 | 0 | 0 | 0 | 0 | 0 | 0 | 0 | 0 | 1 | 0 | 3 | 9 | 4 |
| HG0159.1 | *H. sapiens* | ENSG00000182197 | 0 | 0 | 0 | 0 | 0 | 0 | 0 | 0 | 0 | 0 | 0 | 0 | 0 | 0 | 2 | 0 | 2 | 9 | 3 |
| HG0159.2 | *H. sapiens* | ENSG00000182197 | 0 | 0 | 0 | 0 | 0 | 0 | 0 | 0 | 0 | 0 | 0 | 0 | 0 | 0 | 2 | 0 | 1 | 9 | 3 |
| HG0160 | *D. rerio* | ENSDARG00000060089 | 0 | 0 | 0 | 0 | 0 | 0 | 0 | 0 | 0 | 0 | 12 | 2 | 1 | 1 | 0 | 0 | 0 | 0 | 0 |
| HG0161 | *G. gallus* | ENSGALG00000017191 | 0 | 0 | 0 | 0 | 0 | 0 | 0 | 0 | 0 | 0 | 0 | 0 | 0 | 0 | 1 | 15 | 0 | 0 | 0 |
| HG0162 | *H. sapiens* | ENSG00000138347 | 0 | 0 | 0 | 0 | 0 | 0 | 0 | 0 | 0 | 0 | 0 | 0 | 0 | 1 | 1 | 4 | 0 | 8 | 2 |
| HG0163 | *H. sapiens* | ENSG00000146285 | 0 | 0 | 0 | 0 | 0 | 0 | 0 | 0 | 0 | 0 | 0 | 0 | 0 | 2 | 2 | 2 | 0 | 9 | 1 |
| HG0164 | *H. sapiens* | ENSG00000151292 | 0 | 0 | 0 | 0 | 0 | 0 | 0 | 0 | 0 | 0 | 0 | 0 | 0 | 1 | 1 | 0 | 0 | 10 | 4 |
| HG0165 | *H. sapiens* | ENSG00000177508 | 0 | 0 | 0 | 0 | 0 | 0 | 0 | 0 | 0 | 1 | 3 | 2 | 1 | 0 | 1 | 2 | 0 | 5 | 1 |
| HG0166.1 | *H. sapiens* | ENSG00000148948 | 0 | 0 | 0 | 0 | 0 | 0 | 0 | 0 | 0 | 0 | 0 | 0 | 0 | 0 | 1 | 5 | 0 | 7 | 2 |
| HG0166.2 | *H. sapiens* | ENSG00000148948 | 0 | 0 | 0 | 0 | 0 | 0 | 0 | 0 | 0 | 0 | 0 | 0 | 0 | 0 | 0 | 0 | 0 | 9 | 1 |
| HG0166.3 | *H. sapiens* | ENSG00000148948 | 0 | 0 | 0 | 0 | 0 | 0 | 0 | 0 | 0 | 0 | 0 | 0 | 0 | 0 | 0 | 0 | 0 | 6 | 2 |
| HG0166.4 | *H. sapiens* | ENSG00000148948 | 0 | 0 | 0 | 0 | 0 | 0 | 0 | 0 | 0 | 0 | 0 | 0 | 0 | 0 | 0 | 0 | 0 | 5 | 2 |
| HG0167 | *D. rerio* | ENSDARG00000104005 | 0 | 0 | 0 | 0 | 0 | 0 | 0 | 0 | 0 | 0 | 11 | 3 | 1 | 0 | 0 | 0 | 0 | 0 | 0 |
| HG0168 | *G. gallus* | ENSGALG00000006997 | 0 | 0 | 0 | 0 | 0 | 0 | 0 | 0 | 0 | 0 | 0 | 0 | 0 | 1 | 2 | 10 | 2 | 0 | 0 |
| HG0169 | *H. sapiens* | ENSG00000114933 | 0 | 0 | 0 | 0 | 0 | 0 | 0 | 0 | 0 | 0 | 0 | 0 | 0 | 1 | 0 | 11 | 1 | 2 | 0 |
| HG0170 | *H. sapiens* | ENSG00000180530 | 0 | 0 | 0 | 0 | 0 | 0 | 0 | 0 | 0 | 0 | 0 | 0 | 0 | 0 | 1 | 1 | 0 | 9 | 4 |
| HG0171.1 | *H. sapiens* | ENSG00000177311 | 0 | 0 | 0 | 0 | 0 | 0 | 0 | 0 | 0 | 0 | 0 | 0 | 0 | 0 | 0 | 4 | 1 | 7 | 2 |
| HG0171.2 | *H. sapiens* | ENSG00000177311 | 0 | 0 | 0 | 0 | 0 | 0 | 0 | 0 | 0 | 0 | 0 | 0 | 0 | 0 | 0 | 2 | 1 | 6 | 1 |
| HG0171.3 | *H. sapiens* | ENSG00000177311 | 0 | 0 | 0 | 0 | 0 | 0 | 0 | 0 | 0 | 0 | 0 | 0 | 0 | 0 | 0 | 0 | 0 | 7 | 2 |
| HG0171.4 | *H. sapiens* | ENSG00000177311 | 0 | 0 | 0 | 0 | 0 | 0 | 0 | 0 | 0 | 0 | 0 | 0 | 0 | 0 | 0 | 0 | 0 | 6 | 1 |
| HG0172.1 | *H. sapiens* | ENSG00000196782 | 0 | 0 | 0 | 0 | 0 | 0 | 0 | 0 | 0 | 0 | 0 | 0 | 0 | 0 | 1 | 2 | 1 | 7 | 3 |
| HG0172.2 | *H. sapiens* | ENSG00000196782 | 0 | 0 | 0 | 0 | 0 | 0 | 0 | 0 | 0 | 0 | 0 | 0 | 0 | 0 | 0 | 0 | 0 | 6 | 3 |
| HG0173 | *H. sapiens* | ENSG00000101746 | 0 | 0 | 0 | 0 | 0 | 0 | 0 | 0 | 0 | 0 | 0 | 0 | 0 | 0 | 1 | 4 | 0 | 6 | 3 |
| HG0174 | *H. sapiens* | ENSG00000144355 | 0 | 0 | 0 | 0 | 0 | 0 | 0 | 0 | 0 | 0 | 0 | 0 | 0 | 0 | 1 | 3 | 2 | 4 | 4 |
| HG0175 | *H. sapiens* | ENSG00000151320 | 0 | 0 | 0 | 0 | 0 | 0 | 0 | 0 | 0 | 0 | 0 | 0 | 0 | 0 | 1 | 4 | 1 | 7 | 1 |
| HG0176 | *H. sapiens* | ENSG00000170962 | 0 | 0 | 0 | 0 | 0 | 0 | 0 | 0 | 0 | 0 | 0 | 0 | 0 | 0 | 0 | 2 | 2 | 6 | 4 |
| HG0177 | *H. sapiens* | ENSG00000171843 | 0 | 0 | 0 | 0 | 0 | 0 | 0 | 0 | 0 | 0 | 0 | 0 | 0 | 0 | 2 | 0 | 1 | 8 | 3 |
| HG0178.1 | *G. gallus* | ENSGALG00000029047 | 0 | 0 | 0 | 0 | 0 | 0 | 0 | 0 | 0 | 1 | 0 | 0 | 0 | 0 | 2 | 9 | 0 | 0 | 1 |
| | *H. sapiens* | ENSG00000131634 | 0 | 0 | 0 | 0 | 0 | 0 | 0 | 0 | 0 | 0 | 0 | 0 | 0 | 0 | 0 | 0 | 2 | 4 | 4 |
| HG0178.2 | *G. gallus* | ENSGALG00000029047 | 0 | 0 | 0 | 0 | 0 | 0 | 0 | 0 | 0 | 0 | 0 | 0 | 0 | 0 | 0 | 6 | 0 | 0 | 2 |
| HG0179 | *D. rerio* | ENSDARG00000036549 | 0 | 0 | 0 | 0 | 0 | 0 | 0 | 0 | 0 | 0 | 10 | 1 | 0 | 0 | 1 | 1 | 0 | 0 | 0 |
| | *H. sapiens* | ENSG00000160216 | 0 | 0 | 0 | 0 | 0 | 0 | 0 | 0 | 0 | 0 | 5 | 2 | 0 | 0 | 0 | 0 | 2 | 3 | 1 |
| HG0180.1 | *H. sapiens* | ENSG00000112246 | 0 | 0 | 0 | 0 | 0 | 0 | 0 | 0 | 0 | 0 | 0 | 0 | 0 | 0 | 0 | 1 | 1 | 8 | 3 |
| HG0180.2 | *H. sapiens* | ENSG00000112246 | 0 | 0 | 0 | 0 | 0 | 0 | 0 | 0 | 0 | 0 | 0 | 0 | 0 | 0 | 0 | 0 | 0 | 3 | 1 |
| HG0181.1 | *H. sapiens* | ENSG00000143776 | 0 | 0 | 0 | 0 | 0 | 0 | 0 | 0 | 0 | 0 | 0 | 0 | 0 | 0 | 2 | 1 | 1 | 5 | 4 |
| HG0181.2 | *H. sapiens* | ENSG00000143776 | 0 | 0 | 0 | 0 | 0 | 0 | 0 | 0 | 0 | 0 | 0 | 0 | 0 | 0 | 0 | 0 | 0 | 5 | 3 |
| HG0182 | *G. gallus* | ENSGALG00000031991 | 0 | 0 | 0 | 0 | 0 | 0 | 0 | 0 | 0 | 0 | 0 | 0 | 0 | 0 | 1 | 2 | 2 | 4 | 4 |
| HG0183 | *H. sapiens* | ENSG00000011465 | 0 | 0 | 0 | 0 | 0 | 0 | 0 | 0 | 0 | 0 | 0 | 0 | 0 | 0 | 2 | 0 | 0 | 9 | 2 |
| HG0184 | *H. sapiens* | ENSG00000136542 | 0 | 0 | 0 | 0 | 0 | 0 | 0 | 0 | 0 | 0 | 0 | 0 | 0 | 0 | 1 | 0 | 1 | 9 | 2 |
| HG0185 | *H. sapiens* | ENSG00000143033 | 0 | 0 | 0 | 0 | 0 | 0 | 0 | 0 | 0 | 0 | 0 | 1 | 0 | 0 | 0 | 0 | 0 | 8 | 4 |
| HG0186 | *H. sapiens* | ENSG00000167552 | 0 | 0 | 0 | 0 | 0 | 0 | 0 | 0 | 0 | 0 | 0 | 0 | 0 | 0 | 1 | 0 | 1 | 9 | 2 |
| HG0187.1 | *G. gallus* | ENSGALG00000004058 | 0 | 0 | 0 | 0 | 0 | 0 | 0 | 0 | 0 | 0 | 0 | 0 | 0 | 0 | 3 | 9 | 0 | 0 | 0 |
| HG0187.2 | *D. rerio* | ENSDARG00000059610 | 0 | 0 | 0 | 0 | 0 | 0 | 0 | 0 | 0 | 0 | 2 | 4 | 0 | 0 | 0 | 0 | 0 | 0 | 0 |
| HG0187.3 | *G. gallus* | ENSGALG00000004058 | 0 | 0 | 0 | 0 | 0 | 0 | 0 | 0 | 0 | 0 | 0 | 0 | 0 | 0 | 0 | 6 | 0 | 0 | 0 |
| HG0188 | *G. gallus* | ENSGALG00000016284 | 0 | 0 | 0 | 0 | 0 | 0 | 0 | 0 | 0 | 0 | 0 | 0 | 0 | 1 | 3 | 8 | 0 | 0 | 0 |
| | *H. sapiens* | ENSG00000157625 | 0 | 0 | 0 | 0 | 0 | 0 | 0 | 0 | 0 | 0 | 0 | 0 | 0 | 0 | 0 | 0 | 1 | 9 | 4 |
| HG0189 | *D. rerio* | ENSDARG00000001807 | 0 | 0 | 0 | 0 | 0 | 0 | 0 | 0 | 0 | 2 | 9 | 0 | 1 | 0 | 0 | 0 | 0 | 0 | 0 |
| HG0190 | *G. gallus* | ENSGALG00000006949 | 0 | 0 | 0 | 0 | 0 | 0 | 0 | 0 | 0 | 0 | 0 | 0 | 0 | 0 | 0 | 11 | 1 | 0 | 0 |
| HG0191 | *G. gallus* | ENSGALG00000036097 | 0 | 0 | 0 | 0 | 0 | 0 | 0 | 0 | 0 | 0 | 0 | 0 | 0 | 0 | 3 | 7 | 2 | 0 | 0 |
| HG0192 | *H. sapiens* | ENSG00000125107 | 0 | 0 | 0 | 0 | 0 | 0 | 0 | 0 | 0 | 0 | 0 | 0 | 0 | 0 | 1 | 0 | 2 | 8 | 1 |
| HG0193 | *H. sapiens* | ENSG00000134853 | 0 | 0 | 0 | 0 | 0 | 0 | 0 | 0 | 0 | 0 | 0 | 0 | 0 | 1 | 1 | 2 | 1 | 5 | 2 |
| HG0194 | *H. sapiens* | ENSG00000143507 | 0 | 0 | 0 | 0 | 0 | 0 | 0 | 0 | 0 | 0 | 0 | 0 | 0 | 0 | 1 | 0 | 3 | 6 | 2 |
| HG0195 | *H. sapiens* | ENSG00000165966 | 0 | 0 | 0 | 0 | 0 | 0 | 0 | 0 | 0 | 0 | 0 | 0 | 1 | 0 | 0 | 1 | 0 | 8 | 2 |
| HG0196 | *H. sapiens* | ENSG00000189403 | 0 | 0 | 0 | 0 | 0 | 0 | 0 | 0 | 0 | 0 | 0 | 0 | 0 | 1 | 0 | 0 | 1 | 7 | 3 |
| HG0197 | *H. sapiens* | ENSG00000196132 | 0 | 0 | 0 | 0 | 0 | 0 | 0 | 0 | 0 | 0 | 0 | 0 | 0 | 0 | 1 | 1 | 1 | 5 | 4 |
| HG0198.1 | *D. rerio* | ENSDARG00000018688 | 0 | 0 | 0 | 0 | 0 | 0 | 0 | 0 | 0 | 0 | 2 | 1 | 1 | 0 | 1 | 3 | 2 | 0 | 1 |
| HG0198.2 | *H. sapiens* | ENSG00000111145 | 0 | 0 | 0 | 0 | 0 | 0 | 0 | 0 | 0 | 0 | 0 | 0 | 0 | 0 | 0 | 0 | 0 | 8 | 4 |
| HG0199 | *G. gallus* | ENSGALG00000007989 | 0 | 0 | 0 | 0 | 0 | 0 | 0 | 0 | 0 | 0 | 0 | 0 | 0 | 0 | 2 | 9 | 0 | 0 | 0 |

| | | | | | | | | | | | | | | | | | | | | | |
|---|---|---|---|---|---|---|---|---|---|---|---|---|---|---|---|---|---|---|---|---|---|
| HG0200 | *G. gallus* | ENSGALG00000012015 | 0 | 0 | 0 | 0 | 0 | 0 | 0 | 0 | 0 | 0 | 0 | 0 | 0 | 1 | 2 | 6 | 0 | 1 | 1 |
| HG0201 | *G. gallus* | ENSGALG00000015256 | 0 | 0 | 0 | 0 | 0 | 0 | 0 | 0 | 0 | 0 | 0 | 0 | 0 | 0 | 3 | 7 | 1 | 0 | 0 |
| HG0202 | *H. sapiens* | ENSG00000124766 | 0 | 0 | 0 | 0 | 0 | 0 | 0 | 0 | 0 | 0 | 4 | 1 | 0 | 1 | 1 | 0 | 0 | 3 | 1 |
| HG0203 | *H. sapiens* | ENSG00000128606 | 0 | 0 | 0 | 0 | 0 | 0 | 0 | 0 | 0 | 0 | 0 | 0 | 0 | 1 | 0 | 0 | 0 | 7 | 3 |
| HG0204 | *H. sapiens* | ENSG00000162599 | 0 | 0 | 0 | 0 | 0 | 0 | 0 | 0 | 0 | 0 | 0 | 0 | 0 | 0 | 2 | 0 | 0 | 6 | 3 |
| HG0205 | *H. sapiens* | ENSG00000184611 | 0 | 0 | 0 | 0 | 0 | 0 | 0 | 0 | 0 | 0 | 0 | 0 | 0 | 0 | 1 | 0 | 1 | 7 | 2 |
| HG0206.1 | *G. gallus* | ENSGALG00000002069 | 0 | 0 | 0 | 0 | 0 | 0 | 0 | 0 | 0 | 0 | 0 | 0 | 0 | 0 | 3 | 7 | 0 | 0 | 0 |
| HG0206.2 | *H. sapiens* | ENSG00000005238 | 0 | 0 | 0 | 0 | 0 | 0 | 0 | 0 | 0 | 0 | 0 | 0 | 0 | 0 | 0 | 0 | 2 | 7 | 4 |
| HG0207.1 | *G. gallus* | ENSGALG00000007121 | 0 | 0 | 0 | 0 | 0 | 0 | 0 | 0 | 0 | 0 | 0 | 0 | 0 | 0 | 1 | 2 | 3 | 3 | 1 |
| HG0207.2 | *H. sapiens* | ENSG00000106952 | 0 | 0 | 0 | 0 | 0 | 0 | 0 | 0 | 0 | 0 | 0 | 0 | 0 | 0 | 0 | 0 | 0 | 8 | 3 |
| HG0208 | *G. gallus* | ENSGALG00000013272 | 0 | 0 | 0 | 0 | 0 | 0 | 0 | 0 | 0 | 0 | 0 | 0 | 0 | 0 | 1 | 9 | 0 | 0 | 0 |
| HG0209 | *H. sapiens* | ENSG00000101040 | 0 | 0 | 0 | 0 | 0 | 0 | 0 | 0 | 0 | 0 | 0 | 0 | 0 | 0 | 1 | 0 | 0 | 7 | 2 |
| HG0210 | *H. sapiens* | ENSG00000155849 | 0 | 0 | 0 | 0 | 0 | 0 | 0 | 0 | 0 | 0 | 0 | 0 | 0 | 0 | 1 | 0 | 0 | 7 | 2 |
| HG0211 | *H. sapiens* | ENSG00000176204 | 0 | 0 | 0 | 0 | 0 | 0 | 0 | 0 | 0 | 0 | 0 | 0 | 0 | 0 | 0 | 1 | 0 | 6 | 3 |
| HG0212 | *D. rerio* | ENSDARG00000100244 | 0 | 0 | 0 | 0 | 0 | 0 | 0 | 0 | 0 | 1 | 3 | 1 | 1 | 1 | 2 | 0 | 0 | 0 | 0 |
| HG0213.1 | *G. gallus* | ENSGALG00000001946 | 0 | 0 | 0 | 0 | 0 | 0 | 0 | 0 | 0 | 0 | 0 | 0 | 0 | 2 | 1 | 5 | 0 | 0 | 0 |
| HG0213.2 | *G. gallus* | ENSGALG00000001946 | 0 | 0 | 0 | 0 | 0 | 0 | 0 | 0 | 0 | 0 | 0 | 0 | 0 | 0 | 1 | 4 | 0 | 0 | 0 |
| HG0214.1 | *G. gallus* | ENSGALG00000012917 | 0 | 0 | 0 | 0 | 0 | 0 | 0 | 0 | 0 | 0 | 0 | 0 | 0 | 1 | 2 | 4 | 1 | 0 | 0 |
| HG0214.2 | *H. sapiens* | ENSG00000040731 | 0 | 0 | 0 | 0 | 0 | 0 | 0 | 0 | 0 | 0 | 0 | 0 | 0 | 0 | 0 | 0 | 1 | 3 | 4 |
| HG0215 | *D. rerio* | ENSDARG00000079280 | 0 | 0 | 0 | 0 | 0 | 0 | 0 | 0 | 0 | 0 | 5 | 1 | 0 | 0 | 0 | 0 | 0 | 0 | 0 |
| | *G. gallus* | ENSGALG00000016681 | 0 | 0 | 0 | 0 | 0 | 0 | 0 | 0 | 0 | 0 | 5 | 2 | 0 | 0 | 0 | 1 | 0 | 0 | 0 |
| HG0216.1 | *G. gallus* | ENSGALG00000020922 | 0 | 0 | 0 | 0 | 0 | 0 | 0 | 0 | 0 | 0 | 0 | 0 | 0 | 0 | 3 | 5 | 0 | 0 | 0 |
| HG0216.2 | *H. sapiens* | ENSG00000116667 | 0 | 0 | 0 | 0 | 0 | 0 | 0 | 0 | 0 | 0 | 0 | 0 | 0 | 0 | 0 | 0 | 3 | 10 | 2 |
| HG0217.1 | *H. sapiens* | ENSG00000150394 | 0 | 0 | 0 | 0 | 0 | 0 | 0 | 0 | 0 | 0 | 0 | 0 | 0 | 0 | 0 | 1 | 0 | 5 | 2 |
| HG0217.2 | *G. gallus* | ENSGALG00000005319 | 0 | 0 | 0 | 0 | 0 | 0 | 0 | 0 | 0 | 0 | 0 | 0 | 0 | 0 | 0 | 17 | 0 | 0 | 0 |
| HG0218 | *D. rerio* | ENSDARG00000100149 | 0 | 0 | 0 | 0 | 0 | 0 | 0 | 0 | 0 | 0 | 7 | 0 | 0 | 1 | 0 | 0 | 0 | 0 | 0 |
| HG0219 | *D. rerio* | ENSDARG00000104082 | 0 | 0 | 0 | 0 | 0 | 0 | 0 | 0 | 0 | 0 | 3 | 3 | 0 | 0 | 0 | 0 | 2 | 0 | 0 |
| HG0220 | *G. gallus* | ENSGALG00000008418 | 0 | 0 | 0 | 0 | 0 | 0 | 0 | 0 | 0 | 0 | 0 | 0 | 0 | 0 | 1 | 7 | 0 | 0 | 0 |
| HG0221 | *G. gallus* | ENSGALG00000010859 | 0 | 0 | 0 | 0 | 0 | 0 | 0 | 0 | 0 | 0 | 0 | 0 | 0 | 0 | 0 | 3 | 0 | 2 | 3 |
| HG0222 | *G. gallus* | ENSGALG00000011564 | 0 | 0 | 0 | 0 | 0 | 0 | 0 | 0 | 0 | 0 | 0 | 0 | 0 | 0 | 2 | 6 | 0 | 0 | 0 |
| HG0223 | *H. sapiens* | ENSG00000091656 | 0 | 0 | 0 | 0 | 0 | 0 | 0 | 0 | 0 | 0 | 0 | 0 | 0 | 0 | 0 | 4 | 1 | 1 | 2 |
| HG0224 | *H. sapiens* | ENSG00000118007 | 0 | 0 | 0 | 0 | 0 | 0 | 0 | 0 | 0 | 0 | 0 | 0 | 0 | 0 | 0 | 1 | 0 | 5 | 2 |
| HG0225 | *H. sapiens* | ENSG00000150051 | 0 | 0 | 0 | 0 | 0 | 0 | 0 | 0 | 0 | 0 | 4 | 0 | 0 | 0 | 0 | 0 | 0 | 3 | 1 |
| HG0226.1 | *H. sapiens* | ENSG00000180592 | 0 | 0 | 0 | 0 | 0 | 0 | 0 | 0 | 0 | 0 | 1 | 0 | 0 | 0 | 0 | 1 | 0 | 4 | 1 |
| HG0226.2 | *H. sapiens* | ENSG00000180592 | 0 | 0 | 0 | 0 | 0 | 0 | 0 | 0 | 0 | 0 | 0 | 0 | 0 | 0 | 0 | 0 | 0 | 2 | 1 |
| HG0226.3 | *H. sapiens* | ENSG00000180592 | 0 | 0 | 0 | 0 | 0 | 0 | 0 | 0 | 0 | 0 | 0 | 0 | 0 | 0 | 0 | 0 | 0 | 1 | 1 |
| HG0227.1 | *G. gallus* | ENSGALG00000031929 | 0 | 0 | 0 | 0 | 0 | 0 | 0 | 0 | 0 | 0 | 0 | 0 | 0 | 1 | 1 | 5 | 0 | 0 | 0 |
| HG0227.2 | *H. sapiens* | ENSG00000275163 | 0 | 0 | 0 | 0 | 0 | 0 | 0 | 0 | 0 | 0 | 0 | 0 | 0 | 0 | 0 | 3 | 0 | 0 | 0 |
| HG0228 | *D. rerio* | ENSDARG00000087887 | 0 | 0 | 0 | 0 | 0 | 0 | 0 | 0 | 0 | 0 | 4 | 2 | 1 | 0 | 0 | 0 | 0 | 0 | 0 |
| HG0229 | *G. gallus* | ENSGALG00000005474 | 0 | 0 | 0 | 0 | 0 | 0 | 0 | 0 | 0 | 0 | 0 | 0 | 0 | 0 | 1 | 6 | 0 | 0 | 0 |
| HG0230 | *G. gallus* | ENSGALG00000011007 | 0 | 0 | 0 | 0 | 0 | 0 | 0 | 0 | 0 | 0 | 0 | 0 | 0 | 0 | 2 | 5 | 0 | 0 | 0 |
| HG0231 | *H. sapiens* | ENSG00000150636 | 0 | 0 | 0 | 0 | 0 | 0 | 0 | 0 | 0 | 0 | 0 | 0 | 0 | 0 | 1 | 3 | 1 | 2 | 0 |
| HG0232 | *H. sapiens* | ENSG00000180875 | 0 | 0 | 0 | 0 | 0 | 0 | 0 | 0 | 0 | 0 | 0 | 0 | 0 | 0 | 2 | 0 | 1 | 1 | 3 |
| HG0233 | *H. sapiens* | ENSG00000206432 | 0 | 0 | 0 | 0 | 0 | 0 | 0 | 0 | 0 | 0 | 0 | 0 | 0 | 0 | 1 | 0 | 1 | 4 | 1 |
| HG0234.1 | *G. gallus* | ENSGALG00000011250 | 0 | 0 | 0 | 0 | 0 | 0 | 0 | 0 | 0 | 0 | 0 | 0 | 0 | 0 | 1 | 5 | 0 | 0 | 0 |
| HG0234.2 | *H. sapiens* | ENSG00000108684 | 0 | 0 | 0 | 0 | 0 | 0 | 0 | 0 | 0 | 0 | 0 | 0 | 0 | 0 | 0 | 0 | 1 | 2 | 1 |
| HG0235.1 | *G. gallus* | ENSGALG00000035927 | 0 | 0 | 0 | 0 | 0 | 0 | 0 | 0 | 0 | 0 | 0 | 0 | 0 | 0 | 1 | 5 | 0 | 0 | 0 |
| HG0235.2 | *H. sapiens* | ENSG00000101638 | 0 | 0 | 0 | 0 | 0 | 0 | 0 | 0 | 0 | 0 | 0 | 0 | 0 | 0 | 0 | 0 | 0 | 3 | 1 |
| HG0236 | *G. gallus* | ENSGALG00000004437 | 0 | 0 | 0 | 0 | 0 | 0 | 0 | 0 | 0 | 0 | 0 | 0 | 0 | 0 | 1 | 5 | 0 | 0 | 0 |
| HG0237 | *G. gallus* | ENSGALG00000012123 | 0 | 0 | 0 | 0 | 0 | 0 | 0 | 0 | 0 | 0 | 4 | 0 | 0 | 0 | 1 | 0 | 0 | 0 | 1 |
| HG0238 | *H. sapiens* | ENSG00000049618 | 0 | 0 | 0 | 0 | 0 | 0 | 0 | 0 | 0 | 0 | 4 | 0 | 1 | 0 | 0 | 1 | 0 | 0 | 0 |
| HG0239 | *H. sapiens* | ENSG00000149256 | 0 | 0 | 0 | 0 | 0 | 0 | 0 | 0 | 0 | 0 | 0 | 0 | 0 | 0 | 1 | 0 | 0 | 4 | 1 |
| HG0240 | *H. sapiens* | ENSG00000166342 | 0 | 0 | 0 | 0 | 0 | 0 | 0 | 0 | 0 | 0 | 0 | 0 | 0 | 0 | 1 | 3 | 0 | 1 | 1 |
| HG0241 | *H. sapiens* | ENSG00000198561 | 0 | 0 | 0 | 0 | 0 | 0 | 0 | 0 | 0 | 0 | 0 | 0 | 0 | 0 | 1 | 2 | 0 | 1 | 2 |
| HG0242.1 | *G. gallus* | ENSGALG00000007112 | 0 | 0 | 0 | 0 | 0 | 0 | 0 | 0 | 0 | 0 | 0 | 0 | 0 | 0 | 1 | 4 | 0 | 0 | 0 |
| HG0242.2 | *H. sapiens* | ENSG00000140471 | 0 | 0 | 0 | 0 | 0 | 0 | 0 | 0 | 0 | 0 | 0 | 0 | 0 | 0 | 0 | 0 | 0 | 2 | 0 |
| HG0243.1 | *H. sapiens* | ENSG00000163697 | 0 | 0 | 0 | 0 | 0 | 0 | 0 | 0 | 0 | 0 | 0 | 0 | 0 | 0 | 0 | 1 | 0 | 3 | 1 |
| HG0243.2 | *H. sapiens* | ENSG00000163697 | 0 | 0 | 0 | 0 | 0 | 0 | 0 | 0 | 0 | 0 | 0 | 0 | 0 | 0 | 0 | 0 | 0 | 8 | 3 |
| HG0244 | *G. gallus* | ENSGALG00000010837 | 0 | 0 | 0 | 0 | 0 | 0 | 0 | 0 | 0 | 0 | 0 | 0 | 0 | 1 | 0 | 3 | 0 | 0 | 1 |
| HG0245 | *H. sapiens* | ENSG00000091831 | 0 | 0 | 0 | 0 | 0 | 0 | 0 | 0 | 0 | 0 | 0 | 0 | 0 | 0 | 1 | 0 | 1 | 2 | 1 |
| HG0246 | *H. sapiens* | ENSG00000162676 | 0 | 0 | 0 | 0 | 0 | 0 | 0 | 0 | 0 | 0 | 0 | 0 | 0 | 0 | 0 | 1 | 0 | 2 | 2 |
| HG0247.1 | *D. rerio* | ENSDARG00000014746 | 0 | 0 | 0 | 0 | 0 | 0 | 0 | 0 | 0 | 0 | 0 | 0 | 0 | 0 | 1 | 3 | 0 | 0 | 0 |
| HG0247.2 | *H. sapiens* | ENSG00000100320 | 0 | 0 | 0 | 0 | 0 | 0 | 0 | 0 | 0 | 0 | 0 | 0 | 0 | 0 | 0 | 0 | 1 | 7 | 3 |
| HG0248.1 | *G. gallus* | ENSGALG00000004570 | 0 | 0 | 0 | 0 | 0 | 0 | 0 | 0 | 0 | 0 | 0 | 0 | 0 | 0 | 1 | 3 | 0 | 0 | 0 |
| HG0248.2 | *H. sapiens* | ENSG00000124151 | 0 | 0 | 0 | 0 | 0 | 0 | 0 | 0 | 0 | 0 | 0 | 0 | 0 | 0 | 0 | 0 | 0 | 7 | 3 |
| HG0249 | *D. rerio* | ENSDARG00000062550 | 0 | 0 | 0 | 0 | 0 | 0 | 0 | 0 | 0 | 0 | 1 | 2 | 0 | 0 | 1 | 0 | 0 | 0 | 0 |
| HG0250 | *G. gallus* | ENSGALG00000014861 | 0 | 0 | 0 | 0 | 0 | 0 | 0 | 0 | 0 | 0 | 0 | 0 | 0 | 0 | 0 | 3 | 0 | 1 | 0 |
| HG0251 | *H. sapiens* | ENSG00000185053 | 0 | 0 | 0 | 0 | 0 | 0 | 0 | 0 | 0 | 0 | 0 | 0 | 0 | 0 | 1 | 0 | 0 | 2 | 1 |
| HG0252.1 | *G. gallus* | ENSGALG00000009645 | 0 | 0 | 0 | 0 | 0 | 0 | 0 | 0 | 0 | 0 | 0 | 0 | 0 | 0 | 1 | 1 | 0 | 0 | 1 |
| HG0252.2 | *H. sapiens* | ENSG00000119715 | 0 | 0 | 0 | 0 | 0 | 0 | 0 | 0 | 0 | 0 | 0 | 0 | 0 | 0 | 0 | 1 | 0 | 1 | 1 |
| HG0252.3 | *H. sapiens* | ENSG00000196482 | 0 | 0 | 0 | 0 | 0 | 0 | 0 | 0 | 0 | 0 | 0 | 0 | 0 | 0 | 0 | 0 | 1 | 3 | 1 |
| HG0253.1 | *G. gallus* | ENSGALG00000015184 | 0 | 0 | 0 | 0 | 0 | 0 | 0 | 0 | 0 | 0 | 0 | 0 | 0 | 0 | 1 | 0 | 0 | 2 | 0 |
| HG0253.2 | *H. sapiens* | ENSG00000140332 | 0 | 0 | 0 | 0 | 0 | 0 | 0 | 0 | 0 | 0 | 0 | 0 | 0 | 0 | 0 | 0 | 0 | 8 | 2 |
| HG0253.3 | *H. sapiens* | ENSG00000140332 | 0 | 0 | 0 | 0 | 0 | 0 | 0 | 0 | 0 | 0 | 0 | 0 | 0 | 0 | 0 | 0 | 0 | 3 | 0 |
| HG0254.1 | *G. gallus* | ENSGALG00000035919 | 0 | 0 | 0 | 0 | 0 | 0 | 0 | 0 | 0 | 0 | 0 | 0 | 0 | 0 | 1 | 2 | 0 | 0 | 0 |
| HG0254.2 | *H. sapiens* | ENSG00000072134 | 0 | 0 | 0 | 0 | 0 | 0 | 0 | 0 | 0 | 0 | 0 | 0 | 0 | 0 | 0 | 0 | 0 | 2 | 2 |
| HG0255 | *G. gallus* | ENSGALG00000012139 | 0 | 0 | 0 | 0 | 0 | 0 | 0 | 0 | 0 | 0 | 0 | 0 | 0 | 0 | 2 | 1 | 0 | 0 | 0 |
| HG0256 | *G. gallus* | ENSGALG00000039901 | 0 | 0 | 0 | 0 | 0 | 0 | 0 | 0 | 0 | 0 | 0 | 0 | 0 | 0 | 1 | 2 | 0 | 0 | 0 |
| HG0257 | *H. sapiens* | ENSG00000198791 | 0 | 0 | 0 | 0 | 0 | 0 | 0 | 0 | 0 | 0 | 0 | 0 | 0 | 0 | 0 | 1 | 0 | 1 | 1 |

| | | | | | | | | | | | | | | | | | | | | | |
|---|---|---|---|---|---|---|---|---|---|---|---|---|---|---|---|---|---|---|---|---|---|
| HG0258.1 | *G. gallus* | ENSGALG00000016797 | 0 | 0 | 0 | 0 | 0 | 0 | 0 | 0 | 0 | 0 | 0 | 0 | 0 | 0 | 1 | 1 | 0 | 0 | 0 |
| HG0258.2 | *G. gallus* | ENSGALG00000016797 | 0 | 0 | 0 | 0 | 0 | 0 | 0 | 0 | 0 | 0 | 0 | 0 | 0 | 0 | 0 | 13 | 0 | 0 | 0 |
| HG0259 | *G. gallus* | ENSGALG00000009936 | 0 | 0 | 0 | 0 | 0 | 0 | 0 | 0 | 0 | 0 | 0 | 0 | 0 | 0 | 2 | 0 | 0 | 0 | 0 |
| HG0260.1 | *G. gallus* | ENSGALG00000016890 | 0 | 0 | 0 | 0 | 0 | 0 | 0 | 0 | 0 | 0 | 0 | 0 | 0 | 0 | 0 | 19 | 0 | 0 | 0 |
| HG0260.2 | *H. sapiens* | ENSG00000139793 | 0 | 0 | 0 | 0 | 0 | 0 | 0 | 0 | 0 | 0 | 0 | 0 | 0 | 0 | 0 | 0 | 0 | 10 | 4 |
| HG0260.3 | *H. sapiens* | ENSG00000152601 | 0 | 0 | 0 | 0 | 0 | 0 | 0 | 0 | 0 | 0 | 0 | 0 | 0 | 0 | 0 | 0 | 0 | 6 | 2 |
| HG0260.4 | *H. sapiens* | ENSG00000152601 | 0 | 0 | 0 | 0 | 0 | 0 | 0 | 0 | 0 | 0 | 0 | 0 | 0 | 0 | 0 | 0 | 0 | 6 | 1 |
| HG0260.5 | *H. sapiens* | ENSG00000152601 | 0 | 0 | 0 | 0 | 0 | 0 | 0 | 0 | 0 | 0 | 0 | 0 | 0 | 0 | 0 | 0 | 0 | 3 | 0 |
| HG0260.6 | *H. sapiens* | ENSG00000152601 | 0 | 0 | 0 | 0 | 0 | 0 | 0 | 0 | 0 | 0 | 0 | 0 | 0 | 0 | 0 | 0 | 0 | 3 | 0 |
| HG0261.1 | *G. gallus* | ENSGALG00000005867 | 0 | 0 | 0 | 0 | 0 | 0 | 0 | 0 | 0 | 0 | 0 | 0 | 0 | 0 | 0 | 11 | 0 | 0 | 0 |
| HG0261.2 | *G. gallus* | ENSGALG00000009520 | 0 | 0 | 0 | 0 | 0 | 0 | 0 | 0 | 0 | 0 | 0 | 0 | 0 | 0 | 0 | 4 | 0 | 0 | 0 |
| HG0261.3 | *H. sapiens* | ENSG00000145416 | 0 | 0 | 0 | 0 | 0 | 0 | 0 | 0 | 0 | 0 | 0 | 0 | 0 | 0 | 0 | 0 | 1 | 1 | 2 |
| HG0261.4 | *H. sapiens* | ENSG00000165406 | 0 | 0 | 0 | 0 | 0 | 0 | 0 | 0 | 0 | 0 | 0 | 0 | 0 | 0 | 0 | 0 | 0 | 2 | 0 |
| HG0262.1 | *H. sapiens* | ENSG00000023445 | 0 | 0 | 0 | 0 | 0 | 0 | 0 | 0 | 0 | 0 | 0 | 0 | 0 | 0 | 0 | 0 | 0 | 4 | 2 |
| HG0262.2 | *H. sapiens* | ENSG00000023445 | 0 | 0 | 0 | 0 | 0 | 0 | 0 | 0 | 0 | 0 | 0 | 0 | 0 | 0 | 0 | 0 | 0 | 3 | 1 |
| HG0262.3 | *G. gallus* | ENSGALG00000017186 | 0 | 0 | 0 | 0 | 0 | 0 | 0 | 0 | 0 | 0 | 0 | 0 | 0 | 0 | 0 | 2 | 0 | 0 | 0 |
| HG0262.4 | *H. sapiens* | ENSG00000023445 | 0 | 0 | 0 | 0 | 0 | 0 | 0 | 0 | 0 | 0 | 0 | 0 | 0 | 0 | 0 | 0 | 0 | 2 | 0 |
| HG0263.1 | *H. sapiens* | ENSG00000169826 | 0 | 0 | 0 | 0 | 0 | 0 | 0 | 0 | 0 | 0 | 0 | 0 | 0 | 0 | 0 | 0 | 0 | 5 | 1 |
| HG0263.2 | *G. gallus* | ENSGALG00000002543 | 0 | 0 | 0 | 0 | 0 | 0 | 0 | 0 | 0 | 0 | 0 | 0 | 0 | 0 | 0 | 3 | 0 | 0 | 0 |
| HG0263.3 | *G. gallus* | ENSGALG00000002543 | 0 | 0 | 0 | 0 | 0 | 0 | 0 | 0 | 0 | 0 | 0 | 0 | 0 | 0 | 0 | 3 | 0 | 0 | 0 |
| HG0263.4 | *H. sapiens* | ENSG00000147408 | 0 | 0 | 0 | 0 | 0 | 0 | 0 | 0 | 0 | 0 | 0 | 0 | 0 | 0 | 0 | 0 | 0 | 3 | 0 |
| HG0264.1 | *H. sapiens* | ENSG00000151458 | 0 | 0 | 0 | 0 | 0 | 0 | 0 | 0 | 0 | 0 | 0 | 0 | 0 | 0 | 0 | 0 | 1 | 9 | 3 |
| HG0264.2 | *H. sapiens* | ENSG00000151458 | 0 | 0 | 0 | 0 | 0 | 0 | 0 | 0 | 0 | 0 | 0 | 0 | 0 | 0 | 0 | 0 | 1 | 6 | 3 |
| HG0264.3 | *G. gallus* | ENSGALG00000011831 | 0 | 0 | 0 | 0 | 0 | 0 | 0 | 0 | 0 | 0 | 0 | 0 | 0 | 0 | 0 | 4 | 0 | 0 | 0 |
| HG0264.4 | *H. sapiens* | ENSG00000151458 | 0 | 0 | 0 | 0 | 0 | 0 | 0 | 0 | 0 | 0 | 0 | 0 | 0 | 0 | 0 | 0 | 0 | 3 | 0 |
| HG0265.1 | *H. sapiens* | ENSG00000144285 | 0 | 0 | 0 | 0 | 0 | 0 | 0 | 0 | 0 | 0 | 0 | 0 | 0 | 0 | 0 | 0 | 0 | 5 | 2 |
| HG0265.2 | *H. sapiens* | ENSG00000196876 | 0 | 0 | 0 | 0 | 0 | 0 | 0 | 0 | 0 | 0 | 0 | 0 | 0 | 0 | 0 | 0 | 0 | 4 | 2 |
| HG0265.3 | *D. rerio* | ENSDARG00000018032 | 0 | 0 | 0 | 0 | 0 | 0 | 0 | 0 | 0 | 0 | 3 | 2 | 0 | 0 | 0 | 0 | 0 | 0 | 0 |
| HG0266.1 | *H. sapiens* | ENSG00000184185 | 0 | 0 | 0 | 0 | 0 | 0 | 0 | 0 | 0 | 0 | 0 | 0 | 0 | 0 | 0 | 0 | 0 | 4 | 4 |
| | | ENSG00000260458 | 0 | 0 | 0 | 0 | 0 | 0 | 0 | 0 | 0 | 0 | 0 | 0 | 0 | 0 | 0 | 0 | 0 | 4 | 4 |
| HG0266.2 | *G. gallus* | ENSGALG00000001181 | 0 | 0 | 0 | 0 | 0 | 0 | 0 | 0 | 0 | 0 | 0 | 0 | 0 | 0 | 0 | 4 | 0 | 0 | 0 |
| HG0267 | *D. rerio* | ENSDARG00000028173 | 0 | 0 | 0 | 0 | 0 | 0 | 0 | 0 | 0 | 0 | 2 | 3 | 0 | 0 | 0 | 0 | 0 | 0 | 0 |
| | | ENSDARG00000052330 | 0 | 0 | 0 | 0 | 0 | 0 | 0 | 0 | 0 | 0 | 9 | 2 | 0 | 0 | 0 | 0 | 0 | 0 | 0 |
| | *H. sapiens* | ENSG00000164889 | 0 | 0 | 0 | 0 | 0 | 0 | 0 | 0 | 0 | 0 | 0 | 0 | 0 | 0 | 0 | 0 | 0 | 8 | 3 |
| HG0268.1 | *D. rerio* | ENSDARG00000011703 | 0 | 0 | 0 | 0 | 0 | 0 | 0 | 0 | 0 | 0 | 3 | 2 | 0 | 0 | 0 | 0 | 0 | 0 | 0 |
| HG0268.2 | *H. sapiens* | ENSG00000134852 | 0 | 0 | 0 | 0 | 0 | 0 | 0 | 0 | 0 | 0 | 0 | 0 | 0 | 0 | 0 | 0 | 0 | 3 | 2 |
| HG0268.3 | *H. sapiens* | ENSG00000134852 | 0 | 0 | 0 | 0 | 0 | 0 | 0 | 0 | 0 | 0 | 0 | 0 | 0 | 0 | 0 | 0 | 0 | 3 | 1 |
| HG0269.1 | *G. gallus* | ENSGALG00000028685 | 0 | 0 | 0 | 0 | 0 | 0 | 0 | 0 | 0 | 0 | 0 | 0 | 0 | 0 | 0 | 5 | 0 | 0 | 0 |
| HG0269.2 | *D. rerio* | ENSDARG00000061471 | 0 | 0 | 0 | 0 | 0 | 0 | 0 | 0 | 0 | 0 | 1 | 1 | 0 | 0 | 0 | 0 | 0 | 0 | 0 |
| HG0269.3 | *G. gallus* | ENSGALG00000011686 | 0 | 0 | 0 | 0 | 0 | 0 | 0 | 0 | 0 | 0 | 0 | 0 | 0 | 0 | 0 | 2 | 0 | 0 | 0 |
| HG0270.1 | *G. gallus* | ENSGALG00000015147 | 0 | 0 | 0 | 0 | 0 | 0 | 0 | 0 | 0 | 0 | 0 | 0 | 0 | 0 | 0 | 13 | 0 | 0 | 0 |
| HG0270.2 | *G. gallus* | ENSGALG00000015147 | 0 | 0 | 0 | 0 | 0 | 0 | 0 | 0 | 0 | 0 | 0 | 0 | 0 | 0 | 0 | 12 | 0 | 0 | 0 |
| HG0270.3 | *H. sapiens* | ENSG00000128918 | 0 | 0 | 0 | 0 | 0 | 0 | 0 | 0 | 0 | 0 | 0 | 0 | 0 | 0 | 0 | 0 | 2 | 7 | 1 |
| HG0271.1 | *G. gallus* | ENSGALG00000004064 | 0 | 0 | 0 | 0 | 0 | 0 | 0 | 0 | 0 | 0 | 0 | 0 | 0 | 0 | 0 | 11 | 0 | 0 | 0 |
| HG0271.2 | *H. sapiens* | ENSG00000164850 | 0 | 0 | 0 | 0 | 0 | 0 | 0 | 0 | 0 | 0 | 0 | 0 | 0 | 0 | 0 | 0 | 0 | 2 | 2 |
| HG0271.3 | *D. rerio* | ENSDARG00000074661 | 0 | 0 | 0 | 0 | 0 | 0 | 0 | 0 | 0 | 0 | 1 | 2 | 0 | 0 | 0 | 0 | 0 | 0 | 0 |
| HG0272.1 | *D. melanogaster* | FBgn0034649 | 0 | 0 | 0 | 0 | 0 | 7 | 25 | 0 | 0 | 0 | 0 | 0 | 0 | 0 | 0 | 0 | 0 | 0 | 0 |
| HG0272.2 | *D. melanogaster* | FBgn0034649 | 0 | 0 | 0 | 0 | 0 | 6 | 24 | 0 | 0 | 0 | 0 | 0 | 0 | 0 | 0 | 0 | 0 | 0 | 0 |
| HG0272.3 | *H. sapiens* | ENSG00000143315 | 0 | 0 | 0 | 0 | 0 | 0 | 0 | 0 | 0 | 0 | 0 | 0 | 0 | 0 | 0 | 0 | 0 | 3 | 0 |
| HG0273.1 | *G. gallus* | ENSGALG00000003293 | 0 | 0 | 0 | 0 | 0 | 0 | 0 | 0 | 0 | 0 | 0 | 0 | 0 | 0 | 0 | 16 | 0 | 0 | 0 |
| HG0273.2 | *H. sapiens* | ENSG00000140943 | 0 | 0 | 0 | 0 | 0 | 0 | 0 | 0 | 0 | 0 | 0 | 0 | 0 | 0 | 0 | 0 | 0 | 7 | 1 |
| HG0273.3 | *H. sapiens* | ENSG00000140943 | 0 | 0 | 0 | 0 | 0 | 0 | 0 | 0 | 0 | 0 | 0 | 0 | 0 | 0 | 0 | 0 | 0 | 0 | 2 |
| HG0274.1 | *H. sapiens* | ENSG00000204256 | 0 | 0 | 0 | 0 | 0 | 0 | 0 | 0 | 0 | 0 | 0 | 0 | 0 | 0 | 0 | 0 | 0 | 11 | 2 |
| HG0274.2 | *D. rerio* | ENSDARG00000100129 | 0 | 0 | 0 | 0 | 0 | 0 | 0 | 0 | 0 | 0 | 0 | 2 | 0 | 0 | 0 | 0 | 0 | 0 | 0 |
| HG0275.1 | *H. sapiens* | ENSG00000100239 | 0 | 0 | 0 | 0 | 0 | 0 | 0 | 0 | 0 | 0 | 0 | 0 | 0 | 0 | 0 | 0 | 0 | 5 | 1 |
| HG0275.2 | *D. rerio* | ENSDARG00000045540 | 0 | 0 | 0 | 0 | 0 | 0 | 0 | 0 | 0 | 0 | 0 | 2 | 0 | 0 | 0 | 0 | 0 | 0 | 0 |
| HG0276.1 | *G. gallus* | ENSGALG00000008912 | 0 | 0 | 0 | 0 | 0 | 0 | 0 | 0 | 0 | 0 | 0 | 0 | 0 | 0 | 0 | 4 | 0 | 0 | 0 |
| HG0276.2 | *H. sapiens* | ENSG00000073734 | 0 | 0 | 0 | 0 | 0 | 0 | 0 | 0 | 0 | 0 | 0 | 0 | 0 | 0 | 0 | 0 | 0 | 2 | 0 |
| HG0277.1 | *H. sapiens* | ENSG00000102531 | 0 | 0 | 0 | 0 | 0 | 0 | 0 | 0 | 0 | 0 | 0 | 0 | 0 | 0 | 0 | 0 | 1 | 8 | 2 |
| HG0277.2 | *G. gallus* | ENSGALG00000004169 | 0 | 0 | 0 | 0 | 0 | 0 | 0 | 0 | 0 | 0 | 0 | 0 | 0 | 0 | 0 | 2 | 0 | 0 | 0 |
| HG0278.1 | *H. sapiens* | ENSG00000119946 | 0 | 0 | 0 | 0 | 0 | 0 | 0 | 0 | 0 | 0 | 0 | 0 | 0 | 0 | 0 | 0 | 0 | 4 | 1 |
| HG0278.2 | *D. rerio* | ENSDARG00000078733 | 0 | 0 | 0 | 0 | 0 | 0 | 0 | 0 | 0 | 0 | 3 | 0 | 0 | 0 | 0 | 0 | 0 | 0 | 0 |
| HG0279.1 | *H. sapiens* | ENSG00000175928 | 0 | 0 | 0 | 0 | 0 | 0 | 0 | 0 | 0 | 0 | 0 | 0 | 0 | 0 | 0 | 0 | 0 | 6 | 2 |
| HG0279.2 | *G. gallus* | ENSGALG00000035099 | 0 | 0 | 0 | 0 | 0 | 0 | 0 | 0 | 0 | 0 | 0 | 0 | 0 | 0 | 0 | 2 | 0 | 0 | 0 |
| HG0280.1 | *H. sapiens* | ENSG00000156299 | 0 | 0 | 0 | 0 | 0 | 0 | 0 | 0 | 0 | 0 | 0 | 0 | 0 | 0 | 0 | 0 | 0 | 2 | 3 |
| HG0280.2 | *D. melanogaster* | FBgn0085447 | 0 | 0 | 0 | 0 | 0 | 0 | 4 | 0 | 0 | 0 | 0 | 0 | 0 | 0 | 0 | 0 | 0 | 0 | 0 |
| HG0281.1 | *G. gallus* | ENSGALG00000006766 | 0 | 0 | 0 | 0 | 0 | 0 | 0 | 0 | 0 | 0 | 0 | 0 | 0 | 0 | 0 | 12 | 0 | 0 | 0 |
| HG0281.2 | *H. sapiens* | ENSG00000100150 | 0 | 0 | 0 | 0 | 0 | 0 | 0 | 0 | 0 | 0 | 0 | 0 | 0 | 0 | 0 | 0 | 0 | 7 | 2 |
| HG0282.1 | *D. rerio* | ENSDARG00000061328 | 0 | 0 | 0 | 0 | 0 | 0 | 0 | 0 | 0 | 0 | 9 | 2 | 0 | 0 | 0 | 0 | 0 | 0 | 0 |
| HG0282.2 | *H. sapiens* | ENSG00000144857 | 0 | 0 | 0 | 0 | 0 | 0 | 0 | 0 | 0 | 0 | 0 | 0 | 0 | 0 | 0 | 0 | 0 | 2 | 0 |
| HG0283.1 | *G. gallus* | ENSGALG00000010270 | 0 | 0 | 0 | 0 | 0 | 0 | 0 | 0 | 0 | 0 | 0 | 0 | 0 | 0 | 0 | 13 | 0 | 0 | 0 |
| HG0283.2 | *H. sapiens* | ENSG00000119682 | 0 | 0 | 0 | 0 | 0 | 0 | 0 | 0 | 0 | 0 | 0 | 0 | 0 | 0 | 0 | 0 | 0 | 9 | 4 |
| HG0284.1 | *H. sapiens* | ENSG00000007168 | 0 | 0 | 0 | 0 | 0 | 0 | 0 | 0 | 0 | 0 | 0 | 0 | 0 | 0 | 0 | 0 | 0 | 7 | 4 |
| HG0284.2 | *G. gallus* | ENSGALG00000005834 | 0 | 0 | 0 | 0 | 0 | 0 | 0 | 0 | 0 | 0 | 0 | 0 | 0 | 0 | 0 | 3 | 0 | 0 | 0 |
| HG0285.1 | *G. gallus* | ENSGALG00000012156 | 0 | 0 | 0 | 0 | 0 | 0 | 0 | 0 | 0 | 0 | 0 | 0 | 0 | 0 | 0 | 2 | 0 | 0 | 0 |
| HG0285.2 | *H. sapiens* | ENSG00000175497 | 0 | 0 | 0 | 0 | 0 | 0 | 0 | 0 | 0 | 0 | 0 | 0 | 0 | 0 | 0 | 0 | 0 | 1 | 1 |
| HG0286.1 | *H. sapiens* | ENSG00000116698 | 0 | 0 | 0 | 0 | 0 | 0 | 0 | 0 | 0 | 0 | 0 | 0 | 0 | 0 | 0 | 0 | 0 | 7 | 1 |
| HG0286.2 | *D. rerio* | ENSDARG00000060767 | 0 | 0 | 0 | 0 | 0 | 0 | 0 | 0 | 0 | 0 | 0 | 3 | 0 | 0 | 0 | 0 | 0 | 0 | 0 |
| HG0287.1 | *H. sapiens* | ENSG00000170325 | 0 | 0 | 0 | 0 | 0 | 0 | 0 | 0 | 0 | 0 | 0 | 0 | 0 | 0 | 0 | 0 | 0 | 5 | 3 |

| | | | | | | | | | | | | | | | | | | | | | |
|---|---|---|---|---|---|---|---|---|---|---|---|---|---|---|---|---|---|---|---|---|---|
| HG0287.2 | *D. rerio* | ENSDARG00000104251 | 0 | 0 | 0 | 0 | 0 | 0 | 0 | 0 | 0 | 0 | 5 | 1 | 0 | 0 | 0 | 0 | 0 | 0 | 0 |
| HG0288.1 | *G. gallus* | ENSGALG00000006655 | 0 | 0 | 0 | 0 | 0 | 0 | 0 | 0 | 0 | 0 | 0 | 0 | 0 | 0 | 0 | 4 | 0 | 0 | 0 |
| HG0288.2 | *H. sapiens* | ENSG00000073803 | 0 | 0 | 0 | 0 | 0 | 0 | 0 | 0 | 0 | 0 | 0 | 0 | 0 | 0 | 0 | 0 | 0 | 2 | 2 |
| HG0289 | *G. gallus* | ENSGALG00000009581 | 0 | 0 | 0 | 0 | 0 | 0 | 0 | 0 | 0 | 0 | 0 | 0 | 0 | 0 | 0 | 28 | 0 | 0 | 0 |
| | *H. sapiens* | ENSG00000072415 | 0 | 0 | 0 | 0 | 0 | 0 | 0 | 0 | 0 | 0 | 0 | 0 | 0 | 0 | 0 | 0 | 0 | 10 | 3 |
| HG0290.1 | *H. sapiens* | ENSG00000116539 | 0 | 0 | 0 | 0 | 0 | 0 | 0 | 0 | 0 | 0 | 0 | 0 | 0 | 0 | 0 | 0 | 2 | 10 | 3 |
| HG0290.2 | *D. rerio* | ENSDARG00000070981 | 0 | 0 | 0 | 0 | 0 | 0 | 0 | 0 | 0 | 0 | 4 | 1 | 0 | 0 | 0 | 0 | 0 | 0 | 0 |
| HG0291.1 | *G. gallus* | ENSGALG00000009173 | 0 | 0 | 0 | 0 | 0 | 0 | 0 | 0 | 0 | 0 | 0 | 0 | 0 | 0 | 0 | 14 | 0 | 0 | 0 |
| HG0291.2 | *H. sapiens* | ENSG00000151892 | 0 | 0 | 0 | 0 | 0 | 0 | 0 | 0 | 0 | 0 | 0 | 0 | 0 | 0 | 0 | 0 | 0 | 6 | 3 |
| HG0292.1 | *G. gallus* | ENSGALG00000006502 | 0 | 0 | 0 | 0 | 0 | 0 | 0 | 0 | 0 | 0 | 0 | 0 | 0 | 0 | 0 | 11 | 0 | 0 | 0 |
| HG0292.2 | *H. sapiens* | ENSG00000180628 | 0 | 0 | 0 | 0 | 0 | 0 | 0 | 0 | 0 | 0 | 0 | 0 | 0 | 0 | 0 | 0 | 0 | 4 | 0 |
| HG0293 | *G. gallus* | ENSGALG00000012187 | 0 | 0 | 0 | 0 | 0 | 0 | 0 | 0 | 0 | 0 | 0 | 0 | 0 | 0 | 0 | 16 | 0 | 0 | 0 |
| | *H. sapiens* | ENSG00000152127 | 0 | 0 | 0 | 0 | 0 | 0 | 0 | 0 | 0 | 0 | 0 | 0 | 0 | 0 | 0 | 0 | 2 | 7 | 4 |
| HG0294.1 | *G. gallus* | ENSGALG00000017036 | 0 | 0 | 0 | 0 | 0 | 0 | 0 | 0 | 0 | 0 | 0 | 0 | 0 | 0 | 0 | 8 | 0 | 0 | 0 |
| HG0294.2 | *H. sapiens* | ENSG00000183722 | 0 | 0 | 0 | 0 | 0 | 0 | 0 | 0 | 0 | 0 | 0 | 0 | 0 | 0 | 0 | 0 | 0 | 1 | 4 |
| HG0295.1 | *H. sapiens* | ENSG00000166888 | 0 | 0 | 0 | 0 | 0 | 0 | 0 | 0 | 0 | 0 | 0 | 0 | 0 | 0 | 0 | 0 | 2 | 9 | 3 |
| HG0295.2 | *G. gallus* | ENSGALG00000003282 | 0 | 0 | 0 | 0 | 0 | 0 | 0 | 0 | 0 | 0 | 0 | 0 | 0 | 0 | 0 | 0 | 0 | 6 | 2 |
| HG0296 | *G. gallus* | ENSGALG00000006912 | 0 | 0 | 0 | 0 | 0 | 0 | 0 | 0 | 0 | 0 | 0 | 0 | 0 | 0 | 0 | 0 | 1 | 2 | 2 |
| HG0297 | *G. gallus* | ENSGALG00000031684 | 0 | 0 | 0 | 0 | 0 | 0 | 0 | 0 | 0 | 0 | 0 | 0 | 0 | 0 | 0 | 0 | 0 | 3 | 3 |
| HG0298 | *G. gallus* | ENSGALG00000042308 | 0 | 0 | 0 | 0 | 0 | 0 | 0 | 0 | 0 | 0 | 0 | 0 | 0 | 0 | 0 | 0 | 0 | 2 | 1 |
| HG0299 | *G. gallus* | ENSGALG00000040465 | 0 | 0 | 0 | 0 | 0 | 0 | 0 | 0 | 0 | 0 | 3 | 0 | 0 | 0 | 0 | 0 | 0 | 0 | 0 |
| HG0300 | *D. melanogaster* | FBgn0031668 | 0 | 0 | 0 | 0 | 0 | 14 | 28 | 0 | 0 | 0 | 0 | 0 | 0 | 0 | 0 | 0 | 0 | 0 | 0 |
| HG0301 | *D. melanogaster* | FBgn0036856 | 0 | 0 | 0 | 0 | 0 | 6 | 27 | 0 | 0 | 0 | 0 | 0 | 0 | 0 | 0 | 0 | 0 | 0 | 0 |
| HG0302 | *D. melanogaster* | FBgn0031454 | 0 | 0 | 0 | 0 | 0 | 7 | 24 | 0 | 0 | 0 | 0 | 0 | 0 | 0 | 0 | 0 | 0 | 0 | 0 |
| HG0303 | *D. melanogaster* | FBgn0260464 | 0 | 0 | 0 | 0 | 0 | 7 | 23 | 0 | 0 | 0 | 0 | 0 | 0 | 0 | 0 | 0 | 0 | 0 | 0 |
| HG0304 | *D. melanogaster* | FBgn0260467 | 0 | 0 | 0 | 0 | 0 | 8 | 22 | 0 | 0 | 0 | 0 | 0 | 0 | 0 | 0 | 0 | 0 | 0 | 0 |
| HG0305 | *D. melanogaster* | FBgn0050290 | 0 | 0 | 0 | 0 | 0 | 8 | 21 | 0 | 0 | 0 | 0 | 0 | 0 | 0 | 0 | 0 | 0 | 0 | 0 |
| HG0306 | *D. melanogaster* | FBgn0259726 | 0 | 0 | 0 | 0 | 0 | 5 | 22 | 0 | 0 | 0 | 0 | 0 | 0 | 0 | 0 | 0 | 0 | 0 | 0 |
| HG0307 | *D. melanogaster* | FBgn0037822 | 0 | 0 | 0 | 0 | 0 | 7 | 19 | 0 | 0 | 0 | 0 | 0 | 0 | 0 | 0 | 0 | 0 | 0 | 0 |
| HG0308 | *D. melanogaster* | FBgn0264743 | 0 | 0 | 0 | 0 | 0 | 1 | 24 | 0 | 0 | 0 | 0 | 0 | 0 | 0 | 0 | 0 | 0 | 0 | 0 |
| HG0309 | *D. melanogaster* | FBgn0037689 | 0 | 0 | 0 | 0 | 0 | 8 | 15 | 0 | 0 | 0 | 0 | 0 | 0 | 0 | 0 | 0 | 0 | 0 | 0 |
| HG0310 | *D. melanogaster* | FBgn0039339 | 0 | 0 | 0 | 0 | 0 | 7 | 13 | 0 | 0 | 0 | 0 | 0 | 0 | 0 | 0 | 0 | 0 | 0 | 0 |
| HG0311 | *D. melanogaster* | FBgn0263251 | 0 | 0 | 0 | 0 | 0 | 0 | 20 | 0 | 0 | 0 | 0 | 0 | 0 | 0 | 0 | 0 | 0 | 0 | 0 |
| HG0312 | *D. melanogaster* | FBgn0038641 | 0 | 0 | 0 | 0 | 0 | 11 | 8 | 0 | 0 | 0 | 0 | 0 | 0 | 0 | 0 | 0 | 0 | 0 | 0 |
| HG0313 | *D. melanogaster* | FBgn0259725 | 0 | 0 | 0 | 0 | 0 | 2 | 15 | 0 | 0 | 0 | 0 | 0 | 0 | 0 | 0 | 0 | 0 | 0 | 0 |
| HG0314 | *D. melanogaster* | FBgn0260468 | 0 | 0 | 0 | 0 | 0 | 0 | 17 | 0 | 0 | 0 | 0 | 0 | 0 | 0 | 0 | 0 | 0 | 0 | 0 |
| HG0315 | *D. melanogaster* | FBgn0260234 | 0 | 0 | 0 | 0 | 0 | 1 | 13 | 0 | 0 | 0 | 0 | 0 | 0 | 0 | 0 | 0 | 0 | 0 | 0 |
| HG0316 | *D. melanogaster* | FBgn0041164 | 0 | 0 | 0 | 0 | 0 | 2 | 9 | 0 | 0 | 0 | 0 | 0 | 0 | 0 | 0 | 0 | 0 | 0 | 0 |
| HG0317 | *D. melanogaster* | FBgn0050011 | 0 | 0 | 0 | 0 | 0 | 0 | 7 | 0 | 0 | 0 | 0 | 0 | 0 | 0 | 0 | 0 | 0 | 0 | 0 |
| HG0318 | *D. melanogaster* | FBgn0035293 | 0 | 0 | 0 | 0 | 0 | 0 | 6 | 0 | 0 | 0 | 0 | 0 | 0 | 0 | 0 | 0 | 0 | 0 | 0 |
| HG0319 | *D. melanogaster* | FBgn0026778 | 0 | 0 | 0 | 0 | 0 | 0 | 4 | 0 | 0 | 0 | 0 | 0 | 0 | 0 | 0 | 0 | 0 | 0 | 0 |
| HG0320 | *D. melanogaster* | FBgn0040993 | 0 | 0 | 0 | 0 | 0 | 0 | 2 | 0 | 0 | 0 | 0 | 0 | 0 | 0 | 0 | 0 | 0 | 0 | 0 |
| HG0321 | *D. rerio* | ENSDARG00000070426 | 0 | 0 | 0 | 0 | 0 | 0 | 0 | 0 | 0 | 0 | 20 | 3 | 0 | 0 | 0 | 0 | 0 | 0 | 0 |
| HG0322 | *D. rerio* | ENSDARG00000070917 | 0 | 0 | 0 | 0 | 0 | 0 | 0 | 0 | 0 | 0 | 17 | 3 | 0 | 0 | 0 | 0 | 0 | 0 | 0 |
| HG0323 | *D. rerio* | ENSDARG00000071197 | 0 | 0 | 0 | 0 | 0 | 0 | 0 | 0 | 0 | 0 | 19 | 1 | 0 | 0 | 0 | 0 | 0 | 0 | 0 |
| HG0324 | *D. rerio* | ENSDARG00000094965 | 0 | 0 | 0 | 0 | 0 | 0 | 0 | 0 | 0 | 0 | 18 | 1 | 0 | 0 | 0 | 0 | 0 | 0 | 0 |
| HG0325 | *D. rerio* | ENSDARG00000079738 | 0 | 0 | 0 | 0 | 0 | 0 | 0 | 0 | 0 | 0 | 15 | 3 | 0 | 0 | 0 | 0 | 0 | 0 | 0 |
| HG0326 | *D. rerio* | ENSDARG00000012848 | 0 | 0 | 0 | 0 | 0 | 0 | 0 | 0 | 0 | 0 | 14 | 3 | 0 | 0 | 0 | 0 | 0 | 0 | 0 |
| HG0327 | *D. rerio* | ENSDARG00000022203 | 0 | 0 | 0 | 0 | 0 | 0 | 0 | 0 | 0 | 0 | 14 | 3 | 0 | 0 | 0 | 0 | 0 | 0 | 0 |
| HG0328 | *D. rerio* | ENSDARG00000099999 | 0 | 0 | 0 | 0 | 0 | 0 | 0 | 0 | 0 | 0 | 15 | 2 | 0 | 0 | 0 | 0 | 0 | 0 | 0 |
| HG0329 | *D. rerio* | ENSDARG00000036442 | 0 | 0 | 0 | 0 | 0 | 0 | 0 | 0 | 0 | 0 | 14 | 2 | 0 | 0 | 0 | 0 | 0 | 0 | 0 |
| HG0330 | *D. rerio* | ENSDARG00000039392 | 0 | 0 | 0 | 0 | 0 | 0 | 0 | 0 | 0 | 0 | 13 | 3 | 0 | 0 | 0 | 0 | 0 | 0 | 0 |
| HG0331 | *D. rerio* | ENSDARG00000044899 | 0 | 0 | 0 | 0 | 0 | 0 | 0 | 0 | 0 | 0 | 14 | 2 | 0 | 0 | 0 | 0 | 0 | 0 | 0 |
| HG0332 | *D. rerio* | ENSDARG00000102893 | 0 | 0 | 0 | 0 | 0 | 0 | 0 | 0 | 0 | 0 | 13 | 3 | 0 | 0 | 0 | 0 | 0 | 0 | 0 |
| HG0333 | *D. rerio* | ENSDARG00000042518 | 0 | 0 | 0 | 0 | 0 | 0 | 0 | 0 | 0 | 0 | 12 | 2 | 0 | 0 | 0 | 0 | 0 | 0 | 0 |
| HG0334 | *D. rerio* | ENSDARG00000075349 | 0 | 0 | 0 | 0 | 0 | 0 | 0 | 0 | 0 | 0 | 11 | 3 | 0 | 0 | 0 | 0 | 0 | 0 | 0 |
| HG0335 | *D. rerio* | ENSDARG00000034700 | 0 | 0 | 0 | 0 | 0 | 0 | 0 | 0 | 0 | 0 | 10 | 3 | 0 | 0 | 0 | 0 | 0 | 0 | 0 |
| HG0336 | *D. rerio* | ENSDARG00000028676 | 0 | 0 | 0 | 0 | 0 | 0 | 0 | 0 | 0 | 0 | 11 | 1 | 0 | 0 | 0 | 0 | 0 | 0 | 0 |
| HG0337 | *D. rerio* | ENSDARG00000054906 | 0 | 0 | 0 | 0 | 0 | 0 | 0 | 0 | 0 | 0 | 10 | 1 | 0 | 0 | 0 | 0 | 0 | 0 | 0 |
| HG0338 | *D. rerio* | ENSDARG00000000151 | 0 | 0 | 0 | 0 | 0 | 0 | 0 | 0 | 0 | 0 | 8 | 2 | 0 | 0 | 0 | 0 | 0 | 0 | 0 |
| HG0339 | *D. rerio* | ENSDARG00000063144 | 0 | 0 | 0 | 0 | 0 | 0 | 0 | 0 | 0 | 0 | 9 | 1 | 0 | 0 | 0 | 0 | 0 | 0 | 0 |
| HG0340 | *D. rerio* | ENSDARG00000017591 | 0 | 0 | 0 | 0 | 0 | 0 | 0 | 0 | 0 | 0 | 7 | 1 | 0 | 0 | 0 | 0 | 0 | 0 | 0 |
| HG0341 | *D. rerio* | ENSDARG00000039430 | 0 | 0 | 0 | 0 | 0 | 0 | 0 | 0 | 0 | 0 | 7 | 1 | 0 | 0 | 0 | 0 | 0 | 0 | 0 |
| HG0342 | *D. rerio* | ENSDARG00000045801 | 0 | 0 | 0 | 0 | 0 | 0 | 0 | 0 | 0 | 0 | 6 | 2 | 0 | 0 | 0 | 0 | 0 | 0 | 0 |
| HG0343 | *D. rerio* | ENSDARG00000057062 | 0 | 0 | 0 | 0 | 0 | 0 | 0 | 0 | 0 | 0 | 6 | 2 | 0 | 0 | 0 | 0 | 0 | 0 | 0 |
| HG0344 | *D. rerio* | ENSDARG00000032072 | 0 | 0 | 0 | 0 | 0 | 0 | 0 | 0 | 0 | 0 | 6 | 1 | 0 | 0 | 0 | 0 | 0 | 0 | 0 |
| HG0345 | *D. rerio* | ENSDARG00000035910 | 0 | 0 | 0 | 0 | 0 | 0 | 0 | 0 | 0 | 0 | 5 | 2 | 0 | 0 | 0 | 0 | 0 | 0 | 0 |
| HG0346 | *D. rerio* | ENSDARG00000041982 | 0 | 0 | 0 | 0 | 0 | 0 | 0 | 0 | 0 | 0 | 5 | 2 | 0 | 0 | 0 | 0 | 0 | 0 | 0 |
| HG0347 | *D. rerio* | ENSDARG00000063194 | 0 | 0 | 0 | 0 | 0 | 0 | 0 | 0 | 0 | 0 | 5 | 2 | 0 | 0 | 0 | 0 | 0 | 0 | 0 |
| HG0348 | *D. rerio* | ENSDARG00000036139 | 0 | 0 | 0 | 0 | 0 | 0 | 0 | 0 | 0 | 0 | 5 | 1 | 0 | 0 | 0 | 0 | 0 | 0 | 0 |
| HG0349 | *D. rerio* | ENSDARG00000028699 | 0 | 0 | 0 | 0 | 0 | 0 | 0 | 0 | 0 | 0 | 4 | 1 | 0 | 0 | 0 | 0 | 0 | 0 | 0 |
| HG0350 | *D. rerio* | ENSDARG00000036194 | 0 | 0 | 0 | 0 | 0 | 0 | 0 | 0 | 0 | 0 | 4 | 1 | 0 | 0 | 0 | 0 | 0 | 0 | 0 |
| HG0351 | *D. rerio* | ENSDARG00000068096 | 0 | 0 | 0 | 0 | 0 | 0 | 0 | 0 | 0 | 0 | 2 | 3 | 0 | 0 | 0 | 0 | 0 | 0 | 0 |
| HG0352 | *D. rerio* | ENSDARG00000070214 | 0 | 0 | 0 | 0 | 0 | 0 | 0 | 0 | 0 | 0 | 3 | 2 | 0 | 0 | 0 | 0 | 0 | 0 | 0 |
| HG0353 | *D. rerio* | ENSDARG00000078864 | 0 | 0 | 0 | 0 | 0 | 0 | 0 | 0 | 0 | 0 | 3 | 2 | 0 | 0 | 0 | 0 | 0 | 0 | 0 |
| HG0354 | *D. rerio* | ENSDARG00000019897 | 0 | 0 | 0 | 0 | 0 | 0 | 0 | 0 | 0 | 0 | 1 | 3 | 0 | 0 | 0 | 0 | 0 | 0 | 0 |
| HG0355 | *D. rerio* | ENSDARG00000019930 | 0 | 0 | 0 | 0 | 0 | 0 | 0 | 0 | 0 | 0 | 4 | 0 | 0 | 0 | 0 | 0 | 0 | 0 | 0 |
| HG0356 | *D. rerio* | ENSDARG00000020475 | 0 | 0 | 0 | 0 | 0 | 0 | 0 | 0 | 0 | 0 | 3 | 1 | 0 | 0 | 0 | 0 | 0 | 0 | 0 |
| HG0357 | *D. rerio* | ENSDARG00000037285 | 0 | 0 | 0 | 0 | 0 | 0 | 0 | 0 | 0 | 0 | 4 | 0 | 0 | 0 | 0 | 0 | 0 | 0 | 0 |

| | | | | | | | | | | | | | | | | | | | | | |
|---|---|---|---|---|---|---|---|---|---|---|---|---|---|---|---|---|---|---|---|---|---|
| HG0358 | *D. rerio* | ENSDARG00000062506 | 0 | 0 | 0 | 0 | 0 | 0 | 0 | 0 | 0 | 0 | 2 | 2 | 0 | 0 | 0 | 0 | 0 | 0 | 0 |
| HG0359 | *D. rerio* | ENSDARG00000070348 | 0 | 0 | 0 | 0 | 0 | 0 | 0 | 0 | 0 | 0 | 1 | 3 | 0 | 0 | 0 | 0 | 0 | 0 | 0 |
| HG0360 | *D. rerio* | ENSDARG00000070801 | 0 | 0 | 0 | 0 | 0 | 0 | 0 | 0 | 0 | 0 | 3 | 1 | 0 | 0 | 0 | 0 | 0 | 0 | 0 |
| HG0361 | *D. rerio* | ENSDARG00000077581 | 0 | 0 | 0 | 0 | 0 | 0 | 0 | 0 | 0 | 0 | 2 | 2 | 0 | 0 | 0 | 0 | 0 | 0 | 0 |
| HG0362 | *D. rerio* | ENSDARG00000078355 | 0 | 0 | 0 | 0 | 0 | 0 | 0 | 0 | 0 | 0 | 3 | 1 | 0 | 0 | 0 | 0 | 0 | 0 | 0 |
| HG0363 | *D. rerio* | ENSDARG00000095896 | 0 | 0 | 0 | 0 | 0 | 0 | 0 | 0 | 0 | 0 | 4 | 0 | 0 | 0 | 0 | 0 | 0 | 0 | 0 |
| HG0364 | *D. rerio* | ENSDARG00000004988 | 0 | 0 | 0 | 0 | 0 | 0 | 0 | 0 | 0 | 0 | 3 | 0 | 0 | 0 | 0 | 0 | 0 | 0 | 0 |
| HG0365 | *D. rerio* | ENSDARG00000016132 | 0 | 0 | 0 | 0 | 0 | 0 | 0 | 0 | 0 | 0 | 0 | 3 | 0 | 0 | 0 | 0 | 0 | 0 | 0 |
| HG0366 | *D. rerio* | ENSDARG00000017803 | 0 | 0 | 0 | 0 | 0 | 0 | 0 | 0 | 0 | 0 | 0 | 3 | 0 | 0 | 0 | 0 | 0 | 0 | 0 |
| HG0367 | *D. rerio* | ENSDARG00000034056 | 0 | 0 | 0 | 0 | 0 | 0 | 0 | 0 | 0 | 0 | 1 | 2 | 0 | 0 | 0 | 0 | 0 | 0 | 0 |
| HG0368 | *D. rerio* | ENSDARG00000058606 | 0 | 0 | 0 | 0 | 0 | 0 | 0 | 0 | 0 | 0 | 1 | 2 | 0 | 0 | 0 | 0 | 0 | 0 | 0 |
| HG0369 | *D. rerio* | ENSDARG00000059276 | 0 | 0 | 0 | 0 | 0 | 0 | 0 | 0 | 0 | 0 | 2 | 1 | 0 | 0 | 0 | 0 | 0 | 0 | 0 |
| HG0370 | *D. rerio* | ENSDARG00000062693 | 0 | 0 | 0 | 0 | 0 | 0 | 0 | 0 | 0 | 0 | 2 | 1 | 0 | 0 | 0 | 0 | 0 | 0 | 0 |
| HG0371 | *D. rerio* | ENSDARG00000062765 | 0 | 0 | 0 | 0 | 0 | 0 | 0 | 0 | 0 | 0 | 2 | 1 | 0 | 0 | 0 | 0 | 0 | 0 | 0 |
| HG0372 | *D. rerio* | ENSDARG00000069467 | 0 | 0 | 0 | 0 | 0 | 0 | 0 | 0 | 0 | 0 | 1 | 2 | 0 | 0 | 0 | 0 | 0 | 0 | 0 |
| HG0373 | *D. rerio* | ENSDARG00000074611 | 0 | 0 | 0 | 0 | 0 | 0 | 0 | 0 | 0 | 0 | 0 | 3 | 0 | 0 | 0 | 0 | 0 | 0 | 0 |
| HG0374 | *D. rerio* | ENSDARG00000075147 | 0 | 0 | 0 | 0 | 0 | 0 | 0 | 0 | 0 | 0 | 1 | 2 | 0 | 0 | 0 | 0 | 0 | 0 | 0 |
| HG0375 | *D. rerio* | ENSDARG00000076171 | 0 | 0 | 0 | 0 | 0 | 0 | 0 | 0 | 0 | 0 | 3 | 0 | 0 | 0 | 0 | 0 | 0 | 0 | 0 |
| HG0376 | *D. rerio* | ENSDARG00000077229 | 0 | 0 | 0 | 0 | 0 | 0 | 0 | 0 | 0 | 0 | 2 | 1 | 0 | 0 | 0 | 0 | 0 | 0 | 0 |
| HG0377 | *D. rerio* | ENSDARG00000079549 | 0 | 0 | 0 | 0 | 0 | 0 | 0 | 0 | 0 | 0 | 1 | 2 | 0 | 0 | 0 | 0 | 0 | 0 | 0 |
| HG0378 | *D. rerio* | ENSDARG00000080009 | 0 | 0 | 0 | 0 | 0 | 0 | 0 | 0 | 0 | 0 | 1 | 2 | 0 | 0 | 0 | 0 | 0 | 0 | 0 |
| HG0379 | *D. rerio* | ENSDARG00000104372 | 0 | 0 | 0 | 0 | 0 | 0 | 0 | 0 | 0 | 0 | 1 | 2 | 0 | 0 | 0 | 0 | 0 | 0 | 0 |
| HG0380 | *D. rerio* | ENSDARG00000005350 | 0 | 0 | 0 | 0 | 0 | 0 | 0 | 0 | 0 | 0 | 1 | 1 | 0 | 0 | 0 | 0 | 0 | 0 | 0 |
| HG0381 | *D. rerio* | ENSDARG00000006124 | 0 | 0 | 0 | 0 | 0 | 0 | 0 | 0 | 0 | 0 | 1 | 1 | 0 | 0 | 0 | 0 | 0 | 0 | 0 |
| HG0382 | *D. rerio* | ENSDARG00000017242 | 0 | 0 | 0 | 0 | 0 | 0 | 0 | 0 | 0 | 0 | 1 | 1 | 0 | 0 | 0 | 0 | 0 | 0 | 0 |
| HG0383 | *D. rerio* | ENSDARG00000036501 | 0 | 0 | 0 | 0 | 0 | 0 | 0 | 0 | 0 | 0 | 2 | 0 | 0 | 0 | 0 | 0 | 0 | 0 | 0 |
| HG0384 | *D. rerio* | ENSDARG00000054748 | 0 | 0 | 0 | 0 | 0 | 0 | 0 | 0 | 0 | 0 | 0 | 2 | 0 | 0 | 0 | 0 | 0 | 0 | 0 |
| HG0385 | *D. rerio* | ENSDARG00000059549 | 0 | 0 | 0 | 0 | 0 | 0 | 0 | 0 | 0 | 0 | 1 | 1 | 0 | 0 | 0 | 0 | 0 | 0 | 0 |
| HG0386 | *D. rerio* | ENSDARG00000062084 | 0 | 0 | 0 | 0 | 0 | 0 | 0 | 0 | 0 | 0 | 2 | 0 | 0 | 0 | 0 | 0 | 0 | 0 | 0 |
| HG0387 | *D. rerio* | ENSDARG00000071150 | 0 | 0 | 0 | 0 | 0 | 0 | 0 | 0 | 0 | 0 | 0 | 2 | 0 | 0 | 0 | 0 | 0 | 0 | 0 |
| HG0388 | *D. rerio* | ENSDARG00000077228 | 0 | 0 | 0 | 0 | 0 | 0 | 0 | 0 | 0 | 0 | 2 | 0 | 0 | 0 | 0 | 0 | 0 | 0 | 0 |
| HG0389 | *D. rerio* | ENSDARG00000077361 | 0 | 0 | 0 | 0 | 0 | 0 | 0 | 0 | 0 | 0 | 1 | 1 | 0 | 0 | 0 | 0 | 0 | 0 | 0 |
| HG0390 | *D. rerio* | ENSDARG00000090035 | 0 | 0 | 0 | 0 | 0 | 0 | 0 | 0 | 0 | 0 | 1 | 1 | 0 | 0 | 0 | 0 | 0 | 0 | 0 |
| HG0391 | *D. rerio* | ENSDARG00000099634 | 0 | 0 | 0 | 0 | 0 | 0 | 0 | 0 | 0 | 0 | 0 | 2 | 0 | 0 | 0 | 0 | 0 | 0 | 0 |
| HG0392 | *D. rerio* | ENSDARG00000100003 | 0 | 0 | 0 | 0 | 0 | 0 | 0 | 0 | 0 | 0 | 0 | 2 | 0 | 0 | 0 | 0 | 0 | 0 | 0 |
| HG0393 | *G. gallus* | ENSGALG00000002090 | 0 | 0 | 0 | 0 | 0 | 0 | 0 | 0 | 0 | 0 | 0 | 0 | 0 | 0 | 0 | 28 | 0 | 0 | 0 |
| HG0394 | *G. gallus* | ENSGALG00000002479 | 0 | 0 | 0 | 0 | 0 | 0 | 0 | 0 | 0 | 0 | 0 | 0 | 0 | 0 | 0 | 24 | 0 | 0 | 0 |
| HG0395 | *G. gallus* | ENSGALG00000014936 | 0 | 0 | 0 | 0 | 0 | 0 | 0 | 0 | 0 | 0 | 0 | 0 | 0 | 0 | 0 | 22 | 0 | 0 | 0 |
| HG0396.1 | *G. gallus* | ENSGALG00000034253 | 0 | 0 | 0 | 0 | 0 | 0 | 0 | 0 | 0 | 0 | 0 | 0 | 0 | 0 | 0 | 21 | 0 | 0 | 0 |
| HG0396.2 | *G. gallus* | ENSGALG00000034253 | 0 | 0 | 0 | 0 | 0 | 0 | 0 | 0 | 0 | 0 | 0 | 0 | 0 | 0 | 0 | 8 | 0 | 0 | 0 |
| HG0397 | *G. gallus* | ENSGALG00000009438 | 0 | 0 | 0 | 0 | 0 | 0 | 0 | 0 | 0 | 0 | 0 | 0 | 0 | 0 | 0 | 18 | 0 | 0 | 0 |
| HG0398 | *G. gallus* | ENSGALG00000005369 | 0 | 0 | 0 | 0 | 0 | 0 | 0 | 0 | 0 | 0 | 0 | 0 | 0 | 0 | 0 | 17 | 0 | 0 | 0 |
| HG0399 | *G. gallus* | ENSGALG00000032659 | 0 | 0 | 0 | 0 | 0 | 0 | 0 | 0 | 0 | 0 | 0 | 0 | 0 | 0 | 0 | 17 | 0 | 0 | 0 |
| HG0400 | *G. gallus* | ENSGALG00000004261 | 0 | 0 | 0 | 0 | 0 | 0 | 0 | 0 | 0 | 0 | 0 | 0 | 0 | 0 | 0 | 16 | 0 | 0 | 0 |
| HG0401 | *G. gallus* | ENSGALG00000009057 | 0 | 0 | 0 | 0 | 0 | 0 | 0 | 0 | 0 | 0 | 0 | 0 | 0 | 0 | 0 | 16 | 0 | 0 | 0 |
| HG0402 | *G. gallus* | ENSGALG00000035584 | 0 | 0 | 0 | 0 | 0 | 0 | 0 | 0 | 0 | 0 | 0 | 0 | 0 | 0 | 0 | 16 | 0 | 0 | 0 |
| HG0403.1 | *G. gallus* | ENSGALG00000007717 | 0 | 0 | 0 | 0 | 0 | 0 | 0 | 0 | 0 | 0 | 0 | 0 | 0 | 0 | 0 | 15 | 0 | 0 | 0 |
| HG0403.2 | *G. gallus* | ENSGALG00000007717 | 0 | 0 | 0 | 0 | 0 | 0 | 0 | 0 | 0 | 0 | 0 | 0 | 0 | 0 | 0 | 4 | 0 | 0 | 0 |
| HG0404 | *G. gallus* | ENSGALG00000008700 | 0 | 0 | 0 | 0 | 0 | 0 | 0 | 0 | 0 | 0 | 0 | 0 | 0 | 0 | 0 | 15 | 0 | 0 | 0 |
| HG0405.1 | *G. gallus* | ENSGALG00000009185 | 0 | 0 | 0 | 0 | 0 | 0 | 0 | 0 | 0 | 0 | 0 | 0 | 0 | 0 | 0 | 13 | 0 | 0 | 0 |
| HG0405.2 | *G. gallus* | ENSGALG00000009185 | 0 | 0 | 0 | 0 | 0 | 0 | 0 | 0 | 0 | 0 | 0 | 0 | 0 | 0 | 0 | 11 | 0 | 0 | 0 |
| HG0406 | *G. gallus* | ENSGALG00000004935 | 0 | 0 | 0 | 0 | 0 | 0 | 0 | 0 | 0 | 0 | 0 | 0 | 0 | 0 | 0 | 13 | 0 | 0 | 0 |
| HG0407 | *G. gallus* | ENSGALG00000015027 | 0 | 0 | 0 | 0 | 0 | 0 | 0 | 0 | 0 | 0 | 0 | 0 | 0 | 0 | 0 | 13 | 0 | 0 | 0 |
| HG0408 | *G. gallus* | ENSGALG00000036255 | 0 | 0 | 0 | 0 | 0 | 0 | 0 | 0 | 0 | 0 | 0 | 0 | 0 | 0 | 0 | 13 | 0 | 0 | 0 |
| HG0409 | *G. gallus* | ENSGALG00000002315 | 0 | 0 | 0 | 0 | 0 | 0 | 0 | 0 | 0 | 0 | 0 | 0 | 0 | 0 | 0 | 12 | 0 | 0 | 0 |
| HG0410 | *G. gallus* | ENSGALG00000008148 | 0 | 0 | 0 | 0 | 0 | 0 | 0 | 0 | 0 | 0 | 0 | 0 | 0 | 0 | 0 | 12 | 0 | 0 | 0 |
| HG0411 | *G. gallus* | ENSGALG00000012166 | 0 | 0 | 0 | 0 | 0 | 0 | 0 | 0 | 0 | 0 | 0 | 0 | 0 | 0 | 0 | 12 | 0 | 0 | 0 |
| HG0412 | *G. gallus* | ENSGALG00000003446 | 0 | 0 | 0 | 0 | 0 | 0 | 0 | 0 | 0 | 0 | 0 | 0 | 0 | 0 | 0 | 11 | 0 | 0 | 0 |
| HG0413 | *G. gallus* | ENSGALG00000005475 | 0 | 0 | 0 | 0 | 0 | 0 | 0 | 0 | 0 | 0 | 0 | 0 | 0 | 0 | 0 | 11 | 0 | 0 | 0 |
| HG0414 | *G. gallus* | ENSGALG00000032599 | 0 | 0 | 0 | 0 | 0 | 0 | 0 | 0 | 0 | 0 | 0 | 0 | 0 | 0 | 0 | 11 | 0 | 0 | 0 |
| HG0415 | *G. gallus* | ENSGALG00000034970 | 0 | 0 | 0 | 0 | 0 | 0 | 0 | 0 | 0 | 0 | 0 | 0 | 0 | 0 | 0 | 11 | 0 | 0 | 0 |
| HG0416 | *G. gallus* | ENSGALG00000015874 | 0 | 0 | 0 | 0 | 0 | 0 | 0 | 0 | 0 | 0 | 0 | 0 | 0 | 0 | 0 | 9 | 0 | 0 | 0 |
| HG0417 | *G. gallus* | ENSGALG00000001522 | 0 | 0 | 0 | 0 | 0 | 0 | 0 | 0 | 0 | 0 | 0 | 0 | 0 | 0 | 0 | 8 | 0 | 0 | 0 |
| HG0418 | *G. gallus* | ENSGALG00000010337 | 0 | 0 | 0 | 0 | 0 | 0 | 0 | 0 | 0 | 0 | 0 | 0 | 0 | 0 | 0 | 8 | 0 | 0 | 0 |
| HG0419 | *G. gallus* | ENSGALG00000012542 | 0 | 0 | 0 | 0 | 0 | 0 | 0 | 0 | 0 | 0 | 0 | 0 | 0 | 0 | 0 | 8 | 0 | 0 | 0 |
| HG0420 | *G. gallus* | ENSGALG00000000336 | 0 | 0 | 0 | 0 | 0 | 0 | 0 | 0 | 0 | 0 | 0 | 0 | 0 | 0 | 0 | 7 | 0 | 0 | 0 |
| HG0421 | *G. gallus* | ENSGALG00000003492 | 0 | 0 | 0 | 0 | 0 | 0 | 0 | 0 | 0 | 0 | 0 | 0 | 0 | 0 | 0 | 7 | 0 | 0 | 0 |
| HG0422 | *G. gallus* | ENSGALG00000005772 | 0 | 0 | 0 | 0 | 0 | 0 | 0 | 0 | 0 | 0 | 0 | 0 | 0 | 0 | 0 | 7 | 0 | 0 | 0 |
| HG0423 | *G. gallus* | ENSGALG00000006329 | 0 | 0 | 0 | 0 | 0 | 0 | 0 | 0 | 0 | 0 | 0 | 0 | 0 | 0 | 0 | 7 | 0 | 0 | 0 |
| HG0424 | *G. gallus* | ENSGALG00000039553 | 0 | 0 | 0 | 0 | 0 | 0 | 0 | 0 | 0 | 0 | 0 | 0 | 0 | 0 | 0 | 7 | 0 | 0 | 0 |
| HG0425 | *G. gallus* | ENSGALG00000043044 | 0 | 0 | 0 | 0 | 0 | 0 | 0 | 0 | 0 | 0 | 0 | 0 | 0 | 0 | 0 | 7 | 0 | 0 | 0 |
| HG0426 | *G. gallus* | ENSGALG00000002192 | 0 | 0 | 0 | 0 | 0 | 0 | 0 | 0 | 0 | 0 | 0 | 0 | 0 | 0 | 0 | 6 | 0 | 0 | 0 |
| HG0427 | *G. gallus* | ENSGALG00000005550 | 0 | 0 | 0 | 0 | 0 | 0 | 0 | 0 | 0 | 0 | 0 | 0 | 0 | 0 | 0 | 6 | 0 | 0 | 0 |
| HG0428 | *G. gallus* | ENSGALG00000005976 | 0 | 0 | 0 | 0 | 0 | 0 | 0 | 0 | 0 | 0 | 0 | 0 | 0 | 0 | 0 | 6 | 0 | 0 | 0 |
| HG0429 | *G. gallus* | ENSGALG00000006626 | 0 | 0 | 0 | 0 | 0 | 0 | 0 | 0 | 0 | 0 | 0 | 0 | 0 | 0 | 0 | 6 | 0 | 0 | 0 |
| HG0430 | *G. gallus* | ENSGALG00000015490 | 0 | 0 | 0 | 0 | 0 | 0 | 0 | 0 | 0 | 0 | 0 | 0 | 0 | 0 | 0 | 6 | 0 | 0 | 0 |
| HG0431 | *G. gallus* | ENSGALG00000026542 | 0 | 0 | 0 | 0 | 0 | 0 | 0 | 0 | 0 | 0 | 0 | 0 | 0 | 0 | 0 | 6 | 0 | 0 | 0 |
| HG0432 | *G. gallus* | ENSGALG00000040546 | 0 | 0 | 0 | 0 | 0 | 0 | 0 | 0 | 0 | 0 | 0 | 0 | 0 | 0 | 0 | 6 | 0 | 0 | 0 |
| HG0433 | *G. gallus* | ENSGALG00000002952 | 0 | 0 | 0 | 0 | 0 | 0 | 0 | 0 | 0 | 0 | 0 | 0 | 0 | 0 | 0 | 5 | 0 | 0 | 0 |

| | | | | | | | | | | | | | | | | | | | | |
|---|---|---|---|---|---|---|---|---|---|---|---|---|---|---|---|---|---|---|---|---|---|
| HG0434 | *G. gallus* | ENSGALG00000008357 | 0 | 0 | 0 | 0 | 0 | 0 | 0 | 0 | 0 | 0 | 0 | 0 | 0 | 0 | 0 | 5 | 0 | 0 | 0 |
| HG0435 | *G. gallus* | ENSGALG00000010778 | 0 | 0 | 0 | 0 | 0 | 0 | 0 | 0 | 0 | 0 | 0 | 0 | 0 | 0 | 0 | 5 | 0 | 0 | 0 |
| HG0436 | *G. gallus* | ENSGALG00000026736 | 0 | 0 | 0 | 0 | 0 | 0 | 0 | 0 | 0 | 0 | 0 | 0 | 0 | 0 | 0 | 5 | 0 | 0 | 0 |
| HG0437 | *G. gallus* | ENSGALG00000001728 | 0 | 0 | 0 | 0 | 0 | 0 | 0 | 0 | 0 | 0 | 0 | 0 | 0 | 0 | 0 | 4 | 0 | 0 | 0 |
| HG0438 | *G. gallus* | ENSGALG00000008545 | 0 | 0 | 0 | 0 | 0 | 0 | 0 | 0 | 0 | 0 | 0 | 0 | 0 | 0 | 0 | 4 | 0 | 0 | 0 |
| HG0439 | *G. gallus* | ENSGALG00000009172 | 0 | 0 | 0 | 0 | 0 | 0 | 0 | 0 | 0 | 0 | 0 | 0 | 0 | 0 | 0 | 4 | 0 | 0 | 0 |
| HG0440 | *G. gallus* | ENSGALG00000010090 | 0 | 0 | 0 | 0 | 0 | 0 | 0 | 0 | 0 | 0 | 0 | 0 | 0 | 0 | 0 | 4 | 0 | 0 | 0 |
| HG0441 | *G. gallus* | ENSGALG00000015069 | 0 | 0 | 0 | 0 | 0 | 0 | 0 | 0 | 0 | 0 | 0 | 0 | 0 | 0 | 0 | 4 | 0 | 0 | 0 |
| HG0442 | *G. gallus* | ENSGALG00000030250 | 0 | 0 | 0 | 0 | 0 | 0 | 0 | 0 | 0 | 0 | 0 | 0 | 0 | 0 | 0 | 4 | 0 | 0 | 0 |
| HG0443 | *G. gallus* | ENSGALG00000039454 | 0 | 0 | 0 | 0 | 0 | 0 | 0 | 0 | 0 | 0 | 0 | 0 | 0 | 0 | 0 | 4 | 0 | 0 | 0 |
| HG0444 | *G. gallus* | ENSGALG00000000558 | 0 | 0 | 0 | 0 | 0 | 0 | 0 | 0 | 0 | 0 | 0 | 0 | 0 | 0 | 0 | 3 | 0 | 0 | 0 |
| HG0445 | *G. gallus* | ENSGALG00000004017 | 0 | 0 | 0 | 0 | 0 | 0 | 0 | 0 | 0 | 0 | 0 | 0 | 0 | 0 | 0 | 3 | 0 | 0 | 0 |
| HG0446 | *G. gallus* | ENSGALG00000006807 | 0 | 0 | 0 | 0 | 0 | 0 | 0 | 0 | 0 | 0 | 0 | 0 | 0 | 0 | 0 | 3 | 0 | 0 | 0 |
| HG0447 | *G. gallus* | ENSGALG00000008784 | 0 | 0 | 0 | 0 | 0 | 0 | 0 | 0 | 0 | 0 | 0 | 0 | 0 | 0 | 0 | 3 | 0 | 0 | 0 |
| HG0448 | *G. gallus* | ENSGALG00000011762 | 0 | 0 | 0 | 0 | 0 | 0 | 0 | 0 | 0 | 0 | 0 | 0 | 0 | 0 | 0 | 3 | 0 | 0 | 0 |
| HG0449 | *G. gallus* | ENSGALG00000014106 | 0 | 0 | 0 | 0 | 0 | 0 | 0 | 0 | 0 | 0 | 0 | 0 | 0 | 0 | 0 | 3 | 0 | 0 | 0 |
| HG0450 | *G. gallus* | ENSGALG00000015528 | 0 | 0 | 0 | 0 | 0 | 0 | 0 | 0 | 0 | 0 | 0 | 0 | 0 | 0 | 0 | 3 | 0 | 0 | 0 |
| HG0451 | *G. gallus* | ENSGALG00000019489 | 0 | 0 | 0 | 0 | 0 | 0 | 0 | 0 | 0 | 0 | 0 | 0 | 0 | 0 | 0 | 3 | 0 | 0 | 0 |
| HG0452 | *G. gallus* | ENSGALG00000004227 | 0 | 0 | 0 | 0 | 0 | 0 | 0 | 0 | 0 | 0 | 0 | 0 | 0 | 0 | 0 | 2 | 0 | 0 | 0 |
| HG0453 | *G. gallus* | ENSGALG00000005995 | 0 | 0 | 0 | 0 | 0 | 0 | 0 | 0 | 0 | 0 | 0 | 0 | 0 | 0 | 0 | 2 | 0 | 0 | 0 |
| HG0454 | *G. gallus* | ENSGALG00000007186 | 0 | 0 | 0 | 0 | 0 | 0 | 0 | 0 | 0 | 0 | 0 | 0 | 0 | 0 | 0 | 2 | 0 | 0 | 0 |
| HG0455 | *G. gallus* | ENSGALG00000010718 | 0 | 0 | 0 | 0 | 0 | 0 | 0 | 0 | 0 | 0 | 0 | 0 | 0 | 0 | 0 | 2 | 0 | 0 | 0 |
| HG0456 | *G. gallus* | ENSGALG00000033256 | 0 | 0 | 0 | 0 | 0 | 0 | 0 | 0 | 0 | 0 | 0 | 0 | 0 | 0 | 0 | 2 | 0 | 0 | 0 |
| HG0457 | *G. gallus* | ENSGALG00000039538 | 0 | 0 | 0 | 0 | 0 | 0 | 0 | 0 | 0 | 0 | 0 | 0 | 0 | 0 | 0 | 2 | 0 | 0 | 0 |
| HG0458 | *H. sapiens* | ENSG00000060339 | 0 | 0 | 0 | 0 | 0 | 0 | 0 | 0 | 0 | 0 | 0 | 0 | 0 | 0 | 0 | 0 | 3 | 11 | 4 |
| HG0459.1 | *H. sapiens* | ENSG00000163848 | 0 | 0 | 0 | 0 | 0 | 0 | 0 | 0 | 0 | 0 | 0 | 0 | 0 | 0 | 0 | 0 | 2 | 11 | 4 |
| HG0459.2 | *H. sapiens* | ENSG00000163848 | 0 | 0 | 0 | 0 | 0 | 0 | 0 | 0 | 0 | 0 | 0 | 0 | 0 | 0 | 0 | 0 | 0 | 8 | 4 |
| HG0459.3 | *H. sapiens* | ENSG00000163848 | 0 | 0 | 0 | 0 | 0 | 0 | 0 | 0 | 0 | 0 | 0 | 0 | 0 | 0 | 0 | 0 | 0 | 4 | 1 |
| HG0460 | *H. sapiens* | ENSG00000180008 | 0 | 0 | 0 | 0 | 0 | 0 | 0 | 0 | 0 | 0 | 0 | 0 | 0 | 0 | 0 | 0 | 4 | 10 | 3 |
| HG0461 | *H. sapiens* | ENSG00000064419 | 0 | 0 | 0 | 0 | 0 | 0 | 0 | 0 | 0 | 0 | 0 | 0 | 0 | 0 | 0 | 0 | 2 | 10 | 4 |
| HG0462 | *H. sapiens* | ENSG00000125686 | 0 | 0 | 0 | 0 | 0 | 0 | 0 | 0 | 0 | 0 | 0 | 0 | 0 | 0 | 0 | 0 | 2 | 11 | 3 |
| HG0463 | *H. sapiens* | ENSG00000129473 | 0 | 0 | 0 | 0 | 0 | 0 | 0 | 0 | 0 | 0 | 0 | 0 | 0 | 0 | 0 | 0 | 2 | 11 | 3 |
| HG0464 | *H. sapiens* | ENSG00000188215 | 0 | 0 | 0 | 0 | 0 | 0 | 0 | 0 | 0 | 0 | 0 | 0 | 0 | 0 | 0 | 0 | 3 | 9 | 4 |
| HG0465 | *H. sapiens* | ENSG00000132153 | 0 | 0 | 0 | 0 | 0 | 0 | 0 | 0 | 0 | 0 | 0 | 0 | 0 | 0 | 0 | 0 | 1 | 11 | 3 |
| HG0466 | *H. sapiens* | ENSG00000139679 | 0 | 0 | 0 | 0 | 0 | 0 | 0 | 0 | 0 | 0 | 0 | 0 | 0 | 0 | 0 | 0 | 3 | 9 | 3 |
| HG0467.1 | *H. sapiens* | ENSG00000163320 | 0 | 0 | 0 | 0 | 0 | 0 | 0 | 0 | 0 | 0 | 0 | 0 | 0 | 0 | 0 | 0 | 2 | 9 | 4 |
| HG0467.2 | *H. sapiens* | ENSG00000163320 | 0 | 0 | 0 | 0 | 0 | 0 | 0 | 0 | 0 | 0 | 0 | 0 | 0 | 0 | 0 | 0 | 0 | 3 | 3 |
| HG0468 | *H. sapiens* | ENSG00000187778 | 0 | 0 | 0 | 0 | 0 | 0 | 0 | 0 | 0 | 0 | 0 | 0 | 0 | 0 | 0 | 0 | 1 | 10 | 4 |
| HG0469 | *H. sapiens* | ENSG00000198369 | 0 | 0 | 0 | 0 | 0 | 0 | 0 | 0 | 0 | 0 | 0 | 0 | 0 | 0 | 0 | 0 | 3 | 9 | 3 |
| HG0470 | *H. sapiens* | ENSG00000224470 | 0 | 0 | 0 | 0 | 0 | 0 | 0 | 0 | 0 | 0 | 0 | 0 | 0 | 0 | 0 | 0 | 1 | 11 | 3 |
| HG0471 | *H. sapiens* | ENSG00000023041 | 0 | 0 | 0 | 0 | 0 | 0 | 0 | 0 | 0 | 0 | 0 | 0 | 0 | 0 | 0 | 0 | 1 | 10 | 3 |
| HG0472 | *H. sapiens* | ENSG00000105991 | 0 | 0 | 0 | 0 | 0 | 0 | 0 | 0 | 0 | 0 | 0 | 0 | 0 | 0 | 0 | 0 | 1 | 10 | 3 |
| HG0473.1 | *H. sapiens* | ENSG00000138650 | 0 | 0 | 0 | 0 | 0 | 0 | 0 | 0 | 0 | 0 | 0 | 0 | 0 | 0 | 0 | 0 | 1 | 9 | 4 |
| HG0473.2 | *H. sapiens* | ENSG00000240184 | 0 | 0 | 0 | 0 | 0 | 0 | 0 | 0 | 0 | 0 | 0 | 0 | 0 | 0 | 0 | 0 | 0 | 9 | 2 |
| HG0474 | *H. sapiens* | ENSG00000139083 | 0 | 0 | 0 | 0 | 0 | 0 | 0 | 0 | 0 | 0 | 0 | 0 | 0 | 0 | 0 | 0 | 2 | 9 | 3 |
| HG0475 | *H. sapiens* | ENSG00000109118 | 0 | 0 | 0 | 0 | 0 | 0 | 0 | 0 | 0 | 0 | 0 | 0 | 0 | 0 | 0 | 0 | 1 | 9 | 3 |
| HG0476 | *H. sapiens* | ENSG00000130939 | 0 | 0 | 0 | 0 | 0 | 0 | 0 | 0 | 0 | 0 | 0 | 0 | 0 | 0 | 0 | 0 | 1 | 8 | 4 |
| HG0477 | *H. sapiens* | ENSG00000166925 | 0 | 0 | 0 | 0 | 0 | 0 | 0 | 0 | 0 | 0 | 0 | 0 | 0 | 0 | 0 | 0 | 2 | 8 | 3 |
| HG0478 | *H. sapiens* | ENSG00000168214 | 0 | 0 | 0 | 0 | 0 | 0 | 0 | 0 | 0 | 0 | 0 | 0 | 0 | 0 | 0 | 0 | 3 | 7 | 3 |
| HG0479.1 | *H. sapiens* | ENSG00000168453 | 0 | 0 | 0 | 0 | 0 | 0 | 0 | 0 | 0 | 0 | 0 | 0 | 0 | 0 | 0 | 0 | 2 | 8 | 3 |
| HG0479.2 | *H. sapiens* | ENSG00000168453 | 0 | 0 | 0 | 0 | 0 | 0 | 0 | 0 | 0 | 0 | 0 | 0 | 0 | 0 | 0 | 0 | 1 | 8 | 2 |
| HG0480 | *H. sapiens* | ENSG00000198018 | 0 | 0 | 0 | 0 | 0 | 0 | 0 | 0 | 0 | 0 | 0 | 0 | 0 | 0 | 0 | 0 | 1 | 8 | 4 |
| HG0481 | *H. sapiens* | ENSG00000198963 | 0 | 0 | 0 | 0 | 0 | 0 | 0 | 0 | 0 | 0 | 0 | 0 | 0 | 0 | 0 | 0 | 1 | 10 | 2 |
| HG0482 | *H. sapiens* | ENSG00000067167 | 0 | 0 | 0 | 0 | 0 | 0 | 0 | 0 | 0 | 0 | 0 | 0 | 0 | 0 | 0 | 0 | 1 | 8 | 3 |
| HG0483 | *H. sapiens* | ENSG00000164463 | 0 | 0 | 0 | 0 | 0 | 0 | 0 | 0 | 0 | 0 | 0 | 0 | 0 | 0 | 0 | 0 | 2 | 7 | 3 |
| HG0484 | *H. sapiens* | ENSG00000166444 | 0 | 0 | 0 | 0 | 0 | 0 | 0 | 0 | 0 | 0 | 0 | 0 | 0 | 0 | 0 | 0 | 1 | 8 | 3 |
| HG0485.1 | *H. sapiens* | ENSG00000182667 | 0 | 0 | 0 | 0 | 0 | 0 | 0 | 0 | 0 | 0 | 0 | 0 | 0 | 0 | 0 | 0 | 2 | 7 | 3 |
| HG0485.2 | *H. sapiens* | ENSG00000183715 | 0 | 0 | 0 | 0 | 0 | 0 | 0 | 0 | 0 | 0 | 0 | 0 | 0 | 0 | 0 | 0 | 0 | 3 | 2 |
| HG0486 | *H. sapiens* | ENSG00000067900 | 0 | 0 | 0 | 0 | 0 | 0 | 0 | 0 | 0 | 0 | 0 | 0 | 0 | 0 | 0 | 0 | 1 | 7 | 3 |
| HG0487 | *H. sapiens* | ENSG00000091831 | 0 | 0 | 0 | 0 | 0 | 0 | 0 | 0 | 0 | 0 | 0 | 0 | 0 | 0 | 0 | 0 | 1 | 7 | 3 |
| HG0488.1 | *H. sapiens* | ENSG00000116194 | 0 | 0 | 0 | 0 | 0 | 0 | 0 | 0 | 0 | 0 | 0 | 0 | 0 | 0 | 0 | 0 | 1 | 7 | 3 |
| HG0488.2 | *H. sapiens* | ENSG00000116194 | 0 | 0 | 0 | 0 | 0 | 0 | 0 | 0 | 0 | 0 | 0 | 0 | 0 | 0 | 0 | 0 | 0 | 5 | 4 |
| HG0489 | *H. sapiens* | ENSG00000143847 | 0 | 0 | 0 | 0 | 0 | 0 | 0 | 0 | 0 | 0 | 0 | 0 | 0 | 0 | 0 | 0 | 1 | 7 | 3 |
| HG0490.1 | *H. sapiens* | ENSG00000152413 | 0 | 0 | 0 | 0 | 0 | 0 | 0 | 0 | 0 | 0 | 0 | 0 | 0 | 0 | 0 | 0 | 1 | 7 | 3 |
| HG0490.2 | *H. sapiens* | ENSG00000152413 | 0 | 0 | 0 | 0 | 0 | 0 | 0 | 0 | 0 | 0 | 0 | 0 | 0 | 0 | 0 | 0 | 0 | 6 | 3 |
| HG0491 | *H. sapiens* | ENSG00000154114 | 0 | 0 | 0 | 0 | 0 | 0 | 0 | 0 | 0 | 0 | 0 | 0 | 0 | 0 | 0 | 0 | 1 | 8 | 2 |
| HG0492 | *H. sapiens* | ENSG00000162670 | 0 | 0 | 0 | 0 | 0 | 0 | 0 | 0 | 0 | 0 | 0 | 0 | 0 | 0 | 0 | 0 | 1 | 8 | 2 |
| HG0493 | *H. sapiens* | ENSG00000164707 | 0 | 0 | 0 | 0 | 0 | 0 | 0 | 0 | 0 | 0 | 0 | 0 | 0 | 0 | 0 | 0 | 1 | 6 | 4 |
| HG0494 | *H. sapiens* | ENSG00000184226 | 0 | 0 | 0 | 0 | 0 | 0 | 0 | 0 | 0 | 0 | 0 | 0 | 0 | 0 | 0 | 0 | 1 | 9 | 1 |
| HG0495.1 | *H. sapiens* | ENSG00000196730 | 0 | 0 | 0 | 0 | 0 | 0 | 0 | 0 | 0 | 0 | 0 | 0 | 0 | 0 | 0 | 0 | 0 | 6 | 3 |
| HG0495.2 | *H. sapiens* | ENSG00000196730 | 0 | 0 | 0 | 0 | 0 | 0 | 0 | 0 | 0 | 0 | 0 | 0 | 0 | 0 | 0 | 0 | 1 | 7 | 3 |
| HG0496 | *H. sapiens* | ENSG00000135913 | 0 | 0 | 0 | 0 | 0 | 0 | 0 | 0 | 0 | 0 | 0 | 0 | 0 | 0 | 0 | 0 | 1 | 7 | 2 |
| HG0497 | *H. sapiens* | ENSG00000147571 | 0 | 0 | 0 | 0 | 0 | 0 | 0 | 0 | 0 | 0 | 0 | 0 | 0 | 0 | 0 | 0 | 1 | 5 | 4 |
| HG0498 | *H. sapiens* | ENSG00000005379 | 0 | 0 | 0 | 0 | 0 | 0 | 0 | 0 | 0 | 0 | 0 | 0 | 0 | 0 | 0 | 0 | 1 | 6 | 2 |
| HG0499 | *H. sapiens* | ENSG00000011258 | 0 | 0 | 0 | 0 | 0 | 0 | 0 | 0 | 0 | 0 | 0 | 0 | 0 | 0 | 0 | 0 | 1 | 6 | 2 |
| HG0500 | *H. sapiens* | ENSG00000070778 | 0 | 0 | 0 | 0 | 0 | 0 | 0 | 0 | 0 | 0 | 0 | 0 | 0 | 0 | 0 | 0 | 2 | 6 | 1 |
| HG0501 | *H. sapiens* | ENSG00000110400 | 0 | 0 | 0 | 0 | 0 | 0 | 0 | 0 | 0 | 0 | 0 | 0 | 0 | 0 | 0 | 0 | 2 | 5 | 2 |
| HG0502 | *H. sapiens* | ENSG00000126882 | 0 | 0 | 0 | 0 | 0 | 0 | 0 | 0 | 0 | 0 | 0 | 0 | 0 | 0 | 0 | 0 | 1 | 5 | 3 |
| HG0503 | *H. sapiens* | ENSG00000141564 | 0 | 0 | 0 | 0 | 0 | 0 | 0 | 0 | 0 | 0 | 0 | 0 | 0 | 0 | 0 | 0 | 1 | 6 | 2 |

| | | | | | | | | | | | | | | | | | | | | | |
|---|---|---|---|---|---|---|---|---|---|---|---|---|---|---|---|---|---|---|---|---|---|
| HG0504 | *H. sapiens* | ENSG00000144460 | 0 | 0 | 0 | 0 | 0 | 0 | 0 | 0 | 0 | 0 | 0 | 0 | 0 | 0 | 0 | 0 | 1 | 4 | 4 |
| HG0505 | *H. sapiens* | ENSG00000157470 | 0 | 0 | 0 | 0 | 0 | 0 | 0 | 0 | 0 | 0 | 0 | 0 | 0 | 0 | 0 | 0 | 1 | 6 | 2 |
| HG0506.1 | *H. sapiens* | ENSG00000177853 | 0 | 0 | 0 | 0 | 0 | 0 | 0 | 0 | 0 | 0 | 0 | 0 | 0 | 0 | 0 | 0 | 1 | 5 | 3 |
| HG0506.2 | *H. sapiens* | ENSG00000177853 | 0 | 0 | 0 | 0 | 0 | 0 | 0 | 0 | 0 | 0 | 0 | 0 | 0 | 0 | 0 | 0 | 0 | 7 | 2 |
| HG0507 | *H. sapiens* | ENSG00000060237 | 0 | 0 | 0 | 0 | 0 | 0 | 0 | 0 | 0 | 0 | 0 | 0 | 0 | 0 | 0 | 0 | 1 | 5 | 2 |
| HG0508 | *H. sapiens* | ENSG00000061936 | 0 | 0 | 0 | 0 | 0 | 0 | 0 | 0 | 0 | 0 | 0 | 0 | 0 | 0 | 0 | 0 | 1 | 5 | 2 |
| HG0509 | *H. sapiens* | ENSG00000068024 | 0 | 0 | 0 | 0 | 0 | 0 | 0 | 0 | 0 | 0 | 0 | 0 | 0 | 0 | 0 | 0 | 1 | 5 | 2 |
| HG0510 | *H. sapiens* | ENSG00000134769 | 0 | 0 | 0 | 0 | 0 | 0 | 0 | 0 | 0 | 0 | 0 | 0 | 0 | 0 | 0 | 0 | 1 | 6 | 1 |
| HG0511 | *H. sapiens* | ENSG00000143337 | 0 | 0 | 0 | 0 | 0 | 0 | 0 | 0 | 0 | 0 | 0 | 0 | 0 | 0 | 0 | 0 | 1 | 5 | 2 |
| HG0512.1 | *H. sapiens* | ENSG00000112319 | 0 | 0 | 0 | 0 | 0 | 0 | 0 | 0 | 0 | 0 | 0 | 0 | 0 | 0 | 0 | 0 | 1 | 5 | 1 |
| HG0512.2 | *H. sapiens* | ENSG00000104313 | 0 | 0 | 0 | 0 | 0 | 0 | 0 | 0 | 0 | 0 | 0 | 0 | 0 | 0 | 0 | 0 | 0 | 6 | 3 |
| HG0512.3 | *H. sapiens* | ENSG00000104313 | 0 | 0 | 0 | 0 | 0 | 0 | 0 | 0 | 0 | 0 | 0 | 0 | 0 | 0 | 0 | 0 | 0 | 7 | 1 |
| HG0513 | *H. sapiens* | ENSG00000126860 | 0 | 0 | 0 | 0 | 0 | 0 | 0 | 0 | 0 | 0 | 0 | 0 | 0 | 0 | 0 | 0 | 1 | 5 | 1 |
| HG0514 | *H. sapiens* | ENSG00000147570 | 0 | 0 | 0 | 0 | 0 | 0 | 0 | 0 | 0 | 0 | 0 | 0 | 0 | 0 | 0 | 0 | 1 | 4 | 2 |
| HG0515 | *H. sapiens* | ENSG00000170471 | 0 | 0 | 0 | 0 | 0 | 0 | 0 | 0 | 0 | 0 | 0 | 0 | 0 | 0 | 0 | 0 | 1 | 4 | 2 |
| HG0516 | *H. sapiens* | ENSG00000041515 | 0 | 0 | 0 | 0 | 0 | 0 | 0 | 0 | 0 | 0 | 0 | 0 | 0 | 0 | 0 | 0 | 1 | 3 | 2 |
| HG0517 | *H. sapiens* | ENSG00000086717 | 0 | 0 | 0 | 0 | 0 | 0 | 0 | 0 | 0 | 0 | 0 | 0 | 0 | 0 | 0 | 0 | 1 | 2 | 3 |
| HG0518 | *H. sapiens* | ENSG00000111254 | 0 | 0 | 0 | 0 | 0 | 0 | 0 | 0 | 0 | 0 | 0 | 0 | 0 | 0 | 0 | 0 | 1 | 3 | 2 |
| HG0519 | *H. sapiens* | ENSG00000127074 | 0 | 0 | 0 | 0 | 0 | 0 | 0 | 0 | 0 | 0 | 0 | 0 | 0 | 0 | 0 | 0 | 1 | 4 | 1 |
| HG0520 | *H. sapiens* | ENSG00000197415 | 0 | 0 | 0 | 0 | 0 | 0 | 0 | 0 | 0 | 0 | 0 | 0 | 0 | 0 | 0 | 0 | 1 | 4 | 1 |
| HG0521 | *H. sapiens* | ENSG00000038274 | 0 | 0 | 0 | 0 | 0 | 0 | 0 | 0 | 0 | 0 | 0 | 0 | 0 | 0 | 0 | 0 | 1 | 2 | 1 |
| HG0522 | *H. sapiens* | ENSG00000140396 | 0 | 0 | 0 | 0 | 0 | 0 | 0 | 0 | 0 | 0 | 0 | 0 | 0 | 0 | 0 | 0 | 1 | 1 | 2 |
| HG0523 | *H. sapiens* | ENSG00000075213 | 0 | 0 | 0 | 0 | 0 | 0 | 0 | 0 | 0 | 0 | 0 | 0 | 0 | 0 | 0 | 0 | 2 | 0 | 1 |
| HG0524 | *H. sapiens* | ENSG00000107929 | 0 | 0 | 0 | 0 | 0 | 0 | 0 | 0 | 0 | 0 | 0 | 0 | 0 | 0 | 0 | 0 | 1 | 1 | 1 |
| HG0525 | *H. sapiens* | ENSG00000136167 | 0 | 0 | 0 | 0 | 0 | 0 | 0 | 0 | 0 | 0 | 0 | 0 | 0 | 0 | 0 | 0 | 1 | 1 | 1 |
| HG0526 | *H. sapiens* | ENSG00000143569 | 0 | 0 | 0 | 0 | 0 | 0 | 0 | 0 | 0 | 0 | 0 | 0 | 0 | 0 | 0 | 0 | 1 | 1 | 1 |
| HG0527 | *H. sapiens* | ENSG00000198382 | 0 | 0 | 0 | 0 | 0 | 0 | 0 | 0 | 0 | 0 | 0 | 0 | 0 | 0 | 0 | 0 | 1 | 0 | 2 |
| HG0528 | *H. sapiens* | ENSG00000204767 | 0 | 0 | 0 | 0 | 0 | 0 | 0 | 0 | 0 | 0 | 0 | 0 | 0 | 0 | 0 | 0 | 1 | 1 | 0 |
| HG0529 | *H. sapiens* | ENSG00000112851 | 0 | 0 | 0 | 0 | 0 | 0 | 0 | 0 | 0 | 0 | 0 | 0 | 0 | 0 | 0 | 0 | 0 | 11 | 4 |
| HG0530 | *H. sapiens* | ENSG00000128594 | 0 | 0 | 0 | 0 | 0 | 0 | 0 | 0 | 0 | 0 | 0 | 0 | 0 | 0 | 0 | 0 | 0 | 11 | 4 |
| HG0531 | *H. sapiens* | ENSG00000146834 | 0 | 0 | 0 | 0 | 0 | 0 | 0 | 0 | 0 | 0 | 0 | 0 | 0 | 0 | 0 | 0 | 0 | 11 | 4 |
| HG0532 | *H. sapiens* | ENSG00000105221 | 0 | 0 | 0 | 0 | 0 | 0 | 0 | 0 | 0 | 0 | 0 | 0 | 0 | 0 | 0 | 0 | 0 | 11 | 3 |
| HG0533 | *H. sapiens* | ENSG00000114209 | 0 | 0 | 0 | 0 | 0 | 0 | 0 | 0 | 0 | 0 | 0 | 0 | 0 | 0 | 0 | 0 | 0 | 10 | 4 |
| HG0534 | *H. sapiens* | ENSG00000117408 | 0 | 0 | 0 | 0 | 0 | 0 | 0 | 0 | 0 | 0 | 0 | 0 | 0 | 0 | 0 | 0 | 0 | 11 | 3 |
| HG0535 | *H. sapiens* | ENSG00000118407 | 0 | 0 | 0 | 0 | 0 | 0 | 0 | 0 | 0 | 0 | 0 | 0 | 0 | 0 | 0 | 0 | 0 | 10 | 4 |
| HG0536 | *H. sapiens* | ENSG00000181751 | 0 | 0 | 0 | 0 | 0 | 0 | 0 | 0 | 0 | 0 | 0 | 0 | 0 | 0 | 0 | 0 | 0 | 10 | 4 |
| HG0537 | *H. sapiens* | ENSG00000184481 | 0 | 0 | 0 | 0 | 0 | 0 | 0 | 0 | 0 | 0 | 0 | 0 | 0 | 0 | 0 | 0 | 0 | 10 | 4 |
| HG0538 | *H. sapiens* | ENSG00000006116 | 0 | 0 | 0 | 0 | 0 | 0 | 0 | 0 | 0 | 0 | 0 | 0 | 0 | 0 | 0 | 0 | 0 | 9 | 4 |
| HG0539 | *H. sapiens* | ENSG00000115966 | 0 | 0 | 0 | 0 | 0 | 0 | 0 | 0 | 0 | 0 | 0 | 0 | 0 | 0 | 0 | 0 | 0 | 10 | 3 |
| HG0540 | *H. sapiens* | ENSG00000120519 | 0 | 0 | 0 | 0 | 0 | 0 | 0 | 0 | 0 | 0 | 0 | 0 | 0 | 0 | 0 | 0 | 0 | 10 | 3 |
| HG0541.1 | *H. sapiens* | ENSG00000122584 | 0 | 0 | 0 | 0 | 0 | 0 | 0 | 0 | 0 | 0 | 0 | 0 | 0 | 0 | 0 | 0 | 0 | 10 | 3 |
| HG0541.2 | *H. sapiens* | ENSG00000122584 | 0 | 0 | 0 | 0 | 0 | 0 | 0 | 0 | 0 | 0 | 0 | 0 | 0 | 0 | 0 | 0 | 0 | 9 | 2 |
| HG0542 | *H. sapiens* | ENSG00000126581 | 0 | 0 | 0 | 0 | 0 | 0 | 0 | 0 | 0 | 0 | 0 | 0 | 0 | 0 | 0 | 0 | 0 | 9 | 4 |
| HG0543 | *H. sapiens* | ENSG00000131931 | 0 | 0 | 0 | 0 | 0 | 0 | 0 | 0 | 0 | 0 | 0 | 0 | 0 | 0 | 0 | 0 | 0 | 10 | 3 |
| HG0544 | *H. sapiens* | ENSG00000136802 | 0 | 0 | 0 | 0 | 0 | 0 | 0 | 0 | 0 | 0 | 0 | 0 | 0 | 0 | 0 | 0 | 0 | 9 | 4 |
| HG0545 | *H. sapiens* | ENSG00000142784 | 0 | 0 | 0 | 0 | 0 | 0 | 0 | 0 | 0 | 0 | 0 | 0 | 0 | 0 | 0 | 0 | 0 | 9 | 4 |
| HG0546 | *H. sapiens* | ENSG00000148143 | 0 | 0 | 0 | 0 | 0 | 0 | 0 | 0 | 0 | 0 | 0 | 0 | 0 | 0 | 0 | 0 | 0 | 10 | 3 |
| HG0547 | *H. sapiens* | ENSG00000149596 | 0 | 0 | 0 | 0 | 0 | 0 | 0 | 0 | 0 | 0 | 0 | 0 | 0 | 0 | 0 | 0 | 0 | 9 | 4 |
| HG0548 | *H. sapiens* | ENSG00000155744 | 0 | 0 | 0 | 0 | 0 | 0 | 0 | 0 | 0 | 0 | 0 | 0 | 0 | 0 | 0 | 0 | 0 | 9 | 4 |
| HG0549 | *H. sapiens* | ENSG00000156256 | 0 | 0 | 0 | 0 | 0 | 0 | 0 | 0 | 0 | 0 | 0 | 0 | 0 | 0 | 0 | 0 | 0 | 10 | 3 |
| HG0550.1 | *H. sapiens* | ENSG00000156650 | 0 | 0 | 0 | 0 | 0 | 0 | 0 | 0 | 0 | 0 | 0 | 0 | 0 | 0 | 0 | 0 | 0 | 9 | 4 |
| HG0550.2 | *H. sapiens* | ENSG00000156650 | 0 | 0 | 0 | 0 | 0 | 0 | 0 | 0 | 0 | 0 | 0 | 0 | 0 | 0 | 0 | 0 | 0 | 8 | 0 |
| HG0551 | *H. sapiens* | ENSG00000162951 | 0 | 0 | 0 | 0 | 0 | 0 | 0 | 0 | 0 | 0 | 0 | 0 | 0 | 0 | 0 | 0 | 0 | 9 | 4 |
| HG0552 | *H. sapiens* | ENSG00000169641 | 0 | 0 | 0 | 0 | 0 | 0 | 0 | 0 | 0 | 0 | 0 | 0 | 0 | 0 | 0 | 0 | 0 | 10 | 3 |
| HG0553 | *H. sapiens* | ENSG00000183475 | 0 | 0 | 0 | 0 | 0 | 0 | 0 | 0 | 0 | 0 | 0 | 0 | 0 | 0 | 0 | 0 | 0 | 9 | 4 |
| HG0554 | *H. sapiens* | ENSG00000196914 | 0 | 0 | 0 | 0 | 0 | 0 | 0 | 0 | 0 | 0 | 0 | 0 | 0 | 0 | 0 | 0 | 0 | 9 | 4 |
| HG0555 | *H. sapiens* | ENSG00000204310 | 0 | 0 | 0 | 0 | 0 | 0 | 0 | 0 | 0 | 0 | 0 | 0 | 0 | 0 | 0 | 0 | 0 | 10 | 3 |
| HG0556 | *H. sapiens* | ENSG00000273841 | 0 | 0 | 0 | 0 | 0 | 0 | 0 | 0 | 0 | 0 | 0 | 0 | 0 | 0 | 0 | 0 | 0 | 10 | 3 |
| HG0557 | *H. sapiens* | ENSG00000012232 | 0 | 0 | 0 | 0 | 0 | 0 | 0 | 0 | 0 | 0 | 0 | 0 | 0 | 0 | 0 | 0 | 0 | 9 | 3 |
| HG0558 | *H. sapiens* | ENSG00000029363 | 0 | 0 | 0 | 0 | 0 | 0 | 0 | 0 | 0 | 0 | 0 | 0 | 0 | 0 | 0 | 0 | 0 | 8 | 4 |
| HG0559 | *H. sapiens* | ENSG00000078747 | 0 | 0 | 0 | 0 | 0 | 0 | 0 | 0 | 0 | 0 | 0 | 0 | 0 | 0 | 0 | 0 | 0 | 8 | 4 |
| HG0560.1 | *H. sapiens* | ENSG00000092051 | 0 | 0 | 0 | 0 | 0 | 0 | 0 | 0 | 0 | 0 | 0 | 0 | 0 | 0 | 0 | 0 | 0 | 9 | 3 |
| HG0560.2 | *H. sapiens* | ENSG00000092051 | 0 | 0 | 0 | 0 | 0 | 0 | 0 | 0 | 0 | 0 | 0 | 0 | 0 | 0 | 0 | 0 | 0 | 8 | 3 |
| HG0561 | *H. sapiens* | ENSG00000100393 | 0 | 0 | 0 | 0 | 0 | 0 | 0 | 0 | 0 | 0 | 0 | 0 | 0 | 0 | 0 | 0 | 0 | 10 | 2 |
| HG0562 | *H. sapiens* | ENSG00000105821 | 0 | 0 | 0 | 0 | 0 | 0 | 0 | 0 | 0 | 0 | 0 | 0 | 0 | 0 | 0 | 0 | 0 | 8 | 4 |
| HG0563 | *H. sapiens* | ENSG00000107249 | 0 | 0 | 0 | 0 | 0 | 0 | 0 | 0 | 0 | 0 | 0 | 0 | 0 | 0 | 0 | 0 | 0 | 8 | 4 |
| HG0564 | *H. sapiens* | ENSG00000109171 | 0 | 0 | 0 | 0 | 0 | 0 | 0 | 0 | 0 | 0 | 0 | 0 | 0 | 0 | 0 | 0 | 0 | 9 | 3 |
| HG0565 | *H. sapiens* | ENSG00000120533 | 0 | 0 | 0 | 0 | 0 | 0 | 0 | 0 | 0 | 0 | 0 | 0 | 0 | 0 | 0 | 0 | 0 | 9 | 3 |
| HG0566 | *H. sapiens* | ENSG00000124496 | 0 | 0 | 0 | 0 | 0 | 0 | 0 | 0 | 0 | 0 | 0 | 0 | 0 | 0 | 0 | 0 | 0 | 9 | 3 |
| HG0567 | *H. sapiens* | ENSG00000132155 | 0 | 0 | 0 | 0 | 0 | 0 | 0 | 0 | 0 | 0 | 0 | 0 | 0 | 0 | 0 | 0 | 0 | 9 | 3 |
| HG0568.1 | *H. sapiens* | ENSG00000138271 | 0 | 0 | 0 | 0 | 0 | 0 | 0 | 0 | 0 | 0 | 0 | 0 | 0 | 0 | 0 | 0 | 0 | 8 | 4 |
| HG0568.2 | *H. sapiens* | ENSG00000138271 | 0 | 0 | 0 | 0 | 0 | 0 | 0 | 0 | 0 | 0 | 0 | 0 | 0 | 0 | 0 | 0 | 0 | 4 | 1 |
| HG0569 | *H. sapiens* | ENSG00000139651 | 0 | 0 | 0 | 0 | 0 | 0 | 0 | 0 | 0 | 0 | 0 | 0 | 0 | 0 | 0 | 0 | 0 | 10 | 2 |
| HG0570 | *H. sapiens* | ENSG00000152214 | 0 | 0 | 0 | 0 | 0 | 0 | 0 | 0 | 0 | 0 | 0 | 0 | 0 | 0 | 0 | 0 | 0 | 9 | 3 |
| HG0571 | *H. sapiens* | ENSG00000153266 | 0 | 0 | 0 | 0 | 0 | 0 | 0 | 0 | 0 | 0 | 0 | 0 | 0 | 0 | 0 | 0 | 0 | 9 | 3 |
| HG0572 | *H. sapiens* | ENSG00000158161 | 0 | 0 | 0 | 0 | 0 | 0 | 0 | 0 | 0 | 0 | 0 | 0 | 0 | 0 | 0 | 0 | 0 | 8 | 4 |
| HG0573.1 | *H. sapiens* | ENSG00000162526 | 0 | 0 | 0 | 0 | 0 | 0 | 0 | 0 | 0 | 0 | 0 | 0 | 0 | 0 | 0 | 0 | 0 | 9 | 3 |
| HG0573.2 | *H. sapiens* | ENSG00000206203 | 0 | 0 | 0 | 0 | 0 | 0 | 0 | 0 | 0 | 0 | 0 | 0 | 0 | 0 | 0 | 0 | 0 | 5 | 3 |
| HG0574 | *H. sapiens* | ENSG00000164663 | 0 | 0 | 0 | 0 | 0 | 0 | 0 | 0 | 0 | 0 | 0 | 0 | 0 | 0 | 0 | 0 | 0 | 8 | 4 |

| | | | | | | | | | | | | | | | | | | | | |
|---|---|---|---|---|---|---|---|---|---|---|---|---|---|---|---|---|---|---|---|---|---|
| HG0575 | *H. sapiens* | ENSG00000164663 | 0 | 0 | 0 | 0 | 0 | 0 | 0 | 0 | 0 | 0 | 0 | 0 | 0 | 0 | 0 | 0 | 0 | 8 | 4 |
| HG0576 | *H. sapiens* | ENSG00000175029 | 0 | 0 | 0 | 0 | 0 | 0 | 0 | 0 | 0 | 0 | 0 | 0 | 0 | 0 | 0 | 0 | 0 | 9 | 3 |
| HG0577 | *H. sapiens* | ENSG00000250510 | 0 | 0 | 0 | 0 | 0 | 0 | 0 | 0 | 0 | 0 | 0 | 0 | 0 | 0 | 0 | 0 | 0 | 9 | 3 |
| HG0578 | *H. sapiens* | ENSG00000004897 | 0 | 0 | 0 | 0 | 0 | 0 | 0 | 0 | 0 | 0 | 0 | 0 | 0 | 0 | 0 | 0 | 0 | 8 | 3 |
| HG0579 | *H. sapiens* | ENSG00000010818 | 0 | 0 | 0 | 0 | 0 | 0 | 0 | 0 | 0 | 0 | 0 | 0 | 0 | 0 | 0 | 0 | 0 | 7 | 4 |
| HG0580 | *H. sapiens* | ENSG00000073711 | 0 | 0 | 0 | 0 | 0 | 0 | 0 | 0 | 0 | 0 | 0 | 0 | 0 | 0 | 0 | 0 | 0 | 8 | 3 |
| HG0581 | *H. sapiens* | ENSG00000087074 | 0 | 0 | 0 | 0 | 0 | 0 | 0 | 0 | 0 | 0 | 0 | 0 | 0 | 0 | 0 | 0 | 0 | 7 | 4 |
| HG0582 | *H. sapiens* | ENSG00000087338 | 0 | 0 | 0 | 0 | 0 | 0 | 0 | 0 | 0 | 0 | 0 | 0 | 0 | 0 | 0 | 0 | 0 | 8 | 3 |
| HG0583 | *H. sapiens* | ENSG00000087502 | 0 | 0 | 0 | 0 | 0 | 0 | 0 | 0 | 0 | 0 | 0 | 0 | 0 | 0 | 0 | 0 | 0 | 7 | 4 |
| HG0584 | *H. sapiens* | ENSG00000092051 | 0 | 0 | 0 | 0 | 0 | 0 | 0 | 0 | 0 | 0 | 0 | 0 | 0 | 0 | 0 | 0 | 0 | 8 | 3 |
| HG0585 | *H. sapiens* | ENSG00000106038 | 0 | 0 | 0 | 0 | 0 | 0 | 0 | 0 | 0 | 0 | 0 | 0 | 0 | 0 | 0 | 0 | 0 | 8 | 3 |
| HG0586.1 | *H. sapiens* | ENSG00000108375 | 0 | 0 | 0 | 0 | 0 | 0 | 0 | 0 | 0 | 0 | 0 | 0 | 0 | 0 | 0 | 0 | 0 | 8 | 3 |
| HG0586.2 | *H. sapiens* | ENSG00000108375 | 0 | 0 | 0 | 0 | 0 | 0 | 0 | 0 | 0 | 0 | 0 | 0 | 0 | 0 | 0 | 0 | 0 | 4 | 1 |
| HG0587 | *H. sapiens* | ENSG00000119138 | 0 | 0 | 0 | 0 | 0 | 0 | 0 | 0 | 0 | 0 | 0 | 0 | 0 | 0 | 0 | 0 | 0 | 10 | 1 |
| HG0588 | *H. sapiens* | ENSG00000120162 | 0 | 0 | 0 | 0 | 0 | 0 | 0 | 0 | 0 | 0 | 0 | 0 | 0 | 0 | 0 | 0 | 0 | 7 | 4 |
| HG0589.1 | *H. sapiens* | ENSG00000121871 | 0 | 0 | 0 | 0 | 0 | 0 | 0 | 0 | 0 | 0 | 0 | 0 | 0 | 0 | 0 | 0 | 0 | 7 | 4 |
| HG0589.2 | *H. sapiens* | ENSG00000121871 | 0 | 0 | 0 | 0 | 0 | 0 | 0 | 0 | 0 | 0 | 0 | 0 | 0 | 0 | 0 | 0 | 0 | 7 | 3 |
| HG0589.3 | *H. sapiens* | ENSG00000121871 | 0 | 0 | 0 | 0 | 0 | 0 | 0 | 0 | 0 | 0 | 0 | 0 | 0 | 0 | 0 | 0 | 0 | 5 | 3 |
| HG0590 | *H. sapiens* | ENSG00000122557 | 0 | 0 | 0 | 0 | 0 | 0 | 0 | 0 | 0 | 0 | 0 | 0 | 0 | 0 | 0 | 0 | 0 | 9 | 2 |
| HG0591.1 | *H. sapiens* | ENSG00000127124 | 0 | 0 | 0 | 0 | 0 | 0 | 0 | 0 | 0 | 0 | 0 | 0 | 0 | 0 | 0 | 0 | 0 | 7 | 4 |
| HG0591.2 | *H. sapiens* | ENSG00000127124 | 0 | 0 | 0 | 0 | 0 | 0 | 0 | 0 | 0 | 0 | 0 | 0 | 0 | 0 | 0 | 0 | 0 | 7 | 3 |
| HG0591.3 | *H. sapiens* | ENSG00000127124 | 0 | 0 | 0 | 0 | 0 | 0 | 0 | 0 | 0 | 0 | 0 | 0 | 0 | 0 | 0 | 0 | 0 | 4 | 2 |
| HG0591.4 | *H. sapiens* | ENSG00000127124 | 0 | 0 | 0 | 0 | 0 | 0 | 0 | 0 | 0 | 0 | 0 | 0 | 0 | 0 | 0 | 0 | 0 | 1 | 2 |
| HG0592 | *H. sapiens* | ENSG00000133401 | 0 | 0 | 0 | 0 | 0 | 0 | 0 | 0 | 0 | 0 | 0 | 0 | 0 | 0 | 0 | 0 | 0 | 8 | 3 |
| HG0593 | *H. sapiens* | ENSG00000135048 | 0 | 0 | 0 | 0 | 0 | 0 | 0 | 0 | 0 | 0 | 0 | 0 | 0 | 0 | 0 | 0 | 0 | 10 | 1 |
| HG0594 | *H. sapiens* | ENSG00000136169 | 0 | 0 | 0 | 0 | 0 | 0 | 0 | 0 | 0 | 0 | 0 | 0 | 0 | 0 | 0 | 0 | 0 | 7 | 4 |
| HG0595 | *H. sapiens* | ENSG00000141098 | 0 | 0 | 0 | 0 | 0 | 0 | 0 | 0 | 0 | 0 | 0 | 0 | 0 | 0 | 0 | 0 | 0 | 8 | 3 |
| HG0596 | *H. sapiens* | ENSG00000157483 | 0 | 0 | 0 | 0 | 0 | 0 | 0 | 0 | 0 | 0 | 0 | 0 | 0 | 0 | 0 | 0 | 0 | 7 | 4 |
| HG0597 | *H. sapiens* | ENSG00000163032 | 0 | 0 | 0 | 0 | 0 | 0 | 0 | 0 | 0 | 0 | 0 | 0 | 0 | 0 | 0 | 0 | 0 | 8 | 3 |
| HG0598 | *H. sapiens* | ENSG00000163788 | 0 | 0 | 0 | 0 | 0 | 0 | 0 | 0 | 0 | 0 | 0 | 0 | 0 | 0 | 0 | 0 | 0 | 9 | 2 |
| HG0599 | *H. sapiens* | ENSG00000164930 | 0 | 0 | 0 | 0 | 0 | 0 | 0 | 0 | 0 | 0 | 0 | 0 | 0 | 0 | 0 | 0 | 0 | 8 | 3 |
| HG0600 | *H. sapiens* | ENSG00000173065 | 0 | 0 | 0 | 0 | 0 | 0 | 0 | 0 | 0 | 0 | 0 | 0 | 0 | 0 | 0 | 0 | 0 | 8 | 3 |
| HG0601 | *H. sapiens* | ENSG00000175893 | 0 | 0 | 0 | 0 | 0 | 0 | 0 | 0 | 0 | 0 | 0 | 0 | 0 | 0 | 0 | 0 | 0 | 7 | 4 |
| HG0602 | *H. sapiens* | ENSG00000179195 | 0 | 0 | 0 | 0 | 0 | 0 | 0 | 0 | 0 | 0 | 0 | 0 | 0 | 0 | 0 | 0 | 0 | 8 | 3 |
| HG0603 | *H. sapiens* | ENSG00000197329 | 0 | 0 | 0 | 0 | 0 | 0 | 0 | 0 | 0 | 0 | 0 | 0 | 0 | 0 | 0 | 0 | 0 | 9 | 2 |
| HG0604.1 | *H. sapiens* | ENSG00000003147 | 0 | 0 | 0 | 0 | 0 | 0 | 0 | 0 | 0 | 0 | 0 | 0 | 0 | 0 | 0 | 0 | 0 | 7 | 3 |
| HG0604.2 | *H. sapiens* | ENSG00000003147 | 0 | 0 | 0 | 0 | 0 | 0 | 0 | 0 | 0 | 0 | 0 | 0 | 0 | 0 | 0 | 0 | 0 | 5 | 1 |
| HG0604.3 | *H. sapiens* | ENSG00000003147 | 0 | 0 | 0 | 0 | 0 | 0 | 0 | 0 | 0 | 0 | 0 | 0 | 0 | 0 | 0 | 0 | 0 | 2 | 2 |
| HG0605 | *H. sapiens* | ENSG00000053254 | 0 | 0 | 0 | 0 | 0 | 0 | 0 | 0 | 0 | 0 | 0 | 0 | 0 | 0 | 0 | 0 | 0 | 7 | 3 |
| HG0606 | *H. sapiens* | ENSG00000055130 | 0 | 0 | 0 | 0 | 0 | 0 | 0 | 0 | 0 | 0 | 0 | 0 | 0 | 0 | 0 | 0 | 0 | 7 | 3 |
| HG0607 | *H. sapiens* | ENSG00000074657 | 0 | 0 | 0 | 0 | 0 | 0 | 0 | 0 | 0 | 0 | 0 | 0 | 0 | 0 | 0 | 0 | 0 | 7 | 3 |
| HG0608 | *H. sapiens* | ENSG00000084731 | 0 | 0 | 0 | 0 | 0 | 0 | 0 | 0 | 0 | 0 | 0 | 0 | 0 | 0 | 0 | 0 | 0 | 8 | 2 |
| HG0609 | *H. sapiens* | ENSG00000095539 | 0 | 0 | 0 | 0 | 0 | 0 | 0 | 0 | 0 | 0 | 0 | 0 | 0 | 0 | 0 | 0 | 0 | 6 | 4 |
| HG0610 | *H. sapiens* | ENSG00000106069 | 0 | 0 | 0 | 0 | 0 | 0 | 0 | 0 | 0 | 0 | 0 | 0 | 0 | 0 | 0 | 0 | 0 | 8 | 2 |
| HG0611 | *H. sapiens* | ENSG00000108309 | 0 | 0 | 0 | 0 | 0 | 0 | 0 | 0 | 0 | 0 | 0 | 0 | 0 | 0 | 0 | 0 | 0 | 7 | 3 |
| HG0612.1 | *H. sapiens* | ENSG00000110422 | 0 | 0 | 0 | 0 | 0 | 0 | 0 | 0 | 0 | 0 | 0 | 0 | 0 | 0 | 0 | 0 | 0 | 8 | 2 |
| HG0612.2 | *H. sapiens* | ENSG00000110422 | 0 | 0 | 0 | 0 | 0 | 0 | 0 | 0 | 0 | 0 | 0 | 0 | 0 | 0 | 0 | 0 | 0 | 6 | 3 |
| HG0612.3 | *H. sapiens* | ENSG00000110422 | 0 | 0 | 0 | 0 | 0 | 0 | 0 | 0 | 0 | 0 | 0 | 0 | 0 | 0 | 0 | 0 | 0 | 3 | 1 |
| HG0613 | *H. sapiens* | ENSG00000114861 | 0 | 0 | 0 | 0 | 0 | 0 | 0 | 0 | 0 | 0 | 0 | 0 | 0 | 0 | 0 | 0 | 0 | 6 | 4 |
| HG0614 | *H. sapiens* | ENSG00000122203 | 0 | 0 | 0 | 0 | 0 | 0 | 0 | 0 | 0 | 0 | 0 | 0 | 0 | 0 | 0 | 0 | 0 | 6 | 4 |
| HG0615 | *H. sapiens* | ENSG00000128191 | 0 | 0 | 0 | 0 | 0 | 0 | 0 | 0 | 0 | 0 | 0 | 0 | 0 | 0 | 0 | 0 | 0 | 7 | 3 |
| HG0616 | *H. sapiens* | ENSG00000128739 | 0 | 0 | 0 | 0 | 0 | 0 | 0 | 0 | 0 | 0 | 0 | 0 | 0 | 0 | 0 | 0 | 0 | 7 | 3 |
| HG0617 | *H. sapiens* | ENSG00000135164 | 0 | 0 | 0 | 0 | 0 | 0 | 0 | 0 | 0 | 0 | 0 | 0 | 0 | 0 | 0 | 0 | 0 | 6 | 4 |
| HG0618 | *H. sapiens* | ENSG00000137817 | 0 | 0 | 0 | 0 | 0 | 0 | 0 | 0 | 0 | 0 | 0 | 0 | 0 | 0 | 0 | 0 | 0 | 8 | 2 |
| HG0619 | *H. sapiens* | ENSG00000138593 | 0 | 0 | 0 | 0 | 0 | 0 | 0 | 0 | 0 | 0 | 0 | 0 | 0 | 0 | 0 | 0 | 0 | 8 | 2 |
| HG0620 | *H. sapiens* | ENSG00000145555 | 0 | 0 | 0 | 0 | 0 | 0 | 0 | 0 | 0 | 0 | 0 | 0 | 0 | 0 | 0 | 0 | 0 | 6 | 4 |
| HG0621.1 | *H. sapiens* | ENSG00000146352 | 0 | 0 | 0 | 0 | 0 | 0 | 0 | 0 | 0 | 0 | 0 | 0 | 0 | 0 | 0 | 0 | 0 | 7 | 3 |
| HG0621.2 | *H. sapiens* | ENSG00000146352 | 0 | 0 | 0 | 0 | 0 | 0 | 0 | 0 | 0 | 0 | 0 | 0 | 0 | 0 | 0 | 0 | 0 | 5 | 3 |
| HG0621.3 | *H. sapiens* | ENSG00000146352 | 0 | 0 | 0 | 0 | 0 | 0 | 0 | 0 | 0 | 0 | 0 | 0 | 0 | 0 | 0 | 0 | 0 | 2 | 2 |
| HG0622 | *H. sapiens* | ENSG00000148677 | 0 | 0 | 0 | 0 | 0 | 0 | 0 | 0 | 0 | 0 | 0 | 0 | 0 | 0 | 0 | 0 | 0 | 7 | 3 |
| HG0623 | *H. sapiens* | ENSG00000151422 | 0 | 0 | 0 | 0 | 0 | 0 | 0 | 0 | 0 | 0 | 0 | 0 | 0 | 0 | 0 | 0 | 0 | 7 | 3 |
| HG0624 | *H. sapiens* | ENSG00000156853 | 0 | 0 | 0 | 0 | 0 | 0 | 0 | 0 | 0 | 0 | 0 | 0 | 0 | 0 | 0 | 0 | 0 | 6 | 4 |
| HG0625 | *H. sapiens* | ENSG00000164050 | 0 | 0 | 0 | 0 | 0 | 0 | 0 | 0 | 0 | 0 | 0 | 0 | 0 | 0 | 0 | 0 | 0 | 7 | 3 |
| HG0626 | *H. sapiens* | ENSG00000168939 | 0 | 0 | 0 | 0 | 0 | 0 | 0 | 0 | 0 | 0 | 0 | 0 | 0 | 0 | 0 | 0 | 0 | 7 | 3 |
| HG0627 | *H. sapiens* | ENSG00000169760 | 0 | 0 | 0 | 0 | 0 | 0 | 0 | 0 | 0 | 0 | 0 | 0 | 0 | 0 | 0 | 0 | 0 | 8 | 2 |
| HG0628 | *H. sapiens* | ENSG00000170677 | 0 | 0 | 0 | 0 | 0 | 0 | 0 | 0 | 0 | 0 | 0 | 0 | 0 | 0 | 0 | 0 | 0 | 6 | 4 |
| HG0629 | *H. sapiens* | ENSG00000170961 | 0 | 0 | 0 | 0 | 0 | 0 | 0 | 0 | 0 | 0 | 0 | 0 | 0 | 0 | 0 | 0 | 0 | 7 | 3 |
| HG0630 | *H. sapiens* | ENSG00000171552 | 0 | 0 | 0 | 0 | 0 | 0 | 0 | 0 | 0 | 0 | 0 | 0 | 0 | 0 | 0 | 0 | 0 | 6 | 4 |
| HG0631 | *H. sapiens* | ENSG00000173548 | 0 | 0 | 0 | 0 | 0 | 0 | 0 | 0 | 0 | 0 | 0 | 0 | 0 | 0 | 0 | 0 | 0 | 8 | 2 |
| HG0632.1 | *H. sapiens* | ENSG00000181315 | 0 | 0 | 0 | 0 | 0 | 0 | 0 | 0 | 0 | 0 | 0 | 0 | 0 | 0 | 0 | 0 | 0 | 7 | 3 |
| HG0632.2 | *H. sapiens* | ENSG00000181315 | 0 | 0 | 0 | 0 | 0 | 0 | 0 | 0 | 0 | 0 | 0 | 0 | 0 | 0 | 0 | 0 | 0 | 3 | 0 |
| HG0633 | *H. sapiens* | ENSG00000186174 | 0 | 0 | 0 | 0 | 0 | 0 | 0 | 0 | 0 | 0 | 0 | 0 | 0 | 0 | 0 | 0 | 0 | 7 | 3 |
| HG0634 | *H. sapiens* | ENSG00000198026 | 0 | 0 | 0 | 0 | 0 | 0 | 0 | 0 | 0 | 0 | 0 | 0 | 0 | 0 | 0 | 0 | 0 | 7 | 3 |
| HG0635 | *H. sapiens* | ENSG00000259332 | 0 | 0 | 0 | 0 | 0 | 0 | 0 | 0 | 0 | 0 | 0 | 0 | 0 | 0 | 0 | 0 | 0 | 8 | 2 |
| HG0636 | *H. sapiens* | ENSG00000002587 | 0 | 0 | 0 | 0 | 0 | 0 | 0 | 0 | 0 | 0 | 0 | 0 | 0 | 0 | 0 | 0 | 0 | 7 | 2 |
| HG0637 | *H. sapiens* | ENSG00000005102 | 0 | 0 | 0 | 0 | 0 | 0 | 0 | 0 | 0 | 0 | 0 | 0 | 0 | 0 | 0 | 0 | 0 | 6 | 3 |
| HG0638 | *H. sapiens* | ENSG00000013561 | 0 | 0 | 0 | 0 | 0 | 0 | 0 | 0 | 0 | 0 | 0 | 0 | 0 | 0 | 0 | 0 | 0 | 7 | 2 |
| HG0639 | *H. sapiens* | ENSG00000018236 | 0 | 0 | 0 | 0 | 0 | 0 | 0 | 0 | 0 | 0 | 0 | 0 | 0 | 0 | 0 | 0 | 0 | 6 | 3 |
| HG0640 | *H. sapiens* | ENSG00000062194 | 0 | 0 | 0 | 0 | 0 | 0 | 0 | 0 | 0 | 0 | 0 | 0 | 0 | 0 | 0 | 0 | 0 | 7 | 2 |

| | | | | | | | | | | | | | | | | | | | | | |
|---|---|---|---|---|---|---|---|---|---|---|---|---|---|---|---|---|---|---|---|---|---|
| HG0641 | *H. sapiens* | ENSG00000100359 | 0 | 0 | 0 | 0 | 0 | 0 | 0 | 0 | 0 | 0 | 0 | 0 | 0 | 0 | 0 | 0 | 0 | 7 | 2 |
| HG0642 | *H. sapiens* | ENSG00000100815 | 0 | 0 | 0 | 0 | 0 | 0 | 0 | 0 | 0 | 0 | 0 | 0 | 0 | 0 | 0 | 0 | 0 | 8 | 1 |
| HG0643 | *H. sapiens* | ENSG00000103653 | 0 | 0 | 0 | 0 | 0 | 0 | 0 | 0 | 0 | 0 | 0 | 0 | 0 | 0 | 0 | 0 | 0 | 7 | 2 |
| HG0644 | *H. sapiens* | ENSG00000104299 | 0 | 0 | 0 | 0 | 0 | 0 | 0 | 0 | 0 | 0 | 0 | 0 | 0 | 0 | 0 | 0 | 0 | 6 | 3 |
| HG0645 | *H. sapiens* | ENSG00000107282 | 0 | 0 | 0 | 0 | 0 | 0 | 0 | 0 | 0 | 0 | 0 | 0 | 0 | 0 | 0 | 0 | 0 | 8 | 1 |
| HG0646 | *H. sapiens* | ENSG00000108312 | 0 | 0 | 0 | 0 | 0 | 0 | 0 | 0 | 0 | 0 | 0 | 0 | 0 | 0 | 0 | 0 | 0 | 7 | 2 |
| HG0647 | *H. sapiens* | ENSG00000108395 | 0 | 0 | 0 | 0 | 0 | 0 | 0 | 0 | 0 | 0 | 0 | 0 | 0 | 0 | 0 | 0 | 0 | 6 | 3 |
| HG0648 | *H. sapiens* | ENSG00000114331 | 0 | 0 | 0 | 0 | 0 | 0 | 0 | 0 | 0 | 0 | 0 | 0 | 0 | 0 | 0 | 0 | 0 | 7 | 2 |
| HG0649.1 | *H. sapiens* | ENSG00000114439 | 0 | 0 | 0 | 0 | 0 | 0 | 0 | 0 | 0 | 0 | 0 | 0 | 0 | 0 | 0 | 0 | 0 | 7 | 2 |
| HG0649.2 | *H. sapiens* | ENSG00000114439 | 0 | 0 | 0 | 0 | 0 | 0 | 0 | 0 | 0 | 0 | 0 | 0 | 0 | 0 | 0 | 0 | 0 | 7 | 2 |
| HG0650 | *H. sapiens* | ENSG00000115226 | 0 | 0 | 0 | 0 | 0 | 0 | 0 | 0 | 0 | 0 | 0 | 0 | 0 | 0 | 0 | 0 | 0 | 7 | 2 |
| HG0651 | *H. sapiens* | ENSG00000115935 | 0 | 0 | 0 | 0 | 0 | 0 | 0 | 0 | 0 | 0 | 0 | 0 | 0 | 0 | 0 | 0 | 0 | 7 | 2 |
| HG0652 | *H. sapiens* | ENSG00000118515 | 0 | 0 | 0 | 0 | 0 | 0 | 0 | 0 | 0 | 0 | 0 | 0 | 0 | 0 | 0 | 0 | 0 | 6 | 3 |
| HG0653 | *H. sapiens* | ENSG00000120334 | 0 | 0 | 0 | 0 | 0 | 0 | 0 | 0 | 0 | 0 | 0 | 0 | 0 | 0 | 0 | 0 | 0 | 7 | 2 |
| HG0654 | *H. sapiens* | ENSG00000133794 | 0 | 0 | 0 | 0 | 0 | 0 | 0 | 0 | 0 | 0 | 0 | 0 | 0 | 0 | 0 | 0 | 0 | 6 | 3 |
| HG0655 | *H. sapiens* | ENSG00000143061 | 0 | 0 | 0 | 0 | 0 | 0 | 0 | 0 | 0 | 0 | 0 | 0 | 0 | 0 | 0 | 0 | 0 | 7 | 2 |
| HG0656 | *H. sapiens* | ENSG00000150776 | 0 | 0 | 0 | 0 | 0 | 0 | 0 | 0 | 0 | 0 | 0 | 0 | 0 | 0 | 0 | 0 | 0 | 5 | 4 |
| HG0657 | *H. sapiens* | ENSG00000151276 | 0 | 0 | 0 | 0 | 0 | 0 | 0 | 0 | 0 | 0 | 0 | 0 | 0 | 0 | 0 | 0 | 0 | 6 | 3 |
| HG0658 | *H. sapiens* | ENSG00000153786 | 0 | 0 | 0 | 0 | 0 | 0 | 0 | 0 | 0 | 0 | 0 | 0 | 0 | 0 | 0 | 0 | 0 | 7 | 2 |
| HG0659.1 | *H. sapiens* | ENSG00000158856 | 0 | 0 | 0 | 0 | 0 | 0 | 0 | 0 | 0 | 0 | 0 | 0 | 0 | 0 | 0 | 0 | 0 | 8 | 1 |
| HG0659.2 | *H. sapiens* | ENSG00000158856 | 0 | 0 | 0 | 0 | 0 | 0 | 0 | 0 | 0 | 0 | 0 | 0 | 0 | 0 | 0 | 0 | 0 | 7 | 0 |
| HG0660 | *H. sapiens* | ENSG00000165861 | 0 | 0 | 0 | 0 | 0 | 0 | 0 | 0 | 0 | 0 | 0 | 0 | 0 | 0 | 0 | 0 | 0 | 7 | 2 |
| HG0661 | *H. sapiens* | ENSG00000166845 | 0 | 0 | 0 | 0 | 0 | 0 | 0 | 0 | 0 | 0 | 0 | 0 | 0 | 0 | 0 | 0 | 0 | 6 | 3 |
| HG0662 | *H. sapiens* | ENSG00000166897 | 0 | 0 | 0 | 0 | 0 | 0 | 0 | 0 | 0 | 0 | 0 | 0 | 0 | 0 | 0 | 0 | 0 | 5 | 4 |
| HG0663 | *H. sapiens* | ENSG00000168615 | 0 | 0 | 0 | 0 | 0 | 0 | 0 | 0 | 0 | 0 | 0 | 0 | 0 | 0 | 0 | 0 | 0 | 7 | 2 |
| HG0664 | *H. sapiens* | ENSG00000169085 | 0 | 0 | 0 | 0 | 0 | 0 | 0 | 0 | 0 | 0 | 0 | 0 | 0 | 0 | 0 | 0 | 0 | 5 | 4 |
| HG0665 | *H. sapiens* | ENSG00000169515 | 0 | 0 | 0 | 0 | 0 | 0 | 0 | 0 | 0 | 0 | 0 | 0 | 0 | 0 | 0 | 0 | 0 | 6 | 3 |
| HG0666 | *H. sapiens* | ENSG00000171943 | 0 | 0 | 0 | 0 | 0 | 0 | 0 | 0 | 0 | 0 | 0 | 0 | 0 | 0 | 0 | 0 | 0 | 7 | 2 |
| HG0667.1 | *H. sapiens* | ENSG00000182700 | 0 | 0 | 0 | 0 | 0 | 0 | 0 | 0 | 0 | 0 | 0 | 0 | 0 | 0 | 0 | 0 | 0 | 6 | 3 |
| HG0667.2 | *H. sapiens* | ENSG00000182700 | 0 | 0 | 0 | 0 | 0 | 0 | 0 | 0 | 0 | 0 | 0 | 0 | 0 | 0 | 0 | 0 | 0 | 6 | 2 |
| HG0668 | *H. sapiens* | ENSG00000182944 | 0 | 0 | 0 | 0 | 0 | 0 | 0 | 0 | 0 | 0 | 0 | 0 | 0 | 0 | 0 | 0 | 0 | 7 | 2 |
| HG0669 | *H. sapiens* | ENSG00000183876 | 0 | 0 | 0 | 0 | 0 | 0 | 0 | 0 | 0 | 0 | 0 | 0 | 0 | 0 | 0 | 0 | 0 | 6 | 3 |
| HG0670 | *H. sapiens* | ENSG00000187164 | 0 | 0 | 0 | 0 | 0 | 0 | 0 | 0 | 0 | 0 | 0 | 0 | 0 | 0 | 0 | 0 | 0 | 6 | 3 |
| HG0671 | *H. sapiens* | ENSG00000189184 | 0 | 0 | 0 | 0 | 0 | 0 | 0 | 0 | 0 | 0 | 0 | 0 | 0 | 0 | 0 | 0 | 0 | 7 | 2 |
| HG0672 | *H. sapiens* | ENSG00000196369 | 0 | 0 | 0 | 0 | 0 | 0 | 0 | 0 | 0 | 0 | 0 | 0 | 0 | 0 | 0 | 0 | 0 | 7 | 2 |
| HG0673 | *H. sapiens* | ENSG00000266028 | 0 | 0 | 0 | 0 | 0 | 0 | 0 | 0 | 0 | 0 | 0 | 0 | 0 | 0 | 0 | 0 | 0 | 7 | 2 |
| HG0674.1 | *H. sapiens* | ENSG00000006576 | 0 | 0 | 0 | 0 | 0 | 0 | 0 | 0 | 0 | 0 | 0 | 0 | 0 | 0 | 0 | 0 | 0 | 7 | 1 |
| HG0674.2 | *H. sapiens* | ENSG00000116793 | 0 | 0 | 0 | 0 | 0 | 0 | 0 | 0 | 0 | 0 | 0 | 0 | 0 | 0 | 0 | 0 | 0 | 1 | 2 |
| HG0675 | *H. sapiens* | ENSG00000048991 | 0 | 0 | 0 | 0 | 0 | 0 | 0 | 0 | 0 | 0 | 0 | 0 | 0 | 0 | 0 | 0 | 0 | 5 | 3 |
| HG0676 | *H. sapiens* | ENSG00000058673 | 0 | 0 | 0 | 0 | 0 | 0 | 0 | 0 | 0 | 0 | 0 | 0 | 0 | 0 | 0 | 0 | 0 | 6 | 2 |
| HG0677 | *H. sapiens* | ENSG00000064989 | 0 | 0 | 0 | 0 | 0 | 0 | 0 | 0 | 0 | 0 | 0 | 0 | 0 | 0 | 0 | 0 | 0 | 5 | 3 |
| HG0678.1 | *H. sapiens* | ENSG00000090776 | 0 | 0 | 0 | 0 | 0 | 0 | 0 | 0 | 0 | 0 | 0 | 0 | 0 | 0 | 0 | 0 | 0 | 4 | 4 |
| HG0678.2 | *H. sapiens* | ENSG00000108947 | 0 | 0 | 0 | 0 | 0 | 0 | 0 | 0 | 0 | 0 | 0 | 0 | 0 | 0 | 0 | 0 | 0 | 3 | 2 |
| HG0679 | *H. sapiens* | ENSG00000100281 | 0 | 0 | 0 | 0 | 0 | 0 | 0 | 0 | 0 | 0 | 0 | 0 | 0 | 0 | 0 | 0 | 0 | 6 | 2 |
| HG0680 | *H. sapiens* | ENSG00000100678 | 0 | 0 | 0 | 0 | 0 | 0 | 0 | 0 | 0 | 0 | 0 | 0 | 0 | 0 | 0 | 0 | 0 | 6 | 2 |
| HG0681 | *H. sapiens* | ENSG00000101349 | 0 | 0 | 0 | 0 | 0 | 0 | 0 | 0 | 0 | 0 | 0 | 0 | 0 | 0 | 0 | 0 | 0 | 7 | 1 |
| HG0682 | *H. sapiens* | ENSG00000105829 | 0 | 0 | 0 | 0 | 0 | 0 | 0 | 0 | 0 | 0 | 0 | 0 | 0 | 0 | 0 | 0 | 0 | 5 | 3 |
| HG0683 | *H. sapiens* | ENSG00000111837 | 0 | 0 | 0 | 0 | 0 | 0 | 0 | 0 | 0 | 0 | 0 | 0 | 0 | 0 | 0 | 0 | 0 | 6 | 2 |
| HG0684 | *H. sapiens* | ENSG00000113916 | 0 | 0 | 0 | 0 | 0 | 0 | 0 | 0 | 0 | 0 | 0 | 0 | 0 | 0 | 0 | 0 | 0 | 5 | 3 |
| HG0685 | *H. sapiens* | ENSG00000117262 | 0 | 0 | 0 | 0 | 0 | 0 | 0 | 0 | 0 | 0 | 0 | 0 | 0 | 0 | 0 | 0 | 0 | 5 | 3 |
| HG0686 | *H. sapiens* | ENSG00000118985 | 0 | 0 | 0 | 0 | 0 | 0 | 0 | 0 | 0 | 0 | 0 | 0 | 0 | 0 | 0 | 0 | 0 | 6 | 2 |
| HG0687.1 | *H. sapiens* | ENSG00000120549 | 0 | 0 | 0 | 0 | 0 | 0 | 0 | 0 | 0 | 0 | 0 | 0 | 0 | 0 | 0 | 0 | 0 | 6 | 2 |
| HG0687.2 | *H. sapiens* | ENSG00000120549 | 0 | 0 | 0 | 0 | 0 | 0 | 0 | 0 | 0 | 0 | 0 | 0 | 0 | 0 | 0 | 0 | 0 | 1 | 1 |
| HG0688 | *H. sapiens* | ENSG00000122420 | 0 | 0 | 0 | 0 | 0 | 0 | 0 | 0 | 0 | 0 | 0 | 0 | 0 | 0 | 0 | 0 | 0 | 7 | 1 |
| HG0689 | *H. sapiens* | ENSG00000123352 | 0 | 0 | 0 | 0 | 0 | 0 | 0 | 0 | 0 | 0 | 0 | 0 | 0 | 0 | 0 | 0 | 0 | 5 | 3 |
| HG0690 | *H. sapiens* | ENSG00000125931 | 0 | 0 | 0 | 0 | 0 | 0 | 0 | 0 | 0 | 0 | 0 | 0 | 0 | 0 | 0 | 0 | 0 | 5 | 3 |
| HG0691 | *H. sapiens* | ENSG00000129535 | 0 | 0 | 0 | 0 | 0 | 0 | 0 | 0 | 0 | 0 | 0 | 0 | 0 | 0 | 0 | 0 | 0 | 5 | 3 |
| HG0692.1 | *H. sapiens* | ENSG00000130338 | 0 | 0 | 0 | 0 | 0 | 0 | 0 | 0 | 0 | 0 | 0 | 0 | 0 | 0 | 0 | 0 | 0 | 5 | 3 |
| HG0692.2 | *H. sapiens* | ENSG00000130338 | 0 | 0 | 0 | 0 | 0 | 0 | 0 | 0 | 0 | 0 | 0 | 0 | 0 | 0 | 0 | 0 | 0 | 2 | 1 |
| HG0693 | *H. sapiens* | ENSG00000131653 | 0 | 0 | 0 | 0 | 0 | 0 | 0 | 0 | 0 | 0 | 0 | 0 | 0 | 0 | 0 | 0 | 0 | 6 | 2 |
| HG0694 | *H. sapiens* | ENSG00000133265 | 0 | 0 | 0 | 0 | 0 | 0 | 0 | 0 | 0 | 0 | 0 | 0 | 0 | 0 | 0 | 0 | 0 | 6 | 2 |
| HG0695 | *H. sapiens* | ENSG00000133704 | 0 | 0 | 0 | 0 | 0 | 0 | 0 | 0 | 0 | 0 | 0 | 0 | 0 | 0 | 0 | 0 | 0 | 4 | 4 |
| HG0696 | *H. sapiens* | ENSG00000138380 | 0 | 0 | 0 | 0 | 0 | 0 | 0 | 0 | 0 | 0 | 0 | 0 | 0 | 0 | 0 | 0 | 0 | 6 | 2 |
| HG0697 | *H. sapiens* | ENSG00000143344 | 0 | 0 | 0 | 0 | 0 | 0 | 0 | 0 | 0 | 0 | 0 | 0 | 0 | 0 | 0 | 0 | 0 | 6 | 2 |
| HG0698 | *H. sapiens* | ENSG00000157554 | 0 | 0 | 0 | 0 | 0 | 0 | 0 | 0 | 0 | 0 | 0 | 0 | 0 | 0 | 0 | 0 | 0 | 7 | 1 |
| HG0699 | *H. sapiens* | ENSG00000160785 | 0 | 0 | 0 | 0 | 0 | 0 | 0 | 0 | 0 | 0 | 0 | 0 | 0 | 0 | 0 | 0 | 0 | 4 | 4 |
| HG0700 | *H. sapiens* | ENSG00000165156 | 0 | 0 | 0 | 0 | 0 | 0 | 0 | 0 | 0 | 0 | 0 | 0 | 0 | 0 | 0 | 0 | 0 | 7 | 1 |
| HG0701 | *H. sapiens* | ENSG00000166292 | 0 | 0 | 0 | 0 | 0 | 0 | 0 | 0 | 0 | 0 | 0 | 0 | 0 | 0 | 0 | 0 | 0 | 6 | 2 |
| HG0702 | *H. sapiens* | ENSG00000167377 | 0 | 0 | 0 | 0 | 0 | 0 | 0 | 0 | 0 | 0 | 0 | 0 | 0 | 0 | 0 | 0 | 0 | 5 | 3 |
| HG0703 | *H. sapiens* | ENSG00000171862 | 0 | 0 | 0 | 0 | 0 | 0 | 0 | 0 | 0 | 0 | 0 | 0 | 0 | 0 | 0 | 0 | 0 | 7 | 1 |
| HG0704.1 | *H. sapiens* | ENSG00000173598 | 0 | 0 | 0 | 0 | 0 | 0 | 0 | 0 | 0 | 0 | 0 | 0 | 0 | 0 | 0 | 0 | 0 | 6 | 2 |
| HG0704.2 | *H. sapiens* | ENSG00000272325 | 0 | 0 | 0 | 0 | 0 | 0 | 0 | 0 | 0 | 0 | 0 | 0 | 0 | 0 | 0 | 0 | 0 | 4 | 2 |
| HG0705.1 | *H. sapiens* | ENSG00000175224 | 0 | 0 | 0 | 0 | 0 | 0 | 0 | 0 | 0 | 0 | 0 | 0 | 0 | 0 | 0 | 0 | 0 | 6 | 2 |
| HG0705.2 | *H. sapiens* | ENSG00000175224 | 0 | 0 | 0 | 0 | 0 | 0 | 0 | 0 | 0 | 0 | 0 | 0 | 0 | 0 | 0 | 0 | 0 | 6 | 1 |
| HG0706 | *H. sapiens* | ENSG00000178498 | 0 | 0 | 0 | 0 | 0 | 0 | 0 | 0 | 0 | 0 | 0 | 0 | 0 | 0 | 0 | 0 | 0 | 5 | 3 |
| HG0707 | *H. sapiens* | ENSG00000179796 | 0 | 0 | 0 | 0 | 0 | 0 | 0 | 0 | 0 | 0 | 0 | 0 | 0 | 0 | 0 | 0 | 0 | 5 | 3 |
| HG0708 | *H. sapiens* | ENSG00000184613 | 0 | 0 | 0 | 0 | 0 | 0 | 0 | 0 | 0 | 0 | 0 | 0 | 0 | 0 | 0 | 0 | 0 | 7 | 1 |
| HG0709 | *H. sapiens* | ENSG00000186439 | 0 | 0 | 0 | 0 | 0 | 0 | 0 | 0 | 0 | 0 | 0 | 0 | 0 | 0 | 0 | 0 | 0 | 5 | 3 |
| HG0710 | *H. sapiens* | ENSG00000196505 | 0 | 0 | 0 | 0 | 0 | 0 | 0 | 0 | 0 | 0 | 0 | 0 | 0 | 0 | 0 | 0 | 0 | 5 | 3 |

| | | | | | | | | | | | | | | | | | | | | | |
|---|---|---|---|---|---|---|---|---|---|---|---|---|---|---|---|---|---|---|---|---|---|
| HG0711 | *H. sapiens* | ENSG00000198300 | 0 | 0 | 0 | 0 | 0 | 0 | 0 | 0 | 0 | 0 | 0 | 0 | 0 | 0 | 0 | 0 | 0 | 6 | 2 |
| HG0712 | *H. sapiens* | ENSG00000213949 | 0 | 0 | 0 | 0 | 0 | 0 | 0 | 0 | 0 | 0 | 0 | 0 | 0 | 0 | 0 | 0 | 0 | 6 | 2 |
| HG0713 | *H. sapiens* | ENSG00000008513 | 0 | 0 | 0 | 0 | 0 | 0 | 0 | 0 | 0 | 0 | 0 | 0 | 0 | 0 | 0 | 0 | 0 | 5 | 2 |
| HG0714 | *H. sapiens* | ENSG00000051825 | 0 | 0 | 0 | 0 | 0 | 0 | 0 | 0 | 0 | 0 | 0 | 0 | 0 | 0 | 0 | 0 | 0 | 4 | 3 |
| HG0715 | *H. sapiens* | ENSG00000071246 | 0 | 0 | 0 | 0 | 0 | 0 | 0 | 0 | 0 | 0 | 0 | 0 | 0 | 0 | 0 | 0 | 0 | 5 | 2 |
| HG0716 | *H. sapiens* | ENSG00000075413 | 0 | 0 | 0 | 0 | 0 | 0 | 0 | 0 | 0 | 0 | 0 | 0 | 0 | 0 | 0 | 0 | 0 | 3 | 4 |
| HG0717 | *H. sapiens* | ENSG00000083857 | 0 | 0 | 0 | 0 | 0 | 0 | 0 | 0 | 0 | 0 | 0 | 0 | 0 | 0 | 0 | 0 | 0 | 5 | 2 |
| HG0718 | *H. sapiens* | ENSG00000085274 | 0 | 0 | 0 | 0 | 0 | 0 | 0 | 0 | 0 | 0 | 0 | 0 | 0 | 0 | 0 | 0 | 0 | 5 | 2 |
| HG0719 | *H. sapiens* | ENSG00000085998 | 0 | 0 | 0 | 0 | 0 | 0 | 0 | 0 | 0 | 0 | 0 | 0 | 0 | 0 | 0 | 0 | 0 | 6 | 1 |
| HG0720 | *H. sapiens* | ENSG00000087053 | 0 | 0 | 0 | 0 | 0 | 0 | 0 | 0 | 0 | 0 | 0 | 0 | 0 | 0 | 0 | 0 | 0 | 5 | 2 |
| HG0721 | *H. sapiens* | ENSG00000095951 | 0 | 0 | 0 | 0 | 0 | 0 | 0 | 0 | 0 | 0 | 0 | 0 | 0 | 0 | 0 | 0 | 0 | 5 | 2 |
| HG0722 | *H. sapiens* | ENSG00000101445 | 0 | 0 | 0 | 0 | 0 | 0 | 0 | 0 | 0 | 0 | 0 | 0 | 0 | 0 | 0 | 0 | 0 | 5 | 2 |
| HG0723 | *H. sapiens* | ENSG00000102974 | 0 | 0 | 0 | 0 | 0 | 0 | 0 | 0 | 0 | 0 | 0 | 0 | 0 | 0 | 0 | 0 | 0 | 6 | 1 |
| HG0724 | *H. sapiens* | ENSG00000104783 | 0 | 0 | 0 | 0 | 0 | 0 | 0 | 0 | 0 | 0 | 0 | 0 | 0 | 0 | 0 | 0 | 0 | 5 | 2 |
| HG0725 | *H. sapiens* | ENSG00000112218 | 0 | 0 | 0 | 0 | 0 | 0 | 0 | 0 | 0 | 0 | 0 | 0 | 0 | 0 | 0 | 0 | 0 | 4 | 3 |
| HG0726.1 | *H. sapiens* | ENSG00000115504 | 0 | 0 | 0 | 0 | 0 | 0 | 0 | 0 | 0 | 0 | 0 | 0 | 0 | 0 | 0 | 0 | 0 | 5 | 2 |
| HG0726.2 | *H. sapiens* | ENSG00000115504 | 0 | 0 | 0 | 0 | 0 | 0 | 0 | 0 | 0 | 0 | 0 | 0 | 0 | 0 | 0 | 0 | 0 | 1 | 2 |
| HG0727 | *H. sapiens* | ENSG00000115825 | 0 | 0 | 0 | 0 | 0 | 0 | 0 | 0 | 0 | 0 | 0 | 0 | 0 | 0 | 0 | 0 | 0 | 6 | 1 |
| HG0728 | *H. sapiens* | ENSG00000116991 | 0 | 0 | 0 | 0 | 0 | 0 | 0 | 0 | 0 | 0 | 0 | 0 | 0 | 0 | 0 | 0 | 0 | 6 | 1 |
| HG0729 | *H. sapiens* | ENSG00000119383 | 0 | 0 | 0 | 0 | 0 | 0 | 0 | 0 | 0 | 0 | 0 | 0 | 0 | 0 | 0 | 0 | 0 | 6 | 1 |
| HG0730 | *H. sapiens* | ENSG00000125170 | 0 | 0 | 0 | 0 | 0 | 0 | 0 | 0 | 0 | 0 | 0 | 0 | 0 | 0 | 0 | 0 | 0 | 4 | 3 |
| HG0731 | *H. sapiens* | ENSG00000129255 | 0 | 0 | 0 | 0 | 0 | 0 | 0 | 0 | 0 | 0 | 0 | 0 | 0 | 0 | 0 | 0 | 0 | 5 | 2 |
| HG0732 | *H. sapiens* | ENSG00000135914 | 0 | 0 | 0 | 0 | 0 | 0 | 0 | 0 | 0 | 0 | 0 | 0 | 0 | 0 | 0 | 0 | 0 | 6 | 1 |
| HG0733 | *H. sapiens* | ENSG00000136247 | 0 | 0 | 0 | 0 | 0 | 0 | 0 | 0 | 0 | 0 | 0 | 0 | 0 | 0 | 0 | 0 | 0 | 4 | 3 |
| HG0734 | *H. sapiens* | ENSG00000139318 | 0 | 0 | 0 | 0 | 0 | 0 | 0 | 0 | 0 | 0 | 0 | 0 | 0 | 0 | 0 | 0 | 0 | 6 | 1 |
| HG0735 | *H. sapiens* | ENSG00000139579 | 0 | 0 | 0 | 0 | 0 | 0 | 0 | 0 | 0 | 0 | 0 | 0 | 0 | 0 | 0 | 0 | 0 | 5 | 2 |
| HG0736 | *H. sapiens* | ENSG00000139826 | 0 | 0 | 0 | 0 | 0 | 0 | 0 | 0 | 0 | 0 | 0 | 0 | 0 | 0 | 0 | 0 | 0 | 5 | 2 |
| HG0737 | *H. sapiens* | ENSG00000141639 | 0 | 0 | 0 | 0 | 0 | 0 | 0 | 0 | 0 | 0 | 0 | 0 | 0 | 0 | 0 | 0 | 0 | 5 | 2 |
| HG0738 | *H. sapiens* | ENSG00000143393 | 0 | 0 | 0 | 0 | 0 | 0 | 0 | 0 | 0 | 0 | 0 | 0 | 0 | 0 | 0 | 0 | 0 | 4 | 3 |
| HG0739 | *H. sapiens* | ENSG00000144331 | 0 | 0 | 0 | 0 | 0 | 0 | 0 | 0 | 0 | 0 | 0 | 0 | 0 | 0 | 0 | 0 | 0 | 5 | 2 |
| HG0740 | *H. sapiens* | ENSG00000144331 | 0 | 0 | 0 | 0 | 0 | 0 | 0 | 0 | 0 | 0 | 0 | 0 | 0 | 0 | 0 | 0 | 0 | 5 | 2 |
| HG0741 | *H. sapiens* | ENSG00000144834 | 0 | 0 | 0 | 0 | 0 | 0 | 0 | 0 | 0 | 0 | 0 | 0 | 0 | 0 | 0 | 0 | 0 | 6 | 1 |
| HG0742 | *H. sapiens* | ENSG00000154556 | 0 | 0 | 0 | 0 | 0 | 0 | 0 | 0 | 0 | 0 | 0 | 0 | 0 | 0 | 0 | 0 | 0 | 6 | 1 |
| HG0743 | *H. sapiens* | ENSG00000154814 | 0 | 0 | 0 | 0 | 0 | 0 | 0 | 0 | 0 | 0 | 0 | 0 | 0 | 0 | 0 | 0 | 0 | 5 | 2 |
| HG0744 | *H. sapiens* | ENSG00000163053 | 0 | 0 | 0 | 0 | 0 | 0 | 0 | 0 | 0 | 0 | 0 | 0 | 0 | 0 | 0 | 0 | 0 | 4 | 3 |
| HG0745 | *H. sapiens* | ENSG00000163145 | 0 | 0 | 0 | 0 | 0 | 0 | 0 | 0 | 0 | 0 | 0 | 0 | 0 | 0 | 0 | 0 | 0 | 5 | 2 |
| HG0746 | *H. sapiens* | ENSG00000164741 | 0 | 0 | 0 | 0 | 0 | 0 | 0 | 0 | 0 | 0 | 0 | 0 | 0 | 0 | 0 | 0 | 0 | 6 | 1 |
| HG0747 | *H. sapiens* | ENSG00000168036 | 0 | 0 | 0 | 0 | 0 | 0 | 0 | 0 | 0 | 0 | 0 | 0 | 0 | 0 | 0 | 0 | 0 | 5 | 2 |
| HG0748 | *H. sapiens* | ENSG00000168830 | 0 | 0 | 0 | 0 | 0 | 0 | 0 | 0 | 0 | 0 | 0 | 0 | 0 | 0 | 0 | 0 | 0 | 4 | 3 |
| HG0749 | *H. sapiens* | ENSG00000169760 | 0 | 0 | 0 | 0 | 0 | 0 | 0 | 0 | 0 | 0 | 0 | 0 | 0 | 0 | 0 | 0 | 0 | 5 | 2 |
| HG0750 | *H. sapiens* | ENSG00000170881 | 0 | 0 | 0 | 0 | 0 | 0 | 0 | 0 | 0 | 0 | 0 | 0 | 0 | 0 | 0 | 0 | 0 | 5 | 2 |
| HG0751 | *H. sapiens* | ENSG00000171791 | 0 | 0 | 0 | 0 | 0 | 0 | 0 | 0 | 0 | 0 | 0 | 0 | 0 | 0 | 0 | 0 | 0 | 4 | 3 |
| HG0752 | *H. sapiens* | ENSG00000172269 | 0 | 0 | 0 | 0 | 0 | 0 | 0 | 0 | 0 | 0 | 0 | 0 | 0 | 0 | 0 | 0 | 0 | 5 | 2 |
| HG0753 | *H. sapiens* | ENSG00000173406 | 0 | 0 | 0 | 0 | 0 | 0 | 0 | 0 | 0 | 0 | 0 | 0 | 0 | 0 | 0 | 0 | 0 | 4 | 3 |
| HG0754 | *H. sapiens* | ENSG00000175182 | 0 | 0 | 0 | 0 | 0 | 0 | 0 | 0 | 0 | 0 | 0 | 0 | 0 | 0 | 0 | 0 | 0 | 5 | 2 |
| HG0755 | *H. sapiens* | ENSG00000176597 | 0 | 0 | 0 | 0 | 0 | 0 | 0 | 0 | 0 | 0 | 0 | 0 | 0 | 0 | 0 | 0 | 0 | 5 | 2 |
| HG0756 | *H. sapiens* | ENSG00000177051 | 0 | 0 | 0 | 0 | 0 | 0 | 0 | 0 | 0 | 0 | 0 | 0 | 0 | 0 | 0 | 0 | 0 | 5 | 2 |
| HG0757.1 | *H. sapiens* | ENSG00000177301 | 0 | 0 | 0 | 0 | 0 | 0 | 0 | 0 | 0 | 0 | 0 | 0 | 0 | 0 | 0 | 0 | 0 | 5 | 2 |
| HG0757.2 | *H. sapiens* | ENSG00000182255 | 0 | 0 | 0 | 0 | 0 | 0 | 0 | 0 | 0 | 0 | 0 | 0 | 0 | 0 | 0 | 0 | 0 | 2 | 0 |
| HG0758 | *H. sapiens* | ENSG00000179299 | 0 | 0 | 0 | 0 | 0 | 0 | 0 | 0 | 0 | 0 | 0 | 0 | 0 | 0 | 0 | 0 | 0 | 5 | 2 |
| HG0759 | *H. sapiens* | ENSG00000180929 | 0 | 0 | 0 | 0 | 0 | 0 | 0 | 0 | 0 | 0 | 0 | 0 | 0 | 0 | 0 | 0 | 0 | 4 | 3 |
| HG0760 | *H. sapiens* | ENSG00000182463 | 0 | 0 | 0 | 0 | 0 | 0 | 0 | 0 | 0 | 0 | 0 | 0 | 0 | 0 | 0 | 0 | 0 | 4 | 3 |
| HG0761 | *H. sapiens* | ENSG00000185985 | 0 | 0 | 0 | 0 | 0 | 0 | 0 | 0 | 0 | 0 | 0 | 0 | 0 | 0 | 0 | 0 | 0 | 5 | 2 |
| HG0762 | *H. sapiens* | ENSG00000187147 | 0 | 0 | 0 | 0 | 0 | 0 | 0 | 0 | 0 | 0 | 0 | 0 | 0 | 0 | 0 | 0 | 0 | 6 | 1 |
| HG0763 | *H. sapiens* | ENSG00000188050 | 0 | 0 | 0 | 0 | 0 | 0 | 0 | 0 | 0 | 0 | 0 | 0 | 0 | 0 | 0 | 0 | 0 | 4 | 3 |
| HG0764 | *H. sapiens* | ENSG00000188725 | 0 | 0 | 0 | 0 | 0 | 0 | 0 | 0 | 0 | 0 | 0 | 0 | 0 | 0 | 0 | 0 | 0 | 5 | 2 |
| HG0765 | *H. sapiens* | ENSG00000196123 | 0 | 0 | 0 | 0 | 0 | 0 | 0 | 0 | 0 | 0 | 0 | 0 | 0 | 0 | 0 | 0 | 0 | 6 | 1 |
| HG0766 | *H. sapiens* | ENSG00000196189 | 0 | 0 | 0 | 0 | 0 | 0 | 0 | 0 | 0 | 0 | 0 | 0 | 0 | 0 | 0 | 0 | 0 | 5 | 2 |
| HG0767.1 | *H. sapiens* | ENSG00000197555 | 0 | 0 | 0 | 0 | 0 | 0 | 0 | 0 | 0 | 0 | 0 | 0 | 0 | 0 | 0 | 0 | 0 | 6 | 1 |
| HG0767.2 | *H. sapiens* | ENSG00000197555 | 0 | 0 | 0 | 0 | 0 | 0 | 0 | 0 | 0 | 0 | 0 | 0 | 0 | 0 | 0 | 0 | 0 | 0 | 2 |
| HG0768 | *H. sapiens* | ENSG00000213023 | 0 | 0 | 0 | 0 | 0 | 0 | 0 | 0 | 0 | 0 | 0 | 0 | 0 | 0 | 0 | 0 | 0 | 4 | 3 |
| HG0769.1 | *H. sapiens* | ENSG00000280165 | 0 | 0 | 0 | 0 | 0 | 0 | 0 | 0 | 0 | 0 | 0 | 0 | 0 | 0 | 0 | 0 | 0 | 4 | 3 |
| HG0769.2 | *H. sapiens* | ENSG00000197991 | 0 | 0 | 0 | 0 | 0 | 0 | 0 | 0 | 0 | 0 | 0 | 0 | 0 | 0 | 0 | 0 | 0 | 2 | 1 |
| HG0770 | *H. sapiens* | ENSG00000010165 | 0 | 0 | 0 | 0 | 0 | 0 | 0 | 0 | 0 | 0 | 0 | 0 | 0 | 0 | 0 | 0 | 0 | 6 | 0 |
| HG0771 | *H. sapiens* | ENSG00000013306 | 0 | 0 | 0 | 0 | 0 | 0 | 0 | 0 | 0 | 0 | 0 | 0 | 0 | 0 | 0 | 0 | 0 | 4 | 2 |
| HG0772 | *H. sapiens* | ENSG00000064309 | 0 | 0 | 0 | 0 | 0 | 0 | 0 | 0 | 0 | 0 | 0 | 0 | 0 | 0 | 0 | 0 | 0 | 4 | 2 |
| HG0773 | *H. sapiens* | ENSG00000065548 | 0 | 0 | 0 | 0 | 0 | 0 | 0 | 0 | 0 | 0 | 0 | 0 | 0 | 0 | 0 | 0 | 0 | 4 | 2 |
| HG0774 | *H. sapiens* | ENSG00000066933 | 0 | 0 | 0 | 0 | 0 | 0 | 0 | 0 | 0 | 0 | 0 | 0 | 0 | 0 | 0 | 0 | 0 | 5 | 1 |
| HG0775 | *H. sapiens* | ENSG00000068001 | 0 | 0 | 0 | 0 | 0 | 0 | 0 | 0 | 0 | 0 | 0 | 0 | 0 | 0 | 0 | 0 | 0 | 4 | 2 |
| HG0776 | *H. sapiens* | ENSG00000082014 | 0 | 0 | 0 | 0 | 0 | 0 | 0 | 0 | 0 | 0 | 0 | 0 | 0 | 0 | 0 | 0 | 0 | 5 | 1 |
| HG0777 | *H. sapiens* | ENSG00000085644 | 0 | 0 | 0 | 0 | 0 | 0 | 0 | 0 | 0 | 0 | 0 | 0 | 0 | 0 | 0 | 0 | 0 | 5 | 1 |
| HG0778 | *H. sapiens* | ENSG00000089116 | 0 | 0 | 0 | 0 | 0 | 0 | 0 | 0 | 0 | 0 | 0 | 0 | 0 | 0 | 0 | 0 | 0 | 3 | 3 |
| HG0779 | *H. sapiens* | ENSG00000091140 | 0 | 0 | 0 | 0 | 0 | 0 | 0 | 0 | 0 | 0 | 0 | 0 | 0 | 0 | 0 | 0 | 0 | 3 | 3 |
| HG0780 | *H. sapiens* | ENSG00000092199 | 0 | 0 | 0 | 0 | 0 | 0 | 0 | 0 | 0 | 0 | 0 | 0 | 0 | 0 | 0 | 0 | 0 | 5 | 1 |
| HG0781 | *H. sapiens* | ENSG00000100099 | 0 | 0 | 0 | 0 | 0 | 0 | 0 | 0 | 0 | 0 | 0 | 0 | 0 | 0 | 0 | 0 | 0 | 4 | 2 |
| HG0782 | *H. sapiens* | ENSG00000100368 | 0 | 0 | 0 | 0 | 0 | 0 | 0 | 0 | 0 | 0 | 0 | 0 | 0 | 0 | 0 | 0 | 0 | 4 | 2 |
| HG0783 | *H. sapiens* | ENSG00000105053 | 0 | 0 | 0 | 0 | 0 | 0 | 0 | 0 | 0 | 0 | 0 | 0 | 0 | 0 | 0 | 0 | 0 | 5 | 1 |
| HG0784 | *H. sapiens* | ENSG00000105851 | 0 | 0 | 0 | 0 | 0 | 0 | 0 | 0 | 0 | 0 | 0 | 0 | 0 | 0 | 0 | 0 | 0 | 3 | 3 |
| HG0785 | *H. sapiens* | ENSG00000107242 | 0 | 0 | 0 | 0 | 0 | 0 | 0 | 0 | 0 | 0 | 0 | 0 | 0 | 0 | 0 | 0 | 0 | 2 | 4 |

| | | | | | | | | | | | | | | | | | | | | |
|---|---|---|---|---|---|---|---|---|---|---|---|---|---|---|---|---|---|---|---|---|
| HG0786 | *H. sapiens* | ENSG00000107862 | 0 | 0 | 0 | 0 | 0 | 0 | 0 | 0 | 0 | 0 | 0 | 0 | 0 | 0 | 0 | 0 | 0 | 4 | 2 |
| HG0787 | *H. sapiens* | ENSG00000108592 | 0 | 0 | 0 | 0 | 0 | 0 | 0 | 0 | 0 | 0 | 0 | 0 | 0 | 0 | 0 | 0 | 0 | 3 | 3 |
| HG0788 | *H. sapiens* | ENSG00000113739 | 0 | 0 | 0 | 0 | 0 | 0 | 0 | 0 | 0 | 0 | 0 | 0 | 0 | 0 | 0 | 0 | 0 | 4 | 2 |
| HG0789 | *H. sapiens* | ENSG00000115355 | 0 | 0 | 0 | 0 | 0 | 0 | 0 | 0 | 0 | 0 | 0 | 0 | 0 | 0 | 0 | 0 | 0 | 4 | 2 |
| HG0790.1 | *H. sapiens* | ENSG00000115568 | 0 | 0 | 0 | 0 | 0 | 0 | 0 | 0 | 0 | 0 | 0 | 0 | 0 | 0 | 0 | 0 | 0 | 5 | 1 |
| HG0790.2 | *H. sapiens* | ENSG00000115568 | 0 | 0 | 0 | 0 | 0 | 0 | 0 | 0 | 0 | 0 | 0 | 0 | 0 | 0 | 0 | 0 | 0 | 3 | 2 |
| HG0791 | *H. sapiens* | ENSG00000131323 | 0 | 0 | 0 | 0 | 0 | 0 | 0 | 0 | 0 | 0 | 0 | 0 | 0 | 0 | 0 | 0 | 0 | 3 | 3 |
| HG0792 | *H. sapiens* | ENSG00000132424 | 0 | 0 | 0 | 0 | 0 | 0 | 0 | 0 | 0 | 0 | 0 | 0 | 0 | 0 | 0 | 0 | 0 | 4 | 2 |
| HG0793 | *H. sapiens* | ENSG00000132698 | 0 | 0 | 0 | 0 | 0 | 0 | 0 | 0 | 0 | 0 | 0 | 0 | 0 | 0 | 0 | 0 | 0 | 5 | 1 |
| HG0794.1 | *H. sapiens* | ENSG00000135218 | 0 | 0 | 0 | 0 | 0 | 0 | 0 | 0 | 0 | 0 | 0 | 0 | 0 | 0 | 0 | 0 | 0 | 4 | 2 |
| HG0794.2 | *H. sapiens* | ENSG00000135218 | 0 | 0 | 0 | 0 | 0 | 0 | 0 | 0 | 0 | 0 | 0 | 0 | 0 | 0 | 0 | 0 | 0 | 3 | 1 |
| HG0795 | *H. sapiens* | ENSG00000137601 | 0 | 0 | 0 | 0 | 0 | 0 | 0 | 0 | 0 | 0 | 0 | 0 | 0 | 0 | 0 | 0 | 0 | 5 | 1 |
| HG0796 | *H. sapiens* | ENSG00000138074 | 0 | 0 | 0 | 0 | 0 | 0 | 0 | 0 | 0 | 0 | 0 | 0 | 0 | 0 | 0 | 0 | 0 | 4 | 2 |
| HG0797 | *H. sapiens* | ENSG00000140526 | 0 | 0 | 0 | 0 | 0 | 0 | 0 | 0 | 0 | 0 | 0 | 0 | 0 | 0 | 0 | 0 | 0 | 4 | 2 |
| HG0798.1 | *H. sapiens* | ENSG00000142949 | 0 | 0 | 0 | 0 | 0 | 0 | 0 | 0 | 0 | 0 | 0 | 0 | 0 | 0 | 0 | 0 | 0 | 4 | 2 |
| HG0798.2 | *H. sapiens* | ENSG00000153707 | 0 | 0 | 0 | 0 | 0 | 0 | 0 | 0 | 0 | 0 | 0 | 0 | 0 | 0 | 0 | 0 | 0 | 2 | 1 |
| HG0799 | *H. sapiens* | ENSG00000143398 | 0 | 0 | 0 | 0 | 0 | 0 | 0 | 0 | 0 | 0 | 0 | 0 | 0 | 0 | 0 | 0 | 0 | 4 | 2 |
| HG0800 | *H. sapiens* | ENSG00000143612 | 0 | 0 | 0 | 0 | 0 | 0 | 0 | 0 | 0 | 0 | 0 | 0 | 0 | 0 | 0 | 0 | 0 | 4 | 2 |
| HG0801 | *H. sapiens* | ENSG00000144278 | 0 | 0 | 0 | 0 | 0 | 0 | 0 | 0 | 0 | 0 | 0 | 0 | 0 | 0 | 0 | 0 | 0 | 5 | 1 |
| HG0802 | *H. sapiens* | ENSG00000144283 | 0 | 0 | 0 | 0 | 0 | 0 | 0 | 0 | 0 | 0 | 0 | 0 | 0 | 0 | 0 | 0 | 0 | 4 | 2 |
| HG0803 | *H. sapiens* | ENSG00000144802 | 0 | 0 | 0 | 0 | 0 | 0 | 0 | 0 | 0 | 0 | 0 | 0 | 0 | 0 | 0 | 0 | 0 | 5 | 1 |
| HG0804 | *H. sapiens* | ENSG00000144815 | 0 | 0 | 0 | 0 | 0 | 0 | 0 | 0 | 0 | 0 | 0 | 0 | 0 | 0 | 0 | 0 | 0 | 4 | 2 |
| HG0805.1 | *H. sapiens* | ENSG00000145936 | 0 | 0 | 0 | 0 | 0 | 0 | 0 | 0 | 0 | 0 | 0 | 0 | 0 | 0 | 0 | 0 | 0 | 4 | 2 |
| HG0805.2 | *H. sapiens* | ENSG00000145936 | 0 | 0 | 0 | 0 | 0 | 0 | 0 | 0 | 0 | 0 | 0 | 0 | 0 | 0 | 0 | 0 | 0 | 3 | 2 |
| HG0806 | *H. sapiens* | ENSG00000147481 | 0 | 0 | 0 | 0 | 0 | 0 | 0 | 0 | 0 | 0 | 0 | 0 | 0 | 0 | 0 | 0 | 0 | 5 | 1 |
| HG0807.1 | *H. sapiens* | ENSG00000147488 | 0 | 0 | 0 | 0 | 0 | 0 | 0 | 0 | 0 | 0 | 0 | 0 | 0 | 0 | 0 | 0 | 0 | 4 | 2 |
| HG0807.2 | *H. sapiens* | ENSG00000147488 | 0 | 0 | 0 | 0 | 0 | 0 | 0 | 0 | 0 | 0 | 0 | 0 | 0 | 0 | 0 | 0 | 0 | 4 | 1 |
| HG0807.3 | *H. sapiens* | ENSG00000186487 | 0 | 0 | 0 | 0 | 0 | 0 | 0 | 0 | 0 | 0 | 0 | 0 | 0 | 0 | 0 | 0 | 0 | 2 | 1 |
| HG0808 | *H. sapiens* | ENSG00000149269 | 0 | 0 | 0 | 0 | 0 | 0 | 0 | 0 | 0 | 0 | 0 | 0 | 0 | 0 | 0 | 0 | 0 | 5 | 1 |
| HG0809 | *H. sapiens* | ENSG00000151287 | 0 | 0 | 0 | 0 | 0 | 0 | 0 | 0 | 0 | 0 | 0 | 0 | 0 | 0 | 0 | 0 | 0 | 4 | 2 |
| HG0810 | *H. sapiens* | ENSG00000156504 | 0 | 0 | 0 | 0 | 0 | 0 | 0 | 0 | 0 | 0 | 0 | 0 | 0 | 0 | 0 | 0 | 0 | 4 | 2 |
| HG0811 | *H. sapiens* | ENSG00000157350 | 0 | 0 | 0 | 0 | 0 | 0 | 0 | 0 | 0 | 0 | 0 | 0 | 0 | 0 | 0 | 0 | 0 | 4 | 2 |
| HG0812 | *H. sapiens* | ENSG00000157368 | 0 | 0 | 0 | 0 | 0 | 0 | 0 | 0 | 0 | 0 | 0 | 0 | 0 | 0 | 0 | 0 | 0 | 3 | 3 |
| HG0813 | *H. sapiens* | ENSG00000160199 | 0 | 0 | 0 | 0 | 0 | 0 | 0 | 0 | 0 | 0 | 0 | 0 | 0 | 0 | 0 | 0 | 0 | 5 | 1 |
| HG0814 | *H. sapiens* | ENSG00000163071 | 0 | 0 | 0 | 0 | 0 | 0 | 0 | 0 | 0 | 0 | 0 | 0 | 0 | 0 | 0 | 0 | 0 | 5 | 1 |
| HG0815 | *H. sapiens* | ENSG00000163072 | 0 | 0 | 0 | 0 | 0 | 0 | 0 | 0 | 0 | 0 | 0 | 0 | 0 | 0 | 0 | 0 | 0 | 4 | 2 |
| HG0816 | *H. sapiens* | ENSG00000163794 | 0 | 0 | 0 | 0 | 0 | 0 | 0 | 0 | 0 | 0 | 0 | 0 | 0 | 0 | 0 | 0 | 0 | 5 | 1 |
| HG0817 | *H. sapiens* | ENSG00000163888 | 0 | 0 | 0 | 0 | 0 | 0 | 0 | 0 | 0 | 0 | 0 | 0 | 0 | 0 | 0 | 0 | 0 | 4 | 2 |
| HG0818 | *H. sapiens* | ENSG00000164125 | 0 | 0 | 0 | 0 | 0 | 0 | 0 | 0 | 0 | 0 | 0 | 0 | 0 | 0 | 0 | 0 | 0 | 4 | 2 |
| HG0819 | *H. sapiens* | ENSG00000164188 | 0 | 0 | 0 | 0 | 0 | 0 | 0 | 0 | 0 | 0 | 0 | 0 | 0 | 0 | 0 | 0 | 0 | 5 | 1 |
| HG0820 | *H. sapiens* | ENSG00000164946 | 0 | 0 | 0 | 0 | 0 | 0 | 0 | 0 | 0 | 0 | 0 | 0 | 0 | 0 | 0 | 0 | 0 | 4 | 2 |
| HG0821 | *H. sapiens* | ENSG00000165300 | 0 | 0 | 0 | 0 | 0 | 0 | 0 | 0 | 0 | 0 | 0 | 0 | 0 | 0 | 0 | 0 | 0 | 5 | 1 |
| HG0822 | *H. sapiens* | ENSG00000166170 | 0 | 0 | 0 | 0 | 0 | 0 | 0 | 0 | 0 | 0 | 0 | 0 | 0 | 0 | 0 | 0 | 0 | 5 | 1 |
| HG0823 | *H. sapiens* | ENSG00000166268 | 0 | 0 | 0 | 0 | 0 | 0 | 0 | 0 | 0 | 0 | 0 | 0 | 0 | 0 | 0 | 0 | 0 | 4 | 2 |
| HG0824 | *H. sapiens* | ENSG00000167100 | 0 | 0 | 0 | 0 | 0 | 0 | 0 | 0 | 0 | 0 | 0 | 0 | 0 | 0 | 0 | 0 | 0 | 4 | 2 |
| HG0825 | *H. sapiens* | ENSG00000169180 | 0 | 0 | 0 | 0 | 0 | 0 | 0 | 0 | 0 | 0 | 0 | 0 | 0 | 0 | 0 | 0 | 0 | 4 | 2 |
| HG0826 | *H. sapiens* | ENSG00000169851 | 0 | 0 | 0 | 0 | 0 | 0 | 0 | 0 | 0 | 0 | 0 | 0 | 0 | 0 | 0 | 0 | 0 | 5 | 1 |
| HG0827 | *H. sapiens* | ENSG00000170365 | 0 | 0 | 0 | 0 | 0 | 0 | 0 | 0 | 0 | 0 | 0 | 0 | 0 | 0 | 0 | 0 | 0 | 6 | 0 |
| HG0828 | *H. sapiens* | ENSG00000170390 | 0 | 0 | 0 | 0 | 0 | 0 | 0 | 0 | 0 | 0 | 0 | 0 | 0 | 0 | 0 | 0 | 0 | 6 | 0 |
| HG0829 | *H. sapiens* | ENSG00000171502 | 0 | 0 | 0 | 0 | 0 | 0 | 0 | 0 | 0 | 0 | 0 | 0 | 0 | 0 | 0 | 0 | 0 | 5 | 1 |
| HG0830 | *H. sapiens* | ENSG00000171631 | 0 | 0 | 0 | 0 | 0 | 0 | 0 | 0 | 0 | 0 | 0 | 0 | 0 | 0 | 0 | 0 | 0 | 5 | 1 |
| HG0831.1 | *H. sapiens* | ENSG00000174482 | 0 | 0 | 0 | 0 | 0 | 0 | 0 | 0 | 0 | 0 | 0 | 0 | 0 | 0 | 0 | 0 | 0 | 3 | 3 |
| HG0831.2 | *H. sapiens* | ENSG00000174482 | 0 | 0 | 0 | 0 | 0 | 0 | 0 | 0 | 0 | 0 | 0 | 0 | 0 | 0 | 0 | 0 | 0 | 1 | 3 |
| HG0832 | *H. sapiens* | ENSG00000175048 | 0 | 0 | 0 | 0 | 0 | 0 | 0 | 0 | 0 | 0 | 0 | 0 | 0 | 0 | 0 | 0 | 0 | 3 | 3 |
| HG0833 | *H. sapiens* | ENSG00000176641 | 0 | 0 | 0 | 0 | 0 | 0 | 0 | 0 | 0 | 0 | 0 | 0 | 0 | 0 | 0 | 0 | 0 | 5 | 1 |
| HG0834 | *H. sapiens* | ENSG00000182489 | 0 | 0 | 0 | 0 | 0 | 0 | 0 | 0 | 0 | 0 | 0 | 0 | 0 | 0 | 0 | 0 | 0 | 4 | 2 |
| HG0835 | *H. sapiens* | ENSG00000182901 | 0 | 0 | 0 | 0 | 0 | 0 | 0 | 0 | 0 | 0 | 0 | 0 | 0 | 0 | 0 | 0 | 0 | 3 | 3 |
| HG0836 | *H. sapiens* | ENSG00000197461 | 0 | 0 | 0 | 0 | 0 | 0 | 0 | 0 | 0 | 0 | 0 | 0 | 0 | 0 | 0 | 0 | 0 | 5 | 1 |
| HG0837 | *H. sapiens* | ENSG00000198690 | 0 | 0 | 0 | 0 | 0 | 0 | 0 | 0 | 0 | 0 | 0 | 0 | 0 | 0 | 0 | 0 | 0 | 6 | 0 |
| HG0838 | *H. sapiens* | ENSG00000198707 | 0 | 0 | 0 | 0 | 0 | 0 | 0 | 0 | 0 | 0 | 0 | 0 | 0 | 0 | 0 | 0 | 0 | 4 | 2 |
| HG0839 | *H. sapiens* | ENSG00000198791 | 0 | 0 | 0 | 0 | 0 | 0 | 0 | 0 | 0 | 0 | 0 | 0 | 0 | 0 | 0 | 0 | 0 | 3 | 3 |
| HG0840 | *H. sapiens* | ENSG00000278195 | 0 | 0 | 0 | 0 | 0 | 0 | 0 | 0 | 0 | 0 | 0 | 0 | 0 | 0 | 0 | 0 | 0 | 6 | 0 |
| HG0841 | *H. sapiens* | ENSG00000278259 | 0 | 0 | 0 | 0 | 0 | 0 | 0 | 0 | 0 | 0 | 0 | 0 | 0 | 0 | 0 | 0 | 0 | 4 | 2 |
| HG0842 | *H. sapiens* | ENSG00000002746 | 0 | 0 | 0 | 0 | 0 | 0 | 0 | 0 | 0 | 0 | 0 | 0 | 0 | 0 | 0 | 0 | 0 | 3 | 2 |
| HG0843 | *H. sapiens* | ENSG00000043093 | 0 | 0 | 0 | 0 | 0 | 0 | 0 | 0 | 0 | 0 | 0 | 0 | 0 | 0 | 0 | 0 | 0 | 2 | 3 |
| HG0844 | *H. sapiens* | ENSG00000091009 | 0 | 0 | 0 | 0 | 0 | 0 | 0 | 0 | 0 | 0 | 0 | 0 | 0 | 0 | 0 | 0 | 0 | 3 | 2 |
| HG0845 | *H. sapiens* | ENSG00000099260 | 0 | 0 | 0 | 0 | 0 | 0 | 0 | 0 | 0 | 0 | 0 | 0 | 0 | 0 | 0 | 0 | 0 | 3 | 2 |
| HG0846 | *H. sapiens* | ENSG00000100852 | 0 | 0 | 0 | 0 | 0 | 0 | 0 | 0 | 0 | 0 | 0 | 0 | 0 | 0 | 0 | 0 | 0 | 4 | 1 |
| HG0847 | *H. sapiens* | ENSG00000106086 | 0 | 0 | 0 | 0 | 0 | 0 | 0 | 0 | 0 | 0 | 0 | 0 | 0 | 0 | 0 | 0 | 0 | 5 | 0 |
| HG0848 | *H. sapiens* | ENSG00000111817 | 0 | 0 | 0 | 0 | 0 | 0 | 0 | 0 | 0 | 0 | 0 | 0 | 0 | 0 | 0 | 0 | 0 | 4 | 1 |
| HG0849 | *H. sapiens* | ENSG00000111846 | 0 | 0 | 0 | 0 | 0 | 0 | 0 | 0 | 0 | 0 | 0 | 0 | 0 | 0 | 0 | 0 | 0 | 2 | 3 |
| HG0850 | *H. sapiens* | ENSG00000113916 | 0 | 0 | 0 | 0 | 0 | 0 | 0 | 0 | 0 | 0 | 0 | 0 | 0 | 0 | 0 | 0 | 0 | 3 | 2 |
| HG0851 | *H. sapiens* | ENSG00000114279 | 0 | 0 | 0 | 0 | 0 | 0 | 0 | 0 | 0 | 0 | 0 | 0 | 0 | 0 | 0 | 0 | 0 | 3 | 2 |
| HG0852 | *H. sapiens* | ENSG00000114942 | 0 | 0 | 0 | 0 | 0 | 0 | 0 | 0 | 0 | 0 | 0 | 0 | 0 | 0 | 0 | 0 | 0 | 4 | 1 |
| HG0853 | *H. sapiens* | ENSG00000121807 | 0 | 0 | 0 | 0 | 0 | 0 | 0 | 0 | 0 | 0 | 0 | 0 | 0 | 0 | 0 | 0 | 0 | 3 | 2 |
| HG0854 | *H. sapiens* | ENSG00000127249 | 0 | 0 | 0 | 0 | 0 | 0 | 0 | 0 | 0 | 0 | 0 | 0 | 0 | 0 | 0 | 0 | 0 | 4 | 1 |
| HG0855 | *H. sapiens* | ENSG00000127804 | 0 | 0 | 0 | 0 | 0 | 0 | 0 | 0 | 0 | 0 | 0 | 0 | 0 | 0 | 0 | 0 | 0 | 4 | 1 |
| HG0856 | *H. sapiens* | ENSG00000130055 | 0 | 0 | 0 | 0 | 0 | 0 | 0 | 0 | 0 | 0 | 0 | 0 | 0 | 0 | 0 | 0 | 0 | 4 | 1 |
| HG0857 | *H. sapiens* | ENSG00000130347 | 0 | 0 | 0 | 0 | 0 | 0 | 0 | 0 | 0 | 0 | 0 | 0 | 0 | 0 | 0 | 0 | 0 | 4 | 1 |

| | | | | | | | | | | | | | | | | | | | | | |
|---|---|---|---|---|---|---|---|---|---|---|---|---|---|---|---|---|---|---|---|---|---|
| HG0858 | *H. sapiens* | ENSG00000132702 | 0 | 0 | 0 | 0 | 0 | 0 | 0 | 0 | 0 | 0 | 0 | 0 | 0 | 0 | 0 | 0 | 0 | 2 | 3 |
| HG0859 | *H. sapiens* | ENSG00000132842 | 0 | 0 | 0 | 0 | 0 | 0 | 0 | 0 | 0 | 0 | 0 | 0 | 0 | 0 | 0 | 0 | 0 | 4 | 1 |
| HG0860 | *H. sapiens* | ENSG00000133056 | 0 | 0 | 0 | 0 | 0 | 0 | 0 | 0 | 0 | 0 | 0 | 0 | 0 | 0 | 0 | 0 | 0 | 2 | 3 |
| HG0861 | *H. sapiens* | ENSG00000134001 | 0 | 0 | 0 | 0 | 0 | 0 | 0 | 0 | 0 | 0 | 0 | 0 | 0 | 0 | 0 | 0 | 0 | 3 | 2 |
| HG0862 | *H. sapiens* | ENSG00000134245 | 0 | 0 | 0 | 0 | 0 | 0 | 0 | 0 | 0 | 0 | 0 | 0 | 0 | 0 | 0 | 0 | 0 | 4 | 1 |
| HG0863 | *H. sapiens* | ENSG00000134313 | 0 | 0 | 0 | 0 | 0 | 0 | 0 | 0 | 0 | 0 | 0 | 0 | 0 | 0 | 0 | 0 | 0 | 3 | 2 |
| HG0864 | *H. sapiens* | ENSG00000134533 | 0 | 0 | 0 | 0 | 0 | 0 | 0 | 0 | 0 | 0 | 0 | 0 | 0 | 0 | 0 | 0 | 0 | 4 | 1 |
| HG0865 | *H. sapiens* | ENSG00000135540 | 0 | 0 | 0 | 0 | 0 | 0 | 0 | 0 | 0 | 0 | 0 | 0 | 0 | 0 | 0 | 0 | 0 | 3 | 2 |
| HG0866 | *H. sapiens* | ENSG00000136011 | 0 | 0 | 0 | 0 | 0 | 0 | 0 | 0 | 0 | 0 | 0 | 0 | 0 | 0 | 0 | 0 | 0 | 3 | 2 |
| HG0867 | *H. sapiens* | ENSG00000137203 | 0 | 0 | 0 | 0 | 0 | 0 | 0 | 0 | 0 | 0 | 0 | 0 | 0 | 0 | 0 | 0 | 0 | 3 | 2 |
| HG0868 | *H. sapiens* | ENSG00000137843 | 0 | 0 | 0 | 0 | 0 | 0 | 0 | 0 | 0 | 0 | 0 | 0 | 0 | 0 | 0 | 0 | 0 | 4 | 1 |
| HG0869 | *H. sapiens* | ENSG00000139910 | 0 | 0 | 0 | 0 | 0 | 0 | 0 | 0 | 0 | 0 | 0 | 0 | 0 | 0 | 0 | 0 | 0 | 4 | 1 |
| HG0870 | *H. sapiens* | ENSG00000140993 | 0 | 0 | 0 | 0 | 0 | 0 | 0 | 0 | 0 | 0 | 0 | 0 | 0 | 0 | 0 | 0 | 0 | 3 | 2 |
| HG0871 | *H. sapiens* | ENSG00000141127 | 0 | 0 | 0 | 0 | 0 | 0 | 0 | 0 | 0 | 0 | 0 | 0 | 0 | 0 | 0 | 0 | 0 | 3 | 2 |
| HG0872 | *H. sapiens* | ENSG00000142583 | 0 | 0 | 0 | 0 | 0 | 0 | 0 | 0 | 0 | 0 | 0 | 0 | 0 | 0 | 0 | 0 | 0 | 5 | 0 |
| HG0873 | *H. sapiens* | ENSG00000142875 | 0 | 0 | 0 | 0 | 0 | 0 | 0 | 0 | 0 | 0 | 0 | 0 | 0 | 0 | 0 | 0 | 0 | 3 | 2 |
| HG0874 | *H. sapiens* | ENSG00000143515 | 0 | 0 | 0 | 0 | 0 | 0 | 0 | 0 | 0 | 0 | 0 | 0 | 0 | 0 | 0 | 0 | 0 | 3 | 2 |
| HG0875 | *H. sapiens* | ENSG00000145284 | 0 | 0 | 0 | 0 | 0 | 0 | 0 | 0 | 0 | 0 | 0 | 0 | 0 | 0 | 0 | 0 | 0 | 5 | 0 |
| HG0876 | *H. sapiens* | ENSG00000145908 | 0 | 0 | 0 | 0 | 0 | 0 | 0 | 0 | 0 | 0 | 0 | 0 | 0 | 0 | 0 | 0 | 0 | 4 | 1 |
| HG0877 | *H. sapiens* | ENSG00000146426 | 0 | 0 | 0 | 0 | 0 | 0 | 0 | 0 | 0 | 0 | 0 | 0 | 0 | 0 | 0 | 0 | 0 | 4 | 1 |
| HG0878 | *H. sapiens* | ENSG00000146802 | 0 | 0 | 0 | 0 | 0 | 0 | 0 | 0 | 0 | 0 | 0 | 0 | 0 | 0 | 0 | 0 | 0 | 4 | 1 |
| HG0879 | *H. sapiens* | ENSG00000147145 | 0 | 0 | 0 | 0 | 0 | 0 | 0 | 0 | 0 | 0 | 0 | 0 | 0 | 0 | 0 | 0 | 0 | 2 | 3 |
| HG0880 | *H. sapiens* | ENSG00000151789 | 0 | 0 | 0 | 0 | 0 | 0 | 0 | 0 | 0 | 0 | 0 | 0 | 0 | 0 | 0 | 0 | 0 | 4 | 1 |
| HG0881 | *H. sapiens* | ENSG00000153822 | 0 | 0 | 0 | 0 | 0 | 0 | 0 | 0 | 0 | 0 | 0 | 0 | 0 | 0 | 0 | 0 | 0 | 4 | 1 |
| HG0882.1 | *H. sapiens* | ENSG00000154832 | 0 | 0 | 0 | 0 | 0 | 0 | 0 | 0 | 0 | 0 | 0 | 0 | 0 | 0 | 0 | 0 | 0 | 3 | 2 |
| HG0882.2 | *H. sapiens* | ENSG00000154832 | 0 | 0 | 0 | 0 | 0 | 0 | 0 | 0 | 0 | 0 | 0 | 0 | 0 | 0 | 0 | 0 | 0 | 2 | 0 |
| HG0883 | *H. sapiens* | ENSG00000156011 | 0 | 0 | 0 | 0 | 0 | 0 | 0 | 0 | 0 | 0 | 0 | 0 | 0 | 0 | 0 | 0 | 0 | 2 | 3 |
| HG0884 | *H. sapiens* | ENSG00000156463 | 0 | 0 | 0 | 0 | 0 | 0 | 0 | 0 | 0 | 0 | 0 | 0 | 0 | 0 | 0 | 0 | 0 | 3 | 2 |
| HG0885.1 | *H. sapiens* | ENSG00000157502 | 0 | 0 | 0 | 0 | 0 | 0 | 0 | 0 | 0 | 0 | 0 | 0 | 0 | 0 | 0 | 0 | 0 | 3 | 2 |
| HG0885.2 | *H. sapiens* | ENSG00000157502 | 0 | 0 | 0 | 0 | 0 | 0 | 0 | 0 | 0 | 0 | 0 | 0 | 0 | 0 | 0 | 0 | 0 | 2 | 1 |
| HG0886 | *H. sapiens* | ENSG00000158850 | 0 | 0 | 0 | 0 | 0 | 0 | 0 | 0 | 0 | 0 | 0 | 0 | 0 | 0 | 0 | 0 | 0 | 2 | 3 |
| HG0887 | *H. sapiens* | ENSG00000160959 | 0 | 0 | 0 | 0 | 0 | 0 | 0 | 0 | 0 | 0 | 0 | 0 | 0 | 0 | 0 | 0 | 0 | 4 | 1 |
| HG0888 | *H. sapiens* | ENSG00000164023 | 0 | 0 | 0 | 0 | 0 | 0 | 0 | 0 | 0 | 0 | 0 | 0 | 0 | 0 | 0 | 0 | 0 | 3 | 2 |
| HG0889 | *H. sapiens* | ENSG00000164091 | 0 | 0 | 0 | 0 | 0 | 0 | 0 | 0 | 0 | 0 | 0 | 0 | 0 | 0 | 0 | 0 | 0 | 2 | 3 |
| HG0890 | *H. sapiens* | ENSG00000165813 | 0 | 0 | 0 | 0 | 0 | 0 | 0 | 0 | 0 | 0 | 0 | 0 | 0 | 0 | 0 | 0 | 0 | 4 | 1 |
| HG0891 | *H. sapiens* | ENSG00000166211 | 0 | 0 | 0 | 0 | 0 | 0 | 0 | 0 | 0 | 0 | 0 | 0 | 0 | 0 | 0 | 0 | 0 | 3 | 2 |
| HG0892 | *H. sapiens* | ENSG00000168418 | 0 | 0 | 0 | 0 | 0 | 0 | 0 | 0 | 0 | 0 | 0 | 0 | 0 | 0 | 0 | 0 | 0 | 4 | 1 |
| HG0893 | *H. sapiens* | ENSG00000168769 | 0 | 0 | 0 | 0 | 0 | 0 | 0 | 0 | 0 | 0 | 0 | 0 | 0 | 0 | 0 | 0 | 0 | 4 | 1 |
| HG0894.1 | *H. sapiens* | ENSG00000169282 | 0 | 0 | 0 | 0 | 0 | 0 | 0 | 0 | 0 | 0 | 0 | 0 | 0 | 0 | 0 | 0 | 0 | 4 | 1 |
| HG0894.2 | *H. sapiens* | ENSG00000069424 | 0 | 0 | 0 | 0 | 0 | 0 | 0 | 0 | 0 | 0 | 0 | 0 | 0 | 0 | 0 | 0 | 0 | 3 | 1 |
| HG0895 | *H. sapiens* | ENSG00000170035 | 0 | 0 | 0 | 0 | 0 | 0 | 0 | 0 | 0 | 0 | 0 | 0 | 0 | 0 | 0 | 0 | 0 | 5 | 0 |
| HG0896 | *H. sapiens* | ENSG00000170365 | 0 | 0 | 0 | 0 | 0 | 0 | 0 | 0 | 0 | 0 | 0 | 0 | 0 | 0 | 0 | 0 | 0 | 4 | 1 |
| HG0897 | *H. sapiens* | ENSG00000171435 | 0 | 0 | 0 | 0 | 0 | 0 | 0 | 0 | 0 | 0 | 0 | 0 | 0 | 0 | 0 | 0 | 0 | 4 | 1 |
| HG0898 | *H. sapiens* | ENSG00000171729 | 0 | 0 | 0 | 0 | 0 | 0 | 0 | 0 | 0 | 0 | 0 | 0 | 0 | 0 | 0 | 0 | 0 | 4 | 1 |
| HG0899 | *H. sapiens* | ENSG00000173320 | 0 | 0 | 0 | 0 | 0 | 0 | 0 | 0 | 0 | 0 | 0 | 0 | 0 | 0 | 0 | 0 | 0 | 5 | 0 |
| HG0900 | *H. sapiens* | ENSG00000173517 | 0 | 0 | 0 | 0 | 0 | 0 | 0 | 0 | 0 | 0 | 0 | 0 | 0 | 0 | 0 | 0 | 0 | 4 | 1 |
| HG0901 | *H. sapiens* | ENSG00000175697 | 0 | 0 | 0 | 0 | 0 | 0 | 0 | 0 | 0 | 0 | 0 | 0 | 0 | 0 | 0 | 0 | 0 | 4 | 1 |
| HG0902 | *H. sapiens* | ENSG00000177182 | 0 | 0 | 0 | 0 | 0 | 0 | 0 | 0 | 0 | 0 | 0 | 0 | 0 | 0 | 0 | 0 | 0 | 4 | 1 |
| HG0903 | *H. sapiens* | ENSG00000180616 | 0 | 0 | 0 | 0 | 0 | 0 | 0 | 0 | 0 | 0 | 0 | 0 | 0 | 0 | 0 | 0 | 0 | 2 | 3 |
| HG0904 | *H. sapiens* | ENSG00000182329 | 0 | 0 | 0 | 0 | 0 | 0 | 0 | 0 | 0 | 0 | 0 | 0 | 0 | 0 | 0 | 0 | 0 | 4 | 1 |
| HG0905 | *H. sapiens* | ENSG00000183337 | 0 | 0 | 0 | 0 | 0 | 0 | 0 | 0 | 0 | 0 | 0 | 0 | 0 | 0 | 0 | 0 | 0 | 4 | 1 |
| HG0906 | *H. sapiens* | ENSG00000183773 | 0 | 0 | 0 | 0 | 0 | 0 | 0 | 0 | 0 | 0 | 0 | 0 | 0 | 0 | 0 | 0 | 0 | 2 | 3 |
| HG0907 | *H. sapiens* | ENSG00000186862 | 0 | 0 | 0 | 0 | 0 | 0 | 0 | 0 | 0 | 0 | 0 | 0 | 0 | 0 | 0 | 0 | 0 | 4 | 1 |
| HG0908.1 | *H. sapiens* | ENSG00000188554 | 0 | 0 | 0 | 0 | 0 | 0 | 0 | 0 | 0 | 0 | 0 | 0 | 0 | 0 | 0 | 0 | 0 | 3 | 2 |
| HG0908.2 | *H. sapiens* | ENSG00000188554 | 0 | 0 | 0 | 0 | 0 | 0 | 0 | 0 | 0 | 0 | 0 | 0 | 0 | 0 | 0 | 0 | 0 | 3 | 1 |
| HG0909 | *H. sapiens* | ENSG00000189079 | 0 | 0 | 0 | 0 | 0 | 0 | 0 | 0 | 0 | 0 | 0 | 0 | 0 | 0 | 0 | 0 | 0 | 4 | 1 |
| HG0910 | *H. sapiens* | ENSG00000197935 | 0 | 0 | 0 | 0 | 0 | 0 | 0 | 0 | 0 | 0 | 0 | 0 | 0 | 0 | 0 | 0 | 0 | 3 | 2 |
| HG0911 | *H. sapiens* | ENSG00000198673 | 0 | 0 | 0 | 0 | 0 | 0 | 0 | 0 | 0 | 0 | 0 | 0 | 0 | 0 | 0 | 0 | 0 | 3 | 2 |
| HG0912 | *H. sapiens* | ENSG00000198822 | 0 | 0 | 0 | 0 | 0 | 0 | 0 | 0 | 0 | 0 | 0 | 0 | 0 | 0 | 0 | 0 | 0 | 4 | 1 |
| HG0913.1 | *H. sapiens* | ENSG00000204599 | 0 | 0 | 0 | 0 | 0 | 0 | 0 | 0 | 0 | 0 | 0 | 0 | 0 | 0 | 0 | 0 | 0 | 4 | 1 |
| HG0913.2 | *H. sapiens* | ENSG00000204599 | 0 | 0 | 0 | 0 | 0 | 0 | 0 | 0 | 0 | 0 | 0 | 0 | 0 | 0 | 0 | 0 | 0 | 3 | 1 |
| HG0914 | *H. sapiens* | ENSG00000227877 | 0 | 0 | 0 | 0 | 0 | 0 | 0 | 0 | 0 | 0 | 0 | 0 | 0 | 0 | 0 | 0 | 0 | 4 | 1 |
| HG0915 | *H. sapiens* | ENSG00000254469 | 0 | 0 | 0 | 0 | 0 | 0 | 0 | 0 | 0 | 0 | 0 | 0 | 0 | 0 | 0 | 0 | 0 | 4 | 1 |
| HG0916 | *H. sapiens* | ENSG00000020633 | 0 | 0 | 0 | 0 | 0 | 0 | 0 | 0 | 0 | 0 | 0 | 0 | 0 | 0 | 0 | 0 | 0 | 3 | 1 |
| HG0917 | *H. sapiens* | ENSG00000042980 | 0 | 0 | 0 | 0 | 0 | 0 | 0 | 0 | 0 | 0 | 0 | 0 | 0 | 0 | 0 | 0 | 0 | 3 | 1 |
| HG0918 | *H. sapiens* | ENSG00000047648 | 0 | 0 | 0 | 0 | 0 | 0 | 0 | 0 | 0 | 0 | 0 | 0 | 0 | 0 | 0 | 0 | 0 | 3 | 1 |
| HG0919 | *H. sapiens* | ENSG00000051009 | 0 | 0 | 0 | 0 | 0 | 0 | 0 | 0 | 0 | 0 | 0 | 0 | 0 | 0 | 0 | 0 | 0 | 3 | 1 |
| HG0920 | *H. sapiens* | ENSG00000051382 | 0 | 0 | 0 | 0 | 0 | 0 | 0 | 0 | 0 | 0 | 0 | 0 | 0 | 0 | 0 | 0 | 0 | 3 | 1 |
| HG0921 | *H. sapiens* | ENSG00000058799 | 0 | 0 | 0 | 0 | 0 | 0 | 0 | 0 | 0 | 0 | 0 | 0 | 0 | 0 | 0 | 0 | 0 | 3 | 1 |
| HG0922 | *H. sapiens* | ENSG00000079841 | 0 | 0 | 0 | 0 | 0 | 0 | 0 | 0 | 0 | 0 | 0 | 0 | 0 | 0 | 0 | 0 | 0 | 3 | 1 |
| HG0923 | *H. sapiens* | ENSG00000081138 | 0 | 0 | 0 | 0 | 0 | 0 | 0 | 0 | 0 | 0 | 0 | 0 | 0 | 0 | 0 | 0 | 0 | 3 | 1 |
| HG0924 | *H. sapiens* | ENSG00000081307 | 0 | 0 | 0 | 0 | 0 | 0 | 0 | 0 | 0 | 0 | 0 | 0 | 0 | 0 | 0 | 0 | 0 | 3 | 1 |
| HG0925 | *H. sapiens* | ENSG00000088766 | 0 | 0 | 0 | 0 | 0 | 0 | 0 | 0 | 0 | 0 | 0 | 0 | 0 | 0 | 0 | 0 | 0 | 3 | 1 |
| HG0926 | *H. sapiens* | ENSG00000095397 | 0 | 0 | 0 | 0 | 0 | 0 | 0 | 0 | 0 | 0 | 0 | 0 | 0 | 0 | 0 | 0 | 0 | 4 | 0 |
| HG0927 | *H. sapiens* | ENSG00000095951 | 0 | 0 | 0 | 0 | 0 | 0 | 0 | 0 | 0 | 0 | 0 | 0 | 0 | 0 | 0 | 0 | 0 | 2 | 2 |
| HG0928 | *H. sapiens* | ENSG00000102057 | 0 | 0 | 0 | 0 | 0 | 0 | 0 | 0 | 0 | 0 | 0 | 0 | 0 | 0 | 0 | 0 | 0 | 2 | 2 |
| HG0929 | *H. sapiens* | ENSG00000102239 | 0 | 0 | 0 | 0 | 0 | 0 | 0 | 0 | 0 | 0 | 0 | 0 | 0 | 0 | 0 | 0 | 0 | 1 | 3 |
| HG0930 | *H. sapiens* | ENSG00000102524 | 0 | 0 | 0 | 0 | 0 | 0 | 0 | 0 | 0 | 0 | 0 | 0 | 0 | 0 | 0 | 0 | 0 | 3 | 1 |
| HG0931 | *H. sapiens* | ENSG00000104951 | 0 | 0 | 0 | 0 | 0 | 0 | 0 | 0 | 0 | 0 | 0 | 0 | 0 | 0 | 0 | 0 | 0 | 3 | 1 |

| | | | | | | | | | | | | | | | | | | | | | |
|---|---|---|---|---|---|---|---|---|---|---|---|---|---|---|---|---|---|---|---|---|---|
| HG0932 | *H. sapiens* | ENSG00000105997 | 0 | 0 | 0 | 0 | 0 | 0 | 0 | 0 | 0 | 0 | 0 | 0 | 0 | 0 | 0 | 0 | 0 | 3 | 1 |
| HG0933 | *H. sapiens* | ENSG00000106330 | 0 | 0 | 0 | 0 | 0 | 0 | 0 | 0 | 0 | 0 | 0 | 0 | 0 | 0 | 0 | 0 | 0 | 2 | 2 |
| HG0934 | *H. sapiens* | ENSG00000107443 | 0 | 0 | 0 | 0 | 0 | 0 | 0 | 0 | 0 | 0 | 0 | 0 | 0 | 0 | 0 | 0 | 0 | 3 | 1 |
| HG0935 | *H. sapiens* | ENSG00000108518 | 0 | 0 | 0 | 0 | 0 | 0 | 0 | 0 | 0 | 0 | 0 | 0 | 0 | 0 | 0 | 0 | 0 | 2 | 2 |
| HG0936 | *H. sapiens* | ENSG00000108799 | 0 | 0 | 0 | 0 | 0 | 0 | 0 | 0 | 0 | 0 | 0 | 0 | 0 | 0 | 0 | 0 | 0 | 3 | 1 |
| HG0937 | *H. sapiens* | ENSG00000109846 | 0 | 0 | 0 | 0 | 0 | 0 | 0 | 0 | 0 | 0 | 0 | 0 | 0 | 0 | 0 | 0 | 0 | 2 | 2 |
| HG0938 | *H. sapiens* | ENSG00000111665 | 0 | 0 | 0 | 0 | 0 | 0 | 0 | 0 | 0 | 0 | 0 | 0 | 0 | 0 | 0 | 0 | 0 | 2 | 2 |
| HG0939 | *H. sapiens* | ENSG00000111674 | 0 | 0 | 0 | 0 | 0 | 0 | 0 | 0 | 0 | 0 | 0 | 0 | 0 | 0 | 0 | 0 | 0 | 4 | 0 |
| HG0940 | *H. sapiens* | ENSG00000115556 | 0 | 0 | 0 | 0 | 0 | 0 | 0 | 0 | 0 | 0 | 0 | 0 | 0 | 0 | 0 | 0 | 0 | 3 | 1 |
| HG0941 | *H. sapiens* | ENSG00000116285 | 0 | 0 | 0 | 0 | 0 | 0 | 0 | 0 | 0 | 0 | 0 | 0 | 0 | 0 | 0 | 0 | 0 | 3 | 1 |
| HG0942 | *H. sapiens* | ENSG00000117477 | 0 | 0 | 0 | 0 | 0 | 0 | 0 | 0 | 0 | 0 | 0 | 0 | 0 | 0 | 0 | 0 | 0 | 3 | 1 |
| HG0943 | *H. sapiens* | ENSG00000120690 | 0 | 0 | 0 | 0 | 0 | 0 | 0 | 0 | 0 | 0 | 0 | 0 | 0 | 0 | 0 | 0 | 0 | 2 | 2 |
| HG0944 | *H. sapiens* | ENSG00000120833 | 0 | 0 | 0 | 0 | 0 | 0 | 0 | 0 | 0 | 0 | 0 | 0 | 0 | 0 | 0 | 0 | 0 | 3 | 1 |
| HG0945 | *H. sapiens* | ENSG00000124782 | 0 | 0 | 0 | 0 | 0 | 0 | 0 | 0 | 0 | 0 | 0 | 0 | 0 | 0 | 0 | 0 | 0 | 4 | 0 |
| HG0946.1 | *H. sapiens* | ENSG00000127324 | 0 | 0 | 0 | 0 | 0 | 0 | 0 | 0 | 0 | 0 | 0 | 0 | 0 | 0 | 0 | 0 | 0 | 3 | 1 |
| HG0946.2 | *H. sapiens* | ENSG00000127324 | 0 | 0 | 0 | 0 | 0 | 0 | 0 | 0 | 0 | 0 | 0 | 0 | 0 | 0 | 0 | 0 | 0 | 3 | 1 |
| HG0947 | *H. sapiens* | ENSG00000129226 | 0 | 0 | 0 | 0 | 0 | 0 | 0 | 0 | 0 | 0 | 0 | 0 | 0 | 0 | 0 | 0 | 0 | 3 | 1 |
| HG0948 | *H. sapiens* | ENSG00000129595 | 0 | 0 | 0 | 0 | 0 | 0 | 0 | 0 | 0 | 0 | 0 | 0 | 0 | 0 | 0 | 0 | 0 | 3 | 1 |
| HG0949 | *H. sapiens* | ENSG00000131381 | 0 | 0 | 0 | 0 | 0 | 0 | 0 | 0 | 0 | 0 | 0 | 0 | 0 | 0 | 0 | 0 | 0 | 4 | 0 |
| HG0950 | *H. sapiens* | ENSG00000132964 | 0 | 0 | 0 | 0 | 0 | 0 | 0 | 0 | 0 | 0 | 0 | 0 | 0 | 0 | 0 | 0 | 0 | 2 | 2 |
| HG0951 | *H. sapiens* | ENSG00000134061 | 0 | 0 | 0 | 0 | 0 | 0 | 0 | 0 | 0 | 0 | 0 | 0 | 0 | 0 | 0 | 0 | 0 | 3 | 1 |
| HG0952 | *H. sapiens* | ENSG00000134453 | 0 | 0 | 0 | 0 | 0 | 0 | 0 | 0 | 0 | 0 | 0 | 0 | 0 | 0 | 0 | 0 | 0 | 4 | 0 |
| HG0953 | *H. sapiens* | ENSG00000134815 | 0 | 0 | 0 | 0 | 0 | 0 | 0 | 0 | 0 | 0 | 0 | 0 | 0 | 0 | 0 | 0 | 0 | 3 | 1 |
| HG0954 | *H. sapiens* | ENSG00000134874 | 0 | 0 | 0 | 0 | 0 | 0 | 0 | 0 | 0 | 0 | 0 | 0 | 0 | 0 | 0 | 0 | 0 | 4 | 0 |
| HG0955 | *H. sapiens* | ENSG00000134982 | 0 | 0 | 0 | 0 | 0 | 0 | 0 | 0 | 0 | 0 | 0 | 0 | 0 | 0 | 0 | 0 | 0 | 3 | 1 |
| HG0956 | *H. sapiens* | ENSG00000136235 | 0 | 0 | 0 | 0 | 0 | 0 | 0 | 0 | 0 | 0 | 0 | 0 | 0 | 0 | 0 | 0 | 0 | 4 | 0 |
| HG0957 | *H. sapiens* | ENSG00000136874 | 0 | 0 | 0 | 0 | 0 | 0 | 0 | 0 | 0 | 0 | 0 | 0 | 0 | 0 | 0 | 0 | 0 | 3 | 1 |
| HG0958 | *H. sapiens* | ENSG00000137497 | 0 | 0 | 0 | 0 | 0 | 0 | 0 | 0 | 0 | 0 | 0 | 0 | 0 | 0 | 0 | 0 | 0 | 2 | 2 |
| HG0959 | *H. sapiens* | ENSG00000137558 | 0 | 0 | 0 | 0 | 0 | 0 | 0 | 0 | 0 | 0 | 0 | 0 | 0 | 0 | 0 | 0 | 0 | 3 | 1 |
| HG0960 | *H. sapiens* | ENSG00000140297 | 0 | 0 | 0 | 0 | 0 | 0 | 0 | 0 | 0 | 0 | 0 | 0 | 0 | 0 | 0 | 0 | 0 | 3 | 1 |
| HG0961 | *H. sapiens* | ENSG00000140853 | 0 | 0 | 0 | 0 | 0 | 0 | 0 | 0 | 0 | 0 | 0 | 0 | 0 | 0 | 0 | 0 | 0 | 2 | 2 |
| HG0962.1 | *H. sapiens* | ENSG00000143319 | 0 | 0 | 0 | 0 | 0 | 0 | 0 | 0 | 0 | 0 | 0 | 0 | 0 | 0 | 0 | 0 | 0 | 3 | 1 |
| HG0962.2 | *H. sapiens* | ENSG00000143319 | 0 | 0 | 0 | 0 | 0 | 0 | 0 | 0 | 0 | 0 | 0 | 0 | 0 | 0 | 0 | 0 | 0 | 2 | 2 |
| HG0963 | *H. sapiens* | ENSG00000143867 | 0 | 0 | 0 | 0 | 0 | 0 | 0 | 0 | 0 | 0 | 0 | 0 | 0 | 0 | 0 | 0 | 0 | 2 | 2 |
| HG0964 | *H. sapiens* | ENSG00000145246 | 0 | 0 | 0 | 0 | 0 | 0 | 0 | 0 | 0 | 0 | 0 | 0 | 0 | 0 | 0 | 0 | 0 | 4 | 0 |
| HG0965 | *H. sapiens* | ENSG00000146006 | 0 | 0 | 0 | 0 | 0 | 0 | 0 | 0 | 0 | 0 | 0 | 0 | 0 | 0 | 0 | 0 | 0 | 4 | 0 |
| HG0966 | *H. sapiens* | ENSG00000148356 | 0 | 0 | 0 | 0 | 0 | 0 | 0 | 0 | 0 | 0 | 0 | 0 | 0 | 0 | 0 | 0 | 0 | 2 | 2 |
| HG0967 | *H. sapiens* | ENSG00000149925 | 0 | 0 | 0 | 0 | 0 | 0 | 0 | 0 | 0 | 0 | 0 | 0 | 0 | 0 | 0 | 0 | 0 | 3 | 1 |
| HG0968 | *H. sapiens* | ENSG00000154319 | 0 | 0 | 0 | 0 | 0 | 0 | 0 | 0 | 0 | 0 | 0 | 0 | 0 | 0 | 0 | 0 | 0 | 2 | 2 |
| HG0969 | *H. sapiens* | ENSG00000154645 | 0 | 0 | 0 | 0 | 0 | 0 | 0 | 0 | 0 | 0 | 0 | 0 | 0 | 0 | 0 | 0 | 0 | 3 | 1 |
| HG0970 | *H. sapiens* | ENSG00000154727 | 0 | 0 | 0 | 0 | 0 | 0 | 0 | 0 | 0 | 0 | 0 | 0 | 0 | 0 | 0 | 0 | 0 | 3 | 1 |
| HG0971 | *H. sapiens* | ENSG00000155592 | 0 | 0 | 0 | 0 | 0 | 0 | 0 | 0 | 0 | 0 | 0 | 0 | 0 | 0 | 0 | 0 | 0 | 2 | 2 |
| HG0972 | *H. sapiens* | ENSG00000155926 | 0 | 0 | 0 | 0 | 0 | 0 | 0 | 0 | 0 | 0 | 0 | 0 | 0 | 0 | 0 | 0 | 0 | 2 | 2 |
| HG0973 | *H. sapiens* | ENSG00000156535 | 0 | 0 | 0 | 0 | 0 | 0 | 0 | 0 | 0 | 0 | 0 | 0 | 0 | 0 | 0 | 0 | 0 | 2 | 2 |
| HG0974 | *H. sapiens* | ENSG00000157578 | 0 | 0 | 0 | 0 | 0 | 0 | 0 | 0 | 0 | 0 | 0 | 0 | 0 | 0 | 0 | 0 | 0 | 3 | 1 |
| HG0975.1 | *H. sapiens* | ENSG00000182158 | 0 | 0 | 0 | 0 | 0 | 0 | 0 | 0 | 0 | 0 | 0 | 0 | 0 | 0 | 0 | 0 | 0 | 2 | 2 |
| HG0975.2 | *H. sapiens* | ENSG00000157613 | 0 | 0 | 0 | 0 | 0 | 0 | 0 | 0 | 0 | 0 | 0 | 0 | 0 | 0 | 0 | 0 | 0 | 4 | 0 |
| HG0976 | *H. sapiens* | ENSG00000158008 | 0 | 0 | 0 | 0 | 0 | 0 | 0 | 0 | 0 | 0 | 0 | 0 | 0 | 0 | 0 | 0 | 0 | 2 | 2 |
| HG0977 | *H. sapiens* | ENSG00000158321 | 0 | 0 | 0 | 0 | 0 | 0 | 0 | 0 | 0 | 0 | 0 | 0 | 0 | 0 | 0 | 0 | 0 | 2 | 2 |
| HG0978 | *H. sapiens* | ENSG00000159023 | 0 | 0 | 0 | 0 | 0 | 0 | 0 | 0 | 0 | 0 | 0 | 0 | 0 | 0 | 0 | 0 | 0 | 3 | 1 |
| HG0979 | *H. sapiens* | ENSG00000159202 | 0 | 0 | 0 | 0 | 0 | 0 | 0 | 0 | 0 | 0 | 0 | 0 | 0 | 0 | 0 | 0 | 0 | 3 | 1 |
| HG0980 | *H. sapiens* | ENSG00000162670 | 0 | 0 | 0 | 0 | 0 | 0 | 0 | 0 | 0 | 0 | 0 | 0 | 0 | 0 | 0 | 0 | 0 | 2 | 2 |
| HG0981 | *H. sapiens* | ENSG00000163629 | 0 | 0 | 0 | 0 | 0 | 0 | 0 | 0 | 0 | 0 | 0 | 0 | 0 | 0 | 0 | 0 | 0 | 2 | 2 |
| HG0982 | *H. sapiens* | ENSG00000164116 | 0 | 0 | 0 | 0 | 0 | 0 | 0 | 0 | 0 | 0 | 0 | 0 | 0 | 0 | 0 | 0 | 0 | 2 | 2 |
| HG0983 | *H. sapiens* | ENSG00000164128 | 0 | 0 | 0 | 0 | 0 | 0 | 0 | 0 | 0 | 0 | 0 | 0 | 0 | 0 | 0 | 0 | 0 | 2 | 2 |
| HG0984 | *H. sapiens* | ENSG00000164185 | 0 | 0 | 0 | 0 | 0 | 0 | 0 | 0 | 0 | 0 | 0 | 0 | 0 | 0 | 0 | 0 | 0 | 2 | 2 |
| HG0985 | *H. sapiens* | ENSG00000164604 | 0 | 0 | 0 | 0 | 0 | 0 | 0 | 0 | 0 | 0 | 0 | 0 | 0 | 0 | 0 | 0 | 0 | 3 | 1 |
| HG0986 | *H. sapiens* | ENSG00000164663 | 0 | 0 | 0 | 0 | 0 | 0 | 0 | 0 | 0 | 0 | 0 | 0 | 0 | 0 | 0 | 0 | 0 | 2 | 2 |
| HG0987 | *H. sapiens* | ENSG00000164932 | 0 | 0 | 0 | 0 | 0 | 0 | 0 | 0 | 0 | 0 | 0 | 0 | 0 | 0 | 0 | 0 | 0 | 2 | 2 |
| HG0988 | *H. sapiens* | ENSG00000166130 | 0 | 0 | 0 | 0 | 0 | 0 | 0 | 0 | 0 | 0 | 0 | 0 | 0 | 0 | 0 | 0 | 0 | 3 | 1 |
| HG0989 | *H. sapiens* | ENSG00000166603 | 0 | 0 | 0 | 0 | 0 | 0 | 0 | 0 | 0 | 0 | 0 | 0 | 0 | 0 | 0 | 0 | 0 | 4 | 0 |
| HG0990 | *H. sapiens* | ENSG00000166866 | 0 | 0 | 0 | 0 | 0 | 0 | 0 | 0 | 0 | 0 | 0 | 0 | 0 | 0 | 0 | 0 | 0 | 3 | 1 |
| HG0991 | *H. sapiens* | ENSG00000167654 | 0 | 0 | 0 | 0 | 0 | 0 | 0 | 0 | 0 | 0 | 0 | 0 | 0 | 0 | 0 | 0 | 0 | 4 | 0 |
| HG0992 | *H. sapiens* | ENSG00000168000 | 0 | 0 | 0 | 0 | 0 | 0 | 0 | 0 | 0 | 0 | 0 | 0 | 0 | 0 | 0 | 0 | 0 | 3 | 1 |
| HG0993 | *H. sapiens* | ENSG00000169306 | 0 | 0 | 0 | 0 | 0 | 0 | 0 | 0 | 0 | 0 | 0 | 0 | 0 | 0 | 0 | 0 | 0 | 2 | 2 |
| HG0994 | *H. sapiens* | ENSG00000170852 | 0 | 0 | 0 | 0 | 0 | 0 | 0 | 0 | 0 | 0 | 0 | 0 | 0 | 0 | 0 | 0 | 0 | 4 | 0 |
| HG0995 | *H. sapiens* | ENSG00000171307 | 0 | 0 | 0 | 0 | 0 | 0 | 0 | 0 | 0 | 0 | 0 | 0 | 0 | 0 | 0 | 0 | 0 | 1 | 3 |
| HG0996 | *H. sapiens* | ENSG00000173890 | 0 | 0 | 0 | 0 | 0 | 0 | 0 | 0 | 0 | 0 | 0 | 0 | 0 | 0 | 0 | 0 | 0 | 1 | 3 |
| HG0997 | *H. sapiens* | ENSG00000177728 | 0 | 0 | 0 | 0 | 0 | 0 | 0 | 0 | 0 | 0 | 0 | 0 | 0 | 0 | 0 | 0 | 0 | 2 | 2 |
| HG0998 | *H. sapiens* | ENSG00000178172 | 0 | 0 | 0 | 0 | 0 | 0 | 0 | 0 | 0 | 0 | 0 | 0 | 0 | 0 | 0 | 0 | 0 | 3 | 1 |
| HG0999 | *H. sapiens* | ENSG00000178401 | 0 | 0 | 0 | 0 | 0 | 0 | 0 | 0 | 0 | 0 | 0 | 0 | 0 | 0 | 0 | 0 | 0 | 3 | 1 |
| HG1000 | *H. sapiens* | ENSG00000179083 | 0 | 0 | 0 | 0 | 0 | 0 | 0 | 0 | 0 | 0 | 0 | 0 | 0 | 0 | 0 | 0 | 0 | 2 | 2 |
| HG1001 | *H. sapiens* | ENSG00000179761 | 0 | 0 | 0 | 0 | 0 | 0 | 0 | 0 | 0 | 0 | 0 | 0 | 0 | 0 | 0 | 0 | 0 | 2 | 2 |
| HG1002 | *H. sapiens* | ENSG00000181027 | 0 | 0 | 0 | 0 | 0 | 0 | 0 | 0 | 0 | 0 | 0 | 0 | 0 | 0 | 0 | 0 | 0 | 3 | 1 |
| HG1003 | *H. sapiens* | ENSG00000184867 | 0 | 0 | 0 | 0 | 0 | 0 | 0 | 0 | 0 | 0 | 0 | 0 | 0 | 0 | 0 | 0 | 0 | 2 | 2 |
| HG1004 | *H. sapiens* | ENSG00000185875 | 0 | 0 | 0 | 0 | 0 | 0 | 0 | 0 | 0 | 0 | 0 | 0 | 0 | 0 | 0 | 0 | 0 | 3 | 1 |
| HG1005 | *H. sapiens* | ENSG00000186472 | 0 | 0 | 0 | 0 | 0 | 0 | 0 | 0 | 0 | 0 | 0 | 0 | 0 | 0 | 0 | 0 | 0 | 3 | 1 |
| HG1006.1 | *H. sapiens* | ENSG00000188227 | 0 | 0 | 0 | 0 | 0 | 0 | 0 | 0 | 0 | 0 | 0 | 0 | 0 | 0 | 0 | 0 | 0 | 3 | 1 |
| HG1006.2 | *H. sapiens* | ENSG00000188227 | 0 | 0 | 0 | 0 | 0 | 0 | 0 | 0 | 0 | 0 | 0 | 0 | 0 | 0 | 0 | 0 | 0 | 2 | 1 |

| | | | | | | | | | | | | | | | | | | | | |
|---|---|---|---|---|---|---|---|---|---|---|---|---|---|---|---|---|---|---|---|---|---|
| HG1007 | *H. sapiens* | ENSG00000188910 | 0 | 0 | 0 | 0 | 0 | 0 | 0 | 0 | 0 | 0 | 0 | 0 | 0 | 0 | 0 | 0 | 0 | 3 | 1 |
| HG1008 | *H. sapiens* | ENSG00000198252 | 0 | 0 | 0 | 0 | 0 | 0 | 0 | 0 | 0 | 0 | 0 | 0 | 0 | 0 | 0 | 0 | 0 | 4 | 0 |
| HG1009 | *H. sapiens* | ENSG00000198909 | 0 | 0 | 0 | 0 | 0 | 0 | 0 | 0 | 0 | 0 | 0 | 0 | 0 | 0 | 0 | 0 | 0 | 2 | 2 |
| HG1010 | *H. sapiens* | ENSG00000203740 | 0 | 0 | 0 | 0 | 0 | 0 | 0 | 0 | 0 | 0 | 0 | 0 | 0 | 0 | 0 | 0 | 0 | 1 | 3 |
| HG1011 | *H. sapiens* | ENSG00000204439 | 0 | 0 | 0 | 0 | 0 | 0 | 0 | 0 | 0 | 0 | 0 | 0 | 0 | 0 | 0 | 0 | 0 | 3 | 1 |
| HG1012 | *H. sapiens* | ENSG00000213694 | 0 | 0 | 0 | 0 | 0 | 0 | 0 | 0 | 0 | 0 | 0 | 0 | 0 | 0 | 0 | 0 | 0 | 3 | 1 |
| HG1013 | *H. sapiens* | ENSG00000214022 | 0 | 0 | 0 | 0 | 0 | 0 | 0 | 0 | 0 | 0 | 0 | 0 | 0 | 0 | 0 | 0 | 0 | 3 | 1 |
| HG1014.1 | *H. sapiens* | ENSG00000214595 | 0 | 0 | 0 | 0 | 0 | 0 | 0 | 0 | 0 | 0 | 0 | 0 | 0 | 0 | 0 | 0 | 0 | 3 | 1 |
| HG1014.2 | *H. sapiens* | ENSG00000214595 | 0 | 0 | 0 | 0 | 0 | 0 | 0 | 0 | 0 | 0 | 0 | 0 | 0 | 0 | 0 | 0 | 0 | 3 | 1 |
| HG1015 | *H. sapiens* | ENSG00000240720 | 0 | 0 | 0 | 0 | 0 | 0 | 0 | 0 | 0 | 0 | 0 | 0 | 0 | 0 | 0 | 0 | 0 | 2 | 2 |
| HG1016 | *H. sapiens* | ENSG00000254004 | 0 | 0 | 0 | 0 | 0 | 0 | 0 | 0 | 0 | 0 | 0 | 0 | 0 | 0 | 0 | 0 | 0 | 3 | 1 |
| HG1017 | *H. sapiens* | ENSG00000255154 | 0 | 0 | 0 | 0 | 0 | 0 | 0 | 0 | 0 | 0 | 0 | 0 | 0 | 0 | 0 | 0 | 0 | 4 | 0 |
| HG1018 | *H. sapiens* | ENSG00000259120 | 0 | 0 | 0 | 0 | 0 | 0 | 0 | 0 | 0 | 0 | 0 | 0 | 0 | 0 | 0 | 0 | 0 | 3 | 1 |
| HG1019 | *H. sapiens* | ENSG00000019995 | 0 | 0 | 0 | 0 | 0 | 0 | 0 | 0 | 0 | 0 | 0 | 0 | 0 | 0 | 0 | 0 | 0 | 3 | 0 |
| HG1020 | *H. sapiens* | ENSG00000023902 | 0 | 0 | 0 | 0 | 0 | 0 | 0 | 0 | 0 | 0 | 0 | 0 | 0 | 0 | 0 | 0 | 0 | 2 | 1 |
| HG1021 | *H. sapiens* | ENSG00000064692 | 0 | 0 | 0 | 0 | 0 | 0 | 0 | 0 | 0 | 0 | 0 | 0 | 0 | 0 | 0 | 0 | 0 | 2 | 1 |
| HG1022 | *H. sapiens* | ENSG00000065357 | 0 | 0 | 0 | 0 | 0 | 0 | 0 | 0 | 0 | 0 | 0 | 0 | 0 | 0 | 0 | 0 | 0 | 2 | 1 |
| HG1023 | *H. sapiens* | ENSG00000068001 | 0 | 0 | 0 | 0 | 0 | 0 | 0 | 0 | 0 | 0 | 0 | 0 | 0 | 0 | 0 | 0 | 0 | 3 | 0 |
| HG1024 | *H. sapiens* | ENSG00000073464 | 0 | 0 | 0 | 0 | 0 | 0 | 0 | 0 | 0 | 0 | 0 | 0 | 0 | 0 | 0 | 0 | 0 | 2 | 1 |
| HG1025 | *H. sapiens* | ENSG00000074621 | 0 | 0 | 0 | 0 | 0 | 0 | 0 | 0 | 0 | 0 | 0 | 0 | 0 | 0 | 0 | 0 | 0 | 2 | 1 |
| HG1026.1 | *H. sapiens* | ENSG00000075035 | 0 | 0 | 0 | 0 | 0 | 0 | 0 | 0 | 0 | 0 | 0 | 0 | 0 | 0 | 0 | 0 | 0 | 3 | 0 |
| HG1026.2 | *H. sapiens* | ENSG00000179314 | 0 | 0 | 0 | 0 | 0 | 0 | 0 | 0 | 0 | 0 | 0 | 0 | 0 | 0 | 0 | 0 | 0 | 1 | 2 |
| HG1026.3 | *H. sapiens* | ENSG00000179314 | 0 | 0 | 0 | 0 | 0 | 0 | 0 | 0 | 0 | 0 | 0 | 0 | 0 | 0 | 0 | 0 | 0 | 3 | 0 |
| HG1027 | *H. sapiens* | ENSG00000075223 | 0 | 0 | 0 | 0 | 0 | 0 | 0 | 0 | 0 | 0 | 0 | 0 | 0 | 0 | 0 | 0 | 0 | 2 | 1 |
| HG1028 | *H. sapiens* | ENSG00000076356 | 0 | 0 | 0 | 0 | 0 | 0 | 0 | 0 | 0 | 0 | 0 | 0 | 0 | 0 | 0 | 0 | 0 | 3 | 0 |
| HG1029 | *H. sapiens* | ENSG00000079482 | 0 | 0 | 0 | 0 | 0 | 0 | 0 | 0 | 0 | 0 | 0 | 0 | 0 | 0 | 0 | 0 | 0 | 2 | 1 |
| HG1030 | *H. sapiens* | ENSG00000082175 | 0 | 0 | 0 | 0 | 0 | 0 | 0 | 0 | 0 | 0 | 0 | 0 | 0 | 0 | 0 | 0 | 0 | 2 | 1 |
| HG1031 | *H. sapiens* | ENSG00000082269 | 0 | 0 | 0 | 0 | 0 | 0 | 0 | 0 | 0 | 0 | 0 | 0 | 0 | 0 | 0 | 0 | 0 | 3 | 0 |
| HG1032 | *H. sapiens* | ENSG00000084453 | 0 | 0 | 0 | 0 | 0 | 0 | 0 | 0 | 0 | 0 | 0 | 0 | 0 | 0 | 0 | 0 | 0 | 2 | 1 |
| HG1033 | *H. sapiens* | ENSG00000089225 | 0 | 0 | 0 | 0 | 0 | 0 | 0 | 0 | 0 | 0 | 0 | 0 | 0 | 0 | 0 | 0 | 0 | 2 | 1 |
| HG1034 | *H. sapiens* | ENSG00000090487 | 0 | 0 | 0 | 0 | 0 | 0 | 0 | 0 | 0 | 0 | 0 | 0 | 0 | 0 | 0 | 0 | 0 | 3 | 0 |
| HG1035 | *H. sapiens* | ENSG00000093000 | 0 | 0 | 0 | 0 | 0 | 0 | 0 | 0 | 0 | 0 | 0 | 0 | 0 | 0 | 0 | 0 | 0 | 3 | 0 |
| HG1036 | *H. sapiens* | ENSG00000100813 | 0 | 0 | 0 | 0 | 0 | 0 | 0 | 0 | 0 | 0 | 0 | 0 | 0 | 0 | 0 | 0 | 0 | 2 | 1 |
| HG1037 | *H. sapiens* | ENSG00000102710 | 0 | 0 | 0 | 0 | 0 | 0 | 0 | 0 | 0 | 0 | 0 | 0 | 0 | 0 | 0 | 0 | 0 | 2 | 1 |
| HG1038 | *H. sapiens* | ENSG00000104331 | 0 | 0 | 0 | 0 | 0 | 0 | 0 | 0 | 0 | 0 | 0 | 0 | 0 | 0 | 0 | 0 | 0 | 2 | 1 |
| HG1039 | *H. sapiens* | ENSG00000104804 | 0 | 0 | 0 | 0 | 0 | 0 | 0 | 0 | 0 | 0 | 0 | 0 | 0 | 0 | 0 | 0 | 0 | 3 | 0 |
| HG1040 | *H. sapiens* | ENSG00000105223 | 0 | 0 | 0 | 0 | 0 | 0 | 0 | 0 | 0 | 0 | 0 | 0 | 0 | 0 | 0 | 0 | 0 | 1 | 2 |
| HG1041 | *H. sapiens* | ENSG00000106536 | 0 | 0 | 0 | 0 | 0 | 0 | 0 | 0 | 0 | 0 | 0 | 0 | 0 | 0 | 0 | 0 | 0 | 3 | 0 |
| HG1042 | *H. sapiens* | ENSG00000107249 | 0 | 0 | 0 | 0 | 0 | 0 | 0 | 0 | 0 | 0 | 0 | 0 | 0 | 0 | 0 | 0 | 0 | 3 | 0 |
| HG1043 | *H. sapiens* | ENSG00000107736 | 0 | 0 | 0 | 0 | 0 | 0 | 0 | 0 | 0 | 0 | 0 | 0 | 0 | 0 | 0 | 0 | 0 | 2 | 1 |
| HG1044 | *H. sapiens* | ENSG00000109381 | 0 | 0 | 0 | 0 | 0 | 0 | 0 | 0 | 0 | 0 | 0 | 0 | 0 | 0 | 0 | 0 | 0 | 2 | 1 |
| HG1045 | *H. sapiens* | ENSG00000110925 | 0 | 0 | 0 | 0 | 0 | 0 | 0 | 0 | 0 | 0 | 0 | 0 | 0 | 0 | 0 | 0 | 0 | 2 | 1 |
| HG1046 | *H. sapiens* | ENSG00000112701 | 0 | 0 | 0 | 0 | 0 | 0 | 0 | 0 | 0 | 0 | 0 | 0 | 0 | 0 | 0 | 0 | 0 | 2 | 1 |
| HG1047 | *H. sapiens* | ENSG00000112769 | 0 | 0 | 0 | 0 | 0 | 0 | 0 | 0 | 0 | 0 | 0 | 0 | 0 | 0 | 0 | 0 | 0 | 3 | 0 |
| HG1048 | *H. sapiens* | ENSG00000115295 | 0 | 0 | 0 | 0 | 0 | 0 | 0 | 0 | 0 | 0 | 0 | 0 | 0 | 0 | 0 | 0 | 0 | 2 | 1 |
| HG1049 | *H. sapiens* | ENSG00000116212 | 0 | 0 | 0 | 0 | 0 | 0 | 0 | 0 | 0 | 0 | 0 | 0 | 0 | 0 | 0 | 0 | 0 | 3 | 0 |
| HG1050 | *H. sapiens* | ENSG00000116489 | 0 | 0 | 0 | 0 | 0 | 0 | 0 | 0 | 0 | 0 | 0 | 0 | 0 | 0 | 0 | 0 | 0 | 1 | 2 |
| HG1051 | *H. sapiens* | ENSG00000116852 | 0 | 0 | 0 | 0 | 0 | 0 | 0 | 0 | 0 | 0 | 0 | 0 | 0 | 0 | 0 | 0 | 0 | 2 | 1 |
| HG1052 | *H. sapiens* | ENSG00000118640 | 0 | 0 | 0 | 0 | 0 | 0 | 0 | 0 | 0 | 0 | 0 | 0 | 0 | 0 | 0 | 0 | 0 | 2 | 1 |
| HG1053 | *H. sapiens* | ENSG00000119522 | 0 | 0 | 0 | 0 | 0 | 0 | 0 | 0 | 0 | 0 | 0 | 0 | 0 | 0 | 0 | 0 | 0 | 3 | 0 |
| HG1054 | *H. sapiens* | ENSG00000120832 | 0 | 0 | 0 | 0 | 0 | 0 | 0 | 0 | 0 | 0 | 0 | 0 | 0 | 0 | 0 | 0 | 0 | 2 | 1 |
| HG1055 | *H. sapiens* | ENSG00000122299 | 0 | 0 | 0 | 0 | 0 | 0 | 0 | 0 | 0 | 0 | 0 | 0 | 0 | 0 | 0 | 0 | 0 | 1 | 2 |
| HG1056 | *H. sapiens* | ENSG00000128271 | 0 | 0 | 0 | 0 | 0 | 0 | 0 | 0 | 0 | 0 | 0 | 0 | 0 | 0 | 0 | 0 | 0 | 2 | 1 |
| HG1057 | *H. sapiens* | ENSG00000131097 | 0 | 0 | 0 | 0 | 0 | 0 | 0 | 0 | 0 | 0 | 0 | 0 | 0 | 0 | 0 | 0 | 0 | 1 | 2 |
| HG1058 | *H. sapiens* | ENSG00000131386 | 0 | 0 | 0 | 0 | 0 | 0 | 0 | 0 | 0 | 0 | 0 | 0 | 0 | 0 | 0 | 0 | 0 | 1 | 2 |
| HG1059.1 | *H. sapiens* | ENSG00000132970 | 0 | 0 | 0 | 0 | 0 | 0 | 0 | 0 | 0 | 0 | 0 | 0 | 0 | 0 | 0 | 0 | 0 | 3 | 0 |
| HG1059.2 | *H. sapiens* | ENSG00000112290 | 0 | 0 | 0 | 0 | 0 | 0 | 0 | 0 | 0 | 0 | 0 | 0 | 0 | 0 | 0 | 0 | 0 | 1 | 1 |
| HG1060 | *H. sapiens* | ENSG00000134242 | 0 | 0 | 0 | 0 | 0 | 0 | 0 | 0 | 0 | 0 | 0 | 0 | 0 | 0 | 0 | 0 | 0 | 2 | 1 |
| HG1061 | *H. sapiens* | ENSG00000134802 | 0 | 0 | 0 | 0 | 0 | 0 | 0 | 0 | 0 | 0 | 0 | 0 | 0 | 0 | 0 | 0 | 0 | 1 | 2 |
| HG1062 | *H. sapiens* | ENSG00000137337 | 0 | 0 | 0 | 0 | 0 | 0 | 0 | 0 | 0 | 0 | 0 | 0 | 0 | 0 | 0 | 0 | 0 | 2 | 1 |
| HG1063 | *H. sapiens* | ENSG00000138162 | 0 | 0 | 0 | 0 | 0 | 0 | 0 | 0 | 0 | 0 | 0 | 0 | 0 | 0 | 0 | 0 | 0 | 3 | 0 |
| HG1064 | *H. sapiens* | ENSG00000139734 | 0 | 0 | 0 | 0 | 0 | 0 | 0 | 0 | 0 | 0 | 0 | 0 | 0 | 0 | 0 | 0 | 0 | 2 | 1 |
| HG1065 | *H. sapiens* | ENSG00000140030 | 0 | 0 | 0 | 0 | 0 | 0 | 0 | 0 | 0 | 0 | 0 | 0 | 0 | 0 | 0 | 0 | 0 | 1 | 2 |
| HG1066 | *H. sapiens* | ENSG00000140553 | 0 | 0 | 0 | 0 | 0 | 0 | 0 | 0 | 0 | 0 | 0 | 0 | 0 | 0 | 0 | 0 | 0 | 2 | 1 |
| HG1067 | *H. sapiens* | ENSG00000143340 | 0 | 0 | 0 | 0 | 0 | 0 | 0 | 0 | 0 | 0 | 0 | 0 | 0 | 0 | 0 | 0 | 0 | 3 | 0 |
| HG1068 | *H. sapiens* | ENSG00000144791 | 0 | 0 | 0 | 0 | 0 | 0 | 0 | 0 | 0 | 0 | 0 | 0 | 0 | 0 | 0 | 0 | 0 | 2 | 1 |
| HG1069 | *H. sapiens* | ENSG00000146414 | 0 | 0 | 0 | 0 | 0 | 0 | 0 | 0 | 0 | 0 | 0 | 0 | 0 | 0 | 0 | 0 | 0 | 3 | 0 |
| HG1070 | *H. sapiens* | ENSG00000147459 | 0 | 0 | 0 | 0 | 0 | 0 | 0 | 0 | 0 | 0 | 0 | 0 | 0 | 0 | 0 | 0 | 0 | 2 | 1 |
| HG1071 | *H. sapiens* | ENSG00000147573 | 0 | 0 | 0 | 0 | 0 | 0 | 0 | 0 | 0 | 0 | 0 | 0 | 0 | 0 | 0 | 0 | 0 | 0 | 3 |
| HG1072 | *H. sapiens* | ENSG00000147874 | 0 | 0 | 0 | 0 | 0 | 0 | 0 | 0 | 0 | 0 | 0 | 0 | 0 | 0 | 0 | 0 | 0 | 2 | 1 |
| HG1073 | *H. sapiens* | ENSG00000149187 | 0 | 0 | 0 | 0 | 0 | 0 | 0 | 0 | 0 | 0 | 0 | 0 | 0 | 0 | 0 | 0 | 0 | 2 | 1 |
| HG1074 | *H. sapiens* | ENSG00000150275 | 0 | 0 | 0 | 0 | 0 | 0 | 0 | 0 | 0 | 0 | 0 | 0 | 0 | 0 | 0 | 0 | 0 | 3 | 0 |
| HG1075 | *H. sapiens* | ENSG00000150347 | 0 | 0 | 0 | 0 | 0 | 0 | 0 | 0 | 0 | 0 | 0 | 0 | 0 | 0 | 0 | 0 | 0 | 2 | 1 |
| HG1076 | *H. sapiens* | ENSG00000150967 | 0 | 0 | 0 | 0 | 0 | 0 | 0 | 0 | 0 | 0 | 0 | 0 | 0 | 0 | 0 | 0 | 0 | 1 | 2 |
| HG1077 | *H. sapiens* | ENSG00000151474 | 0 | 0 | 0 | 0 | 0 | 0 | 0 | 0 | 0 | 0 | 0 | 0 | 0 | 0 | 0 | 0 | 0 | 1 | 2 |
| HG1078 | *H. sapiens* | ENSG00000162607 | 0 | 0 | 0 | 0 | 0 | 0 | 0 | 0 | 0 | 0 | 0 | 0 | 0 | 0 | 0 | 0 | 0 | 2 | 1 |
| HG1079 | *H. sapiens* | ENSG00000162975 | 0 | 0 | 0 | 0 | 0 | 0 | 0 | 0 | 0 | 0 | 0 | 0 | 0 | 0 | 0 | 0 | 0 | 1 | 2 |
| HG1080 | *H. sapiens* | ENSG00000164168 | 0 | 0 | 0 | 0 | 0 | 0 | 0 | 0 | 0 | 0 | 0 | 0 | 0 | 0 | 0 | 0 | 0 | 2 | 1 |
| HG1081 | *H. sapiens* | ENSG00000165392 | 0 | 0 | 0 | 0 | 0 | 0 | 0 | 0 | 0 | 0 | 0 | 0 | 0 | 0 | 0 | 0 | 0 | 2 | 1 |

| | | | | | | | | | | | | | | | | | | | | | |
|---|---|---|---|---|---|---|---|---|---|---|---|---|---|---|---|---|---|---|---|---|---|
| HG1082 | *H. sapiens* | ENSG00000165434 | 0 | 0 | 0 | 0 | 0 | 0 | 0 | 0 | 0 | 0 | 0 | 0 | 0 | 0 | 0 | 0 | 0 | 1 | 2 |
| HG1083 | *H. sapiens* | ENSG00000165804 | 0 | 0 | 0 | 0 | 0 | 0 | 0 | 0 | 0 | 0 | 0 | 0 | 0 | 0 | 0 | 0 | 0 | 2 | 1 |
| HG1084 | *H. sapiens* | ENSG00000165923 | 0 | 0 | 0 | 0 | 0 | 0 | 0 | 0 | 0 | 0 | 0 | 0 | 0 | 0 | 0 | 0 | 0 | 3 | 0 |
| HG1085 | *H. sapiens* | ENSG00000166532 | 0 | 0 | 0 | 0 | 0 | 0 | 0 | 0 | 0 | 0 | 0 | 0 | 0 | 0 | 0 | 0 | 0 | 2 | 1 |
| HG1086 | *H. sapiens* | ENSG00000167377 | 0 | 0 | 0 | 0 | 0 | 0 | 0 | 0 | 0 | 0 | 0 | 0 | 0 | 0 | 0 | 0 | 0 | 2 | 1 |
| HG1087 | *H. sapiens* | ENSG00000168675 | 0 | 0 | 0 | 0 | 0 | 0 | 0 | 0 | 0 | 0 | 0 | 0 | 0 | 0 | 0 | 0 | 0 | 0 | 3 |
| HG1088 | *H. sapiens* | ENSG00000169740 | 0 | 0 | 0 | 0 | 0 | 0 | 0 | 0 | 0 | 0 | 0 | 0 | 0 | 0 | 0 | 0 | 0 | 2 | 1 |
| HG1089 | *H. sapiens* | ENSG00000169813 | 0 | 0 | 0 | 0 | 0 | 0 | 0 | 0 | 0 | 0 | 0 | 0 | 0 | 0 | 0 | 0 | 0 | 2 | 1 |
| HG1090.1 | *H. sapiens* | ENSG00000170458 | 0 | 0 | 0 | 0 | 0 | 0 | 0 | 0 | 0 | 0 | 0 | 0 | 0 | 0 | 0 | 0 | 0 | 1 | 2 |
| HG1090.2 | *H. sapiens* | ENSG00000170458 | 0 | 0 | 0 | 0 | 0 | 0 | 0 | 0 | 0 | 0 | 0 | 0 | 0 | 0 | 0 | 0 | 0 | 3 | 0 |
| HG1091 | *H. sapiens* | ENSG00000172183 | 0 | 0 | 0 | 0 | 0 | 0 | 0 | 0 | 0 | 0 | 0 | 0 | 0 | 0 | 0 | 0 | 0 | 3 | 0 |
| HG1092 | *H. sapiens* | ENSG00000173327 | 0 | 0 | 0 | 0 | 0 | 0 | 0 | 0 | 0 | 0 | 0 | 0 | 0 | 0 | 0 | 0 | 0 | 2 | 1 |
| HG1093 | *H. sapiens* | ENSG00000174371 | 0 | 0 | 0 | 0 | 0 | 0 | 0 | 0 | 0 | 0 | 0 | 0 | 0 | 0 | 0 | 0 | 0 | 2 | 1 |
| HG1094 | *H. sapiens* | ENSG00000177917 | 0 | 0 | 0 | 0 | 0 | 0 | 0 | 0 | 0 | 0 | 0 | 0 | 0 | 0 | 0 | 0 | 0 | 2 | 1 |
| HG1095 | *H. sapiens* | ENSG00000179921 | 0 | 0 | 0 | 0 | 0 | 0 | 0 | 0 | 0 | 0 | 0 | 0 | 0 | 0 | 0 | 0 | 0 | 3 | 0 |
| HG1096 | *H. sapiens* | ENSG00000181544 | 0 | 0 | 0 | 0 | 0 | 0 | 0 | 0 | 0 | 0 | 0 | 0 | 0 | 0 | 0 | 0 | 0 | 2 | 1 |
| HG1097 | *H. sapiens* | ENSG00000182481 | 0 | 0 | 0 | 0 | 0 | 0 | 0 | 0 | 0 | 0 | 0 | 0 | 0 | 0 | 0 | 0 | 0 | 2 | 1 |
| HG1098 | *H. sapiens* | ENSG00000183775 | 0 | 0 | 0 | 0 | 0 | 0 | 0 | 0 | 0 | 0 | 0 | 0 | 0 | 0 | 0 | 0 | 0 | 2 | 1 |
| HG1099 | *H. sapiens* | ENSG00000186334 | 0 | 0 | 0 | 0 | 0 | 0 | 0 | 0 | 0 | 0 | 0 | 0 | 0 | 0 | 0 | 0 | 0 | 1 | 2 |
| HG1100 | *H. sapiens* | ENSG00000187815 | 0 | 0 | 0 | 0 | 0 | 0 | 0 | 0 | 0 | 0 | 0 | 0 | 0 | 0 | 0 | 0 | 0 | 2 | 1 |
| HG1101 | *H. sapiens* | ENSG00000189334 | 0 | 0 | 0 | 0 | 0 | 0 | 0 | 0 | 0 | 0 | 0 | 0 | 0 | 0 | 0 | 0 | 0 | 1 | 2 |
| HG1102 | *H. sapiens* | ENSG00000196704 | 0 | 0 | 0 | 0 | 0 | 0 | 0 | 0 | 0 | 0 | 0 | 0 | 0 | 0 | 0 | 0 | 0 | 1 | 2 |
| HG1103 | *H. sapiens* | ENSG00000197620 | 0 | 0 | 0 | 0 | 0 | 0 | 0 | 0 | 0 | 0 | 0 | 0 | 0 | 0 | 0 | 0 | 0 | 3 | 0 |
| HG1104 | *H. sapiens* | ENSG00000205155 | 0 | 0 | 0 | 0 | 0 | 0 | 0 | 0 | 0 | 0 | 0 | 0 | 0 | 0 | 0 | 0 | 0 | 1 | 2 |
| HG1105 | *H. sapiens* | ENSG00000213654 | 0 | 0 | 0 | 0 | 0 | 0 | 0 | 0 | 0 | 0 | 0 | 0 | 0 | 0 | 0 | 0 | 0 | 2 | 1 |
| HG1106 | *H. sapiens* | ENSG00000215021 | 0 | 0 | 0 | 0 | 0 | 0 | 0 | 0 | 0 | 0 | 0 | 0 | 0 | 0 | 0 | 0 | 0 | 2 | 1 |
| HG1107 | *H. sapiens* | ENSG00000217930 | 0 | 0 | 0 | 0 | 0 | 0 | 0 | 0 | 0 | 0 | 0 | 0 | 0 | 0 | 0 | 0 | 0 | 2 | 1 |
| HG1108 | *H. sapiens* | ENSG00000251247 | 0 | 0 | 0 | 0 | 0 | 0 | 0 | 0 | 0 | 0 | 0 | 0 | 0 | 0 | 0 | 0 | 0 | 3 | 0 |
| HG1109 | *H. sapiens* | ENSG00000278311 | 0 | 0 | 0 | 0 | 0 | 0 | 0 | 0 | 0 | 0 | 0 | 0 | 0 | 0 | 0 | 0 | 0 | 2 | 1 |
| HG1110 | *H. sapiens* | ENSG00000034152 | 0 | 0 | 0 | 0 | 0 | 0 | 0 | 0 | 0 | 0 | 0 | 0 | 0 | 0 | 0 | 0 | 0 | 1 | 1 |
| HG1111 | *H. sapiens* | ENSG00000040531 | 0 | 0 | 0 | 0 | 0 | 0 | 0 | 0 | 0 | 0 | 0 | 0 | 0 | 0 | 0 | 0 | 0 | 1 | 1 |
| HG1112 | *H. sapiens* | ENSG00000068885 | 0 | 0 | 0 | 0 | 0 | 0 | 0 | 0 | 0 | 0 | 0 | 0 | 0 | 0 | 0 | 0 | 0 | 1 | 1 |
| HG1113 | *H. sapiens* | ENSG00000072201 | 0 | 0 | 0 | 0 | 0 | 0 | 0 | 0 | 0 | 0 | 0 | 0 | 0 | 0 | 0 | 0 | 0 | 0 | 2 |
| HG1114 | *H. sapiens* | ENSG00000073737 | 0 | 0 | 0 | 0 | 0 | 0 | 0 | 0 | 0 | 0 | 0 | 0 | 0 | 0 | 0 | 0 | 0 | 1 | 1 |
| HG1115 | *H. sapiens* | ENSG00000074219 | 0 | 0 | 0 | 0 | 0 | 0 | 0 | 0 | 0 | 0 | 0 | 0 | 0 | 0 | 0 | 0 | 0 | 2 | 0 |
| HG1116 | *H. sapiens* | ENSG00000084090 | 0 | 0 | 0 | 0 | 0 | 0 | 0 | 0 | 0 | 0 | 0 | 0 | 0 | 0 | 0 | 0 | 0 | 0 | 2 |
| HG1117 | *H. sapiens* | ENSG00000089902 | 0 | 0 | 0 | 0 | 0 | 0 | 0 | 0 | 0 | 0 | 0 | 0 | 0 | 0 | 0 | 0 | 0 | 1 | 1 |
| HG1118 | *H. sapiens* | ENSG00000090975 | 0 | 0 | 0 | 0 | 0 | 0 | 0 | 0 | 0 | 0 | 0 | 0 | 0 | 0 | 0 | 0 | 0 | 1 | 1 |
| HG1119 | *H. sapiens* | ENSG00000092020 | 0 | 0 | 0 | 0 | 0 | 0 | 0 | 0 | 0 | 0 | 0 | 0 | 0 | 0 | 0 | 0 | 0 | 2 | 0 |
| HG1120 | *H. sapiens* | ENSG00000095585 | 0 | 0 | 0 | 0 | 0 | 0 | 0 | 0 | 0 | 0 | 0 | 0 | 0 | 0 | 0 | 0 | 0 | 1 | 1 |
| HG1121 | *H. sapiens* | ENSG00000100350 | 0 | 0 | 0 | 0 | 0 | 0 | 0 | 0 | 0 | 0 | 0 | 0 | 0 | 0 | 0 | 0 | 0 | 2 | 0 |
| HG1122 | *H. sapiens* | ENSG00000100410 | 0 | 0 | 0 | 0 | 0 | 0 | 0 | 0 | 0 | 0 | 0 | 0 | 0 | 0 | 0 | 0 | 0 | 1 | 1 |
| HG1123 | *H. sapiens* | ENSG00000100884 | 0 | 0 | 0 | 0 | 0 | 0 | 0 | 0 | 0 | 0 | 0 | 0 | 0 | 0 | 0 | 0 | 0 | 1 | 1 |
| HG1124 | *H. sapiens* | ENSG00000100934 | 0 | 0 | 0 | 0 | 0 | 0 | 0 | 0 | 0 | 0 | 0 | 0 | 0 | 0 | 0 | 0 | 0 | 0 | 2 |
| HG1125 | *H. sapiens* | ENSG00000101574 | 0 | 0 | 0 | 0 | 0 | 0 | 0 | 0 | 0 | 0 | 0 | 0 | 0 | 0 | 0 | 0 | 0 | 2 | 0 |
| HG1126 | *H. sapiens* | ENSG00000101890 | 0 | 0 | 0 | 0 | 0 | 0 | 0 | 0 | 0 | 0 | 0 | 0 | 0 | 0 | 0 | 0 | 0 | 1 | 1 |
| HG1127 | *H. sapiens* | ENSG00000103064 | 0 | 0 | 0 | 0 | 0 | 0 | 0 | 0 | 0 | 0 | 0 | 0 | 0 | 0 | 0 | 0 | 0 | 1 | 1 |
| HG1128 | *H. sapiens* | ENSG00000103852 | 0 | 0 | 0 | 0 | 0 | 0 | 0 | 0 | 0 | 0 | 0 | 0 | 0 | 0 | 0 | 0 | 0 | 1 | 1 |
| HG1129 | *H. sapiens* | ENSG00000104432 | 0 | 0 | 0 | 0 | 0 | 0 | 0 | 0 | 0 | 0 | 0 | 0 | 0 | 0 | 0 | 0 | 0 | 1 | 1 |
| HG1130 | *H. sapiens* | ENSG00000105559 | 0 | 0 | 0 | 0 | 0 | 0 | 0 | 0 | 0 | 0 | 0 | 0 | 0 | 0 | 0 | 0 | 0 | 2 | 0 |
| HG1131 | *H. sapiens* | ENSG00000106780 | 0 | 0 | 0 | 0 | 0 | 0 | 0 | 0 | 0 | 0 | 0 | 0 | 0 | 0 | 0 | 0 | 0 | 1 | 1 |
| HG1132 | *H. sapiens* | ENSG00000109771 | 0 | 0 | 0 | 0 | 0 | 0 | 0 | 0 | 0 | 0 | 0 | 0 | 0 | 0 | 0 | 0 | 0 | 1 | 1 |
| HG1133 | *H. sapiens* | ENSG00000111328 | 0 | 0 | 0 | 0 | 0 | 0 | 0 | 0 | 0 | 0 | 0 | 0 | 0 | 0 | 0 | 0 | 0 | 1 | 1 |
| HG1134 | *H. sapiens* | ENSG00000112984 | 0 | 0 | 0 | 0 | 0 | 0 | 0 | 0 | 0 | 0 | 0 | 0 | 0 | 0 | 0 | 0 | 0 | 1 | 1 |
| HG1135 | *H. sapiens* | ENSG00000113597 | 0 | 0 | 0 | 0 | 0 | 0 | 0 | 0 | 0 | 0 | 0 | 0 | 0 | 0 | 0 | 0 | 0 | 1 | 1 |
| HG1136 | *H. sapiens* | ENSG00000113749 | 0 | 0 | 0 | 0 | 0 | 0 | 0 | 0 | 0 | 0 | 0 | 0 | 0 | 0 | 0 | 0 | 0 | 2 | 0 |
| HG1137 | *H. sapiens* | ENSG00000115307 | 0 | 0 | 0 | 0 | 0 | 0 | 0 | 0 | 0 | 0 | 0 | 0 | 0 | 0 | 0 | 0 | 0 | 1 | 1 |
| HG1138 | *H. sapiens* | ENSG00000116871 | 0 | 0 | 0 | 0 | 0 | 0 | 0 | 0 | 0 | 0 | 0 | 0 | 0 | 0 | 0 | 0 | 0 | 1 | 1 |
| HG1139.1 | *H. sapiens* | ENSG00000118308 | 0 | 0 | 0 | 0 | 0 | 0 | 0 | 0 | 0 | 0 | 0 | 0 | 0 | 0 | 0 | 0 | 0 | 5 | 0 |
| HG1139.2 | *H. sapiens* | ENSG00000118308 | 0 | 0 | 0 | 0 | 0 | 0 | 0 | 0 | 0 | 0 | 0 | 0 | 0 | 0 | 0 | 0 | 0 | 2 | 0 |
| HG1140 | *H. sapiens* | ENSG00000118495 | 0 | 0 | 0 | 0 | 0 | 0 | 0 | 0 | 0 | 0 | 0 | 0 | 0 | 0 | 0 | 0 | 0 | 1 | 1 |
| HG1141 | *H. sapiens* | ENSG00000121964 | 0 | 0 | 0 | 0 | 0 | 0 | 0 | 0 | 0 | 0 | 0 | 0 | 0 | 0 | 0 | 0 | 0 | 1 | 1 |
| HG1142 | *H. sapiens* | ENSG00000124613 | 0 | 0 | 0 | 0 | 0 | 0 | 0 | 0 | 0 | 0 | 0 | 0 | 0 | 0 | 0 | 0 | 0 | 1 | 1 |
| HG1143 | *H. sapiens* | ENSG00000125245 | 0 | 0 | 0 | 0 | 0 | 0 | 0 | 0 | 0 | 0 | 0 | 0 | 0 | 0 | 0 | 0 | 0 | 1 | 1 |
| HG1144 | *H. sapiens* | ENSG00000128203 | 0 | 0 | 0 | 0 | 0 | 0 | 0 | 0 | 0 | 0 | 0 | 0 | 0 | 0 | 0 | 0 | 0 | 2 | 0 |
| HG1145 | *H. sapiens* | ENSG00000128585 | 0 | 0 | 0 | 0 | 0 | 0 | 0 | 0 | 0 | 0 | 0 | 0 | 0 | 0 | 0 | 0 | 0 | 2 | 0 |
| HG1146 | *H. sapiens* | ENSG00000130413 | 0 | 0 | 0 | 0 | 0 | 0 | 0 | 0 | 0 | 0 | 0 | 0 | 0 | 0 | 0 | 0 | 0 | 1 | 1 |
| HG1147 | *H. sapiens* | ENSG00000131781 | 0 | 0 | 0 | 0 | 0 | 0 | 0 | 0 | 0 | 0 | 0 | 0 | 0 | 0 | 0 | 0 | 0 | 2 | 0 |
| HG1148 | *H. sapiens* | ENSG00000132938 | 0 | 0 | 0 | 0 | 0 | 0 | 0 | 0 | 0 | 0 | 0 | 0 | 0 | 0 | 0 | 0 | 0 | 2 | 0 |
| HG1149 | *H. sapiens* | ENSG00000132975 | 0 | 0 | 0 | 0 | 0 | 0 | 0 | 0 | 0 | 0 | 0 | 0 | 0 | 0 | 0 | 0 | 0 | 1 | 1 |
| HG1150 | *H. sapiens* | ENSG00000135299 | 0 | 0 | 0 | 0 | 0 | 0 | 0 | 0 | 0 | 0 | 0 | 0 | 0 | 0 | 0 | 0 | 0 | 1 | 1 |
| HG1151 | *H. sapiens* | ENSG00000137168 | 0 | 0 | 0 | 0 | 0 | 0 | 0 | 0 | 0 | 0 | 0 | 0 | 0 | 0 | 0 | 0 | 0 | 1 | 1 |
| HG1152 | *H. sapiens* | ENSG00000138658 | 0 | 0 | 0 | 0 | 0 | 0 | 0 | 0 | 0 | 0 | 0 | 0 | 0 | 0 | 0 | 0 | 0 | 1 | 1 |
| HG1153 | *H. sapiens* | ENSG00000138738 | 0 | 0 | 0 | 0 | 0 | 0 | 0 | 0 | 0 | 0 | 0 | 0 | 0 | 0 | 0 | 0 | 0 | 1 | 1 |
| HG1154 | *H. sapiens* | ENSG00000139725 | 0 | 0 | 0 | 0 | 0 | 0 | 0 | 0 | 0 | 0 | 0 | 0 | 0 | 0 | 0 | 0 | 0 | 1 | 1 |
| HG1155 | *H. sapiens* | ENSG00000141668 | 0 | 0 | 0 | 0 | 0 | 0 | 0 | 0 | 0 | 0 | 0 | 0 | 0 | 0 | 0 | 0 | 0 | 1 | 1 |
| HG1156 | *H. sapiens* | ENSG00000144741 | 0 | 0 | 0 | 0 | 0 | 0 | 0 | 0 | 0 | 0 | 0 | 0 | 0 | 0 | 0 | 0 | 0 | 2 | 0 |
| HG1157 | *H. sapiens* | ENSG00000148123 | 0 | 0 | 0 | 0 | 0 | 0 | 0 | 0 | 0 | 0 | 0 | 0 | 0 | 0 | 0 | 0 | 0 | 2 | 0 |
| HG1158 | *H. sapiens* | ENSG00000149571 | 0 | 0 | 0 | 0 | 0 | 0 | 0 | 0 | 0 | 0 | 0 | 0 | 0 | 0 | 0 | 0 | 0 | 1 | 1 |

| | | | | | | | | | | | | | | | | | | | | | |
|---|---|---|---|---|---|---|---|---|---|---|---|---|---|---|---|---|---|---|---|---|---|
| HG1159 | *H. sapiens* | ENSG00000152822 | 0 | 0 | 0 | 0 | 0 | 0 | 0 | 0 | 0 | 0 | 0 | 0 | 0 | 0 | 0 | 0 | 0 | 1 | 1 |
| HG1160 | *H. sapiens* | ENSG00000154162 | 0 | 0 | 0 | 0 | 0 | 0 | 0 | 0 | 0 | 0 | 0 | 0 | 0 | 0 | 0 | 0 | 0 | 1 | 1 |
| HG1161 | *H. sapiens* | ENSG00000154227 | 0 | 0 | 0 | 0 | 0 | 0 | 0 | 0 | 0 | 0 | 0 | 0 | 0 | 0 | 0 | 0 | 0 | 1 | 1 |
| HG1162 | *H. sapiens* | ENSG00000155380 | 0 | 0 | 0 | 0 | 0 | 0 | 0 | 0 | 0 | 0 | 0 | 0 | 0 | 0 | 0 | 0 | 0 | 2 | 0 |
| HG1163 | *H. sapiens* | ENSG00000157637 | 0 | 0 | 0 | 0 | 0 | 0 | 0 | 0 | 0 | 0 | 0 | 0 | 0 | 0 | 0 | 0 | 0 | 1 | 1 |
| HG1164 | *H. sapiens* | ENSG00000162639 | 0 | 0 | 0 | 0 | 0 | 0 | 0 | 0 | 0 | 0 | 0 | 0 | 0 | 0 | 0 | 0 | 0 | 1 | 1 |
| HG1165 | *H. sapiens* | ENSG00000162694 | 0 | 0 | 0 | 0 | 0 | 0 | 0 | 0 | 0 | 0 | 0 | 0 | 0 | 0 | 0 | 0 | 0 | 2 | 0 |
| HG1166 | *H. sapiens* | ENSG00000165071 | 0 | 0 | 0 | 0 | 0 | 0 | 0 | 0 | 0 | 0 | 0 | 0 | 0 | 0 | 0 | 0 | 0 | 1 | 1 |
| HG1167 | *H. sapiens* | ENSG00000165934 | 0 | 0 | 0 | 0 | 0 | 0 | 0 | 0 | 0 | 0 | 0 | 0 | 0 | 0 | 0 | 0 | 0 | 2 | 0 |
| HG1168 | *H. sapiens* | ENSG00000166669 | 0 | 0 | 0 | 0 | 0 | 0 | 0 | 0 | 0 | 0 | 0 | 0 | 0 | 0 | 0 | 0 | 0 | 1 | 1 |
| HG1169 | *H. sapiens* | ENSG00000167637 | 0 | 0 | 0 | 0 | 0 | 0 | 0 | 0 | 0 | 0 | 0 | 0 | 0 | 0 | 0 | 0 | 0 | 2 | 0 |
| HG1170 | *H. sapiens* | ENSG00000167858 | 0 | 0 | 0 | 0 | 0 | 0 | 0 | 0 | 0 | 0 | 0 | 0 | 0 | 0 | 0 | 0 | 0 | 2 | 0 |
| HG1171 | *H. sapiens* | ENSG00000167968 | 0 | 0 | 0 | 0 | 0 | 0 | 0 | 0 | 0 | 0 | 0 | 0 | 0 | 0 | 0 | 0 | 0 | 1 | 1 |
| HG1172 | *H. sapiens* | ENSG00000170381 | 0 | 0 | 0 | 0 | 0 | 0 | 0 | 0 | 0 | 0 | 0 | 0 | 0 | 0 | 0 | 0 | 0 | 2 | 0 |
| HG1173 | *H. sapiens* | ENSG00000170464 | 0 | 0 | 0 | 0 | 0 | 0 | 0 | 0 | 0 | 0 | 0 | 0 | 0 | 0 | 0 | 0 | 0 | 2 | 0 |
| HG1174 | *H. sapiens* | ENSG00000173918 | 0 | 0 | 0 | 0 | 0 | 0 | 0 | 0 | 0 | 0 | 0 | 0 | 0 | 0 | 0 | 0 | 0 | 2 | 0 |
| HG1175 | *H. sapiens* | ENSG00000174473 | 0 | 0 | 0 | 0 | 0 | 0 | 0 | 0 | 0 | 0 | 0 | 0 | 0 | 0 | 0 | 0 | 0 | 1 | 1 |
| HG1176 | *H. sapiens* | ENSG00000176472 | 0 | 0 | 0 | 0 | 0 | 0 | 0 | 0 | 0 | 0 | 0 | 0 | 0 | 0 | 0 | 0 | 0 | 1 | 1 |
| HG1177 | *H. sapiens* | ENSG00000178295 | 0 | 0 | 0 | 0 | 0 | 0 | 0 | 0 | 0 | 0 | 0 | 0 | 0 | 0 | 0 | 0 | 0 | 2 | 0 |
| HG1178 | *H. sapiens* | ENSG00000182670 | 0 | 0 | 0 | 0 | 0 | 0 | 0 | 0 | 0 | 0 | 0 | 0 | 0 | 0 | 0 | 0 | 0 | 2 | 0 |
| HG1179 | *H. sapiens* | ENSG00000185933 | 0 | 0 | 0 | 0 | 0 | 0 | 0 | 0 | 0 | 0 | 0 | 0 | 0 | 0 | 0 | 0 | 0 | 1 | 1 |
| HG1180 | *H. sapiens* | ENSG00000188674 | 0 | 0 | 0 | 0 | 0 | 0 | 0 | 0 | 0 | 0 | 0 | 0 | 0 | 0 | 0 | 0 | 0 | 2 | 0 |
| HG1181 | *H. sapiens* | ENSG00000189058 | 0 | 0 | 0 | 0 | 0 | 0 | 0 | 0 | 0 | 0 | 0 | 0 | 0 | 0 | 0 | 0 | 0 | 2 | 0 |
| HG1182 | *H. sapiens* | ENSG00000196155 | 0 | 0 | 0 | 0 | 0 | 0 | 0 | 0 | 0 | 0 | 0 | 0 | 0 | 0 | 0 | 0 | 0 | 2 | 0 |
| HG1183 | *H. sapiens* | ENSG00000198133 | 0 | 0 | 0 | 0 | 0 | 0 | 0 | 0 | 0 | 0 | 0 | 0 | 0 | 0 | 0 | 0 | 0 | 1 | 1 |
| HG1184 | *H. sapiens* | ENSG00000198894 | 0 | 0 | 0 | 0 | 0 | 0 | 0 | 0 | 0 | 0 | 0 | 0 | 0 | 0 | 0 | 0 | 0 | 1 | 1 |
| HG1185 | *H. sapiens* | ENSG00000198920 | 0 | 0 | 0 | 0 | 0 | 0 | 0 | 0 | 0 | 0 | 0 | 0 | 0 | 0 | 0 | 0 | 0 | 2 | 0 |
| HG1186 | *H. sapiens* | ENSG00000204923 | 0 | 0 | 0 | 0 | 0 | 0 | 0 | 0 | 0 | 0 | 0 | 0 | 0 | 0 | 0 | 0 | 0 | 2 | 0 |
| HG1187 | *H. sapiens* | ENSG00000213762 | 0 | 0 | 0 | 0 | 0 | 0 | 0 | 0 | 0 | 0 | 0 | 0 | 0 | 0 | 0 | 0 | 0 | 2 | 0 |
| HG1188 | *H. sapiens* | ENSG00000214415 | 0 | 0 | 0 | 0 | 0 | 0 | 0 | 0 | 0 | 0 | 0 | 0 | 0 | 0 | 0 | 0 | 0 | 2 | 0 |
| HG1189 | *H. sapiens* | ENSG00000279909 | 0 | 0 | 0 | 0 | 0 | 0 | 0 | 0 | 0 | 0 | 0 | 0 | 0 | 0 | 0 | 0 | 0 | 1 | 1 |
| HG1190 | *H. sapiens* | ENSG00000123684 | 0 | 0 | 0 | 0 | 0 | 0 | 0 | 0 | 0 | 0 | 0 | 0 | 0 | 0 | 0 | 0 | 0 | 0 | 2 |
| HG1191 | *H. sapiens* | ENSG00000164331 | 0 | 0 | 0 | 0 | 0 | 0 | 0 | 0 | 0 | 0 | 0 | 0 | 0 | 0 | 0 | 0 | 0 | 0 | 2 |
| HG1192 | *H. sapiens* | ENSG00000168758 | 0 | 0 | 0 | 0 | 0 | 0 | 0 | 0 | 0 | 0 | 0 | 0 | 0 | 0 | 0 | 0 | 0 | 0 | 2 |
| HG1193 | *H. sapiens* | ENSG00000177706 | 0 | 0 | 0 | 0 | 0 | 0 | 0 | 0 | 0 | 0 | 0 | 0 | 0 | 0 | 0 | 0 | 0 | 0 | 2 |
| HG1194 | *H. sapiens* | ENSG00000214694 | 0 | 0 | 0 | 0 | 0 | 0 | 0 | 0 | 0 | 0 | 0 | 0 | 0 | 0 | 0 | 0 | 0 | 0 | 2 |

** Numbers of orders from which uORF-tBLASTn and mORF-tBLASTn hits were extracted are indicated.

* Orders included in lower taxonomic categories were excluded.

***The category 'Ctenophora' was omitted from animal taxonomic categories because no sequences were classified to this category.
