## Supplementary Table S3 for "Exhaustive identification of conserved upstream open reading frames with potential translational regulatory functions from animal genomes"

Supplementary Table S3. Numbers of bases and sequences of EST/TSA and RefSeq and their assembly.

| | Species | Sequecnes | Bases |
|---|---|---|---|
| Plant | | | |
| Original EST/TSA | 1,134 | 72,369,635 | 53,793,063,962 |
| Assembled EST/TSA | 1,134 | 44,000,517 | 22,309,227,524 |
| RefSeq RNA | 74 | 3,218,211 | 5,490,948,218 |
| Assembled EST/TSA/RefSeq | 1,143 | 47,218,728 | 27,800,175,742 |
| Animal | | | |
| Original EST/TSA | 1,765 | 174,758,651 | 142,647,705,501 |
| Assembled EST/TSA | 1,765 | 94,795,989 | 49,197,476,441 |
| RefSeq RNA | 332 | 10,679,878 | 28,758,793,440 |
| Assembled EST/TSA/RefSeq | 1,920 | 105,475,867 | 77,956,269,881 |
| Eukaryota other than animals and plants | | | |
| Original EST/TSA | 619 | 14,932,588 | 11,598,150,341 |
| RefSeq RNA | 303 | 3,047,192 | 4,604,373,881 |
| Organism other than Eukaryota | | | |
| Original EST/TSA | 37 | 21,275 | 19,198,153 |
| RefSeq RNA | 12,812 | 21,415 | 31,025,423 |
