## Supplementary Table S4 for "Exhaustive identification of conserved upstream open reading frames with potential translational regulatory functions from animal genomes"

Supplementary Table S4. Estimation of the number of potential “spurious” CPuORFs for each species.

| Species | (1): all uORF* | (2): uORFs that overlap with mORFs** | (3): (1)-(2) | (4): (2) & uORFs with FR ≥ 0.3 |
|---|---|---|---|---|
| *D. melanogaster* | 17,035 | 6,320 | 10,715 | 2,493 |
| *D. rerio* | 39,616 | 6,037 | 33,579 | 4,086 |
| *G. gallus* | 8,929 | 3,246 | 5,683 | 1,511 |
| *H. sapiens* | 44,085 | 23,380 | 20,705 | 3,814 |

* When multiple uORFs in a transcript shared the same stop or start codon, they were counted as one.

** uORFs that overlapped with an mORF between splice variants according to genomic infomation of the original species.

The fusion ratio (FR) was calculated as shown in Supplementary Fig. S3b
