## Supplementary Table S5 for "Exhaustive identification of conserved upstream open reading frames with potential translational regulatory functions from animal genomes"

Supplementary Table S5. Information regarding the constructs.

| Gene symbol | Gene ID | Refseq ID | Construct |
|---|---|---|---|
| *PTP4A1* | ENSG00000112245 | NM_003463.4 | *pSV40:5'UTR (PTP4A1) ::luc2* |
| *MKKS* | ENSG00000125863 | NM_170784.2 | *pSV40:5'UTR (MKKS) ::luc2* |
| *SLC6A8* | ENSG00000130821 | NM_005629.3 | *pSV40:5'UTR (SLC6A8) ::luc2* |
| *FAM13B* | ENSG00000031003 | NM_016603.3 | *pSV40:5'UTR (FAM13B) ::luc2* |
| *MIEF1* | ENSG00000100335 | NM_019008.5 | *pSV40:5'UTR (MIEF1) ::luc2* |
| *EIF5* | ENSG00000100664 | NM_001969.4 | *pSV40:5'UTR (EIF5) ::luc2* |
| *MAPK6* | ENSG00000069956 | NM_002748.3 | *pSV40:5'UTR (MAPK6) ::luc2* |
| *MEIS2* | ENSG00000134138 | NM_170674.4 | *pSV40:5'UTR (MEIS2) ::luc2* |
| *KAT6A* | ENSG00000083168 | NM_006766.5 | *pSV40:5'UTR (KAT6A) ::luc2* |
| *SLC35A4* | ENSG00000176087 | NM_080670.3 | *pSV40:5'UTR (SLC35A4) ::luc2* |
| *LRRC8B* | ENSG00000197147 | NM_015350.2 | *pSV40:5'UTR (LRRC8B) ::luc2* |
| *CDH11* | ENSG00000140937 | NM_001797.3 | *pSV40:5'UTR (CDH11) ::luc2* |
| *PNRC2* | ENSG00000189266 | NM_017761.3 | *pSV40:5'UTR (PNRC2) ::luc2* |
| *BACH2* | ENSG00000112182 | NM_021813.3 | *pSV40:5'UTR (BACH2) ::luc2* |
| *FGF9* | ENSG00000102678 | NM_002010.2 | *pSV40:5'UTR (FGF9) ::luc2* |
| *PNISR* | ENSG00000132424 | NM_001322405.2 | *pSV40:5'UTR (PNISR) ::luc2* |
| *TMEM184C* | ENSG00000164168 | NM_018241.3 | *pSV40:5'UTR (TMEM184C) ::luc2* |
